# Genomic regions insertion and deletion in Monkeypox virus causing multi-country outbreak-2022

**DOI:** 10.1101/2022.06.28.497936

**Authors:** Perumal Arumugam Desingu, K. Nagarajan

**Author notes:** Corresponding authors, Perumal Arumugam Desingu, MVSc, Ph.D., DST-INSPIRE Faculty, Division of Biological Science, Department of Microbiology and Cell Biology, Indian Institute of Science (IISc), Bangaluru – 560012, India. **Email addresses:** Perumal Arumugam Desingu;, K. Nagarajan.

## Abstract

The genetic diversity and evolutionary origin of the Monkeypox virus (MPXV) that is currently creating a multi-country outbreak-2022 is not fully understood. Here we report that the MPXVs that cause outbreak-2022 (MPXVs-2022) have deletion/insertion of ∼500 to 2000bp nucleotide in multiple genomic regions. Our analyses revealed that MPXVs-2022 are very close to the West African Clade of MPXVs (WA-MPXVs) that caused the Outbreak in Nigeria in 2017-2018. Furthermore, we classified the WA-MPXVs detected before 2017 that could not be transmitted from human-to-human as WA-MPXVs-I and WA-MPXVs detected after 2017 that could be transmitted from human-to-human as WA-MPXVs-II (including MPXVs-2022), and human-to-human transmissible Central African MPXVs (CA-MPXV) remained as a separate clade. Overall our results suggest that although WA-MPXVs-II are almost identical to WA-MPXVs-I throughout the genome and two large genomic insertions (∼500, 2000bp size insertion), they differ from WA-MPXVs-I in 5’-inverted terminal repeat (5’-ITR) (deletion of the last-2000bp-5’-ITR) and 13 proteins that of CA-MPXVs, and the presence of seven unique proteins in WA-MPXVs-II is likely to be a significant cause of outbreak-2022. This study shed light on the genetic diversity and evolutionary origin of MPXVs causing outbreak-2022.

---

The zoonotic infection of human monkeypox is caused by the monkeypox virus (MPXV) of Orthopoxvirus, which is closely related to the variola virus (which causes smallpox), vaccinia virus, and cowpox virus (CPXV) (*1*). Historically, from 1958 to 1964, MPXVs were detected in monkeys in European countries such as Denmark (*2*) and the Netherlands (*3, 4*), and the USA (*5*). After that, MPXV was first detected in humans in the Congo in 1970 (*6*) and, until 2003, spread only in African countries (*7-9*) as two distinct clades, the central African (CA-MPXVs) and west African clades (WA-MPXVs) of MPXV (*9-12*). Of the two clades of MPXV viruses, CA-MPXVs viruses cause periodic outbreaks in central African countries between 1981 to 1986, 1996–97 (*13*), 2003 (*14*), 2005 to 2007 (*15, 16*), and July-December 2013 (*17*). Further, the CA-MPXVs viruses are highly transmitted from human-to-human, causing severe illness in children, resulting in increased periodic outbreaks (*10-12, 18*), while WA-MPXVs produced short outbreaks in Cote d’Ivoire, Liberia, Nigeria, and Sierra Leone between 1970 and 1981 due to a lack of human-to-human transmission (*7, 19*). However, it is essential to note that CA-MPXV viruses, which cause transmission from human-to-human and cause serious illness, have not yet been identified in countries other than Africa. On the other hand, WA-MPXV viruses, which are not transmitted from human-to-human, were transmitted to people in close contact with infected prairie dogs through rodents imported from Ghana in the USA in 2003 (*7-9*). Moreover, the WA-MPXV virus, detected primarily in adults in the USA in 2003, was quickly controlled due to its lack of human-to-human transmission (*7-9*).

Remarkably, the Outbreak of 2017-18 in west Africa created a turning point in the human Outbreak of WA-MPXV viruses (*7, 9*). Because during the Outbreak of WA-MPXV viruses in Nigeria in 2017-18, the virus became human-to-human transmission, primarily affecting young adults (*7, 9, 20*) and spread to countries such as the United Kingdom (*21*), Israel (*9*), and Singapore (*22*). Furthermore, it is essential to note that the MPXVs-2022 viruses are related to the human-to-human transmission of WA-MPXV viruses that caused the Nigeria-2017-18 outbreak (*23*). In this case, it is not entirely clear how the WA-MPXV virus acquired human-to-human transmission status. The WA-MPXV viruses must be genetically altered in order to be transmitted from human-to-human. As the MPXVs-2022 virus spread to multiple countries and rocking the world healthcare system, in this situation, it is urgently needed to understand the genetic alteration that happened in the MPXVs-2022. It is essential to identify genetic alterations in MPXVs-2022 viruses in order to design future antivirals, immunotherapy, and vaccines and predict future outbreaks by conducting future studies tailored to those genetic alterations. In this concern, we have systematically analyzed this study’s genetic alterations in MPXVs-2022 viruses. Furthermore, this study highlights the deletion and insertion specific to genomic regions in MPXVs-2022 viruses and unique proteins.

In this study, we first retrieved the complete genome sequences of MPXVs detected in the USA this year (2022) from the GISAID database and explored the genetic diversity through the SimPlot analysis. This analysis revealed that more than 99.4% of the nucleotide sequence identities throughout the complete genome sequence were found among MPXVs detected in the USA this year (2022) (**Supplementary Figure 1A**). From this, it is clear that the MPXVs circulating in the USA this year are identical and of a single origin. Following this, we retrieved the complete genome sequences of viruses associated with MPXVs-2022 from the NCBI public database and subjected them to phylogenetic analysis to detect genetic relationships between the MPXVs-2022 and other MPXVs previously reported and other poxviruses. In this analysis, we noted that all MPXVs-2022 detected this year in many parts of the world developed monophylogeny (**Figure 1A; Supplementary Figure 1B**). More interestingly, MPXVs-2022 grouped closely with the viruses that caused the Outbreak in Nigeria in 2017-2018 and spread from Nigeria to countries such as the UK, Israel, and Singapore in 2018 (**Figure 1A; Supplementary Figure 1B**). Furthermore, as in previous reports (*9-12*), in our phylogenetic analysis, the MPXVs split into two groups called WA-MPXVs and CA-MPXVs (**Figure 1A-1B; Supplementary Figure 1B**). Interestingly, we noted two distinct sub-groups within WA-MPXVs (**Figure 1A-1B; Supplementary Figure 1B**). Except for the Nigeria-SE-1971 and W-Nigeria (1978) viruses, other WA-MPXVs detected before 2017 formed the WA-MPXVs-Clade-I (**Figure 1A-1B; Supplementary Figure 1B**). On the other hand, Nigeria outbreak-2017-2018 viruses and MPXVs-2022 viruses combined with Nigeria-SE-1971 and W-Nigeria (1978) viruses to form WA-MPXVs-Clade-II (**Figure 1A-1B; Supplementary Figure 1B**).

**Figure 1.**
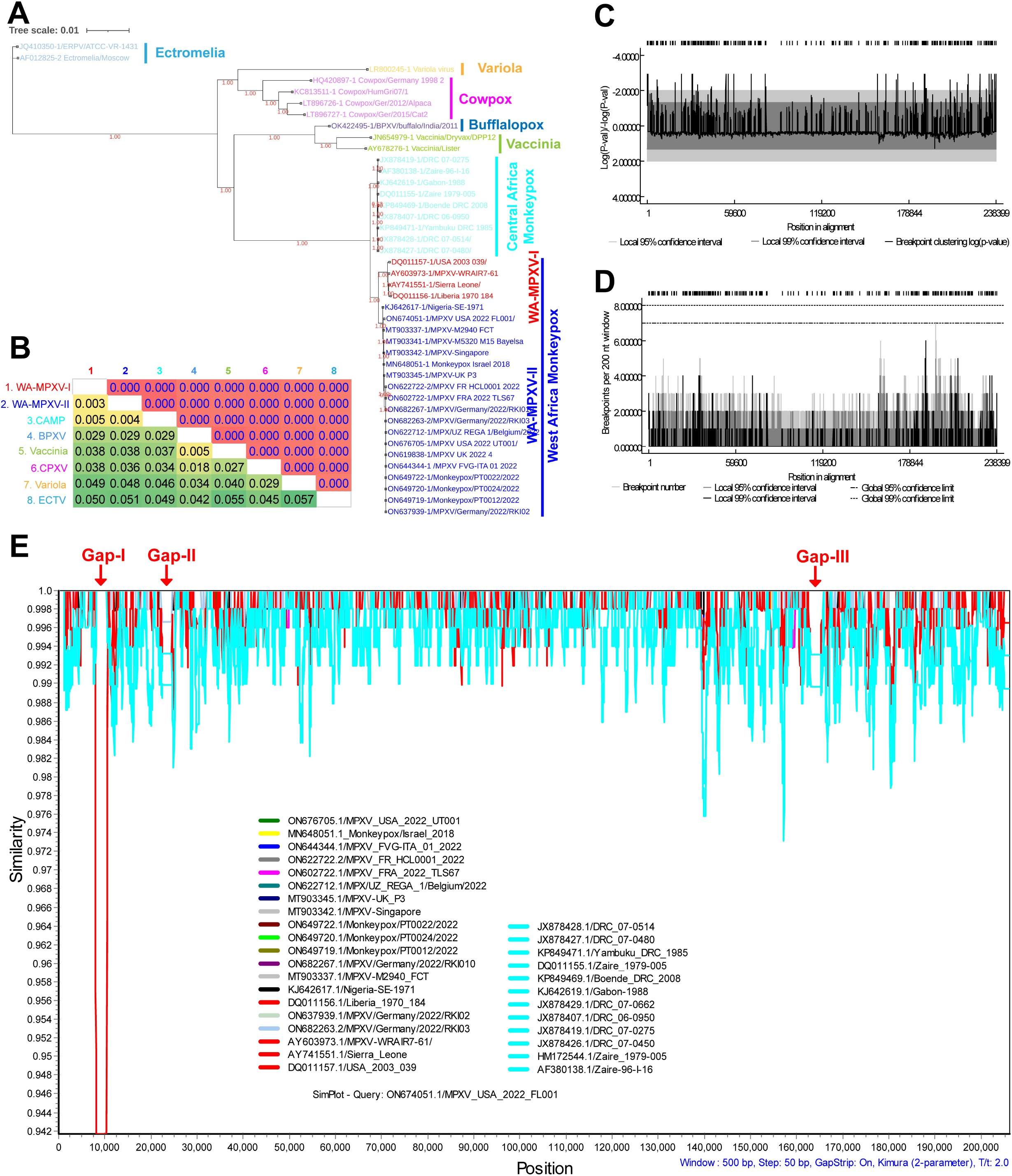
Complete genome-level genetic diversity in the monkeypox viruses. (**A-B**) WA-MPXVs are subdivided into distinct WA-MPXVs-I and WA-MPXVs-II in the complete genome sequence size phylogenetic tree (PhyML 3.3_1) (**A**); and Net Between-Group Mean Distance (NBGMD) (**B**). (**C**) The breakpoint p-value distribution plot and (**D**) breakpoint distribution plot analysis revealed the presence of recombination throughout the genome of poxviruses. (**E**) SimPlot analysis revealed the presence of three distinct genomic gaps between the MPXVs.

Curiously, WA-MPXVs (WA-MPXVs-Clade-I) detected before Nigeria outbreak-2017-2018 are not transmitted from human-to-human (*7, 9*), and WA-MPXV-Clade-II viruses are transmitted from human-to-human, with a genetic diversity within these two clades likely to cause outbreak-2022. Furthermore, CA-MPXVs viruses are transmitted from human-to-human, so it is likely that WA-MPXV-Clade-II viruses differed from WA-MPXVs-Clade-I and acquired genomic regions associated with CA-MPXVs viruses. Since genetic recombination is high among poxviruses (*24-26*), it can be speculated that this genetic diversity may have also been caused by recombination. To this end, we first analyze the breakpoint p-value distribution plot and breakpoint distribution plot using different poxvirus complete genome sequences to determine in which genomic regions the genetic recombination between poxviruses is most likely to occur. This analysis reveals that there are recombination events almost throughout the genome between poxviruses (**Figure 1C-1D**). Next, we performed a SimPlot analysis to determine which genomic areas have the most genetic diversity between the CA-MPXVs, WA-MPXVs-I, and WA-MPXVs-II viruses. This analysis revealed three distinct sequence alignment gaps between CA-MPXVs, WA-MPXVs-I, and WA-MPXVs-II viruses (**Figure 1E**). These three gaps’ first sequence alignment gap was found to be in the 8099bp-10445bp (7047-9357bp as per DQ011156.1/Liberia-1970-184) region, and the WA-MPXVs-I viruses in this region were found to be different from the WA-MPXVs-II and CA-MPXVs viruses (**Figure 1E**). In particular, region 7047-9357bp of the Liberia-1970-184 (WA-MPXVs-I) virus represents the final part of the 5’-ITR. WA-MPXVs-I (DQ011156.1/Liberia-1970-184) viruses contain 1-9357bp 5’-ITR and 190900-200256bp (size 9357bp) 3’-ITR. It should be noted, however, that the WA-MPXVs-II and CA-MPXVs viruses contain ∼6400bp of 5’-ITR and 3’-ITR (WA-MPXVs-II-ON676705.1/MPXV/USA/2022/UT001, 1-6420bp 5’-ITR, and 190745-197154bp 3’-ITR; and CA-MPXVs-DQ011155.1/Zaire/1979-005, 1-6389bp 5’-ITR, and 190579-196967 3’-ITR). Of these, it seems that 5’-ITR and 3’-ITR in WA-MPXVs-II and CA-MPXVs viruses are 3000bp shorter than in WA-MPXVs-I viruses. However, the SimPlot analysis revealed a gap of 8099bp-10445bp with WA-MPXVs-I viruses, corresponding to the narrowing of WA-MPXVs-II and CA-MPXVs viruses in the 5’-ITR region; it is important to note, however, that such a gap is not found in the 3’-ITR region (**Figure 1E**). Of these, it seems that only the 5’-ITR fraction of WA-MPXVs-II and CA-MPXVs viruses is ∼3000bp shorter than that of WA-MPXVs-I viruses.

After that, to ensure that the WA-MPXVs-II and CA-MPXVs viruses are shorter in size than the WA-MPXVs-I viruses only in the 5’-ITR region; we first analyzed the sequences of MPXVs aligning with 5’-ITR (1-9357bp) in WA-MPXVs-I (DQ011156.1/Liberia-1970-184), subject to MAFFT alignment, phylogenetic and NBGMD analysis. The MAFFT alignment revealed gaps in the WA-MPXVs-II and CA-MPXVs viruses at the end of the 5’-ITR compared to WA-MPXVs-I (**Supplementary Data 1**). Next, in the phylogenetic analysis, the WA-MPXVs-I, WA-MPXVs-II, and CA-MPXVs viruses were subdivided into three distinct groups (**Figure 2A**; **Supplementary Data 1**). Interestingly, in the NBGMD analysis, the WA-MPXVs-II viruses revealed the closest genetic identity to the CA-MPXVs viruses (**Figure 2B; Supplementary Data 1**). Furthermore, the sequences of MPXVs align with the additional 5’-ITR (7047-9357bp, gap containing region) region in WA-MPXVs-I (Liberia-1970-184) viruses were subjected to phylogenetic and NBGMD analysis (**Supplementary Data 2**). In this phylogenetic analysis, the WA-MPXVs-II and CA-MPXVs viruses were grouped, and the WA-MPXVs-I viruses as a separate group (**Figure 2C**). In support of this, the NBGMD analysis revealed 0.3% and 81.8% genetic diversity of WA-MPXVs-II viruses with CA-MPXVs and WA-MPXVs-I viruses, respectively (**Figure 2D**). Collectively, the 5’-ITR fraction of WA-MPXVs-II and CA-MPXVs viruses is shorter than that of WA-MPXVs-I viruses, or the WA-MPXVs-II and CA-MPXVs viruses have lost an additional 5’-ITR (7047-9357bp) of WA-MPXVs-I viruses (Liberia-1970-184). The question now is whether the corresponding 3’-ITR sequences of the 5’-ITR (7047-9357bp) sequences are, in addition to the WA-MPXVs-I (Liberia-1970-184) viruses, are present only in the WA-MPXVs-I viruses. To detect this, MAFFT alignment, phylogenetic and NBGMD analysis was performed using sequences of MPXVs that are aligned with 3’-ITR (190900-200256bp) in WA-MPXVs-I (DQ011156.1/Liberia-1970-184) viruses. Surprisingly, gaps in the WA-MPXVs-II and CA-MPXVs viruses in the 5’-ITR of the MAFFT alignment were not found in the corresponding 3’-ITR region (**Supplementary Data 3**). Furthermore, it is noteworthy that WA-MPXVs-II viruses in the 3’-ITR region (full 9357bp size) expressed a closer relationship with WA-MPXVs-I viruses than CA-MPXVs viruses (**Figure 2E-2F**). Overall, the 9357bp length 3’-ITR regions are present in WA-MPXVs-II and CA-MPXVs viruses as they are in WA-MPXVs-I viruses, and the WA-MPXVs-II and CA-MPXVs viruses have lost (∼2000bp) the final portion of the 5’-ITR (has only first ∼6400bp length).

**Figure 2:**
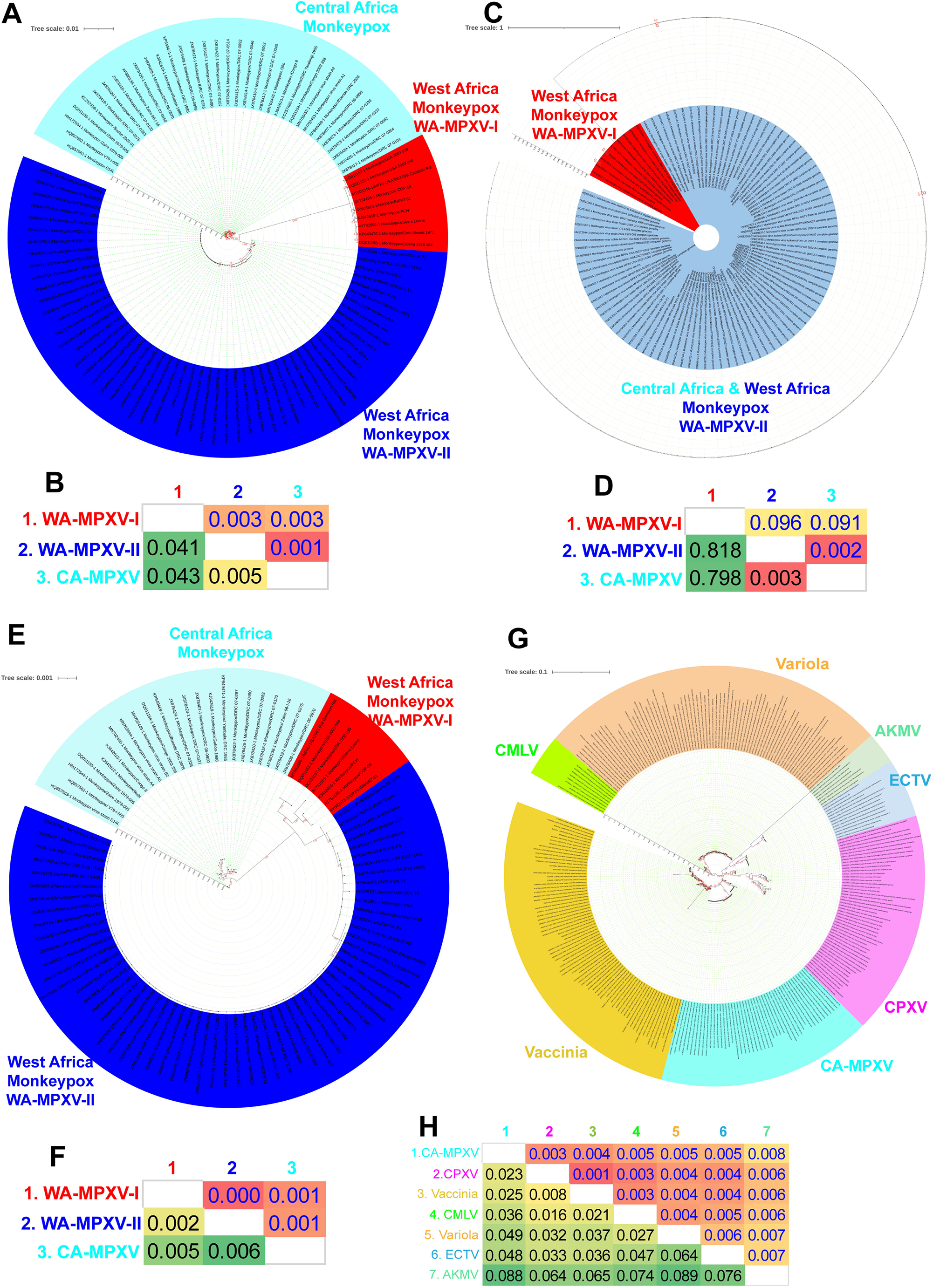
Genomic region-specific deletions in the WA-MPXVs-II. **(A)** Phylogenetic tree and (**B**) NBGMD analysis for MPXVs viruses region related with 5’-ITR (1-9357bp) in WA-MPXVs-I (DQ011156.1/Liberia-1970-184) viruses. WA-MPXVs-II viruses show a closer genetic identity to CA-MPXVs viruses than WA-MPXVs-I viruses. **(C)** Phylogenetic tree and (**D**) NBGMD analysis for MPXVs viruses region related with the additional 5’-ITR (7047-9357bp) region in WA-MPXVs-I (Liberia-1970-184) viruses. The WA-MPXVs-II and CA-MPXVs viruses in this region are almost identical, but the WA-MPXVs-I viruses exhibit enormous genetic diversity. **(E)** Phylogenetic tree and (**F**) NBGMD analysis for MPXVs viruses region related with 3’-ITR (190900-200256bp) in WA-MPXVs-I (DQ011156.1 /Liberia-1970-184) viruses. WA-MPXVs-II viruses in the 3’-ITR region (full 9357bp size) expressed a closer relationship with WA-MPXVs-I viruses than CA-MPXVs viruses. **(G)** Phylogenetic tree and (**H**) NBGMD analysis for poxviruses region related with genomic gap 18875-20830bp (KC257460.1/DRC/Yandongi/1985) CA-MPXVs viruses. The CA-MPXVs viruses in this area express close genetic identity with vaccinia/CPXV viruses. It should be noted, however, that this genomic component is nowhere to be found in the complete genome of WA-MPXVs-I & II viruses.

Next, we focus on the second genomic gap in the SimPlot analysis (**Figure 1E**). The second sequence alignment gap was found to be in the 22550-24503bp (18875-20830bp as per KC257460.1/DRC Yandongi 1985) region (**Figure 1E**), and among MPXVs viruses, these region-specific nucleotide sequences are found only in CA-MPXVs viruses and not in WA-MPXVs-I & II (**Supplementary Data 4**). Next, we are interested in finding out where from CA-MPXVs viruses got this particular genetic component. To this end, phylogenetic and NBGMD analysis was performed by retrieving the aligned sequences that revealed the close relationship of this 18875-20830bp (KC257460.1/DRC/Yandongi/1985) nucleotide sequences (**Supplementary Data 5**) in the NCBI BLAST analysis. These analyzes show that the genomic region of 18875-20830bp (KC257460.1/DRC Yandongi 1985) in CA-MPXVs viruses is derived from vaccinia/CPXV viruses (**Figure 2G-2H**).

Further, we focus on the third genomic gap in the SimPlot analysis (**Figure 1E**). The third sequence alignment gap was found to be in the 163099-165378bp (156341-158604bp as per ON676705.1/MPXV/USA/2022/UT001) region (**Figure 1E**), and among MPXVs viruses, these region-specific nucleotide sequences are found only in WA-MPXVs-I & II viruses and not in CA-MPXVs (**Supplementary Data 6**). To find out where the WA-MPXVs-I & II viruses got this specific genomic region, the phylogenetic and NBGMD analysis was performed by retrieving the aligned sequences that revealed a close relationship with this 156341-158604bp (ON676705.1/MPXV/USA/2022/UT001) sequences (**Supplementary Data 7**) in the NCBI BLAST analysis. These analyzes show that the genomic region of 156341-158604bp (ON676705.1/MPXV/USA/2022/UT001) in WA-MPXVs-I & II viruses is derived from vaccinia/CPXV viruses (**Figure 3A-3B**). Also, the in-depth analysis revealed another genomic gap between MPXVs viruses 139987-140434bp (133523-133970bp as per ON676705.1/MPXV/USA/2022/UT001), and among MPXVs viruses, these region-specific nucleotide sequences are found only in WA-MPXVs-I & II viruses and not in CA-MPXVs. (**Supplementary Data 8**). Further, phylogenetic and NBGMD analysis for this genomic region revealed that WA-MPXVs-I & II viruses may have inherited this genomic region from the vaccinia virus (**Figure 3C-3D; Supplementary Data 9**).

**Figure 3:**
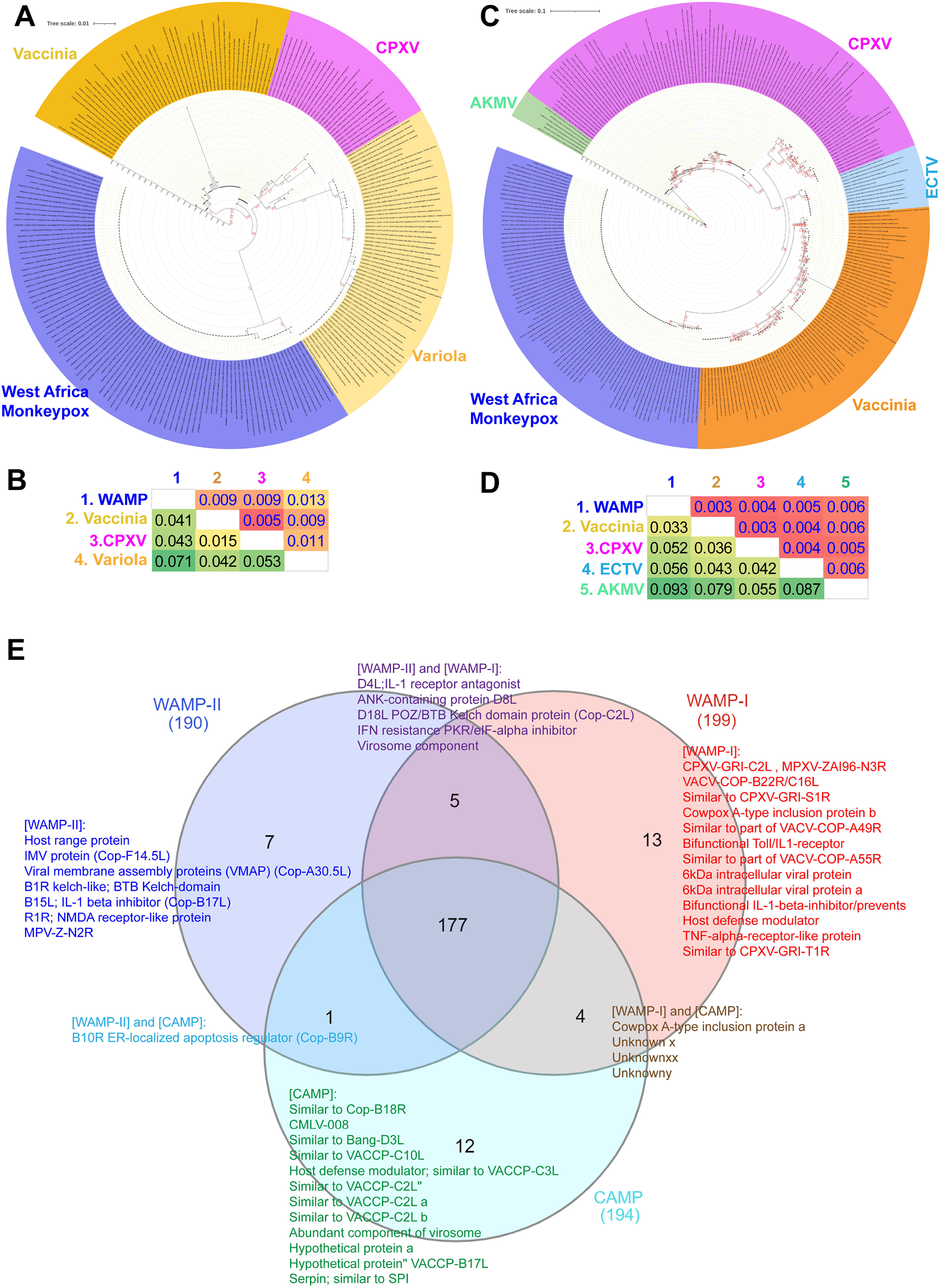
Genomic region-specific insertions in the monkeypox viruses. **(A)** Phylogenetic tree and (**B**) NBGMD analysis for poxviruses region related with genomic gap 156341-158604bp (ON676705.1/MPXV/USA/2022/UT001) WA-MPXVs. This region’s WA-MPXVs-I and WA-MPXVs-II viruses are almost identical and express close genetic identity with vaccinia/CPXV viruses. It should be noted, however, that this genomic component is nowhere to be found in the complete genome of CA-MPXVs viruses. **(C)** Phylogenetic tree and (**D**) NBGMD analysis for poxviruses region related with genomic gap 133523-133970bp (ON676705.1/MPXV/USA/2022/UT001) WA-MPXVs. This region’s WA-MPXVs-I and WA-MPXVs-II viruses are almost identical and express close genetic identity with vaccinia viruses. It should be noted, however, that this genomic component is nowhere to be found in the complete genome of CA-MPXVs viruses. (**E**) The Venn diagram depicts the similarities and differences in proteins between WA-MPXVs-I, WA-MPXVs-II, and CA-MPXVs viruses. The details of each protein are given in the **Supplementary Data 1**, and the details of the unique proteins in the WA-MPXVs-I, WA-MPXVs-II and CA-MPXVs viruses are given in the **Supplementary Data 2**

Finally, since there are differences in genomic regions between WA-MPXVs-I, WA-MPXVs-II, and CA-MPXVs viruses, we examined whether there are any differences in the protein in these viruses. This analysis revealed the presence of 13, 7, and 12 unique proteins in the WA-MPXVs-I, WA-MPXVs-II, and CA-MPXVs viruses, respectively. The WA-MPXVs-II and CA-MPXVs viruses may also have been transmitted from human-to-human because of the loss of the last part of the 5’-ITR and 13 proteins from the WA-MPXVs-I viruses; however, it warrants future studies in this direction. Furthermore, it is speculated that these viruses may infect the adult population with specific two genomic insertions and five proteins typically present between WA-MPXVs-I and WA-MPXVs-II viruses and absent in CA-MPXVs viruses, though it needs future studies in this direction. Finally, this study sheds light on the genomic region-specific deletion and insertion in MPXVs viruses and explains the genetic diversity in MPXVs-2022 viruses causing outbreak-2022. This novel clue is expected to support future research on vaccines, immunotherapy, antiviral therapy design, and the prediction of future outbreaks.

## Author’s Contributions

All the authors contributed significantly to this manuscript. PAD analyzed and wrote the first draft, and KN reviewed the manuscript. All the authors reviewed and approved the final submission.

## Data Availability Statement

Not applicable.

## References

1. H. Adler et al., Clinical features and management of human monkeypox: a retrospective observational study in the UK. The Lancet. Infectious diseases, (May 24, 2022).

2. A. E. Von Magnus P, Petersen KB, Birch-Andersen A., A pox-like disease in cynomolgus monkeys. Acta Path. Micro. Scand. 46(2):156–176, (1956).

3. R. Gispen, J. D. Verlinde, P. Zwart, Histopathological and virological studies on monkeypox. Arch Gesamte Virusforsch 21, 205 (1967).

4. S. Parker, R. M. Buller, A review of experimental and natural infections of animals with monkeypox virus between 1958 and 2012. Future Virol 8, 129 (Feb 1, 2013).

5. H. Y. Mcconnell SJ, Mattson DE, Erickson L., Monkey pox disease in irradiated cynomologous monkeys.. Nature 195:1128–1129., (1962).

6. I. D. Ladnyj, P. Ziegler, E. Kima, A human infection caused by monkeypox virus in Basankusu Territory, Democratic Republic of the Congo. Bull World Health Organ 46, 593 (1972).

7. G. Rezza, Emergence of human monkeypox in west Africa. Lancet Infect Dis 19, 797 (Aug, 2019).

8. A. M. Likos et al., A tale of two clades: monkeypox viruses. J Gen Virol 86, 2661 (Oct, 2005).

9. A. Yinka-Ogunleye et al., Outbreak of human monkeypox in Nigeria in 2017-18: a clinical and epidemiological report. Lancet Infect Dis 19, 872 (Aug, 2019).

10. K. Brown, P. A. Leggat, Human Monkeypox: Current State of Knowledge and Implications for the Future. Trop Med Infect Dis 1, (Dec 20, 2016).

11. M. G. Reynolds et al., Spectrum of infection and risk factors for human monkeypox, United States, 2003. Emerg Infect Dis 13, 1332 (Sep, 2007).

12. Y. J. Hutin et al., Outbreak of human monkeypox, Democratic Republic of Congo, 1996 to 1997. Emerg Infect Dis 7, 434 (May-Jun, 2001).

13. D. L. Heymann, M. Szczeniowski, K. Esteves, Re-emergence of monkeypox in Africa: a review of the past six years. Br Med Bull 54, 693 (1998).

14. L. A. Learned et al., Extended interhuman transmission of monkeypox in a hospital community in the Republic of the Congo, 2003. Am J Trop Med Hyg 73, 428 (Aug, 2005).

15. A. W. Rimoin et al., Major increase in human monkeypox incidence 30 years after smallpox vaccination campaigns cease in the Democratic Republic of Congo. Proc Natl Acad Sci U S A 107, 16262 (Sep 14, 2010).

16. P. Formenty et al., Human monkeypox outbreak caused by novel virus belonging to Congo Basin clade, Sudan, 2005. Emerg Infect Dis 16, 1539 (Oct, 2010).

17. L. D. Nolen et al., Extended Human-to-Human Transmission during a Monkeypox Outbreak in the Democratic Republic of the Congo. Emerg Infect Dis 22, 1014 (Jun, 2016).

18. J. G. Breman et al., Human monkeypox, 1970-79. Bulletin of the World Health Organization 58, 165 (1980).

19. M. G. Reynolds, I. K. Damon, Outbreaks of human monkeypox after cessation of smallpox vaccination. Trends Microbiol 20, 80 (Feb, 2012).

20. Z. Jezek et al., Human monkeypox: a study of 2,510 contacts of 214 patients. J Infect Dis 154, 551 (Oct, 1986).

21. M. R. Mauldin et al., Exportation of Monkeypox Virus From the African Continent. J Infect Dis 225, 1367 (Apr 19, 2022).

22. O. T. Ng et al., A case of imported Monkeypox in Singapore. Lancet Infect Dis 19, 1166 (Nov, 2019).

23. https://www.who.int/emergencies/disease-outbreak-news/item/2022-DON385.

24. L. Qin, D. H. Evans, Genome scale patterns of recombination between coinfecting vaccinia viruses. Journal of virology 88, 5277 (May, 2014).

25. X. D. Yao, D. H. Evans, Effects of DNA structure and homology length on vaccinia virus recombination. Journal of virology 75, 6923 (Aug, 2001).

26. P. D. Gershon, R. P. Kitching, J. M. Hammond, D. N. Black, Poxvirus genetic recombination during natural virus transmission. The Journal of general virology 70 (Pt 2), 485 (Feb, 1989).

