## Supplementary Data 1 for "Genomic regions insertion and deletion in Monkeypox virus causing multi-country outbreak-2022"

>ON674051.1_Monkeypox_virus_isolate_MPXV_USA_2022_FL001_complete_genome

--- --- --- --- --- --- --- --- --- --- --- --- --- --- --- --- --- --- --- --- --- --- --- --- --- --- --- --t att ata ttt tac tat ttt att tag tgt cta gaa aaa aat gtg tga ccc acg acc gta gga aac tct aga ggg taa gaa aaa tca atc gtt tat aga gac cat cag aaa gag gtt taa tat ttt tgt gag acc tat cga a-- --- --- --- --- --- --- --- --- --- --- --- --- --- --- --- --- --- --- --- --- --- --- --- --- --- --- --- --- --- --- --- --- --- --- --- --- --- --- --- --- --- --- --- --- --- --- --- --- --- --- --- --- --- --- --- --- --- --- --- --- --- --- --- --- --- --- --- --- --- --- --- --- --- --- --- --- --- --- --- --- --- --- --- --- --- --- --- --- --- --- --- --- --- --- --- --- --- --- --- --- --- --- --- --- --- --- --- --- --- --- --- --- --- --- --- --- --- --- --- --- --- --- --- --- --- --- --- --- --- --- --- --- --- --- --- --- --- --- --- --- --- --- --- --- --- --- --- --- --- --- --- --- --- --- --- --- --- --- --- --- --- --- --- --- --- --- --- --- --- --- --- --- --- --- --- --- --- --- --- --- --- --- --- --- --- --- --- --- --- --- --- --- --- --- --- --- --- --- --- --- --- --- --- --- --- --- --- --- --- --- --- --- --- --- --- --- --- --- --- --- --- --- --- --- --- --- --- --- --- --- --- --- --- --- --- --- --- --- --- --- --- --- --- --- --- --- --- --- --- --- --- --- --- --- --- --- --- --- --- --- --- --- --- --- --- --- --- --- --- --- --- --- --- --- --- --- --- --- --- -ga gag aaa gga taa aaa ctt ttt acg act cca tca gaa aga ggt tta ata ttt ttg tga gac cca tcg aag aga gaa aga gat ggt tag tca aga tat ttt tct tag tac aaa agt caa tgt ttt aaa ata tat gga cga gaa tta att tgt ctg tat aaa aac ttg tgt gaa att atg tac tag aga aaa aac gtg agc agt gtc ccc tac atg gat ttt aca gat cat tta tat tcc aaa aat att aac tat ata cgt tta tta tat gat gtt aac gtg taa att ata aac att att tta tga tgc aat tgt ctg aca acc tag att ggt ata agg atg ttg ata agc tct acg aga ata tat tgt tgg acg tta tcg ttt acg aaa tag ttg aga cat cag aaa gag gtt taa tat ttt tgt gag a-c cat cga aga gag aaa gag aat aaa aat a-- -?t ttt ttt ttt ttt tgt aaa act ttt tta tga gac --- --- -ca aga gaa tac gaa tag tga tca tat cgt atc aca tat tga aac aga aag aag aag taa cga gag gta act ttt tgt gaa tgt agt taa ata ttt ttg ttt tgc aaa ccg gaa tat agt gcc cgg t-- -ct ttt tta att cgt ggt gcg gtg tct gaa tcg ttc gat taa ccc aac tca tcc att ttc aga tga ata gag tta tcg att cag aca cat gct ttg agt ttt gtt gaa tcg atg agt gaa gta tca tcg gtt gca cct tca gat gcc gat ccg tcg aca tac ttg aat cca tcc ttg act tca agt tca gat gat tcc tca cac atg tct ccg ata cgt acg cta aac tct agg ttc ttg aca cat ttt gta tca acg atc gtt gaa ccg atg ata tct ttg taa ctc act ttc tta tgt gag atg tta gac cca agt act gga tgg gtc ttg atg tca ctg tct ttc tct tct tcg cta cat ctg atg tcg ata gac atc tca cag tct ttg atc ata gcc aga gct tct tca cgc gtg atc gcg gga gag tcc tta cct tgt ccc ggt gac acg ctg gac aat cta gta ttc aca gtg ttt cca tca gag gat tcg gag atg gat gaa atc ttt ggg cat ttg gtg aat cca aag ttc atg tta aga ccc gca ccg acg ata gtg taa taa gtg gtg gga tct cct ttt aca act --- --- --- --- --- --- tct tcg gat acc tca tca tct tcg gtc tct gta act tcc gtt acg gat tga caa atc tta tca ttg gtc ggt gtt tgg tct tgc ttt gtg act ttg ata ata aca tcg att ccc ata tga tgt ttg ttt tct tct tca gta cac gag gat gaa gat tgt tga aga cta gta ggc ata gca gct gcc act agg cac atg cat gcc agg aca ata tat tgt ttc atg att gct att gat tga tta ctg ttc tag atg att cta ctt tct tac cat ata ata aat tag aat ata ttt tct act ttt acg aga aat taa tta ttg tat tta tta ttt ata ggt --a aaa aaa ctt act ata agt ggg tgg gat tct ggg aat tag tga tca gtt tat gta tat cgc aac tag cgg gca tat ggc tat tga cat cga gaa cat tac cca tat gat aag aga ttg tat cat ttt cgt agt ctt gag tat tgg tat tac tat ata gta tgt aga tgt cga cgc tag ata gac agt cgc cca cta gag tta ccg tct ctg aat gcg gca tga tag tat cat tct ttg ttt tcg tta act gtt tgg aag atg aat ctt tgt tgt tac att taa tct cga aat tca gag tac ata tct ttg aag tat tct gat atc tat ttt ctc ctg taa aga atc ctg aag ttg cta cat tat taa gga cag aga agt att ctg cac gaa aga ctg gat cac aat ctt tat gat tca tgg taa tag tta gtt ccg acg ttg aga tgg att cgc tga gac cgg tag tgg tcg tcc gag tac acg atg tgt cgt tga ctg gat aca ggt taa ttt cca cat cga tat agt taa atg tat tgc tgg tta cga cgg gtt cgc att tat ctg tgg aag aga cgg tgt gag aat atg ttc cgg gac cac acg gag aac aga tga cgt ctc cgg tag acg tgt atc cgg ata ctc cgt atc cta ttc cac act ttg ttt tag aaa tac atg ttc tac acc ctg atg ctc ctt tga gaa gac aat aat atc ctg gag agc att cac aga ttc tat tgt gag tcg tgt tac acg atc gcg tct cta cct gat tac tat cac atc ttc cgt tac aac tta gac aag cct gta aat gat tat tgt gag atg taa agg tat ccg aac cac acg gtg tac att gtg tat tag tct tgc tat cac ata atc tgg aag cgt aag ttc ccg gag gac acg ata gac aac ata gat tac ggc ttc tgt att cgt tgt ctt tac act ttc cat tgg atg gtg cat gtg gtg cta tat ctc ttc cgt tta tta tta tac atg aga gaa aca ata tat acg agt ata ata cgg acc tca tga ttt aat aat -gt agt aat cgt cgt ctt gtt act gtt tgt ttc cta ctt ctc caa tca tat aga tta ttt ttt aaa tat ttt ctt tct atc atg gat aat att tgt aat ggt tct ttc cgt aca aca tac tgt tta gat ggt agt cgc tta gct tgg tta tga tat tgc gca taa ttt ccg gag gca aat acg ata gtc tag att gac tat cga tgg tag act cta att tat tga gtg ctt tgt cga cga gtt tac ttt tat gct cca tcg ata gat gac act gtt cta tga gat cgt cgt aca tgg gaa atg aaa tgc gtt tgt ctg aat gta tgg ctt cga gat atg tgt gat acc gga tgt ctt ctg ttc tca ata ccg tat aca agt cgg tgt ctg aga ttc gaa tct ctt tga gga gac tta tgt cac gac tac att ttt cga tga tgg aat cta tct tat cga atg ata tat ttt tca taa ata cac ttt tat agt cct cgt tta aac aga att tag tat gta gtt ccg caa atg act cgt ccc tta ata ggc agt agg cta tta tct tct tta cgt agt gat cgt cgt agg gag aga act ccg aca tct tgt aga aca acg att taa tca tag gta gag ata ctt tca gtc tgt ggt gga taa tgt cat taa cga cat ccg cct tgt ata taa tgt ttc tgt ttt caa aca cca agt cga ata ctg tct tat cgt ctt tag tcg gaa ggt tga tgt cgt atc cga tgt ata cga ggt atg agg caa cat tgt tat tgc aat tct gga agg cgg tat gaa gag gag tca ttg tat tat agt att cgt ctt tct gaa tgt cga atc tat cta gta gat acc gta gta tat tga gag agc gac ttc cat atc ctt gat tat gtt tta tga ata gat aaa gta gat gtt gtc ctt ctt cct ttt gta att tcc cgt att ttt gtt cgt gcc aat tga gta aca tta tga gaa tat gac ctg ttg cac aat cgt tct tta tgt att cca tga tgg gtg tac aat caa gat tat tac gta tcc tcg tat cgg ctc ctc gag ata aaa gag cat aca cca cac gag gac tat gtt tgg tat act gtt gaa ggt aag tgt gta acg gcg tat ttc cga ttt tcg taa ccg cgt taa tgt ttg ctc cat gat cta tta tcg cgt aga tga atc gct tct cag ctc gca tct tag tgt gac tct ttg act tgt aat aat tgc ttt cgt gga acg cgg ata tgt gtt tac agt agt aat gaa gag aag tga gtc cat cct cat cga cgc aat tag ggt cag atc ctt tag tca ata att tgt aca gaa cgt aat agt tta agc tcc cat tga att tat atc taa gat aac aca gca ata gat cgg atg att tac taa agt cat caa tgg ggt ccg tta gta tat caa aga tct tgt tat cga ttg ata gtg aat gaa tca gat agt ggt gta gag gaa tat gtc ctt ttt tat cct tgc tat caa agt tac gca tgc cgt ggc gta aca ata tct tta ata cag atg gat taa atc gtg tat tca tcg tat agc aat gta atg gag agt tac cac att tta gtc gtt tat tca gat cgc agt gtt taa taa cta att taa aca gat gag atg ata tat cca cat caa aga atg cga gat aca tat gac aga cat tat tga cag aaa tgt gac ctt cat tat cac cgt cgt cca taa atg cgt tag gta cgt acc aca tac tat cgt taa cga tgc gca caa tct cgt cca -tt tca tcc atc ttc ata atg att tac ttt ttc ata att aga gaa aaa gat caa ggt ata aaa att aga agt gtt aga cta ta- --- --- --- --- --- --- --- --- --- --- -aa cta act tat gac tta act aac tta tga --- --- --- --- --- --- --- --- --- --- --- --- --- --- --- --- --- --- --- --- --- --- --- --- --- --- --- --- --- --- --- --- --- --- --- --- --- --- --- --- --- --- --- --- --- --- --- --- ctt aac ta- --- --- --- --- --- --- --- --- --- --- -ac tta tga ctt aac taa ctt atg act taa cta act ta- tga ctt aac taa ctt atg act taa cta act tat gac tta act aac tta tga cta tta act cat ttc aag aaa ggt ggg tgg ata gaa ctc tat atg aca gct tgt gaa aca att aga tcc cta att tct aat gga agt ttt gat agg aga ttg tca tca gtt gat aca ttg ttt att atc tca tct att aga gca cgt ctg ttt aga gct tta gtg acc tgc tcg gtt act tct gtg taa atc ttg aat cct tta gtg ata cac tgt gtc aaa act gga tgt tta gaa tac cta tgt aga ata tgg gaa gca tgc ttg ttt ttg tct cta tta tag att aac tca tac atg gtt gta tta tga att ttc atc tgc cta atg tac tcc aat tct tgt tta caa tca att ata taa tca aag agt gat gat gca tac aca tta caa agt gaa taa tct acc atc ata aaa tac ttg ata cag agc ttt atc aca tca tgg ttt tca att gta tta tta agt ata gct aat ttt ata cag tca ata gac aat ggt tct cta agc aat att tct aat att tta aga tgt gct tcc cta cgg gcg atg aca gat ccc cta tcc acg gcc acg tca aga cat gta tat cca tta ctc att act gcg ttg aca ttt gct cca ttt tct aat agc cat gat act aaa tct ata taa cct gca tag ata gcg cga taa agc aag gtc ctt cca cca gca tct agt tga ttg ata tct tca ata tat ggg ata caa agc tta taa att tct aat act gtg ggt tca tct aca agg aat ccc cta gta tac tga att att tta tat aga tct aat tta aca tca ttt tca tct ggg ata cca caa ttc aaa ata aac tca aca aca cta ctt tcc ttt tta cat att ccc cta aaa tag gca ttc aag cat tct att tta tat att aca gcc cca tga tct acc ata aga tca aca atg tct att tct aca tat gca tta gat aga tag taa agt aag aga tct tgc aca gaa tta caa ttc tta ata att ata gag aaa ata tct tcc ata taa ttc ttt gac act aat gca gat ata ata tct tta tat gta ata tat gca aac agt cta tct act ata tac tga tca ata tta tct cta tga atc cta aaa taa tca tac aga aca tct aca gga tca caa att ggt tca agg aga aat cta tca aat att ttc ctg tca aca act ggt tct aga aca taa cag tca aca cct aat cca tgt ttt tta taa tca tct acc aaa gat aat gac caa aga tcg agg tcg tcg tga aac tgc tca tcg aca gcc atg aaa tct acc gac tcc atg gtg cga atc gca ctg tct tat tcg cca ttg att ttc a-t ttt tta taa tta tgt aca tgt ttt cct tct att ctc aag agt cta caa aaa tat a-- -tt ttt tcg ata tct aag tac taa gtt ttt tta ctg ttt ttg tta ctg tct tcc att ctt cta act aaa gat ctg aga taa att ata caa tct tcg cta tcg aac cat ttt tgt agt cta aag cct gaa gta att aac caa ctg ttt tta tta gtg gct ttt ttc gat cta tcc tcg tc- --- ctc tca tca tcc tta ta- --- --- --- --- --- --- tta tta tca tta tta tca tag tct att aaa cac aaa tca tct --- --- --- --- --- --- --- --- --- --- --- --- --- --- --- --- --- --- --- --- --- --- --- --- --- --- --a cgt tta taa c-- --- --- --- --- --- --- --- --a aca ttc --- --- --- --- -tc att att aat tag ttc tgt ag- aat atc ttt aat aat ttg gct ata --- --- -ca tct gtt --- --- --- --- --- --- --- --- --- --- --- --- --- --- --- --- --- --- --c aat act --- --- --- --- --- --- --- --a tct att gat gat ttc ttt t-- --- --- --- --- --- --- --- --- --- --- --t taa gac t-- --- --- --- --- --- --- --- --- --- --- --- --- --- --- --- --- --- --- --- --- --- --- --- --- --- --- --- --- --- --- --- --- --- --- --- --- --- --- --- taa act agt --- --- --- --- --- --- --- --- --- --- --- --- --- --- --- --- --- --- --- --- --- --- --- --- --- --- --- --- --- --- --- --- --- --- --- --- --- --- --- --- --- --- --- --- --- --- --- --- --- --- --- --- --- --- --- --- --- --- --- --- --- --- --- --- --- --- --- --- --- --- --- --- --- --- --- --- --- --- --- --- --- --- --- --- --- --- --- --- --- --- --- --- --- --- --- --- --- --- --- --- --- --- --- --- --- --- --- --- --- --t atg gta atg acg atg aaa --- -tc gag tag ta- --- --a ctt cta ata aag act tga ta- --- --t cat tat cat atg ttt gat cg- --- --- --- --- --- --- --t cat agt taa tag tgt g-- --- --- --- --- --- --- --- --- --- --- --- --- --- --- --- --- --- --- --- -gc taa atg gta ctg tta ata agt tta t-- --- --- --- --- --- --- --- --- --- --- --- --- --- --- --- --- --a gac aat atc ata gta ttt tct ttc cag aat --- --- --- --t aga tta ttt ttt taa ata ctg atc ctc aca att ccg tga tgt agc agt agt tgg t-- --- --- --- --- --- --- --- --- --- --- --- --- --- --- --- --- --- --- --- --- --- --- --- --- --- --- --- -gc atg gtc tat atc gt- --- --- --- --- --- --- --- --- --- --- --- --- --- --- --- --- --- --- --- --- --- --- --- --- --- --- --- --- --- --- --- --- --- --- --- --- --- --- --- --- --- --- --- taa aat gta tca tat ata ata gtt ttc tga cgt gga gta cag aat ttt cga --- --- --- --- --- --- --- --- --- --- --- --- --- --- --- --- --- --- --- --- --- --- --- --- --- --- --- --- --- --- --- --- --- --- --- --- --- --- --- --- --- --- --- --t taa tga gtt cat ggt aag gaa ggg caa atg cct --- gta tat aat ata cat aag tta a-- --- --- --- --- --- --- --- --- --- --- --- --- --t agt ttt tta tca tat tt- --- --- --- --- --- --- --- --t cta ata cca taa taa aaa t-- --- --- --- --- --- --- --- --- --- --- --- --- --- --- --- --- --- --- --- --- --- --- --- --- --t atc att att gcg ttt g-g tag tt- --- --- --- --- ctg ccc ta- --- --- --- --- --- --- --- --t cat cta tat cac tgt cac tct c-- --- --g ctc tca cta tat ctt cta aaa tta caa --- -ac aac --- --- --- --- --- --- --- --- --- --- --- --- --- --- --- --- --- --- --- --- --- --- --- --- --- --- --- --- --- --- --- --- --- --- --- --- tgg ata ttc gat --- --- --- --- --- --- --- --- --- --- --- --- --- --- --- --- --- --- --- --- --- --- --- --- --- --- --a aca gca ttt gtg t-- --- --- --- --- --- --- --- --- --- --- --- --- --- --- --- --- -

>NC_063383.1_Monkeypox_virus_complete_genome

--- --- --- --- --- --- --- --- --- --- --- --- --- --- --- --- --- --- --- --- --- --- --- --- --- --- --- --- --- --a ttt tac tat ttt att tag tgt cta gaa aaa aat gtg tga ccc acg acc gta gga aac tct aga ggg taa gaa aaa tca atc gtt tat aga gac cat cag aaa gag gtt taa tat ttt tgt gag acc tat cga a-- --- --- --- --- --- --- --- --- --- --- --- --- --- --- --- --- --- --- --- --- --- --- --- --- --- --- --- --- --- --- --- --- --- --- --- --- --- --- --- --- --- --- --- --- --- --- --- --- --- --- --- --- --- --- --- --- --- --- --- --- --- --- --- --- --- --- --- --- --- --- --- --- --- --- --- --- --- --- --- --- --- --- --- --- --- --- --- --- --- --- --- --- --- --- --- --- --- --- --- --- --- --- --- --- --- --- --- --- --- --- --- --- --- --- --- --- --- --- --- --- --- --- --- --- --- --- --- --- --- --- --- --- --- --- --- --- --- --- --- --- --- --- --- --- --- --- --- --- --- --- --- --- --- --- --- --- --- --- --- --- --- --- --- --- --- --- --- --- --- --- --- --- --- --- --- --- --- --- --- --- --- --- --- --- --- --- --- --- --- --- --- --- --- --- --- --- --- --- --- --- --- --- --- --- --- --- --- --- --- --- --- --- --- --- --- --- --- --- --- --- --- --- --- --- --- --- --- --- --- --- --- --- --- --- --- --- --- --- --- --- --- --- --- --- --- --- --- --- --- --- --- --- --- --- --- --- --- --- --- --- --- --- --- --- --- --- --- --- --- --- --- --- --- --- --- --- --- --- --- -ga gag aaa gga taa aaa ctt ttt acg act cca tca gaa aga ggt tta ata ttt ttg tga gac cca tcg aag aga gaa aga gat ggt tag tca aga tat ttt tct tag tac aaa agt caa tgt ttt aaa ata tat gga cga gaa tta att tgt ctg tat aaa aac ttg tgt gaa att atg tac tag aga aaa aac gtg agc agt gtc ccc tac atg gat ttt aca gat cat tta tat tcc aaa aat att aac tat ata cgt tta tta tat gat gtt aac gtg taa att ata aac att att tta tga tgc aat tgt ctg aca acc tag att ggt ata agg atg ttg ata agc tct acg aga ata tat tgt tgg acg tta tcg ttt acg aaa tag ttg aga cat cag aaa gag gtt taa tat ttt tgt gag a-c cat cga aga gag aaa gag aat aaa aat at- ttt ttt ttt ttt ttt tgt aaa act ttt tta tga gac --- --- -ca aga gaa tac gaa tag tga tca tat cgt atc aca tat tga aac aga aag aag aag taa cga gag gta act ttt tgt gaa tgt agt taa ata ttt ttg ttt tgc aaa ccg gaa tat agt gcc cgg t-- -ct ttt tta att cgt ggt gcg gtg tct gaa tcg ttc gat taa ccc aac tca tcc att ttc aga tga ata gag tta tcg att cag aca cat gct ttg agt ttt gtt gaa tcg atg agt gaa gta tca tcg gtt gca cct tca gat gcc gat ccg tcg aca tac ttg aat cca tcc ttg act tca agt tca gat gat tcc tca cac atg tct ccg ata cgt acg cta aac tct agg ttc ttg aca cat ttt gta tca acg atc gtt gaa ccg atg ata tct ttg taa ctc act ttc tta tgt gag atg tta gac cca agt act gga tgg gtc ttg atg tca ctg tct ttc tct tct tcg cta cat ctg atg tcg ata gac atc tca cag tct ttg atc ata gcc aga gct tct tca cgc gtg atc gcg gga gag tcc tta cct tgt ccc ggt gac acg ctg gac aat cta gta ttc aca gtg ttt cca tca gag gat tcg gag atg gat gaa atc ttt ggg cat ttg gtg aat cca aag ttc atg tta aga ccc gca ccg acg ata gtg taa taa gtg gtg gga tct cct ttt aca act --- --- --- --- --- --- tct tcg gat acc tca tca tct tcg gtc tct gta act tcc gtt acg gat tga caa atc tta tca ttg gtc ggt gtt tgg tct tgc ttt gtg act ttg ata ata aca tcg att ccc ata tga tgt ttg ttt tct tct tca gta cac gag gat gaa gat tgt tga aga cta gta ggc ata gca gct gcc act agg cac atg cat gcc agg aca ata tat tgt ttc atg att gct att gat tga tta ctg ttc tag atg att cta ctt tct tac cat ata ata aat tag aat ata ttt tct act ttt acg aga aat taa tta ttg tat tta tta ttt ata ggt --a aaa aaa ctt act ata agt ggg tgg gat tct ggg aat tag tga tca gtt tat gta tat cgc aac tag cgg gca tat ggc tat tga cat cga gaa cat tac cca tat gat aag aga ttg tat cat ttt cgt agt ctt gag tat tgg tat tac tat ata gta tgt aga tgt cga cgc tag ata gac agt cgc cca cta gag tta ccg tct ctg aat gcg gca tga tag tat cat tct ttg ttt tcg tta act gtt tgg aag atg aat ctt tgt tgt tac att taa tct cga aat tca gag tac ata tct ttg aag tat tct gat atc tat ttt ctc ctg taa aga atc ctg aag ttg cta cat tat taa gga cag aga agt att ctg cac gaa aga ctg gat cac aat ctt tat gat tca tgg taa tag tta gtt ccg acg ttg aga tgg att cgc tga gac cgg tag tgg tcg tcc gag tac acg atg tgt cgt tga ctg gat aca ggt taa ttt cca cat cga tat agt taa atg tat tgc tgg tta cga cgg gtt cgc att tat ctg tgg aag aga cgg tgt gag aat atg ttc cgg gac cac acg gag aac aga tga cgt ctc cgg tag acg tgt atc cgg ata ctc cgt atc cta ttc cac act ttg ttt tag aaa tac atg ttc tac acc ctg atg ctc ctt tga gaa gac aat aat atc ctg gag agc att cac aga ttc tat tgt gag tcg tgt tac acg atc gcg tct cta cct gat tac tat cac atc ttc cgt tac aac tta gac aag cct gta aat gat tat tgt gag atg taa agg tat ccg aac cac acg gtg tac att gtg tat tag tct tgc tat cac ata atc tgg aag cgt aag ttc ccg gag gac acg ata gac aac ata gat tac ggc ttc tgt att cgt tgt ctt tac act ttc cat tgg atg gtg cat gtg gtg cta tat ctc ttc cgt tta tta tta tac atg aga gaa aca ata tat acg agt ata ata cgg acc tca tga ttt aat aat -gt agt aat cgt cgt ctt gtt act gtt tgt ttc cta ctt ctc caa tca tat aga tta ttt ttt aaa tat ttt ctt tct atc atg gat aat att tgt aat ggt tct ttc cgt aca aca tac tgt tta gat ggt agt cgc tta gct tgg tta tga tat tgc gca taa ttt ccg gag gca aat acg ata gtc tag att gac tat cga tgg tag act cta att tat tga gtg ctt tgt cga cga gtt tac ttt tat gct cca tcg ata gat gac act gtt cta tga gat cgt cgt aca tgg gaa atg aaa tgc gtt tgt ctg aat gta tgg ctt cga gat atg tgt gat acc gga tgt ctt ctg ttc tca ata ccg tat aca agt cgg tgt ctg aga ttc gaa tct ctt tga gga gac tta tgt cac gac tac att ttt cga tga tgg aat cta tct tat cga atg ata tat ttt tca taa ata cac ttt tat agt cct cgt tta aac aga att tag tat gta gtt ccg caa atg act cgt ccc tta ata ggc agt agg cta tta tct tct tta cgt agt gat cgt cgt agg gag aga act ccg aca tct tgt aga aca acg att taa tca tag gta gag ata ctt tca gtc tgt ggt gga taa tgt cat taa cga cat ccg cct tgt ata taa tgt ttc tgt ttt caa aca cca agt cga ata ctg tct tat cgt ctt tag tcg gaa ggt tga tgt cgt atc cga tgt ata cga ggt atg agg caa cat tgt tat tgc aat tct gga agg cgg tat gaa gag gag tca ttg tat tat agt att cgt ctt tct gaa tgt cga atc tat cta gta gat acc gta gta tat tga gag agc gac ttc cat atc ctt gat tat gtt tta tga ata gat aaa gta gat gtt gtc ctt ctt cct ttt gta att tcc cgt att ttt gtt cgt gcc aat tga gta aca tta tga gaa tat gac ctg ttg cac aat cgt tct tta tgt att cca tga tgg gtg tac aat caa gat tat tac gta tcc tcg tat cgg ctc ctc gag ata aaa gag cat aca cca cac gag gac tat gtt tgg tat act gtt gaa ggt aag tgt gta acg gcg tat ttc cga ttt tcg taa ccg cgt taa tgt ttg ctc cat gat cta tta tcg cgt aga tga atc gct tct cag ctc gca tct tag tgt gac tct ttg act tgt aat aat tgc ttt cgt gga acg cgg ata tgt gtt tac agt agt aat gaa gag aag tga gtc cat cct cat cga cgc aat tag ggt cag atc ctt tag tca ata att tgt aca gaa cgt aat agt tta agc tcc cat tga att tat atc taa gat aac aca gca ata gat cgg atg att tac taa agt cat caa tgg ggt ccg tta gta tat caa aga tct tgt tat cga ttg ata gtg aat gaa tca gat agt ggt gta gag gaa tat gtc ctt ttt tat cct tgc tat caa agt tac gca tgc cgt ggc gta aca ata tct tta ata cag atg gat taa atc gtg tat tca tcg tat agc aat gta atg gag agt tac cac att tta gtc gtt tat tca gat cgc agt gtt taa taa cta att taa aca gat gag atg ata tat cca cat caa aga atg cga gat aca tat gac aga cat tat tga cag aaa tgt gac ctt cat tat cac cgt cgt cca taa atg cgt tag gta cgt acc aca tac tat cgt taa cga tgc gca caa tct cgt cca -tt tca tcc atc ttc ata atg att tac ttt ttc ata att aga gaa aaa gat caa ggt ata aaa att aga agt gtt aga cta ta- --- --- --- --- --- --- --- --- --- --- -aa cta act tat gac tta act aac tta tga --- --- --- --- --- --- --- --- --- --- --- --- --- --- --- --- --- --- --- --- --- --- --- --- --- --- --- --- --- --- --- --- --- --- --- --- --- --- --- --- --- --- --- --- --- --- --- --- ctt aac ta- --- --- --- --- --- --- --- --- --- --- -ac tta tga ctt aac taa ctt atg act taa cta act ta- tga ctt aac taa ctt atg act taa cta act tat gac tta act aac tta tga cta tta act cat ttc aag aaa ggt ggg tgg ata gaa ctc tat atg aca gct tgt gaa aca att aga tcc cta att tct aat gga agt ttt gat agg aga ttg tca tca gtt gat aca ttg ttt att atc tca tct att aga gca cgt ctg ttt aga gct tta gtg acc tgc tcg gtt act tct gtg taa atc ttg aat cct tta gtg ata cac tgt gtc aaa act gga tgt tta gaa tac cta tgt aga ata tgg gaa gca tgc ttg ttt ttg tct cta tta tag att aac tca tac atg gtt gta tta tga att ttc atc tgc cta atg tac tcc aat tct tgt tta caa tca att ata taa tca aag agt gat gat gca tac aca tta caa agt gaa taa tct acc atc ata aaa tac ttg ata cag agc ttt atc aca tca tgg ttt tca att gta tta tta agt ata gct aat ttt ata cag tca ata gac aat ggt tct cta agc aat att tct aat att tta aga tgt gct tcc cta cgg gcg atg aca gat ccc cta tcc acg gcc acg tca aga cat gta tat cca tta ctc att act gcg ttg aca ttt gct cca ttt tct aat agc cat gat act aaa tct ata taa cct gca tag ata gcg cga taa agc aag gtc ctt cca cca gca tct agt tga ttg ata tct tca ata tat ggg ata caa agc tta taa att tct aat act gtg ggt tca tct aca agg aat ccc cta gta tac tga att att tta tat aga tct aat tta aca tca ttt tca tct ggg ata cca caa ttc aaa ata aac tca aca aca cta ctt tcc ttt tta cat att ccc cta aaa tag gca ttc aag cat tct att tta tat att aca gcc cca tga tct acc ata aga tca aca atg tct att tct aca tat gca tta gat aga tag taa agt aag aga tct tgc aca gaa tta caa ttc tta ata att ata gag aaa ata tct tcc ata taa ttc ttt gac act aat gca gat ata ata tct tta tat gta ata tat gca aac agt cta tct act ata tac tga tca ata tta tct cta tga atc cta aaa taa tca tac aga aca tct aca gga tca caa att ggt tca agg aga aat cta tca aat att ttc ctg tca aca act ggt tct aga aca taa cag tca aca cct aat cca tgt ttt tta taa tca tct acc aaa gat aat gac caa aga tcg agg tcg tcg tga aac tgc tca tcg aca gcc atg aaa tct acc gac tcc atg gtg cga atc gca ctg tct tat tcg cca ttg att ttc a-t ttt tta taa tta tgt aca tgt ttt cct tct att ctc aag agt cta caa aaa tat a-- -tt ttt tcg ata tct aag tac taa gtt ttt tta ctg ttt ttg tta ctg tct tcc att ctt cta act aaa gat ctg aga taa att ata caa tct tcg cta tcg aac cat ttt tgt agt cta aag cct gaa gta att aac caa ctg ttt tta tta gtg gct ttt ttc gat cta tcc tcg tc- --- ctc tca tca tcc tta ta- --- --- --- --- --- --- tta tta tca tta tta tca tag tct att aaa cac aaa tca tct --- --- --- --- --- --- --- --- --- --- --- --- --- --- --- --- --- --- --- --- --- --- --- --- --- --- --a cgt tta taa c-- --- --- --- --- --- --- --- --a aca ttc --- --- --- --- -tc att att aat tag ttc tgt ag- aat atc ttt aat aat ttg gct ata --- --- -ca tct gtt --- --- --- --- --- --- --- --- --- --- --- --- --- --- --- --- --- --- --c aat act --- --- --- --- --- --- --- --a tct att gat gat ttc ttt t-- --- --- --- --- --- --- --- --- --- --- --t taa gac t-- --- --- --- --- --- --- --- --- --- --- --- --- --- --- --- --- --- --- --- --- --- --- --- --- --- --- --- --- --- --- --- --- --- --- --- --- --- --- --- taa act agt --- --- --- --- --- --- --- --- --- --- --- --- --- --- --- --- --- --- --- --- --- --- --- --- --- --- --- --- --- --- --- --- --- --- --- --- --- --- --- --- --- --- --- --- --- --- --- --- --- --- --- --- --- --- --- --- --- --- --- --- --- --- --- --- --- --- --- --- --- --- --- --- --- --- --- --- --- --- --- --- --- --- --- --- --- --- --- --- --- --- --- --- --- --- --- --- --- --- --- --- --- --- --- --- --- --- --- --- --- --t atg gta atg acg atg aaa --- -tc gag tag ta- --- --a ctt cta ata aag act tga ta- --- --t cat tat cat atg ttt gat cg- --- --- --- --- --- --- --t cat agt taa tag tgt g-- --- --- --- --- --- --- --- --- --- --- --- --- --- --- --- --- --- --- --- -gc taa atg gta ctg tta ata agt tta t-- --- --- --- --- --- --- --- --- --- --- --- --- --- --- --- --- --a gac aat atc ata gta ttt tct ttc cag aat --- --- --- --t aga tta ttt ttt taa ata ctg atc ctc aca att ccg tga tgt agc agt agt tgg t-- --- --- --- --- --- --- --- --- --- --- --- --- --- --- --- --- --- --- --- --- --- --- --- --- --- --- --- -gc atg gtc tat atc gt- --- --- --- --- --- --- --- --- --- --- --- --- --- --- --- --- --- --- --- --- --- --- --- --- --- --- --- --- --- --- --- --- --- --- --- --- --- --- --- --- --- --- --- taa aat gta tca tat ata ata gtt ttc tga cgt gga gta cag aat ttt cga --- --- --- --- --- --- --- --- --- --- --- --- --- --- --- --- --- --- --- --- --- --- --- --- --- --- --- --- --- --- --- --- --- --- --- --- --- --- --- --- --- --- --- --t taa tga gtt cat ggt aag gaa ggg caa atg cct --- gta tat aat ata cat aag tta a-- --- --- --- --- --- --- --- --- --- --- --- --- --t agt ttt tta tca tat tt- --- --- --- --- --- --- --- --t cta ata cca taa taa aaa t-- --- --- --- --- --- --- --- --- --- --- --- --- --- --- --- --- --- --- --- --- --- --- --- --- --t atc att att gcg ttt g-g tag tt- --- --- --- --- ctg ccc ta- --- --- --- --- --- --- --- --t cat cta tat cac tgt cac tct c-- --- --g ctc tca cta tat ctt cta aaa tta caa --- -ac aac --- --- --- --- --- --- --- --- --- --- --- --- --- --- --- --- --- --- --- --- --- --- --- --- --- --- --- --- --- --- --- --- --- --- --- --- tgg ata ttc gat --- --- --- --- --- --- --- --- --- --- --- --- --- --- --- --- --- --- --- --- --- --- --- --- --- --- --a aca gca ttt gtg t-- --- --- --- --- --- --- --- --- --- --- --- --- --- --- --- --- -

>ON676704.1_Monkeypox_virus_isolate_MPXV_USA_2022_FL002_complete_genome

--- --- --- --- --- --- --- --- --- --- --- --- --- --- --- --- --- --- --- --- --- --- --- --- --- --- --- --t att ata ttt tac tat ttt att tag tgt cta gaa aaa aat gtg tga ccc acg acc gta gga aac tct aga ggg taa gaa aaa tca atc gtt tat aga gac cat cag aaa gag gtt taa tat ttt tgt gag acc tat cga a-- --- --- --- --- --- --- --- --- --- --- --- --- --- --- --- --- --- --- --- --- --- --- --- --- --- --- --- --- --- --- --- --- --- --- --- --- --- --- --- --- --- --- --- --- --- --- --- --- --- --- --- --- --- --- --- --- --- --- --- --- --- --- --- --- --- --- --- --- --- --- --- --- --- --- --- --- --- --- --- --- --- --- --- --- --- --- --- --- --- --- --- --- --- --- --- --- --- --- --- --- --- --- --- --- --- --- --- --- --- --- --- --- --- --- --- --- --- --- --- --- --- --- --- --- --- --- --- --- --- --- --- --- --- --- --- --- --- --- --- --- --- --- --- --- --- --- --- --- --- --- --- --- --- --- --- --- --- --- --- --- --- --- --- --- --- --- --- --- --- --- --- --- --- --- --- --- --- --- --- --- --- --- --- --- --- --- --- --- --- --- --- --- --- --- --- --- --- --- --- --- --- --- --- --- --- --- --- --- --- --- --- --- --- --- --- --- --- --- --- --- --- --- --- --- --- --- --- --- --- --- --- --- --- --- --- --- --- --- --- --- --- --- --- --- --- --- --- --- --- --- --- --- --- --- --- --- --- --- --- --- --- --- --- --- --- --- --- --- --- --- --- --- --- --- --- --- --- --- --- -ga gag aaa gga taa aaa ctt ttt acg act cca tca gaa aga ggt tta ata ttt ttg tga gac cca tcg aag aga gaa aga gat ggt tag tca aga tat ttt tct tag tac aaa agt caa tgt ttt aaa ata tat gga cga gaa tta att tgt ctg tat aaa aac ttg tgt gaa att atg tac tag aga aaa aac gtg agc agt gtc ccc tac atg gat ttt aca gat cat tta tat tcc aaa aat att aac tat ata cgt tta tta tat gat gtt aac gtg taa att ata aac att att tta tga tgc aat tgt ctg aca acc tag att ggt ata agg atg ttg ata agc tct acg aga ata tat tgt tgg acg tta tcg ttt acg aaa tag ttg aga cat cag aaa gag gtt taa tat ttt tgt gag a-c cat cga aga gag aaa gag aat aaa aat a-- -?t ttt ttt ttt ttt tgt aaa act ttt tta tga gac --- --- -ca aga gaa tac gaa tag tga tca tat cgt atc aca tat tga aac aga aag aag aag taa cga gag gta act ttt tgt gaa tgt agt taa ata ttt ttg ttt tgc aaa ccg gaa tat agt gcc cgg t-- -ct ttt tta att cgt ggt gcg gtg tct gaa tcg ttc gat taa ccc aac tca tcc att ttc aga tga ata gag tta tcg att cag aca cat gct ttg agt ttt gtt gaa tcg atg agt gaa gta tca tcg gtt gca cct tca gat gcc gat ccg tcg aca tac ttg aat cca tcc ttg act tca agt tca gat gat tcc tca cac atg tct ccg ata cgt acg cta aac tct agg ttc ttg aca cat ttt gta tca acg atc gtt gaa ccg atg ata tct ttg taa ctc act ttc tta tgt gag atg tta gac cca agt act gga tgg gtc ttg atg tca ctg tct ttc tct tct tcg cta cat ctg atg tcg ata gac atc tca cag tct ttg atc ata gcc aga gct tct tca cgc gtg atc gcg gga gag tcc tta cct tgt ccc ggt gac acg ctg gac aat cta gta ttc aca gtg ttt cca tca gag gat tcg gag atg gat aaa atc ttt ggg cat ttg gtg aat cca aag ttc atg tta aga ccc gca ccg acg ata gtg taa taa gtg gtg gga tct cct ttt aca act --- --- --- --- --- --- tct tcg gat acc tca tca tct tcg gtc tct gta act tcc gtt acg gat tga caa atc tta tca ttg gtc ggt gtt tgg tct tgc ttt gtg act ttg ata ata aca tcg att ccc ata tga tgt ttg ttt tct tct tca gta cac gag gat gaa gat tgt tga aga cta gta ggc ata gca gct gcc act agg cac atg cat gcc agg aca ata tat tgt ttc atg att gct att gat tga tta ctg ttc tag atg att cta ctt tct tac cat ata ata aat tag aat ata ttt tct act ttt acg aga aat taa tta ttg tat tta tta ttt ata ggt --a aaa aaa ctt act ata agt ggg tgg gat tct ggg aat tag tga tca gtt tat gta tat cgc aac tag cgg gca tat ggc tat tga cat cga gaa cat tac cca tat gat aag aga ttg tat cat ttt cgt agt ctt gag tat tgg tat tac tat ata gta tgt aga tgt cga cgc tag ata gac agt cgc cca cta gag tta ccg tct ctg aat gcg gca tga tag tat cat tct ttg ttt tcg tta act gtt tgg aag atg aat ctt tgt tgt tac att taa tct cga aat tca gag tac ata tct ttg aag tat tct gat atc tat ttt ctc ctg taa aga atc ctg aag ttg cta cat tat taa gga cag aga agt att ctg cac gaa aga ctg gat cac aat ctt tat gat tca tgg taa tag tta gtt ccg acg ttg aga tgg att cgc tga gac cgg tag tgg tcg tcc gag tac acg atg tgt cgt tga ctg gat aca ggt taa ttt cca cat cga tat agt taa atg tat tgc tgg tta cga cgg gtt cgc att tat ctg tgg aag aga cgg tgt gag aat atg ttc cgg gac cac acg gag aac aga tga cgt ctc cgg tag acg tgt atc cgg ata ctc cgt atc cta ttc cac act ttg ttt tag aaa tac atg ttc tac acc ctg atg ctc ctt tga gaa gac aat aat atc ctg gag agc att cac aga ttc tat tgt gag tcg tgt tac acg atc gcg tct cta cct gat tac tat cac atc ttc cgt tac aac tta gac aag cct gta aat gat tat tgt gag atg taa agg tat ccg aac cac acg gtg tac att gtg tat tag tct tgc tat cac ata atc tga aag cgt aag ttc ccg gag gac acg ata gac aac ata gat tac ggc ttc tgt att cgt tgt ctt tac act ttc cat tgg atg gtg cat gtg gtg cta tat ctc ttc cgt tta tta tta tac atg aga gaa aca ata tat acg agt ata ata cgg acc tca tga ttt aat aat -gt agt aat cgt cgt ctt gtt act gtt tgt ttc cta ctt ctc caa tca tat aga tta ttt ttt aaa tat ttt ctt tct atc atg gat aat att tgt aat ggt tct ttc cgt aca aca tac tgt tta gat ggt agt cgc tta gct tgg tta tga tat tgc gca taa ttt ccg gag gca aat acg ata gtc tag att gac tat cga tgg tag act cta att tat tga gtg ctt tgt cga cga gtt tac ttt tat gct cca tcg ata gat gac act gtt cta tga gat cgt cgt aca tgg gaa atg aaa tgc gtt tgt ctg aat gta tgg ctt cga gat atg tgt gat acc gaa tgt ctt ctg ttc tca ata ccg tat aca agt cgg tgt ctg aga ttc gaa tct ctt tga gga gac tta tgt cac gac tac att ttt cga tga tgg aat cta tct tat cga atg ata tat ttt tca taa ata cac ttt tat agt cct cgt tta aac aga att tag tat gta gtt ccg caa atg act cgt ccc tta ata ggc agt agg cta tta tct tct tta cgt agt gat cgt cgt agg gag aga act ccg aca tct tgt aga aca acg att taa tca tag gta gag ata ctt tca gtc tgt ggt gga taa tgt cat taa cga cat ccg cct tgt ata taa tgt ttc tgt ttt caa aca cca agt cga ata ctg tct tat cgt ctt tag tcg gaa ggt tga tgt cgt atc caa tgt ata cga ggt atg agg caa cat tgt tat tgc aat tct gga agg cgg tat gaa gag gag tca ttg tat tat agt att cgt ctt tct gaa tgt cga atc tat cta gta gat acc gta gta tat tga gag agc gac ttc cat atc ctt gat tat gtt tta tga ata gat aaa gta gat gtt gtc ctt ctt cct ttt gta att tcc cgt att ttt gtt cgt gcc aat tga gta aca tta tga gaa tat gac ctg ttg cac aat cgt tct tta tgt att cca tga tgg gtg tac aat taa gat tat tac gta tcc tcg tat cgg ctc ctc gag ata aaa gag cat aca cca cac gag gac tat gtt tgg tat act gtt gaa ggt aag tgt gta acg gcg tat ttc cga ttt tcg taa ccg cgt taa tgt ttg ctc cat gat cta tta tcg cgt aga tga atc gct tct cag ctc gca tct tag tgt gac tct ttg act tgt aat aat tgc ttt cgt gga acg cgg ata tgt gtt tac agt agt aat gaa gag aag tga gtc cat cct cat cga cgc aat tag ggt cag atc ctt tag tca ata att tgt aca gaa cgt aat agt tta agc tcc cat tga att tat atc taa gat aac aca gca ata gat cgg atg att tac taa agt cat caa tgg ggt ccg tta gta tat caa aga tct tgt tat cga ttg ata gtg aat gaa tca gat agt ggt gta gag gaa tat gtc ctt ttt tat cct tgc tat caa agt tac gca tgc cgt ggc gta aca ata tct tta ata cag atg gat taa atc gtg tat tca tcg tat agc aat gta atg gag agt tac cac att tta gtc gtt tat tca gat cgc agt gtt taa taa cta att taa aca gat gag atg ata tat cca cat caa aga atg cga gat aca tat gac aga cat tat tga cag aaa tgt gac ctt cat tat cac cgt cgt cca taa atg cgt tag gta cgt acc aca tac tat cgt taa cga tgc gca caa tct cgt cca -tt tca tcc atc ttc ata atg att tac ttt ttc ata att aga gaa aaa gat caa ggt ata aaa att aga agt gtt aga cta ta- --- --- --- --- --- --- --- --- --- --- -aa cta act tat gac tta act aac tta tga --- --- --- --- --- --- --- --- --- --- --- --- --- --- --- --- --- --- --- --- --- --- --- --- --- --- --- --- --- --- --- --- --- --- --- --- --- --- --- --- --- --- --- --- --- --- --- --- ctt aac ta- --- --- --- --- --- --- --- --- --- --- -ac tta tga ctt aac taa ctt atg act taa cta act ta- tga ctt aac taa ctt atg act taa cta act tat gac tta act aac tta tga cta tta act cat ttc aag aaa ggt ggg tgg ata gaa ctc tat atg aca gct tgt gaa aca att aga tcc cta att tct aat gga agt ttt gat agg aga ttg tca tca gtt gat aca ttg ttt att atc tca tct att aga gca cgt ctg ttt aga gct tta gtg acc tgc tcg gtt act tct gtg taa atc ttg aat cct tta gtg ata cac tgt gtc aaa act gga tgt tta gaa tac cta tgt aga ata tgg gaa gca tgc ttg ttt ttg tct cta tta tag att aac tca tac atg gtt gta tta tga att ttc atc tgc cta atg tac tcc aat tct tgt tta caa tca att ata taa tca aag agt gat gat gca tac aca tta caa agt gaa taa tct acc atc ata aaa tac ttg ata cag agc ttt atc aca tca tgg ttt tca att gta tta tta agt ata gct aat ttt ata cag tca ata gac aat ggt tct cta agc aat att tct aat att tta aga tgt gct tcc cta cgg gcg atg aca gat ccc cta tcc acg gcc acg tca aga cat gta tat cca tta ctc att act gcg ttg aca ttt gct cca ttt tct aat agc cat gat act aaa tct ata taa cct gca tag ata gcg cga taa agc aag gtc ctt cca cca gca tct agt tga ttg ata tct tca ata tat ggg ata caa agc tta taa att tct aat act gtg ggt tca tct aca agg aat ccc cta gta tac tga att att tta tat aga tct aat tta aca tca ttt tca tct ggg ata cca caa ttc aaa ata aac tca aca aca cta ctt tcc ttt tta cat att ccc cta aaa tag gca ttc aag cat tct att tta tat att aca gcc cca tga tct acc ata aga tca aca atg tct att tct aca tat gca tta gat aga tag taa agt aag aga tct tgc aca gaa tta caa ttc tta ata att ata gag aaa ata tct tcc ata taa ttc ttt gac act aat gca gat ata ata tct tta tat gta ata tat gca aac agt cta tct act ata tac tga tca ata tta tct cta tga atc cta aaa taa tca tac aga aca tct aca gga tca caa att ggt tca agg aga aat cta tca aat att ttc ctg tca aca act ggt tct aga aca taa cag tca aca cct aat cca tgt ttt tta taa tca tct acc aaa gat aat gac caa aga tcg agg tcg tcg tga aac tgc tca tcg aca gcc atg aaa tct acc gac tcc atg gtg cga atc gca ctg tct tat tcg cca ttg att ttc a-t ttt tta taa tta tgt aca tgt ttt cct tct att ctc aag agt cta caa aaa tat a-- -tt ttt tcg ata tct aag tac taa gtt ttt tta ctg ttt ttg tta ctg tct tcc att ctt cta act aaa gat ctg aga taa att ata caa tct tcg cta tcg aac cat ttt tgt agt cta aag cct gaa gta att aac caa ctg ttt tta tta gtg gct ttt ttc gat cta tcc tcg tc- --- ctc tca tca tcc tta ta- --- --- --- --- --- --- tta tta tca tta tta tca tag tct att aaa cac aaa tca tct --- --- --- --- --- --- --- --- --- --- --- --- --- --- --- --- --- --- --- --- --- --- --- --- --- --- --a cgt tta taa c-- --- --- --- --- --- --- --- --a aca ttc --- --- --- --- -tc att att aat tag ttc tgt ag- aat atc ttt aat aat ttg gct ata --- --- -ca tct gtt --- --- --- --- --- --- --- --- --- --- --- --- --- --- --- --- --- --- --c aat act --- --- --- --- --- --- --- --a tct att gat gat ttc ttt t-- --- --- --- --- --- --- --- --- --- --- --t taa gac t-- --- --- --- --- --- --- --- --- --- --- --- --- --- --- --- --- --- --- --- --- --- --- --- --- --- --- --- --- --- --- --- --- --- --- --- --- --- --- --- taa act agt --- --- --- --- --- --- --- --- --- --- --- --- --- --- --- --- --- --- --- --- --- --- --- --- --- --- --- --- --- --- --- --- --- --- --- --- --- --- --- --- --- --- --- --- --- --- --- --- --- --- --- --- --- --- --- --- --- --- --- --- --- --- --- --- --- --- --- --- --- --- --- --- --- --- --- --- --- --- --- --- --- --- --- --- --- --- --- --- --- --- --- --- --- --- --- --- --- --- --- --- --- --- --- --- --- --- --- --- --- --t atg gta atg acg atg aaa --- -tc gag tag ta- --- --a ctt cta ata aag act tga ta- --- --t cat tat cat atg ttt gat cg- --- --- --- --- --- --- --t cat agt taa tag tgt g-- --- --- --- --- --- --- --- --- --- --- --- --- --- --- --- --- --- --- --- -gc taa atg gta ctg tta ata agt tta t-- --- --- --- --- --- --- --- --- --- --- --- --- --- --- --- --- --a gac aat atc ata gta ttt tct ttc cag aat --- --- --- --t aga tta ttt ttt taa ata ctg atc ctc aca att ccg tga tgt agc agt agt tgg t-- --- --- --- --- --- --- --- --- --- --- --- --- --- --- --- --- --- --- --- --- --- --- --- --- --- --- --- -gc atg gtc tat atc gt- --- --- --- --- --- --- --- --- --- --- --- --- --- --- --- --- --- --- --- --- --- --- --- --- --- --- --- --- --- --- --- --- --- --- --- --- --- --- --- --- --- --- --- taa aat gta tca tat ata ata gtt ttc tga cgt gga gta cag aat ttt cga --- --- --- --- --- --- --- --- --- --- --- --- --- --- --- --- --- --- --- --- --- --- --- --- --- --- --- --- --- --- --- --- --- --- --- --- --- --- --- --- --- --- --- --t taa tga gtt cat ggt aag gaa ggg caa atg cct --- gta tat aat ata cat aag tta a-- --- --- --- --- --- --- --- --- --- --- --- --- --t agt ttt tta tca tat tt- --- --- --- --- --- --- --- --t cta ata cca taa taa aaa t-- --- --- --- --- --- --- --- --- --- --- --- --- --- --- --- --- --- --- --- --- --- --- --- --- --t atc att att gcg ttt g-g tag tt- --- --- --- --- ctg ccc ta- --- --- --- --- --- --- --- --t cat cta tat cac tgt cac tct c-- --- --g ctc tca cta tat ctt cta aaa tta caa --- -ac aac --- --- --- --- --- --- --- --- --- --- --- --- --- --- --- --- --- --- --- --- --- --- --- --- --- --- --- --- --- --- --- --- --- --- --- --- tgg ata ttc gat --- --- --- --- --- --- --- --- --- --- --- --- --- --- --- --- --- --- --- --- --- --- --- --- --- --- --a aca gca ttt gtg t-- --- --- --- --- --- --- --- --- --- --- --- --- --- --- --- --- -

>ON563414.3_Monkeypox_virus_isolate_MPXV_USA_2022_MA001_complete_genome

--- --- --- --- --- --- --- --- --- --- --- --- --- --- --- --- --- --t ttt agt a-- --- --- --- --- --c att aat att ata ttt tac tat ttt att tag tgt cta gaa aaa aat gtg tga ccc acg acc gta gga aac tct aga ggg taa gaa aaa tca atc gtt tat aga gac cat cag aaa gag gtt taa tat ttt tgt gag acc tat cga a-- --- --- --- --- --- --- --- --- --- --- --- --- --- --- --- --- --- --- --- --- --- --- --- --- --- --- --- --- --- --- --- --- --- --- --- --- --- --- --- --- --- --- --- --- --- --- --- --- --- --- --- --- --- --- --- --- --- --- --- --- --- --- --- --- --- --- --- --- --- --- --- --- --- --- --- --- --- --- --- --- --- --- --- --- --- --- --- --- --- --- --- --- --- --- --- --- --- --- --- --- --- --- --- --- --- --- --- --- --- --- --- --- --- --- --- --- --- --- --- --- --- --- --- --- --- --- --- --- --- --- --- --- --- --- --- --- --- --- --- --- --- --- --- --- --- --- --- --- --- --- --- --- --- --- --- --- --- --- --- --- --- --- --- --- --- --- --- --- --- --- --- --- --- --- --- --- --- --- --- --- --- --- --- --- --- --- --- --- --- --- --- --- --- --- --- --- --- --- --- --- --- --- --- --- --- --- --- --- --- --- --- --- --- --- --- --- --- --- --- --- --- --- --- --- --- --- --- --- --- --- --- --- --- --- --- --- --- --- --- --- --- --- --- --- --- --- --- --- --- --- --- --- --- --- --- --- --- --- --- --- --- --- --- --- --- --- --- --- --- --- --- --- --- --- --- --- --- --- --- -ga gag aaa gga taa aaa ctt ttt acg act cca tca gaa aga ggt tta ata ttt ttg tga gac cca tcg aag aga gaa aga gat ggt tag tca aga tat ttt tct tag tac aaa agt caa tgt ttt aaa ata tat gga cga gaa tta att tgt ctg tat aaa aac ttg tgt gaa att atg tac tag aga aaa aac gtg agc agt gtc ccc tac atg gat ttt aca gat cat tta tat tcc aaa aat att aac tat ata cgt tta tta tat gat gtt aac gtg taa att ata aac att att tta tga tgc aat tgt ctg aca acc tag att ggt ata agg atg ttg ata agc tct acg aga ata tat tgt tgg acg tta tcg ttt acg aaa tag ttg aga cat cag aaa gag gtt taa tat ttt tgt gag a-c cat cga aga gag aaa gag aat aaa aat a-- --t ttt ttt ttt ttt tgt aaa act ttt tta tga gac --- --- -ca aga gaa tac gaa tag tga tca tat cgt atc aca tat tga aac aga aag aag aag taa cga gag gta act ttt tgt gaa tgt agt taa ata ttt ttg ttt tgc aaa ccg gaa tat agt gcc cgg t-- -ct ttt tta att cgt ggt gcg gtg tct gaa tcg ttc gat taa ccc aac tca tcc att ttc aga tga ata gag tta tcg att cag aca cat gct ttg agt ttt gtt gaa tcg atg agt gaa gta tca tcg gtt gca cct tca gat gcc gat ccg tcg aca tac ttg aat cca tcc ttg act tca agt tca gat gat tcc tca cac atg tct ccg ata cgt acg cta aac tct agg ttc ttg aca cat ttt gta tca acg atc gtt gaa ccg atg ata tct ttg taa ctc act ttc tta tgt gag atg tta gac cca agt act gga tgg gtc ttg atg tca ctg tct ttc tct tct tcg cta cat ctg atg tcg ata gac atc tca cag tct ttg atc ata gcc aga gct tct tca cgc gtg atc gcg gga gag tcc tta cct tgt ccc ggt gac acg ctg gac aat cta gta ttc aca gtg ttt cca tca gag gat tcg gag atg gat aaa atc ttt ggg cat ttg gtg aat cca aag ttc atg tta aga ccc gca ccg acg ata gtg taa taa gtg gtg gga tct cct ttt aca act --- --- --- --- --- --- tct tcg gat acc tca tca tct tcg gtc tct gta act tcc gtt acg gat tga caa atc tta tca ttg gtc ggt gtt tgg tct tgc ttt gtg act ttg ata ata aca tcg att ccc ata tga tgt ttg ttt tct tct tca gta cac gag gat gaa gat tgt tga aga cta gta ggc ata gca gct gcc act agg cac atg cat gcc agg aca ata tat tgt ttc atg att gct att gat tga tta ctg ttc tag atg att cta ctt tct tac cat ata ata aat tag aat ata ttt tct act ttt acg aga aat taa tta ttg tat tta tta ttt ata ggt --a aaa aaa ctt act ata agt ggg tgg gat tct ggg aat tag tga tca gtt tat gta tat cgc aac tag cgg gca tat ggc tat tga cat cga gaa cat tac cca tat gat aag aga ttg tat cat ttt cgt agt ctt gag tat tgg tat tac tat ata gta tgt aga tgt cga cgc tag ata gac agt cgc cca cta gag tta ccg tct ctg aat gcg gca tga tag tat cat tct ttg ttt tcg tta act gtt tgg aag atg aat ctt tgt tgt tac att taa tct cga aat tca gag tac ata tct ttg aag tat tct gat atc tat ttt ctc ctg taa aga atc ctg aag ttg cta cat tat taa gga cag aga agt att ctg cac gaa aga ctg gat cac aat ctt tat gat tca tgg taa tag tta gtt ccg acg ttg aga tgg att cgc tga gac cgg tag tgg tcg tcc gag tac acg atg tgt cgt tga ctg gat aca ggt taa ttt cca cat cga tat agt taa atg tat tgc tgg tta cga cgg gtt cgc att tat ctg tgg aag aga cgg tgt gag aat atg ttc cgg gac cac acg gag aac aga tga cgt ctc cgg tag acg tgt atc cgg ata ctc cgt atc cta ttc cac act ttg ttt tag aaa tac atg ttc tac acc ctg atg ctc ctt tga gaa gac aat aat atc ctg gag agc att cac aga ttc tat tgt gag tcg tgt tac acg atc gcg tct cta cct gat tac tat cac atc ttc cgt tac aac tta gac aag cct gta aat gat tat tgt gag atg taa agg tat ccg aac cac acg gtg tac att gtg tat tag tct tgc tat cac ata atc tga aag cgt aag ttc ccg gag gac acg ata gac aac ata gat tac ggc ttc tgt att cgt tgt ctt tac act ttc cat tgg atg gtg cat gtg gtg cta tat ctc ttc cgt tta tta tta tac atg aga gaa aca ata tat acg agt ata ata cgg acc tca tga ttt aat aat -gt agt aat cgt cgt ctt gtt act gtt tgt ttc cta ctt ctc caa tca tat aga tta ttt ttt aaa tat ttt ctt tct atc atg gat aat att tgt aat ggt tct ttc cgt aca aca tac tgt tta gat ggt agt cgc tta gct tgg tta tga tat tgc gca taa ttt ccg gag gca aat acg ata gtc tag att gac tat cga tgg tag act cta att tat tga gtg ctt tgt cga cga gtt tac ttt tat gct cca tcg ata gat gac act gtt cta tga gat cgt cgt aca tgg gaa atg aaa tgc gtt tgt ctg aat gta tgg ctt cga gat atg tgt gat acc gaa tgt ctt ctg ttc tca ata ccg tat aca agt cgg tgt ctg aga ttc gaa tct ctt tga gga gac tta tgt cac gac tac att ttt cga tga tgg aat cta tct tat cga atg ata tat ttt tca taa ata cac ttt tat agt cct cgt tta aac aga att tag tat gta gtt ccg caa atg act cgt ccc tta ata ggc agt agg cta tta tct tct tta cgt agt gat cgt cgt agg gag aga act ccg aca tct tgt aga aca acg att taa tca tag gta gag ata ctt tca gtc tgt ggt gga taa tgt cat taa cga cat ccg cct tgt ata taa tgt ttc tgt ttt caa aca cca agt cga ata ctg tct tat cgt ctt tag tcg gaa ggt tga tgt cgt atc caa tgt ata cga ggt atg agg caa cat tgt tat tgc aat tct gga agg cgg tat gaa gag gag tca ttg tat tat agt att cgt ctt tct gaa tgt cga atc tat cta gta gat acc gta gta tat tga gag agc gac ttc cat atc ctt gat tat gtt tta tga ata gat aaa gta gat gtt gtc ctt ctt cct ttt gta att tcc cgt att ttt gtt cgt gcc aat tga gta aca tta tga gaa tat gac ctg ttg cac aat cgt tct tta tgt att cca tga tgg gtg tac aat taa gat tat tac gta tcc tcg tat cgg ctc ctc gag ata aaa gag cat aca cca cac gag gac tat gtt tgg tat act gtt gaa ggt aag tgt gta acg gcg tat ttc cga ttt tcg taa ccg cgt taa tgt ttg ctc cat gat cta tta tcg cgt aga tga atc gct tct cag ctc gca tct tag tgt gac tct ttg act tgt aat aat tgc ttt cgt gga acg cgg ata tgt gtt tac agt agt aat gaa gag aag tga gtc cat cct cat cga cgc aat tag ggt cag atc ctt tag tca ata att tgt aca gaa cgt aat agt tta agc tcc cat tga att tat atc taa gat aac aca gca ata gat cgg atg att tac taa agt cat caa tgg ggt ccg tta gta tat caa aga tct tgt tat cga ttg ata gtg aat gaa tca gat agt ggt gta gag gaa tat gtc ctt ttt tat cct tgc tat caa agt tac gca tgc cgt ggc gta aca ata tct tta ata cag atg gat taa atc gtg tat tca tcg tat agc aat gta atg gag agt tac cac att tta gtc gtt tat tca gat cgc agt gtt taa taa cta att taa aca gat gag atg ata tat cca cat caa aga atg cga gat aca tat gac aga cat tat tga cag aaa tgt gac ctt cat tat cac cgt cgt cca taa atg cgt tag gta cgt acc aca tac tat cgt taa cga tgc gca caa tct cgt cca -tt tca tcc atc ttc ata atg att tac ttt ttc ata att aga gaa aaa gat caa ggt ata aaa att aga agt gtt aga cta ta- --- --- --- --- --- --- --- --- --- --- -aa cta act tat gac tta act aac tta tga --- --- --- --- --- --- --- --- --- --- --- --- --- --- --- --- --- --- --- --- --- --- --- --- --- --- --- --- --- --- --- --- --- --- --- --- --- --- --- --- --- --- --- --- --- --- --- --- ctt aac ta- --- --- --- --- --- --- --- --- --- --- -ac tta tga ctt aac taa ctt atg act taa cta act ta- tga ctt aac taa ctt atg act taa cta act tat gac tta act aac tta tga cta tta act cat ttc aag aaa ggt ggg tgg ata gaa ctc tat atg aca gct tgt gaa aca att aga tcc cta att tct aat gga agt ttt gat agg aga ttg tca tca gtt gat aca ttg ttt att atc tca tct att aga gca cgt ctg ttt aga gct tta gtg acc tgc tcg gtt act tct gtg taa atc ttg aat cct tta gtg ata cac tgt gtc aaa act gga tgt tta gaa tac cta tgt aga ata tgg gaa gca tgc ttg ttt ttg tct cta tta tag att aac tca tac atg gtt gta tta tga att ttc atc tgc cta atg tac tcc aat tct tgt tta caa tca att ata taa tca aag agt gat gat gca tac aca tta caa agt gaa taa tct acc atc ata aaa tac ttg ata cag agc ttt atc aca tca tgg ttt tca att gta tta tta agt ata gct aat ttt ata cag tca ata gac aat ggt tct cta agc aat att tct aat att tta aga tgt gct tcc cta cgg gcg atg aca gat ccc cta tcc acg gcc acg tca aga cat gta tat cca tta ctc att act gcg ttg aca ttt gct cca ttt tct aat agc cat gat act aaa tct ata taa cct gca tag ata gcg cga taa agc aag gtc ctt cca cca gca tct agt tga ttg ata tct tca ata tat ggg ata caa agc tta taa att tct aat act gtg ggt tca tct aca agg aat ccc cta gta tac tga att att tta tat aga tct aat tta aca tca ttt tca tct ggg ata cca caa ttc aaa ata aac tca aca aca cta ctt tcc ttt tta cat att ccc cta aaa tag gca ttc aag cat tct att tta tat att aca gcc cca tga tct acc ata aga tca aca atg tct att tct aca tat gca tta gat aga tag taa agt aag aga tct tgc aca gaa tta caa ttc tta ata att ata gag aaa ata tct tcc ata taa ttc ttt gac act aat gca gat ata ata tct tta tat gta ata tat gca aac agt cta tct act ata tac tga tca ata tta tct cta tga atc cta aaa taa tca tac aga aca tct aca gga tca caa att ggt tca agg aga aat cta tca aat att ttc ctg tca aca act ggt tct aga aca taa cag tca aca cct aat cca tgt ttt tta taa tca tct acc aaa gat aat gac caa aga tcg agg tcg tcg tga aac tgc tca tcg aca gcc atg aaa tct acc gac tcc atg gtg cga atc gca ctg tct tat tcg cca ttg att ttc a-t ttt tta taa tta tgt aca tgt ttt cct tct att ctc aag agt cta caa aaa tat a-- -tt ttt tcg ata tct aag tac taa gtt ttt tta ctg ttt ttg tta ctg tct tcc att ctt cta act aaa gat ctg aga taa att ata caa tct tcg cta tcg aac cat ttt tgt agt cta aag cct gaa gta att aac caa ctg ttt tta tta gtg gct ttt ttc gat cta tcc tcg tc- --- ctc tca tca tcc tta ta- --- --- --- --- --- --- tta tta tca tta tta tca tag tct att aaa cac aaa tca tct --- --- --- --- --- --- --- --- --- --- --- --- --- --- --- --- --- --- --- --- --- --- --- --- --- --- --a cgt tta taa c-- --- --- --- --- --- --- --- --a aca ttc --- --- --- --- -tc att att aat tag ttc tgt ag- aat atc ttt aat aat ttg gct ata --- --- -ca tct gtt --- --- --- --- --- --- --- --- --- --- --- --- --- --- --- --- --- --- --c aat act --- --- --- --- --- --- --- --a tct att gat gat ttc ttt t-- --- --- --- --- --- --- --- --- --- --- --t taa gac t-- --- --- --- --- --- --- --- --- --- --- --- --- --- --- --- --- --- --- --- --- --- --- --- --- --- --- --- --- --- --- --- --- --- --- --- --- --- --- --- taa act agt --- --- --- --- --- --- --- --- --- --- --- --- --- --- --- --- --- --- --- --- --- --- --- --- --- --- --- --- --- --- --- --- --- --- --- --- --- --- --- --- --- --- --- --- --- --- --- --- --- --- --- --- --- --- --- --- --- --- --- --- --- --- --- --- --- --- --- --- --- --- --- --- --- --- --- --- --- --- --- --- --- --- --- --- --- --- --- --- --- --- --- --- --- --- --- --- --- --- --- --- --- --- --- --- --- --- --- --- --- --t atg gta atg acg atg aaa --- -tc gag tag ta- --- --a ctt cta ata aag act tga ta- --- --t cat tat cat atg ttt gat cg- --- --- --- --- --- --- --t cat agt taa tag tgt g-- --- --- --- --- --- --- --- --- --- --- --- --- --- --- --- --- --- --- --- -gc taa atg gta ctg tta ata agt tta t-- --- --- --- --- --- --- --- --- --- --- --- --- --- --- --- --- --a gac aat atc ata gta ttt tct ttc cag aat --- --- --- --t aga tta ttt ttt taa ata ctg atc ctc aca att ccg tga tgt agc agt agt tgg t-- --- --- --- --- --- --- --- --- --- --- --- --- --- --- --- --- --- --- --- --- --- --- --- --- --- --- --- -gc atg gtc tat atc gt- --- --- --- --- --- --- --- --- --- --- --- --- --- --- --- --- --- --- --- --- --- --- --- --- --- --- --- --- --- --- --- --- --- --- --- --- --- --- --- --- --- --- --- taa aat gta tca tat ata ata gtt ttc tga cgt gga gta cag aat ttt cga --- --- --- --- --- --- --- --- --- --- --- --- --- --- --- --- --- --- --- --- --- --- --- --- --- --- --- --- --- --- --- --- --- --- --- --- --- --- --- --- --- --- --- --t taa tga gtt cat ggt aag gaa ggg caa atg cct --- gta tat aat ata cat aag tta a-- --- --- --- --- --- --- --- --- --- --- --- --- --t agt ttt tta tca tat tt- --- --- --- --- --- --- --- --t cta ata cca taa taa aaa t-- --- --- --- --- --- --- --- --- --- --- --- --- --- --- --- --- --- --- --- --- --- --- --- --- --t atc att att gcg ttt g-g tag tt- --- --- --- --- ctg ccc ta- --- --- --- --- --- --- --- --t cat cta tat cac tgt cac tct c-- --- --g ctc tca cta tat ctt cta aaa tta caa --- -ac aac --- --- --- --- --- --- --- --- --- --- --- --- --- --- --- --- --- --- --- --- --- --- --- --- --- --- --- --- --- --- --- --- --- --- --- --- tgg ata ttc gat --- --- --- --- --- --- --- --- --- --- --- --- --- --- --- --- --- --- --- --- --- --- --- --- --- --- --a aca gca ttt gtg t-- --- --- --- --- --- --- --- --- --- --- --- --- --- --- --- --- -

>ON676705.1_Monkeypox_virus_isolate_MPXV_USA_2022_UT001_complete_genome

--- --- --- --- --- --- --- --- --- --- --- --- --- --- --- --- --- --- --- --- --- --- --- --- --- --- --- --- --- --a ttt tac tat ttt att tag tgt cta gaa aaa aat gtg tga ccc acg acc gta gga aac tct aga ggg taa gaa aaa tca atc gtt tat aga gac cat cag aaa gag gtt taa tat ttt tgt gag acc tat cga a-- --- --- --- --- --- --- --- --- --- --- --- --- --- --- --- --- --- --- --- --- --- --- --- --- --- --- --- --- --- --- --- --- --- --- --- --- --- --- --- --- --- --- --- --- --- --- --- --- --- --- --- --- --- --- --- --- --- --- --- --- --- --- --- --- --- --- --- --- --- --- --- --- --- --- --- --- --- --- --- --- --- --- --- --- --- --- --- --- --- --- --- --- --- --- --- --- --- --- --- --- --- --- --- --- --- --- --- --- --- --- --- --- --- --- --- --- --- --- --- --- --- --- --- --- --- --- --- --- --- --- --- --- --- --- --- --- --- --- --- --- --- --- --- --- --- --- --- --- --- --- --- --- --- --- --- --- --- --- --- --- --- --- --- --- --- --- --- --- --- --- --- --- --- --- --- --- --- --- --- --- --- --- --- --- --- --- --- --- --- --- --- --- --- --- --- --- --- --- --- --- --- --- --- --- --- --- --- --- --- --- --- --- --- --- --- --- --- --- --- --- --- --- --- --- --- --- --- --- --- --- --- --- --- --- --- --- --- --- --- --- --- --- --- --- --- --- --- --- --- --- --- --- --- --- --- --- --- --- --- --- --- --- --- --- --- --- --- --- --- --- --- --- --- --- --- --- --- --- --- -ga gag aaa gga taa aaa ctt ttt acg act cca tca gaa aga ggt tta ata ttt ttg tga gac cca tcg aag aga gaa aga gat ggt tag tca aga tat ttt tct tag tac aaa agt caa tgt ttt aaa ata tat gga cga gaa tta att tgt ctg tat aaa aac ttg tgt gaa att atg tac tag aga aaa aac gtg agc agt gtc ccc tac atg gat ttt aca gat cat tta tat tcc aaa aat att aac tat ata cgt tta tta tat gat gtt aac gtg taa att ata aac att att tta tga tgc aat tgt ctg aca acc tag att ggt ata agg atg ttg ata agc tct acg aga ata tat tgt tgg acg tta tcg ttt acg aaa tag ttg aga cat cag aaa gag gtt taa tat ttt tgt gag a-c cat cga aga gag aaa gag aat aaa aat a-- -?t ttt ttt ttt ttt tgt aaa act ttt tta tga gac --- --- -ca aga gaa tac gaa tag tga tca tat cgt atc aca tat tga aac aga aag aag aag taa cga gag gta act ttt tgt gaa tgt agt taa ata ttt ttg ttt tgc aaa ccg gaa tat agt gcc cgg t-- -ct ttt tta att cgt ggt gcg gtg tct gaa tcg ttc gat taa ccc aac tca tcc att ttc aga tga ata gag tta tcg att cag aca cat gct ttg agt ttt gtt gaa tcg atg agt gaa gta tca tcg gtt gca cct tca gat gcc gat ccg tcg aca tac ttg aat cca tcc ttg act tca agt tca gat gat tcc tca cac atg tct ccg ata cgt acg cta aac tct agg ttc ttg aca cat ttt gta tca acg atc gtt gaa ccg atg ata tct ttg taa ctc act ttc tta tgt gag atg tta gac cca agt act gga tgg gtc ttg atg tca ctg tct ttc tct tct tcg cta cat ctg atg tcg ata gac atc tca cag tct ttg atc ata gcc aga gct tct tca cgc gtg atc gcg gga gag tcc tta cct tgt ccc ggt gac acg ctg gac aat cta gta ttc aca gtg ttt cca tca gag gat tcg gag atg gat aaa atc ttt ggg cat ttg gtg aat cca aag ttc atg tta aga ccc gca ccg acg ata gtg taa taa gtg gtg gga tct cct ttt aca act --- --- --- --- --- --- tct tcg gat acc tca tca tct tcg gtc tct gta act tcc gtt acg gat tga caa atc tta tca ttg gtc ggt gtt tgg tct tgc ttt gtg act ttg ata ata aca tcg att ccc ata tga tgt ttg ttt tct tct tca gta cac gag gat gaa gat tgt tga aga cta gta ggc ata gca gct gcc act agg cac atg cat gcc agg aca ata tat tgt ttc atg att gct att gat tga tta ctg ttc tag atg att cta ctt tct tac cat ata ata aat tag aat ata ttt tct act ttt acg aga aat taa tta ttg tat tta tta ttt ata ggt --a aaa aaa ctt act ata agt ggg tgg gat tct ggg aat tag tga tca gtt tat gta tat cgc aac tag cgg gca tat ggc tat tga cat cga gaa cat tac cca tat gat aag aga ttg tat cat ttt cgt agt ctt gag tat tgg tat tac tat ata gta tgt aga tgt cga cgc tag ata gac agt cgc cca cta gag tta ccg tct ctg aat gcg gca tga tag tat cat tct ttg ttt tcg tta act gtt tgg aag atg aat ctt tgt tgt tac att taa tct cga aat tca gag tac ata tct ttg aag tat tct gat atc tat ttt ctc ctg taa aga atc ctg aag ttg cta cat tat taa gga cag aga agt att ctg cac gaa aga ctg gat cac aat ctt tat gat tca tgg taa tag tta gtt ccg acg ttg aga tgg att cgc tga gac cgg tag tgg tcg tcc gag tac acg atg tgt cgt tga ctg gat aca ggt taa ttt cca cat cga tat agt taa atg tat tgc tgg tta cga cgg gtt cgc att tat ctg tgg aag aga cgg tgt gag aat atg ttc cgg gac cac acg gag aac aga tga cgt ctc cgg tag acg tgt atc cgg ata ctc cgt atc cta ttc cac act ttg ttt tag aaa tac atg ttc tac acc ctg atg ctc ctt tga gaa gac aat aat atc ctg gag agc att cac aga ttc tat tgt gag tcg tgt tac acg atc gcg tct cta cct gat tac tat cac atc ttc cgt tac aac tta gac aag cct gta aat gat tat tgt gag atg taa agg tat ccg aac cac acg gtg tac att gtg tat tag tct tgc tat cac ata atc tga aag cgt aag ttc ccg gag gac acg ata gac aac ata gat tac ggc ttc tgt att cgt tgt ctt tac act ttc cat tgg atg gtg cat gtg gtg cta tat ctc ttc cgt tta tta tta tac atg aga gaa aca ata tat acg agt ata ata cgg acc tca tga ttt aat aat -gt agt aat cgt cgt ctt gtt act gtt tgt ttc cta ctt ctc caa tca tat aga tta ttt ttt aaa tat ttt ctt tct atc atg gat aat att tgt aat ggt tct ttc cgt aca aca tac tgt tta gat ggt agt cgc tta gct tgg tta tga tat tgc gca taa ttt ccg gag gca aat acg ata gtc tag att gac tat cga tgg tag act cta att tat tga gtg ctt tgt cga cga gtt tac ttt tat gct cca tcg ata gat gac act gtt cta tga gat cgt cgt aca tgg gaa atg aaa tgc gtt tgt ctg aat gta tgg ctt cga gat atg tgt gat acc gaa tgt ctt ctg ttc tca ata ccg tat aca agt cgg tgt ctg aga ttc gaa tct ctt tga gga gac tta tgt cac gac tac att ttt cga tga tgg aat cta tct tat cga atg ata tat ttt tca taa ata cac ttt tat agt cct cgt tta aac aga att tag tat gta gtt ccg caa atg act cgt ccc tta ata ggc agt agg cta tta tct tct tta cgt agt gat cgt cgt agg gag aga act ccg aca tct tgt aga aca acg att taa tca tag gta gag ata ctt tca gtc tgt ggt gga taa tgt cat taa cga cat ccg cct tgt ata taa tgt ttc tgt ttt caa aca cca agt cga ata ctg tct tat cgt ctt tag tcg gaa ggt tga tgt cgt atc caa tgt ata cga ggt atg agg caa cat tgt tat tgc aat tct gga agg cgg tat gaa gag gag tca ttg tat tat agt att cgt ctt tct gaa tgt cga atc tat cta gta gat acc gta gta tat tga gag agc gac ttc cat atc ctt gat tat gtt tta tga ata gat aaa gta gat gtt gtc ctt ctt cct ttt gta att tcc cgt att ttt gtt cgt gcc aat tga gta aca tta tga gaa tat gac ctg ttg cac aat cgt tct tta tgt att cca tga tgg gtg tac aat taa gat tat tac gta tcc tcg tat cgg ctc ctc gag ata aaa gag cat aca cca cac gag gac tat gtt tgg tat act gtt gaa ggt aag tgt gta acg gcg tat ttc cga ttt tcg taa ccg cgt taa tgt ttg ctc cat gat cta tta tcg cgt aga tga atc gct tct cag ctc gca tct tag tgt gac tct ttg act tgt aat aat tgc ttt cgt gga acg cgg ata tgt gtt tac agt agt aat gaa gag aag tga gtc cat cct cat cga cgc aat tag ggt cag atc ctt tag tca ata att tgt aca gaa cgt aat agt tta agc tcc cat tga att tat atc taa gat aac aca gca ata gat cgg atg att tac taa agt cat caa tgg ggt ccg tta gta tat caa aga tct tgt tat cga ttg ata gtg aat gaa tca gat agt ggt gta gag gaa tat gtc ctt ttt tat cct tgc tat caa agt tac gca tgc cgt ggc gta aca ata tct tta ata cag atg gat taa atc gtg tat tca tcg tat agc aat gta atg gag agt tac cac att tta gtc gtt tat tca gat cgc agt gtt taa taa cta att taa aca gat gag atg ata tat cca cat caa aga atg cga gat aca tat gac aga cat tat tga cag aaa tgt gac ctt cat tat cac cgt cgt cca taa atg cgt tag gta cgt acc aca tac tat cgt taa cga tgc gca caa tct cgt cca -tt tca tcc atc ttc ata atg att tac ttt ttc ata att aga gaa aaa gat caa ggt ata aaa att aga agt gtt aga cta ta- --- --- --- --- --- --- --- --- --- --- -aa cta act tat gac tta act aac tta tga --- --- --- --- --- --- --- --- --- --- --- --- --- --- --- --- --- --- --- --- --- --- --- --- --- --- --- --- --- --- --- --- --- --- --- --- --- --- --- --- --- --- --- --- --- --- --- --- ctt aac ta- --- --- --- --- --- --- --- --- --- --- -ac tta tga ctt aac taa ctt atg act taa cta act ta- tga ctt aac taa ctt atg act taa cta act tat gac tta act aac tta tga cta tta act cat ttc aag aaa ggt ggg tgg ata gaa ctc tat atg aca gct tgt gaa aca att aga tcc cta att tct aat gga agt ttt gat agg aga ttg tca tca gtt gat aca ttg ttt att atc tca tct att aga gca cgt ctg ttt aga gct tta gtg acc tgc tcg gtt act tct gtg taa atc ttg aat cct tta gtg ata cac tgt gtc aaa act gga tgt tta gaa tac cta tgt aga ata tgg gaa gca tgc ttg ttt ttg tct cta tta tag att aac tca tac atg gtt gta tta tga att ttc atc tgc cta atg tac tcc aat tct tgt tta caa tca att ata taa tca aag agt gat gat gca tac aca tta caa agt gaa taa tct acc atc ata aaa tac ttg ata cag agc ttt atc aca tca tgg ttt tca att gta tta tta agt ata gct aat ttt ata cag tca ata gac aat ggt tct cta agc aat att tct aat att tta aga tgt gct tcc cta cgg gcg atg aca gat ccc cta tcc acg gcc acg tca aga cat gta tat cca tta ctc att act gcg ttg aca ttt gct cca ttt tct aat agc cat gat act aaa tct ata taa cct gca tag ata gcg cga taa agc aag gtc ctt cca cca gca tct agt tga ttg ata tct tca ata tat ggg ata caa agc tta taa att tct aat act gtg ggt tca tct aca agg aat ccc cta gta tac tga att att tta tat aga tct aat tta aca tca ttt tca tct ggg ata cca caa ttc aaa ata aac tca aca aca cta ctt tcc ttt tta cat att ccc cta aaa tag gca ttc aag cat tct att tta tat att aca gcc cca tga tct acc ata aga tca aca atg tct att tct aca tat gca tta gat aga tag taa agt aag aga tct tgc aca gaa tta caa ttc tta ata att ata gag aaa ata tct tcc ata taa ttc ttt gac act aat gca gat ata ata tct tta tat gta ata tat gca aac agt cta tct act ata tac tga tca ata tta tct cta tga atc cta aaa taa tca tac aga aca tct aca gga tca caa att ggt tca agg aga aat cta tca aat att ttc ctg tca aca act ggt tct aga aca taa cag tca aca cct aat cca tgt ttt tta taa tca tct acc aaa gat aat gac caa aga tcg agg tcg tcg tga aac tgc tca tcg aca gcc atg aaa tct acc gac tcc atg gtg cga atc gca ctg tct tat tcg cca ttg att ttc a-t ttt tta taa tta tgt aca tgt ttt cct tct att ctc aag agt cta caa aaa tat a-- -tt ttt tcg ata tct aag tac taa gtt ttt tta ctg ttt ttg tta ctg tct tcc att ctt cta act aaa gat ctg aga taa att ata caa tct tcg cta tcg aac cat ttt tgt agt cta aag cct gaa gta att aac caa ctg ttt tta tta gtg gct ttt ttc gat cta tcc tcg tc- --- ctc tca tca tcc tta ta- --- --- --- --- --- --- tta tta tca tta tta tca tag tct att aaa cac aaa tca tct --- --- --- --- --- --- --- --- --- --- --- --- --- --- --- --- --- --- --- --- --- --- --- --- --- --- --a cgt tta taa c-- --- --- --- --- --- --- --- --a aca ttc --- --- --- --- -tc att att aat tag ttc tgt ag- aat atc ttt aat aat ttg gct ata --- --- -ca tct gtt --- --- --- --- --- --- --- --- --- --- --- --- --- --- --- --- --- --- --c aat act --- --- --- --- --- --- --- --a tct att gat gat ttc ttt t-- --- --- --- --- --- --- --- --- --- --- --t taa gac t-- --- --- --- --- --- --- --- --- --- --- --- --- --- --- --- --- --- --- --- --- --- --- --- --- --- --- --- --- --- --- --- --- --- --- --- --- --- --- --- taa act agt --- --- --- --- --- --- --- --- --- --- --- --- --- --- --- --- --- --- --- --- --- --- --- --- --- --- --- --- --- --- --- --- --- --- --- --- --- --- --- --- --- --- --- --- --- --- --- --- --- --- --- --- --- --- --- --- --- --- --- --- --- --- --- --- --- --- --- --- --- --- --- --- --- --- --- --- --- --- --- --- --- --- --- --- --- --- --- --- --- --- --- --- --- --- --- --- --- --- --- --- --- --- --- --- --- --- --- --- --- --t atg gta atg acg atg aaa --- -tc gag tag ta- --- --a ctt cta ata aag act tga ta- --- --t cat tat cat atg ttt gat cg- --- --- --- --- --- --- --t cat agt taa tag tgt g-- --- --- --- --- --- --- --- --- --- --- --- --- --- --- --- --- --- --- --- -gc taa atg gta ctg tta ata agt tta t-- --- --- --- --- --- --- --- --- --- --- --- --- --- --- --- --- --a gac aat atc ata gta ttt tct ttc cag aat --- --- --- --t aga tta ttt ttt taa ata ctg atc ctc aca att ccg tga tgt agc agt agt tgg t-- --- --- --- --- --- --- --- --- --- --- --- --- --- --- --- --- --- --- --- --- --- --- --- --- --- --- --- -gc atg gtc tat atc gt- --- --- --- --- --- --- --- --- --- --- --- --- --- --- --- --- --- --- --- --- --- --- --- --- --- --- --- --- --- --- --- --- --- --- --- --- --- --- --- --- --- --- --- taa aat gta tca tat ata ata gtt ttc tga cgt gga gta cag aat ttt cga --- --- --- --- --- --- --- --- --- --- --- --- --- --- --- --- --- --- --- --- --- --- --- --- --- --- --- --- --- --- --- --- --- --- --- --- --- --- --- --- --- --- --- --t taa tga gtt cat ggt aag gaa ggg caa atg cct --- gta tat aat ata cat aag tta a-- --- --- --- --- --- --- --- --- --- --- --- --- --t agt ttt tta tca tat tt- --- --- --- --- --- --- --- --t cta ata cca taa taa aaa t-- --- --- --- --- --- --- --- --- --- --- --- --- --- --- --- --- --- --- --- --- --- --- --- --- --t atc att att gcg ttt g-g tag tt- --- --- --- --- ctg ccc ta- --- --- --- --- --- --- --- --t cat cta tat cac tgt cac tct c-- --- --g ctc tca cta tat ctt cta aaa tta caa --- -ac aac --- --- --- --- --- --- --- --- --- --- --- --- --- --- --- --- --- --- --- --- --- --- --- --- --- --- --- --- --- --- --- --- --- --- --- --- tgg ata ttc gat --- --- --- --- --- --- --- --- --- --- --- --- --- --- --- --- --- --- --- --- --- --- --- --- --- --- --a aca gca ttt gtg t-- --- --- --- --- --- --- --- --- --- --- --- --- --- --- --- --- -

>MT903343.1_Monkeypox_virus_isolate_MPXV-UK_P1

--- --- --- --- --- --- --- --- --- --- --- --- --- --- --- --- --- --- --- --- --- --- --- --- --- --- --- --t att ata ttt tac tat ttt att tag tgt cta gaa aaa aat gtg tga ccc acg acc gta gga aac tct aga ggg taa gaa aaa tca atc gtt tat aga gac cat cag aaa gag gtt taa tat ttt tgt gag acc tat cga a-- --- --- --- --- --- --- --- --- --- --- --- --- --- --- --- --- --- --- --- --- --- --- --- --- --- --- --- --- --- --- --- --- --- --- --- --- --- --- --- --- --- --- --- --- --- --- --- --- --- --- --- --- --- --- --- --- --- --- --- --- --- --- --- --- --- --- --- --- --- --- --- --- --- --- --- --- --- --- --- --- --- --- --- --- --- --- --- --- --- --- --- --- --- --- --- --- --- --- --- --- --- --- --- --- --- --- --- --- --- --- --- --- --- --- --- --- --- --- --- --- --- --- --- --- --- --- --- --- --- --- --- --- --- --- --- --- --- --- --- --- --- --- --- --- --- --- --- --- --- --- --- --- --- --- --- --- --- --- --- --- --- --- --- --- --- --- --- --- --- --- --- --- --- --- --- --- --- --- --- --- --- --- --- --- --- --- --- --- --- --- --- --- --- --- --- --- --- --- --- --- --- --- --- --- --- --- --- --- --- --- --- --- --- --- --- --- --- --- --- --- --- --- --- --- --- --- --- --- --- --- --- --- --- --- --- --- --- --- --- --- --- --- --- --- --- --- --- --- --- --- --- --- --- --- --- --- --- --- --- --- --- --- --- --- --- --- --- --- --- --- --- --- --- --- --- --- --- --- --- -ga gag aaa gga taa aaa ctt ttt acg act cca tca gaa aga ggt tta ata ttt ttg tga gac cca tcg aag aga gaa aga gat ggt tag tca aga tat ttt tct tag tac aaa agt caa tgt ttt aaa ata tat gga cga gaa tta att tgt ctg tat aaa aac ttg tgt gaa att atg tac tag aga aaa aac gtg agc agt gtc ccc tac atg gat ttt aca gat cat tta tat tcc aaa aat att aac tat ata cgt tta tta tat gat gtt aac gtg taa att ata aac att att tta tga tgc aat tgt ctg aca acc tag att ggt ata agg atg ttg ata agc tct acg aga ata tat tgt tgg acg tta tcg ttt acg aaa tag ttg aga cat cag aaa gag gtt taa tat ttt tgt gag a-c cat cga aga gag aaa gag aat aaa aat a-- ttt ttt ttt ttt ttt tgt aaa act ttt tta tga gac --- --- -ca aga gaa tac gaa tag tga tca tat cgt atc aca tat tga aac aga aag aag aag taa cga gag gta act ttt tgt gaa tgt agt taa ata ttt ttg ttt tgc aaa ccg gaa tat agt gcc cgg t-- -ct ttt tta att cgt ggt gcg gtg tct gaa tcg ttc gat taa ccc aac tca tcc att ttc aga tga ata gag tta tcg att cag aca cat gct ttg agt ttt gtt gaa tcg atg agt gaa gta tca tcg gtt gca cct tca gat gcc gat ccg tcg aca tac ttg aat cca tcc ttg act tca agt tca gat gat tcc tca cac atg tct ccg ata cgt acg cta aac tct agg ttc ttg aca cat ttt gta tca acg atc gtt gaa ccg atg ata tct ttg taa ctc act ttc tta tgt gag atg tta gac cca agt act gga tgg gtc ttg atg tca ctg tct ttc tct tct tcg cta cat ctg atg tcg ata gac atc tca cag tct ttg atc ata gcc aga gct tct tca cgc gtg atc gcg gga gag tcc tta cct tgt ccc ggt gac acg ctg gac aat cta gta ttc aca gtg ttt cca tca gag gat tcg gag atg gat gaa atc ttt ggg cat ttg gtg aat cca aag ttc atg tta aga ccc gca ccg acg ata gtg taa taa gtg gtg gga tct cct ttt aca act --- --- --- --- --- --- tct tcg gat acc tca tca tct tcg gtc tct gta act tcc gtt acg gat tga caa atc tta tca ttg gtc ggt gtt tgg tct tgc ttt gtg act ttg ata ata aca tcg att ccc ata tga tgt ttg ttt tct tct tca gta cac gag gat gaa gat tgt tga aga cta gta ggc ata gca gct gcc act agg cac atg cat gcc agg aca ata tat tgt ttc atg att gct att gat tga tta ctg ttc tag atg att cta ctt tct tac cat ata ata aat tag aat ata ttt tct act ttt acg aga aat taa tta ttg tat tta tta ttt ata ggt --a aaa aaa ctt act ata agt ggg tgg gat tct ggg aat tag tga tca gtt tat gta tat cgc aac tag cgg gca tat ggc tat tga cat cga gaa cat tac cca tat gat aag aga ttg tat cat ttt cgt agt ctt gag tat tgg tat tac tat ata gta tgt aga tgt cga cgc tag ata gac agt cgc cca cta gag tta ccg tct ctg aat gcg gca tga tag tat cat tct ttg ttt tcg tta act gtt tgg aag atg aat ctt tgt tgt tac att taa tct cga aat tca gag tac ata tct ttg aag tat tct gat atc tat ttt ctc ctg taa aga atc ctg aag ttg cta cat tat taa gga cag aga agt att ctg cac gaa aga ctg gat cac aat ctt tat gat tca tgg taa tag tta gtt ccg acg ttg aga tgg att cgc tga gac cgg tag tgg tcg tcc gag tac acg atg tgt cgt tga ctg gat aca ggt taa ttt cca cat cga tat agt taa atg tat tgc tgg tta cga cgg gtt cgc att tat ctg tgg aag aga cgg tgt gag aat atg ttc cgg gac cac acg gag aac aga tga cgt ctc cgg tag acg tgt atc cgg ata ctc cgt atc cta ttc cac act ttg ttt tag aaa tac atg ttc tac acc ctg atg ctc ctt tga gaa gac aat aat atc ctg gag agc att cac aga ttc tat tgt gag tcg tgt tac acg atc gcg tct cta cct gat tac tat cac atc ttc cgt tac aac tta gac aag cct gta aat gat tat tgt gag atg taa agg tat ccg aac cac acg gtg tac att gtg tat tag tct tgc tat cac ata atc tgg aag cgt aag ttc ccg gag gac acg ata gac aac ata gat tac ggc ttc tgt att cgt tgt ctt tac act ttc cat tgg atg gtg cat gtg gtg cta tat ctc ttc cgt tta tta tta tac atg aga gaa aca ata tat acg agt ata ata cgg acc tca tga ttt aat aat -gt agt aat cgt cgt ctt gtt act gtt tgt ttc cta ctt ctc caa tca tat aga tta ttt ttt aaa tat ttt ctt tct atc atg gat aat att tgt aat ggt tct ttc cgt aca aca tac tgt tta gat ggt agt cgc tta gct tgg tta tga tat tgc gca taa ttt ccg gag gca aat acg ata gtc tag att gac tat cga tgg tag act cta att tat tga gtg ctt tgt cga cga gtt tac ttt tat gct cca tcg ata gat gac act gtt cta tga gat cgt cgt aca tgg gaa atg aaa tgc gtt tgt ctg aat gta tgg ctt cga gat atg tgt gat acc gga tgt ctt ctg ttc tca ata ccg tat aca agt cgg tgt ctg aga ttc gaa tct ctt tga gga gac tta tgt cac gac tac att ttt cga tga tgg aat cta tct tat cga atg ata tat ttt tca taa ata cac ttt tat agt cct cgt tta aac aga att tag tat gta gtt ccg caa atg act cgt ccc tta ata ggc agt agg cta tta tct tct tta cgt agt gat cgt cgt agg gag aga act ccg aca tct tgt aga aca acg att taa tca tag gta gag ata ctt tca gtc tgt ggt gga taa tgt cat taa cga cat ccg cct tgt ata taa tgt ttc tgt ttt caa aca cca agt cga ata ctg tct tat cgt ctt tag tcg gaa ggt tga tgt cgt atc cga tgt ata cga ggt atg agg caa cat tgt tat tgc aat tct gga agg cgg tat gaa gag gag tca ttg tat tat agt att cgt ctt tct gaa tgt cga atc tat cta gta gat acc gta gta tat tga gag agc gac ttc cat atc ctt gat tat gtt tta tga ata gat aaa gta gat gtt gtc ctt ctt cct ttt gta att tcc cgt att ttt gtt cgt gcc aat tga gta aca tta tga gaa tat gac ctg ttg cac aat cgt tct tta tgt att cca tga tgg gtg tac aat caa gat tat tac gta tcc tcg tat cgg ctc ctc gag ata aaa gag cat aca cca cac gag gac tat gtt tgg tat act gtt gaa ggt aag tgt gta acg gcg tat ttc cga ttt tcg taa ccg cgt taa tgt ttg ctc cat gat cta tta tcg cgt aga tga atc gct tct cag ctc gca tct tag tgt gac tct ttg act tgt aat aat tgc ttt cgt gga acg cgg ata tgt gtt tac agt agt aat gaa gag aag tga gtc cat cct cat cga cgc aat tag ggt cag atc ctt tag tca ata att tgt aca gaa cgt aat agt tta agc tcc cat tga att tat atc taa gat aac aca gca ata gat cgg atg att tac taa agt cat caa tgg ggt ccg tta gta tat caa aga tct tgt tat cga ttg ata gtg aat gaa tca gat agt ggt gta gag gaa tat gtc ctt ttt tat cct tgc tat caa agt tac gca tgc cgt ggc gta aca ata tct tta ata cag atg gat taa atc gtg tat tca tcg tat agc aat gta atg gag agt tac cac att tta gtc gtt tat tca gat cgc agt gtt taa taa cta att taa aca gat gag atg ata tat cca cat caa aga atg cga gat aca tat gac aga cat tat tga cag aaa tgt gac ctt cat tat cac cgt cgt cca taa atg cgt tag gta cgt acc aca tac tat cgt taa cga tgc gca caa tct cgt cca -tt tca tcc atc ttc ata atg att tac ttt ttc ata att aga gaa aaa gat caa ggt ata aaa att aga agt gtt aga cta ta- --- --- --- --- --- --- --- --- --- --- -aa cta act tat gac tta act aac tta tga --- --- --- --- --- --- --- --- --- --- --- --- --- --- --- --- --- --- --- --- --- --- --- --- --- --- --- --- --- --- --- --- --- --- --- --- --- --- --- --- --- --- --- --- --- --- --- --- ctt aac ta- --- --- --- --- --- --- --- --- --- --- -ac tta tga ctt aac taa ctt atg act taa cta act ta- tga ctt aac taa ctt atg act taa cta act tat gac tta act aac tta tga cta tta act cat ttc aag aaa ggt ggg tgg ata gaa ctc tat atg aca gct tgt gaa aca att aga tcc cta att tct aat gga agt ttt gat agg aga ttg tca tca gtt gat aca ttg ttt att atc tca tct att aga gca cgt ctg ttt aga gct tta gtg acc tgc tcg gtt act tct gtg taa atc ttg aat cct tta gtg ata cac tgt gtc aaa act gga tgt tta gaa tac cta tgt aga ata tgg gaa gca tgc ttg ttt ttg tct cta tta tag att aac tca tac atg gtt gta tta tga att ttc atc tgc cta atg tac tcc aat tct tgt tta caa tca att ata taa tca aag agt gat gat gca tac aca tta caa agt gaa taa tct acc atc ata aaa tac ttg ata cag agc ttt atc aca tca tgg ttt tca att gta tta tta agt ata gct aat ttt ata cag tca ata gac aat ggt tct cta agc aat att tct aat att tta aga tgt gct tcc cta cgg gcg atg aca gat ccc cta tcc acg gcc acg tca aga cat gta tat cca tta ctc att act gcg ttg aca ttt gct cca ttt tct aat agc cat gat act aaa tct ata taa cct gca tag ata gcg cga taa agc aag gtc ctt cca cca gca tct agt tga ttg ata tct tca ata tat ggg ata caa agc tta taa att tct aat act gtg ggt tca tct aca agg aat ccc cta gta tac tga att att tta tat aga tct aat tta aca tca ttt tca tct ggg ata cca caa ttc aaa ata aac tca aca aca cta ctt tcc ttt tta cat att ccc cta aaa tag gca ttc aag cat tct att tta tat att aca gcc cca tga tct acc ata aga tca aca atg tct att tct aca tat gca tta gat aga tag taa agt aag aga tct tgc aca gaa tta caa ttc tta ata att ata gag aaa ata tct tcc ata taa ttc ttt gac act aat gca gat ata ata tct tta tat gta ata tat gca aac agt cta tct act ata tac tga tca ata tta tct cta tga atc cta aaa taa tca tac aga aca tct aca gga tca caa att ggt tca agg aga aat cta tca aat att ttc ctg tca aca act ggt tct aga aca taa cag tca aca cct aat cca tgt ttt tta taa tca tct acc aaa gat aat gac caa aga tcg agg tcg tcg tga aac tgc tca tcg aca gcc atg aaa tct acc gac tcc atg gtg cga atc gca ctg tct tat tcg cca ttg att ttc a-t ttt tta taa tta tgt aca tgt ttt cct tct att ctc aag agt cta caa aaa tat a-- -tt ttt tcg ata tct aag tac taa gtt ttt tta ctg ttt ttg tta ctg tct tcc att ctt cta act aaa gat ctg aga taa att ata caa tct tcg cta tcg aac cat ttt tgt agt cta aag cct gaa gta att aac caa ctg ttt tta tta gtg gct ttt ttc gat cta tcc tcg tc- --- ctc tca tca tcc tta ta- --- --- --- --- --- --- tta tta tca tta tta tca tag tct att aaa cac aaa tca tct --- --- --- --- --- --- --- --- --- --- --- --- --- --- --- --- --- --- --- --- --- --- --- --- --- --- --a cgt tta taa c-- --- --- --- --- --- --- --- --a aca ttc --- --- --- --- -tc att att aat tag ttc tgt ag- aat atc ttt aat aat ttg gct ata --- --- -ca tct gtt --- --- --- --- --- --- --- --- --- --- --- --- --- --- --- --- --- --- --c aat act --- --- --- --- --- --- --- --a tct att gat gat ttc ttt t-- --- --- --- --- --- --- --- --- --- --- --t taa gac t-- --- --- --- --- --- --- --- --- --- --- --- --- --- --- --- --- --- --- --- --- --- --- --- --- --- --- --- --- --- --- --- --- --- --- --- --- --- --- --- taa act agt --- --- --- --- --- --- --- --- --- --- --- --- --- --- --- --- --- --- --- --- --- --- --- --- --- --- --- --- --- --- --- --- --- --- --- --- --- --- --- --- --- --- --- --- --- --- --- --- --- --- --- --- --- --- --- --- --- --- --- --- --- --- --- --- --- --- --- --- --- --- --- --- --- --- --- --- --- --- --- --- --- --- --- --- --- --- --- --- --- --- --- --- --- --- --- --- --- --- --- --- --- --- --- --- --- --- --- --- --- --t atg gta atg acg atg aaa --- -tc gag tag ta- --- --a ctt cta ata aag act tga ta- --- --t cat tat cat atg ttt gat cg- --- --- --- --- --- --- --t cat agt taa tag tgt g-- --- --- --- --- --- --- --- --- --- --- --- --- --- --- --- --- --- --- --- -gc taa atg gta ctg tta ata agt tta t-- --- --- --- --- --- --- --- --- --- --- --- --- --- --- --- --- --a gac aat atc ata gta ttt tct ttc cag aat --- --- --- --t aga tta ttt ttt taa ata ctg atc ctc aca att ccg tga tgt agc agt agt tgg t-- --- --- --- --- --- --- --- --- --- --- --- --- --- --- --- --- --- --- --- --- --- --- --- --- --- --- --- -gc atg gtc tat atc gt- --- --- --- --- --- --- --- --- --- --- --- --- --- --- --- --- --- --- --- --- --- --- --- --- --- --- --- --- --- --- --- --- --- --- --- --- --- --- --- --- --- --- --- taa aat gta tca tat ata ata gtt ttc tga cgt gga gta cag aat ttt cga --- --- --- --- --- --- --- --- --- --- --- --- --- --- --- --- --- --- --- --- --- --- --- --- --- --- --- --- --- --- --- --- --- --- --- --- --- --- --- --- --- --- --- --t taa tga gtt cat ggt aag gaa ggg caa atg cct --- gta tat aat ata cat aag tta a-- --- --- --- --- --- --- --- --- --- --- --- --- --t agt ttt tta tca tat tt- --- --- --- --- --- --- --- --t cta ata cca taa taa aaa t-- --- --- --- --- --- --- --- --- --- --- --- --- --- --- --- --- --- --- --- --- --- --- --- --- --t atc att att gcg ttt g-g tag tt- --- --- --- --- ctg ccc ta- --- --- --- --- --- --- --- --t cat cta tat cac tgt cac tct c-- --- --g ctc tca cta tat ctt cta aaa tta caa --- -ac aac --- --- --- --- --- --- --- --- --- --- --- --- --- --- --- --- --- --- --- --- --- --- --- --- --- --- --- --- --- --- --- --- --- --- --- --- tgg ata ttc gat --- --- --- --- --- --- --- --- --- --- --- --- --- --- --- --- --- --- --- --- --- --- --- --- --- --- --a aca gca ttt gtg t-- --- --- --- --- --- --- --- --- --- --- --- --- --- --- --- --- -

>ON631963.1_Monkeypox_virus_isolate_MPxV/VIDRL01/2022_complete_genome

--- --- --- --- --- --- --- --- --- --- --- --- --- --- --- --- --- --- --- --- --- -at aag ttt tag tac att aat att ata ttt tac tat ttt att tag tgt cta gaa aaa aat gtg tga ccc acg acc gta gga aac tct aga ggg taa gaa aaa tca atc gtt tat aga gac cat cag aaa gag gtt taa tat ttt tgt gag acc tat cga a-- --- --- --- --- --- --- --- --- --- --- --- --- --- --- --- --- --- --- --- --- --- --- --- --- --- --- --- --- --- --- --- --- --- --- --- --- --- --- --- --- --- --- --- --- --- --- --- --- --- --- --- --- --- --- --- --- --- --- --- --- --- --- --- --- --- --- --- --- --- --- --- --- --- --- --- --- --- --- --- --- --- --- --- --- --- --- --- --- --- --- --- --- --- --- --- --- --- --- --- --- --- --- --- --- --- --- --- --- --- --- --- --- --- --- --- --- --- --- --- --- --- --- --- --- --- --- --- --- --- --- --- --- --- --- --- --- --- --- --- --- --- --- --- --- --- --- --- --- --- --- --- --- --- --- --- --- --- --- --- --- --- --- --- --- --- --- --- --- --- --- --- --- --- --- --- --- --- --- --- --- --- --- --- --- --- --- --- --- --- --- --- --- --- --- --- --- --- --- --- --- --- --- --- --- --- --- --- --- --- --- --- --- --- --- --- --- --- --- --- --- --- --- --- --- --- --- --- --- --- --- --- --- --- --- --- --- --- --- --- --- --- --- --- --- --- --- --- --- --- --- --- --- --- --- --- --- --- --- --- --- --- --- --- --- --- --- --- --- --- --- --- --- --- --- --- --- --- --- --- -ga gag aaa gga taa aaa ctt ttt acg act cca tca gaa aga ggt tta ata ttt ttg tga gac cca tcg aag aga gaa aga gat ggt tag tca aga tat ttt tct tag tac aaa agt caa tgt ttt aaa ata tat gga cga gaa tta att tgt ctg tat aaa aac ttg tgt gaa att atg tac tag aga aaa aac gtg agc agt gtc ccc tac atg gat ttt aca gat cat tta tat tcc aaa aat att aac tat ata cgt tta tta tat gat gtt aac gtg taa att ata aac att att tta tga tgc aat tgt ctg aca acc tag att ggt ata agg atg ttg ata agc tct acg aga ata tat tgt tgg acg tta tcg ttt acg aaa tag ttg aga cat cag aaa gag gtt taa tat ttt tgt gag a-c cat cga aga gag aaa gag aat aaa aat a-- -tt ttt ttt ttt ttt tgt aaa act ttt tta tga gac --- --- -ca aga gaa tac gaa tag tga tca tat cgt atc aca tat tga aac aga aag aag aag taa cga gag gta act ttt tgt gaa tgt agt taa ata ttt ttg ttt tgc aaa ccg gaa tat agt gcc cgg t-- -ct ttt tta att cgt ggt gcg gtg tct gaa tcg ttc gat taa ccc aac tca tcc att ttc aga tga ata gag tta tcg att cag aca cat gct ttg agt ttt gtt gaa tcg atg agt gaa gta tca tcg gtt gca cct tca gat gcc gat ccg tcg aca tac ttg aat cca tcc ttg act tca agt tca gat gat tcc tca cac atg tct ccg ata cgt acg cta aac tct agg ttc ttg aca cat ttt gta tca acg atc gtt gaa ccg atg ata tct ttg taa ctc act ttc tta tgt gag atg tta gac cca agt act gga tgg gtc ttg atg tca ctg tct ttc tct tct tcg cta cat ctg atg tcg ata gac atc tca cag tct ttg atc ata gcc aga gct tct tca cgc gtg atc gcg gga gag tcc tta cct tgt ccc ggt gac acg ctg gac aat cta gta ttc aca gtg ttt cca tca gag gat tcg gag atg gat aaa atc ttt ggg cat ttg gtg aat cca aag ttc atg tta aga ccc gca ccg acg ata gtg taa taa gtg gtg gga tct cct ttt aca act --- --- --- --- --- --- tct tcg gat acc tca tca tct tcg gtc tct gta act tcc gtt acg gat tga caa atc tta tca ttg gtc ggt gtt tgg tct tgc ttt gtg act ttg ata ata aca tcg att ccc ata tga tgt ttg ttt tct tct tca gta cac gag gat gaa gat tgt tga aga cta gta ggc ata gca gct gcc act agg cac atg cat gcc agg aca ata tat tgt ttc atg att gct att gat tga tta ctg ttc tag atg att cta ctt tct tac cat ata ata aat tag aat ata ttt tct act ttt acg aga aat taa tta ttg tat tta tta ttt ata ggt --a aaa aaa ctt act ata agt ggg tgg gat tct ggg aat tag tga tca gtt tat gta tat cgc aac tag cgg gca tat ggc tat tga cat cga gaa cat tac cca tat gat aag aga ttg tat cat ttt cgt agt ctt gag tat tgg tat tac tat ata gta tgt aga tgt cga cgc tag ata gac agt cgc cca cta gag tta ccg tct ctg aat gcg gca tga tag tat cat tct ttg ttt tcg tta act gtt tgg aag atg aat ctt tgt tgt tac att taa tct cga aat tca gag tac ata tct ttg aag tat tct gat atc tat ttt ctc ctg taa aga atc ctg aag ttg cta cat tat taa gga cag aga agt att ctg cac gaa aga ctg gat cac aat ctt tat gat tca tgg taa tag tta gtt ccg acg ttg aga tgg att cgc tga gac cgg tag tgg tcg tcc gag tac acg atg tgt cgt tga ctg gat aca ggt taa ttt cca cat cga tat agt taa atg tat tgc tgg tta cga cgg gtt cgc att tat ctg tgg aag aga cgg tgt gag aat atg ttc cgg gac cac acg gag aac aga tga cgt ctc cgg tag acg tgt atc cgg ata ctc cgt atc cta ttc cac act ttg ttt tag aaa tac atg ttc tac acc ctg atg ctc ctt tga gaa gac aat aat atc ctg gag agc att cac aga ttc tat tgt gag tcg tgt tac acg atc gcg tct cta cct gat tac tat cac atc ttc cgt tac aac tta gac aag cct gta aat gat tat tgt gag atg taa agg tat ccg aac cac acg gtg tac att gtg tat tag tct tgc tat cac ata atc tga aag cgt aag ttc ccg gag gac acg ata gac aac ata gat tac ggc ttc tgt att cgt tgt ctt tac act ttc cat tgg atg gtg cat gtg gtg cta tat ctc ttc cgt tta tta tta tac atg aga gaa aca ata tat acg agt ata ata cgg acc tca tga ttt aat aat -gt agt aat cgt cgt ctt gtt act gtt tgt ttc cta ctt ctc caa tca tat aga tta ttt ttt aaa tat ttt ctt tct atc atg gat aat att tgt aat ggt tct ttc cgt aca aca tac tgt tta gat ggt agt cgc tta gct tgg tta tga tat tgc gca taa ttt ccg gag gca aat acg ata gtc tag att gac tat cga tgg tag act cta att tat tga gtg ctt tgt cga cga gtt tac ttt tat gct cca tcg ata gat gac act gtt cta tga gat cgt cgt aca tgg gaa atg aaa tgc gtt tgt ctg aat gta tgg ctt cga gat atg tgt gat acc gaa tgt ctt ctg ttc tca ata ccg tat aca agt cgg tgt ctg aga ttc gaa tct ctt tga gga gac tta tgt cac gac tac att ttt cga tga tgg aat cta tct tat cga atg ata tat ttt tca taa ata cac ttt tat agt cct cgt tta aac aga att tag tat gta gtt ccg caa atg act cgt ccc tta ata ggc agt agg cta tta tct tct tta cgt agt gat cgt cgt agg gag aga act ccg aca tct tgt aga aca acg att taa tca tag gta gag ata ctt tca gtc tgt ggt gga taa tgt cat taa cga cat ccg cct tgt ata taa tgt ttc tgt ttt caa aca cca agt cga ata ctg tct tat cgt ctt tag tcg gaa ggt tga tgt cgt atc caa tgt ata cga ggt atg agg caa cat tgt tat tgc aat tct gga agg cgg tat gaa gag gag tca ttg tat tat agt att cgt ctt tct gaa tgt cga atc tat cta gta gat acc gta gta tat tga gag agc gac ttc cat atc ctt gat tat gtt tta tga ata gat aaa gta gat gtt gtc ctt ctt cct ttt gta att tcc cgt att ttt gtt cgt gcc aat tga gta aca tta tga gaa tat gac ctg ttg cac aat cgt tct tta tgt att cca tga tgg gtg tac aat taa gat tat tac gta tcc tcg tat cgg ctc ctc gag ata aaa gag cat aca cca cac gag gac tat gtt tgg tat act gtt gaa ggt aag tgt gta acg gcg tat ttc cga ttt tcg taa ccg cgt taa tgt ttg ctc cat gat cta tta tcg cgt aga tga atc gct tct cag ctc gca tct tag tgt gac tct ttg act tgt aat aat tgc ttt cgt gga acg cgg ata tgt gtt tac agt agt aat gaa gag aag tga gtc cat cct cat cga cgc aat tag ggt cag atc ctt tag tca ata att tgt aca gaa cgt aat agt tta agc tcc cat tga att tat atc taa gat aac aca gca ata gat cgg atg att tac taa agt cat caa tgg ggt ccg tta gta tat caa aga tct tgt tat cga ttg ata gtg aat gaa tca gat agt ggt gta gag gaa tat gtc ctt ttt tat cct tgc tat caa agt tac gca tgc cgt ggc gta aca ata tct tta ata cag atg gat taa atc gtg tat tca tcg tat agc aat gta atg gag agt tac cac att tta gtc gtt tat tca gat cgc agt gtt taa taa cta att taa aca gat gag atg ata tat cca cat caa aga atg cga gat aca tat gac aga cat tat tga cag aaa tgt gac ctt cat tat cac cgt cgt cca taa atg cgt tag gta cgt acc aca tac tat cgt taa cga tgc gca caa tct cgt cca -tt tca tcc atc ttc ata atg att tac ttt ttc ata att aga gaa aaa gat caa ggt ata aaa att aga agt gtt aga cta ta- --- --- --- --- --- --- --- --- --- --- -aa cta act tat gac tta act aac tta tga --- --- --- --- --- --- --- --- --- --- --- --- --- --- --- --- --- --- --- --- --- --- --- --- --- --- --- --- --- --- --- --- --- --- --- --- --- --- --- --- --- --- --- --- --- --- --- --- ctt aac ta- --- --- --- --- --- --- --- --- --- --- -ac tta tga ctt aac taa ctt atg act taa cta act ta- tga ctt aac taa ctt atg act taa cta act tat gac tta act aac tta tga cta tta act cat ttc aag aaa ggt ggg tgg ata gaa ctc tat atg aca gct tgt gaa aca att aga tcc cta att tct aat gga agt ttt gat agg aga ttg tca tca gtt gat aca ttg ttt att atc tca tct att aga gca cgt ctg ttt aga gct tta gtg acc tgc tcg gtt act tct gtg taa atc ttg aat cct tta gtg ata cac tgt gtc aaa act gga tgt tta gaa tac cta tgt aga ata tgg gaa gca tgc ttg ttt ttg tct cta tta tag att aac tca tac atg gtt gta tta tga att ttc atc tgc cta atg tac tcc aat tct tgt tta caa tca att ata taa tca aag agt gat gat gca tac aca tta caa agt gaa taa tct acc atc ata aaa tac ttg ata cag agc ttt atc aca tca tgg ttt tca att gta tta tta agt ata gct aat ttt ata cag tca ata gac aat ggt tct cta agc aat att tct aat att tta aga tgt gct tcc cta cgg gcg atg aca gat ccc cta tcc acg gcc acg tca aga cat gta tat cca tta ctc att act gcg ttg aca ttt gct cca ttt tct aat agc cat gat act aaa tct ata taa cct gca tag ata gcg cga taa agc aag gtc ctt cca cca gca tct agt tga ttg ata tct tca ata tat ggg ata caa agc tta taa att tct aat act gtg ggt tca tct aca agg aat ccc cta gta tac tga att att tta tat aga tct aat tta aca tca ttt tca tct ggg ata cca caa ttc aaa ata aac tca aca aca cta ctt tcc ttt tta cat att ccc cta aaa tag gca ttc aag cat tct att tta tat att aca gcc cca tga tct acc ata aga tca aca atg tct att tct aca tat gca tta gat aga tag taa agt aag aga tct tgc aca gaa tta caa ttc tta ata att ata gag aaa ata tct tcc ata taa ttc ttt gac act aat gca gat ata ata tct tta tat gta ata tat gca aac agt cta tct act ata tac tga tca ata tta tct cta tga atc cta aaa taa tca tac aga aca tct aca gga tca caa att ggt tca agg aga aat cta tca aat att ttc ctg tca aca act ggt tct aga aca taa cag tca aca cct aat cca tgt ttt tta taa tca tct acc aaa gat aat gac caa aga tcg agg tcg tcg tga aac tgc tca tcg aca gcc atg aaa tct acc gac tcc atg gtg cga atc gca ctg tct tat tcg cca ttg att ttc a-t ttt tta taa tta tgt aca tgt ttt cct tct att ctc aag agt cta caa aaa tat a-- -tt ttt tcg ata tct aag tac taa gtt ttt tta ctg ttt ttg tta ctg tct tcc att ctt cta act aaa gat ctg aga taa att ata caa tct tcg cta tcg aac cat ttt tgt agt cta aag cct gaa gta att aac caa ctg ttt tta tta gtg gct ttt ttc gat cta tcc tcg tc- --- ctc tca tca tcc tta ta- --- --- --- --- --- --- tta tta tca tta tta tca tag tct att aaa cac aaa tca tct --- --- --- --- --- --- --- --- --- --- --- --- --- --- --- --- --- --- --- --- --- --- --- --- --- --- --a cgt tta taa c-- --- --- --- --- --- --- --- --a aca ttc --- --- --- --- -tc att att aat tag ttc tgt ag- aat atc ttt aat aat ttg gct ata --- --- -ca tct gtt --- --- --- --- --- --- --- --- --- --- --- --- --- --- --- --- --- --- --c aat act --- --- --- --- --- --- --- --a tct att gat gat ttc ttt t-- --- --- --- --- --- --- --- --- --- --- --t taa gac t-- --- --- --- --- --- --- --- --- --- --- --- --- --- --- --- --- --- --- --- --- --- --- --- --- --- --- --- --- --- --- --- --- --- --- --- --- --- --- --- taa act agt --- --- --- --- --- --- --- --- --- --- --- --- --- --- --- --- --- --- --- --- --- --- --- --- --- --- --- --- --- --- --- --- --- --- --- --- --- --- --- --- --- --- --- --- --- --- --- --- --- --- --- --- --- --- --- --- --- --- --- --- --- --- --- --- --- --- --- --- --- --- --- --- --- --- --- --- --- --- --- --- --- --- --- --- --- --- --- --- --- --- --- --- --- --- --- --- --- --- --- --- --- --- --- --- --- --- --- --- --- --t atg gta atg acg atg aaa --- -tc gag tag ta- --- --a ctt cta ata aag act tga ta- --- --t cat tat cat atg ttt gat cg- --- --- --- --- --- --- --t cat agt taa tag tgt g-- --- --- --- --- --- --- --- --- --- --- --- --- --- --- --- --- --- --- --- -gc taa atg gta ctg tta ata agt tta t-- --- --- --- --- --- --- --- --- --- --- --- --- --- --- --- --- --a gac aat atc ata gta ttt tct ttc cag aat --- --- --- --t aga tta ttt ttt taa ata ctg atc ctc aca att ccg tga tgt agc agt agt tgg t-- --- --- --- --- --- --- --- --- --- --- --- --- --- --- --- --- --- --- --- --- --- --- --- --- --- --- --- -gc atg gtc tat atc gt- --- --- --- --- --- --- --- --- --- --- --- --- --- --- --- --- --- --- --- --- --- --- --- --- --- --- --- --- --- --- --- --- --- --- --- --- --- --- --- --- --- --- --- taa aat gta tca tat ata ata gtt ttc tga cgt gga gta cag aat ttt cga --- --- --- --- --- --- --- --- --- --- --- --- --- --- --- --- --- --- --- --- --- --- --- --- --- --- --- --- --- --- --- --- --- --- --- --- --- --- --- --- --- --- --- --t taa tga gtt cat ggt aag gaa ggg caa atg cct --- gta tat aat ata cat aag tta a-- --- --- --- --- --- --- --- --- --- --- --- --- --t agt ttt tta tca tat tt- --- --- --- --- --- --- --- --t cta ata cca taa taa aaa t-- --- --- --- --- --- --- --- --- --- --- --- --- --- --- --- --- --- --- --- --- --- --- --- --- --t atc att att gcg ttt g-g tag tt- --- --- --- --- ctg ccc ta- --- --- --- --- --- --- --- --t cat cta tat cac tgt cac tct c-- --- --g ctc tca cta tat ctt cta aaa tta caa --- -ac aac --- --- --- --- --- --- --- --- --- --- --- --- --- --- --- --- --- --- --- --- --- --- --- --- --- --- --- --- --- --- --- --- --- --- --- --- tgg ata ttc gat --- --- --- --- --- --- --- --- --- --- --- --- --- --- --- --- --- --- --- --- --- --- --- --- --- --- --a aca gca ttt gtg t-- --- --- --- --- --- --- --- --- --- --- --- --- --- --- --- --- -

>ON622722.2_Monkeypox_virus_isolate_MPXV_FR_HCL0001_2022_complete_genome

--- --- --- --- --- --- --- --- --- --- --- --- --- --- --- --- --- --- --- --- --- --- --- --- --- --- --- --t att ata ttt tac tat ttt att tag tgt cta gaa aaa aat gtg tga ccc acg acc gta gga aac tct aga ggg taa gaa aaa tca atc gtt tat aga gac cat cag aaa gag gtt taa tat ttt tgt gag acc tat cga a-- --- --- --- --- --- --- --- --- --- --- --- --- --- --- --- --- --- --- --- --- --- --- --- --- --- --- --- --- --- --- --- --- --- --- --- --- --- --- --- --- --- --- --- --- --- --- --- --- --- --- --- --- --- --- --- --- --- --- --- --- --- --- --- --- --- --- --- --- --- --- --- --- --- --- --- --- --- --- --- --- --- --- --- --- --- --- --- --- --- --- --- --- --- --- --- --- --- --- --- --- --- --- --- --- --- --- --- --- --- --- --- --- --- --- --- --- --- --- --- --- --- --- --- --- --- --- --- --- --- --- --- --- --- --- --- --- --- --- --- --- --- --- --- --- --- --- --- --- --- --- --- --- --- --- --- --- --- --- --- --- --- --- --- --- --- --- --- --- --- --- --- --- --- --- --- --- --- --- --- --- --- --- --- --- --- --- --- --- --- --- --- --- --- --- --- --- --- --- --- --- --- --- --- --- --- --- --- --- --- --- --- --- --- --- --- --- --- --- --- --- --- --- --- --- --- --- --- --- --- --- --- --- --- --- --- --- --- --- --- --- --- --- --- --- --- --- --- --- --- --- --- --- --- --- --- --- --- --- --- --- --- --- --- --- --- --- --- --- --- --- --- --- --- --- --- --- --- --- --- -ga gag aaa gga taa aaa ctt ttt acg act cca tca gaa aga ggt tta ata ttt ttg tga gac cca tcg aag aga gaa aga gat ggt tag tca aga tat ttt tct tag tac aaa agt caa tgt ttt aaa ata tat gga cga gaa tta att tgt ctg tat aaa aac ttg tgt gaa att atg tac tag aga aaa aac gtg agc agt gtc ccc tac atg gat ttt aca gat cat tta tat tcc aaa aat att aac tat ata cgt tta tta tat gat gtt aac gtg taa att ata aac att att tta tga tgc aat tgt ctg aca acc tag att ggt ata agg atg ttg ata agc tct acg aga ata tat tgt tgg acg tta tcg ttt acg aaa tag ttg aga cat cag aaa gag gtt taa tat ttt tgt gag a-c cat cga aga gag aaa gag aat aaa aat a-- --t ttt ttt ttt ttt tgt aaa act ttt tta tga gac --- --- -ca aga gaa tac gaa tag tga tca tat cgt atc aca tat tga aac aga aag aag aag taa cga gag gta act ttt tgt gaa tgt agt taa ata ttt ttg ttt tgc aaa ccg gaa tat agt gcc cgg t-- -ct ttt tta att cgt ggt gcg gtg tct gaa tcg ttc gat taa ccc aac tca tcc att ttc aga tga ata gag tta tcg att cag aca cat gct ttg agt ttt gtt gaa tcg atg agt gaa gta tca tcg gtt gca cct tca gat gcc gat ccg tcg aca tac ttg aat cca tcc ttg act tca agt tca gat gat tcc tca cac atg tct ccg ata cgt acg cta aac tct agg ttc ttg aca cat ttt gta tca acg atc gtt gaa ccg atg ata tct ttg taa ctc act ttc tta tgt gag atg tta gac cca agt act gga tgg gtc ttg atg tca ctg tct ttc tct tct tcg cta cat ctg atg tcg ata gac atc tca cag tct ttg atc ata gcc aga gct tct tca cgc gtg atc gcg gga gag tcc tta cct tgt ccc ggt gac acg ctg gac aat cta gta ttc aca gtg ttt cca tca gag gat tcg gag atg gat aaa atc ttt ggg cat ttg gtg aat cca aag ttc atg tta aga ccc gca ccg acg ata gtg taa taa gtg gtg gga tct cct ttt aca act --- --- --- --- --- --- tct tcg gat acc tca tca tct tcg gtc tct gta act tcc gtt acg gat tga caa atc tta tca ttg gtc ggt gtt tgg tct tgc ttt gtg act ttg ata ata aca tcg att ccc ata tga tgt ttg ttt tct tct tca gta cac gag gat gaa gat tgt tga aga cta gta ggc ata gca gct gcc act agg cac atg cat gcc agg aca ata tat tgt ttc atg att gct att gat tga tta ctg ttc tag atg att cta ctt tct tac cat ata ata aat tag aat ata ttt tct act ttt acg aga aat taa tta ttg tat tta tta ttt ata ggt --a aaa aaa ctt act ata agt ggg tgg gat tct ggg aat tag tga tca gtt tat gta tat cgc aac tag cgg gca tat ggc tat tga cat cga gaa cat tac cca tat gat aag aga ttg tat cat ttt cgt agt ctt gag tat tgg tat tac tat ata gta tgt aga tgt cga cgc tag ata gac agt cgc cca cta gag tta ccg tct ctg aat gcg gca tga tag tat cat tct ttg ttt tcg tta act gtt tgg aag atg aat ctt tgt tgt tac att taa tct cga aat tca gag tac ata tct ttg aag tat tct gat atc tat ttt ctc ctg taa aga atc ctg aag ttg cta cat tat taa gga cag aga agt att ctg cac gaa aga ctg gat cac aat ctt tat gat tca tgg taa tag tta gtt ccg acg ttg aga tgg att cgc tga gac cgg tag tgg tcg tcc gag tac acg atg tgt cgt tga ctg gat aca ggt taa ttt cca cat cga tat agt taa atg tat tgc tgg tta cga cgg gtt cgc att tat ctg tgg aag aga cgg tgt gag aat atg ttc cgg gac cac acg gag aac aga tga cgt ctc cgg tag acg tgt atc cgg ata ctc cgt atc cta ttc cac act ttg ttt tag aaa tac atg ttc tac acc ctg atg ctc ctt tga gaa gac aat aat atc ctg gag agc att cac aga ttc tat tgt gag tcg tgt tac acg atc gcg tct cta cct gat tac tat cac atc ttc cgt tac aac tta gac aag cct gta aat gat tat tgt gag atg taa agg tat ccg aac cac acg gtg tac att gtg tat tag tct tgc tat cac ata atc tga aag cgt aag ttc ccg gag gac acg ata gac aac ata gat tac ggc ttc tgt att cgt tgt ctt tac act ttc cat tgg atg gtg cat gtg gtg cta tat ctc ttc cgt tta tta tta tac atg aga gaa aca ata tat acg agt ata ata cgg acc tca tga ttt aat aat -gt agt aat cgt cgt ctt gtt act gtt tgt ttc cta ctt ctc caa tca tat aga tta ttt ttt aaa tat ttt ctt tct atc atg gat aat att tgt aat ggt tct ttc cgt aca aca tac tgt tta gat ggt agt cgc tta gct tgg tta tga tat tgc gca taa ttt ccg gag gca aat acg ata gtc tag att gac tat cga tgg tag act cta att tat tga gtg ctt tgt cga cga gtt tac ttt tat gct cca tcg ata gat gac act gtt cta tga gat cgt cgt aca tgg gaa atg aaa tgc gtt tgt ctg aat gta tgg ctt cga gat atg tgt gat acc gaa tgt ctt ctg ttc tca ata ccg tat aca agt cgg tgt ctg aga ttc gaa tct ctt tga gga gac tta tgt cac gac tac att ttt cga tga tgg aat cta tct tat cga atg ata tat ttt tca taa ata cac ttt tat agt cct cgt tta aac aga att tag tat gta gtt ccg caa atg act cgt ccc tta ata ggc agt agg cta tta tct tct tta cgt agt gat cgt cgt agg gag aga act ccg aca tct tgt aga aca acg att taa tca tag gta gag ata ctt tca gtc tgt ggt gga taa tgt cat taa cga cat ccg cct tgt ata taa tgt ttc tgt ttt caa aca cca agt cga ata ctg tct tat cgt ctt tag tcg gaa ggt tga tgt cgt atc caa tgt ata cga ggt atg agg caa cat tgt tat tgc aat tct gga agg cgg tat gaa gag gag tca ttg tat tat agt att cgt ctt tct gaa tgt cga atc tat cta gta gat acc gta gta tat tga gag agc gac ttc cat atc ctt gat tat gtt tta tga ata gat aaa gta gat gtt gtc ctt ctt cct ttt gta att tcc cgt att ttt gtt cgt gcc aat tga gta aca tta tga gaa tat gac ctg ttg cac aat cgt tct tta tgt att cca tga tgg gtg tac aat taa gat tat tac gta tcc tcg tat cgg ctc ctc gag ata aaa gag cat aca cca cac gag gac tat gtt tgg tat act gtt gaa ggt aag tgt gta acg gcg tat ttc cga ttt tcg taa ccg cgt taa tgt ttg ctc cat gat cta tta tcg cgt aga tga atc gct tct cag ctc gca tct tag tgt gac tct ttg act tgt aat aat tgc ttt cgt gga acg cgg ata tgt gtt tac agt agt aat gaa gag aag tga gtc cat cct cat cga cgc aat tag ggt cag atc ctt tag tca ata att tgt aca gaa cgt aat agt tta agc tcc cat tga att tat atc taa gat aac aca gca ata gat cgg atg att tac taa agt cat caa tgg ggt ccg tta gta tat caa aga tct tgt tat cga ttg ata gtg aat gaa tca gat agt ggt gta gag gaa tat gtc ctt ttt tat cct tgc tat caa agt tac gca tgc cgt ggc gta aca ata tct tta ata cag atg gat taa atc gtg tat tca tcg tat agc aat gta atg gag agt tac cac att tta gtc gtt tat tca gat cgc agt gtt taa taa cta att taa aca gat gag atg ata tat cca cat caa aga atg cga gat aca tat gac aga cat tat tga cag aaa tgt gac ctt cat tat cac cgt cgt cca taa atg cgt tag gta cgt acc aca tac tat cgt taa cga tgc gca caa tct cgt cca -tt tca tcc atc ttc ata atg att tac ttt ttc ata att aga gaa aaa gat caa ggt ata aaa att aga agt gtt aga cta ta- --- --- --- --- --- --- --- --- --- --- -aa cta act tat gac tta act aac tta tga --- --- --- --- --- --- --- --- --- --- --- --- --- --- --- --- --- --- --- --- --- --- --- --- --- --- --- --- --- --- --- --- --- --- --- --- --- --- --- --- --- --- --- --- --- --- --- --- ctt aac ta- --- --- --- --- --- --- --- --- --- --- -ac tta tga ctt aac taa ctt atg act taa cta act ta- tga ctt aac taa ctt atg act taa cta act tat gac tta act aac tta tga cta tta act cat ttc aag aaa ggt ggg tgg ata gaa ctc tat atg aca gct tgt gaa aca att aga tcc cta att tct aat gga agt ttt gat agg aga ttg tca tca gtt gat aca ttg ttt att atc tca tct att aga gca cgt ctg ttt aga gct tta gtg acc tgc tcg gtt act tct gtg taa atc ttg aat cct tta gtg ata cac tgt gtc aaa act gga tgt tta gaa tac cta tgt aga ata tgg gaa gca tgc ttg ttt ttg tct cta tta tag att aac tca tac atg gtt gta tta tga att ttc atc tgc cta atg tac tcc aat tct tgt tta caa tca att ata taa tca aag agt gat gat gca tac aca tta caa agt gaa taa tct acc atc ata aaa tac ttg ata cag agc ttt atc aca tca tgg ttt tca att gta tta tta agt ata gct aat ttt ata cag tca ata gac aat ggt tct cta agc aat att tct aat att tta aga tgt gct tcc cta cgg gcg atg aca gat ccc cta tcc acg gcc acg tca aga cat gta tat cca tta ctc att act gcg ttg aca ttt gct cca ttt tct aat agc cat gat act aaa tct ata taa cct gca tag ata gcg cga taa agc aag gtc ctt cca cca gca tct agt tga ttg ata tct tca ata tat ggg ata caa agc tta taa att tct aat act gtg ggt tca tct aca agg aat ccc cta gta tac tga att att tta tat aga tct aat tta aca tca ttt tca tct ggg ata cca caa ttc aaa ata aac tca aca aca cta ctt tcc ttt tta cat att ccc cta aaa tag gca ttc aag cat tct att tta tat att aca gcc cca tga tct acc ata aga tca aca atg tct att tct aca tat gca tta gat aga tag taa agt aag aga tct tgc aca gaa tta caa ttc tta ata att ata gag aaa ata tct tcc ata taa ttc ttt gac act aat gca gat ata ata tct tta tat gta ata tat gca aac agt cta tct act ata tac tga tca ata tta tct cta tga atc cta aaa taa tca tac aga aca tct aca gga tca caa att ggt tca agg aga aat cta tca aat att ttc ctg tca aca act ggt tct aga aca taa cag tca aca cct aat cca tgt ttt tta taa tca tct acc aaa gat aat gac caa aga tcg agg tcg tcg tga aac tgc tca tcg aca gcc atg aaa tct acc gac tcc atg gtg cga atc gca ctg tct tat tcg cca ttg att ttc a-t ttt tta taa tta tgt aca tgt ttt cct tct att ctc aag agt cta caa aaa tat a-- -tt ttt tcg ata tct aag tac taa gtt ttt tta ctg ttt ttg tta ctg tct tcc att ctt cta act aaa gat ctg aga taa att ata caa tct tcg cta tcg aac cat ttt tgt agt cta aag cct gaa gta att aac caa ctg ttt tta tta gtg gct ttt ttc gat cta tcc tcg tc- --- ctc tca tca tcc tta ta- --- --- --- --- --- --- tta tta tca tta tta tca tag tct att aaa cac aaa tca tct --- --- --- --- --- --- --- --- --- --- --- --- --- --- --- --- --- --- --- --- --- --- --- --- --- --- --a cgt tta taa c-- --- --- --- --- --- --- --- --a aca ttc --- --- --- --- -tc att att aat tag ttc tgt ag- aat atc ttt aat aat ttg gct ata --- --- -ca tct gtt --- --- --- --- --- --- --- --- --- --- --- --- --- --- --- --- --- --- --c aat act --- --- --- --- --- --- --- --a tct att gat gat ttc ttt t-- --- --- --- --- --- --- --- --- --- --- --t taa gac t-- --- --- --- --- --- --- --- --- --- --- --- --- --- --- --- --- --- --- --- --- --- --- --- --- --- --- --- --- --- --- --- --- --- --- --- --- --- --- --- taa act agt --- --- --- --- --- --- --- --- --- --- --- --- --- --- --- --- --- --- --- --- --- --- --- --- --- --- --- --- --- --- --- --- --- --- --- --- --- --- --- --- --- --- --- --- --- --- --- --- --- --- --- --- --- --- --- --- --- --- --- --- --- --- --- --- --- --- --- --- --- --- --- --- --- --- --- --- --- --- --- --- --- --- --- --- --- --- --- --- --- --- --- --- --- --- --- --- --- --- --- --- --- --- --- --- --- --- --- --- --- --t atg gta atg acg atg aaa --- -tc gag tag ta- --- --a ctt cta ata aag act tga ta- --- --t cat tat cat atg ttt gat cg- --- --- --- --- --- --- --t cat agt taa tag tgt g-- --- --- --- --- --- --- --- --- --- --- --- --- --- --- --- --- --- --- --- -gc taa atg gta ctg tta ata agt tta t-- --- --- --- --- --- --- --- --- --- --- --- --- --- --- --- --- --a gac aat atc ata gta ttt tct ttc cag aat --- --- --- --t aga tta ttt ttt taa ata ctg atc ctc aca att ccg tga tgt agc agt agt tgg t-- --- --- --- --- --- --- --- --- --- --- --- --- --- --- --- --- --- --- --- --- --- --- --- --- --- --- --- -gc atg gtc tat atc gt- --- --- --- --- --- --- --- --- --- --- --- --- --- --- --- --- --- --- --- --- --- --- --- --- --- --- --- --- --- --- --- --- --- --- --- --- --- --- --- --- --- --- --- taa aat gta tca tat ata ata gtt ttc tga cgt gga gta cag aat ttt cga --- --- --- --- --- --- --- --- --- --- --- --- --- --- --- --- --- --- --- --- --- --- --- --- --- --- --- --- --- --- --- --- --- --- --- --- --- --- --- --- --- --- --- --t taa tga gtt cat ggt aag gaa ggg caa atg cct --- gta tat aat ata cat aag tta a-- --- --- --- --- --- --- --- --- --- --- --- --- --t agt ttt tta tca tat tt- --- --- --- --- --- --- --- --t cta ata cca taa taa aaa t-- --- --- --- --- --- --- --- --- --- --- --- --- --- --- --- --- --- --- --- --- --- --- --- --- --t atc att att gcg ttt g-g tag tt- --- --- --- --- ctg ccc ta- --- --- --- --- --- --- --- --t cat cta tat cac tgt cac tct c-- --- --g ctc tca cta tat ctt cta aaa tta caa --- -ac aac --- --- --- --- --- --- --- --- --- --- --- --- --- --- --- --- --- --- --- --- --- --- --- --- --- --- --- --- --- --- --- --- --- --- --- --- tgg ata ttc gat --- --- --- --- --- --- --- --- --- --- --- --- --- --- --- --- --- --- --- --- --- --- --- --- --- --- --a aca gca ttt gtg t-- --- --- --- --- --- --- --- --- --- --- --- --- --- --- --- --- -

>ON627808.1_Monkeypox_virus_isolate_MPX/human/USA/UT-UPHL-82200022/2022_complete_genome

--- --- --- --- --- --- --- --- --- --- --- --- --- --- --- --- --- --- --- --- --- --- --- --- --- --- --- --- att ata ttt tac tat ttt att tag tgt cta gaa aaa aat gtg tga ccc acg acc gta gga aac tct aga ggg taa gaa aaa tca atc gtt tat aga gac cat cag aaa gag gtt taa tat ttt tgt gag acc tat cga a-- --- --- --- --- --- --- --- --- --- --- --- --- --- --- --- --- --- --- --- --- --- --- --- --- --- --- --- --- --- --- --- --- --- --- --- --- --- --- --- --- --- --- --- --- --- --- --- --- --- --- --- --- --- --- --- --- --- --- --- --- --- --- --- --- --- --- --- --- --- --- --- --- --- --- --- --- --- --- --- --- --- --- --- --- --- --- --- --- --- --- --- --- --- --- --- --- --- --- --- --- --- --- --- --- --- --- --- --- --- --- --- --- --- --- --- --- --- --- --- --- --- --- --- --- --- --- --- --- --- --- --- --- --- --- --- --- --- --- --- --- --- --- --- --- --- --- --- --- --- --- --- --- --- --- --- --- --- --- --- --- --- --- --- --- --- --- --- --- --- --- --- --- --- --- --- --- --- --- --- --- --- --- --- --- --- --- --- --- --- --- --- --- --- --- --- --- --- --- --- --- --- --- --- --- --- --- --- --- --- --- --- --- --- --- --- --- --- --- --- --- --- --- --- --- --- --- --- --- --- --- --- --- --- --- --- --- --- --- --- --- --- --- --- --- --- --- --- --- --- --- --- --- --- --- --- --- --- --- --- --- --- --- --- --- --- --- --- --- --- --- --- --- --- --- --- --- --- --- --- -ga gag aaa gga taa aaa ctt ttt acg act cca tca gaa aga ggt tta ata ttt ttg tga gac cca tcg aag aga gaa aga gat ggt tag tca aga tat ttt tct tag tac aaa agt caa tgt ttt aaa ata tat gga cga gaa tta att tgt ctg tat aaa aac ttg tgt gaa att atg tac tag aga aaa aac gtg agc agt gtc ccc tac atg gat ttt aca gat cat tta tat tcc aaa aat att aac tat ata cgt tta tta tat gat gtt aac gtg taa att ata aac att att tta tga tgc aat tgt ctg aca acc tag att ggt ata agg atg ttg ata agc tct acg aga ata tat tgt tgg acg tta tcg ttt acg aaa tag ttg aga cat cag aaa gag gtt taa tat ttt tgt gag a-c cat cga aga gag aaa gag aat aaa aat a-- --t ttt ttt ttt ttt tgt aaa act ttt tta tga gac --- --- -ca aga gaa tac gaa tag tga tca tat cgt atc aca tat tga aac aga aag aag aag taa cga gag gta act ttt tgt gaa tgt agt taa ata ttt ttg ttt tgc aaa ccg gaa tat agt gcc cgg t-- -ct ttt tta att cgt ggt gcg gtg tct gaa tcg ttc gat taa ccc aac tca tcc att ttc aga tga ata gag tta tcg att cag aca cat gct ttg agt ttt gtt gaa tcg atg agt gaa gta tca tcg gtt gca cct tca gat gcc gat ccg tcg aca tac ttg aat cca tcc ttg act tca agt tca gat gat tcc tca cac atg tct ccg ata cgt acg cta aac tct agg ttc ttg aca cat ttt gta tca acg atc gtt gaa ccg atg ata tct ttg taa ctc act ttc tta tgt gag atg tta gac cca agt act gga tgg gtc ttg atg tca ctg tct ttc tct tct tcg cta cat ctg atg tcg ata gac atc tca cag tct ttg atc ata gcc aga gct tct tca cgc gtg atc gcg gga gag tcc tta cct tgt ccc ggt gac acg ctg gac aat cta gta ttc aca gtg ttt cca tca gag gat tcg gag atg gat aaa atc ttt ggg cat ttg gtg aat cca aag ttc atg tta aga ccc gca ccg acg ata gtg taa taa gtg gtg gga tct cct ttt aca act --- --- --- --- --- --- tct tcg gat acc tca tca tct tcg gtc tct gta act tcc gtt acg gat tga caa atc tta tca ttg gtc ggt gtt tgg tct tgc ttt gtg act ttg ata ata aca tcg att ccc ata tga tgt ttg ttt tct tct tca gta cac gag gat gaa gat tgt tga aga cta gta ggc ata gca gct gcc act agg cac atg cat gcc agg aca ata tat tgt ttc atg att gct att gat tga tta ctg ttc tag atg att cta ctt tct tac cat ata ata aat tag aat ata ttt tct act ttt acg aga aat taa tta ttg tat tta tta ttt ata ggt --a aaa aaa ctt act ata agt ggg tgg gat tct ggg aat tag tga tca gtt tat gta tat cgc aac tag cgg gca tat ggc tat tga cat cga gaa cat tac cca tat gat aag aga ttg tat cat ttt cgt agt ctt gag tat tgg tat tac tat ata gta tgt aga tgt cga cgc tag ata gac agt cgc cca cta gag tta ccg tct ctg aat gcg gca tga tag tat cat tct ttg ttt tcg tta act gtt tgg aag atg aat ctt tgt tgt tac att taa tct cga aat tca gag tac ata tct ttg aag tat tct gat atc tat ttt ctc ctg taa aga atc ctg aag ttg cta cat tat taa gga cag aga agt att ctg cac gaa aga ctg gat cac aat ctt tat gat tca tgg taa tag tta gtt ccg acg ttg aga tgg att cgc tga gac cgg tag tgg tcg tcc gag tac acg atg tgt cgt tga ctg gat aca ggt taa ttt cca cat cga tat agt taa atg tat tgc tgg tta cga cgg gtt cgc att tat ctg tgg aag aga cgg tgt gag aat atg ttc cgg gac cac acg gag aac aga tga cgt ctc cgg tag acg tgt atc cgg ata ctc cgt atc cta ttc cac act ttg ttt tag aaa tac atg ttc tac acc ctg atg ctc ctt tga gaa gac aat aat atc ctg gag agc att cac aga ttc tat tgt gag tcg tgt tac acg atc gcg tct cta cct gat tac tat cac atc ttc cgt tac aac tta gac aag cct gta aat gat tat tgt gag atg taa agg tat ccg aac cac acg gtg tac att gtg tat tag tct tgc tat cac ata atc tga aag cgt aag ttc ccg gag gac acg ata gac aac ata gat tac ggc ttc tgt att cgt tgt ctt tac act ttc cat tgg atg gtg cat gtg gtg cta tat ctc ttc cgt tta tta tta tac atg aga gaa aca ata tat acg agt ata ata cgg acc tca tga ttt aat aat -gt agt aat cgt cgt ctt gtt act gtt tgt ttc cta ctt ctc caa tca tat aga tta ttt ttt aaa tat ttt ctt tct atc atg gat aat att tgt aat ggt tct ttc cgt aca aca tac tgt tta gat ggt agt cgc tta gct tgg tta tga tat tgc gca taa ttt ccg gag gca aat acg ata gtc tag att gac tat cga tgg tag act cta att tat tga gtg ctt tgt cga cga gtt tac ttt tat gct cca tcg ata gat gac act gtt cta tga gat cgt cgt aca tgg gaa atg aaa tgc gtt tgt ctg aat gta tgg ctt cga gat atg tgt gat acc gaa tgt ctt ctg ttc tca ata ccg tat aca agt cgg tgt ctg aga ttc gaa tct ctt tga gga gac tta tgt cac gac tac att ttt cga tga tgg aat cta tct tat cga atg ata tat ttt tca taa ata cac ttt tat agt cct cgt tta aac aga att tag tat gta gtt ccg caa atg act cgt ccc tta ata ggc agt agg cta tta tct tct tta cgt agt gat cgt cgt agg gag aga act ccg aca tct tgt aga aca acg att taa tca tag gta gag ata ctt tca gtc tgt ggt gga taa tgt cat taa cga cat ccg cct tgt ata taa tgt ttc tgt ttt caa aca cca agt cga ata ctg tct tat cgt ctt tag tcg gaa ggt tga tgt cgt atc caa tgt ata cga ggt atg agg caa cat tgt tat tgc aat tct gga agg cgg tat gaa gag gag tca ttg tat tat agt att cgt ctt tct gaa tgt cga atc tat cta gta gat acc gta gta tat tga gag agc gac ttc cat atc ctt gat tat gtt tta tga ata gat aaa gta gat gtt gtc ctt ctt cct ttt gta att tcc cgt att ttt gtt cgt gcc aat tga gta aca tta tga gaa tat gac ctg ttg cac aat cgt tct tta tgt att cca tga tgg gtg tac aat taa gat tat tac gta tcc tcg tat cgg ctc ctc gag ata aaa gag cat aca cca cac gag gac tat gtt tgg tat act gtt gaa ggt aag tgt gta acg gcg tat ttc cga ttt tcg taa ccg cgt taa tgt ttg ctc cat gat cta tta tcg cgt aga tga atc gct tct cag ctc gca tct tag tgt gac tct ttg act tgt aat aat tgc ttt cgt gga acg cgg ata tgt gtt tac agt agt aat gaa gag aag tga gtc cat cct cat cga cgc aat tag ggt cag atc ctt tag tca ata att tgt aca gaa cgt aat agt tta agc tcc cat tga att tat atc taa gat aac aca gca ata gat cgg atg att tac taa agt cat caa tgg ggt ccg tta gta tat caa aga tct tgt tat cga ttg ata gtg aat gaa tca gat agt ggt gta gag gaa tat gtc ctt ttt tat cct tgc tat caa agt tac gca tgc cgt ggc gta aca ata tct tta ata cag atg gat taa atc gtg tat tca tcg tat agc aat gta atg gag agt tac cac att tta gtc gtt tat tca gat cgc agt gtt taa taa cta att taa aca gat gag atg ata tat cca cat caa aga atg cga gat aca tat gac aga cat tat tga cag aaa tgt gac ctt cat tat cac cgt cgt cca taa atg cgt tag gta cgt acc aca tac tat cgt taa cga tgc gca caa tct cgt cca -tt tca tcc atc ttc ata atg att tac ttt ttc ata att aga gaa aaa gat caa ggt ata aaa att aga agt gtt aga cta ta- --- --- --- --- --- --- --- --- --- --- -aa cta act tat gac tta act aac tta tga --- --- --- --- --- --- --- --- --- --- --- --- --- --- --- --- --- --- --- --- --- --- --- --- --- --- --- --- --- --- --- --- --- --- --- --- --- --- --- --- --- --- --- --- --- --- --- --- ctt aac ta- --- --- --- --- --- --- --- --- --- --- -ac tta tga ctt aac taa ctt atg act taa cta act ta- tga ctt aac taa ctt atg act taa cta act tat gac tta act aac tta tga cta tta act cat ttc aag aaa ggt ggg tgg ata gaa ctc tat atg aca gct tgt gaa aca att aga tcc cta att tct aat gga agt ttt gat agg aga ttg tca tca gtt gat aca ttg ttt att atc tca tct att aga gca cgt ctg ttt aga gct tta gtg acc tgc tcg gtt act tct gtg taa atc ttg aat cct tta gtg ata cac tgt gtc aaa act gga tgt tta gaa tac cta tgt aga ata tgg gaa gca tgc ttg ttt ttg tct cta tta tag att aac tca tac atg gtt gta tta tga att ttc atc tgc cta atg tac tcc aat tct tgt tta caa tca att ata taa tca aag agt gat gat gca tac aca tta caa agt gaa taa tct acc atc ata aaa tac ttg ata cag agc ttt atc aca tca tgg ttt tca att gta tta tta agt ata gct aat ttt ata cag tca ata gac aat ggt tct cta agc aat att tct aat att tta aga tgt gct tcc cta cgg gcg atg aca gat ccc cta tcc acg gcc acg tca aga cat gta tat cca tta ctc att act gcg ttg aca ttt gct cca ttt tct aat agc cat gat act aaa tct ata taa cct gca tag ata gcg cga taa agc aag gtc ctt cca cca gca tct agt tga ttg ata tct tca ata tat ggg ata caa agc tta taa att tct aat act gtg ggt tca tct aca agg aat ccc cta gta tac tga att att tta tat aga tct aat tta aca tca ttt tca tct ggg ata cca caa ttc aaa ata aac tca aca aca cta ctt tcc ttt tta cat att ccc cta aaa tag gca ttc aag cat tct att tta tat att aca gcc cca tga tct acc ata aga tca aca atg tct att tct aca tat gca tta gat aga tag taa agt aag aga tct tgc aca gaa tta caa ttc tta ata att ata gag aaa ata tct tcc ata taa ttc ttt gac act aat gca gat ata ata tct tta tat gta ata tat gca aac agt cta tct act ata tac tga tca ata tta tct cta tga atc cta aaa taa tca tac aga aca tct aca gga tca caa att ggt tca agg aga aat cta tca aat att ttc ctg tca aca act ggt tct aga aca taa cag tca aca cct aat cca tgt ttt tta taa tca tct acc aaa gat aat gac caa aga tcg agg tcg tcg tga aac tgc tca tcg aca gcc atg aaa tct acc gac tcc atg gtg cga atc gca ctg tct tat tcg cca ttg att ttc a-t ttt tta taa tta tgt aca tgt ttt cct tct att ctc aag agt cta caa aaa tat a-- -tt ttt tcg ata tct aag tac taa gtt ttt tta ctg ttt ttg tta ctg tct tcc att ctt cta act aaa gat ctg aga taa att ata caa tct tcg cta tcg aac cat ttt tgt agt cta aag cct gaa gta att aac caa ctg ttt tta tta gtg gct ttt ttc gat cta tcc tcg tc- --- ctc tca tca tcc tta ta- --- --- --- --- --- --- tta tta tca tta tta tca tag tct att aaa cac aaa tca tct --- --- --- --- --- --- --- --- --- --- --- --- --- --- --- --- --- --- --- --- --- --- --- --- --- --- --a cgt tta taa c-- --- --- --- --- --- --- --- --a aca ttc --- --- --- --- -tc att att aat tag ttc tgt ag- aat atc ttt aat aat ttg gct ata --- --- -ca tct gtt --- --- --- --- --- --- --- --- --- --- --- --- --- --- --- --- --- --- --c aat act --- --- --- --- --- --- --- --a tct att gat gat ttc ttt t-- --- --- --- --- --- --- --- --- --- --- --t taa gac t-- --- --- --- --- --- --- --- --- --- --- --- --- --- --- --- --- --- --- --- --- --- --- --- --- --- --- --- --- --- --- --- --- --- --- --- --- --- --- --- taa act agt --- --- --- --- --- --- --- --- --- --- --- --- --- --- --- --- --- --- --- --- --- --- --- --- --- --- --- --- --- --- --- --- --- --- --- --- --- --- --- --- --- --- --- --- --- --- --- --- --- --- --- --- --- --- --- --- --- --- --- --- --- --- --- --- --- --- --- --- --- --- --- --- --- --- --- --- --- --- --- --- --- --- --- --- --- --- --- --- --- --- --- --- --- --- --- --- --- --- --- --- --- --- --- --- --- --- --- --- --- --t atg gta atg acg atg aaa --- -tc gag tag ta- --- --a ctt cta ata aag act tga ta- --- --t cat tat cat atg ttt gat cg- --- --- --- --- --- --- --t cat agt taa tag tgt g-- --- --- --- --- --- --- --- --- --- --- --- --- --- --- --- --- --- --- --- -gc taa atg gta ctg tta ata agt tta t-- --- --- --- --- --- --- --- --- --- --- --- --- --- --- --- --- --a gac aat atc ata gta ttt tct ttc cag aat --- --- --- --t aga tta ttt ttt taa ata ctg atc ctc aca att ccg tga tgt agc agt agt tgg t-- --- --- --- --- --- --- --- --- --- --- --- --- --- --- --- --- --- --- --- --- --- --- --- --- --- --- --- -gc atg gtc tat atc gt- --- --- --- --- --- --- --- --- --- --- --- --- --- --- --- --- --- --- --- --- --- --- --- --- --- --- --- --- --- --- --- --- --- --- --- --- --- --- --- --- --- --- --- taa aat gta tca tat ata ata gtt ttc tga cgt gga gta cag aat ttt cga --- --- --- --- --- --- --- --- --- --- --- --- --- --- --- --- --- --- --- --- --- --- --- --- --- --- --- --- --- --- --- --- --- --- --- --- --- --- --- --- --- --- --- --t taa tga gtt cat ggt aag gaa ggg caa atg cct --- gta tat aat ata cat aag tta a-- --- --- --- --- --- --- --- --- --- --- --- --- --t agt ttt tta tca tat tt- --- --- --- --- --- --- --- --t cta ata cca taa taa aaa t-- --- --- --- --- --- --- --- --- --- --- --- --- --- --- --- --- --- --- --- --- --- --- --- --- --t atc att att gcg ttt g-g tag tt- --- --- --- --- ctg ccc ta- --- --- --- --- --- --- --- --t cat cta tat cac tgt cac tct c-- --- --g ctc tca cta tat ctt cta aaa tta caa --- -ac aac --- --- --- --- --- --- --- --- --- --- --- --- --- --- --- --- --- --- --- --- --- --- --- --- --- --- --- --- --- --- --- --- --- --- --- --- tgg ata ttc gat --- --- --- --- --- --- --- --- --- --- --- --- --- --- --- --- --- --- --- --- --- --- --- --- --- --- --a aca gca ttt gtg t-- --- --- --- --- --- --- --- --- --- --- --- --- --- --- --- --- -

>ON676703.1_Monkeypox_virus_isolate_MPXV_USA_2022_CA001_complete_genome

--- --- --- --- --- --- --- --- --- --- --- --- --- --- --- --- --- --- --- --- --- --- --- --- --- --- --- --t att ata ttt tac tat ttt att tag tgt cta gaa aaa aat gtg tga ccc acg acc gta gga aac tct aga ggg taa gaa aaa tca atc gtt tat aga gac cat cag aaa gag gtt taa tat ttt tgt gag acc tat cga a-- --- --- --- --- --- --- --- --- --- --- --- --- --- --- --- --- --- --- --- --- --- --- --- --- --- --- --- --- --- --- --- --- --- --- --- --- --- --- --- --- --- --- --- --- --- --- --- --- --- --- --- --- --- --- --- --- --- --- --- --- --- --- --- --- --- --- --- --- --- --- --- --- --- --- --- --- --- --- --- --- --- --- --- --- --- --- --- --- --- --- --- --- --- --- --- --- --- --- --- --- --- --- --- --- --- --- --- --- --- --- --- --- --- --- --- --- --- --- --- --- --- --- --- --- --- --- --- --- --- --- --- --- --- --- --- --- --- --- --- --- --- --- --- --- --- --- --- --- --- --- --- --- --- --- --- --- --- --- --- --- --- --- --- --- --- --- --- --- --- --- --- --- --- --- --- --- --- --- --- --- --- --- --- --- --- --- --- --- --- --- --- --- --- --- --- --- --- --- --- --- --- --- --- --- --- --- --- --- --- --- --- --- --- --- --- --- --- --- --- --- --- --- --- --- --- --- --- --- --- --- --- --- --- --- --- --- --- --- --- --- --- --- --- --- --- --- --- --- --- --- --- --- --- --- --- --- --- --- --- --- --- --- --- --- --- --- --- --- --- --- --- --- --- --- --- --- --- --- --- -ga gag aaa gga taa aaa ctt ttt acg act cca tca gaa aga ggt tta ata ttt ttg tga gac cca tcg aag aga gaa aga gat ggt tag tca aga tat ttt tct tag tac aaa agt caa tgt ttt aaa ata tat gga cga gaa tta att tgt ctg tat aaa aac ttg tgt gaa att atg tac tag aga aaa aac gtg agc agt gtc ccc tac atg gat ttt aca gat cat tta tat tcc aaa aat att aac tat ata cgt tta tta tat gat gtt aac gtg taa att ata aac att att tta tga tgc aat tgt ctg aca acc tag att ggt ata agg atg ttg ata agc tct acg aga ata tat tgt tgg acg tta tcg ttt acg aaa tag ttg aga cat cag aaa gag gtt taa tat ttt tgt gag a-c cat cga aga gag aaa gag aat aaa aat a-- -?t ttt ttt ttt ttt tgt aaa act ttt tta tga gac --- --- -ca aga gaa tac gaa tag tga tca tat cgt atc aca tat tga aac aga aag aag aag taa cga gag gta act ttt tgt gaa tgt agt taa ata ttt ttg ttt tgc aaa ccg gaa tat agt gcc cgg t-- -ct ttt tta att cgt ggt gcg gtg tct gaa tcg ttc gat taa ccc aac tca tcc att ttc aga tga ata gag tta tcg att cag aca cat gct ttg agt ttt gtt gaa tcg atg agt gaa gta tca tcg gtt gca cct tca gat gcc gat ccg tcg aca tac ttg aat cca tcc ttg act tca agt tca gat gat tcc tca cac atg tct ccg ata cgt acg cta aac tct agg ttc ttg aca cat ttt gta tca acg atc gtt gaa ccg atg ata tct ttg taa ctc act ttc tta tgt gag atg tta gac cca agt act gga tgg gtc ttg atg tca ctg tct ttc tct tct tcg cta cat ctg atg tcg ata gac atc tca cag tct ttg atc ata gcc aga gct tct tca cgc gtg atc gcg gga gag tcc tta cct tgt ccc ggt gac acg ctg gac aat cta gta ttc aca gtg ttt cca tca gag gat tcg gag atg gat aaa atc ttt ggg cat ttg gtg aat cca aag ttc atg tta aga ccc gca ccg acg ata gtg taa taa gtg gtg gga tct cct ttt aca act --- --- --- --- --- --- tct tcg gat acc tca tca tct tcg gtc tct gta act tcc gtt acg gat tga caa atc tta tca ttg gtc ggt gtt tgg tct tgc ttt gtg act ttg ata ata aca tcg att ccc ata tga tgt ttg ttt tct tct tca gta cac gag gat gaa gat tgt tga aga cta gta ggc ata gca gct gcc act agg cac atg cat gcc agg aca ata tat tgt ttc atg att gct att gat tga tta ctg ttc tag atg att cta ctt tct tac cat ata ata aat tag aat ata ttt tct act ttt acg aga aat taa tta ttg tat tta tta ttt ata ggt --a aaa aaa ctt act ata agt ggg tgg gat tct ggg aat tag tga tca gtt tat gta tat cgc aac tag cgg gca tat ggc tat tga cat cga gaa cat tac cca tat gat aag aga ttg tat cat ttt cgt agt ctt gag tat tgg tat tac tat ata gta tgt aga tgt cga cgc tag ata gac agt cgc cca cta gag tta ccg tct ctg aat gcg gca tga tag tat cat tct ttg ttt tcg tta act gtt tgg aag atg aat ctt tgt tgt tac att taa tct cga aat tca gag tac ata tct ttg aag tat tct gat atc tat ttt ctc ctg taa aga atc ctg aag ttg cta cat tat taa gga cag aga agt att ctg cac gaa aga ctg gat cac aat ctt tat gat tca tgg taa tag tta gtt ccg acg ttg aga tgg att cgc tga gac cgg tag tgg tcg tcc gag tac acg atg tgt cgt tga ctg gat aca ggt taa ttt cca cat cga tat agt taa atg tat tgc tgg tta cga cgg gtt cgc att tat ctg tgg aag aga cgg tgt gag aat atg ttc cgg gac cac acg gag aac aga tga cgt ctc cgg tag acg tgt atc cgg ata ctc cgt atc cta ttc cac act ttg ttt tag aaa tac atg ttc tac acc ctg atg ctc ctt tga gaa gac aat aat atc ctg gag agc att cac aga ttc tat tgt gag tcg tgt tac acg atc gcg tct cta cct gat tac tat cac atc ttc cgt tac aac tta gac aag cct gta aat gat tat tgt gag atg taa agg tat ccg aac cac acg gtg tac att gtg tat tag tct tgc tat cac ata atc tga aag cgt aag ttc ccg gag gac acg ata gac aac ata gat tac ggc ttc tgt att cgt tgt ctt tac act ttc cat tgg atg gtg cat gtg gtg cta tat ctc ttc cgt tta tta tta tac atg aga gaa aca ata tat acg agt ata ata cgg acc tca tga ttt aat aat -gt agt aat cgt cgt ctt gtt act gtt tgt ttc cta ctt ctc caa tca tat aga tta ttt ttt aaa tat ttt ctt tct atc atg gat aat att tgt aat ggt tct ttc cgt aca aca tac tgt tta gat ggt agt cgc tta gct tgg tta tga tat tgc gca taa ttt ccg gag gca aat acg ata gtc tag att gac tat cga tgg tag act cta att tat tga gtg ctt tgt cga cga gtt tac ttt tat gct cca tcg ata gat gac act gtt cta tga gat cgt cgt aca tgg gaa atg aaa tgc gtt tgt ctg aat gta tgg ctt cga gat atg tgt gat acc gaa tgt ctt ctg ttc tca ata ccg tat aca agt cgg tgt ctg aga ttc gaa tct ctt tga gga gac tta tgt cac gac tac att ttt cga tga tgg aat cta tct tat cga atg ata tat ttt tca taa ata cac ttt tat agt cct cgt tta aac aga att tag tat gta gtt ccg caa atg act cgt ccc tta ata ggc agt agg cta tta tct tct tta cgt agt gat cgt cgt agg gag aga act ccg aca tct tgt aga aca acg att taa tca tag gta gag ata ctt tca gtc tgt ggt gga taa tgt cat taa cga cat ccg cct tgt ata taa tgt ttc tgt ttt caa aca cca agt cga ata ctg tct tat cgt ctt tag tcg gaa ggt tga tgt cgt atc caa tgt ata cga ggt atg agg caa cat tgt tat tgc aat tct gga agg cgg tat gaa gag gag tca ttg tat tat agt att cgt ctt tct gaa tgt cga atc tat cta gta gat acc gta gta tat tga gag agc gac ttc cat atc ctt gat tat gtt tta tga ata gat aaa gta gat gtt gtc ctt ctt cct ttt gta att tcc cgt att ttt gtt cgt gcc aat tga gta aca tta tga gaa tat gac ctg ttg cac aat cgt tct tta tgt att cca tga tgg gtg tac aat taa gat tat tac gta tcc tcg tat cgg ctc ctc gag ata aaa gag cat aca cca cac gag gac tat gtt tgg tat act gtt gaa ggt aag tgt gta acg gcg tat ttc cga ttt tcg taa ccg cgt taa tgt ttg ctc cat gat cta tta tcg cgt aga tga atc gct tct cag ctc gca tct tag tgt gac tct ttg act tgt aat aat tgc ttt cgt gga acg cgg ata tgt gtt tac agt agt aat gaa gag aag tga gtc cat cct cat cga cgc aat tag ggt cag atc ctt tag tca ata att tgt aca gaa cgt aat agt tta agc tcc cat tga att tat atc taa gat aac aca gca ata gat cgg atg att tac taa agt cat caa tgg ggt ccg tta gta tat caa aga tct tgt tat cga ttg ata gtg aat gaa tca gat agt ggt gta gag gaa tat gtc ctt ttt tat cct tgc tat caa agt tac gca tgc cgt ggc gta aca ata tct tta ata cag atg gat taa atc gtg tat tca tcg tat agc aat gta atg gag agt tac cac att tta gtc gtt tat tca gat cgc agt gtt taa taa cta att taa aca gat gag atg ata tat cca cat caa aga atg cga gat aca tat gac aga cat tat tga cag aaa tgt gac ctt cat tat cac cgt cgt cca taa atg cgt tag gta cgt acc aca tac tat cgt taa cga tgc gca caa tct cgt cca -tt tca tcc atc ttc ata atg att tac ttt ttc ata att aga gaa aaa gat caa ggt ata aaa att aga agt gtt aga cta ta- --- --- --- --- --- --- --- --- --- --- -aa cta act tat gac tta act aac tta tga --- --- --- --- --- --- --- --- --- --- --- --- --- --- --- --- --- --- --- --- --- --- --- --- --- --- --- --- --- --- --- --- --- --- --- --- --- --- --- --- --- --- --- --- --- --- --- --- ctt aac ta- --- --- --- --- --- --- --- --- --- --- -ac tta tga ctt aac taa ctt atg act taa cta act ta- tga ctt aac taa ctt atg act taa cta act tat gac tta act aac tta tga cta tta act cat ttc aag aaa ggt ggg tgg ata gaa ctc tat atg aca gct tgt gaa aca att aga tcc cta att tct aat gga agt ttt gat agg aga ttg tca tca gtt gat aca ttg ttt att atc tca tct att aga gca cgt ctg ttt aga gct tta gtg acc tgc tcg gtt act tct gtg taa atc ttg aat cct tta gtg ata cac tgt gtc aaa act gga tgt tta gaa tac cta tgt aga ata tgg gaa gca tgc ttg ttt ttg tct cta tta tag att aac tca tac atg gtt gta tta tga att ttc atc tgc cta atg tac tcc aat tct tgt tta caa tca att ata taa tca aag agt gat gat gca tac aca tta caa agt gaa taa tct acc atc ata aaa tac ttg ata cag agc ttt atc aca tca tgg ttt tca att gta tta tta agt ata gct aat ttt ata cag tca ata gac aat ggt tct cta agc aat att tct aat att tta aga tgt gct tcc cta cgg gcg atg aca gat ccc cta tcc acg gcc acg tca aga cat gta tat cca tta ctc att act gcg ttg aca ttt gct cca ttt tct aat agc cat gat act aaa tct ata taa cct gca tag ata gcg cga taa agc aag gtc ctt cca cca gca tct agt tga ttg ata tct tca ata tat ggg ata caa agc tta taa att tct aat act gtg ggt tca tct aca agg aat ccc cta gta tac tga att att tta tat aga tct aat tta aca tca ttt tca tct ggg ata cca caa ttc aaa ata aac tca aca aca cta ctt tcc ttt tta cat att ccc cta aaa tag gca ttc aag cat tct att tta tat att aca gcc cca tga tct acc ata aga tca aca atg tct att tct aca tat gca tta gat aga tag taa agt aag aga tct tgc aca gaa tta caa ttc tta ata att ata gag aaa ata tct tcc ata taa ttc ttt gac act aat gca gat ata ata tct tta tat gta ata tat gca aac agt cta tct act ata tac tga tca ata tta tct cta tga atc cta aaa taa tca tac aga aca tct aca gga tca caa att ggt tca agg aga aat cta tca aat att ttc ctg tca aca act ggt tct aga aca taa cag tca aca cct aat cca tgt ttt tta taa tca tct acc aaa gat aat gac caa aga tcg agg tcg tcg tga aac tgc tca tcg aca gcc atg aaa tct acc gac tcc atg gtg cga atc gca ctg tct tat tcg cca ttg att ttc a-t ttt tta taa tta tgt aca tgt ttt cct tct att ctc aag agt cta caa aaa tat a-- -tt ttt tcg ata tct aag tac taa gtt ttt tta ctg ttt ttg tta ctg tct tcc att ctt cta act aaa gat ctg aga taa att ata caa tct tcg cta tcg aac cat ttt tgt agt cta aag cct gaa gta att aac caa ctg ttt tta tta gtg gct ttt ttc gat cta tcc tcg tc- --- ctc tca tca tcc tta ta- --- --- --- --- --- --- tta tta tca tta tta tca tag tct att aaa cac aaa tca tct --- --- --- --- --- --- --- --- --- --- --- --- --- --- --- --- --- --- --- --- --- --- --- --- --- --- --a cgt tta taa c-- --- --- --- --- --- --- --- --a aca ttc --- --- --- --- -tc att att aat tag ttc tgt ag- aat atc ttt aat aat ttg gct ata --- --- -ca tct gtt --- --- --- --- --- --- --- --- --- --- --- --- --- --- --- --- --- --- --c aat act --- --- --- --- --- --- --- --a tct att gat gat ttc ttt t-- --- --- --- --- --- --- --- --- --- --- --t taa gac t-- --- --- --- --- --- --- --- --- --- --- --- --- --- --- --- --- --- --- --- --- --- --- --- --- --- --- --- --- --- --- --- --- --- --- --- --- --- --- --- taa act agt --- --- --- --- --- --- --- --- --- --- --- --- --- --- --- --- --- --- --- --- --- --- --- --- --- --- --- --- --- --- --- --- --- --- --- --- --- --- --- --- --- --- --- --- --- --- --- --- --- --- --- --- --- --- --- --- --- --- --- --- --- --- --- --- --- --- --- --- --- --- --- --- --- --- --- --- --- --- --- --- --- --- --- --- --- --- --- --- --- --- --- --- --- --- --- --- --- --- --- --- --- --- --- --- --- --- --- --- --- --t atg gta atg acg atg aaa --- -tc gag tag ta- --- --a ctt cta ata aag act tga ta- --- --t cat tat cat atg ttt gat cg- --- --- --- --- --- --- --t cat agt taa tag tgt g-- --- --- --- --- --- --- --- --- --- --- --- --- --- --- --- --- --- --- --- -gc taa atg gta ctg tta ata agt tta t-- --- --- --- --- --- --- --- --- --- --- --- --- --- --- --- --- --a gac aat atc ata gta ttt tct ttc cag aat --- --- --- --t aga tta ttt ttt taa ata ctg atc ctc aca att ccg tga tgt agc agt agt tgg t-- --- --- --- --- --- --- --- --- --- --- --- --- --- --- --- --- --- --- --- --- --- --- --- --- --- --- --- -gc atg gtc tat atc gt- --- --- --- --- --- --- --- --- --- --- --- --- --- --- --- --- --- --- --- --- --- --- --- --- --- --- --- --- --- --- --- --- --- --- --- --- --- --- --- --- --- --- --- taa aat gta tca tat ata ata gtt ttc tga cgt gga gta cag aat ttt cga --- --- --- --- --- --- --- --- --- --- --- --- --- --- --- --- --- --- --- --- --- --- --- --- --- --- --- --- --- --- --- --- --- --- --- --- --- --- --- --- --- --- --- --t taa tga gtt cat ggt aag gaa ggg caa atg cct --- gta tat aat ata cat aag tta a-- --- --- --- --- --- --- --- --- --- --- --- --- --t agt ttt tta tca tat tt- --- --- --- --- --- --- --- --t cta ata cca taa taa aaa t-- --- --- --- --- --- --- --- --- --- --- --- --- --- --- --- --- --- --- --- --- --- --- --- --- --t atc att att gcg ttt g-g tag tt- --- --- --- --- ctg ccc ta- --- --- --- --- --- --- --- --t cat cta tat cac tgt cac tct c-- --- --g ctc tca cta tat ctt cta aaa tta caa --- -ac aac --- --- --- --- --- --- --- --- --- --- --- --- --- --- --- --- --- --- --- --- --- --- --- --- --- --- --- --- --- --- --- --- --- --- --- --- tgg ata ttc gat --- --- --- --- --- --- --- --- --- --- --- --- --- --- --- --- --- --- --- --- --- --- --- --- --- --- --a aca gca ttt gtg t-- --- --- --- --- --- --- --- --- --- --- --- --- --- --- --- --- -

>ON676706.1_Monkeypox_virus_isolate_MPXV_USA_2022_UT002_complete_genome

--- --- --- --- --- --- --- --- --- --- --- --- --- --- --- --- --- --- --- --- --- --- --- --- --- --- --- --- --- --- --- --- --- --- --- --- --- --- --- --- --- --- --- --- --g acc gta gga aac tct aga ggg taa gaa aaa tca atc gtt tat aga gac cat cag aaa gag gtt taa tat ttt tgt gag acc tat cga a-- --- --- --- --- --- --- --- --- --- --- --- --- --- --- --- --- --- --- --- --- --- --- --- --- --- --- --- --- --- --- --- --- --- --- --- --- --- --- --- --- --- --- --- --- --- --- --- --- --- --- --- --- --- --- --- --- --- --- --- --- --- --- --- --- --- --- --- --- --- --- --- --- --- --- --- --- --- --- --- --- --- --- --- --- --- --- --- --- --- --- --- --- --- --- --- --- --- --- --- --- --- --- --- --- --- --- --- --- --- --- --- --- --- --- --- --- --- --- --- --- --- --- --- --- --- --- --- --- --- --- --- --- --- --- --- --- --- --- --- --- --- --- --- --- --- --- --- --- --- --- --- --- --- --- --- --- --- --- --- --- --- --- --- --- --- --- --- --- --- --- --- --- --- --- --- --- --- --- --- --- --- --- --- --- --- --- --- --- --- --- --- --- --- --- --- --- --- --- --- --- --- --- --- --- --- --- --- --- --- --- --- --- --- --- --- --- --- --- --- --- --- --- --- --- --- --- --- --- --- --- --- --- --- --- --- --- --- --- --- --- --- --- --- --- --- --- --- --- --- --- --- --- --- --- --- --- --- --- --- --- --- --- --- --- --- --- --- --- --- --- --- --- --- --- --- --- --- --- --- -ga gag aaa gga taa aaa ctt ttt acg act cca tca gaa aga ggt tta ata ttt ttg tga gac cca tcg aag aga gaa aga gat ggt tag tca aga tat ttt tct tag tac aaa agt caa tgt ttt aaa ata tat gga cga gaa tta att tgt ctg tat aaa aac ttg tgt gaa att atg tac tag aga aaa aac gtg agc agt gtc ccc tac atg gat ttt aca gat cat tta tat tcc aaa aat att aac tat ata cgt tta tta tat gat gtt aac gtg taa att ata aac att att tta tga tgc aat tgt ctg aca acc tag att ggt ata agg atg ttg ata agc tct acg aga ata tat tgt tgg acg tta tcg ttt acg aaa tag ttg aga cat cag aaa gag gtt taa tat ttt tgt gag a-c cat cga aga gag aaa gag aat aaa aat a-- -?t ttt ttt ttt ttt tgt aaa act ttt tta tga gac --- --- -ca aga gaa tac gaa tag tga tca tat cgt atc aca tat tga aac aga aag aag aag taa cga gag gta act ttt tgt gaa tgt agt taa ata ttt ttg ttt tgc aaa ccg gaa tat agt gcc cgg t-- -ct ttt tta att cgt ggt gcg gtg tct gaa tcg ttc gat taa ccc aac tca tcc att ttc aga tga ata gag tta tcg att cag aca cat gct ttg agt ttt gtt gaa tcg atg agt gaa gta tca tcg gtt gca cct tca gat gcc gat ccg tcg aca tac ttg aat cca tcc ttg act tca agt tca gat gat tcc tca cac atg tct ccg ata cgt acg cta aac tct agg ttc ttg aca cat ttt gta tca acg atc gtt gaa ccg atg ata tct ttg taa ctc act ttc tta tgt gag atg tta gac cca agt act gga tgg gtc ttg atg tca ctg tct ttc tct tct tcg cta cat ctg atg tcg ata gac atc tca cag tct ttg atc ata gcc aga gct tct tca cgc gtg atc gcg gga gag tcc tta cct tgt ccc ggt gac acg ctg gac aat cta gta ttc aca gtg ttt cca tca gag gat tcg gag atg gat aaa atc ttt ggg cat ttg gtg aat cca aag ttc atg tta aga ccc gca ccg acg ata gtg taa taa gtg gtg gga tct cct ttt aca act --- --- --- --- --- --- tct tcg gat acc tca tca tct tcg gtc tct gta act tcc gtt acg gat tga caa atc tta tca ttg gtc ggt gtt tgg tct tgc ttt gtg act ttg ata ata aca tcg att ccc ata tga tgt ttg ttt tct tct tca gta cac gag gat gaa gat tgt tga aga cta gta ggc ata gca gct gcc act agg cac atg cat gcc agg aca ata tat tgt ttc atg att gct att gat tga tta ctg ttc tag atg att cta ctt tct tac cat ata ata aat tag aat ata ttt tct act ttt acg aga aat taa tta ttg tat tta tta ttt ata ggt --a aaa aaa ctt act ata agt ggg tgg gat tct ggg aat tag tga tca gtt tat gta tat cgc aac tag cgg gca tat ggc tat tga cat cga gaa cat tac cca tat gat aag aga ttg tat cat ttt cgt agt ctt gag tat tgg tat tac tat ata gta tgt aga tgt cga cgc tag ata gac agt cgc cca cta gag tta ccg tct ctg aat gcg gca tga tag tat cat tct ttg ttt tcg tta act gtt tgg aag atg aat ctt tgt tgt tac att taa tct cga aat tca gag tac ata tct ttg aag tat tct gat atc tat ttt ctc ctg taa aga atc ctg aag ttg cta cat tat taa gga cag aga agt att ctg cac gaa aga ctg gat cac aat ctt tat gat tca tgg taa tag tta gtt ccg acg ttg aga tgg att cgc tga gac cgg tag tgg tcg tcc gag tac acg atg tgt cgt tga ctg gat aca ggt taa ttt cca cat cga tat agt taa atg tat tgc tgg tta cga cgg gtt cgc att tat ctg tgg aag aga cgg tgt gag aat atg ttc cgg gac cac acg gag aac aga tga cgt ctc cgg tag acg tgt atc cgg ata ctc cgt atc cta ttc cac act ttg ttt tag aaa tac atg ttc tac acc ctg atg ctc ctt tga gaa gac aat aat atc ctg gag agc att cac aga ttc tat tgt gag tcg tgt tac acg atc gcg tct cta cct gat tac tat cac atc ttc cgt tac aac tta gac aag cct gta aat gat tat tgt gag atg taa agg tat ccg aac cac acg gtg tac att gtg tat tag tct tgc tat cac ata atc tga aag cgt aag ttc ccg gag gac acg ata gac aac ata gat tac ggc ttc tgt att cgt tgt ctt tac act ttc cat tgg atg gtg cat gtg gtg cta tat ctc ttc cgt tta tta tta tac atg aga gaa aca ata tat acg agt ata ata cgg acc tca tga ttt aat aat -gt agt aat cgt cgt ctt gtt act gtt tgt ttc cta ctt ctc caa tca tat aga tta ttt ttt aaa tat ttt ctt tct atc atg gat aat att tgt aat ggt tct ttc cgt aca aca tac tgt tta gat ggt agt cgc tta gct tgg tta tga tat tgc gca taa ttt ccg gag gca aat acg ata gtc tag att gac tat cga tgg tag act cta att tat tga gtg ctt tgt cga cga gtt tac ttt tat gct cca tcg ata gat gac act gtt cta tga gat cgt cgt aca tgg gaa atg aaa tgc gtt tgt ctg aat gta tgg ctt cga gat atg tgt gat acc gaa tgt ctt ctg ttc tca ata ccg tat aca agt cgg tgt ctg aga ttc gaa tct ctt tga gga gac tta tgt cac gac tac att ttt cga tga tgg aat cta tct tat cga atg ata tat ttt tca taa ata cac ttt tat agt cct cgt tta aac aga att tag tat gta gtt ccg caa atg act cgt ccc tta ata ggc agt agg cta tta tct tct tta cgt agt gat cgt cgt agg gag aga act ccg aca tct tgt aga aca acg att taa tca tag gta gag ata ctt tca gtc tgt ggt gga taa tgt cat taa cga cat ccg cct tgt ata taa tgt ttc tgt ttt caa aca cca agt cga ata ctg tct tat cgt ctt tag tcg gaa ggt tga tgt cgt atc caa tgt ata cga ggt atg agg caa cat tgt tat tgc aat tct gga agg cgg tat gaa gag gag tca ttg tat tat agt att cgt ctt tct gaa tgt cga atc tat cta gta gat acc gta gta tat tga gag agc gac ttc cat atc ctt gat tat gtt tta tga ata gat aaa gta gat gtt gtc ctt ctt cct ttt gta att tcc cgt att ttt gtt cgt gcc aat tga gta aca tta tga gaa tat gac ctg ttg cac aat cgt tct tta tgt att cca tga tgg gtg tac aat taa gat tat tac gta tcc tcg tat cgg ctc ctc gag ata aaa gag cat aca cca cac gag gac tat gtt tgg tat act gtt gaa ggt aag tgt gta acg gcg tat ttc cga ttt tcg taa ccg cgt taa tgt ttg ctc cat gat cta tta tcg cgt aga tga atc gct tct cag ctc gca tct tag tgt gac tct ttg act tgt aat aat tgc ttt cgt gga acg cgg ata tgt gtt tac agt agt aat gaa gag aag tga gtc cat cct cat cga cgc aat tag ggt cag atc ctt tag tca ata att tgt aca gaa cgt aat agt tta agc tcc cat tga att tat atc taa gat aac aca gca ata gat cgg atg att tac taa agt cat caa tgg ggt ccg tta gta tat caa aga tct tgt tat cga ttg ata gtg aat gaa tca gat agt ggt gta gag gaa tat gtc ctt ttt tat cct tgc tat caa agt tac gca tgc cgt ggc gta aca ata tct tta ata cag atg gat taa atc gtg tat tca tcg tat agc aat gta atg gag agt tac cac att tta gtc gtt tat tca gat cgc agt gtt taa taa cta att taa aca gat gag atg ata tat cca cat caa aga atg cga gat aca tat gac aga cat tat tga cag aaa tgt gac ctt cat tat cac cgt cgt cca taa atg cgt tag gta cgt acc aca tac tat cgt taa cga tgc gca caa tct cgt cca -tt tca tcc atc ttc ata atg att tac ttt ttc ata att aga gaa aaa gat caa ggt ata aaa att aga agt gtt aga cta ta- --- --- --- --- --- --- --- --- --- --- -aa cta act tat gac tta act aac tta tga --- --- --- --- --- --- --- --- --- --- --- --- --- --- --- --- --- --- --- --- --- --- --- --- --- --- --- --- --- --- --- --- --- --- --- --- --- --- --- --- --- --- --- --- --- --- --- --- ctt aac ta- --- --- --- --- --- --- --- --- --- --- -ac tta tga ctt aac taa ctt atg act taa cta act ta- tga ctt aac taa ctt atg act taa cta act tat gac tta act aac tta tga cta tta act cat ttc aag aaa ggt ggg tgg ata gaa ctc tat atg aca gct tgt gaa aca att aga tcc cta att tct aat gga agt ttt gat agg aga ttg tca tca gtt gat aca ttg ttt att atc tca tct att aga gca cgt ctg ttt aga gct tta gtg acc tgc tcg gtt act tct gtg taa atc ttg aat cct tta gtg ata cac tgt gtc aaa act gga tgt tta gaa tac cta tgt aga ata tgg gaa gca tgc ttg ttt ttg tct cta tta tag att aac tca tac atg gtt gta tta tga att ttc atc tgc cta atg tac tcc aat tct tgt tta caa tca att ata taa tca aag agt gat gat gca tac aca tta caa agt gaa taa tct acc atc ata aaa tac ttg ata cag agc ttt atc aca tca tgg ttt tca att gta tta tta agt ata gct aat ttt ata cag tca ata gac aat ggt tct cta agc aat att tct aat att tta aga tgt gct tcc cta cgg gcg atg aca gat ccc cta tcc acg gcc acg tca aga cat gta tat cca tta ctc att act gcg ttg aca ttt gct cca ttt tct aat agc cat gat act aaa tct ata taa cct gca tag ata gcg cga taa agc aag gtc ctt cca cca gca tct agt tga ttg ata tct tca ata tat ggg ata caa agc tta taa att tct aat act gtg ggt tca tct aca agg aat ccc cta gta tac tga att att tta tat aga tct aat tta aca tca ttt tca tct ggg ata cca caa ttc aaa ata aac tca aca aca cta ctt tcc ttt tta cat att ccc cta aaa tag gca ttc aag cat tct att tta tat att aca gcc cca tga tct acc ata aga tca aca atg tct att tct aca tat gca tta gat aga tag taa agt aag aga tct tgc aca gaa tta caa ttc tta ata att ata gag aaa ata tct tcc ata taa ttc ttt gac act aat gca gat ata ata tct tta tat gta ata tat gca aac agt cta tct act ata tac tga tca ata tta tct cta tga atc cta aaa taa tca tac aga aca tct aca gga tca caa att ggt tca agg aga aat cta tca aat att ttc ctg tca aca act ggt tct aga aca taa cag tca aca cct aat cca tgt ttt tta taa tca tct acc aaa gat aat gac caa aga tcg agg tcg tcg tga aac tgc tca tcg aca gcc atg aaa tct acc gac tcc atg gtg cga atc gca ctg tct tat tcg cca ttg att ttc a-t ttt tta taa tta tgt aca tgt ttt cct tct att ctc aag agt cta caa aaa tat a-- -tt ttt tcg ata tct aag tac taa gtt ttt tta ctg ttt ttg tta ctg tct tcc att ctt cta act aaa gat ctg aga taa att ata caa tct tcg cta tcg aac cat ttt tgt agt cta aag cct gaa gta att aac caa ctg ttt tta tta gtg gct ttt ttc gat cta tcc tcg tc- --- ctc tca tca tcc tta ta- --- --- --- --- --- --- tta tta tca tta tta tca tag tct att aaa cac aaa tca tct --- --- --- --- --- --- --- --- --- --- --- --- --- --- --- --- --- --- --- --- --- --- --- --- --- --- --a cgt tta taa c-- --- --- --- --- --- --- --- --a aca ttc --- --- --- --- -tc att att aat tag ttc tgt ag- aat atc ttt aat aat ttg gct ata --- --- -ca tct gtt --- --- --- --- --- --- --- --- --- --- --- --- --- --- --- --- --- --- --c aat act --- --- --- --- --- --- --- --a tct att gat gat ttc ttt t-- --- --- --- --- --- --- --- --- --- --- --t taa gac t-- --- --- --- --- --- --- --- --- --- --- --- --- --- --- --- --- --- --- --- --- --- --- --- --- --- --- --- --- --- --- --- --- --- --- --- --- --- --- --- taa act agt --- --- --- --- --- --- --- --- --- --- --- --- --- --- --- --- --- --- --- --- --- --- --- --- --- --- --- --- --- --- --- --- --- --- --- --- --- --- --- --- --- --- --- --- --- --- --- --- --- --- --- --- --- --- --- --- --- --- --- --- --- --- --- --- --- --- --- --- --- --- --- --- --- --- --- --- --- --- --- --- --- --- --- --- --- --- --- --- --- --- --- --- --- --- --- --- --- --- --- --- --- --- --- --- --- --- --- --- --- --t atg gta atg acg atg aaa --- -tc gag tag ta- --- --a ctt cta ata aag act tga ta- --- --t cat tat cat atg ttt gat cg- --- --- --- --- --- --- --t cat agt taa tag tgt g-- --- --- --- --- --- --- --- --- --- --- --- --- --- --- --- --- --- --- --- -gc taa atg gta ctg tta ata agt tta t-- --- --- --- --- --- --- --- --- --- --- --- --- --- --- --- --- --a gac aat atc ata gta ttt tct ttc cag aat --- --- --- --t aga tta ttt ttt taa ata ctg atc ctc aca att ccg tga tgt agc agt agt tgg t-- --- --- --- --- --- --- --- --- --- --- --- --- --- --- --- --- --- --- --- --- --- --- --- --- --- --- --- -gc atg gtc tat atc gt- --- --- --- --- --- --- --- --- --- --- --- --- --- --- --- --- --- --- --- --- --- --- --- --- --- --- --- --- --- --- --- --- --- --- --- --- --- --- --- --- --- --- --- taa aat gta tca tat ata ata gtt ttc tga cgt gga gta cag aat ttt cga --- --- --- --- --- --- --- --- --- --- --- --- --- --- --- --- --- --- --- --- --- --- --- --- --- --- --- --- --- --- --- --- --- --- --- --- --- --- --- --- --- --- --- --t taa tga gtt cat ggt aag gaa ggg caa atg cct --- gta tat aat ata cat aag tta a-- --- --- --- --- --- --- --- --- --- --- --- --- --t agt ttt tta tca tat tt- --- --- --- --- --- --- --- --t cta ata cca taa taa aaa t-- --- --- --- --- --- --- --- --- --- --- --- --- --- --- --- --- --- --- --- --- --- --- --- --- --t atc att att gcg ttt g-g tag tt- --- --- --- --- ctg ccc ta- --- --- --- --- --- --- --- --t cat cta tat cac tgt cac tct c-- --- --g ctc tca cta tat ctt cta aaa tta caa --- -ac aac --- --- --- --- --- --- --- --- --- --- --- --- --- --- --- --- --- --- --- --- --- --- --- --- --- --- --- --- --- --- --- --- --- --- --- --- tgg ata ttc gat --- --- --- --- --- --- --- --- --- --- --- --- --- --- --- --- --- --- --- --- --- --- --- --- --- --- --a aca gca ttt gtg t-- --- --- --- --- --- --- --- --- --- --- --- --- --- --- --- --- -

>ON649879.1_Monkeypox_virus_isolate_MPXV_ISR001_2022_complete_genome

--- --- --- --- --- --- --- --- --- --- --- --- --- --- --- --- --- --- --- --- --- --- --- --- --- --- --- --- --- --- --- --- --- --- --- --- --- --- --- --- --- --- --- --- --- --- --- --- --- --- --- --- --- --- --- --- --- --- --- --- --- --- --- --- --- --- --- --- --- --- --- --- --- --- --- --- --- --- --- --- --- --- --- --- --- --- --- --- --- --- --- --- --- --- --- --- --- --- --- --- --- --- --- --- --- --- --- --- --- --- --- --- --- --- --- --- --- --- --- --- --- --- --- --- --- --- --- --- --- --- --- --- --- --- --- --- --- --- --- --- --- --- --- --- --- --- --- --- --- --- --- --- --- --- --- --- --- --- --- --- --- --- --- --- --- --- --- --- --- --- --- --- --- --- --- --- --- --- --- --- --- --- --- --- --- --- --- --- --- --- --- --- --- --- --- --- --- --- --- --- --- --- --- --- --- --- --- --- --- --- --- --- --- --- --- --- --- --- --- --- --- --- --- --- --- --- --- --- --- --- --- --- --- --- --- --- --- --- --- --- --- --- --- --- --- --- --- --- --- --- --- --- --- --- --- --- --- --- --- --- --- --- --- --- --- --- --- --- --- --- --- --- --- --- --- --- --- --- --- --- --- --- --- --- --- --- --- --- --- --- --- --- --- --- --- --- --- --- --- --- --- --- --- --- --- --- --- --- --- --- --- --- --- --- --- --- --- --- --- --- --- --- --- --- --- --- --- --- --- --- --- --- --- --- --- --- --- --- --- --- --- --- --- --- --- --- --- --- --- --- --- --- --- --- --- --- --- --- --- --- --- --- --- --- -ca tca gaa aga ggt tta ata ttt ttg tga gac cca tcg aag aga gaa aga gat ggt tag tca aga tat ttt tct tag tac aaa agt caa tgt ttt aaa ata tat gga cga gaa tta att tgt ctg tat aaa aac ttg tgt gaa att atg tac tag aga aaa aac gtg agc agt gtc ccc tac atg gat ttt aca gat cat tta tat tcc aaa aat att aac tat ata cgt tta tta tat gat gtt aac gtg taa att ata aac att att tta tga tgc aat tgt ctg aca acc tag att ggt ata agg atg ttg ata agc tct acg aga ata tat tgt tgg acg tta tcg ttt acg aaa tag ttg aga cat cag aaa gag gtt taa tat ttt tgt gag a-c cat cga aga gag aaa gag aat aaa aat a-- --t ttt ttt ttt ttt tgt aaa act ttt tta tga gac --- --- -ca aga gaa tac gaa tag tga tca tat cgt atc aca tat tga aac aga aag aag aag taa cga gag gta act ttt tgt gaa tgt agt taa ata ttt ttg ttt tgc aaa ccg gaa tat agt gcc cgg t-- -ct ttt tta att cgt ggt gcg gtg tct gaa tcg ttc gat taa ccc aac tca tcc att ttc aga tga ata gag tta tcg att cag aca cat gct ttg agt ttt gtt gaa tcg atg agt gaa gta tca tcg gtt gca cct tca gat gcc gat ccg tcg aca tac ttg aat cca tcc ttg act tca agt tca gat gat tcc tca cac atg tct ccg ata cgt acg cta aac tct agg ttc ttg aca cat ttt gta tca acg atc gtt gaa ccg atg ata tct ttg taa ctc act ttc tta tgt gag atg tta gac cca agt act gga tgg gtc ttg atg tca ctg tct ttc tct tct tcg cta cat ctg atg tcg ata gac atc tca cag tct ttg atc ata gcc aga gct tct tca cgc gtg atc gcg gga gag tcc tta cct tgt ccc ggt gac acg ctg gac aat cta gta ttc aca gtg ttt cca tca gag gat tcg gag atg gat aaa atc ttt ggg cat ttg gtg aat cca aag ttc atg tta aga ccc gca ccg acg ata gtg taa taa gtg gtg gga tct cct ttt aca act --- --- --- --- --- --- tct tcg gat acc tca tca tct tcg gtc tct gta act tcc gtt acg gat tga caa atc tta tca ttg gtc ggt gtt tgg tct tgc ttt gtg act ttg ata ata aca tcg att ccc ata tga tgt ttg ttt tct tct tca gta cac gag gat gaa gat tgt tga aga cta gta ggc ata gca gct gcc act agg cac atg cat gcc agg aca ata tat tgt ttc atg att gct att gat tga tta ctg ttc tag atg att cta ctt tct tac cat ata ata aat tag aat ata ttt tct act ttt acg aga aat taa tta ttg tat tta tta ttt ata ggt --a aaa aaa ctt act ata agt ggg tgg gat tct ggg aat tag tga tca gtt tat gta tat cgc aac tag cgg gca tat ggc tat tga cat cga gaa cat tac cca tat gat aag aga ttg tat cat ttt cgt agt ctt gag tat tgg tat tac tat ata gta tgt aga tgt cga cgc tag ata gac agt cgc cca cta gag tta ccg tct ctg aat gcg gca tga tag tat cat tct ttg ttt tcg tta act gtt tgg aag atg aat ctt tgt tgt tac att taa tct cga aat tca gag tac ata tct ttg aag tat tct gat atc tat ttt ctc ctg taa aga atc ctg aag ttg cta cat tat taa gga cag aga agt att ctg cac gaa aga ctg gat cac aat ctt tat gat tca tgg taa tag tta gtt ccg acg ttg aga tgg att cgc tga gac cgg tag tgg tcg tcc gag tac acg atg tgt cgt tga ctg gat aca ggt taa ttt cca cat cga tat agt taa atg tat tgc tgg tta cga cgg gtt cgc att tat ctg tgg aag aga cgg tgt gag aat atg ttc cgg gac cac acg gag aac aga tga cgt ctc cgg tag acg tgt atc cgg ata ctc cgt atc cta ttc cac act ttg ttt tag aaa tac atg ttc tac acc ctg atg ctc ctt tga gaa gac aat aat atc ctg gag agc att cac aga ttc tat tgt gag tcg tgt tac acg atc gcg tct cta cct gat tac tat cac atc ttc cgt tac aac tta gac aag cct gta aat gat tat tgt gag atg taa agg tat ccg aac cac acg gtg tac att gtg tat tag tct tgc tat cac ata atc tga aag cgt aag ttc ccg gag gac acg ata gac aac ata gat tac ggc ttc tgt att cgt tgt ctt tac act ttc cat tgg atg gtg cat gtg gtg cta tat ctc ttc cgt tta tta tta tac atg aga gaa aca ata tat acg agt ata ata cgg acc tca tga ttt aat aat -gt agt aat cgt cgt ctt gtt act gtt tgt ttc cta ctt ctc caa tca tat aga tta ttt ttt aaa tat ttt ctt tct atc atg gat aat att tgt aat ggt tct ttc cgt aca aca tac tgt tta gat ggt agt cgc tta gct tgg tta tga tat tgc gca taa ttt ccg gag gca aat acg ata gtc tag att gac tat cga tgg tag act cta att tat tga gtg ctt tgt cga cga gtt tac ttt tat gct cca tcg ata gat gac act gtt cta tga gat cgt cgt aca tgg gaa atg aaa tgc gtt tgt ctg aat gta tgg ctt cga gat atg tgt gat acc gaa tgt ctt ctg ttc tca ata ccg tat aca agt cgg tgt ctg aga ttc gaa tct ctt tga gga gac tta tgt cac gac tac att ttt cga tga tgg aat cta tct tat cga atg ata tat ttt tca taa ata cac ttt tat agt cct cgt tta aac aga att tag tat gta gtt ccg caa atg act cgt ccc tta ata ggc agt agg cta tta tct tct tta cgt agt gat cgt cgt agg gag aga act ccg aca tct tgt aga aca acg att taa tca tag gta gag ata ctt tca gtc tgt ggt gga taa tgt cat taa cga cat ccg cct tgt ata taa tgt ttc tgt ttt caa aca cca agt cga ata ctg tct tat cgt ctt tag tcg gaa ggt tga tgt cgt atc caa tgt ata cga ggt atg agg caa cat tgt tat tgc aat tct gga agg cgg tat gaa gag gag tca ttg tat tat agt att cgt ctt tct gaa tgt cga atc tat cta gta gat acc gta gta tat tga gag agc gac ttc cat atc ctt gat tat gtt tta tga ata gat aaa gta gat gtt gtc ctt ctt cct ttt gta att tcc cgt att ttt gtt cgt gcc aat tga gta aca tta tga gaa tat gac ctg ttg cac aat cgt tct tta tgt att cca tga tgg gtg tac aat taa gat tat tac gta tcc tcg tat cgg ctc ctc gag ata aaa gag cat aca cca cac gag gac tat gtt tgg tat act gtt gaa ggt aag tgt gta acg gcg tat ttc cga ttt tcg taa ccg cgt taa tgt ttg ctc cat gat cta tta tcg cgt aga tga atc gct tct cag ctc gca tct tag tgt gac tct ttg act tgt aat aat tgc ttt cgt gga acg cgg ata tgt gtt tac agt agt aat gaa gag aag tga gtc cat cct cat cga cgc aat tag ggt cag atc ctt tag tca ata att tgt aca gaa cgt aat agt tta agc tcc cat tga att tat atc taa gat aac aca gca ata gat cgg atg att tac taa agt cat caa tgg ggt ccg tta gta tat caa aga tct tgt tat cga ttg ata gtg aat gaa tca gat agt ggt gta gag gaa tat gtc ctt ttt tat cct tgc tat caa agt tac gca tgc cgt ggc gta aca ata tct tta ata cag atg gat taa atc gtg tat tca tcg tat agc aat gta atg gag agt tac cac att tta gtc gtt tat tca gat cgc agt gtt taa taa cta att taa aca gat gag atg ata tat cca cat caa aga atg cga gat aca tat gac aga cat tat tga cag aaa tgt gac ctt cat tat cac cgt cgt cca taa atg cgt tag gta cgt acc aca tac tat cgt taa cga tgc gca caa tct cgt cca -tt tca tcc atc ttc ata atg att tac ttt ttc ata att aga gaa aaa gat caa ggt ata aaa att aga agt gtt aga cta ta- --- --- --- --- --- --- --- --- --- --- -aa cta act tat gac tta act aac tta tga --- --- --- --- --- --- --- --- --- --- --- --- --- --- --- --- --- --- --- --- --- --- --- --- --- --- --- --- --- --- --- --- --- --- --- --- --- --- --- --- --- --- --- --- --- --- --- --- ctt aac ta- --- --- --- --- --- --- --- --- --- --- -ac tta tga ctt aac taa ctt atg act taa cta act ta- tga ctt aac taa ctt atg act taa cta act tat gac tta act aac tta tga cta tta act cat ttc aag aaa ggt ggg tgg ata gaa ctc tat atg aca gct tgt gaa aca att aga tcc cta att tct aat gga agt ttt gat agg aga ttg tca tca gtt gat aca ttg ttt att atc tca tct att aga gca cgt ctg ttt aga gct tta gtg acc tgc tcg gtt act tct gtg taa atc ttg aat cct tta gtg ata cac tgt gtc aaa act gga tgt tta gaa tac cta tgt aga ata tgg gaa gca tgc ttg ttt ttg tct cta tta tag att aac tca tac atg gtt gta tta tga att ttc atc tgc cta atg tac tcc aat tct tgt tta caa tca att ata taa tca aag agt gat gat gca tac aca tta caa agt gaa taa tct acc atc ata aaa tac ttg ata cag agc ttt atc aca tca tgg ttt tca att gta tta tta agt ata gct aat ttt ata cag tca ata gac aat ggt tct cta agc aat att tct aat att tta aga tgt gct tcc cta cgg gcg atg aca gat ccc cta tcc acg gcc acg tca aga cat gta tat cca tta ctc att act gcg ttg aca ttt gct cca ttt tct aat agc cat gat act aaa tct ata taa cct gca tag ata gcg cga taa agc aag gtc ctt cca cca gca tct agt tga ttg ata tct tca ata tat ggg ata caa agc tta taa att tct aat act gtg ggt tca tct aca agg aat ccc cta gta tac tga att att tta tat aga tct aat tta aca tca ttt tca tct ggg ata cca caa ttc aaa ata aac tca aca aca cta ctt tcc ttt tta cat att ccc cta aaa tag gca ttc aag cat tct att tta tat att aca gcc cca tga tct acc ata aga tca aca atg tct att tct aca tat gca tta gat aga tag taa agt aag aga tct tgc aca gaa tta caa ttc tta ata att ata gag aaa ata tct tcc ata taa ttc ttt gac act aat gca gat ata ata tct tta tat gta ata tat gca aac agt cta tct act ata tac tga tca ata tta tct cta tga atc cta aaa taa tca tac aga aca tct aca gga tca caa att ggt tca agg aga aat cta tca aat att ttc ctg tca aca act ggt tct aga aca taa cag tca aca cct aat cca tgt ttt tta taa tca tct acc aaa gat aat gac caa aga tcg agg tcg tcg tga aac tgc tca tcg aca gcc atg aaa tct acc gac tcc atg gtg cga atc gca ctg tct tat tcg cca ttg att ttc a-t ttt tta taa tta tgt aca tgt ttt cct tct att ctc aag agt cta caa aaa tat a-- -tt ttt tcg ata tct aag tac taa gtt ttt tta ctg ttt ttg tta ctg tct tcc att ctt cta act aaa gat ctg aga taa att ata caa tct tcg cta tcg aac cat ttt tgt agt cta aag cct gaa gta att aac caa ctg ttt tta tta gtg gct ttt ttc gat cta tcc tcg tc- --- ctc tca tca tcc tta ta- --- --- --- --- --- --- tta tta tca tta tta tca tag tct att aaa cac aaa tca tct --- --- --- --- --- --- --- --- --- --- --- --- --- --- --- --- --- --- --- --- --- --- --- --- --- --- --a cgt tta taa c-- --- --- --- --- --- --- --- --a aca ttc --- --- --- --- -tc att att aat tag ttc tgt ag- aat atc ttt aat aat ttg gct ata --- --- -ca tct gtt --- --- --- --- --- --- --- --- --- --- --- --- --- --- --- --- --- --- --c aat act --- --- --- --- --- --- --- --a tct att gat gat ttc ttt t-- --- --- --- --- --- --- --- --- --- --- --t taa gac t-- --- --- --- --- --- --- --- --- --- --- --- --- --- --- --- --- --- --- --- --- --- --- --- --- --- --- --- --- --- --- --- --- --- --- --- --- --- --- --- taa act agt --- --- --- --- --- --- --- --- --- --- --- --- --- --- --- --- --- --- --- --- --- --- --- --- --- --- --- --- --- --- --- --- --- --- --- --- --- --- --- --- --- --- --- --- --- --- --- --- --- --- --- --- --- --- --- --- --- --- --- --- --- --- --- --- --- --- --- --- --- --- --- --- --- --- --- --- --- --- --- --- --- --- --- --- --- --- --- --- --- --- --- --- --- --- --- --- --- --- --- --- --- --- --- --- --- --- --- --- --- --t atg gta atg acg atg aaa --- -tc gag tag ta- --- --a ctt cta ata aag act tga ta- --- --t cat tat cat atg ttt gat cg- --- --- --- --- --- --- --t cat agt taa tag tgt g-- --- --- --- --- --- --- --- --- --- --- --- --- --- --- --- --- --- --- --- -gc taa atg gta ctg tta ata agt tta t-- --- --- --- --- --- --- --- --- --- --- --- --- --- --- --- --- --a gac aat atc ata gta ttt tct ttc cag aat --- --- --- --t aga tta ttt ttt taa ata ctg atc ctc aca att ccg tga tgt agc agt agt tgg t-- --- --- --- --- --- --- --- --- --- --- --- --- --- --- --- --- --- --- --- --- --- --- --- --- --- --- --- -gc atg gtc tat atc gt- --- --- --- --- --- --- --- --- --- --- --- --- --- --- --- --- --- --- --- --- --- --- --- --- --- --- --- --- --- --- --- --- --- --- --- --- --- --- --- --- --- --- --- taa aat gta tca tat ata ata gtt ttc tga cgt gga gta cag aat ttt cga --- --- --- --- --- --- --- --- --- --- --- --- --- --- --- --- --- --- --- --- --- --- --- --- --- --- --- --- --- --- --- --- --- --- --- --- --- --- --- --- --- --- --- --t taa tga gtt cat ggt aag gaa ggg caa atg cct --- gta tat aat ata cat aag tta a-- --- --- --- --- --- --- --- --- --- --- --- --- --t agt ttt tta tca tat tt- --- --- --- --- --- --- --- --t cta ata cca taa taa aaa t-- --- --- --- --- --- --- --- --- --- --- --- --- --- --- --- --- --- --- --- --- --- --- --- --- --t atc att att gcg ttt g-g tag tt- --- --- --- --- ctg ccc ta- --- --- --- --- --- --- --- --t cat cta tat cac tgt cac tct c-- --- --g ctc tca cta tat ctt cta aaa tta caa --- -ac aac --- --- --- --- --- --- --- --- --- --- --- --- --- --- --- --- --- --- --- --- --- --- --- --- --- --- --- --- --- --- --- --- --- --- --- --- tgg ata ttc gat --- --- --- --- --- --- --- --- --- --- --- --- --- --- --- --- --- --- --- --- --- --- --- --- --- --- --a aca gca ttt gtg t-- --- --- --- --- --- --- --- --- --- --- --- --- --- --- --- --- -

>ON602722.1_Monkeypox_virus_isolate_MPXV_FRA_2022_TLS67_complete_genome

--- --- --- --- --- --- --- --- --- --- --- --- --- --- --- --- --- --- --- --- --- --- --- --- --- --- --- --t att ata ttt tac tat ttt att tag tgt cta g-a aaa aat gtg tga ccc acg acc gta gga aac tct aga ggg taa gaa aaa tca atc gtt tat aga gac cat cag aaa gag gtt taa tat ttt tgt gag acc tat cga a-- --- --- --- --- --- --- --- --- --- --- --- --- --- --- --- --- --- --- --- --- --- --- --- --- --- --- --- --- --- --- --- --- --- --- --- --- --- --- --- --- --- --- --- --- --- --- --- --- --- --- --- --- --- --- --- --- --- --- --- --- --- --- --- --- --- --- --- --- --- --- --- --- --- --- --- --- --- --- --- --- --- --- --- --- --- --- --- --- --- --- --- --- --- --- --- --- --- --- --- --- --- --- --- --- --- --- --- --- --- --- --- --- --- --- --- --- --- --- --- --- --- --- --- --- --- --- --- --- --- --- --- --- --- --- --- --- --- --- --- --- --- --- --- --- --- --- --- --- --- --- --- --- --- --- --- --- --- --- --- --- --- --- --- --- --- --- --- --- --- --- --- --- --- --- --- --- --- --- --- --- --- --- --- --- --- --- --- --- --- --- --- --- --- --- --- --- --- --- --- --- --- --- --- --- --- --- --- --- --- --- --- --- --- --- --- --- --- --- --- --- --- --- --- --- --- --- --- --- --- --- --- --- --- --- --- --- --- --- --- --- --- --- --- --- --- --- --- --- --- --- --- --- --- --- --- --- --- --- --- --- --- --- --- --- --- --- --- --- --- --- --- --- --- --- --- --- --- --- --- -ga gag aaa gga taa aaa c-t ttt acg act cca tca gaa aga ggt tta ata ttt ttg tga gac cca tcg aag aga gaa aga gat ggt tag tca aga tat ttt tct tag tac aaa agt caa tgt ttt aaa ata tat gga cga gaa tta att tgt ctg tat aaa aac ttg tgt gaa att atg tac tag aga aaa aac gtg agc agt gtc ccc tac atg gat ttt aca gat cat tta tat tcc aaa aat att aac tat ata cgt tta tta tat gat gtt aac gtg taa att ata aac att att tta tga tgc aat tgt ctg aca acc tag att ggt ata agg atg ttg ata agc tct acg aga ata tat tgt tgg acg tta tcg ttt acg aaa tag ttg aga cat cag aaa gag gtt taa tat ttt tgt gag a-c cat cga aga gag aaa gag aat aaa aat a-- --- -tt ttt ttt ttt tgt aaa ac- ttt tta tga gac --- --- -ca aga gaa tac gaa tag tga tca tat cgt atc aca tat tga aac aga aag aag aag taa cga gag gta act ttt tgt gaa tgt agt taa ata ttt ttg ttt tgc aaa ccg gaa tat agt gcc cgg t-- --c ttt tta att cgt ggt gcg gtg tct gaa tcg ttc gat taa ccc aac tca tcc att ttc aga tga ata gag tta tcg att cag aca cat gct ttg agt ttt gtt gaa tcg atg agt gaa gta tca tcg gtt gca cct tca gat gcc gat ccg tcg aca tac ttg aat cca tcc ttg act tca agt tca gat gat tcc tca cac atg tct ccg ata cgt acg cta aac tct agg ttc ttg aca cat ttt gta tca acg atc gtt gaa ccg atg ata tct ttg taa ctc act ttc tta tgt gag atg tta gac cca agt act gga tgg gtc ttg atg tca ctg tct ttc tct tct tcg cta cat ctg atg tcg ata gac atc tca cag tct ttg atc ata gcc aga gct tct tca cgc gtg atc gcg gga gag tcc tta cct tgt ccc ggt gac acg ctg gac aat cta gta ttc aca gtg ttt cca tca gag gat tcg gag atg gat aaa atc ttt ggg cat ttg gtg aat cca aag ttc atg tta aga ccc gca ccg acg ata gtg taa taa gtg gtg gga tct cct ttt aca act --- --- --- --- --- --- tct tcg gat acc tca tca tct tcg gtc tct gta act tcc gtt acg gat tga caa atc tta tca ttg gtc ggt gtt tgg tct tgc ttt gtg act ttg ata ata aca tcg att ccc ata tga tgt ttg ttt tct tct tca gta cac gag gat gaa gat tgt tga aga cta gta ggc ata gca gct gcc act agg cac atg cat gcc agg aca ata tat tgt ttc atg att gct att gat tga tta ctg ttc tag atg att cta ctt tct tac cat ata ata aat tag aat ata ttt tct act ttt acg aga aat taa tta ttg tat tta tta ttt ata ggt --- aaa aaa ctt act ata agt ggg tgg gat tct ggg aat tag tga tca gtt tat gta tat cgc aac tag cgg gca tat ggc tat tga cat cga gaa cat tac cca tat gat aag aga ttg tat cat ttt cgt agt ctt gag tat tgg tat tac tat ata gta tgt aga tgt cga cgc tag ata gac agt cgc cca cta gag tta ccg tct ctg aat gcg gca tga tag tat cat tct ttg ttt tcg tta act gtt tgg aag atg aat ctt tgt tgt tac att taa tct cga aat tca gag tac ata tct ttg aag tat tct gat atc tat ttt ctc ctg taa aga atc ctg aag ttg cta cat tat taa gga cag aga agt att ctg cac gaa aga ctg gat cac aat ctt tat gat tca tgg taa tag tta gtt ccg acg ttg aga tgg att cgc tga gac cgg tag tgg tcg tcc gag tac acg atg tgt cgt tga ctg gat aca ggt taa ttt cca cat cga tat agt taa atg tat tgc tgg tta cga cgg gtt cgc att tat ctg tgg aag aga cgg tgt gag aat atg ttc cgg gac cac acg gag aac aga tga cgt ctc cgg tag acg tgt atc cgg ata ctc cgt atc cta ttc cac act ttg ttt tag aaa tac atg ttc tac acc ctg atg ctc ctt tga gaa gac aat aat atc ctg gag agc att cac aga ttc tat tgt gag tcg tgt tac acg atc gcg tct cta cct gat tac tat cac atc ttc cgt tac aac tta gac aag cct gta aat gat tat tgt gag atg taa agg tat ccg aac cac acg gtg tac att gtg tat tag tct tgc tat cac ata atc tga aag cgt aag ttc ccg gag gac acg ata gac aac ata gat tac ggc ttc tgt att cgt tgt ctt tac act ttc cat tgg atg gtg cat gtg gtg cta tat ctc ttc cgt tta tta tta tac atg aga gaa aca ata tat acg agt ata ata cgg acc tca tga ttt aat aat -gt agt aat cgt cgt ctt gtt act gtt tgt ttc cta ctt ctc caa tca tat aga tta ttt ttt aaa tat ttt ctt tct atc atg gat aat att tgt aat ggt tct ttc cgt aca aca tac tgt tta gat ggt agt cgc tta gct tgg tta tga tat tgc gca taa ttt ccg gag gca aat acg ata gtc tag att gac tat cga tgg tag act cta att tat tga gtg ctt tgt cga cga gtt tac ttt tat gct cca tcg ata gat gac act gtt cta tga gat cgt cgt aca tgg gaa atg aaa tgc gtt tgt ctg aat gta tgg ctt cga gat atg tgt gat acc gaa tgt ctt ctg ttc tca ata ccg tat aca agt cgg tgt ctg aga ttc gaa tct ctt tga gga gac tta tgt cac gac tac att ttt cga tga tgg aat cta tct tat cga atg ata tat ttt tca taa ata cac ttt tat agt cct cgt tta aac aga att tag tat gta gtt ccg caa atg act cgt ccc tta ata ggc agt agg cta tta tct tct tta cgt agt gat cgt cgt agg gag aga act ccg aca tct tgt aga aca acg att taa tca tag gta gag ata ctt tca gtc tgt ggt gga taa tgt cat taa cga cat ccg cct tgt ata taa tgt ttc tgt ttt caa aca cca agt cga ata ctg tct tat cgt ctt tag tcg gaa ggt tga tgt cgt atc caa tgt ata cga ggt atg agg caa cat tgt tat tgc aat tct gga agg cgg tat gaa gag gag tca ttg tat tat agt att cgt ctt tct gaa tgt cga atc tat cta gta gat acc gta gta tat tga gag agc gac ttc cat atc ctt gat tat gtt tta tga ata gat aaa gta gat gtt gtc ctt ctt cct ttt gta att tcc cgt att ttt gtt cgt gcc aat tga gta aca tta tga gaa tat gac ctg ttg cac aat cgt tct tta tgt att cca tga tgg gtg tac aat taa gat tat tac gta tcc tcg tat cgg ctc ctc gag ata aaa gag cat aca cca cac gag gac tat gtt tgg tat act gtt gaa ggt aag tgt gta acg gcg tat ttc cga ttt tcg taa ccg cgt taa tgt ttg ctc cat gat cta tta tcg cgt aga tga atc gct tct cag ctc gca tct tag tgt gac tct ttg act tgt aat aat tgc ttt cgt gga acg cgg ata tgt gtt tac agt agt aat gaa gag aag tga gtc cat cct cat cga cgc aat tag ggt cag atc ctt tag tca ata att tgt aca gaa cgt aat agt tta agc tcc cat tga att tat atc taa gat aac aca gca ata gat cgg atg att tac taa agt cat caa tgg ggt ccg tta gta tat caa aga tct tgt tat cga ttg ata gtg aat gaa tca gat agt ggt gta gag gaa tat gtc c-t ttt tat cct tgc tat caa agt tac gca tgc cgt ggc gta aca ata tct tta ata cag atg gat taa atc gtg tat tca tcg tat agc aat gta atg gag agt tac cac att tta gtc gtt tat tca gat cgc agt gtt taa taa cta att taa aca gat gag atg ata tat cca cat caa aga atg cga gat aca tat gac aga cat tat tga cag aaa tgt gac ctt cat tat cac cgt cgt cca taa atg cgt tag gta cgt acc aca tac tat cgt taa cga tgc gca caa tct cgt cca -tt tca tcc atc ttc ata atg att tac ttt ttc ata att aga gaa aaa gat caa ggt ata aaa att aga agt gtt aga cta ta- --- --- --- --- --- --- --- --- --- --- -aa cta act tat gac tta act aac tta tga --- --- --- --- --- --- --- --- --- --- --- --- --- --- --- --- --- --- --- --- --- --- --- --- --- --- --- --- --- --- --- --- --- --- --- --- --- --- --- --- --- --- --- --- --- --- --- --- ctt aac ta- --- --- --- --- --- --- --- --- --- --- -ac tta tga ctt aac taa ctt atg act taa cta act ta- tga ctt aac taa ctt atg act taa cta act tat gac tta act aac tta tga cta tta act cat ttc aag aaa ggt ggg tgg ata gaa ctc tat atg aca gct tgt gaa aca att aga tcc cta att tct aat gga agt ttt gat agg aga ttg tca tca gtt gat aca ttg ttt att atc tca tct att aga gca cgt ctg ttt aga gct tta gtg acc tgc tcg gtt act tct gtg taa atc ttg aat cct tta gtg ata cac tgt gtc aaa act gga tgt tta gaa tac cta tgt aga ata tgg gaa gca tgc ttg ttt ttg tct cta tta tag att aac tca tac atg gtt gta tta tga att ttc atc tgc cta atg tac tcc aat tct tgt tta caa tca att ata taa tca aag agt gat gat gca tac aca tta caa agt gaa taa tct acc atc ata aaa tac ttg ata cag agc ttt atc aca tca tgg ttt tca att gta tta tta agt ata gct aat ttt ata cag tca ata gac aat ggt tct cta agc aat att tct aat att tta aga tgt gct tcc cta cgg gcg atg aca gat ccc cta tcc acg gcc acg tca aga cat gta tat cca tta ctc att act gcg ttg aca ttt gct cca ttt tct aat agc cat gat act aaa tct ata taa cct gca tag ata gcg cga taa agc aag gtc ctt cca cca gca tct agt tga ttg ata tct tca ata tat ggg ata caa agc tta taa att tct aat act gtg ggt tca tct aca agg aat ccc cta gta tac tga att att tta tat aga tct aat tta aca tca ttt tca tct ggg ata cca caa ttc aaa ata aac tca aca aca cta ctt tcc ttt tta cat att -cc cta aaa tag gca ttc aag cat tct att tta tat att aca gcc cca tga tct acc ata aga tca aca atg tct att tct aca tat gca tta gat aga tag taa agt aag aga tct tgc aca gaa tta caa ttc tta ata att ata gag aaa ata tct tcc ata taa ttc ttt gac act aat gca gat ata ata tct tta tat gta ata tat gca aac agt cta tct act ata tac tga tca ata tta tct cta tga atc cta aaa taa tca tac aga aca tct aca gga tca caa att ggt tca agg aga aat cta tca aat att ttc ctg tca aca act ggt tct aga aca taa cag tca aca cct aat cca tgt ttt tta taa tca tct acc aaa gat aat gac caa aga tcg agg tcg tcg tga aac tgc tca tcg aca gcc atg aaa tct acc gac tcc atg gtg cga atc gca ctg tct tat tcg cca ttg att ttc a-t ttt tta taa tta tgt aca tgt ttt cct tct att ctc aag agt cta caa aaa tat a-- -tt ttt tcg ata tct aag tac taa gtt ttt tta ctg ttt ttg tta ctg tct tcc att ctt cta act aaa gat ctg aga taa att ata caa tct tcg cta tcg aac cat ttt tgt agt cta aag cct gaa gta att aac caa ctg ttt tta tta gtg gct ttt ttc gat cta tcc tcg tc- --- ctc tca tca tcc tta ta- --- --- --- --- --- --- tta tta tca tta tta tca tag tct att aaa cac aaa tca tct --- --- --- --- --- --- --- --- --- --- --- --- --- --- --- --- --- --- --- --- --- --- --- --- --- --- --a cgt tta taa c-- --- --- --- --- --- --- --- --a aca ttc --- --- --- --- -tc att att aat tag ttc tgt ag- aat atc ttt aat aat ttg gct ata --- --- -ca tct gtt --- --- --- --- --- --- --- --- --- --- --- --- --- --- --- --- --- --- --c aat act --- --- --- --- --- --- --- --a tct att gat gat ttc ttt t-- --- --- --- --- --- --- --- --- --- --- --t taa gac t-- --- --- --- --- --- --- --- --- --- --- --- --- --- --- --- --- --- --- --- --- --- --- --- --- --- --- --- --- --- --- --- --- --- --- --- --- --- --- --- taa act agt --- --- --- --- --- --- --- --- --- --- --- --- --- --- --- --- --- --- --- --- --- --- --- --- --- --- --- --- --- --- --- --- --- --- --- --- --- --- --- --- --- --- --- --- --- --- --- --- --- --- --- --- --- --- --- --- --- --- --- --- --- --- --- --- --- --- --- --- --- --- --- --- --- --- --- --- --- --- --- --- --- --- --- --- --- --- --- --- --- --- --- --- --- --- --- --- --- --- --- --- --- --- --- --- --- --- --- --- --- --t atg gta atg acg atg aaa --- -tc gag tag ta- --- --a ctt cta ata aag act tga ta- --- --t cat tat cat atg ttt gat cg- --- --- --- --- --- --- --t cat agt taa tag tgt g-- --- --- --- --- --- --- --- --- --- --- --- --- --- --- --- --- --- --- --- -gc taa atg gta ctg tta ata agt tta t-- --- --- --- --- --- --- --- --- --- --- --- --- --- --- --- --- --a gac aat atc ata gta ttt tct ttc cag aat --- --- --- --t aga tta ttt ttt -aa ata ctg atc ctc aca att ccg tga tgt agc agt agt tgg t-- --- --- --- --- --- --- --- --- --- --- --- --- --- --- --- --- --- --- --- --- --- --- --- --- --- --- --- -gc atg gtc tat atc gt- --- --- --- --- --- --- --- --- --- --- --- --- --- --- --- --- --- --- --- --- --- --- --- --- --- --- --- --- --- --- --- --- --- --- --- --- --- --- --- --- --- --- --- taa aat gta tca tat ata ata gtt ttc tga cgt gga gta cag aat ttt cga --- --- --- --- --- --- --- --- --- --- --- --- --- --- --- --- --- --- --- --- --- --- --- --- --- --- --- --- --- --- --- --- --- --- --- --- --- --- --- --- --- --- --- --t taa tga gtt cat ggt aag gaa ggg caa atg cct --- gta tat aat ata cat aag tta a-- --- --- --- --- --- --- --- --- --- --- --- --- --t agt ttt tta tca tat tt- --- --- --- --- --- --- --- --t cta ata cca taa taa aaa t-- --- --- --- --- --- --- --- --- --- --- --- --- --- --- --- --- --- --- --- --- --- --- --- --- --t atc att att gcg ttt g-g tag tt- --- --- --- --- ctg ccc ta- --- --- --- --- --- --- --- --t cat cta tat cac tgt cac tct c-- --- --g ctc tca cta tat ctt cta aaa tta caa --- -ac aac --- --- --- --- --- --- --- --- --- --- --- --- --- --- --- --- --- --- --- --- --- --- --- --- --- --- --- --- --- --- --- --- --- --- --- --- tgg ata ttc gat --- --- --- --- --- --- --- --- --- --- --- --- --- --- --- --- --- --- --- --- --- --- --- --- --- --- --a aca gca ttt gtg t-- --- --- --- --- --- --- --- --- --- --- --- --- --- --- --- --- -

>ON675438.1_Monkeypox_virus_isolate_MPXV_USA_2022_VA001_complete_genome

--- --- --- --- --- --- --- --- --- --- --- --- --- --- --- --- --- --- --- --- --- --- --- --- --- --- --- --- --- --a ttt tac tat ttt att tag tgt cta gaa aaa aat gtg tga ccc acg acc gta gga aac tct aga ggg taa gaa aaa tca atc gtt tat aga gac cat cag aaa gag gtt taa tat ttt tgt gag acc tat cga a-- --- --- --- --- --- --- --- --- --- --- --- --- --- --- --- --- --- --- --- --- --- --- --- --- --- --- --- --- --- --- --- --- --- --- --- --- --- --- --- --- --- --- --- --- --- --- --- --- --- --- --- --- --- --- --- --- --- --- --- --- --- --- --- --- --- --- --- --- --- --- --- --- --- --- --- --- --- --- --- --- --- --- --- --- --- --- --- --- --- --- --- --- --- --- --- --- --- --- --- --- --- --- --- --- --- --- --- --- --- --- --- --- --- --- --- --- --- --- --- --- --- --- --- --- --- --- --- --- --- --- --- --- --- --- --- --- --- --- --- --- --- --- --- --- --- --- --- --- --- --- --- --- --- --- --- --- --- --- --- --- --- --- --- --- --- --- --- --- --- --- --- --- --- --- --- --- --- --- --- --- --- --- --- --- --- --- --- --- --- --- --- --- --- --- --- --- --- --- --- --- --- --- --- --- --- --- --- --- --- --- --- --- --- --- --- --- --- --- --- --- --- --- --- --- --- --- --- --- --- --- --- --- --- --- --- --- --- --- --- --- --- --- --- --- --- --- --- --- --- --- --- --- --- --- --- --- --- --- --- --- --- --- --- --- --- --- --- --- --- --- --- --- --- --- --- --- --- --- --- -ga gag aaa gga taa aaa ctt ttt acg act cca tca gaa aga ggt tta ata ttt ttg tga gac cca tcg aag aga gaa aga gat ggt tag tca aga tat ttt tct tag tac aaa agt caa tgt ttt aaa ata tat gga cga gaa tta att tgt ctg tat aaa aac ttg tgt gaa att atg tac tag aga aaa aac gtg agc agt gtc ccc tac atg gat ttt aca gat cat tta tat tcc aaa aat att aac tat ata cgt tta tta tat gat gtt aac gtg taa att ata aac att att tta tga tgc aat tgt ctg aca acc tag att ggt ata agg atg ttg ata agc tct acg aga ata tat tgt tgg acg tta tcg ttt acg aaa tag ttg aga cat cag aaa gag gtt taa tat ttt tgt gag a-c cat cga aga gag aaa gag aat aaa aat a-- -?t ttt ttt ttt ttt tgt aaa act ttt tta tga gac --- --- -ca aga gaa tac gaa tag tga tca tat cgt atc aca tat tga aac aga aag aag aag taa cga gag gta act ttt tgt gaa tgt agt taa ata ttt ttg ttt tgc aaa ccg gaa tat agt gcc cgg t-- -ct ttt tta att cgt ggt gcg gtg tct gaa tcg ttc gat taa ccc aac tca tcc att ttc aga tga ata gag tta tcg att cag aca cat gct ttg agt ttt gtt gaa tcg atg agt gaa gta tca tcg gtt gca cct tca gat gcc gat ccg tcg aca tac ttg aat cca tcc ttg act tca agt tca gat gat tcc tca cac atg tct ccg ata cgt acg cta aac tct agg ttc ttg aca cat ttt gta tca acg atc gtt gaa ccg atg ata tct ttg taa ctc act ttc tta tgt gag atg tta gac cca agt act gga tgg gtc ttg atg tca ctg tct ttc tct tct tcg cta cat ctg atg tcg ata gac atc tca cag tct ttg atc ata gcc aga gct tct tca cgc gtg atc gcg gga gag tcc tta cct tgt ccc ggt gac acg ctg gac aat cta gta ttc aca gtg ttt cca tca gag gat tcg gag atg gat gaa atc ttt ggg cat ttg gtg aat cca aag ttc atg tta aga ccc gca ccg acg ata gtg taa taa gtg gtg gga tct cct ttt aca act --- --- --- --- --- --- tct tcg gat acc tca tca tct tcg gtc tct gta act tcc gtt acg gat tga caa atc tta tca ttg gtc ggt gtt tgg tct tgc ttt gtg act ttg ata ata aca tcg att ccc ata tga tgt ttg ttt tct tct tca gta cac gag gat gaa gat tgt tga aga cta gta ggc ata gca gct gcc act agg cac atg cat gcc agg aca ata tat tgt ttc atg att gct att gat tga tta ctg ttc tag atg att cta ctt tct tac cat ata ata aat tag aat ata ttt tct act ttt acg aga aat taa tta ttg tat tta tta ttt ata ggt --a aaa aaa ctt act ata agt ggg tgg gat tct ggg aat tag tga tca gtt tat gta tat cgc aac tag cgg gca tat ggc tat tga cat cga gaa cat tac cca tat gat aag aga ttg tat cat ttt cgt agt ctt gag tat tgg tat tac tat ata gta tgt aga tgt cga cgc tag ata gac agt cgc cca cta gag tta ccg tct ctg aat gcg gca tga tag tat cat tct ttg ttt tcg tta act gtt tgg aag atg aat ctt tgt tgt tac att taa tct cga aat tca gag tac ata tct ttg aag tat tct gat atc tat ttt ctc ctg taa aga atc ctg aag ttg cta cat tat taa gga cag aga agt att ctg cac gaa aga ctg gat cac aat ctt tat gat tca tgg taa tag tta gtt ccg acg ttg aga tgg att cgc tga gac cgg tag tgg tcg tcc gag tac acg atg tgt cgt tga ctg gat aca ggt taa ttt cca cat cga tat agt taa atg tat tgc tgg tta cga cgg gtt cgc att tat ctg tgg aag aga cgg tgt gag aat atg ttc cgg gac cac acg gag aac aga tga cgt ctc cgg tag acg tgt atc cgg ata ctc cgt atc cta ttc cac act ttg ttt tag aaa tac atg ttc tac acc ctg atg ctc ctt tga gaa gac aat aat atc ctg gag agc att cac aga ttc tat tgt gag tcg tgt tac acg atc gcg tct cta cct gat tac tat cac atc ttc cgt tac aac tta gac aag cct gta aat gat tat tgt gag atg taa agg tat ccg aac cac acg gtg tac att gtg tat tag tct tgc tat cac ata atc tgg aag cgt aag ttc ccg gag gac acg ata gac aac ata gat tac ggc ttc tgt att cgt tgt ctt tac act ttc cat tgg atg gtg cat gtg gtg cta tat ctc ttc cgt tta tta tta tac atg aga gaa aca ata tat acg agt ata ata cgg acc tca tga ttt aat aat -gt agt aat cgt cgt ctt gtt act gtt tgt ttc cta ctt ctc caa tca tat aga tta ttt ttt aaa tat ttt ctt tct atc atg gat aat att tgt aat ggt tct ttc cgt aca aca tac tgt tta gat ggt agt cgc tta gct tgg tta tga tat tgc gca taa ttt ccg gag gca aat acg ata gtc tag att gac tat cga tgg tag act cta att tat tga gtg ctt tgt cga cga gtt tac ttt tat gct cca tcg ata gat gac act gtt cta tga gat cgt cgt aca tgg gaa atg aaa tgc gtt tgt ctg aat gta tgg ctt cga gat atg tgt gat acc gga tgt ctt ctg ttc tca ata ccg tat aca agt cgg tgt ctg aga ttc gaa tct ctt tga gga gac tta tgt cac gac tac att ttt cga tga tgg aat cta tct tat cga atg ata tat ttt tca taa ata cac ttt tat agt cct cgt tta aac aga att tag tat gta gtt ccg caa atg act cgt ccc tta ata ggc agt agg cta tta tct tct tta cgt agt gat cgt cgt agg gag aga act ccg aca tct tgt aga aca acg att taa tca tag gta gag ata ctt tca gtc tgt ggt gga taa tgt cat taa cga cat ccg cct tgt ata taa tgt ttc tgt ttt caa aca cca agt cga ata ctg tct tat cgt ctt tag tcg gaa ggt tga tgt cgt atc cga tgt ata cga ggt atg agg caa cat tgt tat tgc aat tct gga agg cgg tat gaa gag gag tca ttg tat tat agt att cgt ctt tct gaa tgt cga atc tat cta gta gat acc gta gta tat tga gag agc gac ttc cat atc ctt gat tat gtt tta tga ata gat aaa gta gat gtt gtc ctt ctt cct ttt gta att tcc cgt att ttt gtt cgt gcc aat tga gta aca tta tga gaa tat gac ctg ttg cac aat cgt tct tta tgt att cca tga tgg gtg tac aat caa gat tat tac gta tcc tcg tat cgg ctc ctc gag ata aaa gag cat aca cca cac gag gac tat gtt tgg tat act gtt gaa ggt aag tgt gta acg gcg tat ttc cga ttt tcg taa ccg cgt taa tgt ttg ctc cat gat cta tta tcg cgt aga tga atc gct tct cag ctc gca tct tag tgt gac tct ttg act tgt aat aat tgc ttt cgt gga acg cgg ata tgt gtt tac agt agt aat gaa gag aag tga gtc cat cct cat cga cgc aat tag ggt cag atc ctt tag tca ata att tgt aca gaa cgt aat agt tta agc tcc cat tga att tat atc taa gat aac aca gca ata gat cgg atg att tac taa agt cat caa tgg ggt ccg tta gta tat caa aga tct tgt tat cga ttg ata gtg aat gaa tca gat agt ggt gta gag gaa tat gtc ctt ttt tat cct tgc tat caa agt tac gca tgc cgt ggc gta aca ata tct tta ata cag atg gat taa atc gtg tat tca tcg tat agc aat gta atg gag agt tac cac att tta gtc gtt tat tca gat cgc agt gtt taa taa cta att taa aca gat gag atg ata tat cca cat caa aga atg cga gat aca tat gac aga cat tat tga cag aaa tgt gac ctt cat tat cac cgt cgt cca taa atg cgt tag gta cgt acc aca tac tat cgt taa cga tgc gca caa tct cgt cca -tt tca tcc atc ttc ata atg att tac ttt ttc ata att aga gaa aaa gat caa ggt ata aaa att aga agt gtt aga cta ta- --- --- --- --- --- --- --- --- --- --- -aa cta act tat gac tta act aac tta tga --- --- --- --- --- --- --- --- --- --- --- --- --- --- --- --- --- --- --- --- --- --- --- --- --- --- --- --- --- --- --- --- --- --- --- --- --- --- --- --- --- --- --- --- --- --- --- --- ctt aac ta- --- --- --- --- --- --- --- --- --- --- -ac tta tga ctt aac taa ctt atg act taa cta act ta- tga ctt aac taa ctt atg act taa cta act tat gac tta act aac tta tga cta tta act cat ttc aag aaa ggt ggg tgg ata gaa ctc tat atg aca gct tgt gaa aca att aga tcc cta att tct aat gga agt ttt gat agg aga ttg tca tca gtt gat aca ttg ttt att atc tca tct att aga gca cgt ctg ttt aga gct tta gtg acc tgc tcg gtt act tct gtg taa atc ttg aat cct tta gtg ata cac tgt gtc aaa act gga tgt tta gaa tac cta tgt aga ata tgg gaa gca tgc ttg ttt ttg tct cta tta tag att aac tca tac atg gtt gta tta tga att ttc atc tgc cta atg tac tcc aat tct tgt tta caa tca att ata taa tca aag agt gat gat gca tac aca tta caa agt gaa taa tct acc atc ata aaa tac ttg ata cag agc ttt atc aca tca tgg ttt tca att gta tta tta agt ata gct aat ttt ata cag tca ata gac aat ggt tct cta agc aat att tct aat att tta aga tgt gct tcc cta cgg gcg atg aca gat ccc cta tcc acg gcc acg tca aga cat gta tat cca tta ctc att act gcg ttg aca ttt gct cca ttt tct aat agc cat gat act aaa tct ata taa cct gca tag ata gcg cga taa agc aag gtc ctt cca cca gca tct agt tga ttg ata tct tca ata tat ggg ata caa agc tta taa att tct aat act gtg ggt tca tct aca agg aat ccc cta gta tac tga att att tta tat aga tct aat tta aca tca ttt tca tct ggg ata cca caa ttc aaa ata aac tca aca aca cta ctt tcc ttt tta cat att ccc cta aaa tag gca ttc aag cat tct att tta tat att aca gcc cca tga tct acc ata aga tca aca atg tct att tct aca tat gca tta gat aga tag taa agt aag aga tct tgc aca gaa tta caa ttc tta ata att ata gag aaa ata tct tcc ata taa ttc ttt gac act aat gca gat ata ata tct tta tat gta ata tat gca aac agt cta tct act ata tac tga tca ata tta tct cta tga atc cta aaa taa tca tac aga aca tct aca gga tca caa att ggt tca agg aga aat cta tca aat att ttc ctg tca aca act ggt tct aga aca taa cag tca aca cct aat cca tgt ttt tta taa tca tct acc aaa gat aat gac caa aga tcg agg tcg tcg tga aac tgc tca tcg aca gcc atg aaa tct acc gac tcc atg gtg cga atc gca ctg tct tat tcg cca ttg att ttc a-t ttt tta taa tta tgt aca tgt ttt cct tct att ctc aag agt cta caa aaa tat a-- -tt ttt tcg ata tct aag tac taa gtt ttt tta ctg ttt ttg tta ctg tct tcc att ctt cta act aaa gat ctg aga taa att ata caa tct tcg cta tcg aac cat ttt tgt agt cta aag cct gaa gta att aac caa ctg ttt tta tta gtg gct ttt ttc gat cta tcc tcg tc- --- ctc tca tca tcc tta ta- --- --- --- --- --- --- tta tta tca tta tta tca tag tct att aaa cac aaa tca tct --- --- --- --- --- --- --- --- --- --- --- --- --- --- --- --- --- --- --- --- --- --- --- --- --- --- --a cgt tta taa c-- --- --- --- --- --- --- --- --a aca ttc --- --- --- --- -tc att att aat tag ttc tgt ag- aat atc ttt aat aat ttg gct ata --- --- -ca tct gtt --- --- --- --- --- --- --- --- --- --- --- --- --- --- --- --- --- --- --c aat act --- --- --- --- --- --- --- --a tct att gat gat ttc ttt t-- --- --- --- --- --- --- --- --- --- --- --t taa gac t-- --- --- --- --- --- --- --- --- --- --- --- --- --- --- --- --- --- --- --- --- --- --- --- --- --- --- --- --- --- --- --- --- --- --- --- --- --- --- --- taa act agt --- --- --- --- --- --- --- --- --- --- --- --- --- --- --- --- --- --- --- --- --- --- --- --- --- --- --- --- --- --- --- --- --- --- --- --- --- --- --- --- --- --- --- --- --- --- --- --- --- --- --- --- --- --- --- --- --- --- --- --- --- --- --- --- --- --- --- --- --- --- --- --- --- --- --- --- --- --- --- --- --- --- --- --- --- --- --- --- --- --- --- --- --- --- --- --- --- --- --- --- --- --- --- --- --- --- --- --- --- --t atg gta atg acg atg aaa --- -tc gag tag ta- --- --a ctt cta ata aag act tga ta- --- --t cat tat cat atg ttt gat cg- --- --- --- --- --- --- --t cat agt taa tag tgt g-- --- --- --- --- --- --- --- --- --- --- --- --- --- --- --- --- --- --- --- -gc taa atg gta ctg tta ata agt tta t-- --- --- --- --- --- --- --- --- --- --- --- --- --- --- --- --- --a gac aat atc ata gta ttt tct ttc cag aat --- --- --- --t aga tta ttt ttt taa ata ctg atc ctc aca att ccg tga tgt agc agt agt tgg t-- --- --- --- --- --- --- --- --- --- --- --- --- --- --- --- --- --- --- --- --- --- --- --- --- --- --- --- -gc atg gtc tat atc gt- --- --- --- --- --- --- --- --- --- --- --- --- --- --- --- --- --- --- --- --- --- --- --- --- --- --- --- --- --- --- --- --- --- --- --- --- --- --- --- --- --- --- --- taa aat gta tca tat ata ata gtt ttc tga cgt gga gta cag aat ttt cga --- --- --- --- --- --- --- --- --- --- --- --- --- --- --- --- --- --- --- --- --- --- --- --- --- --- --- --- --- --- --- --- --- --- --- --- --- --- --- --- --- --- --- --t taa tga gtt cat ggt aag gaa ggg caa atg cct --- gta tat aat ata cat aag tta a-- --- --- --- --- --- --- --- --- --- --- --- --- --t agt ttt tta tca tat tt- --- --- --- --- --- --- --- --t cta ata cca taa taa aaa t-- --- --- --- --- --- --- --- --- --- --- --- --- --- --- --- --- --- --- --- --- --- --- --- --- --t atc att att gcg ttt g-g tag tt- --- --- --- --- ctg ccc ta- --- --- --- --- --- --- --- --t cat cta tat cac tgt cac tct c-- --- --g ctc tca cta tat ctt cta aaa tta caa --- -ac aac --- --- --- --- --- --- --- --- --- --- --- --- --- --- --- --- --- --- --- --- --- --- --- --- --- --- --- --- --- --- --- --- --- --- --- --- tgg ata ttc gat --- --- --- --- --- --- --- --- --- --- --- --- --- --- --- --- --- --- --- --- --- --- --- --- --- --- --a aca gca ttt gtg t-- --- --- --- --- --- --- --- --- --- --- --- --- --- --- --- --- -

>ON676707.1_Monkeypox_virus_isolate_MPXV_USA_2021_TX_complete_genome

--- --- --- --- --- --- --- --- --- --- --- tac taa ctt tag tta gat aaa tta ata ata cat aag ttt tag tac att aat att ata ttt tac tat ttt att tag tgt cta gaa aaa aat gtg tga ccc acg acc gta gga aac tct aga ggg taa gaa aaa tca atc gtt tat aga gac cat cag aaa gag gtt taa tat ttt tgt gag acc tat cga a-- --- --- --- --- --- --- --- --- --- --- --- --- --- --- --- --- --- --- --- --- --- --- --- --- --- --- --- --- --- --- --- --- --- --- --- --- --- --- --- --- --- --- --- --- --- --- --- --- --- --- --- --- --- --- --- --- --- --- --- --- --- --- --- --- --- --- --- --- --- --- --- --- --- --- --- --- --- --- --- --- --- --- --- --- --- --- --- --- --- --- --- --- --- --- --- --- --- --- --- --- --- --- --- --- --- --- --- --- --- --- --- --- --- --- --- --- --- --- --- --- --- --- --- --- --- --- --- --- --- --- --- --- --- --- --- --- --- --- --- --- --- --- --- --- --- --- --- --- --- --- --- --- --- --- --- --- --- --- --- --- --- --- --- --- --- --- --- --- --- --- --- --- --- --- --- --- --- --- --- --- --- --- --- --- --- --- --- --- --- --- --- --- --- --- --- --- --- --- --- --- --- --- --- --- --- --- --- --- --- --- --- --- --- --- --- --- --- --- --- --- --- --- --- --- --- --- --- --- --- --- --- --- --- --- --- --- --- --- --- --- --- --- --- --- --- --- --- --- --- --- --- --- --- --- --- --- --- --- --- --- --- --- --- --- --- --- --- --- --- --- --- --- --- --- --- --- --- --- --- -ga gag aaa gga taa aaa ctt ttt acg act cca tca gaa aga ggt tta ata ttt ttg tga gac cca tcg aag aga gaa aga gat ggt tag tca aga tat ttt tct tag tac aaa agt caa tgt ttt aaa ata tat gga cga gaa tta att tgt ctg tat aaa aac ttg tgt gaa att atg tac tag aga aaa aac gtg agc agt gtc ccc tac atg gat ttt aca gat cat tta tat tcc aaa aat att aac tat ata cgt tta tta tat gat gtt aac gtg taa att ata aac att att tta tga tgc aat tgt ctg aca acc tag att ggt ata agg atg ttg ata agc tct acg aga ata tat tgt tgg acg tta tcg ttt acg aaa tag ttg aga cat cag aaa gag gtt taa tat ttt tgt gag a-c cat cga aga gag aaa gag aat aaa aat a-- --t ttt ttt ttt ttt tgt aaa act ttt tta tga gac --- --- -ca aga gaa tac gaa tag tga tca tat cgt atc aca tat tga aac aga aag aag aag taa cga gag gta act ttt tgt gaa tgt agt taa ata ttt ttg ttt tgc aaa ccg gaa tat agt gcc cgg t-- -ct ttt tta att cgt ggt gcg gtg tct gaa tcg ttc gat taa ccc aac tca tcc att ttc aga tga ata gag tta tcg att cag aca cat gct ttg agt ttt gtt gaa tcg atg agt gaa gta tca tcg gtt gca cct tca gat gcc gat ccg tcg aca tac ttg aat cca tcc ttg act tca agt tca gat gat tcc tca cac atg tct ccg ata cgt acg cta aac tct agg ttc ttg aca cat ttt gta tca acg atc gtt gaa ccg atg ata tct ttg taa ctc act ttc tta tgt gag atg tta gac cca agt act gga tgg gtc ttg atg tca ctg tct ttc tct tct tcg cta cat ctg atg tcg ata gac atc tca cag tct ttg atc ata gcc aga gct tct tca cgc gtg atc gcg gga gag tcc tta cct tgt ccc ggt gac acg ctg gac aat cta gta ttc aca gtg ttt cca tca gag gat tcg gag atg gat gaa atc ttt ggg cat ttg gtg aat cca aag ttc atg tta aga ccc gca ccg acg ata gtg taa taa gtg gtg gga tct cct ttt aca act --- --- --- --- --- --- tct tcg gat acc tca tca tct tcg gtc tct gta act tcc gtt acg gat tga caa atc tta tca ttg gtc ggt gtt tgg tct tgc ttt gtg act ttg ata ata aca tcg att ccc ata tga tgt ttg ttt tct tct tca gta cac gag gat gaa gat tgt tga aga cta gta ggc ata gca gct gcc act agg cac atg cat gcc agg aca ata tat tgt ttc atg att gct att gat tga tta ctg ttc tag atg att cta ctt tct tac cat ata ata aat tag aat ata ttt tct act ttt acg aga aat taa tta ttg tat tta tta ttt ata ggt --a aaa aaa ctt act ata agt ggg tgg gat tct ggg aat tag tga tca gtt tat gta tat cgc aac tag cgg gca tat ggc tat tga cat cga gaa cat tac cca tat gat aag aga ttg tat cat ttt cgt agt ctt gag tat tgg tat tac tat ata gta tgt aga tgt cga cgc tag ata gac agt cgc cca cta gag tta ccg tct ctg aat gcg gca tga tag tat cat tct ttg ttt tcg tta act gtt tgg aag atg aat ctt tgt tgt tac att taa tct cga aat tca gag tac ata tct ttg aag tat tct gat atc tat ttt ctc ctg taa aga atc ctg aag ttg cta cat tat taa gga cag aga agt att ctg cac gaa aga ctg gat cac aat ctt tat gat tca tgg taa tag tta gtt ccg acg ttg aga tgg att cgc tga gac cgg tag tgg tcg tcc gag tac acg atg tgt cgt tga ctg gat aca ggt taa ttt cca cat cga tat agt taa atg tat tgc tgg tta cga cgg gtt cgc att tat ctg tgg aag aga cgg tgt gag aat atg ttc cgg gac cac acg gag aac aga tga cgt ctc cgg tag acg tgt atc cgg ata ctc cgt atc cta ttc cac act ttg ttt tag aaa tac atg ttc tac acc ctg atg ctc ctt tga gaa gac aat aat atc ctg gag agc att cac aga ttc tat tgt gag tcg tgt tac acg atc gcg tct cta cct gat tac tat cac atc ttc cgt tac aac tta gac aag cct gta aat gat tat tgt gag atg taa agg tat ccg aac cac acg gtg tac att gtg tat tag tct tgc tat cac ata atc tgg aag cgt aag ttc ccg gag gac acg ata gac aac ata gat tac ggc ttc tgt att cgt tgt ctt tac act ttc cat tgg atg gtg cat gtg gtg cta tat ctc ttc cgt tta tta tta tac atg aga gaa aca ata tat acg agt ata ata cgg acc tca taa ttt aat aat -gt agt aat cgt cgt ctt gtt act gtt tgt ttc cta ctt ctc caa tca tat aga tta ttt ttt aaa tat ttt ctt tct atc atg gat aat att tgt aat ggt tct ttc cgt aca aca tac tgt tta gat ggt agt cgc tta gct tgg tta tga tat tgc gca taa ttt ccg gag gca aat acg ata gtc tag att gac tat cga tgg tag act cta att tat tga gtg ctt tgt cga cga gtt tac ttt tat gct cca tcg ata gat gac act gtt cta tga gat cgt cgt aca tgg gaa atg aaa tgc gtt tgt ctg aat gta tgg ctt cga gat atg tgt gat acc gga tgt ctt ctg ttc tca ata ccg tat aca agt cgg tgt ctg aga ttc gaa tct ctt tga gga gac tta tgt cac gac tac att ttt cga tga tgg aat cta tct tat cga atg ata tat ttt tca taa ata cac ttt tat agt cct cgt tta aac aga att tag tat gta gtt ccg caa atg act cgt ccc tta ata ggc agt agg cta tta tct tct tta cgt agt gat cgt cgt agg gag aga act ccg aca tct tgt aga aca acg att taa tca tag gta gag ata ctt tca gtc tgt ggt gga taa tgt cat taa cga cat ccg cct tgt ata taa tgt ttc tgt ttt caa aca cca agt cga ata ctg tct tat cgt ctt tag tcg gaa ggt tga tgt cgt atc cga tgt ata cga ggt atg agg caa cat tgt tat tgc aat tct gga agg cgg tat gaa gag gag tca ttg tat tat agt att cgt ctt tct gaa tgt cga atc tat cta gta gat acc gta gta tat tga gag agc gac ttc cat atc ctt gat tat gtt tta tga ata gat aaa gta gat gtt gtc ctt ctt cct ttt gta att tcc cgt att ttt gtt cgt gcc aat tga gta aca tta tga gaa tat gac ctg ttg cac aat cgt tct tta tgt att cca tga tgg gtg tac aat caa gat tat tac gta tcc tcg tat cgg ctc ctc gag ata aaa gag cat aca cca cac gag gac tat gtt tgg tat act gtt gaa ggt aag tgt gta acg gcg tat ttc cga ttt tcg taa ccg cgt taa tgt ttg ctc cat gat cta tta tcg cgt aga tga atc gct tct cag ctc gca tct tag tgt gac tct ttg act tgt aat aat tgc ttt cgt gga acg cgg ata tgt gtt tac agt agt aat gaa gag aag tga gtc cat cct cat cga cgc aat tag ggt cag atc ctt tag tca ata att tgt aca gaa cgt aat agt tta agc tcc cat tga att tat atc taa gat aac aca gca ata gat cgg atg att tac taa agt cat caa tgg ggt ccg tta gta tat caa aga tct tgt tat cga ttg ata gtg aat gaa tca gat agt ggt gta gag gaa tat gtc ctt ttt tat cct tgc tat caa agt tac gca tgc cgt ggc gta aca ata tct tta ata cag atg gat taa atc gtg tat tca tcg tat agc aat gta atg gag agt tac cac att tta gtc gtt tat tca gat cgc agt gtt taa taa cta att taa aca gat gag atg ata tat cca cat caa aga atg cga gat aca tat gac aga cat tat tga cag aaa tgt gac ctt cat tat cac cgt cgt cca taa atg cgt tag gta cgt acc aca tac tat cgt taa cga tgc gca caa tct cgt cca -tt tca tcc atc ttc ata atg att tac ttt ttc ata att aga gaa aaa gat caa ggt ata aaa att aga agt gtt aga cta ta- --- --- --- --- --- --- --- --- --- --- -aa cta act tat gac tta act aac tta tga --- --- --- --- --- --- --- --- --- --- --- --- --- --- --- --- --- --- --- --- --- --- --- --- --- --- --- --- --- --- --- --- --- --- --- --- --- --- --- --- --- --- --- --- --- --- --- --- ctt aac ta- --- --- --- --- --- --- --- --- --- --- -ac tta tga ctt aac taa ctt atg act taa cta act ta- tga ctt aac taa ctt atg act taa cta act tat gac tta act aac tta tga cta tta act cat ttc aag aaa ggt ggg tgg ata gaa ctc tat atg aca gct tgt gaa aca att aga tcc cta att tct aat gga agt ttt gat agg aga ttg tca tca gtt gat aca ttg ttt att atc tca tct att aga gca cgt ctg ttt aga gct tta gtg acc tgc tcg gtt act tct gtg taa atc ttg aat cct tta gtg ata cac tgt gtc aaa act gga tgt tta gaa tac cta tgt aga ata tgg gaa gca tgc ttg ttt ttg tct cta tta tag att aac tca tac atg gtt gta tta tga att ttc atc tgc cta atg tac tcc aat tct tgt tta caa tca att ata taa tca aag agt gat gat gca tac aca tta caa agt gaa taa tct acc atc ata aaa tac ttg ata cag agc ttt atc aca tca tgg ttt tca att gta tta tta agt ata gct aat ttt ata cag tca ata gac aat ggt tct cta agc aat att tct aat att tta aga tgt gct tcc cta cgg gcg atg aca gat ccc cta tcc acg gcc acg tca aga cat gta tat cca tta ctc att act gcg ttg aca ttt gct cca ttt tct aat agc cat gat act aaa tct ata taa cct gca tag ata gcg cga taa agc aag gtc ctt cca cca gca tct agt tga ttg ata tct tca ata tat ggg ata caa agc tta taa att tct aat act gtg ggt tca tct aca agg aat ccc cta gta tac tga att att tta tat aga tct aat tta aca tca ttt tca tct ggg ata cca caa ttc aaa ata aac tca aca aca cta ctt tcc ttt tta cat att ccc cta aaa tag gca ttc aag cat tct att tta tat att aca gcc cca tga tct acc ata aga tca aca atg tct att tct aca tat gca tta gat aga tag taa agt aag aga tct tgc aca gaa tta caa ttc tta ata att ata gag aaa ata tct tcc ata taa ttc ttt gac act aat gca gat ata ata tct tta tat gta ata tat gca aac agt cta tct act ata tac tga tca ata tta tct cta tga atc cta aaa taa tca tac aga aca tct aca gga tca caa att ggt tca agg aga aat cta tca aat att ttc ctg tca aca act ggt tct aga aca taa cag tca aca cct aat cca tgt ttt tta taa tca tct acc aaa gat aat gac caa aga tcg agg tcg tcg tga aac tgc tca tcg aca gcc atg aaa tct acc gac tcc atg gtg cga atc gca ctg tct tat tcg cca ttg att ttc a-t ttt tta taa tta tgt aca tgt ttt cct tct att ctc aag agt cta caa aaa tat a-- -tt ttt tcg ata tct aag tac taa gtt ttt tta ctg ttt ttg tta ctg tct tcc att ctt cta act aaa gat ctg aga taa att ata caa tct tcg cta tcg aac cat ttt tgt agt cta aag cct gaa gta att aac caa ctg ttt tta tta gtg gct ttt ttc gat cta tcc tcg tc- --- ctc tca tca tcc tta ta- --- --- --- --- --- --- tta tta tca tta tta tca tag tct att aaa cac aaa tca tct --- --- --- --- --- --- --- --- --- --- --- --- --- --- --- --- --- --- --- --- --- --- --- --- --- --- --a cgt tta taa c-- --- --- --- --- --- --- --- --a aca ttc --- --- --- --- -tc att att aat tag ttc tgt ag- aat atc ttt aat aat ttg gct ata --- --- -ca tct gtt --- --- --- --- --- --- --- --- --- --- --- --- --- --- --- --- --- --- --c aat act --- --- --- --- --- --- --- --a tct att gat gat ttc ttt t-- --- --- --- --- --- --- --- --- --- --- --t taa gac t-- --- --- --- --- --- --- --- --- --- --- --- --- --- --- --- --- --- --- --- --- --- --- --- --- --- --- --- --- --- --- --- --- --- --- --- --- --- --- --- taa act agt --- --- --- --- --- --- --- --- --- --- --- --- --- --- --- --- --- --- --- --- --- --- --- --- --- --- --- --- --- --- --- --- --- --- --- --- --- --- --- --- --- --- --- --- --- --- --- --- --- --- --- --- --- --- --- --- --- --- --- --- --- --- --- --- --- --- --- --- --- --- --- --- --- --- --- --- --- --- --- --- --- --- --- --- --- --- --- --- --- --- --- --- --- --- --- --- --- --- --- --- --- --- --- --- --- --- --- --- --- --t atg gta atg acg atg aaa --- -tc gag tag ta- --- --a ctt cta ata aag act tga ta- --- --t cat tat cat atg ttt gat cg- --- --- --- --- --- --- --t cat agt taa tag tgt g-- --- --- --- --- --- --- --- --- --- --- --- --- --- --- --- --- --- --- --- -gc taa atg gta ctg tta ata agt tta t-- --- --- --- --- --- --- --- --- --- --- --- --- --- --- --- --- --a gac aat atc ata gta ttt tct ttc cag aat --- --- --- --t aga tta ttt ttt taa ata ctg atc ctc aca att ccg tga tgt agc agt agt tgg t-- --- --- --- --- --- --- --- --- --- --- --- --- --- --- --- --- --- --- --- --- --- --- --- --- --- --- --- -gc atg gtc tat atc gt- --- --- --- --- --- --- --- --- --- --- --- --- --- --- --- --- --- --- --- --- --- --- --- --- --- --- --- --- --- --- --- --- --- --- --- --- --- --- --- --- --- --- --- taa aat gta tca tat ata ata gtt ttc tga cgt gga gta cag aat ttt cga --- --- --- --- --- --- --- --- --- --- --- --- --- --- --- --- --- --- --- --- --- --- --- --- --- --- --- --- --- --- --- --- --- --- --- --- --- --- --- --- --- --- --- --t taa tga gtt cat ggt aag gaa ggg caa atg cct --- gta tat aat ata cat aag tta a-- --- --- --- --- --- --- --- --- --- --- --- --- --t agt ttt tta tca tat tt- --- --- --- --- --- --- --- --t cta ata cca taa taa aaa t-- --- --- --- --- --- --- --- --- --- --- --- --- --- --- --- --- --- --- --- --- --- --- --- --- --t atc att att gcg ttt g-g tag tt- --- --- --- --- ctg ccc ta- --- --- --- --- --- --- --- --t cat cta tat cac tgt cac tct c-- --- --g ctc tca cta tat ctt cta aaa tta caa --- -ac aac --- --- --- --- --- --- --- --- --- --- --- --- --- --- --- --- --- --- --- --- --- --- --- --- --- --- --- --- --- --- --- --- --- --- --- --- tgg ata ttc gat --- --- --- --- --- --- --- --- --- --- --- --- --- --- --- --- --- --- --- --- --- --- --- --- --- --- --a aca gca ttt gtg t-- --- --- --- --- --- --- --- --- --- --- --- --- --- --- --- --- -

>MT903345.1_Monkeypox_virus_isolate_MPXV-UK_P3

--- --- --- --- --- --- --- --- --- --- --- --- --- --- --- --- --- --- --- --- --- -at aag ttt tag tac att aat att ata ttt tac tat ttt att tag tgt cta gaa aaa aat gtg tga ccc acg acc gta gga aac tct aga ggg taa gaa aaa tca atc gtt tat aga gac cat cag aaa gag gtt taa tat ttt tgt gag acc tat cga a-- --- --- --- --- --- --- --- --- --- --- --- --- --- --- --- --- --- --- --- --- --- --- --- --- --- --- --- --- --- --- --- --- --- --- --- --- --- --- --- --- --- --- --- --- --- --- --- --- --- --- --- --- --- --- --- --- --- --- --- --- --- --- --- --- --- --- --- --- --- --- --- --- --- --- --- --- --- --- --- --- --- --- --- --- --- --- --- --- --- --- --- --- --- --- --- --- --- --- --- --- --- --- --- --- --- --- --- --- --- --- --- --- --- --- --- --- --- --- --- --- --- --- --- --- --- --- --- --- --- --- --- --- --- --- --- --- --- --- --- --- --- --- --- --- --- --- --- --- --- --- --- --- --- --- --- --- --- --- --- --- --- --- --- --- --- --- --- --- --- --- --- --- --- --- --- --- --- --- --- --- --- --- --- --- --- --- --- --- --- --- --- --- --- --- --- --- --- --- --- --- --- --- --- --- --- --- --- --- --- --- --- --- --- --- --- --- --- --- --- --- --- --- --- --- --- --- --- --- --- --- --- --- --- --- --- --- --- --- --- --- --- --- --- --- --- --- --- --- --- --- --- --- --- --- --- --- --- --- --- --- --- --- --- --- --- --- --- --- --- --- --- --- --- --- --- --- --- --- --- -ga gag aaa gga taa aaa ctt ttt acg act cca tca gaa aga ggt tta ata ttt ttg tga gac cca tcg aag aga gaa aga gat ggt tag tca aga tat ttt tct tag tac aaa agt caa tgt ttt aaa ata tat gga cga gaa tta att tgt ctg tat aaa aac ttg tgt gaa att atg tac tag aga aaa aac gtg agc agt gtc ccc tac atg gat ttt aca gat cat tta tat tcc aaa aat att aac tat ata cgt tta tta tat gat gtt aac gtg taa att ata aac att att tta tga tgc aat tgt ctg aca acc tag att ggt ata agg atg ttg ata agc tct acg aga ata tat tgt tgg acg tta tcg ttt acg aaa tag ttg aga cat cag aaa gag gtt taa tat ttt tgt gag a-c cat cga aga gag aaa gag aat aaa aat a-- --t ttt ttt ttt ttt tgt aaa act ttt tta tga gac --- --- -ca aga gaa tac gaa tag tga tca tat cgt atc aca tat tga aac aga aag aag aag taa cga gag gta act ttt tgt gaa tgt agt taa ata ttt ttg ttt tgc aaa ccg gaa tat agt gcc cgg t-- -ct ttt tta att cgt ggt gcg gtg tct gaa tcg ttc gat taa ccc aac tca tcc att ttc aga tga ata gag tta tcg att cag aca cat gct ttg agt ttt gtt gaa tcg atg agt gaa gta tca tcg gtt gca cct tca gat gcc gat ccg tcg aca tac ttg aat cca tcc ttg act tca agt tca gat gat tcc tca cac atg tct ccg ata cgt acg cta aac tct agg ttc ttg aca cat ttt gta tca acg atc gtt gaa ccg atg ata tct ttg taa ctc act ttc tta tgt gag atg tta gac cca agt act gga tgg gtc ttg atg tca ctg tct ttc tct tct tcg cta cat ctg atg tcg ata gac atc tca cag tct ttg atc ata gcc aga gct tct tca cgc gtg atc gcg gga gag tcc tta cct tgt ccc ggt gac acg ctg gac aat cta gta ttc aca gtg ttt cca tca gag gat tcg gag atg gat gaa atc ttt ggg cat ttg gtg aat cca aag ttc atg tta aga ccc gca ccg acg ata gtg taa taa gtg gtg gga tct cct ttt aca act --- --- --- --- --- --- tct tcg gat acc tca tca tct tcg gtc tct gta act tcc gtt acg gat tga caa atc tta tca ttg gtc ggt gtt tgg tct tgc ttt gtg act ttg ata ata aca tcg att ccc ata tga tgt ttg ttt tct tct tca gta cac gag gat gaa gat tgt tga aga cta gta ggc ata gca gct gcc act agg cac atg cat gcc agg aca ata tat tgt ttc atg att gct att gat tga tta ctg ttc tag atg att cta ctt tct tac cat ata ata aat tag aat ata ttt tct act ttt acg aga aat taa tta ttg tat tta tta ttt ata ggt --a aaa aaa ctt act ata agt ggg tgg gat tct ggg aat tag tga tca gtt tat gta tat cgc aac tag cgg gca tat ggc tat tga cat cga gaa cat tac cca tat gat aag aga ttg tat cat ttt cgt agt ctt gag tat tgg tat tac tat ata gta tgt aga tgt cga cgc tag ata gac agt cgc cca cta gag tta ccg tct ctg aat gcg gca tga tag tat cat tct ttg ttt tcg tta act gtt tgg aag atg aat ctt tgt tgt tac att taa tct cga aat tca gag tac ata tct ttg aag tat tct gat atc tat ttt ctc ctg taa aga atc ctg aag ttg cta cat tat taa gga cag aga agt att ctg cac gaa aga ctg gat cac aat ctt tat gat tca tgg taa tag tta gtt ccg acg ttg aga tgg att cgc tga gac cgg tag tgg tcg tcc gag tac acg atg tgt cgt tga ctg gat aca ggt taa ttt cca cat cga tat agt taa atg tat tgc tgg tta cga cgg gtt cgc att tat ctg tgg aag aga cgg tgt gag aat atg ttc cgg gac cac acg gag aac aga tga cgt ctc cgg tag acg tgt atc cgg ata ctc cgt atc cta ttc cac act ttg ttt tag aaa tac atg ttc tac acc ctg atg ctc ctt tga gaa gac aat aat atc ctg gag agc att cac aga ttc tat tgt gag tcg tgt tac acg atc gcg tct cta cct gat tac tat cac atc ttc cgt tac aac tta gac aag cct gta aat gat tat tgt gag atg taa agg tat ccg aac cac acg gtg tac att gtg tat tag tct tgc tat cac ata atc tgg aag cgt aag ttc ccg gag gac acg ata gac aac ata gat tac ggc ttc tgt att cgt tgt ctt tac act ttc cat tgg atg gtg cat gtg gtg cta tat ctc ttc cgt tta tta tta tac atg aga gaa aca ata tat acg agt ata ata cgg acc tca tga ttt aat aat -gt agt aat cgt cgt ctt gtt act gtt tgt ttc cta ctt ctc caa tca tat aga tta ttt ttt aaa tat ttt ctt tct atc atg gat aat att tgt aat ggt tct ttc cgt aca aca tac tgt tta gat ggt agt cgc tta gct tgg tta tga tat tgc gca taa ttt ccg gag gca aat acg ata gtc tag att gac tat cga tgg tag act cta att tat tga gtg ctt tgt cga cga gtt tac ttt tat gct cca tcg ata gat gac act gtt cta tga gat cgt cgt aca tgg gaa atg aaa tgc gtt tgt ctg aat gta tgg ctt cga gat atg tgt gat acc gga tgt ctt ctg ttc tca ata ccg tat aca agt cgg tgt ctg aga ttc gaa tct ctt tga gga gac tta tgt cac gac tac att ttt cga tga tgg aat cta tct tat cga atg ata tat ttt tca taa ata cac ttt tat agt cct cgt tta aac aga att tag tat gta gtt ccg caa atg act cgt ccc tta ata ggc agt agg cta tta tct tct tta cgt agt gat cgt cgt agg gag aga act ccg aca tct tgt aga aca acg att taa tca tag gta gag ata ctt tca gtc tgt ggt gga taa tgt cat taa cga cat ccg cct tgt ata taa tgt ttc tgt ttt caa aca cca agt cga ata ctg tct tat cgt ctt tag tcg gaa ggt tga tgt cgt atc cga tgt ata cga ggt atg agg caa cat tgt tat tgc aat tct gga agg cgg tat gaa gag gag tca ttg tat tat agt att cgt ctt tct gaa tgt cga atc tat cta gta gat acc gta gta tat tga gag agc gac ttc cat atc ctt gat tat gtt tta tga ata gat aaa gta gat gtt gtc ctt ctt cct ttt gta att tcc cgt att ttt gtt cgt gcc aat tga gta aca tta tga gaa tat gac ctg ttg cac aat cgt tct tta tgt att cca tga tgg gtg tac aat caa gat tat tac gta tcc tcg tat cgg ctc ctc gag ata aaa gag cat aca cca cac gag gac tat gtt tgg tat act gtt gaa ggt aag tgt gta acg gcg tat ttc cga ttt tcg taa ccg cgt taa tgt ttg ctc cat gat cta tta tcg cgt aga tga atc gct tct cag ctc gca tct tag tgt gac tct ttg act tgt aat aat tgc ttt cgt gga acg cgg ata tgt gtt tac agt agt aat gaa gag aag tga gtc cat cct cat cga cgc aat tag ggt cag atc ctt tag tca ata att tgt aca gaa cgt aat agt tta agc tcc cat tga att tat atc taa gat aac aca gca ata gat cgg atg att tac taa agt cat caa tgg ggt ccg tta gta tat caa aga tct tgt tat cga ttg ata gtg aat gaa tca gat agt ggt gta gag gaa tat gtc ctt ttt tat cct tgc tat caa agt tac gca tgc cgt ggc gta aca ata tct tta ata cag atg gat taa atc gtg tat tca tcg tat agc aat gta atg gag agt tac cac att tta gtc gtt tat tca gat cgc agt gtt taa taa cta att taa aca gat gag atg ata tat cca cat caa aga atg cga gat aca tat gac aga cat tat tga cag aaa tgt gac ctt cat tat cac cgt cgt cca taa atg cgt tag gta cgt acc aca tac tat cgt taa cga tgc gca caa tct cgt cca -tt tca tcc atc ttc ata atg att tac ttt ttc ata att aga gaa aaa gat caa ggt ata aaa att aga agt gtt aga cta ta- --- --- --- --- --- --- --- --- --- --- -aa cta act tat gac tta act aac tta tga --- --- --- --- --- --- --- --- --- --- --- --- --- --- --- --- --- --- --- --- --- --- --- --- --- --- --- --- --- --- --- --- --- --- --- --- --- --- --- --- --- --- --- --- --- --- --- --- ctt aac ta- --- --- --- --- --- --- --- --- --- --- -ac tta tga ctt aac taa ctt atg act taa cta act ta- tga ctt aac taa ctt atg act taa cta act tat gac tta act aac tta tga cta tta act cat ttc aag aaa ggt ggg tgg ata gaa ctc tat atg aca gct tgt gaa aca att aga tcc cta att tct aat gga agt ttt gat agg aga ttg tca tca gtt gat aca ttg ttt att atc tca tct att aga gca cgt ctg ttt aga gct tta gtg acc tgc tcg gtt act tct gtg taa atc ttg aat cct tta gtg ata cac tgt gtc aaa act gga tgt tta gaa tac cta tgt aga ata tgg gaa gca tgc ttg ttt ttg tct cta tta tag att aac tca tac atg gtt gta tta tga att ttc atc tgc cta atg tac tcc aat tct tgt tta caa tca att ata taa tca aag agt gat gat gca tac aca tta caa agt gaa taa tct acc atc ata aaa tac ttg ata cag agc ttt atc aca tca tgg ttt tca att gta tta tta agt ata gct aat ttt ata cag tca ata gac aat ggt tct cta agc aat att tct aat att tta aga tgt gct tcc cta cgg gcg atg aca gat ccc cta tcc acg gcc acg tca aga cat gta tat cca tta ctc att act gcg ttg aca ttt gct cca ttt tct aat agc cat gat act aaa tct ata taa cct gca tag ata gcg cga taa agc aag gtc ctt cca cca gca tct agt tga ttg ata tct tca ata tat ggg ata caa agc tta taa att tct aat act gtg ggt tca tct aca agg aat ccc cta gta tac tga att att tta tat aga tct aat tta aca tca ttt tca tct ggg ata cca caa ttc aaa ata aac tca aca aca cta ctt tcc ttt tta cat att ccc cta aaa tag gca ttc aag cat tct att tta tat att aca gcc cca tga tct acc ata aga tca aca atg tct att tct aca tat gca tta gat aga tag taa agt aag aga tct tgc aca gaa tta caa ttc tta ata att ata gag aaa ata tct tcc ata taa ttc ttt gac act aat gca gat ata ata tct tta tat gta ata tat gca aac agt cta tct act ata tac tga tca ata tta tct cta tga atc cta aaa taa tca tac aga aca tct aca gga tca caa att ggt tca agg aga aat cta tca aat att ttc ctg tca aca act ggt tct aga aca taa cag tca aca cct aat cca tgt ttt tta taa tca tct acc aaa gat aat gac caa aga tcg agg tcg tcg tga aac tgc tca tcg aca gcc atg aaa tct acc gac tcc atg gtg cga atc gca ctg tct tat tcg cca ttg att ttc a-t ttt tta taa tta tgt aca tgt ttt cct tct att ctc aag agt cta caa aaa tat a-- -tt ttt tcg ata tct aag tac taa gtt ttt tta ctg ttt ttg tta ctg tct tcc att ctt cta act aaa gat ctg aga taa att ata caa tct tcg cta tcg aac cat ttt tgt agt cta aag cct gaa gta att aac caa ctg ttt tta tta gtg gct ttt ttc gat cta tcc tcg tc- --- ctc tca tca tcc tta ta- --- --- --- --- --- --- tta tta tca tta tta tca tag tct att aaa cac aaa tca tct --- --- --- --- --- --- --- --- --- --- --- --- --- --- --- --- --- --- --- --- --- --- --- --- --- --- --a cgt tta taa c-- --- --- --- --- --- --- --- --a aca ttc --- --- --- --- -tc att att aat tag ttc tgt ag- aat atc ttt aat aat ttg gct ata --- --- -ca tct gtt --- --- --- --- --- --- --- --- --- --- --- --- --- --- --- --- --- --- --c aat act --- --- --- --- --- --- --- --a tct att gat gat ttc ttt t-- --- --- --- --- --- --- --- --- --- --- --t taa gac t-- --- --- --- --- --- --- --- --- --- --- --- --- --- --- --- --- --- --- --- --- --- --- --- --- --- --- --- --- --- --- --- --- --- --- --- --- --- --- --- taa act agt --- --- --- --- --- --- --- --- --- --- --- --- --- --- --- --- --- --- --- --- --- --- --- --- --- --- --- --- --- --- --- --- --- --- --- --- --- --- --- --- --- --- --- --- --- --- --- --- --- --- --- --- --- --- --- --- --- --- --- --- --- --- --- --- --- --- --- --- --- --- --- --- --- --- --- --- --- --- --- --- --- --- --- --- --- --- --- --- --- --- --- --- --- --- --- --- --- --- --- --- --- --- --- --- --- --- --- --- --- --t atg gta atg acg atg aaa --- -tc gag tag ta- --- --a ctt cta ata aag act tga ta- --- --t cat tat cat atg ttt gat cg- --- --- --- --- --- --- --t cat agt taa tag tgt g-- --- --- --- --- --- --- --- --- --- --- --- --- --- --- --- --- --- --- --- -gc taa atg gta ctg tta ata agt tta t-- --- --- --- --- --- --- --- --- --- --- --- --- --- --- --- --- --a gac aat atc ata gta ttt tct ttc cag aat --- --- --- --t aga tta ttt ttt taa ata ctg atc ctc aca att ccg tga tgt agc agt agt tgg t-- --- --- --- --- --- --- --- --- --- --- --- --- --- --- --- --- --- --- --- --- --- --- --- --- --- --- --- -gc atg gtc tat atc gt- --- --- --- --- --- --- --- --- --- --- --- --- --- --- --- --- --- --- --- --- --- --- --- --- --- --- --- --- --- --- --- --- --- --- --- --- --- --- --- --- --- --- --- taa aat gta tca tat ata ata gtt ttc tga cgt gga gta cag aat ttt cga --- --- --- --- --- --- --- --- --- --- --- --- --- --- --- --- --- --- --- --- --- --- --- --- --- --- --- --- --- --- --- --- --- --- --- --- --- --- --- --- --- --- --- --t taa tga gtt cat ggt aag gaa ggg caa atg cct --- gta tat aat ata cat aag tta a-- --- --- --- --- --- --- --- --- --- --- --- --- --t agt ttt tta tca tat tt- --- --- --- --- --- --- --- --t cta ata cca taa taa aaa t-- --- --- --- --- --- --- --- --- --- --- --- --- --- --- --- --- --- --- --- --- --- --- --- --- --t atc att att gcg ttt g-g tag tt- --- --- --- --- ctg ccc ta- --- --- --- --- --- --- --- --t cat cta tat cac tgt cac tct c-- --- --g ctc tca cta tat ctt cta aaa tta caa --- -ac aac --- --- --- --- --- --- --- --- --- --- --- --- --- --- --- --- --- --- --- --- --- --- --- --- --- --- --- --- --- --- --- --- --- --- --- --- tgg ata ttc gat --- --- --- --- --- --- --- --- --- --- --- --- --- --- --- --- --- --- --- --- --- --- --- --- --- --- --a aca gca ttt gtg t-- --- --- --- --- --- --- --- --- --- --- --- --- --- --- --- --- -

>MT903344.1_Monkeypox_virus_isolate_MPXV-UK_P2

--- --- --- --- --- --- --- --- --- --- --- --- --- --- --- --- --- --- -ta att --- --- --- --- --- --- --- --a att taa ttt tac tat ttt att tag tgt cta gaa aaa aat gtg tga ccc acg acc gta gga aac tct aga ggg taa gaa aaa tca atc gtt tat aga gac cat cag aaa gag gtt taa tat ttt tgt gag acc tat cga a-- --- --- --- --- --- --- --- --- --- --- --- --- --- --- --- --- --- --- --- --- --- --- --- --- --- --- --- --- --- --- --- --- --- --- --- --- --- --- --- --- --- --- --- --- --- --- --- --- --- --- --- --- --- --- --- --- --- --- --- --- --- --- --- --- --- --- --- --- --- --- --- --- --- --- --- --- --- --- --- --- --- --- --- --- --- --- --- --- --- --- --- --- --- --- --- --- --- --- --- --- --- --- --- --- --- --- --- --- --- --- --- --- --- --- --- --- --- --- --- --- --- --- --- --- --- --- --- --- --- --- --- --- --- --- --- --- --- --- --- --- --- --- --- --- --- --- --- --- --- --- --- --- --- --- --- --- --- --- --- --- --- --- --- --- --- --- --- --- --- --- --- --- --- --- --- --- --- --- --- --- --- --- --- --- --- --- --- --- --- --- --- --- --- --- --- --- --- --- --- --- --- --- --- --- --- --- --- --- --- --- --- --- --- --- --- --- --- --- --- --- --- --- --- --- --- --- --- --- --- --- --- --- --- --- --- --- --- --- --- --- --- --- --- --- --- --- --- --- --- --- --- --- --- --- --- --- --- --- --- --- --- --- --- --- --- --- --- --- --- --- --- --- --- --- --- --- --- --- --- -ga gag aaa gga taa aaa ctt ttt acg act cca tca gaa aga ggt tta ata ttt ttg tga gac cca tcg aag aga gaa aga gat ggt tag tca aga tat ttt tct tag tac aaa agt caa tgt ttt aaa ata tat gga cga gaa tta att tgt ctg tat aaa aac ttg tgt gaa att atg tac tag aga aaa aac gtg agc agt gtc ccc tac atg gat ttt aca gat cat tta tat tcc aaa aat att aac tat ata cgt tta tta tat gat gtt aac gtg taa att ata aac att att tta tga tgc aat tgt ctg aca acc tag att ggt ata agg atg ttg ata agc tct acg aga ata tat tgt tgg acg tta tcg ttt acg aaa tag ttg aga cat cag aaa gag gtt taa tat ttt tgt gag a-c cat cga aga gag aaa gag aat aaa aat a-- -tt ttt ttt ttt ttt tgt aaa act ttt tta tga gac --- --- -ca aga gaa tac gaa tag tga tca tat cgt atc aca tat tga aac aga aag aag aag taa cga gag gta act ttt tgt gaa tgt agt taa ata ttt ttg ttt tgc aaa ccg gaa tat agt gcc cgg t-- -ct ttt tta att cgt ggt gcg gtg tct gaa tcg ttc gat taa ccc aac tca tcc att ttc aga tga ata gag tta tcg att cag aca cat gct ttg agt ttt gtt gaa tcg atg agt gaa gta tca tcg gtt gca cct tca gat gcc gat ccg tcg aca tac ttg aat cca tcc ttg act tca agt tca gat gat tcc tca cac atg tct ccg ata cgt acg cta aac tct agg ttc ttg aca cat ttt gta tca acg atc gtt gaa ccg atg ata tct ttg taa ctc act ttc tta tgt gag atg tta gac cca agt act gga tgg gtc ttg atg tca ctg tct ttc tct tct tcg cta cat ctg atg tcg ata gac atc tca cag tct ttg atc ata gcc aga gct tct tca cgc gtg atc gcg gga gag tcc tta cct tgt ccc ggt gac acg ctg gac aat cta gta ttc aca gtg ttt cca tca gag gat tcg gag atg gat gaa atc ttt ggg cat ttg gtg aat cca aag ttc atg tta aga ccc gca ccg acg ata gtg taa taa gtg gtg gga tct cct ttt aca act --- --- --- --- --- --- tct tcg gat acc tca tca tct tcg gtc tct gta act tcc gtt acg gat tga caa atc tta tca ttg gtc ggt gtt tgg tct tgc ttt gtg act ttg ata ata aca tcg att ccc ata tga tgt ttg ttt tct tct tca gta cac gag gat gaa gat tgt tga aga cta gta ggc ata gca gct gcc act agg cac atg cat gcc agg aca ata tat tgt ttc atg att gct att gat tga tta ctg ttc tag atg att cta ctt tct tac cat ata ata aat tag aat ata ttt tct act ttt acg aga aat taa tta ttg tat tta tta ttt ata ggt --a aaa aaa ctt act ata agt ggg tgg gat tct ggg aat tag tga tca gtt tat gta tat cgc aac tag cgg gca tat ggc tat tga cat cga gaa cat tac cca tat gat aag aga ttg tat cat ttt cgt agt ctt gag tat tgg tat tac tat ata gta tgt aga tgt cga cgc tag ata gac agt cgc cca cta gag tta ccg tct ctg aat gcg gca tga tag tat cat tct ttg ttt tcg tta act gtt tgg aag atg aat ctt tgt tgt tac att taa tct cga aat tca gag tac ata tct ttg aag tat tct gat atc tat ttt ctc ctg taa aga atc ctg aag ttg cta cat tat taa gga cag aga agt att ctg cac gaa aga ctg gat cac aat ctt tat gat tca tgg taa tag tta gtt ccg acg ttg aga tgg att cgc tga gac cgg tag tgg tcg tcc gag tac acg atg tgt cgt tga ctg gat aca ggt taa ttt cca cat cga tat agt taa atg tat tgc tgg tta cga cgg gtt cgc att tat ctg tgg aag aga cgg tgt gag aat atg ttc cgg gac cac acg gag aac aga tga cgt ctc cgg tag acg tgt atc cgg ata ctc cgt atc cta ttc cac act ttg ttt tag aaa tac atg ttc tac acc ctg atg ctc ctt tga gaa gac aat aat atc ctg gag agc att cac aga ttc tat tgt gag tcg tgt tac acg atc gcg tct cta cct gat tac tat cac atc ttc cgt tac aac tta gac aag cct gta aat gat tat tgt gag atg taa agg tat ccg aac cac acg gtg tac att gtg tat tag tct tgc tat cac ata atc tgg aag cgt aag ttc ccg gag gac acg ata gac aac ata gat tac ggc ttc tgt att cgt tgt ctt tac act ttc cat tgg atg gtg cat gtg gtg cta tat ctc ttc cgt tta tta tta tac atg aga gaa aca ata tat acg agt ata ata cgg acc tca tga ttt aat aat -gt agt aat cgt cgt ctt gtt act gtt tgt ttc cta ctt ctc caa tca tat aga tta ttt ttt aaa tat ttt ctt tct atc atg gat aat att tgt aat ggt tct ttc cgt aca aca tac tgt tta gat ggt agt cgc tta gct tgg tta tga tat tgc gca taa ttt ccg gag gca aat acg ata gtc tag att gac tat cga tgg tag act cta att tat tga gtg ctt tgt cga cga gtt tac ttt tat gct cca tcg ata gat gac act gtt cta tga gat cgt cgt aca tgg gaa atg aaa tgc gtt tgt ctg aat gta tgg ctt cga gat atg tgt gat acc gga tgt ctt ctg ttc tca ata ccg tat aca agt cgg tgt ctg aga ttc gaa tct ctt tga gga gac tta tgt cac gac tac att ttt cga tga tgg aat cta tct tat cga atg ata tat ttt tca taa ata cac ttt tat agt cct cgt tta aac aga att tag tat gta gtt ccg caa atg act cgt ccc tta ata ggc agt agg cta tta tct tct tta cgt agt gat cgt cgt agg gag aga act ccg aca tct tgt aga aca acg att taa tca tag gta gag ata ctt tca gtc tgt ggt gga taa tgt cat taa cga cat ccg cct tgt ata taa tgt ttc tgt ttt caa aca cca agt cga ata ctg tct tat cgt ctt tag tcg gaa ggt tga tgt cgt atc cga tgt ata cga ggt atg agg caa cat tgt tat tgc aat tct gga agg cgg tat gaa gag gag tca ttg tat tat agt att cgt ctt tct gaa tgt cga atc tat cta gta gat acc gta gta tat tga gag agc gac ttc cat atc ctt gat tat gtt tta tga ata gat aaa gta gat gtt gtc ctt ctt cct ttt gta att tcc cgt att ttt gtt cgt gcc aat tga gta aca tta tga gaa tat gac ctg ttg cac aat cgt tct tta tgt att cca tga tgg gtg tac aat caa gat tat tac gta tcc tcg tat cgg ctc ctc gag ata aaa gag cat aca cca cac gag gac tat gtt tgg tat act gtt gaa ggt aag tgt gta acg gcg tat ttc cga ttt tcg taa ccg cgt taa tgt ttg ctc cat gat cta tta tcg cgt aga tga atc gct tct cag ctc gca tct tag tgt gac tct ttg act tgt aat aat tgc ttt cgt gga acg cgg ata tgt gtt tac agt agt aat gaa gag aag tga gtc cat cct cat cga cgc aat tag ggt cag atc ctt tag tca ata att tgt aca gaa cgt aat agt tta agc tcc cat tga att tat atc taa gat aac aca gca ata gat cgg atg att tac taa agt cat caa tgg ggt ccg tta gta tat caa aga tct tgt tat cga ttg ata gtg aat gaa tca gat agt ggt gta gag gaa tat gtc ctt ttt tat cct tgc tat caa agt tac gca tgc cgt ggc gta aca ata tct tta ata cag atg gat taa atc gtg tat tca tcg tat agc aat gta atg gag agt tac cac att tta gtc gtt tat tca gat cgc agt gtt taa taa cta att taa aca gat gag atg ata tat cca cat caa aga atg cga gat aca tat gac aga cat tat tga cag aaa tgt gac ctt cat tat cac cgt cgt cca taa atg cgt tag gta cgt acc aca tac tat cgt taa cga tgc gca caa tct cgt cca -tt tca tcc atc ttc ata atg att tac ttt ttc ata att aga gaa aaa gat caa ggt ata aaa att aga agt gtt aga cta ta- --- --- --- --- --- --- --- --- --- --- -aa cta act tat gac tta act aac tta tga --- --- --- --- --- --- --- --- --- --- --- --- --- --- --- --- --- --- --- --- --- --- --- --- --- --- --- --- --- --- --- --- --- --- --- --- --- --- --- --- --- --- --- --- --- --- --- --- ctt aac ta- --- --- --- --- --- --- --- --- --- --- -ac tta tga ctt aac taa ctt atg act taa cta act ta- tga ctt aac taa ctt atg act taa cta act tat gac tta act aac tta tga cta tta act cat ttc aag aaa ggt ggg tgg ata gaa ctc tat atg aca gct tgt gaa aca att aga tcc cta att tct aat gga agt ttt gat agg aga ttg tca tca gtt gat aca ttg ttt att atc tca tct att aga gca cgt ctg ttt aga gct tta gtg acc tgc tcg gtt act tct gtg taa atc ttg aat cct tta gtg ata cac tgt gtc aaa act gga tgt tta gaa tac cta tgt aga ata tgg gaa gca tgc ttg ttt ttg tct cta tta tag att aac tca tac atg gtt gta tta tga att ttc atc tgc cta atg tac tcc aat tct tgt tta caa tca att ata taa tca aag agt gat gat gca tac aca tta caa agt gaa taa tct acc atc ata aaa tac ttg ata cag agc ttt atc aca tca tgg ttt tca att gta tta tta agt ata gct aat ttt ata cag tca ata gac aat ggt tct cta agc aat att tct aat att tta aga tgt gct tcc cta cgg gcg atg aca gat ccc cta tcc acg gcc acg tca aga cat gta tat cca tta ctc att act gcg ttg aca ttt gct cca ttt tct aat agc cat gat act aaa tct ata taa cct gca tag ata gcg cga taa agc aag gtc ctt cca cca gca tct agt tga ttg ata tct tca ata tat ggg ata caa agc tta taa att tct aat act gtg ggt tca tct aca agg aat ccc cta gta tac tga att att tta tat aga tct aat tta aca tca ttt tca tct ggg ata cca caa ttc aaa ata aac tca aca aca cta ctt tcc ttt tta cat att ccc cta aaa tag gca ttc aag cat tct att tta tat att aca gcc cca tga tct acc ata aga tca aca atg tct att tct aca tat gca tta gat aga tag taa agt aag aga tct tgc aca gaa tta caa ttc tta ata att ata gag aaa ata tct tcc ata taa ttc ttt gac act aat gca gat ata ata tct tta tat gta ata tat gca aac agt cta tct act ata tac tga tca ata tta tct cta tga atc cta aaa taa tca tac aga aca tct aca gga tca caa att ggt tca agg aga aat cta tca aat att ttc ctg tca aca act ggt tct aga aca taa cag tca aca cct aat cca tgt ttt tta taa tca tct acc aaa gat aat gac caa aga tcg agg tcg tcg tga aac tgc tca tcg aca gcc atg aaa tct acc gac tcc atg gtg cga atc gca ctg tct tat tcg cca ttg att ttc a-t ttt tta taa tta tgt aca tgt ttt cct tct att ctc aag agt cta caa aaa tat a-- -tt ttt tcg ata tct aag tac taa gtt ttt tta ctg ttt ttg tta ctg tct tcc att ctt cta act aaa gat ctg aga taa att ata caa tct tcg cta tcg aac cat ttt tgt agt cta aag cct gaa gta att aac caa ctg ttt tta tta gtg gct ttt ttc gat cta tcc tcg tc- --- ctc tca tca tcc tta ta- --- --- --- --- --- --- tta tta tca tta tta tca tag tct att aaa cac aaa tca tct --- --- --- --- --- --- --- --- --- --- --- --- --- --- --- --- --- --- --- --- --- --- --- --- --- --- --a cgt tta taa c-- --- --- --- --- --- --- --- --a aca ttc --- --- --- --- -tc att att aat tag ttc tgt ag- aat atc ttt aat aat ttg gct ata --- --- -ca tct gtt --- --- --- --- --- --- --- --- --- --- --- --- --- --- --- --- --- --- --c aat act --- --- --- --- --- --- --- --a tct att gat gat ttc ttt t-- --- --- --- --- --- --- --- --- --- --- --t taa gac t-- --- --- --- --- --- --- --- --- --- --- --- --- --- --- --- --- --- --- --- --- --- --- --- --- --- --- --- --- --- --- --- --- --- --- --- --- --- --- --- taa act agt --- --- --- --- --- --- --- --- --- --- --- --- --- --- --- --- --- --- --- --- --- --- --- --- --- --- --- --- --- --- --- --- --- --- --- --- --- --- --- --- --- --- --- --- --- --- --- --- --- --- --- --- --- --- --- --- --- --- --- --- --- --- --- --- --- --- --- --- --- --- --- --- --- --- --- --- --- --- --- --- --- --- --- --- --- --- --- --- --- --- --- --- --- --- --- --- --- --- --- --- --- --- --- --- --- --- --- --- --- --t atg gta atg acg atg aaa --- -tc gag tag ta- --- --a ctt cta ata aag act tga ta- --- --t cat tat cat atg ttt gat cg- --- --- --- --- --- --- --t cat agt taa tag tgt g-- --- --- --- --- --- --- --- --- --- --- --- --- --- --- --- --- --- --- --- -gc taa atg gta ctg tta ata agt tta t-- --- --- --- --- --- --- --- --- --- --- --- --- --- --- --- --- --a gac aat atc ata gta ttt tct ttc cag aat --- --- --- --t aga tta ttt ttt taa ata ctg atc ctc aca att ccg tga tgt agc agt agt tgg t-- --- --- --- --- --- --- --- --- --- --- --- --- --- --- --- --- --- --- --- --- --- --- --- --- --- --- --- -gc atg gtc tat atc gt- --- --- --- --- --- --- --- --- --- --- --- --- --- --- --- --- --- --- --- --- --- --- --- --- --- --- --- --- --- --- --- --- --- --- --- --- --- --- --- --- --- --- --- taa aat gta tca tat ata ata gtt ttc tga cgt gga gta cag aat ttt cga --- --- --- --- --- --- --- --- --- --- --- --- --- --- --- --- --- --- --- --- --- --- --- --- --- --- --- --- --- --- --- --- --- --- --- --- --- --- --- --- --- --- --- --t taa tga gtt cat ggt aag gaa ggg caa atg cct --- gta tat aat ata cat aag tta a-- --- --- --- --- --- --- --- --- --- --- --- --- --t agt ttt tta tca tat tt- --- --- --- --- --- --- --- --t cta ata cca taa taa aaa t-- --- --- --- --- --- --- --- --- --- --- --- --- --- --- --- --- --- --- --- --- --- --- --- --- --t atc att att gcg ttt g-g tag tt- --- --- --- --- ctg ccc ta- --- --- --- --- --- --- --- --t cat cta tat cac tgt cac tct c-- --- --g ctc tca cta tat ctt cta aaa tta caa --- -ac aac --- --- --- --- --- --- --- --- --- --- --- --- --- --- --- --- --- --- --- --- --- --- --- --- --- --- --- --- --- --- --- --- --- --- --- --- tgg ata ttc gat --- --- --- --- --- --- --- --- --- --- --- --- --- --- --- --- --- --- --- --- --- --- --- --- --- --- --a aca gca ttt gtg t-- --- --- --- --- --- --- --- --- --- --- --- --- --- --- --- --- -

>MT903342.1_Monkeypox_virus_isolate_MPXV-Singapore

--- --- --- --- --- --- --- --- --- --- --- --- --- --- --- --- --- --- --- --- --- --- --- --- --- --- --- --- --- --a ttt tac tat ttt att tag tgt cta gaa aaa aat gtg tga ccc acg acc gta gga aac tct aga ggg taa gaa aaa tca atc gtt tat aga gac cat cag aaa gag gtt taa tat ttt tgt gag acc tat cga a-- --- --- --- --- --- --- --- --- --- --- --- --- --- --- --- --- --- --- --- --- --- --- --- --- --- --- --- --- --- --- --- --- --- --- --- --- --- --- --- --- --- --- --- --- --- --- --- --- --- --- --- --- --- --- --- --- --- --- --- --- --- --- --- --- --- --- --- --- --- --- --- --- --- --- --- --- --- --- --- --- --- --- --- --- --- --- --- --- --- --- --- --- --- --- --- --- --- --- --- --- --- --- --- --- --- --- --- --- --- --- --- --- --- --- --- --- --- --- --- --- --- --- --- --- --- --- --- --- --- --- --- --- --- --- --- --- --- --- --- --- --- --- --- --- --- --- --- --- --- --- --- --- --- --- --- --- --- --- --- --- --- --- --- --- --- --- --- --- --- --- --- --- --- --- --- --- --- --- --- --- --- --- --- --- --- --- --- --- --- --- --- --- --- --- --- --- --- --- --- --- --- --- --- --- --- --- --- --- --- --- --- --- --- --- --- --- --- --- --- --- --- --- --- --- --- --- --- --- --- --- --- --- --- --- --- --- --- --- --- --- --- --- --- --- --- --- --- --- --- --- --- --- --- --- --- --- --- --- --- --- --- --- --- --- --- --- --- --- --- --- --- --- --- --- --- --- --- --- --- -ga gag aaa gga taa aaa ctt ttt acg act cca tca gaa aga ggt tta ata ttt ttg tga gac cca tcg aag aga gaa aga gat ggt tag tca aga tat ttt tct tag tac aaa agt caa tgt ttt aaa ata tat gga cga gaa tta att tgt ctg tat aaa aac ttg tgt gaa att atg tac tag aga aaa aac gtg agc agt gtc ccc tac atg gat ttt aca gat cat tta tat tcc aaa aat att aac tat ata cgt tta tta tat gat gtt aac gtg taa att ata aac att att tta tga tgc aat tgt ctg aca acc tag att ggt ata agg atg ttg ata agc tct acg aga ata tat tgt tgg acg tta tcg ttt acg aaa tag ttg aga cat cag aaa gag gtt taa tat ttt tgt gag a-c cat cga aga gag aaa gag aat aaa aat a-- ttt ttt ttt ttt ttt tgt aaa act ttt tta tga gac --- --- -ca aga gaa tac gaa tag tga tca tat cgt atc aca tat tga aac aga aag aag aag taa cga gag gta act ttt tgt gaa tgt agt taa ata ttt ttg ttt tgc aaa ccg gaa tat agt gcc cgg t-- -ct ttt tta att cgt ggt gcg gtg tct gaa tcg ttc gat taa ccc aac tca tcc att ttc aga tga ata gag tta tcg att cag aca cat gct ttg agt ttt gtt gaa tcg atg agt gaa gta tca tcg gtt gca cct tca gat gcc gat ccg tcg aca tac ttg aat cca tcc ttg act tca agt tca gat gat tcc tca cac atg tct ccg ata cgt acg cta aac tct agg ttc ttg aca cat ttt gta tca acg atc gtt gaa ccg atg ata tct ttg taa ctc act ttc tta tgt gag atg tta gac cca agt act gga tgg gtc ttg atg tca ctg tct ttc tct tct tcg cta cat ctg atg tcg ata gac atc tca cag tct ttg atc ata gcc aga gct tct tca cgc gtg atc gcg gga gag tcc tta cct tgt ccc ggt gac acg ctg gac aat cta gta ttc aca gtg ttt cca tca gag gat tcg gag atg gat gaa atc ttt ggg cat ttg gtg aat cca aag ttc atg tta aga ccc gca ccg acg ata gtg taa taa gtg gtg gga tct cct ttt aca act --- --- --- --- --- --- tct tcg gat acc tca tca tct tcg gtc tct gta act tcc gtt acg gat tga caa atc tta tca ttg gtc ggt gtt tgg tct tgc ttt gtg act ttg ata ata aca tcg att ccc ata tga tgt ttg ttt tct tct tca gta cac gag gat gaa gat tgt tga aga cta gta ggc ata gca gct gcc act agg cac atg cat gcc agg aca ata tat tgt ttc atg att gct att gat tga tta ctg ttc tag atg att cta ctt tct tac cat ata ata aat tag aat ata ttt tct act ttt acg aga aat taa tta ttg tat tta tta ttt ata ggt --a aaa aaa ctt act ata agt ggg tgg gat tct ggg aat tag tga tca gtt tat gta tat cgc aac tag cgg gca tat ggc tat tga cat cga gaa cat tac cca tat gat aag aga ttg tat cat ttt cgt agt ctt gag tat tgg tat tac tat ata gta tgt aga tgt cga cgc tag ata gac agt cgc cca cta gag tta ccg tct ctg aat gcg gca tga tag tat cat tct ttg ttt tcg tta act gtt tgg aag atg aat ctt tgt tgt tac att taa tct cga aat tca gag tac ata tct ttg aag tat tct gat atc tat ttt ctc ctg taa aga atc ctg aag ttg cta cat tat taa gga cag aga agt att ctg cac gaa aga ctg gat cac aat ctt tat gat tca tgg taa tag tta gtt ccg acg ttg aga tgg att cgc tga gac cgg tag tgg tcg tcc gag tac acg atg tgt cgt tga ctg gat aca ggt taa ttt cca cat cga tat agt taa atg tat tgc tgg tta cga cgg gtt cgc att tat ctg tgg aag aga cgg tgt gag aat atg ttc cgg gac cac acg gag aac aga tga cgt ctc cgg tag acg tgt atc cgg ata ctc cgt atc cta ttc cac act ttg ttt tag aaa tac atg ttc tac acc ctg atg ctc ctt tga gaa gac aat aat atc ctg gag agc att cac aga ttc tat tgt gag tcg tgt tac acg atc gcg tct cta cct gat tac tat cac atc ttc cgt tac aac tta gac aag cct gta aat gat tat tgt gag atg taa agg tat ccg aac cac acg gtg tac att gtg tat tag tct tgc tat cac ata atc tgg aag cgt aag ttc ccg gag gac acg ata gac aac ata gat tac ggc ttc tgt att cgt tgt ctt tac act ttc cat tgg atg gtg cat gtg gtg cta tat ctc ttc cgt tta tta tta tac atg aga gaa aca ata tat acg agt ata ata cgg acc tca tga ttt aat aat -gt agt aat cgt cgt ctt gtt act gtt tgt ttc cta ctt ctc caa tca tat aga tta ttt ttt aaa tat ttt ctt tct atc atg gat aat att tgt aat ggt tct ttc cgt aca aca tac tgt tta gat ggt agt cgc tta gct tgg tta tga tat tgc gca taa ttt ccg gag gca aat acg ata gtc tag att gac tat cga tgg tag act cta att tat tga gtg ctt tgt cga cga gtt tac ttt tat gct cca tcg ata gat gac act gtt cta tga gat cgt cgt aca tgg gaa atg aaa tgc gtt tgt ctg aat gta tgg ctt cga gat atg tgt gat acc gga tgt ctt ctg ttc tca ata ccg tat aca agt cgg tgt ctg aga ttc gaa tct ctt tga gga gac tta tgt cac gac tac att ttt cga tga tgg aat cta tct tat cga atg ata tat ttt tca taa ata cac ttt tat agt cct cgt tta aac aga att tag tat gta gtt ccg caa atg act cgt ccc tta ata ggc agt agg cta tta tct tct tta cgt agt gat cgt cgt agg gag aga act ccg aca tct tgt aga aca acg att taa tca tag gta gag ata ctt tca gtc tgt ggt gga taa tgt cat taa cga cat ccg cct tgt ata taa tgt ttc tgt ttt caa aca cca agt cga ata ctg tct tat cgt ctt tag tcg gaa ggt tga tgt cgt atc cga tgt ata cga ggt atg agg caa cat tgt tat tgc aat tct gga agg cgg tat gaa gag gag tca ttg tat tat agt att cgt ctt tct gaa tgt cga atc tat cta gta gat acc gta gta tat tga gag agc gac ttc cat atc ctt gat tat gtt tta tga ata gat aaa gta gat gtt gtc ctt ctt cct ttt gta att tcc cgt att ttt gtt cgt gcc aat tga gta aca tta tga gaa tat gac ctg ttg cac aat cgt tct tta tgt att cca tga tgg gtg tac aat caa gat tat tac gta tcc tcg tat cgg ctc ctc gag ata aaa gag cat aca cca cac gag gac tat gtt tgg tat act gtt gaa ggt aag tgt gta acg gcg tat ttc cga ttt tcg taa ccg cgt taa tgt ttg ctc cat gat cta tta tcg cgt aga tga atc gct tct cag ctc gca tct tag tgt gac tct ttg act tgt aat aat tgc ttt cgt gga acg cgg ata tgt gtt tac agt agt aat gaa gag aag tga gtc cat cct cat cga cgc aat tag ggt cag atc ctt tag tca ata att tgt aca gaa cgt aat agt tta agc tcc cat tga att tat atc taa gat aac aca gca ata gat cgg atg att tac taa agt cat caa tgg ggt ccg tta gta tat caa aga tct tgt tat cga ttg ata gtg aat gaa tca gat agt ggt gta gag gaa tat gtc ctt ttt tat cct tgc tat caa agt tac gca tgc cgt ggc gta aca ata tct tta ata cag atg gat taa atc gtg tat tca tcg tat agc aat gta atg gag agt tac cac att tta gtc gtt tat tca gat cgc agt gtt taa taa cta att taa aca gat gag atg ata tat cca cat caa aga atg cga gat aca tat gac aga cat tat tga cag aaa tgt gac ctt cat tat cac cgt cgt cca taa atg cgt tag gta cgt acc aca tac tat cgt taa cga tgc gca caa tct cgt cca -tt tca tcc atc ttc ata atg att tac ttt ttc ata att aga gaa aaa gat caa ggt ata aaa att aga agt gtt aga cta ta- --- --- --- --- --- aac taa ctt atg act taa cta act tat gac tta act aac tta tga --- --- --- --- --- --- --- --- --- --- --- --- --- --- --- --- --- --- --- --- --- --- --- --- --- --- --- --- --- --- --- --- --- --- --- --- --- --- --- --- --- --- --- --- --- --- --- --- ctt aac ta- --- --- --- --- --- --- --- --- --- --- -ac tta tga ctt aac taa ctt atg act taa cta act ta- tga ctt aac taa ctt atg act taa cta act tat gac tta act aac tta tga cta tta act cat ttc aag aaa ggt ggg tgg ata gaa ctc tat atg aca gct tgt gaa aca att aga tcc cta att tct aat gga agt ttt gat agg aga ttg tca tca gtt gat aca ttg ttt att atc tca tct att aga gca cgt ctg ttt aga gct tta gtg acc tgc tcg gtt act tct gtg taa atc ttg aat cct tta gtg ata cac tgt gtc aaa act gga tgt tta gaa tac cta tgt aga ata tgg gaa gca tgc ttg ttt ttg tct cta tta tag att aac tca tac atg gtt gta tta tga att ttc atc tgc cta atg tac tcc aat tct tgt tta caa tca att ata taa tca aag agt gat gat gca tac aca tta caa agt gaa taa tct acc atc ata aaa tac ttg ata cag agc ttt atc aca tca tgg ttt tca att gta tta tta agt ata gct aat ttt ata cag tca ata gac aat ggt tct cta agc aat att tct aat att tta aga tgt gct tcc cta cgg gcg atg aca gat ccc cta tcc acg gcc acg tca aga cat gta tat cca tta ctc att act gcg ttg aca ttt gct cca ttt tct aat agc cat gat act aaa tct ata taa cct gca tag ata gcg cga taa agc aag gtc ctt cca cca gca tct agt tga ttg ata tct tca ata tat ggg ata caa agc tta taa att tct aat act gtg ggt tca tct aca agg aat ccc cta gta tac tga att att tta tat aga tct aat tta aca tca ttt tca tct ggg ata cca caa ttc aaa ata aac tca aca aca cta ctt tcc ttt tta cat att ccc cta aaa tag gca ttc aag cat tct att tta tat att aca gcc cca tga tct acc ata aga tca aca atg tct att tct aca tat gca tta gat aga tag taa agt aag aga tct tgc aca gaa tta caa ttc tta ata att ata gag aaa ata tct tcc ata taa ttc ttt gac act aat gca gat ata ata tct tta tat gta ata tat gca aac agt cta tct act ata tac tga tca ata tta tct cta tga atc cta aaa taa tca tac aga aca tct aca gga tca caa att ggt tca agg aga aat cta tca aat att ttc ctg tca aca act ggt tct aga aca taa cag tca aca cct aat cca tgt ttt tta taa tca tct acc aaa gat aat gac caa aga tcg agg tcg tcg tga aac tgc tca tcg aca gcc atg aaa tct acc gac tcc atg gtg cga atc gca ctg tct tat tcg cca ttg att ttc a-t ttt tta taa tta tgt aca tgt ttt cct tct att ctc aag agt cta caa aaa tat a-- -tt ttt tcg ata tct aag tac taa gtt ttt tta ctg ttt ttg tta ctg tct tcc att ctt cta act aaa gat ctg aga taa att ata caa tct tcg cta tcg aac cat ttt tgt agt cta aag cct gaa gta att aac caa ctg ttt tta tta gtg gct ttt ttc gat cta tcc tcg tc- --- ctc tca tca tcc tta ta- --- --- --- --- --- --- tta tta tca tta tta tca tag tct att aaa cac aaa tca tct --- --- --- --- --- --- --- --- --- --- --- --- --- --- --- --- --- --- --- --- --- --- --- --- --- --- --a cgt tta taa c-- --- --- --- --- --- --- --- --a aca ttc --- --- --- --- -tc att att aat tag ttc tgt ag- aat atc ttt aat aat ttg gct ata --- --- -ca tct gtt --- --- --- --- --- --- --- --- --- --- --- --- --- --- --- --- --- --- --c aat act --- --- --- --- --- --- --- --a tct att gat gat ttc ttt t-- --- --- --- --- --- --- --- --- --- --- --t taa gac t-- --- --- --- --- --- --- --- --- --- --- --- --- --- --- --- --- --- --- --- --- --- --- --- --- --- --- --- --- --- --- --- --- --- --- --- --- --- --- --- taa act agt --- --- --- --- --- --- --- --- --- --- --- --- --- --- --- --- --- --- --- --- --- --- --- --- --- --- --- --- --- --- --- --- --- --- --- --- --- --- --- --- --- --- --- --- --- --- --- --- --- --- --- --- --- --- --- --- --- --- --- --- --- --- --- --- --- --- --- --- --- --- --- --- --- --- --- --- --- --- --- --- --- --- --- --- --- --- --- --- --- --- --- --- --- --- --- --- --- --- --- --- --- --- --- --- --- --- --- --- --- --t atg gta atg acg atg aaa --- -tc gag tag ta- --- --a ctt cta ata aag act tga ta- --- --t cat tat cat atg ttt gat cg- --- --- --- --- --- --- --t cat agt taa tag tgt g-- --- --- --- --- --- --- --- --- --- --- --- --- --- --- --- --- --- --- --- -gc taa atg gta ctg tta ata agt tta t-- --- --- --- --- --- --- --- --- --- --- --- --- --- --- --- --- --a gac aat atc ata gta ttt tct ttc cag aat --- --- --- --t aga tta ttt ttt taa ata ctg atc ctc aca att ccg tga tgt agc agt agt tgg t-- --- --- --- --- --- --- --- --- --- --- --- --- --- --- --- --- --- --- --- --- --- --- --- --- --- --- --- -gc atg gtc tat atc gt- --- --- --- --- --- --- --- --- --- --- --- --- --- --- --- --- --- --- --- --- --- --- --- --- --- --- --- --- --- --- --- --- --- --- --- --- --- --- --- --- --- --- --- taa aat gta tca tat ata ata gtt ttc tga cgt gga gta cag aat ttt cga --- --- --- --- --- --- --- --- --- --- --- --- --- --- --- --- --- --- --- --- --- --- --- --- --- --- --- --- --- --- --- --- --- --- --- --- --- --- --- --- --- --- --- --t taa tga gtt cat ggt aag gaa ggg caa atg cct --- gta tat aat ata cat aag tta a-- --- --- --- --- --- --- --- --- --- --- --- --- --t agt ttt tta tca tat tt- --- --- --- --- --- --- --- --t cta ata cca taa taa aaa t-- --- --- --- --- --- --- --- --- --- --- --- --- --- --- --- --- --- --- --- --- --- --- --- --- --t atc att att gcg ttt g-g tag tt- --- --- --- --- ctg ccc ta- --- --- --- --- --- --- --- --t cat cta tat cac tgt cac tct c-- --- --g ctc tca cta tat ctt cta aaa tta caa --- -ac aac --- --- --- --- --- --- --- --- --- --- --- --- --- --- --- --- --- --- --- --- --- --- --- --- --- --- --- --- --- --- --- --- --- --- --- --- tgg ata ttc gat --- --- --- --- --- --- --- --- --- --- --- --- --- --- --- --- --- --- --- --- --- --- --- --- --- --- --a aca gca ttt gtg t-- --- --- --- --- --- --- --- --- --- --- --- --- --- --- --- --- -

>ON676708.1_Monkeypox_virus_isolate_MPXV_USA_2021_MD_complete_genome

--- --- --- --- --- --- --- --- --- --- --- --- --- --- --- --- --- --- --- --- --- --- --- --- --- --- --- --- --- --a ttt tac tat ttt att tag tgt cta gaa aaa aat gtg tga ccc acg acc gta gga aac tct aga ggg taa gaa aaa tca atc gtt tat aga gac cat cag aaa gag gtt taa tat ttt tgt gag acc tat cga a-- --- --- --- --- --- --- --- --- --- --- --- --- --- --- --- --- --- --- --- --- --- --- --- --- --- --- --- --- --- --- --- --- --- --- --- --- --- --- --- --- --- --- --- --- --- --- --- --- --- --- --- --- --- --- --- --- --- --- --- --- --- --- --- --- --- --- --- --- --- --- --- --- --- --- --- --- --- --- --- --- --- --- --- --- --- --- --- --- --- --- --- --- --- --- --- --- --- --- --- --- --- --- --- --- --- --- --- --- --- --- --- --- --- --- --- --- --- --- --- --- --- --- --- --- --- --- --- --- --- --- --- --- --- --- --- --- --- --- --- --- --- --- --- --- --- --- --- --- --- --- --- --- --- --- --- --- --- --- --- --- --- --- --- --- --- --- --- --- --- --- --- --- --- --- --- --- --- --- --- --- --- --- --- --- --- --- --- --- --- --- --- --- --- --- --- --- --- --- --- --- --- --- --- --- --- --- --- --- --- --- --- --- --- --- --- --- --- --- --- --- --- --- --- --- --- --- --- --- --- --- --- --- --- --- --- --- --- --- --- --- --- --- --- --- --- --- --- --- --- --- --- --- --- --- --- --- --- --- --- --- --- --- --- --- --- --- --- --- --- --- --- --- --- --- --- --- --- --- --- -ga gag aaa gga taa aaa ctt ttt acg act cca tca gaa aga ggt tta ata ttt ttg tga gac cca tcg aag aga gaa aga gat ggt tag tca aga tat ttt tct tag tac aaa agt caa tgt ttt aaa ata tat gga cga gaa tta att tgt ctg tat aaa aac ttg tgt gaa att atg tac tag aga aaa aac gtg agc agt gtc ccc tac atg gat ttt aca gat cat tta tat tcc aaa aat att aac tat ata cgt tta tta tat gat gtt aac gtg taa att ata aac att att tta tga tgc aat tgt ctg aca acc tag att ggt ata agg atg ttg ata agc tct acg aga ata tat tgt tgg acg tta tcg ttt acg aaa tag ttg aga cat cag aaa gag gtt taa tat ttt tgt gag a-c cat cga aga gag aaa gag aat aaa aat a-- --t ttt ttt ttt ttt tgt aaa act ttt tta tga gac --- --- -ca aga gaa tac gaa tag tga tca tat cgt atc aca tat tga aac aga aag aag aag taa cga gag gta act ttt tgt gaa tgt agt taa ata ttt ttg ttt tgc aaa ccg gaa tat agt gcc cgg t-- -ct ttt tta att cgt ggt gcg gtg tct gaa tcg ttc gat taa ccc aac tca tcc att ttc aga tga ata gag tta tcg att cag aca cat gct ttg agt ttt gtt gaa tcg atg agt gaa gta tca tcg gtt gca cct tca gat gcc gat ccg tcg aca tac ttg aat cca tcc ttg act tca agt tca gat gat tcc tca cac atg tct ccg ata cgt acg cta aac tct agg ttc ttg aca cat ttt gta tca acg atc gtt gaa ccg atg ata tct ttg taa ctc act ttc tta tgt gag atg tta gac cca agt act gga tgg gtc ttg atg tca ctg tct ttc tct tct tcg cta cat ctg atg tcg ata gac atc tca cag tct ttg atc ata gcc aga gct tct tca cgc gtg atc gcg gga gag tcc tta cct tgt ccc ggt gac acg ctg gac aat cta gta ttc aca gtg ttt cca tca gag gat tcg gag atg gat aaa atc ttt ggg cat ttg gtg aat cca aag ttc atg tta aga ccc gca ccg acg ata gtg taa taa gtg gtg gga tct cct ttt aca act --- --- --- --- --- --- tct tcg gat acc tca tca tct tcg gtc tct gta act tcc gtt acg gat tga caa atc tta tca ttg gtc ggt gtt tgg tct tgc ttt gtg act ttg ata ata aca tcg att ccc ata tga tgt ttg ttt tct tct tca gta cac gag gat gaa gat tgt tga aga cta gta ggc ata gca gct gcc act agg cac atg cat gcc agg aca ata tat tgt ttc atg att gct att gat tga tta ctg ttc tag atg att cta ctt tct tac cat ata ata aat tag aat ata ttt tct act ttt acg aga aat taa tta ttg tat tta tta ttt ata ggt --a aaa aaa ctt act ata agt ggg tgg gat tct ggg aat tag tga tca gtt tat gta tat cgc aac tag cgg gca tat ggc tat tga cat cga gaa cat tac cca tat gat aag aga ttg tat cat ttt cgt agt ctt gag tat tgg tat tac tat ata gta tgt aga tgt cga cgc tag ata gac agt cgc cca cta gag tta ccg tct ctg aat gcg gca tga tag tat cat tct ttg ttt tcg tta act gtt tgg aag atg aat ctt tgt tgt tac att taa tct cga aat tca gag tac ata tct ttg aag tat tct gat atc tat ttt ctc ctg taa aga atc ctg aag ttg cta cat tat taa gga cag aga agt att ctg cac gaa aga ctg gat cac aat ctt tat gat tca tgg taa tag tta gtt ccg acg ttg aga tgg att cgc tga gac cgg tag tgg tcg tcc gag tac acg atg tgt cgt tga ctg gat aca ggt taa ttt cca cat cga tat agt taa atg tat tgc tgg tta cga cgg gtt cgc att tat ctg tgg aag aga cgg tgt gag aat atg ttc cgg gac cac acg gag aac aga tga cgt ctc cgg tag acg tgt atc cgg ata ctc cgt atc cta ttc cac act ttg ttt tag aaa tac atg ttc tac acc ctg atg ctc ctt tga gaa gac aat aat atc ctg gag agc att cac aga ttc tat tgt gag tcg tgt tac acg atc gcg tct cta cct gat tac tat cac atc ttc cgt tac aac tta gac aag cct gta aat gat tat tgt gag atg taa agg tat ccg aac cac acg gtg tac att gtg tat tag tct tgc tat cac ata atc tga aag cgt aag ttc ccg gag gac acg ata gac aac ata gat tac ggc ttc tgt att cgt tgt ctt tac act ttc cat tgg atg gtg cat gtg gtg cta tat ctc ttc cgt tta tta tta tac atg aga gaa aca ata tat acg agt ata ata cgg acc tca tga ttt aat aat -gt agt aat cgt cgt ctt gtt act gtt tgt ttc cta ctt ctc caa tca tat aga tta ttt ttt aaa tat ttt ctt tct atc atg gat aat att tgt aat ggt tct ttc cgt aca aca tac tgt tta gat ggt agt cgc tta gct tgg tta tga tat tgc gca taa ttt ccg gag gca aat acg ata gtc tag att gac tat cga tgg tag act cta att tat tga gtg ctt tgt cga cga gtt tac ttt tat gct cca tcg ata gat gac act gtt cta tga gat cgt cgt aca tgg gaa atg aaa tgc gtt tgt ctg aat gta tgg ctt cga gat atg tgt gat acc gga tgt ctt ctg ttc tca ata ccg tat aca agt cgg tgt ctg aga ttc gaa tct ctt tga gga gac tta tgt cac gac tac att ttt cga tga tgg aat cta tct tat cga atg ata tat ttt tca taa ata cac ttt tat agt cct cgt tta aac aga att tag tat gta gtt ccg caa atg act cgt ccc tta ata ggc agt agg cta tta tct tct tta cgt agt gat cgt cgt agg gag aga act ccg aca tct tgt aga aca acg att taa tca tag gta gag ata ctt tca gtc tgt ggt gga taa tgt cat taa cga cat ccg cct tgt ata taa tgt ttc tgt ttt caa aca cca agt cga ata ctg tct tat cgt ctt tag tcg gaa ggt tga tgt cgt atc caa tgt ata cga ggt atg agg caa cat tgt tat tgc aat tct gga agg cgg tat gaa gag gag tca ttg tat tat agt att cgt ctt tct gaa tgt cga atc tat cta gta gat acc gta gta tat tga gag agc gac ttc cat atc ctt gat tat gtt tta tga ata gat aaa gta gat gtt gtc ctt ctt cct ttt gta att tcc cgt att ttt gtt cgt gcc aat tga gta aca tta tga gaa tat gac ctg ttg cac aat cgt tct tta tgt att cca tga tgg gtg tac aat taa gat tat tac gta tcc tcg tat cgg ctc ctc gag ata aaa gag cat aca cca cac gag gac tat gtt tgg tat act gtt gaa ggt aag tgt gta acg gcg tat ttc cga ttt tcg taa ccg cgt taa tgt ttg ctc cat gat cta tta tcg cgt aga tga atc gct tct cag ctc gca tct tag tgt gac tct ttg act tgt aat aat tgc ttt cgt gga acg cgg ata tgt gtt tac agt agt aat gaa gag aag tga gtc cat cct cat cga cgc aat tag ggt cag atc ctt tag tca ata att tgt aca gaa cgt aat agt tta agc tcc cat tga att tat atc taa gat aac aca gca ata gat cgg atg att tac taa agt cat caa tgg ggt ccg tta gta tat caa aga tct tgt tat cga ttg ata gtg aat gaa tca gat agt ggt gta gag gaa tat gtc ctt ttt tat cct tgc tat caa agt tac gca tgc cgt ggc gta aca ata tct tta ata cag atg gat taa atc gtg tat tca tcg tat agc aat gta atg gag agt tac cac att tta gtc gtt tat tca gat cgc agt gtt taa taa cta att taa aca gat gag atg ata tat cca cat caa aga atg cga gat aca tat gac aga cat tat tga cag aaa tgt gac ctt cat tat cac cgt cgt cca taa atg cgt tag gta cgt acc aca tac tat cgt taa cga tgc gca caa tct cgt cca -tt tca tcc atc ttc ata atg att tac ttt ttc ata att aga gaa aaa gat caa ggt ata aaa att aga agt gtt aga cta ta- --- --- --- --- --- --- --- --- --- --- -aa cta act tat gac tta act aac tta tga --- --- --- --- --- --- --- --- --- --- --- --- --- --- --- --- --- --- --- --- --- --- --- --- --- --- --- --- --- --- --- --- --- --- --- --- --- --- --- --- --- --- --- --- --- --- --- --- ctt aac ta- --- --- --- --- --- --- --- --- --- --- -ac tta tga ctt aac taa ctt atg act taa cta act ta- tga ctt aac taa ctt atg act taa cta act tat gac tta act aac tta tga cta tta act cat ttc aag aaa ggt ggg tgg ata gaa ctc tat atg aca gct tgt gaa aca att aga tcc cta att tct aat gga agt ttt gat agg aga ttg tca tca gtt gat aca ttg ttt att atc tca tct att aga gca cgt ctg ttt aga gct tta gtg acc tgc tcg gtt act tct gtg taa atc ttg aat cct tta gtg ata cac tgt gtc aaa act gga tgt tta gaa tac cta tgt aga ata tgg gaa gca tgc ttg ttt ttg tct cta tta tag att aac tca tac atg gtt gta tta tga att ttc atc tgc cta atg tac tcc aat tct tgt tta caa tca att ata taa tca aag agt gat gat gca tac aca tta caa agt gaa taa tct acc atc ata aaa tac ttg ata cag agc ttt atc aca tca tgg ttt tca att gta tta tta agt ata gct aat ttt ata cag tca ata gac aat ggt tct cta agc aat att tct aat att tta aga tgt gct tcc cta cgg gcg atg aca gat ccc cta tcc acg gcc acg tca aga cat gta tat cca tta ctc att act gcg ttg aca ttt gct cca ttt tct aat agc cat gat act aaa tct ata taa cct gca tag ata gcg cga taa agc aag gtc ctt cca cca gca tct agt tga ttg ata tct tca ata tat ggg ata caa agc tta taa att tct aat act gtg ggt tca tct aca agg aat ccc cta gta tac tga att att tta tat aga tct aat tta aca tca ttt tca tct ggg ata cca caa ttc aaa ata aac tca aca aca cta ctt tcc ttt tta cat att ccc cta aaa tag gca ttc aag cat tct att tta tat att aca gcc cca tga tct acc ata aga tca aca atg tct att tct aca tat gca tta gat aga tag taa agt aag aga tct tgc aca gaa tta caa ttc tta ata att ata gag aaa ata tct tcc ata taa ttc ttt gac act aat gca gat ata ata tct tta tat gta ata tat gca aac agt cta tct act ata tac tga tca ata tta tct cta tga atc cta aaa taa tca tac aga aca tct aca gga tca caa att ggt tca agg aga aat cta tca aat att ttc ctg tca aca act ggt tct aga aca taa cag tca aca cct aat cca tgt ttt tta taa tca tct acc aaa gat aat gac caa aga tcg agg tcg tcg tga aac tgc tca tcg aca gcc atg aaa tct acc gac tcc atg gtg cga atc gca ctg tct tat tcg cca ttg att ttc a-t ttt tta taa tta tgt aca tgt ttt cct tct att ctc aag agt cta caa aaa tat a-- -tt ttt tcg ata tct aag tac taa gtt ttt tta ctg ttt ttg tta ctg tct tcc att ctt cta act aaa gat ctg aga taa att ata caa tct tcg cta tcg aac cat ttt tgt agt cta aag cct gaa gta att aac caa ctg ttt tta tta gtg gct ttt ttc gat cta tcc tcg tc- --- ctc tca tca tcc tta ta- --- --- --- --- --- --- tta tta tca tta tta tca tag tct att aaa cac aaa tca tct --- --- --- --- --- --- --- --- --- --- --- --- --- --- --- --- --- --- --- --- --- --- --- --- --- --- --a cgt tta taa c-- --- --- --- --- --- --- --- --a aca ttc --- --- --- --- -tc att att aat tag ttc tgt ag- aat atc ttt aat aat ttg gct ata --- --- -ca tct gtt --- --- --- --- --- --- --- --- --- --- --- --- --- --- --- --- --- --- --c aat act --- --- --- --- --- --- --- --a tct att gat gat ttc ttt t-- --- --- --- --- --- --- --- --- --- --- --t taa gac t-- --- --- --- --- --- --- --- --- --- --- --- --- --- --- --- --- --- --- --- --- --- --- --- --- --- --- --- --- --- --- --- --- --- --- --- --- --- --- --- taa act agt --- --- --- --- --- --- --- --- --- --- --- --- --- --- --- --- --- --- --- --- --- --- --- --- --- --- --- --- --- --- --- --- --- --- --- --- --- --- --- --- --- --- --- --- --- --- --- --- --- --- --- --- --- --- --- --- --- --- --- --- --- --- --- --- --- --- --- --- --- --- --- --- --- --- --- --- --- --- --- --- --- --- --- --- --- --- --- --- --- --- --- --- --- --- --- --- --- --- --- --- --- --- --- --- --- --- --- --- --- --t atg gta atg acg atg aaa --- -tc gag tag ta- --- --a ctt cta ata aag act tga ta- --- --t cat tat cat atg ttt gat cg- --- --- --- --- --- --- --t cat agt taa tag tgt g-- --- --- --- --- --- --- --- --- --- --- --- --- --- --- --- --- --- --- --- -gc taa atg gta ctg tta ata agt tta t-- --- --- --- --- --- --- --- --- --- --- --- --- --- --- --- --- --a gac aat atc ata gta ttt tct ttc cag aat --- --- --- --t aga tta ttt ttt taa ata ctg atc ctc aca att ccg tga tgt agc agt agt tgg t-- --- --- --- --- --- --- --- --- --- --- --- --- --- --- --- --- --- --- --- --- --- --- --- --- --- --- --- -gc atg gtc tat atc gt- --- --- --- --- --- --- --- --- --- --- --- --- --- --- --- --- --- --- --- --- --- --- --- --- --- --- --- --- --- --- --- --- --- --- --- --- --- --- --- --- --- --- --- taa aat gta tca tat ata ata gtt ttc tga cgt gga gta cag aat ttt cga --- --- --- --- --- --- --- --- --- --- --- --- --- --- --- --- --- --- --- --- --- --- --- --- --- --- --- --- --- --- --- --- --- --- --- --- --- --- --- --- --- --- --- --t taa tga gtt cat ggt aag gaa ggg caa atg cct --- gta tat aat ata cat aag tta a-- --- --- --- --- --- --- --- --- --- --- --- --- --t agt ttt tta tca tat tt- --- --- --- --- --- --- --- --t cta ata cca taa taa aaa t-- --- --- --- --- --- --- --- --- --- --- --- --- --- --- --- --- --- --- --- --- --- --- --- --- --t atc att att gcg ttt g-g tag tt- --- --- --- --- ctg ccc ta- --- --- --- --- --- --- --- --t cat cta tat cac tgt cac tct c-- --- --g ctc tca cta tat ctt cta aaa tta caa --- -ac aac --- --- --- --- --- --- --- --- --- --- --- --- --- --- --- --- --- --- --- --- --- --- --- --- --- --- --- --- --- --- --- --- --- --- --- --- tgg ata ttc gat --- --- --- --- --- --- --- --- --- --- --- --- --- --- --- --- --- --- --- --- --- --- --- --- --- --- --a aca gca ttt gtg t-- --- --- --- --- --- --- --- --- --- --- --- --- --- --- --- --- -

>ON585033.1_Monkeypox_virus_isolate_Monkeypox/PT0006/2022_complete_genome

--- --- --- --- --- --- --- --- --- --- --- --- --- --- --- --- --- --- -ta att a-- --- --- --- --- --- --- --- att taa ttt tac tat ttt att tag tgt cta gaa aaa aat gtg tga ccc acg acc gta gga aac tct aga ggg taa gaa aaa tca atc gtt tat aga gac cat cag aaa gag gtt taa tat ttt tgt gag acc tat cga a-- --- --- --- --- --- --- --- --- --- --- --- --- --- --- --- --- --- --- --- --- --- --- --- --- --- --- --- --- --- --- --- --- --- --- --- --- --- --- --- --- --- --- --- --- --- --- --- --- --- --- --- --- --- --- --- --- --- --- --- --- --- --- --- --- --- --- --- --- --- --- --- --- --- --- --- --- --- --- --- --- --- --- --- --- --- --- --- --- --- --- --- --- --- --- --- --- --- --- --- --- --- --- --- --- --- --- --- --- --- --- --- --- --- --- --- --- --- --- --- --- --- --- --- --- --- --- --- --- --- --- --- --- --- --- --- --- --- --- --- --- --- --- --- --- --- --- --- --- --- --- --- --- --- --- --- --- --- --- --- --- --- --- --- --- --- --- --- --- --- --- --- --- --- --- --- --- --- --- --- --- --- --- --- --- --- --- --- --- --- --- --- --- --- --- --- --- --- --- --- --- --- --- --- --- --- --- --- --- --- --- --- --- --- --- --- --- --- --- --- --- --- --- --- --- --- --- --- --- --- --- --- --- --- --- --- --- --- --- --- --- --- --- --- --- --- --- --- --- --- --- --- --- --- --- --- --- --- --- --- --- --- --- --- --- --- --- --- --- --- --- --- --- --- --- --- --- --- --- --- -ga gag aaa gga taa aaa ctt ttt acg act cca tca gaa aga ggt tta ata ttt ttg tga gac cca tcg aag aga gaa aga gat ggt tag tca aga tat ttt tct tag tac aaa agt caa tgt ttt aaa ata tat gga cga gaa tta att tgt ctg tat aaa aac ttg tgt gaa att atg tac tag aga aaa aac gtg agc agt gtc ccc tac atg gat ttt aca gat cat tta tat tcc aaa aat att aac tat ata cgt tta tta tat gat gtt aac gtg taa att ata aac att att tta tga tgc aat tgt ctg aca acc tag att ggt ata agg atg ttg ata agc tct acg aga ata tat tgt tgg acg tta tcg ttt acg aaa tag ttg aga cat cag aaa gag gtt taa tat ttt tgt gag a-c cat cga aga gag aaa gag aat aaa aat a-- --t ttt ttt ttt ttt tgt aaa act ttt tta tga gac --- --- -ca aga gaa tac gaa tag tga tca tat cgt atc aca tat tga aac aga aag aag aag taa cga gag gta act ttt tgt gaa tgt agt taa ata ttt ttg ttt tgc aaa ccg gaa tat agt gcc cgg t-- -ct ttt tta att cgt ggt gcg gtg tct gaa tcg ttc gat taa ccc aac tca tcc att ttc aga tga ata gag tta tcg att cag aca cat gct ttg agt ttt gtt gaa tcg atg agt gaa gta tca tcg gtt gca cct tca gat gcc gat ccg tcg aca tac ttg aat cca tcc ttg act tca agt tca gat gat tcc tca cac atg tct ccg ata cgt acg cta aac tct agg ttc ttg aca cat ttt gta tca acg atc gtt gaa ccg atg ata tct ttg taa ctc act ttc tta tgt gag atg tta gac cca agt act gga tgg gtc ttg atg tca ctg tct ttc tct tct tcg cta cat ctg atg tcg ata gac atc tca cag tct ttg atc ata gcc aga gct tct tca cgc gtg atc gcg gga gag tcc tta cct tgt ccc ggt gac acg ctg gac aat cta gta ttc aca gtg ttt cca tca gag gat tcg gag atg gat aaa atc ttt ggg cat ttg gtg aat cca aag ttc atg tta aga ccc gca ccg acg ata gtg taa taa gtg gtg gga tct cct ttt aca act --- --- --- --- --- --- tct tcg gat acc tca tca tct tcg gtc tct gta act tcc gtt acg gat tga caa atc tta tca ttg gtc ggt gtt tgg tct tgc ttt gtg act ttg ata ata aca tcg att ccc ata tga tgt ttg ttt tct tct tca gta cac gag gat gaa gat tgt tga aga cta gta ggc ata gca gct gcc act agg cac atg cat gcc agg aca ata tat tgt ttc atg att gct att gat tga tta ctg ttc tag atg att cta ctt tct tac cat ata ata aat tag aat ata ttt tct act ttt acg aga aat taa tta ttg tat tta tta ttt ata ggt --a aaa aaa ctt act ata agt ggg tgg gat tct ggg aat tag tga tca gtt tat gta tat cgc aac tag cgg gca tat ggc tat tga cat cga gaa cat tac cca tat gat aag aga ttg tat cat ttt cgt agt ctt gag tat tgg tat tac tat ata gta tgt aga tgt cga cgc tag ata gac agt cgc cca cta gag tta ccg tct ctg aat gcg gca tga tag tat cat tct ttg ttt tcg tta act gtt tgg aag atg aat ctt tgt tgt tac att taa tct cga aat tca gag tac ata tct ttg aag tat tct gat atc tat ttt ctc ctg taa aga atc ctg aag ttg cta cat tat taa gga cag aga agt att ctg cac gaa aga ctg gat cac aat ctt tat gat tca tgg taa tag tta gtt ccg acg ttg aga tgg att cgc tga gac cgg tag tgg tcg tcc gag tac acg atg tgt cgt tga ctg gat aca ggt taa ttt cca cat cga tat agt taa atg tat tgc tgg tta cga cgg gtt cgc att tat ctg tgg aag aga cgg tgt gag aat atg ttc cgg gac cac acg gag aac aga tga cgt ctc cgg tag acg tgt atc cgg ata ctc cgt atc cta ttc cac act ttg ttt tag aaa tac atg ttc tac acc ctg atg ctc ctt tga gaa gac aat aat atc ctg gag agc att cac aga ttc tat tgt gag tcg tgt tac acg atc gcg tct cta cct gat tac tat cac atc ttc cgt tac aac tta gac aag cct gta aat gat tat tgt gag atg taa agg tat ccg aac cac acg gtg tac att gtg tat tag tct tgc tat cac ata atc tga aag cgt aag ttc ccg gag gac acg ata gac aac ata gat tac ggc ttc tgt att cgt tgt ctt tac act ttc cat tgg atg gtg cat gtg gtg cta tat ctc ttc cgt tta tta tta tac atg aga gaa aca ata tat acg agt ata ata cgg acc tca tga ttt aat aat -gt agt aat cgt cgt ctt gtt act gtt tgt ttc cta ctt ctc caa tca tat aga tta ttt ttt aaa tat ttt ctt tct atc atg gat aat att tgt aat ggt tct ttc cgt aca aca tac tgt tta gat ggt agt cgc tta gct tgg tta tga tat tgc gca taa ttt ccg gag gca aat acg ata gtc tag att gac tat cga tgg tag act cta att tat tga gtg ctt tgt cga cga gtt tac ttt tat gct cca tcg ata gat gac act gtt cta tga gat cgt cgt aca tgg gaa atg aaa tgc gtt tgt ctg aat gta tgg ctt cga gat atg tgt gat acc gaa tgt ctt ctg ttc tca ata ccg tat aca agt cgg tgt ctg aga ttc gaa tct ctt tga gga gac tta tgt cac gac tac att ttt cga tga tgg aat cta tct tat cga atg ata tat ttt tca taa ata cac ttt tat agt cct cgt tta aac aga att tag tat gta gtt ccg caa atg act cgt ccc tta ata ggc agt agg cta tta tct tct tta cgt agt gat cgt cgt agg gag aga act ccg aca tct tgt aga aca acg att taa tca tag gta gag ata ctt tca gtc tgt ggt gga taa tgt cat taa cga cat ccg cct tgt ata taa tgt ttc tgt ttt caa aca cca agt cga ata ctg tct tat cgt ctt tag tcg gaa ggt tga tgt cgt atc caa tgt ata cga ggt atg agg caa cat tgt tat tgc aat tct gga agg cgg tat gaa gag gag tca ttg tat tat agt att cgt ctt tct gaa tgt cga atc tat cta gta gat acc gta gta tat tga gag agc gac ttc cat atc ctt gat tat gtt tta tga ata gat aaa gta gat gtt gtc ctt ctt cct ttt gta att tcc cgt att ttt gtt cgt gcc aat tga gta aca tta tga gaa tat gac ctg ttg cac aat cgt tct tta tgt att cca tga tgg gtg tac aat taa gat tat tac gta tcc tcg tat cgg ctc ctc gag ata aaa gag cat aca cca cac gag gac tat gtt tgg tat act gtt gaa ggt aag tgt gta acg gcg tat ttc cga ttt tcg taa ccg cgt taa tgt ttg ctc cat gat cta tta tcg cgt aga tga atc gct tct cag ctc gca tct tag tgt gac tct ttg act tgt aat aat tgc ttt cgt gga acg cgg ata tgt gtt tac agt agt aat gaa gag aag tga gtc cat cct cat cga cgc aat tag ggt cag atc ctt tag tca ata att tgt aca gaa cgt aat agt tta agc tcc cat tga att tat atc taa gat aac aca gca ata gat cgg atg att tac taa agt cat caa tgg ggt ccg tta gta tat caa aga tct tgt tat cga ttg ata gtg aat gaa tca gat agt ggt gta gag gaa tat gtc ctt ttt tat cct tgc tat caa agt tac gca tgc cgt ggc gta aca ata tct tta ata cag atg gat taa atc gtg tat tca tcg tat agc aat gta atg gag agt tac cac att tta gtc gtt tat tca gat cgc agt gtt taa taa cta att taa aca gat gag atg ata tat cca cat caa aga atg cga gat aca tat gac aga cat tat tga cag aaa tgt gac ctt cat tat cac cgt cgt cca taa atg cgt tag gta cgt acc aca tac tat cgt taa cga tgc gca caa tct cgt cca -tt tca tcc atc ttc ata atg att tac ttt ttc ata att aga gaa aaa gat caa ggt ata aaa att aga agt gtt aga cta ta- --- --- --- --- --- --- --- --- --- --- -aa cta act tat gac tta act aac tta tga --- --- --- --- --- --- --- --- --- --- --- --- --- --- --- --- --- --- --- --- --- --- --- --- --- --- --- --- --- --- --- --- --- --- --- --- --- --- --- --- --- --- --- --- --- --- --- --- ctt aac ta- --- --- --- --- --- --- --- --- --- --- -ac tta tga ctt aac taa ctt atg act taa cta act ta- tga ctt aac taa ctt atg act taa cta act tat gac tta act aac tta tga cta tta act cat ttc aag aaa ggt ggg tgg ata gaa ctc tat atg aca gct tgt gaa aca att aga tcc cta att tct aat gga agt ttt gat agg aga ttg tca tca gtt gat aca ttg ttt att atc tca tct att aga gca cgt ctg ttt aga gct tta gtg acc tgc tcg gtt act tct gtg taa atc ttg aat cct tta gtg ata cac tgt gtc aaa act gga tgt tta gaa tac cta tgt aga ata tgg gaa gca tgc ttg ttt ttg tct cta tta tag att aac tca tac atg gtt gta tta tga att ttc atc tgc cta atg tac tcc aat tct tgt tta caa tca att ata taa tca aag agt gat gat gca tac aca tta caa agt gaa taa tct acc atc ata aaa tac ttg ata cag agc ttt atc aca tca tgg ttt tca att gta tta tta agt ata gct aat ttt ata cag tca ata gac aat ggt tct cta agc aat att tct aat att tta aga tgt gct tcc cta cgg gcg atg aca gat ccc cta tcc acg gcc acg tca aga cat gta tat cca tta ctc att act gcg ttg aca ttt gct cca ttt tct aat agc cat gat act aaa tct ata taa cct gca tag ata gcg cga taa agc aag gtc ctt cca cca gca tct agt tga ttg ata tct tca ata tat ggg ata caa agc tta taa att tct aat act gtg ggt tca tct aca agg aat ccc cta gta tac tga att att tta tat aga tct aat tta aca tca ttt tca tct ggg ata cca caa ttc aaa ata aac tca aca aca cta ctt tcc ttt tta cat att ccc cta aaa tag gca ttc aag cat tct att tta tat att aca gcc cca tga tct acc ata aga tca aca atg tct att tct aca tat gca tta gat aga tag taa agt aag aga tct tgc aca gaa tta caa ttc tta ata att ata gag aaa ata tct tcc ata taa ttc ttt gac act aat gca gat ata ata tct tta tat gta ata tat gca aac agt cta tct act ata tac tga tca ata tta tct cta tga atc cta aaa taa tca tac aga aca tct aca gga tca caa att ggt tca agg aga aat cta tca aat att ttc ctg tca aca act ggt tct aga aca taa cag tca aca cct aat cca tgt ttt tta taa tca tct acc aaa gat aat gac caa aga tcg agg tcg tcg tga aac tgc tca tcg aca gcc atg aaa tct acc gac tcc atg gtg cga atc gca ctg tct tat tcg cca ttg att ttc a-t ttt tta taa tta tgt aca tgt ttt cct tct att ctc aag agt cta caa aaa tat a-- -tt ttt tcg ata tct aag tac taa gtt ttt tta ctg ttt ttg tta ctg tct tcc att ctt cta act aaa gat ctg aga taa att ata caa tct tcg cta tcg aac cat ttt tgt agt cta aag cct gaa gta att aac caa ctg ttt tta tta gtg gct ttt ttc gat cta tcc tcg tc- --- ctc tca tca tcc tta ta- --- --- --- --- --- --- tta tta tca tta tta tca tag tct att aaa cac aaa tca tct --- --- --- --- --- --- --- --- --- --- --- --- --- --- --- --- --- --- --- --- --- --- --- --- --- --- --a cgt tta taa c-- --- --- --- --- --- --- --- --a aca ttc --- --- --- --- -tc att att aat tag ttc tgt ag- aat atc ttt aat aat ttg gct ata --- --- -ca tct gtt --- --- --- --- --- --- --- --- --- --- --- --- --- --- --- --- --- --- --c aat act --- --- --- --- --- --- --- --a tct att gat gat ttc ttt t-- --- --- --- --- --- --- --- --- --- --- --t taa gac t-- --- --- --- --- --- --- --- --- --- --- --- --- --- --- --- --- --- --- --- --- --- --- --- --- --- --- --- --- --- --- --- --- --- --- --- --- --- --- --- taa act agt --- --- --- --- --- --- --- --- --- --- --- --- --- --- --- --- --- --- --- --- --- --- --- --- --- --- --- --- --- --- --- --- --- --- --- --- --- --- --- --- --- --- --- --- --- --- --- --- --- --- --- --- --- --- --- --- --- --- --- --- --- --- --- --- --- --- --- --- --- --- --- --- --- --- --- --- --- --- --- --- --- --- --- --- --- --- --- --- --- --- --- --- --- --- --- --- --- --- --- --- --- --- --- --- --- --- --- --- --- --t atg gta atg acg atg aaa --- -tc gag tag ta- --- --a ctt cta ata aag act tga ta- --- --t cat tat cat atg ttt gat cg- --- --- --- --- --- --- --t cat agt taa tag tgt g-- --- --- --- --- --- --- --- --- --- --- --- --- --- --- --- --- --- --- --- -gc taa atg gta ctg tta ata agt tta t-- --- --- --- --- --- --- --- --- --- --- --- --- --- --- --- --- --a gac aat atc ata gta ttt tct ttc cag aat --- --- --- --t aga tta ttt ttt taa ata ctg atc ctc aca att ccg tga tgt agc agt agt tgg t-- --- --- --- --- --- --- --- --- --- --- --- --- --- --- --- --- --- --- --- --- --- --- --- --- --- --- --- -gc atg gtc tat atc gt- --- --- --- --- --- --- --- --- --- --- --- --- --- --- --- --- --- --- --- --- --- --- --- --- --- --- --- --- --- --- --- --- --- --- --- --- --- --- --- --- --- --- --- taa aat gta tca tat ata ata gtt ttc tga cgt gga gta cag aat ttt cga --- --- --- --- --- --- --- --- --- --- --- --- --- --- --- --- --- --- --- --- --- --- --- --- --- --- --- --- --- --- --- --- --- --- --- --- --- --- --- --- --- --- --- --t taa tga gtt cat ggt aag gaa ggg caa atg cct --- gta tat aat ata cat aag tta a-- --- --- --- --- --- --- --- --- --- --- --- --- --t agt ttt tta tca tat tt- --- --- --- --- --- --- --- --t cta ata cca taa taa aaa t-- --- --- --- --- --- --- --- --- --- --- --- --- --- --- --- --- --- --- --- --- --- --- --- --- --t atc att att gcg ttt g-g tag tt- --- --- --- --- ctg ccc ta- --- --- --- --- --- --- --- --t cat cta tat cac tgt cac tct c-- --- --g ctc tca cta tat ctt cta aaa tta caa --- -ac aac --- --- --- --- --- --- --- --- --- --- --- --- --- --- --- --- --- --- --- --- --- --- --- --- --- --- --- --- --- --- --- --- --- --- --- --- tgg ata ttc gat --- --- --- --- --- --- --- --- --- --- --- --- --- --- --- --- --- --- --- --- --- --- --- --- --- --- --a aca gca ttt gtg t-- --- --- --- --- --- --- --- --- --- --- --- --- --- --- --- --- -

>ON585035.1_Monkeypox_virus_isolate_Monkeypox/PT0009/2022_complete_genome

--- --- --- --- --- --- --- --- --- --- --- --- --- --- --- --- --- --- -ta att a-- --- --- --- --- --- --- --- att taa ttt tac tat ttt att tag tgt cta gaa aaa aat gtg tga ccc acg acc gta gga aac tct aga ggg taa gaa aaa tca atc gtt tat aga gac cat cag aaa gag gtt taa tat ttt tgt gag acc tat cga a-- --- --- --- --- --- --- --- --- --- --- --- --- --- --- --- --- --- --- --- --- --- --- --- --- --- --- --- --- --- --- --- --- --- --- --- --- --- --- --- --- --- --- --- --- --- --- --- --- --- --- --- --- --- --- --- --- --- --- --- --- --- --- --- --- --- --- --- --- --- --- --- --- --- --- --- --- --- --- --- --- --- --- --- --- --- --- --- --- --- --- --- --- --- --- --- --- --- --- --- --- --- --- --- --- --- --- --- --- --- --- --- --- --- --- --- --- --- --- --- --- --- --- --- --- --- --- --- --- --- --- --- --- --- --- --- --- --- --- --- --- --- --- --- --- --- --- --- --- --- --- --- --- --- --- --- --- --- --- --- --- --- --- --- --- --- --- --- --- --- --- --- --- --- --- --- --- --- --- --- --- --- --- --- --- --- --- --- --- --- --- --- --- --- --- --- --- --- --- --- --- --- --- --- --- --- --- --- --- --- --- --- --- --- --- --- --- --- --- --- --- --- --- --- --- --- --- --- --- --- --- --- --- --- --- --- --- --- --- --- --- --- --- --- --- --- --- --- --- --- --- --- --- --- --- --- --- --- --- --- --- --- --- --- --- --- --- --- --- --- --- --- --- --- --- --- --- --- --- --- -ga gag aaa gga taa aaa ctt ttt acg act cca tca gaa aga ggt tta ata ttt ttg tga gac cca tcg aag aga gaa aga gat ggt tag tca aga tat ttt tct tag tac aaa agt caa tgt ttt aaa ata tat gga cga gaa tta att tgt ctg tat aaa aac ttg tgt gaa att atg tac tag aga aaa aac gtg agc agt gtc ccc tac atg gat ttt aca gat cat tta tat tcc aaa aat att aac tat ata cgt tta tta tat gat gtt aac gtg taa att ata aac att att tta tga tgc aat tgt ctg aca acc tag att ggt ata agg atg ttg ata agc tct acg aga ata tat tgt tgg acg tta tcg ttt acg aaa tag ttg aga cat cag aaa gag gtt taa tat ttt tgt gag a-c cat cga aga gag aaa gag aat aaa aat a-- --t ttt ttt ttt ttt tgt aaa act ttt tta tga gac --- --- -ca aga gaa tac gaa tag tga tca tat cgt atc aca tat tga aac aga aag aag aag taa cga gag gta act ttt tgt gaa tgt agt taa ata ttt ttg ttt tgc aaa ccg gaa tat agt gcc cgg t-- -ct ttt tta att cgt ggt gcg gtg tct gaa tcg ttc gat taa ccc aac tca tcc att ttc aga tga ata gag tta tcg att cag aca cat gct ttg agt ttt gtt gaa tcg atg agt gaa gta tca tcg gtt gca cct tca gat gcc gat ccg tcg aca tac ttg aat cca tcc ttg act tca agt tca gat gat tcc tca cac atg tct ccg ata cgt acg cta aac tct agg ttc ttg aca cat ttt gta tca acg atc gtt gaa ccg atg ata tct ttg taa ctc act ttc tta tgt gag atg tta gac cca agt act gga tgg gtc ttg atg tca ctg tct ttc tct tct tcg cta cat ctg atg tcg ata gac atc tca cag tct ttg atc ata gcc aga gct tct tca cgc gtg atc gcg gga gag tcc tta cct tgt ccc ggt gac acg ctg gac aat cta gta ttc aca gtg ttt cca tca gag gat tcg gag atg gat aaa atc ttt ggg cat ttg gtg aat cca aag ttc atg tta aga ccc gca ccg acg ata gtg taa taa gtg gtg gga tct cct ttt aca act --- --- --- --- --- --- tct tcg gat acc tca tca tct tcg gtc tct gta act tcc gtt acg gat tga caa atc tta tca ttg gtc ggt gtt tgg tct tgc ttt gtg act ttg ata ata aca tcg att ccc ata tga tgt ttg ttt tct tct tca gta cac gag gat gaa gat tgt tga aga cta gta ggc ata gca gct gcc act agg cac atg cat gcc agg aca ata tat tgt ttc atg att gct att gat tga tta ctg ttc tag atg att cta ctt tct tac cat ata ata aat tag aat ata ttt tct act ttt acg aga aat taa tta ttg tat tta tta ttt ata ggt --a aaa aaa ctt act ata agt ggg tgg gat tct ggg aat tag tga tca gtt tat gta tat cgc aac tag cgg gca tat ggc tat tga cat cga gaa cat tac cca tat gat aag aga ttg tat cat ttt cgt agt ctt gag tat tgg tat tac tat ata gta tgt aga tgt cga cgc tag ata gac agt cgc cca cta gag tta ccg tct ctg aat gcg gca tga tag tat cat tct ttg ttt tcg tta act gtt tgg aag atg aat ctt tgt tgt tac att taa tct cga aat tca gag tac ata tct ttg aag tat tct gat atc tat ttt ctc ctg taa aga atc ctg aag ttg cta cat tat taa gga cag aga agt att ctg cac gaa aga ctg gat cac aat ctt tat gat tca tgg taa tag tta gtt ccg acg ttg aga tgg att cgc tga gac cgg tag tgg tcg tcc gag tac acg atg tgt cgt tga ctg gat aca ggt taa ttt cca cat cga tat agt taa atg tat tgc tgg tta cga cgg gtt cgc att tat ctg tgg aag aga cgg tgt gag aat atg ttc cgg gac cac acg gag aac aga tga cgt ctc cgg tag acg tgt atc cgg ata ctc cgt atc cta ttc cac act ttg ttt tag aaa tac atg ttc tac acc ctg atg ctc ctt tga gaa gac aat aat atc ctg gag agc att cac aga ttc tat tgt gag tcg tgt tac acg atc gcg tct cta cct gat tac tat cac atc ttc cgt tac aac tta gac aag cct gta aat gat tat tgt gag atg taa agg tat ccg aac cac acg gtg tac att gtg tat tag tct tgc tat cac ata atc tga aag cgt aag ttc ccg gag gac acg ata gac aac ata gat tac ggc ttc tgt att cgt tgt ctt tac act ttc cat tgg atg gtg cat gtg gtg cta tat ctc ttc cgt tta tta tta tac atg aga gaa aca ata tat acg agt ata ata cgg acc tca tga ttt aat aat -gt agt aat cgt cgt ctt gtt act gtt tgt ttc cta ctt ctc caa tca tat aga tta ttt ttt aaa tat ttt ctt tct atc atg gat aat att tgt aat ggt tct ttc cgt aca aca tac tgt tta gat ggt agt cgc tta gct tgg tta tga tat tgc gca taa ttt ccg gag gca aat acg ata gtc tag att gac tat cga tgg tag act cta att tat tga gtg ctt tgt cga cga gtt tac ttt tat gct cca tcg ata gat gac act gtt cta tga gat cgt cgt aca tgg gaa atg aaa tgc gtt tgt ctg aat gta tgg ctt cga gat atg tgt gat acc gaa tgt ctt ctg ttc tca ata ccg tat aca agt cgg tgt ctg aga ttc gaa tct ctt tga gga gac tta tgt cac gac tac att ttt cga tga tgg aat cta tct tat cga atg ata tat ttt tca taa ata cac ttt tat agt cct cgt tta aac aga att tag tat gta gtt ccg caa atg act cgt ccc tta ata ggc agt agg cta tta tct tct tta cgt agt gat cgt cgt agg gag aga act ccg aca tct tgt aga aca acg att taa tca tag gta gag ata ctt tca gtc tgt ggt gga taa tgt cat taa cga cat ccg cct tgt ata taa tgt ttc tgt ttt caa aca cca agt cga ata ctg tct tat cgt ctt tag tcg gaa ggt tga tgt cgt atc caa tgt ata cga ggt atg agg caa cat tgt tat tgc aat tct gga agg cgg tat gaa gag gag tca ttg tat tat agt att cgt ctt tct gaa tgt cga atc tat cta gta gat acc gta gta tat tga gag agc gac ttc cat atc ctt gat tat gtt tta tga ata gat aaa gta gat gtt gtc ctt ctt cct ttt gta att tcc cgt att ttt gtt cgt gcc aat tga gta aca tta tga gaa tat gac ctg ttg cac aat cgt tct tta tgt att cca tga tgg gtg tac aat taa gat tat tac gta tcc tcg tat cgg ctc ctc gag ata aaa gag cat aca cca cac gag gac tat gtt tgg tat act gtt gaa ggt aag tgt gta acg gcg tat ttc cga ttt tcg taa ccg cgt taa tgt ttg ctc cat gat cta tta tcg cgt aga tga atc gct tct cag ctc gca tct tag tgt gac tct ttg act tgt aat aat tgc ttt cgt gga acg cgg ata tgt gtt tac agt agt aat gaa gag aag tga gtc cat cct cat cga cgc aat tag ggt cag atc ctt tag tca ata att tgt aca gaa cgt aat agt tta agc tcc cat tga att tat atc taa gat aac aca gca ata gat cgg atg att tac taa agt cat caa tgg ggt ccg tta gta tat caa aga tct tgt tat cga ttg ata gtg aat gaa tca gat agt ggt gta gag gaa tat gtc ctt ttt tat cct tgc tat caa agt tac gca tgc cgt ggc gta aca ata tct tta ata cag atg gat taa atc gtg tat tca tcg tat agc aat gta atg gag agt tac cac att tta gtc gtt tat tca gat cgc agt gtt taa taa cta att taa aca gat gag atg ata tat cca cat caa aga atg cga gat aca tat gac aga cat tat tga cag aaa tgt gac ctt cat tat cac cgt cgt cca taa atg cgt tag gta cgt acc aca tac tat cgt taa cga tgc gca caa tct cgt cca -tt tca tcc atc ttc ata atg att tac ttt ttc ata att aga gaa aaa gat caa ggt ata aaa att aga agt gtt aga cta ta- --- --- --- --- --- --- --- --- --- --- -aa cta act tat gac tta act aac tta tga --- --- --- --- --- --- --- --- --- --- --- --- --- --- --- --- --- --- --- --- --- --- --- --- --- --- --- --- --- --- --- --- --- --- --- --- --- --- --- --- --- --- --- --- --- --- --- --- ctt aac ta- --- --- --- --- --- --- --- --- --- --- -ac tta tga ctt aac taa ctt atg act taa cta act ta- tga ctt aac taa ctt atg act taa cta act tat gac tta act aac tta tga cta tta act cat ttc aag aaa ggt ggg tgg ata gaa ctc tat atg aca gct tgt gaa aca att aga tcc cta att tct aat gga agt ttt gat agg aga ttg tca tca gtt gat aca ttg ttt att atc tca tct att aga gca cgt ctg ttt aga gct tta gtg acc tgc tcg gtt act tct gtg taa atc ttg aat cct tta gtg ata cac tgt gtc aaa act gga tgt tta gaa tac cta tgt aga ata tgg gaa gca tgc ttg ttt ttg tct cta tta tag att aac tca tac atg gtt gta tta tga att ttc atc tgc cta atg tac tcc aat tct tgt tta caa tca att ata taa tca aag agt gat gat gca tac aca tta caa agt gaa taa tct acc atc ata aaa tac ttg ata cag agc ttt atc aca tca tgg ttt tca att gta tta tta agt ata gct aat ttt ata cag tca ata gac aat ggt tct cta agc aat att tct aat att tta aga tgt gct tcc cta cgg gcg atg aca gat ccc cta tcc acg gcc acg tca aga cat gta tat cca tta ctc att act gcg ttg aca ttt gct cca ttt tct aat agc cat gat act aaa tct ata taa cct gca tag ata gcg cga taa agc aag gtc ctt cca cca gca tct agt tga ttg ata tct tca ata tat ggg ata caa agc tta taa att tct aat act gtg ggt tca tct aca agg aat ccc cta gta tac tga att att tta tat aga tct aat tta aca tca ttt tca tct ggg ata cca caa ttc aaa ata aac tca aca aca cta ctt tcc ttt tta cat att ccc cta aaa tag gca ttc aag cat tct att tta tat att aca gcc cca tga tct acc ata aga tca aca atg tct att tct aca tat gca tta gat aga tag taa agt aag aga tct tgc aca gaa tta caa ttc tta ata att ata gag aaa ata tct tcc ata taa ttc ttt gac act aat gca gat ata ata tct tta tat gta ata tat gca aac agt cta tct act ata tac tga tca ata tta tct cta tga atc cta aaa taa tca tac aga aca tct aca gga tca caa att ggt tca agg aga aat cta tca aat att ttc ctg tca aca act ggt tct aga aca taa cag tca aca cct aat cca tgt ttt tta taa tca tct acc aaa gat aat gac caa aga tcg agg tcg tcg tga aac tgc tca tcg aca gcc atg aaa tct acc gac tcc atg gtg cga atc gca ctg tct tat tcg cca ttg att ttc a-t ttt tta taa tta tgt aca tgt ttt cct tct att ctc aag agt cta caa aaa tat a-- -tt ttt tcg ata tct aag tac taa gtt ttt tta ctg ttt ttg tta ctg tct tcc att ctt cta act aaa gat ctg aga taa att ata caa tct tcg cta tcg aac cat ttt tgt agt cta aag cct gaa gta att aac caa ctg ttt tta tta gtg gct ttt ttc gat cta tcc tcg tc- --- ctc tca tca tcc tta ta- --- --- --- --- --- --- tta tta tca tta tta tca tag tct att aaa cac aaa tca tct --- --- --- --- --- --- --- --- --- --- --- --- --- --- --- --- --- --- --- --- --- --- --- --- --- --- --a cgt tta taa c-- --- --- --- --- --- --- --- --a aca ttc --- --- --- --- -tc att att aat tag ttc tgt ag- aat atc ttt aat aat ttg gct ata --- --- -ca tct gtt --- --- --- --- --- --- --- --- --- --- --- --- --- --- --- --- --- --- --c aat act --- --- --- --- --- --- --- --a tct att gat gat ttc ttt t-- --- --- --- --- --- --- --- --- --- --- --t taa gac t-- --- --- --- --- --- --- --- --- --- --- --- --- --- --- --- --- --- --- --- --- --- --- --- --- --- --- --- --- --- --- --- --- --- --- --- --- --- --- --- taa act agt --- --- --- --- --- --- --- --- --- --- --- --- --- --- --- --- --- --- --- --- --- --- --- --- --- --- --- --- --- --- --- --- --- --- --- --- --- --- --- --- --- --- --- --- --- --- --- --- --- --- --- --- --- --- --- --- --- --- --- --- --- --- --- --- --- --- --- --- --- --- --- --- --- --- --- --- --- --- --- --- --- --- --- --- --- --- --- --- --- --- --- --- --- --- --- --- --- --- --- --- --- --- --- --- --- --- --- --- --- --t atg gta atg acg atg aaa --- -tc gag tag ta- --- --a ctt cta ata aag act tga ta- --- --t cat tat cat atg ttt gat cg- --- --- --- --- --- --- --t cat agt taa tag tgt g-- --- --- --- --- --- --- --- --- --- --- --- --- --- --- --- --- --- --- --- -gc taa atg gta ctg tta ata agt tta t-- --- --- --- --- --- --- --- --- --- --- --- --- --- --- --- --- --a gac aat atc ata gta ttt tct ttc cag aat --- --- --- --t aga tta ttt ttt taa ata ctg atc ctc aca att ccg tga tgt agc agt agt tgg t-- --- --- --- --- --- --- --- --- --- --- --- --- --- --- --- --- --- --- --- --- --- --- --- --- --- --- --- -gc atg gtc tat atc gt- --- --- --- --- --- --- --- --- --- --- --- --- --- --- --- --- --- --- --- --- --- --- --- --- --- --- --- --- --- --- --- --- --- --- --- --- --- --- --- --- --- --- --- taa aat gta tca tat ata ata gtt ttc tga cgt gga gta cag aat ttt cga --- --- --- --- --- --- --- --- --- --- --- --- --- --- --- --- --- --- --- --- --- --- --- --- --- --- --- --- --- --- --- --- --- --- --- --- --- --- --- --- --- --- --- --t taa tga gtt cat ggt aag gaa ggg caa atg cct --- gta tat aat ata cat aag tta a-- --- --- --- --- --- --- --- --- --- --- --- --- --t agt ttt tta tca tat tt- --- --- --- --- --- --- --- --t cta ata cca taa taa aaa t-- --- --- --- --- --- --- --- --- --- --- --- --- --- --- --- --- --- --- --- --- --- --- --- --- --t atc att att gcg ttt g-g tag tt- --- --- --- --- ctg ccc ta- --- --- --- --- --- --- --- --t cat cta tat cac tgt cac tct c-- --- --g ctc tca cta tat ctt cta aaa tta caa --- -ac aac --- --- --- --- --- --- --- --- --- --- --- --- --- --- --- --- --- --- --- --- --- --- --- --- --- --- --- --- --- --- --- --- --- --- --- --- tgg ata ttc gat --- --- --- --- --- --- --- --- --- --- --- --- --- --- --- --- --- --- --- --- --- --- --- --- --- --- --a aca gca ttt gtg t-- --- --- --- --- --- --- --- --- --- --- --- --- --- --- --- --- -

>ON649725.1_Monkeypox_virus_isolate_Monkeypox/PT0011/2022_complete_genome

--- --- --- --- --- --- --- --- --- --- --- --- --- --- --- --- --- --- -ta att a-- --- --- --- --- --- --- --- att taa ttt tac tat ttt att tag tgt cta gaa aaa aat gtg tga ccc acg acc gta gga aac tct aga ggg taa gaa aaa tca atc gtt tat aga gac cat cag aaa gag gtt taa tat ttt tgt gag acc tat cga a-- --- --- --- --- --- --- --- --- --- --- --- --- --- --- --- --- --- --- --- --- --- --- --- --- --- --- --- --- --- --- --- --- --- --- --- --- --- --- --- --- --- --- --- --- --- --- --- --- --- --- --- --- --- --- --- --- --- --- --- --- --- --- --- --- --- --- --- --- --- --- --- --- --- --- --- --- --- --- --- --- --- --- --- --- --- --- --- --- --- --- --- --- --- --- --- --- --- --- --- --- --- --- --- --- --- --- --- --- --- --- --- --- --- --- --- --- --- --- --- --- --- --- --- --- --- --- --- --- --- --- --- --- --- --- --- --- --- --- --- --- --- --- --- --- --- --- --- --- --- --- --- --- --- --- --- --- --- --- --- --- --- --- --- --- --- --- --- --- --- --- --- --- --- --- --- --- --- --- --- --- --- --- --- --- --- --- --- --- --- --- --- --- --- --- --- --- --- --- --- --- --- --- --- --- --- --- --- --- --- --- --- --- --- --- --- --- --- --- --- --- --- --- --- --- --- --- --- --- --- --- --- --- --- --- --- --- --- --- --- --- --- --- --- --- --- --- --- --- --- --- --- --- --- --- --- --- --- --- --- --- --- --- --- --- --- --- --- --- --- --- --- --- --- --- --- --- --- --- --- -ga gag aaa gga taa aaa ctt ttt acg act cca tca gaa aga ggt tta ata ttt ttg tga gac cca tcg aag aga gaa aga gat ggt tag tca aga tat ttt tct tag tac aaa agt caa tgt ttt aaa ata tat gga cga gaa tta att tgt ctg tat aaa aac ttg tgt gaa att atg tac tag aga aaa aac gtg agc agt gtc ccc tac atg gat ttt aca gat cat tta tat tcc aaa aat att aac tat ata cgt tta tta tat gat gtt aac gtg taa att ata aac att att tta tga tgc aat tgt ctg aca acc tag att ggt ata agg atg ttg ata agc tct acg aga ata tat tgt tgg acg tta tcg ttt acg aaa tag ttg aga cat cag aaa gag gtt taa tat ttt tgt gag a-c cat cga aga gag aaa gag aat aaa aat a-- --t ttt ttt ttt ttt tgt aaa act ttt tta tga gac --- --- -ca aga gaa tac gaa tag tga tca tat cgt atc aca tat tga aac aga aag aag aag taa cga gag gta act ttt tgt gaa tgt agt taa ata ttt ttg ttt tgc aaa ccg gaa tat agt gcc cgg t-- -ct ttt tta att cgt ggt gcg gtg tct gaa tcg ttc gat taa ccc aac tca tcc att ttc aga tga ata gag tta tcg att cag aca cat gct ttg agt ttt gtt gaa tcg atg agt gaa gta tca tcg gtt gca cct tca gat gcc gat ccg tcg aca tac ttg aat cca tcc ttg act tca agt tca gat gat tcc tca cac atg tct ccg ata cgt acg cta aac tct agg ttc ttg aca cat ttt gta tca acg atc gtt gaa ccg atg ata tct ttg taa ctc act ttc tta tgt gag atg tta gac cca agt act gga tgg gtc ttg atg tca ctg tct ttc tct tct tcg cta cat ctg atg tcg ata gac atc tca cag tct ttg atc ata gcc aga gct tct tca cgc gtg atc gcg gga gag tcc tta cct tgt ccc ggt gac acg ctg gac aat cta gta ttc aca gtg ttt cca tca gag gat tcg gag atg gat aaa atc ttt ggg cat ttg gtg aat cca aag ttc atg tta aga ccc gca ccg acg ata gtg taa taa gtg gtg gga tct cct ttt aca act --- --- --- --- --- --- tct tcg gat acc tca tca tct tcg gtc tct gta act tcc gtt acg gat tga caa atc tta tca ttg gtc ggt gtt tgg tct tgc ttt gtg act ttg ata ata aca tcg att ccc ata tga tgt ttg ttt tct tct tca gta cac gag gat gaa gat tgt tga aga cta gta ggc ata gca gct gcc act agg cac atg cat gcc agg aca ata tat tgt ttc atg att gct att gat tga tta ctg ttc tag atg att cta ctt tct tac cat ata ata aat tag aat ata ttt tct act ttt acg aga aat taa tta ttg tat tta tta ttt ata ggt --a aaa aaa ctt act ata agt ggg tgg gat tct ggg aat tag tga tca gtt tat gta tat cgc aac tag cgg gca tat ggc tat tga cat cga gaa cat tac cca tat gat aag aga ttg tat cat ttt cgt agt ctt gag tat tgg tat tac tat ata gta tgt aga tgt cga cgc tag ata gac agt cgc cca cta gag tta ccg tct ctg aat gcg gca tga tag tat cat tct ttg ttt tcg tta act gtt tgg aag atg aat ctt tgt tgt tac att taa tct cga aat tca gag tac ata tct ttg aag tat tct gat atc tat ttt ctc ctg taa aga atc ctg aag ttg cta cat tat taa gga cag aga agt att ctg cac gaa aga ctg gat cac aat ctt tat gat tca tgg taa tag tta gtt ccg acg ttg aga tgg att cgc tga gac cgg tag tgg tcg tcc gag tac acg atg tgt cgt tga ctg gat aca ggt taa ttt cca cat cga tat agt taa atg tat tgc tgg tta cga cgg gtt cgc att tat ctg tgg aag aga cgg tgt gag aat atg ttc cgg gac cac acg gag aac aga tga cgt ctc cgg tag acg tgt atc cgg ata ctc cgt atc cta ttc cac act ttg ttt tag aaa tac atg ttc tac acc ctg atg ctc ctt tga gaa gac aat aat atc ctg gag agc att cac aga ttc tat tgt gag tcg tgt tac acg atc gcg tct cta cct gat tac tat cac atc ttc cgt tac aac tta gac aag cct gta aat gat tat tgt gag atg taa agg tat ccg aac cac acg gtg tac att gtg tat tag tct tgc tat cac ata atc tga aag cgt aag ttc ccg gag gac acg ata gac aac ata gat tac ggc ttc tgt att cgt tgt ctt tac act ttc cat tgg atg gtg cat gtg gtg cta tat ctc ttc cgt tta tta tta tac atg aga gaa aca ata tat acg agt ata ata cgg acc tca tga ttt aat aat -gt agt aat cgt cgt ctt gtt act gtt tgt ttc cta ctt ctc caa tca tat aga tta ttt ttt aaa tat ttt ctt tct atc atg gat aat att tgt aat ggt tct ttc cgt aca aca tac tgt tta gat ggt agt cgc tta gct tgg tta tga tat tgc gca taa ttt ccg gag gca aat acg ata gtc tag att gac tat cga tgg tag act cta att tat tga gtg ctt tgt cga cga gtt tac ttt tat gct cca tcg ata gat gac act gtt cta tga gat cgt cgt aca tgg gaa atg aaa tgc gtt tgt ctg aat gta tgg ctt cga gat atg tgt gat acc gaa tgt ctt ctg ttc tca ata ccg tat aca agt cgg tgt ctg aga ttc gaa tct ctt tga gga gac tta tgt cac gac tac att ttt cga tga tgg aat cta tct tat cga atg ata tat ttt tca taa ata cac ttt tat agt cct cgt tta aac aga att tag tat gta gtt ccg caa atg act cgt ccc tta ata ggc agt agg cta tta tct tct tta cgt agt gat cgt cgt agg gag aga act ccg aca tct tgt aga aca acg att taa tca tag gta gag ata ctt tca gtc tgt ggt gga taa tgt cat taa cga cat ccg cct tgt ata taa tgt ttc tgt ttt caa aca cca agt cga ata ctg tct tat cgt ctt tag tcg gaa ggt tga tgt cgt atc caa tgt ata cga ggt atg agg caa cat tgt tat tgc aat tct gga agg cgg tat gaa gag gag tca ttg tat tat agt att cgt ctt tct gaa tgt cga atc tat cta gta gat acc gta gta tat tga gag agc gac ttc cat atc ctt gat tat gtt tta tga ata gat aaa gta gat gtt gtc ctt ctt cct ttt gta att tcc cgt att ttt gtt cgt gcc aat tga gta aca tta tga gaa tat gac ctg ttg cac aat cgt tct tta tgt att cca tga tgg gtg tac aat taa gat tat tac gta tcc tcg tat cgg ctc ctc gag ata aaa gag cat aca cca cac gag gac tat gtt tgg tat act gtt gaa ggt aag tgt gta acg gcg tat ttc cga ttt tcg taa ccg cgt taa tgt ttg ctc cat gat cta tta tcg cgt aga tga atc gct tct cag ctc gca tct tag tgt gac tct ttg act tgt aat aat tgc ttt cgt gga acg cgg ata tgt gtt tac agt agt aat gaa gag aag tga gtc cat cct cat cga cgc aat tag ggt cag atc ctt tag tca ata att tgt aca gaa cgt aat agt tta agc tcc cat tga att tat atc taa gat aac aca gca ata gat cgg atg att tac taa agt cat caa tgg ggt ccg tta gta tat caa aga tct tgt tat cga ttg ata gtg aat gaa tca gat agt ggt gta gag gaa tat gtc ctt ttt tat cct tgc tat caa agt tac gca tgc cgt ggc gta aca ata tct tta ata cag atg gat taa atc gtg tat tca tcg tat agc aat gta atg gag agt tac cac att tta gtc gtt tat tca gat cgc agt gtt taa taa cta att taa aca gat gag atg ata tat cca cat caa aga atg cga gat aca tat gac aga cat tat tga cag aaa tgt gac ctt cat tat cac cgt cgt cca taa atg cgt tag gta cgt acc aca tac tat cgt taa cga tgc gca caa tct cgt cca -tt tca tcc atc ttc ata atg att tac ttt ttc ata att aga gaa aaa gat caa ggt ata aaa att aga agt gtt aga cta ta- --- --- --- --- --- --- --- --- --- --- -aa cta act tat gac tta act aac tta tga --- --- --- --- --- --- --- --- --- --- --- --- --- --- --- --- --- --- --- --- --- --- --- --- --- --- --- --- --- --- --- --- --- --- --- --- --- --- --- --- --- --- --- --- --- --- --- --- ctt aac ta- --- --- --- --- --- --- --- --- --- --- -ac tta tga ctt aac taa ctt atg act taa cta act ta- tga ctt aac taa ctt atg act taa cta act tat gac tta act aac tta tga cta tta act cat ttc aag aaa ggt ggg tgg ata gaa ctc tat atg aca gct tgt gaa aca att aga tcc cta att tct aat gga agt ttt gat agg aga ttg tca tca gtt gat aca ttg ttt att atc tca tct att aga gca cgt ctg ttt aga gct tta gtg acc tgc tcg gtt act tct gtg taa atc ttg aat cct tta gtg ata cac tgt gtc aaa act gga tgt tta gaa tac cta tgt aga ata tgg gaa gca tgc ttg ttt ttg tct cta tta tag att aac tca tac atg gtt gta tta tga att ttc atc tgc cta atg tac tcc aat tct tgt tta caa tca att ata taa tca aag agt gat gat gca tac aca tta caa agt gaa taa tct acc atc ata aaa tac ttg ata cag agc ttt atc aca tca tgg ttt tca att gta tta tta agt ata gct aat ttt ata cag tca ata gac aat ggt tct cta agc aat att tct aat att tta aga tgt gct tcc cta cgg gcg atg aca gat ccc cta tcc acg gcc acg tca aga cat gta tat cca tta ctc att act gcg ttg aca ttt gct cca ttt tct aat agc cat gat act aaa tct ata taa cct gca tag ata gcg cga taa agc aag gtc ctt cca cca gca tct agt tga ttg ata tct tca ata tat ggg ata caa agc tta taa att tct aat act gtg ggt tca tct aca agg aat ccc cta gta tac tga att att tta tat aga tct aat tta aca tca ttt tca tct ggg ata cca caa ttc aaa ata aac tca aca aca cta ctt tcc ttt tta cat att ccc cta aaa tag gca ttc aag cat tct att tta tat att aca gcc cca tga tct acc ata aga tca aca atg tct att tct aca tat gca tta gat aga tag taa agt aag aga tct tgc aca gaa tta caa ttc tta ata att ata gag aaa ata tct tcc ata taa ttc ttt gac act aat gca gat ata ata tct tta tat gta ata tat gca aac agt cta tct act ata tac tga tca ata tta tct cta tga atc cta aaa taa tca tac aga aca tct aca gga tca caa att ggt tca agg aga aat cta tca aat att ttc ctg tca aca act ggt tct aga aca taa cag tca aca cct aat cca tgt ttt tta taa tca tct acc aaa gat aat gac caa aga tcg agg tcg tcg tga aac tgc tca tcg aca gcc atg aaa tct acc gac tcc atg gtg cga atc gca ctg tct tat tcg cca ttg att ttc a-t ttt tta taa tta tgt aca tgt ttt cct tct att ctc aag agt cta caa aaa tat a-- -tt ttt tcg ata tct aag tac taa gtt ttt tta ctg ttt ttg tta ctg tct tcc att ctt cta act aaa gat ctg aga taa att ata caa tct tcg cta tcg aac cat ttt tgt agt cta aag cct gaa gta att aac caa ctg ttt tta tta gtg gct ttt ttc gat cta tcc tcg tc- --- ctc tca tca tcc tta ta- --- --- --- --- --- --- tta tta tca tta tta tca tag tct att aaa cac aaa tca tct --- --- --- --- --- --- --- --- --- --- --- --- --- --- --- --- --- --- --- --- --- --- --- --- --- --- --a cgt tta taa c-- --- --- --- --- --- --- --- --a aca ttc --- --- --- --- -tc att att aat tag ttc tgt ag- aat atc ttt aat aat ttg gct ata --- --- -ca tct gtt --- --- --- --- --- --- --- --- --- --- --- --- --- --- --- --- --- --- --c aat act --- --- --- --- --- --- --- --a tct att gat gat ttc ttt t-- --- --- --- --- --- --- --- --- --- --- --t taa gac t-- --- --- --- --- --- --- --- --- --- --- --- --- --- --- --- --- --- --- --- --- --- --- --- --- --- --- --- --- --- --- --- --- --- --- --- --- --- --- --- taa act agt --- --- --- --- --- --- --- --- --- --- --- --- --- --- --- --- --- --- --- --- --- --- --- --- --- --- --- --- --- --- --- --- --- --- --- --- --- --- --- --- --- --- --- --- --- --- --- --- --- --- --- --- --- --- --- --- --- --- --- --- --- --- --- --- --- --- --- --- --- --- --- --- --- --- --- --- --- --- --- --- --- --- --- --- --- --- --- --- --- --- --- --- --- --- --- --- --- --- --- --- --- --- --- --- --- --- --- --- --- --t atg gta atg acg atg aaa --- -tc gag tag ta- --- --a ctt cta ata aag act tga ta- --- --t cat tat cat atg ttt gat cg- --- --- --- --- --- --- --t cat agt taa tag tgt g-- --- --- --- --- --- --- --- --- --- --- --- --- --- --- --- --- --- --- --- -gc taa atg gta ctg tta ata agt tta t-- --- --- --- --- --- --- --- --- --- --- --- --- --- --- --- --- --a gac aat atc ata gta ttt tct ttc cag aat --- --- --- --t aga tta ttt ttt taa ata ctg atc ctc aca att ccg tga tgt agc agt agt tgg t-- --- --- --- --- --- --- --- --- --- --- --- --- --- --- --- --- --- --- --- --- --- --- --- --- --- --- --- -gc atg gtc tat atc gt- --- --- --- --- --- --- --- --- --- --- --- --- --- --- --- --- --- --- --- --- --- --- --- --- --- --- --- --- --- --- --- --- --- --- --- --- --- --- --- --- --- --- --- taa aat gta tca tat ata ata gtt ttc tga cgt gga gta cag aat ttt cga --- --- --- --- --- --- --- --- --- --- --- --- --- --- --- --- --- --- --- --- --- --- --- --- --- --- --- --- --- --- --- --- --- --- --- --- --- --- --- --- --- --- --- --t taa tga gtt cat ggt aag gaa ggg caa atg cct --- gta tat aat ata cat aag tta a-- --- --- --- --- --- --- --- --- --- --- --- --- --t agt ttt tta tca tat tt- --- --- --- --- --- --- --- --t cta ata cca taa taa aaa t-- --- --- --- --- --- --- --- --- --- --- --- --- --- --- --- --- --- --- --- --- --- --- --- --- --t atc att att gcg ttt g-g tag tt- --- --- --- --- ctg ccc ta- --- --- --- --- --- --- --- --t cat cta tat cac tgt cac tct c-- --- --g ctc tca cta tat ctt cta aaa tta caa --- -ac aac --- --- --- --- --- --- --- --- --- --- --- --- --- --- --- --- --- --- --- --- --- --- --- --- --- --- --- --- --- --- --- --- --- --- --- --- tgg ata ttc gat --- --- --- --- --- --- --- --- --- --- --- --- --- --- --- --- --- --- --- --- --- --- --- --- --- --- --a aca gca ttt gtg t-- --- --- --- --- --- --- --- --- --- --- --- --- --- --- --- --- -

>ON649724.1_Monkeypox_virus_isolate_Monkeypox/PT0014/2022_complete_genome

--- --- --- --- --- --- --- --- --- --- --- --- --- --- --- --- --- --- -ta att a-- --- --- --- --- --- --- --- att taa ttt tac tat ttt att tag tgt cta gaa aaa aat gtg tga ccc acg acc gta gga aac tct aga ggg taa gaa aaa tca atc gtt tat aga gac cat cag aaa gag gtt taa tat ttt tgt gag acc tat cga a-- --- --- --- --- --- --- --- --- --- --- --- --- --- --- --- --- --- --- --- --- --- --- --- --- --- --- --- --- --- --- --- --- --- --- --- --- --- --- --- --- --- --- --- --- --- --- --- --- --- --- --- --- --- --- --- --- --- --- --- --- --- --- --- --- --- --- --- --- --- --- --- --- --- --- --- --- --- --- --- --- --- --- --- --- --- --- --- --- --- --- --- --- --- --- --- --- --- --- --- --- --- --- --- --- --- --- --- --- --- --- --- --- --- --- --- --- --- --- --- --- --- --- --- --- --- --- --- --- --- --- --- --- --- --- --- --- --- --- --- --- --- --- --- --- --- --- --- --- --- --- --- --- --- --- --- --- --- --- --- --- --- --- --- --- --- --- --- --- --- --- --- --- --- --- --- --- --- --- --- --- --- --- --- --- --- --- --- --- --- --- --- --- --- --- --- --- --- --- --- --- --- --- --- --- --- --- --- --- --- --- --- --- --- --- --- --- --- --- --- --- --- --- --- --- --- --- --- --- --- --- --- --- --- --- --- --- --- --- --- --- --- --- --- --- --- --- --- --- --- --- --- --- --- --- --- --- --- --- --- --- --- --- --- --- --- --- --- --- --- --- --- --- --- --- --- --- --- --- --- -ga gag aaa gga taa aaa ctt ttt acg act cca tca gaa aga ggt tta ata ttt ttg tga gac cca tcg aag aga gaa aga gat ggt tag tca aga tat ttt tct tag tac aaa agt caa tgt ttt aaa ata tat gga cga gaa tta att tgt ctg tat aaa aac ttg tgt gaa att atg tac tag aga aaa aac gtg agc agt gtc ccc tac atg gat ttt aca gat cat tta tat tcc aaa aat att aac tat ata cgt tta tta tat gat gtt aac gtg taa att ata aac att att tta tga tgc aat tgt ctg aca acc tag att ggt ata agg atg ttg ata agc tct acg aga ata tat tgt tgg acg tta tcg ttt acg aaa tag ttg aga cat cag aaa gag gtt taa tat ttt tgt gag a-c cat cga aga gag aaa gag aat aaa aat a-- --t ttt ttt ttt ttt tgt aaa act ttt tta tga gac --- --- -ca aga gaa tac gaa tag tga tca tat cgt atc aca tat tga aac aga aag aag aag taa cga gag gta act ttt tgt gaa tgt agt taa ata ttt ttg ttt tgc aaa ccg gaa tat agt gcc cgg t-- -ct ttt tta att cgt ggt gcg gtg tct gaa tcg ttc gat taa ccc aac tca tcc att ttc aga tga ata gag tta tcg att cag aca cat gct ttg agt ttt gtt gaa tcg atg agt gaa gta tca tcg gtt gca cct tca gat gcc gat ccg tcg aca tac ttg aat cca tcc ttg act tca agt tca gat gat tcc tca cac atg tct ccg ata cgt acg cta aac tct agg ttc ttg aca cat ttt gta tca acg atc gtt gaa ccg atg ata tct ttg taa ctc act ttc tta tgt gag atg tta gac cca agt act gga tgg gtc ttg atg tca ctg tct ttc tct tct tcg cta cat ctg atg tcg ata gac atc tca cag tct ttg atc ata gcc aga gct tct tca cgc gtg atc gcg gga gag tcc tta cct tgt ccc ggt gac acg ctg gac aat cta gta ttc aca gtg ttt cca tca gag gat tcg gag atg gat aaa atc ttt ggg cat ttg gtg aat cca aag ttc atg tta aga ccc gca ccg acg ata gtg taa taa gtg gtg gga tct cct ttt aca act --- --- --- --- --- --- tct tcg gat acc tca tca tct tcg gtc tct gta act tcc gtt acg gat tga caa atc tta tca ttg gtc ggt gtt tgg tct tgc ttt gtg act ttg ata ata aca tcg att ccc ata tga tgt ttg ttt tct tct tca gta cac gag gat gaa gat tgt tga aga cta gta ggc ata gca gct gcc act agg cac atg cat gcc agg aca ata tat tgt ttc atg att gct att gat tga tta ctg ttc tag atg att cta ctt tct tac cat ata ata aat tag aat ata ttt tct act ttt acg aga aat taa tta ttg tat tta tta ttt ata ggt --a aaa aaa ctt act ata agt ggg tgg gat tct ggg aat tag tga tca gtt tat gta tat cgc aac tag cgg gca tat ggc tat tga cat cga gaa cat tac cca tat gat aag aga ttg tat cat ttt cgt agt ctt gag tat tgg tat tac tat ata gta tgt aga tgt cga cgc tag ata gac agt cgc cca cta gag tta ccg tct ctg aat gcg gca tga tag tat cat tct ttg ttt tcg tta act gtt tgg aag atg aat ctt tgt tgt tac att taa tct cga aat tca gag tac ata tct ttg aag tat tct gat atc tat ttt ctc ctg taa aga atc ctg aag ttg cta cat tat taa gga cag aga agt att ctg cac gaa aga ctg gat cac aat ctt tat gat tca tgg taa tag tta gtt ccg acg ttg aga tgg att cgc tga gac cgg tag tgg tcg tcc gag tac acg atg tgt cgt tga ctg gat aca ggt taa ttt cca cat cga tat agt taa atg tat tgc tgg tta cga cgg gtt cgc att tat ctg tgg aag aga cgg tgt gag aat atg ttc cgg gac cac acg gag aac aga tga cgt ctc cgg tag acg tgt atc cgg ata ctc cgt atc cta ttc cac act ttg ttt tag aaa tac atg ttc tac acc ctg atg ctc ctt tga gaa gac aat aat atc ctg gag agc att cac aga ttc tat tgt gag tcg tgt tac acg atc gcg tct cta cct gat tac tat cac atc ttc cgt tac aac tta gac aag cct gta aat gat tat tgt gag atg taa agg tat ccg aac cac acg gtg tac att gtg tat tag tct tgc tat cac ata atc tga aag cgt aag ttc ccg gag gac acg ata gac aac ata gat tac ggc ttc tgt att cgt tgt ctt tac act ttc cat tgg atg gtg cat gtg gtg cta tat ctc ttc cgt tta tta tta tac atg aga gaa aca ata tat acg agt ata ata cgg acc tca tga ttt aat aat -gt agt aat cgt cgt ctt gtt act gtt tgt ttc cta ctt ctc caa tca tat aga tta ttt ttt aaa tat ttt ctt tct atc atg gat aat att tgt aat ggt tct ttc cgt aca aca tac tgt tta gat ggt agt cgc tta gct tgg tta tga tat tgc gca taa ttt ccg gag gca aat acg ata gtc tag att gac tat cga tgg tag act cta att tat tga gtg ctt tgt cga cga gtt tac ttt tat gct cca tcg ata gat gac act gtt cta tga gat cgt cgt aca tgg gaa atg aaa tgc gtt tgt ctg aat gta tgg ctt cga gat atg tgt gat acc gaa tgt ctt ctg ttc tca ata ccg tat aca agt cgg tgt ctg aga ttc gaa tct ctt tga gga gac tta tgt cac gac tac att ttt cga tga tgg aat cta tct tat cga atg ata tat ttt tca taa ata cac ttt tat agt cct cgt tta aac aga att tag tat gta gtt ccg caa atg act cgt ccc tta ata ggc agt agg cta tta tct tct tta cgt agt gat cgt cgt agg gag aga act ccg aca tct tgt aga aca acg att taa tca tag gta gag ata ctt tca gtc tgt ggt gga taa tgt cat taa cga cat ccg cct tgt ata taa tgt ttc tgt ttt caa aca cca agt cga ata ctg tct tat cgt ctt tag tcg gaa ggt tga tgt cgt atc caa tgt ata cga ggt atg agg caa cat tgt tat tgc aat tct gga agg cgg tat gaa gag gag tca ttg tat tat agt att cgt ctt tct gaa tgt cga atc tat cta gta gat acc gta gta tat tga gag agc gac ttc cat atc ctt gat tat gtt tta tga ata gat aaa gta gat gtt gtc ctt ctt cct ttt gta att tcc cgt att ttt gtt cgt gcc aat tga gta aca tta tga gaa tat gac ctg ttg cac aat cgt tct tta tgt att cca tga tgg gtg tac aat taa gat tat tac gta tcc tcg tat cgg ctc ctc gag ata aaa gag cat aca cca cac gag gac tat gtt tgg tat act gtt gaa ggt aag tgt gta acg gcg tat ttc cga ttt tcg taa ccg cgt taa tgt ttg ctc cat gat cta tta tcg cgt aga tga atc gct tct cag ctc gca tct tag tgt gac tct ttg act tgt aat aat tgc ttt cgt gga acg cgg ata tgt gtt tac agt agt aat gaa gag aag tga gtc cat cct cat cga cgc aat tag ggt cag atc ctt tag tca ata att tgt aca gaa cgt aat agt tta agc tcc cat tga att tat atc taa gat aac aca gca ata gat cgg atg att tac taa agt cat caa tgg ggt ccg tta gta tat caa aga tct tgt tat cga ttg ata gtg aat gaa tca gat agt ggt gta gag gaa tat gtc ctt ttt tat cct tgc tat caa agt tac gca tgc cgt ggc gta aca ata tct tta ata cag atg gat taa atc gtg tat tca tcg tat agc aat gta atg gag agt tac cac att tta gtc gtt tat tca gat cgc agt gtt taa taa cta att taa aca gat gag atg ata tat cca cat caa aga atg cga gat aca tat gac aga cat tat tga cag aaa tgt gac ctt cat tat cac cgt cgt cca taa atg cgt tag gta cgt acc aca tac tat cgt taa cga tgc gca caa tct cgt cca -tt tca tcc atc ttc ata atg att tac ttt ttc ata att aga gaa aaa gat caa ggt ata aaa att aga agt gtt aga cta ta- --- --- --- --- --- --- --- --- --- --- -aa cta act tat gac tta act aac tta tga --- --- --- --- --- --- --- --- --- --- --- --- --- --- --- --- --- --- --- --- --- --- --- --- --- --- --- --- --- --- --- --- --- --- --- --- --- --- --- --- --- --- --- --- --- --- --- --- ctt aac ta- --- --- --- --- --- --- --- --- --- --- -ac tta tga ctt aac taa ctt atg act taa cta act ta- tga ctt aac taa ctt atg act taa cta act tat gac tta act aac tta tga cta tta act cat ttc aag aaa ggt ggg tgg ata gaa ctc tat atg aca gct tgt gaa aca att aga tcc cta att tct aat gga agt ttt gat agg aga ttg tca tca gtt gat aca ttg ttt att atc tca tct att aga gca cgt ctg ttt aga gct tta gtg acc tgc tcg gtt act tct gtg taa atc ttg aat cct tta gtg ata cac tgt gtc aaa act gga tgt tta gaa tac cta tgt aga ata tgg gaa gca tgc ttg ttt ttg tct cta tta tag att aac tca tac atg gtt gta tta tga att ttc atc tgc cta atg tac tcc aat tct tgt tta caa tca att ata taa tca aag agt gat gat gca tac aca tta caa agt gaa taa tct acc atc ata aaa tac ttg ata cag agc ttt atc aca tca tgg ttt tca att gta tta tta agt ata gct aat ttt ata cag tca ata gac aat ggt tct cta agc aat att tct aat att tta aga tgt gct tcc cta cgg gcg atg aca gat ccc cta tcc acg gcc acg tca aga cat gta tat cca tta ctc att act gcg ttg aca ttt gct cca ttt tct aat agc cat gat act aaa tct ata taa cct gca tag ata gcg cga taa agc aag gtc ctt cca cca gca tct agt tga ttg ata tct tca ata tat ggg ata caa agc tta taa att tct aat act gtg ggt tca tct aca agg aat ccc cta gta tac tga att att tta tat aga tct aat tta aca tca ttt tca tct ggg ata cca caa ttc aaa ata aac tca aca aca cta ctt tcc ttt tta cat att ccc cta aaa tag gca ttc aag cat tct att tta tat att aca gcc cca tga tct acc ata aga tca aca atg tct att tct aca tat gca tta gat aga tag taa agt aag aga tct tgc aca gaa tta caa ttc tta ata att ata gag aaa ata tct tcc ata taa ttc ttt gac act aat gca gat ata ata tct tta tat gta ata tat gca aac agt cta tct act ata tac tga tca ata tta tct cta tga atc cta aaa taa tca tac aga aca tct aca gga tca caa att ggt tca agg aga aat cta tca aat att ttc ctg tca aca act ggt tct aga aca taa cag tca aca cct aat cca tgt ttt tta taa tca tct acc aaa gat aat gac caa aga tcg agg tcg tcg tga aac tgc tca tcg aca gcc atg aaa tct acc gac tcc atg gtg cga atc gca ctg tct tat tcg cca ttg att ttc a-t ttt tta taa tta tgt aca tgt ttt cct tct att ctc aag agt cta caa aaa tat a-- -tt ttt tcg ata tct aag tac taa gtt ttt tta ctg ttt ttg tta ctg tct tcc att ctt cta act aaa gat ctg aga taa att ata caa tct tcg cta tcg aac cat ttt tgt agt cta aag cct gaa gta att aac caa ctg ttt tta tta gtg gct ttt ttc gat cta tcc tcg tc- --- ctc tca tca tcc tta ta- --- --- --- --- --- --- tta tta tca tta tta tca tag tct att aaa cac aaa tca tct --- --- --- --- --- --- --- --- --- --- --- --- --- --- --- --- --- --- --- --- --- --- --- --- --- --- --a cgt tta taa c-- --- --- --- --- --- --- --- --a aca ttc --- --- --- --- -tc att att aat tag ttc tgt ag- aat atc ttt aat aat ttg gct ata --- --- -ca tct gtt --- --- --- --- --- --- --- --- --- --- --- --- --- --- --- --- --- --- --c aat act --- --- --- --- --- --- --- --a tct att gat gat ttc ttt t-- --- --- --- --- --- --- --- --- --- --- --t taa gac t-- --- --- --- --- --- --- --- --- --- --- --- --- --- --- --- --- --- --- --- --- --- --- --- --- --- --- --- --- --- --- --- --- --- --- --- --- --- --- --- taa act agt --- --- --- --- --- --- --- --- --- --- --- --- --- --- --- --- --- --- --- --- --- --- --- --- --- --- --- --- --- --- --- --- --- --- --- --- --- --- --- --- --- --- --- --- --- --- --- --- --- --- --- --- --- --- --- --- --- --- --- --- --- --- --- --- --- --- --- --- --- --- --- --- --- --- --- --- --- --- --- --- --- --- --- --- --- --- --- --- --- --- --- --- --- --- --- --- --- --- --- --- --- --- --- --- --- --- --- --- --- --t atg gta atg acg atg aaa --- -tc gag tag ta- --- --a ctt cta ata aag act tga ta- --- --t cat tat cat atg ttt gat cg- --- --- --- --- --- --- --t cat agt taa tag tgt g-- --- --- --- --- --- --- --- --- --- --- --- --- --- --- --- --- --- --- --- -gc taa atg gta ctg tta ata agt tta t-- --- --- --- --- --- --- --- --- --- --- --- --- --- --- --- --- --a gac aat atc ata gta ttt tct ttc cag aat --- --- --- --t aga tta ttt ttt taa ata ctg atc ctc aca att ccg tga tgt agc agt agt tgg t-- --- --- --- --- --- --- --- --- --- --- --- --- --- --- --- --- --- --- --- --- --- --- --- --- --- --- --- -gc atg gtc tat atc gt- --- --- --- --- --- --- --- --- --- --- --- --- --- --- --- --- --- --- --- --- --- --- --- --- --- --- --- --- --- --- --- --- --- --- --- --- --- --- --- --- --- --- --- taa aat gta tca tat ata ata gtt ttc tga cgt gga gta cag aat ttt cga --- --- --- --- --- --- --- --- --- --- --- --- --- --- --- --- --- --- --- --- --- --- --- --- --- --- --- --- --- --- --- --- --- --- --- --- --- --- --- --- --- --- --- --t taa tga gtt cat ggt aag gaa ggg caa atg cct --- gta tat aat ata cat aag tta a-- --- --- --- --- --- --- --- --- --- --- --- --- --t agt ttt tta tca tat tt- --- --- --- --- --- --- --- --t cta ata cca taa taa aaa t-- --- --- --- --- --- --- --- --- --- --- --- --- --- --- --- --- --- --- --- --- --- --- --- --- --t atc att att gcg ttt g-g tag tt- --- --- --- --- ctg ccc ta- --- --- --- --- --- --- --- --t cat cta tat cac tgt cac tct c-- --- --g ctc tca cta tat ctt cta aaa tta caa --- -ac aac --- --- --- --- --- --- --- --- --- --- --- --- --- --- --- --- --- --- --- --- --- --- --- --- --- --- --- --- --- --- --- --- --- --- --- --- tgg ata ttc gat --- --- --- --- --- --- --- --- --- --- --- --- --- --- --- --- --- --- --- --- --- --- --- --- --- --- --a aca gca ttt gtg t-- --- --- --- --- --- --- --- --- --- --- --- --- --- --- --- --- -

>ON649722.1_Monkeypox_virus_isolate_Monkeypox/PT0022/2022_complete_genome

--- --- --- --- --- --- --- --- --- --- --- --- --- --- --- --- --- --- -ta att a-- --- --- --- --- --- --- --- att taa ttt tac tat ttt att tag tgt cta gaa aaa aat gtg tga ccc acg acc gta gga aac tct aga ggg taa gaa aaa tca atc gtt tat aga gac cat cag aaa gag gtt taa tat ttt tgt gag acc tat cga a-- --- --- --- --- --- --- --- --- --- --- --- --- --- --- --- --- --- --- --- --- --- --- --- --- --- --- --- --- --- --- --- --- --- --- --- --- --- --- --- --- --- --- --- --- --- --- --- --- --- --- --- --- --- --- --- --- --- --- --- --- --- --- --- --- --- --- --- --- --- --- --- --- --- --- --- --- --- --- --- --- --- --- --- --- --- --- --- --- --- --- --- --- --- --- --- --- --- --- --- --- --- --- --- --- --- --- --- --- --- --- --- --- --- --- --- --- --- --- --- --- --- --- --- --- --- --- --- --- --- --- --- --- --- --- --- --- --- --- --- --- --- --- --- --- --- --- --- --- --- --- --- --- --- --- --- --- --- --- --- --- --- --- --- --- --- --- --- --- --- --- --- --- --- --- --- --- --- --- --- --- --- --- --- --- --- --- --- --- --- --- --- --- --- --- --- --- --- --- --- --- --- --- --- --- --- --- --- --- --- --- --- --- --- --- --- --- --- --- --- --- --- --- --- --- --- --- --- --- --- --- --- --- --- --- --- --- --- --- --- --- --- --- --- --- --- --- --- --- --- --- --- --- --- --- --- --- --- --- --- --- --- --- --- --- --- --- --- --- --- --- --- --- --- --- --- --- --- --- --- -ga gag aaa gga taa aaa ctt ttt acg act cca tca gaa aga ggt tta ata ttt ttg tga gac cca tcg aag aga gaa aga gat ggt tag tca aga tat ttt tct tag tac aaa agt caa tgt ttt aaa ata tat gga cga gaa tta att tgt ctg tat aaa aac ttg tgt gaa att atg tac tag aga aaa aac gtg agc agt gtc ccc tac atg gat ttt aca gat cat tta tat tcc aaa aat att aac tat ata cgt tta tta tat gat gtt aac gtg taa att ata aac att att tta tga tgc aat tgt ctg aca acc tag att ggt ata agg atg ttg ata agc tct acg aga ata tat tgt tgg acg tta tcg ttt acg aaa tag ttg aga cat cag aaa gag gtt taa tat ttt tgt gag a-c cat cga aga gag aaa gag aat aaa aat a-- --t ttt ttt ttt ttt tgt aaa act ttt tta tga gac --- --- -ca aga gaa tac gaa tag tga tca tat cgt atc aca tat tga aac aga aag aag aag taa cga gag gta act ttt tgt gaa tgt agt taa ata ttt ttg ttt tgc aaa ccg gaa tat agt gcc cgg t-- -ct ttt tta att cgt ggt gcg gtg tct gaa tcg ttc gat taa ccc aac tca tcc att ttc aga tga ata gag tta tcg att cag aca cat gct ttg agt ttt gtt gaa tcg atg agt gaa gta tca tcg gtt gca cct tca gat gcc gat ccg tcg aca tac ttg aat cca tcc ttg act tca agt tca gat gat tcc tca cac atg tct ccg ata cgt acg cta aac tct agg ttc ttg aca cat ttt gta tca acg atc gtt gaa ccg atg ata tct ttg taa ctc act ttc tta tgt gag atg tta gac cca agt act gga tgg gtc ttg atg tca ctg tct ttc tct tct tcg cta cat ctg atg tcg ata gac atc tca cag tct ttg atc ata gcc aga gct tct tca cgc gtg atc gcg gga gag tcc tta cct tgt ccc ggt gac acg ctg gac aat cta gta ttc aca gtg ttt cca tca gag gat tcg gag atg gat aaa atc ttt ggg cat ttg gtg aat cca aag ttc atg tta aga ccc gca ccg acg ata gtg taa taa gtg gtg gga tct cct ttt aca act --- --- --- --- --- --- tct tcg gat acc tca tca tct tcg gtc tct gta act tcc gtt acg gat tga caa atc tta tca ttg gtc ggt gtt tgg tct tgc ttt gtg act ttg ata ata aca tcg att ccc ata tga tgt ttg ttt tct tct tca gta cac gag gat gaa gat tgt tga aga cta gta ggc ata gca gct gcc act agg cac atg cat gcc agg aca ata tat tgt ttc atg att gct att gat tga tta ctg ttc tag atg att cta ctt tct tac cat ata ata aat tag aat ata ttt tct act ttt acg aga aat taa tta ttg tat tta tta ttt ata ggt --a aaa aaa ctt act ata agt ggg tgg gat tct ggg aat tag tga tca gtt tat gta tat cgc aac tag cgg gca tat ggc tat tga cat cga gaa cat tac cca tat gat aag aga ttg tat cat ttt cgt agt ctt gag tat tgg tat tac tat ata gta tgt aga tgt cga cgc tag ata gac agt cgc cca cta gag tta ccg tct ctg aat gcg gca tga tag tat cat tct ttg ttt tcg tta act gtt tgg aag atg aat ctt tgt tgt tac att taa tct cga aat tca gag tac ata tct ttg aag tat tct gat atc tat ttt ctc ctg taa aga atc ctg aag ttg cta cat tat taa gga cag aga agt att ctg cac gaa aga ctg gat cac aat ctt tat gat tca tgg taa tag tta gtt ccg acg ttg aga tgg att cgc tga gac cgg tag tgg tcg tcc gag tac acg atg tgt cgt tga ctg gat aca ggt taa ttt cca cat cga tat agt taa atg tat tgc tgg tta cga cgg gtt cgc att tat ctg tgg aag aga cgg tgt gag aat atg ttc cgg gac cac acg gag aac aga tga cgt ctc cgg tag acg tgt atc cgg ata ctc cgt atc cta ttc cac act ttg ttt tag aaa tac atg ttc tac acc ctg atg ctc ctt tga gaa gac aat aat atc ctg gag agc att cac aga ttc tat tgt gag tcg tgt tac acg atc gcg tct cta cct gat tac tat cac atc ttc cgt tac aac tta gac aag cct gta aat gat tat tgt gag atg taa agg tat ccg aac cac acg gtg tac att gtg tat tag tct tgc tat cac ata atc tga aag cgt aag ttc ccg gag gac acg ata gac aac ata gat tac ggc ttc tgt att cgt tgt ctt tac act ttc cat tgg atg gtg cat gtg gtg cta tat ctc ttc cgt tta tta tta tac atg aga gaa aca ata tat acg agt ata ata cgg acc tca tga ttt aat aat -gt agt aat cgt cgt ctt gtt act gtt tgt ttc cta ctt ctc caa tca tat aga tta ttt ttt aaa tat ttt ctt tct atc atg gat aat att tgt aat ggt tct ttc cgt aca aca tac tgt tta gat ggt agt cgc tta gct tgg tta tga tat tgc gca taa ttt ccg gag gca aat acg ata gtc tag att gac tat cga tgg tag act cta att tat tga gtg ctt tgt cga cga gtt tac ttt tat gct cca tcg ata gat gac act gtt cta tga gat cgt cgt aca tgg gaa atg aaa tgc gtt tgt ctg aat gta tgg ctt cga gat atg tgt gat acc gaa tgt ctt ctg ttc tca ata ccg tat aca agt cgg tgt ctg aga ttc gaa tct ctt tga gga gac tta tgt cac gac tac att ttt cga tga tgg aat cta tct tat cga atg ata tat ttt tca taa ata cac ttt tat agt cct cgt tta aac aga att tag tat gta gtt ccg caa atg act cgt ccc tta ata ggc agt agg cta tta tct tct tta cgt agt gat cgt cgt agg gag aga act ccg aca tct tgt aga aca acg att taa tca tag gta gag ata ctt tca gtc tgt ggt gga taa tgt cat taa cga cat ccg cct tgt ata taa tgt ttc tgt ttt caa aca cca agt cga ata ctg tct tat cgt ctt tag tcg gaa ggt tga tgt cgt atc caa tgt ata cga ggt atg agg caa cat tgt tat tgc aat tct gga agg cgg tat gaa gag gag tca ttg tat tat agt att cgt ctt tct gaa tgt cga atc tat cta gta gat acc gta gta tat tga gag agc gac ttc cat atc ctt gat tat gtt tta tga ata gat aaa gta gat gtt gtc ctt ctt cct ttt gta att tcc cgt att ttt gtt cgt gcc aat tga gta aca tta tga gaa tat gac ctg ttg cac aat cgt tct tta tgt att cca tga tgg gtg tac aat taa gat tat tac gta tcc tcg tat cgg ctc ctc gag ata aaa gag cat aca cca cac gag gac tat gtt tgg tat act gtt gaa ggt aag tgt gta acg gcg tat ttc cga ttt tcg taa ccg cgt taa tgt ttg ctc cat gat cta tta tcg cgt aga tga atc gct tct cag ctc gca tct tag tgt gac tct ttg act tgt aat aat tgc ttt cgt gga acg cgg ata tgt gtt tac agt agt aat gaa gag aag tga gtc cat cct cat cga cgc aat tag ggt cag atc ctt tag tca ata att tgt aca gaa cgt aat agt tta agc tcc cat tga att tat atc taa gat aac aca gca ata gat cgg atg att tac taa agt cat caa tgg ggt ccg tta gta tat caa aga tct tgt tat cga ttg ata gtg aat gaa tca gat agt ggt gta gag gaa tat gtc ctt ttt tat cct tgc tat caa agt tac gca tgc cgt ggc gta aca ata tct tta ata cag atg gat taa atc gtg tat tca tcg tat agc aat gta atg gag agt tac cac att tta gtc gtt tat tca gat cgc agt gtt taa taa cta att taa aca gat gag atg ata tat cca cat caa aga atg cga gat aca tat gac aga cat tat tga cag aaa tgt gac ctt cat tat cac cgt cgt cca taa atg cgt tag gta cgt acc aca tac tat cgt taa cga tgc gca caa tct cgt cca -tt tca tcc atc ttc ata atg att tac ttt ttc ata att aga gaa aaa gat caa ggt ata aaa att aga agt gtt aga cta ta- --- --- --- --- --- --- --- --- --- --- -aa cta act tat gac tta act aac tta tga --- --- --- --- --- --- --- --- --- --- --- --- --- --- --- --- --- --- --- --- --- --- --- --- --- --- --- --- --- --- --- --- --- --- --- --- --- --- --- --- --- --- --- --- --- --- --- --- ctt aac ta- --- --- --- --- --- --- --- --- --- --- -ac tta tga ctt aac taa ctt atg act taa cta act ta- tga ctt aac taa ctt atg act taa cta act tat gac tta act aac tta tga cta tta act cat ttc aag aaa ggt ggg tgg ata gaa ctc tat atg aca gct tgt gaa aca att aga tcc cta att tct aat gga agt ttt gat agg aga ttg tca tca gtt gat aca ttg ttt att atc tca tct att aga gca cgt ctg ttt aga gct tta gtg acc tgc tcg gtt act tct gtg taa atc ttg aat cct tta gtg ata cac tgt gtc aaa act gga tgt tta gaa tac cta tgt aga ata tgg gaa gca tgc ttg ttt ttg tct cta tta tag att aac tca tac atg gtt gta tta tga att ttc atc tgc cta atg tac tcc aat tct tgt tta caa tca att ata taa tca aag agt gat gat gca tac aca tta caa agt gaa taa tct acc atc ata aaa tac ttg ata cag agc ttt atc aca tca tgg ttt tca att gta tta tta agt ata gct aat ttt ata cag tca ata gac aat ggt tct cta agc aat att tct aat att tta aga tgt gct tcc cta cgg gcg atg aca gat ccc cta tcc acg gcc acg tca aga cat gta tat cca tta ctc att act gcg ttg aca ttt gct cca ttt tct aat agc cat gat act aaa tct ata taa cct gca tag ata gcg cga taa agc aag gtc ctt cca cca gca tct agt tga ttg ata tct tca ata tat ggg ata caa agc tta taa att tct aat act gtg ggt tca tct aca agg aat ccc cta gta tac tga att att tta tat aga tct aat tta aca tca ttt tca tct ggg ata cca caa ttc aaa ata aac tca aca aca cta ctt tcc ttt tta cat att ccc cta aaa tag gca ttc aag cat tct att tta tat att aca gcc cca tga tct acc ata aga tca aca atg tct att tct aca tat gca tta gat aga tag taa agt aag aga tct tgc aca gaa tta caa ttc tta ata att ata gag aaa ata tct tcc ata taa ttc ttt gac act aat gca gat ata ata tct tta tat gta ata tat gca aac agt cta tct act ata tac tga tca ata tta tct cta tga atc cta aaa taa tca tac aga aca tct aca gga tca caa att ggt tca agg aga aat cta tca aat att ttc ctg tca aca act ggt tct aga aca taa cag tca aca cct aat cca tgt ttt tta taa tca tct acc aaa gat aat gac caa aga tcg agg tcg tcg tga aac tgc tca tcg aca gcc atg aaa tct acc gac tcc atg gtg cga atc gca ctg tct tat tcg cca ttg att ttc a-t ttt tta taa tta tgt aca tgt ttt cct tct att ctc aag agt cta caa aaa tat a-- -tt ttt tcg ata tct aag tac taa gtt ttt tta ctg ttt ttg tta ctg tct tcc att ctt cta act aaa gat ctg aga taa att ata caa tct tcg cta tcg aac cat ttt tgt agt cta aag cct gaa gta att aac caa ctg ttt tta tta gtg gct ttt ttc gat cta tcc tcg tc- --- ctc tca tca tcc tta ta- --- --- --- --- --- --- tta tta tca tta tta tca tag tct att aaa cac aaa tca tct --- --- --- --- --- --- --- --- --- --- --- --- --- --- --- --- --- --- --- --- --- --- --- --- --- --- --a cgt tta taa c-- --- --- --- --- --- --- --- --a aca ttc --- --- --- --- -tc att att aat tag ttc tgt ag- aat atc ttt aat aat ttg gct ata --- --- -ca tct gtt --- --- --- --- --- --- --- --- --- --- --- --- --- --- --- --- --- --- --c aat act --- --- --- --- --- --- --- --a tct att gat gat ttc ttt t-- --- --- --- --- --- --- --- --- --- --- --t taa gac t-- --- --- --- --- --- --- --- --- --- --- --- --- --- --- --- --- --- --- --- --- --- --- --- --- --- --- --- --- --- --- --- --- --- --- --- --- --- --- --- taa act agt --- --- --- --- --- --- --- --- --- --- --- --- --- --- --- --- --- --- --- --- --- --- --- --- --- --- --- --- --- --- --- --- --- --- --- --- --- --- --- --- --- --- --- --- --- --- --- --- --- --- --- --- --- --- --- --- --- --- --- --- --- --- --- --- --- --- --- --- --- --- --- --- --- --- --- --- --- --- --- --- --- --- --- --- --- --- --- --- --- --- --- --- --- --- --- --- --- --- --- --- --- --- --- --- --- --- --- --- --- --t atg gta atg acg atg aaa --- -tc gag tag ta- --- --a ctt cta ata aag act tga ta- --- --t cat tat cat atg ttt gat cg- --- --- --- --- --- --- --t cat agt taa tag tgt g-- --- --- --- --- --- --- --- --- --- --- --- --- --- --- --- --- --- --- --- -gc taa atg gta ctg tta ata agt tta t-- --- --- --- --- --- --- --- --- --- --- --- --- --- --- --- --- --a gac aat atc ata gta ttt tct ttc cag aat --- --- --- --t aga tta ttt ttt taa ata ctg atc ctc aca att ccg tga tgt agc agt agt tgg t-- --- --- --- --- --- --- --- --- --- --- --- --- --- --- --- --- --- --- --- --- --- --- --- --- --- --- --- -gc atg gtc tat atc gt- --- --- --- --- --- --- --- --- --- --- --- --- --- --- --- --- --- --- --- --- --- --- --- --- --- --- --- --- --- --- --- --- --- --- --- --- --- --- --- --- --- --- --- taa aat gta tca tat ata ata gtt ttc tga cgt gga gta cag aat ttt cga --- --- --- --- --- --- --- --- --- --- --- --- --- --- --- --- --- --- --- --- --- --- --- --- --- --- --- --- --- --- --- --- --- --- --- --- --- --- --- --- --- --- --- --t taa tga gtt cat ggt aag gaa ggg caa atg cct --- gta tat aat ata cat aag tta a-- --- --- --- --- --- --- --- --- --- --- --- --- --t agt ttt tta tca tat tt- --- --- --- --- --- --- --- --t cta ata cca taa taa aaa t-- --- --- --- --- --- --- --- --- --- --- --- --- --- --- --- --- --- --- --- --- --- --- --- --- --t atc att att gcg ttt g-g tag tt- --- --- --- --- ctg ccc ta- --- --- --- --- --- --- --- --t cat cta tat cac tgt cac tct c-- --- --g ctc tca cta tat ctt cta aaa tta caa --- -ac aac --- --- --- --- --- --- --- --- --- --- --- --- --- --- --- --- --- --- --- --- --- --- --- --- --- --- --- --- --- --- --- --- --- --- --- --- tgg ata ttc gat --- --- --- --- --- --- --- --- --- --- --- --- --- --- --- --- --- --- --- --- --- --- --- --- --- --- --a aca gca ttt gtg t-- --- --- --- --- --- --- --- --- --- --- --- --- --- --- --- --- -

>ON649721.1_Monkeypox_virus_isolate_Monkeypox/PT0021/2022_complete_genome

--- --- --- --- --- --- --- --- --- --- --- --- --- --- --- --- --- --- -ta att a-- --- --- --- --- --- --- --- att taa ttt tac tat ttt att tag tgt cta gaa aaa aat gtg tga ccc acg acc gta gga aac tct aga ggg taa gaa aaa tca atc gtt tat aga gac cat cag aaa gag gtt taa tat ttt tgt gag acc tat cga a-- --- --- --- --- --- --- --- --- --- --- --- --- --- --- --- --- --- --- --- --- --- --- --- --- --- --- --- --- --- --- --- --- --- --- --- --- --- --- --- --- --- --- --- --- --- --- --- --- --- --- --- --- --- --- --- --- --- --- --- --- --- --- --- --- --- --- --- --- --- --- --- --- --- --- --- --- --- --- --- --- --- --- --- --- --- --- --- --- --- --- --- --- --- --- --- --- --- --- --- --- --- --- --- --- --- --- --- --- --- --- --- --- --- --- --- --- --- --- --- --- --- --- --- --- --- --- --- --- --- --- --- --- --- --- --- --- --- --- --- --- --- --- --- --- --- --- --- --- --- --- --- --- --- --- --- --- --- --- --- --- --- --- --- --- --- --- --- --- --- --- --- --- --- --- --- --- --- --- --- --- --- --- --- --- --- --- --- --- --- --- --- --- --- --- --- --- --- --- --- --- --- --- --- --- --- --- --- --- --- --- --- --- --- --- --- --- --- --- --- --- --- --- --- --- --- --- --- --- --- --- --- --- --- --- --- --- --- --- --- --- --- --- --- --- --- --- --- --- --- --- --- --- --- --- --- --- --- --- --- --- --- --- --- --- --- --- --- --- --- --- --- --- --- --- --- --- --- --- --- -ga gag aaa gga taa aaa ctt ttt acg act cca tca gaa aga ggt tta ata ttt ttg tga gac cca tcg aag aga gaa aga gat ggt tag tca aga tat ttt tct tag tac aaa agt caa tgt ttt aaa ata tat gga cga gaa tta att tgt ctg tat aaa aac ttg tgt gaa att atg tac tag aga aaa aac gtg agc agt gtc ccc tac atg gat ttt aca gat cat tta tat tcc aaa aat att aac tat ata cgt tta tta tat gat gtt aac gtg taa att ata aac att att tta tga tgc aat tgt ctg aca acc tag att ggt ata agg atg ttg ata agc tct acg aga ata tat tgt tgg acg tta tcg ttt acg aaa tag ttg aga cat cag aaa gag gtt taa tat ttt tgt gag a-c cat cga aga gag aaa gag aat aaa aat a-- --t ttt ttt ttt ttt tgt aaa act ttt tta tga gac --- --- -ca aga gaa tac gaa tag tga tca tat cgt atc aca tat tga aac aga aag aag aag taa cga gag gta act ttt tgt gaa tgt agt taa ata ttt ttg ttt tgc aaa ccg gaa tat agt gcc cgg t-- -ct ttt tta att cgt ggt gcg gtg tct gaa tcg ttc gat taa ccc aac tca tcc att ttc aga tga ata gag tta tcg att cag aca cat gct ttg agt ttt gtt gaa tcg atg agt gaa gta tca tcg gtt gca cct tca gat gcc gat ccg tcg aca tac ttg aat cca tcc ttg act tca agt tca gat gat tcc tca cac atg tct ccg ata cgt acg cta aac tct agg ttc ttg aca cat ttt gta tca acg atc gtt gaa ccg atg ata tct ttg taa ctc act ttc tta tgt gag atg tta gac cca agt act gga tgg gtc ttg atg tca ctg tct ttc tct tct tcg cta cat ctg atg tcg ata gac atc tca cag tct ttg atc ata gcc aga gct tct tca cgc gtg atc gcg gga gag tcc tta cct tgt ccc ggt gac acg ctg gac aat cta gta ttc aca gtg ttt cca tca gag gat tcg gag atg gat aaa atc ttt ggg cat ttg gtg aat cca aag ttc atg tta aga ccc gca ccg acg ata gtg taa taa gtg gtg gga tct cct ttt aca act --- --- --- --- --- --- tct tcg gat acc tca tca tct tcg gtc tct gta act tcc gtt acg gat tga caa atc tta tca ttg gtc ggt gtt tgg tct tgc ttt gtg act ttg ata ata aca tcg att ccc ata tga tgt ttg ttt tct tct tca gta cac gag gat gaa gat tgt tga aga cta gta ggc ata gca gct gcc act agg cac atg cat gcc agg aca ata tat tgt ttc atg att gct att gat tga tta ctg ttc tag atg att cta ctt tct tac cat ata ata aat tag aat ata ttt tct act ttt acg aga aat taa tta ttg tat tta tta ttt ata ggt --a aaa aaa ctt act ata agt ggg tgg gat tct ggg aat tag tga tca gtt tat gta tat cgc aac tag cgg gca tat ggc tat tga cat cga gaa cat tac cca tat gat aag aga ttg tat cat ttt cgt agt ctt gag tat tgg tat tac tat ata gta tgt aga tgt cga cgc tag ata gac agt cgc cca cta gag tta ccg tct ctg aat gcg gca tga tag tat cat tct ttg ttt tcg tta act gtt tgg aag atg aat ctt tgt tgt tac att taa tct cga aat tca gag tac ata tct ttg aag tat tct gat atc tat ttt ctc ctg taa aga atc ctg aag ttg cta cat tat taa gga cag aga agt att ctg cac gaa aga ctg gat cac aat ctt tat gat tca tgg taa tag tta gtt ccg acg ttg aga tgg att cgc tga gac cgg tag tgg tcg tcc gag tac acg atg tgt cgt tga ctg gat aca ggt taa ttt cca cat cga tat agt taa atg tat tgc tgg tta cga cgg gtt cgc att tat ctg tgg aag aga cgg tgt gag aat atg ttc cgg gac cac acg gag aac aga tga cgt ctc cgg tag acg tgt atc cgg ata ctc cgt atc cta ttc cac act ttg ttt tag aaa tac atg ttc tac acc ctg atg ctc ctt tga gaa gac aat aat atc ctg gag agc att cac aga ttc tat tgt gag tcg tgt tac acg atc gcg tct cta cct gat tac tat cac atc ttc cgt tac aac tta gac aag cct gta aat gat tat tgt gag atg taa agg tat ccg aac cac acg gtg tac att gtg tat tag tct tgc tat cac ata atc tga aag cgt aag ttc ccg gag gac acg ata gac aac ata gat tac ggc ttc tgt att cgt tgt ctt tac act ttc cat tgg atg gtg cat gtg gtg cta tat ctc ttc cgt tta tta tta tac atg aga gaa aca ata tat acg agt ata ata cgg acc tca tga ttt aat aat -gt agt aat cgt cgt ctt gtt act gtt tgt ttc cta ctt ctc caa tca tat aga tta ttt ttt aaa tat ttt ctt tct atc atg gat aat att tgt aat ggt tct ttc cgt aca aca tac tgt tta gat ggt agt cgc tta gct tgg tta tga tat tgc gca taa ttt ccg gag gca aat acg ata gtc tag att gac tat cga tgg tag act cta att tat tga gtg ctt tgt cga cga gtt tac ttt tat gct cca tcg ata gat gac act gtt cta tga gat cgt cgt aca tgg gaa atg aaa tgc gtt tgt ctg aat gta tgg ctt cga gat atg tgt gat acc gaa tgt ctt ctg ttc tca ata ccg tat aca agt cgg tgt ctg aga ttc gaa tct ctt tga gga gac tta tgt cac gac tac att ttt cga tga tgg aat cta tct tat cga atg ata tat ttt tca taa ata cac ttt tat agt cct cgt tta aac aga att tag tat gta gtt ccg caa atg act cgt ccc tta ata ggc agt agg cta tta tct tct tta cgt agt gat cgt cgt agg gag aga act ccg aca tct tgt aga aca acg att taa tca tag gta gag ata ctt tca gtc tgt ggt gga taa tgt cat taa cga cat ccg cct tgt ata taa tgt ttc tgt ttt caa aca cca agt cga ata ctg tct tat cgt ctt tag tcg gaa ggt tga tgt cgt atc caa tgt ata cga ggt atg agg caa cat tgt tat tgc aat tct gga agg cgg tat gaa gag gag tca ttg tat tat agt att cgt ctt tct gaa tgt cga atc tat cta gta gat acc gta gta tat tga gag agc gac ttc cat atc ctt gat tat gtt tta tga ata gat aaa gta gat gtt gtc ctt ctt cct ttt gta att tcc cgt att ttt gtt cgt gcc aat tga gta aca tta tga gaa tat gac ctg ttg cac aat cgt tct tta tgt att cca tga tgg gtg tac aat taa gat tat tac gta tcc tcg tat cgg ctc ctc gag ata aaa gag cat aca cca cac gag gac tat gtt tgg tat act gtt gaa ggt aag tgt gta acg gcg tat ttc cga ttt tcg taa ccg cgt taa tgt ttg ctc cat gat cta tta tcg cgt aga tga atc gct tct cag ctc gca tct tag tgt gac tct ttg act tgt aat aat tgc ttt cgt gga acg cgg ata tgt gtt tac agt agt aat gaa gag aag tga gtc cat cct cat cga cgc aat tag ggt cag atc ctt tag tca ata att tgt aca gaa cgt aat agt tta agc tcc cat tga att tat atc taa gat aac aca gca ata gat cgg atg att tac taa agt cat caa tgg ggt ccg tta gta tat caa aga tct tgt tat cga ttg ata gtg aat gaa tca gat agt ggt gta gag gaa tat gtc ctt ttt tat cct tgc tat caa agt tac gca tgc cgt ggc gta aca ata tct tta ata cag atg gat taa atc gtg tat tca tcg tat agc aat gta atg gag agt tac cac att tta gtc gtt tat tca gat cgc agt gtt taa taa cta att taa aca gat gag atg ata tat cca cat caa aga atg cga gat aca tat gac aga cat tat tga cag aaa tgt gac ctt cat tat cac cgt cgt cca taa atg cgt tag gta cgt acc aca tac tat cgt taa cga tgc gca caa tct cgt cca -tt tca tcc atc ttc ata atg att tac ttt ttc ata att aga gaa aaa gat caa ggt ata aaa att aga agt gtt aga cta ta- --- --- --- --- --- --- --- --- --- --- -aa cta act tat gac tta act aac tta tga --- --- --- --- --- --- --- --- --- --- --- --- --- --- --- --- --- --- --- --- --- --- --- --- --- --- --- --- --- --- --- --- --- --- --- --- --- --- --- --- --- --- --- --- --- --- --- --- ctt aac ta- --- --- --- --- --- --- --- --- --- --- -ac tta tga ctt aac taa ctt atg act taa cta act ta- tga ctt aac taa ctt atg act taa cta act tat gac tta act aac tta tga cta tta act cat ttc aag aaa ggt ggg tgg ata gaa ctc tat atg aca gct tgt gaa aca att aga tcc cta att tct aat gga agt ttt gat agg aga ttg tca tca gtt gat aca ttg ttt att atc tca tct att aga gca cgt ctg ttt aga gct tta gtg acc tgc tcg gtt act tct gtg taa atc ttg aat cct tta gtg ata cac tgt gtc aaa act gga tgt tta gaa tac cta tgt aga ata tgg gaa gca tgc ttg ttt ttg tct cta tta tag att aac tca tac atg gtt gta tta tga att ttc atc tgc cta atg tac tcc aat tct tgt tta caa tca att ata taa tca aag agt gat gat gca tac aca tta caa agt gaa taa tct acc atc ata aaa tac ttg ata cag agc ttt atc aca tca tgg ttt tca att gta tta tta agt ata gct aat ttt ata cag tca ata gac aat ggt tct cta agc aat att tct aat att tta aga tgt gct tcc cta cgg gcg atg aca gat ccc cta tcc acg gcc acg tca aga cat gta tat cca tta ctc att act gcg ttg aca ttt gct cca ttt tct aat agc cat gat act aaa tct ata taa cct gca tag ata gcg cga taa agc aag gtc ctt cca cca gca tct agt tga ttg ata tct tca ata tat ggg ata caa agc tta taa att tct aat act gtg ggt tca tct aca agg aat ccc cta gta tac tga att att tta tat aga tct aat tta aca tca ttt tca tct ggg ata cca caa ttc aaa ata aac tca aca aca cta ctt tcc ttt tta cat att ccc cta aaa tag gca ttc aag cat tct att tta tat att aca gcc cca tga tct acc ata aga tca aca atg tct att tct aca tat gca tta gat aga tag taa agt aag aga tct tgc aca gaa tta caa ttc tta ata att ata gag aaa ata tct tcc ata taa ttc ttt gac act aat gca gat ata ata tct tta tat gta ata tat gca aac agt cta tct act ata tac tga tca ata tta tct cta tga atc cta aaa taa tca tac aga aca tct aca gga tca caa att ggt tca agg aga aat cta tca aat att ttc ctg tca aca act ggt tct aga aca taa cag tca aca cct aat cca tgt ttt tta taa tca tct acc aaa gat aat gac caa aga tcg agg tcg tcg tga aac tgc tca tcg aca gcc atg aaa tct acc gac tcc atg gtg cga atc gca ctg tct tat tcg cca ttg att ttc a-t ttt tta taa tta tgt aca tgt ttt cct tct att ctc aag agt cta caa aaa tat a-- -tt ttt tcg ata tct aag tac taa gtt ttt tta ctg ttt ttg tta ctg tct tcc att ctt cta act aaa gat ctg aga taa att ata caa tct tcg cta tcg aac cat ttt tgt agt cta aag cct gaa gta att aac caa ctg ttt tta tta gtg gct ttt ttc gat cta tcc tcg tc- --- ctc tca tca tcc tta ta- --- --- --- --- --- --- tta tta tca tta tta tca tag tct att aaa cac aaa tca tct --- --- --- --- --- --- --- --- --- --- --- --- --- --- --- --- --- --- --- --- --- --- --- --- --- --- --a cgt tta taa c-- --- --- --- --- --- --- --- --a aca ttc --- --- --- --- -tc att att aat tag ttc tgt ag- aat atc ttt aat aat ttg gct ata --- --- -ca tct gtt --- --- --- --- --- --- --- --- --- --- --- --- --- --- --- --- --- --- --c aat act --- --- --- --- --- --- --- --a tct att gat gat ttc ttt t-- --- --- --- --- --- --- --- --- --- --- --t taa gac t-- --- --- --- --- --- --- --- --- --- --- --- --- --- --- --- --- --- --- --- --- --- --- --- --- --- --- --- --- --- --- --- --- --- --- --- --- --- --- --- taa act agt --- --- --- --- --- --- --- --- --- --- --- --- --- --- --- --- --- --- --- --- --- --- --- --- --- --- --- --- --- --- --- --- --- --- --- --- --- --- --- --- --- --- --- --- --- --- --- --- --- --- --- --- --- --- --- --- --- --- --- --- --- --- --- --- --- --- --- --- --- --- --- --- --- --- --- --- --- --- --- --- --- --- --- --- --- --- --- --- --- --- --- --- --- --- --- --- --- --- --- --- --- --- --- --- --- --- --- --- --- --t atg gta atg acg atg aaa --- -tc gag tag ta- --- --a ctt cta ata aag act tga ta- --- --t cat tat cat atg ttt gat cg- --- --- --- --- --- --- --t cat agt taa tag tgt g-- --- --- --- --- --- --- --- --- --- --- --- --- --- --- --- --- --- --- --- -gc taa atg gta ctg tta ata agt tta t-- --- --- --- --- --- --- --- --- --- --- --- --- --- --- --- --- --a gac aat atc ata gta ttt tct ttc cag aat --- --- --- --t aga tta ttt ttt taa ata ctg atc ctc aca att ccg tga tgt agc agt agt tgg t-- --- --- --- --- --- --- --- --- --- --- --- --- --- --- --- --- --- --- --- --- --- --- --- --- --- --- --- -gc atg gtc tat atc gt- --- --- --- --- --- --- --- --- --- --- --- --- --- --- --- --- --- --- --- --- --- --- --- --- --- --- --- --- --- --- --- --- --- --- --- --- --- --- --- --- --- --- --- taa aat gta tca tat ata ata gtt ttc tga cgt gga gta cag aat ttt cga --- --- --- --- --- --- --- --- --- --- --- --- --- --- --- --- --- --- --- --- --- --- --- --- --- --- --- --- --- --- --- --- --- --- --- --- --- --- --- --- --- --- --- --t taa tga gtt cat ggt aag gaa ggg caa atg cct --- gta tat aat ata cat aag tta a-- --- --- --- --- --- --- --- --- --- --- --- --- --t agt ttt tta tca tat tt- --- --- --- --- --- --- --- --t cta ata cca taa taa aaa t-- --- --- --- --- --- --- --- --- --- --- --- --- --- --- --- --- --- --- --- --- --- --- --- --- --t atc att att gcg ttt g-g tag tt- --- --- --- --- ctg ccc ta- --- --- --- --- --- --- --- --t cat cta tat cac tgt cac tct c-- --- --g ctc tca cta tat ctt cta aaa tta caa --- -ac aac --- --- --- --- --- --- --- --- --- --- --- --- --- --- --- --- --- --- --- --- --- --- --- --- --- --- --- --- --- --- --- --- --- --- --- --- tgg ata ttc gat --- --- --- --- --- --- --- --- --- --- --- --- --- --- --- --- --- --- --- --- --- --- --- --- --- --- --a aca gca ttt gtg t-- --- --- --- --- --- --- --- --- --- --- --- --- --- --- --- --- -

>ON649720.1_Monkeypox_virus_isolate_Monkeypox/PT0024/2022_complete_genome

--- --- --- --- --- --- --- --- --- --- --- --- --- --- --- --- --- --- -ta att a-- --- --- --- --- --- --- --- att taa ttt tac tat ttt att tag tgt cta gaa aaa aat gtg tga ccc acg acc gta gga aac tct aga ggg taa gaa aaa tca atc gtt tat aga gac cat cag aaa gag gtt taa tat ttt tgt gag acc tat cga a-- --- --- --- --- --- --- --- --- --- --- --- --- --- --- --- --- --- --- --- --- --- --- --- --- --- --- --- --- --- --- --- --- --- --- --- --- --- --- --- --- --- --- --- --- --- --- --- --- --- --- --- --- --- --- --- --- --- --- --- --- --- --- --- --- --- --- --- --- --- --- --- --- --- --- --- --- --- --- --- --- --- --- --- --- --- --- --- --- --- --- --- --- --- --- --- --- --- --- --- --- --- --- --- --- --- --- --- --- --- --- --- --- --- --- --- --- --- --- --- --- --- --- --- --- --- --- --- --- --- --- --- --- --- --- --- --- --- --- --- --- --- --- --- --- --- --- --- --- --- --- --- --- --- --- --- --- --- --- --- --- --- --- --- --- --- --- --- --- --- --- --- --- --- --- --- --- --- --- --- --- --- --- --- --- --- --- --- --- --- --- --- --- --- --- --- --- --- --- --- --- --- --- --- --- --- --- --- --- --- --- --- --- --- --- --- --- --- --- --- --- --- --- --- --- --- --- --- --- --- --- --- --- --- --- --- --- --- --- --- --- --- --- --- --- --- --- --- --- --- --- --- --- --- --- --- --- --- --- --- --- --- --- --- --- --- --- --- --- --- --- --- --- --- --- --- --- --- --- --- -ga gag aaa gga taa aaa ctt ttt acg act cca tca gaa aga ggt tta ata ttt ttg tga gac cca tcg aag aga gaa aga gat ggt tag tca aga tat ttt tct tag tac aaa agt caa tgt ttt aaa ata tat gga cga gaa tta att tgt ctg tat aaa aac ttg tgt gaa att atg tac tag aga aaa aac gtg agc agt gtc ccc tac atg gat ttt aca gat cat tta tat tcc aaa aat att aac tat ata cgt tta tta tat gat gtt aac gtg taa att ata aac att att tta tga tgc aat tgt ctg aca acc tag att ggt ata agg atg ttg ata agc tct acg aga ata tat tgt tgg acg tta tcg ttt acg aaa tag ttg aga cat cag aaa gag gtt taa tat ttt tgt gag a-c cat cga aga gag aaa gag aat aaa aat a-- --t ttt ttt ttt ttt tgt aaa act ttt tta tga gac --- --- -ca aga gaa tac gaa tag tga tca tat cgt atc aca tat tga aac aga aag aag aag taa cga gag gta act ttt tgt gaa tgt agt taa ata ttt ttg ttt tgc aaa ccg gaa tat agt gcc cgg t-- -ct ttt tta att cgt ggt gcg gtg tct gaa tcg ttc gat taa ccc aac tca tcc att ttc aga tga ata gag tta tcg att cag aca cat gct ttg agt ttt gtt gaa tcg atg agt gaa gta tca tcg gtt gca cct tca gat gcc gat ccg tcg aca tac ttg aat cca tcc ttg act tca agt tca gat gat tcc tca cac atg tct ccg ata cgt acg cta aac tct agg ttc ttg aca cat ttt gta tca acg atc gtt gaa ccg atg ata tct ttg taa ctc act ttc tta tgt gag atg tta gac cca agt act gga tgg gtc ttg atg tca ctg tct ttc tct tct tcg cta cat ctg atg tcg ata gac atc tca cag tct ttg atc ata gcc aga gct tct tca cgc gtg atc gcg gga gag tcc tta cct tgt ccc ggt gac acg ctg gac aat cta gta ttc aca gtg ttt cca tca gag gat tcg gag atg gat aaa atc ttt ggg cat ttg gtg aat cca aag ttc atg tta aga ccc gca ccg acg ata gtg taa taa gtg gtg gga tct cct ttt aca act --- --- --- --- --- --- tct tcg gat acc tca tca tct tcg gtc tct gta act tcc gtt acg gat tga caa atc tta tca ttg gtc ggt gtt tgg tct tgc ttt gtg act ttg ata ata aca tcg att ccc ata tga tgt ttg ttt tct tct tca gta cac gag gat gaa gat tgt tga aga cta gta ggc ata gca gct gcc act agg cac atg cat gcc agg aca ata tat tgt ttc atg att gct att gat tga tta ctg ttc tag atg att cta ctt tct tac cat ata ata aat tag aat ata ttt tct act ttt acg aga aat taa tta ttg tat tta tta ttt ata ggt --a aaa aaa ctt act ata agt ggg tgg gat tct ggg aat tag tga tca gtt tat gta tat cgc aac tag cgg gca tat ggc tat tga cat cga gaa cat tac cca tat gat aag aga ttg tat cat ttt cgt agt ctt gag tat tgg tat tac tat ata gta tgt aga tgt cga cgc tag ata gac agt cgc cca cta gag tta ccg tct ctg aat gcg gca tga tag tat cat tct ttg ttt tcg tta act gtt tgg aag atg aat ctt tgt tgt tac att taa tct cga aat tca gag tac ata tct ttg aag tat tct gat atc tat ttt ctc ctg taa aga atc ctg aag ttg cta cat tat taa gga cag aga agt att ctg cac gaa aga ctg gat cac aat ctt tat gat tca tgg taa tag tta gtt ccg acg ttg aga tgg att cgc tga gac cgg tag tgg tcg tcc gag tac acg atg tgt cgt tga ctg gat aca ggt taa ttt cca cat cga tat agt taa atg tat tgc tgg tta cga cgg gtt cgc att tat ctg tgg aag aga cgg tgt gag aat atg ttc cgg gac cac acg gag aac aga tga cgt ctc cgg tag acg tgt atc cgg ata ctc cgt atc cta ttc cac act ttg ttt tag aaa tac atg ttc tac acc ctg atg ctc ctt tga gaa gac aat aat atc ctg gag agc att cac aga ttc tat tgt gag tcg tgt tac acg atc gcg tct cta cct gat tac tat cac atc ttc cgt tac aac tta gac aag cct gta aat gat tat tgt gag atg taa agg tat ccg aac cac acg gtg tac att gtg tat tag tct tgc tat cac ata atc tga aag cgt aag ttc ccg gag gac acg ata gac aac ata gat tac ggc ttc tgt att cgt tgt ctt tac act ttc cat tgg atg gtg cat gtg gtg cta tat ctc ttc cgt tta tta tta tac atg aga gaa aca ata tat acg agt ata ata cgg acc tca tga ttt aat aat -gt agt aat cgt cgt ctt gtt act gtt tgt ttc cta ctt ctc caa tca tat aga tta ttt ttt aaa tat ttt ctt tct atc atg gat aat att tgt aat ggt tct ttc cgt aca aca tac tgt tta gat ggt agt cgc tta gct tgg tta tga tat tgc gca taa ttt ccg gag gca aat acg ata gtc tag att gac tat cga tgg tag act cta att tat tga gtg ctt tgt cga cga gtt tac ttt tat gct cca tcg ata gat gac act gtt cta tga gat cgt cgt aca tgg gaa atg aaa tgc gtt tgt ctg aat gta tgg ctt cga gat atg tgt gat acc gaa tgt ctt ctg ttc tca ata ccg tat aca agt cgg tgt ctg aga ttc gaa tct ctt tga gga gac tta tgt cac gac tac att ttt cga tga tgg aat cta tct tat cga atg ata tat ttt tca taa ata cac ttt tat agt cct cgt tta aac aga att tag tat gta gtt ccg caa atg act cgt ccc tta ata ggc agt agg cta tta tct tct tta cgt agt gat cgt cgt agg gag aga act ccg aca tct tgt aga aca acg att taa tca tag gta gag ata ctt tca gtc tgt ggt gga taa tgt cat taa cga cat ccg cct tgt ata taa tgt ttc tgt ttt caa aca cca agt cga ata ctg tct tat cgt ctt tag tcg gaa ggt tga tgt cgt atc caa tgt ata cga ggt atg agg caa cat tgt tat tgc aat tct gga agg cgg tat gaa gag gag tca ttg tat tat agt att cgt ctt tct gaa tgt cga atc tat cta gta gat acc gta gta tat tga gag agc gac ttc cat atc ctt gat tat gtt tta tga ata gat aaa gta gat gtt gtc ctt ctt cct ttt gta att tcc cgt att ttt gtt cgt gcc aat tga gta aca tta tga gaa tat gac ctg ttg cac aat cgt tct tta tgt att cca tga tgg gtg tac aat taa gat tat tac gta tcc tcg tat cgg ctc ctc gag ata aaa gag cat aca cca cac gag gac tat gtt tgg tat act gtt gaa ggt aag tgt gta acg gcg tat ttc cga ttt tcg taa ccg cgt taa tgt ttg ctc cat gat cta tta tcg cgt aga tga atc gct tct cag ctc gca tct tag tgt gac tct ttg act tgt aat aat tgc ttt cgt gga acg cgg ata tgt gtt tac agt agt aat gaa gag aag tga gtc cat cct cat cga cgc aat tag ggt cag atc ctt tag tca ata att tgt aca gaa cgt aat agt tta agc tcc cat tga att tat atc taa gat aac aca gca ata gat cgg atg att tac taa agt cat caa tgg ggt ccg tta gta tat caa aga tct tgt tat cga ttg ata gtg aat gaa tca gat agt ggt gta gag gaa tat gtc ctt ttt tat cct tgc tat caa agt tac gca tgc cgt ggc gta aca ata tct tta ata cag atg gat taa atc gtg tat tca tcg tat agc aat gta atg gag agt tac cac att tta gtc gtt tat tca gat cgc agt gtt taa taa cta att taa aca gat gag atg ata tat cca cat caa aga atg cga gat aca tat gac aga cat tat tga cag aaa tgt gac ctt cat tat cac cgt cgt cca taa atg cgt tag gta cgt acc aca tac tat cgt taa cga tgc gca caa tct cgt cca -tt tca tcc atc ttc ata atg att tac ttt ttc ata att aga gaa aaa gat caa ggt ata aaa att aga agt gtt aga cta ta- --- --- --- --- --- --- --- --- --- --- -aa cta act tat gac tta act aac tta tga --- --- --- --- --- --- --- --- --- --- --- --- --- --- --- --- --- --- --- --- --- --- --- --- --- --- --- --- --- --- --- --- --- --- --- --- --- --- --- --- --- --- --- --- --- --- --- --- ctt aac ta- --- --- --- --- --- --- --- --- --- --- -ac tta tga ctt aac taa ctt atg act taa cta act ta- tga ctt aac taa ctt atg act taa cta act tat gac tta act aac tta tga cta tta act cat ttc aag aaa ggt ggg tgg ata gaa ctc tat atg aca gct tgt gaa aca att aga tcc cta att tct aat gga agt ttt gat agg aga ttg tca tca gtt gat aca ttg ttt att atc tca tct att aga gca cgt ctg ttt aga gct tta gtg acc tgc tcg gtt act tct gtg taa atc ttg aat cct tta gtg ata cac tgt gtc aaa act gga tgt tta gaa tac cta tgt aga ata tgg gaa gca tgc ttg ttt ttg tct cta tta tag att aac tca tac atg gtt gta tta tga att ttc atc tgc cta atg tac tcc aat tct tgt tta caa tca att ata taa tca aag agt gat gat gca tac aca tta caa agt gaa taa tct acc atc ata aaa tac ttg ata cag agc ttt atc aca tca tgg ttt tca att gta tta tta agt ata gct aat ttt ata cag tca ata gac aat ggt tct cta agc aat att tct aat att tta aga tgt gct tcc cta cgg gcg atg aca gat ccc cta tcc acg gcc acg tca aga cat gta tat cca tta ctc att act gcg ttg aca ttt gct cca ttt tct aat agc cat gat act aaa tct ata taa cct gca tag ata gcg cga taa agc aag gtc ctt cca cca gca tct agt tga ttg ata tct tca ata tat ggg ata caa agc tta taa att tct aat act gtg ggt tca tct aca agg aat ccc cta gta tac tga att att tta tat aga tct aat tta aca tca ttt tca tct ggg ata cca caa ttc aaa ata aac tca aca aca cta ctt tcc ttt tta cat att ccc cta aaa tag gca ttc aag cat tct att tta tat att aca gcc cca tga tct acc ata aga tca aca atg tct att tct aca tat gca tta gat aga tag taa agt aag aga tct tgc aca gaa tta caa ttc tta ata att ata gag aaa ata tct tcc ata taa ttc ttt gac act aat gca gat ata ata tct tta tat gta ata tat gca aac agt cta tct act ata tac tga tca ata tta tct cta tga atc cta aaa taa tca tac aga aca tct aca gga tca caa att ggt tca agg aga aat cta tca aat att ttc ctg tca aca act ggt tct aga aca taa cag tca aca cct aat cca tgt ttt tta taa tca tct acc aaa gat aat gac caa aga tcg agg tcg tcg tga aac tgc tca tcg aca gcc atg aaa tct acc gac tcc atg gtg cga atc gca ctg tct tat tcg cca ttg att ttc a-t ttt tta taa tta tgt aca tgt ttt cct tct att ctc aag agt cta caa aaa tat a-- -tt ttt tcg ata tct aag tac taa gtt ttt tta ctg ttt ttg tta ctg tct tcc att ctt cta act aaa gat ctg aga taa att ata caa tct tcg cta tcg aac cat ttt tgt agt cta aag cct gaa gta att aac caa ctg ttt tta tta gtg gct ttt ttc gat cta tcc tcg tc- --- ctc tca tca tcc tta ta- --- --- --- --- --- --- tta tta tca tta tta tca tag tct att aaa cac aaa tca tct --- --- --- --- --- --- --- --- --- --- --- --- --- --- --- --- --- --- --- --- --- --- --- --- --- --- --a cgt tta taa c-- --- --- --- --- --- --- --- --a aca ttc --- --- --- --- -tc att att aat tag ttc tgt ag- aat atc ttt aat aat ttg gct ata --- --- -ca tct gtt --- --- --- --- --- --- --- --- --- --- --- --- --- --- --- --- --- --- --c aat act --- --- --- --- --- --- --- --a tct att gat gat ttc ttt t-- --- --- --- --- --- --- --- --- --- --- --t taa gac t-- --- --- --- --- --- --- --- --- --- --- --- --- --- --- --- --- --- --- --- --- --- --- --- --- --- --- --- --- --- --- --- --- --- --- --- --- --- --- --- taa act agt --- --- --- --- --- --- --- --- --- --- --- --- --- --- --- --- --- --- --- --- --- --- --- --- --- --- --- --- --- --- --- --- --- --- --- --- --- --- --- --- --- --- --- --- --- --- --- --- --- --- --- --- --- --- --- --- --- --- --- --- --- --- --- --- --- --- --- --- --- --- --- --- --- --- --- --- --- --- --- --- --- --- --- --- --- --- --- --- --- --- --- --- --- --- --- --- --- --- --- --- --- --- --- --- --- --- --- --- --- --t atg gta atg acg atg aaa --- -tc gag tag ta- --- --a ctt cta ata aag act tga ta- --- --t cat tat cat atg ttt gat cg- --- --- --- --- --- --- --t cat agt taa tag tgt g-- --- --- --- --- --- --- --- --- --- --- --- --- --- --- --- --- --- --- --- -gc taa atg gta ctg tta ata agt tta t-- --- --- --- --- --- --- --- --- --- --- --- --- --- --- --- --- --a gac aat atc ata gta ttt tct ttc cag aat --- --- --- --t aga tta ttt ttt taa ata ctg atc ctc aca att ccg tga tgt agc agt agt tgg t-- --- --- --- --- --- --- --- --- --- --- --- --- --- --- --- --- --- --- --- --- --- --- --- --- --- --- --- -gc atg gtc tat atc gt- --- --- --- --- --- --- --- --- --- --- --- --- --- --- --- --- --- --- --- --- --- --- --- --- --- --- --- --- --- --- --- --- --- --- --- --- --- --- --- --- --- --- --- taa aat gta tca tat ata ata gtt ttc tga cgt gga gta cag aat ttt cga --- --- --- --- --- --- --- --- --- --- --- --- --- --- --- --- --- --- --- --- --- --- --- --- --- --- --- --- --- --- --- --- --- --- --- --- --- --- --- --- --- --- --- --t taa tga gtt cat ggt aag gaa ggg caa atg cct --- gta tat aat ata cat aag tta a-- --- --- --- --- --- --- --- --- --- --- --- --- --t agt ttt tta tca tat tt- --- --- --- --- --- --- --- --t cta ata cca taa taa aaa t-- --- --- --- --- --- --- --- --- --- --- --- --- --- --- --- --- --- --- --- --- --- --- --- --- --t atc att att gcg ttt g-g tag tt- --- --- --- --- ctg ccc ta- --- --- --- --- --- --- --- --t cat cta tat cac tgt cac tct c-- --- --g ctc tca cta tat ctt cta aaa tta caa --- -ac aac --- --- --- --- --- --- --- --- --- --- --- --- --- --- --- --- --- --- --- --- --- --- --- --- --- --- --- --- --- --- --- --- --- --- --- --- tgg ata ttc gat --- --- --- --- --- --- --- --- --- --- --- --- --- --- --- --- --- --- --- --- --- --- --- --- --- --- --a aca gca ttt gtg t-- --- --- --- --- --- --- --- --- --- --- --- --- --- --- --- --- -

>ON649723.1_Monkeypox_virus_isolate_Monkeypox/PT0013/2022_complete_genome

--- --- --- --- --- --- --- --- --- --- --- --- --- --- --- --- --- --- -ta att a-- --- --- --- --- --- --- --- att taa ttt tac tat ttt att tag tgt cta gaa aaa aat gtg tga ccc acg acc gta gga aac tct aga ggg taa gaa aaa tca atc gtt tat aga gac cat cag aaa gag gtt taa tat ttt tgt gag acc tat cga a-- --- --- --- --- --- --- --- --- --- --- --- --- --- --- --- --- --- --- --- --- --- --- --- --- --- --- --- --- --- --- --- --- --- --- --- --- --- --- --- --- --- --- --- --- --- --- --- --- --- --- --- --- --- --- --- --- --- --- --- --- --- --- --- --- --- --- --- --- --- --- --- --- --- --- --- --- --- --- --- --- --- --- --- --- --- --- --- --- --- --- --- --- --- --- --- --- --- --- --- --- --- --- --- --- --- --- --- --- --- --- --- --- --- --- --- --- --- --- --- --- --- --- --- --- --- --- --- --- --- --- --- --- --- --- --- --- --- --- --- --- --- --- --- --- --- --- --- --- --- --- --- --- --- --- --- --- --- --- --- --- --- --- --- --- --- --- --- --- --- --- --- --- --- --- --- --- --- --- --- --- --- --- --- --- --- --- --- --- --- --- --- --- --- --- --- --- --- --- --- --- --- --- --- --- --- --- --- --- --- --- --- --- --- --- --- --- --- --- --- --- --- --- --- --- --- --- --- --- --- --- --- --- --- --- --- --- --- --- --- --- --- --- --- --- --- --- --- --- --- --- --- --- --- --- --- --- --- --- --- --- --- --- --- --- --- --- --- --- --- --- --- --- --- --- --- --- --- --- --- -ga gag aaa gga taa aaa ctt ttt acg act cca tca gaa aga ggt tta ata ttt ttg tga gac cca tcg aag aga gaa aga gat ggt tag tca aga tat ttt tct tag tac aaa agt caa tgt ttt aaa ata tat gga cga gaa tta att tgt ctg tat aaa aac ttg tgt gaa att atg tac tag aga aaa aac gtg agc agt gtc ccc tac atg gat ttt aca gat cat tta tat tcc aaa aat att aac tat ata cgt tta tta tat gat gtt aac gtg taa att ata aac att att tta tga tgc aat tgt ctg aca acc tag att ggt ata agg atg ttg ata agc tct acg aga ata tat tgt tgg acg tta tcg ttt acg aaa tag ttg aga cat cag aaa gag gtt taa tat ttt tgt gag a-c cat cga aga gag aaa gag aat aaa aat a-- --t ttt ttt ttt ttt tgt aaa act ttt tta tga gac --- --- -ca aga gaa tac gaa tag tga tca tat cgt atc aca tat tga aac aga aag aag aag taa cga gag gta act ttt tgt gaa tgt agt taa ata ttt ttg ttt tgc aaa ccg gaa tat agt gcc cgg t-- -ct ttt tta att cgt ggt gcg gtg tct gaa tcg ttc gat taa ccc aac tca tcc att ttc aga tga ata gag tta tcg att cag aca cat gct ttg agt ttt gtt gaa tcg atg agt gaa gta tca tcg gtt gca cct tca gat gcc gat ccg tcg aca tac ttg aat cca tcc ttg act tca agt tca gat gat tcc tca cac atg tct ccg ata cgt acg cta aac tct agg ttc ttg aca cat ttt gta tca acg atc gtt gaa ccg atg ata tct ttg taa ctc act ttc tta tgt gag atg tta gac cca agt act gga tgg gtc ttg atg tca ctg tct ttc tct tct tcg cta cat ctg atg tcg ata gac atc tca cag tct ttg atc ata gcc aga gct tct tca cgc gtg atc gcg gga gag tcc tta cct tgt ccc ggt gac acg ctg gac aat cta gta ttc aca gtg ttt cca tca gag gat tcg gag atg gat aaa atc ttt ggg cat ttg gtg aat cca aag ttc atg tta aga ccc gca ccg acg ata gtg taa taa gtg gtg gga tct cct ttt aca act --- --- --- --- --- --- tct tcg gat acc tca tca tct tcg gtc tct gta act tcc gtt acg gat tga caa atc tta tca ttg gtc ggt gtt tgg tct tgc ttt gtg act ttg ata ata aca tcg att ccc ata tga tgt ttg ttt tct tct tca gta cac gag gat gaa gat tgt tga aga cta gta ggc ata gca gct gcc act agg cac atg cat gcc agg aca ata tat tgt ttc atg att gct att gat tga tta ctg ttc tag atg att cta ctt tct tac cat ata ata aat tag aat ata ttt tct act ttt acg aga aat taa tta ttg tat tta tta ttt ata ggt --a aaa aaa ctt act ata agt ggg tgg gat tct ggg aat tag tga tca gtt tat gta tat cgc aac tag cgg gca tat ggc tat tga cat cga gaa cat tac cca tat gat aag aga ttg tat cat ttt cgt agt ctt gag tat tgg tat tac tat ata gta tgt aga tgt cga cgc tag ata gac agt cgc cca cta gag tta ccg tct ctg aat gcg gca tga tag tat cat tct ttg ttt tcg tta act gtt tgg aag atg aat ctt tgt tgt tac att taa tct cga aat tca gag tac ata tct ttg aag tat tct gat atc tat ttt ctc ctg taa aga atc ctg aag ttg cta cat tat taa gga cag aga agt att ctg cac gaa aga ctg gat cac aat ctt tat gat tca tgg taa tag tta gtt ccg acg ttg aga tgg att cgc tga gac cgg tag tgg tcg tcc gag tac acg atg tgt cgt tga ctg gat aca ggt taa ttt cca cat cga tat agt taa atg tat tgc tgg tta cga cgg gtt cgc att tat ctg tgg aag aga cgg tgt gag aat atg ttc cgg gac cac acg gag aac aga tga cgt ctc cgg tag acg tgt atc cgg ata ctc cgt atc cta ttc cac act ttg ttt tag aaa tac atg ttc tac acc ctg atg ctc ctt tga gaa gac aat aat atc ctg gag agc att cac aga ttc tat tgt gag tcg tgt tac acg atc gcg tct cta cct gat tac tat cac atc ttc cgt tac aac tta gac aag cct gta aat gat tat tgt gag atg taa agg tat ccg aac cac acg gtg tac att gtg tat tag tct tgc tat cac ata atc tga aag cgt aag ttc ccg gag gac acg ata gac aac ata gat tac ggc ttc tgt att cgt tgt ctt tac act ttc cat tgg atg gtg cat gtg gtg cta tat ctc ttc cgt tta tta tta tac atg aga gaa aca ata tat acg agt ata ata cgg acc tca tga ttt aat aat -gt agt aat cgt cgt ctt gtt act gtt tgt ttc cta ctt ctc caa tca tat aga tta ttt ttt aaa tat ttt ctt tct atc atg gat aat att tgt aat ggt tct ttc cgt aca aca tac tgt tta gat ggt agt cgc tta gct tgg tta tga tat tgc gca taa ttt ccg gag gca aat acg ata gtc tag att gac tat cga tgg tag act cta att tat tga gtg ctt tgt cga cga gtt tac ttt tat gct cca tcg ata gat gac act gtt cta tga gat cgt cgt aca tgg gaa atg aaa tgc gtt tgt ctg aat gta tgg ctt cga gat atg tgt gat acc gaa tgt ctt ctg ttc tca ata ccg tat aca agt cgg tgt ctg aga ttc gaa tct ctt tga gga gac tta tgt cac gac tac att ttt cga tga tgg aat cta tct tat cga atg ata tat ttt tca taa ata cac ttt tat agt cct cgt tta aac aga att tag tat gta gtt ccg caa atg act cgt ccc tta ata ggc agt agg cta tta tct tct tta cgt agt gat cgt cgt agg gag aga act ccg aca tct tgt aga aca acg att taa tca tag gta gag ata ctt tca gtc tgt ggt gga taa tgt cat taa cga cat ccg cct tgt ata taa tgt ttc tgt ttt caa aca cca agt cga ata ctg tct tat cgt ctt tag tcg gaa ggt tga tgt cgt atc caa tgt ata cga ggt atg agg caa cat tgt tat tgc aat tct gga agg cgg tat gaa gag gag tca ttg tat tat agt att cgt ctt tct gaa tgt cga atc tat cta gta gat acc gta gta tat tga gag agc gac ttc cat atc ctt gat tat gtt tta tga ata gat aaa gta gat gtt gtc ctt ctt cct ttt gta att tcc cgt att ttt gtt cgt gcc aat tga gta aca tta tga gaa tat gac ctg ttg cac aat cgt tct tta tgt att cca tga tgg gtg tac aat taa gat tat tac gta tcc tcg tat cgg ctc ctc gag ata aaa gag cat aca cca cac gag gac tat gtt tgg tat act gtt gaa ggt aag tgt gta acg gcg tat ttc cga ttt tcg taa ccg cgt taa tgt ttg ctc cat gat cta tta tcg cgt aga tga atc gct tct cag ctc gca tct tag tgt gac tct ttg act tgt aat aat tgc ttt cgt gga acg cgg ata tgt gtt tac agt agt aat gaa gag aag tga gtc cat cct cat cga cgc aat tag ggt cag atc ctt tag tca ata att tgt aca gaa cgt aat agt tta agc tcc cat tga att tat atc taa gat aac aca gca ata gat cgg atg att tac taa agt cat caa tgg ggt ccg tta gta tat caa aga tct tgt tat cga ttg ata gtg aat gaa tca gat agt ggt gta gag gaa tat gtc ctt ttt tat cct tgc tat caa agt tac gca tgc cgt ggc gta aca ata tct tta ata cag atg gat taa atc gtg tat tca tcg tat agc aat gta atg gag agt tac cac att tta gtc gtt tat tca gat cgc agt gtt taa taa cta att taa aca gat gag atg ata tat cca cat caa aga atg cga gat aca tat gac aga cat tat tga cag aaa tgt gac ctt cat tat cac cgt cgt cca taa atg cgt tag gta cgt acc aca tac tat cgt taa cga tgc gca caa tct cgt cca -tt tca tcc atc ttc ata atg att tac ttt ttc ata att aga gaa aaa gat caa ggt ata aaa att aga agt gtt aga cta ta- --- --- --- --- --- --- --- --- --- --- -aa cta act tat gac tta act aac tta tga --- --- --- --- --- --- --- --- --- --- --- --- --- --- --- --- --- --- --- --- --- --- --- --- --- --- --- --- --- --- --- --- --- --- --- --- --- --- --- --- --- --- --- --- --- --- --- --- ctt aac ta- --- --- --- --- --- --- --- --- --- --- -ac tta tga ctt aac taa ctt atg act taa cta act ta- tga ctt aac taa ctt atg act taa cta act tat gac tta act aac tta tga cta tta act cat ttc aag aaa ggt ggg tgg ata gaa ctc tat atg aca gct tgt gaa aca att aga tcc cta att tct aat gga agt ttt gat agg aga ttg tca tca gtt gat aca ttg ttt att atc tca tct att aga gca cgt ctg ttt aga gct tta gtg acc tgc tcg gtt act tct gtg taa atc ttg aat cct tta gtg ata cac tgt gtc aaa act gga tgt tta gaa tac cta tgt aga ata tgg gaa gca tgc ttg ttt ttg tct cta tta tag att aac tca tac atg gtt gta tta tga att ttc atc tgc cta atg tac tcc aat tct tgt tta caa tca att ata taa tca aag agt gat gat gca tac aca tta caa agt gaa taa tct acc atc ata aaa tac ttg ata cag agc ttt atc aca tca tgg ttt tca att gta tta tta agt ata gct aat ttt ata cag tca ata gac aat ggt tct cta agc aat att tct aat att tta aga tgt gct tcc cta cgg gcg atg aca gat ccc cta tcc acg gcc acg tca aga cat gta tat cca tta ctc att act gcg ttg aca ttt gct cca ttt tct aat agc cat gat act aaa tct ata taa cct gca tag ata gcg cga taa agc aag gtc ctt cca cca gca tct agt tga ttg ata tct tca ata tat ggg ata caa agc tta taa att tct aat act gtg ggt tca tct aca agg aat ccc cta gta tac tga att att tta tat aga tct aat tta aca tca ttt tca tct ggg ata cca caa ttc aaa ata aac tca aca aca cta ctt tcc ttt tta cat att ccc cta aaa tag gca ttc aag cat tct att tta tat att aca gcc cca tga tct acc ata aga tca aca atg tct att tct aca tat gca tta gat aga tag taa agt aag aga tct tgc aca gaa tta caa ttc tta ata att ata gag aaa ata tct tcc ata taa ttc ttt gac act aat gca gat ata ata tct tta tat gta ata tat gca aac agt cta tct act ata tac tga tca ata tta tct cta tga atc cta aaa taa tca tac aga aca tct aca gga tca caa att ggt tca agg aga aat cta tca aat att ttc ctg tca aca act ggt tct aga aca taa cag tca aca cct aat cca tgt ttt tta taa tca tct acc aaa gat aat gac caa aga tcg agg tcg tcg tga aac tgc tca tcg aca gcc atg aaa tct acc gac tcc atg gtg cga atc gca ctg tct tat tcg cca ttg att ttc a-t ttt tta taa tta tgt aca tgt ttt cct tct att ctc aag agt cta caa aaa tat a-- -tt ttt tcg ata tct aag tac taa gtt ttt tta ctg ttt ttg tta ctg tct tcc att ctt cta act aaa gat ctg aga taa att ata caa tct tcg cta tcg aac cat ttt tgt agt cta aag cct gaa gta att aac caa ctg ttt tta tta gtg gct ttt ttc gat cta tcc tcg tc- --- ctc tca tca tcc tta ta- --- --- --- --- --- --- tta tta tca tta tta tca tag tct att aaa cac aaa tca tct --- --- --- --- --- --- --- --- --- --- --- --- --- --- --- --- --- --- --- --- --- --- --- --- --- --- --a cgt tta taa c-- --- --- --- --- --- --- --- --a aca ttc --- --- --- --- -tc att att aat tag ttc tgt ag- aat atc ttt aat aat ttg gct ata --- --- -ca tct gtt --- --- --- --- --- --- --- --- --- --- --- --- --- --- --- --- --- --- --c aat act --- --- --- --- --- --- --- --a tct att gat gat ttc ttt t-- --- --- --- --- --- --- --- --- --- --- --t taa gac t-- --- --- --- --- --- --- --- --- --- --- --- --- --- --- --- --- --- --- --- --- --- --- --- --- --- --- --- --- --- --- --- --- --- --- --- --- --- --- --- taa act agt --- --- --- --- --- --- --- --- --- --- --- --- --- --- --- --- --- --- --- --- --- --- --- --- --- --- --- --- --- --- --- --- --- --- --- --- --- --- --- --- --- --- --- --- --- --- --- --- --- --- --- --- --- --- --- --- --- --- --- --- --- --- --- --- --- --- --- --- --- --- --- --- --- --- --- --- --- --- --- --- --- --- --- --- --- --- --- --- --- --- --- --- --- --- --- --- --- --- --- --- --- --- --- --- --- --- --- --- --- --t atg gta atg acg atg aaa --- -tc gag tag ta- --- --a ctt cta ata aag act tga ta- --- --t cat tat cat atg ttt gat cg- --- --- --- --- --- --- --t cat agt taa tag tgt g-- --- --- --- --- --- --- --- --- --- --- --- --- --- --- --- --- --- --- --- -gc taa atg gta ctg tta ata agt tta t-- --- --- --- --- --- --- --- --- --- --- --- --- --- --- --- --- --a gac aat atc ata gta ttt tct ttc cag aat --- --- --- --t aga tta ttt ttt taa ata ctg atc ctc aca att ccg tga tgt agc agt agt tgg t-- --- --- --- --- --- --- --- --- --- --- --- --- --- --- --- --- --- --- --- --- --- --- --- --- --- --- --- -gc atg gtc tat atc gt- --- --- --- --- --- --- --- --- --- --- --- --- --- --- --- --- --- --- --- --- --- --- --- --- --- --- --- --- --- --- --- --- --- --- --- --- --- --- --- --- --- --- --- taa aat gta tca tat ata ata gtt ttc tga cgt gga gta cag aat ttt cga --- --- --- --- --- --- --- --- --- --- --- --- --- --- --- --- --- --- --- --- --- --- --- --- --- --- --- --- --- --- --- --- --- --- --- --- --- --- --- --- --- --- --- --t taa tga gtt cat ggt aag gaa ggg caa atg cct --- gta tat aat ata cat aag tta a-- --- --- --- --- --- --- --- --- --- --- --- --- --t agt ttt tta tca tat tt- --- --- --- --- --- --- --- --t cta ata cca taa taa aaa t-- --- --- --- --- --- --- --- --- --- --- --- --- --- --- --- --- --- --- --- --- --- --- --- --- --t atc att att gcg ttt g-g tag tt- --- --- --- --- ctg ccc ta- --- --- --- --- --- --- --- --t cat cta tat cac tgt cac tct c-- --- --g ctc tca cta tat ctt cta aaa tta caa --- -ac aac --- --- --- --- --- --- --- --- --- --- --- --- --- --- --- --- --- --- --- --- --- --- --- --- --- --- --- --- --- --- --- --- --- --- --- --- tgg ata ttc gat --- --- --- --- --- --- --- --- --- --- --- --- --- --- --- --- --- --- --- --- --- --- --- --- --- --- --a aca gca ttt gtg t-- --- --- --- --- --- --- --- --- --- --- --- --- --- --- --- --- -

>ON649718.1_Monkeypox_virus_isolate_Monkeypox/PT0018/2022_complete_genome

--- --- --- --- --- --- --- --- --- --- --- --- --- --- --- --- --- --- -ta att a-- --- --- --- --- --- --- --- att taa ttt tac tat ttt att tag tgt cta gaa aaa aat gtg tga ccc acg acc gta gga aac tct aga ggg taa gaa aaa tca atc gtt tat aga gac cat cag aaa gag gtt taa tat ttt tgt gag acc tat cga a-- --- --- --- --- --- --- --- --- --- --- --- --- --- --- --- --- --- --- --- --- --- --- --- --- --- --- --- --- --- --- --- --- --- --- --- --- --- --- --- --- --- --- --- --- --- --- --- --- --- --- --- --- --- --- --- --- --- --- --- --- --- --- --- --- --- --- --- --- --- --- --- --- --- --- --- --- --- --- --- --- --- --- --- --- --- --- --- --- --- --- --- --- --- --- --- --- --- --- --- --- --- --- --- --- --- --- --- --- --- --- --- --- --- --- --- --- --- --- --- --- --- --- --- --- --- --- --- --- --- --- --- --- --- --- --- --- --- --- --- --- --- --- --- --- --- --- --- --- --- --- --- --- --- --- --- --- --- --- --- --- --- --- --- --- --- --- --- --- --- --- --- --- --- --- --- --- --- --- --- --- --- --- --- --- --- --- --- --- --- --- --- --- --- --- --- --- --- --- --- --- --- --- --- --- --- --- --- --- --- --- --- --- --- --- --- --- --- --- --- --- --- --- --- --- --- --- --- --- --- --- --- --- --- --- --- --- --- --- --- --- --- --- --- --- --- --- --- --- --- --- --- --- --- --- --- --- --- --- --- --- --- --- --- --- --- --- --- --- --- --- --- --- --- --- --- --- --- --- --- -ga gag aaa gga taa aaa ctt ttt acg act cca tca gaa aga ggt tta ata ttt ttg tga gac cca tcg aag aga gaa aga gat ggt tag tca aga tat ttt tct tag tac aaa agt caa tgt ttt aaa ata tat gga cga gaa tta att tgt ctg tat aaa aac ttg tgt gaa att atg tac tag aga aaa aac gtg agc agt gtc ccc tac atg gat ttt aca gat cat tta tat tcc aaa aat att aac tat ata cgt tta tta tat gat gtt aac gtg taa att ata aac att att tta tga tgc aat tgt ctg aca acc tag att ggt ata agg atg ttg ata agc tct acg aga ata tat tgt tgg acg tta tcg ttt acg aaa tag ttg aga cat cag aaa gag gtt taa tat ttt tgt gag a-c cat cga aga gag aaa gag aat aaa aat a-- --t ttt ttt ttt ttt tgt aaa act ttt tta tga gac --- --- -ca aga gaa tac gaa tag tga tca tat cgt atc aca tat tga aac aga aag aag aag taa cga gag gta act ttt tgt gaa tgt agt taa ata ttt ttg ttt tgc aaa ccg gaa tat agt gcc cgg t-- -ct ttt tta att cgt ggt gcg gtg tct gaa tcg ttc gat taa ccc aac tca tcc att ttc aga tga ata gag tta tcg att cag aca cat gct ttg agt ttt gtt gaa tcg atg agt gaa gta tca tcg gtt gca cct tca gat gcc gat ccg tcg aca tac ttg aat cca tcc ttg act tca agt tca gat gat tcc tca cac atg tct ccg ata cgt acg cta aac tct agg ttc ttg aca cat ttt gta tca acg atc gtt gaa ccg atg ata tct ttg taa ctc act ttc tta tgt gag atg tta gac cca agt act gga tgg gtc ttg atg tca ctg tct ttc tct tct tcg cta cat ctg atg tcg ata gac atc tca cag tct ttg atc ata gcc aga gct tct tca cgc gtg atc gcg gga gag tcc tta cct tgt ccc ggt gac acg ctg gac aat cta gta ttc aca gtg ttt cca tca gag gat tcg gag atg gat aaa atc ttt ggg cat ttg gtg aat cca aag ttc atg tta aga ccc gca ccg acg ata gtg taa taa gtg gtg gga tct cct ttt aca act --- --- --- --- --- --- tct tcg gat acc tca tca tct tcg gtc tct gta act tcc gtt acg gat tga caa atc tta tca ttg gtc ggt gtt tgg tct tgc ttt gtg act ttg ata ata aca tcg att ccc ata tga tgt ttg ttt tct tct tca gta cac gag gat gaa gat tgt tga aga cta gta ggc ata gca gct gcc act agg cac atg cat gcc agg aca ata tat tgt ttc atg att gct att gat tga tta ctg ttc tag atg att cta ctt tct tac cat ata ata aat tag aat ata ttt tct act ttt acg aga aat taa tta ttg tat tta tta ttt ata ggt --a aaa aaa ctt act ata agt ggg tgg gat tct ggg aat tag tga tca gtt tat gta tat cgc aac tag cgg gca tat ggc tat tga cat cga gaa cat tac cca tat gat aag aga ttg tat cat ttt cgt agt ctt gag tat tgg tat tac tat ata gta tgt aga tgt cga cgc tag ata gac agt cgc cca cta gag tta ccg tct ctg aat gcg gca tga tag tat cat tct ttg ttt tcg tta act gtt tgg aag atg aat ctt tgt tgt tac att taa tct cga aat tca gag tac ata tct ttg aag tat tct gat atc tat ttt ctc ctg taa aga atc ctg aag ttg cta cat tat taa gga cag aga agt att ctg cac gaa aga ctg gat cac aat ctt tat gat tca tgg taa tag tta gtt ccg acg ttg aga tgg att cgc tga gac cgg tag tgg tcg tcc gag tac acg atg tgt cgt tga ctg gat aca ggt taa ttt cca cat cga tat agt taa atg tat tgc tgg tta cga cgg gtt cgc att tat ctg tgg aag aga cgg tgt gag aat atg ttc cgg gac cac acg gag aac aga tga cgt ctc cgg tag acg tgt atc cgg ata ctc cgt atc cta ttc cac act ttg ttt tag aaa tac atg ttc tac acc ctg atg ctc ctt tga gaa gac aat aat atc ctg gag agc att cac aga ttc tat tgt gag tcg tgt tac acg atc gcg tct cta cct gat tac tat cac atc ttc cgt tac aac tta gac aag cct gta aat gat tat tgt gag atg taa agg tat ccg aac cac acg gtg tac att gtg tat tag tct tgc tat cac ata atc tga aag cgt aag ttc ccg gag gac acg ata gac aac ata gat tac ggc ttc tgt att cgt tgt ctt tac act ttc cat tgg atg gtg cat gtg gtg cta tat ctc ttc cgt tta tta tta tac atg aga gaa aca ata tat acg agt ata ata cgg acc tca tga ttt aat aat -gt agt aat cgt cgt ctt gtt act gtt tgt ttc cta ctt ctc caa tca tat aga tta ttt ttt aaa tat ttt ctt tct atc atg gat aat att tgt aat ggt tct ttc cgt aca aca tac tgt tta gat ggt agt cgc tta gct tgg tta tga tat tgc gca taa ttt ccg gag gca aat acg ata gtc tag att gac tat cga tgg tag act cta att tat tga gtg ctt tgt cga cga gtt tac ttt tat gct cca tcg ata gat gac act gtt cta tga gat cgt cgt aca tgg gaa atg aaa tgc gtt tgt ctg aat gta tgg ctt cga gat atg tgt gat acc gaa tgt ctt ctg ttc tca ata ccg tat aca agt cgg tgt ctg aga ttc gaa tct ctt tga gga gac tta tgt cac gac tac att ttt cga tga tgg aat cta tct tat cga atg ata tat ttt tca taa ata cac ttt tat agt cct cgt tta aac aga att tag tat gta gtt ccg caa atg act cgt ccc tta ata ggc agt agg cta tta tct tct tta cgt agt gat cgt cgt agg gag aga act ccg aca tct tgt aga aca acg att taa tca tag gta gag ata ctt tca gtc tgt ggt gga taa tgt cat taa cga cat ccg cct tgt ata taa tgt ttc tgt ttt caa aca cca agt cga ata ctg tct tat cgt ctt tag tcg gaa ggt tga tgt cgt atc caa tgt ata cga ggt atg agg caa cat tgt tat tgc aat tct gga agg cgg tat gaa gag gag tca ttg tat tat agt att cgt ctt tct gaa tgt cga atc tat cta gta gat acc gta gta tat tga gag agc gac ttc cat atc ctt gat tat gtt tta tga ata gat aaa gta gat gtt gtc ctt ctt cct ttt gta att tcc cgt att ttt gtt cgt gcc aat tga gta aca tta tga gaa tat gac ctg ttg cac aat cgt tct tta tgt att cca tga tgg gtg tac aat taa gat tat tac gta tcc tcg tat cgg ctc ctc gag ata aaa gag cat aca cca cac gag gac tat gtt tgg tat act gtt gaa ggt aag tgt gta acg gcg tat ttc cga ttt tcg taa ccg cgt taa tgt ttg ctc cat gat cta tta tcg cgt aga tga atc gct tct cag ctc gca tct tag tgt gac tct ttg act tgt aat aat tgc ttt cgt gga acg cgg ata tgt gtt tac agt agt aat gaa gag aag tga gtc cat cct cat cga cgc aat tag ggt cag atc ctt tag tca ata att tgt aca gaa cgt aat agt tta agc tcc cat tga att tat atc taa gat aac aca gca ata gat cgg atg att tac taa agt cat caa tgg ggt ccg tta gta tat caa aga tct tgt tat cga ttg ata gtg aat gaa tca gat agt ggt gta gag gaa tat gtc ctt ttt tat cct tgc tat caa agt tac gca tgc cgt ggc gta aca ata tct tta ata cag atg gat taa atc gtg tat tca tcg tat agc aat gta atg gag agt tac cac att tta gtc gtt tat tca gat cgc agt gtt taa taa cta att taa aca gat gag atg ata tat cca cat caa aga atg cga gat aca tat gac aga cat tat tga cag aaa tgt gac ctt cat tat cac cgt cgt cca taa atg cgt tag gta cgt acc aca tac tat cgt taa cga tgc gca caa tct cgt cca -tt tca tcc atc ttc ata atg att tac ttt ttc ata att aga gaa aaa gat caa ggt ata aaa att aga agt gtt aga cta ta- --- --- --- --- --- --- --- --- --- --- -aa cta act tat gac tta act aac tta tga --- --- --- --- --- --- --- --- --- --- --- --- --- --- --- --- --- --- --- --- --- --- --- --- --- --- --- --- --- --- --- --- --- --- --- --- --- --- --- --- --- --- --- --- --- --- --- --- ctt aac ta- --- --- --- --- --- --- --- --- --- --- -ac tta tga ctt aac taa ctt atg act taa cta act ta- tga ctt aac taa ctt atg act taa cta act tat gac tta act aac tta tga cta tta act cat ttc aag aaa ggt ggg tgg ata gaa ctc tat atg aca gct tgt gaa aca att aga tcc cta att tct aat gga agt ttt gat agg aga ttg tca tca gtt gat aca ttg ttt att atc tca tct att aga gca cgt ctg ttt aga gct tta gtg acc tgc tcg gtt act tct gtg taa atc ttg aat cct tta gtg ata cac tgt gtc aaa act gga tgt tta gaa tac cta tgt aga ata tgg gaa gca tgc ttg ttt ttg tct cta tta tag att aac tca tac atg gtt gta tta tga att ttc atc tgc cta atg tac tcc aat tct tgt tta caa tca att ata taa tca aag agt gat gat gca tac aca tta caa agt gaa taa tct acc atc ata aaa tac ttg ata cag agc ttt atc aca tca tgg ttt tca att gta tta tta agt ata gct aat ttt ata cag tca ata gac aat ggt tct cta agc aat att tct aat att tta aga tgt gct tcc cta cgg gcg atg aca gat ccc cta tcc acg gcc acg tca aga cat gta tat cca tta ctc att act gcg ttg aca ttt gct cca ttt tct aat agc cat gat act aaa tct ata taa cct gca tag ata gcg cga taa agc aag gtc ctt cca cca gca tct agt tga ttg ata tct tca ata tat ggg ata caa agc tta taa att tct aat act gtg ggt tca tct aca agg aat ccc cta gta tac tga att att tta tat aga tct aat tta aca tca ttt tca tct ggg ata cca caa ttc aaa ata aac tca aca aca cta ctt tcc ttt tta cat att ccc cta aaa tag gca ttc aag cat tct att tta tat att aca gcc cca tga tct acc ata aga tca aca atg tct att tct aca tat gca tta gat aga tag taa agt aag aga tct tgc aca gaa tta caa ttc tta ata att ata gag aaa ata tct tcc ata taa ttc ttt gac act aat gca gat ata ata tct tta tat gta ata tat gca aac agt cta tct act ata tac tga tca ata tta tct cta tga atc cta aaa taa tca tac aga aca tct aca gga tca caa att ggt tca agg aga aat cta tca aat att ttc ctg tca aca act ggt tct aga aca taa cag tca aca cct aat cca tgt ttt tta taa tca tct acc aaa gat aat gac caa aga tcg agg tcg tcg tga aac tgc tca tcg aca gcc atg aaa tct acc gac tcc atg gtg cga atc gca ctg tct tat tcg cca ttg att ttc a-t ttt tta taa tta tgt aca tgt ttt cct tct att ctc aag agt cta caa aaa tat a-- -tt ttt tcg ata tct aag tac taa gtt ttt tta ctg ttt ttg tta ctg tct tcc att ctt cta act aaa gat ctg aga taa att ata caa tct tcg cta tcg aac cat ttt tgt agt cta aag cct gaa gta att aac caa ctg ttt tta tta gtg gct ttt ttc gat cta tcc tcg tc- --- ctc tca tca tcc tta ta- --- --- --- --- --- --- tta tta tca tta tta tca tag tct att aaa cac aaa tca tct --- --- --- --- --- --- --- --- --- --- --- --- --- --- --- --- --- --- --- --- --- --- --- --- --- --- --a cgt tta taa c-- --- --- --- --- --- --- --- --a aca ttc --- --- --- --- -tc att att aat tag ttc tgt ag- aat atc ttt aat aat ttg gct ata --- --- -ca tct gtt --- --- --- --- --- --- --- --- --- --- --- --- --- --- --- --- --- --- --c aat act --- --- --- --- --- --- --- --a tct att gat gat ttc ttt t-- --- --- --- --- --- --- --- --- --- --- --t taa gac t-- --- --- --- --- --- --- --- --- --- --- --- --- --- --- --- --- --- --- --- --- --- --- --- --- --- --- --- --- --- --- --- --- --- --- --- --- --- --- --- taa act agt --- --- --- --- --- --- --- --- --- --- --- --- --- --- --- --- --- --- --- --- --- --- --- --- --- --- --- --- --- --- --- --- --- --- --- --- --- --- --- --- --- --- --- --- --- --- --- --- --- --- --- --- --- --- --- --- --- --- --- --- --- --- --- --- --- --- --- --- --- --- --- --- --- --- --- --- --- --- --- --- --- --- --- --- --- --- --- --- --- --- --- --- --- --- --- --- --- --- --- --- --- --- --- --- --- --- --- --- --- --t atg gta atg acg atg aaa --- -tc gag tag ta- --- --a ctt cta ata aag act tga ta- --- --t cat tat cat atg ttt gat cg- --- --- --- --- --- --- --t cat agt taa tag tgt g-- --- --- --- --- --- --- --- --- --- --- --- --- --- --- --- --- --- --- --- -gc taa atg gta ctg tta ata agt tta t-- --- --- --- --- --- --- --- --- --- --- --- --- --- --- --- --- --a gac aat atc ata gta ttt tct ttc cag aat --- --- --- --t aga tta ttt ttt taa ata ctg atc ctc aca att ccg tga tgt agc agt agt tgg t-- --- --- --- --- --- --- --- --- --- --- --- --- --- --- --- --- --- --- --- --- --- --- --- --- --- --- --- -gc atg gtc tat atc gt- --- --- --- --- --- --- --- --- --- --- --- --- --- --- --- --- --- --- --- --- --- --- --- --- --- --- --- --- --- --- --- --- --- --- --- --- --- --- --- --- --- --- --- taa aat gta tca tat ata ata gtt ttc tga cgt gga gta cag aat ttt cga --- --- --- --- --- --- --- --- --- --- --- --- --- --- --- --- --- --- --- --- --- --- --- --- --- --- --- --- --- --- --- --- --- --- --- --- --- --- --- --- --- --- --- --t taa tga gtt cat ggt aag gaa ggg caa atg cct --- gta tat aat ata cat aag tta a-- --- --- --- --- --- --- --- --- --- --- --- --- --t agt ttt tta tca tat tt- --- --- --- --- --- --- --- --t cta ata cca taa taa aaa t-- --- --- --- --- --- --- --- --- --- --- --- --- --- --- --- --- --- --- --- --- --- --- --- --- --t atc att att gcg ttt g-g tag tt- --- --- --- --- ctg ccc ta- --- --- --- --- --- --- --- --t cat cta tat cac tgt cac tct c-- --- --g ctc tca cta tat ctt cta aaa tta caa --- -ac aac --- --- --- --- --- --- --- --- --- --- --- --- --- --- --- --- --- --- --- --- --- --- --- --- --- --- --- --- --- --- --- --- --- --- --- --- tgg ata ttc gat --- --- --- --- --- --- --- --- --- --- --- --- --- --- --- --- --- --- --- --- --- --- --- --- --- --- --a aca gca ttt gtg t-- --- --- --- --- --- --- --- --- --- --- --- --- --- --- --- --- -

>ON649719.1_Monkeypox_virus_isolate_Monkeypox/PT0012/2022_complete_genome

--- --- --- --- --- --- --- --- --- --- --- --- --- --- --- --- --- --- -ta att a-- --- --- --- --- --- --- --- att taa ttt tac tat ttt att tag tgt cta gaa aaa aat gtg tga ccc acg acc gta gga aac tct aga ggg taa gaa aaa tca atc gtt tat aga gac cat cag aaa gag gtt taa tat ttt tgt gag acc tat cga a-- --- --- --- --- --- --- --- --- --- --- --- --- --- --- --- --- --- --- --- --- --- --- --- --- --- --- --- --- --- --- --- --- --- --- --- --- --- --- --- --- --- --- --- --- --- --- --- --- --- --- --- --- --- --- --- --- --- --- --- --- --- --- --- --- --- --- --- --- --- --- --- --- --- --- --- --- --- --- --- --- --- --- --- --- --- --- --- --- --- --- --- --- --- --- --- --- --- --- --- --- --- --- --- --- --- --- --- --- --- --- --- --- --- --- --- --- --- --- --- --- --- --- --- --- --- --- --- --- --- --- --- --- --- --- --- --- --- --- --- --- --- --- --- --- --- --- --- --- --- --- --- --- --- --- --- --- --- --- --- --- --- --- --- --- --- --- --- --- --- --- --- --- --- --- --- --- --- --- --- --- --- --- --- --- --- --- --- --- --- --- --- --- --- --- --- --- --- --- --- --- --- --- --- --- --- --- --- --- --- --- --- --- --- --- --- --- --- --- --- --- --- --- --- --- --- --- --- --- --- --- --- --- --- --- --- --- --- --- --- --- --- --- --- --- --- --- --- --- --- --- --- --- --- --- --- --- --- --- --- --- --- --- --- --- --- --- --- --- --- --- --- --- --- --- --- --- --- --- --- -ga gag aaa gga taa aaa ctt ttt acg act cca tca gaa aga ggt tta ata ttt ttg tga gac cca tcg aag aga gaa aga gat ggt tag tca aga tat ttt tct tag tac aaa agt caa tgt ttt aaa ata tat gga cga gaa tta att tgt ctg tat aaa aac ttg tgt gaa att atg tac tag aga aaa aac gtg agc agt gtc ccc tac atg gat ttt aca gat cat tta tat tcc aaa aat att aac tat ata cgt tta tta tat gat gtt aac gtg taa att ata aac att att tta tga tgc aat tgt ctg aca acc tag att ggt ata agg atg ttg ata agc tct acg aga ata tat tgt tgg acg tta tcg ttt acg aaa tag ttg aga cat cag aaa gag gtt taa tat ttt tgt gag a-c cat cga aga gag aaa gag aat aaa aat a-- --t ttt ttt ttt ttt tgt aaa act ttt tta tga gac --- --- -ca aga gaa tac gaa tag tga tca tat cgt atc aca tat tga aac aga aag aag aag taa cga gag gta act ttt tgt gaa tgt agt taa ata ttt ttg ttt tgc aaa ccg gaa tat agt gcc cgg t-- -ct ttt tta att cgt ggt gcg gtg tct gaa tcg ttc gat taa ccc aac tca tcc att ttc aga tga ata gag tta tcg att cag aca cat gct ttg agt ttt gtt gaa tcg atg agt gaa gta tca tcg gtt gca cct tca gat gcc gat ccg tcg aca tac ttg aat cca tcc ttg act tca agt tca gat gat tcc tca cac atg tct ccg ata cgt acg cta aac tct agg ttc ttg aca cat ttt gta tca acg atc gtt gaa ccg atg ata tct ttg taa ctc act ttc tta tgt gag atg tta gac cca agt act gga tgg gtc ttg atg tca ctg tct ttc tct tct tcg cta cat ctg atg tcg ata gac atc tca cag tct ttg atc ata gcc aga gct tct tca cgc gtg atc gcg gga gag tcc tta cct tgt ccc ggt gac acg ctg gac aat cta gta ttc aca gtg ttt cca tca gag gat tcg gag atg gat aaa atc ttt ggg cat ttg gtg aat cca aag ttc atg tta aga ccc gca ccg acg ata gtg taa taa gtg gtg gga tct cct ttt aca act --- --- --- --- --- --- tct tcg gat acc tca tca tct tcg gtc tct gta act tcc gtt acg gat tga caa atc tta tca ttg gtc ggt gtt tgg tct tgc ttt gtg act ttg ata ata aca tcg att ccc ata tga tgt ttg ttt tct tct tca gta cac gag gat gaa gat tgt tga aga cta gta ggc ata gca gct gcc act agg cac atg cat gcc agg aca ata tat tgt ttc atg att gct att gat tga tta ctg ttc tag atg att cta ctt tct tac cat ata ata aat tag aat ata ttt tct act ttt acg aga aat taa tta ttg tat tta tta ttt ata ggt --a aaa aaa ctt act ata agt ggg tgg gat tct ggg aat tag tga tca gtt tat gta tat cgc aac tag cgg gca tat ggc tat tga cat cga gaa cat tac cca tat gat aag aga ttg tat cat ttt cgt agt ctt gag tat tgg tat tac tat ata gta tgt aga tgt cga cgc tag ata gac agt cgc cca cta gag tta ccg tct ctg aat gcg gca tga tag tat cat tct ttg ttt tcg tta act gtt tgg aag atg aat ctt tgt tgt tac att taa tct cga aat tca gag tac ata tct ttg aag tat tct gat atc tat ttt ctc ctg taa aga atc ctg aag ttg cta cat tat taa gga cag aga agt att ctg cac gaa aga ctg gat cac aat ctt tat gat tca tgg taa tag tta gtt ccg acg ttg aga tgg att cgc tga gac cgg tag tgg tcg tcc gag tac acg atg tgt cgt tga ctg gat aca ggt taa ttt cca cat cga tat agt taa atg tat tgc tgg tta cga cgg gtt cgc att tat ctg tgg aag aga cgg tgt gag aat atg ttc cgg gac cac acg gag aac aga tga cgt ctc cgg tag acg tgt atc cgg ata ctc cgt atc cta ttc cac act ttg ttt tag aaa tac atg ttc tac acc ctg atg ctc ctt tga gaa gac aat aat atc ctg gag agc att cac aga ttc tat tgt gag tcg tgt tac acg atc gcg tct cta cct gat tac tat cac atc ttc cgt tac aac tta gac aag cct gta aat gat tat tgt gag atg taa agg tat ccg aac cac acg gtg tac att gtg tat tag tct tgc tat cac ata atc tga aag cgt aag ttc ccg gag gac acg ata gac aac ata gat tac ggc ttc tgt att cgt tgt ctt tac act ttc cat tgg atg gtg cat gtg gtg cta tat ctc ttc cgt tta tta tta tac atg aga gaa aca ata tat acg agt ata ata cgg acc tca tga ttt aat aat -gt agt aat cgt cgt ctt gtt act gtt tgt ttc cta ctt ctc caa tca tat aga tta ttt ttt aaa tat ttt ctt tct atc atg gat aat att tgt aat ggt tct ttc cgt aca aca tac tgt tta gat ggt agt cgc tta gct tgg tta tga tat tgc gca taa ttt ccg gag gca aat acg ata gtc tag att gac tat cga tgg tag act cta att tat tga gtg ctt tgt cga cga gtt tac ttt tat gct cca tcg ata gat gac act gtt cta tga gat cgt cgt aca tgg gaa atg aaa tgc gtt tgt ctg aat gta tgg ctt cga gat atg tgt gat acc gaa tgt ctt ctg ttc tca ata ccg tat aca agt cgg tgt ctg aga ttc gaa tct ctt tga gga gac tta tgt cac gac tac att ttt cga tga tgg aat cta tct tat cga atg ata tat ttt tca taa ata cac ttt tat agt cct cgt tta aac aga att tag tat gta gtt ccg caa atg act cgt ccc tta ata ggc agt agg cta tta tct tct tta cgt agt gat cgt cgt agg gag aga act ccg aca tct tgt aga aca acg att taa tca tag gta gag ata ctt tca gtc tgt ggt gga taa tgt cat taa cga cat ccg cct tgt ata taa tgt ttc tgt ttt caa aca cca agt cga ata ctg tct tat cgt ctt tag tcg gaa ggt tga tgt cgt atc caa tgt ata cga ggt atg agg caa cat tgt tat tgc aat tct gga agg cgg tat gaa gag gag tca ttg tat tat agt att cgt ctt tct gaa tgt cga atc tat cta gta gat acc gta gta tat tga gag agc gac ttc cat atc ctt gat tat gtt tta tga ata gat aaa gta gat gtt gtc ctt ctt cct ttt gta att tcc cgt att ttt gtt cgt gcc aat tga gta aca tta tga gaa tat gac ctg ttg cac aat cgt tct tta tgt att cca tga tgg gtg tac aat taa gat tat tac gta tcc tcg tat cgg ctc ctc gag ata aaa gag cat aca cca cac gag gac tat gtt tgg tat act gtt gaa ggt aag tgt gta acg gcg tat ttc cga ttt tcg taa ccg cgt taa tgt ttg ctc cat gat cta tta tcg cgt aga tga atc gct tct cag ctc gca tct tag tgt gac tct ttg act tgt aat aat tgc ttt cgt gga acg cgg ata tgt gtt tac agt agt aat gaa gag aag tga gtc cat cct cat cga cgc aat tag ggt cag atc ctt tag tca ata att tgt aca gaa cgt aat agt tta agc tcc cat tga att tat atc taa gat aac aca gca ata gat cgg atg att tac taa agt cat caa tgg ggt ccg tta gta tat caa aga tct tgt tat cga ttg ata gtg aat gaa tca gat agt ggt gta gag gaa tat gtc ctt ttt tat cct tgc tat caa agt tac gca tgc cgt ggc gta aca ata tct tta ata cag atg gat taa atc gtg tat tca tcg tat agc aat gta atg gag agt tac cac att tta gtc gtt tat tca gat cgc agt gtt taa taa cta att taa aca gat gag atg ata tat cca cat caa aga atg cga gat aca tat gac aga cat tat tga cag aaa tgt gac ctt cat tat cac cgt cgt cca taa atg cgt tag gta cgt acc aca tac tat cgt taa cga tgc gca caa tct cgt cca -tt tca tcc atc ttc ata atg att tac ttt ttc ata att aga gaa aaa gat caa ggt ata aaa att aga agt gtt aga cta ta- --- --- --- --- --- --- --- --- --- --- -aa cta act tat gac tta act aac tta tga --- --- --- --- --- --- --- --- --- --- --- --- --- --- --- --- --- --- --- --- --- --- --- --- --- --- --- --- --- --- --- --- --- --- --- --- --- --- --- --- --- --- --- --- --- --- --- --- ctt aac ta- --- --- --- --- --- --- --- --- --- --- -ac tta tga ctt aac taa ctt atg act taa cta act ta- tga ctt aac taa ctt atg act taa cta act tat gac tta act aac tta tga cta tta act cat ttc aag aaa ggt ggg tgg ata gaa ctc tat atg aca gct tgt gaa aca att aga tcc cta att tct aat gga agt ttt gat agg aga ttg tca tca gtt gat aca ttg ttt att atc tca tct att aga gca cgt ctg ttt aga gct tta gtg acc tgc tcg gtt act tct gtg taa atc ttg aat cct tta gtg ata cac tgt gtc aaa act gga tgt tta gaa tac cta tgt aga ata tgg gaa gca tgc ttg ttt ttg tct cta tta tag att aac tca tac atg gtt gta tta tga att ttc atc tgc cta atg tac tcc aat tct tgt tta caa tca att ata taa tca aag agt gat gat gca tac aca tta caa agt gaa taa tct acc atc ata aaa tac ttg ata cag agc ttt atc aca tca tgg ttt tca att gta tta tta agt ata gct aat ttt ata cag tca ata gac aat ggt tct cta agc aat att tct aat att tta aga tgt gct tcc cta cgg gcg atg aca gat ccc cta tcc acg gcc acg tca aga cat gta tat cca tta ctc att act gcg ttg aca ttt gct cca ttt tct aat agc cat gat act aaa tct ata taa cct gca tag ata gcg cga taa agc aag gtc ctt cca cca gca tct agt tga ttg ata tct tca ata tat ggg ata caa agc tta taa att tct aat act gtg ggt tca tct aca agg aat ccc cta gta tac tga att att tta tat aga tct aat tta aca tca ttt tca tct ggg ata cca caa ttc aaa ata aac tca aca aca cta ctt tcc ttt tta cat att ccc cta aaa tag gca ttc aag cat tct att tta tat att aca gcc cca tga tct acc ata aga tca aca atg tct att tct aca tat gca tta gat aga tag taa agt aag aga tct tgc aca gaa tta caa ttc tta ata att ata gag aaa ata tct tcc ata taa ttc ttt gac act aat gca gat ata ata tct tta tat gta ata tat gca aac agt cta tct act ata tac tga tca ata tta tct cta tga atc cta aaa taa tca tac aga aca tct aca gga tca caa att ggt tca agg aga aat cta tca aat att ttc ctg tca aca act ggt tct aga aca taa cag tca aca cct aat cca tgt ttt tta taa tca tct acc aaa gat aat gac caa aga tcg agg tcg tcg tga aac tgc tca tcg aca gcc atg aaa tct acc gac tcc atg gtg cga atc gca ctg tct tat tcg cca ttg att ttc a-t ttt tta taa tta tgt aca tgt ttt cct tct att ctc aag agt cta caa aaa tat a-- -tt ttt tcg ata tct aag tac taa gtt ttt tta ctg ttt ttg tta ctg tct tcc att ctt cta act aaa gat ctg aga taa att ata caa tct tcg cta tcg aac cat ttt tgt agt cta aag cct gaa gta att aac caa ctg ttt tta tta gtg gct ttt ttc gat cta tcc tcg tc- --- ctc tca tca tcc tta ta- --- --- --- --- --- --- tta tta tca tta tta tca tag tct att aaa cac aaa tca tct --- --- --- --- --- --- --- --- --- --- --- --- --- --- --- --- --- --- --- --- --- --- --- --- --- --- --a cgt tta taa c-- --- --- --- --- --- --- --- --a aca ttc --- --- --- --- -tc att att aat tag ttc tgt ag- aat atc ttt aat aat ttg gct ata --- --- -ca tct gtt --- --- --- --- --- --- --- --- --- --- --- --- --- --- --- --- --- --- --c aat act --- --- --- --- --- --- --- --a tct att gat gat ttc ttt t-- --- --- --- --- --- --- --- --- --- --- --t taa gac t-- --- --- --- --- --- --- --- --- --- --- --- --- --- --- --- --- --- --- --- --- --- --- --- --- --- --- --- --- --- --- --- --- --- --- --- --- --- --- --- taa act agt --- --- --- --- --- --- --- --- --- --- --- --- --- --- --- --- --- --- --- --- --- --- --- --- --- --- --- --- --- --- --- --- --- --- --- --- --- --- --- --- --- --- --- --- --- --- --- --- --- --- --- --- --- --- --- --- --- --- --- --- --- --- --- --- --- --- --- --- --- --- --- --- --- --- --- --- --- --- --- --- --- --- --- --- --- --- --- --- --- --- --- --- --- --- --- --- --- --- --- --- --- --- --- --- --- --- --- --- --- --t atg gta atg acg atg aaa --- -tc gag tag ta- --- --a ctt cta ata aag act tga ta- --- --t cat tat cat atg ttt gat cg- --- --- --- --- --- --- --t cat agt taa tag tgt g-- --- --- --- --- --- --- --- --- --- --- --- --- --- --- --- --- --- --- --- -gc taa atg gta ctg tta ata agt tta t-- --- --- --- --- --- --- --- --- --- --- --- --- --- --- --- --- --a gac aat atc ata gta ttt tct ttc cag aat --- --- --- --t aga tta ttt ttt taa ata ctg atc ctc aca att ccg tga tgt agc agt agt tgg t-- --- --- --- --- --- --- --- --- --- --- --- --- --- --- --- --- --- --- --- --- --- --- --- --- --- --- --- -gc atg gtc tat atc gt- --- --- --- --- --- --- --- --- --- --- --- --- --- --- --- --- --- --- --- --- --- --- --- --- --- --- --- --- --- --- --- --- --- --- --- --- --- --- --- --- --- --- --- taa aat gta tca tat ata ata gtt ttc tga cgt gga gta cag aat ttt cga --- --- --- --- --- --- --- --- --- --- --- --- --- --- --- --- --- --- --- --- --- --- --- --- --- --- --- --- --- --- --- --- --- --- --- --- --- --- --- --- --- --- --- --t taa tga gtt cat ggt aag gaa ggg caa atg cct --- gta tat aat ata cat aag tta a-- --- --- --- --- --- --- --- --- --- --- --- --- --t agt ttt tta tca tat tt- --- --- --- --- --- --- --- --t cta ata cca taa taa aaa t-- --- --- --- --- --- --- --- --- --- --- --- --- --- --- --- --- --- --- --- --- --- --- --- --- --t atc att att gcg ttt g-g tag tt- --- --- --- --- ctg ccc ta- --- --- --- --- --- --- --- --t cat cta tat cac tgt cac tct c-- --- --g ctc tca cta tat ctt cta aaa tta caa --- -ac aac --- --- --- --- --- --- --- --- --- --- --- --- --- --- --- --- --- --- --- --- --- --- --- --- --- --- --- --- --- --- --- --- --- --- --- --- tgg ata ttc gat --- --- --- --- --- --- --- --- --- --- --- --- --- --- --- --- --- --- --- --- --- --- --- --- --- --- --a aca gca ttt gtg t-- --- --- --- --- --- --- --- --- --- --- --- --- --- --- --- --- -

>ON682267.1_Monkeypox_virus_isolate_MPXV/Germany/2022/RKI010_complete_genome

--- --- --- --- --- --- --- --- --- --- --- --- --- --- --- --- --- --- --- --- --- --- --- --- --- --- --- --- --- --a ttt tac tat ttt att tag tgt cta gaa aaa aat gtg tga ccc acg acc gta gga aac tct aga ggg taa gaa aaa tca atc gtt tat aga gac cat cag aaa gag gtt taa tat ttt tgt gag acc tat cga a-- --- --- --- --- --- --- --- --- --- --- --- --- --- --- --- --- --- --- --- --- --- --- --- --- --- --- --- --- --- --- --- --- --- --- --- --- --- --- --- --- --- --- --- --- --- --- --- --- --- --- --- --- --- --- --- --- --- --- --- --- --- --- --- --- --- --- --- --- --- --- --- --- --- --- --- --- --- --- --- --- --- --- --- --- --- --- --- --- --- --- --- --- --- --- --- --- --- --- --- --- --- --- --- --- --- --- --- --- --- --- --- --- --- --- --- --- --- --- --- --- --- --- --- --- --- --- --- --- --- --- --- --- --- --- --- --- --- --- --- --- --- --- --- --- --- --- --- --- --- --- --- --- --- --- --- --- --- --- --- --- --- --- --- --- --- --- --- --- --- --- --- --- --- --- --- --- --- --- --- --- --- --- --- --- --- --- --- --- --- --- --- --- --- --- --- --- --- --- --- --- --- --- --- --- --- --- --- --- --- --- --- --- --- --- --- --- --- --- --- --- --- --- --- --- --- --- --- --- --- --- --- --- --- --- --- --- --- --- --- --- --- --- --- --- --- --- --- --- --- --- --- --- --- --- --- --- --- --- --- --- --- --- --- --- --- --- --- --- --- --- --- --- --- --- --- --- --- --- --- -ga gag aaa gga taa aaa ctt ttt acg act cca tca gaa aga ggt tta ata ttt ttg tga gac cca tcg aag aga gaa aga gat ggt tag tca aga tat ttt tct tag tac aaa agt caa tgt ttt aaa ata tat gga cga gaa tta att tgt ctg tat aaa aac ttg tgt gaa att atg tac tag aga aaa aac gtg agc agt gtc ccc tac atg gat ttt aca gat cat tta tat tcc aaa aat att aac tat ata cgt tta tta tat gat gtt aac gtg taa att ata aac att att tta tga tgc aat tgt ctg aca acc tag att ggt ata agg atg ttg ata agc tct acg aga ata tat tgt tgg acg tta tcg ttt acg aaa tag ttg aga cat cag aaa gag gtt taa tat ttt tgt gag a-c cat cga aga gag aaa gag aat aaa aat a-- --t ttt ttt ttt ttt tgt aaa act ttt tta tga gac --- --- -ca aga gaa tac gaa tag tga tca tat cgt atc aca tat tga aac aga aag aag aag taa cga gag gta act ttt tgt gaa tgt agt taa ata ttt ttg ttt tgc aaa ccg gaa tat agt gcc cgg t-- -ct ttt tta att cgt ggt gcg gtg tct gaa tcg ttc gat taa ccc aac tca tcc att ttc aga tga ata gag tta tcg att cag aca cat gct ttg agt ttt gtt gaa tcg atg agt gaa gta tca tcg gtt gca cct tca gat gcc gat ccg tcg aca tac ttg aat cca tcc ttg act tca agt tca gat gat tcc tca cac atg tct ccg ata cgt acg cta aac tct agg ttc ttg aca cat ttt gta tca acg atc gtt gaa ccg atg ata tct ttg taa ctc act ttc tta tgt gag atg tta gac cca agt act gga tgg gtc ttg atg tca ctg tct ttc tct tct tcg cta cat ctg atg tcg ata gac atc tca cag tct ttg atc ata gcc aga gct tct tca cgc gtg atc gcg gga gag tcc tta cct tgt ccc ggt gac acg ctg gac aat cta gta ttc aca gtg ttt cca tca gag gat tcg gag atg gat aaa atc ttt ggg cat ttg gtg aat cca aag ttc atg tta aga ccc gca ccg acg ata gtg taa taa gtg gtg gga tct cct ttt aca act --- --- --- --- --- --- tct tcg gat acc tca tca tct tcg gtc tct gta act tcc gtt acg gat tga caa atc tta tca ttg gtc ggt gtt tgg tct tgc ttt gtg act ttg ata ata aca tcg att ccc ata tga tgt ttg ttt tct tct tca gta cac gag gat gaa gat tgt tga aga cta gta ggc ata gca gct gcc act agg cac atg cat gcc agg aca ata tat tgt ttc atg att gct att gat tga tta ctg ttc tag atg att cta ctt tct tac cat ata ata aat tag aat ata ttt tct act ttt acg aga aat taa tta ttg tat tta tta ttt ata ggt --a aaa aaa ctt act ata agt ggg tgg gat tct ggg aat tag tga tca gtt tat gta tat cgc aac tag cgg gca tat ggc tat tga cat cga gaa cat tac cca tat gat aag aga ttg tat cat ttt cgt agt ctt gag tat tgg tat tac tat ata gta tgt aga tgt cga cgc tag ata gac agt cgc cca cta gag tta ccg tct ctg aat gcg gca tga tag tat cat tct ttg ttt tcg tta act gtt tgg aag atg aat ctt tgt tgt tac att taa tct cga aat tca gag tac ata tct ttg aag tat tct gat atc tat ttt ctc ctg taa aga atc ctg aag ttg cta cat tat taa gga cag aga agt att ctg cac gaa aga ctg gat cac aat ctt tat gat tca tgg taa tag tta gtt ccg acg ttg aga tgg att cgc tga gac cgg tag tgg tcg tcc gag tac acg atg tgt cgt tga ctg gat aca ggt taa ttt cca cat cga tat agt taa atg tat tgc tgg tta cga cgg gtt cgc att tat ctg tgg aag aga cgg tgt gag aat atg ttc cgg gac cac acg gag aac aga tga cgt ctc cgg tag acg tgt atc cgg ata ctc cgt atc cta ttc cac act ttg ttt tag aaa tac atg ttc tac acc ctg atg ctc ctt tga gaa gac aat aat atc ctg gag agc att cac aga ttc tat tgt gag tcg tgt tac acg atc gcg tct cta cct gat tac tat cac atc ttc cgt tac aac tta gac aag cct gta aat gat tat tgt gag atg taa agg tat ccg aac cac acg gtg tac att gtg tat tag tct tgc tat cac ata atc tga aag cgt aag ttc ccg gag gac acg ata gac aac ata gat tac ggc ttc tgt att cgt tgt ctt tac act ttc cat tgg atg gtg cat gtg gtg cta tat ctc ttc cgt tta tta tta tac atg aga gaa aca ata tat acg agt ata ata cgg acc tca tga ttt aat aat -gt agt aat cgt cgt ctt gtt act gtt tgt ttc cta ctt ctc caa tca tat aga tta ttt ttt aaa tat ttt ctt tct atc atg gat aat att tgt aat ggt tct ttc cgt aca aca tac tgt tta gat ggt agt cgc tta gct tgg tta tga tat tgc gca taa ttt ccg gag gca aat acg ata gtc tag att gac tat cga tgg tag act cta att tat tga gtg ctt tgt cga cga gtt tac ttt tat gct cca tcg ata gat gac act gtt cta tga gat cgt cgt aca tgg gaa atg aaa tgc gtt tgt ctg aat gta tgg ctt cga gat atg tgt gat acc gaa tgt ctt ctg ttc tca ata ccg tat aca agt cgg tgt ctg aga ttc gaa tct ctt tga gga gac tta tgt cac gac tac att ttt cga tga tgg aat cta tct tat cga atg ata tat ttt tca taa ata cac ttt tat agt cct cgt tta aac aga att tag tat gta gtt ccg caa atg act cgt ccc tta ata ggc agt agg cta tta tct tct tta cgt agt gat cgt cgt agg gag aga act ccg aca tct tgt aga aca acg att taa tca tag gta gag ata ctt tca gtc tgt ggt gga taa tgt cat taa cga cat ccg cct tgt ata taa tgt ttc tgt ttt caa aca cca agt cga ata ctg tct tat cgt ctt tag tcg gaa ggt tga tgt cgt atc caa tgt ata cga ggt atg agg caa cat tgt tat tgc aat tct gga agg cgg tat gaa gag gag tca ttg tat tat agt att cgt ctt tct gaa tgt cga atc tat cta gta gat acc gta gta tat tga gag agc gac ttc cat atc ctt gat tat gtt tta tga ata gat aaa gta gat gtt gtc ctt ctt cct ttt gta att tcc cgt att ttt gtt cgt gcc aat tga gta aca tta tga gaa tat gac ctg ttg cac aat cgt tct tta tgt att cca tga tgg gtg tac aat taa gat tat tac gta tcc tcg tat cgg ctc ctc gag ata aaa gag cat aca cca cac gag gac tat gtt tgg tat act gtt gaa ggt aag tgt gta acg gcg tat ttc cga ttt tcg taa ccg cgt taa tgt ttg ctc cat gat cta tta tcg cgt aga tga atc gct tct cag ctc gca tct tag tgt gac tct ttg act tgt aat aat tgc ttt cgt gga acg cgg ata tgt gtt tac agt agt aat gaa gag aag tga gtc cat cct cat cga cgc aat tag ggt cag atc ctt tag tca ata att tgt aca gaa cgt aat agt tta agc tcc cat tga att tat atc taa gat aac aca gca ata gat cgg atg att tac taa agt cat caa tgg ggt ccg tta gta tat caa aga tct tgt tat cga ttg ata gtg aat gaa tca gat agt ggt gta gag gaa tat gtc ctt ttt tat cct tgc tat caa agt tac gca tgc cgt ggc gta aca ata tct tta ata cag atg gat taa atc gtg tat tca tcg tat agc aat gta atg gag agt tac cac att tta gtc gtt tat tca gat cgc agt gtt taa taa cta att taa aca gat gag atg ata tat cca cat caa aga atg cga gat aca tat gac aga cat tat tga cag aaa tgt gac ctt cat tat cac cgt cgt cca taa atg cgt tag gta cgt acc aca tac tat cgt taa cga tgc gca caa tct cgt cca -tt tca tcc atc ttc ata atg att tac ttt ttc ata att aga gaa aaa gat caa ggt ata aaa att aga agt gtt aga cta ta- --- --- --- --- --- --- --- --- --- --- -aa cta act tat gac tta act aac tta tga --- --- --- --- --- --- --- --- --- --- --- --- --- --- --- --- --- --- --- --- --- --- --- --- --- --- --- --- --- --- --- --- --- --- --- --- --- --- --- --- --- --- --- --- --- --- --- --- ctt aac ta- --- --- --- --- --- --- --- --- --- --- -ac tta tga ctt aac taa ctt atg act taa cta act ta- tga ctt aac taa ctt atg act taa cta act tat gac tta act aac tta tga cta tta act cat ttc aag aaa ggt ggg tgg ata gaa ctc tat atg aca gct tgt gaa aca att aga tcc cta att tct aat gga agt ttt gat agg aga ttg tca tca gtt gat aca ttg ttt att atc tca tct att aga gca cgt ctg ttt aga gct tta gtg acc tgc tcg gtt act tct gtg taa atc ttg aat cct tta gtg ata cac tgt gtc aaa act gga tgt tta gaa tac cta tgt aga ata tgg gaa gca tgc ttg ttt ttg tct cta tta tag att aac tca tac atg gtt gta tta tga att ttc atc tgc cta atg tac tcc aat tct tgt tta caa tca att ata taa tca aag agt gat gat gca tac aca tta caa agt gaa taa tct acc atc ata aaa tac ttg ata cag agc ttt atc aca tca tgg ttt tca att gta tta tta agt ata gct aat ttt ata cag tca ata gac aat ggt tct cta agc aat att tct aat att tta aga tgt gct tcc cta cgg gcg atg aca gat ccc cta tcc acg gcc acg tca aga cat gta tat cca tta ctc att act gcg ttg aca ttt gct cca ttt tct aat agc cat gat act aaa tct ata taa cct gca tag ata gcg cga taa agc aag gtc ctt cca cca gca tct agt tga ttg ata tct tca ata tat ggg ata caa agc tta taa att tct aat act gtg ggt tca tct aca agg aat ccc cta gta tac tga att att tta tat aga tct aat tta aca tca ttt tca tct ggg ata cca caa ttc aaa ata aac tca aca aca cta ctt tcc ttt tta cat att ccc cta aaa tag gca ttc aag cat tct att tta tat att aca gcc cca tga tct acc ata aga tca aca atg tct att tct aca tat gca tta gat aga tag taa agt aag aga tct tgc aca gaa tta caa ttc tta ata att ata gag aaa ata tct tcc ata taa ttc ttt gac act aat gca gat ata ata tct tta tat gta ata tat gca aac agt cta tct act ata tac tga tca ata tta tct cta tga atc cta aaa taa tca tac aga aca tct aca gga tca caa att ggt tca agg aga aat cta tca aat att ttc ctg tca aca act ggt tct aga aca taa cag tca aca cct aat cca tgt ttt tta taa tca tct acc aaa gat aat gac caa aga tcg agg tcg tcg tga aac tgc tca tcg aca gcc atg aaa tct acc gac tcc atg gtg cga atc gca ctg tct tat tcg cca ttg att ttc a-t ttt tta taa tta tgt aca tgt ttt cct tct att ctc aag agt cta caa aaa tat a-- -tt ttt tcg ata tct aag tac taa gtt ttt tta ctg ttt ttg tta ctg tct tcc att ctt cta act aaa gat ctg aga taa att ata caa tct tcg cta tcg aac cat ttt tgt agt cta aag cct gaa gta att aac caa ctg ttt tta tta gtg gct ttt ttc gat cta tcc tcg tc- --- ctc tca tca tcc tta ta- --- --- --- --- --- --- tta tta tca tta tta tca tag tct att aaa cac aaa tca tct --- --- --- --- --- --- --- --- --- --- --- --- --- --- --- --- --- --- --- --- --- --- --- --- --- --- --a cgt tta taa c-- --- --- --- --- --- --- --- --a aca ttc --- --- --- --- -tc att att aat tag ttc tgt ag- aat atc ttt aat aat ttg gct ata --- --- -ca tct gtt --- --- --- --- --- --- --- --- --- --- --- --- --- --- --- --- --- --- --c aat act --- --- --- --- --- --- --- --a tct att gat gat ttc ttt t-- --- --- --- --- --- --- --- --- --- --- --t taa gac t-- --- --- --- --- --- --- --- --- --- --- --- --- --- --- --- --- --- --- --- --- --- --- --- --- --- --- --- --- --- --- --- --- --- --- --- --- --- --- --- taa act agt --- --- --- --- --- --- --- --- --- --- --- --- --- --- --- --- --- --- --- --- --- --- --- --- --- --- --- --- --- --- --- --- --- --- --- --- --- --- --- --- --- --- --- --- --- --- --- --- --- --- --- --- --- --- --- --- --- --- --- --- --- --- --- --- --- --- --- --- --- --- --- --- --- --- --- --- --- --- --- --- --- --- --- --- --- --- --- --- --- --- --- --- --- --- --- --- --- --- --- --- --- --- --- --- --- --- --- --- --- --t atg gta atg acg atg aaa --- -tc gag tag ta- --- --a ctt cta ata aag act tga ta- --- --t cat tat cat atg ttt gat cg- --- --- --- --- --- --- --t cat agt taa tag tgt g-- --- --- --- --- --- --- --- --- --- --- --- --- --- --- --- --- --- --- --- -gc taa atg gta ctg tta ata agt tta t-- --- --- --- --- --- --- --- --- --- --- --- --- --- --- --- --- --a gac aat atc ata gta ttt tct ttc cag aat --- --- --- --t aga tta ttt ttt taa ata ctg atc ctc aca att ccg tga tgt agc agt agt tgg t-- --- --- --- --- --- --- --- --- --- --- --- --- --- --- --- --- --- --- --- --- --- --- --- --- --- --- --- -gc atg gtc tat atc gt- --- --- --- --- --- --- --- --- --- --- --- --- --- --- --- --- --- --- --- --- --- --- --- --- --- --- --- --- --- --- --- --- --- --- --- --- --- --- --- --- --- --- --- taa aat gta tca tat ata ata gtt ttc tga cgt gga gta cag aat ttt cga --- --- --- --- --- --- --- --- --- --- --- --- --- --- --- --- --- --- --- --- --- --- --- --- --- --- --- --- --- --- --- --- --- --- --- --- --- --- --- --- --- --- --- --t taa tga gtt cat ggt aag gaa ggg caa atg cct --- gta tat aat ata cat aag tta a-- --- --- --- --- --- --- --- --- --- --- --- --- --t agt ttt tta tca tat tt- --- --- --- --- --- --- --- --t cta ata cca taa taa aaa t-- --- --- --- --- --- --- --- --- --- --- --- --- --- --- --- --- --- --- --- --- --- --- --- --- --t atc att att gcg ttt g-g tag tt- --- --- --- --- ctg ccc ta- --- --- --- --- --- --- --- --t cat cta tat cac tgt cac tct c-- --- --g ctc tca cta tat ctt cta aaa tta caa --- -ac aac --- --- --- --- --- --- --- --- --- --- --- --- --- --- --- --- --- --- --- --- --- --- --- --- --- --- --- --- --- --- --- --- --- --- --- --- tgg ata ttc gat --- --- --- --- --- --- --- --- --- --- --- --- --- --- --- --- --- --- --- --- --- --- --- --- --- --- --a aca gca ttt gtg t-- --- --- --- --- --- --- --- --- --- --- --- --- --- --- --- --- -

>ON649717.1_Monkeypox_virus_isolate_Monkeypox/PT0019/2022_partial_genome

--- --- --- --- --- --- --- --- --- --- --- --- --- --- --- --- --- --- --- --- --- --- --- --- --- --- --- --- att taa ttt tac tat ttt att tag tgt cta gaa aaa aat gtg tga ccc acg acc gta gga aac tct aga ggg taa gaa aaa tca atc gtt tat aga gac cat cag aaa gag gtt taa tat ttt tgt gag acc tat cga a-- --- --- --- --- --- --- --- --- --- --- --- --- --- --- --- --- --- --- --- --- --- --- --- --- --- --- --- --- --- --- --- --- --- --- --- --- --- --- --- --- --- --- --- --- --- --- --- --- --- --- --- --- --- --- --- --- --- --- --- --- --- --- --- --- --- --- --- --- --- --- --- --- --- --- --- --- --- --- --- --- --- --- --- --- --- --- --- --- --- --- --- --- --- --- --- --- --- --- --- --- --- --- --- --- --- --- --- --- --- --- --- --- --- --- --- --- --- --- --- --- --- --- --- --- --- --- --- --- --- --- --- --- --- --- --- --- --- --- --- --- --- --- --- --- --- --- --- --- --- --- --- --- --- --- --- --- --- --- --- --- --- --- --- --- --- --- --- --- --- --- --- --- --- --- --- --- --- --- --- --- --- --- --- --- --- --- --- --- --- --- --- --- --- --- --- --- --- --- --- --- --- --- --- --- --- --- --- --- --- --- --- --- --- --- --- --- --- --- --- --- --- --- --- --- --- --- --- --- --- --- --- --- --- --- --- --- --- --- --- --- --- --- --- --- --- --- --- --- --- --- --- --- --- --- --- --- --- --- --- --- --- --- --- --- --- --- --- --- --- --- --- --- --- --- --- --- --- --- --- -ga gag aaa gga taa aaa ctt ttt acg act cca tca gaa aga ggt tta ata ttt ttg tga gac cca tcg aag aga gaa aga gat ggt tag tca aga tat ttt tct tag tac aaa agt caa tgt ttt aaa ata tat gga cga gaa tta att tgt ctg tat aaa aac ttg tgt gaa att atg tac tag aga aaa aac gtg agc agt gtc ccc tac atg gat ttt aca gat cat tta tat tcc aaa aat att aac tat ata cgt tta tta tat gat gtt aac gtg taa att ata aac att att tta tga tgc aat tgt ctg aca acc tag att ggt ata agg atg ttg ata agc tct acg aga ata tat tgt tgg acg tta tcg ttt acg aaa tag ttg aga cat cag aaa gag gtt taa tat ttt tgt gag a-c cat cga aga gag aaa gag aat aaa aat a-- --t ttt ttt ttt ttt tgt aaa act ttt tta tga gac --- --- -ca aga gaa tac gaa tag tga tca tat cgt atc aca tat tga aac aga aag aag aag taa cga gag gta act ttt tgt gaa tgt agt taa ata ttt ttg ttt tgc aaa ccg gaa tat agt gcc cgg t-- -ct ttt tta att cgt ggt gcg gtg tct gaa tcg ttc gat taa ccc aac tca tcc att ttc aga tga ata gag tta tcg att cag aca cat gct ttg agt ttt gtt gaa tcg atg agt gaa gta tca tcg gtt gca cct tca gat gcc gat ccg tcg aca tac ttg aat cca tcc ttg act tca agt tca gat gat tcc tca cac atg tct ccg ata cgt acg cta aac tct agg ttc ttg aca cat ttt gta tca acg atc gtt gaa ccg atg ata tct ttg taa ctc act ttc tta tgt gag atg tta gac cca agt act gga tgg gtc ttg atg tca ctg tct ttc tct tct tcg cta cat ctg atg tcg ata gac atc tca cag tct ttg atc ata gcc aga gct tct tca cgc gtg atc gcg gga gag tcc tta cct tgt ccc ggt gac acg ctg gac aat cta gta ttc aca gtg ttt cca tca gag gat tcg gag atg gat aaa atc ttt ggg cat ttg gtg aat cca aag ttc atg tta aga ccc gca ccg acg ata gtg taa taa gtg gtg gga tct cct ttt aca act --- --- --- --- --- --- tct tcg gat acc tca tca tct tcg gtc tct gta act tcc gtt acg gat tga caa atc tta tca ttg gtc ggt gtt tgg tct tgc ttt gtg act ttg ata ata aca tcg att ccc ata tga tgt ttg ttt tct tct tca gta cac gag gat gaa gat tgt tga aga cta gta ggc ata gca gct gcc act agg cac atg cat gcc agg aca ata tat tgt ttc atg att gct att gat tga tta ctg ttc tag atg att cta ctt tct tac cat ata ata aat tag aat ata ttt tct act ttt acg aga aat taa tta ttg tat tta tta ttt ata ggt --a aaa aaa ctt act ata agt ggg tgg gat tct ggg aat tag tga tca gtt tat gta tat cgc aac tag cgg gca tat ggc tat tga cat cga gaa cat tac cca tat gat aag aga ttg tat cat ttt cgt agt ctt gag tat tgg tat tac tat ata gta tgt aga tgt cga cgc tag ata gac agt cgc cca cta gag tta ccg tct ctg aat gcg gca tga tag tat cat tct ttg ttt tcg tta act gtt tgg aag atg aat ctt tgt tgt tac att taa tct cga aat tca gag tac ata tct ttg aag tat tct gat atc tat ttt ctc ctg taa aga atc ctg aag ttg cta cat tat taa gga cag aga agt att ctg cac gaa aga ctg gat cac aat ctt tat gat tca tgg taa tag tta gtt ccg acg ttg aga tgg att cgc tga gac cgg tag tgg tcg tcc gag tac acg atg tgt cgt tga ctg gat aca ggt taa ttt cca cat cga tat agt taa atg tat tgc tgg tta cga cgg gtt cgc att tat ctg tgg aag aga cgg tgt gag aat atg ttc cgg gac cac acg gag aac aga tga cgt ctc cgg tag acg tgt atc cgg ata ctc cgt atc cta ttc cac act ttg ttt tag aaa tac atg ttc tac acc ctg atg ctc ctt tga gaa gac aat aat atc ctg gag agc att cac aga ttc tat tgt gag tcg tgt tac acg atc gcg tct cta cct gat tac tat cac atc ttc cgt tac aac tta gac aag cct gta aat gat tat tgt gag atg taa agg tat ccg aac cac acg gtg tac att gtg tat tag tct tgc tat cac ata atc tga aag cgt aag ttc ccg gag gac acg ata gac aac ata gat tac ggc ttc tgt att cgt tgt ctt tac act ttc cat tgg atg gtg cat gtg gtg cta tat ctc ttc cgt tta tta tta tac atg aga gaa aca ata tat acg agt ata ata cgg acc tca tga ttt aat aat -gt agt aat cgt cgt ctt gtt act gtt tgt ttc cta ctt ctc caa tca tat aga tta ttt ttt aaa tat ttt ctt tct atc atg gat aat att tgt aat ggt tct ttc cgt aca aca tac tgt tta gat ggt agt cgc tta gct tgg tta tga tat tgc gca taa ttt ccg gag gca aat acg ata gtc tag att gac tat cga tgg tag act cta att tat tga gtg ctt tgt cga cga gtt tac ttt tat gct cca tcg ata gat gac act gtt cta tga gat cgt cgt aca tgg gaa atg aaa tgc gtt tgt ctg aat gta tgg ctt cga gat atg tgt gat acc gaa tgt ctt ctg ttc tca ata ccg tat aca agt cgg tgt ctg aga ttc gaa tct ctt tga gga gac tta tgt cac gac tac att ttt cga tga tgg aat cta tct tat cga atg ata tat ttt tca taa ata cac ttt tat agt cct cgt tta aac aga att tag tat gta gtt ccg caa atg act cgt ccc tta ata ggc agt agg cta tta tct tct tta cgt agt gat cgt cgt agg gag aga act ccg aca tct tgt aga aca acg att taa tca tag gta gag ata ctt tca gtc tgt ggt gga taa tgt cat taa cga cat ccg cct tgt ata taa tgt ttc tgt ttt caa aca cca agt cga ata ctg tct tat cgt ctt tag tcg gaa ggt tga tgt cgt atc caa tgt ata cga ggt atg agg caa cat tgt tat tgc aat tct gga agg cgg tat gaa gag gag tca ttg tat tat agt att cgt ctt tct gaa tgt cga atc tat cta gta gat acc gta gta tat tga gag agc gac ttc cat atc ctt gat tat gtt tta tga ata gat aaa gta gat gtt gtc ctt ctt cct ttt gta att tcc cgt att ttt gtt cgt gcc aat tga gta aca tta tga gaa tat gac ctg ttg cac aat cgt tct tta tgt att cca tga tgg gtg tac aat taa gat tat tac gta tcc tcg tat cgg ctc ctc gag ata aaa gag cat aca cca cac gag gac tat gtt tgg tat act gtt gaa ggt aag tgt gta acg gcg tat ttc cga ttt tcg taa ccg cgt taa tgt ttg ctc cat gat cta tta tcg cgt aga tga atc gct tct cag ctc gca tct tag tgt gac tct ttg act tgt aat aat tgc ttt cgt gga acg cgg ata tgt gtt tac agt agt aat gaa gag aag tga gtc cat cct cat cga cgc aat tag ggt cag atc ctt tag tca ata att tgt aca gaa cgt aat agt tta agc tcc cat tga att tat atc taa gat aac aca gca ata gat cgg atg att tac taa agt cat caa tgg ggt ccg tta gta tat caa aga tct tgt tat cga ttg ata gtg aat gaa tca gat agt ggt gta gag gaa tat gtc ctt ttt tat cct tgc tat caa agt tac gca tgc cgt ggc gta aca ata tct tta ata cag atg gat taa atc gtg tat tca tcg tat agc aat gta atg gag agt tac cac att tta gtc gtt tat tca gat cgc agt gtt taa taa cta att taa aca gat gag atg ata tat cca cat caa aga atg cga gat aca tat gac aga cat tat tga cag aaa tgt gac ctt cat tat cac cgt cgt cca taa atg cgt tag gta cgt acc aca tac tat cgt taa cga tgc gca caa tct cgt cca -tt tca tcc atc ttc ata atg att tac ttt ttc ata att aga gaa aaa gat caa ggt ata aaa att aga agt gtt aga cta ta- --- --- --- --- --- --- --- --- --- --- -aa cta act tat gac tta act aac tta tga --- --- --- --- --- --- --- --- --- --- --- --- --- --- --- --- --- --- --- --- --- --- --- --- --- --- --- --- --- --- --- --- --- --- --- --- --- --- --- --- --- --- --- --- --- --- --- --- ctt aac ta- --- --- --- --- --- --- --- --- --- --- -ac tta tga ctt aac taa ctt atg act taa cta act ta- tga ctt aac taa ctt atg act taa cta act tat gac tta act aac tta tga cta tta act cat ttc aag aaa ggt ggg tgg ata gaa ctc tat atg aca gct tgt gaa aca att aga tcc cta att tct aat gga agt ttt gat agg aga ttg tca tca gtt gat aca ttg ttt att atc tca tct att aga gca cgt ctg ttt aga gct tta gtg acc tgc tcg gtt act tct gtg taa atc ttg aat cct tta gtg ata cac tgt gtc aaa act gga tgt tta gaa tac cta tgt aga ata tgg gaa gca tgc ttg ttt ttg tct cta tta tag att aac tca tac atg gtt gta tta tga att ttc atc tgc cta atg tac tcc aat tct tgt tta caa tca att ata taa tca aag agt gat gat gca tac aca tta caa agt gaa taa tct acc atc ata aaa tac ttg ata cag agc ttt atc aca tca tgg ttt tca att gta tta tta agt ata gct aat ttt ata cag tca ata gac aat ggt tct cta agc aat att tct aat att tta aga tgt gct tcc cta cgg gcg atg aca gat ccc cta tcc acg gcc acg tca aga cat gta tat cca tta ctc att act gcg ttg aca ttt gct cca ttt tct aat agc cat gat act aaa tct ata taa cct gca tag ata gcg cga taa agc aag gtc ctt cca cca gca tct agt tga ttg ata tct tca ata tat ggg ata caa agc tta taa att tct aat act gtg ggt tca tct aca agg aat ccc cta gta tac tga att att tta tat aga tct aat tta aca tca ttt tca tct ggg ata cca caa ttc aaa ata aac tca aca aca cta ctt tcc ttt tta cat att ccc cta aaa tag gca ttc aag cat tct att tta tat att aca gcc cca tga tct acc ata aga tca aca atg tct att tct aca tat gca tta gat aga tag taa agt aag aga tct tgc aca gaa tta caa ttc tta ata att ata gag aaa ata tct tcc ata taa ttc ttt gac act aat gca gat ata ata tct tta tat gta ata tat gca aac agt cta tct act ata tac tga tca ata tta tct cta tga atc cta aaa taa tca tac aga aca tct aca gga tca caa att ggt tca agg aga aat cta tca aat att ttc ctg tca aca act ggt tct aga aca taa cag tca aca cct aat cca tgt ttt tta taa tca tct acc aaa gat aat gac caa aga tcg agg tcg tcg tga aac tgc tca tcg aca gcc atg aaa tct acc gac tcc atg gtg cga atc gca ctg tct tat tcg cca ttg att ttc a-t ttt tta taa tta tgt aca tgt ttt cct tct att ctc aag agt cta caa aaa tat a-- -tt ttt tcg ata tct aag tac taa gtt ttt tta ctg ttt ttg tta ctg tct tcc att ctt cta act aaa gat ctg aga taa att ata caa tct tcg cta tcg aac cat ttt tgt agt cta aag cct gaa gta att aac caa ctg ttt tta tta gtg gct ttt ttc gat cta tcc tcg tc- --- ctc tca tca tcc tta ta- --- --- --- --- --- --- tta tta tca tta tta tca tag tct att aaa cac aaa tca tct --- --- --- --- --- --- --- --- --- --- --- --- --- --- --- --- --- --- --- --- --- --- --- --- --- --- --a cgt tta taa c-- --- --- --- --- --- --- --- --a aca ttc --- --- --- --- -tc att att aat tag ttc tgt ag- aat atc ttt aat aat ttg gct ata --- --- -ca tct gtt --- --- --- --- --- --- --- --- --- --- --- --- --- --- --- --- --- --- --c aat act --- --- --- --- --- --- --- --a tct att gat gat ttc ttt t-- --- --- --- --- --- --- --- --- --- --- --t taa gac t-- --- --- --- --- --- --- --- --- --- --- --- --- --- --- --- --- --- --- --- --- --- --- --- --- --- --- --- --- --- --- --- --- --- --- --- --- --- --- --- taa act agt --- --- --- --- --- --- --- --- --- --- --- --- --- --- --- --- --- --- --- --- --- --- --- --- --- --- --- --- --- --- --- --- --- --- --- --- --- --- --- --- --- --- --- --- --- --- --- --- --- --- --- --- --- --- --- --- --- --- --- --- --- --- --- --- --- --- --- --- --- --- --- --- --- --- --- --- --- --- --- --- --- --- --- --- --- --- --- --- --- --- --- --- --- --- --- --- --- --- --- --- --- --- --- --- --- --- --- --- --- --t atg gta atg acg atg aaa --- -tc gag tag ta- --- --a ctt cta ata aag act tga ta- --- --t cat tat cat atg ttt gat cg- --- --- --- --- --- --- --t cat agt taa tag tgt g-- --- --- --- --- --- --- --- --- --- --- --- --- --- --- --- --- --- --- --- -gc taa atg gta ctg tta ata agt tta t-- --- --- --- --- --- --- --- --- --- --- --- --- --- --- --- --- --a gac aat atc ata gta ttt tct ttc cag aat --- --- --- --t aga tta ttt ttt taa ata ctg atc ctc aca att ccg tga tgt agc agt agt tgg t-- --- --- --- --- --- --- --- --- --- --- --- --- --- --- --- --- --- --- --- --- --- --- --- --- --- --- --- -gc atg gtc tat atc gt- --- --- --- --- --- --- --- --- --- --- --- --- --- --- --- --- --- --- --- --- --- --- --- --- --- --- --- --- --- --- --- --- --- --- --- --- --- --- --- --- --- --- --- taa aat gta tca tat ata ata gtt ttc tga cgt gga gta cag aat ttt cga --- --- --- --- --- --- --- --- --- --- --- --- --- --- --- --- --- --- --- --- --- --- --- --- --- --- --- --- --- --- --- --- --- --- --- --- --- --- --- --- --- --- --- --t taa tga gtt cat ggt aag gaa ggg caa atg cct --- gta tat aat ata cat aag tta a-- --- --- --- --- --- --- --- --- --- --- --- --- --t agt ttt tta tca tat tt- --- --- --- --- --- --- --- --t cta ata cca taa taa aaa t-- --- --- --- --- --- --- --- --- --- --- --- --- --- --- --- --- --- --- --- --- --- --- --- --- --t atc att att gcg ttt g-g tag tt- --- --- --- --- ctg ccc ta- --- --- --- --- --- --- --- --t cat cta tat cac tgt cac tct c-- --- --g ctc tca cta tat ctt cta aaa tta caa --- -ac aac --- --- --- --- --- --- --- --- --- --- --- --- --- --- --- --- --- --- --- --- --- --- --- --- --- --- --- --- --- --- --- --- --- --- --- --- tgg ata ttc gat --- --- --- --- --- --- --- --- --- --- --- --- --- --- --- --- --- --- --- --- --- --- --- --- --- --- --a aca gca ttt gtg t-- --- --- --- --- --- --- --- --- --- --- --- --- --- --- --- --- -

>ON649708.1_Monkeypox_virus_isolate_Monkeypox/PT0023/2022_partial_genome

--- --- --- --- --- --- --- --- --- --- --- --- --- --- --- --- --- --- --- --- --- --- --- --- --- --- --- --- att taa ttt tac tat ttt att tag tgt cta gaa aaa aat gtg tga ccc acg acc gta gga aac tct aga ggg taa gaa aaa tca atc gtt tat aga gac cat cag aaa gag gtt taa tat ttt tgt gag acc tat cga a-- --- --- --- --- --- --- --- --- --- --- --- --- --- --- --- --- --- --- --- --- --- --- --- --- --- --- --- --- --- --- --- --- --- --- --- --- --- --- --- --- --- --- --- --- --- --- --- --- --- --- --- --- --- --- --- --- --- --- --- --- --- --- --- --- --- --- --- --- --- --- --- --- --- --- --- --- --- --- --- --- --- --- --- --- --- --- --- --- --- --- --- --- --- --- --- --- --- --- --- --- --- --- --- --- --- --- --- --- --- --- --- --- --- --- --- --- --- --- --- --- --- --- --- --- --- --- --- --- --- --- --- --- --- --- --- --- --- --- --- --- --- --- --- --- --- --- --- --- --- --- --- --- --- --- --- --- --- --- --- --- --- --- --- --- --- --- --- --- --- --- --- --- --- --- --- --- --- --- --- --- --- --- --- --- --- --- --- --- --- --- --- --- --- --- --- --- --- --- --- --- --- --- --- --- --- --- --- --- --- --- --- --- --- --- --- --- --- --- --- --- --- --- --- --- --- --- --- --- --- --- --- --- --- --- --- --- --- --- --- --- --- --- --- --- --- --- --- --- --- --- --- --- --- --- --- --- --- --- --- --- --- --- --- --- --- --- --- --- --- --- --- --- --- --- --- --- --- --- --- -ga gag aaa gga taa aaa ctt ttt acg act cca tca gaa aga ggt tta ata ttt ttg tga gac cca tcg aag aga gaa aga gat ggt tag tca aga tat ttt tct tag tac aaa agt caa tgt ttt aaa ata tat gga cga gaa tta att tgt ctg tat aaa aac ttg tgt gaa att atg tac tag aga aaa aac gtg agc agt gtc ccc tac atg gat ttt aca gat cat tta tat tcc aaa aat att aac tat ata cgt tta tta tat gat gtt aac gtg taa att ata aac att att tta tga tgc aat tgt ctg aca acc tag att ggt ata agg atg ttg ata agc tct acg aga ata tat tgt tgg acg tta tcg ttt acg aaa tag ttg aga cat cag aaa gag gtt taa tat ttt tgt gag a-c cat cga aga gag aaa gag aat aaa aat a-- --t ttt ttt ttt ttt tgt aaa act ttt tta tga gac --- --- -ca aga gaa tac gaa tag tga tca tat cgt atc aca tat tga aac aga aag aag aag taa cga gag gta act ttt tgt gaa tgt agt taa ata ttt ttg ttt tgc aaa ccg gaa tat agt gcc cgg t-- -ct ttt tta att cgt ggt gcg gtg tct gaa tcg ttc gat taa ccc aac tca tcc att ttc aga tga ata gag tta tcg att cag aca cat gct ttg agt ttt gtt gaa tcg atg agt gaa gta tca tcg gtt gca cct tca gat gcc gat ccg tcg aca tac ttg aat cca tcc ttg act tca agt tca gat gat tcc tca cac atg tct ccg ata cgt acg cta aac tct agg ttc ttg aca cat ttt gta tca acg atc gtt gaa ccg atg ata tct ttg taa ctc act ttc tta tgt gag atg tta gac cca agt act gga tgg gtc ttg atg tca ctg tct ttc tct tct tcg cta cat ctg atg tcg ata gac atc tca cag tct ttg atc ata gcc aga gct tct tca cgc gtg atc gcg gga gag tcc tta cct tgt ccc ggt gac acg ctg gac aat cta gta ttc aca gtg ttt cca tca gag gat tcg gag atg gat aaa atc ttt ggg cat ttg gtg aat cca aag ttc atg tta aga ccc gca ccg acg ata gtg taa taa gtg gtg gga tct cct ttt aca act --- --- --- --- --- --- tct tcg gat acc tca tca tct tcg gtc tct gta act tcc gtt acg gat tga caa atc tta tca ttg gtc ggt gtt tgg tct tgc ttt gtg act ttg ata ata aca tcg att ccc ata tga tgt ttg ttt tct tct tca gta cac gag gat gaa gat tgt tga aga cta gta ggc ata gca gct gcc act agg cac atg cat gcc agg aca ata tat tgt ttc atg att gct att gat tga tta ctg ttc tag atg att cta ctt tct tac cat ata ata aat tag aat ata ttt tct act ttt acg aga aat taa tta ttg tat tta tta ttt ata ggt --a aaa aaa ctt act ata agt ggg tgg gat tct ggg aat tag tga tca gtt tat gta tat cgc aac tag cgg gca tat ggc tat tga cat cga gaa cat tac cca tat gat aag aga ttg tat cat ttt cgt agt ctt gag tat tgg tat tac tat ata gta tgt aga tgt cga cgc tag ata gac agt cgc cca cta gag tta ccg tct ctg aat gcg gca tga tag tat cat tct ttg ttt tcg tta act gtt tgg aag atg aat ctt tgt tgt tac att taa tct cga aat tca gag tac ata tct ttg aag tat tct gat atc tat ttt ctc ctg taa aga atc ctg aag ttg cta cat tat taa gga cag aga agt att ctg cac gaa aga ctg gat cac aat ctt tat gat tca tgg taa tag tta gtt ccg acg ttg aga tgg att cgc tga gac cgg tag tgg tcg tcc gag tac acg atg tgt cgt tga ctg gat aca ggt taa ttt cca cat cga tat agt taa atg tat tgc tgg tta cga cgg gtt cgc att tat ctg tgg aag aga cgg tgt gag aat atg ttc cgg gac cac acg gag aac aga tga cgt ctc cgg tag acg tgt atc cgg ata ctc cgt atc cta ttc cac act ttg ttt tag aaa tac atg ttc tac acc ctg atg ctc ctt tga gaa gac aat aat atc ctg gag agc att cac aga ttc tat tgt gag tcg tgt tac acg atc gcg tct cta cct gat tac tat cac atc ttc cgt tac aac tta gac aag cct gta aat gat tat tgt gag atg taa agg tat ccg aac cac acg gtg tac att gtg tat tag tct tgc tat cac ata atc tga aag cgt aag ttc ccg gag gac acg ata gac aac ata gat tac ggc ttc tgt att cgt tgt ctt tac act ttc cat tgg atg gtg cat gtg gtg cta tat ctc ttc cgt tta tta tta tac atg aga gaa aca ata tat acg agt ata ata cgg acc tca tga ttt aat aat -gt agt aat cgt cgt ctt gtt act gtt tgt ttc cta ctt ctc caa tca tat aga tta ttt ttt aaa tat ttt ctt tct atc atg gat aat att tgt aat ggt tct ttc cgt aca aca tac tgt tta gat ggt agt cgc tta gct tgg tta tga tat tgc gca taa ttt ccg gag gca aat acg ata gtc tag att gac tat cga tgg tag act cta att tat tga gtg ctt tgt cga cga gtt tac ttt tat gct cca tcg ata gat gac act gtt cta tga gat cgt cgt aca tgg gaa atg aaa tgc gtt tgt ctg aat gta tgg ctt cga gat atg tgt gat acc gaa tgt ctt ctg ttc tca ata ccg tat aca agt cgg tgt ctg aga ttc gaa tct ctt tga gga gac tta tgt cac gac tac att ttt cga tga tgg aat cta tct tat cga atg ata tat ttt tca taa ata cac ttt tat agt cct cgt tta aac aga att tag tat gta gtt ccg caa atg act cgt ccc tta ata ggc agt agg cta tta tct tct tta cgt agt gat cgt cgt agg gag aga act ccg aca tct tgt aga aca acg att taa tca tag gta gag ata ctt tca gtc tgt ggt gga taa tgt cat taa cga cat ccg cct tgt ata taa tgt ttc tgt ttt caa aca cca agt cga ata ctg tct tat cgt ctt tag tcg gaa ggt tga tgt cgt atc caa tgt ata cga ggt atg agg caa cat tgt tat tgc aat tct gga agg cgg tat gaa gag gag tca ttg tat tat agt att cgt ctt tct gaa tgt cga atc tat cta gta gat acc gta gta tat tga gag agc gac ttc cat atc ctt gat tat gtt tta tga ata gat aaa gta gat gtt gtc ctt ctt cct ttt gta att tcc cgt att ttt gtt cgt gcc aat tga gta aca tta tga gaa tat gac ctg ttg cac aat cgt tct tta tgt att cca tga tgg gtg tac aat taa gat tat tac gta tcc tcg tat cgg ctc ctc gag ata aaa gag cat aca cca cac gag gac tat gtt tgg tat act gtt gaa ggt aag tgt gta acg gcg tat ttc cga ttt tcg taa ccg cgt taa tgt ttg ctc cat gat cta tta tcg cgt aga tga atc gct tct cag ctc gca tct tag tgt gac tct ttg act tgt aat aat tgc ttt cgt gga acg cgg ata tgt gtt tac agt agt aat gaa gag aag tga gtc cat cct cat cga cgc aat tag ggt cag atc ctt tag tca ata att tgt aca gaa cgt aat agt tta agc tcc cat tga att tat atc taa gat aac aca gca ata gat cgg atg att tac taa agt cat caa tgg ggt ccg tta gta tat caa aga tct tgt tat cga ttg ata gtg aat gaa tca gat agt ggt gta gag gaa tat gtc ctt ttt tat cct tgc tat caa agt tac gca tgc cgt ggc gta aca ata tct tta ata cag atg gat taa atc gtg tat tca tcg tat agc aat gta atg gag agt tac cac att tta gtc gtt tat tca gat cgc agt gtt taa taa cta att taa aca gat gag atg ata tat cca cat caa aga atg cga gat aca tat gac aga cat tat tga cag aaa tgt gac ctt cat tat cac cgt cgt cca taa atg cgt tag gta cgt acc aca tac tat cgt taa cga tgc gca caa tct cgt cca -tt tca tcc atc ttc ata atg att tac ttt ttc ata att aga gaa aaa gat caa ggt ata aaa att aga agt gtt aga cta ta- --- --- --- --- --- --- --- --- --- --- -aa cta act tat gac tta act aac tta tga --- --- --- --- --- --- --- --- --- --- --- --- --- --- --- --- --- --- --- --- --- --- --- --- --- --- --- --- --- --- --- --- --- --- --- --- --- --- --- --- --- --- --- --- --- --- --- --- ctt aac ta- --- --- --- --- --- --- --- --- --- --- -ac tta tga ctt aac taa ctt atg act taa cta act ta- tga ctt aac taa ctt atg act taa cta act tat gac tta act aac tta tga cta tta act cat ttc aag aaa ggt ggg tgg ata gaa ctc tat atg aca gct tgt gaa aca att aga tcc cta att tct aat gga agt ttt gat agg aga ttg tca tca gtt gat aca ttg ttt att atc tca tct att aga gca cgt ctg ttt aga gct tta gtg acc tgc tcg gtt act tct gtg taa atc ttg aat cct tta gtg ata cac tgt gtc aaa act gga tgt tta gaa tac cta tgt aga ata tgg gaa gca tgc ttg ttt ttg tct cta tta tag att aac tca tac atg gtt gta tta tga att ttc atc tgc cta atg tac tcc aat tct tgt tta caa tca att ata taa tca aag agt gat gat gca tac aca tta caa agt gaa taa tct acc atc ata aaa tac ttg ata cag agc ttt atc aca tca tgg ttt tca att gta tta tta agt ata gct aat ttt ata cag tca ata gac aat ggt tct cta agc aat att tct aat att tta aga tgt gct tcc cta cgg gcg atg aca gat ccc cta tcc acg gcc acg tca aga cat gta tat cca tta ctc att act gcg ttg aca ttt gct cca ttt tct aat agc cat gat act aaa tct ata taa cct gca tag ata gcg cga taa agc aag gtc ctt cca cca gca tct agt tga ttg ata tct tca ata tat ggg ata caa agc tta taa att tct aat act gtg ggt tca tct aca agg aat ccc cta gta tac tga att att tta tat aga tct aat tta aca tca ttt tca tct ggg ata cca caa ttc aaa ata aac tca aca aca cta ctt tcc ttt tta cat att ccc cta aaa tag gca ttc aag cat tct att tta tat att aca gcc cca tga tct acc ata aga tca aca atg tct att tct aca tat gca tta gat aga tag taa agt aag aga tct tgc aca gaa tta caa ttc tta ata att ata gag aaa ata tct tcc ata taa ttc ttt gac act aat gca gat ata ata tct tta tat gta ata tat gca aac agt cta tct act ata tac tga tca ata tta tct cta tga atc cta aaa taa tca tac aga aca tct aca gga tca caa att ggt tca agg aga aat cta tca aat att ttc ctg tca aca act ggt tct aga aca taa cag tca aca cct aat cca tgt ttt tta taa tca tct acc aaa gat aat gac caa aga tcg agg tcg tcg tga aac tgc tca tcg aca gcc atg aaa tct acc gac tcc atg gtg cga atc gca ctg tct tat tcg cca ttg att ttc a-t ttt tta taa tta tgt aca tgt ttt cct tct att ctc aag agt cta caa aaa tat a-- -tt ttt tcg ata tct aag tac taa gtt ttt tta ctg ttt ttg tta ctg tct tcc att ctt cta act aaa gat ctg aga taa att ata caa tct tcg cta tcg aac cat ttt tgt agt cta aag cct gaa gta att aac caa ctg ttt tta tta gtg gct ttt ttc gat cta tcc tcg tc- --- ctc tca tca tcc tta ta- --- --- --- --- --- --- tta tta tca tta tta tca tag tct att aaa cac aaa tca tct --- --- --- --- --- --- --- --- --- --- --- --- --- --- --- --- --- --- --- --- --- --- --- --- --- --- --a cgt tta taa c-- --- --- --- --- --- --- --- --a aca ttc --- --- --- --- -tc att att aat tag ttc tgt ag- aat atc ttt aat aat ttg gct ata --- --- -ca tct gtt --- --- --- --- --- --- --- --- --- --- --- --- --- --- --- --- --- --- --c aat act --- --- --- --- --- --- --- --a tct att gat gat ttc ttt t-- --- --- --- --- --- --- --- --- --- --- --t taa gac t-- --- --- --- --- --- --- --- --- --- --- --- --- --- --- --- --- --- --- --- --- --- --- --- --- --- --- --- --- --- --- --- --- --- --- --- --- --- --- --- taa act agt --- --- --- --- --- --- --- --- --- --- --- --- --- --- --- --- --- --- --- --- --- --- --- --- --- --- --- --- --- --- --- --- --- --- --- --- --- --- --- --- --- --- --- --- --- --- --- --- --- --- --- --- --- --- --- --- --- --- --- --- --- --- --- --- --- --- --- --- --- --- --- --- --- --- --- --- --- --- --- --- --- --- --- --- --- --- --- --- --- --- --- --- --- --- --- --- --- --- --- --- --- --- --- --- --- --- --- --- --- --t atg gta atg acg atg aaa --- -tc gag tag ta- --- --a ctt cta ata aag act tga ta- --- --t cat tat cat atg ttt gat cg- --- --- --- --- --- --- --t cat agt taa tag tgt g-- --- --- --- --- --- --- --- --- --- --- --- --- --- --- --- --- --- --- --- -gc taa atg gta ctg tta ata agt tta t-- --- --- --- --- --- --- --- --- --- --- --- --- --- --- --- --- --a gac aat atc ata gta ttt tct ttc cag aat --- --- --- --t aga tta ttt ttt taa ata ctg atc ctc aca att ccg tga tgt agc agt agt tgg t-- --- --- --- --- --- --- --- --- --- --- --- --- --- --- --- --- --- --- --- --- --- --- --- --- --- --- --- -gc atg gtc tat atc gt- --- --- --- --- --- --- --- --- --- --- --- --- --- --- --- --- --- --- --- --- --- --- --- --- --- --- --- --- --- --- --- --- --- --- --- --- --- --- --- --- --- --- --- taa aat gta tca tat ata ata gtt ttc tga cgt gga gta cag aat ttt cga --- --- --- --- --- --- --- --- --- --- --- --- --- --- --- --- --- --- --- --- --- --- --- --- --- --- --- --- --- --- --- --- --- --- --- --- --- --- --- --- --- --- --- --t taa tga gtt cat ggt aag gaa ggg caa atg cct --- gta tat aat ata cat aag tta a-- --- --- --- --- --- --- --- --- --- --- --- --- --t agt ttt tta tca tat tt- --- --- --- --- --- --- --- --t cta ata cca taa taa aaa t-- --- --- --- --- --- --- --- --- --- --- --- --- --- --- --- --- --- --- --- --- --- --- --- --- --t atc att att gcg ttt g-g tag tt- --- --- --- --- ctg ccc ta- --- --- --- --- --- --- --- --t cat cta tat cac tgt cac tct c-- --- --g ctc tca cta tat ctt cta aaa tta caa --- -ac aac --- --- --- --- --- --- --- --- --- --- --- --- --- --- --- --- --- --- --- --- --- --- --- --- --- --- --- --- --- --- --- --- --- --- --- --- tgg ata ttc gat --- --- --- --- --- --- --- --- --- --- --- --- --- --- --- --- --- --- --- --- --- --- --- --- --- --- --a aca gca ttt gtg t-- --- --- --- --- --- --- --- --- --- --- --- --- --- --- --- --- -

>ON649709.1_Monkeypox_virus_isolate_Monkeypox/PT0028/2022_complete_genome

--- --- --- --- --- --- --- --- --- --- --- --- --- --- --- --- --- --- -?? ??? ?-- --- --- --- --- --- --- --- att taa ttt tac tat ttt att tag tgt cta gaa aaa aat gtg tga ccc acg acc gta gga aac tct aga ggg taa gaa aaa tca atc gtt tat aga gac cat cag aaa gag gtt taa tat ttt tgt gag acc tat cga a-- --- --- --- --- --- --- --- --- --- --- --- --- --- --- --- --- --- --- --- --- --- --- --- --- --- --- --- --- --- --- --- --- --- --- --- --- --- --- --- --- --- --- --- --- --- --- --- --- --- --- --- --- --- --- --- --- --- --- --- --- --- --- --- --- --- --- --- --- --- --- --- --- --- --- --- --- --- --- --- --- --- --- --- --- --- --- --- --- --- --- --- --- --- --- --- --- --- --- --- --- --- --- --- --- --- --- --- --- --- --- --- --- --- --- --- --- --- --- --- --- --- --- --- --- --- --- --- --- --- --- --- --- --- --- --- --- --- --- --- --- --- --- --- --- --- --- --- --- --- --- --- --- --- --- --- --- --- --- --- --- --- --- --- --- --- --- --- --- --- --- --- --- --- --- --- --- --- --- --- --- --- --- --- --- --- --- --- --- --- --- --- --- --- --- --- --- --- --- --- --- --- --- --- --- --- --- --- --- --- --- --- --- --- --- --- --- --- --- --- --- --- --- --- --- --- --- --- --- --- --- --- --- --- --- --- --- --- --- --- --- --- --- --- --- --- --- --- --- --- --- --- --- --- --- --- --- --- --- --- --- --- --- --- --- --- --- --- --- --- --- --- --- --- --- --- --- --- --- --- -ga gag aaa gga taa aaa ctt ttt acg act cca tca gaa aga ggt tta ata ttt ttg tga gac cca tcg aag aga gaa aga gat ggt tag tca aga tat ttt tct tag tac aaa agt caa tgt ttt aaa ata tat gga cga gaa tta att tgt ctg tat aaa aac ttg tgt gaa att atg tac tag aga aaa aac gtg agc agt gtc ccc tac atg gat ttt aca gat cat tta tat tcc aaa aat att aac tat ata cgt tta tta tat gat gtt aac gtg taa att ata aac att att tta tga tgc aat tgt ctg aca acc tag att ggt ata agg atg ttg ata agc tct acg aga ata tat tgt tgg acg tta tcg ttt acg aaa tag ttg aga cat cag aaa gag gtt taa tat ttt tgt gag a-c cat cga aga gag aaa gag aat aaa aat a-- --t ttt ttt ttt ttt tgt aaa act ttt tta tga gac --- --- -ca aga gaa tac gaa tag tga tca tat cgt atc aca tat tga aac aga aag aag aag taa cga gag gta act ttt tgt gaa tgt agt taa ata ttt ttg ttt tgc aaa ccg gaa tat agt gcc cgg t-- -ct ttt tta att cgt ggt gcg gtg tct gaa tcg ttc gat taa ccc aac tca tcc att ttc aga tga ata gag tta tcg att cag aca cat gct ttg agt ttt gtt gaa tcg atg agt gaa gta tca tcg gtt gca cct tca gat gcc gat ccg tcg aca tac ttg aat cca tcc ttg act tca agt tca gat gat tcc tca cac atg tct ccg ata cgt acg cta aac tct agg ttc ttg aca cat ttt gta tca acg atc gtt gaa ccg atg ata tct ttg taa ctc act ttc tta tgt gag atg tta gac cca agt act gga tgg gtc ttg atg tca ctg tct ttc tct tct tcg cta cat ctg atg tcg ata gac atc tca cag tct ttg atc ata gcc aga gct tct tca cgc gtg atc gcg gga gag tcc tta cct tgt ccc ggt gac acg ctg gac aat cta gta ttc aca gtg ttt cca tca gag gat tcg gag atg gat aaa atc ttt ggg cat ttg gtg aat cca aag ttc atg tta aga ccc gca ccg acg ata gtg taa taa gtg gtg gga tct cct ttt aca act --- --- --- --- --- --- tct tcg gat acc tca tca tct tcg gtc tct gta act tcc gtt acg gat tga caa atc tta tca ttg gtc ggt gtt tgg tct tgc ttt gtg act ttg ata ata aca tcg att ccc ata tga tgt ttg ttt tct tct tca gta cac gag gat gaa gat tgt tga aga cta gta ggc ata gca gct gcc act agg cac atg cat gcc agg aca ata tat tgt ttc atg att gct att gat tga tta ctg ttc tag atg att cta ctt tct tac cat ata ata aat tag aat ata ttt tct act ttt acg aga aat taa tta ttg tat tta tta ttt ata ggt --a aaa aaa ctt act ata agt ggg tgg gat tct ggg aat tag tga tca gtt tat gta tat cgc aac tag cgg gca tat ggc tat tga cat cga gaa cat tac cca tat gat aag aga ttg tat cat ttt cgt agt ctt gag tat tgg tat tac tat ata gta tgt aga tgt cga cgc tag ata gac agt cgc cca cta gag tta ccg tct ctg aat gcg gca tga tag tat cat tct ttg ttt tcg tta act gtt tgg aag atg aat ctt tgt tgt tac att taa tct cga aat tca gag tac ata tct ttg aag tat tct gat atc tat ttt ctc ctg taa aga atc ctg aag ttg cta cat tat taa gga cag aga agt att ctg cac gaa aga ctg gat cac aat ctt tat gat tca tgg taa tag tta gtt ccg acg ttg aga tgg att cgc tga gac cgg tag tgg tcg tcc gag tac acg atg tgt cgt tga ctg gat aca ggt taa ttt cca cat cga tat agt taa atg tat tgc tgg tta cga cgg gtt cgc att tat ctg tgg aag aga cgg tgt gag aat atg ttc cgg gac cac acg gag aac aga tga cgt ctc cgg tag acg tgt atc cgg ata ctc cgt atc cta ttc cac act ttg ttt tag aaa tac atg ttc tac acc ctg atg ctc ctt tga gaa gac aat aat atc ctg gag agc att cac aga ttc tat tgt gag tcg tgt tac acg atc gcg tct cta cct gat tac tat cac atc ttc cgt tac aac tta gac aag cct gta aat gat tat tgt gag atg taa agg tat ccg aac cac acg gtg tac att gtg tat tag tct tgc tat cac ata atc tga aag cgt aag ttc ccg gag gac acg ata gac aac ata gat tac ggc ttc tgt att cgt tgt ctt tac act ttc cat tgg atg gtg cat gtg gtg cta tat ctc ttc cgt tta tta tta tac atg aga gaa aca ata tat acg agt ata ata cgg acc tca tga ttt aat aat -gt agt aat cgt cgt ctt gtt act gtt tgt ttc cta ctt ctc caa tca tat aga tta ttt ttt aaa tat ttt ctt tct atc atg gat aat att tgt aat ggt tct ttc cgt aca aca tac tgt tta gat ggt agt cgc tta gct tgg tta tga tat tgc gca taa ttt ccg gag gca aat acg ata gtc tag att gac tat cga tgg tag act cta att tat tga gtg ctt tgt cga cga gtt tac ttt tat gct cca tcg ata gat gac act gtt cta tga gat cgt cgt aca tgg gaa atg aaa tgc gtt tgt ctg aat gta tgg ctt cga gat atg tgt gat acc gaa tgt ctt ctg ttc tca ata ccg tat aca agt cgg tgt ctg aga ttc gaa tct ctt tga gga gac tta tgt cac gac tac att ttt cga tga tgg aat cta tct tat cga atg ata tat ttt tca taa ata cac ttt tat agt cct cgt tta aac aga att tag tat gta gtt ccg caa atg act cgt ccc tta ata ggc agt agg cta tta tct tct tta cgt agt gat cgt cgt agg gag aga act ccg aca tct tgt aga aca acg att taa tca tag gta gag ata ctt tca gtc tgt ggt gga taa tgt cat taa cga cat ccg cct tgt ata taa tgt ttc tgt ttt caa aca cca agt cga ata ctg tct tat cgt ctt tag tcg gaa ggt tga tgt cgt atc caa tgt ata cga ggt atg agg caa cat tgt tat tgc aat tct gga agg cgg tat gaa gag gag tca ttg tat tat agt att cgt ctt tct gaa tgt cga atc tat cta gta gat acc gta gta tat tga gag agc gac ttc cat atc ctt gat tat gtt tta tga ata gat aaa gta gat gtt gtc ctt ctt cct ttt gta att tcc cgt att ttt gtt cgt gcc aat tga gta aca tta tga gaa tat gac ctg ttg cac aat cgt tct tta tgt att cca tga tgg gtg tac aat taa gat tat tac gta tcc tcg tat cgg ctc ctc gag ata aaa gag cat aca cca cac gag gac tat gtt tgg tat act gtt gaa ggt aag tgt gta acg gcg tat ttc cga ttt tcg taa ccg cgt taa tgt ttg ctc cat gat cta tta tcg cgt aga tga atc gct tct cag ctc gca tct tag tgt gac tct ttg act tgt aat aat tgc ttt cgt gga acg cgg ata tgt gtt tac agt agt aat gaa gag aag tga gtc cat cct cat cga cgc aat tag ggt cag atc ctt tag tca ata att tgt aca gaa cgt aat agt tta agc tcc cat tga att tat atc taa gat aac aca gca ata gat cgg atg att tac taa agt cat caa tgg ggt ccg tta gta tat caa aga tct tgt tat cga ttg ata gtg aat gaa tca gat agt ggt gta gag gaa tat gtc ctt ttt tat cct tgc tat caa agt tac gca tgc cgt ggc gta aca ata tct tta ata cag atg gat taa atc gtg tat tca tcg tat agc aat gta atg gag agt tac cac att tta gtc gtt tat tca gat cgc agt gtt taa taa cta att taa aca gat gag atg ata tat cca cat caa aga atg cga gat aca tat gac aga cat tat tga cag aaa tgt gac ctt cat tat cac cgt cgt cca taa atg cgt tag gta cgt acc aca tac tat cgt taa cga tgc gca caa tct cgt cca -tt tca tcc atc ttc ata atg att tac ttt ttc ata att aga gaa aaa gat caa ggt ata aaa att aga agt gtt aga cta ta- --- --- --- --- --- --- --- --- --- --- -aa cta act tat gac tta act aac tta tga --- --- --- --- --- --- --- --- --- --- --- --- --- --- --- --- --- --- --- --- --- --- --- --- --- --- --- --- --- --- --- --- --- --- --- --- --- --- --- --- --- --- --- --- --- --- --- --- ctt aac ta- --- --- --- --- --- --- --- --- --- --- -ac tta tga ctt aac taa ctt atg act taa cta act ta- tga ctt aac taa ctt atg act taa cta act tat gac tta act aac tta tga cta tta act cat ttc aag aaa ggt ggg tgg ata gaa ctc tat atg aca gct tgt gaa aca att aga tcc cta att tct aat gga agt ttt gat agg aga ttg tca tca gtt gat aca ttg ttt att atc tca tct att aga gca cgt ctg ttt aga gct tta gtg acc tgc tcg gtt act tct gtg taa atc ttg aat cct tta gtg ata cac tgt gtc aaa act gga tgt tta gaa tac cta tgt aga ata tgg gaa gca tgc ttg ttt ttg tct cta tta tag att aac tca tac atg gtt gta tta tga att ttc atc tgc cta atg tac tcc aat tct tgt tta caa tca att ata taa tca aag agt gat gat gca tac aca tta caa agt gaa taa tct acc atc ata aaa tac ttg ata cag agc ttt atc aca tca tgg ttt tca att gta tta tta agt ata gct aat ttt ata cag tca ata gac aat ggt tct cta agc aat att tct aat att tta aga tgt gct tcc cta cgg gcg atg aca gat ccc cta tcc acg gcc acg tca aga cat gta tat cca tta ctc att act gcg ttg aca ttt gct cca ttt tct aat agc cat gat act aaa tct ata taa cct gca tag ata gcg cga taa agc aag gtc ctt cca cca gca tct agt tga ttg ata tct tca ata tat ggg ata caa agc tta taa att tct aat act gtg ggt tca tct aca agg aat ccc cta gta tac tga att att tta tat aga tct aat tta aca tca ttt tca tct ggg ata cca caa ttc aaa ata aac tca aca aca cta ctt tcc ttt tta cat att ccc cta aaa tag gca ttc aag cat tct att tta tat att aca gcc cca tga tct acc ata aga tca aca atg tct att tct aca tat gca tta gat aga tag taa agt aag aga tct tgc aca gaa tta caa ttc tta ata att ata gag aaa ata tct tcc ata taa ttc ttt gac act aat gca gat ata ata tct tta tat gta ata tat gca aac agt cta tct act ata tac tga tca ata tta tct cta tga atc cta aaa taa tca tac aga aca tct aca gga tca caa att ggt tca agg aga aat cta tca aat att ttc ctg tca aca act ggt tct aga aca taa cag tca aca cct aat cca tgt ttt tta taa tca tct acc aaa gat aat gac caa aga tcg agg tcg tcg tga aac tgc tca tcg aca gcc atg aaa tct acc gac tcc atg gtg cga atc gca ctg tct tat tcg cca ttg att ttc a-t ttt tta taa tta tgt aca tgt ttt cct tct att ctc aag agt cta caa aaa tat a-- -tt ttt tcg ata tct aag tac taa gtt ttt tta ctg ttt ttg tta ctg tct tcc att ctt cta act aaa gat ctg aga taa att ata caa tct tcg cta tcg aac cat ttt tgt agt cta aag cct gaa gta att aac caa ctg ttt tta tta gtg gct ttt ttc gat cta tcc tcg tc- --- ctc tca tca tcc tta ta- --- --- --- --- --- --- tta tta tca tta tta tca tag tct att aaa cac aaa tca tct --- --- --- --- --- --- --- --- --- --- --- --- --- --- --- --- --- --- --- --- --- --- --- --- --- --- --a cgt tta taa c-- --- --- --- --- --- --- --- --a aca ttc --- --- --- --- -tc att att aat tag ttc tgt ag- aat atc ttt aat aat ttg gct ata --- --- -ca tct gtt --- --- --- --- --- --- --- --- --- --- --- --- --- --- --- --- --- --- --c aat act --- --- --- --- --- --- --- --a tct att gat gat ttc ttt t-- --- --- --- --- --- --- --- --- --- --- --t taa gac t-- --- --- --- --- --- --- --- --- --- --- --- --- --- --- --- --- --- --- --- --- --- --- --- --- --- --- --- --- --- --- --- --- --- --- --- --- --- --- --- taa act agt --- --- --- --- --- --- --- --- --- --- --- --- --- --- --- --- --- --- --- --- --- --- --- --- --- --- --- --- --- --- --- --- --- --- --- --- --- --- --- --- --- --- --- --- --- --- --- --- --- --- --- --- --- --- --- --- --- --- --- --- --- --- --- --- --- --- --- --- --- --- --- --- --- --- --- --- --- --- --- --- --- --- --- --- --- --- --- --- --- --- --- --- --- --- --- --- --- --- --- --- --- --- --- --- --- --- --- --- --- --t atg gta atg acg atg aaa --- -tc gag tag ta- --- --a ctt cta ata aag act tga ta- --- --t cat tat cat atg ttt gat cg- --- --- --- --- --- --- --t cat agt taa tag tgt g-- --- --- --- --- --- --- --- --- --- --- --- --- --- --- --- --- --- --- --- -gc taa atg gta ctg tta ata agt tta t-- --- --- --- --- --- --- --- --- --- --- --- --- --- --- --- --- --a gac aat atc ata gta ttt tct ttc cag aat --- --- --- --t aga tta ttt ttt taa ata ctg atc ctc aca att ccg tga tgt agc agt agt tgg t-- --- --- --- --- --- --- --- --- --- --- --- --- --- --- --- --- --- --- --- --- --- --- --- --- --- --- --- -gc atg gtc tat atc gt- --- --- --- --- --- --- --- --- --- --- --- --- --- --- --- --- --- --- --- --- --- --- --- --- --- --- --- --- --- --- --- --- --- --- --- --- --- --- --- --- --- --- --- taa aat gta tca tat ata ata gtt ttc tga cgt gga gta cag aat ttt cga --- --- --- --- --- --- --- --- --- --- --- --- --- --- --- --- --- --- --- --- --- --- --- --- --- --- --- --- --- --- --- --- --- --- --- --- --- --- --- --- --- --- --- --t taa tga gtt cat ggt aag gaa ggg caa atg cct --- gta tat aat ata cat aag tta a-- --- --- --- --- --- --- --- --- --- --- --- --- --t agt ttt tta tca tat tt- --- --- --- --- --- --- --- --t cta ata cca taa taa aaa t-- --- --- --- --- --- --- --- --- --- --- --- --- --- --- --- --- --- --- --- --- --- --- --- --- --t atc att att gcg ttt g-g tag tt- --- --- --- --- ctg ccc ta- --- --- --- --- --- --- --- --t cat cta tat cac tgt cac tct c-- --- --g ctc tca cta tat ctt cta aaa tta caa --- -ac aac --- --- --- --- --- --- --- --- --- --- --- --- --- --- --- --- --- --- --- --- --- --- --- --- --- --- --- --- --- --- --- --- --- --- --- --- tgg ata ttc gat --- --- --- --- --- --- --- --- --- --- --- --- --- --- --- --- --- --- --- --- --- --- --- --- --- --- --a aca gca ttt gtg t-- --- --- --- --- --- --- --- --- --- --- --- --- --- --- --- --- -

>ON614676.1_Monkeypox_virus_isolate_INMI-Pt1_partial_genome

--- --- --- --- --- --- --- --- --- --- --- --- --- --- --- --- --- --- --- --- --- --- --- --- --- --- --- --- --- --a ttt tac tat ttt att tag tgt cta gaa aaa aat gtg tga ccc acg acc gta gga aac tct aga ggg taa gaa aaa tca atc gtt tat aga gac cat cag aaa gag gtt taa tat ttt tgt gag acc tat cga a-- --- --- --- --- --- --- --- --- --- --- --- --- --- --- --- --- --- --- --- --- --- --- --- --- --- --- --- --- --- --- --- --- --- --- --- --- --- --- --- --- --- --- --- --- --- --- --- --- --- --- --- --- --- --- --- --- --- --- --- --- --- --- --- --- --- --- --- --- --- --- --- --- --- --- --- --- --- --- --- --- --- --- --- --- --- --- --- --- --- --- --- --- --- --- --- --- --- --- --- --- --- --- --- --- --- --- --- --- --- --- --- --- --- --- --- --- --- --- --- --- --- --- --- --- --- --- --- --- --- --- --- --- --- --- --- --- --- --- --- --- --- --- --- --- --- --- --- --- --- --- --- --- --- --- --- --- --- --- --- --- --- --- --- --- --- --- --- --- --- --- --- --- --- --- --- --- --- --- --- --- --- --- --- --- --- --- --- --- --- --- --- --- --- --- --- --- --- --- --- --- --- --- --- --- --- --- --- --- --- --- --- --- --- --- --- --- --- --- --- --- --- --- --- --- --- --- --- --- --- --- --- --- --- --- --- --- --- --- --- --- --- --- --- --- --- --- --- --- --- --- --- --- --- --- --- --- --- --- --- --- --- --- --- --- --- --- --- --- --- --- --- --- --- --- --- --- --- --- --- -ga gag aaa gga taa aaa ctt ttt acg act cca tca gaa aga ggt tta ata ttt ttg tga gac cca tcg aag aga gaa aga gat ggt tag tca aga tat ttt tct tag tac aaa agt caa tgt ttt aaa ata tat gga cga gaa tta att tgt ctg tat aaa aac ttg tgt gaa att atg tac tag aga aaa aac gtg agc agt gtc ccc tac atg gat ttt aca gat cat tta tat tcc aaa aat att aac tat ata cgt tta tta tat gat gtt aac gtg taa att ata aac att att tta tga tgc aat tgt ctg aca acc tag att ggt ata agg atg ttg ata agc tct acg aga ata tat tgt tgg acg tta tcg ttt acg aaa tag ttg aga cat cag aaa gag gtt taa tat ttt tgt gag a-c cat cga aga gag aaa gag aat aaa aat a-- --? ttt ttt ttt ttt tgt aaa act ttt tta tga gac --- --- -ca aga gaa tac gaa tag tga tca tat cgt atc aca tat tga aac aga aag aag aag taa cga gag gta act ttt tgt gaa tgt agt taa ata ttt ttg ttt tgc aaa ccg gaa tat agt gcc cgg t-- -ct ttt tta att cgt ggt gcg gtg tct gaa tcg ttc gat taa ccc aac tca tcc att ttc aga tga ata gag tta tcg att cag aca cat gct ttg agt ttt gtt gaa tcg atg agt gaa gta tca tcg gtt gca cct tca gat gcc gat ccg tcg aca tac ttg aat cca tcc ttg act tca agt tca gat gat tcc tca cac atg tct ccg ata cgt acg cta aac tct agg ttc ttg aca cat ttt gta tca acg atc gtt gaa ccg atg ata tct ttg taa ctc act ttc tta tgt gag atg tta gac cca agt act gga tgg gtc ttg atg tca ctg tct ttc tct tct tcg cta cat ctg atg tcg ata gac atc tca cag tct ttg atc ata gcc aga gct tct tca cgc gtg atc gcg gga gag tcc tta cct tgt ccc ggt gac acg ctg gac aat cta gta ttc aca gtg ttt cca tca gag gat tcg gag atg gat aaa atc ttt ggg cat ttg gtg aat cca aag ttc atg tta aga ccc gca ccg acg ata gtg taa taa gtg gtg gga tct cct ttt aca act --- --- --- --- --- --- tct tcg gat acc tca tca tct tcg gtc tct gta act tcc gtt acg gat tga caa atc tta tca ttg gtc ggt gtt tgg tct tgc ttt gtg act ttg ata ata aca tcg att ccc ata tga tgt ttg ttt tct tct tca gta cac gag gat gaa gat tgt tga aga cta gta ggc ata gca gct gcc act agg cac atg cat gcc agg aca ata tat tgt ttc atg att gct att gat tga tta ctg ttc tag atg att cta ctt tct tac cat ata ata aat tag aat ata ttt tct act ttt acg aga aat taa tta ttg tat tta tta ttt ata ggt --a aaa aaa ctt act ata agt ggg tgg gat tct ggg aat tag tga tca gtt tat gta tat cgc aac tag cgg gca tat ggc tat tga cat cga gaa cat tac cca tat gat aag aga ttg tat cat ttt cgt agt ctt gag tat tgg tat tac tat ata gta tgt aga tgt cga cgc tag ata gac agt cgc cca cta gag tta ccg tct ctg aat gcg gca tga tag tat cat tct ttg ttt tcg tta act gtt tgg aag atg aat ctt tgt tgt tac att taa tct cga aat tca gag tac ata tct ttg aag tat tct gat atc tat ttt ctc ctg taa aga atc ctg aag ttg cta cat tat taa gga cag aga agt att ctg cac gaa aga ctg gat cac aat ctt tat gat tca tgg taa tag tta gtt ccg acg ttg aga tgg att cgc tga gac cgg tag tgg tcg tcc gag tac acg atg tgt cgt tga ctg gat aca ggt taa ttt cca cat cga tat agt taa atg tat tgc tgg tta cga cgg gtt cgc att tat ctg tgg aag aga cgg tgt gag aat atg ttc cgg gac cac acg gag aac aga tga cgt ctc cgg tag acg tgt atc cgg ata ctc cgt atc cta ttc cac act ttg ttt tag aaa tac atg ttc tac acc ctg atg ctc ctt tga gaa gac aat aat atc ctg gag agc att cac aga ttc tat tgt gag tcg tgt tac acg atc gcg tct cta cct gat tac tat cac atc ttc cgt tac aac tta gac aag cct gta aat gat tat tgt gag atg taa agg tat ccg aac cac acg gtg tac att gtg tat tag tct tgc tat cac ata atc tga aag cgt aag ttc ccg gag gac acg ata gac aac ata gat tac ggc ttc tgt att cgt tgt ctt tac act ttc cat tgg atg gtg cat gtg gtg cta tat ctc ttc cgt tta tta tta tac atg aga gaa aca ata tat acg agt ata ata cgg acc tca tga ttt aat aat -gt agt aat cgt cgt ctt gtt act gtt tgt ttc cta ctt ctc caa tca tat aga tta ttt ttt aaa tat ttt ctt tct atc atg gat aat att tgt aat ggt tct ttc cgt aca aca tac tgt tta gat ggt agt cgc tta gct tgg tta tga tat tgc gca taa ttt ccg gag gca aat acg ata gtc tag att gac tat cga tgg tag act cta att tat tga gtg ctt tgt cga cga gtt tac ttt tat gct cca tcg ata gat gac act gtt cta tga gat cgt cgt aca tgg gaa atg aaa tgc gtt tgt ctg aat gta tgg ctt cga gat atg tgt gat acc gaa tgt ctt ctg ttc tca ata ccg tat aca agt cgg tgt ctg aga ttc gaa tct ctt tga gga gac tta tgt cac gac tac att ttt cga tga tgg aat cta tct tat cga atg ata tat ttt tca taa ata cac ttt tat agt cct cgt tta aac aga att tag tat gta gtt ccg caa atg act cgt ccc tta ata ggc agt agg cta tta tct tct tta cgt agt gat cgt cgt agg gag aga act ccg aca tct tgt aga aca acg att taa tca tag gta gag ata ctt tca gtc tgt ggt gga taa tgt cat taa cga cat ccg cct tgt ata taa tgt ttc tgt ttt caa aca cca agt cga ata ctg tct tat cgt ctt tag tcg gaa ggt tga tgt cgt atc caa tgt ata cga ggt atg agg caa cat tgt tat tgc aat tct gga agg cgg tat gaa gag gag tca ttg tat tat agt att cgt ctt tct gaa tgt cga atc tat cta gta gat acc gta gta tat tga gag agc gac ttc cat atc ctt gat tat gtt tta tga ata gat aaa gta gat gtt gtc ctt ctt cct ttt gta att tcc cgt att ttt gtt cgt gcc aat tga gta aca tta tga gaa tat gac ctg ttg cac aat cgt tct tta tgt att cca tga tgg gtg tac aat taa gat tat tac gta tcc tcg tat cgg ctc ctc gag ata aaa gag cat aca cca cac gag gac tat gtt tgg tat act gtt gaa ggt aag tgt gta acg gcg tat ttc cga ttt tcg taa ccg cgt taa tgt ttg ctc cat gat cta tta tcg cgt aga tga atc gct tct cag ctc gca tct tag tgt gac tct ttg act tgt aat aat tgc ttt cgt gga acg cgg ata tgt gtt tac agt agt aat gaa gag aag tga gtc cat cct cat cga cgc aat tag ggt cag atc ctt tag tca ata att tgt aca gaa cgt aat agt tta agc tcc cat tga att tat atc taa gat aac aca gca ata gat cgg atg att tac taa agt cat caa tgg ggt ccg tta gta tat caa aga tct tgt tat cga ttg ata gtg aat gaa tca gat agt ggt gta gag gaa tat gtc ctt ttt tat cct tgc tat caa agt tac gca tgc cgt ggc gta aca ata tct tta ata cag atg gat taa atc gtg tat tca tcg tat agc aat gta atg gag agt tac cac att tta gtc gtt tat tca gat cgc agt gtt taa taa cta att taa aca gat gag atg ata tat cca cat caa aga atg cga gat aca tat gac aga cat tat tga cag aaa tgt gac ctt cat tat cac cgt cgt cca taa atg cgt tag gta cgt acc aca tac tat cgt taa cga tgc gca caa tct cgt cca -tt tca tcc atc ttc ata atg att tac ttt ttc ata att aga gaa aaa gat caa ggt ata aaa att aga agt gtt aga cta ta- --- --- --- --- --- --- --- --- --- --- -aa cta act tat gac tta act aac tta tga --- --- --- --- --- --- --- --- --- --- --- --- --- --- --- --- --- --- --- --- --- --- --- --- --- --- --- --- --- --- --- --- --- --- --- --- --- --- --- --- --- --- --- --- --- --- --- --- ctt aac ta- --- --- --- --- --- --- --- --- --- --- -ac tta tga ctt aac taa ctt atg act taa cta act ta- tga ctt aac taa ctt atg act taa cta act tat gac tta act aac tta tga cta tta act cat ttc aag aaa ggt ggg tgg ata gaa ctc tat atg aca gct tgt gaa aca att aga tcc cta att tct aat gga agt ttt gat agg aga ttg tca tca gtt gat aca ttg ttt att atc tca tct att aga gca cgt ctg ttt aga gct tta gtg acc tgc tcg gtt act tct gtg taa atc ttg aat cct tta gtg ata cac tgt gtc aaa act gga tgt tta gaa tac cta tgt aga ata tgg gaa gca tgc ttg ttt ttg tct cta tta tag att aac tca tac atg gtt gta tta tga att ttc atc tgc cta atg tac tcc aat tct tgt tta caa tca att ata taa tca aag agt gat gat gca tac aca tta caa agt gaa taa tct acc atc ata aaa tac ttg ata cag agc ttt atc aca tca tgg ttt tca att gta tta tta agt ata gct aat ttt ata cag tca ata gac aat ggt tct cta agc aat att tct aat att tta aga tgt gct tcc cta cgg gcg atg aca gat ccc cta tcc acg gcc acg tca aga cat gta tat cca tta ctc att act gcg ttg aca ttt gct cca ttt tct aat agc cat gat act aaa tct ata taa cct gca tag ata gcg cga taa agc aag gtc ctt cca cca gca tct agt tga ttg ata tct tca ata tat ggg ata caa agc tta taa att tct aat act gtg ggt tca tct aca agg aat ccc cta gta tac tga att att tta tat aga tct aat tta aca tca ttt tca tct ggg ata cca caa ttc aaa ata aac tca aca aca cta ctt tcc ttt tta cat att ccc cta aaa tag gca ttc aag cat tct att tta tat att aca gcc cca tga tct acc ata aga tca aca atg tct att tct aca tat gca tta gat aga tag taa agt aag aga tct tgc aca gaa tta caa ttc tta ata att ata gag aaa ata tct tcc ata taa ttc ttt gac act aat gca gat ata ata tct tta tat gta ata tat gca aac agt cta tct act ata tac tga tca ata tta tct cta tga atc cta aaa taa tca tac aga aca tct aca gga tca caa att ggt tca agg aga aat cta tca aat att ttc ctg tca aca act ggt tct aga aca taa cag tca aca cct aat cca tgt ttt tta taa tca tct acc aaa gat aat gac caa aga tcg agg tcg tcg tga aac tgc tca tcg aca gcc atg aaa tct acc gac tcc atg gtg cga atc gca ctg tct tat tcg cca ttg att ttc a-t ttt tta taa tta tgt aca tgt ttt cct tct att ctc aag agt cta caa aaa tat a-- -tt ttt tcg ata tct aag tac taa gtt ttt tta ctg ttt ttg tta ctg tct tcc att ctt cta act aaa gat ctg aga taa att ata caa tct tcg cta tcg aac cat ttt tgt agt cta aag cct gaa gta att aac caa ctg ttt tta tta gtg gct ttt ttc gat cta tcc tcg tc- --- ctc tca tca tcc tta ta- --- --- --- --- --- --- tta tta tca tta tta tca tag tct att aaa cac aaa tca tct --- --- --- --- --- --- --- --- --- --- --- --- --- --- --- --- --- --- --- --- --- --- --- --- --- --- --a cgt tta taa c-- --- --- --- --- --- --- --- --a aca ttc --- --- --- --- -tc att att aat tag ttc tgt ag- aat atc ttt aat aat ttg gct ata --- --- -ca tct gtt --- --- --- --- --- --- --- --- --- --- --- --- --- --- --- --- --- --- --c aat act --- --- --- --- --- --- --- --a tct att gat gat ttc ttt t-- --- --- --- --- --- --- --- --- --- --- --t taa gac t-- --- --- --- --- --- --- --- --- --- --- --- --- --- --- --- --- --- --- --- --- --- --- --- --- --- --- --- --- --- --- --- --- --- --- --- --- --- --- --- taa act agt --- --- --- --- --- --- --- --- --- --- --- --- --- --- --- --- --- --- --- --- --- --- --- --- --- --- --- --- --- --- --- --- --- --- --- --- --- --- --- --- --- --- --- --- --- --- --- --- --- --- --- --- --- --- --- --- --- --- --- --- --- --- --- --- --- --- --- --- --- --- --- --- --- --- --- --- --- --- --- --- --- --- --- --- --- --- --- --- --- --- --- --- --- --- --- --- --- --- --- --- --- --- --- --- --- --- --- --- --- --t atg gta atg acg atg aaa --- -tc gag tag ta- --- --a ctt cta ata aag act tga ta- --- --t cat tat cat atg ttt gat cg- --- --- --- --- --- --- --t cat agt taa tag tgt g-- --- --- --- --- --- --- --- --- --- --- --- --- --- --- --- --- --- --- --- -gc taa atg gta ctg tta ata agt tta t-- --- --- --- --- --- --- --- --- --- --- --- --- --- --- --- --- --a gac aat atc ata gta ttt tct ttc cag aat --- --- --- --t aga tta ttt ttt taa ata ctg atc ctc aca att ccg tga tgt agc agt agt tgg t-- --- --- --- --- --- --- --- --- --- --- --- --- --- --- --- --- --- --- --- --- --- --- --- --- --- --- --- -gc atg gtc tat atc gt- --- --- --- --- --- --- --- --- --- --- --- --- --- --- --- --- --- --- --- --- --- --- --- --- --- --- --- --- --- --- --- --- --- --- --- --- --- --- --- --- --- --- --- taa aat gta tca tat ata ata gtt ttc tga cgt gga gta cag aat ttt cga --- --- --- --- --- --- --- --- --- --- --- --- --- --- --- --- --- --- --- --- --- --- --- --- --- --- --- --- --- --- --- --- --- --- --- --- --- --- --- --- --- --- --- --t taa tga gtt cat ggt aag gaa ggg caa atg cct --- gta tat aat ata cat aag tta a-- --- --- --- --- --- --- --- --- --- --- --- --- --t agt ttt tta tca tat tt- --- --- --- --- --- --- --- --t cta ata cca taa taa aaa t-- --- --- --- --- --- --- --- --- --- --- --- --- --- --- --- --- --- --- --- --- --- --- --- --- --t atc att att gcg ttt g-g tag tt- --- --- --- --- ctg ccc ta- --- --- --- --- --- --- --- --t cat cta tat cac tgt cac tct c-- --- --g ctc tca cta tat ctt cta aaa tta caa --- -ac aac --- --- --- --- --- --- --- --- --- --- --- --- --- --- --- --- --- --- --- --- --- --- --- --- --- --- --- --- --- --- --- --- --- --- --- --- tgg ata ttc gat --- --- --- --- --- --- --- --- --- --- --- --- --- --- --- --- --- --- --- --- --- --- --- --- --- --- --a aca gca ttt gtg t-- --- --- --- --- --- --- --- --- --- --- --- --- --- --- --- --- -

>ON649712.1_Monkeypox_virus_isolate_Monkeypox/PT0025/2022_partial_genome

--- --- --- --- --- --- --- --- --- --- --- --- --- --- --- --- --- --- --- --- --- --- --- --- --- --- --- --- att taa ttt tac tat ttt att tag tgt cta gaa aaa aat gtg tga ccc acg acc gta gga aac tct aga ggg taa gaa aaa tca atc gtt tat aga gac cat cag aaa gag gtt taa tat ttt tgt gag acc tat cga a-- --- --- --- --- --- --- --- --- --- --- --- --- --- --- --- --- --- --- --- --- --- --- --- --- --- --- --- --- --- --- --- --- --- --- --- --- --- --- --- --- --- --- --- --- --- --- --- --- --- --- --- --- --- --- --- --- --- --- --- --- --- --- --- --- --- --- --- --- --- --- --- --- --- --- --- --- --- --- --- --- --- --- --- --- --- --- --- --- --- --- --- --- --- --- --- --- --- --- --- --- --- --- --- --- --- --- --- --- --- --- --- --- --- --- --- --- --- --- --- --- --- --- --- --- --- --- --- --- --- --- --- --- --- --- --- --- --- --- --- --- --- --- --- --- --- --- --- --- --- --- --- --- --- --- --- --- --- --- --- --- --- --- --- --- --- --- --- --- --- --- --- --- --- --- --- --- --- --- --- --- --- --- --- --- --- --- --- --- --- --- --- --- --- --- --- --- --- --- --- --- --- --- --- --- --- --- --- --- --- --- --- --- --- --- --- --- --- --- --- --- --- --- --- --- --- --- --- --- --- --- --- --- --- --- --- --- --- --- --- --- --- --- --- --- --- --- --- --- --- --- --- --- --- --- --- --- --- --- --- --- --- --- --- --- --- --- --- --- --- --- --- --- --- --- --- --- --- --- --- -ga gag aaa gga taa aaa ctt ttt acg act cca tca gaa aga ggt tta ata ttt ttg tga gac cca tcg aag aga gaa aga gat ggt tag tca aga tat ttt tct tag tac aaa agt caa tgt ttt aaa ata tat gga cga gaa tta att tgt ctg tat aaa aac ttg tgt gaa att atg tac tag aga aaa aac gtg agc agt gtc ccc tac atg gat ttt aca gat cat tta tat tcc aaa aat att aac tat ata cgt tta tta tat gat gtt aac gtg taa att ata aac att att tta tga tgc aat tgt ctg aca acc tag att ggt ata agg atg ttg ata agc tct acg aga ata tat tgt tgg acg tta tcg ttt acg aaa tag ttg aga cat cag aaa gag gtt taa tat ttt tgt gag a-c cat cga aga gag aaa gag aat aaa aat a-- --t ttt ttt ttt ttt tgt aaa act ttt tta tga gac --- --- -ca aga gaa tac gaa tag tga tca tat cgt atc aca tat tga aac aga aag aag aag taa cga gag gta act ttt tgt gaa tgt agt taa ata ttt ttg ttt tgc aaa ccg gaa tat agt gcc cgg t-- -ct ttt tta att cgt ggt gcg gtg tct gaa tcg ttc gat taa ccc aac tca tcc att ttc aga tga ata gag tta tcg att cag aca cat gct ttg agt ttt gtt gaa tcg atg agt gaa gta tca tcg gtt gca cct tca gat gcc gat ccg tcg aca tac ttg aat cca tcc ttg act tca agt tca gat gat tcc tca cac atg tct ccg ata cgt acg cta aac tct agg ttc ttg aca cat ttt gta tca acg atc gtt gaa ccg atg ata tct ttg taa ctc act ttc tta tgt gag atg tta gac cca agt act gga tgg gtc ttg atg tca ctg tct ttc tct tct tcg cta cat ctg atg tcg ata gac atc tca cag tct ttg atc ata gcc aga gct tct tca cgc gtg atc gcg gga gag tcc tta cct tgt ccc ggt gac acg ctg gac aat cta gta ttc aca gtg ttt cca tca gag gat tcg gag atg gat aaa atc ttt ggg cat ttg gtg aat cca aag ttc atg tta aga ccc gca ccg acg ata gtg taa taa gtg gtg gga tct cct ttt aca act --- --- --- --- --- --- tct tcg gat acc tca tca tct tcg gtc tct gta act tcc gtt acg gat tga caa atc tta tca ttg gtc ggt gtt tgg tct tgc ttt gtg act ttg ata ata aca tcg att ccc ata tga tgt ttg ttt tct tct tca gta cac gag gat gaa gat tgt tga aga cta gta ggc ata gca gct gcc act agg cac atg cat gcc agg aca ata tat tgt ttc atg att gct att gat tga tta ctg ttc tag atg att cta ctt tct tac cat ata ata aat tag aat ata ttt tct act ttt acg aga aat taa tta ttg tat tta tta ttt ata ggt --a aaa aaa ctt act ata agt ggg tgg gat tct ggg aat tag tga tca gtt tat gta tat cgc aac tag cgg gca tat ggc tat tga cat cga gaa cat tac cca tat gat aag aga ttg tat cat ttt cgt agt ctt gag tat tgg tat tac tat ata gta tgt aga tgt cga cgc tag ata gac agt cgc cca cta gag tta ccg tct ctg aat gcg gca tga tag tat cat tct ttg ttt tcg tta act gtt tgg aag atg aat ctt tgt tgt tac att taa tct cga aat tca gag tac ata tct ttg aag tat tct gat atc tat ttt ctc ctg taa aga atc ctg aag ttg cta cat tat taa gga cag aga agt att ctg cac gaa aga ctg gat cac aat ctt tat gat tca tgg taa tag tta gtt ccg acg ttg aga tgg att cgc tga gac cgg tag tgg tcg tcc gag tac acg atg tgt cgt tga ctg gat aca ggt taa ttt cca cat cga tat agt taa atg tat tgc tgg tta cga cgg gtt cgc att tat ctg tgg aag aga cgg tgt gag aat atg ttc cgg gac cac acg gag aac aga tga cgt ctc cgg tag acg tgt atc cgg ata ctc cgt atc cta ttc cac act ttg ttt tag aaa tac atg ttc tac acc ctg atg ctc ctt tga gaa gac aat aat atc ctg gag agc att cac aga ttc tat tgt gag tcg tgt tac acg atc gcg tct cta cct gat tac tat cac atc ttc cgt tac aac tta gac aag cct gta aat gat tat tgt gag atg taa agg tat ccg aac cac acg gtg tac att gtg tat tag tct tgc tat cac ata atc tga aag cgt aag ttc ccg gag gac acg ata gac aac ata gat tac ggc ttc tgt att cgt tgt ctt tac act ttc cat tgg atg gtg cat gtg gtg cta tat ctc ttc cgt tta tta tta tac atg aga gaa aca ata tat acg agt ata ata cgg acc tca tga ttt aat aat -gt agt aat cgt cgt ctt gtt act gtt tgt ttc cta ctt ctc caa tca tat aga tta ttt ttt aaa tat ttt ctt tct atc atg gat aat att tgt aat ggt tct ttc cgt aca aca tac tgt tta gat ggt agt cgc tta gct tgg tta tga tat tgc gca taa ttt ccg gag gca aat acg ata gtc tag att gac tat cga tgg tag act cta att tat tga gtg ctt tgt cga cga gtt tac ttt tat gct cca tcg ata gat gac act gtt cta tga gat cgt cgt aca tgg gaa atg aaa tgc gtt tgt ctg aat gta tgg ctt cga gat atg tgt gat acc gaa tgt ctt ctg ttc tca ata ccg tat aca agt cgg tgt ctg aga ttc gaa tct ctt tga gga gac tta tgt cac gac tac att ttt cga tga tgg aat cta tct tat cga atg ata tat ttt tca taa ata cac ttt tat agt cct cgt tta aac aga att tag tat gta gtt ccg caa atg act cgt ccc tta ata ggc agt agg cta tta tct tct tta cgt agt gat cgt cgt agg gag aga act ccg aca tct tgt aga aca acg att taa tca tag gta gag ata ctt tca gtc tgt ggt gga taa tgt cat taa cga cat ccg cct tgt ata taa tgt ttc tgt ttt caa aca cca agt cga ata ctg tct tat cgt ctt tag tcg gaa ggt tga tgt cgt atc caa tgt ata cga ggt atg agg caa cat tgt tat tgc aat tct gga agg cgg tat gaa gag gag tca ttg tat tat agt att cgt ctt tct gaa tgt cga atc tat cta gta gat acc gta gta tat tga gag agc gac ttc cat atc ctt gat tat gtt tta tga ata gat aaa gta gat gtt gtc ctt ctt cct ttt gta att tcc cgt att ttt gtt cgt gcc aat tga gta aca tta tga gaa tat gac ctg ttg cac aat cgt tct tta tgt att cca tga tgg gtg tac aat taa gat tat tac gta tcc tcg tat cgg ctc ctc gag ata aaa gag cat aca cca cac gag gac tat gtt tgg tat act gtt gaa ggt aag tgt gta acg gcg tat ttc cga ttt tcg taa ccg cgt taa tgt ttg ctc cat gat cta tta tcg cgt aga tga atc gct tct cag ctc gca tct tag tgt gac tct ttg act tgt aat aat tgc ttt cgt gga acg cgg ata tgt gtt tac agt agt aat gaa gag aag tga gtc cat cct cat cga cgc aat tag ggt cag atc ctt tag tca ata att tgt aca gaa cgt aat agt tta agc tcc cat tga att tat atc taa gat aac aca gca ata gat cgg atg att tac taa agt cat caa tgg ggt ccg tta gta tat caa aga tct tgt tat cga ttg ata gtg aat gaa tca gat agt ggt gta gag gaa tat gtc ctt ttt tat cct tgc tat caa agt tac gca tgc cgt ggc gta aca ata tct tta ata cag atg gat taa atc gtg tat tca tcg tat agc aat gta atg gag agt tac cac att tta gtc gtt tat tca gat cgc agt gtt taa taa cta att taa aca gat gag atg ata tat cca cat caa aga atg cga gat aca tat gac aga cat tat tga cag aaa tgt gac ctt cat tat cac cgt cgt cca taa atg cgt tag gta cgt acc aca tac tat cgt taa cga tgc gca caa tct cgt cca -tt tca tcc atc ttc ata atg att tac ttt ttc ata att aga gaa aaa gat caa ggt ata aaa att aga agt gtt aga cta ta- --- --- --- --- --- --- --- --- --- --- -aa cta act tat gac tta act aac tta tga --- --- --- --- --- --- --- --- --- --- --- --- --- --- --- --- --- --- --- --- --- --- --- --- --- --- --- --- --- --- --- --- --- --- --- --- --- --- --- --- --- --- --- --- --- --- --- --- ctt aac ta- --- --- --- --- --- --- --- --- --- --- -ac tta tga ctt aac taa ctt atg act taa cta act ta- tga ctt aac taa ctt atg act taa cta act tat gac tta act aac tta tga cta tta act cat ttc aag aaa ggt ggg tgg ata gaa ctc tat atg aca gct tgt gaa aca att aga tcc cta att tct aat gga agt ttt gat agg aga ttg tca tca gtt gat aca ttg ttt att atc tca tct att aga gca cgt ctg ttt aga gct tta gtg acc tgc tcg gtt act tct gtg taa atc ttg aat cct tta gtg ata cac tgt gtc aaa act gga tgt tta gaa tac cta tgt aga ata tgg gaa gca tgc ttg ttt ttg tct cta tta tag att aac tca tac atg gtt gta tta tga att ttc atc tgc cta atg tac tcc aat tct tgt tta caa tca att ata taa tca aag agt gat gat gca tac aca tta caa agt gaa taa tct acc atc ata aaa tac ttg ata cag agc ttt atc aca tca tgg ttt tca att gta tta tta agt ata gct aat ttt ata cag tca ata gac aat ggt tct cta agc aat att tct aat att tta aga tgt gct tcc cta cgg gcg atg aca gat ccc cta tcc acg gcc acg tca aga cat gta tat cca tta ctc att act gcg ttg aca ttt gct cca ttt tct aat agc cat gat act aaa tct ata taa cct gca tag ata gcg cga taa agc aag gtc ctt cca cca gca tct agt tga ttg ata tct tca ata tat ggg ata caa agc tta taa att tct aat act gtg ggt tca tct aca agg aat ccc cta gta tac tga att att tta tat aga tct aat tta aca tca ttt tca tct ggg ata cca caa ttc aaa ata aac tca aca aca cta ctt tcc ttt tta cat att ccc cta aaa tag gca ttc aag cat tct att tta tat att aca gcc cca tga tct acc ata aga tca aca atg tct att tct aca tat gca tta gat aga tag taa agt aag aga tct tgc aca gaa tta caa ttc tta ata att ata gag aaa ata tct tcc ata taa ttc ttt gac act aat gca gat ata ata tct tta tat gta ata tat gca aac agt cta tct act ata tac tga tca ata tta tct cta tga atc cta aaa taa tca tac aga aca tct aca gga tca caa att ggt tca agg aga aat cta tca aat att ttc ctg tca aca act ggt tct aga aca taa cag tca aca cct aat cca tgt ttt tta taa tca tct acc aaa gat aat gac caa aga tcg agg tcg tcg tga aac tgc tca tcg aca gcc atg aaa tct acc gac tcc atg gtg cga atc gca ctg tct tat tcg cca ttg att ttc a-t ttt tta taa tta tgt aca tgt ttt cct tct att ctc aag agt cta caa aaa tat a-- -tt ttt tcg ata tct aag tac taa gtt ttt tta ctg ttt ttg tta ctg tct tcc att ctt cta act aaa gat ctg aga taa att ata caa tct tcg cta tcg aac cat ttt tgt agt cta aag cct gaa gta att aac caa ctg ttt tta tta gtg gct ttt ttc gat cta tcc tcg tc- --- ctc tca tca tcc tta ta- --- --- --- --- --- --- tta tta tca tta tta tca tag tct att aaa cac aaa tca tct --- --- --- --- --- --- --- --- --- --- --- --- --- --- --- --- --- --- --- --- --- --- --- --- --- --- --a cgt tta taa c-- --- --- --- --- --- --- --- --a aca ttc --- --- --- --- -tc att att aat tag ttc tgt ag- aat atc ttt aat aat ttg gct ata --- --- -ca tct gtt --- --- --- --- --- --- --- --- --- --- --- --- --- --- --- --- --- --- --c aat act --- --- --- --- --- --- --- --a tct att gat gat ttc ttt t-- --- --- --- --- --- --- --- --- --- --- --t taa gac t-- --- --- --- --- --- --- --- --- --- --- --- --- --- --- --- --- --- --- --- --- --- --- --- --- --- --- --- --- --- --- --- --- --- --- --- --- --- --- --- taa act agt --- --- --- --- --- --- --- --- --- --- --- --- --- --- --- --- --- --- --- --- --- --- --- --- --- --- --- --- --- --- --- --- --- --- --- --- --- --- --- --- --- --- --- --- --- --- --- --- --- --- --- --- --- --- --- --- --- --- --- --- --- --- --- --- --- --- --- --- --- --- --- --- --- --- --- --- --- --- --- --- --- --- --- --- --- --- --- --- --- --- --- --- --- --- --- --- --- --- --- --- --- --- --- --- --- --- --- --- --- --t atg gta atg acg atg aaa --- -tc gag tag ta- --- --a ctt cta ata aag act tga ta- --- --t cat tat cat atg ttt gat cg- --- --- --- --- --- --- --t cat agt taa tag tgt g-- --- --- --- --- --- --- --- --- --- --- --- --- --- --- --- --- --- --- --- -gc taa atg gta ctg tta ata agt tta t-- --- --- --- --- --- --- --- --- --- --- --- --- --- --- --- --- --a gac aat atc ata gta ttt tct ttc cag aat --- --- --- --t aga tta ttt ttt taa ata ctg atc ctc aca att ccg tga tgt agc agt agt tgg t-- --- --- --- --- --- --- --- --- --- --- --- --- --- --- --- --- --- --- --- --- --- --- --- --- --- --- --- -gc atg gtc tat atc gt- --- --- --- --- --- --- --- --- --- --- --- --- --- --- --- --- --- --- --- --- --- --- --- --- --- --- --- --- --- --- --- --- --- --- --- --- --- --- --- --- --- --- --- taa aat gta tca tat ata ata gtt ttc tga cgt gga gta cag aat ttt cga --- --- --- --- --- --- --- --- --- --- --- --- --- --- --- --- --- --- --- --- --- --- --- --- --- --- --- --- --- --- --- --- --- --- --- --- --- --- --- --- --- --- --- --t taa tga gtt cat ggt aag gaa ggg caa atg cct --- gta tat aat ata cat aag tta a-- --- --- --- --- --- --- --- --- --- --- --- --- --t agt ttt tta tca tat tt- --- --- --- --- --- --- --- --t cta ata cca taa taa aaa t-- --- --- --- --- --- --- --- --- --- --- --- --- --- --- --- --- --- --- --- --- --- --- --- --- --t atc att att gcg ttt g-g tag tt- --- --- --- --- ctg ccc ta- --- --- --- --- --- --- --- --t cat cta tat cac tgt cac tct c-- --- --g ctc tca cta tat ctt cta aaa tta caa --- -ac aac --- --- --- --- --- --- --- --- --- --- --- --- --- --- --- --- --- --- --- --- --- --- --- --- --- --- --- --- --- --- --- --- --- --- --- --- tgg ata ttc gat --- --- --- --- --- --- --- --- --- --- --- --- --- --- --- --- --- --- --- --- --- --- --- --- --- --- --a aca gca ttt gtg t-- --- --- --- --- --- --- --- --- --- --- --- --- --- --- --- --- -

>ON595760.2_Monkeypox_virus_isolate_MPXV-CH-38134631/2022_partial_genome

--- --- --- --- --- --- --- --- --- --- --- --- --- --- --- --- --- --- --- --- --- --- --- --- --- --- --- --- --- --- --- --- --- --- --- --- --- --- --- --- --- --- --- --- --- --- --- --- --- --- --- --- --- --- --- --- --- -tt tat aga gac cat cag aaa gag gtt taa tat ttt tgt gag acc tat cga a-- --- --- --- --- --- --- --- --- --- --- --- --- --- --- --- --- --- --- --- --- --- --- --- --- --- --- --- --- --- --- --- --- --- --- --- --- --- --- --- --- --- --- --- --- --- --- --- --- --- --- --- --- --- --- --- --- --- --- --- --- --- --- --- --- --- --- --- --- --- --- --- --- --- --- --- --- --- --- --- --- --- --- --- --- --- --- --- --- --- --- --- --- --- --- --- --- --- --- --- --- --- --- --- --- --- --- --- --- --- --- --- --- --- --- --- --- --- --- --- --- --- --- --- --- --- --- --- --- --- --- --- --- --- --- --- --- --- --- --- --- --- --- --- --- --- --- --- --- --- --- --- --- --- --- --- --- --- --- --- --- --- --- --- --- --- --- --- --- --- --- --- --- --- --- --- --- --- --- --- --- --- --- --- --- --- --- --- --- --- --- --- --- --- --- --- --- --- --- --- --- --- --- --- --- --- --- --- --- --- --- --- --- --- --- --- --- --- --- --- --- --- --- --- --- --- --- --- --- --- --- --- --- --- --- --- --- --- --- --- --- --- --- --- --- --- --- --- --- --- --- --- --- --- --- --- --- --- --- --- --- --- --- --- --- --- --- --- --- --- --- --- --- --- --- --- --- --- --- --- -ga gag aaa gga t?a aaa c?t ttt acg act cca tca gaa aga ggt tta ata ttt ttg tga gac cca tcg aag aga gaa aga gat ggt tag tca aga tat ttt tct tag tac aaa agt caa tgt ttt aaa ata tat gga cga gaa tta att tgt ctg tat aaa aac ttg tgt gaa att atg tac tag aga aaa aac gtg agc agt gtc ccc tac atg gat ttt aca gat cat tta tat tcc aaa aat att aac tat ata cgt tta tta tat gat gtt aac gtg taa att ata aac att att tta tga tgc aat tgt ctg aca acc tag att ggt ata agg atg ttg ata agc tct acg aga ata tat tgt tgg acg tta tcg ttt acg aaa tag ttg aga cat cag aaa gag gtt taa tat ttt tgt gag a-c cat cga aga gag aaa gag aat aaa aat a-- --- -tt ttt ttt ttt tgt aaa act ttt tta tga gac --- --- -ca aga gaa tac gaa tag tga tca tat cgt atc aca tat tga aac aga aag aag aag taa cga gag gta act ttt tgt gaa tgt agt taa ata ttt ttg ttt tgc aaa ccg gaa tat agt gcc cgg t-- -ct ttt tta att cgt ggt gcg gtg tct gaa tcg ttc gat taa ccc aac tca tcc att ttc aga tga ata gag tta tcg att cag aca cat gct ttg agt ttt gtt gaa tcg atg agt gaa gta tca tcg gtt gca cct tca gat gcc gat ccg tcg aca tac ttg aat cca tcc ttg act tca agt tca gat gat tcc tca cac atg tct ccg ata cgt acg cta aac tct agg ttc ttg aca cat ttt gta tca acg atc gtt gaa ccg atg ata tct ttg taa ctc act ttc tta tgt gag atg tta gac cca agt act gga tgg gtc ttg atg tca ctg tct ttc tct tct tcg cta cat ctg atg tcg ata gac atc tca cag tct ttg atc ata gcc aga gct tct tca cgc gtg atc gcg gga gag tcc tta cct tgt ccc ggt gac acg ctg gac aat cta gta ttc aca gtg ttt cca tca gag gat tcg gag atg gat aaa atc ttt ggg cat ttg gtg aat cca aag ttc atg tta aga ccc gca ccg acg ata gtg taa taa gtg gtg gga tct cct ttt aca act --- --- --- --- --- --- tct tcg gat acc tca tca tct tcg gtc tct gta act tcc gtt acg gat tga caa atc tta tca ttg gtc ggt gtt tgg tct tgc ttt gtg act ttg ata ata aca tcg att ccc ata tga tgt ttg ttt tct tct tca gta cac gag gat gaa gat tgt tga aga cta gta ggc ata gca gct gcc act agg cac atg cat gcc agg aca ata tat tgt ttc atg att gct att gat tga tta ctg ttc tag atg att cta ctt tct tac cat ata ata aat tag aat ata ttt tct act ttt acg aga aat taa tta ttg tat tta tta ttt ata ggt --a aaa aaa ctt act ata agt ggg tgg gat tct ggg aat tag tga tca gtt tat gta tat cgc aac tag cgg gca tat ggc tat tga cat cga gaa cat tac cca tat gat aag aga ttg tat cat ttt cgt agt ctt gag tat tgg tat tac tat ata gta tgt aga tgt cga cgc tag ata gac agt cgc cca cta gag tta ccg tct ctg aat gcg gca tga tag tat cat tct ttg ttt tcg tta act gtt tgg aag atg aat ctt tgt tgt tac att taa tct cga aat tca gag tac ata tct ttg aag tat tct gat atc tat ttt ctc ctg taa aga atc ctg aag ttg cta cat tat taa gga cag aga agt att ctg cac gaa aga ctg gat cac aat ctt tat gat tca tgg taa tag tta gtt ccg acg ttg aga tgg att cgc tga gac cgg tag tgg tcg tcc gag tac acg atg tgt cgt tga ctg gat aca ggt taa ttt cca cat cga tat agt taa atg tat tgc tgg tta cga cgg gtt cgc att tat ctg tgg aag aga cgg tgt gag aat atg ttc cgg gac cac acg gag aac aga tga cgt ctc cgg tag acg tgt atc cgg ata ctc cgt atc cta ttc cac act ttg ttt tag aaa tac atg ttc tac acc ctg atg ctc ctt tga gaa gac aat aat atc ctg gag agc att cac aga ttc tat tgt gag tcg tgt tac acg atc gcg tct cta cct gat tac tat cac atc ttc cgt tac aac tta gac aag cct gta aat gat tat tgt gag atg taa agg tat ccg aac cac acg gtg tac att gtg tat tag tct tgc tat cac ata atc tga aag cgt aag ttc ccg gag gac acg ata gac aac ata gat tac ggc ttc tgt att cgt tgt ctt tac act ttc cat tgg atg gtg cat gtg gtg cta tat ctc ttc cgt tta tta tta tac atg aga gaa aca ata tat acg agt ata ata cgg acc tca tga ttt aat aat -gt agt aat cgt cgt ctt gtt act gtt tgt ttc cta ctt ctc caa tca tat aga tta ttt ttt aaa tat ttt ctt tct atc atg gat aat att tgt aat ggt tct ttc cgt aca aca tac tgt tta gat ggt agt cgc tta gct tgg tta tga tat tgc gca taa ttt ccg gag gca aat acg ata gtc tag att gac tat cga tgg tag act cta att tat tga gtg ctt tgt cga cga gtt tac ttt tat gct cca tcg ata gat gac act gtt cta tga gat cgt cgt aca tgg gaa atg aaa tgc gtt tgt ctg aat gta tgg ctt cga gat atg tgt gat acc gaa tgt ctt ctg ttc tca ata ccg tat aca agt cgg tgt ctg aga ttc gaa tct ctt tga gga gac tta tgt cac gac tac att ttt cga tga tgg aat cta tct tat cga atg ata tat ttt tca taa ata cac ttt tat agt cct cgt tta aac aga att tag tat gta gtt ccg caa atg act cgt ccc tta ata ggc agt agg cta tta tct tct tta cgt agt gat cgt cgt agg gag aga act ccg aca tct tgt aga aca acg att taa tca tag gta gag ata ctt tca gtc tgt ggt gga taa tgt cat taa cga cat ccg cct tgt ata taa tgt ttc tgt ttt caa aca cca agt cga ata ctg tct tat cgt ctt tag tcg gaa ggt tga tgt cgt atc caa tgt ata cga ggt atg agg caa cat tgt tat tgc aat tct gga agg cgg tat gaa gag gag tca ttg tat tat agt att cgt ctt tct gaa tgt cga atc tat cta gta gat acc gta gta tat tga gag agc gac ttc cat atc ctt gat tat gtt tta tga ata gat aaa gta gat gtt gtc ctt ctt cct ttt gta att tcc cgt att ttt gtt cgt gcc aat tga gta aca tta tga gaa tat gac ctg ttg cac aat cgt tct tta tgt att cca tga tgg gtg tac aat taa gat tat tac gta tcc tcg tat cgg ctc ctc gag ata aaa gag cat aca cca cac gag gac tat gtt tgg tat act gtt gaa ggt aag tgt gta acg gcg tat ttc cga ttt tcg taa ccg cgt taa tgt ttg ctc cat gat cta tta tcg cgt aga tga atc gct tct cag ctc gca tct tag tgt gac tct ttg act tgt aat aat tgc ttt cgt gga acg cgg ata tgt gtt tac agt agt aat gaa gag aag tga gtc cat cct cat cga cgc aat tag ggt cag atc ctt tag tca ata att tgt aca gaa cgt aat agt tta agc tcc cat tga att tat atc taa gat aac aca gca ata gat cgg atg att tac taa agt cat caa tgg ggt ccg tta gta tat caa aga tct tgt tat cga ttg ata gtg aat gaa tca gat agt ggt gta gag gaa tat gtc ctt ttt tat cct tgc tat caa agt tac gca tgc cgt ggc gta aca ata tct tta ata cag atg gat taa atc gtg tat tca tcg tat agc aat gta atg gag agt tac cac att tta gtc gtt tat tca gat cgc agt gtt taa taa cta att taa aca gat gag atg ata tat cca cat caa aga atg cga gat aca tat gac aga cat tat tga cag aaa tgt gac ctt cat tat cac cgt cgt cca taa atg cgt tag gta cgt acc aca tac tat cgt taa cga tgc gca caa tct cgt cca -tt tca tcc atc ttc ata atg att tac ttt ttc ata att aga gaa aaa gat caa ggt ata aaa att aga agt gtt aga cta ta- --- --- --- --- --- --- --- --- --- --- -aa cta act tat gac tta act aac tta tga --- --- --- --- --- --- --- --- --- --- --- --- --- --- --- --- --- --- --- --- --- --- --- --- --- --- --- --- --- --- --- --- --- --- --- --- --- --- --- --- --- --- --- --- --- --- --- --- ctt aac ta- --- --- --- --- --- --- --- --- --- --- -ac tta tga ctt aac taa ctt atg act taa cta act ta- tga ctt aac taa ctt atg act taa cta act tat gac tta act aac tta tga cta tta act cat ttc aag aaa ggt ggg tgg ata gaa ctc tat atg aca gct tgt gaa aca att aga tcc cta att tct aat gga agt ttt gat agg aga ttg tca tca gtt gat aca ttg ttt att atc tca tct att aga gca cgt ctg ttt aga gct tta gtg acc tgc tcg gtt act tct gtg taa atc ttg aat cct tta gtg ata cac tgt gtc aaa act gga tgt tta gaa tac cta tgt aga ata tgg gaa gca tgc ttg ttt ttg tct cta tta tag att aac tca tac atg gtt gta tta tga att ttc atc tgc cta atg tac tcc aat tct tgt tta caa tca att ata taa tca aag agt gat gat gca tac aca tta caa agt gaa taa tct acc atc ata aaa tac ttg ata cag agc ttt atc aca tca tgg ttt tca att gta tta tta agt ata gct aat ttt ata cag tca ata gac aat ggt tct cta agc aat att tct aat att tta aga tgt gct tcc cta cgg gcg atg aca gat ccc cta tcc acg gcc acg tca aga cat gta tat cca tta ctc att act gcg ttg aca ttt gct cca ttt tct aat agc cat gat act aaa tct ata taa cct gca tag ata gcg cga taa agc aag gtc ctt cca cca gca tct agt tga ttg ata tct tca ata tat ggg ata caa agc tta taa att tct aat act gtg ggt tca tct aca agg aat ccc cta gta tac tga att att tta tat aga tct aat tta aca tca ttt tca tct ggg ata cca caa ttc aaa ata aac tca aca aca cta ctt tcc ttt tta cat att ccc cta aaa tag gca ttc aag cat tct att tta tat att aca gcc cca tga tct acc ata aga tca aca atg tct att tct aca tat gca tta gat aga tag taa agt aag aga tct tgc aca gaa tta caa ttc tta ata att ata gag aaa ata tct tcc ata taa ttc ttt gac act aat gca gat ata ata tct tta tat gta ata tat gca aac agt cta tct act ata tac tga tca ata tta tct cta tga atc cta aaa taa tca tac aga aca tct aca gga tca caa att ggt tca agg aga aat cta tca aat att ttc ctg tca aca act ggt tct aga aca taa cag tca aca cct aat cca tgt ttt tta taa tca tct acc aaa gat aat gac caa aga tcg agg tcg tcg tga aac tgc tca tcg aca gcc atg aaa tct acc gac tcc atg gtg cga atc gca ctg tct tat tcg cca ttg att ttc a-t ttt tta taa tta tgt aca tgt ttt cct tct att ctc aag agt cta caa aaa tat a-- -tt ttt tcg ata tct aag tac taa gtt ttt tta ctg ttt ttg tta ctg tct tcc att ctt cta act aaa gat ctg aga taa att ata caa tct tcg cta tcg aac cat ttt tgt agt cta aag cct gaa gta att aac caa ctg ttt tta tta gtg gct ttt ttc gat cta tcc tcg tc- --- ctc tca tca tcc tta ta- --- --- --- --- --- --- tta tta tca tta tta tca tag tct att aaa cac aaa tca tct --- --- --- --- --- --- --- --- --- --- --- --- --- --- --- --- --- --- --- --- --- --- --- --- --- --- --a cgt tta taa c-- --- --- --- --- --- --- --- --a aca ttc --- --- --- --- -tc att att aat tag ttc tgt ag- aat atc ttt aat aat ttg gct ata --- --- -ca tct gtt --- --- --- --- --- --- --- --- --- --- --- --- --- --- --- --- --- --- --c aat act --- --- --- --- --- --- --- --a tct att gat gat ttc ttt t-- --- --- --- --- --- --- --- --- --- --- --t taa gac t-- --- --- --- --- --- --- --- --- --- --- --- --- --- --- --- --- --- --- --- --- --- --- --- --- --- --- --- --- --- --- --- --- --- --- --- --- --- --- --- taa act agt --- --- --- --- --- --- --- --- --- --- --- --- --- --- --- --- --- --- --- --- --- --- --- --- --- --- --- --- --- --- --- --- --- --- --- --- --- --- --- --- --- --- --- --- --- --- --- --- --- --- --- --- --- --- --- --- --- --- --- --- --- --- --- --- --- --- --- --- --- --- --- --- --- --- --- --- --- --- --- --- --- --- --- --- --- --- --- --- --- --- --- --- --- --- --- --- --- --- --- --- --- --- --- --- --- --- --- --- --- --t atg gta atg acg atg aaa --- -tc gag tag ta- --- --a ctt cta ata aag act tga ta- --- --t cat tat cat atg ttt gat cg- --- --- --- --- --- --- --t cat agt taa tag tgt g-- --- --- --- --- --- --- --- --- --- --- --- --- --- --- --- --- --- --- --- -gc taa atg gta ctg tta ata agt tta t-- --- --- --- --- --- --- --- --- --- --- --- --- --- --- --- --- --a gac aat atc ata gta ttt tct ttc cag aat --- --- --- --t aga tta ttt ttt taa ata ctg atc ctc aca att ccg tga tgt agc agt agt tgg t-- --- --- --- --- --- --- --- --- --- --- --- --- --- --- --- --- --- --- --- --- --- --- --- --- --- --- --- -gc atg gtc tat atc gt- --- --- --- --- --- --- --- --- --- --- --- --- --- --- --- --- --- --- --- --- --- --- --- --- --- --- --- --- --- --- --- --- --- --- --- --- --- --- --- --- --- --- --- taa aat gta tca tat ata ata gtt ttc tga cgt gga gta cag aat ttt cga --- --- --- --- --- --- --- --- --- --- --- --- --- --- --- --- --- --- --- --- --- --- --- --- --- --- --- --- --- --- --- --- --- --- --- --- --- --- --- --- --- --- --- --t taa tga gtt cat ggt aag gaa ggg caa atg cct --- gta tat aat ata cat aag tta a-- --- --- --- --- --- --- --- --- --- --- --- --- --t agt ttt tta tca tat tt- --- --- --- --- --- --- --- --t cta ata cca taa taa aaa t-- --- --- --- --- --- --- --- --- --- --- --- --- --- --- --- --- --- --- --- --- --- --- --- --- --t atc att att gcg ttt g-g tag tt- --- --- --- --- ctg ccc ta- --- --- --- --- --- --- --- --t cat cta tat cac tgt cac tct c-- --- --g ctc tca cta tat ctt cta aaa tta caa --- -ac aac --- --- --- --- --- --- --- --- --- --- --- --- --- --- --- --- --- --- --- --- --- --- --- --- --- --- --- --- --- --- --- --- --- --- --- --- tgg ata ttc gat --- --- --- --- --- --- --- --- --- --- --- --- --- --- --- --- --- --- --- --- --- --- --- --- --- --- --a aca gca ttt gtg t-- --- --- --- --- --- --- --- --- --- --- --- --- --- --- --- --- -

>ON615424.1_Monkeypox_virus_isolate_MPXV_2022_NL001_partial_genome

--- --- --- --- --- --- --- --- --- --- --- --- --- --- --- --- --- --- --- --- --- --- --- --- --- --- --- --- --- --- --- --c tat ttt att tag tgt cta gaa aaa aat gtg tga ccc acg acc gta gga aac tct aga ggg taa gaa aaa tca atc gtt tat aga gac cat cag aaa gag gtt taa tat ttt tgt gag acc tat cga a-- --- --- --- --- --- --- --- --- --- --- --- --- --- --- --- --- --- --- --- --- --- --- --- --- --- --- --- --- --- --- --- --- --- --- --- --- --- --- --- --- --- --- --- --- --- --- --- --- --- --- --- --- --- --- --- --- --- --- --- --- --- --- --- --- --- --- --- --- --- --- --- --- --- --- --- --- --- --- --- --- --- --- --- --- --- --- --- --- --- --- --- --- --- --- --- --- --- --- --- --- --- --- --- --- --- --- --- --- --- --- --- --- --- --- --- --- --- --- --- --- --- --- --- --- --- --- --- --- --- --- --- --- --- --- --- --- --- --- --- --- --- --- --- --- --- --- --- --- --- --- --- --- --- --- --- --- --- --- --- --- --- --- --- --- --- --- --- --- --- --- --- --- --- --- --- --- --- --- --- --- --- --- --- --- --- --- --- --- --- --- --- --- --- --- --- --- --- --- --- --- --- --- --- --- --- --- --- --- --- --- --- --- --- --- --- --- --- --- --- --- --- --- --- --- --- --- --- --- --- --- --- --- --- --- --- --- --- --- --- --- --- --- --- --- --- --- --- --- --- --- --- --- --- --- --- --- --- --- --- --- --- --- --- --- --- --- --- --- --- --- --- --- --- --- --- --- --- --- --- -ga gag aaa gga taa aaa ctt ttt acg act cca tca gaa aga ggt tta ata ttt ttg tga gac cca tcg aag aga gaa aga gat ggt tag tca aga tat ttt tct tag tac aaa agt caa tgt ttt aaa ata tat gga cga gaa tta att tgt ctg tat aaa aac ttg tgt gaa att atg tac tag aga aaa aac gtg agc agt gtc ccc tac atg gat ttt aca gat cat tta tat tcc aaa aat att aac tat ata cgt tta tta tat gat gtt aac gtg taa att ata aac att att tta tga tgc aat tgt ctg aca acc tag att ggt ata agg atg ttg ata agc tct acg aga ata tat tgt tgg acg tta tcg ttt acg aaa tag ttg aga cat cag aaa gag gtt taa tat ttt tgt gag a-c cat cga aga gag aaa gag aat aaa aat a-- --t ttt ttt ttt ttt tgt aaa act ttt tta tga gac --- --- -ca aga gaa tac gaa tag tga tca tat cgt atc aca tat tga aac aga aag aag aag taa cga gag gta act ttt tgt gaa tgt agt taa ata ttt ttg ttt tgc aaa ccg gaa tat agt gcc cgg t-- -ct ttt tta att cgt ggt gcg gtg tct gaa tcg ttc gat taa ccc aac tca tcc att ttc aga tga ata gag tta tcg att cag aca cat gct ttg agt ttt gtt gaa tcg atg agt gaa gta tca tcg gtt gca cct tca gat gcc gat ccg tcg aca tac ttg aat cca tcc ttg act tca agt tca gat gat tcc tca cac atg tct ccg ata cgt acg cta aac tct agg ttc ttg aca cat ttt gta tca acg atc gtt gaa ccg atg ata tct ttg taa ctc act ttc tta tgt gag atg tta gac cca agt act gga tgg gtc ttg atg tca ctg tct ttc tct tct tcg cta cat ctg atg tcg ata gac atc tca cag tct ttg atc ata gcc aga gct tct tca cgc gtg atc gcg gga gag tcc tta cct tgt ccc ggt gac acg ctg gac aat cta gta ttc aca gtg ttt cca tca gag gat tcg gag atg gat aaa atc ttt ggg cat ttg gtg aat cca aag ttc atg tta aga ccc gca ccg acg ata gtg taa taa gtg gtg gga tct cct ttt aca act --- --- --- --- --- --- tct tcg gat acc tca tca tct tcg gtc tct gta act tcc gtt acg gat tga caa atc tta tca ttg gtc ggt gtt tgg tct tgc ttt gtg act ttg ata ata aca tcg att ccc ata tga tgt ttg ttt tct tct tca gta cac gag gat gaa gat tgt tga aga cta gta ggc ata gca gct gcc act agg cac atg cat gcc agg aca ata tat tgt ttc atg att gct att gat tga tta ctg ttc tag atg att cta ctt tct tac cat ata ata aat tag aat ata ttt tct act ttt acg aga aat taa tta ttg tat tta tta ttt ata ggt --a aaa aaa ctt act ata agt ggg tgg gat tct ggg aat tag tga tca gtt tat gta tat cgc aac tag cgg gca tat ggc tat tga cat cga gaa cat tac cca tat gat aag aga ttg tat cat ttt cgt agt ctt gag tat tgg tat tac tat ata gta tgt aga tgt cga cgc tag ata gac agt cgc cca cta gag tta ccg tct ctg aat gcg gca tga tag tat cat tct ttg ttt tcg tta act gtt tgg aag atg aat ctt tgt tgt tac att taa tct cga aat tca gag tac ata tct ttg aag tat tct gat atc tat ttt ctc ctg taa aga atc ctg aag ttg cta cat tat taa gga cag aga agt att ctg cac gaa aga ctg gat cac aat ctt tat gat tca tgg taa tag tta gtt ccg acg ttg aga tgg att cgc tga gac cgg tag tgg tcg tcc gag tac acg atg tgt cgt tga ctg gat aca ggt taa ttt cca cat cga tat agt taa atg tat tgc tgg tta cga cgg gtt cgc att tat ctg tgg aag aga cgg tgt gag aat atg ttc cgg gac cac acg gag aac aga tga cgt ctc cgg tag acg tgt atc cgg ata ctc cgt atc cta ttc cac act ttg ttt tag aaa tac atg ttc tac acc ctg atg ctc ctt tga gaa gac aat aat atc ctg gag agc att cac aga ttc tat tgt gag tcg tgt tac acg atc gcg tct cta cct gat tac tat cac atc ttc cgt tac aac tta gac aag cct gta aat gat tat tgt gag atg taa agg tat ccg aac cac acg gtg tac att gtg tat tag tct tgc tat cac ata atc tga aag cgt aag ttc ccg gag gac acg ata gac aac ata gat tac ggc ttc tgt att cgt tgt ctt tac act ttc cat tgg atg gtg cat gtg gtg cta tat ctc ttc cgt tta tta tta tac atg aga gaa aca ata tat acg agt ata ata cgg acc tca tga ttt aat aat -gt agt aat cgt cgt ctt gtt act gtt tgt ttc cta ctt ctc caa tca tat aga tta ttt ttt aaa tat ttt ctt tct atc atg gat aat att tgt aat ggt tct ttc cgt aca aca tac tgt tta gat ggt agt cgc tta gct tgg tta tga tat tgc gca taa ttt ccg gag gca aat acg ata gtc tag att gac tat cga tgg tag act cta att tat tga gtg ctt tgt cga cga gtt tac ttt tat gct cca tcg ata gat gac act gtt cta tga gat cgt cgt aca tgg gaa atg aaa tgc gtt tgt ctg aat gta tgg ctt cga gat atg tgt gat acc gaa tgt ctt ctg ttc tca ata ccg tat aca agt cgg tgt ctg aga ttc gaa tct ctt tga gga gac tta tgt cac gac tac att ttt cga tga tgg aat cta tct tat cga atg ata tat ttt tca taa ata cac ttt tat agt cct cgt tta aac aga att tag tat gta gtt ccg caa atg act cgt ccc tta ata ggc agt agg cta tta tct tct tta cgt agt gat cgt cgt agg gag aga act ccg aca tct tgt aga aca acg att taa tca tag gta gag ata ctt tca gtc tgt ggt gga taa tgt cat taa cga cat ccg cct tgt ata taa tgt ttc tgt ttt caa aca cca agt cga ata ctg tct tat cgt ctt tag tcg gaa ggt tga tgt cgt atc caa tgt ata cga ggt atg agg caa cat tgt tat tgc aat tct gga agg cgg tat gaa gag gag tca ttg tat tat agt att cgt ctt tct gaa tgt cga atc tat cta gta gat acc gta gta tat tga gag agc gac ttc cat atc ctt gat tat gtt tta tga ata gat aaa gta gat gtt gtc ctt ctt cct ttt gta att tcc cgt att ttt gtt cgt gcc aat tga gta aca tta tga gaa tat gac ctg ttg cac aat cgt tct tta tgt att cca tga tgg gtg tac aat taa gat tat tac gta tcc tcg tat cgg ctc ctc gag ata aaa gag cat aca cca cac gag gac tat gtt tgg tat act gtt gaa ggt aag tgt gta acg gcg tat ttc cga ttt tcg taa ccg cgt taa tgt ttg ctc cat gat cta tta tcg cgt aga tga atc gct tct cag ctc gca tct tag tgt gac tct ttg act tgt aat aat tgc ttt cgt gga acg cgg ata tgt gtt tac agt agt aat gaa gag aag tga gtc cat cct cat cga cgc aat tag ggt cag atc ctt tag tca ata att tgt aca gaa cgt aat agt tta agc tcc cat tga att tat atc taa gat aac aca gca ata gat cgg atg att tac taa agt cat caa tgg ggt ccg tta gta tat caa aga tct tgt tat cga ttg ata gtg aat gaa tca gat agt ggt gta gag gaa tat gtc ctt ttt tat cct tgc tat caa agt tac gca tgc cgt ggc gta aca ata tct tta ata cag atg gat taa atc gtg tat tca tcg tat agc aat gta atg gag agt tac cac att tta gtc gtt tat tca gat cgc agt gtt taa taa cta att taa aca gat gag atg ata tat cca cat caa aga atg cga gat aca tat gac aga cat tat tga cag aaa tgt gac ctt cat tat cac cgt cgt cca taa atg cgt tag gta cgt acc aca tac tat cgt taa cga tgc gca caa tct cgt cca -tt tca tcc atc ttc ata atg att tac ttt ttc ata att aga gaa aaa gat caa ggt ata aaa att aga agt gtt aga cta ta- --- --- --- --- --- --- --- --- --- --- -aa cta act tat gac tta act aac tta tga --- --- --- --- --- --- --- --- --- --- --- --- --- --- --- --- --- --- --- --- --- --- --- --- --- --- --- --- --- --- --- --- --- --- --- --- --- --- --- --- --- --- --- --- --- --- --- --- ctt aac ta- --- --- --- --- --- --- --- --- --- --- -ac tta tga ctt aac taa ctt atg act taa cta act ta- tga ctt aac taa ctt atg act taa cta act tat gac tta act aac tta tga ct? ??? ??? ??? ??? ??? ??? ??? ??? ??? ??? ??? ??? ??? ??? ??? ??? ??? ??? ??? ??? ??? ??? ?ta att tct aat gga agt ttt gat agg aga ttg tca tca gtt gat aca ttg ttt att atc tca tct att aga gca cgt ctg ttt aga gct tta gtg acc tgc tcg gtt act tct gtg taa atc ttg aat cct tta gtg ata cac tgt gtc aaa act gga tgt tta gaa tac cta tgt aga ata tgg gaa gca tgc ttg ttt ttg tct cta tta tag att aac tca tac atg gtt gta tta tga att ttc atc tgc cta atg tac tcc aat tct tgt tta caa tca att ata taa tca aag agt gat gat gca tac aca tta caa agt gaa taa tct acc atc ata aaa tac ttg ata cag agc ttt atc aca tca tgg ttt tca att gta tta tta agt ata gct aat ttt ata cag tca ata gac aat ggt tct cta agc aat att tct aat att tta aga tgt gct tcc cta cgg gcg atg aca gat ccc cta tcc acg gcc acg tca aga cat gta tat cca tta ctc att act gcg ttg aca ttt gct cca ttt tct aat agc cat gat act aaa tct ata taa cct gca tag ata gcg cga taa agc aag gtc ctt cca cca gca tct agt tga ttg ata tct tca ata tat ggg ata caa agc tta taa att tct aat act gtg ggt tca tct aca agg aat ccc cta gta tac tga att att tta tat aga tct aat tta aca tca ttt tca tct ggg ata cca caa ttc aaa ata aac tca aca aca cta ctt tcc ttt tta cat att ccc cta aaa tag gca ttc aag cat tct att tta tat att aca gcc cca tga tct acc ata aga tca aca atg tct att tct aca tat gca tta gat aga tag taa agt aag aga tct tgc aca gaa tta caa ttc tta ata att ata gag aaa ata tct tcc ata taa ttc ttt gac act aat gca gat ata ata tct tta tat gta ata tat gca aac agt cta tct act ata tac tga tca ata tta tct cta tga atc cta aaa taa tca tac aga aca tct aca gga tca caa att ggt tca agg aga aat cta tca aat att ttc ctg tca aca act ggt tct aga aca taa cag tca aca cct aat cca tgt ttt tta taa tca tct acc aaa gat aat gac caa aga tcg agg tcg tcg tga aac tgc tca tcg aca gcc atg aaa tct acc gac tcc atg gtg cga atc gca ctg tct tat tcg cca ttg att ttc a-t ttt tta taa tta tgt aca tgt ttt cct tct att ctc aag agt cta caa aaa tat a-- -tt ttt tcg ata tct aag tac taa gtt ttt tta ctg ttt ttg tta ctg tct tcc att ctt cta act aaa gat ctg aga taa att ata caa tct tcg cta tcg aac cat ttt tgt agt cta aag cct gaa gta att aac caa ctg ttt tta tta gtg gct ttt ttc gat cta tcc tcg tc- --- ctc tca tca tcc tta ta- --- --- --- --- --- --- tta tta tca tta tta tca tag tct att aaa cac aaa tca tct --- --- --- --- --- --- --- --- --- --- --- --- --- --- --- --- --- --- --- --- --- --- --- --- --- --- --a cgt tta taa c-- --- --- --- --- --- --- --- --a aca ttc --- --- --- --- -tc att att aat tag ttc tgt ag- aat atc ttt aat aat ttg gct ata --- --- -ca tct gtt --- --- --- --- --- --- --- --- --- --- --- --- --- --- --- --- --- --- --c aat act --- --- --- --- --- --- --- --a tct att gat gat ttc ttt t-- --- --- --- --- --- --- --- --- --- --- --t taa gac t-- --- --- --- --- --- --- --- --- --- --- --- --- --- --- --- --- --- --- --- --- --- --- --- --- --- --- --- --- --- --- --- --- --- --- --- --- --- --- --- taa act agt --- --- --- --- --- --- --- --- --- --- --- --- --- --- --- --- --- --- --- --- --- --- --- --- --- --- --- --- --- --- --- --- --- --- --- --- --- --- --- --- --- --- --- --- --- --- --- --- --- --- --- --- --- --- --- --- --- --- --- --- --- --- --- --- --- --- --- --- --- --- --- --- --- --- --- --- --- --- --- --- --- --- --- --- --- --- --- --- --- --- --- --- --- --- --- --- --- --- --- --- --- --- --- --- --- --- --- --- --- --t atg gta atg acg atg aaa --- -tc gag tag ta- --- --a ctt cta ata aag act tga ta- --- --t cat tat cat atg ttt gat cg- --- --- --- --- --- --- --t cat agt taa tag tgt g-- --- --- --- --- --- --- --- --- --- --- --- --- --- --- --- --- --- --- --- -gc taa atg gta ctg tta ata agt tta t-- --- --- --- --- --- --- --- --- --- --- --- --- --- --- --- --- --a gac aat atc ata gta ttt tct ttc cag aat --- --- --- --t aga tta ttt ttt taa ata ctg atc ctc aca att ccg tga tgt agc agt agt tgg t-- --- --- --- --- --- --- --- --- --- --- --- --- --- --- --- --- --- --- --- --- --- --- --- --- --- --- --- -gc atg gtc tat atc gt- --- --- --- --- --- --- --- --- --- --- --- --- --- --- --- --- --- --- --- --- --- --- --- --- --- --- --- --- --- --- --- --- --- --- --- --- --- --- --- --- --- --- --- taa aat gta tca tat ata ata gtt ttc tga cgt gga gta cag aat ttt cga --- --- --- --- --- --- --- --- --- --- --- --- --- --- --- --- --- --- --- --- --- --- --- --- --- --- --- --- --- --- --- --- --- --- --- --- --- --- --- --- --- --- --- --t taa tga gtt cat ggt aag gaa ggg caa atg cct --- gta tat aat ata cat aag tta a-- --- --- --- --- --- --- --- --- --- --- --- --- --t agt ttt tta tca tat tt- --- --- --- --- --- --- --- --t cta ata cca taa taa aaa t-- --- --- --- --- --- --- --- --- --- --- --- --- --- --- --- --- --- --- --- --- --- --- --- --- --t atc att att gcg ttt g-g tag tt- --- --- --- --- ctg ccc ta- --- --- --- --- --- --- --- --t cat cta tat cac tgt cac tct c-- --- --g ctc tca cta tat ctt cta aaa tta caa --- -ac aac --- --- --- --- --- --- --- --- --- --- --- --- --- --- --- --- --- --- --- --- --- --- --- --- --- --- --- --- --- --- --- --- --- --- --- --- tgg ata ttc gat --- --- --- --- --- --- --- --- --- --- --- --- --- --- --- --- --- --- --- --- --- --- --- --- --- --- --a aca gca ttt gtg t-- --- --- --- --- --- --- --- --- --- --- --- --- --- --- --- --- -

>ON585034.1_Monkeypox_virus_isolate_Monkeypox/PT0007/2022_complete_genome

--- --- --- --- --- --- --- --- --- --- --- --- --- --- --- --- --- --- --- --- --- --- --- --- --- --- --- --- att taa ttt tac tat ttt att tag tgt cta gaa aaa aat gtg tga ccc acg acc gta gga aac tct aga ggg taa gaa aaa tca atc gtt tat aga gac cat cag aaa gag gtt taa tat ttt tgt gag acc tat cga a-- --- --- --- --- --- --- --- --- --- --- --- --- --- --- --- --- --- --- --- --- --- --- --- --- --- --- --- --- --- --- --- --- --- --- --- --- --- --- --- --- --- --- --- --- --- --- --- --- --- --- --- --- --- --- --- --- --- --- --- --- --- --- --- --- --- --- --- --- --- --- --- --- --- --- --- --- --- --- --- --- --- --- --- --- --- --- --- --- --- --- --- --- --- --- --- --- --- --- --- --- --- --- --- --- --- --- --- --- --- --- --- --- --- --- --- --- --- --- --- --- --- --- --- --- --- --- --- --- --- --- --- --- --- --- --- --- --- --- --- --- --- --- --- --- --- --- --- --- --- --- --- --- --- --- --- --- --- --- --- --- --- --- --- --- --- --- --- --- --- --- --- --- --- --- --- --- --- --- --- --- --- --- --- --- --- --- --- --- --- --- --- --- --- --- --- --- --- --- --- --- --- --- --- --- --- --- --- --- --- --- --- --- --- --- --- --- --- --- --- --- --- --- --- --- --- --- --- --- --- --- --- --- --- --- --- --- --- --- --- --- --- --- --- --- --- --- --- --- --- --- --- --- --- --- --- --- --- --- --- --- --- --- --- --- --- --- --- --- --- --- --- --- --- --- --- --- --- --- --- -ga gag aaa gga taa aaa ctt ttt acg act cca tca gaa aga ggt tta ata ttt ttg tga gac cca tcg aag aga gaa aga gat ggt tag tca aga tat ttt tct tag tac aaa agt caa tgt ttt aaa ata tat gga cga gaa tta att tgt ctg tat aaa aac ttg tgt gaa att atg tac tag aga aaa aac gtg agc agt gtc ccc tac atg gat ttt aca gat cat tta tat tcc aaa aat att aac tat ata cgt tta tta tat gat gtt aac gtg taa att ata aac att att tta tga tgc aat tgt ctg aca acc tag att ggt ata agg atg ttg ata agc tct acg aga ata tat tgt tgg acg tta tcg ttt acg aaa tag ttg aga cat cag aaa gag gtt taa tat ttt tgt gag a-c cat cga aga gag aaa gag aat aaa aat a-- --t ttt ttt ttt ttt tgt aaa act ttt tta tga gac --- --- -ca aga gaa tac gaa tag tga tca tat cgt atc aca tat tga aac aga aag aag aag taa cga gag gta act ttt tgt gaa tgt agt taa ata ttt ttg ttt tgc aaa ccg gaa tat agt gcc cgg t-- -ct ttt tta att cgt ggt gcg gtg tct gaa tcg ttc gat taa ccc aac tca tcc att ttc aga tga ata gag tta tcg att cag aca cat gct ttg agt ttt gtt gaa tcg atg agt gaa gta tca tcg gtt gca cct tca gat gcc gat ccg tcg aca tac ttg aat cca tcc ttg act tca agt tca gat gat tcc tca cac atg tct ccg ata cgt acg cta aac tct agg ttc ttg aca cat ttt gta tca acg atc gtt gaa ccg atg ata tct ttg taa ctc act ttc tta tgt gag atg tta gac cca agt act gga tgg gtc ttg atg tca ctg tct ttc tct tct tcg cta cat ctg atg tcg ata gac atc tca cag tct ttg atc ata gcc aga gct tct tca cgc gtg atc gcg gga gag tcc tta cct tgt ccc ggt gac acg ctg gac aat cta gta ttc aca gtg ttt cca tca gag gat tcg gag atg gat aaa atc ttt ggg cat ttg gtg aat cca aag ttc atg tta aga ccc gca ccg acg ata gtg taa taa gtg gtg gga tct cct ttt aca act --- --- --- --- --- --- tct tcg gat acc tca tca tct tcg gtc tct gta act tcc gtt acg gat tga caa atc tta tca ttg gtc ggt gtt tgg tct tgc ttt gtg act ttg ata ata aca tcg att ccc ata tga tgt ttg ttt tct tct tca gta cac gag gat gaa gat tgt tga aga cta gta ggc ata gca gct gcc act agg cac atg cat gcc agg aca ata tat tgt ttc atg att gct att gat tga tta ctg ttc tag atg att cta ctt tct tac cat ata ata aat tag aat ata ttt tct act ttt acg aga aat taa tta ttg tat tta tta ttt ata ggt --a aaa aaa ctt act ata agt ggg tgg gat tct ggg aat tag tga tca gtt tat gta tat cgc aac tag cgg gca tat ggc tat tga cat cga gaa cat tac cca tat gat aag aga ttg tat cat ttt cgt agt ctt gag tat tgg tat tac tat ata gta tgt aga tgt cga cgc tag ata gac agt cgc cca cta gag tta ccg tct ctg aat gcg gca tga tag tat cat tct ttg ttt tcg tta act gtt tgg aag atg aat ctt tgt tgt tac att taa tct cga aat tca gag tac ata tct ttg aag tat tct gat atc tat ttt ctc ctg taa aga atc ctg aag ttg cta cat tat taa gga cag aga agt att ctg cac gaa aga ctg gat cac aat ctt tat gat tca tgg taa tag tta gtt ccg acg ttg aga tgg att cgc tga gac cgg tag tgg tcg tcc gag tac acg atg tgt cgt tga ctg gat aca ggt taa ttt cca cat cga tat agt taa atg tat tgc tgg tta cga cgg gtt cgc att tat ctg tgg aag aga cgg tgt gag aat atg ttc cgg gac cac acg gag aac aga tga cgt ctc cgg tag acg tgt atc cgg ata ctc cgt atc cta ttc cac act ttg ttt tag aaa tac atg ttc tac acc ctg atg ctc ctt tga gaa gac aat aat atc ctg gag agc att cac aga ttc tat tgt gag tcg tgt tac acg atc gcg tct cta cct gat tac tat cac atc ttc cgt tac aac tta gac aag cct gta aat gat tat tgt gag atg taa agg tat ccg aac cac acg gtg tac att gtg tat tag tct tgc tat cac ata atc tga aag cgt aag ttc ccg gag gac acg ata gac aac ata gat tac ggc ttc tgt att cgt tgt ctt tac act ttc cat tgg atg gtg cat gtg gtg cta tat ctc ttc cgt tta tta tta tac atg aga gaa aca ata tat acg agt ata ata cgg acc tca tga ttt aat aat -gt agt aat cgt cgt ctt gtt act gtt tgt ttc cta ctt ctc caa tca tat aga tta ttt ttt aaa tat ttt ctt tct atc atg gat aat att tgt aat ggt tct ttc cgt aca aca tac tgt tta gat ggt agt cgc tta gct tgg tta tga tat tgc gca taa ttt ccg gag gca aat acg ata gtc tag att gac tat cga tgg tag act cta att tat tga gtg ctt tgt cga cga gtt tac ttt tat gct cca tcg ata gat gac act gtt cta tga gat cgt cgt aca tgg gaa atg aaa tgc gtt tgt ctg aat gta tgg ctt cga gat atg tgt gat acc gaa tgt ctt ctg ttc tca ata ccg tat aca agt cgg tgt ctg aga ttc gaa tct ctt tga gga gac tta tgt cac gac tac att ttt cga tga tgg aat cta tct tat cga atg ata tat ttt tca taa ata cac ttt tat agt cct cgt tta aac aga att tag tat gta gtt ccg caa atg act cgt ccc tta ata ggc agt agg cta tta tct tct tta cgt agt gat cgt cgt agg gag aga act ccg aca tct tgt aga aca acg att taa tca tag gta gag ata ctt tca gtc tgt ggt gga taa tgt cat taa cga cat ccg cct tgt ata taa tgt ttc tgt ttt caa aca cca agt cga ata ctg tct tat cgt ctt tag tcg gaa ggt tga tgt cgt atc caa tgt ata cga ggt atg agg caa cat tgt tat tgc aat tct gga agg cgg tat gaa gag gag tca ttg tat tat agt att cgt ctt tct gaa tgt cga atc tat cta gta gat acc gta gta tat tga gag agc gac ttc cat atc ctt gat tat gtt tta tga ata gat aaa gta gat gtt gtc ctt ctt cct ttt gta att tcc cgt att ttt gtt cgt gcc aat tga gta aca tta tga gaa tat gac ctg ttg cac aat cgt tct tta tgt att cca tga tgg gtg tac aat taa gat tat tac gta tcc tcg tat cgg ctc ctc gag ata aaa gag cat aca cca cac gag gac tat gtt tgg tat act gtt gaa ggt aag tgt gta acg gcg tat ttc cga ttt tcg taa ccg cgt taa tgt ttg ctc cat gat cta tta tcg cgt aga tga atc gct tct cag ctc gca tct tag tgt gac tct ttg act tgt aat aat tgc ttt cgt gga acg cgg ata tgt gtt tac agt agt aat gaa gag aag tga gtc cat cct cat cga cgc aat tag ggt cag atc ctt tag tca ata att tgt aca gaa cgt aat agt tta agc tcc cat tga att tat atc taa gat aac aca gca ata gat cgg atg att tac taa agt cat caa tgg ggt ccg tta gta tat caa aga tct tgt tat cga ttg ata gtg aat gaa tca gat agt ggt gta gag gaa tat gtc ctt ttt tat cct tgc tat caa agt tac gca tgc cgt ggc gta aca ata tct tta ata cag atg gat taa atc gtg tat tca tcg tat agc aat gta atg gag agt tac cac att tta gtc gtt tat tca gat cgc agt gtt taa taa cta att taa aca gat gag atg ata tat cca cat caa aga atg cga gat aca tat gac aga cat tat tga cag aaa tgt gac ctt cat tat cac cgt cgt cca taa atg cgt tag gta cgt acc aca tac tat cgt taa cga tgc gca caa tct cgt cca -tt tca tcc atc ttc ata atg att tac ttt ttc ata att aga gaa aaa gat caa ggt ata aaa att aga agt gtt aga cta ta- --- --- --- --- --- --- --- --- --- --- -aa cta act tat gac tta act aac tta tga --- --- --- --- --- --- --- --- --- --- --- --- --- --- --- --- --- --- --- --- --- --- --- --- --- --- --- --- --- --- --- --- --- --- --- --- --- --- --- --- --- --- --- --- --- --- --- --- ctt aac ta- --- --- --- --- --- --- --- --- --- --- -ac tta tga ctt aac taa ctt atg act taa cta act ta- tga ctt aac taa ctt atg act taa cta act tat gac tta act aac tta tga cta tta act cat ttc aag aaa ggt ggg tgg ata gaa ctc tat atg aca gct tgt gaa aca att aga tcc cta att tct aat gga agt ttt gat agg aga ttg tca tca gtt gat aca ttg ttt att atc tca tct att aga gca cgt ctg ttt aga gct tta gtg acc tgc tcg gtt act tct gtg taa atc ttg aat cct tta gtg ata cac tgt gtc aaa act gga tgt tta gaa tac cta tgt aga ata tgg gaa gca tgc ttg ttt ttg tct cta tta tag att aac tca tac atg gtt gta tta tga att ttc atc tgc cta atg tac tcc aat tct tgt tta caa tca att ata taa tca aag agt gat gat gca tac aca tta caa agt gaa taa tct acc atc ata aaa tac ttg ata cag agc ttt atc aca tca tgg ttt tca att gta tta tta agt ata gct aat ttt ata cag tca ata gac aat ggt tct cta agc aat att tct aat att tta aga tgt gct tcc cta cgg gcg atg aca gat ccc cta tcc acg gcc acg tca aga cat gta tat cca tta ctc att act gcg ttg aca ttt gct cca ttt tct aat agc cat gat act aaa tct ata taa cct gca tag ata gcg cga taa agc aag gtc ctt cca cca gca tct agt tga ttg ata tct tca ata tat ggg ata caa agc tta taa att tct aat act gtg ggt tca tct aca agg aat ccc cta gta tac tga att att tta tat aga tct aat tta aca tca ttt tca tct ggg ata cca caa ttc aaa ata aac tca aca aca cta ctt tcc ttt tta cat att ccc cta aaa tag gca ttc aag cat tct att tta tat att aca gcc cca tga tct acc ata aga tca aca atg tct att tct aca tat gca tta gat aga tag taa agt aag aga tct tgc aca gaa tta caa ttc tta ata att ata gag aaa ata tct tcc ata taa ttc ttt gac act aat gca gat ata ata tct tta tat gta ata tat gca aac agt cta tct act ata tac tga tca ata tta tct cta tga atc cta aaa taa tca tac aga aca tct aca gga tca caa att ggt tca agg aga aat cta tca aat att ttc ctg tca aca act ggt tct aga aca taa cag tca aca cct aat cca tgt ttt tta taa tca tct acc aaa gat aat gac caa aga tcg agg tcg tcg tga aac tgc tca tcg aca gcc atg aaa tct acc gac tcc atg gtg cga atc gca ctg tct tat tcg cca ttg att ttc a-t ttt tta taa tta tgt aca tgt ttt cct tct att ctc aag ?gt cta caa aaa tat a-- -tt ttt tcg at? tct aag tac taa gtt ttt tta ctg ttt ttg tta ctg tct tcc att ctt cta act aaa gat ctg aga taa att ata caa tct tcg cta tcg aac cat ttt tgt agt cta aag cct gaa gta att aac caa ctg ttt tta tta gtg gct ttt ttc gat cta tcc tcg tc- --- ctc tca tca tcc tta ta- --- --- --- --- --- --- tta tta tca tta tta tca tag tct att aaa cac aaa tca tct --- --- --- --- --- --- --- --- --- --- --- --- --- --- --- --- --- --- --- --- --- --- --- --- --- --- --a cgt tta taa c-- --- --- --- --- --- --- --- --a aca ttc --- --- --- --- -tc att att aat tag ttc tgt ag- aat atc ttt aat aat ttg gct ata --- --- -ca tct gtt --- --- --- --- --- --- --- --- --- --- --- --- --- --- --- --- --- --- --c aat act --- --- --- --- --- --- --- --a tct att gat gat ttc ttt t-- --- --- --- --- --- --- --- --- --- --- --t taa gac t-- --- --- --- --- --- --- --- --- --- --- --- --- --- --- --- --- --- --- --- --- --- --- --- --- --- --- --- --- --- --- --- --- --- --- --- --- --- --- --- taa act agt --- --- --- --- --- --- --- --- --- --- --- --- --- --- --- --- --- --- --- --- --- --- --- --- --- --- --- --- --- --- --- --- --- --- --- --- --- --- --- --- --- --- --- --- --- --- --- --- --- --- --- --- --- --- --- --- --- --- --- --- --- --- --- --- --- --- --- --- --- --- --- --- --- --- --- --- --- --- --- --- --- --- --- --- --- --- --- --- --- --- --- --- --- --- --- --- --- --- --- --- --- --- --- --- --- --- --- --- --- --t atg gta atg acg atg aaa --- -tc gag tag ta- --- --a ctt cta ata aag act tga ta- --- --t cat tat cat atg ttt gat cg- --- --- --- --- --- --- --t cat agt taa tag tgt g-- --- --- --- --- --- --- --- --- --- --- --- --- --- --- --- --- --- --- --- -gc taa atg gta ctg tta ata agt tta t-- --- --- --- --- --- --- --- --- --- --- --- --- --- --- --- --- --a gac aat atc ata gta ttt tct ttc cag aat --- --- --- --t aga tta ttt ttt taa ata ctg atc ctc aca att ccg tga tgt agc agt agt tgg t-- --- --- --- --- --- --- --- --- --- --- --- --- --- --- --- --- --- --- --- --- --- --- --- --- --- --- --- -gc atg gtc tat atc gt- --- --- --- --- --- --- --- --- --- --- --- --- --- --- --- --- --- --- --- --- --- --- --- --- --- --- --- --- --- --- --- --- --- --- --- --- --- --- --- --- --- --- --- taa aat gta tca tat ata ata gtt ttc tga cgt gga gta cag aat ttt cga --- --- --- --- --- --- --- --- --- --- --- --- --- --- --- --- --- --- --- --- --- --- --- --- --- --- --- --- --- --- --- --- --- --- --- --- --- --- --- --- --- --- --- --t taa tga gtt cat ggt aag gaa ggg caa atg cct --- gta tat aat ata cat aag tta a-- --- --- --- --- --- --- --- --- --- --- --- --- --t agt ttt tta tca tat tt- --- --- --- --- --- --- --- --t cta ata cca taa taa aaa t-- --- --- --- --- --- --- --- --- --- --- --- --- --- --- --- --- --- --- --- --- --- --- --- --- --t atc att att gcg ttt g-g tag tt- --- --- --- --- ctg ccc ta- --- --- --- --- --- --- --- --t cat cta tat cac tgt cac tct c-- --- --g ctc tca cta tat ctt cta aaa tta caa --- -ac aac --- --- --- --- --- --- --- --- --- --- --- --- --- --- --- --- --- --- --- --- --- --- --- --- --- --- --- --- --- --- --- --- --- --- --- --- tgg ata ttc gat --- --- --- --- --- --- --- --- --- --- --- --- --- --- --- --- --- --- --- --- --- --- --- --- --- --- --a aca gca ttt gtg t-- --- --- --- --- --- --- --- --- --- --- --- --- --- --- --- --- -

>ON649713.1_Monkeypox_virus_isolate_Monkeypox/PT0020/2022_complete_genome

--- --- --- --- --- --- --- --- --- --- --- --- --- --- --- --- --- --- -ta att a-- --- --- --- --- --- --- --- att taa ttt tac tat ttt att tag tgt cta gaa aaa aat gtg tga ccc acg acc gta gga aac tct aga ggg taa gaa aaa tca atc gtt tat aga gac cat cag aaa gag gtt taa tat ttt tgt gag acc tat cga a-- --- --- --- --- --- --- --- --- --- --- --- --- --- --- --- --- --- --- --- --- --- --- --- --- --- --- --- --- --- --- --- --- --- --- --- --- --- --- --- --- --- --- --- --- --- --- --- --- --- --- --- --- --- --- --- --- --- --- --- --- --- --- --- --- --- --- --- --- --- --- --- --- --- --- --- --- --- --- --- --- --- --- --- --- --- --- --- --- --- --- --- --- --- --- --- --- --- --- --- --- --- --- --- --- --- --- --- --- --- --- --- --- --- --- --- --- --- --- --- --- --- --- --- --- --- --- --- --- --- --- --- --- --- --- --- --- --- --- --- --- --- --- --- --- --- --- --- --- --- --- --- --- --- --- --- --- --- --- --- --- --- --- --- --- --- --- --- --- --- --- --- --- --- --- --- --- --- --- --- --- --- --- --- --- --- --- --- --- --- --- --- --- --- --- --- --- --- --- --- --- --- --- --- --- --- --- --- --- --- --- --- --- --- --- --- --- --- --- --- --- --- --- --- --- --- --- --- --- --- --- --- --- --- --- --- --- --- --- --- --- --- --- --- --- --- --- --- --- --- --- --- --- --- --- --- --- --- --- --- --- --- --- --- --- --- --- --- --- --- --- --- --- --- --- --- --- --- --- --- -ga gag aaa gga taa aaa ctt ttt acg act cca tca gaa aga ggt tta ata ttt ttg tga gac cca tcg aag aga gaa aga gat ggt tag tca aga tat ttt tct tag tac aaa agt caa tgt ttt aaa ata tat gga cga gaa tta att tgt ctg tat aaa aac ttg tgt gaa att atg tac tag aga aaa aac gtg agc agt gtc ccc tac atg gat ttt aca gat cat tta tat tcc aaa aat att aac tat ata cgt tta tta tat gat gtt aac gtg taa att ata aac att att tta tga tgc aat tgt ctg aca acc tag att ggt ata agg atg ttg ata agc tct acg aga ata tat tgt tgg acg tta tcg ttt acg aaa tag ttg aga cat cag aaa gag gtt taa tat ttt tgt gag a-c cat cga aga gag aaa gag aat aaa aat a-- --t ttt ttt ttt ttt tgt aaa act ttt tta tga gac --- --- -ca aga gaa tac gaa tag tga tca tat cgt atc aca tat tga aac aga aag aag aag taa cga gag gta act ttt tgt gaa tgt agt taa ata ttt ttg ttt tgc aaa ccg gaa tat agt gcc cgg t-- -ct ttt tta att cgt ggt gcg gtg tct gaa tcg ttc gat taa ccc aac tca tcc att ttc aga tga ata gag tta tcg att cag aca cat gct ttg agt ttt gtt gaa tcg atg agt gaa gta tca tcg gtt gca cct tca gat gcc gat ccg tcg aca tac ttg aat cca tcc ttg act tca agt tca gat gat tcc tca cac atg tct ccg ata cgt acg cta aac tct agg ttc ttg aca cat ttt gta tca acg atc gtt gaa ccg atg ata tct ttg taa ctc act ttc tta tgt gag atg tta gac cca agt act gga tgg gtc ttg atg tca ctg tct ttc tct tct tcg cta cat ctg atg tcg ata gac atc tca cag tct ttg atc ata gcc aga gct tct tca cgc gtg atc gcg gga gag tcc tta cct tgt ccc ggt gac acg ctg gac aat cta gta ttc aca gtg ttt cca tca gag gat tcg gag atg gat aaa atc ttt ggg cat ttg gtg aat cca aag ttc atg tta aga ccc gca ccg acg ata gtg taa taa gtg gtg gga tct cct ttt aca act --- --- --- --- --- --- tct tcg gat acc tca tca tct tcg gtc tct gta act tcc gtt acg gat tga caa atc tta tca ttg gtc ggt gtt tgg tct tgc ttt gtg act ttg ata ata aca tcg att ccc ata tga tgt ttg ttt tct tct tca gta cac gag gat gaa gat tgt tga aga cta gta ggc ata gca gct gcc act agg cac atg cat gcc agg aca ata tat tgt ttc atg att gct att gat tga tta ctg ttc tag atg att cta ctt tct tac cat ata ata aat tag aat ata ttt tct act ttt acg aga aat taa tta ttg tat tta tta ttt ata ggt --a aaa aaa ctt act ata agt ggg tgg gat tct ggg aat tag tga tca gtt tat gta tat cgc aac tag cgg gca tat ggc tat tga cat cga gaa cat tac cca tat gat aag aga ttg tat cat ttt cgt agt ctt gag tat tgg tat tac tat ata gta tgt aga tgt cga cgc tag ata gac agt cgc cca cta gag tta ccg tct ctg aat gcg gca tga tag tat cat tct ttg ttt tcg tta act gtt tgg aag atg aat ctt tgt tgt tac att taa tct cga aat tca gag tac ata tct ttg aag tat tct gat atc tat ttt ctc ctg taa aga atc ctg aag ttg cta cat tat taa gga cag aga agt att ctg cac gaa aga ctg gat cac aat ctt tat gat tca tgg taa tag tta gtt ccg acg ttg aga tgg att cgc tga gac cgg tag tgg tcg tcc gag tac acg atg tgt cgt tga ctg gat aca ggt taa ttt cca cat cga tat agt taa atg tat tgc tgg tta cga cgg gtt cgc att tat ctg tgg aag aga cgg tgt gag aat atg ttc cgg gac cac acg gag aac aga tga cgt ctc cgg tag acg tgt atc cgg ata ctc cgt atc cta ttc cac act ttg ttt tag aaa tac atg ttc tac acc ctg atg ctc ctt tga gaa gac aat aat atc ctg gag agc att cac aga ttc tat tgt gag tcg tgt tac acg atc gcg tct cta cct gat tac tat cac atc ttc cgt tac aac tta gac aag cct gta aat gat tat tgt gag atg taa agg tat ccg aac cac acg gtg tac att gtg tat tag tct tgc tat cac ata atc tga aag cgt aag ttc ccg gag gac acg ata gac aac ata gat tac ggc ttc tgt att cgt tgt ctt tac act ttc cat tgg atg gtg cat gtg gtg cta tat ctc ttc cgt tta tta tta tac atg aga gaa aca ata tat acg agt ata ata cgg acc tca tga ttt aat aat -gt agt aat cgt cgt ctt gtt act gtt tgt ttc cta ctt ctc caa tca tat aga tta ttt ttt aaa tat ttt ctt tct atc atg gat aat att tgt aat ggt tct ttc cgt aca aca tac tgt tta gat ggt agt cgc tta gct tgg tta tga tat tgc gca taa ttt ccg gag gca aat acg ata gtc tag att gac tat cga tgg tag act cta att tat tga gtg ctt tgt cga cga gtt tac ttt tat gct cca tcg ata gat gac act gtt cta tga gat cgt cgt aca tgg gaa atg aaa tgc gtt tgt ctg aat gta tgg ctt cga gat atg tgt gat acc gaa tgt ctt ctg ttc tca ata ccg tat aca agt cgg tgt ctg aga ttc gaa tct ctt tga gga gac tta tgt cac gac tac att ttt cga tga tgg aat cta tct tat cga atg ata tat ttt tca taa ata cac ttt tat agt cct cgt tta aac aga att tag tat gta gtt ccg caa atg act cgt ccc tta ata ggc agt agg cta tta tct tct tta cgt agt gat cgt cgt agg gag aga act ccg aca tct tgt aga aca acg att taa tca tag gta gag ata ctt tca gtc tgt ggt gga taa tgt cat taa cga cat ccg cct tgt ata taa tgt ttc tgt ttt caa aca cca agt cga ata ctg tct tat cgt ctt tag tcg gaa ggt tga tgt cgt atc caa tgt ata cga ggt atg agg caa cat tgt tat tgc aat tct gga agg cgg tat gaa gag gag tca ttg tat tat agt att cgt ctt tct gaa tgt cga atc tat cta gta gat acc gta gta tat tga gag agc gac ttc cat atc ctt gat tat gtt tta tga ata gat aaa gta gat gtt gtc ctt ctt cct ttt gta att tcc cgt att ttt gtt cgt gcc aat tga gta aca tta tga gaa tat gac ctg ttg cac aat cgt tct tta tgt att cca tga tgg gtg tac aat taa gat tat tac gta tcc tcg tat cgg ctc ctc gag ata aaa gag cat aca cca cac gag gac tat gtt tgg tat act gtt gaa ggt aag tgt gta acg gcg tat ttc cga ttt tcg taa ccg cgt taa tgt ttg ctc cat gat cta tta tcg cgt aga tga atc gct tct cag ctc gca tct tag tgt gac tct ttg act tgt aat aat tgc ttt cgt gga acg cgg ata tgt gtt tac agt agt aat gaa gag aag tga gtc cat cct cat cga cgc aat tag ggt cag atc ctt tag tca ata att tgt aca gaa cgt aat agt tta agc tcc cat tga att tat atc taa gat aac aca gca ata gat cgg atg att tac taa agt cat caa tgg ggt ccg tta gta tat caa aga tct tgt tat cga ttg ata gtg aat gaa tca gat agt ggt gta gag gaa tat gtc ctt ttt tat cct tgc tat caa agt tac gca tgc cgt ggc gta aca ata tct tta ata cag atg gat taa atc gtg tat tca tcg tat agc aat gta atg gag agt tac cac att tta gtc gtt tat tca gat cgc agt gtt taa taa cta att taa aca gat gag atg ata tat cca cat caa aga atg cga gat aca tat gac aga cat tat tga cag aaa tgt gac ctt cat tat cac cgt cgt cca taa atg cgt tag gta cgt acc aca tac tat cgt taa cga tgc gca caa tct cgt cca -tt tca tcc atc ttc ata atg att tac ttt ttc ata att aga gaa aaa gat caa ggt ata aaa att aga agt gtt aga cta ta- --- --- --- --- --- --- --- --- --- --- -aa cta act tat gac tta act aac tta tga --- --- --- --- --- --- --- --- --- --- --- --- --- --- --- --- --- --- --- --- --- --- --- --- --- --- --- --- --- --- --- --- --- --- --- --- --- --- --- --- --- --- --- --- --- --- --- --- ctt aac ta- --- --- --- --- --- --- --- --- --- --- -ac tta tga ctt aac taa ctt atg act taa cta act ta- tga ctt aac taa ctt atg act taa cta act tat gac tta act aac tta tga cta tta act cat ttc aag aaa ggt ggg tgg ata gaa ctc tat atg aca gct tgt gaa aca att aga tcc cta att tct aat gga agt ttt gat agg aga ttg tca tca gtt gat aca ttg ttt att atc tca tct att aga gca cgt ctg ttt aga gct tta gtg acc tgc tcg gtt act tct gtg taa atc ttg aat cct tta gtg ata cac tgt gtc aaa act gga tgt tta gaa tac cta tgt aga ata tgg gaa gca tgc ttg ttt ttg tct cta tta tag att aac tca tac atg gtt gta tta tga att ttc atc tgc cta atg tac tcc aat tct tgt tta caa tca att ata taa tca aag agt gat gat gca tac aca tta caa agt gaa taa tct acc atc ata aaa tac ttg ata cag agc ttt atc aca tca tgg ttt tca att gta tta tta agt ata gct aat ttt ata cag tca ata gac aat ggt tct cta agc aat att tct aat att tta aga tgt gct tcc cta cgg gcg atg aca gat ccc cta tcc acg gcc acg tca aga cat gta tat cca tta ctc att act gcg ttg aca ttt gct cca ttt tct aat agc cat gat act aaa tct ata taa cct gca tag ata gcg cga taa agc aag gtc ctt cca cca gca tct agt tga ttg ata tct tca ata tat ggg ata caa agc tta taa att tct aat act gtg ggt tca tct aca agg aat ccc cta gta tac tga att att tta tat aga tct aat tta aca tca ttt tca tct ggg ata cca caa ttc aaa ata aac tca aca aca cta ctt tcc ttt tta cat att ccc cta aaa tag gca ttc aag cat tct att tta tat att aca gcc cca tga tct acc ata aga tca aca atg tct att tct aca tat gca tta gat aga tag taa agt aag aga tct tgc aca gaa tta caa ttc tta ata att ata gag aaa ata tct tcc ata taa ttc ttt gac act aat gca gat ata ata tct tta tat gta ata tat gca aac agt cta tct act ata tac tga tca ata tta tct cta tga atc cta aaa taa tca tac aga aca tct aca gga tca caa att ggt tca agg aga aat cta tca aat att ttc ctg tca aca act ggt tct aga aca taa cag tca aca cct aat cca tgt ttt tta taa tca tct acc aaa gat aat gac caa aga tcg agg tcg tcg tga aac tgc tca tcg aca gcc atg aaa tct acc gac tcc atg gtg cga atc gca ctg tct tat tcg cca ttg att ttc a-t ttt tta taa tta tgt aca tgt ttt cct tct att ctc aag agt cta caa aaa tat a-- -tt ttt tcg ata tct aag tac taa gtt ttt tta ctg ttt ttg tta ctg tct tcc att ctt cta act aaa gat ctg aga taa att ata caa tct tcg cta tcg aac cat ttt tgt agt cta aag cct gaa gta att aac caa ctg ttt tta tta gtg gct ttt ttc gat cta tcc tcg tc- --- ctc tca tca tcc tta ta- --- --- --- --- --- --- tta tta tca tta tta tca tag tct att aaa cac aaa tca tct --- --- --- --- --- --- --- --- --- --- --- --- --- --- --- --- --- --- --- --- --- --- --- --- --- --- --a cgt tta taa c-- --- --- --- --- --- --- --- --a aca ttc --- --- --- --- -tc att att aat tag ttc tgt ag- aat atc ttt aat aat ttg gct ata --- --- -ca tct gtt --- --- --- --- --- --- --- --- --- --- --- --- --- --- --- --- --- --- --c aat act --- --- --- --- --- --- --- --a tct att gat gat ttc ttt t-- --- --- --- --- --- --- --- --- --- --- --t taa gac t-- --- --- --- --- --- --- --- --- --- --- --- --- --- --- --- --- --- --- --- --- --- --- --- --- --- --- --- --- --- --- --- --- --- --- --- --- --- --- --- taa act agt --- --- --- --- --- --- --- --- --- --- --- --- --- --- --- --- --- --- --- --- --- --- --- --- --- --- --- --- --- --- --- --- --- --- --- --- --- --- --- --- --- --- --- --- --- --- --- --- --- --- --- --- --- --- --- --- --- --- --- --- --- --- --- --- --- --- --- --- --- --- --- --- --- --- --- --- --- --- --- --- --- --- --- --- --- --- --- --- --- --- --- --- --- --- --- --- --- --- --- --- --- --- --- --- --- --- --- --- --- --t atg gta atg acg atg aaa --- -tc gag tag ta- --- --a ctt cta ata aag act tga ta- --- --t cat tat cat atg ttt gat cg- --- --- --- --- --- --- --t cat agt taa tag tgt g-- --- --- --- --- --- --- --- --- --- --- --- --- --- --- --- --- --- --- --- -gc taa atg gta ctg tta ata agt tta t-- --- --- --- --- --- --- --- --- --- --- --- --- --- --- --- --- --a gac aat atc ata gta ttt tct ttc cag aat --- --- --- --t aga tta ttt ttt taa ata ctg atc ctc aca att ccg tga tgt agc agt agt tgg t-- --- --- --- --- --- --- --- --- --- --- --- --- --- --- --- --- --- --- --- --- --- --- --- --- --- --- --- -gc atg gtc tat atc gt- --- --- --- --- --- --- --- --- --- --- --- --- --- --- --- --- --- --- --- --- --- --- --- --- --- --- --- --- --- --- --- --- --- --- --- --- --- --- --- --- --- --- --- taa aat gta tca tat ata ata gtt ttc tga cgt gga gta cag aat ttt cga --- --- --- --- --- --- --- --- --- --- --- --- --- --- --- --- --- --- --- --- --- --- --- --- --- --- --- --- --- --- --- --- --- --- --- --- --- --- --- --- --- --- --- --t taa tga gtt cat ggt aag gaa ggg caa atg cct --- gta tat aat ata cat aag tta a-- --- --- --- --- --- --- --- --- --- --- --- --- --t agt ttt tta tca tat tt- --- --- --- --- --- --- --- --t cta ata cca taa taa aaa t-- --- --- --- --- --- --- --- --- --- --- --- --- --- --- --- --- --- --- --- --- --- --- --- --- --t atc att att gcg ttt g-g tag tt- --- --- --- --- ctg ccc ta- --- --- --- --- --- --- --- --t cat cta tat cac tgt cac tct c-- --- --g ctc tca cta tat ctt cta aaa tta caa --- -ac aac --- --- --- --- --- --- --- --- --- --- --- --- --- --- --- --- --- --- --- --- --- --- --- --- --- --- --- --- --- --- --- --- --- --- --- --- tgg ata ttc gat --- --- --- --- --- --- --- --- --- --- --- --- --- --- --- --- --- --- --- --- --- --- --- --- --- --- --a aca gca ttt gtg t-- --- --- --- --- --- --- --- --- --- --- --- --- --- --- --- --- -

>ON585031.1_Monkeypox_virus_isolate_Monkeypox/PT0003/2022_complete_genome

--- --- --- --- --- --- --- --- --- --- --- --- --- --- --- --- --- --- --- --- --- --- --- --- --- --- --- --- att taa ttt tac tat ttt att tag tgt cta gaa aaa aat gtg tga ccc acg acc gta gga aac tct aga ggg taa gaa aaa tca atc gtt tat aga gac cat cag aaa gag gtt taa tat ttt tgt gag acc tat cga a-- --- --- --- --- --- --- --- --- --- --- --- --- --- --- --- --- --- --- --- --- --- --- --- --- --- --- --- --- --- --- --- --- --- --- --- --- --- --- --- --- --- --- --- --- --- --- --- --- --- --- --- --- --- --- --- --- --- --- --- --- --- --- --- --- --- --- --- --- --- --- --- --- --- --- --- --- --- --- --- --- --- --- --- --- --- --- --- --- --- --- --- --- --- --- --- --- --- --- --- --- --- --- --- --- --- --- --- --- --- --- --- --- --- --- --- --- --- --- --- --- --- --- --- --- --- --- --- --- --- --- --- --- --- --- --- --- --- --- --- --- --- --- --- --- --- --- --- --- --- --- --- --- --- --- --- --- --- --- --- --- --- --- --- --- --- --- --- --- --- --- --- --- --- --- --- --- --- --- --- --- --- --- --- --- --- --- --- --- --- --- --- --- --- --- --- --- --- --- --- --- --- --- --- --- --- --- --- --- --- --- --- --- --- --- --- --- --- --- --- --- --- --- --- --- --- --- --- --- --- --- --- --- --- --- --- --- --- --- --- --- --- --- --- --- --- --- --- --- --- --- --- --- --- --- --- --- --- --- --- --- --- --- --- --- --- --- --- --- --- --- --- --- --- --- --- --- --- --- --- -ga gag aaa gga taa aaa ctt ttt acg act cca tca gaa aga ggt tta ata ttt ttg tga gac cca tcg aag aga gaa aga gat ggt tag tca aga tat ttt tct tag tac aaa agt caa tgt ttt aaa ata tat gga cga gaa tta att tgt ctg tat aaa aac ttg tgt gaa att atg tac tag aga aaa aac gtg agc agt gtc ccc tac atg gat ttt aca gat cat tta tat tcc aaa aat att aac tat ata cgt tta tta tat gat gtt aac gtg taa att ata aac att att tta tga tgc aat tgt ctg aca acc tag att ggt ata agg atg ttg ata agc tct acg aga ata tat tgt tgg acg tta tcg ttt acg aaa tag ttg aga cat cag aaa gag gtt taa tat ttt tgt gag a-c cat cga aga gag aaa gag aat aaa aat a-- --t ttt ttt ttt ttt tgt aaa act ttt tta tga gac --- --- -ca aga gaa tac gaa tag tga tca tat cgt atc aca tat tga aac aga aag aag aag taa cga gag gta act ttt tgt gaa tgt agt taa ata ttt ttg ttt tgc aaa ccg gaa tat agt gcc cgg t-- -ct ttt tta att cgt ggt gcg gtg tct gaa tcg ttc gat taa ccc aac tca tcc att ttc aga tga ata gag tta tcg att cag aca cat gct ttg agt ttt gtt gaa tcg atg agt gaa gta tca tcg gtt gca cct tca gat gcc gat ccg tcg aca tac ttg aat cca tcc ttg act tca agt tca gat gat tcc tca cac atg tct ccg ata cgt acg cta aac tct agg ttc ttg aca cat ttt gta tca acg atc gtt gaa ccg atg ata tct ttg taa ctc act ttc tta tgt gag atg tta gac cca agt act gga tgg gtc ttg atg tca ctg tct ttc tct tct tcg cta cat ctg atg tcg ata gac atc tca cag tct ttg atc ata gcc aga gct tct tca cgc gtg atc gcg gga gag tcc tta cct tgt ccc ggt gac acg ctg gac aat cta gta ttc aca gtg ttt cca tca gag gat tcg gag atg gat aaa atc ttt ggg cat ttg gtg aat cca aag ttc atg tta aga ccc gca ccg acg ata gtg taa taa gtg gtg gga tct cct ttt aca act --- --- --- --- --- --- tct tcg gat acc tca tca tct tcg gtc tct gta act tcc gtt acg gat tga caa atc tta tca ttg gtc ggt gtt tgg tct tgc ttt gtg act ttg ata ata aca tcg att ccc ata tga tgt ttg ttt tct tct tca gta cac gag gat gaa gat tgt tga aga cta gta ggc ata gca gct gcc act agg cac atg cat gcc agg aca ata tat tgt ttc atg att gct att gat tga tta ctg ttc tag atg att cta ctt tct tac cat ata ata aat tag aat ata ttt tct act ttt acg aga aat taa tta ttg tat tta tta ttt ata ggt --a aaa aaa ctt act ata agt ggg tgg gat tct ggg aat tag tga tca gtt tat gta tat cgc aac tag cgg gca tat ggc tat tga cat cga gaa cat tac cca tat gat aag aga ttg tat cat ttt cgt agt ctt gag tat tgg tat tac tat ata gta tgt aga tgt cga cgc tag ata gac agt cgc cca cta gag tta ccg tct ctg aat gcg gca tga tag tat cat tct ttg ttt tcg tta act gtt tgg aag atg aat ctt tgt tgt tac att taa tct cga aat tca gag tac ata tct ttg aag tat tct gat atc tat ttt ctc ctg taa aga atc ctg aag ttg cta cat tat taa gga cag aga agt att ctg cac gaa aga ctg gat cac aat ctt tat gat tca tgg taa tag tta gtt ccg acg ttg aga tgg att cgc tga gac cgg tag tgg tcg tcc gag tac acg atg tgt cgt tga ctg gat aca ggt taa ttt cca cat cga tat agt taa atg tat tgc tgg tta cga cgg gtt cgc att tat ctg tgg aag aga cgg tgt gag aat atg ttc cgg gac cac acg gag aac aga tga cgt ctc cgg tag acg tgt atc cgg ata ctc cgt atc cta ttc cac act ttg ttt tag aaa tac atg ttc tac acc ctg atg ctc ctt tga gaa gac aat aat atc ctg gag agc att cac aga ttc tat tgt gag tcg tgt tac acg atc gcg tct cta cct gat tac tat cac atc ttc cgt tac aac tta gac aag cct gta aat gat tat tgt gag atg taa agg tat ccg aac cac acg gtg tac att gtg tat tag tct tgc tat cac ata atc tga aag cgt aag ttc ccg gag gac acg ata gac aac ata gat tac ggc ttc tgt att cgt tgt ctt tac act ttc cat tgg atg gtg cat gtg gtg cta tat ctc ttc cgt tta tta tta tac atg aga gaa aca ata tat acg agt ata ata cgg acc tca tga ttt aat aat -gt agt aat cgt cgt ctt gtt act gtt tgt ttc cta ctt ctc caa tca tat aga tta ttt ttt aaa tat ttt ctt tct atc atg gat aat att tgt aat ggt tct ttc cgt aca aca tac tgt tta gat ggt agt cgc tta gct tgg tta tga tat tgc gca taa ttt ccg gag gca aat acg ata gtc tag att gac tat cga tgg tag act cta att tat tga gtg ctt tgt cga cga gtt tac ttt tat gct cca tcg ata gat gac act gtt cta tga gat cgt cgt aca tgg gaa atg aaa tgc gtt tgt ctg aat gta tgg ctt cga gat atg tgt gat acc gaa tgt ctt ctg ttc tca ata ccg tat aca agt cgg tgt ctg aga ttc gaa tct ctt tga gga gac tta tgt cac gac tac att ttt cga tga tgg aat cta tct tat cga atg ata tat ttt tca taa ata cac ttt tat agt cct cgt tta aac aga att tag tat gta gtt ccg caa atg act cgt ccc tta ata ggc agt agg cta tta tct tct tta cgt agt gat cgt cgt agg gag aga act ccg aca tct tgt aga aca acg att taa tca tag gta gag ata ctt tca gtc tgt ggt gga taa tgt cat taa cga cat ccg cct tgt ata taa tgt ttc tgt ttt caa aca cca agt cga ata ctg tct tat cgt ctt tag tcg gaa ggt tga tgt cgt atc caa tgt ata cga ggt atg agg caa cat tgt tat tgc aat tct gga agg cgg tat gaa gag gag tca ttg tat tat agt att cgt ctt tct gaa tgt cga atc tat cta gta gat acc gta gta tat tga gag agc gac ttc cat atc ctt gat tat gtt tta tga ata gat aaa gta gat gtt gtc ctt ctt cct ttt gta att tcc cgt att ttt gtt cgt gcc aat tga gta aca tta tga gaa tat gac ctg ttg cac aat cgt tct tta tgt att cca tga tgg gtg tac aat taa gat tat tac gta tcc tcg tat cgg ctc ctc gag ata aaa gag cat aca cca cac gag gac tat gtt tgg tat act gtt gaa ggt aag tgt gta acg gcg tat ttc cga ttt tcg taa ccg cgt taa tgt ttg ctc cat gat cta tta tcg cgt aga tga atc gct tct cag ctc gca tct tag tgt gac tct ttg act tgt aat aat tgc ttt cgt gga acg cgg ata tgt gtt tac agt agt aat gaa gag aag tga gtc cat cct cat cga cgc aat tag ggt cag atc ctt tag tca ata att tgt aca gaa cgt aat agt tta agc tcc cat tga att tat atc taa gat aac aca gca ata gat cgg atg att tac taa agt cat caa tgg ggt ccg tta gta tat caa aga tct tgt tat cga ttg ata gtg aat gaa tca gat agt ggt gta gag gaa tat gtc ctt ttt tat cct tgc tat caa agt tac gca tgc cgt ggc gta aca ata tct tta ata cag atg gat taa atc gtg tat tca tcg tat agc aat gta atg gag agt tac cac att tta gtc gtt tat tca gat cgc agt gtt taa taa cta att taa aca gat gag atg ata tat cca cat caa aga atg cga gat aca tat gac aga cat tat tga cag aaa tgt gac ctt cat tat cac cgt cgt cca taa atg cgt tag gta cgt acc aca tac tat cgt taa cga tgc gca caa tct cgt cca -tt tca tcc atc ttc ata atg att tac ttt ttc ata att aga gaa aaa gat caa ggt ata aaa att aga agt gtt aga cta ta- --- --- --- --- --- --- --- --- --- --- -aa cta act tat gac tta act aac tta tga --- --- --- --- --- --- --- --- --- --- --- --- --- --- --- --- --- --- --- --- --- --- --- --- --- --- --- --- --- --- --- --- --- --- --- --- --- --- --- --- --- --- --- --- --- --- --- --- ctt aac ta- --- --- --- --- --- --- --- --- --- --- -ac tta tga ctt aac taa ctt atg act taa cta act ta- tga ctt aac taa ctt atg act taa cta act tat gac tta act aac tta tga cta tta act cat ttc aag aaa ggt ggg tgg ata gaa ctc tat atg aca gct tgt gaa aca att aga tcc cta att tct aat gga agt ttt gat agg aga ttg tca tca gtt gat aca ttg ttt att atc tca tct att aga gca cgt ctg ttt aga gct tta gtg acc tgc tcg gtt act tct gtg taa atc ttg aat cct tta gtg ata cac tgt gtc aaa act gga tgt tta gaa tac cta tgt aga ata tgg gaa gca tgc ttg ttt ttg tct cta tta tag att aac tca tac atg gtt gta tta tga att ttc atc tgc cta atg tac tcc aat tct tgt tta caa tca att ata taa tca aag agt gat gat gca tac aca tta caa agt gaa taa tct acc atc ata aaa tac ttg ata cag agc ttt atc aca tca tgg ttt tca att gta tta tta agt ata gct aat ttt ata cag tca ata gac aat ggt tct cta agc aat att tct aat att tta aga tgt gct tcc cta cgg gcg atg aca gat ccc cta tcc acg gcc acg tca aga cat gta tat cca tta ctc att act gcg ttg aca ttt gct cca ttt tct aat agc cat gat act aaa tct ata taa cct gca tag ata gcg cga taa agc aag gtc ctt cca cca gca tct agt tga ttg ata tct tca ata tat ggg ata caa agc tta taa att tct aat act gtg ggt tca tct aca agg aat ccc cta gta tac tga att att tta tat aga tct aat tta aca tca ttt tca tct ggg ata cca caa ttc aaa ata aac tca aca aca cta ctt tcc ttt tta cat att ccc cta aaa tag gca ttc aag cat tct att tta tat att aca gcc cca tga tct acc ata aga tca aca atg tct att tct aca tat gca tta gat aga tag taa agt aag aga tct tgc aca gaa tta caa ttc tta ata att ata gag aaa ata tct tcc ata taa ttc ttt gac act aat gca gat ata ata tct tta tat gta ata tat gca aac agt cta tct act ata tac tga tca ata tta tct cta tga atc cta aaa taa tca tac aga aca tct aca gga tca caa att ggt tca agg aga aat cta tca aat att ttc ctg tca aca act ggt tct aga aca taa cag tca aca cct aat cca tgt ttt tta taa tca tct acc aaa gat aat gac caa aga tcg agg tcg tcg tga aac tgc tca tcg aca gcc atg aaa tct acc gac tcc atg gtg cga atc gca ctg tct tat tcg cca ttg att ttc a-t ttt tta taa tta tgt aca tgt ttt cct tct att ctc aag agt cta caa aaa tat a-- -tt ttt tcg ata tct aag tac taa gtt ttt tta ctg ttt ttg tta ctg tct tcc att ctt cta act aaa gat ctg aga taa att ata caa tct tcg cta tcg aac cat ttt tgt agt cta aag cct gaa gta att aac caa ctg ttt tta tta gtg gct ttt ttc gat cta tcc tcg tc- --- ctc tca tca tcc tta ta- --- --- --- --- --- --- tta tta tca tta tta tca tag tct att aaa cac aaa tca tct --- --- --- --- --- --- --- --- --- --- --- --- --- --- --- --- --- --- --- --- --- --- --- --- --- --- --a cgt tta taa c-- --- --- --- --- --- --- --- --a aca ttc --- --- --- --- -tc att att aat tag ttc tgt ag- aat atc ttt aat aat ttg gct ata --- --- -ca tct gtt --- --- --- --- --- --- --- --- --- --- --- --- --- --- --- --- --- --- --c aat act --- --- --- --- --- --- --- --a tct att gat gat ttc ttt t-- --- --- --- --- --- --- --- --- --- --- --t taa gac t-- --- --- --- --- --- --- --- --- --- --- --- --- --- --- --- --- --- --- --- --- --- --- --- --- --- --- --- --- --- --- --- --- --- --- --- --- --- --- --- taa act agt --- --- --- --- --- --- --- --- --- --- --- --- --- --- --- --- --- --- --- --- --- --- --- --- --- --- --- --- --- --- --- --- --- --- --- --- --- --- --- --- --- --- --- --- --- --- --- --- --- --- --- --- --- --- --- --- --- --- --- --- --- --- --- --- --- --- --- --- --- --- --- --- --- --- --- --- --- --- --- --- --- --- --- --- --- --- --- --- --- --- --- --- --- --- --- --- --- --- --- --- --- --- --- --- --- --- --- --- --- --t atg gta atg acg atg aaa --- -tc gag tag ta- --- --a ctt cta ata aag act tga ta- --- --t cat tat cat atg ttt gat cg- --- --- --- --- --- --- --t cat agt taa tag tgt g-- --- --- --- --- --- --- --- --- --- --- --- --- --- --- --- --- --- --- --- -gc taa atg gta ctg tta ata agt tta t-- --- --- --- --- --- --- --- --- --- --- --- --- --- --- --- --- --a gac aat atc ata gta ttt tct ttc cag aat --- --- --- --t aga tta ttt ttt taa ata ctg atc ctc aca att ccg tga tgt agc agt agt tgg t-- --- --- --- --- --- --- --- --- --- --- --- --- --- --- --- --- --- --- --- --- --- --- --- --- --- --- --- -gc atg gtc tat atc gt- --- --- --- --- --- --- --- --- --- --- --- --- --- --- --- --- --- --- --- --- --- --- --- --- --- --- --- --- --- --- --- --- --- --- --- --- --- --- --- --- --- --- --- taa aat gta tca tat ata ata gtt ttc tga cgt gga gta cag aat ttt cga --- --- --- --- --- --- --- --- --- --- --- --- --- --- --- --- --- --- --- --- --- --- --- --- --- --- --- --- --- --- --- --- --- --- --- --- --- --- --- --- --- --- --- --t taa tga gtt cat ggt aag gaa ggg caa atg cct --- gta tat aat ata cat aag tta a-- --- --- --- --- --- --- --- --- --- --- --- --- --t agt ttt tta tca tat tt- --- --- --- --- --- --- --- --t cta ata cca taa taa aaa t-- --- --- --- --- --- --- --- --- --- --- --- --- --- --- --- --- --- --- --- --- --- --- --- --- --t atc att att gcg ttt g-g tag tt- --- --- --- --- ctg ccc ta- --- --- --- --- --- --- --- --t cat cta tat cac tgt cac tct c-- --- --g ctc tca cta tat ctt cta aaa tta caa --- -ac aac --- --- --- --- --- --- --- --- --- --- --- --- --- --- --- --- --- --- --- --- --- --- --- --- --- --- --- --- --- --- --- --- --- --- --- --- tgg ata ttc gat --- --- --- --- --- --- --- --- --- --- --- --- --- --- --- --- --- --- --- --- --- --- --- --- --- --- --a aca gca ttt gtg t-- --- --- --- --- --- --- --- --- --- --- --- --- --- --- --- --- -

>ON585038.1_Monkeypox_virus_isolate_Monkeypox/PT0008/2022_complete_genome

--- --- --- --- --- --- --- --- --- --- --- --- --- --- --- --- --- --- -ta att a-- --- --- --- --- --- --- --- att taa ttt tac tat ttt att tag tgt cta gaa aaa aat gtg tga ccc acg acc gta gga aac tct aga ggg taa gaa aaa tca atc gtt tat aga gac cat cag aaa gag gtt taa tat ttt tgt gag acc tat cga a-- --- --- --- --- --- --- --- --- --- --- --- --- --- --- --- --- --- --- --- --- --- --- --- --- --- --- --- --- --- --- --- --- --- --- --- --- --- --- --- --- --- --- --- --- --- --- --- --- --- --- --- --- --- --- --- --- --- --- --- --- --- --- --- --- --- --- --- --- --- --- --- --- --- --- --- --- --- --- --- --- --- --- --- --- --- --- --- --- --- --- --- --- --- --- --- --- --- --- --- --- --- --- --- --- --- --- --- --- --- --- --- --- --- --- --- --- --- --- --- --- --- --- --- --- --- --- --- --- --- --- --- --- --- --- --- --- --- --- --- --- --- --- --- --- --- --- --- --- --- --- --- --- --- --- --- --- --- --- --- --- --- --- --- --- --- --- --- --- --- --- --- --- --- --- --- --- --- --- --- --- --- --- --- --- --- --- --- --- --- --- --- --- --- --- --- --- --- --- --- --- --- --- --- --- --- --- --- --- --- --- --- --- --- --- --- --- --- --- --- --- --- --- --- --- --- --- --- --- --- --- --- --- --- --- --- --- --- --- --- --- --- --- --- --- --- --- --- --- --- --- --- --- --- --- --- --- --- --- --- --- --- --- --- --- --- --- --- --- --- --- --- --- --- --- --- --- --- --- --- -ga gag aaa gga taa aaa ctt ttt acg act cca tca gaa aga ggt tta ata ttt ttg tga gac cca tcg aag aga gaa aga gat ggt tag tca aga tat ttt tct tag tac aaa agt caa tgt ttt aaa ata tat gga cga gaa tta att tgt ctg tat aaa aac ttg tgt gaa att atg tac tag aga aaa aac gtg agc agt gtc ccc tac atg gat ttt aca gat cat tta tat tcc aaa aat att aac tat ata cgt tta tta tat gat gtt aac gtg taa att ata aac att att tta tga tgc aat tgt ctg aca acc tag att ggt ata agg atg ttg ata agc tct acg aga ata tat tgt tgg acg tta tcg ttt acg aaa tag ttg aga cat cag aaa gag gtt taa tat ttt tgt gag a-c cat cga aga gag aaa gag aat aaa aat a-- --- ttt ttt ttt ttt tgt aaa act ttt tta tga gac --- --- -ca aga gaa tac gaa tag tga tca tat cgt atc aca tat tga aac aga aag aag aag taa cga gag gta act ttt tgt gaa tgt agt taa ata ttt ttg ttt tgc aaa ccg gaa tat agt gcc cgg t-- -ct ttt tta att cgt ggt gcg gtg tct gaa tcg ttc gat taa ccc aac tca tcc att ttc aga tga ata gag tta tcg att cag aca cat gct ttg agt ttt gtt gaa tcg atg agt gaa gta tca tcg gtt gca cct tca gat gcc gat ccg tcg aca tac ttg aat cca tcc ttg act tca agt tca gat gat tcc tca cac atg tct ccg ata cgt acg cta aac tct agg ttc ttg aca cat ttt gta tca acg atc gtt gaa ccg atg ata tct ttg taa ctc act ttc tta tgt gag atg tta gac cca agt act gga tgg gtc ttg atg tca ctg tct ttc tct tct tcg cta cat ctg atg tcg ata gac atc tca cag tct ttg atc ata gcc aga gct tct tca cgc gtg atc gcg gga gag tcc tta cct tgt ccc ggt gac acg ctg gac aat cta gta ttc aca gtg ttt cca tca gag gat tcg gag atg gat aaa atc ttt ggg cat ttg gtg aat cca aag ttc atg tta aga ccc gca ccg acg ata gtg taa taa gtg gtg gga tct cct ttt aca act --- --- --- --- --- --- tct tcg gat acc tca tca tct tcg gtc tct gta act tcc gtt acg gat tga caa atc tta tca ttg gtc ggt gtt tgg tct tgc ttt gtg act ttg ata ata aca tcg att ccc ata tga tgt ttg ttt tct tct tca gta cac gag gat gaa gat tgt tga aga cta gta ggc ata gca gct gcc act agg cac atg cat gcc agg aca ata tat tgt ttc atg att gct att gat tga tta ctg ttc tag atg att cta ctt tct tac cat ata ata aat tag aat ata ttt tct act ttt acg aga aat taa tta ttg tat tta tta ttt ata ggt --a aaa aaa ctt act ata agt ggg tgg gat tct ggg aat tag tga tca gtt tat gta tat cgc aac tag cgg gca tat ggc tat tga cat cga gaa cat tac cca tat gat aag aga ttg tat cat ttt cgt agt ctt gag tat tgg tat tac tat ata gta tgt aga tgt cga cgc tag ata gac agt cgc cca cta gag tta ccg tct ctg aat gcg gca tga tag tat cat tct ttg ttt tcg tta act gtt tgg aag atg aat ctt tgt tgt tac att taa tct cga aat tca gag tac ata tct ttg aag tat tct gat atc tat ttt ctc ctg taa aga atc ctg aag ttg cta cat tat taa gga cag aga agt att ctg cac gaa aga ctg gat cac aat ctt tat gat tca tgg taa tag tta gtt ccg acg ttg aga tgg att cgc tga gac cgg tag tgg tcg tcc gag tac acg atg tgt cgt tga ctg gat aca ggt taa ttt cca cat cga tat agt taa atg tat tgc tgg tta cga cgg gtt cgc att tat ctg tgg aag aga cgg tgt gag aat atg ttc cgg gac cac acg gag aac aga tga cgt ctc cgg tag acg tgt atc cgg ata ctc cgt atc cta ttc cac act ttg ttt tag aaa tac atg ttc tac acc ctg atg ctc ctt tga gaa gac aat aat atc ctg gag agc att cac aga ttc tat tgt gag tcg tgt tac acg atc gcg tct cta cct gat tac tat cac atc ttc cgt tac aac tta gac aag cct gta aat gat tat tgt gag atg taa agg tat ccg aac cac acg gtg tac att gtg tat tag tct tgc tat cac ata atc tga aag cgt aag ttc ccg gag gac acg ata gac aac ata gat tac ggc ttc tgt att cgt tgt ctt tac act ttc cat tgg atg gtg cat gtg gtg cta tat ctc ttc cgt tta tta tta tac atg aga gaa aca ata tat acg agt ata ata cgg acc tca tga ttt aat aat -gt agt aat cgt cgt ctt gtt act gtt tgt ttc cta ctt ctc caa tca tat aga tta ttt ttt aaa tat ttt ctt tct atc atg gat aat att tgt aat ggt tct ttc cgt aca aca tac tgt tta gat ggt agt cgc tta gct tgg tta tga tat tgc gca taa ttt ccg gag gca aat acg ata gtc tag att gac tat cga tgg tag act cta att tat tga gtg ctt tgt cga cga gtt tac ttt tat gct cca tcg ata gat gac act gtt cta tga gat cgt cgt aca tgg gaa atg aaa tgc gtt tgt ctg aat gta tgg ctt cga gat atg tgt gat acc gaa tgt ctt ctg ttc tca ata ccg tat aca agt cgg tgt ctg aga ttc gaa tct ctt tga gga gac tta tgt cac gac tac att ttt cga tga tgg aat cta tct tat cga atg ata tat ttt tca taa ata cac ttt tat agt cct cgt tta aac aga att tag tat gta gtt ccg caa atg act cgt ccc tta ata ggc agt agg cta tta tct tct tta cgt agt gat cgt cgt agg gag aga act ccg aca tct tgt aga aca acg att taa tca tag gta gag ata ctt tca gtc tgt ggt gga taa tgt cat taa cga cat ccg cct tgt ata taa tgt ttc tgt ttt caa aca cca agt cga ata ctg tct tat cgt ctt tag tcg gaa ggt tga tgt cgt atc caa tgt ata cga ggt atg agg caa cat tgt tat tgc aat tct gga agg cgg tat gaa gag gag tca ttg tat tat agt att cgt ctt tct gaa tgt cga atc tat cta gta gat acc gta gta tat tga gag agc gac ttc cat atc ctt gat tat gtt tta tga ata gat aaa gta gat gtt gtc ctt ctt cct ttt gta att tcc cgt att ttt gtt cgt gcc aat tga gta aca tta tga gaa tat gac ctg ttg cac aat cgt tct tta tgt att cca tga tgg gtg tac aat taa gat tat tac gta tcc tcg tat cgg ctc ctc gag ata aaa gag cat aca cca cac gag gac tat gtt tgg tat act gtt gaa ggt aag tgt gta acg gcg tat ttc cga ttt tcg taa ccg cgt taa tgt ttg ctc cat gat cta tta tcg cgt aga tga atc gct tct cag ctc gca tct tag tgt gac tct ttg act tgt aat aat tgc ttt cgt gga acg cgg ata tgt gtt tac agt agt aat gaa gag aag tga gtc cat cct cat cga cgc aat tag ggt cag atc ctt tag tca ata att tgt aca gaa cgt aat agt tta agc tcc cat tga att tat atc taa gat aac aca gca ata gat cgg atg att tac taa agt cat caa tgg ggt ccg tta gta tat caa aga tct tgt tat cga ttg ata gtg aat gaa tca gat agt ggt gta gag gaa tat gtc ctt ttt tat cct tgc tat caa agt tac gca tgc cgt ggc gta aca ata tct tta ata cag atg gat taa atc gtg tat tca tcg tat agc aat gta atg gag agt tac cac att tta gtc gtt tat tca gat cgc agt gtt taa taa cta att taa aca gat gag atg ata tat cca cat caa aga atg cga gat aca tat gac aga cat tat tga cag aaa tgt gac ctt cat tat cac cgt cgt cca taa atg cgt tag gta cgt acc aca tac tat cgt taa cga tgc gca caa tct cgt cca -tt tca tcc atc ttc ata atg att tac ttt ttc ata att aga gaa aaa gat caa ggt ata aaa att aga agt gtt aga cta ta- --- --- --- --- --- --- --- --- --- --- -aa cta act tat gac tta act aac tta tga --- --- --- --- --- --- --- --- --- --- --- --- --- --- --- --- --- --- --- --- --- --- --- --- --- --- --- --- --- --- --- --- --- --- --- --- --- --- --- --- --- --- --- --- --- --- --- --- ctt aac ta- --- --- --- --- --- --- --- --- --- --- -ac tta tga ctt aac taa ctt atg act taa cta act ta- tga ctt aac taa ctt atg act taa cta act tat gac tta act aac tta tga cta tta act cat ttc aag aaa ggt ggg tgg ata gaa ctc tat atg aca gct tgt gaa aca att aga tcc cta att tct aat gga agt ttt gat agg aga ttg tca tca gtt gat aca ttg ttt att atc tca tct att aga gca cgt ctg ttt aga gct tta gtg acc tgc tcg gtt act tct gtg taa atc ttg aat cct tta gtg ata cac tgt gtc aaa act gga tgt tta gaa tac cta tgt aga ata tgg gaa gca tgc ttg ttt ttg tct cta tta tag att aac tca tac atg gtt gta tta tga att ttc atc tgc cta atg tac tcc aat tct tgt tta caa tca att ata taa tca aag agt gat gat gca tac aca tta caa agt gaa taa tct acc atc ata aaa tac ttg ata cag agc ttt atc aca tca tgg ttt tca att gta tta tta agt ata gct aat ttt ata cag tca ata gac aat ggt tct cta agc aat att tct aat att tta aga tgt gct tcc cta cgg gcg atg aca gat ccc cta tcc acg gcc acg tca aga cat gta tat cca tta ctc att act gcg ttg aca ttt gct cca ttt tct aat agc cat gat act aaa tct ata taa cct gca tag ata gcg cga taa agc aag gtc ctt cca cca gca tct agt tga ttg ata tct tca ata tat ggg ata caa agc tta taa att tct aat act gtg ggt tca tct aca agg aat ccc cta gta tac tga att att tta tat aga tct aat tta aca tca ttt tca tct ggg ata cca caa ttc aaa ata aac tca aca aca cta ctt tcc ttt tta cat att ccc cta aaa tag gca ttc aag cat tct att tta tat att aca gcc cca tga tct acc ata aga tca aca atg tct att tct aca tat gca tta gat aga tag taa agt aag aga tct tgc aca gaa tta caa ttc tta ata att ata gag aaa ata tct tcc ata taa ttc ttt gac act aat gca gat ata ata tct tta tat gta ata tat gca aac agt cta tct act ata tac tga tca ata tta tct cta tga atc cta aaa taa tca tac aga aca tct aca gga tca caa att ggt tca agg aga aat cta tca aat att ttc ctg tca aca act ggt tct aga aca taa cag tca aca cct aat cca tgt ttt tta taa tca tct acc aaa gat aat gac caa aga tcg agg tcg tcg tga aac tgc tca tcg aca gcc atg aaa tct acc gac tcc atg gtg cga atc gca ctg tct tat tcg cca ttg att ttc a-t ttt tta taa tta tgt aca tgt ttt cct tct att ctc aag agt cta caa aaa tat a-- -tt ttt tcg ata tct aag tac taa gtt ttt tta ctg ttt ttg tta ctg tct tcc att ctt cta act aaa gat ctg aga taa att ata caa tct tcg cta tcg aac cat ttt tgt agt cta aag cct gaa gta att aac caa ctg ttt tta tta gtg gct ttt ttc gat cta tcc tcg tc- --- ctc tca tca tcc tta ta- --- --- --- --- --- --- tta tta tca tta tta tca tag tct att aaa cac aaa tca tct --- --- --- --- --- --- --- --- --- --- --- --- --- --- --- --- --- --- --- --- --- --- --- --- --- --- --a cgt tta taa c-- --- --- --- --- --- --- --- --a aca ttc --- --- --- --- -tc att att aat tag ttc tgt ag- aat atc ttt aat aat ttg gct ata --- --- -ca tct gtt --- --- --- --- --- --- --- --- --- --- --- --- --- --- --- --- --- --- --c aat act --- --- --- --- --- --- --- --a tct att gat gat ttc ttt t-- --- --- --- --- --- --- --- --- --- --- --t taa gac t-- --- --- --- --- --- --- --- --- --- --- --- --- --- --- --- --- --- --- --- --- --- --- --- --- --- --- --- --- --- --- --- --- --- --- --- --- --- --- --- taa act agt --- --- --- --- --- --- --- --- --- --- --- --- --- --- --- --- --- --- --- --- --- --- --- --- --- --- --- --- --- --- --- --- --- --- --- --- --- --- --- --- --- --- --- --- --- --- --- --- --- --- --- --- --- --- --- --- --- --- --- --- --- --- --- --- --- --- --- --- --- --- --- --- --- --- --- --- --- --- --- --- --- --- --- --- --- --- --- --- --- --- --- --- --- --- --- --- --- --- --- --- --- --- --- --- --- --- --- --- --- --t atg gta atg acg atg aaa --- -tc gag tag ta- --- --a ctt cta ata aag act tga ta- --- --t cat tat cat atg ttt gat cg- --- --- --- --- --- --- --t cat agt taa tag tgt g-- --- --- --- --- --- --- --- --- --- --- --- --- --- --- --- --- --- --- --- -gc taa atg gta ctg tta ata agt tta t-- --- --- --- --- --- --- --- --- --- --- --- --- --- --- --- --- --a gac aat atc ata gta ttt tct ttc cag aat --- --- --- --t aga tta ttt ttt taa ata ctg atc ctc aca att ccg tga tgt agc agt agt tgg t-- --- --- --- --- --- --- --- --- --- --- --- --- --- --- --- --- --- --- --- --- --- --- --- --- --- --- --- -gc atg gtc tat atc gt- --- --- --- --- --- --- --- --- --- --- --- --- --- --- --- --- --- --- --- --- --- --- --- --- --- --- --- --- --- --- --- --- --- --- --- --- --- --- --- --- --- --- --- taa aat gta tca tat ata ata gtt ttc tga cgt gga gta cag aat ttt cga --- --- --- --- --- --- --- --- --- --- --- --- --- --- --- --- --- --- --- --- --- --- --- --- --- --- --- --- --- --- --- --- --- --- --- --- --- --- --- --- --- --- --- --t taa tga gtt cat ggt aag gaa ggg caa atg cct --- gta tat aat ata cat aag tta a-- --- --- --- --- --- --- --- --- --- --- --- --- --t agt ttt tta tca tat tt- --- --- --- --- --- --- --- --t cta ata cca taa taa aaa t-- --- --- --- --- --- --- --- --- --- --- --- --- --- --- --- --- --- --- --- --- --- --- --- --- --t atc att att gcg ttt g-g tag tt- --- --- --- --- ctg ccc ta- --- --- --- --- --- --- --- --t cat cta tat cac tgt cac tct c-- --- --g ctc tca cta tat ctt cta aaa tta caa --- -ac aac --- --- --- --- --- --- --- --- --- --- --- --- --- --- --- --- --- --- --- --- --- --- --- --- --- --- --- --- --- --- --- --- --- --- --- --- tgg ata ttc gat --- --- --- --- --- --- --- --- --- --- --- --- --- --- --- --- --- --- --- --- --- --- --- --- --- --- --a aca gca ttt gtg t-- --- --- --- --- --- --- --- --- --- --- --- --- --- --- --- --- -

>DQ011156.1_Monkeypox_virus_strain_Liberia_1970_184_complete_genome

--- --- --- --- --- --- --- --- --- --- --- --- --- --- --- --- --- --- --- --- --- --- --- --- --- --- --- --- --- --- --- --- --- --- --- --- --- --- --- --- --- --- --- --- --- --- --- --- --- --- --- --- --- --- --- --- --- -tt tat aga gac cat cag aaa gag gtt taa tat ttt tgt gag acc cat cga aga gag aaa gga taa aaa ctt ttt acg act cca tca gaa aga ggt tta ata ttt ttg tga gac cca tcg aag aga gaa agg ata aaa act ttt tac gac tcc atc aga aag agg ttt aat att ttt gtg aga ccc atc gaa gag aga aag gat aaa aac ttt tta cga ctc cat cag aaa gag gtt taa tat ttt tgt gag acc cat cga aga gag aaa gga taa aaa ctt ttt acg act cca tca gaa aga ggt tta ata ttt ttg tga gac cca tcg aag aga gaa agg ata aaa act ttt tac gac tcc atc aga aag agg ttt aat att ttt gtg aga ccc atc gaa gag aga aag gat aaa aac ttt tta cga ctc cat cag aaa gag gtt taa tat ttt tgt gag acc cat cga aga gag aaa gga taa aaa ctt ttt acg act cca tca gaa aga ggt tta ata ttt ttg tga gac cca tcg aag aga gaa agg ata aaa act ttt tac gac tcc atc aga aag agg ttt aat att ttt gtg aga ccc atc gaa gag aga aag gat aaa aac ttt tta cga ctc cat cag aaa gag gtt taa tat ttt tgt gag acc cat cga aga gag aaa gga taa aaa ctt ttt acg act cca tca gaa aga ggt tta --- --- --- --- --- --- --- --- --- --- --- --- --- --- --- --- --- --- --- --- --- --- --- --- --- --- --- --- --- --- --- --- --- --- --- --- --- --- --- --- --- --- --- --- --- --- --a tat ttt tgt gag acc cat cga aga gag aaa gga taa aaa ctt ttt acg act cca tca gaa aga ggt tta ata ttt ttg tga gac cca tcg aag aga gaa aga gat ggt tag tca aga tat ttt tct tag tac aaa agt caa tgt ttt aaa ata tat gga cga gaa tta att tgt ctg tat aaa aac ttg tgt gaa att atg tac tag aga aaa aac gtg agc agt gtc ccc tac atg gat ttt aca gat cat tta tat tcc aaa aat att aac tat ata cgt tta tta tat gat gtt aac gtg taa att ata aac att att tta tga tgc aat tgt ctg aca acc tag att ggt ata agg atg ttg ata agc tct acg aga ata tat tgt tgg acg tta tcg ttt acg aaa tag ttg aga cat cag aaa gag gtt taa tat ttt tgt gag a-c cat cga aga gag aaa tag aat aaa aat a-- --- --- --- ttt ttt tgt aaa act ttt tta tga gac --- --- -ca aga gaa tac gaa tag tga tca tat cgt atc aca tat tga aac aga aag aag aag taa cga gag gta act ttt tgt aaa tgt agt taa ata ttt ttg ttt tgc aaa ccg gaa tat agt gcc cgg t-- -ct ttt tta att cgt ggt gcg gtg tct gaa tcg ttc gat taa ccc aac tca tcc att ttc aga tga ata gag tta tcg att cag aca cat gct ttg agt ttt gtt gaa tcg atg agt gaa gta tca tcg gtt gca cct tca gat gcc gat ccg tcg aca tac ttg aat cca tcc ttg act tca agt tca gat gat tcc tca cac atg tct ccg ata cgt acg cta aac tct agg ttc ttg aca cat ttt gta tca acg atc gtt gaa ccg atg ata tct ttg taa ctc act ttc tta tgt gag atg tta gac cca agt act gga tgg gtc ttg atg tca ctg tct ttc tct tct tcg cta cat ctg atg tcg ata gac atc tca cag tct ttg atc ata gcc aga gct tct tca cgc gtg atc gcg gga gag tcc tta cct tgt ccc ggt gac acg ctg gac aat cta gta ttc aca gtg ttt cca tca gag gat tcg gag atg gat gaa atc ttt ggg cat ttg gtg aat cca aag ttc atg tta aga ccc gca ccg acg ata gtg taa taa gtg gtg gga tct cct ttt aca act tct tcg gat acc tca tca tct tcg gat acc tca tca tct tcg gtc tct gta act tcc gtt acg gat tga caa atc tta tca ttg gtc ggt gtt tgg tct tgc ttt gtg act ttg ata ata aca tcg att ccc ata tga tgt ttg ttt tct tct tca gta cac gag gat gaa gat tgt tga aga cta gta ggc ata gca gct gcc act agg cac atg cat gcc agg aca ata tat tgt ttc atg att gct att gat tga tta ctg ttc tag atg att cta ctt tct tac cat ata ata aat tag aat ata ttt tct act ttt acg aga aat taa tta ttg tat tta tta ttt atg ggt --a aaa aaa ctt act ata agt ggg tgg gat tct ggg aat tag tga tca gtt tat gta tat cgc aac tag cgg gca tat ggc tat tga cat cga gaa cat tac cca tat gat aag aga ttg tat ccg ttt cgt agt ctt gag tat tgg tat tac tat ata gta tgt aga tgt cga cgc tag ata gac agt cgc cca cta gag tta ccg tct ctg aat gcg gca tga tag tat cat tct ttg ttt tcg tta act gtt tgg aag atg aat ctt tgt tgt tac att taa tct cga aat tca gag tac ata tct ttg aag tat tct gat atc tat ttt ctc ctg taa aga atc ctg aag ttg cta cat tat taa gga cag aga agt att ctg cac gaa aga ctg gat cac aat ctt tat gat tca tgg taa tag tta gtt ccg acg ttg aga tgg att cgc tga gac cgg tag tgg tcg tcc gag tac acg atg tgt cgt tga ctg gat aca ggt taa ttt cca cat cga tat agt taa atg tat tgc tgg tta cga cgg gtt cgc att tat ctg tgg aag aga cgg tgt gag aat atg ttc cgg gac cac acg gag aac aga tga cgt ctc cgg tag acg tgt atc cgg ata ctc cgt atc cta ttc cac act ttg ttt tag aaa tac atg ttc tac acc ctg atg ctc ctt tga gaa gac aat aat atc ctg gag agc att cac aga ttc tat tgt gag tcg tgt tac acg atc gcg tct cta cct gat tac tat cac atc ttc cgt tac aac tta gac aag cct gta aat gat tat tgt gag atg taa agg tat ccg aac cac acg gtg tac att gtg tat tag tct tgc tat cac ata atc tgg aag cgt aag ttc ccg gag gac acg ata gac aac ata gat tac ggc ttc tgt att cgt tgt ctt tac act ttc cat tgg atg gtg cat gtg gtg cta tat ctc ttc cgt tta tta tta tac atg aga gaa aca ata tat acg agt ata ata cgg acc tca tga ttt aat aat -gt agt aat cgt cgt ctt gtt act gtt tgt ttc cta ctt ctc caa tca tat aga tta ttt ttt aaa tat ttt ctt tct atc atg gat aat att tgt aat ggt tct ttc cgt aca aca tac tgt tta gat ggt agt cgc tta gct tgg tta tga tat tgc gca taa ttt ccg gag gca aat acg ata gtc tag att gat tat cga tgg tag act cta att tat tga gtg ctt tgt cga cga gtt tac ttt tat gct cca tcg ata gat gac act gtt cta tga gat cgt cgt aca tgg gaa atg aaa tgc gtt tgt ctg aat gta tgg ctt cga gat agg tgt gat acc gga tgt ctt ctg ttc tca ata ccg tat aca agt cgg tgt ctg aga ttc gaa tct ctt tga gga gac tta tgt cac gac tac att ttt cga tga tgg aat cta tct tat cga atg ata tat ttt tca taa ata cac ttt tat agt cct cgt tta aac aga att tag tat gta gtt ccg caa atg act cgt ccc tta ata ggc agt agg cta tta tct tct tta cgt agt gat cgt cgt agg gag aga act ccg aca tct tgt aga aca acg att taa tca tag gta gag ata ctt tca gtc tgt ggt gga tga tgt cat taa cga cat ccg cct tgt ata tga tgt ttc tgt ttt caa aca cca agt cga ata ctg tct tat cgt ctt tag tcg gaa ggt tga tgt cgt atc cga tgt ata cga ggt atg agg caa cat tgt tat tgc aat tct gga agg cgg tat gaa gag gag tca ttg tat tat agt att cgt ctt tct gaa tgt cga atc tat cta gta gat acc gta gta tat tga gag agc gac ttc cat atc ctt gat tat gtt tta tga ata gat aaa gta gat gtt gtc ctt ctt cct ttt gta att tcc cgt att ttt gtt cgt gcc aat tga gta aca tta tga gaa tat gac ctg ttg cac aat cgt tct tta tgt att cca tga tgg gtg tac aat caa gat tat tac gta tcc tcg tat cag ctc ctc gag ata aaa gag cat aca cca cac gag gac tat gtt tgg tat act gtt gaa ggt aag tgt gta acg gcg tat ttc cga ttt tcg taa ccg cgt taa tgt ttg ctc cat gat cta tta tcg cgt aga tga atc gct tct cag ctc gca tct tag tgt gac tct ttg act tgt aat aat tgc ttt cgt gga acg cgg ata tgt gtt tac agt agt aat gaa gag aag tga gtc cat ccg cat cga cgc aat tag ggt cag atc ctt tag tca ata att tgt aca gaa cgt aat agt tta agc tcc cat tga att tat atc taa gat aac aca gca ata gat cgg atg att tac taa agt cat caa tgg ggt ccg tta gta tat caa aga tct tgt tat cga ttg ata gtg aat gaa tca gat agt ggt gta gag gaa tat gtc ctt ttt tat cct tgc tat caa agt tac gca tgc cgt ggc gta aca ata tct tta ata cag atg gat taa atc gtg tat tca tcg tat agc aat gta atg gag agt tac cac att tta gtc gtt tat tca gat cgc agt gtt taa taa cta att taa aca gat gag atg ata tat cca cat caa aga atg cga gat aca tat gac aga cat tat tga cag aaa tgt gac ctt cat tat cac cgt cgt cca taa atg cgt tag gta cgt acc aca tac tat cgt taa cga tgc gca caa tct cgt cca -tt tca tcc atc ttc ata atg att tac ttt ttc ata att aga gaa aaa gat caa ggt ata aaa att aga agt gtt aga cta ta- --- --- --- --- --- aac taa ctt ata a-- -aa cta act tat gac tta act aac tta tga --- --- --- --- --- --- --- --- --- --- --- --- --- --- --- --- --- --- --- --- --- --- --- --- --- --- --- --- --- --- --- --- --- --- --- --- --- --- --- --- --- --- --- --- --- --- --- --- ctt aac taa ctt --- --- --- --- --- --- --- --- --- -ac tta taa --- aac taa ctt atg act taa cta act ta- tga ctt aac taa ctt atg act taa cta act tat gac tta act aac tta tga cta tta act cat ttc aag aaa ggt ggg tgg ata gaa ctc tat atg aca gct tgt gaa acg att aga tcc cta att tct aat gga agt ttt gat agg aga ttg tca tca gtt gat aca ttg ttt att atc tca tct att aga gca cgt ctg ttt aga gct tta gtg acc tgc tcg gtt act tct gtg taa atc ttg aat cct tta gtg ata cac tgt gtc aaa act gga tgt tta gaa tac cta tgt aga ata tgg gaa gca tgc ttg ttt ttg tct cta tta tag att aac tca tac atg gtt gta tta tga att ttc atc tgc cta atg tac tcc aat tct tgt tta caa tca att ata taa tca aag agt gat gat gca tac aca tta caa agt gaa taa tct acc atc ata aaa tac ttg ata cag agc ttt atc aca tca tgg ttt tca att gta tta tta agt ata gct aat ttt ata cag tca ata gac aat ggt tct cta agc aat att tct aat att tta aga tgt gct tcc cta cgg gcg atg aca gat ccc cta tcc acg gcc acg tca aga cat gta tat cca tta ctc att act gcg ttg aca ttt gct cca ttt tct aat agc aat gat act aaa tct ata taa cct gca tag ata gcg cga taa agc aag gtc ctt cca cca gca tct agt tga ttg ata tct tca ata tat ggg ata caa agc tta taa att tct aat act gtg ggt tca tct aca agg aat ccc cta gta tac tga att att tta tat aga tct aat tta aca tca ttt tca tct ggg ata cca caa ttc aaa ata aac tca aca aca cta ctt tcc ttt tta cat att ccc cta aaa tag gca ttc aag cat tct att tta tat att aca gcc cca tga tct acc ata aga tca aca atg tct att tct aca tat gca tta gat aga tag taa agt aag aga tct tgc aca gaa tta caa ttc tta ata att ata gag aaa ata tct tcc ata taa ttc ttt gac act aat gca gat ata ata tct tta tat gta ata tat gca aac agt cta tct act ata tac tga tca ata tta tct cta tga atc cta aaa taa tca tac aga aca tct aca gga tca caa att ggt tca agg aga aat cta tca aat att ttc ctg tca aca act ggt tct aga aca taa cag tca aca cct aat cca tgt ttt tta taa tca tct acc aaa gat aat gac caa aga tcg agg tcg tcg tga aac tgc tca tcg aca gcc atg aaa tct acc gac tcc atg gtg cga atc gca ctg tct tat tcg cca ttg att ttc a-t ttt tta taa tta tgt aca tgt ttt cct tct att ctc aag agt cta caa aaa tat a-- ttt ttt tcg ata tct aag tac taa gtt ttt tta ctg ttt ttg tta ctg tct tcc att ctt cta act aaa gat ctg aga taa att ata caa tct tcg cta tcg aac cat ttt tgt agt cta aag cct gaa gta att aac caa ctg ttt tta tta gtg gct ttt ttc gat cta tca att tca gta tat tct tcg ccg tta taa aag taa tgt tgt tta att gta gga cgg ttg tta gta taa tca cat gaa taa taa tat tct aat tcc tcg tat tga cta ctt aca gat act cga aat agt ctg aaa aat tct tca aag ata ttt tta taa aga tct agg aaa agt tta tta ccg acc atg aac gag ata gat gga taa ata tcc ttt cca tca aag gtc ata att gga taa ttg tcc agc aat ata tct gct gta tta gtt ata tca ctt cca ttt att ttc aga ttg aag taa tgt act agt ttg tga caa tta aca aga tac aaa aga gat gcc gat act aat acg taa ata gct ata cgc gaa tcc att gtt acc ttt ttt tat ttc ata ggt cta tta ata aat ata tgt att act taa gac tag aaa aat caa aag taa gtt ttt gat att tga ttc tta ctt att gtg gga ttg tag ttt act tag taa ttc atc tct gaa tcc tga taa atc atg cat atc aat gat gca act acg cag caa act agt agg aat ata gat atc tgg ata tgt acg taa ata gtc gat tat atc ttt tac aat act att agt ccc tat tgc gtt atc tat ata tcc att aat aat att aca cag tgg ata ctt atg aga aat ata cct ctt aca gat ttt taa cga tat ata atc tag aaa gat atg tgt gta gta ctg tat tac cta aat tat cag tct cat tca aat att gca tga cta tta tcg aga att gct ata tcc ctc tat act cga tgc att tat tac agt aat tca atc cag cta aca taa gag cca atc tca atg ttg gtt taa tta tat cat ctt cat gta ata ata acg atg gaa act ttc cag tag cgt aac act tat cta aga agg ata taa taa cta tgt cta cat tat gtc tct tat cga gaa tat tct taa cga gat atc cat agc tat tct ggt gct aat tat tcc tat att ata ttc cac gaa aaa tga tga agg c-- -ca ttc atc ata aga tga taa aaa gtg tag tga gta aga gta tta gtg aga gag cat gaa gga gat tta gta ttt agc agt gag gat atg atc caa gag ggt gag ata gtc gtt ctc gtt cag aat ctt tcg cag cat aag tag tat gtc gat ata ctt atc att gaa gac tct tcc ggt gac aat agc tga ttg agt aca aag tcc aat tat tgc aca aag ttc ttt ggc ggt ttt cat gga gtc att tct gat gaa aca ttt aat gat ctc cac gca att gtc gat att gtc cca cgg aag tga atc cga gaa ctc ctt caa ctc gct acc aaa tag ctc cat tgc atc aat tct gaa aga gat gag aag cct gta gag agg ccc tgc gct ttc tct atg ggt cca tct atg aga aac cca cag gat gta ttc agt cag aca a-t gtc tga cat cag tca cgg tat tca ggg agt cct tag tag cgt ggc aat gac agg gac tga act ggg cac aag gag agg cca ttg tga agg tag acg aag gta acc tga tgg tag acc tgt agc cgt ctg tgc tta ata gag ggc ttt aat ttc cat ttt taa tgg tgt cgt gaa tga gga atg aga gtg tct ctc gtc ctt ggt tta cat gga tca gag tga gaa aaa ata tct tgt ata tta tta act aac aac ctt ggt ttc tat cca tgt tt- aaa aaa tga cct ata tgt tct tta tta att cta ttt taa act tta tcc tca aga ctc ctg aca aaa tta aaa tcc aga aag cag caa aca atc ctg tta caa gtt tac tga aat ctc tct ttg att gta gag tat atg tag tca gag caa gaa aca ctg cag tag tca aca tga aag ctt gca taa cga tac gtg cat cat aga aag taa caa cag agg cca gcg tta gag att cta aca gtg taa atc cac aa- --a gta tgt aca gat tca ggg gat gtt cat gtc tgt gta aag tca atg cga aaa tca agc cta tag atc cga aca ttg atg cca ata tta gaa cag gac tcc ctt gta taa atg tcc gat gca ttc aaa gta taa aaa tac tgc agc tgt tgc cgt tgt taa agg aaa ttg tag aaa gga tac cgt aga ctt ttc tta gaa atg cca ttc gta tgt aca cgc tgg cag acg cca ccg agc tgt cat agt tga agt cgt cct cga t

>KP849470.1_Monkeypox_virus_isolate_Cote_dIvoire_1971_complete_genome

ata ata tta atg tac taa aac tta tgt att att aat tta tct aac taa agt tag taa att ata tac ata att tta taa tt- --a att taa ttt tac tat ttt att tag tgt cta gaa aaa aat gtg tga ccc acg acc gta gga aac tct aga ggg taa gaa aaa tca atc gtt tat aga gac cat cag aaa gag gtt taa tat ttt tgt gag acc cat cga aga gag aaa gga taa aaa ctt ttt acg act cca tca gaa aga ggt tta ata ttt ttg tga gac cca tcg aag aga gaa agg ata aaa act ttt tac gac tcc atc aga aag agg ttt aat att ttt gtg aga ccc atc gaa gag aga aag gat aaa aac ttt tta cga ctc cat cag aaa gag gtt taa tat ttt tgt gag acc cat cga aga gag aaa gga taa aaa ctt ttt acg act cca tca gaa aga ggt tta ata ttt ttg tga gac cca tcg aag aga gaa agg ata aaa act ttt tac gac tcc atc aga aag agg ttt aat att ttt gtg aga ccc atc gaa gag aga aag gat aaa aac ttt tta cga ctc cat cag aaa gag gtt taa tat ttt tgt gag acc cat cga aga gag aaa gga taa aaa ctt ttt acg act cca tca gaa aga ggt tta ata ttt ttg tga gac cca tcg aag aga gaa agg ata aaa act ttt tac gac tcc atc aga aag agg ttt aat att ttt gtg aga ccc atc gaa gag aga aag gat aaa aac ttt tta cga ctc cat cag aaa gag gtt taa tat ttt tgt gag acc cat cga aga gag aaa gga taa aaa ctt ttt acg act cca tca gaa aga ggt tta --- --- --- --- --- --- --- --- --- --- --- --- --- --- --- --- --- --- --- --- --- --- --- --- --- --- --- --- --- --- --- --- --- --- --- --- --- --- --- --- --- --- --- --- --- --- --a tat ttt tgt gag acc cat cga aga gag aaa gga taa aaa ctt ttt acg act cca tca gaa aga ggt tta ata ttt ttg tga gac cca tcg aag aga gaa aga gat ggt tag tca aga tat ttt tct tag tac aaa agt caa tgt ttt aaa ata tat gga cga gaa tta att tgt ctg tat aaa aac ttg tgt gaa att atg tac tag aga aaa aac gtg agc agt gtc ccc tac atg gat ttt aca gat cat tta tat tcc aaa aat att aac tat ata cgt tta tta tat gat gtt aac gtg taa att ata aac att att tta tga tgc aat tgt ctg aca acc tag att ggt ata agg atg ttg ata agc tct acg aga ata tat tgt tgg acg tta tcg ttt acg aaa tag ttg aga cat cag aaa gag gtt taa tat ttt tgt gag a-c cat cga aga gag aaa tag aat aaa aat a-- --- --- --- ttt ttt tgt aaa act ttt tta tga gac --- --- -ca aga gaa tac gaa tag tga tca tat cgt atc aca tat tga aac aga aag aag aag taa cga gag gta act ttt tgt aaa tgt agt taa ata ttt ttg ttt tgc aaa ccg gaa tat agt gcc cgg t-- -ct ttt tta att cgt ggt gcg gtg tct gaa tcg ttc gat taa ccc aac tca tcc att ttc aga tga ata gag tta tcg att cag aca cat gct ttg agt ttt gtt gaa tcg atg agt gaa gta tca tcg gtt gca cct tca gat gcc gat ccg tcg aca tac ttg aat cca tcc ttg act tca agt tca gat gat tcc tca cac atg tct ccg ata cgt acg cta aac tct agg ttc ttg aca cat ttt gta tca acg atc gtt gaa ccg atg ata tct ttg taa ctc act ttc tta tgt gag atg tta gac cca agt act gga tgg gtc ttg atg tca ctg tct ttc tct tct tcg cta cat ctg atg tcg ata gac atc tca cag tct ttg atc ata gcc aga gct tct tca cgc gtg atc gcg gga gag tcc tta cct tgt ccc ggt gac acg ctg gac aat cta gta ttc aca gtg ttt cca tca gag gat tcg gag atg gat gaa atc ttt ggg cat ttg gtg aat cca aag ttc atg tta aga ccc gca ccg acg ata gtg taa taa gtg gtg gga tct cct ttt aca act tct tcg gat acc tca tca tct tcg gat acc tca tca tct tcg gtc tct gta act tcc gtt acg gat tga caa atc tta tca ttg gtc ggt gtt tgg tct tgc ttt gtg act ttg ata ata aca tcg att ccc ata tga tgt ttg ttt tct tct tca gta cac gag gat gaa gat tgt tga aga cta gta ggc ata gca gct gcc act agg cac atg cat gcc agg aca ata tat tgt ttc atg att gct att gat tga tta ctg ttc tag atg att cta ctt tct tac cat ata ata aat tag aat ata ttt tct act ttt acg aga aat taa tta ttg tat tta tta ttt atg ggt --a aaa aaa ctt act ata agt ggg tgg gat tct ggg aat tag tga tca gtt tat gta tat cgc aac tag cgg gca tat ggc tat tga cat cga gaa cat tac cca tat gat aag aga ttg tat ccg ttt cgt agt ctt gag tat tgg tat tac tat ata gta tgt aga tgt cga cgc tag ata gac agt cgc cca cta gag tta ccg tct ctg aat gcg gca tga tag tat cat tct ttg ttt tcg tta act gtt tgg aag atg aat ctt tgt tgt tac att taa tct cga aat tca gag tac ata tct ttg aag tat tct gat atc tat ttt ctc ctg taa aga atc ctg aag ttg cta cat tat taa gga cag aga agt att ctg cac gaa aga ctg gat cac aat ctt tat gat tca tgg taa tag tta gtt ccg acg ttg aga tgg att cgc tga gac cgg tag tgg tcg tcc gag tac acg atg tgt cgt tga ctg gat aca ggt taa ttt cca cat cga tat agt taa atg tat tgc tgg tta cga cgg gtt cgc att tat ctg tgg aag aga cgg tgt gag aat atg ttc cgg gac cac acg gag aac aga tga cgt ctc cgg tag acg tgt atc cgg ata ctc cgt atc cta ttc cac act ttg ttt tag aaa tac atg ttc tac acc ctg atg ctc ctt tga gaa gac aat aat atc ctg gag agc att cac aga ttc tat tgt gag tcg tgt tac acg atc gcg tct cta cct gat tac tat cac atc ttc cgt tac aac tta gac aag cct gta aat gat tat tgt gag atg taa agg tat ccg aac cac acg gtg tac att gtg tat tag tct tgc tat cac ata atc tgg aag cgt aag ttc ccg gag gac acg ata gac aac ata gat tac ggc ttc tgt att cgt tgt ctt tac act ttc cat tgg atg gtg cat gtg gtg cta tat ctc ttc cgt tta tta tta tac atg aga gaa aca ata tat acg agt ata ata cgg acc tca tga ttt aat aat -gt agt aat cgt cgt ctt gtt act gtt tgt ttc cta ctt ctc caa tca tat aga tta ttt ttt aaa tat ttt ctt tct atc atg gat aat att tgt aat ggt tct ttc cgt aca aca tac tgt tta gat ggt agt cgc tta gct tgg tta tga tat tgc gca taa ttt ccg gag gca aat acg ata gtc tag att gat tat cga tgg tag act cta att tat tga gtg ctt tgt cga cga gtt tac ttt tat gct cca tcg ata gat gac act gtt cta tga gat cgt cgt aca tgg gaa atg aaa tgc gtt tgt ctg aat gta tgg ctt cga gat agg tgt gat acc gga tgt ctt ctg ttc tca ata ccg tat aca agt cgg tgt ctg aga ttc gaa tct ctt tga gga gac tta tgt cac gac tac att ttt cga tga tgg aat cta tct tat cga atg ata tat ttt tca taa ata cac ttt tat agt cct cgt tta aac aga att tag tat gta gtt ccg caa atg act cgt ccc tta ata ggc agt agg cta tta tct tct tta cgt agt gat cgt cgt agg gag aga act ccg aca tct tgt aga aca acg att taa tca tag gta gag ata ctt tca gtc tgt ggt gga tga tgt cat taa cga cat ccg cct tgt ata tga tgt ttc tgt ttt caa aca cca agt cga ata ctg tct tat cgt ctt tag tcg gaa ggt tga tgt cgt atc cga tgt ata cga ggt atg agg caa cat tgt tat tgc aat tct gga agg cgg tat gaa gag gag tca ttg tat tat agt att cgt ctt tct gaa tgt cga atc tat cta gta gat acc gta gta tat tga gag agc gac ttc cat atc ctt gat tat gtt tta tga ata gat aaa gta gat gtt gtc ctt ctt cct ttt gta att tcc cgt att ttt gtt cgt gcc aat tga gta aca tta tga gaa tat gac ctg ttg cac aat cgt tct tta tgt att cca tga tgg gtg tac aat caa gat tat tac gta tcc tcg tat cag ctc ctc gag ata aaa gag cat aca cca cac gag gac tat gtt tgg tat act gtt gaa ggt aag tgt gta acg gcg tat ttc cga ttt tcg taa ccg cgt taa tgt ttg ctc cat gat cta tta tcg cgt aga tga atc gct tct cag ctc gca tct tag tgt gac tct ttg act tgt aat aat tgc ttt cgt gga acg cgg ata tgt gtt tac agt agt aat gaa gag aag tga gtc cat ccg cat cga cgc aat tag ggt cag atc ctt tag tca ata att tgt aca gaa cgt aat agt tta agc tcc cat tga att tat atc taa gat aac aca gca ata gat cgg atg att tac taa agt cat caa tgg ggt ccg tta gta tat caa aga tct tgt tat cga ttg ata gtg aat gaa tca gat agt ggt gta gag gaa tat gtc ctt ttt tat cct tgc tat caa agt tac gca tgc cgt ggc gta aca ata tct tta ata cag atg gat taa atc gtg tat tca tcg tat agc aat gta atg gag agt tac cac att tta gtc gtt tat tca gat cgc agt gtt taa taa cta att taa aca gat gag atg ata tat cca cat caa aga atg cga gat aca tat gac aga cat tat tga cag aaa tgt gac ctt cat tat cac cgt cgt cca taa atg cgt tag gta cgt acc aca tac tat cgt taa cga tgc gca caa tct cgt cca -tt tca tcc atc ttc ata atg att tac ttt ttc ata att aga gaa aaa gat caa ggt ata aaa att aga agt gtt aga cta ta- --- --- --- --- --- aac taa ctt ata a-- -aa cta act tat gac tta act aac tta tga --- --- --- --- --- --- --- --- --- --- --- --- --- --- --- --- --- --- --- --- --- --- --- --- --- --- --- --- --- --- --- --- --- --- --- --- --- --- --- --- --- --- --- --- --- --- --- --- ctt aac taa ctt --- --- --- --- --- --- --- --- --- -ac tta taa --- aac taa ctt atg act taa cta act ta- tga ctt aac taa ctt atg act taa cta act tat gac tta act aac tta tga cta tta act cat ttc aag aaa ggt ggg tgg ata gaa ctc tat atg aca gct tgt gaa acg att aga tcc cta att tct aat gga agt ttt gat agg aga ttg tca tca gtt gat aca ttg ttt att atc tca tct att aga gca cgt ctg ttt aga gct tta gtg acc tgc tcg gtt act tct gtg taa atc ttg aat cct tta gtg ata cac tgt gtc aaa act gga tgt tta gaa tac cta tgt aga ata tgg gaa gca tgc ttg ttt ttg tct cta tta tag att aac tca tac atg gtt gta tta tga att ttc atc tgc cta atg tac tcc aat tct tgt tta caa tca att ata taa tca aag agt gat gat gca tac aca tta caa agt gaa taa tct acc atc ata aaa tac ttg ata cag agc ttt atc aca tca tgg ttt tca att gta tta tta agt ata gct aat ttt ata cag tca ata gac aat ggt tct cta agc aat att tct aat att tta aga tgt gct tcc cta cgg gcg atg aca gat ccc cta tcc acg gcc acg tca aga cat gta tat cca tta ctc att act gcg ttg aca ttt gct cca ttt tct aat agc aat gat act aaa tct ata taa cct gca tag ata gcg cga taa agc aag gtc ctt cca cca gca tct agt tga ttg ata tct tca ata tat ggg ata caa agc tta taa att tct aat act gtg ggt tca tct aca agg aat ccc cta gta tac tga att att tta tat aga tct aat tta aca tca ttt tca tct ggg ata cca caa ttc aaa ata aac tca aca aca cta ctt tcc ttt tta cat att ccc cta aaa tag gca ttc aag cat tct att tta tat att aca gcc cca tga tct acc ata aga tca aca atg tct att tct aca tat gca tta gat aga tag taa agt aag aga tct tgc aca gaa tta caa ttc tta ata att ata gag aaa ata tct tcc ata taa ttc ttt gac act aat gca gat ata ata tct tta tat gta ata tat gca aac agt cta tct act ata tac tga tca ata tta tct cta tga atc cta aaa taa tca tac aga aca tct aca gga tca caa att ggt tca agg aga aat cta tca aat att ttc ctg tca aca act ggt tct aga aca taa cag tca aca cct aat cca tgt ttt tta taa tca tct acc aaa gat aat gac caa aga tcg agg tcg tcg tga aac tgc tca tcg aca gcc atg aaa tct acc gac tcc atg gtg cga atc gca ctg tct tat tcg cca ttg att ttc a-t ttt tta taa tta tgt aca tgt ttt cct tct att ctc aag agt cta caa aaa tat a-- ttt ttt tcg ata tct aag tac taa gtt ttt tta ctg ttt ttg tta ctg tct tcc att ctt cta act aaa gat ctg aga taa att ata caa tct tcg cta tcg aac cat ttt tgt agt cta aag cct gaa gta att aac caa ctg ttt tta tta gtg gct ttt ttc gat cta tca att tca gta tat tct tcg ccg tta taa aag taa tgt tgt tta att gta gga cgg ttg tta gta taa tca cat gaa taa taa tat tct aat tcc tcg tat tga cta ctt aca gat act cga aat agt ctg aaa aat tct tca aag ata ttt tta taa aga tct agg aaa agt tta tta ccg acc atg aac gag ata gat gga taa ata tcc ttt cca tca aag gtc ata att gga taa ttg tcc agc aat ata tct gct gta tta gtt ata tca ctt cca ttt att ttc aga ttg aag taa tgt act agt ttg tga caa tta aca aga tac aaa aga gat gcc gat act aat acg taa ata gct ata cgc gaa tcc att gtt acc ttt ttt tat ttc ata ggt cta tta ata aat ata tgt att act taa gac tag aaa aat caa aag taa gtt ttt gat att tga ttc tta ctt att gtg gga ttg tag ttt act tag taa ttc atc tct gaa tcc tga taa atc atg cat atc aat gat gca act acg cag caa act agt agg aat ata gat atc tgg ata tgt acg taa ata gtc gat tat atc ttt tac aat act att agt ccc tat tgc gtt atc tat ata tcc att aat aat att aca cag tgg ata ctt atg aga aat ata cct ctt aca gat ttt taa cga tat ata atc tag aaa gat atg tgt gta gta ctg tat tac cta aat tat cag tct cat tca aat att gca tga cta tta tcg aga att gct ata tcc ctc tat act cga tgc att tat tac agt aat tca atc cag cta aca taa gag cca atc tca atg ttg gtt taa tta tat cat ctt cat gta ata ata acg atg gaa act ttc cag tag cgt aac act tat cta aga agg ata taa taa cta tgt cta cat tat gtc tct tat cga gaa tat tct taa cga gat atc cat agc tat tct ggt gct aat tat tcc tat att ata ttc cac gaa aaa tga tga agg c-- -ca ttc atc ata aga tga taa aaa gtg tag tga gta aga gta tta gtg aga gag cat gaa gga gat tta gta ttt agc agt gag gat atg atc caa gag ggt gag ata gtc gtt ctc gtt cag aat ctt tcg cag cat aag tag tat gtc gat ata ctt atc att gaa gac tct tcc ggt gac aat agc tga ttg agt aca aag tcc aat tat tgc aca aag ttc ttt ggc ggt ttt cat gga gtc att tct gat gaa aca ttt aat gat ctc cac gca att gtc gat att gtc cca cgg aag tga atc cga gaa ctc ctt caa ctc gct acc aaa tag ctc cat tgc atc aat tct gaa aga gat gag aag cct gta gag agg ccc tgc gct ttc tct atg ggt cca tct atg aga aac cca cag gat gta ttc agt cag aca a-t gtc tga cat cag tca cgg tat tca ggg agt cct tag tag cgt ggc aat gac agg gac tga act ggg cac aag gag agg cca ttg tga agg tag acg aag gta acc tga tgg tag acc tgt agc cgt ctg tgc tta ata gag ggc ttt aat ttc cat ttt taa tgg tgt cgt gaa tga gga atg aga gtg tct ctc gtc ctt ggt tta cat gga tca gag tga gaa aaa ata tct tgt ata tta tta act aac aac ctt ggt ttc tat cca tgt tt- aaa aaa tga cct ata tgt tct tta tta att cta ttt taa act tta tcc tca aga ctc ctg aca aaa tta aaa tcc aga aag cag caa aca atc ctg tta caa gtt tac tga aat ctc tct ttg att gta gag tat atg tag tca gag caa gaa aca ctg cag tag tca aca tga aag ctt gca taa cga tac gtg cat cat aga aag taa caa cag agg cca gcg tta gag att cta aca gtg taa atc cac aa- --a gta tgt aca gat tca ggg gat gtt cat gtc tgt gta aag tca atg cga aaa tca agc cta tag atc cga aca ttg atg cca ata tta gaa cag gac tcc ctt gta taa atg tcc gat gca ttc aaa gta taa aaa tac tgc agc tgt tgc cgt tgt taa agg aaa ttg tag aaa gga tac cgt aga ctt ttc tta gaa atg cca ttc gta tgt aca cgc tgg cag acg cca ccg agc tgt cat agt tga agt cgt cct cga t

>ON682264.2_Monkeypox_virus_isolate_MPXV/Germany/2022/RKI05_complete_genome

--- --- --- --- --- --- --- --- --- --- --- --- --- --- --- --- --- --- --- --- --- --- --- --- --- --- --- --- --- --a ttt tac tat ttt att tag tgt cta gaa aaa aat gtg tga ccc acg acc gta gga aac tct aga ggg taa gaa aaa tca atc gtt tat aga gac cat cag aaa gag gtt taa tat ttt tgt gag acc tat cga a-- --- --- --- --- --- --- --- --- --- --- --- --- --- --- --- --- --- --- --- --- --- --- --- --- --- --- --- --- --- --- --- --- --- --- --- --- --- --- --- --- --- --- --- --- --- --- --- --- --- --- --- --- --- --- --- --- --- --- --- --- --- --- --- --- --- --- --- --- --- --- --- --- --- --- --- --- --- --- --- --- --- --- --- --- --- --- --- --- --- --- --- --- --- --- --- --- --- --- --- --- --- --- --- --- --- --- --- --- --- --- --- --- --- --- --- --- --- --- --- --- --- --- --- --- --- --- --- --- --- --- --- --- --- --- --- --- --- --- --- --- --- --- --- --- --- --- --- --- --- --- --- --- --- --- --- --- --- --- --- --- --- --- --- --- --- --- --- --- --- --- --- --- --- --- --- --- --- --- --- --- --- --- --- --- --- --- --- --- --- --- --- --- --- --- --- --- --- --- --- --- --- --- --- --- --- --- --- --- --- --- --- --- --- --- --- --- --- --- --- --- --- --- --- --- --- --- --- --- --- --- --- --- --- --- --- --- --- --- --- --- --- --- --- --- --- --- --- --- --- --- --- --- --- --- --- --- --- --- --- --- --- --- --- --- --- --- --- --- --- --- --- --- --- --- --- --- --- --- --- -ga gag aaa gga taa aaa ctt ttt acg act cca tca gaa aga ggt tta ata ttt ttg tga gac cca tcg aag aga gaa aga gat ggt tag tca aga tat ttt tct tag tac aaa agt caa tgt ttt aaa ata tat gga cga gaa tta att tgt ctg tat aaa aac ttg tgt gaa att atg tac tag aga aaa aac gtg agc agt gtc ccc tac atg gat ttt aca gat cat tta tat tcc aaa aat att aac tat ata cgt tta tta tat gat gtt aac gtg taa att ata aac att att tta tga tgc aat tgt ctg aca acc tag att ggt ata agg atg ttg ata agc tct acg aga ata tat tgt tgg acg tta tcg ttt acg aaa tag ttg aga cat cag aaa gag gtt taa tat ttt tgt gag a-c cat cga aga gag aaa gag aat aaa aat a-- --t ttt ttt ttt ttt tgt aaa act ttt tta tga gac --- --- -ca aga gaa tac gaa tag tga tca tat cgt atc aca tat tga aac aga aag aag aag taa cga gag gta act ttt tgt gaa tgt agt taa ata ttt ttg ttt tgc aaa ccg gaa tat agt gcc cgg t-- -ct ttt tta att cgt ggt gcg gtg tct gaa tcg ttc gat taa ccc aac tca tcc att ttc aga tga ata gag tta tcg att cag aca cat gct ttg agt ttt gtt gaa tcg atg agt gaa gta tca tcg gtt gca cct tca gat gcc gat ccg tcg aca tac ttg aat cca tcc ttg act tca agt tca gat gat tcc tca cac atg tct ccg ata cgt acg cta aac tct agg ttc ttg aca cat ttt gta tca acg atc gtt gaa ccg atg ata tct ttg taa ctc act ttc tta tgt gag atg tta gac cca agt act gga tgg gtc ttg atg tca ctg tct ttc tct tct tcg cta cat ctg atg tcg ata gac atc tca cag tct ttg atc ata gcc aga gct tct tca cgc gtg atc gcg gga gag tcc tta cct tgt ccc ggt gac acg ctg gac aat cta gta ttc aca gtg ttt cca tca gag gat tcg gag atg gat aaa atc ttt ggg cat ttg gtg aat cca aag ttc atg tta aga ccc gca ccg acg ata gtg taa taa gtg gtg gga tct cct ttt aca act --- --- --- --- --- --- tct tcg gat acc tca tca tct tcg gtc tct gta act tcc gtt acg gat tga caa atc tta tca ttg gtc ggt gtt tgg tct tgc ttt gtg act ttg ata ata aca tcg att ccc ata tga tgt ttg ttt tct tct tca gta cac gag gat gaa gat tgt tga aga cta gta ggc ata gca gct gcc act agg cac atg cat gcc agg aca ata tat tgt ttc atg att gct att gat tga tta ctg ttc tag atg att cta ctt tct tac cat ata ata aat tag aat ata ttt tct act ttt acg aga aat taa tta ttg tat tta tta ttt ata ggt --a aaa aaa ctt act ata agt ggg tgg gat tct ggg aat tag tga tca gtt tat gta tat cgc aac tag cgg gca tat ggc tat tga cat cga gaa cat tac cca tat gat aag aga ttg tat cat ttt cgt agt ctt gag tat tgg tat tac tat ata gta tgt aga tgt cga cgc tag ata gac agt cgc cca cta gag tta ccg tct ctg aat gcg gca tga tag tat cat tct ttg ttt tcg tta act gtt tgg aag atg aat ctt tgt tgt tac att taa tct cga aat tca gag tac ata tct ttg aag tat tct gat atc tat ttt ctc ctg taa aga atc ctg aag ttg cta cat tat taa gga cag aga agt att ctg cac gaa aga ctg gat cac aat ctt tat gat tca tgg taa tag tta gtt ccg acg ttg aga tgg att cgc tga gac cgg tag tgg tcg tcc gag tac acg atg tgt cgt tga ctg gat aca ggt taa ttt cca cat cga tat agt taa atg tat tgc tgg tta cga cgg gtt cgc att tat ctg tgg aag aga cgg tgt gag aat atg ttc cgg gac cac acg gag aac aga tga cgt ctc cgg tag acg tgt atc cgg ata ctc cgt atc cta ttc cac act ttg ttt tag aaa tac atg ttc tac acc ctg atg ctc ctt tga gaa gac aat aat atc ctg gag agc att cac aga ttc tat tgt gag tcg tgt tac acg atc gcg tct cta cct gat tac tat cac atc ttc cgt tac aac tta gac aag cct gta aat gat tat tgt gag atg taa agg tat ccg aac cac acg gtg tac att gtg tat tag tct tgc tat cac ata atc tga aag cgt aag ttc ccg gag gac acg ata gac aac ata gat tac ggc ttc tgt att cgt tgt ctt tac act ttc cat tgg atg gtg cat gtg gtg cta tat ctc ttc cgt tta tta tta tac atg aga gaa aca ata tat acg agt ata ata cgg acc tca tga ttt aat aat -gt agt aat cgt cgt ctt gtt act gtt tgt ttc cta ctt ctc caa tca tat aga tta ttt ttt aaa tat ttt ctt tct atc atg gat aat att tgt aat ggt tct ttc cgt aca aca tac tgt tta gat ggt agt cgc tta gct tgg tta tga tat tgc gca taa ttt ccg gag gca aat acg ata gtc tag att gac tat cga tgg tag act cta att tat tga gtg ctt tgt cga cga gtt tac ttt tat gct cca tcg ata gat gac act gtt cta tga gat cgt cgt aca tgg gaa atg aaa tgc gtt tgt ctg aat gta tgg ctt cga gat atg tgt gat acc gaa tgt ctt ctg ttc tca ata ccg tat aca agt cgg tgt ctg aga ttc gaa tct ctt tga gga gac tta tgt cac gac tac att ttt cga tga tgg aat cta tct tat cga atg ata tat ttt tca taa ata cac ttt tat agt cct cgt tta aac aga att tag tat gta gtt ccg caa atg act cgt ccc tta ata ggc agt agg cta tta tct tct tta cgt agt gat cgt cgt agg gag aga act ccg aca tct tgt aga aca acg att taa tca tag gta gag ata ctt tca gtc tgt ggt gga taa tgt cat taa cga cat ccg cct tgt ata taa tgt ttc tgt ttt caa aca cca agt cga ata ctg tct tat cgt ctt tag tcg gaa ggt tga tgt cgt atc caa tgt ata cga ggt atg agg caa cat tgt tat tgc aat tct gga agg cgg tat gaa gag gag tca ttg tat tat agt att cgt ctt tct gaa tgt cga atc tat cta gta gat acc gta gta tat tga gag agc gac ttc cat atc ctt gat tat gtt tta tga ata gat aaa gta gat gtt gtc ctt ctt cct ttt gta att tcc cgt att ttt gtt cgt gcc aat tga gta aca tta tga gaa tat gac ctg ttg cac aat cgt tct tta tgt att cca tga tgg gtg tac aat taa gat tat tac gta tcc tcg tat cgg ctc ctc gag ata aaa gag cat aca cca cac gag gac tat gtt tgg tat act gtt gaa ggt aag tgt gta acg gcg tat ttc cga ttt tcg taa ccg cgt taa tgt ttg ctc cat gat cta tta tcg cgt aga tga atc gct tct cag ctc gca tct tag tgt gac tct ttg act tgt aat aat tgc ttt cgt gga acg cgg ata tgt gtt tac agt agt aat gaa gag aag tga gtc cat cct cat cga cgc aat tag ggt cag atc ctt tag tca ata att tgt aca gaa cgt aat agt tta agc tcc cat tga att tat atc taa gat aac aca gca ata gat cgg atg att tac taa agt cat caa tgg ggt ccg tta gta tat caa aga tct tgt tat cga ttg ata gtg aat gaa tca gat agt ggt gta gag gaa tat gtc ctt ttt tat cct tgc tat caa agt tac gca tgc cgt ggc gta aca ata tct tta ata cag atg gat taa atc gtg tat tca tcg tat agc aat gta atg gag agt tac cac att tta gtc gtt tat tca gat cgc agt gtt taa taa cta att taa aca gat gag atg ata tat cca cat caa aga atg cga gat aca tat gac aga cat tat tga cag aaa tgt gac ctt cat tat cac cgt cgt cca taa atg cgt tag gta cgt acc aca tac tat cgt taa cga tgc gca caa tct cgt cca -tt tca tcc atc ttc ata atg att tac ttt ttc ata att aga gaa aaa gat caa ggt ata aaa att aga agt gtt aga cta ta- --- --- --- --- --- --- --- --- --- --- -aa cta act tat gac tta act aac tta tga --- --- --- --- --- --- --- --- --- --- --- --- --- --- --- --- --- --- --- --- --- --- --- --- --- --- --- --- --- --- --- --- --- --- --- --- --- --- --- --- --- --- --- --- --- --- --- --- ctt aac ta- --- --- --- --- --- --- --- --- --- --- -ac tta tga ctt aac taa ctt atg act taa cta act ta- tga ctt aac taa ctt atg act taa cta act tat gac tta act aac tta tga cta tta act cat ttc aag aaa ggt ggg tgg ata gaa ctc tat atg aca gct tgt gaa aca att aga tcc cta att tct aat gga agt ttt gat agg aga ttg tca tca gtt gat aca ttg ttt att atc tca tct att aga gca cgt ctg ttt aga gct tta gtg acc tgc tcg gtt act tct gtg taa atc ttg aat cct tta gtg ata cac tgt gtc aaa act gga tgt tta gaa tac cta tgt aga ata tgg gaa gca tgc ttg ttt ttg tct cta tta tag att aac tca tac atg gtt gta tta tga att ttc atc tgc cta atg tac tcc aat tct tgt tta caa tca att ata taa tca aag agt gat gat gca tac aca tta caa agt gaa taa tct acc atc ata aaa tac ttg ata cag agc ttt atc aca tca tgg ttt tca att gta tta tta agt ata gct aat ttt ata cag tca ata gac aat ggt tct cta agc aat att tct aat att tta aga tgt gct tcc cta cgg gcg atg aca gat ccc cta tcc acg gcc acg tca aga cat gta tat cca tta ctc att act gcg ttg aca ttt gct cca ttt tct aat agc cat gat act aaa tct ata taa cct gca tag ata gcg cga taa agc aag gtc ctt cca cca gca tct agt tga ttg ata tct tca ata tat ggg ata caa agc tta taa att tct aat act gtg ggt tca tct aca agg aat ccc cta gta tac tga att att tta tat aga tct aat tta aca tca ttt tca tct ggg ata cca caa ttc aaa ata aac tca aca aca cta ctt tcc ttt tta cat att ccc cta aaa tag gca ttc aag cat tct att tta tat att aca gcc cca tga tct acc ata aga tca aca atg tct att tct aca tat gca tta gat aga tag taa agt aag aga tct tgc aca gaa tta caa ttc tta ata att ata gag aaa ata tct tcc ata taa ttc ttt gac act aat gca gat ata ata tct tta tat gta ata tat gca aac agt cta tct act ata tac tga tca ata tta tct cta tga atc cta aaa taa tca tac aga aca tct aca gga tca caa att ggt tca agg aga aat cta tca aat att ttc ctg tca aca act ggt tct aga aca taa cag tca aca cct aat cca tgt ttt tta taa tca tct acc aaa gat aat gac caa aga tcg agg tcg tcg tga aac tgc tca tcg aca gcc atg aaa tct acc gac tcc atg gtg cga atc gca ctg tct tat tcg cca ttg att ttc a-t ttt tta taa tta tgt aca tgt ttt cct tct att ctc aag agt cta caa aaa tat a-- -tt ttt tcg ata tct aag tac taa gtt ttt tta ctg ttt ttg tta ctg tct tcc att ctt cta act aaa gat ctg aga taa att ata caa tct tcg cta tcg aac cat ttt tgt agt cta aag cct gaa gta att aac caa ctg ttt tta tta gtg gct ttt ttc gat cta tcc tcg tc- --- ctc tca tca tcc tta ta- --- --- --- --- --- --- tta tta tca tta tta tca tag tct att aaa cac aaa tca tct --- --- --- --- --- --- --- --- --- --- --- --- --- --- --- --- --- --- --- --- --- --- --- --- --- --- --a cgt tta taa c-- --- --- --- --- --- --- --- --a aca ttc --- --- --- --- -tc att att aat tag ttc tgt ag- aat atc ttt aat aat ttg gct ata --- --- -ca tct gtt --- --- --- --- --- --- --- --- --- --- --- --- --- --- --- --- --- --- --c aat act --- --- --- --- --- --- --- --a tct att gat gat ttc ttt t-- --- --- --- --- --- --- --- --- --- --- --t taa gac t-- --- --- --- --- --- --- --- --- --- --- --- --- --- --- --- --- --- --- --- --- --- --- --- --- --- --- --- --- --- --- --- --- --- --- --- --- --- --- --- taa act agt --- --- --- --- --- --- --- --- --- --- --- --- --- --- --- --- --- --- --- --- --- --- --- --- --- --- --- --- --- --- --- --- --- --- --- --- --- --- --- --- --- --- --- --- --- --- --- --- --- --- --- --- --- --- --- --- --- --- --- --- --- --- --- --- --- --- --- --- --- --- --- --- --- --- --- --- --- --- --- --- --- --- --- --- --- --- --- --- --- --- --- --- --- --- --- --- --- --- --- --- --- --- --- --- --- --- --- --- --- --t atg gta atg acg atg aaa --- -tc gag tag ta- --- --a ctt cta ata aag act tga ta- --- --t cat tat cat atg ttt gat cg- --- --- --- --- --- --- --t cat agt taa tag tgt g-- --- --- --- --- --- --- --- --- --- --- --- --- --- --- --- --- --- --- --- -gc taa atg gta ctg tta ata agt tta t-- --- --- --- --- --- --- --- --- --- --- --- --- --- --- --- --- --a gac aat atc ata gta ttt tct ttc cag aat --- --- --- --t aga tta ttt ttt taa ata ctg atc ctc aca att ccg tga tgt agc agt agt tgg t-- --- --- --- --- --- --- --- --- --- --- --- --- --- --- --- --- --- --- --- --- --- --- --- --- --- --- --- -gc atg gtc tat atc gt- --- --- --- --- --- --- --- --- --- --- --- --- --- --- --- --- --- --- --- --- --- --- --- --- --- --- --- --- --- --- --- --- --- --- --- --- --- --- --- --- --- --- --- taa aat gta tca tat ata ata gtt ttc tga cgt gga gta cag aat ttt cga --- --- --- --- --- --- --- --- --- --- --- --- --- --- --- --- --- --- --- --- --- --- --- --- --- --- --- --- --- --- --- --- --- --- --- --- --- --- --- --- --- --- --- --t taa tga gtt cat ggt aag gaa ggg caa atg cct --- gta tat aat ata cat aag tta a-- --- --- --- --- --- --- --- --- --- --- --- --- --t agt ttt tta tca tat tt- --- --- --- --- --- --- --- --t cta ata cca taa taa aaa t-- --- --- --- --- --- --- --- --- --- --- --- --- --- --- --- --- --- --- --- --- --- --- --- --- --t atc att att gcg ttt g-g tag tt- --- --- --- --- ctg ccc ta- --- --- --- --- --- --- --- --t cat cta tat cac tgt cac tct c-- --- --g ctc tca cta tat ctt cta aaa tta caa --- -ac aac --- --- --- --- --- --- --- --- --- --- --- --- --- --- --- --- --- --- --- --- --- --- --- --- --- --- --- --- --- --- --- --- --- --- --- --- tgg ata ttc gat --- --- --- --- --- --- --- --- --- --- --- --- --- --- --- --- --- --- --- --- --- --- --- --- --- --- --a aca gca ttt gtg t-- --- --- --- --- --- --- --- --- --- --- --- --- --- --- --- --- -

>ON682268.1_Monkeypox_virus_isolate_MPXV/Germany/2022/RKI08_complete_genome

--- --- --- --- --- --- --- --- --- --- --- --- --- --- --- --- --- --- --- --- --- --- --- --- --- --- --- --- --- --a ttt tac tat ttt att tag tgt cta gaa aaa aat gtg tga ccc acg acc gta gga aac tct aga ggg taa gaa aaa tca atc gtt tat aga gac cat cag aaa gag gtt taa tat ttt tgt gag acc tat cga a-- --- --- --- --- --- --- --- --- --- --- --- --- --- --- --- --- --- --- --- --- --- --- --- --- --- --- --- --- --- --- --- --- --- --- --- --- --- --- --- --- --- --- --- --- --- --- --- --- --- --- --- --- --- --- --- --- --- --- --- --- --- --- --- --- --- --- --- --- --- --- --- --- --- --- --- --- --- --- --- --- --- --- --- --- --- --- --- --- --- --- --- --- --- --- --- --- --- --- --- --- --- --- --- --- --- --- --- --- --- --- --- --- --- --- --- --- --- --- --- --- --- --- --- --- --- --- --- --- --- --- --- --- --- --- --- --- --- --- --- --- --- --- --- --- --- --- --- --- --- --- --- --- --- --- --- --- --- --- --- --- --- --- --- --- --- --- --- --- --- --- --- --- --- --- --- --- --- --- --- --- --- --- --- --- --- --- --- --- --- --- --- --- --- --- --- --- --- --- --- --- --- --- --- --- --- --- --- --- --- --- --- --- --- --- --- --- --- --- --- --- --- --- --- --- --- --- --- --- --- --- --- --- --- --- --- --- --- --- --- --- --- --- --- --- --- --- --- --- --- --- --- --- --- --- --- --- --- --- --- --- --- --- --- --- --- --- --- --- --- --- --- --- --- --- --- --- --- --- --- -ga gag aaa gga taa aaa ctt ttt acg act cca tca gaa aga ggt tta ata ttt ttg tga gac cca tcg aag aga gaa aga gat ggt tag tca aga tat ttt tct tag tac aaa agt caa tgt ttt aaa ata tat gga cga gaa tta att tgt ctg tat aaa aac ttg tgt gaa att atg tac tag aga aaa aac gtg agc agt gtc ccc tac atg gat ttt aca gat cat tta tat tcc aaa aat att aac tat ata cgt tta tta tat gat gtt aac gtg taa att ata aac att att tta tga tgc aat tgt ctg aca acc tag att ggt ata agg atg ttg ata agc tct acg aga ata tat tgt tgg acg tta tcg ttt acg aaa tag ttg aga cat cag aaa gag gtt taa tat ttt tgt gag a-c cat cga aga gag aaa gag aat aaa aat a-- --t ttt ttt ttt ttt tgt aaa act ttt tta tga gac --- --- -ca aga gaa tac gaa tag tga tca tat cgt atc aca tat tga aac aga aag aag aag taa cga gag gta act ttt tgt gaa tgt agt taa ata ttt ttg ttt tgc aaa ccg gaa tat agt gcc cgg t-- -ct ttt tta att cgt ggt gcg gtg tct gaa tcg ttc gat taa ccc aac tca tcc att ttc aga tga ata gag tta tcg att cag aca cat gct ttg agt ttt gtt gaa tcg atg agt gaa gta tca tcg gtt gca cct tca gat gcc gat ccg tcg aca tac ttg aat cca tcc ttg act tca agt tca gat gat tcc tca cac atg tct ccg ata cgt acg cta aac tct agg ttc ttg aca cat ttt gta tca acg atc gtt gaa ccg atg ata tct ttg taa ctc act ttc tta tgt gag atg tta gac cca agt act gga tgg gtc ttg atg tca ctg tct ttc tct tct tcg cta cat ctg atg tcg ata gac atc tca cag tct ttg atc ata gcc aga gct tct tca cgc gtg atc gcg gga gag tcc tta cct tgt ccc ggt gac acg ctg gac aat cta gta ttc aca gtg ttt cca tca gag gat tcg gag atg gat aaa atc ttt ggg cat ttg gtg aat cca aag ttc atg tta aga ccc gca ccg acg ata gtg taa taa gtg gtg gga tct cct ttt aca act --- --- --- --- --- --- tct tcg gat acc tca tca tct tcg gtc tct gta act tcc gtt acg gat tga caa atc tta tca ttg gtc ggt gtt tgg tct tgc ttt gtg act ttg ata ata aca tcg att ccc ata tga tgt ttg ttt tct tct tca gta cac gag gat gaa gat tgt tga aga cta gta ggc ata gca gct gcc act agg cac atg cat gcc agg aca ata tat tgt ttc atg att gct att gat tga tta ctg ttc tag atg att cta ctt tct tac cat ata ata aat tag aat ata ttt tct act ttt acg aga aat taa tta ttg tat tta tta ttt ata ggt --a aaa aaa ctt act ata agt ggg tgg gat tct ggg aat tag tga tca gtt tat gta tat cgc aac tag cgg gca tat ggc tat tga cat cga gaa cat tac cca tat gat aag aga ttg tat cat ttt cgt agt ctt gag tat tgg tat tac tat ata gta tgt aga tgt cga cgc tag ata gac agt cgc cca cta gag tta ccg tct ctg aat gcg gca tga tag tat cat tct ttg ttt tcg tta act gtt tgg aag atg aat ctt tgt tgt tac att taa tct cga aat tca gag tac ata tct ttg aag tat tct gat atc tat ttt ctc ctg taa aga atc ctg aag ttg cta cat tat taa gga cag aga agt att ctg cac gaa aga ctg gat cac aat ctt tat gat tca tgg taa tag tta gtt ccg acg ttg aga tgg att cgc tga gac cgg tag tgg tcg tcc gag tac acg atg tgt cgt tga ctg gat aca ggt taa ttt cca cat cga tat agt taa atg tat tgc tgg tta cga cgg gtt cgc att tat ctg tgg aag aga cgg tgt gag aat atg ttc cgg gac cac acg gag aac aga tga cgt ctc cgg tag acg tgt atc cgg ata ctc cgt atc cta ttc cac act ttg ttt tag aaa tac atg ttc tac acc ctg atg ctc ctt tga gaa gac aat aat atc ctg gag agc att cac aga ttc tat tgt gag tcg tgt tac acg atc gcg tct cta cct gat tac tat cac atc ttc cgt tac aac tta gac aag cct gta aat gat tat tgt gag atg taa agg tat ccg aac cac acg gtg tac att gtg tat tag tct tgc tat cac ata atc tga aag cgt aag ttc ccg gag gac acg ata gac aac ata gat tac ggc ttc tgt att cgt tgt ctt tac act ttc cat tgg atg gtg cat gtg gtg cta tat ctc ttc cgt tta tta tta tac atg aga gaa aca ata tat acg agt ata ata cgg acc tca tga ttt aat aat -gt agt aat cgt cgt ctt gtt act gtt tgt ttc cta ctt ctc caa tca tat aga tta ttt ttt aaa tat ttt ctt tct atc atg gat aat att tgt aat ggt tct ttc cgt aca aca tac tgt tta gat ggt agt cgc tta gct tgg tta tga tat tgc gca taa ttt ccg gag gca aat acg ata gtc tag att gac tat cga tgg tag act cta att tat tga gtg ctt tgt cga cga gtt tac ttt tat gct cca tcg ata gat gac act gtt cta tga gat cgt cgt aca tgg gaa atg aaa tgc gtt tgt ctg aat gta tgg ctt cga gat atg tgt gat acc gaa tgt ctt ctg ttc tca ata ccg tat aca agt cgg tgt ctg aga ttc gaa tct ctt tga gga gac tta tgt cac gac tac att ttt cga tga tgg aat cta tct tat cga atg ata tat ttt tca taa ata cac ttt tat agt cct cgt tta aac aga att tag tat gta gtt ccg caa atg act cgt ccc tta ata ggc agt agg cta tta tct tct tta cgt agt gat cgt cgt agg gag aga act ccg aca tct tgt aga aca acg att taa tca tag gta gag ata ctt tca gtc tgt ggt gga taa tgt cat taa cga cat ccg cct tgt ata taa tgt ttc tgt ttt caa aca cca agt cga ata ctg tct tat cgt ctt tag tcg gaa ggt tga tgt cgt atc caa tgt ata cga ggt atg agg caa cat tgt tat tgc aat tct gga agg cgg tat gaa gag gag tca ttg tat tat agt att cgt ctt tct gaa tgt cga atc tat cta gta gat acc gta gta tat tga gag agc gac ttc cat atc ctt gat tat gtt tta tga ata gat aaa gta gat gtt gtc ctt ctt cct ttt gta att tcc cgt att ttt gtt cgt gcc aat tga gta aca tta tga gaa tat gac ctg ttg cac aat cgt tct tta tgt att cca tga tgg gtg tac aat taa gat tat tac gta tcc tcg tat cgg ctc ctc gag ata aaa gag cat aca cca cac gag gac tat gtt tgg tat act gtt gaa ggt aag tgt gta acg gcg tat ttc cga ttt tcg taa ccg cgt taa tgt ttg ctc cat gat cta tta tcg cgt aga tga atc gct tct cag ctc gca tct tag tgt gac tct ttg act tgt aat aat tgc ttt cgt gga acg cgg ata tgt gtt tac agt agt aat gaa gag aag tga gtc cat cct cat cga cgc aat tag ggt cag atc ctt tag tca ata att tgt aca gaa cgt aat agt tta agc tcc cat tga att tat atc taa gat aac aca gca ata gat cgg atg att tac taa agt cat caa tgg ggt ccg tta gta tat caa aga tct tgt tat cga ttg ata gtg aat gaa tca gat agt ggt gta gag gaa tat gtc ctt ttt tat cct tgc tat caa agt tac gca tgc cgt ggc gta aca ata tct tta ata cag atg gat taa atc gtg tat tca tcg tat agc aat gta atg gag agt tac cac att tta gtc gtt tat tca gat cgc agt gtt taa taa cta att taa aca gat aag atg ata tat cca cat caa aga atg cga gat aca tat gac aga cat tat tga cag aaa tgt gac ctt cat tat cac cgt cgt cca taa atg cgt tag gta cgt acc aca tac tat cgt taa cga tgc gca caa tct cgt cca -tt tca tcc atc ttc ata atg att tac ttt ttc ata att aga gaa aaa gat caa ggt ata aaa att aga agt gtt aga cta ta- --- --- --- --- --- --- --- --- --- --- -aa cta act tat gac tta act aac tta tga --- --- --- --- --- --- --- --- --- --- --- --- --- --- --- --- --- --- --- --- --- --- --- --- --- --- --- --- --- --- --- --- --- --- --- --- --- --- --- --- --- --- --- --- --- --- --- --- ctt aac ta- --- --- --- --- --- --- --- --- --- --- -ac tta tga ctt aac taa ctt atg act taa cta act ta- tga ctt aac taa ctt atg act taa cta act tat gac tta act aac tta tga cta tta act cat ttc aag aaa ggt ggg tgg ata gaa ctc tat atg aca gct tgt gaa aca att aga tcc cta att tct aat gga agt ttt gat agg aga ttg tca tca gtt gat aca ttg ttt att atc tca tct att aga gca cgt ctg ttt aga gct tta gtg acc tgc tcg gtt act tct gtg taa atc ttg aat cct tta gtg ata cac tgt gtc aaa act gga tgt tta gaa tac cta tgt aga ata tgg gaa gca tgc ttg ttt ttg tct cta tta tag att aac tca tac atg gtt gta tta tga att ttc atc tgc cta atg tac tcc aat tct tgt tta caa tca att ata taa tca aag agt gat gat gca tac aca tta caa agt gaa taa tct acc atc ata aaa tac ttg ata cag agc ttt atc aca tca tgg ttt tca att gta tta tta agt ata gct aat ttt ata cag tca ata gac aat ggt tct cta agc aat att tct aat att tta aga tgt gct tcc cta cgg gcg atg aca gat ccc cta tcc acg gcc acg tca aga cat gta tat cca tta ctc att act gcg ttg aca ttt gct cca ttt tct aat agc cat gat act aaa tct ata taa cct gca tag ata gcg cga taa agc aag gtc ctt cca cca gca tct agt tga ttg ata tct tca ata tat ggg ata caa agc tta taa att tct aat act gtg ggt tca tct aca agg aat ccc cta gta tac tga att att tta tat aga tct aat tta aca tca ttt tca tct ggg ata cca caa ttc aaa ata aac tca aca aca cta ctt tcc ttt tta cat att ccc cta aaa tag gca ttc aag cat tct att tta tat att aca gcc cca tga tct acc ata aga tca aca atg tct att tct aca tat gca tta gat aga tag taa agt aag aga tct tgc aca gaa tta caa ttc tta ata att ata gag aaa ata tct tcc ata taa ttc ttt gac act aat gca gat ata ata tct tta tat gta ata tat gca aac agt cta tct act ata tac tga tca ata tta tct cta tga atc cta aaa taa tca tac aga aca tct aca gga tca caa att ggt tca agg aga aat cta tca aat att ttc ctg tca aca act ggt tct aga aca taa cag tca aca cct aat cca tgt ttt tta taa tca tct acc aaa gat aat gac caa aga tcg agg tcg tcg tga aac tgc tca tcg aca gcc atg aaa tct acc gac tcc atg gtg cga atc gca ctg tct tat tcg cca ttg att ttc a-t ttt tta taa tta tgt aca tgt ttt cct tct att ctc aag agt cta caa aaa tat a-- -tt ttt tcg ata tct aag tac taa gtt ttt tta ctg ttt ttg tta ctg tct tcc att ctt cta act aaa gat ctg aga taa att ata caa tct tcg cta tcg aac cat ttt tgt agt cta aag cct gaa gta att aac caa ctg ttt tta tta gtg gct ttt ttc gat cta tcc tcg tc- --- ctc tca tca tcc tta ta- --- --- --- --- --- --- tta tta tca tta tta tca tag tct att aaa cac aaa tca tct --- --- --- --- --- --- --- --- --- --- --- --- --- --- --- --- --- --- --- --- --- --- --- --- --- --- --a cgt tta taa c-- --- --- --- --- --- --- --- --a aca ttc --- --- --- --- -tc att att aat tag ttc tgt ag- aat atc ttt aat aat ttg gct ata --- --- -ca tct gtt --- --- --- --- --- --- --- --- --- --- --- --- --- --- --- --- --- --- --c aat act --- --- --- --- --- --- --- --a tct att gat gat ttc ttt t-- --- --- --- --- --- --- --- --- --- --- --t taa gac t-- --- --- --- --- --- --- --- --- --- --- --- --- --- --- --- --- --- --- --- --- --- --- --- --- --- --- --- --- --- --- --- --- --- --- --- --- --- --- --- taa act agt --- --- --- --- --- --- --- --- --- --- --- --- --- --- --- --- --- --- --- --- --- --- --- --- --- --- --- --- --- --- --- --- --- --- --- --- --- --- --- --- --- --- --- --- --- --- --- --- --- --- --- --- --- --- --- --- --- --- --- --- --- --- --- --- --- --- --- --- --- --- --- --- --- --- --- --- --- --- --- --- --- --- --- --- --- --- --- --- --- --- --- --- --- --- --- --- --- --- --- --- --- --- --- --- --- --- --- --- --- --t atg gta atg acg atg aaa --- -tc gag tag ta- --- --a ctt cta ata aag act tga ta- --- --t cat tat cat atg ttt gat cg- --- --- --- --- --- --- --t cat agt taa tag tgt g-- --- --- --- --- --- --- --- --- --- --- --- --- --- --- --- --- --- --- --- -gc taa atg gta ctg tta ata agt tta t-- --- --- --- --- --- --- --- --- --- --- --- --- --- --- --- --- --a gac aat atc ata gta ttt tct ttc cag aat --- --- --- --t aga tta ttt ttt taa ata ctg atc ctc aca att ccg tga tgt agc agt agt tgg t-- --- --- --- --- --- --- --- --- --- --- --- --- --- --- --- --- --- --- --- --- --- --- --- --- --- --- --- -gc atg gtc tat atc gt- --- --- --- --- --- --- --- --- --- --- --- --- --- --- --- --- --- --- --- --- --- --- --- --- --- --- --- --- --- --- --- --- --- --- --- --- --- --- --- --- --- --- --- taa aat gta tca tat ata ata gtt ttc tga cgt gga gta cag aat ttt cga --- --- --- --- --- --- --- --- --- --- --- --- --- --- --- --- --- --- --- --- --- --- --- --- --- --- --- --- --- --- --- --- --- --- --- --- --- --- --- --- --- --- --- --t taa tga gtt cat ggt aag gaa ggg caa atg cct --- gta tat aat ata cat aag tta a-- --- --- --- --- --- --- --- --- --- --- --- --- --t agt ttt tta tca tat tt- --- --- --- --- --- --- --- --t cta ata cca taa taa aaa t-- --- --- --- --- --- --- --- --- --- --- --- --- --- --- --- --- --- --- --- --- --- --- --- --- --t atc att att gcg ttt g-g tag tt- --- --- --- --- ctg ccc ta- --- --- --- --- --- --- --- --t cat cta tat cac tgt cac tct c-- --- --g ctc tca cta tat ctt cta aaa tta caa --- -ac aac --- --- --- --- --- --- --- --- --- --- --- --- --- --- --- --- --- --- --- --- --- --- --- --- --- --- --- --- --- --- --- --- --- --- --- --- tgg ata ttc gat --- --- --- --- --- --- --- --- --- --- --- --- --- --- --- --- --- --- --- --- --- --- --- --- --- --- --a aca gca ttt gtg t-- --- --- --- --- --- --- --- --- --- --- --- --- --- --- --- --- -

>ON682270.1_Monkeypox_virus_isolate_MPXV/Germany/2022/RKI06_complete_genome

--- --- --- --- --- --- --- --- --- --- --- --- --- --- --- --- --- --- --- --- --- --- --- --- --- --- --- --- --- --a ttt tac tat ttt att tag tgt cta gaa aaa aat gtg tga ccc acg acc gta gga aac tct aga ggg taa gaa aaa tca atc gtt tat aga gac cat cag aaa gag gtt taa tat ttt tgt gag acc tat cga a-- --- --- --- --- --- --- --- --- --- --- --- --- --- --- --- --- --- --- --- --- --- --- --- --- --- --- --- --- --- --- --- --- --- --- --- --- --- --- --- --- --- --- --- --- --- --- --- --- --- --- --- --- --- --- --- --- --- --- --- --- --- --- --- --- --- --- --- --- --- --- --- --- --- --- --- --- --- --- --- --- --- --- --- --- --- --- --- --- --- --- --- --- --- --- --- --- --- --- --- --- --- --- --- --- --- --- --- --- --- --- --- --- --- --- --- --- --- --- --- --- --- --- --- --- --- --- --- --- --- --- --- --- --- --- --- --- --- --- --- --- --- --- --- --- --- --- --- --- --- --- --- --- --- --- --- --- --- --- --- --- --- --- --- --- --- --- --- --- --- --- --- --- --- --- --- --- --- --- --- --- --- --- --- --- --- --- --- --- --- --- --- --- --- --- --- --- --- --- --- --- --- --- --- --- --- --- --- --- --- --- --- --- --- --- --- --- --- --- --- --- --- --- --- --- --- --- --- --- --- --- --- --- --- --- --- --- --- --- --- --- --- --- --- --- --- --- --- --- --- --- --- --- --- --- --- --- --- --- --- --- --- --- --- --- --- --- --- --- --- --- --- --- --- --- --- --- --- --- --- -ga gag aaa gga taa aaa ctt ttt acg act cca tca gaa aga ggt tta ata ttt ttg tga gac cca tcg aag aga gaa aga gat ggt tag tca aga tat ttt tct tag tac aaa agt caa tgt ttt aaa ata tat gga cga gaa tta att tgt ctg tat aaa aac ttg tgt gaa att atg tac tag aga aaa aac gtg agc agt gtc ccc tac atg gat ttt aca gat cat tta tat tcc aaa aat att aac tat ata cgt tta tta tat gat gtt aac gtg taa att ata aac att att tta tga tgc aat tgt ctg aca acc tag att ggt ata agg atg ttg ata agc tct acg aga ata tat tgt tgg acg tta tcg ttt acg aaa tag ttg aga cat cag aaa gag gtt taa tat ttt tgt gag a-c cat cga aga gag aaa gag aat aaa aat a-- --t ttt ttt ttt ttt tgt aaa act ttt tta tga gac --- --- -ca aga gaa tac gaa tag tga tca tat cgt atc aca tat tga aac aga aag aag aag taa cga gag gta act ttt tgt gaa tgt agt taa ata ttt ttg ttt tgc aaa ccg gaa tat agt gcc cgg t-- -ct ttt tta att cgt ggt gcg gtg tct gaa tcg ttc gat taa ccc aac tca tcc att ttc aga tga ata gag tta tcg att cag aca cat gct ttg agt ttt gtt gaa tcg atg agt gaa gta tca tcg gtt gca cct tca gat gcc gat ccg tcg aca tac ttg aat cca tcc ttg act tca agt tca gat gat tcc tca cac atg tct ccg ata cgt acg cta aac tct agg ttc ttg aca cat ttt gta tca acg atc gtt gaa ccg atg ata tct ttg taa ctc act ttc tta tgt gag atg tta gac cca agt act gga tgg gtc ttg atg tca ctg tct ttc tct tct tcg cta cat ctg atg tcg ata gac atc tca cag tct ttg atc ata gcc aga gct tct tca cgc gtg atc gcg gga gag tcc tta cct tgt ccc ggt gac acg ctg gac aat cta gta ttc aca gtg ttt cca tca gag gat tcg gag atg gat aaa atc ttt ggg cat ttg gtg aat cca aag ttc atg tta aga ccc gca ccg acg ata gtg taa taa gtg gtg gga tct cct ttt aca act --- --- --- --- --- --- tct tcg gat acc tca tca tct tcg gtc tct gta act tcc gtt acg gat tga caa atc tta tca ttg gtc ggt gtt tgg tct tgc ttt gtg act ttg ata ata aca tcg att ccc ata tga tgt ttg ttt tct tct tca gta cac gag gat gaa gat tgt tga aga cta gta ggc ata gca gct gcc act agg cac atg cat gcc agg aca ata tat tgt ttc atg att gct att gat tga tta ctg ttc tag atg att cta ctt tct tac cat ata ata aat tag aat ata ttt tct act ttt acg aga aat taa tta ttg tat tta tta ttt ata ggt --a aaa aaa ctt act ata agt ggg tgg gat tct ggg aat tag tga tca gtt tat gta tat cgc aac tag cgg gca tat ggc tat tga cat cga gaa cat tac cca tat gat aag aga ttg tat cat ttt cgt agt ctt gag tat tgg tat tac tat ata gta tgt aga tgt cga cgc tag ata gac agt cgc cca cta gag tta ccg tct ctg aat gcg gca tga tag tat cat tct ttg ttt tcg tta act gtt tgg aag atg aat ctt tgt tgt tac att taa tct cga aat tca gag tac ata tct ttg aag tat tct gat atc tat ttt ctc ctg taa aga atc ctg aag ttg cta cat tat taa gga cag aga agt att ctg cac gaa aga ctg gat cac aat ctt tat gat tca tgg taa tag tta gtt ccg acg ttg aga tgg att cgc tga gac cgg tag tgg tcg tcc gag tac acg atg tgt cgt tga ctg gat aca ggt taa ttt cca cat cga tat agt taa atg tat tgc tgg tta cga cgg gtt cgc att tat ctg tgg aag aga cgg tgt gag aat atg ttc cgg gac cac acg gag aac aga tga cgt ctc cgg tag acg tgt atc cgg ata ctc cgt atc cta ttc cac act ttg ttt tag aaa tac atg ttc tac acc ctg atg ctc ctt tga gaa gac aat aat atc ctg gag agc att cac aga ttc tat tgt gag tcg tgt tac acg atc gcg tct cta cct gat tac tat cac atc ttc cgt tac aac tta gac aag cct gta aat gat tat tgt gag atg taa agg tat ccg aac cac acg gtg tac att gtg tat tag tct tgc tat cac ata atc tga aag cgt aag ttc ccg gag gac acg ata gac aac ata gat tac ggc ttc tgt att cgt tgt ctt tac act ttc cat tgg atg gtg cat gtg gtg cta tat ctc ttc cgt tta tta tta tac atg aga gaa aca ata tat acg agt ata ata cgg acc tca tga ttt aat aat -gt agt aat cgt cgt ctt gtt act gtt tgt ttc cta ctt ctc caa tca tat aga tta ttt ttt aaa tat ttt ctt tct atc atg gat aat att tgt aat ggt tct ttc cgt aca aca tac tgt tta gat ggt agt cgc tta gct tgg tta tga tat tgc gca taa ttt ccg gag gca aat acg ata gtc tag att gac tat cga tgg tag act cta att tat tga gtg ctt tgt cga cga gtt tac ttt tat gct cca tcg ata gat gac act gtt cta tga gat cgt cgt aca tgg gaa atg aaa tgc gtt tgt ctg aat gta tgg ctt cga gat atg tgt gat acc gaa tgt ctt ctg ttc tca ata ccg tat aca agt cgg tgt ctg aga ttc gaa tct ctt tga gga gac tta tgt cac gac tac att ttt cga tga tgg aat cta tct tat cga atg ata tat ttt tca taa ata cac ttt tat agt cct cgt tta aac aga att tag tat gta gtt ccg caa atg act cgt ccc tta ata ggc agt agg cta tta tct tct tta cgt agt gat cgt cgt agg gag aga act ccg aca tct tgt aga aca acg att taa tca tag gta gag ata ctt tca gtc tgt ggt gga taa tgt cat taa cga cat ccg cct tgt ata taa tgt ttc tgt ttt caa aca cca agt cga ata ctg tct tat cgt ctt tag tcg gaa ggt tga tgt cgt atc caa tgt ata cga ggt atg agg caa cat tgt tat tgc aat tct gga agg cgg tat gaa gag gag tca ttg tat tat agt att cgt ctt tct gaa tgt cga atc tat cta gta gat acc gta gta tat tga gag agc gac ttc cat atc ctt gat tat gtt tta tga ata gat aaa gta gat gtt gtc ctt ctt cct ttt gta att tcc cgt att ttt gtt cgt gcc aat tga gta aca tta tga gaa tat gac ctg ttg cac aat cgt tct tta tgt att cca tga tgg gtg tac aat taa gat tat tac gta tcc tcg tat cgg ctc ctc gag ata aaa gag cat aca cca cac gag gac tat gtt tgg tat act gtt gaa ggt aag tgt gta acg gcg tat ttc cga ttt tcg taa ccg cgt taa tgt ttg ctc cat gat cta tta tcg cgt aga tga atc gct tct cag ctc gca tct tag tgt gac tct ttg act tgt aat aat tgc ttt cgt gga acg cgg ata tgt gtt tac agt agt aat gaa gag aag tga gtc cat cct cat cga cgc aat tag ggt cag atc ctt tag tca ata att tgt aca gaa cgt aat agt tta agc tcc cat tga att tat atc taa gat aac aca gca ata gat cgg atg att tac taa agt cat caa tgg ggt ccg tta gta tat caa aga tct tgt tat cga ttg ata gtg aat gaa tca gat agt ggt gta gag gaa tat gtc ctt ttt tat cct tgc tat caa agt tac gca tgc cgt ggc gta aca ata tct tta ata cag atg gat taa atc gtg tat tca tcg tat agc aat gta atg gag agt tac cac att tta gtc gtt tat tca gat cgc agt gtt taa taa cta att taa aca gat gag atg ata tat cca cat caa aga atg cga gat aca tat gac aga cat tat tga cag aaa tgt gac ctt cat tat cac cgt cgt cca taa atg cgt tag gta cgt acc aca tac tat cgt taa cga tgc gca caa tct cgt cca -tt tca tcc atc ttc ata atg att tac ttt ttc ata att aga gaa aaa gat caa ggt ata aaa att aga agt gtt aga cta ta- --- --- --- --- --- --- --- --- --- --- -aa cta act tat gac tta act aac tta tga --- --- --- --- --- --- --- --- --- --- --- --- --- --- --- --- --- --- --- --- --- --- --- --- --- --- --- --- --- --- --- --- --- --- --- --- --- --- --- --- --- --- --- --- --- --- --- --- ctt aac ta- --- --- --- --- --- --- --- --- --- --- -ac tta tga ctt aac taa ctt atg act taa cta act ta- tga ctt aac taa ctt atg act taa cta act tat gac tta act aac tta tga cta tta act cat ttc aag aaa ggt ggg tgg ata gaa ctc tat atg aca gct tgt gaa aca att aga tcc cta att tct aat gga agt ttt gat agg aga ttg tca tca gtt gat aca ttg ttt att atc tca tct att aga gca cgt ctg ttt aga gct tta gtg acc tgc tcg gtt act tct gtg taa atc ttg aat cct tta gtg ata cac tgt gtc aaa act gga tgt tta gaa tac cta tgt aga ata tgg gaa gca tgc ttg ttt ttg tct cta tta tag att aac tca tac atg gtt gta tta tga att ttc atc tgc cta atg tac tcc aat tct tgt tta caa tca att ata taa tca aag agt gat gat gca tac aca tta caa agt gaa taa tct acc atc ata aaa tac ttg ata cag agc ttt atc aca tca tgg ttt tca att gta tta tta agt ata gct aat ttt ata cag tca ata gac aat ggt tct cta agc aat att tct aat att tta aga tgt gct tcc cta cgg gcg atg aca gat ccc cta tcc acg gcc acg tca aga cat gta tat cca tta ctc att act gcg ttg aca ttt gct cca ttt tct aat agc cat gat act aaa tct ata taa cct gca tag ata gcg cga taa agc aag gtc ctt cca cca gca tct agt tga ttg ata tct tca ata tat ggg ata caa agc tta taa att tct aat act gtg ggt tca tct aca agg aat ccc cta gta tac tga att att tta tat aga tct aat tta aca tca ttt tca tct ggg ata cca caa ttc aaa ata aac tca aca aca cta ctt tcc ttt tta cat att ccc cta aaa tag gca ttc aag cat tct att tta tat att aca gcc cca tga tct acc ata aga tca aca atg tct att tct aca tat gca tta gat aga tag taa agt aag aga tct tgc aca gaa tta caa ttc tta ata att ata gag aaa ata tct tcc ata taa ttc ttt gac act aat gca gat ata ata tct tta tat gta ata tat gca aac agt cta tct act ata tac tga tca ata tta tct cta tga atc cta aaa taa tca tac aga aca tct aca gga tca caa att ggt tca agg aga aat cta tca aat att ttc ctg tca aca act ggt tct aga aca taa cag tca aca cct aat cca tgt ttt tta taa tca tct acc aaa gat aat gac caa aga tcg agg tcg tcg tga aac tgc tca tcg aca gcc atg aaa tct acc gac tcc atg gtg cga atc gca ctg tct tat tcg cca ttg att ttc a-t ttt tta taa tta tgt aca tgt ttt cct tct att ctc aag agt cta caa aaa tat a-- -tt ttt tcg ata tct aag tac taa gtt ttt tta ctg ttt ttg tta ctg tct tcc att ctt cta act aaa gat ctg aga taa att ata caa tct tcg cta tcg aac cat ttt tgt agt cta aag cct gaa gta att aac caa ctg ttt tta tta gtg gct ttt ttc gat cta tcc tcg tc- --- ctc tca tca tcc tta ta- --- --- --- --- --- --- tta tta tca tta tta tca tag tct att aaa cac aaa tca tct --- --- --- --- --- --- --- --- --- --- --- --- --- --- --- --- --- --- --- --- --- --- --- --- --- --- --a cgt tta taa c-- --- --- --- --- --- --- --- --a aca ttc --- --- --- --- -tc att att aat tag ttc tgt ag- aat atc ttt aat aat ttg gct ata --- --- -ca tct gtt --- --- --- --- --- --- --- --- --- --- --- --- --- --- --- --- --- --- --c aat act --- --- --- --- --- --- --- --a tct att gat gat ttc ttt t-- --- --- --- --- --- --- --- --- --- --- --t taa gac t-- --- --- --- --- --- --- --- --- --- --- --- --- --- --- --- --- --- --- --- --- --- --- --- --- --- --- --- --- --- --- --- --- --- --- --- --- --- --- --- taa act agt --- --- --- --- --- --- --- --- --- --- --- --- --- --- --- --- --- --- --- --- --- --- --- --- --- --- --- --- --- --- --- --- --- --- --- --- --- --- --- --- --- --- --- --- --- --- --- --- --- --- --- --- --- --- --- --- --- --- --- --- --- --- --- --- --- --- --- --- --- --- --- --- --- --- --- --- --- --- --- --- --- --- --- --- --- --- --- --- --- --- --- --- --- --- --- --- --- --- --- --- --- --- --- --- --- --- --- --- --- --t atg gta atg acg atg aaa --- -tc gag tag ta- --- --a ctt cta ata aag act tga ta- --- --t cat tat cat atg ttt gat cg- --- --- --- --- --- --- --t cat agt taa tag tgt g-- --- --- --- --- --- --- --- --- --- --- --- --- --- --- --- --- --- --- --- -gc taa atg gta ctg tta ata agt tta t-- --- --- --- --- --- --- --- --- --- --- --- --- --- --- --- --- --a gac aat atc ata gta ttt tct ttc cag aat --- --- --- --t aga tta ttt ttt taa ata ctg atc ctc aca att ccg tga tgt agc agt agt tgg t-- --- --- --- --- --- --- --- --- --- --- --- --- --- --- --- --- --- --- --- --- --- --- --- --- --- --- --- -gc atg gtc tat atc gt- --- --- --- --- --- --- --- --- --- --- --- --- --- --- --- --- --- --- --- --- --- --- --- --- --- --- --- --- --- --- --- --- --- --- --- --- --- --- --- --- --- --- --- taa aat gta tca tat ata ata gtt ttc tga cgt gga gta cag aat ttt cga --- --- --- --- --- --- --- --- --- --- --- --- --- --- --- --- --- --- --- --- --- --- --- --- --- --- --- --- --- --- --- --- --- --- --- --- --- --- --- --- --- --- --- --t taa tga gtt cat ggt aag gaa ggg caa atg cct --- gta tat aat ata cat aag tta a-- --- --- --- --- --- --- --- --- --- --- --- --- --t agt ttt tta tca tat tt- --- --- --- --- --- --- --- --t cta ata cca taa taa aaa t-- --- --- --- --- --- --- --- --- --- --- --- --- --- --- --- --- --- --- --- --- --- --- --- --- --t atc att att gcg ttt g-g tag tt- --- --- --- --- ctg ccc ta- --- --- --- --- --- --- --- --t cat cta tat cac tgt cac tct c-- --- --g ctc tca cta tat ctt cta aaa tta caa --- -ac aac --- --- --- --- --- --- --- --- --- --- --- --- --- --- --- --- --- --- --- --- --- --- --- --- --- --- --- --- --- --- --- --- --- --- --- --- tgg ata ttc gat --- --- --- --- --- --- --- --- --- --- --- --- --- --- --- --- --- --- --- --- --- --- --- --- --- --- --a aca gca ttt gtg t-- --- --- --- --- --- --- --- --- --- --- --- --- --- --- --- --- -

>ON637938.1_Monkeypox_virus_isolate_MPXV/Germany/2022/RKI01_complete_genome

--- --- --- --- --- --- --- --- --- --- --- --- --- --- --- --- --- --a tta ata ata cat aag ttt tag tac att aat att taa ttt tac tat ttt att tag tgt cta gaa aaa aat gtg tga ccc acg acc gta gga aac tct aga ggg taa gaa aaa tca atc gtt tat aga gac cat cag aaa gag gtt taa tat ttt tgt gag acc tat cga a-- --- --- --- --- --- --- --- --- --- --- --- --- --- --- --- --- --- --- --- --- --- --- --- --- --- --- --- --- --- --- --- --- --- --- --- --- --- --- --- --- --- --- --- --- --- --- --- --- --- --- --- --- --- --- --- --- --- --- --- --- --- --- --- --- --- --- --- --- --- --- --- --- --- --- --- --- --- --- --- --- --- --- --- --- --- --- --- --- --- --- --- --- --- --- --- --- --- --- --- --- --- --- --- --- --- --- --- --- --- --- --- --- --- --- --- --- --- --- --- --- --- --- --- --- --- --- --- --- --- --- --- --- --- --- --- --- --- --- --- --- --- --- --- --- --- --- --- --- --- --- --- --- --- --- --- --- --- --- --- --- --- --- --- --- --- --- --- --- --- --- --- --- --- --- --- --- --- --- --- --- --- --- --- --- --- --- --- --- --- --- --- --- --- --- --- --- --- --- --- --- --- --- --- --- --- --- --- --- --- --- --- --- --- --- --- --- --- --- --- --- --- --- --- --- --- --- --- --- --- --- --- --- --- --- --- --- --- --- --- --- --- --- --- --- --- --- --- --- --- --- --- --- --- --- --- --- --- --- --- --- --- --- --- --- --- --- --- --- --- --- --- --- --- --- --- --- --- --- --- -ga gag aaa gga taa aaa ctt ttt acg act cca tca gaa aga ggt tta ata ttt ttg tga gac cca tcg aag aga gaa aga gat ggt tag tca aga tat ttt tct tag tac aaa agt caa tgt ttt aaa ata tat gga cga gaa tta att tgt ctg tat aaa aac ttg tgt gaa att atg tac tag aga aaa aac gtg agc agt gtc ccc tac atg gat ttt aca gat cat tta tat tcc aaa aat att aac tat ata cgt tta tta tat gat gtt aac gtg taa att ata aac att att tta tga tgc aat tgt ctg aca acc tag att ggt ata agg atg ttg ata agc tct acg aga ata tat tgt tgg acg tta tcg ttt acg aaa tag ttg aga cat cag aaa gag gtt taa tat ttt tgt gag a-c cat cga aga gag aaa gag aat aaa aat a-- --t ttt ttt ttt ttt tgt aaa act ttt tta tga gac --- --- -ca aga gaa tac gaa tag tga tca tat cgt atc aca tat tga aac aga aag aag aag taa cga gag gta act ttt tgt gaa tgt agt taa ata ttt ttg ttt tgc aaa ccg gaa tat agt gcc cgg t-- -ct ttt tta att cgt ggt gcg gtg tct gaa tcg ttc gat taa ccc aac tca tcc att ttc aga tga ata gag tta tcg att cag aca cat gct ttg agt ttt gtt gaa tcg atg agt gaa gta tca tcg gtt gca cct tca gat gcc gat ccg tcg aca tac ttg aat cca tcc ttg act tca agt tca gat gat tcc tca cac atg tct ccg ata cgt acg cta aac tct agg ttc ttg aca cat ttt gta tca acg atc gtt gaa ccg atg ata tct ttg taa ctc act ttc tta tgt gag atg tta gac cca agt act gga tgg gtc ttg atg tca ctg tct ttc tct tct tcg cta cat ctg atg tcg ata gac atc tca cag tct ttg atc ata gcc aga gct tct tca cgc gtg atc gcg gga gag tcc tta cct tgt ccc ggt gac acg ctg gac aat cta gta ttc aca gtg ttt cca tca gag gat tcg gag atg gat aaa atc ttt ggg cat ttg gtg aat cca aag ttc atg tta aga ccc gca ccg acg ata gtg taa taa gtg gtg gga tct cct ttt aca act --- --- --- --- --- --- tct tcg gat acc tca tca tct tcg gtc tct gta act tcc gtt acg gat tga caa atc tta tca ttg gtc ggt gtt tgg tct tgc ttt gtg act ttg ata ata aca tcg att ccc ata tga tgt ttg ttt tct tct tca gta cac gag gat gaa gat tgt tga aga cta gta ggc ata gca gct gcc act agg cac atg cat gcc agg aca ata tat tgt ttc atg att gct att gat tga tta ctg ttc tag atg att cta ctt tct tac cat ata ata aat tag aat ata ttt tct act ttt acg aga aat taa tta ttg tat tta tta ttt ata ggt --a aaa aaa ctt act ata agt ggg tgg gat tct ggg aat tag tga tca gtt tat gta tat cgc aac tag cgg gca tat ggc tat tga cat cga gaa cat tac cca tat gat aag aga ttg tat cat ttt cgt agt ctt gag tat tgg tat tac tat ata gta tgt aga tgt cga cgc tag ata gac agt cgc cca cta gag tta ccg tct ctg aat gcg gca tga tag tat cat tct ttg ttt tcg tta act gtt tgg aag atg aat ctt tgt tgt tac att taa tct cga aat tca gag tac ata tct ttg aag tat tct gat atc tat ttt ctc ctg taa aga atc ctg aag ttg cta cat tat taa gga cag aga agt att ctg cac gaa aga ctg gat cac aat ctt tat gat tca tgg taa tag tta gtt ccg acg ttg aga tgg att cgc tga gac cgg tag tgg tcg tcc gag tac acg atg tgt cgt tga ctg gat aca ggt taa ttt cca cat cga tat agt taa atg tat tgc tgg tta cga cgg gtt cgc att tat ctg tgg aag aga cgg tgt gag aat atg ttc cgg gac cac acg gag aac aga tga cgt ctc cgg tag acg tgt atc cgg ata ctc cgt atc cta ttc cac act ttg ttt tag aaa tac atg ttc tac acc ctg atg ctc ctt tga gaa gac aat aat atc ctg gag agc att cac aga ttc tat tgt gag tcg tgt tac acg atc gcg tct cta cct gat tac tat cac atc ttc cgt tac aac tta gac aag cct gta aat gat tat tgt gag atg taa agg tat ccg aac cac acg gtg tac att gtg tat tag tct tgc tat cac ata atc tga aag cgt aag ttc ccg gag gac acg ata gac aac ata gat tac ggc ttc tgt att cgt tgt ctt tac act ttc cat tgg atg gtg cat gtg gtg cta tat ctc ttc cgt tta tta tta tac atg aga gaa aca ata tat acg agt ata ata cgg acc tca tga ttt aat aat -gt agt aat cgt cgt ctt gtt act gtt tgt ttc cta ctt ctc caa tca tat aga tta ttt ttt aaa tat ttt ctt tct atc atg gat aat att tgt aat ggt tct ttc cgt aca aca tac tgt tta gat ggt agt cgc tta gct tgg tta tga tat tgc gca taa ttt ccg gag gca aat acg ata gtc tag att gac tat cga tgg tag act cta att tat tga gtg ctt tgt cga cga gtt tac ttt tat gct cca tcg ata gat gac act gtt cta tga gat cgt cgt aca tgg gaa atg aaa tgc gtt tgt ctg aat gta tgg ctt cga gat atg tgt gat acc gaa tgt ctt ctg ttc tca ata ccg tat aca agt cgg tgt ctg aga ttc gaa tct ctt tga gga gac tta tgt cac gac tac att ttt cga tga tgg aat cta tct tat cga atg ata tat ttt tca taa ata cac ttt tat agt cct cgt tta aac aga att tag tat gta gtt ccg caa atg act cgt ccc tta ata ggc agt agg cta tta tct tct tta cgt agt gat cgt cgt agg gag aga act ccg aca tct tgt aga aca acg att taa tca tag gta gag ata ctt tca gtc tgt ggt gga taa tgt cat taa cga cat ccg cct tgt ata taa tgt ttc tgt ttt caa aca cca agt cga ata ctg tct tat cgt ctt tag tcg gaa ggt tga tgt cgt atc caa tgt ata cga ggt atg agg caa cat tgt tat tgc aat tct gga agg cgg tat gaa gag gag tca ttg tat tat agt att cgt ctt tct gaa tgt cga atc tat cta gta gat acc gta gta tat tga gag agc gac ttc cat atc ctt gat tat gtt tta tga ata gat aaa gta gat gtt gtc ctt ctt cct ttt gta att tcc cgt att ttt gtt cgt gcc aat tga gta aca tta tga gaa tat gac ctg ttg cac aat cgt tct tta tgt att cca tga tgg gtg tac aat taa gat tat tac gta tcc tcg tat cgg ctc ctc gag ata aaa gag cat aca cca cac gag gac tat gtt tgg tat act gtt gaa ggt aag tgt gta acg gcg tat ttc cga ttt tcg taa ccg cgt taa tgt ttg ctc cat gat cta tta tcg cgt aga tga atc gct tct cag ctc gca tct tag tgt gac tct ttg act tgt aat aat tgc ttt cgt gga acg cgg ata tgt gtt tac agt agt aat gaa gag aag tga gtc cat cct cat cga cgc aat tag ggt cag atc ctt tag tca ata att tgt aca gaa cgt aat agt tta agc tcc cat tga att tat atc taa gat aac aca gca ata gat cgg atg att tac taa agt cat caa tgg ggt ccg tta gta tat caa aga tct tgt tat cga ttg ata gtg aat gaa tca gat agt ggt gta gag gaa tat gtc ctt ttt tat cct tgc tat caa agt tac gca tgc cgt ggc gta aca ata tct tta ata cag atg gat taa atc gtg tat tca tcg tat agc aat gta atg gag agt tac cac att tta gtc gtt tat tca gat cgc agt gtt taa taa cta att taa aca gat gag atg ata tat cca cat caa aga atg cga gat aca tat gac aga cat tat tga cag aaa tgt gac ctt cat tat cac cgt cgt cca taa atg cgt tag gta cgt acc aca tac tat cgt taa cga tgc gca caa tct cgt cca -tt tca tcc atc ttc ata atg att tac ttt ttc ata att aga gaa aaa gat caa ggt ata aaa att aga agt gtt aga cta taa act aac tta tga ctt aac taa ctt atg act taa cta act tat gac tta act aac tta tga --- --- --- --- --- --- --- --- --- --- --- --- --- --- --- --- --- --- --- --- --- --- --- --- --- --- --- --- --- --- --- --- --- --- --- --- --- --- --- --- --- --- --- --- --- --- --- --- ctt aac ta- --- --- --- --- --- --- --- --- --- --- -ac tta tga ctt aac taa ctt atg act taa cta act ta- tga ctt aac taa ctt atg act taa cta act tat gac tta act aac tta tga cta tta act cat ttc aag aaa ggt ggg tgg ata gaa ctc tat atg aca gct tgt gaa aca att aga tcc cta att tct aat gga agt ttt gat agg aga ttg tca tca gtt gat aca ttg ttt att atc tca tct att aga gca cgt ctg ttt aga gct tta gtg acc tgc tcg gtt act tct gtg taa atc ttg aat cct tta gtg ata cac tgt gtc aaa act gga tgt tta gaa tac cta tgt aga ata tgg gaa gca tgc ttg ttt ttg tct cta tta tag att aac tca tac atg gtt gta tta tga att ttc atc tgc cta atg tac tcc aat tct tgt tta caa tca att ata taa tca aag agt gat gat gca tac aca tta caa agt gaa taa tct acc atc ata aaa tac ttg ata cag agc ttt atc aca tca tgg ttt tca att gta tta tta agt ata gct aat ttt ata cag tca ata gac aat ggt tct cta agc aat att tct aat att tta aga tgt gct tcc cta cgg gcg atg aca gat ccc cta tcc acg gcc acg tca aga cat gta tat cca tta ctc att act gcg ttg aca ttt gct cca ttt tct aat agc cat gat act aaa tct ata taa cct gca tag ata gcg cga taa agc aag gtc ctt cca cca gca tct agt tga ttg ata tct tca ata tat ggg ata caa agc tta taa att tct aat act gtg ggt tca tct aca agg aat ccc cta gta tac tga att att tta tat aga tct aat tta aca tca ttt tca tct ggg ata cca caa ttc aaa ata aac tca aca aca cta ctt tcc ttt tta cat att ccc cta aaa tag gca ttc aag cat tct att tta tat att aca gcc cca tga tct acc ata aga tca aca atg tct att tct aca tat gca tta gat aga tag taa agt aag aga tct tgc aca gaa tta caa ttc tta ata att ata gag aaa ata tct tcc ata taa ttc ttt gac act aat gca gat ata ata tct tta tat gta ata tat gca aac agt cta tct act ata tac tga tca ata tta tct cta tga atc cta aaa taa tca tac aga aca tct aca gga tca caa att ggt tca agg aga aat cta tca aat att ttc ctg tca aca act ggt tct aga aca taa cag tca aca cct aat cca tgt ttt tta taa tca tct acc aaa gat aat gac caa aga tcg agg tcg tcg tga aac tgc tca tcg aca gcc atg aaa tct acc gac tcc atg gtg cga atc gca ctg tct tat tcg cca ttg att ttc a-t ttt tta taa tta tgt aca tgt ttt cct tct att ctc aag agt cta caa aaa tat a-- -tt ttt tcg ata tct aag tac taa gtt ttt tta ctg ttt ttg tta ctg tct tcc att ctt cta act aaa gat ctg aga taa att ata caa tct tcg cta tcg aac cat ttt tgt agt cta aag cct gaa gta att aac caa ctg ttt tta tta gtg gct ttt ttc gat cta tcc tcg tc- --- ctc tca tca tcc tta ta- --- --- --- --- --- --- tta tta tca tta tta tca tag tct att aaa cac aaa tca tct --- --- --- --- --- --- --- --- --- --- --- --- --- --- --- --- --- --- --- --- --- --- --- --- --- --- --a cgt tta taa c-- --- --- --- --- --- --- --- --a aca ttc --- --- --- --- -tc att att aat tag ttc tgt ag- aat atc ttt aat aat ttg gct ata --- --- -ca tct gtt --- --- --- --- --- --- --- --- --- --- --- --- --- --- --- --- --- --- --c aat act --- --- --- --- --- --- --- --a tct att gat gat ttc ttt t-- --- --- --- --- --- --- --- --- --- --- --t taa gac t-- --- --- --- --- --- --- --- --- --- --- --- --- --- --- --- --- --- --- --- --- --- --- --- --- --- --- --- --- --- --- --- --- --- --- --- --- --- --- --- taa act agt --- --- --- --- --- --- --- --- --- --- --- --- --- --- --- --- --- --- --- --- --- --- --- --- --- --- --- --- --- --- --- --- --- --- --- --- --- --- --- --- --- --- --- --- --- --- --- --- --- --- --- --- --- --- --- --- --- --- --- --- --- --- --- --- --- --- --- --- --- --- --- --- --- --- --- --- --- --- --- --- --- --- --- --- --- --- --- --- --- --- --- --- --- --- --- --- --- --- --- --- --- --- --- --- --- --- --- --- --- --t atg gta atg acg atg aaa --- -tc gag tag ta- --- --a ctt cta ata aag act tga ta- --- --t cat tat cat atg ttt gat cg- --- --- --- --- --- --- --t cat agt taa tag tgt g-- --- --- --- --- --- --- --- --- --- --- --- --- --- --- --- --- --- --- --- -gc taa atg gta ctg tta ata agt tta t-- --- --- --- --- --- --- --- --- --- --- --- --- --- --- --- --- --a gac aat atc ata gta ttt tct ttc cag aat --- --- --- --t aga tta ttt ttt taa ata ctg atc ctc aca att ccg tga tgt agc agt agt tgg t-- --- --- --- --- --- --- --- --- --- --- --- --- --- --- --- --- --- --- --- --- --- --- --- --- --- --- --- -gc atg gtc tat atc gt- --- --- --- --- --- --- --- --- --- --- --- --- --- --- --- --- --- --- --- --- --- --- --- --- --- --- --- --- --- --- --- --- --- --- --- --- --- --- --- --- --- --- --- taa aat gta tca tat ata ata gtt ttc tga cgt gga gta cag aat ttt cga --- --- --- --- --- --- --- --- --- --- --- --- --- --- --- --- --- --- --- --- --- --- --- --- --- --- --- --- --- --- --- --- --- --- --- --- --- --- --- --- --- --- --- --t taa tga gtt cat ggt aag gaa ggg caa atg cct --- gta tat aat ata cat aag tta a-- --- --- --- --- --- --- --- --- --- --- --- --- --t agt ttt tta tca tat tt- --- --- --- --- --- --- --- --t cta ata cca taa taa aaa t-- --- --- --- --- --- --- --- --- --- --- --- --- --- --- --- --- --- --- --- --- --- --- --- --- --t atc att att gcg ttt g-g tag tt- --- --- --- --- ctg ccc ta- --- --- --- --- --- --- --- --t cat cta tat cac tgt cac tct c-- --- --g ctc tca cta tat ctt cta aaa tta caa --- -ac aac --- --- --- --- --- --- --- --- --- --- --- --- --- --- --- --- --- --- --- --- --- --- --- --- --- --- --- --- --- --- --- --- --- --- --- --- tgg ata ttc gat --- --- --- --- --- --- --- --- --- --- --- --- --- --- --- --- --- --- --- --- --- --- --- --- --- --- --a aca gca ttt gtg t-- --- --- --- --- --- --- --- --- --- --- --- --- --- --- --- --- -

>ON637939.1_Monkeypox_virus_isolate_MPXV/Germany/2022/RKI02_complete_genome

--- --- --- --- --- --- --- --- --- --- --- --- --- --- --- --- --- --a tta ata ata cat aag ttt tag tac att aat att taa ttt tac tat ttt att tag tgt cta gaa aaa aat gtg tga ccc acg acc gta gga aac tct aga ggg taa gaa aaa tca atc gtt tat aga gac cat cag aaa gag gtt taa tat ttt tgt gag acc tat cga a-- --- --- --- --- --- --- --- --- --- --- --- --- --- --- --- --- --- --- --- --- --- --- --- --- --- --- --- --- --- --- --- --- --- --- --- --- --- --- --- --- --- --- --- --- --- --- --- --- --- --- --- --- --- --- --- --- --- --- --- --- --- --- --- --- --- --- --- --- --- --- --- --- --- --- --- --- --- --- --- --- --- --- --- --- --- --- --- --- --- --- --- --- --- --- --- --- --- --- --- --- --- --- --- --- --- --- --- --- --- --- --- --- --- --- --- --- --- --- --- --- --- --- --- --- --- --- --- --- --- --- --- --- --- --- --- --- --- --- --- --- --- --- --- --- --- --- --- --- --- --- --- --- --- --- --- --- --- --- --- --- --- --- --- --- --- --- --- --- --- --- --- --- --- --- --- --- --- --- --- --- --- --- --- --- --- --- --- --- --- --- --- --- --- --- --- --- --- --- --- --- --- --- --- --- --- --- --- --- --- --- --- --- --- --- --- --- --- --- --- --- --- --- --- --- --- --- --- --- --- --- --- --- --- --- --- --- --- --- --- --- --- --- --- --- --- --- --- --- --- --- --- --- --- --- --- --- --- --- --- --- --- --- --- --- --- --- --- --- --- --- --- --- --- --- --- --- --- --- --- -ga gag aaa gga taa aaa ctt ttt acg act cca tca gaa aga ggt tta ata ttt ttg tga gac cca tcg aag aga gaa aga gat ggt tag tca aga tat ttt tct tag tac aaa agt caa tgt ttt aaa ata tat gga cga gaa tta att tgt ctg tat aaa aac ttg tgt gaa att atg tac tag aga aaa aac gtg agc agt gtc ccc tac atg gat ttt aca gat cat tta tat tcc aaa aat att aac tat ata cgt tta tta tat gat gtt aac gtg taa att ata aac att att tta tga tgc aat tgt ctg aca acc tag att ggt ata agg atg ttg ata agc tct acg aga ata tat tgt tgg acg tta tcg ttt acg aaa tag ttg aga cat cag aaa gag gtt taa tat ttt tgt gag a-c cat cga aga gag aaa gag aat aaa aat a-- --t ttt ttt ttt ttt tgt aaa act ttt tta tga gac --- --- -ca aga gaa tac gaa tag tga tca tat cgt atc aca tat tga aac aga aag aag aag taa cga gag gta act ttt tgt gaa tgt agt taa ata ttt ttg ttt tgc aaa ccg gaa tat agt gcc cgg t-- -ct ttt tta att cgt ggt gcg gtg tct gaa tcg ttc gat taa ccc aac tca tcc att ttc aga tga ata gag tta tcg att cag aca cat gct ttg agt ttt gtt gaa tcg atg agt gaa gta tca tcg gtt gca cct tca gat gcc gat ccg tcg aca tac ttg aat cca tcc ttg act tca agt tca gat gat tcc tca cac atg tct ccg ata cgt acg cta aac tct agg ttc ttg aca cat ttt gta tca acg atc gtt gaa ccg atg ata tct ttg taa ctc act ttc tta tgt gag atg tta gac cca agt act gga tgg gtc ttg atg tca ctg tct ttc tct tct tcg cta cat ctg atg tcg ata gac atc tca cag tct ttg atc ata gcc aga gct tct tca cgc gtg atc gcg gga gag tcc tta cct tgt ccc ggt gac acg ctg gac aat cta gta ttc aca gtg ttt cca tca gag gat tcg gag atg gat aaa atc ttt ggg cat ttg gtg aat cca aag ttc atg tta aga ccc gca ccg acg ata gtg taa taa gtg gtg gga tct cct ttt aca act --- --- --- --- --- --- tct tcg gat acc tca tca tct tcg gtc tct gta act tcc gtt acg gat tga caa atc tta tca ttg gtc ggt gtt tgg tct tgc ttt gtg act ttg ata ata aca tcg att ccc ata tga tgt ttg ttt tct tct tca gta cac gag gat gaa gat tgt tga aga cta gta ggc ata gca gct gcc act agg cac atg cat gcc agg aca ata tat tgt ttc atg att gct att gat tga tta ctg ttc tag atg att cta ctt tct tac cat ata ata aat tag aat ata ttt tct act ttt acg aga aat taa tta ttg tat tta tta ttt ata ggt --a aaa aaa ctt act ata agt ggg tgg gat tct ggg aat tag tga tca gtt tat gta tat cgc aac tag cgg gca tat ggc tat tga cat cga gaa cat tac cca tat gat aag aga ttg tat cat ttt cgt agt ctt gag tat tgg tat tac tat ata gta tgt aga tgt cga cgc tag ata gac agt cgc cca cta gag tta ccg tct ctg aat gcg gca tga tag tat cat tct ttg ttt tcg tta act gtt tgg aag atg aat ctt tgt tgt tac att taa tct cga aat tca gag tac ata tct ttg aag tat tct gat atc tat ttt ctc ctg taa aga atc ctg aag ttg cta cat tat taa gga cag aga agt att ctg cac gaa aga ctg gat cac aat ctt tat gat tca tgg taa tag tta gtt ccg acg ttg aga tgg att cgc tga gac cgg tag tgg tcg tcc gag tac acg atg tgt cgt tga ctg gat aca ggt taa ttt cca cat cga tat agt taa atg tat tgc tgg tta cga cgg gtt cgc att tat ctg tgg aag aga cgg tgt gag aat atg ttc cgg gac cac acg gag aac aga tga cgt ctc cgg tag acg tgt atc cgg ata ctc cgt atc cta ttc cac act ttg ttt tag aaa tac atg ttc tac acc ctg atg ctc ctt tga gaa gac aat aat atc ctg gag agc att cac aga ttc tat tgt gag tcg tgt tac acg atc gcg tct cta cct gat tac tat cac atc ttc cgt tac aac tta gac aag cct gta aat gat tat tgt gag atg taa agg tat ccg aac cac acg gtg tac att gtg tat tag tct tgc tat cac ata atc tga aag cgt aag ttc ccg gag gac acg ata gac aac ata gat tac ggc ttc tgt att cgt tgt ctt tac act ttc cat tgg atg gtg cat gtg gtg cta tat ctc ttc cgt tta tta tta tac atg aga gaa aca ata tat acg agt ata ata cgg acc tca tga ttt aat aat -gt agt aat cgt cgt ctt gtt act gtt tgt ttc cta ctt ctc caa tca tat aga tta ttt ttt aaa tat ttt ctt tct atc atg gat aat att tgt aat ggt tct ttc cgt aca aca tac tgt tta gat ggt agt cgc tta gct tgg tta tga tat tgc gca taa ttt ccg gag gca aat acg ata gtc tag att gac tat cga tgg tag act cta att tat tga gtg ctt tgt cga cga gtt tac ttt tat gct cca tcg ata gat gac act gtt cta tga gat cgt cgt aca tgg gaa atg aaa tgc gtt tgt ctg aat gta tgg ctt cga gat atg tgt gat acc gaa tgt ctt ctg ttc tca ata ccg tat aca agt cgg tgt ctg aga ttc gaa tct ctt tga gga gac tta tgt cac gac tac att ttt cga tga tgg aat cta tct tat cga atg ata tat ttt tca taa ata cac ttt tat agt cct cgt tta aac aga att tag tat gta gtt ccg caa atg act cgt ccc tta ata ggc agt agg cta tta tct tct tta cgt agt gat cgt cgt agg gag aga act ccg aca tct tgt aga aca acg att taa tca tag gta gag ata ctt tca gtc tgt ggt gga taa tgt cat taa cga cat ccg cct tgt ata taa tgt ttc tgt ttt caa aca cca agt cga ata ctg tct tat cgt ctt tag tcg gaa ggt tga tgt cgt atc caa tgt ata cga ggt atg agg caa cat tgt tat tgc aat tct gga agg cgg tat gaa gag gag tca ttg tat tat agt att cgt ctt tct gaa tgt cga atc tat cta gta gat acc gta gta tat tga gag agc gac ttc cat atc ctt gat tat gtt tta tga ata gat aaa gta gat gtt gtc ctt ctt cct ttt gta att tcc cgt att ttt gtt cgt gcc aat tga gta aca tta tga gaa tat gac ctg ttg cac aat cgt tct tta tgt att cca tga tgg gtg tac aat taa gat tat tac gta tcc tcg tat cgg ctc ctc gag ata aaa gag cat aca cca cac gag gac tat gtt tgg tat act gtt gaa ggt aag tgt gta acg gcg tat ttc cga ttt tcg taa ccg cgt taa tgt ttg ctc cat gat cta tta tcg cgt aga tga atc gct tct cag ctc gca tct tag tgt gac tct ttg act tgt aat aat tgc ttt cgt gga acg cgg ata tgt gtt tac agt agt aat gaa gag aag tga gtc cat cct cat cga cgc aat tag ggt cag atc ctt tag tca ata att tgt aca gaa cgt aat agt tta agc tcc cat tga att tat atc taa gat aac aca gca ata gat cgg atg att tac taa agt cat caa tgg ggt ccg tta gta tat caa aga tct tgt tat cga ttg ata gtg aat gaa tca gat agt ggt gta gag gaa tat gtc ctt ttt tat cct tgc tat caa agt tac gca tgc cgt ggc gta aca ata tct tta ata cag atg gat taa atc gtg tat tca tcg tat agc aat gta atg gag agt tac cac att tta gtc gtt tat tca gat cgc agt gtt taa taa cta att taa aca gat gag atg ata tat cca cat caa aga atg cga gat aca tat gac aga cat tat tga cag aaa tgt gac ctt cat tat cac cgt cgt cca taa atg cgt tag gta cgt acc aca tac tat cgt taa cga tgc gca caa tct cgt cca -tt tca tcc atc ttc ata atg att tac ttt ttc ata att aga gaa aaa gat caa ggt ata aaa att aga agt gtt aga cta taa act aac tta tga ctt aac taa ctt atg act taa cta act tat gac tta act aac tta tga --- --- --- --- --- --- --- --- --- --- --- --- --- --- --- --- --- --- --- --- --- --- --- --- --- --- --- --- --- --- --- --- --- --- --- --- --- --- --- --- --- --- --- --- --- --- --- --- ctt aac ta- --- --- --- --- --- --- --- --- --- --- -ac tta tga ctt aac taa ctt atg act taa cta act ta- tga ctt aac taa ctt atg act taa cta act tat gac tta act aac tta tga cta tta act cat ttc aag aaa ggt ggg tgg ata gaa ctc tat atg aca gct tgt gaa aca att aga tcc cta att tct aat gga agt ttt gat agg aga ttg tca tca gtt gat aca ttg ttt att atc tca tct att aga gca cgt ctg ttt aga gct tta gtg acc tgc tcg gtt act tct gtg taa atc ttg aat cct tta gtg ata cac tgt gtc aaa act gga tgt tta gaa tac cta tgt aga ata tgg gaa gca tgc ttg ttt ttg tct cta tta tag att aac tca tac atg gtt gta tta tga att ttc atc tgc cta atg tac tcc aat tct tgt tta caa tca att ata taa tca aag agt gat gat gca tac aca tta caa agt gaa taa tct acc atc ata aaa tac ttg ata cag agc ttt atc aca tca tgg ttt tca att gta tta tta agt ata gct aat ttt ata cag tca ata gac aat ggt tct cta agc aat att tct aat att tta aga tgt gct tcc cta cgg gcg atg aca gat ccc cta tcc acg gcc acg tca aga cat gta tat cca tta ctc att act gcg ttg aca ttt gct cca ttt tct aat agc cat gat act aaa tct ata taa cct gca tag ata gcg cga taa agc aag gtc ctt cca cca gca tct agt tga ttg ata tct tca ata tat ggg ata caa agc tta taa att tct aat act gtg ggt tca tct aca agg aat ccc cta gta tac tga att att tta tat aga tct aat tta aca tca ttt tca tct ggg ata cca caa ttc aaa ata aac tca aca aca cta ctt tcc ttt tta cat att ccc cta aaa tag gca ttc aag cat tct att tta tat att aca gcc cca tga tct acc ata aga tca aca atg tct att tct aca tat gca tta gat aga tag taa agt aag aga tct tgc aca gaa tta caa ttc tta ata att ata gag aaa ata tct tcc ata taa ttc ttt gac act aat gca gat ata ata tct tta tat gta ata tat gca aac agt cta tct act ata tac tga tca ata tta tct cta tga atc cta aaa taa tca tac aga aca tct aca gga tca caa att ggt tca agg aga aat cta tca aat att ttc ctg tca aca act ggt tct aga aca taa cag tca aca cct aat cca tgt ttt tta taa tca tct acc aaa gat aat gac caa aga tcg agg tcg tcg tga aac tgc tca tcg aca gcc atg aaa tct acc gac tcc atg gtg cga atc gca ctg tct tat tcg cca ttg att ttc a-t ttt tta taa tta tgt aca tgt ttt cct tct att ctc aag agt cta caa aaa tat a-- -tt ttt tcg ata tct aag tac taa gtt ttt tta ctg ttt ttg tta ctg tct tcc att ctt cta act aaa gat ctg aga taa att ata caa tct tcg cta tcg aac cat ttt tgt agt cta aag cct gaa gta att aac caa ctg ttt tta tta gtg gct ttt ttc gat cta tcc tcg tc- --- ctc tca tca tcc tta ta- --- --- --- --- --- --- tta tta tca tta tta tca tag tct att aaa cac aaa tca tct --- --- --- --- --- --- --- --- --- --- --- --- --- --- --- --- --- --- --- --- --- --- --- --- --- --- --a cgt tta taa c-- --- --- --- --- --- --- --- --a aca ttc --- --- --- --- -tc att att aat tag ttc tgt ag- aat atc ttt aat aat ttg gct ata --- --- -ca tct gtt --- --- --- --- --- --- --- --- --- --- --- --- --- --- --- --- --- --- --c aat act --- --- --- --- --- --- --- --a tct att gat gat ttc ttt t-- --- --- --- --- --- --- --- --- --- --- --t taa gac t-- --- --- --- --- --- --- --- --- --- --- --- --- --- --- --- --- --- --- --- --- --- --- --- --- --- --- --- --- --- --- --- --- --- --- --- --- --- --- --- taa act agt --- --- --- --- --- --- --- --- --- --- --- --- --- --- --- --- --- --- --- --- --- --- --- --- --- --- --- --- --- --- --- --- --- --- --- --- --- --- --- --- --- --- --- --- --- --- --- --- --- --- --- --- --- --- --- --- --- --- --- --- --- --- --- --- --- --- --- --- --- --- --- --- --- --- --- --- --- --- --- --- --- --- --- --- --- --- --- --- --- --- --- --- --- --- --- --- --- --- --- --- --- --- --- --- --- --- --- --- --- --t atg gta atg acg atg aaa --- -tc gag tag ta- --- --a ctt cta ata aag act tga ta- --- --t cat tat cat atg ttt gat cg- --- --- --- --- --- --- --t cat agt taa tag tgt g-- --- --- --- --- --- --- --- --- --- --- --- --- --- --- --- --- --- --- --- -gc taa atg gta ctg tta ata agt tta t-- --- --- --- --- --- --- --- --- --- --- --- --- --- --- --- --- --a gac aat atc ata gta ttt tct ttc cag aat --- --- --- --t aga tta ttt ttt taa ata ctg atc ctc aca att ccg tga tgt agc agt agt tgg t-- --- --- --- --- --- --- --- --- --- --- --- --- --- --- --- --- --- --- --- --- --- --- --- --- --- --- --- -gc atg gtc tat atc gt- --- --- --- --- --- --- --- --- --- --- --- --- --- --- --- --- --- --- --- --- --- --- --- --- --- --- --- --- --- --- --- --- --- --- --- --- --- --- --- --- --- --- --- taa aat gta tca tat ata ata gtt ttc tga cgt gga gta cag aat ttt cga --- --- --- --- --- --- --- --- --- --- --- --- --- --- --- --- --- --- --- --- --- --- --- --- --- --- --- --- --- --- --- --- --- --- --- --- --- --- --- --- --- --- --- --t taa tga gtt cat ggt aag gaa ggg caa atg cct --- gta tat aat ata cat aag tta a-- --- --- --- --- --- --- --- --- --- --- --- --- --t agt ttt tta tca tat tt- --- --- --- --- --- --- --- --t cta ata cca taa taa aaa t-- --- --- --- --- --- --- --- --- --- --- --- --- --- --- --- --- --- --- --- --- --- --- --- --- --t atc att att gcg ttt g-g tag tt- --- --- --- --- ctg ccc ta- --- --- --- --- --- --- --- --t cat cta tat cac tgt cac tct c-- --- --g ctc tca cta tat ctt cta aaa tta caa --- -ac aac --- --- --- --- --- --- --- --- --- --- --- --- --- --- --- --- --- --- --- --- --- --- --- --- --- --- --- --- --- --- --- --- --- --- --- --- tgg ata ttc gat --- --- --- --- --- --- --- --- --- --- --- --- --- --- --- --- --- --- --- --- --- --- --- --- --- --- --a aca gca ttt gtg t-- --- --- --- --- --- --- --- --- --- --- --- --- --- --- --- --- -

>ON682263.2_Monkeypox_virus_isolate_MPXV/Germany/2022/RKI03_complete_genome

--- --- --- --- --- --- --- --- --- --- --- --- --- --- --- --- --- --- --- --- --- --- --- --- --- --- --- --- --- --a ttt tac tat ttt att tag tgt cta gaa aaa aat gtg tga ccc acg acc gta gga aac tct aga ggg taa gaa aaa tca atc gtt tat aga gac cat cag aaa gag gtt taa tat ttt tgt gag acc tat cga a-- --- --- --- --- --- --- --- --- --- --- --- --- --- --- --- --- --- --- --- --- --- --- --- --- --- --- --- --- --- --- --- --- --- --- --- --- --- --- --- --- --- --- --- --- --- --- --- --- --- --- --- --- --- --- --- --- --- --- --- --- --- --- --- --- --- --- --- --- --- --- --- --- --- --- --- --- --- --- --- --- --- --- --- --- --- --- --- --- --- --- --- --- --- --- --- --- --- --- --- --- --- --- --- --- --- --- --- --- --- --- --- --- --- --- --- --- --- --- --- --- --- --- --- --- --- --- --- --- --- --- --- --- --- --- --- --- --- --- --- --- --- --- --- --- --- --- --- --- --- --- --- --- --- --- --- --- --- --- --- --- --- --- --- --- --- --- --- --- --- --- --- --- --- --- --- --- --- --- --- --- --- --- --- --- --- --- --- --- --- --- --- --- --- --- --- --- --- --- --- --- --- --- --- --- --- --- --- --- --- --- --- --- --- --- --- --- --- --- --- --- --- --- --- --- --- --- --- --- --- --- --- --- --- --- --- --- --- --- --- --- --- --- --- --- --- --- --- --- --- --- --- --- --- --- --- --- --- --- --- --- --- --- --- --- --- --- --- --- --- --- --- --- --- --- --- --- --- --- --- -ga gag aaa gga taa aaa ctt ttt acg act cca tca gaa aga ggt tta ata ttt ttg tga gac cca tcg aag aga gaa aga gat ggt tag tca aga tat ttt tct tag tac aaa agt caa tgt ttt aaa ata tat gga cga gaa tta att tgt ctg tat aaa aac ttg tgt gaa att atg tac tag aga aaa aac gtg agc agt gtc ccc tac atg gat ttt aca gat cat tta tat tcc aaa aat att aac tat ata cgt tta tta tat gat gtt aac gtg taa att ata aac att att tta tga tgc aat tgt ctg aca acc tag att ggt ata agg atg ttg ata agc tct acg aga ata tat tgt tgg acg tta tcg ttt acg aaa tag ttg aga cat cag aaa gag gtt taa tat ttt tgt gag a-c cat cga aga gag aaa gag aat aaa aat a-- --t ttt ttt ttt ttt tgt aaa act ttt tta tga gac --- --- -ca aga gaa tac gaa tag tga tca tat cgt atc aca tat tga aac aga aag aag aag taa cga gag gta act ttt tgt gaa tgt agt taa ata ttt ttg ttt tgc aaa ccg gaa tat agt gcc cgg t-- -ct ttt tta att cgt ggt gcg gtg tct gaa tcg ttc gat taa ccc aac tca tcc att ttc aga tga ata gag tta tcg att cag aca cat gct ttg agt ttt gtt gaa tcg atg agt gaa gta tca tcg gtt gca cct tca gat gcc gat ccg tcg aca tac ttg aat cca tcc ttg act tca agt tca gat gat tcc tca cac atg tct ccg ata cgt acg cta aac tct agg ttc ttg aca cat ttt gta tca acg atc gtt gaa ccg atg ata tct ttg taa ctc act ttc tta tgt gag atg tta gac cca agt act gga tgg gtc ttg atg tca ctg tct ttc tct tct tcg cta cat ctg atg tcg ata gac atc tca cag tct ttg atc ata gcc aga gct tct tca cgc gtg atc gcg gga gag tcc tta cct tgt ccc ggt gac acg ctg gac aat cta gta ttc aca gtg ttt cca tca gag gat tcg gag atg gat aaa atc ttt ggg cat ttg gtg aat cca aag ttc atg tta aga ccc gca ccg acg ata gtg taa taa gtg gtg gga tct cct ttt aca act --- --- --- --- --- --- tct tcg gat acc tca tca tct tcg gtc tct gta act tcc gtt acg gat tga caa atc tta tca ttg gtc ggt gtt tgg tct tgc ttt gtg act ttg ata ata aca tcg att ccc ata tga tgt ttg ttt tct tct tca gta cac gag gat gaa gat tgt tga aga cta gta ggc ata gca gct gcc act agg cac atg cat gcc agg aca ata tat tgt ttc atg att gct att gat tga tta ctg ttc tag atg att cta ctt tct tac cat ata ata aat tag aat ata ttt tct act ttt acg aga aat taa tta ttg tat tta tta ttt ata ggt --a aaa aaa ctt act ata agt ggg tgg gat tct ggg aat tag tga tca gtt tat gta tat cgc aac tag cgg gca tat ggc tat tga cat cga gaa cat tac cca tat gat aag aga ttg tat cat ttt cgt agt ctt gag tat tgg tat tac tat ata gta tgt aga tgt cga cgc tag ata gac agt cgc cca cta gag tta ccg tct ctg aat gcg gca tga tag tat cat tct ttg ttt tcg tta act gtt tgg aag atg aat ctt tgt tgt tac att taa tct cga aat tca gag tac ata tct ttg aag tat tct gat atc tat ttt ctc ctg taa aga atc ctg aag ttg cta cat tat taa gga cag aga agt att ctg cac gaa aga ctg gat cac aat ctt tat gat tca tgg taa tag tta gtt ccg acg ttg aga tgg att cgc tga gac cgg tag tgg tcg tcc gag tac acg atg tgt cgt tga ctg gat aca ggt taa ttt cca cat cga tat agt taa atg tat tgc tgg tta cga cgg gtt cgc att tat ctg tgg aag aga cgg tgt gag aat atg ttc cgg gac cac acg gag aac aga tga cgt ctc cgg tag acg tgt atc cgg ata ctc cgt atc cta ttc cac act ttg ttt tag aaa tac atg ttc tac acc ctg atg ctc ctt tga gaa gac aat aat atc ctg gag agc att cac aga ttc tat tgt gag tcg tgt tac acg atc gcg tct cta cct gat tac tat cac atc ttc cgt tac aac tta gac aag cct gta aat gat tat tgt gag atg taa agg tat ccg aac cac acg gtg tac att gtg tat tag tct tgc tat cac ata atc tga aag cgt aag ttc ccg gag gac acg ata gac aac ata gat tac ggc ttc tgt att cgt tgt ctt tac act ttc cat tgg atg gtg cat gtg gtg cta tat ctc ttc cgt tta tta tta tac atg aga gaa aca ata tat acg agt ata ata cgg acc tca tga ttt aat aat -gt agt aat cgt cgt ctt gtt act gtt tgt ttc cta ctt ctc caa tca tat aga tta ttt ttt aaa tat ttt ctt tct atc atg gat aat att tgt aat ggt tct ttc cgt aca aca tac tgt tta gat ggt agt cgc tta gct tgg tta tga tat tgc gca taa ttt ccg gag gca aat acg ata gtc tag att gac tat cga tgg tag act cta att tat tga gtg ctt tgt cga cga gtt tac ttt tat gct cca tcg ata gat gac act gtt cta tga gat cgt cgt aca tgg gaa atg aaa tgc gtt tgt ctg aat gta tgg ctt cga gat atg tgt gat acc gaa tgt ctt ctg ttc tca ata ccg tat aca agt cgg tgt ctg aga ttc gaa tct ctt tga gga gac tta tgt cac gac tac att ttt cga tga tgg aat cta tct tat cga atg ata tat ttt tca taa ata cac ttt tat agt cct cgt tta aac aga att tag tat gta gtt ccg caa atg act cgt ccc tta ata ggc agt agg cta tta tct tct tta cgt agt gat cgt cgt agg gag aga act ccg aca tct tgt aga aca acg att taa tca tag gta gag ata ctt tca gtc tgt ggt gga taa tgt cat taa cga cat ccg cct tgt ata taa tgt ttc tgt ttt caa aca cca agt cga ata ctg tct tat cgt ctt tag tcg gaa ggt tga tgt cgt atc caa tgt ata cga ggt atg agg caa cat tgt tat tgc aat tct gga agg cgg tat gaa gag gag tca ttg tat tat agt att cgt ctt tct gaa tgt cga atc tat cta gta gat acc gta gta tat tga gag agc gac ttc cat atc ctt gat tat gtt tta tga ata gat aaa gta gat gtt gtc ctt ctt cct ttt gta att tcc cgt att ttt gtt cgt gcc aat tga gta aca tta tga gaa tat gac ctg ttg cac aat cgt tct tta tgt att cca tga tgg gtg tac aat taa gat tat tac gta tcc tcg tat cgg ctc ctc gag ata aaa gag cat aca cca cac gag gac tat gtt tgg tat act gtt gaa ggt aag tgt gta acg gcg tat ttc cga ttt tcg taa ccg cgt taa tgt ttg ctc cat gat cta tta tcg cgt aga tga atc gct tct cag ctc gca tct tag tgt gac tct ttg act tgt aat aat tgc ttt cgt gga acg cgg ata tgt gtt tac agt agt aat gaa gag aag tga gtc cat cct cat cga cgc aat tag ggt cag atc ctt tag tca ata att tgt aca gaa cgt aat agt tta agc tcc cat tga att tat atc taa gat aac aca gca ata gat cgg atg att tac taa agt cat caa tgg ggt ccg tta gta tat caa aga tct tgt tat cga ttg ata gtg aat gaa tca gat agt ggt gta gag gaa tat gtc ctt ttt tat cct tgc tat caa agt tac gca tgc cgt ggc gta aca ata tct tta ata cag atg gat taa atc gtg tat tca tcg tat agc aat gta atg gag agt tac cac att tta gtc gtt tat tca gat cgc agt gtt taa taa cta att taa aca gat gag atg ata tat cca cat caa aga atg cga gat aca tat gac aga cat tat tga cag aaa tgt gac ctt cat tat cac cgt cgt cca taa atg cgt tag gta cgt acc aca tac tat cgt taa cga tgc gca caa tct cgt cca -tt tca tcc atc ttc ata atg att tac ttt ttc ata att aga gaa aaa gat caa ggt ata aaa att aga agt gtt aga cta taa act aac tta tga ctt aac taa ctt atg act taa cta act tat gac tta act aac tta tga --- --- --- --- --- --- --- --- --- --- --- --- --- --- --- --- --- --- --- --- --- --- --- --- --- --- --- --- --- --- --- --- --- --- --- --- --- --- --- --- --- --- --- --- --- --- --- --- ctt aac ta- --- --- --- --- --- --- --- --- --- --- -ac tta tga ctt aac taa ctt atg act taa cta act ta- tga ctt aac taa ctt atg act taa cta act tat gac tta act aac tta tga cta tta act cat ttc aag aaa ggt ggg tgg ata gaa ctc tat atg aca gct tgt gaa aca att aga tcc cta att tct aat gga agt ttt gat agg aga ttg tca tca gtt gat aca ttg ttt att atc tca tct att aga gca cgt ctg ttt aga gct tta gtg acc tgc tcg gtt act tct gtg taa atc ttg aat cct tta gtg ata cac tgt gtc aaa act gga tgt tta gaa tac cta tgt aga ata tgg gaa gca tgc ttg ttt ttg tct cta tta tag att aac tca tac atg gtt gta tta tga att ttc atc tgc cta atg tac tcc aat tct tgt tta caa tca att ata taa tca aag agt gat gat gca tac aca tta caa agt gaa taa tct acc atc ata aaa tac ttg ata cag agc ttt atc aca tca tgg ttt tca att gta tta tta agt ata gct aat ttt ata cag tca ata gac aat ggt tct cta agc aat att tct aat att tta aga tgt gct tcc cta cgg gcg atg aca gat ccc cta tcc acg gcc acg tca aga cat gta tat cca tta ctc att act gcg ttg aca ttt gct cca ttt tct aat agc cat gat act aaa tct ata taa cct gca tag ata gcg cga taa agc aag gtc ctt cca cca gca tct agt tga ttg ata tct tca ata tat ggg ata caa agc tta taa att tct aat act gtg ggt tca tct aca agg aat ccc cta gta tac tga att att tta tat aga tct aat tta aca tca ttt tca tct ggg ata cca caa ttc aaa ata aac tca aca aca cta ctt tcc ttt tta cat att ccc cta aaa tag gca ttc aag cat tct att tta tat att aca gcc cca tga tct acc ata aga tca aca atg tct att tct aca tat gca tta gat aga tag taa agt aag aga tct tgc aca gaa tta caa ttc tta ata att ata gag aaa ata tct tcc ata taa ttc ttt gac act aat gca gat ata ata tct tta tat gta ata tat gca aac agt cta tct act ata tac tga tca ata tta tct cta tga atc cta aaa taa tca tac aga aca tct aca gga tca caa att ggt tca agg aga aat cta tca aat att ttc ctg tca aca act ggt tct aga aca taa cag tca aca cct aat cca tgt ttt tta taa tca tct acc aaa gat aat gac caa aga tcg agg tcg tcg tga aac tgc tca tcg aca gcc atg aaa tct acc gac tcc atg gtg cga atc gca ctg tct tat tcg cca ttg att ttc a-t ttt tta taa tta tgt aca tgt ttt cct tct att ctc aag agt cta caa aaa tat a-- -tt ttt tcg ata tct aag tac taa gtt ttt tta ctg ttt ttg tta ctg tct tcc att ctt cta act aaa gat ctg aga taa att ata caa tct tcg cta tcg aac cat ttt tgt agt cta aag cct gaa gta att aac caa ctg ttt tta tta gtg gct ttt ttc gat cta tcc tcg tc- --- ctc tca tca tcc tta ta- --- --- --- --- --- --- tta tta tca tta tta tca tag tct att aaa cac aaa tca tct --- --- --- --- --- --- --- --- --- --- --- --- --- --- --- --- --- --- --- --- --- --- --- --- --- --- --a cgt tta taa c-- --- --- --- --- --- --- --- --a aca ttc --- --- --- --- -tc att att aat tag ttc tgt ag- aat atc ttt aat aat ttg gct ata --- --- -ca tct gtt --- --- --- --- --- --- --- --- --- --- --- --- --- --- --- --- --- --- --c aat act --- --- --- --- --- --- --- --a tct att gat gat ttc ttt t-- --- --- --- --- --- --- --- --- --- --- --t taa gac t-- --- --- --- --- --- --- --- --- --- --- --- --- --- --- --- --- --- --- --- --- --- --- --- --- --- --- --- --- --- --- --- --- --- --- --- --- --- --- --- taa act agt --- --- --- --- --- --- --- --- --- --- --- --- --- --- --- --- --- --- --- --- --- --- --- --- --- --- --- --- --- --- --- --- --- --- --- --- --- --- --- --- --- --- --- --- --- --- --- --- --- --- --- --- --- --- --- --- --- --- --- --- --- --- --- --- --- --- --- --- --- --- --- --- --- --- --- --- --- --- --- --- --- --- --- --- --- --- --- --- --- --- --- --- --- --- --- --- --- --- --- --- --- --- --- --- --- --- --- --- --- --t atg gta atg acg atg aaa --- -tc gag tag ta- --- --a ctt cta ata aag act tga ta- --- --t cat tat cat atg ttt gat cg- --- --- --- --- --- --- --t cat agt taa tag tgt g-- --- --- --- --- --- --- --- --- --- --- --- --- --- --- --- --- --- --- --- -gc taa atg gta ctg tta ata agt tta t-- --- --- --- --- --- --- --- --- --- --- --- --- --- --- --- --- --a gac aat atc ata gta ttt tct ttc cag aat --- --- --- --t aga tta ttt ttt taa ata ctg atc ctc aca att ccg tga tgt agc agt agt tgg t-- --- --- --- --- --- --- --- --- --- --- --- --- --- --- --- --- --- --- --- --- --- --- --- --- --- --- --- -gc atg gtc tat atc gt- --- --- --- --- --- --- --- --- --- --- --- --- --- --- --- --- --- --- --- --- --- --- --- --- --- --- --- --- --- --- --- --- --- --- --- --- --- --- --- --- --- --- --- taa aat gta tca tat ata ata gtt ttc tga cgt gga gta cag aat ttt cga --- --- --- --- --- --- --- --- --- --- --- --- --- --- --- --- --- --- --- --- --- --- --- --- --- --- --- --- --- --- --- --- --- --- --- --- --- --- --- --- --- --- --- --t taa tga gtt cat ggt aag gaa ggg caa atg cct --- gta tat aat ata cat aag tta a-- --- --- --- --- --- --- --- --- --- --- --- --- --t agt ttt tta tca tat tt- --- --- --- --- --- --- --- --t cta ata cca taa taa aaa t-- --- --- --- --- --- --- --- --- --- --- --- --- --- --- --- --- --- --- --- --- --- --- --- --- --t atc att att gcg ttt g-g tag tt- --- --- --- --- ctg ccc ta- --- --- --- --- --- --- --- --t cat cta tat cac tgt cac tct c-- --- --g ctc tca cta tat ctt cta aaa tta caa --- -ac aac --- --- --- --- --- --- --- --- --- --- --- --- --- --- --- --- --- --- --- --- --- --- --- --- --- --- --- --- --- --- --- --- --- --- --- --- tgg ata ttc gat --- --- --- --- --- --- --- --- --- --- --- --- --- --- --- --- --- --- --- --- --- --- --- --- --- --- --a aca gca ttt gtg t-- --- --- --- --- --- --- --- --- --- --- --- --- --- --- --- --- -

>ON682266.1_Monkeypox_virus_isolate_MPXV/Germany/2022/RKI09_complete_genome

--- --- --- --- --- --- --- --- --- --- --- --- --- --- --- --- --- --- --- --- --- --- --- --- --- --- --- --- --- --a ttt tac tat ttt att tag tgt cta gaa aaa aat gtg tga ccc acg acc gta gga aac tct aga ggg taa gaa aaa tca atc gtt tat aga gac cat cag aaa gag gtt taa tat ttt tgt gag acc tat cga a-- --- --- --- --- --- --- --- --- --- --- --- --- --- --- --- --- --- --- --- --- --- --- --- --- --- --- --- --- --- --- --- --- --- --- --- --- --- --- --- --- --- --- --- --- --- --- --- --- --- --- --- --- --- --- --- --- --- --- --- --- --- --- --- --- --- --- --- --- --- --- --- --- --- --- --- --- --- --- --- --- --- --- --- --- --- --- --- --- --- --- --- --- --- --- --- --- --- --- --- --- --- --- --- --- --- --- --- --- --- --- --- --- --- --- --- --- --- --- --- --- --- --- --- --- --- --- --- --- --- --- --- --- --- --- --- --- --- --- --- --- --- --- --- --- --- --- --- --- --- --- --- --- --- --- --- --- --- --- --- --- --- --- --- --- --- --- --- --- --- --- --- --- --- --- --- --- --- --- --- --- --- --- --- --- --- --- --- --- --- --- --- --- --- --- --- --- --- --- --- --- --- --- --- --- --- --- --- --- --- --- --- --- --- --- --- --- --- --- --- --- --- --- --- --- --- --- --- --- --- --- --- --- --- --- --- --- --- --- --- --- --- --- --- --- --- --- --- --- --- --- --- --- --- --- --- --- --- --- --- --- --- --- --- --- --- --- --- --- --- --- --- --- --- --- --- --- --- --- --- -ga gag aaa gga taa aaa ctt ttt acg act cca tca gaa aga ggt tta ata ttt ttg tga gac cca tcg aag aga gaa aga gat ggt tag tca aga tat ttt tct tag tac aaa agt caa tgt ttt aaa ata tat gga cga gaa tta att tgt ctg tat aaa aac ttg tgt gaa att atg tac tag aga aaa aac gtg agc agt gtc ccc tac atg gat ttt aca gat cat tta tat tcc aaa aat att aac tat ata cgt tta tta tat gat gtt aac gtg taa att ata aac att att tta tga tgc aat tgt ctg aca acc tag att ggt ata agg atg ttg ata agc tct acg aga ata tat tgt tgg acg tta tcg ttt acg aaa tag ttg aga cat cag aaa gag gtt taa tat ttt tgt gag a-c cat cga aga gag aaa gag aat aaa aat a-- --t ttt ttt ttt ttt tgt aaa act ttt tta tga gac --- --- -ca aga gaa tac gaa tag tga tca tat cgt atc aca tat tga aac aga aag aag aag taa cga gag gta act ttt tgt gaa tgt agt taa ata ttt ttg ttt tgc aaa ccg gaa tat agt gcc cgg t-- -ct ttt tta att cgt ggt gcg gtg tct gaa tcg ttc gat taa ccc aac tca tcc att ttc aga tga ata gag tta tcg att cag aca cat gct ttg agt ttt gtt gaa tcg atg agt gaa gta tca tcg gtt gca cct tca gat gcc gat ccg tcg aca tac ttg aat cca tcc ttg act tca agt tca gat gat tcc tca cac atg tct ccg ata cgt acg cta aac tct agg ttc ttg aca cat ttt gta tca acg atc gtt gaa ccg atg ata tct ttg taa ctc act ttc tta tgt gag atg tta gac cca agt act gga tgg gtc ttg atg tca ctg tct ttc tct tct tcg cta cat ctg atg tcg ata gac atc tca cag tct ttg atc ata gcc aga gct tct tca cgc gtg atc gcg gga gag tcc tta cct tgt ccc ggt gac acg ctg gac aat cta gta ttc aca gtg ttt cca tca gag gat tcg gag atg gat aaa atc ttt ggg cat ttg gtg aat cca aag ttc atg tta aga ccc gca ccg acg ata gtg taa taa gtg gtg gga tct cct ttt aca act --- --- --- --- --- --- tct tcg gat acc tca tca tct tcg gtc tct gta act tcc gtt acg gat tga caa atc tta tca ttg gtc ggt gtt tgg tct tgc ttt gtg act ttg ata ata aca tcg att ccc ata tga tgt ttg ttt tct tct tca gta cac gag gat gaa gat tgt tga aga cta gta ggc ata gca gct gcc act agg cac atg cat gcc agg aca ata tat tgt ttc atg att gct att gat tga tta ctg ttc tag atg att cta ctt tct tac cat ata ata aat tag aat ata ttt tct act ttt acg aga aat taa tta ttg tat tta tta ttt ata ggt --a aaa aaa ctt act ata agt ggg tgg gat tct ggg aat tag tga tca gtt tat gta tat cgc aac tag cgg gca tat ggc tat tga cat cga gaa cat tac cca tat gat aag aga ttg tat cat ttt cgt agt ctt gag tat tgg tat tac tat ata gta tgt aga tgt cga cgc tag ata gac agt cgc cca cta gag tta ccg tct ctg aat gcg gca tga tag tat cat tct ttg ttt tcg tta act gtt tgg aag atg aat ctt tgt tgt tac att taa tct cga aat tca gag tac ata tct ttg aag tat tct gat atc tat ttt ctc ctg taa aga atc ctg aag ttg cta cat tat taa gga cag aga agt att ctg cac gaa aga ctg gat cac aat ctt tat gat tca tgg taa tag tta gtt ccg acg ttg aga tgg att cgc tga gac cgg tag tgg tcg tcc gag tac acg atg tgt cgt tga ctg gat aca ggt taa ttt cca cat cga tat agt taa atg tat tgc tgg tta cga cgg gtt cgc att tat ctg tgg aag aga cgg tgt gag aat atg ttc cgg gac cac acg gag aac aga tga cgt ctc cgg tag acg tgt atc cgg ata ctc cgt atc cta ttc cac act ttg ttt tag aaa tac atg ttc tac acc ctg atg ctc ctt tga gaa gac aat aat atc ctg gag agc att cac aga ttc tat tgt gag tcg tgt tac acg atc gcg tct cta cct gat tac tat cac atc ttc cgt tac aac tta gac aag cct gta aat gat tat tgt gag atg taa agg tat ccg aac cac acg gtg tac att gtg tat tag tct tgc tat cac ata atc tga aag cgt aag ttc ccg gag gac acg ata gac aac ata gat tac ggc ttc tgt att cgt tgt ctt tac act ttc cat tgg atg gtg cat gtg gtg cta tat ctc ttc cgt tta tta tta tac atg aga gaa aca ata tat acg agt ata ata cgg acc tca tga ttt aat aat -gt agt aat cgt cgt ctt gtt act gtt tgt ttc cta ctt ctc caa tca tat aga tta ttt ttt aaa tat ttt ctt tct atc atg gat aat att tgt aat ggt tct ttc cgt aca aca tac tgt tta gat ggt agt cgc tta gct tgg tta tga tat tgc gca taa ttt ccg gag gca aat acg ata gtc tag att gac tat cga tgg tag act cta att tat tga gtg ctt tgt cga cga gtt tac ttt tat gct cca tcg ata gat gac act gtt cta tga gat cgt cgt aca tgg gaa atg aaa tgc gtt tgt ctg aat gta tgg ctt cga gat atg tgt gat acc gaa tgt ctt ctg ttc tca ata ccg tat aca agt cgg tgt ctg aga ttc gaa tct ctt tga gga gac tta tgt cac gac tac att ttt cga tga tgg aat cta tct tat cga atg ata tat ttt tca taa ata cac ttt tat agt cct cgt tta aac aga att tag tat gta gtt ccg caa atg act cgt ccc tta ata ggc agt agg cta tta tct tct tta cgt agt gat cgt cgt agg gag aga act ccg aca tct tgt aga aca acg att taa tca tag gta gag ata ctt tca gtc tgt ggt gga taa tgt cat taa cga cat ccg cct tgt ata taa tgt ttc tgt ttt caa aca cca agt cga ata ctg tct tat cgt ctt tag tcg gaa ggt tga tgt cgt atc caa tgt ata cga ggt atg agg caa cat tgt tat tgc aat tct gga agg cgg tat gaa gag gag tca ttg tat tat agt att cgt ctt tct gaa tgt cga atc tat cta gta gat acc gta gta tat tga gag agc gac ttc cat atc ctt gat tat gtt tta tga ata gat aaa gta gat gtt gtc ctt ctt cct ttt gta att tcc cgt att ttt gtt cgt gcc aat tga gta aca tta tga gaa tat gac ctg ttg cac aat cgt tct tta tgt att cca tga tgg gtg tac aat taa gat tat tac gta tcc tcg tat cgg ctc ctc gag ata aaa gag cat aca cca cac gag gac tat gtt tgg tat act gtt gaa ggt aag tgt gta acg gcg tat ttc cga ttt tcg taa ccg cgt taa tgt ttg ctc cat gat cta tta tcg cgt aga tga atc gct tct cag ctc gca tct tag tgt gac tct ttg act tgt aat aat tgc ttt cgt gga acg cgg ata tgt gtt tac agt agt aat gaa gag aag tga gtc cat cct cat cga cgc aat tag ggt cag atc ctt tag tca ata att tgt aca gaa cgt aat agt tta agc tcc cat tga att tat atc taa gat aac aca gca ata gat cgg atg att tac taa agt cat caa tgg ggt ccg tta gta tat caa aga tct tgt tat cga ttg ata gtg aat gaa tca gat agt ggt gta gag gaa tat gtc ctt ttt tat cct tgc tat caa agt tac gca tgc cgt ggc gta aca ata tct tta ata cag atg gat taa atc gtg tat tca tcg tat agc aat gta atg gag agt tac cac att tta gtc gtt tat tca gat cgc agt gtt taa taa cta att taa aca gat aag atg ata tat cca cat caa aga atg cga gat aca tat gac aga cat tat tga cag aaa tgt gac ctt cat tat cac cgt cgt cca taa atg cgt tag gta cgt acc aca tac tat cgt taa cga tgc gca caa tct cgt cca -tt tca tcc atc ttc ata atg att tac ttt ttc ata att aga gaa aaa gat caa ggt ata aaa att aga agt gtt aga cta ta- --- --- --- --- --- --- --- --- --- --- -aa cta act tat gac tta act aac tta tga --- --- --- --- --- --- --- --- --- --- --- --- --- --- --- --- --- --- --- --- --- --- --- --- --- --- --- --- --- --- --- --- --- --- --- --- --- --- --- --- --- --- --- --- --- --- --- --- ctt aac ta- --- --- --- --- --- --- --- --- --- --- -ac tta tga ctt aac taa ctt atg act taa cta act ta- tga ctt aac taa ctt atg act taa cta act tat gac tta act aac tta tga cta tta act cat ttc aag aaa ggt ggg tgg ata gaa ctc tat atg aca gct tgt gaa aca att aga tcc cta att tct aat gga agt ttt gat agg aga ttg tca tca gtt gat aca ttg ttt att atc tca tct att aga gca cgt ctg ttt aga gct tta gtg acc tgc tcg gtt act tct gtg taa atc ttg aat cct tta gtg ata cac tgt gtc aaa act gga tgt tta gaa tac cta tgt aga ata tgg gaa gca tgc ttg ttt ttg tct cta tta tag att aac tca tac atg gtt gta tta tga att ttc atc tgc cta atg tac tcc aat tct tgt tta caa tca att ata taa tca aag agt gat gat gca tac aca tta caa agt gaa taa tct acc atc ata aaa tac ttg ata cag agc ttt atc aca tca tgg ttt tca att gta tta tta agt ata gct aat ttt ata cag tca ata gac aat ggt tct cta agc aat att tct aat att tta aga tgt gct tcc cta cgg gcg atg aca gat ccc cta tcc acg gcc acg tca aga cat gta tat cca tta ctc att act gcg ttg aca ttt gct cca ttt tct aat agc cat gat act aaa tct ata taa cct gca tag ata gcg cga taa agc aag gtc ctt cca cca gca tct agt tga ttg ata tct tca ata tat ggg ata caa agc tta taa att tct aat act gtg ggt tca tct aca agg aat ccc cta gta tac tga att att tta tat aga tct aat tta aca tca ttt tca tct ggg ata cca caa ttc aaa ata aac tca aca aca cta ctt tcc ttt tta cat att ccc cta aaa tag gca ttc aag cat tct att tta tat att aca gcc cca tga tct acc ata aga tca aca atg tct att tct aca tat gca tta gat aga tag taa agt aag aga tct tgc aca gaa tta caa ttc tta ata att ata gag aaa ata tct tcc ata taa ttc ttt gac act aat gca gat ata ata tct tta tat gta ata tat gca aac agt cta tct act ata tac tga tca ata tta tct cta tga atc cta aaa taa tca tac aga aca tct aca gga tca caa att ggt tca agg aga aat cta tca aat att ttc ctg tca aca act ggt tct aga aca taa cag tca aca cct aat cca tgt ttt tta taa tca tct acc aaa gat aat gac caa aga tcg agg tcg tcg tga aac tgc tca tcg aca gcc atg aaa tct acc gac tcc atg gtg cga atc gca ctg tct tat tcg cca ttg att ttc a-t ttt tta taa tta tgt aca tgt ttt cct tct att ctc aag agt cta caa aaa tat a-- -tt ttt tcg ata tct aag tac taa gtt ttt tta ctg ttt ttg tta ctg tct tcc att ctt cta act aaa gat ctg aga taa att ata caa tct tcg cta tcg aac cat ttt tgt agt cta aag cct gaa gta att aac caa ctg ttt tta tta gtg gct ttt ttc gat cta tcc tcg tc- --- ctc tca tca tcc tta ta- --- --- --- --- --- --- tta tta tca tta tta tca tag tct att aaa cac aaa tca tct --- --- --- --- --- --- --- --- --- --- --- --- --- --- --- --- --- --- --- --- --- --- --- --- --- --- --a cgt tta taa c-- --- --- --- --- --- --- --- --a aca ttc --- --- --- --- -tc att att aat tag ttc tgt ag- aat atc ttt aat aat ttg gct ata --- --- -ca tct gtt --- --- --- --- --- --- --- --- --- --- --- --- --- --- --- --- --- --- --c aat act --- --- --- --- --- --- --- --a tct att gat gat ttc ttt t-- --- --- --- --- --- --- --- --- --- --- --t taa gac t-- --- --- --- --- --- --- --- --- --- --- --- --- --- --- --- --- --- --- --- --- --- --- --- --- --- --- --- --- --- --- --- --- --- --- --- --- --- --- --- taa act agt --- --- --- --- --- --- --- --- --- --- --- --- --- --- --- --- --- --- --- --- --- --- --- --- --- --- --- --- --- --- --- --- --- --- --- --- --- --- --- --- --- --- --- --- --- --- --- --- --- --- --- --- --- --- --- --- --- --- --- --- --- --- --- --- --- --- --- --- --- --- --- --- --- --- --- --- --- --- --- --- --- --- --- --- --- --- --- --- --- --- --- --- --- --- --- --- --- --- --- --- --- --- --- --- --- --- --- --- --- --t atg gta atg acg atg aaa --- -tc gag tag ta- --- --a ctt cta ata aag act tga ta- --- --t cat tat cat atg ttt gat cg- --- --- --- --- --- --- --t cat agt taa tag tgt g-- --- --- --- --- --- --- --- --- --- --- --- --- --- --- --- --- --- --- --- -gc taa atg gta ctg tta ata agt tta t-- --- --- --- --- --- --- --- --- --- --- --- --- --- --- --- --- --a gac aat atc ata gta ttt tct ttc cag aat --- --- --- --t aga tta ttt ttt taa ata ctg atc ctc aca att ccg tga tgt agc agt agt tgg t-- --- --- --- --- --- --- --- --- --- --- --- --- --- --- --- --- --- --- --- --- --- --- --- --- --- --- --- -gc atg gtc tat atc gt- --- --- --- --- --- --- --- --- --- --- --- --- --- --- --- --- --- --- --- --- --- --- --- --- --- --- --- --- --- --- --- --- --- --- --- --- --- --- --- --- --- --- --- taa aat gta tca tat ata ata gtt ttc tga cgt gga gta cag aat ttt cga --- --- --- --- --- --- --- --- --- --- --- --- --- --- --- --- --- --- --- --- --- --- --- --- --- --- --- --- --- --- --- --- --- --- --- --- --- --- --- --- --- --- --- --t taa tga gtt cat ggt aag gaa ggg caa atg cct --- gta tat aat ata cat aag tta a-- --- --- --- --- --- --- --- --- --- --- --- --- --t agt ttt tta tca tat tt- --- --- --- --- --- --- --- --t cta ata cca taa taa aaa t-- --- --- --- --- --- --- --- --- --- --- --- --- --- --- --- --- --- --- --- --- --- --- --- --- --t atc att att gcg ttt g-g tag tt- --- --- --- --- ctg ccc ta- --- --- --- --- --- --- --- --t cat cta tat cac tgt cac tct c-- --- --g ctc tca cta tat ctt cta aaa tta caa --- -ac aac --- --- --- --- --- --- --- --- --- --- --- --- --- --- --- --- --- --- --- --- --- --- --- --- --- --- --- --- --- --- --- --- --- --- --- --- tgg ata ttc gat --- --- --- --- --- --- --- --- --- --- --- --- --- --- --- --- --- --- --- --- --- --- --- --- --- --- --a aca gca ttt gtg t-- --- --- --- --- --- --- --- --- --- --- --- --- --- --- --- --- -

>ON619836.1_Monkeypox_virus_isolate_MPXV_UK_2022_2_complete_genome

--- --- --- --- --- --- --- --- --- --- --- --- --- --- --- --- --- --- --- --- --- --- --- --- --- --- --- --t att ata ttt tac tat ttt att tag tgt cta gaa aaa aat gtg tga ccc acg acc gta gga aac tct aga ggg taa gaa aaa tca atc gtt tat aga gac cat cag aaa gag gtt taa tat ttt tgt gag acc tat cga a-- --- --- --- --- --- --- --- --- --- --- --- --- --- --- --- --- --- --- --- --- --- --- --- --- --- --- --- --- --- --- --- --- --- --- --- --- --- --- --- --- --- --- --- --- --- --- --- --- --- --- --- --- --- --- --- --- --- --- --- --- --- --- --- --- --- --- --- --- --- --- --- --- --- --- --- --- --- --- --- --- --- --- --- --- --- --- --- --- --- --- --- --- --- --- --- --- --- --- --- --- --- --- --- --- --- --- --- --- --- --- --- --- --- --- --- --- --- --- --- --- --- --- --- --- --- --- --- --- --- --- --- --- --- --- --- --- --- --- --- --- --- --- --- --- --- --- --- --- --- --- --- --- --- --- --- --- --- --- --- --- --- --- --- --- --- --- --- --- --- --- --- --- --- --- --- --- --- --- --- --- --- --- --- --- --- --- --- --- --- --- --- --- --- --- --- --- --- --- --- --- --- --- --- --- --- --- --- --- --- --- --- --- --- --- --- --- --- --- --- --- --- --- --- --- --- --- --- --- --- --- --- --- --- --- --- --- --- --- --- --- --- --- --- --- --- --- --- --- --- --- --- --- --- --- --- --- --- --- --- --- --- --- --- --- --- --- --- --- --- --- --- --- --- --- --- --- --- --- --- -ga gag aaa gga taa aaa ctt ttt acg act cca tca gaa aga ggt tta ata ttt ttg tga gac cca tcg aag aga gaa aga gat ggt tag tca aga tat ttt tct tag tac aaa agt caa tgt ttt aaa ata tat gga cga gaa tta att tgt ctg tat aaa aac ttg tgt gaa att atg tac tag aga aaa aac gtg agc agt gtc ccc tac atg gat ttt aca gat cat tta tat tcc aaa aat att aac tat ata cgt tta tta tat gat gtt aac gtg taa att ata aac att att tta tga tgc aat tgt ctg aca acc tag att ggt ata agg atg ttg ata agc tct acg aga ata tat tgt tgg acg tta tcg ttt acg aaa tag ttg aga cat cag aaa gag gtt taa tat ttt tgt gag a-c cat cga aga gag aaa gag aat aaa aat a-- --? ttt ttt ttt ttt tgt aaa act ttt tta tga gac --- --- -ca aga gaa tac gaa tag tga tca tat cgt atc aca tat tga aac aga aag aag aag taa cga gag gta act ttt tgt gaa tgt agt taa ata ttt ttg ttt tgc aaa ccg gaa tat agt gcc cgg t-- -ct ttt tta att cgt ggt gcg gtg tct gaa tcg ttc gat taa ccc aac tca tcc att ttc aga tga ata gag tta tcg att cag aca cat gct ttg agt ttt gtt gaa tcg atg agt gaa gta tca tcg gtt gca cct tca gat gcc gat ccg tcg aca tac ttg aat cca tcc ttg act tca agt tca gat gat tcc tca cac atg tct ccg ata cgt acg cta aac tct agg ttc ttg aca cat ttt gta tca acg atc gtt gaa ccg atg ata tct ttg taa ctc act ttc tta tgt gag atg tta gac cca agt act gga tgg gtc ttg atg tca ctg tct ttc tct tct tcg cta cat ctg atg tcg ata gac atc tca cag tct ttg atc ata gcc aga gct tct tca cgc gtg atc gcg gga gag tcc tta cct tgt ccc ggt gac acg ctg gac aat cta gta ttc aca gtg ttt cca tca gag gat tcg gag atg gat aaa atc ttt ggg cat ttg gtg aat cca aag ttc atg tta aga ccc gca ccg acg ata gtg taa taa gtg gtg gga tct cct ttt aca act --- --- --- --- --- --- tct tcg gat acc tca tca tct tcg gtc tct gta act tcc gtt acg gat tga caa atc tta tca ttg gtc ggt gtt tgg tct tgc ttt gtg act ttg ata ata aca tcg att ccc ata tga tgt ttg ttt tct tct tca gta cac gag gat gaa gat tgt tga aga cta gta ggc ata gca gct gcc act agg cac atg cat gcc agg aca ata tat tgt ttc atg att gct att gat tga tta ctg ttc tag atg att cta ctt tct tac cat ata ata aat tag aat ata ttt tct act ttt acg aga aat taa tta ttg tat tta tta ttt ata ggt --a aaa aaa ctt act ata agt ggg tgg gat tct ggg aat tag tga tca gtt tat gta tat cgc aac tag cgg gca tat ggc tat tga cat cga gaa cat tac cca tat gat aag aga ttg tat cat ttt cgt agt ctt gag tat tgg tat tac tat ata gta tgt aga tgt cga cgc tag ata gac agt cgc cca cta gag tta ccg tct ctg aat gcg gca tga tag tat cat tct ttg ttt tcg tta act gtt tgg aag atg aat ctt tgt tgt tac att taa tct cga aat tca gag tac ata tct ttg aag tat tct gat atc tat ttt ctc ctg taa aga atc ctg aag ttg cta cat tat taa gga cag aga agt att ctg cac gaa aga ctg gat cac aat ctt tat gat tca tgg taa tag tta gtt ccg acg ttg aga tgg att cgc tga gac cgg tag tgg tcg tcc gag tac acg atg tgt cgt tga ctg gat aca ggt taa ttt cca cat cga tat agt taa atg tat tgc tgg tta cga cgg gtt cgc att tat ctg tgg aag aga cgg tgt gag aat atg ttc cgg gac cac acg gag aac aga tga cgt ctc cgg tag acg tgt atc cgg ata ctc cgt atc cta ttc cac act ttg ttt tag aaa tac atg ttc tac acc ctg atg ctc ctt tga gaa gac aat aat atc ctg gag agc att cac aga ttc tat tgt gag tcg tgt tac acg atc gcg tct cta cct gat tac tat cac atc ttc cgt tac aac tta gac aag cct gta aat gat tat tgt gag atg taa agg tat ccg aac cac acg gtg tac att gtg tat tag tct tgc tat cac ata atc tga aag cgt aag ttc ccg gag gac acg ata gac aac ata gat tac ggc ttc tgt att cgt tgt ctt tac act ttc cat tgg atg gtg cat gtg gtg cta tat ctc ttc cgt tta tta tta tac atg aga gaa aca ata tat acg agt ata ata cgg acc tca tga ttt aat aat -gt agt aat cgt cgt ctt gtt act gtt tgt ttc cta ctt ctc caa tca tat aga tta ttt ttt aaa tat ttt ctt tct atc atg gat aat att tgt aat ggt tct ttc cgt aca aca tac tgt tta gat ggt agt cgc tta gct tgg tta tga tat tgc gca taa ttt ccg gag gca aat acg ata gtc tag att gac tat cga tgg tag act cta att tat tga gtg ctt tgt cga cga gtt tac ttt tat gct cca tcg ata gat gac act gtt cta tga gat cgt cgt aca tgg gaa atg aaa tgc gtt tgt ctg aat gta tgg ctt cga gat atg tgt gat acc gaa tgt ctt ctg ttc tca ata ccg tat aca agt cgg tgt ctg aga ttc gaa tct ctt tga gga gac tta tgt cac gac tac att ttt cga tga tgg aat cta tct tat cga atg ata tat ttt tca taa ata cac ttt tat agt cct cgt tta aac aga att tag tat gta gtt ccg caa atg act cgt ccc tta ata ggc agt agg cta tta tct tct tta cgt agt gat cgt cgt agg gag aga act ccg aca tct tgt aga aca acg att taa tca tag gta gag ata ctt tca gtc tgt ggt gga taa tgt cat taa cga cat ccg cct tgt ata taa tgt ttc tgt ttt caa aca cca agt cga ata ctg tct tat cgt ctt tag tcg gaa ggt tga tgt cgt atc caa tgt ata cga ggt atg agg caa cat tgt tat tgc aat tct gga agg cgg tat gaa gag gag tca ttg tat tat agt att cgt ctt tct gaa tgt cga atc tat cta gta gat acc gta gta tat tga gag agc gac ttc cat atc ctt gat tat gtt tta tga ata gat aaa gta gat gtt gtc ctt ctt cct ttt gta att tcc cgt att ttt gtt cgt gcc aat tga gta aca tta tga gaa tat gac ctg ttg cac aat cgt tct tta tgt att cca tga tgg gtg tac aat taa gat tat tac gta tcc tcg tat cgg ctc ctc gag ata aaa gag cat aca cca cac gag gac tat gtt tgg tat act gtt gaa ggt aag tgt gta acg gcg tat ttc cga ttt tcg taa ccg cgt taa tgt ttg ctc cat gat cta tta tcg cgt aga tga atc gct tct cag ctc gca tct tag tgt gac tct ttg act tgt aat aat tgc ttt cgt gga acg cgg ata tgt gtt tac agt agt aat gaa gag aag tga gtc cat cct cat cga cgc aat tag ggt cag atc ctt tag tca ata att tgt aca gaa cgt aat agt tta agc tcc cat tga att tat atc taa gat aac aca gca ata gat cgg atg att tac taa agt cat caa tgg ggt ccg tta gta tat caa aga tct tgt tat cga ttg ata gtg aat gaa tca gat agt ggt gta gag gaa tat gtc ctt ttt tat cct tgc tat caa agt tac gca tgc cgt ggc gta aca ata tct tta ata cag atg gat taa atc gtg tat tca tcg tat agc aat gta atg gag agt tac cac att tta gtc gtt tat tca gat cgc agt gtt taa taa cta att taa aca gat gag atg ata tat cca cat caa aga atg cga gat aca tat gac aga cat tat tga cag aaa tgt gac ctt cat tat cac cgt cgt cca taa atg cgt tag gta cgt acc aca tac tat cgt taa cga tgc gca caa tct cgt cca -tt tca tcc atc ttc ata atg att tac ttt ttc ata att aga gaa aaa gat caa ggt ata aaa att aga agt gtt aga cta ta- --- --- --- --- --- --- --- --- --- --- -aa cta act tat gac tta act aac tta tga --- --- --- --- --- --- --- --- --- --- --- --- --- --- --- --- --- --- --- --- --- --- --- --- --- --- --- --- --- --- --- --- --- --- --- --- --- --- --- --- --- --- --- --- --- --- --- --- ctt aac ta- --- --- --- --- --- --- --- --- --- --- -ac tta tga ctt aac taa ctt atg act taa cta act ta- tga ctt aac taa ctt atg act taa cta act tat gac tta act aac tta tga cta tta act cat ttc aag aaa ggt ggg tgg ata gaa ctc tat atg aca gct tgt gaa aca att aga tcc cta att tct aat gga agt ttt gat agg aga ttg tca tca gtt gat aca ttg ttt att atc tca tct att aga gca cgt ctg ttt aga gct tta gtg acc tgc tcg gtt act tct gtg taa atc ttg aat cct tta gtg ata cac tgt gtc aaa act gga tgt tta gaa tac cta tgt aga ata tgg gaa gca tgc ttg ttt ttg tct cta tta tag att aac tca tac atg gtt gta tta tga att ttc atc tgc cta atg tac tcc aat tct tgt tta caa tca att ata taa tca aag agt gat gat gca tac aca tta caa agt gaa taa tct acc atc ata aaa tac ttg ata cag agc ttt atc aca tca tgg ttt tca att gta tta tta agt ata gct aat ttt ata cag tca ata gac aat ggt tct cta agc aat att tct aat att tta aga tgt gct tcc cta cgg gcg atg aca gat ccc cta tcc acg gcc acg tca aga cat gta tat cca tta ctc att act gcg ttg aca ttt gct cca ttt tct aat agc cat gat act aaa tct ata taa cct gca tag ata gcg cga taa agc aag gtc ctt cca cca gca tct agt tga ttg ata tct tca ata tat ggg ata caa agc tta taa att tct aat act gtg ggt tca tct aca agg aat ccc cta gta tac tga att att tta tat aga tct aat tta aca tca ttt tca tct ggg ata cca caa ttc aaa ata aac tca aca aca cta ctt tcc ttt tta cat att ccc cta aaa tag gca ttc aag cat tct att tta tat att aca gcc cca tga tct acc ata aga tca aca atg tct att tct aca tat gca tta gat aga tag taa agt aag aga tct tgc aca gaa tta caa ttc tta ata att ata gag aaa ata tct tcc ata taa ttc ttt gac act aat gca gat ata ata tct tta tat gta ata tat gca aac agt cta tct act ata tac tga tca ata tta tct cta tga atc cta aaa taa tca tac aga aca tct aca gga tca caa att ggt tca agg aga aat cta tca aat att ttc ctg tca aca act ggt tct aga aca taa cag tca aca cct aat cca tgt ttt tta taa tca tct acc aaa gat aat gac caa aga tcg agg tcg tcg tga aac tgc tca tcg aca gcc atg aaa tct acc gac tcc atg gtg cga atc gca ctg tct tat tcg cca ttg att ttc a-t ttt tta taa tta tgt aca tgt ttt cct tct att ctc aag agt cta caa aaa tat a-- -tt ttt tcg ata tct aag tac taa gtt ttt tta ctg ttt ttg tta ctg tct tcc att ctt cta act aaa gat ctg aga taa att ata caa tct tcg cta tcg aac cat ttt tgt agt cta aag cct gaa gta att aac caa ctg ttt tta tta gtg gct ttt ttc gat cta tcc tcg tc- --- ctc tca tca tcc tta ta- --- --- --- --- --- --- tta tta tca tta tta tca tag tct att aaa cac aaa tca tct --- --- --- --- --- --- --- --- --- --- --- --- --- --- --- --- --- --- --- --- --- --- --- --- --- --- --a cgt tta taa c-- --- --- --- --- --- --- --- --a aca ttc --- --- --- --- -tc att att aat tag ttc tgt ag- aat atc ttt aat aat ttg gct ata --- --- -ca tct gtt --- --- --- --- --- --- --- --- --- --- --- --- --- --- --- --- --- --- --c aat act --- --- --- --- --- --- --- --a tct att gat gat ttc ttt t-- --- --- --- --- --- --- --- --- --- --- --t taa gac t-- --- --- --- --- --- --- --- --- --- --- --- --- --- --- --- --- --- --- --- --- --- --- --- --- --- --- --- --- --- --- --- --- --- --- --- --- --- --- --- taa act agt --- --- --- --- --- --- --- --- --- --- --- --- --- --- --- --- --- --- --- --- --- --- --- --- --- --- --- --- --- --- --- --- --- --- --- --- --- --- --- --- --- --- --- --- --- --- --- --- --- --- --- --- --- --- --- --- --- --- --- --- --- --- --- --- --- --- --- --- --- --- --- --- --- --- --- --- --- --- --- --- --- --- --- --- --- --- --- --- --- --- --- --- --- --- --- --- --- --- --- --- --- --- --- --- --- --- --- --- --- --t atg gta atg acg atg aaa --- -tc gag tag ta- --- --a ctt cta ata aag act tga ta- --- --t cat tat cat atg ttt gat cg- --- --- --- --- --- --- --t cat agt taa tag tgt g-- --- --- --- --- --- --- --- --- --- --- --- --- --- --- --- --- --- --- --- -gc taa atg gta ctg tta ata agt tta t-- --- --- --- --- --- --- --- --- --- --- --- --- --- --- --- --- --a gac aat atc ata gta ttt tct ttc cag aat --- --- --- --t aga tta ttt ttt taa ata ctg atc ctc aca att ccg tga tgt agc agt agt tgg t-- --- --- --- --- --- --- --- --- --- --- --- --- --- --- --- --- --- --- --- --- --- --- --- --- --- --- --- -gc atg gtc tat atc gt- --- --- --- --- --- --- --- --- --- --- --- --- --- --- --- --- --- --- --- --- --- --- --- --- --- --- --- --- --- --- --- --- --- --- --- --- --- --- --- --- --- --- --- taa aat gta tca tat ata ata gtt ttc tga cgt gga gta cag aat ttt cga --- --- --- --- --- --- --- --- --- --- --- --- --- --- --- --- --- --- --- --- --- --- --- --- --- --- --- --- --- --- --- --- --- --- --- --- --- --- --- --- --- --- --- --t taa tga gtt cat ggt aag gaa ggg caa atg cct --- gta tat aat ata cat aag tta a-- --- --- --- --- --- --- --- --- --- --- --- --- --t agt ttt tta tca tat tt- --- --- --- --- --- --- --- --t cta ata cca taa taa aaa t-- --- --- --- --- --- --- --- --- --- --- --- --- --- --- --- --- --- --- --- --- --- --- --- --- --t atc att att gcg ttt g-g tag tt- --- --- --- --- ctg ccc ta- --- --- --- --- --- --- --- --t cat cta tat cac tgt cac tct c-- --- --g ctc tca cta tat ctt cta aaa tta caa --- -ac aac --- --- --- --- --- --- --- --- --- --- --- --- --- --- --- --- --- --- --- --- --- --- --- --- --- --- --- --- --- --- --- --- --- --- --- --- tgg ata ttc gat --- --- --- --- --- --- --- --- --- --- --- --- --- --- --- --- --- --- --- --- --- --- --- --- --- --- --a aca gca ttt gtg t-- --- --- --- --- --- --- --- --- --- --- --- --- --- --- --- --- -

>ON619838.1_Monkeypox_virus_isolate_MPXV_UK_2022_4_complete_genome

--- --- --- --- --- --- --- --- --- --- --- --- --- --- --- --- --- --- --- --- --- --- --- --- --- --- --- --- --- --a ttt tac tat ttt att tag tgt cta gaa aaa aat gtg tga ccc acg acc gta gga aac tct aga ggg taa gaa aaa tca atc gtt tat aga gac cat cag aaa gag gtt taa tat ttt tgt gag acc tat cga a-- --- --- --- --- --- --- --- --- --- --- --- --- --- --- --- --- --- --- --- --- --- --- --- --- --- --- --- --- --- --- --- --- --- --- --- --- --- --- --- --- --- --- --- --- --- --- --- --- --- --- --- --- --- --- --- --- --- --- --- --- --- --- --- --- --- --- --- --- --- --- --- --- --- --- --- --- --- --- --- --- --- --- --- --- --- --- --- --- --- --- --- --- --- --- --- --- --- --- --- --- --- --- --- --- --- --- --- --- --- --- --- --- --- --- --- --- --- --- --- --- --- --- --- --- --- --- --- --- --- --- --- --- --- --- --- --- --- --- --- --- --- --- --- --- --- --- --- --- --- --- --- --- --- --- --- --- --- --- --- --- --- --- --- --- --- --- --- --- --- --- --- --- --- --- --- --- --- --- --- --- --- --- --- --- --- --- --- --- --- --- --- --- --- --- --- --- --- --- --- --- --- --- --- --- --- --- --- --- --- --- --- --- --- --- --- --- --- --- --- --- --- --- --- --- --- --- --- --- --- --- --- --- --- --- --- --- --- --- --- --- --- --- --- --- --- --- --- --- --- --- --- --- --- --- --- --- --- --- --- --- --- --- --- --- --- --- --- --- --- --- --- --- --- --- --- --- --- --- --- -ga gag aaa gga taa aaa ctt ttt acg act cca tca gaa aga ggt tta ata ttt ttg tga gac cca tcg aag aga gaa aga gat ggt tag tca aga tat ttt tct tag tac aaa agt caa tgt ttt aaa ata tat gga cga gaa tta att tgt ctg tat aaa aac ttg tgt gaa att atg tac tag aga aaa aac gtg agc agt gtc ccc tac atg gat ttt aca gat cat tta tat tcc aaa aat att aac tat ata cgt tta tta tat gat gtt aac gtg taa att ata aac att att tta tga tgc aat tgt ctg aca acc tag att ggt ata agg atg ttg ata agc tct acg aga ata tat tgt tgg acg tta tcg ttt acg aaa tag ttg aga cat cag aaa gag gtt taa tat ttt tgt gag a-c cat cga aga gag aaa gag aat aaa aat a-- --? ttt ttt ttt ttt tgt aaa act ttt tta tga gac --- --- -ca aga gaa tac gaa tag tga tca tat cgt atc aca tat tga aac aga aag aag aag taa cga gag gta act ttt tgt gaa tgt agt taa ata ttt ttg ttt tgc aaa ccg gaa tat agt gcc cgg t-- -ct ttt tta att cgt ggt gcg gtg tct gaa tcg ttc gat taa ccc aac tca tcc att ttc aga tga ata gag tta tcg att cag aca cat gct ttg agt ttt gtt gaa tcg atg agt gaa gta tca tcg gtt gca cct tca gat gcc gat ccg tcg aca tac ttg aat cca tcc ttg act tca agt tca gat gat tcc tca cac atg tct ccg ata cgt acg cta aac tct agg ttc ttg aca cat ttt gta tca acg atc gtt gaa ccg atg ata tct ttg taa ctc act ttc tta tgt gag atg tta gac cca agt act gga tgg gtc ttg atg tca ctg tct ttc tct tct tcg cta cat ctg atg tcg ata gac atc tca cag tct ttg atc ata gcc aga gct tct tca cgc gtg atc gcg gga gag tcc tta cct tgt ccc ggt gac acg ctg gac aat cta gta ttc aca gtg ttt cca tca gag gat tcg gag atg gat aaa atc ttt ggg cat ttg gtg aat cca aag ttc atg tta aga ccc gca ccg acg ata gtg taa taa gtg gtg gga tct cct ttt aca act --- --- --- --- --- --- tct tcg gat acc tca tca tct tcg gtc tct gta act tcc gtt acg gat tga caa atc tta tca ttg gtc ggt gtt tgg tct tgc ttt gtg act ttg ata ata aca tcg att ccc ata tga tgt ttg ttt tct tct tca gta cac gag gat gaa gat tgt tga aga cta gta ggc ata gca gct gcc act agg cac atg cat gcc agg aca ata tat tgt ttc atg att gct att gat tga tta ctg ttc tag atg att cta ctt tct tac cat ata ata aat tag aat ata ttt tct act ttt acg aga aat taa tta ttg tat tta tta ttt ata ggt --a aaa aaa ctt act ata agt ggg tgg gat tct ggg aat tag tga tca gtt tat gta tat cgc aac tag cgg gca tat ggc tat tga cat cga gaa cat tac cca tat gat aag aga ttg tat cat ttt cgt agt ctt gag tat tgg tat tac tat ata gta tgt aga tgt cga cgc tag ata gac agt cgc cca cta gag tta ccg tct ctg aat gcg gca tga tag tat cat tct ttg ttt tcg tta act gtt tgg aag atg aat ctt tgt tgt tac att taa tct cga aat tca gag tac ata tct ttg aag tat tct gat atc tat ttt ctc ctg taa aga atc ctg aag ttg cta cat tat taa gga cag aga agt att ctg cac gaa aga ctg gat cac aat ctt tat gat tca tgg taa tag tta gtt ccg acg ttg aga tgg att cgc tga gac cgg tag tgg tcg tcc gag tac acg atg tgt cgt tga ctg gat aca ggt taa ttt cca cat cga tat agt taa atg tat tgc tgg tta cga cgg gtt cgc att tat ctg tgg aag aga cgg tgt gag aat atg ttc cgg gac cac acg gag aac aga tga cgt ctc cgg tag acg tgt atc cgg ata ctc cgt atc cta ttc cac act ttg ttt tag aaa tac atg ttc tac acc ctg atg ctc ctt tga gaa gac aat aat atc ctg gag agc att cac aga ttc tat tgt gag tcg tgt tac acg atc gcg tct cta cct gat tac tat cac atc ttc cgt tac aac tta gac aag cct gta aat gat tat tgt gag atg taa agg tat ccg aac cac acg gtg tac att gtg tat tag tct tgc tat cac ata atc tga aag cgt aag ttc ccg gag gac acg ata gac aac ata gat tac ggc ttc tgt att cgt tgt ctt tac act ttc cat tgg atg gtg cat gtg gtg cta tat ctc ttc cgt tta tta tta tac atg aga gaa aca ata tat acg agt ata ata cgg acc tca tga ttt aat aat -gt agt aat cgt cgt ctt gtt act gtt tgt ttc cta ctt ctc caa tca tat aga tta ttt ttt aaa tat ttt ctt tct atc atg gat aat att tgt aat ggt tct ttc cgt aca aca tac tgt tta gat ggt agt cgc tta gct tgg tta tga tat tgc gca taa ttt ccg gag gca aat acg ata gtc tag att gac tat cga tgg tag act cta att tat tga gtg ctt tgt cga cga gtt tac ttt tat gct cca tcg ata gat gac act gtt cta tga gat cgt cgt aca tgg gaa atg aaa tgc gtt tgt ctg aat gta tgg ctt cga gat atg tgt gat acc gaa tgt ctt ctg ttc tca ata ccg tat aca agt cgg tgt ctg aga ttc gaa tct ctt tga gga gac tta tgt cac gac tac att ttt cga tga tgg aat cta tct tat cga atg ata tat ttt tca taa ata cac ttt tat agt cct cgt tta aac aga att tag tat gta gtt ccg caa atg act cgt ccc tta ata ggc agt agg cta tta tct tct tta cgt agt gat cgt cgt agg gag aga act ccg aca tct tgt aga aca acg att taa tca tag gta gag ata ctt tca gtc tgt ggt gga taa tgt cat taa cga cat ccg cct tgt ata taa tgt ttc tgt ttt caa aca cca agt cga ata ctg tct tat cgt ctt tag tcg gaa ggt tga tgt cgt atc caa tgt ata cga ggt atg agg caa cat tgt tat tgc aat tct gga agg cgg tat gaa gag gag tca ttg tat tat agt att cgt ctt tct gaa tgt cga atc tat cta gta gat acc gta gta tat tga gag agc gac ttc cat atc ctt gat tat gtt tta tga ata gat aaa gta gat gtt gtc ctt ctt cct ttt gta att tcc cgt att ttt gtt cgt gcc aat tga gta aca tta tga gaa tat gac ctg ttg cac aat cgt tct tta tgt att cca tga tgg gtg tac aat taa gat tat tac gta tcc tcg tat cgg ctc ctc gag ata aaa gag cat aca cca cac gag gac tat gtt tgg tat act gtt gaa ggt aag tgt gta acg gcg tat ttc cga ttt tcg taa ccg cgt taa tgt ttg ctc cat gat cta tta tcg cgt aga tga atc gct tct cag ctc gca tct tag tgt gac tct ttg act tgt aat aat tgc ttt cgt gga acg cgg ata tgt gtt tac agt agt aat gaa gag aag tga gtc cat cct cat cga cgc aat tag ggt cag atc ctt tag tca ata att tgt aca gaa cgt aat agt tta agc tcc cat tga att tat atc taa gat aac aca gca ata gat cgg atg att tac taa agt cat caa tgg ggt ccg tta gta tat caa aga tct tgt tat cga ttg ata gtg aat gaa tca gat agt ggt gta gag gaa tat gtc ctt ttt tat cct tgc tat caa agt tac gca tgc cgt ggc gta aca ata tct tta ata cag atg gat taa atc gtg tat tca tcg tat agc aat gta atg gag agt tac cac att tta gtc gtt tat tca gat cgc agt gtt taa taa cta att taa aca gat gag atg ata tat cca cat caa aga atg cga gat aca tat gac aga cat tat tga cag aaa tgt gac ctt cat tat cac cgt cgt cca taa atg cgt tag gta cgt acc aca tac tat cgt taa cga tgc gca caa tct cgt cca -tt tca tcc atc ttc ata atg att tac ttt ttc ata att aga gaa aaa gat caa ggt ata aaa att aga agt gtt aga cta ta- --- --- --- --- --- --- --- --- --- --- -aa cta act tat gac tta act aac tta tga --- --- --- --- --- --- --- --- --- --- --- --- --- --- --- --- --- --- --- --- --- --- --- --- --- --- --- --- --- --- --- --- --- --- --- --- --- --- --- --- --- --- --- --- --- --- --- --- ctt aac ta- --- --- --- --- --- --- --- --- --- --- -ac tta tga ctt aac taa ctt atg act taa cta act ta- tga ctt aac taa ctt atg act taa cta act tat gac tta act aac tta tga cta tta act cat ttc aag aaa ggt ggg tgg ata gaa ctc tat atg aca gct tgt gaa aca att aga tcc cta att tct aat gga agt ttt gat agg aga ttg tca tca gtt gat aca ttg ttt att atc tca tct att aga gca cgt ctg ttt aga gct tta gtg acc tgc tcg gtt act tct gtg taa atc ttg aat cct tta gtg ata cac tgt gtc aaa act gga tgt tta gaa tac cta tgt aga ata tgg gaa gca tgc ttg ttt ttg tct cta tta tag att aac tca tac atg gtt gta tta tga att ttc atc tgc cta atg tac tcc aat tct tgt tta caa tca att ata taa tca aag agt gat gat gca tac aca tta caa agt gaa taa tct acc atc ata aaa tac ttg ata cag agc ttt atc aca tca tgg ttt tca att gta tta tta agt ata gct aat ttt ata cag tca ata gac aat ggt tct cta agc aat att tct aat att tta aga tgt gct tcc cta cgg gcg atg aca gat ccc cta tcc acg gcc acg tca aga cat gta tat cca tta ctc att act gcg ttg aca ttt gct cca ttt tct aat agc cat gat act aaa tct ata taa cct gca tag ata gcg cga taa agc aag gtc ctt cca cca gca tct agt tga ttg ata tct tca ata tat ggg ata caa agc tta taa att tct aat act gtg ggt tca tct aca agg aat ccc cta gta tac tga att att tta tat aga tct aat tta aca tca ttt tca tct ggg ata cca caa ttc aaa ata aac tca aca aca cta ctt tcc ttt tta cat att ccc cta aaa tag gca ttc aag cat tct att tta tat att aca gcc cca tga tct acc ata aga tca aca atg tct att tct aca tat gca tta gat aga tag taa agt aag aga tct tgc aca gaa tta caa ttc tta ata att ata gag aaa ata tct tcc ata taa ttc ttt gac act aat gca gat ata ata tct tta tat gta ata tat gca aac agt cta tct act ata tac tga tca ata tta tct cta tga atc cta aaa taa tca tac aga aca tct aca gga tca caa att ggt tca agg aga aat cta tca aat att ttc ctg tca aca act ggt tct aga aca taa cag tca aca cct aat cca tgt ttt tta taa tca tct acc aaa gat aat gac caa aga tcg agg tcg tcg tga aac tgc tca tcg aca gcc atg aaa tct acc gac tcc atg gtg cga atc gca ctg tct tat tcg cca ttg att ttc a-t ttt tta taa tta tgt aca tgt ttt cct tct att ctc aag agt cta caa aaa tat a-- -tt ttt tcg ata tct aag tac taa gtt ttt tta ctg ttt ttg tta ctg tct tcc att ctt cta act aaa gat ctg aga taa att ata caa tct tcg cta tcg aac cat ttt tgt agt cta aag cct gaa gta att aac caa ctg ttt tta tta gtg gct ttt ttc gat cta tcc tcg tc- --- ctc tca tca tcc tta ta- --- --- --- --- --- --- tta tta tca tta tta tca tag tct att aaa cac aaa tca tct --- --- --- --- --- --- --- --- --- --- --- --- --- --- --- --- --- --- --- --- --- --- --- --- --- --- --a cgt tta taa c-- --- --- --- --- --- --- --- --a aca ttc --- --- --- --- -tc att att aat tag ttc tgt ag- aat atc ttt aat aat ttg gct ata --- --- -ca tct gtt --- --- --- --- --- --- --- --- --- --- --- --- --- --- --- --- --- --- --c aat act --- --- --- --- --- --- --- --a tct att gat gat ttc ttt t-- --- --- --- --- --- --- --- --- --- --- --t taa gac t-- --- --- --- --- --- --- --- --- --- --- --- --- --- --- --- --- --- --- --- --- --- --- --- --- --- --- --- --- --- --- --- --- --- --- --- --- --- --- --- taa act agt --- --- --- --- --- --- --- --- --- --- --- --- --- --- --- --- --- --- --- --- --- --- --- --- --- --- --- --- --- --- --- --- --- --- --- --- --- --- --- --- --- --- --- --- --- --- --- --- --- --- --- --- --- --- --- --- --- --- --- --- --- --- --- --- --- --- --- --- --- --- --- --- --- --- --- --- --- --- --- --- --- --- --- --- --- --- --- --- --- --- --- --- --- --- --- --- --- --- --- --- --- --- --- --- --- --- --- --- --- --t atg gta atg acg atg aaa --- -tc gag tag ta- --- --a ctt cta ata aag act tga ta- --- --t cat tat cat atg ttt gat cg- --- --- --- --- --- --- --t cat agt taa tag tgt g-- --- --- --- --- --- --- --- --- --- --- --- --- --- --- --- --- --- --- --- -gc taa atg gta ctg tta ata agt tta t-- --- --- --- --- --- --- --- --- --- --- --- --- --- --- --- --- --a gac aat atc ata gta ttt tct ttc cag aat --- --- --- --t aga tta ttt ttt taa ata ctg atc ctc aca att ccg tga tgt agc agt agt tgg t-- --- --- --- --- --- --- --- --- --- --- --- --- --- --- --- --- --- --- --- --- --- --- --- --- --- --- --- -gc atg gtc tat atc gt- --- --- --- --- --- --- --- --- --- --- --- --- --- --- --- --- --- --- --- --- --- --- --- --- --- --- --- --- --- --- --- --- --- --- --- --- --- --- --- --- --- --- --- taa aat gta tca tat ata ata gtt ttc tga cgt gga gta cag aat ttt cga --- --- --- --- --- --- --- --- --- --- --- --- --- --- --- --- --- --- --- --- --- --- --- --- --- --- --- --- --- --- --- --- --- --- --- --- --- --- --- --- --- --- --- --t taa tga gtt cat ggt aag gaa ggg caa atg cct --- gta tat aat ata cat aag tta a-- --- --- --- --- --- --- --- --- --- --- --- --- --t agt ttt tta tca tat tt- --- --- --- --- --- --- --- --t cta ata cca taa taa aaa t-- --- --- --- --- --- --- --- --- --- --- --- --- --- --- --- --- --- --- --- --- --- --- --- --- --t atc att att gcg ttt g-g tag tt- --- --- --- --- ctg ccc ta- --- --- --- --- --- --- --- --t cat cta tat cac tgt cac tct c-- --- --g ctc tca cta tat ctt cta aaa tta caa --- -ac aac --- --- --- --- --- --- --- --- --- --- --- --- --- --- --- --- --- --- --- --- --- --- --- --- --- --- --- --- --- --- --- --- --- --- --- --- tgg ata ttc gat --- --- --- --- --- --- --- --- --- --- --- --- --- --- --- --- --- --- --- --- --- --- --- --- --- --- --a aca gca ttt gtg t-- --- --- --- --- --- --- --- --- --- --- --- --- --- --- --- --- -

>ON619835.1_Monkeypox_virus_isolate_MPXV_UK_2022_1_complete_genome

--- --- --- --- --- --- --- --- --- --- --- --- --- --- --- --- --- --- --- --- --- --- --- --- --- --- --- --- --- --a ttt tac tat ttt att tag tgt cta gaa aaa aat gtg tga ccc acg acc gta gga aac tct aga ggg taa gaa aaa tca atc gtt tat aga gac cat cag aaa gag gtt taa tat ttt tgt gag acc tat cga a-- --- --- --- --- --- --- --- --- --- --- --- --- --- --- --- --- --- --- --- --- --- --- --- --- --- --- --- --- --- --- --- --- --- --- --- --- --- --- --- --- --- --- --- --- --- --- --- --- --- --- --- --- --- --- --- --- --- --- --- --- --- --- --- --- --- --- --- --- --- --- --- --- --- --- --- --- --- --- --- --- --- --- --- --- --- --- --- --- --- --- --- --- --- --- --- --- --- --- --- --- --- --- --- --- --- --- --- --- --- --- --- --- --- --- --- --- --- --- --- --- --- --- --- --- --- --- --- --- --- --- --- --- --- --- --- --- --- --- --- --- --- --- --- --- --- --- --- --- --- --- --- --- --- --- --- --- --- --- --- --- --- --- --- --- --- --- --- --- --- --- --- --- --- --- --- --- --- --- --- --- --- --- --- --- --- --- --- --- --- --- --- --- --- --- --- --- --- --- --- --- --- --- --- --- --- --- --- --- --- --- --- --- --- --- --- --- --- --- --- --- --- --- --- --- --- --- --- --- --- --- --- --- --- --- --- --- --- --- --- --- --- --- --- --- --- --- --- --- --- --- --- --- --- --- --- --- --- --- --- --- --- --- --- --- --- --- --- --- --- --- --- --- --- --- --- --- --- --- --- -ga gag aaa gga taa aaa ctt ttt acg act cca tca gaa aga ggt tta ata ttt ttg tga gac cca tcg aag aga gaa aga gat ggt tag tca aga tat ttt tct tag tac aaa agt caa tgt ttt aaa ata tat gga cga gaa tta att tgt ctg tat aaa aac ttg tgt gaa att atg tac tag aga aaa aac gtg agc agt gtc ccc tac atg gat ttt aca gat cat tta tat tcc aaa aat att aac tat ata cgt tta tta tat gat gtt aac gtg taa att ata aac att att tta tga tgc aat tgt ctg aca acc tag att ggt ata agg atg ttg ata agc tct acg aga ata tat tgt tgg acg tta tcg ttt acg aaa tag ttg aga cat cag aaa gag gtt taa tat ttt tgt gag a-c cat cga aga gag aaa gag aat aaa aat a-- --? ttt ttt ttt ttt tgt aaa act ttt tta tga gac --- --- -ca aga gaa tac gaa tag tga tca tat cgt atc aca tat tga aac aga aag aag aag taa cga gag gta act ttt tgt gaa tgt agt taa ata ttt ttg ttt tgc aaa ccg gaa tat agt gcc cgg t-- -ct ttt tta att cgt ggt gcg gtg tct gaa tcg ttc gat taa ccc aac tca tcc att ttc aga tga ata gag tta tcg att cag aca cat gct ttg agt ttt gtt gaa tcg atg agt gaa gta tca tcg gtt gca cct tca gat gcc gat ccg tcg aca tac ttg aat cca tcc ttg act tca agt tca gat gat tcc tca cac atg tct ccg ata cgt acg cta aac tct agg ttc ttg aca cat ttt gta tca acg atc gtt gaa ccg atg ata tct ttg taa ctc act ttc tta tgt gag atg tta gac cca agt act gga tgg gtc ttg atg tca ctg tct ttc tct tct tcg cta cat ctg atg tcg ata gac atc tca cag tct ttg atc ata gcc aga gct tct tca cgc gtg atc gcg gga gag tcc tta cct tgt ccc ggt gac acg ctg gac aat cta gta ttc aca gtg ttt cca tca gag gat tcg gag atg gat aaa atc ttt ggg cat ttg gtg aat cca aag ttc atg tta aga ccc gca ccg acg ata gtg taa taa gtg gtg gga tct cct ttt aca act --- --- --- --- --- --- tct tcg gat acc tca tca tct tcg gtc tct gta act tcc gtt acg gat tga caa atc tta tca ttg gtc ggt gtt tgg tct tgc ttt gtg act ttg ata ata aca tcg att ccc ata tga tgt ttg ttt tct tct tca gta cac gag gat gaa gat tgt tga aga cta gta ggc ata gca gct gcc act agg cac atg cat gcc agg aca ata tat tgt ttc atg att gct att gat tga tta ctg ttc tag atg att cta ctt tct tac cat ata ata aat tag aat ata ttt tct act ttt acg aga aat taa tta ttg tat tta tta ttt ata ggt --a aaa aaa ctt act ata agt ggg tgg gat tct ggg aat tag tga tca gtt tat gta tat cgc aac tag cgg gca tat ggc tat tga cat cga gaa cat tac cca tat gat aag aga ttg tat cat ttt cgt agt ctt gag tat tgg tat tac tat ata gta tgt aga tgt cga cgc tag ata gac agt cgc cca cta gag tta ccg tct ctg aat gcg gca tga tag tat cat tct ttg ttt tcg tta act gtt tgg aag atg aat ctt tgt tgt tac att taa tct cga aat tca gag tac ata tct ttg aag tat tct gat atc tat ttt ctc ctg taa aga atc ctg aag ttg cta cat tat taa gga cag aga agt att ctg cac gaa aga ctg gat cac aat ctt tat gat tca tgg taa tag tta gtt ccg acg ttg aga tgg att cgc tga gac cgg tag tgg tcg tcc gag tac acg atg tgt cgt tga ctg gat aca ggt taa ttt cca cat cga tat agt taa atg tat tgc tgg tta cga cgg gtt cgc att tat ctg tgg aag aga cgg tgt gag aat atg ttc cgg gac cac acg gag aac aga tga cgt ctc cgg tag acg tgt atc cgg ata ctc cgt atc cta ttc cac act ttg ttt tag aaa tac atg ttc tac acc ctg atg ctc ctt tga gaa gac aat aat atc ctg gag agc att cac aga ttc tat tgt gag tcg tgt tac acg atc gcg tct cta cct gat tac tat cac atc ttc cgt tac aac tta gac aag cct gta aat gat tat tgt gag atg taa agg tat ccg aac cac acg gtg tac att gtg tat tag tct tgc tat cac ata atc tga aag cgt aag ttc ccg gag gac acg ata gac aac ata gat tac ggc ttc tgt att cgt tgt ctt tac act ttc cat tgg atg gtg cat gtg gtg cta tat ctc ttc cgt tta tta tta tac atg aga gaa aca ata tat acg agt ata ata cgg acc tca tga ttt aat aat -gt agt aat cgt cgt ctt gtt act gtt tgt ttc cta ctt ctc caa tca tat aga tta ttt ttt aaa tat ttt ctt tct atc atg gat aat att tgt aat ggt tct ttc cgt aca aca tac tgt tta gat ggt agt cgc tta gct tgg tta tga tat tgc gca taa ttt ccg gag gca aat acg ata gtc tag att gac tat cga tgg tag act cta att tat tga gtg ctt tgt cga cga gtt tac ttt tat gct cca tcg ata gat gac act gtt cta tga gat cgt cgt aca tgg gaa atg aaa tgc gtt tgt ctg aat gta tgg ctt cga gat atg tgt gat acc gaa tgt ctt ctg ttc tca ata ccg tat aca agt cgg tgt ctg aga ttc gaa tct ctt tga gga gac tta tgt cac gac tac att ttt cga tga tgg aat cta tct tat cga atg ata tat ttt tca taa ata cac ttt tat agt cct cgt tta aac aga att tag tat gta gtt ccg caa atg act cgt ccc tta ata ggc agt agg cta tta tct tct tta cgt agt gat cgt cgt agg gag aga act ccg aca tct tgt aga aca acg att taa tca tag gta gag ata ctt tca gtc tgt ggt gga taa tgt cat taa cga cat ccg cct tgt ata taa tgt ttc tgt ttt caa aca cca agt cga ata ctg tct tat cgt ctt tag tcg gaa ggt tga tgt cgt atc caa tgt ata cga ggt atg agg caa cat tgt tat tgc aat tct gga agg cgg tat gaa gag gag tca ttg tat tat agt att cgt ctt tct gaa tgt cga atc tat cta gta gat acc gta gta tat tga gag agc gac ttc cat atc ctt gat tat gtt tta tga ata gat aaa gta gat gtt gtc ctt ctt cct ttt gta att tcc cgt att ttt gtt cgt gcc aat tga gta aca tta tga gaa tat gac ctg ttg cac aat cgt tct tta tgt att cca tga tgg gtg tac aat taa gat tat tac gta tcc tcg tat cgg ctc ctc gag ata aaa gag cat aca cca cac gag gac tat gtt tgg tat act gtt gaa ggt aag tgt gta acg gcg tat ttc cga ttt tcg taa ccg cgt taa tgt ttg ctc cat gat cta tta tcg cgt aga tga atc gct tct cag ctc gca tct tag tgt gac tct ttg act tgt aat aat tgc ttt cgt gga acg cgg ata tgt gtt tac agt agt aat gaa gag aag tga gtc cat cct cat cga cgc aat tag ggt cag atc ctt tag tca ata att tgt aca gaa cgt aat agt tta agc tcc cat tga att tat atc taa gat aac aca gca ata gat cgg atg att tac taa agt cat caa tgg ggt ccg tta gta tat caa aga tct tgt tat cga ttg ata gtg aat gaa tca gat agt ggt gta gag gaa tat gtc ctt ttt tat cct tgc tat caa agt tac gca tgc cgt ggc gta aca ata tct tta ata cag atg gat taa atc gtg tat tca tcg tat agc aat gta atg gag agt tac cac att tta gtc gtt tat tca gat cgc agt gtt taa taa cta att taa aca gat gag atg ata tat cca cat caa aga atg cga gat aca tat gac aga cat tat tga cag aaa tgt gac ctt cat tat cac cgt cgt cca taa atg cgt tag gta cgt acc aca tac tat cgt taa cga tgc gca caa tct cgt cca -tt tca tcc atc ttc ata atg att tac ttt ttc ata att aga gaa aaa gat caa ggt ata aaa att aga agt gtt aga cta ta- --- --- --- --- --- --- --- --- --- --- -aa cta act tat gac tta act aac tta tga --- --- --- --- --- --- --- --- --- --- --- --- --- --- --- --- --- --- --- --- --- --- --- --- --- --- --- --- --- --- --- --- --- --- --- --- --- --- --- --- --- --- --- --- --- --- --- --- ctt aac ta- --- --- --- --- --- --- --- --- --- --- -ac tta tga ctt aac taa ctt atg act taa cta act ta- tga ctt aac taa ctt atg act taa cta act tat gac tta act aac tta tga cta tta act cat ttc aag aaa ggt ggg tgg ata gaa ctc tat atg aca gct tgt gaa aca att aga tcc cta att tct aat gga agt ttt gat agg aga ttg tca tca gtt gat aca ttg ttt att atc tca tct att aga gca cgt ctg ttt aga gct tta gtg acc tgc tcg gtt act tct gtg taa atc ttg aat cct tta gtg ata cac tgt gtc aaa act gga tgt tta gaa tac cta tgt aga ata tgg gaa gca tgc ttg ttt ttg tct cta tta tag att aac tca tac atg gtt gta tta tga att ttc atc tgc cta atg tac tcc aat tct tgt tta caa tca att ata taa tca aag agt gat gat gca tac aca tta caa agt gaa taa tct acc atc ata aaa tac ttg ata cag agc ttt atc aca tca tgg ttt tca att gta tta tta agt ata gct aat ttt ata cag tca ata gac aat ggt tct cta agc aat att tct aat att tta aga tgt gct tcc cta cgg gcg atg aca gat ccc cta tcc acg gcc acg tca aga cat gta tat cca tta ctc att act gcg ttg aca ttt gct cca ttt tct aat agc cat gat act aaa tct ata taa cct gca tag ata gcg cga taa agc aag gtc ctt cca cca gca tct agt tga ttg ata tct tca ata tat ggg ata caa agc tta taa att tct aat act gtg ggt tca tct aca agg aat ccc cta gta tac tga att att tta tat aga tct aat tta aca tca ttt tca tct ggg ata cca caa ttc aaa ata aac tca aca aca cta ctt tcc ttt tta cat att ccc cta aaa tag gca ttc aag cat tct att tta tat att aca gcc cca tga tct acc ata aga tca aca atg tct att tct aca tat gca tta gat aga tag taa agt aag aga tct tgc aca gaa tta caa ttc tta ata att ata gag aaa ata tct tcc ata taa ttc ttt gac act aat gca gat ata ata tct tta tat gta ata tat gca aac agt cta tct act ata tac tga tca ata tta tct cta tga atc cta aaa taa tca tac aga aca tct aca gga tca caa att ggt tca agg aga aat cta tca aat att ttc ctg tca aca act ggt tct aga aca taa cag tca aca cct aat cca tgt ttt tta taa tca tct acc aaa gat aat gac caa aga tcg agg tcg tcg tga aac tgc tca tcg aca gcc atg aaa tct acc gac tcc atg gtg cga atc gca ctg tct tat tcg cca ttg att ttc a-t ttt tta taa tta tgt aca tgt ttt cct tct att ctc aag agt cta caa aaa tat a-- -tt ttt tcg ata tct aag tac taa gtt ttt tta ctg ttt ttg tta ctg tct tcc att ctt cta act aaa gat ctg aga taa att ata caa tct tcg cta tcg aac cat ttt tgt agt cta aag cct gaa gta att aac caa ctg ttt tta tta gtg gct ttt ttc gat cta tcc tcg tc- --- ctc tca tca tcc tta ta- --- --- --- --- --- --- tta tta tca tta tta tca tag tct att aaa cac aaa tca tct --- --- --- --- --- --- --- --- --- --- --- --- --- --- --- --- --- --- --- --- --- --- --- --- --- --- --a cgt tta taa c-- --- --- --- --- --- --- --- --a aca ttc --- --- --- --- -tc att att aat tag ttc tgt ag- aat atc ttt aat aat ttg gct ata --- --- -ca tct gtt --- --- --- --- --- --- --- --- --- --- --- --- --- --- --- --- --- --- --c aat act --- --- --- --- --- --- --- --a tct att gat gat ttc ttt t-- --- --- --- --- --- --- --- --- --- --- --t taa gac t-- --- --- --- --- --- --- --- --- --- --- --- --- --- --- --- --- --- --- --- --- --- --- --- --- --- --- --- --- --- --- --- --- --- --- --- --- --- --- --- taa act agt --- --- --- --- --- --- --- --- --- --- --- --- --- --- --- --- --- --- --- --- --- --- --- --- --- --- --- --- --- --- --- --- --- --- --- --- --- --- --- --- --- --- --- --- --- --- --- --- --- --- --- --- --- --- --- --- --- --- --- --- --- --- --- --- --- --- --- --- --- --- --- --- --- --- --- --- --- --- --- --- --- --- --- --- --- --- --- --- --- --- --- --- --- --- --- --- --- --- --- --- --- --- --- --- --- --- --- --- --- --t atg gta atg acg atg aaa --- -tc gag tag ta- --- --a ctt cta ata aag act tga ta- --- --t cat tat cat atg ttt gat cg- --- --- --- --- --- --- --t cat agt taa tag tgt g-- --- --- --- --- --- --- --- --- --- --- --- --- --- --- --- --- --- --- --- -gc taa atg gta ctg tta ata agt tta t-- --- --- --- --- --- --- --- --- --- --- --- --- --- --- --- --- --a gac aat atc ata gta ttt tct ttc cag aat --- --- --- --t aga tta ttt ttt taa ata ctg atc ctc aca att ccg tga tgt agc agt agt tgg t-- --- --- --- --- --- --- --- --- --- --- --- --- --- --- --- --- --- --- --- --- --- --- --- --- --- --- --- -gc atg gtc tat atc gt- --- --- --- --- --- --- --- --- --- --- --- --- --- --- --- --- --- --- --- --- --- --- --- --- --- --- --- --- --- --- --- --- --- --- --- --- --- --- --- --- --- --- --- taa aat gta tca tat ata ata gtt ttc tga cgt gga gta cag aat ttt cga --- --- --- --- --- --- --- --- --- --- --- --- --- --- --- --- --- --- --- --- --- --- --- --- --- --- --- --- --- --- --- --- --- --- --- --- --- --- --- --- --- --- --- --t taa tga gtt cat ggt aag gaa ggg caa atg cct --- gta tat aat ata cat aag tta a-- --- --- --- --- --- --- --- --- --- --- --- --- --t agt ttt tta tca tat tt- --- --- --- --- --- --- --- --t cta ata cca taa taa aaa t-- --- --- --- --- --- --- --- --- --- --- --- --- --- --- --- --- --- --- --- --- --- --- --- --- --t atc att att gcg ttt g-g tag tt- --- --- --- --- ctg ccc ta- --- --- --- --- --- --- --- --t cat cta tat cac tgt cac tct c-- --- --g ctc tca cta tat ctt cta aaa tta caa --- -ac aac --- --- --- --- --- --- --- --- --- --- --- --- --- --- --- --- --- --- --- --- --- --- --- --- --- --- --- --- --- --- --- --- --- --- --- --- tgg ata ttc gat --- --- --- --- --- --- --- --- --- --- --- --- --- --- --- --- --- --- --- --- --- --- --- --- --- --- --a aca gca ttt gtg t-- --- --- --- --- --- --- --- --- --- --- --- --- --- --- --- --- -

>ON645312.1_Monkeypox_virus_isolate_MPXV_GSTT_Patient1_partial_genome

--- --- --- --- --- --- --- --- --- --- --- --- --- --- --- --- --- --- --- --- --- --- --- --- --- --- --- --- --t taa ttt tac tat ttt att tag tgt cta gaa aaa aat gtg tga ccc acg acc gta gga aac tct aga ggg taa gaa aaa tca atc gtt tat aga gac cat cag aaa gag gtt taa tat ttt tgt gag acc tat cga a-- --- --- --- --- --- --- --- --- --- --- --- --- --- --- --- --- --- --- --- --- --- --- --- --- --- --- --- --- --- --- --- --- --- --- --- --- --- --- --- --- --- --- --- --- --- --- --- --- --- --- --- --- --- --- --- --- --- --- --- --- --- --- --- --- --- --- --- --- --- --- --- --- --- --- --- --- --- --- --- --- --- --- --- --- --- --- --- --- --- --- --- --- --- --- --- --- --- --- --- --- --- --- --- --- --- --- --- --- --- --- --- --- --- --- --- --- --- --- --- --- --- --- --- --- --- --- --- --- --- --- --- --- --- --- --- --- --- --- --- --- --- --- --- --- --- --- --- --- --- --- --- --- --- --- --- --- --- --- --- --- --- --- --- --- --- --- --- --- --- --- --- --- --- --- --- --- --- --- --- --- --- --- --- --- --- --- --- --- --- --- --- --- --- --- --- --- --- --- --- --- --- --- --- --- --- --- --- --- --- --- --- --- --- --- --- --- --- --- --- --- --- --- --- --- --- --- --- --- --- --- --- --- --- --- --- --- --- --- --- --- --- --- --- --- --- --- --- --- --- --- --- --- --- --- --- --- --- --- --- --- --- --- --- --- --- --- --- --- --- --- --- --- --- --- --- --- --- --- --- -ga gag aaa gga taa aaa ctt ttt acg act cca tca gaa aga ggt tta ata ttt ttg tga gac cca tcg aag aga gaa aga gat ggt tag tca aga tat ttt tct tag tac aaa agt caa tgt ttt aaa ata tat gga cga gaa tta att tgt ctg tat aaa aac ttg tgt gaa att atg tac tag aga aaa aac gtg agc agt gtc ccc tac atg gat ttt aca gat cat tta tat tcc aaa aat att aac tat ata cgt tta tta tat gat gtt aac gtg taa att ata aac att att tta tga tgc aat tgt ctg aca acc tag att ggt ata agg atg ttg ata agc tct acg aga ata tat tgt tgg acg tta tcg ttt acg aaa tag ttg aga cat cag aaa gag gtt taa tat ttt tgt gag a-c cat cga aga gag aaa gag aat aaa aat a-- --- --- --- ttt ttt tgt aaa ac- ttt tta tga gac --- --- -ca aga gaa tac gaa tag tga tca tat cgt atc aca tat tga aac aga aag aag aag taa cga gag gta act ttt tgt gaa tgt agt taa ata ttt ttg ttt tgc aaa ccg gaa tat agt gcc cgg t-- -ct ttt tta att cgt ggt gcg gtg tct gaa tcg ttc gat taa ccc aac tca tcc att ttc aga tga ata gag tta tcg att cag aca cat gct ttg agt ttt gtt gaa tcg atg agt gaa gta tca tcg gtt gca cct tca gat gcc gat ccg tcg aca tac ttg aat cca tcc ttg act tca agt tca gat gat tcc tca cac atg tct ccg ata cgt acg cta aac tct agg ttc ttg aca cat ttt gta tca acg atc gtt gaa ccg atg ata tct ttg taa ctc act ttc tta tgt gag atg tta gac cca agt act gga tgg gtc ttg atg tca ctg tct ttc tct tct tcg cta cat ctg atg tcg ata gac atc tca cag tct ttg atc ata gcc aga gct tct tca cgc gtg atc gcg gga gag tcc tta cct tgt ccc ggt gac acg ctg gac aat cta gta ttc aca gtg ttt cca tca gag gat tcg gag atg gat aaa atc ttt ggg cat ttg gtg aat cca aag ttc atg tta aga ccc gca ccg acg ata gtg taa taa gtg gtg gga tct cct ttt aca act --- --- --- --- --- --- tct tcg gat acc tca tca tct tcg gtc tct gta act tcc gtt acg gat tga caa atc tta tca ttg gtc ggt gtt tgg tct tgc ttt gtg act ttg ata ata aca tcg att ccc ata tga tgt ttg ttt tct tct tca gta cac gag gat gaa gat tgt tga aga cta gta ggc ata gca gct gcc act agg cac atg cat gcc agg aca ata tat tgt ttc atg att gct att gat tga tta ctg ttc tag atg att cta ctt tct tac cat ata ata aat tag aat ata ttt tct act ttt acg aga aat taa tta ttg tat tta tta ttt ata ggt --a aaa aaa ctt act ata agt ggg tgg gat tct ggg aat tag tga tca gtt tat gta tat cgc aac tag cgg gca tat ggc tat tga cat cga gaa cat tac cca tat gat aag aga ttg tat cat ttt cgt agt ctt gag tat tgg tat tac tat ata gta tgt aga tgt cga cgc tag ata gac agt cgc cca cta gag tta ccg tct ctg aat gcg gca tga tag tat cat tct ttg ttt tcg tta act gtt tgg aag atg aat ctt tgt tgt tac att taa tct cga aat tca gag tac ata tct ttg aag tat tct gat atc tat ttt ctc ctg taa aga atc ctg aag ttg cta cat tat taa gga cag aga agt att ctg cac gaa aga ctg gat cac aat ctt tat gat tca tgg taa tag tta gtt ccg acg ttg aga tgg att cgc tga gac cgg tag tgg tcg tcc gag tac acg atg tgt cgt tga ctg gat aca ggt taa ttt cca cat cga tat agt taa atg tat tgc tgg tt- --a cgg gtt cgc att tat ctg tgg aag aga cgg tgt gag aat atg ttc cgg gac cac acg gag aac aga tga cgt ctc cgg tag acg tgt atc cgg ata ctc cgt atc cta ttc cac act ttg ttt tag aaa tac atg ttc tac acc ctg atg ctc ctt tga gaa gac aat aat atc ctg gag agc att cac aga ttc tat tgt gag tcg tgt tac acg atc gcg tct cta cct gat tac tat cac atc ttc cgt tac aac tta gac aag cct gta aat gat tat tgt gag atg taa agg tat ccg aac cac acg gtg tac att gtg tat tag tct tgc tat cac ata atc tga aag cgt aag ttc ccg gag gac acg ata gac aac ata gat tac ggc ttc tgt att cgt tgt ctt tac act ttc cat tgg atg gtg cat gtg gtg cta tat ctc ttc cgt tta tta tta tac atg aga gaa aca ata tat acg agt ata ata cgg acc tca tga ttt aat aat -gt agt aat cgt cgt ctt gtt act gtt tgt ttc cta ctt ctc caa tca tat aga tta ttt ttt aaa tat ttt ctt tct atc atg gat aat att tgt aat ggt tct ttc cgt aca aca tac tgt tta gat ggt agt cgc tta gct tgg tta tga tat tgc gca taa ttt ccg gag gca aat acg ata gtc tag att gac tat cga tgg tag act cta att tat tga gtg ctt tgt cga cga gtt tac ttt tat gct cca tcg ata gat gac act gtt cta tga gat cgt cgt aca tgg gaa atg aaa tgc gtt tgt ctg aat gta tgg ctt cga gat atg tgt gat acc gaa tgt ctt ctg ttc tca ata ccg tat aca agt cgg tgt ctg aga ttc gaa tct ctt tga gga gac tta tgt cac gac tac att ttt cga tga tgg aat cta tct tat cga atg ata tat ttt tca taa ata cac ttt tat agt cct cgt tta aac aga att tag tat gta gtt ccg caa atg act cgt ccc tta ata ggc agt agg cta tta tct tct tta cgt agt gat cgt cgt agg gag aga act ccg aca tct tgt aga aca acg att taa tca tag gta gag ata ctt tca gtc tgt ggt gga taa tgt cat taa cga cat ccg cct tgt ata taa tgt ttc tgt ttt caa aca cca agt cga ata ctg tct tat cgt ctt tag tcg gaa ggt tga tgt cgt atc caa tgt ata cga ggt atg agg caa cat tgt tat tgc aat tct gga agg cgg tat gaa gag gag tca ttg tat tat agt att cgt ctt tct gaa tgt cga atc tat cta gta gat acc gta gta tat tga gag agc gac ttc cat atc ctt gat tat gtt tta tga ata gat aaa gta gat gtt gtc ctt ctt cct ttt gta att tcc cgt att ttt gtt cgt gcc aat tga gta aca tta tga gaa tat gac ctg ttg cac aat cgt tct tta tgt att cca tga tgg gtg tac aat taa gat tat tac gta tcc tcg tat cgg ctc ctc gag ata aaa gag cat aca cca cac gag gac tat gtt tgg tat act gtt gaa ggt aag tgt gta acg gcg tat ttc cga ttt tcg taa ccg cgt taa tgt ttg ctc cat gat cta tta tcg cgt aga tga atc gct tct cag ctc gca tct tag tgt gac tct ttg act tgt aat aat tgc ttt cgt gga acg cgg ata tgt gtt tac agt agt aat gaa gag aag tga gtc cat cct cat cga cgc aat tag ggt cag atc ctt tag tca ata att tgt aca gaa cgt aat agt tta agc tcc cat tga att tat atc taa gat aac aca gca ata gat cgg atg att tac taa agt cat caa tgg ggt ccg tta gta tat caa aga tct tgt tat cga ttg ata gtg aat gaa tca gat agt ggt gta gag gaa tat gtc ctt ttt tat cct tgc tat caa agt tac gca tgc cgt ggc gta aca ata tct tta ata cag atg gat taa atc gtg tat tca tcg tat agc aat gta atg gag agt tac cac att tta gtc gtt tat tca gat cgc agt gtt taa taa cta att taa aca gat gag atg ata tat cca cat caa aga atg cga gat aca tat gac aga cat tat tga cag aaa tgt gac ctt cat tat cac cgt cgt cca taa atg cgt tag gta cgt acc aca tac tat cgt taa cga tgc gca caa tct cgt cca -tt tca tcc atc ttc ata atg att tac ttt ttc ata att aga gaa aaa gat caa ggt ata aaa att aga agt gtt aga cta ta- --- --- --- --- --- --- --- --- --- --- -aa cta act tat ga- --- --- --- --- --- --- --- --- --- --- --- --- --- --- --- --- --- --- --- --- --- --- --- --- --- --- --- --- --- --- --- --- --- --- --- --- --- --- --- --- --- --- --- --- --- --- --- --- --- --- --- --- --- --- --- --- --- --- --- --- --- --- --- --- --- --- --- --- --- --- --- --- --- --- --- --- --- --- --- --- --- --- --- --- --- --- -aa cta act tat gac tta act aac tta tga cta tta act cat ttc aag aaa ggt ggg tgg ata gaa ctc tat atg aca gct tgt gaa aca att aga tcc cta att tct aat gga agt ttt gat agg aga ttg tca tca gtt gat aca ttg ttt att atc tca tct att aga gca cgt ctg ttt aga gct tta gtg acc tgc tcg gtt act tct gtg taa atc ttg aat cct tta gtg ata cac tgt gtc aaa act gga tgt tta gaa tac cta tgt aga ata tgg gaa gca tgc ttg ttt ttg tct cta tta tag att aac tca tac atg gtt gta tta tga att ttc atc tgc cta atg tac tcc aat tct tgt tta caa tca att ata taa tca aag agt gat gat gca tac aca tta caa agt gaa taa tct acc atc ata aaa tac ttg ata cag agc ttt atc aca tca tgg ttt tca att gta tta tta agt ata gct aat ttt ata cag tca ata gac aat ggt tct cta agc aat att tct aat att tta aga tgt gct tcc cta cgg gcg atg aca gat ccc cta tcc acg gcc acg tca aga cat gta tat cca tta ctc att act gcg ttg aca ttt gct cca ttt tct aat agc cat gat act aaa tct ata taa cct gca tag ata gcg cga taa agc aag gtc ctt cca cca gca tct agt tga ttg ata tct tca ata tat ggg ata caa agc tta taa att tct aat act gtg ggt tca tct aca agg aat ccc cta gta tac tga att att tta tat aga tct aat tta aca tca ttt tca tct ggg ata cca caa ttc aaa ata aac tca aca aca cta ctt tcc ttt tta cat att ccc cta aaa tag gca ttc aag cat tct att tta tat att aca gcc cca tga tct acc ata aga tca aca atg tct att tct aca tat gca tta gat aga tag taa agt aag aga tct tgc aca gaa tta caa ttc tta ata att ata gag aaa ata tct tcc ata taa ttc ttt gac act aat gca gat ata ata tct tta tat gta ata tat gca aac agt cta tct act ata tac tga tca ata tta tct cta tga atc cta aaa taa tca tac aga aca tct aca gga tca caa att ggt tca agg aga aat cta tca aat att ttc ctg tca aca act ggt tct aga aca taa cag tca aca cct aat cca tgt ttt tta taa tca tct acc aaa gat aat gac caa aga tcg agg tcg tcg tga aac tgc tca tcg aca gcc atg aaa tct acc gac tcc atg gtg cga atc gca ctg tct tat tcg cca ttg att ttc a-t ttt tta taa tta tgt aca tgt ttt cct tct att ctc aag agt cta caa aaa tat a-- -tt ttt tcg ata tct aag tac taa gtt ttt tta ctg ttt ttg tta ctg tct tcc att ctt cta act aaa gat ctg aga taa att ata caa tct tcg cta tcg aac cat ttt tgt agt cta aag cct gaa gta att aac caa ctg ttt tta tta gtg gct ttt ttc gat cta tcc tcg tc- --- --c tca tca tcc tta ta- --- --- --- --- --- --- tta tta tca tta tta tca tag tct att aaa cac aaa tca tct --- --- --- --- --- --- --- --- --- --- --- --- --- --- --- --- --- --- --- --- --- --- --- --- --- --- --a cgt tta taa c-- --- --- --- --- --- --- --- --a aca ttc --- --- --- --- -tc att att aat tag ttc tgt ag- aat atc ttt aat aat ttg gct ata --- --- -ca tct gtt --- --- --- --- --- --- --- --- --- --- --- --- --- --- --- --- --- --- --c aat act --- --- --- --- --- --- --- --a tct att gat gat ttc ttt t-- --- --- --- --- --- --- --- --- --- --- --t taa gac t-- --- --- --- --- --- --- --- --- --- --- --- --- --- --- --- --- --- --- --- --- --- --- --- --- --- --- --- --- --- --- --- --- --- --- --- --- --- --- --- taa act agt --- --- --- --- --- --- --- --- --- --- --- --- --- --- --- --- --- --- --- --- --- --- --- --- --- --- --- --- --- --- --- --- --- --- --- --- --- --- --- --- --- --- --- --- --- --- --- --- --- --- --- --- --- --- --- --- --- --- --- --- --- --- --- --- --- --- --- --- --- --- --- --- --- --- --- --- --- --- --- --- --- --- --- --- --- --- --- --- --- --- --- --- --- --- --- --- --- --- --- --- --- --- --- --- --- --- --- --- --- --t atg gta atg acg atg aaa --- -tc gag tag ta- --- --a ctt cta ata aag act tga ta- --- --t cat tat cat atg ttt gat cg- --- --- --- --- --- --- --t cat agt taa tag tgt g-- --- --- --- --- --- --- --- --- --- --- --- --- --- --- --- --- --- --- --- -gc taa atg gta ctg tta ata agt tta t-- --- --- --- --- --- --- --- --- --- --- --- --- --- --- --- --- --a gac aat atc ata gta ttt tct ttc cag aat --- --- --- --t aga tta ttt ttt taa ata ctg atc ctc aca att ccg tga tgt agc agt agt tgg t-- --- --- --- --- --- --- --- --- --- --- --- --- --- --- --- --- --- --- --- --- --- --- --- --- --- --- --- -gc atg gtc tat atc gt- --- --- --- --- --- --- --- --- --- --- --- --- --- --- --- --- --- --- --- --- --- --- --- --- --- --- --- --- --- --- --- --- --- --- --- --- --- --- --- --- --- --- --- taa aat gta tca tat ata ata gtt ttc tga cgt gga gta cag aat ttt cga --- --- --- --- --- --- --- --- --- --- --- --- --- --- --- --- --- --- --- --- --- --- --- --- --- --- --- --- --- --- --- --- --- --- --- --- --- --- --- --- --- --- --- --t taa tga gtt cat ggt aag gaa ggg caa atg cct --- gta tat aat ata cat aag tta a-- --- --- --- --- --- --- --- --- --- --- --- --- --t agt ttt tta tca tat tt- --- --- --- --- --- --- --- --t cta ata cca taa taa aaa t-- --- --- --- --- --- --- --- --- --- --- --- --- --- --- --- --- --- --- --- --- --- --- --- --- --t atc att att gcg ttt g-g tag tt- --- --- --- --- ctg ccc ta- --- --- --- --- --- --- --- --t cat cta tat cac tgt ca- --- --- --- --- ctc tca cta tat ctt cta aaa tta caa --- -ac aac --- --- --- --- --- --- --- --- --- --- --- --- --- --- --- --- --- --- --- --- --- --- --- --- --- --- --- --- --- --- --- --- --- --- --- --- tgg ata ttc gat --- --- --- --- --- --- --- --- --- --- --- --- --- --- --- --- --- --- --- --- --- --- --- --- --- --- --a aca gca ttt gtg t-- --- --- --- --- --- --- --- --- --- --- --- --- --- --- --- --- -

>ON682269.2_Monkeypox_virus_isolate_MPXV/Germany/2022/RKI07_complete_genome

--- --- --- --- --- --- --- --- --- --- --- --- --- --- --- --- --- --- --- --- --- --- --- --- --- --- --- --- --- --a ttt tac tat ttt att tag tgt cta gaa aaa aat gtg tga ccc acg acc gta gga aac tct aga ggg taa gaa aaa tca atc gtt tat aga gac cat cag aaa gag gtt taa tat ttt tgt gag acc tat cga a-- --- --- --- --- --- --- --- --- --- --- --- --- --- --- --- --- --- --- --- --- --- --- --- --- --- --- --- --- --- --- --- --- --- --- --- --- --- --- --- --- --- --- --- --- --- --- --- --- --- --- --- --- --- --- --- --- --- --- --- --- --- --- --- --- --- --- --- --- --- --- --- --- --- --- --- --- --- --- --- --- --- --- --- --- --- --- --- --- --- --- --- --- --- --- --- --- --- --- --- --- --- --- --- --- --- --- --- --- --- --- --- --- --- --- --- --- --- --- --- --- --- --- --- --- --- --- --- --- --- --- --- --- --- --- --- --- --- --- --- --- --- --- --- --- --- --- --- --- --- --- --- --- --- --- --- --- --- --- --- --- --- --- --- --- --- --- --- --- --- --- --- --- --- --- --- --- --- --- --- --- --- --- --- --- --- --- --- --- --- --- --- --- --- --- --- --- --- --- --- --- --- --- --- --- --- --- --- --- --- --- --- --- --- --- --- --- --- --- --- --- --- --- --- --- --- --- --- --- --- --- --- --- --- --- --- --- --- --- --- --- --- --- --- --- --- --- --- --- --- --- --- --- --- --- --- --- --- --- --- --- --- --- --- --- --- --- --- --- --- --- --- --- --- --- --- --- --- --- --- -ga gag aaa gga taa aaa ctt ttt acg act cca tca gaa aga ggt tta ata ttt ttg tga gac cca tcg aag aga gaa aga gat ggt tag tca aga tat ttt tct tag tac aaa agt caa tgt ttt aaa ata tat gga cga gaa tta att tgt ctg tat aaa aac ttg tgt gaa att atg tac tag aga aaa aac gtg agc agt gtc ccc tac atg gat ttt aca gat cat tta tat tcc aaa aat att aac tat ata cgt tta tta tat gat gtt aac gtg taa att ata aac att att tta tga tgc aat tgt ctg aca acc tag att ggt ata agg atg ttg ata agc tct acg aga ata tat tgt tgg acg tta tcg ttt acg aaa tag ttg aga cat cag aaa gag gtt taa tat ttt tgt gag a-c cat cga aga gag aaa gag aat aaa aat a-- --t ttt ttt ttt ttt tgt aaa act ttt tta tga gac --- --- -ca aga gaa tac gaa tag tga tca tat cgt atc aca tat tga aac aga aag aag aag taa cga gag gta act ttt tgt gaa tgt agt taa ata ttt ttg ttt tgc aaa ccg gaa tat agt gcc cgg t-- -ct ttt tta att cgt ggt gcg gtg tct gaa tcg ttc gat taa ccc aac tca tcc att ttc aga tga ata gag tta tcg att cag aca cat gct ttg agt ttt gtt gaa tcg atg agt gaa gta tca tcg gtt gca cct tca gat gcc gat ccg tcg aca tac ttg aat cca tcc ttg act tca agt tca gat gat tcc tca cac atg tct ccg ata cgt acg cta aac tct agg ttc ttg aca cat ttt gta tca acg atc gtt gaa ccg atg ata tct ttg taa ctc act ttc tta tgt gag atg tta gac cca agt act gga tgg gtc ttg atg tca ctg tct ttc tct tct tcg cta cat ctg atg tcg ata gac atc tca cag tct ttg atc ata gcc aga gct tct tca cgc gtg atc gcg gga gag tcc tta cct tgt ccc ggt gac acg ctg gac aat cta gta ttc aca gtg ttt cca tca gag gat tcg gag atg gat aaa atc ttt ggg cat ttg gtg aat cca aag ttc atg tta aga ccc gca ccg acg ata gtg taa taa gtg gtg gga tct cct ttt aca act --- --- --- --- --- --- tct tcg gat acc tca tca tct tcg gtc tct gta act tcc gtt acg gat tga caa atc tta tca ttg gtc ggt gtt tgg tct tgc ttt gtg act ttg ata ata aca tcg att ccc ata tga tgt ttg ttt tct tct tca gta cac gag gat gaa gat tgt tga aga cta gta ggc ata gca gct gcc act agg cac atg cat gcc agg aca ata tat tgt ttc atg att gct att gat tga tta ctg ttc tag atg att cta ctt tct tac cat ata ata aat tag aat ata ttt tct act ttt acg aga aat taa tta ttg tat tta tta ttt ata ggt --a aaa aaa ctt act ata agt ggg tgg gat tct ggg aat tag tga tca gtt tat gta tat cgc aac tag cgg gca tat ggc tat tga cat cga gaa cat tac cca tat gat aag aga ttg tat cat ttt cgt agt ctt gag tat tgg tat tac tat ata gta tgt aga tgt cga cgc tag ata gac agt cgc cca cta gag tta ccg tct ctg aat gcg gca tga tag tat cat tct ttg ttt tcg tta act gtt tgg aag atg aat ctt tgt tgt tac att taa tct cga aat tca gag tac ata tct ttg aag tat tct gat atc tat ttt ctc ctg taa aga atc ctg aag ttg cta cat tat taa gga cag aga agt att ctg cac gaa aga ctg gat cac aat ctt tat gat tca tgg taa tag tta gtt ccg acg ttg aga tgg att cgc tga gac cgg tag tgg tcg tcc gag tac acg atg tgt cgt tga ctg gat aca ggt taa ttt cca cat cga tat agt taa atg tat tgc tgg tta cga cgg gtt cgc att tat ctg tgg aag aga cgg tgt gag aat atg ttc cgg gac cac acg gag aac aga tga cgt ctc cgg tag acg tgt atc cgg ata ctc cgt atc cta ttc cac act ttg ttt tag aaa tac atg ttc tac acc ctg atg ctc ctt tga gaa gac aat aat atc ctg gag agc att cac aga ttc tat tgt gag tcg tgt tac acg atc gcg tct cta cct gat tac tat cac atc ttc cgt tac aac tta gac aag cct gta aat gat tat tgt gag atg taa agg tat ccg aac cac acg gtg tac att gtg tat tag tct tgc tat cac ata atc tga aag cgt aag ttc ccg gag gac acg ata gac aac ata gat tac ggc ttc tgt att cgt tgt ctt tac act ttc cat tgg atg gtg cat gtg gtg cta tat ctc ttc cgt tta tta tta tac atg aga gaa aca ata tat acg agt ata ata cgg acc tca tga ttt aat aat -gt agt aat cgt cgt ctt gtt act gtt tgt ttc cta ctt ctc caa tca tat aga tta ttt ttt aaa tat ttt ctt tct atc atg gat aat att tgt aat ggt tct ttc cgt aca aca tac tgt tta gat ggt agt cgc tta gct tgg tta tga tat tgc gca taa ttt ccg gag gca aat acg ata gtc tag att gac tat cga tgg tag act cta att tat tga gtg ctt tgt cga cga gtt tac ttt tat gct cca tcg ata gat gac act gtt cta tga gat cgt cgt aca tgg gaa atg aaa tgc gtt tgt ctg aat gta tgg ctt cga gat atg tgt gat acc gaa tgt ctt ctg ttc tca ata ccg tat aca agt cgg tgt ctg aga ttc gaa tct ctt tga gga gac tta tgt cac gac tac att ttt cga tga tgg aat cta tct tat cga atg ata tat ttt tca taa ata cac ttt tat agt cct cgt tta aac aga att tag tat gta gtt ccg caa atg act cgt ccc tta ata ggc agt agg cta tta tct tct tta cgt agt gat cgt cgt agg gag aga act ccg aca tct tgt aga aca acg att taa tca tag gta gag ata ctt tca gtc tgt ggt gga taa tgt cat taa cga cat ccg cct tgt ata taa tgt ttc tgt ttt caa aca cca agt cga ata ctg tct tat cgt ctt tag tcg gaa ggt tga tgt cgt atc caa tgt ata cga ggt atg agg caa cat tgt tat tgc aat tct gga agg cgg tat gaa gag gag tca ttg tat tat agt att cgt ctt tct gaa tgt cga atc tat cta gta gat acc gta gta tat tga gag agc gac ttc cat atc ctt gat tat gtt tta tga ata gat aaa gta gat gtt gtc ctt ctt cct ttt gta att tcc cgt att ttt gtt cgt gcc aat tga gta aca tta tga gaa tat gac ctg ttg cac aat cgt tct tta tgt att cca tga tgg gtg tac aat taa gat tat tac gta tcc tcg tat cgg ctc ctc gag ata aaa gag cat aca cca cac gag gac tat gtt tgg tat act gtt gaa ggt aag tgt gta acg gcg tat ttc cga ttt tcg taa ccg cgt taa tgt ttg ctc cat gat cta tta tcg cgt aga tga atc gct tct cag ctc gca tct tag tgt gac tct ttg act tgt aat aat tgc ttt cgt gga acg cgg ata tgt gtt tac agt agt aat gaa gag aag tga gtc cat cct cat cga cgc aat tag ggt cag atc ctt tag tca ata att tgt aca gaa cgt aat agt tta agc tcc cat tga att tat atc taa gat aac aca gca ata gat cgg atg att tac taa agt cat caa tgg ggt ccg tta gta tat caa aga tct tgt tat cga ttg ata gtg aat gaa tca gat agt ggt gta gag gaa tat gtc ctt ttt tat cct tgc tat caa agt tac gca tgc cgt ggc gta aca ata tct tta ata cag atg gat taa atc gtg tat tca tcg tat agc aat gta atg gag agt tac cac att tta gtc gtt tat tca gat cgc agt gtt taa taa cta att taa aca gat gag atg ata tat cca cat caa aga atg cga gat aca tat gac aga cat tat tga cag aaa tgt gac ctt cat tat cac cgt cgt cca taa atg cgt tag gta cgt acc aca tac tat cgt taa cga tgc gca caa tct cgt cca -tt tca tcc atc ttc ata atg att tac ttt ttc ata att aga gaa aaa gat caa ggt ata aaa att aga agt gtt aga cta ta- --- --- --- --- --- --- --- --- --- --- -aa cta act tat gac tta act aac tta tga --- --- --- --- --- --- --- --- --- --- --- --- --- --- --- --- --- --- --- --- --- --- --- --- --- --- --- --- --- --- --- --- --- --- --- --- --- --- --- --- --- --- --- --- --- --- --- --- ctt aac ta- --- --- --- --- --- --- --- --- --- --- -ac tta tga ctt aac taa ctt atg act taa cta act ta- tga ctt aac taa ctt atg act taa cta act tat gac tta act aac tta tga cta tta act cat ttc aag aaa ggt ggg tgg ata gaa ctc tat atg aca gct tgt gaa aca att aga tcc cta att tct aat gga agt ttt gat agg aga ttg tca tca gtt gat aca ttg ttt att atc tca tct att aga gca cgt ctg ttt aga gct tta gtg acc tgc tcg gtt act tct gtg taa atc ttg aat cct tta gtg ata cac tgt gtc aaa act gga tgt tta gaa tac cta tgt aga ata tgg gaa gca tgc ttg ttt ttg tct cta tta tag att aac tca tac atg gtt gta tta tga att ttc atc tgc cta atg tac tcc aat tct tgt tta caa tca att ata taa tca aag agt gat gat gca tac aca tta caa agt gaa taa tct acc atc ata aaa tac ttg ata cag agc ttt atc aca tca tgg ttt tca att gta tta tta agt ata gct aat ttt ata cag tca ata gac aat ggt tct cta agc aat att tct aat att tta aga tgt gct tcc cta cgg gcg atg aca gat ccc cta tcc acg gcc acg tca aga cat gta tat cca tta ctc att act gcg ttg aca ttt gct cca ttt tct aat agc cat gat act aaa tct ata taa cct gca tag ata gcg cga taa agc aag gtc ctt cca cca gca tct agt tga ttg ata tct tca ata tat ggg ata caa agc tta taa att tct aat act gtg ggt tca tct aca agg aat ccc cta gta tac tga att att tta tat aga tct aat tta aca tca ttt tca tct ggg ata cca caa ttc aaa ata aac tca aca aca cta ctt tcc ttt tta cat att ccc cta aaa tag gca ttc aag cat tct att tta tat att aca gcc cca tga tct acc ata aga tca aca atg tct att tct aca tat gca tta gat aga tag taa agt aag aga tct tgc aca gaa tta caa ttc tta ata att ata gag aaa ata tct tcc ata taa ttc ttt gac act aat gca gat ata ata tct tta tat gta ata tat gca aac agt cta tct act ata tac tga tca ata tta tct cta tga atc cta aaa taa tca tac aga aca tct aca gga tca caa att ggt tca agg aga aat cta tca aat att ttc ctg tca aca act ggt tct aga aca taa cag tca aca cct aat cca tgt ttt tta taa tca tct acc aaa gat aat gac caa aga tcg agg tcg tcg tga aac tgc tca tcg aca gcc atg aaa tct acc gac tcc atg gtg cga atc gca ctg tct tat tcg cca ttg att ttc a-t ttt tta taa tta tgt aca tgt ttt cct tct att ctc aag agt cta caa aaa tat a-- -tt ttt tcg ata tct aag tac taa gtt ttt tta ctg ttt ttg tta ctg tct tcc att ctt cta act aaa gat ctg aga taa att ata caa tct tcg cta tcg aac cat ttt tgt agt cta aag cct gaa gta att aac caa ctg ttt tta tta gtg gct ttt ttc gat cta tcc tcg tc- --- ctc tca tca tcc tta ta- --- --- --- --- --- --- tta tta tca tta tta tca tag tct att aaa cac aaa tca tct --- --- --- --- --- --- --- --- --- --- --- --- --- --- --- --- --- --- --- --- --- --- --- --- --- --- --a cgt tta taa c-- --- --- --- --- --- --- --- --a aca ttc --- --- --- --- -tc att att aat tag ttc tgt ag- aat atc ttt aat aat ttg gct ata --- --- -ca tct gtt --- --- --- --- --- --- --- --- --- --- --- --- --- --- --- --- --- --- --c aat act --- --- --- --- --- --- --- --a tct att gat gat ttc ttt t-- --- --- --- --- --- --- --- --- --- --- --t taa gac t-- --- --- --- --- --- --- --- --- --- --- --- --- --- --- --- --- --- --- --- --- --- --- --- --- --- --- --- --- --- --- --- --- --- --- --- --- --- --- --- taa act agt --- --- --- --- --- --- --- --- --- --- --- --- --- --- --- --- --- --- --- --- --- --- --- --- --- --- --- --- --- --- --- --- --- --- --- --- --- --- --- --- --- --- --- --- --- --- --- --- --- --- --- --- --- --- --- --- --- --- --- --- --- --- --- --- --- --- --- --- --- --- --- --- --- --- --- --- --- --- --- --- --- --- --- --- --- --- --- --- --- --- --- --- --- --- --- --- --- --- --- --- --- --- --- --- --- --- --- --- --- --t atg gta atg acg atg aaa --- -tc gag tag ta- --- --a ctt cta ata aag act tga ta- --- --t cat tat cat atg ttt gat cg- --- --- --- --- --- --- --t cat agt taa tag tgt g-- --- --- --- --- --- --- --- --- --- --- --- --- --- --- --- --- --- --- --- -gc taa atg gta ctg tta ata agt tta t-- --- --- --- --- --- --- --- --- --- --- --- --- --- --- --- --- --a gac aat atc ata gta ttt tct ttc cag aat --- --- --- --t aga tta ttt ttt taa ata ctg atc ctc aca att ccg tga tgt agc agt agt tgg t-- --- --- --- --- --- --- --- --- --- --- --- --- --- --- --- --- --- --- --- --- --- --- --- --- --- --- --- -gc atg gtc tat atc gt- --- --- --- --- --- --- --- --- --- --- --- --- --- --- --- --- --- --- --- --- --- --- --- --- --- --- --- --- --- --- --- --- --- --- --- --- --- --- --- --- --- --- --- taa aat gta tca tat ata ata gtt ttc tga cgt gga gta cag aat ttt cga --- --- --- --- --- --- --- --- --- --- --- --- --- --- --- --- --- --- --- --- --- --- --- --- --- --- --- --- --- --- --- --- --- --- --- --- --- --- --- --- --- --- --- --t taa tga gtt cat ggt aag gaa ggg caa atg cct --- gta tat aat ata cat aag tta a-- --- --- --- --- --- --- --- --- --- --- --- --- --t agt ttt tta tca tat tt- --- --- --- --- --- --- --- --t cta ata cca taa taa aaa t-- --- --- --- --- --- --- --- --- --- --- --- --- --- --- --- --- --- --- --- --- --- --- --- --- --t atc att att gcg ttt g-g tag tt- --- --- --- --- ctg ccc ta- --- --- --- --- --- --- --- --t cat cta tat cac tgt cac tct c-- --- --g ctc tca cta tat ctt cta aaa tta caa --- -ac aac --- --- --- --- --- --- --- --- --- --- --- --- --- --- --- --- --- --- --- --- --- --- --- --- --- --- --- --- --- --- --- --- --- --- --- --- tgg ata ttc gat --- --- --- --- --- --- --- --- --- --- --- --- --- --- --- --- --- --- --- --- --- --- --- --- --- --- --a aca gca ttt gtg t-- --- --- --- --- --- --- --- --- --- --- --- --- --- --- --- --- -

>KJ642616.1_Monkeypox_virus_strain_PCH_complete_genome

--- --- --- --- --- --- --- --- --- --- --- --- --- --- --- --- --- --- --- --- --- --- --- --- --- --- --- --- --- --- --- --- --- --- --- --- --- --- --- --- --- --- --- --- --- --- --- --- --- --- --- --- --- --- --- --- --- --- --- --- --- --- --- --- --- --- --- --- --- --- --- --- --- --- --- --- --- --- --- --- --- --- --- --- --- --- --- --- --- --- --- --- --- --- --- --- --- --- --- --- --- --- --- --- --- --- --- --- --- --- --- --- --- --- --- --- --- --- --- --- --- --- --- --- --- --- --- --- --- --- --- --- --- --- --- --- --- --- --- --- --- --- --- --- --- --- --- --- --- --- --- --- --- --- --- --- --- --- --- --- --- --- --- --- --- --- --- --- --- --- --- --- --- --- --- --- --- --- --- --- --- --- --- --- --- --- --- --- --- --- --- --- --- --- --- --- --- --- --- --- --- --- --- --- --- --- --- --- --- --- --- --- --- --- --- --- --- --- --- --- --- --- --- --- --- --- --- --- --- --- --- --- --- --- --- --- --- --- --- --- --- --- --- --- --- --- --- --- --- --- --- --- --- --- --- --- --- --- --- --- --- --- --- --- --- --- --- --- --- --- --- --- --- --- --- --- --- --- --- --- --- --- --- --- --- --- --- --- --- --- --- --- --- --- --- --- --- --- --- --- --- --- --- --- --- --- --- --- --- --- --- --- --- --- --- --- --- --- --- --- --- --- --- --- --- --- --- --- --- --- --- --- --- --- --- --- --- --- --- --- --- --- --- --- --- --- --- --- --- --- --- --- --- --- --- --- --- --- --- --- --- --- --- --- --- --- --- --- --- --- --- --- --- --- --- --a tcg aag aga gaa aga gat ggt tag tca aga tat ttt tct tag tac aaa agt caa tgt ttt aaa ata tat gga cga gaa tta att tgt ctg tat aaa aac ttg tgt gaa att atg tac tag aga aaa aac gtg agc agt gtc ccc tac atg gat ttt aca gat cat tta tat tcc aaa aat att aac tat ata cgt tta tta tat gat gtt aac gtg taa att ata aac att att tta tga tgc aat tgt ctg aca acc tag att ggt ata agg atg ttg ata agc tct acg aga ata tat tgt tgg acg tta tcg ttt acg aaa tag ttg aga cat cag aaa gag gtt taa tat ttt tgt gag a-c cat cga aga gag aaa gag aat aaa aat a-- --t ttt ttt ttt ttt tgt aaa act ttt tta tga gac --- --- -ca aga gaa tac gaa tag tga tca tat cgt atc aca tat tga aac aga aag aag aag taa cga gag gta act ttt tgt aaa tgt agt taa ata ttt ttg ttt tgc aaa ccg gaa tat agt gcc cgg t-- -ct ttt tta att cgt ggt gcg gtg tct gaa tcg ttc gat taa ccc aac tca tcc att ttc aga tga ata gag tta tcg att cag aca cat gct ttg agt ttt gtt gaa tcg atg agt gaa gta tca tcg gtt gca cct tca gat gcc gat ccg tcg aca tac ttg aat cca tcc ttg act tca agt tca gat gat tcc tca cac atg tct ccg ata cgt acg cta aac tct agg ttc ttg aca cat ttt gta tca acg atc gtt gaa ccg atg ata tct ttg taa ctc act ttc tta tgt gag atg tta gac cca agt act gga tgg gtc ttg atg tca ctg tct ttc tct tct tcg cta cat ctg atg tcg ata gac atc tca cag tct ttg atc ata gcc aga gct tct tca cgc gtg atc gcg gga gag tcc tta cct tgt ccc ggt gac acg ctg gac aat cta gta ttc aca gtg ttt cca tca gag gat tcg gag atg gat gaa atc ttt ggg cat ttg gtg aat cca aag ttc atg tta aga ccc gca ccg acg ata gtg taa taa gtg gtg gga tct cct ttt aca act --- --- --- --- --- --- tct tcg gat acc tca tca tct tcg gtc tct gta act tcc gtt acg gat tga caa atc tta tca ttg gtc ggt gtt tgg tct tgc ttt gtg act ttg ata ata aca tcg att ccc ata tga tgt ttg ttt tct tct tca gta cac gag gat gaa gat tgt tga aga cta gta ggc ata gca gct gcc act agg cac atg cat gcc agg aca ata tat tgt ttc atg att gct att gat tga tta ctg ttc tag atg att cta ctt tct tac cat ata ata aat tag aat ata ttt tct act ttt acg aga aat taa tta ttg tat tta tta ttt atg ggt --a aaa aaa ctt act ata agt ggg tgg gat tct ggg aat tag tga tca gtt tat gta tat cgc aac tag cgg gca tat ggc tat tga cat cga gaa cat tac cca tat gat aag aga ttg tat ccg ttt cgt agt ctt gag tat tgg tat tac tat ata gta tgt aga tgt cga cgc tag ata gac agt cgc cca cta gag tta ccg tct ctg aat gcg gca tga tag tat cat tct ttg ttt tcg tta act gtt tgg aag atg aat ctt tgt tgt tac att taa tct cga aat tca gag tac ata tct ttg aag tat tct gat atc tat ttt ctc ctg taa aga atc ctg aag ttg cta cat tat taa gga cag aga agt att ctg cac gaa aga ctg gat cac aat ctt tat gat tca tgg taa tag tta gtt ccg acg ttg aga tgg att cgc tga gac cgg tag tgg tcg tcc gag tac acg atg tgt cgt tga ctg gat aca ggt taa ttt cca cat cga tat agt taa atg tat tgc tgg tta cga cgg gtt cgc att tat ctg tgg aag aga cgg tgt gag aat atg ttc cgg gac cac acg gag aac aga tga cgt ctc cgg tag acg tgt atc cgg ata ctc cgt atc cta ttc cac act ttg ttt tag aaa tac atg ttc tac acc ctg atg ctc ctt tga gaa gac aat aat atc ctg gag agc att cac aga ttc tat tgt gag tcg tgt tac acg atc gcg tct cta cct gat tac tat cac atc ttc cgt tac aac tta gac aag cct gta aat gat tat tgt gag atg taa agg tat ccg aac cac acg gtg tac att gtg tat tag tct tgc tat cac ata atc tgg aag cgt aag ttc ccg gag gac acg ata gac aac ata gat tac ggc ttc tgt att cgt tgt ctt tac act ttc cat tgg atg gtg cat gtg gtg cta tat ctc ttc cgt tta tta tta tac atg aga gaa aca ata tat acg agt ata ata cgg acc tca tga ttt aat --t -gt agt aat cgt cgt ctt gtt act gtt tgt ttc cta ctt ctc caa tca tat aga tta ttt ttt aaa tat ttt ctt tct atc atg gat aat att tgt aat ggt tct ttc cgt aca aca tac tgt tta gat ggt agt cgc tta gct tgg tta tga tat tgc gca taa ttt ccg gag gca aat acg ata gtc tag att gat tat cga tgg tag act cta att tat tga gtg ctt tgt cga cga gtt tac ttt tat gct cca tcg ata gat gac act gtt cta tga gat cgt cat aca tgg gaa atg aaa tgc gtt tgt ctg aat gta tgg ctt cga gat agg tgt gat acc gga tgt ctt ctg ttc tca ata ccg tat aca agt cgg tgt ctg aga ttc gaa tct ctt tga gga gac tta tgt cac gac tac att ttt cga tga tgg aat cta tct tat cga atg ata tat ttt tca taa ata cac ttt tat agt cct cgt tta aac aga att tag tat gta gtt ccg caa atg act cgt ccc tta ata ggc agt agg cta tta tct tct tta cgt agt gat cgt cgt agg gag aga act ccg aca tct tgt aga aca acg att taa tca tag gta gag ata ctt tca gtc tgt ggt gga tga tgt cat taa cga cat ccg cct tgt ata tga tgt ttc tgt ttt caa aca cca agt cga ata ctg tct tat cgt ctt tag tcg gaa ggt tga tgt cgt atc cga tgt ata cga ggt atg agg caa cat tgt tat tgc aat tct gga agg cgg tat gaa gag gag tca ttg tat tat agt att cgt ctt tct gaa tgt cga atc tat cta gta gat acc gta gta tat tga gag agc gac ttc cat atc ctt gat tat gtt tta tga ata gat aaa gta gat gtt gtc ctt ctt cct ttt gta att tcc cgt att ttt gtt cgt gcc aat tga gta aca tta tga gaa tat gac ctg ttg cac aat cgt tct tta tgt att cca tga tgg gtg tac aat caa gat tat tac gta tcc tcg tat cag ctc ctc gag ata aaa gag cat aca cca cac gag gac tat gtt tgg tat act gtt gaa ggt aag tgt gta acg gcg tat ttc cga ttt tcg taa ccg cgt taa tgt ttg ctc cat gat cta tta tcg cgt aga tga atc gct tct cag ctc gca tct tag tgt gac tct ttg act tgt aat aat tgc ttt cgt gga acg cgg ata tgt gtt tac agt agt aat gaa gag aag tga gtc cat cct cat cga cgc aat tag ggt cag atc ctt tag tca ata att tgt aca gaa cgt aat agt tta agc tcc cat tga att tat atc taa gat aac aca gca ata gat cgg atg att tac taa agt cat caa tgg ggt ccg tta gta tat caa aga tct tgt tat cga ttg ata gtg aat gaa tca gat agt ggt gta gag gaa tat gtc ctt ttt tat cct tgc tat caa agt tac gca tgc cgt ggc gta aca ata tct tta ata cag atg gat taa atc gtg tat tca tcg tat agc aat gta atg gag agt tac cac att tta gtc gtt tat tca gat cgc agt gtt taa taa cta att taa aca gat gag atg ata tat cca cat caa aga atg cga gat aca tat gac aga cat tat tga cag aaa tgt gac ctt cat tat cac cgt cgt cca taa atg cgt tag gta cgt acc aca tac tat cgt taa cga tgc gca caa tct cgt cca -tt tca tcc atc ttc ata atg att tac ttt ttc ata att aga gaa aaa gat caa ggt ata aaa att aga agt gtt aga cta ta- --- --- --- --- --- aac taa ctt ata a-- -aa cta act tat gac tta act aac tta tga --- --- --- --- --- --- --- --- --- --- --- --- --- --- --- --- --- --- --- --- --- --- --- --- --- --- --- --- --- --- --- --- --- --- --- --- --- --- --- --- --- --- --- --- --- --- --- --- ctt aac taa ctt atg act taa cta act tat gac tta act aac tta tga ctt aac taa ctt atg act taa cta act ta- tga ctt aac taa ctt atg act taa cta act tat gac tta act aac tta tga cta tta act cat ttc aag aaa ggt ggg tgg ata gaa ctc tat atg aca gct tgt gaa aca att aga tcc cta att tct aat gga agt ttt gat agg aga ttg tca tca gtt gat aca ttg ttt att atc tca tct att aga gca cgt ctg ttt aga gct tta gtg acc tgc tcg gtt act tct gtg taa atc ttg aat cct tta gtg ata cac tgt gtc aaa act gga tgt tta gaa tac cta tgt aga ata tgg gaa gca tgc ttg ttt ttg tct cta tta tag att aac tca tac atg gtt gta tta tga att ttc atc tgc cta atg tac tcc aat tct tgt tta caa tca att ata taa tca aag agt gat gat gca tac aca tta caa agt gaa taa tct acc atc ata aaa tac ttg ata cag agc ttt atc aca tca tgg ttt tca att gta tta tta agt ata gct aat ttt ata cag tca ata gac aat ggt tct cta agc aat att tct aat att tta aga tgt gct tcc cta cgg gcg atg aca gat ccc cta tcc acg gcc acg tca aga cat gta tat cca tta ctc att act gcg ttg aca ttt gct cca ttt tct aat agc aat gat act aaa tct ata taa cct gca tag ata gcg cga taa agc aag gtc ctt cca cca gca tct agt tga ttg ata tct tca ata tat ggg ata caa agc tta taa att tct aat act gtg ggt tca tct aca agg aat ccc cta gta tac tga att att tta tat aga tct aat tta aca tca ttt tca tct ggg ata cca caa ttc aaa ata aac tca aca aca cta ctt tcc ttt tta cat att ccc cta aaa tag gca ttc aag cat tct att tta tat att aca gcc cca tga tct acc ata aga tca aca atg tct att tct aca tat gca tta gat aga tag taa agt aag aga tct tgt aca gaa tta caa ttc tta ata att ata gag aaa ata tct tcc ata taa ttc ttt gac act aat gca gat ata ata tct tta tat gta ata tat gca aac agt cta tct act ata tac tga tca ata tta tct cta tga atc cta aaa taa tca tac aga aca tct aca gga tca caa att ggt tca agg aga aat cta tca aat att ttc ctg tca aca act ggt tct aga aca taa cag tca aca cct aat cca tgt ttt tta taa tca tct acc aaa gat aat gac caa aga tcg agg tcg tcg tga aac tgc tca tcg aca gcc atg aaa tct acc gac tcc atg gtg cga atc gca ctg tct tat tcg cca ttg att ttc a-t ttt tta taa tta tgt aca tgt ttt cct tct att ctc aag agt cta caa aaa tat a-- ttt ttt tcg ata tct aag tac taa gtt ttt tta ctg ttt ttg tta ctg tct tcc att ctt cta act aaa gat ctg aga taa att ata caa tct tcg cta tcg aac cat ttt tgt agt cta aag cct gaa gta att aac caa ctg ttt tta tta gtg gct ttt ttc gat cta tca att tca gta tat tct tcg ccg tta taa aag taa tgt tgt tta att gta gga cgg ttg tta gta taa tca cat gaa taa taa tat tct aat tcc tcg tat tga cta ctt aca gat act cga aat agt ctg aaa aat tct tca aag ata ttt tta taa aga tct agg aaa agt tta tta ccg acc atg aac gag ata gat gga taa ata tcc ttt cca tca aag gtc ata att gga taa ttg tcc agc aat ata tct gct gta tta gtt ata tca ctt cca ttt att ttc aga ttg aag taa tgt act agt ttg tga caa tta aca aga tac aaa aga gat gcc gat act aat acg taa ata gct ata cgc gaa tcc att gtt acc ttt ttt tat ttc ata ggt cta tta ata aat ata tgt att act taa gac tag aaa aat caa aag tga gtt ttt gat att tga ttc tta ctt att gtg gga ttg tag ttt act tag taa ttc atc tct gaa tcc tga taa atc atg cat atc aat gat gca act acg cag caa act agt agg aat ata gat atc tgg ata tgt acg taa ata gtc gat tat atc ttt tac aat act att agt ccc tat tgc gtt atc tat ata tcc att aat aat att aca cag tgg ata ctt atg aga aat ata cct ctt aca gat ttt taa cga tat ata atc tag aaa gat atg tgt gta gta ctg tat tac cta aat tat cag tct cat tca aat att gca tga cta tta tcg aga att gct ata tcc ctc tat act cga tgc att tat tac agt aat tca atc cag cta aca taa gag cca atc tca atg ttg gtt taa tta tat cat ctt cat gta ata ata acg atg gaa act ttc cag tag cgt aac act tat cta aga agg ata taa taa cta tgt cta cat tat gtc tct tat cga gaa tat tct taa cga gat atc cat agc tat tct ggt gct aat tat tcc tat att ata ttc cac gaa aaa tga tga agg caa tca ttc atc ata aga tga taa aaa gtg tag tga gta aga gta tta gtg aga gag cat gaa gga gat tta gta ttt agc agt gag gat atg atc caa gag ggt gag ata gtc gtt ctc gtt cag aat ctt tcg cag cat aag tag tat gtc gat ata ctt atc att gaa gac tct tcc agt gac aat agc tga ttg agt aca aag tcc aat tat tgc aca aag ttc ttt ggc ggt ttt cat gga gtc att tct gat gaa aca ttt aat gat ctc cac gca att gtc gat att gtc cca cgg aag tga atc cga gaa ctc ctt caa ctc gct acc aaa tag ctc cat tgc atc aat tct gaa aga gat gag aag cct gta gag agg ccc tgc gct ttc tct atg ggt cca tct atg aga aac cca cag gat gta ttc agt cag aca a-t gtc tga cat cag tca cgg tat tca ggg agt cct tag tag cgt ggc aat gac agg gac tga act ggg cac aag gag agg cca ttg tga agg tag acg aag gta acc tga tgg tag acc tgt agc cgt ctg tgc tta ata gag ggc ttt aat ttc cat ttt taa tgg tgt cgt gaa tga gga atg aga gtg tct ctc gtc ctt ggt tta cat gga tca gag tga gaa aaa ata tct tgt ata tta tta act aac aac ctt ggt ttc tat cca tgt tt- aaa aaa tga cct ata tgt tct tta tta att cta ttt taa act tta tcc tca aga ctc ctg aca aaa tta aaa tcc aga aag cag caa aca atc ctg tta caa gtt tac tga aat ctc tct ttg att gta gag tat atg tag tca gag caa gaa aca ctg cag tag tca aca tga aag ctt gca taa cga tac gtg cat cat aga aag taa caa cag agg cca gcg tta gag att cta aca gtg taa atc cac aat ggt gta tgt aca gat tca ggg gat gtt cat gtc tgt gta aag tca atg cga aaa tca agc cta tag atc cga aca ttg atg cca ata tta gaa cag gac tcc ctt gta taa atg tcc gat gca ttc aaa gta taa aaa tac tgc agc tgt tgc cgt tgt taa agg aaa ttg tag aaa gga tac cgt aga ctt ttc tta gaa atg cca ttc gta tgt aca cgc tgg cag acg cca ccg agc tgt cat agt tga agt cgt cct cga t

>AY753185.1_Monkeypox_virus_strain_COP-58_complete_genome

--- --- --- --- --- --- --- --- --- --- --- --- --- --- --- --- --- --- --- --- --- --- --- --- --- --- --- --- --- --- --- --- --- --- --- --- --- --- --- --- --- --- --- --- --- --- --- --- aac tct aga ggg taa gaa aaa tca atc gtt tat aga gac cat cag aaa gag gtt taa tat ttt tgt gag acc cat cga aga gag aaa g-- --- --- --- --- --- --- --- --- --- --- --- --- --- --- --- --- --- --- --- --- --- --- --- --- --- --- --- --- --- --- --- --- --- --- --- --- --- --- --- --- --- --- --- --- --- --- --- --- --- --- --- --- --- --- --- --- --- --- --- --- --- --- --- --- --- --- --- --- --- --- --- --- --- --- --- --- --- --- --- --- --- --- --- --- --- --- --- --- --- --- --- --- --- --- --- --- --- --- --- --- --- --- --- --- --- --- --- --- --- --- --- --- --- --- --- --- --- --- --- --- --- --- --- --- --- --- --- --- --- --- --- --- --- --- --- --- --- --- --- --- --- --- --- --- --- --- --- --- --- --- --- --- --- --- --- --- --- --- --- --- --- --- --- --- --- --- --- --- --- --- --- --- --- --- --- --- --- --- --- --- --- --- --- --- --- --- --- --- --- --- --- --- --- --- --- --- --- --- --- --- --- --- --- --- --- --- --- --- --- --- --- --- --- --- --- --- --- --- --- --- --- --- --- --- --- --- --- --- --- --- --- --- --- --- --- --- --- --- --- --- --- --- --- --- --- --- --- --- --- --- --- --- --- --- --- --- --- --- --- --- --- --- --- --- --- --- --- --- --- --- --- --- --- --- --- --- --- --- --- -ga taa aaa ctt ttt acg act cca tca gaa aga ggt tta ata ttt ttg tga gac cca tcg aag aga gaa aga gat ggt tag tca aga tat ttt tct tag tac aaa agt caa tgt ttt aaa ata tat gga cga gaa tta att tgt ctg tat aaa aac ttg tgt gaa att atg tac tag aga aaa aac gtg agc agt gtc ccc tac atg gat ttt aca gat cat tta tat tcc aaa aat att aac tat ata cgt tta tta tat gat gtt aac gtg taa att ata aac att att tta tga tgc aat tgt ctg aca acc tag att ggt ata agg atg ttg ata agc tct acg aga ata tat tgt tgg acg tta tcg ttt acg aaa tag ttg aga cat cag aaa gag gtt taa tat ttt tgt gag a-c cat cga aga gag aaa tag aat aaa aat a-- -tt ttt ttt ttt ttt tgt aaa act ttt tta tga gac --- --- -ca aga gaa tac gaa tag tga tca tat cgt atc aca tat tga aac aga aag aag aag taa cga gag gta act ttt tgt aaa tgt agt taa ata ttt ttg ttt tgc aaa ccg gaa tat agt gcc cgg t-- -ct ttt tta att cgt ggt gcg gtg tct gaa tcg ttc gat taa ccc aac tca tcc att ttc aga tga ata gag tta tcg att cag aca cat gct ttg agt ttt gtt gaa tcg atg agt gaa gta tca tcg gtt gca cct tca gat gcc gat ccg tcg aca tac ttg aat cca tcc ttg act tca agt tca gat gat tcc tca cac atg tct ccg ata cgt acg cta aac tct agg ttc ttg aca cat ttt gta tca acg atc gtt gaa ccg atg ata tct ttg taa ctc act ttc tta tgt gag atg tta gac cca agt act gga tgg gtc ttg atg tca ctg tct ttc tct tct tcg cta cat ctg atg tcg ata gac atc tca cag tct ttg atc ata gcc aga gct tct tca cgc gtg atc gcg gga gag tcc tta cct tgt ccc ggt gac acg ctg gac aat cta gta ttc aca gtg ttt cca tca gag gat tcg gag atg gat gaa atc ttt ggg cat ttg gtg aat cca aag ttc atg tta aga ccc gca ccg acg ata gtg taa taa gtg gtg gga tct cct ttt aca act --- --- --- --- --- --- tct tcg gat acc tca tca tct tcg gtc tct gta act tcc gtt acg gat tga caa atc tta tca ttg gtc ggt gtt tgg tct tgc ttt gtg act ttg ata ata aca tcg att ccc ata tga tgt ttg ttt tct tct tca gta cac gag gat gaa gat tgt tga aga cta gta ggc ata gca gct gcc act agg cac atg cat gcc agg aca ata tat tgt ttc atg att gct att gat tga tta ctg ttc tag atg att cta ctt tct tac cat ata ata aat tag aat ata ttt tct act ttt acg aga aat taa tta ttg tat tta tta ttt atg ggt --a aaa aaa ctt act ata agt ggg tgg gat tct ggg aat tag tga tca gtt tat gta tat cgc aac tag cgg gca tat ggc tat tga cat cga gaa cat tac cca tat gat aag aga ttg tat ccg ttt cgt agt ctt gag tat tgg tat tac tat ata gta tgt aga tgt cga cgc tag ata gac agt cgc cca cta gag tta ccg tct ctg aat gcg gca tga tag tat cat tct ttg ttt tcg tta act gtt tgg aag atg aat ctt tgt tgt tac att taa tct cga aat tca gag tac ata tct ttg aag tat tct gat atc tat ttt ctc ctg taa aga atc ctg aag ttg cta cat tat taa gga cag aga agt att ctg cac gaa aga ctg gat cac aat ctt tat gat tca tgg taa tag tta gtt ccg acg ttg aga tgg att cgc tga gac cgg tag tgg tcg tcc gag tac acg atg tgt cgt tga ctg gat aca ggt taa ttt cca cat cga tat agt taa atg tat tgc tgg tta cga cgg gtt cgc att tat ctg tgg aag aga cgg tgt gag aat atg ttc cgg gac cac acg gag aac aga tga cgt ctc cgg tag acg tgt atc cgg ata ctc cgt atc cta ttc cac act ttg ttt tag aaa tac atg ttc tac acc ctg atg ctc ctt tga gaa gac aat aat atc ctg gag agc att cac aga ttc tat tgt gag tcg tgt tac acg atc gcg tct cta cct gat tac tat cac atc ttc cgt tac aac tta gac aag cct gta aat gat tat tgt gag atg taa agg tat ccg aac cac acg gtg tac att gtg tat tag tct tgc tat cac ata atc tgg aag cgt aag ttc ccg gag gac acg ata gac aac ata gat tac ggc ttc tgt att cgt tgt ctt tac act ttc cat tgg atg gtg cat gtg gtg cta tat ctc ttc cgt tta tta tta tac atg aga gaa aca ata tat acg agt ata ata cgg acc tca tga ttt aat --t -gt agt aat cgt cgt ctt gtt act gtt tgt ttc cta ctt ctc caa tca tat aga tta ttt ttt aaa tat ttt ctt tct atc atg gat aat att tgt aat ggt tct ttc cgt aca aca tac tgt tta gat ggt agt cgc tta gct tgg tta tga tat tgc gca taa ttt ccg gag gca aat acg ata gtc tag att gat tat cga tgg tag act cta att tat tga gtg ctt tgt cga cga gtt tac ttt tat gct cca tcg ata gat gac act gtt cta tga gat cgt cat aca tgg gaa atg aaa tgc gtt tgt ctg aat gta tgg ctt cga gat agg tgt gat acc gga tgt ctt ctg ttc tca ata ccg tat aca agt cgg tgt ctg aga ttc gaa tct ctt tga gga gac tta tgt cac gac tac att ttt cga tga tgg aat cta tct tat cga atg ata tat ttt tca taa ata cac ttt tat agt cct cgt tta aac aga att tag tat gta gtt ccg caa atg act cgt ccc tta ata ggc agt agg cta tta tct tct tta cgt agt gat cgt cgt agg gag aga act ccg aca tct tgt aga aca acg att taa tca tag gta gag ata ctt tca gtc tgt ggt gga tga tgt cat taa cga cat ccg cct tgt ata tga tgt ttc tgt ttt caa aca cca agt cga ata ctg tct tat cgt ctt tag tcg gaa ggt tga tgt cgt atc cga tgt ata cga ggt atg agg caa cat tgt tat tgc aat tct gga agg cgg tat gaa gag gag tca ttg tat tat agt att cgt ctt tct gaa tgt cga atc tat cta gta gat acc gta gta tat tga gag agc gac ttc cat atc ctt gat tat gtt tta tga ata gat aaa gta gat gtt gtc ctt ctt cct ttt gta att tcc cgt att ttt gtt cgt gcc aat tga gta aca tta tga gaa tat gac ctg ttg cac aat cgt tct tta tgt att cca tga tgg gtg tac aat caa gat tat tac gta tcc tcg tat cag ctc ctc gag ata aaa gag cat aca cca cac gag gac tat gtt tgg tat act gtt gaa ggt aag tgt gta acg gcg tat ttc cga ttt tcg taa ccg cgt taa tgt ttg ctc cat gat cta tta tcg cgt aga tga atc gct tct cag ctc gca tct tag tgt gac tct ttg act tgt aat aat tgc ttt cgt gga acg cgg ata tgt gtt tac agt agt aat gaa gag aag tga gtc cat cct cat cga cgc aat tag ggt cag atc ctt tag tca ata att tgt aca gaa cgt aat agt tta agc tcc cat tga att tat atc taa gat aac aca gca ata gat cgg atg att tac taa agt cat caa tgg ggt ccg tta gta tat caa aga tct tgt tat cga ttg ata gtg aat gaa tca gat agt ggt gta gag gaa tat gtc ctt ttt tat cct tgc tat caa agt tac gca tgc cgt ggc gta aca ata tct tta ata cag atg gat taa atc gtg tat tca tcg tat agc aat gta atg gag agt tac cac att tta gtc gtt tat tca gat cgc agt gtt taa taa cta att taa aca gat gag atg ata tat cca cat caa aga atg cga gat aca tat gac aga cat tat tga cag aaa tgt gac ctt cat tat cac cgt cgt cca taa atg cgt tag gta cgt acc aca tac tat cgt taa cga tgc gca caa tct cgt cca -tt tca tcc atc ttc ata atg att tac ttt ttc ata att aga gaa aaa gat caa ggt ata aaa att aga agt gtt aga cta ta- --- --- --- --- --- aac taa ctt ata a-- -aa cta act tat gac tta act aac tta tga ctt aac taa ctt atg act taa cta act tat gac tta act aac tta tga ctt aac taa ctt atg act taa cta act tat gac tta act aac tta tga ctt aac taa ctt atg act taa cta act tat gac tta act aac tta tga ctt aac taa ctt atg act taa cta act tat gac tta act aac tta tga ctt aac taa ctt atg act taa cta act ta- tga ctt aac taa ctt atg act taa cta act tat gac tta act aac tta tga cta tta act cat ttc aag aaa ggt ggg tgg ata gaa ctc tat atg aca gct tgt gaa aca att aga tcc cta att tct aat gga agt ttt gat agg aga ttg tca tca gtt gat aca ttg ttt att atc tca tct att aga gca cgt ctg ttt aga gct tta gtg acc tgc tcg gtt act tct gtg taa atc ttg aat cct tta gtg ata cac tgt gtc aaa act gga tgt tta gaa tac cta tgt aga ata tgg gaa gca tgc ttg ttt ttg tct cta tta tag att aac tca tac atg gtt gta tta tga att ttc atc tgc cta atg tac tcc aat tct tgt tta caa tca att ata taa tca aag agt gat gat gca tac aca tta caa agt gaa taa tct acc atc ata aaa tac ttg ata cag agc ttt atc aca tca tgg ttt tca att gta tta tta agt ata gct aat ttt ata cag tca ata gac aat ggt tct cta agc aat att tct aat att tta aga tgt gct tcc cta cgg gcg atg aca gat ccc cta tcc acg gcc acg tca aga cat gta tat cca tta ctc att act gcg ttg aca ttt gct cca ttt tct aat agc aat gat act aaa tct ata taa cct gca tag ata gcg cga taa agc aag gtc ctt cca cca gca tct agt tga ttg ata tct tca ata tat ggg ata caa agc tta taa att tct aat act gtg ggt tca tct aca agg aat ccc cta gta tac tga att att tta tat aga tct aat tta aca tca ttt tca tct ggg ata cca caa ttc aaa ata aac tca aca aca cta ctt tcc ttt tta cat att ccc cta aaa tag gca ttc aag cat tct att tta tat att aca gcc cca tga tct acc ata aga tca aca atg tct att tct aca tat gca tta gat aga tag taa agt aag aga tct tgt aca gaa tta caa ttc tta ata att ata gag aaa ata tct tcc ata taa ttc ttt gac act aat gca gat ata ata tct tta tat gta ata tat gca aac agt cta tct act ata tac tga tca ata tta tct cta tga atc cta aaa taa tca tac aga aca tct aca gga tca caa att ggt tca agg aga aat cta tca aat att ttc ctg tca aca act ggt tct aga aca taa cag tca aca cct aat cca tgt ttt tta taa tca tct acc aaa gat aat gac caa aga tcg agg tcg tcg tga aac tgc tca tcg aca gcc atg aaa tct acc gac tcc atg gtg cga atc gca ctg tct tat tcg cca ttg att ttc a-t ttt tta taa tta tgt aca tgt ttt cct tct att ctc aag agt cta caa aaa tat a-- ttt ttt tcg ata tct aag tac taa gtt ttt tta ctg ttt ttg tta ctg tct tcc att ctt cta act aaa gat ctg aga taa att ata caa tct tcg cta tcg aac cat ttt tgt agt cta aag cct gaa gta att aac caa ctg ttt tta tta gtg gct ttt ttc gat cta tca att tca gta tat tct tcg ccg tta taa aag taa tgt tgt tta att gta gga cgg ttg tta gta taa tca cat gaa taa taa tat tct aat tcc tcg tat tga cta ctt aca gat act cga aat agt ctg aaa aat tct tca aag ata ttt tta taa aga tct agg aaa agt tta tta ccg acc atg aac gag ata gat gga taa ata tcc ttt cca tca aag gtc ata att gga taa ttg tcc agc aat ata tct gct gta tta gtt ata tca ctt cca ttt att ttc aga ttg aag taa tgt act agt ttg tga caa tta aca aga tac aaa aga gat gcc gat act aat acg taa ata gct ata cgc gaa tcc att gtt acc ttt ttt tat ttc ata ggt cta tta ata aat ata tgt att act taa gac tag aaa aat caa aag tga gtt ttt gat att tga ttc tta ctt att gtg gga ttg tag ttt act tag taa ttc atc tct gaa tcc tga taa atc atg cat atc aat gat gca act acg cag caa act agt agg aat ata gat atc tgg ata tgt acg taa ata gtc gat tat atc ttt tac aat act att agt ccc tat tgc gtt atc tat ata tcc att aat aat att aca cag tgg ata ctt atg aga aat ata cct ctt aca gat ttt taa cga tat ata atc tag aaa gat atg tgt gta gta ctg tat tac cta aat tat cag tct cat tca aat att gca tga cta tta tcg aga att gct ata tcc ctc tat act cga tgc att tat tac agt aat tca atc cag cta aca taa gag cca atc tca atg ttg gtt taa tta tat cat ctt cat gta ata ata acg atg gaa act ttc cag tag cgt aac act tat cta aga agg ata taa taa cta tgt cta cat tat gtc tct tat cga gaa tat tct taa cga gat atc cat agc tat tct ggt gct aat tat tcc tat att ata ttc cac gaa aaa tga tga agg caa tca ttc atc ata aga tga taa aaa gtg tag tga gta aga gta tta gtg aga gag cat gaa gga gat tta gta ttt agc agt gag gat atg atc caa gag ggt gag ata gtc gtt ctc gtt cag aat ctt tcg cag cat aag tag tat gtc gat ata ctt atc att gaa gac tct tcc agt gac aat agc tga ttg agt aca aag tcc aat tat tgc aca aag ttc ttt ggc ggt ttt cat gga gtc att tct gat gaa aca ttt aat gat ctc cac gca att gtc gat att gtc cca cgg aag tga atc cga gaa ctc ctt caa ctc gct acc aaa tag ctc cat tgc atc aat tct gaa aga gat gag aag cct gta gag agg ccc tgc gct ttc tct atg ggt cca tct atg aga aac cca cag gat gta ttc agt cag aca a-t gtc tga cat cag tca cgg tat tca ggg agt cct tag tag cgt ggc aat gac agg gac tga act ggg cac aag gag agg cca ttg tga agg tag acg aag gta acc tga tgg tag acc tgt agc cgt ctg tgc tta ata gag ggc ttt aat ttc cat ttt taa tgg tgt cgt gaa tga gga atg aga gtg tct ctc gtc ctt ggt tta cat gga tca gag tga gaa aaa ata tct tgt ata tta tta act aac aac ctt ggt ttc tat cca tgt tt- aaa aaa tga cct ata tgt tct tta tta att cta ttt taa act tta tcc tca aga ctc ctg aca aaa tta aaa tcc aga aag cag caa aca atc ctg tta caa gtt tac tga aat ctc tct ttg att gta gag tat atg tag tca gag caa gaa aca ctg cag tag tca aca tga aag ctt gca taa cga tac gtg cat cat aga aag taa caa cag agg cca gcg tta gag att cta aca gtg taa atc cac aa- --a gta tgt aca gat tca ggg gat gtt cat gtc tgt gta aag tca atg cga aaa tca agc cta tag atc cga aca ttg atg cca ata tta gaa cag gac tcc ctt gta taa atg tcc gat gca ttc aaa gta taa aaa tac tgc agc tgt tgc cgt tgt taa agg aaa ttg tag aaa gga tac cgt aga ctt ttc tta gaa atg cca ttc gta tgt aca cgc tgg cag acg cca ccg agc tgt cat agt tga agt cgt cct cga t

>AY603973.1_Monkeypox_virus_strain_MPXV-WRAIR7-61_complete_genome

--- --- --- --- --- --- --- --- --- --- --- --- --- --- --- --- --- --- --- --- --- --- --- --- --- --- --- --- --- --- --- --- --- --- --- --- --- --- --- --- --- --- --- --- --- --- --- --- --- --- aga ggg taa gaa aaa tca atc gtt tat aga gac cat cag aaa gag gtt taa tat ttt tgt gag acc cat cga aga gag aaa g-- --- --- --- --- --- --- --- --- --- --- --- --- --- --- --- --- --- --- --- --- --- --- --- --- --- --- --- --- --- --- --- --- --- --- --- --- --- --- --- --- --- --- --- --- --- --- --- --- --- --- --- --- --- --- --- --- --- --- --- --- --- --- --- --- --- --- --- --- --- --- --- --- --- --- --- --- --- --- --- --- --- --- --- --- --- --- --- --- --- --- --- --- --- --- --- --- --- --- --- --- --- --- --- --- --- --- --- --- --- --- --- --- --- --- --- --- --- --- --- --- --- --- --- --- --- --- --- --- --- --- --- --- --- --- --- --- --- --- --- --- --- --- --- --- --- --- --- --- --- --- --- --- --- --- --- --- --- --- --- --- --- --- --- --- --- --- --- --- --- --- --- --- --- --- --- --- --- --- --- --- --- --- --- --- --- --- --- --- --- --- --- --- --- --- --- --- --- --- --- --- --- --- --- --- --- --- --- --- --- --- --- --- --- --- --- --- --- --- --- --- --- --- --- --- --- --- --- --- --- --- --- --- --- --- --- --- --- --- --- --- --- --- --- --- --- --- --- --- --- --- --- --- --- --- --- --- --- --- --- --- --- --- --- --- --- --- --- --- --- --- --- --- --- --- --- --- --- --- --- -ga taa aaa ctt ttt acg act cca tca gaa aga ggt tta ata ttt ttg tga gac cca tcg aag aga gaa aga gat ggt tag tca aga tat ttt tct tag tac aaa agt caa tgt ttt aaa ata tat gga cga gaa tta att tgt ctg tat aaa aac ttg tgt gaa att atg tac tag aga aaa aac gtg agc agt gtc ccc tac atg gat ttt aca gat cat tta tat tcc aaa aat att aac tat ata cgt tta tta tat gat gtt aac gtg taa att ata aac att att tta tga tgc aat tgt ctg aca acc tag att ggt ata agg atg ttg ata agc tct acg aga ata tat tgt tgg acg tta tcg ttt acg aaa tag ttg aga cat cag aaa gag gtt taa tat ttt tgt gag a-c cat cga aga gag aaa tag aat aaa aat a-- -tt ttt ttt ttt ttt tgt aaa act ttt tta tga gac --- --- -ca aga gaa tac gaa tag tga tca tat cgt atc aca tat tga aac aga aag aag aag taa cga gag gta act ttt tgt aaa tgt agt taa ata ttt ttg ttt tgc aaa ccg gaa tat agt gcc cgg t-- -ct ttt tta att cgt ggt gca gtg tct gaa tcg ttc gat taa ccc aac tca tcc att ttc aga tga ata gag tta tcg att cag aca cat gct ttg agt ttt gtt gaa tcg atg agt gaa gta tca tcg gtt gca cct tca gat gcc gat ccg tcg aca tac ttg aat cca tcc ttg act tca agt tca gat gat tcc tca cac atg tct ccg ata cgt acg cta aac tct agg ttc ttg aca cat ttt gta tca acg atc gtt gaa ccg atg ata tct ttg taa ctc act ttc tta tgt gag atg tta gac cca agt act gga tgg gtc ttg atg tca ctg tct ttc tct tct tcg cta cat ctg atg tcg ata gac atc tca cag tct ttg atc ata gcc aga gct tct tca cgc gtg atc gcg gga gag tcc tta cct tgt ccc ggt gac acg ctg gac aat cta gta ttc aca gtg ttt cca tca gag gat tcg gag atg gat gaa atc ttt ggg cat ttg gtg aat cca aag ttc atg tta aga ccc gca ccg acg ata gtg taa taa gtg gtg gga tct cct ttt aca act --- --- --- --- --- --- tct tcg gat acc tca tca tct tcg gtc tct gta act tcc gtt acg gat tga caa atc tta tca ttg gtc ggt gtt tgg tct tgc ttt gtg act ttg ata ata aca tcg att ccc ata tga tgt ttg ttt tct tct tca gta cac gag gat gaa gat tgt tga aga cta gta ggc ata gca gct gcc act agg cac atg cat gcc agg aca ata tat tgt ttc atg att gct att gat tga tta ctg ttc tag atg att cta ctt tct tac cat ata ata aat tag aat ata ttt tct act ttt acg aga aat taa tta ttg tat tta tta ttt atg ggt --a aaa aaa ctt act ata agt ggg tgg gat tct ggg aat tag tga tca gtt tat gta tat cgc aac tag cgg gca tat ggc tat tga cat cga gaa cat tac cca tat gat aag aga ttg tat ccg ttt cgt agt ctt gag tat tgg tat tac tat ata gta tgt aga tgt cga cgc tag ata gac agt cgc cca cta gag tta ccg tct ctg aat gcg gca tga tag tat cat tct ttg ttt tcg tta act gtt tgg aag atg aat ctt tgt tgt tac att taa tct cga aat tca gag tac ata tct ttg aag tat tct gat atc tat ttt ctc ctg taa aga atc ctg aag ttg cta cat tat taa gga cag aga agt att ctg cac gaa aga ctg gat cac aat ctt tat gat tca tgg taa tag tta gtt ccg acg ttg aga tgg att cgc tga gac cgg tag tgg tcg tcc gag tac acg atg tgt cgt tga ctg gat aca ggt taa ttt cca cat cga tat agt taa atg tat tgc tgg tta cga cgg gtt cgc att tat ctg tgg aag aga cgg tgt gag aat atg ttc cgg gac cac acg gag aac aga tga cgt ctc cgg tag acg tgt atc cgg ata ctc cgt atc cta ttc cac act ttg ttt tag aaa tac atg ttc tac acc ctg atg ctc ctt tga gaa gac aat aat atc ctg gag agc att cac aga ttc tat tgt gag tcg tgt tac acg atc gcg tct cta cct gat tac tat cac atc ttc cgt tac aac tta gac aag cct gta aat gat tat tgt gag atg taa agg tat ccg aac cac acg gtg tac att gtg tat tag tct tgc tat cac ata atc tgg aag cgt aag ttc ccg gag gac acg ata gac aac ata gat tac ggc ttc tgt att cgt tgt ctt tac act ttc cat tgg atg gtg cat gtg gtg cta tat ctc ttc cgt tta tta tta tac atg aga gaa aca ata tat acg agt ata ata cgg acc tca tga ttt aat --t -gt agt aat cgt cgt ctt gtt act gtt tgt ttc cta ctt ctc caa tca tat aga tta ttt ttt aaa tat ttt ctt tct atc atg gat aat att tgt aat ggt tct ttc cgt aca aca tac tgt tta gat ggt agt cgc tta gct tgg tta tga tat tgc gca taa ttt ccg gag gca aat acg ata gtc tag att gat tat cga tgg tag act cta att tat tga gtg ctt tgt cga cga gtt tac ttt tat gct cca tcg ata gat gac act gtt cta tga gat cgt cat aca tgg gaa atg aaa tgc gtt tgt ctg aat gta tgg ctt cga gat agg tgt gat acc gga tgt ctt ctg ttc tca ata ccg tat aca agt cgg tgt ctg aga ttc gaa tct ctt tga gga gac tta tgt cac gac tac att ttt cga tga tgg aat cta tct tat cga atg ata tat ttt tca taa ata cac ttt tat agt cct cgt tta aac aga att tag tat gta gtt ccg caa atg act cgt ccc tta ata ggc agt agg cta tta tct tct tta cgt agt gat cgt cgt agg gag aga act ccg aca tct tgt aga aca acg att taa tca tag gta gag ata ctt tca gtc tgt ggt gga tga tgt cat taa cga cat ccg cct tgt ata tga tgt ttc tgt ttt caa aca cca agt cga ata ctg tct tat cgt ctt tag tcg gaa ggt tga tgt cgt atc cga tgt ata cga ggt atg agg caa cat tgt tat tgc aat tct gga agg cgg tat gaa gag gag tca ttg tat tat agt att cgt ctt tct gaa tgt cga atc tat cta gta gat acc gta gta tat tga gag agc gac ttc cat atc ctt gat tat gtt tta tga ata gat aaa gta gat gtt gtc ctt ctt cct ttt gta att tcc cgt att ttt gtt cgt gcc aat tga gta aca tta tga gaa tat gac ctg ttg cac aat cgt tct tta tgt att cca tga tgg gtg tac aat caa gat tat tac gta tcc tcg tat cag ctc ctc gag ata aaa gag cat aca cca cac gag gac tat gtt tgg tat act gtt gaa ggt aag tgt gta acg gcg tat ttc cga ttt tcg taa ccg cgt taa tgt ttg ctc cat gat cta tta tcg cgt aga tga atc gct tct cag ctc gca tct tag tgt gac tct ttg act tgt aat aat tgc ttt cgt gga acg cgg ata tgt gtt tac agt agt aat gaa gag aag tga gtc cat cct cat cga cgc aat tag ggt cag atc ctt tag tca ata att tgt aca gaa cgt aat agt tta agc tcc cat tga att tat atc taa gat aac aca gca ata gat cgg atg att tac taa agt cat caa tgg ggt ccg tta gta tat caa aga tct tgt tat cga ttg ata gtg aat gaa tca gat agt ggt gta gag gaa tat gtc ctt ttt tat cct tgc tat caa agt tac gca tgc cgt ggc gta aca ata tct tta ata cag atg gat taa atc gtg tat tca tcg tat agc aat gta atg gag agt tac cac att tta gtc gtt tat tca gat cgc agt gtt taa taa cta att taa aca gat gag atg ata tat cca cat caa aga atg cga gat aca tat gac aga cat tat tga cag aaa tgt gac ctt cat tat cac cgt cgt cca taa atg cgt tag gta cgt acc aca tac tat cgt taa cga tgc gca caa tct cgt cca -tt tca tcc atc ttc ata atg att tac ttt ttc ata att aga gaa aaa gat caa ggt ata aaa att aga agt gtt aga cta ta- --- --- --- --- --- aac taa ctt ata a-- -aa cta act tat gac tta act aac tta tga --- --- --- --- --- --- --- --- --- --- --- --- --- --- --- --- --- --- --- --- --- --- --- --- --- --- --- --- --- --- --- --- ctt aac taa ctt atg act taa cta act tat gac tta act aac tta tga ctt aac taa ctt atg act taa cta act tat gac tta act aac tta tga ctt aac taa ctt atg act taa cta act ta- tga ctt aac taa ctt atg act taa cta act tat gac tta act aac tta tga cta tta act cat ttc aag aaa ggt ggg tgg ata gaa ctc tat atg aca gct tgt gaa aca att aga tcc cta att tct aat gga agt ttt gat agg aga ttg tca tca gtt gat aca ttg ttt att atc tca tct att aga gca cgt ctg ttt aga gct tta gtg acc tgc tcg gtt act tct gtg taa atc ttg aat cct tta gtg ata cac tgt gtc aaa act gga tgt tta gaa tac cta tgt aga ata tgg gaa gca tgc ttg ttt ttg tct cta tta tag att aac tca tac atg gtt gta tta tga att ttc atc tgc cta atg tac tcc aat tct tgt tta caa tca att ata taa tca aag agt gat gat gca tac aca tta caa agt gaa taa tct acc atc ata aaa tac ttg ata cag agc ttt atc aca tca tgg ttt tca att gta tta tta agt ata gct aat ttt ata cag tca ata gac aat ggt tct cta agc aat att tct aat att tta aga tgt gct tcc cta cgg gcg atg aca gat ccc cta tcc acg gcc acg tca aga cat gta tat cca tta ctc att act gcg ttg aca ttt gct cca ttt tct aat agc aat gat act aaa tct ata taa cct gca tag ata gcg cga taa agc aag gtc ctt cca cca gca tct agt tga ttg ata tct tca ata tat ggg ata caa agc tta taa att tct aat act gtg ggt tca tct aca agg aat ccc cta gta tac tga att att tta tat aga tct aat tta aca tca ttt tca tct ggg ata cca caa ttc aaa ata aac tca aca aca cta ctt tcc ttt tta cat att ccc cta aaa tag gca ttc aag cat tct att tta tat att aca gcc cca tga tct acc ata aga tca aca atg tct att tct aca tat gca tta gat aga tag taa agt aag aga tct tgt aca gaa tta caa ttc tta ata att ata gag aaa ata tct tcc ata taa ttc ttt gac act aat gca gat ata ata tct tta tat gta ata tat gca aac agt cta tct act ata tac tga tca ata tta tct cta tga atc cta aaa taa tca tac aga aca tct aca gga tca caa att ggt tca agg aga aat cta tca aat att ttc ctg tca aca act ggt tct aga aca taa cag tca aca cct aat cca tgt ttt tta taa tca tct acc aaa gat aat gac caa aga tcg agg tcg tcg tga aac tgc tca tcg aca gcc atg aaa tct acc gac tcc atg gtg cga atc gca ctg tct tat tcg cca ttg att ttc a-t ttt tta taa tta tgt aca tgt ttt cct tct att ctc aag agt cta caa aaa tat a-- ttt ttt tcg ata tct aag tac taa gtt ttt tta ctg ttt ttg tta ctg tct tcc att ctt cta act aaa gat ctg aga taa att ata caa tct tcg cta tcg aac cat ttt tgt agt cta aag cct gaa gta att aac caa ctg ttt tta tta gtg gct ttt ttc gat cta tca att tca gta tat tct tcg ccg tta taa aag taa tgt tgt tta att gta gga cgg ttg tta gta taa tca cat gaa taa taa tat tct aat tcc tcg tat tga cta ctt aca gat act cga aat agt ctg aaa aat tct tca aag ata ttt tta taa aga tct agg aaa agt tta tta ccg acc atg aac gag ata gat gga taa ata tcc ttt cca tca aag gtc ata att gga taa ttg tcc agc aat ata tct gct gta tta gtt ata tca ctt cca ttt att ttc aga ttg aag taa tgt act agt ttg tga caa tta aca aga tac aaa aga gat gcc gat act aat acg taa ata gct ata cgc gaa tcc att gtt acc ttt ttt tat ttc ata ggt cta tta ata aat ata tgt att act taa gac tag aaa aat caa aag tga gtt ttt gat att tga ttc tta ctt att gtg gga ttg tag ttt act tag taa ttc atc tct gaa tcc tga taa atc atg cat atc aat gat gca act acg cag caa act agt agg aat ata gat atc tgg ata tgt acg taa ata gtc gat tat atc ttt tac aat act att agt ccc tat tgc gtt atc tat ata tcc att aat aat att aca cag tgg ata ctt atg aga aat ata cct ctt aca gat ttt taa cga tat ata atc tag aaa gat atg tgt gta gta ctg tat tac cta aat tat cag tct cat tca aat att gca tga cta tta tcg aga att gct ata tcc ctc tat act cga tgc att tat tac agt aat tca atc cag cta aca taa gag cca atc tca atg ttg gtt taa tta tat cat ctt cat gta ata ata acg atg gaa act ttc cag tag cgt aac act tat cta aga agg ata taa taa cta tgt cta cat tat gtc tct tat cga gaa tat tct taa cga gat atc cat agc tat tct ggt gct aat tat tcc tat att ata ttc cac gaa aaa tga tga agg caa tca ttc atc ata aga tga taa aaa gtg tag tga gta aga gta tta gtg aga gag cat gaa gga gat tta gta ttt agc agt gag gat atg atc caa gag ggt gag ata gtc gtt ctc gtt cag aat ctt tcg cag cat aag tag tat gtc gat ata ctt atc att gaa gac tct tcc agt gac aat agc tga ttg agt aca aag tcc aat tat tgc aca aag ttc ttt ggc ggt ttt cat gga gtc att tct gat gaa aca ttt aat gat ctc cac gca att gtc gat att gtc cca cgg aag tga atc cga gaa ctc ctt caa ctc gct acc aaa tag ctc cat tgc atc aat tct gaa aga gat gag aag cct gta gag agg ccc tgc gct ttc tct atg ggt cca tct atg aga aac cca cag gat gta ttc agt cag aca a-t gtc tga cat cag tca cgg tat tca ggg agt cct tag tag cgt ggc aat gac agg gac tga act ggg cac aag gag agg cca ttg tga agg tag acg aag gta acc tga tgg tag acc tgt agc cgt ctg tgc tta ata gag ggc ttt aat ttc cat ttt taa tgg tgt cgt gaa tga gga atg aga gtg tct ctc gtc ctt ggt tta cat gga tca gag tga gaa aaa ata tct tgt ata tta tta act aac aac ctt ggt ttc tat cca tgt tt- aaa aaa tga cct ata tgt tct tta tta att cta ttt taa act tta tcc tca aga ctc ctg aca aaa tta aaa tcc aga aag cag caa aca atc ctg tta caa gtt tac tga aat ctc tct ttg att gta gag tat atg tag tca gag caa gaa aca ctg cag tag tca aca tga aag ctt gca taa cga tac gtg cat cat aga aag taa caa cag agg cca gcg tta gag att cta aca gtg taa atc cac aa- --a gta tgt aca gat tca ggg gat gtt cat gtc tgt gta aag tca atg cga aaa tca agc cta tag atc cga aca ttg atg cca ata tta gaa cag gac tcc ctt gta taa atg tcc gat gca ttc aaa gta taa aaa tac tgc agc tgt tgc cgt tgt taa agg aaa ttg tag aaa gga tac cgt aga ctt ttc tta gaa atg cca ttc gta tgt aca cgc tgg cag acg cca ccg agc tgt cat agt tga agt cgt cct cga t

>ON619837.1_Monkeypox_virus_isolate_MPXV_UK_2022_3_complete_genome

--- --- --- --- --- --- --- --- --- --- --- --- --- --- --- --- --- --- --- --- --- --- --- --- --- --- --- --- --- --a ttt tac tat ttt att tag tgt cta gaa aaa aat gtg tga ccc acg acc gta gga aac tct aga ggg taa gaa aaa tca atc gtt tat aga gac cat cag aaa gag gtt taa tat ttt tgt gag acc tat cga a-- --- --- --- --- --- --- --- --- --- --- --- --- --- --- --- --- --- --- --- --- --- --- --- --- --- --- --- --- --- --- --- --- --- --- --- --- --- --- --- --- --- --- --- --- --- --- --- --- --- --- --- --- --- --- --- --- --- --- --- --- --- --- --- --- --- --- --- --- --- --- --- --- --- --- --- --- --- --- --- --- --- --- --- --- --- --- --- --- --- --- --- --- --- --- --- --- --- --- --- --- --- --- --- --- --- --- --- --- --- --- --- --- --- --- --- --- --- --- --- --- --- --- --- --- --- --- --- --- --- --- --- --- --- --- --- --- --- --- --- --- --- --- --- --- --- --- --- --- --- --- --- --- --- --- --- --- --- --- --- --- --- --- --- --- --- --- --- --- --- --- --- --- --- --- --- --- --- --- --- --- --- --- --- --- --- --- --- --- --- --- --- --- --- --- --- --- --- --- --- --- --- --- --- --- --- --- --- --- --- --- --- --- --- --- --- --- --- --- --- --- --- --- --- --- --- --- --- --- --- --- --- --- --- --- --- --- --- --- --- --- --- --- --- --- --- --- --- --- --- --- --- --- --- --- --- --- --- --- --- --- --- --- --- --- --- --- --- --- --- --- --- --- --- --- --- --- --- --- --- -ga gag aaa gga taa aaa ctt ttt acg act cca tca gaa aga ggt tta ata ttt ttg tga gac cca tcg aag aga gaa aga gat ggt tag tca aga tat ttt tct tag tac aaa agt caa tgt ttt aaa ata tat gga cga gaa tta att tgt ctg tat aaa aac ttg tgt gaa att atg tac tag aga aaa aac gtg agc agt gtc ccc tac atg gat ttt aca gat cat tta tat tcc aaa aat att aac tat ata cgt tta tta tat gat gtt aac gtg taa att ata aac att att tta tga tgc aat tgt ctg aca acc tag att ggt ata agg atg ttg ata agc tct acg aga ata tat tgt tgg acg tta tcg ttt acg aaa tag ttg aga cat cag aaa gag gtt taa tat ttt tgt gag a-c cat cga aga gag aaa gag aat aaa aat a-- --- ttt ttt ttt ttt tgt aaa act ttt tta tga gac --- --- -ca aga gaa tac gaa tag tga tca tat cgt atc aca tat tga aac aga aag aag aag taa cga gag gta act ttt tgt gaa tgt agt taa ata ttt ttg ttt tgc aaa ccg gaa tat agt gcc cgg t-- -ct ttt tta att cgt ggt gcg gtg tct gaa tcg ttc gat taa ccc aac tca tcc att ttc aga tga ata gag tta tcg att cag aca cat gct ttg agt ttt gtt gaa tcg atg agt gaa gta tca tcg gtt gca cct tca gat gcc gat ccg tcg aca tac ttg aat cca tcc ttg act tca agt tca gat gat tcc tca cac atg tct ccg ata cgt acg cta aac tct agg ttc ttg aca cat ttt gta tca acg atc gtt gaa ccg atg ata tct ttg taa ctc act ttc tta tgt gag atg tta gac cca agt act gga tgg gtc ttg atg tca ctg tct ttc tct tct tcg cta cat ctg atg tcg ata gac atc tca cag tct ttg atc ata gcc aga gct tct tca cgc gtg atc gcg gga gag tcc tta cct tgt ccc ggt gac acg ctg gac aat cta gta ttc aca gtg ttt cca tca gag gat tcg gag atg gat aaa atc ttt ggg cat ttg gtg aat cca aag ttc atg tta aga ccc gca ccg acg ata gtg taa taa gtg gtg gga tct cct ttt aca act --- --- --- --- --- --- tct tcg gat acc tca tca tct tcg gtc tct gta act tcc gtt acg gat tga caa atc tta tca ttg gtc ggt gtt tgg tct tgc ttt gtg act ttg ata ata aca tcg att ccc ata tga tgt ttg ttt tct tct tca gta cac gag gat gaa gat tgt tga aga cta gta ggc ata gca gct gcc act agg cac atg cat gcc agg aca ata tat tgt ttc atg att gct att gat tga tta ctg ttc tag atg att cta ctt tct tac cat ata ata aat tag aat ata ttt tct act ttt acg aga aat taa tta ttg tat tta tta ttt ata ggt --a aaa aaa ctt act ata agt ggg tgg gat tct ggg aat tag tga tca gtt tat gta tat cgc aac tag cgg gca tat ggc tat tga cat cga gaa cat tac cca tat gat aag aga ttg tat cat ttt cgt agt ctt gag tat tgg tat tac tat ata gta tgt aga tgt cga cgc tag ata gac agt cgc cca cta gag tta ccg tct ctg aat gcg gca tga tag tat cat tct ttg ttt tcg tta act gtt tgg aag atg aat ctt tgt tgt tac att taa tct cga aat tca gag tac ata tct ttg aag tat tct gat atc tat ttt ctc ctg taa aga atc ctg aag ttg cta cat tat taa gga cag aga agt att ctg cac gaa aga ctg gat cac aat ctt tat gat tca tgg taa tag tta gtt ccg acg ttg aga tgg att cgc tga gac cgg tag tgg tcg tcc gag tac acg atg tgt cgt tga ctg gat aca ggt taa ttt cca cat cga tat agt taa atg tat tgc tgg tta cga cgg gtt cgc att tat ctg tgg aag aga cgg tgt gag aat atg ttc cgg gac cac acg gag aac aga tga cgt ctc cgg tag acg tgt atc cgg ata ctc cgt atc cta ttc cac act ttg ttt tag aaa tac atg ttc tac acc ctg atg ctc ctt tga gaa gac aat aat atc ctg gag agc att cac aga ttc tat tgt gag tcg tgt tac acg atc gcg tct cta cct gat tac tat cac atc ttc cgt tac aac tta gac aag cct gta aat gat tat tgt gag atg taa agg tat ccg aac cac acg gtg tac att gtg tat tag tct tgc tat cac ata atc tga aag cgt aag ttc ccg gag gac acg ata gac aac ata gat tac ggc ttc tgt att cgt tgt ctt tac act ttc cat tgg atg gtg cat gtg gtg cta tat ctc ttc cgt tta tta tta tac atg aga gaa aca ata tat acg agt ata ata cgg acc tca tga ttt aat aat -gt agt aat cgt cgt ctt gtt act gtt tgt ttc cta ctt ctc caa tca tat aga tta ttt ttt aaa tat ttt ctt tct atc atg gat aat att tgt aat ggt tct ttc cgt aca aca tac tgt tta gat ggt agt cgc tta gct tgg tta tga tat tgc gca taa ttt ccg gag gca aat acg ata gtc tag att gac tat cga tgg tag act cta att tat tga gtg ctt tgt cga cga gtt tac ttt tat gct cca tcg ata gat gac act gtt cta tga gat cgt cgt aca tgg gaa atg aaa tgc gtt tgt ctg aat gta tgg ctt cga gat atg tgt gat acc gaa tgt ctt ctg ttc tca ata ccg tat aca agt cgg tgt ctg aga ttc gaa tct ctt tga gga gac tta tgt cac gac tac att ttt cga tga tgg aat cta tct tat cga atg ata tat ttt tca taa ata cac ttt tat agt cct cgt tta aac aga att tag tat gta gtt ccg caa atg act cgt ccc tta ata ggc agt agg cta tta tct tct tta cgt agt gat cgt cgt agg gag aga act ccg aca tct tgt aga aca acg att taa tca tag gta gag ata ctt tca gtc tgt ggt gga taa tgt cat taa cga cat ccg cct tgt ata taa tgt ttc tgt ttt caa aca cca agt cga ata ctg tct tat cgt ctt tag tcg gaa ggt tga tgt cgt atc caa tgt ata cga ggt atg agg caa cat tgt tat tgc aat tct gga agg cgg tat gaa gag gag tca ttg tat tat agt att cgt ctt tct gaa tgt cga atc tat cta gta gat acc gta gta tat tga gag agc gac ttc cat atc ctt gat tat gtt tta tga ata gat aaa gta gat gtt gtc ctt ctt cct ttt gta att tcc cgt att ttt gtt cgt gcc aat tga gta aca tta tga gaa tat gac ctg ttg cac aat cgt tct tta tgt att cca tga tgg gtg tac aat taa gat tat tac gta tcc tcg tat cgg ctc ctc gag ata aaa gag cat aca cca cac gag gac tat gtt tgg tat act gtt gaa ggt aag tgt gta acg gcg tat ttc cga ttt tcg taa ccg cgt taa tgt ttg ctc cat gat cta tta tcg cgt aga tga atc gct tct cag ctc gca tct tag tgt gac tct ttg act tgt aat aat tgc ttt cgt gga acg cgg ata tgt gtt tac agt agt aat gaa gag aag tga gtc cat cct cat cga cgc aat tag ggt cag atc ctt tag tca ata att tgt aca gaa cgt aat agt tta agc tcc cat tga att tat atc taa gat aac aca gca ata gat cgg atg att tac taa agt cat caa tgg ggt ccg tta gta tat caa aga tct tgt tat cga ttg ata gtg aat gaa tca gat agt ggt gta gag gaa tat gtc ctt ttt tat cct tgc tat caa agt tac gca tgc cgt ggc gta aca ata tct tta ata cag atg gat taa atc gtg tat tca tcg tat agc aat gta atg gag agt tac cac att tta gtc gtt tat tca gat cgc agt gtt taa taa cta att taa aca gat gag atg ata tat cca cat caa aga atg cga gat aca tat gac aga cat tat tga cag aaa tgt gac ctt cat tat cac cgt cgt cca taa atg cgt tag gta cgt acc aca tac tat cgt taa cga tgc gca caa tct cgt cca -tt tca tcc atc ttc ata atg att tac ttt ttc ata att aga gaa aaa gat caa ggt ata aaa att aga agt gtt aga cta ta- --- --- --- --- --- --- --- --- --- --- -aa cta act tat gac tta act aac tta tga --- --- --- --- --- --- --- --- --- --- --- --- --- --- --- --- --- --- --- --- --- --- --- --- --- --- --- --- --- --- --- --- --- --- --- --- --- --- --- --- --- --- --- --- --- --- --- --- ctt aac ta- --- --- --- --- --- --- --- --- --- --- -ac tta tga ctt aac taa ctt atg act taa cta act ta- tga ctt aac taa ctt atg act taa cta act tat gac tta act aac tta tga cta tta act cat ttc aag aaa ggt ggg tgg ata gaa ctc tat atg aca gct tgt gaa aca att aga tcc cta att tct aat gga agt ttt gat agg aga ttg tca tca gtt gat aca ttg ttt att atc tca tct att aga gca cgt ctg ttt aga gct tta gtg acc tgc tcg gtt act tct gtg taa atc ttg aat cct tta gtg ata cac tgt gtc aaa act gga tgt tta gaa tac cta tgt aga ata tgg gaa gca tgc ttg ttt ttg tct cta tta tag att aac tca tac atg gtt gta tta tga att ttc atc tgc cta atg tac tcc aat tct tgt tta caa tca att ata taa tca aag agt gat gat gca tac aca tta caa agt gaa taa tct acc atc ata aaa tac ttg ata cag agc ttt atc aca tca tgg ttt tca att gta tta tta agt ata gct aat ttt ata cag tca ata gac aat ggt tct cta agc aat att tct aat att tta aga tgt gct tcc cta cgg gcg atg aca gat ccc cta tcc acg gcc acg tca aga cat gta tat cca tta ctc att act gcg ttg aca ttt gct cca ttt tct aat agc cat gat act aaa tct ata taa cct gca tag ata gcg cga taa agc aag gtc ctt cca cca gca tct agt tga ttg ata tct tca ata tat ggg ata caa agc tta taa att tct aat act gtg ggt tca tct aca agg aat ccc cta gta tac tga att att tta tat aga tct aat tta aca tca ttt tca tct ggg ata cca caa ttc aaa ata aac tca aca aca cta ctt tcc ttt tta cat att ccc cta aaa tag gca ttc aag cat tct att tta tat att aca gcc cca tga tct acc ata aga tca aca atg tct att tct aca tat gca tta gat aga tag taa agt aag aga tct tgc aca gaa tta caa ttc tta ata att ata gag aaa ata tct tcc ata taa ttc ttt gac act aat gca gat ata ata tct tta tat gta ata tat gca aac agt cta tct act ata tac tga tca ata tta tct cta tga atc cta aaa taa tca tac aga aca tct aca gga tca caa att ggt tca agg aga aat cta tca aat att ttc ctg tca aca act ggt tct aga aca taa cag tca aca cct aat cca tgt ttt tta taa tca tct acc aaa gat aat gac caa aga tcg agg tcg tcg tga aac tgc tca tcg aca gcc atg aaa tct acc gac tcc atg gtg cga atc gca ctg tct tat tcg cca ttg att ttc a-t ttt tta taa tta tgt aca tgt ttt cct tct att ctc aag agt cta caa aaa tat a-- -tt ttt tcg ata tct aag tac taa gtt ttt tta ctg ttt ttg tta ctg tct tcc att ctt cta act aaa gat ctg aga taa att ata caa tct tcg cta tcg aac cat ttt tgt agt cta aag cct gaa gta att aac caa ctg ttt tta tta gtg gct ttt ttc gat cta tcc tcg tc- --- ctc tca tca tcc tta ta- --- --- --- --- --- --- tta tta tca tta tta tca tag tct att aaa cac aaa tca tct --- --- --- --- --- --- --- --- --- --- --- --- --- --- --- --- --- --- --- --- --- --- --- --- --- --- --a cgt tta taa c-- --- --- --- --- --- --- --- --a aca ttc --- --- --- --- -tc att att aat tag ttc tgt ag- aat atc ttt aat aat ttg gct ata --- --- -ca tct gtt --- --- --- --- --- --- --- --- --- --- --- --- --- --- --- --- --- --- --c aat act --- --- --- --- --- --- --- --a tct att gat gat ttc ttt t-- --- --- --- --- --- --- --- --- --- --- --t taa gac t-- --- --- --- --- --- --- --- --- --- --- --- --- --- --- --- --- --- --- --- --- --- --- --- --- --- --- --- --- --- --- --- --- --- --- --- --- --- --- --- taa act agt --- --- --- --- --- --- --- --- --- --- --- --- --- --- --- --- --- --- --- --- --- --- --- --- --- --- --- --- --- --- --- --- --- --- --- --- --- --- --- --- --- --- --- --- --- --- --- --- --- --- --- --- --- --- --- --- --- --- --- --- --- --- --- --- --- --- --- --- --- --- --- --- --- --- --- --- --- --- --- --- --- --- --- --- --- --- --- --- --- --- --- --- --- --- --- --- --- --- --- --- --- --- --- --- --- --- --- --- --- --t atg gta atg acg atg aaa --- -tc gag tag ta- --- --a ctt cta ata aag act tga ta- --- --t cat tat cat atg ttt gat cg- --- --- --- --- --- --- --t cat agt taa tag tgt g-- --- --- --- --- --- --- --- --- --- --- --- --- --- --- --- --- --- --- --- -gc taa atg gta ctg tta ata agt tta t-- --- --- --- --- --- --- --- --- --- --- --- --- --- --- --- --- --a gac aat atc ata gta ttt tct ttc cag aat --- --- --- --t aga tta ttt ttt taa ata ctg atc ctc aca att ccg tga tgt agc agt agt tgg t-- --- --- --- --- --- --- --- --- --- --- --- --- --- --- --- --- --- --- --- --- --- --- --- --- --- --- --- -gc atg gtc tat atc gt- --- --- --- --- --- --- --- --- --- --- --- --- --- --- --- --- --- --- --- --- --- --- --- --- --- --- --- --- --- --- --- --- --- --- --- --- --- --- --- --- --- --- --- taa aat gta tca tat ata ata gtt ttc tga cgt gga gta cag aat ttt cga --- --- --- --- --- --- --- --- --- --- --- --- --- --- --- --- --- --- --- --- --- --- --- --- --- --- --- --- --- --- --- --- --- --- --- --- --- --- --- --- --- --- --- --t taa tga gtt cat ggt aag gaa ggg caa atg cct --- gta tat aat ata cat aag tta a-- --- --- --- --- --- --- --- --- --- --- --- --- --t agt ttt tta tca tat tt- --- --- --- --- --- --- --- --t cta ata cca taa taa aaa t-- --- --- --- --- --- --- --- --- --- --- --- --- --- --- --- --- --- --- --- --- --- --- --- --- --t atc att att gcg ttt g-g tag tt- --- --- --- --- ctg ccc ta- --- --- --- --- --- --- --- --t cat cta tat cac tgt cac tct c-- --- --g ctc tca cta tat ctt cta aaa tta caa --- -ac aac --- --- --- --- --- --- --- --- --- --- --- --- --- --- --- --- --- --- --- --- --- --- --- --- --- --- --- --- --- --- --- --- --- --- --- --- tgg ata ttc gat --- --- --- --- --- --- --- --- --- --- --- --- --- --- --- --- --- --- --- --- --- --- --- --- --- --- --a aca gca ttt gtg t-- --- --- --- --- --- --- --- --- --- --- --- --- --- --- --- --- -

>ON585037.1_Monkeypox_virus_isolate_Monkeypox/PT0005/2022_complete_genome

--- --- --- --- --- --- --- --- --- --- --- --- --- --- --- --- --- --- --- --- --- --- --- --- --- --- --- --- att taa ttt tac tat ttt att tag tgt cta gaa aaa aat gtg tga ccc acg acc gta gga aac tct aga ggg taa gaa aaa tca atc gtt tat aga gac cat cag aaa gag gtt taa tat ttt tgt gag acc tat cga a-- --- --- --- --- --- --- --- --- --- --- --- --- --- --- --- --- --- --- --- --- --- --- --- --- --- --- --- --- --- --- --- --- --- --- --- --- --- --- --- --- --- --- --- --- --- --- --- --- --- --- --- --- --- --- --- --- --- --- --- --- --- --- --- --- --- --- --- --- --- --- --- --- --- --- --- --- --- --- --- --- --- --- --- --- --- --- --- --- --- --- --- --- --- --- --- --- --- --- --- --- --- --- --- --- --- --- --- --- --- --- --- --- --- --- --- --- --- --- --- --- --- --- --- --- --- --- --- --- --- --- --- --- --- --- --- --- --- --- --- --- --- --- --- --- --- --- --- --- --- --- --- --- --- --- --- --- --- --- --- --- --- --- --- --- --- --- --- --- --- --- --- --- --- --- --- --- --- --- --- --- --- --- --- --- --- --- --- --- --- --- --- --- --- --- --- --- --- --- --- --- --- --- --- --- --- --- --- --- --- --- --- --- --- --- --- --- --- --- --- --- --- --- --- --- --- --- --- --- --- --- --- --- --- --- --- --- --- --- --- --- --- --- --- --- --- --- --- --- --- --- --- --- --- --- --- --- --- --- --- --- --- --- --- --- --- --- --- --- --- --- --- --- --- --- --- --- --- --- --- -ga gag aaa gga taa aaa ctt ttt acg act cca tca gaa aga ggt tta ata ttt ttg tga gac cca tcg aag aga gaa aga gat ggt tag tca aga tat ttt tct tag tac aaa agt caa tgt ttt aaa ata tat gga cga gaa tta att tgt ctg tat aaa aac ttg tgt gaa att atg tac tag aga aaa aac gtg agc agt gtc ccc tac atg gat ttt aca gat cat tta tat tcc aaa aat att aac tat ata cgt tta tta tat gat gtt aac gtg taa att ata aac att att tta tga tgc aat tgt ctg aca acc tag att ggt ata agg atg ttg ata agc tct acg aga ata tat tgt tgg acg tta tcg ttt acg aaa tag ttg aga cat cag aaa gag gtt taa tat ttt tgt gag a-c cat cga aga gag aaa gag aat aaa aat a-- --t ttt ttt ttt ttt tgt aaa act ttt tta tga gac --- --- -ca aga gaa tac gaa tag tga tca tat cgt atc aca tat tga aac aga aag aag aag taa cga gag gta act ttt tgt gaa tgt agt taa ata ttt ttg ttt tgc aaa ccg gaa tat agt gcc cgg t-- -ct ttt tta att cgt ggt gcg gtg tct gaa tcg ttc gat taa ccc aac tca tcc att ttc aga tga ata gag tta tcg att cag aca cat gct ttg agt ttt gtt gaa tcg atg agt gaa gta tca tcg gtt gca cct tca gat gcc gat ccg tcg aca tac ttg aat cca tcc ttg act tca agt tca gat gat tcc tca cac atg tct ccg ata cgt acg cta aac tct agg ttc ttg aca cat ttt gta tca acg atc gtt gaa ccg atg ata tct ttg taa ctc act ttc tta tgt gag atg tta gac cca agt act gga tgg gtc ttg atg tca ctg tct ttc tct tct tcg cta cat ctg atg tcg ata gac atc tca cag tct ttg atc ata gcc aga gct tct tca cgc gtg atc gcg gga gag tcc tta cct tgt ccc ggt gac acg ctg gac aat cta gta ttc aca gtg ttt cca tca gag gat tcg gag atg gat aaa atc ttt ggg cat ttg gtg aat cca aag ttc atg tta aga ccc gca ccg acg ata gtg taa taa gtg gtg gga tct cct ttt aca act --- --- --- --- --- --- tct tcg gat acc tca tca tct tcg gtc tct gta act tcc gtt acg gat tga caa atc tta tca ttg gtc ggt gtt tgg tct tgc ttt gtg act ttg ata ata aca tcg att ccc ata tga tgt ttg ttt tct tct tca gta cac gag gat gaa gat tgt tga aga cta gta ggc ata gca gct gcc act agg cac atg cat gcc agg aca ata tat tgt ttc atg att gct att gat tga tta ctg ttc tag atg att cta ctt tct tac cat ata ata aat tag aat ata ttt tct act ttt acg aga aat taa tta ttg tat tta tta ttt ata ggt --a aaa aaa ctt act ata agt ggg tgg gat tct ggg aat tag tga tca gtt tat gta tat cgc aac tag cgg gca tat ggc tat tga cat cga gaa cat tac cca tat gat aag aga ttg tat cat ttt cgt agt ctt gag tat tgg tat tac tat ata gta tgt aga tgt cga cgc tag ata gac agt cgc cca cta gag tta ccg tct ctg aat gcg gca tga tag tat cat tct ttg ttt tcg tta act gtt tgg aag atg aat ctt tgt tgt tac att taa tct cga aat tca gag tac ata tct ttg aag tat tct gat atc tat ttt ctc ctg taa aga atc ctg aag ttg cta cat tat taa gga cag aga agt att ctg cac gaa aga ctg gat cac aat ctt tat gat tca tgg taa tag tta gtt ccg acg ttg aga tgg att cgc tga gac cgg tag tgg tcg tcc gag tac acg atg tgt cgt tga ctg gat aca ggt taa ttt cca cat cga tat agt taa atg tat tgc tgg tta cga cgg gtt cgc att tat ctg tgg aag aga cgg tgt gag aat atg ttc cgg gac cac acg gag aac aga tga cgt ctc cgg tag acg tgt atc cgg ata ctc cgt atc cta ttc cac act ttg ttt tag aaa tac atg ttc tac acc ctg atg ctc ctt tga gaa gac aat aat atc ctg gag agc att cac aga ttc tat tgt gag tcg tgt tac acg atc gcg tct cta cct gat tac tat cac atc ttc cgt tac aac tta gac aag cct gta aat gat tat tgt gag atg taa agg tat ccg aac cac acg gtg tac att gtg tat tag tct tgc tat cac ata atc tga aag cgt aag ttc ccg gag gac acg ata gac aac ata gat tac ggc ttc tgt att cgt tgt ctt tac act ttc cat tgg atg gtg cat gtg gtg cta tat ctc ttc cgt tta tta tta tac atg aga gaa aca ata tat acg agt ata ata cgg acc tca tga ttt aat aat -gt agt aat cgt cgt ctt gtt act gtt tgt ttc cta ctt ctc caa tca tat aga tta ttt ttt aaa tat ttt ctt tct atc atg gat aat att tgt aat ggt tct ttc cgt aca aca tac tgt tta gat ggt agt cgc tta gct tgg tta tga tat tgc gca taa ttt ccg gag gca aat acg ata gtc tag att gac tat cga tgg tag act cta att tat tga gtg ctt tgt cga cga gtt tac ttt tat gct cca tcg ata gat gac act gtt cta tga gat cgt cgt aca tgg gaa atg aaa tgc gtt tgt ctg aat gta tgg ctt cga gat atg tgt gat acc gaa tgt ctt ctg ttc tca ata ccg tat aca agt cgg tgt ctg aga ttc gaa tct ctt tga gga gac tta tgt cac gac tac att ttt cga tga tgg aat cta tct tat cga atg ata tat ttt tca taa ata cac ttt tat agt cct cgt tta aac aga att tag tat gta gtt ccg caa atg act cgt ccc tta ata ggc agt agg cta tta tct tct tta cgt agt gat cgt cgt agg gag aga act ccg aca tct tgt aga aca acg att taa tca tag gta gag ata ctt tca gtc tgt ggt gga taa tgt cat taa cga cat ccg cct tgt ata taa tgt ttc tgt ttt caa aca cca agt cga ata ctg tct tat cgt ctt tag tcg gaa ggt tga tgt cgt atc caa tgt ata cga ggt atg agg caa cat tgt tat tgc aat tct gga agg cgg tat gaa gag gag tca ttg tat tat agt att cgt ctt tct gaa tgt cga atc tat cta gta gat acc gta gta tat tga gag agc gac ttc cat atc ctt gat tat gtt tta tga ata gat aaa gta gat gtt gtc ctt ctt cct ttt gta att tcc cgt att ttt gtt cgt gcc aat tga gta aca tta tga gaa tat gac ctg ttg cac aat cgt tct tta tgt att cca tga tgg gtg tac aat taa gat tat tac gta tcc tcg tat cgg ctc ctc gag ata aaa gag cat aca cca cac gag gac tat gtt tgg tat act gtt gaa ggt aag tgt gta acg gcg tat ttc cga ttt tcg taa ccg cgt taa tgt ttg ctc cat gat cta tta tcg cgt aga tga atc gct tct cag ctc gca tct tag tgt gac tct ttg act tgt aat aat tgc ttt cgt gga acg cgg ata tgt gtt tac agt agt aat gaa gag aag tga gtc cat cct cat cga cgc aat tag ggt cag atc ctt tag tca ata att tgt aca gaa cgt aat agt tta agc tcc cat tga att tat atc taa gat aac aca gca ata gat cgg atg att tac taa agt cat caa tgg ggt ccg tta gta tat caa aga tct tgt tat cga ttg ata gtg aat gaa tca gat agt ggt gta gag gaa tat gtc ctt ttt tat cct tgc tat caa agt tac gca tgc cgt ggc gta aca ata tct tta ata cag atg gat taa atc gtg tat tca tcg tat agc aat gta atg gag agt tac cac att tta gtc gtt tat tca gat cgc agt gtt taa taa cta att taa aca gat gag atg ata tat cca cat caa aga atg cga gat aca tat gac aga cat tat tga cag aaa tgt gac ctt cat tat cac cgt cgt cca taa atg cgt tag gta cgt acc aca tac tat cgt taa cga tgc gca caa tct cgt cca -tt tca tcc atc ttc ata atg att tac ttt ttc ata att aga gaa aaa gat caa ggt ata aaa att aga agt gtt aga cta ta- --- --- --- --- --- --- --- --- --- --- -aa cta act tat gac tta act aac tta tga --- --- --- --- --- --- --- --- --- --- --- --- --- --- --- --- --- --- --- --- --- --- --- --- --- --- --- --- --- --- --- --- --- --- --- --- --- --- --- --- --- --- --- --- --- --- --- --- ctt aac ta- --- --- --- --- --- --- --- --- --- --- -ac tta tga ctt aac taa ctt atg act taa cta act ta- tga ctt aac taa ctt atg act taa cta act tat gac tta act aac tta tga cta tta act cat ttc aag aaa ggt ggg tgg ata gaa ctc tat atg aca gct tgt gaa aca att aga tcc cta att tct aat gga agt ttt gat agg aga ttg tca tca gtt gat aca ttg ttt att atc tca tct att aga gca cgt ctg ttt aga gct tta gtg acc tgc tcg gtt act tct gtg taa atc ttg aat cct tta gtg ata cac tgt gtc aaa act gga tgt tta gaa tac cta tgt aga ata tgg gaa gca tgc ttg ttt ttg tct cta tta tag att aac tca tac atg gtt gta tta tga att ttc atc tgc cta atg tac tcc aat tct tgt tta caa tca att ata taa tca aag agt gat gat gca tac aca tta caa agt gaa taa tct acc atc ata aaa tac ttg ata cag agc ttt atc aca tca tgg ttt tca att gta tta tta agt ata gct aat ttt ata cag tca ata gac aat ggt tct cta agc aat att tct aat att tta aga tgt gct tcc cta cgg gcg atg aca gat ccc cta tcc acg gcc acg tca aga cat gta tat cca tta ctc att act gcg ttg aca ttt gct cca ttt tct aat agc cat gat act aaa tct ata taa cct gca tag ata gcg cga taa agc aag gtc ctt cca cca gca tct agt tga ttg ata tct tca ata tat ggg ata caa agc tta taa att tct aat act gtg ggt tca tct aca agg aat ccc cta gta tac tga att att tta tat aga tct aat tta aca tca ttt tca tct ggg ata cca caa ttc aaa ata aac tca aca aca cta ctt tcc ttt tta cat att ccc cta aaa tag gca ttc aag cat tct att tta tat att aca gcc cca tga tct acc ata aga tca aca atg tct att tct aca tat gca tta gat aga tag taa agt aag aga tct tgc aca gaa tta caa ttc tta ata att ata gag aaa ata tct tcc ata taa ttc ttt gac act aat gca gat ata ata tct tta tat gta ata tat gca aac agt cta tct act ata tac tga tca ata tta tct cta tga atc cta aaa taa tca tac aga aca tct aca gga tca caa att ggt tca agg aga aat cta tca aat att ttc ctg tca aca act ggt tct aga aca taa cag tca aca cct aat cca tgt ttt tta taa tca tct acc aaa gat aat gac caa aga tcg agg tcg tcg tga aac tgc tca tcg aca gcc atg aaa tct acc gac tcc atg gtg cga atc gca ctg tct tat tcg cca ttg att ttc a-t ttt tta taa tta tgt aca tgt ttt cct tct att ctc aag agt cta caa aaa tat a-- -tt ttt tcg ata tct aag tac taa gtt ttt tta ctg ttt ttg tta ctg tct tcc att ctt cta act aaa gat ctg aga taa att ata caa tct tcg cta tcg aac cat ttt tgt agt cta aag cct gaa gta att aac caa ctg ttt tta tta gtg gct ttt ttc gat cta tcc tcg tc- --- ctc tca tca tcc tta ta- --- --- --- --- --- --- tta tta tca tta tta tca tag tct att aaa cac aaa tca tct --- --- --- --- --- --- --- --- --- --- --- --- --- --- --- --- --- --- --- --- --- --- --- --- --- --- --a cgt tta taa c-- --- --- --- --- --- --- --- --a aca ttc --- --- --- --- -tc att att aat tag ttc tgt ag- aat atc ttt aat aat ttg gct ata --- --- -ca tct gtt --- --- --- --- --- --- --- --- --- --- --- --- --- --- --- --- --- --- --c aat act --- --- --- --- --- --- --- --a tct att gat gat ttc ttt t-- --- --- --- --- --- --- --- --- --- --- --t taa gac t-- --- --- --- --- --- --- --- --- --- --- --- --- --- --- --- --- --- --- --- --- --- --- --- --- --- --- --- --- --- --- --- --- --- --- --- --- --- --- --- taa act agt --- --- --- --- --- --- --- --- --- --- --- --- --- --- --- --- --- --- --- --- --- --- --- --- --- --- --- --- --- --- --- --- --- --- --- --- --- --- --- --- --- --- --- --- --- --- --- --- --- --- --- --- --- --- --- --- --- --- --- --- --- --- --- --- --- --- --- --- --- --- --- --- --- --- --- --- --- --- --- --- --- --- --- --- --- --- --- --- --- --- --- --- --- --- --- --- --- --- --- --- --- --- --- --- --- --- --- --- --- --t atg gta atg acg atg aaa --- -tc gag tag ta- --- --a ctt cta ata aag act tga ta- --- --t cat tat cat atg ttt gat cg- --- --- --- --- --- --- --t cat agt taa tag tgt g-- --- --- --- --- --- --- --- --- --- --- --- --- --- --- --- --- --- --- --- -gc taa atg gta ctg tta ata agt tta t-- --- --- --- --- --- --- --- --- --- --- --- --- --- --- --- --- --a gac aat atc ata gta ttt tct ttc cag aat --- --- --- --t aga tta ttt ttt taa ata ctg atc ctc aca att ccg tga tgt agc agt agt tgg t-- --- --- --- --- --- --- --- --- --- --- --- --- --- --- --- --- --- --- --- --- --- --- --- --- --- --- --- -gc atg gtc tat atc gt- --- --- --- --- --- --- --- --- --- --- --- --- --- --- --- --- --- --- --- --- --- --- --- --- --- --- --- --- --- --- --- --- --- --- --- --- --- --- --- --- --- --- --- taa aat gta tca tat ata ata gtt ttc tga cgt gga gta cag aat ttt cga --- --- --- --- --- --- --- --- --- --- --- --- --- --- --- --- --- --- --- --- --- --- --- --- --- --- --- --- --- --- --- --- --- --- --- --- --- --- --- --- --- --- --- --t taa tga gtt cat ggt aag gaa ggg caa atg cct --- gta tat aat ata cat aag tta a-- --- --- --- --- --- --- --- --- --- --- --- --- --t agt ttt tta tca tat tt- --- --- --- --- --- --- --- --t cta ata cca taa taa aaa t-- --- --- --- --- --- --- --- --- --- --- --- --- --- --- --- --- --- --- --- --- --- --- --- --- --t atc att att gcg ttt g-g tag tt- --- --- --- --- ctg ccc ta- --- --- --- --- --- --- --- --t cat cta tat cac tgt cac tct c-- --- --g ctc tca cta tat ctt cta aaa tta caa --- -ac aac --- --- --- --- --- --- --- --- --- --- --- --- --- --- --- --- --- --- --- --- --- --- --- --- --- --- --- --- --- --- --- --- --- --- --- --- tgg ata ttc gat --- --- --- --- --- --- --- --- --- --- --- --- --- --- --- --- --- --- --- --- --- --- --- --- --- --- --a aca gca ttt gtg t-- --- --- --- --- --- --- --- --- --- --- --- --- --- --- --- --- -

>ON585032.1_Monkeypox_virus_isolate_Monkeypox/PT0004/2022_complete_genome

--- --- --- --- --- --- --- --- --- --- --- --- --- --- --- --- --- --- --- --- --- --- --- --- --- --- --- --- att taa ttt tac tat ttt att tag tgt cta gaa aaa aat gtg tga ccc acg acc gta gga aac tct aga ggg taa gaa aaa tca atc gtt tat aga gac cat cag aaa gag gtt taa tat ttt tgt gag acc tat cga a-- --- --- --- --- --- --- --- --- --- --- --- --- --- --- --- --- --- --- --- --- --- --- --- --- --- --- --- --- --- --- --- --- --- --- --- --- --- --- --- --- --- --- --- --- --- --- --- --- --- --- --- --- --- --- --- --- --- --- --- --- --- --- --- --- --- --- --- --- --- --- --- --- --- --- --- --- --- --- --- --- --- --- --- --- --- --- --- --- --- --- --- --- --- --- --- --- --- --- --- --- --- --- --- --- --- --- --- --- --- --- --- --- --- --- --- --- --- --- --- --- --- --- --- --- --- --- --- --- --- --- --- --- --- --- --- --- --- --- --- --- --- --- --- --- --- --- --- --- --- --- --- --- --- --- --- --- --- --- --- --- --- --- --- --- --- --- --- --- --- --- --- --- --- --- --- --- --- --- --- --- --- --- --- --- --- --- --- --- --- --- --- --- --- --- --- --- --- --- --- --- --- --- --- --- --- --- --- --- --- --- --- --- --- --- --- --- --- --- --- --- --- --- --- --- --- --- --- --- --- --- --- --- --- --- --- --- --- --- --- --- --- --- --- --- --- --- --- --- --- --- --- --- --- --- --- --- --- --- --- --- --- --- --- --- --- --- --- --- --- --- --- --- --- --- --- --- --- --- --- -ga gag aaa gga taa aaa ctt ttt acg act cca tca gaa aga ggt tta ata ttt ttg tga gac cca tcg aag aga gaa aga gat ggt tag tca aga tat ttt tct tag tac aaa agt caa tgt ttt aaa ata tat gga cga gaa tta att tgt ctg tat aaa aac ttg tgt gaa att atg tac tag aga aaa aac gtg agc agt gtc ccc tac atg gat ttt aca gat cat tta tat tcc aaa aat att aac tat ata cgt tta tta tat gat gtt aac gtg taa att ata aac att att tta tga tgc aat tgt ctg aca acc tag att ggt ata agg atg ttg ata agc tct acg aga ata tat tgt tgg acg tta tcg ttt acg aaa tag ttg aga cat cag aaa gag gtt taa tat ttt tgt gag a-c cat cga aga gag aaa gag aat aaa aat a-- --t ttt ttt ttt ttt tgt aaa act ttt tta tga gac --- --- -ca aga gaa tac gaa tag tga tca tat cgt atc aca tat tga aac aga aag aag aag taa cga gag gta act ttt tgt gaa tgt agt taa ata ttt ttg ttt tgc aaa ccg gaa tat agt gcc cgg t-- -ct ttt tta att cgt ggt gcg gtg tct gaa tcg ttc gat taa ccc aac tca tcc att ttc aga tga ata gag tta tcg att cag aca cat gct ttg agt ttt gtt gaa tcg atg agt gaa gta tca tcg gtt gca cct tca gat gcc gat ccg tcg aca tac ttg aat cca tcc ttg act tca agt tca gat gat tcc tca cac atg tct ccg ata cgt acg cta aac tct agg ttc ttg aca cat ttt gta tca acg atc gtt gaa ccg atg ata tct ttg taa ctc act ttc tta tgt gag atg tta gac cca agt act gga tgg gtc ttg atg tca ctg tct ttc tct tct tcg cta cat ctg atg tcg ata gac atc tca cag tct ttg atc ata gcc aga gct tct tca cgc gtg atc gcg gga gag tcc tta cct tgt ccc ggt gac acg ctg gac aat cta gta ttc aca gtg ttt cca tca gag gat tcg gag atg gat aaa atc ttt ggg cat ttg gtg aat cca aag ttc atg tta aga ccc gca ccg acg ata gtg taa taa gtg gtg gga tct cct ttt aca act --- --- --- --- --- --- tct tcg gat acc tca tca tct tcg gtc tct gta act tcc gtt acg gat tga caa atc tta tca ttg gtc ggt gtt tgg tct tgc ttt gtg act ttg ata ata aca tcg att ccc ata tga tgt ttg ttt tct tct tca gta cac gag gat gaa gat tgt tga aga cta gta ggc ata gca gct gcc act agg cac atg cat gcc agg aca ata tat tgt ttc atg att gct att gat tga tta ctg ttc tag atg att cta ctt tct tac cat ata ata aat tag aat ata ttt tct act ttt acg aga aat taa tta ttg tat tta tta ttt ata ggt --a aaa aaa ctt act ata agt ggg tgg gat tct ggg aat tag tga tca gtt tat gta tat cgc aac tag cgg gca tat ggc tat tga cat cga gaa cat tac cca tat gat aag aga ttg tat cat ttt cgt agt ctt gag tat tgg tat tac tat ata gta tgt aga tgt cga cgc tag ata gac agt cgc cca cta gag tta ccg tct ctg aat gcg gca tga tag tat cat tct ttg ttt tcg tta act gtt tgg aag atg aat ctt tgt tgt tac att taa tct cga aat tca gag tac ata tct ttg aag tat tct gat atc tat ttt ctc ctg taa aga atc ctg aag ttg cta cat tat taa gga cag aga agt att ctg cac gaa aga ctg gat cac aat ctt tat gat tca tgg taa tag tta gtt ccg acg ttg aga tgg att cgc tga gac cgg tag tgg tcg tcc gag tac acg atg tgt cgt tga ctg gat aca ggt taa ttt cca cat cga tat agt taa atg tat tgc tgg tta cga cgg gtt cgc att tat ctg tgg aag aga cgg tgt gag aat atg ttc cgg gac cac acg gag aac aga tga cgt ctc cgg tag acg tgt atc cgg ata ctc cgt atc cta ttc cac act ttg ttt tag aaa tac atg ttc tac acc ctg atg ctc ctt tga gaa gac aat aat atc ctg gag agc att cac aga ttc tat tgt gag tcg tgt tac acg atc gcg tct cta cct gat tac tat cac atc ttc cgt tac aac tta gac aag cct gta aat gat tat tgt gag atg taa agg tat ccg aac cac acg gtg tac att gtg tat tag tct tgc tat cac ata atc tga aag cgt aag ttc ccg gag gac acg ata gac aac ata gat tac ggc ttc tgt att cgt tgt ctt tac act ttc cat tgg atg gtg cat gtg gtg cta tat ctc ttc cgt tta tta tta tac atg aga gaa aca ata tat acg agt ata ata cgg acc tca tga ttt aat aat -gt agt aat cgt cgt ctt gtt act gtt tgt ttc cta ctt ctc caa tca tat aga tta ttt ttt aaa tat ttt ctt tct atc atg gat aat att tgt aat ggt tct ttc cgt aca aca tac tgt tta gat ggt agt cgc tta gct tgg tta tga tat tgc gca taa ttt ccg gag gca aat acg ata gtc tag att gac tat cga tgg tag act cta att tat tga gtg ctt tgt cga cga gtt tac ttt tat gct cca tcg ata gat gac act gtt cta tga gat cgt cgt aca tgg gaa atg aaa tgc gtt tgt ctg aat gta tgg ctt cga gat atg tgt gat acc gaa tgt ctt ctg ttc tca ata ccg tat aca agt cgg tgt ctg aga ttc gaa tct ctt tga gga gac tta tgt cac gac tac att ttt cga tga tgg aat cta tct tat cga atg ata tat ttt tca taa ata cac ttt tat agt cct cgt tta aac aga att tag tat gta gtt ccg caa atg act cgt ccc tta ata ggc agt agg cta tta tct tct tta cgt agt gat cgt cgt agg gag aga act ccg aca tct tgt aga aca acg att taa tca tag gta gag ata ctt tca gtc tgt ggt gga taa tgt cat taa cga cat ccg cct tgt ata taa tgt ttc tgt ttt caa aca cca agt cga ata ctg tct tat cgt ctt tag tcg gaa ggt tga tgt cgt atc caa tgt ata cga ggt atg agg caa cat tgt tat tgc aat tct gga agg cgg tat gaa gag gag tca ttg tat tat agt att cgt ctt tct gaa tgt cga atc tat cta gta gat acc gta gta tat tga gag agc gac ttc cat atc ctt gat tat gtt tta tga ata gat aaa gta gat gtt gtc ctt ctt cct ttt gta att tcc cgt att ttt gtt cgt gcc aat tga gta aca tta tga gaa tat gac ctg ttg cac aat cgt tct tta tgt att cca tga tgg gtg tac aat taa gat tat tac gta tcc tcg tat cgg ctc ctc gag ata aaa gag cat aca cca cac gag gac tat gtt tgg tat act gtt gaa ggt aag tgt gta acg gcg tat ttc cga ttt tcg taa ccg cgt taa tgt ttg ctc cat gat cta tta tcg cgt aga tga atc gct tct cag ctc gca tct tag tgt gac tct ttg act tgt aat aat tgc ttt cgt gga acg cgg ata tgt gtt tac agt agt aat gaa gag aag tga gtc cat cct cat cga cgc aat tag ggt cag atc ctt tag tca ata att tgt aca gaa cgt aat agt tta agc tcc cat tga att tat atc taa gat aac aca gca ata gat cgg atg att tac taa agt cat caa tgg ggt ccg tta gta tat caa aga tct tgt tat cga ttg ata gtg aat gaa tca gat agt ggt gta gag gaa tat gtc ctt ttt tat cct tgc tat caa agt tac gca tgc cgt ggc gta aca ata tct tta ata cag atg gat taa atc gtg tat tca tcg tat agc aat gta atg gag agt tac cac att tta gtc gtt tat tca gat cgc agt gtt taa taa cta att taa aca gat gag atg ata tat cca cat caa aga atg cga gat aca tat gac aga cat tat tga cag aaa tgt gac ctt cat tat cac cgt cgt cca taa atg cgt tag gta cgt acc aca tac tat cgt taa cga tgc gca caa tct cgt cca -tt tca tcc atc ttc ata atg att tac ttt ttc ata att aga gaa aaa gat caa ggt ata aaa att aga agt gtt aga cta ta- --- --- --- --- --- --- --- --- --- --- -aa cta act tat gac tta act aac tta tga --- --- --- --- --- --- --- --- --- --- --- --- --- --- --- --- --- --- --- --- --- --- --- --- --- --- --- --- --- --- --- --- --- --- --- --- --- --- --- --- --- --- --- --- --- --- --- --- ctt aac ta- --- --- --- --- --- --- --- --- --- --- -ac tta tga ctt aac taa ctt atg act taa cta act ta- tga ctt aac taa ctt atg act taa cta act tat gac tta act aac tta tga cta tta act cat ttc aag aaa ggt ggg tgg ata gaa ctc tat atg aca gct tgt gaa aca att aga tcc cta att tct aat gga agt ttt gat agg aga ttg tca tca gtt gat aca ttg ttt att atc tca tct att aga gca cgt ctg ttt aga gct tta gtg acc tgc tcg gtt act tct gtg taa atc ttg aat cct tta gtg ata cac tgt gtc aaa act gga tgt tta gaa tac cta tgt aga ata tgg gaa gca tgc ttg ttt ttg tct cta tta tag att aac tca tac atg gtt gta tta tga att ttc atc tgc cta atg tac tcc aat tct tgt tta caa tca att ata taa tca aag agt gat gat gca tac aca tta caa agt gaa taa tct acc atc ata aaa tac ttg ata cag agc ttt atc aca tca tgg ttt tca att gta tta tta agt ata gct aat ttt ata cag tca ata gac aat ggt tct cta agc aat att tct aat att tta aga tgt gct tcc cta cgg gcg atg aca gat ccc cta tcc acg gcc acg tca aga cat gta tat cca tta ctc att act gcg ttg aca ttt gct cca ttt tct aat agc cat gat act aaa tct ata taa cct gca tag ata gcg cga taa agc aag gtc ctt cca cca gca tct agt tga ttg ata tct tca ata tat ggg ata caa agc tta taa att tct aat act gtg ggt tca tct aca agg aat ccc cta gta tac tga att att tta tat aga tct aat tta aca tca ttt tca tct ggg ata cca caa ttc aaa ata aac tca aca aca cta ctt tcc ttt tta cat att ccc cta aaa tag gca ttc aag cat tct att tta tat att aca gcc cca tga tct acc ata aga tca aca atg tct att tct aca tat gca tta gat aga tag taa agt aag aga tct tgc aca gaa tta caa ttc tta ata att ata gag aaa ata tct tcc ata taa ttc ttt gac act aat gca gat ata ata tct tta tat gta ata tat gca aac agt cta tct act ata tac tga tca ata tta tct cta tga atc cta aaa taa tca tac aga aca tct aca gga tca caa att ggt tca agg aga aat cta tca aat att ttc ctg tca aca act ggt tct aga aca taa cag tca aca cct aat cca tgt ttt tta taa tca tct acc aaa gat aat gac caa aga tcg agg tcg tcg tga aac tgc tca tcg aca gcc atg aaa tct acc gac tcc atg gtg cga atc gca ctg tct tat tc? ??? ??? ??? ??? ?-? ??? ??? ??? ??? ??? ??? ??? ??? ??? ??? ??? ??? ??? ??? ??? ??? ??? ??? ?-- -?? ??? ??? ??? tct aag tac taa gtt ttt tta ctg ttt ttg tta ctg tct tcc att ctt cta act aaa gat ctg aga taa att ata caa tct tcg cta tcg aac cat ttt tgt agt cta aag cct gaa gta att aac caa ctg ttt tta tta gtg gct ttt ttc gat cta tcc tcg tc- --- ctc tca tca tcc tta ta- --- --- --- --- --- --- tta tta tca tta tta tca tag tct att aaa cac aaa tca tct --- --- --- --- --- --- --- --- --- --- --- --- --- --- --- --- --- --- --- --- --- --- --- --- --- --- --a cgt tta taa c-- --- --- --- --- --- --- --- --a aca ttc --- --- --- --- -tc att att aat tag ttc tgt ag- aat atc ttt aat aat ttg gct ata --- --- -ca tct gtt --- --- --- --- --- --- --- --- --- --- --- --- --- --- --- --- --- --- --c aat act --- --- --- --- --- --- --- --a tct att gat gat ttc ttt t-- --- --- --- --- --- --- --- --- --- --- --t taa gac t-- --- --- --- --- --- --- --- --- --- --- --- --- --- --- --- --- --- --- --- --- --- --- --- --- --- --- --- --- --- --- --- --- --- --- --- --- --- --- --- taa act agt --- --- --- --- --- --- --- --- --- --- --- --- --- --- --- --- --- --- --- --- --- --- --- --- --- --- --- --- --- --- --- --- --- --- --- --- --- --- --- --- --- --- --- --- --- --- --- --- --- --- --- --- --- --- --- --- --- --- --- --- --- --- --- --- --- --- --- --- --- --- --- --- --- --- --- --- --- --- --- --- --- --- --- --- --- --- --- --- --- --- --- --- --- --- --- --- --- --- --- --- --- --- --- --- --- --- --- --- --- --t atg gta atg acg atg aaa --- -tc gag tag ta- --- --a ctt cta ata aag act tga ta- --- --t cat tat cat atg ttt gat cg- --- --- --- --- --- --- --t cat agt taa tag tgt g-- --- --- --- --- --- --- --- --- --- --- --- --- --- --- --- --- --- --- --- -gc taa atg gta ctg tta ata agt tta t-- --- --- --- --- --- --- --- --- --- --- --- --- --- --- --- --- --a gac aat atc ata gta ttt tct ttc cag aat --- --- --- --t aga tta ttt ttt taa ata ctg atc ctc aca att ccg tga tgt agc agt agt tgg t-- --- --- --- --- --- --- --- --- --- --- --- --- --- --- --- --- --- --- --- --- --- --- --- --- --- --- --- -gc atg gtc tat atc gt- --- --- --- --- --- --- --- --- --- --- --- --- --- --- --- --- --- --- --- --- --- --- --- --- --- --- --- --- --- --- --- --- --- --- --- --- --- --- --- --- --- --- --- taa aat gta tca tat ata ata gtt ttc tga cgt gga gta cag aat ttt cga --- --- --- --- --- --- --- --- --- --- --- --- --- --- --- --- --- --- --- --- --- --- --- --- --- --- --- --- --- --- --- --- --- --- --- --- --- --- --- --- --- --- --- --t taa tga gtt cat ggt aag gaa ggg caa atg cct --- gta tat aat ata cat aag tta a-- --- --- --- --- --- --- --- --- --- --- --- --- --t agt ttt tta tca tat tt- --- --- --- --- --- --- --- --t cta ata cca taa taa aaa t-- --- --- --- --- --- --- --- --- --- --- --- --- --- --- --- --- --- --- --- --- --- --- --- --- --t atc att att gcg ttt g-g tag tt- --- --- --- --- ctg ccc ta- --- --- --- --- --- --- --- --t cat cta tat cac tgt cac tct c-- --- --g ctc tca cta tat ctt cta aaa tta caa --- -ac aac --- --- --- --- --- --- --- --- --- --- --- --- --- --- --- --- --- --- --- --- --- --- --- --- --- --- --- --- --- --- --- --- --- --- --- --- tgg ata ttc gat --- --- --- --- --- --- --- --- --- --- --- --- --- --- --- --- --- --- --- --- --- --- --- --- --- --- --a aca gca ttt gtg t-- --- --- --- --- --- --- --- --- --- --- --- --- --- --- --- --- -

>AY741551.1_Monkeypox_virus_isolate_Sierra_Leone_complete_genome

--- --- --- --- --- --- --- --- --- --- --- --- --- --- --- --- --- --- --- --- --- --- --- --- --- --- --- --- --- --- --- --- --- --- --- --- --- --- --- --- --- --- --- --- --- --- --- --- --- --- --- --g taa gaa aaa tca atc gtt tat aga gac cat cag aaa gag gtt taa tat ttt tgt gag acc cat cga aga gag aaa g-- --- --- --- --- --- --- --- --- --- --- --- --- --- --- --- --- --- --- --- --- --- --- --- --- --- --- --- --- --- --- --- --- --- --- --- --- --- --- --- --- --- --- --- --- --- --- --- --- --- --- --- --- --- --- --- --- --- --- --- --- --- --- --- --- --- --- --- --- --- --- --- --- --- --- --- --- --- --- --- --- --- --- --- --- --- --- --- --- --- --- --- --- --- --- --- --- --- --- --- --- --- --- --- --- --- --- --- --- --- --- --- --- --- --- --- --- --- --- --- --- --- --- --- --- --- --- --- --- --- --- --- --- --- --- --- --- --- --- --- --- --- --- --- --- --- --- --- --- --- --- --- --- --- --- --- --- --- --- --- --- --- --- --- --- --- --- --- --- --- --- --- --- --- --- --- --- --- --- --- --- --- --- --- --- --- --- --- --- --- --- --- --- --- --- --- --- --- --- --- --- --- --- --- --- --- --- --- --- --- --- --- --- --- --- --- --- --- --- --- --- --- --- --- --- --- --- --- --- --- --- --- --- --- --- --- --- --- --- --- --- --- --- --- --- --- --- --- --- --- --- --- --- --- --- --- --- --- --- --- --- --- --- --- --- --- --- --- --- --- --- --- --- --- --- --- --- --- --- --- -ga taa aaa ctt ttt acg act cca tca gaa aga ggt tta ata ttt ttg tga gac cca tcg aag aga gaa aga gat ggt tag tca aga tat ttt tct tag tac aaa agt caa tgt ttt aaa ata tat gga cga gaa tta att tgt ctg tat aaa aac ttg tgt gaa att atg tac tag aga aaa aac gtg agc agt gtc ccc tac atg gat ttt aca gat cat tta tat tcc aaa aat att aac tat ata cgt tta tta tat gat gtt aac gtg taa att ata aac att att tta tga tgc aat tgt ctg aca acc tag att ggt ata agg atg ttg ata agc tct acg aga ata tat tgt tgg acg tta tcg ttt acg aaa tag ttg aga cat cag aaa gag gtt taa tat ttt tgt gag a-c cat cga aga gag aaa tag aat aaa aat att ttt ttt ttt ttt ttt tgt aaa act ttt tta tga gac --- --- -ca aga gaa tac gaa tag tga tca tat cgt atc aca tat tga aac aga aag aag aag taa cga gag gta act ttt tgt aaa tgt agt taa ata ttt ttg ttt tgc aaa ccg gaa tat agt gcc cgg t-- -ct ttt tta att cgt ggt gcg gtg tct gaa tcg ttc gat taa ccc aac tca tcc att ttc aga tga ata gag tta tcg att cag aca cat gct ttg agt ttt gtt gaa tcg atg agt gaa gta tca tcg gtt gca cct tca gat gcc gat ccg tcg aca tac ttg aat cca tcc ttg act tca agt tca gat gat tcc tca cac atg tct ccg ata cgt acg cta aac tct agg ttc ttg aca cat ttt gta tca acg atc gtt gaa ccg atg ata tct ttg taa ctc act ttc tta tgt gag atg tta gac cca agt act gga tgg gtc ttg atg tca ctg tct ttc tct tct tcg cta cat ctg atg tcg ata gac atc tca cag tct ttg atc ata gcc aga gct tct tca cgc gtg atc gcg gga gag tcc tta cct tgt ccc ggt gac acg ctg gac aat cta gta ttc aca gtg ttt cca tca gag gat tcg gag atg gat gaa atc ttt ggg cat ttg gtg aat cca aag ttc atg tta aga ccc gca ccg acg ata gtg taa taa gtg gtg gga tct cct ttt aca act --- --- --- --- --- --- tct tcg gat acc tca tca tct tcg gtc tct gta act tcc gtt acg gat tga caa atc tta tca ttg gtc ggt gtt tgg tct tgc ttt gtg act ttg ata ata aca tcg att ccc ata tga tgt ttg ttt tct tct tca gta cac gag gat gaa gat tgt tga aga cta gta ggc ata gca gct gcc act agg cac atg cat gcc agg aca ata tat tgt ttc atg att gct att gat tga tta ctg ttc tag atg att cta ctt tct tac cat ata ata aat tag aat ata ttt tct act ttt acg aga aat taa tta ttg tat tta tta ttt atg ggt --a aaa aaa ctt act ata agt ggg tgg gat tct ggg aat tag tga tca gtt tat gta tat cgc aac tag cgg gca tat ggc tat tga cat cga gaa cat tac cca tat gat aag aga ttg tat ccg ttt cgt agt ctt gag tat tgg tat tac tat ata gta tgt aga tgt cga cgc tag ata gac agt cgc cca cta gag tta ccg tct ctg aat gcg gca tga tag tat cat tct ttg ttt tcg tta act gtt tgg aag atg aat ctt tgt tgt tac att taa tct cga aat tca gag tac ata tct ttg aag tat tct gat atc tat ttt ctc ctg taa aga atc ctg aag ttg cta cat tat taa gga cag aga agt att ctg cac gaa aga ctg gat cac aat ctt tat gat tca tgg taa tag tta gtt ccg acg ttg aga tgg att cgc tga gac cgg tag tgg tcg tcc gag tac acg atg tgt cgt tga ctg gat aca ggt taa ttt cca cat cga tat agt taa atg tat tgc tgg tta cga cgg gtt cgc att tat ctg tgg aag aga cgg tgt gag aat atg ttc cgg gac cac acg gag aac aga tga cgt ctc cgg tag acg tgt atc cgg ata ctc cgt atc cta ttc cac act ttg ttt tag aaa tac atg ttc tac acc cta atg ctc ctt tga gaa gac aat aat atc ctg gag agc att cac aga ttc tat tgt gag tcg tgt tac acg atc gcg tct cta cct gat tac tat cac atc ttc cgt tac aac tta gac aag cct gta aat gat tat tgt gag atg taa agg tat ccg aac cac acg gtg tac att gtg tat tag tct tgc tat cac ata atc tgg aag cgt aag ttc ccg gag gac acg ata gac aac ata gat tac ggc ttc tgt att cgt tgt ctt tac act ttc cat tgg atg gtg cat gtg gtg cta tat ctc ttc cgt tta tta tta tac atg aga gaa aca ata tat acg agt ata ata cgg acc tca tga ttt aat aat -gt agt aat cgt cgt ctt gtt act gtt tgt ttc cta ctt ctc caa tca tat aga tta ttt ttt aaa tat ttt ctt tct atc atg gat aat att tgt aat ggt tct ttc cgt aca aca tac tgt tta gat ggt agt cgc tta gct tgg tta tga tat tgc gca taa ttt ccg gag gca aat acg ata gtc tag att gat tat cga tgg tag act cta att tat tga gtg ctt tgt cga cga gtt tac ttt tat gct cca tcg ata gat gac act gtt cta tga gat cgt cgt aca tgg gaa atg aaa tgc gtt tgt ctg aat gta tgg ctt cga gat agg tgt gat acc gga tgt ctt ctg ttc tca ata ccg tat aca agt cgg tgt ctg aga ttc gaa tct ctt tga gga gac tta tgt cac gac tac att ttt cga tga tgg aat cta tct tat cga atg ata tat ttt tca taa ata cac ttt tat agt cct cgt tta aac aga att tag tat gta gtt ccg caa atg act cgt ccc tta ata ggc agt agg cta tta tct tct tta cgt agt gat cgt cgt agg gag aga act ccg aca tct tgt aga aca acg att taa tca tag gta gag ata ctt tca gtc tgt ggt gga tga tgt cat taa caa cat ccg cct tgt ata tga tgt ttc tgt ttt caa aca cca agt cga ata ctg tct tat cgt ctt tag tcg gaa ggt tga tgt cgt atc cga tgt ata cga ggt atg agg caa cat tgt tat tgc aat tct gga agg cgg tat gaa gag gag tca ttg tat tat agt att cgt ctt tct gaa tgt cga atc tat cta gta gat acc gta gta tat tga gag agc gac ttc cat atc ctt gat tat gtt tta tga ata gat aaa gta gat gtt gtc ctt ctt cct ttt gta att tcc cgt att ttt gtt cgt gcc aat tga gta aca tta tga gaa tat gac ctg ttg cac aat cgt tct tta tgt att cca tga tgg gtg tac aat caa gat tat tac gta tcc tcg tat cag ctc ctc gag ata aaa gag cat aca cca cac gag gac tat gtt tgg tat act gtt gaa ggt aag tgt gta acg gcg tat ttc cga ttt tcg taa ccg cgt taa tgt ttg ctc cat gat cta tta tcg cgt aga tga atc gct tct cag ctc gca tct tag tgt gac tct ttg act tgt aat aat tgc ttt cgt gga acg cgg ata tgt gtt tac agt agt aat gaa gag aag tga gtc cat ccg cat cga cgc aat tag ggt cag atc ctt tag tca ata att tgt aca gaa cgt aat agt tta agc tcc cat tga att tat atc taa gat aac aca gca ata gat cgg atg att tac taa agt cat caa tgg ggt ccg tta gta tat caa aga tct tgt tat cga ttg ata gtg aat gaa tca gat agt ggt gta gag gaa tat gtc ctt ttt tat cct tgc tat caa agt tac gca tgc cgt ggc gta aca ata tct tta ata cag atg gat taa atc gtg tat tca tcg tat agc aat gta atg gag agt tac cac att tta gtc gtt tat tca gat cgc agt gtt taa taa cta att taa aca gat gag atg ata tat cca cat caa aga atg cga gat aca tat gac aga cat tat tga cag aaa tgt gac ctt cat tat cac cgt cgt cca taa atg cgt tag gta cgt acc aca tac tat cgt taa cga tgc gca caa tct cgt cca -tt tca tcc atc ttc ata atg att tac ttt ttc ata att aga gaa aaa gat caa ggt ata aaa att aga agt gtt aga cta ta- --- --- --- --- --- aac taa ctt ata a-- -aa cta act tat ga- --- --- --- --- --- --- --- --- --- --- --- --- --- --- --- --- --- --- --- --- --- --- --- --- --- --- --- --- --- --- --- --- --- --- --- --- --- --- --- --- --- --- --- --- --- --- --- --- --- --- --- --- --- --- --- --- --- --- --- --- --- --- --- --- --- --- --- --- --- --- --- --- --- --- --- --- --- --- --- --- --- --- --- --- --- --- --- --- --- --- --c tta act aac tta tga cta tta act cat ttc aag aaa ggt ggg tgg ata gaa ctc tat atg aca gct tgt gaa acg att aga tcc cta att tct aat gga agt ttt gat agg aga ttg tca tca gtt gat aca ttg ttt att atc tca tct att aga gca cgt ctg ttt aga gct tta gtg acc tgc tcg gtt act tct gtg taa atc ttg aat cct tta gtg ata cac tgt gtc aaa act gga tgt tta gaa tac cta tgt aga ata tgg gaa gca tgc ttg ttt ttg tct cta tta tag att aac tca tac atg gtt gta tta tga att ttc atc tgc cta atg tac tcc aat tct tgt tta caa tca att ata taa tca aag agt gat gat gca tac aca tta caa agt gaa taa tct acc atc ata aaa tac ttg ata cag agc ttt atc aca tca tgg ttt tca att gta tta tta agt ata gct aat ttt ata cag tca ata gac aat ggt tct cta agc aat att tct aat att tta aga tgt gct tcc cta cgg gcg atg aca gat ccc cta tcc acg gcc acg tca aga cat gta tat cca tta ctc att act gcg ttg aca ttt gct cca ttt tct aat agc aat gat act aaa tct ata taa cct gca tag ata gcg cga taa agc aag gtc ctt cca cca gca tct agt tga ttg ata tct tca ata tat ggg ata caa agc tta taa att tct aat act gtg ggt tca tct aca agg aat ccc cta gta tac tga att att tta tat aga tct aat tta aca tca ttt tca tct ggg ata cca caa ttc aaa ata aac tca aca aca cta ctt tcc ttt tta cat att ccc cta aaa tag gca ttc aag cat tct att tta tat att aca gcc cca tga tct acc ata aga tca aca atg tct att tct aca tat tca tta gat aga tag taa agt aag aga tct tgc aca gaa tta caa ttc tta ata att ata gag aaa ata tct tcc ata taa ttc ttt gac act aat gca gat ata ata tct tta tat gta ata tat gca aac agt cta tct act ata tac tga tca ata tta tct cta tga atc cta aaa taa tca tac aga aca tct aca gga tca caa att ggt tca agg aga aat cta tca aat att ttc ctg tca aca act ggt tct aga aca taa cag tca aca cct aat cca tgt ttt tta taa tca tct acc aaa gat aat gac caa aga tcg agg tcg tcg tga aac tgc tca tcg aca gcc atg aaa tct acc gac tcc atg gtg cga atc gca ctg tct tat tcg cca ttg att ttc a-t ttt tta taa tta tgt aca tgt ttt cct tct att ctc aag agt cta caa aaa tat a-- ttt ttt tcg ata tct aag tac taa gtt ttt tta ctg ttt ttg tta ctg tct tcc att ctt cta act aaa gat ctg aga taa att ata caa tct tcg cta tcg aac cat ttt tgt agt cta aag cct gaa gta att aac caa ctg ttt tta tta gtg gct ttt ttc gat cta tca att tca gta tat tct tcg ccg tta taa aag taa tgt tgt tta att gta gga cgg ttg tta gta taa tca cat gaa taa taa tat tct aat tcc tcg tat tga cta ctt aca gat act cga aat agt ctg aaa aat tct tca aag ata ttt tta taa aga tct agg aaa agt tta tta ccg acc atg aac gag ata gat gga taa ata tcc ttt cca tca aag gtc ata att gga taa ttg tcc agc aat ata tct gct gta tta gtt ata tca ctt cca ttt att ttc aga ttg aag taa tgt act agt ttg tga caa tta aca aga tac aaa aga gat gcc gat act aat acg taa ata gct ata cgc gaa tcc att gtt acc ttt ttt tat ttc ata ggt cta tta ata aat ata tgt att act taa gac tag aaa aat caa aag taa gtt ttt gat att tga ttc tta ctt att gtg gga ttg tag ttt act tag taa ttc atc tct gaa tcc tga taa atc atg cat atc aat gat gca act acg cag caa act agt agg aat ata gat atc tgg ata tgt acg taa ata gtc gat tat atc ttt tac aat act att agt ccc tat tgc gtt atc tat ata tcc att aat aat att aca cag tgg ata ctt atg aga aat ata cct ctt aca gat ttt taa cga tat ata atc tag aaa gat atg tgt gta gta ctg tat tac cta aat tat cag tct cat tca aat att gca tga cta tta tcg aga att gct ata tcc ctc tat act cga tgc att tat tac agt aat tca atc cag cta aca taa gag cca atc tca atg ttg gtt taa tta tat cat ctt cat gca ata ata acg atg gaa act ttc cag tag cgt aac act tat cta aga agg ata taa taa cta tgt cta cat tat gtc tct tat cga gaa tat tct taa cga gat atc cat agc tat tct ggt gct aat tat tcc tat att ata ttc cac gaa aaa tga tga agg caa tca ttc atc ata aga tga taa aaa gtg tag tga gta aga gta tta gtg aga gag cat gaa gga gat tta gta ttt agc agt gag gat atg atc caa gag ggt gag ata gtc gtt ctc gtt cag aat ctt tcg cag cat aag tag tat gtc gat ata ctt atc att gaa gac tct tcc agt gac aat agc tga ttg agt aca aag tcc aat tat tgc aca aag ttc ttt ggc ggt ttt cat gga gtc att tct gat gaa aca ttt aat gat ctc cac gca att gtc gat att gtc cca cgg aag tga atc cga gaa ctc ctt caa ctc gct acc aaa tag ctc cat tgc atc aat tct gaa aga gat gag aag cct gta gag agg ccc tgc gct ttc tct atg ggt cca tct atg aga aac cca cag gat gta ttc agt cag aca a-t gtc tga cat cag tca cgg tat tca ggg agt cct tag tag cgt ggc aat gac agg gac tga act ggg cac aag gag agg cca ttg tga agg tag acg aag gta acc tga tgg tag acc tgt agc cgt ctg tgc tta ata gag ggc ttt aat ttc cat ttt taa tgg tgt cgt gaa tga gga atg aga gtg tct ctc gtc ctt ggt tta cat gga tca gag tga gaa aaa ata tct tgt ata tta tta act aac aac ctt ggt ttc tat cca tgt tt- aaa aaa tga cct ata tgt tct tta tta att cta ttt taa act tta tcc tca aga ctc ctg aca aaa tta aaa tcc aga aag cag caa aca atc ctg tta caa gtt tac tga aat ctc tct ttg att ata gag tat atg tag tca gag caa gaa aca ctg cag tag tca aca tga aag ctt gca taa cga tac gtg cat cat aga aag taa caa cag agg cca gcg tta gag att cta aca gtg taa atc cac aa- --a gta tgt aca gat tca ggg gat gtt cat gtc tgt gta aag tca atg cga aaa tca agc cta tag atc cga aca ttg atg cca ata tta gaa cag gac tcc ctt gta taa atg tcc gat gca ttc aaa gta taa aaa tac tgc agc tgt tgc cgt tgt taa agg aaa ttg tag aaa gga tac cgt aga ctt ttc tta gaa atg cca ttc gta tgt aca cgc tgg cag acg cca ccg agc tgt cat agt tga agt cgt cct cga t

>MT903346.1_Monkeypox_virus_isolate_MPXV-USA2003_099_Gambian_Rat

--- --- --- --- --- --- --- --- --- --- --- --- --- --- --- --- --- --- --- --- --- --- --- --- --- --- --- --- --- --- --- --- --- --- --- --g tgt cta gaa aaa aat gtg tga ccc acg acc gta gga aac tct aga ggg taa gaa aaa tca atc gtt tat aga gac cat cag aaa gag gtt taa tat ttt tgt gag acc cat cga aga gag aaa g-- --- --- --- --- --- --- --- --- --- --- --- --- --- --- --- --- --- --- --- --- --- --- --- --- --- --- --- --- --- --- --- --- --- --- --- --- --- --- --- --- --- --- --- --- --- --- --- --- --- --- --- --- --- --- --- --- --- --- --- --- --- --- --- --- --- --- --- --- --- --- --- --- --- --- --- --- --- --- --- --- --- --- --- --- --- --- --- --- --- --- --- --- --- --- --- --- --- --- --- --- --- --- --- --- --- --- --- --- --- --- --- --- --- --- --- --- --- --- --- --- --- --- --- --- --- --- --- --- --- --- --- --- --- --- --- --- --- --- --- --- --- --- --- --- --- --- --- --- --- --- --- --- --- --- --- --- --- --- --- --- --- --- --- --- --- --- --- --- --- --- --- --- --- --- --- --- --- --- --- --- --- --- --- --- --- --- gat aaa aac ttt tta cga ctc cat cag aaa gag gtt taa tat ttt tgt gag acc cat cga aga gag aaa gga taa aaa ctt ttt acg act cca tca gaa aga ggt tta --- --- --- --- --- --- --- --- --- --- --- --- --- --- --- --- --- --- --- --- --- --- --- --- --- --- --- --- --- --- --- --- --- --- --- --- --- --- --- --- --- --- --- --- --- --- --a tat ttt tgt gag acc cat cga aga gag aaa gga taa aaa ctt ttt acg act cca tca gaa aga ggt tta ata ttt ttg tga gac cca tcg aag aga gaa aga gat ggt tag tca aga tat ttt tct tag tac aaa agt caa tgt ttt aaa ata tat gga cga gaa tta att tgt ctg tat aaa aac ttg tgt gaa att atg tac tag aga aaa aac gtg agt agt gtc ccc tac atg gat ttt aca gat cat tta tat tcc aaa aat att aac tat ata cgt tta tta tat gat gtt aac gtg taa att ata aac att att tta tga tgc aat tgt ctg aca acc tag att ggt ata agg atg ttg ata agc tct acg aga ata tat tgt tgg acg tta tcg ttt acg aaa tag ttg aga cat cag aaa gag gtt taa tat ttt tgt gag a-c cat cga aga gag aaa tag aat aaa aat a-- --- --- -tt ttt ttt tgt aaa act ttt tta tga gac --- --- -ca aga gaa tac gaa tag tga tca tat cgt atc aca tat tga aac aga aag aag aag taa cga gag gta act ttt tgt gaa tgt agt taa ata ttt ttg ttt tgc aaa ccg gaa tat agt gcc cgg t-- -ct ttt tta att cgt ggt gcg gtg tct gaa tcg ttc gat taa ccc aac tca tcc att ttc aga tga ata gag tta tcg att cag aca cat gct ttg agt ttt gtt gaa tcg atg agt gaa gta tca tcg gtt gca cct tca gat gcc gat ccg tcg aca tac ttg aat cca tcc ttg act tca agt tca gat gat tcc tca cac atg tct ccg ata cgt acg cta aac tct agg ttc ttg aca cat ttt gta tca acg atc gtt gaa ccg atg ata tct ttg taa ctc act ttc tta tgt gag atg tta gac cca agt act gga tgg gtc ttg atg tca ctg tct ttc tct tct tcg cta cat ctg atg tcg ata gac atc tca cag tct ttg atc ata gcc aga gct tct tca cgc gtg atc gcg gga gag tcc tta cct tgt ccc ggt gac acg ctg gac aat cta gta ttc aca gtg ttt cca tca gag gat tcg gag atg gat gaa atc ttt ggg cat ttg gtg aat cca aag ttc atg tta aga ccc gca ccg acg ata gtg taa taa gtg gtg gga tct cct ttt aca act --- --- --- --- --- --- tct tcg gat acc tca tca tct tcg gtc tct gta act tcc gtt acg gat tga caa atc tta tca ttg gtc ggt gtt tgg tct tgc ttt gtg act ttg ata ata aca tcg att ccc ata tga tgt ttg ttt tct tct tca gta cac gag gat gaa gat tgt tga aga cta gta ggc ata gca gct gcc act agg cac atg cat gcc agg aca ata tat tgt ttc atg att gct att gat tga tta ctg ttc tag atg att cta ctt tct tac cat ata ata aat tag aat ata ttt tct act ttt acg aga aat taa tta ttg tat tta tta ttt atg ggt --a aaa aaa ctt act ata agt ggg tgg gat tct ggg aat tag tga tca gtt tat gta tat cgc aac tag cgg gca tat ggc tat tga cat cga gaa cat tac cca tat gat aag aga ttg tat ccg ttt cgt agt ctt gag tat tgg tat tac tat ata gta tgt aga tgt cga cgc tag ata gac agt cgc cca cta gag tta ccg tct ctg aat gcg gca tga tag tat cat tct ttg ttt tcg tta act gtt tgg aag atg aat ctt tgt tgt tac att taa tct cga aat tca gag tac ata tct ttg aag tat tct gat atc tat ttt ctc ctg taa aga atc ctg aag ttg cta cat tat taa gga cag aga agt att ctg cac gaa aga ctg gat cac aat ctt tat gat tca tgg taa tag tta gtt ccg acg ttg aga tgg att cgc tga gac cgg tag tgg tcg tcc gag tac acg atg tgt cgt tga ctg gat aca ggt taa ttt cca cat cga tat agt taa atg tat tgc tgg tta cga cgg gtt cgc att tat ctg tgg aag aga cgg tgt gag aat atg ttc cgg gac cac acg gag aac aga tga cgt ctc cgg tag acg tgt atc cgg ata ctc cgt atc cta ttc cac act ttg ttt tag aaa tac atg ttc tac acc ctg atg ctc ctt tga gaa gac aat aat atc ctg gag agc att cac aga ttc tat tgt gag tcg tgt tac acg atc gcg tct cta cct gat tac tat cac atc ttc cgt tac aac tta gac aag cct gta aat gat tat tgt gag atg taa agg tat ccg aac cac acg gtg tac att gtg tat tag tct tgc tat cac ata atc tgg aag cgt aag ttc ccg gag gac acg ata gac aac ata gat tac ggc ttc tgt att cgt tgt ctt tac act ttc cat tgg atg gtg cat gtg gtg cta tat ctc ttc cgt tta tta tta tac atg aga gaa aca ata tat acg agt ata ata cgg acc tca tga ttt aat aat -gt agt aat cgt cgt ctt gtt act gtt tgt ttc cta ctt ctc caa tca tat aga tta ttt ttt aaa tat ttt ctt tct atc atg gat aat att tgt aat ggt tct ttc cgt aca aca tac tgt tta gat ggt agt cgc tta gct tgg tta tga tat tgc gca taa ttt ccg gag gca aat acg ata gtc tag att gac tat cga tgg tag act cta att tat tga gtg ctt tgt cga cga gtt tac ttt tat gct cca tcg ata gat gac act gtt cta tga gat cgt cgt aca tgg gaa atg aaa tgc gtt tgt ctg aat gta tgg ctt cga gat agg tgt gat acc gga tgt ctt ctg ttc tca ata ccg tat aca agt cgg tgt ctg aga ttc gaa tct ctt tga gga gac tta tgt cac gac tac att ttt cga tga tgg aat cta tct tat cga atg ata tat ttt tca taa ata cac ttt tat agt cct cgt tta aac aga att tag tat gta gtt ccg caa atg act cgt acc tta ata ggc agt agg cta tta tct tct tta cgt agt gat cgt cgt agg gag aga act ccg aca tct tgt aga aca acg att taa tca tag gta gag ata ctt tca gtc tgt ggt gga tga tgt cat taa cga cat ccg cct tgt ata tga tgt ttc tgt ttt caa aca cca agt cga ata ctg tct tat cgt ctt tag tcg gaa ggt tga tgt cgt atc cga tgt ata cga ggt atg agg caa cat tgt tat tgc aat tct gga agg cgg tat gaa gag gag tca ttg tat tat agt att cgt ctt tct gaa tgt cga atc tat cta gta gat acc gta gta tat tga gag agc gac ttc cat atc ctt gat tat gtt tta tga ata gat aaa gta gat gtt gtc ctt ctt cct ttt gta att tcc cgt att ttt gtt cgt gcc aat tga gta aca tta tga gaa tat gac ctg ttg cac aat cgt tct tta tgt att cca tga tgg gtg tac aat caa gat tat tac gta tcc tcg tat cgg ctc ctc gag ata aaa gag cat aca cca cac gag gac tat gtt tgg tat act gtt gaa ggt aag tgt gta acg gcg tat ttc cga ttt tcg taa ccg cgt taa tgt ttg ctc cat gat cta tta tcg cgt aga tga atc gct tct cag ctc gca tct tag tgt gac tct ttg act tgt aat aat tgc ttt cgt gga acg cgg ata tgt gtt tac agt agt aat gaa gag aag tga gtc cat cct cat cga cgc aat tag ggt cag atc ctt tag tca ata att tgt aca gaa cgt aat agt tta agc tcc cat tga att tat atc taa gat aac aca gca ata gat cgg atg att tac taa agt cat caa tgg ggt ccg tta gta tat caa aga tct tgt tat cga ttg ata gtg aat gaa tca gat agt ggt gta gag gaa tat gtc ctt ttt tat cct tgc tat caa agt tac gca tgc cgt ggc gta aca ata tct tta ata cag atg gat taa atc gtg tat tca tcg tat agc aat gta atg gag agt tac cac att tta gtc gtt tat tca gat cgc agt gtt taa taa cta att taa aca gat gag atg ata tat cca cat caa aga atg cga gat aca tat gac aga cat tat tga cag aaa tgt gac ctt cat tat cac cgt cgt cca taa atg cgt tag gta cgt acc aca tac tat cgt taa cga tgc gca caa tct cgt cca -tt tca tcc atc ttc ata atg att tac ttt ttc ata att aga gaa aaa gat caa ggt ata aaa att aga agt gtt aga cta ta- --- --- --- --- --- aac taa ctt ata a-- -aa cta act tat gac tta act aac tt- --- --- --- --- --- --- --- --- --- --- --- --- --- --- --- --- --- --- --- --- --- --- --- --- --- --- --- --- --- --- --- --- --- --- --- --- --- --- --- --- --- --- --- --- --- --- --- --- --- --- --- --- --- --- --- --- --- --- --- --- --- --- -ac tta taa --- aac taa ctt atg act taa cta act tac tta taa aac taa ctt atg act taa cta act tat gac tta act aac tta tga cta tta act cat ttc aag aaa ggt ggg tgg ata gaa ctc tat atg aca gct tgt gaa aca att aga tcc cta att tct aat gga agt ttt gat agg aga ttg tca tca gtt gat aca ttg ttt att atc tca tct att aga gca cgt ctg ttt aga gct tta gtg acc tgc tcg gtt act tct gtg taa atc ttg aat cct tta gtg ata cac tgt gtc aaa act gga tgt tta gaa tac cta tgt aga ata tgg gaa gca tgc ttg ttt ttg tct cta tta tag att aac tca tac atg gtt gta tta tga att ttc atc tgc cta atg tac tcc aat tct tgt tta caa tca att ata taa tca aag agt gat gat gca tac aca tta caa agt gaa taa tct acc atc ata aaa tac ttg ata cag agc ttt atc aca tca tgg ttt tca att gta tta tta agt ata gct aat ttt ata cag tca ata gac aat ggt tct cta agc aat att tct aat att tta aga tgt gct tcc cta cgg gcg atg aca gat ccc cta tcc acg gcc acg tca aga cat gta tat cca tta ctc att act gcg ttg aca ttt gct cca ttt tct aat agc cat gat act aaa tct ata taa cct gca tag ata gcg cga taa agc aag gtc ctt cca cca gca tct agt tga ttg ata tct tca ata tat ggg ata caa agc tta taa att tct aat act gtg ggt tca tct aca agg aat ccc cta gta tac tga att att tta tat aga tct aat tta aca tca ttt tca tct ggg ata cca caa ttc aaa ata aac tca aca aca cta ctt tcc ttt tta cat att ccc cta aaa tag gca ttc aag cat tct att tta tat att aca gcc cca tga tct acc ata aga tca aca atg tct att tct aca tat gca tta gat aga tag taa agt aag aga tct tgc aca gaa tta caa ttc tta ata att ata gag aaa ata tct tcc ata taa ttc ttt gac act aat gca gat ata ata tct tta tat gta ata tat gca aac agt cta tct act ata tac tga tca ata tta tct cta tga atc cta aaa taa tca tac aga aca tct aca gga tca caa att ggt tca agg aga aat cta tca aat att ttc ctg tca aca act ggt tct aga aca taa cag tca aca cct aat cca tgt ttt tta taa tca tct acc aaa gat aat gac caa aga tcg agg tcg tcg tga aac tgc tca tcg aca gcc atg aaa tct acc gac tcc atg gtg cga atc gca ctg tct tat tcg cca ttg att ttc a-t ttt tta taa tta tgt aca tgt ttt cct tct att ctc aag agt cta caa aaa tat a-- ttt ttt tcg ata tct aag tac taa gtt ttt tta ctg ttt ttg tta ctg tct tcc att ctt cta act aaa gat ctg aga taa att ata caa tct tcg cta tcg aac cat ttt tgt agt cta aag cct gaa gta att aac caa ctg ttt tta tta gtg gct ttt ttc gat cta tca att tca gta tat tct tcg ccg tta taa aag taa tgt tgt tta att gta gga cgg ttg tta gta taa tca cat gaa taa taa tat tct aat tcc tcg tat tga cta ctt aca gat act cga aat agt ctg aaa aat tct tca aag ata ttt tta taa aga tct agg aaa agt tta tta ccg acc atg aac gag ata gat gga taa ata tcc ttt cca tca aag gtc ata att gga taa ttg tcc agc aat ata tct gct gta tta gtt ata tca ctt cca ttt att ttc aga ttg aag taa tgt act agt ttg tga caa tta aca aga tac aaa aga gat gcc gat act aat acg taa ata gct ata cgc gaa tcc att gtt acc ttt ttt tat ttc ata ggt cta tta ata aat ata tgt att act taa gac tag aaa aat caa aag tga gtt ttt gat att tga ttc tta ctt att gtg gga ttg tag ttt act tag taa ttc atc tct gaa tcc tga taa atc atg cat atc aat gat gca act acg cag caa act agt agg aat ata gat atc tgg ata tgt acg taa ata gtc gat tat atc ttt tac aat act att agt ccc tat tgc gtt atc tat ata tcc att aat aat att aca cag tgg ata ctt atg aga aat ata cct ctt aca gat ttt taa cga tat ata atc tag aaa gat atg tgt gta gta ctg tat tac cta aat tat cag tct cat tca aat att gca tga cta tta tcg aga att gct ata tcc ctc tat act cga tgc att tat tac agt aat tca atc cag cta aca taa gag cca atc tca atg ttg gtt taa tta tat cat ctt cat gta ata ata acg atg gaa act ttc cag tag cgt aac act tat cta aga agg ata taa taa cta tgt cta cat tat gtc tct tat cga gaa tat tct taa cga gat atc cat agc tat tct ggt gct aat tat tcc tat att ata ttc cac gaa aaa tga tga agg caa tca ttc atc ata aga tga taa aaa gtg tag tga gta aga gta tta gtg aga gag cat gaa gga gat tta gta ttt agc agt gag gat atg atc caa gag tgt gag ata gtc gtt ctc gtt cag aat ctt tcg cag cat aag tag tat gtc gat ata ctt atc att gaa gac tct tcc agt gac aat agc tga ttg agt aca aag tcc aat tat tgc aca aag ttc ttt ggc ggt ttt cat gga gtc att tct gat gaa aca ttt aat gat ctc cac gca att gtc gat att gtc cca cgg aag tga atc cga gaa ctc ctt caa ctc gct acc aaa tag ctc cat tgc atc aat tct gaa aga gat gag aag cct gta gag agg ccc tgc gct ttc tct atg ggt cca tct atg aga aac cca cag gat gta ttc agt cag aca a-t gtc tga cat cag tca cgg tat tca ggg agt cct tag tag cgt ggc aat gac agg gac tga act ggg cac aag gag agg cca ttg tga agg tag acg aag gta acc tga tgg tag acc tgt agc cgt ctg tgc tta ata gag ggc ttt aat ttc cat ttt taa tgg tgt cgt gaa tga gga atg aga gtg tct ctc gtc ctt ggt tta cat gga tca gag tga gaa aaa ata tct tgt ata tta tta act aac aac ctt ggt ttc tat cca tgt tta aaa aaa tga cct ata tgt tct tta tta att cta ttt taa act tta tcc tca aga ctc ctg aca aaa tta aaa tcc aga aag cag caa aca atc ctg tta caa gtt tac tga aat ctc tct ttg att gta gag tat atg tag tca gag caa gaa aca ctg cag tag tca aca tga aag ctt gca taa cga tac gtg cat cat aga aag taa caa cag agg cca gcg tta gag att cta aca gtg taa atc cac aa- --a gta tgt aca gat tca ggg gat gtt cat gtc tgt gta aag tca atg cga aaa tca agc cta tag atc cga aca ttg atg cca ata tta gaa cag gac tcc ctt gta taa atg tcc gat gca ttc aaa gta taa aaa tac tgc agc tgt tgc cgt tgt taa agg aaa ttg tag aaa gaa tac cgt aga ctt ttc tta gaa atg cca ttc gta tgt aca cgc tgg cag acg cca ccg agc tgt cat agt tga agt cgt cct cga t

>DQ011157.1_Monkeypox_virus_strain_USA_2003_039_complete_genome

--- --- --- --- --- --- --- --- --- --- --- --- --- --- --- --- --- --- --- --- --- --- --- --- --- --- --- --- --- --- --- --- --- --- --- --g tgt cta gaa aaa aat gtg tga ccc acg acc gta gga aac tct aga ggg taa gaa aaa tca atc gtt tat aga gac cat cag aaa gag gtt taa tat ttt tgt gag acc cat cga aga gag aaa g-- --- --- --- --- --- --- --- --- --- --- --- --- --- --- --- --- --- --- --- --- --- --- --- --- --- --- --- --- --- --- --- --- --- --- --- --- --- --- --- --- --- --- --- --- --- --- --- --- --- --- --- --- --- --- --- --- --- --- --- --- --- --- --- --- --- --- --- --- --- --- --- --- --- --- --- --- --- --- --- --- --- --- --- --- --- --- --- --- --- --- --- --- --- --- --- --- --- --- --- --- --- --- --- --- --- --- --- --- --- --- --- --- --- --- --- --- --- --- --- --- --- --- --- --- --- --- --- --- --- --- --- --- --- --- --- --- --- --- --- --- --- --- --- --- --- --- --- --- --- --- --- --- --- --- --- --- --- --- --- --- --- --- --- --- --- --- --- --- --- --- --- --- --- --- --- --- --- --- --- --- --- --- --- --- --- --- gat aaa aac ttt tta cga ctc cat cag aaa gag gtt taa tat ttt tgt gag acc cat cga aga gag aaa gga taa aaa ctt ttt acg act cca tca gaa aga ggt tta --- --- --- --- --- --- --- --- --- --- --- --- --- --- --- --- --- --- --- --- --- --- --- --- --- --- --- --- --- --- --- --- --- --- --- --- --- --- --- --- --- --- --- --- --- --- --a tat ttt tgt gag acc cat cga aga gag aaa gga taa aaa ctt ttt acg act cca tca gaa aga ggt tta ata ttt ttg tga gac cca tcg aag aga gaa aga gat ggt tag tca aga tat ttt tct tag tac aaa agt caa tgt ttt aaa ata tat gga cga gaa tta att tgt ctg tat aaa aac ttg tgt gaa att atg tac tag aga aaa aac gtg agt agt gtc ccc tac atg gat ttt aca gat cat tta tat tcc aaa aat att aac tat ata cgt tta tta tat gat gtt aac gtg taa att ata aac att att tta tga tgc aat tgt ctg aca acc tag att ggt ata agg atg ttg ata agc tct acg aga ata tat tgt tgg acg tta tcg ttt acg aaa tag ttg aga cat cag aaa gag gtt taa tat ttt tgt gag a-c cat cga aga gag aaa tag aat aaa aat a-- --- --- ttt ttt ttt tgt aaa act ttt tta tga gac --- --- -ca aga gaa tac gaa tag tga tca tat cgt atc aca tat tga aac aga aag aag aag taa cga gag gta act ttt tgt gaa tgt agt taa ata ttt ttg ttt tgc aaa ccg gaa tat agt gcc cgg t-- -ct ttt tta att cgt ggt gcg gtg tct gaa tcg ttc gat taa ccc aac tca tcc att ttc aga tga ata gag tta tcg att cag aca cat gct ttg agt ttt gtt gaa tcg atg agt gaa gta tca tcg gtt gca cct tca gat gcc gat ccg tcg aca tac ttg aat cca tcc ttg act tca agt tca gat gat tcc tca cac atg tct ccg ata cgt acg cta aac tct agg ttc ttg aca cat ttt gta tca acg atc gtt gaa ccg atg ata tct ttg taa ctc act ttc tta tgt gag atg tta gac cca agt act gga tgg gtc ttg atg tca ctg tct ttc tct tct tcg cta cat ctg atg tcg ata gac atc tca cag tct ttg atc ata gcc aga gct tct tca cgc gtg atc gcg gga gag tcc tta cct tgt ccc ggt gac acg ctg gac aat cta gta ttc aca gtg ttt cca tca gag gat tcg gag atg gat gaa atc ttt ggg cat ttg gtg aat cca aag ttc atg tta aga ccc gca ccg acg ata gtg taa taa gtg gtg gga tct cct ttt aca act --- --- --- --- --- --- tct tcg gat acc tca tca tct tcg gtc tct gta act tcc gtt acg gat tga caa atc tta tca ttg gtc ggt gtt tgg tct tgc ttt gtg act ttg ata ata aca tcg att ccc ata tga tgt ttg ttt tct tct tca gta cac gag gat gaa gat tgt tga aga cta gta ggc ata gca gct gcc act agg cac atg cat gcc agg aca ata tat tgt ttc atg att gct att gat tga tta ctg ttc tag atg att cta ctt tct tac cat ata ata aat tag aat ata ttt tct act ttt acg aga aat taa tta ttg tat tta tta ttt atg ggt --a aaa aaa ctt act ata agt ggg tgg gat tct ggg aat tag tga tca gtt tat gta tat cgc aac tag cgg gca tat ggc tat tga cat cga gaa cat tac cca tat gat aag aga ttg tat ccg ttt cgt agt ctt gag tat tgg tat tac tat ata gta tgt aga tgt cga cgc tag ata gac agt cgc cca cta gag tta ccg tct ctg aat gcg gca tga tag tat cat tct ttg ttt tcg tta act gtt tgg aag atg aat ctt tgt tgt tac att taa tct cga aat tca gag tac ata tct ttg aag tat tct gat atc tat ttt ctc ctg taa aga atc ctg aag ttg cta cat tat taa gga cag aga agt att ctg cac gaa aga ctg gat cac aat ctt tat gat tca tgg taa tag tta gtt ccg acg ttg aga tgg att cgc tga gac cgg tag tgg tcg tcc gag tac acg atg tgt cgt tga ctg gat aca ggt taa ttt cca cat cga tat agt taa atg tat tgc tgg tta cga cgg gtt cgc att tat ctg tgg aag aga cgg tgt gag aat atg ttc cgg gac cac acg gag aac aga tga cgt ctc cgg tag acg tgt atc cgg ata ctc cgt atc cta ttc cac act ttg ttt tag aaa tac atg ttc tac acc ctg atg ctc ctt tga gaa gac aat aat atc ctg gag agc att cac aga ttc tat tgt gag tcg tgt tac acg atc gcg tct cta cct gat tac tat cac atc ttc cgt tac aac tta gac aag cct gta aat gat tat tgt gag atg taa agg tat ccg aac cac acg gtg tac att gtg tat tag tct tgc tat cac ata atc tgg aag cgt aag ttc ccg gag gac acg ata gac aac ata gat tac ggc ttc tgt att cgt tgt ctt tac act ttc cat tgg atg gtg cat gtg gtg cta tat ctc ttc cgt tta tta tta tac atg aga gaa aca ata tat acg agt ata ata cgg acc tca tga ttt aat aat -gt agt aat cgt cgt ctt gtt act gtt tgt ttc cta ctt ctc caa tca tat aga tta ttt ttt aaa tat ttt ctt tct atc atg gat aat att tgt aat ggt tct ttc cgt aca aca tac tgt tta gat ggt agt cgc tta gct tgg tta tga tat tgc gca taa ttt ccg gag gca aat acg ata gtc tag att gac tat cga tgg tag act cta att tat tga gtg ctt tgt cga cga gtt tac ttt tat gct cca tcg ata gat gac act gtt cta tga gat cgt cgt aca tgg gaa atg aaa tgc gtt tgt ctg aat gta tgg ctt cga gat agg tgt gat acc gga tgt ctt ctg ttc tca ata ccg tat aca agt cgg tgt ctg aga ttc gaa tct ctt tga gga gac tta tgt cac gac tac att ttt cga tga tgg aat cta tct tat cga atg ata tat ttt tca taa ata cac ttt tat agt cct cgt tta aac aga att tag tat gta gtt ccg caa atg act cgt acc tta ata ggc agt agg cta tta tct tct tta cgt agt gat cgt cgt agg gag aga act ccg aca tct tgt aga aca acg att taa tca tag gta gag ata ctt tca gtc tgt ggt gga tga tgt cat taa cga cat ccg cct tgt ata tga tgt ttc tgt ttt caa aca cca agt cga ata ctg tct tat cgt ctt tag tcg gaa ggt tga tgt cgt atc cga tgt ata cga ggt atg agg caa cat tgt tat tgc aat tct gga agg cgg tat gaa gag gag tca ttg tat tat agt att cgt ctt tct gaa tgt cga atc tat cta gta gat acc gta gta tat tga gag agc gac ttc cat atc ctt gat tat gtt tta tga ata gat aaa gta gat gtt gtc ctt ctt cct ttt gta att tcc cgt att ttt gtt cgt gcc aat tga gta aca tta tga gaa tat gac ctg ttg cac aat cgt tct tta tgt att cca tga tgg gtg tac aat caa gat tat tac gta tcc tcg tat cgg ctc ctc gag ata aaa gag cat aca cca cac gag gac tat gtt tgg tat act gtt gaa ggt aag tgt gta acg gcg tat ttc cga ttt tcg taa ccg cgt taa tgt ttg ctc cat gat cta tta tcg cgt aga tga atc gct tct cag ctc gca tct tag tgt gac tct ttg act tgt aat aat tgc ttt cgt gga acg cgg ata tgt gtt tac agt agt aat gaa gag aag tga gtc cat cct cat cga cgc aat tag ggt cag atc ctt tag tca ata att tgt aca gaa cgt aat agt tta agc tcc cat tga att tat atc taa gat aac aca gca ata gat cgg atg att tac taa agt cat caa tgg ggt ccg tta gta tat caa aga tct tgt tat cga ttg ata gtg aat gaa tca gat agt ggt gta gag gaa tat gtc ctt ttt tat cct tgc tat caa agt tac gca tgc cgt ggc gta aca ata tct tta ata cag atg gat taa atc gtg tat tca tcg tat agc aat gta atg gag agt tac cac att tta gtc gtt tat tca gat cgc agt gtt taa taa cta att taa aca gat gag atg ata tat cca cat caa aga atg cga gat aca tat gac aga cat tat tga cag aaa tgt gac ctt cat tat cac cgt cgt cca taa atg cgt tag gta cgt acc aca tac tat cgt taa cga tgc gca caa tct cgt cca -tt tca tcc atc ttc ata atg att tac ttt ttc ata att aga gaa aaa gat caa ggt ata aaa att aga agt gtt aga cta ta- --- --- --- --- --- aac taa ctt ata a-- -aa cta act tat gac tta act aac tt- --- --- --- --- --- --- --- --- --- --- --- --- --- --- --- --- --- --- --- --- --- --- --- --- --- --- --- --- --- --- --- --- --- --- --- --- --- --- --- --- --- --- --- --- --- --- --- --- --- --- --- --- --- --- --- --- --- --- --- --- --- --- -ac tta taa --- aac taa ctt atg act taa cta act tac tta taa aac taa ctt atg act taa cta act tat gac tta act aac tta tga cta tta act cat ttc aag aaa ggt ggg tgg ata gaa ctc tat atg aca gct tgt gaa aca att aga tcc cta att tct aat gga agt ttt gat agg aga ttg tca tca gtt gat aca ttg ttt att atc tca tct att aga gca cgt ctg ttt aga gct tta gtg acc tgc tcg gtt act tct gtg taa atc ttg aat cct tta gtg ata cac tgt gtc aaa act gga tgt tta gaa tac cta tgt aga ata tgg gaa gca tgc ttg ttt ttg tct cta tta tag att aac tca tac atg gtt gta tta tga att ttc atc tgc cta atg tac tcc aat tct tgt tta caa tca att ata taa tca aag agt gat gat gca tac aca tta caa agt gaa taa tct acc atc ata aaa tac ttg ata cag agc ttt atc aca tca tgg ttt tca att gta tta tta agt ata gct aat ttt ata cag tca ata gac aat ggt tct cta agc aat att tct aat att tta aga tgt gct tcc cta cgg gcg atg aca gat ccc cta tcc acg gcc acg tca aga cat gta tat cca tta ctc att act gcg ttg aca ttt gct cca ttt tct aat agc cat gat act aaa tct ata taa cct gca tag ata gcg cga taa agc aag gtc ctt cca cca gca tct agt tga ttg ata tct tca ata tat ggg ata caa agc tta taa att tct aat act gtg ggt tca tct aca agg aat ccc cta gta tac tga att att tta tat aga tct aat tta aca tca ttt tca tct ggg ata cca caa ttc aaa ata aac tca aca aca cta ctt tcc ttt tta cat att ccc cta aaa tag gca ttc aag cat tct att tta tat att aca gcc cca tga tct acc ata aga tca aca atg tct att tct aca tat gca tta gat aga tag taa agt aag aga tct tgc aca gaa tta caa ttc tta ata att ata gag aaa ata tct tcc ata taa ttc ttt gac act aat gca gat ata ata tct tta tat gta ata tat gca aac agt cta tct act ata tac tga tca ata tta tct cta tga atc cta aaa taa tca tac aga aca tct aca gga tca caa att ggt tca agg aga aat cta tca aat att ttc ctg tca aca act ggt tct aga aca taa cag tca aca cct aat cca tgt ttt tta taa tca tct acc aaa gat aat gac caa aga tcg agg tcg tcg tga aac tgc tca tcg aca gcc atg aaa tct acc gac tcc atg gtg cga atc gca ctg tct tat tcg cca ttg att ttc a-t ttt tta taa tta tgt aca tgt ttt cct tct att ctc aag agt cta caa aaa tat a-- ttt ttt tcg ata tct aag tac taa gtt ttt tta ctg ttt ttg tta ctg tct tcc att ctt cta act aaa gat ctg aga taa att ata caa tct tcg cta tcg aac cat ttt tgt agt cta aag cct gaa gta att aac caa ctg ttt tta tta gtg gct ttt ttc gat cta tca att tca gta tat tct tcg ccg tta taa aag taa tgt tgt tta att gta gga cgg ttg tta gta taa tca cat gaa taa taa tat tct aat tcc tcg tat tga cta ctt aca gat act cga aat agt ctg aaa aat tct tca aag ata ttt tta taa aga tct agg aaa agt tta tta ccg acc atg aac gag ata gat gga taa ata tcc ttt cca tca aag gtc ata att gga taa ttg tcc agc aat ata tct gct gta tta gtt ata tca ctt cca ttt att ttc aga ttg aag taa tgt act agt ttg tga caa tta aca aga tac aaa aga gat gcc gat act aat acg taa ata gct ata cgc gaa tcc att gtt acc ttt ttt tat ttc ata ggt cta tta ata aat ata tgt att act taa gac tag aaa aat caa aag tga gtt ttt gat att tga ttc tta ctt att gtg gga ttg tag ttt act tag taa ttc atc tct gaa tcc tga taa atc atg cat atc aat gat gca act acg cag caa act agt agg aat ata gat atc tgg ata tgt acg taa ata gtc gat tat atc ttt tac aat act att agt ccc tat tgc gtt atc tat ata tcc att aat aat att aca cag tgg ata ctt atg aga aat ata cct ctt aca gat ttt taa cga tat ata atc tag aaa gat atg tgt gta gta ctg tat tac cta aat tat cag tct cat tca aat att gca tga cta tta tcg aga att gct ata tcc ctc tat act cga tgc att tat tac agt aat tca atc cag cta aca taa gag cca atc tca atg ttg gtt taa tta tat cat ctt cat gta ata ata acg atg gaa act ttc cag tag cgt aac act tat cta aga agg ata taa taa cta tgt cta cat tat gtc tct tat cga gaa tat tct taa cga gat atc cat agc tat tct ggt gct aat tat tcc tat att ata ttc cac gaa aaa tga tga agg caa tca ttc atc ata aga tga taa aaa gtg tag tga gta aga gta tta gtg aga gag cat gaa gga gat tta gta ttt agc agt gag gat atg atc caa gag tgt gag ata gtc gtt ctc gtt cag aat ctt tcg cag cat aag tag tat gtc gat ata ctt atc att gaa gac tct tcc agt gac aat agc tga ttg agt aca aag tcc aat tat tgc aca aag ttc ttt ggc ggt ttt cat gga gtc att tct gat gaa aca ttt aat gat ctc cac gca att gtc gat att gtc cca cgg aag tga atc cga gaa ctc ctt caa ctc gct acc aaa tag ctc cat tgc atc aat tct gaa aga gat gag aag cct gta gag agg ccc tgc gct ttc tct atg ggt cca tct atg aga aac cca cag gat gta ttc agt cag aca a-t gtc tga cat cag tca cgg tat tca ggg agt cct tag tag cgt ggc aat gac agg gac tga act ggg cac aag gag agg cca ttg tga agg tag acg aag gta acc tga tgg tag acc tgt agc cgt ctg tgc tta ata gag ggc ttt aat ttc cat ttt taa tgg tgt cgt gaa tga gga atg aga gtg tct ctc gtc ctt ggt tta cat gga tca gag tga gaa aaa ata tct tgt ata tta tta act aac aac ctt ggt ttc tat cca tgt tta aaa aaa tga cct ata tgt tct tta tta att cta ttt taa act tta tcc tca aga ctc ctg aca aaa tta aaa tcc aga aag cag caa aca atc ctg tta caa gtt tac tga aat ctc tct ttg att gta gag tat atg tag tca gag caa gaa aca ctg cag tag tca aca tga aag ctt gca taa cga tac gtg cat cat aga aag taa caa cag agg cca gcg tta gag att cta aca gtg taa atc cac aa- --a gta tgt aca gat tca ggg gat gtt cat gtc tgt gta aag tca atg cga aaa tca agc cta tag atc cga aca ttg atg cca ata tta gaa cag gac tcc ctt gta taa atg tcc gat gca ttc aaa gta taa aaa tac tgc agc tgt tgc cgt tgt taa agg aaa ttg tag aaa gaa tac cgt aga ctt ttc tta gaa atg cca ttc gta tgt aca cgc tgg cag acg cca ccg agc tgt cat agt tga agt cgt cct cga t

>DQ011153.1_Monkeypox_virus_strain_USA_2003_044_complete_genome

--- --- --- --- --- --- --- --- --- --- --- --- --- --- --- --- --- --- --- --- --- --- --- --- --- --- --- --- --- --- --- --- --- --- --- --g tgt cta gaa aaa aat gtg tga ccc acg act gta gga aac tct aga ggg taa gaa aaa tca atc gtt tat aga gac cat cag aaa gag gtt taa tat ttt tgt gag acc cat cga aga gag aaa g-- --- --- --- --- --- --- --- --- --- --- --- --- --- --- --- --- --- --- --- --- --- --- --- --- --- --- --- --- --- --- --- --- --- --- --- --- --- --- --- --- --- --- --- --- --- --- --- --- --- --- --- --- --- --- --- --- --- --- --- --- --- --- --- --- --- --- --- --- --- --- --- --- --- --- --- --- --- --- --- --- --- --- --- --- --- --- --- --- --- --- --- --- --- --- --- --- --- --- --- --- --- --- --- --- --- --- --- --- --- --- --- --- --- --- --- --- --- --- --- --- --- --- --- --- --- --- --- --- --- --- --- --- --- --- --- --- --- --- --- --- --- --- --- --- --- --- --- --- --- --- --- --- --- --- --- --- --- --- --- --- --- --- --- --- --- --- --- --- --- --- --- --- --- --- --- --- --- --- --- --- --- --- --- --- --- --- gat aaa aac ttt tta cga ctc cat cag aaa gag gtt taa tat ttt tgt gag acc cat cga aga gag aaa gga taa aaa ctt ttt acg act cca tca gaa aga ggt tta --- --- --- --- --- --- --- --- --- --- --- --- --- --- --- --- --- --- --- --- --- --- --- --- --- --- --- --- --- --- --- --- --- --- --- --- --- --- --- --- --- --- --- --- --- --- --a tat ttt tgt gag acc cat cga aga gag aaa gga taa aaa ctt ttt acg act cca tca gaa aga ggt tta ata ttt ttg tga gac cca tcg aag aga gaa aga gat ggt tag tca aga tat ttt tct tag tac aaa agt caa tgt ttt aaa ata tat gga cga gaa tta att tgt ctg tat aaa aac ttg tgt gaa att atg tac tag aga aaa aac gtg agt agt gtc ccc tac atg gat ttt aca gat cat tta tat tcc aaa aat att aac tat ata cgt tta tta tat gat gtt aac gtg taa att ata aac att att tta tga tgc aat tgt ctg aca acc tag att ggt ata agg atg ttg ata agc tct acg aga ata tat tgt tgg acg tta tcg ttt acg aaa tag ttg aga cat cag aaa gag gtt taa tat ttt tgt gag a-c cat cga aga gag aaa tag aat aaa aat a-- --- --- ttt ttt ttt tgt aaa act ttt tta tga gac --- --- -ca aga gaa tac gaa tag tga tca tat cgt atc aca tat tga aac aga aag aag aag taa cga gag gta act ttt tgt gaa tgt agt taa ata ttt ttg ttt tgc aaa ccg gaa tat agt gcc cgg t-- -ct ttt tta att cgt ggt gcg gtg tct gaa tcg ttc gat taa ccc aac tca tcc att ttc aga tga ata gag tta tcg att cag aca cat gct ttg agt ttt gtt gaa tcg atg agt gaa gta tca tcg gtt gca cct tca gat gcc gat ccg tcg aca tac ttg aat cca tcc ttg act tca agt tca gat gat tcc tca cac atg tct ccg ata cgt acg cta aac tct agg ttc ttg aca cat ttt gta tca acg atc gtt gaa ccg atg ata tct ttg taa ctc act ttc tta tgt gag atg tta gac cca agt act gga tgg gtc ttg atg tca ctg tct ttc tct tct tcg cta cat ctg atg tcg ata gac atc tca cag tct ttg atc ata gcc aga gct tct tca cgc gtg atc gcg gga gag tcc tta cct tgt ccc ggt gac acg ctg gac aat cta gta ttc aca gtg ttt cca tca gag gat tcg gag atg gat gaa atc ttt ggg cat ttg gtg aat cca aag ttc atg tta aga ccc gca ccg acg ata gtg taa taa gtg gtg gga tct cct ttt aca act --- --- --- --- --- --- tct tcg gat acc tca tca tct tcg gtc tct gta act tcc gtt acg gat tga caa atc tta tca ttg gtc ggt gtt tgg tct tgc ttt gtg act ttg ata ata aca tcg att ccc ata tga tgt ttg ttt tct tct tca gta cac gag gat gaa gat tgt tga aga cta gta ggc ata gca gct gcc act agg cac atg cat gcc agg aca ata tat tgt ttc atg att gct att gat tga tta ctg ttc tag atg att cta ctt tct tac cat ata ata aat tag aat ata ttt tct act ttt acg aga aat taa tta ttg tat tta tta ttt atg ggt --a aaa aaa ctt act ata agt ggg tgg gat tct ggg aat tag tga tca gtt tat gta tat cgc aac tag cgg gca tat ggc tat tga cat cga gaa cat tac cca tat gat aag aga ttg tat ccg ttt cgt agt ctt gag tat tgg tat tac tat ata gta tgt aga tgt cga cgc tag ata gac agt cgc cca cta gag tta ccg tct ctg aat gcg gca tga tag tat cat tct ttg ttt tcg tta act gtt tgg aag atg aat ctt tgt tgt tac att taa tct cga aat tca gag tac ata tct ttg aag tat tct gat atc tat ttt ctc ctg taa aga atc ctg aag ttg cta cat tat taa gga cag aga agt att ctg cac gaa aga ctg gat cac aat ctt tat gat tca tgg taa tag tta gtt ccg acg ttg aga tgg att cgc tga gac cgg tag tgg tcg tcc gag tac acg atg tgt cgt tga ctg gat aca ggt taa ttt cca cat cga tat agt taa atg tat tgc tgg tta cga cgg gtt cgc att tat ctg tgg aag aga cgg tgt gag aat atg ttc cgg gac cac acg gag aac aga tga cgt ctc cgg tag acg tgt atc cgg ata ctc cgt atc cta ttc cac act ttg ttt tag aaa tac atg ttc tac acc ctg atg ctc ctt tga gaa gac aat aat atc ctg gag agc att cac aga ttc tat tgt gag tcg tgt tac acg atc gcg tct cta cct gat tac tat cac atc ttc cgt tac aac tta gac aag cct gta aat gat tat tgt gag atg taa agg tat ccg aac cac acg gtg tac att gtg tat tag tct tgc tat cac ata atc tgg aag cgt aag ttc ccg gag gac acg ata gac aac ata gat tac ggc ttc tgt att cgt tgt ctt tac act ttc cat tgg atg gtg cat gtg gtg cta tat ctc ttc cgt tta tta tta tac atg aga gaa aca ata tat acg agt ata ata cgg acc tca tga ttt aat aat -gt agt aat cgt cgt ctt gtt act gtt tgt ttc cta ctt ctc caa tca tat aga tta ttt ttt aaa tat ttt ctt tct atc atg gat aat att tgt aat ggt tct ttc cgt aca aca tac tgt tta gat ggt agt cgc tta gct tgg tta tga tat tgc gca taa ttt ccg gag gca aat acg ata gtc tag att gac tat cga tgg tag act cta att tat tga gtg ctt tgt cga cga gtt tac ttt tat gct cca tcg ata gat gac act gtt cta tga gat cgt cgt aca tgg gaa atg aaa tgc gtt tgt ctg aat gta tgg ctt cga gat agg tgt gat acc gga tgt ctt ctg ttc tca ata ccg tat aca agt cgg tgt ctg aga ttc gaa tct ctt tga gga gac tta tgt cac gac tac att ttt cga tga tgg aat cta tct tat cga atg ata tat ttt tca taa ata cac ttt tat agt cct cgt tta aac aga att tag tat gta gtt ccg caa atg act cgt acc tta ata ggc agt agg cta tta tct tct tta cgt agt gat cgt cgt agg gag aga act ccg aca tct tgt aga aca acg att taa tca tag gta gag ata ctt tca gtc tgt ggt gga tga tgt cat taa cga cat ccg cct tgt ata tga tgt ttc tgt ttt caa aca cca agt cga ata ctg tct tat cgt ctt tag tcg gaa ggt tga tgt cgt atc cga tgt ata cga ggt atg agg caa cat tgt tat tgc aat tct gga agg cgg tat gaa gag gag tca ttg tat tat agt att cgt ctt tct gaa tgt cga atc tat cta gta gat acc gta gta tat tga gag agc gac ttc cat atc ctt gat tat gtt tta tga ata gat aaa gta gat gtt gtc ctt ctt cct ttt gta att tcc cgt att ttt gtt cgt gcc aat tga gta aca tta tga gaa tat gac ctg ttg cac aat cgt tct tta tgt att cca tga tgg gtg tac aat caa gat tat tac gta tcc tcg tat cgg ctc ctc gag ata aaa gag cat aca cca cac gag gac tat gtt tgg tat act gtt gaa ggt aag tgt gta acg gcg tat ttc cga ttt tcg taa ccg cgt taa tgt ttg ctc cat gat cta tta tcg cgt aga tga atc gct tct cag ctc gca tct tag tgt gac tct ttg act tgt aat aat tgc ttt cgt gga acg cgg ata tgt gtt tac agt agt aat gaa gag aag tga gtc cat cct cat cga cgc aat tag ggt cag atc ctt tag tca ata att tgt aca gaa cgt aat agt tta agc tcc cat tga att tat atc taa gat aac aca gca ata gat cgg atg att tac taa agt cat caa tgg ggt ccg tta gta tat caa aga tct tgt tat cga ttg ata gtg aat gaa tca gat agt ggt gta gag gaa tat gtc ctt ttt tat cct tgc tat caa agt tac gca tgc cgt ggc gta aca ata tct tta ata cag atg gat taa atc gtg tat tca tcg tat agc aat gta atg gag agt tac cac att tta gtc gtt tat tca gat cgc agt gtt taa taa cta att taa aca gat gag atg ata tat cca cat caa aga atg cga gat aca tat gac aga cat tat tga cag aaa tgt gac ctt cat tat cac cgt cgt cca taa atg cgt tag gta cgt acc aca tac tat cgt taa cga tgc gca caa tct cgt cca -tt tca tcc atc ttc ata atg att tac ttt ttc ata att aga gaa aaa gat caa ggt ata aaa att aga agt gtt aga cta ta- --- --- --- --- --- aac taa ctt ata a-- -aa cta act tat gac tta act aac tt- --- --- --- --- --- --- --- --- --- --- --- --- --- --- --- --- --- --- --- --- --- --- --- --- --- --- --- --- --- --- --- --- --- --- --- --- --- --- --- --- --- --- --- --- --- --- --- --- --- --- --- --- --- --- --- --- --- --- --- --- --- --- -ac tta taa --- aac taa ctt atg act taa cta act tac tta taa aac taa ctt atg act taa cta act tat gac tta act aac tta tga cta tta act cat ttc aag aaa ggt ggg tgg ata gaa ctc tat atg aca gct tgt gaa aca att aga tcc cta att tct aat gga agt ttt gat agg aga ttg tca tca gtt gat aca ttg ttt att atc tca tct att aga gca cgt ctg ttt aga gct tta gtg acc tgc tcg gtt act tct gtg taa atc ttg aat cct tta gtg ata cac tgt gtc aaa act gga tgt tta gaa tac cta tgt aga ata tgg gaa gca tgc ttg ttt ttg tct cta tta tag att aac tca tac atg gtt gta tta tga att ttc atc tgc cta atg tac tcc aat tct tgt tta caa tca att ata taa tca aag agt gat gat gca tac aca tta caa agt gaa taa tct acc atc ata aaa tac ttg ata cag agc ttt atc aca tca tgg ttt tca att gta tta tta agt ata gct aat ttt ata cag tca ata gac aat ggt tct cta agc aat att tct aat att tta aga tgt gct tcc cta cgg gcg atg aca gat ccc cta tcc acg gcc acg tca aga cat gta tat cca tta ctc att act gcg ttg aca ttt gct cca ttt tct aat agc cat gat act aaa tct ata taa cct gca tag ata gcg cga taa agc aag gtc ctt cca cca gca tct agt tga ttg ata tct tca ata tat ggg ata caa agc tta taa att tct aat act gtg ggt tca tct aca agg aat ccc cta gta tac tga att att tta tat aga tct aat tta aca tca ttt tca tct ggg ata cca caa ttc aaa ata aac tca aca aca cta ctt tcc ttt tta cat att ccc cta aaa tag gca ttc aag cat tct att tta tat att aca gcc cca tga tct acc ata aga tca aca atg tct att tct aca tat gca tta gat aga tag taa agt aag aga tct tgc aca gaa tta caa ttc tta ata att ata gag aaa ata tct tcc ata taa ttc ttt gac act aat gca gat ata ata tct tta tat gta ata tat gca aac agt cta tct act ata tac tga tca ata tta tct cta tga atc cta aaa taa tca tac aga aca tct aca gga tca caa att ggt tca agg aga aat cta tca aat att ttc ctg tca aca act ggt tct aga aca taa cag tca aca cct aat cca tgt ttt tta taa tca tct acc aaa gat aat gac caa aga tcg agg tcg tcg tga aac tgc tca tcg aca gcc atg aaa tct acc gac tcc atg gtg cga atc gca ctg tct tat tcg cca ttg att ttc a-t ttt tta taa tta tgt aca tgt ttt cct tct att ctc aag agt cta caa aaa tat a-- ttt ttt tcg ata tct aag tac taa gtt ttt tta ctg ttt ttg tta ctg tct tcc att ctt cta act aaa gat ctg aga taa att ata caa tct tcg cta tcg aac cat ttt tgt agt cta aag cct gaa gta att aac caa ctg ttt tta tta gtg gct ttt ttc gat cta tca att tca gta tat tct tcg ccg tta taa aag taa tgt tgt tta att gta gga cgg ttg tta gta taa tca cat gaa taa taa tat tct aat tcc tcg tat tga cta ctt aca gat act cga aat agt ctg aaa aat tct tca aag ata ttt tta taa aga tct agg aaa agt tta tta ccg acc atg aac gag ata gat gga taa ata tcc ttt cca tca aag gtc ata att gga taa ttg tcc agc aat ata tct gct gta tta gtt ata tca ctt cca ttt att ttc aga ttg aag taa tgt act agt ttg tga caa tta aca aga tac aaa aga gat gcc gat act aat acg taa ata gct ata cgc gaa tcc att gtt acc ttt ttt tat ttc ata ggt cta tta ata aat ata tgt att act taa gac tag aaa aat caa aag tga gtt ttt gat att tga ttc tta ctt att gtg gga ttg tag ttt act tag taa ttc atc tct gaa tcc tga taa atc atg cat atc aat gat gca act acg cag caa act agt agg aat ata gat atc tgg ata tgt acg taa ata gtc gat tat atc ttt tac aat act att agt ccc tat tgc gtt atc tat ata tcc att aat aat att aca cag tgg ata ctt atg aga aat ata cct ctt aca gat ttt taa cga tat ata atc tag aaa gat atg tgt gta gta ctg tat tac cta aat tat cag tct cat tca aat att gca tga cta tta tcg aga att gct ata tcc ctc tat act cga tgc att tat tac agt aat tca atc cag cta aca taa gag cca atc tca atg ttg gtt taa tta tat cat ctt cat gta ata ata acg atg gaa act ttc cag tag cgt aac act tat cta aga agg ata taa taa cta tgt cta cat tat gtc tct tat cga gaa tat tct taa cga gat atc cat agc tat tct ggt gct aat tat tcc tat att ata ttc cac gaa aaa tga tga agg caa tca ttc atc ata aga tga taa aaa gtg tag tga gta aga gta tta gtg aga gag cat gaa gga gat tta gta ttt agc agt gag gat atg atc caa gag tgt gag ata gtc gtt ctc gtt cag aat ctt tcg cag cat aag tag tat gtc gat ata ctt atc att gaa gac tct tcc agt gac aat agc tga ttg agt aca aag tcc aat tat tgc aca aag ttc ttt ggc ggt ttt cat gga gtc att tct gat gaa aca ttt aat gat ctc cac gca att gtc gat att gtc cca cgg aag tga atc cga gaa ctc ctt caa ctc gct acc aaa tag ctc cat tgc atc aat tct gaa aga gat gag aag cct gta gag agg ccc tgc gct ttc tct atg ggt cca tct atg aga aac cca cag gat gta ttc agt cag aca a-t gtc tga cat cag tca cgg tat tca ggg agt cct tag tag cgt ggc aat gac agg gac tga act ggg cac aag gag agg cca ttg tga agg tag acg aag gta acc tga tgg tag acc tgt agc cgt ctg tgc tta ata gag ggc ttt aat ttc cat ttt taa tgg tgt cgt gaa tga gga atg aga gtg tct ctc gtc ctt ggt tta cat gga tca gag tga gaa aaa ata tct tgt ata tta tta act aac aac ctt ggt ttc tat cca tgt tta aaa aaa tga cct ata tgt tct tta tta att cta ttt taa act tta tcc tca aga ctc ctg aca aaa tta aaa tcc aga aag cag caa aca atc ctg tta caa gtt tac tga aat ctc tct ttg att gta gag tat atg tag tca gag caa gaa aca ctg cag tag tca aca tga aag ctt gca taa cga tac gtg cat cat aga aag taa caa cag agg cca gcg tta gag att cta aca gtg taa atc cac aa- --a gta tgt aca gat tca ggg gat gtt cat gtc tgt gta aag tca atg cga aaa tca agc cta tag atc cga aca ttg atg cca ata tta gaa cag gac tcc ctt gta taa atg tcc gat gca ttc aaa gta taa aaa tac tgc agc tgt tgc cgt tgt taa agg aaa ttg tag aaa gaa tac cgt aga ctt ttc tta gaa atg cca ttc gta tgt aca cgc tgg cag acg cca ccg agc tgt cat agt tga agt cgt cct cga t

>JX878428.1_Monkeypox_virus_isolate_DRC_07-0514_complete_genome

--- --- --- --- --- --- --- --- --- --- --- --- --- --- --- --- --- --- --- --- --- --- --- --- --- --- --- --- --- --- --- --- --- --- --- --g tgt cta gaa aaa aat gtg tga ccc acg acc gta gga aac tct aga ggg taa gaa aaa tca atc gtt tat aga gac cat cag aaa gag gtt taa tat ttt tgt gag acc cat cga aga gag aaa gga taa aaa ctt ttt acg act cca tca gaa aga ggt tta ata ttt ttg tga gac cca tcg aa- --- --- --- --- --- --- --- --- --- --- --- --- --- --- --- --- --- --- --- --- --- --- --- --- --- --- --- --- --- --- --- --- --- --- --- --- --- --- --- --- --- --- --- --- --- --- --- --- --- --- --- --- --- --- --- --- --- --- --- --- --- --- --- --- --- --- --- --- --- --- --- --- --- --- --- --- --- --- --- --- --- --- --- --- --- --- --- --- --- --- --- --- --- --- --- --- --- --- --- --- --- --- --- --- --- --- --- --- --- --- --- --- --- --- --- --- --- --- --- --- --- --- --- --- --- --- --- --- --- --- --- --- --- --- --- --- --- --- --- --- --- --- --- --- --- --- --- --- --- --- --- --- --- --- --- --- --- --- --- --- --- --- --- gag aga aag gat aaa aac ttt tta cga ctc cat cag aaa gag gtt taa tat ttt tgt gag acc cat cga aga gag aaa gga taa aaa ctt ttt acg act cca tca gaa aga ggt tta --- --- --- --- --- --- --- --- --- --- --- --- --- --- --- --- --- --- --- --- --- --- --- --- --- --- --- --- --- --- --- --- --- --- --- --- --- --- --- --- --- --- --- --- --- --- --a tat ttt tgt gag acc cat cga cga gag aaa gga taa aaa ctt ttt acg act cca tca gaa aga ggt tta ata ttt ttg tga gac cca tcg acg aga gaa aga gat ggt tag tca aga tat ttt tct tag tac aaa agt caa tgt ttt aaa ata tat gga cga gaa tta att tgt ctg tat aaa aac ttg tgt gaa att atg tac tag aga aaa aac gtg agc agt gtc ccc tac atg gat ttt aca gat cat tta tat tcc aaa aat att aac tat ata cgt tta tta tat gat gtt aac gtg taa att ata aac att att tta tga tgc aat tgt ctg aca acc --- att ggc ata agg ata ttg ata agc tct acg aga ata tat tgt tgg acg tta tcg ttt acg aaa tag ttg aga cat cag aaa gag gtt taa tat ttt tgt gag a-c cat cga aga gag aaa gag aat aaa aat a-- --- --- --- ttt ttt tgt aaa act ttt tta tga gac aag aga gaa aga gaa tac gaa tag tga tca tat cgt atc aca tat tga aac aga aag aag aag taa cga gag gta act ttt tgt gaa tgt agt taa aca ttt ttg ttt tgc aaa ccg gaa tat agt acc cgg tac act ttt tta att cgt ggt gcg gtg tct gaa tcg ttt gat taa ccc aac tca tcc att ttc aga tga ata gag tta tcg att cag aca cat gct ttg agt ttt gtt gaa tcg atg agt gaa gta tca tcg gtt gca cct tca gat gcc gat ccg tcg aca tac ttg aat cca tcc ttg act tca agt tca gat gat tcc tca cac atg tct ccg ata cgt acg cta aac tct agg ttc tta aca cat ttt gta tca acg atc gtt gaa ccg atg ata tct ttg taa ctc act ttc tta tgt gag atg tta gac cca agt act gga tgg gtc ttg atg tcg ctg tct ttc tct tct tcg cta cat ctg atg tcg ata gac atc tca cag tct ttg atc ata gcc aga gct tct tca cgc gtg atc gcg gga gag tcc tta cct tgt cct ggt gac acg ctg gac aat cta gta ttc aca gtg ttt cca tca gag gat tcg gag atg gat gaa atc ttt ggg cat ttg gtg aat cca aag ttc atg tta aga ccc gcg ccg acg ata gtg taa taa gtg gtg gga tct cct ttt aca act tct tcg gat acc tca tca tct tcg gat acc tca tca tct tcg gtc tct gta act tcc gtt acg gat tga caa atc tta tca ttg gtc ggt gtt tgg tct tgc ttt gtg act ttg ata ata aca tcg att ccc ata tga tgt ttg ttt tct tct tca gta cac gag gat gag gat tgt tga aga cta gta ggc ata gca gct gcc act agg cac atg cat gcc agg aca ata tat tgt ttc atg att gct att gat tga tta ctg ttc tag atg att cta ctt tct tac cat ata ata aat tag aat ata ttt tct act ttt acg aga aat taa tta ttg tat tta tta ttt atg ggt -aa aaa aaa ctt act ata agt ggg tgg gat tct ggg aat tag tga tca gtt tat gta tat cac aac tag cgg gca tat ggc tat tga cat cga gaa cat tac cca tat gat aag aga ttg tat ccg ttt cgt agt ctt gag tat tgg tat tac tat ata gta tgt aga tgt cga cgc tag ata gac agt cgc cca cta gag tta ccg tct ctg aat gcg gca tga tag tat cat tct ttg ttt tcg tta act gtt tgg aag atg aat ctt tgt tgt tac att taa tct cga aat tca gag tac ata tct ttg aag tat tct gat atc tat ttt ctc ctg taa aga atc ctg aag ttg cta cat tat taa gga cag aga agt att ctg cac gaa aga cgg gat cac aat ctt tat gat tca tgg taa tag tta gtt ccg acg ttg aga tgg att cgc tga gac cgg tag tgg tcg tcc gag tac acg atg tgt cgt tga ctg gat aca ggt taa ttt cca cat cga tat agt taa atg tat tgc tgg tt- --a cgg gtt cgc att tat ctg tgg aag aga cgg tgt gag aat atg ttc cgg gac cac acg gag aac aga tga cgt ctc cgg tag acg tgt atc cgg ata ctc cgt atc cta ttc cac act ttg ttt tag aaa tac atg ttc tac acc ctg atg atc ctt tga gaa gac aat aat atc ctg gag agc att cac aga ttc tat tgt gag tcg tgt tac acg atc gcg tct cta cct gat tac tat cac atc ttc cgt tac aac tta gac aag cct gta aat gat tat tgt gag atg taa agg tat ccg aac cac acg gcg tac att gtg tat tag tct tgc tat cac ata atc tgg aag cgt aag ttc ccg gag gac acg ata gac aac ata gat tac ggc ttc tgt att cgt tgt ctt tac att ttc cat tgg atg gtg cat gtg gtg cta tat ctc ttc cgt tta tta tta tac atg aga gaa aca ata tat acg agt ata ata cgg acc tca tga ttt aat aat -gt agt aat cgt cgt ctt gtt act gtt tgt ttc cta ctt ctc caa tca tat aga tta ttt ttt aaa tat ttt ctt tct atc atg gat aat att tgt aat ggt tct ttc cgt aca aca tac tgt tta gat ggt agt cgc tta gct tgg tta tga tat tgc gca taa ttt ccg gag gca aat acg ata gtc tag att gac tat cga tgg tag act cta att tat tga gtg ctt tgt cga cga gtt tac ttt tat gct cca tcg ata gat gac act gtt cta tga gat cgt cgt aca tgg gaa atg aaa tgc gtt tgt ctg aat gta tgg ctt cga gat agg tgt gat acc gga tgt ctt ctg ttc tca ata ccg tat aca agt tgg tgt ctg aga ttc gaa tct ctt tga gga gac tta tgt cac gac tac att ttt cga tga tgg aat cta tct tat cga atg ata tat ttt tca taa ata cac ttt tat agt cct cgt tta aac aga att tag tat gta gtt ccg caa atg act cgt ccc tta ata ggc agt agg cta tta tct tct tta cgt agt gat cgt cgt agg gag aga act ccg aca tct tgt aga aca acg att taa tca tag gta gag ata ctt tca gtc tgt ggt gga tga tgt cat taa cga cat ccg cct tgt ata tga tgt ttc tgt ttt caa aca cca agt cga ata ctg tct tat cgt ctt tag tcg gaa ggt tga tgt cgt atc cga tgt ata cga ggt atg agg caa cat tgt tat tgc aat tct gga agg cgg tat gaa gag gag tca ttg tat tat agt att cgt ctt tct gaa tgt cga atc tat cta gta gat acc gta gta tat tga gag agc gac ttc cat atc ctt gat tat gtt tta tga ata gat aaa gta gat gtt gtc ctt ctt cct ttt gta att tcc cgt att ttt gtt cgt gcc aat tga gta aca tta tga gaa tat gac ctg ttg cac aat cgt tct tta tgt att cca tga tgg gtg tac aat caa gat tat tac gta tcc tcg tat cgg ctc ctc gag ata aaa gag cat aca cca cac gag gac tat gtt tgg tat act gtt gaa ggt aag tgt gta acg gcg tat ttc cga ttt tcg taa ccg cgt taa tgt ttg ctc cat gat cta tta tcg cgt aga tga atc gct tct cag ctc gca tct tag tgt gac tct ttg act tgt aat aat tgc ttt cgt gga aca cgg ata tgt gtt tac agt agt aat gaa gag aag tga gtc cat cct cat cga cgc aat tag ggt cag atc ctt tag tca ata att tgt aca gaa cgt aat agt tta agc tcc cat tga att tat atc taa gat aac aca gca ata gat cgg atg att tac taa agt cat caa tgg ggt ccg tta gta tat caa aga tct tgt tat cga ttg ata gtg aat gaa tca gat agt ggt gta gag gaa tat gtc ctt ttt tat cct tgc tat caa agt tac gca tgc cgt ggc gta aca ata tct tta ata cag atg gat taa atc gtg tat tca tcg tat agc aat gta atg gag agt tac cac att tta gtc gtt tat tca gat cgc agt gtt taa taa cta att taa aca gat gag atg ata tat cca cat caa aga atg cga gat aca tat gac aga cat tat tga cag aaa tgt gac ctt cat tat cac cgt cgt cca taa atg cgt tag gta cgt acc aca tac tat cgt taa cga tgc gca caa tct cgt cca ttt tca tcc atc ttc ata atg att tac ttt ttc ata att aga gaa aaa gat caa ggt ata aaa att aga agt gtt aga cta ta- --- --- --- --- --- --- --- --- --- --- -aa cta act tat aa- --- --- --- --- --- --- --- --- --- --- --- --- --- --- --- --- --- --- --- --- --- --- --- --- --- --- --- --- --- --- --- --- --- --- --- --- --- --- --- --- --- --- --- --- --- --- --- --- --- --- --- --- --- --- --- --- --- --- --- --- --- --- --- --- --- --- --- --- --- --- --- --- --- --- --- --- --- --- --- --- --- --- --- --- --- --- -aa cta act tat gac tta act aac tta tga cta tta act cat ttc aag aaa ggt ggg tgg aga gaa ctc tat atg aca gct tgt gaa aca att aga tcc cta att tct aat gga agt ttt gat agg aga ttg tca tca gtt gat aca ttg ttt att atc tca tct att aga gca cgt ctg ttt aga gct tta gtg acc tgc tcg gtt act tct gtg taa atc ttg aat cct tta gtg ata cac tgt gtc aaa act gga tgt tta gaa tac cta tgt aga ata tgg gaa gca tgc ttg ttt ttg tct cta tta tag att aac tca tac atg gtt gta tta tga att ttc atc tgc cta atg tac tcc aat tct tgt tta caa tca att ata taa tca aag agt gat gat gca tac aca tta caa agt gaa taa tct acc atc ata aaa tac ttg ata cag agc ttt atc aca tca tgg ttt tca att gta tta tta agt ata gct aat ttt ata cag tca ata gac aat ggt tct cta agc aat att tct aat att tta aga tgt gct tcc cta cgg gcg atg aca gat ccc cta tcc acg gcc acg tca aga cat gta tat cca tta ctc att act gcg ttg aca ttt gct cca ttt tct aat agc cat gat act aaa tct ata taa cct gca tag ata gcg cga taa agc aag gtc ctt cca cca gca tct agt tga ttg ata tct tca ata tat ggg ata caa agc tta taa att tct aat act gtg ggt tca tct aca agg aat ccc cta gta tac tga att att tta tat aga tct aat tta aca tca ttt tca tct ggg ata cca caa ttc aaa ata aac tca aca aca cta ctt tcc ttt tta cat att ccc cta aaa tag gca ttc aag cat tct att tta tat att aca gcc cca tga tct acc ata aaa tca aca atg tct att tct aca tat gca tta gat aga tag taa agt aag aga tct tgt aca gaa tta caa ttc tta ata att ata gag aaa ata tct tcc ata taa ttc ttt gac act aat gca gat ata ata tct tta tat gta ata tat gca aac agt cta tct act ata tac tga tca ata tta tct cta tga atc cta aaa taa tca tac aga aca tct aca gga tca caa att ggt tca agg aga aat cta tca aat att ttc ctg tca aca act ggt tct aga aca taa cag tca aca cct aat cca tgt ttt tta tag tca tct acc aaa gat aat gac caa aga tcg agg tcg tcg tga aac tgc tca tcg aca gcc atg aaa tct gcc gac tcc atg gcg cga atc gca ctg tct tat tcg cca ttg att ttc a-t ttt tta taa tta tgt aca tgt ttt cct tct att ctc aag agt cta caa aaa tat a-t ttt ttt tcg ata tct aag tac taa gtt ttt tta ctg ttt ttg tta ctg tct tcc att ctt cta act aaa gat ctg aga taa att ata caa tct tcg cta tcg aac cat ttt tgt agt cta aag cct gaa gta att aac caa ctg ttt tta tta gtg gct ttt ttc gat cta tcc tcg tc- --- --c tca tca tcc tta ta- --- --- --- --- --- --- tta tta tca tta tta tca tag tct att aaa cac aaa tca tct --- --- --- --- --- --- --- --- --- --- --- --- --- --- --- --- --- --- --- --- --- --- --- --- --- --- --a cgt tta taa c-- --- --- --- --- --- --- --- --a aca ttc --- --- --- --- -tc att att aat tag ttc tgt agt aat atc ttt aat aat ttg gct ata --- --- -ca tct gtt --- --- --- --- --- --- --- --- --- --- --- --- --- --- --- --- --- --- --c aat act --- --- --- --- --- --- --- --a tct att gat gat ttc ttt t-- --- --- --- --- --- --- --- --- --- --- --t taa gac t-- --- --- --- --- --- --- --- --- --- --- --- --- --- --- --- --- --- --- --- --- --- --- --- --- --- --- --- --- --- --- --- --- --- --- --- --- --- --- --- taa act agt --- --- --- --- --- --- --- --- --- --- --- --- --- --- --- --- --- --- --- --- --- --- --- --- --- --- --- --- --- --- --- --- --- --- --- --- --- --- --- --- --- --- --- --- --- --- --- --- --- --- --- --- --- --- --- --- --- --- --- --- --- --- --- --- --- --- --- --- --- --- --- --- --- --- --- --- --- --- --- --- --- --- --- --- --- --- --- --- --- --- --- --- --- --- --- --- --- --- --- --- --- --- --- --- --- --- --- --- --- --t atg gta atg acg atg aaa --- -tc gag tag ta- --- --a ctt cta ata aag act tga ta- --- --t cat tat cat atg ttt gat cg- --- --- --- --- --- --- --t cat agt taa tag tgt g-- --- --- --- --- --- --- --- --- --- --- --- --- --- --- --- --- --- --- --- -gc taa atg gta ctg tta ata agt tta t-- --- --- --- --- --- --- --- --- --- --- --- --- --- --- --- --- --a gac aat atc ata gta ttt tct ttc cag aat --- --- --- --t aga tta ttt ttt taa ata ctg atc ctc aca att ccg tga tgt agc agt agt tgg t-- --- --- --- --- --- --- --- --- --- --- --- --- --- --- --- --- --- --- --- --- --- --- --- --- --- --- --- -gc atg gtc tat atc gt- --- --- --- --- --- --- --- --- --- --- --- --- --- --- --- --- --- --- --- --- --- --- --- --- --- --- --- --- --- --- --- --- --- --- --- --- --- --- --- --- --- --- --- taa aat gta tca tat ata ata gtt ttc tga cgt gga gta cag aat ttt cga --- --- --- --- --- --- --- --- --- --- --- --- --- --- --- --- --- --- --- --- --- --- --- --- --- --- --- --- --- --- --- --- --- --- --- --- --- --- --- --- --- --- --- --t taa tga gtt cat ggt aag gaa ggg caa atg tct --- gta tat aat ata cat aag tta a-- --- --- --- --- --- --- --- --- --- --- --- --- --t agt ttt tta tca tat tt- --- --- --- --- --- --- --- --t cta ata cca taa taa aaa tta tca tta tg- --- --- -ta taa tca --- --- --- --- --- --- --- --- --- --- --- --t cac tg- --- -tc gct atc att att gcg ttt gtg tag tt- --- --- --- --- ctg ccc ta- --- --- --- --- --- --- --- --t cat cta cat cac tgt ca- --- --- --- --- ctc tca cta tat ctt cta aaa tta caa --- -ac aac --- --- --- --- --- --- --- --- --- --- --- --- --- --- --- --- --- --- --- --- --- --- --- --- --- --- --- --- --- --- --- --- --- --- --- --- tgg ata ttc gat --- --- --- --- --- --- --- --- --- --- --- --- --- --- --- --- --- --- --- --- --- --- --- --- --- --- --a aca gca ttt gtg t-- --- --- --- --- --- --- --- --- --- --- --- --- --- --- --- --- -

>JX878427.1_Monkeypox_virus_isolate_DRC_07-0480_complete_genome

--- --- --- --- --- --- --- --- --- --- --- --- --- --- --- --- --- --- --- --- --- --- --- --- --- --- --- --- --- --- --- --- --- --- --- --g tgt cta gaa aaa aat gtg tga ccc acg acc gta gga aac tct aga ggg taa gaa aaa tca atc gtt tat aga gac cat cag aaa gag gtt taa tat ttt tgt gag acc cat cga aga gag aaa gga taa aaa ctt ttt acg act cca tca gaa aga ggt tta ata ttt ttg tga gac cca tcg aa- --- --- --- --- --- --- --- --- --- --- --- --- --- --- --- --- --- --- --- --- --- --- --- --- --- --- --- --- --- --- --- --- --- --- --- --- --- --- --- --- --- --- --- --- --- --- --- --- --- --- --- --- --- --- --- --- --- --- --- --- --- --- --- --- --- --- --- --- --- --- --- --- --- --- --- --- --- --- --- --- --- --- --- --- --- --- --- --- --- --- --- --- --- --- --- --- --- --- --- --- --- --- --- --- --- --- --- --- --- --- --- --- --- --- --- --- --- --- --- --- --- --- --- --- --- --- --- --- --- --- --- --- --- --- --- --- --- --- --- --- --- --- --- --- --- --- --- --- --- --- --- --- --- --- --- --- --- --- --- --- --- --- --- gag aga aag gat aaa aac ttt tta cga ctc cat cag aaa gag gtt taa tat ttt tgt gag acc cat cga aga gag aaa gga taa aaa ctt ttt acg act cca tca gaa aga ggt tta --- --- --- --- --- --- --- --- --- --- --- --- --- --- --- --- --- --- --- --- --- --- --- --- --- --- --- --- --- --- --- --- --- --- --- --- --- --- --- --- --- --- --- --- --- --- --a tat ttt tgt gag acc cat cga cga gag aaa gga taa aaa ctt ttt acg act cca tca gaa aga ggt tta ata ttt ttg tga gac cca tcg acg aga gaa aga gat ggt tag tca aga tat ttt tct tag tac aaa agt caa tgt ttt aaa ata tat gga cga gaa tta att tgt ctg tat aaa aac ttg tgt gaa att atg tac tag aga aaa aac gtg agc agt gtc ccc tac atg gat ttt aca gat cat tta tat tcc aaa aat att aac tat ata cgt tta tta tat gat gtt aac gtg taa att ata aac att att tta tga tgc aat tgt ctg aca acc --- att ggc ata agg ata ttg ata agc tct acg aga ata tat tgt tgg acg tta tcg ttt acg aaa tag ttg aga cat cag aaa gag gtt taa tat ttt tgt gag a-c cat cga aga gag aaa gag aat aaa aat a-- --- --- --- ttt ttt tgt aaa act ttt tta tga gac aag aga gaa aga gaa tac gaa tag tga tca tat cgt atc aca tat tga aac aga aag aag aag taa cga gag gta act ttt tgt gaa tgt agt taa aca ttt ttg ttt tgc aaa ccg gaa tat agt acc cgg tac act ttt tta att cgt ggt gcg gtg tct gaa tcg ttt gat taa ccc aac tca tcc att ttc aga tga ata gag tta tcg att cag aca cat gct ttg agt ttt gtt gaa tcg atg agt gaa gta tca tcg gtt gca cct tca gat gcc gat ccg tcg aca tac ttg aat cca tcc ttg act tca agt tca gat gat tcc tca cac atg tct ccg ata cgt acg cta aac tct agg ttc tta aca cat ttt gta tca acg atc gtt gaa ccg atg ata tct ttg taa ctc act ttc tta tgt gag atg tta gac cca agt act gga tgg gtc ttg atg tcg ctg tct ttc tct tct tcg cta cat ctg atg tcg ata gac atc tca cag tct ttg atc ata gcc aga gct tct tca cgc gtg atc gcg gga gag tcc tta cct tgt cct ggt gac acg ctg gac aat cta gta ttc aca gtg ttt cca tca gag gat tcg gag atg gat gaa atc ttt ggg cat ttg gtg aat cca aag ttc atg tta aga ccc gcg ccg acg ata gtg taa taa gtg gtg gga tct cct ttt aca act tct tcg gat acc tca tca tct tcg gat acc tca tca tct tcg gtc tct gta act tcc gtt acg gat tga caa atc tta tca ttg gtc ggt gtt tgg tct tgc ttt gtg act ttg ata ata aca tcg att ccc ata tga tgt ttg ttt tct tct tca gta cac gag gat gag gat tgt tga aga cta gta ggc ata gca gct gcc act agg cac atg cat gcc agg aca ata tat tgt ttc atg att gct att gat tga tta ctg ttc tag atg att cta ctt tct tac cat ata ata aat tag aat ata ttt tct act ttt acg aga aat taa tta ttg tat tta tta ttt atg ggt -aa aaa aaa ctt act ata agt ggg tgg gat tct ggg aat tag tga tca gtt tat gta tat cac aac tag cgg gca tat ggc tat tga cat cga gaa cat tac cca tat gat aag aga ttg tat ccg ttt cgt agt ctt gag tat tgg tat tac tat ata gta tgt aga tgt cga cgc tag ata gac agt cgc cca cta gag tta ccg tct ctg aat gcg gca tga tag tat cat tct ttg ttt tcg tta act gtt tgg aag atg aat ctt tgt tgt tac att taa tct cga aat tca gag tac ata tct ttg aag tat tct gat atc tat ttt ctc ctg taa aga atc ctg aag ttg cta cat tat taa gga cag aga agt att ctg cac gaa aga cgg gat cac aat ctt tat gat tca tgg taa tag tta gtt ccg acg ttg aga tgg att cgc tga gac cgg tag tgg tcg tcc gag tac acg atg tgt cgt tga ctg gat aca ggt taa ttt cca cat cga tat agt taa atg tat tgc tgg tt- --a cgg gtt cgc att tat ctg tgg aag aga cgg tgt gag aat atg ttc cgg gac cac acg gag aac aga tga cgt ctc cgg tag acg tgt atc cgg ata ctc cgt atc cta ttc cac act ttg ttt tag aaa tac atg ttc tac acc ctg atg atc ctt tga gaa gac aat aat atc ctg gag agc att cac aga ttc tat tgt gag tcg tgt tac acg atc gcg tct cta cct gat tac tat cac atc ttc cgt tac aac tta gac aag cct gta aat gat tat tgt gag atg taa agg tat ccg aac cac acg gcg tac att gtg tat tag tct tgc tat cac ata atc tgg aag cgt aag ttc ccg gag gac acg ata gac aac ata gat tac ggc ttc tgt att cgt tgt ctt tac att ttc cat tgg atg gtg cat gtg gtg cta tat ctc ttc cgt tta tta tta tac atg aga gaa aca ata tat acg agt ata ata cgg acc tca tga ttt aat aat -gt agt aat cgt cgt ctt gtt act gtt tgt ttc cta ctt ctc caa tca tat aga tta ttt ttt aaa tat ttt ctt tct atc atg gat aat att tgt aat ggt tct ttc cgt aca aca tac tgt tta gat ggt agt cgc tta gct tgg tta tga tat tgc gca taa ttt ccg gag gca aat acg ata gtc tag att gac tat cga tgg tag act cta att tat tga gtg ctt tgt cga cga gtt tac ttt tat gct cca tcg ata gat gac act gtt cta tga gat cgt cgt aca tgg gaa atg aaa tgc gtt tgt ctg aat gta tgg ctt cga gat agg tgt gat acc gga tgt ctt ctg ttc tca ata ccg tat aca agt tgg tgt ctg aga ttc gaa tct ctt tga gga gac tta tgt cac gac tac att ttt cga tga tgg aat cta tct tat cga atg ata tat ttt tca taa ata cac ttt tat agt cct cgt tta aac aga att tag tat gta gtt ccg caa atg act cgt ccc tta ata ggc agt agg cta tta tct tct tta cgt agt gat cgt cgt agg gag aga act ccg aca tct tgt aga aca acg att taa tca tag gta gag ata ctt tca gtc tgt ggt gga tga tgt cat taa cga cat ccg cct tgt ata tga tgt ttc tgt ttt caa aca cca agt cga ata ctg tct tat cgt ctt tag tcg gaa ggt tga tgt cgt atc cga tgt ata cga ggt atg agg caa cat tgt tat tgc aat tct gga agg cgg tat gaa gag gag tca ttg tat tat agt att cgt ctt tct gaa tgt cga atc tat cta gta gat acc gta gta tat tga gag agc gac ttc cat atc ctt gat tat gtt tta tga ata gat aaa gta gat gtt gtc ctt ctt cct ttt gta att tcc cgt att ttt gtt cgt gcc aat tga gta aca tta tga gaa tat gac ctg ttg cac aat cgt tct tta tgt att cca tga tgg gtg tac aat caa gat tat tac gta tcc tcg tat cgg ctc ctc gag ata aaa gag cat aca cca cac gag gac tat gtt tgg tat act gtt gaa ggt aag tgt gta acg gcg tat ttc cga ttt tcg taa ccg cgt taa tgt ttg ctc cat gat cta tta tcg cgt aga tga atc gct tct cag ctc gca tct tag tgt gac tct ttg act tgt aat aat tgc ttt cgt gga aca cgg ata tgt gtt tac agt agt aat gaa gag aag tga gtc cat cct cat cga cgc aat tag ggt cag atc ctt tag tca ata att tgt aca gaa cgt aat agt tta agc tcc cat tga att tat atc taa gat aac aca gca ata gat cgg atg att tac taa agt cat caa tgg ggt ccg tta gta tat caa aga tct tgt tat cga ttg ata gtg aat gaa tca gat agt ggt gta gag gaa tat gtc ctt ttt tat cct tgc tat caa agt tac gca tgc cgt ggc gta aca ata tct tta ata cag atg gat taa atc gtg tat tca tcg tat agc aat gta atg gag agt tac cac att tta gtc gtt tat tca gat cgc agt gtt taa taa cta att taa aca gat gag atg ata tat cca cat caa aga atg cga gat aca tat gac aga cat tat tga cag aaa tgt gac ctt cat tat cac cgt cgt cca taa atg cgt tag gta cgt acc aca tac tat cgt taa cga tgc gca caa tct cgt cca ttt tca tcc atc ttc ata atg att tac ttt ttc ata att aga gaa aaa gat caa ggt ata aaa att aga agt gtt aga cta ta- --- --- --- --- --- --- --- --- --- --- -aa cta act tat aa- --- --- --- --- --- --- --- --- --- --- --- --- --- --- --- --- --- --- --- --- --- --- --- --- --- --- --- --- --- --- --- --- --- --- --- --- --- --- --- --- --- --- --- --- --- --- --- --- --- --- --- --- --- --- --- --- --- --- --- --- --- --- --- --- --- --- --- --- --- --- --- --- --- --- --- --- --- --- --- --- --- --- --- --- --- --- -aa cta act tat gac tta act aac tta tga cta tta act cat ttc aag aaa ggt ggg tgg aga gaa ctc tat atg aca gct tgt gaa aca att aga tcc cta att tct aat gga agt ttt gat agg aga ttg tca tca gtt gat aca ttg ttt att atc tca tct att aga gca cgt ctg ttt aga gct tta gtg acc tgc tcg gtt act tct gtg taa atc ttg aat cct tta gtg ata cac tgt gtc aaa act gga tgt tta gaa tac cta tgt aga ata tgg gaa gca tgc ttg ttt ttg tct cta tta tag att aac tca tac atg gtt gta tta tga att ttc atc tgc cta atg tac tcc aat tct tgt tta caa tca att ata taa tca aag agt gat gat gca tac aca tta caa agt gaa taa tct acc atc ata aaa tac ttg ata cag agc ttt atc aca tca tgg ttt tca att gta tta tta agt ata gct aat ttt ata cag tca ata gac aat ggt tct cta agc aat att tct aat att tta aga tgt gct tcc cta cgg gcg atg aca gat ccc cta tcc acg gcc acg tca aga cat gta tat cca tta ctc att act gcg ttg aca ttt gct cca ttt tct aat agc cat gat act aaa tct ata taa cct gca tag ata gcg cga taa agc aag gtc ctt cca cca gca tct agt tga ttg ata tct tca ata tat ggg ata caa agc tta taa att tct aat act gtg ggt tca tct aca agg aat ccc cta gta tac tga att att tta tat aga tct aat tta aca tca ttt tca tct ggg ata cca caa ttc aaa ata aac tca aca aca cta ctt tcc ttt tta cat att ccc cta aaa tag gca ttc aag cat tct att tta tat att aca gcc cca tga tct acc ata aaa tca aca atg tct att tct aca tat gca tta gat aga tag taa agt aag aga tct tgt aca gaa tta caa ttc tta ata att ata gag aaa ata tct tcc ata taa ttc ttt gac act aat gca gat ata ata tct tta tat gta ata tat gca aac agt cta tct act ata tac tga tca ata tta tct cta tga atc cta aaa taa tca tac aga aca tct aca gga tca caa att ggt tca agg aga aat cta tca aat att ttc ctg tca aca act ggt tct aga aca taa cag tca aca cct aat cca tgt ttt tta tag tca tct acc aaa gat aat gac caa aga tcg agg tcg tcg tga aac tgc tca tcg aca gcc atg aaa tct gcc gac tcc atg gcg cga atc gca ctg tct tat tcg cca ttg att ttc a-t ttt tta taa tta tgt aca tgt ttt cct tct att ctc aag agt cta caa aaa tat a-t ttt ttt tcg ata tct aag tac taa gtt ttt tta ctg ttt ttg tta ctg tct tcc att ctt cta act aaa gat ctg aga taa att ata caa tct tcg cta tcg aac cat ttt tgt agt cta aag cct gaa gta att aac caa ctg ttt tta tta gtg gct ttt ttc gat cta tcc tcg tc- --- --c tca tca tcc tta ta- --- --- --- --- --- --- tta tta tca tta tta tca tag tct att aaa cac aaa tca tct --- --- --- --- --- --- --- --- --- --- --- --- --- --- --- --- --- --- --- --- --- --- --- --- --- --- --a cgt tta taa c-- --- --- --- --- --- --- --- --a aca ttc --- --- --- --- -tc att att aat tag ttc tgt agt aat atc ttt aat aat ttg gct ata --- --- -ca tct gtt --- --- --- --- --- --- --- --- --- --- --- --- --- --- --- --- --- --- --c aat act --- --- --- --- --- --- --- --a tct att gat gat ttc ttt t-- --- --- --- --- --- --- --- --- --- --- --t taa gac t-- --- --- --- --- --- --- --- --- --- --- --- --- --- --- --- --- --- --- --- --- --- --- --- --- --- --- --- --- --- --- --- --- --- --- --- --- --- --- --- taa act agt --- --- --- --- --- --- --- --- --- --- --- --- --- --- --- --- --- --- --- --- --- --- --- --- --- --- --- --- --- --- --- --- --- --- --- --- --- --- --- --- --- --- --- --- --- --- --- --- --- --- --- --- --- --- --- --- --- --- --- --- --- --- --- --- --- --- --- --- --- --- --- --- --- --- --- --- --- --- --- --- --- --- --- --- --- --- --- --- --- --- --- --- --- --- --- --- --- --- --- --- --- --- --- --- --- --- --- --- --- --t atg gta atg acg atg aaa --- -tc gag tag ta- --- --a ctt cta ata aag act tga ta- --- --t cat tat cat atg ttt gat cg- --- --- --- --- --- --- --t cat agt taa tag tgt g-- --- --- --- --- --- --- --- --- --- --- --- --- --- --- --- --- --- --- --- -gc taa atg gta ctg tta ata agt tta t-- --- --- --- --- --- --- --- --- --- --- --- --- --- --- --- --- --a gac aat atc ata gta ttt tct ttc cag aat --- --- --- --t aga tta ttt ttt taa ata ctg atc ctc aca att ccg tga tgt agc agt agt tgg t-- --- --- --- --- --- --- --- --- --- --- --- --- --- --- --- --- --- --- --- --- --- --- --- --- --- --- --- -gc atg gtc tat atc gt- --- --- --- --- --- --- --- --- --- --- --- --- --- --- --- --- --- --- --- --- --- --- --- --- --- --- --- --- --- --- --- --- --- --- --- --- --- --- --- --- --- --- --- taa aat gta tca tat ata ata gtt ttc tga cgt gga gta cag aat ttt cga --- --- --- --- --- --- --- --- --- --- --- --- --- --- --- --- --- --- --- --- --- --- --- --- --- --- --- --- --- --- --- --- --- --- --- --- --- --- --- --- --- --- --- --t taa tga gtt cat ggt aag gaa ggg caa atg tct --- gta tat aat ata cat aag tta a-- --- --- --- --- --- --- --- --- --- --- --- --- --t agt ttt tta tca tat tt- --- --- --- --- --- --- --- --t cta ata cca taa taa aaa tta tca tta tg- --- --- -ta taa tca --- --- --- --- --- --- --- --- --- --- --- --t cac tg- --- -tc gct atc att att gcg ttt gtg tag tt- --- --- --- --- ctg ccc ta- --- --- --- --- --- --- --- --t cat cta cat cac tgt ca- --- --- --- --- ctc tca cta tat ctt cta aaa tta caa --- -ac aac --- --- --- --- --- --- --- --- --- --- --- --- --- --- --- --- --- --- --- --- --- --- --- --- --- --- --- --- --- --- --- --- --- --- --- --- tgg ata ttc gat --- --- --- --- --- --- --- --- --- --- --- --- --- --- --- --- --- --- --- --- --- --- --- --- --- --- --a aca gca ttt gtg t-- --- --- --- --- --- --- --- --- --- --- --- --- --- --- --- --- -

>JX878421.1_Monkeypox_virus_isolate_DRC_07-0286_complete_genome

--- --- --- --- --- --- --- --- --- --- --- --- --- --- --- --- --- --- --- --- --- --- --- --- --- --- --- --- --- --- --- --- --- --- --- --g tgt cta gaa aaa aat gtg tga ccc acg acc gta gga aac tct aga ggg taa gaa aaa tca atc gtt tat aga gac cat cag aaa gag gtt taa tat ttt tgt gag acc cat cga aga gag aaa gga taa aaa ctt ttt acg act cca tca gaa aga ggt tta ata ttt ttg tga gac cca tcg aag aga gaa agg ata aaa act ttt tac gac tcc atc aga aag agg ttt aat att ttt gtg aga ccc atc gaa --- --- --- --- --- --- --- --- --- --- --- --- --- --- --- --- --- --- --- --- --- --- --- --- --- --- --- --- --- --- --- --- --- --- --- --- --- --- --- --- --- --- --- --- --- --- --- --- --- --- --- --- --- --- --- --- --- --- --- --- --- --- --- --- --- --- --- --- --- --- --- --- --- --- --- --- --- --- --- --- --- --- --- --- --- --- --- --- --- --- --- --- --- -ga gag aaa gga taa aaa ctt ttt acg act cca tca gaa aga ggt tta ata ttt ttg tga gac cca tcg acg aga gaa agg ata aaa act ttt tac gac tcc atc aga aag agg ttt aat att ttt gtg aga ccc atc gaa gag aga aag gat aaa aac ttt tta cga ctc cat cag aaa gag gtt taa tat ttt tgt gag acc cat cga cga gag aaa gga taa aaa ctt ttt acg act cca tca gaa aga ggt tta --- --- --- --- --- --- --- --- --- --- --- --- --- --- --- --- --- --- --- --- --- --- --- --- --- --- --- --- --- --- --- --- --- --- --- --- --- --- --- --- --- --- --- --- --- --- --a tat ttt tgt gag acc cat cga aga gag aaa gga taa aaa ctt ttt acg act cca tca gaa aga ggt tta ata ttt ttg tga gac cca tcg acg aga gaa aga gat ggt tag tca aga tat ttt tct tag tac aaa agt caa tgt ttt aaa ata tat gga cga gaa tta att tgt ctg tat aaa aac ttg tgt gaa att atg tac tag aga aaa aac gtg agc agt gtc ccc tac atg gat ttt aca gat cat tta tat tcc aaa aat att aac tat ata cgt tta tta tat gat gtt aac gtg taa att ata aac att att tta tga tgc aat tgt ctg aca acc --- att ggc ata agg ata ttg ata agc tct acg aga ata tat tgt tgg acg tta tcg ttt acg aaa tag ttg aga cat cag aaa gag gtt taa tat ttt tgt gag a-c cat cga aga gag aaa gag aat aaa aat a-- --- --- --- ttt ttt tgt aaa act ttt tta tga gac aag aga gaa aga gaa tac gaa tag tga tca tat cgt atc aca tat tga aac aga aag aag aag taa cga gag gta act ttt tgt gaa tgt agt taa aca ttt ttg ttt tgc aaa ccg gaa tat agt acc cgg tac act ttt tta att cgt ggt gcg gtg tct gaa tcg ttt gat taa ccc aac tca tcc att ttc aga tga ata gag tta tcg att cag aca cat gct ttg agt ttt gtt gaa tcg atg agt gaa gta tca tcg gtt gca cct tca gat gcc gat ccg tcg aca tac ttg aat cca tcc ttg act tca agt tca gat gat tcc tca cac atg tct ccg ata cgt acg cta aac tct agg ttc tta aca cat ttt gta tca acg atc gtt gaa ccg atg ata tct ttg taa ctc act ttc tta tgt gag atg tta gac cca agt act gga tgg gtc ttg atg tcg ctg tct ttc tct tct tcg cta cat ctg atg tcg ata gac atc tca cag tct ttg atc ata gcc aga gct tct tca cgc gtg atc gcg gga gag tcc tta cct tgt cct ggt gac acg ctg gac aat cta gta ttc aca gtg ttt cca tca gag gat tcg gag atg gat gaa atc ttt ggg cat ttg gtg aat cca aag ttc atg tta aga ccc gcg ccg acg ata gtg taa taa gtg gtg gga tct cct ttt aca act tct tcg gat acc tca tca tct tcg gat acc tca tca tct tcg gtc tct gta act tcc gtt acg gat tga caa atc tta tca ttg gtc ggt gtt tgg tct tgc ttt gtg act ttg ata ata aca tcg att ccc ata tga tgt ttg ttt tct tct tca gta cac gag gat gag gat tgt tga aga cta gta ggc ata gca gct gcc act agg cac atg cat gcc agg aca ata tat tgt ttc atg att gct att gat tga tta ctg ttc tag atg att cta ctt tct tac cat ata ata aat tag aat ata ttt tct act ttt acg aga aat taa tta ttg tat tta tta ttt atg ggt -aa aaa aaa ctt act ata agt ggg tgg gat tct ggg aat tag tga tca gtt tat gta tat cac aac tag cgg gca tat ggc tat tga cat cga gaa cat tac cca tat gat aag aga ttg tat ccg ttt cgt agt ctt gag tat tgg tat tac tat ata gta tgt aga tgt cga cgc tag ata gac agt cgc cca cta gag tta ccg tct ctg aat gcg gca tga tag tat cat tct ttg ttt tcg tta act gtt tgg aag atg aat ctt tgt tgt tac att taa tct cga aat tca gag tac ata tct ttg aag tat tct gat atc tat ttt ctc ctg taa aga atc ctg aag ttg cta cat tat taa gga cag aga agt att ctg cac gaa aga cgg gat cac aat ctt tat gat tca tgg taa tag tta gtt ccg acg ttg aga tgg att cgc tga gac cgg tag tgg tcg tcc gag tac acg atg tgt cgt tga ctg gat aca ggt taa ttt cca cat cga tat agt taa atg tat tgc tgg tt- --a cgg gtt cgc att tat ctg tgg aag aga cgg tgt gag aat atg ttc cgg gac cac acg gag aac aga tga cgt ctc cgg tag acg tgt atc cgg ata ctc cgt atc cta ttc cac act ttg ttt tag aaa tac atg ttc tac acc ctg atg atc ctt tga gaa gac aat aat atc ctg gag agc att cac aga ttc tat tgt gag tcg tgt tac acg atc gcg tct cta cct gat tac tat cac atc ttc cgt tac aac tta gac aag cct gta aat gat tat tgt gag atg taa agg tat ccg aac cac acg gcg tac att gtg tat tag tct tgc tat cac ata atc tgg aag cgt aag ttc ccg gag gac acg ata gac aac ata gat tac ggc ttc tgt att cgt tgt ctt tac att ttc cat tgg atg gtg cat gtg gtg cta tat ctc ttc cgt tta tta tta tac atg aga gaa aca ata tat acg agt ata ata cgg acc tca tga ttt aat aat -gt agt aat cgt cgt ctt gtt act gtt tgt ttc cta ctt ctc caa tca tat aga tta ttt ttt aaa tat ttt ctt tct atc atg gat aat att tgt aat ggt tct ttc cgt aca aca tac tgt tta gat ggt agt cgc tta gct tgg tta tga tat tgc gca taa ttt ccg gag gca aat acg ata gtc tag att gac tat cga tgg tag act cta att tat tga gtg ctt tgt cga cga gtt tac ttt tat gct cca tcg ata gat gac act gtt cta tga gat cgt cgt aca tgg gaa atg aaa tgc gtt tgt ctg aat gta tgg ctt cga gat agg tgt gat acc gga tgt ctt ctg ttc tca ata ccg tat aca agt tgg tgt ctg aga ttc gaa tct ctt tga gga gac tta tgt cac gac tac att ttt cga tga tgg aat cta tct tat cga atg ata tat ttt tca taa ata cac ttt tat agt cct cgt tta aac aga att tag tat gta gtt ccg caa atg act cgt ccc tta ata ggc agt agg cta tta tct tct tta cgt agt gat cgt cgt agg gag aga act ccg aca tct tgt aga aca acg att taa tca tag gta gag ata ctt tca gtc tgt ggt gga tga tgt cat taa cga cat ccg cct tgt ata tga tgt ttc tgt ttt caa aca cca agt cga ata ctg tct tat cgt ctt tag tcg gaa ggt tga tgt cgt atc cga tgt ata cga ggt atg agg caa cat tgt tat tgc aat tct gga agg cgg tat gaa gag gag tca ttg tat tat agt att cgt ctt tct gaa tgt cga atc tat cta gta gat acc gta gta tat tga gag agc gac ttc cat atc ctt gat tat gtt tta tga ata gat aaa gta gat gtt gtc ctt ctt cct ttt gta att tcc cgt att ttt gtt cgt gcc aat tga gta aca tta tga gaa tat gac ctg ttg cac aat cgt tct tta tgt att cca tga tgg gtg tac aat caa gat tat tac gta tcc tcg tat cgg ctc ctc gag ata aaa gag cat aca cca cac gag gac tat gtt tgg tat act gtt gaa ggt aag tgt gta acg gcg tat ttc cga ttt tcg taa ccg cgt taa tgt ttg ctc cat gat cta tta tcg cgt aga tga atc gct tct cag ctc gca tct tag tgt gac tct ttg act tgt aat aat tgc ttt cgt gga aca cgg ata tgt gtt tac agt agt aat gaa gag aag tga gtc cat cct cat cga cgc aat tag ggt cag atc ctt tag tca ata att tgt aca gaa cgt aat agt tta agc tcc cat tga att tat atc taa gat aac aca gca ata gat cgg atg att tac taa agt cat caa tgg ggt ccg tta gta tat caa aga tct tgt tat cga ttg ata gtg aat gaa tca gat agt ggt gta gag gaa tat gtc ctt ttt tat cct tgc tat caa agt tac gca tgc cgt ggc gta aca ata tct tta ata cag atg gat taa atc gtg tat tca tcg tat agc aat gta atg gag agt tac cac att tta gtc gtt tat tca gat cgc agt gtt taa taa cta att taa aca gat gag atg ata tat cca cat caa aga atg cga gat aca tat gac aga cat tat tga cag aaa tgt gac ctt cat tat cac cgt cgt cca taa atg cgt tag gta cgt acc aca tac tat cgt taa cga tgc gca caa tct cgt cca ttt tca tcc atc ttc ata atg att tac ttt ttc ata att aga gaa aaa gat caa ggt ata aaa att aga agt gtt aga cta ta- --- --- --- --- --- --- --- --- --- --- -aa cta act tat aa- --- --- --- --- --- --- --- --- --- --- --- --- --- --- --- --- --- --- --- --- --- --- --- --- --- --- --- --- --- --- --- --- --- --- --- --- --- --- --- --- --- --- --- --- --- --- --- --- --- --- --- --- --- --- --- --- --- --- --- --- --- --- --- --- --- --- --- --- --- --- --- --- --- --- --- --- --- --- --- --- --- --- --- --- --- --- -aa cta act tat gac tta act aac tta tga cta tta act cat ttc aag aaa ggt ggg tgg aga gaa ctc tat atg aca gct tgt gaa aca att aga tcc cta att tct aat gga agt ttt gat agg aga ttg tca tca gtt gat aca ttg ttt att atc tca tct att aga gca cgt ctg ttt aga gct tta gtg acc tgc tcg gtt act tct gtg taa atc ttg aat cct tta gtg ata cac tgt gtc aaa act gga tgt tta gaa tac cta tgt aga ata tgg gaa gca tgc ttg ttt ttg tct cta tta tag att aac tca tac atg gtt gta tta tga att ttc atc tgc cta atg tac tcc aat tct tgt tta caa tca att ata taa tca aag agt gat gat gca tac aca tta caa agt gaa taa tct acc atc ata aaa tac ttg ata cag agc ttt atc aca tca tgg ttt tca att gta tta tta agt ata gct aat ttt ata cag tca ata gac aat ggt tct cta agc aat att tct aat att tta aga tgt gct tcc cta cgg gcg atg aca gat ccc cta tcc acg gcc acg tca aga cat gta tat cca tta ctc att act gcg ttg aca ttt gct cca ttt tct aat agc cat gat act aaa tct ata taa cct gca tag ata gcg cga taa agc aag gtc ctt cca cca gca tct agt tga ttg ata tct tca ata tat ggg ata caa agc tta taa att tct aat act gtg ggt tca tct aca agg aat ccc cta gta tac tga att att tta tat aga tct aat tta aca tca ttt tca tct ggg ata cca caa ttc aaa ata aac tca aca aca cta ctt tcc ttt tta cat att ccc cta aaa tag gca ttc aag cat tct att tta tat att aca gcc cca tga tct acc ata aaa tca aca atg tct att tct aca tat gca tta gat aga tag taa agt aag aga tct tgt aca gaa tta caa ttc tta ata att ata gag aaa ata tct tcc ata taa ttc ttt gac act aat gca gat ata ata tct tta tat gta ata tat gca aac agt cta tct act ata tac tga tca ata tta tct cta tga atc cta aaa taa tca tac aga aca tct aca gga tca caa att ggt tca agg aga aat cta tca aat att ttc ctg tca aca act ggt tct aga aca taa cag tca aca cct aat cca tgt ttt tta tag tca tct acc aaa gat aat gac caa aga tcg agg tcg tcg tga aac tgc tca tcg aca gcc atg aaa tct gcc gac tcc atg gcg cga atc gca ctg tct tat tcg cca ttg att ttc a-t ttt tta taa tta tgt aca tgt ttt cct tct att ctc aag agt cta caa aaa tat a-t ttt ttt tcg ata tct aag tac taa gtt ttt tta ctg ttt ttg tta ctg tct tcc att ctt cta act aaa gat ctg aga taa att ata caa tct tcg cta tcg aac cat ttt tgt agt cta aag cct gaa gta att aac caa ctg ttt tta tta gtg gct ttt ttc gat cta tcc tcg tc- --- --c tca tca tcc tta ta- --- --- --- --- --- --- tta tta tca tta tta tca tag tct att aaa cac aaa tca tct --- --- --- --- --- --- --- --- --- --- --- --- --- --- --- --- --- --- --- --- --- --- --- --- --- --- --a cgt tta taa c-- --- --- --- --- --- --- --- --a aca ttc --- --- --- --- -tc att att aat tag ttc tgt agt aat atc ttt aat aat ttg gct ata --- --- -ca tct gtt --- --- --- --- --- --- --- --- --- --- --- --- --- --- --- --- --- --- --c aat act --- --- --- --- --- --- --- --a tct att gat gat ttc ttt t-- --- --- --- --- --- --- --- --- --- --- --t taa gac t-- --- --- --- --- --- --- --- --- --- --- --- --- --- --- --- --- --- --- --- --- --- --- --- --- --- --- --- --- --- --- --- --- --- --- --- --- --- --- --- taa act agt --- --- --- --- --- --- --- --- --- --- --- --- --- --- --- --- --- --- --- --- --- --- --- --- --- --- --- --- --- --- --- --- --- --- --- --- --- --- --- --- --- --- --- --- --- --- --- --- --- --- --- --- --- --- --- --- --- --- --- --- --- --- --- --- --- --- --- --- --- --- --- --- --- --- --- --- --- --- --- --- --- --- --- --- --- --- --- --- --- --- --- --- --- --- --- --- --- --- --- --- --- --- --- --- --- --- --- --- --- --t atg gta atg acg atg aaa --- -tc gag tag ta- --- --a ctt cta ata aag act tga ta- --- --t cat tat cat atg ttt gat cg- --- --- --- --- --- --- --t cat agt taa tag tgt g-- --- --- --- --- --- --- --- --- --- --- --- --- --- --- --- --- --- --- --- -gc taa atg gta ctg tta ata agt tta t-- --- --- --- --- --- --- --- --- --- --- --- --- --- --- --- --- --a gac aat atc ata gta ttt tct ttc cag aat --- --- --- --t aga tta ttt ttt taa ata ctg atc ctc aca att ccg tga tgt agc agt agt tgg t-- --- --- --- --- --- --- --- --- --- --- --- --- --- --- --- --- --- --- --- --- --- --- --- --- --- --- --- -gc atg gtc tat atc gt- --- --- --- --- --- --- --- --- --- --- --- --- --- --- --- --- --- --- --- --- --- --- --- --- --- --- --- --- --- --- --- --- --- --- --- --- --- --- --- --- --- --- --- taa aat gta tca tat ata ata gtt ttc tga cgt gga gta cag aat ttt cga --- --- --- --- --- --- --- --- --- --- --- --- --- --- --- --- --- --- --- --- --- --- --- --- --- --- --- --- --- --- --- --- --- --- --- --- --- --- --- --- --- --- --- --t taa tga gtt cat ggt aag gaa ggg caa atg tct --- gta tat aat ata cat aag tta a-- --- --- --- --- --- --- --- --- --- --- --- --- --t agt ttt tta tca tat tt- --- --- --- --- --- --- --- --t cta ata cca taa taa aaa tta tca tta tg- --- --- -ta taa tca --- --- --- --- --- --- --- --- --- --- --- --t cac tg- --- -tc gct atc att att gcg ttt gtg tag tt- --- --- --- --- ctg ccc ta- --- --- --- --- --- --- --- --t cat cta cat cac tgt ca- --- --- --- --- ctc tca cta tat ctt cta aaa tta caa --- -ac aac --- --- --- --- --- --- --- --- --- --- --- --- --- --- --- --- --- --- --- --- --- --- --- --- --- --- --- --- --- --- --- --- --- --- --- --- tgg ata ttc gat --- --- --- --- --- --- --- --- --- --- --- --- --- --- --- --- --- --- --- --- --- --- --- --- --- --- --a aca gca ttt gtg t-- --- --- --- --- --- --- --- --- --- --- --- --- --- --- --- --- -

>JX878415.1_Monkeypox_virus_isolate_DRC_07-0092_complete_genome

--- --- --- --- --- --- --- --- --- --- --- --- --- --- --- --- --- --- --- --- --- --- --- --- --- --- --- --- --- --- --- --- --- --- --- --g tgt cta gaa aaa aat gtg tga ccc acg acc gta gga aac tct aga ggg taa gaa aaa tca atc gtt tat aga gac cat cag aaa gag gtt taa tat ttt tgt gag acc cat cga aga gag aaa gga taa aaa ctt ttt acg act cca tca gaa aga ggt tta ata ttt ttg tga gac cca tcg aa- --- --- --- --- --- --- --- --- --- --- --- --- --- --- --- --- --- --- --- --- --- --- --- --- --- --- --- --- --- --- --- --- --- --- --- --- --- --- --- --- --- --- --- --- --- --- --- --- --- --- --- --- --- --- --- --- --- --- --- --- --- --- --- --- --- --- --- --- --- --- --- --- --- --- --- --- --- --- --- --- --- --- --- --- --- --- --- --- --- --- --- --- --- --- --- --- --- --- --- --- --- --- --- --- --- --- --- --- --- --- --- --- --- --- --- --- --- --- --- --- --- --- --- --- --- --- --- --- --- --- --- --- --- --- --- --- --- --- --- --g aga gaa agg ata aaa act ttt tac gac tcc atc aga aag agg ttt aat att ttt gtg aga ccc atc gaa gag aga aag gat aaa aac ttt tta cga ctc cat cag aaa gag gtt taa tat ttt tgt gag acc cat cga cga gag aaa gga taa aaa ctt ttt acg act cca tca gaa aga ggt tta --- --- --- --- --- --- --- --- --- --- --- --- --- --- --- --- --- --- --- --- --- --- --- --- --- --- --- --- --- --- --- --- --- --- --- --- --- --- --- --- --- --- --- --- --- --- --a tat ttt tgt gag acc cat cga cga gag aaa gga taa aaa ctt ttt acg act cca tca gaa aga ggt tta ata ttt ttg tga gac cca tcg acg aga gaa aga gat ggt tag tca aga tat ttt tct tag tac aaa agt caa tgt ttt aaa ata tat gga cga gaa tta att tgt ctg tat aaa aac ttg tgt gaa att atg tac tag aga aaa aac gtg agc agt gtc ccc tac atg gat ttt aca gat cat tta tat tcc aaa aat att aac tat ata cgt tta tta tat gat gtt aac gtg taa att ata aac att att tta tga tgc aat tgt ctg aca acc --- att ggc ata agg ata ttg ata agc tct acg aga ata tat tgt tgg acg tta tcg ttt acg aaa tag ttg aga cat cag aaa gag gtt taa tat ttt tgt gag a-c cat cga aga gag aaa gag aat aaa aat a-- --- --- --- ttt ttt tgt aaa act ttt tta tga gac aag aga gaa aga gaa tac gaa tag tga tca tat cgt atc aca tat tga aac aga aag aag aag taa cga gag gta act ttt tgt gaa tgt agt taa aca ttt ttg ttt tgc aaa ccg gaa tat agt acc cgg tac act ttt tta att cgt ggt gcg gtg tct gaa tcg ttt gat taa ccc aac tca tcc att ttc aga tga ata gag tta tcg att cag aca cat gct ttg agt ttt gtt gaa tcg atg agt gaa gta tca tcg gtt gca cct tca gat gcc gat ccg tcg aca tac ttg aat cca tcc ttg act tca agt tca gat gat tcc tca cac atg tct ccg ata cgt acg cta aac tct agg ttc tta aca cat ttt gta tca acg atc gtt gaa ccg atg ata tct ttg taa ctc act ttc tta tgt gag atg tta gac cca agt act gga tgg gtc ttg atg tcg ctg tct ttc tct tct tcg cta cat ctg atg tcg ata gac atc tca cag tct ttg atc ata gcc aga gct tct tca cgc gtg atc gcg gga gag tcc tta cct tgt cct ggt gac acg ctg gac aat cta gta ttc aca gtg ttt cca tca gag gat tcg gag atg gat gaa atc ttt ggg cat ttg gtg aat cca aag ttc atg tta aga ccc gcg ccg acg ata gtg taa taa gtg gtg gga tct cct ttt aca act tct tcg gat acc tca tca tct tcg gat acc tca tca tct tcg gtc tct gta act tcc gtt acg gat tga caa atc tta tca ttg gtc ggt gtt tgg tct tgc ttt gtg act ttg ata ata aca tcg att ccc ata tga tgt ttg ttt tct tct tca gta cac gag gat gag gat tgt tga aga cta gta ggc ata gca gct gcc act agg cac atg cat gcc agg aca ata tat tgt ttc atg att gct att gat tga tta ctg ttc tag atg att cta ctt tct tac cat ata ata aat tag aat ata ttt tct act ttt acg aga aat taa tta ttg tat tta tta ttt atg ggt -aa aaa aaa ctt act ata agt ggg tgg gat tct ggg aat tag tga tca gtt tat gta tat cac aac tag cgg gca tat ggc tat tga cat cga gaa cat tac cca tat gat aag aga ttg tat ccg ttt cgt agt ctt gag tat tgg tat tac tat ata gta tgt aga tgt cga cgc tag ata gac agt cgc cca cta gag tta ccg tct ctg aat gcg gca tga tag tat cat tct ttg ttt tcg tta act gtt tgg aag atg aat ctt tgt tgt tac att taa tct cga aat tca gag tac ata tct ttg aag tat tct gat atc tat ttt ctc ctg taa aga atc ctg aag ttg cta cat tat taa gga cag aga agt att ctg cac gaa aga cgg gat cac aat ctt tat gat tca tgg taa tag tta gtt ccg acg ttg aga tgg att cgc tga gac cgg tag tgg tcg tcc gag tac acg atg tgt cgt tga ctg gat aca ggt taa ttt cca cat cga tat agt taa atg tat tgc tgg tt- --a cgg gtt cgc att tat ctg tgg aag aga cgg tgt gag aat atg ttc cgg gac cac acg gag aac aga tga cgt ctc cgg tag acg tgt atc cgg ata ctc cgt atc cta ttc cac act ttg ttt tag aaa tac atg ttc tac acc ctg atg atc ctt tga gaa gac aat aat atc ctg gag agc att cac aga ttc tat tgt gag tcg tgt tac acg atc gcg tct cta cct gat tac tat cac atc ttc cgt tac aac tta gac aag cct gta aat gat tat tgt gag atg taa agg tat ccg aac cac acg gcg tac att gtg tat tag tct tgc tat cac ata atc tgg aag cgt aag ttc ccg gag gac acg ata gac aac ata gat tac ggc ttc tgt att cgt tgt ctt tac att ttc cat tgg atg gtg cat gtg gtg cta tat ctc ttc cgt tta tta tta tac atg aga gaa aca ata tat acg agt ata ata cgg acc tca tga ttt aat aat -gt agt aat cgt cgt ctt gtt act gtt tgt ttc cta ctt ctc caa tca tat aga tta ttt ttt aaa tat ttt ctt tct atc atg gat aat att tgt aat ggt tct ttc cgt aca aca tac tgt tta gat ggt agt cgc tta gct tgg tta tga tat tgc gca taa ttt ccg gag gca aat acg ata gtc tag att gac tat cga tgg tag act cta att tat tga gtg ctt tgt cga cga gtt tac ttt tat gct cca tcg ata gat gac act gtt cta tga gat cgt cgt aca tgg gaa atg aaa tgc gtt tgt ctg aat gta tgg ctt cga gat agg tgt gat acc gga tgt ctt ctg ttc tca ata ccg tat aca agt tgg tgt ctg aga ttc gaa tct ctt tga gga gac tta tgt cac gac tac att ttt cga tga tgg aat cta tct tat cga atg ata tat ttt tca taa ata cac ttt tat agt cct cgt tta aac aga att tag tat gta gtt ccg caa atg act cgt ccc tta ata ggc agt agg cta tta tct tct tta cgt agt gat cgt cgt agg gag aga act ccg aca tct tgt aga aca acg att taa tca tag gta gag ata ctt tca gtc tgt ggt gga tga tgt cat taa cga cat ccg cct tgt ata tga tgt ttc tgt ttt caa aca cca agt cga ata ctg tct tat cgt ctt tag tcg gaa ggt tga tgt cgt atc cga tgt ata cga ggt atg agg caa cat tgt tat tgc aat tct gga agg cgg tat gaa gag gag tca ttg tat tat agt att cgt ctt tct gaa tgt cga atc tat cta gta gat acc gta gta tat tga gag agc gac ttc cat atc ctt gat tat gtt tta tga ata gat aaa gta gat gtt gtc ctt ctt cct ttt gta att tcc cgt att ttt gtt cgt gcc aat tga gta aca tta tga gaa tat gac ctg ttg cac aat cgt tct tta tgt att cca tga tgg gtg tac aat caa gat tat tac gta tcc tcg tat cgg ctc ctc gag ata aaa gag cat aca cca cac gag gac tat gtt tgg tat act gtt gaa ggt aag tgt gta acg gcg tat ttc cga ttt tcg taa ccg cgt taa tgt ttg ctc cat gat cta tta tcg cgt aga tga atc gct tct cag ctc gca tct tag tgt gac tct ttg act tgt aat aat tgc ttt cgt gga aca cgg ata tgt gtt tac agt agt aat gaa gag aag tga gtc cat cct cat cga cgc aat tag ggt cag atc ctt tag tca ata att tgt aca gaa cgt aat agt tta agc tcc cat tga att tat atc taa gat aac aca gca ata gat cgg atg att tac taa agt cat caa tgg ggt ccg tta gta tat caa aga tct tgt tat cga ttg ata gtg aat gaa tca gat agt ggt gta gag gaa tat gtc ctt ttt tat cct tgc tat caa agt tac gca tgc cgt ggc gta aca ata tct tta ata cag atg gat taa atc gtg tat tca tcg tat agc aat gta atg gag agt tac cac att tta gtc gtt tat tca gat cgc agt gtt taa taa cta att taa aca gat gag atg ata tat cca cat caa aga atg cga gat aca tat gac aga cat tat tga cag aaa tgt gac ctt cat tat cac cgt cgt cca taa atg cgt tag gta cgt acc aca tac tat cgt taa cga tgc gca caa tct cgt cca ttt tca tcc atc ttc ata atg att tac ttt ttc ata att aga gaa aaa gat caa ggt ata aaa att aga agt gtt aga cta ta- --- --- --- --- --- --- --- --- --- --- -aa cta act tat aa- --- --- --- --- --- --- --- --- --- --- --- --- --- --- --- --- --- --- --- --- --- --- --- --- --- --- --- --- --- --- --- --- --- --- --- --- --- --- --- --- --- --- --- --- --- --- --- --- --- --- --- --- --- --- --- --- --- --- --- --- --- --- --- --- --- --- --- --- --- --- --- --- --- --- --- --- --- --- --- --- --- --- --- --- --- --- -aa cta act tat gac tta act aac tta tga cta tta act cat ttc aag aaa ggt ggg tgg aga gaa ctc tat atg aca gct tgt gaa aca att aga tcc cta att tct aat gga agt ttt gat agg aga ttg tca tca gtt gat aca ttg ttt att atc tca tct att aga gca cgt ctg ttt aga gct tta gtg acc tgc tcg gtt act tct gtg taa atc ttg aat cct tta gtg ata cac tgt gtc aaa act gga tgt tta gaa tac cta tgt aga ata tgg gaa gca tgc ttg ttt ttg tct cta tta tag att aac tca tac atg gtt gta tta tga att ttc atc tgc cta atg tac tcc aat tct tgt tta caa tca att ata taa tca aag agt gat gat gca tac aca tta caa agt gaa taa tct acc atc ata aaa tac ttg ata cag agc ttt atc aca tca tgg ttt tca att gta tta tta agt ata gct aat ttt ata cag tca ata gac aat ggt tct cta agc aat att tct aat att tta aga tgt gct tcc cta cgg gcg atg aca gat ccc cta tcc acg gcc acg tca aga cat gta tat cca tta ctc att act gcg ttg aca ttt gct cca ttt tct aat agc cat gat act aaa tct ata taa cct gca tag ata gcg cga taa agc aag gtc ctt cca cca gca tct agt tga ttg ata tct tca ata tat ggg ata caa agc tta taa att tct aat act gtg ggt tca tct aca agg aat ccc cta gta tac tga att att tta tat aga tct aat tta aca tca ttt tca tct ggg ata cca caa ttc aaa ata aac tca aca aca cta ctt tcc ttt tta cat att ccc cta aaa tag gca ttc aag cat tct att tta tat att aca gcc cca tga tct acc ata aaa tca aca atg tct att tct aca tat gca tta gat aga tag taa agt aag aga tct tgt aca gaa tta caa ttc tta ata att ata gag aaa ata tct tcc ata taa ttc ttt gac act aat gca gat ata ata tct tta tat gta ata tat gca aac agt cta tct act ata tac tga tca ata tta tct cta tga atc cta aaa taa tca tac aga aca tct aca gga tca caa att ggt tca agg aga aat cta tca aat att ttc ctg tca aca act ggt tct aga aca taa cag tca aca cct aat cca tgt ttt tta tag tca tct acc aaa gat aat gac caa aga tcg agg tcg tcg tga aac tgc tca tcg aca gcc atg aaa tct gcc gac tcc atg gcg cga atc gca ctg tct tat tcg cca ttg att ttc a-t ttt tta taa tta tgt aca tgt ttt cct tct att ctc aag agt cta caa aaa tat a-t ttt ttt tcg ata tct aag tac taa gtt ttt tta ctg ttt ttg tta ctg tct tcc att ctt cta act aaa gat ctg aga taa att ata caa tct tcg cta tcg aac cat ttt tgt agt cta aag cct gaa gta att aac caa ctg ttt tta tta gtg gct ttt ttc gat cta tcc tcg tc- --- --c tca tca tcc tta ta- --- --- --- --- --- --- tta tta tca tta tta tca tag tct att aaa cac aaa tca tct --- --- --- --- --- --- --- --- --- --- --- --- --- --- --- --- --- --- --- --- --- --- --- --- --- --- --a cgt tta taa c-- --- --- --- --- --- --- --- --a aca ttc --- --- --- --- -tc att att aat tag ttc tgt agt aat atc ttt aat aat ttg gct ata --- --- -ca tct gtt --- --- --- --- --- --- --- --- --- --- --- --- --- --- --- --- --- --- --c aat act --- --- --- --- --- --- --- --a tct att gat gat ttc ttt t-- --- --- --- --- --- --- --- --- --- --- --t taa gac t-- --- --- --- --- --- --- --- --- --- --- --- --- --- --- --- --- --- --- --- --- --- --- --- --- --- --- --- --- --- --- --- --- --- --- --- --- --- --- --- taa act agt --- --- --- --- --- --- --- --- --- --- --- --- --- --- --- --- --- --- --- --- --- --- --- --- --- --- --- --- --- --- --- --- --- --- --- --- --- --- --- --- --- --- --- --- --- --- --- --- --- --- --- --- --- --- --- --- --- --- --- --- --- --- --- --- --- --- --- --- --- --- --- --- --- --- --- --- --- --- --- --- --- --- --- --- --- --- --- --- --- --- --- --- --- --- --- --- --- --- --- --- --- --- --- --- --- --- --- --- --- --t atg gta atg acg atg aaa --- -tc gag tag ta- --- --a ctt cta ata aag act tga ta- --- --t cat tat cat atg ttt gat cg- --- --- --- --- --- --- --t cat agt taa tag tgt g-- --- --- --- --- --- --- --- --- --- --- --- --- --- --- --- --- --- --- --- -gc taa atg gta ctg tta ata agt tta t-- --- --- --- --- --- --- --- --- --- --- --- --- --- --- --- --- --a gac aat atc ata gta ttt tct ttc cag aat --- --- --- --t aga tta ttt ttt taa ata ctg atc ctc aca att ccg tga tgt agc agt agt tgg t-- --- --- --- --- --- --- --- --- --- --- --- --- --- --- --- --- --- --- --- --- --- --- --- --- --- --- --- -gc atg gtc tat atc gt- --- --- --- --- --- --- --- --- --- --- --- --- --- --- --- --- --- --- --- --- --- --- --- --- --- --- --- --- --- --- --- --- --- --- --- --- --- --- --- --- --- --- --- taa aat gta tca tat ata ata gtt ttc tga cgt gga gta cag aat ttt cga --- --- --- --- --- --- --- --- --- --- --- --- --- --- --- --- --- --- --- --- --- --- --- --- --- --- --- --- --- --- --- --- --- --- --- --- --- --- --- --- --- --- --- --t taa tga gtt cat ggt aag gaa ggg caa atg tct --- gta tat aat ata cat aag tta a-- --- --- --- --- --- --- --- --- --- --- --- --- --t agt ttt tta tca tat tt- --- --- --- --- --- --- --- --t cta ata cca taa taa aaa tta tca tta tg- --- --- -ta taa tca --- --- --- --- --- --- --- --- --- --- --- --t cac tg- --- -tc gct atc att att gcg ttt gtg tag tt- --- --- --- --- ctg ccc ta- --- --- --- --- --- --- --- --t cat cta cat cac tgt ca- --- --- --- --- ctc tca cta tat ctt cta aaa tta caa --- -ac aac --- --- --- --- --- --- --- --- --- --- --- --- --- --- --- --- --- --- --- --- --- --- --- --- --- --- --- --- --- --- --- --- --- --- --- --- tgg ata ttc gat --- --- --- --- --- --- --- --- --- --- --- --- --- --- --- --- --- --- --- --- --- --- --- --- --- --- --a aca gca ttt gtg t-- --- --- --- --- --- --- --- --- --- --- --- --- --- --- --- --- -

>JX878414.1_Monkeypox_virus_isolate_DRC_07-0046_complete_genome

--- --- --- --- --- --- --- --- --- --- --- --- --- --- --- --- --- --- --- --- --- --- --- --- --- --- --- --- --- --- --- --- --- --- --- --g tgt cta gaa aaa aat gtg tga ccc acg acc gta gga aac tct aga ggg taa gaa aaa tca atc gtt tat aga gac cat cag aaa gag gtt taa tat ttt tgt gag acc cat cga aga gag aaa gga taa aaa ctt ttt acg act cca tca gaa aga ggt tta ata ttt ttg tga gac cca tcg aag aga gaa agg ata aaa act ttt tac gac tcc atc aga aag agg ttt aat att ttt gtg aga ccc atc gaa g-- --- --- --- --- --- --- --- --- --- --- --- --- --- --- --- --- --- --- --- --- --- --- --- --- --- --- --- --- --- --- --- --- --- --- --- --- --- --- --- --- --- --- --- --- --- --- --- --- --- --- --- --- --- --- --- --- --- --- --- --- --- --- --- --- --- --- --- --- --- -ag aga aag gat aaa aac ttt tta cga ctc cat cag aaa gag gtt taa tat ttt tgt gag acc cat cga cga gag aaa gga taa aaa ctt ttt acg act cca tca gaa aga ggt tta ata ttt ttg tga gac cca tcg acg aga gaa agg ata aaa act ttt tac gac tcc atc aga aag agg ttt aat att ttt gtg aga ccc atc gac gag aga aag gat aaa aac ttt tta cga ctc cat cag aaa gag gtt taa tat ttt tgt gag acc cat cga cga gag aaa gga taa aaa ctt ttt acg act cca tca gaa aga ggt tta --- --- --- --- --- --- --- --- --- --- --- --- --- --- --- --- --- --- --- --- --- --- --- --- --- --- --- --- --- --- --- --- --- --- --- --- --- --- --- --- --- --- --- --- --- --- --a tat ttt tgt gag acc cat cga cga gag aaa gga taa aaa ctt ttt acg act cca tca gaa aga ggt tta ata ttt ttg tga gac cca tcg acg aga gaa aga gat ggt tag tca aga tat ttt tct tag tac aaa agt caa tgt ttt aaa ata tat gga cga gaa tta att tgt ctg tat aaa aac ttg tgt gaa att atg tac tag aga aaa aac gtg agc agt gtc ccc tac atg gat ttt aca gat cat tta tat tcc aaa aat att aac tat ata cgt tta tta tat gat gtt aac gtg taa att ata aac att att tta tga tgc aat tgt ctg aca acc --- att ggc ata agg ata ttg ata agc tct acg aga ata tat tgt tgg acg tta tcg ttt acg aaa tag ttg aga cat cag aaa gag gtt taa tat ttt tgt gag a-c cat cga aga gag aaa gag aat aaa aat a-- --- --- --- ttt ttt tgt aaa act ttt tta tga gac aag aga gaa aga gaa tac gaa tag tga tca tat cgt atc aca tat tga aac aga aag aag aag taa cga gag gta act ttt tgt gaa tgt agt taa aca ttt ttg ttt tgc aaa ccg gaa tat agt acc cgg tac act ttt tta att cgt ggt gcg gtg tct gaa tcg ttt gat taa ccc aac tca tcc att ttc aga tga ata gag tta tcg att cag aca cat gct ttg agt ttt gtt gaa tcg atg agt gaa gta tca tcg gtt gca cct tca gat gcc gat ccg tcg aca tac ttg aat cca tcc ttg act tca agt tca gat gat tcc tca cac atg tct ccg ata cgt acg cta aac tct agg ttc tta aca cat ttt gta tca acg atc gtt gaa ccg atg ata tct ttg taa ctc act ttc tta tgt gag atg tta gac cca agt act gga tgg gtc ttg atg tcg ctg tct ttc tct tct tcg cta cat ctg atg tcg ata gac atc tca cag tct ttg atc ata gcc aga gct tct tca cgc gtg atc gcg gga gag tcc tta cct tgt cct ggt gac acg ctg gac aat cta gta ttc aca gtg ttt cca tca gag gat tcg gag atg gat gaa atc ttt ggg cat ttg gtg aat cca aag ttc atg tta aga ccc gcg ccg acg ata gtg taa taa gtg gtg gga tct cct ttt aca act tct tcg gat acc tca tca tct tcg gat acc tca tca tct tcg gtc tct gta act tcc gtt acg gat tga caa atc tta tca ttg gtc ggt gtt tgg tct tgc ttt gtg act ttg ata ata aca tcg att ccc ata tga tgt ttg ttt tct tct tca gta cac gag gat gag gat tgt tga aga cta gta ggc ata gca gct gcc act agg cac atg cat gcc agg aca ata tat tgt ttc atg att gct att gat tga tta ctg ttc tag atg att cta ctt tct tac cat ata ata aat tag aat ata ttt tct act ttt acg aga aat taa tta ttg tat tta tta ttt atg ggt -aa aaa aaa ctt act ata agt ggg tgg gat tct ggg aat tag tga tca gtt tat gta tat cac aac tag cgg gca tat ggc tat tga cat cga gaa cat tac cca tat gat aag aga ttg tat ccg ttt cgt agt ctt gag tat tgg tat tac tat ata gta tgt aga tgt cga cgc tag ata gac agt cgc cca cta gag tta ccg tct ctg aat gcg gca tga tag tat cat tct ttg ttt tcg tta act gtt tgg aag atg aat ctt tgt tgt tac att taa tct cga aat tca gag tac ata tct ttg aag tat tct gat atc tat ttt ctc ctg taa aga atc ctg aag ttg cta cat tat taa gga cag aga agt att ctg cac gaa aga cgg gat cac aat ctt tat gat tca tgg taa tag tta gtt ccg acg ttg aga tgg att cgc tga gac cgg tag tgg tcg tcc gag tac acg atg tgt cgt tga ctg gat aca ggt taa ttt cca cat cga tat agt taa atg tat tgc tgg tt- --a cgg gtt cgc att tat ctg tgg aag aga cgg tgt gag aat atg ttc cgg gac cac acg gag aac aga tga cgt ctc cgg tag acg tgt atc cgg ata ctc cgt atc cta ttc cac act ttg ttt tag aaa tac atg ttc tac acc ctg atg atc ctt tga gaa gac aat aat atc ctg gag agc att cac aga ttc tat tgt gag tcg tgt tac acg atc gcg tct cta cct gat tac tat cac atc ttc cgt tac aac tta gac aag cct gta aat gat tat tgt gag atg taa agg tat ccg aac cac acg gcg tac att gtg tat tag tct tgc tat cac ata atc tgg aag cgt aag ttc ccg gag gac acg ata gac aac ata gat tac ggc ttc tgt att cgt tgt ctt tac att ttc cat tgg atg gtg cat gtg gtg cta tat ctc ttc cgt tta tta tta tac atg aga gaa aca ata tat acg agt ata ata cgg acc tca tga ttt aat aat -gt agt aat cgt cgt ctt gtt act gtt tgt ttc cta ctt ctc caa tca tat aga tta ttt ttt aaa tat ttt ctt tct atc atg gat aat att tgt aat ggt tct ttc cgt aca aca tac tgt tta gat ggt agt cgc tta gct tgg tta tga tat tgc gca taa ttt ccg gag gca aat acg ata gtc tag att gac tat cga tgg tag act cta att tat tga gtg ctt tgt cga cga gtt tac ttt tat gct cca tcg ata gat gac act gtt cta tga gat cgt cgt aca tgg gaa atg aaa tgc gtt tgt ctg aat gta tgg ctt cga gat agg tgt gat acc gga tgt ctt ctg ttc tca ata ccg tat aca agt tgg tgt ctg aga ttc gaa tct ctt tga gga gac tta tgt cac gac tac att ttt cga tga tgg aat cta tct tat cga atg ata tat ttt tca taa ata cac ttt tat agt cct cgt tta aac aga att tag tat gta gtt ccg caa atg act cgt ccc tta ata ggc agt agg cta tta tct tct tta cgt agt gat cgt cgt agg gag aga act ccg aca tct tgt aga aca acg att taa tca tag gta gag ata ctt tca gtc tgt ggt gga tga tgt cat taa cga cat ccg cct tgt ata tga tgt ttc tgt ttt caa aca cca agt cga ata ctg tct tat cgt ctt tag tcg gaa ggt tga tgt cgt atc cga tgt ata cga ggt atg agg caa cat tgt tat tgc aat tct gga agg cgg tat gaa gag gag tca ttg tat tat agt att cgt ctt tct gaa tgt cga atc tat cta gta gat acc gta gta tat tga gag agc gac ttc cat atc ctt gat tat gtt tta tga ata gat aaa gta gat gtt gtc ctt ctt cct ttt gta att tcc cgt att ttt gtt cgt gcc aat tga gta aca tta tga gaa tat gac ctg ttg cac aat cgt tct tta tgt att cca tga tgg gtg tac aat caa gat tat tac gta tcc tcg tat cgg ctc ctc gag ata aaa gag cat aca cca cac gag gac tat gtt tgg tat act gtt gaa ggt aag tgt gta acg gcg tat ttc cga ttt tcg taa ccg cgt taa tgt ttg ctc cat gat cta tta tcg cgt aga tga atc gct tct cag ctc gca tct tag tgt gac tct ttg act tgt aat aat tgc ttt cgt gga aca cgg ata tgt gtt tac agt agt aat gaa gag aag tga gtc cat cct cat cga cgc aat tag ggt cag atc ctt tag tca ata att tgt aca gaa cgt aat agt tta agc tcc cat tga att tat atc taa gat aac aca gca ata gat cgg atg att tac taa agt cat caa tgg ggt ccg tta gta tat caa aga tct tgt tat cga ttg ata gtg aat gaa tca gat agt ggt gta gag gaa tat gtc ctt ttt tat cct tgc tat caa agt tac gca tgc cgt ggc gta aca ata tct tta ata cag atg gat taa atc gtg tat tca tcg tat agc aat gta atg gag agt tac cac att tta gtc gtt tat tca gat cgc agt gtt taa taa cta att taa aca gat gag atg ata tat cca cat caa aga atg cga gat aca tat gac aga cat tat tga cag aaa tgt gac ctt cat tat cac cgt cgt cca taa atg cgt tag gta cgt acc aca tac tat cgt taa cga tgc gca caa tct cgt cca ttt tca tcc atc ttc ata atg att tac ttt ttc ata att aga gaa aaa gat caa ggt ata aaa att aga agt gtt aga cta ta- --- --- --- --- --- --- --- --- --- --- -aa cta act tat aa- --- --- --- --- --- --- --- --- --- --- --- --- --- --- --- --- --- --- --- --- --- --- --- --- --- --- --- --- --- --- --- --- --- --- --- --- --- --- --- --- --- --- --- --- --- --- --- --- --- --- --- --- --- --- --- --- --- --- --- --- --- --- --- --- --- --- --- --- --- --- --- --- --- --- --- --- --- --- --- --- --- --- --- --- --- --- -aa cta act tat gac tta act aac tta tga cta tta act cat ttc aag aaa ggt ggg tgg aga gaa ctc tat atg aca gct tgt gaa aca att aga tcc cta att tct aat gga agt ttt gat agg aga ttg tca tca gtt gat aca ttg ttt att atc tca tct att aga gca cgt ctg ttt aga gct tta gtg acc tgc tcg gtt act tct gtg taa atc ttg aat cct tta gtg ata cac tgt gtc aaa act gga tgt tta gaa tac cta tgt aga ata tgg gaa gca tgc ttg ttt ttg tct cta tta tag att aac tca tac atg gtt gta tta tga att ttc atc tgc cta atg tac tcc aat tct tgt tta caa tca att ata taa tca aag agt gat gat gca tac aca tta caa agt gaa taa tct acc atc ata aaa tac ttg ata cag agc ttt atc aca tca tgg ttt tca att gta tta tta agt ata gct aat ttt ata cag tca ata gac aat ggt tct cta agc aat att tct aat att tta aga tgt gct tcc cta cgg gcg atg aca gat ccc cta tcc acg gcc acg tca aga cat gta tat cca tta ctc att act gcg ttg aca ttt gct cca ttt tct aat agc cat gat act aaa tct ata taa cct gca tag ata gcg cga taa agc aag gtc ctt cca cca gca tct agt tga ttg ata tct tca ata tat ggg ata caa agc tta taa att tct aat act gtg ggt tca tct aca agg aat ccc cta gta tac tga att att tta tat aga tct aat tta aca tca ttt tca tct ggg ata cca caa ttc aaa ata aac tca aca aca cta ctt tcc ttt tta cat att ccc cta aaa tag gca ttc aag cat tct att tta tat att aca gcc cca tga tct acc ata aaa tca aca atg tct att tct aca tat gca tta gat aga tag taa agt aag aga tct tgt aca gaa tta caa ttc tta ata att ata gag aaa ata tct tcc ata taa ttc ttt gac act aat gca gat ata ata tct tta tat gta ata tat gca aac agt cta tct act ata tac tga tca ata tta tct cta tga atc cta aaa taa tca tac aga aca tct aca gga tca caa att ggt tca agg aga aat cta tca aat att ttc ctg tca aca act ggt tct aga aca taa cag tca aca cct aat cca tgt ttt tta tag tca tct acc aaa gat aat gac caa aga tcg agg tcg tcg tga aac tgc tca tcg aca gcc atg aaa tct gcc gac tcc atg gcg cga atc gca ctg tct tat tcg cca ttg att ttc a-t ttt tta taa tta tgt aca tgt ttt cct tct att ctc aag agt cta caa aaa tat a-t ttt ttt tcg ata tct aag tac taa gtt ttt tta ctg ttt ttg tta ctg tct tcc att ctt cta act aaa gat ctg aga taa att ata caa tct tcg cta tcg aac cat ttt tgt agt cta aag cct gaa gta att aac caa ctg ttt tta tta gtg gct ttt ttc gat cta tcc tcg tc- --- --c tca tca tcc tta ta- --- --- --- --- --- --- tta tta tca tta tta tca tag tct att aaa cac aaa tca tct --- --- --- --- --- --- --- --- --- --- --- --- --- --- --- --- --- --- --- --- --- --- --- --- --- --- --a cgt tta taa c-- --- --- --- --- --- --- --- --a aca ttc --- --- --- --- -tc att att aat tag ttc tgt agt aat atc ttt aat aat ttg gct ata --- --- -ca tct gtt --- --- --- --- --- --- --- --- --- --- --- --- --- --- --- --- --- --- --c aat act --- --- --- --- --- --- --- --a tct att gat gat ttc ttt t-- --- --- --- --- --- --- --- --- --- --- --t taa gac t-- --- --- --- --- --- --- --- --- --- --- --- --- --- --- --- --- --- --- --- --- --- --- --- --- --- --- --- --- --- --- --- --- --- --- --- --- --- --- --- taa act agt --- --- --- --- --- --- --- --- --- --- --- --- --- --- --- --- --- --- --- --- --- --- --- --- --- --- --- --- --- --- --- --- --- --- --- --- --- --- --- --- --- --- --- --- --- --- --- --- --- --- --- --- --- --- --- --- --- --- --- --- --- --- --- --- --- --- --- --- --- --- --- --- --- --- --- --- --- --- --- --- --- --- --- --- --- --- --- --- --- --- --- --- --- --- --- --- --- --- --- --- --- --- --- --- --- --- --- --- --- --t atg gta atg acg atg aaa --- -tc gag tag ta- --- --a ctt cta ata aag act tga ta- --- --t cat tat cat atg ttt gat cg- --- --- --- --- --- --- --t cat agt taa tag tgt g-- --- --- --- --- --- --- --- --- --- --- --- --- --- --- --- --- --- --- --- -gc taa atg gta ctg tta ata agt tta t-- --- --- --- --- --- --- --- --- --- --- --- --- --- --- --- --- --a gac aat atc ata gta ttt tct ttc cag aat --- --- --- --t aga tta ttt ttt taa ata ctg atc ctc aca att ccg tga tgt agc agt agt tgg t-- --- --- --- --- --- --- --- --- --- --- --- --- --- --- --- --- --- --- --- --- --- --- --- --- --- --- --- -gc atg gtc tat atc gt- --- --- --- --- --- --- --- --- --- --- --- --- --- --- --- --- --- --- --- --- --- --- --- --- --- --- --- --- --- --- --- --- --- --- --- --- --- --- --- --- --- --- --- taa aat gta tca tat ata ata gtt ttc tga cgt gga gta cag aat ttt cga --- --- --- --- --- --- --- --- --- --- --- --- --- --- --- --- --- --- --- --- --- --- --- --- --- --- --- --- --- --- --- --- --- --- --- --- --- --- --- --- --- --- --- --t taa tga gtt cat ggt aag gaa ggg caa atg tct --- gta tat aat ata cat aag tta a-- --- --- --- --- --- --- --- --- --- --- --- --- --t agt ttt tta tca tat tt- --- --- --- --- --- --- --- --t cta ata cca taa taa aaa tta tca tta tg- --- --- -ta taa tca --- --- --- --- --- --- --- --- --- --- --- --t cac tg- --- -tc gct atc att att gcg ttt gtg tag tt- --- --- --- --- ctg ccc ta- --- --- --- --- --- --- --- --t cat cta cat cac tgt ca- --- --- --- --- ctc tca cta tat ctt cta aaa tta caa --- -ac aac --- --- --- --- --- --- --- --- --- --- --- --- --- --- --- --- --- --- --- --- --- --- --- --- --- --- --- --- --- --- --- --- --- --- --- --- tgg ata ttc gat --- --- --- --- --- --- --- --- --- --- --- --- --- --- --- --- --- --- --- --- --- --- --- --- --- --- --a aca gca ttt gtg t-- --- --- --- --- --- --- --- --- --- --- --- --- --- --- --- --- -

>JX878413.1_Monkeypox_virus_isolate_DRC_07-0045_complete_genome

--- --- --- --- --- --- --- --- --- --- --- --- --- --- --- --- --- --- --- --- --- --- --- --- --- --- --- --- --- --- --- --- --- --- --- --g tgt cta gaa aaa aat gtg tga ccc acg acc gta gga aac tct aga ggg taa gaa aaa tca atc gtt tat aga gac cat cag aaa gag gtt taa tat ttt tgt gag acc cat cga aga gag aaa gga taa aaa ctt ttt acg act cca tca gaa aga ggt tta ata ttt ttg tga gac cca tcg aa- --- --- --- --- --- --- --- --- --- --- --- --- --- --- --- --- --- --- --- --- --- --- --- --- --- --- --- --- --- --- --- --- --- --- --- --- --- --- --- --- --- --- --- --- --- --- --- --- --- --- --- --- --- --- --- --- --- --- --- --- --- --- --- --- --- --- --- --- --- --- --- --- --- --- --- --- --- --- --- --- --- --- --- --- --- --- --- --- --- --- --- --- --- --- --- --- --- --- --- --- --- --- --- --- --- --- --- --- --- --- --- --- --- --- --- --- -ga gag aaa gga taa aaa ctt ttt acg act cca tca gaa aga ggt tta ata ttt ttg tga gac cca tcg aag aga gaa agg ata aaa act ttt tac gac tcc atc aga aag agg ttt aat att ttt gtg aga ccc atc gac gag aga aag gat aaa aac ttt tta cga ctc cat cag aaa gag gtt taa tat ttt tgt gag acc cat cga cga gag aaa gga taa aaa ctt ttt acg act cca tca gaa aga ggt tta --- --- --- --- --- --- --- --- --- --- --- --- --- --- --- --- --- --- --- --- --- --- --- --- --- --- --- --- --- --- --- --- --- --- --- --- --- --- --- --- --- --- --- --- --- --- --a tat ttt tgt gag acc cat cga cga gag aaa gga taa aaa ctt ttt acg act cca tca gaa aga ggt tta ata ttt ttg tga gac cca tcg acg aga gaa aga gat ggt tag tca aga tat ttt tct tag tac aaa agt caa tgt ttt aaa ata tat gga cga gaa tta att tgt ctg tat aaa aac ttg tgt gaa att atg tac tag aga aaa aac gtg agc agt gtc ccc tac atg gat ttt aca gat cat tta tat tcc aaa aat att aac tat ata cgt tta tta tat gat gtt aac gtg taa att ata aac att att tta tga tgc aat tgt ctg aca acc --- att ggc ata agg ata ttg ata agc tct acg aga ata tat tgt tgg acg tta tcg ttt acg aaa tag ttg aga cat cag aaa gag gtt taa tat ttt tgt gag a-c cat cga aga gag aaa gag aat aaa aat a-- --- --- --- ttt ttt tgt aaa act ttt tta tga gac aag aga gaa aga gaa tac gaa tag tga tca tat cgt atc aca tat tga aac aga aag aag aag taa cga gag gta act ttt tgt gaa tgt agt taa aca ttt ttg ttt tgc aaa ccg gaa tat agt acc cgg tac act ttt tta att cgt ggt gcg gtg tct gaa tcg ttt gat taa ccc aac tca tcc att ttc aga tga ata gag tta tcg att cag aca cat gct ttg agt ttt gtt gaa tcg atg agt gaa gta tca tcg gtt gca cct tca gat gcc gat ccg tcg aca tac ttg aat cca tcc ttg act tca agt tca gat gat tcc tca cac atg tct ccg ata cgt acg cta aac tct agg ttc tta aca cat ttt gta tca acg atc gtt gaa ccg atg ata tct ttg taa ctc act ttc tta tgt gag atg tta gac cca agt act gga tgg gtc ttg atg tcg ctg tct ttc tct tct tcg cta cat ctg atg tcg ata gac atc tca cag tct ttg atc ata gcc aga gct tct tca cgc gtg atc gcg gga gag tcc tta cct tgt cct ggt gac acg ctg gac aat cta gta ttc aca gtg ttt cca tca gag gat tcg gag atg gat gaa atc ttt ggg cat ttg gtg aat cca aag ttc atg tta aga ccc gcg ccg acg ata gtg taa taa gtg gtg gga tct cct ttt aca act tct tcg gat acc tca tca tct tcg gat acc tca tca tct tcg gtc tct gta act tcc gtt acg gat tga caa atc tta tca ttg gtc ggt gtt tgg tct tgc ttt gtg act ttg ata ata aca tcg att ccc ata tga tgt ttg ttt tct tct tca gta cac gag gat gag gat tgt tga aga cta gta ggc ata gca gct gcc act agg cac atg cat gcc agg aca ata tat tgt ttc atg att gct att gat tga tta ctg ttc tag atg att cta ctt tct tac cat ata ata aat tag aat ata ttt tct act ttt acg aga aat taa tta ttg tat tta tta ttt atg ggt -aa aaa aaa ctt act ata agt ggg tgg gat tct ggg aat tag tga tca gtt tat gta tat cac aac tag cgg gca tat ggc tat tga cat cga gaa cat tac cca tat gat aag aga ttg tat ccg ttt cgt agt ctt gag tat tgg tat tac tat ata gta tgt aga tgt cga cgc tag ata gac agt cgc cca cta gag tta ccg tct ctg aat gcg gca tga tag tat cat tct ttg ttt tcg tta act gtt tgg aag atg aat ctt tgt tgt tac att taa tct cga aat tca gag tac ata tct ttg aag tat tct gat atc tat ttt ctc ctg taa aga atc ctg aag ttg cta cat tat taa gga cag aga agt att ctg cac gaa aga cgg gat cac aat ctt tat gat tca tgg taa tag tta gtt ccg acg ttg aga tgg att cgc tga gac cgg tag tgg tcg tcc gag tac acg atg tgt cgt tga ctg gat aca ggt taa ttt cca cat cga tat agt taa atg tat tgc tgg tt- --a cgg gtt cgc att tat ctg tgg aag aga cgg tgt gag aat atg ttc cgg gac cac acg gag aac aga tga cgt ctc cgg tag acg tgt atc cgg ata ctc cgt atc cta ttc cac act ttg ttt tag aaa tac atg ttc tac acc ctg atg atc ctt tga gaa gac aat aat atc ctg gag agc att cac aga ttc tat tgt gag tcg tgt tac acg atc gcg tct cta cct gat tac tat cac atc ttc cgt tac aac tta gac aag cct gta aat gat tat tgt gag atg taa agg tat ccg aac cac acg gcg tac att gtg tat tag tct tgc tat cac ata atc tgg aag cgt aag ttc ccg gag gac acg ata gac aac ata gat tac ggc ttc tgt att cgt tgt ctt tac att ttc cat tgg atg gtg cat gtg gtg cta tat ctc ttc cgt tta tta tta tac atg aga gaa aca ata tat acg agt ata ata cgg acc tca tga ttt aat aat -gt agt aat cgt cgt ctt gtt act gtt tgt ttc cta ctt ctc caa tca tat aga tta ttt ttt aaa tat ttt ctt tct atc atg gat aat att tgt aat ggt tct ttc cgt aca aca tac tgt tta gat ggt agt cgc tta gct tgg tta tga tat tgc gca taa ttt ccg gag gca aat acg ata gtc tag att gac tat cga tgg tag act cta att tat tga gtg ctt tgt cga cga gtt tac ttt tat gct cca tcg ata gat gac act gtt cta tga gat cgt cgt aca tgg gaa atg aaa tgc gtt tgt ctg aat gta tgg ctt cga gat agg tgt gat acc gga tgt ctt ctg ttc tca ata ccg tat aca agt tgg tgt ctg aga ttc gaa tct ctt tga gga gac tta tgt cac gac tac att ttt cga tga tgg aat cta tct tat cga atg ata tat ttt tca taa ata cac ttt tat agt cct cgt tta aac aga att tag tat gta gtt ccg caa atg act cgt ccc tta ata ggc agt agg cta tta tct tct tta cgt agt gat cgt cgt agg gag aga act ccg aca tct tgt aga aca acg att taa tca tag gta gag ata ctt tca gtc tgt ggt gga tga tgt cat taa cga cat ccg cct tgt ata tga tgt ttc tgt ttt caa aca cca agt cga ata ctg tct tat cgt ctt tag tcg gaa ggt tga tgt cgt atc cga tgt ata cga ggt atg agg caa cat tgt tat tgc aat tct gga agg cgg tat gaa gag gag tca ttg tat tat agt att cgt ctt tct gaa tgt cga atc tat cta gta gat acc gta gta tat tga gag agc gac ttc cat atc ctt gat tat gtt tta tga ata gat aaa gta gat gtt gtc ctt ctt cct ttt gta att tcc cgt att ttt gtt cgt gcc aat tga gta aca tta tga gaa tat gac ctg ttg cac aat cgt tct tta tgt att cca tga tgg gtg tac aat caa gat tat tac gta tcc tcg tat cgg ctc ctc gag ata aaa gag cat aca cca cac gag gac tat gtt tgg tat act gtt gaa ggt aag tgt gta acg gcg tat ttc cga ttt tcg taa ccg cgt taa tgt ttg ctc cat gat cta tta tcg cgt aga tga atc gct tct cag ctc gca tct tag tgt gac tct ttg act tgt aat aat tgc ttt cgt gga aca cgg ata tgt gtt tac agt agt aat gaa gag aag tga gtc cat cct cat cga cgc aat tag ggt cag atc ctt tag tca ata att tgt aca gaa cgt aat agt tta agc tcc cat tga att tat atc taa gat aac aca gca ata gat cgg atg att tac taa agt cat caa tgg ggt ccg tta gta tat caa aga tct tgt tat cga ttg ata gtg aat gaa tca gat agt ggt gta gag gaa tat gtc ctt ttt tat cct tgc tat caa agt tac gca tgc cgt ggc gta aca ata tct tta ata cag atg gat taa atc gtg tat tca tcg tat agc aat gta atg gag agt tac cac att tta gtc gtt tat tca gat cgc agt gtt taa taa cta att taa aca gat gag atg ata tat cca cat caa aga atg cga gat aca tat gac aga cat tat tga cag aaa tgt gac ctt cat tat cac cgt cgt cca taa atg cgt tag gta cgt acc aca tac tat cgt taa cga tgc gca caa tct cgt cca ttt tca tcc atc ttc ata atg att tac ttt ttc ata att aga gaa aaa gat caa ggt ata aaa att aga agt gtt aga cta ta- --- --- --- --- --- --- --- --- --- --- -aa cta act tat aa- --- --- --- --- --- --- --- --- --- --- --- --- --- --- --- --- --- --- --- --- --- --- --- --- --- --- --- --- --- --- --- --- --- --- --- --- --- --- --- --- --- --- --- --- --- --- --- --- --- --- --- --- --- --- --- --- --- --- --- --- --- --- --- --- --- --- --- --- --- --- --- --- --- --- --- --- --- --- --- --- --- --- --- --- --- --- -aa cta act tat gac tta act aac tta tga cta tta act cat ttc aag aaa ggt ggg tgg aga gaa ctc tat atg aca gct tgt gaa aca att aga tcc cta att tct aat gga agt ttt gat agg aga ttg tca tca gtt gat aca ttg ttt att atc tca tct att aga gca cgt ctg ttt aga gct tta gtg acc tgc tcg gtt act tct gtg taa atc ttg aat cct tta gtg ata cac tgt gtc aaa act gga tgt tta gaa tac cta tgt aga ata tgg gaa gca tgc ttg ttt ttg tct cta tta tag att aac tca tac atg gtt gta tta tga att ttc atc tgc cta atg tac tcc aat tct tgt tta caa tca att ata taa tca aag agt gat gat gca tac aca tta caa agt gaa taa tct acc atc ata aaa tac ttg ata cag agc ttt atc aca tca tgg ttt tca att gta tta tta agt ata gct aat ttt ata cag tca ata gac aat ggt tct cta agc aat att tct aat att tta aga tgt gct tcc cta cgg gcg atg aca gat ccc cta tcc acg gcc acg tca aga cat gta tat cca tta ctc att act gcg ttg aca ttt gct cca ttt tct aat agc cat gat act aaa tct ata taa cct gca tag ata gcg cga taa agc aag gtc ctt cca cca gca tct agt tga ttg ata tct tca ata tat ggg ata caa agc tta taa att tct aat act gtg ggt tca tct aca agg aat ccc cta gta tac tga att att tta tat aga tct aat tta aca tca ttt tca tct ggg ata cca caa ttc aaa ata aac tca aca aca cta ctt tcc ttt tta cat att ccc cta aaa tag gca ttc aag cat tct att tta tat att aca gcc cca tga tct acc ata aaa tca aca atg tct att tct aca tat gca tta gat aga tag taa agt aag aga tct tgt aca gaa tta caa ttc tta ata att ata gag aaa ata tct tcc ata taa ttc ttt gac act aat gca gat ata ata tct tta tat gta ata tat gca aac agt cta tct act ata tac tga tca ata tta tct cta tga atc cta aaa taa tca tac aga aca tct aca gga tca caa att ggt tca agg aga aat cta tca aat att ttc ctg tca aca act ggt tct aga aca taa cag tca aca cct aat cca tgt ttt tta tag tca tct acc aaa gat aat gac caa aga tcg agg tcg tcg tga aac tgc tca tcg aca gcc atg aaa tct gcc gac tcc atg gcg cga atc gca ctg tct tat tcg cca ttg att ttc a-t ttt tta taa tta tgt aca tgt ttt cct tct att ctc aag agt cta caa aaa tat a-t ttt ttt tcg ata tct aag tac taa gtt ttt tta ctg ttt ttg tta ctg tct tcc att ctt cta act aaa gat ctg aga taa att ata caa tct tcg cta tcg aac cat ttt tgt agt cta aag cct gaa gta att aac caa ctg ttt tta tta gtg gct ttt ttc gat cta tcc tcg tc- --- --c tca tca tcc tta ta- --- --- --- --- --- --- tta tta tca tta tta tca tag tct att aaa cac aaa tca tct --- --- --- --- --- --- --- --- --- --- --- --- --- --- --- --- --- --- --- --- --- --- --- --- --- --- --a cgt tta taa c-- --- --- --- --- --- --- --- --a aca ttc --- --- --- --- -tc att att aat tag ttc tgt agt aat atc ttt aat aat ttg gct ata --- --- -ca tct gtt --- --- --- --- --- --- --- --- --- --- --- --- --- --- --- --- --- --- --c aat act --- --- --- --- --- --- --- --a tct att gat gat ttc ttt t-- --- --- --- --- --- --- --- --- --- --- --t taa gac t-- --- --- --- --- --- --- --- --- --- --- --- --- --- --- --- --- --- --- --- --- --- --- --- --- --- --- --- --- --- --- --- --- --- --- --- --- --- --- --- taa act agt --- --- --- --- --- --- --- --- --- --- --- --- --- --- --- --- --- --- --- --- --- --- --- --- --- --- --- --- --- --- --- --- --- --- --- --- --- --- --- --- --- --- --- --- --- --- --- --- --- --- --- --- --- --- --- --- --- --- --- --- --- --- --- --- --- --- --- --- --- --- --- --- --- --- --- --- --- --- --- --- --- --- --- --- --- --- --- --- --- --- --- --- --- --- --- --- --- --- --- --- --- --- --- --- --- --- --- --- --- --t atg gta atg acg atg aaa --- -tc gag tag ta- --- --a ctt cta ata aag act tga ta- --- --t cat tat cat atg ttt gat cg- --- --- --- --- --- --- --t cat agt taa tag tgt g-- --- --- --- --- --- --- --- --- --- --- --- --- --- --- --- --- --- --- --- -gc taa atg gta ctg tta ata agt tta t-- --- --- --- --- --- --- --- --- --- --- --- --- --- --- --- --- --a gac aat atc ata gta ttt tct ttc cag aat --- --- --- --t aga tta ttt ttt taa ata ctg atc ctc aca att ccg tga tgt agc agt agt tgg t-- --- --- --- --- --- --- --- --- --- --- --- --- --- --- --- --- --- --- --- --- --- --- --- --- --- --- --- -gc atg gtc tat atc gt- --- --- --- --- --- --- --- --- --- --- --- --- --- --- --- --- --- --- --- --- --- --- --- --- --- --- --- --- --- --- --- --- --- --- --- --- --- --- --- --- --- --- --- taa aat gta tca tat ata ata gtt ttc tga cgt gga gta cag aat ttt cga --- --- --- --- --- --- --- --- --- --- --- --- --- --- --- --- --- --- --- --- --- --- --- --- --- --- --- --- --- --- --- --- --- --- --- --- --- --- --- --- --- --- --- --t taa tga gtt cat ggt aag gaa ggg caa atg tct --- gta tat aat ata cat aag tta a-- --- --- --- --- --- --- --- --- --- --- --- --- --t agt ttt tta tca tat tt- --- --- --- --- --- --- --- --t cta ata cca taa taa aaa tta tca tta tg- --- --- -ta taa tca --- --- --- --- --- --- --- --- --- --- --- --t cac tg- --- -tc gct atc att att gcg ttt gtg tag tt- --- --- --- --- ctg ccc ta- --- --- --- --- --- --- --- --t cat cta cat cac tgt ca- --- --- --- --- ctc tca cta tat ctt cta aaa tta caa --- -ac aac --- --- --- --- --- --- --- --- --- --- --- --- --- --- --- --- --- --- --- --- --- --- --- --- --- --- --- --- --- --- --- --- --- --- --- --- tgg ata ttc gat --- --- --- --- --- --- --- --- --- --- --- --- --- --- --- --- --- --- --- --- --- --- --- --- --- --- --a aca gca ttt gtg t-- --- --- --- --- --- --- --- --- --- --- --- --- --- --- --- --- -

>JX878422.1_Monkeypox_virus_isolate_DRC_07-0287_complete_genome

--- --- --- --- --- --- --- --- --- --- --- --- --- --- --- --- --- --- --- --- --- --- --- --- --- --- --- --- --- --- --- --- --- --- --- --g tgt cta gaa aaa aat gtg tga ccc acg acc gta gga aac tct aga ggg taa gaa aaa tca atc gtt tat aga gac cat cag aaa gag gtt taa tat ttt tgt gag acc cat cga aga gag aaa gga taa aaa ctt ttt acg act cca tca gaa aga ggt tta ata ttt ttg tga gac cca tcg aa- --- --- --- --- --- --- --- --- --- --- --- --- --- --- --- --- --- --- --- --- --- --- --- --- --- --- --- --- --- --- --- --- --- --- --- --- --- --- --- --- --- --- --- --- --- --- --- --- --- --- --- --- --- --- --- --- --- --- --- --- --- --- --- --- --- --- --- --- --- --- --- --- --- --- --- --- --- --- --- --- --- --- --- --- --- --- --- --- --- --- --- --- --- --- --- --- --- --- --- --- --- --- --- --- --- --- --- --- --- --- --- --- --- --- --- --- --- --- --- --- --- --- --- --- --- --- --- --- --- --- --- --- --- --- --- --- --- --- --- --- --- --- --- --- --- --- --- --- --- --- --- --- --- --- --- --- --- --- --- --- --- --- --- gag aga aag gat aaa aac ttt tta cga ctc cat cag aaa gag gtt taa tat ttt tgt gag acc cat cga aga gag aaa gga taa aaa ctt ttt acg act cca tca gaa aga ggt tta --- --- --- --- --- --- --- --- --- --- --- --- --- --- --- --- --- --- --- --- --- --- --- --- --- --- --- --- --- --- --- --- --- --- --- --- --- --- --- --- --- --- --- --- --- --- --a tat ttt tgt gag acc cat cga cga gag aaa gga taa aaa ctt ttt acg act cca tca gaa aga ggt tta ata ttt ttg tga gac cca tcg acg aga gaa aga gat ggt tag tca aga tat ttt tct tag tac aaa agt caa tgt ttt aaa ata tat gga cga gaa tta att tgt ctg tat aaa aac ttg tgt gaa att atg tac tag aga aaa aac gtg agc agt gtc ccc tac atg gat ttt aca gat cat tta tat tcc aaa aat att aac tat ata cgt tta tta tat gat gtt aac gtg taa att ata aac att att tta tga tgc aat tgt ctg aca acc --- att ggc ata agg ata ttg ata agc tct acg aga ata tat tgt tgg acg tta tcg ttt acg aaa tag ttg aga cat cag aaa gag gtt taa tat ttt tgt gag a-c cat cga aga gag aaa gag aat aaa aat a-- --- --- --- ttt ttt tgt aaa act ttt tta tga gac aag aga gaa aga gaa tac gaa tag tga tca tat cgt atc aca tat tga aac aga aag aag aag taa cga gag gta act ttt tgt gaa tgt agt taa aca ttt ttg ttt tgc aaa ccg gaa tat agt acc cgg tac act ttt tta att cgt ggt gcg gtg tct gaa tcg ttt gat taa ccc aac tca tcc att ttc aga tga ata gag tta tcg att cag aca cat gct ttg agt ttt gtt gaa tcg atg agt gaa gta tca tcg gtt gca cct tca gat gcc gat ccg tcg aca tac ttg aat cca tcc ttg act tca agt tca gat gat tcc tca cac atg tct ccg ata cgt acg cta aac tct agg ttc tta aca cat ttt gta tca acg atc gtt gaa ccg atg ata tct ttg taa ctc act ttc tta tgt gag atg tta gac cca agt act gga tgg gtc ttg atg tcg ctg tct ttc tct tct tcg cta cat ctg atg tcg ata gac atc tca cag tct ttg atc ata gcc aga gct tct tca cgc gtg atc gcg gga gag tcc tta cct tgt cct ggt gac acg ctg gac aat cta gta ttc aca gtg ttt cca tca gag gat tcg gag atg gat gaa atc ttt ggg cat ttg gtg aat cca aag ttc atg tta aga ccc gcg ccg acg ata gtg taa taa gtg gtg gga tct cct ttt aca act tct tcg gat acc tca tca tct tcg gat acc tca tca tct tcg gtc tct gta act tcc gtt acg gat tga caa atc tta tca ttg gtc ggt gtt tgg tct tgc ttt gtg act ttg ata ata aca tcg att ccc ata tga tgt ttg ttt tct tct tca gta cac gag gat gag gat tgt tga aga cta gta ggc ata gca gct gcc act agg cac atg cat gcc agg aca ata tat tgt ttc atg att gct att gat tga tta ctg ttc tag atg att cta ctt tct tac cat ata ata aat tag aat ata ttt tct act ttt acg aga aat taa tta ttg tat tta tta ttt atg ggt -aa aaa aaa ctt act ata agt ggg tgg gat tct ggg aat tag tga tca gtt tat gta tat cac aac tag cgg gca tat ggc tat tga cat cga gaa cat tac cca tat gat aag aga ttg tat ccg ttt cgt agt ctt gag tat tgg tat tac tat ata gta tgt aga tgt cga cgc tag ata gac agt cgc cca cta gag tta ccg tct ctg aat gcg gca tga tag tat cat tct ttg ttt tcg tta act gtt tgg aag atg aat ctt tgt tgt tac att taa tct cga aat tca gag tac ata tct ttg aag tat tct gat atc tat ttt ctc ctg taa aga atc ctg aag ttg cta cat tat taa gga cag aga agt att ctg cac gaa aga cgg gat cac aat ctt tat gat tca tgg taa tag tta gtt ccg acg ttg aga tgg att cgc tga gac cgg tag tgg tcg tcc gag tac acg atg tgt cgt tga ctg gat aca ggt taa ttt cca cat cga tat agt taa atg tat tgc tgg tt- --a cgg gtt cgc att tat ctg tgg aag aga cgg tgt gag aat atg ttc cgg gac cac acg gag aac aga tga cgt ctc cgg tag acg tgt atc cgg ata ctc cgt atc cta ttc cac act ttg ttt tag aaa tac atg ttc tac acc ctg atg atc ctt tga gaa gac aat aat atc ctg gag agc att cac aga ttc tat tgt gag tcg tgt tac acg atc gcg tct cta cct gat tac tat cac atc ttc cgt tac aac tta gac aag cct gta aat gat tat tgt gag atg taa agg tat ccg aac cac acg gcg tac att gtg tat tag tct tgc tat cac ata atc tgg aag cgt aag ttc ccg gag gac acg ata gac aac ata gat tac ggc ttc tgt att cgt tgt ctt tac att ttc cat tgg atg gtg cat gtg gtg cta tat ctc ttc cgt tta tta tta tac atg aga gaa aca ata tat acg agt ata ata cgg acc tca tga ttt aat aat -gt agt aat cgt cgt ctt gtt act gtt tgt ttc cta ctt ctc caa tca tat aga tta ttt ttt aaa tat ttt ctt tct atc atg gat aat att tgt aat ggt tct ttc cgt aca aca tac tgt tta gat ggt agt cgc tta gct tgg tta tga tat tgc gca taa ttt ccg gag gca aat acg ata gtc tag att gac tat cga tgg tag act cta att tat tga gtg ctt tgt cga cga gtt tac ttt tat gct cca tcg ata gat gac act gtt cta tga gat cgt cgt aca tgg gaa atg aaa tgc gtt tgt ctg aat gta tgg ctt cga gat agg tgt gat acc gga tgt ctt ctg ttc tca ata ccg tat aca agt tgg tgt ctg aga ttc gaa tct ctt tga gga gac tta tgt cac gac tac att ttt cga tga tgg aat cta tct tat cga atg ata tat ttt tca taa ata cac ttt tat agt cct cgt tta aac aga att tag tat gta gtt ccg caa atg act cgt ccc tta ata ggc agt agg cta tta tct tct tta cgt agt gat cgt cgt agg gag aga act ccg aca tct tgt aga aca acg att taa tca tag gta gag ata ctt tca gtc tgt ggt gga tga tgt cat taa cga cat ccg cct tgt ata tga tgt ttc tgt ttt caa aca cca agt cga ata ctg tct tat cgt ctt tag tcg gaa ggt tga tgt cgt atc cga tgt ata cga ggt atg agg caa cat tgt tat tgc aat tct gga agg cgg tat gaa gag gag tca ttg tat tat agt att cgt ctt tct gaa tgt cga atc tat cta gta gat acc gta gta tat tga gag agc gac ttc cat atc ctt gat tat gtt tta tga ata gat aaa gta gat gtt gtc ctt ctt cct ttt gta att tcc cgt att ttt gtt cgt gcc aat tga gta aca tta tga gaa tat gac ctg ttg cac aat cgt tct tta tgt att cca tga tgg gtg tac aat caa gat tat tac gta tcc tcg tat cgg ctc ctc gag ata aaa gag cat aca cca cac gag gac tat gtt tgg tat act gtt gaa ggt aag tgt gta acg gcg tat ttc cga ttt tcg taa ccg cgt taa tgt ttg ctc cat gat cta tta tcg cgt aga tga atc gct tct cag ctc gca tct tag tgt gac tct ttg act tgt aat aat tgc ttt cgt gga aca cgg ata tgt gtt tac agt agt aat gaa gag aag tga gtc cat cct cat cga cgc aat tag ggt cag atc ctt tag tca ata att tgt aca gaa cgt aat agt tta agc tcc cat tga att tat atc taa gat aac aca gca ata gat cgg atg att tac taa agt cat caa tgg ggt ccg tta gta tat caa aga tct tgt tat cga ttg ata gtg aat gaa tca gat agt ggt gta gag gaa tat gtc ctt ttt tat cct tgc tat caa agt tac gca tgc cgt ggc gta aca ata tct tta ata cag atg gat taa atc gtg tat tca tcg tat agc aat gta atg gag agt tac cac att tta gtc gtt tat tca gat cgc agt gtt taa taa cta att taa aca gat gag atg ata tat cca cat caa aga atg cga gat aca tat gac aga cat tat tga cag aaa tgt gac ctt cat tat cac cgt cgt cca taa atg cgt tag gta cgt acc aca tac tat cgt taa cga tgc gca caa tct cgt cca ttt tca tcc atc ttc ata atg att tac ttt ttc ata att aga gaa aaa gat caa ggt ata aaa att aga agt gtt aga cta ta- --- --- --- --- --- --- --- --- --- --- -aa cta act tat aa- --- --- --- --- --- --- --- --- --- --- --- --- --- --- --- --- --- --- --- --- --- --- --- --- --- --- --- --- --- --- --- --- --- --- --- --- --- --- --- --- --- --- --- --- --- --- --- --- --- --- --- --- --- --- --- --- --- --- --- --- --- --- --- --- --- --- --- --- --- --- --- --- --- --- --- --- --- --- --- --- --- --- --- --- --- --- -aa cta act tat gac tta act aac tta tga cta tta act cat ttc aag aaa ggt ggg tgg aga gaa ctc tat atg aca gct tgt gaa aca att aga tcc cta att tct aat gga agt ttt gat agg aga ttg tca tca gtt gat aca ttg ttt att atc tca tct att aga gca cgt ctg ttt aga gct tta gtg acc tgc tcg gtt act tct gtg taa atc ttg aat cct tta gtg ata cac tgt gtc aaa act gga tgt tta gaa tac cta tgt aga ata tgg gaa gca tgc ttg ttt ttg tct cta tta tag att aac tca tac atg gtt gta tta tga att ttc atc tgc cta atg tac tcc aat tct tgt tta caa tca att ata taa tca aag agt gat gat gca tac aca tta caa agt gaa taa tct acc atc ata aaa tac ttg ata cag agc ttt atc aca tca tgg ttt tca att gta tta tta agt ata gct aat ttt ata cag tca ata gac aat ggt tct cta agc aat att tct aat att tta aga tgt gct tcc cta cgg gcg atg aca gat ccc cta tcc acg gcc acg tca aga cat gta tat cca tta ctc att act gcg ttg aca ttt gct cca ttt tct aat agc cat gat act aaa tct ata taa cct gca tag ata gcg cga taa agc aag gtc ctt cca cca gca tct agt tga ttg ata tct tca ata tat ggg ata caa agc tta taa att tct aat act gtg ggt tca tct aca agg aat ccc cta gta tac tga att att tta tat aga tct aat tta aca tca ttt tca tct ggg ata cca caa ttc aaa ata aac tca aca aca cta ctt tcc ttt tta cat att ccc cta aaa tag gca ttc aag cat tct att tta tat att aca gcc cca tga tct acc ata aaa tca aca atg tct att tct aca tat gca tta gat aga tag taa agt aag aga tct tgt aca gaa tta caa ttc tta ata att ata gag aaa ata tct tcc ata taa ttc ttt gac act aat gca gat ata ata tct tta tat gta ata tat gca aac agt cta tct act ata tac tga tca ata tta tct cta tga atc cta aaa taa tca tac aga aca tct aca gga tca caa att ggt tca agg aga aat cta tca aat att ttc ctg tca aca act ggt tct aga aca taa cag tca aca cct aat cca tgt ttt tta tag tca tct acc aaa gat aat gac caa aga tcg agg tcg tcg tga aac tgc tca tcg aca gcc atg aaa tct gcc gac tcc atg gcg cga atc gca ctg tct tat tcg cca ttg att ttc a-t ttt tta taa tta tgt aca tgt ttt cct tct att ctc aag agt cta caa aaa tat a-t ttt ttt tcg ata tct aag tac taa gtt ttt tta ctg ttt ttg tta ctg tct tcc att ctt cta act aaa gat ctg aga taa att ata caa tct tcg cta tcg aac cat ttt tgt agt cta aag cct gaa gta att aac caa ctg ttt tta tta gtg gct ttt ttc gat cta tcc tcg tc- --- --c tca tca tcc tta ta- --- --- --- --- --- --- tta tta tca tta tta tca tag tct att aaa cac aaa tca tct --- --- --- --- --- --- --- --- --- --- --- --- --- --- --- --- --- --- --- --- --- --- --- --- --- --- --a cgt tta taa c-- --- --- --- --- --- --- --- --a aca ttc --- --- --- --- -tc att att aat tag ttc tgt agt aat atc ttt aat aat ttg gct ata --- --- -ca tct gtt --- --- --- --- --- --- --- --- --- --- --- --- --- --- --- --- --- --- --c aat act --- --- --- --- --- --- --- --a tct att gat gat ttc ttt t-- --- --- --- --- --- --- --- --- --- --- --t taa gac t-- --- --- --- --- --- --- --- --- --- --- --- --- --- --- --- --- --- --- --- --- --- --- --- --- --- --- --- --- --- --- --- --- --- --- --- --- --- --- --- taa act agt --- --- --- --- --- --- --- --- --- --- --- --- --- --- --- --- --- --- --- --- --- --- --- --- --- --- --- --- --- --- --- --- --- --- --- --- --- --- --- --- --- --- --- --- --- --- --- --- --- --- --- --- --- --- --- --- --- --- --- --- --- --- --- --- --- --- --- --- --- --- --- --- --- --- --- --- --- --- --- --- --- --- --- --- --- --- --- --- --- --- --- --- --- --- --- --- --- --- --- --- --- --- --- --- --- --- --- --- --- --t atg gta atg acg atg aaa --- -tc gag tag ta- --- --a ctt cta ata aag act tga ta- --- --t cat tat cat atg ttt gat cg- --- --- --- --- --- --- --t cat agt taa tag tgt g-- --- --- --- --- --- --- --- --- --- --- --- --- --- --- --- --- --- --- --- -gc taa atg gta ctg tta ata agt tta t-- --- --- --- --- --- --- --- --- --- --- --- --- --- --- --- --- --a gac aat atc ata gta ttt tct ttc cag aat --- --- --- --t aga tta ttt ttt taa ata ctg atc ctc aca att ccg tga tgt agc agt agt tgg t-- --- --- --- --- --- --- --- --- --- --- --- --- --- --- --- --- --- --- --- --- --- --- --- --- --- --- --- -gc atg gtc tat atc gt- --- --- --- --- --- --- --- --- --- --- --- --- --- --- --- --- --- --- --- --- --- --- --- --- --- --- --- --- --- --- --- --- --- --- --- --- --- --- --- --- --- --- --- taa aat gta tca tat ata ata gtt ttc tga cgt gga gta cag aat ttt cga --- --- --- --- --- --- --- --- --- --- --- --- --- --- --- --- --- --- --- --- --- --- --- --- --- --- --- --- --- --- --- --- --- --- --- --- --- --- --- --- --- --- --- --t taa tga gtt cat ggt aag gaa ggg caa atg tct --- gta tat aat ata cat aag tta a-- --- --- --- --- --- --- --- --- --- --- --- --- --t agt ttt tta tca tat tt- --- --- --- --- --- --- --- --t cta ata cca taa taa aaa tta tca tta tg- --- --- -ta taa tca --- --- --- --- --- --- --- --- --- --- --- --t cac tg- --- -tc gct atc att att gcg ttt gtg tag tt- --- --- --- --- ctg ccc ta- --- --- --- --- --- --- --- --t cat cta cat cac tgt ca- --- --- --- --- ctc tca cta tat ctt cta aaa tta caa --- -ac aac --- --- --- --- --- --- --- --- --- --- --- --- --- --- --- --- --- --- --- --- --- --- --- --- --- --- --- --- --- --- --- --- --- --- --- --- tgg ata ttc gat --- --- --- --- --- --- --- --- --- --- --- --- --- --- --- --- --- --- --- --- --- --- --- --- --- --- --a aca gca ttt gtg t-- --- --- --- --- --- --- --- --- --- --- --- --- --- --- --- --- -

>JX878416.1_Monkeypox_virus_isolate_DRC_07-0093_complete_genome

--- --- --- --- --- --- --- --- --- --- --- --- --- --- --- --- --- --- --- --- --- --- --- --- --- --- --- --- --- --- --- --- --- --- --- --g tgt cta gaa aaa aat gtg tga ccc acg acc gta gga aac tct aga ggg taa gaa aaa tca atc gtt tat aga gac cat cag aaa gag gtt taa tat ttt tgt gag acc cat cga aga gag aaa gga taa aaa ctt ttt acg act cca tca gaa aga ggt tta ata ttt ttg tga gac cca tcg aa- --- --- --- --- --- --- --- --- --- --- --- --- --- --- --- --- --- --- --- --- --- --- --- --- --- --- --- --- --- --- --- --- --- --- --- --- --- --- --- --- --- --- --- --- --- --- --- --- --- --- --- --- --- --- --- --- --- --- --- --- --- --- --- --- --- --- --- --- --- --- --- --- --- --- --- --- --- --- --- --- --- --- --- --- --- --- --- --- --- --- --- --- --- --- --- --- --- --- --- --- --- --- --- --- --- --- --- --- --- --- --- --- --- --- --- --- -ga gag aaa gga taa aaa ctt ttt acg act cca tca gaa aga ggt tta ata ttt ttg tga gac cca tcg aag aga gaa agg ata aaa act ttt tac gac tcc atc aga aag agg ttt aat att ttt gtg aga ccc atc gac gag aga aag gat aaa aac ttt tta cga ctc cat cag aaa gag gtt taa tat ttt tgt gag acc cat cga cga gag aaa gga taa aaa ctt ttt acg act cca tca gaa aga ggt tta --- --- --- --- --- --- --- --- --- --- --- --- --- --- --- --- --- --- --- --- --- --- --- --- --- --- --- --- --- --- --- --- --- --- --- --- --- --- --- --- --- --- --- --- --- --- --a tat ttt tgt gag acc cat cga cga gag aaa gga taa aaa ctt ttt acg act cca tca gaa aga ggt tta ata ttt ttg tga gac cca tcg acg aga gaa aga gat ggt tag tca aga tat ttt tct tag tac aaa agt caa tgt ttt aaa ata tat gga cga gaa tta att tgt ctg tat aaa aac ttg tgt gaa att atg tac tag aga aaa aac gtg agc agt gtc ccc tac atg gat ttt aca gat cat tta tat tcc aaa aat att aac tat ata cgt tta tta tat gat gtt aac gtg taa att ata aac att att tta tga tgc aat tgt ctg aca acc --- att ggc ata agg ata ttg ata agc tct acg aga ata tat tgt tgg acg tta tcg ttt acg aaa tag ttg aga cat cag aaa gag gtt taa tat ttt tgt gag a-c cat cga aga gag aaa gag aat aaa aat a-- --- --- --- ttt ttt tgt aaa act ttt tta tga gac aag aga gaa aga gaa tac gaa tag tga tca tat cgt atc aca tat tga aac aga aag aag aag taa cga gag gta act ttt tgt gaa tgt agt taa aca ttt ttg ttt tgc aaa ccg gaa tat agt acc cgg tac act ttt tta att cgt ggt gcg gtg tct gaa tcg ttt gat taa ccc aac tca tcc att ttc aga tga ata gag tta tcg att cag aca cat gct ttg agt ttt gtt gaa tcg atg agt gaa gta tca tcg gtt gca cct tca gat gcc gat ccg tcg aca tac ttg aat cca tcc ttg act tca agt tca gat gat tcc tca cac atg tct ccg ata cgt acg cta aac tct agg ttc tta aca cat ttt gta tca acg atc gtt gaa ccg atg ata tct ttg taa ctc act ttc tta tgt gag atg tta gac cca agt act gga tgg gtc ttg atg tcg ctg tct ttc tct tct tcg cta cat ctg atg tcg ata gac atc tca cag tct ttg atc ata gcc aga gct tct tca cgc gtg atc gcg gga gag tcc tta cct tgt cct ggt gac acg ctg gac aat cta gta ttc aca gtg ttt cca tca gag gat tcg gag atg gat gaa atc ttt ggg cat ttg gtg aat cca aag ttc atg tta aga ccc gcg ccg acg ata gtg taa taa gtg gtg gga tct cct ttt aca act tct tcg gat acc tca tca tct tcg gat acc tca tca tct tcg gtc tct gta act tcc gtt acg gat tga caa atc tta tca ttg gtc ggt gtt tgg tct tgc ttt gtg act ttg ata ata aca tcg att ccc ata tga tgt ttg ttt tct tct tca gta cac gag gat gag gat tgt tga aga cta gta ggc ata gca gct gcc act agg cac atg cat gcc agg aca ata tat tgt ttc atg att gct att gat tga tta ctg ttc tag atg att cta ctt tct tac cat ata ata aat tag aat ata ttt tct act ttt acg aga aat taa tta ttg tat tta tta ttt atg ggt -aa aaa aaa ctt act ata agt ggg tgg gat tct ggg aat tag tga tca gtt tat gta tat cac aac tag cgg gca tat ggc tat tga cat cga gaa cat tac cca tat gat aag aga ttg tat ccg ttt cgt agt ctt gag tat tgg tat tac tat ata gta tgt aga tgt cga cgc tag ata gac agt cgc cca cta gag tta ccg tct ctg aat gcg gca tga tag tat cat tct ttg ttt tcg tta act gtt tgg aag atg aat ctt tgt tgt tac att taa tct cga aat tca gag tac ata tct ttg aag tat tct gat atc tat ttt ctc ctg taa aga atc ctg aag ttg cta cat tat taa gga cag aga agt att ctg cac gaa aga cgg gat cac aat ctt tat gat tca tgg taa tag tta gtt ccg acg ttg aga tgg att cgc tga gac cgg tag tgg tcg tcc gag tac acg atg tgt cgt tga ctg gat aca ggt taa ttt cca cat cga tat agt taa atg tat tgc tgg tt- --a cgg gtt cgc att tat ctg tgg aag aga cgg tgt gag aat atg ttc cgg gac cac acg gag aac aga tga cgt ctc cgg tag acg tgt atc cgg ata ctc cgt atc cta ttc cac act ttg ttt tag aaa tac atg ttc tac acc ctg atg atc ctt tga gaa gac aat aat atc ctg gag agc att cac aga ttc tat tgt gag tcg tgt tac acg atc gcg tct cta cct gat tac tat cac atc ttc cgt tac aac tta gac aag cct gta aat gat tat tgt gag atg taa agg tat ccg aac cac acg gcg tac att gtg tat tag tct tgc tat cac ata atc tgg aag cgt aag ttc ccg gag gac acg ata gac aac ata gat tac ggc ttc tgt att cgt tgt ctt tac att ttc cat tgg atg gtg cat gtg gtg cta tat ctc ttc cgt tta tta tta tac atg aga gaa aca ata tat acg agt ata ata cgg acc tca tga ttt aat aat -gt agt aat cgt cgt ctt gtt act gtt tgt ttc cta ctt ctc caa tca tat aga tta ttt ttt aaa tat ttt ctt tct atc atg gat aat att tgt aat ggt tct ttc cgt aca aca tac tgt tta gat ggt agt cgc tta gct tgg tta tga tat tgc gca taa ttt ccg gag gca aat acg ata gtc tag att gac tat cga tgg tag act cta att tat tga gtg ctt tgt cga cga gtt tac ttt tat gct cca tcg ata gat gac act gtt cta tga gat cgt cgt aca tgg gaa atg aaa tgc gtt tgt ctg aat gta tgg ctt cga gat agg tgt gat acc gga tgt ctt ctg ttc tca ata ccg tat aca agt tgg tgt ctg aga ttc gaa tct ctt tga gga gac tta tgt cac gac tac att ttt cga tga tgg aat cta tct tat cga atg ata tat ttt tca taa ata cac ttt tat agt cct cgt tta aac aga att tag tat gta gtt ccg caa atg act cgt ccc tta ata ggc agt agg cta tta tct tct tta cgt agt gat cgt cgt agg gag aga act ccg aca tct tgt aga aca acg att taa tca tag gta gag ata ctt tca gtc tgt ggt gga tga tgt cat taa cga cat ccg cct tgt ata tga tgt ttc tgt ttt caa aca cca agt cga ata ctg tct tat cgt ctt tag tcg gaa ggt tga tgt cgt atc cga tgt ata cga ggt atg agg caa cat tgt tat tgc aat tct gga agg cgg tat gaa gag gag tca ttg tat tat agt att cgt ctt tct gaa tgt cga atc tat cta gta gat acc gta gta tat tga gag agc gac ttc cat atc ctt gat tat gtt tta tga ata gat aaa gta gat gtt gtc ctt ctt cct ttt gta att tcc cgt att ttt gtt cgt gcc aat tga gta aca tta tga gaa tat gac ctg ttg cac aat cgt tct tta tgt att cca tga tgg gtg tac aat caa gat tat tac gta tcc tcg tat cgg ctc ctc gag ata aaa gag cat aca cca cac gag gac tat gtt tgg tat act gtt gaa ggt aag tgt gta acg gcg tat ttc cga ttt tcg taa ccg cgt taa tgt ttg ctc cat gat cta tta tcg cgt aga tga atc gct tct cag ctc gca tct tag tgt gac tct ttg act tgt aat aat tgc ttt cgt gga aca cgg ata tgt gtt tac agt agt aat gaa gag aag tga gtc cat cct cat cga cgc aat tag ggt cag atc ctt tag tca ata att tgt aca gaa cgt aat agt tta agc tcc cat tga att tat atc taa gat aac aca gca ata gat cgg atg att tac taa agt cat caa tgg ggt ccg tta gta tat caa aga tct tgt tat cga ttg ata gtg aat gaa tca gat agt ggt gta gag gaa tat gtc ctt ttt tat cct tgc tat caa agt tac gca tgc cgt ggc gta aca ata tct tta ata cag atg gat taa atc gtg tat tca tcg tat agc aat gta atg gag agt tac cac att tta gtc gtt tat tca gat cgc agt gtt taa taa cta att taa aca gat gag atg ata tat cca cat caa aga atg cga gat aca tat gac aga cat tat tga cag aaa tgt gac ctt cat tat cac cgt cgt cca taa atg cgt tag gta cgt acc aca tac tat cgt taa cga tgc gca caa tct cgt cca ttt tca tcc atc ttc ata atg att tac ttt ttc ata att aga gaa aaa gat caa ggt ata aaa att aga agt gtt aga cta ta- --- --- --- --- --- --- --- --- --- --- -aa cta act tat aa- --- --- --- --- --- --- --- --- --- --- --- --- --- --- --- --- --- --- --- --- --- --- --- --- --- --- --- --- --- --- --- --- --- --- --- --- --- --- --- --- --- --- --- --- --- --- --- --- --- --- --- --- --- --- --- --- --- --- --- --- --- --- --- --- --- --- --- --- --- --- --- --- --- --- --- --- --- --- --- --- --- --- --- --- --- --- -aa cta act tat gac tta act aac tta tga cta tta act cat ttc aag aaa ggt ggg tgg aga gaa ctc tat atg aca gct tgt gaa aca att aga tcc cta att tct aat gga agt ttt gat agg aga ttg tca tca gtt gat aca ttg ttt att atc tca tct att aga gca cgt ctg ttt aga gct tta gtg acc tgc tcg gtt act tct gtg taa atc ttg aat cct tta gtg ata cac tgt gtc aaa act gga tgt tta gaa tac cta tgt aga ata tgg gaa gca tgc ttg ttt ttg tct cta tta tag att aac tca tac atg gtt gta tta tga att ttc atc tgc cta atg tac tcc aat tct tgt tta caa tca att ata taa tca aag agt gat gat gca tac aca tta caa agt gaa taa tct acc atc ata aaa tac ttg ata cag agc ttt atc aca tca tgg ttt tca att gta tta tta agt ata gct aat ttt ata cag tca ata gac aat ggt tct cta agc aat att tct aat att tta aga tgt gct tcc cta cgg gcg atg aca gat ccc cta tcc acg gcc acg tca aga cat gta tat cca tta ctc att act gcg ttg aca ttt gct cca ttt tct aat agc cat gat act aaa tct ata taa cct gca tag ata gcg cga taa agc aag gtc ctt cca cca gca tct agt tga ttg ata tct tca ata tat ggg ata caa agc tta taa att tct aat act gtg ggt tca tct aca agg aat ccc cta gta tac tga att att tta tat aga tct aat tta aca tca ttt tca tct ggg ata cca caa ttc aaa ata aac tca aca aca cta ctt tcc ttt tta cat att ccc cta aaa tag gca ttc aag cat tct att tta tat att aca gcc cca tga tct acc ata aaa tca aca atg tct att tct aca tat gca tta gat aga tag taa agt aag aga tct tgt aca gaa tta caa ttc tta ata att ata gag aaa ata tct tcc ata taa ttc ttt gac act aat gca gat ata ata tct tta tat gta ata tat gca aac agt cta tct act ata tac tga tca ata tta tct cta tga atc cta aaa taa tca tac aga aca tct aca gga tca caa att ggt tca agg aga aat cta tca aat att ttc ctg tca aca act ggt tct aga aca taa cag tca aca cct aat cca tgt ttt tta tag tca tct acc aaa gat aat gac caa aga tcg agg tcg tcg tga aac tgc tca tcg aca gcc atg aaa tct gcc gac tcc atg gcg cga atc gca ctg tct tat tcg cca ttg att ttc a-t ttt tta taa tta tgt aca tgt ttt cct tct att ctc aag agt cta caa aaa tat a-t ttt ttt tcg ata tct aag tac taa gtt ttt tta ctg ttt ttg tta ctg tct tcc att ctt cta act aaa gat ctg aga taa att ata caa tct tcg cta tcg aac cat ttt tgt agt cta aag cct gaa gta att aac caa ctg ttt tta tta gtg gct ttt ttc gat cta tcc tcg tc- --- --c tca tca tcc tta ta- --- --- --- --- --- --- tta tta tca tta tta tca tag tct att aaa cac aaa tca tct --- --- --- --- --- --- --- --- --- --- --- --- --- --- --- --- --- --- --- --- --- --- --- --- --- --- --a cgt tta taa c-- --- --- --- --- --- --- --- --a aca ttc --- --- --- --- -tc att att aat tag ttc tgt agt aat atc ttt aat aat ttg gct ata --- --- -ca tct gtt --- --- --- --- --- --- --- --- --- --- --- --- --- --- --- --- --- --- --c aat act --- --- --- --- --- --- --- --a tct att gat gat ttc ttt t-- --- --- --- --- --- --- --- --- --- --- --t taa gac t-- --- --- --- --- --- --- --- --- --- --- --- --- --- --- --- --- --- --- --- --- --- --- --- --- --- --- --- --- --- --- --- --- --- --- --- --- --- --- --- taa act agt --- --- --- --- --- --- --- --- --- --- --- --- --- --- --- --- --- --- --- --- --- --- --- --- --- --- --- --- --- --- --- --- --- --- --- --- --- --- --- --- --- --- --- --- --- --- --- --- --- --- --- --- --- --- --- --- --- --- --- --- --- --- --- --- --- --- --- --- --- --- --- --- --- --- --- --- --- --- --- --- --- --- --- --- --- --- --- --- --- --- --- --- --- --- --- --- --- --- --- --- --- --- --- --- --- --- --- --- --- --t atg gta atg acg atg aaa --- -tc gag tag ta- --- --a ctt cta ata aag act tga ta- --- --t cat tat cat atg ttt gat cg- --- --- --- --- --- --- --t cat agt taa tag tgt g-- --- --- --- --- --- --- --- --- --- --- --- --- --- --- --- --- --- --- --- -gc taa atg gta ctg tta ata agt tta t-- --- --- --- --- --- --- --- --- --- --- --- --- --- --- --- --- --a gac aat atc ata gta ttt tct ttc cag aat --- --- --- --t aga tta ttt ttt taa ata ctg atc ctc aca att ccg tga tgt agc agt agt tgg t-- --- --- --- --- --- --- --- --- --- --- --- --- --- --- --- --- --- --- --- --- --- --- --- --- --- --- --- -gc atg gtc tat atc gt- --- --- --- --- --- --- --- --- --- --- --- --- --- --- --- --- --- --- --- --- --- --- --- --- --- --- --- --- --- --- --- --- --- --- --- --- --- --- --- --- --- --- --- taa aat gta tca tat ata ata gtt ttc tga cgt gga gta cag aat ttt cga --- --- --- --- --- --- --- --- --- --- --- --- --- --- --- --- --- --- --- --- --- --- --- --- --- --- --- --- --- --- --- --- --- --- --- --- --- --- --- --- --- --- --- --t taa tga gtt cat ggt aag gaa ggg caa atg tct --- gta tat aat ata cat aag tta a-- --- --- --- --- --- --- --- --- --- --- --- --- --t agt ttt tta tca tat tt- --- --- --- --- --- --- --- --t cta ata cca taa taa aaa tta tca tta tg- --- --- -ta taa tca --- --- --- --- --- --- --- --- --- --- --- --t cac tg- --- -tc gct atc att att gcg ttt gtg tag tt- --- --- --- --- ctg ccc ta- --- --- --- --- --- --- --- --t cat cta cat cac tgt ca- --- --- --- --- ctc tca cta tat ctt cta aaa tta caa --- -ac aac --- --- --- --- --- --- --- --- --- --- --- --- --- --- --- --- --- --- --- --- --- --- --- --- --- --- --- --- --- --- --- --- --- --- --- --- tgg ata ttc gat --- --- --- --- --- --- --- --- --- --- --- --- --- --- --- --- --- --- --- --- --- --- --- --- --- --- --a aca gca ttt gtg t-- --- --- --- --- --- --- --- --- --- --- --- --- --- --- --- --- -

>KP849471.1_Monkeypox_virus_isolate_Yambuku_DRC_1985_complete_genome

ata ata tta atg tac taa aac tta tgt att att aat tta tct aac taa agt tag taa att ata tac ata att tta taa tt- --a att taa ttt tac tat ttt att tag tgt cta gaa aaa aat gtg tga ccc acg acc gta gga aac tct aga ggg taa gaa aaa tca atc gtt tat aga gac cat cag aaa gag gtt taa tat ttt tgt gag acc cat cga aga gag aaa gg- --- --- --- --- --- --- --- --- --- --- --- --- --- --- --- --- --- --- --- --- --- --- --- --- --- --- --- --- --- --- --- --- --- --- --- --- --- --- --- --- --- --- --- --- --- --- --- --- --- --- --- --- --- --- --- --- --- --- --- --- --- --- --- --- --- --- --- --- --- --- --- --- --- --- --- --- --- --- --- --- --- --- --- --- --- --- --- --- --- --- --- --- --- --- --- --- --- --- --- --- --- --- --- --- --- --- --- --- --- --- --- --- --- --- --- --- --- --- --- --- --- --- --- --- --- --- --- --- --- --- --- --- --- --- --- --- --- --- --- --- --- --- --- --- --- --- --- --- --- --- --- --- --- --- --- --- --- --- --- --- --- --- --- --- --- --- --- --- --- --- --- --- --- --- --- --- --- --- --- --- --- --- --- --- --- --- -at aaa aac ttt tta cga ctc cat cag aaa gag gtt taa tat ttt tgt gag acc cat cga aga gag aaa gga taa aaa ctt ttt acg act cca tca gaa aga ggt tta --- --- --- --- --- --- --- --- --- --- --- --- --- --- --- --- --- --- --- --- --- --- --- --- --- --- --- --- --- --- --- --- --- --- --- --- --- --- --- --- --- --- --- --- --- --- --a tat ttt tgt gag acc cat cga aga gag aaa gga taa aaa ctt ttt acg act cca tca gaa aga ggt tta ata ttt ttg tga gac cca tcg acg aga gaa aga gat ggt tag tca aga tat ttt tct tag tac aaa agt caa tgt ttt aaa ata tat gga cga gaa tta att tgt ctg tat aaa aac ttg tgt gaa att atg tac tag aga aaa aac gtg agc agt gtc ccc tac atg gat ttt aca gat cat tta tat tcc aaa aat att aac tat ata cgt tta tta tat gat gtt aac gtg taa att ata aac att att tta tga tgc aat tgt ctg aca acc --- att ggc ata agg ata ttg ata agc tct acg aga ata tat tgt tgg acg tta tcg ttt acg aaa tag ttg aga cat cag aaa gag gtt taa tat ttt tgt gag a-c cat cga aga gag aaa gag aat aaa aat a-- --- --- --- ttt ttt tgt aaa act ttt tta tga gac aag aga gaa aga gaa tac gaa tag tga tca tat cgt atc aca tat tga aac aga aag aag aag taa cga gag gta act ttt tgt gaa tgt agt taa aca ttt ttg ttt tgc aaa ccg gaa tat agt acc cgg tac act ttt tta att cgt ggt gcg gtg tct gaa tcg ttt gat taa ccc aac tca tcc att ttc aga tga ata gag tta tcg att cag aca cat gct ttg agt ttt gtt gaa tcg atg agt gaa gta tca tcg gtt gca cct tca gat gcc gat ccg tcg aca tac ttg aat cca tcc ttg act tca agt tca gat gat tcc tca cac atg tct ccg ata cgt acg cta aac tct agg ttc tta aca cat ttt gta tca acg atc gtt gaa ccg atg ata tct ttg taa ctc act ttc tta tgt gag atg tta gac cca agt act gga tgg gtc ttg atg tcg ctg tct ttc tct tct tcg cta cat ctg atg tcg ata gac atc tca cag tct ttg atc ata gcc aga gct tct tca cgc gtg atc gcg gga gag tcc tta cct tgt cct ggt gac acg ctg gac aat cta gta ttc aca gtg ttt cca tca gag gat tcg gag atg gat gaa atc ttt ggg cat ttg gtg aat cca aag ttc atg tta aga ccc gcg ccg acg ata gtg taa taa gtg gtg gga tct cct ttt aca act --- --- --- --- --- --- tct tcg gat acc tca tca tct tcg gtc tct gta act tcc gtt acg gat tga caa atc tta tca ttg gtc ggt gtt tgg tct tgc ttt gtg act ttg ata ata aca tcg att ccc ata tga tgt ttg ttt tct tct tca gta cac gag gat gag gat tgt tga aga cta gta ggc ata gca gct gcc act agg cac atg cat gcc agg aca ata tat tgt ttc atg att gct att gat tga tta ctg ttc tag atg att cta ctt tct tac cat ata ata aat tag aat ata ttt tct act ttt acg aga aat taa tta ttg tat tta tta ttt atg ggt -aa aaa aaa ctt act ata agt ggg tgg gat tct ggg aat tag tga tca gtt tat gta tat cgc aac tag cgg gca tat ggc tat tga cat cga gaa cat tac cca tat gat aag aga ttg tat ccg ttt cgt agt ctt gag tat tgg tat tac tat ata gta tgt aga tgt cga cgc tag ata gac agt cgc cca cta gag tta ccg tct ctg aat gcg gca tga tag tat cat tct ttg ttt tcg tta act gtt tgg aag atg aat ctt tgt tgt tac att taa tct cga aat tca gag tac ata tct ttg aag tat tct gat atc tat ttt ctc ctg taa aga atc ctg aag ttg cta cat tat taa gga cag aga agt att ctg cac gaa aga cgg gat cac aat ctt tat gat tca tgg taa tag tta gtt ccg acg ttg aga tgg att cgc tga gac cgg tag tgg tcg tcc gag tac acg atg tgt cgt tga ctg gat aca ggt taa ttt cca cat cga tat agt taa atg tat tgc tgg tt- --a cgg gtt cgc att tat ctg tgg aag aga cgg tgt gag aat atg ttc cgg gac cac acg gag aac aga tga cgt ctc cgg tag acg tgt atc cgg ata ctc cgt atc cta ttc cac act ttg ttt tag aaa tac atg ttc tac acc ctg atg atc ctt tga gaa gac aat aat atc ctg gag agc att cac aga ttc tat tgt gag tcg tgt tac acg atc gcg tct cta cct gat tac tat cac atc ttc cgt tac aac tta gac aag cct gta aat gat tat tgt gag atg taa agg tat ccg aac cac acg gcg tac att gtg tat tag tct tgc tat cac ata atc tgg aag cgt aag ttc ccg gag gac acg ata gac aac ata gat tac ggc ttc tgt att cgt tgt ctt tac att ttc cat tgg atg gtg cat gtg gtg cta tat ctc ttc cgt tta tta tta tac atg aga gaa aca ata tat acg agt ata ata cgg acc tca tga ttt aat aat -gt agt aat cgt cgt ctt gtt act gtt tgt ttc cta ctt ctc caa tca tat aga tta ttt ttt aaa tat ttt ctt tct atc atg gat aat att tgt aat ggt tct ttc cgt aca aca tac tgt tta gat ggt agt cgc tta gct tgg tta tga tat tgc gca taa ttt ccg gag gca aat acg ata gtc tag att gac tat cga tgg tag act cta att tat tga gtg ctt tgt cga cga gtt tac ttt tat gct cca tcg ata gat gac act gtt cta tga gat cgt cgt aca tgg gaa atg aaa tgc gtt tgt ctg aat gta tgg ctt cga gat agg tgt gat acc gga tgt ctt ctg ttc tca ata ccg tat aca agt tgg tgt ctg aga ttc gaa tct ctt tga gga gac tta tgt cac gac tac att ttt cga tga tgg aat cta tct tat cga atg ata tat ttt tca taa ata cac ttt tat agt cct cgt tta aac aga att tag tat gta gtt ccg caa atg act cgt ccc tta ata ggc agt agg cta tta tct tct tta cgt agt gat cgt cgt agg gag aga act ccg aca tct tgt aga aca acg att taa tca tag gta gag ata ctt tca gtc tgt ggt gga tga tgt cat taa cga cat ccg cct tgt ata tga tgt ttc tgt ttt caa aca cca agt cga ata ctg tct tat cgt ctt tag tcg gaa ggt tga tgt cgt atc cga tgt ata cga ggt atg agg caa cat tgt tat tgc aat tct gga agg cgg tat gaa gag gag tca ttg tat tat agt att cgt ctt tct gaa tgt cga atc tat cta gta gat acc gta gta tat tga gag agc gac ttc cat atc ctt gat tat gtt tta tga ata gat aaa gta gat gtt gtc ctt ctt cct ttt gta att tcc cgt att ttt gtt cgt gcc aat tga gta aca tta tga gaa tat gac ctg ttg cac aat cgt tct tta tgt att cca tga tgg gtg tac aat caa gat tat tac gta tcc tcg tat cgg ctc ctc gag ata aaa gag cat aca cca cac gag gac tat gtt tgg tat act gtt gaa ggt aag tgt gta acg gcg tat ttc cga ttt tcg taa ccg cgt taa tgt ttg ctc cat gat cta tta tcg cgt aaa tga atc gct tct cag ctc gca tct tag tgt gac tct ttg act tgt aat aat tgc ttt cgt gga aca cgg ata tgt gtt tac agt agt aat gaa gag aag tga gtc cat cct cat cga cgc aat tag ggt cag atc ctt tag tca ata att tgt aca gaa cgt aat agt tta agc tcc cat tga att tat atc taa gat aac aca gca ata gat cgg atg att tac taa agt cat caa tgg ggt ccg tta gta tat caa aga tct tgt tat cga ttg ata gtg aat gaa tca gat agt ggt gta gag gaa tat gtc ctt ttt tat cct tgc tat caa agt tac gca tgc cgt ggc gta aca ata tct tta ata cag atg gat taa atc gtg tat tca tcg tat agc aat gta atg gag agt tac cac att tta gtc gtt tat tca gat cgc agt gtt taa taa cta att taa aca gat gag atg ata tat cca cat caa aga atg cga gat aca tat gac aga cat tat tga cag aaa tgt gac ctt cat tat cac cgt cgt cca taa atg cgt tag gta cgt acc aca tac tat cgt taa cga tgc gca caa tct cgt cca ttt tca tcc atc ttc ata atg att tac ttt ttc ata att aga gaa aaa gat caa ggt ata aaa att aga agt gtt aga cta ta- --- --- --- --- --- --- --- --- --- --- -aa cta act tat aa- --- --- --- --- --- --- --- --- --- --- --- --- --- --- --- --- --- --- --- --- --- --- --- --- --- --- --- --- --- --- --- --- --- --- --- --- --- --- --- --- --- --- --- --- --- --- --- --- --- --- --- --- --- --- --- --- --- --- --- --- --- --- --- --- --- --- --- --- --- --- --- --- --- --- --- --- --- --- --- --- --- --- --- --- --- --- -aa cta act tat gac tta act aac tta tga cta tta act cat ttc aag aaa ggt ggg tgg aga gaa ctc tat atg aca gct tgt gaa aca att aga tcc cta att tct aat gga agt ttt gat agg aga ttg tca tca gtt gat aca ttg ttt att atc tca tct att aga gca cgt ctg ttt aga gct tta gtg acc tgc tcg gtt act tct gtg taa atc ttg aat cct tta gtg ata cac tgt gtc aaa act gga tgt tta gaa tac cta tgt aga ata tgg gaa gca tgc ttg ttt ttg tct cta tta tag att aac tca tac atg gtt gta tta tga att ttc atc tgc cta atg tac tcc aat tct tgt tta caa tca att ata taa tca aag agt gat gat gca tac aca tta caa agt gaa taa tct acc atc ata aaa tac ttg ata cag agc ttt atc aca tca tgg ttt tca att gta tta tta agt ata gct aat ttt ata cag tca ata gac aat ggt tct cta agc aat att tct aat att tta aga tgt gct tcc cta cgg gcg atg aca gat ccc cta tcc acg gcc acg tca aga cat gta tat cca tta ctc att act gcg ttg aca ttt gct cca ttt tct aat agc cat gat act aaa tct ata taa cct gca tag ata gcg cga taa agc aag gtc ctt cca cca gca tct agt tga ttg ata tct tca ata tat ggg ata caa agc tta taa att tct aat act gtg ggt tca tct aca agg aat ccc cta gta tac tga att att tta tat aga tct aat tta aca tca ttt tca tct ggg ata cca caa ttc aaa ata aac tca aca aca cta ctt tcc ttt tta cat att ccc cta aaa tag gca ttc aag cat tct att tta tat att aca gcc cca tga tct acc ata aaa tca aca atg tct att tct aca tat gca tta gat aga tag taa agt aag aga tct tgt aca gaa tta caa ttc tta ata att ata gag aaa ata tct tcc ata taa ttc ttt gac act aat gca gat ata ata tct tta tat gta ata tat gca aac agt cta tct act ata tac tga tca ata tta tct cta tga atc cta aaa taa tca tac aga aca tct aca gga tca caa att ggt tca agg aga aat cta tca aat att ttc ctg tca aca act ggt tct aga aca taa cag tca aca cct aat cca tgt ttt tta tag tca tct acc aaa gat aat gac caa aga tcg agg tcg tcg tga aac tgc tca tcg aca gcc atg aaa tct gcc gac tcc atg gcg cga atc gca ctg tct tat tcg cca ttg att ttc a-t ttt tta taa tta tgt aca tgt ttt cct tct att ctc aag agt cta caa aaa tat a-t ttt ttt tcg ata tct aag tac taa gtt ttt tta ctg ttt ttg tta ctg tct tcc att ctt cta act aaa gat ctg aga taa att ata caa tct tcg cta tcg aac cat ttt tgt agt cta aag cct gaa gta att aac caa ctg ttt tta tta gtg gct ttt ttc gat cta tcc tcg tc- --- --c tca tca tcc tta ta- --- --- --- --- --- --- tta tta tca tta tta tca tag tct att aaa cac aaa tca tct --- --- --- --- --- --- --- --- --- --- --- --- --- --- --- --- --- --- --- --- --- --- --- --- --- --- --a cgt tta taa c-- --- --- --- --- --- --- --- --a aca ttc --- --- --- --- -tc att att aat tag ttc tgt agt aat atc ttt aat aat ttg gct ata --- --- -ca tct gtt --- --- --- --- --- --- --- --- --- --- --- --- --- --- --- --- --- --- --c aat act --- --- --- --- --- --- --- --a tct att gat gat ttc ttt t-- --- --- --- --- --- --- --- --- --- --- --t taa gac t-- --- --- --- --- --- --- --- --- --- --- --- --- --- --- --- --- --- --- --- --- --- --- --- --- --- --- --- --- --- --- --- --- --- --- --- --- --- --- --- taa act agt --- --- --- --- --- --- --- --- --- --- --- --- --- --- --- --- --- --- --- --- --- --- --- --- --- --- --- --- --- --- --- --- --- --- --- --- --- --- --- --- --- --- --- --- --- --- --- --- --- --- --- --- --- --- --- --- --- --- --- --- --- --- --- --- --- --- --- --- --- --- --- --- --- --- --- --- --- --- --- --- --- --- --- --- --- --- --- --- --- --- --- --- --- --- --- --- --- --- --- --- --- --- --- --- --- --- --- --- --- --t atg gta atg acg atg aaa --- -tc gag tag ta- --- --a ctt cta ata aag act tga ta- --- --t cat tat cat atg ttt gat cg- --- --- --- --- --- --- --t cat agt taa tag tgt g-- --- --- --- --- --- --- --- --- --- --- --- --- --- --- --- --- --- --- --- -gc taa atg gta ctg tta ata agt tta t-- --- --- --- --- --- --- --- --- --- --- --- --- --- --- --- --- --a gac aat atc ata gta ttt tct ttc cag aat --- --- --- --t aga tta ttt ttt taa ata ctg atc ctc aca att ccg tga tgt agc agt agt tgg t-- --- --- --- --- --- --- --- --- --- --- --- --- --- --- --- --- --- --- --- --- --- --- --- --- --- --- --- -gc atg gtc tat atc gt- --- --- --- --- --- --- --- --- --- --- --- --- --- --- --- --- --- --- --- --- --- --- --- --- --- --- --- --- --- --- --- --- --- --- --- --- --- --- --- --- --- --- --- taa aat gta tca tat ata ata gtt ttc tga cgt gga gta cag aat ttt cga --- --- --- --- --- --- --- --- --- --- --- --- --- --- --- --- --- --- --- --- --- --- --- --- --- --- --- --- --- --- --- --- --- --- --- --- --- --- --- --- --- --- --- --t taa tga gtt cat ggt aag gaa ggg caa atg tct --- gta tat aat ata cat aag tta a-- --- --- --- --- --- --- --- --- --- --- --- --- --t agt ttt tta tca tat tt- --- --- --- --- --- --- --- --t cta ata cca taa taa aaa tta tca tta tg- --- --- -ta taa tca --- --- --- --- --- --- --- --- --- --- --- --t cac tg- --- -tc gct atc att att gcg ttt gtg tag tt- --- --- --- --- ctg ccc ta- --- --- --- --- --- --- --- --t cat cta cat cac tgt ca- --- --- --- --- ctc tca cta tat ctt cta aaa tta caa --- -ac aac --- --- --- --- --- --- --- --- --- --- --- --- --- --- --- --- --- --- --- --- --- --- --- --- --- --- --- --- --- --- --- --- --- --- --- --- tgg ata ttc gat --- --- --- --- --- --- --- --- --- --- --- --- --- --- --- --- --- --- --- --- --- --- --- --- --- --- --a aca gca ttt gtg t-- --- --- --- --- --- --- --- --- --- --- --- --- --- --- --- --- -

>DQ011155.1_Monkeypox_virus_strain_Zaire_1979-005_complete_genome

--- --- --- --- --- --- --- --- --- --- --- --- --- --- --- --- --- --- --- --- --- --- --- --- --- --- --- --- --- --- --- --- --- --- --- --g tgt cta gaa aaa aat gtg tga ccc acg acc gta gga aac tct aga ggg taa gaa aaa tca atc gtt tat aga gac cat cag aaa gag gtt taa tat ttt tgt gag acc cat cga aga gag aaa gg- --- --- --- --- --- --- --- --- --- --- --- --- --- --- --- --- --- --- --- --- --- --- --- --- --- --- --- --- --- --- --- --- --- --- --- --- --- --- --- --- --- --- --- --- --- --- --- --- --- --- --- --- --- --- --- --- --- --- --- --- --- --- --- --- --- --- --- --- --- --- --- --- --- --- --- --- --- --- --- --- --- --- --- --- --- --- --- --- --- --- --- --- --- --- --- --- --- --- --- --- --- --- --- --- --- --- --- --- --- --- --- --- --- --- --- --- --- --- --- --- --- --- --- --- --- --- --- --- --- --- --- --- --- --- --- --- --- --- --- --- --- --- --- --- --- --- --- --- --- --- --- --- --- --- --- --- --- --- --- --- --- --- --- --- --- --- --- --- --- --- --- --- --- --- --- --- --- --- --- --- --- --- --- --- --- --- --- --- --- --- --- --- --- --- --- --- --- --- --- --- --- --- --- --- --- --- --- --- --- --a taa aaa ctt ttt acg act cca tca gaa aga ggt tta --- --- --- --- --- --- --- --- --- --- --- --- --- --- --- --- --- --- --- --- --- --- --- --- --- --- --- --- --- --- --- --- --- --- --- --- --- --- --- --- --- --- --- --- --- --- --a tat ttt tgt gag acc cat cga aga gag aaa gga taa aaa ctt ttt acg act cca tca gaa aga ggt tta ata ttt ttg tga gac cca tcg aag aga gaa aga gat ggt tag tca aga tat ttt tct tag tac aaa agt caa tgt ttt aaa ata tat gga cga gaa tta att tgt ctg tat aaa aac ttg tgt gaa att atg tac tag aga aaa aac gtg agc agt gtc ccc tac atg gat ttt aca gat cat tta tat tcc aaa aat att aac tat ata cgt tta tta tat gat gtt aac gtg taa att ata aac att att tta tga tgc aat tgt ctg aca acc --- att ggc ata agg ata ttg ata agc tct acg aga ata tat tgt tgg acg tta tcg ttt acg aaa tag ttg aga cat cag aaa gag gtt taa tat ttt tgt gag a-c cat cga aga gag aaa gag aat aaa aat a-- --- --- --- ttt ttt tgt aaa act ttt tta tga gac aag aga gaa aga gaa tac gaa tag tga tca tat cgt atc aca tat tga aac aga aag aag aag taa cga gag gta act ttt tgt gaa tgt agt taa aca ttt ttg ttt tgc aaa ccg gaa tat agt acc cgg tac act ttt tta att cgt ggt gcg gtg tct gaa tcg ttc gat taa ccc aac tca tcc att ttc aga tga ata gag tta tcg att cag aca cat gct ttg agt ttt gtt gaa tcg atg agt gaa gta tca tcg gtt gca cct tca gat gcc gat ccg tcg aca tac ttg aat cca tcc ttg act tca agt tca gat gat tcc tca cac atg tct ccg ata cgt acg cta aac tct agg ttc tta aca cat ttt gta tca acg atc gtt gaa ccg atg ata tct ttg taa ctc act ttc tta tgt gag atg tta gac cca agt act gga tgg gtc ttg atg tcg ctg tct ttc tct tct tcg cta cat ctg atg tcg ata gac atc tca cag tct ttg atc ata gcc aga gct tct tca cgc gtg atc gcg gga gag tcc tta cct tgt cct ggt gac acg ctg gac aat cta gta ttc aca gtg ttt cca tca gag gat tcg gag atg gat gaa atc ttt ggg cat ttg gtg aat cca aag ttc atg tta aga ccc gcg ccg acg ata ttg taa taa gtg gtg gga tct cct ttt aca act --- --- --- --- --- --- tct tcg gat acc tca tca tct tcg gtc tct gta act tcc gtt acg gat tga caa atc tta tca ttg gtc ggt gtt tgg tct tgc ttt gtg act ttg ata ata aca tcg att ccc ata tga tgt ttg ttt tct tct tca gta cac gag gat gag gat tgt tga aga cta gta ggc ata gca gct gcc act agg cac atg cat gcc agg aca ata tat tgt ttc atg att gct att gat tga tta ctg ttc tag atg att cta ctt tct tac cat ata ata aat tag aat ata ttt tct act ttt acg aga aat taa tta ttg tat tta tta ttt atg ggt -aa aaa aaa ctt act ata agt ggg tgg gat tct ggg aat tag tga tca gtt tat gta tat cgc aac tag cgg gca tat ggc tat tga cat cga gaa cat tac cca tat gat aag aga ttg tat ccg ttt cgt agt ctt gag tat tgg tat tac tat ata gta tgt aga tgt cga cgc tag ata gac agt cgc cca cta gag tta ccg tct ctg aat gcg gca tga tag tat cat tct ttg ttt tcg tta act gtt tgg aag atg aat ctt tgt tgt tac att taa tct cga aat tca gag tac ata tct ttg aag tat tct gat atc tat ttt ctc ctg taa aga atc ctg aag ttg cta cat tat taa gga cag aga agt att ctg cac gaa aga cgg gat cac aat ctt tat gat tca tgg taa tag tta gtt ccg acg ttg aga tgg att cgc tga gac cgg tag tgg tcg tcc gag tac acg atg tgt cgt tga ctg gat aca ggt taa ttt cca cat cga tat agt taa atg tat tgc tgg tt- --a cgg gtt cgc att tat ctg tgg aag aga cgg tgt gag aat atg ttc cgg gac cac acg gag aac aga tga cgt ctc cgg tag acg tgt atc cgg ata ctc cgt atc cta ttc cac act ttg ttt tag aaa tac atg ttc tac acc ctg atg atc ctt tga gaa gac aat aat atc ctg gag agc att cac aga ttc tat tgt gag tcg tgt tac acg atc gcg tct cta cct gat tac tat cac atc ttc cgt tac aac tta gac aag cct gta aat gat tat tgt gag atg taa agg tat ccg aac cac acg gcg tac att gtg tat tag tct tgc tat cac ata atc tgg aag cgt aag ttc ccg gag gac acg ata gac aac ata gat tac ggc ttc tgt att cgt tgt ctt tac att ttc cat tgg atg gtg cat gtg gtg cta tat ctc ttc cgt tta tta tta tac atg aga gaa aca ata tat acg agt ata ata cgg acc tca tga ttt aat aat -gt agt aat cgt cgt ctt gtt act gtt tgt ttc cta ctt ctc caa tca tat aga tta ttt ttt aaa tat ttt ctt tct atc atg gat aat att tgt aat ggt tct ttc cgt aca aca tac tgt tta gat ggt agt cgc tta gct tgg tta tga tat tgc gca taa ttt ccg gag gca aat acg ata gtc tag att gac tat cga tgg tag act cta att tat tga gtg ctt tgt cga cga gtt tac ttt tat gct cca tcg ata gat gac act gtt cta tga gat cgt cgt aca tgg gaa atg aaa tgc gtt tgt ctg aat gta tgg ctt cga gat agg tgt gat acc gga tgt ctt ctg ttc tca ata ccg tat aca agt tgg tgt ctg aga ttc gaa tct ctt tga gga gac tta tgt cac gac tac att ttt cga tga tgg aat cta tct tat cga atg ata tat ttt tca taa ata cac ttt tat agt cct cgt tta aac aga att tag tat gta gtt ccg caa atg act cgt ccc tta ata ggc agt agg cta tta tct tct tta cgt agt gat cgt cgt agg gag aga act ccg aca tct tgt aga aca acg att taa tca tag gta gag ata ctt tca gtc tgt ggt gga tga tgt cat taa cga cat ccg cct tgt ata tga tgt ttc tgt ttt caa aca cca agt cga ata ctg tct tat cgt ctt tag tcg gaa ggt tga tgt cgt atc cga tgt ata cga ggt atg agg caa cat tgt tat tgc aat tct gga agg cgg tat gaa gag gag tca ttg tat tat agt att cgt ctt tct gaa tgt cga atc tat cta gta gat acc gta gta tat tga gag agc gac ttc cat atc ctt gat tat gtt tta tga ata gat aaa gta gat gtt gtc ctt ctt cct ttt gta att tcc cgt att ttt gtt cgt gcc aat tga gta aca tta tga gaa tat gac ctg ttg cac aat cgt tct tta tgt att cca tga tgg gtg tac aat caa gat tat tac gta tcc tcg tat cgg ctc ctc gag ata aaa gag cat aca cca cac gag gac tat gtt tgg tat act gtt gaa ggt aag tgt gta acg gcg tat ttc cga ttt tcg taa ccg cgt taa tgt ttg ctc cat gat cta tta tcg cgt aga tga atc gct tct cag ctc gca tct tag tgt gac tct ttg act tgt aat aat tgc ttt cgt gga aca cgg ata tgt gtt tac agt agt aat gaa gag aag tga gtc cat cct cat cga cgc aat tag ggt cag atc ctt tag tca ata att tgt aca gaa cgt aat agt tta agc tcc cat tga att tat atc taa gat aac aca gca ata gat cgg atg att tac taa agt cat caa tgg ggt ccg tta gta tat caa aga tct tgt tat cga ttg ata gtg aat gaa tca gat agt ggt gta gag gaa tat gtc ctt ttt tat cct tgc tat caa agt tac gca tgc cgt ggc gta aca ata tct tta ata cag atg gat taa atc gtg tat tca tcg tat agc aat gta atg gag agt tac cac att tta gtc gtt tat tca gat cgc agt gtt taa taa cta att taa aca gat gag atg ata tat cca cat caa aga atg cga gat aca tat gac aga cat tat tga cag aaa tgt gac ctt cat tat cac cgt cgt cca taa atg cgt tag gta cgt acc aca tac tat cgt taa cga tgc gca caa tct cgt cca ttt tca tcc atc ttc ata atg att tac ttt ttc ata att aga gaa aaa gat caa ggt ata aaa att aga agt gtt aga cta ta- --- --- --- --- --- --- --- --- --- --- -aa cta act tat aa- --- --- --- --- --- --- --- --- --- --- --- --- --- --- --- --- --- --- --- --- --- --- --- --- --- --- --- --- --- --- --- --- --- --- --- --- --- --- --- --- --- --- --- --- --- --- --- --- --- --- --- --- --- --- --- --- --- --- --- --- --- --- --- --- --- --- --- --- --- --- --- --- --- --- --- --- --- --- --- --- --- --- --- --- --- --- -aa cta act tat gac tta act aac tta tga cta tta act cat ttc aag aaa ggt ggg tgg aga gaa ctc tat atg aca gct tgt gaa aca att aga tcc cta att tct aat gga agt ttt gat agg aga ttg tca tca gtt gat aca ttg ttt att atc tca tct att aga gca cgt ctg ttt aga gct tta gtg acc tgc tcg gtt act tct gtg taa atc ttg aat cct tta gtg ata cac tgt gtc aaa act gga tgt tta gaa tac cta tgt aga ata tgg gaa gca tgc ttg ttt ttg tct cta tta tag att aac tca tac atg gtt gta tta tga att ttc atc tgc cta atg tac tcc aat tct tgt tta caa tca att ata taa tca aag agt gat gat gca tac aca tta caa agt gaa taa tct acc atc ata aaa tac ttg ata cag agc ttt atc aca tca tgg ttt tca att gta tta tta agt ata gct aat ttt ata cag tca ata gac aat ggt tct cta agc aat att tct aat att tta aga tgt gct tcc cta cgg gcg atg aca gat ccc cta tcc acg gcc acg tca aga cat gta tat cca tta ctc att act gcg ttg aca ttt gct cca ttt tct aat agc cat gat act aaa tct ata taa cct gca tag ata gcg cga taa agc aag gtc ctt cca cca gca tct agt tga ttg ata tct tca ata tat ggg ata caa agc tta taa att tct aat act gtg ggt tca tct aca agg aat ccc cta gta tac tga att att tta tat aga tct aat tta aca tca ttt tca tct ggg ata cca caa ttc aaa ata aac tca aca aca cta ctt tcc ttt tta cat att ccc cta aaa tag gca ttc aag cat tct att tta tat att aca gcc cca tga tct acc ata aaa tca aca atg tct att tct aca tat gca tta gat aga tag taa agt aag aga tct tgt aca gaa tta caa ttc tta ata att ata gag aaa ata tct tcc ata taa ttc ttt gac act aat gca gat ata ata tct tta tat gta ata tat gca aac agt cta tct act ata tac tga tca ata tta tct cta tga atc cta aaa taa tca tac aga aca tct aca gga tca caa att ggt tca agg aga aat cta tca aat att ttc ctg tca aca act ggt tct aga aca taa cag tca aca cct aat cca tgt ttt tta tag tca tct acc aaa gat aat gac caa aga tcg agg tcg tcg tga aac tgc tca tcg aca gcc atg aaa tct acc gac tcc atg gcg cga atc gca ctg tct tat tcg cca ttg att ttc a-t ttt tta taa tta tgt aca tgt ttt cct tct att ctc aag agt cta caa aaa tat a-t ttt ttt tcg ata tct aag tac taa gtt ttt tta ctg ttt ttg tta ctg tct tcc att ctt cta act aaa gat ctg aga taa att ata caa tct tcg cta tcg aac cat ttt tgt agt cta aag cct gaa gta att aac caa ctg ttt tta tta gtg gct ttt ttc gat cta tcc tcg tc- --- --c tca tca tcc tta ta- --- --- --- --- --- --- tta tta tca tta tta tca tag tct att aaa cac aaa tca tct --- --- --- --- --- --- --- --- --- --- --- --- --- --- --- --- --- --- --- --- --- --- --- --- --- --- --a cgt tta taa c-- --- --- --- --- --- --- --- --a aca ttc --- --- --- --- -tc att att aat tag ttc tgt agt aat atc ttt aat aat ttg gct ata --- --- -ca tct gtt --- --- --- --- --- --- --- --- --- --- --- --- --- --- --- --- --- --- --c aat act --- --- --- --- --- --- --- --a tct att gat gat ttc ttt t-- --- --- --- --- --- --- --- --- --- --- --t taa gac t-- --- --- --- --- --- --- --- --- --- --- --- --- --- --- --- --- --- --- --- --- --- --- --- --- --- --- --- --- --- --- --- --- --- --- --- --- --- --- --- taa act agt --- --- --- --- --- --- --- --- --- --- --- --- --- --- --- --- --- --- --- --- --- --- --- --- --- --- --- --- --- --- --- --- --- --- --- --- --- --- --- --- --- --- --- --- --- --- --- --- --- --- --- --- --- --- --- --- --- --- --- --- --- --- --- --- --- --- --- --- --- --- --- --- --- --- --- --- --- --- --- --- --- --- --- --- --- --- --- --- --- --- --- --- --- --- --- --- --- --- --- --- --- --- --- --- --- --- --- --- --- --t atg gta atg acg atg aaa --- -tc gag tag ta- --- --a ctt cta ata aat act tga ta- --- --t cat tat cat atg ttt gat cg- --- --- --- --- --- --- --t cat agt taa tag tgt g-- --- --- --- --- --- --- --- --- --- --- --- --- --- --- --- --- --- --- --- -gc taa atg gta ctg tta ata agt tta t-- --- --- --- --- --- --- --- --- --- --- --- --- --- --- --- --- --a gac aat atc ata gta ttt tct ttc cag aat --- --- --- --t aga tta ttt ttt taa ata ctg atc ctc aca att ccg tga tgt agc agt agt tgg t-- --- --- --- --- --- --- --- --- --- --- --- --- --- --- --- --- --- --- --- --- --- --- --- --- --- --- --- -gc atg gtc tat atc gt- --- --- --- --- --- --- --- --- --- --- --- --- --- --- --- --- --- --- --- --- --- --- --- --- --- --- --- --- --- --- --- --- --- --- --- --- --- --- --- --- --- --- --- taa aat gta tca tat ata ata gtt ttc tga cgt gga gta cag aat ttt cga --- --- --- --- --- --- --- --- --- --- --- --- --- --- --- --- --- --- --- --- --- --- --- --- --- --- --- --- --- --- --- --- --- --- --- --- --- --- --- --- --- --- --- --t taa tga gtt cat ggt aag gaa ggg caa atg tct --- gta tat aat ata cat aag tta a-- --- --- --- --- --- --- --- --- --- --- --- --- --t agt ttt tta tca tat tt- --- --- --- --- --- --- --- --t cta ata cca taa taa aaa tta tca tta tg- --- --- -ta taa tca --- --- --- --- --- --- --- --- --- --- --- --t cac tg- --- -tc gct atc att att gcg ttt gtg tag tt- --- --- --- --- ctg ccc ta- --- --- --- --- --- --- --- --t cat cta cat cac tgt ca- --- --- --- --- ctc tca cta tat ctt cta aaa tta caa --- -ac aac --- --- --- --- --- --- --- --- --- --- --- --- --- --- --- --- --- --- --- --- --- --- --- --- --- --- --- --- --- --- --- --- --- --- --- --- tgg ata ttc gat --- --- --- --- --- --- --- --- --- --- --- --- --- --- --- --- --- --- --- --- --- --- --- --- --- --- --a aca gca ttt gtg t-- --- --- --- --- --- --- --- --- --- --- --- --- --- --- --- --- -

>KP849469.1_Monkeypox_virus_isolate_Boende_DRC_2008_complete_genome

ata ata tta atg tac taa aac tta tgt att att aat tta tct aac taa agt tag taa att ata tac ata att tta taa tt- --a att taa ttt tac tat ttt att tag tgt cta gaa aaa aat gtg tga ccc acg acc gta gga aac tct aga ggg taa gaa aaa tca atc gtt tat aga gac cat cag aaa gag gtt taa tat ttt tgt gag acc cat cga aga gag aaa gg- --- --- --- --- --- --- --- --- --- --- --- --- --- --- --- --- --- --- --- --- --- --- --- --- --- --- --- --- --- --- --- --- --- --- --- --- --- --- --- --- --- --- --- --- --- --- --- --- --- --- --- --- --- --- --- --- --- --- --- --- --- --- --- --- --- --- --- --- --- --- --- --- --- --- --- --- --- --- --- --- --- --- --- --- --- --- --- --- --- --- --- --- --- --- --- --- --- --- --- --- --- --- --- --- --- --- --- --- --- --- --- --- --- --- --- --- --- --- --- --- --- --- --- --- --- --- --- --- --- --- --- --- --- --- --- --- --- --- --- --- --- --- --- --- --- --- --- --- --- --- --- --- --- --- --- --- --- --- --- --- --- --- --- --- --- --- --- --- --- --- --- --- --- --- --- --- --- --- --- --- --- --- --- --- --- --- -at aaa aac ttt tta cga ctc cat cag aaa gag gtt taa tat ttt tgt gag acc cat cga cga gag aaa gga taa aaa ctt ttt acg act cca tca gaa aga ggt tta --- --- --- --- --- --- --- --- --- --- --- --- --- --- --- --- --- --- --- --- --- --- --- --- --- --- --- --- --- --- --- --- --- --- --- --- --- --- --- --- --- --- --- --- --- --- --a tat ttt tgt gag acc cat cga cga gag aaa gga taa aaa ctt ttt acg act cca tca gaa aga ggt tta ata ttt ttg tga gac cca tcg acg aga gaa aga gat ggt tag tca aga tat ttt tct tag tac aaa agt caa tgt ttt aaa ata tat gga cga gaa tta att tgt ctg tat aaa aac ttg tgt gaa att atg tac tag aga aaa aac gtg agc agt gtc ccc tac atg gat ttt aca gat cat tta tat tcc aaa aat att aac tat ata cgt tta tta tat gat gtt aac gtg taa att ata aac att att tta tga tgc aat tgt ctg aca acc --- att ggc ata agg ata ttg ata agc tct acg aga ata tat tgt tgg acg tta tcg ttt acg aaa tag ttg aga cat cag aaa gag gtt taa tat ttt tgt gag a-c cat cga aga gag aaa gag aat aaa aat a-- --- --- --- ttt ttt tgt aaa act ttt tta tga gac aag aga gaa aga gaa tac gaa tag tga tca tat cgt atc aca tat tga aac aga aag aag aag taa cga gag gta act ttt tgt gaa tgt agt taa aca ttt ttg ttt tgc aaa ccg gaa tat agt acc cgg tac act ttt tta att cgt ggt gcg gtg tct gaa tcg ttc gat taa ccc aac tca tcc att ttc aga tga ata gag tta tcg att cag aca cat gct ttg agt ttt gtt gaa tcg atg agt gaa gta tca tcg gtt gca cct tca gat gcc gat cca tcg aca tac ttg aat cca tcc ttg act tca agt tca gat gat tcc tca cac atg tct ccg ata cgt acg cta aac tct agg ttc tta aca cat ttt gta tca acg atc gtt gaa ccg atg ata tct ttg taa ctc act ttc tta tgt gag atg tta gac cca agt act gga tgg gtc ttg atg tcg ctg tct ttc tct tct tcg cta cat ctg atg tcg ata gac atc tca cag tct ttg atc ata gcc aga gct tct tca cgc gtg atc gcg gga gag tcc tta cct tgt cct ggt gac acg ctg gac aat cta gta ttc aca gtg ttt cca tca gag gat tcg gag atg gat gaa atc ttt ggg cat ttg gtg aat cca aag ttc atg tta aga ccc gcg ccg acg ata ttg taa taa gtg gtg gga tct cct ttt aca act tct tcg gat acc tca tca tct tcg gat acc tca tca tct tcg gtc tct gta act tcc gtt acg gat tga caa atc tta tca ttg gtc ggt gtt tgg tct tgc ttt gtg act ttg ata ata aca tcg att ccc ata tga tgt ttg ttt tct tct tca gta cac gag gat gag gat tgt tga aga cta gta ggc ata gca gct gcc act agg cac atg cat gcc agg aca ata tat tgt ttc atg att gct att gat tga tta ctg ttc tag atg att cta ctt tct tac cat ata ata aat tag aat ata ttt tct act ttt acg aga aat taa tta ttg tat tta tta ttt atg ggt -aa aaa aaa ctt act ata agt ggg tgg gat tct ggg aat tag tga tca gtt tat gta tat cgc aac tag cgg gca tat ggc tat tga cat cga gaa cat tac cca tat gat aag aga ttg tat ccg ttt cgt agt ctt gag tat tgg tat tac tat ata gta tgt aga tgt cga cgc tag ata gac agt cgc cca cta gag tta ccg tct ctg aat gcg gca tga tag tat cat tct ttg ttt tcg tta act gtt tgg aag atg aat ctt tgt tgt tac att taa tct cga aat tca gag tac ata tct ttg aag tat tct gat atc tat ttt ctc ctg taa aga atc ctg aag ttg cta cat tat taa gga cag aga agt att ctg cac gaa aga cgg gat cac aat ctt tat gat tca tgg taa tag tta gtt ccg acg ttg aga tgg att cgc tga gac cgg tag tgg tcg tcc gag tac acg atg tgt cgt tga ctg gat aca ggt taa ttt cca cat cga tat agt taa atg tat tgc tgg tt- --a cgg gtt cgc att tat ctg tgg aag aga cgg tgt gag aat atg ttc cgg gac cac acg gag aac aga tga cgt ctc cgg tag acg tgt atc cgg ata ctc cgt atc cta ttc cac act ttg ttt tag aaa tac atg ttc tac acc ctg atg atc ctt tga gaa gac aat aat atc ctg gag agc att cac aga ttc tat tgt gag tcg tgt tac acg atc gcg tct cta cct gat tac tat cac atc ttc cgt tac aac tta gac aag cct gta aat gat tat tgt gag atg taa agg tat ccg aac cac acg gcg tac att gtg tat tag tct tgc tat cac ata atc tgg aag cgt aag ttc ccg gag gac acg ata gac aac ata gat tac ggc ttc tgt att cgt tgt ctt tac att ttc cat tgg atg gtg cat gtg gtg cta tat ctc ttc cgt tta tta tta tac atg aga gaa aca ata tat acg agt ata ata cgg acc tca tga ttt aat aat -gt agt aat cgt cgt ctt gtt act gtt tgt ttc cta ctt ctc caa tca tat aga tta ttt ttt aaa tat ttt ctt tct atc atg gat aat att tgt aat ggt tct ttc cgt aca aca tac tgt tta gat ggt agt cgc tta gct tgg tta tga tat tgc gca taa ttt ccg gag gca aat acg ata gtc tag att gac tat cga tgg tag act cta att tat tga gtg ctt tgt cga cga gtt tac ttt tat gct cca tcg ata gat gac act gtt cta tga gat cgt cgt aca tgg gaa atg aaa tgc gtt tgt ctg aat gta tgg ctt cga gat agg tgt gat acc gga tgt ctt ctg ttc tca ata ccg tat aca agt tgg tgt ctg aga ttc gaa tct ctt tga gga gac tta tgt cac gac tac att ttt cga tga tgg aat cta tct tat cga atg ata tat ttt tca taa ata cac ttt tat agt cct cgt tta aac aga att tag tat gta gtt ccg caa atg act cgt ccc tta ata tgc agt agg cta tta tct tct tta cgt agt gat cgt cgt agg gag aga act ccg aca tct tgt aga aca acg att taa tca tag gta gag ata ctt tca gtc tgt ggt gga tga tgt cat taa cga cat ccg cct tgt ata tga tgt ttc tgt ttt caa aca cca agt cga ata ctg tct tat cgt ctt tag tcg gaa ggt tga tgt cgt atc cga tgt ata cga ggt atg agg caa cat tgt tat tgc aat tct gga agg cgg tat gaa gag gag tca ttg tat tat agt att cgt ctt tct gaa tgt cga atc tat cta gta gat acc gta gta tat tga gag agc gac ttc cat atc ctt gat tat gtt tta tga ata gat aaa gta gat gtt gtc ctt ctt cct ttt gta att tcc cgt att ttt gtt cgt gcc aat tga gta aca tta tga gaa tat gac ctg ttg cac aat cgt tct tta tgt att cca tga tgg gtg tac aat caa gat tat tac gta tcc tcg tat cgg ctc ctc gag ata aaa gag cat aca cca cac gag gac tat gtt tgg tat act gtt gaa ggt aag tgt gta acg gcg tat ttc cga ttt tcg taa ccg cgt taa tgt ttg ctc cat gat cta tta tcg cgt aga tga atc gct tct cag ctc gca tct tag tgt gac tct ttg act tgt aat aat tgc ttt cgt gga aca cgg ata tgt gtt tac agt agt aat gaa gag aag tga gtc cat cct cat cga cgc aat tag ggt cag atc ctt tag tca ata att tgt aca gaa cgt aat agt tta agc tcc cat tga att tat atc taa gat aac aca gca ata gat cgg atg att tac taa agt cat caa tgg ggt ccg tta gta tat caa aga tct tgt tat cga ttg ata gtg aat gaa tca gat agt ggt gta gag gaa tat gtc ctt ttt tat cct tgc tat caa agt tac gca tgc cgt ggc gta aca ata tct tta ata cag atg gat taa atc gtg tat tca tcg tat agc aat gta atg gag agt tac cac att tta gtc gtt tat tca gat cgc agt gtt taa taa cta att taa aca gat gag atg ata tat cca cat caa aga atg cga gat aca tat gac aga cat tat tga cag aaa tgt gac ctt cat tat cac cgt cgt cca taa atg cgt tag gta cgt acc aca tac tat cgt taa cga tgc gca caa tct cgt cca ttt tca tcc atc ttc ata atg att tac ttt ttc ata att aga gaa aaa gat caa ggt ata aaa att aga agt gtt aga cta ta- --- --- --- --- --- --- --- --- --- --- -aa cta act tat aa- --- --- --- --- --- --- --- --- --- --- --- --- --- --- --- --- --- --- --- --- --- --- --- --- --- --- --- --- --- --- --- --- --- --- --- --- --- --- --- --- --- --- --- --- --- --- --- --- --- --- --- --- --- --- --- --- --- --- --- --- --- --- --- --- --- --- --- --- --- --- --- --- --- --- --- --- --- --- --- --- --- --- --- --- --- --- -aa cta act tat gac tta act aac tta tga cta tta act cat ttc aag aaa ggt ggg tgg aga gaa ctc tat atg aca gct tgt gaa aca att aga tcc cta att tct aat gga agt ttt gat agg aga ttg tca tca gtt gat aca ttg ttt att atc tca tct att aga gca cgt ctg ttt aga gct tta gtg acc tgc tcg gtt act tct gtg taa atc ttg aat cct tta gtg ata cac tgt gtc aaa act gga tgt tta gaa tac cta tgt aga ata tgg gaa gca tgc ttg ttt ttg tct cta tta tag att aac tca tac atg gtt gta tta tga att ttc atc tgc cta atg tac tcc aat tct tgt tta caa tca att ata taa tca aag agt gat gat gca tac aca tta caa agt gaa taa tct acc atc ata aaa tac ttg ata cag agc ttt atc aca tca tgg ttt tca att gta tta tta agt ata gct aat ttt ata cag tca ata gac aat ggt tct cta agc aat att tct aat att tta aga tgt gct tcc cta cgg gcg atg aca gat ccc cta tcc acg gcc acg tca aga cat gta tat cca tta ctc att act gcg ttg aca ttt gct cca ttt tct aat agc cat gat act aaa tct ata taa cct gca tag ata gcg cga taa agc aag gtc ctt cca cca gca tct agt tga ttg ata tct tca ata tat ggg ata caa agc tta taa att tct aat act gtg ggt tca tct aca agg aat ccc cta gta tac tga att att tta tat aga tct aat tta aca tca ttt tca tct ggg ata cca cga ttc aaa ata aac tca aca aca cta ctt tcc ttt tta cat att ccc cta aaa tag gca ttc aag cat tct att tta tat att aca gcc cca tga tct acc ata aaa tca aca atg tct att tct aca tat gca tta gat aga tag taa agt aag aga tct tgt aca gaa tta caa ttc tta ata att ata gag aaa ata tct tcc ata taa ttc ttt gac act aat gca gat ata ata tct tta tat gta ata tat gca aac agt cta tct act ata tac tga tca ata tta tct cta tga atc cta aaa taa tca tac aga aca tct aca gga tca caa att ggt tca agg aga aat cta tca aat att ttc ctg tca aca act ggt tct aga aca taa cag tca aca cct aat cca tgt ttt tta tag tca tct acc aaa gat aat gac caa aga tcg agg tcg tcg tga aac tgc tca tcg aca gcc atg aaa tct gcc gac tcc atg gcg cga atc gca ctg tct tat tcg cca ttg att ttc a-t ttt tta taa tta tgt aca tg- ttt cct tct att ctc aag agt cta caa aaa tat a-t ttt ttt tcg ata tct aag tac taa gtt ttt tta ctg ttt ttg tta ctg tct tcc att ctt cta act aaa gat ctg aga taa att ata caa tct tcg cta tcg aac cat ttt tgt agt cta aag cct gaa gta att aac caa ctg ttt tta tta gtg gct ttt ttc gat cta tcc tcg tc- --- --c tca tca tcc tta ta- --- --- --- --- --- --- tta tta tca tta tta tca tag tct att aaa cac aaa tca tct --- --- --- --- --- --- --- --- --- --- --- --- --- --- --- --- --- --- --- --- --- --- --- --- --- --- --a cgt tta taa c-- --- --- --- --- --- --- --- --a aca ttc --- --- --- --- -tc att att aat tag ttc tgt agt aat atc ttt aat aat ttg gct ata --- --- -ca tct gtt --- --- --- --- --- --- --- --- --- --- --- --- --- --- --- --- --- --- --c aat act --- --- --- --- --- --- --- --a tct att gat gat ttc ttt t-- --- --- --- --- --- --- --- --- --- --- --t taa gac t-- --- --- --- --- --- --- --- --- --- --- --- --- --- --- --- --- --- --- --- --- --- --- --- --- --- --- --- --- --- --- --- --- --- --- --- --- --- --- --- taa act agt --- --- --- --- --- --- --- --- --- --- --- --- --- --- --- --- --- --- --- --- --- --- --- --- --- --- --- --- --- --- --- --- --- --- --- --- --- --- --- --- --- --- --- --- --- --- --- --- --- --- --- --- --- --- --- --- --- --- --- --- --- --- --- --- --- --- --- --- --- --- --- --- --- --- --- --- --- --- --- --- --- --- --- --- --- --- --- --- --- --- --- --- --- --- --- --- --- --- --- --- --- --- --- --- --- --- --- --- --- --t atg gta atg acg atg aaa --- -tc gag tag ta- --- --a ctt cta ata aag act tga ta- --- --t cat tat cat atg ttt gat cg- --- --- --- --- --- --- --t cat agt taa tag tgt g-- --- --- --- --- --- --- --- --- --- --- --- --- --- --- --- --- --- --- --- -gc taa atg gta ctg tta ata agt tta t-- --- --- --- --- --- --- --- --- --- --- --- --- --- --- --- --- --a gac aat atc ata gta ttt tct ttc cag aat --- --- --- --t aga tta ttt ttt taa ata ctg atc ctc aca att ccg tga tgt agc agt agt tgg t-- --- --- --- --- --- --- --- --- --- --- --- --- --- --- --- --- --- --- --- --- --- --- --- --- --- --- --- -gc atg gtc tat atc gt- --- --- --- --- --- --- --- --- --- --- --- --- --- --- --- --- --- --- --- --- --- --- --- --- --- --- --- --- --- --- --- --- --- --- --- --- --- --- --- --- --- --- --- taa aat gta tca tat ata ata gtt ttc tga cgt gga gta cag aat ttt cga --- --- --- --- --- --- --- --- --- --- --- --- --- --- --- --- --- --- --- --- --- --- --- --- --- --- --- --- --- --- --- --- --- --- --- --- --- --- --- --- --- --- --- --t taa tga gtt cat ggt aag gaa ggg caa atg tct --- gta tat aat ata cat aag tta a-- --- --- --- --- --- --- --- --- --- --- --- --- --t agt ttt tta tca tat tt- --- --- --- --- --- --- --- --t cta ata cca taa taa aaa tta tca tta tg- --- --- -ta taa tca --- --- --- --- --- --- --- --- --- --- --- --t cac tg- --- -tc gct atc att att gcg ttt gtg tag tt- --- --- --- --- ctg ccc ta- --- --- --- --- --- --- --- --t cat cta cat cac tgt ca- --- --- --- --- ctc tca cta tat ctt cta aaa tta caa --- -ac aac --- --- --- --- --- --- --- --- --- --- --- --- --- --- --- --- --- --- --- --- --- --- --- --- --- --- --- --- --- --- --- --- --- --- --- --- tgg ata ttc gat --- --- --- --- --- --- --- --- --- --- --- --- --- --- --- --- --- --- --- --- --- --- --- --- --- --- --a aca gca ttt gtg t-- --- --- --- --- --- --- --- --- --- --- --- --- --- --- --- --- -

>KJ642619.1_Monkeypox_virus_strain_Gabon-1988_complete_genome

--- --- --- --- --- --- --- --- --- --- --- --- --- --- --- --- --- --- --- --- --- --- --- --- --- --- --- --- --- --- --- --- --- --- --- --g tgt cta gaa aaa aat gtg tga ccc acg acc gta gga aac tct aga ggg taa gaa aaa tca atc gtt tat aga gac cat cag aaa gag gtt taa tat ttt tgt gag acc cat cga a-- --- --- --- --- --- --- --- --- --- --- --- --- --- --- --- --- --- --- --- --- --- --- --- --- --- --- --- --- --- --- --- --- --- --- --- --- --- --- --- --- --- --- --- --- --- --- --- --- --- --- --- --- --- --- --- --- --- --- --- --- --- --- --- --- --- --- --- --- --- --- --- --- --- --- --- --- --- --- --- --- --- --- --- --- --- --- --- --- --- --- --- --- --- --- --- --- --- --- --- --- --- --- --- --- --- --- --- --- --- --- --- --- --- --- --- --- --- --- --- --- --- --- --- --- --- --- --- --- --- --- --- --- --- --- --- --- --- --- --- --- --- --- --- --- --- --- --- --- --- --- --- --- --- --- --- --- --- --- --- --- --- --- --- --- --- --- --- --- --- --- --- --- --- --- --- --- --- --- --- --- --- --- --- --- --- --- --- --- --- --- --- --- --- --- --- --- --- --- --- --- --- --- --- --- --- --- --- --- --- --- --- --- --- --- --- --- --- --- --- --- --- --- --- --- --- --- --- --- --- --- --- --- --- --- --- --- --- --- --- --- --- --- --- --- --- --- --- --- --- --- --- --- --- --- --- --- --- --- --- --- --- --- --- --- --- --- --- --- --- --- --- --- --- --- --- --- --- --- --- -ga gag aaa gga taa aaa ctt ttt acg act cca tca gaa aga ggt tta ata ttt ttg tga gac cca tcg acg aga gaa aga gat ggt tag tca aga tat ttt tct tag tac aaa agt caa tgt ttt aaa ata tat gga cga gaa tta att tgt ctg tat aaa aac ttg tgt gaa att atg tac tag aga aaa aac gtg agc agt gtc ccc tac atg gat ttt aca gat cat tta tat tcc aaa aat att aac tat ata cgt tta tta tat gat gtt aac gtg taa att ata aac att att tta tga tgc aat tgt ctg aca acc tag att ggc ata agg ata ttg ata agc tct acg aga ata tat tgt tgg acg tta tcg ttt acg aaa tag ttg aga cat cag aaa gag gtt taa tat ttt tgt gag a-c cat cga aga gag aaa gag aat aaa aat a-- --- --- --- ttt ttt tgt aaa act ttt tta tga gac aag aga gaa aga gaa tac gaa tag tga tca tat cgt atc aca tat tga aac aga aag aag aag taa cga gag gta act ttt tgt gaa tgt agt taa aca ttt ttg ttt tgc aaa ccg gaa tat agt acc cgg tac act ttt tta att cgt ggt gcg gtg tct gaa tcg ttc gat taa ccc aac tca tcc att ttc aga tga ata gag tta tcg att cag aca cat gct ttg agt ttt gtt gaa tcg atg agt gaa gta tca tcg gtt gca cct tca gat gcc gat ccg tcg aca tac ttg aat cca tcc ttg act tca agt tca gat gat tcc tca cac atg tct ccg ata cgt acg cta aac tct agg ttc tta aca cat ttt gta tca acg atc gtt gaa ccg atg ata tct ttg taa ctc act ttc tta tgt gag atg tta gac cca agt act gga tgg gtc ttg atg tcg ctg tct ttc tct tct tcg cta cat ctg atg tcg ata gac atc tca cag tct ttg atc ata gcc aga gct tct tca cgc gtg atc gcg gga gag tcc tta cct tgt cct ggt gac acg ctg gac aat cta gta ttc aca gtg ttt cca tca gag gat tcg gag atg gat gaa atc ttt ggg cat ttg gtg aat cca aag ttc atg tta aga ccc gcg ccg acg ata gtg taa taa gtg gtg gga tct cct ttt aca act --- --- --- --- --- --- tct tcg gat acc tca tca tct tcg gtc tct gta act tcc gtt acg gat tga caa atc tta tca ttg gtc ggt gtt tgg tct tgc ttt gtg act ttg ata ata aca tcg att ccc ata tga tgt ttg ttt tct tct tca gta cac gag gat gag gat tgt tga aga cta gta ggc ata gca gct gcc act agg cac atg cat gcc agg aca ata tat tgt ttc atg att gct att gat tga tta ctg ttc tag atg att cta ctt tct tac cat ata ata aat tag aat ata ttt tct act ttt acg aga aat taa tta ttg tat tta tta ttt atg ggt -aa aaa aaa ctt act ata agt ggg tgg gat tct ggg aat tag tga tca gtt tat gta tat cgc aac tag cgg gca tat ggc tat tga cat cga gaa cat tac cca tat gat aag aga ttg tat ccg ttt cgt agt ctt gag tat tgg tat tac tat ata gta tgt aga tgt caa cgc tag ata gac agt cgc cca cta gag tta ccg tct ctg aat gcg gca tga tag tat cat tct ttg ttt tcg tta act gtt tgg aag atg aat ctt tgt tgt tac att taa tct cga aat tca gag tac ata tct ttg aag tat tct gat atc tat ttt ctc ctg taa aga atc ctg aag ttg cta cat tat taa gga cag aga agt att ctg cac gaa aga cgg gat cac aat ctt tat gat tca tgg taa tag tta gtt ccg acg ttg aga tgg att cgc tga gac cgg tag tgg tcg tcc gag tac acg atg tgt cgt tga ctg gat aca ggt taa ttt cca cat cga tat agt taa atg tat tgc tgg tt- --a cgg gtt cgc att tat ctg tgg aag aga cgg tgt gag aat atg ttc cgg gac cac acg gag aac aga tga cgt ctc cgg tag acg tgt atc cgg ata ctc cgt atc cta ttc cac act ttg ttt tag aaa tac atg ttc tac acc ctg atg atc ctt tga gaa gac aat aat atc ctg gag agc att cac aga ttc tat tgt gag tcg tgt tac acg atc gcg tct cta cct gat tac tat cac atc ttc cgt tac aac tta gac aag cct gta aat gat tat tgt gag atg taa agg tat ccg aac cac acg gcg tac att gtg tat tag tct tgc tat cac ata atc tgg aag cgt aag ttc ccg gag gac acg ata gac aac ata gat tac ggc ttc tgt att cgt tgt ctt tac att ttc cat tgg atg gtg cat gtg gtg cta tat ctc ttc cgt tta tta tta tac atg aga gaa aca ata tat acg agt ata ata cgg acc tca tga ttt aat aat -gt agt aat cgt cgt ctt gtt act gtt tgt ttc cta ctt ctc caa tca tat aga tta ttt ttt aaa tat ttt ctt tct atc atg gat aat att tgt aat ggt tct ttc cgt aca aca tac tgt tta gat ggt agt cgc tta gct tgg tta tga tat tgc gca taa ttt ccg gag gca aat acg ata gtc tag att gac tat cga tgg tag act cta att tat tga gtg ctt tgt cga cga gtt tac ttt tat gct cca tcg ata gat gac act gtt cta tga gat cgt cgt aca tgg gaa atg aaa tgc gtt tgt ctg aat gta tgg ctt cga gat agg tgt gat acc gga tgt ctt ctg ttc tca ata ccg tat aca agt tgg tgt ctg aga ttc gaa tct ctt tga gga gac tta tgt cac gac tac att ttt cga tga tgg aat cta tct tat cga atg ata tat ttt tca taa ata cac ttt tat agt cct cgt tta aac aga att tag tat gta gtt ccg caa atg act cgt ccc tta ata ggc agt agg cta tta tct tct tta cgt agt gat cgt cgt agg gag aga act ccg aca tct tgt aga aca acg att taa tca tag gta gag ata ctt tca gtc tgt ggt gga tga tgt cat taa cga cat ccg cct tgt ata tga tgt ttc tgt ttt caa aca cca agt cga ata ctg tct tat cgt ctt tag tcg gaa ggt tga tgt cgt atc cga tgt ata cga ggt atg agg caa cat tgt tat tgc aat tct gga agg cgg tat gaa gag gag tca ttg tat tat agt att cgt ctt tct gaa tgt cga atc tat cta gta gat acc gta gta tat tga gag agc gac ttc cat atc ctt gat tat gtt tta tga ata gat aaa gta gat gtt gtc ctt ctt cct ttt gta att tcc cgt att ttt gtt cgt gcc aat tga gta aca tta tga gaa tat gac ctg ttg cac aat cgt tct tta tgt att cca tga tgg gtg tac aat caa gat tat tac gta tcc tcg tat cgg ctc ctc gag ata aaa gag cat aca cca cac gag gac tat gtt tgg tat act gtt gaa ggt aag tgt gta acg gcg tat ttc cga ttt tcg taa ccg cgt taa tgt ttg ctc cat gat cta tta tcg cgt aga tga atc gct tct cag ctc gca tct tag tgt gac tct ttg act tgt aat aat tgc ttt cgt gga aca cgg ata tgt gtt tac agt agt aat gaa gag aag tga gtc cat cct cat cga cgc aat tag ggt cag atc ctt tag tca ata att tgt aca gaa cgt aat agt tta agc tcc cat tga att tat atc taa gat aac aca gca ata gat cgg atg att tac taa agt cat caa tgg ggt ccg tta gta tat caa aga tct tgt tat cga ttg ata gtg aat gaa tca gat agt ggt gta gag gaa tat gtc ctt ttt tat cct tgc tat caa agt tac gca tgc cgt ggc gta aca ata tct tta ata cag atg gat taa atc gtg tat tca tcg tat agc aat gta atg gag agt tac cac att tta gtc gtt tat tca gat cgc agt gtt taa taa cta att taa aca gat gag atg ata tat cca cat caa aga atg cga gat aca tat gac aga cat tat tga cag aaa tgt gac ctt cat tat cac cgt cgt cca taa atg cgt tag gta cgt acc aca tac tat cgt taa cga tgc gca caa tct cgt cca ttt tca tcc atc ttc ata atg att tac ttt ttc ata att aga gaa aaa gat caa ggt ata aaa att aga agt gtt aga cta ta- --- --- --- --- --- --- --- --- --- --- -aa cta act tat aa- --- --- --- --- --- --- --- --- --- --- --- --- --- --- --- --- --- --- --- --- --- --- --- --- --- --- --- --- --- --- --- --- --- --- --- --- --- --- --- --- --- --- --- --- --- --- --- --- --- --- --- --- --- --- --- --- --- --- --- --- --- --- --- --- --- --- --- --- --- --- --- --- --- --- --- --- --- --- --- --- --- --- --- --- --- --- --- --- --- --- --- --a act aac tta tga cta tta act cat ttc aag aaa ggt ggg tgg aga gaa ctc tat atg aca gct tgt gaa aca att aga tcc cta att tct aat gga agt ttt gat agg aga ttg tca tca gtt gat aca ttg ttt att atc tca tct att aga gca cgt ctg ttt aga gct tta gtg acc tgc tcg gtt act tct gtg taa atc ttg aat cct tta gtg ata cac tgt gtc aaa act gga tgt tta gaa tac cta tgt aga ata tgg gaa gca tgc ttg ttt ttg tct cta tta tag att aac tca tac atg gtt gta tta tga att ttc atc tgc cta atg tac tcc aat tct tgt tta caa tca att ata taa tca aag agt gat gat gca tac aca tta caa agt gaa taa tct acc atc ata aaa tac ttg ata cag agc ttt atc aca tca tgg ttt tca att gta tta tta agt ata gct aat ttt ata cag tca ata gac aat ggt tct cta agc aat att tct aat att tta aga tgt gct tcc cta cgg gcg atg aca gat ccc cta tcc acg gcc acg tca aga cat gta tat cca tta ctc att act gcg ttg aca ttt gct cca ttt tct aat agc cat gat act aaa tct ata taa cct gca tag ata gcg cga taa agc aag gtc ctt cca cca gca tct agt tga ttg ata tct tca ata tat ggg ata caa agc tta taa att tct aat act gtg ggt tca tct aca agg aat ccc cta gta tac tga att att tta tat aga tct aat tta aca tca ttt tca tct ggg ata cca caa ttc aaa ata aac tca aca aca cta ctt tcc ttt tta cat att ccc cta aaa tag gca ttc aag cat tct att tta tat att aca gcc cca tga tct acc ata aaa tca aca atg tct att tct aca tat gca tta gat aga tag taa agt aag aga tct tgt aca gaa tta caa ttc tta ata att ata gag aaa ata tct tcc ata taa ttc ttt gac act aat gca gat ata ata tct tta tat gta ata tat gca aac agt cta tct act ata tac tga tca ata tta tct cta tga atc cta aaa taa tca tac aga aca tct aca gga tca caa att ggt tca agg aga aat cta tca aat att ttc ctg tca aca act ggt tct aga aca taa cag tca aca cct aat cca tgt ttt tta tag tca tct acc aaa gat aat gac caa aga tcg agg tcg tcg tga aac tgc tca tcg aca gcc atg aaa tct gcc gac tcc atg gcg cga atc gca ctg tct tat tcg cca ttg att ttc a-t ttt tta taa tta tgt aca tgt ttt cc- --- -tt ctc aag agt cta caa aaa tat a-t ttt ttt tcg ata tct aag tac taa g-t ttt tta ctg ttt ttg tta ctg tct tcc att ctt cta act aaa gat ctg aga taa att ata caa tct tcg cta tcg aac cat ttt tgt agt cta aag cct gaa gta att aac caa ctg ttt tta tta gtg gct ttt ttc gat cta tcc tcg tc- --- --c tca tca tcc tta ta- --- --- --- --- --- --- tta tta tca tta tta tca tag tct att aaa cac aaa tca tct --- --- --- --- --- --- --- --- --- --- --- --- --- --- --- --- --- --- --- --- --- --- --- --- --- --- --a cgt tta taa c-- --- --- --- --- --- --- --- --a aca ttc --- --- --- --- -tc att att aat tag ttc tgt agt aat atc ttt aat aat ttg gct ata --- --- -ca tct gtt --- --- --- --- --- --- --- --- --- --- --- --- --- --- --- --- --- --- --c aat act --- --- --- --- --- --- --- --a tct att gat gat ttc ttt t-- --- --- --- --- --- --- --- --- --- --- --t taa gac t-- --- --- --- --- --- --- --- --- --- --- --- --- --- --- --- --- --- --- --- --- --- --- --- --- --- --- --- --- --- --- --- --- --- --- --- --- --- --- --- taa act agt --- --- --- --- --- --- --- --- --- --- --- --- --- --- --- --- --- --- --- --- --- --- --- --- --- --- --- --- --- --- --- --- --- --- --- --- --- --- --- --- --- --- --- --- --- --- --- --- --- --- --- --- --- --- --- --- --- --- --- --- --- --- --- --- --- --- --- --- --- --- --- --- --- --- --- --- --- --- --- --- --- --- --- --- --- --- --- --- --- --- --- --- --- --- --- --- --- --- --- --- --- --- --- --- --- --- --- --- --- --t atg gta atg acg atg aaa --- -tc gag tag ta- --- --a ctt cta ata aag act tga ta- --- --t cat tat cat atg ttt gat cg- --- --- --- --- --- --- --t cat agt taa tag tgt g-- --- --- --- --- --- --- --- --- --- --- --- --- --- --- --- --- --- --- --- -gc taa atg gta ctg tta ata agt tta t-- --- --- --- --- --- --- --- --- --- --- --- --- --- --- --- --- --a gac aat atc ata gta ttt tct ttc cag aat --- --- --- --t aga tta ttt ttt taa ata ctg atc ctc aca att ccg tga tgt agc agt agt tgg t-- --- --- --- --- --- --- --- --- --- --- --- --- --- --- --- --- --- --- --- --- --- --- --- --- --- --- --- -gc atg gtc tat atc gt- --- --- --- --- --- --- --- --- --- --- --- --- --- --- --- --- --- --- --- --- --- --- --- --- --- --- --- --- --- --- --- --- --- --- --- --- --- --- --- --- --- --- --- taa aat gta tca tat ata ata gtt ttc tga cgt gga gta cag aat ttt cga --- --- --- --- --- --- --- --- --- --- --- --- --- --- --- --- --- --- --- --- --- --- --- --- --- --- --- --- --- --- --- --- --- --- --- --- --- --- --- --- --- --- --- --t taa tga gtt cat ggt aag gaa ggg caa atg tct --- gta tat aat ata cat aag tta a-- --- --- --- --- --- --- --- --- --- --- --- --- --t ag- ttt tta tca tat tt- --- --- --- --- --- --- --- --t cta ata cca taa taa aaa tta tca tta tg- --- --- -ta taa tca --- --- --- --- --- --- --- --- --- --- --- --t cac tg- --- -tc gct atc att att gcg ttt gtg tag tt- --- --- --- --- ctg ccc ta- --- --- --- --- --- --- --- --t cat cta cat cac tgt ca- --- --- --- --- ctc tca cta tat ctt cta aaa tta caa --- -ac aac --- --- --- --- --- --- --- --- --- --- --- --- --- --- --- --- --- --- --- --- --- --- --- --- --- --- --- --- --- --- --- --- --- --- --- --- tgg ata ttc gat --- --- --- --- --- --- --- --- --- --- --- --- --- --- --- --- --- --- --- --- --- --- --- --- --- --- --a aca gca ttt gtg t-- --- --- --- --- --- --- --- --- --- --- --- --- --- --- --- --- -

>JX878409.1_Monkeypox_virus_isolate_DRC_06-0999_complete_genome

--- --- --- --- --- --- --- --- --- --- --- --- --- --- --- --- --- --- --- --- --- --- --- --- --- --- --- --- --- --- --- --- --- --- --- --g tgt cta gaa aaa aat gtg tga ccc acg acc gta gga aac tct aga ggg taa gaa aaa tca atc gtt tat aga gac cat cag aaa gag gtt taa tat ttt tgt gag acc cat cga aga gag aaa gga taa aaa ctt ttt acg act cca tca gaa aga ggt tta ata ttt ttg tga gac cca tcg aag aga gaa agg ata aaa act ttt tac gac tcc atc aga aag agg ttt aat att ttt gtg aga ccc atc gaa gag aga aag gat aaa aac ttt tta cga ctc cat cag aaa gag gtt taa tat ttt tgt gag acc cat cga cga gag aaa gga taa aaa ctt ttt acg act cca tca gaa aga ggt tta ata ttt ttg tga gac cca tcg aag aga gaa agg ata aaa act ttt tac gac tcc atc aga aag agg ttt aat att ttt gtg aga ccc atc gac gag aga aag gat aaa aac ttt tta cga ctc cat cag aaa gag gtt taa tat ttt tgt gag acc cat cga aga gag aaa gga taa aaa ctt ttt acg act cca tca gaa aga ggt tta ata ttt ttg tga gac cca tcg acg aga gaa agg ata aaa act ttt tac gac tcc atc aga aag agg ttt aat att ttt gtg aga ccc atc gaa gag aga aag gat aaa aac ttt tta cga ctc cat cag aaa gag gtt taa tat ttt tgt gag acc cat cga cga gag aaa gga taa aaa ctt ttt acg act cca tca gaa aga ggt tta ata ttt ttg tga gac cca tcg aag aga gaa agg ata aaa act ttt tac gac tcc atc aga aag agg ttt aat att ttt gtg aga ccc atc gac gag aga aag gat aaa aac ttt tta cga ctc cat cag aaa gag gtt taa tat ttt tgt gag acc cat cga aga gag aaa gga taa aaa ctt ttt acg act cca tca gaa aga ggt tta ata ttt ttg tga gac cca tcg acg aga gaa aga gat ggt tag tca aga tat ttt tct tag tac aaa agt caa tgt ttt aaa ata tat gga cga gaa tta att tgt ctg tat aaa aac ttg tgt gaa att atg tac tag aga aaa aac gtg agc agt gtc ccc tac atg gat ttt aca gat cat tta tat tcc aaa aat att aac tat ata cgt tta tta tat gat gtt aac gtg taa att ata aac att att tta tga tgc aat tgt ctg aca acc --- att ggc ata agg ata ttg ata agc tct acg aga ata tat tgt tgg acg tta tcg ttt acg aaa tag ttg aga cat cag aaa gag gtt taa tat ttt tgt gag a-c cat cga aga gag aaa gag aat aaa aat a-- --- --- --- ttt ttt tgt aaa act ttt tta tga gac aag aga gaa aga gaa tac gaa tag tga tca tat cgt atc aca tat tga aac aga aag aag aag taa cga gag gta act ttt tgt gaa tgt agt taa aca ttt ttg ttt tgc aaa ccg gaa tat agt acc cgg tac act ttt tta att cgt ggt gcg gtg tct gaa tcg ttt gat taa ccc aac tca tcc att ttc aga tga ata gag tta tcg att cag aca cat gct ttg agt ttt gtt gaa tcg atg agt gaa gta tca tcg gtt gca cct tca gat gcc gat ccg tcg aca tac ttg aat cca tcc ttg act tca agt tca gat gat tcc tca cac atg tct ccg ata cgt acg cta aac tct agg ttc tta aca cat ttt gta tca acg atc gtt gaa ccg atg ata tct ttg taa ctc act ttc tta tgt gag atg tta gac cca agt act gga tgg gtc ttg atg tcg ctg tct ttc tct tct tcg cta cat ctg atg tcg ata gac atc tca cag tct ttg atc ata gcc aga gct tct tca cgc gtg atc gcg gga gag tcc tta cct tgt cct ggt gac acg ctg gac aat cta gta ttc aca gtg ttt cca tca gag gat tcg gag atg gat gaa atc ttt ggg cat ttg gtg aat cca aag ttc atg tta aga ccc gcg ccg acg ata gtg taa taa gtg gtg gga tct cct ttt aca act tct tcg gat acc tca tca tct tcg gat acc tca tca tct tcg gtc tct gta act tcc gtt acg gat tga caa atc tta tca ttg gtc ggt gtt tgg tct tgc ttt gtg act ttg ata ata aca tcg att ccc ata tga tgt ttg ttt tct tct tca gta cac gag gat gag gat tgt tga aga cta gta ggc ata gca gct gcc act agg cac atg cat gcc agg aca ata tat tgt ttc atg att gct att gat tga tta ctg ttc tag atg att cta ctt tct tac cat ata ata aat tag aat ata ttt tct act ttt acg aga aat taa tta ttg tat tta tta ttt atg ggt -aa aaa aaa ctt act ata agt ggg tgg gat tct ggg aat tag tga tca gtt tat gta tat cgc aac tag cgg gca tat ggc tat tga cat cga gaa cat tac cca tat gat aag aga ttg tat ccg ttt cgt agt ctt gag tat tgg tat tac tat ata gta tgt aga tgt cga cgc tag ata gac agt cgc cca cta gag tta ccg tct ctg aat gcg gca tga tag tat cat tct ttg ttt tcg tta act gtt tgg aag atg aat ctt tgt tgt tac att taa tct cga aat tca gag tac ata tct ttg aag tat tct gat atc tat ttt ctc ctg taa aga atc ctg aag ttg cta cat tat taa gga cag aga agt att ctg cac gaa aga cgg gat cac aat ctt tat gat tca tgg taa tag tta gtt ccg acg ttg aga tgg att cgc tga gac cgg tag tgg tcg tcc gag tac acg atg tgt cgt tga ctg gat aca ggt taa ttt cca cat cga tat agt taa atg tat tgc tgg tt- --a cgg gtt cgc att tat ctg tgg aag aga cgg tgt gag aat atg ttc cgg gac cac acg gag aac aga tga cgt ctc cgg tag acg tgt atc cgg ata ctc cgt atc cta ttc cac act ttg ttt tag aaa tac atg ttc tac acc ctg atg atc ctt tga gaa gac aat aat atc ctg gag agc att cac aga ttc tat tgt gag tcg tgt tac acg atc gcg tct cta cct gat tac tat cac atc ttc cgt tac aac tta gac aag cct gta aat gat tat tgt gag atg taa agg tat ccg aac cac acg gcg tac att gtg tat tag tct tgc tat cac ata atc tgg aag cgt aag ttc ccg gag gac acg ata gac aac ata gat tac ggc ttc tgt att cgt tgt ctt tac att ttc cat tgg atg gtg cat gtg gtg cta tat ctc ttc cgt tta tta tta tac atg aga gaa aca ata tat acg agt ata ata cgg acc tca tga ttt aat aat -gt agt aat cgt cgt ctt gtt act gtt tgt ttc cta ctt ctc caa tca tat aga tta ttt ttt aaa tat ttt ctt tct atc atg gat aat att tgt aat ggt tct ttc cgt aca aca tac tgt tta gat ggt agt cgc tta gct tgg tta tga tat tgc gca taa ttt ccg gag gca aat acg ata gtc tag att gac tat cga tgg tag act cta att tat tga gtg ctt tgt cga cga gtt tac ttt tat gct cca tcg ata gat gac act gtt cta tga gat cgt cgt aca tgg gaa atg aaa tgc gtt tgt ctg aat gta tgg ctt cga gat agg tgt gat acc gga tgt ctt ctg ttc tca ata ccg tat aca agt tgg tgt ctg aga ttc gaa tct ctt tga gga gac tta tgt cac gac tac att ttt cga tga tgg aat cta tct tat cga atg ata tat ttt tca taa ata cac ttt tat agt cct cgt tta aac aga att tag tat gta gtt ccg caa atg act cgt ccc tta ata ggc agt agg cta tta tct tct tta cgt agt gat cgt cgt agg gag aga act ccg aca tct tgt aga aca acg att taa tca tag gta gag ata ctt tca gtc tgt ggt gga tga tgt cat taa cga cat ccg cct tgt ata tga tgt ttc tgt ttt caa aca cca agt cga ata ctg tct tat cgt ctt tag tcg gaa ggt tga tgt cgt atc cga tgt ata cga ggt atg agg caa cat tgt tat tgc aat tct gga agg cgg tat gaa gag gag tca ttg tat tat agt att cgt ctt tct gaa tgt cga atc tat cta gta gat acc gta gta tat tga gag agc gac ttc cat atc ctt gat tat gtt tta tga ata gat aaa gta gat gtt gtc ctt ctt cct ttt gta att tcc cgt att ttt gtt cgt gcc aat tga gta aca tta tga gaa tat gac ctg ttg cac aat cgt tct tta tgt att cca tga tgg gtg tac aat caa gat tat tac gta tcc tcg tat cgg ctc ctc gag ata aaa gag cat aca cca cac gag gac tat gtt tgg tat act gtt gaa ggt aag tgt gta acg gcg tat ttc cga ttt tcg taa ccg cgt taa tgt ttg ctc cat gat cta tta tcg cgt aga tga atc gct tct cag ctc gca tct tag tgt gac tct ttg act tgt aat aat tgc ttt cgt gga aca cgg ata tgt gtt tac agt agt aat gaa gag aag tga gtc cat cct cat cga cgc aat tag ggt cag atc ctt tag tca ata att tgt aca gaa cgt aat agt tta agc tcc cat tga att tat atc taa gat aac aca gca ata gat cgg atg att tac taa agt cat caa tgg ggt ccg tta gta tat caa aga tct tgt tat cga ttg ata gtg aat gaa tca gat agt ggt gta gag gaa tat gtc ctt ttt tat cct tgc tat caa agt tac gca tgc cgt ggc gta aca ata tct tta ata cag atg gat taa atc gtg tat tca tcg tat agc aat gta atg gag agt tac cac att tta gtc gtt tat tca gat cgc agt gtt taa taa cta att taa aca gat gag atg ata tat cca cat caa aga atg cga gat aca tat gac aga cat tat tga cag aaa tgt gac ctt cat tat cac cgt cgt cca taa atg cgt tag gta cgt acc aca tac tat cgt taa cga tgc gca caa tct cgt cca ttt tca tcc atc ttc ata atg att tac ttt ttc ata att aga gaa aaa gat caa ggt ata aaa att aga agt gtt aga cta ta- --- --- --- --- --- --- --- --- --- --- -aa cta act tat aa- --- --- --- --- --- --- --- --- --- --- --- --- --- --- --- --- --- --- --- --- --- --- --- --- --- --- --- --- --- --- --- --- --- --- --- --- --- --- --- --- --- --- --- --- --- --- --- --- --- --- --- --- --- --- --- --- --- --- --- --- --- --- --- --- --- --- --- --- --- --- --- --- --- --- --- --- --- --- --- --- --- --- --- --- --- --- -aa cta act tat gac tta act aac tta tga cta tta act cat ttc aag aaa ggt ggg tgg aga gaa ctc tat atg aca gct tgt gaa aca att aga tcc cta att tct aat gga agt ttt gat agg aga ttg tca tca gtt gat aca ttg ttt att atc tca tct att aga gca cgt ctg ttt aga gct tta gtg acc tgc tcg gtt act tct gtg taa atc ttg aat cct tta gtg ata cac tgt gtc aaa act gga tgt tta gaa tac cta tgt aga ata tgg gaa gca tgc ttg ttt ttg tct cta tta tag att aac tca tac atg gtt gta tta tga att ttc atc tgc cta atg tac tcc aat tct tgt tta caa tca att ata taa tca aag agt gat gat gca tac aca tta caa agt gaa taa tct acc atc ata aaa tac ttg ata cag agc ttt atc aca tca tgg ttt tca att gta tta tta agt ata gct aat ttt ata cag tca ata gac aat ggt tct cta agc aat att tct aat att tta aga tgt gct tcc cta cgg gcg atg aca gat ccc cta tcc acg gcc acg tca aga cat gta tat cca tta ctc att act gcg ttg aca ttt gct cca ttt tct aat agc cat gat act aaa tct ata taa cct gca tag ata gcg cga taa agc aag gtc ctt cca cca gca tct agt tga ttg ata tct tca ata tat ggg ata caa agc tta taa att tct aat act gtg ggt tca tct aca agg aat ccc cta gta tac tga att att tta tat aga tct aat tta aca tca ttt tca tct ggg ata cca caa ttc aaa ata aac tca aca aca cta ctt tcc ttt tta cat att ccc cta aaa tag gca ttc aag cat tct att tta tat att aca gcc cca tga tct acc ata aaa tca aca atg tct att tct aca tat gca tta gat aga tag taa agt aag aga tct tgt aca gaa tta caa ttc tta ata att ata gag aaa ata tct tcc ata taa ttc ttt gac act aat gca gat ata ata tct tta tat gta ata tat gca aac agt cta tct act ata tac tga tca ata tta tct cta tga atc cta aaa taa tca tac aga aca tct aca gga tca caa att ggt tca agg aga aat cta tca aat att ttc ctg tca aca act ggt tct aga aca taa cag tca aca cct aat cca tgt ttt tta tag tca tct acc aaa gat aat gac caa aga tcg agg tcg tcg tga aac tgc tca tcg aca gcc atg aaa tct gcc gac tcc atg gcg cga atc gca ctg tct tat tcg cca ttg att ttc a-t ttt tta taa tta tgt aca tgt ttt cct tct att ctc aag agt cta caa aaa tat a-t ttt ttt tcg ata tct aag tac taa gtt ttt tta ctg ttt ttg tta ctg tct tcc att ctt cta act aaa gat ctg aga taa att ata caa tct tcg cta tcg aac cat ttt tgt agt cta aag cct gaa gta att aac caa ctg ttt tta tta gtg gct ttt ttc gat cta tcc tcg tc- --- --c tca tca tcc tta ta- --- --- --- --- --- --- tta tta tca tta tta tca tag tct att aaa cac aaa tca tct --- --- --- --- --- --- --- --- --- --- --- --- --- --- --- --- --- --- --- --- --- --- --- --- --- --- --a cgt tta taa c-- --- --- --- --- --- --- --- --a aca ttc --- --- --- --- -tc att att aat tag ttc tgt agt aat atc ttt aat aat ttg gct ata --- --- -ca tct gtt --- --- --- --- --- --- --- --- --- --- --- --- --- --- --- --- --- --- --c aat act --- --- --- --- --- --- --- --a tct att gat gat ttc ttt t-- --- --- --- --- --- --- --- --- --- --- --t taa gac t-- --- --- --- --- --- --- --- --- --- --- --- --- --- --- --- --- --- --- --- --- --- --- --- --- --- --- --- --- --- --- --- --- --- --- --- --- --- --- --- taa act agt --- --- --- --- --- --- --- --- --- --- --- --- --- --- --- --- --- --- --- --- --- --- --- --- --- --- --- --- --- --- --- --- --- --- --- --- --- --- --- --- --- --- --- --- --- --- --- --- --- --- --- --- --- --- --- --- --- --- --- --- --- --- --- --- --- --- --- --- --- --- --- --- --- --- --- --- --- --- --- --- --- --- --- --- --- --- --- --- --- --- --- --- --- --- --- --- --- --- --- --- --- --- --- --- --- --- --- --- --- --t atg gta atg acg atg aaa --- -tc gag tag ta- --- --a ctt cta ata aag act tga ta- --- --t cat tat cat atg ttt gat cg- --- --- --- --- --- --- --t cat agt taa tag tgt g-- --- --- --- --- --- --- --- --- --- --- --- --- --- --- --- --- --- --- --- -gc taa atg gta ctg tta ata agt tta t-- --- --- --- --- --- --- --- --- --- --- --- --- --- --- --- --- --a gac aat atc ata gta ttt tct ttc cag aat --- --- --- --t aga tta ttt ttt taa ata ctg atc ctc aca att ccg tga tgt agc agt agt tgg t-- --- --- --- --- --- --- --- --- --- --- --- --- --- --- --- --- --- --- --- --- --- --- --- --- --- --- --- -gc atg gtc tat atc gt- --- --- --- --- --- --- --- --- --- --- --- --- --- --- --- --- --- --- --- --- --- --- --- --- --- --- --- --- --- --- --- --- --- --- --- --- --- --- --- --- --- --- --- taa aat gta tca tat ata ata gtt ttc tga cgt gga gta cag aat ttt cga --- --- --- --- --- --- --- --- --- --- --- --- --- --- --- --- --- --- --- --- --- --- --- --- --- --- --- --- --- --- --- --- --- --- --- --- --- --- --- --- --- --- --- --t taa tga gtt cat ggt aag gaa ggg caa atg tct --- gta tat aat ata cat aag tta a-- --- --- --- --- --- --- --- --- --- --- --- --- --t ag- ttt tta tca tat tt- --- --- --- --- --- --- --- --t cta ata cca taa taa aaa tta tca tta tg- --- --- -ta taa tca --- --- --- --- --- --- --- --- --- --- --- --t cac tg- --- -tc gct atc att att gcg ttt gtg tag tt- --- --- --- --- ctg ccc ta- --- --- --- --- --- --- --- --t cat cta cat cac tgt ca- --- --- --- --- ctc tca cta tat ctt cta aaa tta caa --- -ac aac --- --- --- --- --- --- --- --- --- --- --- --- --- --- --- --- --- --- --- --- --- --- --- --- --- --- --- --- --- --- --- --- --- --- --- --- tgg aga ttc gat --- --- --- --- --- --- --- --- --- --- --- --- --- --- --- --- --- --- --- --- --- --- --- --- --- --- --a aca gca ttt gtg t-- --- --- --- --- --- --- --- --- --- --- --- --- --- --- --- --- -

>DQ011154.1_Monkeypox_virus_strain_Congo_2003_358_complete_genome

--- --- --- --- --- --- --- --- --- --- --- --- --- --- --- --- --- --- --- --- --- --- --- --- --- --- --- --- --- --- --- --- --- --- --- --g tgt cta gaa aaa aat gtg tga ccc acg acc gta gga aac tct aga ggg taa gaa aaa tca atc gtt tat aga gac cat cag aaa gag gtt taa tat ttt tgt gag acc cat cga aga gag aaa gg- --- --- --- --- --- --- --- --- --- --- --- --- --- --- --- --- --- --- --- --- --- --- --- --- --- --- --- --- --- --- --- --- --- --- --- --- --- --- --- --- --- --- --- --- --- --- --- --- --- --- --- --- --- --- --- --- --- --- --- --- --- --- --- --- --- --- --- --- --- --- --- --- --- --- --- --- --- --- --- --- --- --- --- --- --- --- --- --- --- --- --- --- --- --- --- --- --- --- --- --- --- --- --- --- --- --- --- --- --- --- --- --- --- --- --- --- --- --- --- --- --- --- --- --- --- --- --- --- --- --- --- --- --- --- --- --- --- --- --- --- --- --- --- --- --- --- --- --- --- --- --- --- --- --- --- --- --- --- --- --- --- --- --- --- --- --- --- --- --- --- --- --- --- --- --- --- --- --- --- --- --- --- --- --- --- --- -at aaa aac ttt tta cga ctc cat cag aaa gag gtt taa tat ttt tgt gag acc cat cga aga gag aaa gga taa aaa ctt ttt acg act cca tca gaa aga ggt tta --- --- --- --- --- --- --- --- --- --- --- --- --- --- --- --- --- --- --- --- --- --- --- --- --- --- --- --- --- --- --- --- --- --- --- --- --- --- --- --- --- --- --- --- --- --- --a tat ttt tgt gag acc cat cga aga gag aaa gga taa aaa ctt ttt acg act cca tca gaa aga ggt tta ata ttt ttg tga gac cca tcg acg aga gaa aga gat ggt tag tca aga tat ttt tct tag tac aaa agt caa tgt ttt aaa ata tat gga cga gaa tta att tgt ctg tat aaa aac ttg tgt gaa att atg tac tag aga aaa aac gtg agc agt gtc ccc tac atg gat ttt aca gat cat tta tat tcc aaa aat att aac tat ata cgt tta tta tat gat gtt aac gtg taa att ata aac att att tta tga tgc aat tgt ctg aca acc --- att ggc ata agg ata ttg ata agc tct acg aga ata tat tgt tgg acg tta tcg ttt acg aaa tag ttg aga cat cag aaa gag gtt taa tat ttt tgt gag a-c cat cga aga gag aaa gag aat aaa aat a-- --- --- --- ttt ttt tgt aaa act ttt tta tga gac aag aga gaa aga gaa tac gaa tag tga tca tat cgt atc aca tat tga aac aga aag aag aag taa cga gag gta act ttt tgt gaa tgt agt taa aca ttt ttg ttt tgc aaa ccg gaa tat agt acc cgg tac act ttt tta att cgt ggt gcg gtg tct gaa tcg ttc gat taa ccc aac tca tcc att ttc aga tga ata gag tta tcg att cag aca cat gct ttg agt ttt gtt gaa tcg atg agt gaa gta tca tcg gtt gca cct tca gat gcc gat cca tcg aca tac ttg aat cca tcc ttg act tca agt tca gat gat tcc tca cac atg tct ccg ata cgt acg cta aac tct agg ttc tta aca cat ttt gta tca acg atc gtt gaa ccg atg ata tct ttg taa ctc act ttc tta tgt gag atg tta gac cca agt act gga tgg gtc ttg atg tcg ctg tct ttc tct tct tcg cta cat ctg atg tcg ata gac atc tca cag tct ttg atc ata gcc aga gct tct tca cgc gtg atc gcg gga gag tcc tta cct tgt cct ggt gac acg ctg gac aat cta gta ttc aca gtg ttt cca tca gag gat tcg gag atg gat gaa atc ttt ggg cat ttg gtg aat cca aag ttc atg tta aga ccc gcg ccg acg ata ttg taa taa gtg gtg gga tct cct ttt aca act tct tcg gat acc tca tca tct tcg gat acc tca tca tct tcg gtc tct gta act tcc gtt acg gat tga caa atc tta tca ttg gtc ggt gtt tgg tct tgc ttt gtg act ttg ata ata aca tcg att ccc ata tga tgt ttg ttt tct tct tca gta cac gag gat gag gat tgt tga aga cta gta ggc ata gca gct gcc act agg cac atg cat gcc agg aca ata tat tgt ttc atg att gct att gat tga tta ctg ttc tag atg att cta ctt tct tac cat ata ata aat tag aat ata ttt tct act ttt acg aga aat taa tta ttg tat tta tta ttt atg ggt -aa aaa aaa ctt act ata agt ggg tgg gat tct ggg aat tag tga tca gtt tat gta tat cgc aac tag cgg gca tat ggc tat tga cat cga gaa cat tac cca tat gat aag aga ttg tat ccg ttt cgt agt ctt gag tat tgg tat tac tat ata gta tgt aga tgt cga cgc tag ata gac agt cgc cca cta gag tta ccg tct ctg aat gcg gca tga tag tat cat tct ttg ttt tcg tta act gtt tgg aag atg aat ctt tgt tgt tac att taa tct cga aat tca gag tac ata tct ttg aag tat tct gat atc tat ttt ctc ctg taa aga atc ctg aag ttg cta cat tat taa gga cag aga agt att ctg cac gaa aga cgg gat cac aat ctt tat gat tca tgg taa tag tta gtt ccg acg ttg aga tgg att cgc tga gac cgg tag tgg tcg tcc gag tac acg atg tgt cgt tga ctg gat aca ggt taa ttt cca cat cga tat agt taa atg tat tgc tgg tt- --a cgg gtt cgc att tat ctg tgg aag aga cgg tgt gag aat atg ttc cgg gac cac acg gag aac aga tga cgt ctc cgg tag acg tgt atc cgg ata ctc cgt atc cta ttc cac act ttg ttt tag aaa tac atg ttc tac acc ctg atg atc ctt tga gaa gac aat aat atc ctg gag agc att cac aga ttc tat tgt gag tcg tgt tac acg atc gcg tct cta cct gat tac tat cac atc ttc cgt tac aac tta gac aag cct gta aat gat tat tgt gag atg taa agg tat ccg aac cac acg gcg tac att gtg tat tag tct tgc tat cac ata atc tgg aag cgt aag ttc ccg gag gac acg ata gac aac ata gat tac ggc ttc tgt att cgt tgt ctt tac att ttc cat tgg atg gtg cat gtg gtg cta tat ctc ttc cgt tta tta tta tac atg aga gaa aca ata tat acg agt ata ata cgg acc tca tga ttt aat aat -gt agt aat cgt cgt ctt gtt act gtt tgt ttc cta ctt ctc caa tca tat aga tta ttt ttt aaa tat ttt ctt tct atc atg gat aat att tgt aat ggt tct ttc cgt aca aca tac tgt tta gat ggt agt cgc tta gct tgg tta tga tat tgc gca taa ttt ccg gag gca aat acg ata gtc tag att gac tat cga tgg tag act cta att tat tga gtg ctt tgt cga cga gtt tac ttt tat gct cca tcg ata gat gac act gtt cta tga gat cgt cgt aca tgg gaa atg aaa tgc gtt tgt ctg aat gta tgg ctt cga gat agg tgt gat acc gga tgt ctt ctg ttc tca ata ccg tat aca agt tgg tgt ctg aga ttc gaa tct ctt tga gga gac tta tgt cac gac tac att ttt cga tga tgg aat cta tct tat cga atg ata tat ttt tca taa ata cac ttt tat agt cct cgt tta aac aga att tag tat gta gtt ccg caa atg act cgt ccc tta ata tgc agt agg cta tta tct tct tta cgt agt gat cgt cgt agg gag aga act ccg aca tct tgt aga aca acg att taa tca tag gta gag ata ctt tca gtc tgt ggt gga tga tgt cat taa cga cat ccg cct tgt ata tga tgt ttc tgt ttt caa aca cca agt cga ata ctg tct tat cgt ctt tag tcg gaa ggt tga tgt cgt atc cga tgt ata cga ggt atg agg caa cat tgt tat tgc aat tct gga agg cgg tat gaa gag gag tca ttg tat tat agt att cgt ctt tct gaa tgt cga atc tat cta gta gat acc gta gta tat tga gag agc gac ttc cat atc ctt gat tat gtt tta tga ata gat aaa gta gat gtt gtc ctt ctt cct ttt gta att tcc cgt att ttt gtt cgt gcc aat tga gta aca tta tga gaa tat gac ctg ttg cac aat cgt tct tta tgt att cca tga tgg gtg tac aat caa gat tat tac gta tcc tcg tat cgg ctc ctc gag ata aaa gag cat aca cca cac gag gac tat gtt tgg tat act gtt gaa ggt aag tgt gta acg gcg tat ttc cga ttt tcg taa ccg cgt taa tgt ttg ctc cat gat cta tta tcg cgt aga tga atc gct tct cag ctc gca tct tag tgt gac tct ttg act tgt aat aat tgc ttt cgt gga aca cgg ata tgt gtt tac agt agt aat gaa gag aag tga gtc cat cct cat cga cgc aat tag ggt cag atc ctt tag tca ata att tgt aca gaa cgt aat agt tta agc tcc cat tga att tat atc taa gat aac aca gca ata gat cgg atg att tac taa agt cat caa tgg ggt ccg tta gta tat caa aga tct tgt tat cga ttg ata gtg aat gaa tca gat agt ggt gta gag gaa tat gtc ctt ttt tat cct tgc tat caa agt tac gca tgc cgt ggc gta aca ata tct tta ata cag atg gat taa atc gtg tat tca tcg tat agc aat gta atg gag agt tac cac att tta gtc gtt tat tca gat cgc agt gtt taa taa cta att taa aca gat gag atg ata tat cca cat caa aga atg cga gat aca tat gac aga cat tat tga cag aaa tgt gac ctt cat tat cac cgt cgt cca taa atg cgt tag gta cgt acc aca tac tat cgt taa cga tgc gca caa tct cgt cca ttt tca tcc atc ttc ata atg att tac ttt ttc ata att aga gaa aaa gat caa ggt ata aaa att aga agt gtt aga cta ta- --- --- --- --- --- --- --- --- --- --- -aa cta act tat aa- --- --- --- --- --- --- --- --- --- --- --- --- --- --- --- --- --- --- --- --- --- --- --- --- --- --- --- --- --- --- --- --- --- --- --- --- --- --- --- --- --- --- --- --- --- --- --- --- --- --- --- --- --- --- --- --- --- --- --- --- --- --- --- --- --- --- --- --- --- --- --- --- --- --- --- --- --- --- --- --- --- --- --- --- --- --- -aa cta act tat gac tta act aac tta tga cta tta act cat ttc aag aaa ggt ggg tgg aga gaa ctc tat atg aca gct tgt gaa aca att aga tcc cta att tct aat gga agt ttt gat agg aga ttg tca tca gtt gat aca ttg ttt att atc tca tct att aga gca cgt ctg ttt aga gct tta gtg acc tgc tcg gtt act tct gtg taa atc ttg aat cct tta gtg ata cac tgt gtc aaa act gga tgt tta gaa tac cta tgt aga ata tgg gaa gca tgc ttg ttt ttg tct cta tta tag att aac tca tac atg gtt gta tta tga att ttc atc tgc cta atg tac tcc aat tct tgt tta caa tca att ata taa tca aag agt gat gat gca tac aca tta caa agt gaa taa tct acc atc ata aaa tac ttg ata cag agc ttt atc aca tca tgg ttt tca att gta tta tta agt ata gct aat ttt ata cag tca ata gac aat ggt tct cta agc aat att tct aat att tta aga tgt gct tcc cta cgg gcg atg aca gat ccc cta tcc acg gcc acg tca aga cat gta tat cca tta ctc att act gcg ttg aca ttt gct cca ttt tct aat agc cat gat act aaa tct ata taa cct gca tag ata gcg cga taa agc aag gtc ctt cca cca gca tct agt tga ttg ata tct tca ata tat ggg ata caa agc tta taa att tct aat act gtg ggt tca tct aca agg aat ccc cta gta tac tga att att tta tat aga tct aat tta aca tca ttt tca tct ggg ata cca caa ttc aaa ata aac tca aca aca cta ctt tcc ttt tta cat att ccc cta aaa tag gca ttc aag cat tct att tta tat att aca gcc cca tga tct acc ata aaa tca aca atg tct att tct aca tat gca tta gat aga tag taa agt aag aga tct tgt aca gaa tta caa ttc tta ata att ata gag aaa ata tct tcc ata taa ttc ttt gac act aat gca gat ata ata tct tta tat gta ata tat gca aac agt cta tct act ata tac tga tca ata tta tct cta tga atc cta aaa taa tca tac aga aca tct aca gga tca caa att ggt tca agg aga aat cta tca aat att ttc ctg tca aca act ggt tct aga aca taa cag tca aca cct aat cca tgt ttt tta tag tca tct acc aaa gat aat gac caa aga tcg agg tcg tcg tga aac tgc tca tcg aca gcc atg aaa tct gcc gac tcc atg gcg cga atc gca ctg tct tat tcg cca ttg att ttc a-t ttt tta taa tta tgt aca tgt ttt cct tct att ctc aag agt cta caa aaa tat a-t ttt ttt tcg ata tct aag tac taa gtt ttt tta ctg ttt ttg tta ctg tct tcc att ctt cta act aaa gat ctg aga taa att ata caa tct tcg cta tcg aac cat ttt tgt agt cta aag cct gaa gta att aac caa ctg ttt tta tta gtg gct ttt ttc gat cta tcc tcg tc- --- --c tca tca tcc tta ta- --- --- --- --- --- --- tta tta tca tta tta tca tag tct att aaa cac aaa tca tct --- --- --- --- --- --- --- --- --- --- --- --- --- --- --- --- --- --- --- --- --- --- --- --- --- --- --a cgt tta taa c-- --- --- --- --- --- --- --- --a aca ttc --- --- --- --- -tc att att aat tag ttc tgt agt aat atc ttt aat aat ttg gct ata --- --- -ca tct gtt --- --- --- --- --- --- --- --- --- --- --- --- --- --- --- --- --- --- --c aat act --- --- --- --- --- --- --- --a tct att gat gat ttc ttt t-- --- --- --- --- --- --- --- --- --- --- --t taa gac t-- --- --- --- --- --- --- --- --- --- --- --- --- --- --- --- --- --- --- --- --- --- --- --- --- --- --- --- --- --- --- --- --- --- --- --- --- --- --- --- taa act agt --- --- --- --- --- --- --- --- --- --- --- --- --- --- --- --- --- --- --- --- --- --- --- --- --- --- --- --- --- --- --- --- --- --- --- --- --- --- --- --- --- --- --- --- --- --- --- --- --- --- --- --- --- --- --- --- --- --- --- --- --- --- --- --- --- --- --- --- --- --- --- --- --- --- --- --- --- --- --- --- --- --- --- --- --- --- --- --- --- --- --- --- --- --- --- --- --- --- --- --- --- --- --- --- --- --- --- --- --- --t atg gta atg acg atg aaa --- -tc gag tag ta- --- --a ctt cta ata aag act tga ta- --- --t cat tat cat atg ttt gat cg- --- --- --- --- --- --- --t cat agt taa tag tgt g-- --- --- --- --- --- --- --- --- --- --- --- --- --- --- --- --- --- --- --- -gc taa atg gta ctg tta ata agt tta t-- --- --- --- --- --- --- --- --- --- --- --- --- --- --- --- --- --a gac aat atc ata gta ttt tct ttc cag aat --- --- --- --t aga tta ttt ttt taa ata ctg atc ctc aca att ccg tga tgt agc agt agt tgg t-- --- --- --- --- --- --- --- --- --- --- --- --- --- --- --- --- --- --- --- --- --- --- --- --- --- --- --- -gc atg gtc tat atc gt- --- --- --- --- --- --- --- --- --- --- --- --- --- --- --- --- --- --- --- --- --- --- --- --- --- --- --- --- --- --- --- --- --- --- --- --- --- --- --- --- --- --- --- taa aat gta tca tat ata ata gtt ttc tga cgt gga gta cag aat ttt cga --- --- --- --- --- --- --- --- --- --- --- --- --- --- --- --- --- --- --- --- --- --- --- --- --- --- --- --- --- --- --- --- --- --- --- --- --- --- --- --- --- --- --- --t taa tga gtt cat ggt aag gaa ggg caa atg tct --- gta tat aat ata cat aag tta a-- --- --- --- --- --- --- --- --- --- --- --- --- --t agt ttt tta tca tat tt- --- --- --- --- --- --- --- --t cta ata cca taa taa aaa tta tca tta tg- --- --- -ta taa tca --- --- --- --- --- --- --- --- --- --- --- --t cac tg- --- -tc gct atc att att gcg ttt gtg tag tt- --- --- --- --- ctg ccc ta- --- --- --- --- --- --- --- --t cat cta cat cac tgt ca- --- --- --- --- ctc tca cta tat ctt cta aaa tta caa --- -ac aac --- --- --- --- --- --- --- --- --- --- --- --- --- --- --- --- --- --- --- --- --- --- --- --- --- --- --- --- --- --- --- --- --- --- --- --- tgg ata ttc gat --- --- --- --- --- --- --- --- --- --- --- --- --- --- --- --- --- --- --- --- --- --- --- --- --- --- --a aca gca ttt gtg t-- --- --- --- --- --- --- --- --- --- --- --- --- --- --- --- --- -

>MN702453.1_Monkeypox_virus_strain_A1_contig_SPADES

--- --- --- --- --- --- --- --- --- --- --- --- --- --- --- --- --- --- --- --- --- --- --- --- --- --- --- --- --- --- --- --- --- --- --- --- --- --- --- --- --- --- --- --- --- --- --- --- --- --- --- --- --- --- --- --- --- --- --- --- --- --- --- --- --- --- --- --- --- --- --- --- --- --- --- --- --- --- --- --- --- --- --- --- --- --- --- --- --- --- --- --- --- --- --- --- --- --- --- --- --- --- --- --- --- --- --- --- --- --- --- --- --- --- --- --- --- --- --- --- --- --- --- --- --- --- --- --- --- --- --- --- --- --- --- --- --- --- --- --- --- --- --- --- --- --- --- --- --- --- --- --- --- --- --- --- --- --- --- --- --- --- --- --- --- --- --- --- --- --- --- --- --- --- --- --- --- --- --- --- --- --- --- --- --- --- --- --- --- --- --- --- --- --- --- --- --- --- --- --- --- --- --- --- --- --- --- --- --- --- --- --- --- --- --- --- --- --- --- --- --- --- --- --- --- --- --- --- --- --- --- --- --- --- --- --- --- --- --- --- --- --- --- --- --- --- --- --- --- --- --- --- --- --- --- --- --- --- --- --- --- --- --- --- --- --- --- --- --- --- --- --- --- --- --- --- --- --- --- --- --- --- --- --- --- --- --- --- --- --- --- --- --- --- --- --- --- --- --- --- --- --- --- --- --- --- --- --- --- --- --- --- --- --- --- --- --- --- --- --- --- --- --- --- --- --- --- --- --- --- --- --- --- --- --- --- --- --- --- --- --- --- --- --- --- --- --- --- --- --- --- --c cat cga aga gag aaa gga taa aaa ctt ttt acg act cca tca gaa aga ggt tta ata ttt ttg tga gac cca tcg acg aga gaa aga gat ggt tag tca aga tat ttt tct tag tac aaa agt caa tgt ttt aaa ata tat gga cga gaa tta att tgt ctg tat aaa aac ttg tgt gaa att atg tac tag aga aaa aac gtg agc agt gtc ccc tac atg gat ttt aca gat cat tta tat tcc aaa aat att aac tat ata cgt tta tta tat gat gtt aac gtg taa att ata aac att att tta tga tgc aat tgt ctg aca acc --- att ggc ata agg ata ttg ata agc tct acg aga ata tat tgt tgg acg tta tcg ttt acg aaa tag ttg aga cat cag aaa gag gtt taa tat ttt tgt gag a-c cat cga aga gag aaa gag aat aaa aat a-- --- --- --- ttt ttt tgt aaa act ttt tta tga gac aag aga gaa aga gaa tac gaa tag tga tca tat cgt atc aca tat tga aac aga aag aag aag taa cga gag gta act ttt tgt gaa tgt agt taa aca ttt ttg ttt tgc aaa ccg gaa tat agt acc cgg tac act ttt tta att cgt ggt gcg gtg tct gaa tcg ttc gat taa ccc aac tca tcc att ttc aga tga ata gag tta tcg att cag aca cat gct ttg agt ttt gtt gaa tcg atg agt gaa gta tca tcg gtt gca cct tca gat gcc gat cca tcg aca tac ttg aat cca tcc ttg act tca agt tca gat gat tcc tca cac atg tct ccg ata cgt acg cta aac tct agg ttc tta aca cat ttt gta tca acg atc gtt gaa ccg atg ata tct ttg taa ctc act ttc tta tgt gag atg tta gac cca agt act gga tgg gtc ttg atg tcg ctg tct ttc tct tct tcg cta cat ctg atg tcg ata gac atc tca cag tct ttg atc ata gcc aga gct tct tca cgc gtg atc gcg gga gag tcc tta cct tgt cct ggt gac acg ctg gac aat cta gta ttc aca gtg ttt cca tca gag gat tcg gag atg gat gaa atc ttt ggg cat ttg gtg aat cca aag ttc atg tta aga ccc gcg ccg acg ata ttg taa taa gtg gtg gga tct cct ttt aca act tct tcg gat acc tca tca tct tcg gat acc tca tca tct tcg gtc tct gta act tcc gtt acg gat tga caa atc tta tca ttg gtc ggt gtt tgg tct tgc ttt gtg act ttg ata ata aca tcg att ccc ata tga tgt ttg ttt tct tct tca gta cac gag gat gag gat tgt tga aga cta gta ggc ata gca gct gcc act agg cac atg cat gcc agg aca ata tat tgt ttc atg att gct att gat tga tta ctg ttc tag atg att cta ctt tct tac cat ata ata aat tag aat ata ttt tct act ttt acg aga aat taa tta ttg tat tta tta ttt atg ggt -aa aaa aaa ctt act ata agt ggg tgg gat tct ggg aat tag tga tca gtt tat gta tat cgc aac tag cgg gca tat ggc tat tga cat cga gaa cat tac cca tat gat aag aga ttg tat ccg ttt cgt agt ctt gag tat tgg tat tac tat ata gta tgt aga tgt cga cgc tag ata gac agt cgc cca cta gag tta ccg tct ctg aat gcg gca tga tag tat cat tct ttg ttt tcg tta act gtt tgg aag atg aat ctt tgt tgt tac att taa tct cga aat tca gag tac ata tct ttg aag tat tct gat atc tat ttt ctc ctg taa aga atc ctg aag ttg cta cat tat taa gga cag aga agt att ctg cac gaa aga cgg gat cac aat ctt tat gat tca tgg taa tag tta gtt ccg acg ttg aga tgg att cgc tga gac cgg tag tgg tcg tcc gag tac acg atg tgt cgt tga ctg gat aca ggt taa ttt cca cat cga tat agt taa atg tat tgc tgg tt- --a cgg gtt cgc att tat ctg tgg aag aga cgg tgt gag aat atg ttc cgg gac cac acg gag aac aga tga cgt ctc cgg tag acg tgt atc cgg ata ctc cgt atc cta ttc cac act ttg ttt tag aaa tac atg ttc tac acc ctg atg atc ctt tga gaa gac aat aat atc ctg gag agc att cac aga ttc tat tgt gag tcg tgt tac acg atc gcg tct cta cct gat tac tat cac atc ttc cgt tac aac tta gac aag cct gta aat gat tat tgt gag atg taa agg tat ccg aac cac acg gcg tac att gtg tat tag tct tgc tat cac ata atc tgg aag cgt aag ttc ccg gag gac acg ata gac aac ata gat tac ggc ttc tgt att cgt tgt ctt tac att ttc cat tgg atg gtg cat gtg gtg cta tat ctc ttc cgt tta tta tta tac atg aga gaa aca ata tat acg agt ata ata cgg acc tca tga ttt aat aat -gt agt aat cgt cgt ctt gtt act gtt tgt ttc cta ctt ctc caa tca tat aga tta ttt ttt aaa tat ttt ctt tct atc atg gat aat att tgt aat ggt tct ttc cgt aca aca tac tgt tta gat ggt agt cgc tta gct tgg tta tga tat tgc gca taa ttt ccg gag gca aat acg ata gtc tag att gac tat cga tgg tag act cta att tat tga gtg ctt tgt cga cga gtt tac ttt tat gct cca tcg ata gat gac act gtt cta tga gat cgt cgt aca tgg gaa atg aaa tgc gtt tgt ctg aat gta tgg ctt cga gat agg tgt gat acc gga tgt ctt ctg ttc tca ata ccg tat aca agt tgg tgt ctg aga ttc gaa tct ctt tga gga gac tta tgt cac gac tac att ttt cga tga tgg aat cta tct tat cga atg ata tat ttt tca taa ata cac ttt tat agt cct cgt tta aac aga att tag tat gta gtt ccg caa atg act cgt ccc tta ata tgc agt agg cta tta tct tct tta cgt agt gat cgt cgt agg gag aga act ccg aca tct tgt aga aca acg att taa tca tag gta gag ata ctt tca gtc tgt ggt gga tga tgt cat taa cga cat ccg cct tgt ata tga tgt ttc tgt ttt caa aca cca agt cga ata ctg tct tat cgt ctt tag tcg gaa ggt tga tgt cgt atc cga tgt ata cga ggt atg agg caa cat tgt tat tgc aat tct gga agg cgg tat gaa gag gag tca ttg tat tat agt att cgt ctt tct gaa tgt cga atc tat cta gta gat acc gta gta tat tga gag agc gac ttc cat atc ctt gat tat gtt tta tga ata gat aaa gta gat gtt gtc ctt ctt cct ttt gta att tcc cgt att ttt gtt cgt gcc aat tga gta aca tta tga gaa tat gac ctg ttg cac aat cgt tct tta tgt att cca tga tgg gtg tac aat caa gat tat tac gta tcc tcg tat cgg ctc ctc gag ata aaa gag cat aca cca cac gag gac tat gtt tgg tat act gtt gaa ggt aag tgt gta acg gcg tat ttc cga ttt tcg taa ccg cgt taa tgt ttg ctc cat gat cta tta tcg cgt aga tga atc gct tct cag ctc gca tct tag tgt gac tct ttg act tgt aat aat tgc ttt cgt gga aca cgg ata tgt gtt tac agt agt aat gaa gag aag tga gtc cat cct cat cga cgc aat tag ggt cag atc ctt tag tca ata att tgt aca gaa cgt aat agt tta agc tcc cat tga att tat atc taa gat aac aca gca ata gat cgg atg att tac taa agt cat caa tgg ggt ccg tta gta tat caa aga tct tgt tat cga ttg ata gtg aat gaa tca gat agt ggt gta gag gaa tat gtc ctt ttt tat cct tgc tat caa agt tac gca tgc cgt ggc gta aca ata tct tta ata cag atg gat taa atc gtg tat tca tcg tat agc aat gta atg gag agt tac cac att tta gtc gtt tat tca gat cgc agt gtt taa taa cta att taa aca gat gag atg ata tat cca cat caa aga atg cga gat aca tat gac aga cat tat tga cag aaa tgt gac ctt cat tat cac cgt cgt cca taa atg cgt tag gta cgt acc aca tac tat cgt taa cga tgc gta caa tct cgt cca ttt tca tcc atc ttc ata atg att tac ttt ttc ata att aga gaa aaa gat caa ggt ata aaa att aga agt gtt aga cta ta- --- --- --- --- --- --- --- --- --- --- -aa cta act tat aa- --- --- --- --- --- --- --- --- --- --- --- --- --- --- --- --- --- --- --- --- --- --- --- --- --- --- --- --- --- --- --- --- --- --- --- --- --- --- --- --- --- --- --- --- --- --- --- --- --- --- --- --- --- --- --- --- --- --- --- --- --- --- --- --- --- --- --- --- --- --- --- --- --- --- --- --- --- --- --- --- --- --- --- --- --- --- -aa cta act tat gac tta act aac tta tga cta tta act cat ttc aag aaa ggt ggg tgg aga gaa ctc tat atg aca gct tgt gaa aca att aga tcc cta att tct aat gga agt ttt gat agg aga ttg tca tca gtt gat aca ttg ttt att atc tca tct att aga gca cgt ctg ttt aga gct tta gtg acc tgc tcg gtt act tct gtg taa atc ttg aat cct tta gtg ata cac tgt gtc aaa act gga tgt tta gaa tac cta tgt aga ata tgg gaa gca tgc ttg ttt ttg tct cta tta tag att aac tca tac atg gtt gta tta tga att ttc atc tgc cta atg tac tcc aat tct tgt tta caa tca att ata taa tca aag agt gat gat gca tac aca tta caa agt gaa taa tct acc atc ata aaa tac ttg ata cag agc ttt atc aca tca tgg ttt tca att gta tta tta agt ata gct aat ttt ata cag tca ata gac aat ggt tct cta agc aat att tct aat att tta aga tgt gct tcc cta cgg gcg atg aca gat ccc cta tcc acg gcc acg tca aga cat gta tat cca tta ctc att act gcg ttg aca ttt gct cca ttt tct aat agc cat gat act aaa tct ata taa cct gca tag ata gcg cga taa agc aag gtc ctt cca cca gca tct agt tga ttg ata tct tca ata tat ggg ata caa agc tta taa att tct aat act gtg ggt tca tct aca agg aat ccc cta gta tac tga att att tta tat aga tct aat tta aca tca ttt tca tct ggg ata cca caa ttc aaa ata aac tca aca aca cta ctt tcc ttt tta cat att ccc cta aaa tag gca ttc aag cat tct att tta tat att aca gcc cca tga tct acc ata aaa tca aca atg tct att tct aca tat gca tta gat aga tag taa agt aag aga tct tgt aca gaa tta caa ttc tta ata att ata gag aaa ata tct tcc ata taa ttc ttt gac act aat gca gat ata ata tct tta tat gta ata tat gca aac agt cta tct act ata tac tga tca ata tta tct cta tga atc cta aaa taa tca tac aga aca tct aca gga tca caa att ggt tca agg aga aat cta tca aat att ttc ctg tca aca act ggt tct aga aca taa cag tca aca cct aat cca tgt ttt tta tag tca tct acc aaa gat aat gac caa aga tcg agg tcg tcg tga aac tgc tca tcg aca gcc atg aaa tct gcc gac tcc atg gcg cga atc gca ctg tct tat tcg cca ttg att ttc a-t ttt tta taa tta tgt aca tgt ttt cct tct att ctc aag agt cta caa aaa tat a-t ttt ttt tcg ata tct aag tac taa gtt ttt tta ctg ttt ttg tta ctg tct tcc att ctt cta act aaa gat ctg aga taa att ata caa tct tcg cta tcg aac cat ttt tgt agt cta aag cct gaa gta att aac caa ctg ttt tta tta gtg gct ttt ttc gat cta tcc tcg tc- --- --c tca tca tcc tta ta- --- --- --- --- --- --- tta tta tca tta tta tca tag tct att aaa cac aaa tca tct --- --- --- --- --- --- --- --- --- --- --- --- --- --- --- --- --- --- --- --- --- --- --- --- --- --- --a cgt tta taa c-- --- --- --- --- --- --- --- --a aca ttc --- --- --- --- -tc att att aat tag ttc tgt agt aat atc ttt aat aat ttg gct ata --- --- -ca tct gtt --- --- --- --- --- --- --- --- --- --- --- --- --- --- --- --- --- --- --c aat act --- --- --- --- --- --- --- --a tct att gat gat ttc ttt t-- --- --- --- --- --- --- --- --- --- --- --t taa gac t-- --- --- --- --- --- --- --- --- --- --- --- --- --- --- --- --- --- --- --- --- --- --- --- --- --- --- --- --- --- --- --- --- --- --- --- --- --- --- --- taa act agt --- --- --- --- --- --- --- --- --- --- --- --- --- --- --- --- --- --- --- --- --- --- --- --- --- --- --- --- --- --- --- --- --- --- --- --- --- --- --- --- --- --- --- --- --- --- --- --- --- --- --- --- --- --- --- --- --- --- --- --- --- --- --- --- --- --- --- --- --- --- --- --- --- --- --- --- --- --- --- --- --- --- --- --- --- --- --- --- --- --- --- --- --- --- --- --- --- --- --- --- --- --- --- --- --- --- --- --- --- --t atg gta atg acg atg aaa --- -tc gag tag ta- --- --a ctt cta ata aag act tga ta- --- --t cat tat cat atg ttt gat cg- --- --- --- --- --- --- --t cat agt taa tag tgt g-- --- --- --- --- --- --- --- --- --- --- --- --- --- --- --- --- --- --- --- -gc taa atg gta ctg tta ata agt tta t-- --- --- --- --- --- --- --- --- --- --- --- --- --- --- --- --- --a gac aat atc ata gta ttt tct ttc cag aat --- --- --- --t aga tta ttt ttt taa ata ctg atc ctc aca att ccg tga tgt agc agt agt tgg t-- --- --- --- --- --- --- --- --- --- --- --- --- --- --- --- --- --- --- --- --- --- --- --- --- --- --- --- -gc atg gtc tat atc gt- --- --- --- --- --- --- --- --- --- --- --- --- --- --- --- --- --- --- --- --- --- --- --- --- --- --- --- --- --- --- --- --- --- --- --- --- --- --- --- --- --- --- --- taa aat gta tca tat ata ata gtt ttc tga cgt gga gta cag aat ttt cga --- --- --- --- --- --- --- --- --- --- --- --- --- --- --- --- --- --- --- --- --- --- --- --- --- --- --- --- --- --- --- --- --- --- --- --- --- --- --- --- --- --- --- --t taa tga gtt cat ggt aag gaa ggg caa atg tct --- gta tat aat ata cat aag tta a-- --- --- --- --- --- --- --- --- --- --- --- --- --t agt ttt tta tca tat tt- --- --- --- --- --- --- --- --t cta ata cca taa taa aaa tta tca tta tg- --- --- -ta taa tca --- --- --- --- --- --- --- --- --- --- --- --t cac tg- --- -tc gct atc att att gcg ttt gtg tag tt- --- --- --- --- ctg ccc ta- --- --- --- --- --- --- --- --t cat cta cat cac tgt ca- --- --- --- --- ctc tca cta tat ctt cta aaa tta caa --- -ac aac --- --- --- --- --- --- --- --- --- --- --- --- --- --- --- --- --- --- --- --- --- --- --- --- --- --- --- --- --- --- --- --- --- --- --- --- tgg ata ttc gat --- --- --- --- --- --- --- --- --- --- --- --- --- --- --- --- --- --- --- --- --- --- --- --- --- --- --a aca gca ttt gtg t-- --- --- --- --- --- --- --- --- --- --- --- --- --- --- --- --- -

>MN702452.1_Monkeypox_virus_strain_A2_contig_SPADES

--- --- --- --- --- --- --- --- --- --- --- --- --- --- --- --- --- --- --- --- --- --- --- --- --- --- --- --- --- --- --- --- --- --- --- --- --- --- --- --- --- --- --- --- --- --- --- --- --- --- --- --- --- --- --- --- --- --- --- --- --- --- --- --- --- --- --- --- --- --- --- --- --- --- --- --- --- --- --- --- --- --- --- --- --- --- --- --- --- --- --- --- --- --- --- --- --- --- --- --- --- --- --- --- --- --- --- --- --- --- --- --- --- --- --- --- --- --- --- --- --- --- --- --- --- --- --- --- --- --- --- --- --- --- --- --- --- --- --- --- --- --- --- --- --- --- --- --- --- --- --- --- --- --- --- --- --- --- --- --- --- --- --- --- --- --- --- --- --- --- --- --- --- --- --- --- --- --- --- --- --- --- --- --- --- --- --- --- --- --- --- --- --- --- --- --- --- --- --- --- --- --- --- --- --- --- --- --- --- --- --- --- --- --- --- --- --- --- --- --- --- --- --- --- --- --- --- --- --- --- --- --- --- --- --- --- --- --- --- --- --- --- --- --- --- --- --- --- --- --- --- --- --- --- --- --- --- --- --- --- --- --- --- --- --- --- --- --- --- --- --- --- --- --- --- --- --- --- --- --- --- --- --- --- --- --- --- --- --- --- --- --- --- --- --- --- --- --- --- --- --- --- --- --- --- --- --- --- --- --- --- --- --- --- --- --- --- --- --- --- --- --- --- --- --- --- --- --- --- --- --- --- --- --- --- --- --- --- --- --- --- --- --- --- --- --- --- --- --- --- --- --c cat cga aga gag aaa gga taa aaa ctt ttt acg act cca tca gaa aga ggt tta ata ttt ttg tga gac cca tcg acg aga gaa aga gat ggt tag tca aga tat ttt tct tag tac aaa agt caa tgt ttt aaa ata tat gga cga gaa tta att tgt ctg tat aaa aac ttg tgt gaa att atg tac tag aga aaa aac gtg agc agt gtc ccc tac atg gat ttt aca gat cat tta tat tcc aaa aat att aac tat ata cgt tta tta tat gat gtt aac gtg taa att ata aac att att tta tga tgc aat tgt ctg aca acc --- att ggc ata agg ata ttg ata agc tct acg aga ata tat tgt tgg acg tta tcg ttt acg aaa tag ttg aga cat cag aaa gag gtt taa tat ttt tgt gag a-c cat cga aga gag aaa gag aat aaa aat a-- --- --- --- ttt ttt tgt aaa act ttt tta tga gac aag aga gaa aga gaa tac gaa tag tga tca tat cgt atc aca tat tga aac aga aag aag aag taa cga gag gta act ttt tgt gaa tgt agt taa aca ttt ttg ttt tgc aaa ccg gaa tat agt acc cgg tac act ttt tta att cgt ggt gcg gtg tct gaa tcg ttc gat taa ccc aac tca tcc att ttc aga tga ata gag tta tcg att cag aca cat gct ttg agt ttt gtt gaa tcg atg agt gaa gta tca tcg gtt gca cct tca gat gcc gat cca tcg aca tac ttg aat cca tcc ttg act tca agt tca gat gat tcc tca cac atg tct ccg ata cgt acg cta aac tct agg ttc tta aca cat ttt gta tca acg atc gtt gaa ccg atg ata tct ttg taa ctc act ttc tta tgt gag atg tta gac cca agt act gga tgg gtc ttg atg tcg ctg tct ttc tct tct tcg cta cat ctg atg tcg ata gac atc tca cag tct ttg atc ata gcc aga gct tct tca cgc gtg atc gcg gga gag tcc tta cct tgt cct ggt gac acg ctg gac aat cta gta ttc aca gtg ttt cca tca gag gat tcg gag atg gat gaa atc ttt ggg cat ttg gtg aat cca aag ttc atg tta aga ccc gcg ccg acg ata ttg taa taa gtg gtg gga tct cct ttt aca act tct tcg gat acc tca tca tct tcg gat acc tca tca tct tcg gtc tct gta act tcc gtt acg gat tga caa atc tta tca ttg gtc ggt gtt tgg tct tgc ttt gtg act ttg ata ata aca tcg att ccc ata tga tgt ttg ttt tct tct tca gta cac gag gat gag gat tgt tga aga cta gta ggc ata gca gct gcc act agg cac atg cat gcc agg aca ata tat tgt ttc atg att gct att gat tga tta ctg ttc tag atg att cta ctt tct tac cat ata ata aat tag aat ata ttt tct act ttt acg aga aat taa tta ttg tat tta tta ttt atg ggt -aa aaa aaa ctt act ata agt ggg tgg gat tct ggg aat tag tga tca gtt tat gta tat cgc aac tag cgg gca tat ggc tat tga cat cga gaa cat tac cca tat gat aag aga ttg tat ccg ttt cgt agt ctt gag tat tgg tat tac tat ata gta tgt aga tgt cga cgc tag ata gac agt cgc cca cta gag tta ccg tct ctg aat gcg gca tga tag tat cat tct ttg ttt tcg tta act gtt tgg aag atg aat ctt tgt tgt tac att taa tct cga aat tca gag tac ata tct ttg aag tat tct gat atc tat ttt ctc ctg taa aga atc ctg aag ttg cta cat tat taa gga cag aga agt att ctg cac gaa aga cgg gat cac aat ctt tat gat tca tgg taa tag tta gtt ccg acg ttg aga tgg att cgc tga gac cgg tag tgg tcg tcc gag tac acg atg tgt cgt tga ctg gat aca ggt taa ttt cca cat cga tat agt taa atg tat tgc tgg tt- --a cgg gtt cgc att tat ctg tgg aag aga cgg tgt gag aat atg ttc cgg gac cac acg gag aac aga tga cgt ctc cgg tag acg tgt atc cgg ata ctc cgt atc cta ttc cac act ttg ttt tag aaa tac atg ttc tac acc ctg atg atc ctt tga gaa gac aat aat atc ctg gag agc att cac aga ttc tat tgt gag tcg tgt tac acg atc gcg tct cta cct gat tac tat cac atc ttc cgt tac aac tta gac aag cct gta aat gat tat tgt gag atg taa agg tat ccg aac cac acg gcg tac att gtg tat tag tct tgc tat cac ata atc tgg aag cgt aag ttc ccg gag gac acg ata gac aac ata gat tac ggc ttc tgt att cgt tgt ctt tac att ttc cat tgg atg gtg cat gtg gtg cta tat ctc ttc cgt tta tta tta tac atg aga gaa aca ata tat acg agt ata ata cgg acc tca tga ttt aat aat -gt agt aat cgt cgt ctt gtt act gtt tgt ttc cta ctt ctc caa tca tat aga tta ttt ttt aaa tat ttt ctt tct atc atg gat aat att tgt aat ggt tct ttc cgt aca aca tac tgt tta gat ggt agt cgc tta gct tgg tta tga tat tgc gca taa ttt ccg gag gca aat acg ata gtc tag att gac tat cga tgg tag act cta att tat tga gtg ctt tgt cga cga gtt tac ttt tat gct cca tcg ata gat gac act gtt cta tga gat cgt cgt aca tgg gaa atg aaa tgc gtt tgt ctg aat gta tgg ctt cga gat agg tgt gat acc gga tgt ctt ctg ttc tca ata ccg tat aca agt tgg tgt ctg aga ttc gaa tct ctt tga gga gac tta tgt cac gac tac att ttt cga tga tgg aat cta tct tat cga atg ata tat ttt tca taa ata cac ttt tat agt cct cgt tta aac aga att tag tat gta gtt ccg caa atg act cgt ccc tta ata tgc agt agg cta tta tct tct tta cgt agt gat cgt cgt agg gag aga act ccg aca tct tgt aga aca acg att taa tca tag gta gag ata ctt tca gtc tgt ggt gga tga tgt cat taa cga cat ccg cct tgt ata tga tgt ttc tgt ttt caa aca cca agt cga ata ctg tct tat cgt ctt tag tcg gaa ggt tga tgt cgt atc cga tgt ata cga ggt atg agg caa cat tgt tat tgc aat tct gga agg cgg tat gaa gag gag tca ttg tat tat agt att cgt ctt tct gaa tgt cga atc tat cta gta gat acc gta gta tat tga gag agc gac ttc cat atc ctt gat tat gtt tta tga ata gat aaa gta gat gtt gtc ctt ctt cct ttt gta att tcc cgt att ttt gtt cgt gcc aat tga gta aca tta tga gaa tat gac ctg ttg cac aat cgt tct tta tgt att cca tga tgg gtg tac aat caa gat tat tac gta tcc tcg tat cgg ctc ctc gag ata aaa gag cat aca cca cac gag gac tat gtt tgg tat act gtt gaa ggt aag tgt gta acg gcg tat ttc cga ttt tcg taa ccg cgt taa tgt ttg ctc cat gat cta tta tcg cgt aga tga atc gct tct cag ctc gca tct tag tgt gac tct ttg act tgt aat aat tgc ttt cgt gga aca cgg ata tgt gtt tac agt agt aat gaa gag aag tga gtc cat cct cat cga cgc aat tag ggt cag atc ctt tag tca ata att tgt aca gaa cgt aat agt tta agc tcc cat tga att tat atc taa gat aac aca gca ata gat cgg atg att tac taa agt cat caa tgg ggt ccg tta gta tat caa aga tct tgt tat cga ttg ata gtg aat gaa tca gat agt ggt gta gag gaa tat gtc ctt ttt tat cct tgc tat caa agt tac gca tgc cgt ggc gta aca ata tct tta ata cag atg gat taa atc gtg tat tca tcg tat agc aat gta atg gag agt tac cac att tta gtc gtt tat tca gat cgc agt gtt taa taa cta att taa aca gat gag atg ata tat cca cat caa aga atg cga gat aca tat gac aga cat tat tga cag aaa tgt gac ctt cat tat cac cgt cgt cca taa atg cgt tag gta cgt acc aca tac tat cgt taa cga tgc gta caa tct cgt cca ttt tca tcc atc ttc ata atg att tac ttt ttc ata att aga gaa aaa gat caa ggt ata aaa att aga agt gtt aga cta ta- --- --- --- --- --- --- --- --- --- --- -aa cta act tat aa- --- --- --- --- --- --- --- --- --- --- --- --- --- --- --- --- --- --- --- --- --- --- --- --- --- --- --- --- --- --- --- --- --- --- --- --- --- --- --- --- --- --- --- --- --- --- --- --- --- --- --- --- --- --- --- --- --- --- --- --- --- --- --- --- --- --- --- --- --- --- --- --- --- --- --- --- --- --- --- --- --- --- --- --- --- --- -aa cta act tat gac tta act aac tta tga cta tta act cat ttc aag aaa ggt ggg tgg aga gaa ctc tat atg aca gct tgt gaa aca att aga tcc cta att tct aat gga agt ttt gat agg aga ttg tca tca gtt gat aca ttg ttt att atc tca tct att aga gca cgt ctg ttt aga gct tta gtg acc tgc tcg gtt act tct gtg taa atc ttg aat cct tta gtg ata cac tgt gtc aaa act gga tgt tta gaa tac cta tgt aga ata tgg gaa gca tgc ttg ttt ttg tct cta tta tag att aac tca tac atg gtt gta tta tga att ttc atc tgc cta atg tac tcc aat tct tgt tta caa tca att ata taa tca aag agt gat gat gca tac aca tta caa agt gaa taa tct acc atc ata aaa tac ttg ata cag agc ttt atc aca tca tgg ttt tca att gta tta tta agt ata gct aat ttt ata cag tca ata gac aat ggt tct cta agc aat att tct aat att tta aga tgt gct tcc cta cgg gcg atg aca gat ccc cta tcc acg gcc acg tca aga cat gta tat cca tta ctc att act gcg ttg aca ttt gct cca ttt tct aat agc cat gat act aaa tct ata taa cct gca tag ata gcg cga taa agc aag gtc ctt cca cca gca tct agt tga ttg ata tct tca ata tat ggg ata caa agc tta taa att tct aat act gtg ggt tca tct aca agg aat ccc cta gta tac tga att att tta tat aga tct aat tta aca tca ttt tca tct ggg ata cca caa ttc aaa ata aac tca aca aca cta ctt tcc ttt tta cat att ccc cta aaa tag gca ttc aag cat tct att tta tat att aca gcc cca tga tct acc ata aaa tca aca atg tct att tct aca tat gca tta gat aga tag taa agt aag aga tct tgt aca gaa tta caa ttc tta ata att ata gag aaa ata tct tcc ata taa ttc ttt gac act aat gca gat ata ata tct tta tat gta ata tat gca aac agt cta tct act ata tac tga tca ata tta tct cta tga atc cta aaa taa tca tac aga aca tct aca gga tca caa att ggt tca agg aga aat cta tca aat att ttc ctg tca aca act ggt tct aga aca taa cag tca aca cct aat cca tgt ttt tta tag tca tct acc aaa gat aat gac caa aga tcg agg tcg tcg tga aac tgc tca tcg aca gcc atg aaa tct gcc gac tcc atg gcg cga atc gca ctg tct tat tcg cca ttg att ttc a-t ttt tta taa tta tgt aca tgt ttt cct tct att ctc aag agt cta caa aaa tat a-t ttt ttt tcg ata tct aag tac taa gtt ttt tta ctg ttt ttg tta ctg tct tcc att ctt cta act aaa gat ctg aga taa att ata caa tct tcg cta tcg aac cat ttt tgt agt cta aag cct gaa gta att aac caa ctg ttt tta tta gtg gct ttt ttc gat cta tcc tcg tc- --- --c tca tca tcc tta ta- --- --- --- --- --- --- tta tta tca tta tta tca tag tct att aaa cac aaa tca tct --- --- --- --- --- --- --- --- --- --- --- --- --- --- --- --- --- --- --- --- --- --- --- --- --- --- --a cgt tta taa c-- --- --- --- --- --- --- --- --a aca ttc --- --- --- --- -tc att att aat tag ttc tgt agt aat atc ttt aat aat ttg gct ata --- --- -ca tct gtt --- --- --- --- --- --- --- --- --- --- --- --- --- --- --- --- --- --- --c aat act --- --- --- --- --- --- --- --a tct att gat gat ttc ttt t-- --- --- --- --- --- --- --- --- --- --- --t taa gac t-- --- --- --- --- --- --- --- --- --- --- --- --- --- --- --- --- --- --- --- --- --- --- --- --- --- --- --- --- --- --- --- --- --- --- --- --- --- --- --- taa act agt --- --- --- --- --- --- --- --- --- --- --- --- --- --- --- --- --- --- --- --- --- --- --- --- --- --- --- --- --- --- --- --- --- --- --- --- --- --- --- --- --- --- --- --- --- --- --- --- --- --- --- --- --- --- --- --- --- --- --- --- --- --- --- --- --- --- --- --- --- --- --- --- --- --- --- --- --- --- --- --- --- --- --- --- --- --- --- --- --- --- --- --- --- --- --- --- --- --- --- --- --- --- --- --- --- --- --- --- --- --t atg gta atg acg atg aaa --- -tc gag tag ta- --- --a ctt cta ata aag act tga ta- --- --t cat tat cat atg ttt gat cg- --- --- --- --- --- --- --t cat agt taa tag tgt g-- --- --- --- --- --- --- --- --- --- --- --- --- --- --- --- --- --- --- --- -gc taa atg gta ctg tta ata agt tta t-- --- --- --- --- --- --- --- --- --- --- --- --- --- --- --- --- --a gac aat atc ata gta ttt tct ttc cag aat --- --- --- --t aga tta ttt ttt taa ata ctg atc ctc aca att ccg tga tgt agc agt agt tgg t-- --- --- --- --- --- --- --- --- --- --- --- --- --- --- --- --- --- --- --- --- --- --- --- --- --- --- --- -gc atg gtc tat atc gt- --- --- --- --- --- --- --- --- --- --- --- --- --- --- --- --- --- --- --- --- --- --- --- --- --- --- --- --- --- --- --- --- --- --- --- --- --- --- --- --- --- --- --- taa aat gta tca tat ata ata gtt ttc tga cgt gga gta cag aat ttt cga --- --- --- --- --- --- --- --- --- --- --- --- --- --- --- --- --- --- --- --- --- --- --- --- --- --- --- --- --- --- --- --- --- --- --- --- --- --- --- --- --- --- --- --t taa tga gtt cat ggt aag gaa ggg caa atg tct --- gta tat aat ata cat aag tta a-- --- --- --- --- --- --- --- --- --- --- --- --- --t agt ttt tta tca tat tt- --- --- --- --- --- --- --- --t cta ata cca taa taa aaa tta tca tta tg- --- --- -ta taa tca --- --- --- --- --- --- --- --- --- --- --- --t cac tg- --- -tc gct atc att att gcg ttt gtg tag tt- --- --- --- --- ctg ccc ta- --- --- --- --- --- --- --- --t cat cta cat cac tgt ca- --- --- --- --- ctc tca cta tat ctt cta aaa tta caa --- -ac aac --- --- --- --- --- --- --- --- --- --- --- --- --- --- --- --- --- --- --- --- --- --- --- --- --- --- --- --- --- --- --- --- --- --- --- --- tgg ata ttc gat --- --- --- --- --- --- --- --- --- --- --- --- --- --- --- --- --- --- --- --- --- --- --- --- --- --- --a aca gca ttt gtg t-- --- --- --- --- --- --- --- --- --- --- --- --- --- --- --- --- -

>JX878424.1_Monkeypox_virus_isolate_DRC_07-0338_complete_genome

--- --- --- --- --- --- --- --- --- --- --- --- --- --- --- --- --- --- --- --- --- --- --- --- --- --- --- --- --- --- --- --- --- --- --- --g tgt cta gaa aaa aat gtg tga ccc acg acc gta gga aac tct aga ggg taa gaa aaa tca atc gtt tat aga gac cat cag aaa gag gtt taa tat ttt tgt gag acc cat cga cga gag aaa gg- --- --- --- --- --- --- --- --- --- --- --- --- --- --- --- --- --- --- --- --- --- --- --- --- --- --- --- --- --- --- --- --- --- --- --- --- --- --- --- --- --- --- --- --- --- --- --- --- --- --- --- --- --- --- --- --- --- --- --- --- --- --- --- --- --- --- --- --- --- --- --- --- --- --- --- --- --- --- --- --- --- --- --- --- --- --- --- --- --- --- --- --- --- --- --- --- --- --- --- --- --- --- --- --- --- --- --- --- --- --- --- --- --- --- --- --- --- --- --- --- --- --- --- --- --- --- --- --- --- --- --- --- --- --- --- --- --- --- --- --- --- --- --- --- --- --- --- --- --- --- --- --- --- --- --- --- --- --- --- --- --- --- --- --- --- --- --- --- --- --- --- --- --- --- --- --- --- --- --- --- --- --- --- --- --- --- --- --- --- --- --- --- --- --- --- --- --- --- --- --- --- --- --- --- --- --- --- --- --- --a taa aaa ctt ttt acg act cca tca gaa aga ggt tta --- --- --- --- --- --- --- --- --- --- --- --- --- --- --- --- --- --- --- --- --- --- --- --- --- --- --- --- --- --- --- --- --- --- --- --- --- --- --- --- --- --- --- --- --- --- --a tat ttt tgt gag acc cat cga cga gag aaa gga taa aaa ctt ttt acg act cca tca gaa aga ggt tta ata ttt ttg tga gac cca tcg acg aga gaa aga gat ggt tag tca aga tat ttt tct tag tac aaa agt caa tgt ttt aaa ata tat gga cga gaa tta att tgt ctg tat aaa aac ttg tgt gaa att atg tac tag aga aaa aac gtg agc agt gtc ccc tac atg gat ttt aca gat cat tta tat tcc aaa aat att aac tat ata cgt tta tta tat gat gtt aac gtg taa att ata aac att att tta tga tgc aat tgt ctg aca acc --- att ggc ata agg ata ttg ata agc tct acg aga ata tat tgt tgg acg tta tcg ttt acg aaa tag ttg aga cat cag aaa gag gtt taa tat ttt tgt gag a-c cat cga aga gag aaa gag aat aaa aat a-- --- --- --- ttt ttt tgt aaa act ttt tta tga gac aag aga gaa aga gaa tac gaa tag tga tca tat cgt atc aca tat tga aac aga aag aag aag taa cga gag gta act ttt tgt gaa tgt agt taa aca ttt ttg ttt tgc aaa ccg gaa tat agt acc cgg tac act ttt tta att cgt ggt gcg gtg tct gaa tcg ttc gat taa ccc aac tca tcc att ttc aga tga ata gag tta tcg att cag aca cat gct ttg agt ttt gtt gaa tcg atg agt gaa gta tca tcg gtt gca cct tca gat gcc gat cca tcg aca tac ttg aat cca tcc ttg act tca agt tca gat gat tcc tca cac atg tct ccg ata cgt acg cta aac tct agg ttc tta aca cat ttt gta tca acg atc gtt gaa ccg atg ata tct ttg taa ctc act ttc tta tgt gag atg tta gac cca agt act gga tgg gtc ttg atg tcg ctg tct ttc tct tct tcg cta cat ctg atg tcg ata gac atc tca cag tct ttg atc ata gcc aga gct tct tca cgc gtg atc gcg gga gag tcc tta cct tgt cct ggt gac acg ctg gac aat cta gta ttc aca gtg ttt cca tca gag gat tcg gag atg gat gaa atc ttt ggg cat ttg gtg aat cca aag ttc atg tta aga ccc gcg ccg acg ata ttg taa taa gtg gtg gga tct cct ttt aca act tct tcg gat acc tca tca tct tcg gat acc tca tca tct tcg gtc tct gta act tcc gtt acg gat tga caa atc tta tca ttg gtc ggt gtt tgg tct tgc ttt gtg act ttg ata ata aca tcg att ccc ata tga tgt ttg ttt tct tct tca gta cac gag gat gag gat tgt tga aga cta gta ggc ata gca gct gcc act agg cac atg cat gcc agg aca ata tat tgt ttc atg att gct att gat tga tta ctg ttc tag atg att cta ctt tct tac cat ata ata aat tag aat ata ttt tct act ttt acg aga aat taa tta ttg tat tta tta ttt atg ggt -aa aaa aaa ctt act ata agt ggg tgg gat tct ggg aat tag tga tca gtt tat gta tat cgc aac tag cgg gca tat ggc tat tga cat cga gaa cat tac cca tat gat aag aga ttg tat ccg ttt cgt agt ctt gag tat tgg tat tac tat ata gta tgt aga tgt cga cgc tag ata gac agt cgc cca cta gag tta ccg tct ctg aat gcg gca tga tag tat cat tct ttg ttt tcg tta act gtt tgg aag atg aat ctt tgt tgt tac att taa tct cga aat tca gag tac ata tct ttg aag tat tct gat atc tat ttt ctc ctg taa aga atc ctg aag ttg cta cat tat taa gga cag aga agt att ctg cac gaa aga cgg gat cac aat ctt tat gat tca tgg taa tag tta gtt ccg acg ttg aga tgg att cgc tga gac cgg tag tgg tcg tcc gag tac acg atg tgt cgt tga ctg gat aca ggt taa ttt cca cat cga tat agt taa atg tat tgc tgg tt- --a cgg gtt cgc att tat ctg tgg aag aga cgg tgt gag aat atg ttc cgg gac cac acg gag aac aga tga cgt ctc cgg tag acg tgt atc cgg ata ctc cgt atc cta ttc cac act ttg ttt tag aaa tac atg ttc tac acc ctg atg atc ctt tga gaa gac aat aat atc ctg gag agc att cac aga ttc tat tgt gag tcg tgt tac acg atc gcg tct cta cct gat tac tat cac atc ttc cgt tac aac tta gac aag cct gta aat gat tat tgt gag atg taa agg tat ccg aac cac acg gcg tac att gtg tat tag tct tgc tat cac ata atc tgg aag cgt aag ttc ccg gag gac acg ata gac aac ata gat tac ggc ttc tgt att cgt tgt ctt tac att ttc cat tgg atg gtg cat gtg gtg cta tat ctc ttc cgt tta tta tta tac atg aga gaa aca ata tat acg agt ata ata cgg acc tca tga ttt aat aat -gt agt aat cgt cgt ctt gtt act gtt tgt ttc cta ctt ctc caa tca tat aga tta ttt ttt aaa tat ttt ctt tct atc atg gat aat att tgt aat ggt tct ttc cgt aca aca tac tgt tta gat ggt agt cgc tta gct tgg tta tga tat tgc gca taa ttt ccg gag gca aat acg ata gtc tag att gac tat cga tgg tag act cta att tat tga gtg ctt tgt cga cga gtt tac ttt tat gct cca tcg ata gat gac act gtt cta tga gat cgt cgt aca tgg gaa atg aaa tgc gtt tgt ctg aat gta tgg ctt cga gat agg tgt gat acc gga tgt ctt ctg ttc tca ata ccg tat aca agt tgg tgt ctg aga ttc gaa tct ctt tga gga gac tta tgt cac gac tac att ttt cga tga tgg aat cta tct tat cga atg ata tat ttt tca taa ata cac ttt tat agt cct cgt tta aac aga att tag tat gta gtt ccg caa atg act cgt ccc tta ata tgc agt agg cta tta tct tct tta cgt agt gat cgt cgt agg gag aga act ccg aca tct tgt aga aca acg att taa tca tag gta gag ata ctt tca gtc tgt ggt gga tga tgt cat taa cga cat ccg cct tgt ata tga tgt ttc tgt ttt caa aca cca agt cga ata ctg tct tat cgt ctt tag tcg gaa ggt tga tgt cgt atc cga tgt ata cga ggt atg agg caa cat tgt tat tgc aat tct gga agg cgg tat gaa gag gag tca ttg tat tat agt att cgt ctt tct gaa tgt cga atc tat cta gta gat acc gta gta tat tga gag agc gac ttc cat atc ctt gat tat gtt tta tga ata gat aaa gta gat gtt gtc ctt ctt cct ttt gta att tcc cgt att ttt gtt cgt gcc aat tga gta aca tta tga gaa tat gac ctg ttg cac aat cgt tct tta tgt att cca tga tgg gtg tac aat caa gat tat tac gta tcc tcg tat cgg ctc ctc gag ata aaa gag cat aca cca cac gag gac tat gtt tgg tat act gtt gaa ggt aag tgt gta acg gcg tat ttc cga ttt tcg taa ccg cgt taa tgt ttg ctc cat gat cta tta tcg cgt aga tga atc gct tct cag ctc gca tct tag tgt gac tct ttg act tgt aat aat tgc ttt cgt gga aca cgg ata tgt gtt tac agt agt aat gaa gag aag tga gtc cat cct cat cga cgc aat tag ggt cag atc ctt tag tca ata att tgt aca gaa cgt aat agt tta agc tcc cat tga att tat atc taa gat aac aca gca ata gat cgg atg att tac taa agt cat caa tgg ggt ccg tta gta tat caa aga tct tgt tat cga ttg ata gtg aat gaa tca gat agt ggt gta gag gaa tat gtc ctt ttt tat cct tgc tat caa agt tac gca tgc cgt ggc gta aca ata tct tta ata cag atg gat taa atc gtg tat tca tcg tat agc aat gta atg gag agt tac cac att tta gtc gtt tat tca gat cgc agt gtt taa taa cta att taa aca gat gag atg ata tat cca cat caa aga atg cga gat aca tat gac aga cat tat tga cag aaa tgt gac ctt cat tat cac cgt cgt cca taa atg cgt tag gta cgt acc aca tac tat cgt taa cga tgc gca caa tct cgt cca ttt tca tcc atc ttc ata atg att tac ttt ttc ata att aga gaa aaa gat caa ggt ata aaa att aga agt gtt aga cta ta- --- --- --- --- --- --- --- --- --- --- -aa cta act tat aa- --- --- --- --- --- --- --- --- --- --- --- --- --- --- --- --- --- --- --- --- --- --- --- --- --- --- --- --- --- --- --- --- --- --- --- --- --- --- --- --- --- --- --- --- --- --- --- --- --- --- --- --- --- --- --- --- --- --- --- --- --- --- --- --- --- --- --- --- --- --- --- --- --- --- --- --- --- --- --- --- --- --- --- --- --- --- -aa cta act tat gac tta act aac tta tga cta tta act cat ttc aag aaa ggt ggg tgg aga gaa ctc tat atg aca gct tgt gaa aca att aga tcc cta att tct aat gga agt ttt gat agg aga ttg tca tca gtt gat aca ttg ttt att atc tca tct att aga gca cgt ctg ttt aga gct tta gtg acc tgc tcg gtt act tct gtg taa atc ttg aat cct tta gtg ata cac tgt gtc aaa act gga tgt tta gaa tac cta tgt aga ata tgg gaa gca tgc ttg ttt ttg tct cta tta tag att aac tca tac atg gtt gta tta tga att ttc atc tgc cta atg tac tcc aat tct tgt tta caa tca att ata taa tca aag agt gat gat gca tac aca tta caa agt gaa taa tct acc atc ata aaa tac ttg ata cag agc ttt atc aca tca tgg ttt tca att gta tta tta agt ata gct aat ttt ata cag tca ata gac aat ggt tct cta agc aat att tct aat att tta aga tgt gct tcc cta cgg gcg atg aca gat ccc cta tcc acg gcc acg tca aga cat gta tat cca tta ctc att act gcg ttg aca ttt gct cca ttt tct aat agc cat gat act aaa tct ata taa cct gca tag ata gcg cga taa agc aag gtc ctt cca cca gca tct agt tga ttg ata tct tca ata tat ggg ata caa agc tta taa att tct aat act gtg ggt tca tct aca agg aat ccc cta gta tac tga att att tta tat aga tct aat tta aca tca ttt tca tct ggg ata cca caa ttc aaa ata aac tca aca aca cta ctt tcc ttt tta cat att ccc cta aaa tag gca ttc aag cat tct att tta tat att aca gcc cca tga tct acc ata aaa tca aca atg tct att tct aca tat gca tta gat aga tag taa agt aag aga tct tgt aca gaa tta caa ttc tta ata att ata gag aaa ata tct tcc ata taa ttc ttt gac act aat gca gat ata ata tct tta tat gta ata tat gca aac agt cta tct act ata tac tga tca ata tta tct cta tga atc cta aaa taa tca tac aga aca tct aca gga tca caa att ggt tca agg aga aat cta tca aat att ttc ctg tca aca act ggt tct aga aca taa cag tca aca cct aat cca tgt ttt tta tag tca tct acc aaa gat aat gac caa aga tcg agg tcg tcg tga aac tgc tca tcg aca gcc atg aaa tct gcc gac tcc atg gcg cga atc gca ctg tct tat tcg cca ttg att ttc a-t ttt tta taa tta tgt aca tgt ttt cct tct att ctc aag agt cta caa aaa tat a-t ttt ttt tcg ata tct aag tac taa gtt ttt tta ctg ttt ttg tta ctg tct tcc att ctt cta act aaa gat ctg aga taa att ata caa tct tcg cta tcg aac cat ttt tgt agt cta aag cct gaa gta att aac caa ctg ttt tta tta gtg gct ttt ttc gat cta tcc tcg tc- --- --c tca tca tcc tta ta- --- --- --- --- --- --- tta tta tca tta tta tca tag tct att aaa cac aaa tca tct --- --- --- --- --- --- --- --- --- --- --- --- --- --- --- --- --- --- --- --- --- --- --- --- --- --- --a cgt tta taa c-- --- --- --- --- --- --- --- --a aca ttc --- --- --- --- -tc att att aat tag ttc tgt agt aat atc ttt aat aat ttg gct ata --- --- -ca tct gtt --- --- --- --- --- --- --- --- --- --- --- --- --- --- --- --- --- --- --c aat act --- --- --- --- --- --- --- --a tct att gat gat ttc ttt t-- --- --- --- --- --- --- --- --- --- --- --t taa gac t-- --- --- --- --- --- --- --- --- --- --- --- --- --- --- --- --- --- --- --- --- --- --- --- --- --- --- --- --- --- --- --- --- --- --- --- --- --- --- --- taa act agt --- --- --- --- --- --- --- --- --- --- --- --- --- --- --- --- --- --- --- --- --- --- --- --- --- --- --- --- --- --- --- --- --- --- --- --- --- --- --- --- --- --- --- --- --- --- --- --- --- --- --- --- --- --- --- --- --- --- --- --- --- --- --- --- --- --- --- --- --- --- --- --- --- --- --- --- --- --- --- --- --- --- --- --- --- --- --- --- --- --- --- --- --- --- --- --- --- --- --- --- --- --- --- --- --- --- --- --- --- --t atg gta atg acg atg aaa --- -tc gag tag ta- --- --a ctt cta ata aag act tga ta- --- --t cat tat cat atg ttt gat cg- --- --- --- --- --- --- --t cat agt taa tag tgt g-- --- --- --- --- --- --- --- --- --- --- --- --- --- --- --- --- --- --- --- -gc taa atg gta ctg tta ata agt tta t-- --- --- --- --- --- --- --- --- --- --- --- --- --- --- --- --- --a gac aat atc ata gta ttt tct ttc cag aat --- --- --- --t aga tta ttt ttt taa ata ctg atc ctc aca att ccg tga tgt agc agt agt tgg t-- --- --- --- --- --- --- --- --- --- --- --- --- --- --- --- --- --- --- --- --- --- --- --- --- --- --- --- -gc atg gtc tat atc gt- --- --- --- --- --- --- --- --- --- --- --- --- --- --- --- --- --- --- --- --- --- --- --- --- --- --- --- --- --- --- --- --- --- --- --- --- --- --- --- --- --- --- --- taa aat gta tca tat ata ata gtt ttc tga cgt gga gta cag aat ttt cga --- --- --- --- --- --- --- --- --- --- --- --- --- --- --- --- --- --- --- --- --- --- --- --- --- --- --- --- --- --- --- --- --- --- --- --- --- --- --- --- --- --- --- --t taa tga gtt cat ggt aag gaa ggg caa atg tct --- gta tat aat ata cat aag tta a-- --- --- --- --- --- --- --- --- --- --- --- --- --t agt ttt tta tca tat tt- --- --- --- --- --- --- --- --t cta ata cca taa taa aaa tta tca tta tg- --- --- -ta taa tca --- --- --- --- --- --- --- --- --- --- --- --t cac tg- --- -tc gct atc att att gcg ttt gtg tag tt- --- --- --- --- ctg ccc ta- --- --- --- --- --- --- --- --t cat cta cat cac tgt ca- --- --- --- --- ctc tca cta tat ctt cta aaa tta caa --- -ac aac --- --- --- --- --- --- --- --- --- --- --- --- --- --- --- --- --- --- --- --- --- --- --- --- --- --- --- --- --- --- --- --- --- --- --- --- tgg ata ttc gat --- --- --- --- --- --- --- --- --- --- --- --- --- --- --- --- --- --- --- --- --- --- --- --- --- --- --a aca gca ttt gtg t-- --- --- --- --- --- --- --- --- --- --- --- --- --- --- --- --- -

>KC257460.1_Monkeypox_virus_strain_DRC_Yandongi_1985_complete_genome

--- --- --- --- --- --- --- --- --- --- --- --- --- --- --- --- --- --- --- --- --- --- --- --- --- --- --- --- --- --- --- --- --- --- --- --- --- --- --- --- --- --- --- --- --- --- --- --- --- --- --- --- --- --- --- --- --- --- --- --- --- --- --- --- --- --- --- --- --- --- --- --- --- --- --- --- --- --- --- --- --- --- --- --- --- --- --- --- --- --- --- --- --- --- --- --- --- --- --- --- --- --- --- --- --- --- --- --- --- --- --- --- --- --- --- --- --- --- --- --- --- --- --- --- --- --- --- --- --- --- --- --- --- --- --- --- --- --- --- --- --- --- --- --- --- --- --- --- --- --- --- --- --- --- --- --- --- --- --- --- --- --- --- --- --- --- --- --- --- --- --- --- --- --- --- --- --- --- --- --- --- --- --- --- --- --- --- --- --- --- --- --- --- --- --- --- --- --- --- --- --- --- --- --- --- --- --- --- --- --- --- --- --- --- --- --- --- --- --- --- --- --- --- --- --- --- --- --- --- --- --- --- --- --- --- --- --- --- --- --- --- --- --- --- --- --- --- --- --- --- --- --- --- --- --- --- --- --- --- --- --- --- --- --- --- --- --- --- --- --- --- --- --- --- --- --- --- --- --- --- --- --- --- --- --- --- --- --- --- --- --- --- --- --- --- --- --- --- --- --- --- --- --- --- --- --- --- --- --- --- --- --- --- --- --- --- --- --- --- --- --- --- --- --- --- --- --- --- --- --- --- --- --- --- --- --- --- --- --- --- --- --- --- --- --- --- --- --- --- --- --- --- --- --- --- --- --- --- --- --- --- --- --- --- --- --- --- --- --- --- --- --- --- --- gac cca tcg acg aga gaa aga gat ggt tag tca aga tat ttt tct tag tac aaa agt caa tgt ttt aaa ata tat gga cga gaa tta att tgt ctg tat aaa aac ttg tgt gaa att atg tac tag aga aaa aac gtg agc agt gtc ccc tac atg gat ttt aca gat cat tta tat tcc aaa aat att aac tat ata cgt tta tta tat gat gtt aac gtg taa att ata aac att att tta tga tgc aat tgt ctg aca ac- --- --- --c ata agg ata ttg ata agc tct acg aga ata tat tgt tgg acg tta tcg ttt acg aaa tag ttg aga cat cag aaa gag gtt taa tat ttt tgt gag acc cat cga aga gag aa- -ag gat aaa aat a-- --- --- --- ttt ttt tgt aaa act ttt tta tga gac aag aga gaa aga gaa tac gaa tag tga tca tat cgt atc aca tat tga aac aga aag aag aag taa cga gag gta act ttt tgt gaa tgt agt taa aca ttt ttg ttt tgc aaa ccg gaa tat agt acc cgg tac act ttt tta att cgt ggt gcg gtg tct gaa tcg ttc gat taa ccc aac tca tcc att ttc aga tga ata gag tta tcg att cag aca cat gct ttg agt ttt gtt gaa tcg atg agt gaa gta tca tcg gtt gca cct tca gat gcc gat ccg tcg aca tac ttg aat cca tcc ttg act tca agt tca gat gat tcc tca cac atg tct ccg ata cgt acg cta aac tct agg ttc tta aca cat ttt gta tca acg atc gtt gaa ccg atg ata tct ttg taa ctc act ttc tta tgt gag atg tta gac cca agt act gga tgg gtc ttg atg tcg ctg tct ttc tct tct tcg cta cat ctg atg tcg ata gac atc tca cag tct ttg atc ata gcc aga gct tct tca cgc gtg atc gcg gga gag tcc tta cct tgt cct ggt gac acg ctg gac aat cta gta ttc aca gtg ttt cca tca gag gat tcg gag atg gat gaa atc ttt ggg cat ttg gtg aat cca aag ttc atg tta aga ccc gcg ccg acg ata ttg taa taa gtg gtg gga tct cct ttt aca act tct tcg gat acc tca tca tct tcg gat acc tca tca tct tcg gtc tct gta act tcc gtt acg gat tga caa atc tta tca ttg gtc ggt gtt tgg tct tgc ttt gtg act ttg ata ata aca tcg att ccc ata tga tgt ttg ttt tct tct tca gta cac gag gat gag gat tgt tga aga cta gta ggc ata gca gct gcc act agg cac atg cat gcc agg aca ata tat tgt ttc atg att gct att gat tga tta ctg ttc tag atg att cta ctt tct tac cat ata ata aat tag aat ata ttt tct act ttt acg aga aat taa tta ttg tat tta tta ttt atg ggt -aa aaa aaa ctt act ata agt ggg tgg gat tct ggg aat tag tga tca gtt tat gta tat cgc aac tag cgg gca tat ggc tat tga cat cga gaa cat tac cca tat gat aag aga ttg tat ccg ttt cgt agt ctt gag tat tgg tat tac tat ata gta tgt aga tgt cga cgc tag ata gac agt cgc cca cta gag tta ccg tct ctg aat gcg gca tga tag tat cat tct ttg ttt tcg tta act gtt tgg aag atg aat ctt tgt tgt tac att taa tct cga aat tca gag tac ata tct ttg aag tat tct gat atc tat ttt ctc ctg taa aga atc ctg aag ttg cta cat tat taa gga cag aga agt att ctg cac gaa aga cgg gat cac aat ctt tat gat tca tgg taa tag tta gtt ccg acg ttg aga tgg att cgc tga gac cgg tag tgg tcg tcc gag tac acg atg tgt cgt tga ctg gat aca ggt taa ttt cca cat cga tat agt taa atg tat tgc tgg tt- --a cgg gtt cgc att tat ctg tgg aag aga cgg tgt gag aat atg ttc cgg gac cac acg gag aac aga tga cgt ctc cgg tag acg tgt atc cgg ata ctc cgt atc cta ttc cac act ttg ttt tag aaa tac atg ttc tac acc ctg atg atc ctt tga gaa gac aat aat atc ctg gag agc att cac aga ttc tat tgt gag tcg tgt tac acg atc gcg tct cta cct gat tac tat cac atc ttc cgt tac aac tta gac aag cct gta aat gat tat tgt gag atg taa agg tat ccg aac cac acg gcg tac att gtg tat tag tct tgc tat cac ata atc tgg aag cgt aag ttc ccg gag gac acg ata gac aac ata gat tac ggc ttc tgt att cgt tgt ctt tac att ttc cat tgg atg gtg cat gtg gtg cta tat ctc ttc cgt tta tta tta tac atg aga gaa aca ata tat acg agt ata ata cgg acc tca tga ttt aat aat -gt agt aat cgt cgt ctt gtt act gtt tgt ttc cta ctt ctc caa tca tat aga tta ttt ttt aaa tat ttt ctt tct atc atg gat aat att tgt aat ggt tct ttc cgt aca aca tac tgt tta gat ggt agt cgc tta gct tgg tta tga tat tgc gca taa ttt ccg gag gca aat acg ata gtc tag att gac tat cga tgg tag act cta att tat tga gtg ctt tgt cga cga gtt tac ttt tat gct cca tcg ata gat gac act gtt cta tga gat cgt cgt aca tgg gaa atg aaa tgc gtt tgt ctg aat gta tgg ctt cga gat agg tgt gat acc gga tgt ctt ctg ttc tca ata ccg tat aca agt tgg tgt ctg aga ttc gaa tct ctt tga gga gac tta tgt cac gac tac att ttt cga tga tgg aat cta tct tat cga atg ata tat ttt tca taa ata cac ttt tat agt cct cgt tta aac aga att tag tat gta gtt ccg caa atg act cgt ccc tta ata ggc agt agg cta tta tct tct tta cgt agt gat cgt cgt agg gag aga act ccg aca tct tgt aga aca acg att taa tca tag gta gag ata ctt tca gtc tgt ggt gga tga tgt cat taa cga cat ccg cct tgt ata tga tgt ttc tgt ttt caa aca cca agt cga ata ctg tct tat cgt ctt tag tcg gaa ggt tga tgt cgt atc cga tgt ata cga ggt atg agg caa cat tgt tat tgc aat tct gga agg cgg tat gaa gag gag tca ttg tat tat agt att cgt ctt tct gaa tgt cga atc tat cta gta gat acc gta gta tat tga gag agc gac ttc cat atc ctt gat tat gtt tta tga ata gat aaa gta gat gtt gtc ctt ctt cct ttt gta att tcc cgt att ttt gtt cgt gcc aat tga gta aca tta tga gaa tat gac ctg ttg cac aat cgt tct tta tgt att cca tga tgg gtg tac aat caa gat tat tac gta tcc tcg tat cgg ctc ctc gag ata aaa gag cat aca cca cac gag gac tat gtt tgg tat act gtt gaa ggt aag tgt gta acg gcg tat ttc cga ttt tcg taa ccg cgt taa tgt ttg ctc cat gat cta tta tcg cgt aga tga atc gct tct cag ctc gca tct tag tgt gac tct ttg act tgt aat aat tgc ttt cgt gga aca cgg ata tgt gtt tac agt agt aat gaa gag aag tga gtc cat cct cat cga cgc aat tag ggt cag atc ctt tag tca ata att tgt aca gaa cgt aat agt tta agc tcc cat tga att tat atc taa gat aac aca gca ata gat cgg atg att tac taa agt cat caa tgg ggt ccg tta gta tat caa aga tct tgt tat cga ttg ata gtg aat gaa tca gat agt ggt gta gag gaa tat gtc ctt ttt tat cct tgc tat caa agt tac gca tgc cgt ggc gta aca ata tct tta ata cag atg gat taa atc gtg tat tca tcg tat agc aat gta atg gag agt tac cac att tta gtc gtt tat tca gat cgc agt gtt taa taa cta att taa aca gat gag atg ata tat cca cat caa aga atg cga gat aca tat gac aga cat tat tga cag aaa tgt gac ctt cat tat cac cgt cgt cca taa atg cgt tag gta cgt acc aca tac tat cgt taa cga tgc gca caa tct cgt cca ttt tca tcc atc ttc ata atg att tac ttt ttc ata att aga gaa aaa gat caa ggt ata aaa att aga agt gtt aga cta ta- --- --- --- --- --- --- --- --- --- --- -aa cta act tat aa- --- --- --- --- --- --- --- --- --- --- --- --- --- --- --- --- --- --- --- --- --- --- --- --- --- --- --- --- --- --- --- --- --- --- --- --- --- --- --- --- --- --- --- --- --- --- --- --- --- --- --- --- --- --- --- --- --- --- --- --- --- --- --- --- --- --- --- --- --- --- --- --- --- --- --- --- --- --- --- --- --- --- --- --- --- --- -aa cta act tat gac tta act aac tta tga cta tta act cat ttc aag aaa ggt ggg tgg aga gaa ctc tat atg aca gct tgt gaa aca att aga tcc cta att tct aat gga agt ttt gat agg aga ttg tca tca gtt gat aca ttg ttt att atc tca tct att aga gca cgt ctg ttt aga gct tta gtg acc tgc tcg gtt act tct gtg taa atc ttg aat cct tta gtg ata cac tgt gtc aaa act gga tgt tta gaa tac cta tgt aga ata tgg gaa gca tgc ttg ttt ttg tct cta tta tag att aac tca tac atg gtt gta tta tga att ttc atc tgc cta atg tac tcc aat tct tgt tta caa tca att ata taa tca aag agt gat gat gca tac aca tta caa agt gaa taa tct acc atc ata aaa tac ttg ata cag agc ttt atc aca tca tgg ttt tca att gta tta tta agt ata gct aat ttt ata cag tca ata gac aat ggt tct cta agc aat att tct aat att tta aga tgt gct tcc cta cgg gcg atg aca gat ccc cta tcc acg gcc acg tca aga cat gta tat cca tta ctc att act gcg ttg aca ttt gct cca ttt tct aat agc cat gat act aaa tct ata taa cct gca tag ata gcg cga taa agc aag gtc ctt cca cca gca tct agt tga ttg ata tct tca ata tat ggg ata caa agc tta taa att tct aat act gtg ggt tca tct aca agg aat ccc cta gta tac tga att att tta tat aga tct aat tta aca tca ttt tca tct ggg ata cca caa ttc aaa ata aac tca aca aca cta ctt tcc ttt tta cat att ccc cta aaa tag gca ttc aag cat tct att tta tat att aca gcc cca tga tct acc ata aaa tca aca atg tct att tct aca tat gca tta gat aga tag taa agt aag aga tct tgt aca gaa tta caa ttc tta ata att ata gag aaa ata tct tcc ata taa ttc ttt gac act aat gca gat ata ata tct tta tat gta ata tat gca aac agt cta tct act ata tac tga tca ata tta tct cta tga atc cta aaa taa tca tac aga aca tct aca gga tca caa att ggt tca agg aga aat cta tca aat att ttc ctg tca aca act ggt tct aga aca taa cag tca aca cct aat cca tgt ttt tta tag tca tct acc aaa gat aat gac caa aga tcg agg tcg tcg tga aac tgc tca tcg aca gcc atg aaa tct gcc gac tcc atg gcg cga atc gca ctg tct tat tcg cca ttg att ttc a-t ttt tta taa tta tgt aca tgt ttt cct tct att ctc aag agt cta caa aaa tat a-t ttt ttt tcg ata tct aag tac taa gtt ttt tta ctg ttt ttg tta ctg tct tcc att ctt cta act aaa gat ctg aga taa att ata caa tct tcg cta tcg aac cat ttt tgt agt cta aag cct gaa gta att aac caa ctg ttt tta tta gtg gct ttt ttc gat cta tcc tcg tc- --- --c tca tca tcc tta ta- --- --- --- --- --- --- tta tta tca tta tta tca tag tct att aaa cac aaa tca tct --- --- --- --- --- --- --- --- --- --- --- --- --- --- --- --- --- --- --- --- --- --- --- --- --- --- --a cgt tta taa c-- --- --- --- --- --- --- --- --a aca ttc --- --- --- --- -tc att att aat tag ttc tgt agt aat atc ttt aat aat ttg gct ata --- --- -ca tct gtt --- --- --- --- --- --- --- --- --- --- --- --- --- --- --- --- --- --- --c aat act --- --- --- --- --- --- --- --a tct att gat gat ttc ttt t-- --- --- --- --- --- --- --- --- --- --- --t taa gac t-- --- --- --- --- --- --- --- --- --- --- --- --- --- --- --- --- --- --- --- --- --- --- --- --- --- --- --- --- --- --- --- --- --- --- --- --- --- --- --- taa act agt --- --- --- --- --- --- --- --- --- --- --- --- --- --- --- --- --- --- --- --- --- --- --- --- --- --- --- --- --- --- --- --- --- --- --- --- --- --- --- --- --- --- --- --- --- --- --- --- --- --- --- --- --- --- --- --- --- --- --- --- --- --- --- --- --- --- --- --- --- --- --- --- --- --- --- --- --- --- --- --- --- --- --- --- --- --- --- --- --- --- --- --- --- --- --- --- --- --- --- --- --- --- --- --- --- --- --- --- --- --t atg gta atg acg atg aaa --- -tc gag tag ta- --- --a ctt cta ata aat act tga ta- --- --t cat tat cat atg ttt gat cg- --- --- --- --- --- --- --t cat agt taa tag tgt g-- --- --- --- --- --- --- --- --- --- --- --- --- --- --- --- --- --- --- --- -gc taa atg gta ctg tta ata agt tta t-- --- --- --- --- --- --- --- --- --- --- --- --- --- --- --- --- --a gac aat atc ata gta ttt tct ttc cag aat --- --- --- --t aga tta ttt ttt taa ata ctg atc ctc aca att ccg tga tgt agc agt agt tgg t-- --- --- --- --- --- --- --- --- --- --- --- --- --- --- --- --- --- --- --- --- --- --- --- --- --- --- --- -gc atg gtc tat atc gt- --- --- --- --- --- --- --- --- --- --- --- --- --- --- --- --- --- --- --- --- --- --- --- --- --- --- --- --- --- --- --- --- --- --- --- --- --- --- --- --- --- --- --- taa aat gta tca tat ata ata gtt ttc tga cgt gga gta cag aat ttt cga --- --- --- --- --- --- --- --- --- --- --- --- --- --- --- --- --- --- --- --- --- --- --- --- --- --- --- --- --- --- --- --- --- --- --- --- --- --- --- --- --- --- --- --t taa tga gtt cat ggt aag gaa ggg caa atg tct --- gta tat aat ata cat aag tta a-- --- --- --- --- --- --- --- --- --- --- --- --- --t agt ttt tta tca tat tt- --- --- --- --- --- --- --- --t cta ata cca taa taa aaa tta tca tta tg- --- --- -ta taa tca --- --- --- --- --- --- --- --- --- --- --- --t cac tg- --- -tc gct atc att att gcg ttt gtg tag tt- --- --- --- --- ctg ccc ta- --- --- --- --- --- --- --- --t cat cta cat cac tgt ca- --- --- --- --- ctc tca cta tat ctt cta aaa tta caa --- -ac aac --- --- --- --- --- --- --- --- --- --- --- --- --- --- --- --- --- --- --- --- --- --- --- --- --- --- --- --- --- --- --- --- --- --- --- --- tgg ata ttc gat --- --- --- --- --- --- --- --- --- --- --- --- --- --- --- --- --- --- --- --- --- --- --- --- --- --- --a aca gca ttt gtg t-- --- --- --- --- --- --- --- --- --- --- --- --- --- --- --- --- -

>JX878429.1_Monkeypox_virus_isolate_DRC_07-0662_complete_genome

--- --- --- --- --- --- --- --- --- --- --- --- --- --- --- --- --- --- --- --- --- --- --- --- --- --- --- --- --- --- --- --- --- --- --- --g tgt cta gaa aaa aat gtg tga ccc acg acc gta gga aac tct aga ggg taa gaa aaa tca atc gtt tat aga gac cat cag aaa gag gtt taa tat ttt tgt gag acc cat cga cga gag aaa gga taa aaa ctt ttt acg act cca tca gaa aga ggt tta ata ttt ttg tga gac cca tcg ac- --- --- --- --- --- --- --- --- --- --- --- --- --- --- --- --- --- --- --- --- --- --- --- --- --- --- --- --- --- --- --- --- --- --- --- --- --- --- --- --- --- --- --- --- --- --- --- --- --- --- --- --- --- --- --- --- --- --- --- --- --- --- --- --- --- --- --- --- --- --- --- --- --- --- --- --- --- --- --- --- --- --- --- --- --- --- --- --- --- --- --- --- --- --- --- --- --- --- --- --- --- --- --- --- --- --- --- --- --- --- --- --- --- --- --- --- --- --- --- --- --- --- --- --- --- --- --- --- --- --- --- --- --- --- --- --- --- --- --- --g aga gaa agg ata aaa act ttt tac gac tcc atc aga aag agg ttt aat att ttt gtg aga ccc atc gac gag aga aag gat aaa aac ttt tta cga ctc cat cag aaa gag gtt taa tat ttt tgt gag acc cat cga cga gag aaa gga taa aaa ctt ttt acg act cca tca gaa aga ggt tta --- --- --- --- --- --- --- --- --- --- --- --- --- --- --- --- --- --- --- --- --- --- --- --- --- --- --- --- --- --- --- --- --- --- --- --- --- --- --- --- --- --- --- --- --- --- --a tat ttt tgt gag acc cat cga cga gag aaa gga taa aaa ctt ttt acg act cca tca gaa aga ggt tta ata ttt ttg tga gac cca tcg acg aga gaa aga gat ggt tag tca aga tat ttt tct tag tac aaa agt caa tgt ttt aaa ata tat gga cga gaa tta att tgt ctg tat aaa aac ttg tgt gaa att atg tac tag aga aaa aac gtg agc agt gtc ccc tac atg gat ttt aca gat cat tta tat tcc aaa aat att aac tat ata cgt tta tta tat gat gtt aac gtg taa att ata aac att att tta tga tgc aat tgt ctg aca acc --- att ggc ata agg ata ttg ata agc tct acg aga ata tat tgt tgg acg tta tcg ttt acg aaa tag ttg aga cat cag aaa gag gtt taa tat ttt tgt gag a-c cat cga aga gag aaa gag aat aaa aat a-- --- --- --- ttt ttt tgt aaa act ttt tta tga gac aag aga gaa aga gaa tac gaa tag tga tca tat cgt atc aca tat tga aac aga aag aag aag taa cga gag gta act ttt tgt gaa tgt agt taa aca ttt ttg ttt tgc aaa ccg gaa tat agt acc cgg tac act ttt tta att cgt ggt gcg gtg tct gaa tcg ttc gat taa ccc aac tca tcc att ttc aga tga ata gag tta tcg att cag aca cat gct ttg agt ttt gtt gaa tcg atg agt gaa gta tca tcg gtt gca cct tca gat gcc gat cca tcg aca tac ttg aat cca tcc ttg act tca agt tca gat gat tcc tca cac atg tct ccg ata cgt acg cta aac tct agg ttc tta aca cat ttt gta tca acg atc gtt gaa ccg atg ata tct ttg taa ctc act ttc tta tgt gag atg tta gac cca agt act gga tgg gtc ttg atg tcg ctg tct ttc tct tct tcg cta cat ctg atg tcg ata gac atc tca cag tct ttg atc ata gcc aga gct tct tca cgc gtg atc gcg gga gag tcc tta cct tgt cct ggt gac acg ctg gac aat cta gta ttc aca gtg ttt cca tca gag gat tcg gag atg gat gaa atc ttt ggg cat ttg gtg aat cca aag ttc atg tta aga ccc gcg ccg acg ata ttg taa taa gtg gtg gga tct cct ttt aca act tct tcg gat acc tca tca tct tcg gat acc tca tca tct tcg gtc tct gta act tcc gtt acg gat tga caa atc tta tca ttg gtc ggt gtt tgg tct tgc ttt gtg act ttg ata ata aca tcg att ccc ata tga tgt ttg ttt tct tct tca gta cac gag gat gag gat tgt tga aga cta gta ggc ata gca gct gcc act agg cac atg cat gcc agg aca ata tat tgt ttc atg att gct att gat tga tta ctg ttc tag atg att cta ctt tct tac cat ata ata aat tag aat ata ttt tct act ttt acg aga aat taa tta ttg tat tta tta ttt atg ggt -aa aaa aaa ctt act ata agt ggg tgg gat tct ggg aat tag tga tca gtt tat gta tat cgc aac tag cgg gca tat ggc tat tga cat cga gaa cat tac cca tat gat aag aga ttg tat ccg ttt cgt agt ctt gag tat tgg tat tac tat ata gta tgt aga tgt cga cgc tag ata gac agt cgc cca cta gag tta ccg tct ctg aat gcg gca tga tag tat cat tct ttg ttt tcg tta act gtt tgg aag atg aat ctt tgt tgt tac att taa tct cga aat tca gag tac ata tct ttg aag tat tct gat atc tat ttt ctc ctg taa aga atc ctg aag ttg cta cat tat taa gga cag aga agt att ctg cac gaa aga cgg gat cac aat ctt tat gat tca tgg taa tag tta gtt ccg acg ttg aga tgg att cgc tga gac cgg tag tgg tcg tcc gag tac acg atg tgt cgt tga ctg gat aca ggt taa ttt cca cat cga tat agt taa atg tat tgc tgg tt- --a cgg gtt cgc att tat ctg tgg aag aga cgg tgt gag aat atg ttc cgg gac cac acg gag aac aga tga cgt ctc cgg tag acg tgt atc cgg ata ctc cgt atc cta ttc cac act ttg ttt tag aaa tac atg ttc tac acc ctg atg atc ctt tga gaa gac aat aat atc ctg gag agc att cac aga ttc tat tgt gag tcg tgt tac acg atc gcg tct cta cct gat tac tat cac atc ttc cgt tac aac tta gac aag cct gta aat gat tat tgt gag atg taa agg tat ccg aac cac acg gcg tac att gtg tat tag tct tgc tat cac ata atc tgg aag cgt aag ttc ccg gag gac acg ata gac aac ata gat tac ggc ttc tgt att cgt tgt ctt tac att ttc cat tgg atg gtg cat gtg gtg cta tat ctc ttc cgt tta tta tta tac atg aga gaa aca ata tat acg agt ata ata cgg acc tca tga ttt aat aat -gt agt aat cgt cgt ctt gtt act gtt tgt ttc cta ctt ctc caa tca tat aga tta ttt ttt aaa tat ttt ctt tct atc atg gat aat att tgt aat ggt tct ttc cgt aca aca tac tgt tta gat ggt agt cgc tta gct tgg tta tga tat tgc gca taa ttt ccg gag gca aat acg ata gtc tag att gac tat cga tgg tag act cta att tat tga gtg ctt tgt cga cga gtt tac ttt tat gct cca tcg ata gat gac act gtt cta tga gat cgt cgt aca tgg gaa atg aaa tgc gtt tgt ctg aat gta tgg ctt cga gat agg tgt gat acc gga tgt ctt ctg ttc tca ata ccg tat aca agt tgg tgt ctg aga ttc gaa tct ctt tga gga gac tta tgt cac gac tac att ttt cga tga tgg aat cta tct tat cga atg ata tat ttt tca taa ata cac ttt tat agt cct cgt tta aac aga att tag tat gta gtt ccg caa atg act cgt ccc tta ata tgc agt agg cta tta tct tct tta cgt agt gat cgt cgt agg gag aga act ccg aca tct tgt aga aca acg att taa tca tag gta gag ata ctt tca gtc tgt ggt gga tga tgt cat taa cga cat ccg cct tgt ata tga tgt ttc tgt ttt caa aca cca agt cga ata ctg tct tat cgt ctt tag tcg gaa ggt tga tgt cgt atc cga tgt ata cga ggt atg agg caa cat tgt tat tgc aat tct gga agg cgg tat gaa gag gag tca ttg tat tat agt att cgt ctt tct gaa tgt cga atc tat cta gta gat acc gta gta tat tga gag agc gac ttc cat atc ctt gat tat gtt tta tga ata gat aaa gta gat gtt gtc ctt ctt cct ttt gta att tcc cgt att ttt gtt cgt gcc aat tga gta aca tta tga gaa tat gac ctg ttg cac aat cgt tct tta tgt att cca tga tgg gtg tac aat caa gat tat tac gta tcc tcg tat cgg ctc ctc gag ata aaa gag cat aca cca cac gag gac tat gtt tgg tat act gtt gaa ggt aag tgt gta acg gcg tat ttc cga ttt tcg taa ccg cgt taa tgt ttg ctc cat gat cta tta tcg cgt aga tga atc gct tct cag ctc gca tct tag tgt gac tct ttg act tgt aat aat tgc ttt cgt gga aca cgg ata tgt gtt tac agt agt aat gaa gag aag tga gtc cat cct cat cga cgc aat tag ggt cag atc ctt tag tca ata att tgt aca gaa cgt aat agt tta agc tcc cat tga att tat atc taa gat aac aca gca ata gat cgg atg att tac taa agt cat caa tgg ggt ccg tta gta tat caa aga tct tgt tat cga ttg ata gtg aat gaa tca gat agt ggt gta gag gaa tat gtc ctt ttt tat cct tgc tat caa agt tac gca tgc cgt ggc gta aca ata tct tta ata cag atg gat taa atc gtg tat tca tcg tat agc aat gta atg gag agt tac cac att tta gtc gtt tat tca gat cgc agt gtt taa taa cta att taa aca gat gag atg ata tat cca cat caa aga atg cga gat aca tat gac aga cat tat tga cag aaa tgt gac ctt cat tat cac cgt cgt cca taa atg cgt tag gta cgt acc aca tac tat cgt taa cga tgc gca caa tct cgt cca ttt tca tcc atc ttc ata atg att tac ttt ttc ata att aga gaa aaa gat caa ggt ata aaa att aga agt gtt aga cta ta- --- --- --- --- --- --- --- --- --- --- -aa cta act tat aa- --- --- --- --- --- --- --- --- --- --- --- --- --- --- --- --- --- --- --- --- --- --- --- --- --- --- --- --- --- --- --- --- --- --- --- --- --- --- --- --- --- --- --- --- --- --- --- --- --- --- --- --- --- --- --- --- --- --- --- --- --- --- --- --- --- --- --- --- --- --- --- --- --- --- --- --- --- --- --- --- --- --- --- --- --- --- -aa cta act tat gac tta act aac tta tga cta tta act cat ttc aag aaa ggt ggg tgg aga gaa ctc tat atg aca gct tgt gaa aca att aga tcc cta att tct aat gga agt ttt gat agg aga ttg tca tca gtt gat aca ttg ttt att atc tca tct att aga gca cgt ctg ttt aga gct tta gtg acc tgc tcg gtt act tct gtg taa atc ttg aat cct tta gtg ata cac tgt gtc aaa act gga tgt tta gaa tac cta tgt aga ata tgg gaa gca tgc ttg ttt ttg tct cta tta tag att aac tca tac atg gtt gta tta tga att ttc atc tgc cta atg tac tcc aat tct tgt tta caa tca att ata taa tca aag agt gat gat gca tac aca tta caa agt gaa taa tct acc atc ata aaa tac ttg ata cag agc ttt atc aca tca tgg ttt tca att gta tta tta agt ata gct aat ttt ata cag tca ata gac aat ggt tct cta agc aat att tct aat att tta aga tgt gct tcc cta cgg gcg atg aca gat ccc cta tcc acg gcc acg tca aga cat gta tat cca tta ctc att act gcg ttg aca ttt gct cca ttt tct aat agc cat gat act aaa tct ata taa cct gca tag ata gcg cga taa agc aag gtc ctt cca cca gca tct agt tga ttg ata tct tca ata tat ggg ata caa agc tta taa att tct aat act gtg ggt tca tct aca agg aat ccc cta gta tac tga att att tta tat aga tct aat tta aca tca ttt tca tct ggg ata cca caa ttc aaa ata aac tca aca aca cta ctt tcc ttt tta cat att ccc cta aaa tag gca ttc aag cat tct att tta tat att aca gcc cca tga tct acc ata aaa tca aca atg tct att tct aca tat gca tta gat aga tag taa agt aag aga tct tgt aca gaa tta caa ttc tta ata att ata gag aaa ata tct tcc ata taa ttc ttt gac act aat gca gat ata ata tct tta tat gta ata tat gca aac agt cta tct act ata tac tga tca ata tta tct cta tga atc cta aaa taa tca tac aga aca tct aca gga tca caa att ggt tca agg aga aat cta tca aat att ttc ctg tca aca act ggt tct aga aca taa cag tca aca cct aat cca tgt ttt tta tag tca tct acc aaa gat aat gac caa aga tcg agg tcg tcg tga aac tgc tca tcg aca gcc atg aaa tct gcc gac tcc atg gcg cga atc gca ctg tct tat tcg cca ttg att ttc a-t ttt tta taa tta tgt aca tgt ttt cct tct att ctc aag agt cta caa aaa tat a-t ttt ttt tcg ata tct aag tac taa gtt ttt tta ctg ttt ttg tta ctg tct tcc att ctt cta act aaa gat ctg aga taa att ata caa tct tcg cta tcg aac cat ttt tgt agt cta aag cct gaa gta att aac caa ctg ttt tta tta gtg gct ttt ttc gat cta tcc tcg tc- --- --c tca tca tcc tta ta- --- --- --- --- --- --- tta tta tca tta tta tca tag tct att aaa cac aaa tca tct --- --- --- --- --- --- --- --- --- --- --- --- --- --- --- --- --- --- --- --- --- --- --- --- --- --- --a cgt tta taa c-- --- --- --- --- --- --- --- --a aca ttc --- --- --- --- -tc att att aat tag ttc tgt agt aat atc ttt aat aat ttg gct ata --- --- -ca tct gtt --- --- --- --- --- --- --- --- --- --- --- --- --- --- --- --- --- --- --c aat act --- --- --- --- --- --- --- --a tct att gat gat ttc ttt t-- --- --- --- --- --- --- --- --- --- --- --t taa gac t-- --- --- --- --- --- --- --- --- --- --- --- --- --- --- --- --- --- --- --- --- --- --- --- --- --- --- --- --- --- --- --- --- --- --- --- --- --- --- --- taa act agt --- --- --- --- --- --- --- --- --- --- --- --- --- --- --- --- --- --- --- --- --- --- --- --- --- --- --- --- --- --- --- --- --- --- --- --- --- --- --- --- --- --- --- --- --- --- --- --- --- --- --- --- --- --- --- --- --- --- --- --- --- --- --- --- --- --- --- --- --- --- --- --- --- --- --- --- --- --- --- --- --- --- --- --- --- --- --- --- --- --- --- --- --- --- --- --- --- --- --- --- --- --- --- --- --- --- --- --- --- --t atg gta atg acg atg aaa --- -tc gag tag ta- --- --a ctt cta ata aag act tga ta- --- --t cat tat cat atg ttt gat cg- --- --- --- --- --- --- --t cat agt taa tag tgt g-- --- --- --- --- --- --- --- --- --- --- --- --- --- --- --- --- --- --- --- -gc taa atg gta ctg tta ata agt tta t-- --- --- --- --- --- --- --- --- --- --- --- --- --- --- --- --- --a gac aat atc ata gta ttt tct ttc cag aat --- --- --- --t aga tta ttt ttt taa ata ctg atc ctc aca att ccg tga tgt agc agt agt tgg t-- --- --- --- --- --- --- --- --- --- --- --- --- --- --- --- --- --- --- --- --- --- --- --- --- --- --- --- -gc atg gtc tat atc gt- --- --- --- --- --- --- --- --- --- --- --- --- --- --- --- --- --- --- --- --- --- --- --- --- --- --- --- --- --- --- --- --- --- --- --- --- --- --- --- --- --- --- --- taa aat gta tca tat ata ata gtt ttc tga cgt gga gta cag aat ttt cga --- --- --- --- --- --- --- --- --- --- --- --- --- --- --- --- --- --- --- --- --- --- --- --- --- --- --- --- --- --- --- --- --- --- --- --- --- --- --- --- --- --- --- --t taa tga gtt cat ggt aag gaa ggg caa atg tct --- gta tat aat ata cat aag tta a-- --- --- --- --- --- --- --- --- --- --- --- --- --t agt ttt tta tca tat tt- --- --- --- --- --- --- --- --t cta ata cca taa taa aaa tta tca tta tg- --- --- -ta taa tca --- --- --- --- --- --- --- --- --- --- --- --t cac tg- --- -tc gct atc att att gcg ttt gtg tag tt- --- --- --- --- ctg ccc ta- --- --- --- --- --- --- --- --t cat cta cat cac tgt ca- --- --- --- --- ctc tca cta tat ctt cta aaa tta caa --- -ac aac --- --- --- --- --- --- --- --- --- --- --- --- --- --- --- --- --- --- --- --- --- --- --- --- --- --- --- --- --- --- --- --- --- --- --- --- tgg ata ttc gat --- --- --- --- --- --- --- --- --- --- --- --- --- --- --- --- --- --- --- --- --- --- --- --- --- --- --a aca gca ttt gtg t-- --- --- --- --- --- --- --- --- --- --- --- --- --- --- --- --- -

>JX878425.1_Monkeypox_virus_isolate_DRC_07-0354_complete_genome

--- --- --- --- --- --- --- --- --- --- --- --- --- --- --- --- --- --- --- --- --- --- --- --- --- --- --- --- --- --- --- --- --- --- --- --g tgt cta gaa aaa aat gtg tga ccc acg acc gta gga aac tct aga ggg taa gaa aaa tca atc gtt tat aga gac cat cag aaa gag gtt taa tat ttt tgt gag acc cat cga cga gag aaa gga taa aaa ctt ttt acg act cca tca gaa aga ggt tta ata ttt ttg tga gac cca tcg acg aga gaa agg ata aaa act ttt tac gac tcc atc aga aag agg ttt aat att ttt gtg aga ccc atc gac --- --- --- --- --- --- --- --- --- --- --- --- --- --- --- --- --- --- --- --- --- --- --- --- --- --- --- --- --- --- --- --- --- --- --- --- --- --- --- --- --- --- --- --- --- --- --- --- --- --- --- --- --- --- --- --- --- --- --- --- --- --- --- --- --- --- --- --- --- --- --- --- --- --- --- --- --- --- --- --- --- --- --- --- --- --- --- --- --- --- --- --- --- -ga gag aaa gga taa aaa ctt ttt acg act cca tca gaa aga ggt tta ata ttt ttg tga gac cca tcg acg aga gaa agg ata aaa act ttt tac gac tcc atc aga aag agg ttt aat att ttt gtg aga ccc atc gac gag aga aag gat aaa aac ttt tta cga ctc cat cag aaa gag gtt taa tat ttt tgt gag acc cat cga cga gag aaa gga taa aaa ctt ttt acg act cca tca gaa aga ggt tta --- --- --- --- --- --- --- --- --- --- --- --- --- --- --- --- --- --- --- --- --- --- --- --- --- --- --- --- --- --- --- --- --- --- --- --- --- --- --- --- --- --- --- --- --- --- --a tat ttt tgt gag acc cat cga cga gag aaa gga taa aaa ctt ttt acg act cca tca gaa aga ggt tta ata ttt ttg tga gac cca tcg acg aga gaa aga gat ggt tag tca aga tat ttt tct tag tac aaa agt caa tgt ttt aaa ata tat gga cga gaa tta att tgt ctg tat aaa aac ttg tgt gaa att atg tac tag aga aaa aac gtg agc agt gtc ccc tac atg gat ttt aca gat cat tta tat tcc aaa aat att aac tat ata cgt tta tta tat gat gtt aac gtg taa att ata aac att att tta tga tgc aat tgt ctg aca acc --- att ggc ata agg ata ttg ata agc tct acg aga ata tat tgt tgg acg tta tcg ttt acg aaa tag ttg aga cat cag aaa gag gtt taa tat ttt tgt gag a-c cat cga aga gag aaa gag aat aaa aat a-- --- --- --- ttt ttt tgt aaa act ttt tta tga gac aag aga gaa aga gaa tac gaa tag tga tca tat cgt atc aca tat tga aac aga aag aag aag taa cga gag gta act ttt tgt gaa tgt agt taa aca ttt ttg ttt tgc aaa ccg gaa tat agt acc cgg tac act ttt tta att cgt ggt gcg gtg tct gaa tcg ttc gat taa ccc aac tca tcc att ttc aga tga ata gag tta tcg att cag aca cat gct ttg agt ttt gtt gaa tcg atg agt gaa gta tca tcg gtt gca cct tca gat gcc gat cca tcg aca tac ttg aat cca tcc ttg act tca agt tca gat gat tcc tca cac atg tct ccg ata cgt acg cta aac tct agg ttc tta aca cat ttt gta tca acg atc gtt gaa ccg atg ata tct ttg taa ctc act ttc tta tgt gag atg tta gac cca agt act gga tgg gtc ttg atg tcg ctg tct ttc tct tct tcg cta cat ctg atg tcg ata gac atc tca cag tct ttg atc ata gcc aga gct tct tca cgc gtg atc gcg gga gag tcc tta cct tgt cct ggt gac acg ctg gac aat cta gta ttc aca gtg ttt cca tca gag gat tcg gag atg gat gaa atc ttt ggg cat ttg gtg aat cca aag ttc atg tta aga ccc gcg ccg acg ata ttg taa taa gtg gtg gga tct cct ttt aca act tct tcg gat acc tca tca tct tcg gat acc tca tca tct tcg gtc tct gta act tcc gtt acg gat tga caa atc tta tca ttg gtc ggt gtt tgg tct tgc ttt gtg act ttg ata ata aca tcg att ccc ata tga tgt ttg ttt tct tct tca gta cac gag gat gag gat tgt tga aga cta gta ggc ata gca gct gcc act agg cac atg cat gcc agg aca ata tat tgt ttc atg att gct att gat tga tta ctg ttc tag atg att cta ctt tct tac cat ata ata aat tag aat ata ttt tct act ttt acg aga aat taa tta ttg tat tta tta ttt atg ggt -aa aaa aaa ctt act ata agt ggg tgg gat tct ggg aat tag tga tca gtt tat gta tat cgc aac tag cgg gca tat ggc tat tga cat cga gaa cat tac cca tat gat aag aga ttg tat ccg ttt cgt agt ctt gag tat tgg tat tac tat ata gta tgt aga tgt cga cgc tag ata gac agt cgc cca cta gag tta ccg tct ctg aat gcg gca tga tag tat cat tct ttg ttt tcg tta act gtt tgg aag atg aat ctt tgt tgt tac att taa tct cga aat tca gag tac ata tct ttg aag tat tct gat atc tat ttt ctc ctg taa aga atc ctg aag ttg cta cat tat taa gga cag aga agt att ctg cac gaa aga cgg gat cac aat ctt tat gat tca tgg taa tag tta gtt ccg acg ttg aga tgg att cgc tga gac cgg tag tgg tcg tcc gag tac acg atg tgt cgt tga ctg gat aca ggt taa ttt cca cat cga tat agt taa atg tat tgc tgg tt- --a cgg gtt cgc att tat ctg tgg aag aga cgg tgt gag aat atg ttc cgg gac cac acg gag aac aga tga cgt ctc cgg tag acg tgt atc cgg ata ctc cgt atc cta ttc cac act ttg ttt tag aaa tac atg ttc tac acc ctg atg atc ctt tga gaa gac aat aat atc ctg gag agc att cac aga ttc tat tgt gag tcg tgt tac acg atc gcg tct cta cct gat tac tat cac atc ttc cgt tac aac tta gac aag cct gta aat gat tat tgt gag atg taa agg tat ccg aac cac acg gcg tac att gtg tat tag tct tgc tat cac ata atc tgg aag cgt aag ttc ccg gag gac acg ata gac aac ata gat tac ggc ttc tgt att cgt tgt ctt tac att ttc cat tgg atg gtg cat gtg gtg cta tat ctc ttc cgt tta tta tta tac atg aga gaa aca ata tat acg agt ata ata cgg acc tca tga ttt aat aat -gt agt aat cgt cgt ctt gtt act gtt tgt ttc cta ctt ctc caa tca tat aga tta ttt ttt aaa tat ttt ctt tct atc atg gat aat att tgt aat ggt tct ttc cgt aca aca tac tgt tta gat ggt agt cgc tta gct tgg tta tga tat tgc gca taa ttt ccg gag gca aat acg ata gtc tag att gac tat cga tgg tag act cta att tat tga gtg ctt tgt cga cga gtt tac ttt tat gct cca tcg ata gat gac act gtt cta tga gat cgt cgt aca tgg gaa atg aaa tgc gtt tgt ctg aat gta tgg ctt cga gat agg tgt gat acc gga tgt ctt ctg ttc tca ata ccg tat aca agt tgg tgt ctg aga ttc gaa tct ctt tga gga gac tta tgt cac gac tac att ttt cga tga tgg aat cta tct tat cga atg ata tat ttt tca taa ata cac ttt tat agt cct cgt tta aac aga att tag tat gta gtt ccg caa atg act cgt ccc tta ata tgc agt agg cta tta tct tct tta cgt agt gat cgt cgt agg gag aga act ccg aca tct tgt aga aca acg att taa tca tag gta gag ata ctt tca gtc tgt ggt gga tga tgt cat taa cga cat ccg cct tgt ata tga tgt ttc tgt ttt caa aca cca agt cga ata ctg tct tat cgt ctt tag tcg gaa ggt tga tgt cgt atc cga tgt ata cga ggt atg agg caa cat tgt tat tgc aat tct gga agg cgg tat gaa gag gag tca ttg tat tat agt att cgt ctt tct gaa tgt cga atc tat cta gta gat acc gta gta tat tga gag agc gac ttc cat atc ctt gat tat gtt tta tga ata gat aaa gta gat gtt gtc ctt ctt cct ttt gta att tcc cgt att ttt gtt cgt gcc aat tga gta aca tta tga gaa tat gac ctg ttg cac aat cgt tct tta tgt att cca tga tgg gtg tac aat caa gat tat tac gta tcc tcg tat cgg ctc ctc gag ata aaa gag cat aca cca cac gag gac tat gtt tgg tat act gtt gaa ggt aag tgt gta acg gcg tat ttc cga ttt tcg taa ccg cgt taa tgt ttg ctc cat gat cta tta tcg cgt aga tga atc gct tct cag ctc gca tct tag tgt gac tct ttg act tgt aat aat tgc ttt cgt gga aca cgg ata tgt gtt tac agt agt aat gaa gag aag tga gtc cat cct cat cga cgc aat tag ggt cag atc ctt tag tca ata att tgt aca gaa cgt aat agt tta agc tcc cat tga att tat atc taa gat aac aca gca ata gat cgg atg att tac taa agt cat caa tgg ggt ccg tta gta tat caa aga tct tgt tat cga ttg ata gtg aat gaa tca gat agt ggt gta gag gaa tat gtc ctt ttt tat cct tgc tat caa agt tac gca tgc cgt ggc gta aca ata tct tta ata cag atg gat taa atc gtg tat tca tcg tat agc aat gta atg gag agt tac cac att tta gtc gtt tat tca gat cgc agt gtt taa taa cta att taa aca gat gag atg ata tat cca cat caa aga atg cga gat aca tat gac aga cat tat tga cag aaa tgt gac ctt cat tat cac cgt cgt cca taa atg cgt tag gta cgt acc aca tac tat cgt taa cga tgc gca caa tct cgt cca ttt tca tcc atc ttc ata atg att tac ttt ttc ata att aga gaa aaa gat caa ggt ata aaa att aga agt gtt aga cta ta- --- --- --- --- --- --- --- --- --- --- -aa cta act tat aa- --- --- --- --- --- --- --- --- --- --- --- --- --- --- --- --- --- --- --- --- --- --- --- --- --- --- --- --- --- --- --- --- --- --- --- --- --- --- --- --- --- --- --- --- --- --- --- --- --- --- --- --- --- --- --- --- --- --- --- --- --- --- --- --- --- --- --- --- --- --- --- --- --- --- --- --- --- --- --- --- --- --- --- --- --- --- -aa cta act tat gac tta act aac tta tga cta tta act cat ttc aag aaa ggt ggg tgg aga gaa ctc tat atg aca gct tgt gaa aca att aga tcc cta att tct aat gga agt ttt gat agg aga ttg tca tca gtt gat aca ttg ttt att atc tca tct att aga gca cgt ctg ttt aga gct tta gtg acc tgc tcg gtt act tct gtg taa atc ttg aat cct tta gtg ata cac tgt gtc aaa act gga tgt tta gaa tac cta tgt aga ata tgg gaa gca tgc ttg ttt ttg tct cta tta tag att aac tca tac atg gtt gta tta tga att ttc atc tgc cta atg tac tcc aat tct tgt tta caa tca att ata taa tca aag agt gat gat gca tac aca tta caa agt gaa taa tct acc atc ata aaa tac ttg ata cag agc ttt atc aca tca tgg ttt tca att gta tta tta agt ata gct aat ttt ata cag tca ata gac aat ggt tct cta agc aat att tct aat att tta aga tgt gct tcc cta cgg gcg atg aca gat ccc cta tcc acg gcc acg tca aga cat gta tat cca tta ctc att act gcg ttg aca ttt gct cca ttt tct aat agc cat gat act aaa tct ata taa cct gca tag ata gcg cga taa agc aag gtc ctt cca cca gca tct agt tga ttg ata tct tca ata tat ggg ata caa agc tta taa att tct aat act gtg ggt tca tct aca agg aat ccc cta gta tac tga att att tta tat aga tct aat tta aca tca ttt tca tct ggg ata cca caa ttc aaa ata aac tca aca aca cta ctt tcc ttt tta cat att ccc cta aaa tag gca ttc aag cat tct att tta tat att aca gcc cca tga tct acc ata aaa tca aca atg tct att tct aca tat gca tta gat aga tag taa agt aag aga tct tgt aca gaa tta caa ttc tta ata att ata gag aaa ata tct tcc ata taa ttc ttt gac act aat gca gat ata ata tct tta tat gta ata tat gca aac agt cta tct act ata tac tga tca ata tta tct cta tga atc cta aaa taa tca tac aga aca tct aca gga tca caa att ggt tca agg aga aat cta tca aat att ttc ctg tca aca act ggt tct aga aca taa cag tca aca cct aat cca tgt ttt tta tag tca tct acc aaa gat aat gac caa aga tcg agg tcg tcg tga aac tgc tca tcg aca gcc atg aaa tct gcc gac tcc atg gcg cga atc gca ctg tct tat tcg cca ttg att ttc a-t ttt tta taa tta tgt aca tgt ttt cct tct att ctc aag agt cta caa aaa tat a-t ttt ttt tcg ata tct aag tac taa gtt ttt tta ctg ttt ttg tta ctg tct tcc att ctt cta act aaa gat ctg aga taa att ata caa tct tcg cta tcg aac cat ttt tgt agt cta aag cct gaa gta att aac caa ctg ttt tta tta gtg gct ttt ttc gat cta tcc tcg tc- --- --c tca tca tcc tta ta- --- --- --- --- --- --- tta tta tca tta tta tca tag tct att aaa cac aaa tca tct --- --- --- --- --- --- --- --- --- --- --- --- --- --- --- --- --- --- --- --- --- --- --- --- --- --- --a cgt tta taa c-- --- --- --- --- --- --- --- --a aca ttc --- --- --- --- -tc att att aat tag ttc tgt agt aat atc ttt aat aat ttg gct ata --- --- -ca tct gtt --- --- --- --- --- --- --- --- --- --- --- --- --- --- --- --- --- --- --c aat act --- --- --- --- --- --- --- --a tct att gat gat ttc ttt t-- --- --- --- --- --- --- --- --- --- --- --t taa gac t-- --- --- --- --- --- --- --- --- --- --- --- --- --- --- --- --- --- --- --- --- --- --- --- --- --- --- --- --- --- --- --- --- --- --- --- --- --- --- --- taa act agt --- --- --- --- --- --- --- --- --- --- --- --- --- --- --- --- --- --- --- --- --- --- --- --- --- --- --- --- --- --- --- --- --- --- --- --- --- --- --- --- --- --- --- --- --- --- --- --- --- --- --- --- --- --- --- --- --- --- --- --- --- --- --- --- --- --- --- --- --- --- --- --- --- --- --- --- --- --- --- --- --- --- --- --- --- --- --- --- --- --- --- --- --- --- --- --- --- --- --- --- --- --- --- --- --- --- --- --- --- --t atg gta atg acg atg aaa --- -tc gag tag ta- --- --a ctt cta ata aag act tga ta- --- --t cat tat cat atg ttt gat cg- --- --- --- --- --- --- --t cat agt taa tag tgt g-- --- --- --- --- --- --- --- --- --- --- --- --- --- --- --- --- --- --- --- -gc taa atg gta ctg tta ata agt tta t-- --- --- --- --- --- --- --- --- --- --- --- --- --- --- --- --- --a gac aat atc ata gta ttt tct ttc cag aat --- --- --- --t aga tta ttt ttt taa ata ctg atc ctc aca att ccg tga tgt agc agt agt tgg t-- --- --- --- --- --- --- --- --- --- --- --- --- --- --- --- --- --- --- --- --- --- --- --- --- --- --- --- -gc atg gtc tat atc gt- --- --- --- --- --- --- --- --- --- --- --- --- --- --- --- --- --- --- --- --- --- --- --- --- --- --- --- --- --- --- --- --- --- --- --- --- --- --- --- --- --- --- --- taa aat gta tca tat ata ata gtt ttc tga cgt gga gta cag aat ttt cga --- --- --- --- --- --- --- --- --- --- --- --- --- --- --- --- --- --- --- --- --- --- --- --- --- --- --- --- --- --- --- --- --- --- --- --- --- --- --- --- --- --- --- --t taa tga gtt cat ggt aag gaa ggg caa atg tct --- gta tat aat ata cat aag tta a-- --- --- --- --- --- --- --- --- --- --- --- --- --t agt ttt tta tca tat tt- --- --- --- --- --- --- --- --t cta ata cca taa taa aaa tta tca tta tg- --- --- -ta taa tca --- --- --- --- --- --- --- --- --- --- --- --t cac tg- --- -tc gct atc att att gcg ttt gtg tag tt- --- --- --- --- ctg ccc ta- --- --- --- --- --- --- --- --t cat cta cat cac tgt ca- --- --- --- --- ctc tca cta tat ctt cta aaa tta caa --- -ac aac --- --- --- --- --- --- --- --- --- --- --- --- --- --- --- --- --- --- --- --- --- --- --- --- --- --- --- --- --- --- --- --- --- --- --- --- tgg ata ttc gat --- --- --- --- --- --- --- --- --- --- --- --- --- --- --- --- --- --- --- --- --- --- --- --- --- --- --a aca gca ttt gtg t-- --- --- --- --- --- --- --- --- --- --- --- --- --- --- --- --- -

>JX878407.1_Monkeypox_virus_isolate_DRC_06-0950_complete_genome

--- --- --- --- --- --- --- --- --- --- --- --- --- --- --- --- --- --- --- --- --- --- --- --- --- --- --- --- --- --- --- --- --- --- --- --g tgt cta gaa aaa aat gtg tga ccc acg acc gta gga aac tct aga ggg taa gaa aaa tca atc gtt tat aga gac cat cag aaa gag gtt taa tat ttt tgt gag acc cat cga cga gag aaa gg- --- --- --- --- --- --- --- --- --- --- --- --- --- --- --- --- --- --- --- --- --- --- --- --- --- --- --- --- --- --- --- --- --- --- --- --- --- --- --- --- --- --- --- --- --- --- --- --- --- --- --- --- --- --- --- --- --- --- --- --- --- --- --- --- --- --- --- --- --- --- --- --- --- --- --- --- --- --- --- --- --- --- --- --- --- --- --- --- --- --- --- --- --- --- --- --- --- --- --- --- --- --- --- --- --- --- --- --- --- --- --- --- --- --- --- --- --- --- --- --- --- --- --- --- --- --- --- --- --- --- --- --- --- --- --- --- --- --- --- --- --- --- --- --- --- --- --- --- --- --- --- --- --- --- --- --- --- --- --- --- --- --- --- --- --- --- --- --- --- --- --- --- --- --- --- --- --- --- --- --- --- --- --- --- --- --- --- --- --- --- --- --- --- --- --- --- --- --- --- --- --- --- --- --- --- --- --- --- --- --a taa aaa ctt ttt acg act cca tca gaa aga ggt tta --- --- --- --- --- --- --- --- --- --- --- --- --- --- --- --- --- --- --- --- --- --- --- --- --- --- --- --- --- --- --- --- --- --- --- --- --- --- --- --- --- --- --- --- --- --- --a tat ttt tgt gag acc cat cga cga gag aaa gga taa aaa ctt ttt acg act cca tca gaa aga ggt tta ata ttt ttg tga gac cca tcg acg aga gaa aga gat ggt tag tca aga tat ttt tct tag tac aaa agt caa tgt ttt aaa ata tat gga cga gaa tta att tgt ctg tat aaa aac ttg tgt gaa att atg tac tag aga aaa aac gtg agc agt gtc ccc tac atg gat ttt aca gat cat tta tat tcc aaa aat att aac tat ata cgt tta tta tat gat gtt aac gtg taa att ata aac att att tta tga tgc aat tgt ctg aca acc --- att ggc ata agg ata ttg ata agc tct acg aga ata tat tgt tgg acg tta tcg ttt acg aaa tag ttg aga cat cag aaa gag gtt taa tat ttt tgt gag a-c cat cga aga gag aaa gag aat aaa aat a-- --- --- --- ttt ttt tgt aaa act ttt tta tga gac aag aga gaa aga gaa tac gaa tag tga tca tat cgt atc aca tat tga aac aga aag aag aag taa cga gag gta act ttt tgt gaa tgt agt taa aca ttt ttg ttt tgc aaa ccg gaa tat agt acc cgg tac act ttt tta att cgt ggt gcg gtg tct gaa tcg ttc gat taa ccc aac tca tcc att ttc aga tga ata gag tta tcg att cag aca cat gct ttg agt ttt gtt gaa tcg atg agt gaa gta tca tcg gtt gca cct tca gat gcc gat cca tcg aca tac ttg aat cca tcc ttg act tca agt tca gat gat tcc tca cac atg tct ccg ata cgt acg cta aac tct agg ttc tta aca cat ttt gta tca acg atc gtt gaa ccg atg ata tct ttg taa ctc act ttc tta tgt gag atg tta gac cca agt act gga tgg gtc ttg atg tcg ctg tct ttc tct tct tcg cta cat ctg atg tcg ata gac atc tca cag tct ttg atc ata gcc aga gct tct tca cgc gtg atc gcg gga gag tcc tta cct tgt cct ggt gac acg ctg gac aat cta gta ttc aca gtg ttt cca tca gag gat tcg gag atg gat gaa atc ttt ggg cat ttg gtg aat cca aag ttc atg tta aga ccc gcg ccg acg ata ttg taa taa gtg gtg gga tct cct ttt aca act tct tcg gat acc tca tca tct tcg gat acc tca tca tct tcg gtc tct gta act tcc gtt acg gat tga caa atc tta tca ttg gtc ggt gtt tgg tct tgc ttt gtg act ttg ata ata aca tcg att ccc ata tga tgt ttg ttt tct tct tca gta cac gag gat gag gat tgt tga aga cta gta ggc ata gca gct gcc act agg cac atg cat gcc agg aca ata tat tgt ttc atg att gct att gat tga tta ctg ttc tag atg att cta ctt tct tac cat ata ata aat tag aat ata ttt tct act ttt acg aga aat taa tta ttg tat tta tta ttt atg ggt -aa aaa aaa ctt act ata agt ggg tgg gat tct ggg aat tag tga tca gtt tat gta tat cgc aac tag cgg gca tat ggc tat tga cat cga gaa cat tac cca tat gat aag aga ttg tat ccg ttt cgt agt ctt gag tat tgg tat tac tat ata gta tgt aga tgt cga cgc tag ata gac agt cgc cca cta gag tta ccg tct ctg aat gcg gca tga tag tat cat tct ttg ttt tcg tta act gtt tgg aag atg aat ctt tgt tgt tac att taa tct cga aat tca gag tac ata tct ttg aag tat tct gat atc tat ttt ctc ctg taa aga atc ctg aag ttg cta cat tat taa gga cag aga agt att ctg cac gaa aga cgg gat cac aat ctt tat gat tca tgg taa tag tta gtt ccg acg ttg aga tgg att cgc tga gac cgg tag tgg tcg tcc gag tac acg atg tgt cgt tga ctg gat aca ggt taa ttt cca cat cga tat agt taa atg tat tgc tgg tt- --a cgg gtt cgc att tat ctg tgg aag aga cgg tgt gag aat atg ttc cgg gac cac acg gag aac aga tga cgt ctc cgg tag acg tgt atc cgg ata ctc cgt atc cta ttc cac act ttg ttt tag aaa tac atg ttc tac acc ctg atg atc ctt tga gaa gac aat aat atc ctg gag agc att cac aga ttc tat tgt gag tcg tgt tac acg atc gcg tct cta cct gat tac tat cac atc ttc cgt tac aac tta gac aag cct gta aat gat tat tgt gag atg taa agg tat ccg aac cac acg gcg tac att gtg tat tag tct tgc tat cac ata atc tgg aag cgt aag ttc ccg gag gac acg ata gac aac ata gat tac ggc ttc tgt att cgt tgt ctt tac att ttc cat tgg atg gtg cat gtg gtg cta tat ctc ttc cgt tta tta tta tac atg aga gaa aca ata tat acg agt ata ata cgg acc tca tga ttt aat aat -gt agt aat cgt cgt ctt gtt act gtt tgt ttc cta ctt ctc caa tca tat aga tta ttt ttt aaa tat ttt ctt tct atc atg gat aat att tgt aat ggt tct ttc cgt aca aca tac tgt tta gat ggt agt cgc tta gct tgg tta tga tat tgc gca taa ttt ccg gag gca aat acg ata gtc tag att gac tat cga tgg tag act cta att tat tga gtg ctt tgt cga cga gtt tac ttt tat gct cca tcg ata gat gac act gtt cta tga gat cgt cgt aca tgg gaa atg aaa tgc gtt tgt ctg aat gta tgg ctt cga gat agg tgt gat acc gga tgt ctt ctg ttc tca ata ccg tat aca agt tgg tgt ctg aga ttc gaa tct ctt tga gga gac tta tgt cac gac tac att ttt cga tga tgg aat cta tct tat cga atg ata tat ttt tca taa ata cac ttt tat agt cct cgt tta aac aga att tag tat gta gtt ccg caa atg act cgt ccc tta ata tgc agt agg cta tta tct tct tta cgt agt gat cgt cgt agg gag aga act ccg aca tct tgt aga aca acg att taa tca tag gta gag ata ctt tca gtc tgt ggt gga tga tgt cat taa cga cat ccg cct tgt ata tga tgt ttc tgt ttt caa aca cca agt cga ata ctg tct tat cgt ctt tag tcg gaa ggt tga tgt cgt atc cga tgt ata cga ggt atg agg caa cat tgt tat tgc aat tct gga agg cgg tat gaa gag gag tca ttg tat tat agt att cgt ctt tct gaa tgt cga atc tat cta gta gat acc gta gta tat tga gag agc gac ttc cat atc ctt gat tat gtt tta tga ata gat aaa gta gat gtt gtc ctt ctt cct ttt gta att tcc cgt att ttt gtt cgt gcc aat tga gta aca tta tga gaa tat gac ctg ttg cac aat cgt tct tta tgt att cca tga tgg gtg tac aat caa gat tat tac gta tcc tcg tat cgg ctc ctc gag ata aaa gag cat aca cca cac gag gac tat gtt tgg tat act gtt gaa ggt aag tgt gta acg gcg tat ttc cga ttt tcg taa ccg cgt taa tgt ttg ctc cat gat cta tta tcg cgt aga tga atc gct tct cag ctc gca tct tag tgt gac tct ttg act tgt aat aat tgc ttt cgt gga aca cgg ata tgt gtt tac agt agt aat gaa gag aag tga gtc cat cct cat cga cgc aat tag ggt cag atc ctt tag tca ata att tgt aca gaa cgt aat agt tta agc tcc cat tga att tat atc taa gat aac aca gca ata gat cgg atg att tac taa agt cat caa tgg ggt ccg tta gta tat caa aga tct tgt tat cga ttg ata gtg aat gaa tca gat agt ggt gta gag gaa tat gtc ctt ttt tat cct tgc tat caa agt tac gca tgc cgt ggc gta aca ata tct tta ata cag atg gat taa atc gtg tat tca tcg tat agc aat gta atg gag agt tac cac att tta gtc gtt tat tca gat cgc agt gtt taa taa cta att taa aca gat gag atg ata tat cca cat caa aga atg cga gat aca tat gac aga cat tat tga cag aaa tgt gac ctt cat tat cac cgt cgt cca taa atg cgt tag gta cgt acc aca tac tat cgt taa cga tgc gca caa tct cgt cca ttt tca tcc atc ttc ata atg att tac ttt ttc ata att aga gaa aaa gat caa ggt ata aaa att aga agt gtt aga cta ta- --- --- --- --- --- --- --- --- --- --- -aa cta act tat aa- --- --- --- --- --- --- --- --- --- --- --- --- --- --- --- --- --- --- --- --- --- --- --- --- --- --- --- --- --- --- --- --- --- --- --- --- --- --- --- --- --- --- --- --- --- --- --- --- --- --- --- --- --- --- --- --- --- --- --- --- --- --- --- --- --- --- --- --- --- --- --- --- --- --- --- --- --- --- --- --- --- --- --- --- --- --- -aa cta act tat gac tta act aac tta tga cta tta act cat ttc aag aaa ggt ggg tgg aga gaa ctc tat atg aca gct tgt gaa aca att aga tcc cta att tct aat gga agt ttt gat agg aga ttg tca tca gtt gat aca ttg ttt att atc tca tct att aga gca cgt ctg ttt aga gct tta gtg acc tgc tcg gtt act tct gtg taa atc ttg aat cct tta gtg ata cac tgt gtc aaa act gga tgt tta gaa tac cta tgt aga ata tgg gaa gca tgc ttg ttt ttg tct cta tta tag att aac tca tac atg gtt gta tta tga att ttc atc tgc cta atg tac tcc aat tct tgt tta caa tca att ata taa tca aag agt gat gat gca tac aca tta caa agt gaa taa tct acc atc ata aaa tac ttg ata cag agc ttt atc aca tca tgg ttt tca att gta tta tta agt ata gct aat ttt ata cag tca ata gac aat ggt tct cta agc aat att tct aat att tta aga tgt gct tcc cta cgg gcg atg aca gat ccc cta tcc acg gcc acg tca aga cat gta tat cca tta ctc att act gcg ttg aca ttt gct cca ttt tct aat agc cat gat act aaa tct ata taa cct gca tag ata gcg cga taa agc aag gtc ctt cca cca gca tct agt tga ttg ata tct tca ata tat ggg ata caa agc tta taa att tct aat act gtg ggt tca tct aca agg aat ccc cta gta tac tga att att tta tat aga tct aat tta aca tca ttt tca tct ggg ata cca caa ttc aaa ata aac tca aca aca cta ctt tcc ttt tta cat att ccc cta aaa tag gca ttc aag cat tct att tta tat att aca gcc cca tga tct acc ata aaa tca aca atg tct att tct aca tat gca tta gat aga tag taa agt aag aga tct tgt aca gaa tta caa ttc tta ata att ata gag aaa ata tct tcc ata taa ttc ttt gac act aat gca gat ata ata tct tta tat gta ata tat gca aac agt cta tct act ata tac tga tca ata tta tct cta tga atc cta aaa taa tca tac aga aca tct aca gga tca caa att ggt tca agg aga aat cta tca aat att ttc ctg tca aca act ggt tct aga aca taa cag tca aca cct aat cca tgt ttt tta tag tca tct acc aaa gat aat gac caa aga tcg agg tcg tcg tga aac tgc tca tcg aca gcc atg aaa tct gcc gac tcc atg gcg cga atc gca ctg tct tat tcg cca ttg att ttc a-t ttt tta taa tta tgt aca tgt ttt cct tct att ctc aag agt cta caa aaa tat a-t ttt ttt tcg ata tct aag tac taa gtt ttt tta ctg ttt ttg tta ctg tct tcc att ctt cta act aaa gat ctg aga taa att ata caa tct tcg cta tcg aac cat ttt tgt agt cta aag cct gaa gta att aac caa ctg ttt tta tta gtg gct ttt ttc gat cta tcc tcg tc- --- --c tca tca tcc tta ta- --- --- --- --- --- --- tta tta tca tta tta tca tag tct att aaa cac aaa tca tct --- --- --- --- --- --- --- --- --- --- --- --- --- --- --- --- --- --- --- --- --- --- --- --- --- --- --a cgt tta taa c-- --- --- --- --- --- --- --- --a aca ttc --- --- --- --- -tc att att aat tag ttc tgt agt aat atc ttt aat aat ttg gct ata --- --- -ca tct gtt --- --- --- --- --- --- --- --- --- --- --- --- --- --- --- --- --- --- --c aat act --- --- --- --- --- --- --- --a tct att gat gat ttc ttt t-- --- --- --- --- --- --- --- --- --- --- --t taa gac t-- --- --- --- --- --- --- --- --- --- --- --- --- --- --- --- --- --- --- --- --- --- --- --- --- --- --- --- --- --- --- --- --- --- --- --- --- --- --- --- taa act agt --- --- --- --- --- --- --- --- --- --- --- --- --- --- --- --- --- --- --- --- --- --- --- --- --- --- --- --- --- --- --- --- --- --- --- --- --- --- --- --- --- --- --- --- --- --- --- --- --- --- --- --- --- --- --- --- --- --- --- --- --- --- --- --- --- --- --- --- --- --- --- --- --- --- --- --- --- --- --- --- --- --- --- --- --- --- --- --- --- --- --- --- --- --- --- --- --- --- --- --- --- --- --- --- --- --- --- --- --- --t atg gta atg acg atg aaa --- -tc gag tag ta- --- --a ctt cta ata aag act tga ta- --- --t cat tat cat atg ttt gat cg- --- --- --- --- --- --- --t cat agt taa tag tgt g-- --- --- --- --- --- --- --- --- --- --- --- --- --- --- --- --- --- --- --- -gc taa atg gta ctg tta ata agt tta t-- --- --- --- --- --- --- --- --- --- --- --- --- --- --- --- --- --a gac aat atc ata gta ttt tct ttc cag aat --- --- --- --t aga tta ttt ttt taa ata ctg atc ctc aca att ccg tga tgt agc agt agt tgg t-- --- --- --- --- --- --- --- --- --- --- --- --- --- --- --- --- --- --- --- --- --- --- --- --- --- --- --- -gc atg gtc tat atc gt- --- --- --- --- --- --- --- --- --- --- --- --- --- --- --- --- --- --- --- --- --- --- --- --- --- --- --- --- --- --- --- --- --- --- --- --- --- --- --- --- --- --- --- taa aat gta tca tat ata ata gtt ttc tga cgt gga gta cag aat ttt cga --- --- --- --- --- --- --- --- --- --- --- --- --- --- --- --- --- --- --- --- --- --- --- --- --- --- --- --- --- --- --- --- --- --- --- --- --- --- --- --- --- --- --- --t taa tga gtt cat ggt aag gaa ggg caa atg tct --- gta tat aat ata cat aag tta a-- --- --- --- --- --- --- --- --- --- --- --- --- --t agt ttt tta tca tat tt- --- --- --- --- --- --- --- --t cta ata cca taa taa aaa tta tca tta tg- --- --- -ta taa tca --- --- --- --- --- --- --- --- --- --- --- --t cac tg- --- -tc gct atc att att gcg ttt gtg tag tt- --- --- --- --- ctg ccc ta- --- --- --- --- --- --- --- --t cat cta cat cac tgt ca- --- --- --- --- ctc tca cta tat ctt cta aaa tta caa --- -ac aac --- --- --- --- --- --- --- --- --- --- --- --- --- --- --- --- --- --- --- --- --- --- --- --- --- --- --- --- --- --- --- --- --- --- --- --- tgg ata ttc gat --- --- --- --- --- --- --- --- --- --- --- --- --- --- --- --- --- --- --- --- --- --- --- --- --- --- --a aca gca ttt gtg t-- --- --- --- --- --- --- --- --- --- --- --- --- --- --- --- --- -

>JX878423.1_Monkeypox_virus_isolate_DRC_07-0337_complete_genome

--- --- --- --- --- --- --- --- --- --- --- --- --- --- --- --- --- --- --- --- --- --- --- --- --- --- --- --- --- --- --- --- --- --- --- --g tgt cta gaa aaa aat gtg tga ccc acg acc gta gga aac tct aga ggg taa gaa aaa tca atc gtt tat aga gac cat cag aaa gag gtt taa tat ttt tgt gag acc cat cga cga gag aaa gg- --- --- --- --- --- --- --- --- --- --- --- --- --- --- --- --- --- --- --- --- --- --- --- --- --- --- --- --- --- --- --- --- --- --- --- --- --- --- --- --- --- --- --- --- --- --- --- --- --- --- --- --- --- --- --- --- --- --- --- --- --- --- --- --- --- --- --- --- --- --- --- --- --- --- --- --- --- --- --- --- --- --- --- --- --- --- --- --- --- --- --- --- --- --- --- --- --- --- --- --- --- --- --- --- --- --- --- --- --- --- --- --- --- --- --- --- --- --- --- --- --- --- --- --- --- --- --- --- --- --- --- --- --- --- --- --- --- --- --- --- --- --- --- --- --- --- --- --- --- --- --- --- --- --- --- --- --- --- --- --- --- --- --- --- --- --- --- --- --- --- --- --- --- --- --- --- --- --- --- --- --- --- --- --- --- --- -at aaa aac ttt tta cga ctc cat cag aaa gag gtt taa tat ttt tgt gag acc cat cga cga gag aaa gga taa aaa ctt ttt acg act cca tca gaa aga ggt tta --- --- --- --- --- --- --- --- --- --- --- --- --- --- --- --- --- --- --- --- --- --- --- --- --- --- --- --- --- --- --- --- --- --- --- --- --- --- --- --- --- --- --- --- --- --- --a tat ttt tgt gag acc cat cga cga gag aaa gga taa aaa ctt ttt acg act cca tca gaa aga ggt tta ata ttt ttg tga gac cca tcg acg aga gaa aga gat ggt tag tca aga tat ttt tct tag tac aaa agt caa tgt ttt aaa ata tat gga cga gaa tta att tgt ctg tat aaa aac ttg tgt gaa att atg tac tag aga aaa aac gtg agc agt gtc ccc tac atg gat ttt aca gat cat tta tat tcc aaa aat att aac tat ata cgt tta tta tat gat gtt aac gtg taa att ata aac att att tta tga tgc aat tgt ctg aca acc --- att ggc ata agg ata ttg ata agc tct acg aga ata tat tgt tgg acg tta tcg ttt acg aaa tag ttg aga cat cag aaa gag gtt taa tat ttt tgt gag a-c cat cga aga gag aaa gag aat aaa aat a-- --- --- --- ttt ttt tgt aaa act ttt tta tga gac aag aga gaa aga gaa tac gaa tag tga tca tat cgt atc aca tat tga aac aga aag aag aag taa cga gag gta act ttt tgt gaa tgt agt taa aca ttt ttg ttt tgc aaa ccg gaa tat agt acc cgg tac act ttt tta att cgt ggt gcg gtg tct gaa tcg ttc gat taa ccc aac tca tcc att ttc aga tga ata gag tta tcg att cag aca cat gct ttg agt ttt gtt gaa tcg atg agt gaa gta tca tcg gtt gca cct tca gat gcc gat cca tcg aca tac ttg aat cca tcc ttg act tca agt tca gat gat tcc tca cac atg tct ccg ata cgt acg cta aac tct agg ttc tta aca cat ttt gta tca acg atc gtt gaa ccg atg ata tct ttg taa ctc act ttc tta tgt gag atg tta gac cca agt act gga tgg gtc ttg atg tcg ctg tct ttc tct tct tcg cta cat ctg atg tcg ata gac atc tca cag tct ttg atc ata gcc aga gct tct tca cgc gtg atc gcg gga gag tcc tta cct tgt cct ggt gac acg ctg gac aat cta gta ttc aca gtg ttt cca tca gag gat tcg gag atg gat gaa atc ttt ggg cat ttg gtg aat cca aag ttc atg tta aga ccc gcg ccg acg ata ttg taa taa gtg gtg gga tct cct ttt aca act tct tcg gat acc tca tca tct tcg gat acc tca tca tct tcg gtc tct gta act tcc gtt acg gat tga caa atc tta tca ttg gtc ggt gtt tgg tct tgc ttt gtg act ttg ata ata aca tcg att ccc ata tga tgt ttg ttt tct tct tca gta cac gag gat gag gat tgt tga aga cta gta ggc ata gca gct gcc act agg cac atg cat gcc agg aca ata tat tgt ttc atg att gct att gat tga tta ctg ttc tag atg att cta ctt tct tac cat ata ata aat tag aat ata ttt tct act ttt acg aga aat taa tta ttg tat tta tta ttt atg ggt -aa aaa aaa ctt act ata agt ggg tgg gat tct ggg aat tag tga tca gtt tat gta tat cgc aac tag cgg gca tat ggc tat tga cat cga gaa cat tac cca tat gat aag aga ttg tat ccg ttt cgt agt ctt gag tat tgg tat tac tat ata gta tgt aga tgt cga cgc tag ata gac agt cgc cca cta gag tta ccg tct ctg aat gcg gca tga tag tat cat tct ttg ttt tcg tta act gtt tgg aag atg aat ctt tgt tgt tac att taa tct cga aat tca gag tac ata tct ttg aag tat tct gat atc tat ttt ctc ctg taa aga atc ctg aag ttg cta cat tat taa gga cag aga agt att ctg cac gaa aga cgg gat cac aat ctt tat gat tca tgg taa tag tta gtt ccg acg ttg aga tgg att cgc tga gac cgg tag tgg tcg tcc gag tac acg atg tgt cgt tga ctg gat aca ggt taa ttt cca cat cga tat agt taa atg tat tgc tgg tt- --a cgg gtt cgc att tat ctg tgg aag aga cgg tgt gag aat atg ttc cgg gac cac acg gag aac aga tga cgt ctc cgg tag acg tgt atc cgg ata ctc cgt atc cta ttc cac act ttg ttt tag aaa tac atg ttc tac acc ctg atg atc ctt tga gaa gac aat aat atc ctg gag agc att cac aga ttc tat tgt gag tcg tgt tac acg atc gcg tct cta cct gat tac tat cac atc ttc cgt tac aac tta gac aag cct gta aat gat tat tgt gag atg taa agg tat ccg aac cac acg gcg tac att gtg tat tag tct tgc tat cac ata atc tgg aag cgt aag ttc ccg gag gac acg ata gac aac ata gat tac ggc ttc tgt att cgt tgt ctt tac att ttc cat tgg atg gtg cat gtg gtg cta tat ctc ttc cgt tta tta tta tac atg aga gaa aca ata tat acg agt ata ata cgg acc tca tga ttt aat aat -gt agt aat cgt cgt ctt gtt act gtt tgt ttc cta ctt ctc caa tca tat aga tta ttt ttt aaa tat ttt ctt tct atc atg gat aat att tgt aat ggt tct ttc cgt aca aca tac tgt tta gat ggt agt cgc tta gct tgg tta tga tat tgc gca taa ttt ccg gag gca aat acg ata gtc tag att gac tat cga tgg tag act cta att tat tga gtg ctt tgt cga cga gtt tac ttt tat gct cca tcg ata gat gac act gtt cta tga gat cgt cgt aca tgg gaa atg aaa tgc gtt tgt ctg aat gta tgg ctt cga gat agg tgt gat acc gga tgt ctt ctg ttc tca ata ccg tat aca agt tgg tgt ctg aga ttc gaa tct ctt tga gga gac tta tgt cac gac tac att ttt cga tga tgg aat cta tct tat cga atg ata tat ttt tca taa ata cac ttt tat agt cct cgt tta aac aga att tag tat gta gtt ccg caa atg act cgt ccc tta ata tgc agt agg cta tta tct tct tta cgt agt gat cgt cgt agg gag aga act ccg aca tct tgt aga aca acg att taa tca tag gta gag ata ctt tca gtc tgt ggt gga tga tgt cat taa cga cat ccg cct tgt ata tga tgt ttc tgt ttt caa aca cca agt cga ata ctg tct tat cgt ctt tag tcg gaa ggt tga tgt cgt atc cga tgt ata cga ggt atg agg caa cat tgt tat tgc aat tct gga agg cgg tat gaa gag gag tca ttg tat tat agt att cgt ctt tct gaa tgt cga atc tat cta gta gat acc gta gta tat tga gag agc gac ttc cat atc ctt gat tat gtt tta tga ata gat aaa gta gat gtt gtc ctt ctt cct ttt gta att tcc cgt att ttt gtt cgt gcc aat tga gta aca tta tga gaa tat gac ctg ttg cac aat cgt tct tta tgt att cca tga tgg gtg tac aat caa gat tat tac gta tcc tcg tat cgg ctc ctc gag ata aaa gag cat aca cca cac gag gac tat gtt tgg tat act gtt gaa ggt aag tgt gta acg gcg tat ttc cga ttt tcg taa ccg cgt taa tgt ttg ctc cat gat cta tta tcg cgt aga tga atc gct tct cag ctc gca tct tag tgt gac tct ttg act tgt aat aat tgc ttt cgt gga aca cgg ata tgt gtt tac agt agt aat gaa gag aag tga gtc cat cct cat cga cgc aat tag ggt cag atc ctt tag tca ata att tgt aca gaa cgt aat agt tta agc tcc cat tga att tat atc taa gat aac aca gca ata gat cgg atg att tac taa agt cat caa tgg ggt ccg tta gta tat caa aga tct tgt tat cga ttg ata gtg aat gaa tca gat agt ggt gta gag gaa tat gtc ctt ttt tat cct tgc tat caa agt tac gca tgc cgt ggc gta aca ata tct tta ata cag atg gat taa atc gtg tat tca tcg tat agc aat gta atg gag agt tac cac att tta gtc gtt tat tca gat cgc agt gtt taa taa cta att taa aca gat gag atg ata tat cca cat caa aga atg cga gat aca tat gac aga cat tat tga cag aaa tgt gac ctt cat tat cac cgt cgt cca taa atg cgt tag gta cgt acc aca tac tat cgt taa cga tgc gca caa tct cgt cca ttt tca tcc atc ttc ata atg att tac ttt ttc ata att aga gaa aaa gat caa ggt ata aaa att aga agt gtt aga cta ta- --- --- --- --- --- --- --- --- --- --- -aa cta act tat aa- --- --- --- --- --- --- --- --- --- --- --- --- --- --- --- --- --- --- --- --- --- --- --- --- --- --- --- --- --- --- --- --- --- --- --- --- --- --- --- --- --- --- --- --- --- --- --- --- --- --- --- --- --- --- --- --- --- --- --- --- --- --- --- --- --- --- --- --- --- --- --- --- --- --- --- --- --- --- --- --- --- --- --- --- --- --- -aa cta act tat gac tta act aac tta tga cta tta act cat ttc aag aaa ggt ggg tgg aga gaa ctc tat atg aca gct tgt gaa aca att aga tcc cta att tct aat gga agt ttt gat agg aga ttg tca tca gtt gat aca ttg ttt att atc tca tct att aga gca cgt ctg ttt aga gct tta gtg acc tgc tcg gtt act tct gtg taa atc ttg aat cct tta gtg ata cac tgt gtc aaa act gga tgt tta gaa tac cta tgt aga ata tgg gaa gca tgc ttg ttt ttg tct cta tta tag att aac tca tac atg gtt gta tta tga att ttc atc tgc cta atg tac tcc aat tct tgt tta caa tca att ata taa tca aag agt gat gat gca tac aca tta caa agt gaa taa tct acc atc ata aaa tac ttg ata cag agc ttt atc aca tca tgg ttt tca att gta tta tta agt ata gct aat ttt ata cag tca ata gac aat ggt tct cta agc aat att tct aat att tta aga tgt gct tcc cta cgg gcg atg aca gat ccc cta tcc acg gcc acg tca aga cat gta tat cca tta ctc att act gcg ttg aca ttt gct cca ttt tct aat agc cat gat act aaa tct ata taa cct gca tag ata gcg cga taa agc aag gtc ctt cca cca gca tct agt tga ttg ata tct tca ata tat ggg ata caa agc tta taa att tct aat act gtg ggt tca tct aca agg aat ccc cta gta tac tga att att tta tat aga tct aat tta aca tca ttt tca tct ggg ata cca caa ttc aaa ata aac tca aca aca cta ctt tcc ttt tta cat att ccc cta aaa tag gca ttc aag cat tct att tta tat att aca gcc cca tga tct acc ata aaa tca aca atg tct att tct aca tat gca tta gat aga tag taa agt aag aga tct tgt aca gaa tta caa ttc tta ata att ata gag aaa ata tct tcc ata taa ttc ttt gac act aat gca gat ata ata tct tta tat gta ata tat gca aac agt cta tct act ata tac tga tca ata tta tct cta tga atc cta aaa taa tca tac aga aca tct aca gga tca caa att ggt tca agg aga aat cta tca aat att ttc ctg tca aca act ggt tct aga aca taa cag tca aca cct aat cca tgt ttt tta tag tca tct acc aaa gat aat gac caa aga tcg agg tcg tcg tga aac tgc tca tcg aca gcc atg aaa tct gcc gac tcc atg gcg cga atc gca ctg tct tat tcg cca ttg att ttc a-t ttt tta taa tta tgt aca tgt ttt cct tct att ctc aag agt cta caa aaa tat a-t ttt ttt tcg ata tct aag tac taa gtt ttt tta ctg ttt ttg tta ctg tct tcc att ctt cta act aaa gat ctg aga taa att ata caa tct tcg cta tcg aac cat ttt tgt agt cta aag cct gaa gta att aac caa ctg ttt tta tta gtg gct ttt ttc gat cta tcc tcg tc- --- --c tca tca tcc tta ta- --- --- --- --- --- --- tta tta tca tta tta tca tag tct att aaa cac aaa tca tct --- --- --- --- --- --- --- --- --- --- --- --- --- --- --- --- --- --- --- --- --- --- --- --- --- --- --a cgt tta taa c-- --- --- --- --- --- --- --- --a aca ttc --- --- --- --- -tc att att aat tag ttc tgt agt aat atc ttt aat aat ttg gct ata --- --- -ca tct gtt --- --- --- --- --- --- --- --- --- --- --- --- --- --- --- --- --- --- --c aat act --- --- --- --- --- --- --- --a tct att gat gat ttc ttt t-- --- --- --- --- --- --- --- --- --- --- --t taa gac t-- --- --- --- --- --- --- --- --- --- --- --- --- --- --- --- --- --- --- --- --- --- --- --- --- --- --- --- --- --- --- --- --- --- --- --- --- --- --- --- taa act agt --- --- --- --- --- --- --- --- --- --- --- --- --- --- --- --- --- --- --- --- --- --- --- --- --- --- --- --- --- --- --- --- --- --- --- --- --- --- --- --- --- --- --- --- --- --- --- --- --- --- --- --- --- --- --- --- --- --- --- --- --- --- --- --- --- --- --- --- --- --- --- --- --- --- --- --- --- --- --- --- --- --- --- --- --- --- --- --- --- --- --- --- --- --- --- --- --- --- --- --- --- --- --- --- --- --- --- --- --- --t atg gta atg acg atg aaa --- -tc gag tag ta- --- --a ctt cta ata aag act tga ta- --- --t cat tat cat atg ttt gat cg- --- --- --- --- --- --- --t cat agt taa tag tgt g-- --- --- --- --- --- --- --- --- --- --- --- --- --- --- --- --- --- --- --- -gc taa atg gta ctg tta ata agt tta t-- --- --- --- --- --- --- --- --- --- --- --- --- --- --- --- --- --a gac aat atc ata gta ttt tct ttc cag aat --- --- --- --t aga tta ttt ttt taa ata ctg atc ctc aca att ccg tga tgt agc agt agt tgg t-- --- --- --- --- --- --- --- --- --- --- --- --- --- --- --- --- --- --- --- --- --- --- --- --- --- --- --- -gc atg gtc tat atc gt- --- --- --- --- --- --- --- --- --- --- --- --- --- --- --- --- --- --- --- --- --- --- --- --- --- --- --- --- --- --- --- --- --- --- --- --- --- --- --- --- --- --- --- taa aat gta tca tat ata ata gtt ttc tga cgt gga gta cag aat ttt cga --- --- --- --- --- --- --- --- --- --- --- --- --- --- --- --- --- --- --- --- --- --- --- --- --- --- --- --- --- --- --- --- --- --- --- --- --- --- --- --- --- --- --- --t taa tga gtt cat ggt aag gaa ggg caa atg tct --- gta tat aat ata cat aag tta a-- --- --- --- --- --- --- --- --- --- --- --- --- --t agt ttt tta tca tat tt- --- --- --- --- --- --- --- --t cta ata cca taa taa aaa tta tca tta tg- --- --- -ta taa tca --- --- --- --- --- --- --- --- --- --- --- --t cac tg- --- -tc gct atc att att gcg ttt gtg tag tt- --- --- --- --- ctg ccc ta- --- --- --- --- --- --- --- --t cat cta cat cac tgt ca- --- --- --- --- ctc tca cta tat ctt cta aaa tta caa --- -ac aac --- --- --- --- --- --- --- --- --- --- --- --- --- --- --- --- --- --- --- --- --- --- --- --- --- --- --- --- --- --- --- --- --- --- --- --- tgg ata ttc gat --- --- --- --- --- --- --- --- --- --- --- --- --- --- --- --- --- --- --- --- --- --- --- --- --- --- --a aca gca ttt gtg t-- --- --- --- --- --- --- --- --- --- --- --- --- --- --- --- --- -

>JX878408.1_Monkeypox_virus_isolate_DRC_06-0970_complete_genome

--- --- --- --- --- --- --- --- --- --- --- --- --- --- --- --- --- --- --- --- --- --- --- --- --- --- --- --- --- --- --- --- --- --- --- --g tgt cta gaa aaa aat gtg tga ccc acg acc gta gga aac tct aga ggg taa gaa aaa tca atc gtt tat aga gac cat cag aaa gag gtt taa tat ttt tgt gag acc cat cga a-- --- --- --- --- --- --- --- --- --- --- --- --- --- --- --- --- --- --- --- --- --- --- --- --- --- --- --- --- --- --- --- --- --- --- --- --- --- --- --- --- --- --- --- --- --- --- --- --- --- --- --- --- --- --- --- --- --- --- --- --- --- --- --- --- --- --- --- --- --- --- --- --- --- --- --- --- --- --- --- --- --- --- --- --- --- --- --- --- --- --- --- --- --- --- --- --- --- --- --- --- --- --- --- --- --- --- --- --- --- --- --- --- --- --- --- --- --- --- --- --- --- --- --- --- --- --- --- --- --- --- --- --- --- --- --- --- --- --- --- --- --- --- --- --- --- --- --- --- --- --- --- --- --- --- --- --- --- --- --- --- --- --- --- --- --- --- --- --- --- --- --- --- --- --- --- --- --- --- --- --- --- --- --- --- --- --- --- --- --- --- --- --- --- --- --- --- --- --- --- --- --- --- --- --- --- --- --- --- --- --- --- --- --- --- --- --- --- --- --- --- --- --- --- --- --- --- --- --- --- --- --- --- --- --- --- --- --- --- --- --- --- --- --- --- --- --- --- --- --- --- --- --- --- --- --- --- --- --- --- --- --- --- --- --- --- --- --- --- --- --- --- --- --- --- --- --- --- --- --- -ga gag aaa gga taa aaa ctt ttt acg act cca tca gaa aga ggt tta ata ttt ttg tga gac cca tcg acg aga gaa aga gat ggt tag tca aga tat ttt tct tag tac aaa agt caa tgt ttt aaa ata tat gga cga gaa tta att tgt ctg tat aaa aac ttg tgt gaa att atg tac tag aga aaa aac gtg agc agt gtc ccc tac atg gat ttt aca gat cat tta tat tcc aaa aat att aac tat ata cgt tta tta tat gat gtt aac gtg taa att ata aac att att tta tga tgc aat tgt ctg aca acc tag att ggc ata agg ata ttg ata agc tct acg aga ata tat tgt tgg acg tta tcg ttt acg aaa tag ttg aga cat cag aaa gag gtt taa tat ttt tgt gag a-c cat cga aga gag aaa gag aat aaa aat a-- --- --- --- ttt ttt tgt aaa ac- ttt tta tga gac aag aga gaa aga gaa tac gaa tag tga tca tat cgt atc aca tat tga aac aga aag aag aag taa cga gag gta act ttt tgt gaa tgt agt taa aca ttt ttg ttt tgc aaa ccg gaa tat agt acc cgg tac act ttt tta att cgt ggt gcg gtg tct gaa tcg ttc gat taa ccc aac tca tcc att ttc aga tga ata gag tta tcg att cag aca cat gct ttg agt ttt gtt gaa tcg atg agt gaa gta tca tcg gtt gca cct tca gat gcc gat ccg tcg aca tac ttg aat cca tcc ttg act tca agt tca gat gat tcc tca cac atg tct ccg ata cgt acg cta aac tct agg ttc tta aca cat ttt gta tca acg atc gtt gaa ccg atg ata tct ttg taa ctc act ttc tta tgt gag atg tta gac cca agt act gga tgg gtc ttg atg tcg ctg tct ttc tct tct tcg cta cat ctg atg tcg ata gac atc tca cag tct ttg atc ata gcc aga gct tct tca cgc gtg atc gcg gga gag tcc tta cct tgt cct ggt gac acg ctg gac aat cta gta ttc aca gtg ttt cca tca gag gat tcg gag atg gat gaa atc ttt ggg cat ttg gtg aat cca aag ttc atg tta aga ccc gcg ccg acg ata gtg taa taa gtg gtg gga tct cct ttt aca act --- --- --- --- --- --- tct tcg gat acc tca tca tct tcg gtc tct gta act tcc gtt acg gat tga caa atc tta tca ttg gtc ggt gtt tgg tct tgc ttt gtg act ttg ata ata aca tcg att ccc ata tga tgt ttg ttt tct tct tca gta cac gag gat gag gat tgt tga aga cta gta ggc ata gca gct gcc act agg cac atg cat gcc agg aca ata tat tgt ttc atg att gct att gat tga tta ctg ttc tag atg att cta ctt tct tac cat ata ata aat tag aat ata ttt tct act ttt acg aga aat taa tta ttg tat tta tta ttt atg ggt aaa aaa aaa ctt act ata agt ggg tgg gat tct ggg aat tag tga tca gtt tat gta tat cgc aac tag cgg gca tat ggc tat tga cat cga gaa cat tac cca tat gat aag aga ttg tat ccg ttt cgt agt ctt gag tat tgg tat tac tat ata gta tgt aga tgt cga cgc tag ata gac agt cgc cca cta gag tta ccg tct ctg aat gcg gca tga tag tat cat tct ttg ttt tcg tta act gtt tgg aag atg aat ctt tgt tgt tac att taa tct cga aat tca gag tac ata tct ttg aag tat tct gat atc tat ttt ctc ctg taa aga atc ctg aag ttg cta cat tat taa gga cag aga agt att ctg cac gaa aga cgg gat cac aat ctt tat gat tca tgg taa tag tta gtt ccg acg ttg aga tgg att cgc tga gac cgg tag tgg tcg tcc gag tac acg atg tgt cgt tga ctg gat aca ggt taa ttt cca cat cga tat agt taa atg tat tgc tgg tt- --a cgg gtt cgc att tat ctg tgg aag aga cgg tgt gag aat atg ttc cgg gac cac acg gag aac aga tga cgt ctc cgg tag acg tgt atc cgg ata ctc cgt atc cta ttc cac act ttg ttt tag aaa tac atg ttc tac acc ctg atg atc ctt tga gaa gac aat aat atc ctg gag agc att cac aga ttc tat tgt gag tcg tgt tac acg atc gcg tct cta cct gat tac tat cac atc ttc cgt tac aac tta gac aag cct gta aat gat tat tgt gag atg taa agg tat ccg aac cac acg gcg tac att gtg tat tag tct tgc tat cac ata atc tgg aag cgt aag ttc ccg gag gac acg ata gac aac ata gat tac ggc ttc tgt att cgt tgt ctt tac att ttc cat tgg atg gtg cat gtg gtg cta tat ctc ttc cgt tta tta tta tac atg aga gaa aca ata tat acg agt ata ata cgg acc tca tga ttt aat aat -gt agt aat cgt cgt ctt gtt act gtt tgt ttc cta ctt ctc caa tca tat aga tta ttt ttt aaa tat ttt ctt tct atc atg gat aat att tgt aat ggt tct ttc cgt aca aca tac tgt tta gat ggt agt cgc tta gct tgg tta tga tat tgc gca taa ttt ccg gag gca aat acg ata gtc tag att gac tat cga tgg tag act cta att tat tga gtg ctt tgt cga cga gtt tac ttt tat gct cca tcg ata gat gac act gtt cta tga gat cgt cgt aca tgg gaa atg aaa tgc gtt tgt ctg aat gta tgg ctt cga gat agg tgt gat acc gga tgt ctt ctg ttc tca ata ccg tat aca agt tgg tgt ctg aga ttc gaa tct ctt tga gga gac tta tgt cac gac tac att ttt cga tga tgg aat cta tct tat cga atg ata tat ttt tca taa ata cac ttt tat agt cct cgt tta aac aga att tag tat gta gtt ccg caa atg act cgt ccc tta ata ggc agt agg cta tta tct tct tta cgt agt gat cgt cgt agg gag aga act ccg aca tct tgt aga aca acg att taa tca tag gta gag ata ctt tca gtc tgt ggt gga tga tgt cat taa cga cat ccg cct tgt ata tga tgt ttc tgt ttt caa aca cca agt cga ata ctg tct tat cgt ctt tag tcg gaa ggt tga tgt cgt atc cga tgt ata cga ggt atg agg caa cat tgt tat tgc aat tct gga agg cgg tat gaa gag gag tca ttg tat tat agt att cgt ctt tct gaa tgt cga atc tat cta gta gat acc gta gta tat tga gag agc gac ttc cat atc ctt gat tat gtt tta tga ata gat aaa gta gat gtt gtc ctt ctt cct ttt gta att tcc cgt att ttt gtt cgt gcc aat tga gta aca tta tga gaa tat gac ctg ttg cac aat cgt tct tta tgt att cca tga tgg gtg tac aat caa gat tat tac gta tcc tcg tat cgg ctc ctc gag ata aaa gag cat aca cca cac gag gac tat gtt tgg tat act gtt gaa ggt aag tgt gta acg gcg tat ttc cga ttt tcg taa ccg cgt taa tgt ttg ctc cat gat cta tta tcg cgt aga tga atc gct tct cag ctc gca tct tag tgt gac tct ttg act tgt aat aat tgc ttt cgt gga aca cgg ata tgt gtt tac agt agt aat gaa gag aag tga gtc cat cct cat cga cgc aat tag ggt cag atc ctt tag tca ata att tgt aca gaa cgt aat agt tta agc tcc cat tga att tat atc taa gat aac aca gca ata gat cgg atg att tac taa agt cat caa tgg ggt ccg tta gta tat caa aga tct tgt tat cga ttg ata gtg aat gaa tca gat agt ggt gta gag gaa tat gtc ctt ttt tat cct tgc tat caa agt tac gca tgc cgt ggc gta aca ata tct tta ata cag atg gat taa atc gtg tat tca tcg tat agc aat gta atg gag agt tac cac att tta gtc gtt tat tca gat cgc agt gtt taa taa cta att taa aca gat gag atg ata tat cca cat caa aga atg cga gat aca tat gac aga cat tat tga cag aaa tgt gac ctt cat tat cac cgt cgt cca taa atg cgt tag gta cgt acc aca tac tat cgt taa cga tgc gca caa tct cgt cca ttt tca tcc atc ttc ata atg att tac ttt ttc ata att aga gaa aaa gat caa ggt ata aaa att aga agt gtt aga cta ta- --- --- --- --- --- --- --- --- --- --- -aa cta act tat aa- --- --- --- --- --- --- --- --- --- --- --- --- --- --- --- --- --- --- --- --- --- --- --- --- --- --- --- --- --- --- --- --- --- --- --- --- --- --- --- --- --- --- --- --- --- --- --- --- --- --- --- --- --- --- --- --- --- --- --- --- --- --- --- --- --- --- --- --- --- --- --- --- --- --- --- --- --- --- --- --- --- --- --- --- --- --- -aa cta act tat gac tta act aac tta tga cta tta act cat ttc aag aaa ggt ggg tgg aga gaa ctc tat atg aca gct tgt gaa aca att aga tcc cta att tct aat gga agt ttt gat agg aga ttg tca tca gtt gat aca ttg ttt att atc tca tct att aga gca cgt ctg ttt aga gct tta gtg acc tgc tcg gtt act tct gtg taa atc ttg aat cct tta gtg ata cac tgt gtc aaa act gga tgt tta gaa tac cta tgt aga ata tgg gaa gca tgc ttg ttt ttg tct cta tta tag att aac tca tac atg gtt gta tta tga att ttc atc tgc cta atg tac tcc aat tct tgt tta caa tca att ata taa tca aag agt gat gat gca tac aca tta caa agt gaa taa tct acc atc ata aaa tac ttg ata cag agc ttt atc aca tca tgg ttt tca att gta tta tta agt ata gct aat ttt ata cag tca ata gac aat ggt tct cta agc aat att tct aat att tta aga tgt gtt tcc cta cgg gcg atg aca gat ccc cta tcc acg gcc acg tca aga cat gta tat cca tta ctc att act gcg ttg aca ttt gct cca ttt tct aat agc cat gat act aaa tct ata taa cct gca tag ata gcg cga taa agc aag gtc ctt cca cca gca tct agt tga ttg ata tct tca ata tat ggg ata caa agc tta taa att tct aat act gtg ggt tca tct aca agg aat ccc cta gta tac tga att att tta tat aga tct aat tta aca tca ttt tca tct ggg ata cca caa ttc aaa ata aac tca aca aca cta ctt tcc ttt tta cat att ccc cta aaa tag gca ttc aag cat tct att tta tat att aca gcc cca tga tct acc ata aaa tca aca atg tct att tct aca tat gca tta gat aga tag taa agt aag aga tct tgt aca gaa tta caa ttc tta ata att ata gag aaa ata tct tcc ata taa ttc ttt gac act aat gca gat ata ata tct tta tat gta ata tat gca aac agt cta tct act ata tac tga tca ata tta tct cta tga atc cta aaa taa tca tac aga aca tct aca gga tca caa att ggt tca agg aga aat cta tca aat att ttc ctg tca aca act ggt tct aga aca taa cag tca aca cct aat cca tgt ttt tta tag tca tct acc aaa gat aat gac caa aga tcg agg tcg tcg tga aac tgc tca tcg aca gcc atg aaa tct gcc gac tcc atg gcg cga atc gca ctg tct tat tcg cca ttg att ttc a-t ttt tta taa tta tgt aca tgt ttt cct tct att ctc aag agt cta caa aaa tat a-t ttt ttt tcg ata tct aag tac taa gtt ttt tta ctg ttt ttg tta ctg tct tcc att ctt cta act aaa gat ctg aga taa att ata caa tct tcg cta tcg aac cat ttt tgt agt cta aag cct gaa gta att aac caa ctg ttt tta tta gtg gct ttt ttc gat cta tcc tcg tc- --- --c tca tca tcc tta ta- --- --- --- --- --- --- tta tta tca tta tta tca tag tct att aaa cac aaa tca tct --- --- --- --- --- --- --- --- --- --- --- --- --- --- --- --- --- --- --- --- --- --- --- --- --- --- --a cgt tta taa c-- --- --- --- --- --- --- --- --a aca ttc --- --- --- --- -tc att att aat tag ttc tgt agt aat atc ttt aat aat ttg gct ata --- --- -ca tct gtt --- --- --- --- --- --- --- --- --- --- --- --- --- --- --- --- --- --- --c aat act --- --- --- --- --- --- --- --a tct att gat gat ttc ttt t-- --- --- --- --- --- --- --- --- --- --- --t taa gac t-- --- --- --- --- --- --- --- --- --- --- --- --- --- --- --- --- --- --- --- --- --- --- --- --- --- --- --- --- --- --- --- --- --- --- --- --- --- --- --- taa act agt --- --- --- --- --- --- --- --- --- --- --- --- --- --- --- --- --- --- --- --- --- --- --- --- --- --- --- --- --- --- --- --- --- --- --- --- --- --- --- --- --- --- --- --- --- --- --- --- --- --- --- --- --- --- --- --- --- --- --- --- --- --- --- --- --- --- --- --- --- --- --- --- --- --- --- --- --- --- --- --- --- --- --- --- --- --- --- --- --- --- --- --- --- --- --- --- --- --- --- --- --- --- --- --- --- --- --- --- --- --t atg gta atg acg atg aaa --- -tc gag tag ta- --- --a ctt cta ata aag act tga ta- --- --t cat tat cat atg ttt gat cg- --- --- --- --- --- --- --t cat agt taa tag tgt g-- --- --- --- --- --- --- --- --- --- --- --- --- --- --- --- --- --- --- --- -gc taa atg gta ctg tta ata agt tta t-- --- --- --- --- --- --- --- --- --- --- --- --- --- --- --- --- --a gac aat atc ata gta ttt tct ttc cag aat --- --- --- --t aga tta ttt ttt taa ata ctg atc ctc aca att ccg tga tgt agc agt agt tgg t-- --- --- --- --- --- --- --- --- --- --- --- --- --- --- --- --- --- --- --- --- --- --- --- --- --- --- --- -gc atg gtc tat atc gt- --- --- --- --- --- --- --- --- --- --- --- --- --- --- --- --- --- --- --- --- --- --- --- --- --- --- --- --- --- --- --- --- --- --- --- --- --- --- --- --- --- --- --- taa aat gta tca tat ata ata gtt ttc tga cgt gga gta cag aat ttt cga --- --- --- --- --- --- --- --- --- --- --- --- --- --- --- --- --- --- --- --- --- --- --- --- --- --- --- --- --- --- --- --- --- --- --- --- --- --- --- --- --- --- --- --t taa tga gtt cat ggt aag gaa ggg caa atg tct --- gta tat aat ata cat aag tta a-- --- --- --- --- --- --- --- --- --- --- --- --- --t agt ttt tta tca tat tt- --- --- --- --- --- --- --- --t cta ata cca taa taa aaa tta tca tta tg- --- --- -ta taa tca --- --- --- --- --- --- --- --- --- --- --- --t cac tg- --- -tc gct atc att att gcg ttt gtg tag tt- --- --- --- --- ctg ccc ta- --- --- --- --- --- --- --- --t cat cta cat cac tgt ca- --- --- --- --- ctc tca cta tat ctt cta aaa tta caa --- -ac aac --- --- --- --- --- --- --- --- --- --- --- --- --- --- --- --- --- --- --- --- --- --- --- --- --- --- --- --- --- --- --- --- --- --- --- --- tgg ata ttc gat --- --- --- --- --- --- --- --- --- --- --- --- --- --- --- --- --- --- --- --- --- --- --- --- --- --- --a aca gca ttt gtg t-- --- --- --- --- --- --- --- --- --- --- --- --- --- --- --- --- -

>JX878420.1_Monkeypox_virus_isolate_DRC_07-0283_complete_genome

--- --- --- --- --- --- --- --- --- --- --- --- --- --- --- --- --- --- --- --- --- --- --- --- --- --- --- --- --- --- --- --- --- --- --- --g tgt cta gaa aaa aat gtg tga ccc acg acc gta gga aac tct aga ggg taa gaa aaa tca atc gtt tat aga gac cat cag aaa gag gtt taa tat ttt tgt gag acc cat cga a-- --- --- --- --- --- --- --- --- --- --- --- --- --- --- --- --- --- --- --- --- --- --- --- --- --- --- --- --- --- --- --- --- --- --- --- --- --- --- --- --- --- --- --- --- --- --- --- --- --- --- --- --- --- --- --- --- --- --- --- --- --- --- --- --- --- --- --- --- --- --- --- --- --- --- --- --- --- --- --- --- --- --- --- --- --- --- --- --- --- --- --- --- --- --- --- --- --- --- --- --- --- --- --- --- --- --- --- --- --- --- --- --- --- --- --- --- --- --- --- --- --- --- --- --- --- --- --- --- --- --- --- --- --- --- --- --- --- --- --- --- --- --- --- --- --- --- --- --- --- --- --- --- --- --- --- --- --- --- --- --- --- --- --- --- --- --- --- --- --- --- --- --- --- --- --- --- --- --- --- --- --- --- --- --- --- --- --- --- --- --- --- --- --- --- --- --- --- --- --- --- --- --- --- --- --- --- --- --- --- --- --- --- --- --- --- --- --- --- --- --- --- --- --- --- --- --- --- --- --- --- --- --- --- --- --- --- --- --- --- --- --- --- --- --- --- --- --- --- --- --- --- --- --- --- --- --- --- --- --- --- --- --- --- --- --- --- --- --- --- --- --- --- --- --- --- --- --- --- --- -ga gag aaa gga taa aaa ctt ttt acg act cca tca gaa aga ggt tta ata ttt ttg tga gac cca tcg acg aga gaa aga gat ggt tag tca aga tat ttt tct tag tac aaa agt caa tgt ttt aaa ata tat gga cga gaa tta att tgt ctg tat aaa aac ttg tgt gaa att atg tac tag aga aaa aac gtg agc agt gtc ccc tac atg gat ttt aca gat cat tta tat tcc aaa aat att aac tat ata cgt tta tta tat gat gtt aac gtg taa att ata aac att att tta tga tgc aat tgt ctg aca acc tag att ggc ata agg ata ttg ata agc tct acg aga ata tat tgt tgg acg tta tcg ttt acg aaa tag ttg aga cat cag aaa gag gtt taa tat ttt tgt gag a-c cat cga aga gag aaa gag aat aaa aat a-- --- --- --- ttt ttt tgt aaa ac- ttt tta tga gac aag aga gaa aga gaa tac gaa tag tga tca tat cgt atc aca tat tga aac aga aag aag aag taa cga gag gta act ttt tgt gaa tgt agt taa aca ttt ttg ttt tgc aaa ccg gaa tat agt acc cgg tac act ttt tta att cgt ggt gcg gtg tct gaa tcg ttc gat taa ccc aac tca tcc att ttc aga tga ata gag tta tcg att cag aca cat gct ttg agt ttt gtt gaa tcg atg agt gaa gta tca tcg gtt gca cct tca gat gcc gat ccg tcg aca tac ttg aat cca tcc ttg act tca agt tca gat gat tcc tca cac atg tct ccg ata cgt acg cta aac tct agg ttc tta aca cat ttt gta tca acg atc gtt gaa ccg atg ata tct ttg taa ctc act ttc tta tgt gag atg tta gac cca agt act gga tgg gtc ttg atg tcg ctg tct ttc tct tct tcg cta cat ctg atg tcg ata gac atc tca cag tct ttg atc ata gcc aga gct tct tca cgc gtg atc gcg gga gag tcc tta cct tgt cct ggt gac acg ctg gac aat cta gta ttc aca gtg ttt cca tca gag gat tcg gag atg gat gaa atc ttt ggg cat ttg gtg aat cca aag ttc atg tta aga ccc gcg ccg acg ata gtg taa taa gtg gtg gga tct cct ttt aca act --- --- --- --- --- --- tct tcg gat acc tca tca tct tcg gtc tct gta act tcc gtt acg gat tga caa atc tta tca ttg gtc ggt gtt tgg tct tgc ttt gtg act ttg ata ata aca tcg att ccc ata tga tgt ttg ttt tct tct tca gta cac gag gat gag gat tgt tga aga cta gta ggc ata gca gct gcc act agg cac atg cat gcc agg aca ata tat tgt ttc atg att gct att gat tga tta ctg ttc tag atg att cta ctt tct tac cat ata ata aat tag aat ata ttt tct act ttt acg aga aat taa tta ttg tat tta tta ttt atg ggt -aa aaa aaa ctt act ata agt ggg tgg gat tct ggg aat tag tga tca gtt tat gta tat cgc aac tag cgg gca tat ggc tat tga cat cga gaa cat tac cca tat gat aag aga ttg tat ccg ttt cgt agt ctt gag tat tgg tat tac tat ata gta tgt aga tgt cga cgc tag ata gac agt cgc cca cta gag tta ccg tct ctg aat gcg gca tga tag tat cat tct ttg ttt tcg tta act gtt tgg aag atg aat ctt tgt tgt tac att taa tct cga aat tca gag tac ata tct ttg aag tat tct gat atc tat ttt ctc ctg taa aga atc ctg aag ttg cta cat tat taa gga cag aga agt att ctg cac gaa aga cgg gat cac aat ctt tat gat tca tgg taa tag tta gtt ccg acg ttg aga tgg att cgc tga gac cgg tag tgg tcg tcc gag tac acg atg tgt cgt tga ctg gat aca ggt taa ttt cca cat cga tat agt taa atg tat tgc tgg tt- --a cgg gtt cgc att tat ctg tgg aag aga cgg tgt gag aat atg ttc cgg gac cac acg gag aac aga tga cgt ctc cgg tag acg tgt atc cgg ata ctc cgt atc cta ttc cac act ttg ttt tag aaa tac atg ttc tac acc ctg atg atc ctt tga gaa gac aat aat atc ctg gag agc att cac aga ttc tat tgt gag tcg tgt tac acg atc gcg tct cta cct gat tac tat cac atc ttc cgt tac aac tta gac aag cct gta aat gat tat tgt gag atg taa agg tat ccg aac cac acg gcg tac att gtg tat tag tct tgc tat cac ata atc tgg aag cgt aag ttc ccg gag gac acg ata gac aac ata gat tac ggc ttc tgt att cgt tgt ctt tac att ttc cat tgg atg gtg cat gtg gtg cta tat ctc ttc cgt tta tta tta tac atg aga gaa aca ata tat acg agt ata ata cgg acc tca tga ttt aat aat -gt agt aat cgt cgt ctt gtt act gtt tgt ttc cta ctt ctc caa tca tat aga tta ttt ttt aaa tat ttt ctt tct atc atg gat aat att tgt aat ggt tct ttc cgt aca aca tac tgt tta gat ggt agt cgc tta gct tgg tta tga tat tgc gca taa ttt ccg gag gca aat acg ata gtc tag att gac tat cga tgg tag act cta att tat tga gtg ctt tgt cga cga gtt tac ttt tat gct cca tcg ata gat gac act gtt cta tga gat cgt cgt aca tgg gaa atg aaa tgc gtt tgt ctg aat gta tgg ctt cga gat agg tgt gat acc gga tgt ctt ctg ttc tca ata ccg tat aca agt tgg tgt ctg aga ttc gaa tct ctt tga gga gac tta tgt cac gac tac att ttt cga tga tgg aat cta tct tat cga atg ata tat ttt tca taa ata cac ttt tat agt cct cgt tta aac aga att tag tat gta gtt ccg caa atg act cgt ccc tta ata ggc agt agg cta tta tct tct tta cgt agt gat cgt cgt agg gag aga act ccg aca tct tgt aga aca acg att taa tca tag gta gag ata ctt tca gtc tgt ggt gga tga tgt cat taa cga cat ccg cct tgt ata tga tgt ttc tgt ttt caa aca cca agt cga ata ctg tct tat cgt ctt tag tcg gaa ggt tga tgt cgt atc cga tgt ata cga ggt atg agg caa cat tgt tat tgc aat tct gga agg cgg tat gaa gag gag tca ttg tat tat agt att cgt ctt tct gaa tgt cga atc tat cta gta gat acc gta gta tat tga gag agc gac ttc cat atc ctt gat tat gtt tta tga ata gat aaa gta gat gtt gtc ctt ctt cct ttt gta att tcc cgt att ttt gtt cgt gcc aat tga gta aca tta tga gaa tat gac ctg ttg cac aat cgt tct tta tgt att cca tga tgg gtg tac aat caa gat tat tac gta tcc tcg tat cgg ctc ctc gag ata aaa gag cat aca cca cac gag gac tat gtt tgg tat act gtt gaa ggt aag tgt gta acg gcg tat ttc cga ttt tcg taa ccg cgt taa tgt ttg ctc cat gat cta tta tcg cgt aga tga atc gct tct cag ctc gca tct tag tgt gac tct ttg act tgt aat aat tgc ttt cgt gga aca cgg ata tgt gtt tac agt agt aat gaa gag aag tga gtc cat cct cat cga cgc aat tag ggt cag atc ctt tag tca ata att tgt aca gaa cgt aat agt tta agc tcc cat tga att tat atc taa gat aac aca gca ata gat cgg atg att tac taa agt cat caa tgg ggt ccg tta gta tat caa aga tct tgt tat cga ttg ata gtg aat gaa tca gat agt ggt gta gag gaa tat gtc ctt ttt tat cct tgc tat caa agt tac gca tgc cgt ggc gta aca ata tct tta ata cag atg gat taa atc gtg tat tca tcg tat agc aat gta atg gag agt tac cac att tta gtc gtt tat tca gat cgc agt gtt taa taa cta att taa aca gat gag atg ata tat cca cat caa aga atg cga gat aca tat gac aga cat tat tga cag aaa tgt gac ctt cat tat cac cgt cgt cca taa atg cgt tag gta cgt acc aca tac tat cgt taa cga tgc gca caa tct cgt cca ttt tca tcc atc ttc ata atg att tac ttt ttc ata att aga gaa aaa gat caa ggt ata aaa att aga agt gtt aga cta ta- --- --- --- --- --- --- --- --- --- --- -aa cta act tat aa- --- --- --- --- --- --- --- --- --- --- --- --- --- --- --- --- --- --- --- --- --- --- --- --- --- --- --- --- --- --- --- --- --- --- --- --- --- --- --- --- --- --- --- --- --- --- --- --- --- --- --- --- --- --- --- --- --- --- --- --- --- --- --- --- --- --- --- --- --- --- --- --- --- --- --- --- --- --- --- --- --- --- --- --- --- --- -aa cta act tat gac tta act aac tta tga cta tta act cat ttc aag aaa ggt ggg tgg aga gaa ctc tat atg aca gct tgt gaa aca att aga tcc cta att tct aat gga agt ttt gat agg aga ttg tca tca gtt gat aca ttg ttt att atc tca tct att aga gca cgt ctg ttt aga gct tta gtg acc tgc tcg gtt act tct gtg taa atc ttg aat cct tta gtg ata cac tgt gtc aaa act gga tgt tta gaa tac cta tgt aga ata tgg gaa gca tgc ttg ttt ttg tct cta tta tag att aac tca tac atg gtt gta tta tga att ttc atc tgc cta atg tac tcc aat tct tgt tta caa tca att ata taa tca aag agt gat gat gca tac aca tta caa agt gaa taa tct acc atc ata aaa tac ttg ata cag agc ttt atc aca tca tgg ttt tca att gta tta tta agt ata gct aat ttt ata cag tca ata gac aat ggt tct cta agc aat att tct aat att tta aga tgt gtt tcc cta cgg gcg atg aca gat ccc cta tcc acg gcc acg tca aga cat gta tat cca tta ctc att act gcg ttg aca ttt gct cca ttt tct aat agc cat gat act aaa tct ata taa cct gca tag ata gcg cga taa agc aag gtc ctt cca cca gca tct agt tga ttg ata tct tca ata tat ggg ata caa agc tta taa att tct aat act gtg ggt tca tct aca agg aat ccc cta gta tac tga att att tta tat aga tct aat tta aca tca ttt tca tct ggg ata cca caa ttc aaa ata aac tca aca aca cta ctt tcc ttt tta cat att ccc cta aaa tag gca ttc aag cat tct att tta tat att aca gcc cca tga tct acc ata aaa tca aca atg tct att tct aca tat gca tta gat aga tag taa agt aag aga tct tgt aca gaa tta caa ttc tta ata att ata gag aaa ata tct tcc ata taa ttc ttt gac act aat gca gat ata ata tct tta tat gta ata tat gca aac agt cta tct act ata tac tga tca ata tta tct cta tga atc cta aaa taa tca tac aga aca tct aca gga tca caa att ggt tca agg aga aat cta tca aat att ttc ctg tca aca act ggt tct aga aca taa cag tca aca cct aat cca tgt ttt tta tag tca tct acc aaa gat aat gac caa aga tcg agg tcg tcg tga aac tgc tca tcg aca gcc atg aaa tct gcc gac tcc atg gcg cga atc gca ctg tct tat tcg cca ttg att ttc a-t ttt tta taa tta tgt aca tgt ttt cct tct att ctc aag agt cta caa aaa tat a-t ttt ttt tcg ata tct aag tac taa gtt ttt tta ctg ttt ttg tta ctg tct tcc att ctt cta act aaa gat ctg aga taa att ata caa tct tcg cta tcg aac cat ttt tgt agt cta aag cct gaa gta att aac caa ctg ttt tta tta gtg gct ttt ttc gat cta tcc tcg tc- --- --c tca tca tcc tta ta- --- --- --- --- --- --- tta tta tca tta tta tca tag tct att aaa cac aaa tca tct --- --- --- --- --- --- --- --- --- --- --- --- --- --- --- --- --- --- --- --- --- --- --- --- --- --- --a cgt tta taa c-- --- --- --- --- --- --- --- --a aca ttc --- --- --- --- -tc att att aat tag ttc tgt agt aat atc ttt aat aat ttg gct ata --- --- -ca tct gtt --- --- --- --- --- --- --- --- --- --- --- --- --- --- --- --- --- --- --c aat act --- --- --- --- --- --- --- --a tct att gat gat ttc ttt t-- --- --- --- --- --- --- --- --- --- --- --t taa gac t-- --- --- --- --- --- --- --- --- --- --- --- --- --- --- --- --- --- --- --- --- --- --- --- --- --- --- --- --- --- --- --- --- --- --- --- --- --- --- --- taa act agt --- --- --- --- --- --- --- --- --- --- --- --- --- --- --- --- --- --- --- --- --- --- --- --- --- --- --- --- --- --- --- --- --- --- --- --- --- --- --- --- --- --- --- --- --- --- --- --- --- --- --- --- --- --- --- --- --- --- --- --- --- --- --- --- --- --- --- --- --- --- --- --- --- --- --- --- --- --- --- --- --- --- --- --- --- --- --- --- --- --- --- --- --- --- --- --- --- --- --- --- --- --- --- --- --- --- --- --- --- --t atg gta atg acg atg aaa --- -tc gag tag ta- --- --a ctt cta ata aag act tga ta- --- --t cat tat cat atg ttt gat cg- --- --- --- --- --- --- --t cat agt taa tag tgt g-- --- --- --- --- --- --- --- --- --- --- --- --- --- --- --- --- --- --- --- -gc taa atg gta ctg tta ata agt tta t-- --- --- --- --- --- --- --- --- --- --- --- --- --- --- --- --- --a gac aat atc ata gta ttt tct ttc cag aat --- --- --- --t aga tta ttt ttt taa ata ctg atc ctc aca att ccg tga tgt agc agt agt tgg t-- --- --- --- --- --- --- --- --- --- --- --- --- --- --- --- --- --- --- --- --- --- --- --- --- --- --- --- -gc atg gtc tat atc gt- --- --- --- --- --- --- --- --- --- --- --- --- --- --- --- --- --- --- --- --- --- --- --- --- --- --- --- --- --- --- --- --- --- --- --- --- --- --- --- --- --- --- --- taa aat gta tca tat ata ata gtt ttc tga cgt gga gta cag aat ttt cga --- --- --- --- --- --- --- --- --- --- --- --- --- --- --- --- --- --- --- --- --- --- --- --- --- --- --- --- --- --- --- --- --- --- --- --- --- --- --- --- --- --- --- --t taa tga gtt cat ggt aag gaa ggg caa atg tct --- gta tat aat ata cat aag tta a-- --- --- --- --- --- --- --- --- --- --- --- --- --t agt ttt tta tca tat tt- --- --- --- --- --- --- --- --t cta ata cca taa taa aaa tta tca tta tg- --- --- -ta taa tca --- --- --- --- --- --- --- --- --- --- --- --t cac tg- --- -tc gct atc att att gcg ttt gtg tag tt- --- --- --- --- ctg ccc ta- --- --- --- --- --- --- --- --t cat cta cat cac tgt ca- --- --- --- --- ctc tca cta tat ctt cta aaa tta caa --- -ac aac --- --- --- --- --- --- --- --- --- --- --- --- --- --- --- --- --- --- --- --- --- --- --- --- --- --- --- --- --- --- --- --- --- --- --- --- tgg ata ttc gat --- --- --- --- --- --- --- --- --- --- --- --- --- --- --- --- --- --- --- --- --- --- --- --- --- --- --a aca gca ttt gtg t-- --- --- --- --- --- --- --- --- --- --- --- --- --- --- --- --- -

>JX878419.1_Monkeypox_virus_isolate_DRC_07-0275_complete_genome

--- --- --- --- --- --- --- --- --- --- --- --- --- --- --- --- --- --- --- --- --- --- --- --- --- --- --- --- --- --- --- --- --- --- --- --g tgt cta gaa aaa aat gtg tga ccc acg acc gta gga aac tct aga ggg taa gaa aaa tca atc gtt tat aga gac cat cag aaa gag gtt taa tat ttt tgt gag acc cat cga a-- --- --- --- --- --- --- --- --- --- --- --- --- --- --- --- --- --- --- --- --- --- --- --- --- --- --- --- --- --- --- --- --- --- --- --- --- --- --- --- --- --- --- --- --- --- --- --- --- --- --- --- --- --- --- --- --- --- --- --- --- --- --- --- --- --- --- --- --- --- --- --- --- --- --- --- --- --- --- --- --- --- --- --- --- --- --- --- --- --- --- --- --- --- --- --- --- --- --- --- --- --- --- --- --- --- --- --- --- --- --- --- --- --- --- --- --- --- --- --- --- --- --- --- --- --- --- --- --- --- --- --- --- --- --- --- --- --- --- --- --- --- --- --- --- --- --- --- --- --- --- --- --- --- --- --- --- --- --- --- --- --- --- --- --- --- --- --- --- --- --- --- --- --- --- --- --- --- --- --- --- --- --- --- --- --- --- --- --- --- --- --- --- --- --- --- --- --- --- --- --- --- --- --- --- --- --- --- --- --- --- --- --- --- --- --- --- --- --- --- --- --- --- --- --- --- --- --- --- --- --- --- --- --- --- --- --- --- --- --- --- --- --- --- --- --- --- --- --- --- --- --- --- --- --- --- --- --- --- --- --- --- --- --- --- --- --- --- --- --- --- --- --- --- --- --- --- --- --- --- -ga gag aaa gga taa aaa ctt ttt acg act cca tca gaa aga ggt tta ata ttt ttg tga gac cca tcg acg aga gaa aga gat ggt tag tca aga tat ttt tct tag tac aaa agt caa tgt ttt aaa ata tat gga cga gaa tta att tgt ctg tat aaa aac ttg tgt gaa att atg tac tag aga aaa aac gtg agc agt gtc ccc tac atg gat ttt aca gat cat tta tat tcc aaa aat att aac tat ata cgt tta tta tat gat gtt aac gtg taa att ata aac att att tta tga tgc aat tgt ctg aca acc tag att ggc ata agg ata ttg ata agc tct acg aga ata tat tgt tgg acg tta tcg ttt acg aaa tag ttg aga cat cag aaa gag gtt taa tat ttt tgt gag a-c cat cga aga gag aaa gag aat aaa aat a-- --- --- --- ttt ttt tgt aaa ac- ttt tta tga gac aag aga gaa aga gaa tac gaa tag tga tca tat cgt atc aca tat tga aac aga aag aag aag taa cga gag gta act ttt tgt gaa tgt agt taa aca ttt ttg ttt tgc aaa ccg gaa tat agt acc cgg tac act ttt tta att cgt ggt gcg gtg tct gaa tcg ttc gat taa ccc aac tca tcc att ttc aga tga ata gag tta tcg att cag aca cat gct ttg agt ttt gtt gaa tcg atg agt gaa gta tca tcg gtt gca cct tca gat gcc gat ccg tcg aca tac ttg aat cca tcc ttg act tca agt tca gat gat tcc tca cac atg tct ccg ata cgt acg cta aac tct agg ttc tta aca cat ttt gta tca acg atc gtt gaa ccg atg ata tct ttg taa ctc act ttc tta tgt gag atg tta gac cca agt act gga tgg gtc ttg atg tcg ctg tct ttc tct tct tcg cta cat ctg atg tcg ata gac atc tca cag tct ttg atc ata gcc aga gct tct tca cgc gtg atc gcg gga gag tcc tta cct tgt cct ggt gac acg ctg gac aat cta gta ttc aca gtg ttt cca tca gag gat tcg gag atg gat gaa atc ttt ggg cat ttg gtg aat cca aag ttc atg tta aga ccc gcg ccg acg ata gtg taa taa gtg gtg gga tct cct ttt aca act --- --- --- --- --- --- tct tcg gat acc tca tca tct tcg gtc tct gta act tcc gtt acg gat tga caa atc tta tca ttg gtc ggt gtt tgg tct tgc ttt gtg act ttg ata ata aca tcg att ccc ata tga tgt ttg ttt tct tct tca gta cac gag gat gag gat tgt tga aga cta gta ggc ata gca gct gcc act agg cac atg cat gcc agg aca ata tat tgt ttc atg att gct att gat tga tta ctg ttc tag atg att cta ctt tct tac cat ata ata aat tag aat ata ttt tct act ttt acg aga aat taa tta ttg tat tta tta ttt atg ggt -aa aaa aaa ctt act ata agt ggg tgg gat tct ggg aat tag tga tca gtt tat gta tat cgc aac tag cgg gca tat ggc tat tga cat cga gaa cat tac cca tat gat aag aga ttg tat ccg ttt cgt agt ctt gag tat tgg tat tac tat ata gta tgt aga tgt cga cgc tag ata gac agt cgc cca cta gag tta ccg tct ctg aat gcg gca tga tag tat cat tct ttg ttt tcg tta act gtt tgg aag atg aat ctt tgt tgt tac att taa tct cga aat tca gag tac ata tct ttg aag tat tct gat atc tat ttt ctc ctg taa aga atc ctg aag ttg cta cat tat taa gga cag aga agt att ctg cac gaa aga cgg gat cac aat ctt tat gat tca tgg taa tag tta gtt ccg acg ttg aga tgg att cgc tga gac cgg tag tgg tcg tcc gag tac acg atg tgt cgt tga ctg gat aca ggt taa ttt cca cat cga tat agt taa atg tat tgc tgg tt- --a cgg gtt cgc att tat ctg tgg aag aga cgg tgt gag aat atg ttc cgg gac cac acg gag aac aga tga cgt ctc cgg tag acg tgt atc cgg ata ctc cgt atc cta ttc cac act ttg ttt tag aaa tac atg ttc tac acc ctg atg atc ctt tga gaa gac aat aat atc ctg gag agc att cac aga ttc tat tgt gag tcg tgt tac acg atc gcg tct cta cct gat tac tat cac atc ttc cgt tac aac tta gac aag cct gta aat gat tat tgt gag atg taa agg tat ccg aac cac acg gcg tac att gtg tat tag tct tgc tat cac ata atc tgg aag cgt aag ttc ccg gag gac acg ata gac aac ata gat tac ggc ttc tgt att cgt tgt ctt tac att ttc cat tgg atg gtg cat gtg gtg cta tat ctc ttc cgt tta tta tta tac atg aga gaa aca ata tat acg agt ata ata cgg acc tca tga ttt aat aat -gt agt aat cgt cgt ctt gtt act gtt tgt ttc cta ctt ctc caa tca tat aga tta ttt ttt aaa tat ttt ctt tct atc atg gat aat att tgt aat ggt tct ttc cgt aca aca tac tgt tta gat ggt agt cgc tta gct tgg tta tga tat tgc gca taa ttt ccg gag gca aat acg ata gtc tag att gac tat cga tgg tag act cta att tat tga gtg ctt tgt cga cga gtt tac ttt tat gct cca tcg ata gat gac act gtt cta tga gat cgt cgt aca tgg gaa atg aaa tgc gtt tgt ctg aat gta tgg ctt cga gat agg tgt gat acc gga tgt ctt ctg ttc tca ata ccg tat aca agt tgg tgt ctg aga ttc gaa tct ctt tga gga gac tta tgt cac gac tac att ttt cga tga tgg aat cta tct tat cga atg ata tat ttt tca taa ata cac ttt tat agt cct cgt tta aac aga att tag tat gta gtt ccg caa atg act cgt ccc tta ata ggc agt agg cta tta tct tct tta cgt agt gat cgt cgt agg gag aga act ccg aca tct tgt aga aca acg att taa tca tag gta gag ata ctt tca gtc tgt ggt gga tga tgt cat taa cga cat ccg cct tgt ata tga tgt ttc tgt ttt caa aca cca agt cga ata ctg tct tat cgt ctt tag tcg gaa ggt tga tgt cgt atc cga tgt ata cga ggt atg agg caa cat tgt tat tgc aat tct gga agg cgg tat gaa gag gag tca ttg tat tat agt att cgt ctt tct gaa tgt cga atc tat cta gta gat acc gta gta tat tga gag agc gac ttc cat atc ctt gat tat gtt tta tga ata gat aaa gta gat gtt gtc ctt ctt cct ttt gta att tcc cgt att ttt gtt cgt gcc aat tga gta aca tta tga gaa tat gac ctg ttg cac aat cgt tct tta tgt att cca tga tgg gtg tac aat caa gat tat tac gta tcc tcg tat cgg ctc ctc gag ata aaa gag cat aca cca cac gag gac tat gtt tgg tat act gtt gaa ggt aag tgt gta acg gcg tat ttc cga ttt tcg taa ccg cgt taa tgt ttg ctc cat gat cta tta tcg cgt aga tga atc gct tct cag ctc gca tct tag tgt gac tct ttg act tgt aat aat tgc ttt cgt gga aca cgg ata tgt gtt tac agt agt aat gaa gag aag tga gtc cat cct cat cga cgc aat tag ggt cag atc ctt tag tca ata att tgt aca gaa cgt aat agt tta agc tcc cat tga att tat atc taa gat aac aca gca ata gat cgg atg att tac taa agt cat caa tgg ggt ccg tta gta tat caa aga tct tgt tat cga ttg ata gtg aat gaa tca gat agt ggt gta gag gaa tat gtc ctt ttt tat cct tgc tat caa agt tac gca tgc cgt ggc gta aca ata tct tta ata cag atg gat taa atc gtg tat tca tcg tat agc aat gta atg gag agt tac cac att tta gtc gtt tat tca gat cgc agt gtt taa taa cta att taa aca gat gag atg ata tat cca cat caa aga atg cga gat aca tat gac aga cat tat tga cag aaa tgt gac ctt cat tat cac cgt cgt cca taa atg cgt tag gta cgt acc aca tac tat cgt taa cga tgc gca caa tct cgt cca ttt tca tcc atc ttc ata atg att tac ttt ttc ata att aga gaa aaa gat caa ggt ata aaa att aga agt gtt aga cta ta- --- --- --- --- --- --- --- --- --- --- -aa cta act tat aa- --- --- --- --- --- --- --- --- --- --- --- --- --- --- --- --- --- --- --- --- --- --- --- --- --- --- --- --- --- --- --- --- --- --- --- --- --- --- --- --- --- --- --- --- --- --- --- --- --- --- --- --- --- --- --- --- --- --- --- --- --- --- --- --- --- --- --- --- --- --- --- --- --- --- --- --- --- --- --- --- --- --- --- --- --- --- -aa cta act tat gac tta act aac tta tga cta tta act cat ttc aag aaa ggt ggg tgg aga gaa ctc tat atg aca gct tgt gaa aca att aga tcc cta att tct aat gga agt ttt gat agg aga ttg tca tca gtt gat aca ttg ttt att atc tca tct att aga gca cgt ctg ttt aga gct tta gtg acc tgc tcg gtt act tct gtg taa atc ttg aat cct tta gtg ata cac tgt gtc aaa act gga tgt tta gaa tac cta tgt aga ata tgg gaa gca tgc ttg ttt ttg tct cta tta tag att aac tca tac atg gtt gta tta tga att ttc atc tgc cta atg tac tcc aat tct tgt tta caa tca att ata taa tca aag agt gat gat gca tac aca tta caa agt gaa taa tct acc atc ata aaa tac ttg ata cag agc ttt atc aca tca tgg ttt tca att gta tta tta agt ata gct aat ttt ata cag tca ata gac aat ggt tct cta agc aat att tct aat att tta aga tgt gtt tcc cta cgg gcg atg aca gat ccc cta tcc acg gcc acg tca aga cat gta tat cca tta ctc att act gcg ttg aca ttt gct cca ttt tct aat agc cat gat act aaa tct ata taa cct gca tag ata gcg cga taa agc aag gtc ctt cca cca gca tct agt tga ttg ata tct tca ata tat ggg ata caa agc tta taa att tct aat act gtg ggt tca tct aca agg aat ccc cta gta tac tga att att tta tat aga tct aat tta aca tca ttt tca tct ggg ata cca caa ttc aaa ata aac tca aca aca cta ctt tcc ttt tta cat att ccc cta aaa tag gca ttc aag cat tct att tta tat att aca gcc cca tga tct acc ata aaa tca aca atg tct att tct aca tat gca tta gat aga tag taa agt aag aga tct tgt aca gaa tta caa ttc tta ata att ata gag aaa ata tct tcc ata taa ttc ttt gac act aat gca gat ata ata tct tta tat gta ata tat gca aac agt cta tct act ata tac tga tca ata tta tct cta tga atc cta aaa taa tca tac aga aca tct aca gga tca caa att ggt tca agg aga aat cta tca aat att ttc ctg tca aca act ggt tct aga aca taa cag tca aca cct aat cca tgt ttt tta tag tca tct acc aaa gat aat gac caa aga tcg agg tcg tcg tga aac tgc tca tcg aca gcc atg aaa tct gcc gac tcc atg gcg cga atc gca ctg tct tat tcg cca ttg att ttc a-t ttt tta taa tta tgt aca tgt ttt cct tct att ctc aag agt cta caa aaa tat a-t ttt ttt tcg ata tct aag tac taa gtt ttt tta ctg ttt ttg tta ctg tct tcc att ctt cta act aaa gat ctg aga taa att ata caa tct tcg cta tcg aac cat ttt tgt agt cta aag cct gaa gta att aac caa ctg ttt tta tta gtg gct ttt ttc gat cta tcc tcg tc- --- --c tca tca tcc tta ta- --- --- --- --- --- --- tta tta tca tta tta tca tag tct att aaa cac aaa tca tct --- --- --- --- --- --- --- --- --- --- --- --- --- --- --- --- --- --- --- --- --- --- --- --- --- --- --a cgt tta taa c-- --- --- --- --- --- --- --- --a aca ttc --- --- --- --- -tc att att aat tag ttc tgt agt aat atc ttt aat aat ttg gct ata --- --- -ca tct gtt --- --- --- --- --- --- --- --- --- --- --- --- --- --- --- --- --- --- --c aat act --- --- --- --- --- --- --- --a tct att gat gat ttc ttt t-- --- --- --- --- --- --- --- --- --- --- --t taa gac t-- --- --- --- --- --- --- --- --- --- --- --- --- --- --- --- --- --- --- --- --- --- --- --- --- --- --- --- --- --- --- --- --- --- --- --- --- --- --- --- taa act agt --- --- --- --- --- --- --- --- --- --- --- --- --- --- --- --- --- --- --- --- --- --- --- --- --- --- --- --- --- --- --- --- --- --- --- --- --- --- --- --- --- --- --- --- --- --- --- --- --- --- --- --- --- --- --- --- --- --- --- --- --- --- --- --- --- --- --- --- --- --- --- --- --- --- --- --- --- --- --- --- --- --- --- --- --- --- --- --- --- --- --- --- --- --- --- --- --- --- --- --- --- --- --- --- --- --- --- --- --- --t atg gta atg acg atg aaa --- -tc gag tag ta- --- --a ctt cta ata aag act tga ta- --- --t cat tat cat atg ttt gat cg- --- --- --- --- --- --- --t cat agt taa tag tgt g-- --- --- --- --- --- --- --- --- --- --- --- --- --- --- --- --- --- --- --- -gc taa atg gta ctg tta ata agt tta t-- --- --- --- --- --- --- --- --- --- --- --- --- --- --- --- --- --a gac aat atc ata gta ttt tct ttc cag aat --- --- --- --t aga tta ttt ttt taa ata ctg atc ctc aca att ccg tga tgt agc agt agt tgg t-- --- --- --- --- --- --- --- --- --- --- --- --- --- --- --- --- --- --- --- --- --- --- --- --- --- --- --- -gc atg gtc tat atc gt- --- --- --- --- --- --- --- --- --- --- --- --- --- --- --- --- --- --- --- --- --- --- --- --- --- --- --- --- --- --- --- --- --- --- --- --- --- --- --- --- --- --- --- taa aat gta tca tat ata ata gtt ttc tga cgt gga gta cag aat ttt cga --- --- --- --- --- --- --- --- --- --- --- --- --- --- --- --- --- --- --- --- --- --- --- --- --- --- --- --- --- --- --- --- --- --- --- --- --- --- --- --- --- --- --- --t taa tga gtt cat ggt aag gaa ggg caa atg tct --- gta tat aat ata cat aag tta a-- --- --- --- --- --- --- --- --- --- --- --- --- --t agt ttt tta tca tat tt- --- --- --- --- --- --- --- --t cta ata cca taa taa aaa tta tca tta tg- --- --- -ta taa tca --- --- --- --- --- --- --- --- --- --- --- --t cac tg- --- -tc gct atc att att gcg ttt gtg tag tt- --- --- --- --- ctg ccc ta- --- --- --- --- --- --- --- --t cat cta cat cac tgt ca- --- --- --- --- ctc tca cta tat ctt cta aaa tta caa --- -ac aac --- --- --- --- --- --- --- --- --- --- --- --- --- --- --- --- --- --- --- --- --- --- --- --- --- --- --- --- --- --- --- --- --- --- --- --- tgg ata ttc gat --- --- --- --- --- --- --- --- --- --- --- --- --- --- --- --- --- --- --- --- --- --- --- --- --- --- --a aca gca ttt gtg t-- --- --- --- --- --- --- --- --- --- --- --- --- --- --- --- --- -

>JX878417.1_Monkeypox_virus_isolate_DRC_07-0104_complete_genome

--- --- --- --- --- --- --- --- --- --- --- --- --- --- --- --- --- --- --- --- --- --- --- --- --- --- --- --- --- --- --- --- --- --- --- --g tgt cta gaa aaa aat gtg tga ccc acg acc gta gga aac tct aga ggg taa gaa aaa tca atc gtt tat aga gac cat cag aaa gag gtt taa tat ttt tgt gag acc cat cga aga gag aaa gga taa aaa ctt ttt acg act cca tca gaa aga ggt tta ata ttt ttg tga gac cca tcg aag aga gaa agg ata aaa act ttt tac gac tcc atc aga aag agg ttt aat att ttt gtg aga ccc atc gaa gag aga aag gat aaa aac ttt tta cga ctc cat cag aaa gag gtt taa tat ttt tgt gag acc cat cga cga gag aaa gga taa aaa ctt ttt acg act cca tca gaa aga ggt tta ata ttt ttg tga gac cca tcg aag --- --- --- --- --- --- --- --- --- --- --- --- --- --- --- --- --- --- --- --- --- --- --- -ag aga aag gat aaa aac ttt tta cga ctc cat cag aaa gag gtt taa tat ttt tgt gag acc cat cga cga gag aaa gga taa aaa ctt ttt acg act cca tca gaa aga ggt tta ata ttt ttg tga gac cca tcg aag aga gaa agg ata aaa act ttt tac gac tcc atc aga aag agg ttt aat att ttt gtg aga ccc atc gac gag aga aag gat aaa aac ttt tta cga ctc cat cag aaa gag gtt taa tat ttt tgt gag acc cat cga aga gag aaa gga taa aaa ctt ttt acg act cca tca gaa aga ggt tta --- --- --- --- --- --- --- --- --- --- --- --- --- --- --- --- --- --- --- --- --- --- --- --- --- --- --- --- --- --- --- --- --- --- --- --- --- --- --- --- --- --- --- --- --- --- --a tat ttt tgt gag acc cat cga cga gag aaa gga taa aaa ctt ttt acg act cca tca gaa aga ggt tta ata ttt ttg tga gac cca tcg aag aga gaa aga gat ggt tag tca aga tat ttt tct tag tac aaa agt caa tgt ttt aaa ata tat gga cga gaa tta att tgt ctg tat aaa aac ttg tgt gaa att atg tac tag aga aaa aac gtg agc agt gtc ccc tac atg gat ttt aca gat cat tta tat tcc aaa aat att aac tat ata cgt tta tta tat gat gtt aac gtg taa att ata aac att att tta tga tgc aat tgt ctg aca acc tag att ggc ata agg ata ttg ata agc tct acg aga ata tat tgt tgg acg tta tcg ttt acg aaa tag ttg aga cat cag aaa gag gtt taa tat ttt tgt gag a-c cat cga aga gag aaa gag aat aaa aat a-- --- --- --- ttt ttt tgt aaa ac- ttt tta tga gac aag aga gaa aga gaa tac gaa tag tga tca tat cgt atc aca tat tga aac aga aag aag aag taa cga gag gta act ttt tgt gaa tgt agt taa aca ttt ttg ttt tgc aaa ccg gaa tat agt acc cgg tac act ttt tta att cgt ggt gcg gtg tct gaa tcg ttc gat taa ccc aac tca tcc att ttc aga tga ata gag tta tcg att cag aca cat gct ttg agt ttt gtt gaa tcg atg agt gaa gta tca tcg gtt gca cct tca gat gcc gat ccg tcg aca tac ttg aat cca tcc ttg act tca agt tca gat gat tcc tca cac atg tct ccg ata cgt acg cta aac tct agg ttc ttg aca cat ttt gta tca acg atc gtt gaa ccg atg ata tct ttg taa ctc act ttc tta tgt gag atg tta gac cca agt act gga tgg gtc ttg atg tcg ctg tct ttc tct tct tcg cta cat ctg atg tcg ata gac atc tca cag tct ttg atc ata gcc aga gct tct tca cgc gtg atc gcg gga gag tcc tta cct tgt cct ggt gac acg ctg gac aat cta gta ttc aca gtg ttt cca tca gag gat tcg gag atg gat gaa atc ttt ggg cat ttg gtg aat cca aag ttc atg tta aga ccc gcg ccg acg ata gtg taa taa gtg gtg gga tct cct ttt aca act --- --- --- --- --- --- tct tcg gat acc tca tca tct tcg gtc tct gta act tcc gtt acg gat tga caa atc tta tca ttg gtc ggt gtt tgg tct tgc ttt gtg act ttg ata ata aca tcg att ccc ata tga tgt ttg ttt tct tct tca gta cac gag gat gag gat tgt tga aga cta gta ggc ata gca gct gcc act agg cac atg cat gcc agg aca ata tat tgt ttc atg att gct att gat tga tta ctg ttc tag atg att cta ctt tct tac cat ata ata aat tag aat ata ttt tct act ttt acg aga aat taa tta ttg tat tta tta ttt atg ggt -aa aaa aaa ctt act ata agt ggg tgg gat tct ggg aat tag tga tca gtt tat gta tat cgc aac tag cgg gca tat ggc tat tga cat cga gaa cat tac cca tat gat aag aga ttg tat ccg ttt cgt agt ctt gag tat tgg tat tac tat ata gta tgt aga tgt cga cgc tag ata gac agt cgc cca cta gag tta ccg tct ctg aat gcg gca tga tag tat cat tct ttg ttt tcg tta act gtt tgg aag atg aat ctt tgt tgt tac att taa tct cga aat tca gag tac ata tct ttg aag tat tct gat atc tat ttt ctc ctg taa aga atc ctg aag ttg cta cat tat taa gga cag aga agt att ctg cac gaa aga cgg gat cac aat ctt tat gat tca tgg taa tag tta gtt ccg acg ttg aga tgg att cgc tga gac cgg tag tgg tcg tcc gag tac acg atg tgt cgt tga ctg gat aca ggt taa ttt cca cat cga tat agt taa atg tat tgc tgg tt- --a cgg gtt cgc att tat ctg tgg aag aga cgg tgt gag aat atg ttc cgg gac cac acg gag aac aga tga cgt ctc cgg tag acg tgt atc cgg ata ctc cgt atc cta ttc cac act ttg ttt tag aaa tac atg ttc tac acc ctg atg atc ctt tga gaa gac aat aat atc ctg gag agc att cac aga ttc tat tgt gag tcg tgt tac acg atc gcg tct cta cct gat tac tat cac atc ttc cgt tac aac tta gac aag cct gta aat gat tat tgt gag atg taa agg tat ccg aac cac acg gcg tac att gtg tat tag tct tgc tat cac ata atc tgg aag cgt aag ttc ccg gag gac acg ata gac aac ata gat tac ggc ttc tgt att cgt tgt ctt tac att ttc cat tgg atg gtg cat gtg gtg cta tat ctc ttc cgt tta tta tta tac atg aga gaa aca ata tat acg agt ata ata cgg acc tca tga ttt aat aat -gt agt aat cgt cgt ctt gtt act gtt tgt ttc cta ctt ctc caa tca tat aga tta ttt ttt aaa tat ttt ctt tct atc atg gat aat att tgt aat ggt tct ttc cgt aca aca tac tgt tta gat ggt agt cgc tta gct tgg tta tga tat tgc gca taa ttt ccg gag gca aat acg ata gtc tag att gac tat cga tgg tag act cta att tat tga gtg ctt tgt cga cga gtt tac ttt tat gct cca tcg ata gat gac act gtt cta tga gat cgt cgt aca tgg gaa atg aaa tgc gtt tgt ctg aat gta tgg ctt cga gat agg tgt gat acc gga tgt ctt ctg ttc tca ata ccg tat aca agt tgg tgt ctg aga ttc gaa tct ctt tga gga gac tta tgt cac gac tac att ttt cga tga tgg aat cta tct tat cga atg ata tat ttt tca taa ata cac ttt tat agt cct cgt tta aac aga att tag tat gta gtt ccg caa atg act cgt ccc tta ata ggc agt agg cta tta tct tct tta cgt agt gat cgt cgt agg gag aga act ccg aca tct tgt aga aca acg att taa tca tag gta gag ata ctt tca gtc tgt ggt gga tga tgt cat taa cga cat ccg cct tgt ata tga tgt ttc tgt ttt caa aca cca agt cga ata ctg tct tat cgt ctt tag tcg gaa ggt tga tgt cgt atc cga tgt ata cga ggt atg agg caa cat tgt tat tgc aat tct gga agg cgg tat gaa gag gag tca ttg tat tat agt att cgt ctt tct gaa tgt cga atc tat cta gta gat acc gta gta tat tga gag agc gac ttc cat atc ctt gat tat gtt tta tga ata gat aaa gta gat gtt gtc ctt ctt cct ttt gta att tcc cgt att ttt gtt cgt gcc aat tga gta aca tta tga gaa tat gac ctg ttg cac aat cgt tct tta tgt att cca tga tgg gtg tac aat caa gat tat tac gta tcc tcg tat cgg ctc ctc gag ata aaa gag cat aca cca cac gag gac tat gtt tgg tat act gtt gaa ggt aag tgt gta acg gcg tat ttc cga ttt tcg taa ccg cgt taa tgt ttg ctc cat gat cta tta tcg cgt aga tga atc gct tct cag ctc gca tct tag tgt gac tct ttg act tgt aat aat tgc ttt cgt gga aca cgg ata tgt gtt tac agt agt aat gaa gag aag tga gtc cat cct cat cga cgc aat tag ggt cag atc ctt tag tca ata att tgt aca gaa cgt aat agt tta agc tcc cat tga att tat atc taa gat aac aca gca ata gat cgg atg att tac taa agt cat caa tgg ggt ccg tta gta tat caa aga tct tgt tat cga ttg ata gtg aat gaa tca gat agt ggt gta gag gaa tat gtc ctt ttt tat cct tgc tat caa agt tac gca tgc cgt ggc gta aca ata tct tta ata cag atg gat taa atc gtg tat tca tcg tat agc aat gta atg gag agt tac cac att tta gtc gtt tat tca gat cgc agt gtt taa taa cta att taa aca gat gag atg ata tat cca cat caa aga atg cga gat aca tat gac aga cat tat tga cag aaa tgt gac ctt cat tat cac cgt cgt cca taa atg cgt tag gta cgt acc aca tac tat cgt taa cga tgc gca caa tct cgt cca ttt tca tcc atc ttc ata atg att tac ttt ttc ata att aga gaa aaa gat caa ggt ata aaa att aga agt gtt aga cta ta- --- --- --- --- --- --- --- --- --- --- -aa cta act tat aa- --- --- --- --- --- --- --- --- --- --- --- --- --- --- --- --- --- --- --- --- --- --- --- --- --- --- --- --- --- --- --- --- --- --- --- --- --- --- --- --- --- --- --- --- --- --- --- --- --- --- --- --- --- --- --- --- --- --- --- --- --- --- --- --- --- --- --- --- --- --- --- --- --- --- --- --- --- --- --- --- --- --- --- --- --- --- -aa cta act tat gac tta act aac tta tga cta tta act cat ttc aag aaa ggt ggg tgg aga gaa ctc tat atg aca gct tgt gaa aca att aga tcc cta att tct aat gga agt ttt gat agg aga ttg tca tca gtt gat aca ttg ttt att atc tca tct att aga gca cgt ctg ttt aga gct tta gtg acc tgc tcg gtt act tct gtg taa atc ttg aat cct tta gtg ata cac tgt gtc aaa act gga tgt tta gaa tac cta tgt aga ata tgg gaa gca tgc ttg ttt ttg tct cta tta tag att aac tca tac atg gtt gta tta tga att ttc atc tgc cta atg tac tcc aat tct tgt tta caa tca att ata taa tca aag agt gat gat gca tac aca tta caa agt gaa taa tct acc atc ata aaa tac ttg ata cag agc ttt atc aca tca tgg ttt tca att gta tta tta agt ata gct aat ttt ata cag tca ata gac aat ggt tct cta agc aat att tct aat att tta aga tgt gct tcc cta cgg gcg atg aca gat ccc cta tcc acg gcc acg tca aga cat gta tat cca tta ctc att act gcg ttg aca ttt gct cca ttt tct aat agc cat gat act aaa tct ata taa cct gca tag ata gcg cga taa agc aag gtc ctt cca cca gca tct agt tga ttg ata tct tca ata tat ggg ata caa agc tta taa att tct aat act gtg ggt tca tct aca agg aat ccc cta gta tac tga att att tta tat aga tct aat tta aca tca ttt tca tct ggg ata cca caa ttc aaa ata aac tca aca aca cta ctt tcc ttt tta cat att ccc cta aaa tag gca ttc aag cat tct att tta tat att aca gcc cca tga tct acc ata aaa tca aca atg tct att tct aca tat gca tta gat aga tag taa agt aag aga tct tgt aca gaa tta caa ttc tta ata att ata gag aaa ata tct tcc ata taa ttc ttt gac act aat gca gat ata ata tct tta tat gta ata tat gca aac agt cta tct act ata tac tga tca ata tta tct cta tga atc cta aaa taa tca tac aga aca tct aca gga tca caa att ggt tca agg aga aat cta tca aat att ttc ctg tca aca act ggt tct aga aca taa cag tca aca cct aat cca tgt ttt tta tag tca tct acc aaa gat aat gac caa aga tcg agg tcg tcg tga aac tgc tca tcg aca gcc atg aaa tct gcc gac tcc atg gcg cga atc gca ctg tct tat tcg cca ttg att ttc a-t ttt tta taa tta tgt aca tgt ttt cct tct att ctc aag agt cta caa aaa tat a-t ttt ttt tcg ata tct aag tac taa gtt ttt tta ctg ttt ttg tta ctg tct tcc att ctt cta act aaa gat ctg aga taa att ata caa tct tcg cta tcg aac cat ttt tgt agt cta aag cct gaa gta att aac caa ctg ttt tta tta gtg gct ttt ttc gat cta tcc tcg tc- --- --c tca tca tcc tta ta- --- --- --- --- --- --- tta tta tca tta tta tca tag tct att aaa cac aaa tca tct --- --- --- --- --- --- --- --- --- --- --- --- --- --- --- --- --- --- --- --- --- --- --- --- --- --- --a cgt tta taa c-- --- --- --- --- --- --- --- --a aca ttc --- --- --- --- -tc att att aat tag ttc tgt agt aat atc ttt aat aat ttg gct ata --- --- -ca tct gtt --- --- --- --- --- --- --- --- --- --- --- --- --- --- --- --- --- --- --c aat act --- --- --- --- --- --- --- --a tct att gat gat ttc ttt t-- --- --- --- --- --- --- --- --- --- --- --t taa gac t-- --- --- --- --- --- --- --- --- --- --- --- --- --- --- --- --- --- --- --- --- --- --- --- --- --- --- --- --- --- --- --- --- --- --- --- --- --- --- --- taa act agt --- --- --- --- --- --- --- --- --- --- --- --- --- --- --- --- --- --- --- --- --- --- --- --- --- --- --- --- --- --- --- --- --- --- --- --- --- --- --- --- --- --- --- --- --- --- --- --- --- --- --- --- --- --- --- --- --- --- --- --- --- --- --- --- --- --- --- --- --- --- --- --- --- --- --- --- --- --- --- --- --- --- --- --- --- --- --- --- --- --- --- --- --- --- --- --- --- --- --- --- --- --- --- --- --- --- --- --- --- --t atg gta atg a-g atg aaa --- -tc gag tag ta- --- --a ctt cta ata aag act tga ta- --- --t cat tat cat atg ttt gat cg- --- --- --- --- --- --- --t cat agt taa tag tgt g-- --- --- --- --- --- --- --- --- --- --- --- --- --- --- --- --- --- --- --- -gc taa atg gta ctg tta ata agt tta t-- --- --- --- --- --- --- --- --- --- --- --- --- --- --- --- --- --a gac aat atc ata gta ttt tct ttc cag aat --- --- --- --t aga tta ttt ttt taa ata ctg atc ctc aca att ccg tga tgt agc agt agt tgg t-- --- --- --- --- --- --- --- --- --- --- --- --- --- --- --- --- --- --- --- --- --- --- --- --- --- --- --- -gc atg gtc tat atc gt- --- --- --- --- --- --- --- --- --- --- --- --- --- --- --- --- --- --- --- --- --- --- --- --- --- --- --- --- --- --- --- --- --- --- --- --- --- --- --- --- --- --- --- taa aat gta tca tat ata ata gtt ttc tga cgt gga gta cag aat ttt cga --- --- --- --- --- --- --- --- --- --- --- --- --- --- --- --- --- --- --- --- --- --- --- --- --- --- --- --- --- --- --- --- --- --- --- --- --- --- --- --- --- --- --- --t taa tga gtt cat ggt aag gaa ggg caa atg tct --- gta tat aat ata cat aag tta a-- --- --- --- --- --- --- --- --- --- --- --- --- --t agt ttt tta tca tat tt- --- --- --- --- --- --- --- --t cta ata cca taa taa aaa tta tca tta tg- --- --- -ta taa tca --- --- --- --- --- --- --- --- --- --- --- --t cac tg- --- -tc gct atc att att gcg ttt gtg tag tt- --- --- --- --- ctg ccc ta- --- --- --- --- --- --- --- --t cat cta cat cac tgt ca- --- --- --- --- ctc tca cta tat ctt cta aaa tta caa --- -ac aac --- --- --- --- --- --- --- --- --- --- --- --- --- --- --- --- --- --- --- --- --- --- --- --- --- --- --- --- --- --- --- --- --- --- --- --- tgg ata ttc gat --- --- --- --- --- --- --- --- --- --- --- --- --- --- --- --- --- --- --- --- --- --- --- --- --- --- --a aca gca ttt gtg t-- --- --- --- --- --- --- --- --- --- --- --- --- --- --- --- --- -

>JX878426.1_Monkeypox_virus_isolate_DRC_07-0450_complete_genome

--- --- --- --- --- --- --- --- --- --- --- --- --- --- --- --- --- --- --- --- --- --- --- --- --- --- --- --- --- --- --- --- --- --- --- --g tgt cta gaa aaa aat gtg tga ccc acg acc gta gga aac tct aga ggg taa gaa aaa tca atc gtt tat aga gac cat cag aaa gag gtt taa tat ttt tgt gag acc cat cga a-- --- --- --- --- --- --- --- --- --- --- --- --- --- --- --- --- --- --- --- --- --- --- --- --- --- --- --- --- --- --- --- --- --- --- --- --- --- --- --- --- --- --- --- --- --- --- --- --- --- --- --- --- --- --- --- --- --- --- --- --- --- --- --- --- --- --- --- --- --- --- --- --- --- --- --- --- --- --- --- --- --- --- --- --- --- --- --- --- --- --- --- --- --- --- --- --- --- --- --- --- --- --- --- --- --- --- --- --- --- --- --- --- --- --- --- --- --- --- --- --- --- --- --- --- --- --- --- --- --- --- --- --- --- --- --- --- --- --- --- --- --- --- --- --- --- --- --- --- --- --- --- --- --- --- --- --- --- --- --- --- --- --- --- --- --- --- --- --- --- --- --- --- --- --- --- --- --- --- --- --- --- --- --- --- --- --- --- --- --- --- --- --- --- --- --- --- --- --- --- --- --- --- --- --- --- --- --- --- --- --- --- --- --- --- --- --- --- --- --- --- --- --- --- --- --- --- --- --- --- --- --- --- --- --- --- --- --- --- --- --- --- --- --- --- --- --- --- --- --- --- --- --- --- --- --- --- --- --- --- --- --- --- --- --- --- --- --- --- --- --- --- --- --- --- --- --- --- --- --- -ga gag aaa gga taa aaa ctt ttt acg act cca tca gaa aga ggt tta ata ttt ttg tga gac cca tcg acg aga gaa aga gat ggt tag tca aga tat ttt tct tag tac aaa agt caa tgt ttt aaa ata tat gga cga gaa tta att tgt ctg tat aaa aac ttg tgt gaa att atg tac tag aga aaa aac gtg agc agt gtc ccc tac atg gat ttt aca gat cat tta tat tcc aaa aat att aac tat ata cgt tta tta tat gat gtt aac gtg taa att ata aac att att tta tga tgc aat tgt ctg aca acc tag att ggc ata agg ata ttg ata agc tct acg aga ata tat tgt tgg acg tta tcg ttt acg aaa tag ttg aga cat cag aaa gag gtt taa tat ttt tgt gag a-c cat cga aga gag aaa gag aat aaa aat a-- --- --- --- ttt ttt tgt aaa ac- ttt tta tga gac aag aga gaa aga gaa tac gaa tag tga tca tat cgt atc aca tat tga aac aga aag aag aag taa cga gag gta act ttt tgt gaa tgt agt taa aca ttt ttg ttt tgc aaa ccg gaa tat agt acc cgg tac act ttt tta att cgt ggt gcg gtg tct gaa tcg ttc gat taa ccc aac tca tcc att ttc aga tga ata gag tta tcg att cag aca cat gct ttg agt ttt gtt gaa tcg atg agt gaa gta tca tcg gtt gca cct tca gat gcc gat ccg tcg aca tac ttg aat cca tcc ttg act tca agt tca gat gat tcc tca cac atg tct ccg ata cgt acg cta aac tct agg ttc tta aca cat ttt gta tca acg atc gtt gaa ccg atg ata tct ttg taa ctc act ttc tta tgt gag atg tta gac cca agt act gga tgg gtc ttg atg tcg ctg tct ttc tct tct tcg cta cat ctg atg tcg ata gac atc tca cag tct ttg atc ata gcc aga gct tct tca cgc gtg atc gcg gga gag tcc tta cct tgt cct ggt gac acg ctg gac aat cta gta ttc aca gtg ttt cca tca gag gat tcg gag atg gat gaa atc ttt ggg cat ttg gtg aat cca aag ttc atg tta aga ccc gcg ccg acg ata gtg taa taa gtg gtg gga tct cct ttt aca act --- --- --- --- --- --- tct tcg gat acc tca tca tct tcg gtc tct gta act tcc gtt acg gat tga caa atc tta tca ttg gtc ggt gtt tgg tct tgc ttt gtg act ttg ata ata aca tcg att ccc ata tga tgt ttg ttt tct tct tca gta cac gag gat gag gat tgt tga aga cta gta ggc ata gca gct gcc act agg cac atg cat gcc agg aca ata tat tgt ttc atg att gct att gat tga tta ctg ttc tag atg att cta ctt tct tac cat ata ata aat tag aat ata ttt tct act ttt acg aga aat taa tta ttg tat tta tta ttt atg ggt aaa aaa aaa ctt act ata agt ggg tgg gat tct ggg aat tag tga tca gtt tat gta tat cgc aac tag cgg gca tat ggc tat tga cat cga gaa cat tac cca tat gat aag aga ttg tat ccg ttt cgt agt ctt gag tat tgg tat tac tat ata gta tgt aga tgt cga cgc tag ata gac agt cgc cca cta gag tta ccg tct ctg aat gcg gca tga tag tat cat tct ttg ttt tcg tta act gtt tgg aag atg aat ctt tgt tgt tac att taa tct cga aat tca gag tac ata tct ttg aag tat tct gat atc tat ttt ctc ctg taa aga atc ctg aag ttg cta cat tat taa gga cag aga agt att ctg cac gaa aga cgg gat cac aat ctt tat gat tca tgg taa tag tta gtt ccg acg ttg aga tgg att cgc tga gac cgg tag tgg tcg tcc gag tac acg atg tgt cgt tga ctg gat aca ggt taa ttt cca cat cga tat agt taa atg tat tgc tgg tt- --a cgg gtt cgc att tat ctg tgg aag aga cgg tgt gag aat atg ttc cgg gac cac acg gag aac aga tga cgt ctc cgg tag acg tgt atc cgg ata ctc cgt atc cta ttc cac act ttg ttt tag aaa tac atg ttc tac acc ctg atg atc ctt tga gaa gac aat aat atc ctg gag agc att cac aga ttc tat tgt gag tcg tgt tac acg atc gcg tct cta cct gat tac tat cac atc ttc cgt tac aac tta gac aag cct gta aat gat tat tgt gag atg taa agg tat ccg aac cac acg gcg tac att gtg tat tag tct tgc tat cac ata atc tgg aag cgt aag ttc ccg gag gac acg ata gac aac ata gat tac ggc ttc tgt att cgt tgt ctt tac att ttc cat tgg atg gtg cat gtg gtg cta tat ctc ttc cgt tta tta tta tac atg aga gaa aca ata tat acg agt ata ata cgg acc tca tga ttt aat aat -gt agt aat cgt cgt ctt gtt act gtt tgt ttc cta ctt ctc caa tca tat aga tta ttt ttt aaa tat ttt ctt tct atc atg gat aat att tgt aat ggt tct ttc cgt aca aca tac tgt tta gat ggt agt cgc tta gct tgg tta tga tat tgc gca taa ttt ccg gag gca aat acg ata gtc tag att gac tat cga tgg tag act cta att tat tga gtg ctt tgt cga cga gtt tac ttt tat gct cca tcg ata gat gac act gtt cta tga gat cgt cgt aca tgg gaa atg aaa tgc gtt tgt ctg aat gta tgg ctt cga gat agg tgt gat acc gga tgt ctt ctg ttc tca ata ccg tat aca agt tgg tgt ctg aga ttc gaa tct ctt tga gga gac tta tgt cac gac tac att ttt cga tga tgg aat cta tct tat cga atg ata tat ttt tca taa ata cac ttt tat agt cct cgt tta aac aga att tag tat gta gtt ccg caa atg act cgt ccc tta ata ggc agt agg cta tta tct tct tta cgt agt gat cgt cgt agg gag aga act ccg aca tct tgt aga aca acg att taa tca tag gta gag ata ctt tca gtc tgt ggt gga tga tgt cat taa cga cat ccg cct tgt ata tga tgt ttc tgt ttt caa aca cca agt cga ata ctg tct tat cgt ctt tag tcg gaa ggt tga tgt cgt atc cga tgt ata cga ggt atg agg caa cat tgt tat tgc aat tct gga agg cgg tat gaa gag gag tca ttg tat tat agt att cgt ctt tct gaa tgt cga atc tat cta gta gat acc gta gta tat tga gag agc gac ttc cat atc ctt gat tat gtt tta tga ata gat aaa gta gat gtt gtc ctt ctt cct ttt gta att tcc cgt att ttt gtt cgt gcc aat tga gta aca tta tga gaa tat gac ctg ttg cac aat cgt tct tta tgt att cca tga tgg gtg tac aat caa gat tat tac gta tcc tcg tat cgg ctc ctc gag ata aaa gag cat aca cca cac gag gac tat gtt tgg tat act gtt gaa ggt aag tgt gta acg gcg tat ttc cga ttt tcg taa ccg cgt taa tgt ttg ctc cat gat cta tta tcg cgt aga tga atc gct tct cag ctc gca tct tag tgt gac tct ttg act tgt aat aat tgc ttt cgt gga aca cgg ata tgt gtt tac agt agt aat gaa gag aag tga gtc cat cct cat cga cgc aat tag ggt cag atc ctt tag tca ata att tgt aca gaa cgt aat agt tta agc tcc cat tga att tat atc taa gat aac aca gca ata gat cgg atg att tac taa agt cat caa tgg ggt ccg tta gta tat caa aga tct tgt tat cga ttg ata gtg aat gaa tca gat agt ggt gta gag gaa tat gtc ctt ttt tat cct tgc tat caa agt tac gca tgc cgt ggc gta aca ata tct tta ata cag atg gat taa atc gtg tat tca tcg tat agc aat gta atg gag agt tac cac att tta gtc gtt tat tca gat cgc agt gtt taa taa cta att taa aca gat gag atg ata tat cca cat caa aga atg cga gat aca tat gac aga cat tat tga cag aaa tgt gac ctt cat tat cac cgt cgt cca taa atg cgt tag gta cgt acc aca tac tat cgt taa cga tgc gca caa tct cgt cca ttt tca tcc atc ttc ata atg att tac ttt ttc ata att aga gaa aaa gat caa ggt ata aaa att aga agt gtt aga cta ta- --- --- --- --- --- --- --- --- --- --- -aa cta act tat aa- --- --- --- --- --- --- --- --- --- --- --- --- --- --- --- --- --- --- --- --- --- --- --- --- --- --- --- --- --- --- --- --- --- --- --- --- --- --- --- --- --- --- --- --- --- --- --- --- --- --- --- --- --- --- --- --- --- --- --- --- --- --- --- --- --- --- --- --- --- --- --- --- --- --- --- --- --- --- --- --- --- --- --- --- --- --- -aa cta act tat gac tta act aac tta tga cta tta act cat ttc aag aaa ggt ggg tgg aga gaa ctc tat atg aca gct tgt gaa aca att aga tcc cta att tct aat gga agt ttt gat agg aga ttg tca tca gtt gat aca ttg ttt att atc tca tct att aga gca cgt ctg ttt aga gct tta gtg acc tgc tcg gtt act tct gtg taa atc ttg aat cct tta gtg ata cac tgt gtc aaa act gga tgt tta gaa tac cta tgt aga ata tgg gaa gca tgc ttg ttt ttg tct cta tta tag att aac tca tac atg gtt gta tta tga att ttc atc tgc cta atg tac tcc aat tct tgt tta caa tca att ata taa tca aag agt gat gat gca tac aca tta caa agt gaa taa tct acc atc ata aaa tac ttg ata cag agc ttt atc aca tca tgg ttt tca att gta tta tta agt ata gct aat ttt ata cag tca ata gac aat ggt tct cta agc aat att tct aat att tta aga tgt gtt tcc cta cgg gcg atg aca gat ccc cta tcc acg gcc acg tca aga cat gta tat cca tta ctc att act gcg ttg aca ttt gct cca ttt tct aat agc cat gat act aaa tct ata taa cct gca tag ata gcg cga taa agc aag gtc ctt cca cca gca tct agt tga ttg ata tct tca ata tat ggg ata caa agc tta taa att tct aat act gtg ggt tca tct aca agg aat ccc cta gta tac tga att att tta tat aga tct aat tta aca tca ttt tca tct ggg ata cca caa ttc aaa ata aac tca aca aca cta ctt tcc ttt tta cat att ccc cta aaa tag gca ttc aag cat tct att tta tat att aca gcc cca tga tct acc ata aaa tca aca atg tct att tct aca tat gca tta gat aga tag taa agt aag aga tct tgt aca gaa tta caa ttc tta ata att ata gag aaa ata tct tcc ata taa ttc ttt gac act aat gca gat ata ata tct tta tat gta ata tat gca aac agt cta tct act ata tac tga tca ata tta tct cta tga atc cta aaa taa tca tac aga aca tct aca gga tca caa att ggt tca agg aga aat cta tca aat att ttc ctg tca aca act ggt tct aga aca taa cag tca aca cct aat cca tgt ttt tta tag tca tct acc aaa gat aat gac caa aga tcg agg tcg tcg tga aac tgc tca tcg aca gcc atg aaa tct gcc gac tcc atg gcg cga atc gca ctg tct tat tcg cca ttg att ttc a-t ttt tta taa tta tgt aca tgt ttt cct tct att ctc aag agt cta caa aaa tat a-t ttt ttt tcg ata tct aag tac taa gtt ttt tta ctg ttt ttg tta ctg tct tcc att ctt cta act aaa gat ctg aga taa att ata caa tct tcg cta tcg aac cat ttt tgt agt cta aag cct gaa gta att aac caa ctg ttt tta tta gtg gct ttt ttc gat cta tcc tcg tc- --- --c tca tca tcc tta ta- --- --- --- --- --- --- tta tta tca tta tta tca tag tct att aaa cac aaa tca tct --- --- --- --- --- --- --- --- --- --- --- --- --- --- --- --- --- --- --- --- --- --- --- --- --- --- --a cgt tta taa c-- --- --- --- --- --- --- --- --a aca ttc --- --- --- --- -tc att att aat tag ttc tgt agt aat atc ttt aat aat ttg gct ata --- --- -ca tct gtt --- --- --- --- --- --- --- --- --- --- --- --- --- --- --- --- --- --- --c aat act --- --- --- --- --- --- --- --a tct att gat gat ttc ttt t-- --- --- --- --- --- --- --- --- --- --- --t taa gac t-- --- --- --- --- --- --- --- --- --- --- --- --- --- --- --- --- --- --- --- --- --- --- --- --- --- --- --- --- --- --- --- --- --- --- --- --- --- --- --- taa act agt --- --- --- --- --- --- --- --- --- --- --- --- --- --- --- --- --- --- --- --- --- --- --- --- --- --- --- --- --- --- --- --- --- --- --- --- --- --- --- --- --- --- --- --- --- --- --- --- --- --- --- --- --- --- --- --- --- --- --- --- --- --- --- --- --- --- --- --- --- --- --- --- --- --- --- --- --- --- --- --- --- --- --- --- --- --- --- --- --- --- --- --- --- --- --- --- --- --- --- --- --- --- --- --- --- --- --- --- --- --t atg gta atg acg atg aaa --- -tc gag tag ta- --- --a ctt cta ata aag act tga ta- --- --t cat tat cat atg ttt gat cg- --- --- --- --- --- --- --t cat agt taa tag tgt g-- --- --- --- --- --- --- --- --- --- --- --- --- --- --- --- --- --- --- --- -gc taa atg gta ctg tta ata agt tta t-- --- --- --- --- --- --- --- --- --- --- --- --- --- --- --- --- --a gac aat atc ata gta ttt tct ttc cag aat --- --- --- --t aga tta ttt ttt taa ata ctg atc ctc aca att ccg tga tgt agc agt agt tgg t-- --- --- --- --- --- --- --- --- --- --- --- --- --- --- --- --- --- --- --- --- --- --- --- --- --- --- --- -gc atg gtc tat atc gt- --- --- --- --- --- --- --- --- --- --- --- --- --- --- --- --- --- --- --- --- --- --- --- --- --- --- --- --- --- --- --- --- --- --- --- --- --- --- --- --- --- --- --- taa aat gta tca tat ata ata gtt ttc tga cgt gga gta cag aat ttt cga --- --- --- --- --- --- --- --- --- --- --- --- --- --- --- --- --- --- --- --- --- --- --- --- --- --- --- --- --- --- --- --- --- --- --- --- --- --- --- --- --- --- --- --t taa tga gtt cat ggt aag gaa ggg caa atg tct --- gta tat aat ata cat aag tta a-- --- --- --- --- --- --- --- --- --- --- --- --- --t agt ttt tta tca tat tt- --- --- --- --- --- --- --- --t cta ata cca taa taa aaa tta tca tta tg- --- --- -ta taa tca --- --- --- --- --- --- --- --- --- --- --- --t cac tg- --- -tc gct atc att att gcg ttt gtg tag tt- --- --- --- --- ctg ccc ta- --- --- --- --- --- --- --- --t cat cta cat cac tgt ca- --- --- --- --- ctc tca cta tat ctt cta aaa tta caa --- -ac aac --- --- --- --- --- --- --- --- --- --- --- --- --- --- --- --- --- --- --- --- --- --- --- --- --- --- --- --- --- --- --- --- --- --- --- --- tgg ata ttc gat --- --- --- --- --- --- --- --- --- --- --- --- --- --- --- --- --- --- --- --- --- --- --- --- --- --- --a aca gca ttt gtg t-- --- --- --- --- --- --- --- --- --- --- --- --- --- --- --- --- -

>KC257459.1_Monkeypox_virus_strain_Sudan_2005_01_complete_genome

--- --- --- --- --- --- --- --- --- --- --- --- --- --- --- --- --- --- --- --- --- --- --- --- --- --- --- --- --- --- --- --- --- --- --- --g tgt cta gaa aaa aat gtg tga ccc acg acc gta gga aac tct aga ggg taa gaa aaa tca atc gtt tat aga gac cat cag aaa gag gtt taa tat ttt tgt gag acc cat cga aga gag aaa gga taa aaa ctt ttt acg act cca tca gaa aga ggt tta ata ttt ttg tga gac cca tcg aag aga gaa agg ata aaa act ttt tac gac tcc atc aga aag agg ttt aat att ttt gtg aga ccc atc gaa g-- --- --- --- --- --- --- --- --- --- --- --- --- --- --- --- --- --- --- --- --- --- --- --- --- --- --- --- --- --- --- --- --- --- --- --- --- --- --- --- --- --- --- --- --- --- --- --- --- --- --- --- --- --- --- --- --- --- --- --- --- --- --- --- --- --- --- --- --- --- -ag aga aag gat aaa aac ttt tta cga ctc cat cag aaa gag gtt taa tat ttt tgt gag acc cat cga aga gag aaa gga taa aaa ctt ttt acg act cca tca gaa aga ggt tta ata ttt ttg tga gac cca tcg aag aga gaa agg ata aaa act ttt tac gac tcc atc aga aag agg ttt aat att ttt gtg aga ccc atc gaa gag aga aag gat aaa aac ttt tta cga ctc cat cag aaa gag gtt taa tat ttt tgt gag acc cat cga aga gag aaa gga taa aaa ctt ttt acg act cca tca gaa aga ggt tta --- --- --- --- --- --- --- --- --- --- --- --- --- --- --- --- --- --- --- --- --- --- --- --- --- --- --- --- --- --- --- --- --- --- --- --- --- --- --- --- --- --- --- --- --- --- --a tat ttt tgt gag acc cat cga aga gag aaa gga taa aaa ctt ttt acg act cca tca gaa aga ggt tta ata ttt ttg tga gac cca tcg aag aga gaa aga gat ggt tag tca aga tat ttt tct tag tac aaa agt caa tgt ttt aaa ata tat gga cga gaa tta att tgt ctg tat aaa aac ttg tgt gaa att atg tac tag aga aaa aac gtg agc agt gtc ccc tac atg gat ttt aca gat cat tta tat tcc aaa aat att aac tat ata cgt tta tta tat gat gtt aac gtg taa att ata aac att att tta tga tgc aat tgt ctg aca ac- --- --- --c ata agg ata ttg ata agc tct acg aga ata tat tgt tgg acg tta tcg ttt acg aaa tag ttg aga cat cag aaa gag gtt taa tat ttt tgt gag a-c cat cga aga gag aaa gag aat aaa aat a-- --- --- --- ttt ttt tgt aaa act ttt tta tga gac aag aga gaa aga gaa tac gaa tag tga tca tat cgt atc aca tat tga aac aga aag aag aag taa cga gag gta act ttt tgt gaa tgt agt taa aca ttt ttg ttt tgc aaa ccg gaa tat agt acc cgg tac act ttt tta att cgt ggt gcg gtg tct gaa tcg ttc gat taa ccc aac tca tcc att ttc aga tga ata gag tta tcg att cag aca cat gct ttg agt ttt gtt gaa tcg atg agt gaa gta tca tcg gtt gca cct tca gat gcc gat ccg tcg aca tac ttg aat cca tcc ttg act tca agt tca gat gat tcc tca cac atg tct ccg ata cgt acg cta aac tct agg ttc tta aca cat ttt gta tca acg atc gtt gaa ccg atg ata tct ttg taa ctc act ttc tta tgt gag atg tta gac cca agt act gga tgg gtc ttg atg tcg ctg tct ttc tct tct tcg cta cat ctg atg tcg ata gac atc tca cag tct ttg atc ata gcc aga gct tct tca cgc gtg atc gcg gga gag tcc tta cct tgt cct ggt gac acg ctg gac aat cta gta ttc aca gtg ttt cca tca gag gat tcg gag atg gat gaa atc ttt ggg cat ttg gtg aat cca aag ttc atg tta aga ccc gcg ccg acg ata ttg taa taa gtg gtg gga tct cct ttt aca act tct tcg gat acc tca tca tct tcg gat acc tca tca tct tcg gtc tct gta act tcc gtt acg gat tga caa atc tta tca ttg gtc ggt gtt tgg tct tgc ttt gtg act ttg ata ata aca tcg att ccc ata tga tgt ttg ttt tct tct tca gta cac gag gat gag gat tgt tga aga cta gta ggc ata gca gct gcc act agg cac atg cat gcc agg aca ata tat tgt ttc atg att gct att gat tga tta ctg ttc tag atg att cta ctt tct tac cat ata ata aat tag aat ata ttt tct act ttt acg aga aat taa tta ttg tat tta tta ttt atg ggt -aa aaa aaa ctt act ata agt ggg tgg gat tct ggg aat tag tga tca gtt tat gta tat cgc aac tag cgg gca tat ggc tat tga cat cga gaa cat tac cca tat gat aag aga ttg tat ccg ttt cgt agt ctt gag tat tgg tat tac tat ata gta tgt aga tgt cga cgc tag ata gac agt cgc cca cta gag tta ccg tct ctg aat gcg gca tga tag tat cat tct ttg ttt tcg tta act gtt tgg aag atg aat ctt tgt tgt tac att taa tct cga aat tca gag tac ata tct ttg aag tat tct gat atc tat ttt ctc ctg taa aga atc ctg aag ttg cta cat tat taa gga cag aga agt att ctg cac gaa aga cgg gat cac aat ctt tat gat tca tgg taa tag tta gtt ccg acg ttg aga tgg att cgc tga gac cgg tag tgg tcg tcc gag tac acg atg tgt cgt tga ctg gat aca ggt taa ttt cca cat cga tat agt taa atg tat tgc tgg tt- --a cgg gtt cgc att tat ctg tgg aag aga cgg tgt gag aat atg ttc cgg gac cac acg gag aac aga tga cgt ctc cgg tag acg tgt atc cgg ata ctc cgt atc cta ttc cac act ttg ttt tag aaa tac atg ttc tac acc ctg atg atc ctt tga gaa gac aat aat atc ctg gag agc att cac aga ttc tat tgt gag tcg tgt tac acg atc gcg tct cta cct gat tac tat cac atc ttc cgt tac aac tta gac aag cct gta aat gat tat tgt gag atg taa agg tat ccg aac cac acg gcg tac att gtg tat tag tct tgc tat cac ata atc tgg aag cgt aag ttc ccg gag gac acg ata gac aac ata gat tac ggc ttc tgt att cgt tgt ctt tac att ttc cat tgg atg gtg cat gtg gtg cta tat ctc ttc cgt tta tta tta tac atg aga gaa aca ata tat acg agt ata ata cgg acc tca tga ttt aat aat -gt agt aat cgt cgt ctt gtt act gtt tgt ttc cta ctt ctc caa tca tat aga tta ttt ttt aaa tat ttt ctt tct atc atg gat aat att tgt aat ggt tct ttc cgt aca aca tac tgt tta gat ggt agt cgc tta gct tgg tta tga tat tgc gca taa ttt ccg gag gca aat acg ata gtc tag att gac tat cga tgg tag act cta att tat tga gtg ctt tgt cga cga gtt tac ttt tat gct cca tcg ata gat gac act gtt cta tga gat cgt cgt aca tgg gaa atg aaa tgc gtt tgt ctg aat gta tgg ctt cga gat agg tgt gat acc gga tgt ctt ctg ttc tca ata ccg tat aca agt tgg tgt ctg aga ttc gaa tct ctt tga gga gac tta tgt cac gac tac att ttt cga tga tgg aat cta tct tat cga atg ata tat ttt tca taa ata cac ttt tat agt cct cgt tta aac aga att tag tat gta gtt ccg caa atg act cgt ccc tta ata ggc agt agg cta tta tct tct tta cgt agt gat cgt cgt agg gag aga act ccg aca tct tgt aga aca acg att taa tca tag gta gag ata ctt tca gtc tgt ggt gga tga tgt cat taa cga cat ccg cct tgt ata tga tgt ttc tgt ttt caa aca cca agt cga ata ctg tct tat cgt ctt tag tcg gaa ggt tga tgt cgt atc cga tgt ata cga ggt atg agg caa cat tgt tat tgc aat tct gga agg cgg tat gaa gag gag tca ttg tat tat agt att cgt ctt tct gaa tgt cga atc tat cta gta gat acc gta gta tat tga gag agc gac ttc cat atc ctt gat tat gtt tta tga ata gat aaa gta gat gtt gtc ctt ctt cct ttt gta att tcc cgt att ttt gtt cgt gcc aat tga gta aca tta tga gaa tat gac ctg ttg cac aat cgt tct tta tgt att cca tga tgg gtg tac aat caa gat tat tac gta tcc tcg tat cgg ctc ctc gag ata aaa gag cat aca cca cac gag gac tat gtt tgg tat act gtt gaa ggt aag tgt gta acg gcg tat ttc cga ttt tcg taa ccg cgt taa tgt ttg ctc cat gat cta tta tcg cgt aga tga atc gct tct cag ctc gca tct tag tgt gac tct ttg act tgt aat aat tgc ttt cgt gga aca cgg ata tgt gtt tac agt agt aat gaa gag aag tga gtc cat cct cat cga cgc aat tag ggt cag atc ctt tag tca ata att tgt aca gaa cgt aat agt tta agc tcc cat tga att tat atc taa gat aac aca gca ata gat cgg atg att tac taa agt tat caa tgg ggt ccg tta gta tat caa aga tct tgt tat cga ttg ata gtg aat gaa tca gat agt ggt gta gag gaa tat gtc ctt ttt tat cct tgc tat caa agt tac gca tgc cgt ggc gta aca ata tct tta ata cag atg gat taa atc gtg tat tca tcg tat agc aat gta atg gag agt tac cac att tta gtc gtt tat tca gat cgc agt gtt taa taa cta att taa aca gat gag atg ata tat cca cat caa aga atg cga gat aca tat gac aga cat tat tga cag aaa tgt gac ctt cat tat cac cgt cgt cca taa atg cgt tag gta cgt acc aca tac tat cgt taa cga tgc gca caa tct cgt cca ttt tca tcc atc ttc ata atg att tac ttt ttc ata att aga gaa aaa gat caa ggt ata aaa att aga agt gtt aga cta ta- --- --- --- --- --- --- --- --- --- --- -aa cta act tat aa- --- --- --- --- --- --- --- --- --- --- --- --- --- --- --- --- --- --- --- --- --- --- --- --- --- --- --- --- --- --- --- --- --- --- --- --- --- --- --- --- --- --- --- --- --- --- --- --- --- --- --- --- --- --- --- --- --- --- --- --- --- --- --- --- --- --- --- --- --- --- --- --- --- --- --- --- --- --- --- --- --- --- --- --- --- --- -aa cta act tat gac tta act aac tta tga cta tta act cat ttc aag aaa ggt ggg tgg aga gaa ctc tat atg aca gct tgt gaa aca att aga tcc cta att tct aat gga agt ttt gat agg aga ttg tca tca gtt gat aca ttg ttt att atc tca tct att aga gca cgt ctg ttt aga gct tta gtg acc tgc tcg gtt act tct gtg taa atc ttg aat cct tta gtg ata cac tgt gtc aaa act gga tgt tta gaa tac cta tgt aga ata tgg gaa gca tgc ttg ttt ttg tct cta tta tag att aac tca tac atg gtt gta tta tga att ttc atc tgc cta atg tac tcc aat tct tgt tta caa tca att ata taa tca aag agt gat gat gca tac aca tta caa agt gaa taa tct acc atc ata aaa tac ttg ata cag agc ttt atc aca tca tgg ttt tca att gta tta tta agt ata gct aat ttt ata cag tca ata gac aat ggt tct cta agc aat att tct aat att tta aga tgt gct tcc cta cgg gcg atg aca gat ccc cta tcc acg gcc acg tca aga cat gta tat cca tta ctc att act gcg ttg aca ttt gct cca ttt tct aat agc cat gat act aaa tct ata taa cct gca tag ata gcg cga taa agc aag gtc ctt cca cca gca tct agt tga ttg ata tct tca ata tat ggg ata caa agc tta taa att tct aat act gtg ggt tca tct aca agg aat ccc cta gta tac tga att att tta tat aga tct aat tta aca tca ttt tca tct ggg ata cca caa ttc aaa ata aac tca aca aca cta ctt tcc ttt tta cat att ccc cta aaa tag gca ttc aag cat tct att tta tat att aca gcc cca tga tct acc ata aaa tca aca atg tct att tct aca tat gca tta gat aga tag taa agt aag aga tct tgt aca gaa tta caa ttc tta ata att ata gag aaa ata tct tcc ata taa ttc ttt gac act aat gca gat ata ata tct tta tat gta ata tat gca aac agt cta tct act ata tac tga tca ata tta tct cta tga atc cta aaa taa tca tac aga aca tct aca gga tca caa att ggt tca agg aga aat cta tca aat att ttc ctg tca aca act ggt tct aga aca taa cag tca aca cct aat cca tgt ttt tta tag tca tct acc aaa gat aat gac caa aga tcg agg tcg tcg tga aac tgc tca tcg aca gcc atg aaa tct gcc gac tcc atg gcg cga atc gca ctg tct tat tcg cca ttg att ttc a-t ttt tta taa tta tgt aca tgt ttt cct tct att ctc aag agt cta caa aaa tat a-t ttt ttt tcg ata tct aag tac taa gtt ttt tta ctg ttt ttg tta ctg tct tcc att ctt cta act aaa gat ctg aga taa att ata caa tct tcg cta tcg aac cat ttt tgt agt cta aag cct gaa gta att aac caa ctg ttt tta tta gtg gct ttt ttc gat cta tcc tcg tc- --- --c tca tca tcc tta ta- --- --- --- --- --- --- tta tta tca tta tta tca tag tct att aaa cac aaa tca tct --- --- --- --- --- --- --- --- --- --- --- --- --- --- --- --- --- --- --- --- --- --- --- --- --- --- --a cgt tta taa c-- --- --- --- --- --- --- --- --a aca ttc --- --- --- --- -tc att att aat tag ttc tgt agt aat atc ttt aat aat ttg gct ata --- --- -ca tct gtt --- --- --- --- --- --- --- --- --- --- --- --- --- --- --- --- --- --- --c aat act --- --- --- --- --- --- --- --a tct att gat gat ttc ttt t-- --- --- --- --- --- --- --- --- --- --- --t taa gac t-- --- --- --- --- --- --- --- --- --- --- --- --- --- --- --- --- --- --- --- --- --- --- --- --- --- --- --- --- --- --- --- --- --- --- --- --- --- --- --- taa act agt --- --- --- --- --- --- --- --- --- --- --- --- --- --- --- --- --- --- --- --- --- --- --- --- --- --- --- --- --- --- --- --- --- --- --- --- --- --- --- --- --- --- --- --- --- --- --- --- --- --- --- --- --- --- --- --- --- --- --- --- --- --- --- --- --- --- --- --- --- --- --- --- --- --- --- --- --- --- --- --- --- --- --- --- --- --- --- --- --- --- --- --- --- --- --- --- --- --- --- --- --- --- --- --- --- --- --- --- --- --t atg gta atg acg atg aaa --- -tc gag tag ta- --- --a ctt cta ata aat act tga ta- --- --t cat tat cat atg ttt gat cg- --- --- --- --- --- --- --t cat agt taa tag tgt g-- --- --- --- --- --- --- --- --- --- --- --- --- --- --- --- --- --- --- --- -gc taa atg gta ctg tta ata agt tta t-- --- --- --- --- --- --- --- --- --- --- --- --- --- --- --- --- --a gac aat atc ata gta ttt tct ttc cag aat --- --- --- --t aga tta ttt ttt taa ata ctg atc ctc aca att ccg tga tgt agc agt agt tgg t-- --- --- --- --- --- --- --- --- --- --- --- --- --- --- --- --- --- --- --- --- --- --- --- --- --- --- --- -gc atg gtc tat atc gt- --- --- --- --- --- --- --- --- --- --- --- --- --- --- --- --- --- --- --- --- --- --- --- --- --- --- --- --- --- --- --- --- --- --- --- --- --- --- --- --- --- --- --- taa aat gta tca tat ata ata gtt ttc tga cgt gga gta cag aat ttt cga --- --- --- --- --- --- --- --- --- --- --- --- --- --- --- --- --- --- --- --- --- --- --- --- --- --- --- --- --- --- --- --- --- --- --- --- --- --- --- --- --- --- --- --t taa tga gtt cat ggt aag gaa ggg caa atg tct --- gta tat aat ata cat aag tta a-- --- --- --- --- --- --- --- --- --- --- --- --- --t agt ttt tta tca tat tt- --- --- --- --- --- --- --- --t cta ata cca taa taa aaa tta tca tta tg- --- --- -ta taa tca --- --- --- --- --- --- --- --- --- --- --- --t cac tg- --- -tc gct atc att att gcg ttt gtg tag tt- --- --- --- --- ctg ccc ta- --- --- --- --- --- --- --- --t cat cta cat cac tgt ca- --- --- --- --- ctc tca cta tat ctt cta aaa tta caa --- -ac aac --- --- --- --- --- --- --- --- --- --- --- --- --- --- --- --- --- --- --- --- --- --- --- --- --- --- --- --- --- --- --- --- --- --- --- --- tgg ata ttc gat --- --- --- --- --- --- --- --- --- --- --- --- --- --- --- --- --- --- --- --- --- --- --- --- --- --- --a aca gca ttt gtg t-- --- --- --- --- --- --- --- --- --- --- --- --- --- --- --- --- -

>MN702446.1_Monkeypox_virus_strain_38c_contig_SPADES

--- --- --- --- --- --- --- --- --- --- --- --- --- --- --- --- --- --- --- --- --- --- --- --- --- --- --- --- --- --- --- --- --- --- --- --- --- --- --- --- --- --- --- --- --- --- --- --- --- --- --- --- --- --- --- --- --- --- --- --- --- --- --- --- --- --- --- --- --- --- --- --- --- --- --- --- --- --- --- --- --- --- --- --- --- --- --- --- --- --- --- --- --- --- --- --- --- --- --- --- --- --- --- --- --- --- --- --- --- --- --- --- --- --- --- --- --- --- --- --- --- --- --- --- --- --- --- --- --- --- --- --- --- --- --- --- --- --- --- --- --- --- --- --- --- --- --- --- --- --- --- --- --- --- --- --- --- --- --- --- --- --- --- --- --- --- --- --- --- --- --- --- --- --- --- --- --- --- --- --- --- --- --- --- --- --- --- --- --- --- --- --- --- --- --- --- --- --- --- --- --- --- --- --- --- --- --- --- --- --- --- --- --- --- --- --- --- --- --- --- --- --- --- --- --- --- --- --- --- --- --- --- --- --- --- --- --- --- --- --- --- --- --- --- --- --- --- --- --- --- --- --- --- --- --- --- --- --- --- --- --- --- --- --- --- --- --- --- --- --- --- --- --- --- --- --- --- --- --- --- --- --- --- --- --- --- --- --- --- --- --- --- --- --- --- --- --- --- --- --- --- --- --- --- --- --- --- --- --- --- --- --- --- --- --- --- --- --- --- --- --- --- --- --- --- --- --- --- --- --- --- --- --- --- --- --- --- --- --- --- --- --- --- --- --- --- --- --- --- --- --- --c cat cga aga gag aaa gga taa aaa ctt ttt acg act cca tca gaa aga ggt tta ata ttt ttg tga gac cca tcg acg aga gaa aga gat ggt tag tca aga tat ttt tct tag tac aaa agt caa tgt ttt aaa ata tat gga cga gaa tta att tgt ctg tat aaa aac ttg tgt gaa att atg tac tag aga aaa aac gtg agc agt gtc ccc tac atg gat ttt aca gat cat tta tat tcc aaa aat att aac tat ata cgt tta tta tat gat gtt aac gtg taa att ata aac att att tta tga tgc aat tgt ctg aca acc --- att ggc ata agg ata ttg ata agc tct acg aga ata tat tgt tgg acg tta tcg ttt acg aaa tag ttg aga cat cag aaa gag gtt taa tat ttt tgt gag a-c cat cga aga gag aaa gag aat aaa aat a-- --- --- --- ttt ttt tgt aaa act ttt tta tga gac aag aga gaa aga gaa tac gaa tag tga tca tat cgt atc aca tat tga aac aga aag aag aag taa cga gag gta act ttt tgt gaa tgt agt taa aca ttt ttg ttt tgc aaa ccg gaa tat agt acc cgg tac act ttt tta att cgt ggt gcg gtg tct gaa tcg ttc gat taa ccc aac tca tcc att ttc aga tga ata gag tta tcg att cag aca cat gct ttg agt ttt gtt gaa tcg atg agt gaa gta tca tcg gtt gca cct tca gat gcc gat ccg tcg aca tac ttg aat cca tcc ttg act tca agt tca gat gat tcc tca cac atg tct ccg ata cgt acg cta aac tct agg ttc tta aca cat ttt gta tca acg atc gtt gaa ccg atg ata tct ttg taa ctc act ttc tta tgt gag atg tta gac cca agt act gga tgg gtc ttg atg tcg ctg tct ttc tct tct tcg cta cat ctg atg tcg ata gac atc tca cag tct ttg atc ata gcc aga gct tct tca cgc gtg atc gcg gga gag tcc tta cct tgt cct ggt gac acg ctg gac aat cta gta ttc aca gtg ttt cca tca gag gat tcg gag atg gat gaa atc ttt ggg cat ttg gtg aat cca aag ttc atg tta aga ccc gcg ccg acg ata ttg taa taa gtg gtg gga tct cct ttt aca act --- --- --- --- --- --- tct tcg gat acc tca tca tct tcg gtc tct gta act tcc gtt acg gat tga caa atc tta tca ttg gtc ggt gtt tgg tct tgc ttt gtg act ttg ata ata aca tcg att ccc ata tga tgt ttg ttt tct tct tca gta cac gag gat gag gat tgt tga aga cta gta ggc ata gca gct gcc act agg cac atg cat gcc agg aca ata tat tgt ttc atg att gct att gat tga tta ctg ttc tag atg att cta ctt tct tac cat ata ata aat tag aat ata ttt tct act ttt acg aga aat taa tta ttg tat tta tta ttt atg ggt --a aaa aaa ctt act ata agt ggg tgg gat tct ggg aat tag tga tca gtt tat gta tat cgc aac tag cgg gca tat ggc tat tga cat cga gaa cat tac cca tat gat aag aga ttg tat ccg ttt cgt agt ctt gag tat tgg tat tac tat ata gta tgt aga tgt cga cgc tag ata gac agt cgc cca cta gag tta ccg tct ctg aat gcg gca tga tag tat cat tct ttg ttt tcg tta act gtt tgg aag atg aat ctt tgt tgt tac att taa tct cga aat tca gag tac ata tct ttg aag tat tct gat atc tat ttt ctc ctg taa aga atc ctg aag ttg cta cat tat taa gga cag aga agt att ctg cac gaa aga cgg gat cac aat ctt tat gat tca tgg taa tag tta gtt ccg acg ttg aga tgg att cgc tga gac cgg tag tgg tcg tcc gag tac acg atg tgt cgt tga ctg gat aca ggt taa ttt cca cat cga tat agt taa atg tat tgc tgg tt- --a cgg gtt cgc att tat ctg tgg aag aga cgg tgt gag aat atg ttc cgg gac cac acg gag aac aga tga cgt ctc cgg tag acg tgt atc cgg ata ctc cgt atc cta ttc cac act ttg ttt tag aaa tac atg ttc tac acc ctg atg atc ctt tga gaa gac aat aat atc ctg gag agc att cac aga ttc tat tgt gag tcg tgt tac acg atc gcg tct cta cct gat tac tat cac atc ttc cgt tac aac tta gac aag cct gta aat gat tat tgt gag atg taa agg tat ccg aac cac acg gcg tac att gtg tat tag tct tgc tat cac ata atc tgg aag cgt aag ttc ccg gag gac acg ata gac aac ata gat tac ggc ttc tgt att cgt tgt ctt tac att ttc cat tgg atg gtg cat gtg gtg cta tat ctc ttc cgt tta tta tta tac atg aga gaa aca ata tat acg agt ata ata cgg acc tca tga ttt aat aat -gt agt aat cgt cgt ctt gtt act gtt tgt ttc cta ctt ctc caa tca tat aga tta ttt ttt aaa tat ttt ctt tct atc atg gat aat att tgt aat ggt tct ttc cgt aca aca tac tgt tta gat ggt agt cgc tta gct tgg tta tga tat tgc gca taa ttt ccg gag gca aat acg ata gtc tag att gac tat cga tgg tag act cta att tat tga gtg ctt tgt cga cga gtt tac ttt tat gct cca tcg ata gat gac act gtt cta tga gat cgt cgt aca tgg gaa atg aaa tgc gtt tgt ctg aat gta tgg ctt cga gat agg tgt gat acc gga tgt ctt ctg ttc tca ata ccg tat aca agt tgg tgt ctg aga ttc gaa tct ctt tga gga gac tta tgt cac gac tac att ttt cga tga tgg aat cta tct tat cga atg ata tat ttt tca taa ata cac ttt tat agt cct cgt tta aac aga att tag tat gta gtt ccg caa atg act cgt ccc tta ata ggc agt agg cta tta tct tct tta cgt agt gat cgt cgt agg gag aga act ccg aca tct tgt aga aca acg att taa tca tag gta gag ata ctt tca gtc tgt ggt gga tga tgt cat taa cga cat ccg cct tgt ata tga tgt ttc tgt ttt caa aca cca agt cga ata ctg tct tat cgt ctt tag tcg gaa ggt tga tgt cgt atc cga tgt ata cga ggt atg agg caa cat tgt tat tgc aat tct gga agg cgg tat gaa gag gag tca ttg tat tat agt att cgt ctt tct gaa tgt cga atc tat cta gta gat acc gta gta tat tga gag agc gac ttc cat atc ctt gat tat gtt tta tga ata gat aaa gta gat gtt gtc ctt ctt cct ttt gta att tcc cgt att ttt gtt cgt gcc aat tga gta aca tta tga gaa tat gac ctg ttg cac aat cgt tct tta tgt att cca tga tgg gtg tac aat caa gat tat tac gta tcc tcg tat cgg ctc ctc gag ata aaa gag cat aca cca cac gag gac tat gtt tgg tat act gtt gaa ggt aag tgt gta acg gcg tat ttc cga ttt tcg taa ccg cgt taa tgt ttg ctc cat gat cta tta tcg cgt aga tga atc gct tct cag ctc gca tct tag tgt gac tct ttg act tgt aat aat tgc ttt cgt gga aca cgg ata tgt gtt tac agt agt aat gaa gag aag tga gtc cat cct cat cga cgc aat tag ggt cag atc ctt tag tca ata att tgt aca gaa cgt aat agt tta agc tcc cat tga att tat atc taa gat aac aca gca ata gat cgg atg att tac taa agt cat caa tgg ggt ccg tta gta tat caa aga tct tgt tat cga ttg ata gtg aat gaa tca gat agt ggt gta gag gaa tat gtc ctt ttt tat cct tgc tat caa agt tac gca tgc cgt ggc gta aca ata tct tta ata cag atg gat taa atc gtg tat tca tcg tat agc aat gta atg gag agt tac cac att tta gtc gtt tat tca gat cgc agt gtt taa taa cta att taa aca gat gag atg ata tat cca cat caa aga atg cga gat aca tat gac aga cat tat tga cag aaa tgt gac ctt cat tat cac cgt cgt cca taa atg cgt tag gta cgt acc aca tac tat cgt taa cga tgc gca caa tct cgt cca ttt tca tcc atc ttc ata atg att tac ttt ttc ata att aga gaa aaa gat caa ggt ata aaa att aga agt gtt aga cta ta- --- --- --- --- --- --- --- --- --- --- -aa cta act tat aa- --- --- --- --- --- --- --- --- --- --- --- --- --- --- --- --- --- --- --- --- --- --- --- --- --- --- --- --- --- --- --- --- --- --- --- --- --- --- --- --- --- --- --- --- --- --- --- --- --- --- --- --- --- --- --- --- --- --- --- --- --- --- --- --- --- --- --- --- --- --- --- --- --- --- --- --- --- --- --- --- --- --- --- --- --- --- -aa cta act tat gac tta act aac tta tga cta tta act cat ttc aag aaa ggt ggg tgg aga gaa ctc tat atg aca gct tgt gaa aca att aga tcc cta att tct aat gga agt ttt gat agg aga ttg tca tca gtt gat aca ttg ttt att atc tca tct att aga gca cgt ctg ttt aga gct tta gtg acc tgc tcg gtt act tct gtg taa atc ttg aat cct tta gtg ata cac tgt gtc aaa act gga tgt tta gaa tac cta tgt aga ata tgg gaa gca tgc ttg ttt ttg tct cta tta tag att aac tca tac atg gtt gta tta tga att ttc atc tgc cta atg tac tcc aat tct tgt tta caa tca att ata taa tca aag agt gat gat gca tac aca tta caa agt gaa taa tct acc atc ata aaa tac ttg ata cag agc ttt atc aca tca tgg ttt tca att gta tta tta agt ata gct aat ttt ata cag tca ata gac aat ggt tct cta agc aat att tct aat att tta aga tgt gct tcc cta cgg gcg atg aca gat ccc cta tcc acg gcc acg tca aga cat gta tat cca tta ctc att act gcg ttg aca ttt gct cca ttt tct aat agc cat gat act aaa tct ata taa cct gca tag ata gcg cga taa agc aag gtc ctt cca cca gca tct agt tga ttg ata tct tca ata tat ggg ata caa agc tta taa att tct aat act gtg ggt tca tct aca agg aat ccc cta gta tac tga att att tta tat aga tct aat tta aca tca ttt tca tct ggg ata cca caa ttc aaa ata aac tca aca aca cta ctt tcc ttt tta cat att ccc cta aaa tag gca ttc aag cat tct att tta tat att aca gcc cca tga tct acc ata aaa tca aca atg tct att tct aca tat gca tta gat aga tag taa agt aag aga tct tgt aca gaa tta caa ttc tta ata att ata gag aaa ata tct tcc ata taa ttc ttt gac act aat gca gat ata ata tct tta tat gta ata tat gca aac agt cta tct act ata tac tga tca ata tta tct cta tga atc cta aaa taa tca tac aga aca tct aca gga tca caa att ggt tca agg aga aat cta tca aat att ttc ctg tca aca act ggt tct aga aca taa cag tca aca cct aat cca tgt ttt tta tag tca tct acc aaa gat aat gac caa aga tcg agg tcg tcg tga aac tgc tca tcg aca gcc atg aaa tct gcc gac tcc atg gcg cga atc gca ctg tct tat tcg cca ttg att ttc a-t ttt tta taa tta tgt aca tgt ttt cct tct att ctc aag agt cta caa aaa tat att ttt ttt tcg ata tct aag tac taa gtt ttt tta ctg ttt ttg tta ctg tct tcc att ctt cta act aaa gat ctg aga taa att ata caa tct tcg cta tcg aac cat ttt tgt agt cta aag cct gaa gta att aac caa ctg ttt tta tta gtg gct ttt ttc gat cta tcc tcg tc- --- --c tca tca tcc tta ta- --- --- --- --- --- --- tta tta tca tta tta tca tag tct att aaa cac aaa tca tct --- --- --- --- --- --- --- --- --- --- --- --- --- --- --- --- --- --- --- --- --- --- --- --- --- --- --a cgt tta taa c-- --- --- --- --- --- --- --- --a aca ttc --- --- --- --- -tc att att aat tag ttc tgt agt aat atc ttt aat aat ttg gct ata --- --- -ca tct gtt --- --- --- --- --- --- --- --- --- --- --- --- --- --- --- --- --- --- --c aat act --- --- --- --- --- --- --- --a tct att gat gat ttc ttt t-- --- --- --- --- --- --- --- --- --- --- --t taa gac t-- --- --- --- --- --- --- --- --- --- --- --- --- --- --- --- --- --- --- --- --- --- --- --- --- --- --- --- --- --- --- --- --- --- --- --- --- --- --- --- taa act agt --- --- --- --- --- --- --- --- --- --- --- --- --- --- --- --- --- --- --- --- --- --- --- --- --- --- --- --- --- --- --- --- --- --- --- --- --- --- --- --- --- --- --- --- --- --- --- --- --- --- --- --- --- --- --- --- --- --- --- --- --- --- --- --- --- --- --- --- --- --- --- --- --- --- --- --- --- --- --- --- --- --- --- --- --- --- --- --- --- --- --- --- --- --- --- --- --- --- --- --- --- --- --- --- --- --- --- --- --- --t atg gta atg acg atg aaa --- -tc gag tag ta- --- --a ctt cta ata aag act tga ta- --- --t cat tat cat atg ttt gat cg- --- --- --- --- --- --- --t cat agt taa tag tgt g-- --- --- --- --- --- --- --- --- --- --- --- --- --- --- --- --- --- --- --- -gc taa atg gta ctg tta ata agt tta t-- --- --- --- --- --- --- --- --- --- --- --- --- --- --- --- --- --a gac aat atc ata gta ttt tct ttc cag aat --- --- --- --t aga tta ttt ttt taa ata ctg atc ctc aca att ccg tga tgt agc agt agt tgg t-- --- --- --- --- --- --- --- --- --- --- --- --- --- --- --- --- --- --- --- --- --- --- --- --- --- --- --- -gc atg gtc tat atc gt- --- --- --- --- --- --- --- --- --- --- --- --- --- --- --- --- --- --- --- --- --- --- --- --- --- --- --- --- --- --- --- --- --- --- --- --- --- --- --- --- --- --- --- taa aat gta tca tat ata ata gtt ttc tga cgt gga gta cag aat ttt cga --- --- --- --- --- --- --- --- --- --- --- --- --- --- --- --- --- --- --- --- --- --- --- --- --- --- --- --- --- --- --- --- --- --- --- --- --- --- --- --- --- --- --- --t taa tga gtt cat ggt aag gaa ggg caa atg tct --- gta tat aat ata cat aag tta a-- --- --- --- --- --- --- --- --- --- --- --- --- --t agt ttt tta tca tat tt- --- --- --- --- --- --- --- --t cta ata cca taa taa aaa tta tca tta tg- --- --- -ta taa tca --- --- --- --- --- --- --- --- --- --- --- --t cac tg- --- -tc gct atc att att gcg ttt gtg tag tt- --- --- --- --- ctg ccc ta- --- --- --- --- --- --- --- --t cat cta cat cac tgt ca- --- --- --- --- ctc tca cta tat ctt cta aaa tta caa --- -ac aac --- --- --- --- --- --- --- --- --- --- --- --- --- --- --- --- --- --- --- --- --- --- --- --- --- --- --- --- --- --- --- --- --- --- --- --- tgg ata ttc gat --- --- --- --- --- --- --- --- --- --- --- --- --- --- --- --- --- --- --- --- --- --- --- --- --- --- --a aca gca ttt gtg t-- --- --- --- --- --- --- --- --- --- --- --- --- --- --- --- --- -

>JX878418.1_Monkeypox_virus_isolate_DRC_07-0120_complete_genome

--- --- --- --- --- --- --- --- --- --- --- --- --- --- --- --- --- --- --- --- --- --- --- --- --- --- --- --- --- --- --- --- --- --- --- --g tgt cta gaa aaa aat gtg tga ccc acg acc gta gga aac tct aga ggg taa gaa aaa tca atc gtt tat aga gac cat cag aaa gag gtt taa tat ttt tgt gag acc cat cga a-- --- --- --- --- --- --- --- --- --- --- --- --- --- --- --- --- --- --- --- --- --- --- --- --- --- --- --- --- --- --- --- --- --- --- --- --- --- --- --- --- --- --- --- --- --- --- --- --- --- --- --- --- --- --- --- --- --- --- --- --- --- --- --- --- --- --- --- --- --- --- --- --- --- --- --- --- --- --- --- --- --- --- --- --- --- --- --- --- --- --- --- --- --- --- --- --- --- --- --- --- --- --- --- --- --- --- --- --- --- --- --- --- --- --- --- --- --- --- --- --- --- --- --- --- --- --- --- --- --- --- --- --- --- --- --- --- --- --- --- --- --- --- --- --- --- --- --- --- --- --- --- --- --- --- --- --- --- --- --- --- --- --- --- --- --- --- --- --- --- --- --- --- --- --- --- --- --- --- --- --- --- --- --- --- --- --- --- --- --- --- --- --- --- --- --- --- --- --- --- --- --- --- --- --- --- --- --- --- --- --- --- --- --- --- --- --- --- --- --- --- --- --- --- --- --- --- --- --- --- --- --- --- --- --- --- --- --- --- --- --- --- --- --- --- --- --- --- --- --- --- --- --- --- --- --- --- --- --- --- --- --- --- --- --- --- --- --- --- --- --- --- --- --- --- --- --- --- --- --- -ga gag aaa gga taa aaa ctt ttt acg act cca tca gaa aga ggt tta ata ttt ttg tga gac cca tcg acg aga gaa aga gat ggt tag tca aga tat ttt tct tag tac aaa agt caa tgt ttt aaa ata tat gga cga gaa tta att tgt ctg tat aaa aac ttg tgt gaa att atg tac tag aga aaa aac gtg agc agt gtc ccc tac atg gat ttt aca gat cat tta tat tcc aaa aat att aac tat ata cgt tta tta tat gat gtt aac gtg taa att ata aac att att tta tga tgc aat tgt ctg aca acc tag att ggc ata agg ata ttg ata agc tct acg aga ata tat tgt tgg acg tta tcg ttt acg aaa tag ttg aga cat cag aaa gag gtt taa tat ttt tgt gag a-c cat cga aga gag aaa gag aat aaa aat a-- --- --- --- ttt ttt tgt aaa ac- ttt tta tga gac aag aga gaa aga gaa tac gaa tag tga tca tat cgt atc aca tat tga aac aga aag aag aag taa cga gag gta act ttt tgt gaa tgt agt taa aca ttt ttg ttt tgc aaa ccg gaa tat agt acc cgg tac act ttt tta att cgt ggt gcg gtg tct gaa tcg ttc gat taa ccc aac tca tcc att ttc aga tga ata gag tta tcg att cag aca cat gct ttg agt ttt gtt gaa tcg atg agt gaa gta tca tcg gtt gca cct tca gat gcc gat ccg tcg aca tac ttg aat cca tcc ttg act tca agt tca gat gat tcc tca cac atg tct ccg ata cgt acg cta aac tct agg ttc tta aca cat ttt gta tca acg atc gtt gaa ccg atg ata tct ttg taa ctc act ttc tta tgt gag atg tta gac cca agt act gga tgg gtc ttg atg tcg ctg tct ttc tct tct tcg cta cat ctg atg tcg ata gac atc tca cag tct ttg atc ata gcc aga gct tct tca cgc gtg atc gcg gga gag tcc tta cct tgt cct ggt gac acg ctg gac aat cta gta ttc aca gtg ttt cca tca gag gat tcg gag atg gat gaa atc ttt ggg cat ttg gtg aat cca aag ttc atg tta aga ccc gcg ccg acg ata gtg taa taa gtg gtg gga tct cct ttt aca act --- --- --- --- --- --- tct tcg gat acc tca tca tct tcg gtc tct gta act tcc gtt acg gat tga caa atc tta tca ttg gtc ggt gtt tgg tct tgc ttt gtg act ttg ata ata aca tcg att ccc ata tga tgt ttg ttt tct tct tca gta cac gag gat gag gat tgt tga aga cta gta ggc ata gca gct gcc act agg cac atg cat gcc agg aca ata tat tgt ttc atg att gct att gat tga tta ctg ttc tag atg att cta ctt tct tac cat ata ata aat tag aat ata ttt tct act ttt acg aga aat taa tta ttg tat tta tta ttt atg ggt aaa aaa aaa ctt act ata agt ggg tgg gat tct ggg aat tag tga tca gtt tat gta tat cgc aac tag cgg gca tat ggc tat tga cat cga gaa cat tac cca tat gat aag aga ttg tat ccg ttt cgt agt ctt gag tat tgg tat tac tat ata gta tgt aga tgt cga cgc tag ata gac agt cgc cca cta gag tta ccg tct ctg aat gcg gca tga tag tat cat tct ttg ttt tca tta act gtt tgg aag atg aat ctt tgt tgt tac att taa tct cga aat tca gag tac ata tct ttg aag tat tct gat atc tat ttt ctc ctg taa aga atc ctg aag ttg cta cat tat taa gga cag aga agt att ctg cac gaa aga cgg gat cac aat ctt tat gat tca tgg taa tag tta gtt ccg acg ttg aga tgg att cgc tga gac cgg tag tgg tcg tcc gag tac acg atg tgt cgt tga ctg gat aca ggt taa ttt cca cat cga tat agt taa atg tat tgc tgg tt- --a cgg gtt cgc att tat ctg tgg aag aga cgg tgt gag aat atg ttc cgg gac cac acg gag aac aga tga cgt ctc cgg tag acg tgt atc cgg ata ctc cgt atc cta ttc cac act ttg ttt tag aaa tac atg ttc tac acc ctg atg atc ctt tga gaa gac aat aat atc ctg gag agc att cac aga ttc tat tgt gag tcg tgt tac acg atc gcg tct cta cct gat tac tat cac atc ttc cgt tac aac tta gac aag cct gta aat gat tat tgt gag atg taa agg tat ccg aac cac acg gcg tac att gtg tat tag tct tgc tat cac ata atc tgg aag cgt aag ttc ccg gag gac acg ata gac aac ata gat tac ggc ttc tgt att cgt tgt ctt tac att ttc cat tgg atg gtg cat gtg gtg cta tat ctc ttc cgt tta tta tta tac atg aga gaa aca ata tat acg agt ata ata cgg acc tca tga ttt aat aat -gt agt aat cgt cgt ctt gtt act gtt tgt ttc cta ctt ctc caa tca tat aga tta ttt ttt aaa tat ttt ctt tct atc atg gat aat att tgt aat ggt tct ttc cgt aca aca tac tgt tta gat ggt agt cgc tta gct tgg tta tga tat tgc gca taa ttt ccg gag gca aat acg ata gtc tag att gac tat cga tgg tag act cta att tat tga gtg ctt tgt cga cga gtt tac ttt tat gct cca tcg ata gat gac act gtt cta tga gat cgt cgt aca tgg gaa atg aaa tgc gtt tgt ctg aat gta tgg ctt cga gat agg tgt gat acc gga tgt ctt ctg ttc tca ata ccg tat aca agt tgg tgt ctg aga ttc gaa tct ctt tga gga gac tta tgt cac gac tac att ttt cga tga tgg aat cta tct tat cga atg ata tat ttt tca taa ata cac ttt tat agt cct cgt tta aac aga att tag tat gta gtt ccg caa atg act cgt ccc tta ata ggc agt agg cta tta tct tct tta cgt agt gat cgt cgt agg gag aga act ccg aca tct tgt aga aca acg att taa tca tag gta gag ata ctt tca gtc tgt ggt gga tga tgt cat taa cga cat ccg cct tgt ata tga tgt ttc tgt ttt caa aca cca agt cga ata ctg tct tat cgt ctt tag tcg gaa ggt tga tgt cgt atc cga tgt ata cga ggt atg agg caa cat tgt tat tgc aat tct gga agg cgg tat gaa gag gag tca ttg tat tat agt att cgt ctt tct gaa tgt cga atc tat cta gta gat acc gta gta tat tga gag agc gac ttc cat atc ctt gat tat gtt tta tga ata gat aaa gta gat gtt gtc ctt ctt cct ttt gta att tcc cgt att ttt gtt cgt gcc aat tga gta aca tta tga gaa tat gac ctg ttg cac aat cgt tct tta tgt att cca tga tgg gtg tac aat caa gat tat tac gta tcc tcg tat cgg ctc ctc gag ata aaa gag cat aca cca cac gag gac tat gtt tgg tat act gtt gaa ggt aag tgt gta acg gcg tat ttc cga ttt tcg taa ccg cgt taa tgt ttg ctc cat gat cta tta tcg cgt aga tga atc gct tct cag ctc gca tct tag tgt gac tct ttg act tgt aat aat tgc ttt cgt gga aca cgg ata tgt gtt tac agt agt aat gaa gag aag tga gtc cat cct cat cga cgc aat tag ggt cag atc ctt tag tca ata att tgt aca gaa cgt aat agt tta agc tcc cat tga att tat atc taa gat aac aca gca ata gat cgg atg att tac taa agt cat caa tgg ggt ccg tta gta tat caa aga tct tgt tat cga ttg ata gtg aat gaa tca gat agt ggt gta gag gaa tat gtc ctt ttt tat cct tgc tat caa agt tac gca tgc cgt ggc gta aca ata tct tta ata cag atg gat taa atc gtg tat tca tcg tat agc aat gta atg gag agt tac cac att tta gtc gtt tat tca gat cgc agt gtt taa taa cta att taa aca gat gag atg ata tat cca cat caa aga atg cga gat aca tat gac aga cat tat tga cag aaa tgt gac ctt cat tat cac cgt cgt cca taa atg cgt tag gta cgt acc aca tac tat cgt taa cga tgc gca caa tct cgt cca ttt tca tcc atc ttc ata atg att tac ttt ttc ata att aga gaa aaa gat caa ggt ata aaa att aga agt gtt aga cta ta- --- --- --- --- --- --- --- --- --- --- -aa cta act tat aa- --- --- --- --- --- --- --- --- --- --- --- --- --- --- --- --- --- --- --- --- --- --- --- --- --- --- --- --- --- --- --- --- --- --- --- --- --- --- --- --- --- --- --- --- --- --- --- --- --- --- --- --- --- --- --- --- --- --- --- --- --- --- --- --- --- --- --- --- --- --- --- --- --- --- --- --- --- --- --- --- --- --- --- --- --- --- -aa cta act tat gac tta act aac tta tga cta tta act cat ttc aag aaa ggt ggg tgg aga gaa ctc tat atg aca gct tgt gaa aca att aga tcc cta att tct aat gga agt ttt gat agg aga ttg tca tca gtt gat aca ttg ttt att atc tca tct att aga gca cgt ctg ttt aga gct tta gtg acc tgc tcg gtt act tct gtg taa atc ttg aat cct tta gtg ata cac tgt gtc aaa act gga tgt tta gaa tac cta tgt aga ata tgg gaa gca tgc ttg ttt ttg tct cta tta tag att aac tca tac atg gtt gta tta tga att ttc atc tgc cta atg tac tcc aat tct tgt tta caa tca att ata taa tca aag agt gat gat gca tac aca tta caa agt gaa taa tct acc atc ata aaa tac ttg ata cag agc ttt atc aca tca tgg ttt tca att gta tta tta agt ata gct aat ttt ata cag tca ata gac aat ggt tct cta agc aat att tct aat att tta aga tgt gtt tcc cta cgg gcg atg aca gat ccc cta tcc acg gcc acg tca aga cat gta tat cca tta ctc att act gcg ttg aca ttt gct cca ttt tct aat agc cat gat act aaa tct ata taa cct gca tag ata gcg cga taa agc aag gtc ctt cca cca gca tct agt tga ttg ata tct tca ata tat ggg ata caa agc tta taa att tct aat act gtg ggt tca tct aca agg aat ccc cta gta tac tga att att tta tat aga tct aat tta aca tca ttt tca tct ggg ata cca caa ttc aaa ata aac tca aca aca cta ctt tcc ttt tta cat att ccc cta aaa tag gca ttc aag cat tct att tta tat att aca gcc cca tga tct acc ata aaa tca aca atg tct att tct aca tat gca tta gat aga tag taa agt aag aga tct tgt aca gaa tta caa ttc tta ata att ata gag aaa ata tct tcc ata taa ttc ttt gac act aat gca gat ata ata tct tta tat gta ata tat gca aac agt cta tct act ata tac tga tca ata tta tct cta tga atc cta aaa taa tca tac aga aca tct aca gga tca caa att ggt tca agg aga aat cta tca aat att ttc ctg tca aca act ggt tct aga aca taa cag tca aca cct aat cca tgt ttt tta tag tca tct acc aaa gat aat gac caa aga tcg agg tcg tcg tga aac tgc tca tcg aca gcc atg aaa tct gcc gac tcc atg gcg cga atc gca ctg tct tat tcg cca ttg att ttc a-t ttt tta taa tta tgt aca tgt ttt cct tct att ctc aag agt cta caa aaa tat a-t ttt ttt tcg ata tct aag tac taa gtt ttt tta ctg ttt ttg tta ctg tct tcc att ctt cta act aaa gat ctg aga taa att ata caa tct tcg cta tcg aac cat ttt tgt agt cta aag cct gaa gta att aac caa ctg ttt tta tta gtg gct ttt ttc gat cta tcc tcg tc- --- --c tca tca tcc tta ta- --- --- --- --- --- --- tta tta tca tta tta tca tag tct att aaa cac aaa tca tct --- --- --- --- --- --- --- --- --- --- --- --- --- --- --- --- --- --- --- --- --- --- --- --- --- --- --a cgt tta taa c-- --- --- --- --- --- --- --- --a aca ttc --- --- --- --- -tc att att aat tag ttc tgt agt aat atc ttt aat aat ttg gct ata --- --- -ca tct gtt --- --- --- --- --- --- --- --- --- --- --- --- --- --- --- --- --- --- --c aat act --- --- --- --- --- --- --- --a tct att gat gat ttc ttt t-- --- --- --- --- --- --- --- --- --- --- --t taa gac t-- --- --- --- --- --- --- --- --- --- --- --- --- --- --- --- --- --- --- --- --- --- --- --- --- --- --- --- --- --- --- --- --- --- --- --- --- --- --- --- taa act agt --- --- --- --- --- --- --- --- --- --- --- --- --- --- --- --- --- --- --- --- --- --- --- --- --- --- --- --- --- --- --- --- --- --- --- --- --- --- --- --- --- --- --- --- --- --- --- --- --- --- --- --- --- --- --- --- --- --- --- --- --- --- --- --- --- --- --- --- --- --- --- --- --- --- --- --- --- --- --- --- --- --- --- --- --- --- --- --- --- --- --- --- --- --- --- --- --- --- --- --- --- --- --- --- --- --- --- --- --- --t atg gta atg acg atg aaa --- -tc gag tag ta- --- --a ctt cta ata aag act tga ta- --- --t cat tat cat atg ttt gat cg- --- --- --- --- --- --- --t cat agt taa tag tgt g-- --- --- --- --- --- --- --- --- --- --- --- --- --- --- --- --- --- --- --- -gc taa atg gta ctg tta ata agt tta t-- --- --- --- --- --- --- --- --- --- --- --- --- --- --- --- --- --a gac aat atc ata gta ttt tct ttc cag aat --- --- --- --t aga tta ttt ttt taa ata ctg atc ctc aca att ccg tga tgt agc agt agt tgg t-- --- --- --- --- --- --- --- --- --- --- --- --- --- --- --- --- --- --- --- --- --- --- --- --- --- --- --- -gc atg gtc tat atc gt- --- --- --- --- --- --- --- --- --- --- --- --- --- --- --- --- --- --- --- --- --- --- --- --- --- --- --- --- --- --- --- --- --- --- --- --- --- --- --- --- --- --- --- taa aat gta tca tat ata ata gtt ttc tga cgt gga gta cag aat ttt cga --- --- --- --- --- --- --- --- --- --- --- --- --- --- --- --- --- --- --- --- --- --- --- --- --- --- --- --- --- --- --- --- --- --- --- --- --- --- --- --- --- --- --- --t taa tga gtt cat ggt aag gaa ggg caa atg tct --- gta tat aat ata cat aag tta a-- --- --- --- --- --- --- --- --- --- --- --- --- --t agt ttt tta tca tat tt- --- --- --- --- --- --- --- --t cta ata cca taa taa aaa tta tca tta tg- --- --- -ta taa tca --- --- --- --- --- --- --- --- --- --- --- --t cac tg- --- -tc gct atc att att gcg ttt gtg tag tt- --- --- --- --- ctg ccc ta- --- --- --- --- --- --- --- --t cat cta cat cac tgt ca- --- --- --- --- ctc tca cta tat ctt cta aaa tta caa --- -ac aac --- --- --- --- --- --- --- --- --- --- --- --- --- --- --- --- --- --- --- --- --- --- --- --- --- --- --- --- --- --- --- --- --- --- --- --- tgg ata ttc gat --- --- --- --- --- --- --- --- --- --- --- --- --- --- --- --- --- --- --- --- --- --- --- --- --- --- --a aca gca ttt gtg t-- --- --- --- --- --- --- --- --- --- --- --- --- --- --- --- --- -

>KJ642613.1_Monkeypox_virus_strain_Congo_8_complete_genome

--- --- --- --- --- --- --- --- --- --- --- --- --- --- --- --- --- --- --- --- --- --- --- --- --- --- --- --- --- --- --- --- --- --- --- --- --- --- --- --- --- --- --- --- --- --- --- --- --- --- --- --- --- --- --- --- --- --- --- --- --- --- --- --- --- --- --- --- --- --- --- --- --- --- --- --- --- --- --- --- --- --- --- --- --- --- --- --- --- --- --- --- --- --- --- --- --- --- --- --- --- --- --- --- --- --- --- --- --- --- --- --- --- --- --- --- --- --- --- --- --- --- --- --- --- --- --- --- --- --- --- --- --- --- --- --- --- --- --- --- --- --- --- --- --- --- --- --- --- --- --- --- --- --- --- --- --- --- --- --- --- --- --- --- --- --- --- --- --- --- --- --- --- --- --- --- --- --- --- --- --- --- --- --- --- --- --- --- --- --- --- --- --- --- --- --- --- --- --- --- --- --- --- --- --- --- --- --- --- --- --- --- --- --- --- --- --- --- --- --- --- --- --- --- --- --- --- --- --- --- --- --- --- --- --- --- --- --- --- --- --- --- --- --- --- --- --- --- --- --- --- --- --- --- --- --- --- --- --- --- --- --- --- --- --- --- --- --- --- --- --- --- --- --- --- --- --- --- --- --- --- --- --- --- --- --- --- --- --- --- --- --- --- --- --- --- --- --- --- --- --- --- --- --- --- --- --- --- --- --- --- --- --- --- --- --- --- --- --- --- --- --- --- --- --- --- --- --- --- --- --- --- --- --- --- --- --- --- --- --- --- --- --- --- --- --- --- --- --- --- --- --- --- --- --- --- --- --- --- --- --- --- --- --- --- --- --- --- --- --- --- --- --- --- -ac cca tcg acg aga gaa aga gat ggt tag tca aga tat ttt tct tag tac aaa agt caa tgt ttt aaa ata tat gga cga gaa tta att tgt ctg tat aaa aac ttg tgt gaa att atg tac tag aga aaa aac gtg agc agt gtc ccc tac atg gat ttt aca gat cat tta tat tcc aaa aat att aac tat ata cgt tta tta tat gat gtt aac gtg taa att ata aac att att tta tga tgc aat tgt ctg aca acc --- att ggc ata agg ata ttg ata agc tct acg aga ata tat tgt tgg acg tta tcg ttt acg aaa tag ttg aga cat cag aaa gag gtt taa tat ttt tgt gag acc cat cga aga gag aa- -ag gat aaa aat a-- --- --- --- ttt ttt tgt aaa act ttt tta tga gac aag aga gaa aga gaa tac gaa tag tga tca tat cgt atc aca tat tga aac aga aag aag aag taa cga gag gta act ttt tgt gaa tgt agt taa aca ttt ttg ttt tgc aaa ccg gaa tat agt acc cgg tac act ttt tta att cgt ggt gcg gtg tct gaa tcg ttc gat taa ccc aac tca tcc att ttc aga tga ata gag tta tcg att cag aca cat gct ttg agt ttt gtt gaa tcg atg agt gaa gta tca tcg gtt gca cct tca gat gcc gat ccg tcg aca tac ttg aat cca tcc ttg act tca agt tca gat gat tcc tca cac atg tct ccg ata cgt acg cta aac tct agg ttc tta aca cat ttt gta tca acg atc gtt gaa ccg atg ata tct ttg taa ctc act ttc tta tgt gag atg tta gac cca agt act gga tgg gtc ttg atg tcg ctg tct ttc tct tct tcg cta cat ctg atg tcg ata gac atc tca cag tct ttg atc ata gcc aga gct tct tca cgc gtg atc gcg gga gag tcc tta cct tgt cct ggt gac acg ctg gac aat cta gta ttc aca gtg ttt cca tca gag gat tcg gag atg gat gaa atc ttt ggg cat ttg gtg aat cca aag ttc atg tta aga ccc gcg ccg acg ata ttg taa taa gtg gtg gga tct cct ttt aca act tct tcg gat acc tca tca tct tcg gat acc tca tca tct tcg gtc tct gta act tcc gtt acg gat tga caa atc tta tca ttg gtc ggt gtt tgg tct tgc ttt gtg act ttg ata ata aca tcg att ccc ata tga tgt ttg ttt tct tct tca gta cac gag gat gag gat tgt tga aga cta gta ggc ata gca gct gcc act agg cac atg cat gcc agg aca ata tat tgt ttc atg att gct att gat tga tta ctg ttc tag atg att cta ctt tct tac cat ata ata aat tag aat ata ttt tct act ttt acg aga aat taa tta ttg tat tta tta ttt atg ggt -aa aaa aaa ctt act ata agt ggg tgg gat tct ggg aat tag tga tca gtt tat gta tat cgc aac tag cgg gca tat ggc tat tga cat cga gaa cat tac cca tat gat aag aga ttg tat ccg ttt cgt agt ctt gag tat tgg tat tac tat ata gta tgt aga tgt cga cgc tag ata gac agt cgc cca cta gag tta ccg tct ctg aat gcg gca tga tag tat cat tct ttg ttt tcg tta act gtt tgg aag atg aat ctt tgt tgt tac att taa tct cga aat tca gag tac ata tct ttg aag tat tct gat atc tat ttt ctc ctg taa aga atc ctg aag ttg cta cat tat taa gga cag aga agt att ctg cac gaa aga cgg gat cac aat ctt tat gat tca tgg taa tag tta gtt ccg acg ttg aga tgg att cgc tga gac cgg tag tgg tcg tcc gag tac acg atg tgt cgt tga ctg gat aca ggt taa ttt cca cat cga tat agt taa atg tat tgc tgg tt- --a cgg gtt cgc att tat ctg tgg aag aga cgg tgt gag aat atg ttc cgg gac cac acg gag aac aga tga cgt ctc cgg tag acg tgt atc cgg ata ctc cgt atc cta ttc cac act ttg ttt tag aaa tac atg ttc tac acc ctg atg atc ctt tga gaa gac aat aat atc ctg gag agc att cac aga ttc tat tgt gag tcg tgt tac acg atc gcg tct cta cct gat tac tat cac atc ttc cgt tac aac tta gac aag cct gta aat gat tat tgt gag atg taa agg tat ccg aac cac acg gcg tac att gtg tat tag tct tgc tat cac ata atc tgg aag cgt aag ttc ccg gag gac acg ata gac aac ata gat tac ggc ttc tgt att cgt tgt ctt tac att ttc cat tgg atg gtg cat gtg gtg cta tat ctc ttc cgt tta tta tta tac atg aga gaa aca ata tat acg agt ata ata cgg acc tca tga ttt aat aat -gt agt aat cgt cgt ctt gtt act gtt tgt ttc cta ctt ctc caa tca tat aga tta ttt ttt aaa tat ttt ctt tct atc atg gat aat att tgt aat ggt tct ttc cgt aca aca tac tgt tta gat ggt agt cgc tta gct tgg tta tga tat tgc gca taa ttt ccg gag gca aat acg ata gtc tag att gac tat cga tgg tag act cta att tat tga gtg ctt tgt cga cga gtt tac ttt tat gct cca tcg ata gat gac act gtt cta tga gat cgt cgt aca tgg gaa atg aaa tgc gtt tgt ctg aat gta tgg ctt cga gat agg tgt gat acc gga tgt ctt ctg ttc tca ata ccg tat aca agt tgg tgt ctg aga ttc gaa tct ctt tga gga gac tta tgt cac gac tac att ttt cga tga tgg aat cta tct tat cga atg ata tat ttt tca taa ata cac ttt tat agt cct cgt tta aac aga att tag tat gta gtt ccg caa atg act cgt ccc tta ata ggc agt agg cta tta tct tct tta cgt agt gat cgt cgt agg gag aga act ccg aca tct tgt aga aca acg att taa tca tag gta gag ata ctt tca gtc tgt ggt gga tga tgt cat taa cga cat ccg cct tgt ata tga tgt ttc tgt ttt caa aca cca agt cga ata ctg tct tat cgt ctt tag tcg gaa ggt tga tgt cgt atc cga tgt ata cga ggt atg agg caa cat tgt tat tgc aat tct gga agg cgg tat gaa gag gag tca ttg tat tat agt att cgt ctt tct gaa tgt cga atc tat cta gta gat acc gta gta tat tga gag agc gac ttc cat atc ctt gat tat gtt tta tga ata gat aaa gta gat gtt gtc ctt ctt cct ttt gta att tcc cgt att ttt gtt cgt gcc aat tga gta aca tta tga gaa tat gac ctg ttg cac aat cgt tct tta tgt att cca tga tgg gtg tac aat caa gat tat tac gta tcc tcg tat cgg ctc ctc gag ata aaa gag cat aca cca cac gag gac tat gtt tgg tat act gtt gaa ggt aag tgt gta acg gcg tat ttc cga ttt tcg taa ccg cgt taa tgt ttg ctc cat gat cta tta tcg cgt aga tga atc gct tct cag ctc gca tct tag tgt gac tct ttg act tgt aat aat tgc ttt cgt gga aca cgg ata tgt gtt tac agt agt aat gaa gag aag tga gtc cat cct cat cga cgc aat tag ggt cag atc ctt tag tca ata att tgt aca gaa cgt aat agt tta agc tcc cat tga att tat atc taa gat aac aca gca ata gat cgg atg att tac taa agt cat caa tgg ggt ccg tta gta tat caa aga tct tgt tat cga ttg ata gtg aat gaa tca gat agt ggt gta gag gaa tat gtc ctt ttt tat cct tgc tat caa agt tac gca tgc cgt ggc gta aca ata tct tta ata cag atg gat taa atc gtg tat tca tcg tat agc aat gta atg gag agt tac cac att tta gtc gtt tat tca gat cgc agt gtt taa taa cta att taa aca gat gag atg ata tat cca cat caa aga atg cga gat aca tat gac aga cat tat tga cag aaa tgt gac ctt cat tat cac cgt cgt cca taa atg cgt tag gta cgt acc aca tac tat cgt taa cga tgc gca caa tct cgt cca ttt tca tcc atc ttc ata atg att tac ttt ttc ata att aga gaa aaa gat caa ggt ata aaa att aga agt gtt aga cta ta- --- --- --- --- --- --- --- --- --- --- -aa cta act tat aa- --- --- --- --- --- --- --- --- --- --- --- --- --- --- --- --- --- --- --- --- --- --- --- --- --- --- --- --- --- --- --- --- --- --- --- --- --- --- --- --- --- --- --- --- --- --- --- --- --- --- --- --- --- --- --- --- --- --- --- --- --- --- --- --- --- --- --- --- --- --- --- --- --- --- --- --- --- --- --- --- --- --- --- --- --- --- -aa cta act tat gac tta act aac tta tga cta tta act cat ttc aag aaa ggt ggg tgg aga gaa ctc tat atg aca gct tgt gaa aca att aga tcc cta att tct aat gga agt ttt gat agg aga ttg tca tca gtt gat aca ttg ttt att atc tca tct att aga gca cgt ctg ttt aga gct tta gtg acc tgc tcg gtt act tct gtg taa atc ttg aat cct tta gtg ata cac tgt gtc aaa act gga tgt tta gaa tac cta tgt aga ata tgg gaa gca tgc ttg ttt ttg tct cta tta tag att aac tca tac atg gtt gta tta tga att ttc atc tgc cta atg tac tcc aat tct tgt tta caa tca att ata taa tca aag agt gat gat gca tac aca tta caa agt gaa taa tct acc atc ata aaa tac ttg ata cag agc ttt atc aca tca tgg ttt tca att gta tta tta agt ata gct aat ttt ata cag tca ata gac aat ggt tct cta agc aat att tct aat att tta aga tgt gct tcc cta cgg gcg atg aca gat ccc cta tcc acg gcc acg tca aga cat gta tat cca tta ctc att act gcg ttg aca ttt gct cca ttt tct aat agc cat gat act aaa tct ata taa cct gca tag ata gcg cga taa agc aag gtc ctt cca cca gca tct agt tga ttg ata tct tca ata tat ggg ata caa agc tta taa att tct aat act gtg ggt tca tct aca agg aat ccc cta gta tac tga att att tta tat aga tct aat tta aca tca ttt tca tct ggg ata cca caa ttc aaa ata aac tca aca aca cta ctt tcc ttt tta cat att ccc cta aaa tag gca ttc aag cat tct att tta tat att aca gcc cca tga tct acc ata aaa tca aca atg tct att tct aca tat gca tta gat aga tag taa agt aag aga tct tgt aca gaa tta caa ttc tta ata att ata gag aaa ata tct tcc ata taa ttc ttt gac act aat gca gat ata ata tct tta tat gta ata tat gca aac agt cta tct act ata tac tga tca ata tta tct cta tga atc cta aaa taa tca tac aga aca tct aca gga tca caa att ggt tca agg aga aat cta tca aat att ttc ctg tca aca act ggt tct aga aca taa cag tca aca cct aat cca tgt ttt tta tag tca tct acc aaa gat aat gac caa aga tcg agg tcg tcg tga aac tgc tca tcg aca gcc atg aaa tct gcc gac tcc atg gcg cga atc gca ctg tct tat tcg cca ttg att ttc a-t ttt tta taa tta tgt aca tgt ttt cct tct att ctc aag agt cta caa aaa tat a-t ttt ttt tcg ata tct aag tac taa gtt ttt tta ctg ttt ttg tta ctg tct tcc att ctt cta act aaa gat ctg aga taa att ata caa tct tcg cta tcg aac cat ttt tgt agt cta aag cct gaa gta att aac caa ctg ttt tta tta gtg gct ttt ttc gat cta tcc tcg tc- --- --c tca tca tcc tta ta- --- --- --- --- --- --- tta tta tca tta tta tca tag tct att aaa cac aaa tca tct --- --- --- --- --- --- --- --- --- --- --- --- --- --- --- --- --- --- --- --- --- --- --- --- --- --- --a cgt tta taa c-- --- --- --- --- --- --- --- --a aca ttc --- --- --- --- -tc att att aat tag ttc tgt agt aat atc ttt aat aat ttg gct ata --- --- -ca tct gtt --- --- --- --- --- --- --- --- --- --- --- --- --- --- --- --- --- --- --c aat act --- --- --- --- --- --- --- --a tct att gat gat ttc ttt t-- --- --- --- --- --- --- --- --- --- --- --t taa gac t-- --- --- --- --- --- --- --- --- --- --- --- --- --- --- --- --- --- --- --- --- --- --- --- --- --- --- --- --- --- --- --- --- --- --- --- --- --- --- --- taa act agt --- --- --- --- --- --- --- --- --- --- --- --- --- --- --- --- --- --- --- --- --- --- --- --- --- --- --- --- --- --- --- --- --- --- --- --- --- --- --- --- --- --- --- --- --- --- --- --- --- --- --- --- --- --- --- --- --- --- --- --- --- --- --- --- --- --- --- --- --- --- --- --- --- --- --- --- --- --- --- --- --- --- --- --- --- --- --- --- --- --- --- --- --- --- --- --- --- --- --- --- --- --- --- --- --- --- --- --- --- --t atg gta atg acg atg aaa --- -tc gag tag ta- --- --a ctt cta ata aag act tga ta- --- --t cat tat cat atg ttt gat cg- --- --- --- --- --- --- --t cat agt taa tag tgt g-- --- --- --- --- --- --- --- --- --- --- --- --- --- --- --- --- --- --- --- -gc taa atg gta ctg tta ata agt tta t-- --- --- --- --- --- --- --- --- --- --- --- --- --- --- --- --- --a gac aat atc ata gta ttt tct ttc cag aat --- --- --- --t aga tta ttt ttt taa ata ctg atc ctc aca att ccg tga tgt agc agt agt tgg t-- --- --- --- --- --- --- --- --- --- --- --- --- --- --- --- --- --- --- --- --- --- --- --- --- --- --- --- -gc atg gtc tat atc gt- --- --- --- --- --- --- --- --- --- --- --- --- --- --- --- --- --- --- --- --- --- --- --- --- --- --- --- --- --- --- --- --- --- --- --- --- --- --- --- --- --- --- --- taa aat gta tca tat ata ata gtt ttc tga cgt gga gta cag aat ttt cga --- --- --- --- --- --- --- --- --- --- --- --- --- --- --- --- --- --- --- --- --- --- --- --- --- --- --- --- --- --- --- --- --- --- --- --- --- --- --- --- --- --- --- --t taa tga gtt cat ggt aag gaa ggg caa atg tct --- gta tat aat ata cat aag tta a-- --- --- --- --- --- --- --- --- --- --- --- --- --t agt ttt tta tca tat tt- --- --- --- --- --- --- --- --t cta ata cca taa taa aaa tta tca tta tg- --- --- -ta taa tca --- --- --- --- --- --- --- --- --- --- --- --t cac tg- --- -tc gct atc att att gcg ttt gtg tag tt- --- --- --- --- ctg ccc ta- --- --- --- --- --- --- --- --t cat cta cat cac tgt ca- --- --- --- --- ctc tca cta tat ctt cta aaa tta caa --- -ac aac --- --- --- --- --- --- --- --- --- --- --- --- --- --- --- --- --- --- --- --- --- --- --- --- --- --- --- --- --- --- --- --- --- --- --- --- tgg ata ttc gat --- --- --- --- --- --- --- --- --- --- --- --- --- --- --- --- --- --- --- --- --- --- --- --- --- --- --a aca gca ttt gtg t-- --- --- --- --- --- --- --- --- --- --- --- --- --- --- --- --- -

>HM172544.1_Monkeypox_virus_strain_Zaire_1979-005_complete_genome

--- --- --- --- --- --- --- --- --- --- --- --- --- --- --- --- --- --- --- --- --- --- --- --- --- --- --- --- --- --- --- --- --- --- --- --g tgt cta gaa aaa aat gtg tga ccc acg acc gta gga aac tct aga ggg taa gaa aaa tca atc gtt tat aga gac cat cag aaa gag gtt taa tat ttt tgt gag acc cat cga aga gag aaa gg- --- --- --- --- --- --- --- --- --- --- --- --- --- --- --- --- --- --- --- --- --- --- --- --- --- --- --- --- --- --- --- --- --- --- --- --- --- --- --- --- --- --- --- --- --- --- --- --- --- --- --- --- --- --- --- --- --- --- --- --- --- --- --- --- --- --- --- --- --- --- --- --- --- --- --- --- --- --- --- --- --- --- --- --- --- --- --- --- --- --- --- --- --- --- --- --- --- --- --- --- --- --- --- --- --- --- --- --- --- --- --- --- --- --- --- --- --- --- --- --- --- --- --- --- --- --- --- --- --- --- --- --- --- --- --- --- --- --- --- --- --- --- --- --- --- --- --- --- --- --- --- --- --- --- --- --- --- --- --- --- --- --- --- --- --- --- --- --- --- --- --- --- --- --- --- --- --- --- --- --- --- --- --- --- --- --- --- --- --- --- --- --- --- --- --- --- --- --- --- --- --- --- --- --- --- --- --- --- --- --a taa aaa ctt ttt acg act cca tca gaa aga ggt tta --- --- --- --- --- --- --- --- --- --- --- --- --- --- --- --- --- --- --- --- --- --- --- --- --- --- --- --- --- --- --- --- --- --- --- --- --- --- --- --- --- --- --- --- --- --- --a tat ttt tgt gag acc cat cga aga gag aaa gga taa aaa ctt ttt acg act cca tca gaa aga ggt tta ata ttt ttg tga gac cca tcg aag aga gaa aga gat ggt tag tca aga tat ttt tct tag tac aaa agt caa tgt ttt aaa ata tat gga cga gaa tta att tgt ctg tat aaa aac ttg tgt gaa att atg tac tag aga aaa aac gtg agc agt gtc ccc tac atg gat ttt aca gat cat tta tat tcc aaa aat att aac tat ata cgt tta tta tat gat gtt aac gtg taa att ata aac att att tta tga tgc aat tgt ctg aca acc --- att ggc ata agg ata ttg ata agc tct acg aga ata tat tgt tgg acg tta tcg ttt acg aaa tag ttg aga cat cag aaa gag gtt taa tat ttt tgt gag a-c cat cga aga gag aaa gag aat aaa aat a-- --- --- --- ttt ttt tgt aaa act ttt tta tga gac aag aga gaa aga gaa tac gaa tag tga tca tat cgt atc aca tat tga aac aga aag aag aag taa cga gag gta act ttt tgt gaa tgt agt taa aca ttt ttg ttt tgc aaa ccg gaa tat agt acc cgg tac act ttt tta att cgt ggt gcg gtg tct gaa tcg ttc gat taa ccc aac tca tcc att ttc aga tga ata gag tta tcg att cag aca cat gct ttg agt ttt gtt gaa tcg atg agt gaa gta tca tcg gtt gca cct tca gat gcc gat ccg tcg aca tac ttg aat cca tcc ttg act tca agt tca gat gat tcc tca cac atg tct ccg ata cgt acg cta aac tct agg ttc tta aca cat ttt gta tca acg atc gtt gaa ccg atg ata tct ttg taa ctc act ttc tta tgt gag atg tta gac cca agt act gga tgg gtc ttg atg tcg ctg tct ttc tct tct tcg cta cat ctg atg tcg ata gac atc tca cag tct ttg atc ata gcc aga gct tct tca cgc gtg atc gcg gga gag tcc tta cct tgt cct ggt gac acg ctg gac aat cta gta ttc aca gtg ttt cca tca gag gat tcg gag atg gat gaa atc ttt ggg cat ttg gtg aat cca aag ttc atg tta aga ccc gcg ccg acg ata ttg taa taa gtg gtg gga tct cct ttt aca act --- --- --- --- --- --- tct tcg gat acc tca tca tct tcg gtc tct gta act tcc gtt acg gat tga caa atc tta tca ttg gtc ggt gtt tgg tct tgc ttt gtg act ttg ata ata aca tcg att ccc ata tga tgt ttg ttt tct tct tca gta cac gag gat gag gat tgt tga aga cta gta ggc ata gca gct gcc act agg cac atg cat gcc agg aca ata tat tgt ttc atg att gct att gat tga tta ctg ttc tag atg att cta ctt tct tac cat ata ata aat tag aat ata ttt tct act ttt acg aga aat taa tta ttg tat tta tta ttt atg ggt -aa aaa aaa ctt act ata agt ggg tgg gat tct ggg aat tag tga tca gtt tat gta tat cgc aac tag cgg gca tat ggc tat tga cat cga gaa cat tac cca tat gat aag aga ttg tat ccg ttt cgt agt ctt gag tat tgg tat tac tat ata gta tgt aga tgt cga cgc tag ata gac agt cgc cca cta gag tta ccg tct ctg aat gcg gca tga tag tat cat tct ttg ttt tcg tta act gtt tgg aag atg aat ctt tgt tgt tac att taa tct cga aat tca gag tac ata tct ttg aag tat tct gat atc tat ttt ctc ctg taa aga atc ctg aag ttg cta cat tat taa gga cag aga agt att ctg cac gaa aga cgg gat cac aat ctt tat gat tca tgg taa tag tta gtt ccg acg ttg aga tgg att cgc tga gac cgg tag tgg tcg tcc gag tac acg atg tgt cgt tga ctg gat aca ggt taa ttt cca cat cga tat agt taa atg tat tgc tgg tt- --a cgg gtt cgc att tat ctg tgg aag aga cgg tgt gag aat atg ttc cgg gac cac acg gag aac aga tga cgt ctc cgg tag acg tgt atc cgg ata ctc cgt atc cta ttc cac act ttg ttt tag aaa tac atg ttc tac acc ctg atg atc ctt tga gaa gac aat aat atc ctg gag agc att cac aga ttc tat tgt gag tcg tgt tac acg atc gcg tct cta cct gat tac tat cac atc ttc cgt tac aac tta gac aag cct gta aat gat tat tgt gag atg taa agg tat ccg aac cac acg gcg tac att gtg tat tag tct tgc tat cac ata atc tgg aag cgt aag ttc ccg gag gac acg ata gac aac ata gat tac ggc ttc tgt att cgt tgt ctt tac att ttc cat tgg atg gtg cat gtg gtg cta tat ctc ttc cgt tta tta tta tac atg aga gaa aca ata tat acg agt ata ata cgg acc tca tga ttt aat aat -gt agt aat cgt cgt ctt gtt act gtt tgt ttc cta ctt ctc caa tca tat aga tta ttt ttt aaa tat ttt ctt tct atc atg gat aat att tgt aat ggt tct ttc cgt aca aca tac tgt tta gat ggt agt cgc tta gct tgg tta tga tat tgc gca taa ttt ccg gag gca aat acg ata gtc tag att gac tat cga tgg tag act cta att tat tga gtg ctt tgt cga cga gtt tac ttt tat gct cca tcg ata gat gac act gtt cta tga gat cgt cgt aca tgg gaa atg aaa tgc gtt tgt ctg aat gta tgg ctt cga gat agg tgt gat acc gga tgt ctt ctg ttc tca ata ccg tat aca agt tgg tgt ctg aga ttc gaa tct ctt tga gga gac tta tgt cac gac tac att ttt cga tga tgg aat cta tct tat cga atg ata tat ttt tca taa ata cac ttt tat agt cct cgt tta aac aga att tag tat gta gtt ccg caa atg act cgt ccc tta ata ggc agt agg cta tta tct tct tta cgt agt gat cgt cgt agg gag aga act ccg aca tct tgt aga aca acg att taa tca tag gta gag ata ctt tca gtc tgt ggt gga tga tgt cat taa cga cat ccg cct tgt ata tga tgt ttc tgt ttt caa aca cca agt cga ata ctg tct tat cgt ctt tag tcg gaa ggt tga tgt cgt atc cga tgt ata cga ggt atg agg caa cat tgt tat tgc aat tct gga agg cgg tat gaa gag gag tca ttg tat tat agt att cgt ctt tct gaa tgt cga atc tat cta gta gat acc gta gta tat tga gag agc gac ttc cat atc ctt gat tat gtt tta tga ata gat aaa gta gat gtt gtc ctt ctt cct ttt gta att tcc cgt att ttt gtt cgt gcc aat tga gta aca tta tga gaa tat gac ctg ttg cac aat cgt tct tta tgt att cca tga tgg gtg tac aat caa gat tat tac gta tcc tcg tat cgg ctc ctc gag ata aaa gag cat aca cca cac gag gac tat gtt tgg tat act gtt gaa ggt aag tgt gta acg gcg tat ttc cga ttt tcg taa ccg cgt taa tgt ttg ctc cat gat cta tta tcg cgt aga tga atc gct tct cag ctc gca tct tag tgt gac tct ttg act tgt aat aat tgc ttt cgt gga aca cgg ata tgt gtt tac agt agt aat gaa gag aag tga gtc cat cct cat cga cgc aat tag ggt cag atc ctt tag tca ata att tgt aca gaa cgt aat agt tta agc tcc cat tga att tat atc taa gat aac aca gca ata gat cgg atg att tac taa agt cat caa tgg ggt ccg tta gta tat caa aga tct tgt tat cga ttg ata gtg aat gaa tca gat agt ggt gta gag gaa tat gtc ctt ttt tat cct tgc tat caa agt tac gca tgc cgt ggc gta aca ata tct tta ata cag atg gat taa atc gtg tat tca tcg tat agc aat gta atg gag agt tac cac att tta gtc gtt tat tca gat cgc agt gtt taa taa cta att taa aca gat gag atg ata tat cca cat caa aga atg cga gat aca tat gac aga cat tat tga cag aaa tgt gac ctt cat tat cac cgt cgt cca taa atg cgt tag gta cgt acc aca tac tat cgt taa cga tgc gca caa tct cgt cca ttt tca tcc atc ttc ata atg att tac ttt ttc ata att aga gaa aaa gat caa ggt ata aaa att aga agt gtt aga cta ta- --- --- --- --- --- --- --- --- --- --- -aa cta act tat aa- --- --- --- --- --- --- --- --- --- --- --- --- --- --- --- --- --- --- --- --- --- --- --- --- --- --- --- --- --- --- --- --- --- --- --- --- --- --- --- --- --- --- --- --- --- --- --- --- --- --- --- --- --- --- --- --- --- --- --- --- --- --- --- --- --- --- --- --- --- --- --- --- --- --- --- --- --- --- --- --- --- --- --- --- --- --- -aa cta act tat gac tta act aac tta tga cta tta act cat ttc aag aaa ggt ggg tgg aga gaa ctc tat atg aca gct tgt gaa aca att aga tcc cta att tct aat gga agt ttt gat agg aga ttg tca tca gtt gat aca ttg ttt att atc tca tct att aga gca cgt ctg ttt aga gct tta gtg acc tgc tcg gtt act tct gtg taa atc ttg aat cct tta gtg ata cac tgt gtc aaa act gga tgt tta gaa tac cta tgt aga ata tgg gaa gca tgc ttg ttt ttg tct cta tta tag att aac tca tac atg gtt gta tta tga att ttc atc tgc cta atg tac tcc aat tct tgt tta caa tca att ata taa tca aag agt gat gat gca tac aca tta caa agt gaa taa tct acc atc ata aaa tac ttg ata cag agc ttt atc aca tca tgg ttt tca att gta tta tta agt ata gct aat ttt ata cag tca ata gac aat ggt tct cta agc aat att tct aat att tta aga tgt gct tcc cta cgg gcg atg aca gat ccc cta tcc acg gcc acg tca aga cat gta tat cca tta ctc att act gcg ttg aca ttt gct cca ttt tct aat agc cat gat act aaa tct ata taa cct gca tag ata gcg cga taa agc aag gtc ctt cca cca gca tct agt tga ttg ata tct tca ata tat ggg ata caa agc tta taa att tct aat act gtg ggt tca tct aca agg aat ccc cta gta tac tga att att tta tat aga tct aat tta aca tca ttt tca tct ggg ata cca caa ttc aaa ata aac tca aca aca cta ctt tcc ttt tta cat att ccc cta aaa tag gca ttc aag cat tct att tta tat att aca gcc cca tga tct acc ata aaa tca aca atg tct att tct aca tat gca tta gat aga tag taa agt aag aga tct tgt aca gaa tta caa ttc tta ata att ata gag aaa ata tct tcc ata taa ttc ttt gac act aat gca gat ata ata tct tta tat gta ata tat gca aac agt cta tct act ata tac tga tca ata tta tct cta tga atc cta aaa taa tca tac aga aca tct aca gga tca caa att ggt tca agg aga aat cta tca aat att ttc ctg tca aca act ggt tct aga aca taa cag tca aca cct aat cca tgt ttt tta tag tca tct acc aaa gat aat gac caa aga tcg agg tcg tcg tga aac tgc tca tcg aca gcc atg aaa tct acc gac tcc atg gcg cga atc gca ctg tct tat tcg cca ttg att ttc att ttt tta taa tta tgt aca tgt ttt cct tct att ctc aag agt cta caa aaa tat a-t ttt ttt tcg ata tct aag tac taa gtt ttt tta ctg ttt ttg tta ctg tct tcc att ctt cta act aaa gat ctg aga taa att ata caa tct tcg cta tcg aac cat ttt tgt agt cta aag cct gaa gta att aac caa ctg ttt tta tta gtg gct ttt ttc gat cta tcc tcg tc- --- --c tca tca tcc tta ta- --- --- --- --- --- --- tta tta tca tta tta tca tag tct att aaa cac aaa tca tct --- --- --- --- --- --- --- --- --- --- --- --- --- --- --- --- --- --- --- --- --- --- --- --- --- --- --a cgt tta taa c-- --- --- --- --- --- --- --- --a aca ttc --- --- --- --- -tc att att aat tag ttc tgt agt aat atc ttt aat aat ttg gct ata --- --- -ca tct gtt --- --- --- --- --- --- --- --- --- --- --- --- --- --- --- --- --- --- --c aat act --- --- --- --- --- --- --- --a tct att gat gat ttc ttt t-- --- --- --- --- --- --- --- --- --- --- --t taa gac t-- --- --- --- --- --- --- --- --- --- --- --- --- --- --- --- --- --- --- --- --- --- --- --- --- --- --- --- --- --- --- --- --- --- --- --- --- --- --- --- taa act agt --- --- --- --- --- --- --- --- --- --- --- --- --- --- --- --- --- --- --- --- --- --- --- --- --- --- --- --- --- --- --- --- --- --- --- --- --- --- --- --- --- --- --- --- --- --- --- --- --- --- --- --- --- --- --- --- --- --- --- --- --- --- --- --- --- --- --- --- --- --- --- --- --- --- --- --- --- --- --- --- --- --- --- --- --- --- --- --- --- --- --- --- --- --- --- --- --- --- --- --- --- --- --- --- --- --- --- --- --- --t atg gta atg acg atg aaa --- -tc gag tag ta- --- --a ctt cta ata aat act tga ta- --- --t cat tat cat atg ttt gat cg- --- --- --- --- --- --- --t cat agt taa tag tgt g-- --- --- --- --- --- --- --- --- --- --- --- --- --- --- --- --- --- --- --- -gc taa atg gta ctg tta ata agt tta t-- --- --- --- --- --- --- --- --- --- --- --- --- --- --- --- --- --a gac aat atc ata gta ttt tct ttc cag aat --- --- --- --t aga tta ttt ttt taa ata ctg atc ctc aca att ccg tga tgt agc agt agt tgg t-- --- --- --- --- --- --- --- --- --- --- --- --- --- --- --- --- --- --- --- --- --- --- --- --- --- --- --- -gc atg gtc tat atc gt- --- --- --- --- --- --- --- --- --- --- --- --- --- --- --- --- --- --- --- --- --- --- --- --- --- --- --- --- --- --- --- --- --- --- --- --- --- --- --- --- --- --- --- taa aat gta tca tat ata ata gtt ttc tga cgt gga gta cag aat ttt cga --- --- --- --- --- --- --- --- --- --- --- --- --- --- --- --- --- --- --- --- --- --- --- --- --- --- --- --- --- --- --- --- --- --- --- --- --- --- --- --- --- --- --- --t taa tga gtt cat ggt aag gaa gga caa atg tct --- gta tat aat ata cat aag tta a-- --- --- --- --- --- --- --- --- --- --- --- --- --t agt ttt tta tca tat tt- --- --- --- --- --- --- --- --t cta ata cca taa taa aaa tta tca tta tg- --- --- -ta taa tca --- --- --- --- --- --- --- --- --- --- --- --t cac tg- --- -tc gct atc att att gcg ttt gtg tag tt- --- --- --- --- ctg ccc ta- --- --- --- --- --- --- --- --t cat cta cat cac tgt ca- --- --- --- --- ctc tca cta tat ctt cta aaa tta caa --- -ac aac --- --- --- --- --- --- --- --- --- --- --- --- --- --- --- --- --- --- --- --- --- --- --- --- --- --- --- --- --- --- --- --- --- --- --- --- tgg ata ttc gat --- --- --- --- --- --- --- --- --- --- --- --- --- --- --- --- --- --- --- --- --- --- --- --- --- --- --a aca gca ttt gtg t-- --- --- --- --- --- --- --- --- --- --- --- --- --- --- --- --- -

>AF380138.1_Monkeypox_virus_strain_Zaire-96-I-16_complete_genome

--- --- --- --- --- --- --- --- --- --- --- --- --- --- --- --- -gt tag taa att ata tac ata att tta taa tt- --a att taa ttt tac tat ttt att tag tgt cta gaa aaa aat gtg tga ccc acg acc gta gga aac tct aga ggg taa gaa aaa tca atc gtt tat aga gac cat cag aaa gag gtt taa tat ttt tgt gag acc cat cga a-- --- --- --- --- --- --- --- --- --- --- --- --- --- --- --- --- --- --- --- --- --- --- --- --- --- --- --- --- --- --- --- --- --- --- --- --- --- --- --- --- --- --- --- --- --- --- --- --- --- --- --- --- --- --- --- --- --- --- --- --- --- --- --- --- --- --- --- --- --- --- --- --- --- --- --- --- --- --- --- --- --- --- --- --- --- --- --- --- --- --- --- --- --- --- --- --- --- --- --- --- --- --- --- --- --- --- --- --- --- --- --- --- --- --- --- --- --- --- --- --- --- --- --- --- --- --- --- --- --- --- --- --- --- --- --- --- --- --- --- --- --- --- --- --- --- --- --- --- --- --- --- --- --- --- --- --- --- --- --- --- --- --- --- --- --- --- --- --- --- --- --- --- --- --- --- --- --- --- --- --- --- --- --- --- --- --- --- --- --- --- --- --- --- --- --- --- --- --- --- --- --- --- --- --- --- --- --- --- --- --- --- --- --- --- --- --- --- --- --- --- --- --- --- --- --- --- --- --- --- --- --- --- --- --- --- --- --- --- --- --- --- --- --- --- --- --- --- --- --- --- --- --- --- --- --- --- --- --- --- --- --- --- --- --- --- --- --- --- --- --- --- --- --- --- --- --- --- --- --- -ga gag aaa gga taa aaa ctt ttt acg act cca tca gaa aga ggt tta ata ttt ttg tga gac cca tcg acg aga gaa aga gat ggt tag tca aga tat ttt tct tag tac aaa agt caa tgt ttt aaa ata tat gga cga gaa tta att tgt ctg tat aaa aac ttg tgt gaa att atg tac tag aga aaa aac gtg agc agt gtc ccc tac atg gat ttt aca gat cat tta tat tcc aaa aat att aac tat ata cgt tta tta tat gat gtt aac gtg taa att ata aac att att tta tga tgc aat tgt ctg aca acc tag att ggc ata agg ata ttg ata agc tct acg aga ata tat tgt tgg acg tta tcg ttt acg aaa tag ttg aga cat cag aaa gag gtt taa tat ttt tgt gag a-c cat cga aga gag aaa gag aat aaa aat a-- --- --- --- ttt ttt tgt aaa ac- ttt tta tga gac aag aga gaa aga gaa tac gaa tag tga tca tat cgt atc aca tat tga aac aga aag aag aag taa cga gag gta act ttt tgt gaa tgt agt taa aca ttt ttg ttt tgc aaa ccg gaa tat agt acc cgg tac act ttt tta att cgt ggt gcg gtg tct gaa tcg ttc gat taa ccc aac tca tcc att ttc aga tga ata gag tta tcg att cag aca cat gct ttg agt ttt gtt gaa tcg atg agt gaa gta tca tcg gtt gca cct tca gat gcc gat ccg tcg aca tac ttg aat cca tcc ttg act tca agt tca gat gat tcc tca cac atg tct ccg ata cgt acg cta aac tct agg ttc tta aca cat ttt gta tca acg atc gtt gaa ccg atg ata tct ttg taa ctc act ttc tta tgt gag atg tta gac cca agt act gga tgg gtc ttg atg tcg ctg tct ttc tct tct tcg cta cat ctg atg tcg ata gac atc tca cag tct ttg atc ata gcc aga gct tct tca cgc gtg atc gcg gga gag tcc tta cct tgt cct ggt gac acg ctg gac aat cta gta ttc aca gtg ttt cca tca gag gat tcg gag atg gat gaa atc ttt ggg cat ttg gtg aat cca aag ttc atg tta aga ccc gcg ccg acg ata gtg taa taa gtg gtg gga tct cct ttt aca act --- --- --- --- --- --- tct tcg gat acc tca tca tct tcg gtc tct gta act tcc gtt acg gat tga caa atc tta tca ttg gtc ggt gtt tgg tct tgc ttt gtg act ttg ata ata aca tcg att ccc ata tga tgt ttg ttt tct tct tca gta cac gag gat gag gat tgt tga aga cta gta ggc ata gca gct gcc act agg cac atg cat gcc agg aca ata tat tgt ttc atg att gct att gat tga tta ctg ttc tag atg att cta ctt tct tac cat ata ata aat tag aat ata ttt tct act ttt acg aga aat taa tta ttg tat tta tta ttt atg ggt aaa aaa aaa ctt act ata agt ggg tgg gat tct ggg aat tag tga tca gtt tat gta tat cgc aac tag cgg gca tat ggc tat tga cat cga gaa cat tac cca tat gat aag aga ttg tat ccg ttt cgt agt ctt gag tat tgg tat tac tat ata gta tgt aga tgt cga cgc tag ata gac agt cgc cca cta gag tta ccg tct ctg aat gcg gca tga tag tat cat tct ttg ttt tcg tta act gtt tgg aag atg aat ctt tgt tgt tac att taa tct cga aat tca gag tac ata tct ttg aag tat tct gat atc tat ttt ctc ctg taa aga atc ctg aag ttg cta cat tat taa gga cag aga agt att ctg cac gaa aga cgg gat cac aat ctt tat gat tca tgg taa tag tta gtt ccg acg ttg aga tgg att cgc tga gac cgg tag tgg tcg tcc gag tac acg atg tgt cgt tga ctg gat aca ggt taa ttt cca cat cga tat agt taa atg tat tgc tgg tt- --a cgg gtt cgc att tat ctg tgg aag aga cgg tgt gag aat atg ttc cgg gac cac acg gag aac aga tga cgt ctc cgg tag acg tgt atc cgg ata ctc cgt atc cta ttc cac act ttg ttt tag aaa tac atg ttc tac acc ctg atg atc ctt tga gaa gac aat aat atc ctg gag agc att cac aga ttc tat tgt gag tcg tgt tac acg atc gcg tct cta cct gat tac tat cac atc ttc cgt tac aac tta gac aag cct gta aat gat tat tgt gag atg taa agg tat ccg aac cac acg gcg tac att gtg tat tag tct tgc tat cac ata atc tgg aag cgt aag ttc ccg gag gac acg ata gac aac ata gat tac ggc ttc tgt att cgt tgt ctt tac att ttc cat tgg atg gtg cat gtg gtg cta aat ctc ttc cgt tta tta tta tac atg aga gaa aca ata tat acg agt ata ata cgg acc tca tga ttt aat aat agt agt aat cgt cgt ctt gtt act gtt tgt ttc cta ctt ctc caa tca tat aga tta ttt ttt aaa tat ttt ctt tct atc atg gat aat att tgt aat ggt tct ttc cgt aca aca tac tgt tta gat ggt agt cgc tta gct tgg tta tga tat tgc gca taa ttt ccg gag gca aat acg ata gtc tag att gac tat cga tgg tag act cta att tat tga gtg ctt tgt cga cga gtt tac ttt tat gct cca tcg ata gat gac act gtt cta tga gat cgt cgt aca tgg gaa atg aaa tgc gtt tgt ctg aat gta tgg ctt cga gat agg tgt gat acc gga tgt ctt ctg ttc tca ata ccg tat aca agt tgg tgt ctg aga ttc gaa tct ctt tga gga gac tta tgt cac gac tac att ttt cga tga tgg aat cta tct tat cga atg ata tat ttt tca taa ata cac ttt tat agt cct cgt tta aac aga att tag tat gta gtt ccg caa atg act cgt ccc tta ata ggc agt agg cta tta tct tct tta cgt agt gat cgt cgt agg gag aga act ccg aca tct tgt aga aca acg att taa tca tag gta gag ata ctt tca gtc tgt ggt gga tga tgt cat taa cga cat ccg cct tgt ata tga tgt ttc tgt ttt caa aca cca agt cga ata ctg tct tat cgt ctt tag tcg gaa ggt tga tgt cgt atc cga tgt ata cga ggt atg agg caa cat tgt tat tgc aat tct gga agg cgg tat gaa gag gag tca ttg tat tat agt att cgt ctt tct gaa tgt cga atc tat cta gta gat acc gta gta tat tga gag agc gac ttc cat atc ctt gat tat gtt tta tga ata gat aaa gta gat gtt gtc ctt ctt cct ttt gta att tcc cgt att ttt gtt cgt gcc aat tga gta aca tta tga gaa tat gac ctg ttg cac aat cgt tct tta tgt att cca tga tgg gtg tac aat caa gat tat tac gta tcc tcg tat cgg ctc ctc gag ata aaa gag cat aca cca cac gag gac tat gtt tgg tat act gtt gaa ggt aag tgt gta acg gcg tat ttc cga ttt tcg taa ccg cgt taa tgt ttg ctc cat gat cta tta tcg cgt aga tga atc gct tct cag ctc gca tct tag tgt gac tct ttg act tgt aat aat tgc ttt cgt gga aca cgg ata tgt gtt tac agt agt aat gaa gag aag tga gtc cat cct cat cga cgc aat tag ggt cag atc ctt tag tca ata att tgt aca gaa cgt aat agt tta agc tcc cat tga att tat atc taa gat aac aca gca ata gat cgg atg att tac taa agt cat caa tgg ggt ccg tta gta tat caa aga tct tgt tat cga ttg ata gtg aat gaa tca gat agt ggt gta gag gaa tat gtc ctt ttt tat cct tgc tat caa agt tac gca tgc cgt ggc gta aca ata tct tta ata cag atg gat taa atc gtg tat tca tcg tat agc aat gta atg gag agt tac cac att tta gtc gtt tat tca gat cgc agt gtt taa taa cta att taa aca gat gag atg ata tat cca cat caa aga atg cga gat aca tat gac aga cat tat tga cag aaa tgt gac ctt cat tat cac cgt cgt cca taa atg cgt tag gta cgt acc aca tac tat cgt taa cga tgc gca caa tct cgt cca ttt tca tcc atc ttc ata atg att tac ttt ttc ata att aga gaa aaa gat caa ggt ata aaa att aga agt gtt aga cta ta- --- --- --- --- --- --- --- --- --- --- -aa cta act tat aa- --- --- --- --- --- --- --- --- --- --- --- --- --- --- --- --- --- --- --- --- --- --- --- --- --- --- --- --- --- --- --- --- --- --- --- --- --- --- --- --- --- --- --- --- --- --- --- --- --- --- --- --- --- --- --- --- --- --- --- --- --- --- --- --- --- --- --- --- --- --- --- --- --- --- --- --- --- --- --- --- --- --- --- --- --- --- -aa cta act tat gac tta act aac tta tga cta tta act cat ttc aag aaa ggt ggg tgg aga gaa ctc tat atg aca gct tgt gaa aca att aga tcc cta att tct aat gga agt ttt gat agg aga ttg tca tca gtt gat aca ttg ttt att atc tca tct att aga gca cgt ctg ttt aga gct tta gtg acc tgc tcg gtt act tct gtg taa atc ttg aat cct tta gtg ata cac tgt gtc aaa act gga tgt tta gaa tac cta tgt aga ata tgg gaa gca tgc ttg ttt ttg tct cta tta tag att aac tca tac atg gtt gta tta tga att ttc atc tgc cta atg tac tcc aat tct tgt tta caa tca att ata taa tca aag agt gat gat gca tac aca tta caa agt gaa taa tct acc atc ata aaa tac ttg ata cag agc ttt atc aca tca tgg ttt tca att gta tta tta agt ata gct aat ttt ata cag tca ata gac aat ggt tct cta agc aat att tct aat att tta aga tgt gtt tcc cta cgg gcg atg aca gat ccc cta tcc acg gcc acg tca aga cat gta tat cca tta ctc att act gcg ttg aca ttt gct cca ttt tct aat agc cat gat act aaa tct ata taa cct gca tag ata gcg cga taa agc aag gtc ctt cca cca gca tct agt tga ttg ata tct tca ata tat ggg ata caa agc tta taa att tct aat act gtg ggt tca tct aca agg aat ccc cta gta tac tga att att tta tat aga tct aat tta aca tca ttt tca tct ggg ata cca caa ttc aaa ata aac tca aca aca cta ctt tcc ttt tta cat att ccc cta aaa tag gca ttc aag cat tct att tta tat att aca gcc cca tga tct acc ata aaa tca aca atg tct att tct aca tat gca tta gat aga tag taa agt aag aga tct tgt aca gaa tta caa ttc tta ata att ata gag aaa ata tct tcc ata taa ttc ttt gac act aat gca gat ata ata tct tta tat gta ata tat gca aac agt cta tct act ata tac tga tca ata tta tct cta tga atc cta aaa taa tca tac aga aca tct aca gga tca caa att ggt tca agg aga aat cta tca aat att ttc ctg tca aca act ggt tct aga aca taa cag tca aca cct aat cca tgt ttt tta tag tca tct acc aaa gat aat gac caa aga tcg agg tcg tcg tga aac tgc tca tcg aca gcc atg aaa tct gcc gac tcc atg gcg cga atc gca ctg tct tat tcg cca ttg att ttc a-t ttt tta taa tta tgt aca tgt ttt cct tct att ctc aag agt cta caa aaa tat a-t ttt ttt tcg ata tct aag tac taa gtt ttt tta ctg ttt ttg tta ctg tct tcc att ctt cta act aaa gat ctg aga taa att ata caa tct tcg cta tcg aac cat ttt tgt agt cta aag cct gaa gta att aac caa ctg ttt tta tta gtg gct ttt ttc gat cta tcc tcg tc- --- --c tca tca tcc tta ta- --- --- --- --- --- --- tta tta tca tta tta tca tag tct att aaa cac aaa tca tct --- --- --- --- --- --- --- --- --- --- --- --- --- --- --- --- --- --- --- --- --- --- --- --- --- --- --a cgt tta taa c-- --- --- --- --- --- --- --- --a aca ttc --- --- --- --- -tc att att aat tag ttc tgt agt aat atc ttt aat aat ttg gct ata --- --- -ca tct gtt --- --- --- --- --- --- --- --- --- --- --- --- --- --- --- --- --- --- --c aat act --- --- --- --- --- --- --- --a tct att gat gat ttc ttt t-- --- --- --- --- --- --- --- --- --- --- --t taa gac t-- --- --- --- --- --- --- --- --- --- --- --- --- --- --- --- --- --- --- --- --- --- --- --- --- --- --- --- --- --- --- --- --- --- --- --- --- --- --- --- taa act agt --- --- --- --- --- --- --- --- --- --- --- --- --- --- --- --- --- --- --- --- --- --- --- --- --- --- --- --- --- --- --- --- --- --- --- --- --- --- --- --- --- --- --- --- --- --- --- --- --- --- --- --- --- --- --- --- --- --- --- --- --- --- --- --- --- --- --- --- --- --- --- --- --- --- --- --- --- --- --- --- --- --- --- --- --- --- --- --- --- --- --- --- --- --- --- --- --- --- --- --- --- --- --- --- --- --- --- --- --- --t atg gta atg acg atg aaa --- -tc gag tag ta- --- --a ctt cta ata aag act tga ta- --- --t cat tat cat atg ttt gat cg- --- --- --- --- --- --- --t cat agt taa tag tgt g-- --- --- --- --- --- --- --- --- --- --- --- --- --- --- --- --- --- --- --- -gc taa atg gta ctg tta ata agt tta t-- --- --- --- --- --- --- --- --- --- --- --- --- --- --- --- --- --a gac aat atc ata gta ttt tct ttc cag aat --- --- --- --t aga tta ttt ttt taa ata ctg atc ctc aca att ccg tga tgt agc agt agt tgg t-- --- --- --- --- --- --- --- --- --- --- --- --- --- --- --- --- --- --- --- --- --- --- --- --- --- --- --- -gc atg gtc tat atc gt- --- --- --- --- --- --- --- --- --- --- --- --- --- --- --- --- --- --- --- --- --- --- --- --- --- --- --- --- --- --- --- --- --- --- --- --- --- --- --- --- --- --- --- taa aat gta tca tat ata ata gtt ttc tga cgt gga gta cag aat ttt cga --- --- --- --- --- --- --- --- --- --- --- --- --- --- --- --- --- --- --- --- --- --- --- --- --- --- --- --- --- --- --- --- --- --- --- --- --- --- --- --- --- --- --- --t taa tga gtt cat ggt aag gaa ggg caa atg tct --- gta tat aat ata cat aag tta a-- --- --- --- --- --- --- --- --- --- --- --- --- --t agt ttt tta tca tat tt- --- --- --- --- --- --- --- --t cta ata cca taa taa aaa tta tca tta tg- --- --- -ta taa tca --- --- --- --- --- --- --- --- --- --- --- --t cac tg- --- -tc gct atc att att gcg ttt gtg tag tt- --- --- --- --- ctg ccc ta- --- --- --- --- --- --- --- --t cat cta cat cac tgt ca- --- --- --- --- ctc tca cta tat ctt cta aaa tta caa --- -ac aac --- --- --- --- --- --- --- --- --- --- --- --- --- --- --- --- --- --- --- --- --- --- --- --- --- --- --- --- --- --- --- --- --- --- --- --- tgg ata ttc gat --- --- --- --- --- --- --- --- --- --- --- --- --- --- --- --- --- --- --- --- --- --- --- --- --- --- --a aca gca ttt gtg t-- --- --- --- --- --- --- --- --- --- --- --- --- --- --- --- --- -

>HQ857562.1_Monkeypox_virus_strain_V79-I-005_complete_genome

--- --- --- --- --- --- --- --- --- --- --- --- --- --- --- --- --- --- --- att ata tac ata att tta taa tt- --a att taa ttt tac tat ttt att tag tgt cta gaa aaa aat gtg tga ccc acg acc gta gga aac tct aga ggg taa gaa aaa tca atc gtt tat aga gac cat cag aaa gag gtt taa tat ttt tgt gag acc cat cga a-- --- --- --- --- --- --- --- --- --- --- --- --- --- --- --- --- --- --- --- --- --- --- --- --- --- --- --- --- --- --- --- --- --- --- --- --- --- --- --- --- --- --- --- --- --- --- --- --- --- --- --- --- --- --- --- --- --- --- --- --- --- --- --- --- --- --- --- --- --- --- --- --- --- --- --- --- --- --- --- --- --- --- --- --- --- --- --- --- --- --- --- --- --- --- --- --- --- --- --- --- --- --- --- --- --- --- --- --- --- --- --- --- --- --- --- --- --- --- --- --- --- --- --- --- --- --- --- --- --- --- --- --- --- --- --- --- --- --- --- --- --- --- --- --- --- --- --- --- --- --- --- --- --- --- --- --- --- --- --- --- --- --- --- --- --- --- --- --- --- --- --- --- --- --- --- --- --- --- --- --- --- --- --- --- --- --- --- --- --- --- --- --- --- --- --- --- --- --- --- --- --- --- --- --- --- --- --- --- --- --- --- --- --- --- --- --- --- --- --- --- --- --- --- --- --- --- --- --- --- --- --- --- --- --- --- --- --- --- --- --- --- --- --- --- --- --- --- --- --- --- --- --- --- --- --- --- --- --- --- --- --- --- --- --- --- --- --- --- --- --- --- --- --- --- --- --- --- --- --- -ga gag aaa gga taa aaa ctt ttt acg act cca tca gaa aga ggt tta ata ttt ttg tga gac cca tcg aag aga gaa aga gat ggt tag tca aga tat ttt tct tag tac aaa agt caa tgt ttt aaa ata tat gga cga gaa tta att tgt ctg tat aaa aac ttg tgt gaa att atg tac tag aga aaa aac gtg agc agt gtc ccc tac atg gat ttt aca gat cat tta tat tcc aaa aat att aac tat ata cgt tta tta tat gat gtt aac gtg taa att ata aac att att tta tga tgc aat tgt ctg aca acc atc att ggc ata agg ata ttg ata agc tct acg aga ata tat tgt tgg acg tta tcg ttt acg aaa tag ttg aga cat cag aaa gag gtt taa tat ttt tgt gag a-c cat cga aga gag aaa gag aat aaa aat a-- --- --- --- ttt ttt tgt aaa ac- ttt tta tga gac aag aga gaa aga gaa tac gaa tag tga tca tat cgt atc aca tat tga aac aga aag aag aag taa cga gag gta act ttt tgt gaa tgt agt taa aca ttt ttg ttt tgc aaa ccg gaa tat agt acc cgg tac act ttt tta att cgt ggt gcg gtg tct gaa tcg ttc gat taa ccc aac tca tcc att ttc aga tga ata gag tta tcg att cag aca cat gct ttg agt ttt gtt gaa tcg atg agt gaa gta tca tcg gtt gca cct tca gat gcc gat ccg tcg aca tac ttg aat cca tcc ttg act tca agt tca gat gat tcc tca cac atg tct ccg ata cgt acg cta aac tct agg ttc tta aca cat ttt gta tca acg atc gtt gaa ccg atg ata tct ttg taa ctc act ttc tta tgt gag atg tta gac cca agt act gga tgg gtc ttg atg tcg ctg tct ttc tct tct tcg cta cat ctg atg tcg ata gac atc tca cag tct ttg atc ata gcc aga gct tct tca cgc gtg atc gcg gga gag tcc tta cct tgt cct ggt gac acg ctg gac aat cta gta ttc aca gtg ttt cca tca gag gat tcg gag atg gat gaa atc ttt ggg cat ttg gtg aat cca aag ttc atg tta aga ccc gcg ccg acg ata ttg taa taa gtg gtg gga tct cct ttt aca act --- --- --- --- --- --- tct tcg gat acc tca tca tct tcg gtc tct gta act tcc gtt acg gat tga caa atc tta tca ttg gtc ggt gtt tgg tct tgc ttt gtg act ttg ata ata aca tcg att ccc ata tga tgt ttg ttt tct tct tca gta cac gag gat gag gat tgt tga aga cta gta ggc ata gca gct gcc act agg cac atg cat gcc agg aca ata tat tgt ttc atg att gct att gat tga tta ctg ttc tag atg att cta ctt tct tac cat ata ata aat tag aat ata ttt tct act ttt acg aga aat taa tta ttg tat tta tta ttt atg ggt aaa aaa aaa ctt act ata agt ggg tgg gat tct ggg aat tag tga tca gtt tat gta tat cgc aac tag cgg gca tat ggc tat tga cat cga gaa cat tac cca tat gat aag aga ttg tat ccg ttt cgt agt ctt gag tat tgg tat tac tat ata gta tgt aga tgt cga cgc tag ata gac agt cgc cca cta gag tta ccg tct ctg aat gcg gca tga tag tat cat tct ttg ttt tcg tta act gtt tgg aag atg aat ctt tgt tgt tac att taa tct cga aat tca gag tac ata tct ttg aag tat tct gat atc tat ttt ctc ctg taa aga atc ctg aag ttg cta cat tat taa gga cag aga agt att ctg cac gaa aga cgg gat cac aat ctt tat gat tca tgg taa tag tta gtt ccg acg ttg aga tgg att cgc tga gac cgg tag tgg tcg tcc gag tac acg atg tgt cgt tga ctg gat aca ggt taa ttt cca cat cga tat agt taa atg tat tgc tgg tt- --a cgg gtt cgc att tat ctg tgg aag aga cgg tgt gag aat atg ttc cgg gac cac acg gag aac aga tga cgt ctc cgg tag acg tgt atc cgg ata ctc cgt atc cta ttc cac act ttg ttt tag aaa tac atg ttc tac acc ctg atg atc ctt tga gaa gac aat aat atc ctg gag agc att cac aga ttc tat tgt gag tcg tgt tac acg atc gcg tct cta cct gat tac tat cac atc ttc cgt tac aac tta gac aag cct gta aat gat tat tgt gag atg taa agg tat ccg aac cac acg gcg tac att gtg tat tag tct tgc tat cac ata atc tgg aag cgt aag ttc ccg gag gac acg ata gac aac ata gat tac ggc ttc tgt att cgt tgt ctt tac att ttc cat tgg atg gtg cat gtg gtg cta tat ctc ttc cgt tta tta tta tac atg aga gaa aca ata tat acg agt ata ata cgg acc tca tga ttt aat aat tgt agt aat cgt cgt ctt gtt act gtt tgt ttc cta ctt ctc caa tca tat aga tta ttt ttt aaa tat ttt ctt tct atc atg gat aat att tgt aat ggt tct ttc cgt aca aca tac tgt tta gat ggt agt cgc tta gct tgg tta tga tat tgc gca taa ttt ccg gag gca aat acg ata gtc tag att gac tat cga tgg tag act cta att tat tga gtg ctt tgt cga cga gtt tac ttt tat gct cca tcg ata gat gac act gtt cta tga gat cgt cgt aca tgg gaa atg aaa tgc gtt tgt ctg aat gta tgg ctt cga gat agg tgt gat acc gga tgt ctt ctg ttc tca ata ccg tat aca agt tgg tgt ctg aga ttc gaa tct ctt tga gga gac tta tgt cac gac tac att ttt cga tga tgg aat cta tct tat cga atg ata tat ttt tca taa ata cac ttt tat agt cct cgt tta aac aga att tag tat gta gtt ccg caa atg act cgt ccc tta ata ggc agt agg cta tta tct tct tta cgt agt gat cgt cgt agg gag aga act ccg aca tct tgt aga aca acg att taa tca tag gta gag ata ctt tca gtc tgt ggt gga tga tgt cat taa cga cat ccg cct tgt ata tga tgt ttc tgt ttt caa aca cca agt cga ata ctg tct tat cgt ctt tag tcg gaa ggt tga tgt cgt atc cga tgt ata cga ggt atg agg caa cat tgt tat tgc aat tct gga agg cgg tat gaa gag gag tca ttg tat tat agt att cgt ctt tct gaa tgt cga atc tat cta gta gat acc gta gta tat tga gag agc gac ttc cat atc ctt gat tat gtt tta tga ata gat aaa gta gat gtt gtc ctt ctt cct ttt gta att tcc cgt att ttt gtt cgt gcc aat tga gta aca tta tga gaa tat gac ctg ttg cac aat cgt tct tta tgt att cca tga tgg gtg tac aat caa gat tat tac gta tcc tcg tat cgg ctc ctc gag ata aaa gag cat aca cca cac gag gac tat gtt tgg tat act gtt gaa ggt aag tgt gta acg gcg tat ttc cga ttt tcg taa ccg cgt taa tgt ttg ctc cat gat cta tta tcg cgt aga tga atc gct tct cag ctc gca tct tag tgt gac tct ttg act tgt aat aat tgc ttt cgt gga aca cgg ata tgt gtt tac agt agt aat gaa gag aag tga gtc cat cct cat cga cgc aat tag ggt cag atc ctt tag tca ata att tgt aca gaa cgt aat agt tta agc tcc cat tga att tat atc taa gat aac aca gca ata gat cgg atg att tac taa agt cat caa tgg ggt ccg tta gta tat caa aga tct tgt tat cga ttg ata gtg aat gaa tca gat agt ggt gta gag gaa tat gtc ctt ttt tat cct tgc tat caa agt tac gca tgc cgt ggc gta aca ata tct tta ata cag atg gat taa atc gtg tat tca tcg tat agc aat gta atg gag agt tac cac att tta gtc gtt tat tca gat cgc agt gtt taa taa cta att taa aca gat gag atg ata tat cca cat caa aga atg cga gat aca tat gac aga cat tat tga cag aaa tgt gac ctt cat tat cac cgt cgt cca taa atg cgt tag gta cgt acc aca tac tat cgt taa cga tgc gca caa tct cgt cca ttt tca tcc atc ttc ata atg att tac ttt ttc ata att aga gaa aaa gat caa ggt ata aaa att aga agt gtt aga cta ta- --- --- --- --- --- --- --- --- --- --- -aa cta act tat aa- --- --- --- --- --- --- --- --- --- --- --- --- --- --- --- --- --- --- --- --- --- --- --- --- --- --- --- --- --- --- --- --- --- --- --- --- --- --- --- --- --- --- --- --- --- --- --- --- --- --- --- --- --- --- --- --- --- --- --- --- --- --- --- --- --- --- --- --- --- --- --- --- --- --- --- --- --- --- --- --- --- --- --- --- --- --- -aa cta act tat gac tta act aac tta tga cta tta act cat ttc aag aaa ggt ggg tgg aga gaa ctc tat atg aca gct tgt gaa aca att aga tcc cta att tct aat gga agt ttt gat agg aga ttg tca tca gtt gat aca ttg ttt att atc tca tct att aga gca cgt ctg ttt aga gct tta gtg acc tgc tcg gtt act tct gtg taa atc ttg aat cct tta gtg ata cac tgt gtc aaa act gga tgt tta gaa tac cta tgt aga ata tgg gaa gca tgc ttg ttt ttg tct cta tta tag att aac tca tac atg gtt gta tta tga att ttc atc tgc cta atg tac tcc aat tct tgt tta caa tca att ata taa tca aag agt gat gat gca tac aca tta caa agt gaa taa tct acc atc ata aaa tac ttg ata cag agc ttt atc aca tca tgg ttt tca att gta tta tta agt ata gct aat ttt ata cag tca ata gac aat ggt tct cta agc aat att tct aat att tta aga tgt gct tcc cta cgg gcg atg aca gat ccc cta tcc acg gcc acg tca aga cat gta tat cca tta ctc att act gcg ttg aca ttt gct cca ttt tct aat agc cat gat act aaa tct ata taa cct gca tag ata gcg cga taa agc aag gtc ctt cca cca gca tct agt tga ttg ata tct tca ata tat ggg ata caa agc tta taa att tct aat act gtg ggt tca tct aca agg aat ccc cta gta tac tga att att tta tat aga tct aat tta aca tca ttt tca tct ggg ata cca caa ttc aaa ata aac tca aca aca cta ctt tcc ttt tta cat att ccc cta aaa tag gca ttc aag cat tct att tta tat att aca gcc cca tga tct acc ata aaa tca aca atg tct att tct aca tat gca tta gat aga tag taa agt aag aga tct tgt aca gaa tta caa ttc tta ata att ata gag aaa ata tct tcc ata taa ttc ttt gac act aat gca gat ata ata tct tta tat gta ata tat gca aac agt cta tct act ata tac tga tca ata tta tct cta tga atc cta aaa taa tca tac aga aca tct aca gga tca caa att ggt tca agg aga aat cta tca aat att ttc ctg tca aca act ggt tct aga aca taa cag tca aca cct aat cca tgt ttt tta tag tca tct acc aaa gat aat gac caa aga tcg agg tcg tcg tga aac tgc tca tcg aca gcc atg aaa tct acc gac tcc atg gcg cga atc gca ctg tct tat tcg cca ttg att ttc a-t ttt tta taa tta tgt aca tgt ttt cct tct att ctc aag agt cta caa aaa tat a-t ttt ttt tcg ata tct aag tac taa gtt ttt tta ctg ttt ttg tta ctg tct tcc att ctt cta act aaa gat ctg aga taa att ata caa tct tcg cta tcg aac cat ttt tgt agt cta aag cct gaa gta att aac caa ctg ttt tta tta gtg gct ttt ttc gat cta tcc tcg tc- --- --c tca tca tcc tta ta- --- --- --- --- --- --- tta tta tca tta tta tca tag tct att aaa cac aaa tca tct --- --- --- --- --- --- --- --- --- --- --- --- --- --- --- --- --- --- --- --- --- --- --- --- --- --- --a cgt tta taa c-- --- --- --- --- --- --- --- --a aca ttc --- --- --- --- -tc att att aat tag ttc tgt agt aat atc ttt aat aat ttg gct ata --- --- -ca tct gtt --- --- --- --- --- --- --- --- --- --- --- --- --- --- --- --- --- --- --c aat act --- --- --- --- --- --- --- --a tct att gat gat ttc ttt t-- --- --- --- --- --- --- --- --- --- --- --t taa gac t-- --- --- --- --- --- --- --- --- --- --- --- --- --- --- --- --- --- --- --- --- --- --- --- --- --- --- --- --- --- --- --- --- --- --- --- --- --- --- --- taa act agt --- --- --- --- --- --- --- --- --- --- --- --- --- --- --- --- --- --- --- --- --- --- --- --- --- --- --- --- --- --- --- --- --- --- --- --- --- --- --- --- --- --- --- --- --- --- --- --- --- --- --- --- --- --- --- --- --- --- --- --- --- --- --- --- --- --- --- --- --- --- --- --- --- --- --- --- --- --- --- --- --- --- --- --- --- --- --- --- --- --- --- --- --- --- --- --- --- --- --- --- --- --- --- --- --- --- --- --- --- --t atg gta atg acg atg aaa --- -tc gag tag ta- --- --a ctt cta ata aat act tga ta- --- --t cat tat cat atg ttt gat cg- --- --- --- --- --- --- --t cat agt taa tag tgt g-- --- --- --- --- --- --- --- --- --- --- --- --- --- --- --- --- --- --- --- -gc taa atg gta ctg tta ata agt tta t-- --- --- --- --- --- --- --- --- --- --- --- --- --- --- --- --- --a gac aat atc ata gta ttt tct ttc cag aat --- --- --- --t aga tta ttt ttt taa ata ctg atc ctc aca att ccg tga tgt agc agt agt tgg t-- --- --- --- --- --- --- --- --- --- --- --- --- --- --- --- --- --- --- --- --- --- --- --- --- --- --- --- -gc atg gtc tat atc gt- --- --- --- --- --- --- --- --- --- --- --- --- --- --- --- --- --- --- --- --- --- --- --- --- --- --- --- --- --- --- --- --- --- --- --- --- --- --- --- --- --- --- --- taa aat gta tca tat ata ata gtt ttc tga cgt gga gta cag aat ttt cga --- --- --- --- --- --- --- --- --- --- --- --- --- --- --- --- --- --- --- --- --- --- --- --- --- --- --- --- --- --- --- --- --- --- --- --- --- --- --- --- --- --- --- --t taa tga gtt cat ggt aag gaa ggg caa atg tct --- gta tat aat ata cat aag tta a-- --- --- --- --- --- --- --- --- --- --- --- --- --t agt ttt tta tca tat tt- --- --- --- --- --- --- --- --t cta ata cca taa taa aaa tta tca tta tg- --- --- -ta taa tca --- --- --- --- --- --- --- --- --- --- --- --t cac tg- --- -tc gct atc att att gcg ttt gtg tag tt- --- --- --- --- ctg ccc ta- --- --- --- --- --- --- --- --t cat cta cat cac tgt ca- --- --- --- --- ctc tca cta tat ctt cta aaa tta caa --- -ac aac --- --- --- --- --- --- --- --- --- --- --- --- --- --- --- --- --- --- --- --- --- --- --- --- --- --- --- --- --- --- --- --- --- --- --- --- tgg ata ttc gat --- --- --- --- --- --- --- --- --- --- --- --- --- --- --- --- --- --- --- --- --- --- --- --- --- --- --a aca gca ttt gtg t-- --- --- --- --- --- --- --- --- --- --- --- --- --- --- --- --- -

>HQ857563.1_Monkeypox_virus_strain_D14L_knockout_complete_genome

--- --- --- --- --- --- --- --- --- --- --- --- --- --- --- --- --- --- --- --- --- --- --- --- --- --- --- --- --- --a ttt tac tat ttt att tag tgt cta gaa aaa aat gtg tga ccc acg acc gta gga aac tct aga ggg taa gaa aaa tca atc gtt tat aga gac cat cag aaa gag gtt taa tat ttt tgt gag acc cat cga a-- --- --- --- --- --- --- --- --- --- --- --- --- --- --- --- --- --- --- --- --- --- --- --- --- --- --- --- --- --- --- --- --- --- --- --- --- --- --- --- --- --- --- --- --- --- --- --- --- --- --- --- --- --- --- --- --- --- --- --- --- --- --- --- --- --- --- --- --- --- --- --- --- --- --- --- --- --- --- --- --- --- --- --- --- --- --- --- --- --- --- --- --- --- --- --- --- --- --- --- --- --- --- --- --- --- --- --- --- --- --- --- --- --- --- --- --- --- --- --- --- --- --- --- --- --- --- --- --- --- --- --- --- --- --- --- --- --- --- --- --- --- --- --- --- --- --- --- --- --- --- --- --- --- --- --- --- --- --- --- --- --- --- --- --- --- --- --- --- --- --- --- --- --- --- --- --- --- --- --- --- --- --- --- --- --- --- --- --- --- --- --- --- --- --- --- --- --- --- --- --- --- --- --- --- --- --- --- --- --- --- --- --- --- --- --- --- --- --- --- --- --- --- --- --- --- --- --- --- --- --- --- --- --- --- --- --- --- --- --- --- --- --- --- --- --- --- --- --- --- --- --- --- --- --- --- --- --- --- --- --- --- --- --- --- --- --- --- --- --- --- --- --- --- --- --- --- --- --- --- -ga gag aaa gga taa aaa ctt ttt acg act cca tca gaa aga ggt tta ata ttt ttg tga gac cca tcg aag aga gaa aga gat ggt tag tca aga tat ttt tct tag tac aaa agt caa tgt ttt aaa ata tat gga cga gaa tta att tgt ctg tat aaa aac ttg tgt gaa att atg tac tag aga aaa aac gtg agc agt gtc ccc tac atg gat ttt aca gat cat tta tat tcc aaa aat att aac tat ata cgt tta tta tat gat gtt aac gtg taa att ata aac att att tta tga tgc aat tgt ctg aca acc atc att ggc ata agg ata ttg ata agc tct acg aga ata tat tgt tgg acg tta tcg ttt acg aaa tag ttg aga cat cag aaa gag gtt taa tat ttt tgt gag a-c cat cga aga gag aaa gag aat aaa aat a-- --- --- --- ttt ttt tgt aaa ac- ttt tta tga gac aag aga gaa aga gaa tac gaa tag tga tca tat cgt atc aca tat tga aac aga aag aag aag taa cga gag gta act ttt tgt gaa tgt agt taa aca ttt ttg ttt tgc aaa ccg gaa tat agt acc cgg tac act ttt tta att cgt ggt gcg gtg tct gaa tcg ttc gat taa ccc aac tca tcc att ttc aga tga ata gag tta tcg att cag aca cat gct ttg agt ttt gtt gaa tcg atg agt gaa gta tca tcg gtt gca cct tca gat gcc gat ccg tcg aca tac ttg aat cca tcc ttg act tca agt tca gat gat tcc tca cac atg tct ccg ata cgt acg cta aac tct agg ttc tta aca cat ttt gta tca acg atc gtt gaa ccg atg ata tct ttg taa ctc act ttc tta tgt gag atg tta gac cca agt act gga tgg gtc ttg atg tcg ctg tct ttc tct tct tcg cta cat ctg atg tcg ata gac atc tca cag tct ttg atc ata gcc aga gct tct tca cgc gtg atc gcg gga gag tcc tta cct tgt cct ggt gac acg ctg gac aat cta gta ttc aca gtg ttt cca tca gag gat tcg gag atg gat gaa atc ttt ggg cat ttg gtg aat cca aag ttc atg tta aga ccc gcg ccg acg ata ttg taa taa gtg gtg gga tct cct ttt aca act --- --- --- --- --- --- tct tcg gat acc tca tca tct tcg gtc tct gta act tcc gtt acg gat tga caa atc tta tca ttg gtc ggt gtt tgg tct tgc ttt gtg act ttg ata ata aca tcg att ccc ata tga tgt ttg ttt tct tct tca gta cac gag gat gag gat tgt tga aga cta gta ggc ata gca gct gcc act agg cac atg cat gcc agg aca ata tat tgt ttc atg att gct att gat tga tta ctg ttc tag atg att cta ctt tct tac cat ata ata aat tag aat ata ttt tct act ttt acg aga aat taa tta ttg tat tta tta ttt atg ggt aaa aaa aaa ctt act ata agt ggg tgg gat tct ggg aat tag tga tca gtt tat gta tat cgc aac tas cgg gca tat ggc tat yga cat cga gaa cat tac cca yat gat aag aga ttg tat cmg ttt cgt agt ctt gag tat tgg tat tac tat ata gta tgt aga tgt cga cgc tag ata gac agt cgc cca cta gag tta ccg tct ctg aat gcg gca tga tag tat cat tct ttg ttt tcg tta act gtt tgg aag atg aat ctt tgt tgt tac att taa tct cga aat tca gag tac ata tct ttg aag tat tct gat atc tat ttt ctc ctg taa aga atc ctg aag ttg cta cat tat taa gga cag aga agt att ctg cac gaa aga cgg gat cac aat ctt tat gat tca tgg taa tag tta gtt ccg acg ttg aga tgg att cgc tga gac cgg tag tgg tcg tcc gag tac acg atg tgt cgt tga ctg gat aca ggt taa ttt cca cat cga tat agt taa atg tat tgc tgg tt- --a cgg gtt cgc att tat ctg tgg aag aga cgg tgt gag aat atg ttc cgg gac cac acg gag aac aga tga cgt ctc cgg tag acg tgt atc cgg ata ctc cgt atc cta ttc cac act ttg ttt tag aaa tac atg ttc tac acc ctg atg atc ctt tga gaa gac aat aat atc ctg gag agc att cac aga ttc tat tgt gag tcg tgt tac acg atc gcg tct cta cct gat tac tat cac atc ttc cgt tac aac tta gac aag cct gta aat gat tat tgt gag atg taa agg tat ccg aac cac acg gcg tac att gtg tat tag tct tgc tat cac ata atc tgg aag cgt aag ttc ccg gag gac acg ata gac aac ata gat tac ggc ttc tgt att cgt tgt ctt tac att ttc cat tgg atg gtg cat gtg gtg cta tat ctc ttc cgt tta tta tta tac atg aga gaa aca ata tat acg agt ata ata cgg acc tca tga ttt aat aat tgt agt aat cgt cgt ctt gtt act gtt tgt ttc cta ctt ctc caa tca tat aga tta ttt ttt aaa tat ttt ctt tct atc atg gat aat att tgt aat ggt tct ttc cgt aca aca tac tgt tta gat ggt agt cgc tta gct tgg tta tga tat tgc gca taa ttt ccg gag gca aat acg ata gtc tag att gac tat cga tgg tag act cta att tat tga gtg ctt tgt cga cga gtt tac ttt tat gct cca tcg ata gat gac act gtt cta tga gat cgt cgt aca tgg gaa atg aaa tgc gtt tgt ctg aat gta tgg ctt cga gat agg tgt gat acc gga tgt ctt ctg ttc tca ata ccg tat aca agt tgg tgt ctg aga ttc gaa tct ctt tga gga gac tta tgt cac gac tac att ttt cga tga tgg aat cta tct tat cga atg ata tat ttt tca taa ata cac ttt tat agt cct cgt tta aac aga att tag tat gta gtt ccg caa atg act cgt ccc tta ata ggc agt agg cta tta tct tct tta cgt agt gat cgt cgt agg gag aga act ccg aca tct tgt aga aca acg att taa tca tag gta gag ata ctt tca gtc tgt ggt gga tga tgt cat taa cga cat ccg cct tgt ata tga tgt ttc tgt ttt caa aca cca agt cga ata ctg tct tat cgt ctt tag tcg gaa ggt tga tgt cgt atc cga tgt ata cga ggt atg agg caa cat tgt tat tgc aat tct gga agg cgg tat gaa gag gag tca ttg tat tat agt att cgt ctt tct gaa tgt cga atc tat cta gta gat acc gta gta tat tga gag agc gac ttc cat atc ctt gat tat gtt tta tga ata gat aaa gta gat gtt gtc ctt ctt cct ttt gta att tcc cgt att ttt gtt cgt gcc aat tga gta aca tta tga gaa tat gac ctg ttg cac aat cgt tct tta tgt att cca tga tgg gtg tac aat caa gat tat tac gta tcc tcg tat cgg ctc ctc gag ata aaa gag cat aca cca cac gag gac tat gtt tgg tat act gtt gaa ggt aag tgt gta acg gcg tat ttc cga ttt tcg taa ccg cgt taa tgt ttg ctc cat gat cta tta tcg cgt aga tga atc gct tct cag ctc gca tct tag tgt gac tct ttg act tgt aat aat tgc ttt cgt gga aca cgg ata tgt gtt tac agt agt aat gaa gag aag tga gtc cat cct cat cga cgc aat tag ggt cag atc ctt tag tca ata att tgt aca gaa cgt aat agt tta agc tcc cat tga att tat atc taa gat aac aca gca ata gat cgg atg att tac taa agt cat caa tgg ggt ccg tta gta tat caa aga tct tgt tat cga ttg ata gtg aat gaa tca gat agt ggt gta gag gaa tat gtc ctt ttt tat cct tgc tat caa agt tac gca tgc cgt ggc gta aca ata tct tta ata cag atg gat taa atc gtg tat tca tcg tat agc aat gta atg gag agt tac cac att tta gtc gtt tat tca gat cgc agt gtt taa taa cta att taa aca gat gag atg ata tat cca cat caa aga atg cga gat aca tat gac aga cat tat tga cag aaa tgt gac ctt cat tat cac cgt cgt cca taa atg cgt tag gta cgt acc aca tac tat cgt taa cga tgc gca caa tct cgt cca ttt tca tcc atc ttc ata atg att tac ttt ttc ata att aga gaa aaa gat caa ggt ata aaa att aga agt gtt aga cta ta- --- --- --- --- --- --- --- --- --- --- -aa cta act tat aa- --- --- --- --- --- --- --- --- --- --- --- --- --- --- --- --- --- --- --- --- --- --- --- --- --- --- --- --- --- --- --- --- --- --- --- --- --- --- --- --- --- --- --- --- --- --- --- --- --- --- --- --- --- --- --- --- --- --- --- --- --- --- --- --- --- --- --- --- --- --- --- --- --- --- --- --- --- --- --- --- --- --- --- --- --- --- -aa cta act tat gac tta act aac tta tga cta tta act cat ttc aag aaa ggt ggg tgg aga gaa ctc tat atg aca gct tgt gaa aca att aga tcc cta att tct aat gga agt ttt gat agg aga ttg tca tca gtt gat aca ttg ttt att atc tca tct att aga gca cgt ctg ttt aga gct tta gtg acc tgc tcg gtt act tct gtg taa atc ttg aat cct tta gtg ata cac tgt gtc aaa act gga tgt tta gaa tac cta tgt aga ata tgg gaa gca tgc ttg ttt ttg tct cta tta tag att aac tca tac atg gtt gta tta tga att ttc atc tgc cta atg tac tcc aat tct tgt tta caa tca att ata taa tca aag agt gat gat gca tac aca tta caa agt gaa taa tct acc atc ata aaa tac ttg ata cag agc ttt atc aca tca tgg ttt tca att gta tta tta agt ata gct aat ttt ata cag tca ata gac aat ggt tct cta agc aat att tct aat att tta aga tgt gct tcc cta cgg gcg atg aca gat ccc cta tcc acg gcc acg tca aga cat gta tat cca tta ctc att act gcg ttg aca ttt gct cca ttt tct aat agc cat gat act aaa tct ata taa cct gca tag ata gcg cga taa agc aag gtc ctt cca cca gca tct agt tga ttg ata tct tca ata tat ggg ata caa agc tta taa att tct aat act gtg ggt tca tct aca agg aat ccc cta gta tac tga att att tta tat aga tct aat tta aca tca ttt tca tct ggg ata cca caa ttc aaa ata aac tca aca aca cta ctt tcc ttt tta cat att ccc cta aaa tag gca ttc aag cat tct att tta tat att aca gcc cca tga tct acc ata aaa tca aca atg tct att tct aca tat gca tta gat aga tag taa agt aag aga tct tgt aca gaa tta caa ttc tta ata att ata gag aaa ata tct tcc ata taa ttc ttt gac act aat gca gat ata ata tct tta tat gta ata tat gca aac agt cta tct act ata tac tga tca ata tta tct cta tga atc cta aaa taa tca tac aga aca tct aca gga tca caa att ggt tca agg aga aat cta tca aat att ttc ctg tca aca act ggt tct aga aca taa cag tca aca cct aat cca tgt ttt tta tag tca tct acc aaa gat aat gac caa aga tcg agg tcg tcg tga aac tgc tca tcg aca gcc atg aaa tct acc gac tcc atg gcg cga atc gca ctg tct tat tcg cca ttg att ttc a-t ttt tta taa tta tgt aca tgt ttt cct tct att ctc aag agt cta caa aaa tat a-t ttt ttt tcg ata tct aag tac taa gtt ttt tta ctg ttt ttg tta ctg tct tcc att ctt cta act aaa gat ctg aga taa att ata caa tct tcg cta tcg aac cat ttt tgt agt cta aag cct gaa gta att aac caa ctg ttt tta tta gtg gct ttt ttc gat cta tcc tcg tc- --- --c tca tca tcc tta ta- --- --- --- --- --- --- tta tta tca tta tta tca tag tct att aaa cac aaa tca tct --- --- --- --- --- --- --- --- --- --- --- --- --- --- --- --- --- --- --- --- --- --- --- --- --- --- --a cgt tta taa c-- --- --- --- --- --- --- --- --a aca ttc --- --- --- --- -tc att att aat tag ttc tgt agt aat atc ttt aat aat ttg gct ata --- --- -ca tct gtt --- --- --- --- --- --- --- --- --- --- --- --- --- --- --- --- --- --- --c aat act --- --- --- --- --- --- --- --a tct att gat gat ttc ttt t-- --- --- --- --- --- --- --- --- --- --- --t taa gac t-- --- --- --- --- --- --- --- --- --- --- --- --- --- --- --- --- --- --- --- --- --- --- --- --- --- --- --- --- --- --- --- --- --- --- --- --- --- --- --- taa act agt --- --- --- --- --- --- --- --- --- --- --- --- --- --- --- --- --- --- --- --- --- --- --- --- --- --- --- --- --- --- --- --- --- --- --- --- --- --- --- --- --- --- --- --- --- --- --- --- --- --- --- --- --- --- --- --- --- --- --- --- --- --- --- --- --- --- --- --- --- --- --- --- --- --- --- --- --- --- --- --- --- --- --- --- --- --- --- --- --- --- --- --- --- --- --- --- --- --- --- --- --- --- --- --- --- --- --- --- --- --t atg gta atg acg atg aaa --- -tc gag tag ta- --- --a ctt cta ata aat act tga ta- --- --t cat tat cat atg ttt gat cg- --- --- --- --- --- --- --t cat agt taa tag tgt g-- --- --- --- --- --- --- --- --- --- --- --- --- --- --- --- --- --- --- --- -gc taa atg gta ctg tta ata agt tta t-- --- --- --- --- --- --- --- --- --- --- --- --- --- --- --- --- --a gac aat atc ata gta ttt tct ttc cag aat --- --- --- --t aga tta ttt ttt taa ata ctg atc ctc aca att ccg tga tgt agc agt agt tgg t-- --- --- --- --- --- --- --- --- --- --- --- --- --- --- --- --- --- --- --- --- --- --- --- --- --- --- --- -gc atg gtc tat atc gt- --- --- --- --- --- --- --- --- --- --- --- --- --- --- --- --- --- --- --- --- --- --- --- --- --- --- --- --- --- --- --- --- --- --- --- --- --- --- --- --- --- --- --- taa aat gta tca tat ata ata gtt ttc tga cgt gga gta cag aat ttt cga --- --- --- --- --- --- --- --- --- --- --- --- --- --- --- --- --- --- --- --- --- --- --- --- --- --- --- --- --- --- --- --- --- --- --- --- --- --- --- --- --- --- --- --t taa tga gtt cat ggt aag gaa ggg caa atg tct --- gta tat aat ata cat aag tta a-- --- --- --- --- --- --- --- --- --- --- --- --- --t agt ttt tta tca tat tt- --- --- --- --- --- --- --- --t cta ata cca taa taa aaa tta tca tta tg- --- --- -ta taa tca --- --- --- --- --- --- --- --- --- --- --- --t cac tg- --- -tc gct atc att att gcg ttt gtg tag tt- --- --- --- --- ctg ccc ta- --- --- --- --- --- --- --- --t cat cta cat cac tgt ca- --- --- --- --- ctc tca cta tat ctt cta aaa tta caa --- -ac aac --- --- --- --- --- --- --- --- --- --- --- --- --- --- --- --- --- --- --- --- --- --- --- --- --- --- --- --- --- --- --- --- --- --- --- --- tgg ata ttc gat --- --- --- --- --- --- --- --- --- --- --- --- --- --- --- --- --- --- --- --- --- --- --- --- --- --- --a aca gca ttt gtg t-- --- --- --- --- --- --- --- --- --- --- --- --- --- --- --- --- -
