## Supplementary Data 2 for "Genomic regions insertion and deletion in Monkeypox virus causing multi-country outbreak-2022"

>ON674051.1_Monkeypox_virus_isolate_MPXV_USA_2022_FL001_complete_genome

ttt ttt cga tct atc ctc gtc c-- -t- ctc atc atc ctt ata --- --- --- --- --- --- -tt att atc att att atc ata gtc tat taa aca caa atc atc t-- --- --- --- --- --- --- --- --- --- --- --- --- --- --- --- --- --- --- --- --- --- --- --- --- --- --- acg ttt ata ac- --- --- --- --- --- --- --- --- aac att c-- --- --- --- --t cat tat taa tta gtt ctg tag -aa tat ctt taa taa ttt ggc tat a-- --- --c atc tgt t-- --- --- --- --- --- --- --- --- --- --- --- --- --- --- --- --- --- --- caa tac t-- --- --- --- --- --- --- --- atc tat tga tga ttt ctt tt- --- --- --- --- --- --- --- --- --- --- --- tta aga ct- --- --- --- --- --- --- --- --- --- --- --- --- --- --- --- --- --- --- --- --- --- --- --- --- --- --- --- --- --- --- --- --- --- --- --- --- --- --- --- -ta aac tag t-- --- --- --- --- --- --- --- --- --- --- --- --- --- --- --- --- --- --- --- --- --- --- --- --- --- --- --- --- --- --- --- --- --- --- --- --- --- --- --- --- --- --- --- --- --- --- --- --- --- --- --- --- --- --- --- --- --- --- --- --- --- --- --- --- --- --- --- --- --- --- --- --- --- --- --- --- --- --- --- --- --- --- --- --- --- --- --- --- --- --- --- --- --- --- --- --- --- --- --- --- --- --- --- --- --- --- --- --- --- tat ggt aat gac gat gaa a-- --t cga gta gta --- --- act tct aat aaa gac ttg ata --- --- tca tta tca tat gtt tga tcg --- --- --- --- --- --- --- tca tag tta ata gtg tg- --- --- --- --- --- --- --- --- --- --- --- --- --- --- --- --- --- --- --- --g cta aat ggt act gtt aat aag ttt at- --- --- --- --- --- --- --- --- --- --- --- --- --- --- --- --- --- aga caa tat cat agt att ttc ttt cca gaa t-- --- --- --- tag att att ttt tta aat act gat cct cac aat tcc gtg atg tag cag tag ttg gt- --- --- --- --- --- --- --- --- --- --- --- --- --- --- --- --- --- --- --- --- --- --- --- --- --- --- --- --g cat ggt cta tat cgt --- --- --- --- --- --- --- --- --- --- --- --- --- --- --- --- --- --- --- --- --- --- --- --- --- --- --- --- --- --- --- --- --- --- --- --- --- --- --- --- --- --- --- -ta aaa tgt atc ata tat aat agt ttt ctg acg tgg agt aca gaa ttt tcg a-- --- --- --- --- --- --- --- --- --- --- --- --- --- --- --- --- --- --- --- --- --- --- --- --- --- --- --- --- --- --- --- --- --- --- --- --- --- --- --- --- --- --- --- tta atg agt tca tgg taa gga agg gca aat gcc t-- -gt ata taa tat aca taa gtt aa- --- --- --- --- --- --- --- --- --- --- --- --- --- tag ttt ttt atc ata ttt --- --- --- --- --- --- --- --- tct aat acc ata ata aaa att atc --- --- --- --- --- --- --- --- --- --- --- --- --- --- --- --- --- --- --- --- --- --- --- --- --- -at tat tgc gtt tg- gta gtt --- --- --- --- -ct gcc cta --- --- --- --- --- --- --- --- tca tct ata tca ctg tca ctc tc- --- --- gct ctc act ata tct tct aaa att aca a-- --a caa c-- --- --- --- --- --- --- --- --- --- --- --- --- --- --- --- --- --- --- --- --- --- --- --- --- --- --- --- --- --- --- --- --- --- --- --- -tg gat att cga t-- --- --- --- --- --- --- --- --- --- --- --- --- --- --- --- --- --- --- --- --- --- --- --- --- --- --- aac agc att tgt gt- --- --- --- --- --- --- --- --- --- --- --- --- --- --- --- --- --

>NC_063383.1_Monkeypox_virus_complete_genome

ttt ttt cga tct atc ctc gtc c-- -t- ctc atc atc ctt ata --- --- --- --- --- --- -tt att atc att att atc ata gtc tat taa aca caa atc atc t-- --- --- --- --- --- --- --- --- --- --- --- --- --- --- --- --- --- --- --- --- --- --- --- --- --- --- acg ttt ata ac- --- --- --- --- --- --- --- --- aac att c-- --- --- --- --t cat tat taa tta gtt ctg tag -aa tat ctt taa taa ttt ggc tat a-- --- --c atc tgt t-- --- --- --- --- --- --- --- --- --- --- --- --- --- --- --- --- --- --- caa tac t-- --- --- --- --- --- --- --- atc tat tga tga ttt ctt tt- --- --- --- --- --- --- --- --- --- --- --- tta aga ct- --- --- --- --- --- --- --- --- --- --- --- --- --- --- --- --- --- --- --- --- --- --- --- --- --- --- --- --- --- --- --- --- --- --- --- --- --- --- --- -ta aac tag t-- --- --- --- --- --- --- --- --- --- --- --- --- --- --- --- --- --- --- --- --- --- --- --- --- --- --- --- --- --- --- --- --- --- --- --- --- --- --- --- --- --- --- --- --- --- --- --- --- --- --- --- --- --- --- --- --- --- --- --- --- --- --- --- --- --- --- --- --- --- --- --- --- --- --- --- --- --- --- --- --- --- --- --- --- --- --- --- --- --- --- --- --- --- --- --- --- --- --- --- --- --- --- --- --- --- --- --- --- --- tat ggt aat gac gat gaa a-- --t cga gta gta --- --- act tct aat aaa gac ttg ata --- --- tca tta tca tat gtt tga tcg --- --- --- --- --- --- --- tca tag tta ata gtg tg- --- --- --- --- --- --- --- --- --- --- --- --- --- --- --- --- --- --- --- --g cta aat ggt act gtt aat aag ttt at- --- --- --- --- --- --- --- --- --- --- --- --- --- --- --- --- --- aga caa tat cat agt att ttc ttt cca gaa t-- --- --- --- tag att att ttt tta aat act gat cct cac aat tcc gtg atg tag cag tag ttg gt- --- --- --- --- --- --- --- --- --- --- --- --- --- --- --- --- --- --- --- --- --- --- --- --- --- --- --- --g cat ggt cta tat cgt --- --- --- --- --- --- --- --- --- --- --- --- --- --- --- --- --- --- --- --- --- --- --- --- --- --- --- --- --- --- --- --- --- --- --- --- --- --- --- --- --- --- --- -ta aaa tgt atc ata tat aat agt ttt ctg acg tgg agt aca gaa ttt tcg a-- --- --- --- --- --- --- --- --- --- --- --- --- --- --- --- --- --- --- --- --- --- --- --- --- --- --- --- --- --- --- --- --- --- --- --- --- --- --- --- --- --- --- --- tta atg agt tca tgg taa gga agg gca aat gcc t-- -gt ata taa tat aca taa gtt aa- --- --- --- --- --- --- --- --- --- --- --- --- --- tag ttt ttt atc ata ttt --- --- --- --- --- --- --- --- tct aat acc ata ata aaa att atc --- --- --- --- --- --- --- --- --- --- --- --- --- --- --- --- --- --- --- --- --- --- --- --- --- -at tat tgc gtt tg- gta gtt --- --- --- --- -ct gcc cta --- --- --- --- --- --- --- --- tca tct ata tca ctg tca ctc tc- --- --- gct ctc act ata tct tct aaa att aca a-- --a caa c-- --- --- --- --- --- --- --- --- --- --- --- --- --- --- --- --- --- --- --- --- --- --- --- --- --- --- --- --- --- --- --- --- --- --- --- -tg gat att cga t-- --- --- --- --- --- --- --- --- --- --- --- --- --- --- --- --- --- --- --- --- --- --- --- --- --- --- aac agc att tgt gt- --- --- --- --- --- --- --- --- --- --- --- --- --- --- --- --- --

>ON676704.1_Monkeypox_virus_isolate_MPXV_USA_2022_FL002_complete_genome

ttt ttt cga tct atc ctc gtc c-- -t- ctc atc atc ctt ata --- --- --- --- --- --- -tt att atc att att atc ata gtc tat taa aca caa atc atc t-- --- --- --- --- --- --- --- --- --- --- --- --- --- --- --- --- --- --- --- --- --- --- --- --- --- --- acg ttt ata ac- --- --- --- --- --- --- --- --- aac att c-- --- --- --- --t cat tat taa tta gtt ctg tag -aa tat ctt taa taa ttt ggc tat a-- --- --c atc tgt t-- --- --- --- --- --- --- --- --- --- --- --- --- --- --- --- --- --- --- caa tac t-- --- --- --- --- --- --- --- atc tat tga tga ttt ctt tt- --- --- --- --- --- --- --- --- --- --- --- tta aga ct- --- --- --- --- --- --- --- --- --- --- --- --- --- --- --- --- --- --- --- --- --- --- --- --- --- --- --- --- --- --- --- --- --- --- --- --- --- --- --- -ta aac tag t-- --- --- --- --- --- --- --- --- --- --- --- --- --- --- --- --- --- --- --- --- --- --- --- --- --- --- --- --- --- --- --- --- --- --- --- --- --- --- --- --- --- --- --- --- --- --- --- --- --- --- --- --- --- --- --- --- --- --- --- --- --- --- --- --- --- --- --- --- --- --- --- --- --- --- --- --- --- --- --- --- --- --- --- --- --- --- --- --- --- --- --- --- --- --- --- --- --- --- --- --- --- --- --- --- --- --- --- --- --- tat ggt aat gac gat gaa a-- --t cga gta gta --- --- act tct aat aaa gac ttg ata --- --- tca tta tca tat gtt tga tcg --- --- --- --- --- --- --- tca tag tta ata gtg tg- --- --- --- --- --- --- --- --- --- --- --- --- --- --- --- --- --- --- --- --g cta aat ggt act gtt aat aag ttt at- --- --- --- --- --- --- --- --- --- --- --- --- --- --- --- --- --- aga caa tat cat agt att ttc ttt cca gaa t-- --- --- --- tag att att ttt tta aat act gat cct cac aat tcc gtg atg tag cag tag ttg gt- --- --- --- --- --- --- --- --- --- --- --- --- --- --- --- --- --- --- --- --- --- --- --- --- --- --- --- --g cat ggt cta tat cgt --- --- --- --- --- --- --- --- --- --- --- --- --- --- --- --- --- --- --- --- --- --- --- --- --- --- --- --- --- --- --- --- --- --- --- --- --- --- --- --- --- --- --- -ta aaa tgt atc ata tat aat agt ttt ctg acg tgg agt aca gaa ttt tcg a-- --- --- --- --- --- --- --- --- --- --- --- --- --- --- --- --- --- --- --- --- --- --- --- --- --- --- --- --- --- --- --- --- --- --- --- --- --- --- --- --- --- --- --- tta atg agt tca tgg taa gga agg gca aat gcc t-- -gt ata taa tat aca taa gtt aa- --- --- --- --- --- --- --- --- --- --- --- --- --- tag ttt ttt atc ata ttt --- --- --- --- --- --- --- --- tct aat acc ata ata aaa att atc --- --- --- --- --- --- --- --- --- --- --- --- --- --- --- --- --- --- --- --- --- --- --- --- --- -at tat tgc gtt tg- gta gtt --- --- --- --- -ct gcc cta --- --- --- --- --- --- --- --- tca tct ata tca ctg tca ctc tc- --- --- gct ctc act ata tct tct aaa att aca a-- --a caa c-- --- --- --- --- --- --- --- --- --- --- --- --- --- --- --- --- --- --- --- --- --- --- --- --- --- --- --- --- --- --- --- --- --- --- --- -tg gat att cga t-- --- --- --- --- --- --- --- --- --- --- --- --- --- --- --- --- --- --- --- --- --- --- --- --- --- --- aac agc att tgt gt- --- --- --- --- --- --- --- --- --- --- --- --- --- --- --- --- --

>ON563414.3_Monkeypox_virus_isolate_MPXV_USA_2022_MA001_complete_genome

ttt ttt cga tct atc ctc gtc c-- -t- ctc atc atc ctt ata --- --- --- --- --- --- -tt att atc att att atc ata gtc tat taa aca caa atc atc t-- --- --- --- --- --- --- --- --- --- --- --- --- --- --- --- --- --- --- --- --- --- --- --- --- --- --- acg ttt ata ac- --- --- --- --- --- --- --- --- aac att c-- --- --- --- --t cat tat taa tta gtt ctg tag -aa tat ctt taa taa ttt ggc tat a-- --- --c atc tgt t-- --- --- --- --- --- --- --- --- --- --- --- --- --- --- --- --- --- --- caa tac t-- --- --- --- --- --- --- --- atc tat tga tga ttt ctt tt- --- --- --- --- --- --- --- --- --- --- --- tta aga ct- --- --- --- --- --- --- --- --- --- --- --- --- --- --- --- --- --- --- --- --- --- --- --- --- --- --- --- --- --- --- --- --- --- --- --- --- --- --- --- -ta aac tag t-- --- --- --- --- --- --- --- --- --- --- --- --- --- --- --- --- --- --- --- --- --- --- --- --- --- --- --- --- --- --- --- --- --- --- --- --- --- --- --- --- --- --- --- --- --- --- --- --- --- --- --- --- --- --- --- --- --- --- --- --- --- --- --- --- --- --- --- --- --- --- --- --- --- --- --- --- --- --- --- --- --- --- --- --- --- --- --- --- --- --- --- --- --- --- --- --- --- --- --- --- --- --- --- --- --- --- --- --- --- tat ggt aat gac gat gaa a-- --t cga gta gta --- --- act tct aat aaa gac ttg ata --- --- tca tta tca tat gtt tga tcg --- --- --- --- --- --- --- tca tag tta ata gtg tg- --- --- --- --- --- --- --- --- --- --- --- --- --- --- --- --- --- --- --- --g cta aat ggt act gtt aat aag ttt at- --- --- --- --- --- --- --- --- --- --- --- --- --- --- --- --- --- aga caa tat cat agt att ttc ttt cca gaa t-- --- --- --- tag att att ttt tta aat act gat cct cac aat tcc gtg atg tag cag tag ttg gt- --- --- --- --- --- --- --- --- --- --- --- --- --- --- --- --- --- --- --- --- --- --- --- --- --- --- --- --g cat ggt cta tat cgt --- --- --- --- --- --- --- --- --- --- --- --- --- --- --- --- --- --- --- --- --- --- --- --- --- --- --- --- --- --- --- --- --- --- --- --- --- --- --- --- --- --- --- -ta aaa tgt atc ata tat aat agt ttt ctg acg tgg agt aca gaa ttt tcg a-- --- --- --- --- --- --- --- --- --- --- --- --- --- --- --- --- --- --- --- --- --- --- --- --- --- --- --- --- --- --- --- --- --- --- --- --- --- --- --- --- --- --- --- tta atg agt tca tgg taa gga agg gca aat gcc t-- -gt ata taa tat aca taa gtt aa- --- --- --- --- --- --- --- --- --- --- --- --- --- tag ttt ttt atc ata ttt --- --- --- --- --- --- --- --- tct aat acc ata ata aaa att atc --- --- --- --- --- --- --- --- --- --- --- --- --- --- --- --- --- --- --- --- --- --- --- --- --- -at tat tgc gtt tg- gta gtt --- --- --- --- -ct gcc cta --- --- --- --- --- --- --- --- tca tct ata tca ctg tca ctc tc- --- --- gct ctc act ata tct tct aaa att aca a-- --a caa c-- --- --- --- --- --- --- --- --- --- --- --- --- --- --- --- --- --- --- --- --- --- --- --- --- --- --- --- --- --- --- --- --- --- --- --- -tg gat att cga t-- --- --- --- --- --- --- --- --- --- --- --- --- --- --- --- --- --- --- --- --- --- --- --- --- --- --- aac agc att tgt gt- --- --- --- --- --- --- --- --- --- --- --- --- --- --- --- --- --

>ON676705.1_Monkeypox_virus_isolate_MPXV_USA_2022_UT001_complete_genome

ttt ttt cga tct atc ctc gtc c-- -t- ctc atc atc ctt ata --- --- --- --- --- --- -tt att atc att att atc ata gtc tat taa aca caa atc atc t-- --- --- --- --- --- --- --- --- --- --- --- --- --- --- --- --- --- --- --- --- --- --- --- --- --- --- acg ttt ata ac- --- --- --- --- --- --- --- --- aac att c-- --- --- --- --t cat tat taa tta gtt ctg tag -aa tat ctt taa taa ttt ggc tat a-- --- --c atc tgt t-- --- --- --- --- --- --- --- --- --- --- --- --- --- --- --- --- --- --- caa tac t-- --- --- --- --- --- --- --- atc tat tga tga ttt ctt tt- --- --- --- --- --- --- --- --- --- --- --- tta aga ct- --- --- --- --- --- --- --- --- --- --- --- --- --- --- --- --- --- --- --- --- --- --- --- --- --- --- --- --- --- --- --- --- --- --- --- --- --- --- --- -ta aac tag t-- --- --- --- --- --- --- --- --- --- --- --- --- --- --- --- --- --- --- --- --- --- --- --- --- --- --- --- --- --- --- --- --- --- --- --- --- --- --- --- --- --- --- --- --- --- --- --- --- --- --- --- --- --- --- --- --- --- --- --- --- --- --- --- --- --- --- --- --- --- --- --- --- --- --- --- --- --- --- --- --- --- --- --- --- --- --- --- --- --- --- --- --- --- --- --- --- --- --- --- --- --- --- --- --- --- --- --- --- --- tat ggt aat gac gat gaa a-- --t cga gta gta --- --- act tct aat aaa gac ttg ata --- --- tca tta tca tat gtt tga tcg --- --- --- --- --- --- --- tca tag tta ata gtg tg- --- --- --- --- --- --- --- --- --- --- --- --- --- --- --- --- --- --- --- --g cta aat ggt act gtt aat aag ttt at- --- --- --- --- --- --- --- --- --- --- --- --- --- --- --- --- --- aga caa tat cat agt att ttc ttt cca gaa t-- --- --- --- tag att att ttt tta aat act gat cct cac aat tcc gtg atg tag cag tag ttg gt- --- --- --- --- --- --- --- --- --- --- --- --- --- --- --- --- --- --- --- --- --- --- --- --- --- --- --- --g cat ggt cta tat cgt --- --- --- --- --- --- --- --- --- --- --- --- --- --- --- --- --- --- --- --- --- --- --- --- --- --- --- --- --- --- --- --- --- --- --- --- --- --- --- --- --- --- --- -ta aaa tgt atc ata tat aat agt ttt ctg acg tgg agt aca gaa ttt tcg a-- --- --- --- --- --- --- --- --- --- --- --- --- --- --- --- --- --- --- --- --- --- --- --- --- --- --- --- --- --- --- --- --- --- --- --- --- --- --- --- --- --- --- --- tta atg agt tca tgg taa gga agg gca aat gcc t-- -gt ata taa tat aca taa gtt aa- --- --- --- --- --- --- --- --- --- --- --- --- --- tag ttt ttt atc ata ttt --- --- --- --- --- --- --- --- tct aat acc ata ata aaa att atc --- --- --- --- --- --- --- --- --- --- --- --- --- --- --- --- --- --- --- --- --- --- --- --- --- -at tat tgc gtt tg- gta gtt --- --- --- --- -ct gcc cta --- --- --- --- --- --- --- --- tca tct ata tca ctg tca ctc tc- --- --- gct ctc act ata tct tct aaa att aca a-- --a caa c-- --- --- --- --- --- --- --- --- --- --- --- --- --- --- --- --- --- --- --- --- --- --- --- --- --- --- --- --- --- --- --- --- --- --- --- -tg gat att cga t-- --- --- --- --- --- --- --- --- --- --- --- --- --- --- --- --- --- --- --- --- --- --- --- --- --- --- aac agc att tgt gt- --- --- --- --- --- --- --- --- --- --- --- --- --- --- --- --- --

>MT903343.1_Monkeypox_virus_isolate_MPXV-UK_P1

ttt ttt cga tct atc ctc gtc c-- -t- ctc atc atc ctt ata --- --- --- --- --- --- -tt att atc att att atc ata gtc tat taa aca caa atc atc t-- --- --- --- --- --- --- --- --- --- --- --- --- --- --- --- --- --- --- --- --- --- --- --- --- --- --- acg ttt ata ac- --- --- --- --- --- --- --- --- aac att c-- --- --- --- --t cat tat taa tta gtt ctg tag -aa tat ctt taa taa ttt ggc tat a-- --- --c atc tgt t-- --- --- --- --- --- --- --- --- --- --- --- --- --- --- --- --- --- --- caa tac t-- --- --- --- --- --- --- --- atc tat tga tga ttt ctt tt- --- --- --- --- --- --- --- --- --- --- --- tta aga ct- --- --- --- --- --- --- --- --- --- --- --- --- --- --- --- --- --- --- --- --- --- --- --- --- --- --- --- --- --- --- --- --- --- --- --- --- --- --- --- -ta aac tag t-- --- --- --- --- --- --- --- --- --- --- --- --- --- --- --- --- --- --- --- --- --- --- --- --- --- --- --- --- --- --- --- --- --- --- --- --- --- --- --- --- --- --- --- --- --- --- --- --- --- --- --- --- --- --- --- --- --- --- --- --- --- --- --- --- --- --- --- --- --- --- --- --- --- --- --- --- --- --- --- --- --- --- --- --- --- --- --- --- --- --- --- --- --- --- --- --- --- --- --- --- --- --- --- --- --- --- --- --- --- tat ggt aat gac gat gaa a-- --t cga gta gta --- --- act tct aat aaa gac ttg ata --- --- tca tta tca tat gtt tga tcg --- --- --- --- --- --- --- tca tag tta ata gtg tg- --- --- --- --- --- --- --- --- --- --- --- --- --- --- --- --- --- --- --- --g cta aat ggt act gtt aat aag ttt at- --- --- --- --- --- --- --- --- --- --- --- --- --- --- --- --- --- aga caa tat cat agt att ttc ttt cca gaa t-- --- --- --- tag att att ttt tta aat act gat cct cac aat tcc gtg atg tag cag tag ttg gt- --- --- --- --- --- --- --- --- --- --- --- --- --- --- --- --- --- --- --- --- --- --- --- --- --- --- --- --g cat ggt cta tat cgt --- --- --- --- --- --- --- --- --- --- --- --- --- --- --- --- --- --- --- --- --- --- --- --- --- --- --- --- --- --- --- --- --- --- --- --- --- --- --- --- --- --- --- -ta aaa tgt atc ata tat aat agt ttt ctg acg tgg agt aca gaa ttt tcg a-- --- --- --- --- --- --- --- --- --- --- --- --- --- --- --- --- --- --- --- --- --- --- --- --- --- --- --- --- --- --- --- --- --- --- --- --- --- --- --- --- --- --- --- tta atg agt tca tgg taa gga agg gca aat gcc t-- -gt ata taa tat aca taa gtt aa- --- --- --- --- --- --- --- --- --- --- --- --- --- tag ttt ttt atc ata ttt --- --- --- --- --- --- --- --- tct aat acc ata ata aaa att atc --- --- --- --- --- --- --- --- --- --- --- --- --- --- --- --- --- --- --- --- --- --- --- --- --- -at tat tgc gtt tg- gta gtt --- --- --- --- -ct gcc cta --- --- --- --- --- --- --- --- tca tct ata tca ctg tca ctc tc- --- --- gct ctc act ata tct tct aaa att aca a-- --a caa c-- --- --- --- --- --- --- --- --- --- --- --- --- --- --- --- --- --- --- --- --- --- --- --- --- --- --- --- --- --- --- --- --- --- --- --- -tg gat att cga t-- --- --- --- --- --- --- --- --- --- --- --- --- --- --- --- --- --- --- --- --- --- --- --- --- --- --- aac agc att tgt gt- --- --- --- --- --- --- --- --- --- --- --- --- --- --- --- --- --

>ON631963.1_Monkeypox_virus_isolate_MPxV/VIDRL01/2022_complete_genome

ttt ttt cga tct atc ctc gtc c-- -t- ctc atc atc ctt ata --- --- --- --- --- --- -tt att atc att att atc ata gtc tat taa aca caa atc atc t-- --- --- --- --- --- --- --- --- --- --- --- --- --- --- --- --- --- --- --- --- --- --- --- --- --- --- acg ttt ata ac- --- --- --- --- --- --- --- --- aac att c-- --- --- --- --t cat tat taa tta gtt ctg tag -aa tat ctt taa taa ttt ggc tat a-- --- --c atc tgt t-- --- --- --- --- --- --- --- --- --- --- --- --- --- --- --- --- --- --- caa tac t-- --- --- --- --- --- --- --- atc tat tga tga ttt ctt tt- --- --- --- --- --- --- --- --- --- --- --- tta aga ct- --- --- --- --- --- --- --- --- --- --- --- --- --- --- --- --- --- --- --- --- --- --- --- --- --- --- --- --- --- --- --- --- --- --- --- --- --- --- --- -ta aac tag t-- --- --- --- --- --- --- --- --- --- --- --- --- --- --- --- --- --- --- --- --- --- --- --- --- --- --- --- --- --- --- --- --- --- --- --- --- --- --- --- --- --- --- --- --- --- --- --- --- --- --- --- --- --- --- --- --- --- --- --- --- --- --- --- --- --- --- --- --- --- --- --- --- --- --- --- --- --- --- --- --- --- --- --- --- --- --- --- --- --- --- --- --- --- --- --- --- --- --- --- --- --- --- --- --- --- --- --- --- --- tat ggt aat gac gat gaa a-- --t cga gta gta --- --- act tct aat aaa gac ttg ata --- --- tca tta tca tat gtt tga tcg --- --- --- --- --- --- --- tca tag tta ata gtg tg- --- --- --- --- --- --- --- --- --- --- --- --- --- --- --- --- --- --- --- --g cta aat ggt act gtt aat aag ttt at- --- --- --- --- --- --- --- --- --- --- --- --- --- --- --- --- --- aga caa tat cat agt att ttc ttt cca gaa t-- --- --- --- tag att att ttt tta aat act gat cct cac aat tcc gtg atg tag cag tag ttg gt- --- --- --- --- --- --- --- --- --- --- --- --- --- --- --- --- --- --- --- --- --- --- --- --- --- --- --- --g cat ggt cta tat cgt --- --- --- --- --- --- --- --- --- --- --- --- --- --- --- --- --- --- --- --- --- --- --- --- --- --- --- --- --- --- --- --- --- --- --- --- --- --- --- --- --- --- --- -ta aaa tgt atc ata tat aat agt ttt ctg acg tgg agt aca gaa ttt tcg a-- --- --- --- --- --- --- --- --- --- --- --- --- --- --- --- --- --- --- --- --- --- --- --- --- --- --- --- --- --- --- --- --- --- --- --- --- --- --- --- --- --- --- --- tta atg agt tca tgg taa gga agg gca aat gcc t-- -gt ata taa tat aca taa gtt aa- --- --- --- --- --- --- --- --- --- --- --- --- --- tag ttt ttt atc ata ttt --- --- --- --- --- --- --- --- tct aat acc ata ata aaa att atc --- --- --- --- --- --- --- --- --- --- --- --- --- --- --- --- --- --- --- --- --- --- --- --- --- -at tat tgc gtt tg- gta gtt --- --- --- --- -ct gcc cta --- --- --- --- --- --- --- --- tca tct ata tca ctg tca ctc tc- --- --- gct ctc act ata tct tct aaa att aca a-- --a caa c-- --- --- --- --- --- --- --- --- --- --- --- --- --- --- --- --- --- --- --- --- --- --- --- --- --- --- --- --- --- --- --- --- --- --- --- -tg gat att cga t-- --- --- --- --- --- --- --- --- --- --- --- --- --- --- --- --- --- --- --- --- --- --- --- --- --- --- aac agc att tgt gt- --- --- --- --- --- --- --- --- --- --- --- --- --- --- --- --- --

>ON622722.2_Monkeypox_virus_isolate_MPXV_FR_HCL0001_2022_complete_genome

ttt ttt cga tct atc ctc gtc c-- -t- ctc atc atc ctt ata --- --- --- --- --- --- -tt att atc att att atc ata gtc tat taa aca caa atc atc t-- --- --- --- --- --- --- --- --- --- --- --- --- --- --- --- --- --- --- --- --- --- --- --- --- --- --- acg ttt ata ac- --- --- --- --- --- --- --- --- aac att c-- --- --- --- --t cat tat taa tta gtt ctg tag -aa tat ctt taa taa ttt ggc tat a-- --- --c atc tgt t-- --- --- --- --- --- --- --- --- --- --- --- --- --- --- --- --- --- --- caa tac t-- --- --- --- --- --- --- --- atc tat tga tga ttt ctt tt- --- --- --- --- --- --- --- --- --- --- --- tta aga ct- --- --- --- --- --- --- --- --- --- --- --- --- --- --- --- --- --- --- --- --- --- --- --- --- --- --- --- --- --- --- --- --- --- --- --- --- --- --- --- -ta aac tag t-- --- --- --- --- --- --- --- --- --- --- --- --- --- --- --- --- --- --- --- --- --- --- --- --- --- --- --- --- --- --- --- --- --- --- --- --- --- --- --- --- --- --- --- --- --- --- --- --- --- --- --- --- --- --- --- --- --- --- --- --- --- --- --- --- --- --- --- --- --- --- --- --- --- --- --- --- --- --- --- --- --- --- --- --- --- --- --- --- --- --- --- --- --- --- --- --- --- --- --- --- --- --- --- --- --- --- --- --- --- tat ggt aat gac gat gaa a-- --t cga gta gta --- --- act tct aat aaa gac ttg ata --- --- tca tta tca tat gtt tga tcg --- --- --- --- --- --- --- tca tag tta ata gtg tg- --- --- --- --- --- --- --- --- --- --- --- --- --- --- --- --- --- --- --- --g cta aat ggt act gtt aat aag ttt at- --- --- --- --- --- --- --- --- --- --- --- --- --- --- --- --- --- aga caa tat cat agt att ttc ttt cca gaa t-- --- --- --- tag att att ttt tta aat act gat cct cac aat tcc gtg atg tag cag tag ttg gt- --- --- --- --- --- --- --- --- --- --- --- --- --- --- --- --- --- --- --- --- --- --- --- --- --- --- --- --g cat ggt cta tat cgt --- --- --- --- --- --- --- --- --- --- --- --- --- --- --- --- --- --- --- --- --- --- --- --- --- --- --- --- --- --- --- --- --- --- --- --- --- --- --- --- --- --- --- -ta aaa tgt atc ata tat aat agt ttt ctg acg tgg agt aca gaa ttt tcg a-- --- --- --- --- --- --- --- --- --- --- --- --- --- --- --- --- --- --- --- --- --- --- --- --- --- --- --- --- --- --- --- --- --- --- --- --- --- --- --- --- --- --- --- tta atg agt tca tgg taa gga agg gca aat gcc t-- -gt ata taa tat aca taa gtt aa- --- --- --- --- --- --- --- --- --- --- --- --- --- tag ttt ttt atc ata ttt --- --- --- --- --- --- --- --- tct aat acc ata ata aaa att atc --- --- --- --- --- --- --- --- --- --- --- --- --- --- --- --- --- --- --- --- --- --- --- --- --- -at tat tgc gtt tg- gta gtt --- --- --- --- -ct gcc cta --- --- --- --- --- --- --- --- tca tct ata tca ctg tca ctc tc- --- --- gct ctc act ata tct tct aaa att aca a-- --a caa c-- --- --- --- --- --- --- --- --- --- --- --- --- --- --- --- --- --- --- --- --- --- --- --- --- --- --- --- --- --- --- --- --- --- --- --- -tg gat att cga t-- --- --- --- --- --- --- --- --- --- --- --- --- --- --- --- --- --- --- --- --- --- --- --- --- --- --- aac agc att tgt gt- --- --- --- --- --- --- --- --- --- --- --- --- --- --- --- --- --

>ON627808.1_Monkeypox_virus_isolate_MPX/human/USA/UT-UPHL-82200022/2022_complete_genome

ttt ttt cga tct atc ctc gtc c-- -t- ctc atc atc ctt ata --- --- --- --- --- --- -tt att atc att att atc ata gtc tat taa aca caa atc atc t-- --- --- --- --- --- --- --- --- --- --- --- --- --- --- --- --- --- --- --- --- --- --- --- --- --- --- acg ttt ata ac- --- --- --- --- --- --- --- --- aac att c-- --- --- --- --t cat tat taa tta gtt ctg tag -aa tat ctt taa taa ttt ggc tat a-- --- --c atc tgt t-- --- --- --- --- --- --- --- --- --- --- --- --- --- --- --- --- --- --- caa tac t-- --- --- --- --- --- --- --- atc tat tga tga ttt ctt tt- --- --- --- --- --- --- --- --- --- --- --- tta aga ct- --- --- --- --- --- --- --- --- --- --- --- --- --- --- --- --- --- --- --- --- --- --- --- --- --- --- --- --- --- --- --- --- --- --- --- --- --- --- --- -ta aac tag t-- --- --- --- --- --- --- --- --- --- --- --- --- --- --- --- --- --- --- --- --- --- --- --- --- --- --- --- --- --- --- --- --- --- --- --- --- --- --- --- --- --- --- --- --- --- --- --- --- --- --- --- --- --- --- --- --- --- --- --- --- --- --- --- --- --- --- --- --- --- --- --- --- --- --- --- --- --- --- --- --- --- --- --- --- --- --- --- --- --- --- --- --- --- --- --- --- --- --- --- --- --- --- --- --- --- --- --- --- --- tat ggt aat gac gat gaa a-- --t cga gta gta --- --- act tct aat aaa gac ttg ata --- --- tca tta tca tat gtt tga tcg --- --- --- --- --- --- --- tca tag tta ata gtg tg- --- --- --- --- --- --- --- --- --- --- --- --- --- --- --- --- --- --- --- --g cta aat ggt act gtt aat aag ttt at- --- --- --- --- --- --- --- --- --- --- --- --- --- --- --- --- --- aga caa tat cat agt att ttc ttt cca gaa t-- --- --- --- tag att att ttt tta aat act gat cct cac aat tcc gtg atg tag cag tag ttg gt- --- --- --- --- --- --- --- --- --- --- --- --- --- --- --- --- --- --- --- --- --- --- --- --- --- --- --- --g cat ggt cta tat cgt --- --- --- --- --- --- --- --- --- --- --- --- --- --- --- --- --- --- --- --- --- --- --- --- --- --- --- --- --- --- --- --- --- --- --- --- --- --- --- --- --- --- --- -ta aaa tgt atc ata tat aat agt ttt ctg acg tgg agt aca gaa ttt tcg a-- --- --- --- --- --- --- --- --- --- --- --- --- --- --- --- --- --- --- --- --- --- --- --- --- --- --- --- --- --- --- --- --- --- --- --- --- --- --- --- --- --- --- --- tta atg agt tca tgg taa gga agg gca aat gcc t-- -gt ata taa tat aca taa gtt aa- --- --- --- --- --- --- --- --- --- --- --- --- --- tag ttt ttt atc ata ttt --- --- --- --- --- --- --- --- tct aat acc ata ata aaa att atc --- --- --- --- --- --- --- --- --- --- --- --- --- --- --- --- --- --- --- --- --- --- --- --- --- -at tat tgc gtt tg- gta gtt --- --- --- --- -ct gcc cta --- --- --- --- --- --- --- --- tca tct ata tca ctg tca ctc tc- --- --- gct ctc act ata tct tct aaa att aca a-- --a caa c-- --- --- --- --- --- --- --- --- --- --- --- --- --- --- --- --- --- --- --- --- --- --- --- --- --- --- --- --- --- --- --- --- --- --- --- -tg gat att cga t-- --- --- --- --- --- --- --- --- --- --- --- --- --- --- --- --- --- --- --- --- --- --- --- --- --- --- aac agc att tgt gt- --- --- --- --- --- --- --- --- --- --- --- --- --- --- --- --- --

>ON676703.1_Monkeypox_virus_isolate_MPXV_USA_2022_CA001_complete_genome

ttt ttt cga tct atc ctc gtc c-- -t- ctc atc atc ctt ata --- --- --- --- --- --- -tt att atc att att atc ata gtc tat taa aca caa atc atc t-- --- --- --- --- --- --- --- --- --- --- --- --- --- --- --- --- --- --- --- --- --- --- --- --- --- --- acg ttt ata ac- --- --- --- --- --- --- --- --- aac att c-- --- --- --- --t cat tat taa tta gtt ctg tag -aa tat ctt taa taa ttt ggc tat a-- --- --c atc tgt t-- --- --- --- --- --- --- --- --- --- --- --- --- --- --- --- --- --- --- caa tac t-- --- --- --- --- --- --- --- atc tat tga tga ttt ctt tt- --- --- --- --- --- --- --- --- --- --- --- tta aga ct- --- --- --- --- --- --- --- --- --- --- --- --- --- --- --- --- --- --- --- --- --- --- --- --- --- --- --- --- --- --- --- --- --- --- --- --- --- --- --- -ta aac tag t-- --- --- --- --- --- --- --- --- --- --- --- --- --- --- --- --- --- --- --- --- --- --- --- --- --- --- --- --- --- --- --- --- --- --- --- --- --- --- --- --- --- --- --- --- --- --- --- --- --- --- --- --- --- --- --- --- --- --- --- --- --- --- --- --- --- --- --- --- --- --- --- --- --- --- --- --- --- --- --- --- --- --- --- --- --- --- --- --- --- --- --- --- --- --- --- --- --- --- --- --- --- --- --- --- --- --- --- --- --- tat ggt aat gac gat gaa a-- --t cga gta gta --- --- act tct aat aaa gac ttg ata --- --- tca tta tca tat gtt tga tcg --- --- --- --- --- --- --- tca tag tta ata gtg tg- --- --- --- --- --- --- --- --- --- --- --- --- --- --- --- --- --- --- --- --g cta aat ggt act gtt aat aag ttt at- --- --- --- --- --- --- --- --- --- --- --- --- --- --- --- --- --- aga caa tat cat agt att ttc ttt cca gaa t-- --- --- --- tag att att ttt tta aat act gat cct cac aat tcc gtg atg tag cag tag ttg gt- --- --- --- --- --- --- --- --- --- --- --- --- --- --- --- --- --- --- --- --- --- --- --- --- --- --- --- --g cat ggt cta tat cgt --- --- --- --- --- --- --- --- --- --- --- --- --- --- --- --- --- --- --- --- --- --- --- --- --- --- --- --- --- --- --- --- --- --- --- --- --- --- --- --- --- --- --- -ta aaa tgt atc ata tat aat agt ttt ctg acg tgg agt aca gaa ttt tcg a-- --- --- --- --- --- --- --- --- --- --- --- --- --- --- --- --- --- --- --- --- --- --- --- --- --- --- --- --- --- --- --- --- --- --- --- --- --- --- --- --- --- --- --- tta atg agt tca tgg taa gga agg gca aat gcc t-- -gt ata taa tat aca taa gtt aa- --- --- --- --- --- --- --- --- --- --- --- --- --- tag ttt ttt atc ata ttt --- --- --- --- --- --- --- --- tct aat acc ata ata aaa att atc --- --- --- --- --- --- --- --- --- --- --- --- --- --- --- --- --- --- --- --- --- --- --- --- --- -at tat tgc gtt tg- gta gtt --- --- --- --- -ct gcc cta --- --- --- --- --- --- --- --- tca tct ata tca ctg tca ctc tc- --- --- gct ctc act ata tct tct aaa att aca a-- --a caa c-- --- --- --- --- --- --- --- --- --- --- --- --- --- --- --- --- --- --- --- --- --- --- --- --- --- --- --- --- --- --- --- --- --- --- --- -tg gat att cga t-- --- --- --- --- --- --- --- --- --- --- --- --- --- --- --- --- --- --- --- --- --- --- --- --- --- --- aac agc att tgt gt- --- --- --- --- --- --- --- --- --- --- --- --- --- --- --- --- --

>ON676706.1_Monkeypox_virus_isolate_MPXV_USA_2022_UT002_complete_genome

ttt ttt cga tct atc ctc gtc c-- -t- ctc atc atc ctt ata --- --- --- --- --- --- -tt att atc att att atc ata gtc tat taa aca caa atc atc t-- --- --- --- --- --- --- --- --- --- --- --- --- --- --- --- --- --- --- --- --- --- --- --- --- --- --- acg ttt ata ac- --- --- --- --- --- --- --- --- aac att c-- --- --- --- --t cat tat taa tta gtt ctg tag -aa tat ctt taa taa ttt ggc tat a-- --- --c atc tgt t-- --- --- --- --- --- --- --- --- --- --- --- --- --- --- --- --- --- --- caa tac t-- --- --- --- --- --- --- --- atc tat tga tga ttt ctt tt- --- --- --- --- --- --- --- --- --- --- --- tta aga ct- --- --- --- --- --- --- --- --- --- --- --- --- --- --- --- --- --- --- --- --- --- --- --- --- --- --- --- --- --- --- --- --- --- --- --- --- --- --- --- -ta aac tag t-- --- --- --- --- --- --- --- --- --- --- --- --- --- --- --- --- --- --- --- --- --- --- --- --- --- --- --- --- --- --- --- --- --- --- --- --- --- --- --- --- --- --- --- --- --- --- --- --- --- --- --- --- --- --- --- --- --- --- --- --- --- --- --- --- --- --- --- --- --- --- --- --- --- --- --- --- --- --- --- --- --- --- --- --- --- --- --- --- --- --- --- --- --- --- --- --- --- --- --- --- --- --- --- --- --- --- --- --- --- tat ggt aat gac gat gaa a-- --t cga gta gta --- --- act tct aat aaa gac ttg ata --- --- tca tta tca tat gtt tga tcg --- --- --- --- --- --- --- tca tag tta ata gtg tg- --- --- --- --- --- --- --- --- --- --- --- --- --- --- --- --- --- --- --- --g cta aat ggt act gtt aat aag ttt at- --- --- --- --- --- --- --- --- --- --- --- --- --- --- --- --- --- aga caa tat cat agt att ttc ttt cca gaa t-- --- --- --- tag att att ttt tta aat act gat cct cac aat tcc gtg atg tag cag tag ttg gt- --- --- --- --- --- --- --- --- --- --- --- --- --- --- --- --- --- --- --- --- --- --- --- --- --- --- --- --g cat ggt cta tat cgt --- --- --- --- --- --- --- --- --- --- --- --- --- --- --- --- --- --- --- --- --- --- --- --- --- --- --- --- --- --- --- --- --- --- --- --- --- --- --- --- --- --- --- -ta aaa tgt atc ata tat aat agt ttt ctg acg tgg agt aca gaa ttt tcg a-- --- --- --- --- --- --- --- --- --- --- --- --- --- --- --- --- --- --- --- --- --- --- --- --- --- --- --- --- --- --- --- --- --- --- --- --- --- --- --- --- --- --- --- tta atg agt tca tgg taa gga agg gca aat gcc t-- -gt ata taa tat aca taa gtt aa- --- --- --- --- --- --- --- --- --- --- --- --- --- tag ttt ttt atc ata ttt --- --- --- --- --- --- --- --- tct aat acc ata ata aaa att atc --- --- --- --- --- --- --- --- --- --- --- --- --- --- --- --- --- --- --- --- --- --- --- --- --- -at tat tgc gtt tg- gta gtt --- --- --- --- -ct gcc cta --- --- --- --- --- --- --- --- tca tct ata tca ctg tca ctc tc- --- --- gct ctc act ata tct tct aaa att aca a-- --a caa c-- --- --- --- --- --- --- --- --- --- --- --- --- --- --- --- --- --- --- --- --- --- --- --- --- --- --- --- --- --- --- --- --- --- --- --- -tg gat att cga t-- --- --- --- --- --- --- --- --- --- --- --- --- --- --- --- --- --- --- --- --- --- --- --- --- --- --- aac agc att tgt gt- --- --- --- --- --- --- --- --- --- --- --- --- --- --- --- --- --

>ON649879.1_Monkeypox_virus_isolate_MPXV_ISR001_2022_complete_genome

ttt ttt cga tct atc ctc gtc c-- -t- ctc atc atc ctt ata --- --- --- --- --- --- -tt att atc att att atc ata gtc tat taa aca caa atc atc t-- --- --- --- --- --- --- --- --- --- --- --- --- --- --- --- --- --- --- --- --- --- --- --- --- --- --- acg ttt ata ac- --- --- --- --- --- --- --- --- aac att c-- --- --- --- --t cat tat taa tta gtt ctg tag -aa tat ctt taa taa ttt ggc tat a-- --- --c atc tgt t-- --- --- --- --- --- --- --- --- --- --- --- --- --- --- --- --- --- --- caa tac t-- --- --- --- --- --- --- --- atc tat tga tga ttt ctt tt- --- --- --- --- --- --- --- --- --- --- --- tta aga ct- --- --- --- --- --- --- --- --- --- --- --- --- --- --- --- --- --- --- --- --- --- --- --- --- --- --- --- --- --- --- --- --- --- --- --- --- --- --- --- -ta aac tag t-- --- --- --- --- --- --- --- --- --- --- --- --- --- --- --- --- --- --- --- --- --- --- --- --- --- --- --- --- --- --- --- --- --- --- --- --- --- --- --- --- --- --- --- --- --- --- --- --- --- --- --- --- --- --- --- --- --- --- --- --- --- --- --- --- --- --- --- --- --- --- --- --- --- --- --- --- --- --- --- --- --- --- --- --- --- --- --- --- --- --- --- --- --- --- --- --- --- --- --- --- --- --- --- --- --- --- --- --- --- tat ggt aat gac gat gaa a-- --t cga gta gta --- --- act tct aat aaa gac ttg ata --- --- tca tta tca tat gtt tga tcg --- --- --- --- --- --- --- tca tag tta ata gtg tg- --- --- --- --- --- --- --- --- --- --- --- --- --- --- --- --- --- --- --- --g cta aat ggt act gtt aat aag ttt at- --- --- --- --- --- --- --- --- --- --- --- --- --- --- --- --- --- aga caa tat cat agt att ttc ttt cca gaa t-- --- --- --- tag att att ttt tta aat act gat cct cac aat tcc gtg atg tag cag tag ttg gt- --- --- --- --- --- --- --- --- --- --- --- --- --- --- --- --- --- --- --- --- --- --- --- --- --- --- --- --g cat ggt cta tat cgt --- --- --- --- --- --- --- --- --- --- --- --- --- --- --- --- --- --- --- --- --- --- --- --- --- --- --- --- --- --- --- --- --- --- --- --- --- --- --- --- --- --- --- -ta aaa tgt atc ata tat aat agt ttt ctg acg tgg agt aca gaa ttt tcg a-- --- --- --- --- --- --- --- --- --- --- --- --- --- --- --- --- --- --- --- --- --- --- --- --- --- --- --- --- --- --- --- --- --- --- --- --- --- --- --- --- --- --- --- tta atg agt tca tgg taa gga agg gca aat gcc t-- -gt ata taa tat aca taa gtt aa- --- --- --- --- --- --- --- --- --- --- --- --- --- tag ttt ttt atc ata ttt --- --- --- --- --- --- --- --- tct aat acc ata ata aaa att atc --- --- --- --- --- --- --- --- --- --- --- --- --- --- --- --- --- --- --- --- --- --- --- --- --- -at tat tgc gtt tg- gta gtt --- --- --- --- -ct gcc cta --- --- --- --- --- --- --- --- tca tct ata tca ctg tca ctc tc- --- --- gct ctc act ata tct tct aaa att aca a-- --a caa c-- --- --- --- --- --- --- --- --- --- --- --- --- --- --- --- --- --- --- --- --- --- --- --- --- --- --- --- --- --- --- --- --- --- --- --- -tg gat att cga t-- --- --- --- --- --- --- --- --- --- --- --- --- --- --- --- --- --- --- --- --- --- --- --- --- --- --- aac agc att tgt gt- --- --- --- --- --- --- --- --- --- --- --- --- --- --- --- --- --

>ON602722.1_Monkeypox_virus_isolate_MPXV_FRA_2022_TLS67_complete_genome

ttt ttt cga tct atc ctc gtc c-- -t- ctc atc atc ctt ata --- --- --- --- --- --- -tt att atc att att atc ata gtc tat taa aca caa atc atc t-- --- --- --- --- --- --- --- --- --- --- --- --- --- --- --- --- --- --- --- --- --- --- --- --- --- --- acg ttt ata ac- --- --- --- --- --- --- --- --- aac att c-- --- --- --- --t cat tat taa tta gtt ctg tag -aa tat ctt taa taa ttt ggc tat a-- --- --c atc tgt t-- --- --- --- --- --- --- --- --- --- --- --- --- --- --- --- --- --- --- caa tac t-- --- --- --- --- --- --- --- atc tat tga tga ttt ctt tt- --- --- --- --- --- --- --- --- --- --- --- tta aga ct- --- --- --- --- --- --- --- --- --- --- --- --- --- --- --- --- --- --- --- --- --- --- --- --- --- --- --- --- --- --- --- --- --- --- --- --- --- --- --- -ta aac tag t-- --- --- --- --- --- --- --- --- --- --- --- --- --- --- --- --- --- --- --- --- --- --- --- --- --- --- --- --- --- --- --- --- --- --- --- --- --- --- --- --- --- --- --- --- --- --- --- --- --- --- --- --- --- --- --- --- --- --- --- --- --- --- --- --- --- --- --- --- --- --- --- --- --- --- --- --- --- --- --- --- --- --- --- --- --- --- --- --- --- --- --- --- --- --- --- --- --- --- --- --- --- --- --- --- --- --- --- --- --- tat ggt aat gac gat gaa a-- --t cga gta gta --- --- act tct aat aaa gac ttg ata --- --- tca tta tca tat gtt tga tcg --- --- --- --- --- --- --- tca tag tta ata gtg tg- --- --- --- --- --- --- --- --- --- --- --- --- --- --- --- --- --- --- --- --g cta aat ggt act gtt aat aag ttt at- --- --- --- --- --- --- --- --- --- --- --- --- --- --- --- --- --- aga caa tat cat agt att ttc ttt cca gaa t-- --- --- --- tag att att ttt t-a aat act gat cct cac aat tcc gtg atg tag cag tag ttg gt- --- --- --- --- --- --- --- --- --- --- --- --- --- --- --- --- --- --- --- --- --- --- --- --- --- --- --- --g cat ggt cta tat cgt --- --- --- --- --- --- --- --- --- --- --- --- --- --- --- --- --- --- --- --- --- --- --- --- --- --- --- --- --- --- --- --- --- --- --- --- --- --- --- --- --- --- --- -ta aaa tgt atc ata tat aat agt ttt ctg acg tgg agt aca gaa ttt tcg a-- --- --- --- --- --- --- --- --- --- --- --- --- --- --- --- --- --- --- --- --- --- --- --- --- --- --- --- --- --- --- --- --- --- --- --- --- --- --- --- --- --- --- --- tta atg agt tca tgg taa gga agg gca aat gcc t-- -gt ata taa tat aca taa gtt aa- --- --- --- --- --- --- --- --- --- --- --- --- --- tag ttt ttt atc ata ttt --- --- --- --- --- --- --- --- tct aat acc ata ata aaa att atc --- --- --- --- --- --- --- --- --- --- --- --- --- --- --- --- --- --- --- --- --- --- --- --- --- -at tat tgc gtt tg- gta gtt --- --- --- --- -ct gcc cta --- --- --- --- --- --- --- --- tca tct ata tca ctg tca ctc tc- --- --- gct ctc act ata tct tct aaa att aca a-- --a caa c-- --- --- --- --- --- --- --- --- --- --- --- --- --- --- --- --- --- --- --- --- --- --- --- --- --- --- --- --- --- --- --- --- --- --- --- -tg gat att cga t-- --- --- --- --- --- --- --- --- --- --- --- --- --- --- --- --- --- --- --- --- --- --- --- --- --- --- aac agc att tgt gt- --- --- --- --- --- --- --- --- --- --- --- --- --- --- --- --- --

>ON675438.1_Monkeypox_virus_isolate_MPXV_USA_2022_VA001_complete_genome

ttt ttt cga tct atc ctc gtc c-- -t- ctc atc atc ctt ata --- --- --- --- --- --- -tt att atc att att atc ata gtc tat taa aca caa atc atc t-- --- --- --- --- --- --- --- --- --- --- --- --- --- --- --- --- --- --- --- --- --- --- --- --- --- --- acg ttt ata ac- --- --- --- --- --- --- --- --- aac att c-- --- --- --- --t cat tat taa tta gtt ctg tag -aa tat ctt taa taa ttt ggc tat a-- --- --c atc tgt t-- --- --- --- --- --- --- --- --- --- --- --- --- --- --- --- --- --- --- caa tac t-- --- --- --- --- --- --- --- atc tat tga tga ttt ctt tt- --- --- --- --- --- --- --- --- --- --- --- tta aga ct- --- --- --- --- --- --- --- --- --- --- --- --- --- --- --- --- --- --- --- --- --- --- --- --- --- --- --- --- --- --- --- --- --- --- --- --- --- --- --- -ta aac tag t-- --- --- --- --- --- --- --- --- --- --- --- --- --- --- --- --- --- --- --- --- --- --- --- --- --- --- --- --- --- --- --- --- --- --- --- --- --- --- --- --- --- --- --- --- --- --- --- --- --- --- --- --- --- --- --- --- --- --- --- --- --- --- --- --- --- --- --- --- --- --- --- --- --- --- --- --- --- --- --- --- --- --- --- --- --- --- --- --- --- --- --- --- --- --- --- --- --- --- --- --- --- --- --- --- --- --- --- --- --- tat ggt aat gac gat gaa a-- --t cga gta gta --- --- act tct aat aaa gac ttg ata --- --- tca tta tca tat gtt tga tcg --- --- --- --- --- --- --- tca tag tta ata gtg tg- --- --- --- --- --- --- --- --- --- --- --- --- --- --- --- --- --- --- --- --g cta aat ggt act gtt aat aag ttt at- --- --- --- --- --- --- --- --- --- --- --- --- --- --- --- --- --- aga caa tat cat agt att ttc ttt cca gaa t-- --- --- --- tag att att ttt tta aat act gat cct cac aat tcc gtg atg tag cag tag ttg gt- --- --- --- --- --- --- --- --- --- --- --- --- --- --- --- --- --- --- --- --- --- --- --- --- --- --- --- --g cat ggt cta tat cgt --- --- --- --- --- --- --- --- --- --- --- --- --- --- --- --- --- --- --- --- --- --- --- --- --- --- --- --- --- --- --- --- --- --- --- --- --- --- --- --- --- --- --- -ta aaa tgt atc ata tat aat agt ttt ctg acg tgg agt aca gaa ttt tcg a-- --- --- --- --- --- --- --- --- --- --- --- --- --- --- --- --- --- --- --- --- --- --- --- --- --- --- --- --- --- --- --- --- --- --- --- --- --- --- --- --- --- --- --- tta atg agt tca tgg taa gga agg gca aat gcc t-- -gt ata taa tat aca taa gtt aa- --- --- --- --- --- --- --- --- --- --- --- --- --- tag ttt ttt atc ata ttt --- --- --- --- --- --- --- --- tct aat acc ata ata aaa att atc --- --- --- --- --- --- --- --- --- --- --- --- --- --- --- --- --- --- --- --- --- --- --- --- --- -at tat tgc gtt tg- gta gtt --- --- --- --- -ct gcc cta --- --- --- --- --- --- --- --- tca tct ata tca ctg tca ctc tc- --- --- gct ctc act ata tct tct aaa att aca a-- --a caa c-- --- --- --- --- --- --- --- --- --- --- --- --- --- --- --- --- --- --- --- --- --- --- --- --- --- --- --- --- --- --- --- --- --- --- --- -tg gat att cga t-- --- --- --- --- --- --- --- --- --- --- --- --- --- --- --- --- --- --- --- --- --- --- --- --- --- --- aac agc att tgt gt- --- --- --- --- --- --- --- --- --- --- --- --- --- --- --- --- --

>ON676707.1_Monkeypox_virus_isolate_MPXV_USA_2021_TX_complete_genome

ttt ttt cga tct atc ctc gtc c-- -t- ctc atc atc ctt ata --- --- --- --- --- --- -tt att atc att att atc ata gtc tat taa aca caa atc atc t-- --- --- --- --- --- --- --- --- --- --- --- --- --- --- --- --- --- --- --- --- --- --- --- --- --- --- acg ttt ata ac- --- --- --- --- --- --- --- --- aac att c-- --- --- --- --t cat tat taa tta gtt ctg tag -aa tat ctt taa taa ttt ggc tat a-- --- --c atc tgt t-- --- --- --- --- --- --- --- --- --- --- --- --- --- --- --- --- --- --- caa tac t-- --- --- --- --- --- --- --- atc tat tga tga ttt ctt tt- --- --- --- --- --- --- --- --- --- --- --- tta aga ct- --- --- --- --- --- --- --- --- --- --- --- --- --- --- --- --- --- --- --- --- --- --- --- --- --- --- --- --- --- --- --- --- --- --- --- --- --- --- --- -ta aac tag t-- --- --- --- --- --- --- --- --- --- --- --- --- --- --- --- --- --- --- --- --- --- --- --- --- --- --- --- --- --- --- --- --- --- --- --- --- --- --- --- --- --- --- --- --- --- --- --- --- --- --- --- --- --- --- --- --- --- --- --- --- --- --- --- --- --- --- --- --- --- --- --- --- --- --- --- --- --- --- --- --- --- --- --- --- --- --- --- --- --- --- --- --- --- --- --- --- --- --- --- --- --- --- --- --- --- --- --- --- --- tat ggt aat gac gat gaa a-- --t cga gta gta --- --- act tct aat aaa gac ttg ata --- --- tca tta tca tat gtt tga tcg --- --- --- --- --- --- --- tca tag tta ata gtg tg- --- --- --- --- --- --- --- --- --- --- --- --- --- --- --- --- --- --- --- --g cta aat ggt act gtt aat aag ttt at- --- --- --- --- --- --- --- --- --- --- --- --- --- --- --- --- --- aga caa tat cat agt att ttc ttt cca gaa t-- --- --- --- tag att att ttt tta aat act gat cct cac aat tcc gtg atg tag cag tag ttg gt- --- --- --- --- --- --- --- --- --- --- --- --- --- --- --- --- --- --- --- --- --- --- --- --- --- --- --- --g cat ggt cta tat cgt --- --- --- --- --- --- --- --- --- --- --- --- --- --- --- --- --- --- --- --- --- --- --- --- --- --- --- --- --- --- --- --- --- --- --- --- --- --- --- --- --- --- --- -ta aaa tgt atc ata tat aat agt ttt ctg acg tgg agt aca gaa ttt tcg a-- --- --- --- --- --- --- --- --- --- --- --- --- --- --- --- --- --- --- --- --- --- --- --- --- --- --- --- --- --- --- --- --- --- --- --- --- --- --- --- --- --- --- --- tta atg agt tca tgg taa gga agg gca aat gcc t-- -gt ata taa tat aca taa gtt aa- --- --- --- --- --- --- --- --- --- --- --- --- --- tag ttt ttt atc ata ttt --- --- --- --- --- --- --- --- tct aat acc ata ata aaa att atc --- --- --- --- --- --- --- --- --- --- --- --- --- --- --- --- --- --- --- --- --- --- --- --- --- -at tat tgc gtt tg- gta gtt --- --- --- --- -ct gcc cta --- --- --- --- --- --- --- --- tca tct ata tca ctg tca ctc tc- --- --- gct ctc act ata tct tct aaa att aca a-- --a caa c-- --- --- --- --- --- --- --- --- --- --- --- --- --- --- --- --- --- --- --- --- --- --- --- --- --- --- --- --- --- --- --- --- --- --- --- -tg gat att cga t-- --- --- --- --- --- --- --- --- --- --- --- --- --- --- --- --- --- --- --- --- --- --- --- --- --- --- aac agc att tgt gt- --- --- --- --- --- --- --- --- --- --- --- --- --- --- --- --- --

>MT903345.1_Monkeypox_virus_isolate_MPXV-UK_P3

ttt ttt cga tct atc ctc gtc c-- -t- ctc atc atc ctt ata --- --- --- --- --- --- -tt att atc att att atc ata gtc tat taa aca caa atc atc t-- --- --- --- --- --- --- --- --- --- --- --- --- --- --- --- --- --- --- --- --- --- --- --- --- --- --- acg ttt ata ac- --- --- --- --- --- --- --- --- aac att c-- --- --- --- --t cat tat taa tta gtt ctg tag -aa tat ctt taa taa ttt ggc tat a-- --- --c atc tgt t-- --- --- --- --- --- --- --- --- --- --- --- --- --- --- --- --- --- --- caa tac t-- --- --- --- --- --- --- --- atc tat tga tga ttt ctt tt- --- --- --- --- --- --- --- --- --- --- --- tta aga ct- --- --- --- --- --- --- --- --- --- --- --- --- --- --- --- --- --- --- --- --- --- --- --- --- --- --- --- --- --- --- --- --- --- --- --- --- --- --- --- -ta aac tag t-- --- --- --- --- --- --- --- --- --- --- --- --- --- --- --- --- --- --- --- --- --- --- --- --- --- --- --- --- --- --- --- --- --- --- --- --- --- --- --- --- --- --- --- --- --- --- --- --- --- --- --- --- --- --- --- --- --- --- --- --- --- --- --- --- --- --- --- --- --- --- --- --- --- --- --- --- --- --- --- --- --- --- --- --- --- --- --- --- --- --- --- --- --- --- --- --- --- --- --- --- --- --- --- --- --- --- --- --- --- tat ggt aat gac gat gaa a-- --t cga gta gta --- --- act tct aat aaa gac ttg ata --- --- tca tta tca tat gtt tga tcg --- --- --- --- --- --- --- tca tag tta ata gtg tg- --- --- --- --- --- --- --- --- --- --- --- --- --- --- --- --- --- --- --- --g cta aat ggt act gtt aat aag ttt at- --- --- --- --- --- --- --- --- --- --- --- --- --- --- --- --- --- aga caa tat cat agt att ttc ttt cca gaa t-- --- --- --- tag att att ttt tta aat act gat cct cac aat tcc gtg atg tag cag tag ttg gt- --- --- --- --- --- --- --- --- --- --- --- --- --- --- --- --- --- --- --- --- --- --- --- --- --- --- --- --g cat ggt cta tat cgt --- --- --- --- --- --- --- --- --- --- --- --- --- --- --- --- --- --- --- --- --- --- --- --- --- --- --- --- --- --- --- --- --- --- --- --- --- --- --- --- --- --- --- -ta aaa tgt atc ata tat aat agt ttt ctg acg tgg agt aca gaa ttt tcg a-- --- --- --- --- --- --- --- --- --- --- --- --- --- --- --- --- --- --- --- --- --- --- --- --- --- --- --- --- --- --- --- --- --- --- --- --- --- --- --- --- --- --- --- tta atg agt tca tgg taa gga agg gca aat gcc t-- -gt ata taa tat aca taa gtt aa- --- --- --- --- --- --- --- --- --- --- --- --- --- tag ttt ttt atc ata ttt --- --- --- --- --- --- --- --- tct aat acc ata ata aaa att atc --- --- --- --- --- --- --- --- --- --- --- --- --- --- --- --- --- --- --- --- --- --- --- --- --- -at tat tgc gtt tg- gta gtt --- --- --- --- -ct gcc cta --- --- --- --- --- --- --- --- tca tct ata tca ctg tca ctc tc- --- --- gct ctc act ata tct tct aaa att aca a-- --a caa c-- --- --- --- --- --- --- --- --- --- --- --- --- --- --- --- --- --- --- --- --- --- --- --- --- --- --- --- --- --- --- --- --- --- --- --- -tg gat att cga t-- --- --- --- --- --- --- --- --- --- --- --- --- --- --- --- --- --- --- --- --- --- --- --- --- --- --- aac agc att tgt gt- --- --- --- --- --- --- --- --- --- --- --- --- --- --- --- --- --

>MT903344.1_Monkeypox_virus_isolate_MPXV-UK_P2

ttt ttt cga tct atc ctc gtc c-- -t- ctc atc atc ctt ata --- --- --- --- --- --- -tt att atc att att atc ata gtc tat taa aca caa atc atc t-- --- --- --- --- --- --- --- --- --- --- --- --- --- --- --- --- --- --- --- --- --- --- --- --- --- --- acg ttt ata ac- --- --- --- --- --- --- --- --- aac att c-- --- --- --- --t cat tat taa tta gtt ctg tag -aa tat ctt taa taa ttt ggc tat a-- --- --c atc tgt t-- --- --- --- --- --- --- --- --- --- --- --- --- --- --- --- --- --- --- caa tac t-- --- --- --- --- --- --- --- atc tat tga tga ttt ctt tt- --- --- --- --- --- --- --- --- --- --- --- tta aga ct- --- --- --- --- --- --- --- --- --- --- --- --- --- --- --- --- --- --- --- --- --- --- --- --- --- --- --- --- --- --- --- --- --- --- --- --- --- --- --- -ta aac tag t-- --- --- --- --- --- --- --- --- --- --- --- --- --- --- --- --- --- --- --- --- --- --- --- --- --- --- --- --- --- --- --- --- --- --- --- --- --- --- --- --- --- --- --- --- --- --- --- --- --- --- --- --- --- --- --- --- --- --- --- --- --- --- --- --- --- --- --- --- --- --- --- --- --- --- --- --- --- --- --- --- --- --- --- --- --- --- --- --- --- --- --- --- --- --- --- --- --- --- --- --- --- --- --- --- --- --- --- --- --- tat ggt aat gac gat gaa a-- --t cga gta gta --- --- act tct aat aaa gac ttg ata --- --- tca tta tca tat gtt tga tcg --- --- --- --- --- --- --- tca tag tta ata gtg tg- --- --- --- --- --- --- --- --- --- --- --- --- --- --- --- --- --- --- --- --g cta aat ggt act gtt aat aag ttt at- --- --- --- --- --- --- --- --- --- --- --- --- --- --- --- --- --- aga caa tat cat agt att ttc ttt cca gaa t-- --- --- --- tag att att ttt tta aat act gat cct cac aat tcc gtg atg tag cag tag ttg gt- --- --- --- --- --- --- --- --- --- --- --- --- --- --- --- --- --- --- --- --- --- --- --- --- --- --- --- --g cat ggt cta tat cgt --- --- --- --- --- --- --- --- --- --- --- --- --- --- --- --- --- --- --- --- --- --- --- --- --- --- --- --- --- --- --- --- --- --- --- --- --- --- --- --- --- --- --- -ta aaa tgt atc ata tat aat agt ttt ctg acg tgg agt aca gaa ttt tcg a-- --- --- --- --- --- --- --- --- --- --- --- --- --- --- --- --- --- --- --- --- --- --- --- --- --- --- --- --- --- --- --- --- --- --- --- --- --- --- --- --- --- --- --- tta atg agt tca tgg taa gga agg gca aat gcc t-- -gt ata taa tat aca taa gtt aa- --- --- --- --- --- --- --- --- --- --- --- --- --- tag ttt ttt atc ata ttt --- --- --- --- --- --- --- --- tct aat acc ata ata aaa att atc --- --- --- --- --- --- --- --- --- --- --- --- --- --- --- --- --- --- --- --- --- --- --- --- --- -at tat tgc gtt tg- gta gtt --- --- --- --- -ct gcc cta --- --- --- --- --- --- --- --- tca tct ata tca ctg tca ctc tc- --- --- gct ctc act ata tct tct aaa att aca a-- --a caa c-- --- --- --- --- --- --- --- --- --- --- --- --- --- --- --- --- --- --- --- --- --- --- --- --- --- --- --- --- --- --- --- --- --- --- --- -tg gat att cga t-- --- --- --- --- --- --- --- --- --- --- --- --- --- --- --- --- --- --- --- --- --- --- --- --- --- --- aac agc att tgt gt- --- --- --- --- --- --- --- --- --- --- --- --- --- --- --- --- --

>MT903342.1_Monkeypox_virus_isolate_MPXV-Singapore

ttt ttt cga tct atc ctc gtc c-- -t- ctc atc atc ctt ata --- --- --- --- --- --- -tt att atc att att atc ata gtc tat taa aca caa atc atc t-- --- --- --- --- --- --- --- --- --- --- --- --- --- --- --- --- --- --- --- --- --- --- --- --- --- --- acg ttt ata ac- --- --- --- --- --- --- --- --- aac att c-- --- --- --- --t cat tat taa tta gtt ctg tag -aa tat ctt taa taa ttt ggc tat a-- --- --c atc tgt t-- --- --- --- --- --- --- --- --- --- --- --- --- --- --- --- --- --- --- caa tac t-- --- --- --- --- --- --- --- atc tat tga tga ttt ctt tt- --- --- --- --- --- --- --- --- --- --- --- tta aga ct- --- --- --- --- --- --- --- --- --- --- --- --- --- --- --- --- --- --- --- --- --- --- --- --- --- --- --- --- --- --- --- --- --- --- --- --- --- --- --- -ta aac tag t-- --- --- --- --- --- --- --- --- --- --- --- --- --- --- --- --- --- --- --- --- --- --- --- --- --- --- --- --- --- --- --- --- --- --- --- --- --- --- --- --- --- --- --- --- --- --- --- --- --- --- --- --- --- --- --- --- --- --- --- --- --- --- --- --- --- --- --- --- --- --- --- --- --- --- --- --- --- --- --- --- --- --- --- --- --- --- --- --- --- --- --- --- --- --- --- --- --- --- --- --- --- --- --- --- --- --- --- --- --- tat ggt aat gac gat gaa a-- --t cga gta gta --- --- act tct aat aaa gac ttg ata --- --- tca tta tca tat gtt tga tcg --- --- --- --- --- --- --- tca tag tta ata gtg tg- --- --- --- --- --- --- --- --- --- --- --- --- --- --- --- --- --- --- --- --g cta aat ggt act gtt aat aag ttt at- --- --- --- --- --- --- --- --- --- --- --- --- --- --- --- --- --- aga caa tat cat agt att ttc ttt cca gaa t-- --- --- --- tag att att ttt tta aat act gat cct cac aat tcc gtg atg tag cag tag ttg gt- --- --- --- --- --- --- --- --- --- --- --- --- --- --- --- --- --- --- --- --- --- --- --- --- --- --- --- --g cat ggt cta tat cgt --- --- --- --- --- --- --- --- --- --- --- --- --- --- --- --- --- --- --- --- --- --- --- --- --- --- --- --- --- --- --- --- --- --- --- --- --- --- --- --- --- --- --- -ta aaa tgt atc ata tat aat agt ttt ctg acg tgg agt aca gaa ttt tcg a-- --- --- --- --- --- --- --- --- --- --- --- --- --- --- --- --- --- --- --- --- --- --- --- --- --- --- --- --- --- --- --- --- --- --- --- --- --- --- --- --- --- --- --- tta atg agt tca tgg taa gga agg gca aat gcc t-- -gt ata taa tat aca taa gtt aa- --- --- --- --- --- --- --- --- --- --- --- --- --- tag ttt ttt atc ata ttt --- --- --- --- --- --- --- --- tct aat acc ata ata aaa att atc --- --- --- --- --- --- --- --- --- --- --- --- --- --- --- --- --- --- --- --- --- --- --- --- --- -at tat tgc gtt tg- gta gtt --- --- --- --- -ct gcc cta --- --- --- --- --- --- --- --- tca tct ata tca ctg tca ctc tc- --- --- gct ctc act ata tct tct aaa att aca a-- --a caa c-- --- --- --- --- --- --- --- --- --- --- --- --- --- --- --- --- --- --- --- --- --- --- --- --- --- --- --- --- --- --- --- --- --- --- --- -tg gat att cga t-- --- --- --- --- --- --- --- --- --- --- --- --- --- --- --- --- --- --- --- --- --- --- --- --- --- --- aac agc att tgt gt- --- --- --- --- --- --- --- --- --- --- --- --- --- --- --- --- --

>ON676708.1_Monkeypox_virus_isolate_MPXV_USA_2021_MD_complete_genome

ttt ttt cga tct atc ctc gtc c-- -t- ctc atc atc ctt ata --- --- --- --- --- --- -tt att atc att att atc ata gtc tat taa aca caa atc atc t-- --- --- --- --- --- --- --- --- --- --- --- --- --- --- --- --- --- --- --- --- --- --- --- --- --- --- acg ttt ata ac- --- --- --- --- --- --- --- --- aac att c-- --- --- --- --t cat tat taa tta gtt ctg tag -aa tat ctt taa taa ttt ggc tat a-- --- --c atc tgt t-- --- --- --- --- --- --- --- --- --- --- --- --- --- --- --- --- --- --- caa tac t-- --- --- --- --- --- --- --- atc tat tga tga ttt ctt tt- --- --- --- --- --- --- --- --- --- --- --- tta aga ct- --- --- --- --- --- --- --- --- --- --- --- --- --- --- --- --- --- --- --- --- --- --- --- --- --- --- --- --- --- --- --- --- --- --- --- --- --- --- --- -ta aac tag t-- --- --- --- --- --- --- --- --- --- --- --- --- --- --- --- --- --- --- --- --- --- --- --- --- --- --- --- --- --- --- --- --- --- --- --- --- --- --- --- --- --- --- --- --- --- --- --- --- --- --- --- --- --- --- --- --- --- --- --- --- --- --- --- --- --- --- --- --- --- --- --- --- --- --- --- --- --- --- --- --- --- --- --- --- --- --- --- --- --- --- --- --- --- --- --- --- --- --- --- --- --- --- --- --- --- --- --- --- --- tat ggt aat gac gat gaa a-- --t cga gta gta --- --- act tct aat aaa gac ttg ata --- --- tca tta tca tat gtt tga tcg --- --- --- --- --- --- --- tca tag tta ata gtg tg- --- --- --- --- --- --- --- --- --- --- --- --- --- --- --- --- --- --- --- --g cta aat ggt act gtt aat aag ttt at- --- --- --- --- --- --- --- --- --- --- --- --- --- --- --- --- --- aga caa tat cat agt att ttc ttt cca gaa t-- --- --- --- tag att att ttt tta aat act gat cct cac aat tcc gtg atg tag cag tag ttg gt- --- --- --- --- --- --- --- --- --- --- --- --- --- --- --- --- --- --- --- --- --- --- --- --- --- --- --- --g cat ggt cta tat cgt --- --- --- --- --- --- --- --- --- --- --- --- --- --- --- --- --- --- --- --- --- --- --- --- --- --- --- --- --- --- --- --- --- --- --- --- --- --- --- --- --- --- --- -ta aaa tgt atc ata tat aat agt ttt ctg acg tgg agt aca gaa ttt tcg a-- --- --- --- --- --- --- --- --- --- --- --- --- --- --- --- --- --- --- --- --- --- --- --- --- --- --- --- --- --- --- --- --- --- --- --- --- --- --- --- --- --- --- --- tta atg agt tca tgg taa gga agg gca aat gcc t-- -gt ata taa tat aca taa gtt aa- --- --- --- --- --- --- --- --- --- --- --- --- --- tag ttt ttt atc ata ttt --- --- --- --- --- --- --- --- tct aat acc ata ata aaa att atc --- --- --- --- --- --- --- --- --- --- --- --- --- --- --- --- --- --- --- --- --- --- --- --- --- -at tat tgc gtt tg- gta gtt --- --- --- --- -ct gcc cta --- --- --- --- --- --- --- --- tca tct ata tca ctg tca ctc tc- --- --- gct ctc act ata tct tct aaa att aca a-- --a caa c-- --- --- --- --- --- --- --- --- --- --- --- --- --- --- --- --- --- --- --- --- --- --- --- --- --- --- --- --- --- --- --- --- --- --- --- -tg gat att cga t-- --- --- --- --- --- --- --- --- --- --- --- --- --- --- --- --- --- --- --- --- --- --- --- --- --- --- aac agc att tgt gt- --- --- --- --- --- --- --- --- --- --- --- --- --- --- --- --- --

>ON585033.1_Monkeypox_virus_isolate_Monkeypox/PT0006/2022_complete_genome

ttt ttt cga tct atc ctc gtc c-- -t- ctc atc atc ctt ata --- --- --- --- --- --- -tt att atc att att atc ata gtc tat taa aca caa atc atc t-- --- --- --- --- --- --- --- --- --- --- --- --- --- --- --- --- --- --- --- --- --- --- --- --- --- --- acg ttt ata ac- --- --- --- --- --- --- --- --- aac att c-- --- --- --- --t cat tat taa tta gtt ctg tag -aa tat ctt taa taa ttt ggc tat a-- --- --c atc tgt t-- --- --- --- --- --- --- --- --- --- --- --- --- --- --- --- --- --- --- caa tac t-- --- --- --- --- --- --- --- atc tat tga tga ttt ctt tt- --- --- --- --- --- --- --- --- --- --- --- tta aga ct- --- --- --- --- --- --- --- --- --- --- --- --- --- --- --- --- --- --- --- --- --- --- --- --- --- --- --- --- --- --- --- --- --- --- --- --- --- --- --- -ta aac tag t-- --- --- --- --- --- --- --- --- --- --- --- --- --- --- --- --- --- --- --- --- --- --- --- --- --- --- --- --- --- --- --- --- --- --- --- --- --- --- --- --- --- --- --- --- --- --- --- --- --- --- --- --- --- --- --- --- --- --- --- --- --- --- --- --- --- --- --- --- --- --- --- --- --- --- --- --- --- --- --- --- --- --- --- --- --- --- --- --- --- --- --- --- --- --- --- --- --- --- --- --- --- --- --- --- --- --- --- --- --- tat ggt aat gac gat gaa a-- --t cga gta gta --- --- act tct aat aaa gac ttg ata --- --- tca tta tca tat gtt tga tcg --- --- --- --- --- --- --- tca tag tta ata gtg tg- --- --- --- --- --- --- --- --- --- --- --- --- --- --- --- --- --- --- --- --g cta aat ggt act gtt aat aag ttt at- --- --- --- --- --- --- --- --- --- --- --- --- --- --- --- --- --- aga caa tat cat agt att ttc ttt cca gaa t-- --- --- --- tag att att ttt tta aat act gat cct cac aat tcc gtg atg tag cag tag ttg gt- --- --- --- --- --- --- --- --- --- --- --- --- --- --- --- --- --- --- --- --- --- --- --- --- --- --- --- --g cat ggt cta tat cgt --- --- --- --- --- --- --- --- --- --- --- --- --- --- --- --- --- --- --- --- --- --- --- --- --- --- --- --- --- --- --- --- --- --- --- --- --- --- --- --- --- --- --- -ta aaa tgt atc ata tat aat agt ttt ctg acg tgg agt aca gaa ttt tcg a-- --- --- --- --- --- --- --- --- --- --- --- --- --- --- --- --- --- --- --- --- --- --- --- --- --- --- --- --- --- --- --- --- --- --- --- --- --- --- --- --- --- --- --- tta atg agt tca tgg taa gga agg gca aat gcc t-- -gt ata taa tat aca taa gtt aa- --- --- --- --- --- --- --- --- --- --- --- --- --- tag ttt ttt atc ata ttt --- --- --- --- --- --- --- --- tct aat acc ata ata aaa att atc --- --- --- --- --- --- --- --- --- --- --- --- --- --- --- --- --- --- --- --- --- --- --- --- --- -at tat tgc gtt tg- gta gtt --- --- --- --- -ct gcc cta --- --- --- --- --- --- --- --- tca tct ata tca ctg tca ctc tc- --- --- gct ctc act ata tct tct aaa att aca a-- --a caa c-- --- --- --- --- --- --- --- --- --- --- --- --- --- --- --- --- --- --- --- --- --- --- --- --- --- --- --- --- --- --- --- --- --- --- --- -tg gat att cga t-- --- --- --- --- --- --- --- --- --- --- --- --- --- --- --- --- --- --- --- --- --- --- --- --- --- --- aac agc att tgt gt- --- --- --- --- --- --- --- --- --- --- --- --- --- --- --- --- --

>ON585035.1_Monkeypox_virus_isolate_Monkeypox/PT0009/2022_complete_genome

ttt ttt cga tct atc ctc gtc c-- -t- ctc atc atc ctt ata --- --- --- --- --- --- -tt att atc att att atc ata gtc tat taa aca caa atc atc t-- --- --- --- --- --- --- --- --- --- --- --- --- --- --- --- --- --- --- --- --- --- --- --- --- --- --- acg ttt ata ac- --- --- --- --- --- --- --- --- aac att c-- --- --- --- --t cat tat taa tta gtt ctg tag -aa tat ctt taa taa ttt ggc tat a-- --- --c atc tgt t-- --- --- --- --- --- --- --- --- --- --- --- --- --- --- --- --- --- --- caa tac t-- --- --- --- --- --- --- --- atc tat tga tga ttt ctt tt- --- --- --- --- --- --- --- --- --- --- --- tta aga ct- --- --- --- --- --- --- --- --- --- --- --- --- --- --- --- --- --- --- --- --- --- --- --- --- --- --- --- --- --- --- --- --- --- --- --- --- --- --- --- -ta aac tag t-- --- --- --- --- --- --- --- --- --- --- --- --- --- --- --- --- --- --- --- --- --- --- --- --- --- --- --- --- --- --- --- --- --- --- --- --- --- --- --- --- --- --- --- --- --- --- --- --- --- --- --- --- --- --- --- --- --- --- --- --- --- --- --- --- --- --- --- --- --- --- --- --- --- --- --- --- --- --- --- --- --- --- --- --- --- --- --- --- --- --- --- --- --- --- --- --- --- --- --- --- --- --- --- --- --- --- --- --- --- tat ggt aat gac gat gaa a-- --t cga gta gta --- --- act tct aat aaa gac ttg ata --- --- tca tta tca tat gtt tga tcg --- --- --- --- --- --- --- tca tag tta ata gtg tg- --- --- --- --- --- --- --- --- --- --- --- --- --- --- --- --- --- --- --- --g cta aat ggt act gtt aat aag ttt at- --- --- --- --- --- --- --- --- --- --- --- --- --- --- --- --- --- aga caa tat cat agt att ttc ttt cca gaa t-- --- --- --- tag att att ttt tta aat act gat cct cac aat tcc gtg atg tag cag tag ttg gt- --- --- --- --- --- --- --- --- --- --- --- --- --- --- --- --- --- --- --- --- --- --- --- --- --- --- --- --g cat ggt cta tat cgt --- --- --- --- --- --- --- --- --- --- --- --- --- --- --- --- --- --- --- --- --- --- --- --- --- --- --- --- --- --- --- --- --- --- --- --- --- --- --- --- --- --- --- -ta aaa tgt atc ata tat aat agt ttt ctg acg tgg agt aca gaa ttt tcg a-- --- --- --- --- --- --- --- --- --- --- --- --- --- --- --- --- --- --- --- --- --- --- --- --- --- --- --- --- --- --- --- --- --- --- --- --- --- --- --- --- --- --- --- tta atg agt tca tgg taa gga agg gca aat gcc t-- -gt ata taa tat aca taa gtt aa- --- --- --- --- --- --- --- --- --- --- --- --- --- tag ttt ttt atc ata ttt --- --- --- --- --- --- --- --- tct aat acc ata ata aaa att atc --- --- --- --- --- --- --- --- --- --- --- --- --- --- --- --- --- --- --- --- --- --- --- --- --- -at tat tgc gtt tg- gta gtt --- --- --- --- -ct gcc cta --- --- --- --- --- --- --- --- tca tct ata tca ctg tca ctc tc- --- --- gct ctc act ata tct tct aaa att aca a-- --a caa c-- --- --- --- --- --- --- --- --- --- --- --- --- --- --- --- --- --- --- --- --- --- --- --- --- --- --- --- --- --- --- --- --- --- --- --- -tg gat att cga t-- --- --- --- --- --- --- --- --- --- --- --- --- --- --- --- --- --- --- --- --- --- --- --- --- --- --- aac agc att tgt gt- --- --- --- --- --- --- --- --- --- --- --- --- --- --- --- --- --

>ON649725.1_Monkeypox_virus_isolate_Monkeypox/PT0011/2022_complete_genome

ttt ttt cga tct atc ctc gtc c-- -t- ctc atc atc ctt ata --- --- --- --- --- --- -tt att atc att att atc ata gtc tat taa aca caa atc atc t-- --- --- --- --- --- --- --- --- --- --- --- --- --- --- --- --- --- --- --- --- --- --- --- --- --- --- acg ttt ata ac- --- --- --- --- --- --- --- --- aac att c-- --- --- --- --t cat tat taa tta gtt ctg tag -aa tat ctt taa taa ttt ggc tat a-- --- --c atc tgt t-- --- --- --- --- --- --- --- --- --- --- --- --- --- --- --- --- --- --- caa tac t-- --- --- --- --- --- --- --- atc tat tga tga ttt ctt tt- --- --- --- --- --- --- --- --- --- --- --- tta aga ct- --- --- --- --- --- --- --- --- --- --- --- --- --- --- --- --- --- --- --- --- --- --- --- --- --- --- --- --- --- --- --- --- --- --- --- --- --- --- --- -ta aac tag t-- --- --- --- --- --- --- --- --- --- --- --- --- --- --- --- --- --- --- --- --- --- --- --- --- --- --- --- --- --- --- --- --- --- --- --- --- --- --- --- --- --- --- --- --- --- --- --- --- --- --- --- --- --- --- --- --- --- --- --- --- --- --- --- --- --- --- --- --- --- --- --- --- --- --- --- --- --- --- --- --- --- --- --- --- --- --- --- --- --- --- --- --- --- --- --- --- --- --- --- --- --- --- --- --- --- --- --- --- --- tat ggt aat gac gat gaa a-- --t cga gta gta --- --- act tct aat aaa gac ttg ata --- --- tca tta tca tat gtt tga tcg --- --- --- --- --- --- --- tca tag tta ata gtg tg- --- --- --- --- --- --- --- --- --- --- --- --- --- --- --- --- --- --- --- --g cta aat ggt act gtt aat aag ttt at- --- --- --- --- --- --- --- --- --- --- --- --- --- --- --- --- --- aga caa tat cat agt att ttc ttt cca gaa t-- --- --- --- tag att att ttt tta aat act gat cct cac aat tcc gtg atg tag cag tag ttg gt- --- --- --- --- --- --- --- --- --- --- --- --- --- --- --- --- --- --- --- --- --- --- --- --- --- --- --- --g cat ggt cta tat cgt --- --- --- --- --- --- --- --- --- --- --- --- --- --- --- --- --- --- --- --- --- --- --- --- --- --- --- --- --- --- --- --- --- --- --- --- --- --- --- --- --- --- --- -ta aaa tgt atc ata tat aat agt ttt ctg acg tgg agt aca gaa ttt tcg a-- --- --- --- --- --- --- --- --- --- --- --- --- --- --- --- --- --- --- --- --- --- --- --- --- --- --- --- --- --- --- --- --- --- --- --- --- --- --- --- --- --- --- --- tta atg agt tca tgg taa gga agg gca aat gcc t-- -gt ata taa tat aca taa gtt aa- --- --- --- --- --- --- --- --- --- --- --- --- --- tag ttt ttt atc ata ttt --- --- --- --- --- --- --- --- tct aat acc ata ata aaa att atc --- --- --- --- --- --- --- --- --- --- --- --- --- --- --- --- --- --- --- --- --- --- --- --- --- -at tat tgc gtt tg- gta gtt --- --- --- --- -ct gcc cta --- --- --- --- --- --- --- --- tca tct ata tca ctg tca ctc tc- --- --- gct ctc act ata tct tct aaa att aca a-- --a caa c-- --- --- --- --- --- --- --- --- --- --- --- --- --- --- --- --- --- --- --- --- --- --- --- --- --- --- --- --- --- --- --- --- --- --- --- -tg gat att cga t-- --- --- --- --- --- --- --- --- --- --- --- --- --- --- --- --- --- --- --- --- --- --- --- --- --- --- aac agc att tgt gt- --- --- --- --- --- --- --- --- --- --- --- --- --- --- --- --- --

>ON649724.1_Monkeypox_virus_isolate_Monkeypox/PT0014/2022_complete_genome

ttt ttt cga tct atc ctc gtc c-- -t- ctc atc atc ctt ata --- --- --- --- --- --- -tt att atc att att atc ata gtc tat taa aca caa atc atc t-- --- --- --- --- --- --- --- --- --- --- --- --- --- --- --- --- --- --- --- --- --- --- --- --- --- --- acg ttt ata ac- --- --- --- --- --- --- --- --- aac att c-- --- --- --- --t cat tat taa tta gtt ctg tag -aa tat ctt taa taa ttt ggc tat a-- --- --c atc tgt t-- --- --- --- --- --- --- --- --- --- --- --- --- --- --- --- --- --- --- caa tac t-- --- --- --- --- --- --- --- atc tat tga tga ttt ctt tt- --- --- --- --- --- --- --- --- --- --- --- tta aga ct- --- --- --- --- --- --- --- --- --- --- --- --- --- --- --- --- --- --- --- --- --- --- --- --- --- --- --- --- --- --- --- --- --- --- --- --- --- --- --- -ta aac tag t-- --- --- --- --- --- --- --- --- --- --- --- --- --- --- --- --- --- --- --- --- --- --- --- --- --- --- --- --- --- --- --- --- --- --- --- --- --- --- --- --- --- --- --- --- --- --- --- --- --- --- --- --- --- --- --- --- --- --- --- --- --- --- --- --- --- --- --- --- --- --- --- --- --- --- --- --- --- --- --- --- --- --- --- --- --- --- --- --- --- --- --- --- --- --- --- --- --- --- --- --- --- --- --- --- --- --- --- --- --- tat ggt aat gac gat gaa a-- --t cga gta gta --- --- act tct aat aaa gac ttg ata --- --- tca tta tca tat gtt tga tcg --- --- --- --- --- --- --- tca tag tta ata gtg tg- --- --- --- --- --- --- --- --- --- --- --- --- --- --- --- --- --- --- --- --g cta aat ggt act gtt aat aag ttt at- --- --- --- --- --- --- --- --- --- --- --- --- --- --- --- --- --- aga caa tat cat agt att ttc ttt cca gaa t-- --- --- --- tag att att ttt tta aat act gat cct cac aat tcc gtg atg tag cag tag ttg gt- --- --- --- --- --- --- --- --- --- --- --- --- --- --- --- --- --- --- --- --- --- --- --- --- --- --- --- --g cat ggt cta tat cgt --- --- --- --- --- --- --- --- --- --- --- --- --- --- --- --- --- --- --- --- --- --- --- --- --- --- --- --- --- --- --- --- --- --- --- --- --- --- --- --- --- --- --- -ta aaa tgt atc ata tat aat agt ttt ctg acg tgg agt aca gaa ttt tcg a-- --- --- --- --- --- --- --- --- --- --- --- --- --- --- --- --- --- --- --- --- --- --- --- --- --- --- --- --- --- --- --- --- --- --- --- --- --- --- --- --- --- --- --- tta atg agt tca tgg taa gga agg gca aat gcc t-- -gt ata taa tat aca taa gtt aa- --- --- --- --- --- --- --- --- --- --- --- --- --- tag ttt ttt atc ata ttt --- --- --- --- --- --- --- --- tct aat acc ata ata aaa att atc --- --- --- --- --- --- --- --- --- --- --- --- --- --- --- --- --- --- --- --- --- --- --- --- --- -at tat tgc gtt tg- gta gtt --- --- --- --- -ct gcc cta --- --- --- --- --- --- --- --- tca tct ata tca ctg tca ctc tc- --- --- gct ctc act ata tct tct aaa att aca a-- --a caa c-- --- --- --- --- --- --- --- --- --- --- --- --- --- --- --- --- --- --- --- --- --- --- --- --- --- --- --- --- --- --- --- --- --- --- --- -tg gat att cga t-- --- --- --- --- --- --- --- --- --- --- --- --- --- --- --- --- --- --- --- --- --- --- --- --- --- --- aac agc att tgt gt- --- --- --- --- --- --- --- --- --- --- --- --- --- --- --- --- --

>ON649722.1_Monkeypox_virus_isolate_Monkeypox/PT0022/2022_complete_genome

ttt ttt cga tct atc ctc gtc c-- -t- ctc atc atc ctt ata --- --- --- --- --- --- -tt att atc att att atc ata gtc tat taa aca caa atc atc t-- --- --- --- --- --- --- --- --- --- --- --- --- --- --- --- --- --- --- --- --- --- --- --- --- --- --- acg ttt ata ac- --- --- --- --- --- --- --- --- aac att c-- --- --- --- --t cat tat taa tta gtt ctg tag -aa tat ctt taa taa ttt ggc tat a-- --- --c atc tgt t-- --- --- --- --- --- --- --- --- --- --- --- --- --- --- --- --- --- --- caa tac t-- --- --- --- --- --- --- --- atc tat tga tga ttt ctt tt- --- --- --- --- --- --- --- --- --- --- --- tta aga ct- --- --- --- --- --- --- --- --- --- --- --- --- --- --- --- --- --- --- --- --- --- --- --- --- --- --- --- --- --- --- --- --- --- --- --- --- --- --- --- -ta aac tag t-- --- --- --- --- --- --- --- --- --- --- --- --- --- --- --- --- --- --- --- --- --- --- --- --- --- --- --- --- --- --- --- --- --- --- --- --- --- --- --- --- --- --- --- --- --- --- --- --- --- --- --- --- --- --- --- --- --- --- --- --- --- --- --- --- --- --- --- --- --- --- --- --- --- --- --- --- --- --- --- --- --- --- --- --- --- --- --- --- --- --- --- --- --- --- --- --- --- --- --- --- --- --- --- --- --- --- --- --- --- tat ggt aat gac gat gaa a-- --t cga gta gta --- --- act tct aat aaa gac ttg ata --- --- tca tta tca tat gtt tga tcg --- --- --- --- --- --- --- tca tag tta ata gtg tg- --- --- --- --- --- --- --- --- --- --- --- --- --- --- --- --- --- --- --- --g cta aat ggt act gtt aat aag ttt at- --- --- --- --- --- --- --- --- --- --- --- --- --- --- --- --- --- aga caa tat cat agt att ttc ttt cca gaa t-- --- --- --- tag att att ttt tta aat act gat cct cac aat tcc gtg atg tag cag tag ttg gt- --- --- --- --- --- --- --- --- --- --- --- --- --- --- --- --- --- --- --- --- --- --- --- --- --- --- --- --g cat ggt cta tat cgt --- --- --- --- --- --- --- --- --- --- --- --- --- --- --- --- --- --- --- --- --- --- --- --- --- --- --- --- --- --- --- --- --- --- --- --- --- --- --- --- --- --- --- -ta aaa tgt atc ata tat aat agt ttt ctg acg tgg agt aca gaa ttt tcg a-- --- --- --- --- --- --- --- --- --- --- --- --- --- --- --- --- --- --- --- --- --- --- --- --- --- --- --- --- --- --- --- --- --- --- --- --- --- --- --- --- --- --- --- tta atg agt tca tgg taa gga agg gca aat gcc t-- -gt ata taa tat aca taa gtt aa- --- --- --- --- --- --- --- --- --- --- --- --- --- tag ttt ttt atc ata ttt --- --- --- --- --- --- --- --- tct aat acc ata ata aaa att atc --- --- --- --- --- --- --- --- --- --- --- --- --- --- --- --- --- --- --- --- --- --- --- --- --- -at tat tgc gtt tg- gta gtt --- --- --- --- -ct gcc cta --- --- --- --- --- --- --- --- tca tct ata tca ctg tca ctc tc- --- --- gct ctc act ata tct tct aaa att aca a-- --a caa c-- --- --- --- --- --- --- --- --- --- --- --- --- --- --- --- --- --- --- --- --- --- --- --- --- --- --- --- --- --- --- --- --- --- --- --- -tg gat att cga t-- --- --- --- --- --- --- --- --- --- --- --- --- --- --- --- --- --- --- --- --- --- --- --- --- --- --- aac agc att tgt gt- --- --- --- --- --- --- --- --- --- --- --- --- --- --- --- --- --

>ON649721.1_Monkeypox_virus_isolate_Monkeypox/PT0021/2022_complete_genome

ttt ttt cga tct atc ctc gtc c-- -t- ctc atc atc ctt ata --- --- --- --- --- --- -tt att atc att att atc ata gtc tat taa aca caa atc atc t-- --- --- --- --- --- --- --- --- --- --- --- --- --- --- --- --- --- --- --- --- --- --- --- --- --- --- acg ttt ata ac- --- --- --- --- --- --- --- --- aac att c-- --- --- --- --t cat tat taa tta gtt ctg tag -aa tat ctt taa taa ttt ggc tat a-- --- --c atc tgt t-- --- --- --- --- --- --- --- --- --- --- --- --- --- --- --- --- --- --- caa tac t-- --- --- --- --- --- --- --- atc tat tga tga ttt ctt tt- --- --- --- --- --- --- --- --- --- --- --- tta aga ct- --- --- --- --- --- --- --- --- --- --- --- --- --- --- --- --- --- --- --- --- --- --- --- --- --- --- --- --- --- --- --- --- --- --- --- --- --- --- --- -ta aac tag t-- --- --- --- --- --- --- --- --- --- --- --- --- --- --- --- --- --- --- --- --- --- --- --- --- --- --- --- --- --- --- --- --- --- --- --- --- --- --- --- --- --- --- --- --- --- --- --- --- --- --- --- --- --- --- --- --- --- --- --- --- --- --- --- --- --- --- --- --- --- --- --- --- --- --- --- --- --- --- --- --- --- --- --- --- --- --- --- --- --- --- --- --- --- --- --- --- --- --- --- --- --- --- --- --- --- --- --- --- --- tat ggt aat gac gat gaa a-- --t cga gta gta --- --- act tct aat aaa gac ttg ata --- --- tca tta tca tat gtt tga tcg --- --- --- --- --- --- --- tca tag tta ata gtg tg- --- --- --- --- --- --- --- --- --- --- --- --- --- --- --- --- --- --- --- --g cta aat ggt act gtt aat aag ttt at- --- --- --- --- --- --- --- --- --- --- --- --- --- --- --- --- --- aga caa tat cat agt att ttc ttt cca gaa t-- --- --- --- tag att att ttt tta aat act gat cct cac aat tcc gtg atg tag cag tag ttg gt- --- --- --- --- --- --- --- --- --- --- --- --- --- --- --- --- --- --- --- --- --- --- --- --- --- --- --- --g cat ggt cta tat cgt --- --- --- --- --- --- --- --- --- --- --- --- --- --- --- --- --- --- --- --- --- --- --- --- --- --- --- --- --- --- --- --- --- --- --- --- --- --- --- --- --- --- --- -ta aaa tgt atc ata tat aat agt ttt ctg acg tgg agt aca gaa ttt tcg a-- --- --- --- --- --- --- --- --- --- --- --- --- --- --- --- --- --- --- --- --- --- --- --- --- --- --- --- --- --- --- --- --- --- --- --- --- --- --- --- --- --- --- --- tta atg agt tca tgg taa gga agg gca aat gcc t-- -gt ata taa tat aca taa gtt aa- --- --- --- --- --- --- --- --- --- --- --- --- --- tag ttt ttt atc ata ttt --- --- --- --- --- --- --- --- tct aat acc ata ata aaa att atc --- --- --- --- --- --- --- --- --- --- --- --- --- --- --- --- --- --- --- --- --- --- --- --- --- -at tat tgc gtt tg- gta gtt --- --- --- --- -ct gcc cta --- --- --- --- --- --- --- --- tca tct ata tca ctg tca ctc tc- --- --- gct ctc act ata tct tct aaa att aca a-- --a caa c-- --- --- --- --- --- --- --- --- --- --- --- --- --- --- --- --- --- --- --- --- --- --- --- --- --- --- --- --- --- --- --- --- --- --- --- -tg gat att cga t-- --- --- --- --- --- --- --- --- --- --- --- --- --- --- --- --- --- --- --- --- --- --- --- --- --- --- aac agc att tgt gt- --- --- --- --- --- --- --- --- --- --- --- --- --- --- --- --- --

>ON649720.1_Monkeypox_virus_isolate_Monkeypox/PT0024/2022_complete_genome

ttt ttt cga tct atc ctc gtc c-- -t- ctc atc atc ctt ata --- --- --- --- --- --- -tt att atc att att atc ata gtc tat taa aca caa atc atc t-- --- --- --- --- --- --- --- --- --- --- --- --- --- --- --- --- --- --- --- --- --- --- --- --- --- --- acg ttt ata ac- --- --- --- --- --- --- --- --- aac att c-- --- --- --- --t cat tat taa tta gtt ctg tag -aa tat ctt taa taa ttt ggc tat a-- --- --c atc tgt t-- --- --- --- --- --- --- --- --- --- --- --- --- --- --- --- --- --- --- caa tac t-- --- --- --- --- --- --- --- atc tat tga tga ttt ctt tt- --- --- --- --- --- --- --- --- --- --- --- tta aga ct- --- --- --- --- --- --- --- --- --- --- --- --- --- --- --- --- --- --- --- --- --- --- --- --- --- --- --- --- --- --- --- --- --- --- --- --- --- --- --- -ta aac tag t-- --- --- --- --- --- --- --- --- --- --- --- --- --- --- --- --- --- --- --- --- --- --- --- --- --- --- --- --- --- --- --- --- --- --- --- --- --- --- --- --- --- --- --- --- --- --- --- --- --- --- --- --- --- --- --- --- --- --- --- --- --- --- --- --- --- --- --- --- --- --- --- --- --- --- --- --- --- --- --- --- --- --- --- --- --- --- --- --- --- --- --- --- --- --- --- --- --- --- --- --- --- --- --- --- --- --- --- --- --- tat ggt aat gac gat gaa a-- --t cga gta gta --- --- act tct aat aaa gac ttg ata --- --- tca tta tca tat gtt tga tcg --- --- --- --- --- --- --- tca tag tta ata gtg tg- --- --- --- --- --- --- --- --- --- --- --- --- --- --- --- --- --- --- --- --g cta aat ggt act gtt aat aag ttt at- --- --- --- --- --- --- --- --- --- --- --- --- --- --- --- --- --- aga caa tat cat agt att ttc ttt cca gaa t-- --- --- --- tag att att ttt tta aat act gat cct cac aat tcc gtg atg tag cag tag ttg gt- --- --- --- --- --- --- --- --- --- --- --- --- --- --- --- --- --- --- --- --- --- --- --- --- --- --- --- --g cat ggt cta tat cgt --- --- --- --- --- --- --- --- --- --- --- --- --- --- --- --- --- --- --- --- --- --- --- --- --- --- --- --- --- --- --- --- --- --- --- --- --- --- --- --- --- --- --- -ta aaa tgt atc ata tat aat agt ttt ctg acg tgg agt aca gaa ttt tcg a-- --- --- --- --- --- --- --- --- --- --- --- --- --- --- --- --- --- --- --- --- --- --- --- --- --- --- --- --- --- --- --- --- --- --- --- --- --- --- --- --- --- --- --- tta atg agt tca tgg taa gga agg gca aat gcc t-- -gt ata taa tat aca taa gtt aa- --- --- --- --- --- --- --- --- --- --- --- --- --- tag ttt ttt atc ata ttt --- --- --- --- --- --- --- --- tct aat acc ata ata aaa att atc --- --- --- --- --- --- --- --- --- --- --- --- --- --- --- --- --- --- --- --- --- --- --- --- --- -at tat tgc gtt tg- gta gtt --- --- --- --- -ct gcc cta --- --- --- --- --- --- --- --- tca tct ata tca ctg tca ctc tc- --- --- gct ctc act ata tct tct aaa att aca a-- --a caa c-- --- --- --- --- --- --- --- --- --- --- --- --- --- --- --- --- --- --- --- --- --- --- --- --- --- --- --- --- --- --- --- --- --- --- --- -tg gat att cga t-- --- --- --- --- --- --- --- --- --- --- --- --- --- --- --- --- --- --- --- --- --- --- --- --- --- --- aac agc att tgt gt- --- --- --- --- --- --- --- --- --- --- --- --- --- --- --- --- --

>ON649723.1_Monkeypox_virus_isolate_Monkeypox/PT0013/2022_complete_genome

ttt ttt cga tct atc ctc gtc c-- -t- ctc atc atc ctt ata --- --- --- --- --- --- -tt att atc att att atc ata gtc tat taa aca caa atc atc t-- --- --- --- --- --- --- --- --- --- --- --- --- --- --- --- --- --- --- --- --- --- --- --- --- --- --- acg ttt ata ac- --- --- --- --- --- --- --- --- aac att c-- --- --- --- --t cat tat taa tta gtt ctg tag -aa tat ctt taa taa ttt ggc tat a-- --- --c atc tgt t-- --- --- --- --- --- --- --- --- --- --- --- --- --- --- --- --- --- --- caa tac t-- --- --- --- --- --- --- --- atc tat tga tga ttt ctt tt- --- --- --- --- --- --- --- --- --- --- --- tta aga ct- --- --- --- --- --- --- --- --- --- --- --- --- --- --- --- --- --- --- --- --- --- --- --- --- --- --- --- --- --- --- --- --- --- --- --- --- --- --- --- -ta aac tag t-- --- --- --- --- --- --- --- --- --- --- --- --- --- --- --- --- --- --- --- --- --- --- --- --- --- --- --- --- --- --- --- --- --- --- --- --- --- --- --- --- --- --- --- --- --- --- --- --- --- --- --- --- --- --- --- --- --- --- --- --- --- --- --- --- --- --- --- --- --- --- --- --- --- --- --- --- --- --- --- --- --- --- --- --- --- --- --- --- --- --- --- --- --- --- --- --- --- --- --- --- --- --- --- --- --- --- --- --- --- tat ggt aat gac gat gaa a-- --t cga gta gta --- --- act tct aat aaa gac ttg ata --- --- tca tta tca tat gtt tga tcg --- --- --- --- --- --- --- tca tag tta ata gtg tg- --- --- --- --- --- --- --- --- --- --- --- --- --- --- --- --- --- --- --- --g cta aat ggt act gtt aat aag ttt at- --- --- --- --- --- --- --- --- --- --- --- --- --- --- --- --- --- aga caa tat cat agt att ttc ttt cca gaa t-- --- --- --- tag att att ttt tta aat act gat cct cac aat tcc gtg atg tag cag tag ttg gt- --- --- --- --- --- --- --- --- --- --- --- --- --- --- --- --- --- --- --- --- --- --- --- --- --- --- --- --g cat ggt cta tat cgt --- --- --- --- --- --- --- --- --- --- --- --- --- --- --- --- --- --- --- --- --- --- --- --- --- --- --- --- --- --- --- --- --- --- --- --- --- --- --- --- --- --- --- -ta aaa tgt atc ata tat aat agt ttt ctg acg tgg agt aca gaa ttt tcg a-- --- --- --- --- --- --- --- --- --- --- --- --- --- --- --- --- --- --- --- --- --- --- --- --- --- --- --- --- --- --- --- --- --- --- --- --- --- --- --- --- --- --- --- tta atg agt tca tgg taa gga agg gca aat gcc t-- -gt ata taa tat aca taa gtt aa- --- --- --- --- --- --- --- --- --- --- --- --- --- tag ttt ttt atc ata ttt --- --- --- --- --- --- --- --- tct aat acc ata ata aaa att atc --- --- --- --- --- --- --- --- --- --- --- --- --- --- --- --- --- --- --- --- --- --- --- --- --- -at tat tgc gtt tg- gta gtt --- --- --- --- -ct gcc cta --- --- --- --- --- --- --- --- tca tct ata tca ctg tca ctc tc- --- --- gct ctc act ata tct tct aaa att aca a-- --a caa c-- --- --- --- --- --- --- --- --- --- --- --- --- --- --- --- --- --- --- --- --- --- --- --- --- --- --- --- --- --- --- --- --- --- --- --- -tg gat att cga t-- --- --- --- --- --- --- --- --- --- --- --- --- --- --- --- --- --- --- --- --- --- --- --- --- --- --- aac agc att tgt gt- --- --- --- --- --- --- --- --- --- --- --- --- --- --- --- --- --

>ON649718.1_Monkeypox_virus_isolate_Monkeypox/PT0018/2022_complete_genome

ttt ttt cga tct atc ctc gtc c-- -t- ctc atc atc ctt ata --- --- --- --- --- --- -tt att atc att att atc ata gtc tat taa aca caa atc atc t-- --- --- --- --- --- --- --- --- --- --- --- --- --- --- --- --- --- --- --- --- --- --- --- --- --- --- acg ttt ata ac- --- --- --- --- --- --- --- --- aac att c-- --- --- --- --t cat tat taa tta gtt ctg tag -aa tat ctt taa taa ttt ggc tat a-- --- --c atc tgt t-- --- --- --- --- --- --- --- --- --- --- --- --- --- --- --- --- --- --- caa tac t-- --- --- --- --- --- --- --- atc tat tga tga ttt ctt tt- --- --- --- --- --- --- --- --- --- --- --- tta aga ct- --- --- --- --- --- --- --- --- --- --- --- --- --- --- --- --- --- --- --- --- --- --- --- --- --- --- --- --- --- --- --- --- --- --- --- --- --- --- --- -ta aac tag t-- --- --- --- --- --- --- --- --- --- --- --- --- --- --- --- --- --- --- --- --- --- --- --- --- --- --- --- --- --- --- --- --- --- --- --- --- --- --- --- --- --- --- --- --- --- --- --- --- --- --- --- --- --- --- --- --- --- --- --- --- --- --- --- --- --- --- --- --- --- --- --- --- --- --- --- --- --- --- --- --- --- --- --- --- --- --- --- --- --- --- --- --- --- --- --- --- --- --- --- --- --- --- --- --- --- --- --- --- --- tat ggt aat gac gat gaa a-- --t cga gta gta --- --- act tct aat aaa gac ttg ata --- --- tca tta tca tat gtt tga tcg --- --- --- --- --- --- --- tca tag tta ata gtg tg- --- --- --- --- --- --- --- --- --- --- --- --- --- --- --- --- --- --- --- --g cta aat ggt act gtt aat aag ttt at- --- --- --- --- --- --- --- --- --- --- --- --- --- --- --- --- --- aga caa tat cat agt att ttc ttt cca gaa t-- --- --- --- tag att att ttt tta aat act gat cct cac aat tcc gtg atg tag cag tag ttg gt- --- --- --- --- --- --- --- --- --- --- --- --- --- --- --- --- --- --- --- --- --- --- --- --- --- --- --- --g cat ggt cta tat cgt --- --- --- --- --- --- --- --- --- --- --- --- --- --- --- --- --- --- --- --- --- --- --- --- --- --- --- --- --- --- --- --- --- --- --- --- --- --- --- --- --- --- --- -ta aaa tgt atc ata tat aat agt ttt ctg acg tgg agt aca gaa ttt tcg a-- --- --- --- --- --- --- --- --- --- --- --- --- --- --- --- --- --- --- --- --- --- --- --- --- --- --- --- --- --- --- --- --- --- --- --- --- --- --- --- --- --- --- --- tta atg agt tca tgg taa gga agg gca aat gcc t-- -gt ata taa tat aca taa gtt aa- --- --- --- --- --- --- --- --- --- --- --- --- --- tag ttt ttt atc ata ttt --- --- --- --- --- --- --- --- tct aat acc ata ata aaa att atc --- --- --- --- --- --- --- --- --- --- --- --- --- --- --- --- --- --- --- --- --- --- --- --- --- -at tat tgc gtt tg- gta gtt --- --- --- --- -ct gcc cta --- --- --- --- --- --- --- --- tca tct ata tca ctg tca ctc tc- --- --- gct ctc act ata tct tct aaa att aca a-- --a caa c-- --- --- --- --- --- --- --- --- --- --- --- --- --- --- --- --- --- --- --- --- --- --- --- --- --- --- --- --- --- --- --- --- --- --- --- -tg gat att cga t-- --- --- --- --- --- --- --- --- --- --- --- --- --- --- --- --- --- --- --- --- --- --- --- --- --- --- aac agc att tgt gt- --- --- --- --- --- --- --- --- --- --- --- --- --- --- --- --- --

>ON649719.1_Monkeypox_virus_isolate_Monkeypox/PT0012/2022_complete_genome

ttt ttt cga tct atc ctc gtc c-- -t- ctc atc atc ctt ata --- --- --- --- --- --- -tt att atc att att atc ata gtc tat taa aca caa atc atc t-- --- --- --- --- --- --- --- --- --- --- --- --- --- --- --- --- --- --- --- --- --- --- --- --- --- --- acg ttt ata ac- --- --- --- --- --- --- --- --- aac att c-- --- --- --- --t cat tat taa tta gtt ctg tag -aa tat ctt taa taa ttt ggc tat a-- --- --c atc tgt t-- --- --- --- --- --- --- --- --- --- --- --- --- --- --- --- --- --- --- caa tac t-- --- --- --- --- --- --- --- atc tat tga tga ttt ctt tt- --- --- --- --- --- --- --- --- --- --- --- tta aga ct- --- --- --- --- --- --- --- --- --- --- --- --- --- --- --- --- --- --- --- --- --- --- --- --- --- --- --- --- --- --- --- --- --- --- --- --- --- --- --- -ta aac tag t-- --- --- --- --- --- --- --- --- --- --- --- --- --- --- --- --- --- --- --- --- --- --- --- --- --- --- --- --- --- --- --- --- --- --- --- --- --- --- --- --- --- --- --- --- --- --- --- --- --- --- --- --- --- --- --- --- --- --- --- --- --- --- --- --- --- --- --- --- --- --- --- --- --- --- --- --- --- --- --- --- --- --- --- --- --- --- --- --- --- --- --- --- --- --- --- --- --- --- --- --- --- --- --- --- --- --- --- --- --- tat ggt aat gac gat gaa a-- --t cga gta gta --- --- act tct aat aaa gac ttg ata --- --- tca tta tca tat gtt tga tcg --- --- --- --- --- --- --- tca tag tta ata gtg tg- --- --- --- --- --- --- --- --- --- --- --- --- --- --- --- --- --- --- --- --g cta aat ggt act gtt aat aag ttt at- --- --- --- --- --- --- --- --- --- --- --- --- --- --- --- --- --- aga caa tat cat agt att ttc ttt cca gaa t-- --- --- --- tag att att ttt tta aat act gat cct cac aat tcc gtg atg tag cag tag ttg gt- --- --- --- --- --- --- --- --- --- --- --- --- --- --- --- --- --- --- --- --- --- --- --- --- --- --- --- --g cat ggt cta tat cgt --- --- --- --- --- --- --- --- --- --- --- --- --- --- --- --- --- --- --- --- --- --- --- --- --- --- --- --- --- --- --- --- --- --- --- --- --- --- --- --- --- --- --- -ta aaa tgt atc ata tat aat agt ttt ctg acg tgg agt aca gaa ttt tcg a-- --- --- --- --- --- --- --- --- --- --- --- --- --- --- --- --- --- --- --- --- --- --- --- --- --- --- --- --- --- --- --- --- --- --- --- --- --- --- --- --- --- --- --- tta atg agt tca tgg taa gga agg gca aat gcc t-- -gt ata taa tat aca taa gtt aa- --- --- --- --- --- --- --- --- --- --- --- --- --- tag ttt ttt atc ata ttt --- --- --- --- --- --- --- --- tct aat acc ata ata aaa att atc --- --- --- --- --- --- --- --- --- --- --- --- --- --- --- --- --- --- --- --- --- --- --- --- --- -at tat tgc gtt tg- gta gtt --- --- --- --- -ct gcc cta --- --- --- --- --- --- --- --- tca tct ata tca ctg tca ctc tc- --- --- gct ctc act ata tct tct aaa att aca a-- --a caa c-- --- --- --- --- --- --- --- --- --- --- --- --- --- --- --- --- --- --- --- --- --- --- --- --- --- --- --- --- --- --- --- --- --- --- --- -tg gat att cga t-- --- --- --- --- --- --- --- --- --- --- --- --- --- --- --- --- --- --- --- --- --- --- --- --- --- --- aac agc att tgt gt- --- --- --- --- --- --- --- --- --- --- --- --- --- --- --- --- --

>ON682267.1_Monkeypox_virus_isolate_MPXV/Germany/2022/RKI010_complete_genome

ttt ttt cga tct atc ctc gtc c-- -t- ctc atc atc ctt ata --- --- --- --- --- --- -tt att atc att att atc ata gtc tat taa aca caa atc atc t-- --- --- --- --- --- --- --- --- --- --- --- --- --- --- --- --- --- --- --- --- --- --- --- --- --- --- acg ttt ata ac- --- --- --- --- --- --- --- --- aac att c-- --- --- --- --t cat tat taa tta gtt ctg tag -aa tat ctt taa taa ttt ggc tat a-- --- --c atc tgt t-- --- --- --- --- --- --- --- --- --- --- --- --- --- --- --- --- --- --- caa tac t-- --- --- --- --- --- --- --- atc tat tga tga ttt ctt tt- --- --- --- --- --- --- --- --- --- --- --- tta aga ct- --- --- --- --- --- --- --- --- --- --- --- --- --- --- --- --- --- --- --- --- --- --- --- --- --- --- --- --- --- --- --- --- --- --- --- --- --- --- --- -ta aac tag t-- --- --- --- --- --- --- --- --- --- --- --- --- --- --- --- --- --- --- --- --- --- --- --- --- --- --- --- --- --- --- --- --- --- --- --- --- --- --- --- --- --- --- --- --- --- --- --- --- --- --- --- --- --- --- --- --- --- --- --- --- --- --- --- --- --- --- --- --- --- --- --- --- --- --- --- --- --- --- --- --- --- --- --- --- --- --- --- --- --- --- --- --- --- --- --- --- --- --- --- --- --- --- --- --- --- --- --- --- --- tat ggt aat gac gat gaa a-- --t cga gta gta --- --- act tct aat aaa gac ttg ata --- --- tca tta tca tat gtt tga tcg --- --- --- --- --- --- --- tca tag tta ata gtg tg- --- --- --- --- --- --- --- --- --- --- --- --- --- --- --- --- --- --- --- --g cta aat ggt act gtt aat aag ttt at- --- --- --- --- --- --- --- --- --- --- --- --- --- --- --- --- --- aga caa tat cat agt att ttc ttt cca gaa t-- --- --- --- tag att att ttt tta aat act gat cct cac aat tcc gtg atg tag cag tag ttg gt- --- --- --- --- --- --- --- --- --- --- --- --- --- --- --- --- --- --- --- --- --- --- --- --- --- --- --- --g cat ggt cta tat cgt --- --- --- --- --- --- --- --- --- --- --- --- --- --- --- --- --- --- --- --- --- --- --- --- --- --- --- --- --- --- --- --- --- --- --- --- --- --- --- --- --- --- --- -ta aaa tgt atc ata tat aat agt ttt ctg acg tgg agt aca gaa ttt tcg a-- --- --- --- --- --- --- --- --- --- --- --- --- --- --- --- --- --- --- --- --- --- --- --- --- --- --- --- --- --- --- --- --- --- --- --- --- --- --- --- --- --- --- --- tta atg agt tca tgg taa gga agg gca aat gcc t-- -gt ata taa tat aca taa gtt aa- --- --- --- --- --- --- --- --- --- --- --- --- --- tag ttt ttt atc ata ttt --- --- --- --- --- --- --- --- tct aat acc ata ata aaa att atc --- --- --- --- --- --- --- --- --- --- --- --- --- --- --- --- --- --- --- --- --- --- --- --- --- -at tat tgc gtt tg- gta gtt --- --- --- --- -ct gcc cta --- --- --- --- --- --- --- --- tca tct ata tca ctg tca ctc tc- --- --- gct ctc act ata tct tct aaa att aca a-- --a caa c-- --- --- --- --- --- --- --- --- --- --- --- --- --- --- --- --- --- --- --- --- --- --- --- --- --- --- --- --- --- --- --- --- --- --- --- -tg gat att cga t-- --- --- --- --- --- --- --- --- --- --- --- --- --- --- --- --- --- --- --- --- --- --- --- --- --- --- aac agc att tgt gt- --- --- --- --- --- --- --- --- --- --- --- --- --- --- --- --- --

>ON649717.1_Monkeypox_virus_isolate_Monkeypox/PT0019/2022_partial_genome

ttt ttt cga tct atc ctc gtc c-- -t- ctc atc atc ctt ata --- --- --- --- --- --- -tt att atc att att atc ata gtc tat taa aca caa atc atc t-- --- --- --- --- --- --- --- --- --- --- --- --- --- --- --- --- --- --- --- --- --- --- --- --- --- --- acg ttt ata ac- --- --- --- --- --- --- --- --- aac att c-- --- --- --- --t cat tat taa tta gtt ctg tag -aa tat ctt taa taa ttt ggc tat a-- --- --c atc tgt t-- --- --- --- --- --- --- --- --- --- --- --- --- --- --- --- --- --- --- caa tac t-- --- --- --- --- --- --- --- atc tat tga tga ttt ctt tt- --- --- --- --- --- --- --- --- --- --- --- tta aga ct- --- --- --- --- --- --- --- --- --- --- --- --- --- --- --- --- --- --- --- --- --- --- --- --- --- --- --- --- --- --- --- --- --- --- --- --- --- --- --- -ta aac tag t-- --- --- --- --- --- --- --- --- --- --- --- --- --- --- --- --- --- --- --- --- --- --- --- --- --- --- --- --- --- --- --- --- --- --- --- --- --- --- --- --- --- --- --- --- --- --- --- --- --- --- --- --- --- --- --- --- --- --- --- --- --- --- --- --- --- --- --- --- --- --- --- --- --- --- --- --- --- --- --- --- --- --- --- --- --- --- --- --- --- --- --- --- --- --- --- --- --- --- --- --- --- --- --- --- --- --- --- --- --- tat ggt aat gac gat gaa a-- --t cga gta gta --- --- act tct aat aaa gac ttg ata --- --- tca tta tca tat gtt tga tcg --- --- --- --- --- --- --- tca tag tta ata gtg tg- --- --- --- --- --- --- --- --- --- --- --- --- --- --- --- --- --- --- --- --g cta aat ggt act gtt aat aag ttt at- --- --- --- --- --- --- --- --- --- --- --- --- --- --- --- --- --- aga caa tat cat agt att ttc ttt cca gaa t-- --- --- --- tag att att ttt tta aat act gat cct cac aat tcc gtg atg tag cag tag ttg gt- --- --- --- --- --- --- --- --- --- --- --- --- --- --- --- --- --- --- --- --- --- --- --- --- --- --- --- --g cat ggt cta tat cgt --- --- --- --- --- --- --- --- --- --- --- --- --- --- --- --- --- --- --- --- --- --- --- --- --- --- --- --- --- --- --- --- --- --- --- --- --- --- --- --- --- --- --- -ta aaa tgt atc ata tat aat agt ttt ctg acg tgg agt aca gaa ttt tcg a-- --- --- --- --- --- --- --- --- --- --- --- --- --- --- --- --- --- --- --- --- --- --- --- --- --- --- --- --- --- --- --- --- --- --- --- --- --- --- --- --- --- --- --- tta atg agt tca tgg taa gga agg gca aat gcc t-- -gt ata taa tat aca taa gtt aa- --- --- --- --- --- --- --- --- --- --- --- --- --- tag ttt ttt atc ata ttt --- --- --- --- --- --- --- --- tct aat acc ata ata aaa att atc --- --- --- --- --- --- --- --- --- --- --- --- --- --- --- --- --- --- --- --- --- --- --- --- --- -at tat tgc gtt tg- gta gtt --- --- --- --- -ct gcc cta --- --- --- --- --- --- --- --- tca tct ata tca ctg tca ctc tc- --- --- gct ctc act ata tct tct aaa att aca a-- --a caa c-- --- --- --- --- --- --- --- --- --- --- --- --- --- --- --- --- --- --- --- --- --- --- --- --- --- --- --- --- --- --- --- --- --- --- --- -tg gat att cga t-- --- --- --- --- --- --- --- --- --- --- --- --- --- --- --- --- --- --- --- --- --- --- --- --- --- --- aac agc att tgt gt- --- --- --- --- --- --- --- --- --- --- --- --- --- --- --- --- --

>ON649708.1_Monkeypox_virus_isolate_Monkeypox/PT0023/2022_partial_genome

ttt ttt cga tct atc ctc gtc c-- -t- ctc atc atc ctt ata --- --- --- --- --- --- -tt att atc att att atc ata gtc tat taa aca caa atc atc t-- --- --- --- --- --- --- --- --- --- --- --- --- --- --- --- --- --- --- --- --- --- --- --- --- --- --- acg ttt ata ac- --- --- --- --- --- --- --- --- aac att c-- --- --- --- --t cat tat taa tta gtt ctg tag -aa tat ctt taa taa ttt ggc tat a-- --- --c atc tgt t-- --- --- --- --- --- --- --- --- --- --- --- --- --- --- --- --- --- --- caa tac t-- --- --- --- --- --- --- --- atc tat tga tga ttt ctt tt- --- --- --- --- --- --- --- --- --- --- --- tta aga ct- --- --- --- --- --- --- --- --- --- --- --- --- --- --- --- --- --- --- --- --- --- --- --- --- --- --- --- --- --- --- --- --- --- --- --- --- --- --- --- -ta aac tag t-- --- --- --- --- --- --- --- --- --- --- --- --- --- --- --- --- --- --- --- --- --- --- --- --- --- --- --- --- --- --- --- --- --- --- --- --- --- --- --- --- --- --- --- --- --- --- --- --- --- --- --- --- --- --- --- --- --- --- --- --- --- --- --- --- --- --- --- --- --- --- --- --- --- --- --- --- --- --- --- --- --- --- --- --- --- --- --- --- --- --- --- --- --- --- --- --- --- --- --- --- --- --- --- --- --- --- --- --- --- tat ggt aat gac gat gaa a-- --t cga gta gta --- --- act tct aat aaa gac ttg ata --- --- tca tta tca tat gtt tga tcg --- --- --- --- --- --- --- tca tag tta ata gtg tg- --- --- --- --- --- --- --- --- --- --- --- --- --- --- --- --- --- --- --- --g cta aat ggt act gtt aat aag ttt at- --- --- --- --- --- --- --- --- --- --- --- --- --- --- --- --- --- aga caa tat cat agt att ttc ttt cca gaa t-- --- --- --- tag att att ttt tta aat act gat cct cac aat tcc gtg atg tag cag tag ttg gt- --- --- --- --- --- --- --- --- --- --- --- --- --- --- --- --- --- --- --- --- --- --- --- --- --- --- --- --g cat ggt cta tat cgt --- --- --- --- --- --- --- --- --- --- --- --- --- --- --- --- --- --- --- --- --- --- --- --- --- --- --- --- --- --- --- --- --- --- --- --- --- --- --- --- --- --- --- -ta aaa tgt atc ata tat aat agt ttt ctg acg tgg agt aca gaa ttt tcg a-- --- --- --- --- --- --- --- --- --- --- --- --- --- --- --- --- --- --- --- --- --- --- --- --- --- --- --- --- --- --- --- --- --- --- --- --- --- --- --- --- --- --- --- tta atg agt tca tgg taa gga agg gca aat gcc t-- -gt ata taa tat aca taa gtt aa- --- --- --- --- --- --- --- --- --- --- --- --- --- tag ttt ttt atc ata ttt --- --- --- --- --- --- --- --- tct aat acc ata ata aaa att atc --- --- --- --- --- --- --- --- --- --- --- --- --- --- --- --- --- --- --- --- --- --- --- --- --- -at tat tgc gtt tg- gta gtt --- --- --- --- -ct gcc cta --- --- --- --- --- --- --- --- tca tct ata tca ctg tca ctc tc- --- --- gct ctc act ata tct tct aaa att aca a-- --a caa c-- --- --- --- --- --- --- --- --- --- --- --- --- --- --- --- --- --- --- --- --- --- --- --- --- --- --- --- --- --- --- --- --- --- --- --- -tg gat att cga t-- --- --- --- --- --- --- --- --- --- --- --- --- --- --- --- --- --- --- --- --- --- --- --- --- --- --- aac agc att tgt gt- --- --- --- --- --- --- --- --- --- --- --- --- --- --- --- --- --

>ON649709.1_Monkeypox_virus_isolate_Monkeypox/PT0028/2022_complete_genome

ttt ttt cga tct atc ctc gtc c-- -t- ctc atc atc ctt ata --- --- --- --- --- --- -tt att atc att att atc ata gtc tat taa aca caa atc atc t-- --- --- --- --- --- --- --- --- --- --- --- --- --- --- --- --- --- --- --- --- --- --- --- --- --- --- acg ttt ata ac- --- --- --- --- --- --- --- --- aac att c-- --- --- --- --t cat tat taa tta gtt ctg tag -aa tat ctt taa taa ttt ggc tat a-- --- --c atc tgt t-- --- --- --- --- --- --- --- --- --- --- --- --- --- --- --- --- --- --- caa tac t-- --- --- --- --- --- --- --- atc tat tga tga ttt ctt tt- --- --- --- --- --- --- --- --- --- --- --- tta aga ct- --- --- --- --- --- --- --- --- --- --- --- --- --- --- --- --- --- --- --- --- --- --- --- --- --- --- --- --- --- --- --- --- --- --- --- --- --- --- --- -ta aac tag t-- --- --- --- --- --- --- --- --- --- --- --- --- --- --- --- --- --- --- --- --- --- --- --- --- --- --- --- --- --- --- --- --- --- --- --- --- --- --- --- --- --- --- --- --- --- --- --- --- --- --- --- --- --- --- --- --- --- --- --- --- --- --- --- --- --- --- --- --- --- --- --- --- --- --- --- --- --- --- --- --- --- --- --- --- --- --- --- --- --- --- --- --- --- --- --- --- --- --- --- --- --- --- --- --- --- --- --- --- --- tat ggt aat gac gat gaa a-- --t cga gta gta --- --- act tct aat aaa gac ttg ata --- --- tca tta tca tat gtt tga tcg --- --- --- --- --- --- --- tca tag tta ata gtg tg- --- --- --- --- --- --- --- --- --- --- --- --- --- --- --- --- --- --- --- --g cta aat ggt act gtt aat aag ttt at- --- --- --- --- --- --- --- --- --- --- --- --- --- --- --- --- --- aga caa tat cat agt att ttc ttt cca gaa t-- --- --- --- tag att att ttt tta aat act gat cct cac aat tcc gtg atg tag cag tag ttg gt- --- --- --- --- --- --- --- --- --- --- --- --- --- --- --- --- --- --- --- --- --- --- --- --- --- --- --- --g cat ggt cta tat cgt --- --- --- --- --- --- --- --- --- --- --- --- --- --- --- --- --- --- --- --- --- --- --- --- --- --- --- --- --- --- --- --- --- --- --- --- --- --- --- --- --- --- --- -ta aaa tgt atc ata tat aat agt ttt ctg acg tgg agt aca gaa ttt tcg a-- --- --- --- --- --- --- --- --- --- --- --- --- --- --- --- --- --- --- --- --- --- --- --- --- --- --- --- --- --- --- --- --- --- --- --- --- --- --- --- --- --- --- --- tta atg agt tca tgg taa gga agg gca aat gcc t-- -gt ata taa tat aca taa gtt aa- --- --- --- --- --- --- --- --- --- --- --- --- --- tag ttt ttt atc ata ttt --- --- --- --- --- --- --- --- tct aat acc ata ata aaa att atc --- --- --- --- --- --- --- --- --- --- --- --- --- --- --- --- --- --- --- --- --- --- --- --- --- -at tat tgc gtt tg- gta gtt --- --- --- --- -ct gcc cta --- --- --- --- --- --- --- --- tca tct ata tca ctg tca ctc tc- --- --- gct ctc act ata tct tct aaa att aca a-- --a caa c-- --- --- --- --- --- --- --- --- --- --- --- --- --- --- --- --- --- --- --- --- --- --- --- --- --- --- --- --- --- --- --- --- --- --- --- -tg gat att cga t-- --- --- --- --- --- --- --- --- --- --- --- --- --- --- --- --- --- --- --- --- --- --- --- --- --- --- aac agc att tgt gt- --- --- --- --- --- --- --- --- --- --- --- --- --- --- --- --- --

>ON614676.1_Monkeypox_virus_isolate_INMI-Pt1_partial_genome

ttt ttt cga tct atc ctc gtc c-- -t- ctc atc atc ctt ata --- --- --- --- --- --- -tt att atc att att atc ata gtc tat taa aca caa atc atc t-- --- --- --- --- --- --- --- --- --- --- --- --- --- --- --- --- --- --- --- --- --- --- --- --- --- --- acg ttt ata ac- --- --- --- --- --- --- --- --- aac att c-- --- --- --- --t cat tat taa tta gtt ctg tag -aa tat ctt taa taa ttt ggc tat a-- --- --c atc tgt t-- --- --- --- --- --- --- --- --- --- --- --- --- --- --- --- --- --- --- caa tac t-- --- --- --- --- --- --- --- atc tat tga tga ttt ctt tt- --- --- --- --- --- --- --- --- --- --- --- tta aga ct- --- --- --- --- --- --- --- --- --- --- --- --- --- --- --- --- --- --- --- --- --- --- --- --- --- --- --- --- --- --- --- --- --- --- --- --- --- --- --- -ta aac tag t-- --- --- --- --- --- --- --- --- --- --- --- --- --- --- --- --- --- --- --- --- --- --- --- --- --- --- --- --- --- --- --- --- --- --- --- --- --- --- --- --- --- --- --- --- --- --- --- --- --- --- --- --- --- --- --- --- --- --- --- --- --- --- --- --- --- --- --- --- --- --- --- --- --- --- --- --- --- --- --- --- --- --- --- --- --- --- --- --- --- --- --- --- --- --- --- --- --- --- --- --- --- --- --- --- --- --- --- --- --- tat ggt aat gac gat gaa a-- --t cga gta gta --- --- act tct aat aaa gac ttg ata --- --- tca tta tca tat gtt tga tcg --- --- --- --- --- --- --- tca tag tta ata gtg tg- --- --- --- --- --- --- --- --- --- --- --- --- --- --- --- --- --- --- --- --g cta aat ggt act gtt aat aag ttt at- --- --- --- --- --- --- --- --- --- --- --- --- --- --- --- --- --- aga caa tat cat agt att ttc ttt cca gaa t-- --- --- --- tag att att ttt tta aat act gat cct cac aat tcc gtg atg tag cag tag ttg gt- --- --- --- --- --- --- --- --- --- --- --- --- --- --- --- --- --- --- --- --- --- --- --- --- --- --- --- --g cat ggt cta tat cgt --- --- --- --- --- --- --- --- --- --- --- --- --- --- --- --- --- --- --- --- --- --- --- --- --- --- --- --- --- --- --- --- --- --- --- --- --- --- --- --- --- --- --- -ta aaa tgt atc ata tat aat agt ttt ctg acg tgg agt aca gaa ttt tcg a-- --- --- --- --- --- --- --- --- --- --- --- --- --- --- --- --- --- --- --- --- --- --- --- --- --- --- --- --- --- --- --- --- --- --- --- --- --- --- --- --- --- --- --- tta atg agt tca tgg taa gga agg gca aat gcc t-- -gt ata taa tat aca taa gtt aa- --- --- --- --- --- --- --- --- --- --- --- --- --- tag ttt ttt atc ata ttt --- --- --- --- --- --- --- --- tct aat acc ata ata aaa att atc --- --- --- --- --- --- --- --- --- --- --- --- --- --- --- --- --- --- --- --- --- --- --- --- --- -at tat tgc gtt tg- gta gtt --- --- --- --- -ct gcc cta --- --- --- --- --- --- --- --- tca tct ata tca ctg tca ctc tc- --- --- gct ctc act ata tct tct aaa att aca a-- --a caa c-- --- --- --- --- --- --- --- --- --- --- --- --- --- --- --- --- --- --- --- --- --- --- --- --- --- --- --- --- --- --- --- --- --- --- --- -tg gat att cga t-- --- --- --- --- --- --- --- --- --- --- --- --- --- --- --- --- --- --- --- --- --- --- --- --- --- --- aac agc att tgt gt- --- --- --- --- --- --- --- --- --- --- --- --- --- --- --- --- --

>ON649712.1_Monkeypox_virus_isolate_Monkeypox/PT0025/2022_partial_genome

ttt ttt cga tct atc ctc gtc c-- -t- ctc atc atc ctt ata --- --- --- --- --- --- -tt att atc att att atc ata gtc tat taa aca caa atc atc t-- --- --- --- --- --- --- --- --- --- --- --- --- --- --- --- --- --- --- --- --- --- --- --- --- --- --- acg ttt ata ac- --- --- --- --- --- --- --- --- aac att c-- --- --- --- --t cat tat taa tta gtt ctg tag -aa tat ctt taa taa ttt ggc tat a-- --- --c atc tgt t-- --- --- --- --- --- --- --- --- --- --- --- --- --- --- --- --- --- --- caa tac t-- --- --- --- --- --- --- --- atc tat tga tga ttt ctt tt- --- --- --- --- --- --- --- --- --- --- --- tta aga ct- --- --- --- --- --- --- --- --- --- --- --- --- --- --- --- --- --- --- --- --- --- --- --- --- --- --- --- --- --- --- --- --- --- --- --- --- --- --- --- -ta aac tag t-- --- --- --- --- --- --- --- --- --- --- --- --- --- --- --- --- --- --- --- --- --- --- --- --- --- --- --- --- --- --- --- --- --- --- --- --- --- --- --- --- --- --- --- --- --- --- --- --- --- --- --- --- --- --- --- --- --- --- --- --- --- --- --- --- --- --- --- --- --- --- --- --- --- --- --- --- --- --- --- --- --- --- --- --- --- --- --- --- --- --- --- --- --- --- --- --- --- --- --- --- --- --- --- --- --- --- --- --- --- tat ggt aat gac gat gaa a-- --t cga gta gta --- --- act tct aat aaa gac ttg ata --- --- tca tta tca tat gtt tga tcg --- --- --- --- --- --- --- tca tag tta ata gtg tg- --- --- --- --- --- --- --- --- --- --- --- --- --- --- --- --- --- --- --- --g cta aat ggt act gtt aat aag ttt at- --- --- --- --- --- --- --- --- --- --- --- --- --- --- --- --- --- aga caa tat cat agt att ttc ttt cca gaa t-- --- --- --- tag att att ttt tta aat act gat cct cac aat tcc gtg atg tag cag tag ttg gt- --- --- --- --- --- --- --- --- --- --- --- --- --- --- --- --- --- --- --- --- --- --- --- --- --- --- --- --g cat ggt cta tat cgt --- --- --- --- --- --- --- --- --- --- --- --- --- --- --- --- --- --- --- --- --- --- --- --- --- --- --- --- --- --- --- --- --- --- --- --- --- --- --- --- --- --- --- -ta aaa tgt atc ata tat aat agt ttt ctg acg tgg agt aca gaa ttt tcg a-- --- --- --- --- --- --- --- --- --- --- --- --- --- --- --- --- --- --- --- --- --- --- --- --- --- --- --- --- --- --- --- --- --- --- --- --- --- --- --- --- --- --- --- tta atg agt tca tgg taa gga agg gca aat gcc t-- -gt ata taa tat aca taa gtt aa- --- --- --- --- --- --- --- --- --- --- --- --- --- tag ttt ttt atc ata ttt --- --- --- --- --- --- --- --- tct aat acc ata ata aaa att atc --- --- --- --- --- --- --- --- --- --- --- --- --- --- --- --- --- --- --- --- --- --- --- --- --- -at tat tgc gtt tg- gta gtt --- --- --- --- -ct gcc cta --- --- --- --- --- --- --- --- tca tct ata tca ctg tca ctc tc- --- --- gct ctc act ata tct tct aaa att aca a-- --a caa c-- --- --- --- --- --- --- --- --- --- --- --- --- --- --- --- --- --- --- --- --- --- --- --- --- --- --- --- --- --- --- --- --- --- --- --- -tg gat att cga t-- --- --- --- --- --- --- --- --- --- --- --- --- --- --- --- --- --- --- --- --- --- --- --- --- --- --- aac agc att tgt gt- --- --- --- --- --- --- --- --- --- --- --- --- --- --- --- --- --

>ON595760.2_Monkeypox_virus_isolate_MPXV-CH-38134631/2022_partial_genome

ttt ttt cga tct atc ctc gtc c-- -t- ctc atc atc ctt ata --- --- --- --- --- --- -tt att atc att att atc ata gtc tat taa aca caa atc atc t-- --- --- --- --- --- --- --- --- --- --- --- --- --- --- --- --- --- --- --- --- --- --- --- --- --- --- acg ttt ata ac- --- --- --- --- --- --- --- --- aac att c-- --- --- --- --t cat tat taa tta gtt ctg tag -aa tat ctt taa taa ttt ggc tat a-- --- --c atc tgt t-- --- --- --- --- --- --- --- --- --- --- --- --- --- --- --- --- --- --- caa tac t-- --- --- --- --- --- --- --- atc tat tga tga ttt ctt tt- --- --- --- --- --- --- --- --- --- --- --- tta aga ct- --- --- --- --- --- --- --- --- --- --- --- --- --- --- --- --- --- --- --- --- --- --- --- --- --- --- --- --- --- --- --- --- --- --- --- --- --- --- --- -ta aac tag t-- --- --- --- --- --- --- --- --- --- --- --- --- --- --- --- --- --- --- --- --- --- --- --- --- --- --- --- --- --- --- --- --- --- --- --- --- --- --- --- --- --- --- --- --- --- --- --- --- --- --- --- --- --- --- --- --- --- --- --- --- --- --- --- --- --- --- --- --- --- --- --- --- --- --- --- --- --- --- --- --- --- --- --- --- --- --- --- --- --- --- --- --- --- --- --- --- --- --- --- --- --- --- --- --- --- --- --- --- --- tat ggt aat gac gat gaa a-- --t cga gta gta --- --- act tct aat aaa gac ttg ata --- --- tca tta tca tat gtt tga tcg --- --- --- --- --- --- --- tca tag tta ata gtg tg- --- --- --- --- --- --- --- --- --- --- --- --- --- --- --- --- --- --- --- --g cta aat ggt act gtt aat aag ttt at- --- --- --- --- --- --- --- --- --- --- --- --- --- --- --- --- --- aga caa tat cat agt att ttc ttt cca gaa t-- --- --- --- tag att att ttt tta aat act gat cct cac aat tcc gtg atg tag cag tag ttg gt- --- --- --- --- --- --- --- --- --- --- --- --- --- --- --- --- --- --- --- --- --- --- --- --- --- --- --- --g cat ggt cta tat cgt --- --- --- --- --- --- --- --- --- --- --- --- --- --- --- --- --- --- --- --- --- --- --- --- --- --- --- --- --- --- --- --- --- --- --- --- --- --- --- --- --- --- --- -ta aaa tgt atc ata tat aat agt ttt ctg acg tgg agt aca gaa ttt tcg a-- --- --- --- --- --- --- --- --- --- --- --- --- --- --- --- --- --- --- --- --- --- --- --- --- --- --- --- --- --- --- --- --- --- --- --- --- --- --- --- --- --- --- --- tta atg agt tca tgg taa gga agg gca aat gcc t-- -gt ata taa tat aca taa gtt aa- --- --- --- --- --- --- --- --- --- --- --- --- --- tag ttt ttt atc ata ttt --- --- --- --- --- --- --- --- tct aat acc ata ata aaa att atc --- --- --- --- --- --- --- --- --- --- --- --- --- --- --- --- --- --- --- --- --- --- --- --- --- -at tat tgc gtt tg- gta gtt --- --- --- --- -ct gcc cta --- --- --- --- --- --- --- --- tca tct ata tca ctg tca ctc tc- --- --- gct ctc act ata tct tct aaa att aca a-- --a caa c-- --- --- --- --- --- --- --- --- --- --- --- --- --- --- --- --- --- --- --- --- --- --- --- --- --- --- --- --- --- --- --- --- --- --- --- -tg gat att cga t-- --- --- --- --- --- --- --- --- --- --- --- --- --- --- --- --- --- --- --- --- --- --- --- --- --- --- aac agc att tgt gt- --- --- --- --- --- --- --- --- --- --- --- --- --- --- --- --- --

>ON615424.1_Monkeypox_virus_isolate_MPXV_2022_NL001_partial_genome

ttt ttt cga tct atc ctc gtc c-- -t- ctc atc atc ctt ata --- --- --- --- --- --- -tt att atc att att atc ata gtc tat taa aca caa atc atc t-- --- --- --- --- --- --- --- --- --- --- --- --- --- --- --- --- --- --- --- --- --- --- --- --- --- --- acg ttt ata ac- --- --- --- --- --- --- --- --- aac att c-- --- --- --- --t cat tat taa tta gtt ctg tag -aa tat ctt taa taa ttt ggc tat a-- --- --c atc tgt t-- --- --- --- --- --- --- --- --- --- --- --- --- --- --- --- --- --- --- caa tac t-- --- --- --- --- --- --- --- atc tat tga tga ttt ctt tt- --- --- --- --- --- --- --- --- --- --- --- tta aga ct- --- --- --- --- --- --- --- --- --- --- --- --- --- --- --- --- --- --- --- --- --- --- --- --- --- --- --- --- --- --- --- --- --- --- --- --- --- --- --- -ta aac tag t-- --- --- --- --- --- --- --- --- --- --- --- --- --- --- --- --- --- --- --- --- --- --- --- --- --- --- --- --- --- --- --- --- --- --- --- --- --- --- --- --- --- --- --- --- --- --- --- --- --- --- --- --- --- --- --- --- --- --- --- --- --- --- --- --- --- --- --- --- --- --- --- --- --- --- --- --- --- --- --- --- --- --- --- --- --- --- --- --- --- --- --- --- --- --- --- --- --- --- --- --- --- --- --- --- --- --- --- --- --- tat ggt aat gac gat gaa a-- --t cga gta gta --- --- act tct aat aaa gac ttg ata --- --- tca tta tca tat gtt tga tcg --- --- --- --- --- --- --- tca tag tta ata gtg tg- --- --- --- --- --- --- --- --- --- --- --- --- --- --- --- --- --- --- --- --g cta aat ggt act gtt aat aag ttt at- --- --- --- --- --- --- --- --- --- --- --- --- --- --- --- --- --- aga caa tat cat agt att ttc ttt cca gaa t-- --- --- --- tag att att ttt tta aat act gat cct cac aat tcc gtg atg tag cag tag ttg gt- --- --- --- --- --- --- --- --- --- --- --- --- --- --- --- --- --- --- --- --- --- --- --- --- --- --- --- --g cat ggt cta tat cgt --- --- --- --- --- --- --- --- --- --- --- --- --- --- --- --- --- --- --- --- --- --- --- --- --- --- --- --- --- --- --- --- --- --- --- --- --- --- --- --- --- --- --- -ta aaa tgt atc ata tat aat agt ttt ctg acg tgg agt aca gaa ttt tcg a-- --- --- --- --- --- --- --- --- --- --- --- --- --- --- --- --- --- --- --- --- --- --- --- --- --- --- --- --- --- --- --- --- --- --- --- --- --- --- --- --- --- --- --- tta atg agt tca tgg taa gga agg gca aat gcc t-- -gt ata taa tat aca taa gtt aa- --- --- --- --- --- --- --- --- --- --- --- --- --- tag ttt ttt atc ata ttt --- --- --- --- --- --- --- --- tct aat acc ata ata aaa att atc --- --- --- --- --- --- --- --- --- --- --- --- --- --- --- --- --- --- --- --- --- --- --- --- --- -at tat tgc gtt tg- gta gtt --- --- --- --- -ct gcc cta --- --- --- --- --- --- --- --- tca tct ata tca ctg tca ctc tc- --- --- gct ctc act ata tct tct aaa att aca a-- --a caa c-- --- --- --- --- --- --- --- --- --- --- --- --- --- --- --- --- --- --- --- --- --- --- --- --- --- --- --- --- --- --- --- --- --- --- --- -tg gat att cga t-- --- --- --- --- --- --- --- --- --- --- --- --- --- --- --- --- --- --- --- --- --- --- --- --- --- --- aac agc att tgt gt- --- --- --- --- --- --- --- --- --- --- --- --- --- --- --- --- --

>ON585034.1_Monkeypox_virus_isolate_Monkeypox/PT0007/2022_complete_genome

ttt ttt cga tct atc ctc gtc c-- -t- ctc atc atc ctt ata --- --- --- --- --- --- -tt att atc att att atc ata gtc tat taa aca caa atc atc t-- --- --- --- --- --- --- --- --- --- --- --- --- --- --- --- --- --- --- --- --- --- --- --- --- --- --- acg ttt ata ac- --- --- --- --- --- --- --- --- aac att c-- --- --- --- --t cat tat taa tta gtt ctg tag -aa tat ctt taa taa ttt ggc tat a-- --- --c atc tgt t-- --- --- --- --- --- --- --- --- --- --- --- --- --- --- --- --- --- --- caa tac t-- --- --- --- --- --- --- --- atc tat tga tga ttt ctt tt- --- --- --- --- --- --- --- --- --- --- --- tta aga ct- --- --- --- --- --- --- --- --- --- --- --- --- --- --- --- --- --- --- --- --- --- --- --- --- --- --- --- --- --- --- --- --- --- --- --- --- --- --- --- -ta aac tag t-- --- --- --- --- --- --- --- --- --- --- --- --- --- --- --- --- --- --- --- --- --- --- --- --- --- --- --- --- --- --- --- --- --- --- --- --- --- --- --- --- --- --- --- --- --- --- --- --- --- --- --- --- --- --- --- --- --- --- --- --- --- --- --- --- --- --- --- --- --- --- --- --- --- --- --- --- --- --- --- --- --- --- --- --- --- --- --- --- --- --- --- --- --- --- --- --- --- --- --- --- --- --- --- --- --- --- --- --- --- tat ggt aat gac gat gaa a-- --t cga gta gta --- --- act tct aat aaa gac ttg ata --- --- tca tta tca tat gtt tga tcg --- --- --- --- --- --- --- tca tag tta ata gtg tg- --- --- --- --- --- --- --- --- --- --- --- --- --- --- --- --- --- --- --- --g cta aat ggt act gtt aat aag ttt at- --- --- --- --- --- --- --- --- --- --- --- --- --- --- --- --- --- aga caa tat cat agt att ttc ttt cca gaa t-- --- --- --- tag att att ttt tta aat act gat cct cac aat tcc gtg atg tag cag tag ttg gt- --- --- --- --- --- --- --- --- --- --- --- --- --- --- --- --- --- --- --- --- --- --- --- --- --- --- --- --g cat ggt cta tat cgt --- --- --- --- --- --- --- --- --- --- --- --- --- --- --- --- --- --- --- --- --- --- --- --- --- --- --- --- --- --- --- --- --- --- --- --- --- --- --- --- --- --- --- -ta aaa tgt atc ata tat aat agt ttt ctg acg tgg agt aca gaa ttt tcg a-- --- --- --- --- --- --- --- --- --- --- --- --- --- --- --- --- --- --- --- --- --- --- --- --- --- --- --- --- --- --- --- --- --- --- --- --- --- --- --- --- --- --- --- tta atg agt tca tgg taa gga agg gca aat gcc t-- -gt ata taa tat aca taa gtt aa- --- --- --- --- --- --- --- --- --- --- --- --- --- tag ttt ttt atc ata ttt --- --- --- --- --- --- --- --- tct aat acc ata ata aaa att atc --- --- --- --- --- --- --- --- --- --- --- --- --- --- --- --- --- --- --- --- --- --- --- --- --- -at tat tgc gtt tg- gta gtt --- --- --- --- -ct gcc cta --- --- --- --- --- --- --- --- tca tct ata tca ctg tca ctc tc- --- --- gct ctc act ata tct tct aaa att aca a-- --a caa c-- --- --- --- --- --- --- --- --- --- --- --- --- --- --- --- --- --- --- --- --- --- --- --- --- --- --- --- --- --- --- --- --- --- --- --- -tg gat att cga t-- --- --- --- --- --- --- --- --- --- --- --- --- --- --- --- --- --- --- --- --- --- --- --- --- --- --- aac agc att tgt gt- --- --- --- --- --- --- --- --- --- --- --- --- --- --- --- --- --

>ON649713.1_Monkeypox_virus_isolate_Monkeypox/PT0020/2022_complete_genome

ttt ttt cga tct atc ctc gtc c-- -t- ctc atc atc ctt ata --- --- --- --- --- --- -tt att atc att att atc ata gtc tat taa aca caa atc atc t-- --- --- --- --- --- --- --- --- --- --- --- --- --- --- --- --- --- --- --- --- --- --- --- --- --- --- acg ttt ata ac- --- --- --- --- --- --- --- --- aac att c-- --- --- --- --t cat tat taa tta gtt ctg tag -aa tat ctt taa taa ttt ggc tat a-- --- --c atc tgt t-- --- --- --- --- --- --- --- --- --- --- --- --- --- --- --- --- --- --- caa tac t-- --- --- --- --- --- --- --- atc tat tga tga ttt ctt tt- --- --- --- --- --- --- --- --- --- --- --- tta aga ct- --- --- --- --- --- --- --- --- --- --- --- --- --- --- --- --- --- --- --- --- --- --- --- --- --- --- --- --- --- --- --- --- --- --- --- --- --- --- --- -ta aac tag t-- --- --- --- --- --- --- --- --- --- --- --- --- --- --- --- --- --- --- --- --- --- --- --- --- --- --- --- --- --- --- --- --- --- --- --- --- --- --- --- --- --- --- --- --- --- --- --- --- --- --- --- --- --- --- --- --- --- --- --- --- --- --- --- --- --- --- --- --- --- --- --- --- --- --- --- --- --- --- --- --- --- --- --- --- --- --- --- --- --- --- --- --- --- --- --- --- --- --- --- --- --- --- --- --- --- --- --- --- --- tat ggt aat gac gat gaa a-- --t cga gta gta --- --- act tct aat aaa gac ttg ata --- --- tca tta tca tat gtt tga tcg --- --- --- --- --- --- --- tca tag tta ata gtg tg- --- --- --- --- --- --- --- --- --- --- --- --- --- --- --- --- --- --- --- --g cta aat ggt act gtt aat aag ttt at- --- --- --- --- --- --- --- --- --- --- --- --- --- --- --- --- --- aga caa tat cat agt att ttc ttt cca gaa t-- --- --- --- tag att att ttt tta aat act gat cct cac aat tcc gtg atg tag cag tag ttg gt- --- --- --- --- --- --- --- --- --- --- --- --- --- --- --- --- --- --- --- --- --- --- --- --- --- --- --- --g cat ggt cta tat cgt --- --- --- --- --- --- --- --- --- --- --- --- --- --- --- --- --- --- --- --- --- --- --- --- --- --- --- --- --- --- --- --- --- --- --- --- --- --- --- --- --- --- --- -ta aaa tgt atc ata tat aat agt ttt ctg acg tgg agt aca gaa ttt tcg a-- --- --- --- --- --- --- --- --- --- --- --- --- --- --- --- --- --- --- --- --- --- --- --- --- --- --- --- --- --- --- --- --- --- --- --- --- --- --- --- --- --- --- --- tta atg agt tca tgg taa gga agg gca aat gcc t-- -gt ata taa tat aca taa gtt aa- --- --- --- --- --- --- --- --- --- --- --- --- --- tag ttt ttt atc ata ttt --- --- --- --- --- --- --- --- tct aat acc ata ata aaa att atc --- --- --- --- --- --- --- --- --- --- --- --- --- --- --- --- --- --- --- --- --- --- --- --- --- -at tat tgc gtt tg- gta gtt --- --- --- --- -ct gcc cta --- --- --- --- --- --- --- --- tca tct ata tca ctg tca ctc tc- --- --- gct ctc act ata tct tct aaa att aca a-- --a caa c-- --- --- --- --- --- --- --- --- --- --- --- --- --- --- --- --- --- --- --- --- --- --- --- --- --- --- --- --- --- --- --- --- --- --- --- -tg gat att cga t-- --- --- --- --- --- --- --- --- --- --- --- --- --- --- --- --- --- --- --- --- --- --- --- --- --- --- aac agc att tgt gt- --- --- --- --- --- --- --- --- --- --- --- --- --- --- --- --- --

>ON585031.1_Monkeypox_virus_isolate_Monkeypox/PT0003/2022_complete_genome

ttt ttt cga tct atc ctc gtc c-- -t- ctc atc atc ctt ata --- --- --- --- --- --- -tt att atc att att atc ata gtc tat taa aca caa atc atc t-- --- --- --- --- --- --- --- --- --- --- --- --- --- --- --- --- --- --- --- --- --- --- --- --- --- --- acg ttt ata ac- --- --- --- --- --- --- --- --- aac att c-- --- --- --- --t cat tat taa tta gtt ctg tag -aa tat ctt taa taa ttt ggc tat a-- --- --c atc tgt t-- --- --- --- --- --- --- --- --- --- --- --- --- --- --- --- --- --- --- caa tac t-- --- --- --- --- --- --- --- atc tat tga tga ttt ctt tt- --- --- --- --- --- --- --- --- --- --- --- tta aga ct- --- --- --- --- --- --- --- --- --- --- --- --- --- --- --- --- --- --- --- --- --- --- --- --- --- --- --- --- --- --- --- --- --- --- --- --- --- --- --- -ta aac tag t-- --- --- --- --- --- --- --- --- --- --- --- --- --- --- --- --- --- --- --- --- --- --- --- --- --- --- --- --- --- --- --- --- --- --- --- --- --- --- --- --- --- --- --- --- --- --- --- --- --- --- --- --- --- --- --- --- --- --- --- --- --- --- --- --- --- --- --- --- --- --- --- --- --- --- --- --- --- --- --- --- --- --- --- --- --- --- --- --- --- --- --- --- --- --- --- --- --- --- --- --- --- --- --- --- --- --- --- --- --- tat ggt aat gac gat gaa a-- --t cga gta gta --- --- act tct aat aaa gac ttg ata --- --- tca tta tca tat gtt tga tcg --- --- --- --- --- --- --- tca tag tta ata gtg tg- --- --- --- --- --- --- --- --- --- --- --- --- --- --- --- --- --- --- --- --g cta aat ggt act gtt aat aag ttt at- --- --- --- --- --- --- --- --- --- --- --- --- --- --- --- --- --- aga caa tat cat agt att ttc ttt cca gaa t-- --- --- --- tag att att ttt tta aat act gat cct cac aat tcc gtg atg tag cag tag ttg gt- --- --- --- --- --- --- --- --- --- --- --- --- --- --- --- --- --- --- --- --- --- --- --- --- --- --- --- --g cat ggt cta tat cgt --- --- --- --- --- --- --- --- --- --- --- --- --- --- --- --- --- --- --- --- --- --- --- --- --- --- --- --- --- --- --- --- --- --- --- --- --- --- --- --- --- --- --- -ta aaa tgt atc ata tat aat agt ttt ctg acg tgg agt aca gaa ttt tcg a-- --- --- --- --- --- --- --- --- --- --- --- --- --- --- --- --- --- --- --- --- --- --- --- --- --- --- --- --- --- --- --- --- --- --- --- --- --- --- --- --- --- --- --- tta atg agt tca tgg taa gga agg gca aat gcc t-- -gt ata taa tat aca taa gtt aa- --- --- --- --- --- --- --- --- --- --- --- --- --- tag ttt ttt atc ata ttt --- --- --- --- --- --- --- --- tct aat acc ata ata aaa att atc --- --- --- --- --- --- --- --- --- --- --- --- --- --- --- --- --- --- --- --- --- --- --- --- --- -at tat tgc gtt tg- gta gtt --- --- --- --- -ct gcc cta --- --- --- --- --- --- --- --- tca tct ata tca ctg tca ctc tc- --- --- gct ctc act ata tct tct aaa att aca a-- --a caa c-- --- --- --- --- --- --- --- --- --- --- --- --- --- --- --- --- --- --- --- --- --- --- --- --- --- --- --- --- --- --- --- --- --- --- --- -tg gat att cga t-- --- --- --- --- --- --- --- --- --- --- --- --- --- --- --- --- --- --- --- --- --- --- --- --- --- --- aac agc att tgt gt- --- --- --- --- --- --- --- --- --- --- --- --- --- --- --- --- --

>ON585038.1_Monkeypox_virus_isolate_Monkeypox/PT0008/2022_complete_genome

ttt ttt cga tct atc ctc gtc c-- -t- ctc atc atc ctt ata --- --- --- --- --- --- -tt att atc att att atc ata gtc tat taa aca caa atc atc t-- --- --- --- --- --- --- --- --- --- --- --- --- --- --- --- --- --- --- --- --- --- --- --- --- --- --- acg ttt ata ac- --- --- --- --- --- --- --- --- aac att c-- --- --- --- --t cat tat taa tta gtt ctg tag -aa tat ctt taa taa ttt ggc tat a-- --- --c atc tgt t-- --- --- --- --- --- --- --- --- --- --- --- --- --- --- --- --- --- --- caa tac t-- --- --- --- --- --- --- --- atc tat tga tga ttt ctt tt- --- --- --- --- --- --- --- --- --- --- --- tta aga ct- --- --- --- --- --- --- --- --- --- --- --- --- --- --- --- --- --- --- --- --- --- --- --- --- --- --- --- --- --- --- --- --- --- --- --- --- --- --- --- -ta aac tag t-- --- --- --- --- --- --- --- --- --- --- --- --- --- --- --- --- --- --- --- --- --- --- --- --- --- --- --- --- --- --- --- --- --- --- --- --- --- --- --- --- --- --- --- --- --- --- --- --- --- --- --- --- --- --- --- --- --- --- --- --- --- --- --- --- --- --- --- --- --- --- --- --- --- --- --- --- --- --- --- --- --- --- --- --- --- --- --- --- --- --- --- --- --- --- --- --- --- --- --- --- --- --- --- --- --- --- --- --- --- tat ggt aat gac gat gaa a-- --t cga gta gta --- --- act tct aat aaa gac ttg ata --- --- tca tta tca tat gtt tga tcg --- --- --- --- --- --- --- tca tag tta ata gtg tg- --- --- --- --- --- --- --- --- --- --- --- --- --- --- --- --- --- --- --- --g cta aat ggt act gtt aat aag ttt at- --- --- --- --- --- --- --- --- --- --- --- --- --- --- --- --- --- aga caa tat cat agt att ttc ttt cca gaa t-- --- --- --- tag att att ttt tta aat act gat cct cac aat tcc gtg atg tag cag tag ttg gt- --- --- --- --- --- --- --- --- --- --- --- --- --- --- --- --- --- --- --- --- --- --- --- --- --- --- --- --g cat ggt cta tat cgt --- --- --- --- --- --- --- --- --- --- --- --- --- --- --- --- --- --- --- --- --- --- --- --- --- --- --- --- --- --- --- --- --- --- --- --- --- --- --- --- --- --- --- -ta aaa tgt atc ata tat aat agt ttt ctg acg tgg agt aca gaa ttt tcg a-- --- --- --- --- --- --- --- --- --- --- --- --- --- --- --- --- --- --- --- --- --- --- --- --- --- --- --- --- --- --- --- --- --- --- --- --- --- --- --- --- --- --- --- tta atg agt tca tgg taa gga agg gca aat gcc t-- -gt ata taa tat aca taa gtt aa- --- --- --- --- --- --- --- --- --- --- --- --- --- tag ttt ttt atc ata ttt --- --- --- --- --- --- --- --- tct aat acc ata ata aaa att atc --- --- --- --- --- --- --- --- --- --- --- --- --- --- --- --- --- --- --- --- --- --- --- --- --- -at tat tgc gtt tg- gta gtt --- --- --- --- -ct gcc cta --- --- --- --- --- --- --- --- tca tct ata tca ctg tca ctc tc- --- --- gct ctc act ata tct tct aaa att aca a-- --a caa c-- --- --- --- --- --- --- --- --- --- --- --- --- --- --- --- --- --- --- --- --- --- --- --- --- --- --- --- --- --- --- --- --- --- --- --- -tg gat att cga t-- --- --- --- --- --- --- --- --- --- --- --- --- --- --- --- --- --- --- --- --- --- --- --- --- --- --- aac agc att tgt gt- --- --- --- --- --- --- --- --- --- --- --- --- --- --- --- --- --

>DQ011156.1_Monkeypox_virus_strain_Liberia_1970_184_complete_genome

ttt ttt cga tct atc aat ttc agt ata ttc ttc gcc gtt ata aaa gta atg ttg ttt aat tgt agg acg gtt gtt agt ata atc aca tga ata ata ata ttc taa ttc ctc gta ttg act act tac aga tac tcg aaa tag tct gaa aaa ttc ttc aaa gat att ttt ata aag atc tag gaa aag ttt att acc gac cat gaa cga gat aga tgg ata aat atc ctt tcc atc aaa ggt cat aat tgg ata att gtc cag caa tat atc tgc tgt att agt tat atc act tcc att tat ttt cag att gaa gta atg tac tag ttt gtg aca att aac aag ata caa aag aga tgc cga tac taa tac gta aat agc tat acg cga atc cat tgt tac ctt ttt tta ttt cat agg tct att aat aaa tat atg tat tac tta aga cta gaa aaa tca aaa gta agt ttt tga tat ttg att ctt act tat tgt ggg att gta gtt tac tta gta att cat ctc tga atc ctg ata aat cat gca tat caa tga tgc aac tac gca gca aac tag tag gaa tat aga tat ctg gat atg tac gta aat agt cga tta tat ctt tta caa tac tat tag tcc cta ttg cgt tat cta tat atc cat taa taa tat tac aca gtg gat act tat gag aaa tat acc tct tac aga ttt tta acg ata tat aat cta gaa aga tat gtg tgt agt act gta tta cct aaa tta tca gtc tca ttc aaa tat tgc atg act att atc gag aat tgc tat atc cct cta tac tcg atg cat tta tta cag taa ttc aat cca gct aac ata aga gcc aat ctc aat gtt ggt tta att ata tca tct tca tgt aat aat aac gat gga aac ttt cca gta gcg taa cac tta tct aag aag gat ata ata act atg tct aca tta tgt ctc tta tcg aga ata ttc tta acg aga tat cca tag cta ttc tgg tgc taa tta ttc cta tat tat att cca cga aaa atg atg aag gc- --c att cat cat aag atg ata aaa agt gta gtg agt aag agt att agt gag aga gca tga agg aga ttt agt att tag cag tga gga tat gat cca aga ggg tga gat agt cgt tct cgt tca gaa tct ttc gca gca taa gta gta tgt cga tat act tat cat tga aga ctc ttc cgg tga caa tag ctg att gag tac aaa gtc caa tta ttg cac aaa gtt ctt tgg cgg ttt tca tgg agt cat ttc tga tga aac att taa tga tct cca cgc aat tgt cga tat tgt ccc acg gaa gtg aat ccg aga act cct tca act cgc tac caa ata gct cca ttg cat caa ttc tga aag aga tga gaa gcc tgt aga gag gcc ctg cgc ttt ctc tat ggg tcc atc tat gag aaa ccc aca gga tgt att cag tca gac aa- tgt ctg aca tca gtc acg gta ttc agg gag tcc tta gta gcg tgg caa tga cag gga ctg aac tgg gca caa gga gag gcc att gtg aag gta gac gaa ggt aac ctg atg gta gac ctg tag ccg tct gtg ctt aat aga ggg ctt taa ttt cca ttt tta atg gtg tcg tga atg agg aat gag agt gtc tct cgt cct tgg ttt aca tgg atc aga gtg aga aaa aat atc ttg tat att att aac taa caa cct tgg ttt cta tcc atg ttt -aa aaa atg acc tat atg ttc ttt att aat tct att tta aac ttt atc ctc aag act cct gac aaa att aaa atc cag aaa gca gca aac aat cct gtt aca agt tta ctg aaa tct ctc ttt gat tgt aga gta tat gta gtc aga gca aga aac act gca gta gtc aac atg aaa gct tgc ata acg ata cgt gca tca tag aaa gta aca aca gag gcc agc gtt aga gat tct aac agt gta aat cca caa --- agt atg tac aga ttc agg gga tgt tca tgt ctg tgt aaa gtc aat gcg aaa atc aag cct ata gat ccg aac att gat gcc aat att aga aca gga ctc cct tgt ata aat gtc cga tgc att caa agt ata aaa ata ctg cag ctg ttg ccg ttg tta aag gaa att gta gaa agg ata ccg tag act ttt ctt aga aat gcc att cgt atg tac acg ctg gca gac gcc acc gag ctg tca tag ttg aag tcg tcc tcg at

>KP849470.1_Monkeypox_virus_isolate_Cote_dIvoire_1971_complete_genome

ttt ttt cga tct atc aat ttc agt ata ttc ttc gcc gtt ata aaa gta atg ttg ttt aat tgt agg acg gtt gtt agt ata atc aca tga ata ata ata ttc taa ttc ctc gta ttg act act tac aga tac tcg aaa tag tct gaa aaa ttc ttc aaa gat att ttt ata aag atc tag gaa aag ttt att acc gac cat gaa cga gat aga tgg ata aat atc ctt tcc atc aaa ggt cat aat tgg ata att gtc cag caa tat atc tgc tgt att agt tat atc act tcc att tat ttt cag att gaa gta atg tac tag ttt gtg aca att aac aag ata caa aag aga tgc cga tac taa tac gta aat agc tat acg cga atc cat tgt tac ctt ttt tta ttt cat agg tct att aat aaa tat atg tat tac tta aga cta gaa aaa tca aaa gta agt ttt tga tat ttg att ctt act tat tgt ggg att gta gtt tac tta gta att cat ctc tga atc ctg ata aat cat gca tat caa tga tgc aac tac gca gca aac tag tag gaa tat aga tat ctg gat atg tac gta aat agt cga tta tat ctt tta caa tac tat tag tcc cta ttg cgt tat cta tat atc cat taa taa tat tac aca gtg gat act tat gag aaa tat acc tct tac aga ttt tta acg ata tat aat cta gaa aga tat gtg tgt agt act gta tta cct aaa tta tca gtc tca ttc aaa tat tgc atg act att atc gag aat tgc tat atc cct cta tac tcg atg cat tta tta cag taa ttc aat cca gct aac ata aga gcc aat ctc aat gtt ggt tta att ata tca tct tca tgt aat aat aac gat gga aac ttt cca gta gcg taa cac tta tct aag aag gat ata ata act atg tct aca tta tgt ctc tta tcg aga ata ttc tta acg aga tat cca tag cta ttc tgg tgc taa tta ttc cta tat tat att cca cga aaa atg atg aag gc- --c att cat cat aag atg ata aaa agt gta gtg agt aag agt att agt gag aga gca tga agg aga ttt agt att tag cag tga gga tat gat cca aga ggg tga gat agt cgt tct cgt tca gaa tct ttc gca gca taa gta gta tgt cga tat act tat cat tga aga ctc ttc cgg tga caa tag ctg att gag tac aaa gtc caa tta ttg cac aaa gtt ctt tgg cgg ttt tca tgg agt cat ttc tga tga aac att taa tga tct cca cgc aat tgt cga tat tgt ccc acg gaa gtg aat ccg aga act cct tca act cgc tac caa ata gct cca ttg cat caa ttc tga aag aga tga gaa gcc tgt aga gag gcc ctg cgc ttt ctc tat ggg tcc atc tat gag aaa ccc aca gga tgt att cag tca gac aa- tgt ctg aca tca gtc acg gta ttc agg gag tcc tta gta gcg tgg caa tga cag gga ctg aac tgg gca caa gga gag gcc att gtg aag gta gac gaa ggt aac ctg atg gta gac ctg tag ccg tct gtg ctt aat aga ggg ctt taa ttt cca ttt tta atg gtg tcg tga atg agg aat gag agt gtc tct cgt cct tgg ttt aca tgg atc aga gtg aga aaa aat atc ttg tat att att aac taa caa cct tgg ttt cta tcc atg ttt -aa aaa atg acc tat atg ttc ttt att aat tct att tta aac ttt atc ctc aag act cct gac aaa att aaa atc cag aaa gca gca aac aat cct gtt aca agt tta ctg aaa tct ctc ttt gat tgt aga gta tat gta gtc aga gca aga aac act gca gta gtc aac atg aaa gct tgc ata acg ata cgt gca tca tag aaa gta aca aca gag gcc agc gtt aga gat tct aac agt gta aat cca caa --- agt atg tac aga ttc agg gga tgt tca tgt ctg tgt aaa gtc aat gcg aaa atc aag cct ata gat ccg aac att gat gcc aat att aga aca gga ctc cct tgt ata aat gtc cga tgc att caa agt ata aaa ata ctg cag ctg ttg ccg ttg tta aag gaa att gta gaa agg ata ccg tag act ttt ctt aga aat gcc att cgt atg tac acg ctg gca gac gcc acc gag ctg tca tag ttg aag tcg tcc tcg at

>ON682264.2_Monkeypox_virus_isolate_MPXV/Germany/2022/RKI05_complete_genome

ttt ttt cga tct atc ctc gtc c-- -t- ctc atc atc ctt ata --- --- --- --- --- --- -tt att atc att att atc ata gtc tat taa aca caa atc atc t-- --- --- --- --- --- --- --- --- --- --- --- --- --- --- --- --- --- --- --- --- --- --- --- --- --- --- acg ttt ata ac- --- --- --- --- --- --- --- --- aac att c-- --- --- --- --t cat tat taa tta gtt ctg tag -aa tat ctt taa taa ttt ggc tat a-- --- --c atc tgt t-- --- --- --- --- --- --- --- --- --- --- --- --- --- --- --- --- --- --- caa tac t-- --- --- --- --- --- --- --- atc tat tga tga ttt ctt tt- --- --- --- --- --- --- --- --- --- --- --- tta aga ct- --- --- --- --- --- --- --- --- --- --- --- --- --- --- --- --- --- --- --- --- --- --- --- --- --- --- --- --- --- --- --- --- --- --- --- --- --- --- --- -ta aac tag t-- --- --- --- --- --- --- --- --- --- --- --- --- --- --- --- --- --- --- --- --- --- --- --- --- --- --- --- --- --- --- --- --- --- --- --- --- --- --- --- --- --- --- --- --- --- --- --- --- --- --- --- --- --- --- --- --- --- --- --- --- --- --- --- --- --- --- --- --- --- --- --- --- --- --- --- --- --- --- --- --- --- --- --- --- --- --- --- --- --- --- --- --- --- --- --- --- --- --- --- --- --- --- --- --- --- --- --- --- --- tat ggt aat gac gat gaa a-- --t cga gta gta --- --- act tct aat aaa gac ttg ata --- --- tca tta tca tat gtt tga tcg --- --- --- --- --- --- --- tca tag tta ata gtg tg- --- --- --- --- --- --- --- --- --- --- --- --- --- --- --- --- --- --- --- --g cta aat ggt act gtt aat aag ttt at- --- --- --- --- --- --- --- --- --- --- --- --- --- --- --- --- --- aga caa tat cat agt att ttc ttt cca gaa t-- --- --- --- tag att att ttt tta aat act gat cct cac aat tcc gtg atg tag cag tag ttg gt- --- --- --- --- --- --- --- --- --- --- --- --- --- --- --- --- --- --- --- --- --- --- --- --- --- --- --- --g cat ggt cta tat cgt --- --- --- --- --- --- --- --- --- --- --- --- --- --- --- --- --- --- --- --- --- --- --- --- --- --- --- --- --- --- --- --- --- --- --- --- --- --- --- --- --- --- --- -ta aaa tgt atc ata tat aat agt ttt ctg acg tgg agt aca gaa ttt tcg a-- --- --- --- --- --- --- --- --- --- --- --- --- --- --- --- --- --- --- --- --- --- --- --- --- --- --- --- --- --- --- --- --- --- --- --- --- --- --- --- --- --- --- --- tta atg agt tca tgg taa gga agg gca aat gcc t-- -gt ata taa tat aca taa gtt aa- --- --- --- --- --- --- --- --- --- --- --- --- --- tag ttt ttt atc ata ttt --- --- --- --- --- --- --- --- tct aat acc ata ata aaa att atc --- --- --- --- --- --- --- --- --- --- --- --- --- --- --- --- --- --- --- --- --- --- --- --- --- -at tat tgc gtt tg- gta gtt --- --- --- --- -ct gcc cta --- --- --- --- --- --- --- --- tca tct ata tca ctg tca ctc tc- --- --- gct ctc act ata tct tct aaa att aca a-- --a caa c-- --- --- --- --- --- --- --- --- --- --- --- --- --- --- --- --- --- --- --- --- --- --- --- --- --- --- --- --- --- --- --- --- --- --- --- -tg gat att cga t-- --- --- --- --- --- --- --- --- --- --- --- --- --- --- --- --- --- --- --- --- --- --- --- --- --- --- aac agc att tgt gt- --- --- --- --- --- --- --- --- --- --- --- --- --- --- --- --- --

>ON682268.1_Monkeypox_virus_isolate_MPXV/Germany/2022/RKI08_complete_genome

ttt ttt cga tct atc ctc gtc c-- -t- ctc atc atc ctt ata --- --- --- --- --- --- -tt att atc att att atc ata gtc tat taa aca caa atc atc t-- --- --- --- --- --- --- --- --- --- --- --- --- --- --- --- --- --- --- --- --- --- --- --- --- --- --- acg ttt ata ac- --- --- --- --- --- --- --- --- aac att c-- --- --- --- --t cat tat taa tta gtt ctg tag -aa tat ctt taa taa ttt ggc tat a-- --- --c atc tgt t-- --- --- --- --- --- --- --- --- --- --- --- --- --- --- --- --- --- --- caa tac t-- --- --- --- --- --- --- --- atc tat tga tga ttt ctt tt- --- --- --- --- --- --- --- --- --- --- --- tta aga ct- --- --- --- --- --- --- --- --- --- --- --- --- --- --- --- --- --- --- --- --- --- --- --- --- --- --- --- --- --- --- --- --- --- --- --- --- --- --- --- -ta aac tag t-- --- --- --- --- --- --- --- --- --- --- --- --- --- --- --- --- --- --- --- --- --- --- --- --- --- --- --- --- --- --- --- --- --- --- --- --- --- --- --- --- --- --- --- --- --- --- --- --- --- --- --- --- --- --- --- --- --- --- --- --- --- --- --- --- --- --- --- --- --- --- --- --- --- --- --- --- --- --- --- --- --- --- --- --- --- --- --- --- --- --- --- --- --- --- --- --- --- --- --- --- --- --- --- --- --- --- --- --- --- tat ggt aat gac gat gaa a-- --t cga gta gta --- --- act tct aat aaa gac ttg ata --- --- tca tta tca tat gtt tga tcg --- --- --- --- --- --- --- tca tag tta ata gtg tg- --- --- --- --- --- --- --- --- --- --- --- --- --- --- --- --- --- --- --- --g cta aat ggt act gtt aat aag ttt at- --- --- --- --- --- --- --- --- --- --- --- --- --- --- --- --- --- aga caa tat cat agt att ttc ttt cca gaa t-- --- --- --- tag att att ttt tta aat act gat cct cac aat tcc gtg atg tag cag tag ttg gt- --- --- --- --- --- --- --- --- --- --- --- --- --- --- --- --- --- --- --- --- --- --- --- --- --- --- --- --g cat ggt cta tat cgt --- --- --- --- --- --- --- --- --- --- --- --- --- --- --- --- --- --- --- --- --- --- --- --- --- --- --- --- --- --- --- --- --- --- --- --- --- --- --- --- --- --- --- -ta aaa tgt atc ata tat aat agt ttt ctg acg tgg agt aca gaa ttt tcg a-- --- --- --- --- --- --- --- --- --- --- --- --- --- --- --- --- --- --- --- --- --- --- --- --- --- --- --- --- --- --- --- --- --- --- --- --- --- --- --- --- --- --- --- tta atg agt tca tgg taa gga agg gca aat gcc t-- -gt ata taa tat aca taa gtt aa- --- --- --- --- --- --- --- --- --- --- --- --- --- tag ttt ttt atc ata ttt --- --- --- --- --- --- --- --- tct aat acc ata ata aaa att atc --- --- --- --- --- --- --- --- --- --- --- --- --- --- --- --- --- --- --- --- --- --- --- --- --- -at tat tgc gtt tg- gta gtt --- --- --- --- -ct gcc cta --- --- --- --- --- --- --- --- tca tct ata tca ctg tca ctc tc- --- --- gct ctc act ata tct tct aaa att aca a-- --a caa c-- --- --- --- --- --- --- --- --- --- --- --- --- --- --- --- --- --- --- --- --- --- --- --- --- --- --- --- --- --- --- --- --- --- --- --- -tg gat att cga t-- --- --- --- --- --- --- --- --- --- --- --- --- --- --- --- --- --- --- --- --- --- --- --- --- --- --- aac agc att tgt gt- --- --- --- --- --- --- --- --- --- --- --- --- --- --- --- --- --

>ON682270.1_Monkeypox_virus_isolate_MPXV/Germany/2022/RKI06_complete_genome

ttt ttt cga tct atc ctc gtc c-- -t- ctc atc atc ctt ata --- --- --- --- --- --- -tt att atc att att atc ata gtc tat taa aca caa atc atc t-- --- --- --- --- --- --- --- --- --- --- --- --- --- --- --- --- --- --- --- --- --- --- --- --- --- --- acg ttt ata ac- --- --- --- --- --- --- --- --- aac att c-- --- --- --- --t cat tat taa tta gtt ctg tag -aa tat ctt taa taa ttt ggc tat a-- --- --c atc tgt t-- --- --- --- --- --- --- --- --- --- --- --- --- --- --- --- --- --- --- caa tac t-- --- --- --- --- --- --- --- atc tat tga tga ttt ctt tt- --- --- --- --- --- --- --- --- --- --- --- tta aga ct- --- --- --- --- --- --- --- --- --- --- --- --- --- --- --- --- --- --- --- --- --- --- --- --- --- --- --- --- --- --- --- --- --- --- --- --- --- --- --- -ta aac tag t-- --- --- --- --- --- --- --- --- --- --- --- --- --- --- --- --- --- --- --- --- --- --- --- --- --- --- --- --- --- --- --- --- --- --- --- --- --- --- --- --- --- --- --- --- --- --- --- --- --- --- --- --- --- --- --- --- --- --- --- --- --- --- --- --- --- --- --- --- --- --- --- --- --- --- --- --- --- --- --- --- --- --- --- --- --- --- --- --- --- --- --- --- --- --- --- --- --- --- --- --- --- --- --- --- --- --- --- --- --- tat ggt aat gac gat gaa a-- --t cga gta gta --- --- act tct aat aaa gac ttg ata --- --- tca tta tca tat gtt tga tcg --- --- --- --- --- --- --- tca tag tta ata gtg tg- --- --- --- --- --- --- --- --- --- --- --- --- --- --- --- --- --- --- --- --g cta aat ggt act gtt aat aag ttt at- --- --- --- --- --- --- --- --- --- --- --- --- --- --- --- --- --- aga caa tat cat agt att ttc ttt cca gaa t-- --- --- --- tag att att ttt tta aat act gat cct cac aat tcc gtg atg tag cag tag ttg gt- --- --- --- --- --- --- --- --- --- --- --- --- --- --- --- --- --- --- --- --- --- --- --- --- --- --- --- --g cat ggt cta tat cgt --- --- --- --- --- --- --- --- --- --- --- --- --- --- --- --- --- --- --- --- --- --- --- --- --- --- --- --- --- --- --- --- --- --- --- --- --- --- --- --- --- --- --- -ta aaa tgt atc ata tat aat agt ttt ctg acg tgg agt aca gaa ttt tcg a-- --- --- --- --- --- --- --- --- --- --- --- --- --- --- --- --- --- --- --- --- --- --- --- --- --- --- --- --- --- --- --- --- --- --- --- --- --- --- --- --- --- --- --- tta atg agt tca tgg taa gga agg gca aat gcc t-- -gt ata taa tat aca taa gtt aa- --- --- --- --- --- --- --- --- --- --- --- --- --- tag ttt ttt atc ata ttt --- --- --- --- --- --- --- --- tct aat acc ata ata aaa att atc --- --- --- --- --- --- --- --- --- --- --- --- --- --- --- --- --- --- --- --- --- --- --- --- --- -at tat tgc gtt tg- gta gtt --- --- --- --- -ct gcc cta --- --- --- --- --- --- --- --- tca tct ata tca ctg tca ctc tc- --- --- gct ctc act ata tct tct aaa att aca a-- --a caa c-- --- --- --- --- --- --- --- --- --- --- --- --- --- --- --- --- --- --- --- --- --- --- --- --- --- --- --- --- --- --- --- --- --- --- --- -tg gat att cga t-- --- --- --- --- --- --- --- --- --- --- --- --- --- --- --- --- --- --- --- --- --- --- --- --- --- --- aac agc att tgt gt- --- --- --- --- --- --- --- --- --- --- --- --- --- --- --- --- --

>ON637938.1_Monkeypox_virus_isolate_MPXV/Germany/2022/RKI01_complete_genome

ttt ttt cga tct atc ctc gtc c-- -t- ctc atc atc ctt ata --- --- --- --- --- --- -tt att atc att att atc ata gtc tat taa aca caa atc atc t-- --- --- --- --- --- --- --- --- --- --- --- --- --- --- --- --- --- --- --- --- --- --- --- --- --- --- acg ttt ata ac- --- --- --- --- --- --- --- --- aac att c-- --- --- --- --t cat tat taa tta gtt ctg tag -aa tat ctt taa taa ttt ggc tat a-- --- --c atc tgt t-- --- --- --- --- --- --- --- --- --- --- --- --- --- --- --- --- --- --- caa tac t-- --- --- --- --- --- --- --- atc tat tga tga ttt ctt tt- --- --- --- --- --- --- --- --- --- --- --- tta aga ct- --- --- --- --- --- --- --- --- --- --- --- --- --- --- --- --- --- --- --- --- --- --- --- --- --- --- --- --- --- --- --- --- --- --- --- --- --- --- --- -ta aac tag t-- --- --- --- --- --- --- --- --- --- --- --- --- --- --- --- --- --- --- --- --- --- --- --- --- --- --- --- --- --- --- --- --- --- --- --- --- --- --- --- --- --- --- --- --- --- --- --- --- --- --- --- --- --- --- --- --- --- --- --- --- --- --- --- --- --- --- --- --- --- --- --- --- --- --- --- --- --- --- --- --- --- --- --- --- --- --- --- --- --- --- --- --- --- --- --- --- --- --- --- --- --- --- --- --- --- --- --- --- --- tat ggt aat gac gat gaa a-- --t cga gta gta --- --- act tct aat aaa gac ttg ata --- --- tca tta tca tat gtt tga tcg --- --- --- --- --- --- --- tca tag tta ata gtg tg- --- --- --- --- --- --- --- --- --- --- --- --- --- --- --- --- --- --- --- --g cta aat ggt act gtt aat aag ttt at- --- --- --- --- --- --- --- --- --- --- --- --- --- --- --- --- --- aga caa tat cat agt att ttc ttt cca gaa t-- --- --- --- tag att att ttt tta aat act gat cct cac aat tcc gtg atg tag cag tag ttg gt- --- --- --- --- --- --- --- --- --- --- --- --- --- --- --- --- --- --- --- --- --- --- --- --- --- --- --- --g cat ggt cta tat cgt --- --- --- --- --- --- --- --- --- --- --- --- --- --- --- --- --- --- --- --- --- --- --- --- --- --- --- --- --- --- --- --- --- --- --- --- --- --- --- --- --- --- --- -ta aaa tgt atc ata tat aat agt ttt ctg acg tgg agt aca gaa ttt tcg a-- --- --- --- --- --- --- --- --- --- --- --- --- --- --- --- --- --- --- --- --- --- --- --- --- --- --- --- --- --- --- --- --- --- --- --- --- --- --- --- --- --- --- --- tta atg agt tca tgg taa gga agg gca aat gcc t-- -gt ata taa tat aca taa gtt aa- --- --- --- --- --- --- --- --- --- --- --- --- --- tag ttt ttt atc ata ttt --- --- --- --- --- --- --- --- tct aat acc ata ata aaa att atc --- --- --- --- --- --- --- --- --- --- --- --- --- --- --- --- --- --- --- --- --- --- --- --- --- -at tat tgc gtt tg- gta gtt --- --- --- --- -ct gcc cta --- --- --- --- --- --- --- --- tca tct ata tca ctg tca ctc tc- --- --- gct ctc act ata tct tct aaa att aca a-- --a caa c-- --- --- --- --- --- --- --- --- --- --- --- --- --- --- --- --- --- --- --- --- --- --- --- --- --- --- --- --- --- --- --- --- --- --- --- -tg gat att cga t-- --- --- --- --- --- --- --- --- --- --- --- --- --- --- --- --- --- --- --- --- --- --- --- --- --- --- aac agc att tgt gt- --- --- --- --- --- --- --- --- --- --- --- --- --- --- --- --- --

>ON637939.1_Monkeypox_virus_isolate_MPXV/Germany/2022/RKI02_complete_genome

ttt ttt cga tct atc ctc gtc c-- -t- ctc atc atc ctt ata --- --- --- --- --- --- -tt att atc att att atc ata gtc tat taa aca caa atc atc t-- --- --- --- --- --- --- --- --- --- --- --- --- --- --- --- --- --- --- --- --- --- --- --- --- --- --- acg ttt ata ac- --- --- --- --- --- --- --- --- aac att c-- --- --- --- --t cat tat taa tta gtt ctg tag -aa tat ctt taa taa ttt ggc tat a-- --- --c atc tgt t-- --- --- --- --- --- --- --- --- --- --- --- --- --- --- --- --- --- --- caa tac t-- --- --- --- --- --- --- --- atc tat tga tga ttt ctt tt- --- --- --- --- --- --- --- --- --- --- --- tta aga ct- --- --- --- --- --- --- --- --- --- --- --- --- --- --- --- --- --- --- --- --- --- --- --- --- --- --- --- --- --- --- --- --- --- --- --- --- --- --- --- -ta aac tag t-- --- --- --- --- --- --- --- --- --- --- --- --- --- --- --- --- --- --- --- --- --- --- --- --- --- --- --- --- --- --- --- --- --- --- --- --- --- --- --- --- --- --- --- --- --- --- --- --- --- --- --- --- --- --- --- --- --- --- --- --- --- --- --- --- --- --- --- --- --- --- --- --- --- --- --- --- --- --- --- --- --- --- --- --- --- --- --- --- --- --- --- --- --- --- --- --- --- --- --- --- --- --- --- --- --- --- --- --- --- tat ggt aat gac gat gaa a-- --t cga gta gta --- --- act tct aat aaa gac ttg ata --- --- tca tta tca tat gtt tga tcg --- --- --- --- --- --- --- tca tag tta ata gtg tg- --- --- --- --- --- --- --- --- --- --- --- --- --- --- --- --- --- --- --- --g cta aat ggt act gtt aat aag ttt at- --- --- --- --- --- --- --- --- --- --- --- --- --- --- --- --- --- aga caa tat cat agt att ttc ttt cca gaa t-- --- --- --- tag att att ttt tta aat act gat cct cac aat tcc gtg atg tag cag tag ttg gt- --- --- --- --- --- --- --- --- --- --- --- --- --- --- --- --- --- --- --- --- --- --- --- --- --- --- --- --g cat ggt cta tat cgt --- --- --- --- --- --- --- --- --- --- --- --- --- --- --- --- --- --- --- --- --- --- --- --- --- --- --- --- --- --- --- --- --- --- --- --- --- --- --- --- --- --- --- -ta aaa tgt atc ata tat aat agt ttt ctg acg tgg agt aca gaa ttt tcg a-- --- --- --- --- --- --- --- --- --- --- --- --- --- --- --- --- --- --- --- --- --- --- --- --- --- --- --- --- --- --- --- --- --- --- --- --- --- --- --- --- --- --- --- tta atg agt tca tgg taa gga agg gca aat gcc t-- -gt ata taa tat aca taa gtt aa- --- --- --- --- --- --- --- --- --- --- --- --- --- tag ttt ttt atc ata ttt --- --- --- --- --- --- --- --- tct aat acc ata ata aaa att atc --- --- --- --- --- --- --- --- --- --- --- --- --- --- --- --- --- --- --- --- --- --- --- --- --- -at tat tgc gtt tg- gta gtt --- --- --- --- -ct gcc cta --- --- --- --- --- --- --- --- tca tct ata tca ctg tca ctc tc- --- --- gct ctc act ata tct tct aaa att aca a-- --a caa c-- --- --- --- --- --- --- --- --- --- --- --- --- --- --- --- --- --- --- --- --- --- --- --- --- --- --- --- --- --- --- --- --- --- --- --- -tg gat att cga t-- --- --- --- --- --- --- --- --- --- --- --- --- --- --- --- --- --- --- --- --- --- --- --- --- --- --- aac agc att tgt gt- --- --- --- --- --- --- --- --- --- --- --- --- --- --- --- --- --

>ON682263.2_Monkeypox_virus_isolate_MPXV/Germany/2022/RKI03_complete_genome

ttt ttt cga tct atc ctc gtc c-- -t- ctc atc atc ctt ata --- --- --- --- --- --- -tt att atc att att atc ata gtc tat taa aca caa atc atc t-- --- --- --- --- --- --- --- --- --- --- --- --- --- --- --- --- --- --- --- --- --- --- --- --- --- --- acg ttt ata ac- --- --- --- --- --- --- --- --- aac att c-- --- --- --- --t cat tat taa tta gtt ctg tag -aa tat ctt taa taa ttt ggc tat a-- --- --c atc tgt t-- --- --- --- --- --- --- --- --- --- --- --- --- --- --- --- --- --- --- caa tac t-- --- --- --- --- --- --- --- atc tat tga tga ttt ctt tt- --- --- --- --- --- --- --- --- --- --- --- tta aga ct- --- --- --- --- --- --- --- --- --- --- --- --- --- --- --- --- --- --- --- --- --- --- --- --- --- --- --- --- --- --- --- --- --- --- --- --- --- --- --- -ta aac tag t-- --- --- --- --- --- --- --- --- --- --- --- --- --- --- --- --- --- --- --- --- --- --- --- --- --- --- --- --- --- --- --- --- --- --- --- --- --- --- --- --- --- --- --- --- --- --- --- --- --- --- --- --- --- --- --- --- --- --- --- --- --- --- --- --- --- --- --- --- --- --- --- --- --- --- --- --- --- --- --- --- --- --- --- --- --- --- --- --- --- --- --- --- --- --- --- --- --- --- --- --- --- --- --- --- --- --- --- --- --- tat ggt aat gac gat gaa a-- --t cga gta gta --- --- act tct aat aaa gac ttg ata --- --- tca tta tca tat gtt tga tcg --- --- --- --- --- --- --- tca tag tta ata gtg tg- --- --- --- --- --- --- --- --- --- --- --- --- --- --- --- --- --- --- --- --g cta aat ggt act gtt aat aag ttt at- --- --- --- --- --- --- --- --- --- --- --- --- --- --- --- --- --- aga caa tat cat agt att ttc ttt cca gaa t-- --- --- --- tag att att ttt tta aat act gat cct cac aat tcc gtg atg tag cag tag ttg gt- --- --- --- --- --- --- --- --- --- --- --- --- --- --- --- --- --- --- --- --- --- --- --- --- --- --- --- --g cat ggt cta tat cgt --- --- --- --- --- --- --- --- --- --- --- --- --- --- --- --- --- --- --- --- --- --- --- --- --- --- --- --- --- --- --- --- --- --- --- --- --- --- --- --- --- --- --- -ta aaa tgt atc ata tat aat agt ttt ctg acg tgg agt aca gaa ttt tcg a-- --- --- --- --- --- --- --- --- --- --- --- --- --- --- --- --- --- --- --- --- --- --- --- --- --- --- --- --- --- --- --- --- --- --- --- --- --- --- --- --- --- --- --- tta atg agt tca tgg taa gga agg gca aat gcc t-- -gt ata taa tat aca taa gtt aa- --- --- --- --- --- --- --- --- --- --- --- --- --- tag ttt ttt atc ata ttt --- --- --- --- --- --- --- --- tct aat acc ata ata aaa att atc --- --- --- --- --- --- --- --- --- --- --- --- --- --- --- --- --- --- --- --- --- --- --- --- --- -at tat tgc gtt tg- gta gtt --- --- --- --- -ct gcc cta --- --- --- --- --- --- --- --- tca tct ata tca ctg tca ctc tc- --- --- gct ctc act ata tct tct aaa att aca a-- --a caa c-- --- --- --- --- --- --- --- --- --- --- --- --- --- --- --- --- --- --- --- --- --- --- --- --- --- --- --- --- --- --- --- --- --- --- --- -tg gat att cga t-- --- --- --- --- --- --- --- --- --- --- --- --- --- --- --- --- --- --- --- --- --- --- --- --- --- --- aac agc att tgt gt- --- --- --- --- --- --- --- --- --- --- --- --- --- --- --- --- --

>ON682266.1_Monkeypox_virus_isolate_MPXV/Germany/2022/RKI09_complete_genome

ttt ttt cga tct atc ctc gtc c-- -t- ctc atc atc ctt ata --- --- --- --- --- --- -tt att atc att att atc ata gtc tat taa aca caa atc atc t-- --- --- --- --- --- --- --- --- --- --- --- --- --- --- --- --- --- --- --- --- --- --- --- --- --- --- acg ttt ata ac- --- --- --- --- --- --- --- --- aac att c-- --- --- --- --t cat tat taa tta gtt ctg tag -aa tat ctt taa taa ttt ggc tat a-- --- --c atc tgt t-- --- --- --- --- --- --- --- --- --- --- --- --- --- --- --- --- --- --- caa tac t-- --- --- --- --- --- --- --- atc tat tga tga ttt ctt tt- --- --- --- --- --- --- --- --- --- --- --- tta aga ct- --- --- --- --- --- --- --- --- --- --- --- --- --- --- --- --- --- --- --- --- --- --- --- --- --- --- --- --- --- --- --- --- --- --- --- --- --- --- --- -ta aac tag t-- --- --- --- --- --- --- --- --- --- --- --- --- --- --- --- --- --- --- --- --- --- --- --- --- --- --- --- --- --- --- --- --- --- --- --- --- --- --- --- --- --- --- --- --- --- --- --- --- --- --- --- --- --- --- --- --- --- --- --- --- --- --- --- --- --- --- --- --- --- --- --- --- --- --- --- --- --- --- --- --- --- --- --- --- --- --- --- --- --- --- --- --- --- --- --- --- --- --- --- --- --- --- --- --- --- --- --- --- --- tat ggt aat gac gat gaa a-- --t cga gta gta --- --- act tct aat aaa gac ttg ata --- --- tca tta tca tat gtt tga tcg --- --- --- --- --- --- --- tca tag tta ata gtg tg- --- --- --- --- --- --- --- --- --- --- --- --- --- --- --- --- --- --- --- --g cta aat ggt act gtt aat aag ttt at- --- --- --- --- --- --- --- --- --- --- --- --- --- --- --- --- --- aga caa tat cat agt att ttc ttt cca gaa t-- --- --- --- tag att att ttt tta aat act gat cct cac aat tcc gtg atg tag cag tag ttg gt- --- --- --- --- --- --- --- --- --- --- --- --- --- --- --- --- --- --- --- --- --- --- --- --- --- --- --- --g cat ggt cta tat cgt --- --- --- --- --- --- --- --- --- --- --- --- --- --- --- --- --- --- --- --- --- --- --- --- --- --- --- --- --- --- --- --- --- --- --- --- --- --- --- --- --- --- --- -ta aaa tgt atc ata tat aat agt ttt ctg acg tgg agt aca gaa ttt tcg a-- --- --- --- --- --- --- --- --- --- --- --- --- --- --- --- --- --- --- --- --- --- --- --- --- --- --- --- --- --- --- --- --- --- --- --- --- --- --- --- --- --- --- --- tta atg agt tca tgg taa gga agg gca aat gcc t-- -gt ata taa tat aca taa gtt aa- --- --- --- --- --- --- --- --- --- --- --- --- --- tag ttt ttt atc ata ttt --- --- --- --- --- --- --- --- tct aat acc ata ata aaa att atc --- --- --- --- --- --- --- --- --- --- --- --- --- --- --- --- --- --- --- --- --- --- --- --- --- -at tat tgc gtt tg- gta gtt --- --- --- --- -ct gcc cta --- --- --- --- --- --- --- --- tca tct ata tca ctg tca ctc tc- --- --- gct ctc act ata tct tct aaa att aca a-- --a caa c-- --- --- --- --- --- --- --- --- --- --- --- --- --- --- --- --- --- --- --- --- --- --- --- --- --- --- --- --- --- --- --- --- --- --- --- -tg gat att cga t-- --- --- --- --- --- --- --- --- --- --- --- --- --- --- --- --- --- --- --- --- --- --- --- --- --- --- aac agc att tgt gt- --- --- --- --- --- --- --- --- --- --- --- --- --- --- --- --- --

>ON619836.1_Monkeypox_virus_isolate_MPXV_UK_2022_2_complete_genome

ttt ttt cga tct atc ctc gtc c-- -t- ctc atc atc ctt ata --- --- --- --- --- --- -tt att atc att att atc ata gtc tat taa aca caa atc atc t-- --- --- --- --- --- --- --- --- --- --- --- --- --- --- --- --- --- --- --- --- --- --- --- --- --- --- acg ttt ata ac- --- --- --- --- --- --- --- --- aac att c-- --- --- --- --t cat tat taa tta gtt ctg tag -aa tat ctt taa taa ttt ggc tat a-- --- --c atc tgt t-- --- --- --- --- --- --- --- --- --- --- --- --- --- --- --- --- --- --- caa tac t-- --- --- --- --- --- --- --- atc tat tga tga ttt ctt tt- --- --- --- --- --- --- --- --- --- --- --- tta aga ct- --- --- --- --- --- --- --- --- --- --- --- --- --- --- --- --- --- --- --- --- --- --- --- --- --- --- --- --- --- --- --- --- --- --- --- --- --- --- --- -ta aac tag t-- --- --- --- --- --- --- --- --- --- --- --- --- --- --- --- --- --- --- --- --- --- --- --- --- --- --- --- --- --- --- --- --- --- --- --- --- --- --- --- --- --- --- --- --- --- --- --- --- --- --- --- --- --- --- --- --- --- --- --- --- --- --- --- --- --- --- --- --- --- --- --- --- --- --- --- --- --- --- --- --- --- --- --- --- --- --- --- --- --- --- --- --- --- --- --- --- --- --- --- --- --- --- --- --- --- --- --- --- --- tat ggt aat gac gat gaa a-- --t cga gta gta --- --- act tct aat aaa gac ttg ata --- --- tca tta tca tat gtt tga tcg --- --- --- --- --- --- --- tca tag tta ata gtg tg- --- --- --- --- --- --- --- --- --- --- --- --- --- --- --- --- --- --- --- --g cta aat ggt act gtt aat aag ttt at- --- --- --- --- --- --- --- --- --- --- --- --- --- --- --- --- --- aga caa tat cat agt att ttc ttt cca gaa t-- --- --- --- tag att att ttt tta aat act gat cct cac aat tcc gtg atg tag cag tag ttg gt- --- --- --- --- --- --- --- --- --- --- --- --- --- --- --- --- --- --- --- --- --- --- --- --- --- --- --- --g cat ggt cta tat cgt --- --- --- --- --- --- --- --- --- --- --- --- --- --- --- --- --- --- --- --- --- --- --- --- --- --- --- --- --- --- --- --- --- --- --- --- --- --- --- --- --- --- --- -ta aaa tgt atc ata tat aat agt ttt ctg acg tgg agt aca gaa ttt tcg a-- --- --- --- --- --- --- --- --- --- --- --- --- --- --- --- --- --- --- --- --- --- --- --- --- --- --- --- --- --- --- --- --- --- --- --- --- --- --- --- --- --- --- --- tta atg agt tca tgg taa gga agg gca aat gcc t-- -gt ata taa tat aca taa gtt aa- --- --- --- --- --- --- --- --- --- --- --- --- --- tag ttt ttt atc ata ttt --- --- --- --- --- --- --- --- tct aat acc ata ata aaa att atc --- --- --- --- --- --- --- --- --- --- --- --- --- --- --- --- --- --- --- --- --- --- --- --- --- -at tat tgc gtt tg- gta gtt --- --- --- --- -ct gcc cta --- --- --- --- --- --- --- --- tca tct ata tca ctg tca ctc tc- --- --- gct ctc act ata tct tct aaa att aca a-- --a caa c-- --- --- --- --- --- --- --- --- --- --- --- --- --- --- --- --- --- --- --- --- --- --- --- --- --- --- --- --- --- --- --- --- --- --- --- -tg gat att cga t-- --- --- --- --- --- --- --- --- --- --- --- --- --- --- --- --- --- --- --- --- --- --- --- --- --- --- aac agc att tgt gt- --- --- --- --- --- --- --- --- --- --- --- --- --- --- --- --- --

>ON619838.1_Monkeypox_virus_isolate_MPXV_UK_2022_4_complete_genome

ttt ttt cga tct atc ctc gtc c-- -t- ctc atc atc ctt ata --- --- --- --- --- --- -tt att atc att att atc ata gtc tat taa aca caa atc atc t-- --- --- --- --- --- --- --- --- --- --- --- --- --- --- --- --- --- --- --- --- --- --- --- --- --- --- acg ttt ata ac- --- --- --- --- --- --- --- --- aac att c-- --- --- --- --t cat tat taa tta gtt ctg tag -aa tat ctt taa taa ttt ggc tat a-- --- --c atc tgt t-- --- --- --- --- --- --- --- --- --- --- --- --- --- --- --- --- --- --- caa tac t-- --- --- --- --- --- --- --- atc tat tga tga ttt ctt tt- --- --- --- --- --- --- --- --- --- --- --- tta aga ct- --- --- --- --- --- --- --- --- --- --- --- --- --- --- --- --- --- --- --- --- --- --- --- --- --- --- --- --- --- --- --- --- --- --- --- --- --- --- --- -ta aac tag t-- --- --- --- --- --- --- --- --- --- --- --- --- --- --- --- --- --- --- --- --- --- --- --- --- --- --- --- --- --- --- --- --- --- --- --- --- --- --- --- --- --- --- --- --- --- --- --- --- --- --- --- --- --- --- --- --- --- --- --- --- --- --- --- --- --- --- --- --- --- --- --- --- --- --- --- --- --- --- --- --- --- --- --- --- --- --- --- --- --- --- --- --- --- --- --- --- --- --- --- --- --- --- --- --- --- --- --- --- --- tat ggt aat gac gat gaa a-- --t cga gta gta --- --- act tct aat aaa gac ttg ata --- --- tca tta tca tat gtt tga tcg --- --- --- --- --- --- --- tca tag tta ata gtg tg- --- --- --- --- --- --- --- --- --- --- --- --- --- --- --- --- --- --- --- --g cta aat ggt act gtt aat aag ttt at- --- --- --- --- --- --- --- --- --- --- --- --- --- --- --- --- --- aga caa tat cat agt att ttc ttt cca gaa t-- --- --- --- tag att att ttt tta aat act gat cct cac aat tcc gtg atg tag cag tag ttg gt- --- --- --- --- --- --- --- --- --- --- --- --- --- --- --- --- --- --- --- --- --- --- --- --- --- --- --- --g cat ggt cta tat cgt --- --- --- --- --- --- --- --- --- --- --- --- --- --- --- --- --- --- --- --- --- --- --- --- --- --- --- --- --- --- --- --- --- --- --- --- --- --- --- --- --- --- --- -ta aaa tgt atc ata tat aat agt ttt ctg acg tgg agt aca gaa ttt tcg a-- --- --- --- --- --- --- --- --- --- --- --- --- --- --- --- --- --- --- --- --- --- --- --- --- --- --- --- --- --- --- --- --- --- --- --- --- --- --- --- --- --- --- --- tta atg agt tca tgg taa gga agg gca aat gcc t-- -gt ata taa tat aca taa gtt aa- --- --- --- --- --- --- --- --- --- --- --- --- --- tag ttt ttt atc ata ttt --- --- --- --- --- --- --- --- tct aat acc ata ata aaa att atc --- --- --- --- --- --- --- --- --- --- --- --- --- --- --- --- --- --- --- --- --- --- --- --- --- -at tat tgc gtt tg- gta gtt --- --- --- --- -ct gcc cta --- --- --- --- --- --- --- --- tca tct ata tca ctg tca ctc tc- --- --- gct ctc act ata tct tct aaa att aca a-- --a caa c-- --- --- --- --- --- --- --- --- --- --- --- --- --- --- --- --- --- --- --- --- --- --- --- --- --- --- --- --- --- --- --- --- --- --- --- -tg gat att cga t-- --- --- --- --- --- --- --- --- --- --- --- --- --- --- --- --- --- --- --- --- --- --- --- --- --- --- aac agc att tgt gt- --- --- --- --- --- --- --- --- --- --- --- --- --- --- --- --- --

>ON619835.1_Monkeypox_virus_isolate_MPXV_UK_2022_1_complete_genome

ttt ttt cga tct atc ctc gtc c-- -t- ctc atc atc ctt ata --- --- --- --- --- --- -tt att atc att att atc ata gtc tat taa aca caa atc atc t-- --- --- --- --- --- --- --- --- --- --- --- --- --- --- --- --- --- --- --- --- --- --- --- --- --- --- acg ttt ata ac- --- --- --- --- --- --- --- --- aac att c-- --- --- --- --t cat tat taa tta gtt ctg tag -aa tat ctt taa taa ttt ggc tat a-- --- --c atc tgt t-- --- --- --- --- --- --- --- --- --- --- --- --- --- --- --- --- --- --- caa tac t-- --- --- --- --- --- --- --- atc tat tga tga ttt ctt tt- --- --- --- --- --- --- --- --- --- --- --- tta aga ct- --- --- --- --- --- --- --- --- --- --- --- --- --- --- --- --- --- --- --- --- --- --- --- --- --- --- --- --- --- --- --- --- --- --- --- --- --- --- --- -ta aac tag t-- --- --- --- --- --- --- --- --- --- --- --- --- --- --- --- --- --- --- --- --- --- --- --- --- --- --- --- --- --- --- --- --- --- --- --- --- --- --- --- --- --- --- --- --- --- --- --- --- --- --- --- --- --- --- --- --- --- --- --- --- --- --- --- --- --- --- --- --- --- --- --- --- --- --- --- --- --- --- --- --- --- --- --- --- --- --- --- --- --- --- --- --- --- --- --- --- --- --- --- --- --- --- --- --- --- --- --- --- --- tat ggt aat gac gat gaa a-- --t cga gta gta --- --- act tct aat aaa gac ttg ata --- --- tca tta tca tat gtt tga tcg --- --- --- --- --- --- --- tca tag tta ata gtg tg- --- --- --- --- --- --- --- --- --- --- --- --- --- --- --- --- --- --- --- --g cta aat ggt act gtt aat aag ttt at- --- --- --- --- --- --- --- --- --- --- --- --- --- --- --- --- --- aga caa tat cat agt att ttc ttt cca gaa t-- --- --- --- tag att att ttt tta aat act gat cct cac aat tcc gtg atg tag cag tag ttg gt- --- --- --- --- --- --- --- --- --- --- --- --- --- --- --- --- --- --- --- --- --- --- --- --- --- --- --- --g cat ggt cta tat cgt --- --- --- --- --- --- --- --- --- --- --- --- --- --- --- --- --- --- --- --- --- --- --- --- --- --- --- --- --- --- --- --- --- --- --- --- --- --- --- --- --- --- --- -ta aaa tgt atc ata tat aat agt ttt ctg acg tgg agt aca gaa ttt tcg a-- --- --- --- --- --- --- --- --- --- --- --- --- --- --- --- --- --- --- --- --- --- --- --- --- --- --- --- --- --- --- --- --- --- --- --- --- --- --- --- --- --- --- --- tta atg agt tca tgg taa gga agg gca aat gcc t-- -gt ata taa tat aca taa gtt aa- --- --- --- --- --- --- --- --- --- --- --- --- --- tag ttt ttt atc ata ttt --- --- --- --- --- --- --- --- tct aat acc ata ata aaa att atc --- --- --- --- --- --- --- --- --- --- --- --- --- --- --- --- --- --- --- --- --- --- --- --- --- -at tat tgc gtt tg- gta gtt --- --- --- --- -ct gcc cta --- --- --- --- --- --- --- --- tca tct ata tca ctg tca ctc tc- --- --- gct ctc act ata tct tct aaa att aca a-- --a caa c-- --- --- --- --- --- --- --- --- --- --- --- --- --- --- --- --- --- --- --- --- --- --- --- --- --- --- --- --- --- --- --- --- --- --- --- -tg gat att cga t-- --- --- --- --- --- --- --- --- --- --- --- --- --- --- --- --- --- --- --- --- --- --- --- --- --- --- aac agc att tgt gt- --- --- --- --- --- --- --- --- --- --- --- --- --- --- --- --- --

>ON645312.1_Monkeypox_virus_isolate_MPXV_GSTT_Patient1_partial_genome

ttt ttt cga tct atc ctc gtc --- --- ctc atc atc ctt ata --- --- --- --- --- --- -tt att atc att att atc ata gtc tat taa aca caa atc atc t-- --- --- --- --- --- --- --- --- --- --- --- --- --- --- --- --- --- --- --- --- --- --- --- --- --- --- acg ttt ata ac- --- --- --- --- --- --- --- --- aac att c-- --- --- --- --t cat tat taa tta gtt ctg tag -aa tat ctt taa taa ttt ggc tat a-- --- --c atc tgt t-- --- --- --- --- --- --- --- --- --- --- --- --- --- --- --- --- --- --- caa tac t-- --- --- --- --- --- --- --- atc tat tga tga ttt ctt tt- --- --- --- --- --- --- --- --- --- --- --- tta aga ct- --- --- --- --- --- --- --- --- --- --- --- --- --- --- --- --- --- --- --- --- --- --- --- --- --- --- --- --- --- --- --- --- --- --- --- --- --- --- --- -ta aac tag t-- --- --- --- --- --- --- --- --- --- --- --- --- --- --- --- --- --- --- --- --- --- --- --- --- --- --- --- --- --- --- --- --- --- --- --- --- --- --- --- --- --- --- --- --- --- --- --- --- --- --- --- --- --- --- --- --- --- --- --- --- --- --- --- --- --- --- --- --- --- --- --- --- --- --- --- --- --- --- --- --- --- --- --- --- --- --- --- --- --- --- --- --- --- --- --- --- --- --- --- --- --- --- --- --- --- --- --- --- --- tat ggt aat gac gat gaa a-- --t cga gta gta --- --- act tct aat aaa gac ttg ata --- --- tca tta tca tat gtt tga tcg --- --- --- --- --- --- --- tca tag tta ata gtg tg- --- --- --- --- --- --- --- --- --- --- --- --- --- --- --- --- --- --- --- --g cta aat ggt act gtt aat aag ttt at- --- --- --- --- --- --- --- --- --- --- --- --- --- --- --- --- --- aga caa tat cat agt att ttc ttt cca gaa t-- --- --- --- tag att att ttt tta aat act gat cct cac aat tcc gtg atg tag cag tag ttg gt- --- --- --- --- --- --- --- --- --- --- --- --- --- --- --- --- --- --- --- --- --- --- --- --- --- --- --- --g cat ggt cta tat cgt --- --- --- --- --- --- --- --- --- --- --- --- --- --- --- --- --- --- --- --- --- --- --- --- --- --- --- --- --- --- --- --- --- --- --- --- --- --- --- --- --- --- --- -ta aaa tgt atc ata tat aat agt ttt ctg acg tgg agt aca gaa ttt tcg a-- --- --- --- --- --- --- --- --- --- --- --- --- --- --- --- --- --- --- --- --- --- --- --- --- --- --- --- --- --- --- --- --- --- --- --- --- --- --- --- --- --- --- --- tta atg agt tca tgg taa gga agg gca aat gcc t-- -gt ata taa tat aca taa gtt aa- --- --- --- --- --- --- --- --- --- --- --- --- --- tag ttt ttt atc ata ttt --- --- --- --- --- --- --- --- tct aat acc ata ata aaa att atc --- --- --- --- --- --- --- --- --- --- --- --- --- --- --- --- --- --- --- --- --- --- --- --- --- -at tat tgc gtt tg- gta gtt --- --- --- --- -ct gcc cta --- --- --- --- --- --- --- --- tca tct ata tca ctg tca --- --- --- --- -ct ctc act ata tct tct aaa att aca a-- --a caa c-- --- --- --- --- --- --- --- --- --- --- --- --- --- --- --- --- --- --- --- --- --- --- --- --- --- --- --- --- --- --- --- --- --- --- --- -tg gat att cga t-- --- --- --- --- --- --- --- --- --- --- --- --- --- --- --- --- --- --- --- --- --- --- --- --- --- --- aac agc att tgt gt- --- --- --- --- --- --- --- --- --- --- --- --- --- --- --- --- --

>ON682269.2_Monkeypox_virus_isolate_MPXV/Germany/2022/RKI07_complete_genome

ttt ttt cga tct atc ctc gtc c-- -t- ctc atc atc ctt ata --- --- --- --- --- --- -tt att atc att att atc ata gtc tat taa aca caa atc atc t-- --- --- --- --- --- --- --- --- --- --- --- --- --- --- --- --- --- --- --- --- --- --- --- --- --- --- acg ttt ata ac- --- --- --- --- --- --- --- --- aac att c-- --- --- --- --t cat tat taa tta gtt ctg tag -aa tat ctt taa taa ttt ggc tat a-- --- --c atc tgt t-- --- --- --- --- --- --- --- --- --- --- --- --- --- --- --- --- --- --- caa tac t-- --- --- --- --- --- --- --- atc tat tga tga ttt ctt tt- --- --- --- --- --- --- --- --- --- --- --- tta aga ct- --- --- --- --- --- --- --- --- --- --- --- --- --- --- --- --- --- --- --- --- --- --- --- --- --- --- --- --- --- --- --- --- --- --- --- --- --- --- --- -ta aac tag t-- --- --- --- --- --- --- --- --- --- --- --- --- --- --- --- --- --- --- --- --- --- --- --- --- --- --- --- --- --- --- --- --- --- --- --- --- --- --- --- --- --- --- --- --- --- --- --- --- --- --- --- --- --- --- --- --- --- --- --- --- --- --- --- --- --- --- --- --- --- --- --- --- --- --- --- --- --- --- --- --- --- --- --- --- --- --- --- --- --- --- --- --- --- --- --- --- --- --- --- --- --- --- --- --- --- --- --- --- --- tat ggt aat gac gat gaa a-- --t cga gta gta --- --- act tct aat aaa gac ttg ata --- --- tca tta tca tat gtt tga tcg --- --- --- --- --- --- --- tca tag tta ata gtg tg- --- --- --- --- --- --- --- --- --- --- --- --- --- --- --- --- --- --- --- --g cta aat ggt act gtt aat aag ttt at- --- --- --- --- --- --- --- --- --- --- --- --- --- --- --- --- --- aga caa tat cat agt att ttc ttt cca gaa t-- --- --- --- tag att att ttt tta aat act gat cct cac aat tcc gtg atg tag cag tag ttg gt- --- --- --- --- --- --- --- --- --- --- --- --- --- --- --- --- --- --- --- --- --- --- --- --- --- --- --- --g cat ggt cta tat cgt --- --- --- --- --- --- --- --- --- --- --- --- --- --- --- --- --- --- --- --- --- --- --- --- --- --- --- --- --- --- --- --- --- --- --- --- --- --- --- --- --- --- --- -ta aaa tgt atc ata tat aat agt ttt ctg acg tgg agt aca gaa ttt tcg a-- --- --- --- --- --- --- --- --- --- --- --- --- --- --- --- --- --- --- --- --- --- --- --- --- --- --- --- --- --- --- --- --- --- --- --- --- --- --- --- --- --- --- --- tta atg agt tca tgg taa gga agg gca aat gcc t-- -gt ata taa tat aca taa gtt aa- --- --- --- --- --- --- --- --- --- --- --- --- --- tag ttt ttt atc ata ttt --- --- --- --- --- --- --- --- tct aat acc ata ata aaa att atc --- --- --- --- --- --- --- --- --- --- --- --- --- --- --- --- --- --- --- --- --- --- --- --- --- -at tat tgc gtt tg- gta gtt --- --- --- --- -ct gcc cta --- --- --- --- --- --- --- --- tca tct ata tca ctg tca ctc tc- --- --- gct ctc act ata tct tct aaa att aca a-- --a caa c-- --- --- --- --- --- --- --- --- --- --- --- --- --- --- --- --- --- --- --- --- --- --- --- --- --- --- --- --- --- --- --- --- --- --- --- -tg gat att cga t-- --- --- --- --- --- --- --- --- --- --- --- --- --- --- --- --- --- --- --- --- --- --- --- --- --- --- aac agc att tgt gt- --- --- --- --- --- --- --- --- --- --- --- --- --- --- --- --- --

>KJ642616.1_Monkeypox_virus_strain_PCH_complete_genome

ttt ttt cga tct atc aat ttc agt ata ttc ttc gcc gtt ata aaa gta atg ttg ttt aat tgt agg acg gtt gtt agt ata atc aca tga ata ata ata ttc taa ttc ctc gta ttg act act tac aga tac tcg aaa tag tct gaa aaa ttc ttc aaa gat att ttt ata aag atc tag gaa aag ttt att acc gac cat gaa cga gat aga tgg ata aat atc ctt tcc atc aaa ggt cat aat tgg ata att gtc cag caa tat atc tgc tgt att agt tat atc act tcc att tat ttt cag att gaa gta atg tac tag ttt gtg aca att aac aag ata caa aag aga tgc cga tac taa tac gta aat agc tat acg cga atc cat tgt tac ctt ttt tta ttt cat agg tct att aat aaa tat atg tat tac tta aga cta gaa aaa tca aaa gtg agt ttt tga tat ttg att ctt act tat tgt ggg att gta gtt tac tta gta att cat ctc tga atc ctg ata aat cat gca tat caa tga tgc aac tac gca gca aac tag tag gaa tat aga tat ctg gat atg tac gta aat agt cga tta tat ctt tta caa tac tat tag tcc cta ttg cgt tat cta tat atc cat taa taa tat tac aca gtg gat act tat gag aaa tat acc tct tac aga ttt tta acg ata tat aat cta gaa aga tat gtg tgt agt act gta tta cct aaa tta tca gtc tca ttc aaa tat tgc atg act att atc gag aat tgc tat atc cct cta tac tcg atg cat tta tta cag taa ttc aat cca gct aac ata aga gcc aat ctc aat gtt ggt tta att ata tca tct tca tgt aat aat aac gat gga aac ttt cca gta gcg taa cac tta tct aag aag gat ata ata act atg tct aca tta tgt ctc tta tcg aga ata ttc tta acg aga tat cca tag cta ttc tgg tgc taa tta ttc cta tat tat att cca cga aaa atg atg aag gca atc att cat cat aag atg ata aaa agt gta gtg agt aag agt att agt gag aga gca tga agg aga ttt agt att tag cag tga gga tat gat cca aga ggg tga gat agt cgt tct cgt tca gaa tct ttc gca gca taa gta gta tgt cga tat act tat cat tga aga ctc ttc cag tga caa tag ctg att gag tac aaa gtc caa tta ttg cac aaa gtt ctt tgg cgg ttt tca tgg agt cat ttc tga tga aac att taa tga tct cca cgc aat tgt cga tat tgt ccc acg gaa gtg aat ccg aga act cct tca act cgc tac caa ata gct cca ttg cat caa ttc tga aag aga tga gaa gcc tgt aga gag gcc ctg cgc ttt ctc tat ggg tcc atc tat gag aaa ccc aca gga tgt att cag tca gac aa- tgt ctg aca tca gtc acg gta ttc agg gag tcc tta gta gcg tgg caa tga cag gga ctg aac tgg gca caa gga gag gcc att gtg aag gta gac gaa ggt aac ctg atg gta gac ctg tag ccg tct gtg ctt aat aga ggg ctt taa ttt cca ttt tta atg gtg tcg tga atg agg aat gag agt gtc tct cgt cct tgg ttt aca tgg atc aga gtg aga aaa aat atc ttg tat att att aac taa caa cct tgg ttt cta tcc atg ttt -aa aaa atg acc tat atg ttc ttt att aat tct att tta aac ttt atc ctc aag act cct gac aaa att aaa atc cag aaa gca gca aac aat cct gtt aca agt tta ctg aaa tct ctc ttt gat tgt aga gta tat gta gtc aga gca aga aac act gca gta gtc aac atg aaa gct tgc ata acg ata cgt gca tca tag aaa gta aca aca gag gcc agc gtt aga gat tct aac agt gta aat cca caa tgg tgt atg tac aga ttc agg gga tgt tca tgt ctg tgt aaa gtc aat gcg aaa atc aag cct ata gat ccg aac att gat gcc aat att aga aca gga ctc cct tgt ata aat gtc cga tgc att caa agt ata aaa ata ctg cag ctg ttg ccg ttg tta aag gaa att gta gaa agg ata ccg tag act ttt ctt aga aat gcc att cgt atg tac acg ctg gca gac gcc acc gag ctg tca tag ttg aag tcg tcc tcg at

>AY753185.1_Monkeypox_virus_strain_COP-58_complete_genome

ttt ttt cga tct atc aat ttc agt ata ttc ttc gcc gtt ata aaa gta atg ttg ttt aat tgt agg acg gtt gtt agt ata atc aca tga ata ata ata ttc taa ttc ctc gta ttg act act tac aga tac tcg aaa tag tct gaa aaa ttc ttc aaa gat att ttt ata aag atc tag gaa aag ttt att acc gac cat gaa cga gat aga tgg ata aat atc ctt tcc atc aaa ggt cat aat tgg ata att gtc cag caa tat atc tgc tgt att agt tat atc act tcc att tat ttt cag att gaa gta atg tac tag ttt gtg aca att aac aag ata caa aag aga tgc cga tac taa tac gta aat agc tat acg cga atc cat tgt tac ctt ttt tta ttt cat agg tct att aat aaa tat atg tat tac tta aga cta gaa aaa tca aaa gtg agt ttt tga tat ttg att ctt act tat tgt ggg att gta gtt tac tta gta att cat ctc tga atc ctg ata aat cat gca tat caa tga tgc aac tac gca gca aac tag tag gaa tat aga tat ctg gat atg tac gta aat agt cga tta tat ctt tta caa tac tat tag tcc cta ttg cgt tat cta tat atc cat taa taa tat tac aca gtg gat act tat gag aaa tat acc tct tac aga ttt tta acg ata tat aat cta gaa aga tat gtg tgt agt act gta tta cct aaa tta tca gtc tca ttc aaa tat tgc atg act att atc gag aat tgc tat atc cct cta tac tcg atg cat tta tta cag taa ttc aat cca gct aac ata aga gcc aat ctc aat gtt ggt tta att ata tca tct tca tgt aat aat aac gat gga aac ttt cca gta gcg taa cac tta tct aag aag gat ata ata act atg tct aca tta tgt ctc tta tcg aga ata ttc tta acg aga tat cca tag cta ttc tgg tgc taa tta ttc cta tat tat att cca cga aaa atg atg aag gca atc att cat cat aag atg ata aaa agt gta gtg agt aag agt att agt gag aga gca tga agg aga ttt agt att tag cag tga gga tat gat cca aga ggg tga gat agt cgt tct cgt tca gaa tct ttc gca gca taa gta gta tgt cga tat act tat cat tga aga ctc ttc cag tga caa tag ctg att gag tac aaa gtc caa tta ttg cac aaa gtt ctt tgg cgg ttt tca tgg agt cat ttc tga tga aac att taa tga tct cca cgc aat tgt cga tat tgt ccc acg gaa gtg aat ccg aga act cct tca act cgc tac caa ata gct cca ttg cat caa ttc tga aag aga tga gaa gcc tgt aga gag gcc ctg cgc ttt ctc tat ggg tcc atc tat gag aaa ccc aca gga tgt att cag tca gac aa- tgt ctg aca tca gtc acg gta ttc agg gag tcc tta gta gcg tgg caa tga cag gga ctg aac tgg gca caa gga gag gcc att gtg aag gta gac gaa ggt aac ctg atg gta gac ctg tag ccg tct gtg ctt aat aga ggg ctt taa ttt cca ttt tta atg gtg tcg tga atg agg aat gag agt gtc tct cgt cct tgg ttt aca tgg atc aga gtg aga aaa aat atc ttg tat att att aac taa caa cct tgg ttt cta tcc atg ttt -aa aaa atg acc tat atg ttc ttt att aat tct att tta aac ttt atc ctc aag act cct gac aaa att aaa atc cag aaa gca gca aac aat cct gtt aca agt tta ctg aaa tct ctc ttt gat tgt aga gta tat gta gtc aga gca aga aac act gca gta gtc aac atg aaa gct tgc ata acg ata cgt gca tca tag aaa gta aca aca gag gcc agc gtt aga gat tct aac agt gta aat cca caa --- agt atg tac aga ttc agg gga tgt tca tgt ctg tgt aaa gtc aat gcg aaa atc aag cct ata gat ccg aac att gat gcc aat att aga aca gga ctc cct tgt ata aat gtc cga tgc att caa agt ata aaa ata ctg cag ctg ttg ccg ttg tta aag gaa att gta gaa agg ata ccg tag act ttt ctt aga aat gcc att cgt atg tac acg ctg gca gac gcc acc gag ctg tca tag ttg aag tcg tcc tcg at

>AY603973.1_Monkeypox_virus_strain_MPXV-WRAIR7-61_complete_genome

ttt ttt cga tct atc aat ttc agt ata ttc ttc gcc gtt ata aaa gta atg ttg ttt aat tgt agg acg gtt gtt agt ata atc aca tga ata ata ata ttc taa ttc ctc gta ttg act act tac aga tac tcg aaa tag tct gaa aaa ttc ttc aaa gat att ttt ata aag atc tag gaa aag ttt att acc gac cat gaa cga gat aga tgg ata aat atc ctt tcc atc aaa ggt cat aat tgg ata att gtc cag caa tat atc tgc tgt att agt tat atc act tcc att tat ttt cag att gaa gta atg tac tag ttt gtg aca att aac aag ata caa aag aga tgc cga tac taa tac gta aat agc tat acg cga atc cat tgt tac ctt ttt tta ttt cat agg tct att aat aaa tat atg tat tac tta aga cta gaa aaa tca aaa gtg agt ttt tga tat ttg att ctt act tat tgt ggg att gta gtt tac tta gta att cat ctc tga atc ctg ata aat cat gca tat caa tga tgc aac tac gca gca aac tag tag gaa tat aga tat ctg gat atg tac gta aat agt cga tta tat ctt tta caa tac tat tag tcc cta ttg cgt tat cta tat atc cat taa taa tat tac aca gtg gat act tat gag aaa tat acc tct tac aga ttt tta acg ata tat aat cta gaa aga tat gtg tgt agt act gta tta cct aaa tta tca gtc tca ttc aaa tat tgc atg act att atc gag aat tgc tat atc cct cta tac tcg atg cat tta tta cag taa ttc aat cca gct aac ata aga gcc aat ctc aat gtt ggt tta att ata tca tct tca tgt aat aat aac gat gga aac ttt cca gta gcg taa cac tta tct aag aag gat ata ata act atg tct aca tta tgt ctc tta tcg aga ata ttc tta acg aga tat cca tag cta ttc tgg tgc taa tta ttc cta tat tat att cca cga aaa atg atg aag gca atc att cat cat aag atg ata aaa agt gta gtg agt aag agt att agt gag aga gca tga agg aga ttt agt att tag cag tga gga tat gat cca aga ggg tga gat agt cgt tct cgt tca gaa tct ttc gca gca taa gta gta tgt cga tat act tat cat tga aga ctc ttc cag tga caa tag ctg att gag tac aaa gtc caa tta ttg cac aaa gtt ctt tgg cgg ttt tca tgg agt cat ttc tga tga aac att taa tga tct cca cgc aat tgt cga tat tgt ccc acg gaa gtg aat ccg aga act cct tca act cgc tac caa ata gct cca ttg cat caa ttc tga aag aga tga gaa gcc tgt aga gag gcc ctg cgc ttt ctc tat ggg tcc atc tat gag aaa ccc aca gga tgt att cag tca gac aa- tgt ctg aca tca gtc acg gta ttc agg gag tcc tta gta gcg tgg caa tga cag gga ctg aac tgg gca caa gga gag gcc att gtg aag gta gac gaa ggt aac ctg atg gta gac ctg tag ccg tct gtg ctt aat aga ggg ctt taa ttt cca ttt tta atg gtg tcg tga atg agg aat gag agt gtc tct cgt cct tgg ttt aca tgg atc aga gtg aga aaa aat atc ttg tat att att aac taa caa cct tgg ttt cta tcc atg ttt -aa aaa atg acc tat atg ttc ttt att aat tct att tta aac ttt atc ctc aag act cct gac aaa att aaa atc cag aaa gca gca aac aat cct gtt aca agt tta ctg aaa tct ctc ttt gat tgt aga gta tat gta gtc aga gca aga aac act gca gta gtc aac atg aaa gct tgc ata acg ata cgt gca tca tag aaa gta aca aca gag gcc agc gtt aga gat tct aac agt gta aat cca caa --- agt atg tac aga ttc agg gga tgt tca tgt ctg tgt aaa gtc aat gcg aaa atc aag cct ata gat ccg aac att gat gcc aat att aga aca gga ctc cct tgt ata aat gtc cga tgc att caa agt ata aaa ata ctg cag ctg ttg ccg ttg tta aag gaa att gta gaa agg ata ccg tag act ttt ctt aga aat gcc att cgt atg tac acg ctg gca gac gcc acc gag ctg tca tag ttg aag tcg tcc tcg at

>ON619837.1_Monkeypox_virus_isolate_MPXV_UK_2022_3_complete_genome

ttt ttt cga tct atc ctc gtc c-- -t- ctc atc atc ctt ata --- --- --- --- --- --- -tt att atc att att atc ata gtc tat taa aca caa atc atc t-- --- --- --- --- --- --- --- --- --- --- --- --- --- --- --- --- --- --- --- --- --- --- --- --- --- --- acg ttt ata ac- --- --- --- --- --- --- --- --- aac att c-- --- --- --- --t cat tat taa tta gtt ctg tag -aa tat ctt taa taa ttt ggc tat a-- --- --c atc tgt t-- --- --- --- --- --- --- --- --- --- --- --- --- --- --- --- --- --- --- caa tac t-- --- --- --- --- --- --- --- atc tat tga tga ttt ctt tt- --- --- --- --- --- --- --- --- --- --- --- tta aga ct- --- --- --- --- --- --- --- --- --- --- --- --- --- --- --- --- --- --- --- --- --- --- --- --- --- --- --- --- --- --- --- --- --- --- --- --- --- --- --- -ta aac tag t-- --- --- --- --- --- --- --- --- --- --- --- --- --- --- --- --- --- --- --- --- --- --- --- --- --- --- --- --- --- --- --- --- --- --- --- --- --- --- --- --- --- --- --- --- --- --- --- --- --- --- --- --- --- --- --- --- --- --- --- --- --- --- --- --- --- --- --- --- --- --- --- --- --- --- --- --- --- --- --- --- --- --- --- --- --- --- --- --- --- --- --- --- --- --- --- --- --- --- --- --- --- --- --- --- --- --- --- --- --- tat ggt aat gac gat gaa a-- --t cga gta gta --- --- act tct aat aaa gac ttg ata --- --- tca tta tca tat gtt tga tcg --- --- --- --- --- --- --- tca tag tta ata gtg tg- --- --- --- --- --- --- --- --- --- --- --- --- --- --- --- --- --- --- --- --g cta aat ggt act gtt aat aag ttt at- --- --- --- --- --- --- --- --- --- --- --- --- --- --- --- --- --- aga caa tat cat agt att ttc ttt cca gaa t-- --- --- --- tag att att ttt tta aat act gat cct cac aat tcc gtg atg tag cag tag ttg gt- --- --- --- --- --- --- --- --- --- --- --- --- --- --- --- --- --- --- --- --- --- --- --- --- --- --- --- --g cat ggt cta tat cgt --- --- --- --- --- --- --- --- --- --- --- --- --- --- --- --- --- --- --- --- --- --- --- --- --- --- --- --- --- --- --- --- --- --- --- --- --- --- --- --- --- --- --- -ta aaa tgt atc ata tat aat agt ttt ctg acg tgg agt aca gaa ttt tcg a-- --- --- --- --- --- --- --- --- --- --- --- --- --- --- --- --- --- --- --- --- --- --- --- --- --- --- --- --- --- --- --- --- --- --- --- --- --- --- --- --- --- --- --- tta atg agt tca tgg taa gga agg gca aat gcc t-- -gt ata taa tat aca taa gtt aa- --- --- --- --- --- --- --- --- --- --- --- --- --- tag ttt ttt atc ata ttt --- --- --- --- --- --- --- --- tct aat acc ata ata aaa att atc --- --- --- --- --- --- --- --- --- --- --- --- --- --- --- --- --- --- --- --- --- --- --- --- --- -at tat tgc gtt tg- gta gtt --- --- --- --- -ct gcc cta --- --- --- --- --- --- --- --- tca tct ata tca ctg tca ctc tc- --- --- gct ctc act ata tct tct aaa att aca a-- --a caa c-- --- --- --- --- --- --- --- --- --- --- --- --- --- --- --- --- --- --- --- --- --- --- --- --- --- --- --- --- --- --- --- --- --- --- --- -tg gat att cga t-- --- --- --- --- --- --- --- --- --- --- --- --- --- --- --- --- --- --- --- --- --- --- --- --- --- --- aac agc att tgt gt- --- --- --- --- --- --- --- --- --- --- --- --- --- --- --- --- --

>ON585037.1_Monkeypox_virus_isolate_Monkeypox/PT0005/2022_complete_genome

ttt ttt cga tct atc ctc gtc c-- -t- ctc atc atc ctt ata --- --- --- --- --- --- -tt att atc att att atc ata gtc tat taa aca caa atc atc t-- --- --- --- --- --- --- --- --- --- --- --- --- --- --- --- --- --- --- --- --- --- --- --- --- --- --- acg ttt ata ac- --- --- --- --- --- --- --- --- aac att c-- --- --- --- --t cat tat taa tta gtt ctg tag -aa tat ctt taa taa ttt ggc tat a-- --- --c atc tgt t-- --- --- --- --- --- --- --- --- --- --- --- --- --- --- --- --- --- --- caa tac t-- --- --- --- --- --- --- --- atc tat tga tga ttt ctt tt- --- --- --- --- --- --- --- --- --- --- --- tta aga ct- --- --- --- --- --- --- --- --- --- --- --- --- --- --- --- --- --- --- --- --- --- --- --- --- --- --- --- --- --- --- --- --- --- --- --- --- --- --- --- -ta aac tag t-- --- --- --- --- --- --- --- --- --- --- --- --- --- --- --- --- --- --- --- --- --- --- --- --- --- --- --- --- --- --- --- --- --- --- --- --- --- --- --- --- --- --- --- --- --- --- --- --- --- --- --- --- --- --- --- --- --- --- --- --- --- --- --- --- --- --- --- --- --- --- --- --- --- --- --- --- --- --- --- --- --- --- --- --- --- --- --- --- --- --- --- --- --- --- --- --- --- --- --- --- --- --- --- --- --- --- --- --- --- tat ggt aat gac gat gaa a-- --t cga gta gta --- --- act tct aat aaa gac ttg ata --- --- tca tta tca tat gtt tga tcg --- --- --- --- --- --- --- tca tag tta ata gtg tg- --- --- --- --- --- --- --- --- --- --- --- --- --- --- --- --- --- --- --- --g cta aat ggt act gtt aat aag ttt at- --- --- --- --- --- --- --- --- --- --- --- --- --- --- --- --- --- aga caa tat cat agt att ttc ttt cca gaa t-- --- --- --- tag att att ttt tta aat act gat cct cac aat tcc gtg atg tag cag tag ttg gt- --- --- --- --- --- --- --- --- --- --- --- --- --- --- --- --- --- --- --- --- --- --- --- --- --- --- --- --g cat ggt cta tat cgt --- --- --- --- --- --- --- --- --- --- --- --- --- --- --- --- --- --- --- --- --- --- --- --- --- --- --- --- --- --- --- --- --- --- --- --- --- --- --- --- --- --- --- -ta aaa tgt atc ata tat aat agt ttt ctg acg tgg agt aca gaa ttt tcg a-- --- --- --- --- --- --- --- --- --- --- --- --- --- --- --- --- --- --- --- --- --- --- --- --- --- --- --- --- --- --- --- --- --- --- --- --- --- --- --- --- --- --- --- tta atg agt tca tgg taa gga agg gca aat gcc t-- -gt ata taa tat aca taa gtt aa- --- --- --- --- --- --- --- --- --- --- --- --- --- tag ttt ttt atc ata ttt --- --- --- --- --- --- --- --- tct aat acc ata ata aaa att atc --- --- --- --- --- --- --- --- --- --- --- --- --- --- --- --- --- --- --- --- --- --- --- --- --- -at tat tgc gtt tg- gta gtt --- --- --- --- -ct gcc cta --- --- --- --- --- --- --- --- tca tct ata tca ctg tca ctc tc- --- --- gct ctc act ata tct tct aaa att aca a-- --a caa c-- --- --- --- --- --- --- --- --- --- --- --- --- --- --- --- --- --- --- --- --- --- --- --- --- --- --- --- --- --- --- --- --- --- --- --- -tg gat att cga t-- --- --- --- --- --- --- --- --- --- --- --- --- --- --- --- --- --- --- --- --- --- --- --- --- --- --- aac agc att tgt gt- --- --- --- --- --- --- --- --- --- --- --- --- --- --- --- --- --

>ON585032.1_Monkeypox_virus_isolate_Monkeypox/PT0004/2022_complete_genome

ttt ttt cga tct atc ctc gtc c-- -t- ctc atc atc ctt ata --- --- --- --- --- --- -tt att atc att att atc ata gtc tat taa aca caa atc atc t-- --- --- --- --- --- --- --- --- --- --- --- --- --- --- --- --- --- --- --- --- --- --- --- --- --- --- acg ttt ata ac- --- --- --- --- --- --- --- --- aac att c-- --- --- --- --t cat tat taa tta gtt ctg tag -aa tat ctt taa taa ttt ggc tat a-- --- --c atc tgt t-- --- --- --- --- --- --- --- --- --- --- --- --- --- --- --- --- --- --- caa tac t-- --- --- --- --- --- --- --- atc tat tga tga ttt ctt tt- --- --- --- --- --- --- --- --- --- --- --- tta aga ct- --- --- --- --- --- --- --- --- --- --- --- --- --- --- --- --- --- --- --- --- --- --- --- --- --- --- --- --- --- --- --- --- --- --- --- --- --- --- --- -ta aac tag t-- --- --- --- --- --- --- --- --- --- --- --- --- --- --- --- --- --- --- --- --- --- --- --- --- --- --- --- --- --- --- --- --- --- --- --- --- --- --- --- --- --- --- --- --- --- --- --- --- --- --- --- --- --- --- --- --- --- --- --- --- --- --- --- --- --- --- --- --- --- --- --- --- --- --- --- --- --- --- --- --- --- --- --- --- --- --- --- --- --- --- --- --- --- --- --- --- --- --- --- --- --- --- --- --- --- --- --- --- --- tat ggt aat gac gat gaa a-- --t cga gta gta --- --- act tct aat aaa gac ttg ata --- --- tca tta tca tat gtt tga tcg --- --- --- --- --- --- --- tca tag tta ata gtg tg- --- --- --- --- --- --- --- --- --- --- --- --- --- --- --- --- --- --- --- --g cta aat ggt act gtt aat aag ttt at- --- --- --- --- --- --- --- --- --- --- --- --- --- --- --- --- --- aga caa tat cat agt att ttc ttt cca gaa t-- --- --- --- tag att att ttt tta aat act gat cct cac aat tcc gtg atg tag cag tag ttg gt- --- --- --- --- --- --- --- --- --- --- --- --- --- --- --- --- --- --- --- --- --- --- --- --- --- --- --- --g cat ggt cta tat cgt --- --- --- --- --- --- --- --- --- --- --- --- --- --- --- --- --- --- --- --- --- --- --- --- --- --- --- --- --- --- --- --- --- --- --- --- --- --- --- --- --- --- --- -ta aaa tgt atc ata tat aat agt ttt ctg acg tgg agt aca gaa ttt tcg a-- --- --- --- --- --- --- --- --- --- --- --- --- --- --- --- --- --- --- --- --- --- --- --- --- --- --- --- --- --- --- --- --- --- --- --- --- --- --- --- --- --- --- --- tta atg agt tca tgg taa gga agg gca aat gcc t-- -gt ata taa tat aca taa gtt aa- --- --- --- --- --- --- --- --- --- --- --- --- --- tag ttt ttt atc ata ttt --- --- --- --- --- --- --- --- tct aat acc ata ata aaa att atc --- --- --- --- --- --- --- --- --- --- --- --- --- --- --- --- --- --- --- --- --- --- --- --- --- -at tat tgc gtt tg- gta gtt --- --- --- --- -ct gcc cta --- --- --- --- --- --- --- --- tca tct ata tca ctg tca ctc tc- --- --- gct ctc act ata tct tct aaa att aca a-- --a caa c-- --- --- --- --- --- --- --- --- --- --- --- --- --- --- --- --- --- --- --- --- --- --- --- --- --- --- --- --- --- --- --- --- --- --- --- -tg gat att cga t-- --- --- --- --- --- --- --- --- --- --- --- --- --- --- --- --- --- --- --- --- --- --- --- --- --- --- aac agc att tgt gt- --- --- --- --- --- --- --- --- --- --- --- --- --- --- --- --- --

>AY741551.1_Monkeypox_virus_isolate_Sierra_Leone_complete_genome

ttt ttt cga tct atc aat ttc agt ata ttc ttc gcc gtt ata aaa gta atg ttg ttt aat tgt agg acg gtt gtt agt ata atc aca tga ata ata ata ttc taa ttc ctc gta ttg act act tac aga tac tcg aaa tag tct gaa aaa ttc ttc aaa gat att ttt ata aag atc tag gaa aag ttt att acc gac cat gaa cga gat aga tgg ata aat atc ctt tcc atc aaa ggt cat aat tgg ata att gtc cag caa tat atc tgc tgt att agt tat atc act tcc att tat ttt cag att gaa gta atg tac tag ttt gtg aca att aac aag ata caa aag aga tgc cga tac taa tac gta aat agc tat acg cga atc cat tgt tac ctt ttt tta ttt cat agg tct att aat aaa tat atg tat tac tta aga cta gaa aaa tca aaa gta agt ttt tga tat ttg att ctt act tat tgt ggg att gta gtt tac tta gta att cat ctc tga atc ctg ata aat cat gca tat caa tga tgc aac tac gca gca aac tag tag gaa tat aga tat ctg gat atg tac gta aat agt cga tta tat ctt tta caa tac tat tag tcc cta ttg cgt tat cta tat atc cat taa taa tat tac aca gtg gat act tat gag aaa tat acc tct tac aga ttt tta acg ata tat aat cta gaa aga tat gtg tgt agt act gta tta cct aaa tta tca gtc tca ttc aaa tat tgc atg act att atc gag aat tgc tat atc cct cta tac tcg atg cat tta tta cag taa ttc aat cca gct aac ata aga gcc aat ctc aat gtt ggt tta att ata tca tct tca tgc aat aat aac gat gga aac ttt cca gta gcg taa cac tta tct aag aag gat ata ata act atg tct aca tta tgt ctc tta tcg aga ata ttc tta acg aga tat cca tag cta ttc tgg tgc taa tta ttc cta tat tat att cca cga aaa atg atg aag gca atc att cat cat aag atg ata aaa agt gta gtg agt aag agt att agt gag aga gca tga agg aga ttt agt att tag cag tga gga tat gat cca aga ggg tga gat agt cgt tct cgt tca gaa tct ttc gca gca taa gta gta tgt cga tat act tat cat tga aga ctc ttc cag tga caa tag ctg att gag tac aaa gtc caa tta ttg cac aaa gtt ctt tgg cgg ttt tca tgg agt cat ttc tga tga aac att taa tga tct cca cgc aat tgt cga tat tgt ccc acg gaa gtg aat ccg aga act cct tca act cgc tac caa ata gct cca ttg cat caa ttc tga aag aga tga gaa gcc tgt aga gag gcc ctg cgc ttt ctc tat ggg tcc atc tat gag aaa ccc aca gga tgt att cag tca gac aa- tgt ctg aca tca gtc acg gta ttc agg gag tcc tta gta gcg tgg caa tga cag gga ctg aac tgg gca caa gga gag gcc att gtg aag gta gac gaa ggt aac ctg atg gta gac ctg tag ccg tct gtg ctt aat aga ggg ctt taa ttt cca ttt tta atg gtg tcg tga atg agg aat gag agt gtc tct cgt cct tgg ttt aca tgg atc aga gtg aga aaa aat atc ttg tat att att aac taa caa cct tgg ttt cta tcc atg ttt -aa aaa atg acc tat atg ttc ttt att aat tct att tta aac ttt atc ctc aag act cct gac aaa att aaa atc cag aaa gca gca aac aat cct gtt aca agt tta ctg aaa tct ctc ttt gat tat aga gta tat gta gtc aga gca aga aac act gca gta gtc aac atg aaa gct tgc ata acg ata cgt gca tca tag aaa gta aca aca gag gcc agc gtt aga gat tct aac agt gta aat cca caa --- agt atg tac aga ttc agg gga tgt tca tgt ctg tgt aaa gtc aat gcg aaa atc aag cct ata gat ccg aac att gat gcc aat att aga aca gga ctc cct tgt ata aat gtc cga tgc att caa agt ata aaa ata ctg cag ctg ttg ccg ttg tta aag gaa att gta gaa agg ata ccg tag act ttt ctt aga aat gcc att cgt atg tac acg ctg gca gac gcc acc gag ctg tca tag ttg aag tcg tcc tcg at

>MT903346.1_Monkeypox_virus_isolate_MPXV-USA2003_099_Gambian_Rat

ttt ttt cga tct atc aat ttc agt ata ttc ttc gcc gtt ata aaa gta atg ttg ttt aat tgt agg acg gtt gtt agt ata atc aca tga ata ata ata ttc taa ttc ctc gta ttg act act tac aga tac tcg aaa tag tct gaa aaa ttc ttc aaa gat att ttt ata aag atc tag gaa aag ttt att acc gac cat gaa cga gat aga tgg ata aat atc ctt tcc atc aaa ggt cat aat tgg ata att gtc cag caa tat atc tgc tgt att agt tat atc act tcc att tat ttt cag att gaa gta atg tac tag ttt gtg aca att aac aag ata caa aag aga tgc cga tac taa tac gta aat agc tat acg cga atc cat tgt tac ctt ttt tta ttt cat agg tct att aat aaa tat atg tat tac tta aga cta gaa aaa tca aaa gtg agt ttt tga tat ttg att ctt act tat tgt ggg att gta gtt tac tta gta att cat ctc tga atc ctg ata aat cat gca tat caa tga tgc aac tac gca gca aac tag tag gaa tat aga tat ctg gat atg tac gta aat agt cga tta tat ctt tta caa tac tat tag tcc cta ttg cgt tat cta tat atc cat taa taa tat tac aca gtg gat act tat gag aaa tat acc tct tac aga ttt tta acg ata tat aat cta gaa aga tat gtg tgt agt act gta tta cct aaa tta tca gtc tca ttc aaa tat tgc atg act att atc gag aat tgc tat atc cct cta tac tcg atg cat tta tta cag taa ttc aat cca gct aac ata aga gcc aat ctc aat gtt ggt tta att ata tca tct tca tgt aat aat aac gat gga aac ttt cca gta gcg taa cac tta tct aag aag gat ata ata act atg tct aca tta tgt ctc tta tcg aga ata ttc tta acg aga tat cca tag cta ttc tgg tgc taa tta ttc cta tat tat att cca cga aaa atg atg aag gca atc att cat cat aag atg ata aaa agt gta gtg agt aag agt att agt gag aga gca tga agg aga ttt agt att tag cag tga gga tat gat cca aga gtg tga gat agt cgt tct cgt tca gaa tct ttc gca gca taa gta gta tgt cga tat act tat cat tga aga ctc ttc cag tga caa tag ctg att gag tac aaa gtc caa tta ttg cac aaa gtt ctt tgg cgg ttt tca tgg agt cat ttc tga tga aac att taa tga tct cca cgc aat tgt cga tat tgt ccc acg gaa gtg aat ccg aga act cct tca act cgc tac caa ata gct cca ttg cat caa ttc tga aag aga tga gaa gcc tgt aga gag gcc ctg cgc ttt ctc tat ggg tcc atc tat gag aaa ccc aca gga tgt att cag tca gac aa- tgt ctg aca tca gtc acg gta ttc agg gag tcc tta gta gcg tgg caa tga cag gga ctg aac tgg gca caa gga gag gcc att gtg aag gta gac gaa ggt aac ctg atg gta gac ctg tag ccg tct gtg ctt aat aga ggg ctt taa ttt cca ttt tta atg gtg tcg tga atg agg aat gag agt gtc tct cgt cct tgg ttt aca tgg atc aga gtg aga aaa aat atc ttg tat att att aac taa caa cct tgg ttt cta tcc atg ttt aaa aaa atg acc tat atg ttc ttt att aat tct att tta aac ttt atc ctc aag act cct gac aaa att aaa atc cag aaa gca gca aac aat cct gtt aca agt tta ctg aaa tct ctc ttt gat tgt aga gta tat gta gtc aga gca aga aac act gca gta gtc aac atg aaa gct tgc ata acg ata cgt gca tca tag aaa gta aca aca gag gcc agc gtt aga gat tct aac agt gta aat cca caa --- agt atg tac aga ttc agg gga tgt tca tgt ctg tgt aaa gtc aat gcg aaa atc aag cct ata gat ccg aac att gat gcc aat att aga aca gga ctc cct tgt ata aat gtc cga tgc att caa agt ata aaa ata ctg cag ctg ttg ccg ttg tta aag gaa att gta gaa aga ata ccg tag act ttt ctt aga aat gcc att cgt atg tac acg ctg gca gac gcc acc gag ctg tca tag ttg aag tcg tcc tcg at

>DQ011157.1_Monkeypox_virus_strain_USA_2003_039_complete_genome

ttt ttt cga tct atc aat ttc agt ata ttc ttc gcc gtt ata aaa gta atg ttg ttt aat tgt agg acg gtt gtt agt ata atc aca tga ata ata ata ttc taa ttc ctc gta ttg act act tac aga tac tcg aaa tag tct gaa aaa ttc ttc aaa gat att ttt ata aag atc tag gaa aag ttt att acc gac cat gaa cga gat aga tgg ata aat atc ctt tcc atc aaa ggt cat aat tgg ata att gtc cag caa tat atc tgc tgt att agt tat atc act tcc att tat ttt cag att gaa gta atg tac tag ttt gtg aca att aac aag ata caa aag aga tgc cga tac taa tac gta aat agc tat acg cga atc cat tgt tac ctt ttt tta ttt cat agg tct att aat aaa tat atg tat tac tta aga cta gaa aaa tca aaa gtg agt ttt tga tat ttg att ctt act tat tgt ggg att gta gtt tac tta gta att cat ctc tga atc ctg ata aat cat gca tat caa tga tgc aac tac gca gca aac tag tag gaa tat aga tat ctg gat atg tac gta aat agt cga tta tat ctt tta caa tac tat tag tcc cta ttg cgt tat cta tat atc cat taa taa tat tac aca gtg gat act tat gag aaa tat acc tct tac aga ttt tta acg ata tat aat cta gaa aga tat gtg tgt agt act gta tta cct aaa tta tca gtc tca ttc aaa tat tgc atg act att atc gag aat tgc tat atc cct cta tac tcg atg cat tta tta cag taa ttc aat cca gct aac ata aga gcc aat ctc aat gtt ggt tta att ata tca tct tca tgt aat aat aac gat gga aac ttt cca gta gcg taa cac tta tct aag aag gat ata ata act atg tct aca tta tgt ctc tta tcg aga ata ttc tta acg aga tat cca tag cta ttc tgg tgc taa tta ttc cta tat tat att cca cga aaa atg atg aag gca atc att cat cat aag atg ata aaa agt gta gtg agt aag agt att agt gag aga gca tga agg aga ttt agt att tag cag tga gga tat gat cca aga gtg tga gat agt cgt tct cgt tca gaa tct ttc gca gca taa gta gta tgt cga tat act tat cat tga aga ctc ttc cag tga caa tag ctg att gag tac aaa gtc caa tta ttg cac aaa gtt ctt tgg cgg ttt tca tgg agt cat ttc tga tga aac att taa tga tct cca cgc aat tgt cga tat tgt ccc acg gaa gtg aat ccg aga act cct tca act cgc tac caa ata gct cca ttg cat caa ttc tga aag aga tga gaa gcc tgt aga gag gcc ctg cgc ttt ctc tat ggg tcc atc tat gag aaa ccc aca gga tgt att cag tca gac aa- tgt ctg aca tca gtc acg gta ttc agg gag tcc tta gta gcg tgg caa tga cag gga ctg aac tgg gca caa gga gag gcc att gtg aag gta gac gaa ggt aac ctg atg gta gac ctg tag ccg tct gtg ctt aat aga ggg ctt taa ttt cca ttt tta atg gtg tcg tga atg agg aat gag agt gtc tct cgt cct tgg ttt aca tgg atc aga gtg aga aaa aat atc ttg tat att att aac taa caa cct tgg ttt cta tcc atg ttt aaa aaa atg acc tat atg ttc ttt att aat tct att tta aac ttt atc ctc aag act cct gac aaa att aaa atc cag aaa gca gca aac aat cct gtt aca agt tta ctg aaa tct ctc ttt gat tgt aga gta tat gta gtc aga gca aga aac act gca gta gtc aac atg aaa gct tgc ata acg ata cgt gca tca tag aaa gta aca aca gag gcc agc gtt aga gat tct aac agt gta aat cca caa --- agt atg tac aga ttc agg gga tgt tca tgt ctg tgt aaa gtc aat gcg aaa atc aag cct ata gat ccg aac att gat gcc aat att aga aca gga ctc cct tgt ata aat gtc cga tgc att caa agt ata aaa ata ctg cag ctg ttg ccg ttg tta aag gaa att gta gaa aga ata ccg tag act ttt ctt aga aat gcc att cgt atg tac acg ctg gca gac gcc acc gag ctg tca tag ttg aag tcg tcc tcg at

>DQ011153.1_Monkeypox_virus_strain_USA_2003_044_complete_genome

ttt ttt cga tct atc aat ttc agt ata ttc ttc gcc gtt ata aaa gta atg ttg ttt aat tgt agg acg gtt gtt agt ata atc aca tga ata ata ata ttc taa ttc ctc gta ttg act act tac aga tac tcg aaa tag tct gaa aaa ttc ttc aaa gat att ttt ata aag atc tag gaa aag ttt att acc gac cat gaa cga gat aga tgg ata aat atc ctt tcc atc aaa ggt cat aat tgg ata att gtc cag caa tat atc tgc tgt att agt tat atc act tcc att tat ttt cag att gaa gta atg tac tag ttt gtg aca att aac aag ata caa aag aga tgc cga tac taa tac gta aat agc tat acg cga atc cat tgt tac ctt ttt tta ttt cat agg tct att aat aaa tat atg tat tac tta aga cta gaa aaa tca aaa gtg agt ttt tga tat ttg att ctt act tat tgt ggg att gta gtt tac tta gta att cat ctc tga atc ctg ata aat cat gca tat caa tga tgc aac tac gca gca aac tag tag gaa tat aga tat ctg gat atg tac gta aat agt cga tta tat ctt tta caa tac tat tag tcc cta ttg cgt tat cta tat atc cat taa taa tat tac aca gtg gat act tat gag aaa tat acc tct tac aga ttt tta acg ata tat aat cta gaa aga tat gtg tgt agt act gta tta cct aaa tta tca gtc tca ttc aaa tat tgc atg act att atc gag aat tgc tat atc cct cta tac tcg atg cat tta tta cag taa ttc aat cca gct aac ata aga gcc aat ctc aat gtt ggt tta att ata tca tct tca tgt aat aat aac gat gga aac ttt cca gta gcg taa cac tta tct aag aag gat ata ata act atg tct aca tta tgt ctc tta tcg aga ata ttc tta acg aga tat cca tag cta ttc tgg tgc taa tta ttc cta tat tat att cca cga aaa atg atg aag gca atc att cat cat aag atg ata aaa agt gta gtg agt aag agt att agt gag aga gca tga agg aga ttt agt att tag cag tga gga tat gat cca aga gtg tga gat agt cgt tct cgt tca gaa tct ttc gca gca taa gta gta tgt cga tat act tat cat tga aga ctc ttc cag tga caa tag ctg att gag tac aaa gtc caa tta ttg cac aaa gtt ctt tgg cgg ttt tca tgg agt cat ttc tga tga aac att taa tga tct cca cgc aat tgt cga tat tgt ccc acg gaa gtg aat ccg aga act cct tca act cgc tac caa ata gct cca ttg cat caa ttc tga aag aga tga gaa gcc tgt aga gag gcc ctg cgc ttt ctc tat ggg tcc atc tat gag aaa ccc aca gga tgt att cag tca gac aa- tgt ctg aca tca gtc acg gta ttc agg gag tcc tta gta gcg tgg caa tga cag gga ctg aac tgg gca caa gga gag gcc att gtg aag gta gac gaa ggt aac ctg atg gta gac ctg tag ccg tct gtg ctt aat aga ggg ctt taa ttt cca ttt tta atg gtg tcg tga atg agg aat gag agt gtc tct cgt cct tgg ttt aca tgg atc aga gtg aga aaa aat atc ttg tat att att aac taa caa cct tgg ttt cta tcc atg ttt aaa aaa atg acc tat atg ttc ttt att aat tct att tta aac ttt atc ctc aag act cct gac aaa att aaa atc cag aaa gca gca aac aat cct gtt aca agt tta ctg aaa tct ctc ttt gat tgt aga gta tat gta gtc aga gca aga aac act gca gta gtc aac atg aaa gct tgc ata acg ata cgt gca tca tag aaa gta aca aca gag gcc agc gtt aga gat tct aac agt gta aat cca caa --- agt atg tac aga ttc agg gga tgt tca tgt ctg tgt aaa gtc aat gcg aaa atc aag cct ata gat ccg aac att gat gcc aat att aga aca gga ctc cct tgt ata aat gtc cga tgc att caa agt ata aaa ata ctg cag ctg ttg ccg ttg tta aag gaa att gta gaa aga ata ccg tag act ttt ctt aga aat gcc att cgt atg tac acg ctg gca gac gcc acc gag ctg tca tag ttg aag tcg tcc tcg at

>JX878428.1_Monkeypox_virus_isolate_DRC_07-0514_complete_genome

ttt ttt cga tct atc ctc gtc --- --- ctc atc atc ctt ata --- --- --- --- --- --- -tt att atc att att atc ata gtc tat taa aca caa atc atc t-- --- --- --- --- --- --- --- --- --- --- --- --- --- --- --- --- --- --- --- --- --- --- --- --- --- --- acg ttt ata ac- --- --- --- --- --- --- --- --- aac att c-- --- --- --- --t cat tat taa tta gtt ctg tag taa tat ctt taa taa ttt ggc tat a-- --- --c atc tgt t-- --- --- --- --- --- --- --- --- --- --- --- --- --- --- --- --- --- --- caa tac t-- --- --- --- --- --- --- --- atc tat tga tga ttt ctt tt- --- --- --- --- --- --- --- --- --- --- --- tta aga ct- --- --- --- --- --- --- --- --- --- --- --- --- --- --- --- --- --- --- --- --- --- --- --- --- --- --- --- --- --- --- --- --- --- --- --- --- --- --- --- -ta aac tag t-- --- --- --- --- --- --- --- --- --- --- --- --- --- --- --- --- --- --- --- --- --- --- --- --- --- --- --- --- --- --- --- --- --- --- --- --- --- --- --- --- --- --- --- --- --- --- --- --- --- --- --- --- --- --- --- --- --- --- --- --- --- --- --- --- --- --- --- --- --- --- --- --- --- --- --- --- --- --- --- --- --- --- --- --- --- --- --- --- --- --- --- --- --- --- --- --- --- --- --- --- --- --- --- --- --- --- --- --- --- tat ggt aat gac gat gaa a-- --t cga gta gta --- --- act tct aat aaa gac ttg ata --- --- tca tta tca tat gtt tga tcg --- --- --- --- --- --- --- tca tag tta ata gtg tg- --- --- --- --- --- --- --- --- --- --- --- --- --- --- --- --- --- --- --- --g cta aat ggt act gtt aat aag ttt at- --- --- --- --- --- --- --- --- --- --- --- --- --- --- --- --- --- aga caa tat cat agt att ttc ttt cca gaa t-- --- --- --- tag att att ttt tta aat act gat cct cac aat tcc gtg atg tag cag tag ttg gt- --- --- --- --- --- --- --- --- --- --- --- --- --- --- --- --- --- --- --- --- --- --- --- --- --- --- --- --g cat ggt cta tat cgt --- --- --- --- --- --- --- --- --- --- --- --- --- --- --- --- --- --- --- --- --- --- --- --- --- --- --- --- --- --- --- --- --- --- --- --- --- --- --- --- --- --- --- -ta aaa tgt atc ata tat aat agt ttt ctg acg tgg agt aca gaa ttt tcg a-- --- --- --- --- --- --- --- --- --- --- --- --- --- --- --- --- --- --- --- --- --- --- --- --- --- --- --- --- --- --- --- --- --- --- --- --- --- --- --- --- --- --- --- tta atg agt tca tgg taa gga agg gca aat gtc t-- -gt ata taa tat aca taa gtt aa- --- --- --- --- --- --- --- --- --- --- --- --- --- tag ttt ttt atc ata ttt --- --- --- --- --- --- --- --- tct aat acc ata ata aaa att atc att atg --- --- --t ata atc a-- --- --- --- --- --- --- --- --- --- --- --- tca ctg --- --t cgc tat cat tat tgc gtt tgt gta gtt --- --- --- --- -ct gcc cta --- --- --- --- --- --- --- --- tca tct aca tca ctg tca --- --- --- --- -ct ctc act ata tct tct aaa att aca a-- --a caa c-- --- --- --- --- --- --- --- --- --- --- --- --- --- --- --- --- --- --- --- --- --- --- --- --- --- --- --- --- --- --- --- --- --- --- --- -tg gat att cga t-- --- --- --- --- --- --- --- --- --- --- --- --- --- --- --- --- --- --- --- --- --- --- --- --- --- --- aac agc att tgt gt- --- --- --- --- --- --- --- --- --- --- --- --- --- --- --- --- --

>JX878427.1_Monkeypox_virus_isolate_DRC_07-0480_complete_genome

ttt ttt cga tct atc ctc gtc --- --- ctc atc atc ctt ata --- --- --- --- --- --- -tt att atc att att atc ata gtc tat taa aca caa atc atc t-- --- --- --- --- --- --- --- --- --- --- --- --- --- --- --- --- --- --- --- --- --- --- --- --- --- --- acg ttt ata ac- --- --- --- --- --- --- --- --- aac att c-- --- --- --- --t cat tat taa tta gtt ctg tag taa tat ctt taa taa ttt ggc tat a-- --- --c atc tgt t-- --- --- --- --- --- --- --- --- --- --- --- --- --- --- --- --- --- --- caa tac t-- --- --- --- --- --- --- --- atc tat tga tga ttt ctt tt- --- --- --- --- --- --- --- --- --- --- --- tta aga ct- --- --- --- --- --- --- --- --- --- --- --- --- --- --- --- --- --- --- --- --- --- --- --- --- --- --- --- --- --- --- --- --- --- --- --- --- --- --- --- -ta aac tag t-- --- --- --- --- --- --- --- --- --- --- --- --- --- --- --- --- --- --- --- --- --- --- --- --- --- --- --- --- --- --- --- --- --- --- --- --- --- --- --- --- --- --- --- --- --- --- --- --- --- --- --- --- --- --- --- --- --- --- --- --- --- --- --- --- --- --- --- --- --- --- --- --- --- --- --- --- --- --- --- --- --- --- --- --- --- --- --- --- --- --- --- --- --- --- --- --- --- --- --- --- --- --- --- --- --- --- --- --- --- tat ggt aat gac gat gaa a-- --t cga gta gta --- --- act tct aat aaa gac ttg ata --- --- tca tta tca tat gtt tga tcg --- --- --- --- --- --- --- tca tag tta ata gtg tg- --- --- --- --- --- --- --- --- --- --- --- --- --- --- --- --- --- --- --- --g cta aat ggt act gtt aat aag ttt at- --- --- --- --- --- --- --- --- --- --- --- --- --- --- --- --- --- aga caa tat cat agt att ttc ttt cca gaa t-- --- --- --- tag att att ttt tta aat act gat cct cac aat tcc gtg atg tag cag tag ttg gt- --- --- --- --- --- --- --- --- --- --- --- --- --- --- --- --- --- --- --- --- --- --- --- --- --- --- --- --g cat ggt cta tat cgt --- --- --- --- --- --- --- --- --- --- --- --- --- --- --- --- --- --- --- --- --- --- --- --- --- --- --- --- --- --- --- --- --- --- --- --- --- --- --- --- --- --- --- -ta aaa tgt atc ata tat aat agt ttt ctg acg tgg agt aca gaa ttt tcg a-- --- --- --- --- --- --- --- --- --- --- --- --- --- --- --- --- --- --- --- --- --- --- --- --- --- --- --- --- --- --- --- --- --- --- --- --- --- --- --- --- --- --- --- tta atg agt tca tgg taa gga agg gca aat gtc t-- -gt ata taa tat aca taa gtt aa- --- --- --- --- --- --- --- --- --- --- --- --- --- tag ttt ttt atc ata ttt --- --- --- --- --- --- --- --- tct aat acc ata ata aaa att atc att atg --- --- --t ata atc a-- --- --- --- --- --- --- --- --- --- --- --- tca ctg --- --t cgc tat cat tat tgc gtt tgt gta gtt --- --- --- --- -ct gcc cta --- --- --- --- --- --- --- --- tca tct aca tca ctg tca --- --- --- --- -ct ctc act ata tct tct aaa att aca a-- --a caa c-- --- --- --- --- --- --- --- --- --- --- --- --- --- --- --- --- --- --- --- --- --- --- --- --- --- --- --- --- --- --- --- --- --- --- --- -tg gat att cga t-- --- --- --- --- --- --- --- --- --- --- --- --- --- --- --- --- --- --- --- --- --- --- --- --- --- --- aac agc att tgt gt- --- --- --- --- --- --- --- --- --- --- --- --- --- --- --- --- --

>JX878421.1_Monkeypox_virus_isolate_DRC_07-0286_complete_genome

ttt ttt cga tct atc ctc gtc --- --- ctc atc atc ctt ata --- --- --- --- --- --- -tt att atc att att atc ata gtc tat taa aca caa atc atc t-- --- --- --- --- --- --- --- --- --- --- --- --- --- --- --- --- --- --- --- --- --- --- --- --- --- --- acg ttt ata ac- --- --- --- --- --- --- --- --- aac att c-- --- --- --- --t cat tat taa tta gtt ctg tag taa tat ctt taa taa ttt ggc tat a-- --- --c atc tgt t-- --- --- --- --- --- --- --- --- --- --- --- --- --- --- --- --- --- --- caa tac t-- --- --- --- --- --- --- --- atc tat tga tga ttt ctt tt- --- --- --- --- --- --- --- --- --- --- --- tta aga ct- --- --- --- --- --- --- --- --- --- --- --- --- --- --- --- --- --- --- --- --- --- --- --- --- --- --- --- --- --- --- --- --- --- --- --- --- --- --- --- -ta aac tag t-- --- --- --- --- --- --- --- --- --- --- --- --- --- --- --- --- --- --- --- --- --- --- --- --- --- --- --- --- --- --- --- --- --- --- --- --- --- --- --- --- --- --- --- --- --- --- --- --- --- --- --- --- --- --- --- --- --- --- --- --- --- --- --- --- --- --- --- --- --- --- --- --- --- --- --- --- --- --- --- --- --- --- --- --- --- --- --- --- --- --- --- --- --- --- --- --- --- --- --- --- --- --- --- --- --- --- --- --- --- tat ggt aat gac gat gaa a-- --t cga gta gta --- --- act tct aat aaa gac ttg ata --- --- tca tta tca tat gtt tga tcg --- --- --- --- --- --- --- tca tag tta ata gtg tg- --- --- --- --- --- --- --- --- --- --- --- --- --- --- --- --- --- --- --- --g cta aat ggt act gtt aat aag ttt at- --- --- --- --- --- --- --- --- --- --- --- --- --- --- --- --- --- aga caa tat cat agt att ttc ttt cca gaa t-- --- --- --- tag att att ttt tta aat act gat cct cac aat tcc gtg atg tag cag tag ttg gt- --- --- --- --- --- --- --- --- --- --- --- --- --- --- --- --- --- --- --- --- --- --- --- --- --- --- --- --g cat ggt cta tat cgt --- --- --- --- --- --- --- --- --- --- --- --- --- --- --- --- --- --- --- --- --- --- --- --- --- --- --- --- --- --- --- --- --- --- --- --- --- --- --- --- --- --- --- -ta aaa tgt atc ata tat aat agt ttt ctg acg tgg agt aca gaa ttt tcg a-- --- --- --- --- --- --- --- --- --- --- --- --- --- --- --- --- --- --- --- --- --- --- --- --- --- --- --- --- --- --- --- --- --- --- --- --- --- --- --- --- --- --- --- tta atg agt tca tgg taa gga agg gca aat gtc t-- -gt ata taa tat aca taa gtt aa- --- --- --- --- --- --- --- --- --- --- --- --- --- tag ttt ttt atc ata ttt --- --- --- --- --- --- --- --- tct aat acc ata ata aaa att atc att atg --- --- --t ata atc a-- --- --- --- --- --- --- --- --- --- --- --- tca ctg --- --t cgc tat cat tat tgc gtt tgt gta gtt --- --- --- --- -ct gcc cta --- --- --- --- --- --- --- --- tca tct aca tca ctg tca --- --- --- --- -ct ctc act ata tct tct aaa att aca a-- --a caa c-- --- --- --- --- --- --- --- --- --- --- --- --- --- --- --- --- --- --- --- --- --- --- --- --- --- --- --- --- --- --- --- --- --- --- --- -tg gat att cga t-- --- --- --- --- --- --- --- --- --- --- --- --- --- --- --- --- --- --- --- --- --- --- --- --- --- --- aac agc att tgt gt- --- --- --- --- --- --- --- --- --- --- --- --- --- --- --- --- --

>JX878415.1_Monkeypox_virus_isolate_DRC_07-0092_complete_genome

ttt ttt cga tct atc ctc gtc --- --- ctc atc atc ctt ata --- --- --- --- --- --- -tt att atc att att atc ata gtc tat taa aca caa atc atc t-- --- --- --- --- --- --- --- --- --- --- --- --- --- --- --- --- --- --- --- --- --- --- --- --- --- --- acg ttt ata ac- --- --- --- --- --- --- --- --- aac att c-- --- --- --- --t cat tat taa tta gtt ctg tag taa tat ctt taa taa ttt ggc tat a-- --- --c atc tgt t-- --- --- --- --- --- --- --- --- --- --- --- --- --- --- --- --- --- --- caa tac t-- --- --- --- --- --- --- --- atc tat tga tga ttt ctt tt- --- --- --- --- --- --- --- --- --- --- --- tta aga ct- --- --- --- --- --- --- --- --- --- --- --- --- --- --- --- --- --- --- --- --- --- --- --- --- --- --- --- --- --- --- --- --- --- --- --- --- --- --- --- -ta aac tag t-- --- --- --- --- --- --- --- --- --- --- --- --- --- --- --- --- --- --- --- --- --- --- --- --- --- --- --- --- --- --- --- --- --- --- --- --- --- --- --- --- --- --- --- --- --- --- --- --- --- --- --- --- --- --- --- --- --- --- --- --- --- --- --- --- --- --- --- --- --- --- --- --- --- --- --- --- --- --- --- --- --- --- --- --- --- --- --- --- --- --- --- --- --- --- --- --- --- --- --- --- --- --- --- --- --- --- --- --- --- tat ggt aat gac gat gaa a-- --t cga gta gta --- --- act tct aat aaa gac ttg ata --- --- tca tta tca tat gtt tga tcg --- --- --- --- --- --- --- tca tag tta ata gtg tg- --- --- --- --- --- --- --- --- --- --- --- --- --- --- --- --- --- --- --- --g cta aat ggt act gtt aat aag ttt at- --- --- --- --- --- --- --- --- --- --- --- --- --- --- --- --- --- aga caa tat cat agt att ttc ttt cca gaa t-- --- --- --- tag att att ttt tta aat act gat cct cac aat tcc gtg atg tag cag tag ttg gt- --- --- --- --- --- --- --- --- --- --- --- --- --- --- --- --- --- --- --- --- --- --- --- --- --- --- --- --g cat ggt cta tat cgt --- --- --- --- --- --- --- --- --- --- --- --- --- --- --- --- --- --- --- --- --- --- --- --- --- --- --- --- --- --- --- --- --- --- --- --- --- --- --- --- --- --- --- -ta aaa tgt atc ata tat aat agt ttt ctg acg tgg agt aca gaa ttt tcg a-- --- --- --- --- --- --- --- --- --- --- --- --- --- --- --- --- --- --- --- --- --- --- --- --- --- --- --- --- --- --- --- --- --- --- --- --- --- --- --- --- --- --- --- tta atg agt tca tgg taa gga agg gca aat gtc t-- -gt ata taa tat aca taa gtt aa- --- --- --- --- --- --- --- --- --- --- --- --- --- tag ttt ttt atc ata ttt --- --- --- --- --- --- --- --- tct aat acc ata ata aaa att atc att atg --- --- --t ata atc a-- --- --- --- --- --- --- --- --- --- --- --- tca ctg --- --t cgc tat cat tat tgc gtt tgt gta gtt --- --- --- --- -ct gcc cta --- --- --- --- --- --- --- --- tca tct aca tca ctg tca --- --- --- --- -ct ctc act ata tct tct aaa att aca a-- --a caa c-- --- --- --- --- --- --- --- --- --- --- --- --- --- --- --- --- --- --- --- --- --- --- --- --- --- --- --- --- --- --- --- --- --- --- --- -tg gat att cga t-- --- --- --- --- --- --- --- --- --- --- --- --- --- --- --- --- --- --- --- --- --- --- --- --- --- --- aac agc att tgt gt- --- --- --- --- --- --- --- --- --- --- --- --- --- --- --- --- --

>JX878414.1_Monkeypox_virus_isolate_DRC_07-0046_complete_genome

ttt ttt cga tct atc ctc gtc --- --- ctc atc atc ctt ata --- --- --- --- --- --- -tt att atc att att atc ata gtc tat taa aca caa atc atc t-- --- --- --- --- --- --- --- --- --- --- --- --- --- --- --- --- --- --- --- --- --- --- --- --- --- --- acg ttt ata ac- --- --- --- --- --- --- --- --- aac att c-- --- --- --- --t cat tat taa tta gtt ctg tag taa tat ctt taa taa ttt ggc tat a-- --- --c atc tgt t-- --- --- --- --- --- --- --- --- --- --- --- --- --- --- --- --- --- --- caa tac t-- --- --- --- --- --- --- --- atc tat tga tga ttt ctt tt- --- --- --- --- --- --- --- --- --- --- --- tta aga ct- --- --- --- --- --- --- --- --- --- --- --- --- --- --- --- --- --- --- --- --- --- --- --- --- --- --- --- --- --- --- --- --- --- --- --- --- --- --- --- -ta aac tag t-- --- --- --- --- --- --- --- --- --- --- --- --- --- --- --- --- --- --- --- --- --- --- --- --- --- --- --- --- --- --- --- --- --- --- --- --- --- --- --- --- --- --- --- --- --- --- --- --- --- --- --- --- --- --- --- --- --- --- --- --- --- --- --- --- --- --- --- --- --- --- --- --- --- --- --- --- --- --- --- --- --- --- --- --- --- --- --- --- --- --- --- --- --- --- --- --- --- --- --- --- --- --- --- --- --- --- --- --- --- tat ggt aat gac gat gaa a-- --t cga gta gta --- --- act tct aat aaa gac ttg ata --- --- tca tta tca tat gtt tga tcg --- --- --- --- --- --- --- tca tag tta ata gtg tg- --- --- --- --- --- --- --- --- --- --- --- --- --- --- --- --- --- --- --- --g cta aat ggt act gtt aat aag ttt at- --- --- --- --- --- --- --- --- --- --- --- --- --- --- --- --- --- aga caa tat cat agt att ttc ttt cca gaa t-- --- --- --- tag att att ttt tta aat act gat cct cac aat tcc gtg atg tag cag tag ttg gt- --- --- --- --- --- --- --- --- --- --- --- --- --- --- --- --- --- --- --- --- --- --- --- --- --- --- --- --g cat ggt cta tat cgt --- --- --- --- --- --- --- --- --- --- --- --- --- --- --- --- --- --- --- --- --- --- --- --- --- --- --- --- --- --- --- --- --- --- --- --- --- --- --- --- --- --- --- -ta aaa tgt atc ata tat aat agt ttt ctg acg tgg agt aca gaa ttt tcg a-- --- --- --- --- --- --- --- --- --- --- --- --- --- --- --- --- --- --- --- --- --- --- --- --- --- --- --- --- --- --- --- --- --- --- --- --- --- --- --- --- --- --- --- tta atg agt tca tgg taa gga agg gca aat gtc t-- -gt ata taa tat aca taa gtt aa- --- --- --- --- --- --- --- --- --- --- --- --- --- tag ttt ttt atc ata ttt --- --- --- --- --- --- --- --- tct aat acc ata ata aaa att atc att atg --- --- --t ata atc a-- --- --- --- --- --- --- --- --- --- --- --- tca ctg --- --t cgc tat cat tat tgc gtt tgt gta gtt --- --- --- --- -ct gcc cta --- --- --- --- --- --- --- --- tca tct aca tca ctg tca --- --- --- --- -ct ctc act ata tct tct aaa att aca a-- --a caa c-- --- --- --- --- --- --- --- --- --- --- --- --- --- --- --- --- --- --- --- --- --- --- --- --- --- --- --- --- --- --- --- --- --- --- --- -tg gat att cga t-- --- --- --- --- --- --- --- --- --- --- --- --- --- --- --- --- --- --- --- --- --- --- --- --- --- --- aac agc att tgt gt- --- --- --- --- --- --- --- --- --- --- --- --- --- --- --- --- --

>JX878413.1_Monkeypox_virus_isolate_DRC_07-0045_complete_genome

ttt ttt cga tct atc ctc gtc --- --- ctc atc atc ctt ata --- --- --- --- --- --- -tt att atc att att atc ata gtc tat taa aca caa atc atc t-- --- --- --- --- --- --- --- --- --- --- --- --- --- --- --- --- --- --- --- --- --- --- --- --- --- --- acg ttt ata ac- --- --- --- --- --- --- --- --- aac att c-- --- --- --- --t cat tat taa tta gtt ctg tag taa tat ctt taa taa ttt ggc tat a-- --- --c atc tgt t-- --- --- --- --- --- --- --- --- --- --- --- --- --- --- --- --- --- --- caa tac t-- --- --- --- --- --- --- --- atc tat tga tga ttt ctt tt- --- --- --- --- --- --- --- --- --- --- --- tta aga ct- --- --- --- --- --- --- --- --- --- --- --- --- --- --- --- --- --- --- --- --- --- --- --- --- --- --- --- --- --- --- --- --- --- --- --- --- --- --- --- -ta aac tag t-- --- --- --- --- --- --- --- --- --- --- --- --- --- --- --- --- --- --- --- --- --- --- --- --- --- --- --- --- --- --- --- --- --- --- --- --- --- --- --- --- --- --- --- --- --- --- --- --- --- --- --- --- --- --- --- --- --- --- --- --- --- --- --- --- --- --- --- --- --- --- --- --- --- --- --- --- --- --- --- --- --- --- --- --- --- --- --- --- --- --- --- --- --- --- --- --- --- --- --- --- --- --- --- --- --- --- --- --- --- tat ggt aat gac gat gaa a-- --t cga gta gta --- --- act tct aat aaa gac ttg ata --- --- tca tta tca tat gtt tga tcg --- --- --- --- --- --- --- tca tag tta ata gtg tg- --- --- --- --- --- --- --- --- --- --- --- --- --- --- --- --- --- --- --- --g cta aat ggt act gtt aat aag ttt at- --- --- --- --- --- --- --- --- --- --- --- --- --- --- --- --- --- aga caa tat cat agt att ttc ttt cca gaa t-- --- --- --- tag att att ttt tta aat act gat cct cac aat tcc gtg atg tag cag tag ttg gt- --- --- --- --- --- --- --- --- --- --- --- --- --- --- --- --- --- --- --- --- --- --- --- --- --- --- --- --g cat ggt cta tat cgt --- --- --- --- --- --- --- --- --- --- --- --- --- --- --- --- --- --- --- --- --- --- --- --- --- --- --- --- --- --- --- --- --- --- --- --- --- --- --- --- --- --- --- -ta aaa tgt atc ata tat aat agt ttt ctg acg tgg agt aca gaa ttt tcg a-- --- --- --- --- --- --- --- --- --- --- --- --- --- --- --- --- --- --- --- --- --- --- --- --- --- --- --- --- --- --- --- --- --- --- --- --- --- --- --- --- --- --- --- tta atg agt tca tgg taa gga agg gca aat gtc t-- -gt ata taa tat aca taa gtt aa- --- --- --- --- --- --- --- --- --- --- --- --- --- tag ttt ttt atc ata ttt --- --- --- --- --- --- --- --- tct aat acc ata ata aaa att atc att atg --- --- --t ata atc a-- --- --- --- --- --- --- --- --- --- --- --- tca ctg --- --t cgc tat cat tat tgc gtt tgt gta gtt --- --- --- --- -ct gcc cta --- --- --- --- --- --- --- --- tca tct aca tca ctg tca --- --- --- --- -ct ctc act ata tct tct aaa att aca a-- --a caa c-- --- --- --- --- --- --- --- --- --- --- --- --- --- --- --- --- --- --- --- --- --- --- --- --- --- --- --- --- --- --- --- --- --- --- --- -tg gat att cga t-- --- --- --- --- --- --- --- --- --- --- --- --- --- --- --- --- --- --- --- --- --- --- --- --- --- --- aac agc att tgt gt- --- --- --- --- --- --- --- --- --- --- --- --- --- --- --- --- --

>JX878422.1_Monkeypox_virus_isolate_DRC_07-0287_complete_genome

ttt ttt cga tct atc ctc gtc --- --- ctc atc atc ctt ata --- --- --- --- --- --- -tt att atc att att atc ata gtc tat taa aca caa atc atc t-- --- --- --- --- --- --- --- --- --- --- --- --- --- --- --- --- --- --- --- --- --- --- --- --- --- --- acg ttt ata ac- --- --- --- --- --- --- --- --- aac att c-- --- --- --- --t cat tat taa tta gtt ctg tag taa tat ctt taa taa ttt ggc tat a-- --- --c atc tgt t-- --- --- --- --- --- --- --- --- --- --- --- --- --- --- --- --- --- --- caa tac t-- --- --- --- --- --- --- --- atc tat tga tga ttt ctt tt- --- --- --- --- --- --- --- --- --- --- --- tta aga ct- --- --- --- --- --- --- --- --- --- --- --- --- --- --- --- --- --- --- --- --- --- --- --- --- --- --- --- --- --- --- --- --- --- --- --- --- --- --- --- -ta aac tag t-- --- --- --- --- --- --- --- --- --- --- --- --- --- --- --- --- --- --- --- --- --- --- --- --- --- --- --- --- --- --- --- --- --- --- --- --- --- --- --- --- --- --- --- --- --- --- --- --- --- --- --- --- --- --- --- --- --- --- --- --- --- --- --- --- --- --- --- --- --- --- --- --- --- --- --- --- --- --- --- --- --- --- --- --- --- --- --- --- --- --- --- --- --- --- --- --- --- --- --- --- --- --- --- --- --- --- --- --- --- tat ggt aat gac gat gaa a-- --t cga gta gta --- --- act tct aat aaa gac ttg ata --- --- tca tta tca tat gtt tga tcg --- --- --- --- --- --- --- tca tag tta ata gtg tg- --- --- --- --- --- --- --- --- --- --- --- --- --- --- --- --- --- --- --- --g cta aat ggt act gtt aat aag ttt at- --- --- --- --- --- --- --- --- --- --- --- --- --- --- --- --- --- aga caa tat cat agt att ttc ttt cca gaa t-- --- --- --- tag att att ttt tta aat act gat cct cac aat tcc gtg atg tag cag tag ttg gt- --- --- --- --- --- --- --- --- --- --- --- --- --- --- --- --- --- --- --- --- --- --- --- --- --- --- --- --g cat ggt cta tat cgt --- --- --- --- --- --- --- --- --- --- --- --- --- --- --- --- --- --- --- --- --- --- --- --- --- --- --- --- --- --- --- --- --- --- --- --- --- --- --- --- --- --- --- -ta aaa tgt atc ata tat aat agt ttt ctg acg tgg agt aca gaa ttt tcg a-- --- --- --- --- --- --- --- --- --- --- --- --- --- --- --- --- --- --- --- --- --- --- --- --- --- --- --- --- --- --- --- --- --- --- --- --- --- --- --- --- --- --- --- tta atg agt tca tgg taa gga agg gca aat gtc t-- -gt ata taa tat aca taa gtt aa- --- --- --- --- --- --- --- --- --- --- --- --- --- tag ttt ttt atc ata ttt --- --- --- --- --- --- --- --- tct aat acc ata ata aaa att atc att atg --- --- --t ata atc a-- --- --- --- --- --- --- --- --- --- --- --- tca ctg --- --t cgc tat cat tat tgc gtt tgt gta gtt --- --- --- --- -ct gcc cta --- --- --- --- --- --- --- --- tca tct aca tca ctg tca --- --- --- --- -ct ctc act ata tct tct aaa att aca a-- --a caa c-- --- --- --- --- --- --- --- --- --- --- --- --- --- --- --- --- --- --- --- --- --- --- --- --- --- --- --- --- --- --- --- --- --- --- --- -tg gat att cga t-- --- --- --- --- --- --- --- --- --- --- --- --- --- --- --- --- --- --- --- --- --- --- --- --- --- --- aac agc att tgt gt- --- --- --- --- --- --- --- --- --- --- --- --- --- --- --- --- --

>JX878416.1_Monkeypox_virus_isolate_DRC_07-0093_complete_genome

ttt ttt cga tct atc ctc gtc --- --- ctc atc atc ctt ata --- --- --- --- --- --- -tt att atc att att atc ata gtc tat taa aca caa atc atc t-- --- --- --- --- --- --- --- --- --- --- --- --- --- --- --- --- --- --- --- --- --- --- --- --- --- --- acg ttt ata ac- --- --- --- --- --- --- --- --- aac att c-- --- --- --- --t cat tat taa tta gtt ctg tag taa tat ctt taa taa ttt ggc tat a-- --- --c atc tgt t-- --- --- --- --- --- --- --- --- --- --- --- --- --- --- --- --- --- --- caa tac t-- --- --- --- --- --- --- --- atc tat tga tga ttt ctt tt- --- --- --- --- --- --- --- --- --- --- --- tta aga ct- --- --- --- --- --- --- --- --- --- --- --- --- --- --- --- --- --- --- --- --- --- --- --- --- --- --- --- --- --- --- --- --- --- --- --- --- --- --- --- -ta aac tag t-- --- --- --- --- --- --- --- --- --- --- --- --- --- --- --- --- --- --- --- --- --- --- --- --- --- --- --- --- --- --- --- --- --- --- --- --- --- --- --- --- --- --- --- --- --- --- --- --- --- --- --- --- --- --- --- --- --- --- --- --- --- --- --- --- --- --- --- --- --- --- --- --- --- --- --- --- --- --- --- --- --- --- --- --- --- --- --- --- --- --- --- --- --- --- --- --- --- --- --- --- --- --- --- --- --- --- --- --- --- tat ggt aat gac gat gaa a-- --t cga gta gta --- --- act tct aat aaa gac ttg ata --- --- tca tta tca tat gtt tga tcg --- --- --- --- --- --- --- tca tag tta ata gtg tg- --- --- --- --- --- --- --- --- --- --- --- --- --- --- --- --- --- --- --- --g cta aat ggt act gtt aat aag ttt at- --- --- --- --- --- --- --- --- --- --- --- --- --- --- --- --- --- aga caa tat cat agt att ttc ttt cca gaa t-- --- --- --- tag att att ttt tta aat act gat cct cac aat tcc gtg atg tag cag tag ttg gt- --- --- --- --- --- --- --- --- --- --- --- --- --- --- --- --- --- --- --- --- --- --- --- --- --- --- --- --g cat ggt cta tat cgt --- --- --- --- --- --- --- --- --- --- --- --- --- --- --- --- --- --- --- --- --- --- --- --- --- --- --- --- --- --- --- --- --- --- --- --- --- --- --- --- --- --- --- -ta aaa tgt atc ata tat aat agt ttt ctg acg tgg agt aca gaa ttt tcg a-- --- --- --- --- --- --- --- --- --- --- --- --- --- --- --- --- --- --- --- --- --- --- --- --- --- --- --- --- --- --- --- --- --- --- --- --- --- --- --- --- --- --- --- tta atg agt tca tgg taa gga agg gca aat gtc t-- -gt ata taa tat aca taa gtt aa- --- --- --- --- --- --- --- --- --- --- --- --- --- tag ttt ttt atc ata ttt --- --- --- --- --- --- --- --- tct aat acc ata ata aaa att atc att atg --- --- --t ata atc a-- --- --- --- --- --- --- --- --- --- --- --- tca ctg --- --t cgc tat cat tat tgc gtt tgt gta gtt --- --- --- --- -ct gcc cta --- --- --- --- --- --- --- --- tca tct aca tca ctg tca --- --- --- --- -ct ctc act ata tct tct aaa att aca a-- --a caa c-- --- --- --- --- --- --- --- --- --- --- --- --- --- --- --- --- --- --- --- --- --- --- --- --- --- --- --- --- --- --- --- --- --- --- --- -tg gat att cga t-- --- --- --- --- --- --- --- --- --- --- --- --- --- --- --- --- --- --- --- --- --- --- --- --- --- --- aac agc att tgt gt- --- --- --- --- --- --- --- --- --- --- --- --- --- --- --- --- --

>KP849471.1_Monkeypox_virus_isolate_Yambuku_DRC_1985_complete_genome

ttt ttt cga tct atc ctc gtc --- --- ctc atc atc ctt ata --- --- --- --- --- --- -tt att atc att att atc ata gtc tat taa aca caa atc atc t-- --- --- --- --- --- --- --- --- --- --- --- --- --- --- --- --- --- --- --- --- --- --- --- --- --- --- acg ttt ata ac- --- --- --- --- --- --- --- --- aac att c-- --- --- --- --t cat tat taa tta gtt ctg tag taa tat ctt taa taa ttt ggc tat a-- --- --c atc tgt t-- --- --- --- --- --- --- --- --- --- --- --- --- --- --- --- --- --- --- caa tac t-- --- --- --- --- --- --- --- atc tat tga tga ttt ctt tt- --- --- --- --- --- --- --- --- --- --- --- tta aga ct- --- --- --- --- --- --- --- --- --- --- --- --- --- --- --- --- --- --- --- --- --- --- --- --- --- --- --- --- --- --- --- --- --- --- --- --- --- --- --- -ta aac tag t-- --- --- --- --- --- --- --- --- --- --- --- --- --- --- --- --- --- --- --- --- --- --- --- --- --- --- --- --- --- --- --- --- --- --- --- --- --- --- --- --- --- --- --- --- --- --- --- --- --- --- --- --- --- --- --- --- --- --- --- --- --- --- --- --- --- --- --- --- --- --- --- --- --- --- --- --- --- --- --- --- --- --- --- --- --- --- --- --- --- --- --- --- --- --- --- --- --- --- --- --- --- --- --- --- --- --- --- --- --- tat ggt aat gac gat gaa a-- --t cga gta gta --- --- act tct aat aaa gac ttg ata --- --- tca tta tca tat gtt tga tcg --- --- --- --- --- --- --- tca tag tta ata gtg tg- --- --- --- --- --- --- --- --- --- --- --- --- --- --- --- --- --- --- --- --g cta aat ggt act gtt aat aag ttt at- --- --- --- --- --- --- --- --- --- --- --- --- --- --- --- --- --- aga caa tat cat agt att ttc ttt cca gaa t-- --- --- --- tag att att ttt tta aat act gat cct cac aat tcc gtg atg tag cag tag ttg gt- --- --- --- --- --- --- --- --- --- --- --- --- --- --- --- --- --- --- --- --- --- --- --- --- --- --- --- --g cat ggt cta tat cgt --- --- --- --- --- --- --- --- --- --- --- --- --- --- --- --- --- --- --- --- --- --- --- --- --- --- --- --- --- --- --- --- --- --- --- --- --- --- --- --- --- --- --- -ta aaa tgt atc ata tat aat agt ttt ctg acg tgg agt aca gaa ttt tcg a-- --- --- --- --- --- --- --- --- --- --- --- --- --- --- --- --- --- --- --- --- --- --- --- --- --- --- --- --- --- --- --- --- --- --- --- --- --- --- --- --- --- --- --- tta atg agt tca tgg taa gga agg gca aat gtc t-- -gt ata taa tat aca taa gtt aa- --- --- --- --- --- --- --- --- --- --- --- --- --- tag ttt ttt atc ata ttt --- --- --- --- --- --- --- --- tct aat acc ata ata aaa att atc att atg --- --- --t ata atc a-- --- --- --- --- --- --- --- --- --- --- --- tca ctg --- --t cgc tat cat tat tgc gtt tgt gta gtt --- --- --- --- -ct gcc cta --- --- --- --- --- --- --- --- tca tct aca tca ctg tca --- --- --- --- -ct ctc act ata tct tct aaa att aca a-- --a caa c-- --- --- --- --- --- --- --- --- --- --- --- --- --- --- --- --- --- --- --- --- --- --- --- --- --- --- --- --- --- --- --- --- --- --- --- -tg gat att cga t-- --- --- --- --- --- --- --- --- --- --- --- --- --- --- --- --- --- --- --- --- --- --- --- --- --- --- aac agc att tgt gt- --- --- --- --- --- --- --- --- --- --- --- --- --- --- --- --- --

>DQ011155.1_Monkeypox_virus_strain_Zaire_1979-005_complete_genome

ttt ttt cga tct atc ctc gtc --- --- ctc atc atc ctt ata --- --- --- --- --- --- -tt att atc att att atc ata gtc tat taa aca caa atc atc t-- --- --- --- --- --- --- --- --- --- --- --- --- --- --- --- --- --- --- --- --- --- --- --- --- --- --- acg ttt ata ac- --- --- --- --- --- --- --- --- aac att c-- --- --- --- --t cat tat taa tta gtt ctg tag taa tat ctt taa taa ttt ggc tat a-- --- --c atc tgt t-- --- --- --- --- --- --- --- --- --- --- --- --- --- --- --- --- --- --- caa tac t-- --- --- --- --- --- --- --- atc tat tga tga ttt ctt tt- --- --- --- --- --- --- --- --- --- --- --- tta aga ct- --- --- --- --- --- --- --- --- --- --- --- --- --- --- --- --- --- --- --- --- --- --- --- --- --- --- --- --- --- --- --- --- --- --- --- --- --- --- --- -ta aac tag t-- --- --- --- --- --- --- --- --- --- --- --- --- --- --- --- --- --- --- --- --- --- --- --- --- --- --- --- --- --- --- --- --- --- --- --- --- --- --- --- --- --- --- --- --- --- --- --- --- --- --- --- --- --- --- --- --- --- --- --- --- --- --- --- --- --- --- --- --- --- --- --- --- --- --- --- --- --- --- --- --- --- --- --- --- --- --- --- --- --- --- --- --- --- --- --- --- --- --- --- --- --- --- --- --- --- --- --- --- --- tat ggt aat gac gat gaa a-- --t cga gta gta --- --- act tct aat aaa tac ttg ata --- --- tca tta tca tat gtt tga tcg --- --- --- --- --- --- --- tca tag tta ata gtg tg- --- --- --- --- --- --- --- --- --- --- --- --- --- --- --- --- --- --- --- --g cta aat ggt act gtt aat aag ttt at- --- --- --- --- --- --- --- --- --- --- --- --- --- --- --- --- --- aga caa tat cat agt att ttc ttt cca gaa t-- --- --- --- tag att att ttt tta aat act gat cct cac aat tcc gtg atg tag cag tag ttg gt- --- --- --- --- --- --- --- --- --- --- --- --- --- --- --- --- --- --- --- --- --- --- --- --- --- --- --- --g cat ggt cta tat cgt --- --- --- --- --- --- --- --- --- --- --- --- --- --- --- --- --- --- --- --- --- --- --- --- --- --- --- --- --- --- --- --- --- --- --- --- --- --- --- --- --- --- --- -ta aaa tgt atc ata tat aat agt ttt ctg acg tgg agt aca gaa ttt tcg a-- --- --- --- --- --- --- --- --- --- --- --- --- --- --- --- --- --- --- --- --- --- --- --- --- --- --- --- --- --- --- --- --- --- --- --- --- --- --- --- --- --- --- --- tta atg agt tca tgg taa gga agg gca aat gtc t-- -gt ata taa tat aca taa gtt aa- --- --- --- --- --- --- --- --- --- --- --- --- --- tag ttt ttt atc ata ttt --- --- --- --- --- --- --- --- tct aat acc ata ata aaa att atc att atg --- --- --t ata atc a-- --- --- --- --- --- --- --- --- --- --- --- tca ctg --- --t cgc tat cat tat tgc gtt tgt gta gtt --- --- --- --- -ct gcc cta --- --- --- --- --- --- --- --- tca tct aca tca ctg tca --- --- --- --- -ct ctc act ata tct tct aaa att aca a-- --a caa c-- --- --- --- --- --- --- --- --- --- --- --- --- --- --- --- --- --- --- --- --- --- --- --- --- --- --- --- --- --- --- --- --- --- --- --- -tg gat att cga t-- --- --- --- --- --- --- --- --- --- --- --- --- --- --- --- --- --- --- --- --- --- --- --- --- --- --- aac agc att tgt gt- --- --- --- --- --- --- --- --- --- --- --- --- --- --- --- --- --

>KP849469.1_Monkeypox_virus_isolate_Boende_DRC_2008_complete_genome

ttt ttt cga tct atc ctc gtc --- --- ctc atc atc ctt ata --- --- --- --- --- --- -tt att atc att att atc ata gtc tat taa aca caa atc atc t-- --- --- --- --- --- --- --- --- --- --- --- --- --- --- --- --- --- --- --- --- --- --- --- --- --- --- acg ttt ata ac- --- --- --- --- --- --- --- --- aac att c-- --- --- --- --t cat tat taa tta gtt ctg tag taa tat ctt taa taa ttt ggc tat a-- --- --c atc tgt t-- --- --- --- --- --- --- --- --- --- --- --- --- --- --- --- --- --- --- caa tac t-- --- --- --- --- --- --- --- atc tat tga tga ttt ctt tt- --- --- --- --- --- --- --- --- --- --- --- tta aga ct- --- --- --- --- --- --- --- --- --- --- --- --- --- --- --- --- --- --- --- --- --- --- --- --- --- --- --- --- --- --- --- --- --- --- --- --- --- --- --- -ta aac tag t-- --- --- --- --- --- --- --- --- --- --- --- --- --- --- --- --- --- --- --- --- --- --- --- --- --- --- --- --- --- --- --- --- --- --- --- --- --- --- --- --- --- --- --- --- --- --- --- --- --- --- --- --- --- --- --- --- --- --- --- --- --- --- --- --- --- --- --- --- --- --- --- --- --- --- --- --- --- --- --- --- --- --- --- --- --- --- --- --- --- --- --- --- --- --- --- --- --- --- --- --- --- --- --- --- --- --- --- --- --- tat ggt aat gac gat gaa a-- --t cga gta gta --- --- act tct aat aaa gac ttg ata --- --- tca tta tca tat gtt tga tcg --- --- --- --- --- --- --- tca tag tta ata gtg tg- --- --- --- --- --- --- --- --- --- --- --- --- --- --- --- --- --- --- --- --g cta aat ggt act gtt aat aag ttt at- --- --- --- --- --- --- --- --- --- --- --- --- --- --- --- --- --- aga caa tat cat agt att ttc ttt cca gaa t-- --- --- --- tag att att ttt tta aat act gat cct cac aat tcc gtg atg tag cag tag ttg gt- --- --- --- --- --- --- --- --- --- --- --- --- --- --- --- --- --- --- --- --- --- --- --- --- --- --- --- --g cat ggt cta tat cgt --- --- --- --- --- --- --- --- --- --- --- --- --- --- --- --- --- --- --- --- --- --- --- --- --- --- --- --- --- --- --- --- --- --- --- --- --- --- --- --- --- --- --- -ta aaa tgt atc ata tat aat agt ttt ctg acg tgg agt aca gaa ttt tcg a-- --- --- --- --- --- --- --- --- --- --- --- --- --- --- --- --- --- --- --- --- --- --- --- --- --- --- --- --- --- --- --- --- --- --- --- --- --- --- --- --- --- --- --- tta atg agt tca tgg taa gga agg gca aat gtc t-- -gt ata taa tat aca taa gtt aa- --- --- --- --- --- --- --- --- --- --- --- --- --- tag ttt ttt atc ata ttt --- --- --- --- --- --- --- --- tct aat acc ata ata aaa att atc att atg --- --- --t ata atc a-- --- --- --- --- --- --- --- --- --- --- --- tca ctg --- --t cgc tat cat tat tgc gtt tgt gta gtt --- --- --- --- -ct gcc cta --- --- --- --- --- --- --- --- tca tct aca tca ctg tca --- --- --- --- -ct ctc act ata tct tct aaa att aca a-- --a caa c-- --- --- --- --- --- --- --- --- --- --- --- --- --- --- --- --- --- --- --- --- --- --- --- --- --- --- --- --- --- --- --- --- --- --- --- -tg gat att cga t-- --- --- --- --- --- --- --- --- --- --- --- --- --- --- --- --- --- --- --- --- --- --- --- --- --- --- aac agc att tgt gt- --- --- --- --- --- --- --- --- --- --- --- --- --- --- --- --- --

>KJ642619.1_Monkeypox_virus_strain_Gabon-1988_complete_genome

ttt ttt cga tct atc ctc gtc --- --- ctc atc atc ctt ata --- --- --- --- --- --- -tt att atc att att atc ata gtc tat taa aca caa atc atc t-- --- --- --- --- --- --- --- --- --- --- --- --- --- --- --- --- --- --- --- --- --- --- --- --- --- --- acg ttt ata ac- --- --- --- --- --- --- --- --- aac att c-- --- --- --- --t cat tat taa tta gtt ctg tag taa tat ctt taa taa ttt ggc tat a-- --- --c atc tgt t-- --- --- --- --- --- --- --- --- --- --- --- --- --- --- --- --- --- --- caa tac t-- --- --- --- --- --- --- --- atc tat tga tga ttt ctt tt- --- --- --- --- --- --- --- --- --- --- --- tta aga ct- --- --- --- --- --- --- --- --- --- --- --- --- --- --- --- --- --- --- --- --- --- --- --- --- --- --- --- --- --- --- --- --- --- --- --- --- --- --- --- -ta aac tag t-- --- --- --- --- --- --- --- --- --- --- --- --- --- --- --- --- --- --- --- --- --- --- --- --- --- --- --- --- --- --- --- --- --- --- --- --- --- --- --- --- --- --- --- --- --- --- --- --- --- --- --- --- --- --- --- --- --- --- --- --- --- --- --- --- --- --- --- --- --- --- --- --- --- --- --- --- --- --- --- --- --- --- --- --- --- --- --- --- --- --- --- --- --- --- --- --- --- --- --- --- --- --- --- --- --- --- --- --- --- tat ggt aat gac gat gaa a-- --t cga gta gta --- --- act tct aat aaa gac ttg ata --- --- tca tta tca tat gtt tga tcg --- --- --- --- --- --- --- tca tag tta ata gtg tg- --- --- --- --- --- --- --- --- --- --- --- --- --- --- --- --- --- --- --- --g cta aat ggt act gtt aat aag ttt at- --- --- --- --- --- --- --- --- --- --- --- --- --- --- --- --- --- aga caa tat cat agt att ttc ttt cca gaa t-- --- --- --- tag att att ttt tta aat act gat cct cac aat tcc gtg atg tag cag tag ttg gt- --- --- --- --- --- --- --- --- --- --- --- --- --- --- --- --- --- --- --- --- --- --- --- --- --- --- --- --g cat ggt cta tat cgt --- --- --- --- --- --- --- --- --- --- --- --- --- --- --- --- --- --- --- --- --- --- --- --- --- --- --- --- --- --- --- --- --- --- --- --- --- --- --- --- --- --- --- -ta aaa tgt atc ata tat aat agt ttt ctg acg tgg agt aca gaa ttt tcg a-- --- --- --- --- --- --- --- --- --- --- --- --- --- --- --- --- --- --- --- --- --- --- --- --- --- --- --- --- --- --- --- --- --- --- --- --- --- --- --- --- --- --- --- tta atg agt tca tgg taa gga agg gca aat gtc t-- -gt ata taa tat aca taa gtt aa- --- --- --- --- --- --- --- --- --- --- --- --- --- tag ttt tt- atc ata ttt --- --- --- --- --- --- --- --- tct aat acc ata ata aaa att atc att atg --- --- --t ata atc a-- --- --- --- --- --- --- --- --- --- --- --- tca ctg --- --t cgc tat cat tat tgc gtt tgt gta gtt --- --- --- --- -ct gcc cta --- --- --- --- --- --- --- --- tca tct aca tca ctg tca --- --- --- --- -ct ctc act ata tct tct aaa att aca a-- --a caa c-- --- --- --- --- --- --- --- --- --- --- --- --- --- --- --- --- --- --- --- --- --- --- --- --- --- --- --- --- --- --- --- --- --- --- --- -tg gat att cga t-- --- --- --- --- --- --- --- --- --- --- --- --- --- --- --- --- --- --- --- --- --- --- --- --- --- --- aac agc att tgt gt- --- --- --- --- --- --- --- --- --- --- --- --- --- --- --- --- --

>JX878409.1_Monkeypox_virus_isolate_DRC_06-0999_complete_genome

ttt ttt cga tct atc ctc gtc --- --- ctc atc atc ctt ata --- --- --- --- --- --- -tt att atc att att atc ata gtc tat taa aca caa atc atc t-- --- --- --- --- --- --- --- --- --- --- --- --- --- --- --- --- --- --- --- --- --- --- --- --- --- --- acg ttt ata ac- --- --- --- --- --- --- --- --- aac att c-- --- --- --- --t cat tat taa tta gtt ctg tag taa tat ctt taa taa ttt ggc tat a-- --- --c atc tgt t-- --- --- --- --- --- --- --- --- --- --- --- --- --- --- --- --- --- --- caa tac t-- --- --- --- --- --- --- --- atc tat tga tga ttt ctt tt- --- --- --- --- --- --- --- --- --- --- --- tta aga ct- --- --- --- --- --- --- --- --- --- --- --- --- --- --- --- --- --- --- --- --- --- --- --- --- --- --- --- --- --- --- --- --- --- --- --- --- --- --- --- -ta aac tag t-- --- --- --- --- --- --- --- --- --- --- --- --- --- --- --- --- --- --- --- --- --- --- --- --- --- --- --- --- --- --- --- --- --- --- --- --- --- --- --- --- --- --- --- --- --- --- --- --- --- --- --- --- --- --- --- --- --- --- --- --- --- --- --- --- --- --- --- --- --- --- --- --- --- --- --- --- --- --- --- --- --- --- --- --- --- --- --- --- --- --- --- --- --- --- --- --- --- --- --- --- --- --- --- --- --- --- --- --- --- tat ggt aat gac gat gaa a-- --t cga gta gta --- --- act tct aat aaa gac ttg ata --- --- tca tta tca tat gtt tga tcg --- --- --- --- --- --- --- tca tag tta ata gtg tg- --- --- --- --- --- --- --- --- --- --- --- --- --- --- --- --- --- --- --- --g cta aat ggt act gtt aat aag ttt at- --- --- --- --- --- --- --- --- --- --- --- --- --- --- --- --- --- aga caa tat cat agt att ttc ttt cca gaa t-- --- --- --- tag att att ttt tta aat act gat cct cac aat tcc gtg atg tag cag tag ttg gt- --- --- --- --- --- --- --- --- --- --- --- --- --- --- --- --- --- --- --- --- --- --- --- --- --- --- --- --g cat ggt cta tat cgt --- --- --- --- --- --- --- --- --- --- --- --- --- --- --- --- --- --- --- --- --- --- --- --- --- --- --- --- --- --- --- --- --- --- --- --- --- --- --- --- --- --- --- -ta aaa tgt atc ata tat aat agt ttt ctg acg tgg agt aca gaa ttt tcg a-- --- --- --- --- --- --- --- --- --- --- --- --- --- --- --- --- --- --- --- --- --- --- --- --- --- --- --- --- --- --- --- --- --- --- --- --- --- --- --- --- --- --- --- tta atg agt tca tgg taa gga agg gca aat gtc t-- -gt ata taa tat aca taa gtt aa- --- --- --- --- --- --- --- --- --- --- --- --- --- tag ttt tt- atc ata ttt --- --- --- --- --- --- --- --- tct aat acc ata ata aaa att atc att atg --- --- --t ata atc a-- --- --- --- --- --- --- --- --- --- --- --- tca ctg --- --t cgc tat cat tat tgc gtt tgt gta gtt --- --- --- --- -ct gcc cta --- --- --- --- --- --- --- --- tca tct aca tca ctg tca --- --- --- --- -ct ctc act ata tct tct aaa att aca a-- --a caa c-- --- --- --- --- --- --- --- --- --- --- --- --- --- --- --- --- --- --- --- --- --- --- --- --- --- --- --- --- --- --- --- --- --- --- --- -tg gag att cga t-- --- --- --- --- --- --- --- --- --- --- --- --- --- --- --- --- --- --- --- --- --- --- --- --- --- --- aac agc att tgt gt- --- --- --- --- --- --- --- --- --- --- --- --- --- --- --- --- --

>DQ011154.1_Monkeypox_virus_strain_Congo_2003_358_complete_genome

ttt ttt cga tct atc ctc gtc --- --- ctc atc atc ctt ata --- --- --- --- --- --- -tt att atc att att atc ata gtc tat taa aca caa atc atc t-- --- --- --- --- --- --- --- --- --- --- --- --- --- --- --- --- --- --- --- --- --- --- --- --- --- --- acg ttt ata ac- --- --- --- --- --- --- --- --- aac att c-- --- --- --- --t cat tat taa tta gtt ctg tag taa tat ctt taa taa ttt ggc tat a-- --- --c atc tgt t-- --- --- --- --- --- --- --- --- --- --- --- --- --- --- --- --- --- --- caa tac t-- --- --- --- --- --- --- --- atc tat tga tga ttt ctt tt- --- --- --- --- --- --- --- --- --- --- --- tta aga ct- --- --- --- --- --- --- --- --- --- --- --- --- --- --- --- --- --- --- --- --- --- --- --- --- --- --- --- --- --- --- --- --- --- --- --- --- --- --- --- -ta aac tag t-- --- --- --- --- --- --- --- --- --- --- --- --- --- --- --- --- --- --- --- --- --- --- --- --- --- --- --- --- --- --- --- --- --- --- --- --- --- --- --- --- --- --- --- --- --- --- --- --- --- --- --- --- --- --- --- --- --- --- --- --- --- --- --- --- --- --- --- --- --- --- --- --- --- --- --- --- --- --- --- --- --- --- --- --- --- --- --- --- --- --- --- --- --- --- --- --- --- --- --- --- --- --- --- --- --- --- --- --- --- tat ggt aat gac gat gaa a-- --t cga gta gta --- --- act tct aat aaa gac ttg ata --- --- tca tta tca tat gtt tga tcg --- --- --- --- --- --- --- tca tag tta ata gtg tg- --- --- --- --- --- --- --- --- --- --- --- --- --- --- --- --- --- --- --- --g cta aat ggt act gtt aat aag ttt at- --- --- --- --- --- --- --- --- --- --- --- --- --- --- --- --- --- aga caa tat cat agt att ttc ttt cca gaa t-- --- --- --- tag att att ttt tta aat act gat cct cac aat tcc gtg atg tag cag tag ttg gt- --- --- --- --- --- --- --- --- --- --- --- --- --- --- --- --- --- --- --- --- --- --- --- --- --- --- --- --g cat ggt cta tat cgt --- --- --- --- --- --- --- --- --- --- --- --- --- --- --- --- --- --- --- --- --- --- --- --- --- --- --- --- --- --- --- --- --- --- --- --- --- --- --- --- --- --- --- -ta aaa tgt atc ata tat aat agt ttt ctg acg tgg agt aca gaa ttt tcg a-- --- --- --- --- --- --- --- --- --- --- --- --- --- --- --- --- --- --- --- --- --- --- --- --- --- --- --- --- --- --- --- --- --- --- --- --- --- --- --- --- --- --- --- tta atg agt tca tgg taa gga agg gca aat gtc t-- -gt ata taa tat aca taa gtt aa- --- --- --- --- --- --- --- --- --- --- --- --- --- tag ttt ttt atc ata ttt --- --- --- --- --- --- --- --- tct aat acc ata ata aaa att atc att atg --- --- --t ata atc a-- --- --- --- --- --- --- --- --- --- --- --- tca ctg --- --t cgc tat cat tat tgc gtt tgt gta gtt --- --- --- --- -ct gcc cta --- --- --- --- --- --- --- --- tca tct aca tca ctg tca --- --- --- --- -ct ctc act ata tct tct aaa att aca a-- --a caa c-- --- --- --- --- --- --- --- --- --- --- --- --- --- --- --- --- --- --- --- --- --- --- --- --- --- --- --- --- --- --- --- --- --- --- --- -tg gat att cga t-- --- --- --- --- --- --- --- --- --- --- --- --- --- --- --- --- --- --- --- --- --- --- --- --- --- --- aac agc att tgt gt- --- --- --- --- --- --- --- --- --- --- --- --- --- --- --- --- --

>MN702453.1_Monkeypox_virus_strain_A1_contig_SPADES

ttt ttt cga tct atc ctc gtc --- --- ctc atc atc ctt ata --- --- --- --- --- --- -tt att atc att att atc ata gtc tat taa aca caa atc atc t-- --- --- --- --- --- --- --- --- --- --- --- --- --- --- --- --- --- --- --- --- --- --- --- --- --- --- acg ttt ata ac- --- --- --- --- --- --- --- --- aac att c-- --- --- --- --t cat tat taa tta gtt ctg tag taa tat ctt taa taa ttt ggc tat a-- --- --c atc tgt t-- --- --- --- --- --- --- --- --- --- --- --- --- --- --- --- --- --- --- caa tac t-- --- --- --- --- --- --- --- atc tat tga tga ttt ctt tt- --- --- --- --- --- --- --- --- --- --- --- tta aga ct- --- --- --- --- --- --- --- --- --- --- --- --- --- --- --- --- --- --- --- --- --- --- --- --- --- --- --- --- --- --- --- --- --- --- --- --- --- --- --- -ta aac tag t-- --- --- --- --- --- --- --- --- --- --- --- --- --- --- --- --- --- --- --- --- --- --- --- --- --- --- --- --- --- --- --- --- --- --- --- --- --- --- --- --- --- --- --- --- --- --- --- --- --- --- --- --- --- --- --- --- --- --- --- --- --- --- --- --- --- --- --- --- --- --- --- --- --- --- --- --- --- --- --- --- --- --- --- --- --- --- --- --- --- --- --- --- --- --- --- --- --- --- --- --- --- --- --- --- --- --- --- --- --- tat ggt aat gac gat gaa a-- --t cga gta gta --- --- act tct aat aaa gac ttg ata --- --- tca tta tca tat gtt tga tcg --- --- --- --- --- --- --- tca tag tta ata gtg tg- --- --- --- --- --- --- --- --- --- --- --- --- --- --- --- --- --- --- --- --g cta aat ggt act gtt aat aag ttt at- --- --- --- --- --- --- --- --- --- --- --- --- --- --- --- --- --- aga caa tat cat agt att ttc ttt cca gaa t-- --- --- --- tag att att ttt tta aat act gat cct cac aat tcc gtg atg tag cag tag ttg gt- --- --- --- --- --- --- --- --- --- --- --- --- --- --- --- --- --- --- --- --- --- --- --- --- --- --- --- --g cat ggt cta tat cgt --- --- --- --- --- --- --- --- --- --- --- --- --- --- --- --- --- --- --- --- --- --- --- --- --- --- --- --- --- --- --- --- --- --- --- --- --- --- --- --- --- --- --- -ta aaa tgt atc ata tat aat agt ttt ctg acg tgg agt aca gaa ttt tcg a-- --- --- --- --- --- --- --- --- --- --- --- --- --- --- --- --- --- --- --- --- --- --- --- --- --- --- --- --- --- --- --- --- --- --- --- --- --- --- --- --- --- --- --- tta atg agt tca tgg taa gga agg gca aat gtc t-- -gt ata taa tat aca taa gtt aa- --- --- --- --- --- --- --- --- --- --- --- --- --- tag ttt ttt atc ata ttt --- --- --- --- --- --- --- --- tct aat acc ata ata aaa att atc att atg --- --- --t ata atc a-- --- --- --- --- --- --- --- --- --- --- --- tca ctg --- --t cgc tat cat tat tgc gtt tgt gta gtt --- --- --- --- -ct gcc cta --- --- --- --- --- --- --- --- tca tct aca tca ctg tca --- --- --- --- -ct ctc act ata tct tct aaa att aca a-- --a caa c-- --- --- --- --- --- --- --- --- --- --- --- --- --- --- --- --- --- --- --- --- --- --- --- --- --- --- --- --- --- --- --- --- --- --- --- -tg gat att cga t-- --- --- --- --- --- --- --- --- --- --- --- --- --- --- --- --- --- --- --- --- --- --- --- --- --- --- aac agc att tgt gt- --- --- --- --- --- --- --- --- --- --- --- --- --- --- --- --- --

>MN702452.1_Monkeypox_virus_strain_A2_contig_SPADES

ttt ttt cga tct atc ctc gtc --- --- ctc atc atc ctt ata --- --- --- --- --- --- -tt att atc att att atc ata gtc tat taa aca caa atc atc t-- --- --- --- --- --- --- --- --- --- --- --- --- --- --- --- --- --- --- --- --- --- --- --- --- --- --- acg ttt ata ac- --- --- --- --- --- --- --- --- aac att c-- --- --- --- --t cat tat taa tta gtt ctg tag taa tat ctt taa taa ttt ggc tat a-- --- --c atc tgt t-- --- --- --- --- --- --- --- --- --- --- --- --- --- --- --- --- --- --- caa tac t-- --- --- --- --- --- --- --- atc tat tga tga ttt ctt tt- --- --- --- --- --- --- --- --- --- --- --- tta aga ct- --- --- --- --- --- --- --- --- --- --- --- --- --- --- --- --- --- --- --- --- --- --- --- --- --- --- --- --- --- --- --- --- --- --- --- --- --- --- --- -ta aac tag t-- --- --- --- --- --- --- --- --- --- --- --- --- --- --- --- --- --- --- --- --- --- --- --- --- --- --- --- --- --- --- --- --- --- --- --- --- --- --- --- --- --- --- --- --- --- --- --- --- --- --- --- --- --- --- --- --- --- --- --- --- --- --- --- --- --- --- --- --- --- --- --- --- --- --- --- --- --- --- --- --- --- --- --- --- --- --- --- --- --- --- --- --- --- --- --- --- --- --- --- --- --- --- --- --- --- --- --- --- --- tat ggt aat gac gat gaa a-- --t cga gta gta --- --- act tct aat aaa gac ttg ata --- --- tca tta tca tat gtt tga tcg --- --- --- --- --- --- --- tca tag tta ata gtg tg- --- --- --- --- --- --- --- --- --- --- --- --- --- --- --- --- --- --- --- --g cta aat ggt act gtt aat aag ttt at- --- --- --- --- --- --- --- --- --- --- --- --- --- --- --- --- --- aga caa tat cat agt att ttc ttt cca gaa t-- --- --- --- tag att att ttt tta aat act gat cct cac aat tcc gtg atg tag cag tag ttg gt- --- --- --- --- --- --- --- --- --- --- --- --- --- --- --- --- --- --- --- --- --- --- --- --- --- --- --- --g cat ggt cta tat cgt --- --- --- --- --- --- --- --- --- --- --- --- --- --- --- --- --- --- --- --- --- --- --- --- --- --- --- --- --- --- --- --- --- --- --- --- --- --- --- --- --- --- --- -ta aaa tgt atc ata tat aat agt ttt ctg acg tgg agt aca gaa ttt tcg a-- --- --- --- --- --- --- --- --- --- --- --- --- --- --- --- --- --- --- --- --- --- --- --- --- --- --- --- --- --- --- --- --- --- --- --- --- --- --- --- --- --- --- --- tta atg agt tca tgg taa gga agg gca aat gtc t-- -gt ata taa tat aca taa gtt aa- --- --- --- --- --- --- --- --- --- --- --- --- --- tag ttt ttt atc ata ttt --- --- --- --- --- --- --- --- tct aat acc ata ata aaa att atc att atg --- --- --t ata atc a-- --- --- --- --- --- --- --- --- --- --- --- tca ctg --- --t cgc tat cat tat tgc gtt tgt gta gtt --- --- --- --- -ct gcc cta --- --- --- --- --- --- --- --- tca tct aca tca ctg tca --- --- --- --- -ct ctc act ata tct tct aaa att aca a-- --a caa c-- --- --- --- --- --- --- --- --- --- --- --- --- --- --- --- --- --- --- --- --- --- --- --- --- --- --- --- --- --- --- --- --- --- --- --- -tg gat att cga t-- --- --- --- --- --- --- --- --- --- --- --- --- --- --- --- --- --- --- --- --- --- --- --- --- --- --- aac agc att tgt gt- --- --- --- --- --- --- --- --- --- --- --- --- --- --- --- --- --

>JX878424.1_Monkeypox_virus_isolate_DRC_07-0338_complete_genome

ttt ttt cga tct atc ctc gtc --- --- ctc atc atc ctt ata --- --- --- --- --- --- -tt att atc att att atc ata gtc tat taa aca caa atc atc t-- --- --- --- --- --- --- --- --- --- --- --- --- --- --- --- --- --- --- --- --- --- --- --- --- --- --- acg ttt ata ac- --- --- --- --- --- --- --- --- aac att c-- --- --- --- --t cat tat taa tta gtt ctg tag taa tat ctt taa taa ttt ggc tat a-- --- --c atc tgt t-- --- --- --- --- --- --- --- --- --- --- --- --- --- --- --- --- --- --- caa tac t-- --- --- --- --- --- --- --- atc tat tga tga ttt ctt tt- --- --- --- --- --- --- --- --- --- --- --- tta aga ct- --- --- --- --- --- --- --- --- --- --- --- --- --- --- --- --- --- --- --- --- --- --- --- --- --- --- --- --- --- --- --- --- --- --- --- --- --- --- --- -ta aac tag t-- --- --- --- --- --- --- --- --- --- --- --- --- --- --- --- --- --- --- --- --- --- --- --- --- --- --- --- --- --- --- --- --- --- --- --- --- --- --- --- --- --- --- --- --- --- --- --- --- --- --- --- --- --- --- --- --- --- --- --- --- --- --- --- --- --- --- --- --- --- --- --- --- --- --- --- --- --- --- --- --- --- --- --- --- --- --- --- --- --- --- --- --- --- --- --- --- --- --- --- --- --- --- --- --- --- --- --- --- --- tat ggt aat gac gat gaa a-- --t cga gta gta --- --- act tct aat aaa gac ttg ata --- --- tca tta tca tat gtt tga tcg --- --- --- --- --- --- --- tca tag tta ata gtg tg- --- --- --- --- --- --- --- --- --- --- --- --- --- --- --- --- --- --- --- --g cta aat ggt act gtt aat aag ttt at- --- --- --- --- --- --- --- --- --- --- --- --- --- --- --- --- --- aga caa tat cat agt att ttc ttt cca gaa t-- --- --- --- tag att att ttt tta aat act gat cct cac aat tcc gtg atg tag cag tag ttg gt- --- --- --- --- --- --- --- --- --- --- --- --- --- --- --- --- --- --- --- --- --- --- --- --- --- --- --- --g cat ggt cta tat cgt --- --- --- --- --- --- --- --- --- --- --- --- --- --- --- --- --- --- --- --- --- --- --- --- --- --- --- --- --- --- --- --- --- --- --- --- --- --- --- --- --- --- --- -ta aaa tgt atc ata tat aat agt ttt ctg acg tgg agt aca gaa ttt tcg a-- --- --- --- --- --- --- --- --- --- --- --- --- --- --- --- --- --- --- --- --- --- --- --- --- --- --- --- --- --- --- --- --- --- --- --- --- --- --- --- --- --- --- --- tta atg agt tca tgg taa gga agg gca aat gtc t-- -gt ata taa tat aca taa gtt aa- --- --- --- --- --- --- --- --- --- --- --- --- --- tag ttt ttt atc ata ttt --- --- --- --- --- --- --- --- tct aat acc ata ata aaa att atc att atg --- --- --t ata atc a-- --- --- --- --- --- --- --- --- --- --- --- tca ctg --- --t cgc tat cat tat tgc gtt tgt gta gtt --- --- --- --- -ct gcc cta --- --- --- --- --- --- --- --- tca tct aca tca ctg tca --- --- --- --- -ct ctc act ata tct tct aaa att aca a-- --a caa c-- --- --- --- --- --- --- --- --- --- --- --- --- --- --- --- --- --- --- --- --- --- --- --- --- --- --- --- --- --- --- --- --- --- --- --- -tg gat att cga t-- --- --- --- --- --- --- --- --- --- --- --- --- --- --- --- --- --- --- --- --- --- --- --- --- --- --- aac agc att tgt gt- --- --- --- --- --- --- --- --- --- --- --- --- --- --- --- --- --

>KC257460.1_Monkeypox_virus_strain_DRC_Yandongi_1985_complete_genome

ttt ttt cga tct atc ctc gtc --- --- ctc atc atc ctt ata --- --- --- --- --- --- -tt att atc att att atc ata gtc tat taa aca caa atc atc t-- --- --- --- --- --- --- --- --- --- --- --- --- --- --- --- --- --- --- --- --- --- --- --- --- --- --- acg ttt ata ac- --- --- --- --- --- --- --- --- aac att c-- --- --- --- --t cat tat taa tta gtt ctg tag taa tat ctt taa taa ttt ggc tat a-- --- --c atc tgt t-- --- --- --- --- --- --- --- --- --- --- --- --- --- --- --- --- --- --- caa tac t-- --- --- --- --- --- --- --- atc tat tga tga ttt ctt tt- --- --- --- --- --- --- --- --- --- --- --- tta aga ct- --- --- --- --- --- --- --- --- --- --- --- --- --- --- --- --- --- --- --- --- --- --- --- --- --- --- --- --- --- --- --- --- --- --- --- --- --- --- --- -ta aac tag t-- --- --- --- --- --- --- --- --- --- --- --- --- --- --- --- --- --- --- --- --- --- --- --- --- --- --- --- --- --- --- --- --- --- --- --- --- --- --- --- --- --- --- --- --- --- --- --- --- --- --- --- --- --- --- --- --- --- --- --- --- --- --- --- --- --- --- --- --- --- --- --- --- --- --- --- --- --- --- --- --- --- --- --- --- --- --- --- --- --- --- --- --- --- --- --- --- --- --- --- --- --- --- --- --- --- --- --- --- --- tat ggt aat gac gat gaa a-- --t cga gta gta --- --- act tct aat aaa tac ttg ata --- --- tca tta tca tat gtt tga tcg --- --- --- --- --- --- --- tca tag tta ata gtg tg- --- --- --- --- --- --- --- --- --- --- --- --- --- --- --- --- --- --- --- --g cta aat ggt act gtt aat aag ttt at- --- --- --- --- --- --- --- --- --- --- --- --- --- --- --- --- --- aga caa tat cat agt att ttc ttt cca gaa t-- --- --- --- tag att att ttt tta aat act gat cct cac aat tcc gtg atg tag cag tag ttg gt- --- --- --- --- --- --- --- --- --- --- --- --- --- --- --- --- --- --- --- --- --- --- --- --- --- --- --- --g cat ggt cta tat cgt --- --- --- --- --- --- --- --- --- --- --- --- --- --- --- --- --- --- --- --- --- --- --- --- --- --- --- --- --- --- --- --- --- --- --- --- --- --- --- --- --- --- --- -ta aaa tgt atc ata tat aat agt ttt ctg acg tgg agt aca gaa ttt tcg a-- --- --- --- --- --- --- --- --- --- --- --- --- --- --- --- --- --- --- --- --- --- --- --- --- --- --- --- --- --- --- --- --- --- --- --- --- --- --- --- --- --- --- --- tta atg agt tca tgg taa gga agg gca aat gtc t-- -gt ata taa tat aca taa gtt aa- --- --- --- --- --- --- --- --- --- --- --- --- --- tag ttt ttt atc ata ttt --- --- --- --- --- --- --- --- tct aat acc ata ata aaa att atc att atg --- --- --t ata atc a-- --- --- --- --- --- --- --- --- --- --- --- tca ctg --- --t cgc tat cat tat tgc gtt tgt gta gtt --- --- --- --- -ct gcc cta --- --- --- --- --- --- --- --- tca tct aca tca ctg tca --- --- --- --- -ct ctc act ata tct tct aaa att aca a-- --a caa c-- --- --- --- --- --- --- --- --- --- --- --- --- --- --- --- --- --- --- --- --- --- --- --- --- --- --- --- --- --- --- --- --- --- --- --- -tg gat att cga t-- --- --- --- --- --- --- --- --- --- --- --- --- --- --- --- --- --- --- --- --- --- --- --- --- --- --- aac agc att tgt gt- --- --- --- --- --- --- --- --- --- --- --- --- --- --- --- --- --

>JX878429.1_Monkeypox_virus_isolate_DRC_07-0662_complete_genome

ttt ttt cga tct atc ctc gtc --- --- ctc atc atc ctt ata --- --- --- --- --- --- -tt att atc att att atc ata gtc tat taa aca caa atc atc t-- --- --- --- --- --- --- --- --- --- --- --- --- --- --- --- --- --- --- --- --- --- --- --- --- --- --- acg ttt ata ac- --- --- --- --- --- --- --- --- aac att c-- --- --- --- --t cat tat taa tta gtt ctg tag taa tat ctt taa taa ttt ggc tat a-- --- --c atc tgt t-- --- --- --- --- --- --- --- --- --- --- --- --- --- --- --- --- --- --- caa tac t-- --- --- --- --- --- --- --- atc tat tga tga ttt ctt tt- --- --- --- --- --- --- --- --- --- --- --- tta aga ct- --- --- --- --- --- --- --- --- --- --- --- --- --- --- --- --- --- --- --- --- --- --- --- --- --- --- --- --- --- --- --- --- --- --- --- --- --- --- --- -ta aac tag t-- --- --- --- --- --- --- --- --- --- --- --- --- --- --- --- --- --- --- --- --- --- --- --- --- --- --- --- --- --- --- --- --- --- --- --- --- --- --- --- --- --- --- --- --- --- --- --- --- --- --- --- --- --- --- --- --- --- --- --- --- --- --- --- --- --- --- --- --- --- --- --- --- --- --- --- --- --- --- --- --- --- --- --- --- --- --- --- --- --- --- --- --- --- --- --- --- --- --- --- --- --- --- --- --- --- --- --- --- --- tat ggt aat gac gat gaa a-- --t cga gta gta --- --- act tct aat aaa gac ttg ata --- --- tca tta tca tat gtt tga tcg --- --- --- --- --- --- --- tca tag tta ata gtg tg- --- --- --- --- --- --- --- --- --- --- --- --- --- --- --- --- --- --- --- --g cta aat ggt act gtt aat aag ttt at- --- --- --- --- --- --- --- --- --- --- --- --- --- --- --- --- --- aga caa tat cat agt att ttc ttt cca gaa t-- --- --- --- tag att att ttt tta aat act gat cct cac aat tcc gtg atg tag cag tag ttg gt- --- --- --- --- --- --- --- --- --- --- --- --- --- --- --- --- --- --- --- --- --- --- --- --- --- --- --- --g cat ggt cta tat cgt --- --- --- --- --- --- --- --- --- --- --- --- --- --- --- --- --- --- --- --- --- --- --- --- --- --- --- --- --- --- --- --- --- --- --- --- --- --- --- --- --- --- --- -ta aaa tgt atc ata tat aat agt ttt ctg acg tgg agt aca gaa ttt tcg a-- --- --- --- --- --- --- --- --- --- --- --- --- --- --- --- --- --- --- --- --- --- --- --- --- --- --- --- --- --- --- --- --- --- --- --- --- --- --- --- --- --- --- --- tta atg agt tca tgg taa gga agg gca aat gtc t-- -gt ata taa tat aca taa gtt aa- --- --- --- --- --- --- --- --- --- --- --- --- --- tag ttt ttt atc ata ttt --- --- --- --- --- --- --- --- tct aat acc ata ata aaa att atc att atg --- --- --t ata atc a-- --- --- --- --- --- --- --- --- --- --- --- tca ctg --- --t cgc tat cat tat tgc gtt tgt gta gtt --- --- --- --- -ct gcc cta --- --- --- --- --- --- --- --- tca tct aca tca ctg tca --- --- --- --- -ct ctc act ata tct tct aaa att aca a-- --a caa c-- --- --- --- --- --- --- --- --- --- --- --- --- --- --- --- --- --- --- --- --- --- --- --- --- --- --- --- --- --- --- --- --- --- --- --- -tg gat att cga t-- --- --- --- --- --- --- --- --- --- --- --- --- --- --- --- --- --- --- --- --- --- --- --- --- --- --- aac agc att tgt gt- --- --- --- --- --- --- --- --- --- --- --- --- --- --- --- --- --

>JX878425.1_Monkeypox_virus_isolate_DRC_07-0354_complete_genome

ttt ttt cga tct atc ctc gtc --- --- ctc atc atc ctt ata --- --- --- --- --- --- -tt att atc att att atc ata gtc tat taa aca caa atc atc t-- --- --- --- --- --- --- --- --- --- --- --- --- --- --- --- --- --- --- --- --- --- --- --- --- --- --- acg ttt ata ac- --- --- --- --- --- --- --- --- aac att c-- --- --- --- --t cat tat taa tta gtt ctg tag taa tat ctt taa taa ttt ggc tat a-- --- --c atc tgt t-- --- --- --- --- --- --- --- --- --- --- --- --- --- --- --- --- --- --- caa tac t-- --- --- --- --- --- --- --- atc tat tga tga ttt ctt tt- --- --- --- --- --- --- --- --- --- --- --- tta aga ct- --- --- --- --- --- --- --- --- --- --- --- --- --- --- --- --- --- --- --- --- --- --- --- --- --- --- --- --- --- --- --- --- --- --- --- --- --- --- --- -ta aac tag t-- --- --- --- --- --- --- --- --- --- --- --- --- --- --- --- --- --- --- --- --- --- --- --- --- --- --- --- --- --- --- --- --- --- --- --- --- --- --- --- --- --- --- --- --- --- --- --- --- --- --- --- --- --- --- --- --- --- --- --- --- --- --- --- --- --- --- --- --- --- --- --- --- --- --- --- --- --- --- --- --- --- --- --- --- --- --- --- --- --- --- --- --- --- --- --- --- --- --- --- --- --- --- --- --- --- --- --- --- --- tat ggt aat gac gat gaa a-- --t cga gta gta --- --- act tct aat aaa gac ttg ata --- --- tca tta tca tat gtt tga tcg --- --- --- --- --- --- --- tca tag tta ata gtg tg- --- --- --- --- --- --- --- --- --- --- --- --- --- --- --- --- --- --- --- --g cta aat ggt act gtt aat aag ttt at- --- --- --- --- --- --- --- --- --- --- --- --- --- --- --- --- --- aga caa tat cat agt att ttc ttt cca gaa t-- --- --- --- tag att att ttt tta aat act gat cct cac aat tcc gtg atg tag cag tag ttg gt- --- --- --- --- --- --- --- --- --- --- --- --- --- --- --- --- --- --- --- --- --- --- --- --- --- --- --- --g cat ggt cta tat cgt --- --- --- --- --- --- --- --- --- --- --- --- --- --- --- --- --- --- --- --- --- --- --- --- --- --- --- --- --- --- --- --- --- --- --- --- --- --- --- --- --- --- --- -ta aaa tgt atc ata tat aat agt ttt ctg acg tgg agt aca gaa ttt tcg a-- --- --- --- --- --- --- --- --- --- --- --- --- --- --- --- --- --- --- --- --- --- --- --- --- --- --- --- --- --- --- --- --- --- --- --- --- --- --- --- --- --- --- --- tta atg agt tca tgg taa gga agg gca aat gtc t-- -gt ata taa tat aca taa gtt aa- --- --- --- --- --- --- --- --- --- --- --- --- --- tag ttt ttt atc ata ttt --- --- --- --- --- --- --- --- tct aat acc ata ata aaa att atc att atg --- --- --t ata atc a-- --- --- --- --- --- --- --- --- --- --- --- tca ctg --- --t cgc tat cat tat tgc gtt tgt gta gtt --- --- --- --- -ct gcc cta --- --- --- --- --- --- --- --- tca tct aca tca ctg tca --- --- --- --- -ct ctc act ata tct tct aaa att aca a-- --a caa c-- --- --- --- --- --- --- --- --- --- --- --- --- --- --- --- --- --- --- --- --- --- --- --- --- --- --- --- --- --- --- --- --- --- --- --- -tg gat att cga t-- --- --- --- --- --- --- --- --- --- --- --- --- --- --- --- --- --- --- --- --- --- --- --- --- --- --- aac agc att tgt gt- --- --- --- --- --- --- --- --- --- --- --- --- --- --- --- --- --

>JX878407.1_Monkeypox_virus_isolate_DRC_06-0950_complete_genome

ttt ttt cga tct atc ctc gtc --- --- ctc atc atc ctt ata --- --- --- --- --- --- -tt att atc att att atc ata gtc tat taa aca caa atc atc t-- --- --- --- --- --- --- --- --- --- --- --- --- --- --- --- --- --- --- --- --- --- --- --- --- --- --- acg ttt ata ac- --- --- --- --- --- --- --- --- aac att c-- --- --- --- --t cat tat taa tta gtt ctg tag taa tat ctt taa taa ttt ggc tat a-- --- --c atc tgt t-- --- --- --- --- --- --- --- --- --- --- --- --- --- --- --- --- --- --- caa tac t-- --- --- --- --- --- --- --- atc tat tga tga ttt ctt tt- --- --- --- --- --- --- --- --- --- --- --- tta aga ct- --- --- --- --- --- --- --- --- --- --- --- --- --- --- --- --- --- --- --- --- --- --- --- --- --- --- --- --- --- --- --- --- --- --- --- --- --- --- --- -ta aac tag t-- --- --- --- --- --- --- --- --- --- --- --- --- --- --- --- --- --- --- --- --- --- --- --- --- --- --- --- --- --- --- --- --- --- --- --- --- --- --- --- --- --- --- --- --- --- --- --- --- --- --- --- --- --- --- --- --- --- --- --- --- --- --- --- --- --- --- --- --- --- --- --- --- --- --- --- --- --- --- --- --- --- --- --- --- --- --- --- --- --- --- --- --- --- --- --- --- --- --- --- --- --- --- --- --- --- --- --- --- --- tat ggt aat gac gat gaa a-- --t cga gta gta --- --- act tct aat aaa gac ttg ata --- --- tca tta tca tat gtt tga tcg --- --- --- --- --- --- --- tca tag tta ata gtg tg- --- --- --- --- --- --- --- --- --- --- --- --- --- --- --- --- --- --- --- --g cta aat ggt act gtt aat aag ttt at- --- --- --- --- --- --- --- --- --- --- --- --- --- --- --- --- --- aga caa tat cat agt att ttc ttt cca gaa t-- --- --- --- tag att att ttt tta aat act gat cct cac aat tcc gtg atg tag cag tag ttg gt- --- --- --- --- --- --- --- --- --- --- --- --- --- --- --- --- --- --- --- --- --- --- --- --- --- --- --- --g cat ggt cta tat cgt --- --- --- --- --- --- --- --- --- --- --- --- --- --- --- --- --- --- --- --- --- --- --- --- --- --- --- --- --- --- --- --- --- --- --- --- --- --- --- --- --- --- --- -ta aaa tgt atc ata tat aat agt ttt ctg acg tgg agt aca gaa ttt tcg a-- --- --- --- --- --- --- --- --- --- --- --- --- --- --- --- --- --- --- --- --- --- --- --- --- --- --- --- --- --- --- --- --- --- --- --- --- --- --- --- --- --- --- --- tta atg agt tca tgg taa gga agg gca aat gtc t-- -gt ata taa tat aca taa gtt aa- --- --- --- --- --- --- --- --- --- --- --- --- --- tag ttt ttt atc ata ttt --- --- --- --- --- --- --- --- tct aat acc ata ata aaa att atc att atg --- --- --t ata atc a-- --- --- --- --- --- --- --- --- --- --- --- tca ctg --- --t cgc tat cat tat tgc gtt tgt gta gtt --- --- --- --- -ct gcc cta --- --- --- --- --- --- --- --- tca tct aca tca ctg tca --- --- --- --- -ct ctc act ata tct tct aaa att aca a-- --a caa c-- --- --- --- --- --- --- --- --- --- --- --- --- --- --- --- --- --- --- --- --- --- --- --- --- --- --- --- --- --- --- --- --- --- --- --- -tg gat att cga t-- --- --- --- --- --- --- --- --- --- --- --- --- --- --- --- --- --- --- --- --- --- --- --- --- --- --- aac agc att tgt gt- --- --- --- --- --- --- --- --- --- --- --- --- --- --- --- --- --

>JX878423.1_Monkeypox_virus_isolate_DRC_07-0337_complete_genome

ttt ttt cga tct atc ctc gtc --- --- ctc atc atc ctt ata --- --- --- --- --- --- -tt att atc att att atc ata gtc tat taa aca caa atc atc t-- --- --- --- --- --- --- --- --- --- --- --- --- --- --- --- --- --- --- --- --- --- --- --- --- --- --- acg ttt ata ac- --- --- --- --- --- --- --- --- aac att c-- --- --- --- --t cat tat taa tta gtt ctg tag taa tat ctt taa taa ttt ggc tat a-- --- --c atc tgt t-- --- --- --- --- --- --- --- --- --- --- --- --- --- --- --- --- --- --- caa tac t-- --- --- --- --- --- --- --- atc tat tga tga ttt ctt tt- --- --- --- --- --- --- --- --- --- --- --- tta aga ct- --- --- --- --- --- --- --- --- --- --- --- --- --- --- --- --- --- --- --- --- --- --- --- --- --- --- --- --- --- --- --- --- --- --- --- --- --- --- --- -ta aac tag t-- --- --- --- --- --- --- --- --- --- --- --- --- --- --- --- --- --- --- --- --- --- --- --- --- --- --- --- --- --- --- --- --- --- --- --- --- --- --- --- --- --- --- --- --- --- --- --- --- --- --- --- --- --- --- --- --- --- --- --- --- --- --- --- --- --- --- --- --- --- --- --- --- --- --- --- --- --- --- --- --- --- --- --- --- --- --- --- --- --- --- --- --- --- --- --- --- --- --- --- --- --- --- --- --- --- --- --- --- --- tat ggt aat gac gat gaa a-- --t cga gta gta --- --- act tct aat aaa gac ttg ata --- --- tca tta tca tat gtt tga tcg --- --- --- --- --- --- --- tca tag tta ata gtg tg- --- --- --- --- --- --- --- --- --- --- --- --- --- --- --- --- --- --- --- --g cta aat ggt act gtt aat aag ttt at- --- --- --- --- --- --- --- --- --- --- --- --- --- --- --- --- --- aga caa tat cat agt att ttc ttt cca gaa t-- --- --- --- tag att att ttt tta aat act gat cct cac aat tcc gtg atg tag cag tag ttg gt- --- --- --- --- --- --- --- --- --- --- --- --- --- --- --- --- --- --- --- --- --- --- --- --- --- --- --- --g cat ggt cta tat cgt --- --- --- --- --- --- --- --- --- --- --- --- --- --- --- --- --- --- --- --- --- --- --- --- --- --- --- --- --- --- --- --- --- --- --- --- --- --- --- --- --- --- --- -ta aaa tgt atc ata tat aat agt ttt ctg acg tgg agt aca gaa ttt tcg a-- --- --- --- --- --- --- --- --- --- --- --- --- --- --- --- --- --- --- --- --- --- --- --- --- --- --- --- --- --- --- --- --- --- --- --- --- --- --- --- --- --- --- --- tta atg agt tca tgg taa gga agg gca aat gtc t-- -gt ata taa tat aca taa gtt aa- --- --- --- --- --- --- --- --- --- --- --- --- --- tag ttt ttt atc ata ttt --- --- --- --- --- --- --- --- tct aat acc ata ata aaa att atc att atg --- --- --t ata atc a-- --- --- --- --- --- --- --- --- --- --- --- tca ctg --- --t cgc tat cat tat tgc gtt tgt gta gtt --- --- --- --- -ct gcc cta --- --- --- --- --- --- --- --- tca tct aca tca ctg tca --- --- --- --- -ct ctc act ata tct tct aaa att aca a-- --a caa c-- --- --- --- --- --- --- --- --- --- --- --- --- --- --- --- --- --- --- --- --- --- --- --- --- --- --- --- --- --- --- --- --- --- --- --- -tg gat att cga t-- --- --- --- --- --- --- --- --- --- --- --- --- --- --- --- --- --- --- --- --- --- --- --- --- --- --- aac agc att tgt gt- --- --- --- --- --- --- --- --- --- --- --- --- --- --- --- --- --

>JX878408.1_Monkeypox_virus_isolate_DRC_06-0970_complete_genome

ttt ttt cga tct atc ctc gtc --- --- ctc atc atc ctt ata --- --- --- --- --- --- -tt att atc att att atc ata gtc tat taa aca caa atc atc t-- --- --- --- --- --- --- --- --- --- --- --- --- --- --- --- --- --- --- --- --- --- --- --- --- --- --- acg ttt ata ac- --- --- --- --- --- --- --- --- aac att c-- --- --- --- --t cat tat taa tta gtt ctg tag taa tat ctt taa taa ttt ggc tat a-- --- --c atc tgt t-- --- --- --- --- --- --- --- --- --- --- --- --- --- --- --- --- --- --- caa tac t-- --- --- --- --- --- --- --- atc tat tga tga ttt ctt tt- --- --- --- --- --- --- --- --- --- --- --- tta aga ct- --- --- --- --- --- --- --- --- --- --- --- --- --- --- --- --- --- --- --- --- --- --- --- --- --- --- --- --- --- --- --- --- --- --- --- --- --- --- --- -ta aac tag t-- --- --- --- --- --- --- --- --- --- --- --- --- --- --- --- --- --- --- --- --- --- --- --- --- --- --- --- --- --- --- --- --- --- --- --- --- --- --- --- --- --- --- --- --- --- --- --- --- --- --- --- --- --- --- --- --- --- --- --- --- --- --- --- --- --- --- --- --- --- --- --- --- --- --- --- --- --- --- --- --- --- --- --- --- --- --- --- --- --- --- --- --- --- --- --- --- --- --- --- --- --- --- --- --- --- --- --- --- --- tat ggt aat gac gat gaa a-- --t cga gta gta --- --- act tct aat aaa gac ttg ata --- --- tca tta tca tat gtt tga tcg --- --- --- --- --- --- --- tca tag tta ata gtg tg- --- --- --- --- --- --- --- --- --- --- --- --- --- --- --- --- --- --- --- --g cta aat ggt act gtt aat aag ttt at- --- --- --- --- --- --- --- --- --- --- --- --- --- --- --- --- --- aga caa tat cat agt att ttc ttt cca gaa t-- --- --- --- tag att att ttt tta aat act gat cct cac aat tcc gtg atg tag cag tag ttg gt- --- --- --- --- --- --- --- --- --- --- --- --- --- --- --- --- --- --- --- --- --- --- --- --- --- --- --- --g cat ggt cta tat cgt --- --- --- --- --- --- --- --- --- --- --- --- --- --- --- --- --- --- --- --- --- --- --- --- --- --- --- --- --- --- --- --- --- --- --- --- --- --- --- --- --- --- --- -ta aaa tgt atc ata tat aat agt ttt ctg acg tgg agt aca gaa ttt tcg a-- --- --- --- --- --- --- --- --- --- --- --- --- --- --- --- --- --- --- --- --- --- --- --- --- --- --- --- --- --- --- --- --- --- --- --- --- --- --- --- --- --- --- --- tta atg agt tca tgg taa gga agg gca aat gtc t-- -gt ata taa tat aca taa gtt aa- --- --- --- --- --- --- --- --- --- --- --- --- --- tag ttt ttt atc ata ttt --- --- --- --- --- --- --- --- tct aat acc ata ata aaa att atc att atg --- --- --t ata atc a-- --- --- --- --- --- --- --- --- --- --- --- tca ctg --- --t cgc tat cat tat tgc gtt tgt gta gtt --- --- --- --- -ct gcc cta --- --- --- --- --- --- --- --- tca tct aca tca ctg tca --- --- --- --- -ct ctc act ata tct tct aaa att aca a-- --a caa c-- --- --- --- --- --- --- --- --- --- --- --- --- --- --- --- --- --- --- --- --- --- --- --- --- --- --- --- --- --- --- --- --- --- --- --- -tg gat att cga t-- --- --- --- --- --- --- --- --- --- --- --- --- --- --- --- --- --- --- --- --- --- --- --- --- --- --- aac agc att tgt gt- --- --- --- --- --- --- --- --- --- --- --- --- --- --- --- --- --

>JX878420.1_Monkeypox_virus_isolate_DRC_07-0283_complete_genome

ttt ttt cga tct atc ctc gtc --- --- ctc atc atc ctt ata --- --- --- --- --- --- -tt att atc att att atc ata gtc tat taa aca caa atc atc t-- --- --- --- --- --- --- --- --- --- --- --- --- --- --- --- --- --- --- --- --- --- --- --- --- --- --- acg ttt ata ac- --- --- --- --- --- --- --- --- aac att c-- --- --- --- --t cat tat taa tta gtt ctg tag taa tat ctt taa taa ttt ggc tat a-- --- --c atc tgt t-- --- --- --- --- --- --- --- --- --- --- --- --- --- --- --- --- --- --- caa tac t-- --- --- --- --- --- --- --- atc tat tga tga ttt ctt tt- --- --- --- --- --- --- --- --- --- --- --- tta aga ct- --- --- --- --- --- --- --- --- --- --- --- --- --- --- --- --- --- --- --- --- --- --- --- --- --- --- --- --- --- --- --- --- --- --- --- --- --- --- --- -ta aac tag t-- --- --- --- --- --- --- --- --- --- --- --- --- --- --- --- --- --- --- --- --- --- --- --- --- --- --- --- --- --- --- --- --- --- --- --- --- --- --- --- --- --- --- --- --- --- --- --- --- --- --- --- --- --- --- --- --- --- --- --- --- --- --- --- --- --- --- --- --- --- --- --- --- --- --- --- --- --- --- --- --- --- --- --- --- --- --- --- --- --- --- --- --- --- --- --- --- --- --- --- --- --- --- --- --- --- --- --- --- --- tat ggt aat gac gat gaa a-- --t cga gta gta --- --- act tct aat aaa gac ttg ata --- --- tca tta tca tat gtt tga tcg --- --- --- --- --- --- --- tca tag tta ata gtg tg- --- --- --- --- --- --- --- --- --- --- --- --- --- --- --- --- --- --- --- --g cta aat ggt act gtt aat aag ttt at- --- --- --- --- --- --- --- --- --- --- --- --- --- --- --- --- --- aga caa tat cat agt att ttc ttt cca gaa t-- --- --- --- tag att att ttt tta aat act gat cct cac aat tcc gtg atg tag cag tag ttg gt- --- --- --- --- --- --- --- --- --- --- --- --- --- --- --- --- --- --- --- --- --- --- --- --- --- --- --- --g cat ggt cta tat cgt --- --- --- --- --- --- --- --- --- --- --- --- --- --- --- --- --- --- --- --- --- --- --- --- --- --- --- --- --- --- --- --- --- --- --- --- --- --- --- --- --- --- --- -ta aaa tgt atc ata tat aat agt ttt ctg acg tgg agt aca gaa ttt tcg a-- --- --- --- --- --- --- --- --- --- --- --- --- --- --- --- --- --- --- --- --- --- --- --- --- --- --- --- --- --- --- --- --- --- --- --- --- --- --- --- --- --- --- --- tta atg agt tca tgg taa gga agg gca aat gtc t-- -gt ata taa tat aca taa gtt aa- --- --- --- --- --- --- --- --- --- --- --- --- --- tag ttt ttt atc ata ttt --- --- --- --- --- --- --- --- tct aat acc ata ata aaa att atc att atg --- --- --t ata atc a-- --- --- --- --- --- --- --- --- --- --- --- tca ctg --- --t cgc tat cat tat tgc gtt tgt gta gtt --- --- --- --- -ct gcc cta --- --- --- --- --- --- --- --- tca tct aca tca ctg tca --- --- --- --- -ct ctc act ata tct tct aaa att aca a-- --a caa c-- --- --- --- --- --- --- --- --- --- --- --- --- --- --- --- --- --- --- --- --- --- --- --- --- --- --- --- --- --- --- --- --- --- --- --- -tg gat att cga t-- --- --- --- --- --- --- --- --- --- --- --- --- --- --- --- --- --- --- --- --- --- --- --- --- --- --- aac agc att tgt gt- --- --- --- --- --- --- --- --- --- --- --- --- --- --- --- --- --

>JX878419.1_Monkeypox_virus_isolate_DRC_07-0275_complete_genome

ttt ttt cga tct atc ctc gtc --- --- ctc atc atc ctt ata --- --- --- --- --- --- -tt att atc att att atc ata gtc tat taa aca caa atc atc t-- --- --- --- --- --- --- --- --- --- --- --- --- --- --- --- --- --- --- --- --- --- --- --- --- --- --- acg ttt ata ac- --- --- --- --- --- --- --- --- aac att c-- --- --- --- --t cat tat taa tta gtt ctg tag taa tat ctt taa taa ttt ggc tat a-- --- --c atc tgt t-- --- --- --- --- --- --- --- --- --- --- --- --- --- --- --- --- --- --- caa tac t-- --- --- --- --- --- --- --- atc tat tga tga ttt ctt tt- --- --- --- --- --- --- --- --- --- --- --- tta aga ct- --- --- --- --- --- --- --- --- --- --- --- --- --- --- --- --- --- --- --- --- --- --- --- --- --- --- --- --- --- --- --- --- --- --- --- --- --- --- --- -ta aac tag t-- --- --- --- --- --- --- --- --- --- --- --- --- --- --- --- --- --- --- --- --- --- --- --- --- --- --- --- --- --- --- --- --- --- --- --- --- --- --- --- --- --- --- --- --- --- --- --- --- --- --- --- --- --- --- --- --- --- --- --- --- --- --- --- --- --- --- --- --- --- --- --- --- --- --- --- --- --- --- --- --- --- --- --- --- --- --- --- --- --- --- --- --- --- --- --- --- --- --- --- --- --- --- --- --- --- --- --- --- --- tat ggt aat gac gat gaa a-- --t cga gta gta --- --- act tct aat aaa gac ttg ata --- --- tca tta tca tat gtt tga tcg --- --- --- --- --- --- --- tca tag tta ata gtg tg- --- --- --- --- --- --- --- --- --- --- --- --- --- --- --- --- --- --- --- --g cta aat ggt act gtt aat aag ttt at- --- --- --- --- --- --- --- --- --- --- --- --- --- --- --- --- --- aga caa tat cat agt att ttc ttt cca gaa t-- --- --- --- tag att att ttt tta aat act gat cct cac aat tcc gtg atg tag cag tag ttg gt- --- --- --- --- --- --- --- --- --- --- --- --- --- --- --- --- --- --- --- --- --- --- --- --- --- --- --- --g cat ggt cta tat cgt --- --- --- --- --- --- --- --- --- --- --- --- --- --- --- --- --- --- --- --- --- --- --- --- --- --- --- --- --- --- --- --- --- --- --- --- --- --- --- --- --- --- --- -ta aaa tgt atc ata tat aat agt ttt ctg acg tgg agt aca gaa ttt tcg a-- --- --- --- --- --- --- --- --- --- --- --- --- --- --- --- --- --- --- --- --- --- --- --- --- --- --- --- --- --- --- --- --- --- --- --- --- --- --- --- --- --- --- --- tta atg agt tca tgg taa gga agg gca aat gtc t-- -gt ata taa tat aca taa gtt aa- --- --- --- --- --- --- --- --- --- --- --- --- --- tag ttt ttt atc ata ttt --- --- --- --- --- --- --- --- tct aat acc ata ata aaa att atc att atg --- --- --t ata atc a-- --- --- --- --- --- --- --- --- --- --- --- tca ctg --- --t cgc tat cat tat tgc gtt tgt gta gtt --- --- --- --- -ct gcc cta --- --- --- --- --- --- --- --- tca tct aca tca ctg tca --- --- --- --- -ct ctc act ata tct tct aaa att aca a-- --a caa c-- --- --- --- --- --- --- --- --- --- --- --- --- --- --- --- --- --- --- --- --- --- --- --- --- --- --- --- --- --- --- --- --- --- --- --- -tg gat att cga t-- --- --- --- --- --- --- --- --- --- --- --- --- --- --- --- --- --- --- --- --- --- --- --- --- --- --- aac agc att tgt gt- --- --- --- --- --- --- --- --- --- --- --- --- --- --- --- --- --

>JX878417.1_Monkeypox_virus_isolate_DRC_07-0104_complete_genome

ttt ttt cga tct atc ctc gtc --- --- ctc atc atc ctt ata --- --- --- --- --- --- -tt att atc att att atc ata gtc tat taa aca caa atc atc t-- --- --- --- --- --- --- --- --- --- --- --- --- --- --- --- --- --- --- --- --- --- --- --- --- --- --- acg ttt ata ac- --- --- --- --- --- --- --- --- aac att c-- --- --- --- --t cat tat taa tta gtt ctg tag taa tat ctt taa taa ttt ggc tat a-- --- --c atc tgt t-- --- --- --- --- --- --- --- --- --- --- --- --- --- --- --- --- --- --- caa tac t-- --- --- --- --- --- --- --- atc tat tga tga ttt ctt tt- --- --- --- --- --- --- --- --- --- --- --- tta aga ct- --- --- --- --- --- --- --- --- --- --- --- --- --- --- --- --- --- --- --- --- --- --- --- --- --- --- --- --- --- --- --- --- --- --- --- --- --- --- --- -ta aac tag t-- --- --- --- --- --- --- --- --- --- --- --- --- --- --- --- --- --- --- --- --- --- --- --- --- --- --- --- --- --- --- --- --- --- --- --- --- --- --- --- --- --- --- --- --- --- --- --- --- --- --- --- --- --- --- --- --- --- --- --- --- --- --- --- --- --- --- --- --- --- --- --- --- --- --- --- --- --- --- --- --- --- --- --- --- --- --- --- --- --- --- --- --- --- --- --- --- --- --- --- --- --- --- --- --- --- --- --- --- --- tat ggt aat ga- gat gaa a-- --t cga gta gta --- --- act tct aat aaa gac ttg ata --- --- tca tta tca tat gtt tga tcg --- --- --- --- --- --- --- tca tag tta ata gtg tg- --- --- --- --- --- --- --- --- --- --- --- --- --- --- --- --- --- --- --- --g cta aat ggt act gtt aat aag ttt at- --- --- --- --- --- --- --- --- --- --- --- --- --- --- --- --- --- aga caa tat cat agt att ttc ttt cca gaa t-- --- --- --- tag att att ttt tta aat act gat cct cac aat tcc gtg atg tag cag tag ttg gt- --- --- --- --- --- --- --- --- --- --- --- --- --- --- --- --- --- --- --- --- --- --- --- --- --- --- --- --g cat ggt cta tat cgt --- --- --- --- --- --- --- --- --- --- --- --- --- --- --- --- --- --- --- --- --- --- --- --- --- --- --- --- --- --- --- --- --- --- --- --- --- --- --- --- --- --- --- -ta aaa tgt atc ata tat aat agt ttt ctg acg tgg agt aca gaa ttt tcg a-- --- --- --- --- --- --- --- --- --- --- --- --- --- --- --- --- --- --- --- --- --- --- --- --- --- --- --- --- --- --- --- --- --- --- --- --- --- --- --- --- --- --- --- tta atg agt tca tgg taa gga agg gca aat gtc t-- -gt ata taa tat aca taa gtt aa- --- --- --- --- --- --- --- --- --- --- --- --- --- tag ttt ttt atc ata ttt --- --- --- --- --- --- --- --- tct aat acc ata ata aaa att atc att atg --- --- --t ata atc a-- --- --- --- --- --- --- --- --- --- --- --- tca ctg --- --t cgc tat cat tat tgc gtt tgt gta gtt --- --- --- --- -ct gcc cta --- --- --- --- --- --- --- --- tca tct aca tca ctg tca --- --- --- --- -ct ctc act ata tct tct aaa att aca a-- --a caa c-- --- --- --- --- --- --- --- --- --- --- --- --- --- --- --- --- --- --- --- --- --- --- --- --- --- --- --- --- --- --- --- --- --- --- --- -tg gat att cga t-- --- --- --- --- --- --- --- --- --- --- --- --- --- --- --- --- --- --- --- --- --- --- --- --- --- --- aac agc att tgt gt- --- --- --- --- --- --- --- --- --- --- --- --- --- --- --- --- --

>JX878426.1_Monkeypox_virus_isolate_DRC_07-0450_complete_genome

ttt ttt cga tct atc ctc gtc --- --- ctc atc atc ctt ata --- --- --- --- --- --- -tt att atc att att atc ata gtc tat taa aca caa atc atc t-- --- --- --- --- --- --- --- --- --- --- --- --- --- --- --- --- --- --- --- --- --- --- --- --- --- --- acg ttt ata ac- --- --- --- --- --- --- --- --- aac att c-- --- --- --- --t cat tat taa tta gtt ctg tag taa tat ctt taa taa ttt ggc tat a-- --- --c atc tgt t-- --- --- --- --- --- --- --- --- --- --- --- --- --- --- --- --- --- --- caa tac t-- --- --- --- --- --- --- --- atc tat tga tga ttt ctt tt- --- --- --- --- --- --- --- --- --- --- --- tta aga ct- --- --- --- --- --- --- --- --- --- --- --- --- --- --- --- --- --- --- --- --- --- --- --- --- --- --- --- --- --- --- --- --- --- --- --- --- --- --- --- -ta aac tag t-- --- --- --- --- --- --- --- --- --- --- --- --- --- --- --- --- --- --- --- --- --- --- --- --- --- --- --- --- --- --- --- --- --- --- --- --- --- --- --- --- --- --- --- --- --- --- --- --- --- --- --- --- --- --- --- --- --- --- --- --- --- --- --- --- --- --- --- --- --- --- --- --- --- --- --- --- --- --- --- --- --- --- --- --- --- --- --- --- --- --- --- --- --- --- --- --- --- --- --- --- --- --- --- --- --- --- --- --- --- tat ggt aat gac gat gaa a-- --t cga gta gta --- --- act tct aat aaa gac ttg ata --- --- tca tta tca tat gtt tga tcg --- --- --- --- --- --- --- tca tag tta ata gtg tg- --- --- --- --- --- --- --- --- --- --- --- --- --- --- --- --- --- --- --- --g cta aat ggt act gtt aat aag ttt at- --- --- --- --- --- --- --- --- --- --- --- --- --- --- --- --- --- aga caa tat cat agt att ttc ttt cca gaa t-- --- --- --- tag att att ttt tta aat act gat cct cac aat tcc gtg atg tag cag tag ttg gt- --- --- --- --- --- --- --- --- --- --- --- --- --- --- --- --- --- --- --- --- --- --- --- --- --- --- --- --g cat ggt cta tat cgt --- --- --- --- --- --- --- --- --- --- --- --- --- --- --- --- --- --- --- --- --- --- --- --- --- --- --- --- --- --- --- --- --- --- --- --- --- --- --- --- --- --- --- -ta aaa tgt atc ata tat aat agt ttt ctg acg tgg agt aca gaa ttt tcg a-- --- --- --- --- --- --- --- --- --- --- --- --- --- --- --- --- --- --- --- --- --- --- --- --- --- --- --- --- --- --- --- --- --- --- --- --- --- --- --- --- --- --- --- tta atg agt tca tgg taa gga agg gca aat gtc t-- -gt ata taa tat aca taa gtt aa- --- --- --- --- --- --- --- --- --- --- --- --- --- tag ttt ttt atc ata ttt --- --- --- --- --- --- --- --- tct aat acc ata ata aaa att atc att atg --- --- --t ata atc a-- --- --- --- --- --- --- --- --- --- --- --- tca ctg --- --t cgc tat cat tat tgc gtt tgt gta gtt --- --- --- --- -ct gcc cta --- --- --- --- --- --- --- --- tca tct aca tca ctg tca --- --- --- --- -ct ctc act ata tct tct aaa att aca a-- --a caa c-- --- --- --- --- --- --- --- --- --- --- --- --- --- --- --- --- --- --- --- --- --- --- --- --- --- --- --- --- --- --- --- --- --- --- --- -tg gat att cga t-- --- --- --- --- --- --- --- --- --- --- --- --- --- --- --- --- --- --- --- --- --- --- --- --- --- --- aac agc att tgt gt- --- --- --- --- --- --- --- --- --- --- --- --- --- --- --- --- --

>KC257459.1_Monkeypox_virus_strain_Sudan_2005_01_complete_genome

ttt ttt cga tct atc ctc gtc --- --- ctc atc atc ctt ata --- --- --- --- --- --- -tt att atc att att atc ata gtc tat taa aca caa atc atc t-- --- --- --- --- --- --- --- --- --- --- --- --- --- --- --- --- --- --- --- --- --- --- --- --- --- --- acg ttt ata ac- --- --- --- --- --- --- --- --- aac att c-- --- --- --- --t cat tat taa tta gtt ctg tag taa tat ctt taa taa ttt ggc tat a-- --- --c atc tgt t-- --- --- --- --- --- --- --- --- --- --- --- --- --- --- --- --- --- --- caa tac t-- --- --- --- --- --- --- --- atc tat tga tga ttt ctt tt- --- --- --- --- --- --- --- --- --- --- --- tta aga ct- --- --- --- --- --- --- --- --- --- --- --- --- --- --- --- --- --- --- --- --- --- --- --- --- --- --- --- --- --- --- --- --- --- --- --- --- --- --- --- -ta aac tag t-- --- --- --- --- --- --- --- --- --- --- --- --- --- --- --- --- --- --- --- --- --- --- --- --- --- --- --- --- --- --- --- --- --- --- --- --- --- --- --- --- --- --- --- --- --- --- --- --- --- --- --- --- --- --- --- --- --- --- --- --- --- --- --- --- --- --- --- --- --- --- --- --- --- --- --- --- --- --- --- --- --- --- --- --- --- --- --- --- --- --- --- --- --- --- --- --- --- --- --- --- --- --- --- --- --- --- --- --- --- tat ggt aat gac gat gaa a-- --t cga gta gta --- --- act tct aat aaa tac ttg ata --- --- tca tta tca tat gtt tga tcg --- --- --- --- --- --- --- tca tag tta ata gtg tg- --- --- --- --- --- --- --- --- --- --- --- --- --- --- --- --- --- --- --- --g cta aat ggt act gtt aat aag ttt at- --- --- --- --- --- --- --- --- --- --- --- --- --- --- --- --- --- aga caa tat cat agt att ttc ttt cca gaa t-- --- --- --- tag att att ttt tta aat act gat cct cac aat tcc gtg atg tag cag tag ttg gt- --- --- --- --- --- --- --- --- --- --- --- --- --- --- --- --- --- --- --- --- --- --- --- --- --- --- --- --g cat ggt cta tat cgt --- --- --- --- --- --- --- --- --- --- --- --- --- --- --- --- --- --- --- --- --- --- --- --- --- --- --- --- --- --- --- --- --- --- --- --- --- --- --- --- --- --- --- -ta aaa tgt atc ata tat aat agt ttt ctg acg tgg agt aca gaa ttt tcg a-- --- --- --- --- --- --- --- --- --- --- --- --- --- --- --- --- --- --- --- --- --- --- --- --- --- --- --- --- --- --- --- --- --- --- --- --- --- --- --- --- --- --- --- tta atg agt tca tgg taa gga agg gca aat gtc t-- -gt ata taa tat aca taa gtt aa- --- --- --- --- --- --- --- --- --- --- --- --- --- tag ttt ttt atc ata ttt --- --- --- --- --- --- --- --- tct aat acc ata ata aaa att atc att atg --- --- --t ata atc a-- --- --- --- --- --- --- --- --- --- --- --- tca ctg --- --t cgc tat cat tat tgc gtt tgt gta gtt --- --- --- --- -ct gcc cta --- --- --- --- --- --- --- --- tca tct aca tca ctg tca --- --- --- --- -ct ctc act ata tct tct aaa att aca a-- --a caa c-- --- --- --- --- --- --- --- --- --- --- --- --- --- --- --- --- --- --- --- --- --- --- --- --- --- --- --- --- --- --- --- --- --- --- --- -tg gat att cga t-- --- --- --- --- --- --- --- --- --- --- --- --- --- --- --- --- --- --- --- --- --- --- --- --- --- --- aac agc att tgt gt- --- --- --- --- --- --- --- --- --- --- --- --- --- --- --- --- --

>MN702446.1_Monkeypox_virus_strain_38c_contig_SPADES

ttt ttt cga tct atc ctc gtc --- --- ctc atc atc ctt ata --- --- --- --- --- --- -tt att atc att att atc ata gtc tat taa aca caa atc atc t-- --- --- --- --- --- --- --- --- --- --- --- --- --- --- --- --- --- --- --- --- --- --- --- --- --- --- acg ttt ata ac- --- --- --- --- --- --- --- --- aac att c-- --- --- --- --t cat tat taa tta gtt ctg tag taa tat ctt taa taa ttt ggc tat a-- --- --c atc tgt t-- --- --- --- --- --- --- --- --- --- --- --- --- --- --- --- --- --- --- caa tac t-- --- --- --- --- --- --- --- atc tat tga tga ttt ctt tt- --- --- --- --- --- --- --- --- --- --- --- tta aga ct- --- --- --- --- --- --- --- --- --- --- --- --- --- --- --- --- --- --- --- --- --- --- --- --- --- --- --- --- --- --- --- --- --- --- --- --- --- --- --- -ta aac tag t-- --- --- --- --- --- --- --- --- --- --- --- --- --- --- --- --- --- --- --- --- --- --- --- --- --- --- --- --- --- --- --- --- --- --- --- --- --- --- --- --- --- --- --- --- --- --- --- --- --- --- --- --- --- --- --- --- --- --- --- --- --- --- --- --- --- --- --- --- --- --- --- --- --- --- --- --- --- --- --- --- --- --- --- --- --- --- --- --- --- --- --- --- --- --- --- --- --- --- --- --- --- --- --- --- --- --- --- --- --- tat ggt aat gac gat gaa a-- --t cga gta gta --- --- act tct aat aaa gac ttg ata --- --- tca tta tca tat gtt tga tcg --- --- --- --- --- --- --- tca tag tta ata gtg tg- --- --- --- --- --- --- --- --- --- --- --- --- --- --- --- --- --- --- --- --g cta aat ggt act gtt aat aag ttt at- --- --- --- --- --- --- --- --- --- --- --- --- --- --- --- --- --- aga caa tat cat agt att ttc ttt cca gaa t-- --- --- --- tag att att ttt tta aat act gat cct cac aat tcc gtg atg tag cag tag ttg gt- --- --- --- --- --- --- --- --- --- --- --- --- --- --- --- --- --- --- --- --- --- --- --- --- --- --- --- --g cat ggt cta tat cgt --- --- --- --- --- --- --- --- --- --- --- --- --- --- --- --- --- --- --- --- --- --- --- --- --- --- --- --- --- --- --- --- --- --- --- --- --- --- --- --- --- --- --- -ta aaa tgt atc ata tat aat agt ttt ctg acg tgg agt aca gaa ttt tcg a-- --- --- --- --- --- --- --- --- --- --- --- --- --- --- --- --- --- --- --- --- --- --- --- --- --- --- --- --- --- --- --- --- --- --- --- --- --- --- --- --- --- --- --- tta atg agt tca tgg taa gga agg gca aat gtc t-- -gt ata taa tat aca taa gtt aa- --- --- --- --- --- --- --- --- --- --- --- --- --- tag ttt ttt atc ata ttt --- --- --- --- --- --- --- --- tct aat acc ata ata aaa att atc att atg --- --- --t ata atc a-- --- --- --- --- --- --- --- --- --- --- --- tca ctg --- --t cgc tat cat tat tgc gtt tgt gta gtt --- --- --- --- -ct gcc cta --- --- --- --- --- --- --- --- tca tct aca tca ctg tca --- --- --- --- -ct ctc act ata tct tct aaa att aca a-- --a caa c-- --- --- --- --- --- --- --- --- --- --- --- --- --- --- --- --- --- --- --- --- --- --- --- --- --- --- --- --- --- --- --- --- --- --- --- -tg gat att cga t-- --- --- --- --- --- --- --- --- --- --- --- --- --- --- --- --- --- --- --- --- --- --- --- --- --- --- aac agc att tgt gt- --- --- --- --- --- --- --- --- --- --- --- --- --- --- --- --- --

>JX878418.1_Monkeypox_virus_isolate_DRC_07-0120_complete_genome

ttt ttt cga tct atc ctc gtc --- --- ctc atc atc ctt ata --- --- --- --- --- --- -tt att atc att att atc ata gtc tat taa aca caa atc atc t-- --- --- --- --- --- --- --- --- --- --- --- --- --- --- --- --- --- --- --- --- --- --- --- --- --- --- acg ttt ata ac- --- --- --- --- --- --- --- --- aac att c-- --- --- --- --t cat tat taa tta gtt ctg tag taa tat ctt taa taa ttt ggc tat a-- --- --c atc tgt t-- --- --- --- --- --- --- --- --- --- --- --- --- --- --- --- --- --- --- caa tac t-- --- --- --- --- --- --- --- atc tat tga tga ttt ctt tt- --- --- --- --- --- --- --- --- --- --- --- tta aga ct- --- --- --- --- --- --- --- --- --- --- --- --- --- --- --- --- --- --- --- --- --- --- --- --- --- --- --- --- --- --- --- --- --- --- --- --- --- --- --- -ta aac tag t-- --- --- --- --- --- --- --- --- --- --- --- --- --- --- --- --- --- --- --- --- --- --- --- --- --- --- --- --- --- --- --- --- --- --- --- --- --- --- --- --- --- --- --- --- --- --- --- --- --- --- --- --- --- --- --- --- --- --- --- --- --- --- --- --- --- --- --- --- --- --- --- --- --- --- --- --- --- --- --- --- --- --- --- --- --- --- --- --- --- --- --- --- --- --- --- --- --- --- --- --- --- --- --- --- --- --- --- --- --- tat ggt aat gac gat gaa a-- --t cga gta gta --- --- act tct aat aaa gac ttg ata --- --- tca tta tca tat gtt tga tcg --- --- --- --- --- --- --- tca tag tta ata gtg tg- --- --- --- --- --- --- --- --- --- --- --- --- --- --- --- --- --- --- --- --g cta aat ggt act gtt aat aag ttt at- --- --- --- --- --- --- --- --- --- --- --- --- --- --- --- --- --- aga caa tat cat agt att ttc ttt cca gaa t-- --- --- --- tag att att ttt tta aat act gat cct cac aat tcc gtg atg tag cag tag ttg gt- --- --- --- --- --- --- --- --- --- --- --- --- --- --- --- --- --- --- --- --- --- --- --- --- --- --- --- --g cat ggt cta tat cgt --- --- --- --- --- --- --- --- --- --- --- --- --- --- --- --- --- --- --- --- --- --- --- --- --- --- --- --- --- --- --- --- --- --- --- --- --- --- --- --- --- --- --- -ta aaa tgt atc ata tat aat agt ttt ctg acg tgg agt aca gaa ttt tcg a-- --- --- --- --- --- --- --- --- --- --- --- --- --- --- --- --- --- --- --- --- --- --- --- --- --- --- --- --- --- --- --- --- --- --- --- --- --- --- --- --- --- --- --- tta atg agt tca tgg taa gga agg gca aat gtc t-- -gt ata taa tat aca taa gtt aa- --- --- --- --- --- --- --- --- --- --- --- --- --- tag ttt ttt atc ata ttt --- --- --- --- --- --- --- --- tct aat acc ata ata aaa att atc att atg --- --- --t ata atc a-- --- --- --- --- --- --- --- --- --- --- --- tca ctg --- --t cgc tat cat tat tgc gtt tgt gta gtt --- --- --- --- -ct gcc cta --- --- --- --- --- --- --- --- tca tct aca tca ctg tca --- --- --- --- -ct ctc act ata tct tct aaa att aca a-- --a caa c-- --- --- --- --- --- --- --- --- --- --- --- --- --- --- --- --- --- --- --- --- --- --- --- --- --- --- --- --- --- --- --- --- --- --- --- -tg gat att cga t-- --- --- --- --- --- --- --- --- --- --- --- --- --- --- --- --- --- --- --- --- --- --- --- --- --- --- aac agc att tgt gt- --- --- --- --- --- --- --- --- --- --- --- --- --- --- --- --- --

>KJ642613.1_Monkeypox_virus_strain_Congo_8_complete_genome

ttt ttt cga tct atc ctc gtc --- --- ctc atc atc ctt ata --- --- --- --- --- --- -tt att atc att att atc ata gtc tat taa aca caa atc atc t-- --- --- --- --- --- --- --- --- --- --- --- --- --- --- --- --- --- --- --- --- --- --- --- --- --- --- acg ttt ata ac- --- --- --- --- --- --- --- --- aac att c-- --- --- --- --t cat tat taa tta gtt ctg tag taa tat ctt taa taa ttt ggc tat a-- --- --c atc tgt t-- --- --- --- --- --- --- --- --- --- --- --- --- --- --- --- --- --- --- caa tac t-- --- --- --- --- --- --- --- atc tat tga tga ttt ctt tt- --- --- --- --- --- --- --- --- --- --- --- tta aga ct- --- --- --- --- --- --- --- --- --- --- --- --- --- --- --- --- --- --- --- --- --- --- --- --- --- --- --- --- --- --- --- --- --- --- --- --- --- --- --- -ta aac tag t-- --- --- --- --- --- --- --- --- --- --- --- --- --- --- --- --- --- --- --- --- --- --- --- --- --- --- --- --- --- --- --- --- --- --- --- --- --- --- --- --- --- --- --- --- --- --- --- --- --- --- --- --- --- --- --- --- --- --- --- --- --- --- --- --- --- --- --- --- --- --- --- --- --- --- --- --- --- --- --- --- --- --- --- --- --- --- --- --- --- --- --- --- --- --- --- --- --- --- --- --- --- --- --- --- --- --- --- --- --- tat ggt aat gac gat gaa a-- --t cga gta gta --- --- act tct aat aaa gac ttg ata --- --- tca tta tca tat gtt tga tcg --- --- --- --- --- --- --- tca tag tta ata gtg tg- --- --- --- --- --- --- --- --- --- --- --- --- --- --- --- --- --- --- --- --g cta aat ggt act gtt aat aag ttt at- --- --- --- --- --- --- --- --- --- --- --- --- --- --- --- --- --- aga caa tat cat agt att ttc ttt cca gaa t-- --- --- --- tag att att ttt tta aat act gat cct cac aat tcc gtg atg tag cag tag ttg gt- --- --- --- --- --- --- --- --- --- --- --- --- --- --- --- --- --- --- --- --- --- --- --- --- --- --- --- --g cat ggt cta tat cgt --- --- --- --- --- --- --- --- --- --- --- --- --- --- --- --- --- --- --- --- --- --- --- --- --- --- --- --- --- --- --- --- --- --- --- --- --- --- --- --- --- --- --- -ta aaa tgt atc ata tat aat agt ttt ctg acg tgg agt aca gaa ttt tcg a-- --- --- --- --- --- --- --- --- --- --- --- --- --- --- --- --- --- --- --- --- --- --- --- --- --- --- --- --- --- --- --- --- --- --- --- --- --- --- --- --- --- --- --- tta atg agt tca tgg taa gga agg gca aat gtc t-- -gt ata taa tat aca taa gtt aa- --- --- --- --- --- --- --- --- --- --- --- --- --- tag ttt ttt atc ata ttt --- --- --- --- --- --- --- --- tct aat acc ata ata aaa att atc att atg --- --- --t ata atc a-- --- --- --- --- --- --- --- --- --- --- --- tca ctg --- --t cgc tat cat tat tgc gtt tgt gta gtt --- --- --- --- -ct gcc cta --- --- --- --- --- --- --- --- tca tct aca tca ctg tca --- --- --- --- -ct ctc act ata tct tct aaa att aca a-- --a caa c-- --- --- --- --- --- --- --- --- --- --- --- --- --- --- --- --- --- --- --- --- --- --- --- --- --- --- --- --- --- --- --- --- --- --- --- -tg gat att cga t-- --- --- --- --- --- --- --- --- --- --- --- --- --- --- --- --- --- --- --- --- --- --- --- --- --- --- aac agc att tgt gt- --- --- --- --- --- --- --- --- --- --- --- --- --- --- --- --- --

>HM172544.1_Monkeypox_virus_strain_Zaire_1979-005_complete_genome

ttt ttt cga tct atc ctc gtc --- --- ctc atc atc ctt ata --- --- --- --- --- --- -tt att atc att att atc ata gtc tat taa aca caa atc atc t-- --- --- --- --- --- --- --- --- --- --- --- --- --- --- --- --- --- --- --- --- --- --- --- --- --- --- acg ttt ata ac- --- --- --- --- --- --- --- --- aac att c-- --- --- --- --t cat tat taa tta gtt ctg tag taa tat ctt taa taa ttt ggc tat a-- --- --c atc tgt t-- --- --- --- --- --- --- --- --- --- --- --- --- --- --- --- --- --- --- caa tac t-- --- --- --- --- --- --- --- atc tat tga tga ttt ctt tt- --- --- --- --- --- --- --- --- --- --- --- tta aga ct- --- --- --- --- --- --- --- --- --- --- --- --- --- --- --- --- --- --- --- --- --- --- --- --- --- --- --- --- --- --- --- --- --- --- --- --- --- --- --- -ta aac tag t-- --- --- --- --- --- --- --- --- --- --- --- --- --- --- --- --- --- --- --- --- --- --- --- --- --- --- --- --- --- --- --- --- --- --- --- --- --- --- --- --- --- --- --- --- --- --- --- --- --- --- --- --- --- --- --- --- --- --- --- --- --- --- --- --- --- --- --- --- --- --- --- --- --- --- --- --- --- --- --- --- --- --- --- --- --- --- --- --- --- --- --- --- --- --- --- --- --- --- --- --- --- --- --- --- --- --- --- --- --- tat ggt aat gac gat gaa a-- --t cga gta gta --- --- act tct aat aaa tac ttg ata --- --- tca tta tca tat gtt tga tcg --- --- --- --- --- --- --- tca tag tta ata gtg tg- --- --- --- --- --- --- --- --- --- --- --- --- --- --- --- --- --- --- --- --g cta aat ggt act gtt aat aag ttt at- --- --- --- --- --- --- --- --- --- --- --- --- --- --- --- --- --- aga caa tat cat agt att ttc ttt cca gaa t-- --- --- --- tag att att ttt tta aat act gat cct cac aat tcc gtg atg tag cag tag ttg gt- --- --- --- --- --- --- --- --- --- --- --- --- --- --- --- --- --- --- --- --- --- --- --- --- --- --- --- --g cat ggt cta tat cgt --- --- --- --- --- --- --- --- --- --- --- --- --- --- --- --- --- --- --- --- --- --- --- --- --- --- --- --- --- --- --- --- --- --- --- --- --- --- --- --- --- --- --- -ta aaa tgt atc ata tat aat agt ttt ctg acg tgg agt aca gaa ttt tcg a-- --- --- --- --- --- --- --- --- --- --- --- --- --- --- --- --- --- --- --- --- --- --- --- --- --- --- --- --- --- --- --- --- --- --- --- --- --- --- --- --- --- --- --- tta atg agt tca tgg taa gga agg aca aat gtc t-- -gt ata taa tat aca taa gtt aa- --- --- --- --- --- --- --- --- --- --- --- --- --- tag ttt ttt atc ata ttt --- --- --- --- --- --- --- --- tct aat acc ata ata aaa att atc att atg --- --- --t ata atc a-- --- --- --- --- --- --- --- --- --- --- --- tca ctg --- --t cgc tat cat tat tgc gtt tgt gta gtt --- --- --- --- -ct gcc cta --- --- --- --- --- --- --- --- tca tct aca tca ctg tca --- --- --- --- -ct ctc act ata tct tct aaa att aca a-- --a caa c-- --- --- --- --- --- --- --- --- --- --- --- --- --- --- --- --- --- --- --- --- --- --- --- --- --- --- --- --- --- --- --- --- --- --- --- -tg gat att cga t-- --- --- --- --- --- --- --- --- --- --- --- --- --- --- --- --- --- --- --- --- --- --- --- --- --- --- aac agc att tgt gt- --- --- --- --- --- --- --- --- --- --- --- --- --- --- --- --- --

>AF380138.1_Monkeypox_virus_strain_Zaire-96-I-16_complete_genome

ttt ttt cga tct atc ctc gtc --- --- ctc atc atc ctt ata --- --- --- --- --- --- -tt att atc att att atc ata gtc tat taa aca caa atc atc t-- --- --- --- --- --- --- --- --- --- --- --- --- --- --- --- --- --- --- --- --- --- --- --- --- --- --- acg ttt ata ac- --- --- --- --- --- --- --- --- aac att c-- --- --- --- --t cat tat taa tta gtt ctg tag taa tat ctt taa taa ttt ggc tat a-- --- --c atc tgt t-- --- --- --- --- --- --- --- --- --- --- --- --- --- --- --- --- --- --- caa tac t-- --- --- --- --- --- --- --- atc tat tga tga ttt ctt tt- --- --- --- --- --- --- --- --- --- --- --- tta aga ct- --- --- --- --- --- --- --- --- --- --- --- --- --- --- --- --- --- --- --- --- --- --- --- --- --- --- --- --- --- --- --- --- --- --- --- --- --- --- --- -ta aac tag t-- --- --- --- --- --- --- --- --- --- --- --- --- --- --- --- --- --- --- --- --- --- --- --- --- --- --- --- --- --- --- --- --- --- --- --- --- --- --- --- --- --- --- --- --- --- --- --- --- --- --- --- --- --- --- --- --- --- --- --- --- --- --- --- --- --- --- --- --- --- --- --- --- --- --- --- --- --- --- --- --- --- --- --- --- --- --- --- --- --- --- --- --- --- --- --- --- --- --- --- --- --- --- --- --- --- --- --- --- --- tat ggt aat gac gat gaa a-- --t cga gta gta --- --- act tct aat aaa gac ttg ata --- --- tca tta tca tat gtt tga tcg --- --- --- --- --- --- --- tca tag tta ata gtg tg- --- --- --- --- --- --- --- --- --- --- --- --- --- --- --- --- --- --- --- --g cta aat ggt act gtt aat aag ttt at- --- --- --- --- --- --- --- --- --- --- --- --- --- --- --- --- --- aga caa tat cat agt att ttc ttt cca gaa t-- --- --- --- tag att att ttt tta aat act gat cct cac aat tcc gtg atg tag cag tag ttg gt- --- --- --- --- --- --- --- --- --- --- --- --- --- --- --- --- --- --- --- --- --- --- --- --- --- --- --- --g cat ggt cta tat cgt --- --- --- --- --- --- --- --- --- --- --- --- --- --- --- --- --- --- --- --- --- --- --- --- --- --- --- --- --- --- --- --- --- --- --- --- --- --- --- --- --- --- --- -ta aaa tgt atc ata tat aat agt ttt ctg acg tgg agt aca gaa ttt tcg a-- --- --- --- --- --- --- --- --- --- --- --- --- --- --- --- --- --- --- --- --- --- --- --- --- --- --- --- --- --- --- --- --- --- --- --- --- --- --- --- --- --- --- --- tta atg agt tca tgg taa gga agg gca aat gtc t-- -gt ata taa tat aca taa gtt aa- --- --- --- --- --- --- --- --- --- --- --- --- --- tag ttt ttt atc ata ttt --- --- --- --- --- --- --- --- tct aat acc ata ata aaa att atc att atg --- --- --t ata atc a-- --- --- --- --- --- --- --- --- --- --- --- tca ctg --- --t cgc tat cat tat tgc gtt tgt gta gtt --- --- --- --- -ct gcc cta --- --- --- --- --- --- --- --- tca tct aca tca ctg tca --- --- --- --- -ct ctc act ata tct tct aaa att aca a-- --a caa c-- --- --- --- --- --- --- --- --- --- --- --- --- --- --- --- --- --- --- --- --- --- --- --- --- --- --- --- --- --- --- --- --- --- --- --- -tg gat att cga t-- --- --- --- --- --- --- --- --- --- --- --- --- --- --- --- --- --- --- --- --- --- --- --- --- --- --- aac agc att tgt gt- --- --- --- --- --- --- --- --- --- --- --- --- --- --- --- --- --

>HQ857562.1_Monkeypox_virus_strain_V79-I-005_complete_genome

ttt ttt cga tct atc ctc gtc --- --- ctc atc atc ctt ata --- --- --- --- --- --- -tt att atc att att atc ata gtc tat taa aca caa atc atc t-- --- --- --- --- --- --- --- --- --- --- --- --- --- --- --- --- --- --- --- --- --- --- --- --- --- --- acg ttt ata ac- --- --- --- --- --- --- --- --- aac att c-- --- --- --- --t cat tat taa tta gtt ctg tag taa tat ctt taa taa ttt ggc tat a-- --- --c atc tgt t-- --- --- --- --- --- --- --- --- --- --- --- --- --- --- --- --- --- --- caa tac t-- --- --- --- --- --- --- --- atc tat tga tga ttt ctt tt- --- --- --- --- --- --- --- --- --- --- --- tta aga ct- --- --- --- --- --- --- --- --- --- --- --- --- --- --- --- --- --- --- --- --- --- --- --- --- --- --- --- --- --- --- --- --- --- --- --- --- --- --- --- -ta aac tag t-- --- --- --- --- --- --- --- --- --- --- --- --- --- --- --- --- --- --- --- --- --- --- --- --- --- --- --- --- --- --- --- --- --- --- --- --- --- --- --- --- --- --- --- --- --- --- --- --- --- --- --- --- --- --- --- --- --- --- --- --- --- --- --- --- --- --- --- --- --- --- --- --- --- --- --- --- --- --- --- --- --- --- --- --- --- --- --- --- --- --- --- --- --- --- --- --- --- --- --- --- --- --- --- --- --- --- --- --- --- tat ggt aat gac gat gaa a-- --t cga gta gta --- --- act tct aat aaa tac ttg ata --- --- tca tta tca tat gtt tga tcg --- --- --- --- --- --- --- tca tag tta ata gtg tg- --- --- --- --- --- --- --- --- --- --- --- --- --- --- --- --- --- --- --- --g cta aat ggt act gtt aat aag ttt at- --- --- --- --- --- --- --- --- --- --- --- --- --- --- --- --- --- aga caa tat cat agt att ttc ttt cca gaa t-- --- --- --- tag att att ttt tta aat act gat cct cac aat tcc gtg atg tag cag tag ttg gt- --- --- --- --- --- --- --- --- --- --- --- --- --- --- --- --- --- --- --- --- --- --- --- --- --- --- --- --g cat ggt cta tat cgt --- --- --- --- --- --- --- --- --- --- --- --- --- --- --- --- --- --- --- --- --- --- --- --- --- --- --- --- --- --- --- --- --- --- --- --- --- --- --- --- --- --- --- -ta aaa tgt atc ata tat aat agt ttt ctg acg tgg agt aca gaa ttt tcg a-- --- --- --- --- --- --- --- --- --- --- --- --- --- --- --- --- --- --- --- --- --- --- --- --- --- --- --- --- --- --- --- --- --- --- --- --- --- --- --- --- --- --- --- tta atg agt tca tgg taa gga agg gca aat gtc t-- -gt ata taa tat aca taa gtt aa- --- --- --- --- --- --- --- --- --- --- --- --- --- tag ttt ttt atc ata ttt --- --- --- --- --- --- --- --- tct aat acc ata ata aaa att atc att atg --- --- --t ata atc a-- --- --- --- --- --- --- --- --- --- --- --- tca ctg --- --t cgc tat cat tat tgc gtt tgt gta gtt --- --- --- --- -ct gcc cta --- --- --- --- --- --- --- --- tca tct aca tca ctg tca --- --- --- --- -ct ctc act ata tct tct aaa att aca a-- --a caa c-- --- --- --- --- --- --- --- --- --- --- --- --- --- --- --- --- --- --- --- --- --- --- --- --- --- --- --- --- --- --- --- --- --- --- --- -tg gat att cga t-- --- --- --- --- --- --- --- --- --- --- --- --- --- --- --- --- --- --- --- --- --- --- --- --- --- --- aac agc att tgt gt- --- --- --- --- --- --- --- --- --- --- --- --- --- --- --- --- --

>HQ857563.1_Monkeypox_virus_strain_D14L_knockout_complete_genome

ttt ttt cga tct atc ctc gtc --- --- ctc atc atc ctt ata --- --- --- --- --- --- -tt att atc att att atc ata gtc tat taa aca caa atc atc t-- --- --- --- --- --- --- --- --- --- --- --- --- --- --- --- --- --- --- --- --- --- --- --- --- --- --- acg ttt ata ac- --- --- --- --- --- --- --- --- aac att c-- --- --- --- --t cat tat taa tta gtt ctg tag taa tat ctt taa taa ttt ggc tat a-- --- --c atc tgt t-- --- --- --- --- --- --- --- --- --- --- --- --- --- --- --- --- --- --- caa tac t-- --- --- --- --- --- --- --- atc tat tga tga ttt ctt tt- --- --- --- --- --- --- --- --- --- --- --- tta aga ct- --- --- --- --- --- --- --- --- --- --- --- --- --- --- --- --- --- --- --- --- --- --- --- --- --- --- --- --- --- --- --- --- --- --- --- --- --- --- --- -ta aac tag t-- --- --- --- --- --- --- --- --- --- --- --- --- --- --- --- --- --- --- --- --- --- --- --- --- --- --- --- --- --- --- --- --- --- --- --- --- --- --- --- --- --- --- --- --- --- --- --- --- --- --- --- --- --- --- --- --- --- --- --- --- --- --- --- --- --- --- --- --- --- --- --- --- --- --- --- --- --- --- --- --- --- --- --- --- --- --- --- --- --- --- --- --- --- --- --- --- --- --- --- --- --- --- --- --- --- --- --- --- --- tat ggt aat gac gat gaa a-- --t cga gta gta --- --- act tct aat aaa tac ttg ata --- --- tca tta tca tat gtt tga tcg --- --- --- --- --- --- --- tca tag tta ata gtg tg- --- --- --- --- --- --- --- --- --- --- --- --- --- --- --- --- --- --- --- --g cta aat ggt act gtt aat aag ttt at- --- --- --- --- --- --- --- --- --- --- --- --- --- --- --- --- --- aga caa tat cat agt att ttc ttt cca gaa t-- --- --- --- tag att att ttt tta aat act gat cct cac aat tcc gtg atg tag cag tag ttg gt- --- --- --- --- --- --- --- --- --- --- --- --- --- --- --- --- --- --- --- --- --- --- --- --- --- --- --- --g cat ggt cta tat cgt --- --- --- --- --- --- --- --- --- --- --- --- --- --- --- --- --- --- --- --- --- --- --- --- --- --- --- --- --- --- --- --- --- --- --- --- --- --- --- --- --- --- --- -ta aaa tgt atc ata tat aat agt ttt ctg acg tgg agt aca gaa ttt tcg a-- --- --- --- --- --- --- --- --- --- --- --- --- --- --- --- --- --- --- --- --- --- --- --- --- --- --- --- --- --- --- --- --- --- --- --- --- --- --- --- --- --- --- --- tta atg agt tca tgg taa gga agg gca aat gtc t-- -gt ata taa tat aca taa gtt aa- --- --- --- --- --- --- --- --- --- --- --- --- --- tag ttt ttt atc ata ttt --- --- --- --- --- --- --- --- tct aat acc ata ata aaa att atc att atg --- --- --t ata atc a-- --- --- --- --- --- --- --- --- --- --- --- tca ctg --- --t cgc tat cat tat tgc gtt tgt gta gtt --- --- --- --- -ct gcc cta --- --- --- --- --- --- --- --- tca tct aca tca ctg tca --- --- --- --- -ct ctc act ata tct tct aaa att aca a-- --a caa c-- --- --- --- --- --- --- --- --- --- --- --- --- --- --- --- --- --- --- --- --- --- --- --- --- --- --- --- --- --- --- --- --- --- --- --- -tg gat att cga t-- --- --- --- --- --- --- --- --- --- --- --- --- --- --- --- --- --- --- --- --- --- --- --- --- --- --- aac agc att tgt gt- --- --- --- --- --- --- --- --- --- --- --- --- --- --- --- --- --
