## Supplementary Data 3 for "Genomic regions insertion and deletion in Monkeypox virus causing multi-country outbreak-2022"

>ON674051.1_Monkeypox_virus_isolate_MPXV_USA_2022_FL001_complete_genome

atc gag gac gac ttc aac tat gac agc tcg gtg gcg tct gcc agc gtg tac ata cga atg gca ttt cta aga aaa gtc tac ggt atc ctt -tc tac aat ttc ctt taa caa cgg caa cag ctg cag tat ttt tat act ttg aat gca -tc gga cat tta tac aag gga gtc ctg ttc taa tat tgg cat caa tgt tcg gat cta tag gct tga ttt tcg cat tga ctt tac aca gac atg aac atc ccc tga atc tgt aca tac ttt gtg gat tta cac tgt tag aat ctc taa cgc tgg cct ctg ttg tta ctt tct atg atg cac gta tcg tta tgc aag ctt tca tgt tga cta ctg cag tgt ttc ttg ctc tga cta cat ata ctc tac aat caa aga gag att tca gta aac ttg taa cag gat tgt ttg ctg ctt tct gga ttt taa ttt tgt cag gag tct tga gga taa agt tta aaa tag aat taa taa aga aca tat agg tca -tt ttt taa aca tgg ata gaa acc aag gtt gtt agt taa taa tat aca aga tat ttt ttc tca ctc tga tcc atg taa acc aag gac gag aga cac tct cat tcc tca ttc acg aca cca tta aaa atg gaa att aaa gcc ctc tat taa gca cag acg gct aca ggt cta cca tca ggt t-a cct tcg tct acc ttc aca atg gcc tct cct tgt gcc cag ttc agt ccc tgt cat tgc cac gct act aag gac tcc ctg aat acc gtg act gac gtc aga cat tgt ctg act gaa tac atc ctg tgg gtt tct cat aga tgg acc cat aga gaa agc gca ggg cct ctc tac agg ctt ctc atc tct ttc aga att gat gca atg gag cta ttt ggt agc gag ttg aag gag ttc tcg aat tca ctt ccg tgg gac aat atc gac aat tgc gtg gag atc att aaa tgt ttc atc aga aat gac tcc atg aaa acc gcc aaa gaa ctt tgt gca ata att gga ctt tgt act caa tca gct att gtc act gga aga gtc ttc aat gat aag tat atc gac ata cta ctt atg ctg cga aag att ctg aac gag aac gac tat ctc acc ctc ttg gat cat atc ctc act gct aaa tac taa atc tcc ttc atg ctc tct cac taa tac tct tac tca cta cac ttt tta tca tct tat gat gaa tga ttg cct tca tca ttt ttt cgt gga ata taa tat agg aat aat tag cac cag aat agc tat gga tat ctc gtt aag aat att ctc tat aag aga cat aat gta gac ata gtt att ata tcc ttc tta gat aag tgt tac gct act gga aag ttt cca tcg tta tta tta cat gaa gat gat ata att aaa cca aca ttg aga ttg gct ctt atg tta gct gga ttg aat tac tgt aat aaa tgc atc gag tat aga ggg ata tag caa ttc tcg ata ata gtc atg caa tat ttg aat gag act gat aat tta ggt aat aca gta ct- --- aca cac ata tct ttc tag att ata tat cgt taa aaa tct gta aga tgt ata ttt ctc ata agt atc cac tgt gta ata tta tta atg gat ata tag ata acg caa tag gga cta ata gta ttg taa aag ata taa tcg act att tac gta cat atc cag ata tct ata ttc cta cta gtt tgc tgc gta gtt gca tca ttg ata tgc atg -at tta tca gga ttc aga gat gaa tta cta agt aaa cta caa tcc cac aat aag taa gaa tca aat atc aaa aac tca ctt ttg att ttt cta gtc tta agt aat aca tat att tat taa tag acc tat gaa ata aaa aaa ggt aac aat gga ttc gcg tat agc tat tta cgt att agt atc ggc atc tct ttt gta tct tgt taa ttg tca caa act agt aca tta ctt caa tct gaa aat aaa tgg aag tga tat aac taa tac agc aga tat att gct gga caa tta tcc aat tat gac ctt tga tgg aaa gga tat tta tcc atc tat ctc gtt cat ggt cgg taa taa act ttt cct aga tct tta taa aaa tat ctt tga aga att ttt cag act att tcg agt atc tgt aag tag tca ata cga gga att aga ata tta tta ttc atg tga tta tac taa caa ccg tcc tac aat taa aca aca tta ctt tta taa cgg cga cga ata tac tga aat tga tag atc gaa aaa agc cac taa taa aaa cag ttg gtt aat tac ttc agg ctt tag act aca aaa atg gtt cga tag cga aga ttg tat aat tta tct cag atc ttt agt tag aag aat gga aga cag taa caa aaa cag taa aaa aac tta gta ctt aga tat cg- --a aaa aat ata ttt ttg tag act ctt gag aat aga agg aaa aca tgt aca taa tta taa aaa atg aaa atc aat ggc gaa taa gac agt gcg att cgc acc atg gag tcg gta gat ttc atg gct gtc gat gag cag ttt cac gac gac ctc gat ctt tgg tca tta tct ttg gta gat gat tat aaa aaa cat gga tta ggt gtt gac tgt tat gtt cta gaa cca gtt gtt gac agg aaa ata ttt gat aga ttt ctc ctt gaa cca att tgt gat cct gta gat gtt ctg tat gat tat ttt agg att cat aga gat aat att gat cag tat ata gta gat aga ctg ttt gca tat att aca tat aaa gat att ata tct gca tta gtg tca aag aat tat atg gaa gat att ttc tct ata att att aag aat tgt aat tct gtg caa gat ctc tta ctt tac tat cta tct aat gca tat gta gaa ata gac att gtt gat ctt atg gta gat cat ggg gct gta ata tat aaa ata gaa tgc ttg aat gcc tat ttt agg gga ata tgt aaa aag gaa agt agt gtt gtt gag ttt att ttg aat tgt ggt atc cca gat gaa aat gat gtt aaa tta gat cta tat aaa ata att cag tat act agg gga ttc ctt gta gat gaa ccc aca gta tta gaa att tat aag ctt tgt atc cca tat att gaa gat atc aat caa cta gat gct ggt gga agg acc ttg ctt tat cgc gct atc tat gca ggt tat ata gat tta gta tca tgg cta tta gaa aat gga gca aat gtc aac gca gta atg agt aat gga tat aca tgt ctt gac gtg gcc gtg gat agg gga tct gtc atc gcc cgt agg gaa gca cat ctt aaa ata tta gaa ata ttg ctt aga gaa cca ttg tct att gac tgt ata aaa tta gct ata ctt aat aat aca att gaa aac cat gat gtg ata aag ctc tgt atc aag tat ttt atg atg gta gat tat tca ctt tgt aat gtg tat gca tca tca ctc ttt gat tat ata att gat tgt aaa caa gaa ttg gag tac att agg cag atg aaa att cat aat aca acc atg tat gag tta atc tat aat aga gac aaa aac aag cat gct tcc cat att cta cat agg tat tct aaa cat cca gtt ttg aca cag tgt atc act aaa gga ttc aag att tac aca gaa gta acc gag cag gtc act aaa gct cta aac aga cgt gct cta ata gat gag ata ata aac aat gta tca act gat gac aat ctc cta tca aaa ctt cca tta gaa att agg gat cta att gtt tca caa gct gtc ata tag agt tct atc cac cca cct ttc ttg aaa tga gtt aat --- --- --- --- --- --- --- --- --- --- --- --- --- --- --- --- --- --- --- --- --- --- --- --- --- --- --- --- --- --- --- --- --- --- --- --- --- --- --- --- --- --- --- --- --- --- --- --- --- --- --- --- --- --- --- --- --- --- --- --- --- --- --- --- --- --- --- --- --- --- --- --- --- --- --- --- --- --- --- --- --- --- --- --- --- --- --- --- --- --- --- --- --- --- --- --- --- --- --- --- --- --- --- --- --- -ag tca taa gtt agt taa --- --- --- --- --- -gt cat aag tta gtt aag tca t-- --- --- --- --- --- --- --- --- --- --- --- --- --- --- --- -aa gtt agt taa gtc ata agt tag tta agt cat aag tta gtt aag tca taa gtt agt taa gtc ata agt tag tta agt cat aag tta gtt tat agt cta aca ctt cta att ttt ata cct tga tct ttt tct cta att atg aaa aag taa atc att atg aag atg gat g-a aat gga cga gat tgt gcg cat cgt taa cga tag tat gtg gta cgt acc taa cgc att tat gga cga cgg tga taa tga agg tca cat ttc tgt caa taa tgt ctg tca tat gta tct cgc att ctt tga tgt gga tat atc atc tca tct gtt taa att agt tat taa aca ctg cga tct gaa taa acg act aaa atg tgg taa ctc tcc att aca ttg cta tac gat gaa tac acg att taa tcc atc tgt att aaa gat att gtt acg cca cgg cat gcg taa ctt tga tag caa gga taa aaa agg aca tat tcc tct aca cca cta tct gat tca ttc act atc aat cga taa caa gat ctt tga tat act aac gga ccc cat tga tga ctt tag taa atc atc cga tct att gct gtg tta tct tag ata taa att caa tgg gag ctt aaa cta tta cgt tct gta caa att att gac taa agg atc tga ccc taa ttg cgt cga tga gga tgg act cac ttc tct tca tta cta ctg taa aca cat atc cgc gtt cca cga aag caa tta tta caa gtc aaa gag tca cac taa gat gcg agc tga gaa gcg att cat cta cgc gat aat aga tca tgg agc aaa cat taa cgc ggt tac gaa aat cgg aaa tac gcc gtt aca cac tta cct tca aca gta tac caa aca tag tcc tcg tgt ggt gta tgc tct ttt atc tcg agg agc cga tac gag gat acg taa taa tct tga ttg tac acc cat cat gga ata cat aaa gaa cga ttg tgc aac agg tca tat tct cat aat gtt act caa ttg gca cga aca aaa ata cgg gaa att aca aaa gga aga agg aca aca tct act tta tct att cat aaa aca taa tca agg ata tgg aag tcg ctc tct caa tat act acg gta tct act aga tag att cga cat tca gaa aga cga ata cta taa tac aat gac tcc tct tca tac cgc ctt cca gaa ttg caa taa caa tgt tgc ctc ata cct cgt ata cat cgg ata cga cat caa cct tcc gac taa aga cga taa gac agt att cga ctt ggt gtt tga aaa cag aaa cat tat ata caa ggc gga tgt cgt taa tga cat tat cca cca cag act gaa agt atc tct acc tat gat taa atc gtt gtt cta caa gat gtc gga gtt ctc tcc cta cga cga tca cta cgt aaa gaa gat aat agc cta ctg cct att aag gga cga gtc att tgc gga act aca tac taa att ctg ttt aaa cga gga cta taa aag tgt att tat gaa aaa tat atc att cga taa gat aga ttc cat cat cga aaa atg tag tcg tga cat aag tct cct caa aga gat tcg aat ctc aga cac cga ctt gta tac ggt att gag aac aga aga cat ccg gta tca cac ata tct cga agc cat aca ttc aga caa acg cat ttc att tcc cat gta cga cga tct cat aga aca gtg tca tct atc gat gga gca taa aag taa act cgt cga caa agc act caa taa att aga gtc tac cat cga tag tca atc tag act atc gta ttt gcc tcc gga aat tat gcg caa tat cat aac caa gct aag cga cta cca tct aaa cag tat gtt gta cgg aaa gaa cca tta caa ata tta tcc atg ata gaa aga aaa tat tta aaa aat aat cta tat gat tgg aga agt agg aaa caa aca gta aca aga cga cga tta cta c-a tta tta aat cat gag gtc cgt att ata ctc gta tat att gtt tct ctc atg tat aat aat aaa cgg aag aga tat agc acc aca tgc acc atc caa tgg aaa gtg taa aga caa cga ata cag aag ccg taa tct atg ttg tct atc gtg tcc tcc ggg aac tta cgc ttc cag att atg tga tag caa gac taa tac aca atg tac acc gtg tgg ttc gga tac ctt tac atc tca caa taa tca ttt aca ggc ttg tct aag ttg taa cgg aag atg tga tag taa tca ggt aga gac gcg atc gtg taa cac gac tca caa tag aat ctg tga atg ctc tcc agg ata tta ttg tct tct caa agg agc atc agg gtg tag aac atg tat ttc taa aac aaa gtg tgg aat agg ata cgg agt atc cgg ata cac gtc tac cgg aga cgt cat ctg ttc tcc gtg tgg tcc cgg aac ata ttc tca cac cgt ctc ttc cac aga taa atg cga acc cgt cgt aac cag caa tac att taa cta tat cga tgt gga aat taa cct gta tcc agt caa cga cac atc gtg tac tcg gac gac cac tac cgg tct cag cga atc cat ctc aac gtc gga act aac tat tac cat gaa tca taa aga ttg tga tcc agt ctt tcg tgc aga ata ctt ctc tgt cct taa taa tgt agc aac ttc agg att ctt tac agg aga aaa tag ata tca gaa tac ttc aaa gat atg tac tct gaa ttt cga gat taa atg taa caa caa aga ttc atc ttc caa aca gtt aac gaa aac aaa gaa tga tac tat cat gcc gca ttc aga gac ggt aac tct agt ggg cga ctg tct atc tag cgt cga cat cta cat act ata tag taa tac caa tac tca aga cta cga aaa tga tac aat ctc tta tca tat ggg taa tgt tct cga tgt caa tag cca tat gcc cgc tag ttg cga tat aca taa act gat cac taa ttc cca gaa tcc cac cca ctt ata gta ag- -tt ttt tta cct ata aat aat aaa tac aat aat taa ttt ctc gta aaa gta gaa aat ata ttc taa ttt att ata tgg taa gaa agt aga atc atc tag aac agt aat caa tca ata gca atc atg aaa caa tat att gtc ctg gca tgc atg tgc cta gtg gca gct gct atg cct act agt ctt caa caa tct tca tcc tcg tgt act gaa gaa gaa aac aaa cat cat atg gga atc gat gtt att atc aaa gtc aca aag caa gac caa aca ccg acc aat gat aag att tgt caa tcc gta acg gaa gtt aca gag a-- --- --- --- --- --- -cc gaa gat gat gag gta tcc gaa gaa gtt gta aaa gga gat ccc acc act tat tac act atc gtc ggt gcg ggt ctt aac atg aac ttt gga ttc acc aaa tgc cca aag att tca tcc atc tcc gaa tcc tct gat gga aac act gtg aat act aga ttg tcc agc gtg tca ccg gga caa ggt aag gac tct ccc gcg atc acg cgt gaa gaa gct ctg gct atg atc aaa gac tgt gag atg tct atc gac atc aga tgt agc gaa gaa gag aaa gac agt gac atc aag acc cat cca gta ctt ggg tct aac atc tca cat aag aaa gtg agt tac aaa gat atc atc ggt tca acg atc gtt gat aca aaa tgt gtc aag aac cta gag ttt agc gta cgt atc gga gac atg tgt gag gaa tca tct gaa ctt gaa gtc aag gat gga ttc aag tat gtc gac gga tcg gca tct gaa ggt gca acc gat gat act tca ctc atc gat tca aca aaa ctc aaa gca tgt gtc tga atc gat aac tct att cat ctg aaa atg gat gag ttg ggt taa tcg aac gat tca gac acc gca cca cga att aaa aaa g-- -ac cgg gca cta tat tcc ggt ttg caa aac aaa aat att taa cta cat tca caa aaa gtt acc tct cgt tac ttc ttc ttt ctg ttt caa tat gtg ata cga tat gat cac tat tcg tat tct ct- --- --- tgg tct cat aaa aaa gtt tta c-- --a aaa aaa aaa aaa a?t att ttt att ctc ttt ctc tct tcg at- ggt ctc aca aaa ata tta aac ctc ttt ctg atg tct caa cta ttt cgt aaa cga taa cgt cca aca ata tat tct cgt aga gct tat caa cat cct tat acc aat cta ggt tgt cag aca att gca tca taa aat aat gtt tat aat tta cac gtt aac atc ata taa taa acg tat ata gtt aat att ttt gga ata taa atg atc tgt aaa atc cat gta ggg gac act gct cac gtt ttt tct cta gta cat aat ttc aca caa gtt ttt ata cag aca aat taa ttc tcg tcc ata ta- ttt taa aac att gac ttt tgt act aag aaa aat atc ttg act aac cat ctc ttt ctc tct tcg atg ggt ctc aca aaa ata tta aac ctc ttt ctg atg g-- --- --- --- --- --- --- --- --- --- --- --- --- --- --- --- --- --- --- --- --- --- --- --- --- --- --- --- --- --- --- --- --- --- --- --- --- --- --- --- --- --- --- --- --- --- --- agt cgt aaa aag ttt tta tcc ttt ctc tct tcg ata ggt ctc --- --- --- --- --- --- --- --- --- --- --- --- --- --- --- --- --- --- --- --- --- --- --- --- --- --- --- --- --- --- --- --- --- --- --- --- --- --- --- --- --- --- --- --- --- --- --- --- --- --- --- --- --- --- --- --- --- --- --- --- --- --- --- --- --- --- --- --- --- --- aca aaa ata tta aac ctc ttt ctg atg gtc tct ata aa

>NC_063383.1_Monkeypox_virus_complete_genome

atc gag gac gac ttc aac tat gac agc tcg gtg gcg tct gcc agc gtg tac ata cga atg gca ttt cta aga aaa gtc tac ggt atc ctt -tc tac aat ttc ctt taa caa cgg caa cag ctg cag tat ttt tat act ttg aat gca -tc gga cat tta tac aag gga gtc ctg ttc taa tat tgg cat caa tgt tcg gat cta tag gct tga ttt tcg cat tga ctt tac aca gac atg aac atc ccc tga atc tgt aca tac ttt gtg gat tta cac tgt tag aat ctc taa cgc tgg cct ctg ttg tta ctt tct atg atg cac gta tcg tta tgc aag ctt tca tgt tga cta ctg cag tgt ttc ttg ctc tga cta cat ata ctc tac aat caa aga gag att tca gta aac ttg taa cag gat tgt ttg ctg ctt tct gga ttt taa ttt tgt cag gag tct tga gga taa agt tta aaa tag aat taa taa aga aca tat agg tca -tt ttt taa aca tgg ata gaa acc aag gtt gtt agt taa taa tat aca aga tat ttt ttc tca ctc tga tcc atg taa acc aag gac gag aga cac tct cat tcc tca ttc acg aca cca tta aaa atg gaa att aaa gcc ctc tat taa gca cag acg gct aca ggt cta cca tca ggt t-a cct tcg tct acc ttc aca atg gcc tct cct tgt gcc cag ttc agt ccc tgt cat tgc cac gct act aag gac tcc ctg aat acc gtg act gac gtc aga cat tgt ctg act gaa tac atc ctg tgg gtt tct cat aga tgg acc cat aga gaa agc gca ggg cct ctc tac agg ctt ctc atc tct ttc aga att gat gca atg gag cta ttt ggt agc gag ttg aag gag ttc tcg aat tca ctt ccg tgg gac aat atc gac aat tgc gtg gag atc att aaa tgt ttc atc aga aat gac tcc atg aaa acc gcc aaa gaa ctt tgt gca ata att gga ctt tgt act caa tca gct att gtc act gga aga gtc ttc aat gat aag tat atc gac ata cta ctt atg ctg cga aag att ctg aac gag aac gac tat ctc acc ctc ttg gat cat atc ctc act gct aaa tac taa atc tcc ttc atg ctc tct cac taa tac tct tac tca cta cac ttt tta tca tct tat gat gaa tga ttg cct tca tca ttt ttt cgt gga ata taa tat agg aat aat tag cac cag aat agc tat gga tat ctc gtt aag aat att ctc tat aag aga cat aat gta gac ata gtt att ata tcc ttc tta gat aag tgt tac gct act gga aag ttt cca tcg tta tta tta cat gaa gat gat ata att aaa cca aca ttg aga ttg gct ctt atg tta gct gga ttg aat tac tgt aat aaa tgc atc gag tat aga ggg ata tag caa ttc tcg ata ata gtc atg caa tat ttg aat gag act gat aat tta ggt aat aca gta ct- --- aca cac ata tct ttc tag att ata tat cgt taa aaa tct gta aga tgt ata ttt ctc ata agt atc cac tgt gta ata tta tta atg gat ata tag ata acg caa tag gga cta ata gta ttg taa aag ata taa tcg act att tac gta cat atc cag ata tct ata ttc cta cta gtt tgc tgc gta gtt gca tca ttg ata tgc atg -at tta tca gga ttc aga gat gaa tta cta agt aaa cta caa tcc cac aat aag taa gaa tca aat atc aaa aac tca ctt ttg att ttt cta gtc tta agt aat aca tat att tat taa tag acc tat gaa ata aaa aaa ggt aac aat gga ttc gcg tat agc tat tta cgt att agt atc ggc atc tct ttt gta tct tgt taa ttg tca caa act agt aca tta ctt caa tct gaa aat aaa tgg aag tga tat aac taa tac agc aga tat att gct gga caa tta tcc aat tat gac ctt tga tgg aaa gga tat tta tcc atc tat ctc gtt cat ggt cgg taa taa act ttt cct aga tct tta taa aaa tat ctt tga aga att ttt cag act att tcg agt atc tgt aag tag tca ata cga gga att aga ata tta tta ttc atg tga tta tac taa caa ccg tcc tac aat taa aca aca tta ctt tta taa cgg cga cga ata tac tga aat tga tag atc gaa aaa agc cac taa taa aaa cag ttg gtt aat tac ttc agg ctt tag act aca aaa atg gtt cga tag cga aga ttg tat aat tta tct cag atc ttt agt tag aag aat gga aga cag taa caa aaa cag taa aaa aac tta gta ctt aga tat cg- --a aaa aat ata ttt ttg tag act ctt gag aat aga agg aaa aca tgt aca taa tta taa aaa atg aaa atc aat ggc gaa taa gac agt gcg att cgc acc atg gag tcg gta gat ttc atg gct gtc gat gag cag ttt cac gac gac ctc gat ctt tgg tca tta tct ttg gta gat gat tat aaa aaa cat gga tta ggt gtt gac tgt tat gtt cta gaa cca gtt gtt gac agg aaa ata ttt gat aga ttt ctc ctt gaa cca att tgt gat cct gta gat gtt ctg tat gat tat ttt agg att cat aga gat aat att gat cag tat ata gta gat aga ctg ttt gca tat att aca tat aaa gat att ata tct gca tta gtg tca aag aat tat atg gaa gat att ttc tct ata att att aag aat tgt aat tct gtg caa gat ctc tta ctt tac tat cta tct aat gca tat gta gaa ata gac att gtt gat ctt atg gta gat cat ggg gct gta ata tat aaa ata gaa tgc ttg aat gcc tat ttt agg gga ata tgt aaa aag gaa agt agt gtt gtt gag ttt att ttg aat tgt ggt atc cca gat gaa aat gat gtt aaa tta gat cta tat aaa ata att cag tat act agg gga ttc ctt gta gat gaa ccc aca gta tta gaa att tat aag ctt tgt atc cca tat att gaa gat atc aat caa cta gat gct ggt gga agg acc ttg ctt tat cgc gct atc tat gca ggt tat ata gat tta gta tca tgg cta tta gaa aat gga gca aat gtc aac gca gta atg agt aat gga tat aca tgt ctt gac gtg gcc gtg gat agg gga tct gtc atc gcc cgt agg gaa gca cat ctt aaa ata tta gaa ata ttg ctt aga gaa cca ttg tct att gac tgt ata aaa tta gct ata ctt aat aat aca att gaa aac cat gat gtg ata aag ctc tgt atc aag tat ttt atg atg gta gat tat tca ctt tgt aat gtg tat gca tca tca ctc ttt gat tat ata att gat tgt aaa caa gaa ttg gag tac att agg cag atg aaa att cat aat aca acc atg tat gag tta atc tat aat aga gac aaa aac aag cat gct tcc cat att cta cat agg tat tct aaa cat cca gtt ttg aca cag tgt atc act aaa gga ttc aag att tac aca gaa gta acc gag cag gtc act aaa gct cta aac aga cgt gct cta ata gat gag ata ata aac aat gta tca act gat gac aat ctc cta tca aaa ctt cca tta gaa att agg gat cta att gtt tca caa gct gtc ata tag agt tct atc cac cca cct ttc ttg aaa tga gtt aat --- --- --- --- --- --- --- --- --- --- --- --- --- --- --- --- --- --- --- --- --- --- --- --- --- --- --- --- --- --- --- --- --- --- --- --- --- --- --- --- --- --- --- --- --- --- --- --- --- --- --- --- --- --- --- --- --- --- --- --- --- --- --- --- --- --- --- --- --- --- --- --- --- --- --- --- --- --- --- --- --- --- --- --- --- --- --- --- --- --- --- --- --- --- --- --- --- --- --- --- --- --- --- --- --- -ag tca taa gtt agt taa --- --- --- --- --- -gt cat aag tta gtt aag tca t-- --- --- --- --- --- --- --- --- --- --- --- --- --- --- --- -aa gtt agt taa gtc ata agt tag tta agt cat aag tta gtt aag tca taa gtt agt taa gtc ata agt tag tta agt cat aag tta gtt tat agt cta aca ctt cta att ttt ata cct tga tct ttt tct cta att atg aaa aag taa atc att atg aag atg gat g-a aat gga cga gat tgt gcg cat cgt taa cga tag tat gtg gta cgt acc taa cgc att tat gga cga cgg tga taa tga agg tca cat ttc tgt caa taa tgt ctg tca tat gta tct cgc att ctt tga tgt gga tat atc atc tca tct gtt taa att agt tat taa aca ctg cga tct gaa taa acg act aaa atg tgg taa ctc tcc att aca ttg cta tac gat gaa tac acg att taa tcc atc tgt att aaa gat att gtt acg cca cgg cat gcg taa ctt tga tag caa gga taa aaa agg aca tat tcc tct aca cca cta tct gat tca ttc act atc aat cga taa caa gat ctt tga tat act aac gga ccc cat tga tga ctt tag taa atc atc cga tct att gct gtg tta tct tag ata taa att caa tgg gag ctt aaa cta tta cgt tct gta caa att att gac taa agg atc tga ccc taa ttg cgt cga tga gga tgg act cac ttc tct tca tta cta ctg taa aca cat atc cgc gtt cca cga aag caa tta tta caa gtc aaa gag tca cac taa gat gcg agc tga gaa gcg att cat cta cgc gat aat aga tca tgg agc aaa cat taa cgc ggt tac gaa aat cgg aaa tac gcc gtt aca cac tta cct tca aca gta tac caa aca tag tcc tcg tgt ggt gta tgc tct ttt atc tcg agg agc cga tac gag gat acg taa taa tct tga ttg tac acc cat cat gga ata cat aaa gaa cga ttg tgc aac agg tca tat tct cat aat gtt act caa ttg gca cga aca aaa ata cgg gaa att aca aaa gga aga agg aca aca tct act tta tct att cat aaa aca taa tca agg ata tgg aag tcg ctc tct caa tat act acg gta tct act aga tag att cga cat tca gaa aga cga ata cta taa tac aat gac tcc tct tca tac cgc ctt cca gaa ttg caa taa caa tgt tgc ctc ata cct cgt ata cat cgg ata cga cat caa cct tcc gac taa aga cga taa gac agt att cga ctt ggt gtt tga aaa cag aaa cat tat ata caa ggc gga tgt cgt taa tga cat tat cca cca cag act gaa agt atc tct acc tat gat taa atc gtt gtt cta caa gat gtc gga gtt ctc tcc cta cga cga tca cta cgt aaa gaa gat aat agc cta ctg cct att aag gga cga gtc att tgc gga act aca tac taa att ctg ttt aaa cga gga cta taa aag tgt att tat gaa aaa tat atc att cga taa gat aga ttc cat cat cga aaa atg tag tcg tga cat aag tct cct caa aga gat tcg aat ctc aga cac cga ctt gta tac ggt att gag aac aga aga cat ccg gta tca cac ata tct cga agc cat aca ttc aga caa acg cat ttc att tcc cat gta cga cga tct cat aga aca gtg tca tct atc gat gga gca taa aag taa act cgt cga caa agc act caa taa att aga gtc tac cat cga tag tca atc tag act atc gta ttt gcc tcc gga aat tat gcg caa tat cat aac caa gct aag cga cta cca tct aaa cag tat gtt gta cgg aaa gaa cca tta caa ata tta tcc atg ata gaa aga aaa tat tta aaa aat aat cta tat gat tgg aga agt agg aaa caa aca gta aca aga cga cga tta cta c-a tta tta aat cat gag gtc cgt att ata ctc gta tat att gtt tct ctc atg tat aat aat aaa cgg aag aga tat agc acc aca tgc acc atc caa tgg aaa gtg taa aga caa cga ata cag aag ccg taa tct atg ttg tct atc gtg tcc tcc ggg aac tta cgc ttc cag att atg tga tag caa gac taa tac aca atg tac acc gtg tgg ttc gga tac ctt tac atc tca caa taa tca ttt aca ggc ttg tct aag ttg taa cgg aag atg tga tag taa tca ggt aga gac gcg atc gtg taa cac gac tca caa tag aat ctg tga atg ctc tcc agg ata tta ttg tct tct caa agg agc atc agg gtg tag aac atg tat ttc taa aac aaa gtg tgg aat agg ata cgg agt atc cgg ata cac gtc tac cgg aga cgt cat ctg ttc tcc gtg tgg tcc cgg aac ata ttc tca cac cgt ctc ttc cac aga taa atg cga acc cgt cgt aac cag caa tac att taa cta tat cga tgt gga aat taa cct gta tcc agt caa cga cac atc gtg tac tcg gac gac cac tac cgg tct cag cga atc cat ctc aac gtc gga act aac tat tac cat gaa tca taa aga ttg tga tcc agt ctt tcg tgc aga ata ctt ctc tgt cct taa taa tgt agc aac ttc agg att ctt tac agg aga aaa tag ata tca gaa tac ttc aaa gat atg tac tct gaa ttt cga gat taa atg taa caa caa aga ttc atc ttc caa aca gtt aac gaa aac aaa gaa tga tac tat cat gcc gca ttc aga gac ggt aac tct agt ggg cga ctg tct atc tag cgt cga cat cta cat act ata tag taa tac caa tac tca aga cta cga aaa tga tac aat ctc tta tca tat ggg taa tgt tct cga tgt caa tag cca tat gcc cgc tag ttg cga tat aca taa act gat cac taa ttc cca gaa tcc cac cca ctt ata gta ag- -tt ttt tta cct ata aat aat aaa tac aat aat taa ttt ctc gta aaa gta gaa aat ata ttc taa ttt att ata tgg taa gaa agt aga atc atc tag aac agt aat caa tca ata gca atc atg aaa caa tat att gtc ctg gca tgc atg tgc cta gtg gca gct gct atg cct act agt ctt caa caa tct tca tcc tcg tgt act gaa gaa gaa aac aaa cat cat atg gga atc gat gtt att atc aaa gtc aca aag caa gac caa aca ccg acc aat gat aag att tgt caa tcc gta acg gaa gtt aca gag a-- --- --- --- --- --- -cc gaa gat gat gag gta tcc gaa gaa gtt gta aaa gga gat ccc acc act tat tac act atc gtc ggt gcg ggt ctt aac atg aac ttt gga ttc acc aaa tgc cca aag att tca tcc atc tcc gaa tcc tct gat gga aac act gtg aat act aga ttg tcc agc gtg tca ccg gga caa ggt aag gac tct ccc gcg atc acg cgt gaa gaa gct ctg gct atg atc aaa gac tgt gag atg tct atc gac atc aga tgt agc gaa gaa gag aaa gac agt gac atc aag acc cat cca gta ctt ggg tct aac atc tca cat aag aaa gtg agt tac aaa gat atc atc ggt tca acg atc gtt gat aca aaa tgt gtc aag aac cta gag ttt agc gta cgt atc gga gac atg tgt gag gaa tca tct gaa ctt gaa gtc aag gat gga ttc aag tat gtc gac gga tcg gca tct gaa ggt gca acc gat gat act tca ctc atc gat tca aca aaa ctc aaa gca tgt gtc tga atc gat aac tct att cat ctg aaa atg gat gag ttg ggt taa tcg aac gat tca gac acc gca cca cga att aaa aaa g-- -ac cgg gca cta tat tcc ggt ttg caa aac aaa aat att taa cta cat tca caa aaa gtt acc tct cgt tac ttc ttc ttt ctg ttt caa tat gtg ata cga tat gat cac tat tcg tat tct ct- --- --- tgg tct cat aaa aaa gtt tta ca- -aa aaa aaa aaa aaa aat att ttt att ctc ttt ctc tct tcg at- ggt ctc aca aaa ata tta aac ctc ttt ctg atg tct caa cta ttt cgt aaa cga taa cgt cca aca ata tat tct cgt aga gct tat caa cat cct tat acc aat cta ggt tgt cag aca att gca tca taa aat aat gtt tat aat tta cac gtt aac atc ata taa taa acg tat ata gtt aat att ttt gga ata taa atg atc tgt aaa atc cat gta ggg gac act gct cac gtt ttt tct cta gta cat aat ttc aca caa gtt ttt ata cag aca aat taa ttc tcg tcc ata ta- ttt taa aac att gac ttt tgt act aag aaa aat atc ttg act aac cat ctc ttt ctc tct tcg atg ggt ctc aca aaa ata tta aac ctc ttt ctg atg g-- --- --- --- --- --- --- --- --- --- --- --- --- --- --- --- --- --- --- --- --- --- --- --- --- --- --- --- --- --- --- --- --- --- --- --- --- --- --- --- --- --- --- --- --- --- --- agt cgt aaa aag ttt tta tcc ttt ctc tct tcg ata ggt ctc --- --- --- --- --- --- --- --- --- --- --- --- --- --- --- --- --- --- --- --- --- --- --- --- --- --- --- --- --- --- --- --- --- --- --- --- --- --- --- --- --- --- --- --- --- --- --- --- --- --- --- --- --- --- --- --- --- --- --- --- --- --- --- --- --- --- --- --- --- --- aca aaa ata tta aac ctc ttt ctg atg gtc tct ata aa

>ON676704.1_Monkeypox_virus_isolate_MPXV_USA_2022_FL002_complete_genome

atc gag gac gac ttc aac tat gac agc tcg gtg gcg tct gcc agc gtg tac ata cga atg gca ttt cta aga aaa gtc tac ggt atc ctt -tc tac aat ttc ctt taa caa cgg caa cag ctg cag tat ttt tat act ttg aat gca -tc gga cat tta tac aag gga gtc ctg ttc taa tat tgg cat caa tgt tcg gat cta tag gct tga ttt tcg cat tga ctt tac aca gac atg aac atc ccc tga atc tgt aca tac ttt gtg gat tta cac tgt tag aat ctc taa cgc tgg cct ctg ttg tta ctt tct atg atg cac gta tcg tta tgc aag ctt tca tgt tga cta ctg cag tgt ttc ttg ctc tga cta cat ata ctc tac aat caa aga gag att tca gta aac ttg taa cag gat tgt ttg ctg ctt tct gga ttt taa ttt tgt cag gag tct tga gga taa agt tta aaa tag aat taa taa aga aca tat agg tca -tt ttt taa aca tgg ata gaa acc aag gtt gtt agt taa taa tat aca aga tat ttt ttc tca ctc tga tcc atg taa acc aag gac gag aga cac tct cat tcc tca ttc acg aca cca tta aaa atg gaa att aaa gcc ctc tat taa gca cag acg gct aca ggt cta cca tca ggt t-a cct tcg tct acc ttc aca atg gcc tct cct tgt gcc cag ttc agt ccc tgt cat tgc cac gct act aag gac tcc ctg aat acc gtg act gac gtc aga cat tgt ctg act gaa tac atc ctg tgg gtt tct cat aga tgg acc cat aga gaa agc gca ggg cct ctc tac agg ctt ctc atc tct ttc aga att gat gca atg gag cta ttt ggt agc gag ttg aag gag ttc tcg aat tca ctt ccg tgg gac aat atc gac aat tgc gtg gag atc att aaa tgt ttc atc aga aat gac tcc atg aaa acc gcc aaa gaa ctt tgt gca ata att gga ctt tgt act caa tca gct att gtc act gga aga gtc ttc aat gat aag tat atc gac ata cta ctt atg ctg cga aag att ctg aac gag aac gac tat ctc acc ctc ttg gat cat atc ctc act gct aaa tac taa atc tcc ttc atg ctc tct cac taa tac tct tac tca cta cac ttt tta tca tct tat gat gaa tga ttg cct tca tca ttt ttt cgt gga ata taa tat agg aat aat tag cac cag aat agc tat gga tat ctc gtt aag aat att ctc tat aag aga cat aat gta gac ata gtt att ata tcc ttc tta gat aag tgt tac gct act gga aag ttt cca tcg tta tta tta cat gaa gat gat ata att aaa cca aca ttg aga ttg gct ctt atg tta gct gga ttg aat tac tgt aat aaa tgc atc gag tat aga ggg ata tag caa ttc tcg ata ata gtc atg caa tat ttg aat gag act gat aat tta ggt aat aca gta ct- --- aca cac ata tct ttc tag att ata tat cgt taa aaa tct gta aga tgt ata ttt ctc ata agt atc cac tgt gta ata tta tta atg gat ata tag ata acg caa tag gga cta ata gta ttg taa aag ata taa tcg act att tac gta cat atc cag ata tct ata ttc cta cta gtt tgc tgc gta gtt gca tca ttg ata tgc atg -at tta tca gga ttc aga gat gaa tta cta agt aaa cta caa tcc cac aat aag taa gaa tca aat atc aaa aac tca ctt ttg att ttt cta gtc tta agt aat aca tat att tat taa tag acc tat gaa ata aaa aaa ggt aac aat gga ttc gcg tat agc tat tta cgt att agt atc ggc atc tct ttt gta tct tgt taa ttg tca caa act agt aca tta ctt caa tct gaa aat aaa tgg aag tga tat aac taa tac agc aga tat att gct gga caa tta tcc aat tat gac ctt tga tgg aaa gga tat tta tcc atc tat ctc gtt cat ggt cgg taa taa act ttt cct aga tct tta taa aaa tat ctt tga aga att ttt cag act att tcg agt atc tgt aag tag tca ata cga gga att aga ata tta tta ttc atg tga tta tac taa caa ccg tcc tac aat taa aca aca tta ctt tta taa cgg cga cga ata tac tga aat tga tag atc gaa aaa agc cac taa taa aaa cag ttg gtt aat tac ttc agg ctt tag act aca aaa atg gtt cga tag cga aga ttg tat aat tta tct cag atc ttt agt tag aag aat gga aga cag taa caa aaa cag taa aaa aac tta gta ctt aga tat cg- --a aaa aat ata ttt ttg tag act ctt gag aat aga agg aaa aca tgt aca taa tta taa aaa atg aaa atc aat ggc gaa taa gac agt gcg att cgc acc atg gag tcg gta gat ttc atg gct gtc gat gag cag ttt cac gac gac ctc gat ctt tgg tca tta tct ttg gta gat gat tat aaa aaa cat gga tta ggt gtt gac tgt tat gtt cta gaa cca gtt gtt gac agg aaa ata ttt gat aga ttt ctc ctt gaa cca att tgt gat cct gta gat gtt ctg tat gat tat ttt agg att cat aga gat aat att gat cag tat ata gta gat aga ctg ttt gca tat att aca tat aaa gat att ata tct gca tta gtg tca aag aat tat atg gaa gat att ttc tct ata att att aag aat tgt aat tct gtg caa gat ctc tta ctt tac tat cta tct aat gca tat gta gaa ata gac att gtt gat ctt atg gta gat cat ggg gct gta ata tat aaa ata gaa tgc ttg aat gcc tat ttt agg gga ata tgt aaa aag gaa agt agt gtt gtt gag ttt att ttg aat tgt ggt atc cca gat gaa aat gat gtt aaa tta gat cta tat aaa ata att cag tat act agg gga ttc ctt gta gat gaa ccc aca gta tta gaa att tat aag ctt tgt atc cca tat att gaa gat atc aat caa cta gat gct ggt gga agg acc ttg ctt tat cgc gct atc tat gca ggt tat ata gat tta gta tca tgg cta tta gaa aat gga gca aat gtc aac gca gta atg agt aat gga tat aca tgt ctt gac gtg gcc gtg gat agg gga tct gtc atc gcc cgt agg gaa gca cat ctt aaa ata tta gaa ata ttg ctt aga gaa cca ttg tct att gac tgt ata aaa tta gct ata ctt aat aat aca att gaa aac cat gat gtg ata aag ctc tgt atc aag tat ttt atg atg gta gat tat tca ctt tgt aat gtg tat gca tca tca ctc ttt gat tat ata att gat tgt aaa caa gaa ttg gag tac att agg cag atg aaa att cat aat aca acc atg tat gag tta atc tat aat aga gac aaa aac aag cat gct tcc cat att cta cat agg tat tct aaa cat cca gtt ttg aca cag tgt atc act aaa gga ttc aag att tac aca gaa gta acc gag cag gtc act aaa gct cta aac aga cgt gct cta ata gat gag ata ata aac aat gta tca act gat gac aat ctc cta tca aaa ctt cca tta gaa att agg gat cta att gtt tca caa gct gtc ata tag agt tct atc cac cca cct ttc ttg aaa tga gtt aat --- --- --- --- --- --- --- --- --- --- --- --- --- --- --- --- --- --- --- --- --- --- --- --- --- --- --- --- --- --- --- --- --- --- --- --- --- --- --- --- --- --- --- --- --- --- --- --- --- --- --- --- --- --- --- --- --- --- --- --- --- --- --- --- --- --- --- --- --- --- --- --- --- --- --- --- --- --- --- --- --- --- --- --- --- --- --- --- --- --- --- --- --- --- --- --- --- --- --- --- --- --- --- --- --- -ag tca taa gtt agt taa --- --- --- --- --- -gt cat aag tta gtt aag tca t-- --- --- --- --- --- --- --- --- --- --- --- --- --- --- --- -aa gtt agt taa gtc ata agt tag tta agt cat aag tta gtt aag tca taa gtt agt taa gtc ata agt tag tta agt cat aag tta gtt tat agt cta aca ctt cta att ttt ata cct tga tct ttt tct cta att atg aaa aag taa atc att atg aag atg gat g-a aat gga cga gat tgt gcg cat cgt taa cga tag tat gtg gta cgt acc taa cgc att tat gga cga cgg tga taa tga agg tca cat ttc tgt caa taa tgt ctg tca tat gta tct cgc att ctt tga tgt gga tat atc atc tca tct gtt taa att agt tat taa aca ctg cga tct gaa taa acg act aaa atg tgg taa ctc tcc att aca ttg cta tac gat gaa tac acg att taa tcc atc tgt att aaa gat att gtt acg cca cgg cat gcg taa ctt tga tag caa gga taa aaa agg aca tat tcc tct aca cca cta tct gat tca ttc act atc aat cga taa caa gat ctt tga tat act aac gga ccc cat tga tga ctt tag taa atc atc cga tct att gct gtg tta tct tag ata taa att caa tgg gag ctt aaa cta tta cgt tct gta caa att att gac taa agg atc tga ccc taa ttg cgt cga tga gga tgg act cac ttc tct tca tta cta ctg taa aca cat atc cgc gtt cca cga aag caa tta tta caa gtc aaa gag tca cac taa gat gcg agc tga gaa gcg att cat cta cgc gat aat aga tca tgg agc aaa cat taa cgc ggt tac gaa aat cgg aaa tac gcc gtt aca cac tta cct tca aca gta tac caa aca tag tcc tcg tgt ggt gta tgc tct ttt atc tcg agg agc cga tac gag gat acg taa taa tct taa ttg tac acc cat cat gga ata cat aaa gaa cga ttg tgc aac agg tca tat tct cat aat gtt act caa ttg gca cga aca aaa ata cgg gaa att aca aaa gga aga agg aca aca tct act tta tct att cat aaa aca taa tca agg ata tgg aag tcg ctc tct caa tat act acg gta tct act aga tag att cga cat tca gaa aga cga ata cta taa tac aat gac tcc tct tca tac cgc ctt cca gaa ttg caa taa caa tgt tgc ctc ata cct cgt ata cat tgg ata cga cat caa cct tcc gac taa aga cga taa gac agt att cga ctt ggt gtt tga aaa cag aaa cat tat ata caa ggc gga tgt cgt taa tga cat tat cca cca cag act gaa agt atc tct acc tat gat taa atc gtt gtt cta caa gat gtc gga gtt ctc tcc cta cga cga tca cta cgt aaa gaa gat aat agc cta ctg cct att aag gga cga gtc att tgc gga act aca tac taa att ctg ttt aaa cga gga cta taa aag tgt att tat gaa aaa tat atc att cga taa gat aga ttc cat cat cga aaa atg tag tcg tga cat aag tct cct caa aga gat tcg aat ctc aga cac cga ctt gta tac ggt att gag aac aga aga cat tcg gta tca cac ata tct cga agc cat aca ttc aga caa acg cat ttc att tcc cat gta cga cga tct cat aga aca gtg tca tct atc gat gga gca taa aag taa act cgt cga caa agc act caa taa att aga gtc tac cat cga tag tca atc tag act atc gta ttt gcc tcc gga aat tat gcg caa tat cat aac caa gct aag cga cta cca tct aaa cag tat gtt gta cgg aaa gaa cca tta caa ata tta tcc atg ata gaa aga aaa tat tta aaa aat aat cta tat gat tgg aga agt agg aaa caa aca gta aca aga cga cga tta cta c-a tta tta aat cat gag gtc cgt att ata ctc gta tat att gtt tct ctc atg tat aat aat aaa cgg aag aga tat agc acc aca tgc acc atc caa tgg aaa gtg taa aga caa cga ata cag aag ccg taa tct atg ttg tct atc gtg tcc tcc ggg aac tta cgc ttt cag att atg tga tag caa gac taa tac aca atg tac acc gtg tgg ttc gga tac ctt tac atc tca caa taa tca ttt aca ggc ttg tct aag ttg taa cgg aag atg tga tag taa tca ggt aga gac gcg atc gtg taa cac gac tca caa tag aat ctg tga atg ctc tcc agg ata tta ttg tct tct caa agg agc atc agg gtg tag aac atg tat ttc taa aac aaa gtg tgg aat agg ata cgg agt atc cgg ata cac gtc tac cgg aga cgt cat ctg ttc tcc gtg tgg tcc cgg aac ata ttc tca cac cgt ctc ttc cac aga taa atg cga acc cgt cgt aac cag caa tac att taa cta tat cga tgt gga aat taa cct gta tcc agt caa cga cac atc gtg tac tcg gac gac cac tac cgg tct cag cga atc cat ctc aac gtc gga act aac tat tac cat gaa tca taa aga ttg tga tcc agt ctt tcg tgc aga ata ctt ctc tgt cct taa taa tgt agc aac ttc agg att ctt tac agg aga aaa tag ata tca gaa tac ttc aaa gat atg tac tct gaa ttt cga gat taa atg taa caa caa aga ttc atc ttc caa aca gtt aac gaa aac aaa gaa tga tac tat cat gcc gca ttc aga gac ggt aac tct agt ggg cga ctg tct atc tag cgt cga cat cta cat act ata tag taa tac caa tac tca aga cta cga aaa tga tac aat ctc tta tca tat ggg taa tgt tct cga tgt caa tag cca tat gcc cgc tag ttg cga tat aca taa act gat cac taa ttc cca gaa tcc cac cca ctt ata gta ag- -tt ttt tta cct ata aat aat aaa tac aat aat taa ttt ctc gta aaa gta gaa aat ata ttc taa ttt att ata tgg taa gaa agt aga atc atc tag aac agt aat caa tca ata gca atc atg aaa caa tat att gtc ctg gca tgc atg tgc cta gtg gca gct gct atg cct act agt ctt caa caa tct tca tcc tcg tgt act gaa gaa gaa aac aaa cat cat atg gga atc gat gtt att atc aaa gtc aca aag caa gac caa aca ccg acc aat gat aag att tgt caa tcc gta acg gaa gtt aca gag a-- --- --- --- --- --- -cc gaa gat gat gag gta tcc gaa gaa gtt gta aaa gga gat ccc acc act tat tac act atc gtc ggt gcg ggt ctt aac atg aac ttt gga ttc acc aaa tgc cca aag att tta tcc atc tcc gaa tcc tct gat gga aac act gtg aat act aga ttg tcc agc gtg tca ccg gga caa ggt aag gac tct ccc gcg atc acg cgt gaa gaa gct ctg gct atg atc aaa gac tgt gag atg tct atc gac atc aga tgt agc gaa gaa gag aaa gac agt gac atc aag acc cat cca gta ctt ggg tct aac atc tca cat aag aaa gtg agt tac aaa gat atc atc ggt tca acg atc gtt gat aca aaa tgt gtc aag aac cta gag ttt agc gta cgt atc gga gac atg tgt gag gaa tca tct gaa ctt gaa gtc aag gat gga ttc aag tat gtc gac gga tcg gca tct gaa ggt gca acc gat gat act tca ctc atc gat tca aca aaa ctc aaa gca tgt gtc tga atc gat aac tct att cat ctg aaa atg gat gag ttg ggt taa tcg aac gat tca gac acc gca cca cga att aaa aaa g-- -ac cgg gca cta tat tcc ggt ttg caa aac aaa aat att taa cta cat tca caa aaa gtt acc tct cgt tac ttc ttc ttt ctg ttt caa tat gtg ata cga tat gat cac tat tcg tat tct ct- --- --- tgg tct cat aaa aaa gtt tta c-- --a aaa aaa aaa aaa a?t att ttt att ctc ttt ctc tct tcg at- ggt ctc aca aaa ata tta aac ctc ttt ctg atg tct caa cta ttt cgt aaa cga taa cgt cca aca ata tat tct cgt aga gct tat caa cat cct tat acc aat cta ggt tgt cag aca att gca tca taa aat aat gtt tat aat tta cac gtt aac atc ata taa taa acg tat ata gtt aat att ttt gga ata taa atg atc tgt aaa atc cat gta ggg gac act gct cac gtt ttt tct cta gta cat aat ttc aca caa gtt ttt ata cag aca aat taa ttc tcg tcc ata ta- ttt taa aac att gac ttt tgt act aag aaa aat atc ttg act aac cat ctc ttt ctc tct tcg atg ggt ctc aca aaa ata tta aac ctc ttt ctg atg g-- --- --- --- --- --- --- --- --- --- --- --- --- --- --- --- --- --- --- --- --- --- --- --- --- --- --- --- --- --- --- --- --- --- --- --- --- --- --- --- --- --- --- --- --- --- --- agt cgt aaa aag ttt tta tcc ttt ctc tct tcg ata ggt ctc --- --- --- --- --- --- --- --- --- --- --- --- --- --- --- --- --- --- --- --- --- --- --- --- --- --- --- --- --- --- --- --- --- --- --- --- --- --- --- --- --- --- --- --- --- --- --- --- --- --- --- --- --- --- --- --- --- --- --- --- --- --- --- --- --- --- --- --- --- --- aca aaa ata tta aac ctc ttt ctg atg gtc tct ata aa

>ON563414.3_Monkeypox_virus_isolate_MPXV_USA_2022_MA001_complete_genome

atc gag gac gac ttc aac tat gac agc tcg gtg gcg tct gcc agc gtg tac ata cga atg gca ttt cta aga aaa gtc tac ggt atc ctt -tc tac aat ttc ctt taa caa cgg caa cag ctg cag tat ttt tat act ttg aat gca -tc gga cat tta tac aag gga gtc ctg ttc taa tat tgg cat caa tgt tcg gat cta tag gct tga ttt tcg cat tga ctt tac aca gac atg aac atc ccc tga atc tgt aca tac ttt gtg gat tta cac tgt tag aat ctc taa cgc tgg cct ctg ttg tta ctt tct atg atg cac gta tcg tta tgc aag ctt tca tgt tga cta ctg cag tgt ttc ttg ctc tga cta cat ata ctc tac aat caa aga gag att tca gta aac ttg taa cag gat tgt ttg ctg ctt tct gga ttt taa ttt tgt cag gag tct tga gga taa agt tta aaa tag aat taa taa aga aca tat agg tca -tt ttt taa aca tgg ata gaa acc aag gtt gtt agt taa taa tat aca aga tat ttt ttc tca ctc tga tcc atg taa acc aag gac gag aga cac tct cat tcc tca ttc acg aca cca tta aaa atg gaa att aaa gcc ctc tat taa gca cag acg gct aca ggt cta cca tca ggt t-a cct tcg tct acc ttc aca atg gcc tct cct tgt gcc cag ttc agt ccc tgt cat tgc cac gct act aag gac tcc ctg aat acc gtg act gac gtc aga cat tgt ctg act gaa tac atc ctg tgg gtt tct cat aga tgg acc cat aga gaa agc gca ggg cct ctc tac agg ctt ctc atc tct ttc aga att gat gca atg gag cta ttt ggt agc gag ttg aag gag ttc tcg aat tca ctt ccg tgg gac aat atc gac aat tgc gtg gag atc att aaa tgt ttc atc aga aat gac tcc atg aaa acc gcc aaa gaa ctt tgt gca ata att gga ctt tgt act caa tca gct att gtc act gga aga gtc ttc aat gat aag tat atc gac ata cta ctt atg ctg cga aag att ctg aac gag aac gac tat ctc acc ctc ttg gat cat atc ctc act gct aaa tac taa atc tcc ttc atg ctc tct cac taa tac tct tac tca cta cac ttt tta tca tct tat gat gaa tga ttg cct tca tca ttt ttt cgt gga ata taa tat agg aat aat tag cac cag aat agc tat gga tat ctc gtt aag aat att ctc tat aag aga cat aat gta gac ata gtt att ata tcc ttc tta gat aag tgt tac gct act gga aag ttt cca tcg tta tta tta cat gaa gat gat ata att aaa cca aca ttg aga ttg gct ctt atg tta gct gga ttg aat tac tgt aat aaa tgc atc gag tat aga ggg ata tag caa ttc tcg ata ata gtc atg caa tat ttg aat gag act gat aat tta ggt aat aca gta ct- --- aca cac ata tct ttc tag att ata tat cgt taa aaa tct gta aga tgt ata ttt ctc ata agt atc cac tgt gta ata tta tta atg gat ata tag ata acg caa tag gga cta ata gta ttg taa aag ata taa tcg act att tac gta cat atc cag ata tct ata ttc cta cta gtt tgc tgc gta gtt gca tca ttg ata tgc atg -at tta tca gga ttc aga gat gaa tta cta agt aaa cta caa tcc cac aat aag taa gaa tca aat atc aaa aac tca ctt ttg att ttt cta gtc tta agt aat aca tat att tat taa tag acc tat gaa ata aaa aaa ggt aac aat gga ttc gcg tat agc tat tta cgt att agt atc ggc atc tct ttt gta tct tgt taa ttg tca caa act agt aca tta ctt caa tct gaa aat aaa tgg aag tga tat aac taa tac agc aga tat att gct gga caa tta tcc aat tat gac ctt tga tgg aaa gga tat tta tcc atc tat ctc gtt cat ggt cgg taa taa act ttt cct aga tct tta taa aaa tat ctt tga aga att ttt cag act att tcg agt atc tgt aag tag tca ata cga gga att aga ata tta tta ttc atg tga tta tac taa caa ccg tcc tac aat taa aca aca tta ctt tta taa cgg cga cga ata tac tga aat tga tag atc gaa aaa agc cac taa taa aaa cag ttg gtt aat tac ttc agg ctt tag act aca aaa atg gtt cga tag cga aga ttg tat aat tta tct cag atc ttt agt tag aag aat gga aga cag taa caa aaa cag taa aaa aac tta gta ctt aga tat cg- --a aaa aat ata ttt ttg tag act ctt gag aat aga agg aaa aca tgt aca taa tta taa aaa atg aaa atc aat ggc gaa taa gac agt gcg att cgc acc atg gag tcg gta gat ttc atg gct gtc gat gag cag ttt cac gac gac ctc gat ctt tgg tca tta tct ttg gta gat gat tat aaa aaa cat gga tta ggt gtt gac tgt tat gtt cta gaa cca gtt gtt gac agg aaa ata ttt gat aga ttt ctc ctt gaa cca att tgt gat cct gta gat gtt ctg tat gat tat ttt agg att cat aga gat aat att gat cag tat ata gta gat aga ctg ttt gca tat att aca tat aaa gat att ata tct gca tta gtg tca aag aat tat atg gaa gat att ttc tct ata att att aag aat tgt aat tct gtg caa gat ctc tta ctt tac tat cta tct aat gca tat gta gaa ata gac att gtt gat ctt atg gta gat cat ggg gct gta ata tat aaa ata gaa tgc ttg aat gcc tat ttt agg gga ata tgt aaa aag gaa agt agt gtt gtt gag ttt att ttg aat tgt ggt atc cca gat gaa aat gat gtt aaa tta gat cta tat aaa ata att cag tat act agg gga ttc ctt gta gat gaa ccc aca gta tta gaa att tat aag ctt tgt atc cca tat att gaa gat atc aat caa cta gat gct ggt gga agg acc ttg ctt tat cgc gct atc tat gca ggt tat ata gat tta gta tca tgg cta tta gaa aat gga gca aat gtc aac gca gta atg agt aat gga tat aca tgt ctt gac gtg gcc gtg gat agg gga tct gtc atc gcc cgt agg gaa gca cat ctt aaa ata tta gaa ata ttg ctt aga gaa cca ttg tct att gac tgt ata aaa tta gct ata ctt aat aat aca att gaa aac cat gat gtg ata aag ctc tgt atc aag tat ttt atg atg gta gat tat tca ctt tgt aat gtg tat gca tca tca ctc ttt gat tat ata att gat tgt aaa caa gaa ttg gag tac att agg cag atg aaa att cat aat aca acc atg tat gag tta atc tat aat aga gac aaa aac aag cat gct tcc cat att cta cat agg tat tct aaa cat cca gtt ttg aca cag tgt atc act aaa gga ttc aag att tac aca gaa gta acc gag cag gtc act aaa gct cta aac aga cgt gct cta ata gat gag ata ata aac aat gta tca act gat gac aat ctc cta tca aaa ctt cca tta gaa att agg gat cta att gtt tca caa gct gtc ata tag agt tct atc cac cca cct ttc ttg aaa tga gtt aat --- --- --- --- --- --- --- --- --- --- --- --- --- --- --- --- --- --- --- --- --- --- --- --- --- --- --- --- --- --- --- --- --- --- --- --- --- --- --- --- --- --- --- --- --- --- --- --- --- --- --- --- --- --- --- --- --- --- --- --- --- --- --- --- --- --- --- --- --- --- --- --- --- --- --- --- --- --- --- --- --- --- --- --- --- --- --- --- --- --- --- --- --- --- --- --- --- --- --- --- --- --- --- --- --- -ag tca taa gtt agt taa --- --- --- --- --- -gt cat aag tta gtt aag tca t-- --- --- --- --- --- --- --- --- --- --- --- --- --- --- --- -aa gtt agt taa gtc ata agt tag tta agt cat aag tta gtt aag tca taa gtt agt taa gtc ata agt tag tta agt cat aag tta gtt tat agt cta aca ctt cta att ttt ata cct tga tct ttt tct cta att atg aaa aag taa atc att atg aag atg gat g-a aat gga cga gat tgt gcg cat cgt taa cga tag tat gtg gta cgt acc taa cgc att tat gga cga cgg tga taa tga agg tca cat ttc tgt caa taa tgt ctg tca tat gta tct cgc att ctt tga tgt gga tat atc atc tca tct gtt taa att agt tat taa aca ctg cga tct gaa taa acg act aaa atg tgg taa ctc tcc att aca ttg cta tac gat gaa tac acg att taa tcc atc tgt att aaa gat att gtt acg cca cgg cat gcg taa ctt tga tag caa gga taa aaa agg aca tat tcc tct aca cca cta tct gat tca ttc act atc aat cga taa caa gat ctt tga tat act aac gga ccc cat tga tga ctt tag taa atc atc cga tct att gct gtg tta tct tag ata taa att caa tgg gag ctt aaa cta tta cgt tct gta caa att att gac taa agg atc tga ccc taa ttg cgt cga tga gga tgg act cac ttc tct tca tta cta ctg taa aca cat atc cgc gtt cca cga aag caa tta tta caa gtc aaa gag tca cac taa gat gcg agc tga gaa gcg att cat cta cgc gat aat aga tca tgg agc aaa cat taa cgc ggt tac gaa aat cgg aaa tac gcc gtt aca cac tta cct tca aca gta tac caa aca tag tcc tcg tgt ggt gta tgc tct ttt atc tcg agg agc cga tac gag gat acg taa taa tct taa ttg tac acc cat cat gga ata cat aaa gaa cga ttg tgc aac agg tca tat tct cat aat gtt act caa ttg gca cga aca aaa ata cgg gaa att aca aaa gga aga agg aca aca tct act tta tct att cat aaa aca taa tca agg ata tgg aag tcg ctc tct caa tat act acg gta tct act aga tag att cga cat tca gaa aga cga ata cta taa tac aat gac tcc tct tca tac cgc ctt cca gaa ttg caa taa caa tgt tgc ctc ata cct cgt ata cat tgg ata cga cat caa cct tcc gac taa aga cga taa gac agt att cga ctt ggt gtt tga aaa cag aaa cat tat ata caa ggc gga tgt cgt taa tga cat tat cca cca cag act gaa agt atc tct acc tat gat taa atc gtt gtt cta caa gat gtc gga gtt ctc tcc cta cga cga tca cta cgt aaa gaa gat aat agc cta ctg cct att aag gga cga gtc att tgc gga act aca tac taa att ctg ttt aaa cga gga cta taa aag tgt att tat gaa aaa tat atc att cga taa gat aga ttc cat cat cga aaa atg tag tcg tga cat aag tct cct caa aga gat tcg aat ctc aga cac cga ctt gta tac ggt att gag aac aga aga cat tcg gta tca cac ata tct cga agc cat aca ttc aga caa acg cat ttc att tcc cat gta cga cga tct cat aga aca gtg tca tct atc gat gga gca taa aag taa act cgt cga caa agc act caa taa att aga gtc tac cat cga tag tca atc tag act atc gta ttt gcc tcc gga aat tat gcg caa tat cat aac caa gct aag cga cta cca tct aaa cag tat gtt gta cgg aaa gaa cca tta caa ata tta tcc atg ata gaa aga aaa tat tta aaa aat aat cta tat gat tgg aga agt agg aaa caa aca gta aca aga cga cga tta cta c-a tta tta aat cat gag gtc cgt att ata ctc gta tat att gtt tct ctc atg tat aat aat aaa cgg aag aga tat agc acc aca tgc acc atc caa tgg aaa gtg taa aga caa cga ata cag aag ccg taa tct atg ttg tct atc gtg tcc tcc ggg aac tta cgc ttt cag att atg tga tag caa gac taa tac aca atg tac acc gtg tgg ttc gga tac ctt tac atc tca caa taa tca ttt aca ggc ttg tct aag ttg taa cgg aag atg tga tag taa tca ggt aga gac gcg atc gtg taa cac gac tca caa tag aat ctg tga atg ctc tcc agg ata tta ttg tct tct caa agg agc atc agg gtg tag aac atg tat ttc taa aac aaa gtg tgg aat agg ata cgg agt atc cgg ata cac gtc tac cgg aga cgt cat ctg ttc tcc gtg tgg tcc cgg aac ata ttc tca cac cgt ctc ttc cac aga taa atg cga acc cgt cgt aac cag caa tac att taa cta tat cga tgt gga aat taa cct gta tcc agt caa cga cac atc gtg tac tcg gac gac cac tac cgg tct cag cga atc cat ctc aac gtc gga act aac tat tac cat gaa tca taa aga ttg tga tcc agt ctt tcg tgc aga ata ctt ctc tgt cct taa taa tgt agc aac ttc agg att ctt tac agg aga aaa tag ata tca gaa tac ttc aaa gat atg tac tct gaa ttt cga gat taa atg taa caa caa aga ttc atc ttc caa aca gtt aac gaa aac aaa gaa tga tac tat cat gcc gca ttc aga gac ggt aac tct agt ggg cga ctg tct atc tag cgt cga cat cta cat act ata tag taa tac caa tac tca aga cta cga aaa tga tac aat ctc tta tca tat ggg taa tgt tct cga tgt caa tag cca tat gcc cgc tag ttg cga tat aca taa act gat cac taa ttc cca gaa tcc cac cca ctt ata gta ag- -tt ttt tta cct ata aat aat aaa tac aat aat taa ttt ctc gta aaa gta gaa aat ata ttc taa ttt att ata tgg taa gaa agt aga atc atc tag aac agt aat caa tca ata gca atc atg aaa caa tat att gtc ctg gca tgc atg tgc cta gtg gca gct gct atg cct act agt ctt caa caa tct tca tcc tcg tgt act gaa gaa gaa aac aaa cat cat atg gga atc gat gtt att atc aaa gtc aca aag caa gac caa aca ccg acc aat gat aag att tgt caa tcc gta acg gaa gtt aca gag a-- --- --- --- --- --- -cc gaa gat gat gag gta tcc gaa gaa gtt gta aaa gga gat ccc acc act tat tac act atc gtc ggt gcg ggt ctt aac atg aac ttt gga ttc acc aaa tgc cca aag att tta tcc atc tcc gaa tcc tct gat gga aac act gtg aat act aga ttg tcc agc gtg tca ccg gga caa ggt aag gac tct ccc gcg atc acg cgt gaa gaa gct ctg gct atg atc aaa gac tgt gag atg tct atc gac atc aga tgt agc gaa gaa gag aaa gac agt gac atc aag acc cat cca gta ctt ggg tct aac atc tca cat aag aaa gtg agt tac aaa gat atc atc ggt tca acg atc gtt gat aca aaa tgt gtc aag aac cta gag ttt agc gta cgt atc gga gac atg tgt gag gaa tca tct gaa ctt gaa gtc aag gat gga ttc aag tat gtc gac gga tcg gca tct gaa ggt gca acc gat gat act tca ctc atc gat tca aca aaa ctc aaa gca tgt gtc tga atc gat aac tct att cat ctg aaa atg gat gag ttg ggt taa tcg aac gat tca gac acc gca cca cga att aaa aaa g-- -ac cgg gca cta tat tcc ggt ttg caa aac aaa aat att taa cta cat tca caa aaa gtt acc tct cgt tac ttc ttc ttt ctg ttt caa tat gtg ata cga tat gat cac tat tcg tat tct ct- --- --- tgg tct cat aaa aaa gtt tta c-- --a aaa aaa aaa aaa a-t att ttt att ctc ttt ctc tct tcg at- ggt ctc aca aaa ata tta aac ctc ttt ctg atg tct caa cta ttt cgt aaa cga taa cgt cca aca ata tat tct cgt aga gct tat caa cat cct tat acc aat cta ggt tgt cag aca att gca tca taa aat aat gtt tat aat tta cac gtt aac atc ata taa taa acg tat ata gtt aat att ttt gga ata taa atg atc tgt aaa atc cat gta ggg gac act gct cac gtt ttt tct cta gta cat aat ttc aca caa gtt ttt ata cag aca aat taa ttc tcg tcc ata ta- ttt taa aac att gac ttt tgt act aag aaa aat atc ttg act aac cat ctc ttt ctc tct tcg atg ggt ctc aca aaa ata tta aac ctc ttt ctg atg g-- --- --- --- --- --- --- --- --- --- --- --- --- --- --- --- --- --- --- --- --- --- --- --- --- --- --- --- --- --- --- --- --- --- --- --- --- --- --- --- --- --- --- --- --- --- --- agt cgt aaa aag ttt tta tcc ttt ctc tct tcg ata ggt ctc --- --- --- --- --- --- --- --- --- --- --- --- --- --- --- --- --- --- --- --- --- --- --- --- --- --- --- --- --- --- --- --- --- --- --- --- --- --- --- --- --- --- --- --- --- --- --- --- --- --- --- --- --- --- --- --- --- --- --- --- --- --- --- --- --- --- --- --- --- --- aca aaa ata tta aac ctc ttt ctg atg gtc tct ata aa

>ON676705.1_Monkeypox_virus_isolate_MPXV_USA_2022_UT001_complete_genome

atc gag gac gac ttc aac tat gac agc tcg gtg gcg tct gcc agc gtg tac ata cga atg gca ttt cta aga aaa gtc tac ggt atc ctt -tc tac aat ttc ctt taa caa cgg caa cag ctg cag tat ttt tat act ttg aat gca -tc gga cat tta tac aag gga gtc ctg ttc taa tat tgg cat caa tgt tcg gat cta tag gct tga ttt tcg cat tga ctt tac aca gac atg aac atc ccc tga atc tgt aca tac ttt gtg gat tta cac tgt tag aat ctc taa cgc tgg cct ctg ttg tta ctt tct atg atg cac gta tcg tta tgc aag ctt tca tgt tga cta ctg cag tgt ttc ttg ctc tga cta cat ata ctc tac aat caa aga gag att tca gta aac ttg taa cag gat tgt ttg ctg ctt tct gga ttt taa ttt tgt cag gag tct tga gga taa agt tta aaa tag aat taa taa aga aca tat agg tca -tt ttt taa aca tgg ata gaa acc aag gtt gtt agt taa taa tat aca aga tat ttt ttc tca ctc tga tcc atg taa acc aag gac gag aga cac tct cat tcc tca ttc acg aca cca tta aaa atg gaa att aaa gcc ctc tat taa gca cag acg gct aca ggt cta cca tca ggt t-a cct tcg tct acc ttc aca atg gcc tct cct tgt gcc cag ttc agt ccc tgt cat tgc cac gct act aag gac tcc ctg aat acc gtg act gac gtc aga cat tgt ctg act gaa tac atc ctg tgg gtt tct cat aga tgg acc cat aga gaa agc gca ggg cct ctc tac agg ctt ctc atc tct ttc aga att gat gca atg gag cta ttt ggt agc gag ttg aag gag ttc tcg aat tca ctt ccg tgg gac aat atc gac aat tgc gtg gag atc att aaa tgt ttc atc aga aat gac tcc atg aaa acc gcc aaa gaa ctt tgt gca ata att gga ctt tgt act caa tca gct att gtc act gga aga gtc ttc aat gat aag tat atc gac ata cta ctt atg ctg cga aag att ctg aac gag aac gac tat ctc acc ctc ttg gat cat atc ctc act gct aaa tac taa atc tcc ttc atg ctc tct cac taa tac tct tac tca cta cac ttt tta tca tct tat gat gaa tga ttg cct tca tca ttt ttt cgt gga ata taa tat agg aat aat tag cac cag aat agc tat gga tat ctc gtt aag aat att ctc tat aag aga cat aat gta gac ata gtt att ata tcc ttc tta gat aag tgt tac gct act gga aag ttt cca tcg tta tta tta cat gaa gat gat ata att aaa cca aca ttg aga ttg gct ctt atg tta gct gga ttg aat tac tgt aat aaa tgc atc gag tat aga ggg ata tag caa ttc tcg ata ata gtc atg caa tat ttg aat gag act gat aat tta ggt aat aca gta ct- --- aca cac ata tct ttc tag att ata tat cgt taa aaa tct gta aga tgt ata ttt ctc ata agt atc cac tgt gta ata tta tta atg gat ata tag ata acg caa tag gga cta ata gta ttg taa aag ata taa tcg act att tac gta cat atc cag ata tct ata ttc cta cta gtt tgc tgc gta gtt gca tca ttg ata tgc atg -at tta tca gga ttc aga gat gaa tta cta agt aaa cta caa tcc cac aat aag taa gaa tca aat atc aaa aac tca ctt ttg att ttt cta gtc tta agt aat aca tat att tat taa tag acc tat gaa ata aaa aaa ggt aac aat gga ttc gcg tat agc tat tta cgt att agt atc ggc atc tct ttt gta tct tgt taa ttg tca caa act agt aca tta ctt caa tct gaa aat aaa tgg aag tga tat aac taa tac agc aga tat att gct gga caa tta tcc aat tat gac ctt tga tgg aaa gga tat tta tcc atc tat ctc gtt cat ggt cgg taa taa act ttt cct aga tct tta taa aaa tat ctt tga aga att ttt cag act att tcg agt atc tgt aag tag tca ata cga gga att aga ata tta tta ttc atg tga tta tac taa caa ccg tcc tac aat taa aca aca tta ctt tta taa cgg cga cga ata tac tga aat tga tag atc gaa aaa agc cac taa taa aaa cag ttg gtt aat tac ttc agg ctt tag act aca aaa atg gtt cga tag cga aga ttg tat aat tta tct cag atc ttt agt tag aag aat gga aga cag taa caa aaa cag taa aaa aac tta gta ctt aga tat cg- --a aaa aat ata ttt ttg tag act ctt gag aat aga agg aaa aca tgt aca taa tta taa aaa atg aaa atc aat ggc gaa taa gac agt gcg att cgc acc atg gag tcg gta gat ttc atg gct gtc gat gag cag ttt cac gac gac ctc gat ctt tgg tca tta tct ttg gta gat gat tat aaa aaa cat gga tta ggt gtt gac tgt tat gtt cta gaa cca gtt gtt gac agg aaa ata ttt gat aga ttt ctc ctt gaa cca att tgt gat cct gta gat gtt ctg tat gat tat ttt agg att cat aga gat aat att gat cag tat ata gta gat aga ctg ttt gca tat att aca tat aaa gat att ata tct gca tta gtg tca aag aat tat atg gaa gat att ttc tct ata att att aag aat tgt aat tct gtg caa gat ctc tta ctt tac tat cta tct aat gca tat gta gaa ata gac att gtt gat ctt atg gta gat cat ggg gct gta ata tat aaa ata gaa tgc ttg aat gcc tat ttt agg gga ata tgt aaa aag gaa agt agt gtt gtt gag ttt att ttg aat tgt ggt atc cca gat gaa aat gat gtt aaa tta gat cta tat aaa ata att cag tat act agg gga ttc ctt gta gat gaa ccc aca gta tta gaa att tat aag ctt tgt atc cca tat att gaa gat atc aat caa cta gat gct ggt gga agg acc ttg ctt tat cgc gct atc tat gca ggt tat ata gat tta gta tca tgg cta tta gaa aat gga gca aat gtc aac gca gta atg agt aat gga tat aca tgt ctt gac gtg gcc gtg gat agg gga tct gtc atc gcc cgt agg gaa gca cat ctt aaa ata tta gaa ata ttg ctt aga gaa cca ttg tct att gac tgt ata aaa tta gct ata ctt aat aat aca att gaa aac cat gat gtg ata aag ctc tgt atc aag tat ttt atg atg gta gat tat tca ctt tgt aat gtg tat gca tca tca ctc ttt gat tat ata att gat tgt aaa caa gaa ttg gag tac att agg cag atg aaa att cat aat aca acc atg tat gag tta atc tat aat aga gac aaa aac aag cat gct tcc cat att cta cat agg tat tct aaa cat cca gtt ttg aca cag tgt atc act aaa gga ttc aag att tac aca gaa gta acc gag cag gtc act aaa gct cta aac aga cgt gct cta ata gat gag ata ata aac aat gta tca act gat gac aat ctc cta tca aaa ctt cca tta gaa att agg gat cta att gtt tca caa gct gtc ata tag agt tct atc cac cca cct ttc ttg aaa tga gtt aat --- --- --- --- --- --- --- --- --- --- --- --- --- --- --- --- --- --- --- --- --- --- --- --- --- --- --- --- --- --- --- --- --- --- --- --- --- --- --- --- --- --- --- --- --- --- --- --- --- --- --- --- --- --- --- --- --- --- --- --- --- --- --- --- --- --- --- --- --- --- --- --- --- --- --- --- --- --- --- --- --- --- --- --- --- --- --- --- --- --- --- --- --- --- --- --- --- --- --- --- --- --- --- --- --- -ag tca taa gtt agt taa --- --- --- --- --- -gt cat aag tta gtt aag tca t-- --- --- --- --- --- --- --- --- --- --- --- --- --- --- --- -aa gtt agt taa gtc ata agt tag tta agt cat aag tta gtt aag tca taa gtt agt taa gtc ata agt tag tta agt cat aag tta gtt tat agt cta aca ctt cta att ttt ata cct tga tct ttt tct cta att atg aaa aag taa atc att atg aag atg gat g-a aat gga cga gat tgt gcg cat cgt taa cga tag tat gtg gta cgt acc taa cgc att tat gga cga cgg tga taa tga agg tca cat ttc tgt caa taa tgt ctg tca tat gta tct cgc att ctt tga tgt gga tat atc atc tca tct gtt taa att agt tat taa aca ctg cga tct gaa taa acg act aaa atg tgg taa ctc tcc att aca ttg cta tac gat gaa tac acg att taa tcc atc tgt att aaa gat att gtt acg cca cgg cat gcg taa ctt tga tag caa gga taa aaa agg aca tat tcc tct aca cca cta tct gat tca ttc act atc aat cga taa caa gat ctt tga tat act aac gga ccc cat tga tga ctt tag taa atc atc cga tct att gct gtg tta tct tag ata taa att caa tgg gag ctt aaa cta tta cgt tct gta caa att att gac taa agg atc tga ccc taa ttg cgt cga tga gga tgg act cac ttc tct tca tta cta ctg taa aca cat atc cgc gtt cca cga aag caa tta tta caa gtc aaa gag tca cac taa gat gcg agc tga gaa gcg att cat cta cgc gat aat aga tca tgg agc aaa cat taa cgc ggt tac gaa aat cgg aaa tac gcc gtt aca cac tta cct tca aca gta tac caa aca tag tcc tcg tgt ggt gta tgc tct ttt atc tcg agg agc cga tac gag gat acg taa taa tct taa ttg tac acc cat cat gga ata cat aaa gaa cga ttg tgc aac agg tca tat tct cat aat gtt act caa ttg gca cga aca aaa ata cgg gaa att aca aaa gga aga agg aca aca tct act tta tct att cat aaa aca taa tca agg ata tgg aag tcg ctc tct caa tat act acg gta tct act aga tag att cga cat tca gaa aga cga ata cta taa tac aat gac tcc tct tca tac cgc ctt cca gaa ttg caa taa caa tgt tgc ctc ata cct cgt ata cat tgg ata cga cat caa cct tcc gac taa aga cga taa gac agt att cga ctt ggt gtt tga aaa cag aaa cat tat ata caa ggc gga tgt cgt taa tga cat tat cca cca cag act gaa agt atc tct acc tat gat taa atc gtt gtt cta caa gat gtc gga gtt ctc tcc cta cga cga tca cta cgt aaa gaa gat aat agc cta ctg cct att aag gga cga gtc att tgc gga act aca tac taa att ctg ttt aaa cga gga cta taa aag tgt att tat gaa aaa tat atc att cga taa gat aga ttc cat cat cga aaa atg tag tcg tga cat aag tct cct caa aga gat tcg aat ctc aga cac cga ctt gta tac ggt att gag aac aga aga cat tcg gta tca cac ata tct cga agc cat aca ttc aga caa acg cat ttc att tcc cat gta cga cga tct cat aga aca gtg tca tct atc gat gga gca taa aag taa act cgt cga caa agc act caa taa att aga gtc tac cat cga tag tca atc tag act atc gta ttt gcc tcc gga aat tat gcg caa tat cat aac caa gct aag cga cta cca tct aaa cag tat gtt gta cgg aaa gaa cca tta caa ata tta tcc atg ata gaa aga aaa tat tta aaa aat aat cta tat gat tgg aga agt agg aaa caa aca gta aca aga cga cga tta cta c-a tta tta aat cat gag gtc cgt att ata ctc gta tat att gtt tct ctc atg tat aat aat aaa cgg aag aga tat agc acc aca tgc acc atc caa tgg aaa gtg taa aga caa cga ata cag aag ccg taa tct atg ttg tct atc gtg tcc tcc ggg aac tta cgc ttt cag att atg tga tag caa gac taa tac aca atg tac acc gtg tgg ttc gga tac ctt tac atc tca caa taa tca ttt aca ggc ttg tct aag ttg taa cgg aag atg tga tag taa tca ggt aga gac gcg atc gtg taa cac gac tca caa tag aat ctg tga atg ctc tcc agg ata tta ttg tct tct caa agg agc atc agg gtg tag aac atg tat ttc taa aac aaa gtg tgg aat agg ata cgg agt atc cgg ata cac gtc tac cgg aga cgt cat ctg ttc tcc gtg tgg tcc cgg aac ata ttc tca cac cgt ctc ttc cac aga taa atg cga acc cgt cgt aac cag caa tac att taa cta tat cga tgt gga aat taa cct gta tcc agt caa cga cac atc gtg tac tcg gac gac cac tac cgg tct cag cga atc cat ctc aac gtc gga act aac tat tac cat gaa tca taa aga ttg tga tcc agt ctt tcg tgc aga ata ctt ctc tgt cct taa taa tgt agc aac ttc agg att ctt tac agg aga aaa tag ata tca gaa tac ttc aaa gat atg tac tct gaa ttt cga gat taa atg taa caa caa aga ttc atc ttc caa aca gtt aac gaa aac aaa gaa tga tac tat cat gcc gca ttc aga gac ggt aac tct agt ggg cga ctg tct atc tag cgt cga cat cta cat act ata tag taa tac caa tac tca aga cta cga aaa tga tac aat ctc tta tca tat ggg taa tgt tct cga tgt caa tag cca tat gcc cgc tag ttg cga tat aca taa act gat cac taa ttc cca gaa tcc cac cca ctt ata gta ag- -tt ttt tta cct ata aat aat aaa tac aat aat taa ttt ctc gta aaa gta gaa aat ata ttc taa ttt att ata tgg taa gaa agt aga atc atc tag aac agt aat caa tca ata gca atc atg aaa caa tat att gtc ctg gca tgc atg tgc cta gtg gca gct gct atg cct act agt ctt caa caa tct tca tcc tcg tgt act gaa gaa gaa aac aaa cat cat atg gga atc gat gtt att atc aaa gtc aca aag caa gac caa aca ccg acc aat gat aag att tgt caa tcc gta acg gaa gtt aca gag a-- --- --- --- --- --- -cc gaa gat gat gag gta tcc gaa gaa gtt gta aaa gga gat ccc acc act tat tac act atc gtc ggt gcg ggt ctt aac atg aac ttt gga ttc acc aaa tgc cca aag att tta tcc atc tcc gaa tcc tct gat gga aac act gtg aat act aga ttg tcc agc gtg tca ccg gga caa ggt aag gac tct ccc gcg atc acg cgt gaa gaa gct ctg gct atg atc aaa gac tgt gag atg tct atc gac atc aga tgt agc gaa gaa gag aaa gac agt gac atc aag acc cat cca gta ctt ggg tct aac atc tca cat aag aaa gtg agt tac aaa gat atc atc ggt tca acg atc gtt gat aca aaa tgt gtc aag aac cta gag ttt agc gta cgt atc gga gac atg tgt gag gaa tca tct gaa ctt gaa gtc aag gat gga ttc aag tat gtc gac gga tcg gca tct gaa ggt gca acc gat gat act tca ctc atc gat tca aca aaa ctc aaa gca tgt gtc tga atc gat aac tct att cat ctg aaa atg gat gag ttg ggt taa tcg aac gat tca gac acc gca cca cga att aaa aaa g-- -ac cgg gca cta tat tcc ggt ttg caa aac aaa aat att taa cta cat tca caa aaa gtt acc tct cgt tac ttc ttc ttt ctg ttt caa tat gtg ata cga tat gat cac tat tcg tat tct ct- --- --- tgg tct cat aaa aaa gtt tta c-- --a aaa aaa aaa aaa a?t att ttt att ctc ttt ctc tct tcg at- ggt ctc aca aaa ata tta aac ctc ttt ctg atg tct caa cta ttt cgt aaa cga taa cgt cca aca ata tat tct cgt aga gct tat caa cat cct tat acc aat cta ggt tgt cag aca att gca tca taa aat aat gtt tat aat tta cac gtt aac atc ata taa taa acg tat ata gtt aat att ttt gga ata taa atg atc tgt aaa atc cat gta ggg gac act gct cac gtt ttt tct cta gta cat aat ttc aca caa gtt ttt ata cag aca aat taa ttc tcg tcc ata ta- ttt taa aac att gac ttt tgt act aag aaa aat atc ttg act aac cat ctc ttt ctc tct tcg atg ggt ctc aca aaa ata tta aac ctc ttt ctg atg g-- --- --- --- --- --- --- --- --- --- --- --- --- --- --- --- --- --- --- --- --- --- --- --- --- --- --- --- --- --- --- --- --- --- --- --- --- --- --- --- --- --- --- --- --- --- --- agt cgt aaa aag ttt tta tcc ttt ctc tct tcg ata ggt ctc --- --- --- --- --- --- --- --- --- --- --- --- --- --- --- --- --- --- --- --- --- --- --- --- --- --- --- --- --- --- --- --- --- --- --- --- --- --- --- --- --- --- --- --- --- --- --- --- --- --- --- --- --- --- --- --- --- --- --- --- --- --- --- --- --- --- --- --- --- --- aca aaa ata tta aac ctc ttt ctg atg gtc tct ata aa

>MN648051.1_Monkeypox_virus_strain_Israel_2018

atc gag gac gac ttc aac tat gac agc tcg gtg gcg tct gcc agc gtg tac ata cga atg gca ttt cta aga aaa gtc tac ggt atc ctt -tc tac aat ttc ctt taa caa cgg caa cag ctg cag tat ttt tat act ttg aat gca -tc gga cat tta tac aag gga gtc ctg ttc taa tat tgg cat caa tgt tcg gat cta tag gct tga ttt tcg cat tga ctt tac aca gac atg aac atc ccc tga atc tgt aca tac ttt gtg gat tta cac tgt tag aat ctc taa cgc tgg cct ctg ttg tta ctt tct atg atg cac gta tcg tta tgc aag ctt tca tgt tga cta ctg cag tgt ttc ttg ctc tga cta cat ata ctc tac aat caa aga gag att tca gta aac ttg taa cag gat tgt ttg ctg ctt tct gga ttt taa ttt tgt cag gag tct tga gga taa agt tta aaa tag aat taa taa aga aca tat agg tca -tt ttt taa aca tgg ata gaa acc aag gtt gtt agt taa taa tat aca aga tat ttt ttc tca ctc tga tcc atg taa acc aag gac gag aga cac tct cat tcc tca ttc acg aca cca tta aaa atg gaa att aaa gcc ctc tat taa gca cag acg gct aca ggt cta cca tca ggt t-a cct tcg tct acc ttc aca atg gcc tct cct tgt gcc cag ttc agt ccc tgt cat tgc cac gct act aag gac tcc ctg aat acc gtg act gac gtc aga cat tgt ctg act gaa tac atc ctg tgg gtt tct cat aga tgg acc cat aga gaa agc gca ggg cct ctc tac agg ctt ctc atc tct ttc aga att gat gca atg gag cta ttt ggt agc gag ttg aag gag ttc tcg aat tca ctt ccg tgg gac aat atc gac aat tgc gtg gag atc att aaa tgt ttc atc aga aat gac tcc atg aaa acc gcc aaa gaa ctt tgt gca ata att gga ctt tgt act caa tca gct att gtc act gga aga gtc ttc aat gat aag tat atc gac ata cta ctt atg ctg cga aag att ctg aac gag aac gac tat ctc acc ctc ttg gat cat atc ctc act gct aaa tac taa atc tcc ttc atg ctc tct cac taa tac tct tac tca cta cac ttt tta tca tct tat gat gaa tga ttg cct tca tca ttt ttt cgt gga ata taa tat agg aat aat tag cac cag aat agc tat gga tat ctc gtt aag aat att ctc tat aag aga cat aat gta gac ata gtt att ata tcc ttc tta gat aag tgt tac gct act gga aag ttt cca tcg tta tta tta cat gaa gat gat ata att aaa cca aca ttg aga ttg gct ctt atg tta gct gga ttg aat tac tgt aat aaa tgc atc gag tat aga ggg ata tag caa ttc tcg ata ata gtc atg caa tat ttg aat gag act gat aat tta ggt aat aca gta ct- --- aca cac ata tct ttc tag att ata tat cgt taa aaa tct gta aga tgt ata ttt ctc ata agt atc cac tgt gta ata tta tta atg gat ata tag ata acg caa tag gga cta ata gta ttg taa aag ata taa tcg act att tac gta cat atc cag ata tct ata ttc cta cta gtt tgc tgc gta gtt gca tca ttg ata tgc atg -at tta tca gga ttc aga gat gaa tta cta agt aaa cta caa tcc cac aat aag taa gaa tca aat atc aaa aac tca ctt ttg att ttt cta gtc tta agt aat aca tat att tat taa tag acc tat gaa ata aaa aaa ggt aac aat gga ttc gcg tat agc tat tta cgt att agt atc ggc atc tct ttt gta tct tgt taa ttg tca caa act agt aca tta ctt caa tct gaa aat aaa tgg aag tga tat aac taa tac agc aga tat att gct gga caa tta tcc aat tat gac ctt tga tgg aaa gga tat tta tcc atc tat ctc gtt cat ggt cgg taa taa act ttt cct aga tct tta taa aaa tat ctt tga aga att ttt cag act att tcg agt atc tgt aag tag tca ata cga gga att aga ata tta tta ttc atg tga tta tac taa caa ccg tcc tac aat taa aca aca tta ctt tta taa cgg cga cga ata tac tga aat tga tag atc gaa aaa agc cac taa taa aaa cag ttg gtt aat tac ttc agg ctt tag act aca aaa atg gtt cga tag cga aga ttg tat aat tta tct cag atc ttt agt tag aag aat gga aga cag taa caa aaa cag taa aaa aac tta gta ctt aga tat cg- --a aaa aat ata ttt ttg tag act ctt gag aat aga agg aaa aca tgt aca taa tta taa aaa atg aaa atc aat ggc gaa taa gac agt gcg att cgc acc atg gag tcg gta gat ttc atg gct gtc gat gag cag ttt cac gac gac ctc gat ctt tgg tca tta tct ttg gta gat gat tat aaa aaa cat gga tta ggt gtt gac tgt tat gtt cta gaa cca gtt gtt gac agg aaa ata ttt gat aga ttt ctc ctt gaa cca att tgt gat cct gta gat gtt ctg tat gat tat ttt agg att cat aga gat aat att gat cag tat ata gta gat aga ctg ttt gca tat att aca tat aaa gat att ata tct gca tta gtg tca aag aat tat atg gaa gat att ttc tct ata att att aag aat tgt aat tct gtg caa gat ctc tta ctt tac tat cta tct aat gca tat gta gaa ata gac att gtt gat ctt atg gta gat cat ggg gct gta ata tat aaa ata gaa tgc ttg aat gcc tat ttt agg gga ata tgt aaa aag gaa agt agt gtt gtt gag ttt att ttg aat tgt ggt atc cca gat gaa aat gat gtt aaa tta gat cta tat aaa ata att cag tat act agg gga ttc ctt gta gat gaa ccc aca gta tta gaa att tat aag ctt tgt atc cca tat att gaa gat atc aat caa cta gat gct ggt gga agg acc ttg ctt tat cgc gct atc tat gca ggt tat ata gat tta gta tca tgg cta tta gaa aat gga gca aat gtc aac gca gta atg agt aat gga tat aca tgt ctt gac gtg gcc gtg gat agg gga tct gtc atc gcc cgt agg gaa gca cat ctt aaa ata tta gaa ata ttg ctt aga gaa cca ttg tct att gac tgt ata aaa tta gct ata ctt aat aat aca att gaa aac cat gat gtg ata aag ctc tgt atc aag tat ttt atg atg gta gat tat tca ctt tgt aat gtg tat gca tca tca ctc ttt gat tat ata att gat tgt aaa caa gaa ttg gag tac att agg cag atg aaa att cat aat aca acc atg tat gag tta atc tat aat aga gac aaa aac aag cat gct tcc cat att cta cat agg tat tct aaa cat cca gtt ttg aca cag tgt atc act aaa gga ttc aag att tac aca gaa gta acc gag cag gtc act aaa gct cta aac aga cgt gct cta ata gat gag ata ata aac aat gta tca act gat gac aat ctc cta tca aaa ctt cca tta gaa att agg gat cta att gtt tca caa gct gtc ata tag agt tct atc cac cca cct ttc ttg aaa tga gtt aat agt cat aag tta gtt aag tca taa gtt agt taa gtc ata agt tag tta agt cat aag tta gtt aag tca taa gtt agt taa g-- --- --- --- --- --- --- --- --- --- --- --- --- --- --- --- --- --- --- --- --- --- --- --- --- --- --- --- --- --- --- --- --- --- --- --- --- --- --- -ag tta at- -ag tca taa gtt --- --- --- --- agt taa gtc ata agt tag tta agt cat aag tta gtt aag tca taa gtt agt taa gtc ata agt tag tta agt cat aag tta gtt aag tca taa gtt agt taa --- --- --- --- --- -gt cat aag tta gtt aag tca taa gtt agt taa gtc ata agt tag tta agt cat aag tta gtt aag tca taa gtt agt taa gtc ata agt tag tta agt cat aag tta gtt aag tca taa gtt agt taa gtc ata agt tag tta agt cat aag tta gtt tat agt cta aca ctt cta att ttt ata cct tga tct ttt tct cta att atg aaa aag taa atc att atg aag atg gat g-a aat gga cga gat tgt gcg cat cgt taa cga tag tat gtg gta cgt acc taa cgc att tat gga cga cgg tga taa tga agg tca cat ttc tgt caa taa tgt ctg tca tat gta tct cgc att ctt tga tgt gga tat atc atc tca tct gtt taa att agt tat taa aca ctg cga tct gaa taa acg act aaa atg tgg taa ctc tcc att aca ttg cta tac gat gaa tac acg att taa tcc atc tgt att aaa gat att gtt acg cca cgg cat gcg taa ctt tga tag caa gga taa aaa agg aca tat tcc tct aca cca cta tct gat tca ttc act atc aat cga taa caa gat ctt tga tat act aac gga ccc cat tga tga ctt tag taa atc atc cga tct att gct gtg tta tct tag ata taa att caa tgg gag ctt aaa cta tta cgt tct gta caa att att gac taa agg atc tga ccc taa ttg cgt cga tga gga tgg act cac ttc tct tca tta cta ctg taa aca cat atc cgc gtt cca cga aag caa tta tta caa gtc aaa gag tca cac taa gat gcg agc tga gaa gcg att cat cta cgc gat aat aga tca tgg agc aaa cat taa cgc ggt tac gaa aat cgg aaa tac gcc gtt aca cac tta cct tca aca gta tac caa aca tag tcc tcg tgt ggt gta tgc tct ttt atc tcg agg agc cga tac gag gat acg taa taa tct tga ttg tac acc cat cat gga ata cat aaa gaa cga ttg tgc aac agg tca tat tct cat aat gtt act caa ttg gca cga aca aaa ata cgg gaa att aca aaa gga aga agg aca aca tct act tta tct att cat aaa aca taa tca agg ata tgg aag tcg ctc tct caa tat act acg gta tct act aga tag att cga cat tca gaa aga cga ata cta taa tac aat gac tcc tct tca tac cgc ctt cca gaa ttg caa taa caa tgt tgc ctc ata cct cgt ata cat cgg ata cga cat caa cct tcc gac taa aga cga taa gac agt att cga ctt ggt gtt tga aaa cag aaa cat tat ata caa ggc gga tgt cgt taa tga cat tat cca cca cag act gaa agt atc tct acc tat gat taa atc gtt gtt cta caa gat gtc gga gtt ctc tcc cta cga cga tca cta cgt aaa gaa gat aat agc cta ctg cct att aag gga cga gtc att tgc gga act aca tac taa att ctg ttt aaa cga gga cta taa aag tgt att tat gaa aaa tat atc att cga taa gat aga ttc cat cat cga aaa atg tag tcg tga cat aag tct cct caa aga gat tcg aat ctc aga cac cga ctt gta tac ggt att gag aac aga aga cat ccg gta tca cac ata tct cga agc cat aca ttc aga caa acg cat ttc att tcc cat gta cga cga tct cat aga aca gtg tca tct atc gat gga gca taa aag taa act cgt cga caa agc act caa taa att aga gtc tac cat cga tag tca atc tag act atc gta ttt gcc tcc gga aat tat gcg caa tat cat aac caa gct aag cga cta cca tct aaa cag tat gtt gta cgg aaa gaa cca tta caa ata tta tcc atg ata gaa aga aaa tat tta aaa aat aat cta tat gat tgg aga agt agg aaa caa aca gta aca aga cga cga tta cta c-a tta tta aat cat gag gtc cgt att ata ctc gta tat att gtt tct ctc atg tat aat aat aaa cgg aag aga tat agc acc aca tgc acc atc caa tgg aaa gtg taa aga caa cga ata cag aag ccg taa tct atg ttg tct atc gtg tcc tcc ggg aac tta cgc ttc cag att atg tga tag caa gac taa tac aca atg tac acc gtg tgg ttc gga tac ctt tac atc tca caa taa tca ttt aca ggc ttg tct aag ttg taa cgg aag atg tga tag taa tca ggt aga gac gcg atc gtg taa cac gac tca caa tag aat ctg tga atg ctc tcc agg ata tta ttg tct tct caa agg agc atc agg gtg tag aac atg tat ttc taa aac aaa gtg tgg aat agg ata cgg agt atc cgg ata cac gtc tac cgg aga cgt cat ctg ttc tcc gtg tgg tcc cgg aac ata ttc tca cac cgt ctc ttc cac aga taa atg cga acc cgt cgt aac cag caa tac att taa cta tat cga tgt gga aat taa cct gta tcc agt caa cga cac atc gtg tac tcg gac gac cac tac cgg tct cag cga atc cat ctc aac gtc gga act aac tat tac cat gaa tca taa aga ttg tga tcc agt ctt tcg tgc aga ata ctt ctc tgt cct taa taa tgt agc aac ttc agg att ctt tac agg aga aaa tag ata tca gaa tac ttc aaa gat atg tac tct gaa ttt cga gat taa atg taa caa caa aga ttc atc ttc caa aca gtt aac gaa aac aaa gaa tga tac tat cat gcc gca ttc aga gac ggt aac tct agt ggg cga ctg tct atc tag cgt cga cat cta cat act ata tag taa tac caa tac tca aga cta cga aaa tga tac aat ctc tta tca tat ggg taa tgt tct cga tgt caa tag cca tat gcc cgc tag ttg cga tat aca taa act gat cac taa ttc cca gaa tcc cac cca ctt ata gta ag- -tt ttt tta cct ata aat aat aaa tac aat aat taa ttt ctc gta aaa gta gaa aat ata ttc taa ttt att ata tgg taa gaa agt aga atc atc tag aac agt aat caa tca ata gca atc atg aaa caa tat att gtc ctg gca tgc atg tgc cta gtg gca gct gct atg cct act agt ctt caa caa tct tca tcc tcg tgt act gaa gaa gaa aac aaa cat cat atg gga atc gat gtt att atc aaa gtc aca aag caa gac caa aca ccg acc aat gat aag att tgt caa tcc gta acg gaa gtt aca gag a-- --- --- --- --- --- -cc gaa gat gat gag gta tcc gaa gaa gtt gta aaa gga gat ccc acc act tat tac act atc gtc ggt gcg ggt ctt aac atg aac ttt gga ttc acc aaa tgc cca aag att tca tcc atc tcc gaa tcc tct gat gga aac act gtg aat act aga ttg tcc agc gtg tca ccg gga caa ggt aag gac tct ccc gcg atc acg cgt gaa gaa gct ctg gct atg atc aaa gac tgt gag atg tct atc gac atc aga tgt agc gaa gaa gag aaa gac agt gac atc aag acc cat cca gta ctt ggg tct aac atc tca cat aag aaa gtg agt tac aaa gat atc atc ggt tca acg atc gtt gat aca aaa tgt gtc aag aac cta gag ttt agc gta cgt atc gga gac atg tgt gag gaa tca tct gaa ctt gaa gtc aag gat gga ttc aag tat gtc gac gga tcg gca tct gaa ggt gca acc gat gat act tca ctc atc gat tca aca aaa ctc aaa gca tgt gtc tga atc gat aac tct att cat ctg aaa atg gat gag ttg ggt taa tcg aac gat tca gac acc gca cca cga att aaa aaa g-- -ac cgg gca cta tat tcc ggt ttg caa aac aaa aat att taa cta cat tca caa aaa gtt acc tct cgt tac ttc ttc ttt ctg ttt caa tat gtg ata cga tat gat cac tat tcg tat tct ct- --- --- tgg tct cat aaa aaa gtt tta c-- --a aaa aaa aaa aaa a-t att ttt att ctc ttt ctc tct tcg at- ggt ctc aca aaa ata tta aac ctc ttt ctg atg tct caa cta ttt cgt aaa cga taa cgt cca aca ata tat tct cgt aga gct tat caa cat cct tat acc aat cta ggt tgt cag aca att gca tca taa aat aat gtt tat aat tta cac gtt aac atc ata taa taa acg tat ata gtt aat att ttt gga ata taa atg atc tgt aaa atc cat gta ggg gac act gct cac gtt ttt tct cta gta cat aat ttc aca caa gtt ttt ata cag aca aat taa ttc tcg tcc ata ta- ttt taa aac att gac ttt tgt act aag aaa aat atc ttg act aac cat ctc ttt ctc tct tcg --- --- --- --- --- --- --- --- --- --- --- --- --- --- --- --- --- --- --- --- --- --- --- --- --- --- --- --- --- --- --- --- --- --- --- --- --- --- --- --- --- --- --- --- --- --- --- --- --- --- --- --- --- --- --- --- --- --- --- --- --- --- --- --- --- --- --- --- --- --- --- --- --- --- --- --- --- --- --- --- --- --- --- --- --- --- --- --- --- --- --- --- --- --- --- --- --- --- --- --- --- --- --- --- --- --- --- --- --- --- --- --- --- --- --- --- --- --- --- --- --- --- --- --- --- --- --- --- --- --- --- --- --- --- --- --- --- --- --- --- --- --- --- --- --- --- --- --- --- --- --- --- --- --- --- --

>ON644344.1_Monkeypox_virus_isolate_MPXV_FVG-ITA_01_2022_complete_genome

atc gag gac gac ttc aac tat gac agc tcg gtg gcg tct gcc agc gtg tac ata cga atg gca ttt cta aga aaa gtc tac ggt atc ctt -tc tac aat ttc ctt taa caa cgg caa cag ctg cag tat ttt tat act ttg aat gca -tc gga cat tta tac aag gga gtc ctg ttc taa tat tgg cat caa tgt tcg gat cta tag gct tga ttt tcg cat tga ctt tac aca gac atg aac atc ccc tga atc tgt aca tac ttt gtg gat tta cac tgt tag aat ctc taa cgc tgg cct ctg ttg tta ctt tct atg atg cac gta tcg tta tgc aag ctt tca tgt tga cta ctg cag tgt ttc ttg ctc tga cta cat ata ctc tac aat caa aga gag att tca gta aac ttg taa cag gat tgt ttg ctg ctt tct gga ttt taa ttt tgt cag gag tct tga gga taa agt tta aaa tag aat taa taa aga aca tat agg tca -tt ttt taa aca tgg ata gaa acc aag gtt gtt agt taa taa tat aca aga tat ttt ttc tca ctc tga tcc atg taa acc aag gac gag aga cac tct cat tcc tca ttc acg aca cca tta aaa atg gaa att aaa gcc ctc tat taa gca cag acg gct aca ggt cta cca tca ggt t-a cct tcg tct acc ttc aca atg gcc tct cct tgt gcc cag ttc agt ccc tgt cat tgc cac gct act aag gac tcc ctg aat acc gtg act gac gtc aga cat tgt ctg act gaa tac atc ctg tgg gtt tct cat aga tgg acc cat aga gaa agc gca ggg cct ctc tac agg ctt ctc atc tct ttc aga att gat gca atg gag cta ttt ggt agc gag ttg aag gag ttc tcg aat tca ctt ccg tgg gac aat atc gac aat tgc gtg gag atc att aaa tgt ttc atc aga aat gac tcc atg aaa acc gcc aaa gaa ctt tgt gca ata att gga ctt tgt act caa tca gct att gtc act gga aga gtc ttc aat gat aag tat atc gac ata cta ctt atg ctg cga aag att ctg aac gag aac gac tat ctc acc ctc ttg gat cat atc ctc act gct aaa tac taa atc tcc ttc atg ctc tct cac taa tac tct tac tca cta cac ttt tta tca tct tat gat gaa tga ttg cct tca tca ttt ttt cgt gga ata taa tat agg aat aat tag cac cag aat agc tat gga tat ctc gtt aag aat att ctc tat aag aga cat aat gta gac ata gtt att ata tcc ttc tta gat aag tgt tac gct act gga aag ttt cca tcg tta tta tta cat gaa gat gat ata att aaa cca aca ttg aga ttg gct ctt atg tta gct gga ttg aat tac tgt aat aaa tgc atc gag tat aga ggg ata tag caa ttc tcg ata ata gtc atg caa tat ttg aat gag act gat aat tta ggt aat aca gta ct- --- aca cac ata tct ttc tag att ata tat cgt taa aaa tct gta aga tgt ata ttt ctc ata agt atc cac tgt gta ata tta tta atg gat ata tag ata acg caa tag gga cta ata gta ttg taa aag ata taa tcg act att tac gta cat atc cag ata tct ata ttc cta cta gtt tgc tgc gta gtt gca tca ttg ata tgc atg -at tta tca gga ttc aga gat gaa tta cta agt aaa cta caa tcc cac aat aag taa gaa tca aat atc aaa aac tca ctt ttg att ttt cta gtc tta agt aat aca tat att tat taa tag acc tat gaa ata aaa aaa ggt aac aat gga ttc gcg tat agc tat tta cgt att agt atc ggc atc tct ttt gta tct tgt taa ttg tca caa act agt aca tta ctt caa tct gaa aat aaa tgg aag tga tat aac taa tac agc aga tat att gct gga caa tta tcc aat tat gac ctt tga tgg aaa gga tat tta tcc atc tat ctc gtt cat ggt cgg taa taa act ttt cct aga tct tta taa aaa tat ctt tga aga att ttt cag act att tcg agt atc tgt aag tag tca ata cga gga att aga ata tta tta ttc atg tga tta tac taa caa ccg tcc tac aat taa aca aca tta ctt tta taa cgg cga cga ata tac tga aat tga tag atc gaa aaa agc cac taa taa aaa cag ttg gtt aat tac ttc agg ctt tag act aca aaa atg gtt cga tag cga aga ttg tat aat tta tct cag atc ttt agt tag aag aat gga aga cag taa caa aaa cag taa aaa aac tta gta ctt aga tat cg- --a aaa aat ata ttt ttg tag act ctt gag aat aga agg aaa aca tgt aca taa tta taa aaa atg aaa atc aat ggc gaa taa gac agt gcg att cgc acc atg gag tcg gta gat ttc atg gct gtc gat gag cag ttt cac gac gac ctc gat ctt tgg tca tta tct ttg gta gat gat tat aaa aaa cat gga tta ggt gtt gac tgt tat gtt cta gaa cca gtt gtt gac agg aaa ata ttt gat aga ttt ctc ctt gaa cca att tgt gat cct gta gat gtt ctg tat gat tat ttt agg att cat aga gat aat att gat cag tat ata gta gat aga ctg ttt gca tat att aca tat aaa gat att ata tct gca tta gtg tca aag aat tat atg gaa gat att ttc tct ata att att aag aat tgt aat tct gtg caa gat ctc tta ctt tac tat cta tct aat gca tat gta gaa ata gac att gtt gat ctt atg gta gat cat ggg gct gta ata tat aaa ata gaa tgc ttg aat gcc tat ttt agg gga ata tgt aaa aag gaa agt agt gtt gtt gag ttt att ttg aat tgt ggt atc cca gat gaa aat gat gtt aaa tta gat cta tat aaa ata att cag tat act agg gga ttc ctt gta gat gaa ccc aca gta tta gaa att tat aag ctt tgt atc cca tat att gaa gat atc aat caa cta gat gct ggt gga agg acc ttg ctt tat cgc gct atc tat gca ggt tat ata gat tta gta tca tgg cta tta gaa aat gga gca aat gtc aac gca gta atg agt aat gga tat aca tgt ctt gac gtg gcc gtg gat agg gga tct gtc atc gcc cgt agg gaa gca cat ctt aaa ata tta gaa ata ttg ctt aga gaa cca ttg tct att gac tgt ata aaa tta gct ata ctt aat aat aca att gaa aac cat gat gtg ata aag ctc tgt atc aag tat ttt atg atg gta gat tat tca ctt tgt aat gtg tat gca tca tca ctc ttt gat tat ata att gat tgt aaa caa gaa ttg gag tac att agg cag atg aaa att cat aat aca acc atg tat gag tta atc tat aat aga gac aaa aac aag cat gct tcc cat att cta cat agg tat tct aaa cat cca gtt ttg aca cag tgt atc act aaa gga ttc aag att tac aca gaa gta acc gag cag gtc act aaa gct cta aac aga cgt gct cta ata gat gag ata ata aac aat gta tca act gat gac aat ctc cta tca aaa ctt cca tta gaa att agg gat cta att gtt tca caa gct gtc ata tag agt tct atc cac cca cct ttc ttg aaa tga gtt aat agt cat aag tta gtt aag tca taa gtt agt taa gtc ata agt tag tta agt cat aag tta gtt aag tca taa gtt agt taa g-- --- --- --- --- --- --- --- --- --- --- --- --- --- --- --- --- --- --- --- --- --- --- --- --- --- --- --- --- --- --- --- --- --- --- --- --- --- --- -ag tta at- -ag tca taa gtt --- --- --- --- agt taa gtc ata agt tag tta agt cat aag tta gtt aag tca taa gtt agt taa gtc ata agt tag tta agt cat aag tta gtt aag tca taa gtt agt taa --- --- --- --- --- -gt cat aag tta gtt aag tca taa gtt agt taa gtc ata agt tag tta agt cat aag tta gtt aag tca taa gtt agt taa gtc ata agt tag tta agt cat aag tta gtt aag tca taa gtt agt taa gtc ata agt tag tta agt cat aag tta gtt tat agt cta aca ctt cta att ttt ata cct tga tct ttt tct cta att atg aaa aag taa atc att atg aag atg gat g-a aat gga cga gat tgt gcg cat cgt taa cga tag tat gtg gta cgt acc taa cgc att tat gga cga cgg tga taa tga agg tca cat ttc tgt caa taa tgt ctg tca tat gta tct cgc att ctt tga tgt gga tat atc atc tca tct gtt taa att agt tat taa aca ctg cga tct gaa taa acg act aaa atg tgg taa ctc tcc att aca ttg cta tac gat gaa tac acg att taa tcc atc tgt att aaa gat att gtt acg cca cgg cat gcg taa ctt tga tag caa gga taa aaa agg aca tat tcc tct aca cca cta tct gat tca ttc act atc aat cga taa caa gat ctt tga tat act aac gga ccc cat tga tga ctt tag taa atc atc cga tct att gct gtg tta tct tag ata taa att caa tgg gag ctt aaa cta tta cgt tct gta caa att att gac taa agg atc tga ccc taa ttg cgt cga tga gga tgg act cac ttc tct tca tta cta ctg taa aca cat atc cgc gtt cca cga aag caa tta tta caa gtc aaa gag tca cac taa gat gcg agc tga gaa gcg att cat cta cgc gat aat aga tca tgg agc aaa cat taa cgc ggt tac gaa aat cgg aaa tac gcc gtt aca cac tta cct tca aca gta tac caa aca tag tcc tcg tgt ggt gta tgc tct ttt atc tcg agg agc cga tac gag gat acg taa taa tct taa ttg tac acc cat cat gga ata cat aaa gaa cga ttg tgc aac agg tca tat tct cat aat gtt act caa ttg gca cga aca aaa ata cgg gaa att aca aaa gga aga agg aca aca tct act tta tct att cat aaa aca taa tca agg ata tgg aag tcg ctc tct caa tat act acg gta tct act aga tag att cga cat tca gaa aga cga ata cta taa tac aat gac tcc tct tca tac cgc ctt cca gaa ttg caa taa caa tgt tgc ctc ata cct cgt ata cat tgg ata cga cat caa cct tcc gac taa aga cga taa gac agt att cga ctt ggt gtt tga aaa cag aaa cat tat ata caa ggc gga tgt cgt taa tga cat tat cca cca cag act gaa agt atc tct acc tat gat taa atc gtt gtt cta caa gat gtc gga gtt ctc tcc cta cga cga tca cta cgt aaa gaa gat aat agc cta ctg cct att aag gga cga gtc att tgc gga act aca tac taa att ctg ttt aaa cga gga cta taa aag tgt att tat gaa aaa tat atc att cga taa gat aga ttc cat cat cga aaa atg tag tcg tga cat aag tct cct caa aga gat tcg aat ctc aga cac cga ctt gta tac ggt att gag aac aga aga cat tcg gta tca cac ata tct cga agc cat aca ttc aga caa acg cat ttc att tcc cat gta cga cga tct cat aga aca gtg tca tct atc gat gga gca taa aag taa act cgt cga caa agc act caa taa att aga gtc tac cat cga tag tca atc tag act atc gta ttt gcc tcc gga aat tat gcg caa tat cat aac caa gct aag cga cta cca tct aaa cag tat gtt gta cgg aaa gaa cca tta caa ata tta tcc atg ata gaa aga aaa tat tta aaa aat aat cta tat gat tgg aga agt agg aaa caa aca gta aca aga cga cga tta cta c-a tta tta aat cat gag gtc cgt att ata ctc gta tat att gtt tct ctc atg tat aat aat aaa cgg aag aga tat agc acc aca tgc acc atc caa tgg aaa gtg taa aga caa cga ata cag aag ccg taa tct atg ttg tct atc gtg tcc tcc ggg aac tta cgc ttt cag att atg tga tag caa gac taa tac aca atg tac acc gtg tgg ttc gga tac ctt tac atc tca caa taa tca ttt aca ggc ttg tct aag ttg taa cgg aag atg tga tag taa tca ggt aga gac gcg atc gtg taa cac gac tca caa tag aat ctg tga atg ctc tcc agg ata tta ttg tct tct caa agg agc atc agg gtg tag aac atg tat ttc taa aac aaa gtg tgg aat agg ata cgg agt atc cgg ata cac gtc tac cgg aga cgt cat ctg ttc tcc gtg tgg tcc cgg aac ata ttc tca cac cgt ctc ttc cac aga taa atg cga acc cgt cgt aac cag caa tac att taa cta tat cga tgt gga aat taa cct gta tcc agt caa cga cac atc gtg tac tcg gac gac cac tac cgg tct cag cga atc cat ctc aac gtc gga act aac tat tac cat gaa tca taa aga ttg tga tcc agt ctt tcg tgc aga ata ctt ctc tgt cct taa taa tgt agc aac ttc agg att ctt tac agg aga aaa tag ata tca gaa tac ttc aaa gat atg tac tct gaa ttt cga gat taa atg taa caa caa aga ttc atc ttc caa aca gtt aac gaa aac aaa gaa tga tac tat cat gcc gca ttc aga gac ggt aac tct agt ggg cga ctg tct atc tag cgt cga cat cta cat act ata tag taa tac caa tac tca aga cta cga aaa tga tac aat ctc tta tca tat ggg taa tgt tct cga tgt caa tag cca tat gcc cgc tag ttg cga tat aca taa act gat cac taa ttc cca gaa tcc cac cca ctt ata gta ag- -tt ttt tta cct ata aat aat aaa tac aat aat taa ttt ctc gta aaa gta gaa aat ata ttc taa ttt att ata tgg taa gaa agt aga atc atc tag aac agt aat caa tca ata gca atc atg aaa caa tat att gtc ctg gca tgc atg tgc cta gtg gca gct gct atg cct act agt ctt caa caa tct tca tcc tcg tgt act gaa gaa gaa aac aaa cat cat atg gga atc gat gtt att atc aaa gtc aca aag caa gac caa aca ccg acc aat gat aag att tgt caa tcc gta acg gaa gtt aca gag a-- --- --- --- --- --- -cc gaa gat gat gag gta tcc gaa gaa gtt gta aaa gga gat ccc acc act tat tac act atc gtc ggt gcg ggt ctt aac atg aac ttt gga ttc acc aaa tgc cca aag att tta tcc atc tcc gaa tcc tct gat gga aac act gtg aat act aga ttg tcc agc gtg tca ccg gga caa ggt aag gac tct ccc gcg atc acg cgt gaa gaa gct ctg gct atg atc aaa gac tgt gag atg tct atc gac atc aga tgt agc gaa gaa gag aaa gac agt gac atc aag acc cat cca gta ctt ggg tct aac atc tca cat aag aaa gtg agt tac aaa gat atc atc ggt tca acg atc gtt gat aca aaa tgt gtc aag aac cta gag ttt agc gta cgt atc gga gac atg tgt gag gaa tca tct gaa ctt gaa gtc aag gat gga ttc aag tat gtc gac gga tcg gca tct gaa ggt gca acc gat gat act tca ctc atc gat tca aca aaa ctc aaa gca tgt gtc tga atc gat aac tct att cat ctg aaa atg gat gag ttg ggt taa tcg aac gat tca gac acc gca cca cga att aaa aaa g-- -ac cgg gca cta tat tcc ggt ttg caa aac aaa aat att taa cta cat tca caa aaa gtt acc tct cgt tac ttc ttc ttt ctg ttt caa tat gtg ata cga tat gat cac tat tcg tat tct ct- --- --- tgg tct cat aaa aaa gtt tta c-- --a aaa aaa aaa aaa a-t att ttt att ctc ttt ctc tct tcg at- ggt ctc aca aaa ata tta aac ctc ttt ctg atg tct caa cta ttt cgt aaa cga taa cgt cca aca ata tat tct cgt aga gct tat caa cat cct tat acc aat cta ggt tgt cag aca att gca tca taa aat aat gtt tat aat tta cac gtt aac atc ata taa taa acg tat ata gtt aat att ttt gga ata taa atg atc tgt aaa atc cat gta ggg gac act gct cac gtt ttt tct cta gta cat aat ttc aca caa gtt ttt ata cag aca aat taa ttc tcg tcc ata ta- ttt taa aac att gac ttt tgt act aag aaa aat atc ttg act aac cat ctc ttt ctc tct tcg --- --- --- --- --- --- --- --- --- --- --- --- --- --- --- --- --- --- --- --- --- --- --- --- --- --- --- --- --- --- --- --- --- --- --- --- --- --- --- --- --- --- --- --- --- --- --- --- --- --- --- --- --- --- --- --- --- --- --- --- --- --- --- --- --- --- --- --- --- --- --- --- --- --- --- --- --- --- --- --- --- --- --- --- --- --- --- --- --- --- --- --- --- --- --- --- --- --- --- --- --- --- --- --- --- --- --- --- --- --- --- --- --- --- --- --- --- --- --- --- --- --- --- --- --- --- --- --- --- --- --- --- --- --- --- --- --- --- --- --- --- --- --- --- --- --- --- --- --- --- --- --- --- --- --- --

>MT903343.1_Monkeypox_virus_isolate_MPXV-UK_P1

atc gaa gac gac ttc aac tat gac agc tcg gtg gcg tct gcc agc gtg tac ata cga atg gca ttt cta aga aaa gtc tac ggt atc ctt -tc tac aat ttc ctt taa caa cgg caa cag ctg cag tat ttt tat act ttg aat gca -tc gga cat tta tac aag gga gtc ctg ttc taa tat tgg cat caa tgt tcg gat cta tag gct tga ttt tcg cat tga ctt tac aca gac atg aac atc ccc tga atc tgt aca tac ttt gtg gat tta cac tgt tag aat ctc taa cgc tgg cct ctg ttg tta ctt tct atg atg cac gta tcg tta tgc aag ctt tca tgt tga cta ctg cag tgt ttc ttg ctc tga cta cat ata ctc tac aat caa aga gag att tca gta aac ttg taa cag gat tgt ttg ctg ctt tct gga ttt taa ttt tgt cag gag tct tga gga taa agt tta aaa tag aat taa taa aga aca tat agg tca -tt ttt taa aca tgg ata gaa acc aag gtt gtt agt taa taa tat aca aga tat ttt ttc tca ctc tga tcc atg taa acc aag gac gag aga cac tct cat tcc tca ttc acg aca cca tta aaa atg gaa att aaa gcc ctc tat taa gca cag acg gct aca ggt cta cca tca ggt t-a cct tcg tct acc ttc aca atg gcc tct cct tgt gcc cag ttc agt ccc tgt cat tgc cac gct act aag gac tcc ctg aat acc gtg act gac gtc aga cat tgt ctg act gaa tac atc ctg tgg gtt tct cat aga tgg acc cat aga gaa agc gca ggg cct ctc tac agg ctt ctc atc tct ttc aga att gat gca atg gag cta ttt ggt agc gag ttg aag gag ttc tcg aat tca ctt ccg tgg gac aat atc gac aat tgc gtg gag atc att aaa tgt ttc atc aga aat gac tcc atg aaa acc gcc aaa gaa ctt tgt gca ata att gga ctt tgt act caa tca gct att gtc act gga aga gtc ttc aat gat aag tat atc gac ata cta ctt atg ctg cga aag att ctg aac gag aac gac tat ctc acc ctc ttg gat cat atc ctc act gct aaa tac taa atc tcc ttc atg ctc tct cac taa tac tct tac tca cta cac ttt tta tca tct tat gat gaa tga ttg cct tca tca ttt ttt cgt gga ata taa tat agg aat aat tag cac cag aat agc tat gga tat ctc gtt aag aat att ctc tat aag aga cat aat gta gac ata gtt att ata tcc ttc tta gat aag tgt tac gct act gga aag ttt cca tcg tta tta tta cat gaa gat gat ata att aaa cca aca ttg aga ttg gct ctt atg tta gct gga ttg aat tac tgt aat aaa tgc atc gag tat aga ggg ata tag caa ttc tcg ata ata gtc atg caa tat ttg aat gag act gat aat tta ggt aat aca gta ct- --- aca cac ata tct ttc tag att ata tat cgt taa aaa tct gta aga tgt ata ttt ctc ata agt atc cac tgt gta ata tta tta atg gat ata tag ata acg caa tag gga cta ata gta ttg taa aag ata taa tcg act att tac gta cat atc cag ata tct ata ttc cta cta gtt tgc tgc gta gtt gca tca ttg ata tgc atg -at tta tca gga ttc aga gat gaa tta cta agt aaa cta caa tcc cac aat aag taa gaa tca aat atc aaa aac tca ctt ttg att ttt cta gtc tta agt aat aca tat att tat taa tag acc tat gaa ata aaa aaa ggt aac aat gga ttc gcg tat agc tat tta cgt att agt atc ggc atc tct ttt gta tct tgt taa ttg tca caa act agt aca tta ctt caa tct gaa aat aaa tgg aag tga tat aac taa tac agc aga tat att gct gga caa tta tcc aat tat gac ctt tga tgg aaa gga tat tta tcc atc tat ctc gtt cat ggt cgg taa taa act ttt cct aga tct tta taa aaa tat ctt tga aga att ttt cag act att tcg agt atc tgt aag tag tca ata cga gga att aga ata tta tta ttc atg tga tta tac taa caa ccg tcc tac aat taa aca aca tta ctt tta taa cgg cga cga ata tac tga aat tga tag atc gaa aaa agc cac taa taa aaa cag ttg gtt aat tac ttc agg ctt tag act aca aaa atg gtt cga tag cga aga ttg tat aat tta tct cag atc ttt agt tag aag aat gga aga cag taa caa aaa cag taa aaa aac tta gta ctt aga tat cg- --a aaa aat ata ttt ttg tag act ctt gag aat aga agg aaa aca tgt aca taa tta taa aaa atg aaa atc aat ggc gaa taa gac agt gcg att cgc acc atg gag tcg gta gat ttc atg gct gtc gat gag cag ttt cac gac gac ctc gat ctt tgg tca tta tct ttg gta gat gat tat aaa aaa cat gga tta ggt gtt gac tgt tat gtt cta gaa cca gtt gtt gac agg aaa ata ttt gat aga ttt ctc ctt gaa cca att tgt gat cct gta gat gtt ctg tat gat tat ttt agg att cat aga gat aat att gat cag tat ata gta gat aga ctg ttt gca tat att aca tat aaa gat att ata tct gca tta gtg tca aag aat tat atg gaa gat att ttc tct ata att att aag aat tgt aat tct gtg caa gat ctc tta ctt tac tat cta tct aat gca tat gta gaa ata gac att gtt gat ctt atg gta gat cat ggg gct gta ata tat aaa ata gaa tgc ttg aat gcc tat ttt agg gga ata tgt aaa aag gaa agt agt gtt gtt gag ttt att ttg aat tgt ggt atc cca gat gaa aat gat gtt aaa tta gat cta tat aaa ata att cag tat act agg gga ttc ctt gta gat gaa ccc aca gta tta gaa att tat aag ctt tgt atc cca tat att gaa gat atc aat caa cta gat gct ggt gga agg acc ttg ctt tat cgc gct atc tat gca ggt tat ata gat tta gta tca tgg cta tta gaa aat gga gca aat gtc aac gca gta atg agt aat gga tat aca tgt ctt gac gtg gcc gtg gat agg gga tct gtc atc gcc cgt agg gaa gca cat ctt aaa ata tta gaa ata ttg ctt aga gaa cca ttg tct att gac tgt ata aaa tta gct ata ctt aat aat aca att gaa aac cat gat gtg ata aag ctc tgt atc aag tat ttt atg atg gta gat tat tca ctt tgt aat gtg tat gca tca tca ctc ttt gat tat ata att gat tgt aaa caa gaa ttg gag tac att agg cag atg aaa att cat aat aca acc atg tat gag tta atc tat aat aga gac aaa aac aag cat gct tcc cat att cta cat agg tat tct aaa cat cca gtt ttg aca cag tgt atc act aaa gga ttc aag att tac aca gaa gta acc gag cag gtc act aaa gct cta aac aga cgt gct cta ata gat gag ata ata aac aat gta tca act gat gac aat ctc cta tca aaa ctt cca tta gaa att agg gat cta att gtt tca caa gct gtc ata tag agt tct atc cac cca cct ttc ttg aaa tga gtt aat --- --- --- --- --- --- --- --- --- --- --- --- --- --- --- --- --- --- --- --- --- --- --- --- --- --- --- --- --- --- --- --- --- --- --- --- --- --- --- --- --- --- --- --- --- --- --- --- --- --- --- --- --- --- --- --- --- --- --- --- --- --- --- --- --- --- --- --- --- --- --- --- --- --- --- --- --- --- --- --- --- --- --- --- --- --- --- --- --- --- --- --- --- --- --- --- --- --- --- --- --- --- --- --- --- -ag tca taa gtt agt taa --- --- --- --- --- -gt cat aag tta gtt aag tca t-- --- --- --- --- --- --- --- --- --- --- --- --- --- --- --- -aa gtt agt taa gtc ata agt tag tta agt cat aag tta gtt aag tca taa gtt agt taa gtc ata agt tag tta agt cat aag tta gtt tat agt cta aca ctt cta att ttt ata cct tga tct ttt tct cta att atg aaa aag taa atc att atg aag atg gat g-a aat gga cga gat tgt gcg cat cgt taa cga tag tat gtg gta cgt acc taa cgc att tat gga cga cgg tga taa tga agg tca cat ttc tgt caa taa tgt ctg tca tat gta tct cgc att ctt tga tgt gga tat atc atc tca tct gtt taa att agt tat taa aca ctg cga tct gaa taa acg act aaa atg tgg taa ctc tcc att aca ttg cta tac gat gaa tac acg att taa tcc atc tgt att aaa gat att gtt acg cca cgg cat gcg taa ctt tga tag caa gga taa aaa agg aca tat tcc tct aca cca cta tct gat tca ttc act atc aat cga taa caa gat ctt tga tat act aac gga ccc cat tga tga ctt tag taa atc atc cga tct att gct gtg tta tct tag ata taa att caa tgg gag ctt aaa cta tta cgt tct gta caa att att gac taa agg atc tga ccc taa ttg cgt cga tga gga tgg act cac ttc tct tca tta cta ctg taa aca cat atc cgc gtt cca cga aag caa tta tta caa gtc aaa gag tca cac taa gat gcg agc tga gaa gcg att cat cta cgc gat aat aga tca tgg agc aaa cat taa cgc ggt tac gaa aat cgg aaa tac gcc gtt aca cac tta cct tca aca gta tac caa aca tag tcc tcg tgt ggt gta tgc tct ttt atc tcg agg agc cga tac gag gat acg taa taa tct tga ttg tac acc cat cat gga ata cat aaa gaa cga ttg tgc aac agg tca tat tct cat aat gtt act caa ttg gca cga aca aaa ata cgg gaa att aca aaa gga aga agg aca aca tct act tta tct att cat aaa aca taa tca agg ata tgg aag tcg ctc tct caa tat act acg gta tct act aga tag att cga cat tca gaa aga cga ata cta taa tac aat gac tcc tct tca tac cgc ctt cca gaa ttg caa taa caa tgt tgc ctc ata cct cgt ata cat cgg ata cga cat caa cct tcc gac taa aga cga taa gac agt att cga ctt ggt gtt tga aaa cag aaa cat tat ata caa ggc gga tgt cgt taa tga cat tat cca cca cag act gaa agt atc tct acc tat gat taa atc gtt gtt cta caa gat gtc gga gtt ctc tcc cta cga cga tca cta cgt aaa gaa gat aat agc cta ctg cct att aag gga cga gtc att tgc gga act aca tac taa att ctg ttt aaa cga gga cta taa aag tgt att tat gaa aaa tat atc att cga taa gat aga ttc cat cat cga aaa atg tag tcg tga cat aag tct cct caa aga gat tcg aat ctc aga cac cga ctt gta tac ggt att gag aac aga aga cat ccg gta tca cac ata tct cga agc cat aca ttc aga caa acg cat ttc att tcc cat gta cga cga tct cat aga aca gtg tca tct atc gat gga gca taa aag taa act cgt cga caa agc act caa taa att aga gtc tac cat cga tag tca atc tag act atc gta ttt gcc tcc gga aat tat gcg caa tat cat aac caa gct aag cga cta cca tct aaa cag tat gtt gta cgg aaa gaa cca tta caa ata tta tcc atg ata gaa aga aaa tat tta aaa aat aat cta tat gat tgg aga agt agg aaa caa aca gta aca aga cga cga tta cta c-a tta tta aat cat gag gtc cgt att ata ctc gta tat att gtt tct ctc atg tat aat aat aaa cgg aag aga tat agc acc aca tgc acc atc caa tgg aaa gtg taa aga caa cga ata cag aag ccg taa tct atg ttg tct atc gtg tcc tcc ggg aac tta cgc ttc cag att atg tga tag caa gac taa tac aca atg tac acc gtg tgg ttc gga tac ctt tac atc tca caa taa tca ttt aca ggc ttg tct aag ttg taa cgg aag atg tga tag taa tca ggt aga gac gcg atc gtg taa cac gac tca caa tag aat ctg tga atg ctc tcc agg ata tta ttg tct tct caa agg agc atc agg gtg tag aac atg tat ttc taa aac aaa gtg tgg aat agg ata cgg agt atc cgg ata cac gtc tac cgg aga cgt cat ctg ttc tcc gtg tgg tcc cgg aac ata ttc tca cac cgt ctc ttc cac aga taa atg cga acc cgt cgt aac cag caa tac att taa cta tat cga tgt gga aat taa cct gta tcc agt caa cga cac atc gtg tac tcg gac gac cac tac cgg tct cag cga atc cat ctc aac gtc gga act aac tat tac cat gaa tca taa aga ttg tga tcc agt ctt tcg tgc aga ata ctt ctc tgt cct taa taa tgt agc aac ttc agg att ctt tac agg aga aaa tag ata tca gaa tac ttc aaa gat atg tac tct gaa ttt cga gat taa atg taa caa caa aga ttc atc ttc caa aca gtt aac gaa aac aaa gaa tga tac tat cat gcc gca ttc aga gac ggt aac tct agt ggg cga ctg tct atc tag cgt cga cat cta cat act ata tag taa tac caa tac tca aga cta cga aaa tga tac aat ctc tta tca tat ggg taa tgt tct cga tgt caa tag cca tat gcc cgc tag ttg cga tat aca taa act gat cac taa ttc cca gaa tcc cac cca ctt ata gta ag- -tt ttt tta cct ata aat aat aaa tac aat aat taa ttt ctc gta aaa gta gaa aat ata ttc taa ttt att ata tgg taa gaa agt aga atc atc tag aac agt aat caa tca ata gca atc atg aaa caa tat att gtc ctg gca tgc atg tgc cta gtg gca gct gct atg cct act agt ctt caa caa tct tca tcc tcg tgt act gaa gaa gaa aac aaa cat cat atg gga atc gat gtt att atc aaa gtc aca aag caa gac caa aca ccg acc aat gat aag att tgt caa tcc gta acg gaa gtt aca gag a-- --- --- --- --- --- -cc gaa gat gat gag gta tcc gaa gaa gtt gta aaa gga gat ccc acc act tat tac act atc gtc ggt gcg ggt ctt aac atg aac ttt gga ttc acc aaa tgc cca aag att tca tcc atc tcc gaa tcc tct gat gga aac act gtg aat act aga ttg tcc agc gtg tca ccg gga caa ggt aag gac tct ccc gcg atc acg cgt gaa gaa gct ctg gct atg atc aaa gac tgt gag atg tct atc gac atc aga tgt agc gaa gaa gag aaa gac agt gac atc aag acc cat cca gta ctt ggg tct aac atc tca cat aag aaa gtg agt tac aaa gat atc atc ggt tca acg atc gtt gat aca aaa tgt gtc aag aac cta gag ttt agc gta cgt atc gga gac atg tgt gag gaa tca tct gaa ctt gaa gtc aag gat gga ttc aag tat gtc gac gga tcg gca tct gaa ggt gca acc gat gat act tca ctc atc gat tca aca aaa ctc aaa gca tgt gtc tga atc gat aac tct att cat ctg aaa atg gat gag ttg ggt taa tcg aac gat tca gac acc gca cca cga att aaa aaa g-- -ac cgg gca cta tat tcc ggt ttg caa aac aaa aat att taa cta cat tca caa aaa gtt acc tct cgt tac ttc ttc ttt ctg ttt caa tat gtg ata cga tat gat cac tat tcg tat tct ct- --- --- tgg tct cat aaa aaa gtt tta c-- -aa aaa aaa aaa aaa aat att ttt att ctc ttt ctc tct tcg at- ggt ctc aca aaa ata tta aac ctc ttt ctg atg tct caa cta ttt cgt aaa cga taa cgt cca aca ata tat tct cgt aga gct tat caa cat cct tat acc aat cta ggt tgt cag aca att gca tca taa aat aat gtt tat aat tta cac gtt aac atc ata taa taa acg tat ata gtt aat att ttt gga ata taa atg atc tgt aaa atc cat gta ggg gac act gct cac gtt ttt tct cta gta cat aat ttc aca caa gtt ttt ata cag aca aat taa ttc tcg tcc ata ta- ttt taa aac att gac ttt tgt act aag aaa aat atc ttg act aac cat ctc ttt ctc tct tcg atg ggt ctc aca aaa ata tta aac ctc ttt ctg atg g-- --- --- --- --- --- --- --- --- --- --- --- --- --- --- --- --- --- --- --- --- --- --- --- --- --- --- --- --- --- --- --- --- --- --- --- --- --- --- --- --- --- --- --- --- --- --- agt cgt aaa aag ttt tta tcc ttt ctc tct tcg ata ggt ctc --- --- --- --- --- --- --- --- --- --- --- --- --- --- --- --- --- --- --- --- --- --- --- --- --- --- --- --- --- --- --- --- --- --- --- --- --- --- --- --- --- --- --- --- --- --- --- --- --- --- --- --- --- --- --- --- --- --- --- --- --- --- --- --- --- --- --- --- --- --- aca aaa ata tta aac ctc ttt ctg atg gtc tct ata aa

>ON631963.1_Monkeypox_virus_isolate_MPxV/VIDRL01/2022_complete_genome

atc gag gac gac ttc aac tat gac agc tcg gtg gcg tct gcc agc gtg tac ata cga atg gca ttt cta aga aaa gtc tac ggt atc ctt -tc tac aat ttc ctt taa caa cgg caa cag ctg cag tat ttt tat act ttg aat gca -tc gga cat tta tac aag gga gtc ctg ttc taa tat tgg cat caa tgt tcg gat cta tag gct tga ttt tcg cat tga ctt tac aca gac atg aac atc ccc tga atc tgt aca tac ttt gtg gat tta cac tgt tag aat ctc taa cgc tgg cct ctg ttg tta ctt tct atg atg cac gta tcg tta tgc aag ctt tca tgt tga cta ctg cag tgt ttc ttg ctc tga cta cat ata ctc tac aat caa aga gag att tca gta aac ttg taa cag gat tgt ttg ctg ctt tct gga ttt taa ttt tgt cag gag tct tga gga taa agt tta aaa tag aat taa taa aga aca tat agg tca -tt ttt taa aca tgg ata gaa acc aag gtt gtt agt taa taa tat aca aga tat ttt ttc tca ctc tga tcc atg taa acc aag gac gag aga cac tct cat tcc tca ttc acg aca cca tta aaa atg gaa att aaa gcc ctc tat taa gca cag acg gct aca ggt cta cca tca ggt t-a cct tcg tct acc ttc aca atg gcc tct cct tgt gcc cag ttc agt ccc tgt cat tgc cac gct act aag gac tcc ctg aat acc gtg act gac gtc aga cat tgt ctg act gaa tac atc ctg tgg gtt tct cat aga tgg acc cat aga gaa agc gca ggg cct ctc tac agg ctt ctc atc tct ttc aga att gat gca atg gag cta ttt ggt agc gag ttg aag gag ttc tcg aat tca ctt ccg tgg gac aat atc gac aat tgc gtg gag atc att aaa tgt ttc atc aga aat gac tcc atg aaa acc gcc aaa gaa ctt tgt gca ata att gga ctt tgt act caa tca gct att gtc act gga aga gtc ttc aat gat aag tat atc gac ata cta ctt atg ctg cga aag att ctg aac gag aac gac tat ctc acc ctc ttg gat cat atc ctc act gct aaa tac taa atc tcc ttc atg ctc tct cac taa tac tct tac tca cta cac ttt tta tca tct tat gat gaa tga ttg cct tca tca ttt ttt cgt gga ata taa tat agg aat aat tag cac cag aat agc tat gga tat ctc gtt aag aat att ctc tat aag aga cat aat gta gac ata gtt att ata tcc ttc tta gat aag tgt tac gct act gga aag ttt cca tcg tta tta tta cat gaa gat gat ata att aaa cca aca ttg aga ttg gct ctt atg tta gct gga ttg aat tac tgt aat aaa tgc atc gag tat aga ggg ata tag caa ttc tcg ata ata gtc atg caa tat ttg aat gag act gat aat tta ggt aat aca gta ct- --- aca cac ata tct ttc tag att ata tat cgt taa aaa tct gta aga tgt ata ttt ctc ata agt atc cac tgt gta ata tta tta atg gat ata tag ata acg caa tag gga cta ata gta ttg taa aag ata taa tcg act att tac gta cat atc cag ata tct ata ttc cta cta gtt tgc tgc gta gtt gca tca ttg ata tgc atg -at tta tca gga ttc aga gat gaa tta cta agt aaa cta caa tcc cac aat aag taa gaa tca aat atc aaa aac tca ctt ttg att ttt cta gtc tta agt aat aca tat att tat taa tag acc tat gaa ata aaa aaa ggt aac aat gga ttc gcg tat agc tat tta cgt att agt atc ggc atc tct ttt gta tct tgt taa ttg tca caa act agt aca tta ctt caa tct gaa aat aaa tgg aag tga tat aac taa tac agc aga tat att gct gga caa tta tcc aat tat gac ctt tga tgg aaa gga tat tta tcc atc tat ctc gtt cat ggt cgg taa taa act ttt cct aga tct tta taa aaa tat ctt tga aga att ttt cag act att tcg agt atc tgt aag tag tca ata cga gga att aga ata tta tta ttc atg tga tta tac taa caa ccg tcc tac aat taa aca aca tta ctt tta taa cgg cga cga ata tac tga aat tga tag atc gaa aaa agc cac taa taa aaa cag ttg gtt aat tac ttc agg ctt tag act aca aaa atg gtt cga tag cga aga ttg tat aat tta tct cag atc ttt agt tag aag aat gga aga cag taa caa aaa cag taa aaa aac tta gta ctt aga tat cg- --a aaa aat ata ttt ttg tag act ctt gag aat aga agg aaa aca tgt aca taa tta taa aaa atg aaa atc aat ggc gaa taa gac agt gcg att cgc acc atg gag tcg gta gat ttc atg gct gtc gat gag cag ttt cac gac gac ctc gat ctt tgg tca tta tct ttg gta gat gat tat aaa aaa cat gga tta ggt gtt gac tgt tat gtt cta gaa cca gtt gtt gac agg aaa ata ttt gat aga ttt ctc ctt gaa cca att tgt gat cct gta gat gtt ctg tat gat tat ttt agg att cat aga gat aat att gat cag tat ata gta gat aga ctg ttt gca tat att aca tat aaa gat att ata tct gca tta gtg tca aag aat tat atg gaa gat att ttc tct ata att att aag aat tgt aat tct gtg caa gat ctc tta ctt tac tat cta tct aat gca tat gta gaa ata gac att gtt gat ctt atg gta gat cat ggg gct gta ata tat aaa ata gaa tgc ttg aat gcc tat ttt agg gga ata tgt aaa aag gaa agt agt gtt gtt gag ttt att ttg aat tgt ggt atc cca gat gaa aat gat gtt aaa tta gat cta tat aaa ata att cag tat act agg gga ttc ctt gta gat gaa ccc aca gta tta gaa att tat aag ctt tgt atc cca tat att gaa gat atc aat caa cta gat gct ggt gga agg acc ttg ctt tat cgc gct atc tat gca ggt tat ata gat tta gta tca tgg cta tta gaa aat gga gca aat gtc aac gca gta atg agt aat gga tat aca tgt ctt gac gtg gcc gtg gat agg gga tct gtc atc gcc cgt agg gaa gca cat ctt aaa ata tta gaa ata ttg ctt aga gaa cca ttg tct att gac tgt ata aaa tta gct ata ctt aat aat aca att gaa aac cat gat gtg ata aag ctc tgt atc aag tat ttt atg atg gta gat tat tca ctt tgt aat gtg tat gca tca tca ctc ttt gat tat ata att gat tgt aaa caa gaa ttg gag tac att agg cag atg aaa att cat aat aca acc atg tat gag tta atc tat aat aga gac aaa aac aag cat gct tcc cat att cta cat agg tat tct aaa cat cca gtt ttg aca cag tgt atc act aaa gga ttc aag att tac aca gaa gta acc gag cag gtc act aaa gct cta aac aga cgt gct cta ata gat gag ata ata aac aat gta tca act gat gac aat ctc cta tca aaa ctt cca tta gaa att agg gat cta att gtt tca caa gct gtc ata tag agt tct atc cac cca cct ttc ttg aaa tga gtt aat --- --- --- --- --- --- --- --- --- --- --- --- --- --- --- --- --- --- --- --- --- --- --- --- --- --- --- --- --- --- --- --- --- --- --- --- --- --- --- --- --- --- --- --- --- --- --- --- --- --- --- --- --- --- --- --- --- --- --- --- --- --- --- --- --- --- --- --- --- --- --- --- --- --- --- --- --- --- --- --- --- --- --- --- --- --- --- --- --- --- --- --- --- --- --- --- --- --- --- --- --- --- --- --- --- -ag tca taa gtt agt taa --- --- --- --- --- -gt cat aag tta gtt aag tca t-- --- --- --- --- --- --- --- --- --- --- --- --- --- --- --- -aa gtt agt taa gtc ata agt tag tta agt cat aag tta gtt aag tca taa gtt agt taa gtc ata agt tag tta agt cat aag tta gtt tat agt cta aca ctt cta att ttt ata cct tga tct ttt tct cta att atg aaa aag taa atc att atg aag atg gat g-a aat gga cga gat tgt gcg cat cgt taa cga tag tat gtg gta cgt acc taa cgc att tat gga cga cgg tga taa tga agg tca cat ttc tgt caa taa tgt ctg tca tat gta tct cgc att ctt tga tgt gga tat atc atc tca tct gtt taa att agt tat taa aca ctg cga tct gaa taa acg act aaa atg tgg taa ctc tcc att aca ttg cta tac gat gaa tac acg att taa tcc atc tgt att aaa gat att gtt acg cca cgg cat gcg taa ctt tga tag caa gga taa aaa agg aca tat tcc tct aca cca cta tct gat tca ttc act atc aat cga taa caa gat ctt tga tat act aac gga ccc cat tga tga ctt tag taa atc atc cga tct att gct gtg tta tct tag ata taa att caa tgg gag ctt aaa cta tta cgt tct gta caa att att gac taa agg atc tga ccc taa ttg cgt cga tga gga tgg act cac ttc tct tca tta cta ctg taa aca cat atc cgc gtt cca cga aag caa tta tta caa gtc aaa gag tca cac taa gat gcg agc tga gaa gcg att cat cta cgc gat aat aga tca tgg agc aaa cat taa cgc ggt tac gaa aat cgg aaa tac gcc gtt aca cac tta cct tca aca gta tac caa aca tag tcc tcg tgt ggt gta tgc tct ttt atc tcg agg agc cga tac gag gat acg taa taa tct taa ttg tac acc cat cat gga ata cat aaa gaa cga ttg tgc aac agg tca tat tct cat aat gtt act caa ttg gca cga aca aaa ata cgg gaa att aca aaa gga aga agg aca aca tct act tta tct att cat aaa aca taa tca agg ata tgg aag tcg ctc tct caa tat act acg gta tct act aga tag att cga cat tca gaa aga cga ata cta taa tac aat gac tcc tct tca tac cgc ctt cca gaa ttg caa taa caa tgt tgc ctc ata cct cgt ata cat tgg ata cga cat caa cct tcc gac taa aga cga taa gac agt att cga ctt ggt gtt tga aaa cag aaa cat tat ata caa ggc gga tgt cgt taa tga cat tat cca cca cag act gaa agt atc tct acc tat gat taa atc gtt gtt cta caa gat gtc gga gtt ctc tcc cta cga cga tca cta cgt aaa gaa gat aat agc cta ctg cct att aag gga cga gtc att tgc gga act aca tac taa att ctg ttt aaa cga gga cta taa aag tgt att tat gaa aaa tat atc att cga taa gat aga ttc cat cat cga aaa atg tag tcg tga cat aag tct cct caa aga gat tcg aat ctc aga cac cga ctt gta tac ggt att gag aac aga aga cat tcg gta tca cac ata tct cga agc cat aca ttc aga caa acg cat ttc att tcc cat gta cga cga tct cat aga aca gtg tca tct atc gat gga gca taa aag taa act cgt cga caa agc act caa taa att aga gtc tac cat cga tag tca atc tag act atc gta ttt gcc tcc gga aat tat gcg caa tat cat aac caa gct aag cga cta cca tct aaa cag tat gtt gta cgg aaa gaa cca tta caa ata tta tcc atg ata gaa aga aaa tat tta aaa aat aat cta tat gat tgg aga agt agg aaa caa aca gta aca aga cga cga tta cta c-a tta tta aat cat gag gtc cgt att ata ctc gta tat att gtt tct ctc atg tat aat aat aaa cgg aag aga tat agc acc aca tgc acc atc caa tgg aaa gtg taa aga caa cga ata cag aag ccg taa tct atg ttg tct atc gtg tcc tcc ggg aac tta cgc ttt cag att atg tga tag caa gac taa tac aca atg tac acc gtg tgg ttc gga tac ctt tac atc tca caa taa tca ttt aca ggc ttg tct aag ttg taa cgg aag atg tga tag taa tca ggt aga gac gcg atc gtg taa cac gac tca caa tag aat ctg tga atg ctc tcc agg ata tta ttg tct tct caa agg agc atc agg gtg tag aac atg tat ttc taa aac aaa gtg tgg aat agg ata cgg agt atc cgg ata cac gtc tac cgg aga cgt cat ctg ttc tcc gtg tgg tcc cgg aac ata ttc tca cac cgt ctc ttc cac aga taa atg cga acc cgt cgt aac cag caa tac att taa cta tat cga tgt gga aat taa cct gta tcc agt caa cga cac atc gtg tac tcg gac gac cac tac cgg tct cag cga atc cat ctc aac gtc gga act aac tat tac cat gaa tca taa aga ttg tga tcc agt ctt tcg tgc aga ata ctt ctc tgt cct taa taa tgt agc aac ttc agg att ctt tac agg aga aaa tag ata tca gaa tac ttc aaa gat atg tac tct gaa ttt cga gat taa atg taa caa caa aga ttc atc ttc caa aca gtt aac gaa aac aaa gaa tga tac tat cat gcc gca ttc aga gac ggt aac tct agt ggg cga ctg tct atc tag cgt cga cat cta cat act ata tag taa tac caa tac tca aga cta cga aaa tga tac aat ctc tta tca tat ggg taa tgt tct cga tgt caa tag cca tat gcc cgc tag ttg cga tat aca taa act gat cac taa ttc cca gaa tcc cac cca ctt ata gta ag- -tt ttt tta cct ata aat aat aaa tac aat aat taa ttt ctc gta aaa gta gaa aat ata ttc taa ttt att ata tgg taa gaa agt aga atc atc tag aac agt aat caa tca ata gca atc atg aaa caa tat att gtc ctg gca tgc atg tgc cta gtg gca gct gct atg cct act agt ctt caa caa tct tca tcc tcg tgt act gaa gaa gaa aac aaa cat cat atg gga atc gat gtt att atc aaa gtc aca aag caa gac caa aca ccg acc aat gat aag att tgt caa tcc gta acg gaa gtt aca gag a-- --- --- --- --- --- -cc gaa gat gat gag gta tcc gaa gaa gtt gta aaa gga gat ccc acc act tat tac act atc gtc ggt gcg ggt ctt aac atg aac ttt gga ttc acc aaa tgc cca aag att tta tcc atc tcc gaa tcc tct gat gga aac act gtg aat act aga ttg tcc agc gtg tca ccg gga caa ggt aag gac tct ccc gcg atc acg cgt gaa gaa gct ctg gct atg atc aaa gac tgt gag atg tct atc gac atc aga tgt agc gaa gaa gag aaa gac agt gac atc aag acc cat cca gta ctt ggg tct aac atc tca cat aag aaa gtg agt tac aaa gat atc atc ggt tca acg atc gtt gat aca aaa tgt gtc aag aac cta gag ttt agc gta cgt atc gga gac atg tgt gag gaa tca tct gaa ctt gaa gtc aag gat gga ttc aag tat gtc gac gga tcg gca tct gaa ggt gca acc gat gat act tca ctc atc gat tca aca aaa ctc aaa gca tgt gtc tga atc gat aac tct att cat ctg aaa atg gat gag ttg ggt taa tcg aac gat tca gac acc gca cca cga att aaa aaa g-- -ac cgg gca cta tat tcc ggt ttg caa aac aaa aat att taa cta cat tca caa aaa gtt acc tct cgt tac ttc ttc ttt ctg ttt caa tat gtg ata cga tat gat cac tat tcg tat tct ct- --- --- tgg tct cat aaa aaa gtt tta c-- --a aaa aaa aaa aaa aat att ttt att ctc ttt ctc tct tcg at- ggt ctc aca aaa ata tta aac ctc ttt ctg atg tct caa cta ttt cgt aaa cga taa cgt cca aca ata tat tct cgt aga gct tat caa cat cct tat acc aat cta ggt tgt cag aca att gca tca taa aat aat gtt tat aat tta cac gtt aac atc ata taa taa acg tat ata gtt aat att ttt gga ata taa atg atc tgt aaa atc cat gta ggg gac act gct cac gtt ttt tct cta gta cat aat ttc aca caa gtt ttt ata cag aca aat taa ttc tcg tcc ata ta- ttt taa aac att gac ttt tgt act aag aaa aat atc ttg act aac cat ctc ttt ctc tct tcg atg ggt ctc aca aaa ata tta aac ctc ttt ctg atg g-- --- --- --- --- --- --- --- --- --- --- --- --- --- --- --- --- --- --- --- --- --- --- --- --- --- --- --- --- --- --- --- --- --- --- --- --- --- --- --- --- --- --- --- --- --- --- agt cgt aaa aag ttt tta tcc ttt ctc tct tcg ata ggt ctc --- --- --- --- --- --- --- --- --- --- --- --- --- --- --- --- --- --- --- --- --- --- --- --- --- --- --- --- --- --- --- --- --- --- --- --- --- --- --- --- --- --- --- --- --- --- --- --- --- --- --- --- --- --- --- --- --- --- --- --- --- --- --- --- --- --- --- --- --- --- aca aaa ata tta aac ctc ttt ctg atg gtc tct ata aa

>ON568298.1_Monkeypox_virus_isolate_MPXV-BY-IMB25241_complete_genome

atc gag gac gac ttc aac tat gac agc tcg gtg gcg tct gcc agc gtg tac ata cga atg gca ttt cta aga aaa gtc tac ggt atc ctt -tc tac aat ttc ctt taa caa cgg caa cag ctg cag tat ttt tat act ttg aat gca -tc gga cat tta tac aag gga gtc ctg ttc taa tat tgg cat caa tgt tcg gat cta tag gct tga ttt tcg cat tga ctt tac aca gac atg aac atc ccc tga atc tgt aca tac ttt gtg gat tta cac tgt tag aat ctc taa cgc tgg cct ctg ttg tta ctt tct atg atg cac gta tcg tta tgc aag ctt tca tgt tga cta ctg cag tgt ttc ttg ctc tga cta cat ata ctc tac aat caa aga gag att tca gta aac ttg taa cag gat tgt ttg ctg ctt tct gga ttt taa ttt tgt cag gag tct tga gga taa agt tta aaa tag aat taa taa aga aca tat agg tca -tt ttt taa aca tgg ata gaa acc aag gtt gtt agt taa taa tat aca aga tat ttt ttc tca ctc tga tcc atg taa acc aag gac gag aga cac tct cat tcc tca ttc acg aca cca tta aaa atg gaa att aaa gcc ctc tat taa gca cag acg gct aca ggt cta cca tca ggt t-a cct tcg tct acc ttc aca atg gcc tct cct tgt gcc cag ttc agt ccc tgt cat tgc cac gct act aag gac tcc ctg aat acc gtg act gac gtc aga cat tgt ctg act gaa tac atc ctg tgg gtt tct cat aga tgg acc cat aga gaa agc gca ggg cct ctc tac agg ctt ctc atc tct ttc aga att gat gca atg gag cta ttt ggt agc gag ttg aag gag ttc tcg aat tca ctt ccg tgg gac aat atc gac aat tgc gtg gag atc att aaa tgt ttc atc aga aat gac tcc atg aaa acc gcc aaa gaa ctt tgt gca ata att gga ctt tgt act caa tca gct att gtc act gga aga gtc ttc aat gat aag tat atc gac ata cta ctt atg ctg cga aag att ctg aac gag aac gac tat ctc acc ctc ttg gat cat atc ctc act gct aaa tac taa atc tcc ttc atg ctc tct cac taa tac tct tac tca cta cac ttt tta tca tct tat gat gaa tga ttg cct tca tca ttt ttt cgt gga ata taa tat agg aat aat tag cac cag aat agc tat gga tat ctc gtt aag aat att ctc tat aag aga cat aat gta gac ata gtt att ata tcc ttc tta gat aag tgt tac gct act gga aag ttt cca tcg tta tta tta cat gaa gat gat ata att aaa cca aca ttg aga ttg gct ctt atg tta gct gga ttg aat tac tgt aat aaa tgc atc gag tat aga ggg ata tag caa ttc tcg ata ata gtc atg caa tat ttg aat gag act gat aat tta ggt aat aca gta ct- --- aca cac ata tct ttc tag att ata tat cgt taa aaa tct gta aga tgt ata ttt ctc ata agt atc cac tgt gta ata tta tta atg gat ata tag ata acg caa tag gga cta ata gta ttg taa aag ata taa tcg act att tac gta cat atc cag ata tct ata ttc cta cta gtt tgc tgc gta gtt gca tca ttg ata tgc atg -at tta tca gga ttc aga gat gaa tta cta agt aaa cta caa tcc cac aat aag taa gaa tca aat atc aaa aac tca ctt ttg att ttt cta gtc tta agt aat aca tat att tat taa tag acc tat gaa ata aaa aaa ggt aac aat gga ttc gcg tat agc tat tta cgt att agt atc ggc atc tct ttt gta tct tgt taa ttg tca caa act agt aca tta ctt caa tct gaa aat aaa tgg aag tga tat aac taa tac agc aga tat att gct gga caa tta tcc aat tat gac ctt tga tgg aaa gga tat tta tcc atc tat ctc gtt cat ggt cgg taa taa act ttt cct aga tct tta taa aaa tat ctt tga aga att ttt cag act att tcg agt atc tgt aag tag tca ata cga gga att aga ata tta tta ttc atg tga tta tac taa caa ccg tcc tac aat taa aca aca tta ctt tta taa cgg cga cga ata tac tga aat tga tag atc gaa aaa agc cac taa taa aaa cag ttg gtt aat tac ttc agg ctt tag act aca aaa atg gtt cga tag cga aga ttg tat aat tta tct cag atc ttt agt tag aag aat gga aga cag taa caa aaa cag taa aaa aac tta gta ctt aga tat cg- --a aaa aat ata ttt ttg tag act ctt gag aat aga agg aaa aca tgt aca taa tta taa aaa atg aaa atc aat ggc gaa taa gac agt gcg att cgc acc atg gag tcg gta gat ttc atg gct gtc gat gag cag ttt cac gac gac ctc gat ctt tgg tca tta tct ttg gta gat gat tat aaa aaa cat gga tta ggt gtt gac tgt tat gtt cta gaa cca gtt gtt gac agg aaa ata ttt gat aga ttt ctc ctt gaa cca att tgt gat cct gta gat gtt ctg tat gat tat ttt agg att cat aga gat aat att gat cag tat ata gta gat aga ctg ttt gca tat att aca tat aaa gat att ata tct gca tta gtg tca aag aat tat atg gaa gat att ttc tct ata att att aag aat tgt aat tct gtg caa gat ctc tta ctt tac tat cta tct aat gca tat gta gaa ata gac att gtt gat ctt atg gta gat cat ggg gct gta ata tat aaa ata gaa tgc ttg aat gcc tat ttt agg gga ata tgt aaa aag gaa agt agt gtt gtt gag ttt att ttg aat tgt ggt atc cca gat gaa aat gat gtt aaa tta gat cta tat aaa ata att cag tat act agg gga ttc ctt gta gat gaa ccc aca gta tta gaa att tat aag ctt tgt atc cca tat att gaa gat atc aat caa cta gat gct ggt gga agg acc ttg ctt tat cgc gct atc tat gca ggt tat ata gat tta gta tca tgg cta tta gaa aat gga gca aat gtc aac gca gta atg agt aat gga tat aca tgt ctt gac gtg gcc gtg gat agg gga tct gtc atc gcc cgt agg gaa gca cat ctt aaa ata tta gaa ata ttg ctt aga gaa cca ttg tct att gac tgt ata aaa tta gct ata ctt aat aat aca att gaa aac cat gat gtg ata aag ctc tgt atc aag tat ttt atg atg gta gat tat tca ctt tgt aat gtg tat gca tca tca ctc ttt gat tat ata att gat tgt aaa caa gaa ttg gag tac att agg cag atg aaa att cat aat aca acc atg tat gag tta atc tat aat aga gac aaa aac aag cat gct tcc cat att cta cat agg tat tct aaa cat cca gtt ttg aca cag tgt atc act aaa gga ttc aag att tac aca gaa gta acc gag cag gtc act aaa gct cta aac aga cgt gct cta ata gat gag ata ata aac aat gta tca act gat gac aat ctc cta tca aaa ctt cca tta gaa att agg gat cta att gtt tca caa gct gtc ata tag agt tct atc cac cca cct ttc ttg aaa tga gtt aat --- --- --- --- --- --- --- --- --- --- --- --- --- --- --- --- --- --- --- --- --- --- --- --- --- --- --- --- --- --- --- --- --- --- --- --- --- --- --- --- --- --- --- --- --- --- --- --- --- --- --- --- --- --- --- --- --- --- --- --- --- --- --- --- --- --- --- --- --- --- --- --- --- --- --- --- --- --- --- --- --- --- --- --- --- --- --- --- --- --- --- --- --- --- --- --- --- --- --- --- --- --- --- --- --- -ag tca taa gtt agt taa --- --- --- --- --- -gt cat aag tta gtt aag tca t-- --- --- --- --- --- --- --- --- --- --- --- --- --- --- --- -aa gtt agt taa gtc ata agt tag tta agt cat aag tta gtt aag tca taa gtt agt taa gtc ata agt tag tta agt cat aag tta gtt tat agt cta aca ctt cta att ttt ata cct tga tct ttt tct cta att atg aaa aag taa atc att atg aag atg gat g-a aat gga cga gat tgt gcg cat cgt taa cga tag tat gtg gta cgt acc taa cgc att tat gga cga cgg tga taa tga agg tca cat ttc tgt caa taa tgt ctg tca tat gta tct cgc att ctt tga tgt gga tat atc atc tca tct gtt taa att agt tat taa aca ctg cga tct gaa taa acg act aaa atg tgg taa ctc tcc att aca ttg cta tac gat gaa tac acg att taa tcc atc tgt att aaa gat att gtt acg cca cgg cat gcg taa ctt tga tag caa gga taa aaa agg aca tat tcc tct aca cca cta tct gat tca ttc act atc aat cga taa caa gat ctt tga tat act aac gga ccc cat tga tga ctt tag taa atc atc cga tct att gct gtg tta tct tag ata taa att caa tgg gag ctt aaa cta tta cgt tct gta caa att att gac taa agg atc tga ccc taa ttg cgt cga tga gga tgg act cac ttc tct tca tta cta ctg taa aca cat atc cgc gtt cca cga aag caa tta tta caa gtc aaa gag tca cac taa gat gcg agc tga gaa gcg att cat cta cgc gat aat aga tca tgg agc aaa cat taa cgc ggt tac gaa aat cgg aaa tac gcc gtt aca cac tta cct tca aca gta tac caa aca tag tcc tcg tgt ggt gta tgc tct ttt atc tcg agg agc cga tac gag gat acg taa taa tct taa ttg tac acc cat cat gga ata cat aaa gaa cga ttg tgc aac agg tca tat tct cat aat gtt act caa ttg gca cga aca aaa ata cgg gaa att aca aaa gga aga agg aca aca tct act tta tct att cat aaa aca taa tca agg ata tgg aag tcg ctc tct caa tat act acg gta tct act aga tag att cga cat tca gaa aga cga ata cta taa tac aat gac tcc tct tca tac cgc ctt cca gaa ttg caa taa caa tgt tgc ctc ata cct cgt ata cat tgg ata cga cat caa cct tcc gac taa aga cga taa gac agt att cga ctt ggt gtt tga aaa cag aaa cat tat ata caa ggc gga tgt cgt taa tga cat tat cca cca cag act gaa agt atc tct acc tat gat taa atc gtt gtt cta caa gat gtc gga gtt ctc tcc cta cga cga tca cta cgt aaa gaa gat aat agc cta ctg cct att aag gga cga gtc att tgc gga act aca tac taa att ctg ttt aaa cga gga cta taa aag tgt att tat gaa aaa tat atc att cga taa gat aga ttc cat cat cga aaa atg tag tcg tga cat aag tct cct caa aga gat tcg aat ctc aga cac cga ctt gta tac ggt att gag aac aga aga cat tcg gta tca cac ata tct cga agc cat aca ttc aga caa acg cat ttc att tcc cat gta cga cga tct cat aga aca gtg tca tct atc gat gga gca taa aag taa act cgt cga caa agc act caa taa att aga gtc tac cat cga tag tca atc tag act atc gta ttt gcc tcc gga aat tat gcg caa tat cat aac caa gct aag cga cta cca tct aaa cag tat gtt gta cgg aaa gaa cca tta caa ata tta tcc atg ata gaa aga aaa tat tta aaa aat aat cta tat gat tgg aga agt agg aaa caa aca gta aca aga cga cga tta cta c-a tta tta aat cat gag gtc cgt att ata ctc gta tat att gtt tct ctc atg tat aat aat aaa cgg aag aga tat agc acc aca tgc acc atc caa tgg aaa gtg taa aga caa cga ata cag aag ccg taa tct atg ttg tct atc gtg tcc tcc ggg aac tta cgc ttt cag att atg tga tag caa gac taa tac aca atg tac acc gtg tgg ttc gga tac ctt tac atc tca caa taa tca ttt aca ggc ttg tct aag ttg taa cgg aag atg tga tag taa tca ggt aga gac gcg atc gtg taa cac gac tca caa tag aat ctg tga atg ctc tcc agg ata tta ttg tct tct caa agg agc atc agg gtg tag aac atg tat ttc taa aac aaa gtg tgg aat agg ata cgg agt atc cgg ata cac gtc tac cgg aga cgt cat ctg ttc tcc gtg tgg tcc cgg aac ata ttc tca cac cgt ctc ttc cac aga taa atg cga acc cgt cgt aac cag caa tac att taa cta tat cga tgt gga aat taa cct gta tcc agt caa cga cac atc gtg tac tcg gac gac cac tac cgg tct cag cga atc cat ctc aac gtc gga act aac tat tac cat gaa tca taa aga ttg tga tcc agt ctt tcg tgc aga ata ctt ctc tgt cct taa taa tgt agc aac ttc agg att ctt tac agg aga aaa tag ata tca gaa tac ttc aaa gat atg tac tct gaa ttt cga gat taa atg taa caa caa aga ttc atc ttc caa aca gtt aac gaa aac aaa gaa tga tac tat cat gcc gca ttc aga gac ggt aac tct agt ggg cga ctg tct atc tag cgt cga cat cta cat act ata tag taa tac caa tac tca aga cta cga aaa tga tac aat ctc tta tca tat ggg taa tgt tct cga tgt caa tag cca tat gcc cgc tag ttg cga tat aca taa act gat cac taa ttc cca gaa tcc cac cca ctt ata gta ag- -tt ttt tta cct ata aat aat aaa tac aat aat taa ttt ctc gta aaa gta gaa aat ata ttc taa ttt att ata tgg taa gaa agt aga atc atc tag aac agt aat caa tca ata gca atc atg aaa caa tat att gtc ctg gca tgc atg tgc cta gtg gca gct gct atg cct act agt ctt caa caa tct tca tcc tcg tgt act gaa gaa gaa aac aaa cat cat atg gga atc gat gtt att atc aaa gtc aca aag caa gac caa aca ccg acc aat gat aag att tgt caa tcc gta acg gaa gtt aca gag a-- --- --- --- --- --- -cc gaa gat gat gag gta tcc gaa gaa gtt gta aaa gga gat ccc acc act tat tac act atc gtc ggt gcg ggt ctt aac atg aac ttt gga ttc acc aaa tgc cca aag att tta tcc atc tcc gaa tcc tct gat gga aac act gtg aat act aga ttg tcc agc gtg tca ccg gga caa ggt aag gac tct ccc gcg atc acg cgt gaa gaa gct ctg gct atg atc aaa gac tgt gag atg tct atc gac atc aga tgt agc gaa gaa gag aaa gac agt gac atc aag acc cat cca gta ctt ggg tct aac atc tca cat aag aaa gtg agt tac aaa gat atc atc ggt tca acg atc gtt gat aca aaa tgt gtc aag aac cta gag ttt agc gta cgt atc gga gac atg tgt gag gaa tca tct gaa ctt gaa gtc aag gat gga ttc aag tat gtc gac gga tcg gca tct gaa ggt gca acc gat gat act tca ctc atc gat tca aca aaa ctc aaa gca tgt gtc tga atc gat aac tct att cat ctg aaa atg gat gag ttg ggt taa tcg aac gat tca gac acc gca cca cga att aaa aaa g-- -ac cgg gca cta tat tcc ggt ttg caa aac aaa aat att taa cta cat tca caa aaa gtt acc tct cgt tac ttc ttc ttt ctg ttt caa tat gtg ata cga tat gat cac tat tcg tat tct ct- --- --- tgg tct cat aaa aaa gtt tta c-- --a aaa aaa aaa aaa aat att ttt att ctc ttt ctc tct tcg at- ggt ctc aca aaa ata tta aac ctc ttt ctg atg tct caa cta ttt cgt aaa cga taa cgt cca aca ata tat tct cgt aga gct tat caa cat cct tat acc aat cta ggt tgt cag aca att gca tca taa aat aat gtt tat aat tta cac gtt aac atc ata taa taa acg tat ata gtt aat att ttt gga ata taa atg atc tgt aaa atc cat gta ggg gac act gct cac gtt ttt tct cta gta cat aat ttc aca caa gtt ttt ata cag aca aat taa ttc tcg tcc ata ta- ttt taa aac att gac ttt tgt act aag aaa aat atc ttg act aac cat ctc ttt ctc tct tcg atg ggt ctc aca aaa ata tta aac ctc ttt ctg atg g-- --- --- --- --- --- --- --- --- --- --- --- --- --- --- --- --- --- --- --- --- --- --- --- --- --- --- --- --- --- --- --- --- --- --- --- --- --- --- --- --- --- --- --- --- --- --- agt cgt aaa aag ttt tta tcc ttt ctc tct tcg ata ggt ctc --- --- --- --- --- --- --- --- --- --- --- --- --- --- --- --- --- --- --- --- --- --- --- --- --- --- --- --- --- --- --- --- --- --- --- --- --- --- --- --- --- --- --- --- --- --- --- --- --- --- --- --- --- --- --- --- --- --- --- --- --- --- --- --- --- --- --- --- --- --- aca aaa ata tta aac ctc ttt ctg atg gtc tct ata aa

>ON622722.2_Monkeypox_virus_isolate_MPXV_FR_HCL0001_2022_complete_genome

atc gag gac gac ttc aac tat gac agc tcg gtg gcg tct gcc agc gtg tac ata cga atg gca ttt cta aga aaa gtc tac ggt atc ctt -tc tac aat ttc ctt taa caa cgg caa cag ctg cag tat ttt tat act ttg aat gca -tc gga cat tta tac aag gga gtc ctg ttc taa tat tgg cat caa tgt tcg gat cta tag gct tga ttt tcg cat tga ctt tac aca gac atg aac atc ccc tga atc tgt aca tac ttt gtg gat tta cac tgt tag aat ctc taa cgc tgg cct ctg ttg tta ctt tct atg atg cac gta tcg tta tgc aag ctt tca tgt tga cta ctg cag tgt ttc ttg ctc tga cta cat ata ctc tac aat caa aga gag att tca gta aac ttg taa cag gat tgt ttg ctg ctt tct gga ttt taa ttt tgt cag gag tct tga gga taa agt tta aaa tag aat taa taa aga aca tat agg tca -tt ttt taa aca tgg ata gaa acc aag gtt gtt agt taa taa tat aca aga tat ttt ttc tca ctc tga tcc atg taa acc aag gac gag aga cac tct cat tcc tca ttc acg aca cca tta aaa atg gaa att aaa gcc ctc tat taa gca cag acg gct aca ggt cta cca tca ggt t-a cct tcg tct acc ttc aca atg gcc tct cct tgt gcc cag ttc agt ccc tgt cat tgc cac gct act aag gac tcc ctg aat acc gtg act gac gtc aga cat tgt ctg act gaa tac atc ctg tgg gtt tct cat aga tgg acc cat aga gaa agc gca ggg cct ctc tac agg ctt ctc atc tct ttc aga att gat gca atg gag cta ttt ggt agc gag ttg aag gag ttc tcg aat tca ctt ccg tgg gac aat atc gac aat tgc gtg gag atc att aaa tgt ttc atc aga aat gac tcc atg aaa acc gcc aaa gaa ctt tgt gca ata att gga ctt tgt act caa tca gct att gtc act gga aga gtc ttc aat gat aag tat atc gac ata cta ctt atg ctg cga aag att ctg aac gag aac gac tat ctc acc ctc ttg gat cat atc ctc act gct aaa tac taa atc tcc ttc atg ctc tct cac taa tac tct tac tca cta cac ttt tta tca tct tat gat gaa tga ttg cct tca tca ttt ttt cgt gga ata taa tat agg aat aat tag cac cag aat agc tat gga tat ctc gtt aag aat att ctc tat aag aga cat aat gta gac ata gtt att ata tcc ttc tta gat aag tgt tac gct act gga aag ttt cca tcg tta tta tta cat gaa gat gat ata att aaa cca aca ttg aga ttg gct ctt atg tta gct gga ttg aat tac tgt aat aaa tgc atc gag tat aga ggg ata tag caa ttc tcg ata ata gtc atg caa tat ttg aat gag act gat aat tta ggt aat aca gta ct- --- aca cac ata tct ttc tag att ata tat cgt taa aaa tct gta aga tgt ata ttt ctc ata agt atc cac tgt gta ata tta tta atg gat ata tag ata acg caa tag gga cta ata gta ttg taa aag ata taa tcg act att tac gta cat atc cag ata tct ata ttc cta cta gtt tgc tgc gta gtt gca tca ttg ata tgc atg -at tta tca gga ttc aga gat gaa tta cta agt aaa cta caa tcc cac aat aag taa gaa tca aat atc aaa aac tca ctt ttg att ttt cta gtc tta agt aat aca tat att tat taa tag acc tat gaa ata aaa aaa ggt aac aat gga ttc gcg tat agc tat tta cgt att agt atc ggc atc tct ttt gta tct tgt taa ttg tca caa act agt aca tta ctt caa tct gaa aat aaa tgg aag tga tat aac taa tac agc aga tat att gct gga caa tta tcc aat tat gac ctt tga tgg aaa gga tat tta tcc atc tat ctc gtt cat ggt cgg taa taa act ttt cct aga tct tta taa aaa tat ctt tga aga att ttt caa act att tcg agt atc tgt aag tag tca ata cga gga att aga ata tta tta ttc atg tga tta tac taa caa ccg tcc tac aat taa aca aca tta ctt tta taa cgg cga cga ata tac tga aat tga tag atc gaa aaa agc cac taa taa aaa cag ttg gtt aat tac ttc agg ctt tag act aca aaa atg gtt cga tag cga aga ttg tat aat tta tct cag atc ttt agt tag aag aat gga aga cag taa caa aaa cag taa aaa aac tta gta ctt aga tat cg- --a aaa aat ata ttt ttg tag act ctt gag aat aga agg aaa aca tgt aca taa tta taa aaa atg aaa atc aat ggc gaa taa gac agt gcg att cgc acc atg gag tcg gta gat ttc atg gct gtc gat gag cag ttt cac gac gac ctc gat ctt tgg tca tta tct ttg gta gat gat tat aaa aaa cat gga tta ggt gtt gac tgt tat gtt cta gaa cca gtt gtt gac agg aaa ata ttt gat aga ttt ctc ctt gaa cca att tgt gat cct gta gat gtt ctg tat gat tat ttt agg att cat aga gat aat att gat cag tat ata gta gat aga ctg ttt gca tat att aca tat aaa gat att ata tct gca tta gtg tca aag aat tat atg gaa gat att ttc tct ata att att aag aat tgt aat tct gtg caa gat ctc tta ctt tac tat cta tct aat gca tat gta gaa ata gac att gtt gat ctt atg gta gat cat ggg gct gta ata tat aaa ata gaa tgc ttg aat gcc tat ttt agg gga ata tgt aaa aag gaa agt agt gtt gtt gag ttt att ttg aat tgt ggt atc cca gat gaa aat gat gtt aaa tta gat cta tat aaa ata att cag tat act agg gga ttc ctt gta gat gaa ccc aca gta tta gaa att tat aag ctt tgt atc cca tat att gaa gat atc aat caa cta gat gct ggt gga agg acc ttg ctt tat cgc gct atc tat gca ggt tat ata gat tta gta tca tgg cta tta gaa aat gga gca aat gtc aac gca gta atg agt aat gga tat aca tgt ctt gac gtg gcc gtg gat agg gga tct gtc atc gcc cgt agg gaa gca cat ctt aaa ata tta gaa ata ttg ctt aga gaa cca ttg tct att gac tgt ata aaa tta gct ata ctt aat aat aca att gaa aac cat gat gtg ata aag ctc tgt atc aag tat ttt atg atg gta gat tat tca ctt tgt aat gtg tat gca tca tca ctc ttt gat tat ata att gat tgt aaa caa gaa ttg gag tac att agg cag atg aaa att cat aat aca acc atg tat gag tta atc tat aat aga gac aaa aac aag cat gct tcc cat att cta cat agg tat tct aaa cat cca gtt ttg aca cag tgt atc act aaa gga ttc aag att tac aca gaa gta acc gag cag gtc act aaa gct cta aac aga cgt gct cta ata gat gag ata ata aac aat gta tca act gat gac aat ctc cta tca aaa ctt cca tta gaa att agg gat cta att gtt tca caa gct gtc ata tag agt tct atc cac cca cct ttc ttg aaa tga gtt aat --- --- --- --- --- --- --- --- --- --- --- --- --- --- --- --- --- --- --- --- --- --- --- --- --- --- --- --- --- --- --- --- --- --- --- --- --- --- --- --- --- --- --- --- --- --- --- --- --- --- --- --- --- --- --- --- --- --- --- --- --- --- --- --- --- --- --- --- --- --- --- --- --- --- --- --- --- --- --- --- --- --- --- --- --- --- --- --- --- --- --- --- --- --- --- --- --- --- --- --- --- --- --- --- --- -ag tca taa gtt agt taa --- --- --- --- --- -gt cat aag tta gtt aag tca t-- --- --- --- --- --- --- --- --- --- --- --- --- --- --- --- -aa gtt agt taa gtc ata agt tag tta agt cat aag tta gtt aag tca taa gtt agt taa gtc ata agt tag tta agt cat aag tta gtt tat agt cta aca ctt cta att ttt ata cct tga tct ttt tct cta att atg aaa aag taa atc att atg aag atg gat g-a aat gga cga gat tgt gcg cat cgt taa cga tag tat gtg gta cgt acc taa cgc att tat gga cga cgg tga taa tga agg tca cat ttc tgt caa taa tgt ctg tca tat gta tct cgc att ctt tga tgt gga tat atc atc tca tct gtt taa att agt tat taa aca ctg cga tct gaa taa acg act aaa atg tgg taa ctc tcc att aca ttg cta tac gat gaa tac acg att taa tcc atc tgt att aaa gat att gtt acg cca cgg cat gcg taa ctt tga tag caa gga taa aaa agg aca tat tcc tct aca cca cta tct gat tca ttc act atc aat cga taa caa gat ctt tga tat act aac gga ccc cat tga tga ctt tag taa atc atc cga tct att gct gtg tta tct tag ata taa att caa tgg gag ctt aaa cta tta cgt tct gta caa att att gac taa agg atc tga ccc taa ttg cgt cga tga gga tgg act cac ttc tct tca tta cta ctg taa aca cat atc cgc gtt cca cga aag caa tta tta caa gtc aaa gag tca cac taa gat gcg agc tga gaa gcg att cat cta cgc gat aat aga tca tgg agc aaa cat taa cgc ggt tac gaa aat cgg aaa tac gcc gtt aca cac tta cct tca aca gta tac caa aca tag tcc tcg tgt ggt gta tgc tct ttt atc tcg agg agc cga tac gag gat acg taa taa tct taa ttg tac acc cat cat gga ata cat aaa gaa cga ttg tgc aac agg tca tat tct cat aat gtt act caa ttg gca cga aca aaa ata cgg gaa att aca aaa gga aga agg aca aca tct act tta tct att cat aaa aca taa tca agg ata tgg aag tcg ctc tct caa tat act acg gta tct act aga tag att cga cat tca gaa aga cga ata cta taa tac aat gac tcc tct tca tac cgc ctt cca gaa ttg caa taa caa tgt tgc ctc ata cct cgt ata cat tgg ata cga cat caa cct tcc gac taa aga cga taa gac agt att cga ctt ggt gtt tga aaa cag aaa cat tat ata caa ggc gga tgt cgt taa tga cat tat cca cca cag act gaa agt atc tct acc tat gat taa atc gtt gtt cta caa gat gtc gga gtt ctc tcc cta cga cga tca cta cgt aaa gaa gat aat agc cta ctg cct att aag gga cga gtc att tgc gga act aca tac taa att ctg ttt aaa cga gga cta taa aag tgt att tat gaa aaa tat atc att cga taa gat aga ttc cat cat cga aaa atg tag tcg tga cat aag tct cct caa aga gat tcg aat ctc aga cac cga ctt gta tac ggt att gag aac aga aga cat tcg gta tca cac ata tct cga agc cat aca ttc aga caa acg cat ttc att tcc cat gta cga cga tct cat aga aca gtg tca tct atc gat gga gca taa aag taa act cgt cga caa agc act caa taa att aga gtc tac cat cga tag tca atc tag act atc gta ttt gcc tcc gga aat tat gcg caa tat cat aac caa gct aag cga cta cca tct aaa cag tat gtt gta cgg aaa gaa cca tta caa ata tta tcc atg ata gaa aga aaa tat tta aaa aat aat cta tat gat tgg aga agt agg aaa caa aca gta aca aga cga cga tta cta c-a tta tta aat cat gag gtc cgt att ata ctc gta tat att gtt tct ctc atg tat aat aat aaa cgg aag aga tat agc acc aca tgc acc atc caa tgg aaa gtg taa aga caa cga ata cag aag ccg taa tct atg ttg tct atc gtg tcc tcc ggg aac tta cgc ttt cag att atg tga tag caa gac taa tac aca atg tac acc gtg tgg ttc gga tac ctt tac atc tca caa taa tca ttt aca ggc ttg tct aag ttg taa cgg aag atg tga tag taa tca ggt aga gac gcg atc gtg taa cac gac tca caa tag aat ctg tga atg ctc tcc agg ata tta ttg tct tct caa agg agc atc agg gtg tag aac atg tat ttc taa aac aaa gtg tgg aat agg ata cgg agt atc cgg ata cac gtc tac cgg aga cgt cat ctg ttc tcc gtg tgg tcc cgg aac ata ttc tca cac cgt ctc ttc cac aga taa atg cga acc cgt cgt aac cag caa tac att taa cta tat cga tgt gga aat taa cct gta tcc agt caa cga cac atc gtg tac tcg gac gac cac tac cgg tct cag cga atc cat ctc aac gtc gga act aac tat tac cat gaa tca taa aga ttg tga tcc agt ctt tcg tgc aga ata ctt ctc tgt cct taa taa tgt agc aac ttc agg att ctt tac agg aga aaa tag ata tca gaa tac ttc aaa gat atg tac tct gaa ttt cga gat taa atg taa caa caa aga ttc atc ttc caa aca gtt aac gaa aac aaa gaa tga tac tat cat gcc gca ttc aga gac ggt aac tct agt ggg cga ctg tct atc tag cgt cga cat cta cat act ata tag taa tac caa tac tca aga cta cga aaa tga tac aat ctc tta tca tat ggg taa tgt tct cga tgt caa tag cca tat gcc cgc tag ttg cga tat aca taa act gat cac taa ttc cca gaa tcc cac cca ctt ata gta ag- -tt ttt tta cct ata aat aat aaa tac aat aat taa ttt ctc gta aaa gta gaa aat ata ttc taa ttt att ata tgg taa gaa agt aga atc atc tag aac agt aat caa tca ata gca atc atg aaa caa tat att gtc ctg gca tgc atg tgc cta gtg gca gct gct atg cct act agt ctt caa caa tct tca tcc tcg tgt act gaa gaa gaa aac aaa cat cat atg gga atc gat gtt att atc aaa gtc aca aag caa gac caa aca ccg acc aat gat aag att tgt caa tcc gta acg gaa gtt aca gag a-- --- --- --- --- --- -cc gaa gat gat gag gta tcc gaa gaa gtt gta aaa gga gat ccc acc act tat tac act atc gtc ggt gcg ggt ctt aac atg aac ttt gga ttc acc aaa tgc cca aag att tta tcc atc tcc gaa tcc tct gat gga aac act gtg aat act aga ttg tcc agc gtg tca ccg gga caa ggt aag gac tct ccc gcg atc acg cgt gaa gaa gct ctg gct atg atc aaa gac tgt gag atg tct atc gac atc aga tgt agc gaa gaa gag aaa gac agt gac atc aag acc cat cca gta ctt ggg tct aac atc tca cat aag aaa gtg agt tac aaa gat atc atc ggt tca acg atc gtt gat aca aaa tgt gtc aag aac cta gag ttt agc gta cgt atc gga gac atg tgt gag gaa tca tct gaa ctt gaa gtc aag gat gga ttc aag tat gtc gac gga tcg gca tct gaa ggt gca acc gat gat act tca ctc atc gat tca aca aaa ctc aaa gca tgt gtc tga atc gat aac tct att cat ctg aaa atg gat gag ttg ggt taa tcg aac gat tca gac acc gca cca cga att aaa aaa g-- -ac cgg gca cta tat tcc ggt ttg caa aac aaa aat att taa cta cat tca caa aaa gtt acc tct cgt tac ttc ttc ttt ctg ttt caa tat gtg ata cga tat gat cac tat tcg tat tct ct- --- --- tgg tct cat aaa aaa gtt tta c-- --a aaa aaa aaa aaa a-t att ttt att ctc ttt ctc tct tcg at- ggt ctc aca aaa ata tta aac ctc ttt ctg atg tct caa cta ttt cgt aaa cga taa cgt cca aca ata tat tct cgt aga gct tat caa cat cct tat acc aat cta ggt tgt cag aca att gca tca taa aat aat gtt tat aat tta cac gtt aac atc ata taa taa acg tat ata gtt aat att ttt gga ata taa atg atc tgt aaa atc cat gta ggg gac act gct cac gtt ttt tct cta gta cat aat ttc aca caa gtt ttt ata cag aca aat taa ttc tcg tcc ata ta- ttt taa aac att gac ttt tgt act aag aaa aat atc ttg act aac cat ctc ttt ctc tct tcg atg ggt ctc aca aaa ata tta aac ctc ttt ctg atg g-- --- --- --- --- --- --- --- --- --- --- --- --- --- --- --- --- --- --- --- --- --- --- --- --- --- --- --- --- --- --- --- --- --- --- --- --- --- --- --- --- --- --- --- --- --- --- agt cgt aaa aag ttt tta tcc ttt ctc tct tcg ata ggt ctc --- --- --- --- --- --- --- --- --- --- --- --- --- --- --- --- --- --- --- --- --- --- --- --- --- --- --- --- --- --- --- --- --- --- --- --- --- --- --- --- --- --- --- --- --- --- --- --- --- --- --- --- --- --- --- --- --- --- --- --- --- --- --- --- --- --- --- --- --- --- aca aaa ata tta aac ctc ttt ctg atg gtc tct ata aa

>ON627808.1_Monkeypox_virus_isolate_MPX/human/USA/UT-UPHL-82200022/2022_complete_genome

atc gag gac gac ttc aac tat gac agc tcg gtg gcg tct gcc agc gtg tac ata cga atg gca ttt cta aga aaa gtc tac ggt atc ctt -tc tac aat ttc ctt taa caa cgg caa cag ctg cag tat ttt tat act ttg aat gca -tc gga cat tta tac aag gga gtc ctg ttc taa tat tgg cat caa tgt tcg gat cta tag gct tga ttt tcg cat tga ctt tac aca gac atg aac atc ccc tga atc tgt aca tac ttt gtg gat tta cac tgt tag aat ctc taa cgc tgg cct ctg ttg tta ctt tct atg atg cac gta tcg tta tgc aag ctt tca tgt tga cta ctg cag tgt ttc ttg ctc tga cta cat ata ctc tac aat caa aga gag att tca gta aac ttg taa cag gat tgt ttg ctg ctt tct gga ttt taa ttt tgt cag gag tct tga gga taa agt tta aaa tag aat taa taa aga aca tat agg tca -tt ttt taa aca tgg ata gaa acc aag gtt gtt agt taa taa tat aca aga tat ttt ttc tca ctc tga tcc atg taa acc aag gac gag aga cac tct cat tcc tca ttc acg aca cca tta aaa atg gaa att aaa gcc ctc tat taa gca cag acg gct aca ggt cta cca tca ggt t-a cct tcg tct acc ttc aca atg gcc tct cct tgt gcc cag ttc agt ccc tgt cat tgc cac gct act aag gac tcc ctg aat acc gtg act gac gtc aga cat tgt ctg act gaa tac atc ctg tgg gtt tct cat aga tgg acc cat aga gaa agc gca ggg cct ctc tac agg ctt ctc atc tct ttc aga att gat gca atg gag cta ttt ggt agc gag ttg aag gag ttc tcg aat tca ctt ccg tgg gac aat atc gac aat tgc gtg gag atc att aaa tgt ttc atc aga aat gac tcc atg aaa acc gcc aaa gaa ctt tgt gca ata att gga ctt tgt act caa tca gct att gtc act gga aga gtc ttc aat gat aag tat atc gac ata cta ctt atg ctg cga aag att ctg aac gag aac gac tat ctc acc ctc ttg gat cat atc ctc act gct aaa tac taa atc tcc ttc atg ctc tct cac taa tac tct tac tca cta cac ttt tta tca tct tat gat gaa tga ttg cct tca tca ttt ttt cgt gga ata taa tat agg aat aat tag cac cag aat agc tat gga tat ctc gtt aag aat att ctc tat aag aga cat aat gta gac ata gtt att ata tcc ttc tta gat aag tgt tac gct act gga aag ttt cca tcg tta tta tta cat gaa gat gat ata att aaa cca aca ttg aga ttg gct ctt atg tta gct gga ttg aat tac tgt aat aaa tgc atc gag tat aga ggg ata tag caa ttc tcg ata ata gtc atg caa tat ttg aat gag act gat aat tta ggt aat aca gta ct- --- aca cac ata tct ttc tag att ata tat cgt taa aaa tct gta aga tgt ata ttt ctc ata agt atc cac tgt gta ata tta tta atg gat ata tag ata acg caa tag gga cta ata gta ttg taa aag ata taa tcg act att tac gta cat atc cag ata tct ata ttc cta cta gtt tgc tgc gta gtt gca tca ttg ata tgc atg -at tta tca gga ttc aga gat gaa tta cta agt aaa cta caa tcc cac aat aag taa gaa tca aat atc aaa aac tca ctt ttg att ttt cta gtc tta agt aat aca tat att tat taa tag acc tat gaa ata aaa aaa ggt aac aat gga ttc gcg tat agc tat tta cgt att agt atc ggc atc tct ttt gta tct tgt taa ttg tca caa act agt aca tta ctt caa tct gaa aat aaa tgg aag tga tat aac taa tac agc aga tat att gct gga caa tta tcc aat tat gac ctt tga tgg aaa gga tat tta tcc atc tat ctc gtt cat ggt cgg taa taa act ttt cct aga tct tta taa aaa tat ctt tga aga att ttt cag act att tcg agt atc tgt aag tag tca ata cga gga att aga ata tta tta ttc atg tga tta tac taa caa ccg tcc tac aat taa aca aca tta ctt tta taa cgg cga cga ata tac tga aat tga tag atc gaa aaa agc cac taa taa aaa cag ttg gtt aat tac ttc agg ctt tag act aca aaa atg gtt cga tag cga aga ttg tat aat tta tct cag atc ttt agt tag aag aat gga aga cag taa caa aaa cag taa aaa aac tta gta ctt aga tat cg- --a aaa aat ata ttt ttg tag act ctt gag aat aga agg aaa aca tgt aca taa tta taa aaa atg aaa atc aat ggc gaa taa gac agt gcg att cgc acc atg gag tcg gta gat ttc atg gct gtc gat gag cag ttt cac gac gac ctc gat ctt tgg tca tta tct ttg gta gat gat tat aaa aaa cat gga tta ggt gtt gac tgt tat gtt cta gaa cca gtt gtt gac agg aaa ata ttt gat aga ttt ctc ctt gaa cca att tgt gat cct gta gat gtt ctg tat gat tat ttt agg att cat aga gat aat att gat cag tat ata gta gat aga ctg ttt gca tat att aca tat aaa gat att ata tct gca tta gtg tca aag aat tat atg gaa gat att ttc tct ata att att aag aat tgt aat tct gtg caa gat ctc tta ctt tac tat cta tct aat gca tat gta gaa ata gac att gtt gat ctt atg gta gat cat ggg gct gta ata tat aaa ata gaa tgc ttg aat gcc tat ttt agg gga ata tgt aaa aag gaa agt agt gtt gtt gag ttt att ttg aat tgt ggt atc cca gat gaa aat gat gtt aaa tta gat cta tat aaa ata att cag tat act agg gga ttc ctt gta gat gaa ccc aca gta tta gaa att tat aag ctt tgt atc cca tat att gaa gat atc aat caa cta gat gct ggt gga agg acc ttg ctt tat cgc gct atc tat gca ggt tat ata gat tta gta tca tgg cta tta gaa aat gga gca aat gtc aac gca gta atg agt aat gga tat aca tgt ctt gac gtg gcc gtg gat agg gga tct gtc atc gcc cgt agg gaa gca cat ctt aaa ata tta gaa ata ttg ctt aga gaa cca ttg tct att gac tgt ata aaa tta gct ata ctt aat aat aca att gaa aac cat gat gtg ata aag ctc tgt atc aag tat ttt atg atg gta gat tat tca ctt tgt aat gtg tat gca tca tca ctc ttt gat tat ata att gat tgt aaa caa gaa ttg gag tac att agg cag atg aaa att cat aat aca acc atg tat gag tta atc tat aat aga gac aaa aac aag cat gct tcc cat att cta cat agg tat tct aaa cat cca gtt ttg aca cag tgt atc act aaa gga ttc aag att tac aca gaa gta acc gag cag gtc act aaa gct cta aac aga cgt gct cta ata gat gag ata ata aac aat gta tca act gat gac aat ctc cta tca aaa ctt cca tta gaa att agg gat cta att gtt tca caa gct gtc ata tag agt tct atc cac cca cct ttc ttg aaa tga gtt aat --- --- --- --- --- --- --- --- --- --- --- --- --- --- --- --- --- --- --- --- --- --- --- --- --- --- --- --- --- --- --- --- --- --- --- --- --- --- --- --- --- --- --- --- --- --- --- --- --- --- --- --- --- --- --- --- --- --- --- --- --- --- --- --- --- --- --- --- --- --- --- --- --- --- --- --- --- --- --- --- --- --- --- --- --- --- --- --- --- --- --- --- --- --- --- --- --- --- --- --- --- --- --- --- --- -ag tca taa gtt agt taa --- --- --- --- --- -gt cat aag tta gtt aag tca t-- --- --- --- --- --- --- --- --- --- --- --- --- --- --- --- -aa gtt agt taa gtc ata agt tag tta agt cat aag tta gtt aag tca taa gtt agt taa gtc ata agt tag tta agt cat aag tta gtt tat agt cta aca ctt cta att ttt ata cct tga tct ttt tct cta att atg aaa aag taa atc att atg aag atg gat g-a aat gga cga gat tgt gcg cat cgt taa cga tag tat gtg gta cgt acc taa cgc att tat gga cga cgg tga taa tga agg tca cat ttc tgt caa taa tgt ctg tca tat gta tct cgc att ctt tga tgt gga tat atc atc tca tct gtt taa att agt tat taa aca ctg cga tct gaa taa acg act aaa atg tgg taa ctc tcc att aca ttg cta tac gat gaa tac acg att taa tcc atc tgt att aaa gat att gtt acg cca cgg cat gcg taa ctt tga tag caa gga taa aaa agg aca tat tcc tct aca cca cta tct gat tca ttc act atc aat cga taa caa gat ctt tga tat act aac gga ccc cat tga tga ctt tag taa atc atc cga tct att gct gtg tta tct tag ata taa att caa tgg gag ctt aaa cta tta cgt tct gta caa att att gac taa agg atc tga ccc taa ttg cgt cga tga gga tgg act cac ttc tct tca tta cta ctg taa aca cat atc cgc gtt cca cga aag caa tta tta caa gtc aaa gag tca cac taa gat gcg agc tga gaa gcg att cat cta cgc gat aat aga tca tgg agc aaa cat taa cgc ggt tac gaa aat cgg aaa tac gcc gtt aca cac tta cct tca aca gta tac caa aca tag tcc tcg tgt ggt gta tgc tct ttt atc tcg agg agc cga tac gag gat acg taa taa tct taa ttg tac acc cat cat gga ata cat aaa gaa cga ttg tgc aac agg tca tat tct cat aat gtt act caa ttg gca cga aca aaa ata cgg gaa att aca aaa gga aga agg aca aca tct act tta tct att cat aaa aca taa tca agg ata tgg aag tcg ctc tct caa tat act acg gta tct act aga tag att cga cat tca gaa aga cga ata cta taa tac aat gac tcc tct tca tac cgc ctt cca gaa ttg caa taa caa tgt tgc ctc ata cct cgt ata cat tgg ata cga cat caa cct tcc gac taa aga cga taa gac agt att cga ctt ggt gtt tga aaa cag aaa cat tat ata caa ggc gga tgt cgt taa tga cat tat cca cca cag act gaa agt atc tct acc tat gat taa atc gtt gtt cta caa gat gtc gga gtt ctc tcc cta cga cga tca cta cgt aaa gaa gat aat agc cta ctg cct att aag gga cga gtc att tgc gga act aca tac taa att ctg ttt aaa cga gga cta taa aag tgt att tat gaa aaa tat atc att cga taa gat aga ttc cat cat cga aaa atg tag tcg tga cat aag tct cct caa aga gat tcg aat ctc aga cac cga ctt gta tac ggt att gag aac aga aga cat tcg gta tca cac ata tct cga agc cat aca ttc aga caa acg cat ttc att tcc cat gta cga cga tct cat aga aca gtg tca tct atc gat gga gca taa aag taa act cgt cga caa agc act caa taa att aga gtc tac cat cga tag tca atc tag act atc gta ttt gcc tcc gga aat tat gcg caa tat cat aac caa gct aag cga cta cca tct aaa cag tat gtt gta cgg aaa gaa cca tta caa ata tta tcc atg ata gaa aga aaa tat tta aaa aat aat cta tat gat tgg aga agt agg aaa caa aca gta aca aga cga cga tta cta c-a tta tta aat cat gag gtc cgt att ata ctc gta tat att gtt tct ctc atg tat aat aat aaa cgg aag aga tat agc acc aca tgc acc atc caa tgg aaa gtg taa aga caa cga ata cag aag ccg taa tct atg ttg tct atc gtg tcc tcc ggg aac tta cgc ttt cag att atg tga tag caa gac taa tac aca atg tac acc gtg tgg ttc gga tac ctt tac atc tca caa taa tca ttt aca ggc ttg tct aag ttg taa cgg aag atg tga tag taa tca ggt aga gac gcg atc gtg taa cac gac tca caa tag aat ctg tga atg ctc tcc agg ata tta ttg tct tct caa agg agc atc agg gtg tag aac atg tat ttc taa aac aaa gtg tgg aat agg ata cgg agt atc cgg ata cac gtc tac cgg aga cgt cat ctg ttc tcc gtg tgg tcc cgg aac ata ttc tca cac cgt ctc ttc cac aga taa atg cga acc cgt cgt aac cag caa tac att taa cta tat cga tgt gga aat taa cct gta tcc agt caa cga cac atc gtg tac tcg gac gac cac tac cgg tct cag cga atc cat ctc aac gtc gga act aac tat tac cat gaa tca taa aga ttg tga tcc agt ctt tcg tgc aga ata ctt ctc tgt cct taa taa tgt agc aac ttc agg att ctt tac agg aga aaa tag ata tca gaa tac ttc aaa gat atg tac tct gaa ttt cga gat taa atg taa caa caa aga ttc atc ttc caa aca gtt aac gaa aac aaa gaa tga tac tat cat gcc gca ttc aga gac ggt aac tct agt ggg cga ctg tct atc tag cgt cga cat cta cat act ata tag taa tac caa tac tca aga cta cga aaa tga tac aat ctc tta tca tat ggg taa tgt tct cga tgt caa tag cca tat gcc cgc tag ttg cga tat aca taa act gat cac taa ttc cca gaa tcc cac cca ctt ata gta ag- -tt ttt tta cct ata aat aat aaa tac aat aat taa ttt ctc gta aaa gta gaa aat ata ttc taa ttt att ata tgg taa gaa agt aga atc atc tag aac agt aat caa tca ata gca atc atg aaa caa tat att gtc ctg gca tgc atg tgc cta gtg gca gct gct atg cct act agt ctt caa caa tct tca tcc tcg tgt act gaa gaa gaa aac aaa cat cat atg gga atc gat gtt att atc aaa gtc aca aag caa gac caa aca ccg acc aat gat aag att tgt caa tcc gta acg gaa gtt aca gag a-- --- --- --- --- --- -cc gaa gat gat gag gta tcc gaa gaa gtt gta aaa gga gat ccc acc act tat tac act atc gtc ggt gcg ggt ctt aac atg aac ttt gga ttc acc aaa tgc cca aag att tta tcc atc tcc gaa tcc tct gat gga aac act gtg aat act aga ttg tcc agc gtg tca ccg gga caa ggt aag gac tct ccc gcg atc acg cgt gaa gaa gct ctg gct atg atc aaa gac tgt gag atg tct atc gac atc aga tgt agc gaa gaa gag aaa gac agt gac atc aag acc cat cca gta ctt ggg tct aac atc tca cat aag aaa gtg agt tac aaa gat atc atc ggt tca acg atc gtt gat aca aaa tgt gtc aag aac cta gag ttt agc gta cgt atc gga gac atg tgt gag gaa tca tct gaa ctt gaa gtc aag gat gga ttc aag tat gtc gac gga tcg gca tct gaa ggt gca acc gat gat act tca ctc atc gat tca aca aaa ctc aaa gca tgt gtc tga atc gat aac tct att cat ctg aaa atg gat gag ttg ggt taa tcg aac gat tca gac acc gca cca cga att aaa aaa g-- -ac cgg gca cta tat tcc ggt ttg caa aac aaa aat att taa cta cat tca caa aaa gtt acc tct cgt tac ttc ttc ttt ctg ttt caa tat gtg ata cga tat gat cac tat tcg tat tct ct- --- --- tgg tct cat aaa aaa gtt tta c-- --? aaa aaa aaa aaa a-t att ttt att ctc ttt ctc tct tcg at- ggt ctc aca aaa ata tta aac ctc ttt ctg atg tct caa cta ttt cgt aaa cga taa cgt cca aca ata tat tct cgt aga gct tat caa cat cct tat acc aat cta ggt tgt cag aca att gca tca taa aat aat gtt tat aat tta cac gtt aac atc ata taa taa acg tat ata gtt aat att ttt gga ata taa atg atc tgt aaa atc cat gta ggg gac act gct cac gtt ttt tct cta gta cat aat ttc aca caa gtt ttt ata cag aca aat taa ttc tcg tcc ata ta- ttt taa aac att gac ttt tgt act aag aaa aat atc ttg act aac cat ctc ttt ctc tct tcg atg ggt ctc aca aaa ata tta aac ctc ttt ctg atg g-- --- --- --- --- --- --- --- --- --- --- --- --- --- --- --- --- --- --- --- --- --- --- --- --- --- --- --- --- --- --- --- --- --- --- --- --- --- --- --- --- --- --- --- --- --- --- agt cgt aaa aag ttt tta tcc ttt ctc tct tcg ata ggt ctc --- --- --- --- --- --- --- --- --- --- --- --- --- --- --- --- --- --- --- --- --- --- --- --- --- --- --- --- --- --- --- --- --- --- --- --- --- --- --- --- --- --- --- --- --- --- --- --- --- --- --- --- --- --- --- --- --- --- --- --- --- --- --- --- --- --- --- --- --- --- aca aaa ata tta aac ctc ttt ctg atg gtc tct ata aa

>ON676703.1_Monkeypox_virus_isolate_MPXV_USA_2022_CA001_complete_genome

atc gag gac gac ttc aac tat gac agc tcg gtg gcg tct gcc agc gtg tac ata cga atg gca ttt cta aga aaa gtc tac ggt atc ctt -tc tac aat ttc ctt taa caa cgg caa cag ctg cag tat ttt tat act ttg aat gca -tc gga cat tta tac aag gga gtc ctg ttc taa tat tgg cat caa tgt tcg gat cta tag gct tga ttt tcg cat tga ctt tac aca gac atg aac atc ccc tga atc tgt aca tac ttt gtg gat tta cac tgt tag aat ctc taa cgc tgg cct ctg ttg tta ctt tct atg atg cac gta tcg tta tgc aag ctt tca tgt tga cta ctg cag tgt ttc ttg ctc tga cta cat ata ctc tac aat caa aga gag att tca gta aac ttg taa cag gat tgt ttg ctg ctt tct gga ttt taa ttt tgt cag gag tct tga gga taa agt tta aaa tag aat taa taa aga aca tat agg tca -tt ttt taa aca tgg ata gaa acc aag gtt gtt agt taa taa tat aca aga tat ttt ttc tca ctc tga tcc atg taa acc aag gac gag aga cac tct cat tcc tca ttc acg aca cca tta aaa atg gaa att aaa gcc ctc tat taa gca cag acg gct aca ggt cta cca tca ggt t-a cct tcg tct acc ttc aca atg gcc tct cct tgt gcc cag ttc agt ccc tgt cat tgc cac gct act aag gac tcc ctg aat acc gtg act gac gtc aga cat tgt ctg act gaa tac atc ctg tgg gtt tct cat aga tgg acc cat aga gaa agc gca ggg cct ctc tac agg ctt ctc atc tct ttc aga att gat gca atg gag cta ttt ggt agc gag ttg aag gag ttc tcg aat tca ctt ccg tgg gac aat atc gac aat tgc gtg gag atc att aaa tgt ttc atc aga aat gac tcc atg aaa acc gcc aaa gaa ctt tgt gca ata att gga ctt tgt act caa tca gct att gtc act gga aga gtc ttc aat gat aag tat atc gac ata cta ctt atg ctg cga aag att ctg aac gag aac gac tat ctc acc ctc ttg gat cat atc ctc act gct aaa tac taa atc tcc ttc atg ctc tct cac taa tac tct tac tca cta cac ttt tta tca tct tat gat gaa tga ttg cct tca tca ttt ttt cgt gga ata taa tat agg aat aat tag cac cag aat agc tat gga tat ctc gtt aag aat att ctc tat aag aga cat aat gta gac ata gtt att ata tcc ttc tta gat aag tgt tac gct act gga aag ttt cca tcg tta tta tta cat gaa gat gat ata att aaa cca aca ttg aga ttg gct ctt atg tta gct gga ttg aat tac tgt aat aaa tgc atc gag tat aga ggg ata tag caa ttc tcg ata ata gtc atg caa tat ttg aat gag act gat aat tta ggt aat aca gta ct- --- aca cac ata tct ttc tag att ata tat cgt taa aaa tct gta aga tgt ata ttt ctc ata agt atc cac tgt gta ata tta tta atg gat ata tag ata acg caa tag gga cta ata gta ttg taa aag ata taa tcg act att tac gta cat atc cag ata tct ata ttc cta cta gtt tgc tgc gta gtt gca tca ttg ata tgc atg -at tta tca gga ttc aga gat gaa tta cta agt aaa cta caa tcc cac aat aag taa gaa tca aat atc aaa aac tca ctt ttg att ttt cta gtc tta agt aat aca tat att tat taa tag acc tat gaa ata aaa aaa ggt aac aat gga ttc gcg tat agc tat tta cgt att agt atc ggc atc tct ttt gta tct tgt taa ttg tca caa act agt aca tta ctt caa tct gaa aat aaa tgg aag tga tat aac taa tac agc aga tat att gct gga caa tta tcc aat tat gac ctt tga tgg aaa gga tat tta tcc atc tat ctc gtt cat ggt cgg taa taa act ttt cct aga tct tta taa aaa tat ctt tga aga att ttt cag act att tcg agt atc tgt aag tag tca ata cga gga att aga ata tta tta ttc atg tga tta tac taa caa ccg tcc tac aat taa aca aca tta ctt tta taa cgg cga cga ata tac tga aat tga tag atc gaa aaa agc cac taa taa aaa cag ttg gtt aat tac ttc agg ctt tag act aca aaa atg gtt cga tag cga aga ttg tat aat tta tct cag atc ttt agt tag aag aat gga aga cag taa caa aaa cag taa aaa aac tta gta ctt aga tat cg- --a aaa aat ata ttt ttg tag act ctt gag aat aga agg aaa aca tgt aca taa tta taa aaa atg aaa atc aat ggc gaa taa gac agt gcg att cgc acc atg gag tcg gta gat ttc atg gct gtc gat gag cag ttt cac gac gac ctc gat ctt tgg tca tta tct ttg gta gat gat tat aaa aaa cat gga tta ggt gtt gac tgt tat gtt cta gaa cca gtt gtt gac agg aaa ata ttt gat aga ttt ctc ctt gaa cca att tgt gat cct gta gat gtt ctg tat gat tat ttt agg att cat aga gat aat att gat cag tat ata gta gat aga ctg ttt gca tat att aca tat aaa gat att ata tct gca tta gtg tca aag aat tat atg gaa gat att ttc tct ata att att aag aat tgt aat tct gtg caa gat ctc tta ctt tac tat cta tct aat gca tat gta gaa ata gac att gtt gat ctt atg gta gat cat ggg gct gta ata tat aaa ata gaa tgc ttg aat gcc tat ttt agg gga ata tgt aaa aag gaa agt agt gtt gtt gag ttt att ttg aat tgt ggt atc cca gat gaa aat gat gtt aaa tta gat cta tat aaa ata att cag tat act agg gga ttc ctt gta gat gaa ccc aca gta tta gaa att tat aag ctt tgt atc cca tat att gaa gat atc aat caa cta gat gct ggt gga agg acc ttg ctt tat cgc gct atc tat gca ggt tat ata gat tta gta tca tgg cta tta gaa aat gga gca aat gtc aac gca gta atg agt aat gga tat aca tgt ctt gac gtg gcc gtg gat agg gga tct gtc atc gcc cgt agg gaa gca cat ctt aaa ata tta gaa ata ttg ctt aga gaa cca ttg tct att gac tgt ata aaa tta gct ata ctt aat aat aca att gaa aac cat gat gtg ata aag ctc tgt atc aag tat ttt atg atg gta gat tat tca ctt tgt aat gtg tat gca tca tca ctc ttt gat tat ata att gat tgt aaa caa gaa ttg gag tac att agg cag atg aaa att cat aat aca acc atg tat gag tta atc tat aat aga gac aaa aac aag cat gct tcc cat att cta cat agg tat tct aaa cat cca gtt ttg aca cag tgt atc act aaa gga ttc aag att tac aca gaa gta acc gag cag gtc act aaa gct cta aac aga cgt gct cta ata gat gag ata ata aac aat gta tca act gat gac aat ctc cta tca aaa ctt cca tta gaa att agg gat cta att gtt tca caa gct gtc ata tag agt tct atc cac cca cct ttc ttg aaa tga gtt aat --- --- --- --- --- --- --- --- --- --- --- --- --- --- --- --- --- --- --- --- --- --- --- --- --- --- --- --- --- --- --- --- --- --- --- --- --- --- --- --- --- --- --- --- --- --- --- --- --- --- --- --- --- --- --- --- --- --- --- --- --- --- --- --- --- --- --- --- --- --- --- --- --- --- --- --- --- --- --- --- --- --- --- --- --- --- --- --- --- --- --- --- --- --- --- --- --- --- --- --- --- --- --- --- --- -ag tca taa gtt agt taa --- --- --- --- --- -gt cat aag tta gtt aag tca t-- --- --- --- --- --- --- --- --- --- --- --- --- --- --- --- -aa gtt agt taa gtc ata agt tag tta agt cat aag tta gtt aag tca taa gtt agt taa gtc ata agt tag tta agt cat aag tta gtt tat agt cta aca ctt cta att ttt ata cct tga tct ttt tct cta att atg aaa aag taa atc att atg aag atg gat g-a aat gga cga gat tgt gcg cat cgt taa cga tag tat gtg gta cgt acc taa cgc att tat gga cga cgg tga taa tga agg tca cat ttc tgt caa taa tgt ctg tca tat gta tct cgc att ctt tga tgt gga tat atc atc tca tct gtt taa att agt tat taa aca ctg cga tct gaa taa acg act aaa atg tgg taa ctc tcc att aca ttg cta tac gat gaa tac acg att taa tcc atc tgt att aaa gat att gtt acg cca cgg cat gcg taa ctt tga tag caa gga taa aaa agg aca tat tcc tct aca cca cta tct gat tca ttc act atc aat cga taa caa gat ctt tga tat act aac gga ccc cat tga tga ctt tag taa atc atc cga tct att gct gtg tta tct tag ata taa att caa tgg gag ctt aaa cta tta cgt tct gta caa att att gac taa agg atc tga ccc taa ttg cgt cga tga gga tgg act cac ttc tct tca tta cta ctg taa aca cat atc cgc gtt cca cga aag caa tta tta caa gtc aaa gag tca cac taa gat gcg agc tga gaa gcg att cat cta cgc gat aat aga tca tgg agc aaa cat taa cgc ggt tac gaa aat cgg aaa tac gcc gtt aca cac tta cct tca aca gta tac caa aca tag tcc tcg tgt ggt gta tgc tct ttt atc tcg agg agc cga tac gag gat acg taa taa tct taa ttg tac acc cat cat gga ata cat aaa gaa cga ttg tgc aac agg tca tat tct cat aat gtt act caa ttg gca cga aca aaa ata cgg gaa att aca aaa gga aga agg aca aca tct act tta tct att cat aaa aca taa tca agg ata tgg aag tcg ctc tct caa tat act acg gta tct act aga tag att cga cat tca gaa aga cga ata cta taa tac aat gac tcc tct tca tac cgc ctt cca gaa ttg caa taa caa tgt tgc ctc ata cct cgt ata cat tgg ata cga cat caa cct tcc gac taa aga cga taa gac agt att cga ctt ggt gtt tga aaa cag aaa cat tat ata caa ggc gga tgt cgt taa tga cat tat cca cca cag act gaa agt atc tct acc tat gat taa atc gtt gtt cta caa gat gtc gga gtt ctc tcc cta cga cga tca cta cgt aaa gaa gat aat agc cta ctg cct att aag gga cga gtc att tgc gga act aca tac taa att ctg ttt aaa cga gga cta taa aag tgt att tat gaa aaa tat atc att cga taa gat aga ttc cat cat cga aaa atg tag tcg tga cat aag tct cct caa aga gat tcg aat ctc aga cac cga ctt gta tac ggt att gag aac aga aga cat tcg gta tca cac ata tct cga agc cat aca ttc aga caa acg cat ttc att tcc cat gta cga cga tct cat aga aca gtg tca tct atc gat gga gca taa aag taa act cgt cga caa agc act caa taa att aga gtc tac cat cga tag tca atc tag act atc gta ttt gcc tcc gga aat tat gcg caa tat cat aac caa gct aag cga cta cca tct aaa cag tat gtt gta cgg aaa gaa cca tta caa ata tta tcc atg ata gaa aga aaa tat tta aaa aat aat cta tat gat tgg aga agt agg aaa caa aca gta aca aga cga cga tta cta c-a tta tta aat cat gag gtc cgt att ata ctc gta tat att gtt tct ctc atg tat aat aat aaa cgg aag aga tat agc acc aca tgc acc atc caa tgg aaa gtg taa aga caa cga ata cag aag ccg taa tct atg ttg tct atc gtg tcc tcc ggg aac tta cgc ttt cag att atg tga tag caa gac taa tac aca atg tac acc gtg tgg ttc gga tac ctt tac atc tca caa taa tca ttt aca ggc ttg tct aag ttg taa cgg aag atg tga tag taa tca ggt aga gac gcg atc gtg taa cac gac tca caa tag aat ctg tga atg ctc tcc agg ata tta ttg tct tct caa agg agc atc agg gtg tag aac atg tat ttc taa aac aaa gtg tgg aat agg ata cgg agt atc cgg ata cac gtc tac cgg aga cgt cat ctg ttc tcc gtg tgg tcc cgg aac ata ttc tca cac cgt ctc ttc cac aga taa atg cga acc cgt cgt aac cag caa tac att taa cta tat cga tgt gga aat taa cct gta tcc agt caa cga cac atc gtg tac tcg gac gac cac tac cgg tct cag cga atc cat ctc aac gtc gga act aac tat tac cat gaa tca taa aga ttg tga tcc agt ctt tcg tgc aga ata ctt ctc tgt cct taa taa tgt agc aac ttc agg att ctt tac agg aga aaa tag ata tca gaa tac ttc aaa gat atg tac tct gaa ttt cga gat taa atg taa caa caa aga ttc atc ttc caa aca gtt aac gaa aac aaa gaa tga tac tat cat gcc gca ttc aga gac ggt aac tct agt ggg cga ctg tct atc tag cgt cga cat cta cat act ata tag taa tac caa tac tca aga cta cga aaa tga tac aat ctc tta tca tat ggg taa tgt tct cga tgt caa tag cca tat gcc cgc tag ttg cga tat aca taa act gat cac taa ttc cca gaa tcc cac cca ctt ata gta ag- -tt ttt tta cct ata aat aat aaa tac aat aat taa ttt ctc gta aaa gta gaa aat ata ttc taa ttt att ata tgg taa gaa agt aga atc atc tag aac agt aat caa tca ata gca atc atg aaa caa tat att gtc ctg gca tgc atg tgc cta gtg gca gct gct atg cct act agt ctt caa caa tct tca tcc tcg tgt act gaa gaa gaa aac aaa cat cat atg gga atc gat gtt att atc aaa gtc aca aag caa gac caa aca ccg acc aat gat aag att tgt caa tcc gta acg gaa gtt aca gag a-- --- --- --- --- --- -cc gaa gat gat gag gta tcc gaa gaa gtt gta aaa gga gat ccc acc act tat tac act atc gtc ggt gcg ggt ctt aac atg aac ttt gga ttc acc aaa tgc cca aag att tta tcc atc tcc gaa tcc tct gat gga aac act gtg aat act aga ttg tcc agc gtg tca ccg gga caa ggt aag gac tct ccc gcg atc acg cgt gaa gaa gct ctg gct atg atc aaa gac tgt gag atg tct atc gac atc aga tgt agc gaa gaa gag aaa gac agt gac atc aag acc cat cca gta ctt ggg tct aac atc tca cat aag aaa gtg agt tac aaa gat atc atc ggt tca acg atc gtt gat aca aaa tgt gtc aag aac cta gag ttt agc gta cgt atc gga gac atg tgt gag gaa tca tct gaa ctt gaa gtc aag gat gga ttc aag tat gtc gac gga tcg gca tct gaa ggt gca acc gat gat act tca ctc atc gat tca aca aaa ctc aaa gca tgt gtc tga atc gat aac tct att cat ctg aaa atg gat gag ttg ggt taa tcg aac gat tca gac acc gca cca cga att aaa aaa g-- -ac cgg gca cta tat tcc ggt ttg caa aac aaa aat att taa cta cat tca caa aaa gtt acc tct cgt tac ttc ttc ttt ctg ttt caa tat gtg ata cga tat gat cac tat tcg tat tct ct- --- --- tgg tct cat aaa aaa gtt tta c-- --a aaa aaa aaa aaa a?t att ttt att ctc ttt ctc tct tcg at- ggt ctc aca aaa ata tta aac ctc ttt ctg atg tct caa cta ttt cgt aaa cga taa cgt cca aca ata tat tct cgt aga gct tat caa cat cct tat acc aat cta ggt tgt cag aca att gca tca taa aat aat gtt tat aat tta cac gtt aac atc ata taa taa acg tat ata gtt aat att ttt gga ata taa atg atc tgt aaa atc cat gta ggg gac act gct cac gtt ttt tct cta gta cat aat ttc aca caa gtt ttt ata cag aca aat taa ttc tcg tcc ata ta- ttt taa aac att gac ttt tgt act aag aaa aat atc ttg act aac cat ctc ttt ctc tct tcg atg ggt ctc aca aaa ata tta aac ctc ttt ctg atg g-- --- --- --- --- --- --- --- --- --- --- --- --- --- --- --- --- --- --- --- --- --- --- --- --- --- --- --- --- --- --- --- --- --- --- --- --- --- --- --- --- --- --- --- --- --- --- agt cgt aaa aag ttt tta tcc ttt ctc tct tcg ata ggt ctc --- --- --- --- --- --- --- --- --- --- --- --- --- --- --- --- --- --- --- --- --- --- --- --- --- --- --- --- --- --- --- --- --- --- --- --- --- --- --- --- --- --- --- --- --- --- --- --- --- --- --- --- --- --- --- --- --- --- --- --- --- --- --- --- --- --- --- --- --- --- aca aaa ata tta aac ctc ttt ctg atg gtc tct ata aa

>ON676706.1_Monkeypox_virus_isolate_MPXV_USA_2022_UT002_complete_genome

atc gag gac gac ttc aac tat gac agc tcg gtg gcg tct gcc agc gtg tac ata cga atg gca ttt cta aga aaa gtc tac ggt atc ctt -tc tac aat ttc ctt taa caa cgg caa cag ctg cag tat ttt tat act ttg aat gca -tc gga cat tta tac aag gga gtc ctg ttc taa tat tgg cat caa tgt tcg gat cta tag gct tga ttt tcg cat tga ctt tac aca gac atg aac atc ccc tga atc tgt aca tac ttt gtg gat tta cac tgt tag aat ctc taa cgc tgg cct ctg ttg tta ctt tct atg atg cac gta tcg tta tgc aag ctt tca tgt tga cta ctg cag tgt ttc ttg ctc tga cta cat ata ctc tac aat caa aga gag att tca gta aac ttg taa cag gat tgt ttg ctg ctt tct gga ttt taa ttt tgt cag gag tct tga gga taa agt tta aaa tag aat taa taa aga aca tat agg tca -tt ttt taa aca tgg ata gaa acc aag gtt gtt agt taa taa tat aca aga tat ttt ttc tca ctc tga tcc atg taa acc aag gac gag aga cac tct cat tcc tca ttc acg aca cca tta aaa atg gaa att aaa gcc ctc tat taa gca cag acg gct aca ggt cta cca tca ggt t-a cct tcg tct acc ttc aca atg gcc tct cct tgt gcc cag ttc agt ccc tgt cat tgc cac gct act aag gac tcc ctg aat acy gtg act gac gtc aga cat tgt ctg act gaa tac atc ctg tgg gtt tct cat aga tgg acc cat aga gaa agc gca ggg cct ctc tac agg ctt ctc atc tct ttc aga att gat gca atg gag cta ttt ggt agc gag ttg aag gag ttc tcg aat tca ctt ccg tgg gac aat atc gac aat tgc gtg gag atc att aaa tgt ttc atc aga aat gac tcc atg aaa acc gcc aaa gaa ctt tgt gca ata att gga ctt tgt act caa tca gct att gtc act gga aga gtc ttc aat gat aag tat atc gac ata cta ctt atg ctg cga aag att ctg aac gag aac gac tat ctc acc ctc ttg gat cat atc ctc act gct aaa tac taa atc tcc ttc atg ctc tct cac taa tac tct tac tca cta cac ttt tta tca tct tat gat gaa tga ttg cct tca tca ttt ttt cgt gga ata taa tat agg aat aat tag cac cag aat agc tat gga tat ctc gtt aag aat att ctc tat aag aga cat aat gta gac ata gtt att ata tcc ttc tta gat aag tgt tac gct act gga aag ttt cca tcg tta tta tta cat gaa gat gat ata att aaa cca aca ttg aga ttg gct ctt atg tta gct gga ttg aat tac tgt aat aaa tgc atc gag tat aga ggg ata tag caa ttc tcg ata ata gtc atg caa tat ttg aat gag act gat aat tta ggt aat aca gta ct- --- aca cac ata tct ttc tag att ata tat cgt taa aaa tct gta aga tgt ata ttt ctc ata agt atc cac tgt gta ata tta tta atg gat ata tag ata acg caa tag gga cta ata gta ttg taa aag ata taa tcg act att tac gta cat atc cag ata tct ata ttc cta cta gtt tgc tgc gta gtt gca tca ttg ata tgc atg -at tta tca gga ttc aga gat gaa tta cta agt aaa cta caa tcc cac aat aag taa gaa tca aat atc aaa aac tca ctt ttg att ttt cta gtc tta agt aat aca tat att tat taa tag acc tat gaa ata aaa aaa ggt aac aat gga ttc gcg tat agc tat tta cgt att agt atc ggc atc tct ttt gta tct tgt taa ttg tca caa act agt aca tta ctt caa tct gaa aat aaa tgg aag tga tat aac taa tac agc aga tat att gct gga caa tta tcc aat tat gac ctt tga tgg aaa gga tat tta tcc atc tat ctc gtt cat ggt cgg taa taa act ttt cct aga tct tta taa aaa tat ctt tga aga att ttt cag act att tcg agt atc tgt aag tag tca ata cga gga att aga ata tta tta ttc atg tga tta tac taa caa ccg tcc tac aat taa aca aca tta ctt tta taa cgg cga cga ata tac tga aat tga tag atc gaa aaa agc cac taa taa aaa cag ttg gtt aat tac ttc agg ctt tag act aca aaa atg gtt cga tag cga aga ttg tat aat tta tct cag atc ttt agt tag aag aat gga aga cag taa caa aaa cag taa aaa aac tta gta ctt aga tat cg- --a aaa aat ata ttt ttg tag act ctt gag aat aga agg aaa aca tgt aca taa tta taa aaa atg aaa atc aat ggc gaa taa gac agt gcg att cgc acc atg gag tcg gta gat ttc atg gct gtc gat gag cag ttt cac gac gac ctc gat ctt tgg tca tta tct ttg gta gat gat tat aaa aaa cat gga tta ggt gtt gac tgt tat gtt cta gaa cca gtt gtt gac agg aaa ata ttt gat aga ttt ctc ctt gaa cca att tgt gat cct gta gat gtt ctg tat gat tat ttt agg att cat aga gat aat att gat cag tat ata gta gat aga ctg ttt gca tat att aca tat aaa gat att ata tct gca tta gtg tca aag aat tat atg gaa gat att ttc tct ata att att aag aat tgt aat tct gtg caa gat ctc tta ctt tac tat cta tct aat gca tat gta gaa ata gac att gtt gat ctt atg gta gat cat ggg gct gta ata tat aaa ata gaa tgc ttg aat gcc tat ttt agg gga ata tgt aaa aag gaa agt agt gtt gtt gag ttt att ttg aat tgt ggt atc cca gat gaa aat gat gtt aaa tta gat cta tat aaa ata att cag tat act agg gga ttc ctt gta gat gaa ccc aca gta tta gaa att tat aag ctt tgt atc cca tat att gaa gat atc aat caa cta gat gct ggt gga agg acc ttg ctt tat cgc gct atc tat gca ggt tat ata gat tta gta tca tgg cta tta gaa aat gga gca aat gtc aac gca gta atg agt aat gga tat aca tgt ctt gac gtg gcc gtg gat agg gga tct gtc atc gcc cgt agg gaa gca cat ctt aaa ata tta gaa ata ttg ctt aga gaa cca ttg tct att gac tgt ata aaa tta gct ata ctt aat aat aca att gaa aac cat gat gtg ata aag ctc tgt atc aag tat ttt atg atg gta gat tat tca ctt tgt aat gtg tat gca tca tca ctc ttt gat tat ata att gat tgt aaa caa gaa ttg gag tac att agg cag atg aaa att cat aat aca acc atg tat gag tta atc tat aat aga gac aaa aac aag cat gct tcc cat att cta cat agg tat tct aaa cat cca gtt ttg aca cag tgt atc act aaa gga ttc aag att tac aca gaa gta acc gag cag gtc act aaa gct cta aac aga cgt gct cta ata gat gag ata ata aac aat gta tca act gat gac aat ctc cta tca aaa ctt cca tta gaa att agg gat cta att gtt tca caa gct gtc ata tag agt tct atc cac cca cct ttc ttg aaa tga gtt aat --- --- --- --- --- --- --- --- --- --- --- --- --- --- --- --- --- --- --- --- --- --- --- --- --- --- --- --- --- --- --- --- --- --- --- --- --- --- --- --- --- --- --- --- --- --- --- --- --- --- --- --- --- --- --- --- --- --- --- --- --- --- --- --- --- --- --- --- --- --- --- --- --- --- --- --- --- --- --- --- --- --- --- --- --- --- --- --- --- --- --- --- --- --- --- --- --- --- --- --- --- --- --- --- --- -ag tca taa gtt agt taa --- --- --- --- --- -gt cat aag tta gtt aag tca t-- --- --- --- --- --- --- --- --- --- --- --- --- --- --- --- -aa gtt agt taa gtc ata agt tag tta agt cat aag tta gtt aag tca taa gtt agt taa gtc ata agt tag tta agt cat aag tta gtt tat agt cta aca ctt cta att ttt ata cct tga tct ttt tct cta att atg aaa aag taa atc att atg aag atg gat g-a aat gga cga gat tgt gcg cat cgt taa cga tag tat gtg gta cgt acc taa cgc att tat gga cga cgg tga taa tga agg tca cat ttc tgt caa taa tgt ctg tca tat gta tct cgc att ctt tga tgt gga tat atc atc tca tct gtt taa att agt tat taa aca ctg cga tct gaa taa acg act aaa atg tgg taa ctc tcc att aca ttg cta tac gat gaa tac acg att taa tcc atc tgt att aaa gat att gtt acg cca cgg cat gcg taa ctt tga tag caa gga taa aaa agg aca tat tcc tct aca cca cta tct gat tca ttc act atc aat cga taa caa gat ctt tga tat act aac gga ccc cat tga tga ctt tag taa atc atc cga tct att gct gtg tta tct tag ata taa att caa tgg gag ctt aaa cta tta cgt tct gta caa att att gac taa agg atc tga ccc taa ttg cgt cga tga gga tgg act cac ttc tct tca tta cta ctg taa aca cat atc cgc gtt cca cga aag caa tta tta caa gtc aaa gag tca cac taa gat gcg agc tga gaa gcg att cat cta cgc gat aat aga tca tgg agc aaa cat taa cgc ggt tac gaa aat cgg aaa tac gcc gtt aca cac tta cct tca aca gta tac caa aca tag tcc tcg tgt ggt gta tgc tct ttt atc tcg agg agc cga tac gag gat acg taa taa tct taa ttg tac acc cat cat gga ata cat aaa gaa cga ttg tgc aac agg tca tat tct cat aat gtt act caa ttg gca cga aca aaa ata cgg gaa att aca aaa gga aga agg aca aca tct act tta tct att cat aaa aca taa tca agg ata tgg aag tcg ctc tct caa tat act acg gta tct act aga tag att cga cat tca gaa aga cga ata cta taa tac aat gac tcc tct tca tac cgc ctt cca gaa ttg caa taa caa tgt tgc ctc ata cct cgt ata cat tgg ata cga cat caa cct tcc gac taa aga cga taa gac agt att cga ctt ggt gtt tga aaa cag aaa cat tat ata caa ggc gga tgt cgt taa tga cat tat cca cca cag act gaa agt atc tct acc tat gat taa atc gtt gtt cta caa gat gtc gga gtt ctc tcc cta cga cga tca cta cgt aaa gaa gat aat agc cta ctg cct att aag gga cga gtc att tgc gga act aca tac taa att ctg ttt aaa cga gga cta taa aag tgt att tat gaa aaa tat atc att cga taa gat aga ttc cat cat cga aaa atg tag tcg tga cat aag tct cct caa aga gat tcg aat ctc aga cac cga ctt gta tac ggt att gag aac aga aga cat tcg gta tca cac ata tct cga agc cat aca ttc aga caa acg cat ttc att tcc cat gta cga cga tct cat aga aca gtg tca tct atc gat gga gca taa aag taa act cgt cga caa agc act caa taa att aga gtc tac cat cga tag tca atc tag act atc gta ttt gcc tcc gga aat tat gcg caa tat cat aac caa gct aag cga cta cca tct aaa cag tat gtt gta cgg aaa gaa cca tta caa ata tta tcc atg ata gaa aga aaa tat tta aaa aat aat cta tat gat tgg aga agt agg aaa caa aca gta aca aga cga cga tta cta c-a tta tta aat cat gag gtc cgt att ata ctc gta tat att gtt tct ctc atg tat aat aat aaa cgg aag aga tat agc acc aca tgc acc atc caa tgg aaa gtg taa aga caa cga ata cag aag ccg taa tct atg ttg tct atc gtg tcc tcc ggg aac tta cgc ttt cag att atg tga tag caa gac taa tac aca atg tac acc gtg tgg ttc gga tac ctt tac atc tca caa taa tca ttt aca ggc ttg tct aag ttg taa cgg aag atg tga tag taa tca ggt aga gac gcg atc gtg taa cac gac tca caa tag aat ctg tga atg ctc tcc agg ata tta ttg tct tct caa agg agc atc agg gtg tag aac atg tat ttc taa aac aaa gtg tgg aat agg ata cgg agt atc cgg ata cac gtc tac cgg aga cgt cat ctg ttc tcc gtg tgg tcc cgg aac ata ttc tca cac cgt ctc ttc cac aga taa atg cga acc cgt cgt aac cag caa tac att taa cta tat cga tgt gga aat taa cct gta tcc agt caa cga cac atc gtg tac tcg gac gac cac tac cgg tct cag cga atc cat ctc aac gtc gga act aac tat tac cat gaa tca taa aga ttg tga tcc agt ctt tcg tgc aga ata ctt ctc tgt cct taa taa tgt agc aac ttc agg att ctt tac agg aga aaa tag ata tca gaa tac ttc aaa gat atg tac tct gaa ttt cga gat taa atg taa caa caa aga ttc atc ttc caa aca gtt aac gaa aac aaa gaa tga tac tat cat gcc gca ttc aga gac ggt aac tct agt ggg cga ctg tct atc tag cgt cga cat cta cat act ata tag taa tac caa tac tca aga cta cga aaa tga tac aat ctc tta tca tat ggg taa tgt tct cga tgt caa tag cca tat gcc cgc tag ttg cga tat aca taa act gat cac taa ttc cca gaa tcc cac cca ctt ata gta ag- -tt ttt tta cct ata aat aat aaa tac aat aat taa ttt ctc gta aaa gta gaa aat ata ttc taa ttt att ata tgg taa gaa agt aga atc atc tag aac agt aat caa tca ata gca atc atg aaa caa tat att gtc ctg gca tgc atg tgc cta gtg gca gct gct atg cct act agt ctt caa caa tct tca tcc tcg tgt act gaa gaa gaa aac aaa cat cat atg gga atc gat gtt att atc aaa gtc aca aag caa gac caa aca ccg acc aat gat aag att tgt caa tcc gta acg gaa gtt aca gag a-- --- --- --- --- --- -cc gaa gat gat gag gta tcc gaa gaa gtt gta aaa gga gat ccc acc act tat tac act atc gtc ggt gcg ggt ctt aac atg aac ttt gga ttc acc aaa tgc cca aag att tta tcc atc tcc gaa tcc tct gat gga aac act gtg aat act aga ttg tcc agc gtg tca ccg gga caa ggt aag gac tct ccc gcg atc acg cgt gaa gaa gct ctg gct atg atc aaa gac tgt gag atg tct atc gac atc aga tgt agc gaa gaa gag aaa gac agt gac atc aag acc cat cca gta ctt ggg tct aac atc tca cat aag aaa gtg agt tac aaa gat atc atc ggt tca acg atc gtt gat aca aaa tgt gtc aag aac cta gag ttt agc gta cgt atc gga gac atg tgt gag gaa tca tct gaa ctt gaa gtc aag gat gga ttc aag tat gtc gac gga tcg gca tct gaa ggt gca acc gat gat act tca ctc atc gat tca aca aaa ctc aaa gca tgt gtc tga atc gat aac tct att cat ctg aaa atg gat gag ttg ggt taa tcg aac gat tca gac acc gca cca cga att aaa aaa g-- -ac cgg gca cta tat tcc ggt ttg caa aac aaa aat att taa cta cat tca caa aaa gtt acc tct cgt tac ttc ttc ttt ctg ttt caa tat gtg ata cga tat gat cac tat tcg tat tct ct- --- --- tgg tct cat aaa aaa gtt tta c-- --a aaa aaa aaa aaa a?t att ttt att ctc ttt ctc tct tcg at- ggt ctc aca aaa ata tta aac ctc ttt ctg atg tct caa cta ttt cgt aaa cga taa cgt cca aca ata tat tct cgt aga gct tat caa cat cct tat acc aat cta ggt tgt cag aca att gca tca taa aat aat gtt tat aat tta cac gtt aac atc ata taa taa acg tat ata gtt aat att ttt gga ata taa atg atc tgt aaa atc cat gta ggg gac act gct cac gtt ttt tct cta gta cat aat ttc aca caa gtt ttt ata cag aca aat taa ttc tcg tcc ata ta- ttt taa aac att gac ttt tgt act aag aaa aat atc ttg act aac cat ctc ttt ctc tct tcg atg ggt ctc aca aaa ata tta aac ctc ttt ctg atg g-- --- --- --- --- --- --- --- --- --- --- --- --- --- --- --- --- --- --- --- --- --- --- --- --- --- --- --- --- --- --- --- --- --- --- --- --- --- --- --- --- --- --- --- --- --- --- agt cgt aaa aag ttt tta tcc ttt ctc tct tcg ata ggt ctc --- --- --- --- --- --- --- --- --- --- --- --- --- --- --- --- --- --- --- --- --- --- --- --- --- --- --- --- --- --- --- --- --- --- --- --- --- --- --- --- --- --- --- --- --- --- --- --- --- --- --- --- --- --- --- --- --- --- --- --- --- --- --- --- --- --- --- --- --- --- aca aaa ata tta aac ctc ttt ctg atg gtc tct ata aa

>ON649879.1_Monkeypox_virus_isolate_MPXV_ISR001_2022_complete_genome

atc gag gac gac ttc aac tat gac agc tcg gtg gcg tct gcc agc gtg tac ata cga atg gca ttt cta aga aaa gtc tac ggt atc ctt -tc tac aat ttc ctt taa caa cgg caa cag ctg cag tat ttt tat act ttg aat gca -tc gga cat tta tac aag gga gtc ctg ttc taa tat tgg cat caa tgt tcg gat cta tag gct tga ttt tcg cat tga ctt tac aca gac atg aac atc ccc tga atc tgt aca tac ttt gtg gat tta cac tgt tag aat ctc taa cgc tgg cct ctg ttg tta ctt tct atg atg cac gta tcg tta tgc aag ctt tca tgt tga cta ctg cag tgt ttc ttg ctc tga cta cat ata ctc tac aat caa aga gag att tca gta aac ttg taa cag gat tgt ttg ctg ctt tct gga ttt taa ttt tgt cag gag tct tga gga taa agt tta aaa tag aat taa taa aga aca tat agg tca -tt ttt taa aca tgg ata gaa acc aag gtt gtt agt taa taa tat aca aga tat ttt ttc tca ctc tga tcc atg taa acc aag gac gag aga cac tct cat tcc tca ttc acg aca cca tta aaa atg gaa att aaa gcc ctc tat taa gca cag acg gct aca ggt cta cca tca ggt t-a cct tcg tct acc ttc aca atg gcc tct cct tgt gcc cag ttc agt ccc tgt cat tgc cac gct act aag gac tcc ctg aat acc gtg act gac gtc aga cat tgt ctg act gaa tac atc ctg tgg gtt tct cat aga tgg acc cat aga gaa agc gca ggg cct ctc tac agg ctt ctc atc tct ttc aga att gat gca atg gag cta ttt ggt agc gag ttg aag gag ttc tcg aat tca ctt ccg tgg gac aat atc gac aat tgc gtg gag atc att aaa tgt ttc atc aga aat gac tcc atg aaa acc gcc aaa gaa ctt tgt gca ata att gga ctt tgt act caa tca gct att gtc act gga aga gtc ttc aat gat aag tat atc gac ata cta ctt atg ctg cga aag att ctg aac gag aac gac tat ctc acc ctc ttg gat cat atc ctc act gct aaa tac taa atc tcc ttc atg ctc tct cac taa tac tct tac tca cta cac ttt tta tca tct tat gat gaa tga ttg cct tca tca ttt ttt cgt gga ata taa tat agg aat aat tag cac cag aat agc tat gga tat ctc gtt aag aat att ctc tat aag aga cat aat gta gac ata gtt att ata tcc ttc tta gat aag tgt tac gct act gga aag ttt cca tcg tta tta tta cat gaa gat gat ata att aaa cca aca ttg aga ttg gct ctt atg tta gct gga ttg aat tac tgt aat aaa tgc atc gag tat aga ggg ata tag caa ttc tcg ata ata gtc atg caa tat ttg aat gag act gat aat tta ggt aat aca gta ct- --- aca cac ata tct ttc tag att ata tat cgt taa aaa tct gta aga tgt ata ttt ctc ata agt atc cac tgt gta ata tta tta atg gat ata tag ata acg caa tag gga cta ata gta ttg taa aag ata taa tcg act att tac gta cat atc cag ata tct ata ttc cta cta gtt tgc tgc gta gtt gca tca ttg ata tgc atg -at tta tca gga ttc aga gat gaa tta cta agt aaa cta caa tcc cac aat aag taa gaa tca aat atc aaa aac tca ctt ttg att ttt cta gtc tta agt aat aca tat att tat taa tag acc tat gaa ata aaa aaa ggt aac aat gga ttc gcg tat agc tat tta cgt att agt atc ggc atc tct ttt gta tct tgt taa ttg tca caa act agt aca tta ctt caa tct gaa aat aaa tgg aag tga tat aac taa tac agc aga tat att gct gga caa tta tcc aat tat gac ctt tga tgg aaa gga tat tta tcc atc tat ctc gtt cat ggt cgg taa taa act ttt cct aga tct tta taa aaa tat ctt tga aga att ttt cag act att tcg agt atc tgt aag tag tca ata cga gga att aga ata tta tta ttc atg tga tta tac taa caa ccg tcc tac aat taa aca aca tta ctt tta taa cgg cga cga ata tac tga aat tga tag atc gaa aaa agc cac taa taa aaa cag ttg gtt aat tac ttc agg ctt tag act aca aaa atg gtt cga tag cga aga ttg tat aat tta tct cag atc ttt agt tag aag aat gga aga cag taa caa aaa cag taa aaa aac tta gta ctt aga tat cg- --a aaa aat ata ttt ttg tag act ctt gag aat aga agg aaa aca tgt aca taa tta taa aaa atg aaa atc aat ggc gaa taa gac agt gcg att cgc acc atg gag tcg gta gat ttc atg gct gtc gat gag cag ttt cac gac gac ctc gat ctt tgg tca tta tct ttg gta gat gat tat aaa aaa cat gga tta ggt gtt gac tgt tat gtt cta gaa cca gtt gtt gac agg aaa ata ttt gat aga ttt ctc ctt gaa cca att tgt gat cct gta gat gtt ctg tat gat tat ttt agg att cat aga gat aat att gat cag tat ata gta gat aga ctg ttt gca tat att aca tat aaa gat att ata tct gca tta gtg tca aag aat tat atg gaa gat att ttc tct ata att att aag aat tgt aat tct gtg caa gat ctc tta ctt tac tat cta tct aat gca tat gta gaa ata gac att gtt gat ctt atg gta gat cat ggg gct gta ata tat aaa ata gaa tgc ttg aat gcc tat ttt agg gga ata tgt aaa aag gaa agt agt gtt gtt gag ttt att ttg aat tgt ggt atc cca gat gaa aat gat gtt aaa tta gat cta tat aaa ata att cag tat act agg gga ttc ctt gta gat gaa ccc aca gta tta gaa att tat aag ctt tgt atc cca tat att gaa gat atc aat caa cta gat gct ggt gga agg acc ttg ctt tat cgc gct atc tat gca ggt tat ata gat tta gta tca tgg cta tta gaa aat gga gca aat gtc aac gca gta atg agt aat gga tat aca tgt ctt gac gtg gcc gtg gat agg gga tct gtc atc gcc cgt agg gaa gca cat ctt aaa ata tta gaa ata ttg ctt aga gaa cca ttg tct att gac tgt ata aaa tta gct ata ctt aat aat aca att gaa aac cat gat gtg ata aag ctc tgt atc aag tat ttt atg atg gta gat tat tca ctt tgt aat gtg tat gca tca tca ctc ttt gat tat ata att gat tgt aaa caa gaa ttg gag tac att agg cag atg aaa att cat aat aca acc atg tat gag tta atc tat aat aga gac aaa aac aag cat gct tcc cat att cta cat agg tat tct aaa cat cca gtt ttg aca cag tgt atc act aaa gga ttc aag att tac aca gaa gta acc gag cag gtc act aaa gct cta aac aga cgt gct cta ata gat gag ata ata aac aat gta tca act gat gac aat ctc cta tca aaa ctt cca tta gaa att agg gat cta att gtt tca caa gct gtc ata tag agt tct atc cac cca cct ttc ttg aaa tga gtt aat --- --- --- --- --- --- --- --- --- --- --- --- --- --- --- --- --- --- --- --- --- --- --- --- --- --- --- --- --- --- --- --- --- --- --- --- --- --- --- --- --- --- --- --- --- --- --- --- --- --- --- --- --- --- --- --- --- --- --- --- --- --- --- --- --- --- --- --- --- --- --- --- --- --- --- --- --- --- --- --- --- --- --- --- --- --- --- --- --- --- --- --- --- --- --- --- --- --- --- --- --- --- --- --- --- -ag tca taa gtt agt taa --- --- --- --- --- -gt cat aag tta gtt aag tca t-- --- --- --- --- --- --- --- --- --- --- --- --- --- --- --- -aa gtt agt taa gtc ata agt tag tta agt cat aag tta gtt aag tca taa gtt agt taa gtc ata agt tag tta agt cat aag tta gtt tat agt cta aca ctt cta att ttt ata cct tga tct ttt tct cta att atg aaa aag taa atc att atg aag atg gat g-a aat gga cga gat tgt gcg cat cgt taa cga tag tat gtg gta cgt acc taa cgc att tat gga cga cgg tga taa tga agg tca cat ttc tgt caa taa tgt ctg tca tat gta tct cgc att ctt tga tgt gga tat atc atc tca tct gtt taa att agt tat taa aca ctg cga tct gaa taa acg act aaa atg tgg taa ctc tcc att aca ttg cta tac gat gaa tac acg att taa tcc atc tgt att aaa gat att gtt acg cca cgg cat gcg taa ctt tga tag caa gga taa aaa agg aca tat tcc tct aca cca cta tct gat tca ttc act atc aat cga taa caa gat ctt tga tat act aac gga ccc cat tga tga ctt tag taa atc atc cga tct att gct gtg tta tct tag ata taa att caa tgg gag ctt aaa cta tta cgt tct gta caa att att gac taa agg atc tga ccc taa ttg cgt cga tga gga tgg act cac ttc tct tca tta cta ctg taa aca cat atc cgc gtt cca cga aag caa tta tta caa gtc aaa gag tca cac taa gat gcg agc tga gaa gcg att cat cta cgc gat aat aga tca tgg agc aaa cat taa cgc ggt tac gaa aat cgg aaa tac gcc gtt aca cac tta cct tca aca gta tac caa aca tag tcc tcg tgt ggt gta tgc tct ttt atc tcg agg agc cga tac gag gat acg taa taa tct taa ttg tac acc cat cat gga ata cat aaa gaa cga ttg tgc aac agg tca tat tct cat aat gtt act caa ttg gca cga aca aaa ata cgg gaa att aca aaa gga aga agg aca aca tct act tta tct att cat aaa aca taa tca agg ata tgg aag tcg ctc tct caa tat act acg gta tct act aga tag att cga cat tca gaa aga cga ata cta taa tac aat gac tcc tct tca tac cgc ctt cca gaa ttg caa taa caa tgt tgc ctc ata cct cgt ata cat tgg ata cga cat caa cct tcc gac taa aga cga taa gac agt att cga ctt ggt gtt tga aaa cag aaa cat tat ata caa ggc gga tgt cgt taa tga cat tat cca cca cag act gaa agt atc tct acc tat gat taa atc gtt gtt cta caa gat gtc gga gtt ctc tcc cta cga cga tca cta cgt aaa gaa gat aat agc cta ctg cct att aag gga cga gtc att tgc gga act aca tac taa att ctg ttt aaa cga gga cta taa aag tgt att tat gaa aaa tat atc att cga taa gat aga ttc cat cat cga aaa atg tag tcg tga cat aag tct cct caa aga gat tcg aat ctc aga cac cga ctt gta tac ggt att gag aac aga aga cat tcg gta tca cac ata tct cga agc cat aca ttc aga caa acg cat ttc att tcc cat gta cga cga tct cat aga aca gtg tca tct atc gat gga gca taa aag taa act cgt cga caa agc act caa taa att aga gtc tac cat cga tag tca atc tag act atc gta ttt gcc tcc gga aat tat gcg caa tat cat aac caa gct aag cga cta cca tct aaa cag tat gtt gta cgg aaa gaa cca tta caa ata tta tcc atg ata gaa aga aaa tat tta aaa aat aat cta tat gat tgg aga agt agg aaa caa aca gta aca aga cga cga tta cta c-a tta tta aat cat gag gtc cgt att ata ctc gta tat att gtt tct ctc atg tat aat aat aaa cgg aag aga tat agc acc aca tgc acc atc caa tgg aaa gtg taa aga caa cga ata cag aag ccg taa tct atg ttg tct atc gtg tcc tcc ggg aac tta cgc ttt cag att atg tga tag caa gac taa tac aca atg tac acc gtg tgg ttc gga tac ctt tac atc tca caa taa tca ttt aca ggc ttg tct aag ttg taa cgg aag atg tga tag taa tca ggt aga gac gcg atc gtg taa cac gac tca caa tag aat ctg tga atg ctc tcc agg ata tta ttg tct tct caa agg agc atc agg gtg tag aac atg tat ttc taa aac aaa gtg tgg aat agg ata cgg agt atc cgg ata cac gtc tac cgg aga cgt cat ctg ttc tcc gtg tgg tcc cgg aac ata ttc tca cac cgt ctc ttc cac aga taa atg cga acc cgt cgt aac cag caa tac att taa cta tat cga tgt gga aat taa cct gta tcc agt caa cga cac atc gtg tac tcg gac gac cac tac cgg tct cag cga atc cat ctc aac gtc gga act aac tat tac cat gaa tca taa aga ttg tga tcc agt ctt tcg tgc aga ata ctt ctc tgt cct taa taa tgt agc aac ttc agg att ctt tac agg aga aaa tag ata tca gaa tac ttc aaa gat atg tac tct gaa ttt cga gat taa atg taa caa caa aga ttc atc ttc caa aca gtt aac gaa aac aaa gaa tga tac tat cat gcc gca ttc aga gac ggt aac tct agt ggg cga ctg tct atc tag cgt cga cat cta cat act ata tag taa tac caa tac tca aga cta cga aaa tga tac aat ctc tta tca tat ggg taa tgt tct cga tgt caa tag cca tat gcc cgc tag ttg cga tat aca taa act gat cac taa ttc cca gaa tcc cac cca ctt ata gta ag- -tt ttt tta cct ata aat aat aaa tac aat aat taa ttt ctc gta aaa gta gaa aat ata ttc taa ttt att ata tgg taa gaa agt aga atc atc tag aac agt aat caa tca ata gca atc atg aaa caa tat att gtc ctg gca tgc atg tgc cta gtg gca gct gct atg cct act agt ctt caa caa tct tca tcc tcg tgt act gaa gaa gaa aac aaa cat cat atg gga atc gat gtt att atc aaa gtc aca aag caa gac caa aca ccg acc aat gat aag att tgt caa tcc gta acg gaa gtt aca gag a-- --- --- --- --- --- -cc gaa gat gat gag gta tcc gaa gaa gtt gta aaa gga gat ccc acc act tat tac act atc gtc ggt gcg ggt ctt aac atg aac ttt gga ttc acc aaa tgc cca aag att tta tcc atc tcc gaa tcc tct gat gga aac act gtg aat act aga ttg tcc agc gtg tca ccg gga caa ggt aag gac tct ccc gcg atc acg cgt gaa gaa gct ctg gct atg atc aaa gac tgt gag atg tct atc gac atc aga tgt agc gaa gaa gag aaa gac agt gac atc aag acc cat cca gta ctt ggg tct aac atc tca cat aag aaa gtg agt tac aaa gat atc atc ggt tca acg atc gtt gat aca aaa tgt gtc aag aac cta gag ttt agc gta cgt atc gga gac atg tgt gag gaa tca tct gaa ctt gaa gtc aag gat gga ttc aag tat gtc gac gga tcg gca tct gaa ggt gca acc gat gat act tca ctc atc gat tca aca aaa ctc aaa gca tgt gtc tga atc gat aac tct att cat ctg aaa atg gat gag ttg ggt taa tcg aac gat tca gac acc gca cca cga att aaa aaa g-- -ac cgg gca cta tat tcc ggt ttg caa aac aaa aat att taa cta cat tca caa aaa gtt acc tct cgt tac ttc ttc ttt ctg ttt caa tat gtg ata cga tat gat cac tat tcg tat tct ct- --- --- tgg tct cat aaa aaa gtt tta c-- --a aaa aaa aaa aaa a-t att ttt att ctc ttt ctc tct tcg at- ggt ctc aca aaa ata tta aac ctc ttt ctg atg tct caa cta ttt cgt aaa cga taa cgt cca aca ata tat tct cgt aga gct tat caa cat cct tat acc aat cta ggt tgt cag aca att gca tca taa aat aat gtt tat aat tta cac gtt aac atc ata taa taa acg tat ata gtt aat att ttt gga ata taa atg atc tgt aaa atc cat gta ggg gac act gct cac gtt ttt tct cta gta cat aat ttc aca caa gtt ttt ata cag aca aat taa ttc tcg tcc ata ta- ttt taa aac att gac ttt tgt act aag aaa aat atc ttg act aac cat ctc ttt ctc tct tcg atg ggt ctc aca aaa ata tta aac ctc ttt ctg atg --- --- --- --- --- --- --- --- --- --- --- --- --- --- --- --- --- --- --- --- --- --- --- --- --- --- --- --- --- --- --- --- --- --- --- --- --- --- --- --- --- --- --- --- --- --- --- --- --- --- --- --- --- --- --- --- --- --- --- --- --- --- --- --- --- --- --- --- --- --- --- --- --- --- --- --- --- --- --- --- --- --- --- --- --- --- --- --- --- --- --- --- --- --- --- --- --- --- --- --- --- --- --- --- --- --- --- --- --- --- --- --- --- --- --- --- --- --- --- --- --- --- --- --- --- --- --- --- --- --- --- --- --- --- --- --- --- --- --- --- --- --- --- --

>ON602722.1_Monkeypox_virus_isolate_MPXV_FRA_2022_TLS67_complete_genome

atc gag gac gac ttc aac tat gac agc tcg gtg gcg tct gcc agc gtg tac ata cga atg gca ttt cta aga aaa gtc tac ggt atc ctt -tc tac aat ttc ctt taa caa cgg caa cag ctg cag tat ttt tat act ttg aat gca -tc gga cat tta tac aag gga gtc ctg ttc taa tat tgg cat caa tgt tcg gat cta tag gct tga ttt tcg cat tga ctt tac aca gac atg aac atc ccc tga atc tgt aca tac ttt gtg gat tta cac tgt tag aat ctc taa cgc tgg cct ctg ttg tta ctt tct atg atg cac gta tcg tta tgc aag ctt tca tgt tga cta ctg cag tgt ttc ttg ctc tga cta cat ata ctc tac aat caa aga gag att tca gta aac ttg taa cag gat tgt ttg ctg ctt tct gga ttt taa ttt tgt cag gag tct tga gga taa agt tta aaa tag aat taa taa aga aca tat agg tca -tt ttt taa aca tgg ata gaa acc aag gtt gtt agt taa taa tat aca aga tat ttt ttc tca ctc tga tcc atg taa acc aag gac gag aga cac tct cat tcc tca ttc acg aca cca tta aaa atg gaa att aaa gcc ctc tat taa gca cag acg gct aca ggt cta cca tca ggt t-a cct tcg tct acc ttc aca atg gcc tct cct tgt gcc cag ttc agt ccc tgt cat tgc cac gct act aag gac tcc ctg aat acc gtg act gac gtc aga cat tgt ctg act gaa tac atc ctg tgg gtt tct cat aga tgg acc cat aga gaa agc gca ggg cct ctc tac agg ctt ctc atc tct ttc aga att gat gca atg gag cta ttt ggt agc gag ttg aag gag ttc tcg aat tca ctt ccg tgg gac aat atc gac aat tgc gtg gag atc att aaa tgt ttc atc aga aat gac tcc atg aaa acc gcc aaa gaa ctt tgt gca ata att gga ctt tgt act caa tca gct att gtc act gga aga gtc ttc aat gat aag tat atc gac ata cta ctt atg ctg cga aag att ctg aac gag aac gac tat ctc acc ctc ttg gat cat atc ctc act gct aaa tac taa atc tcc ttc atg ctc tct cac taa tac tct tac tca cta cac ttt tta tca tct tat gat gaa tga ttg cct tca tca ttt ttt cgt gga ata taa tat agg aat aat tag cac cag aat agc tat gga tat ctc gtt aag aat att ctc tat aag aga cat aat gta gac ata gtt att ata tcc ttc tta gat aag tgt tac gct act gga aag ttt cca tcg tta tta tta cat gaa gat gat ata att aaa cca aca ttg aga ttg gct ctt atg tta gct gga ttg aat tac tgt aat aaa tgc atc gag tat aga ggg ata tag caa ttc tcg ata ata gtc atg caa tat ttg aat gag act gat aat tta ggt aat aca gta ct- --- aca cac ata tct ttc tag att ata tat cgt taa aaa tct gta aga tgt ata ttt ctc ata agt atc cac tgt gta ata tta tta atg gat ata tag ata acg caa tag gga cta ata gta ttg taa aag ata taa tcg act att tac gta cat atc cag ata tct ata ttc cta cta gtt tgc tgc gta gtt gca tca ttg ata tgc atg -at tta tca gga ttc aga gat gaa tta cta agt aaa cta caa tcc cac aat aag taa gaa tca aat atc aaa aac tca ctt ttg att ttt cta gtc tta agt aat aca tat att tat taa tag acc tat gaa at? aaa aaa ggt aac aat gga ttc gcg tat agc tat tta cgt att agt atc ggc atc tct ttt gta tct tgt taa ttg tca caa act agt aca tta ctt caa tct gaa aat aaa tgg aag tga tat aac taa tac agc aga tat att gct gga caa tta tcc aat tat gac ctt tga tgg aaa gga tat tta tcc atc tat ctc gtt cat ggt cgg taa taa act ttt cct aga tct tta taa aaa tat ctt tga aga att ttt cag act att tcg agt atc tgt aag tag tca ata cga gga att aga ata tta tta ttc atg tga tta tac taa caa ccg tcc tac aat taa aca aca tta ctt tta taa cgg cga cga ata tac tga aat tga tag atc gaa aaa agc cac taa taa aaa cag ttg gtt aat tac ttc agg ctt tag act aca aaa atg gtt cga tag cga aga ttg tat aat tta tct cag atc ttt agt tag aag aat gga aga cag taa caa aaa cag t-a aaa aac tta gta ctt aga tat cg- --a aaa aat ata ttt ttg tag act ctt gag aat aga agg aaa aca tgt aca taa tta taa aaa atg aaa atc aat ggc gaa taa gac agt gcg att cgc acc atg gag tcg gta gat ttc atg gct gtc gat gag cag ttt cac gac gac ctc gat ctt tgg tca tta tct ttg gta gat gat tat aaa aaa cat gga tta ggt gtt gac tgt tat gtt cta gaa cca gtt gtt gac agg aaa ata ttt gat aga ttt ctc ctt gaa cca att tgt gat cct gta gat gtt ctg tat gat tat ttt agg att cat aga gat aat att gat cag tat ata gta gat aga ctg ttt gca tat att aca tat aaa gat att ata tct gca tta gtg tca aag aat tat atg gaa gat att ttc tct ata att att aag aat tgt aat tct gtg caa gat ctc tta ctt tac tat cta tct aat gca tat gta gaa ata gac att gtt gat ctt atg gta gat cat ggg gct gta ata tat aaa ata gaa tgc ttg aat gcc tat ttt agg gga ata tgt aaa aag gaa agt agt gtt gtt gag ttt att ttg aat tgt ggt atc cca gat gaa aat gat gtt aaa tta gat cta tat aaa ata att cag tat act agg gga ttc ctt gta gat gaa ccc aca gta tta gaa att tat aag ctt tgt atc cca tat att gaa gat atc aat caa cta gat gct ggt gga agg acc ttg ctt tat cgc gct atc tat gca ggt tat ata gat tta gta tca tgg cta tta gaa aat gga gca aat gtc aac gca gta atg agt aat gga tat aca tgt ctt gac gtg gcc gtg gat agg gga tct gtc atc gcc cgt agg gaa gca cat ctt aaa ata tta gaa ata ttg ctt aga gaa cca ttg tct att gac tgt ata aaa tta gct ata ctt aat aat aca att gaa aac cat gat gtg ata aag ctc tgt atc aag tat ttt atg atg gta gat tat tca ctt tgt aat gtg tat gca tca tca ctc ttt gat tat ata att gat tgt aaa caa gaa ttg gag tac att agg cag atg aaa att cat aat aca acc atg tat gag tta atc tat aat aga gac aaa aac aag cat gct tcc cat att cta cat agg tat tct aaa cat cca gtt ttg aca cag tgt atc act aaa gga ttc aag att tac aca gaa gta acc gag cag gtc act aaa gct cta aac aga cgt gct cta ata gat gag ata ata aac aat gta tca act gat gac aat ctc cta tca aaa ctt cca tta gaa att agg gat cta att gtt tca caa gct gtc ata tag agt tct atc cac cca cct ttc ttg aaa tga gtt aat --- --- --- --- --- --- --- --- --- --- --- --- --- --- --- --- --- --- --- --- --- --- --- --- --- --- --- --- --- --- --- --- --- --- --- --- --- --- --- --- --- --- --- --- --- --- --- --- --- --- --- --- --- --- --- --- --- --- --- --- --- --- --- --- --- --- --- --- --- --- --- --- --- --- --- --- --- --- --- --- --- --- --- --- --- --- --- --- --- --- --- --- --- --- --- --- --- --- --- --- --- --- --- --- --- -ag tca taa gtt agt taa --- --- --- --- --- -gt cat aag tta gtt aag tca t-- --- --- --- --- --- --- --- --- --- --- --- --- --- --- --- -aa gtt agt taa gtc ata agt tag tta agt cat aag tta gtt aag tca taa gtt agt taa gtc ata agt tag tta agt cat aag tta gtt tat agt cta aca ctt cta att ttt ata cct tga tct ttt tct cta att atg aaa aag taa atc att atg aag atg gat g-a aat gga cga gat tgt gcg cat cgt taa cga tag tat gtg gta cgt acc taa cgc att tat gga cga cgg tga taa tga agg tca cat ttc tgt caa taa tgt ctg tca tat gta tct cgc att ctt tga tgt gga tat atc atc tca tct gtt taa att agt tat taa aca ctg cga tct gaa taa acg act aaa atg tgg taa ctc tcc att aca ttg cta tac gat gaa tac acg att taa tcc atc tgt att aaa gat att gtt acg cca cgg cat gcg taa ctt tga tag caa gga t-a aaa agg aca tat tcc tct aca cca cta tct gat tca ttc act atc aat cga taa caa gat ctt tga tat act aac gga ccc cat tga tga ctt tag taa atc atc cga tct att gct gtg tta tct tag ata taa att caa tgg gag ctt aaa cta tta cgt tct gta caa att att gac taa agg atc tga ccc taa ttg cgt cga tga gga tgg act cac ttc tct tca tta cta ctg taa aca cat atc cgc gtt cca cga aag caa tta tta caa gtc aaa gag tca cac taa gat gcg agc tga gaa gcg att cat cta cgc gat aat aga tca tgg agc aaa cat taa cgc ggt tac gaa aat cgg aaa tac gcc gtt aca cac tta cct tca aca gta tac caa aca tag tcc tcg tgt ggt gta tgc tct ttt atc tcg agg agc cga tac gag gat acg taa taa tct taa ttg tac acc cat cat gga ata cat aaa gaa cga ttg tgc aac agg tca tat tct cat aat gtt act caa ttg gca cga aca aaa ata cgg gaa att aca aaa gga aga agg aca aca tct act tta tct att cat aaa aca taa tca agg ata tgg aag tcg ctc tct caa tat act acg gta tct act aga tag att cga cat tca gaa aga cga ata cta taa tac aat gac tcc tct tca tac cgc ctt cca gaa ttg caa taa caa tgt tgc ctc ata cct cgt ata cat tgg ata cga cat caa cct tcc gac taa aga cga taa gac agt att cga ctt ggt gtt tga aaa cag aaa cat tat ata caa ggc gga tgt cgt taa tga cat tat cca cca cag act gaa agt atc tct acc tat gat taa atc gtt gtt cta caa gat gtc gga gtt ctc tcc cta cga cga tca cta cgt aaa gaa gat aat agc cta ctg cct att aag gga cga gtc att tgc gga act aca tac taa att ctg ttt aaa cga gga cta taa aag tgt att tat gaa aaa tat atc att cga taa gat aga ttc cat cat cga aaa atg tag tcg tga cat aag tct cct caa aga gat tcg aat ctc aga cac cga ctt gta tac ggt att gag aac aga aga cat tcg gta tca cac ata tct cga agc cat aca ttc aga caa acg cat ttc att tcc cat gta cga cga tct cat aga aca gtg tca tct atc gat gga gca taa aag taa act cgt cga caa agc act caa taa att aga gtc tac cat cga tag tca atc tag act atc gta ttt gcc tcc gga aat tat gcg caa tat cat aac caa gct aag cga cta cca tct aaa cag tat gtt gta cgg aaa gaa cca tta caa ata tta tcc atg ata gaa aga aaa tat tt- aaa aat aat cta tat gat tgg aga agt agg aaa caa aca gta aca aga cga cga tta cta c-a tta tta aat cat gag gtc cgt att ata ctc gta tat att gtt tct ctc atg tat aat aat aaa cgg aag aga tat agc acc aca tgc acc atc caa tgg aaa gtg taa aga caa cga ata cag aag ccg taa tct atg ttg tct atc gtg tcc tcc ggg aac tta cgc ttt cag att atg tga tag caa gac taa tac aca atg tac acc gtg tgg ttc gga tac ctt tac atc tca caa taa tca ttt aca ggc ttg tct aag ttg taa cgg aag atg tga tag taa tca ggt aga gac gcg atc gtg taa cac gac tca caa tag aat ctg tga atg ctc tcc agg ata tta ttg tct tct caa agg agc atc agg gtg tag aac atg tat ttc taa aac aaa gtg tgg aat agg ata cgg agt atc cgg ata cac gtc tac cgg aga cgt cat ctg ttc tcc gtg tgg tcc cgg aac ata ttc tca cac cgt ctc ttc cac aga taa atg cga acc cgt cgt aac cag caa tac att taa cta tat cga tgt gga aat taa cct gta tcc agt caa cga cac atc gtg tac tcg gac gac cac tac cgg tct cag cga atc cat ctc aac gtc gga act aac tat tac cat gaa tca taa aga ttg tga tcc agt ctt tcg tgc aga ata ctt ctc tgt cct taa taa tgt agc aac ttc agg att ctt tac agg aga aaa tag ata tca gaa tac ttc aaa gat atg tac tct gaa ttt cga gat taa atg taa caa caa aga ttc atc ttc caa aca gtt aac gaa aac aaa gaa tga tac tat cat gcc gca ttc aga gac ggt aac tct agt ggg cga ctg tct atc tag cgt cga cat cta cat act ata tag taa tac caa tac tca aga cta cga aaa tga tac aat ctc tta tca tat ggg taa tgt tct cga tgt caa tag cca tat gcc cgc tag ttg cga tat aca taa act gat cac taa ttc cca gaa tcc cac cca ctt ata gta ag- -tt ttt tta cct ata aat aat aaa tac aat aat taa ttt ctc gta aaa gta gaa aat ata ttc taa ttt att ata tgg taa gaa agt aga atc atc tag aac agt aat caa tca ata gca atc atg aaa caa tat att gtc ctg gca tgc atg tgc cta gtg gca gct gct atg cct act agt ctt caa caa tct tca tcc tcg tgt act gaa gaa gaa aac aaa cat cat atg gga atc gat gtt att atc aaa gtc aca aag caa gac caa aca ccg acc aat gat aag att tgt caa tcc gta acg gaa gtt aca gag a-- --- --- --- --- --- -cc gaa gat gat gag gta tcc gaa gaa gtt gta aaa gga gat ccc acc act tat tac act atc gtc ggt gcg ggt ctt aac atg aac ttt gga ttc acc aaa tgc cca aag att tta tcc atc tcc gaa tcc tct gat gga aac act gtg aat act aga ttg tcc agc gtg tca ccg gga caa ggt aag gac tct ccc gcg atc acg cgt gaa gaa gct ctg gct atg atc aaa gac tgt gag atg tct atc gac atc aga tgt agc gaa gaa gag aaa gac agt gac atc aag acc cat cca gta ctt ggg tct aac atc tca cat aag aaa gtg agt tac aaa gat atc atc ggt tca acg atc gtt gat aca aaa tgt gtc aag aac cta gag ttt agc gta cgt atc gga gac atg tgt gag gaa tca tct gaa ctt gaa gtc aag gat gga ttc aag tat gtc gac gga tcg gca tct gaa ggt gca acc gat gat act tca ctc atc gat tca aca aaa ctc aaa gca tgt gtc tga atc gat aac tct att cat ctg aaa atg gat gag ttg ggt taa tcg aac gat tca gac acc gca cca cga att aaa aaa g-- -ac cgg gca cta tat tcc ggt ttg caa aac aaa aat att taa cta cat tca caa aaa gtt acc tct cgt tac ttc ttc ttt ctg ttt caa tat gtg ata cga tat gat cac tat tcg tat tct ct- --- --- tgg tct cat aaa aaa gtt tta c-- --- --a aaa aaa aaa a-t att ttt att ctc ttt ctc tct tcg at- ggt ctc aca aaa ata tta aac ctc ttt ctg atg tct caa cta ttt cgt aaa cga taa cgt cca aca ata tat tct cgt aga gct tat caa cat cct tat acc aat cta ggt tgt cag aca att gca tca taa aat aat gtt tat aat tta cac gtt aac atc ata taa taa acg tat ata gtt aat att ttt gga ata taa atg atc tgt aaa atc cat gta ggg gac act gct cac gtt ttt tct cta gta cat aat ttc aca caa gtt ttt ata cag aca aat taa ttc tcg tcc ata ta- ttt taa aac att gac ttt tgt act aag aaa aat atc ttg act aac cat ctc ttt ctc tct tcg atg ggt ctc aca aaa ata tta aac ctc ttt ctg atg g-- --- --- --- --- --- --- --- --- --- --- --- --- --- --- --- --- --- --- --- --- --- --- --- --- --- --- --- --- --- --- --- --- --- --- --- --- --- --- --- --- --- --- --- --- --- --- agt cgt aaa aag ttt tta tcc ttt ctc tct tcg ata ggt ctc --- --- --- --- --- --- --- --- --- --- --- --- --- --- --- --- --- --- --- --- --- --- --- --- --- --- --- --- --- --- --- --- --- --- --- --- --- --- --- --- --- --- --- --- --- --- --- --- --- --- --- --- --- --- --- --- --- --- --- --- --- --- --- --- --- --- --- --- --- --- aca aaa ata tta aac ctc ttt ctg atg gtc tct ata aa

>ON675438.1_Monkeypox_virus_isolate_MPXV_USA_2022_VA001_complete_genome

atc gag gac gac ttc aac tat gac agc tcg gtg gcg tct gcc agc gtg tac ata caa atg gca ttt cta aga aaa gtc tac ggt atc ctt -tc tac aat ttc ctt taa caa cgg caa cag ctg cag tat ttt tat act ttg aat gca -tc gga cat tta tac aag gga gtc ctg ttc taa tat tgg cat caa tgt tcg gat cta tag gct tga ttt tcg cat tga ctt tac aca gac atg aac atc ccc tga atc tgt aca tac ttt gtg gat tta cac tgt tag aat ctc taa cgc tgg cct ctg ttg tta ctt tct atg atg cac gta tcg tta tgc aag ctt tca tgt tga cta ctg cag tgt ttc ttg ctc tga cta cat ata ctc tac aat caa aga gag att tca gta aac ttg taa cag gat tgt ttg ctg ctt tct gga ttt taa ttt tgt cag gag tct tga gga taa agt tta aaa tag aat taa taa aga aca tat agg tca -tt ttt taa aca tgg ata gaa acc aag gtt gtt agt taa taa tat aca aga tat ttt ttc tca ctc tga tcc atg taa acc aag gac gag aga cac tct cat tcc tca ttc acg aca cca tta aaa atg gaa att aaa gcc ctc tat taa gca cag acg gct aca ggt cta cca tca ggt t-a cct tcg tct acc ttc aca atg gcc tct cct tgt gcc cag ttc agt ccc tgt cat tgc cac gct act aag gac tcc ctg aat acc gtg act gac gtc aga cat tgt ctg act gaa tac atc ctg tgg gtt tct cat aga tgg acc cat aga gaa agc gca ggg cct ctc tac agg ctt ctc atc tct ttc aga att gat gca atg gag cta ttt ggt agc gag ttg aag gag ttc tcg aat tca ctt ccg tgg gac aat atc gac aat tgc gtg gag atc att aaa tgt ttc atc aga aat gac tcc atg aaa acc gcc aaa gaa ctt tgt gca ata att gga ctt tgt act caa tca gct att gtc act gga aga gtc ttc aat gat aag tat atc gac ata cta ctt atg ctg cga aag att ctg aac gag aac gac tat ctc acc ctc ttg gat cat atc ctc act gct aaa tac taa atc tcc ttc atg ctc tct cac taa tac tct tac tca cta cac ttt tta tca tct tat gat gaa tga ttg cct tca tca ttt ttt cgt gga ata taa tat agg aat aat tag cac cag aat agc tat gga tat ctc gtt aag aat att ctc tat aag aga cat aat gta gac ata gtt att ata tcc ttc tta gat aag tgt tac gct act gga aag ttt cca tcg tta tta tta cat gaa gat gat ata att aaa cca aca ttg aga ttg gct ctt atg tta gct gga ttg aat tac tgt aat aaa tgc atc gag tat aga ggg ata tag caa ttc tcg ata ata gtc atg caa tat ttg aat gag act gat aat tta ggt aat aca gta ct- --- aca cac ata tct ttc tag att ata tat cgt taa aaa tct gta aga tgt ata ttt ctc ata agt atc cac tgt gta ata tta tta atg gat ata tag ata acg caa tag gga cta ata gta ttg taa aag ata taa tcg act att tac gta cat atc cag ata tct ata ttc cta cta gtt tgc tgc gta gtt gca tca ttg ata tgc atg -at tta tca gga ttc aga gat gaa tta cta agt aaa cta caa tcc cac aat aag taa gaa tca aat atc aaa aac tca ctt ttg att ttt cta gtc tta agt aat aca tat att tat taa tag acc tat gaa ata aaa aaa ggt aac aat gga ttc gcg tat agc tat tta cgt att agt atc ggc atc tct ttt gta tct tgt taa ttg tca caa act agt aca tta ctt caa tct gaa aat aaa tgg aag tga tat aac taa tac agc aga tat att gct gga caa tta tcc aat tat gac ctt tga tgg aaa gga tat tta tcc atc tat ctc gtt cat ggt cgg taa taa act ttt cct aga tct tta taa aaa tat ctt tga aga att ttt cag act att tcg agt atc tgt aag tag tca ata cga gga att aga ata tta tta ttc atg tga tta tac taa caa ccg tcc tac aat taa aca aca tta ctt tta taa cgg cga cga ata tac tga aat tga tag atc gaa aaa agc cac taa taa aaa cag ttg gtt aat tac ttc agg ctt tag act aca aaa atg gtt cga tag cga aga ttg tat aat tta tct cag atc ttt agt tag aag aat gga aga cag taa caa aaa cag taa aaa aac tta gta ctt aga tat cg- --a aaa aat ata ttt ttg tag act ctt gag aat aga agg aaa aca tgt aca taa tta taa aaa atg aaa atc aat ggc gaa taa gac agt gcg att cgc acc atg gag tcg gta gat ttc atg gct gtc gat gag cag ttt cac gac gac ctc gat ctt tgg tca tta tct ttg gta gat gat tat aaa aaa cat gga tta ggt gtt gac tgt tat gtt cta gaa cca gtt gtt gac agg aaa ata ttt gat aga ttt ctc ctt gaa cca att tgt gat cct gta gat gtt ctg tat gat tat ttt agg att cat aga gat aat att gat cag tat ata gta gat aga ctg ttt gca tat att aca tat aaa gat att ata tct gca tta gtg tca aag aat tat atg gaa gat att ttc tct ata att att aag aat tgt aat tct gtg caa gat ctc tta ctt tac tat cta tct aat gca tat gta gaa ata gac att gtt gat ctt atg gta gat cat ggg gct gta ata tat aaa ata gaa tgc ttg aat gcc tat ttt agg gga ata tgt aaa aag gaa agt agt gtt gtt gag ttt att ttg aat tgt ggt atc cca gat gaa aat gat gtt aaa tta gat cta tat aaa ata att cag tat act agg gga ttc ctt gta gat gaa ccc aca gta tta gaa att tat aag ctt tgt atc cca tat att gaa gat atc aat caa cta gat gct ggt gga agg acc ttg ctt tat cgc gct atc tat gca ggt tat ata gat tta gta tca tgg cta tta gaa aat gga gca aat gtc aac gca gta atg agt aat gga tat aca tgt ctt gac gtg gcc gtg gat agg gga tct gtc atc gcc cgt agg gaa gca cat ctt aaa ata tta gaa ata ttg ctt aga gaa cca ttg tct att gac tgt ata aaa tta gct ata ctt aat aat aca att gaa aac cat gat gtg ata aag ctc tgt atc aag tat ttt atg atg gta gat tat tca ctt tgt aat gtg tat gca tca tca ctc ttt gat tat ata att gat tgt aaa caa gaa ttg gag tac att agg cag atg aaa att cat aat aca acc atg tat gag tta atc tat aat aga gac aaa aac aag cat gct tcc cat att cta cat agg tat tct aaa cat cca gtt ttg aca cag tgt atc act aaa gga ttc aag att tac aca gaa gta acc gag cag gtc act aaa gct cta aac aga cgt gct cta ata gat gag ata ata aac aat gta tca act gat gac aat ctc cta tca aaa ctt cca tta gaa att agg gat cta att gtt tca caa gct gtc ata tag agt tct atc cac cca cct ttc ttg aaa tga gtt aat --- --- --- --- --- --- --- --- --- --- --- --- --- --- --- --- --- --- --- --- --- --- --- --- --- --- --- --- --- --- --- --- --- --- --- --- --- --- --- --- --- --- --- --- --- --- --- --- --- --- --- --- --- --- --- --- --- --- --- --- --- --- --- --- --- --- --- --- --- --- --- --- --- --- --- --- --- --- --- --- --- --- --- --- --- --- --- --- --- --- --- --- --- --- --- --- --- --- --- --- --- --- --- --- --- -ag tca taa gtt agt taa --- --- --- --- --- -gt cat aag tta gtt aag tca t-- --- --- --- --- --- --- --- --- --- --- --- --- --- --- --- -aa gtt agt taa gtc ata agt tag tta agt cat aag tta gtt aag tca taa gtt agt taa gtc ata agt tag tta agt cat aag tta gtt tat agt cta aca ctt cta att ttt ata cct tga tct ttt tct cta att atg aaa aag taa atc att atg aag atg gat g-a aat gga cga gat tgt gcg cat cgt taa cga tag tat gtg gta cgt acc taa cgc att tat gga cga cgg tga taa tga agg tca cat ttc tgt caa taa tgt ctg tca tat gta tct cgc att ctt tga tgt gga tat atc atc tca tct gtt taa att agt tat taa aca ctg cga tct gaa taa acg act aaa atg tgg taa ctc tcc att aca ttg cta tac gat gaa tac acg att taa tcc atc tgt att aaa gat att gtt acg cca cgg cat gcg taa ctt tga tag caa gga taa aaa agg aca tat tcc tct aca cca cta tct gat tca ttc act atc aat cga taa caa gat ctt tga tat act aac gga ccc cat tga tga ctt tag taa atc atc cga tct att gct gtg tta tct tag ata taa att caa tgg gag ctt aaa cta tta cgt tct gta caa att att gac taa agg atc tga ccc taa ttg cgt cga tga gga tgg act cac ttc tct tca tta cta ctg taa aca cat atc cgc gtt cca cga aag caa tta tta caa gtc aaa gag tca cac taa gat gcg agc tga gaa gcg att cat cta cgc gat aat aga tca tgg agc aaa cat taa cgc ggt tac gaa aat cgg aaa tac gcc gtt aca cac tta cct tca aca gta tac caa aca tag tcc tcg tgt ggt gta tgc tct ttt atc tcg agg agc cga tac gag gat acg taa taa tct tga ttg tac acc cat cat gga ata cat aaa gaa cga ttg tgc aac agg tca tat tct cat aat gtt act caa ttg gca cga aca aaa ata cgg gaa att aca aaa gga aga agg aca aca tct act tta tct att cat aaa aca taa tca agg ata tgg aag tcg ctc tct caa tat act acg gta tct act aga tag att cga cat tca gaa aga cga ata cta taa tac aat gac tcc tct tca tac cgc ctt cca gaa ttg caa taa caa tgt tgc ctc ata cct cgt ata cat cgg ata cga cat caa cct tcc gac taa aga cga taa gac agt att cga ctt ggt gtt tga aaa cag aaa cat tat ata caa ggc gga tgt cgt taa tga cat tat cca cca cag act gaa agt atc tct acc tat gat taa atc gtt gtt cta caa gat gtc gga gtt ctc tcc cta cga cga tca cta cgt aaa gaa gat aat agc cta ctg cct att aag gga cga gtc att tgc gga act aca tac taa att ctg ttt aaa cga gga cta taa aag tgt att tat gaa aaa tat atc att cga taa gat aga ttc cat cat cga aaa atg tag tcg tga cat aag tct cct caa aga gat tcg aat ctc aga cac cga ctt gta tac ggt att gag aac aga aga cat ccg gta tca cac ata tct cga agc cat aca ttc aga caa acg cat ttc att tcc cat gta cga cga tct cat aga aca gtg tca tct atc gat gga gca taa aag taa act cgt cga caa agc act caa taa att aga gtc tac cat cga tag tca atc tag act atc gta ttt gcc tcc gga aat tat gcg caa tat cat aac caa gct aag cga cta cca tct aaa cag tat gtt gta cgg aaa gaa cca tta caa ata tta tcc atg ata gaa aga aaa tat tta aaa aat aat cta tat gat tgg aga agt agg aaa caa aca gta aca aga cga cga tta cta c-a tta tta aat cat gag gtc cgt att ata ctc gta tat att gtt tct ctc atg tat aat aat aaa cgg aag aga tat agc acc aca tgc acc atc caa tgg aaa gtg taa aga caa cga ata cag aag ccg taa tct atg ttg tct atc gtg tcc tcc ggg aac tta cgc ttc cag att atg tga tag caa gac taa tac aca atg tac acc gtg tgg ttc gga tac ctt tac atc tca caa taa tca ttt aca ggc ttg tct aag ttg taa cgg aag atg tga tag taa tca ggt aga gac gcg atc gtg taa cac gac tca caa tag aat ctg tga atg ctc tcc agg ata tta ttg tct tct caa agg agc atc agg gtg tag aac atg tat ttc taa aac aaa gtg tgg aat agg ata cgg agt atc cgg ata cac gtc tac cgg aga cgt cat ctg ttc tcc gtg tgg tcc cgg aac ata ttc tca cac cgt ctc ttc cac aga taa atg cga acc cgt cgt aac cag caa tac att taa cta tat cga tgt gga aat taa cct gta tcc agt caa cga cac atc gtg tac tcg gac gac cac tac cgg tct cag cga atc cat ctc aac gtc gga act aac tat tac cat gaa tca taa aga ttg tga tcc agt ctt tcg tgc aga ata ctt ctc tgt cct taa taa tgt agc aac ttc agg att ctt tac agg aga aaa tag ata tca gaa tac ttc aaa gat atg tac tct gaa ttt cga gat taa atg taa caa caa aga ttc atc ttc caa aca gtt aac gaa aac aaa gaa tga tac tat cat gcc gca ttc aga gac ggt aac tct agt ggg cga ctg tct atc tag cgt cga cat cta cat act ata tag taa tac caa tac tca aga cta cga aaa tga tac aat ctc tta tca tat ggg taa tgt tct cga tgt caa tag cca tat gcc cgc tag ttg cga tat aca taa act gat cac taa ttc cca gaa tcc cac cca ctt ata gta ag- -tt ttt tta cct ata aat aat aaa tac aat aat taa ttt ctc gta aaa gta gaa aat ata ttc taa ttt att ata tgg taa gaa agt aga atc atc tag aac agt aat caa tca ata gca atc atg aaa caa tat att gtc ctg gca tgc atg tgc cta gtg gca gct gct atg cct act agt ctt caa caa tct tca tcc tcg tgt act gaa gaa gaa aac aaa cat cat atg gga atc gat gtt att atc aaa gtc aca aag caa gac caa aca ccg acc aat gat aag att tgt caa tcc gta acg gaa gtt aca gag a-- --- --- --- --- --- -cc gaa gat gat gag gta tcc gaa gaa gtt gta aaa gga gat ccc acc act tat tac act atc gtc ggt gcg ggt ctt aac atg aac ttt gga ttc acc aaa tgc cca aag att tca tcc atc tcc gaa tcc tct gat gga aac act gtg aat act aga ttg tcc agc gtg tca ccg gga caa ggt aag gac tct ccc gcg atc acg cgt gaa gaa gct ctg gct atg atc aaa gac tgt gag atg tct atc gac atc aga tgt agc gaa gaa gag aaa gac agt gac atc aag acc cat cca gta ctt ggg tct aac atc tca cat aag aaa gtg agt tac aaa gat atc atc ggt tca acg atc gtt gat aca aaa tgt gtc aag aac cta gag ttt agc gta cgt atc gga gac atg tgt gag gaa tca tct gaa ctt gaa gtc aag gat gga ttc aag tat gtc gac gga tcg gca tct gaa ggt gca acc gat gat act tca ctc atc gat tca aca aaa ctc aaa gca tgt gtc tga atc gat aac tct att cat ctg aaa atg gat gag ttg ggt taa tcg aac gat tca gac acc gca cca cga att aaa aaa g-- -ac cgg gca cta tat tcc ggt ttg caa aac aaa aat att taa cta cat tca caa aaa gtt acc tct cgt tac ttc ttc ttt ctg ttt caa tat gtg ata cga tat gat cac tat tcg tat tct ct- --- --- tgg tct cat aaa aaa gtt tta c-- --a aaa aaa aaa aaa a?t att ttt att ctc ttt ctc tct tcg at- ggt ctc aca aaa ata tta aac ctc ttt ctg atg tct caa cta ttt cgt aaa cga taa cgt cca aca ata tat tct cgt aga gct tat caa cat cct tat acc aat cta ggt tgt cag aca att gca tca taa aat aat gtt tat aat tta cac gtt aac atc ata taa taa acg tat ata gtt aat att ttt gga ata taa atg atc tgt aaa atc cat gta ggg gac act gct cac gtt ttt tct cta gta cat aat ttc aca caa gtt ttt ata cag aca aat taa ttc tcg tcc ata ta- ttt taa aac att gac ttt tgt act aag aaa aat atc ttg act aac cat ctc ttt ctc tct tcg atg ggt ctc aca aaa ata tta aac ctc ttt ctg atg g-- --- --- --- --- --- --- --- --- --- --- --- --- --- --- --- --- --- --- --- --- --- --- --- --- --- --- --- --- --- --- --- --- --- --- --- --- --- --- --- --- --- --- --- --- --- --- agt cgt aaa aag ttt tta tcc ttt ctc tct tcg ata ggt ctc --- --- --- --- --- --- --- --- --- --- --- --- --- --- --- --- --- --- --- --- --- --- --- --- --- --- --- --- --- --- --- --- --- --- --- --- --- --- --- --- --- --- --- --- --- --- --- --- --- --- --- --- --- --- --- --- --- --- --- --- --- --- --- --- --- --- --- --- --- --- aca aaa ata tta aac ctc ttt ctg atg gtc tct ata aa

>ON631241.1_Monkeypox_virus_isolate_2022/2_SLO_complete_genome

atc gag gac gac ttc aac tat gac agc tcg gtg gcg tct gcc agc gtg tac ata cga atg gca ttt cta aga aaa gtc tac ggt atc ctt -tc tac aat ttc ctt taa caa cgg caa cag ctg cag tat ttt tat act ttg aat gca -tc gga cat tta tac aag gga gtc ctg ttc taa tat tgg cat caa tgt tcg gat cta tag gct tga ttt tcg cat tga ctt tac aca gac atg aac atc ccc tga atc tgt aca tac ttt gtg gat tta cac tgt tag aat ctc taa cgc tgg cct ctg ttg tta ctt tct atg atg cac gta tcg tta tgc aag ctt tca tgt tga cta ctg cag tgt ttc ttg ctc tga cta cat ata ctc tac aat caa aga gag att tca gta aac ttg taa cag gat tgt ttg ctg ctt tct gga ttt taa ttt tgt cag gag tct tga gga taa agt tta aaa tag aat taa taa aga aca tat agg tca -tt ttt taa aca tgg ata gaa acc aag gtt gtt agt taa taa tat aca aga tat ttt ttc tca ctc tga tcc atg taa acc aag gac gag aga cac tct cat tcc tca ttc acg aca cca tta aaa atg gaa att aaa gcc ctc tat taa gca cag acg gct aca ggt cta cca tca ggt t-a cct tcg tct acc ttc aca atg gcc tct cct tgt gcc cag ttc agt ccc tgt cat tgc cac gct act aag gac tcc ctg aat acc gtg act gac gtc aga cat tgt ctg act gaa tac atc ctg tgg gtt tct cat aga tgg acc cat aga gaa agc gca ggg cct ctc tac agg ctt ctc atc tct ttc aga att gat gca atg gag cta ttt ggt agc gag ttg aag gag ttc tcg aat tca ctt ccg tgg gac aat atc gac aat tgc gtg gag atc att aaa tgt ttc atc aga aat gac tcc atg aaa acc gcc aaa gaa ctt tgt gca ata att gga ctt tgt act caa tca gct att gtc act gga aga gtc ttc aat gat aag tat atc gac ata cta ctt atg ctg cga aag att ctg aac gag aac gac tat ctc acc ctc ttg gat cat atc ctc act gct aaa tac taa atc tcc ttc atg ctc tct cac taa tac tct tac tca cta cac ttt tta tca tct tat gat gaa tga ttg cct tca tca ttt ttt cgt gga ata taa tat agg aat aat tag cac cag aat agc tat gga tat ctc gtt aag aat att ctc tat aag aga cat aat gta gac ata gtt att ata tcc ttc tta gat aag tgt tac gct act gga aag ttt cca tcg tta tta tta cat gaa gat gat ata att aaa cca aca ttg aga ttg gct ctt atg tta gct gga ttg aat tac tgt aat aaa tgc atc gag tat aga ggg ata tag caa ttc tcg ata ata gtc atg caa tat ttg aat gag act gat aat tta ggt aat aca gta ct- --- aca cac ata tct ttc tag att ata tat cgt taa aaa tct gta aga tgt ata ttt ctc ata agt atc cac tgt gta ata tta tta atg gat ata tag ata acg caa tag gga cta ata gta ttg taa aag ata taa tcg act att tac gta cat atc cag ata tct ata ttc cta cta gtt tgc tgc gta gtt gca tca ttg ata tgc atg -at tta tca gga ttc aga gat gaa tta cta agt aaa cta caa tcc cac aat aag taa gaa tca aat atc aaa aac tca ctt ttg att ttt cta gtc tta agt aat aca tat att tat taa tag acc tat gaa ata aaa aaa ggt aac aat gga ttc gcg tat agc tat tta cgt att agt atc ggc atc tct ttt gta tct tgt taa ttg tca caa act agt aca tta ctt caa tct gaa aat aaa tgg aag tga tat aac taa tac agc aga tat att gct gga caa tta tcc aat tat gac ctt tga tgg aaa gga tat tta tcc atc tat ctc gtt cat ggt cgg taa taa act ttt cct aga tct tta taa aaa tat ctt tga aga att ttt cag act att tcg agt atc tgt aag tag tca ata cga gga att aga ata tta tta ttc atg tga tta tac taa caa ccg tcc tac aat taa aca aca tta ctt tta taa cgg cga cga ata tac tga aat tga tag atc gaa aaa agc cac taa taa aaa cag ttg gtt aat tac ttc agg ctt tag act aca aaa atg gtt cga tag cga aga ttg tat aat tta tct cag atc ttt agt tag aag aat gga aga cag taa caa aaa cag taa aaa aac tta gta ctt aga tat cg- --a aaa aat ata ttt ttg tag act ctt gag aat aga agg aaa aca tgt aca taa tta taa aaa atg aaa atc aat ggc gaa taa gac agt gcg att cgc acc atg gag tcg gta gat ttc atg gct gtc gat gag cag ttt cac gac gac ctc gat ctt tgg tca tta tct ttg gta gat gat tat aaa aaa cat gga tta ggt gtt gac tgt tat gtt cta gaa cca gtt gtt gac agg aaa ata ttt gat aga ttt ctc ctt gaa cca att tgt gat cct gta gat gtt ctg tat gat tat ttt agg att cat aga gat aat att gat cag tat ata gta gat aga ctg ttt gca tat att aca tat aaa gat att ata tct gca tta gtg tca aag aat tat atg gaa gat att ttc tct ata att att aag aat tgt aat tct gtg caa gat ctc tta ctt tac tat cta tct aat gca tat gta gaa ata gac att gtt gat ctt atg gta gat cat ggg gct gta ata tat aaa ata gaa tgc ttg aat gcc tat ttt agg gga ata tgt aaa aag gaa agt agt gtt gtt gag ttt att ttg aat tgt ggt atc cca gat gaa aat gat gtt aaa tta gat cta tat aaa ata att cag tat act agg gga ttc ctt gta gat gaa ccc aca gta tta gaa att tat aag ctt tgt atc cca tat att gaa gat atc aat caa cta gat gct ggt gga agg acc ttg ctt tat cgc gct atc tat gca ggt tat ata gat tta gta tca tgg cta tta gaa aat gga gca aat gtc aac gca gta atg agt aat gga tat aca tgt ctt gac gtg gcc gtg gat agg gga tct gtc atc gcc cgt agg gaa gca cat ctt aaa ata tta gaa ata ttg ctt aga gaa cca ttg tct att gac tgt ata aaa tta gct ata ctt aat aat aca att gaa aac cat gat gtg ata aag ctc tgt atc aag tat ttt atg atg gta gat tat tca ctt tgt aat gtg tat gca tca tca ctc ttt gat tat ata att gat tgt aaa caa gaa ttg gag tac att agg cag atg aaa att cat aat aca acc atg tat gag tta atc tat aat aga gac aaa aac aag cat gct tcc cat att cta cat agg tat tct aaa cat cca gtt ttg aca cag tgt atc act aaa gga ttc aag att tac aca gaa gta acc gag cag gtc act aaa gct cta aac aga cgt gct cta ata gat gag ata ata aac aat gta tca act gat gac aat ctc cta tca aaa ctt cca tta gaa att agg gat cta att gtt tca caa gct gtc ata tag agt tct atc cac cca cct ttc ttg aaa tga gtt aat a-- --- --- --- --- --- --- --- --- --- --- --- --- --- --- --- --- --- --- --- --- --- --- --- --- --- --- --- --- --- --- --- --- --- --- --- --- --- --- --- --- --- --- --- --- --- --- --- --- --- --- --- --- --- --- --- --- --- --- --- --- --- --- --- --- --- --- --- --- --- --- --- --- --- --- --- --- --- --- --- --- --- --- --- --- --- --- --- --- --- --- --- --- --- --- --- --- --- --- --- --- --- --- --- --- -ag tca taa gtt agt taa --- --- --- --- --- -gt cat aag tta gtt aag tca t-- --- --- --- --- --- --- --- --- --- --- --- --- --- --- --- -aa gtt agt taa gtc ata agt tag tta agt cat aag tta gtt aag tca taa gtt agt taa gtc ata agt tag tta agt cat aag tta gtt tat agt cta aca ctt cta att ttt ata cct tga tct ttt tct cta att atg aaa aag taa atc att atg aag atg gat g-a aat gga cga gat tgt gcg cat cgt taa cga tag tat gtg gta cgt acc taa cgc att tat gga cga cgg tga taa tga agg tca cat ttc tgt caa taa tgt ctg tca tat gta tct cgc att ctt tga tgt gga tat atc atc tca tct gtt taa att agt tat taa aca ctg cga tct gaa taa acg act aaa atg tgg taa ctc tcc att aca ttg cta tac gat gaa tac acg att taa tcc atc tgt att aaa gat att gtt acg cca cgg cat gcg taa ctt tga tag caa gga taa aaa agg aca tat tcc tct aca cca cta tct gat tca ttc act atc aat cga taa caa gat ctt tga tat act aac gga ccc cat tga tga ctt tag taa atc atc cga tct att gct gtg tta tct tag ata taa att caa tgg gag ctt aaa cta tta cgt tct gta caa att att gac taa agg atc tga ccc taa ttg cgt cga tga gga tgg act cac ttc tct tca tta cta ctg taa aca cat atc cgc gtt cca cga aag caa tta tta caa gtc aaa gag tca cac taa gat gcg agc tga gaa gcg att cat cta cgc gat aat aga tca tgg agc aaa cat taa cgc ggt tac gaa aat cgg aaa tac gcc gtt aca cac tta cct tca aca gta tac caa aca tag tcc tcg tgt ggt gta tgc tct ttt atc tcg agg agc cga tac gag gat acg taa taa tct taa ttg tac acc cat cat gga ata cat aaa gaa cga ttg tgc aac agg tca tat tct cat aat gtt act caa ttg gca cga aca aaa ata cgg gaa att aca aaa gga aga agg aca aca tct act tta tct att cat aaa aca taa tca agg ata tgg aag tcg ctc tct caa tat act acg gta tct act aga tag att cga cat tca gaa aga cga ata cta taa tac aat gac tcc tct tca tac cgc ctt cca gaa ttg caa taa caa tgt tgc ctc ata cct cgt ata cat tgg ata cga cat caa cct tcc gac taa aga cga taa gac agt att cga ctt ggt gtt tga aaa cag aaa cat tat ata caa ggc gga tgt cgt taa tga cat tat cca cca cag act gaa agt atc tct acc tat gat taa atc gtt gtt cta caa gat gtc gga gtt ctc tcc cta cga cga tca cta cgt aaa gaa gat aat agc cta ctg cct att aag gga cga gtc att tgc gga act aca tac taa att ctg ttt aaa cga gga cta taa aag tgt att tat gaa aaa tat atc att cga taa gat aga ttc cat cat cga aaa atg tag tcg tga cat aag tct cct caa aga gat tcg aat ctc aga cac cga ctt gta tac ggt att gag aac aga aga cat tcg gta tca cac ata tct cga agc cat aca ttc aga caa acg cat ttc att tcc cat gta cga cga tct cat aga aca gtg tca tct atc gat gga gca taa aag taa act cgt cga caa agc act caa taa att aga gtc tac cat cga tag tca atc tag act atc gta ttt gcc tcc gga aat tat gcg caa tat cat aac caa gct aag cga cta cca tct aaa cag tat gtt gta cgg aaa gaa cca tta caa ata tta tcc atg ata gaa aga aaa tat tta aaa aat aat cta tat gat tgg aga agt agg aaa caa aca gta aca aga cga cga tta cta c-a tta tta aat cat gag gtc cgt att ata ctc gta tat att gtt tct ctc atg tat aat aat aaa cgg aag aga tat agc acc aca tgc acc atc caa tgg aaa gtg taa aga caa cga ata cag aag ccg taa tct atg ttg tct atc gtg tcc tcc ggg aac tta cgc ttt cag att atg tga tag caa gac taa tac aca atg tac acc gtg tgg ttc gga tac ctt tac atc tca caa taa tca ttt aca ggc ttg tct aag ttg taa cgg aag atg tga tag taa tca ggt aga gac gcg atc gtg taa cac gac tca caa tag aat ctg tga atg ctc tcc agg ata tta ttg tct tct caa agg agc atc agg gtg tag aac atg tat ttc taa aac aaa gtg tgg aat agg ata cgg agt atc cgg ata cac gtc tac cgg aga cgt cat ctg ttc tcc gtg tgg tcc cgg aac ata ttc tca cac cgt ctc ttc cac aga taa atg cga acc cgt cgt aac cag caa tac att taa cta tat cga tgt gga aat taa cct gta tcc agt caa cga cac atc gtg tac tcg gac gac cac tac cgg tct cag cga atc cat ctc aac gtc gga act aac tat tac cat gaa tca taa aga ttg tga tcc agt ctt tcg tgc aga ata ctt ctc tgt cct taa taa tgt agc aac ttc agg att ctt tac agg aga aaa tag ata tca gaa tac ttc aaa gat atg tac tct gaa ttt cga gat taa atg taa caa caa aga ttc atc ttc caa aca gtt aac gaa aac aaa gaa tga tac tat cat gcc gca ttc aga gac ggt aac tct agt ggg cga ctg tct atc tag cgt cga cat cta cat act ata tag taa tac caa tac tca aga cta cga aaa tga tac aat ctc tta tca tat ggg taa tgt tct cga tgt caa tag cca tat gcc cgc tag ttg cga tat aca taa act gat cac taa ttc cca gaa tcc cac cca ctt ata gta ag- -tt ttt tta cct ata aat aat aaa tac aat aat taa ttt ctc gta aaa gta gaa aat ata ttc taa ttt att ata tgg taa gaa agt aga atc atc tag aac agt aat caa tca ata gca atc atg aaa caa tat att gtc ctg gca tgc atg tgc cta gtg gca gct gct atg cct act agt ctt caa caa tct tca tcc tcg tgt act gaa gaa gaa aac aaa cat cat atg gga atc gat gtt att atc aaa gtc aca aag caa gac caa aca ccg acc aat gat aag att tgt caa tcc gta acg gaa gtt aca gag a-- --- --- --- --- --- -cc gaa gat gat gag gta tcc gaa gaa gtt gta aaa gga gat ccc acc act tat tac act atc gtc ggt gcg ggt ctt aac atg aac ttt gga ttc acc aaa tgc cca aag att tta tcc atc tcc gaa tcc tct gat gga aac act gtg aat act aga ttg tcc agc gtg tca ccg gga caa ggt aag gac tct ccc gcg atc acg cgt gaa gaa gct ctg gct atg atc aaa gac tgt gag atg tct atc gac atc aga tgt agc gaa gaa gag aaa gac agt gac atc aag acc cat cca gta ctt ggg tct aac atc tca cat aag aaa gtg agt tac aaa gat atc atc ggt tca acg atc gtt gat aca aaa tgt gtc aag aac cta gag ttt agc gta cgt atc gga gac atg tgt gag gaa tca tct gaa ctt gaa gtc aag gat gga ttc aag tat gtc gac gga tcg gca tct gaa ggt gca acc gat gat act tca ctc atc gat tca aca aaa ctc aaa gca tgt gtc tga atc gat aac tct att cat ctg aaa atg gat gag ttg ggt taa tcg aac gat tca gac acc gca cca cga att aaa aaa g-- -ac cgg gca cta tat tcc ggt ttg caa aac aaa aat att taa cta cat tca caa aaa gtt acc tct cgt tac ttc ttc ttt ctg ttt caa tat gtg ata cga tat gat cac tat tcg tat tct ct- --- --- tgg tct cat aaa aaa gtt tta c-- --a aaa aaa aaa aaa a-t att ttt att ctc ttt ctc tct tcg at- ggt ctc aca aaa ata tta aac ctc ttt ctg atg tct caa cta ttt cgt aaa cga taa cgt cca aca ata tat tct cgt aga gct tat caa cat cct tat acc aat cta ggt tgt cag aca att gca tca taa aat aat gtt tat aat tta cac gtt aac atc ata taa taa acg tat ata gtt aat att ttt gga ata taa atg atc tgt aaa atc cat gta ggg gac act gct cac gtt ttt tct cta gta cat aat ttc aca caa gtt ttt ata cag aca aat taa ttc tcg tcc ata ta- ttt taa aac att gac ttt tgt act aag aaa aat atc ttg act aac cat ctc ttt ctc tct tcg atg ggt ctc aca aaa ata tta aac ctc ttt ctg atg g-- --- --- --- --- --- --- --- --- --- --- --- --- --- --- --- --- --- --- --- --- --- --- --- --- --- --- --- --- --- --- --- --- --- --- --- --- --- --- --- --- --- --- --- --- --- --- agt cgt aaa aag ttt tta tcc ttt ctc tct tcg ata ggt ctc --- --- --- --- --- --- --- --- --- --- --- --- --- --- --- --- --- --- --- --- --- --- --- --- --- --- --- --- --- --- --- --- --- --- --- --- --- --- --- --- --- --- --- --- --- --- --- --- --- --- --- --- --- --- --- --- --- --- --- --- --- --- --- --- --- --- --- --- --- --- aca aaa ata tta aac ctc ttt ctg atg gtc tct ata aa

>ON622712.1_Monkeypox_virus_isolate_MPX/UZ_REGA_1/Belgium/2022_complete_genome

atc gag gac gac ttc aac tat gac agc tcg gtg gcg tct gcc agc gtg tac ata cga atg gca ttt cta aga aaa gtc tac ggt atc ctt -tc tac aat ttc ctt taa caa cgg caa cag ctg cag tat ttt tat act ttg aat gca -tc gga cat tta tac aag gga gtc ctg ttc taa tat tgg cat caa tgt tcg gat cta tag gct tga ttt tcg cat tga ctt tac aca gac atg aac atc ccc tga atc tgt aca tac ttt gtg gat tta cac tgt tag aat ctc taa cgc tgg cct ctg ttg tta ctt tct atg atg cac gta tcg tta tgc aag ctt tca tgt tga cta ctg cag tgt ttc ttg ctc tga cta cat ata ctc tac aat caa aga gag att tca gta aac ttg taa cag gat tgt ttg ctg ctt tct gga ttt taa ttt tgt cag gag tct tga gga taa agt tta aaa tag aat taa taa aga aca tat agg tca -tt ttt taa aca tgg ata gaa acc aag gtt gtt agt taa taa tat aca aga tat ttt ttc tca ctc tga tcc atg taa acc aag gac gag aga cac tct cat tcc tca ttc acg aca cca tta aaa atg gaa att aaa gcc ctc tat taa gca cag acg gct aca ggt cta cca tca ggt t-a cct tcg tct acc ttc aca atg gcc tct cct tgt gcc cag ttc agt ccc tgt cat tgc cac gct act aag gac tcc ctg aat acc gtg act gac gtc aga cat tgt ctg act gaa tac atc ctg tgg gtt tct cat aga tgg acc cat aga gaa agc gca ggg cct ctc tac agg ctt ctc atc tct ttc aga att gat gca atg gag cta ttt ggt agc gag ttg aag gag ttc tcg aat tca ctt ccg tgg gac aat atc gac aat tgc gtg gag atc att aaa tgt ttc atc aga aat gac tcc atg aaa acc gcc aaa gaa ctt tgt gca ata att gga ctt tgt act caa tca gct att gtc act gga aga gtc ttc aat gat aag tat atc gac ata cta ctt atg ctg cga aag att ctg aac gag aac gac tat ctc acc ctc ttg gat cat atc ctc act gct aaa tac taa atc tcc ttc atg ctc tct cac taa tac tct tac tca cta cac ttt tta tca tct tat gat gaa tga ttg cct tca tca ttt ttt cgt gga ata taa tat agg aat aat tag cac cag aat agc tat gga tat ctc gtt aag aat att ctc tat aag aga cat aat gta gac ata gtt att ata tcc ttc tta gat aag tgt tac gct act gga aag ttt cca tcg tta tta tta cat gaa gat gat ata att aaa cca aca ttg aga ttg gct ctt atg tta gct gga ttg aat tac tgt aat aaa tgc atc gag tat aga ggg ata tag caa ttc tcg ata ata gtc atg caa tat ttg aat gag act gat aat tta ggt aat aca gta ct- --- aca cac ata tct ttc tag att ata tat cgt taa aaa tct gta aga tgt ata ttt ctc ata agt atc cac tgt gta ata tta tta atg gat ata tag ata acg caa tag gga cta ata gta ttg taa aag ata taa tcg act att tac gta cat atc cag ata tct ata ttc cta cta gtt tgc tgc gta gtt gca tca ttg ata tgc atg -at tta tca gga ttc aga gat gaa tta cta agt aaa cta caa tcc cac aat aag taa gaa tca aat atc aaa aac tca ctt ttg att ttt cta gtc tta agt aat aca tat att tat taa tag acc tat gaa ata aaa aaa ggt aac aat gga ttc gcg tat agc tat tta cgt att agt atc ggc atc tct ttt gta tct tgt taa ttg tca caa act agt aca tta ctt caa tct gaa aat aaa tgg aag tga tat aac taa tac agc aga tat att gct gga caa tta tcc aat tat gac ctt tga tgg aaa gga tat tta tcc atc tat ctc gtt cat ggt cgg taa taa act ttt cct aga tct tta taa aaa tat ctt tga aga att ttt cag act att tcg agt atc tgt aag tag tca ata cga gga att aga ata tta tta ttc atg tga tta tac taa caa ccg tcc tac aat taa aca aca tta ctt tta taa cgg cga cga ata tac tga aat tga tag atc gaa aaa agc cac taa taa aaa cag ttg gtt aat tac ttc agg ctt tag act aca aaa atg gtt cga tag cga aga ttg tat aat tta tct cag atc ttt agt tag aag aat gga aga cag taa caa aaa cag taa aaa aac tta gta ctt aga tat cg- --a aaa aat ata ttt ttg tag act ctt gag aat aga agg aaa aca tgt aca taa tta taa aaa atg aaa atc aat ggc gaa taa gac agt gcg att cgc acc atg gag tcg gta gat ttc atg gct gtc gat gag cag ttt cac gac gac ctc gat ctt tgg tca tta tct ttg gta gat gat tat aaa aaa cat gga tta ggt gtt gac tgt tat gtt cta gaa cca gtt gtt gac agg aaa ata ttt gat aga ttt ctc ctt gaa cca att tgt gat cct gta gat gtt ctg tat gat tat ttt agg att cat aga gat aat att gat cag tat ata gta gat aga ctg ttt gca tat att aca tat aaa gat att ata tct gca tta gtg tca aag aat tat atg gaa gat att ttc tct ata att att aag aat tgt aat tct gtg caa gat ctc tta ctt tac tat cta tct aat gca tat gta gaa ata gac att gtt gat ctt atg gta gat cat ggg gct gta ata tat aaa ata gaa tgc ttg aat gcc tat ttt agg gga ata tgt aaa aag gaa agt agt gtt gtt gag ttt att ttg aat tgt ggt atc cca gat gaa aat gat gtt aaa tta gat cta tat aaa ata att cag tat act agg gga ttc ctt gta gat gaa ccc aca gta tta gaa att tat aag ctt tgt atc cca tat att gaa gat atc aat caa cta gat gct ggt gga agg acc ttg ctt tat cgc gct atc tat gca ggt tat ata gat tta gta tca tgg cta tta gaa aat gga gca aat gtc aac gca gta atg agt aat gga tat aca tgt ctt gac gtg gcc gtg gat agg gga tct gtc atc gcc cgt agg gaa gca cat ctt aaa ata tta gaa ata ttg ctt aga gaa cca ttg tct att gac tgt ata aaa tta gct ata ctt aat aat aca att gaa aac cat gat gtg ata aag ctc tgt atc aag tat ttt atg atg gta gat tat tca ctt tgt aat gtg tat gca tca tca ctc ttt gat tat ata att gat tgt aaa caa gaa ttg gag tac att agg cag atg aaa att cat aat aca acc atg tat gag tta atc tat aat aga gac aaa aac aag cat gct tcc cat att cta cat agg tat tct aaa cat cca gtt ttg aca cag tgt atc act aaa gga ttc aag att tac aca gaa gta acc gag cag gtc act aaa gct cta aac aga cgt gct cta ata gat gag ata ata aac aat gta tca act gat gac aat ctc cta tca aaa ctt cca tta gaa att agg gat cta att gtt tca caa gct gtc ata tag agt tct atc cac cca cct ttc ttg aaa tga gtt aat agt cat aag tta gtt aag tca taa gtt agt taa gtc ata agt tag tta agt cat aag tt- --- --- --- --- --- --- --- --- --- --- --- --- --- --- --- --- --- --- --- --- --- --- --- --- --- --- --- --- --- --- --- --- --- --- --- --- --- --- --- --- --- --- --- --- --- --- --- --- --- --- --- --- --- --- --- --- --- agt taa gtc ata agt tag tta --- --- --- --- --- -ag tca taa gtt agt taa gtc ata agt tag tta agt cat aag tta gtt aag tca taa gtt agt taa --- --- --- --- --- -gt cat aag tta gtt aag tca t-- --- --- --- --- --- --- --- --- --- --- --- --- --- --- --- -aa gtt agt taa gtc ata agt tag tta agt cat aag tta gtt aag tca taa gtt agt taa gtc ata agt tag tta agt cat aag tta gtt tat agt cta aca ctt cta att ttt ata cct tga tct ttt tct cta att atg aaa aag taa atc att atg aag atg gat g-a aat gga cga gat tgt gcg cat cgt taa cga tag tat gtg gta cgt acc taa cgc att tat gga cga cgg tga taa tga agg tca cat ttc tgt caa taa tgt ctg tca tat gta tct cgc att ctt tga tgt gga tat atc atc tca tct gtt taa att agt tat taa aca ctg cga tct gaa taa acg act aaa atg tgg taa ctc tcc att aca ttg cta tac gat gaa tac acg att taa tcc atc tgt att aaa gat att gtt acg cca cgg cat gcg taa ctt tga tag caa gga taa aaa agg aca tat tcc tct aca cca cta tct gat tca ttc act atc aat cga taa caa gat ctt tga tat act aac gga ccc cat tga tga ctt tag taa atc atc cga tct att gct gtg tta tct tag ata taa att caa tgg gag ctt aaa cta tta cgt tct gta caa att att gac taa agg atc tga ccc taa ttg cgt cga tga gga tgg act cac ttc tct tca tta cta ctg taa aca cat atc cgc gtt cca cga aag caa tta tta caa gtc aaa gag tca cac taa gat gcg agc tga gaa gcg att cat cta cgc gat aat aga tca tgg agc aaa cat taa cgc ggt tac gaa aat cgg aaa tac gcc gtt aca cac tta cct tca aca gta tac caa aca tag tcc tcg tgt ggt gta tgc tct ttt atc tcg agg agc cga tac gag gat acg taa taa tct taa ttg tac acc cat cat gga ata cat aaa gaa cga ttg tgc aac agg tca tat tct cat aat gtt act caa ttg gca cga aca aaa ata cgg gaa att aca aaa gga aga agg aca aca tct act tta tct att cat aaa aca taa tca agg ata tgg aag tcg ctc tct caa tat act acg gta tct act aga tag att cga cat tca gaa aga cga ata cta taa tac aat gac tcc tct tca tac cgc ctt cca gaa ttg caa taa caa tgt tgc ctc ata cct cgt ata cat tgg ata cga cat caa cct tcc gac taa aga cga taa gac agt att cga ctt ggt gtt tga aaa cag aaa cat tat ata caa ggc gga tgt cgt taa tga cat tat cca cca cag act gaa agt atc tct acc tat gat taa atc gtt gtt cta caa gat gtc gga gtt ctc tcc cta cga cga tca cta cgt aaa gaa gat aat agc cta ctg cct att aag gga cga gtc att tgc gga act aca tac taa att ctg ttt aaa cga gga cta taa aag tgt att tat gaa aaa tat atc att cga taa gat aga ttc cat cat cga aaa atg tag tcg tga cat aag tct cct caa aga gat tcg aat ctc aga cac cga ctt gta tac ggt att gag aac aga aga cat tcg gta tca cac ata tct cga agc cat aca ttc aga caa acg cat ttc att tcc cat gta cga cga tct cat aga aca gtg tca tct atc gat gga gca taa aag taa act cgt cga caa agc act caa taa att aga gtc tac cat cga tag tca atc tag act atc gta ttt gcc tcc gga aat tat gcg caa tat cat aac caa gct aag cga cta cca tct aaa cag tat gtt gta cgg aaa gaa cca tta caa ata tta tcc atg ata gaa aga aaa tat tta aaa aat aat cta tat gat tgg aga agt agg aaa caa aca gta aca aga cga cga tta cta c-a tta tta aat cat gag gtc cgt att ata ctc gta tat att gtt tct ctc atg tat aat aat aaa cgg aag aga tat agc acc aca tgc acc atc caa tgg aaa gtg taa aga caa cga ata cag aag ccg taa tct atg ttg tct atc gtg tcc tcc ggg aac tta cgc ttt cag att atg tga tag caa gac taa tac aca atg tac acc gtg tgg ttc gga tac ctt tac atc tca caa taa tca ttt aca ggc ttg tct aag ttg taa cgg aag atg tga tag taa tca ggt aga gac gcg atc gtg taa cac gac tca caa tag aat ctg tga atg ctc tcc agg ata tta ttg tct tct caa agg agc atc agg gtg tag aac atg tat ttc taa aac aaa gtg tgg aat agg ata cgg agt atc cgg ata cac gtc tac cgg aga cgt cat ctg ttc tcc gtg tgg tcc cgg aac ata ttc tca cac cgt ctc ttc cac aga taa atg cga acc cgt cgt aac cag caa tac att taa cta tat cga tgt gga aat taa cct gta tcc agt caa cga cac atc gtg tac tcg gac gac cac tac cgg tct cag cga atc cat ctc aac gtc gga act aac tat tac cat gaa tca taa aga ttg tga tcc agt ctt tcg tgc aga ata ctt ctc tgt cct taa taa tgt agc aac ttc agg att ctt tac agg aga aaa tag ata tca gaa tac ttc aaa gat atg tac tct gaa ttt cga gat taa atg taa caa caa aga ttc atc ttc caa aca gtt aac gaa aac aaa gaa tga tac tat cat gcc gca ttc aga gac ggt aac tct agt ggg cga ctg tct atc tag cgt cga cat cta cat act ata tag taa tac caa tac tca aga cta cga aaa tga tac aat ctc tta tca tat ggg taa tgt tct cga tgt caa tag cca tat gcc cgc tag ttg cga tat aca taa act gat cac taa ttc cca gaa tcc cac cca ctt ata gta ag- -tt ttt tta cct ata aat aat aaa tac aat aat taa ttt ctc gta aaa gta gaa aat ata ttc taa ttt att ata tgg taa gaa agt aga atc atc tag aac agt aat caa tca ata gca atc atg aaa caa tat att gtc ctg gca tgc atg tgc cta gtg gca gct gct atg cct act agt ctt caa caa tct tca tcc tcg tgt act gaa gaa gaa aac aaa cat cat atg gga atc gat gtt att atc aaa gtc aca aag caa gac caa aca ccg acc aat gat aag att tgt caa tcc gta acg gaa gtt aca gag a-- --- --- --- --- --- -cc gaa gat gat gag gta tcc gaa gaa gtt gta aaa gga gat ccc acc act tat tac act atc gtc ggt gcg ggt ctt aac atg aac ttt gga ttc acc aaa tgc cca aag att tta tcc atc tcc gaa tcc tct gat gga aac act gtg aat act aga ttg tcc agc gtg tca ccg gga caa ggt aag gac tct ccc gcg atc acg cgt gaa gaa gct ctg gct atg atc aaa gac tgt gag atg tct atc gac atc aga tgt agc gaa gaa gag aaa gac agt gac atc aag acc cat cca gta ctt ggg tct aac atc tca cat aag aaa gtg agt tac aaa gat atc atc ggt tca acg atc gtt gat aca aaa tgt gtc aag aac cta gag ttt agc gta cgt atc gga gac atg tgt gag gaa tca tct gaa ctt gaa gtc aag gat gga ttc aag tat gtc gac gga tcg gca tct gaa ggt gca acc gat gat act tca ctc atc gat tca aca aaa ctc aaa gca tgt gtc tga atc gat aac tct att cat ctg aaa atg gat gag ttg ggt taa tcg aac gat tca gac acc gca cca cga att aaa aaa g-- -ac cgg gca cta tat tcc ggt ttg caa aac aaa aat att taa cta cat tca caa aaa gtt acc tct cgt tac ttc ttc ttt ctg ttt caa tat gtg ata cga tat gat cac tat tcg tat tct ct- --- --- tgg tct cat aaa aaa gtt tta c-- --a aaa aaa aaa aaa a-t att ttt att ctc ttt ctc tct tcg at- ggt ctc aca aaa ata tta aac ctc ttt ctg atg tct caa cta ttt cgt aaa cga taa cgt cca aca ata tat tct cgt aga gct tat caa cat cct tat acc aat cta ggt tgt cag aca att gca tca taa aat aat gtt tat aat tta cac gtt aac atc ata taa taa acg tat ata gtt aat att ttt gga ata taa atg atc tgt aaa atc cat gta ggg gac act gct cac gtt ttt tct cta gta cat aat ttc aca caa gtt ttt ata cag aca aat taa ttc tcg tcc ata ta- ttt taa aac att gac ttt tgt act aag aaa aat atc ttg act aac cat ctc ttt ctc tct tcg atg ggt ctc aca aaa ata tta aac ctc ttt ctg atg g-- --- --- --- --- --- --- --- --- --- --- --- --- --- --- --- --- --- --- --- --- --- --- --- --- --- --- --- --- --- --- --- --- --- --- --- --- --- --- --- --- --- --- --- --- --- --- agt cgt aaa aag ttt tta tcc ttt ctc tct tcg ata ggt ctc --- --- --- --- --- --- --- --- --- --- --- --- --- --- --- --- --- --- --- --- --- --- --- --- --- --- --- --- --- --- --- --- --- --- --- --- --- --- --- --- --- --- --- --- --- --- --- --- --- --- --- --- --- --- --- --- --- --- --- --- --- --- --- --- --- --- --- --- --- --- aca aaa ata tta aac ctc ttt ctg atg gtc tct ata aa

>ON609725.2_Monkeypox_virus_isolate_SLO_complete_genome

atc gag gac gac ttc aac tat gac agc tcg gtg gcg tct gcc agc gtg tac ata cga atg gca ttt cta aga aaa gtc tac ggt atc ctt -tc tac aat ttc ctt taa caa cgg caa cag ctg cag tat ttt tat act ttg aat gca -tc gga cat tta tac aag gga gtc ctg ttc taa tat tgg cat caa tgt tcg gat cta tag gct tga ttt tcg cat tga ctt tac aca gac atg aac atc ccc tga atc tgt aca tac ttt gtg gat tta cac tgt tag aat ctc taa cgc tgg cct ctg ttg tta ctt tct atg atg cac gta tcg tta tgc aag ctt tca tgt tga cta ctg cag tgt ttc ttg ctc tga cta cat ata ctc tac aat caa aga gag att tca gta aac ttg taa cag gat tgt ttg ctg ctt tct gga ttt taa ttt tgt cag gag tct tga gga taa agt tta aaa tag aat taa taa aga aca tat agg tca -tt ttt taa aca tgg ata gaa acc aag gtt gtt agt taa taa tat aca aga tat ttt ttc tca ctc tga tcc atg taa acc aag gac gag aga cac tct cat tcc tca ttc acg aca cca tta aaa atg gaa att aaa gcc ctc tat taa gca cag acg gct aca ggt cta cca tca ggt t-a cct tcg tct acc ttc aca atg gcc tct cct tgt gcc cag ttc agt ccc tgt cat tgc cac gct act aag gac tcc ctg aat acc gtg act gac gtc aga cat tgt ctg act gaa tac atc ctg tgg gtt tct cat aga tgg acc cat aga gaa agc gca ggg cct ctc tac agg ctt ctc atc tct ttc aga att gat gca atg gag cta ttt ggt agc gag ttg aag gag ttc tcg aat tca ctt ccg tgg gac aat atc gac aat tgc gtg gag atc att aaa tgt ttc atc aga aat gac tcc atg aaa acc gcc aaa gaa ctt tgt gca ata att gga ctt tgt act caa tca gct att gtc act gga aga gtc ttc aat gat aag tat atc gac ata cta ctt atg ctg cga aag att ctg aac gag aac gac tat ctc acc ctc ttg gat cat atc ctc act gct aaa tac taa atc tcc ttc atg ctc tct cac taa tac tct tac tca cta cac ttt tta tca tct tat gat gaa tga ttg cct tca tca ttt ttt cgt gga ata taa tat agg aat aat tag cac cag aat agc tat gga tat ctc gtt aag aat att ctc tat aag aga cat aat gta gac ata gtt att ata tcc ttc tta gat aag tgt tac gct act gga aag ttt cca tcg tta tta tta cat gaa gat gat ata att aaa cca aca ttg aga ttg gct ctt atg tta gct gga ttg aat tac tgt aat aaa tgc atc gag tat aga ggg ata tag caa ttc tcg ata ata gtc atg caa tat ttg aat gag act gat aat tta ggt aat aca gta ct- --- aca cac ata tct ttc tag att ata tat cgt taa aaa tct gta aga tgt ata ttt ctc ata agt atc cac tgt gta ata tta tta atg gat ata tag ata acg caa tag gga cta ata gta ttg taa aag ata taa tcg act att tac gta cat atc cag ata tct ata ttc cta cta gtt tgc tgc gta gtt gca tca ttg ata tgc atg -at tta tca gga ttc aga gat gaa tta cta agt aaa cta caa tcc cac aat aag taa gaa tca aat atc aaa aac tca ctt ttg att ttt cta gtc tta agt aat aca tat att tat taa tag acc tat gaa ata aaa aaa ggt aac aat gga ttc gcg tat agc tat tta cgt att agt atc ggc atc tct ttt gta tct tgt taa ttg tca caa act agt aca tta ctt caa tct gaa aat aaa tgg aag tga tat aac taa tac agc aga tat att gct gga caa tta tcc aat tat gac ctt tga tgg aaa gga tat tta tcc atc tat ctc gtt cat ggt cgg taa taa act ttt cct aga tct tta taa aaa tat ctt tga aga att ttt caa act att tcg agt atc tgt aag tag tca ata cga gga att aga ata tta tta ttc atg tga tta tac taa caa ccg tcc tac aat taa aca aca tta ctt tta taa cgg cga cga ata tac tga aat tga tag atc gaa aaa agc cac taa taa aaa cag ttg gtt aat tac ttc agg ctt tag act aca aaa atg gtt cga tag cga aga ttg tat aat tta tct cag atc ttt agt tag aag aat gga aga cag taa caa aaa cag taa aaa aac tta gta ctt aga tat cg- --a aaa aat ata ttt ttg tag act ctt gag aat aga agg aaa aca tgt aca taa tta taa aaa atg aaa atc aat ggc gaa taa gac agt gcg att cgc acc atg gag tcg gta gat ttc atg gct gtc gat gag cag ttt cac gac gac ctc gat ctt tgg tca tta tct ttg gta gat gat tat aaa aaa cat gga tta ggt gtt gac tgt tat gtt cta gaa cca gtt gtt gac agg aaa ata ttt gat aga ttt ctc ctt gaa cca att tgt gat cct gta gat gtt ctg tat gat tat ttt agg att cat aga gat aat att gat cag tat ata gta gat aga ctg ttt gca tat att aca tat aaa gat att ata tct gca tta gtg tca aag aat tat atg gaa gat att ttc tct ata att att aag aat tgt aat tct gtg caa gat ctc tta ctt tac tat cta tct aat gca tat gta gaa ata gac att gtt gat ctt atg gta gat cat ggg gct gta ata tat aaa ata gaa tgc ttg aat gcc tat ttt agg gga ata tgt aaa aag gaa agt agt gtt gtt gag ttt att ttg aat tgt ggt atc cca gat gaa aat gat gtt aaa tta gat cta tat aaa ata att cag tat act agg gga ttc ctt gta gat gaa ccc aca gta tta gaa att tat aag ctt tgt atc cca tat att gaa gat atc aat caa cta gat gct ggt gga agg acc ttg ctt tat cgc gct atc tat gca ggt tat ata gat tta gta tca tgg cta tta gaa aat gga gca aat gtc aac gca gta atg agt aat gga tat aca tgt ctt gac gtg gcc gtg gat agg gga tct gtc atc gcc cgt agg gaa gca cat ctt aaa ata tta gaa ata ttg ctt aga gaa cca ttg tct att gac tgt ata aaa tta gct ata ctt aat aat aca att gaa aac cat gat gtg ata aag ctc tgt atc aag tat ttt atg atg gta gat tat tca ctt tgt aat gtg tat gca tca tca ctc ttt gat tat ata att gat tgt aaa caa gaa ttg gag tac att agg cag atg aaa att cat aat aca acc atg tat gag tta atc tat aat aga gac aaa aac aag cat gct tcc cat att cta cat agg tat tct aaa cat cca gtt ttg aca cag tgt atc act aaa gga ttc aag att tac aca gaa gta acc gag cag gtc act aaa gct cta aac aga cgt gct cta ata gat gag ata ata aac aat gta tca act gat gac aat ctc cta tca aaa ctt cca tta gaa att agg gat cta att gtt tca caa gct gtc ata tag agt tct atc cac cca cct ttc ttg aaa tga gtt aat a-- --- --- --- --- --- --- --- --- --- --- --- --- --- --- --- --- --- --- --- --- --- --- --- --- --- --- --- --- --- --- --- --- --- --- --- --- --- --- --- --- --- --- --- --- --- --- --- --- --- --- --- --- --- --- --- --- --- --- --- --- --- --- --- --- --- --- --- --- --- --- --- --- --- --- --- --- --- --- --- --- --- --- --- --- --- --- --- --- --- --- --- --- --- --- --- --- --- --- --- --- --- --- --- --- -ag tca taa gtt agt taa --- --- --- --- --- -gt cat aag tta gtt aag tca t-- --- --- --- --- --- --- --- --- --- --- --- --- --- --- --- -aa gtt agt taa gtc ata agt tag tta agt cat aag tta gtt aag tca taa gtt agt taa gtc ata agt tag tta agt cat aag tta gtt tat agt cta aca ctt cta att ttt ata cct tga tct ttt tct cta att atg aaa aag taa atc att atg aag atg gat g-a aat gga cga gat tgt gcg cat cgt taa cga tag tat gtg gta cgt acc taa cgc att tat gga cga cgg tga taa tga agg tca cat ttc tgt caa taa tgt ctg tca tat gta tct cgc att ctt tga tgt gga tat atc atc tca tct gtt taa att agt tat taa aca ctg cga tct gaa taa acg act aaa atg tgg taa ctc tcc att aca ttg cta tac gat gaa tac acg att taa tcc atc tgt att aaa gat att gtt acg cca cgg cat gcg taa ctt tga tag caa gga taa aaa agg aca tat tcc tct aca cca cta tct gat tca ttc act atc aat cga taa caa gat ctt tga tat act aac gga ccc cat tga tga ctt tag taa atc atc cga tct att gct gtg tta tct tag ata taa att caa tgg gag ctt aaa cta tta cgt tct gta caa att att gac taa agg atc tga ccc taa ttg cgt cga tga gga tgg act cac ttc tct tca tta cta ctg taa aca cat atc cgc gtt cca cga aag caa tta tta caa gtc aaa gag tca cac taa gat gcg agc tga gaa gcg att cat cta cgc gat aat aga tca tgg agc aaa cat taa cgc ggt tac gaa aat cgg aaa tac gcc gtt aca cac tta cct tca aca gta tac caa aca tag tcc tcg tgt ggt gta tgc tct ttt atc tcg agg agc cga tac gag gat acg taa taa tct taa ttg tac acc cat cat gga ata cat aaa gaa cga ttg tgc aac agg tca tat tct cat aat gtt act caa ttg gca cga aca aaa ata cgg gaa att aca aaa gga aga agg aca aca tct act tta tct att cat aaa aca taa tca agg ata tgg aag tcg ctc tct caa tat act acg gta tct act aga tag att cga cat tca gaa aga cga ata cta taa tac aat gac tcc tct tca tac cgc ctt cca gaa ttg caa taa caa tgt tgc ctc ata cct cgt ata cat tgg ata cga cat caa cct tcc gac taa aga cga taa gac agt att cga ctt ggt gtt tga aaa cag aaa cat tat ata caa ggc gga tgt cgt taa tga cat tat cca cca cag act gaa agt atc tct acc tat gat taa atc gtt gtt cta caa gat gtc gga gtt ctc tcc cta cga cga tca cta cgt aaa gaa gat aat agc cta ctg cct att aag gga cga gtc att tgc gga act aca tac taa att ctg ttt aaa cga gga cta taa aag tgt att tat gaa aaa tat atc att cga taa gat aga ttc cat cat cga aaa atg tag tcg tga cat aag tct cct caa aga gat tcg aat ctc aga cac cga ctt gta tac ggt att gag aac aga aga cat tcg gta tca cac ata tct cga agc cat aca ttc aga caa acg cat ttc att tcc cat gta cga cga tct cat aga aca gtg tca tct atc gat gga gca taa aag taa act cgt cga caa agc act caa taa att aga gtc tac cat cga tag tca atc tag act atc gta ttt gcc tcc gga aat tat gcg caa tat cat aac caa gct aag cga cta cca tct aaa cag tat gtt gta cgg aaa gaa cca tta caa ata tta tcc atg ata gaa aga aaa tat tta aaa aat aat cta tat gat tgg aga agt agg aaa caa aca gta aca aga cga cga tta cta c-a tta tta aat cat gag gtc cgt att ata ctc gta tat att gtt tct ctc atg tat aat aat aaa cgg aag aga tat agc acc aca tgc acc atc caa tgg aaa gtg taa aga caa cga ata cag aag ccg taa tct atg ttg tct atc gtg tcc tcc ggg aac tta cgc ttt cag att atg tga tag caa gac taa tac aca atg tac acc gtg tgg ttc gga tac ctt tac atc tca caa taa tca ttt aca ggc ttg tct aag ttg taa cgg aag atg tga tag taa tca ggt aga gac gcg atc gtg taa cac gac tca caa tag aat ctg tga atg ctc tcc agg ata tta ttg tct tct caa agg agc atc agg gtg tag aac atg tat ttc taa aac aaa gtg tgg aat agg ata cgg agt atc cgg ata cac gtc tac cgg aga cgt cat ctg ttc tcc gtg tgg tcc cgg aac ata ttc tca cac cgt ctc ttc cac aga taa atg cga acc cgt cgt aac cag caa tac att taa cta tat cga tgt gga aat taa cct gta tcc agt caa cga cac atc gtg tac tcg gac gac cac tac cgg tct cag cga atc cat ctc aac gtc gga act aac tat tac cat gaa tca taa aga ttg tga tcc agt ctt tcg tgc aga ata ctt ctc tgt cct taa taa tgt agc aac ttc agg att ctt tac agg aga aaa tag ata tca gaa tac ttc aaa gat atg tac tct gaa ttt cga gat taa atg taa caa caa aga ttc atc ttc caa aca gtt aac gaa aac aaa gaa tga tac tat cat gcc gca ttc aga gac ggt aac tct agt ggg cga ctg tct atc tag cgt cga cat cta cat act ata tag taa tac caa tac tca aga cta cga aaa tga tac aat ctc tta tca tat ggg taa tgt tct cga tgt caa tag cca tat gcc cgc tag ttg cga tat aca taa act gat cac taa ttc cca gaa tcc cac cca ctt ata gta ag- -tt ttt tta cct ata aat aat aaa tac aat aat taa ttt ctc gta aaa gta gaa aat ata ttc taa ttt att ata tgg taa gaa agt aga atc atc tag aac agt aat caa tca ata gca atc atg aaa caa tat att gtc ctg gca tgc atg tgc cta gtg gca gct gct atg cct act agt ctt caa caa tct tca tcc tcg tgt act gaa gaa gaa aac aaa cat cat atg gga atc gat gtt att atc aaa gtc aca aag caa gac caa aca ccg acc aat gat aag att tgt caa tcc gta acg gaa gtt aca gag a-- --- --- --- --- --- -cc gaa gat gat gag gta tcc gaa gaa gtt gta aaa gga gat ccc acc act tat tac act atc gtc ggt gcg ggt ctt aac atg aac ttt gga ttc acc aaa tgc cca aag att tta tcc atc tcc gaa tcc tct gat gga aac act gtg aat act aga ttg tcc agc gtg tca ccg gga caa ggt aag gac tct ccc gcg atc acg cgt gaa gaa gct ctg gct atg atc aaa gac tgt gag atg tct atc gac atc aga tgt agc gaa gaa gag aaa gac agt gac atc aag acc cat cca gta ctt ggg tct aac atc tca cat aag aaa gtg agt tac aaa gat atc atc ggt tca acg atc gtt gat aca aaa tgt gtc aag aac cta gag ttt agc gta cgt atc gga gac atg tgt gag gaa tca tct gaa ctt gaa gtc aag gat gga ttc aag tat gtc gac gga tcg gca tct gaa ggt gca acc gat gat act tca ctc atc gat tca aca aaa ctc aaa gca tgt gtc tga atc gat aac tct att cat ctg aaa atg gat gag ttg ggt taa tcg aac gat tca gac acc gca cca cga att aaa aaa g-- -ac cgg gca cta tat tcc ggt ttg caa aac aaa aat att taa cta cat tca caa aaa gtt acc tct cgt tac ttc ttc ttt ctg ttt caa tat gtg ata cga tat gat cac tat tcg tat tct ct- --- --- tgg tct cat aaa aaa gtt tta c-- --a aaa aaa aaa aaa a-t att ttt att ctc ttt ctc tct tcg at- ggt ctc aca aaa ata tta aac ctc ttt ctg atg tct caa cta ttt cgt aaa cga taa cgt cca aca ata tat tct cgt aga gct tat caa cat cct tat acc aat cta ggt tgt cag aca att gca tca taa aat aat gtt tat aat tta cac gtt aac atc ata taa taa acg tat ata gtt aat att ttt gga ata taa atg atc tgt aaa atc cat gta ggg gac act gct cac gtt ttt tct cta gta cat aat ttc aca caa gtt ttt ata cag aca aat taa ttc tcg tcc ata ta- ttt taa aac att gac ttt tgt act aag aaa aat atc ttg act aac cat ctc ttt ctc tct tcg atg ggt ctc aca aaa ata tta aac ctc ttt ctg atg g-- --- --- --- --- --- --- --- --- --- --- --- --- --- --- --- --- --- --- --- --- --- --- --- --- --- --- --- --- --- --- --- --- --- --- --- --- --- --- --- --- --- --- --- --- --- --- agt cgt aaa aag ttt tta tcc ttt ctc tct tcg ata ggt ctc --- --- --- --- --- --- --- --- --- --- --- --- --- --- --- --- --- --- --- --- --- --- --- --- --- --- --- --- --- --- --- --- --- --- --- --- --- --- --- --- --- --- --- --- --- --- --- --- --- --- --- --- --- --- --- --- --- --- --- --- --- --- --- --- --- --- --- --- --- --- aca aaa ata tta aac ctc ttt ctg atg gtc tct ata aa

>ON622713.1_Monkeypox_virus_isolate_MPX/UZ_REGA_2/Belgium/2022_complete_genome

atc gag gac gac ttc aac tat gac agc tcg gtg gcg tct gcc agc gtg tac ata cga atg gca ttt cta aga aaa gtc tac ggt atc ctt -tc tac aat ttc ctt taa caa cgg caa cag ctg cag tat ttt tat act ttg aat gca -tc gga cat tta tac aag gga gtc ctg ttc taa tat tgg cat caa tgt tcg gat cta tag gct tga ttt tcg cat tga ctt tac aca gac atg aac atc ccc tga atc tgt aca tac ttt gtg gat tta cac tgt tag aat ctc taa cgc tgg cct ctg ttg tta ctt tct atg atg cac gta tcg tta tgc aag ctt tca tgt tga cta ctg cag tgt ttc ttg ctc tga cta cat ata ctc tac aat caa aga gag att tca gta aac ttg taa cag gat tgt ttg ctg ctt tct gga ttt taa ttt tgt cag gag tct tga gga taa agt tta aaa tag aat taa taa aga aca tat agg tca -tt ttt taa aca tgg ata gaa acc aag gtt gtt agt taa taa tat aca aga tat ttt ttc tca ctc tga tcc atg taa acc aag gac gag aga cac tct cat tcc tca ttc acg aca cca tta aaa atg gaa att aaa gcc ctc tat taa gca cag acg gct aca ggt cta cca tca ggt t-a cct tcg tct acc ttc aca atg gcc tct cct tgt gcc cag ttc agt ccc tgt cat tgc cac gct act aag gac tcc ctg aat acc gtg act gac gtc aga cat tgt ctg act gaa tac atc ctg tgg gtt tct cat aga tgg acc cat aga gaa agc gca ggg cct ctc tac agg ctt ctc atc tct ttc aga att gat gca atg gag cta ttt ggt agc gag ttg aag gag ttc tcg aat tca ctt ccg tgg gac aat atc gac aat tgc gtg gag atc att aaa tgt ttc atc aga aat gac tcc atg aaa acc gcc aaa gaa ctt tgt gca ata att gga ctt tgt act caa tca gct att gtc act gga aga gtc ttc aat gat aag tat atc gac ata cta ctt atg ctg cga aag att ctg aac gag aac gac tat ctc acc ctc ttg gat cat atc ctc act gct aaa tac taa atc tcc ttc atg ctc tct cac taa tac tct tac tca cta cac ttt tta tca tct tat gat gaa tga ttg cct tca tca ttt ttt cgt gga ata taa tat agg aat aat tag cac cag aat agc tat gga tat ctc gtt aag aat att ctc tat aag aga cat aat gta gac ata gtt att ata tcc ttc tta gat aag tgt tac gct act gga aag ttt cca tcg tta tta tta cat gaa gat gat ata att aaa cca aca ttg aga ttg gct ctt atg tta gct gga ttg aat tac tgt aat aaa tgc atc gag tat aga ggg ata tag caa ttc tcg ata ata gtc atg caa tat ttg aat gag act gat aat tta ggt aat aca gta ct- --- aca cac ata tct ttc tag att ata tat cgt taa aaa tct gta aga tgt ata ttt ctc ata agt atc cac tgt gta ata tta tta atg gat ata tag ata acg caa tag gga cta ata gta ttg taa aag ata taa tcg act att tac gta cat atc cag ata tct ata ttc cta cta gtt tgc tgc gta gtt gca tca ttg ata tgc atg -at tta tca gga ttc aga gat gaa tta cta agt aaa cta caa tcc cac aat aag taa gaa tca aat atc aaa aac tca ctt ttg att ttt cta gtc tta agt aat aca tat att tat taa tag acc tat gaa ata aaa aaa ggt aac aat gga ttc gcg tat agc tat tta cgt att agt atc ggc atc tct ttt gta tct tgt taa ttg tca caa act agt aca tta ctt caa tct gaa aat aaa tgg aag tga tat aac taa tac agc aga tat att gct gga caa tta tcc aat tat gac ctt tga tgg aaa gga tat tta tcc atc tat ctc gtt cat ggt cgg taa taa act ttt cct aga tct tta taa aaa tat ctt tga aga att ttt cag act att tcg agt atc tgt aag tag tca ata cga gga att aga ata tta tta ttc atg tga tta tac taa caa ccg tcc tac aat taa aca aca tta ctt tta taa cgg cga cga ata tac tga aat tga tag atc gaa aaa agc cac taa taa aaa cag ttg gtt aat tac ttc agg ctt tag act aca aaa atg gtt cga tag cga aga ttg tat aat tta tct cag atc ttt agt tag aag aat gga aga cag taa caa aaa cag taa aaa aac tta gta ctt aga tat cg- --a aaa aat ata ttt ttg tag act ctt gag aat aga agg aaa aca tgt aca taa tta taa aaa atg aaa atc aat ggc gaa taa gac agt gcg att cgc acc atg gag tcg gta gat ttc atg gct gtc gat gag cag ttt cac gac gac ctc gat ctt tgg tca tta tct ttg gta gat gat tat aaa aaa cat gga tta ggt gtt gac tgt tat gtt cta gaa cca gtt gtt gac agg aaa ata ttt gat aga ttt ctc ctt gaa cca att tgt gat cct gta gat gtt ctg tat gat tat ttt agg att cat aga gat aat att gat cag tat ata gta gat aga ctg ttt gca tat att aca tat aaa gat att ata tct gca tta gtg tca aag aat tat atg gaa gat att ttc tct ata att att aag aat tgt aat tct gtg caa gat ctc tta ctt tac tat cta tct aat gca tat gta gaa ata gac att gtt gat ctt atg gta gat cat ggg gct gta ata tat aaa ata gaa tgc ttg aat gcc tat ttt agg gga ata tgt aaa aag gaa agt agt gtt gtt gag ttt att ttg aat tgt ggt atc cca gat gaa aat gat gtt aaa tta gat cta tat aaa ata att cag tat act agg gga ttc ctt gta gat gaa ccc aca gta tta gaa att tat aag ctt tgt atc cca tat att gaa gat atc aat caa cta gat gct ggt gga agg acc ttg ctt tat cgc gct atc tat gca ggt tat ata gat tta gta tca tgg cta tta gaa aat gga gca aat gtc aac gca gta atg agt aat gga tat aca tgt ctt gac gtg gcc gtg gat agg gga tct gtc atc gcc cgt agg gaa gca cat ctt aaa ata tta gaa ata ttg ctt aga gaa cca ttg tct att gac tgt ata aaa tta gct ata ctt aat aat aca att gaa aac cat gat gtg ata aag ctc tgt atc aag tat ttt atg atg gta gat tat tca ctt tgt aat gtg tat gca tca tca ctc ttt gat tat ata att gat tgt aaa caa gaa ttg gag tac att agg cag atg aaa att cat aat aca acc atg tat gag tta atc tat aat aga gac aaa aac aag cat gct tcc cat att cta cat agg tat tct aaa cat cca gtt ttg aca cag tgt atc act aaa gga ttc aag att tac aca gaa gta acc gag cag gtc act aaa gct cta aac aga cgt gct cta ata gat gag ata ata aac aat gta tca act gat gac aat ctc cta tca aaa ctt cca tta gaa att agg gat cta att gtt tca caa gct gtc ata tag agt tct atc cac cca cct ttc ttg aaa tga gtt aat agt cat aag tta gtt aag tca taa gtt agt taa gtc ata agt tag tta agt cat aag tt- --- --- --- --- --- --- --- --- --- --- --- --- --- --- --- --- --- --- --- --- --- --- --- --- --- --- --- --- --- --- --- --- --- --- --- --- --- --- --- --- --- --- --- --- --- --- --- --- --- --- --- --- --- --- --- --- --- agt taa gtc ata agt tag tta --- --- --- --- --- -ag tca taa gtt agt taa gtc ata agt tag tta agt cat aag tta gtt aag tca taa gtt agt taa --- --- --- --- --- -gt cat aag tta gtt aag tca t-- --- --- --- --- --- --- --- --- --- --- --- --- --- --- --- -aa gtt agt taa gtc ata agt tag tta agt cat aag tta gtt aag tca taa gtt agt taa gtc ata agt tag tta agt cat aag tta gtt tat agt cta aca ctt cta att ttt ata cct tga tct ttt tct cta att ata aaa aag taa atc att atg aag atg gat g-a aat gga cga gat tgt gcg cat cgt taa cga tag tat gtg gta cgt acc taa cgc att tat gga cga cgg tga taa tga agg tca cat ttc tgt caa taa tgt ctg tca tat gta tct cgc att ctt tga tgt gga tat atc atc tca tct gtt taa att agt tat taa aca ctg cga tct gaa taa acg act aaa atg tgg taa ctc tcc att aca ttg cta tac gat gaa tac acg att taa tcc atc tgt att aaa gat att gtt acg cca cgg cat gcg taa ctt tga tag caa gga taa aaa agg aca tat tcc tct aca cca cta tct gat tca ttc act atc aat cga taa caa gat ctt tga tat act aac gga ccc cat tga tga ctt tag taa atc atc cga tct att gct gtg tta tct tag ata taa att caa tgg gag ctt aaa cta tta cgt tct gta caa att att gac taa agg atc tga ccc taa ttg cgt cga tga gga tgg act cac ttc tct tca tta cta ctg taa aca cat atc cgc gtt cca cga aag caa tta tta caa gtc aaa gag tca cac taa gat gcg agc tga gaa gcg att cat cta cgc gat aat aga tca tgg agc aaa cat taa cgc ggt tac gaa aat cgg aaa tac gcc gtt aca cac tta cct tca aca gta tac caa aca tag tcc tcg tgt ggt gta tgc tct ttt atc tcg agg agc cga tac gag gat acg taa taa tct taa ttg tac acc cat cat gga ata cat aaa gaa cga ttg tgc aac agg tca tat tct cat aat gtt act caa ttg gca cga aca aaa ata cgg gaa att aca aaa gga aga agg aca aca tct act tta tct att cat aaa aca taa tca agg ata tgg aag tcg ctc tct caa tat act acg gta tct act aga tag att cga cat tca gaa aga cga ata cta taa tac aat gac tcc tct tca tac cgc ctt cca gaa ttg caa taa caa tgt tgc ctc ata cct cgt ata cat tgg ata cga cat caa cct tcc gac taa aga cga taa gac agt att cga ctt ggt gtt tga aaa cag aaa cat tat ata caa ggc gga tgt cgt taa tga cat tat cca cca cag act gaa agt atc tct acc tat gat taa atc gtt gtt cta caa gat gtc gga gtt ctc tcc cta cga cga tca cta cgt aaa gaa gat aat agc cta ctg cct att aag gga cga gtc att tgc gga act aca tac taa att ctg ttt aaa cga gga cta taa aag tgt att tat gaa aaa tat atc att cga taa gat aga ttc cat cat cga aaa atg tag tcg tga cat aag tct cct caa aga gat tcg aat ctc aga cac cga ctt gta tac ggt att gag aac aga aga cat tcg gta tca cac ata tct cga agc cat aca ttc aga caa acg cat ttc att tcc cat gta cga cga tct cat aga aca gtg tca tct atc gat gga gca taa aag taa act cgt cga caa agc act caa taa att aga gtc tac cat cga tag tca atc tag act atc gta ttt gcc tcc gga aat tat gcg caa tat cat aac caa gct aag cga cta cca tct aaa cag tat gtt gta cgg aaa gaa cca tta caa ata tta tcc atg ata gaa aga aaa tat tta aaa aat aat cta tat gat tgg aga agt agg aaa caa aca gta aca aga cga cga tta cta c-a tta tta aat cat gag gtc cgt att ata ctc gta tat att gtt tct ctc atg tat aat aat aaa cgg aag aga tat agc acc aca tgc acc atc caa tgg aaa gtg taa aga caa cga ata cag aag ccg taa tct atg ttg tct atc gtg tcc tcc ggg aac tta cgc ttt cag att atg tga tag caa gac taa tac aca atg tac acc gtg tgg ttc gga tac ctt tac atc tca caa taa tca ttt aca ggc ttg tct aag ttg taa cgg aag atg tga tag taa tca ggt aga gac gcg atc gtg taa cac gac tca caa tag aat ctg tga atg ctc tcc agg ata tta ttg tct tct caa agg agc atc agg gtg tag aac atg tat ttc taa aac aaa gtg tgg aat agg ata cgg agt atc cgg ata cac gtc tac cgg aga cgt cat ctg ttc tcc gtg tgg tcc cgg aac ata ttc tca cac cgt ctc ttc cac aga taa atg cga acc cgt cgt aac cag caa tac att taa cta tat cga tgt gga aat taa cct gta tcc agt caa cga cac atc gtg tac tcg gac gac cac tac cgg tct cag cga atc cat ctc aac gtc gga act aac tat tac cat gaa tca taa aga ttg tga tcc agt ctt tcg tgc aga ata ctt ctc tgt cct taa taa tgt agc aac ttc agg att ctt tac agg aga aaa tag ata tca gaa tac ttc aaa gat atg tac tct gaa ttt cga gat taa atg taa caa caa aga ttc atc ttc caa aca gtt aac gaa aac aaa gaa tga tac tat cat gcc gca ttc aga gac ggt aac tct agt ggg cga ctg tct atc tag cgt cga cat cta cat act ata tag taa tac caa tac tca aga cta cga aaa tga tac aat ctc tta tca tat ggg taa tgt tct cga tgt caa tag cca tat gcc cgc tag ttg cga tat aca taa act gat cac taa ttc cca gaa tcc cac cca ctt ata gta ag- -tt ttt tta cct ata aat aat aaa tac aat aat taa ttt ctc gta aaa gta gaa aat ata ttc taa ttt att ata tgg taa gaa agt aga atc atc tag aac agt aat caa tca ata gca atc atg aaa caa tat att gtc ctg gca tgc atg tgc cta gtg gca gct gct atg cct act agt ctt caa caa tct tca tcc tcg tgt act gaa gaa gaa aac aaa cat cat atg gga atc gat gtt att atc aaa gtc aca aag caa gac caa aca ccg acc aat gat aag att tgt caa tcc gta acg gaa gtt aca gag a-- --- --- --- --- --- -cc gaa gat gat gag gta tcc gaa gaa gtt gta aaa gga gat ccc acc act tat tac act atc gtc ggt gcg ggt ctt aac atg aac ttt gga ttc acc aaa tgc cca aag att tta tcc atc tcc gaa tcc tct gat gga aac act gtg aat act aga ttg tcc agc gtg tca ccg gga caa ggt aag gac tct ccc gcg atc acg cgt gaa gaa gct ctg gct atg atc aaa gac tgt gag atg tct atc gac atc aga tgt agc gaa gaa gag aaa gac agt gac atc aag acc cat cca gta ctt ggg tct aac atc tca cat aag aaa gtg agt tac aaa gat atc atc ggt tca acg atc gtt gat aca aaa tgt gtc aag aac cta gag ttt agc gta cgt atc gga gac atg tgt gag gaa tca tct gaa ctt gaa gtc aag gat gga ttc aag tat gtc gac gga tcg gca tct gaa ggt gca acc gat gat act tca ctc atc gat tca aca aaa ctc aaa gca tgt gtc tga atc gat aac tct att cat ctg aaa atg gat gag ttg ggt taa tcg aac gat tca gac acc gca cca cga att aaa aaa g-- -ac cgg gca cta tat tcc ggt ttg caa aac aaa aat att taa cta cat tca caa aaa gtt acc tct cgt tac ttc ttc ttt ctg ttt caa tat gtg ata cga tat gat cac tat tcg tat tct ct- --- --- tgg tct cat aaa aaa gtt tta c-- --a aaa aaa aaa aaa a-t att ttt att ctc ttt ctc tct tcg at- ggt ctc aca aaa ata tta aac ctc ttt ctg atg tct caa cta ttt cgt aaa cga taa cgt cca aca ata tat tct cgt aga gct tat caa cat cct tat acc aat cta ggt tgt cag aca att gca tca taa aat aat gtt tat aat tta cac gtt aac atc ata taa taa acg tat ata gtt aat att ttt gga ata taa atg atc tgt aaa atc cat gta ggg gac act gct cac gtt ttt tct cta gta cat aat ttc aca caa gtt ttt ata cag aca aat taa ttc tcg tcc ata ta- ttt taa aac att gac ttt tgt act aag aaa aat atc ttg act aac cat ctc ttt ctc tct tcg atg ggt ctc aca aaa ata tta aac ctc ttt ctg atg g-- --- --- --- --- --- --- --- --- --- --- --- --- --- --- --- --- --- --- --- --- --- --- --- --- --- --- --- --- --- --- --- --- --- --- --- --- --- --- --- --- --- --- --- --- --- --- agt cgt aaa aag ttt tta tcc ttt ctc tct tcg ata ggt ctc --- --- --- --- --- --- --- --- --- --- --- --- --- --- --- --- --- --- --- --- --- --- --- --- --- --- --- --- --- --- --- --- --- --- --- --- --- --- --- --- --- --- --- --- --- --- --- --- --- --- --- --- --- --- --- --- --- --- --- --- --- --- --- --- --- --- --- --- --- --- aca aaa ata tta aac ctc ttt ctg atg gtc tct ata aa

>ON676707.1_Monkeypox_virus_isolate_MPXV_USA_2021_TX_complete_genome

atc gag gac gac ttc aac tat gac agc tcg gtg gcg tct gcc agc gtg tac ata cga atg gca ttt cta aga aaa gtc tac ggt atc ctt -tc tac aat ttc ctt taa caa cgg caa cag ctg cag tat ttt tat act ttg aat gca -tc gga cat tta tac aag gga gtc ctg ttc taa tat tgg cat caa tgt tcg gat cta tag gct tga ttt tcg cat tga ctt tac aca gac atg aac atc ccc tga atc tgt aca tac ttt gtg gat tta cac tgt tag aat ctc taa cgc tgg cct ctg ttg tta ctt tct atg atg cac gta tcg tta tgc aag ctt tca tgt tga cta ctg cag tgt ttc ttg ctc tga cta cat ata ctc tac aat caa aga gag att tca gta aac ttg taa cag gat tgt ttg ctg ctt tct gga ttt taa ttt tgt cag gag tct tga gga taa agt tta aaa tag aat taa taa aga aca tat agg tca -tt ttt taa aca tgg ata gaa acc aag gtt gtt agt taa taa tat aca aga tat ttt ttc tca ctc tga tcc atg taa acc aag gac gag aga cac tct cat tcc tca ttc acg aca cca tta aaa atg gaa att aaa gcc ctc tat taa gca cag acg gct aca ggt cta cca tca ggt t-a cct tcg tct acc ttc aca atg gcc tct cct tgt gcc cag ttc agt ccc tgt cat tgc cac gct act aag gac tcc ctg aat acc gtg act gac gtc aga cat tgt ctg act gaa tac atc ctg tgg gtt tct cat aga tgg acc cat aga gaa agc gca ggg cct ctc tac agg ctt ctc atc tct ttc aga att gat gca atg gag cta ttt ggt agc gag ttg aag gag ttc tca aat tca ctt ccg tgg gac aat atc gac aat tgc gtg gag atc att aaa tgt ttc att aga aat gac tcc atg aaa acc gcc aaa gaa ctt tgt gca ata att gga ctt tgt act caa tca gct att gtc act gga aga gtc ttc aat gat aag tat atc gac ata cta ctt atg ctg cga aag att ctg aac gag aac gac tat ctc acc ctc ttg gat cat atc ctc act gct aaa tac taa atc tcc ttc atg ctc tct cac taa tac tct tac tca cta cac ttt tta tca tct tat gat gaa tga ttg cct tca tca ttt ttt cgt gga ata taa tat agg aat aat tag cac cag aat agc tat gga tat ctc gtt aag aat att ctc tat aag aga cat aat gta gac ata gtt att ata tcc ttc tta gat aag tgt tac gct act gga aag ttt cca tcg tta tta tta cat gaa gat gat ata att aaa cca aca ttg aga ttg gct ctt atg tta gct gga ttg aat tac tgt aat aaa tgc atc gag tat aga ggg ata tag caa ttc tcg ata ata gtc atg caa tat ttg aat gag act gat aat tta ggt aat aca gta ct- --- aca cac ata tct ttc tag att ata tat cgt taa aaa tct gta aga tgt ata ttt ctc ata agt atc cac tgt gta ata tta tta atg gat ata tag ata acg caa tag gga cta ata gta ttg taa aag ata taa tcg act att tac gta cat atc cag ata tct ata ttc cta cta gtt tgc tgc gta gtt gca tca ttg ata tgc atg -at tta tca gga ttc aga gat gaa tta cta agt aaa cta caa tcc cac aat aag taa gaa tca aat atc aaa aac tca ctt ttg att ttt cta gtc tta agt aat aca tat att tat taa tag acc tat gaa ata aaa aaa ggt aac aat gga ttc gcg tat agc tat tta cgt att agt atc ggc atc tct ttt gta tct tgt taa ttg tca caa act agt aca tta ctt caa tct gaa aat aaa tgg aag tga tat aac taa tac agc aga tat att gct gga caa tta tcc aat tat gac ctt tga tgg aaa gga tat tta tcc atc tat ctc gtt cat ggt cgg taa taa act ttt cct aga tct tta taa aaa tat ctt tga aga att ttt cag act att tcg agt atc tgt aag tag tca ata cga gga att aga ata tta tta ttc atg tga tta tac taa caa ccg tcc tac aat taa aca aca tta ctt tta taa cgg cga cga ata tac tga aat tga tag atc gaa aaa agc cac taa taa aaa cag ttg gtt aat tac ttc agg ctt tag act aca aaa atg gtt cga tag cga aga ttg tat aat tta tct cag atc ttt agt tag aag aat gga aga cag taa caa aaa cag taa aaa aac tta gta ctt aga tat cg- --a aaa aat ata ttt ttg tag act ctt gag aat aga agg aaa aca tgt aca taa tta taa aaa atg aaa atc aat ggc gaa taa gac agt gcg att cgc acc atg gag tcg gta gat ttc atg gct gtc gat gag cag ttt cac gac gac ctc gat ctt tgg tca tta tct ttg gta gat gat tat aaa aaa cat gga tta ggt gtt gac tgt tat gtt cta gaa cca gtt gtt gac agg aaa ata ttt gat aga ttt ctc ctt gaa cca att tgt gat cct gta gat gtt ctg tat gat tat ttt agg att cat aga gat aat att gat cag tat ata gta gat aga ctg ttt gca tat att aca tat aaa gat att ata tct gca tta gtg tca aag aat tat atg gaa gat att ttc tct ata att att aag aat tgt aat tct gtg caa gat ctc tta ctt tac tat cta tct aat gca tat gta gaa ata gac att gtt gat ctt atg gta gat cat ggg gct gta ata tat aaa ata gaa tgc ttg aat gcc tat ttt agg gga ata tgt aaa aag gaa agt agt gtt gtt gag ttt att ttg aat tgt ggt atc cca gat gaa aat gat gtt aaa tta gat cta tat aaa ata att cag tat act agg gga ttc ctt gta gat gaa ccc aca gta tta gaa att tat aag ctt tgt atc cca tat att gaa gat atc aat caa cta gat gct ggt gga agg acc ttg ctt tat cgc gct atc tat gca ggt tat ata gat tta gta tca tgg cta tta gaa aat gga gca aat gtc aac gca gta atg agt aat gga tat aca tgt ctt gac gtg gcc gtg gat agg gga tct gtc atc gcc cgt agg gaa gca cat ctt aaa ata tta gaa ata ttg ctt aga gaa cca ttg tct att gac tgt ata aaa tta gct ata ctt aat aat aca att gaa aac cat gat gtg ata aag ctc tgt atc aag tat ttt atg atg gta gat tat tca ctt tgt aat gtg tat gca tca tca ctc ttt gat tat ata att gat tgt aaa caa gaa ttg gag tac att agg cag atg aaa att cat aat aca acc atg tat gag tta atc tat aat aga gac aaa aac aag cat gct tcc cat att cta cat agg tat tct aaa cat cca gtt ttg aca cag tgt atc act aaa gga ttc aag att tac aca gaa gta acc gag cag gtc act aaa gct cta aac aga cgt gct cta ata gat gag ata ata aac aat gta tca act gat gac aat ctc cta tca aaa ctt cca tta gaa att agg gat cta att gtt tca caa gct gtc ata tag agt tct atc cac cca cct ttc ttg aaa tga gtt aat --- --- --- --- --- --- --- --- --- --- --- --- --- --- --- --- --- --- --- --- --- --- --- --- --- --- --- --- --- --- --- --- --- --- --- --- --- --- --- --- --- --- --- --- --- --- --- --- --- --- --- --- --- --- --- --- --- --- --- --- --- --- --- --- --- --- --- --- --- --- --- --- --- --- --- --- --- --- --- --- --- --- --- --- --- --- --- --- --- --- --- --- --- --- --- --- --- --- --- --- --- --- --- --- --- -ag tca taa gtt agt taa --- --- --- --- --- -gt cat aag tta gtt aag tca t-- --- --- --- --- --- --- --- --- --- --- --- --- --- --- --- -aa gtt agt taa gtc ata agt tag tta agt cat aag tta gtt aag tca taa gtt agt taa gtc ata agt tag tta agt cat aag tta gtt tat agt cta aca ctt cta att ttt ata cct tga tct ttt tct cta att atg aaa aag taa atc att atg aag atg gat g-a aat gga cga gat tgt gcg cat cgt taa cga tag tat gtg gta cgt acc taa cgc att tat gga cga cgg tga taa tga agg tca cat ttc tgt caa taa tgt ctg tca tat gta tct cgc att ctt tga tgt gga tat atc atc tca tct gtt taa att agt tat taa aca ctg cga tct gaa taa acg act aaa atg tgg taa ctc tcc att aca ttg cta tac gat gaa tac acg att taa tcc atc tgt att aaa gat att gtt acg cca cgg cat gcg taa ctt tga tag caa gga taa aaa agg aca tat tcc tct aca cca cta tct gat tca ttc act atc aat cga taa caa gat ctt tga tat act aac gga ccc cat tga tga ctt tag taa atc atc cga tct att gct gtg tta tct tag ata taa att caa tgg gag ctt aaa cta tta cgt tct gta caa att att gac taa agg atc tga ccc taa ttg cgt cga tga gga tgg act cac ttc tct tca tta cta ctg taa aca cat atc cgc gtt cca cga aag caa tta tta caa gtc aaa gag tca cac taa gat gcg agc tga gaa gcg att cat cta cgc gat aat aga tca tgg agc aaa cat taa cgc ggt tac gaa aat cgg aaa tac gcc gtt aca cac tta cct tca aca gta tac caa aca tag tcc tcg tgt ggt gta tgc tct ttt atc tcg agg agc cga tac gag gat acg taa taa tct tga ttg tac acc cat cat gga ata cat aaa gaa cga ttg tgc aac agg tca tat tct cat aat gtt act caa ttg gca cga aca aaa ata cgg gaa att aca aaa gga aga agg aca aca tct act tta tct att cat aaa aca taa tca agg ata tgg aag tcg ctc tct caa tat act acg gta tct act aga tag att cga cat tca gaa aga cga ata cta taa tac aat gac tcc tct tca tac cgc ctt cca gaa ttg caa taa caa tgt tgc ctc ata cct cgt ata cat cgg ata cga cat caa cct tcc gac taa aga cga taa gac agt att cga ctt ggt gtt tga aaa cag aaa cat tat ata caa ggc gga tgt cgt taa tga cat tat cca cca cag act gaa agt atc tct acc tat gat taa atc gtt gtt cta caa gat gtc gga gtt ctc tcc cta cga cga tca cta cgt aaa gaa gat aat agc cta ctg cct att aag gga cga gtc att tgc gga act aca tac taa att ctg ttt aaa cga gga cta taa aag tgt att tat gaa aaa tat atc att cga taa gat aga ttc cat cat cga aaa atg tag tcg tga cat aag tct cct caa aga gat tcg aat ctc aga cac cga ctt gta tac ggt att gag aac aga aga cat ccg gta tca cac ata tct cga agc cat aca ttc aga caa acg cat ttc att tcc cat gta cga cga tct cat aga aca gtg tca tct atc gat gga gca taa aag taa act cgt cga caa agc act caa taa att aga gtc tac cat cga tag tca atc tag act atc gta ttt gcc tcc gga aat tat gcg caa tat cat aac caa gct aag cga cta cca tct aaa cag tat gtt gta cgg aaa gaa cca tta caa ata tta tcc atg ata gaa aga aaa tat tta aaa aat aat cta tat gat tgg aga agt agg aaa caa aca gta aca aga cga cga tta cta c-a tta tta aat tat gag gtc cgt att ata ctc gta tat att gtt tct ctc atg tat aat aat aaa cgg aag aga tat agc acc aca tgc acc atc caa tgg aaa gtg taa aga caa cga ata cag aag ccg taa tct atg ttg tct atc gtg tcc tcc ggg aac tta cgc ttc cag att atg tga tag caa gac taa tac aca atg tac acc gtg tgg ttc gga tac ctt tac atc tca caa taa tca ttt aca ggc ttg tct aag ttg taa cgg aag atg tga tag taa tca ggt aga gac gcg atc gtg taa cac gac tca caa tag aat ctg tga atg ctc tcc agg ata tta ttg tct tct caa agg agc atc agg gtg tag aac atg tat ttc taa aac aaa gtg tgg aat agg ata cgg agt atc cgg ata cac gtc tac cgg aga cgt cat ctg ttc tcc gtg tgg tcc cgg aac ata ttc tca cac cgt ctc ttc cac aga taa atg cga acc cgt cgt aac cag caa tac att taa cta tat cga tgt gga aat taa cct gta tcc agt caa cga cac atc gtg tac tcg gac gac cac tac cgg tct cag cga atc cat ctc aac gtc gga act aac tat tac cat gaa tca taa aga ttg tga tcc agt ctt tcg tgc aga ata ctt ctc tgt cct taa taa tgt agc aac ttc agg att ctt tac agg aga aaa tag ata tca gaa tac ttc aaa gat atg tac tct gaa ttt cga gat taa atg taa caa caa aga ttc atc ttc caa aca gtt aac gaa aac aaa gaa tga tac tat cat gcc gca ttc aga gac ggt aac tct agt ggg cga ctg tct atc tag cgt cga cat cta cat act ata tag taa tac caa tac tca aga cta cga aaa tga tac aat ctc tta tca tat ggg taa tgt tct cga tgt caa tag cca tat gcc cgc tag ttg cga tat aca taa act gat cac taa ttc cca gaa tcc cac cca ctt ata gta ag- -tt ttt tta cct ata aat aat aaa tac aat aat taa ttt ctc gta aaa gta gaa aat ata ttc taa ttt att ata tgg taa gaa agt aga atc atc tag aac agt aat caa tca ata gca atc atg aaa caa tat att gtc ctg gca tgc atg tgc cta gtg gca gct gct atg cct act agt ctt caa caa tct tca tcc tcg tgt act gaa gaa gaa aac aaa cat cat atg gga atc gat gtt att atc aaa gtc aca aag caa gac caa aca ccg acc aat gat aag att tgt caa tcc gta acg gaa gtt aca gag a-- --- --- --- --- --- -cc gaa gat gat gag gta tcc gaa gaa gtt gta aaa gga gat ccc acc act tat tac act atc gtc ggt gcg ggt ctt aac atg aac ttt gga ttc acc aaa tgc cca aag att tca tcc atc tcc gaa tcc tct gat gga aac act gtg aat act aga ttg tcc agc gtg tca ccg gga caa ggt aag gac tct ccc gcg atc acg cgt gaa gaa gct ctg gct atg atc aaa gac tgt gag atg tct atc gac atc aga tgt agc gaa gaa gag aaa gac agt gac atc aag acc cat cca gta ctt ggg tct aac atc tca cat aag aaa gtg agt tac aaa gat atc atc ggt tca acg atc gtt gat aca aaa tgt gtc aag aac cta gag ttt agc gta cgt atc gga gac atg tgt gag gaa tca tct gaa ctt gaa gtc aag gat gga ttc aag tat gtc gac gga tcg gca tct gaa ggt gca acc gat gat act tca ctc atc gat tca aca aaa ctc aaa gca tgt gtc tga atc gat aac tct att cat ctg aaa atg gat gag ttg ggt taa tcg aac gat tca gac acc gca cca cga att aaa aaa g-- -ac cgg gca cta tat tcc ggt ttg caa aac aaa aat att taa cta cat tca caa aaa gtt acc tct cgt tac ttc ttc ttt ctg ttt caa tat gtg ata cga tat gat cac tat tcg tat tct ct- --- --- tgg tct cat aaa aaa gtt tta c-- --a aaa aaa aaa aaa a-t att ttt att ctc ttt ctc tct tcg at- ggt ctc aca aaa ata tta aac ctc ttt ctg atg tct caa cta ttt cgt aaa cga taa cgt cca aca ata tat tct cgt aga gct tat caa cat cct tat acc aat cta ggt tgt cag aca att gca tca taa aat aat gtt tat aat tta cac gtt aac atc ata taa taa acg tat ata gtt aat att ttt gga ata taa atg atc tgt aaa atc cat gta ggg gac act gct cac gtt ttt tct cta gta cat aat ttc aca caa gtt ttt ata cag aca aat taa ttc tcg tcc ata ta- ttt taa aac att gac ttt tgt act aag aaa aat atc ttg act aac cat ctc ttt ctc tct tcg atg ggt ctc aca aaa ata tta aac ctc ttt ctg atg g-- --- --- --- --- --- --- --- --- --- --- --- --- --- --- --- --- --- --- --- --- --- --- --- --- --- --- --- --- --- --- --- --- --- --- --- --- --- --- --- --- --- --- --- --- --- --- agt cgt aaa aag ttt tta tcc ttt ctc tct tcg ata ggt ctc --- --- --- --- --- --- --- --- --- --- --- --- --- --- --- --- --- --- --- --- --- --- --- --- --- --- --- --- --- --- --- --- --- --- --- --- --- --- --- --- --- --- --- --- --- --- --- --- --- --- --- --- --- --- --- --- --- --- --- --- --- --- --- --- --- --- --- --- --- --- aca aaa ata tta aac ctc ttt ctg atg gtc tct ata aa

>MT903345.1_Monkeypox_virus_isolate_MPXV-UK_P3

atc gag gac gac ttc aac tat gac agc tcg gtg gcg tct gcc agc gtg tac ata cga atg gca ttt cta aga aaa gtc tac ggt atc ctt -tc tac aat ttc ctt taa caa cgg caa cag ctg cag tat ttt tat act ttg aat gca -tc gga cat tta tac aag gga gtc ctg ttc taa tat tgg cat caa tgt tcg gat cta tag gct tga ttt tcg cat tga ctt tac aca gac atg aac atc ccc tga atc tgt aca tac ttt gtg gat tta cac tgt tag aat ctc taa cgc tgg cct ctg ttg tta ctt tct atg atg cac gta tcg tta tgc aag ctt tca tgt tga cta ctg cag tgt ttc ttg ctc tga cta cat ata ctc tac aat caa aga gag att tca gta aac ttg taa cag gat tgt ttg ctg ctt tct gga ttt taa ttt tgt cag gag tct tga gga taa agt tta aaa tag aat taa taa aga aca tat agg tca -tt ttt taa aca tgg ata gaa acc aag gtt gtt agt taa taa tat aca aga tat ttt ttc tca ctc tga tcc atg taa acc aag gac gag aga cac tct cat tcc tca ttc acg aca cca tta aaa atg gaa att aaa gcc ctc tat taa gca cag acg gct aca ggt cta cca tca ggt t-a cct tcg tct acc ttc aca atg gcc tct cct tgt gcc cag ttc agt ccc tgt cat tgc cac gct act aag gac tcc ctg aat acc gtg act gac gtc aga cat tgt ctg act gaa tac atc ctg tgg gtt tct cat aga tgg acc cat aga gaa agc gca ggg cct ctc tac agg ctt ctc atc tct ttc aga att gat gca atg gag cta ttt ggt agc gag ttg aag gag ttc tcg aat tca ctt ccg tgg gac aat atc gac aat tgc gtg gag atc att aaa tgt ttc atc aga aat gac tcc atg aaa acc gcc aaa gaa ctt tgt gca ata att gga ctt tgt act caa tca gct att gtc act gga aga gtc ttc aat gat aag tat atc gac ata cta ctt atg ctg cga aag att ctg aac gag aac gac tat ctc acc ctc ttg gat cat atc ctc act gct aaa tac taa atc tcc ttc atg ctc tct cac taa tac tct tac tca cta cac ttt tta tca tct tat gat gaa tga ttg cct tca tca ttt ttt cgt gga ata taa tat agg aat aat tag cac cag aat agc tat gga tat ctc gtt aag aat att ctc tat aag aga cat aat gta gac ata gtt att ata tcc ttc tta gat aag tgt tac gct act gga aag ttt cca tcg tta tta tta cat gaa gat gat ata att aaa cca aca ttg aga ttg gct ctt atg tta gct gga ttg aat tac tgt aat aaa tgc atc gag tat aga ggg ata tag caa ttc tcg ata ata gtc atg caa tat ttg aat gag act gat aat tta ggt aat aca gta ct- --- aca cac ata tct ttc tag att ata tat cgt taa aaa tct gta aga tgt ata ttt ctc ata agt atc cac tgt gta ata tta tta atg gat ata tag ata acg caa tag gga cta ata gta ttg taa aag ata taa tcg act att tac gta cat atc cag ata tct ata ttc cta cta gtt tgc tgc gta gtt gca tca ttg ata tgc atg -at tta tca gga ttc aga gat gaa tta cta agt aaa cta caa tcc cac aat aag taa gaa tca aat atc aaa aac tca ctt ttg att ttt cta gtc tta agt aat aca tat att tat taa tag acc tat gaa ata aaa aaa ggt aac aat gga ttc gcg tat agc tat tta cgt att agt atc ggc atc tct ttt gta tct tgt taa ttg tca caa act agt aca tta ctt caa tct gaa aat aaa tgg aag tga tat aac taa tac agc aga tat att gct gga caa tta tcc aat tat gac ctt tga tgg aaa gga tat tta tcc atc tat ctc gtt cat ggt cgg taa taa act ttt cct aga tct tta taa aaa tat ctt tga aga att ttt cag act att tcg agt atc tgt aag tag tca ata cga gga att aga ata tta tta ttc atg tga tta tac taa caa ccg tcc tac aat taa aca aca tta ctt tta taa cgg cga cga ata tac tga aat tga tag atc gaa aaa agc cac taa taa aaa cag ttg gtt aat tac ttc agg ctt tag act aca aaa atg gtt cga tag cga aga ttg tat aat tta tct cag atc ttt agt tag aag aat gga aga cag taa caa aaa cag taa aaa aac tta gta ctt aga tat cg- --a aaa aat ata ttt ttg tag act ctt gag aat aga agg aaa aca tgt aca taa tta taa aaa atg aaa atc aat ggc gaa taa gac agt gcg att cgc acc atg gag tcg gta gat ttc atg gct gtc gat gag cag ttt cac gac gac ctc gat ctt tgg tca tta tct ttg gta gat gat tat aaa aaa cat gga tta ggt gtt gac tgt tat gtt cta gaa cca gtt gtt gac agg aaa ata ttt gat aga ttt ctc ctt gaa cca att tgt gat cct gta gat gtt ctg tat gat tat ttt agg att cat aga gat aat att gat cag tat ata gta gat aga ctg ttt gca tat att aca tat aaa gat att ata tct gca tta gtg tca aag aat tat atg gaa gat att ttc tct ata att att aag aat tgt aat tct gtg caa gat ctc tta ctt tac tat cta tct aat gca tat gta gaa ata gac att gtt gat ctt atg gta gat cat ggg gct gta ata tat aaa ata gaa tgc ttg aat gcc tat ttt agg gga ata tgt aaa aag gaa agt agt gtt gtt gag ttt att ttg aat tgt ggt atc cca gat gaa aat gat gtt aaa tta gat cta tat aaa ata att cag tat act agg gga ttc ctt gta gat gaa ccc aca gta tta gaa att tat aag ctt tgt atc cca tat att gaa gat atc aat caa cta gat gct ggt gga agg acc ttg ctt tat cgc gct atc tat gca ggt tat ata gat tta gta tca tgg cta tta gaa aat gga gca aat gtc aac gca gta atg agt aat gga tat aca tgt ctt gac gtg gcc gtg gat agg gga tct gtc atc gcc cgt agg gaa gca cat ctt aaa ata tta gaa ata ttg ctt aga gaa cca ttg tct att gac tgt ata aaa tta gct ata ctt aat aat aca att gaa aac cat gat gtg ata aag ctc tgt atc aag tat ttt atg atg gta gat tat tca ctt tgt aat gtg tat gca tca tca ctc ttt gat tat ata att gat tgt aaa caa gaa ttg gag tac att agg cag atg aaa att cat aat aca acc atg tat gag tta atc tat aat aga gac aaa aac aag cat gct tcc cat att cta cat agg tat tct aaa cat cca gtt ttg aca cag tgt atc act aaa gga ttc aag att tac aca gaa gta acc gag cag gtc act aaa gct cta aac aga cgt gct cta ata gat gag ata ata aac aat gta tca act gat gac aat ctc cta tca aaa ctt cca tta gaa att agg gat cta att gtt tca caa gct gtc ata tag agt tct atc cac cca cct ttc ttg aaa tga gtt aat --- --- --- --- --- --- --- --- --- --- --- --- --- --- --- --- --- --- --- --- --- --- --- --- --- --- --- --- --- --- --- --- --- --- --- --- --- --- --- --- --- --- --- --- --- --- --- --- --- --- --- --- --- --- --- --- --- --- --- --- --- --- --- --- --- --- --- --- --- --- --- --- --- --- --- --- --- --- --- --- --- --- --- --- --- --- --- --- --- --- --- --- --- --- --- --- --- --- --- --- --- --- --- --- --- -ag tca taa gtt agt taa --- --- --- --- --- -gt cat aag tta gtt aag tca t-- --- --- --- --- --- --- --- --- --- --- --- --- --- --- --- -aa gtt agt taa gtc ata agt tag tta agt cat aag tta gtt aag tca taa gtt agt taa gtc ata agt tag tta agt cat aag tta gtt tat agt cta aca ctt cta att ttt ata cct tga tct ttt tct cta att atg aaa aag taa atc att atg aag atg gat g-a aat gga cga gat tgt gcg cat cgt taa cga tag tat gtg gta cgt acc taa cgc att tat gga cga cgg tga taa tga agg tca cat ttc tgt caa taa tgt ctg tca tat gta tct cgc att ctt tga tgt gga tat atc atc tca tct gtt taa att agt tat taa aca ctg cga tct gaa taa acg act aaa atg tgg taa ctc tcc att aca ttg cta tac gat gaa tac acg att taa tcc atc tgt att aaa gat att gtt acg cca cgg cat gcg taa ctt tga tag caa gga taa aaa agg aca tat tcc tct aca cca cta tct gat tca ttc act atc aat cga taa caa gat ctt tga tat act aac gga ccc cat tga tga ctt tag taa atc atc cga tct att gct gtg tta tct tag ata taa att caa tgg gag ctt aaa cta tta cgt tct gta caa att att gac taa agg atc tga ccc taa ttg cgt cga tga gga tgg act cac ttc tct tca tta cta ctg taa aca cat atc cgc gtt cca cga aag caa tta tta caa gtc aaa gag tca cac taa gat gcg agc tga gaa gcg att cat cta cgc gat aat aga tca tgg agc aaa cat taa cgc ggt tac gaa aat cgg aaa tac gcc gtt aca cac tta cct tca aca gta tac caa aca tag tcc tcg tgt ggt gta tgc tct ttt atc tcg agg agc cga tac gag gat acg taa taa tct tga ttg tac acc cat cat gga ata cat aaa gaa cga ttg tgc aac agg tca tat tct cat aat gtt act caa ttg gca cga aca aaa ata cgg gaa att aca aaa gga aga agg aca aca tct act tta tct att cat aaa aca taa tca agg ata tgg aag tcg ctc tct caa tat act acg gta tct act aga tag att cga cat tca gaa aga cga ata cta taa tac aat gac tcc tct tca tac cgc ctt cca gaa ttg caa taa caa tgt tgc ctc ata cct cgt ata cat cgg ata cga cat caa cct tcc gac taa aga cga taa gac agt att cga ctt ggt gtt tga aaa cag aaa cat tat ata caa ggc gga tgt cgt taa tga cat tat cca cca cag act gaa agt atc tct acc tat gat taa atc gtt gtt cta caa gat gtc gga gtt ctc tcc cta cga cga tca cta cgt aaa gaa gat aat agc cta ctg cct att aag gga cga gtc att tgc gga act aca tac taa att ctg ttt aaa cga gga cta taa aag tgt att tat gaa aaa tat atc att cga taa gat aga ttc cat cat cga aaa atg tag tcg tga cat aag tct cct caa aga gat tcg aat ctc aga cac cga ctt gta tac ggt att gag aac aga aga cat ccg gta tca cac ata tct cga agc cat aca ttc aga caa acg cat ttc att tcc cat gta cga cga tct cat aga aca gtg tca tct atc gat gga gca taa aag taa act cgt cga caa agc act caa taa att aga gtc tac cat cga tag tca atc tag act atc gta ttt gcc tcc gga aat tat gcg caa tat cat aac caa gct aag cga cta cca tct aaa cag tat gtt gta cgg aaa gaa cca tta caa ata tta tcc atg ata gaa aga aaa tat tta aaa aat aat cta tat gat tgg aga agt agg aaa caa aca gta aca aga cga cga tta cta c-a tta tta aat cat gag gtc cgt att ata ctc gta tat att gtt tct ctc atg tat aat aat aaa cgg aag aga tat agc acc aca tgc acc atc caa tgg aaa gtg taa aga caa cga ata cag aag ccg taa tct atg ttg tct atc gtg tcc tcc ggg aac tta cgc ttc cag att atg tga tag caa gac taa tac aca atg tac acc gtg tgg ttc gga tac ctt tac atc tca caa taa tca ttt aca ggc ttg tct aag ttg taa cgg aag atg tga tag taa tca ggt aga gac gcg atc gtg taa cac gac tca caa tag aat ctg tga atg ctc tcc agg ata tta ttg tct tct caa agg agc atc agg gtg tag aac atg tat ttc taa aac aaa gtg tgg aat agg ata cgg agt atc cgg ata cac gtc tac cgg aga cgt cat ctg ttc tcc gtg tgg tcc cgg aac ata ttc tca cac cgt ctc ttc cac aga taa atg cga acc cgt cgt aac cag caa tac att taa cta tat cga tgt gga aat taa cct gta tcc agt caa cga cac atc gtg tac tcg gac gac cac tac cgg tct cag cga atc cat ctc aac gtc gga act aac tat tac cat gaa tca taa aga ttg tga tcc agt ctt tcg tgc aga ata ctt ctc tgt cct taa taa tgt agc aac ttc agg att ctt tac agg aga aaa tag ata tca gaa tac ttc aaa gat atg tac tct gaa ttt cga gat taa atg taa caa caa aga ttc atc ttc caa aca gtt aac gaa aac aaa gaa tga tac tat cat gcc gca ttc aga gac ggt aac tct agt ggg cga ctg tct atc tag cgt cga cat cta cat act ata tag taa tac caa tac tca aga cta cga aaa tga tac aat ctc tta tca tat ggg taa tgt tct cga tgt caa tag cca tat gcc cgc tag ttg cga tat aca taa act gat cac taa ttc cca gaa tcc cac cca ctt ata gta ag- -tt ttt tta cct ata aat aat aaa tac aat aat taa ttt ctc gta aaa gta gaa aat ata ttc taa ttt att ata tgg taa gaa agt aga atc atc tag aac agt aat caa tca ata gca atc atg aaa caa tat att gtc ctg gca tgc atg tgc cta gtg gca gct gct atg cct act agt ctt caa caa tct tca tcc tcg tgt act gaa gaa gaa aac aaa cat cat atg gga atc gat gtt att atc aaa gtc aca aag caa gac caa aca ccg acc aat gat aag att tgt caa tcc gta acg gaa gtt aca gag a-- --- --- --- --- --- -cc gaa gat gat gag gta tcc gaa gaa gtt gta aaa gga gat ccc acc act tat tac act atc gtc ggt gcg ggt ctt aac atg aac ttt gga ttc acc aaa tgc cca aag att tca tcc atc tcc gaa tcc tct gat gga aac act gtg aat act aga ttg tcc agc gtg tca ccg gga caa ggt aag gac tct ccc gcg atc acg cgt gaa gaa gct ctg gct atg atc aaa gac tgt gag atg tct atc gac atc aga tgt agc gaa gaa gag aaa gac agt gac atc aag acc cat cca gta ctt ggg tct aac atc tca cat aag aaa gtg agt tac aaa gat atc atc ggt tca acg atc gtt gat aca aaa tgt gtc aag aac cta gag ttt agc gta cgt atc gga gac atg tgt gag gaa tca tct gaa ctt gaa gtc aag gat gga ttc aag tat gtc gac gga tcg gca tct gaa ggt gca acc gat gat act tca ctc atc gat tca aca aaa ctc aaa gca tgt gtc tga atc gat aac tct att cat ctg aaa atg gat gag ttg ggt taa tcg aac gat tca gac acc gca cca cga att aaa aaa g-- -ac cgg gca cta tat tcc ggt ttg caa aac aaa aat att taa cta cat tca caa aaa gtt acc tct cgt tac ttc ttc ttt ctg ttt caa tat gtg ata cga tat gat cac tat tcg tat tct ct- --- --- tgg tct cat aaa aaa gtt tta c-- --a aaa aaa aaa aaa a-t att ttt att ctc ttt ctc tct tcg at- ggt ctc aca aaa ata tta aac ctc ttt ctg atg tct caa cta ttt cgt aaa cga taa cgt cca aca ata tat tct cgt aga gct tat caa cat cct tat acc aat cta ggt tgt cag aca att gca tca taa aat aat gtt tat aat tta cac gtt aac atc ata taa taa acg tat ata gtt aat att ttt gga ata taa atg atc tgt aaa atc cat gta ggg gac act gct cac gtt ttt tct cta gta cat aat ttc aca caa gtt ttt ata cag aca aat taa ttc tcg tcc ata ta- ttt taa aac att gac ttt tgt act aag aaa aat atc ttg act aac cat ctc ttt ctc tct tcg atg ggt ctc aca aaa ata tta aac ctc ttt ctg atg g-- --- --- --- --- --- --- --- --- --- --- --- --- --- --- --- --- --- --- --- --- --- --- --- --- --- --- --- --- --- --- --- --- --- --- --- --- --- --- --- --- --- --- --- --- --- --- agt cgt aaa aag ttt tta tcc ttt ctc tct tcg ata ggt ctc --- --- --- --- --- --- --- --- --- --- --- --- --- --- --- --- --- --- --- --- --- --- --- --- --- --- --- --- --- --- --- --- --- --- --- --- --- --- --- --- --- --- --- --- --- --- --- --- --- --- --- --- --- --- --- --- --- --- --- --- --- --- --- --- --- --- --- --- --- --- aca aaa ata tta aac ctc ttt ctg atg gtc tct ata aa

>MT903344.1_Monkeypox_virus_isolate_MPXV-UK_P2

atc gag gac gac ttc aac tat gac agc tcg gtg gcg tct gcc agc gtg tac ata cga atg gca ttt cta aga aaa gtc tac ggt atc ctt -tc tac aat ttc ctt taa caa cgg caa cag ctg cag tat ttt tat act ttg aat gca -tc gga cat tta tac aag gga gtc ctg ttc taa tat tgg cat caa tgt tcg gat cta tag gct tga ttt tcg cat tga ctt tac aca gac atg aac atc ccc tga atc tgt aca tac ttt gtg gat tta cac tgt tag aat ctc taa cgc tgg cct ctg ttg tta ctt tct atg atg cac gta tcg tta tgc aag ctt tca tgt tga cta ctg cag tgt ttc ttg ctc tga cta cat ata ctc tac aat caa aga gag att tca gta aac ttg taa cag gat tgt ttg ctg ctt tct gga ttt taa ttt tgt cag gag tct tga gga taa agt tta aaa tag aat taa taa aga aca tat agg tca -tt ttt taa aca tgg ata gaa acc aag gtt gtt agt taa taa tat aca aga tat ttt ttc tca ctc tga tcc atg taa acc aag gac gag aga cac tct cat tcc tca ttc acg aca cca tta aaa atg gaa att aaa gcc ctc tat taa gca cag acg gct aca ggt cta cca tca ggt t-a cct tcg tct acc ttc aca atg gcc tct cct tgt gcc cag ttc agt ccc tgt cat tgc cac gct act aag gac tcc ctg aat acc gtg act gac gtc aga cat tgt ctg act gaa tac atc ctg tgg gtt tct cat aga tgg acc cat aga gaa agc gca ggg cct ctc tac agg ctt ctc atc tct ttc aga att gat gca atg gag cta ttt ggt agc gag ttg aag gag ttc tcg aat tca ctt ccg tgg gac aat atc gac aat tgc gtg gag atc att aaa tgt ttc atc aga aat gac tcc atg aaa acc gcc aaa gaa ctt tgt gca ata att gga ctt tgt act caa tca gct att gtc act gga aga gtc ttc aat gat aag tat atc gac ata cta ctt atg ctg cga aag att ctg aac gag aac gac tat ctc acc ctc ttg gat cat atc ctc act gct aaa tac taa atc tcc ttc atg ctc tct cac taa tac tct tac tca cta cac ttt tta tca tct tat gat gaa tga ttg cct tca tca ttt ttt cgt gga ata taa tat agg aat aat tag cac cag aat agc tat gga tat ctc gtt aag aat att ctc tat aag aga cat aat gta gac ata gtt att ata tcc ttc tta gat aag tgt tac gct act gga aag ttt cca tcg tta tta tta cat gaa gat gat ata att aaa cca aca ttg aga ttg gct ctt atg tta gct gga ttg aat tac tgt aat aaa tgc atc gag tat aga ggg ata tag caa ttc tcg ata ata gtc atg caa tat ttg aat gag act gat aat tta ggt aat aca gta ct- --- aca cac ata tct ttc tag att ata tat cgt taa aaa tct gta aga tgt ata ttt ctc ata agt atc cac tgt gta ata tta tta atg gat ata tag ata acg caa tag gga cta ata gta ttg taa aag ata taa tcg act att tac gta cat atc cag ata tct ata ttc cta cta gtt tgc tgc gta gtt gca tca ttg ata tgc atg -at tta tca gga ttc aga gat gaa tta cta agt aaa cta caa tcc cac aat aag taa gaa tca aat atc aaa aac tca ctt ttg att ttt cta gtc tta agt aat aca tat att tat taa tag acc tat gaa ata aaa aaa ggt aac aat gga ttc gcg tat agc tat tta cgt att agt atc ggc atc tct ttt gta tct tgt taa ttg tca caa act agt aca tta ctt caa tct gaa aat aaa tgg aag tga tat aac taa tac agc aga tat att gct gga caa tta tcc aat tat gac ctt tga tgg aaa gga tat tta tcc atc tat ctc gtt cat ggt cgg taa taa act ttt cct aga tct tta taa aaa tat ctt tga aga att ttt cag act att tcg agt atc tgt aag tag tca ata cga gga att aga ata tta tta ttc atg tga tta tac taa caa ccg tcc tac aat taa aca aca tta ctt tta taa cgg cga cga ata tac tga aat tga tag atc gaa aaa agc cac taa taa aaa cag ttg gtt aat tac ttc agg ctt tag act aca aaa atg gtt cga tag cga aga ttg tat aat tta tct cag atc ttt agt tag aag aat gga aga cag taa caa aaa cag taa aaa aac tta gta ctt aga tat cg- --a aaa aat ata ttt ttg tag act ctt gag aat aga agg aaa aca tgt aca taa tta taa aaa atg aaa atc aat ggc gaa taa gac agt gcg att cgc acc atg gag tcg gta gat ttc atg gct gtc gat gag cag ttt cac gac gac ctc gat ctt tgg tca tta tct ttg gta gat gat tat aaa aaa cat gga tta ggt gtt gac tgt tat gtt cta gaa cca gtt gtt gac agg aaa ata ttt gat aga ttt ctc ctt gaa cca att tgt gat cct gta gat gtt ctg tat gat tat ttt agg att cat aga gat aat att gat cag tat ata gta gat aga ctg ttt gca tat att aca tat aaa gat att ata tct gca tta gtg tca aag aat tat atg gaa gat att ttc tct ata att att aag aat tgt aat tct gtg caa gat ctc tta ctt tac tat cta tct aat gca tat gta gaa ata gac att gtt gat ctt atg gta gat cat ggg gct gta ata tat aaa ata gaa tgc ttg aat gcc tat ttt agg gga ata tgt aaa aag gaa agt agt gtt gtt gag ttt att ttg aat tgt ggt atc cca gat gaa aat gat gtt aaa tta gat cta tat aaa ata att cag tat act agg gga ttc ctt gta gat gaa ccc aca gta tta gaa att tat aag ctt tgt atc cca tat att gaa gat atc aat caa cta gat gct ggt gga agg acc ttg ctt tat cgc gct atc tat gca ggt tat ata gat tta gta tca tgg cta tta gaa aat gga gca aat gtc aac gca gta atg agt aat gga tat aca tgt ctt gac gtg gcc gtg gat agg gga tct gtc atc gcc cgt agg gaa gca cat ctt aaa ata tta gaa ata ttg ctt aga gaa cca ttg tct att gac tgt ata aaa tta gct ata ctt aat aat aca att gaa aac cat gat gtg ata aag ctc tgt atc aag tat ttt atg atg gta gat tat tca ctt tgt aat gtg tat gca tca tca ctc ttt gat tat ata att gat tgt aaa caa gaa ttg gag tac att agg cag atg aaa att cat aat aca acc atg tat gag tta atc tat aat aga gac aaa aac aag cat gct tcc cat att cta cat agg tat tct aaa cat cca gtt ttg aca cag tgt atc act aaa gga ttc aag att tac aca gaa gta acc gag cag gtc act aaa gct cta aac aga cgt gct cta ata gat gag ata ata aac aat gta tca act gat gac aat ctc cta tca aaa ctt cca tta gaa att agg gat cta att gtt tca caa gct gtc ata tag agt tct atc cac cca cct ttc ttg aaa tga gtt aat --- --- --- --- --- --- --- --- --- --- --- --- --- --- --- --- --- --- --- --- --- --- --- --- --- --- --- --- --- --- --- --- --- --- --- --- --- --- --- --- --- --- --- --- --- --- --- --- --- --- --- --- --- --- --- --- --- --- --- --- --- --- --- --- --- --- --- --- --- --- --- --- --- --- --- --- --- --- --- --- --- --- --- --- --- --- --- --- --- --- --- --- --- --- --- --- --- --- --- --- --- --- --- --- --- -ag tca taa gtt agt taa --- --- --- --- --- -gt cat aag tta gtt aag tca t-- --- --- --- --- --- --- --- --- --- --- --- --- --- --- --- -aa gtt agt taa gtc ata agt tag tta agt cat aag tta gtt aag tca taa gtt agt taa gtc ata agt tag tta agt cat aag tta gtt tat agt cta aca ctt cta att ttt ata cct tga tct ttt tct cta att atg aaa aag taa atc att atg aag atg gat g-a aat gga cga gat tgt gcg cat cgt taa cga tag tat gtg gta cgt acc taa cgc att tat gga cga cgg tga taa tga agg tca cat ttc tgt caa taa tgt ctg tca tat gta tct cgc att ctt tga tgt gga tat atc atc tca tct gtt taa att agt tat taa aca ctg cga tct gaa taa acg act aaa atg tgg taa ctc tcc att aca ttg cta tac gat gaa tac acg att taa tcc atc tgt att aaa gat att gtt acg cca cgg cat gcg taa ctt tga tag caa gga taa aaa agg aca tat tcc tct aca cca cta tct gat tca ttc act atc aat cga taa caa gat ctt tga tat act aac gga ccc cat tga tga ctt tag taa atc atc cga tct att gct gtg tta tct tag ata taa att caa tgg gag ctt aaa cta tta cgt tct gta caa att att gac taa agg atc tga ccc taa ttg cgt cga tga gga tgg act cac ttc tct tca tta cta ctg taa aca cat atc cgc gtt cca cga aag caa tta tta caa gtc aaa gag tca cac taa gat gcg agc tga gaa gcg att cat cta cgc gat aat aga tca tgg agc aaa cat taa cgc ggt tac gaa aat cgg aaa tac gcc gtt aca cac tta cct tca aca gta tac caa aca tag tcc tcg tgt ggt gta tgc tct ttt atc tcg agg agc cga tac gag gat acg taa taa tct tga ttg tac acc cat cat gga ata cat aaa gaa cga ttg tgc aac agg tca tat tct cat aat gtt act caa ttg gca cga aca aaa ata cgg gaa att aca aaa gga aga agg aca aca tct act tta tct att cat aaa aca taa tca agg ata tgg aag tcg ctc tct caa tat act acg gta tct act aga tag att cga cat tca gaa aga cga ata cta taa tac aat gac tcc tct tca tac cgc ctt cca gaa ttg caa taa caa tgt tgc ctc ata cct cgt ata cat cgg ata cga cat caa cct tcc gac taa aga cga taa gac agt att cga ctt ggt gtt tga aaa cag aaa cat tat ata caa ggc gga tgt cgt taa tga cat tat cca cca cag act gaa agt atc tct acc tat gat taa atc gtt gtt cta caa gat gtc gga gtt ctc tcc cta cga cga tca cta cgt aaa gaa gat aat agc cta ctg cct att aag gga cga gtc att tgc gga act aca tac taa att ctg ttt aaa cga gga cta taa aag tgt att tat gaa aaa tat atc att cga taa gat aga ttc cat cat cga aaa atg tag tcg tga cat aag tct cct caa aga gat tcg aat ctc aga cac cga ctt gta tac ggt att gag aac aga aga cat ccg gta tca cac ata tct cga agc cat aca ttc aga caa acg cat ttc att tcc cat gta cga cga tct cat aga aca gtg tca tct atc gat gga gca taa aag taa act cgt cga caa agc act caa taa att aga gtc tac cat cga tag tca atc tag act atc gta ttt gcc tcc gga aat tat gcg caa tat cat aac caa gct aag cga cta cca tct aaa cag tat gtt gta cgg aaa gaa cca tta caa ata tta tcc atg ata gaa aga aaa tat tta aaa aat aat cta tat gat tgg aga agt agg aaa caa aca gta aca aga cga cga tta cta c-a tta tta aat cat gag gtc cgt att ata ctc gta tat att gtt tct ctc atg tat aat aat aaa cgg aag aga tat agc acc aca tgc acc atc caa tgg aaa gtg taa aga caa cga ata cag aag ccg taa tct atg ttg tct atc gtg tcc tcc ggg aac tta cgc ttc cag att atg tga tag caa gac taa tac aca atg tac acc gtg tgg ttc gga tac ctt tac atc tca caa taa tca ttt aca ggc ttg tct aag ttg taa cgg aag atg tga tag taa tca ggt aga gac gcg atc gtg taa cac gac tca caa tag aat ctg tga atg ctc tcc agg ata tta ttg tct tct caa agg agc atc agg gtg tag aac atg tat ttc taa aac aaa gtg tgg aat agg ata cgg agt atc cgg ata cac gtc tac cgg aga cgt cat ctg ttc tcc gtg tgg tcc cgg aac ata ttc tca cac cgt ctc ttc cac aga taa atg cga acc cgt cgt aac cag caa tac att taa cta tat cga tgt gga aat taa cct gta tcc agt caa cga cac atc gtg tac tcg gac gac cac tac cgg tct cag cga atc cat ctc aac gtc gga act aac tat tac cat gaa tca taa aga ttg tga tcc agt ctt tcg tgc aga ata ctt ctc tgt cct taa taa tgt agc aac ttc agg att ctt tac agg aga aaa tag ata tca gaa tac ttc aaa gat atg tac tct gaa ttt cga gat taa atg taa caa caa aga ttc atc ttc caa aca gtt aac gaa aac aaa gaa tga tac tat cat gcc gca ttc aga gac ggt aac tct agt ggg cga ctg tct atc tag cgt cga cat cta cat act ata tag taa tac caa tac tca aga cta cga aaa tga tac aat ctc tta tca tat ggg taa tgt tct cga tgt caa tag cca tat gcc cgc tag ttg cga tat aca taa act gat cac taa ttc cca gaa tcc cac cca ctt ata gta ag- -tt ttt tta cct ata aat aat aaa tac aat aat taa ttt ctc gta aaa gta gaa aat ata ttc taa ttt att ata tgg taa gaa agt aga atc atc tag aac agt aat caa tca ata gca atc atg aaa caa tat att gtc ctg gca tgc atg tgc cta gtg gca gct gct atg cct act agt ctt caa caa tct tca tcc tcg tgt act gaa gaa gaa aac aaa cat cat atg gga atc gat gtt att atc aaa gtc aca aag caa gac caa aca ccg acc aat gat aag att tgt caa tcc gta acg gaa gtt aca gag a-- --- --- --- --- --- -cc gaa gat gat gag gta tcc gaa gaa gtt gta aaa gga gat ccc acc act tat tac act atc gtc ggt gcg ggt ctt aac atg aac ttt gga ttc acc aaa tgc cca aag att tca tcc atc tcc gaa tcc tct gat gga aac act gtg aat act aga ttg tcc agc gtg tca ccg gga caa ggt aag gac tct ccc gcg atc acg cgt gaa gaa gct ctg gct atg atc aaa gac tgt gag atg tct atc gac atc aga tgt agc gaa gaa gag aaa gac agt gac atc aag acc cat cca gta ctt ggg tct aac atc tca cat aag aaa gtg agt tac aaa gat atc atc ggt tca acg atc gtt gat aca aaa tgt gtc aag aac cta gag ttt agc gta cgt atc gga gac atg tgt gag gaa tca tct gaa ctt gaa gtc aag gat gga ttc aag tat gtc gac gga tcg gca tct gaa ggt gca acc gat gat act tca ctc atc gat tca aca aaa ctc aaa gca tgt gtc tga atc gat aac tct att cat ctg aaa atg gat gag ttg ggt taa tcg aac gat tca gac acc gca cca cga att aaa aaa g-- -ac cgg gca cta tat tcc ggt ttg caa aac aaa aat att taa cta cat tca caa aaa gtt acc tct cgt tac ttc ttc ttt ctg ttt caa tat gtg ata cga tat gat cac tat tcg tat tct ct- --- --- tgg tct cat aaa aaa gtt tta c-- --a aaa aaa aaa aaa aat att ttt att ctc ttt ctc tct tcg at- ggt ctc aca aaa ata tta aac ctc ttt ctg atg tct caa cta ttt cgt aaa cga taa cgt cca aca ata tat tct cgt aga gct tat caa cat cct tat acc aat cta ggt tgt cag aca att gca tca taa aat aat gtt tat aat tta cac gtt aac atc ata taa taa acg tat ata gtt aat att ttt gga ata taa atg atc tgt aaa atc cat gta ggg gac act gct cac gtt ttt tct cta gta cat aat ttc aca caa gtt ttt ata cag aca aat taa ttc tcg tcc ata ta- ttt taa aac att gac ttt tgt act aag aaa aat atc ttg act aac cat ctc ttt ctc tct tcg atg ggt ctc aca aaa ata tta aac ctc ttt ctg atg g-- --- --- --- --- --- --- --- --- --- --- --- --- --- --- --- --- --- --- --- --- --- --- --- --- --- --- --- --- --- --- --- --- --- --- --- --- --- --- --- --- --- --- --- --- --- --- agt cgt aaa aag ttt tta tcc ttt ctc tct tcg ata ggt ctc --- --- --- --- --- --- --- --- --- --- --- --- --- --- --- --- --- --- --- --- --- --- --- --- --- --- --- --- --- --- --- --- --- --- --- --- --- --- --- --- --- --- --- --- --- --- --- --- --- --- --- --- --- --- --- --- --- --- --- --- --- --- --- --- --- --- --- --- --- --- aca aaa ata tta aac ctc ttt ctg atg gtc tct ata aa

>MT903342.1_Monkeypox_virus_isolate_MPXV-Singapore

atc gag gac gac ttc aac tat gac agc tcg gtg gcg tct gcc agc gtg tac ata cga atg gca ttt cta aga aaa gtc tac ggt atc ctt -tc tac aat ttc ctt taa caa cgg caa cag ctg cag tat ttt tat act ttg aat gca -tc gga cat tta tac aag gga gtc ctg ttc taa tat tgg cat caa tgt tcg gat cta tag gct tga ttt tcg cat tga ctt tac aca gac atg aac atc ccc tga atc tgt aca tac ttt gtg gat tta cac tgt tag aat ctc taa cgc tgg cct ctg ttg tta ctt tct atg atg cac gta tcg tta tgc aag ctt tca tgt tga cta ctg cag tgt ttc ttg ctc tga cta cat ata ctc tac aat caa aga gag att tca gta aac ttg taa cag gat tgt ttg ctg ctt tct gga ttt taa ttt tgt cag gag tct tga gga taa agt tta aaa tag aat taa taa aga aca tat agg tca -tt ttt taa aca tgg ata gaa acc aag gtt gtt agt taa taa tat aca aga tat ttt ttc tca ctc tga tcc atg taa acc aag gac gag aga cac tct cat tcc tca ttc acg aca cca tta aaa atg gaa att aaa gcc ctc tat taa gca cag acg gct aca ggt cta cca tca ggt t-a cct tcg tct acc ttc aca atg gcc tct cct tgt gcc cag ttc agt ccc tgt cat tgc cac gct act aag gac tcc ctg aat acc gtg act gac gtc aga cat tgt ctg act gaa tac atc ctg tgg gtt tct cat aga tgg acc cat aga gaa agc gca ggg cct ctc tac agg ctt ctc atc tct ttc aga att gat gca atg gag cta ttt ggt agc gag ttg aag gag ttc tcg aat tca ctt ccg tgg gac aat atc gac aat tgc gtg gag atc att aaa tgt ttc atc aga aat gac tcc atg aaa acc gcc aaa gaa ctt tgt gca ata att gga ctt tgt act caa tca gct att gtc act gga aga gtc ttc aat gat aag tat atc gac ata cta ctt atg ctg cga aag att ctg aac gag aac gac tat ctc acc ctc ttg gat cat atc ctc act gct aaa tac taa atc tcc ttc atg ctc tct cac taa tac tct tac tca cta cac ttt tta tca tct tat gat gaa tga ttg cct tca tca ttt ttt cgt gga ata taa tat agg aat aat tag cac cag aat agc tat gga tat ctc gtt aag aat att ctc tat aag aga cat aat gta gac ata gtt att ata tcc ttc tta gat aag tgt tac gct act gga aag ttt cca tcg tta tta tta cat gaa gat gat ata att aaa cca aca ttg aga ttg gct ctt atg tta gct gga ttg aat tac tgt aat aaa tgc atc gag tat aga ggg ata tag caa ttc tcg ata ata gtc atg caa tat ttg aat gag act gat aat tta ggt aat aca gta ct- --- aca cac ata tct ttc tag att ata tat cgt taa aaa tct gta aga tgt ata ttt ctc ata agt atc cac tgt gta ata tta tta atg gat ata tag ata acg caa tag gga cta ata gta ttg taa aag ata taa tcg act att tac gta cat atc cag ata tct ata ttc cta cta gtt tgc tgc gta gtt gca tca ttg ata tgc atg -at tta tca gga ttc aga gat gaa tta cta agt aaa cta caa tcc cac aat aag taa gaa tca aat atc aaa aac tca ctt ttg att ttt cta gtc tta agt aat aca tat att tat taa tag acc tat gaa ata aaa aaa ggt aac aat gga ttc gcg tat agc tat tta cgt att agt atc ggc atc tct ttt gta tct tgt taa ttg tca caa act agt aca tta ctt caa tct gaa aat aaa tgg aag tga tat aac taa tac agc aga tat att gct gga caa tta tcc aat tat gac ctt tga tgg aaa gga tat tta tcc atc tat ctc gtt cat ggt cgg taa taa act ttt cct aga tct tta taa aaa tat ctt tga aga att ttt cag act att tcg agt atc tgt aag tag tca ata cga gga att aga ata tta tta ttc atg tga tta tac taa caa ccg tcc tac aat taa aca aca tta ctt tta taa cgg cga cga ata tac tga aat tga tag atc gaa aaa agc cac taa taa aaa cag ttg gtt aat tac ttc agg ctt tag act aca aaa atg gtt cga tag cga aga ttg tat aat tta tct cag atc ttt agt tag aag aat gga aga cag taa caa aaa cag taa aaa aac tta gta ctt aga tat cg- --a aaa aat ata ttt ttg tag act ctt gag aat aga agg aaa aca tgt aca taa tta taa aaa atg aaa atc aat ggc gaa taa gac agt gcg att cgc acc atg gag tcg gta gat ttc atg gct gtc gat gag cag ttt cac gac gac ctc gat ctt tgg tca tta tct ttg gta gat gat tat aaa aaa cat gga tta ggt gtt gac tgt tat gtt cta gaa cca gtt gtt gac agg aaa ata ttt gat aga ttt ctc ctt gaa cca att tgt gat cct gta gat gtt ctg tat gat tat ttt agg att cat aga gat aat att gat cag tat ata gta gat aga ctg ttt gca tat att aca tat aaa gat att ata tct gca tta gtg tca aag aat tat atg gaa gat att ttc tct ata att att aag aat tgt aat tct gtg caa gat ctc tta ctt tac tat cta tct aat gca tat gta gaa ata gac att gtt gat ctt atg gta gat cat ggg gct gta ata tat aaa ata gaa tgc ttg aat gcc tat ttt agg gga ata tgt aaa aag gaa agt agt gtt gtt gag ttt att ttg aat tgt ggt atc cca gat gaa aat gat gtt aaa tta gat cta tat aaa ata att cag tat act agg gga ttc ctt gta gat gaa ccc aca gta tta gaa att tat aag ctt tgt atc cca tat att gaa gat atc aat caa cta gat gct ggt gga agg acc ttg ctt tat cgc gct atc tat gca ggt tat ata gat tta gta tca tgg cta tta gaa aat gga gca aat gtc aac gca gta atg agt aat gga tat aca tgt ctt gac gtg gcc gtg gat agg gga tct gtc atc gcc cgt agg gaa gca cat ctt aaa ata tta gaa ata ttg ctt aga gaa cca ttg tct att gac tgt ata aaa tta gct ata ctt aat aat aca att gaa aac cat gat gtg ata aag ctc tgt atc aag tat ttt atg atg gta gat tat tca ctt tgt aat gtg tat gca tca tca ctc ttt gat tat ata att gat tgt aaa caa gaa ttg gag tac att agg cag atg aaa att cat aat aca acc atg tat gag tta atc tat aat aga gac aaa aac aag cat gct tcc cat att cta cat agg tat tct aaa cat cca gtt ttg aca cag tgt atc act aaa gga ttc aag att tac aca gaa gta acc gag cag gtc act aaa gct cta aac aga cgt gct cta ata gat gag ata ata aac aat gta tca act gat gac aat ctc cta tca aaa ctt cca tta gaa att agg gat cta att gtt tca caa gct gtc ata tag agt tct atc cac cca cct ttc ttg aaa tga gtt aat a-- --- --- --- --- --- --- --- --- --- --- --- --- --- --- --- --- --- --- --- --- --- --- --- --- --- --- --- --- --- --- --- --- --- --- --- --- --- --- --- --- --- --- --- --- --- --- --- --- --- --- --- --- --- --- --- --- --- --- --- --- --- --- --- --- --- --- --- --- --- --- --- --- --- --- --- --- --- --- --- --- --- --- --- --- --- --- --- --- --- --- --- --- --- --- --- --- --- --- --- -gt cat aag tta gtt aag tca taa gtt agt taa --- --- --- --- --- -gt cat aag tta gtt aag tca t-- --- --- --- --- --- --- --- --- --- --- --- --- --- --- --- -aa gtt agt taa gtc ata agt tag tta agt cat aag tta gtt aag tca taa gtt agt taa gtc ata agt tag tta agt cat aag tta gtt tat agt cta aca ctt cta att ttt ata cct tga tct ttt tct cta att atg aaa aag taa atc att atg aag atg gat g-a aat gga cga gat tgt gcg cat cgt taa cga tag tat gtg gta cgt acc taa cgc att tat gga cga cgg tga taa tga agg tca cat ttc tgt caa taa tgt ctg tca tat gta tct cgc att ctt tga tgt gga tat atc atc tca tct gtt taa att agt tat taa aca ctg cga tct gaa taa acg act aaa atg tgg taa ctc tcc att aca ttg cta tac gat gaa tac acg att taa tcc atc tgt att aaa gat att gtt acg cca cgg cat gcg taa ctt tga tag caa gga taa aaa agg aca tat tcc tct aca cca cta tct gat tca ttc act atc aat cga taa caa gat ctt tga tat act aac gga ccc cat tga tga ctt tag taa atc atc cga tct att gct gtg tta tct tag ata taa att caa tgg gag ctt aaa cta tta cgt tct gta caa att att gac taa agg atc tga ccc taa ttg cgt cga tga gga tgg act cac ttc tct tca tta cta ctg taa aca cat atc cgc gtt cca cga aag caa tta tta caa gtc aaa gag tca cac taa gat gcg agc tga gaa gcg att cat cta cgc gat aat aga tca tgg agc aaa cat taa cgc ggt tac gaa aat cgg aaa tac gcc gtt aca cac tta cct tca aca gta tac caa aca tag tcc tcg tgt ggt gta tgc tct ttt atc tcg agg agc cga tac gag gat acg taa taa tct tga ttg tac acc cat cat gga ata cat aaa gaa cga ttg tgc aac agg tca tat tct cat aat gtt act caa ttg gca cga aca aaa ata cgg gaa att aca aaa gga aga agg aca aca tct act tta tct att cat aaa aca taa tca agg ata tgg aag tcg ctc tct caa tat act acg gta tct act aga tag att cga cat tca gaa aga cga ata cta taa tac aat gac tcc tct tca tac cgc ctt cca gaa ttg caa taa caa tgt tgc ctc ata cct cgt ata cat cgg ata cga cat caa cct tcc gac taa aga cga taa gac agt att cga ctt ggt gtt tga aaa cag aaa cat tat ata caa ggc gga tgt cgt taa tga cat tat cca cca cag act gaa agt atc tct acc tat gat taa atc gtt gtt cta caa gat gtc gga gtt ctc tcc cta cga cga tca cta cgt aaa gaa gat aat agc cta ctg cct att aag gga cga gtc att tgc gga act aca tac taa att ctg ttt aaa cga gga cta taa aag tgt att tat gaa aaa tat atc att cga taa gat aga ttc cat cat cga aaa atg tag tcg tga cat aag tct cct caa aga gat tcg aat ctc aga cac cga ctt gta tac ggt att gag aac aga aga cat ccg gta tca cac ata tct cga agc cat aca ttc aga caa acg cat ttc att tcc cat gta cga cga tct cat aga aca gtg tca tct atc gat gga gca taa aag taa act cgt cga caa agc act caa taa att aga gtc tac cat cga tag tca atc tag act atc gta ttt gcc tcc gga aat tat gcg caa tat cat aac caa gct aag cga cta cca tct aaa cag tat gtt gta cgg aaa gaa cca tta caa ata tta tcc atg ata gaa aga aaa tat tta aaa aat aat cta tat gat tgg aga agt agg aaa caa aca gta aca aga cga cga tta cta c-a tta tta aat cat gag gtc cgt att ata ctc gta tat att gtt tct ctc atg tat aat aat aaa cgg aag aga tat agc acc aca tgc acc atc caa tgg aaa gtg taa aga caa cga ata cag aag ccg taa tct atg ttg tct atc gtg tcc tcc ggg aac tta cgc ttc cag att atg tga tag caa gac taa tac aca atg tac acc gtg tgg ttc gga tac ctt tac atc tca caa taa tca ttt aca ggc ttg tct aag ttg taa cgg aag atg tga tag taa tca ggt aga gac gcg atc gtg taa cac gac tca caa tag aat ctg tga atg ctc tcc agg ata tta ttg tct tct caa agg agc atc agg gtg tag aac atg tat ttc taa aac aaa gtg tgg aat agg ata cgg agt atc cgg ata cac gtc tac cgg aga cgt cat ctg ttc tcc gtg tgg tcc cgg aac ata ttc tca cac cgt ctc ttc cac aga taa atg cga acc cgt cgt aac cag caa tac att taa cta tat cga tgt gga aat taa cct gta tcc agt caa cga cac atc gtg tac tcg gac gac cac tac cgg tct cag cga atc cat ctc aac gtc gga act aac tat tac cat gaa tca taa aga ttg tga tcc agt ctt tcg tgc aga ata ctt ctc tgt cct taa taa tgt agc aac ttc agg att ctt tac agg aga aaa tag ata tca gaa tac ttc aaa gat atg tac tct gaa ttt cga gat taa atg taa caa caa aga ttc atc ttc caa aca gtt aac gaa aac aaa gaa tga tac tat cat gcc gca ttc aga gac ggt aac tct agt ggg cga ctg tct atc tag cgt cga cat cta cat act ata tag taa tac caa tac tca aga cta cga aaa tga tac aat ctc tta tca tat ggg taa tgt tct cga tgt caa tag cca tat gcc cgc tag ttg cga tat aca taa act gat cac taa ttc cca gaa tcc cac cca ctt ata gta ag- -tt ttt tta cct ata aat aat aaa tac aat aat taa ttt ctc gta aaa gta gaa aat ata ttc taa ttt att ata tgg taa gaa agt aga atc atc tag aac agt aat caa tca ata gca atc atg aaa caa tat att gtc ctg gca tgc atg tgc cta gtg gca gct gct atg cct act agt ctt caa caa tct tca tcc tcg tgt act gaa gaa gaa aac aaa cat cat atg gga atc gat gtt att atc aaa gtc aca aag caa gac caa aca ccg acc aat gat aag att tgt caa tcc gta acg gaa gtt aca gag a-- --- --- --- --- --- -cc gaa gat gat gag gta tcc gaa gaa gtt gta aaa gga gat ccc acc act tat tac act atc gtc ggt gcg ggt ctt aac atg aac ttt gga ttc acc aaa tgc cca aag att tca tcc atc tcc gaa tcc tct gat gga aac act gtg aat act aga ttg tcc agc gtg tca ccg gga caa ggt aag gac tct ccc gcg atc acg cgt gaa gaa gct ctg gct atg atc aaa gac tgt gag atg tct atc gac atc aga tgt agc gaa gaa gag aaa gac agt gac atc aag acc cat cca gta ctt ggg tct aac atc tca cat aag aaa gtg agt tac aaa gat atc atc ggt tca acg atc gtt gat aca aaa tgt gtc aag aac cta gag ttt agc gta cgt atc gga gac atg tgt gag gaa tca tct gaa ctt gaa gtc aag gat gga ttc aag tat gtc gac gga tcg gca tct gaa ggt gca acc gat gat act tca ctc atc gat tca aca aaa ctc aaa gca tgt gtc tga atc gat aac tct att cat ctg aaa atg gat gag ttg ggt taa tcg aac gat tca gac acc gca cca cga att aaa aaa g-- -ac cgg gca cta tat tcc ggt ttg caa aac aaa aat att taa cta cat tca caa aaa gtt acc tct cgt tac ttc ttc ttt ctg ttt caa tat gtg ata cga tat gat cac tat tcg tat tct ct- --- --- tgg tct cat aaa aaa gtt tta c-- -aa aaa aaa aaa aaa aat att ttt att ctc ttt ctc tct tcg at- ggt ctc aca aaa ata tta aac ctc ttt ctg atg tct caa cta ttt cgt aaa cga taa cgt cca aca ata tat tct cgt aga gct tat caa cat cct tat acc aat cta ggt tgt cag aca att gca tca taa aat aat gtt tat aat tta cac gtt aac atc ata taa taa acg tat ata gtt aat att ttt gga ata taa atg atc tgt aaa atc cat gta ggg gac act gct cac gtt ttt tct cta gta cat aat ttc aca caa gtt ttt ata cag aca aat taa ttc tcg tcc ata ta- ttt taa aac att gac ttt tgt act aag aaa aat atc ttg act aac cat ctc ttt ctc tct tcg atg ggt ctc aca aaa ata tta aac ctc ttt ctg atg g-- --- --- --- --- --- --- --- --- --- --- --- --- --- --- --- --- --- --- --- --- --- --- --- --- --- --- --- --- --- --- --- --- --- --- --- --- --- --- --- --- --- --- --- --- --- --- agt cgt aaa aag ttt tta tcc ttt ctc tct tcg ata ggt ctc --- --- --- --- --- --- --- --- --- --- --- --- --- --- --- --- --- --- --- --- --- --- --- --- --- --- --- --- --- --- --- --- --- --- --- --- --- --- --- --- --- --- --- --- --- --- --- --- --- --- --- --- --- --- --- --- --- --- --- --- --- --- --- --- --- --- --- --- --- --- aca aaa ata tta aac ctc ttt ctg atg gtc tct ata aa

>ON676708.1_Monkeypox_virus_isolate_MPXV_USA_2021_MD_complete_genome

atc gag gac gac ttc aac tat gac agc tcg gtg gcg tct gcc agc gtg tac ata cga atg gca ttt cta aga aaa gtc tac ggt atc ctt -tc tac aat ttc ctt taa caa cgg caa cag ctg cag tat ttt tat act ttg aat gca -tc gga cat tta tac aag gga gtc ctg ttc taa tat tgg cat caa tgt tcg gat cta tag gct tga ttt tcg cat tga ctt tac aca gac atg aac atc ccc tga atc tgt aca tac ttt gtg gat tta cac tgt tag aat ctc taa cgc tgg cct ctg ttg tta ctt tct atg atg cac gta tcg tta tgc aag ctt tca tgt tga cta ctg cag tgt ttc ttg ctc tga cta cat ata ctc tac aat caa aga gag att tca gta aac ttg taa cag gat tgt ttg ctg ctt tct gga ttt taa ttt tgt cag gag tct tga gga taa agt tta aaa tag aat taa taa aga aca tat agg tca -tt ttt taa aca tgg ata gaa acc aag gtt gtt agt taa taa tat aca aga tat ttt ttc tca ctc tga tcc atg taa acc aag gac gag aga cac tct cat tcc tca ttc acg aca cca tta aaa atg gaa att aaa gcc ctc tat taa gca cag acg gct aca ggt cta cca tca ggt t-a cct tcg tct acc ttc aca atg gcc tct cct tgt gcc cag ttc agt ccc tgt cat tgc cac gct act aag gac tcc ctg aat acc gtg act gac gtc aga cat tgt ctg act gaa tac atc ctg tgg gtt tct cat aga tgg acc cat aga gaa agc gca ggg cct ctc tac agg ctt ctc atc tct ttc aga att gat gca atg gag cta ttt ggt agc gag ttg aag gag ttc tcg aat tca ctt ccg tgg gac aat atc gac aat tgc gtg gag atc att aaa tgt ttc atc aga aat gac tcc atg aaa acc gcc aaa gaa ctt tgt gca ata att gga ctt tgt act caa tca gct att gtc act gga aga gtc ttc aat gat aag tat atc gac ata cta ctt atg ctg cga aag att ctg aac gag aac gac tat ctc acc ctc ttg gat cat atc ctc act gct aaa tac taa atc tcc ttc atg ctc tct cac taa tac tct tac tca cta cac ttt tta tca tct tat gat gaa tga ttg cct tca tca ttt ttt cgt gga ata taa tat agg aat aat tag cac cag aat agc tat gga tat ctc gtt aag aat att ctc tat aag aga cat aat gta gac ata gtt att ata tcc ttc tta gat aag tgt tac gct act gga aag ttt cca tcg tta tta tta cat gaa gat gat ata att aaa cca aca ttg aga ttg gct ctt atg tta gct gga ttg aat tac tgt aat aaa tgc atc gag tat aga ggg ata tag caa ttc tcg ata ata gtc atg caa tat ttg aat gag act gat aat tta ggt aat aca gta ct- --- aca cac ata tct ttc tag att ata tat cgt taa aaa tct gta aga tgt ata ttt ctc ata agt atc cac tgt gta ata tta tta atg gat ata tag ata acg caa tag gga cta ata gta ttg taa aag ata taa tcg act att tac gta cat atc cag ata tct ata ttc cta cta gtt tgc tgc gta gtt gca tca ttg ata tgc atg -at tta tca gga ttc aga gat gaa tta cta agt aaa cta caa tcc cac aat aag taa gaa tca aat atc aaa aac tca ctt ttg att ttt cta gtc tta agt aat aca tat att tat taa tag acc tat gaa ata aaa aaa ggt aac aat gga ttc gcg tat agc tat tta cgt att agt atc ggc atc tct ttt gta tct tgt taa ttg tca caa act agt aca tta ctt caa tct gaa aat aaa tgg aag tga tat aac taa tac agc aga tat att gct gga caa tta tcc aat tat gac ctt tga tgg aaa gga tat tta tcc atc tat ctc gtt cat ggt cgg taa taa act ttt cct aga tct tta taa aaa tat ctt tga aga att ttt cag act att tcg agt atc tgt aag tag tca ata cga gga att aga ata tta tta ttc atg tga tta tac taa caa ccg tcc tac aat taa aca aca tta ctt tta taa cgg cga cga ata tac tga aat tga tag atc gaa aaa agc cac taa taa aaa cag ttg gtt aat tac ttc agg ctt tag act aca aaa atg gtt cga tag cga aga ttg tat aat tta tct cag atc ttt agt tag aag aat gga aga cag taa caa aaa cag taa aaa aac tta gta ctt aga tat cg- --a aaa aat ata ttt ttg tag act ctt gag aat aga agg aaa aca tgt aca taa tta taa aaa atg aaa atc aat ggc gaa taa gac agt gcg att cgc acc atg gag tcg gta gat ttc atg gct gtc gat gag cag ttt cac gac gac ctc gat ctt tgg tca tta tct ttg gta gat gat tat aaa aaa cat gga tta ggt gtt gac tgt tat gtt cta gaa cca gtt gtt gac agg aaa ata ttt gat aga ttt ctc ctt gaa cca att tgt gat cct gta gat gtt ctg tat gat tat ttt agg att cat aga gat aat att gat cag tat ata gta gat aga ctg ttt gca tat att aca tat aaa gat att ata tct gca tta gtg tca aag aat tat atg gaa gat att ttc tct ata att att aag aat tgt aat tct gtg caa gat ctc tta ctt tac tat cta tct aat gca tat gta gaa ata gac att gtt gat ctt atg gta gat cat ggg gct gta ata tat aaa ata gaa tgc ttg aat gcc tat ttt agg gga ata tgt aaa aag gaa agt agt gtt gtt gag ttt att ttg aat tgt ggt atc cca gat gaa aat gat gtt aaa tta gat cta tat aaa ata att cag tat act agg gga ttc ctt gta gat gaa ccc aca gta tta gaa att tat aag ctt tgt atc cca tat att gaa gat atc aat caa cta gat gct ggt gga agg acc ttg ctt tat cgc gct atc tat gca ggt tat ata gat tta gta tca tgg cta tta gaa aat gga gca aat gtc aac gca gta atg agt aat gga tat aca tgt ctt gac gtg gcc gtg gat agg gga tct gtc atc gcc cgt agg gaa gca cat ctt aaa ata tta gaa ata ttg ctt aga gaa cca ttg tct att gac tgt ata aaa tta gct ata ctt aat aat aca att gaa aac cat gat gtg ata aag ctc tgt atc aag tat ttt atg atg gta gat tat tca ctt tgt aat gtg tat gca tca tca ctc ttt gat tat ata att gat tgt aaa caa gaa ttg gag tac att agg cag atg aaa att cat aat aca acc atg tat gag tta atc tat aat aga gac aaa aac aag cat gct tcc cat att cta cat agg tat tct aaa cat cca gtt ttg aca cag tgt atc act aaa gga ttc aag att tac aca gaa gta acc gag cag gtc act aaa gct cta aac aga cgt gct cta ata gat gag ata ata aac aat gta tca act gat gac aat ctc cta tca aaa ctt cca tta gaa att agg gat cta att gtt tca caa gct gtc ata tag agt tct atc cac cca cct ttc ttg aaa tga gtt aat --- --- --- --- --- --- --- --- --- --- --- --- --- --- --- --- --- --- --- --- --- --- --- --- --- --- --- --- --- --- --- --- --- --- --- --- --- --- --- --- --- --- --- --- --- --- --- --- --- --- --- --- --- --- --- --- --- --- --- --- --- --- --- --- --- --- --- --- --- --- --- --- --- --- --- --- --- --- --- --- --- --- --- --- --- --- --- --- --- --- --- --- --- --- --- --- --- --- --- --- --- --- --- --- --- -ag tca taa gtt agt taa --- --- --- --- --- -gt cat aag tta gtt aag tca t?- --- --- --- --- --- --- --- --- --- --- --- --- --- --- --? ?aa gtt agt taa gtc ata agt tag tta agt cat aag tta gtt aag tca taa gtt agt taa gtc ata agt tag tta agt cat aag tta gtt tat agt cta aca ctt cta att ttt ata cct tga tct ttt tct cta att atg aaa aag taa atc att atg aag atg gat g-a aat gga cga gat tgt gcg cat cgt taa cga tag tat gtg gta cgt acc taa cgc att tat gga cga cgg tga taa tga agg tca cat ttc tgt caa taa tgt ctg tca tat gta tct cgc att ctt tga tgt gga tat atc atc tca tct gtt taa att agt tat taa aca ctg cga tct gaa taa acg act aaa atg tgg taa ctc tcc att aca ttg cta tac gat gaa tac acg att taa tcc atc tgt att aaa gat att gtt acg cca cgg cat gcg taa ctt tga tag caa gga taa aaa agg aca tat tcc tct aca cca cta tct gat tca ttc act atc aat cga taa caa gat ctt tga tat act aac gga ccc cat tga tga ctt tag taa atc atc cga tct att gct gtg tta tct tag ata taa att caa tgg gag ctt aaa cta tta cgt tct gta caa att att gac taa agg atc tga ccc taa ttg cgt cga tga gga tgg act cac ttc tct tca tta cta ctg taa aca cat atc cgc gtt cca cga aag caa tta tta caa gtc aaa gag tca cac taa gat gcg agc tga gaa gcg att cat cta cgc gat aat aga tca tgg agc aaa cat taa cgc ggt tac gaa aat cgg aaa tac gcc gtt aca cac tta cct tca aca gta tac caa aca tag tcc tcg tgt ggt gta tgc tct ttt atc tcg agg agc cga tac gag gat acg taa taa tct taa ttg tac acc cat cat gga ata cat aaa gaa cga ttg tgc aac agg tca tat tct cat aat gtt act caa ttg gca cga aca aaa ata cgg gaa att aca aaa gga aga agg aca aca tct act tta tct att cat aaa aca taa tca agg ata tgg aag tcg ctc tct caa tat act acg gta tct act aga tag att cga cat tca gaa aga cga ata cta taa tac aat gac tcc tct tca tac cgc ctt cca gaa ttg caa taa caa tgt tgc ctc ata cct cgt ata cat tgg ata cga cat caa cct tcc gac taa aga cga taa gac agt att cga ctt ggt gtt tga aaa cag aaa cat tat ata caa ggc gga tgt cgt taa tga cat tat cca cca cag act gaa agt atc tct acc tat gat taa atc gtt gtt cta caa gat gtc gga gtt ctc tcc cta cga cga tca cta cgt aaa gaa gat aat agc cta ctg cct att aag gga cga gtc att tgc gga act aca tac taa att ctg ttt aaa cga gga cta taa aag tgt att tat gaa aaa tat atc att cga taa gat aga ttc cat cat cga aaa atg tag tcg tga cat aag tct cct caa aga gat tcg aat ctc aga cac cga ctt gta tac ggt att gag aac aga aga cat ccg gta tca cac ata tct cga agc cat aca ttc aga caa acg cat ttc att tcc cat gta cga cga tct cat aga aca gtg tca tct atc gat gga gca taa aag taa act cgt cga caa agc act caa taa att aga gtc tac cat cga tag tca atc tag act atc gta ttt gcc tcc gga aat tat gcg caa tat cat aac caa gct aag cga cta cca tct aaa cag tat gtt gta cgg aaa gaa cca tta caa ata tta tcc atg ata gaa aga aaa tat tta aaa aat aat cta tat gat tgg aga agt agg aaa caa aca gta aca aga cga cga tta cta c-a tta tta aat cat gag gtc cgt att ata ctc gta tat att gtt tct ctc atg tat aat aat aaa cgg aag aga tat agc acc aca tgc acc atc caa tgg aaa gtg taa aga caa cga ata cag aag ccg taa tct atg ttg tct atc gtg tcc tcc ggg aac tta cgc ttt cag att atg tga tag caa gac taa tac aca atg tac acc gtg tgg ttc gga tac ctt tac atc tca caa taa tca ttt aca ggc ttg tct aag ttg taa cgg aag atg tga tag taa tca ggt aga gac gcg atc gtg taa cac gac tca caa tag aat ctg tga atg ctc tcc agg ata tta ttg tct tct caa agg agc atc agg gtg tag aac atg tat ttc taa aac aaa gtg tgg aat agg ata cgg agt atc cgg ata cac gtc tac cgg aga cgt cat ctg ttc tcc gtg tgg tcc cgg aac ata ttc tca cac cgt ctc ttc cac aga taa atg cga acc cgt cgt aac cag caa tac att taa cta tat cga tgt gga aat taa cct gta tcc agt caa cga cac atc gtg tac tcg gac gac cac tac cgg tct cag cga atc cat ctc aac gtc gga act aac tat tac cat gaa tca taa aga ttg tga tcc agt ctt tcg tgc aga ata ctt ctc tgt cct taa taa tgt agc aac ttc agg att ctt tac agg aga aaa tag ata tca gaa tac ttc aaa gat atg tac tct gaa ttt cga gat taa atg taa caa caa aga ttc atc ttc caa aca gtt aac gaa aac aaa gaa tga tac tat cat gcc gca ttc aga gac ggt aac tct agt ggg cga ctg tct atc tag cgt cga cat cta cat act ata tag taa tac caa tac tca aga cta cga aaa tga tac aat ctc tta tca tat ggg taa tgt tct cga tgt caa tag cca tat gcc cgc tag ttg cga tat aca taa act gat cac taa ttc cca gaa tcc cac cca ctt ata gta ag- -tt ttt tta cct ata aat aat aaa tac aat aat taa ttt ctc gta aaa gta gaa aat ata ttc taa ttt att ata tgg taa gaa agt aga atc atc tag aac agt aat caa tca ata gca atc atg aaa caa tat att gtc ctg gca tgc atg tgc cta gtg gca gct gct atg cct act agt ctt caa caa tct tca tcc tcg tgt act gaa gaa gaa aac aaa cat cat atg gga atc gat gtt att atc aaa gtc aca aag caa gac caa aca ccg acc aat gat aag att tgt caa tcc gta acg gaa gtt aca gag a-- --- --- --- --- --- -cc gaa gat gat gag gta tcc gaa gaa gtt gta aaa gga gat ccc acc act tat tac act atc gtc ggt gcg ggt ctt aac atg aac ttt gga ttc acc aaa tgc cca aag att tta tcc atc tcc gaa tcc tct gat gga aac act gtg aat act aga ttg tcc agc gtg tca ccg gga caa ggt aag gac tct ccc gcg atc acg cgt gaa gaa gct ctg gct atg atc aaa gac tgt gag atg tct atc gac atc aga tgt agc gaa gaa gag aaa gac agt gac atc aag acc cat cca gta ctt ggg tct aac atc tca cat aag aaa gtg agt tac aaa gat atc atc ggt tca acg atc gtt gat aca aaa tgt gtc aag aac cta gag ttt agc gta cgt atc gga gac atg tgt gag gaa tca tct gaa ctt gaa gtc aag gat gga ttc aag tat gtc gac gga tcg gca tct gaa ggt gca acc gat gat act tca ctc atc gat tca aca aaa ctc aaa gca tgt gtc tga atc gat aac tct att cat ctg aaa atg gat gag ttg ggt taa tcg aac gat tca gac acc gca cca cga att aaa aaa g-- -ac cgg gca cta tat tcc ggt ttg caa aac aaa aat att taa cta cat tca caa aaa gtt acc tct cgt tac ttc ttc ttt ctg ttt caa tat gtg ata cga tat gat cac tat tcg tat tct ct- --- --- tgg tct cat aaa aaa gtt tta c-- --a aaa aaa aaa aaa a-t att ttt att ctc ttt ctc tct tcg at- ggt ctc aca aaa ata tta aac ctc ttt ctg atg tct caa cta ttt cgt aaa cga taa cgt cca aca ata tat tct cgt aga gct tat caa cat cct tat acc aat cta ggt tgt cag aca att gca tca taa aat aat gtt tat aat tta cac gtt aac atc ata taa taa acg tat ata gtt aat att ttt gga ata taa atg atc tgt aaa atc cat gta ggg gac act gct cac gtt ttt tct cta gta cat aat ttc aca caa gtt ttt ata cag aca aat taa ttc tcg tcc ata ta- ttt taa aac att gac ttt tgt act aag aaa aat atc ttg act aac cat ctc ttt ctc tct tcg atg ggt ctc aca aaa ata tta aac ctc ttt ctg atg g-- --- --- --- --- --- --- --- --- --- --- --- --- --- --- --- --- --- --- --- --- --- --- --- --- --- --- --- --- --- --- --- --- --- --- --- --- --- --- --- --- --- --- --- --- --- --- agt cgt aaa aag ttt tta tcc ttt ctc tct tcg ata ggt ctc --- --- --- --- --- --- --- --- --- --- --- --- --- --- --- --- --- --- --- --- --- --- --- --- --- --- --- --- --- --- --- --- --- --- --- --- --- --- --- --- --- --- --- --- --- --- --- --- --- --- --- --- --- --- --- --- --- --- --- --- --- --- --- --- --- --- --- --- --- --- aca aaa ata tta aac ctc ttt ctg atg gtc tct ata aa

>ON585033.1_Monkeypox_virus_isolate_Monkeypox/PT0006/2022_complete_genome

atc gag gac gac ttc aac tat gac agc tcg gtg gcg tct gcc agc gtg tac ata cga atg gca ttt cta aga aaa gtc tac ggt atc ctt -tc tac aat ttc ctt taa caa cgg caa cag ctg cag tat ttt tat act ttg aat gca -tc gga cat tta tac aag gga gtc ctg ttc taa tat tgg cat caa tgt tcg gat cta tag gct tga ttt tcg cat tga ctt tac aca gac atg aac atc ccc tga atc tgt aca tac ttt gtg gat tta cac tgt tag aat ctc taa cgc tgg cct ctg ttg tta ctt tct atg atg cac gta tcg tta tgc aag ctt tca tgt tga cta ctg cag tgt ttc ttg ctc tga cta cat ata ctc tac aat caa aga gag att tca gta aac ttg taa cag gat tgt ttg ctg ctt tct gga ttt taa ttt tgt cag gag tct tga gga taa agt tta aaa tag aat taa taa aga aca tat agg tca -tt ttt taa aca tgg ata gaa acc aag gtt gtt agt taa taa tat aca aga tat ttt ttc tca ctc tga tcc atg taa acc aag gac gag aga cac tct cat tcc tca ttc acg aca cca tta aaa atg gaa att aaa gcc ctc tat taa gca cag acg gct aca ggt cta cca tca ggt t-a cct tcg tct acc ttc aca atg gcc tct cct tgt gcc cag ttc agt ccc tgt cat tgc cac gct act aag gac tcc ctg aat acc gtg act gac gtc aga cat tgt ctg act gaa tac atc ctg tgg gtt tct cat aga tgg acc cat aga gaa agc gca ggg cct ctc tac agg ctt ctc atc tct ttc aga att gat gca atg gag cta ttt ggt agc gag ttg aag gag ttc tcg aat tca ctt ccg tgg gac aat atc gac aat tgc gtg gag atc att aaa tgt ttc atc aga aat gac tcc atg aaa acc gcc aaa gaa ctt tgt gca ata att gga ctt tgt act caa tca gct att gtc act gga aga gtc ttc aat gat aag tat atc gac ata cta ctt atg ctg cga aag att ctg aac gag aac gac tat ctc acc ctc ttg gat cat atc ctc act gct aaa tac taa atc tcc ttc atg ctc tct cac taa tac tct tac tca cta cac ttt tta tca tct tat gat gaa tga ttg cct tca tca ttt ttt cgt gga ata taa tat agg aat aat tag cac cag aat agc tat gga tat ctc gtt aag aat att ctc tat aag aga cat aat gta gac ata gtt att ata tcc ttc tta gat aag tgt tac gct act gga aag ttt cca tcg tta tta tta cat gaa gat gat ata att aaa cca aca ttg aga ttg gct ctt atg tta gct gga ttg aat tac tgt aat aaa tgc atc gag tat aga ggg ata tag caa ttc tcg ata ata gtc atg caa tat ttg aat gag act gat aat tta ggt aat aca gta ct- --- aca cac ata tct ttc tag att ata tat cgt taa aaa tct gta aga tgt ata ttt ctc ata agt atc cac tgt gta ata tta tta atg gat ata tag ata acg caa tag gga cta ata gta ttg taa aag ata taa tcg act att tac gta cat atc cag ata tct ata ttc cta cta gtt tgc tgc gta gtt gca tca ttg ata tgc atg -at tta tca gga ttc aga gat gaa tta cta agt aaa cta caa tcc cac aat aag taa gaa tca aat atc aaa aac tca ctt ttg att ttt cta gtc tta agt aat aca tat att tat taa tag acc tat gaa ata aaa aaa ggt aac aat gga ttc gcg tat agc tat tta cgt att agt atc ggc atc tct ttt gta tct tgt taa ttg tca caa act agt aca tta ctt caa tct gaa aat aaa tgg aag tga tat aac taa tac agc aga tat att gct gga caa tta tcc aat tat gac ctt tga tgg aaa gga tat tta tcc atc tat ctc gtt cat ggt cgg taa taa act ttt cct aga tct tta taa aaa tat ctt tga aga att ttt cag act att tcg agt atc tgt aag tag tca ata cga gga att aga ata tta tta ttc atg tga tta tac taa caa ccg tcc tac aat taa aca aca tta ctt tta taa cgg cga cga ata tac tga aat tga tag atc gaa aaa agc cac taa taa aaa cag ttg gtt aat tac ttc agg ctt tag act aca aaa atg gtt cga tag cga aga ttg tat aat tta tct cag atc ttt agt tag aag aat gga aga cag taa caa aaa cag taa aaa aac tta gta ctt aga tat cg- --a aaa aat ata ttt ttg tag act ctt gag aat aga agg aaa aca tgt aca taa tta taa aaa atg aaa atc aat ggc gaa taa gac agt gcg att cgc acc atg gag tcg gta gat ttc atg gct gtc gat gag cag ttt cac gac gac ctc gat ctt tgg tca tta tct ttg gta gat gat tat aaa aaa cat gga tta ggt gtt gac tgt tat gtt cta gaa cca gtt gtt gac agg aaa ata ttt gat aga ttt ctc ctt gaa cca att tgt gat cct gta gat gtt ctg tat gat tat ttt agg att cat aga gat aat att gat cag tat ata gta gat aga ctg ttt gca tat att aca tat aaa gat att ata tct gca tta gtg tca aag aat tat atg gaa gat att ttc tct ata att att aag aat tgt aat tct gtg caa gat ctc tta ctt tac tat cta tct aat gca tat gta gaa ata gac att gtt gat ctt atg gta gat cat ggg gct gta ata tat aaa ata gaa tgc ttg aat gcc tat ttt agg gga ata tgt aaa aag gaa agt agt gtt gtt gag ttt att ttg aat tgt ggt atc cca gat gaa aat gat gtt aaa tta gat cta tat aaa ata att cag tat act agg gga ttc ctt gta gat gaa ccc aca gta tta gaa att tat aag ctt tgt atc cca tat att gaa gat atc aat caa cta gat gct ggt gga agg acc ttg ctt tat cgc gct atc tat gca ggt tat ata gat tta gta tca tgg cta tta gaa aat gga gca aat gtc aac gca gta atg agt aat gga tat aca tgt ctt gac gtg gcc gtg gat agg gga tct gtc atc gcc cgt agg gaa gca cat ctt aaa ata tta gaa ata ttg ctt aga gaa cca ttg tct att gac tgt ata aaa tta gct ata ctt aat aat aca att gaa aac cat gat gtg ata aag ctc tgt atc aag tat ttt atg atg gta gat tat tca ctt tgt aat gtg tat gca tca tca ctc ttt gat tat ata att gat tgt aaa caa gaa ttg gag tac att agg cag atg aaa att cat aat aca acc atg tat gag tta atc tat aat aga gac aaa aac aag cat gct tcc cat att cta cat agg tat tct aaa cat cca gtt ttg aca cag tgt atc act aaa gga ttc aag att tac aca gaa gta acc gag cag gtc act aaa gct cta aac aga cgt gct cta ata gat gag ata ata aac aat gta tca act gat gac aat ctc cta tca aaa ctt cca tta gaa att agg gat cta att gtt tca caa gct gtc ata tag agt tct atc cac cca cct ttc ttg aaa tga gtt aat --- --- --- --- --- --- --- --- --- --- --- --- --- --- --- --- --- --- --- --- --- --- --- --- --- --- --- --- --- --- --- --- --- --- --- --- --- --- --- --- --- --- --- --- --- --- --- --- --- --- --- --- --- --- --- --- --- --- --- --- --- --- --- --- --- --- --- --- --- --- --- --- --- --- --- --- --- --- --- --- --- --- --- --- --- --- --- --- --- --- --- --- --- --- --- --- --- --- --- --- --- --- --- --- --- -ag tca taa gtt agt taa --- --- --- --- --- -gt cat aag tta gtt aag tca t-- --- --- --- --- --- --- --- --- --- --- --- --- --- --- --- -aa gtt agt taa gtc ata agt tag tta agt cat aag tta gtt aag tca taa gtt agt taa gtc ata agt tag tta agt cat aag tta gtt tat agt cta aca ctt cta att ttt ata cct tga tct ttt tct cta att atg aaa aag taa atc att atg aag atg gat g-a aat gga cga gat tgt gcg cat cgt taa cga tag tat gtg gta cgt acc taa cgc att tat gga cga cgg tga taa tga agg tca cat ttc tgt caa taa tgt ctg tca tat gta tct cgc att ctt tga tgt gga tat atc atc tca tct gtt taa att agt tat taa aca ctg cga tct gaa taa acg act aaa atg tgg taa ctc tcc att aca ttg cta tac gat gaa tac acg att taa tcc atc tgt att aaa gat att gtt acg cca cgg cat gcg taa ctt tga tag caa gga taa aaa agg aca tat tcc tct aca cca cta tct gat tca ttc act atc aat cga taa caa gat ctt tga tat act aac gga ccc cat tga tga ctt tag taa atc atc cga tct att gct gtg tta tct tag ata taa att caa tgg gag ctt aaa cta tta cgt tct gta caa att att gac taa agg atc tga ccc taa ttg cgt cga tga gga tgg act cac ttc tct tca tta cta ctg taa aca cat atc cgc gtt cca cga aag caa tta tta caa gtc aaa gag tca cac taa gat gcg agc tga gaa gcg att cat cta cgc gat aat aga tca tgg agc aaa cat taa cgc ggt tac gaa aat cgg aaa tac gcc gtt aca cac tta cct tca aca gta tac caa aca tag tcc tcg tgt ggt gta tgc tct ttt atc tcg agg agc cga tac gag gat acg taa taa tct taa ttg tac acc cat cat gga ata cat aaa gaa cga ttg tgc aac agg tca tat tct cat aat gtt act caa ttg gca cga aca aaa ata cgg gaa att aca aaa gga aga agg aca aca tct act tta tct att cat aaa aca taa tca agg ata tgg aag tcg ctc tct caa tat act acg gta tct act aga tag att cga cat tca gaa aga cga ata cta taa tac aat gac tcc tct tca tac cgc ctt cca gaa ttg caa taa caa tgt tgc ctc ata cct cgt ata cat tgg ata cga cat caa cct tcc gac taa aga cga taa gac agt att cga ctt ggt gtt tga aaa cag aaa cat tat ata caa ggc gga tgt cgt taa tga cat tat cca cca cag act gaa agt atc tct acc tat gat taa atc gtt gtt cta caa gat gtc gga gtt ctc tcc cta cga cga tca cta cgt aaa gaa gat aat agc cta ctg cct att aag gga cga gtc att tgc gga act aca tac taa att ctg ttt aaa cga gga cta taa aag tgt att tat gaa aaa tat atc att cga taa gat aga ttc cat cat cga aaa atg tag tcg tga cat aag tct cct caa aga gat tcg aat ctc aga cac cga ctt gta tac ggt att gag aac aga aga cat tcg gta tca cac ata tct cga agc cat aca ttc aga caa acg cat ttc att tcc cat gta cga cga tct cat aga aca gtg tca tct atc gat gga gca taa aag taa act cgt cga caa agc act caa taa att aga gtc tac cat cga tag tca atc tag act atc gta ttt gcc tcc gga aat tat gcg caa tat cat aac caa gct aag cga cta cca tct aaa cag tat gtt gta cgg aaa gaa cca tta caa ata tta tcc atg ata gaa aga aaa tat tta aaa aat aat cta tat gat tgg aga agt agg aaa caa aca gta aca aga cga cga tta cta c-a tta tta aat cat gag gtc cgt att ata ctc gta tat att gtt tct ctc atg tat aat aat aaa cgg aag aga tat agc acc aca tgc acc atc caa tgg aaa gtg taa aga caa cga ata cag aag ccg taa tct atg ttg tct atc gtg tcc tcc ggg aac tta cgc ttt cag att atg tga tag caa gac taa tac aca atg tac acc gtg tgg ttc gga tac ctt tac atc tca caa taa tca ttt aca ggc ttg tct aag ttg taa cgg aag atg tga tag taa tca ggt aga gac gcg atc gtg taa cac gac tca caa tag aat ctg tga atg ctc tcc agg ata tta ttg tct tct caa agg agc atc agg gtg tag aac atg tat ttc taa aac aaa gtg tgg aat agg ata cgg agt atc cgg ata cac gtc tac cgg aga cgt cat ctg ttc tcc gtg tgg tcc cgg aac ata ttc tca cac cgt ctc ttc cac aga taa atg cga acc cgt cgt aac cag caa tac att taa cta tat cga tgt gga aat taa cct gta tcc agt caa cga cac atc gtg tac tcg gac gac cac tac cgg tct cag cga atc cat ctc aac gtc gga act aac tat tac cat gaa tca taa aga ttg tga tcc agt ctt tcg tgc aga ata ctt ctc tgt cct taa taa tgt agc aac ttc agg att ctt tac agg aga aaa tag ata tca gaa tac ttc aaa gat atg tac tct gaa ttt cga gat taa atg taa caa caa aga ttc atc ttc caa aca gtt aac gaa aac aaa gaa tga tac tat cat gcc gca ttc aga gac ggt aac tct agt ggg cga ctg tct atc tag cgt cga cat cta cat act ata tag taa tac caa tac tca aga cta cga aaa tga tac aat ctc tta tca tat ggg taa tgt tct cga tgt caa tag cca tat gcc cgc tag ttg cga tat aca taa act gat cac taa ttc cca gaa tcc cac cca ctt ata gta ag- -tt ttt tta cct ata aat aat aaa tac aat aat taa ttt ctc gta aaa gta gaa aat ata ttc taa ttt att ata tgg taa gaa agt aga atc atc tag aac agt aat caa tca ata gca atc atg aaa caa tat att gtc ctg gca tgc atg tgc cta gtg gca gct gct atg cct act agt ctt caa caa tct tca tcc tcg tgt act gaa gaa gaa aac aaa cat cat atg gga atc gat gtt att atc aaa gtc aca aag caa gac caa aca ccg acc aat gat aag att tgt caa tcc gta acg gaa gtt aca gag a-- --- --- --- --- --- -cc gaa gat gat gag gta tcc gaa gaa gtt gta aaa gga gat ccc acc act tat tac act atc gtc ggt gcg ggt ctt aac atg aac ttt gga ttc acc aaa tgc cca aag att tta tcc atc tcc gaa tcc tct gat gga aac act gtg aat act aga ttg tcc agc gtg tca ccg gga caa ggt aag gac tct ccc gcg atc acg cgt gaa gaa gct ctg gct atg atc aaa gac tgt gag atg tct atc gac atc aga tgt agc gaa gaa gag aaa gac agt gac atc aag acc cat cca gta ctt ggg tct aac atc tca cat aag aaa gtg agt tac aaa gat atc atc ggt tca acg atc gtt gat aca aaa tgt gtc aag aac cta gag ttt agc gta cgt atc gga gac atg tgt gag gaa tca tct gaa ctt gaa gtc aag gat gga ttc aag tat gtc gac gga tcg gca tct gaa ggt gca acc gat gat act tca ctc atc gat tca aca aaa ctc aaa gca tgt gtc tga atc gat aac tct att cat ctg aaa atg gat gag ttg ggt taa tcg aac gat tca gac acc gca cca cga att aaa aaa g-- -ac cgg gca cta tat tcc ggt ttg caa aac aaa aat att taa cta cat tca caa aaa gtt acc tct cgt tac ttc ttc ttt ctg ttt caa tat gtg ata cga tat gat cac tat tcg tat tct ct- --- --- tgg tct cat aaa aaa gtt tta c-- --a aaa aaa aaa aaa a-t att ttt att ctc ttt ctc tct tcg at- ggt ctc aca aaa ata tta aac ctc ttt ctg atg tct caa cta ttt cgt aaa cga taa cgt cca aca ata tat tct cgt aga gct tat caa cat cct tat acc aat cta ggt tgt cag aca att gca tca taa aat aat gtt tat aat tta cac gtt aac atc ata taa taa acg tat ata gtt aat att ttt gga ata taa atg atc tgt aaa atc cat gta ggg gac act gct cac gtt ttt tct cta gta cat aat ttc aca caa gtt ttt ata cag aca aat taa ttc tcg tcc ata ta- ttt taa aac att gac ttt tgt act aag aaa aat atc ttg act aac cat ctc ttt ctc tct tcg atg ggt ctc aca aaa ata tta aac ctc ttt ctg atg g-- --- --- --- --- --- --- --- --- --- --- --- --- --- --- --- --- --- --- --- --- --- --- --- --- --- --- --- --- --- --- --- --- --- --- --- --- --- --- --- --- --- --- --- --- --- --- agt cgt aaa aag ttt tta tcc ttt ctc tct tcg ata ggt ctc --- --- --- --- --- --- --- --- --- --- --- --- --- --- --- --- --- --- --- --- --- --- --- --- --- --- --- --- --- --- --- --- --- --- --- --- --- --- --- --- --- --- --- --- --- --- --- --- --- --- --- --- --- --- --- --- --- --- --- --- --- --- --- --- --- --- --- --- --- --- aca aaa ata tta aac ctc ttt ctg atg gtc tct ata aa

>ON585035.1_Monkeypox_virus_isolate_Monkeypox/PT0009/2022_complete_genome

atc gag gac gac ttc aac tat gac agc tcg gtg gcg tct gcc agc gtg tac ata cga atg gca ttt cta aga aaa gtc tac ggt atc ctt -tc tac aat ttc ctt taa caa cgg caa cag ctg cag tat ttt tat act ttg aat gca -tc gga cat tta tac aag gga gtc ctg ttc taa tat tgg cat caa tgt tcg gat cta tag gct tga ttt tcg cat tga ctt tac aca gac atg aac atc ccc tga atc tgt aca tac ttt gtg gat tta cac tgt tag aat ctc taa cgc tgg cct ctg ttg tta ctt tct atg atg cac gta tcg tta tgc aag ctt tca tgt tga cta ctg cag tgt ttc ttg ctc tga cta cat ata ctc tac aat caa aga gag att tca gta aac ttg taa cag gat tgt ttg ctg ctt tct gga ttt taa ttt tgt cag gag tct tga gga taa agt tta aaa tag aat taa taa aga aca tat agg tca -tt ttt taa aca tgg ata gaa acc aag gtt gtt agt taa taa tat aca aga tat ttt ttc tca ctc tga tcc atg taa acc aag gac gag aga cac tct cat tcc tca ttc acg aca cca tta aaa atg gaa att aaa gcc ctc tat taa gca cag acg gct aca ggt cta cca tca ggt t-a cct tcg tct acc ttc aca atg gcc tct cct tgt gcc cag ttc agt ccc tgt cat tgc cac gct act aag gac tcc ctg aat acc gtg act gac gtc aga cat tgt ctg act gaa tac atc ctg tgg gtt tct cat aga tgg acc cat aga gaa agc gca ggg cct ctc tac agg ctt ctc atc tct ttc aga att gat gca atg gag cta ttt ggt agc gag ttg aag gag ttc tcg aat tca ctt ccg tgg gac aat atc gac aat tgc gtg gag atc att aaa tgt ttc atc aga aat gac tcc atg aaa acc gcc aaa gaa ctt tgt gca ata att gga ctt tgt act caa tca gct att gtc act gga aga gtc ttc aat gat aag tat atc gac ata cta ctt atg ctg cga aag att ctg aac gag aac gac tat ctc acc ctc ttg gat cat atc ctc act gct aaa tac taa atc tcc ttc atg ctc tct cac taa tac tct tac tca cta cac ttt tta tca tct tat gat gaa tga ttg cct tca tca ttt ttt cgt gga ata taa tat agg aat aat tag cac cag aat agc tat gga tat ctc gtt aag aat att ctc tat aag aga cat aat gta gac ata gtt att ata tcc ttc tta gat aag tgt tac gct act gga aag ttt cca tcg tta tta tta cat gaa gat gat ata att aaa cca aca ttg aga ttg gct ctt atg tta gct gga ttg aat tac tgt aat aaa tgc atc gag tat aga ggg ata tag caa ttc tcg ata ata gtc atg caa tat ttg aat gag act gat aat tta ggt aat aca gta ct- --- aca cac ata tct ttc tag att ata tat cgt taa aaa tct gta aga tgt ata ttt ctc ata agt atc cac tgt gta ata tta tta atg gat ata tag ata acg caa tag gga cta ata gta ttg taa aag ata taa tcg act att tac gta cat atc cag ata tct ata ttc cta cta gtt tgc tgc gta gtt gca tca ttg ata tgc atg -at tta tca gga ttc aga gat gaa tta cta agt aaa cta caa tcc cac aat aag taa gaa tca aat atc aaa aac tca ctt ttg att ttt cta gtc tta agt aat aca tat att tat taa tag acc tat gaa ata aaa aaa ggt aac aat gga ttc gcg tat agc tat tta cgt att agt atc ggc atc tct ttt gta tct tgt taa ttg tca caa act agt aca tta ctt caa tct gaa aat aaa tgg aag tga tat aac taa tac agc aga tat att gct gga caa tta tcc aat tat gac ctt tga tgg aaa gga tat tta tcc atc tat ctc gtt cat ggt cgg taa taa act ttt cct aga tct tta taa aaa tat ctt tga aga att ttt cag act att tcg agt atc tgt aag tag tca ata cga gga att aga ata tta tta ttc atg tga tta tac taa caa ccg tcc tac aat taa aca aca tta ctt tta taa cgg cga cga ata tac tga aat tga tag atc gaa aaa agc cac taa taa aaa cag ttg gtt aat tac ttc agg ctt tag act aca aaa atg gtt cga tag cga aga ttg tat aat tta tct cag atc ttt agt tag aag aat gga aga cag taa caa aaa cag taa aaa aac tta gta ctt aga tat cg- --a aaa aat ata ttt ttg tag act ctt gag aat aga agg aaa aca tgt aca taa tta taa aaa atg aaa atc aat ggc gaa taa gac agt gcg att cgc acc atg gag tcg gta gat ttc atg gct gtc gat gag cag ttt cac gac gac ctc gat ctt tgg tca tta tct ttg gta gat gat tat aaa aaa cat gga tta ggt gtt gac tgt tat gtt cta gaa cca gtt gtt gac agg aaa ata ttt gat aga ttt ctc ctt gaa cca att tgt gat cct gta gat gtt ctg tat gat tat ttt agg att cat aga gat aat att gat cag tat ata gta gat aga ctg ttt gca tat att aca tat aaa gat att ata tct gca tta gtg tca aag aat tat atg gaa gat att ttc tct ata att att aag aat tgt aat tct gtg caa gat ctc tta ctt tac tat cta tct aat gca tat gta gaa ata gac att gtt gat ctt atg gta gat cat ggg gct gta ata tat aaa ata gaa tgc ttg aat gcc tat ttt agg gga ata tgt aaa aag gaa agt agt gtt gtt gag ttt att ttg aat tgt ggt atc cca gat gaa aat gat gtt aaa tta gat cta tat aaa ata att cag tat act agg gga ttc ctt gta gat gaa ccc aca gta tta gaa att tat aag ctt tgt atc cca tat att gaa gat atc aat caa cta gat gct ggt gga agg acc ttg ctt tat cgc gct atc tat gca ggt tat ata gat tta gta tca tgg cta tta gaa aat gga gca aat gtc aac gca gta atg agt aat gga tat aca tgt ctt gac gtg gcc gtg gat agg gga tct gtc atc gcc cgt agg gaa gca cat ctt aaa ata tta gaa ata ttg ctt aga gaa cca ttg tct att gac tgt ata aaa tta gct ata ctt aat aat aca att gaa aac cat gat gtg ata aag ctc tgt atc aag tat ttt atg atg gta gat tat tca ctt tgt aat gtg tat gca tca tca ctc ttt gat tat ata att gat tgt aaa caa gaa ttg gag tac att agg cag atg aaa att cat aat aca acc atg tat gag tta atc tat aat aga gac aaa aac aag cat gct tcc cat att cta cat agg tat tct aaa cat cca gtt ttg aca cag tgt atc act aaa gga ttc aag att tac aca gaa gta acc gag cag gtc act aaa gct cta aac aga cgt gct cta ata gat gag ata ata aac aat gta tca act gat gac aat ctc cta tca aaa ctt cca tta gaa att agg gat cta att gtt tca caa gct gtc ata tag agt tct atc cac cca cct ttc ttg aaa tga gtt aat --- --- --- --- --- --- --- --- --- --- --- --- --- --- --- --- --- --- --- --- --- --- --- --- --- --- --- --- --- --- --- --- --- --- --- --- --- --- --- --- --- --- --- --- --- --- --- --- --- --- --- --- --- --- --- --- --- --- --- --- --- --- --- --- --- --- --- --- --- --- --- --- --- --- --- --- --- --- --- --- --- --- --- --- --- --- --- --- --- --- --- --- --- --- --- --- --- --- --- --- --- --- --- --- --- -ag tca taa gtt agt taa --- --- --- --- --- -gt cat aag tta gtt aag tca t-- --- --- --- --- --- --- --- --- --- --- --- --- --- --- --- -aa gtt agt taa gtc ata agt tag tta agt cat aag tta gtt aag tca taa gtt agt taa gtc ata agt tag tta agt cat aag tta gtt tat agt cta aca ctt cta att ttt ata cct tga tct ttt tct cta att atg aaa aag taa atc att atg aag atg gat g-a aat gga cga gat tgt gcg cat cgt taa cga tag tat gtg gta cgt acc taa cgc att tat gga cga cgg tga taa tga agg tca cat ttc tgt caa taa tgt ctg tca tat gta tct cgc att ctt tga tgt gga tat atc atc tca tct gtt taa att agt tat taa aca ctg cga tct gaa taa acg act aaa atg tgg taa ctc tcc att aca ttg cta tac gat gaa tac acg att taa tcc atc tgt att aaa gat att gtt acg cca cgg cat gcg taa ctt tga tag caa gga taa aaa agg aca tat tcc tct aca cca cta tct gat tca ttc act atc aat cga taa caa gat ctt tga tat act aac gga ccc cat tga tga ctt tag taa atc atc cga tct att gct gtg tta tct tag ata taa att caa tgg gag ctt aaa cta tta cgt tct gta caa att att gac taa agg atc tga ccc taa ttg cgt cga tga gga tgg act cac ttc tct tca tta cta ctg taa aca cat atc cgc gtt cca cga aag caa tta tta caa gtc aaa gag tca cac taa gat gcg agc tga gaa gcg att cat cta cgc gat aat aga tca tgg agc aaa cat taa cgc ggt tac gaa aat cgg aaa tac gcc gtt aca cac tta cct tca aca gta tac caa aca tag tcc tcg tgt ggt gta tgc tct ttt atc tcg agg agc cga tac gag gat acg taa taa tct taa ttg tac acc cat cat gga ata cat aaa gaa cga ttg tgc aac agg tca tat tct cat aat gtt act caa ttg gca cga aca aaa ata cgg gaa att aca aaa gga aga agg aca aca tct act tta tct att cat aaa aca taa tca agg ata tgg aag tcg ctc tct caa tat act acg gta tct act aga tag att cga cat tca gaa aga cga ata cta taa tac aat gac tcc tct tca tac cgc ctt cca gaa ttg caa taa caa tgt tgc ctc ata cct cgt ata cat tgg ata cga cat caa cct tcc gac taa aga cga taa gac agt att cga ctt ggt gtt tga aaa cag aaa cat tat ata caa ggc gga tgt cgt taa tga cat tat cca cca cag act gaa agt atc tct acc tat gat taa atc gtt gtt cta caa gat gtc gga gtt ctc tcc cta cga cga tca cta cgt aaa gaa gat aat agc cta ctg cct att aag gga cga gtc att tgc gga act aca tac taa att ctg ttt aaa cga gga cta taa aag tgt att tat gaa aaa tat atc att cga taa gat aga ttc cat cat cga aaa atg tag tcg tga cat aag tct cct caa aga gat tcg aat ctc aga cac cga ctt gta tac ggt att gag aac aga aga cat tcg gta tca cac ata tct cga agc cat aca ttc aga caa acg cat ttc att tcc cat gta cga cga tct cat aga aca gtg tca tct atc gat gga gca taa aag taa act cgt cga caa agc act caa taa att aga gtc tac cat cga tag tca atc tag act atc gta ttt gcc tcc gga aat tat gcg caa tat cat aac caa gct aag cga cta cca tct aaa cag tat gtt gta cgg aaa gaa cca tta caa ata tta tcc atg ata gaa aga aaa tat tta aaa aat aat cta tat gat tgg aga agt agg aaa caa aca gta aca aga cga cga tta cta c-a tta tta aat cat gag gtc cgt att ata ctc gta tat att gtt tct ctc atg tat aat aat aaa cgg aag aga tat agc acc aca tgc acc atc caa tgg aaa gtg taa aga caa cga ata cag aag ccg taa tct atg ttg tct atc gtg tcc tcc ggg aac tta cgc ttt cag att atg tga tag caa gac taa tac aca atg tac acc gtg tgg ttc gga tac ctt tac atc tca caa taa tca ttt aca ggc ttg tct aag ttg taa cgg aag atg tga tag taa tca ggt aga gac gcg atc gtg taa cac gac tca caa tag aat ctg tga atg ctc tcc agg ata tta ttg tct tct caa agg agc atc agg gtg tag aac atg tat ttc taa aac aaa gtg tgg aat agg ata cgg agt atc cgg ata cac gtc tac cgg aga cgt cat ctg ttc tcc gtg tgg tcc cgg aac ata ttc tca cac cgt ctc ttc cac aga taa atg cga acc cgt cgt aac cag caa tac att taa cta tat cga tgt gga aat taa cct gta tcc agt caa cga cac atc gtg tac tcg gac gac cac tac cgg tct cag cga atc cat ctc aac gtc gga act aac tat tac cat gaa tca taa aga ttg tga tcc agt ctt tcg tgc aga ata ctt ctc tgt cct taa taa tgt agc aac ttc agg att ctt tac agg aga aaa tag ata tca gaa tac ttc aaa gat atg tac tct gaa ttt cga gat taa atg taa caa caa aga ttc atc ttc caa aca gtt aac gaa aac aaa gaa tga tac tat cat gcc gca ttc aga gac ggt aac tct agt ggg cga ctg tct atc tag cgt cga cat cta cat act ata tag taa tac caa tac tca aga cta cga aaa tga tac aat ctc tta tca tat ggg taa tgt tct cga tgt caa tag cca tat gcc cgc tag ttg cga tat aca taa act gat cac taa ttc cca gaa tcc cac cca ctt ata gta ag- -tt ttt tta cct ata aat aat aaa tac aat aat taa ttt ctc gta aaa gta gaa aat ata ttc taa ttt att ata tgg taa gaa agt aga atc atc tag aac agt aat caa tca ata gca atc atg aaa caa tat att gtc ctg gca tgc atg tgc cta gtg gca gct gct atg cct act agt ctt caa caa tct tca tcc tcg tgt act gaa gaa gaa aac aaa cat cat atg gga atc gat gtt att atc aaa gtc aca aag caa gac caa aca ccg acc aat gat aag att tgt caa tcc gta acg gaa gtt aca gag a-- --- --- --- --- --- -cc gaa gat gat gag gta tcc gaa gaa gtt gta aaa gga gat ccc acc act tat tac act atc gtc ggt gcg ggt ctt aac atg aac ttt gga ttc acc aaa tgc cca aag att tta tcc atc tcc gaa tcc tct gat gga aac act gtg aat act aga ttg tcc agc gtg tca ccg gga caa ggt aag gac tct ccc gcg atc acg cgt gaa gaa gct ctg gct atg atc aaa gac tgt gag atg tct atc gac atc aga tgt agc gaa gaa gag aaa gac agt gac atc aag acc cat cca gta ctt ggg tct aac atc tca cat aag aaa gtg agt tac aaa gat atc atc ggt tca acg atc gtt gat aca aaa tgt gtc aag aac cta gag ttt agc gta cgt atc gga gac atg tgt gag gaa tca tct gaa ctt gaa gtc aag gat gga ttc aag tat gtc gac gga tcg gca tct gaa ggt gca acc gat gat act tca ctc atc gat tca aca aaa ctc aaa gca tgt gtc tga atc gat aac tct att cat ctg aaa atg gat gag ttg ggt taa tcg aac gat tca gac acc gca cca cga att aaa aaa g-- -ac cgg gca cta tat tcc ggt ttg caa aac aaa aat att taa cta cat tca caa aaa gtt acc tct cgt tac ttc ttc ttt ctg ttt caa tat gtg ata cga tat gat cac tat tcg tat tct ct- --- --- tgg tct cat aaa aaa gtt tta c-- --a aaa aaa aaa aaa a-t att ttt att ctc ttt ctc tct tcg at- ggt ctc aca aaa ata tta aac ctc ttt ctg atg tct caa cta ttt cgt aaa cga taa cgt cca aca ata tat tct cgt aga gct tat caa cat cct tat acc aat cta ggt tgt cag aca att gca tca taa aat aat gtt tat aat tta cac gtt aac atc ata taa taa acg tat ata gtt aat att ttt gga ata taa atg atc tgt aaa atc cat gta ggg gac act gct cac gtt ttt tct cta gta cat aat ttc aca caa gtt ttt ata cag aca aat taa ttc tcg tcc ata ta- ttt taa aac att gac ttt tgt act aag aaa aat atc ttg act aac cat ctc ttt ctc tct tcg atg ggt ctc aca aaa ata tta aac ctc ttt ctg atg g-- --- --- --- --- --- --- --- --- --- --- --- --- --- --- --- --- --- --- --- --- --- --- --- --- --- --- --- --- --- --- --- --- --- --- --- --- --- --- --- --- --- --- --- --- --- --- agt cgt aaa aag ttt tta tcc ttt ctc tct tcg ata ggt ctc --- --- --- --- --- --- --- --- --- --- --- --- --- --- --- --- --- --- --- --- --- --- --- --- --- --- --- --- --- --- --- --- --- --- --- --- --- --- --- --- --- --- --- --- --- --- --- --- --- --- --- --- --- --- --- --- --- --- --- --- --- --- --- --- --- --- --- --- --- --- aca aaa ata tta aac ctc ttt ctg atg gtc tct ata aa

>ON649725.1_Monkeypox_virus_isolate_Monkeypox/PT0011/2022_complete_genome

atc gag gac gac ttc aac tat gac agc tcg gtg gcg tct gcc agc gtg tac ata cga atg gca ttt cta aga aaa gtc tac ggt atc ctt -tc tac aat ttc ctt taa caa cgg caa cag ctg cag tat ttt tat act ttg aat gca -tc gga cat tta tac aag gga gtc ctg ttc taa tat tgg cat caa tgt tcg gat cta tag gct tga ttt tcg cat tga ctt tac aca gac atg aac atc ccc tga atc tgt aca tac ttt gtg gat tta cac tgt tag aat ctc taa cgc tgg cct ctg ttg tta ctt tct atg atg cac gta tcg tta tgc aag ctt tca tgt tga cta ctg cag tgt ttc ttg ctc tga cta cat ata ctc tac aat caa aga gag att tca gta aac ttg taa cag gat tgt ttg ctg ctt tct gga ttt taa ttt tgt cag gag tct tga gga taa agt tta aaa tag aat taa taa aga aca tat agg tca -tt ttt taa aca tgg ata gaa acc aag gtt gtt agt taa taa tat aca aga tat ttt ttc tca ctc tga tcc atg taa acc aag gac gag aga cac tct cat tcc tca ttc acg aca cca tta aaa atg gaa att aaa gcc ctc tat taa gca cag acg gct aca ggt cta cca tca ggt t-a cct tcg tct acc ttc aca atg gcc tct cct tgt gcc cag ttc agt ccc tgt cat tgc cac gct act aag gac tcc ctg aat acc gtg act gac gtc aga cat tgt ctg act gaa tac atc ctg tgg gtt tct cat aga tgg acc cat aga gaa agc gca ggg cct ctc tac agg ctt ctc atc tct ttc aga att gat gca atg gag cta ttt ggt agc gag ttg aag gag ttc tcg aat tca ctt ccg tgg gac aat atc gac aat tgc gtg gag atc att aaa tgt ttc atc aga aat gac tcc atg aaa acc gcc aaa gaa ctt tgt gca ata att gga ctt tgt act caa tca gct att gtc act gga aga gtc ttc aat gat aag tat atc gac ata cta ctt atg ctg cga aag att ctg aac gag aac gac tat ctc acc ctc ttg gat cat atc ctc act gct aaa tac taa atc tcc ttc atg ctc tct cac taa tac tct tac tca cta cac ttt tta tca tct tat gat gaa tga ttg cct tca tca ttt ttt cgt gga ata taa tat agg aat aat tag cac cag aat agc tat gga tat ctc gtt aag aat att ctc tat aag aga cat aat gta gac ata gtt att ata tcc ttc tta gat aag tgt tac gct act gga aag ttt cca tcg tta tta tta cat gaa gat gat ata att aaa cca aca ttg aga ttg gct ctt atg tta gct gga ttg aat tac tgt aat aaa tgc atc gag tat aga ggg ata tag caa ttc tcg ata ata gtc atg caa tat ttg aat gag act gat aat tta ggt aat aca gta ct- --- aca cac ata tct ttc tag att ata tat cgt taa aaa tct gta aga tgt ata ttt ctc ata agt atc cac tgt gta ata tta tta atg gat ata tag ata acg caa tag gga cta ata gta ttg taa aag ata taa tcg act att tac gta cat atc cag ata tct ata ttc cta cta gtt tgc tgc gta gtt gca tca ttg ata tgc atg -at tta tca gga ttc aga gat gaa tta cta agt aaa cta caa tcc cac aat aag taa gaa tca aat atc aaa aac tca ctt ttg att ttt cta gtc tta agt aat aca tat att tat taa tag acc tat gaa ata aaa aaa ggt aac aat gga ttc gcg tat agc tat tta cgt att agt atc ggc atc tct ttt gta tct tgt taa ttg tca caa act agt aca tta ctt caa tct gaa aat aaa tgg aag tga tat aac taa tac agc aga tat att gct gga caa tta tcc aat tat gac ctt tga tgg aaa gga tat tta tcc atc tat ctc gtt cat ggt cgg taa taa act ttt cct aga tct tta taa aaa tat ctt tga aga att ttt cag act att tcg agt atc tgt aag tag tca ata cga gga att aga ata tta tta ttc atg tga tta tac taa caa ccg tcc tac aat taa aca aca tta ctt tta taa cgg cga cga ata tac tga aat tga tag atc gaa aaa agc cac taa taa aaa cag ttg gtt aat tac ttc agg ctt tag act aca aaa atg gtt cga tag cga aga ttg tat aat tta tct cag atc ttt agt tag aag aat gga aga cag taa caa aaa cag taa aaa aac tta gta ctt aga tat cg- --a aaa aat ata ttt ttg tag act ctt gag aat aga agg aaa aca tgt aca taa tta taa aaa atg aaa atc aat ggc gaa taa gac agt gcg att cgc acc atg gag tcg gta gat ttc atg gct gtc gat gag cag ttt cac gac gac ctc gat ctt tgg tca tta tct ttg gta gat gat tat aaa aaa cat gga tta ggt gtt gac tgt tat gtt cta gaa cca gtt gtt gac agg aaa ata ttt gat aga ttt ctc ctt gaa cca att tgt gat cct gta gat gtt ctg tat gat tat ttt agg att cat aga gat aat att gat cag tat ata gta gat aga ctg ttt gca tat att aca tat aaa gat att ata tct gca tta gtg tca aag aat tat atg gaa gat att ttc tct ata att att aag aat tgt aat tct gtg caa gat ctc tta ctt tac tat cta tct aat gca tat gta gaa ata gac att gtt gat ctt atg gta gat cat ggg gct gta ata tat aaa ata gaa tgc ttg aat gcc tat ttt agg gga ata tgt aaa aag gaa agt agt gtt gtt gag ttt att ttg aat tgt ggt atc cca gat gaa aat gat gtt aaa tta gat cta tat aaa ata att cag tat act agg gga ttc ctt gta gat gaa ccc aca gta tta gaa att tat aag ctt tgt atc cca tat att gaa gat atc aat caa cta gat gct ggt gga agg acc ttg ctt tat cgc gct atc tat gca ggt tat ata gat tta gta tca tgg cta tta gaa aat gga gca aat gtc aac gca gta atg agt aat gga tat aca tgt ctt gac gtg gcc gtg gat agg gga tct gtc atc gcc cgt agg gaa gca cat ctt aaa ata tta gaa ata ttg ctt aga gaa cca ttg tct att gac tgt ata aaa tta gct ata ctt aat aat aca att gaa aac cat gat gtg ata aag ctc tgt atc aag tat ttt atg atg gta gat tat tca ctt tgt aat gtg tat gca tca tca ctc ttt gat tat ata att gat tgt aaa caa gaa ttg gag tac att agg cag atg aaa att cat aat aca acc atg tat gag tta atc tat aat aga gac aaa aac aag cat gct tcc cat att cta cat agg tat tct aaa cat cca gtt ttg aca cag tgt atc act aaa gga ttc aag att tac aca gaa gta acc gag cag gtc act aaa gct cta aac aga cgt gct cta ata gat gag ata ata aac aat gta tca act gat gac aat ctc cta tca aaa ctt cca tta gaa att agg gat cta att gtt tca caa gct gtc ata tag agt tct atc cac cca cct ttc ttg aaa tga gtt aat --- --- --- --- --- --- --- --- --- --- --- --- --- --- --- --- --- --- --- --- --- --- --- --- --- --- --- --- --- --- --- --- --- --- --- --- --- --- --- --- --- --- --- --- --- --- --- --- --- --- --- --- --- --- --- --- --- --- --- --- --- --- --- --- --- --- --- --- --- --- --- --- --- --- --- --- --- --- --- --- --- --- --- --- --- --- --- --- --- --- --- --- --- --- --- --- --- --- --- --- --- --- --- --- --- -ag tca taa gtt agt taa --- --- --- --- --- -gt cat aag tta gtt aag tca t-- --- --- --- --- --- --- --- --- --- --- --- --- --- --- --- -aa gtt agt taa gtc ata agt tag tta agt cat aag tta gtt aag tca taa gtt agt taa gtc ata agt tag tta agt cat aag tta gtt tat agt cta aca ctt cta att ttt ata cct tga tct ttt tct cta att atg aaa aag taa atc att atg aag atg gat g-a aat gga cga gat tgt gcg cat cgt taa cga tag tat gtg gta cgt acc taa cgc att tat gga cga cgg tga taa tga agg tca cat ttc tgt caa taa tgt ctg tca tat gta tct cgc att ctt tga tgt gga tat atc atc tca tct gtt taa att agt tat taa aca ctg cga tct gaa taa acg act aaa atg tgg taa ctc tcc att aca ttg cta tac gat gaa tac acg att taa tcc atc tgt att aaa gat att gtt acg cca cgg cat gcg taa ctt tga tag caa gga taa aaa agg aca tat tcc tct aca cca cta tct gat tca ttc act atc aat cga taa caa gat ctt tga tat act aac gga ccc cat tga tga ctt tag taa atc atc cga tct att gct gtg tta tct tag ata taa att caa tgg gag ctt aaa cta tta cgt tct gta caa att att gac taa agg atc tga ccc taa ttg cgt cga tga gga tgg act cac ttc tct tca tta cta ctg taa aca cat atc cgc gtt cca cga aag caa tta tta caa gtc aaa gag tca cac taa gat gcg agc tga gaa gcg att cat cta cgc gat aat aga tca tgg agc aaa cat taa cgc ggt tac gaa aat cgg aaa tac gcc gtt aca cac tta cct tca aca gta tac caa aca tag tcc tcg tgt ggt gta tgc tct ttt atc tcg agg agc cga tac gag gat acg taa taa tct taa ttg tac acc cat cat gga ata cat aaa gaa cga ttg tgc aac agg tca tat tct cat aat gtt act caa ttg gca cga aca aaa ata cgg gaa att aca aaa gga aga agg aca aca tct act tta tct att cat aaa aca taa tca agg ata tgg aag tcg ctc tct caa tat act acg gta tct act aga tag att cga cat tca gaa aga cga ata cta taa tac aat gac tcc tct tca tac cgc ctt cca gaa ttg caa taa caa tgt tgc ctc ata cct cgt ata cat tgg ata cga cat caa cct tcc gac taa aga cga taa gac agt att cga ctt ggt gtt tga aaa cag aaa cat tat ata caa ggc gga tgt cgt taa tga cat tat cca cca cag act gaa agt atc tct acc tat gat taa atc gtt gtt cta caa gat gtc gga gtt ctc tcc cta cga cga tca cta cgt aaa gaa gat aat agc cta ctg cct att aag gga cga gtc att tgc gga act aca tac taa att ctg ttt aaa cga gga cta taa aag tgt att tat gaa aaa tat atc att cga taa gat aga ttc cat cat cga aaa atg tag tcg tga cat aag tct cct caa aga gat tcg aat ctc aga cac cga ctt gta tac ggt att gag aac aga aga cat tcg gta tca cac ata tct cga agc cat aca ttc aga caa acg cat ttc att tcc cat gta cga cga tct cat aga aca gtg tca tct atc gat gga gca taa aag taa act cgt cga caa agc act caa taa att aga gtc tac cat cga tag tca atc tag act atc gta ttt gcc tcc gga aat tat gcg caa tat cat aac caa gct aag cga cta cca tct aaa cag tat gtt gta cgg aaa gaa cca tta caa ata tta tcc atg ata gaa aga aaa tat tta aaa aat aat cta tat gat tgg aga agt agg aaa caa aca gta aca aga cga cga tta cta c-a tta tta aat cat gag gtc cgt att ata ctc gta tat att gtt tct ctc atg tat aat aat aaa cgg aag aga tat agc acc aca tgc acc atc caa tgg aaa gtg taa aga caa cga ata cag aag ccg taa tct atg ttg tct atc gtg tcc tcc ggg aac tta cgc ttt cag att atg tga tag caa gac taa tac aca atg tac acc gtg tgg ttc gga tac ctt tac atc tca caa taa tca ttt aca ggc ttg tct aag ttg taa cgg aag atg tga tag taa tca ggt aga gac gcg atc gtg taa cac gac tca caa tag aat ctg tga atg ctc tcc agg ata tta ttg tct tct caa agg agc atc agg gtg tag aac atg tat ttc taa aac aaa gtg tgg aat agg ata cgg agt atc cgg ata cac gtc tac cgg aga cgt cat ctg ttc tcc gtg tgg tcc cgg aac ata ttc tca cac cgt ctc ttc cac aga taa atg cga acc cgt cgt aac cag caa tac att taa cta tat cga tgt gga aat taa cct gta tcc agt caa cga cac atc gtg tac tcg gac gac cac tac cgg tct cag cga atc cat ctc aac gtc gga act aac tat tac cat gaa tca taa aga ttg tga tcc agt ctt tcg tgc aga ata ctt ctc tgt cct taa taa tgt agc aac ttc agg att ctt tac agg aga aaa tag ata tca gaa tac ttc aaa gat atg tac tct gaa ttt cga gat taa atg taa caa caa aga ttc atc ttc caa aca gtt aac gaa aac aaa gaa tga tac tat cat gcc gca ttc aga gac ggt aac tct agt ggg cga ctg tct atc tag cgt cga cat cta cat act ata tag taa tac caa tac tca aga cta cga aaa tga tac aat ctc tta tca tat ggg taa tgt tct cga tgt caa tag cca tat gcc cgc tag ttg cga tat aca taa act gat cac taa ttc cca gaa tcc cac cca ctt ata gta ag- -tt ttt tta cct ata aat aat aaa tac aat aat taa ttt ctc gta aaa gta gaa aat ata ttc taa ttt att ata tgg taa gaa agt aga atc atc tag aac agt aat caa tca ata gca atc atg aaa caa tat att gtc ctg gca tgc atg tgc cta gtg gca gct gct atg cct act agt ctt caa caa tct tca tcc tcg tgt act gaa gaa gaa aac aaa cat cat atg gga atc gat gtt att atc aaa gtc aca aag caa gac caa aca ccg acc aat gat aag att tgt caa tcc gta acg gaa gtt aca gag a-- --- --- --- --- --- -cc gaa gat gat gag gta tcc gaa gaa gtt gta aaa gga gat ccc acc act tat tac act atc gtc ggt gcg ggt ctt aac atg aac ttt gga ttc acc aaa tgc cca aag att tta tcc atc tcc gaa tcc tct gat gga aac act gtg aat act aga ttg tcc agc gtg tca ccg gga caa ggt aag gac tct ccc gcg atc acg cgt gaa gaa gct ctg gct atg atc aaa gac tgt gag atg tct atc gac atc aga tgt agc gaa gaa gag aaa gac agt gac atc aag acc cat cca gta ctt ggg tct aac atc tca cat aag aaa gtg agt tac aaa gat atc atc ggt tca acg atc gtt gat aca aaa tgt gtc aag aac cta gag ttt agc gta cgt atc gga gac atg tgt gag gaa tca tct gaa ctt gaa gtc aag gat gga ttc aag tat gtc gac gga tcg gca tct gaa ggt gca acc gat gat act tca ctc atc gat tca aca aaa ctc aaa gca tgt gtc tga atc gat aac tct att cat ctg aaa atg gat gag ttg ggt taa tcg aac gat tca gac acc gca cca cga att aaa aaa g-- -ac cgg gca cta tat tcc ggt ttg caa aac aaa aat att taa cta cat tca caa aaa gtt acc tct cgt tac ttc ttc ttt ctg ttt caa tat gtg ata cga tat gat cac tat tcg tat tct ct- --- --- tgg tct cat aaa aaa gtt tta c-- --a aaa aaa aaa aaa a-t att ttt att ctc ttt ctc tct tcg at- ggt ctc aca aaa ata tta aac ctc ttt ctg atg tct caa cta ttt cgt aaa cga taa cgt cca aca ata tat tct cgt aga gct tat caa cat cct tat acc aat cta ggt tgt cag aca att gca tca taa aat aat gtt tat aat tta cac gtt aac atc ata taa taa acg tat ata gtt aat att ttt gga ata taa atg atc tgt aaa atc cat gta ggg gac act gct cac gtt ttt tct cta gta cat aat ttc aca caa gtt ttt ata cag aca aat taa ttc tcg tcc ata ta- ttt taa aac att gac ttt tgt act aag aaa aat atc ttg act aac cat ctc ttt ctc tct tcg atg ggt ctc aca aaa ata tta aac ctc ttt ctg atg g-- --- --- --- --- --- --- --- --- --- --- --- --- --- --- --- --- --- --- --- --- --- --- --- --- --- --- --- --- --- --- --- --- --- --- --- --- --- --- --- --- --- --- --- --- --- --- agt cgt aaa aag ttt tta tcc ttt ctc tct tcg ata ggt ctc --- --- --- --- --- --- --- --- --- --- --- --- --- --- --- --- --- --- --- --- --- --- --- --- --- --- --- --- --- --- --- --- --- --- --- --- --- --- --- --- --- --- --- --- --- --- --- --- --- --- --- --- --- --- --- --- --- --- --- --- --- --- --- --- --- --- --- --- --- --- aca aaa ata tta aac ctc ttt ctg atg gtc tct ata aa

>ON649724.1_Monkeypox_virus_isolate_Monkeypox/PT0014/2022_complete_genome

atc gag gac gac ttc aac tat gac agc tcg gtg gcg tct gcc agc gtg tac ata cga atg gca ttt cta aga aaa gtc tac ggt atc ctt -tc tac aat ttc ctt taa caa cgg caa cag ctg cag tat ttt tat act ttg aat gca -tc gga cat tta tac aag gga gtc ctg ttc taa tat tgg cat caa tgt tcg gat cta tag gct tga ttt tcg cat tga ctt tac aca gac atg aac atc ccc tga atc tgt aca tac ttt gtg gat tta cac tgt tag aat ctc taa cgc tgg cct ctg ttg tta ctt tct atg atg cac gta tcg tta tgc aag ctt tca tgt tga cta ctg cag tgt ttc ttg ctc tga cta cat ata ctc tac aat caa aga gag att tca gta aac ttg taa cag gat tgt ttg ctg ctt tct gga ttt taa ttt tgt cag gag tct tga gga taa agt tta aaa tag aat taa taa aga aca tat agg tca -tt ttt taa aca tgg ata gaa acc aag gtt gtt agt taa taa tat aca aga tat ttt ttc tca ctc tga tcc atg taa acc aag gac gag aga cac tct cat tcc tca ttc acg aca cca tta aaa atg gaa att aaa gcc ctc tat taa gca cag acg gct aca ggt cta cca tca ggt t-a cct tcg tct acc ttc aca atg gcc tct cct tgt gcc cag ttc agt ccc tgt cat tgc cac gct act aag gac tcc ctg aat acc gtg act gac gtc aga cat tgt ctg act gaa tac atc ctg tgg gtt tct cat aga tgg acc cat aga gaa agc gca ggg cct ctc tac agg ctt ctc atc tct ttc aga att gat gca atg gag cta ttt ggt agc gag ttg aag gag ttc tcg aat tca ctt ccg tgg gac aat atc gac aat tgc gtg gag atc att aaa tgt ttc atc aga aat gac tcc atg aaa acc gcc aaa gaa ctt tgt gca ata att gga ctt tgt act caa tca gct att gtc act gga aga gtc ttc aat gat aag tat atc gac ata cta ctt atg ctg cga aag att ctg aac gag aac gac tat ctc acc ctc ttg gat cat atc ctc act gct aaa tac taa atc tcc ttc atg ctc tct cac taa tac tct tac tca cta cac ttt tta tca tct tat gat gaa tga ttg cct tca tca ttt ttt cgt gga ata taa tat agg aat aat tag cac cag aat agc tat gga tat ctc gtt aag aat att ctc tat aag aga cat aat gta gac ata gtt att ata tcc ttc tta gat aag tgt tac gct act gga aag ttt cca tcg tta tta tta cat gaa gat gat ata att aaa cca aca ttg aga ttg gct ctt atg tta gct gga ttg aat tac tgt aat aaa tgc atc gag tat aga ggg ata tag caa ttc tcg ata ata gtc atg caa tat ttg aat gag act gat aat tta ggt aat aca gta ct- --- aca cac ata tct ttc tag att ata tat cgt taa aaa tct gta aga tgt ata ttt ctc ata agt atc cac tgt gta ata tta tta atg gat ata tag ata acg caa tag gga cta ata gta ttg taa aag ata taa tcg act att tac gta cat atc cag ata tct ata ttc cta cta gtt tgc tgc gta gtt gca tca ttg ata tgc atg -at tta tca gga ttc aga gat gaa tta cta agt aaa cta caa tcc cac aat aag taa gaa tca aat atc aaa aac tca ctt ttg att ttt cta gtc tta agt aat aca tat att tat taa tag acc tat gaa ata aaa aaa ggt aac aat gga ttc gcg tat agc tat tta cgt att agt atc ggc atc tct ttt gta tct tgt taa ttg tca caa act agt aca tta ctt caa tct gaa aat aaa tgg aag tga tat aac taa tac agc aga tat att gct gga caa tta tcc aat tat gac ctt tga tgg aaa gga tat tta tcc atc tat ctc gtt cat ggt cgg taa taa act ttt cct aga tct tta taa aaa tat ctt tga aga att ttt cag act att tcg agt atc tgt aag tag tca ata cga gga att aga ata tta tta ttc atg tga tta tac taa caa ccg tcc tac aat taa aca aca tta ctt tta taa cgg cga cga ata tac tga aat tga tag atc gaa aaa agc cac taa taa aaa cag ttg gtt aat tac ttc agg ctt tag act aca aaa atg gtt cga tag cga aga ttg tat aat tta tct cag atc ttt agt tag aag aat gga aga cag taa caa aaa cag taa aaa aac tta gta ctt aga tat cg- --a aaa aat ata ttt ttg tag act ctt gag aat aga agg aaa aca tgt aca taa tta taa aaa atg aaa atc aat ggc gaa taa gac agt gcg att cgc acc atg gag tcg gta gat ttc atg gct gtc gat gag cag ttt cac gac gac ctc gat ctt tgg tca tta tct ttg gta gat gat tat aaa aaa cat gga tta ggt gtt gac tgt tat gtt cta gaa cca gtt gtt gac agg aaa ata ttt gat aga ttt ctc ctt gaa cca att tgt gat cct gta gat gtt ctg tat gat tat ttt agg att cat aga gat aat att gat cag tat ata gta gat aga ctg ttt gca tat att aca tat aaa gat att ata tct gca tta gtg tca aag aat tat atg gaa gat att ttc tct ata att att aag aat tgt aat tct gtg caa gat ctc tta ctt tac tat cta tct aat gca tat gta gaa ata gac att gtt gat ctt atg gta gat cat ggg gct gta ata tat aaa ata gaa tgc ttg aat gcc tat ttt agg gga ata tgt aaa aag gaa agt agt gtt gtt gag ttt att ttg aat tgt ggt atc cca gat gaa aat gat gtt aaa tta gat cta tat aaa ata att cag tat act agg gga ttc ctt gta gat gaa ccc aca gta tta gaa att tat aag ctt tgt atc cca tat att gaa gat atc aat caa cta gat gct ggt gga agg acc ttg ctt tat cgc gct atc tat gca ggt tat ata gat tta gta tca tgg cta tta gaa aat gga gca aat gtc aac gca gta atg agt aat gga tat aca tgt ctt gac gtg gcc gtg gat agg gga tct gtc atc gcc cgt agg gaa gca cat ctt aaa ata tta gaa ata ttg ctt aga gaa cca ttg tct att gac tgt ata aaa tta gct ata ctt aat aat aca att gaa aac cat gat gtg ata aag ctc tgt atc aag tat ttt atg atg gta gat tat tca ctt tgt aat gtg tat gca tca tca ctc ttt gat tat ata att gat tgt aaa caa gaa ttg gag tac att agg cag atg aaa att cat aat aca acc atg tat gag tta atc tat aat aga gac aaa aac aag cat gct tcc cat att cta cat agg tat tct aaa cat cca gtt ttg aca cag tgt atc act aaa gga ttc aag att tac aca gaa gta acc gag cag gtc act aaa gct cta aac aga cgt gct cta ata gat gag ata ata aac aat gta tca act gat gac aat ctc cta tca aaa ctt cca tta gaa att agg gat cta att gtt tca caa gct gtc ata tag agt tct atc cac cca cct ttc ttg aaa tga gtt aat --- --- --- --- --- --- --- --- --- --- --- --- --- --- --- --- --- --- --- --- --- --- --- --- --- --- --- --- --- --- --- --- --- --- --- --- --- --- --- --- --- --- --- --- --- --- --- --- --- --- --- --- --- --- --- --- --- --- --- --- --- --- --- --- --- --- --- --- --- --- --- --- --- --- --- --- --- --- --- --- --- --- --- --- --- --- --- --- --- --- --- --- --- --- --- --- --- --- --- --- --- --- --- --- --- -ag tca taa gtt agt taa --- --- --- --- --- -gt cat aag tta gtt aag tca t-- --- --- --- --- --- --- --- --- --- --- --- --- --- --- --- -aa gtt agt taa gtc ata agt tag tta agt cat aag tta gtt aag tca taa gtt agt taa gtc ata agt tag tta agt cat aag tta gtt tat agt cta aca ctt cta att ttt ata cct tga tct ttt tct cta att atg aaa aag taa atc att atg aag atg gat g-a aat gga cga gat tgt gcg cat cgt taa cga tag tat gtg gta cgt acc taa cgc att tat gga cga cgg tga taa tga agg tca cat ttc tgt caa taa tgt ctg tca tat gta tct cgc att ctt tga tgt gga tat atc atc tca tct gtt taa att agt tat taa aca ctg cga tct gaa taa acg act aaa atg tgg taa ctc tcc att aca ttg cta tac gat gaa tac acg att taa tcc atc tgt att aaa gat att gtt acg cca cgg cat gcg taa ctt tga tag caa gga taa aaa agg aca tat tcc tct aca cca cta tct gat tca ttc act atc aat cga taa caa gat ctt tga tat act aac gga ccc cat tga tga ctt tag taa atc atc cga tct att gct gtg tta tct tag ata taa att caa tgg gag ctt aaa cta tta cgt tct gta caa att att gac taa agg atc tga ccc taa ttg cgt cga tga gga tgg act cac ttc tct tca tta cta ctg taa aca cat atc cgc gtt cca cga aag caa tta tta caa gtc aaa gag tca cac taa gat gcg agc tga gaa gcg att cat cta cgc gat aat aga tca tgg agc aaa cat taa cgc ggt tac gaa aat cgg aaa tac gcc gtt aca cac tta cct tca aca gta tac caa aca tag tcc tcg tgt ggt gta tgc tct ttt atc tcg agg agc cga tac gag gat acg taa taa tct taa ttg tac acc cat cat gga ata cat aaa gaa cga ttg tgc aac agg tca tat tct cat aat gtt act caa ttg gca cga aca aaa ata cgg gaa att aca aaa gga aga agg aca aca tct act tta tct att cat aaa aca taa tca agg ata tgg aag tcg ctc tct caa tat act acg gta tct act aga tag att cga cat tca gaa aga cga ata cta taa tac aat gac tcc tct tca tac cgc ctt cca gaa ttg caa taa caa tgt tgc ctc ata cct cgt ata cat tgg ata cga cat caa cct tcc gac taa aga cga taa gac agt att cga ctt ggt gtt tga aaa cag aaa cat tat ata caa ggc gga tgt cgt taa tga cat tat cca cca cag act gaa agt atc tct acc tat gat taa atc gtt gtt cta caa gat gtc gga gtt ctc tcc cta cga cga tca cta cgt aaa gaa gat aat agc cta ctg cct att aag gga cga gtc att tgc gga act aca tac taa att ctg ttt aaa cga gga cta taa aag tgt att tat gaa aaa tat atc att cga taa gat aga ttc cat cat cga aaa atg tag tcg tga cat aag tct cct caa aga gat tcg aat ctc aga cac cga ctt gta tac ggt att gag aac aga aga cat tcg gta tca cac ata tct cga agc cat aca ttc aga caa acg cat ttc att tcc cat gta cga cga tct cat aga aca gtg tca tct atc gat gga gca taa aag taa act cgt cga caa agc act caa taa att aga gtc tac cat cga tag tca atc tag act atc gta ttt gcc tcc gga aat tat gcg caa tat cat aac caa gct aag cga cta cca tct aaa cag tat gtt gta cgg aaa gaa cca tta caa ata tta tcc atg ata gaa aga aaa tat tta aaa aat aat cta tat gat tgg aga agt agg aaa caa aca gta aca aga cga cga tta cta c-a tta tta aat cat gag gtc cgt att ata ctc gta tat att gtt tct ctc atg tat aat aat aaa cgg aag aga tat agc acc aca tgc acc atc caa tgg aaa gtg taa aga caa cga ata cag aag ccg taa tct atg ttg tct atc gtg tcc tcc ggg aac tta cgc ttt cag att atg tga tag caa gac taa tac aca atg tac acc gtg tgg ttc gga tac ctt tac atc tca caa taa tca ttt aca ggc ttg tct aag ttg taa cgg aag atg tga tag taa tca ggt aga gac gcg atc gtg taa cac gac tca caa tag aat ctg tga atg ctc tcc agg ata tta ttg tct tct caa agg agc atc agg gtg tag aac atg tat ttc taa aac aaa gtg tgg aat agg ata cgg agt atc cgg ata cac gtc tac cgg aga cgt cat ctg ttc tcc gtg tgg tcc cgg aac ata ttc tca cac cgt ctc ttc cac aga taa atg cga acc cgt cgt aac cag caa tac att taa cta tat cga tgt gga aat taa cct gta tcc agt caa cga cac atc gtg tac tcg gac gac cac tac cgg tct cag cga atc cat ctc aac gtc gga act aac tat tac cat gaa tca taa aga ttg tga tcc agt ctt tcg tgc aga ata ctt ctc tgt cct taa taa tgt agc aac ttc agg att ctt tac agg aga aaa tag ata tca gaa tac ttc aaa gat atg tac tct gaa ttt cga gat taa atg taa caa caa aga ttc atc ttc caa aca gtt aac gaa aac aaa gaa tga tac tat cat gcc gca ttc aga gac ggt aac tct agt ggg cga ctg tct atc tag cgt cga cat cta cat act ata tag taa tac caa tac tca aga cta cga aaa tga tac aat ctc tta tca tat ggg taa tgt tct cga tgt caa tag cca tat gcc cgc tag ttg cga tat aca taa act gat cac taa ttc cca gaa tcc cac cca ctt ata gta ag- -tt ttt tta cct ata aat aat aaa tac aat aat taa ttt ctc gta aaa gta gaa aat ata ttc taa ttt att ata tgg taa gaa agt aga atc atc tag aac agt aat caa tca ata gca atc atg aaa caa tat att gtc ctg gca tgc atg tgc cta gtg gca gct gct atg cct act agt ctt caa caa tct tca tcc tcg tgt act gaa gaa gaa aac aaa cat cat atg gga atc gat gtt att atc aaa gtc aca aag caa gac caa aca ccg acc aat gat aag att tgt caa tcc gta acg gaa gtt aca gag a-- --- --- --- --- --- -cc gaa gat gat gag gta tcc gaa gaa gtt gta aaa gga gat ccc acc act tat tac act atc gtc ggt gcg ggt ctt aac atg aac ttt gga ttc acc aaa tgc cca aag att tta tcc atc tcc gaa tcc tct gat gga aac act gtg aat act aga ttg tcc agc gtg tca ccg gga caa ggt aag gac tct ccc gcg atc acg cgt gaa gaa gct ctg gct atg atc aaa gac tgt gag atg tct atc gac atc aga tgt agc gaa gaa gag aaa gac agt gac atc aag acc cat cca gta ctt ggg tct aac atc tca cat aag aaa gtg agt tac aaa gat atc atc ggt tca acg atc gtt gat aca aaa tgt gtc aag aac cta gag ttt agc gta cgt atc gga gac atg tgt gag gaa tca tct gaa ctt gaa gtc aag gat gga ttc aag tat gtc gac gga tcg gca tct gaa ggt gca acc gat gat act tca ctc atc gat tca aca aaa ctc aaa gca tgt gtc tga atc gat aac tct att cat ctg aaa atg gat gag ttg ggt taa tcg aac gat tca gac acc gca cca cga att aaa aaa g-- -ac cgg gca cta tat tcc ggt ttg caa aac aaa aat att taa cta cat tca caa aaa gtt acc tct cgt tac ttc ttc ttt ctg ttt caa tat gtg ata cga tat gat cac tat tcg tat tct ct- --- --- tgg tct cat aaa aaa gtt tta c-- --a aaa aaa aaa aaa a-t att ttt att ctc ttt ctc tct tcg at- ggt ctc aca aaa ata tta aac ctc ttt ctg atg tct caa cta ttt cgt aaa cga taa cgt cca aca ata tat tct cgt aga gct tat caa cat cct tat acc aat cta ggt tgt cag aca att gca tca taa aat aat gtt tat aat tta cac gtt aac atc ata taa taa acg tat ata gtt aat att ttt gga ata taa atg atc tgt aaa atc cat gta ggg gac act gct cac gtt ttt tct cta gta cat aat ttc aca caa gtt ttt ata cag aca aat taa ttc tcg tcc ata ta- ttt taa aac att gac ttt tgt act aag aaa aat atc ttg act aac cat ctc ttt ctc tct tcg atg ggt ctc aca aaa ata tta aac ctc ttt ctg atg g-- --- --- --- --- --- --- --- --- --- --- --- --- --- --- --- --- --- --- --- --- --- --- --- --- --- --- --- --- --- --- --- --- --- --- --- --- --- --- --- --- --- --- --- --- --- --- agt cgt aaa aag ttt tta tcc ttt ctc tct tcg ata ggt ctc --- --- --- --- --- --- --- --- --- --- --- --- --- --- --- --- --- --- --- --- --- --- --- --- --- --- --- --- --- --- --- --- --- --- --- --- --- --- --- --- --- --- --- --- --- --- --- --- --- --- --- --- --- --- --- --- --- --- --- --- --- --- --- --- --- --- --- --- --- --- aca aaa ata tta aac ctc ttt ctg atg gtc tct ata aa

>ON649722.1_Monkeypox_virus_isolate_Monkeypox/PT0022/2022_complete_genome

atc gag gac gac ttc aac tat gac agc tcg gtg gcg tct gcc agc gtg tac ata cga atg gca ttt cta aga aaa gtc tac ggt atc ctt -tc tac aat ttc ctt taa caa cgg caa cag ctg cag tat ttt tat act ttg aat gca -tc gga cat tta tac aag gga gtc ctg ttc taa tat tgg cat caa tgt tcg gat cta tag gct tga ttt tcg cat tga ctt tac aca gac atg aac atc ccc tga atc tgt aca tac ttt gtg gat tta cac tgt tag aat ctc taa cgc tgg cct ctg ttg tta ctt tct atg atg cac gta tcg tta tgc aag ctt tca tgt tga cta ctg cag tgt ttc ttg ctc tga cta cat ata ctc tac aat caa aga gag att tca gta aac ttg taa cag gat tgt ttg ctg ctt tct gga ttt taa ttt tgt cag gag tct tga gga taa agt tta aaa tag aat taa taa aga aca tat agg tca -tt ttt taa aca tgg ata gaa acc aag gtt gtt agt taa taa tat aca aga tat ttt ttc tca ctc tga tcc atg taa acc aag gac gag aga cac tct cat tcc tca ttc acg aca cca tta aaa atg gaa att aaa gcc ctc tat taa gca cag acg gct aca ggt cta cca tca ggt t-a cct tcg tct acc ttc aca atg gcc tct cct tgt gcc cag ttc agt ccc tgt cat tgc cac gct act aag gac tcc ctg aat acc gtg act gac gtc aga cat tgt ctg act gaa tac atc ctg tgg gtt tct cat aga tgg acc cat aga gaa agc gca ggg cct ctc tac agg ctt ctc atc tct ttc aga att gat gca atg gag cta ttt ggt agc gag ttg aag gag ttc tcg aat tca ctt ccg tgg gac aat atc gac aat tgc gtg gag atc att aaa tgt ttc atc aga aat gac tcc atg aaa acc gcc aaa gaa ctt tgt gca ata att gga ctt tgt act caa tca gct att gtc act gga aga gtc ttc aat gat aag tat atc gac ata cta ctt atg ctg cga aag att ctg aac gag aac gac tat ctc acc ctc ttg gat cat atc ctc act gct aaa tac taa atc tcc ttc atg ctc tct cac taa tac tct tac tca cta cac ttt tta tca tct tat gat gaa tga ttg cct tca tca ttt ttt cgt gga ata taa tat agg aat aat tag cac cag aat agc tat gga tat ctc gtt aag aat att ctc tat aag aga cat aat gta gac ata gtt att ata tcc ttc tta gat aag tgt tac gct act gga aag ttt cca tcg tta tta tta cat gaa gat gat ata att aaa cca aca ttg aga ttg gct ctt atg tta gct gga ttg aat tac tgt aat aaa tgc atc gag tat aga ggg ata tag caa ttc tcg ata ata gtc atg caa tat ttg aat gag act gat aat tta ggt aat aca gta ct- --- aca cac ata tct ttc tag att ata tat cgt taa aaa tct gta aga tgt ata ttt ctc ata agt atc cac tgt gta ata tta tta atg gat ata tag ata acg caa tag gga cta ata gta ttg taa aag ata taa tcg act att tac gta cat atc cag ata tct ata ttc cta cta gtt tgc tgc gta gtt gca tca ttg ata tgc atg -at tta tca gga ttc aga gat gaa tta cta agt aaa cta caa tcc cac aat aag taa gaa tca aat atc aaa aac tca ctt ttg att ttt cta gtc tta agt aat aca tat att tat taa tag acc tat gaa ata aaa aaa ggt aac aat gga ttc gcg tat agc tat tta cgt att agt atc ggc atc tct ttt gta tct tgt taa ttg tca caa act agt aca tta ctt caa tct gaa aat aaa tgg aag tga tat aac taa tac agc aga tat att gct gga caa tta tcc aat tat gac ctt tga tgg aaa gga tat tta tcc atc tat ctc gtt cat ggt cgg taa taa act ttt cct aga tct tta taa aaa tat ctt tga aga att ttt cag act att tcg agt atc tgt aag tag tca ata cga gga att aga ata tta tta ttc atg tga tta tac taa caa ccg tcc tac aat taa aca aca tta ctt tta taa cgg cga cga ata tac tga aat tga tag atc gaa aaa agc cac taa taa aaa cag ttg gtt aat tac ttc agg ctt tag act aca aaa atg gtt cga tag cga aga ttg tat aat tta tct cag atc ttt agt tag aag aat gga aga cag taa caa aaa cag taa aaa aac tta gta ctt aga tat cg- --a aaa aat ata ttt ttg tag act ctt gag aat aga agg aaa aca tgt aca taa tta taa aaa atg aaa atc aat ggc gaa taa gac agt gcg att cgc acc atg gag tcg gta gat ttc atg gct gtc gat gag cag ttt cac gac gac ctc gat ctt tgg tca tta tct ttg gta gat gat tat aaa aaa cat gga tta ggt gtt gac tgt tat gtt cta gaa cca gtt gtt gac agg aaa ata ttt gat aga ttt ctc ctt gaa cca att tgt gat cct gta gat gtt ctg tat gat tat ttt agg att cat aga gat aat att gat cag tat ata gta gat ??? ?tg ttt gca tat att aca tat aaa gat att ata tct gca tta gtg tca aag aat tat atg gaa gat att ttc tct ata att att aag aat tgt aat tct gtg caa gat ctc tta ctt tac tat cta tct aat gca tat gta gaa ata gac att gtt gat ctt atg gta gat cat ggg gct gta ata tat aaa ata gaa tgc ttg aat gcc tat ttt agg gga ata tgt aaa aag gaa agt agt gtt gtt gag ttt att ttg aat tgt ggt atc cca gat gaa aat gat gtt aaa tta gat cta tat aaa ata att cag tat act agg gga ttc ctt gta gat gaa ccc aca gta tta gaa att tat aag ctt tgt atc cca tat att gaa gat atc aat caa cta gat gct ggt gga agg acc ttg ctt tat cgc gct atc tat gca ggt tat ata gat tta gta tca tgg cta tta gaa aat gga gca aat gtc aac gca gta atg agt aat gga tat aca tgt ctt gac gtg gcc gtg gat agg gga tct gtc atc gcc cgt agg gaa gca cat ctt aaa ata tta gaa ata ttg ctt aga gaa cca ttg tct att gac tgt ata aaa tta gct ata ctt aat aat aca att gaa aac cat gat gtg ata aag ctc tgt atc aag tat ttt atg atg gta gat tat tca ctt tgt aat gtg tat gca tca tca ctc ttt gat tat ata att gat tgt aaa caa gaa ttg gag tac att agg cag atg aaa att cat aat aca acc atg tat gag tta atc tat aat aga gac aaa aac aag cat gct tcc cat att cta cat agg tat tct aaa cat cca gtt ttg aca cag tgt atc act aaa gga ttc aag att tac aca gaa gta acc gag cag gtc act aaa gct cta aac aga cgt gct cta ata gat gag ata ata aac aat gta tca act gat gac aat ctc cta tca aaa ctt cca tta gaa att agg gat cta att gtt tca caa gct gtc ata tag agt tct atc cac cca cct ttc ttg aaa tga gtt aat --- --- --- --- --- --- --- --- --- --- --- --- --- --- --- --- --- --- --- --- --- --- --- --- --- --- --- --- --- --- --- --- --- --- --- --- --- --- --- --- --- --- --- --- --- --- --- --- --- --- --- --- --- --- --- --- --- --- --- --- --- --- --- --- --- --- --- --- --- --- --- --- --- --- --- --- --- --- --- --- --- --- --- --- --- --- --- --- --- --- --- --- --- --- --- --- --- --- --- --- --- --- --- --- --- -ag tca taa gtt agt taa --- --- --- --- --- -gt cat aag tta gtt aag tca t-- --- --- --- --- --- --- --- --- --- --- --- --- --- --- --- -aa gtt agt taa gtc ata agt tag tta agt cat aag tta gtt aag tca taa gtt agt taa gtc ata agt tag tta agt cat aag tta gtt tat agt cta aca ctt cta att ttt ata cct tga tct ttt tct cta att atg aaa aag taa atc att atg aag atg gat g-a aat gga cga gat tgt gcg cat cgt taa cga tag tat gtg gta cgt acc taa cgc att tat gga cga cgg tga taa tga agg tca cat ttc tgt caa taa tgt ctg tca tat gta tct cgc att ctt tga tgt gga tat atc atc tca tct gtt taa att agt tat taa aca ctg cga tct gaa taa acg act aaa atg tgg taa ctc tcc att aca ttg cta tac gat gaa tac acg att taa tcc atc tgt att aaa gat att gtt acg cca cgg cat gcg taa ctt tga tag caa gga taa aaa agg aca tat tcc tct aca cca cta tct gat tca ttc act atc aat cga taa caa gat ctt tga tat act aac gga ccc cat tga tga ctt tag taa atc atc cga tct att gct gtg tta tct tag ata taa att caa tgg gag ctt aaa cta tta cgt tct gta caa att att gac taa agg atc tga ccc taa ttg cgt cga tga gga tgg act cac ttc tct tca tta cta ctg taa aca cat atc cgc gtt cca cga aag caa tta tta caa gtc aaa gag tca cac taa gat gcg agc tga gaa gcg att cat cta cgc gat aat aga tca tgg agc aaa cat taa cgc ggt tac gaa aat cgg aaa tac gcc gtt aca cac tta cct tca aca gta tac caa aca tag tcc tcg tgt ggt gta tgc tct ttt atc tcg agg agc cga tac gag gat acg taa taa tct taa ttg tac acc cat cat gga ata cat aaa gaa cga ttg tgc aac agg tca tat tct cat aat gtt act caa ttg gca cga aca aaa ata cgg gaa att aca aaa gga aga agg aca aca tct act tta tct att cat aaa aca taa tca agg ata tgg aag tcg ctc tct caa tat act acg gta tct act aga tag att cga cat tca gaa aga cga ata cta taa tac aat gac tcc tct tca tac cgc ctt cca gaa ttg caa taa caa tgt tgc ctc ata cct cgt ata cat tgg ata cga cat caa cct tcc gac taa aga cga taa gac agt att cga ctt ggt gtt tga aaa cag aaa cat tat ata caa ggc gga tgt cgt taa tga cat tat cca cca cag act gaa agt atc tct acc tat gat taa atc gtt gtt cta caa gat gtc gga gtt ctc tcc cta cga cga tca cta cgt aaa gaa gat aat agc cta ctg cct att aag gga cga gtc att tgc gga act aca tac taa att ctg ttt aaa cga gga cta taa aag tgt att tat gaa aaa tat atc att cga taa gat aga ttc cat cat cga aaa atg tag tcg tga cat aag tct cct caa aga gat tcg aat ctc aga cac cga ctt gta tac ggt att gag aac aga aga cat tcg gta tca cac ata tct cga agc cat aca ttc aga caa acg cat ttc att tcc cat gta cga cga tct cat aga aca gtg tca tct atc gat gga gca taa aag taa act cgt cga caa agc act caa taa att aga gtc tac cat cga tag tca atc tag act atc gta ttt gcc tcc gga aat tat gcg caa tat cat aac caa gct aag cga cta cca tct aaa cag tat gtt gta cgg aaa gaa cca tta caa ata tta tcc atg ata gaa aga aaa tat tta aaa aat aat cta tat gat tgg aga agt agg aaa caa aca gta aca aga cga cga tta cta c-a tta tta aat cat gag gtc cgt att ata ctc gta tat att gtt tct ctc atg tat aat aat aaa cgg aag aga tat agc acc aca tgc acc atc caa tgg aaa gtg taa aga caa cga ata cag aag ccg taa tct atg ttg tct atc gtg tcc tcc ggg aac tta cgc ttt cag att atg tga tag caa gac taa tac aca atg tac acc gtg tgg ttc gga tac ctt tac atc tca caa taa tca ttt aca ggc ttg tct aag ttg taa cgg aag atg tga tag taa tca ggt aga gac gcg atc gtg taa cac gac tca caa tag aat ctg tga atg ctc tcc agg ata tta ttg tct tct caa agg agc atc agg gtg tag aac atg tat ttc taa aac aaa gtg tgg aat agg ata cgg agt atc cgg ata cac gtc tac cgg aga cgt cat ctg ttc tcc gtg tgg tcc cgg aac ata ttc tca cac cgt ctc ttc cac aga taa atg cga acc cgt cgt aac cag caa tac att taa cta tat cga tgt gga aat taa cct gta tcc agt caa cga cac atc gtg tac tcg gac gac cac tac cgg tct cag cga atc cat ctc aac gtc gga act aac tat tac cat gaa tca taa aga ttg tga tcc agt ctt tcg tgc aga ata ctt ctc tgt cct taa taa tgt agc aac ttc agg att ctt tac agg aga aaa tag ata tca gaa tac ttc aaa gat atg tac tct gaa ttt cga gat taa atg taa caa caa aga ttc atc ttc caa aca gtt aac gaa aac aaa gaa tga tac tat cat gcc gca ttc aga gac ggt aac tct agt ggg cga ctg tct atc tag cgt cga cat cta cat act ata tag taa tac caa tac tca aga cta cga aaa tga tac aat ctc tta tca tat ggg taa tgt tct cga tgt caa tag cca tat gcc cgc tag ttg cga tat aca taa act gat cac taa ttc cca gaa tcc cac cca ctt ata gta ag- -tt ttt tta cct ata aat aat aaa tac aat aat taa ttt ctc gta aaa gta gaa aat ata ttc taa ttt att ata tgg taa gaa agt aga atc atc tag aac agt aat caa tca ata gca atc atg aaa caa tat att gtc ctg gca tgc atg tgc cta gtg gca gct gct atg cct act agt ctt caa caa tct tca tcc tcg tgt act gaa gaa gaa aac aaa cat cat atg gga atc gat gtt att atc aaa gtc aca aag caa gac caa aca ccg acc aat gat aag att tgt caa tcc gta acg gaa gtt aca gag a-- --- --- --- --- --- -cc gaa gat gat gag gta tcc gaa gaa gtt gta aaa gga gat ccc acc act tat tac act atc gtc ggt gcg ggt ctt aac atg aac ttt gga ttc acc aaa tgc cca aag att tta tcc atc tcc gaa tcc tct gat gga aac act gtg aat act aga ttg tcc agc gtg tca ccg gga caa ggt aag gac tct ccc gcg atc acg cgt gaa gaa gct ctg gct atg atc aaa gac tgt gag atg tct atc gac atc aga tgt agc gaa gaa gag aaa gac agt gac atc aag acc cat cca gta ctt ggg tct aac atc tca cat aag aaa gtg agt tac aaa gat atc atc ggt tca acg atc gtt gat aca aaa tgt gtc aag aac cta gag ttt agc gta cgt atc gga gac atg tgt gag gaa tca tct gaa ctt gaa gtc aag gat gga ttc aag tat gtc gac gga tcg gca tct gaa ggt gca acc gat gat act tca ctc atc gat tca aca aaa ctc aaa gca tgt gtc tga atc gat aac tct att cat ctg aaa atg gat gag ttg ggt taa tcg aac gat tca gac acc gca cca cga att aaa aaa g-- -ac cgg gca cta tat tcc ggt ttg caa aac aaa aat att taa cta cat tca caa aaa gtt acc tct cgt tac ttc ttc ttt ctg ttt caa tat gtg ata cga tat gat cac tat tcg tat tct ct- --- --- tgg tct cat aaa aaa gtt tta c-- --a aaa aaa aaa aaa a-t att ttt att ctc ttt ctc tct tcg at- ggt ctc aca aaa ata tta aac ctc ttt ctg atg tct caa cta ttt cgt aaa cga taa cgt cca aca ata tat tct cgt aga gct tat caa cat cct tat acc aat cta ggt tgt cag aca att gca tca taa aat aat gtt tat aat tta cac gtt aac atc ata taa taa acg tat ata gtt aat att ttt gga ata taa atg atc tgt aaa atc cat gta ggg gac act gct cac gtt ttt tct cta gta cat aat ttc aca caa gtt ttt ata cag aca aat taa ttc tcg tcc ata ta- ttt taa aac att gac ttt tgt act aag aaa aat atc ttg act aac cat ctc ttt ctc tct tcg atg ggt ctc aca aaa ata tta aac ctc ttt ctg atg g-- --- --- --- --- --- --- --- --- --- --- --- --- --- --- --- --- --- --- --- --- --- --- --- --- --- --- --- --- --- --- --- --- --- --- --- --- --- --- --- --- --- --- --- --- --- --- agt cgt aaa aag ttt tta tcc ttt ctc tct tcg ata ggt ctc --- --- --- --- --- --- --- --- --- --- --- --- --- --- --- --- --- --- --- --- --- --- --- --- --- --- --- --- --- --- --- --- --- --- --- --- --- --- --- --- --- --- --- --- --- --- --- --- --- --- --- --- --- --- --- --- --- --- --- --- --- --- --- --- --- --- --- --- --- --- aca aaa ata tta aac ctc ttt ctg atg gtc tct ata aa

>ON649721.1_Monkeypox_virus_isolate_Monkeypox/PT0021/2022_complete_genome

atc gag gac gac ttc aac tat gac agc tcg gtg gcg tct gcc agc gtg tac ata cga atg gca ttt cta aga aaa gtc tac ggt atc ctt -tc tac aat ttc ctt taa caa cgg caa cag ctg cag tat ttt tat act ttg aat gca -tc gga cat tta tac aag gga gtc ctg ttc taa tat tgg cat caa tgt tcg gat cta tag gct tga ttt tcg cat tga ctt tac aca gac atg aac atc ccc tga atc tgt aca tac ttt gtg gat tta cac tgt tag aat ctc taa cgc tgg cct ctg ttg tta ctt tct atg atg cac gta tcg tta tgc aag ctt tca tgt tga cta ctg cag tgt ttc ttg ctc tga cta cat ata ctc tac aat caa aga gag att tca gta aac ttg taa cag gat tgt ttg ctg ctt tct gga ttt taa ttt tgt cag gag tct tga gga taa agt tta aaa tag aat taa taa aga aca tat agg tca -tt ttt taa aca tgg ata gaa acc aag gtt gtt agt taa taa tat aca aga tat ttt ttc tca ctc tga tcc atg taa acc aag gac gag aga cac tct cat tcc tca ttc acg aca cca tta aaa atg gaa att aaa gcc ctc tat taa gca cag acg gct aca ggt cta cca tca ggt t-a cct tcg tct acc ttc aca atg gcc tct cct tgt gcc cag ttc agt ccc tgt cat tgc cac gct act aag gac tcc ctg aat acc gtg act gac gtc aga cat tgt ctg act gaa tac atc ctg tgg gtt tct cat aga tgg acc cat aga gaa agc gca ggg cct ctc tac agg ctt ctc atc tct ttc aga att gat gca atg gag cta ttt ggt agc gag ttg aag gag ttc tcg aat tca ctt ccg tgg gac aat atc gac aat tgc gtg gag atc att aaa tgt ttc atc aga aat gac tcc atg aaa acc gcc aaa gaa ctt tgt gca ata att gga ctt tgt act caa tca gct att gtc act gga aga gtc ttc aat gat aag tat atc gac ata cta ctt atg ctg cga aag att ctg aac gag aac gac tat ctc acc ctc ttg gat cat atc ctc act gct aaa tac taa atc tcc ttc atg ctc tct cac taa tac tct tac tca cta cac ttt tta tca tct tat gat gaa tga ttg cct tca tca ttt ttt cgt gga ata taa tat agg aat aat tag cac cag aat agc tat gga tat ctc gtt aag aat att ctc tat aag aga cat aat gta gac ata gtt att ata tcc ttc tta gat aag tgt tac gct act gga aag ttt cca tcg tta tta tta cat gaa gat gat ata att aaa cca aca ttg aga ttg gct ctt atg tta gct gga ttg aat tac tgt aat aaa tgc atc gag tat aga ggg ata tag caa ttc tcg ata ata gtc atg caa tat ttg aat gag act gat aat tta ggt aat aca gta ct- --- aca cac ata tct ttc tag att ata tat cgt taa aaa tct gta aga tgt ata ttt ctc ata agt atc cac tgt gta ata tta tta atg gat ata tag ata acg caa tag gga cta ata gta ttg taa aag ata taa tcg act att tac gta cat atc cag ata tct ata ttc cta cta gtt tgc tgc gta gtt gca tca ttg ata tgc atg -at tta tca gga ttc aga gat gaa tta cta agt aaa cta caa tcc cac aat aag taa gaa tca aat atc aaa aac tca ctt ttg att ttt cta gtc tta agt aat aca tat att tat taa tag acc tat gaa ata aaa aaa ggt aac aat gga ttc gcg tat agc tat tta cgt att agt atc ggc atc tct ttt gta tct tgt taa ttg tca caa act agt aca tta ctt caa tct gaa aat aaa tgg aag tga tat aac taa tac agc aga tat att gct gga caa tta tcc aat tat gac ctt tga tgg aaa gga tat tta tcc atc tat ctc gtt cat ggt cgg taa taa act ttt cct aga tct tta taa aaa tat ctt tga aga att ttt cag act att tcg agt atc tgt aag tag tca ata cga gga att aga ata tta tta ttc atg tga tta tac taa caa ccg tcc tac aat taa aca aca tta ctt tta taa cgg cga cga ata tac tga aat tga tag atc gaa aaa agc cac taa taa aaa cag ttg gtt aat tac ttc agg ctt tag act aca aaa atg gtt cga tag cga aga ttg tat aat tta tct cag atc ttt agt tag aag aat gga aga cag taa caa aaa cag taa aaa aac tta gta ctt aga tat cg- --a aaa aat ata ttt ttg tag act ctt gag aat aga agg aaa aca tgt aca taa tta taa aaa atg aaa atc aat ggc gaa taa gac agt gcg att cgc acc atg gag tcg gta gat ttc atg gct gtc gat gag cag ttt cac gac gac ctc gat ctt tgg tca tta tct ttg gta gat gat tat aaa aaa cat gga tta ggt gtt gac tgt tat gtt cta gaa cca gtt gtt gac agg aaa ata ttt gat aga ttt ctc ctt gaa cca att tgt gat cct gta gat gtt ctg tat gat tat ttt agg att cat aga gat aat att gat cag tat ata gta gat ??? ?tg ttt gca tat att aca tat aaa gat att ata tct gca tta gtg tca aag aat tat atg gaa gat att ttc tct ata att att aag aat tgt aat tct gtg caa gat ctc tta ctt tac tat cta tct aat gca tat gta gaa ata gac att gtt gat ctt atg gta gat cat ggg gct gta ata tat aaa ata gaa tgc ttg aat gcc tat ttt agg gga ata tgt aaa aag gaa agt agt gtt gtt gag ttt att ttg aat tgt ggt atc cca gat gaa aat gat gtt aaa tta gat cta tat aaa ata att cag tat act agg gga ttc ctt gta gat gaa ccc aca gta tta gaa att tat aag ctt tgt atc cca tat att gaa gat atc aat caa cta gat gct ggt gga agg acc ttg ctt tat cgc gct atc tat gca ggt tat ata gat tta gta tca tgg cta tta gaa aat gga gca aat gtc aac gca gta atg agt aat gga tat aca tgt ctt gac gtg gcc gtg gat agg gga tct gtc atc gcc cgt agg gaa gca cat ctt aaa ata tta gaa ata ttg ctt aga gaa cca ttg tct att gac tgt ata aaa tta gct ata ctt aat aat aca att gaa aac cat gat gtg ata aag ctc tgt atc aag tat ttt atg atg gta gat tat tca ctt tgt aat gtg tat gca tca tca ctc ttt gat tat ata att gat tgt aaa caa gaa ttg gag tac att agg cag atg aaa att cat aat aca acc atg tat gag tta atc tat aat aga gac aaa aac aag cat gct tcc cat att cta cat agg tat tct aaa cat cca gtt ttg aca cag tgt atc act aaa gga ttc aag att tac aca gaa gta acc gag cag gtc act aaa gct cta aac aga cgt gct cta ata gat gag ata ata aac aat gta tca act gat gac aat ctc cta tca aaa ctt cca tta gaa att agg gat cta att gtt tca caa gct gtc ata tag agt tct atc cac cca cct ttc ttg aaa tga gtt aat --- --- --- --- --- --- --- --- --- --- --- --- --- --- --- --- --- --- --- --- --- --- --- --- --- --- --- --- --- --- --- --- --- --- --- --- --- --- --- --- --- --- --- --- --- --- --- --- --- --- --- --- --- --- --- --- --- --- --- --- --- --- --- --- --- --- --- --- --- --- --- --- --- --- --- --- --- --- --- --- --- --- --- --- --- --- --- --- --- --- --- --- --- --- --- --- --- --- --- --- --- --- --- --- --- -ag tca taa gtt agt taa --- --- --- --- --- -gt cat aag tta gtt aag tca t-- --- --- --- --- --- --- --- --- --- --- --- --- --- --- --- -aa gtt agt taa gtc ata agt tag tta agt cat aag tta gtt aag tca taa gtt agt taa gtc ata agt tag tta agt cat aag tta gtt tat agt cta aca ctt cta att ttt ata cct tga tct ttt tct cta att atg aaa aag taa atc att atg aag atg gat g-a aat gga cga gat tgt gcg cat cgt taa cga tag tat gtg gta cgt acc taa cgc att tat gga cga cgg tga taa tga agg tca cat ttc tgt caa taa tgt ctg tca tat gta tct cgc att ctt tga tgt gga tat atc atc tca tct gtt taa att agt tat taa aca ctg cga tct gaa taa acg act aaa atg tgg taa ctc tcc att aca ttg cta tac gat gaa tac acg att taa tcc atc tgt att aaa gat att gtt acg cca cgg cat gcg taa ctt tga tag caa gga taa aaa agg aca tat tcc tct aca cca cta tct gat tca ttc act atc aat cga taa caa gat ctt tga tat act aac gga ccc cat tga tga ctt tag taa atc atc cga tct att gct gtg tta tct tag ata taa att caa tgg gag ctt aaa cta tta cgt tct gta caa att att gac taa agg atc tga ccc taa ttg cgt cga tga gga tgg act cac ttc tct tca tta cta ctg taa aca cat atc cgc gtt cca cga aag caa tta tta caa gtc aaa gag tca cac taa gat gcg agc tga gaa gcg att cat cta cgc gat aat aga tca tgg agc aaa cat taa cgc ggt tac gaa aat cgg aaa tac gcc gtt aca cac tta cct tca aca gta tac caa aca tag tcc tcg tgt ggt gta tgc tct ttt atc tcg agg agc cga tac gag gat acg taa taa tct taa ttg tac acc cat cat gga ata cat aaa gaa cga ttg tgc aac agg tca tat tct cat aat gtt act caa ttg gca cga aca aaa ata cgg gaa att aca aaa gga aga agg aca aca tct act tta tct att cat aaa aca taa tca agg ata tgg aag tcg ctc tct caa tat act acg gta tct act aga tag att cga cat tca gaa aga cga ata cta taa tac aat gac tcc tct tca tac cgc ctt cca gaa ttg caa taa caa tgt tgc ctc ata cct cgt ata cat tgg ata cga cat caa cct tcc gac taa aga cga taa gac agt att cga ctt ggt gtt tga aaa cag aaa cat tat ata caa ggc gga tgt cgt taa tga cat tat cca cca cag act gaa agt atc tct acc tat gat taa atc gtt gtt cta caa gat gtc gga gtt ctc tcc cta cga cga tca cta cgt aaa gaa gat aat agc cta ctg cct att aag gga cga gtc att tgc gga act aca tac taa att ctg ttt aaa cga gga cta taa aag tgt att tat gaa aaa tat atc att cga taa gat aga ttc cat cat cga aaa atg tag tcg tga cat aag tct cct caa aga gat tcg aat ctc aga cac cga ctt gta tac ggt att gag aac aga aga cat tcg gta tca cac ata tct cga agc cat aca ttc aga caa acg cat ttc att tcc cat gta cga cga tct cat aga aca gtg tca tct atc gat gga gca taa aag taa act cgt cga caa agc act caa taa att aga gtc tac cat cga tag tca atc tag act atc gta ttt gcc tcc gga aat tat gcg caa tat cat aac caa gct aag cga cta cca tct aaa cag tat gtt gta cgg aaa gaa cca tta caa ata tta tcc atg ata gaa aga aaa tat tta aaa aat aat cta tat gat tgg aga agt agg aaa caa aca gta aca aga cga cga tta cta c-a tta tta aat cat gag gtc cgt att ata ctc gta tat att gtt tct ctc atg tat aat aat aaa cgg aag aga tat agc acc aca tgc acc atc caa tgg aaa gtg taa aga caa cga ata cag aag ccg taa tct atg ttg tct atc gtg tcc tcc ggg aac tta cgc ttt cag att atg tga tag caa gac taa tac aca atg tac acc gtg tgg ttc gga tac ctt tac atc tca caa taa tca ttt aca ggc ttg tct aag ttg taa cgg aag atg tga tag taa tca ggt aga gac gcg atc gtg taa cac gac tca caa tag aat ctg tga atg ctc tcc agg ata tta ttg tct tct caa agg agc atc agg gtg tag aac atg tat ttc taa aac aaa gtg tgg aat agg ata cgg agt atc cgg ata cac gtc tac cgg aga cgt cat ctg ttc tcc gtg tgg tcc cgg aac ata ttc tca cac cgt ctc ttc cac aga taa atg cga acc cgt cgt aac cag caa tac att taa cta tat cga tgt gga aat taa cct gta tcc agt caa cga cac atc gtg tac tcg gac gac cac tac cgg tct cag cga atc cat ctc aac gtc gga act aac tat tac cat gaa tca taa aga ttg tga tcc agt ctt tcg tgc aga ata ctt ctc tgt cct taa taa tgt agc aac ttc agg att ctt tac agg aga aaa tag ata tca gaa tac ttc aaa gat atg tac tct gaa ttt cga gat taa atg taa caa caa aga ttc atc ttc caa aca gtt aac gaa aac aaa gaa tga tac tat cat gcc gca ttc aga gac ggt aac tct agt ggg cga ctg tct atc tag cgt cga cat cta cat act ata tag taa tac caa tac tca aga cta cga aaa tga tac aat ctc tta tca tat ggg taa tgt tct cga tgt caa tag cca tat gcc cgc tag ttg cga tat aca taa act gat cac taa ttc cca gaa tcc cac cca ctt ata gta ag- -tt ttt tta cct ata aat aat aaa tac aat aat taa ttt ctc gta aaa gta gaa aat ata ttc taa ttt att ata tgg taa gaa agt aga atc atc tag aac agt aat caa tca ata gca atc atg aaa caa tat att gtc ctg gca tgc atg tgc cta gtg gca gct gct atg cct act agt ctt caa caa tct tca tcc tcg tgt act gaa gaa gaa aac aaa cat cat atg gga atc gat gtt att atc aaa gtc aca aag caa gac caa aca ccg acc aat gat aag att tgt caa tcc gta acg gaa gtt aca gag a-- --- --- --- --- --- -cc gaa gat gat gag gta tcc gaa gaa gtt gta aaa gga gat ccc acc act tat tac act atc gtc ggt gcg ggt ctt aac atg aac ttt gga ttc acc aaa tgc cca aag att tta tcc atc tcc gaa tcc tct gat gga aac act gtg aat act aga ttg tcc agc gtg tca ccg gga caa ggt aag gac tct ccc gcg atc acg cgt gaa gaa gct ctg gct atg atc aaa gac tgt gag atg tct atc gac atc aga tgt agc gaa gaa gag aaa gac agt gac atc aag acc cat cca gta ctt ggg tct aac atc tca cat aag aaa gtg agt tac aaa gat atc atc ggt tca acg atc gtt gat aca aaa tgt gtc aag aac cta gag ttt agc gta cgt atc gga gac atg tgt gag gaa tca tct gaa ctt gaa gtc aag gat gga ttc aag tat gtc gac gga tcg gca tct gaa ggt gca acc gat gat act tca ctc atc gat tca aca aaa ctc aaa gca tgt gtc tga atc gat aac tct att cat ctg aaa atg gat gag ttg ggt taa tcg aac gat tca gac acc gca cca cga att aaa aaa g-- -ac cgg gca cta tat tcc ggt ttg caa aac aaa aat att taa cta cat tca caa aaa gtt acc tct cgt tac ttc ttc ttt ctg ttt caa tat gtg ata cga tat gat cac tat tcg tat tct ct- --- --- tgg tct cat aaa aaa gtt tta c-- --a aaa aaa aaa aaa a-t att ttt att ctc ttt ctc tct tcg at- ggt ctc aca aaa ata tta aac ctc ttt ctg atg tct caa cta ttt cgt aaa cga taa cgt cca aca ata tat tct cgt aga gct tat caa cat cct tat acc aat cta ggt tgt cag aca att gca tca taa aat aat gtt tat aat tta cac gtt aac atc ata taa taa acg tat ata gtt aat att ttt gga ata taa atg atc tgt aaa atc cat gta ggg gac act gct cac gtt ttt tct cta gta cat aat ttc aca caa gtt ttt ata cag aca aat taa ttc tcg tcc ata ta- ttt taa aac att gac ttt tgt act aag aaa aat atc ttg act aac cat ctc ttt ctc tct tcg atg ggt ctc aca aaa ata tta aac ctc ttt ctg atg g-- --- --- --- --- --- --- --- --- --- --- --- --- --- --- --- --- --- --- --- --- --- --- --- --- --- --- --- --- --- --- --- --- --- --- --- --- --- --- --- --- --- --- --- --- --- --- agt cgt aaa aag ttt tta tcc ttt ctc tct tcg ata ggt ctc --- --- --- --- --- --- --- --- --- --- --- --- --- --- --- --- --- --- --- --- --- --- --- --- --- --- --- --- --- --- --- --- --- --- --- --- --- --- --- --- --- --- --- --- --- --- --- --- --- --- --- --- --- --- --- --- --- --- --- --- --- --- --- --- --- --- --- --- --- --- aca aaa ata tta aac ctc ttt ctg atg gtc tct ata aa

>ON649720.1_Monkeypox_virus_isolate_Monkeypox/PT0024/2022_complete_genome

atc gag gac gac ttc aac tat gac agc tcg gtg gcg tct gcc agc gtg tac ata cga atg gca ttt cta aga aaa gtc tac ggt atc ctt -tc tac aat ttc ctt taa caa cgg caa cag ctg cag tat ttt tat act ttg aat gca -tc gga cat tta tac aag gga gtc ctg ttc taa tat tgg cat caa tgt tcg gat cta tag gct tga ttt tcg cat tga ctt tac aca gac atg aac atc ccc tga atc tgt aca tac ttt gtg gat tta cac tgt tag aat ctc taa cgc tgg cct ctg ttg tta ctt tct atg atg cac gta tcg tta tgc aag ctt tca tgt tga cta ctg cag tgt ttc ttg ctc tga cta cat ata ctc tac aat caa aga gag att tca gta aac ttg taa cag gat tgt ttg ctg ctt tct gga ttt taa ttt tgt cag gag tct tga gga taa agt tta aaa tag aat taa taa aga aca tat agg tca -tt ttt taa aca tgg ata gaa acc aag gtt gtt agt taa taa tat aca aga tat ttt ttc tca ctc tga tcc atg taa acc aag gac gag aga cac tct cat tcc tca ttc acg aca cca tta aaa atg gaa att aaa gcc ctc tat taa gca cag acg gct aca ggt cta cca tca ggt t-a cct tcg tct acc ttc aca atg gcc tct cct tgt gcc cag ttc agt ccc tgt cat tgc cac gct act aag gac tcc ctg aat acc gtg act gac gtc aga cat tgt ctg act gaa tac atc ctg tgg gtt tct cat aga tgg acc cat aga gaa agc gca ggg cct ctc tac agg ctt ctc atc tct ttc aga att gat gca atg gag cta ttt ggt agc gag ttg aag gag ttc tcg aat tca ctt ccg tgg gac aat atc gac aat tgc gtg gag atc att aaa tgt ttc atc aga aat gac tcc atg aaa acc gcc aaa gaa ctt tgt gca ata att gga ctt tgt act caa tca gct att gtc act gga aga gtc ttc aat gat aag tat atc gac ata cta ctt atg ctg cga aag att ctg aac gag aac gac tat ctc acc ctc ttg gat cat atc ctc act gct aaa tac taa atc tcc ttc atg ctc tct cac taa tac tct tac tca cta cac ttt tta tca tct tat gat gaa tga ttg cct tca tca ttt ttt cgt gga ata taa tat agg aat aat tag cac cag aat agc tat gga tat ctc gtt aag aat att ctc tat aag aga cat aat gta gac ata gtt att ata tcc ttc tta gat aag tgt tac gct act gga aag ttt cca tcg tta tta tta cat gaa gat gat ata att aaa cca aca ttg aga ttg gct ctt atg tta gct gga ttg aat tac tgt aat aaa tgc atc gag tat aga ggg ata tag caa ttc tcg ata ata gtc atg caa tat ttg aat gag act gat aat tta ggt aat aca gta ct- --- aca cac ata tct ttc tag att ata tat cgt taa aaa tct gta aga tgt ata ttt ctc ata agt atc cac tgt gta ata tta tta atg gat ata tag ata acg caa tag gga cta ata gta ttg taa aag ata taa tcg act att tac gta cat atc cag ata tct ata ttc cta cta gtt tgc tgc gta gtt gca tca ttg ata tgc atg -at tta tca gga ttc aga gat gaa tta cta agt aaa cta caa tcc cac aat aag taa gaa tca aat atc aaa aac tca ctt ttg att ttt cta gtc tta agt aat aca tat att tat taa tag acc tat gaa ata aaa aaa ggt aac aat gga ttc gcg tat agc tat tta cgt att agt atc ggc atc tct ttt gta tct tgt taa ttg tca caa act agt aca tta ctt caa tct gaa aat aaa tgg aag tga tat aac taa tac agc aga tat att gct gga caa tta tcc aat tat gac ctt tga tgg aaa gga tat tta tcc atc tat ctc gtt cat ggt cgg taa taa act ttt cct aga tct tta taa aaa tat ctt tga aga att ttt cag act att tcg agt atc tgt aag tag tca ata cga gga att aga ata tta tta ttc atg tga tta tac taa caa ccg tcc tac aat taa aca aca tta ctt tta taa cgg cga cga ata tac tga aat tga tag atc gaa aaa agc cac taa taa aaa cag ttg gtt aat tac ttc agg ctt tag act aca aaa atg gtt cga tag cga aga ttg tat aat tta tct cag atc ttt agt tag aag aat gga aga cag taa caa aaa cag taa aaa aac tta gta ctt aga tat cg- --a aaa aat ata ttt ttg tag act ctt gag aat aga agg aaa aca tgt aca taa tta taa aaa atg aaa atc aat ggc gaa taa gac agt gcg att cgc acc atg gag tcg gta gat ttc atg gct gtc gat gag cag ttt cac gac gac ctc gat ctt tgg tca tta tct ttg gta gat gat tat aaa aaa cat gga tta ggt gtt gac tgt tat gtt cta gaa cca gtt gtt gac agg aaa ata ttt gat aga ttt ctc ctt gaa cca att tgt gat cct gta gat gtt ctg tat gat tat ttt agg att cat aga gat aat att gat cag tat ata gta gat aga ctg ttt gca tat att aca tat aaa gat att ata tct gca tta gtg tca aag aat tat atg gaa gat att ttc tct ata att att aag aat tgt aat tct gtg caa gat ctc tta ctt tac tat cta tct aat gca tat gta gaa ata gac att gtt gat ctt atg gta gat cat ggg gct gta ata tat aaa ata gaa tgc ttg aat gcc tat ttt agg gga ata tgt aaa aag gaa agt agt gtt gtt gag ttt att ttg aat tgt ggt atc cca gat gaa aat gat gtt aaa tta gat cta tat aaa ata att cag tat act agg gga ttc ctt gta gat gaa ccc aca gta tta gaa att tat aag ctt tgt atc cca tat att gaa gat atc aat caa cta gat gct ggt gga agg acc ttg ctt tat cgc gct atc tat gca ggt tat ata gat tta gta tca tgg cta tta gaa aat gga gca aat gtc aac gca gta atg agt aat gga tat aca tgt ctt gac gtg gcc gtg gat agg gga tct gtc atc gcc cgt agg gaa gca cat ctt aaa ata tta gaa ata ttg ctt aga gaa cca ttg tct att gac tgt ata aaa tta gct ata ctt aat aat aca att gaa aac cat gat gtg ata aag ctc tgt atc aag tat ttt atg atg gta gat tat tca ctt tgt aat gtg tat gca tca tca ctc ttt gat tat ata att gat tgt aaa caa gaa ttg gag tac att agg cag atg aaa att cat aat aca acc atg tat gag tta atc tat aat aga gac aaa aac aag cat gct tcc cat att cta cat agg tat tct aaa cat cca gtt ttg aca cag tgt atc act aaa gga ttc aag att tac aca gaa gta acc gag cag gtc act aaa gct cta aac aga cgt gct cta ata gat gag ata ata aac aat gta tca act gat gac aat ctc cta tca aaa ctt cca tta gaa att agg gat cta att gtt tca caa gct gtc ata tag agt tct atc cac cca cct ttc ttg aaa tga gtt aat --- --- --- --- --- --- --- --- --- --- --- --- --- --- --- --- --- --- --- --- --- --- --- --- --- --- --- --- --- --- --- --- --- --- --- --- --- --- --- --- --- --- --- --- --- --- --- --- --- --- --- --- --- --- --- --- --- --- --- --- --- --- --- --- --- --- --- --- --- --- --- --- --- --- --- --- --- --- --- --- --- --- --- --- --- --- --- --- --- --- --- --- --- --- --- --- --- --- --- --- --- --- --- --- --- -ag tca taa gtt agt taa --- --- --- --- --- -gt cat aag tta gtt aag tca t-- --- --- --- --- --- --- --- --- --- --- --- --- --- --- --- -aa gtt agt taa gtc ata agt tag tta agt cat aag tta gtt aag tca taa gtt agt taa gtc ata agt tag tta agt cat aag tta gtt tat agt cta aca ctt cta att ttt ata cct tga tct ttt tct cta att atg aaa aag taa atc att atg aag atg gat g-a aat gga cga gat tgt gcg cat cgt taa cga tag tat gtg gta cgt acc taa cgc att tat gga cga cgg tga taa tga agg tca cat ttc tgt caa taa tgt ctg tca tat gta tct cgc att ctt tga tgt gga tat atc atc tca tct gtt taa att agt tat taa aca ctg cga tct gaa taa acg act aaa atg tgg taa ctc tcc att aca ttg cta tac gat gaa tac acg att taa tcc atc tgt att aaa gat att gtt acg cca cgg cat gcg taa ctt tga tag caa gga taa aaa agg aca tat tcc tct aca cca cta tct gat tca ttc act atc aat cga taa caa gat ctt tga tat act aac gga ccc cat tga tga ctt tag taa atc atc cga tct att gct gtg tta tct tag ata taa att caa tgg gag ctt aaa cta tta cgt tct gta caa att att gac taa agg atc tga ccc taa ttg cgt cga tga gga tgg act cac ttc tct tca tta cta ctg taa aca cat atc cgc gtt cca cga aag caa tta tta caa gtc aaa gag tca cac taa gat gcg agc tga gaa gcg att cat cta cgc gat aat aga tca tgg agc aaa cat taa cgc ggt tac gaa aat cgg aaa tac gcc gtt aca cac tta cct tca aca gta tac caa aca tag tcc tcg tgt ggt gta tgc tct ttt atc tcg agg agc cga tac gag gat acg taa taa tct taa ttg tac acc cat cat gga ata cat aaa gaa cga ttg tgc aac agg tca tat tct cat aat gtt act caa ttg gca cga aca aaa ata cgg gaa att aca aaa gga aga agg aca aca tct act tta tct att cat aaa aca taa tca agg ata tgg aag tcg ctc tct caa tat act acg gta tct act aga tag att cga cat tca gaa aga cga ata cta taa tac aat gac tcc tct tca tac cgc ctt cca gaa ttg caa taa caa tgt tgc ctc ata cct cgt ata cat tgg ata cga cat caa cct tcc gac taa aga cga taa gac agt att cga ctt ggt gtt tga aaa cag aaa cat tat ata caa ggc gga tgt cgt taa tga cat tat cca cca cag act gaa agt atc tct acc tat gat taa atc gtt gtt cta caa gat gtc gga gtt ctc tcc cta cga cga tca cta cgt aaa gaa gat aat agc cta ctg cct att aag gga cga gtc att tgc gga act aca tac taa att ctg ttt aaa cga gga cta taa aag tgt att tat gaa aaa tat atc att cga taa gat aga ttc cat cat cga aaa atg tag tcg tga cat aag tct cct caa aga gat tcg aat ctc aga cac cga ctt gta tac ggt att gag aac aga aga cat tcg gta tca cac ata tct cga agc cat aca ttc aga caa acg cat ttc att tcc cat gta cga cga tct cat aga aca gtg tca tct atc gat gga gca taa aag taa act cgt cga caa agc act caa taa att aga gtc tac cat cga tag tca atc tag act atc gta ttt gcc tcc gga aat tat gcg caa tat cat aac caa gct aag cga cta cca tct aaa cag tat gtt gta cgg aaa gaa cca tta caa ata tta tcc atg ata gaa aga aaa tat tta aaa aat aat cta tat gat tgg aga agt agg aaa caa aca gta aca aga cga cga tta cta c-a tta tta aat cat gag gtc cgt att ata ctc gta tat att gtt tct ctc atg tat aat aat aaa cgg aag aga tat agc acc aca tgc acc atc caa tgg aaa gtg taa aga caa cga ata cag aag ccg taa tct atg ttg tct atc gtg tcc tcc ggg aac tta cgc ttt cag att atg tga tag caa gac taa tac aca atg tac acc gtg tgg ttc gga tac ctt tac atc tca caa taa tca ttt aca ggc ttg tct aag ttg taa cgg aag atg tga tag taa tca ggt aga gac gcg atc gtg taa cac gac tca caa tag aat ctg tga atg ctc tcc agg ata tta ttg tct tct caa agg agc atc agg gtg tag aac atg tat ttc taa aac aaa gtg tgg aat agg ata cgg agt atc cgg ata cac gtc tac cgg aga cgt cat ctg ttc tcc gtg tgg tcc cgg aac ata ttc tca cac cgt ctc ttc cac aga taa atg cga acc cgt cgt aac cag caa tac att taa cta tat cga tgt gga aat taa cct gta tcc agt caa cga cac atc gtg tac tcg gac gac cac tac cgg tct cag cga atc cat ctc aac gtc gga act aac tat tac cat gaa tca taa aga ttg tga tcc agt ctt tcg tgc aga ata ctt ctc tgt cct taa taa tgt agc aac ttc agg att ctt tac agg aga aaa tag ata tca gaa tac ttc aaa gat atg tac tct gaa ttt cga gat taa atg taa caa caa aga ttc atc ttc caa aca gtt aac gaa aac aaa gaa tga tac tat cat gcc gca ttc aga gac ggt aac tct agt ggg cga ctg tct atc tag cgt cga cat cta cat act ata tag taa tac caa tac tca aga cta cga aaa tga tac aat ctc tta tca tat ggg taa tgt tct cga tgt caa tag cca tat gcc cgc tag ttg cga tat aca taa act gat cac taa ttc cca gaa tcc cac cca ctt ata gta ag- -tt ttt tta cct ata aat aat aaa tac aat aat taa ttt ctc gta aaa gta gaa aat ata ttc taa ttt att ata tgg taa gaa agt aga atc atc tag aac agt aat caa tca ata gca atc atg aaa caa tat att gtc ctg gca tgc atg tgc cta gtg gca gct gct atg cct act agt ctt caa caa tct tca tcc tcg tgt act gaa gaa gaa aac aaa cat cat atg gga atc gat gtt att atc aaa gtc aca aag caa gac caa aca ccg acc aat gat aag att tgt caa tcc gta acg gaa gtt aca gag a-- --- --- --- --- --- -cc gaa gat gat gag gta tcc gaa gaa gtt gta aaa gga gat ccc acc act tat tac act atc gtc ggt gcg ggt ctt aac atg aac ttt gga ttc acc aaa tgc cca aag att tta tcc atc tcc gaa tcc tct gat gga aac act gtg aat act aga ttg tcc agc gtg tca ccg gga caa ggt aag gac tct ccc gcg atc acg cgt gaa gaa gct ctg gct atg atc aaa gac tgt gag atg tct atc gac atc aga tgt agc gaa gaa gag aaa gac agt gac atc aag acc cat cca gta ctt ggg tct aac atc tca cat aag aaa gtg agt tac aaa gat atc atc ggt tca acg atc gtt gat aca aaa tgt gtc aag aac cta gag ttt agc gta cgt atc gga gac atg tgt gag gaa tca tct gaa ctt gaa gtc aag gat gga ttc aag tat gtc gac gga tcg gca tct gaa ggt gca acc gat gat act tca ctc atc gat tca aca aaa ctc aaa gca tgt gtc tga atc gat aac tct att cat ctg aaa atg gat gag ttg ggt taa tcg aac gat tca gac acc gca cca cga att aaa aaa g-- -ac cgg gca cta tat tcc ggt ttg caa aac aaa aat att taa cta cat tca caa aaa gtt acc tct cgt tac ttc ttc ttt ctg ttt caa tat gtg ata cga tat gat cac tat tcg tat tct ct- --- --- tgg tct cat aaa aaa gtt tta c-- --a aaa aaa aaa aaa a-t att ttt att ctc ttt ctc tct tcg at- ggt ctc aca aaa ata tta aac ctc ttt ctg atg tct caa cta ttt cgt aaa cga taa cgt cca aca ata tat tct cgt aga gct tat caa cat cct tat acc aat cta ggt tgt cag aca att gca tca taa aat aat gtt tat aat tta cac gtt aac atc ata taa taa acg tat ata gtt aat att ttt gga ata taa atg atc tgt aaa atc cat gta ggg gac act gct cac gtt ttt tct cta gta cat aat ttc aca caa gtt ttt ata cag aca aat taa ttc tcg tcc ata ta- ttt taa aac att gac ttt tgt act aag aaa aat atc ttg act aac cat ctc ttt ctc tct tcg atg ggt ctc aca aaa ata tta aac ctc ttt ctg atg g-- --- --- --- --- --- --- --- --- --- --- --- --- --- --- --- --- --- --- --- --- --- --- --- --- --- --- --- --- --- --- --- --- --- --- --- --- --- --- --- --- --- --- --- --- --- --- agt cgt aaa aag ttt tta tcc ttt ctc tct tcg ata ggt ctc --- --- --- --- --- --- --- --- --- --- --- --- --- --- --- --- --- --- --- --- --- --- --- --- --- --- --- --- --- --- --- --- --- --- --- --- --- --- --- --- --- --- --- --- --- --- --- --- --- --- --- --- --- --- --- --- --- --- --- --- --- --- --- --- --- --- --- --- --- --- aca aaa ata tta aac ctc ttt ctg atg gtc tct ata aa

>ON649723.1_Monkeypox_virus_isolate_Monkeypox/PT0013/2022_complete_genome

atc gag gac gac ttc aac tat gac agc tcg gtg gcg tct gcc agc gtg tac ata cga atg gca ttt cta aga aaa gtc tac ggt atc ctt -tc tac aat ttc ctt taa caa cgg caa cag ctg cag tat ttt tat act ttg aat gca -tc gga cat tta tac aag gga gtc ctg ttc taa tat tgg cat caa tgt tcg gat cta tag gct tga ttt tcg cat tga ctt tac aca gac atg aac atc ccc tga atc tgt aca tac ttt gtg gat tta cac tgt tag aat ctc taa cgc tgg cct ctg ttg tta ctt tct atg atg cac gta tcg tta tgc aag ctt tca tgt tga cta ctg cag tgt ttc ttg ctc tga cta cat ata ctc tac aat caa aga gag att tca gta aac ttg taa cag gat tgt ttg ctg ctt tct gga ttt taa ttt tgt cag gag tct tga gga taa agt tta aaa tag aat taa taa aga aca tat agg tca -tt ttt taa aca tgg ata gaa acc aag gtt gtt agt taa taa tat aca aga tat ttt ttc tca ctc tga tcc atg taa acc aag gac gag aga cac tct cat tcc tca ttc acg aca cca tta aaa atg gaa att aaa gcc ctc tat taa gca cag acg gct aca ggt cta cca tca ggt t-a cct tcg tct acc ttc aca atg gcc tct cct tgt gcc cag ttc agt ccc tgt cat tgc cac gct act aag gac tcc ctg aat acc gtg act gac gtc aga cat tgt ctg act gaa tac atc ctg tgg gtt tct cat aga tgg acc cat aga gaa agc gca ggg cct ctc tac agg ctt ctc atc tct ttc aga att gat gca atg gag cta ttt ggt agc gag ttg aag gag ttc tcg aat tca ctt ccg tgg gac aat atc gac aat tgc gtg gag atc att aaa tgt ttc atc aga aat gac tcc atg aaa acc gcc aaa gaa ctt tgt gca ata att gga ctt tgt act caa tca gct att gtc act gga aga gtc ttc aat gat aag tat atc gac ata cta ctt atg ctg cga aag att ctg aac gag aac gac tat ctc acc ctc ttg gat cat atc ctc act gct aaa tac taa atc tcc ttc atg ctc tct cac taa tac tct tac tca cta cac ttt tta tca tct tat gat gaa tga ttg cct tca tca ttt ttt cgt gga ata taa tat agg aat aat tag cac cag aat agc tat gga tat ctc gtt aag aat att ctc tat aag aga cat aat gta gac ata gtt att ata tcc ttc tta gat aag tgt tac gct act gga aag ttt cca tcg tta tta tta cat gaa gat gat ata att aaa cca aca ttg aga ttg gct ctt atg tta gct gga ttg aat tac tgt aat aaa tgc atc gag tat aga ggg ata tag caa ttc tcg ata ata gtc atg caa tat ttg aat gag act gat aat tta ggt aat aca gta ct- --- aca cac ata tct ttc tag att ata tat cgt taa aaa tct gta aga tgt ata ttt ctc ata agt atc cac tgt gta ata tta tta atg gat ata tag ata acg caa tag gga cta ata gta ttg taa aag ata taa tcg act att tac gta cat atc cag ata tct ata ttc cta cta gtt tgc tgc gta gtt gca tca ttg ata tgc atg -at tta tca gga ttc aga gat gaa tta cta agt aaa cta caa tcc cac aat aag taa gaa tca aat atc aaa aac tca ctt ttg att ttt cta gtc tta agt aat aca tat att tat taa tag acc tat gaa ata aaa aaa ggt aac aat gga ttc gcg tat agc tat tta cgt att agt atc ggc atc tct ttt gta tct tgt taa ttg tca caa act agt aca tta ctt caa tct gaa aat aaa tgg aag tga tat aac taa tac agc aga tat att gct gga caa tta tcc aat tat gac ctt tga tgg aaa gga tat tta tcc atc tat ctc gtt cat ggt cgg taa taa act ttt cct aga tct tta taa aaa tat ctt tga aga att ttt cag act att tcg agt atc tgt aag tag tca ata cga gga att aga ata tta tta ttc atg tga tta tac taa caa ccg tcc tac aat taa aca aca tta ctt tta taa cgg cga cga ata tac tga aat tga tag atc gaa aaa agc cac taa taa aaa cag ttg gtt aat tac ttc agg ctt tag act aca aaa atg gtt cga tag cga aga ttg tat aat tta tct cag atc ttt agt tag aag aat gga aga cag taa caa aaa cag taa aaa aac tta gta ctt aga tat cg- --a aaa aat ata ttt ttg tag act ctt gag aat aga agg aaa aca tgt aca taa tta taa aaa atg aaa atc aat ggc gaa taa gac agt gcg att cgc acc atg gag tcg gta gat ttc atg gct gtc gat gag cag ttt cac gac gac ctc gat ctt tgg tca tta tct ttg gta gat gat tat aaa aaa cat gga tta ggt gtt gac tgt tat gtt cta gaa cca gtt gtt gac agg aaa ata ttt gat aga ttt ctc ctt gaa cca att tgt gat cct gta gat gtt ctg tat gat tat ttt agg att cat aga gat aat att gat cag tat ata gt? ??? ??? ?tg ttt gca tat att aca tat aaa gat att ata tct gca tta gtg tca aag aat tat atg gaa gat att ttc tct ata att att aag aat tgt aat tct gtg caa gat ctc tta ctt tac tat cta tct aat gca tat gta gaa ata gac att gtt gat ctt atg gta gat cat ggg gct gta ata tat aaa ata gaa tgc ttg aat gcc tat ttt agg gga ata tgt aaa aag gaa agt agt gtt gtt gag ttt att ttg aat tgt ggt atc cca gat gaa aat gat gtt aaa tta gat cta tat aaa ata att cag tat act agg gga ttc ctt gta gat gaa ccc aca gta tta gaa att tat aag ctt tgt atc cca tat att gaa gat atc aat caa cta gat gct ggt gga agg acc ttg ctt tat cgc gct atc tat gca ggt tat ata gat tta gta tca tgg cta tta gaa aat gga gca aat gtc aac gca gta atg agt aat gga tat aca tgt ctt gac gtg gcc gtg gat agg gga tct gtc atc gcc cgt agg gaa gca cat ctt aaa ata tta gaa ata ttg ctt aga gaa cca ttg tct att gac tgt ata aaa tta gct ata ctt aat aat aca att gaa aac cat gat gtg ata aag ctc tgt atc aag tat ttt atg atg gta gat tat tca ctt tgt aat gtg tat gca tca tca ctc ttt gat tat ata att gat tgt aaa caa gaa ttg gag tac att agg cag atg aaa att cat aat aca acc atg tat gag tta atc tat aat aga gac aaa aac aag cat gct tcc cat att cta cat agg tat tct aaa cat cca gtt ttg aca cag tgt atc act aaa gga ttc aag att tac aca gaa gta acc gag cag gtc act aaa gct cta aac aga cgt gct cta ata gat gag ata ata aac aat gta tca act gat gac aat ctc cta tca aaa ctt cca tta gaa att agg gat cta att gtt tca caa gct gtc ata tag agt tct atc cac cca cct ttc ttg aaa tga gtt aat --- --- --- --- --- --- --- --- --- --- --- --- --- --- --- --- --- --- --- --- --- --- --- --- --- --- --- --- --- --- --- --- --- --- --- --- --- --- --- --- --- --- --- --- --- --- --- --- --- --- --- --- --- --- --- --- --- --- --- --- --- --- --- --- --- --- --- --- --- --- --- --- --- --- --- --- --- --- --- --- --- --- --- --- --- --- --- --- --- --- --- --- --- --- --- --- --- --- --- --- --- --- --- --- --- -ag tca taa gtt agt taa --- --- --- --- --- -gt cat aag tta gtt aag tca t-- --- --- --- --- --- --- --- --- --- --- --- --- --- --- --- -aa gtt agt taa gtc ata agt tag tta agt cat aag tta gtt aag tca taa gtt agt taa gtc ata agt tag tta agt cat aag tta gtt tat agt cta aca ctt cta att ttt ata cct tga tct ttt tct cta att atg aaa aag taa atc att atg aag atg gat g-a aat gga cga gat tgt gcg cat cgt taa cga tag tat gtg gta cgt acc taa cgc att tat gga cga cgg tga taa tga agg tca cat ttc tgt caa taa tgt ctg tca tat gta tct cgc att ctt tga tgt gga tat atc atc tca tct gtt taa att agt tat taa aca ctg cga tct gaa taa acg act aaa atg tgg taa ctc tcc att aca ttg cta tac gat gaa tac acg att taa tcc atc tgt att aaa gat att gtt acg cca cgg cat gcg taa ctt tga tag caa gga taa aaa agg aca tat tcc tct aca cca cta tct gat tca ttc act atc aat cga taa caa gat ctt tga tat act aac gga ccc cat tga tga ctt tag taa atc atc cga tct att gct gtg tta tct tag ata taa att caa tgg gag ctt aaa cta tta cgt tct gta caa att att gac taa agg atc tga ccc taa ttg cgt cga tga gga tgg act cac ttc tct tca tta cta ctg taa aca cat atc cgc gtt cca cga aag caa tta tta caa gtc aaa gag tca cac taa gat gcg agc tga gaa gcg att cat cta cgc gat aat aga tca tgg agc aaa cat taa cgc ggt tac gaa aat cgg aaa tac gcc gtt aca cac tta cct tca aca gta tac caa aca tag tcc tcg tgt ggt gta tgc tct ttt atc tcg agg agc cga tac gag gat acg taa taa tct taa ttg tac acc cat cat gga ata cat aaa gaa cga ttg tgc aac agg tca tat tct cat aat gtt act caa ttg gca cga aca aaa ata cgg gaa att aca aaa gga aga agg aca aca tct act tta tct att cat aaa aca taa tca agg ata tgg aag tcg ctc tct caa tat act acg gta tct act aga tag att cga cat tca gaa aga cga ata cta taa tac aat gac tcc tct tca tac cgc ctt cca gaa ttg caa taa caa tgt tgc ctc ata cct cgt ata cat tgg ata cga cat caa cct tcc gac taa aga cga taa gac agt att cga ctt ggt gtt tga aaa cag aaa cat tat ata caa ggc gga tgt cgt taa tga cat tat cca cca cag act gaa agt atc tct acc tat gat taa atc gtt gtt cta caa gat gtc gga gtt ctc tcc cta cga cga tca cta cgt aaa gaa gat aat agc cta ctg cct att aag gga cga gtc att tgc gga act aca tac taa att ctg ttt aaa cga gga cta taa aag tgt att tat gaa aaa tat atc att cga taa gat aga ttc cat cat cga aaa atg tag tcg tga cat aag tct cct caa aga gat tcg aat ctc aga cac cga ctt gta tac ggt att gag aac aga aga cat tcg gta tca cac ata tct cga agc cat aca ttc aga caa acg cat ttc att tcc cat gta cga cga tct cat aga aca gtg tca tct atc gat gga gca taa aag taa act cgt cga caa agc act caa taa att aga gtc tac cat cga tag tca atc tag act atc gta ttt gcc tcc gga aat tat gcg caa tat cat aac caa gct aag cga cta cca tct aaa cag tat gtt gta cgg aaa gaa cca tta caa ata tta tcc atg ata gaa aga aaa tat tta aaa aat aat cta tat gat tgg aga agt agg aaa caa aca gta aca aga cga cga tta cta c-a tta tta aat cat gag gtc cgt att ata ctc gta tat att gtt tct ctc atg tat aat aat aaa cgg aag aga tat agc acc aca tgc acc atc caa tgg aaa gtg taa aga caa cga ata cag aag ccg taa tct atg ttg tct atc gtg tcc tcc ggg aac tta cgc ttt cag att atg tga tag caa gac taa tac aca atg tac acc gtg tgg ttc gga tac ctt tac atc tca caa taa tca ttt aca ggc ttg tct aag ttg taa cgg aag atg tga tag taa tca ggt aga gac gcg atc gtg taa cac gac tca caa tag aat ctg tga atg ctc tcc agg ata tta ttg tct tct caa agg agc atc agg gtg tag aac atg tat ttc taa aac aaa gtg tgg aat agg ata cgg agt atc cgg ata cac gtc tac cgg aga cgt cat ctg ttc tcc gtg tgg tcc cgg aac ata ttc tca cac cgt ctc ttc cac aga taa atg cga acc cgt cgt aac cag caa tac att taa cta tat cga tgt gga aat taa cct gta tcc agt caa cga cac atc gtg tac tcg gac gac cac tac cgg tct cag cga atc cat ctc aac gtc gga act aac tat tac cat gaa tca taa aga ttg tga tcc agt ctt tcg tgc aga ata ctt ctc tgt cct taa taa tgt agc aac ttc agg att ctt tac agg aga aaa tag ata tca gaa tac ttc aaa gat atg tac tct gaa ttt cga gat taa atg taa caa caa aga ttc atc ttc caa aca gtt aac gaa aac aaa gaa tga tac tat cat gcc gca ttc aga gac ggt aac tct agt ggg cga ctg tct atc tag cgt cga cat cta cat act ata tag taa tac caa tac tca aga cta cga aaa tga tac aat ctc tta tca tat ggg taa tgt tct cga tgt caa tag cca tat gcc cgc tag ttg cga tat aca taa act gat cac taa ttc cca gaa tcc cac cca ctt ata gta ag- -tt ttt tta cct ata aat aat aaa tac aat aat taa ttt ctc gta aaa gta gaa aat ata ttc taa ttt att ata tgg taa gaa agt aga atc atc tag aac agt aat caa tca ata gca atc atg aaa caa tat att gtc ctg gca tgc atg tgc cta gtg gca gct gct atg cct act agt ctt caa caa tct tca tcc tcg tgt act gaa gaa gaa aac aaa cat cat atg gga atc gat gtt att atc aaa gtc aca aag caa gac caa aca ccg acc aat gat aag att tgt caa tcc gta acg gaa gtt aca gag a-- --- --- --- --- --- -cc gaa gat gat gag gta tcc gaa gaa gtt gta aaa gga gat ccc acc act tat tac act atc gtc ggt gcg ggt ctt aac atg aac ttt gga ttc acc aaa tgc cca aag att tta tcc atc tcc gaa tcc tct gat gga aac act gtg aat act aga ttg tcc agc gtg tca ccg gga caa ggt aag gac tct ccc gcg atc acg cgt gaa gaa gct ctg gct atg atc aaa gac tgt gag atg tct atc gac atc aga tgt agc gaa gaa gag aaa gac agt gac atc aag acc cat cca gta ctt ggg tct aac atc tca cat aag aaa gtg agt tac aaa gat atc atc ggt tca acg atc gtt gat aca aaa tgt gtc aag aac cta gag ttt agc gta cgt atc gga gac atg tgt gag gaa tca tct gaa ctt gaa gtc aag gat gga ttc aag tat gtc gac gga tcg gca tct gaa ggt gca acc gat gat act tca ctc atc gat tca aca aaa ctc aaa gca tgt gtc tga atc gat aac tct att cat ctg aaa atg gat gag ttg ggt taa tcg aac gat tca gac acc gca cca cga att aaa aaa g-- -ac cgg gca cta tat tcc ggt ttg caa aac aaa aat att taa cta cat tca caa aaa gtt acc tct cgt tac ttc ttc ttt ctg ttt caa tat gtg ata cga tat gat cac tat tcg tat tct ct- --- --- tgg tct cat aaa aaa gtt tta c-- --a aaa aaa aaa aaa a-t att ttt att ctc ttt ctc tct tcg at- ggt ctc aca aaa ata tta aac ctc ttt ctg atg tct caa cta ttt cgt aaa cga taa cgt cca aca ata tat tct cgt aga gct tat caa cat cct tat acc aat cta ggt tgt cag aca att gca tca taa aat aat gtt tat aat tta cac gtt aac atc ata taa taa acg tat ata gtt aat att ttt gga ata taa atg atc tgt aaa atc cat gta ggg gac act gct cac gtt ttt tct cta gta cat aat ttc aca caa gtt ttt ata cag aca aat taa ttc tcg tcc ata ta- ttt taa aac att gac ttt tgt act aag aaa aat atc ttg act aac cat ctc ttt ctc tct tcg atg ggt ctc aca aaa ata tta aac ctc ttt ctg atg g-- --- --- --- --- --- --- --- --- --- --- --- --- --- --- --- --- --- --- --- --- --- --- --- --- --- --- --- --- --- --- --- --- --- --- --- --- --- --- --- --- --- --- --- --- --- --- agt cgt aaa aag ttt tta tcc ttt ctc tct tcg ata ggt ctc --- --- --- --- --- --- --- --- --- --- --- --- --- --- --- --- --- --- --- --- --- --- --- --- --- --- --- --- --- --- --- --- --- --- --- --- --- --- --- --- --- --- --- --- --- --- --- --- --- --- --- --- --- --- --- --- --- --- --- --- --- --- --- --- --- --- --- --- --- --- aca aaa ata tta aac ctc ttt ctg atg gtc tct ata aa

>ON649718.1_Monkeypox_virus_isolate_Monkeypox/PT0018/2022_complete_genome

atc gag gac gac ttc aac tat gac agc tcg gtg gcg tct gcc agc gtg tac ata cga atg gca ttt cta aga aaa gtc tac ggt atc ctt -tc tac aat ttc ctt taa caa cgg caa cag ctg cag tat ttt tat act ttg aat gca -tc gga cat tta tac aag gga gtc ctg ttc taa tat tgg cat caa tgt tcg gat cta tag gct tga ttt tcg cat tga ctt tac aca gac atg aac atc ccc tga atc tgt aca tac ttt gtg gat tta cac tgt tag aat ctc taa cgc tgg cct ctg ttg tta ctt tct atg atg cac gta tcg tta tgc aag ctt tca tgt tga cta ctg cag tgt ttc ttg ctc tga cta cat ata ctc tac aat caa aga gag att tca gta aac ttg taa cag gat tgt ttg ctg ctt tct gga ttt taa ttt tgt cag gag tct tga gga taa agt tta aaa tag aat taa taa aga aca tat agg tca -tt ttt taa aca tgg ata gaa acc aag gtt gtt agt taa taa tat aca aga tat ttt ttc tca ctc tga tcc atg taa acc aag gac gag aga cac tct cat tcc tca ttc acg aca cca tta aaa atg gaa att aaa gcc ctc tat taa gca cag acg gct aca ggt cta cca tca ggt t-a cct tcg tct acc ttc aca atg gcc tct cct tgt gcc cag ttc agt ccc tgt cat tgc cac gct act aag gac tcc ctg aat acc gtg act gac gtc aga cat tgt ctg act gaa tac atc ctg tgg gtt tct cat aga tgg acc cat aga gaa agc gca ggg cct ctc tac agg ctt ctc atc tct ttc aga att gat gca atg gag cta ttt ggt agc gag ttg aag gag ttc tcg aat tca ctt ccg tgg gac aat atc gac aat tgc gtg gag atc att aaa tgt ttc atc aga aat gac tcc atg aaa acc gcc aaa gaa ctt tgt gca ata att gga ctt tgt act caa tca gct att gtc act gga aga gtc ttc aat gat aag tat atc gac ata cta ctt atg ctg cga aag att ctg aac gag aac gac tat ctc acc ctc ttg gat cat atc ctc act gct aaa tac taa atc tcc ttc atg ctc tct cac taa tac tct tac tca cta cac ttt tta tca tct tat gat gaa tga ttg cct tca tca ttt ttt cgt gga ata taa tat agg aat aat tag cac cag aat agc tat gga tat ctc gtt aag aat att ctc tat aag aga cat aat gta gac ata gtt att ata tcc ttc tta gat aag tgt tac gct act gga aag ttt cca tcg tta tta tta cat gaa gat gat ata att aaa cca aca ttg aga ttg gct ctt atg tta gct gga ttg aat tac tgt aat aaa tgc atc gag tat aga ggg ata tag caa ttc tcg ata ata gtc atg caa tat ttg aat gag act gat aat tta ggt aat aca gta ct- --- aca cac ata tct ttc tag att ata tat cgt taa aaa tct gta aga tgt ata ttt ctc ata agt atc cac tgt gta ata tta tta atg gat ata tag ata acg caa tag gga cta ata gta ttg taa aag ata taa tcg act att tac gta cat atc cag ata tct ata ttc cta cta gtt tgc tgc gta gtt gca tca ttg ata tgc atg -at tta tca gga ttc aga gat gaa tta cta agt aaa cta caa tcc cac aat aag taa gaa tca aat atc aaa aac tca ctt ttg att ttt cta gtc tta agt aat aca tat att tat taa tag acc tat gaa ata aaa aaa ggt aac aat gga ttc gcg tat agc tat tta cgt att agt atc ggc atc tct ttt gta tct tgt taa ttg tca caa act agt aca tta ctt caa tct gaa aat aaa tgg aag tga tat aac taa tac agc aga tat att gct gga caa tta tcc aat tat gac ctt tga tgg aaa gga tat tta tcc atc tat ctc gtt cat ggt cgg taa taa act ttt cct aga tct tta taa aaa tat ctt tga aga att ttt cag act att tcg agt atc tgt aag tag tca ata cga gga att aga ata tta tta ttc atg tga tta tac taa caa ccg tcc tac aat taa aca aca tta ctt tta taa cgg cga cga ata tac tga aat tga tag atc gaa aaa agc cac taa taa aaa cag ttg gtt aat tac ttc agg ctt tag act aca aaa atg gtt cga tag cga aga ttg tat aat tta tct cag atc ttt agt tag aag aat gga aga cag taa caa aaa cag taa aaa aac tta gta ctt aga tat cg- --a aaa aat ata ttt ttg tag act ctt gag aat aga agg aaa aca tgt aca taa tta taa aaa atg aaa atc aat ggc gaa taa gac agt gcg att cgc acc atg gag tcg gta gat ttc atg gct gtc gat gag cag ttt cac gac gac ctc gat ctt tgg tca tta tct ttg gta gat gat tat aaa aaa cat gga tta ggt gtt gac tgt tat gtt cta gaa cca gtt gtt gac agg aaa ata ttt gat aga ttt ctc ctt gaa cca att tgt gat cct gta gat gtt ctg tat gat tat ttt agg att cat aga gat aat att gat cag tat at? ??? ??? ??? ?tg ttt gca tat att aca tat aaa gat att ata tct gca tta gtg tca aag aat tat atg gaa gat att ttc tct ata att att aag aat tgt aat tct gtg caa gat ctc tta ctt tac tat cta tct aat gca tat gta gaa ata gac att gtt gat ctt atg gta gat cat ggg gct gta ata tat aaa ata gaa tgc ttg aat gcc tat ttt agg gga ata tgt aaa aag gaa agt agt gtt gtt gag ttt att ttg aat tgt ggt atc cca gat gaa aat gat gtt aaa tta gat cta tat aaa ata att cag tat act agg gga ttc ctt gta gat gaa ccc aca gta tta gaa att tat aag ctt tgt atc cca tat att gaa gat atc aat caa cta gat gct ggt gga agg acc ttg ctt tat cgc gct atc tat gca ggt tat ata gat tta gta tca tgg cta tta gaa aat gga gca aat gtc aac gca gta atg agt aat gga tat aca tgt ctt gac gtg gcc gtg gat agg gga tct gtc atc gcc cgt agg gaa gca cat ctt aaa ata tta gaa ata ttg ctt aga gaa cca ttg tct att gac tgt ata aaa tta gct ata ctt aat aat aca att gaa aac cat gat gtg ata aag ctc tgt atc aag tat ttt atg atg gta gat tat tca ctt tgt aat gtg tat gca tca tca ctc ttt gat tat ata att gat tgt aaa caa gaa ttg gag tac att agg cag atg aaa att cat aat aca acc atg tat gag tta atc tat aat aga gac aaa aac aag cat gct tcc cat att cta cat agg tat tct aaa cat cca gtt ttg aca cag tgt atc act aaa gga ttc aag att tac aca gaa gta acc gag cag gtc act aaa gct cta aac aga cgt gct cta ata gat gag ata ata aac aat gta tca act gat gac aat ctc cta tca aaa ctt cca tta gaa att agg gat cta att gtt tca caa gct gtc ata tag agt tct atc cac cca cct ttc ttg aaa tga gtt aat --- --- --- --- --- --- --- --- --- --- --- --- --- --- --- --- --- --- --- --- --- --- --- --- --- --- --- --- --- --- --- --- --- --- --- --- --- --- --- --- --- --- --- --- --- --- --- --- --- --- --- --- --- --- --- --- --- --- --- --- --- --- --- --- --- --- --- --- --- --- --- --- --- --- --- --- --- --- --- --- --- --- --- --- --- --- --- --- --- --- --- --- --- --- --- --- --- --- --- --- --- --- --- --- --- -ag tca taa gtt agt taa --- --- --- --- --- -gt cat aag tta gtt aag tca t-- --- --- --- --- --- --- --- --- --- --- --- --- --- --- --- -aa gtt agt taa gtc ata agt tag tta agt cat aag tta gtt aag tca taa gtt agt taa gtc ata agt tag tta agt cat aag tta gtt tat agt cta aca ctt cta att ttt ata cct tga tct ttt tct cta att atg aaa aag taa atc att atg aag atg gat g-a aat gga cga gat tgt gcg cat cgt taa cga tag tat gtg gta cgt acc taa cgc att tat gga cga cgg tga taa tga agg tca cat ttc tgt caa taa tgt ctg tca tat gta tct cgc att ctt tga tgt gga tat atc atc tca tct gtt taa att agt tat taa aca ctg cga tct gaa taa acg act aaa atg tgg taa ctc tcc att aca ttg cta tac gat gaa tac acg att taa tcc atc tgt att aaa gat att gtt acg cca cgg cat gcg taa ctt tga tag caa gga taa aaa agg aca tat tcc tct aca cca cta tct gat tca ttc act atc aat cga taa caa gat ctt tga tat act aac gga ccc cat tga tga ctt tag taa atc atc cga tct att gct gtg tta tct tag ata taa att caa tgg gag ctt aaa cta tta cgt tct gta caa att att gac taa agg atc tga ccc taa ttg cgt cga tga gga tgg act cac ttc tct tca tta cta ctg taa aca cat atc cgc gtt cca cga aag caa tta tta caa gtc aaa gag tca cac taa gat gcg agc tga gaa gcg att cat cta cgc gat aat aga tca tgg agc aaa cat taa cgc ggt tac gaa aat cgg aaa tac gcc gtt aca cac tta cct tca aca gta tac caa aca tag tcc tcg tgt ggt gta tgc tct ttt atc tcg agg agc cga tac gag gat acg taa taa tct taa ttg tac acc cat cat gga ata cat aaa gaa cga ttg tgc aac agg tca tat tct cat aat gtt act caa ttg gca cga aca aaa ata cgg gaa att aca aaa gga aga agg aca aca tct act tta tct att cat aaa aca taa tca agg ata tgg aag tcg ctc tct caa tat act acg gta tct act aga tag att cga cat tca gaa aga cga ata cta taa tac aat gac tcc tct tca tac cgc ctt cca gaa ttg caa taa caa tgt tgc ctc ata cct cgt ata cat tgg ata cga cat caa cct tcc gac taa aga cga taa gac agt att cga ctt ggt gtt tga aaa cag aaa cat tat ata caa ggc gga tgt cgt taa tga cat tat cca cca cag act gaa agt atc tct acc tat gat taa atc gtt gtt cta caa gat gtc gga gtt ctc tcc cta cga cga tca cta cgt aaa gaa gat aat agc cta ctg cct att aag gga cga gtc att tgc gga act aca tac taa att ctg ttt aaa cga gga cta taa aag tgt att tat gaa aaa tat atc att cga taa gat aga ttc cat cat cga aaa atg tag tcg tga cat aag tct cct caa aga gat tcg aat ctc aga cac cga ctt gta tac ggt att gag aac aga aga cat tcg gta tca cac ata tct cga agc cat aca ttc aga caa acg cat ttc att tcc cat gta cga cga tct cat aga aca gtg tca tct atc gat gga gca taa aag taa act cgt cga caa agc act caa taa att aga gtc tac cat cga tag tca atc tag act atc gta ttt gcc tcc gga aat tat gcg caa tat cat aac caa gct aag cga cta cca tct aaa cag tat gtt gta cgg aaa gaa cca tta caa ata tta tcc atg ata gaa aga aaa tat tta aaa aat aat cta tat gat tgg aga agt agg aaa caa aca gta aca aga cga cga tta cta c-a tta tta aat cat gag gtc cgt att ata ctc gta tat att gtt tct ctc atg tat aat aat aaa cgg aag aga tat agc acc aca tgc acc atc caa tgg aaa gtg taa aga caa cga ata cag aag ccg taa tct atg ttg tct atc gtg tcc tcc ggg aac tta cgc ttt cag att atg tga tag caa gac taa tac aca atg tac acc gtg tgg ttc gga tac ctt tac atc tca caa taa tca ttt aca ggc ttg tct aag ttg taa cgg aag atg tga tag taa tca ggt aga gac gcg atc gtg taa cac gac tca caa tag aat ctg tga atg ctc tcc agg ata tta ttg tct tct caa agg agc atc agg gtg tag aac atg tat ttc taa aac aaa gtg tgg aat agg ata cgg agt atc cgg ata cac gtc tac cgg aga cgt cat ctg ttc tcc gtg tgg tcc cgg aac ata ttc tca cac cgt ctc ttc cac aga taa atg cga acc cgt cgt aac cag caa tac att taa cta tat cga tgt gga aat taa cct gta tcc agt caa cga cac atc gtg tac tcg gac gac cac tac cgg tct cag cga atc cat ctc aac gtc gga act aac tat tac cat gaa tca taa aga ttg tga tcc agt ctt tcg tgc aga ata ctt ctc tgt cct taa taa tgt agc aac ttc agg att ctt tac agg aga aaa tag ata tca gaa tac ttc aaa gat atg tac tct gaa ttt cga gat taa atg taa caa caa aga ttc atc ttc caa aca gtt aac gaa aac aaa gaa tga tac tat cat gcc gca ttc aga gac ggt aac tct agt ggg cga ctg tct atc tag cgt cga cat cta cat act ata tag taa tac caa tac tca aga cta cga aaa tga tac aat ctc tta tca tat ggg taa tgt tct cga tgt caa tag cca tat gcc cgc tag ttg cga tat aca taa act gat cac taa ttc cca gaa tcc cac cca ctt ata gta ag- -tt ttt tta cct ata aat aat aaa tac aat aat taa ttt ctc gta aaa gta gaa aat ata ttc taa ttt att ata tgg taa gaa agt aga atc atc tag aac agt aat caa tca ata gca atc atg aaa caa tat att gtc ctg gca tgc atg tgc cta gtg gca gct gct atg cct act agt ctt caa caa tct tca tcc tcg tgt act gaa gaa gaa aac aaa cat cat atg gga atc gat gtt att atc aaa gtc aca aag caa gac caa aca ccg acc aat gat aag att tgt caa tcc gta acg gaa gtt aca gag a-- --- --- --- --- --- -cc gaa gat gat gag gta tcc gaa gaa gtt gta aaa gga gat ccc acc act tat tac act atc gtc ggt gcg ggt ctt aac atg aac ttt gga ttc acc aaa tgc cca aag att tta tcc atc tcc gaa tcc tct gat gga aac act gtg aat act aga ttg tcc agc gtg tca ccg gga caa ggt aag gac tct ccc gcg atc acg cgt gaa gaa gct ctg gct atg atc aaa gac tgt gag atg tct atc gac atc aga tgt agc gaa gaa gag aaa gac agt gac atc aag acc cat cca gta ctt ggg tct aac atc tca cat aag aaa gtg agt tac aaa gat atc atc ggt tca acg atc gtt gat aca aaa tgt gtc aag aac cta gag ttt agc gta cgt atc gga gac atg tgt gag gaa tca tct gaa ctt gaa gtc aag gat gga ttc aag tat gtc gac gga tcg gca tct gaa ggt gca acc gat gat act tca ctc atc gat tca aca aaa ctc aaa gca tgt gtc tga atc gat aac tct att cat ctg aaa atg gat gag ttg ggt taa tcg aac gat tca gac acc gca cca cga att aaa aaa g-- -ac cgg gca cta tat tcc ggt ttg caa aac aaa aat att taa cta cat tca caa aaa gtt acc tct cgt tac ttc ttc ttt ctg ttt caa tat gtg ata cga tat gat cac tat tcg tat tct ct- --- --- tgg tct cat aaa aaa gtt tta c-- --a aaa aaa aaa aaa a-t att ttt att ctc ttt ctc tct tcg at- ggt ctc aca aaa ata tta aac ctc ttt ctg atg tct caa cta ttt cgt aaa cga taa cgt cca aca ata tat tct cgt aga gct tat caa cat cct tat acc aat cta ggt tgt cag aca att gca tca taa aat aat gtt tat aat tta cac gtt aac atc ata taa taa acg tat ata gtt aat att ttt gga ata taa atg atc tgt aaa atc cat gta ggg gac act gct cac gtt ttt tct cta gta cat aat ttc aca caa gtt ttt ata cag aca aat taa ttc tcg tcc ata ta- ttt taa aac att gac ttt tgt act aag aaa aat atc ttg act aac cat ctc ttt ctc tct tcg atg ggt ctc aca aaa ata tta aac ctc ttt ctg atg g-- --- --- --- --- --- --- --- --- --- --- --- --- --- --- --- --- --- --- --- --- --- --- --- --- --- --- --- --- --- --- --- --- --- --- --- --- --- --- --- --- --- --- --- --- --- --- agt cgt aaa aag ttt tta tcc ttt ctc tct tcg ata ggt ctc --- --- --- --- --- --- --- --- --- --- --- --- --- --- --- --- --- --- --- --- --- --- --- --- --- --- --- --- --- --- --- --- --- --- --- --- --- --- --- --- --- --- --- --- --- --- --- --- --- --- --- --- --- --- --- --- --- --- --- --- --- --- --- --- --- --- --- --- --- --- aca aaa ata tta aac ctc ttt ctg atg gtc tct ata aa

>ON649719.1_Monkeypox_virus_isolate_Monkeypox/PT0012/2022_complete_genome

atc gag gac gac ttc aac tat gac agc tcg gtg gcg tct gcc agc gtg tac ata cga atg gca ttt cta aga aaa gtc tac ggt atc ctt -tc tac aat ttc ctt taa caa cgg caa cag ctg cag tat ttt tat act ttg aat gca -tc gga cat tta tac aag gga gtc ctg ttc taa tat tgg cat caa tgt tcg gat cta tag gct tga ttt tcg cat tga ctt tac aca gac atg aac atc ccc tga atc tgt aca tac ttt gtg gat tta cac tgt tag aat ctc taa cgc tgg cct ctg ttg tta ctt tct atg atg cac gta tcg tta tgc aag ctt tca tgt tga cta ctg cag tgt ttc ttg ctc tga cta cat ata ctc tac aat caa aga gag att tca gta aac ttg taa cag gat tgt ttg ctg ctt tct gga ttt taa ttt tgt cag gag tct tga gga taa agt tta aaa tag aat taa taa aga aca tat agg tca -tt ttt taa aca tgg ata gaa acc aag gtt gtt agt taa taa tat aca aga tat ttt ttc tca ctc tga tcc atg taa acc aag gac gag aga cac tct cat tcc tca ttc acg aca cca tta aaa atg gaa att aaa gcc ctc tat taa gca cag acg gct aca ggt cta cca tca ggt t-a cct tcg tct acc ttc aca atg gcc tct cct tgt gcc cag ttc agt ccc tgt cat tgc cac gct act aag gac tcc ctg aat acc gtg act gac gtc aga cat tgt ctg act gaa tac atc ctg tgg gtt tct cat aga tgg acc cat aga gaa agc gca ggg cct ctc tac agg ctt ctc atc tct ttc aga att gat gca atg gag cta ttt ggt agc gag ttg aag gag ttc tcg aat tca ctt ccg tgg gac aat atc gac aat tgc gtg gag atc att aaa tgt ttc atc aga aat gac tcc atg aaa acc gcc aaa gaa ctt tgt gca ata att gga ctt tgt act caa tca gct att gtc act gga aga gtc ttc aat gat aag tat atc gac ata cta ctt atg ctg cga aag att ctg aac gag aac gac tat ctc acc ctc ttg gat cat atc ctc act gct aaa tac taa atc tcc ttc atg ctc tct cac taa tac tct tac tca cta cac ttt tta tca tct tat gat gaa tga ttg cct tca tca ttt ttt cgt gga ata taa tat agg aat aat tag cac cag aat agc tat gga tat ctc gtt aag aat att ctc tat aag aga cat aat gta gac ata gtt att ata tcc ttc tta gat aag tgt tac gct act gga aag ttt cca tcg tta tta tta cat gaa gat gat ata att aaa cca aca ttg aga ttg gct ctt atg tta gct gga ttg aat tac tgt aat aaa tgc atc gag tat aga ggg ata tag caa ttc tcg ata ata gtc atg caa tat ttg aat gag act gat aat tta ggt aat aca gta ct- --- aca cac ata tct ttc tag att ata tat cgt taa aaa tct gta aga tgt ata ttt ctc ata agt atc cac tgt gta ata tta tta atg gat ata tag ata acg caa tag gga cta ata gta ttg taa aag ata taa tcg act att tac gta cat atc cag ata tct ata ttc cta cta gtt tgc tgc gta gtt gca tca ttg ata tgc atg -at tta tca gga ttc aga gat gaa tta cta agt aaa cta caa tcc cac aat aag taa gaa tca aat atc aaa aac tca ctt ttg att ttt cta gtc tta agt aat aca tat att tat taa tag acc tat gaa ata aaa aaa ggt aac aat gga ttc gcg tat agc tat tta cgt att agt atc ggc atc tct ttt gta tct tgt taa ttg tca caa act agt aca tta ctt caa tct gaa aat aaa tgg aag tga tat aac taa tac agc aga tat att gct gga caa tta tcc aat tat gac ctt tga tgg aaa gga tat tta tcc atc tat ctc gtt cat ggt cgg taa taa act ttt cct aga tct tta taa aaa tat ctt tga aga att ttt cag act att tcg agt atc tgt aag tag tca ata cga gga att aga ata tta tta ttc atg tga tta tac taa caa ccg tcc tac aat taa aca aca tta ctt tta taa cgg cga cga ata tac tga aat tga tag atc gaa aaa agc cac taa taa aaa cag ttg gtt aat tac ttc agg ctt tag act aca aaa atg gtt cga tag cga aga ttg tat aat tta tct cag atc ttt agt tag aag aat gga aga cag taa caa aaa cag taa aaa aac tta gta ctt aga tat cg- --a aaa aat ata ttt ttg tag act ctt gag aat aga agg aaa aca tgt aca taa tta taa aaa atg aaa atc aat ggc gaa taa gac agt gcg att cgc acc atg gag tcg gta gat ttc atg gct gtc gat gag cag ttt cac gac gac ctc gat ctt tgg tca tta tct ttg gta gat gat tat aaa aaa cat gga tta ggt gtt gac tgt tat gtt cta gaa cca gtt gtt gac agg aaa ata ttt gat aga ttt ctc ctt gaa cca att tgt gat cct gta gat gtt ctg tat gat tat ttt agg att cat aga gat aat att gat cag tat ata gt? ??? ??? ?tg ttt gca tat att aca tat aaa gat att ata tct gca tta gtg tca aag aat tat atg gaa gat att ttc tct ata att att aag aat tgt aat tct gtg caa gat ctc tta ctt tac tat cta tct aat gca tat gta gaa ata gac att gtt gat ctt atg gta gat cat ggg gct gta ata tat aaa ata gaa tgc ttg aat gcc tat ttt agg gga ata tgt aaa aag gaa agt agt gtt gtt gag ttt att ttg aat tgt ggt atc cca gat gaa aat gat gtt aaa tta gat cta tat aaa ata att cag tat act agg gga ttc ctt gta gat gaa ccc aca gta tta gaa att tat aag ctt tgt atc cca tat att gaa gat atc aat caa cta gat gct ggt gga agg acc ttg ctt tat cgc gct atc tat gca ggt tat ata gat tta gta tca tgg cta tta gaa aat gga gca aat gtc aac gca gta atg agt aat gga tat aca tgt ctt gac gtg gcc gtg gat agg gga tct gtc atc gcc cgt agg gaa gca cat ctt aaa ata tta gaa ata ttg ctt aga gaa cca ttg tct att gac tgt ata aaa tta gct ata ctt aat aat aca att gaa aac cat gat gtg ata aag ctc tgt atc aag tat ttt atg atg gta gat tat tca ctt tgt aat gtg tat gca tca tca ctc ttt gat tat ata att gat tgt aaa caa gaa ttg gag tac att agg cag atg aaa att cat aat aca acc atg tat gag tta atc tat aat aga gac aaa aac aag cat gct tcc cat att cta cat agg tat tct aaa cat cca gtt ttg aca cag tgt atc act aaa gga ttc aag att tac aca gaa gta acc gag cag gtc act aaa gct cta aac aga cgt gct cta ata gat gag ata ata aac aat gta tca act gat gac aat ctc cta tca aaa ctt cca tta gaa att agg gat cta att gtt tca caa gct gtc ata tag agt tct atc cac cca cct ttc ttg aaa tga gtt aat --- --- --- --- --- --- --- --- --- --- --- --- --- --- --- --- --- --- --- --- --- --- --- --- --- --- --- --- --- --- --- --- --- --- --- --- --- --- --- --- --- --- --- --- --- --- --- --- --- --- --- --- --- --- --- --- --- --- --- --- --- --- --- --- --- --- --- --- --- --- --- --- --- --- --- --- --- --- --- --- --- --- --- --- --- --- --- --- --- --- --- --- --- --- --- --- --- --- --- --- --- --- --- --- --- -ag tca taa gtt agt taa --- --- --- --- --- -gt cat aag tta gtt aag tca t-- --- --- --- --- --- --- --- --- --- --- --- --- --- --- --- -aa gtt agt taa gtc ata agt tag tta agt cat aag tta gtt aag tca taa gtt agt taa gtc ata agt tag tta agt cat aag tta gtt tat agt cta aca ctt cta att ttt ata cct tga tct ttt tct cta att atg aaa aag taa atc att atg aag atg gat g-a aat gga cga gat tgt gcg cat cgt taa cga tag tat gtg gta cgt acc taa cgc att tat gga cga cgg tga taa tga agg tca cat ttc tgt caa taa tgt ctg tca tat gta tct cgc att ctt tga tgt gga tat atc atc tca tct gtt taa att agt tat taa aca ctg cga tct gaa taa acg act aaa atg tgg taa ctc tcc att aca ttg cta tac gat gaa tac acg att taa tcc atc tgt att aaa gat att gtt acg cca cgg cat gcg taa ctt tga tag caa gga taa aaa agg aca tat tcc tct aca cca cta tct gat tca ttc act atc aat cga taa caa gat ctt tga tat act aac gga ccc cat tga tga ctt tag taa atc atc cga tct att gct gtg tta tct tag ata taa att caa tgg gag ctt aaa cta tta cgt tct gta caa att att gac taa agg atc tga ccc taa ttg cgt cga tga gga tgg act cac ttc tct tca tta cta ctg taa aca cat atc cgc gtt cca cga aag caa tta tta caa gtc aaa gag tca cac taa gat gcg agc tga gaa gcg att cat cta cgc gat aat aga tca tgg agc aaa cat taa cgc ggt tac gaa aat cgg aaa tac gcc gtt aca cac tta cct tca aca gta tac caa aca tag tcc tcg tgt ggt gta tgc tct ttt atc tcg agg agc cga tac gag gat acg taa taa tct taa ttg tac acc cat cat gga ata cat aaa gaa cga ttg tgc aac agg tca tat tct cat aat gtt act caa ttg gca cga aca aaa ata cgg gaa att aca aaa gga aga agg aca aca tct act tta tct att cat aaa aca taa tca agg ata tgg aag tcg ctc tct caa tat act acg gta tct act aga tag att cga cat tca gaa aga cga ata cta taa tac aat gac tcc tct tca tac cgc ctt cca gaa ttg caa taa caa tgt tgc ctc ata cct cgt ata cat tgg ata cga cat caa cct tcc gac taa aga cga taa gac agt att cga ctt ggt gtt tga aaa cag aaa cat tat ata caa ggc gga tgt cgt taa tga cat tat cca cca cag act gaa agt atc tct acc tat gat taa atc gtt gtt cta caa gat gtc gga gtt ctc tcc cta cga cga tca cta cgt aaa gaa gat aat agc cta ctg cct att aag gga cga gtc att tgc gga act aca tac taa att ctg ttt aaa cga gga cta taa aag tgt att tat gaa aaa tat atc att cga taa gat aga ttc cat cat cga aaa atg tag tcg tga cat aag tct cct caa aga gat tcg aat ctc aga cac cga ctt gta tac ggt att gag aac aga aga cat tcg gta tca cac ata tct cga agc cat aca ttc aga caa acg cat ttc att tcc cat gta cga cga tct cat aga aca gtg tca tct atc gat gga gca taa aag taa act cgt cga caa agc act caa taa att aga gtc tac cat cga tag tca atc tag act atc gta ttt gcc tcc gga aat tat gcg caa tat cat aac caa gct aag cga cta cca tct aaa cag tat gtt gta cgg aaa gaa cca tta caa ata tta tcc atg ata gaa aga aaa tat tta aaa aat aat cta tat gat tgg aga agt agg aaa caa aca gta aca aga cga cga tta cta c-a tta tta aat cat gag gtc cgt att ata ctc gta tat att gtt tct ctc atg tat aat aat aaa cgg aag aga tat agc acc aca tgc acc atc caa tgg aaa gtg taa aga caa cga ata cag aag ccg taa tct atg ttg tct atc gtg tcc tcc ggg aac tta cgc ttt cag att atg tga tag caa gac taa tac aca atg tac acc gtg tgg ttc gga tac ctt tac atc tca caa taa tca ttt aca ggc ttg tct aag ttg taa cgg aag atg tga tag taa tca ggt aga gac gcg atc gtg taa cac gac tca caa tag aat ctg tga atg ctc tcc agg ata tta ttg tct tct caa agg agc atc agg gtg tag aac atg tat ttc taa aac aaa gtg tgg aat agg ata cgg agt atc cgg ata cac gtc tac cgg aga cgt cat ctg ttc tcc gtg tgg tcc cgg aac ata ttc tca cac cgt ctc ttc cac aga taa atg cga acc cgt cgt aac cag caa tac att taa cta tat cga tgt gga aat taa cct gta tcc agt caa cga cac atc gtg tac tcg gac gac cac tac cgg tct cag cga atc cat ctc aac gtc gga act aac tat tac cat gaa tca taa aga ttg tga tcc agt ctt tcg tgc aga ata ctt ctc tgt cct taa taa tgt agc aac ttc agg att ctt tac agg aga aaa tag ata tca gaa tac ttc aaa gat atg tac tct gaa ttt cga gat taa atg taa caa caa aga ttc atc ttc caa aca gtt aac gaa aac aaa gaa tga tac tat cat gcc gca ttc aga gac ggt aac tct agt ggg cga ctg tct atc tag cgt cga cat cta cat act ata tag taa tac caa tac tca aga cta cga aaa tga tac aat ctc tta tca tat ggg taa tgt tct cga tgt caa tag cca tat gcc cgc tag ttg cga tat aca taa act gat cac taa ttc cca gaa tcc cac cca ctt ata gta ag- -tt ttt tta cct ata aat aat aaa tac aat aat taa ttt ctc gta aaa gta gaa aat ata ttc taa ttt att ata tgg taa gaa agt aga atc atc tag aac agt aat caa tca ata gca atc atg aaa caa tat att gtc ctg gca tgc atg tgc cta gtg gca gct gct atg cct act agt ctt caa caa tct tca tcc tcg tgt act gaa gaa gaa aac aaa cat cat atg gga atc gat gtt att atc aaa gtc aca aag caa gac caa aca ccg acc aat gat aag att tgt caa tcc gta acg gaa gtt aca gag a-- --- --- --- --- --- -cc gaa gat gat gag gta tcc gaa gaa gtt gta aaa gga gat ccc acc act tat tac act atc gtc ggt gcg ggt ctt aac atg aac ttt gga ttc acc aaa tgc cca aag att tta tcc atc tcc gaa tcc tct gat gga aac act gtg aat act aga ttg tcc agc gtg tca ccg gga caa ggt aag gac tct ccc gcg atc acg cgt gaa gaa gct ctg gct atg atc aaa gac tgt gag atg tct atc gac atc aga tgt agc gaa gaa gag aaa gac agt gac atc aag acc cat cca gta ctt ggg tct aac atc tca cat aag aaa gtg agt tac aaa gat atc atc ggt tca acg atc gtt gat aca aaa tgt gtc aag aac cta gag ttt agc gta cgt atc gga gac atg tgt gag gaa tca tct gaa ctt gaa gtc aag gat gga ttc aag tat gtc gac gga tcg gca tct gaa ggt gca acc gat gat act tca ctc atc gat tca aca aaa ctc aaa gca tgt gtc tga atc gat aac tct att cat ctg aaa atg gat gag ttg ggt taa tcg aac gat tca gac acc gca cca cga att aaa aaa g-- -ac cgg gca cta tat tcc ggt ttg caa aac aaa aat att taa cta cat tca caa aaa gtt acc tct cgt tac ttc ttc ttt ctg ttt caa tat gtg ata cga tat gat cac tat tcg tat tct ct- --- --- tgg tct cat aaa aaa gtt tta c-- --a aaa aaa aaa aaa a-t att ttt att ctc ttt ctc tct tcg at- ggt ctc aca aaa ata tta aac ctc ttt ctg atg tct caa cta ttt cgt aaa cga taa cgt cca aca ata tat tct cgt aga gct tat caa cat cct tat acc aat cta ggt tgt cag aca att gca tca taa aat aat gtt tat aat tta cac gtt aac atc ata taa taa acg tat ata gtt aat att ttt gga ata taa atg atc tgt aaa atc cat gta ggg gac act gct cac gtt ttt tct cta gta cat aat ttc aca caa gtt ttt ata cag aca aat taa ttc tcg tcc ata ta- ttt taa aac att gac ttt tgt act aag aaa aat atc ttg act aac cat ctc ttt ctc tct tcg atg ggt ctc aca aaa ata tta aac ctc ttt ctg atg g-- --- --- --- --- --- --- --- --- --- --- --- --- --- --- --- --- --- --- --- --- --- --- --- --- --- --- --- --- --- --- --- --- --- --- --- --- --- --- --- --- --- --- --- --- --- --- agt cgt aaa aag ttt tta tcc ttt ctc tct tcg ata ggt ctc --- --- --- --- --- --- --- --- --- --- --- --- --- --- --- --- --- --- --- --- --- --- --- --- --- --- --- --- --- --- --- --- --- --- --- --- --- --- --- --- --- --- --- --- --- --- --- --- --- --- --- --- --- --- --- --- --- --- --- --- --- --- --- --- --- --- --- --- --- --- aca aaa ata tta aac ctc ttt ctg atg gtc tct ata aa

>ON682267.1_Monkeypox_virus_isolate_MPXV/Germany/2022/RKI010_complete_genome

atc gag gac gac ttc aac tat gac agc tcg gtg gcg tct gcc agc gtg tac ata cga atg gca ttt cta aga aaa gtc tac ggt atc ctt -tc tac aat ttc ctt taa caa cgg caa cag ctg cag tat ttt tat act ttg aat gca -tc gga cat tta tac aag gga gtc ctg ttc taa tat tgg cat caa tgt tcg gat cta tag gct tga ttt tcg cat tga ctt tac aca gac atg aac atc ccc tga atc tgt aca tac ttt gtg gat tta cac tgt tag aat ctc taa cgc tgg cct ctg ttg tta ctt tct atg atg cac gta tcg tta tgc aag ctt tca tgt tga cta ctg cag tgt ttc ttg ctc tga cta cat ata ctc tac aat caa aga gag att tca gta aac ttg taa cag gat tgt ttg ctg ctt tct gga ttt taa ttt tgt cag gag tct tga gga taa agt tta aaa tag aat taa taa aga aca tat agg tca -tt ttt taa aca tgg ata gaa acc aag gtt gtt agt taa taa tat aca aga tat ttt ttc tca ctc tga tcc atg taa acc aag gac gag aga cac tct cat tcc tca ttc acg aca cca tta aaa atg gaa att aaa gcc ctc tat taa gca cag acg gct aca ggt cta cca tca ggt t-a cct tcg tct acc ttc aca atg gcc tct cct tgt gcc cag ttc agt ccc tgt cat tgc cac gct act aag gac tcc ctg aat acc gtg act gac gtc aga cat tgt ctg act gaa tac atc ctg tgg gtt tct cat aga tgg acc cat aga gaa agc gca ggg cct ctc tac agg ctt ctc atc tct ttc aga att gat gca atg gag cta ttt ggt agc gag ttg aag gag ttc tcg aat tca ctt ccg tgg gac aat atc gac aat tgc gtg gag atc att aaa tgt ttc atc aga aat gac tcc atg aaa acc gcc aaa gaa ctt tgt gca ata att gga ctt tgt act caa tca gct att gtc act gga aga gtc ttc aat gat aag tat atc gac ata cta ctt atg ctg cga aag att ctg aac gag aac gac tat ctc acc ctc ttg gat cat atc ctc act gct aaa tac taa atc tcc ttc atg ctc tct cac taa tac tct tac tca cta cac ttt tta tca tct tat gat gaa tga ttg cct tca tca ttt ttt cgt gga ata taa tat agg aat aat tag cac cag aat agc tat gga tat ctc gtt aag aat att ctc tat aag aga cat aat gta gac ata gtt att ata tcc ttc tta gat aag tgt tac gct act gga aag ttt cca tcg tta tta tta cat gaa gat gat ata att aaa cca aca ttg aga ttg gct ctt atg tta gct gga ttg aat tac tgt aat aaa tgc atc gag tat aga ggg ata tag caa ttc tcg ata ata gtc atg caa tat ttg aat gag act gat aat tta ggt aat aca gta ct- --- aca cac ata tct ttc tag att ata tat cgt taa aaa tct gta aga tgt ata ttt ctc ata agt atc cac tgt gta ata tta tta atg gat ata tag ata acg caa tag gga cta ata gta ttg taa aag ata taa tcg act att tac gta cat atc cag ata tct ata ttc cta cta gtt tgc tgc gta gtt gca tca ttg ata tgc atg -at tta tca gga ttc aga gat gaa tta cta agt aaa cta caa tcc cac aat aag taa gaa tca aat atc aaa aac tca ctt ttg att ttt cta gtc tta agt aat aca tat att tat taa tag acc tat gaa ata aaa aaa ggt aac aat gga ttc gcg tat agc tat tta cgt att agt atc ggc atc tct ttt gta tct tgt taa ttg tca caa act agt aca tta ctt caa tct gaa aat aaa tgg aag tga tat aac taa tac agc aga tat att gct gga caa tta tcc aat tat gac ctt tga tgg aaa gga tat tta tcc atc tat ctc gtt cat ggt cgg taa taa act ttt cct aga tct tta taa aaa tat ctt tga aga att ttt cag act att tcg agt atc tgt aag tag tca ata cga gga att aga ata tta tta ttc atg tga tta tac taa caa ccg tcc tac aat taa aca aca tta ctt tta taa cgg cga cga ata tac tga aat tga tag atc gaa aaa agc cac taa taa aaa cag ttg gtt aat tac ttc agg ctt tag act aca aaa atg gtt cga tag cga aga ttg tat aat tta tct cag atc ttt agt tag aag aat gga aga cag taa caa aaa cag taa aaa aac tta gta ctt aga tat cg- --a aaa aat ata ttt ttg tag act ctt gag aat aga agg aaa aca tgt aca taa tta taa aaa atg aaa atc aat ggc gaa taa gac agt gcg att cgc acc atg gag tcg gta gat ttc atg gct gtc gat gag cag ttt cac gac gac ctc gat ctt tgg tca tta tct ttg gta gat gat tat aaa aaa cat gga tta ggt gtt gac tgt tat gtt cta gaa cca gtt gtt gac agg aaa ata ttt gat aga ttt ctc ctt gaa cca att tgt gat cct gta gat gtt ctg tat gat tat ttt agg att cat aga gat aat att gat cag tat ata gta gat aga ctg ttt gca tat att aca tat aaa gat att ata tct gca tta gtg tca aag aat tat atg gaa gat att ttc tct ata att att aag aat tgt aat tct gtg caa gat ctc tta ctt tac tat cta tct aat gca tat gta gaa ata gac att gtt gat ctt atg gta gat cat ggg gct gta ata tat aaa ata gaa tgc ttg aat gcc tat ttt agg gga ata tgt aaa aag gaa agt agt gtt gtt gag ttt att ttg aat tgt ggt atc cca gat gaa aat gat gtt aaa tta gat cta tat aaa ata att cag tat act agg gga ttc ctt gta gat gaa ccc aca gta tta gaa att tat aag ctt tgt atc cca tat att gaa gat atc aat caa cta gat gct ggt gga agg acc ttg ctt tat cgc gct atc tat gca ggt tat ata gat tta gta tca tgg cta tta gaa aat gga gca aat gtc aac gca gta atg agt aat gga tat aca tgt ctt gac gtg gcc gtg gat agg gga tct gtc atc gcc cgt agg gaa gca cat ctt aaa ata tta gaa ata ttg ctt aga gaa cca ttg tct att gac tgt ata aaa tta gct ata ctt aat aat aca att gaa aac cat gat gtg ata aag ctc tgt atc aag tat ttt atg atg gta gat tat tca ctt tgt aat gtg tat gca tca tca ctc ttt gat tat ata att gat tgt aaa caa gaa ttg gag tac att agg cag atg aaa att cat aat aca acc atg tat gag tta atc tat aat aga gac aaa aac aag cat gct tcc cat att cta cat agg tat tct aaa cat cca gtt ttg aca cag tgt atc act aaa gga ttc aag att tac aca gaa gta acc gag cag gtc act aaa gct cta aac aga cgt gct cta ata gat gag ata ata aac aat gta tca act gat gac aat ctc cta tca aaa ctt cca tta gaa att agg gat cta att gtt tca caa gct gtc ata tag agt tct atc cac cca cct ttc ttg aaa tga gtt aat a-- --- --- --- --- --- --- --- --- --- --- --- --- --- --- --- --- --- --- --- --- --- --- --- --- --- --- --- --- --- --- --- --- --- --- --- --- --- --- --- --- --- --- --- --- --- --- --- --- --- --- --- --- --- --- --- --- --- --- --- --- --- --- --- --- --- --- --- --- --- --- --- --- --- --- --- --- --- --- --- --- --- --- --- --- --- --- --- --- --- --- --- --- --- --- --- --- --- --- --- --- --- --- --- --- --- --- --- --- --- --- --- --- --- --- --- --- --- --- --- --- --g tca t-- --- --- --- --- --- --- --- --- --- --- --- --- --- --- --- -aa gtt agt taa gtc ata agt tag tta agt cat aag tta gtt aag tca taa gtt agt taa gtc ata agt tag tta agt cat aag tta gtt tat agt cta aca ctt cta att ttt ata cct tga tct ttt tct cta att atg aaa aag taa atc att atg aag atg gat g-a aat gga cga gat tgt gcg cat cgt taa cga tag tat gtg gta cgt acc taa cgc att tat gga cga cgg tga taa tga agg tca cat ttc tgt caa taa tgt ctg tca tat gta tct cgc att ctt tga tgt gga tat atc atc tca tct gtt taa att agt tat taa aca ctg cga tct gaa taa acg act aaa atg tgg taa ctc tcc att aca ttg cta tac gat gaa tac acg att taa tcc atc tgt att aaa gat att gtt acg cca cgg cat gcg taa ctt tga tag caa gga taa aaa agg aca tat tcc tct aca cca cta tct gat tca ttc act atc aat cga taa caa gat ctt tga tat act aac gga ccc cat tga tga ctt tag taa atc atc cga tct att gct gtg tta tct tag ata taa att caa tgg gag ctt aaa cta tta cgt tct gta caa att att gac taa agg atc tga ccc taa ttg cgt cga tga gga tgg act cac ttc tct tca tta cta ctg taa aca cat atc cgc gtt cca cga aag caa tta tta caa gtc aaa gag tca cac taa gat gcg agc tga gaa gcg att cat cta cgc gat aat aga tca tgg agc aaa cat taa cgc ggt tac gaa aat cgg aaa tac gcc gtt aca cac tta cct tca aca gta tac caa aca tag tcc tcg tgt ggt gta tgc tct ttt atc tcg agg agc cga tac gag gat acg taa taa tct taa ttg tac acc cat cat gga ata cat aaa gaa cga ttg tgc aac agg tca tat tct cat aat gtt act caa ttg gca cga aca aaa ata cgg gaa att aca aaa gga aga agg aca aca tct act tta tct att cat aaa aca taa tca agg ata tgg aag tcg ctc tct caa tat act acg gta tct act aga tag att cga cat tca gaa aga cga ata cta taa tac aat gac tcc tct tca tac cgc ctt cca gaa ttg caa taa caa tgt tgc ctc ata cct cgt ata cat tgg ata cga cat caa cct tcc gac taa aga cga taa gac agt att cga ctt ggt gtt tga aaa cag aaa cat tat ata caa ggc gga tgt cgt taa tga cat tat cca cca cag act gaa agt atc tct acc tat gat taa atc gtt gtt cta caa gat gtc gga gtt ctc tcc cta cga cga tca cta cgt aaa gaa gat aat agc cta ctg cct att aag gga cga gtc att tgc gga act aca tac taa att ctg ttt aaa cga gga cta taa aag tgt att tat gaa aaa tat atc att cga taa gat aga ttc cat cat cga aaa atg tag tcg tga cat aag tct cct caa aga gat tcg aat ctc aga cac cga ctt gta tac ggt att gag aac aga aga cat tcg gta tca cac ata tct cga agc cat aca ttc aga caa acg cat ttc att tcc cat gta cga cga tct cat aga aca gtg tca tct atc gat gga gca taa aag taa act cgt cga caa agc act caa taa att aga gtc tac cat cga tag tca atc tag act atc gta ttt gcc tcc gga aat tat gcg caa tat cat aac caa gct aag cga cta cca tct aaa cag tat gtt gta cgg aaa gaa cca tta caa ata tta tcc atg ata gaa aga aaa tat tta aaa aat aat cta tat gat tgg aga agt agg aaa caa aca gta aca aga cga cga tta cta c-a tta tta aat cat gag gtc cgt att ata ctc gta tat att gtt tct ctc atg tat aat aat aaa cgg aag aga tat agc acc aca tgc acc atc caa tgg aaa gtg taa aga caa cga ata cag aag ccg taa tct atg ttg tct atc gtg tcc tcc ggg aac tta cgc ttt cag att atg tga tag caa gac taa tac aca atg tac acc gtg tgg ttc gga tac ctt tac atc tca caa taa tca ttt aca ggc ttg tct aag ttg taa cgg aag atg tga tag taa tca ggt aga gac gcg atc gtg taa cac gac tca caa tag aat ctg tga atg ctc tcc agg ata tta ttg tct tct caa agg agc atc agg gtg tag aac atg tat ttc taa aac aaa gtg tgg aat agg ata cgg agt atc cgg ata cac gtc tac cgg aga cgt cat ctg ttc tcc gtg tgg tcc cgg aac ata ttc tca cac cgt ctc ttc cac aga taa atg cga acc cgt cgt aac cag caa tac att taa cta tat cga tgt gga aat taa cct gta tcc agt caa cga cac atc gtg tac tcg gac gac cac tac cgg tct cag cga atc cat ctc aac gtc gga act aac tat tac cat gaa tca taa aga ttg tga tcc agt ctt tcg tgc aga ata ctt ctc tgt cct taa taa tgt agc aac ttc agg att ctt tac agg aga aaa tag ata tca gaa tac ttc aaa gat atg tac tct gaa ttt cga gat taa atg taa caa caa aga ttc atc ttc caa aca gtt aac gaa aac aaa gaa tga tac tat cat gcc gca ttc aga gac ggt aac tct agt ggg cga ctg tct atc tag cgt cga cat cta cat act ata tag taa tac caa tac tca aga cta cga aaa tga tac aat ctc tta tca tat ggg taa tgt tct cga tgt caa tag cca tat gcc cgc tag ttg cga tat aca taa act gat cac taa ttc cca gaa tcc cac cca ctt ata gta ag- -tt ttt tta cct ata aat aat aaa tac aat aat taa ttt ctc gta aaa gta gaa aat ata ttc taa ttt att ata tgg taa gaa agt aga atc atc tag aac agt aat caa tca ata gca atc atg aaa caa tat att gtc ctg gca tgc atg tgc cta gtg gca gct gct atg cct act agt ctt caa caa tct tca tcc tcg tgt act gaa gaa gaa aac aaa cat cat atg gga atc gat gtt att atc aaa gtc aca aag caa gac caa aca ccg acc aat gat aag att tgt caa tcc gta acg gaa gtt aca gag a-- --- --- --- --- --- -cc gaa gat gat gag gta tcc gaa gaa gtt gta aaa gga gat ccc acc act tat tac act atc gtc ggt gcg ggt ctt aac atg aac ttt gga ttc acc aaa tgc cca aag att tta tcc atc tcc gaa tcc tct gat gga aac act gtg aat act aga ttg tcc agc gtg tca ccg gga caa ggt aag gac tct ccc gcg atc acg cgt gaa gaa gct ctg gct atg atc aaa gac tgt gag atg tct atc gac atc aga tgt agc gaa gaa gag aaa gac agt gac atc aag acc cat cca gta ctt ggg tct aac atc tca cat aag aaa gtg agt tac aaa gat atc atc ggt tca acg atc gtt gat aca aaa tgt gtc aag aac cta gag ttt agc gta cgt atc gga gac atg tgt gag gaa tca tct gaa ctt gaa gtc aag gat gga ttc aag tat gtc gac gga tcg gca tct gaa ggt gca acc gat gat act tca ctc atc gat tca aca aaa ctc aaa gca tgt gtc tga atc gat aac tct att cat ctg aaa atg gat gag ttg ggt taa tcg aac gat tca gac acc gca cca cga att aaa aaa g-- -ac cgg gca cta tat tcc ggt ttg caa aac aaa aat att taa cta cat tca caa aaa gtt acc tct cgt tac ttc ttc ttt ctg ttt caa tat gtg ata cga tat gat cac tat tcg tat tct ct- --- --- tgg tct cat aaa aaa gtt tta c-- --a aaa aaa aaa aaa a-t att ttt att ctc ttt ctc tct tcg at- ggt ctc aca aaa ata tta aac ctc ttt ctg atg tct caa cta ttt cgt aaa cga taa cgt cca aca ata tat tct cgt aga gct tat caa cat cct tat acc aat cta ggt tgt cag aca att gca tca taa aat aat gtt tat aat tta cac gtt aac atc ata taa taa acg tat ata gtt aat att ttt gga ata taa atg atc tgt aaa atc cat gta ggg gac act gct cac gtt ttt tct cta gta cat aat ttc aca caa gtt ttt ata cag aca aat taa ttc tcg tcc ata ta- ttt taa aac att gac ttt tgt act aag aaa aat atc ttg act aac cat ctc ttt ctc tct tcg atg ggt ctc aca aaa ata tta aac ctc ttt ctg atg g-- --- --- --- --- --- --- --- --- --- --- --- --- --- --- --- --- --- --- --- --- --- --- --- --- --- --- --- --- --- --- --- --- --- --- --- --- --- --- --- --- --- --- --- --- --- --- agt cgt aaa aag ttt tta tcc ttt ctc tct tcg ata ggt ctc --- --- --- --- --- --- --- --- --- --- --- --- --- --- --- --- --- --- --- --- --- --- --- --- --- --- --- --- --- --- --- --- --- --- --- --- --- --- --- --- --- --- --- --- --- --- --- --- --- --- --- --- --- --- --- --- --- --- --- --- --- --- --- --- --- --- --- --- --- --- aca aaa ata tta aac ctc ttt ctg atg gtc tct ata aa

>ON649709.1_Monkeypox_virus_isolate_Monkeypox/PT0028/2022_complete_genome

atc gag gac gac ttc aac tat gac agc tcg gtg gcg tct gcc agc gtg tac ata cga atg gca ttt cta aga aaa gtc tac ggt atc ctt -tc tac aat ttc ctt taa caa cgg caa cag ctg cag tat ttt tat act ttg aat gca -tc gga cat tta tac aag gga gtc ctg ttc taa tat tgg cat caa tgt tcg gat cta tag gct tga ttt tcg cat tga ctt tac aca gac atg aac atc ccc tga atc tgt aca tac ttt gtg gat tta cac tgt tag aat ctc taa cgc tgg cct ctg ttg tta ctt tct atg atg cac gta tcg tta tgc aag ctt tca tgt tga cta ctg cag tgt ttc ttg ctc tga cta cat ata ctc tac aat caa aga gag att tca gta aac ttg taa cag gat tgt ttg ctg ctt tct gga ttt taa ttt tgt cag gag tct tga gga taa agt tta aaa tag aat taa taa aga aca tat agg tca -tt ttt taa aca tgg ata gaa acc aag gtt gtt agt taa taa tat aca aga tat ttt ttc tca ctc tga tcc atg taa acc aag gac gag aga cac tct cat tcc tca ttc acg aca cca tta aaa atg gaa att aaa gcc ctc tat taa gca cag acg gct aca ggt cta cca tca ggt t-a cct tcg tct acc ttc aca atg gcc tct cct tgt gcc cag ttc agt ccc tgt cat tgc cac gct act aag gac tcc ctg aat acc gtg act gac gtc aga cat tgt ctg act gaa tac atc ctg tgg gtt tct cat aga tgg acc cat aga gaa agc gca ggg cct ctc tac agg ctt ctc atc tct ttc aga att gat gca atg gag cta ttt ggt agc gag ttg aag gag ttc tcg aat tca ctt ccg tgg gac aat atc gac aat tgc gtg gag atc att aaa tgt ttc atc aga aat gac tcc atg aaa acc gcc aaa gaa ctt tgt gca ata att gga ctt tgt act caa tca gct att gtc act gga aga gtc ttc aat gat aag tat atc gac ata cta ctt atg ctg cga aag att ctg aac gag aac gac tat ctc acc ctc ttg gat cat atc ctc act gct aaa tac taa atc tcc ttc atg ctc tct cac taa tac tct tac tca cta cac ttt tta tca tct tat gat gaa tga ttg cct tca tca ttt ttt cgt gga ata taa tat agg aat aat tag cac cag aat agc tat gga tat ctc gtt aag aat att ctc tat aag aga cat aat gta gac ata gtt att ata tcc ttc tta gat aag tgt tac gct act gga aag ttt cca tcg tta tta tta cat gaa gat gat ata att aaa cca aca ttg aga ttg gct ctt atg tta gct gga ttg aat tac tgt aat aaa tgc atc gag tat aga ggg ata tag caa ttc tcg ata ata gtc atg caa tat ttg aat gag act gat aat tta ggt aat aca gta ct- --- aca cac ata tct ttc tag att ata tat cgt taa aaa tct gta aga tgt ata ttt ctc ata agt atc cac tgt gta ata tta tta atg gat ata tag ata acg caa tag gga cta ata gta ttg taa aag ata taa tcg act att tac gta cat atc cag ata tct ata ttc cta cta gtt tgc tgc gta gtt gca tca ttg ata tgc atg -at tta tca gga ttc aga gat gaa tta cta agt aaa cta caa tcc cac aat aag taa gaa tca aat atc aaa aac tca ctt ttg att ttt cta gtc tta agt aat aca tat att tat taa tag acc tat gaa ata aaa aaa ggt aac aat gga ttc gcg tat agc tat tta cgt att agt atc ggc atc tct ttt gta tct tgt taa ttg tca caa act agt aca tta ctt caa tct gaa aat aaa tgg aag tga tat aac taa tac agc aga tat att gct gga caa tta tcc aat tat gac ctt tga tgg aaa gga tat tta tcc atc tat ctc gtt cat ggt cgg taa taa act ttt cct aga tct tta taa aaa tat ctt tga aga att ttt cag act att tcg agt atc tgt aag tag tca ata cga gga att aga ata tta tta ttc atg tga tta tac taa caa ccg tcc tac aat taa aca aca tta ctt tta taa cgg cga cga ata tac tga aat tga tag atc gaa aaa agc cac taa taa aaa cag ttg gtt aat tac ttc agg ctt tag act aca aaa atg gtt cga tag cga aga ttg tat aat tta tct cag atc ttt agt tag aag aat gga aga cag taa caa aaa cag taa aaa aac tta gta ctt aga tat cg- --a aaa aat ata ttt ttg tag act ctt gag aat aga agg aaa aca tgt aca taa tta taa aaa atg aaa atc aat ggc gaa taa gac agt gcg att cgc acc atg gag tcg gta gat ttc atg gct gtc gat gag cag ttt cac gac gac ctc gat ctt tgg tca tta tct ttg gta gat gat tat aaa aaa cat gga tta ggt gtt gac tgt tat gtt cta gaa cca gtt gtt gac agg aaa ata ttt gat aga ttt ctc ctt gaa cca att tgt gat cct gta gat gtt ctg tat gat tat ttt agg att cat aga gat aat at? ??? ??? ??? ??? ??? ??? ??? ?tg ttt gca tat att aca tat aaa gat att ata tct gca tta gtg tca aag aat tat atg gaa gat att ttc tct ata att att aag aat tgt aat tct gtg caa gat ctc tta ctt tac tat cta tct aat gca tat gta gaa ata gac att gtt gat ctt atg gta gat cat ggg gct gta ata tat aaa ata gaa tgc ttg aat gcc tat ttt agg gga ata tgt aaa aag gaa agt agt gtt gtt gag ttt att ttg aat tgt ggt atc cca gat gaa aat gat gtt aaa tta gat cta tat aaa ata att cag tat act agg gga ttc ctt gta gat gaa ccc aca gta tta gaa att tat aag ctt tgt atc cca tat att gaa gat atc aat caa cta gat gct ggt gga agg acc ttg ctt tat cgc gct atc tat gca ggt tat ata gat tta gta tca tgg cta tta gaa aat gga gca aat gtc aac gca gta atg agt aat gga tat aca tgt ctt gac gtg gcc gtg gat agg gga tct gtc atc gcc cgt agg gaa gca cat ctt aaa ata tta gaa ata ttg ctt aga gaa cca ttg tct att gac tgt ata aaa tta gct ata ctt aat aat aca att gaa aac cat gat gtg ata aag ctc tgt atc aag tat ttt atg atg gta gat tat tca ctt tgt aat gtg tat gca tca tca ctc ttt gat tat ata att gat tgt aaa caa gaa ttg gag tac att agg cag atg aaa att cat aat aca acc atg tat gag tta atc tat aat aga gac aaa aac aag cat gct tcc cat att cta cat agg tat tct aaa cat cca gtt ttg aca cag tgt atc act aaa gga ttc aag att tac aca gaa gta acc gag cag gtc act aaa gct cta aac aga cgt gct cta ata gat gag ata ata aac aat gta tca act gat gac aat ctc cta tca aaa ctt cca tta gaa att agg gat cta att gtt tca caa gct gtc ata tag agt tct atc cac cca cct ttc ttg aaa tga gtt aat --- --- --- --- --- --- --- --- --- --- --- --- --- --- --- --- --- --- --- --- --- --- --- --- --- --- --- --- --- --- --- --- --- --- --- --- --- --- --- --- --- --- --- --- --- --- --- --- --- --- --- --- --- --- --- --- --- --- --- --- --- --- --- --- --- --- --- --- --- --- --- --- --- --- --- --- --- --- --- --- --- --- --- --- --- --- --- --- --- --- --- --- --- --- --- --- --- --- --- --- --- --- --- --- --- -ag tca taa gtt agt taa --- --- --- --- --- -gt cat aag tta gtt aag tca t-- --- --- --- --- --- --- --- --- --- --- --- --- --- --- --- -aa gtt agt taa gtc ata agt tag tta agt cat aag tta gtt aag tca taa gtt agt taa gtc ata agt tag tta agt cat aag tta gtt tat agt cta aca ctt cta att ttt ata cct tga tct ttt tct cta att atg aaa aag taa atc att atg aag atg gat g-a aat gga cga gat tgt gcg cat cgt taa cga tag tat gtg gta cgt acc taa cgc att tat gga cga cgg tga taa tga agg tca cat ttc tgt caa taa tgt ctg tca tat gta tct cgc att ctt tga tgt gga tat atc atc tca tct gtt taa att agt tat taa aca ctg cga tct gaa taa acg act aaa atg tgg taa ctc tcc att aca ttg cta tac gat gaa tac acg att taa tcc atc tgt att aaa gat att gtt acg cca cgg cat gcg taa ctt tga tag caa gga taa aaa agg aca tat tcc tct aca cca cta tct gat tca ttc act atc aat cga taa caa gat ctt tga tat act aac gga ccc cat tga tga ctt tag taa atc atc cga tct att gct gtg tta tct tag ata taa att caa tgg gag ctt aaa cta tta cgt tct gta caa att att gac taa agg atc tga ccc taa ttg cgt cga tga gga tgg act cac ttc tct tca tta cta ctg taa aca cat atc cgc gtt cca cga aag caa tta tta caa gtc aaa gag tca cac taa gat gcg agc tga gaa gcg att cat cta cgc gat aat aga tca tgg agc aaa cat taa cgc ggt tac gaa aat cgg aaa tac gcc gtt aca cac tta cct tca aca gta tac caa aca tag tcc tcg tgt ggt gta tgc tct ttt atc tcg agg agc cga tac gag gat acg taa taa tct taa ttg tac acc cat cat gga ata cat aaa gaa cga ttg tgc aac agg tca tat tct cat aat gtt act caa ttg gca cga aca aaa ata cgg gaa att aca aaa gga aga agg aca aca tct act tta tct att cat aaa aca taa tca agg ata tgg aag tcg ctc tct caa tat act acg gta tct act aga tag att cga cat tca gaa aga cga ata cta taa tac aat gac tcc tct tca tac cgc ctt cca gaa ttg caa taa caa tgt tgc ctc ata cct cgt ata cat tgg ata cga cat caa cct tcc gac taa aga cga taa gac agt att cga ctt ggt gtt tga aaa cag aaa cat tat ata caa ggc gga tgt cgt taa tga cat tat cca cca cag act gaa agt atc tct acc tat gat taa atc gtt gtt cta caa gat gtc gga gtt ctc tcc cta cga cga tca cta cgt aaa gaa gat aat agc cta ctg cct att aag gga cga gtc att tgc gga act aca tac taa att ctg ttt aaa cga gga cta taa aag tgt att tat gaa aaa tat atc att cga taa gat aga ttc cat cat cga aaa atg tag tcg tga cat aag tct cct caa aga gat tcg aat ctc aga cac cga ctt gta tac ggt att gag aac aga aga cat tcg gta tca cac ata tct cga agc cat aca ttc aga caa acg cat ttc att tcc cat gta cga cga tct cat aga aca gtg tca tct atc gat gga gca taa aag taa act cgt cga caa agc act caa taa att aga gtc tac cat cga tag tca atc tag act atc gta ttt gcc tcc gga aat tat gcg caa tat cat aac caa gct aag cga cta cca tct aaa cag tat gtt gta cgg aaa gaa cca tta caa ata tta tcc atg ata gaa aga aaa tat tta aaa aat aat cta tat gat tgg aga agt agg aaa caa aca gta aca aga cga cga tta cta c-a tta tta aat cat gag gtc cgt att ata ctc gta tat att gtt tct ctc atg tat aat aat aaa cgg aag aga tat agc acc aca tgc acc atc caa tgg aaa gtg taa aga caa cga ata cag aag ccg taa tct atg ttg tct atc gtg tcc tcc ggg aac tta cgc ttt cag att atg tga tag caa gac taa tac aca atg tac acc gtg tgg ttc gga tac ctt tac atc tca caa taa tca ttt aca ggc ttg tct aag ttg taa cgg aag atg tga tag taa tca ggt aga gac gcg atc gtg taa cac gac tca caa tag aat ctg tga atg ctc tcc agg ata tta ttg tct tct caa agg agc atc agg gtg tag aac atg tat ttc taa aac aaa gtg tgg aat agg ata cgg agt atc cgg ata cac gtc tac cgg aga cgt cat ctg ttc tcc gtg tgg tcc cgg aac ata ttc tca cac cgt ctc ttc cac aga taa atg cga acc cgt cgt aac cag caa tac att taa cta tat cga tgt gga aat taa cct gta tcc agt caa cga cac atc gtg tac tcg gac gac cac tac cgg tct cag cga atc cat ctc aac gtc gga act aac tat tac cat gaa tca taa aga ttg tga tcc agt ctt tcg tgc aga ata ctt ctc tgt cct taa taa tgt agc aac ttc agg att ctt tac agg aga aaa tag ata tca gaa tac ttc aaa gat atg tac tct gaa ttt cga gat taa atg taa caa caa aga ttc atc ttc caa aca gtt aac gaa aac aaa gaa tga tac tat cat gcc gca ttc aga gac ggt aac tct agt ggg cga ctg tct atc tag cgt cga cat cta cat act ata tag taa tac caa tac tca aga cta cga aaa tga tac aat ctc tta tca tat ggg taa tgt tct cga tgt caa tag cca tat gcc cgc tag ttg cga tat aca taa act gat cac taa ttc cca gaa tcc cac cca ctt ata gta ag- -tt ttt tta cct ata aat aat aaa tac aat aat taa ttt ctc gta aaa gta gaa aat ata ttc taa ttt att ata tgg taa gaa agt aga atc atc tag aac agt aat caa tca ata gca atc atg aaa caa tat att gtc ctg gca tgc atg tgc cta gtg gca gct gct atg cct act agt ctt caa caa tct tca tcc tcg tgt act gaa gaa gaa aac aaa cat cat atg gga atc gat gtt att atc aaa gtc aca aag caa gac caa aca ccg acc aat gat aag att tgt caa tcc gta acg gaa gtt aca gag a-- --- --- --- --- --- -cc gaa gat gat gag gta tcc gaa gaa gtt gta aaa gga gat ccc acc act tat tac act atc gtc ggt gcg ggt ctt aac atg aac ttt gga ttc acc aaa tgc cca aag att tta tcc atc tcc gaa tcc tct gat gga aac act gtg aat act aga ttg tcc agc gtg tca ccg gga caa ggt aag gac tct ccc gcg atc acg cgt gaa gaa gct ctg gct atg atc aaa gac tgt gag atg tct atc gac atc aga tgt agc gaa gaa gag aaa gac agt gac atc aag acc cat cca gta ctt ggg tct aac atc tca cat aag aaa gtg agt tac aaa gat atc atc ggt tca acg atc gtt gat aca aaa tgt gtc aag aac cta gag ttt agc gta cgt atc gga gac atg tgt gag gaa tca tct gaa ctt gaa gtc aag gat gga ttc aag tat gtc gac gga tcg gca tct gaa ggt gca acc gat gat act tca ctc atc gat tca aca aaa ctc aaa gca tgt gtc tga atc gat aac tct att cat ctg aaa atg gat gag ttg ggt taa tcg aac gat tca gac acc gca cca cga att aaa aaa g-- -ac cgg gca cta tat tcc ggt ttg caa aac aaa aat att taa cta cat tca caa aaa gtt acc tct cgt tac ttc ttc ttt ctg ttt caa tat gtg ata cga tat gat cac tat tcg tat tct ct- --- --- tgg tct cat aaa aaa gtt tta c-- --a aaa aaa aaa aaa a-t att ttt att ctc ttt ctc tct tcg at- ggt ctc aca aaa ata tta aac ctc ttt ctg atg tct caa cta ttt cgt aaa cga taa cgt cca aca ata tat tct cgt aga gct tat caa cat cct tat acc aat cta ggt tgt cag aca att gca tca taa aat aat gtt tat aat tta cac gtt aac atc ata taa taa acg tat ata gtt aat att ttt gga ata taa atg atc tgt aaa atc cat gta ggg gac act gct cac gtt ttt tct cta gta cat aat ttc aca caa gtt ttt ata cag aca aat taa ttc tcg tcc ata ta- ttt taa aac att gac ttt tgt act aag aaa aat atc ttg act aac cat ctc ttt ctc tct tcg atg ggt ctc aca aaa ata tta aac ctc ttt ctg atg g-- --- --- --- --- --- --- --- --- --- --- --- --- --- --- --- --- --- --- --- --- --- --- --- --- --- --- --- --- --- --- --- --- --- --- --- --- --- --- --- --- --- --- --- --- --- --- agt cgt aaa aag ttt tta tcc ttt ctc tct tcg ata ggt ctc --- --- --- --- --- --- --- --- --- --- --- --- --- --- --- --- --- --- --- --- --- --- --- --- --- --- --- --- --- --- --- --- --- --- --- --- --- --- --- --- --- --- --- --- --- --- --- --- --- --- --- --- --- --- --- --- --- --- --- --- --- --- --- --- --- --- --- --- --- --- aca aaa ata tta aac ctc ttt ctg atg gtc tct ata aa

>ON595760.2_Monkeypox_virus_isolate_MPXV-CH-38134631/2022_partial_genome

atc gag gac gac ttc aac tat gac agc tcg gtg gcg tct gcc agc gtg tac ata cga atg gca ttt cta aga aaa gtc tac ggt atc ctt -tc tac aat ttc ctt taa caa cgg caa cag ctg cag tat ttt tat act ttg aat gca -tc gga cat tta tac aag gga gtc ctg ttc taa tat tgg cat caa tgt tcg gat cta tag gct tga ttt tcg cat tga ctt tac aca gac atg aac atc ccc tga atc tgt aca tac ttt gtg gat tta cac tgt tag aat ctc taa cgc tgg cct ctg ttg tta ctt tct atg atg cac gta tcg tta tgc aag ctt tca tgt tga cta ctg cag tgt ttc ttg ctc tga cta cat ata ctc tac aat caa aga gag att tca gta aac ttg taa cag gat tgt ttg ctg ctt tct gga ttt taa ttt tgt cag gag tct tga gga taa agt tta aaa tag aat taa taa aga aca tat agg tca -tt ttt taa aca tgg ata gaa acc aag gtt gtt agt taa taa tat aca aga tat ttt ttc tca ctc tga tcc atg taa acc aag gac gag aga cac tct cat tcc tca ttc acg aca cca tta aaa atg gaa att aaa gcc ctc tat taa gca cag acg gct aca ggt cta cca tca ggt t-a cct tcg tct acc ttc aca atg gcc tct cct tgt gcc cag ttc agt ccc tgt cat tgc cac gct act aag gac tcc ctg aat acc gtg act gac gtc aga cat tgt ctg act gaa tac atc ctg tgg gtt tct cat aga tgg acc cat aga gaa agc gca ggg cct ctc tac agg ctt ctc atc tct ttc aga att gat gca atg gag cta ttt ggt agc gag ttg aag gag ttc tcg aat tca ctt ccg tgg gac aat atc gac aat tgc gtg gag atc att aaa tgt ttc atc aga aat gac tcc atg aaa acc gcc aaa gaa ctt tgt gca ata att gga ctt tgt act caa tca gct att gtc act gga aga gtc ttc aat gat aag tat atc gac ata cta ctt atg ctg cga aag att ctg aac gag aac gac tat ctc acc ctc ttg gat cat atc ctc act gct aaa tac taa atc tcc ttc atg ctc tct cac taa tac tct tac tca cta cac ttt tta tca tct tat gat gaa tga ttg cct tca tca ttt ttt cgt gga ata taa tat agg aat aat tag cac cag aat agc tat gga tat ctc gtt aag aat att ctc tat aag aga cat aat gta gac ata gtt att ata tcc ttc tta gat aag tgt tac gct act gga aag ttt cca tcg tta tta tta cat gaa gat gat ata att aaa cca aca ttg aga ttg gct ctt atg tta gct gga ttg aat tac tgt aat aaa tgc atc gag tat aga ggg ata tag caa ttc tcg ata ata gtc atg caa tat ttg aat gag act gat aat tta ggt aat aca gta ct- --- aca cac ata tct ttc tag att ata tat cgt taa aaa tct gta aga tgt ata ttt ctc ata agt atc cac tgt gta ata tta tta atg gat ata tag ata acg caa tag gga cta ata gta ttg taa aag ata taa tcg act att tac gta cat atc cag ata tct ata ttc cta cta gtt tgc tgc gta gtt gca tca ttg ata tgc atg -at tta tca gga ttc aga gat gaa tta cta agt aaa cta caa tcc cac aat aag taa gaa tca aat atc aaa aac tca ctt ttg att ttt cta gtc tta agt aat aca tat att tat taa tag acc tat gaa ata aaa aaa ggt aac aat gga ttc gcg tat agc tat tta cgt att agt atc ggc atc tct ttt gta tct tgt taa ttg tca caa act agt aca tta ctt caa tct gaa aat aaa tgg aag tga tat aac taa tac agc aga tat att gct gga caa tta tcc aat tat gac ctt tga tgg aaa gga tat tta tcc atc tat ctc gtt cat ggt cgg taa taa act ttt cct aga tct tta taa aaa tat ctt tga aga att ttt cag act att tcg agt atc tgt aag tag tca ata cga gga att aga ata tta tta ttc atg tga tta tac taa caa ccg tcc tac aat taa aca aca tta ctt tta taa cgg cga cga ata tac tga aat tga tag atc gaa aaa agc cac taa taa aaa cag ttg gtt aat tac ttc agg ctt tag act aca aaa atg gtt cga tag cga aga ttg tat aat tta tct cag atc ttt agt tag aag aat gga aga cag taa caa aaa cag taa aaa aac tta gta ctt aga tat cg- --a aaa aat ata ttt ttg tag act ctt gag aat aga agg aaa aca tgt aca taa tta taa aaa atg aaa atc aat ggc gaa taa gac agt gcg att cgc acc atg gag tcg gta gat ttc atg gct gtc gat gag cag ttt cac gac gac ctc gat ctt tgg tca tta tct ttg gta gat gat tat aaa aaa cat gga tta ggt gtt gac tgt tat gtt cta gaa cca gtt gtt gac agg aaa ata ttt gat aga ttt ctc ctt gaa cca att tgt gat cct gta gat gtt ctg tat gat tat ttt agg att cat aga gat aat att gat cag tat ata gta gat aga ctg ttt gca tat att aca tat aaa gat att ata tct gca tta gtg tca aag aat tat atg gaa gat att ttc tct ata att att aag aat tgt aat tct gtg caa gat ctc tta ctt tac tat cta tct aat gca tat gta gaa ata gac att gtt gat ctt atg gta gat cat ggg gct gta ata tat aaa ata gaa tgc ttg aat gcc tat ttt agg gga ata tgt aaa aag gaa agt agt gtt gtt gag ttt att ttg aat tgt ggt atc cca gat gaa aat gat gtt aaa tta gat cta tat aaa ata att cag tat act agg gga ttc ctt gta gat gaa ccc aca gta tta gaa att tat aag ctt tgt atc cca tat att gaa gat atc aat caa cta gat gct ggt gga agg acc ttg ctt tat cgc gct atc tat gca ggt tat ata gat tta gta tca tgg cta tta gaa aat gga gca aat gtc aac gca gta atg agt aat gga tat aca tgt ctt gac gtg gcc gtg gat agg gga tct gtc atc gcc cgt agg gaa gca cat ctt aaa ata tta gaa ata ttg ctt aga gaa cca ttg tct att gac tgt ata aaa tta gct ata ctt aat aat aca att gaa aac cat gat gtg ata aag ctc tgt atc aag tat ttt atg atg gta gat tat tca ctt tgt aat gtg tat gca tca tca ctc ttt gat tat ata att gat tgt aaa caa gaa ttg gag tac att agg cag atg aaa att cat aat aca acc atg tat gag tta atc tat aat aga gac aaa aac aag cat gct tcc cat att cta cat agg tat tct aaa cat cca gtt ttg aca cag tgt atc act aaa gga ttc aag att tac aca gaa gta acc gag cag gtc act aaa gct cta aac aga cgt gct cta ata gat gag ata ata aac aat gta tca act gat gac aat ctc cta tca aaa ctt cca tta gaa att agg gat cta att gtt tca caa gct gtc ata tag agt tct atc cac cca cct ttc ttg aaa tga gtt aat --- --- --- --- --- --- --- --- --- --- --- --- --- --- --- --- --- --- --- --- --- --- --- --- --- --- --- --- --- --- --- --- --- --- --- --- --- --- --- --- --- --- --- --- --- --- --- --- --- --- --- --- --- --- --- --- --- --- --- --- --- --- --- --- --- --- --- --- --- --- --- --- --- --- --- --- --- --- --- --- --- --- --- --- --- --- --- --- --- --- --- --- --- --- --- --- --- --- --- --- --- --- --- --- --- -ag tca taa gtt agt taa --- --- --- --- --- -gt cat aag tta gtt aag tca t-- --- --- --- --- --- --- --- --- --- --- --- --- --- --- --- -aa gtt agt taa gtc ata agt tag tta agt cat aag tta gtt aag tca taa gtt agt taa gtc ata agt tag tta agt cat aag tta gtt tat agt cta aca ctt cta att ttt ata cct tga tct ttt tct cta att atg aaa aag taa atc att atg aag atg gat g-a aat gga cga gat tgt gcg cat cgt taa cga tag tat gtg gta cgt acc taa cgc att tat gga cga cgg tga taa tga agg tca cat ttc tgt caa taa tgt ctg tca tat gta tct cgc att ctt tga tgt gga tat atc atc tca tct gtt taa att agt tat taa aca ctg cga tct gaa taa acg act aaa atg tgg taa ctc tcc att aca ttg cta tac gat gaa tac acg att taa tcc atc tgt att aaa gat att gtt acg cca cgg cat gcg taa ctt tga tag caa gga taa aaa agg aca tat tcc tct aca cca cta tct gat tca ttc act atc aat cga taa caa gat ctt tga tat act aac gga ccc cat tga tga ctt tag taa atc atc cga tct att gct gtg tta tct tag ata taa att caa tgg gag ctt aaa cta tta cgt tct gta caa att att gac taa agg atc tga ccc taa ttg cgt cga tga gga tgg act cac ttc tct tca tta cta ctg taa aca cat atc cgc gtt cca cga aag caa tta tta caa gtc aaa gag tca cac taa gat gcg agc tga gaa gcg att cat cta cgc gat aat aga tca tgg agc aaa cat taa cgc ggt tac gaa aat cgg aaa tac gcc gtt aca cac tta cct tca aca gta tac caa aca tag tcc tcg tgt ggt gta tgc tct ttt atc tcg agg agc cga tac gag gat acg taa taa tct taa ttg tac acc cat cat gga ata cat aaa gaa cga ttg tgc aac agg tca tat tct cat aat gtt act caa ttg gca cga aca aaa ata cgg gaa att aca aaa gga aga agg aca aca tct act tta tct att cat aaa aca taa tca agg ata tgg aag tcg ctc tct caa tat act acg gta tct act aga tag att cga cat tca gaa aga cga ata cta taa tac aat gac tcc tct tca tac cgc ctt cca gaa ttg caa taa caa tgt tgc ctc ata cct cgt ata cat tgg ata cga cat caa cct tcc gac taa aga cga taa gac agt att cga ctt ggt gtt tga aaa cag aaa cat tat ata caa ggc gga tgt cgt taa tga cat tat cca cca cag act gaa agt atc tct acc tat gat taa atc gtt gtt cta caa gat gtc gga gtt ctc tcc cta cga cga tca cta cgt aaa gaa gat aat agc cta ctg cct att aag gga cga gtc att tgc gga act aca tac taa att ctg ttt aaa cga gga cta taa aag tgt att tat gaa aaa tat atc att cga taa gat aga ttc cat cat cga aaa atg tag tcg tga cat aag tct cct caa aga gat tcg aat ctc aga cac cga ctt gta tac ggt att gag aac aga aga cat tcg gta tca cac ata tct cga agc cat aca ttc aga caa acg cat ttc att tcc cat gta cga cga tct cat aga aca gtg tca tct atc gat gga gca taa aag taa act cgt cga caa agc act caa taa att aga gtc tac cat cga tag tca atc tag act atc gta ttt gcc tcc gga aat tat gcg caa tat cat aac caa gct aag cga cta cca tct aaa cag tat gtt gta cgg aaa gaa cca tta caa ata tta tcc atg ata gaa aga aaa tat tta aaa aat aat cta tat gat tgg aga agt agg aaa caa aca gta aca aga cga cga tta cta c-a tta tta aat cat gag gtc cgt att ata ctc gta tat att gtt tct ctc atg tat aat aat aaa cgg aag aga tat agc acc aca tgc acc atc caa tgg aaa gtg taa aga caa cga ata cag aag ccg taa tct atg ttg tct atc gtg tcc tcc ggg aac tta cgc ttt cag att atg tga tag caa gac taa tac aca atg tac acc gtg tgg ttc gga tac ctt tac atc tca caa taa tca ttt aca ggc ttg tct aag ttg taa cgg aag atg tga tag taa tca ggt aga gac gcg atc gtg taa cac gac tca caa tag aat ctg tga atg ctc tcc agg ata tta ttg tct tct caa agg agc atc agg gtg tag aac atg tat ttc taa aac aaa gtg tgg aat agg ata cgg agt atc cgg ata cac gtc tac cgg aga cgt cat ctg ttc tcc gtg tgg tcc cgg aac ata ttc tca cac cgt ctc ttc cac aga taa atg cga acc cgt cgt aac cag caa tac att taa cta tat cga tgt gga aat taa cct gta tcc agt caa cga cac atc gtg tac tcg gac gac cac tac cgg tct cag cga atc cat ctc aac gtc gga act aac tat tac cat gaa tca taa aga ttg tga tcc agt ctt tcg tgc aga ata ctt ctc tgt cct taa taa tgt agc aac ttc agg att ctt tac agg aga aaa tag ata tca gaa tac ttc aaa gat atg tac tct gaa ttt cga gat taa atg taa caa caa aga ttc atc ttc caa aca gtt aac gaa aac aaa gaa tga tac tat cat gcc gca ttc aga gac ggt aac tct agt ggg cga ctg tct atc tag cgt cga cat cta cat act ata tag taa tac caa tac tca aga cta cga aaa tga tac aat ctc tta tca tat ggg taa tgt tct cga tgt caa tag cca tat gcc cgc tag ttg cga tat aca taa act gat cac taa ttc cca gaa tcc cac cca ctt ata gta ag- -tt ttt tta cct ata aat aat aaa tac aat aat taa ttt ctc gta aaa gta gaa aat ata ttc taa ttt att ata tgg taa gaa agt aga atc atc tag aac agt aat caa tca ata gca atc atg aaa caa tat att gtc ctg gca tgc atg tgc cta gtg gca gct gct atg cct act agt ctt caa caa tct tca tcc tcg tgt act gaa gaa gaa aac aaa cat cat atg gga atc gat gtt att atc aaa gtc aca aag caa gac caa aca ccg acc aat gat aag att tgt caa tcc gta acg gaa gtt aca gag a-- --- --- --- --- --- -cc gaa gat gat gag gta tcc gaa gaa gtt gta aaa gga gat ccc acc act tat tac act atc gtc ggt gcg ggt ctt aac atg aac ttt gga ttc acc aaa tgc cca aag att tta tcc atc tcc gaa tcc tct gat gga aac act gtg aat act aga ttg tcc agc gtg tca ccg gga caa ggt aag gac tct ccc gcg atc acg cgt gaa gaa gct ctg gct atg atc aaa gac tgt gag atg tct atc gac atc aga tgt agc gaa gaa gag aaa gac agt gac atc aag acc cat cca gta ctt ggg tct aac atc tca cat aag aaa gtg agt tac aaa gat atc atc ggt tca acg atc gtt gat aca aaa tgt gtc aag aac cta gag ttt agc gta cgt atc gga gac atg tgt gag gaa tca tct gaa ctt gaa gtc aag gat gga ttc aag tat gtc gac gga tcg gca tct gaa ggt gca acc gat gat act tca ctc atc gat tca aca aaa ctc aaa gca tgt gtc tga atc gat aac tct att cat ctg aaa atg gat gag ttg ggt taa tcg aac gat tca gac acc gca cca cga att aaa aaa g-- -ac cgg gca cta tat tcc ggt ttg caa aac aaa aat att taa cta cat tca caa aaa gtt acc tct cgt tac ttc ttc ttt ctg ttt caa tat gtg ata cga tat gat cac tat tcg tat tct ct- --- --- tgg tct cat aaa aaa gtt tta c-- --a aaa aaa aaa aaa --t att ttt att ctc ttt ctc tct tcg at- ggt ctc aca aaa ata tta aac ctc ttt ctg atg tct caa cta ttt cgt aaa cga taa cgt cca aca ata tat tct cgt aga gct tat caa cat cct tat acc aat cta ggt tgt cag aca att gca tca taa aat aat gtt tat aat tta cac gtt aac atc ata taa taa acg tat ata gtt aat att ttt gga ata taa atg atc tgt aaa atc cat gta ggg gac act gct cac gtt ttt tct cta gta cat aat ttc aca caa gtt ttt ata cag aca aat taa ttc tcg tcc ata ta- ttt taa aac att gac ttt tgt act aag aaa aat atc ttg act aac cat ctc tt? ?tc tct tcg atg ggt ctc aca aaa ata tta aac ctc ttt ctg atg g-- --- --- --- --- --- --- --- --- --- --- --- --- --- --- --- --- --- --- --- --- --- --- --- --- --- --- --- --- --- --- --- --- --- --- --- --- --- --- --- --- --- --- --- --- --- --- agt cgt ?aa aag ttt tta tcc tt? ?tc tct tcg ata ggt ctc --- --- --- --- --- --- --- --- --- --- --- --- --- --- --- --- --- --- --- --- --- --- --- --- --- --- --- --- --- --- --- --- --- --- --- --- --- --- --- --- --- --- --- --- --- --- --- --- --- --- --- --- --- --- --- --- --- --- --- --- --- --- --- --- --- --- --- --- --- --- aca aaa ata tta aac ctc ttt ctg atg gtc tct ata aa

>MT903337.1_Monkeypox_virus_isolate_MPXV-M2940_FCT

atc gag gac gac ttc aac tat gac agc tcg gtg gcg tct gcc agc gtg tac ata cga atg gca ttt cta aga aaa gtc tac ggt atc ctt -tc tac aat ttc ctt taa caa cgg caa cag ctg cag tat ttt tat act ttg aat gca -tc gga cat tta tac aag gga gtc ctg ttc taa tat tgg cat caa tgt tcg gat cta tag gct tga ttt tcg cat tga ctt tac aca gac atg aac atc ccc tga atc tgt aca tac ttt gtg gat tta cac tgt tag aat ctc taa cgc tgg cct ctg ttg tta ctt tct atg atg cac gta tcg tta tgc aag ctt tca tgt tga cta ctg cag tgt ttc ttg ctc tga cta cat ata ctc tac aat caa aga gag att tca gta aac ttg taa cag gat tgt ttg ctg ctt tct gga ttt taa ttt tgt cag gag tct tga gga taa agt tta aaa tag aat taa taa aga aca tat agg tca -tt ttt taa aca tgg ata gaa acc aag gtt gtt agt taa taa tat aca aga tat ttt ttc tca ctc tga tcc atg taa acc aag gac gag aga cac tct cat tcc tca ttc acg aca cca tta aaa atg gaa att aaa gcc ctc tat taa gca cag acg gct aca ggt cta cca tca ggt t-a cct tcg tct acc ttc aca atg gcc tct cct tgt gcc cag ttc agt ccc tgt cat tgc cac gct act aag gac tcc ctg aat acc gtg act gac gtc aga cat tgt ctg act gaa tac atc ctg tgg gtt tct cat aga tgg acc cat aga gaa agc gca ggg cct ctc tac agg ctt ctc atc tct ttc aga att gat gca atg gag cta ttt ggt agc gag ttg aag gag ttc tcg aat tca ctt ccg tgg gac aat atc gac aat tgc gtg gag atc att aaa tgt ttc atc aga aat gac tcc atg aaa acc gcc aaa gaa ctt tgt gca ata att gga ctt tgt act caa tca gct att gtc act gga aga gtc ttc aat gat aag tat atc gac ata cta ctt atg ctg cga aag att ctg aac gag aac gac tat ctc acc ctc ttg gat cat atc ctc act gct aaa tac taa atc tcc ttc atg ctc tct cac taa tac tct tac tca cta cac ttt tta tca tct tat gat gaa tga ttg cct tca tca ttt ttt cgt gga ata taa tat agg aat aat tag cac cag aat agc tat gga tat ctc gtt aag aat att ctc tat aag aga cat aat gta gac ata gtt att ata tcc ttc tta gat aag tgt tac gct act gga aag ttt cca tcg tta tta tta cat gaa gat gat ata att aaa cca aca ttg aga ttg gct ctt atg tta gct gga ttg aat tac tgt aat aaa tgc atc gag tat aga ggg ata tag caa ttc tcg ata ata gtc atg caa tat ttg aat gag act gat aat tta ggt aat aca gta ct- --- aca cac ata tct ttc tag att ata tat cgt taa aaa tct gta aga tgt ata ttt ctc ata agt atc cac tgt gta ata tta tta atg gat ata tag ata acg caa tag gga cta ata gta ttg taa aag ata taa tcg act att tac gta cat atc cag ata tct ata ttc cta cta gtt tgc tgc gta gtt gca tca ttg ata tgc atg -at tta tca gga ttc aga gat gaa tta cta agt aaa cta caa tcc cac aat aag taa gaa tca aat atc aaa aac tca ctt ttg att ttt cta gtc tta agt aat aca tat att tat taa tag acc tat gaa ata aaa aaa ggt aac aat gga ttc gcg tat agc tat tta cgt att agt atc ggc atc tct ttt gta tct tgt taa ttg tca caa act agt aca tta ctt caa tct gaa aat aaa tgg aag tga tat aac taa tac agc aga tat att gct gga caa tta tcc aat tat gac ctt tga tgg aaa gga tat tta tcc atc tat ctc gtt cat ggt cgg taa taa act ttt cct aga tct tta taa aaa tat ctt tga aga att ttt cag act att tcg agt atc tgt aag tag tca ata cga gga att aga ata tta tta ttc atg tga tta tac taa caa ccg tcc tac aat taa aca aca tta ctt tta taa cgg cga cga ata tac tga aat tga tag atc gaa aaa agc cac taa taa aaa cag ttg gtt aat tac ttc agg ctt tag act aca aaa atg gtt cga tag cga aga ttg tat aat tta tct cag atc ttt agt tag aag aat gga aga cag taa caa aaa cag taa aaa aac tta gta ctt aga tat cg- --a aaa aat ata ttt ttg tag act ctt gag aat aga agg aaa aca tgt aca taa tta taa aaa atg aaa atc aat ggc gaa taa gac agt gcg att cgc acc atg gag tcg gta gat ttc atg gct gtc gat gag cag ttt cac gac gac ctc gat ctt tgg tca tta tct ttg gta gat gat tat aaa aaa cat gga tta ggt gtt gac tgt tat gtt cta gaa cca gtt gtt gac agg aaa ata ttt gat aga ttt ctc ctt gaa cca att tgt gat cct gta gat gtt ctg tat gat tat ttt agg att cat aga gat aat att gat cag tat ata gta gat aga ctg ttt gca tat att aca tat aaa gat att ata tct gca tta gtg tca aag aat tat atg gaa gat att ttc tct ata att att aag aat tgt aat tct gtg caa gat ctc tta ctt tac tat cta tct aat gca tat gta gaa ata gac att gtt gat ctt atg gta gat cat ggg gct gta ata tat aaa ata gaa tgc ttg aat gcc tat ttt agg gga ata tgt aaa aag gaa agt agt gtt gtt gag ttt att ttg aat tgt ggt atc cca gat gaa aat gat gtt aaa tta gat cta tat aaa ata att cag tat act agg gga ttc ctt gta gat gaa ccc aca gta tta gaa att tat aag ctt tgt atc cca tat att gaa gat atc aat caa cta gat gct ggt gga agg acc ttg ctt tat cgc gct atc tat gca ggt tat ata gat tta gta tca tgg cta tta gaa aat gga gca aat gtc aac gca gta atg agt aat gga tat aca tgt ctt gac gtg gcc gtg gat agg gga tct gtc atc gcc cgt agg gaa gca cat ctt aaa ata tta gaa ata ttg ctt aga gaa cca ttg tct att gac tgt ata aaa tta gct ata ctt aat aat aca att gaa aac cat gat gtg ata aag ctc tgt atc aag tat ttt atg atg gta gat tat tca ctt tgt aat gtg tat gca tca tca ctc ttt gat tat ata att gat tgt aaa caa gaa ttg gag tac att agg cag atg aaa att cat aat aca acc atg tat gag tta atc tat aat aga gac aaa aac aag cat gct tcc cat att cta cat agg tat tct aaa cat cca gtt ttg aca cag tgt atc act aaa gga ttc aag att tac aca gaa gta acc gag cag gtc act aaa gct cta aac aga cgt gct cta ata gat gag ata ata aac aat gta tca act gat gac aat ctc cta tca aaa ctt cca tta gaa att agg gat cta att gtt tca caa gct gtc ata tag agt tct atc cac cca cct ttc ttg aaa tga gtt aat agt cat aag tta gtt aag tca taa gtt agt taa gtc ata agt tag tta agt cat aag tta gtt aag tca taa gtt agt taa gtc ata agt tag tta agt cat aag tta gtt aag tca taa gtt agt taa gtc ata agt tag tta agt cat aag tta gtt aag tca taa gtt agt taa gtc ata agt tag tta agt cat aag tta gtt aag tca taa gtt agt taa gtc ata agt taa gtc ata agt tag tta agt cat aag tta gtt aag tca taa gtt agt taa gtc ata agt tag tta agt cat aag tta gtt aag tca taa gtt agt taa --- --- --- --- --- -gt cat aag tta gtt aag tca taa gtt agt taa gtc ata agt tag tta agt cat aag tta gtt aag tca taa gtt agt taa gtc ata agt tag tta agt cat aag tta gtt aag tca taa gtt agt taa gtc ata agt tag tta agt cat aag tta gtt tat agt cta aca ctt cta att ttt ata cct tga tct ttt tct cta att atg aaa aag taa atc att atg aag atg gat g-a aat gga cga gat tgt gcg cat cgt taa cga tag tat gtg gta cgt acc taa cgc att tat gga cga cgg tga taa tga agg tca cat ttc tgt caa taa tgt ctg tca tat gta tct cgc att ctt tga tgt gga tat atc atc tca tct gtt taa att agt tat taa aca ctg cga tct gaa taa acg act aaa atg tgg taa ctc tcc att aca ttg cta tac gat gaa tac acg att taa tcc atc tgt att aaa gat att gtt acg cca cgg cat gcg taa ctt tga tag caa gga taa aaa agg aca tat tcc tct aca cca cta tct gat tca ttc act atc aat cga taa caa gat ctt tga tat act aac gga ccc cat tga tga ctt tag taa atc atc cga tct att gct gtg tta tct tag ata taa att caa tgg gag ctt aaa cta tta cgt tct gta caa att att gac taa agg atc tga ccc taa ttg cgt cga tga gga tgg act cac ttc tct tca tta cta ctg taa aca cat atc cgc gtt cca cga aag caa tta tta caa gtc aaa gag tca cac taa gat gcg agc tga gaa gcg att cat cta cgc gat aat aga tca tgg agc aaa cat taa cgc ggt tac gaa aat cgg aaa tac gcc gtt aca cac tta cct tca aca gta tac caa aca tag tcc tcg tgt ggt gta tgc tct ttt atc tcg agg agc cga tac gag gat acg taa taa tct tga ttg tac acc cat cat gga ata cat aaa gaa cga ttg tgc aac agg tca tat tct cat aat gtt act caa ttg gca cga aca aaa ata cgg gaa att aca aaa gga aga agg aca aca tct act tta tct att cat aaa aca taa tca agg ata tgg aag tcg ctc tct caa tat act acg gta tct act aga tag att cga cat tca gaa aga cga ata cta taa tac aat gac tcc tct tca tac cgc ctt cca gaa ttg caa taa caa tgt tgc ctc ata cct cgt ata cat cgg ata cga cat caa cct tcc gac taa aga cga taa gac agt att cga ctt ggt gtt tga aaa cag aaa cat tat ata caa ggc gga tgt cgt taa tga cat tat cca cca cag act gaa agt atc tct acc tat gat taa atc gtt gtt cta caa gat gtc gga gtt ctc tcc cta cga cga tca cta cgt aaa gaa gat aat agc cta ctg cct att aag gga cga gtc att tgc gga act aca tac taa att ctg ttt aaa cga gga cta taa aag tgt att tat gaa aaa tat atc att cga taa gat aga ttc cat cat cga aaa atg tag tcg tga cat aag tct cct caa aga gat tcg aat ctc aga cac cga ctt gta tac ggt att gag aac aga aga cat ccg gta tca cac ata tct cga agc cat aca ttc aga caa acg cat ttc att tcc cat gta cga cga tct cat aga aca gtg tca tct atc gat gga gca taa aag taa act cgt cga caa agc act caa taa att aga gtc tac cat cga tag tca atc tag act atc gta ttt gcc tcc gga aat tat gcg caa tat cat aac caa gct aag cga cta cca tct aaa cag tat gtt gta cgg aaa gaa cca tta caa ata tta tcc atg ata gaa aga aaa tat tta aaa aat aat cta tat gat tgg aga agt agg aaa caa aca gta aca aga cga cga tta cta c-a tta tta aat cat gag gtc cgt att ata ctc gta tat att gtt tct ctc atg tat aat aat aaa cgg aag aga tat agc acc aca tgc acc atc caa tgg aaa gtg taa aga caa cga ata cag aag ccg taa tct atg ttg tct atc gtg tcc tcc ggg aac tta cgc ttc cag att atg tga tag caa gac taa tac aca atg tac acc gtg tgg ttc gga tac ctt tac atc tca caa taa tca ttt aca ggc ttg tct aag ttg taa cgg aag atg tga tag taa tca ggt aga gac gcg atc gtg taa cac gac tca caa tag aat ctg tga atg ctc tcc agg ata tta ttg tct tct caa agg agc atc agg gtg tag aac atg tat ttc taa aac aaa gtg tgg aat agg ata cgg agt atc cgg ata cac gtc tac cgg aga cgt cat ctg ttc tcc gtg tgg tcc cgg aac ata ttc tca cac cgt ctc ttc cac aga taa atg cga acc cgt cgt aac cag caa tac att taa cta tat cga tgt gga aat taa cct gta tcc agt caa cga cac atc gtg tac tcg gac gac cac tac cgg tct cag cga atc cat ctc aac gtc gga act aac tat tac cat gaa tca taa aga ttg tga tcc agt ctt tcg tgc aga ata ctt ctc tgt cct taa taa tgt agc aac ttc agg att ctt tac agg aga aaa tag ata tca gaa tac ttc aaa gat atg tac tct gaa ttt cga gat taa atg taa caa caa aga ttc atc ttc caa aca gtt aac gaa aac aaa gaa tga tac tat cat gcc gca ttc aga gac ggt aac tct agt ggg cga ctg tct atc tag cgt cga cat cta cat act ata tag taa tac caa tac tca aga cta cga aaa tga tac aat ctc tta tca tat ggg taa tgt tct cga tgt caa tag cca tat gcc cgc tag ttg cga tat aca taa act gat cac taa ttc cca gaa tcc cac cca ctt ata gta ag- -tt ttt tta cct ata aat aat aaa tac aat aat taa ttt ctc gta aaa gta gaa aat ata ttc taa ttt att ata tgg taa gaa agt aga atc atc tag aac agt aat caa tca ata gca atc atg aaa caa tat att gtc ctg gca tgc atg tgc cta gtg gca gct gct atg cct act agt ctt caa caa tct tca tcc tcg tgt act gaa gaa gaa aac aaa cat cat atg gga atc gat gtt att atc aaa gtc aca aag caa gac caa aca ccg acc aat gat aag att tgt caa tcc gta acg gaa gtt aca gag a-- --- --- --- --- --- -cc gaa gat gat gag gta tcc gaa gaa gtt gta aaa gga gat ccc acc act tat tac act atc gtc ggt gcg ggt ctt aac atg aac ttt gga ttc acc aaa tgc cca aag att tca tcc atc tcc gaa tcc tct gat gga aac act gtg aat act aga ttg tcc agc gtg tca ccg gga caa ggt aag gac tct ccc gcg atc acg cgt gaa gaa gct ctg gct atg atc aaa gac tgt gag atg tct atc gac atc aga tgt agc gaa gaa gag aaa gac agt gac atc aag acc cat cca gta ctt ggg tct aac atc tca cat aag aaa gtg agt tac aaa gat atc atc ggt tca acg atc gtt gat aca aaa tgt gtc aag aac cta gag ttt agc gta cgt atc gga gac atg tgt gag gaa tca tct gaa ctt gaa gtc aag gat gga ttc aag tat gtc gac gga tcg gca tct gaa ggt gca acc gat gat act tca ctc atc gat tca aca aaa ctc aaa gca tgt gtc tga atc gat aac tct att cat ctg aaa atg gat gag ttg ggt taa tcg aac gat tca gac acc gca cca cga att aaa aaa g-- -ac cgg gca cta tat tcc ggt ttg caa aac aaa aat att taa cta cat tca caa aaa gtt acc tct cgt tac ttc ttc ttt ctg ttt caa tat gtg ata cga tat gat cac tat tcg tat tct ct- --- --- tgg tct cat aaa aaa gtt tta c-- -aa aaa aaa aaa aaa a-t att ttt att ctc ttt ctc tct tcg at- ggt ctc aca aaa ata tta aac ctc ttt ctg atg tct caa cta ttt cgt aaa cga taa cgt cca aca ata tat tct cgt aga gct tat caa cat cct tat acc aat cta ggt tgt cag aca att gca tca taa aat aat gtt tat aat tta cac gtt aac atc ata taa taa acg tat ata gtt aat att ttt gga ata taa atg atc tgt aaa atc cat gta ggg gac act gct cac gtt ttt tct cta gta cat aat ttc aca caa gtt ttt ata cag aca aat taa ttc tcg tcc ata ta- ttt taa aac att gac ttt tgt act aag aaa aat atc ttg act aac cat ctc ttt ctc tct tcg --- --- --- --- --- --- --- --- --- --- --- --- --- --- --- --- --- --- --- --- --- --- --- --- --- --- --- --- --- --- --- --- --- --- --- --- --- --- --- --- --- --- --- --- --- --- --- --- --- --- --- --- --- --- --- --- --- --- --- --- --- --- --- --- --- --- --- --- --- --- --- --- --- --- --- --- --- --- --- --- --- --- --- --- --- --- --- --- --- --- --- --- --- --- --- --- --- --- --- --- --- --- --- --- --- --- --- --- --- --- --- --- --- --- --- --- --- --- --- --- --- --- --- --- --- --- --- --- --- --- --- --- --- --- --- --- --- --- --- --- --- --- --- --- --- --- --- --- --- --- --- --- --- --- --- --

>MT903339.1_Monkeypox_virus_isolate_MPXV-M3021_Delta

atc gag gac gac ttc aac tat gac agc tcg gtg gcg tct gcc agc gtg tac ata cga atg gca ttt cta aga aaa gtc tac ggt atc ctt -tc tac aat ttc ctt taa caa cgg caa cag ctg cag tat ttt tat act ttg aat gca -tc gga cat tta tac aag gga gtc ctg ttc taa tat tgg cat caa tgt tcg gat cta tag gct tga ttt tcg cat tga ctt tac aca gac atg aac atc ccc tga atc tgt aca tac ttt gtg gat tta cac tgt tag aat ctc taa cgc tgg cct ctg ttg tta ctt tct atg atg cac gta tcg tta tgc aag ctt tca tgt tga cta ctg cag tgt ttc ttg ctc tga cta cat ata ctc tac aat caa aga gag att tca gta aac ttg taa cag gat tgt ttg ctg ctt tct gga ttt taa ttt tgt cag gag tct tga gga taa agt tta aaa tag aat taa taa aga aca tat agg tca -tt ttt taa aca tgg ata gaa acc aag gtt gtt agt taa taa tat aca aga tat ttt ttc tca ctc tga tcc atg taa acc aag gac gag aga cac tct cat tcc tca ttc acg aca cca tta aaa atg gaa att aaa gcc ctc tat taa gca cag acg gct aca ggt cta cca tca ggt t-a cct tcg tct acc ttc aca atg gcc tct cct tgt gcc cag ttc agt ccc tgt cat tgc cac gct act aag gac tcc ctg aat acc gtg act gac gtc aga cat tgt ctg act gaa tac atc ctg tgg gtt tct cat aga tgg acc cat aga gaa agc gca ggg cct ctc tac agg ctt ctc atc tct ttc aga att gat gca atg gag cta ttt ggt agc gag ttg aag gag ttc tcg aat tca ctt ccg tgg gac aat atc gac aat tgc gtg gag atc att aaa tgt ttc atc aga aat gac tcc atg aaa acc gcc aaa gaa ctt tgt gca ata att gga ctt tgt act caa tca gct att gtc act gga aga gtc ttc aat gat aag tat atc gac ata cta ctt atg ctg cga aag att ctg aac gag aac gac tat ctc acc ctc ttg gat cat atc ctc act gct aaa tac taa atc tcc ttc atg ctc tct cac taa tac tct tac tca cta cac ttt tta tca tct tat gat gaa tga ttg cct tca tca ttt ttt cgt gga ata taa tat agg aat aat tag cac cag aat agc tat gga tat ctc gtt aag aat att ctc tat aag aga cat aat gta gac ata gtt att ata tcc ttc tta gat aag tgt tac gct act gga aag ttt cca tcg tta tta tta cat gaa gat gat ata att aaa cca aca ttg aga ttg gct ctt atg tta gct gga ttg aat tac tgt aat aaa tgc atc gag tat aga ggg ata tag caa ttc tcg ata ata gtc atg caa tat ttg aat gag act gat aat tta ggt aat aca gta ct- --- aca cac ata tct ttc tag att ata tat cgt taa aaa tct gta aga tgt ata ttt ctc ata agt atc cac tgt gta ata tta tta atg gat ata tag ata acg caa tag gga cta ata gta ttg taa aag ata taa tcg act att tac gta cat atc cag ata tct ata ttc cta cta gtt tgc tgc gta gtt gca tca ttg ata tgc atg -at tta tca gga ttc aga gat gaa tta cta agt aaa cta caa tcc cac aat aag taa gaa tca aat atc aaa aac tca ctt ttg att ttt cta gtc tta agt aat aca tat att tat taa tag acc tat gaa ata aaa aaa ggt aac aat gga ttc gcg tat agc tat tta cgt att agt atc ggc atc tct ttt gta tct tgt taa ttg tca caa act agt aca tta ctt caa tct gaa aat aaa tgg aag tga tat aac taa tac agc aga tat att gct gga caa tta tcc aat tat gac ctt tga tgg aaa gga tat tta tcc atc tat ctc gtt cat ggt cgg taa taa act ttt cct aga tct tta taa aaa tat ctt tga aga att ttt cag act att tcg agt atc tgt aag tag tca ata cga gga att aga ata tta tta ttc atg tga tta tac taa caa ccg tcc tac aat taa aca aca tta ctt tta taa cgg cga cga ata tac tga aat tga tag atc gaa aaa agc cac taa taa aaa cag ttg gtt aat tac ttc agg ctt tag act aca aaa atg gtt cga tag cga aga ttg tat aat tta tct cag atc ttt agt tag aag aat gga aga cag taa caa aaa cag taa aaa aac tta gta ctt aga tat cg- --a aaa aat ata ttt ttg tag act ctt gag aat aga agg aaa aca tgt aca taa tta taa aaa atg aaa atc aat ggc gaa taa gac agt gcg att cgc acc atg gag tcg gta gat ttc atg gct gtc gat gag cag ttt cac gac gac ctc gat ctt tgg tca tta tct ttg gta gat gat tat aaa aaa cat gga tta ggt gtt gac tgt tat gtt cta gaa cca gtt gtt gac agg aaa ata ttt gat aga ttt ctc ctt gaa cca att tgt gat cct gta gat gtt ctg tat gat tat ttt agg att cat aga gat aat att gat cag tat ata gta gat aga ctg ttt gca tat att aca tat aaa gat att ata tct gca tta gtg tca aag aat tat atg gaa gat att ttc tct ata att att aag aat tgt aat tct gtg caa gat ctc tta ctt tac tat cta tct aat gca tat gta gaa ata gac att gtt gat ctt atg gta gat cat ggg gct gta ata tat aaa ata gaa tgc ttg aat gcc tat ttt agg gga ata tgt aaa aag gaa agt agt gtt gtt gag ttt att ttg aat tgt ggt atc cca gat gaa aat gat gtt aaa tta gat cta tat aaa ata att cag tat act agg gga ttc ctt gta gat gaa ccc aca gta tta gaa att tat aag ctt tgt atc cca tat att gaa gat atc aat caa cta gat gct ggt gga agg acc ttg ctt tat cgc gct atc tat gca ggt tat ata gat tta gta tca tgg cta tta gaa aat gga gca aat gtc aac gca gta atg agt aat gga tat aca tgt ctt gac gtg gcc gtg gat agg gga tct gtc atc gcc cgt agg gaa gca cat ctt aaa ata tta gaa ata ttg ctt aga gaa cca ttg tct att gac tgt ata aaa tta gct ata ctt aat aat aca att gaa aac cat gat gtg ata aag ctc tgt atc aag tat ttt atg atg gta gat tat tca ctt tgt aat gtg tat gca tca tca ctc ttt gat tat ata att gat tgt aaa caa gaa ttg gag tac att agg cag atg aaa att cat aat aca acc atg tat gag tta atc tat aat aga gac aaa aac aag cat gct tcc cat att cta cat agg tat tct aaa cat cca gtt ttg aca cag tgt atc act aaa gga ttc aag att tac aca gaa gta acc gag cag gtc act aaa gct cta aac aga cgt gct cta ata gat gag ata ata aac aat gta tca act gat gac aat ctc cta tca aaa ctt cca tta gaa att agg gat cta att gtt tca caa gct gtc ata tag agt tct atc cac cca cct ttc ttg aaa tga gtt aat agt cat aag tta gtt aag tca taa gtt agt taa gtc ata agt tag tta agt cat aag tta gtt aag tca taa gtt agt taa gtc ata agt tag tta agt cat aag tta gtt aag tca taa gtt agt taa gtc ata agt tag tta agt cat aag tta gtt aag tca taa gtt agt taa gtc ata agt tag tta agt cat aag tta gtt aag tca taa gtt agt taa gtc ata agt taa gtc ata agt tag tta agt cat aag tta gtt aag tca taa gtt agt taa gtc ata agt tag tta agt cat aag tta gtt aag tca taa gtt agt taa --- --- --- --- --- -gt cat aag tta gtt aag tca taa gtt agt taa gtc ata agt tag tta agt cat aag tta gtt aag tca taa gtt agt taa gtc ata agt tag tta agt cat aag tta gtt aag tca taa gtt agt taa gtc ata agt tag tta agt cat aag tta gtt tat agt cta aca ctt cta att ttt ata cct tga tct ttt tct cta att atg aaa aag taa atc att atg aag atg gat g-a aat gga cga gat tgt gcg cat cgt taa cga tag tat gtg gta cgt acc taa cgc att tat gga cga cgg tga taa tga agg tca cat ttc tgt caa taa tgt ctg tca tat gta tct cgc att ctt tga tgt gga tat atc atc tca tct gtt taa att agt tat taa aca ctg cga tct gaa taa acg act aaa atg tgg taa ctc tcc att aca ttg cta tac gat gaa tac acg att taa tcc atc tgt att aaa gat att gtt acg cca cgg cat gcg taa ctt tga tag caa gga taa aaa agg aca tat tcc tct aca cca cta tct gat tca ttc act atc aat cga taa caa gat ctt tga tat act aac gga ccc cat tga tga ctt tag taa atc atc cga tct att gct gtg tta tct tag ata taa att caa tgg gag ctt aaa cta tta cgt tct gta caa att att gac taa agg atc tga ccc taa ttg cgt cga tga gga tgg act cac ttc tct tca tta cta ctg taa aca cat atc cgc gtt cca cga aag caa tta tta caa gtc aaa gag tca cac taa gat gcg agc tga gaa gcg att cat cta cgc gat aat aga tca tgg agc aaa cat taa cgc ggt tac gaa aat cgg aaa tac gcc gtt aca cac tta cct tca aca gta tac caa aca tag tcc tcg tgt ggt gta tgc tct ttt atc tcg agg agc cga tac gag gat acg taa taa tct tga ttg tac acc cat cat gga ata cat aaa gaa cga ttg tgc aac agg tca tat tct cat aat gtt act caa ttg gca cga aca aaa ata cgg gaa att aca aaa gga aga agg aca aca tct act tta tct att cat aaa aca taa tca agg ata tgg aag tcg ctc tct caa tat act acg gta tct act aga tag att cga cat tca gaa aga cga ata cta taa tac aat gac tcc tct tca tac cgc ctt cca gaa ttg caa taa caa tgt tgc ctc ata cct cgt ata cat cgg ata cga cat caa cct tcc gac taa aga cga taa gac agt att cga ctt ggt gtt tga aaa cag aaa cat tat ata caa ggc gga tgt cgt taa tga cat tat cca cca cag act gaa agt atc tct acc tat gat taa atc gtt gtt cta caa gat gtc gga gtt ctc tcc cta cga cga tca cta cgt aaa gaa gat aat agc cta ctg cct att aag gga cga gtc att tgc gga act aca tac taa att ctg ttt aaa cga gga cta taa aag tgt att tat gaa aaa tat atc att cga taa gat aga ttc cat cat cga aaa atg tag tcg tga cat aag tct cct caa aga gat tcg aat ctc aga cac cga ctt gta tac ggt att gag aac aga aga cat ccg gta tca cac ata tct cga agc cat aca ttc aga caa acg cat ttc att tcc cat gta cga cga tct cat aga aca gtg tca tct atc gat gga gca taa aag taa act cgt cga caa agc act caa taa att aga gtc tac cat cga tag tca atc tag act atc gta ttt gcc tcc gga aat tat gcg caa tat cat aac caa gct aag cga cta cca tct aaa cag tat gtt gta cgg aaa gaa cca tta caa ata tta tcc atg ata gaa aga aaa tat tta aaa aat aat cta tat gat tgg aga agt agg aaa caa aca gta aca aga cga cga tta cta c-a tta tta aat cat gag gtc cgt att ata ctc gta tat att gtt tct ctc atg tat aat aat aaa cgg aag aga tat agc acc aca tgc acc atc caa tgg aaa gtg taa aga caa cga ata cag aag ccg taa tct atg ttg tct atc gtg tcc tcc ggg aac tta cgc ttc cag att atg tga tag caa gac taa tac aca atg tac acc gtg tgg ttc gga tac ctt tac atc tca caa taa tca ttt aca ggc ttg tct aag ttg taa cgg aag atg tga tag taa tca ggt aga gac gcg atc gtg taa cac gac tca caa tag aat ctg tga atg ctc tcc agg ata tta ttg tct tct caa agg agc atc agg gtg tag aac atg tat ttc taa aac aaa gtg tgg aat agg ata cgg agt atc cgg ata cac gtc tac cgg aga cgt cat ctg ttc tcc gtg tgg tcc cgg aac ata ttc tca cac cgt ctc ttc cac aga taa atg cga acc cgt cgt aac cag caa tac att taa cta tat cga tgt gga aat taa cct gta tcc agt caa cga cac atc gtg tac tcg gac gac cac tac cgg tct cag cga atc cat ctc aac gtc gga act aac tat tac cat gaa tca taa aga ttg tga tcc agt ctt tcg tgc aga ata ctt ctc tgt cct taa taa tgt agc aac ttc agg att ctt tac agg aga aaa tag ata tca gaa tac ttc aaa gat atg tac tct gaa ttt cga gat taa atg taa caa caa aga ttc atc ttc caa aca gtt aac gaa aac aaa gaa tga tac tat cat gcc gca ttc aga gac ggt aac tct agt ggg cga ctg tct atc tag cgt cga cat cta cat act ata tag taa tac caa tac tca aga cta cga aaa tga tac aat ctc tta tca tat ggg taa tgt tct cga tgt caa tag cca tat gcc cgc tag ttg cga tat aca taa act gat cac taa ttc cca gaa tcc cac cca ctt ata gta ag- -tt ttt tta cct ata aat aat aaa tac aat aat taa ttt ctc gta aaa gta gaa aat ata ttc taa ttt att ata tgg taa gaa agt aga atc atc tag aac agt aat caa tca ata gca atc atg aaa caa tat att gtc ctg gca tgc atg tgc cta gtg gca gct gct atg cct act agt ctt caa caa tct tca tcc tcg tgt act gaa gaa gaa aac aaa cat cat atg gga atc gat gtt att atc aaa gtc aca aag caa gac caa aca ccg acc aat gat aag att tgt caa tcc gta acg gaa gtt aca gag a-- --- --- --- --- --- -cc gaa gat gat gag gta tcc gaa gaa gtt gta aaa gga gat ccc acc act tat tac act atc gtc ggt gcg ggt ctt aac atg aac ttt gga ttc acc aaa tgc cca aag att tca tcc atc tcc gaa tcc tct gat gga aac act gtg aat act aga ttg tcc agc gtg tca ccg gga caa ggt aag gac tct ccc gcg atc acg cgt gaa gaa gct ctg gct atg atc aaa gac tgt gag atg tct atc gac atc aga tgt agc gaa gaa gag aaa gac agt gac atc aag acc cat cca gta ctt ggg tct aac atc tca cat aag aaa gtg agt tac aaa gat atc atc ggt tca acg atc gtt gat aca aaa tgt gtc aag aac cta gag ttt agc gta cgt atc gga gac atg tgt gag gaa tca tct gaa ctt gaa gtc aag gat gga ttc aag tat gtc gac gga tcg gca tct gaa ggt gca acc gat gat act tca ctc atc gat tca aca aaa ctc aaa gca tgt gtc tga atc gat aac tct att cat ctg aaa atg gat gag ttg ggt taa tcg aac gat tca gac acc gca cca cga att aaa aaa g-- -ac cgg gca cta tat tcc ggt ttg caa aac aaa aat att taa cta cat tca caa aaa gtt acc tct cgt tac ttc ttc ttt ctg ttt caa tat gtg ata cga tat gat cac tat tcg tat tct ct- --- --- tgg tct cat aaa aaa gtt tta ca- -aa aaa aaa aaa aaa a-t att ttt att ctc ttt ctc tct tcg at- ggt ctc aca aaa ata tta aac ctc ttt ctg atg tct caa cta ttt cgt aaa cga taa cgt cca aca ata tat tct cgt aga gct tat caa cat cct tat acc aat cta ggt tgt cag aca att gca tca taa aat aat gtt tat aat tta cac gtt aac atc ata taa taa acg tat ata gtt aat att ttt gga ata taa atg atc tgt aaa atc cat gta ggg gac act gct cac gtt ttt tct cta gta cat aat ttc aca caa gtt ttt ata cag aca aat taa ttc tcg tcc ata ta- ttt taa aac att gac ttt tgt act aag aaa aat atc ttg act aac cat ctc ttt ctc tct tcg --- --- --- --- --- --- --- --- --- --- --- --- --- --- --- --- --- --- --- --- --- --- --- --- --- --- --- --- --- --- --- --- --- --- --- --- --- --- --- --- --- --- --- --- --- --- --- --- --- --- --- --- --- --- --- --- --- --- --- --- --- --- --- --- --- --- --- --- --- --- --- --- --- --- --- --- --- --- --- --- --- --- --- --- --- --- --- --- --- --- --- --- --- --- --- --- --- --- --- --- --- --- --- --- --- --- --- --- --- --- --- --- --- --- --- --- --- --- --- --- --- --- --- --- --- --- --- --- --- --- --- --- --- --- --- --- --- --- --- --- --- --- --- --- --- --- --- --- --- --- --- --- --- --- --- --

>MT903338.1_Monkeypox_virus_isolate_MPXV-M2957_Lagos

atc gag gac gac ttc aac tat gac agc tcg gtg gcg tct gcc agc gtg tac ata cga atg gca ttt cta aga aaa gtc tac ggt atc ctt -tc tac aat ttc ctt taa caa cgg caa cag ctg cag tat ttt tat act ttg aat gca -tc gga cat tta tac aag gga gtc ctg ttc taa tat tgg cat caa tgt tcg gat cta tag gct tga ttt tcg cat tga ctt tac aca gac atg aac atc ccc tga atc tgt aca tac ttt gtg gat tta cac tgt tag aat ctc taa cgc tgg cct ctg ttg tta ctt tct atg atg cac gta tcg tta tgc aag ctt tca tgt tga cta ctg cag tgt ttc ttg ctc tga cta cat ata ctc tac aat caa aga gag att tca gta aac ttg taa cag gat tgt ttg ctg ctt tct gga ttt taa ttt tgt cag gag tct tga gga taa agt tta aaa tag aat taa taa aga aca tat agg tca -tt ttt taa aca tgg ata gaa acc aag gtt gtt agt taa taa tat aca aga tat ttt ttc tca ctc tga tcc atg taa acc aag gac gag aga cac tct cat tcc tca ttc acg aca cca tta aaa atg gaa att aaa gcc ctc tat taa gca cag acg gct aca ggt cta cca tca ggt t-a cct tcg tct acc ttc aca atg gcc tct cct tgt gcc cag ttc agt ccc tgt cat tgc cac gct act aag gac tcc ctg aat acc gtg act gac gtc aga cat tgt ctg act gaa tac atc ctg tgg gtt tct cat aga tgg acc cat aga gaa agc gca ggg cct ctc tac agg ctt ctc atc tct ttc aga att gat gca atg gag cta ttt ggt agc gag ttg aag gag ttc tcg aat tca ctt ccg tgg gac aat atc gac aat tgc gtg gag atc att aaa tgt ttc atc aga aat gac tcc atg aaa acc gcc aaa gaa ctt tgt gca ata att gga ctt tgt act caa tca gct att gtc act gga aga gtc ttc aat gat aag tat atc gac ata cta ctt atg ctg cga aag att ctg aac gag aac gac tat ctc acc ctc ttg gat cat atc ctc act gct aaa tac taa atc tcc ttc atg ctc tct cac taa tac tct tac tca cta cac ttt tta tca tct tat gat gaa tga ttg cct tca tca ttt ttt cgt gga ata taa tat agg aat aat tag cac cag aat agc tat gga tat ctc gtt aag aat att ctc tat aag aga cat aat gta gac ata gtt att ata tcc ttc tta gat aag tgt tac gct act gga aag ttt cca tcg tta tta tta cat gaa gat gat ata att aaa cca aca ttg aga ttg gct ctt atg tta gct gga ttg aat tac tgt aat aaa tgc atc gag tat aga ggg ata tag caa ttc tcg ata ata gtc atg caa tat ttg aat gag act gat aat tta ggt aat aca gta ct- --- aca cac ata tct ttc tag att ata tat cgt taa aaa tct gta aga tgt ata ttt ctc ata agt atc cac tgt gta ata tta tta atg gat ata tag ata acg caa tag gga cta ata gta ttg taa aag ata taa tcg act att tac gta cat atc cag ata tct ata ttc cta cta gtt tgc tgc gta gtt gca tca ttg ata tgc atg -at tta tca gga ttc aga gat gaa tta cta agt aaa cta caa tcc cac aat aag taa gaa tca aat atc aaa aac tca ctt ttg att ttt cta gtc tta agt aat aca tat att tat taa tag acc tat gaa ata aaa aaa ggt aac aat gga ttc gcg tat agc tat tta cgt att agt atc ggc atc tct ttt gta tct tgt taa ttg tca caa act agt aca tta ctt caa tct gaa aat aaa tgg aag tga tat aac taa tac agc aga tat att gct gga caa tta tcc aat tat gac ctt tga tgg aaa gga tat tta tcc atc tat ctc gtt cat ggt cgg taa taa act ttt cct aga tct tta taa aaa tat ctt tga aga att ttt cag act att tcg agt atc tgt aag tag tca ata cga gga att aga ata tta tta ttc atg tga tta tac taa caa ccg tcc tac aat taa aca aca tta ctt tta taa cgg cga cga ata tac tga aat tga tag atc gaa aaa agc cac taa taa aaa cag ttg gtt aat tac ttc agg ctt tag act aca aaa atg gtt cga tag cga aga ttg tat aat tta tct cag atc ttt agt tag aag aat gga aga cag taa caa aaa cag taa aaa aac tta gta ctt aga tat cg- --a aaa aat ata ttt ttg tag act ctt gag aat aga agg aaa aca tgt aca taa tta taa aaa atg aaa atc aat ggc gaa taa gac agt gcg att cgc acc atg gag tcg gta gat ttc atg gct gtc gat gag cag ttt cac gac gac ctc gat ctt tgg tca tta tct ttg gta gat gat tat aaa aaa cat gga tta ggt gtt gac tgt tat gtt cta gaa cca gtt gtt gac agg aaa ata ttt gat aga ttt ctc ctt gaa cca att tgt gat cct gta gat gtt ctg tat gat tat ttt agg att cat aga gat aat att gat cag tat ata gta gat aga ctg ttt gca tat att aca tat aaa gat att ata tct gca tta gtg tca aag aat tat atg gaa gat att ttc tct ata att att aag aat tgt aat tct gtg caa gat ctc tta ctt tac tat cta tct aat gca tat gta gaa ata gac att gtt gat ctt atg gta gat cat ggg gct gta ata tat aaa ata gaa tgc ttg aat gcc tat ttt agg gga ata tgt aaa aag gaa agt agt gtt gtt gag ttt att ttg aat tgt ggt atc cca gat gaa aat gat gtt aaa tta gat cta tat aaa ata att cag tat act agg gga ttc ctt gta gat gaa ccc aca gta tta gaa att tat aag ctt tgt atc cca tat att gaa gat atc aat caa cta gat gct ggt gga agg acc ttg ctt tat cgc gct atc tat gca ggt tat ata gat tta gta tca tgg cta tta gaa aat gga gca aat gtc aac gca gta atg agt aat gga tat aca tgt ctt gac gtg gcc gtg gat agg gga tct gtc atc gcc cgt agg gaa gca cat ctt aaa ata tta gaa ata ttg ctt aga gaa cca ttg tct att gac tgt ata aaa tta gct ata ctt aat aat aca att gaa aac cat gat gtg ata aag ctc tgt atc aag tat ttt atg atg gta gat tat tca ctt tgt aat gtg tat gca tca tca ctc ttt gat tat ata att gat tgt aaa caa gaa ttg gag tac att agg cag atg aaa att cat aat aca acc atg tat gag tta atc tat aat aga gac aaa aac aag cat gct tcc cat att cta cat agg tat tct aaa cat cca gtt ttg aca cag tgt atc act aaa gga ttc aag att tac aca gaa gta acc gag cag gtc act aaa gct cta aac aga cgt gct cta ata gat gag ata ata aac aat gta tca act gat gac aat ctc cta tca aaa ctt cca tta gaa att agg gat cta att gtt tca caa gct gtc ata tag agt tct atc cac cca cct ttc ttg aaa tga gtt aat agt cat aag tta gtt aag tca taa gtt agt taa gtc ata agt tag tta agt cat aag tta gtt aag tca taa gtt agt taa gtc ata agt tag tta agt cat aag tta gtt aag tca taa gtt agt taa gtc ata agt tag tta agt cat aag tta gtt aag tca taa gtt agt taa gtc ata agt tag tta agt cat aag tta gtt aag tca taa gtt --- --- --- --- agt taa gtc ata agt tag tta agt cat aag tta gtt aag tca taa gtt agt taa gtc ata agt tag tta agt cat aag tta gtt aag tca taa gtt agt taa gtc ata agt tag tta agt cat aag tta gtt aag tca taa gtt agt taa gtc ata agt tag tta agt cat aag tta gtt aag tca taa gtt agt taa gtc ata agt tag tta agt cat aag tta gtt aag tca taa gtt agt taa gtc ata agt tag tta agt cat aag tta gtt tat agt cta aca ctt cta att ttt ata cct tga tct ttt tct cta att atg aaa aag taa atc att atg aag atg gat g-a aat gga cga gat tgt gcg cat cgt taa cga tag tat gtg gta cgt acc taa cgc att tat gga cga cgg tga taa tga agg tca cat ttc tgt caa taa tgt ctg tca tat gta tct cgc att ctt tga tgt gga tat atc atc tca tct gtt taa att agt tat taa aca ctg cga tct gaa taa acg act aaa atg tgg taa ctc tcc att aca ttg cta tac gat gaa tac acg att taa tcc atc tgt att aaa gat att gtt acg cca cgg cat gcg taa ctt tga tag caa gga taa aaa agg aca tat tcc tct aca cca cta tct gat tca ttc act atc aat cga taa caa gat ctt tga tat act aac gga ccc cat tga tga ctt tag taa atc atc cga tct att gct gtg tta tct tag ata taa att caa tgg gag ctt aaa cta tta cgt tct gta caa att att gac taa agg atc tga ccc taa ttg cgt cga tga gga tgg act cac ttc tct tca tta cta ctg taa aca cat atc cgc gtt cca cga aag caa tta tta caa gtc aaa gag tca cac taa gat gcg agc tga gaa gcg att cat cta cgc gat aat aga tca tgg agc aaa cat taa cgc ggt tac gaa aat cgg aaa tac gcc gtt aca cac tta cct tca aca gta tac caa aca tag tcc tcg tgt ggt gta tgc tct ttt atc tcg agg agc cga tac gag gat acg taa taa tct tga ttg tac acc cat cat gga ata cat aaa gaa cga ttg tgc aac agg tca tat tct cat aat gtt act caa ttg gca cga aca aaa ata cgg gaa att aca aaa gga aga agg aca aca tct act tta tct att cat aaa aca taa tca agg ata tgg aag tcg ctc tct caa tat act acg gta tct act aga tag att cga cat tca gaa aga cga ata cta taa tac aat gac tcc tct tca tac cgc ctt cca gaa ttg caa taa caa tgt tgc ctc ata cct cgt ata cat cgg ata cga cat caa cct tcc gac taa aga cga taa gac agt att cga ctt ggt gtt tga aaa cag aaa cat tat ata caa ggc gga tgt cgt taa tga cat tat cca cca cag act gaa agt atc tct acc tat gat taa atc gtt gtt cta caa gat gtc gga gtt ctc tcc cta cga cga tca cta cgt aaa gaa gat aat agc cta ctg cct att aag gga cga gtc att tgc gga act aca tac taa att ctg ttt aaa cga gga cta taa aag tgt att tat gaa aaa tat atc att cga taa gat aga ttc cat cat cga aaa atg tag tcg tga cat aag tct cct caa aga gat tcg aat ctc aga cac cga ctt gta tac ggt att gag aac aga aga cat ccg gta tca cac ata tct cga agc cat aca ttc aga caa acg cat ttc att tcc cat gta cga cga tct cat aga aca gtg tca tct atc gat gga gca taa aag taa act cgt cga caa agc act caa taa att aga gtc tac cat cga tag tca atc tag act atc gta ttt gcc tcc gga aat tat gcg caa tat cat aac caa gct aag cga cta cca tct aaa cag tat gtt gta cgg aaa gaa cca tta caa ata tta tcc atg ata gaa aga aaa tat tta aaa aat aat cta tat gat tgg aga agt agg aaa caa aca gta aca aga cga cga tta cta c-a tta tta aat cat gag gtc cgt att ata ctc gta tat att gtt tct ctc atg tat aat aat aaa cgg aag aga tat agc acc aca tgc acc atc caa tgg aaa gtg taa aga caa cga ata cag aag ccg taa tct atg ttg tct atc gtg tcc tcc ggg aac tta cgc ttc cag att atg tga tag caa gac taa tac aca atg tac acc gtg tgg ttc gga tac ctt tac atc tca caa taa tca ttt aca ggc ttg tct aag ttg taa cgg aag atg tga tag taa tca ggt aga gac gcg atc gtg taa cac gac tca caa tag aat ctg tga atg ctc tcc agg ata tta ttg tct tct caa agg agc atc agg gtg tag aac atg tat ttc taa aac aaa gtg tgg aat agg ata cgg agt atc cgg ata cac gtc tac cgg aga cgt cat ctg ttc tcc gtg tgg tcc cgg aac ata ttc tca cac cgt ctc ttc cac aga taa atg cga acc cgt cgt aac cag caa tac att taa cta tat cga tgt gga aat taa cct gta tcc agt caa cga cac atc gtg tac tcg gac gac cac tac cgg tct cag cga atc cat ctc aac gtc gga act aac tat tac cat gaa tca taa aga ttg tga tcc agt ctt tcg tgc aga ata ctt ctc tgt cct taa taa tgt agc aac ttc agg att ctt tac agg aga aaa tag ata tca gaa tac ttc aaa gat atg tac tct gaa ttt cga gat taa atg taa caa caa aga ttc atc ttc caa aca gtt aac gaa aac aaa gaa tga tac tat cat gcc gca ttc aga gac ggt aac tct agt ggg cga ctg tct atc tag cgt cga cat cta cat act ata tag taa tac caa tac tca aga cta cga aaa tga tac aat ctc tta tca tat ggg taa tgt tct cga tgt caa tag cca tat gcc cgc tag ttg cga tat aca taa act gat cac taa ttc cca gaa tcc cac cca ctt ata gta ag- -tt ttt tta cct ata aat aat aaa tac aat aat taa ttt ctc gta aaa gta gaa aat ata ttc taa ttt att ata tgg taa gaa agt aga atc atc tag aac agt aat caa tca ata gca atc atg aaa caa tat att gtc ctg gca tgc atg tgc cta gtg gca gct gct atg cct act agt ctt caa caa tct tca tcc tcg tgt act gaa gaa gaa aac aaa cat cat atg gga atc gat gtt att atc aaa gtc aca aag caa gac caa aca ccg acc aat gat aag att tgt caa tcc gta acg gaa gtt aca gag a-- --- --- --- --- --- -cc gaa gat gat gag gta tcc gaa gaa gtt gta aaa gga gat ccc acc act tat tac act atc gtc ggt gcg ggt ctt aac atg aac ttt gga ttc acc aaa tgc cca aag att tca tcc atc tcc gaa tcc tct gat gga aac act gtg aat act aga ttg tcc agc gtg tca ccg gga caa ggt aag gac tct ccc gcg atc acg cgt gaa gaa gct ctg gct atg atc aaa gac tgt gag atg tct atc gac atc aga tgt agc gaa gaa gag aaa gac agt gac atc aag acc cat cca gta ctt ggg tct aac atc tca cat aag aaa gtg agt tac aaa gat atc atc ggt tca acg atc gtt gat aca aaa tgt gtc aag aac cta gag ttt agc gta cgt atc gga gac atg tgt gag gaa tca tct gaa ctt gaa gtc aag gat gga ttc aag tat gtc gac gga tcg gca tct gaa ggt gca acc gat gat act tca ctc atc gat tca aca aaa ctc aaa gca tgt gtc tga atc gat aac tct att cat ctg aaa atg gat gag ttg ggt taa tcg aac gat tca gac acc gca cca cga att aaa aaa g-- -ac cgg gca cta tat tcc ggt ttg caa aac aaa aat att taa cta cat tca caa aaa gtt acc tct cgt tac ttc ttc ttt ctg ttt caa tat gtg ata cga tat gat cac tat tcg tat tct ct- --- --- tgg tct cat aaa aaa gtt tta ca- -aa aaa aaa aaa aaa a-t att ttt att ctc ttt ctc tct tcg at- ggt ctc aca aaa ata tta aac ctc ttt ctg atg tct caa cta ttt cgt aaa cga taa cgt cca aca ata tat tct cgt aga gct tat caa cat cct tat acc aat cta ggt tgt cag aca att gca tca taa aat aat gtt tat aat tta cac gtt aac atc ata taa taa acg tat ata gtt aat att ttt gga ata taa atg atc tgt aaa atc cat gta ggg gac act gct cac gtt ttt tct cta gta cat aat ttc aca caa gtt ttt ata cag aca aat taa ttc tcg tcc ata ta- ttt taa aac att gac ttt tgt act aag aaa aat atc ttg act aac cat ctc ttt ctc tct tcg --- --- --- --- --- --- --- --- --- --- --- --- --- --- --- --- --- --- --- --- --- --- --- --- --- --- --- --- --- --- --- --- --- --- --- --- --- --- --- --- --- --- --- --- --- --- --- --- --- --- --- --- --- --- --- --- --- --- --- --- --- --- --- --- --- --- --- --- --- --- --- --- --- --- --- --- --- --- --- --- --- --- --- --- --- --- --- --- --- --- --- --- --- --- --- --- --- --- --- --- --- --- --- --- --- --- --- --- --- --- --- --- --- --- --- --- --- --- --- --- --- --- --- --- --- --- --- --- --- --- --- --- --- --- --- --- --- --- --- --- --- --- --- --- --- --- --- --- --- --- --- --- --- --- --- --

>KJ642617.1_Monkeypox_virus_strain_Nigeria-SE-1971_complete_genome

atc gag gac gac ttc aac tat gac agc tcg gtg gcg tct gcc agc gtg tac ata cga atg gca ttt cta aga aaa gtc tac ggt atc ctt -tc tac aat ttc ctt taa caa cgg caa cag ctg cag tat ttt tat act ttg aat gca -tc gga cat tta tac aag gga gtc ctg ttc taa tat tgg cat caa tgt tcg gat cta tag gct tga ttt tcg cat tga ctt tac aca gac atg aac atc ccc tga atc tgt aca tac ttt gtg gat tta cac tgt tag aat ctc taa cgc tgg cct ctg ttg tta ctt tct atg atg cac gta tcg tta tgc aag ctt tca tgt tga cta ctg cag tgt ttc ttg ctc tga cta cat ata ctc tac aat caa aga gag att tca gta aac ttg taa cag gat tgt ttg ctg ctt tct gga ttt taa ttt tgt cag gag tct tga gga taa agt tta aaa tag aat taa taa aga aca tat agg tca -tt ttt taa aca tgg ata gaa acc aag gtt gtt agt taa taa tat aca aga tat ttt ttc tca ctc tga tcc atg taa acc aag gac gag aga cac tct cat tcc tca ttc acg aca cca tta aaa atg gaa att aaa gcc ctc tat taa gca cag acg gct aca ggt cta cca tca ggt t-a cct tcg tct acc ttc aca atg gcc tct cct tgt gcc cag ttc agt ccc tgt cat tgc cac gct act aag gac tcc ctg aat acc gtg act gac gtc aga cat tgt ctg act gaa tac atc ctg tgg gtt tct cat aga tgg acc cat aga gaa agc gca ggg cct ctc tac agg ctt ctc atc tct ttc aga att gat gca atg gag cta ttt ggt agc gag ttg aag gag ttc tcg aat tca ctt ccg tgg gac aat atc gac aat tgc gtg gag atc att aaa tgt ttc atc aga aat gac tcc atg aaa acc gcc aaa gaa ctt tgt gca ata att gga ctt tgt act caa tca gct att gtc act gga aga gtc ttc aat gat aag tat atc gac ata cta ctt atg ctg cga aag att ctg aac gag aac gac tat ctc acc ctc ttg gat cat atc ctc act gct aaa tac taa atc tcc ttc atg ctc tct cac taa tac tct tac tca cta cac ttt tta tca tct tat gat gaa tga ttg cct tca tca ttt ttt cgt gga ata taa tat agg aat aat tag cac cag aat agc tat gga tat ctc gtt aag aat att ctc tat aag aga cat aat gta gac ata gtt att ata tcc ttc tta gat aag tgt tac gct act gga aag ttt cca tcg tta tta tta cat gaa gat gat ata att aaa cca aca ttg aga ttg gct ctt atg tta gct gga ttg aat tac tgt aat aaa tgc atc gag tat aga ggg ata tag caa ttc tcg ata ata gtc atg caa tat ttg aat gag act gat aat tta ggt aat aca gta ct- --- aca cac ata tct ttc tag att ata tat cgt taa aaa tct gta aga tgt ata ttt ctc ata agt atc cac tgt gta ata tta tta atg gat ata tag ata acg caa tag gga cta ata gta ttg taa aag ata taa tcg act att tac gta cat atc cag ata tct ata ttc cta cta gtt tgc tgc gta gtt gca tca ttg ata tgc atg -at tta tca gga ttc aga gat gaa tta cta agt aaa cta caa tcc cac aat aag taa gaa tca aat atc aaa aac tca ctt ttg att ttt cta gtc tta agt aat aca tat att tat taa tag acc tat gaa ata aaa aaa ggt aac aat gga ttc gcg tat agc tat tta cgt att agt atc ggc atc tct ttt gta tct tgt taa ttg tca caa act agt aca tta ctt caa tct gaa aat aaa tgg aag tga tat aac taa tac agc aga tat att gct gga caa tta tcc aat tat gac ctt tga tgg aaa gga tat tta tcc atc tat ctc gtt cat ggt cgg taa taa act ttt cct aga tct tta taa aaa tat ctt tga aga att ttt cag act att tcg agt atc tgt aag tag tca ata cga gga att aga ata tta tta ttc atg tga tta tac taa caa ccg tcc tac aat taa aca aca tta ctt tta taa cgg cga cga ata tac tga aat tga tag atc gaa aaa agc cac taa taa aaa cag ttg gtt aat tac ttc agg ctt tag act aca aaa atg gtt cga tag cga aga ttg tat aat tta tct cag atc ttt agt tag aag aat gga aga cag taa caa aaa cag taa aaa aac tta gta ctt aga tat cg- --a aaa aat ata ttt ttg tag act ctt gag aat aga agg aaa aca tgt aca taa tta taa aaa atg aaa atc aat ggc gaa taa gac agt gcg att cgc acc atg gag tcg gta gat ttc atg gct gtc gat gag cag ttt cac gac gac ctc gat ctt tgg tca tta tct ttg gta gat gat tat aaa aaa cat gga tta ggt gtt gac tgt tat gtt cta gaa cca gtt gtt gac agg aaa ata ttt gat aga ttt ctc ctt gaa cca att tgt gat cct gta gat gtt ctg tat gat tat ttt agg att cat aga gat aat att gat cag tat ata gta gat aga ctg ttt gca tat att aca tat aaa gat att ata tct gca tta gtg tca aag aat tat atg gaa gat att ttc tct ata att att aag aat tgt aat tct gtg caa gat ctc tta ctt tac tat cta tct aat gca tat gta gaa ata gac att gtt gat ctt atg gta gat cat ggg gct gta ata tat aaa ata gaa tgc ttg aat gcc tat ttt agg gga ata tgt aaa aag gaa agt agt gtt gtt gag ttt att ttg aat tgt ggt atc cca gat gaa aat gat gtt aaa tta gat cta tat aaa ata att cag tat act agg gga ttc ctt gta gat gaa ccc aca gta tta gaa att tat aag ctt tgt atc cca tat att gaa gat atc aat caa cta gat gct ggt gga agg acc ttg ctt tat cgc gct atc tat gca ggt tat ata gat tta gta tca tgg cta tta gaa aat gga gca aat gtc aac gca gta atg agt aat gga tat aca tgt ctt gac gtg gcc gtg gat agg gga tct gtc atc gcc cgt agg gaa gca cat ctt aaa ata tta gaa ata ttg ctt aga gaa cca ttg tct att gac tgt ata aaa tta gct ata ctt aat aat aca att gaa aac cat gat gtg ata aag ctc tgt atc aag tat ttt atg atg gta gat tat tca ctt tgt aat gtg tat gca tca tca ctc ttt gat tat ata att gat tgt aaa caa gaa ttg gag tac att agg cag atg aaa att cat aat aca acc atg tat gag tta atc tat aat aga gac aaa aac aag cat gct tcc cat att cta cat agg tat tct aaa cat cca gtt ttg aca cag tgt atc act aaa gga ttc aag att tac aca gaa gta acc gag cag gtc act aaa gct cta aac aga cgt gct cta ata gat gag ata ata aac aat gta tca act gat gac aat ctc cta tca aaa ctt cca tta gaa att agg gat cta att gtt tca caa gct gtc ata tag agt tct atc cac cca cct ttc ttg aaa tga gtt aat agt cat aag tta gtt aag tca taa gtt agt taa gtc ata agt tag tta agt cat aag tta gtt aag tca taa gtt agt taa gtc ata agt tag tta agt cat aag tta gtt aag tca taa gtt agt taa gtc ata agt tag tta agt cat aag tta gtt aag tca taa gtt agt taa gtc ata agt tag tta agt cat aag tta gtt aag tca taa gtt agt taa gtc ata agt taa gtc ata agt tag tta agt cat aag tta gtt aag tca taa gtt agt taa gtc ata agt tag tta agt cat aag tta gtt aag tca taa gtt agt taa --- --- --- --- --- -gt cat aag tta gtt aag tca taa gtt agt taa gtc ata agt tag tta agt cat aag tta gtt aag tca taa gtt agt taa gtc ata agt tag tta agt cat aag tta gtt aag tca taa gtt agt taa gtc ata agt tag tta agt cat aag tta gtt tat agt cta aca ctt cta att ttt ata cct tga tct ttt tct cta att atg aaa aag taa atc att atg aag atg gat g-a aat gga cga gat tgt gcg cat cgt taa cga tag tat gtg gta cgt acc taa cgc att tat gga cga cgg tga taa tga agg tca cat ttc tgt caa taa tgt ctg tca tat gta tct cgc att ctt tga tgt gga tat atc atc tca tct gtt taa att agt tat taa aca ctg cga tct gaa taa acg act aaa atg tgg taa ctc tcc att aca ttg cta tac gat gaa tac acg att taa tcc atc tgt att aaa gat att gtt acg cca cgg cat gcg taa ctt tga tag caa gga taa aaa agg aca tat tcc tct aca cca cta tct gat tca ttc act atc aat cga taa caa gat ctt tga tat act aac gga ccc cat tga tga ctt tag taa atc atc cga tct att gct gtg tta tct tag ata taa att caa tgg gag ctt aaa cta tta cgt tct gta caa att att gac taa agg atc tga ccc taa ttg cgt cga tga gga tgg act cac ttc tct tca tta cta ctg taa aca cat atc cgc gtt cca cga aag caa tta tta caa gtc aaa gag tca cac taa gat gcg agc tga gaa gcg att cat cta cgc gat aat aga tca tgg agc aaa cat taa cgc ggt tac gaa aat cgg aaa tac gcc gtt aca cac tta cct tca aca gta tac caa aca tag tcc tcg tgt ggt gta tgc tct ttt atc tcg agg agc cga tac gag gat acg taa taa tct tga ttg tac acc cat cat gga ata cat aaa gaa cga ttg tgc aac agg tca tat tct cat aat gtt act caa ttg gca cga aca aaa ata cgg gaa att aca aaa gga aga agg aca aca tct act tta tct att cat aaa aca taa tca agg ata tgg aag tcg ctc tct caa tat act acg gta tct act aga tag att cga cat tca gaa aga cga ata cta taa tac aat gac tcc tct tca tac cgc ctt cca gaa ttg caa taa caa tgt tgc ctc ata cct cgt ata cat cgg ata cga cat caa cct tcc gac taa aga cga taa gac agt att cga ctt ggt gtt tga aaa cag aaa cat cat ata caa ggc gga tgt cgt taa tga cat tat cca cca cag act gaa agt atc tct acc tat gat taa atc gtt gtt cta caa gat gtc gga gtt ctc tcc cta cga cga tca cta cgt aaa gaa gat aat agc cta ctg cct att aag gga cga gtc att tgc gga act aca tac taa att ctg ttt aaa cga gga cta taa aag tgt att tat gaa aaa tat atc att cga taa gat aga ttc cat cat cga aaa atg tag tcg tga cat aag tct cct caa aga gat tcg aat ctc aga cac cga ctt gta tac ggt att gag aac aga aga cat ccg gta tca cac ata tct cga agc cat aca ttc aga caa acg cat ttc att tcc cat gta cga cga tct cat aga aca gtg tca tct atc gat gga gca taa aag taa act cgt cga caa agc act caa taa att aga gtc tac cat cga tag tca atc tag act atc gta ttt gcc tcc gga aat tat gcg caa tat cat aac caa gct aag cga cta cca tct aaa cag tat gtt gta cgg aaa gaa cca tta caa ata tta tcc atg ata gaa aga aaa tat tta aaa aat aat cta tat gat tgg aga agt agg aaa caa aca gta aca aga cga cga tta cta c-a tta tta aat cat gag gtc cgt att ata ctc gta tat att gtt tct ctc atg tat aat aat aaa cgg aag aga tat agc acc aca tgc acc atc caa tgg aaa gtg taa aga caa cga ata cag aag ccg taa tct atg ttg tct atc gtg tcc tcc ggg aac tta cgc ttc cag att atg tga tag caa gac taa tac aca atg tac acc gtg tgg ttc gga tac ctt tac atc tca caa taa tca ttt aca ggc ttg tct aag ttg taa cgg aag atg tga tag taa tca ggt aga gac gcg atc gtg taa cac gac tca caa tag aat ctg tga atg ctc tcc agg ata tta ttg tct tct caa agg agc atc agg gtg tag aac atg tat ttc taa aac aaa gtg tgg aat agg ata cgg agt atc cgg ata cac gtc tac cgg aga cgt cat ctg ttc tcc gtg tgg tcc cgg aac ata ttc tca cac cgt ctc ttc cac aga taa atg cga acc cgt cgt aac cag caa tac att taa cta tat cga tgt gga aat taa cct gta tcc agt caa cga cac atc gtg tac tcg gac gac cac tac cgg tct cag cga atc cat ctc aac gtc gga act aac tat tac cat gaa tca taa aga ttg tga tcc agt ctt tcg tgc aga ata ctt ctc tgt cct taa taa tgt agc aac ttc agg att ctt tac agg aga aaa tag ata tca gaa tac ttc aaa gat atg tac tct gaa ttt cga gat taa atg taa caa caa aga ttc atc ttc caa aca gtt aac gaa aac aaa gaa tga tac tat cat gcc gca ttc aga gac ggt aac tct agt ggg cga ctg tct atc tag cgt cga cat cta cat act ata tag taa tac caa tac tca aga cta cga aaa tga tac aat ctc tta tca tat ggg taa tgt tct cga tgt caa tag cca tat gcc cgc tag ttg cga tat aca taa act gat cac taa ttc cca gaa tcc cac cca ctt ata gta ag- ttt ttt tta cct ata aat aat aaa tac aat aat taa ttt ctc gta aaa gta gaa aat ata ttc taa ttt att ata tgg taa gaa agt aga atc atc tag aac agt aat caa tca ata gca atc atg aaa caa tat att gtc ctg gca tgc atg tgc cta gtg gca gct gct atg cct act agt ctt caa caa tct tca tcc tcg tgt act gaa gaa gaa aac aaa cat cat atg gga atc gat gtt att atc aaa gtc aca aag caa gac caa aca ccg acc aat gat aag att tgt caa tcc gta acg gaa gtt aca gag a-- --- --- --- --- --- -cc gaa gat gat gag gta tcc gaa gaa gtt gta aaa gga gat ccc acc act tat tac act atc gtc ggt gcg ggt ctt aac atg aac ttt gga ttc acc aaa tgc cca aag att tca tcc atc tcc gaa tcc tct gat gga aac act gtg aat act aga ttg tcc agc gtg tca ccg gga caa ggt aag gac tct ccc gcg atc acg cgt gaa gaa gct ctg gct atg atc aaa gac tgt gag atg tct atc gac atc aga tgt agc gaa gaa gag aaa gac agt gac atc aag acc cat cca gta ctt ggg tct aac atc tca cat aag aaa gtg agt tac aaa gat atc atc ggt tca acg atc gtt gat aca aaa tgt gtc aag aac cta gag ttt agc gta cgt atc gga gac atg tgt gag gaa tca tct gaa ctt gaa gtc aag gat gga ttc aag tat gtc gac gga tcg gca tct gaa ggt gca acc gat gat act tca ctc atc gat tca aca aaa ctc aaa gca tgt gtc tga atc gat aac tct att cat ctg aaa atg gat gag ttg ggt taa tcg aac gat tca gac acc gca cca cga att aaa aaa g-- -ac cgg gca cta tat tcc ggt ttg caa aac aaa aat att taa cta cat tca caa aaa gtt acc tct cgt tac ttc ttc ttt ctg ttt caa tat gtg ata cga tat gat cac tat tcg tat tct ct- --- --- tgg tct cat aaa aaa gtt tta c-- --a aaa aaa aaa aaa a-t att ttt att ctc ttt ctc tct tcg at- ggt ctc aca aaa ata tta aac ctc ttt ctg atg tct caa cta ttt cgt aaa cga taa cgt cca aca ata tat tct cgt aga gct tat caa cat cct tat acc aat cta ggt tgt cag aca att gca tca taa aat aat gtt tat aat tta cac gtt aac atc ata taa taa acg tat ata gtt aat att ttt gga ata taa atg atc tgt aaa atc cat gta ggg gac act gct cac gtt ttt tct cta gta cat aat ttc aca caa gtt ttt ata cag aca aat taa ttc tcg tcc ata ta- ttt taa aac att gac ttt tgt act aag aaa aat atc ttg act aac cat ctc ttt ctc tct tcg --- --- --- --- --- --- --- --- --- --- --- --- --- --- --- --- --- --- --- --- --- --- --- --- --- --- --- --- --- --- --- --- --- --- --- --- --- --- --- --- --- --- --- --- --- --- --- --- --- --- --- --- --- --- --- --- --- --- --- --- --- --- --- --- --- --- --- --- --- --- --- --- --- --- --- --- --- --- --- --- --- --- --- --- --- --- --- --- --- --- --- --- --- --- --- --- --- --- --- --- --- --- --- --- --- --- --- --- --- --- --- --- --- --- --- --- --- --- --- --- --- --- --- --- --- --- --- --- --- --- --- --- --- --- --- --- --- --- --- --- --- --- --- --- --- --- --- --- --- --- --- --- --- --- --- --

>ON585034.1_Monkeypox_virus_isolate_Monkeypox/PT0007/2022_complete_genome

atc gag gac gac ttc aac tat gac agc tcg gtg gcg tct gcc agc gtg tac ata cga atg gca ttt cta aga aaa gtc tac ggt atc ctt -tc tac aat ttc ctt taa caa cgg caa cag ctg cag tat ttt tat act ttg aat gca -tc gga cat tta tac aag gga gtc ctg ttc taa tat tgg cat caa tgt tcg gat cta tag gct tga ttt tcg cat tga ctt tac aca gac atg aac atc ccc tga atc tgt aca tac ttt gtg gat tta cac tgt tag aat ctc taa cgc tgg cct ctg ttg tta ctt tct atg atg cac gta tcg tta tgc aag ctt tca tgt tga cta ctg cag tgt ttc ttg ctc tga cta cat ata ctc tac aat caa aga gag att tca gta aac ttg taa cag gat tgt ttg ctg ctt tct gga ttt taa ttt tgt cag gag tct tga gga taa agt tta aaa tag aat taa taa aga aca tat agg tca -tt ttt taa aca tgg ata gaa acc aag gtt gtt agt taa taa tat aca aga tat ttt ttc tca ctc tga tcc atg taa acc aag gac gag aga cac tct cat tcc tca ttc acg aca cca tta aaa atg gaa att aaa gcc ctc tat taa gca cag acg gct aca ggt cta cca tca ggt t-a cct tcg tct acc ttc aca atg gcc tct cct tgt gcc cag ttc agt ccc tgt cat tgc cac gct act aag gac tcc ctg aat acc gtg act gac gtc aga cat tgt ctg act gaa tac atc ctg tgg gtt tct cat aga tgg acc cat aga gaa agc gca ggg cct ctc tac agg ctt ctc atc tct ttc aga att gat gca atg gag cta ttt ggt agc gag ttg aag gag ttc tcg aat tca ctt ccg tgg gac aat atc gac aat tgc gtg gag atc att aaa tgt ttc atc aga aat gac tcc atg aaa acc gcc aaa gaa ctt tgt gca ata att gga ctt tgt act caa tca gct att gtc act gga aga gtc ttc aat gat aag tat atc gac ata cta ctt atg ctg cga aag att ctg aac gag aac gac tat ctc acc ctc ttg gat cat atc ctc act gct aaa tac taa atc tcc ttc atg ctc tct cac taa tac tct tac tca cta cac ttt tta tca tct tat gat gaa tga ttg cct tca tca ttt ttt cgt gga ata taa tat agg aat aat tag cac cag aat agc tat gga tat ctc gtt aag aat att ctc tat aag aga cat aat gta gac ata gtt att ata tcc ttc tta gat aag tgt tac gct act gga aag ttt cca tcg tta tta tta cat gaa gat gat ata att aaa cca aca ttg aga ttg gct ctt atg tta gct gga ttg aat tac tgt aat aaa tgc atc gag tat aga ggg ata tag caa ttc tcg ata ata gtc atg caa tat ttg aat gag act gat aat tta ggt aat aca gta ct- --- aca cac ata tct ttc tag att ata tat cgt taa aaa tct gta aga tgt ata ttt ctc ata agt atc cac tgt gta ata tta tta atg gat ata tag ata acg caa tag gga cta ata gta ttg taa aag ata taa tcg act att tac gta cat atc cag ata tct ata ttc cta cta gtt tgc tgc gta gtt gca tca ttg ata tgc atg -at tta tca gga ttc aga gat gaa tta cta agt aaa cta caa tcc cac aat aag taa gaa tca aat atc aaa aac tca ctt ttg att ttt cta gtc tta agt aat aca tat att tat taa tag acc tat gaa ata aaa aaa ggt aac aat gga ttc gcg tat agc tat tta cgt att agt atc ggc atc tct ttt gta tct tgt taa ttg tca caa act agt aca tta ctt caa tct gaa aat aaa tgg aag tga tat aac taa tac agc aga tat att gct gga caa tta tcc aat tat gac ctt tga tgg aaa gga tat tta tcc atc tat ctc gtt cat ggt cgg taa taa act ttt cct aga tct tta taa aaa tat ctt tga aga att ttt cag act att tcg agt atc tgt aag tag tca ata cga gga att aga ata tta tta ttc atg tga tta tac taa caa ccg tcc tac aat taa aca aca tta ctt tta taa cgg cga cga ata tac tga aat tga tag atc gaa aaa agc cac taa taa aaa cag ttg gtt aat tac ttc agg ctt tag act aca aaa atg gtt cga tag cga aga ttg tat aat tta tct cag atc ttt agt tag aag aat gga aga cag taa caa aaa cag taa aaa aac tta gta ctt aga tat cg- --a aaa aat ata ttt ttg tag act ctt gag aat aga agg aaa aca tgt aca taa tta taa aaa atg aaa atc aat ggc gaa taa gac agt gcg att cgc acc atg gag tcg gta gat ttc atg gct gtc gat gag cag ttt cac gac gac ctc gat ctt tgg tca tta tct ttg gta gat gat tat aaa aaa cat gga tta ggt gtt gac tgt tat gtt cta gaa cca gtt gtt gac agg aaa ata ttt gat aga ttt ctc ctt gaa cca att tgt gat cct gta gat gtt ctg tat gat tat ttt agg att cat aga gat aat att gat cag tat ata gta gat aga ctg ttt gca tat att aca tat aaa gat att ata tct gca tta gtg tca aag aat tat atg gaa gat att ttc tct ata att att aag aat tgt aat tct gtg caa gat ctc tta ctt tac tat cta tct aat gca tat gta gaa ata gac att gtt gat ctt atg gta gat cat ggg gct gta ata tat aaa ata gaa tgc ttg aat gcc tat ttt agg gga ata tgt aaa aag gaa agt agt gtt gtt gag ttt att ttg aat tgt ggt atc cca gat gaa aat gat gtt aaa tta gat cta tat aaa ata att cag tat act agg gga ttc ctt gta gat gaa ccc aca gta tta gaa att tat aag ctt tgt atc cca tat att gaa gat atc aat caa cta gat gct ggt gga agg acc ttg ctt tat cgc gct atc tat gca ggt tat ata gat tta gta tca tgg cta tta gaa aat gga gca aat gtc aac gca gta atg agt aat gga tat aca tgt ctt gac gtg gcc gtg gat agg gga tct gtc atc gcc cgt agg gaa gca cat ctt aaa ata tta gaa ata ttg ctt aga gaa cca ttg tct att gac tgt ata aaa tta gct ata ctt aat aat aca att gaa aac cat gat gtg ata aag ctc tgt atc aag tat ttt atg atg gta gat tat tca ctt tgt aat gtg tat gca tca tca ctc ttt gat tat ata att gat tgt aaa caa gaa ttg gag tac att agg cag atg aaa att cat aat aca acc atg tat gag tta atc tat aat aga gac aaa aac aag cat gct tcc cat att cta cat agg tat tct aaa cat cca gtt ttg aca cag tgt atc act aaa gga ttc aag att tac aca gaa gta acc gag cag gtc act aaa gct cta aac aga cgt gct cta ata gat gag ata ata aac aat gta tca act gat gac aat ctc cta tca aaa ctt cca tta gaa att agg gat cta att gtt tca caa gct gtc ata tag agt tct atc cac cca cct ttc ttg aaa tga gtt aat --- --- --- --- --- --- --- --- --- --- --- --- --- --- --- --- --- --- --- --- --- --- --- --- --- --- --- --- --- --- --- --- --- --- --- --- --- --- --- --- --- --- --- --- --- --- --- --- --- --- --- --- --- --- --- --- --- --- --- --- --- --- --- --- --- --- --- --- --- --- --- --- --- --- --- --- --- --- --- --- --- --- --- --- --- --- --- --- --- --- --- --- --- --- --- --- --- --- --- --- --- --- --- --- --- -ag tca taa gtt agt taa --- --- --- --- --- -gt cat aag tta gtt aag tca t-- --- --- --- --- --- --- --- --- --- --- --- --- --- --- --- -aa gtt agt taa gtc ata agt tag tta agt cat aag tta gtt aag tca taa gtt agt taa gtc ata agt tag tta agt cat aag tta gtt tat agt cta aca ctt cta att ttt ata cct tga tct ttt tct cta att atg aaa aag taa atc att atg aag atg gat g-a aat gga cga gat tgt gcg cat cgt taa cga tag tat gtg gta cgt acc taa cgc att tat gga cga cgg tga taa tga agg tca cat ttc tgt caa taa tgt ctg tca tat gta tct cgc att ctt tga tgt gga tat atc atc tca tct gtt taa att agt tat taa aca ctg cga tct gaa taa acg act aaa atg tgg taa ctc tcc att aca ttg cta tac gat gaa tac acg att taa tcc atc tgt att aaa gat att gtt acg cca cgg cat gcg taa ctt tga tag caa gga taa aaa agg aca tat tcc tct aca cca cta tct gat tca ttc act atc aat cga taa caa gat ctt tga tat act aac gga ccc cat tga tga ctt tag taa atc atc cga tct att gct gtg tta tct tag ata taa att caa tgg gag ctt aaa cta tta cgt tct gta caa att att gac taa agg atc tga ccc taa ttg cgt cga tga gga tgg act cac ttc tct tca tta cta ctg taa aca cat atc cgc gtt cca cga aag caa tta tta caa gtc aaa gag tca cac taa gat gcg agc tga gaa gcg att cat cta cgc gat aat aga tca tgg agc aaa cat taa cgc ggt tac gaa aat cgg aaa tac gcc gtt aca cac tta cct tca aca gta tac caa aca tag tcc tcg tgt ggt gta tgc tct ttt atc tcg agg agc cga tac gag gat acg taa taa tct taa ttg tac acc cat cat gga ata cat aaa gaa cga ttg tgc aac agg tca tat tct cat aat gtt act caa ttg gca cga aca aaa ata cgg gaa att aca aaa gga aga agg aca aca tct act tta tct att cat aaa aca taa tca agg ata tgg aag tcg ctc tct caa tat act acg gta tct act aga tag att cga cat tca gaa aga cga ata cta taa tac aat gac tcc tct tca tac cgc ctt cca gaa ttg caa taa caa tgt tgc ctc ata cct cgt ata cat tgg ata cga cat caa cct tcc gac taa aga cga taa gac agt att cga ctt ggt gtt tga aaa cag aaa cat tat ata caa ggc gga tgt cgt taa tga cat tat cca cca cag act gaa agt atc tct acc tat gat taa atc gtt gtt cta caa gat gtc gga gtt ctc tcc cta cga cga tca cta cgt aaa gaa gat aat agc cta ctg cct att aag gga cga gtc att tgc gga act aca tac taa att ctg ttt aaa cga gga cta taa aag tgt att tat gaa aaa tat atc att cga taa gat aga ttc cat cat cga aaa atg tag tcg tga cat aag tct cct caa aga gat tcg aat ctc aga cac cga ctt gta tac ggt att gag aac aga aga cat tcg gta tca cac ata tct cga agc cat aca ttc aga caa acg cat ttc att tcc cat gta cga cga tct cat aga aca gtg tca tct atc gat gga gca taa aag taa act cgt cga caa agc act caa taa att aga gtc tac cat cga tag tca atc tag act atc gta ttt gcc tcc gga aat tat gcg caa tat cat aac caa gct aag cga cta cca tct aaa cag tat gtt gta cgg aaa gaa cca tta caa ata tta tcc atg ata gaa aga aaa tat tta aaa aat aat cta tat gat tgg aga agt agg aaa caa aca gta aca aga cga cga tta cta c-a tta tta aat cat gag gtc cgt att ata ctc gta tat att gtt tct ctc atg tat aat aat aaa cgg aag aga tat agc acc aca tgc acc atc caa tgg aaa gtg taa aga caa cga ata cag aag ccg taa tct atg ttg tct atc gtg tcc tcc ggg aac tta cgc ttt cag att atg tga tag caa gac taa tac aca atg tac acc gtg tgg ttc gga tac ctt tac atc tca caa taa tca ttt aca ggc ttg tct aag ttg taa cgg aag atg tga tag taa tca ggt aga gac gcg atc gtg taa cac gac tca caa tag aat ctg tga atg ctc tcc agg ata tta ttg tct tct caa agg agc atc agg gtg tag aac atg tat ttc taa aac aaa gtg tgg aat agg ata cgg agt atc cgg ata cac gtc tac cgg aga cgt cat ctg ttc tcc gtg tgg tcc cgg aac ata ttc tca cac cgt ctc ttc cac aga taa atg cga acc cgt cgt aac cag caa tac att taa cta tat cga tgt gga aat taa cct gta tcc agt caa cga cac atc gtg tac tcg gac gac cac tac cgg tct cag cga atc cat ctc aac gtc gga act aac tat tac cat gaa tca taa aga ttg tga tcc agt ctt tcg tgc aga ata ctt ctc tgt cct taa taa tgt agc aac ttc agg att ctt tac agg aga aaa tag ata tca gaa tac ttc aaa gat atg tac tct gaa ttt cga gat taa atg taa caa caa aga ttc atc ttc caa aca gtt aac gaa aac aaa gaa tga tac tat cat gcc gca ttc aga gac ggt aac tct agt ggg cga ctg tct atc tag cgt cga cat cta cat act ata tag taa tac caa tac tca aga cta cga aaa tga tac aat ctc tta tca tat ggg taa tgt tct cga tgt caa tag cca tat gcc cgc tag ttg cga tat aca taa act gat cac taa ttc cca gaa tcc cac cca ctt ata gta ag- -tt ttt tta cct ata aat aat aaa tac aat aat taa ttt ctc gta aaa gta gaa aat ata ttc taa ttt att ata tgg taa gaa agt aga atc atc tag aac agt aat caa tca ata gca atc atg aaa caa tat att gtc ctg gca tgc atg tgc cta gtg gca gct gct atg cct act agt ctt caa caa tct tca tcc tcg tgt act gaa gaa gaa aac aaa cat cat atg gga atc gat gtt att atc aaa gtc aca aag caa gac caa aca ccg acc aat gat aag att tgt caa tcc gta acg gaa gtt aca gag a-- --- --- --- --- --- -cc gaa gat gat gag gta tcc gaa gaa gtt gta aaa gga gat ccc acc act tat tac act atc gtc ggt gcg ggt ctt aac atg aac ttt gga ttc acc aaa tgc cca aag att tta tcc atc tcc gaa tcc tct gat gga aac act gtg aat act aga ttg tcc agc gtg tca ccg gga caa ggt aag gac tct ccc gcg atc acg cgt gaa gaa gct ctg gct atg atc aaa gac tgt gag atg tct atc gac atc aga tgt agc gaa gaa gag aaa gac agt gac atc aag acc cat cca gta ctt ggg tct aac atc tca cat aag aaa gtg agt tac aaa gat atc atc ggt tca acg atc gtt gat aca aaa tgt gtc aag aac cta gag ttt agc gta cgt atc gga gac atg tgt gag gaa tca tct gaa ctt gaa gtc aag gat gga ttc aag tat gtc gac gga tcg gca tct gaa ggt gca acc gat gat act tca ctc atc gat tca aca aaa ctc aaa gca tgt gtc tga atc gat aac tct att cat ctg aaa atg gat gag ttg ggt taa tcg aac gat tca gac acc gca cca cga att aaa aaa g-- -ac cgg gca cta tat tcc ggt ttg caa aac aaa aat att taa cta cat tca caa aaa gtt acc tct cgt tac ttc ttc ttt ctg ttt caa tat gtg ata cga tat gat cac tat tcg tat tct ct- --- --- tgg tct cat aaa aaa gtt tta c-- --a aaa aaa aaa aaa a-t att ttt att ctc ttt ctc tct tcg at- ggt ctc aca aaa ata tta aac ctc ttt ctg atg tct caa cta ttt cgt aaa cga taa cgt cca aca ata tat tct cgt aga gct tat caa cat cct tat acc aat cta ggt tgt cag aca att gca tca taa aat aat gtt tat aat tta cac gtt aac atc ata taa taa acg tat ata gtt aat att ttt gga ata taa atg atc tgt aaa atc cat gta ggg gac act gct cac gtt ttt tct cta gta cat aat ttc aca caa gtt ttt ata cag aca aat taa ttc tcg tcc ata ta- ttt taa aac att gac ttt tgt act aag aaa aat atc ttg act aac cat ctc ttt ctc tct tcg atg ggt ctc aca aaa ata tta aac ctc ttt ctg atg g-- --- --- --- --- --- --- --- --- --- --- --- --- --- --- --- --- --- --- --- --- --- --- --- --- --- --- --- --- --- --- --- --- --- --- --- --- --- --- --- --- --- --- --- --- --- --- agt cgt aaa aag ttt tta tcc ttt ctc tct tcg ata ggt ctc --- --- --- --- --- --- --- --- --- --- --- --- --- --- --- --- --- --- --- --- --- --- --- --- --- --- --- --- --- --- --- --- --- --- --- --- --- --- --- --- --- --- --- --- --- --- --- --- --- --- --- --- --- --- --- --- --- --- --- --- --- --- --- --- --- --- --- --- --- --- aca aaa ata tta aac ctc ttt ctg atg gtc tct ata aa

>ON649713.1_Monkeypox_virus_isolate_Monkeypox/PT0020/2022_complete_genome

atc gag gac gac ttc aac tat gac agc tcg gtg gcg tct gcc agc gtg tac ata cga atg gca ttt cta aga aaa gtc tac ggt atc ctt -tc tac aat ttc ctt taa caa cgg caa cag ctg cag tat ttt tat act ttg aat gca -tc gga cat tta tac aag gga gtc ctg ttc taa tat tgg cat caa tgt tcg gat cta tag gct tga ttt tcg cat tga ctt tac aca gac atg aac atc ccc tga atc tgt aca tac ttt gtg gat tta cac tgt tag aat ctc taa cgc tgg cct ctg ttg tta ctt tct atg atg cac gta tcg tta tgc aag ctt tca tgt tga cta ctg cag tgt ttc ttg ctc tga cta cat ata ctc tac aat caa aga gag att tca gta aac ttg taa cag gat tgt ttg ctg ctt tct gga ttt taa ttt tgt cag gag tct tga gga taa agt tta aaa tag aat taa taa aga aca tat agg tca -tt ttt taa aca tgg ata gaa acc aag gtt gtt agt taa taa tat aca aga tat ttt ttc tca ctc tga tcc atg taa acc aag gac gag aga cac tct cat tcc tca ttc acg aca cca tta aaa atg gaa att aaa gcc ctc tat taa gca cag acg gct aca ggt cta cca tca ggt t-a cct tcg tct acc ttc aca atg gcc tct cct tgt gcc cag ttc agt ccc tgt cat tgc cac gct act aag gac tcc ctg aat acc gtg act gac gtc aga cat tgt ctg act gaa tac atc ctg tgg gtt tct cat aga tgg acc cat aga gaa agc gca ggg cct ctc tac agg ctt ctc atc tct ttc aga att gat gca atg gag cta ttt ggt agc gag ttg aag gag ttc tcg aat tca ctt ccg tgg gac aat atc gac aat tgc gtg gag atc att aaa tgt ttc atc aga aat gac tcc atg aaa acc gcc aaa gaa ctt tgt gca ata att gga ctt tgt act caa tca gct att gtc act gga aga gtc ttc aat gat aag tat atc gac ata cta ctt atg ctg cga aag att ctg aac gag aac gac tat ctc acc ctc ttg gat cat atc ctc act gct aaa tac taa atc tcc ttc atg ctc tct cac taa tac tct tac tca cta cac ttt tta tca tct tat gat gaa tga ttg cct tca tca ttt ttt cgt gga ata taa tat agg aat aat tag cac cag aat agc tat gga tat ctc gtt aag aat att ctc tat aag aga cat aat gta gac ata gtt att ata tcc ttc tta gat aag tgt tac gct act gga aag ttt cca tcg tta tta tta cat gaa gat gat ata att aaa cca aca ttg aga ttg gct ctt atg tta gct gga ttg aat tac tgt aat aaa tgc atc gag tat aga ggg ata tag caa ttc tcg ata ata gtc atg caa tat ttg aat gag act gat aat tta ggt aat aca gta ct- --- aca cac ata tct ttc tag att ata tat cgt taa aaa tct gta aga tgt ata ttt ctc ata agt atc cac tgt gta ata tta tta atg gat ata tag ata acg caa tag gga cta ata gta ttg taa aag ata taa tcg act att tac gta cat atc cag ata tct ata ttc cta cta gtt tgc tgc gta gtt gca tca ttg ata tgc atg -at tta tca gga ttc aga gat gaa tta cta agt aaa cta caa tcc cac aat aag taa gaa tca aat atc aaa aac tca ctt ttg att ttt cta gtc tta agt aat aca tat att tat taa tag acc tat gaa ata aaa aaa ggt aac aat gga ttc gcg tat agc tat tta cgt att agt atc ggc atc tct ttt gta tct tgt taa ttg tca caa act agt aca tta ctt caa tct gaa aat aaa tgg aag tga tat aac taa tac agc aga tat att gct gga caa tta tcc aat tat gac ctt tga tgg aaa gga tat tta tcc atc tat ctc gtt cat ggt cgg taa taa act ttt cct aga tct tta taa aaa tat ctt tga aga att ttt cag act att tcg agt atc tgt aag tag tca ata cga gga att aga ata tta tta ttc atg tga tta tac taa caa ccg tcc tac aat taa aca aca tta ctt tta taa cgg cga cga ata tac tga aat tga tag atc gaa aaa agc cac taa taa aaa cag ttg gtt aat tac ttc agg ctt tag act aca aaa atg gtt cga tag cga aga ttg tat aat tta tct cag atc ttt agt tag aag aat gga aga cag taa caa aaa cag taa aaa aac tta gta ctt aga tat cg- --a aaa aat ata ttt ttg tag act ctt gag aat aga agg aaa aca tgt aca taa tta taa aaa atg aaa atc aat ggc gaa taa gac agt gcg att cgc acc atg gag tcg gta gat ttc atg gct gtc gat gag cag ttt cac gac gac ctc gat ctt tgg tca tta tct ttg gta gat gat tat aaa aaa cat gga tta ggt gtt gac tgt tat gtt cta gaa cca gtt gtt gac agg aaa ata ttt gat aga ttt ctc ctt gaa cca att tgt gat cct gta gat gtt ctg tat gat tat ttt agg att cat aga gat aat att gat cag tat ata gta g?? ??? ?tg ttt gca tat att aca tat aaa gat att ata tct gca tta gtg tca aag aat tat atg gaa gat att ttc tct ata att att aag aat tgt aat tct gtg caa gat ctc tta ctt tac tat cta tct aat gca tat gta gaa ata gac att gtt gat ctt atg gta gat cat ggg gct gta ata tat aaa ata gaa tgc ttg aat gcc tat ttt agg gga ata tgt aaa aag gaa agt agt gtt gtt gag ttt att ttg aat tgt ggt atc cca gat gaa aat gat gtt aaa tta gat cta tat aaa ata att cag tat act agg gga ttc ctt gta gat gaa ccc aca gta tta gaa att tat aag ctt tgt atc cca tat att gaa gat atc aat caa cta gat gct ggt gga agg acc ttg ctt tat cgc gct atc tat gca ggt tat ata gat tta gta tca tgg cta tta gaa aat gga gca aat gtc aac gca gta atg agt aat gga tat aca tgt ctt gac gtg gcc gtg gat agg gga tct gtc atc gcc cgt agg gaa gca cat ctt aaa ata tta gaa ata ttg ctt aga gaa cca ttg tct att gac tgt ata aaa tta gct ata ctt aat aat aca att gaa aac cat gat gtg ata aag ctc tgt atc aag tat ttt atg atg gta gat tat tca ctt tgt aat gtg tat gca tca tca ctc ttt gat tat ata att gat tgt aaa caa gaa ttg gag tac att agg cag atg aaa att cat aat aca acc atg tat gag tta atc tat aat aga gac aaa aac aag cat gct tcc cat att cta cat agg tat tct aaa cat cca gtt ttg aca cag tgt atc act aaa gga ttc aag att tac aca gaa gta acc gag cag gtc act aaa gct cta aac aga cgt gct cta ata gat gag ata ata aac aat gta tca act gat gac aat ctc cta tca aaa ctt cca tta gaa att agg gat cta att gtt tca caa gct gtc ata tag agt tct atc cac cca cct ttc ttg aaa tga gtt aat --- --- --- --- --- --- --- --- --- --- --- --- --- --- --- --- --- --- --- --- --- --- --- --- --- --- --- --- --- --- --- --- --- --- --- --- --- --- --- --- --- --- --- --- --- --- --- --- --- --- --- --- --- --- --- --- --- --- --- --- --- --- --- --- --- --- --- --- --- --- --- --- --- --- --- --- --- --- --- --- --- --- --- --- --- --- --- --- --- --- --- --- --- --- --- --- --- --- --- --- --- --- --- --- --- -ag tca taa gtt agt taa --- --- --- --- --- -gt cat aag tta gtt aag tca t-- --- --- --- --- --- --- --- --- --- --- --- --- --- --- --- -aa gtt agt taa gtc ata agt tag tta agt cat aag tta gtt aag tca taa gtt agt taa gtc ata agt tag tta agt cat aag tta gtt tat agt cta aca ctt cta att ttt ata cct tga tct ttt tct cta att atg aaa aag taa atc att atg aag atg gat g-a aat gga cga gat tgt gcg cat cgt taa cga tag tat gtg gta cgt acc taa cgc att tat gga cga cgg tga taa tga agg tca cat ttc tgt caa taa tgt ctg tca tat gta tct cgc att ctt tga tgt gga tat atc atc tca tct gtt taa att agt tat taa aca ctg cga tct gaa taa acg act aaa atg tgg taa ctc tcc att aca ttg cta tac gat gaa tac acg att taa tcc atc tgt att aaa gat att gtt acg cca cgg cat gcg taa ctt tga tag caa gga taa aaa agg aca tat tcc tct aca cca cta tct gat tca ttc act atc aat cga taa caa gat ctt tga tat act aac gga ccc cat tga tga ctt tag taa atc atc cga tct att gct gtg tta tct tag ata taa att caa tgg gag ctt aaa cta tta cgt tct gta caa att att gac taa agg atc tga ccc taa ttg cgt cga tga gga tgg act cac ttc tct tca tta cta ctg taa aca cat atc cgc gtt cca cga aag caa tta tta caa gtc aaa gag tca cac taa gat gcg agc tga gaa gcg att cat cta cgc gat aat aga tca tgg agc aaa cat taa cgc ggt tac gaa aat cgg aaa tac gcc gtt aca cac tta cct tca aca gta tac caa aca tag tcc tcg tgt ggt gta tgc tct ttt atc tcg agg agc cga tac gag gat acg taa taa tct taa ttg tac acc cat cat gga ata cat aaa gaa cga ttg tgc aac agg tca tat tct cat aat gtt act caa ttg gca cga aca aaa ata cgg gaa att aca aaa gga aga agg aca aca tct act tta tct att cat aaa aca taa tca agg ata tgg aag tcg ctc tct caa tat act acg gta tct act aga tag att cga cat tca gaa aga cga ata cta taa tac aat gac tcc tct tca tac cgc ctt cca gaa ttg caa taa caa tgt tgc ctc ata cct cgt ata cat tgg ata cga cat caa cct tcc gac taa aga cga taa gac agt att cga ctt ggt gtt tga aaa cag aaa cat tat ata caa ggc gga tgt cgt taa tga cat tat cca cca cag act gaa agt atc tct acc tat gat taa atc gtt gtt cta caa gat gtc gga gtt ctc tcc cta cga cga tca cta cgt aaa gaa gat aat agc cta ctg cct att aag gga cga gtc att tgc gga act aca tac taa att ctg ttt aaa cga gga cta taa aag tgt att tat gaa aaa tat atc att cga taa gat aga ttc cat cat cga aaa atg tag tcg tga cat aag tct cct caa aga gat tcg aat ctc aga cac cga ctt gta tac ggt att gag aac aga aga cat tcg gta tca cac ata tct cga agc cat aca ttc aga caa acg cat ttc att tcc cat gta cga cga tct cat aga aca gtg tca tct atc gat gga gca taa aag taa act cgt cga caa agc act caa taa att aga gtc tac cat cga tag tca atc tag act atc gta ttt gcc tcc gga aat tat gcg caa tat cat aac caa gct aag cga cta cca tct aaa cag tat gtt gta cgg aaa gaa cca tta caa ata tta tcc atg ata gaa aga aaa tat tta aaa aat aat cta tat gat tgg aga agt agg aaa caa aca gta aca aga cga cga tta cta c-a tta tta aat cat gag gtc cgt att ata ctc gta tat att gtt tct ctc atg tat aat aat aaa cgg aag aga tat agc acc aca tgc acc atc caa tgg aaa gtg taa aga caa cga ata cag aag ccg taa tct atg ttg tct atc gtg tcc tcc ggg aac tta cgc ttt cag att atg tga tag caa gac taa tac aca atg tac acc gtg tgg ttc gga tac ctt tac atc tca caa taa tca ttt aca ggc ttg tct aag ttg taa cgg aag atg tga tag taa tca ggt aga gac gcg atc gtg taa cac gac tca caa tag aat ctg tga atg ctc tcc agg ata tta ttg tct tct caa agg agc atc agg gtg tag aac atg tat ttc taa aac aaa gtg tgg aat agg ata cgg agt atc cgg ata cac gtc tac cgg aga cgt cat ctg ttc tcc gtg tgg tcc cgg aac ata ttc tca cac cgt ctc ttc cac aga taa atg cga acc cgt cgt aac cag caa tac att taa cta tat cga tgt gga aat taa cct gta tcc agt caa cga cac atc gtg tac tcg gac gac cac tac cgg tct cag cga atc cat ctc aac gtc gga act aac tat tac cat gaa tca taa aga ttg tga tcc agt ctt tcg tgc aga ata ctt ctc tgt cct taa taa tgt agc aac ttc agg att ctt tac agg aga aaa tag ata tca gaa tac ttc aaa gat atg tac tct gaa ttt cga gat taa atg taa caa caa aga ttc atc ttc caa aca gtt aac gaa aac aaa gaa tga tac tat cat gcc gca ttc aga gac ggt aac tct agt ggg cga ctg tct atc tag cgt cga cat cta cat act ata tag taa tac caa tac tca aga cta cga aaa tga tac aat ctc tta tca tat ggg taa tgt tct cga tgt caa tag cca tat gcc cgc tag ttg cga tat aca taa act gat cac taa ttc cca gaa tcc cac cca ctt ata gta ag- -tt ttt tta cct ata aat aat aaa tac aat aat taa ttt ctc gta aaa gta gaa aat ata ttc taa ttt att ata tgg taa gaa agt aga atc atc tag aac agt aat caa tca ata gca atc atg aaa caa tat att gtc ctg gca tgc atg tgc cta gtg gca gct gct atg cct act agt ctt caa caa tct tca tcc tcg tgt act gaa gaa gaa aac aaa cat cat atg gga atc gat gtt att atc aaa gtc aca aag caa gac caa aca ccg acc aat gat aag att tgt caa tcc gta acg gaa gtt aca gag a-- --- --- --- --- --- -cc gaa gat gat gag gta tcc gaa gaa gtt gta aaa gga gat ccc acc act tat tac act atc gtc ggt gcg ggt ctt aac atg aac ttt gga ttc acc aaa tgc cca aag att tta tcc atc tcc gaa tcc tct gat gga aac act gtg aat act aga ttg tcc agc gtg tca ccg gga caa ggt aag gac tct ccc gcg atc acg cgt gaa gaa gct ctg gct atg atc aaa gac tgt gag atg tct atc gac atc aga tgt agc gaa gaa gag aaa gac agt gac atc aag acc cat cca gta ctt ggg tct aac atc tca cat aag aaa gtg agt tac aaa gat atc atc ggt tca acg atc gtt gat aca aaa tgt gtc aag aac cta gag ttt agc gta cgt atc gga gac atg tgt gag gaa tca tct gaa ctt gaa gtc aag gat gga ttc aag tat gtc gac gga tcg gca tct gaa ggt gca acc gat gat act tca ctc atc gat tca aca aaa ctc aaa gca tgt gtc tga atc gat aac tct att cat ctg aaa atg gat gag ttg ggt taa tcg aac gat tca gac acc gca cca cga att aaa aaa g-- -ac cgg gca cta tat tcc ggt ttg caa aac aaa aat att taa cta cat tca caa aaa gtt acc tct cgt tac ttc ttc ttt ctg ttt caa tat gtg ata cga tat gat cac tat tcg tat tct ct- --- --- tgg tct cat aaa aaa gtt tta c-- --a aaa aaa aaa aaa a-t att ttt att ctc ttt ctc tct tcg at- ggt ctc aca aaa ata tta aac ctc ttt ctg atg tct caa cta ttt cgt aaa cga taa cgt cca aca ata tat tct cgt aga gct tat caa cat cct tat acc aat cta ggt tgt cag aca att gca tca taa aat aat gtt tat aat tta cac gtt aac atc ata taa taa acg tat ata gtt aat att ttt gga ata taa atg atc tgt aaa atc cat gta ggg gac act gct cac gtt ttt tct cta gta cat aat ttc aca caa gtt ttt ata cag aca aat taa ttc tcg tcc ata ta- ttt taa aac att gac ttt tgt act aag aaa aat atc ttg act aac cat ctc ttt ctc tct tcg atg ggt ctc aca aaa ata tta aac ctc ttt ctg atg g-- --- --- --- --- --- --- --- --- --- --- --- --- --- --- --- --- --- --- --- --- --- --- --- --- --- --- --- --- --- --- --- --- --- --- --- --- --- --- --- --- --- --- --- --- --- --- agt cgt aaa aag ttt tta tcc ttt ctc tct tcg ata ggt ctc --- --- --- --- --- --- --- --- --- --- --- --- --- --- --- --- --- --- --- --- --- --- --- --- --- --- --- --- --- --- --- --- --- --- --- --- --- --- --- --- --- --- --- --- --- --- --- --- --- --- --- --- --- --- --- --- --- --- --- --- --- --- --- --- --- --- --- --- --- --- aca aaa ata tta aac ctc ttt ctg atg gtc tct ata aa

>ON585031.1_Monkeypox_virus_isolate_Monkeypox/PT0003/2022_complete_genome

atc gag gac gac ttc aac tat gac agc tcg gtg gcg tct gcc agc gtg tac ata cga atg gca ttt cta aga aaa gtc tac ggt atc ctt -tc tac aat ttc ctt taa caa cgg caa cag ctg cag tat ttt tat act ttg aat gca -tc gga cat tta tac aag gga gtc ctg ttc taa tat tgg cat caa tgt tcg gat cta tag gct tga ttt tcg cat tga ctt tac aca gac atg aac atc ccc tga atc tgt aca tac ttt gtg gat tta cac tgt tag aat ctc taa cgc tgg cct ctg ttg tta ctt tct atg atg cac gta tcg tta tgc aag ctt tca tgt tga cta ctg cag tgt ttc ttg ctc tga cta cat ata ctc tac aat caa aga gag att tca gta aac ttg taa cag gat tgt ttg ctg ctt tct gga ttt taa ttt tgt cag gag tct tga gga taa agt tta aaa tag aat taa taa aga aca tat agg tca -tt ttt taa aca tgg ata gaa acc aag gtt gtt agt taa taa tat aca aga tat ttt ttc tca ctc tga tcc atg taa acc aag gac gag aga cac tct cat tcc tca ttc acg aca cca tta aaa atg gaa att aaa gcc ctc tat taa gca cag acg gct aca ggt cta cca tca ggt t-a cct tcg tct acc ttc aca atg gcc tct cct tgt gcc cag ttc agt ccc tgt cat tgc cac gct act aag gac tcc ctg aat acc gtg act gac gtc aga cat tgt ctg act gaa tac atc ctg tgg gtt tct cat aga tgg acc cat aga gaa agc gca ggg cct ctc tac agg ctt ctc atc tct ttc aga att gat gca atg gag cta ttt ggt agc gag ttg aag gag ttc tcg aat tca ctt ccg tgg gac aat atc gac aat tgc gtg gag atc att aaa tgt ttc atc aga aat gac tcc atg aaa acc gcc aaa gaa ctt tgt gca ata att gga ctt tgt act caa tca gct att gtc act gga aga gtc ttc aat gat aag tat atc gac ata cta ctt atg ctg cga aag att ctg aac gag aac gac tat ctc acc ctc ttg gat cat atc ctc act gct aaa tac taa atc tcc ttc atg ctc tct cac taa tac tct tac tca cta cac ttt tta tca tct tat gat gaa tga ttg cct tca tca ttt ttt cgt gga ata taa tat agg aat aat tag cac cag aat agc tat gga tat ctc gtt aag aat att ctc tat aag aga cat aat gta gac ata gtt att ata tcc ttc tta gat aag tgt tac gct act gga aag ttt cca tcg tta tta tta cat gaa gat gat ata att aaa cca aca ttg aga ttg gct ctt atg tta gct gga ttg aat tac tgt aat aaa tgc atc gag tat aga ggg ata tag caa ttc tcg ata ata gtc atg caa tat ttg aat gag act gat aat tta ggt aat aca gta ct- --- aca cac ata tct ttc tag att ata tat cgt taa aaa tct gta aga tgt ata ttt ctc ata agt atc cac tgt gta ata tta tta atg gat ata tag ata acg caa tag gga cta ata gta ttg taa aag ata taa tcg act att tac gta cat atc cag ata tct ata ttc cta cta gtt tgc tgc gta gtt gca tca ttg ata tgc atg -at tta tca gga ttc aga gat gaa tta cta agt aaa cta caa tcc cac aat aag taa gaa tca aat atc aaa aac tca ctt ttg att ttt cta gtc tta agt aat aca tat att tat taa tag acc tat gaa ata aaa aaa ggt aac aat gga ttc gcg tat agc tat tta cgt att agt atc ggc atc tct ttt gta tct tgt taa ttg tca caa act agt aca tta ctt caa tct gaa aat aaa tgg aag tga tat aac taa tac agc aga tat att gct gga caa tta tcc aat tat gac ctt tga tgg aaa gga tat tta tcc atc tat ctc gtt cat ggt cgg taa taa act ttt cct aga tct tta taa aaa tat ctt tga aga att ttt cag act att tcg agt atc tgt aag tag tca ata cga gga att aga ata tta tta ttc atg tga tta tac taa caa ccg tcc tac aat taa aca aca tta ctt tta taa cgg cga cga ata tac tga aat tga tag atc gaa aaa agc cac taa taa aaa cag ttg gtt aat tac ttc agg ctt tag act aca aaa atg gtt cga tag cga aga ttg tat aat tta tct cag atc ttt agt tag aag aat gga aga cag taa caa aaa cag taa aaa aac tta gta ctt aga tat cg- --a aaa aat ata ttt ttg tag act ctt gag aat aga agg aaa aca tgt aca taa tta taa aaa atg aaa atc aat ggc gaa taa gac agt gcg att cgc acc atg gag tcg gta gat ttc atg gct gtc gat gag cag ttt cac gac gac ctc gat ctt tgg tca tta tct ttg gta gat gat tat aaa aaa cat gga tta ggt gtt gac tgt tat gtt cta gaa cca gtt gtt gac agg aaa ata ttt gat aga ttt ctc ctt gaa cca att tgt gat cct gta gat gtt ctg tat gat tat ttt agg att cat aga gat aat att gat cag tat ata gta gat aga ctg ttt gca tat att aca tat aaa gat att ata tct gca tta gtg tca aag aat tat atg gaa gat att ttc tct ata att att aag aat tgt aat tct gtg caa gat ctc tta ctt tac tat cta tct aat gca tat gta gaa ata gac att gtt gat ctt atg gta gat cat ggg gct gta ata tat aaa ata gaa tgc ttg aat gcc tat ttt agg gga ata tgt aaa aag gaa agt agt gtt gtt gag ttt att ttg aat tgt ggt atc cca gat gaa aat gat gtt aaa tta gat cta tat aaa ata att cag tat act agg gga ttc ctt gta gat gaa ccc aca gta tta gaa att tat aag ctt tgt atc cca tat att gaa gat atc aat caa cta gat gct ggt gga agg acc ttg ctt tat cgc gct atc tat gca ggt tat ata gat tta gta tca tgg cta tta gaa aat gga gca aat gtc aac gca gta atg agt aat gga tat aca tgt ctt gac gtg gcc gtg gat agg gga tct gtc atc gcc cgt agg gaa gca cat ctt aaa ata tta gaa ata ttg ctt aga gaa cca ttg tct att gac tgt ata aaa tta gct ata ctt aat aat aca att gaa aac cat gat gtg ata aag ctc tgt atc aag tat ttt atg atg gta gat tat tca ctt tgt aat gtg tat gca tca tca ctc ttt gat tat ata att gat tgt aaa caa gaa ttg gag tac att agg cag atg aaa att cat aat aca acc atg tat gag tta atc tat aat aga gac aaa aac aag cat gct tcc cat att cta cat agg tat tct aaa cat cca gtt ttg aca cag tgt atc act aaa gga ttc aag att tac aca gaa gta acc gag cag gtc act aaa gct cta aac aga cgt gct cta ata gat gag ata ata aac aat gta tca act gat gac aat ctc cta tca aaa ctt cca tta gaa att agg gat cta att gtt tca caa gct gtc ata tag agt tct atc cac cca cct ttc ttg aaa tga gtt aat --- --- --- --- --- --- --- --- --- --- --- --- --- --- --- --- --- --- --- --- --- --- --- --- --- --- --- --- --- --- --- --- --- --- --- --- --- --- --- --- --- --- --- --- --- --- --- --- --- --- --- --- --- --- --- --- --- --- --- --- --- --- --- --- --- --- --- --- --- --- --- --- --- --- --- --- --- --- --- --- --- --- --- --- --- --- --- --- --- --- --- --- --- --- --- --- --- --- --- --- --- --- --- --- --- -ag tca taa gtt agt taa --- --- --- --- --- -gt cat aag tta gtt aag tca t-- --- --- --- --- --- --- --- --- --- --- --- --- --- --- --- -aa gtt agt taa gtc ata agt tag tta agt cat aag tta gtt aag tca taa gtt agt taa gtc ata agt tag tta agt cat aag tta gtt tat agt cta aca ctt cta att ttt ata cct tga tct ttt tct cta att atg aaa aag taa atc att atg aag atg gat g-a aat gga cga gat tgt gcg cat cgt taa cga tag tat gtg gta cgt acc taa cgc att tat gga cga cgg tga taa tga agg tca cat ttc tgt caa taa tgt ctg tca tat gta tct cgc att ctt tga tgt gga tat atc atc tca tct gtt taa att agt tat taa aca ctg cga tct gaa taa acg act aaa atg tgg taa ctc tcc att aca ttg cta tac gat gaa tac acg att taa tcc atc tgt att aaa gat att gtt acg cca cgg cat gcg taa ctt tga tag caa gga taa aaa agg aca tat tcc tct aca cca cta tct gat tca ttc act atc aat cga taa caa gat ctt tga tat act aac gga ccc cat tga tga ctt tag taa atc atc cga tct att gct gtg tta tct tag ata taa att caa tgg gag ctt aaa cta tta cgt tct gta caa att att gac taa agg atc tga ccc taa ttg cgt cga tga gga tgg act cac ttc tct tca tta cta ctg taa aca cat atc cgc gtt cca cga aag caa tta tta caa gtc aaa gag tca cac taa gat gcg agc tga gaa gcg att cat cta cgc gat aat aga tca tgg agc aaa cat taa cgc ggt tac gaa aat cgg aaa tac gcc gtt aca cac tta cct tca aca gta tac caa aca tag tcc tcg tgt ggt gta tgc tct ttt atc tcg agg agc cga tac gag gat acg taa taa tct taa ttg tac acc cat cat gga ata cat aaa gaa cga ttg tgc aac agg tca tat tct cat aat gtt act caa ttg gca cga aca aaa ata cgg gaa att aca aaa gga aga agg aca aca tct act tta tct att cat aaa aca taa tca agg ata tgg aag tcg ctc tct caa tat act acg gta tct act aga tag att cga cat tca gaa aga cga ata cta taa tac aat gac tcc tct tca tac cgc ctt cca gaa ttg caa taa caa tgt tgc ctc ata cct cgt ata cat tgg ata cga cat caa cct tcc gac taa aga cga taa gac agt att cga ctt ggt gtt tga aaa cag aaa cat tat ata caa ggc gga tgt cgt taa tga cat tat cca cca cag act gaa agt atc tct acc tat gat taa atc gtt gtt cta caa gat gtc gga gtt ctc tcc cta cga cga tca cta cgt aaa gaa gat aat agc cta ctg cct att aag gga cga gtc att tgc gga act aca tac taa att ctg ttt aaa cga gga cta taa aag tgt att tat gaa aaa tat atc att cga taa gat aga ttc cat cat cga aaa atg tag tcg tga cat aag tct cct caa aga gat tcg aat ctc aga cac cga ctt gta tac ggt att gag aac aga aga cat tcg gta tca cac ata tct cga agc cat aca ttc aga caa acg cat ttc att tcc cat gta cga cga tct cat aga aca gtg tca tct atc gat gga gca taa aag taa act cgt cga caa agc act caa taa att aga gtc tac cat cga tag tca atc tag act atc gta ttt gcc tcc gga aat tat gcg caa tat cat aac caa gct aag cga cta cca tct aaa cag tat gtt gta cgg aaa gaa cca tta caa ata tta tcc atg ata gaa aga aaa tat tta aaa aat aat cta tat gat tgg aga agt agg aaa caa aca gta aca aga cga cga tta cta c-a tta tta aat cat gag gtc cgt att ata ctc gta tat att gtt tct ctc atg tat aat aat aaa cgg aag aga tat agc acc aca tgc acc atc caa tgg aaa gtg taa aga caa cga ata cag aag ccg taa tct atg ttg tct atc gtg tcc tcc ggg aac tta cgc ttt cag att atg tga tag caa gac taa tac aca atg tac acc gtg tgg ttc gga tac ctt tac atc tca caa taa tca ttt aca ggc ttg tct aag ttg taa cgg aag atg tga tag taa tca ggt aga gac gcg atc gtg taa cac gac tca caa tag aat ctg tga atg ctc tcc agg ata tta ttg tct tct caa agg agc atc agg gtg tag aac atg tat ttc taa aac aaa gtg tgg aat agg ata cgg agt atc cgg ata cac gtc tac cgg aga cgt cat ctg ttc tcc gtg tgg tcc cgg aac ata ttc tca cac cgt ctc ttc cac aga taa atg cga acc cgt cgt aac cag caa tac att taa cta tat cga tgt gga aat taa cct gta tcc agt caa cga cac atc gtg tac tcg gac gac cac tac cgg tct cag cga atc cat ctc aac gtc gga act aac tat tac cat gaa tca taa aga ttg tga tcc agt ctt tcg tgc aga ata ctt ctc tgt cct taa taa tgt agc aac ttc agg att ctt tac agg aga aaa tag ata tca gaa tac ttc aaa gat atg tac tct gaa ttt cga gat taa atg taa caa caa aga ttc atc ttc caa aca gtt aac gaa aac aaa gaa tga tac tat cat gcc gca ttc aga gac ggt aac tct agt ggg cga ctg tct atc tag cgt cga cat cta cat act ata tag taa tac caa tac tca aga cta cga aaa tga tac aat ctc tta tca tat ggg taa tgt tct cga tgt caa tag cca tat gcc cgc tag ttg cga tat aca taa act gat cac taa ttc cca gaa tcc cac cca ctt ata gta ag- -tt ttt tta cct ata aat aat aaa tac aat aat taa ttt ctc gta aaa gta gaa aat ata ttc taa ttt att ata tgg taa gaa agt aga atc atc tag aac agt aat caa tca ata gca atc atg aaa caa tat att gtc ctg gca tgc atg tgc cta gtg gca gct gct atg cct act agt ctt caa caa tct tca tcc tcg tgt act gaa gaa gaa aac aaa cat cat atg gga atc gat gtt att atc aaa gtc aca aag caa gac caa aca ccg acc aat gat aag att tgt caa tcc gta acg gaa gtt aca gag a-- --- --- --- --- --- -cc gaa gat gat gag gta tcc gaa gaa gtt gta aaa gga gat ccc acc act tat tac act atc gtc ggt gcg ggt ctt aac atg aac ttt gga ttc acc aaa tgc cca aag att tta tcc atc tcc gaa tcc tct gat gga aac act gtg aat act aga ttg tcc agc gtg tca ccg gga caa ggt aag gac tct ccc gcg atc acg cgt gaa gaa gct ctg gct atg atc aaa gac tgt gag atg tct atc gac atc aga tgt agc gaa gaa gag aaa gac agt gac atc aag acc cat cca gta ctt ggg tct aac atc tca cat aag aaa gtg agt tac aaa gat atc atc ggt tca acg atc gtt gat aca aaa tgt gtc aag aac cta gag ttt agc gta cgt atc gga gac atg tgt gag gaa tca tct gaa ctt gaa gtc aag gat gga ttc aag tat gtc gac gga tcg gca tct gaa ggt gca acc gat gat act tca ctc atc gat tca aca aaa ctc aaa gca tgt gtc tga atc gat aac tct att cat ctg aaa atg gat gag ttg ggt taa tcg aac gat tca gac acc gca cca cga att aaa aaa g-- -ac cgg gca cta tat tcc ggt ttg caa aac aaa aat att taa cta cat tca caa aaa gtt acc tct cgt tac ttc ttc ttt ctg ttt caa tat gtg ata cga tat gat cac tat tcg tat tct ct- --- --- tgg tct cat aaa aaa gtt tta c-- --a aaa aaa aaa aaa a-t att ttt att ctc ttt ctc tct tcg at- ggt ctc aca aaa ata tta aac ctc ttt ctg atg tct caa cta ttt cgt aaa cga taa cgt cca aca ata tat tct cgt aga gct tat caa cat cct tat acc aat cta ggt tgt cag aca att gca tca taa aat aat gtt tat aat tta cac gtt aac atc ata taa taa acg tat ata gtt aat att ttt gga ata taa atg atc tgt aaa atc cat gta ggg gac act gct cac gtt ttt tct cta gta cat aat ttc aca caa gtt ttt ata cag aca aat taa ttc tcg tcc ata ta- ttt taa aac att gac ttt tgt act aag aaa aat atc ttg act aac cat ctc ttt ctc tct tcg atg ggt ctc aca aaa ata tta aac ctc ttt ctg atg g-- --- --- --- --- --- --- --- --- --- --- --- --- --- --- --- --- --- --- --- --- --- --- --- --- --- --- --- --- --- --- --- --- --- --- --- --- --- --- --- --- --- --- --- --- --- --- agt cgt aaa aag ttt tta tcc ttt ctc tct tcg ata ggt ctc --- --- --- --- --- --- --- --- --- --- --- --- --- --- --- --- --- --- --- --- --- --- --- --- --- --- --- --- --- --- --- --- --- --- --- --- --- --- --- --- --- --- --- --- --- --- --- --- --- --- --- --- --- --- --- --- --- --- --- --- --- --- --- --- --- --- --- --- --- --- aca aaa ata tta aac ctc ttt ctg atg gtc tct ata aa

>ON585038.1_Monkeypox_virus_isolate_Monkeypox/PT0008/2022_complete_genome

atc gag gac gac ttc aac tat gac agc tcg gtg gcg tct gcc agc gtg tac ata cga atg gca ttt cta aga aaa gtc tac ggt atc ctt -tc tac aat ttc ctt taa caa cgg caa cag ctg cag tat ttt tat act ttg aat gca -tc gga cat tta tac aag gga gtc ctg ttc taa tat tgg cat caa tgt tcg gat cta tag gct tga ttt tcg cat tga ctt tac aca gac atg aac atc ccc tga atc tgt aca tac ttt gtg gat tta cac tgt tag aat ctc taa cgc tgg cct ctg ttg tta ctt tct atg atg cac gta tcg tta tgc aag ctt tca tgt tga cta ctg cag tgt ttc ttg ctc tga cta cat ata ctc tac aat caa aga gag att tca gta aac ttg taa cag gat tgt ttg ctg ctt tct gga ttt taa ttt tgt cag gag tct tga gga taa agt tta aaa tag aat taa taa aga aca tat agg tca -tt ttt taa aca tgg ata gaa acc aag gtt gtt agt taa taa tat aca aga tat ttt ttc tca ctc tga tcc atg taa acc aag gac gag aga cac tct cat tcc tca ttc acg aca cca tta aaa atg gaa att aaa gcc ctc tat taa gca cag acg gct aca ggt cta cca tca ggt t-a cct tcg tct acc ttc aca atg gcc tct cct tgt gcc cag ttc agt ccc tgt cat tgc cac gct act aag gac tcc ctg aat acc gtg act gac gtc aga cat tgt ctg act gaa tac atc ctg tgg gtt tct cat aga tgg acc cat aga gaa agc gca ggg cct ctc tac agg ctt ctc atc tct ttc aga att gat gca atg gag cta ttt ggt agc gag ttg aag gag ttc tcg aat tca ctt ccg tgg gac aat atc gac aat tgc gtg gag atc att aaa tgt ttc atc aga aat gac tcc atg aaa acc gcc aaa gaa ctt tgt gca ata att gga ctt tgt act caa tca gct att gtc act gga aga gtc ttc aat gat aag tat atc gac ata cta ctt atg ctg cga aag att ctg aac gag aac gac tat ctc acc ctc ttg gat cat atc ctc act gct aaa tac taa atc tcc ttc atg ctc tct cac taa tac tct tac tca cta cac ttt tta tca tct tat gat gaa tga ttg cct tca tca ttt ttt cgt gga ata taa tat agg aat aat tag cac cag aat agc tat gga tat ctc gtt aag aat att ctc tat aag aga cat aat gta gac ata gtt att ata tcc ttc tta gat aag tgt tac gct act gga aag ttt cca tcg tta tta tta cat gaa gat gat ata att aaa cca aca ttg aga ttg gct ctt atg tta gct gga ttg aat tac tgt aat aaa tgc atc gag tat aga ggg ata tag caa ttc tcg ata ata gtc atg caa tat ttg aat gag act gat aat tta ggt aat aca gta ct- --- aca cac ata tct ttc tag att ata tat cgt taa aaa tct gta aga tgt ata ttt ctc ata agt atc cac tgt gta ata tta tta atg gat ata tag ata acg caa tag gga cta ata gta ttg taa aag ata taa tcg act att tac gta cat atc cag ata tct ata ttc cta cta gtt tgc tgc gta gtt gca tca ttg ata tgc atg -at tta tca gga ttc aga gat gaa tta cta agt aaa cta caa tcc cac aat aag taa gaa tca aat atc aaa aac tca ctt ttg att ttt cta gtc tta agt aat aca tat att tat taa tag acc tat gaa ata aaa aaa ggt aac aat gga ttc gcg tat agc tat tta cgt att agt atc ggc atc tct ttt gta tct tgt taa ttg tca caa act agt aca tta ctt caa tct gaa aat aaa tgg aag tga tat aac taa tac agc aga tat att gct gga caa tta tcc aat tat gac ctt tga tgg aaa gga tat tta tcc atc tat ctc gtt cat ggt cgg taa taa act ttt cct aga tct tta taa aaa tat ctt tga aga att ttt cag act att tcg agt atc tgt aag tag tca ata cga gga att aga ata tta tta ttc atg tga tta tac taa caa ccg tcc tac aat taa aca aca tta ctt tta taa cgg cga cga ata tac tga aat tga tag atc gaa aaa agc cac taa taa aaa cag ttg gtt aat tac ttc agg ctt tag act aca aaa atg gtt cga tag cga aga ttg tat aat tta tct cag atc ttt agt tag aag aat gga aga cag taa caa aaa cag taa aaa aac tta gta ctt aga tat cg- --a aaa aat ata ttt ttg tag act ctt gag aat aga agg aaa aca tgt aca taa tta taa aaa atg aaa atc aat ggc gaa taa gac agt gcg att cgc acc atg gag tcg gta gat ttc atg gct gtc gat gag cag ttt cac gac gac ctc gat ctt tgg tca tta tct ttg gta gat gat tat aaa aaa cat gga tta ggt gtt gac tgt tat gtt cta gaa cca gtt gtt gac agg aaa ata ttt gat aga ttt ctc ctt gaa cca att tgt gat cct gta gat gtt ctg tat gat tat ttt agg att cat aga gat aat att gat cag tat ata gta gat aga ctg ttt gca tat att aca tat aaa gat att ata tct gca tta gtg tca aag aat tat atg gaa gat att ttc tct ata att att aag aat tgt aat tct gtg caa gat ctc tta ctt tac tat cta tct aat gca tat gta gaa ata gac att gtt gat ctt atg gta gat cat ggg gct gta ata tat aaa ata gaa tgc ttg aat gcc tat ttt agg gga ata tgt aaa aag gaa agt agt gtt gtt gag ttt att ttg aat tgt ggt atc cca gat gaa aat gat gtt aaa tta gat cta tat aaa ata att cag tat act agg gga ttc ctt gta gat gaa ccc aca gta tta gaa att tat aag ctt tgt atc cca tat att gaa gat atc aat caa cta gat gct ggt gga agg acc ttg ctt tat cgc gct atc tat gca ggt tat ata gat tta gta tca tgg cta tta gaa aat gga gca aat gtc aac gca gta atg agt aat gga tat aca tgt ctt gac gtg gcc gtg gat agg gga tct gtc atc gcc cgt agg gaa gca cat ctt aaa ata tta gaa ata ttg ctt aga gaa cca ttg tct att gac tgt ata aaa tta gct ata ctt aat aat aca att gaa aac cat gat gtg ata aag ctc tgt atc aag tat ttt atg atg gta gat tat tca ctt tgt aat gtg tat gca tca tca ctc ttt gat tat ata att gat tgt aaa caa gaa ttg gag tac att agg cag atg aaa att cat aat aca acc atg tat gag tta atc tat aat aga gac aaa aac aag cat gct tcc cat att cta cat agg tat tct aaa cat cca gtt ttg aca cag tgt atc act aaa gga ttc aag att tac aca gaa gta acc gag cag gtc act aaa gct cta aac aga cgt gct cta ata gat gag ata ata aac aat gta tca act gat gac aat ctc cta tca aaa ctt cca tta gaa att agg gat cta att gtt tca caa gct gtc ata tag agt tct atc cac cca cct ttc ttg aaa tga gtt aat --- --- --- --- --- --- --- --- --- --- --- --- --- --- --- --- --- --- --- --- --- --- --- --- --- --- --- --- --- --- --- --- --- --- --- --- --- --- --- --- --- --- --- --- --- --- --- --- --- --- --- --- --- --- --- --- --- --- --- --- --- --- --- --- --- --- --- --- --- --- --- --- --- --- --- --- --- --- --- --- --- --- --- --- --- --- --- --- --- --- --- --- --- --- --- --- --- --- --- --- --- --- --- --- --- -ag tca taa gtt agt taa --- --- --- --- --- -gt cat aag tta gtt aag tca t-- --- --- --- --- --- --- --- --- --- --- --- --- --- --- --- -aa gtt agt taa gtc ata agt tag tta agt cat aag tta gtt aag tca taa gtt agt taa gtc ata agt tag tta agt cat aag tta gtt tat agt cta aca ctt cta att ttt ata cct tga tct ttt tct cta att atg aaa aag taa atc att atg aag atg gat g-a aat gga cga gat tgt gcg cat cgt taa cga tag tat gtg gta cgt acc taa cgc att tat gga cga cgg tga taa tga agg tca cat ttc tgt caa taa tgt ctg tca tat gta tct cgc att ctt tga tgt gga tat atc atc tca tct gtt taa att agt tat taa aca ctg cga tct gaa taa acg act aaa atg tgg taa ctc tcc att aca ttg cta tac gat gaa tac acg att taa tcc atc tgt att aaa gat att gtt acg cca cgg cat gcg taa ctt tga tag caa gga taa aaa agg aca tat tcc tct aca cca cta tct gat tca ttc act atc aat cga taa caa gat ctt tga tat act aac gga ccc cat tga tga ctt tag taa atc atc cga tct att gct gtg tta tct tag ata taa att caa tgg gag ctt aaa cta tta cgt tct gta caa att att gac taa agg atc tga ccc taa ttg cgt cga tga gga tgg act cac ttc tct tca tta cta ctg taa aca cat atc cgc gtt cca cga aag caa tta tta caa gtc aaa gag tca cac taa gat gcg agc tga gaa gcg att cat cta cgc gat aat aga tca tgg agc aaa cat taa cgc ggt tac gaa aat cgg aaa tac gcc gtt aca cac tta cct tca aca gta tac caa aca tag tcc tcg tgt ggt gta tgc tct ttt atc tcg agg agc cga tac gag gat acg taa taa tct taa ttg tac acc cat cat gga ata cat aaa gaa cga ttg tgc aac agg tca tat tct cat aat gtt act caa ttg gca cga aca aaa ata cgg gaa att aca aaa gga aga agg aca aca tct act tta tct att cat aaa aca taa tca agg ata tgg aag tcg ctc tct caa tat act acg gta tct act aga tag att cga cat tca gaa aga cga ata cta taa tac aat gac tcc tct tca tac cgc ctt cca gaa ttg caa taa caa tgt tgc ctc ata cct cgt ata cat tgg ata cga cat caa cct tcc gac taa aga cga taa gac agt att cga ctt ggt gtt tga aaa cag aaa cat tat ata caa ggc gga tgt cgt taa tga cat tat cca cca cag act gaa agt atc tct acc tat gat taa atc gtt gtt cta caa gat gtc gga gtt ctc tcc cta cga cga tca cta cgt aaa gaa gat aat agc cta ctg cct att aag gga cga gtc att tgc gga act aca tac taa att ctg ttt aaa cga gga cta taa aag tgt att tat gaa aaa tat atc att cga taa gat aga ttc cat cat cga aaa atg tag tcg tga cat aag tct cct caa aga gat tcg aat ctc aga cac cga ctt gta tac ggt att gag aac aga aga cat tcg gta tca cac ata tct cga agc cat aca ttc aga caa acg cat ttc att tcc cat gta cga cga tct cat aga aca gtg tca tct atc gat gga gca taa aag taa act cgt cga caa agc act caa taa att aga gtc tac cat cga tag tca atc tag act atc gta ttt gcc tcc gga aat tat gcg caa tat cat aac caa gct aag cga cta cca tct aaa cag tat gtt gta cgg aaa gaa cca tta caa ata tta tcc atg ata gaa aga aaa tat tta aaa aat aat cta tat gat tgg aga agt agg aaa caa aca gta aca aga cga cga tta cta c-a tta tta aat cat gag gtc cgt att ata ctc gta tat att gtt tct ctc atg tat aat aat aaa cgg aag aga tat agc acc aca tgc acc atc caa tgg aaa gtg taa aga caa cga ata cag aag ccg taa tct atg ttg tct atc gtg tcc tcc ggg aac tta cgc ttt cag att atg tga tag caa gac taa tac aca atg tac acc gtg tgg ttc gga tac ctt tac atc tca caa taa tca ttt aca ggc ttg tct aag ttg taa cgg aag atg tga tag taa tca ggt aga gac gcg atc gtg taa cac gac tca caa tag aat ctg tga atg ctc tcc agg ata tta ttg tct tct caa agg agc atc agg gtg tag aac atg tat ttc taa aac aaa gtg tgg aat agg ata cgg agt atc cgg ata cac gtc tac cgg aga cgt cat ctg ttc tcc gtg tgg tcc cgg aac ata ttc tca cac cgt ctc ttc cac aga taa atg cga acc cgt cgt aac cag caa tac att taa cta tat cga tgt gga aat taa cct gta tcc agt caa cga cac atc gtg tac tcg gac gac cac tac cgg tct cag cga atc cat ctc aac gtc gga act aac tat tac cat gaa tca taa aga ttg tga tcc agt ctt tcg tgc aga ata ctt ctc tgt cct taa taa tgt agc aac ttc agg att ctt tac agg aga aaa tag ata tca gaa tac ttc aaa gat atg tac tct gaa ttt cga gat taa atg taa caa caa aga ttc atc ttc caa aca gtt aac gaa aac aaa gaa tga tac tat cat gcc gca ttc aga gac ggt aac tct agt ggg cga ctg tct atc tag cgt cga cat cta cat act ata tag taa tac caa tac tca aga cta cga aaa tga tac aat ctc tta tca tat ggg taa tgt tct cga tgt caa tag cca tat gcc cgc tag ttg cga tat aca taa act gat cac taa ttc cca gaa tcc cac cca ctt ata gta ag- -tt ttt tta cct ata aat aat aaa tac aat aat taa ttt ctc gta aaa gta gaa aat ata ttc taa ttt att ata tgg taa gaa agt aga atc atc tag aac agt aat caa tca ata gca atc atg aaa caa tat att gtc ctg gca tgc atg tgc cta gtg gca gct gct atg cct act agt ctt caa caa tct tca tcc tcg tgt act gaa gaa gaa aac aaa cat cat atg gga atc gat gtt att atc aaa gtc aca aag caa gac caa aca ccg acc aat gat aag att tgt caa tcc gta acg gaa gtt aca gag a-- --- --- --- --- --- -cc gaa gat gat gag gta tcc gaa gaa gtt gta aaa gga gat ccc acc act tat tac act atc gtc ggt gcg ggt ctt aac atg aac ttt gga ttc acc aaa tgc cca aag att tta tcc atc tcc gaa tcc tct gat gga aac act gtg aat act aga ttg tcc agc gtg tca ccg gga caa ggt aag gac tct ccc gcg atc acg cgt gaa gaa gct ctg gct atg atc aaa gac tgt gag atg tct atc gac atc aga tgt agc gaa gaa gag aaa gac agt gac atc aag acc cat cca gta ctt ggg tct aac atc tca cat aag aaa gtg agt tac aaa gat atc atc ggt tca acg atc gtt gat aca aaa tgt gtc aag aac cta gag ttt agc gta cgt atc gga gac atg tgt gag gaa tca tct gaa ctt gaa gtc aag gat gga ttc aag tat gtc gac gga tcg gca tct gaa ggt gca acc gat gat act tca ctc atc gat tca aca aaa ctc aaa gca tgt gtc tga atc gat aac tct att cat ctg aaa atg gat gag ttg ggt taa tcg aac gat tca gac acc gca cca cga att aaa aaa g-- -ac cgg gca cta tat tcc ggt ttg caa aac aaa aat att taa cta cat tca caa aaa gtt acc tct cgt tac ttc ttc ttt ctg ttt caa tat gtg ata cga tat gat cac tat tcg tat tct ct- --- --- tgg tct cat aaa aaa gtt tta c-- --a aaa aaa aaa aaa --t att ttt att ctc ttt ctc tct tcg at- ggt ctc aca aaa ata tta aac ctc ttt ctg atg tct caa cta ttt cgt aaa cga taa cgt cca aca ata tat tct cgt aga gct tat caa cat cct tat acc aat cta ggt tgt cag aca att gca tca taa aat aat gtt tat aat tta cac gtt aac atc ata taa taa acg tat ata gtt aat att ttt gga ata taa atg atc tgt aaa atc cat gta ggg gac act gct cac gtt ttt tct cta gta cat aat ttc aca caa gtt ttt ata cag aca aat taa ttc tcg tcc ata ta- ttt taa aac att gac ttt tgt act aag aaa aat atc ttg act aac cat ctc ttt ctc tct tcg atg ggt ctc aca aaa ata tta aac ctc ttt ctg atg g-- --- --- --- --- --- --- --- --- --- --- --- --- --- --- --- --- --- --- --- --- --- --- --- --- --- --- --- --- --- --- --- --- --- --- --- --- --- --- --- --- --- --- --- --- --- --- agt cgt aaa aag ttt tta tcc ttt ctc tct tcg ata ggt ctc --- --- --- --- --- --- --- --- --- --- --- --- --- --- --- --- --- --- --- --- --- --- --- --- --- --- --- --- --- --- --- --- --- --- --- --- --- --- --- --- --- --- --- --- --- --- --- --- --- --- --- --- --- --- --- --- --- --- --- --- --- --- --- --- --- --- --- --- --- --- aca aaa ata tta aac ctc ttt ctg atg gtc tct ata aa

>ON682264.2_Monkeypox_virus_isolate_MPXV/Germany/2022/RKI05_complete_genome

atc gag gac gac ttc aac tat gac agc tcg gtg gcg tct gcc agc gtg tac ata cga atg gca ttt cta aga aaa gtc tac ggt atc ctt -tc tac aat ttc ctt taa caa cgg caa cag ctg cag tat ttt tat act ttg aat gca -tc gga cat tta tac aag gga gtc ctg ttc taa tat tgg cat caa tgt tcg gat cta tag gct tga ttt tcg cat tga ctt tac aca gac atg aac atc ccc tga atc tgt aca tac ttt gtg gat tta cac tgt tag aat ctc taa cgc tgg cct ctg ttg tta ctt tct atg atg cac gta tcg tta tgc aag ctt tca tgt tga cta ctg cag tgt ttc ttg ctc tga cta cat ata ctc tac aat caa aga gag att tca gta aac ttg taa cag gat tgt ttg ctg ctt tct gga ttt taa ttt tgt cag gag tct tga gga taa agt tta aaa tag aat taa taa aga aca tat agg tca -tt ttt taa aca tgg ata gaa acc aag gtt gtt agt taa taa tat aca aga tat ttt ttc tca ctc tga tcc atg taa acc aag gac gag aga cac tct cat tcc tca ttc acg aca cca tta aaa atg gaa att aaa gcc ctc tat taa gca cag acg gct aca ggt cta cca tca ggt t-a cct tcg tct acc ttc aca atg gcc tct cct tgt gcc cag ttc agt ccc tgt cat tgc cac gct act aag gac tcc ctg aat acc gtg act gac gtc aga cat tgt ctg act gaa tac atc ctg tgg gtt tct cat aga tgg acc cat aga gaa agc gca ggg cct ctc tac agg ctt ctc atc tct ttc aga att gat gca atg gag cta ttt ggt agc gag ttg aag gag ttc tcg aat tca ctt ccg tgg gac aat atc gac aat tgc gtg gag atc att aaa tgt ttc atc aga aat gac tcc atg aaa acc gcc aaa gaa ctt tgt gca ata att gga ctt tgt act caa tca gct att gtc act gga aga gtc ttc aat gat aag tat atc gac ata cta ctt atg ctg cga aag att ctg aac gag aac gac tat ctc acc ctc ttg gat cat atc ctc act gct aaa tac taa atc tcc ttc atg ctc tct cac taa tac tct tac tca cta cac ttt tta tca tct tat gat gaa tga ttg cct tca tca ttt ttt cgt gga ata taa tat agg aat aat tag cac cag aat agc tat gga tat ctc gtt aag aat att ctc tat aag aga cat aat gta gac ata gtt att ata tcc ttc tta gat aag tgt tac gct act gga aag ttt cca tcg tta tta tta cat gaa gat gat ata att aaa cca aca ttg aga ttg gct ctt atg tta gct gga ttg aat tac tgt aat aaa tgc atc gag tat aga ggg ata tag caa ttc tcg ata ata gtc atg caa tat ttg aat gag act gat aat tta ggt aat aca gta ct- --- aca cac ata tct ttc tag att ata tat cgt taa aaa tct gta aga tgt ata ttt ctc ata agt atc cac tgt gta ata tta tta atg gat ata tag ata acg caa tag gga cta ata gta ttg taa aag ata taa tcg act att tac gta cat atc cag ata tct ata ttc cta cta gtt tgc tgc gta gtt gca tca ttg ata tgc atg -at tta tca gga ttc aga gat gaa tta cta agt aaa cta caa tcc cac aat aag taa gaa tca aat atc aaa aac tca ctt ttg att ttt cta gtc tta agt aat aca tat att tat taa tag acc tat gaa ata aaa aaa ggt aac aat gga ttc gcg tat agc tat tta cgt att agt atc ggc atc tct ttt gta tct tgt taa ttg tca caa act agt aca tta ctt caa tct gaa aat aaa tgg aag tga tat aac taa tac agc aga tat att gct gga caa tta tcc aat tat gac ctt tga tgg aaa gga tat tta tcc atc tat ctc gtt cat ggt cgg taa taa act ttt cct aga tct tta taa aaa tat ctt tga aga att ttt cag act att tcg agt atc tgt aag tag tca ata cga gga att aga ata tta tta ttc atg tga tta tac taa caa ccg tcc tac aat taa aca aca tta ctt tta taa cgg cga cga ata tac tga aat tga tag atc gaa aaa agc cac taa taa aaa cag ttg gtt aat tac ttc agg ctt tag act aca aaa atg gtt cga tag cga aga ttg tat aat tta tct cag atc ttt agt tag aag aat gga aga cag taa caa aaa cag taa aaa aac tta gta ctt aga tat cg- --a aaa aat ata ttt ttg tag act ctt gag aat aga agg aaa aca tgt aca taa tta taa aaa atg aaa atc aat ggc gaa taa gac agt gcg att cgc acc atg gag tcg gta gat ttc atg gct gtc gat gag cag ttt cac gac gac ctc gat ctt tgg tca tta tct ttg gta gat gat tat aaa aaa cat gga tta ggt gtt gac tgt tat gtt cta gaa cca gtt gtt gac agg aaa ata ttt gat aga ttt ctc ctt gaa cca att tgt gat cct gta gat gtt ctg tat gat tat ttt agg att cat aga gat aat att gat cag tat ata gta gat aga ctg ttt gca tat att aca tat aaa gat att ata tct gca tta gtg tca aag aat tat atg gaa gat att ttc tct ata att att aag aat tgt aat tct gtg caa gat ctc tta ctt tac tat cta tct aat gca tat gta gaa ata gac att gtt gat ctt atg gta gat cat ggg gct gta ata tat aaa ata gaa tgc ttg aat gcc tat ttt agg gga ata tgt aaa aag gaa agt agt gtt gtt gag ttt att ttg aat tgt ggt atc cca gat gaa aat gat gtt aaa tta gat cta tat aaa ata att cag tat act agg gga ttc ctt gta gat gaa ccc aca gta tta gaa att tat aag ctt tgt atc cca tat att gaa gat atc aat caa cta gat gct ggt gga agg acc ttg ctt tat cgc gct atc tat gca ggt tat ata gat tta gta tca tgg cta tta gaa aat gga gca aat gtc aac gca gta atg agt aat gga tat aca tgt ctt gac gtg gcc gtg gat agg gga tct gtc atc gcc cgt agg gaa gca cat ctt aaa ata tta gaa ata ttg ctt aga gaa cca ttg tct att gac tgt ata aaa tta gct ata ctt aat aat aca att gaa aac cat gat gtg ata aag ctc tgt atc aag tat ttt atg atg gta gat tat tca ctt tgt aat gtg tat gca tca tca ctc ttt gat tat ata att gat tgt aaa caa gaa ttg gag tac att agg cag atg aaa att cat aat aca acc atg tat gag tta atc tat aat aga gac aaa aac aag cat gct tcc cat att cta cat agg tat tct aaa cat cca gtt ttg aca cag tgt atc act aaa gga ttc aag att tac aca gaa gta acc gag cag gtc act aaa gct cta aac aga cgt gct cta ata gat gag ata ata aac aat gta tca act gat gac aat ctc cta tca aaa ctt cca tta gaa att agg gat cta att gtt tca caa gct gtc ata tag agt tct atc cac cca cct ttc ttg aaa tga gtt aat a-- --- --- --- --- --- --- --- --- --- --- --- --- --- --- --- --- --- --- --- --- --- --- --- --- --- --- --- --- --- --- --- --- --- --- --- --- --- --- --- --- --- --- --- --- --- --- --- --- --- --- --- --- --- --- --- --- --- --- --- --- --- --- --- --- --- --- --- --- --- --- --- --- --- --- --- --- --- --- --- --- --- --- --- --- --- --- --- --- --- --- --- --- --- --- --- --- --- --- --- -gt cat aag tta gtt aag tca taa gtt agt taa --- --- --- --- --- -gt cat aag tta gtt aag tca t-- --- --- --- --- --- --- --- --- --- --- --- --- --- --- --- -aa gtt agt taa gtc ata agt tag tta agt cat aag tta gtt aag tca taa gtt agt taa gtc ata agt tag tta agt cat aag tta gtt tat agt cta aca ctt cta att ttt ata cct tga tct ttt tct cta att atg aaa aag taa atc att atg aag atg gat g-a aat gga cga gat tgt gcg cat cgt taa cga tag tat gtg gta cgt acc taa cgc att tat gga cga cgg tga taa tga agg tca cat ttc tgt caa taa tgt ctg tca tat gta tct cgc att ctt tga tgt gga tat atc atc tca tct gtt taa att agt tat taa aca ctg cga tct gaa taa acg act aaa atg tgg taa ctc tcc att aca ttg cta tac gat gaa tac acg att taa tcc atc tgt att aaa gat att gtt acg cca cgg cat gcg taa ctt tga tag caa gga taa aaa agg aca tat tcc tct aca cca cta tct gat tca ttc act atc aat cga taa caa gat ctt tga tat act aac gga ccc cat tga tga ctt tag taa atc atc cga tct att gct gtg tta tct tag ata taa att caa tgg gag ctt aaa cta tta cgt tct gta caa att att gac taa agg atc tga ccc taa ttg cgt cga tga gga tgg act cac ttc tct tca tta cta ctg taa aca cat atc cgc gtt cca cga aag caa tta tta caa gtc aaa gag tca cac taa gat gcg agc tga gaa gcg att cat cta cgc gat aat aga tca tgg agc aaa cat taa cgc ggt tac gaa aat cgg aaa tac gcc gtt aca cac tta cct tca aca gta tac caa aca tag tcc tcg tgt ggt gta tgc tct ttt atc tcg agg agc cga tac gag gat acg taa taa tct taa ttg tac acc cat cat gga ata cat aaa gaa cga ttg tgc aac agg tca tat tct cat aat gtt act caa ttg gca cga aca aaa ata cgg gaa att aca aaa gga aga agg aca aca tct act tta tct att cat aaa aca taa tca agg ata tgg aag tcg ctc tct caa tat act acg gta tct act aga tag att cga cat tca gaa aga cga ata cta taa tac aat gac tcc tct tca tac cgc ctt cca gaa ttg caa taa caa tgt tgc ctc ata cct cgt ata cat tgg ata cga cat caa cct tcc gac taa aga cga taa gac agt att cga ctt ggt gtt tga aaa cag aaa cat tat ata caa ggc gga tgt cgt taa tga cat tat cca cca cag act gaa agt atc tct acc tat gat taa atc gtt gtt cta caa gat gtc gga gtt ctc tcc cta cga cga tca cta cgt aaa gaa gat aat agc cta ctg cct att aag gga cga gtc att tgc gga act aca tac taa att ctg ttt aaa cga gga cta taa aag tgt att tat gaa aaa tat atc att cga taa gat aga ttc cat cat cga aaa atg tag tcg tga cat aag tct cct caa aga gat tcg aat ctc aga cac cga ctt gta tac ggt att gag aac aga aga cat tcg gta tca cac ata tct cga agc cat aca ttc aga caa acg cat ttc att tcc cat gta cga cga tct cat aga aca gtg tca tct atc gat gga gca taa aag taa act cgt cga caa agc act caa taa att aga gtc tac cat cga tag tca atc tag act atc gta ttt gcc tcc gga aat tat gcg caa tat cat aac caa gct aag cga cta cca tct aaa cag tat gtt gta cgg aaa gaa cca tta caa ata tta tcc atg ata gaa aga aaa tat tta aaa aat aat cta tat gat tgg aga agt agg aaa caa aca gta aca aga cga cga tta cta c-a tta tta aat cat gag gtc cgt att ata ctc gta tat att gtt tct ctc atg tat aat aat aaa cgg aag aga tat agc acc aca tgc acc atc caa tgg aaa gtg taa aga caa cga ata cag aag ccg taa tct atg ttg tct atc gtg tcc tcc ggg aac tta cgc ttt cag att atg tga tag caa gac taa tac aca atg tac acc gtg tgg ttc gga tac ctt tac atc tca caa taa tca ttt aca ggc ttg tct aag ttg taa cgg aag atg tga tag taa tca ggt aga gac gcg atc gtg taa cac gac tca caa tag aat ctg tga atg ctc tcc agg ata tta ttg tct tct caa agg agc atc agg gtg tag aac atg tat ttc taa aac aaa gtg tgg aat agg ata cgg agt atc cgg ata cac gtc tac cgg aga cgt cat ctg ttc tcc gtg tgg tcc cgg aac ata ttc tca cac cgt ctc ttc cac aga taa atg cga acc cgt cgt aac cag caa tac att taa cta tat cga tgt gga aat taa cct gta tcc agt caa cga cac atc gtg tac tcg gac gac cac tac cgg tct cag cga atc cat ctc aac gtc gga act aac tat tac cat gaa tca taa aga ttg tga tcc agt ctt tcg tgc aga ata ctt ctc tgt cct taa taa tgt agc aac ttc agg att ctt tac agg aga aaa tag ata tca gaa tac ttc aaa gat atg tac tct gaa ttt cga gat taa atg taa caa caa aga ttc atc ttc caa aca gtt aac gaa aac aaa gaa tga tac tat cat gcc gca ttc aga gac ggt aac tct agt ggg cga ctg tct atc tag cgt cga cat cta cat act ata tag taa tac caa tac tca aga cta cga aaa tga tac aat ctc tta tca tat ggg taa tgt tct cga tgt caa tag cca tat gcc cgc tag ttg cga tat aca taa act gat cac taa ttc cca gaa tcc cac cca ctt ata gta ag- -tt ttt tta cct ata aat aat aaa tac aat aat taa ttt ctc gta aaa gta gaa aat ata ttc taa ttt att ata tgg taa gaa agt aga atc atc tag aac agt aat caa tca ata gca atc atg aaa caa tat att gtc ctg gca tgc atg tgc cta gtg gca gct gct atg cct act agt ctt caa caa tct tca tcc tcg tgt act gaa gaa gaa aac aaa cat cat atg gga atc gat gtt att atc aaa gtc aca aag caa gac caa aca ccg acc aat gat aag att tgt caa tcc gta acg gaa gtt aca gag a-- --- --- --- --- --- -cc gaa gat gat gag gta tcc gaa gaa gtt gta aaa gga gat ccc acc act tat tac act atc gtc ggt gcg ggt ctt aac atg aac ttt gga ttc acc aaa tgc cca aag att tta tcc atc tcc gaa tcc tct gat gga aac act gtg aat act aga ttg tcc agc gtg tca ccg gga caa ggt aag gac tct ccc gcg atc acg cgt gaa gaa gct ctg gct atg atc aaa gac tgt gag atg tct atc gac atc aga tgt agc gaa gaa gag aaa gac agt gac atc aag acc cat cca gta ctt ggg tct aac atc tca cat aag aaa gtg agt tac aaa gat atc atc ggt tca acg atc gtt gat aca aaa tgt gtc aag aac cta gag ttt agc gta cgt atc gga gac atg tgt gag gaa tca tct gaa ctt gaa gtc aag gat gga ttc aag tat gtc gac gga tcg gca tct gaa ggt gca acc gat gat act tca ctc atc gat tca aca aaa ctc aaa gca tgt gtc tga atc gat aac tct att cat ctg aaa atg gat gag ttg ggt taa tcg aac gat tca gac acc gca cca cga att aaa aaa g-- -ac cgg gca cta tat tcc ggt ttg caa aac aaa aat att taa cta cat tca caa aaa gtt acc tct cgt tac ttc ttc ttt ctg ttt caa tat gtg ata cga tat gat cac tat tcg tat tct ct- --- --- tgg tct cat aaa aaa gtt tta c-- --a aaa aaa aaa aaa a-t att ttt att ctc ttt ctc tct tcg at- ggt ctc aca aaa ata tta aac ctc ttt ctg atg tct caa cta ttt cgt aaa cga taa cgt cca aca ata tat tct cgt aga gct tat caa cat cct tat acc aat cta ggt tgt cag aca att gca tca taa aat aat gtt tat aat tta cac gtt aac atc ata taa taa acg tat ata gtt aat att ttt gga ata taa atg atc tgt aaa atc cat gta ggg gac act gct cac gtt ttt tct cta gta cat aat ttc aca caa gtt ttt ata cag aca aat taa ttc tcg tcc ata ta- ttt taa aac att gac ttt tgt act aag aaa aat atc ttg act aac cat ctc ttt ctc tct tcg atg ggt ctc aca aaa ata tta aac ctc ttt ctg atg g-- --- --- --- --- --- --- --- --- --- --- --- --- --- --- --- --- --- --- --- --- --- --- --- --- --- --- --- --- --- --- --- --- --- --- --- --- --- --- --- --- --- --- --- --- --- --- agt cgt aaa aag ttt tta tcc ttt ctc tct tcg ata ggt ctc --- --- --- --- --- --- --- --- --- --- --- --- --- --- --- --- --- --- --- --- --- --- --- --- --- --- --- --- --- --- --- --- --- --- --- --- --- --- --- --- --- --- --- --- --- --- --- --- --- --- --- --- --- --- --- --- --- --- --- --- --- --- --- --- --- --- --- --- --- --- aca aaa ata tta aac ctc ttt ctg atg gtc tct ata aa

>ON682268.1_Monkeypox_virus_isolate_MPXV/Germany/2022/RKI08_complete_genome

atc gag gac gac ttc aac tat gac agc tcg gtg gcg tct gcc agc gtg tac ata cga atg gca ttt cta aga aaa gtc tac ggt atc ctt -tc tac aat ttc ctt taa caa cgg caa cag ctg cag tat ttt tat act ttg aat gca -tc gga cat tta tac aag gga gtc ctg ttc taa tat tgg cat caa tgt tcg gat cta tag gct tga ttt tcg cat tga ctt tac aca gac atg aac atc ccc tga atc tgt aca tac ttt gtg gat tta cac tgt tag aat ctc taa cgc tgg cct ctg ttg tta ctt tct atg atg cac gta tcg tta tgc aag ctt tca tgt tga cta ctg cag tgt ttc ttg ctc tga cta cat ata ctc tac aat caa aga gag att tca gta aac ttg taa cag gat tgt ttg ctg ctt tct gga ttt taa ttt tgt cag gag tct tga gga taa agt tta aaa tag aat taa taa aga aca tat agg tca -tt ttt taa aca tgg ata gaa acc aag gtt gtt agt taa taa tat aca aga tat ttt ttc tca ctc tga tcc atg taa acc aag gac gag aga cac tct cat tcc tca ttc acg aca cca tta aaa atg gaa att aaa gcc ctc tat taa gca cag acg gct aca ggt cta cca tca ggt t-a cct tcg tct acc ttc aca atg gcc tct cct tgt gcc cag ttc agt ccc tgt cat tgc cac gct act aag gac tcc ctg aat acc gtg act gac gtc aga cat tgt ctg act gaa tac atc ctg tgg gtt tct cat aga tgg acc cat aga gaa agc gca ggg cct ctc tac agg ctt ctc atc tct ttc aga att gat gca atg gag cta ttt ggt agc gag ttg aag gag ttc tcg aat tca ctt ccg tgg gac aat atc gac aat tgc gtg gag atc att aaa tgt ttc atc aga aat gac tcc atg aaa acc gcc aaa gaa ctt tgt gca ata att gga ctt tgt act caa tca gct att gtc act gga aga gtc ttc aat gat aag tat atc gac ata cta ctt atg ctg cga aag att ctg aac gag aac gac tat ctc acc ctc ttg gat cat atc ctc act gct aaa tac taa atc tcc ttc atg ctc tct cac taa tac tct tac tca cta cac ttt tta tca tct tat gat gaa tga ttg cct tca tca ttt ttt cgt gga ata taa tat agg aat aat tag cac cag aat agc tat gga tat ctc gtt aag aat att ctc tat aag aga cat aat gta gac ata gtt att ata tcc ttc tta gat aag tgt tac gct act gga aag ttt cca tcg tta tta tta cat gaa gat gat ata att aaa cca aca ttg aga ttg gct ctt atg tta gct gga ttg aat tac tgt aat aaa tgc atc gag tat aga ggg ata tag caa ttc tcg ata ata gtc atg caa tat ttg aat gag act gat aat tta ggt aat aca gta ct- --- aca cac ata tct ttc tag att ata tat cgt taa aaa tct gta aga tgt ata ttt ctc ata agt atc cac tgt gta ata tta tta atg gat ata tag ata acg caa tag gga cta ata gta ttg taa aag ata taa tcg act att tac gta cat atc cag ata tct ata ttc cta cta gtt tgc tgc gta gtt gca tca ttg ata tgc atg -at tta tca gga ttc aga gat gaa tta cta agt aaa cta caa tcc cac aat aag taa gaa tca aat atc aaa aac tca ctt ttg att ttt cta gtc tta agt aat aca tat att tat taa tag acc tat gaa ata aaa aaa ggt aac aat gga ttc gcg tat agc tat tta cgt att agt atc ggc atc tct ttt gta tct tgt taa ttg tca caa act agt aca tta ctt caa tct gaa aat aaa tgg aag tga tat aac taa tac agc aga tat att gct gga caa tta tcc aat tat gac ctt tga tgg aaa gga tat tta tcc atc tat ctc gtt cat ggt cgg taa taa act ttt cct aga tct tta taa aaa tat ctt tga aga att ttt cag act att tcg agt atc tgt aag tag tca ata cga gga att aga ata tta tta ttc atg tga tta tac taa caa ccg tcc tac aat taa aca aca tta ctt tta taa cgg cga cga ata tac tga aat tga tag atc gaa aaa agc cac taa taa aaa cag ttg gtt aat tac ttc agg ctt tag act aca aaa atg gtt cga tag cga aga ttg tat aat tta tct cag atc ttt agt tag aag aat gga aga cag taa caa aaa cag taa aaa aac tta gta ctt aga tat cg- --a aaa aat ata ttt ttg tag act ctt gag aat aga agg aaa aca tgt aca taa tta taa aaa atg aaa atc aat ggc gaa taa gac agt gcg att cgc acc atg gag tcg gta gat ttc atg gct gtc gat gag cag ttt cac gac gac ctc gat ctt tgg tca tta tct ttg gta gat gat tat aaa aaa cat gga tta ggt gtt gac tgt tat gtt cta gaa cca gtt gtt gac agg aaa ata ttt gat aga ttt ctc ctt gaa cca att tgt gat cct gta gat gtt ctg tat gat tat ttt agg att cat aga gat aat att gat cag tat ata gta gat aga ctg ttt gca tat att aca tat aaa gat att ata tct gca tta gtg tca aag aat tat atg gaa gat att ttc tct ata att att aag aat tgt aat tct gtg caa gat ctc tta ctt tac tat cta tct aat gca tat gta gaa ata gac att gtt gat ctt atg gta gat cat ggg gct gta ata tat aaa ata gaa tgc ttg aat gcc tat ttt agg gga ata tgt aaa aag gaa agt agt gtt gtt gag ttt att ttg aat tgt ggt atc cca gat gaa aat gat gtt aaa tta gat cta tat aaa ata att cag tat act agg gga ttc ctt gta gat gaa ccc aca gta tta gaa att tat aag ctt tgt atc cca tat att gaa gat atc aat caa cta gat gct ggt gga agg acc ttg ctt tat cgc gct atc tat gca ggt tat ata gat tta gta tca tgg cta tta gaa aat gga gca aat gtc aac gca gta atg agt aat gga tat aca tgt ctt gac gtg gcc gtg gat agg gga tct gtc atc gcc cgt agg gaa gca cat ctt aaa ata tta gaa ata ttg ctt aga gaa cca ttg tct att gac tgt ata aaa tta gct ata ctt aat aat aca att gaa aac cat gat gtg ata aag ctc tgt atc aag tat ttt atg atg gta gat tat tca ctt tgt aat gtg tat gca tca tca ctc ttt gat tat ata att gat tgt aaa caa gaa ttg gag tac att agg cag atg aaa att cat aat aca acc atg tat gag tta atc tat aat aga gac aaa aac aag cat gct tcc cat att cta cat agg tat tct aaa cat cca gtt ttg aca cag tgt atc act aaa gga ttc aag att tac aca gaa gta acc gag cag gtc act aaa gct cta aac aga cgt gct cta ata gat gag ata ata aac aat gta tca act gat gac aat ctc cta tca aaa ctt cca tta gaa att agg gat cta att gtt tca caa gct gtc ata tag agt tct atc cac cca cct ttc ttg aaa tga gtt aat agt cat aag tt- --- --- --- --- --- --- --- --- --- --- --- --- --- --- --- --- --- --- --- --- --- --- --- --- --- --- --- --- --- --- --- --- --- --- --- --- --- --- --- --- --- --- --- --- --- --- --- --- --- --- --- --- --- --- --- --- --- --- --- --- --- --- --- --- --- --- --- --- --- --- --- --- --- --- --- --- --- --- --- --- --- --- --- --- --- --- --- --- --- agt taa gtc ata agt tag tta agt cat aag tta gtt aag tca taa gtt agt taa --- --- --- --- --- -gt cat aag tta gtt aag tca t-- --- --- --- --- --- --- --- --- --- --- --- --- --- --- --- -aa gtt agt taa gtc ata agt tag tta agt cat aag tta gtt aag tca taa gtt agt taa gtc ata agt tag tta agt cat aag tta gtt tat agt cta aca ctt cta att ttt ata cct tga tct ttt tct cta att atg aaa aag taa atc att atg aag atg gat g-a aat gga cga gat tgt gcg cat cgt taa cga tag tat gtg gta cgt acc taa cgc att tat gga cga cgg tga taa tga agg tca cat ttc tgt caa taa tgt ctg tca tat gta tct cgc att ctt tga tgt gga tat atc atc tta tct gtt taa att agt tat taa aca ctg cga tct gaa taa acg act aaa atg tgg taa ctc tcc att aca ttg cta tac gat gaa tac acg att taa tcc atc tgt att aaa gat att gtt acg cca cgg cat gcg taa ctt tga tag caa gga taa aaa agg aca tat tcc tct aca cca cta tct gat tca ttc act atc aat cga taa caa gat ctt tga tat act aac gga ccc cat tga tga ctt tag taa atc atc cga tct att gct gtg tta tct tag ata taa att caa tgg gag ctt aaa cta tta cgt tct gta caa att att gac taa agg atc tga ccc taa ttg cgt cga tga gga tgg act cac ttc tct tca tta cta ctg taa aca cat atc cgc gtt cca cga aag caa tta tta caa gtc aaa gag tca cac taa gat gcg agc tga gaa gcg att cat cta cgc gat aat aga tca tgg agc aaa cat taa cgc ggt tac gaa aat cgg aaa tac gcc gtt aca cac tta cct tca aca gta tac caa aca tag tcc tcg tgt ggt gta tgc tct ttt atc tcg agg agc cga tac gag gat acg taa taa tct taa ttg tac acc cat cat gga ata cat aaa gaa cga ttg tgc aac agg tca tat tct cat aat gtt act caa ttg gca cga aca aaa ata cgg gaa att aca aaa gga aga agg aca aca tct act tta tct att cat aaa aca taa tca agg ata tgg aag tcg ctc tct caa tat act acg gta tct act aga tag att cga cat tca gaa aga cga ata cta taa tac aat gac tcc tct tca tac cgc ctt cca gaa ttg caa taa caa tgt tgc ctc ata cct cgt ata cat tgg ata cga cat caa cct tcc gac taa aga cga taa gac agt att cga ctt ggt gtt tga aaa cag aaa cat tat ata caa ggc gga tgt cgt taa tga cat tat cca cca cag act gaa agt atc tct acc tat gat taa atc gtt gtt cta caa gat gtc gga gtt ctc tcc cta cga cga tca cta cgt aaa gaa gat aat agc cta ctg cct att aag gga cga gtc att tgc gga act aca tac taa att ctg ttt aaa cga gga cta taa aag tgt att tat gaa aaa tat atc att cga taa gat aga ttc cat cat cga aaa atg tag tcg tga cat aag tct cct caa aga gat tcg aat ctc aga cac cga ctt gta tac ggt att gag aac aga aga cat tcg gta tca cac ata tct cga agc cat aca ttc aga caa acg cat ttc att tcc cat gta cga cga tct cat aga aca gtg tca tct atc gat gga gca taa aag taa act cgt cga caa agc act caa taa att aga gtc tac cat cga tag tca atc tag act atc gta ttt gcc tcc gga aat tat gcg caa tat cat aac caa gct aag cga cta cca tct aaa cag tat gtt gta cgg aaa gaa cca tta caa ata tta tcc atg ata gaa aga aaa tat tta aaa aat aat cta tat gat tgg aga agt agg aaa caa aca gta aca aga cga cga tta cta c-a tta tta aat cat gag gtc cgt att ata ctc gta tat att gtt tct ctc atg tat aat aat aaa cgg aag aga tat agc acc aca tgc acc atc caa tgg aaa gtg taa aga caa cga ata cag aag ccg taa tct atg ttg tct atc gtg tcc tcc ggg aac tta cgc ttt cag att atg tga tag caa gac taa tac aca atg tac acc gtg tgg ttc gga tac ctt tac atc tca caa taa tca ttt aca ggc ttg tct aag ttg taa cgg aag atg tga tag taa tca ggt aga gac gcg atc gtg taa cac gac tca caa tag aat ctg tga atg ctc tcc agg ata tta ttg tct tct caa agg agc atc agg gtg tag aac atg tat ttc taa aac aaa gtg tgg aat agg ata cgg agt atc cgg ata cac gtc tac cgg aga cgt cat ctg ttc tcc gtg tgg tcc cgg aac ata ttc tca cac cgt ctc ttc cac aga taa atg cga acc cgt cgt aac cag caa tac att taa cta tat cga tgt gga aat taa cct gta tcc agt caa cga cac atc gtg tac tcg gac gac cac tac cgg tct cag cga atc cat ctc aac gtc gga act aac tat tac cat gaa tca taa aga ttg tga tcc agt ctt tcg tgc aga ata ctt ctc tgt cct taa taa tgt agc aac ttc agg att ctt tac agg aga aaa tag ata tca gaa tac ttc aaa gat atg tac tct gaa ttt cga gat taa atg taa caa caa aga ttc atc ttc caa aca gtt aac gaa aac aaa gaa tga tac tat cat gcc gca ttc aga gac ggt aac tct agt ggg cga ctg tct atc tag cgt cga cat cta cat act ata tag taa tac caa tac tca aga cta cga aaa tga tac aat ctc tta tca tat ggg taa tgt tct cga tgt caa tag cca tat gcc cgc tag ttg cga tat aca taa act gat cac taa ttc cca gaa tcc cac cca ctt ata gta ag- -tt ttt tta cct ata aat aat aaa tac aat aat taa ttt ctc gta aaa gta gaa aat ata ttc taa ttt att ata tgg taa gaa agt aga atc atc tag aac agt aat caa tca ata gca atc atg aaa caa tat att gtc ctg gca tgc atg tgc cta gtg gca gct gct atg cct act agt ctt caa caa tct tca tcc tcg tgt act gaa gaa gaa aac aaa cat cat atg gga atc gat gtt att atc aaa gtc aca aag caa gac caa aca ccg acc aat gat aag att tgt caa tcc gta acg gaa gtt aca gag a-- --- --- --- --- --- -cc gaa gat gat gag gta tcc gaa gaa gtt gta aaa gga gat ccc acc act tat tac act atc gtc ggt gcg ggt ctt aac atg aac ttt gga ttc acc aaa tgc cca aag att tta tcc atc tcc gaa tcc tct gat gga aac act gtg aat act aga ttg tcc agc gtg tca ccg gga caa ggt aag gac tct ccc gcg atc acg cgt gaa gaa gct ctg gct atg atc aaa gac tgt gag atg tct atc gac atc aga tgt agc gaa gaa gag aaa gac agt gac atc aag acc cat cca gta ctt ggg tct aac atc tca cat aag aaa gtg agt tac aaa gat atc atc ggt tca acg atc gtt gat aca aaa tgt gtc aag aac cta gag ttt agc gta cgt atc gga gac atg tgt gag gaa tca tct gaa ctt gaa gtc aag gat gga ttc aag tat gtc gac gga tcg gca tct gaa ggt gca acc gat gat act tca ctc atc gat tca aca aaa ctc aaa gca tgt gtc tga atc gat aac tct att cat ctg aaa atg gat gag ttg ggt taa tcg aac gat tca gac acc gca cca cga att aaa aaa g-- -ac cgg gca cta tat tcc ggt ttg caa aac aaa aat att taa cta cat tca caa aaa gtt acc tct cgt tac ttc ttc ttt ctg ttt caa tat gtg ata cga tat gat cac tat tcg tat tct ct- --- --- tgg tct cat aaa aaa gtt tta c-- --a aaa aaa aaa aaa a-t att ttt att ctc ttt ctc tct tcg at- ggt ctc aca aaa ata tta aac ctc ttt ctg atg tct caa cta ttt cgt aaa cga taa cgt cca aca ata tat tct cgt aga gct tat caa cat cct tat acc aat cta ggt tgt cag aca att gca tca taa aat aat gtt tat aat tta cac gtt aac atc ata taa taa acg tat ata gtt aat att ttt gga ata taa atg atc tgt aaa atc cat gta ggg gac act gct cac gtt ttt tct cta gta cat aat ttc aca caa gtt ttt ata cag aca aat taa ttc tcg tcc ata ta- ttt taa aac att gac ttt tgt act aag aaa aat atc ttg act aac cat ctc ttt ctc tct tcg atg ggt ctc aca aaa ata tta aac ctc ttt ctg atg g-- --- --- --- --- --- --- --- --- --- --- --- --- --- --- --- --- --- --- --- --- --- --- --- --- --- --- --- --- --- --- --- --- --- --- --- --- --- --- --- --- --- --- --- --- --- --- agt cgt aaa aag ttt tta tcc ttt ctc tct tcg ata ggt ctc --- --- --- --- --- --- --- --- --- --- --- --- --- --- --- --- --- --- --- --- --- --- --- --- --- --- --- --- --- --- --- --- --- --- --- --- --- --- --- --- --- --- --- --- --- --- --- --- --- --- --- --- --- --- --- --- --- --- --- --- --- --- --- --- --- --- --- --- --- --- aca aaa ata tta aac ctc ttt ctg atg gtc tct ata aa

>ON682270.1_Monkeypox_virus_isolate_MPXV/Germany/2022/RKI06_complete_genome

atc gag gac gac ttc aac tat gac agc tcg gtg gcg tct gcc agc gtg tac ata cga atg gca ttt cta aga aaa gtc tac ggt atc ctt -tc tac aat ttc ctt taa caa cgg caa cag ctg cag tat ttt tat act ttg aat gca -tc gga cat tta tac aag gga gtc ctg ttc taa tat tgg cat caa tgt tcg gat cta tag gct tga ttt tcg cat tga ctt tac aca gac atg aac atc ccc tga atc tgt aca tac ttt gtg gat tta cac tgt tag aat ctc taa cgc tgg cct ctg ttg tta ctt tct atg atg cac gta tcg tta tgc aag ctt tca tgt tga cta ctg cag tgt ttc ttg ctc tga cta cat ata ctc tac aat caa aga gag att tca gta aac ttg taa cag gat tgt ttg ctg ctt tct gga ttt taa ttt tgt cag gag tct tga gga taa agt tta aaa tag aat taa taa aga aca tat agg tca -tt ttt taa aca tgg ata gaa acc aag gtt gtt agt taa taa tat aca aga tat ttt ttc tca ctc tga tcc atg taa acc aag gac gag aga cac tct cat tcc tca ttc acg aca cca tta aaa atg gaa att aaa gcc ctc tat taa gca cag acg gct aca ggt cta cca tca ggt t-a cct tcg tct acc ttc aca atg gcc tct cct tgt gcc cag ttc agt ccc tgt cat tgc cac gct act aag gac tcc ctg aat acc gtg act gac gtc aga cat tgt ctg act gaa tac atc ctg tgg gtt tct cat aga tgg acc cat aga gaa agc gca ggg cct ctc tac agg ctt ctc atc tct ttc aga att gat gca atg gag cta ttt ggt agc gag ttg aag gag ttc tcg aat tca ctt ccg tgg gac aat atc gac aat tgc gtg gag atc att aaa tgt ttc atc aga aat gac tcc atg aaa acc gcc aaa gaa ctt tgt gca ata att gga ctt tgt act caa tca gct att gtc act gga aga gtc ttc aat gat aag tat atc gac ata cta ctt atg ctg cga aag att ctg aac gag aac gac tat ctc acc ctc ttg gat cat atc ctc act gct aaa tac taa atc tcc ttc atg ctc tct cac taa tac tct tac tca cta cac ttt tta tca tct tat gat gaa tga ttg cct tca tca ttt ttt cgt gga ata taa tat agg aat aat tag cac cag aat agc tat gga tat ctc gtt aag aat att ctc tat aag aga cat aat gta gac ata gtt att ata tcc ttc tta gat aag tgt tac gct act gga aag ttt cca tcg tta tta tta cat gaa gat gat ata att aaa cca aca ttg aga ttg gct ctt atg tta gct gga ttg aat tac tgt aat aaa tgc atc gag tat aga ggg ata tag caa ttc tcg ata ata gtc atg caa tat ttg aat gag act gat aat tta ggt aat aca gta ct- --- aca cac ata tct ttc tag att ata tat cgt taa aaa tct gta aga tgt ata ttt ctc ata agt atc cac tgt gta ata tta tta atg gat ata tag ata acg caa tag gga cta ata gta ttg taa aag ata taa tcg act att tac gta cat atc cag ata tct ata ttc cta cta gtt tgc tgc gta gtt gca tca ttg ata tgc atg -at tta tca gga ttc aga gat gaa tta cta agt aaa cta caa tcc cac aat aag taa gaa tca aat atc aaa aac tca ctt ttg att ttt cta gtc tta agt aat aca tat att tat taa tag acc tat gaa ata aaa aaa ggt aac aat gga ttc gcg tat agc tat tta cgt att agt atc ggc atc tct ttt gta tct tgt taa ttg tca caa act agt aca tta ctt caa tct gaa aat aaa tgg aag tga tat aac taa tac agc aga tat att gct gga caa tta tcc aat tat gac ctt tga tgg aaa gga tat tta tcc atc tat ctc gtt cat ggt cgg taa taa act ttt cct aga tct tta taa aaa tat ctt tga aga att ttt cag act att tcg agt atc tgt aag tag tca ata cga gga att aga ata tta tta ttc atg tga tta tac taa caa ccg tcc tac aat taa aca aca tta ctt tta taa cgg cga cga ata tac tga aat tga tag atc gaa aaa agc cac taa taa aaa cag ttg gtt aat tac ttc agg ctt tag act aca aaa atg gtt cga tag cga aga ttg tat aat tta tct cag atc ttt agt tag aag aat gga aga cag taa caa aaa cag taa aaa aac tta gta ctt aga tat cg- --a aaa aat ata ttt ttg tag act ctt gag aat aga agg aaa aca tgt aca taa tta taa aaa atg aaa atc aat ggc gaa taa gac agt gcg att cgc acc atg gag tcg gta gat ttc atg gct gtc gat gag cag ttt cac gac gac ctc gat ctt tgg tca tta tct ttg gta gat gat tat aaa aaa cat gga tta ggt gtt gac tgt tat gtt cta gaa cca gtt gtt gac agg aaa ata ttt gat aga ttt ctc ctt gaa cca att tgt gat cct gta gat gtt ctg tat gat tat ttt agg att cat aga gat aat att gat cag tat ata gta gat aga ctg ttt gca tat att aca tat aaa gat att ata tct gca tta gtg tca aag aat tat atg gaa gat att ttc tct ata att att aag aat tgt aat tct gtg caa gat ctc tta ctt tac tat cta tct aat gca tat gta gaa ata gac att gtt gat ctt atg gta gat cat ggg gct gta ata tat aaa ata gaa tgc ttg aat gcc tat ttt agg gga ata tgt aaa aag gaa agt agt gtt gtt gag ttt att ttg aat tgt ggt atc cca gat gaa aat gat gtt aaa tta gat cta tat aaa ata att cag tat act agg gga ttc ctt gta gat gaa ccc aca gta tta gaa att tat aag ctt tgt atc cca tat att gaa gat atc aat caa cta gat gct ggt gga agg acc ttg ctt tat cgc gct atc tat gca ggt tat ata gat tta gta tca tgg cta tta gaa aat gga gca aat gtc aac gca gta atg agt aat gga tat aca tgt ctt gac gtg gcc gtg gat agg gga tct gtc atc gcc cgt agg gaa gca cat ctt aaa ata tta gaa ata ttg ctt aga gaa cca ttg tct att gac tgt ata aaa tta gct ata ctt aat aat aca att gaa aac cat gat gtg ata aag ctc tgt atc aag tat ttt atg atg gta gat tat tca ctt tgt aat gtg tat gca tca tca ctc ttt gat tat ata att gat tgt aaa caa gaa ttg gag tac att agg cag atg aaa att cat aat aca acc atg tat gag tta atc tat aat aga gac aaa aac aag cat gct tcc cat att cta cat agg tat tct aaa cat cca gtt ttg aca cag tgt atc act aaa gga ttc aag att tac aca gaa gta acc gag cag gtc act aaa gct cta aac aga cgt gct cta ata gat gag ata ata aac aat gta tca act gat gac aat ctc cta tca aaa ctt cca tta gaa att agg gat cta att gtt tca caa gct gtc ata tag agt tct atc cac cca cct ttc ttg aaa tga gtt aat agt cat aag tt- --- --- --- --- --- --- --- --- --- --- --- --- --- --- --- --- --- --- --- --- --- --- --- --- --- --- --- --- --- --- --- --- --- --- --- --- --- --- --- --- --- --- --- --- --- --- --- --- --- --- --- --- --- --- --- --- --- --- --- --- --- --- --- --- --- --- --- --- --- --- --- --- --- --- --- --- --- --- --- --- --- --- --- --- --- --- --- --- --- agt taa gtc ata agt tag tta agt cat aag tta gtt aag tca taa gtt agt taa --- --- --- --- --- -gt cat aag tta gtt aag tca t-- --- --- --- --- --- --- --- --- --- --- --- --- --- --- --- -aa gtt agt taa gtc ata agt tag tta agt cat aag tta gtt aag tca taa gtt agt taa gtc ata agt tag tta agt cat aag tta gtt tat agt cta aca ctt cta att ttt ata cct tga tct ttt tct cta att atg aaa aag taa atc att atg aag atg gat g-a aat gga cga gat tgt gcg cat cgt taa cga tag tat gtg gta cgt acc taa cgc att tat gga cga cgg tga taa tga agg tca cat ttc tgt caa taa tgt ctg tca tat gta tct cgc att ctt tga tgt gga tat atc atc tca tct gtt taa att agt tat taa aca ctg cga tct gaa taa acg act aaa atg tgg taa ctc tcc att aca ttg cta tac gat gaa tac acg att taa tcc atc tgt att aaa gat att gtt acg cca cgg cat gcg taa ctt tga tag caa gga taa aaa agg aca tat tcc tct aca cca cta tct gat tca ttc act atc aat cga taa caa gat ctt tga tat act aac gga ccc cat tga tga ctt tag taa atc atc cga tct att gct gtg tta tct tag ata taa att caa tgg gag ctt aaa cta tta cgt tct gta caa att att gac taa agg atc tga ccc taa ttg cgt cga tga gga tgg act cac ttc tct tca tta cta ctg taa aca cat atc cgc gtt cca cga aag caa tta tta caa gtc aaa gag tca cac taa gat gcg agc tga gaa gcg att cat cta cgc gat aat aga tca tgg agc aaa cat taa cgc ggt tac gaa aat cgg aaa tac gcc gtt aca cac tta cct tca aca gta tac caa aca tag tcc tcg tgt ggt gta tgc tct ttt atc tcg agg agc cga tac gag gat acg taa taa tct taa ttg tac acc cat cat gga ata cat aaa gaa cga ttg tgc aac agg tca tat tct cat aat gtt act caa ttg gca cga aca aaa ata cgg gaa att aca aaa gga aga agg aca aca tct act tta tct att cat aaa aca taa tca agg ata tgg aag tcg ctc tct caa tat act acg gta tct act aga tag att cga cat tca gaa aga cga ata cta taa tac aat gac tcc tct tca tac cgc ctt cca gaa ttg caa taa caa tgt tgc ctc ata cct cgt ata cat tgg ata cga cat caa cct tcc gac taa aga cga taa gac agt att cga ctt ggt gtt tga aaa cag aaa cat tat ata caa ggc gga tgt cgt taa tga cat tat cca cca cag act gaa agt atc tct acc tat gat taa atc gtt gtt cta caa gat gtc gga gtt ctc tcc cta cga cga tca cta cgt aaa gaa gat aat agc cta ctg cct att aag gga cga gtc att tgc gga act aca tac taa att ctg ttt aaa cga gga cta taa aag tgt att tat gaa aaa tat atc att cga taa gat aga ttc cat cat cga aaa atg tag tcg tga cat aag tct cct caa aga gat tcg aat ctc aga cac cga ctt gta tac ggt att gag aac aga aga cat tcg gta tca cac ata tct cga agc cat aca ttc aga caa acg cat ttc att tcc cat gta cga cga tct cat aga aca gtg tca tct atc gat gga gca taa aag taa act cgt cga caa agc act caa taa att aga gtc tac cat cga tag tca atc tag act atc gta ttt gcc tcc gga aat tat gcg caa tat cat aac caa gct aag cga cta cca tct aaa cag tat gtt gta cgg aaa gaa cca tta caa ata tta tcc atg ata gaa aga aaa tat tta aaa aat aat cta tat gat tgg aga agt agg aaa caa aca gta aca aga cga cga tta cta c-a tta tta aat cat gag gtc cgt att ata ctc gta tat att gtt tct ctc atg tat aat aat aaa cgg aag aga tat agc acc aca tgc acc atc caa tgg aaa gtg taa aga caa cga ata cag aag ccg taa tct atg ttg tct atc gtg tcc tcc ggg aac tta cgc ttt cag att atg tga tag caa gac taa tac aca atg tac acc gtg tgg ttc gga tac ctt tac atc tca caa taa tca ttt aca ggc ttg tct aag ttg taa cgg aag atg tga tag taa tca ggt aga gac gcg atc gtg taa cac gac tca caa tag aat ctg tga atg ctc tcc agg ata tta ttg tct tct caa agg agc atc agg gtg tag aac atg tat ttc taa aac aaa gtg tgg aat agg ata cgg agt atc cgg ata cac gtc tac cgg aga cgt cat ctg ttc tcc gtg tgg tcc cgg aac ata ttc tca cac cgt ctc ttc cac aga taa atg cga acc cgt cgt aac cag caa tac att taa cta tat cga tgt gga aat taa cct gta tcc agt caa cga cac atc gtg tac tcg gac gac cac tac cgg tct cag cga atc cat ctc aac gtc gga act aac tat tac cat gaa tca taa aga ttg tga tcc agt ctt tcg tgc aga ata ctt ctc tgt cct taa taa tgt agc aac ttc agg att ctt tac agg aga aaa tag ata tca gaa tac ttc aaa gat atg tac tct gaa ttt cga gat taa atg taa caa caa aga ttc atc ttc caa aca gtt aac gaa aac aaa gaa tga tac tat cat gcc gca ttc aga gac ggt aac tct agt ggg cga ctg tct atc tag cgt cga cat cta cat act ata tag taa tac caa tac tca aga cta cga aaa tga tac aat ctc tta tca tat ggg taa tgt tct cga tgt caa tag cca tat gcc cgc tag ttg cga tat aca taa act gat cac taa ttc cca gaa tcc cac cca ctt ata gta ag- -tt ttt tta cct ata aat aat aaa tac aat aat taa ttt ctc gta aaa gta gaa aat ata ttc taa ttt att ata tgg taa gaa agt aga atc atc tag aac agt aat caa tca ata gca atc atg aaa caa tat att gtc ctg gca tgc atg tgc cta gtg gca gct gct atg cct act agt ctt caa caa tct tca tcc tcg tgt act gaa gaa gaa aac aaa cat cat atg gga atc gat gtt att atc aaa gtc aca aag caa gac caa aca ccg acc aat gat aag att tgt caa tcc gta acg gaa gtt aca gag a-- --- --- --- --- --- -cc gaa gat gat gag gta tcc gaa gaa gtt gta aaa gga gat ccc acc act tat tac act atc gtc ggt gcg ggt ctt aac atg aac ttt gga ttc acc aaa tgc cca aag att tta tcc atc tcc gaa tcc tct gat gga aac act gtg aat act aga ttg tcc agc gtg tca ccg gga caa ggt aag gac tct ccc gcg atc acg cgt gaa gaa gct ctg gct atg atc aaa gac tgt gag atg tct atc gac atc aga tgt agc gaa gaa gag aaa gac agt gac atc aag acc cat cca gta ctt ggg tct aac atc tca cat aag aaa gtg agt tac aaa gat atc atc ggt tca acg atc gtt gat aca aaa tgt gtc aag aac cta gag ttt agc gta cgt atc gga gac atg tgt gag gaa tca tct gaa ctt gaa gtc aag gat gga ttc aag tat gtc gac gga tcg gca tct gaa ggt gca acc gat gat act tca ctc atc gat tca aca aaa ctc aaa gca tgt gtc tga atc gat aac tct att cat ctg aaa atg gat gag ttg ggt taa tcg aac gat tca gac acc gca cca cga att aaa aaa g-- -ac cgg gca cta tat tcc ggt ttg caa aac aaa aat att taa cta cat tca caa aaa gtt acc tct cgt tac ttc ttc ttt ctg ttt caa tat gtg ata cga tat gat cac tat tcg tat tct ct- --- --- tgg tct cat aaa aaa gtt tta c-- --a aaa aaa aaa aaa a-t att ttt att ctc ttt ctc tct tcg at- ggt ctc aca aaa ata tta aac ctc ttt ctg atg tct caa cta ttt cgt aaa cga taa cgt cca aca ata tat tct cgt aga gct tat caa cat cct tat acc aat cta ggt tgt cag aca att gca tca taa aat aat gtt tat aat tta cac gtt aac atc ata taa taa acg tat ata gtt aat att ttt gga ata taa atg atc tgt aaa atc cat gta ggg gac act gct cac gtt ttt tct cta gta cat aat ttc aca caa gtt ttt ata cag aca aat taa ttc tcg tcc ata ta- ttt taa aac att gac ttt tgt act aag aaa aat atc ttg act aac cat ctc ttt ctc tct tcg atg ggt ctc aca aaa ata tta aac ctc ttt ctg atg g-- --- --- --- --- --- --- --- --- --- --- --- --- --- --- --- --- --- --- --- --- --- --- --- --- --- --- --- --- --- --- --- --- --- --- --- --- --- --- --- --- --- --- --- --- --- --- agt cgt aaa aag ttt tta tcc ttt ctc tct tcg ata ggt ctc --- --- --- --- --- --- --- --- --- --- --- --- --- --- --- --- --- --- --- --- --- --- --- --- --- --- --- --- --- --- --- --- --- --- --- --- --- --- --- --- --- --- --- --- --- --- --- --- --- --- --- --- --- --- --- --- --- --- --- --- --- --- --- --- --- --- --- --- --- --- aca aaa ata tta aac ctc ttt ctg atg gtc tct ata aa

>ON637938.1_Monkeypox_virus_isolate_MPXV/Germany/2022/RKI01_complete_genome

atc gag gac gac ttc aac tat gac agc tcg gtg gcg tct gcc agc gtg tac ata cga atg gca ttt cta aga aaa gtc tac ggt atc ctt -tc tac aat ttc ctt taa caa cgg caa cag ctg cag tat ttt tat act ttg aat gca -tc gga cat tta tac aag gga gtc ctg ttc taa tat tgg cat caa tgt tcg gat cta tag gct tga ttt tcg cat tga ctt tac aca gac atg aac atc ccc tga atc tgt aca tac ttt gtg gat tta cac tgt tag aat ctc taa cgc tgg cct ctg ttg tta ctt tct atg atg cac gta tcg tta tgc aag ctt tca tgt tga cta ctg cag tgt ttc ttg ctc tga cta cat ata ctc tac aat caa aga gag att tca gta aac ttg taa cag gat tgt ttg ctg ctt tct gga ttt taa ttt tgt cag gag tct tga gga taa agt tta aaa tag aat taa taa aga aca tat agg tca -tt ttt taa aca tgg ata gaa acc aag gtt gtt agt taa taa tat aca aga tat ttt ttc tca ctc tga tcc atg taa acc aag gac gag aga cac tct cat tcc tca ttc acg aca cca tta aaa atg gaa att aaa gcc ctc tat taa gca cag acg gct aca ggt cta cca tca ggt t-a cct tcg tct acc ttc aca atg gcc tct cct tgt gcc cag ttc agt ccc tgt cat tgc cac gct act aag gac tcc ctg aat acc gtg act gac gtc aga cat tgt ctg act gaa tac atc ctg tgg gtt tct cat aga tgg acc cat aga gaa agc gca ggg cct ctc tac agg ctt ctc atc tct ttc aga att gat gca atg gag cta ttt ggt agc gag ttg aag gag ttc tcg aat tca ctt ccg tgg gac aat atc gac aat tgc gtg gag atc att aaa tgt ttc atc aga aat gac tcc atg aaa acc gcc aaa gaa ctt tgt gca ata att gga ctt tgt act caa tca gct att gtc act gga aga gtc ttc aat gat aag tat atc gac ata cta ctt atg ctg cga aag att ctg aac gag aac gac tat ctc acc ctc ttg gat cat atc ctc act gct aaa tac taa atc tcc ttc atg ctc tct cac taa tac tct tac tca cta cac ttt tta tca tct tat gat gaa tga ttg cct tca tca ttt ttt cgt gga ata taa tat agg aat aat tag cac cag aat agc tat gga tat ctc gtt aag aat att ctc tat aag aga cat aat gta gac ata gtt att ata tcc ttc tta gat aag tgt tac gct act gga aag ttt cca tcg tta tta tta cat gaa gat gat ata att aaa cca aca ttg aga ttg gct ctt atg tta gct gga ttg aat tac tgt aat aaa tgc atc gag tat aga ggg ata tag caa ttc tcg ata ata gtc atg caa tat ttg aat gag act gat aat tta ggt aat aca gta ct- --- aca cac ata tct ttc tag att ata tat cgt taa aaa tct gta aga tgt ata ttt ctc ata agt atc cac tgt gta ata tta tta atg gat ata tag ata acg caa tag gga cta ata gta ttg taa aag ata taa tcg act att tac gta cat atc cag ata tct ata ttc cta cta gtt tgc tgc gta gtt gca tca ttg ata tgc atg -at tta tca gga ttc aga gat gaa tta cta agt aaa cta caa tcc cac aat aag taa gaa tca aat atc aaa aac tca ctt ttg att ttt cta gtc tta agt aat aca tat att tat taa tag acc tat gaa ata aaa aaa ggt aac aat gga ttc gcg tat agc tat tta cgt att agt atc ggc atc tct ttt gta tct tgt taa ttg tca caa act agt aca tta ctt caa tct gaa aat aaa tgg aag tga tat aac taa tac agc aga tat att gct gga caa tta tcc aat tat gac ctt tga tgg aaa gga tat tta tcc atc tat ctc gtt cat ggt cgg taa taa act ttt cct aga tct tta taa aaa tat ctt tga aga att ttt cag act att tcg agt atc tgt aag tag tca ata cga gga att aga ata tta tta ttc atg tga tta tac taa caa ccg tcc tac aat taa aca aca tta ctt tta taa cgg cga cga ata tac tga aat tga tag atc gaa aaa agc cac taa taa aaa cag ttg gtt aat tac ttc agg ctt tag act aca aaa atg gtt cga tag cga aga ttg tat aat tta tct cag atc ttt agt tag aag aat gga aga cag taa caa aaa cag taa aaa aac tta gta ctt aga tat cg- --a aaa aat ata ttt ttg tag act ctt gag aat aga agg aaa aca tgt aca taa tta taa aaa atg aaa atc aat ggc gaa taa gac agt gcg att cgc acc atg gag tcg gta gat ttc atg gct gtc gat gag cag ttt cac gac gac ctc gat ctt tgg tca tta tct ttg gta gat gat tat aaa aaa cat gga tta ggt gtt gac tgt tat gtt cta gaa cca gtt gtt gac agg aaa ata ttt gat aga ttt ctc ctt gaa cca att tgt gat cct gta gat gtt ctg tat gat tat ttt agg att cat aga gat aat att gat cag tat ata gta gat aga ctg ttt gca tat att aca tat aaa gat att ata tct gca tta gtg tca aag aat tat atg gaa gat att ttc tct ata att att aag aat tgt aat tct gtg caa gat ctc tta ctt tac tat cta tct aat gca tat gta gaa ata gac att gtt gat ctt atg gta gat cat ggg gct gta ata tat aaa ata gaa tgc ttg aat gcc tat ttt agg gga ata tgt aaa aag gaa agt agt gtt gtt gag ttt att ttg aat tgt ggt atc cca gat gaa aat gat gtt aaa tta gat cta tat aaa ata att cag tat act agg gga ttc ctt gta gat gaa ccc aca gta tta gaa att tat aag ctt tgt atc cca tat att gaa gat atc aat caa cta gat gct ggt gga agg acc ttg ctt tat cgc gct atc tat gca ggt tat ata gat tta gta tca tgg cta tta gaa aat gga gca aat gtc aac gca gta atg agt aat gga tat aca tgt ctt gac gtg gcc gtg gat agg gga tct gtc atc gcc cgt agg gaa gca cat ctt aaa ata tta gaa ata ttg ctt aga gaa cca ttg tct att gac tgt ata aaa tta gct ata ctt aat aat aca att gaa aac cat gat gtg ata aag ctc tgt atc aag tat ttt atg atg gta gat tat tca ctt tgt aat gtg tat gca tca tca ctc ttt gat tat ata att gat tgt aaa caa gaa ttg gag tac att agg cag atg aaa att cat aat aca acc atg tat gag tta atc tat aat aga gac aaa aac aag cat gct tcc cat att cta cat agg tat tct aaa cat cca gtt ttg aca cag tgt atc act aaa gga ttc aag att tac aca gaa gta acc gag cag gtc act aaa gct cta aac aga cgt gct cta ata gat gag ata ata aac aat gta tca act gat gac aat ctc cta tca aaa ctt cca tta gaa att agg gat cta att gtt tca caa gct gtc ata tag agt tct atc cac cca cct ttc ttg aaa tga gtt aat a-- --- --- --- --- --- --- --- --- --- --- --- --- --- --- --- --- --- --- --- --- --- --- --- --- --- --- --- --- --- --- --- --- --- --- --- --- --- --- --- --- --- --- --- --- --- --- --- --- --- --- --- --- --- --- --- --- --- --- --- --- --- --- --- --- --- --- --- --- --- --- --- --- --- --- --- --- --- --- --- --- --- --- --- --- --- --- --- --- --- --- --- --- --- --- gtc ata agt tag tta agt cat aag tta gtt aag tca taa gtt agt taa --- --- --- --- --- -gt cat aag tta gtt aag tca t-- --- --- --- --- --- --- --- --- --- --- --- --- --- --- --- -aa gtt agt taa gtc ata agt tag tta agt cat aag tta gtt aag tca taa gtt agt taa gtc ata agt tag tta agt cat aag tta gtt tat agt cta aca ctt cta att ttt ata cct tga tct ttt tct cta att atg aaa aag taa atc att atg aag atg gat g-a aat gga cga gat tgt gcg cat cgt taa cga tag tat gtg gta cgt acc taa cgc att tat gga cga cgg tga taa tga agg tca cat ttc tgt caa taa tgt ctg tca tat gta tct cgc att ctt tga tgt gga tat atc atc tca tct gtt taa att agt tat taa aca ctg cga tct gaa taa acg act aaa atg tgg taa ctc tcc att aca ttg cta tac gat gaa tac acg att taa tcc atc tgt att aaa gat att gtt acg cca cgg cat gcg taa ctt tga tag caa gga taa aaa agg aca tat tcc tct aca cca cta tct gat tca ttc act atc aat cga taa caa gat ctt tga tat act aac gga ccc cat tga tga ctt tag taa atc atc cga tct att gct gtg tta tct tag ata taa att caa tgg gag ctt aaa cta tta cgt tct gta caa att att gac taa agg atc tga ccc taa ttg cgt cga tga gga tgg act cac ttc tct tca tta cta ctg taa aca cat atc cgc gtt cca cga aag caa tta tta caa gtc aaa gag tca cac taa gat gcg agc tga gaa gcg att cat cta cgc gat aat aga tca tgg agc aaa cat taa cgc ggt tac gaa aat cgg aaa tac gcc gtt aca cac tta cct tca aca gta tac caa aca tag tcc tcg tgt ggt gta tgc tct ttt atc tcg agg agc cga tac gag gat acg taa taa tct taa ttg tac acc cat cat gga ata cat aaa gaa cga ttg tgc aac agg tca tat tct cat aat gtt act caa ttg gca cga aca aaa ata cgg gaa att aca aaa gga aga agg aca aca tct act tta tct att cat aaa aca taa tca agg ata tgg aag tcg ctc tct caa tat act acg gta tct act aga tag att cga cat tca gaa aga cga ata cta taa tac aat gac tcc tct tca tac cgc ctt cca gaa ttg caa taa caa tgt tgc ctc ata cct cgt ata cat tgg ata cga cat caa cct tcc gac taa aga cga taa gac agt att cga ctt ggt gtt tga aaa cag aaa cat tat ata caa ggc gga tgt cgt taa tga cat tat cca cca cag act gaa agt atc tct acc tat gat taa atc gtt gtt cta caa gat gtc gga gtt ctc tcc cta cga cga tca cta cgt aaa gaa gat aat agc cta ctg cct att aag gga cga gtc att tgc gga act aca tac taa att ctg ttt aaa cga gga cta taa aag tgt att tat gaa aaa tat atc att cga taa gat aga ttc cat cat cga aaa atg tag tcg tga cat aag tct cct caa aga gat tcg aat ctc aga cac cga ctt gta tac ggt att gag aac aga aga cat tcg gta tca cac ata tct cga agc cat aca ttc aga caa acg cat ttc att tcc cat gta cga cga tct cat aga aca gtg tca tct atc gat gga gca taa aag taa act cgt cga caa agc act caa taa att aga gtc tac cat cga tag tca atc tag act atc gta ttt gcc tcc gga aat tat gcg caa tat cat aac caa gct aag cga cta cca tct aaa cag tat gtt gta cgg aaa gaa cca tta caa ata tta tcc atg ata gaa aga aaa tat tta aaa aat aat cta tat gat tgg aga agt agg aaa caa aca gta aca aga cga cga tta cta c-a tta tta aat cat gag gtc cgt att ata ctc gta tat att gtt tct ctc atg tat aat aat aaa cgg aag aga tat agc acc aca tgc acc atc caa tgg aaa gtg taa aga caa cga ata cag aag ccg taa tct atg ttg tct atc gtg tcc tcc ggg aac tta cgc ttt cag att atg tga tag caa gac taa tac aca atg tac acc gtg tgg ttc gga tac ctt tac atc tca caa taa tca ttt aca ggc ttg tct aag ttg taa cgg aag atg tga tag taa tca ggt aga gac gcg atc gtg taa cac gac tca caa tag aat ctg tga atg ctc tcc agg ata tta ttg tct tct caa agg agc atc agg gtg tag aac atg tat ttc taa aac aaa gtg tgg aat agg ata cgg agt atc cgg ata cac gtc tac cgg aga cgt cat ctg ttc tcc gtg tgg tcc cgg aac ata ttc tca cac cgt ctc ttc cac aga taa atg cga acc cgt cgt aac cag caa tac att taa cta tat cga tgt gga aat taa cct gta tcc agt caa cga cac atc gtg tac tcg gac gac cac tac cgg tct cag cga atc cat ctc aac gtc gga act aac tat tac cat gaa tca taa aga ttg tga tcc agt ctt tcg tgc aga ata ctt ctc tgt cct taa taa tgt agc aac ttc agg att ctt tac agg aga aaa tag ata tca gaa tac ttc aaa gat atg tac tct gaa ttt cga gat taa atg taa caa caa aga ttc atc ttc caa aca gtt aac gaa aac aaa gaa tga tac tat cat gcc gca ttc aga gac ggt aac tct agt ggg cga ctg tct atc tag cgt cga cat cta cat act ata tag taa tac caa tac tca aga cta cga aaa tga tac aat ctc tta tca tat ggg taa tgt tct cga tgt caa tag cca tat gcc cgc tag ttg cga tat aca taa act gat cac taa ttc cca gaa tcc cac cca ctt ata gta ag- -tt ttt tta cct ata aat aat aaa tac aat aat taa ttt ctc gta aaa gta gaa aat ata ttc taa ttt att ata tgg taa gaa agt aga atc atc tag aac agt aat caa tca ata gca atc atg aaa caa tat att gtc ctg gca tgc atg tgc cta gtg gca gct gct atg cct act agt ctt caa caa tct tca tcc tcg tgt act gaa gaa gaa aac aaa cat cat atg gga atc gat gtt att atc aaa gtc aca aag caa gac caa aca ccg acc aat gat aag att tgt caa tcc gta acg gaa gtt aca gag a-- --- --- --- --- --- -cc gaa gat gat gag gta tcc gaa gaa gtt gta aaa gga gat ccc acc act tat tac act atc gtc ggt gcg ggt ctt aac atg aac ttt gga ttc acc aaa tgc cca aag att tta tcc atc tcc gaa tcc tct gat gga aac act gtg aat act aga ttg tcc agc gtg tca ccg gga caa ggt aag gac tct ccc gcg atc acg cgt gaa gaa gct ctg gct atg atc aaa gac tgt gag atg tct atc gac atc aga tgt agc gaa gaa gag aaa gac agt gac atc aag acc cat cca gta ctt ggg tct aac atc tca cat aag aaa gtg agt tac aaa gat atc atc ggt tca acg atc gtt gat aca aaa tgt gtc aag aac cta gag ttt agc gta cgt atc gga gac atg tgt gag gaa tca tct gaa ctt gaa gtc aag gat gga ttc aag tat gtc gac gga tcg gca tct gaa ggt gca acc gat gat act tca ctc atc gat tca aca aaa ctc aaa gca tgt gtc tga atc gat aac tct att cat ctg aaa atg gat gag ttg ggt taa tcg aac gat tca gac acc gca cca cga att aaa aaa g-- -ac cgg gca cta tat tcc ggt ttg caa aac aaa aat att taa cta cat tca caa aaa gtt acc tct cgt tac ttc ttc ttt ctg ttt caa tat gtg ata cga tat gat cac tat tcg tat tct ct- --- --- tgg tct cat aaa aaa gtt tta c-- --a aaa aaa aaa aaa a-t att ttt att ctc ttt ctc tct tcg at- ggt ctc aca aaa ata tta aac ctc ttt ctg atg tct caa cta ttt cgt aaa cga taa cgt cca aca ata tat tct cgt aga gct tat caa cat cct tat acc aat cta ggt tgt cag aca att gca tca taa aat aat gtt tat aat tta cac gtt aac atc ata taa taa acg tat ata gtt aat att ttt gga ata taa atg atc tgt aaa atc cat gta ggg gac act gct cac gtt ttt tct cta gta cat aat ttc aca caa gtt ttt ata cag aca aat taa ttc tcg tcc ata ta- ttt taa aac att gac ttt tgt act aag aaa aat atc ttg act aac cat ctc ttt ctc tct tcg atg ggt ctc aca aaa ata tta aac ctc ttt ctg atg g-- --- --- --- --- --- --- --- --- --- --- --- --- --- --- --- --- --- --- --- --- --- --- --- --- --- --- --- --- --- --- --- --- --- --- --- --- --- --- --- --- --- --- --- --- --- --- agt cgt aaa aag ttt tta tcc ttt ctc tct tcg ata ggt ctc --- --- --- --- --- --- --- --- --- --- --- --- --- --- --- --- --- --- --- --- --- --- --- --- --- --- --- --- --- --- --- --- --- --- --- --- --- --- --- --- --- --- --- --- --- --- --- --- --- --- --- --- --- --- --- --- --- --- --- --- --- --- --- --- --- --- --- --- --- --- aca aaa ata tta aac ctc ttt ctg atg gtc tct ata aa

>ON637939.1_Monkeypox_virus_isolate_MPXV/Germany/2022/RKI02_complete_genome

atc gag gac gac ttc aac tat gac agc tcg gtg gcg tct gcc agc gtg tac ata cga atg gca ttt cta aga aaa gtc tac ggt atc ctt -tc tac aat ttc ctt taa caa cgg caa cag ctg cag tat ttt tat act ttg aat gca -tc gga cat tta tac aag gga gtc ctg ttc taa tat tgg cat caa tgt tcg gat cta tag gct tga ttt tcg cat tga ctt tac aca gac atg aac atc ccc tga atc tgt aca tac ttt gtg gat tta cac tgt tag aat ctc taa cgc tgg cct ctg ttg tta ctt tct atg atg cac gta tcg tta tgc aag ctt tca tgt tga cta ctg cag tgt ttc ttg ctc tga cta cat ata ctc tac aat caa aga gag att tca gta aac ttg taa cag gat tgt ttg ctg ctt tct gga ttt taa ttt tgt cag gag tct tga gga taa agt tta aaa tag aat taa taa aga aca tat agg tca -tt ttt taa aca tgg ata gaa acc aag gtt gtt agt taa taa tat aca aga tat ttt ttc tca ctc tga tcc atg taa acc aag gac gag aga cac tct cat tcc tca ttc acg aca cca tta aaa atg gaa att aaa gcc ctc tat taa gca cag acg gct aca ggt cta cca tca ggt t-a cct tcg tct acc ttc aca atg gcc tct cct tgt gcc cag ttc agt ccc tgt cat tgc cac gct act aag gac tcc ctg aat acc gtg act gac gtc aga cat tgt ctg act gaa tac atc ctg tgg gtt tct cat aga tgg acc cat aga gaa agc gca ggg cct ctc tac agg ctt ctc atc tct ttc aga att gat gca atg gag cta ttt ggt agc gag ttg aag gag ttc tcg aat tca ctt ccg tgg gac aat atc gac aat tgc gtg gag atc att aaa tgt ttc atc aga aat gac tcc atg aaa acc gcc aaa gaa ctt tgt gca ata att gga ctt tgt act caa tca gct att gtc act gga aga gtc ttc aat gat aag tat atc gac ata cta ctt atg ctg cga aag att ctg aac gag aac gac tat ctc acc ctc ttg gat cat atc ctc act gct aaa tac taa atc tcc ttc atg ctc tct cac taa tac tct tac tca cta cac ttt tta tca tct tat gat gaa tga ttg cct tca tca ttt ttt cgt gga ata taa tat agg aat aat tag cac cag aat agc tat gga tat ctc gtt aag aat att ctc tat aag aga cat aat gta gac ata gtt att ata tcc ttc tta gat aag tgt tac gct act gga aag ttt cca tcg tta tta tta cat gaa gat gat ata att aaa cca aca ttg aga ttg gct ctt atg tta gct gga ttg aat tac tgt aat aaa tgc atc gag tat aga ggg ata tag caa ttc tcg ata ata gtc atg caa tat ttg aat gag act gat aat tta ggt aat aca gta ct- --- aca cac ata tct ttc tag att ata tat cgt taa aaa tct gta aga tgt ata ttt ctc ata agt atc cac tgt gta ata tta tta atg gat ata tag ata acg caa tag gga cta ata gta ttg taa aag ata taa tcg act att tac gta cat atc cag ata tct ata ttc cta cta gtt tgc tgc gta gtt gca tca ttg ata tgc atg -at tta tca gga ttc aga gat gaa tta cta agt aaa cta caa tcc cac aat aag taa gaa tca aat atc aaa aac tca ctt ttg att ttt cta gtc tta agt aat aca tat att tat taa tag acc tat gaa ata aaa aaa ggt aac aat gga ttc gcg tat agc tat tta cgt att agt atc ggc atc tct ttt gta tct tgt taa ttg tca caa act agt aca tta ctt caa tct gaa aat aaa tgg aag tga tat aac taa tac agc aga tat att gct gga caa tta tcc aat tat gac ctt tga tgg aaa gga tat tta tcc atc tat ctc gtt cat ggt cgg taa taa act ttt cct aga tct tta taa aaa tat ctt tga aga att ttt cag act att tcg agt atc tgt aag tag tca ata cga gga att aga ata tta tta ttc atg tga tta tac taa caa ccg tcc tac aat taa aca aca tta ctt tta taa cgg cga cga ata tac tga aat tga tag atc gaa aaa agc cac taa taa aaa cag ttg gtt aat tac ttc agg ctt tag act aca aaa atg gtt cga tag cga aga ttg tat aat tta tct cag atc ttt agt tag aag aat gga aga cag taa caa aaa cag taa aaa aac tta gta ctt aga tat cg- --a aaa aat ata ttt ttg tag act ctt gag aat aga agg aaa aca tgt aca taa tta taa aaa atg aaa atc aat ggc gaa taa gac agt gcg att cgc acc atg gag tcg gta gat ttc atg gct gtc gat gag cag ttt cac gac gac ctc gat ctt tgg tca tta tct ttg gta gat gat tat aaa aaa cat gga tta ggt gtt gac tgt tat gtt cta gaa cca gtt gtt gac agg aaa ata ttt gat aga ttt ctc ctt gaa cca att tgt gat cct gta gat gtt ctg tat gat tat ttt agg att cat aga gat aat att gat cag tat ata gta gat aga ctg ttt gca tat att aca tat aaa gat att ata tct gca tta gtg tca aag aat tat atg gaa gat att ttc tct ata att att aag aat tgt aat tct gtg caa gat ctc tta ctt tac tat cta tct aat gca tat gta gaa ata gac att gtt gat ctt atg gta gat cat ggg gct gta ata tat aaa ata gaa tgc ttg aat gcc tat ttt agg gga ata tgt aaa aag gaa agt agt gtt gtt gag ttt att ttg aat tgt ggt atc cca gat gaa aat gat gtt aaa tta gat cta tat aaa ata att cag tat act agg gga ttc ctt gta gat gaa ccc aca gta tta gaa att tat aag ctt tgt atc cca tat att gaa gat atc aat caa cta gat gct ggt gga agg acc ttg ctt tat cgc gct atc tat gca ggt tat ata gat tta gta tca tgg cta tta gaa aat gga gca aat gtc aac gca gta atg agt aat gga tat aca tgt ctt gac gtg gcc gtg gat agg gga tct gtc atc gcc cgt agg gaa gca cat ctt aaa ata tta gaa ata ttg ctt aga gaa cca ttg tct att gac tgt ata aaa tta gct ata ctt aat aat aca att gaa aac cat gat gtg ata aag ctc tgt atc aag tat ttt atg atg gta gat tat tca ctt tgt aat gtg tat gca tca tca ctc ttt gat tat ata att gat tgt aaa caa gaa ttg gag tac att agg cag atg aaa att cat aat aca acc atg tat gag tta atc tat aat aga gac aaa aac aag cat gct tcc cat att cta cat agg tat tct aaa cat cca gtt ttg aca cag tgt atc act aaa gga ttc aag att tac aca gaa gta acc gag cag gtc act aaa gct cta aac aga cgt gct cta ata gat gag ata ata aac aat gta tca act gat gac aat ctc cta tca aaa ctt cca tta gaa att agg gat cta att gtt tca caa gct gtc ata tag agt tct atc cac cca cct ttc ttg aaa tga gtt aat a-- --- --- --- --- --- --- --- --- --- --- --- --- --- --- --- --- --- --- --- --- --- --- --- --- --- --- --- --- --- --- --- --- --- --- --- --- --- --- --- --- --- --- --- --- --- --- --- --- --- --- --- --- --- --- --- --- --- --- --- --- --- --- --- --- --- --- --- --- --- --- --- --- --- --- --- --- --- --- --- --- --- --- --- --- --- --- --- --- --- --- --- --- --- --- gtc ata agt tag tta agt cat aag tta gtt aag tca taa gtt agt taa --- --- --- --- --- -gt cat aag tta gtt aag tca t-- --- --- --- --- --- --- --- --- --- --- --- --- --- --- --- -aa gtt agt taa gtc ata agt tag tta agt cat aag tta gtt aag tca taa gtt agt taa gtc ata agt tag tta agt cat aag tta gtt tat agt cta aca ctt cta att ttt ata cct tga tct ttt tct cta att atg aaa aag taa atc att atg aag atg gat g-a aat gga cga gat tgt gcg cat cgt taa cga tag tat gtg gta cgt acc taa cgc att tat gga cga cgg tga taa tga agg tca cat ttc tgt caa taa tgt ctg tca tat gta tct cgc att ctt tga tgt gga tat atc atc tca tct gtt taa att agt tat taa aca ctg cga tct gaa taa acg act aaa atg tgg taa ctc tcc att aca ttg cta tac gat gaa tac acg att taa tcc atc tgt att aaa gat att gtt acg cca cgg cat gcg taa ctt tga tag caa gga taa aaa agg aca tat tcc tct aca cca cta tct gat tca ttc act atc aat cga taa caa gat ctt tga tat act aac gga ccc cat tga tga ctt tag taa atc atc cga tct att gct gtg tta tct tag ata taa att caa tgg gag ctt aaa cta tta cgt tct gta caa att att gac taa agg atc tga ccc taa ttg cgt cga tga gga tgg act cac ttc tct tca tta cta ctg taa aca cat atc cgc gtt cca cga aag caa tta tta caa gtc aaa gag tca cac taa gat gcg agc tga gaa gcg att cat cta cgc gat aat aga tca tgg agc aaa cat taa cgc ggt tac gaa aat cgg aaa tac gcc gtt aca cac tta cct tca aca gta tac caa aca tag tcc tcg tgt ggt gta tgc tct ttt atc tcg agg agc cga tac gag gat acg taa taa tct taa ttg tac acc cat cat gga ata cat aaa gaa cga ttg tgc aac agg tca tat tct cat aat gtt act caa ttg gca cga aca aaa ata cgg gaa att aca aaa gga aga agg aca aca tct act tta tct att cat aaa aca taa tca agg ata tgg aag tcg ctc tct caa tat act acg gta tct act aga tag att cga cat tca gaa aga cga ata cta taa tac aat gac tcc tct tca tac cgc ctt cca gaa ttg caa taa caa tgt tgc ctc ata cct cgt ata cat tgg ata cga cat caa cct tcc gac taa aga cga taa gac agt att cga ctt ggt gtt tga aaa cag aaa cat tat ata caa ggc gga tgt cgt taa tga cat tat cca cca cag act gaa agt atc tct acc tat gat taa atc gtt gtt cta caa gat gtc gga gtt ctc tcc cta cga cga tca cta cgt aaa gaa gat aat agc cta ctg cct att aag gga cga gtc att tgc gga act aca tac taa att ctg ttt aaa cga gga cta taa aag tgt att tat gaa aaa tat atc att cga taa gat aga ttc cat cat cga aaa atg tag tcg tga cat aag tct cct caa aga gat tcg aat ctc aga cac cga ctt gta tac ggt att gag aac aga aga cat tcg gta tca cac ata tct cga agc cat aca ttc aga caa acg cat ttc att tcc cat gta cga cga tct cat aga aca gtg tca tct atc gat gga gca taa aag taa act cgt cga caa agc act caa taa att aga gtc tac cat cga tag tca atc tag act atc gta ttt gcc tcc gga aat tat gcg caa tat cat aac caa gct aag cga cta cca tct aaa cag tat gtt gta cgg aaa gaa cca tta caa ata tta tcc atg ata gaa aga aaa tat tta aaa aat aat cta tat gat tgg aga agt agg aaa caa aca gta aca aga cga cga tta cta c-a tta tta aat cat gag gtc cgt att ata ctc gta tat att gtt tct ctc atg tat aat aat aaa cgg aag aga tat agc acc aca tgc acc atc caa tgg aaa gtg taa aga caa cga ata cag aag ccg taa tct atg ttg tct atc gtg tcc tcc ggg aac tta cgc ttt cag att atg tga tag caa gac taa tac aca atg tac acc gtg tgg ttc gga tac ctt tac atc tca caa taa tca ttt aca ggc ttg tct aag ttg taa cgg aag atg tga tag taa tca ggt aga gac gcg atc gtg taa cac gac tca caa tag aat ctg tga atg ctc tcc agg ata tta ttg tct tct caa agg agc atc agg gtg tag aac atg tat ttc taa aac aaa gtg tgg aat agg ata cgg agt atc cgg ata cac gtc tac cgg aga cgt cat ctg ttc tcc gtg tgg tcc cgg aac ata ttc tca cac cgt ctc ttc cac aga taa atg cga acc cgt cgt aac cag caa tac att taa cta tat cga tgt gga aat taa cct gta tcc agt caa cga cac atc gtg tac tcg gac gac cac tac cgg tct cag cga atc cat ctc aac gtc gga act aac tat tac cat gaa tca taa aga ttg tga tcc agt ctt tcg tgc aga ata ctt ctc tgt cct taa taa tgt agc aac ttc agg att ctt tac agg aga aaa tag ata tca gaa tac ttc aaa gat atg tac tct gaa ttt cga gat taa atg taa caa caa aga ttc atc ttc caa aca gtt aac gaa aac aaa gaa tga tac tat cat gcc gca ttc aga gac ggt aac tct agt ggg cga ctg tct atc tag cgt cga cat cta cat act ata tag taa tac caa tac tca aga cta cga aaa tga tac aat ctc tta tca tat ggg taa tgt tct cga tgt caa tag cca tat gcc cgc tag ttg cga tat aca taa act gat cac taa ttc cca gaa tcc cac cca ctt ata gta ag- -tt ttt tta cct ata aat aat aaa tac aat aat taa ttt ctc gta aaa gta gaa aat ata ttc taa ttt att ata tgg taa gaa agt aga atc atc tag aac agt aat caa tca ata gca atc atg aaa caa tat att gtc ctg gca tgc atg tgc cta gtg gca gct gct atg cct act agt ctt caa caa tct tca tcc tcg tgt act gaa gaa gaa aac aaa cat cat atg gga atc gat gtt att atc aaa gtc aca aag caa gac caa aca ccg acc aat gat aag att tgt caa tcc gta acg gaa gtt aca gag a-- --- --- --- --- --- -cc gaa gat gat gag gta tcc gaa gaa gtt gta aaa gga gat ccc acc act tat tac act atc gtc ggt gcg ggt ctt aac atg aac ttt gga ttc acc aaa tgc cca aag att tta tcc atc tcc gaa tcc tct gat gga aac act gtg aat act aga ttg tcc agc gtg tca ccg gga caa ggt aag gac tct ccc gcg atc acg cgt gaa gaa gct ctg gct atg atc aaa gac tgt gag atg tct atc gac atc aga tgt agc gaa gaa gag aaa gac agt gac atc aag acc cat cca gta ctt ggg tct aac atc tca cat aag aaa gtg agt tac aaa gat atc atc ggt tca acg atc gtt gat aca aaa tgt gtc aag aac cta gag ttt agc gta cgt atc gga gac atg tgt gag gaa tca tct gaa ctt gaa gtc aag gat gga ttc aag tat gtc gac gga tcg gca tct gaa ggt gca acc gat gat act tca ctc atc gat tca aca aaa ctc aaa gca tgt gtc tga atc gat aac tct att cat ctg aaa atg gat gag ttg ggt taa tcg aac gat tca gac acc gca cca cga att aaa aaa g-- -ac cgg gca cta tat tcc ggt ttg caa aac aaa aat att taa cta cat tca caa aaa gtt acc tct cgt tac ttc ttc ttt ctg ttt caa tat gtg ata cga tat gat cac tat tcg tat tct ct- --- --- tgg tct cat aaa aaa gtt tta c-- --a aaa aaa aaa aaa a-t att ttt att ctc ttt ctc tct tcg at- ggt ctc aca aaa ata tta aac ctc ttt ctg atg tct caa cta ttt cgt aaa cga taa cgt cca aca ata tat tct cgt aga gct tat caa cat cct tat acc aat cta ggt tgt cag aca att gca tca taa aat aat gtt tat aat tta cac gtt aac atc ata taa taa acg tat ata gtt aat att ttt gga ata taa atg atc tgt aaa atc cat gta ggg gac act gct cac gtt ttt tct cta gta cat aat ttc aca caa gtt ttt ata cag aca aat taa ttc tcg tcc ata ta- ttt taa aac att gac ttt tgt act aag aaa aat atc ttg act aac cat ctc ttt ctc tct tcg atg ggt ctc aca aaa ata tta aac ctc ttt ctg atg g-- --- --- --- --- --- --- --- --- --- --- --- --- --- --- --- --- --- --- --- --- --- --- --- --- --- --- --- --- --- --- --- --- --- --- --- --- --- --- --- --- --- --- --- --- --- --- agt cgt aaa aag ttt tta tcc ttt ctc tct tcg ata ggt ctc --- --- --- --- --- --- --- --- --- --- --- --- --- --- --- --- --- --- --- --- --- --- --- --- --- --- --- --- --- --- --- --- --- --- --- --- --- --- --- --- --- --- --- --- --- --- --- --- --- --- --- --- --- --- --- --- --- --- --- --- --- --- --- --- --- --- --- --- --- --- aca aaa ata tta aac ctc ttt ctg atg gtc tct ata aa

>ON682263.2_Monkeypox_virus_isolate_MPXV/Germany/2022/RKI03_complete_genome

atc gag gac gac ttc aac tat gac agc tcg gtg gcg tct gcc agc gtg tac ata cga atg gca ttt cta aga aaa gtc tac ggt atc ctt -tc tac aat ttc ctt taa caa cgg caa cag ctg cag tat ttt tat act ttg aat gca -tc gga cat tta tac aag gga gtc ctg ttc taa tat tgg cat caa tgt tcg gat cta tag gct tga ttt tcg cat tga ctt tac aca gac atg aac atc ccc tga atc tgt aca tac ttt gtg gat tta cac tgt tag aat ctc taa cgc tgg cct ctg ttg tta ctt tct atg atg cac gta tcg tta tgc aag ctt tca tgt tga cta ctg cag tgt ttc ttg ctc tga cta cat ata ctc tac aat caa aga gag att tca gta aac ttg taa cag gat tgt ttg ctg ctt tct gga ttt taa ttt tgt cag gag tct tga gga taa agt tta aaa tag aat taa taa aga aca tat agg tca -tt ttt taa aca tgg ata gaa acc aag gtt gtt agt taa taa tat aca aga tat ttt ttc tca ctc tga tcc atg taa acc aag gac gag aga cac tct cat tcc tca ttc acg aca cca tta aaa atg gaa att aaa gcc ctc tat taa gca cag acg gct aca ggt cta cca tca ggt t-a cct tcg tct acc ttc aca atg gcc tct cct tgt gcc cag ttc agt ccc tgt cat tgc cac gct act aag gac tcc ctg aat acc gtg act gac gtc aga cat tgt ctg act gaa tac atc ctg tgg gtt tct cat aga tgg acc cat aga gaa agc gca ggg cct ctc tac agg ctt ctc atc tct ttc aga att gat gca atg gag cta ttt ggt agc gag ttg aag gag ttc tcg aat tca ctt ccg tgg gac aat atc gac aat tgc gtg gag atc att aaa tgt ttc atc aga aat gac tcc atg aaa acc gcc aaa gaa ctt tgt gca ata att gga ctt tgt act caa tca gct att gtc act gga aga gtc ttc aat gat aag tat atc gac ata cta ctt atg ctg cga aag att ctg aac gag aac gac tat ctc acc ctc ttg gat cat atc ctc act gct aaa tac taa atc tcc ttc atg ctc tct cac taa tac tct tac tca cta cac ttt tta tca tct tat gat gaa tga ttg cct tca tca ttt ttt cgt gga ata taa tat agg aat aat tag cac cag aat agc tat gga tat ctc gtt aag aat att ctc tat aag aga cat aat gta gac ata gtt att ata tcc ttc tta gat aag tgt tac gct act gga aag ttt cca tcg tta tta tta cat gaa gat gat ata att aaa cca aca ttg aga ttg gct ctt atg tta gct gga ttg aat tac tgt aat aaa tgc atc gag tat aga ggg ata tag caa ttc tcg ata ata gtc atg caa tat ttg aat gag act gat aat tta ggt aat aca gta ct- --- aca cac ata tct ttc tag att ata tat cgt taa aaa tct gta aga tgt ata ttt ctc ata agt atc cac tgt gta ata tta tta atg gat ata tag ata acg caa tag gga cta ata gta ttg taa aag ata taa tcg act att tac gta cat atc cag ata tct ata ttc cta cta gtt tgc tgc gta gtt gca tca ttg ata tgc atg -at tta tca gga ttc aga gat gaa tta cta agt aaa cta caa tcc cac aat aag taa gaa tca aat atc aaa aac tca ctt ttg att ttt cta gtc tta agt aat aca tat att tat taa tag acc tat gaa ata aaa aaa ggt aac aat gga ttc gcg tat agc tat tta cgt att agt atc ggc atc tct ttt gta tct tgt taa ttg tca caa act agt aca tta ctt caa tct gaa aat aaa tgg aag tga tat aac taa tac agc aga tat att gct gga caa tta tcc aat tat gac ctt tga tgg aaa gga tat tta tcc atc tat ctc gtt cat ggt cgg taa taa act ttt cct aga tct tta taa aaa tat ctt tga aga att ttt cag act att tcg agt atc tgt aag tag tca ata cga gga att aga ata tta tta ttc atg tga tta tac taa caa ccg tcc tac aat taa aca aca tta ctt tta taa cgg cga cga ata tac tga aat tga tag atc gaa aaa agc cac taa taa aaa cag ttg gtt aat tac ttc agg ctt tag act aca aaa atg gtt cga tag cga aga ttg tat aat tta tct cag atc ttt agt tag aag aat gga aga cag taa caa aaa cag taa aaa aac tta gta ctt aga tat cg- --a aaa aat ata ttt ttg tag act ctt gag aat aga agg aaa aca tgt aca taa tta taa aaa atg aaa atc aat ggc gaa taa gac agt gcg att cgc acc atg gag tcg gta gat ttc atg gct gtc gat gag cag ttt cac gac gac ctc gat ctt tgg tca tta tct ttg gta gat gat tat aaa aaa cat gga tta ggt gtt gac tgt tat gtt cta gaa cca gtt gtt gac agg aaa ata ttt gat aga ttt ctc ctt gaa cca att tgt gat cct gta gat gtt ctg tat gat tat ttt agg att cat aga gat aat att gat cag tat ata gta gat aga ctg ttt gca tat att aca tat aaa gat att ata tct gca tta gtg tca aag aat tat atg gaa gat att ttc tct ata att att aag aat tgt aat tct gtg caa gat ctc tta ctt tac tat cta tct aat gca tat gta gaa ata gac att gtt gat ctt atg gta gat cat ggg gct gta ata tat aaa ata gaa tgc ttg aat gcc tat ttt agg gga ata tgt aaa aag gaa agt agt gtt gtt gag ttt att ttg aat tgt ggt atc cca gat gaa aat gat gtt aaa tta gat cta tat aaa ata att cag tat act agg gga ttc ctt gta gat gaa ccc aca gta tta gaa att tat aag ctt tgt atc cca tat att gaa gat atc aat caa cta gat gct ggt gga agg acc ttg ctt tat cgc gct atc tat gca ggt tat ata gat tta gta tca tgg cta tta gaa aat gga gca aat gtc aac gca gta atg agt aat gga tat aca tgt ctt gac gtg gcc gtg gat agg gga tct gtc atc gcc cgt agg gaa gca cat ctt aaa ata tta gaa ata ttg ctt aga gaa cca ttg tct att gac tgt ata aaa tta gct ata ctt aat aat aca att gaa aac cat gat gtg ata aag ctc tgt atc aag tat ttt atg atg gta gat tat tca ctt tgt aat gtg tat gca tca tca ctc ttt gat tat ata att gat tgt aaa caa gaa ttg gag tac att agg cag atg aaa att cat aat aca acc atg tat gag tta atc tat aat aga gac aaa aac aag cat gct tcc cat att cta cat agg tat tct aaa cat cca gtt ttg aca cag tgt atc act aaa gga ttc aag att tac aca gaa gta acc gag cag gtc act aaa gct cta aac aga cgt gct cta ata gat gag ata ata aac aat gta tca act gat gac aat ctc cta tca aaa ctt cca tta gaa att agg gat cta att gtt tca caa gct gtc ata tag agt tct atc cac cca cct ttc ttg aaa tga gtt aat a-- --- --- --- --- --- --- --- --- --- --- --- --- --- --- --- --- --- --- --- --- --- --- --- --- --- --- --- --- --- --- --- --- --- --- --- --- --- --- --- --- --- --- --- --- --- --- --- --- --- --- --- --- --- --- --- --- --- --- --- --- --- --- --- --- --- --- --- --- --- --- --- --- --- --- --- --- --- --- --- --- --- --- --- --- --- --- --- --- --- --- --- --- --- --- gtc ata agt tag tta agt cat aag tta gtt aag tca taa gtt agt taa --- --- --- --- --- -gt cat aag tta gtt aag tca t-- --- --- --- --- --- --- --- --- --- --- --- --- --- --- --- -aa gtt agt taa gtc ata agt tag tta agt cat aag tta gtt aag tca taa gtt agt taa gtc ata agt tag tta agt cat aag tta gtt tat agt cta aca ctt cta att ttt ata cct tga tct ttt tct cta att atg aaa aag taa atc att atg aag atg gat g-a aat gga cga gat tgt gcg cat cgt taa cga tag tat gtg gta cgt acc taa cgc att tat gga cga cgg tga taa tga agg tca cat ttc tgt caa taa tgt ctg tca tat gta tct cgc att ctt tga tgt gga tat atc atc tca tct gtt taa att agt tat taa aca ctg cga tct gaa taa acg act aaa atg tgg taa ctc tcc att aca ttg cta tac gat gaa tac acg att taa tcc atc tgt att aaa gat att gtt acg cca cgg cat gcg taa ctt tga tag caa gga taa aaa agg aca tat tcc tct aca cca cta tct gat tca ttc act atc aat cga taa caa gat ctt tga tat act aac gga ccc cat tga tga ctt tag taa atc atc cga tct att gct gtg tta tct tag ata taa att caa tgg gag ctt aaa cta tta cgt tct gta caa att att gac taa agg atc tga ccc taa ttg cgt cga tga gga tgg act cac ttc tct tca tta cta ctg taa aca cat atc cgc gtt cca cga aag caa tta tta caa gtc aaa gag tca cac taa gat gcg agc tga gaa gcg att cat cta cgc gat aat aga tca tgg agc aaa cat taa cgc ggt tac gaa aat cgg aaa tac gcc gtt aca cac tta cct tca aca gta tac caa aca tag tcc tcg tgt ggt gta tgc tct ttt atc tcg agg agc cga tac gag gat acg taa taa tct taa ttg tac acc cat cat gga ata cat aaa gaa cga ttg tgc aac agg tca tat tct cat aat gtt act caa ttg gca cga aca aaa ata cgg gaa att aca aaa gga aga agg aca aca tct act tta tct att cat aaa aca taa tca agg ata tgg aag tcg ctc tct caa tat act acg gta tct act aga tag att cga cat tca gaa aga cga ata cta taa tac aat gac tcc tct tca tac cgc ctt cca gaa ttg caa taa caa tgt tgc ctc ata cct cgt ata cat tgg ata cga cat caa cct tcc gac taa aga cga taa gac agt att cga ctt ggt gtt tga aaa cag aaa cat tat ata caa ggc gga tgt cgt taa tga cat tat cca cca cag act gaa agt atc tct acc tat gat taa atc gtt gtt cta caa gat gtc gga gtt ctc tcc cta cga cga tca cta cgt aaa gaa gat aat agc cta ctg cct att aag gga cga gtc att tgc gga act aca tac taa att ctg ttt aaa cga gga cta taa aag tgt att tat gaa aaa tat atc att cga taa gat aga ttc cat cat cga aaa atg tag tcg tga cat aag tct cct caa aga gat tcg aat ctc aga cac cga ctt gta tac ggt att gag aac aga aga cat tcg gta tca cac ata tct cga agc cat aca ttc aga caa acg cat ttc att tcc cat gta cga cga tct cat aga aca gtg tca tct atc gat gga gca taa aag taa act cgt cga caa agc act caa taa att aga gtc tac cat cga tag tca atc tag act atc gta ttt gcc tcc gga aat tat gcg caa tat cat aac caa gct aag cga cta cca tct aaa cag tat gtt gta cgg aaa gaa cca tta caa ata tta tcc atg ata gaa aga aaa tat tta aaa aat aat cta tat gat tgg aga agt agg aaa caa aca gta aca aga cga cga tta cta c-a tta tta aat cat gag gtc cgt att ata ctc gta tat att gtt tct ctc atg tat aat aat aaa cgg aag aga tat agc acc aca tgc acc atc caa tgg aaa gtg taa aga caa cga ata cag aag ccg taa tct atg ttg tct atc gtg tcc tcc ggg aac tta cgc ttt cag att atg tga tag caa gac taa tac aca atg tac acc gtg tgg ttc gga tac ctt tac atc tca caa taa tca ttt aca ggc ttg tct aag ttg taa cgg aag atg tga tag taa tca ggt aga gac gcg atc gtg taa cac gac tca caa tag aat ctg tga atg ctc tcc agg ata tta ttg tct tct caa agg agc atc agg gtg tag aac atg tat ttc taa aac aaa gtg tgg aat agg ata cgg agt atc cgg ata cac gtc tac cgg aga cgt cat ctg ttc tcc gtg tgg tcc cgg aac ata ttc tca cac cgt ctc ttc cac aga taa atg cga acc cgt cgt aac cag caa tac att taa cta tat cga tgt gga aat taa cct gta tcc agt caa cga cac atc gtg tac tcg gac gac cac tac cgg tct cag cga atc cat ctc aac gtc gga act aac tat tac cat gaa tca taa aga ttg tga tcc agt ctt tcg tgc aga ata ctt ctc tgt cct taa taa tgt agc aac ttc agg att ctt tac agg aga aaa tag ata tca gaa tac ttc aaa gat atg tac tct gaa ttt cga gat taa atg taa caa caa aga ttc atc ttc caa aca gtt aac gaa aac aaa gaa tga tac tat cat gcc gca ttc aga gac ggt aac tct agt ggg cga ctg tct atc tag cgt cga cat cta cat act ata tag taa tac caa tac tca aga cta cga aaa tga tac aat ctc tta tca tat ggg taa tgt tct cga tgt caa tag cca tat gcc cgc tag ttg cga tat aca taa act gat cac taa ttc cca gaa tcc cac cca ctt ata gta ag- -tt ttt tta cct ata aat aat aaa tac aat aat taa ttt ctc gta aaa gta gaa aat ata ttc taa ttt att ata tgg taa gaa agt aga atc atc tag aac agt aat caa tca ata gca atc atg aaa caa tat att gtc ctg gca tgc atg tgc cta gtg gca gct gct atg cct act agt ctt caa caa tct tca tcc tcg tgt act gaa gaa gaa aac aaa cat cat atg gga atc gat gtt att atc aaa gtc aca aag caa gac caa aca ccg acc aat gat aag att tgt caa tcc gta acg gaa gtt aca gag a-- --- --- --- --- --- -cc gaa gat gat gag gta tcc gaa gaa gtt gta aaa gga gat ccc acc act tat tac act atc gtc ggt gcg ggt ctt aac atg aac ttt gga ttc acc aaa tgc cca aag att tta tcc atc tcc gaa tcc tct gat gga aac act gtg aat act aga ttg tcc agc gtg tca ccg gga caa ggt aag gac tct ccc gcg atc acg cgt gaa gaa gct ctg gct atg atc aaa gac tgt gag atg tct atc gac atc aga tgt agc gaa gaa gag aaa gac agt gac atc aag acc cat cca gta ctt ggg tct aac atc tca cat aag aaa gtg agt tac aaa gat atc atc ggt tca acg atc gtt gat aca aaa tgt gtc aag aac cta gag ttt agc gta cgt atc gga gac atg tgt gag gaa tca tct gaa ctt gaa gtc aag gat gga ttc aag tat gtc gac gga tcg gca tct gaa ggt gca acc gat gat act tca ctc atc gat tca aca aaa ctc aaa gca tgt gtc tga atc gat aac tct att cat ctg aaa atg gat gag ttg ggt taa tcg aac gat tca gac acc gca cca cga att aaa aaa g-- -ac cgg gca cta tat tcc ggt ttg caa aac aaa aat att taa cta cat tca caa aaa gtt acc tct cgt tac ttc ttc ttt ctg ttt caa tat gtg ata cga tat gat cac tat tcg tat tct ct- --- --- tgg tct cat aaa aaa gtt tta c-- --a aaa aaa aaa aaa a-t att ttt att ctc ttt ctc tct tcg at- ggt ctc aca aaa ata tta aac ctc ttt ctg atg tct caa cta ttt cgt aaa cga taa cgt cca aca ata tat tct cgt aga gct tat caa cat cct tat acc aat cta ggt tgt cag aca att gca tca taa aat aat gtt tat aat tta cac gtt aac atc ata taa taa acg tat ata gtt aat att ttt gga ata taa atg atc tgt aaa atc cat gta ggg gac act gct cac gtt ttt tct cta gta cat aat ttc aca caa gtt ttt ata cag aca aat taa ttc tcg tcc ata ta- ttt taa aac att gac ttt tgt act aag aaa aat atc ttg act aac cat ctc ttt ctc tct tcg atg ggt ctc aca aaa ata tta aac ctc ttt ctg atg g-- --- --- --- --- --- --- --- --- --- --- --- --- --- --- --- --- --- --- --- --- --- --- --- --- --- --- --- --- --- --- --- --- --- --- --- --- --- --- --- --- --- --- --- --- --- --- agt cgt aaa aag ttt tta tcc ttt ctc tct tcg ata ggt ctc --- --- --- --- --- --- --- --- --- --- --- --- --- --- --- --- --- --- --- --- --- --- --- --- --- --- --- --- --- --- --- --- --- --- --- --- --- --- --- --- --- --- --- --- --- --- --- --- --- --- --- --- --- --- --- --- --- --- --- --- --- --- --- --- --- --- --- --- --- --- aca aaa ata tta aac ctc ttt ctg atg gtc tct ata aa

>ON682266.1_Monkeypox_virus_isolate_MPXV/Germany/2022/RKI09_complete_genome

atc gag gac gac ttc aac tat gac agc tcg gtg gcg tct gcc agc gtg tac ata cga atg gca ttt cta aga aaa gtc tac ggt atc ctt -tc tac aat ttc ctt taa caa cgg caa cag ctg cag tat ttt tat act ttg aat gca -tc gga cat tta tac aag gga gtc ctg ttc taa tat tgg cat caa tgt tcg gat cta tag gct tga ttt tcg cat tga ctt tac aca gac atg aac atc ccc tga atc tgt aca tac ttt gtg gat tta cac tgt tag aat ctc taa cgc tgg cct ctg ttg tta ctt tct atg atg cac gta tcg tta tgc aag ctt tca tgt tga cta ctg cag tgt ttc ttg ctc tga cta cat ata ctc tac aat caa aga gag att tca gta aac ttg taa cag gat tgt ttg ctg ctt tct gga ttt taa ttt tgt cag gag tct tga gga taa agt tta aaa tag aat taa taa aga aca tat agg tca -tt ttt taa aca tgg ata gaa acc aag gtt gtt agt taa taa tat aca aga tat ttt ttc tca ctc tga tcc atg taa acc aag gac gag aga cac tct cat tcc tca ttc acg aca cca tta aaa atg gaa att aaa gcc ctc tat taa gca cag acg gct aca ggt cta cca tca ggt t-a cct tcg tct acc ttc aca atg gcc tct cct tgt gcc cag ttc agt ccc tgt cat tgc cac gct act aag gac tcc ctg aat acc gtg act gac gtc aga cat tgt ctg act gaa tac atc ctg tgg gtt tct cat aga tgg acc cat aga gaa agc gca ggg cct ctc tac agg ctt ctc atc tct ttc aga att gat gca atg gag cta ttt ggt agc gag ttg aag gag ttc tcg aat tca ctt ccg tgg gac aat atc gac aat tgc gtg gag atc att aaa tgt ttc atc aga aat gac tcc atg aaa acc gcc aaa gaa ctt tgt gca ata att gga ctt tgt act caa tca gct att gtc act gga aga gtc ttc aat gat aag tat atc gac ata cta ctt atg ctg cga aag att ctg aac gag aac gac tat ctc acc ctc ttg gat cat atc ctc act gct aaa tac taa atc tcc ttc atg ctc tct cac taa tac tct tac tca cta cac ttt tta tca tct tat gat gaa tga ttg cct tca tca ttt ttt cgt gga ata taa tat agg aat aat tag cac cag aat agc tat gga tat ctc gtt aag aat att ctc tat aag aga cat aat gta gac ata gtt att ata tcc ttc tta gat aag tgt tac gct act gga aag ttt cca tcg tta tta tta cat gaa gat gat ata att aaa cca aca ttg aga ttg gct ctt atg tta gct gga ttg aat tac tgt aat aaa tgc atc gag tat aga ggg ata tag caa ttc tcg ata ata gtc atg caa tat ttg aat gag act gat aat tta ggt aat aca gta ct- --- aca cac ata tct ttc tag att ata tat cgt taa aaa tct gta aga tgt ata ttt ctc ata agt atc cac tgt gta ata tta tta atg gat ata tag ata acg caa tag gga cta ata gta ttg taa aag ata taa tcg act att tac gta cat atc cag ata tct ata ttc cta cta gtt tgc tgc gta gtt gca tca ttg ata tgc atg -at tta tca gga ttc aga gat gaa tta cta agt aaa cta caa tcc cac aat aag taa gaa tca aat atc aaa aac tca ctt ttg att ttt cta gtc tta agt aat aca tat att tat taa tag acc tat gaa ata aaa aaa ggt aac aat gga ttc gcg tat agc tat tta cgt att agt atc ggc atc tct ttt gta tct tgt taa ttg tca caa act agt aca tta ctt caa tct gaa aat aaa tgg aag tga tat aac taa tac agc aga tat att gct gga caa tta tcc aat tat gac ctt tga tgg aaa gga tat tta tcc atc tat ctc gtt cat ggt cgg taa taa act ttt cct aga tct tta taa aaa tat ctt tga aga att ttt cag act att tcg agt atc tgt aag tag tca ata cga gga att aga ata tta tta ttc atg tga tta tac taa caa ccg tcc tac aat taa aca aca tta ctt tta taa cgg cga cga ata tac tga aat tga tag atc gaa aaa agc cac taa taa aaa cag ttg gtt aat tac ttc agg ctt tag act aca aaa atg gtt cga tag cga aga ttg tat aat tta tct cag atc ttt agt tag aag aat gga aga cag taa caa aaa cag taa aaa aac tta gta ctt aga tat cg- --a aaa aat ata ttt ttg tag act ctt gag aat aga agg aaa aca tgt aca taa tta taa aaa atg aaa atc aat ggc gaa taa gac agt gcg att cgc acc atg gag tcg gta gat ttc atg gct gtc gat gag cag ttt cac gac gac ctc gat ctt tgg tca tta tct ttg gta gat gat tat aaa aaa cat gga tta ggt gtt gac tgt tat gtt cta gaa cca gtt gtt gac agg aaa ata ttt gat aga ttt ctc ctt gaa cca att tgt gat cct gta gat gtt ctg tat gat tat ttt agg att cat aga gat aat att gat cag tat ata gta gat aga ctg ttt gca tat att aca tat aaa gat att ata tct gca tta gtg tca aag aat tat atg gaa gat att ttc tct ata att att aag aat tgt aat tct gtg caa gat ctc tta ctt tac tat cta tct aat gca tat gta gaa ata gac att gtt gat ctt atg gta gat cat ggg gct gta ata tat aaa ata gaa tgc ttg aat gcc tat ttt agg gga ata tgt aaa aag gaa agt agt gtt gtt gag ttt att ttg aat tgt ggt atc cca gat gaa aat gat gtt aaa tta gat cta tat aaa ata att cag tat act agg gga ttc ctt gta gat gaa ccc aca gta tta gaa att tat aag ctt tgt atc cca tat att gaa gat atc aat caa cta gat gct ggt gga agg acc ttg ctt tat cgc gct atc tat gca ggt tat ata gat tta gta tca tgg cta tta gaa aat gga gca aat gtc aac gca gta atg agt aat gga tat aca tgt ctt gac gtg gcc gtg gat agg gga tct gtc atc gcc cgt agg gaa gca cat ctt aaa ata tta gaa ata ttg ctt aga gaa cca ttg tct att gac tgt ata aaa tta gct ata ctt aat aat aca att gaa aac cat gat gtg ata aag ctc tgt atc aag tat ttt atg atg gta gat tat tca ctt tgt aat gtg tat gca tca tca ctc ttt gat tat ata att gat tgt aaa caa gaa ttg gag tac att agg cag atg aaa att cat aat aca acc atg tat gag tta atc tat aat aga gac aaa aac aag cat gct tcc cat att cta cat agg tat tct aaa cat cca gtt ttg aca cag tgt atc act aaa gga ttc aag att tac aca gaa gta acc gag cag gtc act aaa gct cta aac aga cgt gct cta ata gat gag ata ata aac aat gta tca act gat gac aat ctc cta tca aaa ctt cca tta gaa att agg gat cta att gtt tca caa gct gtc ata tag agt tct atc cac cca cct ttc ttg aaa tga gtt aat a-- --- --- --- --- --- --- --- --- --- --- --- --- --- --- --- --- --- --- --- --- --- --- --- --- --- --- --- --- --- --- --- --- --- --- --- --- --- --- --- --- --- --- --- --- --- --- --- --- --- --- --- --- --- --- --- --- --- --- --- --- --- --- --- --- --- --- --- --- --- --- --- --- --- --- --- --- --- --- --- --- --- --- --- --- --- --- --- --- --- --- --- --- --- --- --- --- --- --- --- --- --- --- --- --- --- --- --- --- --- --- --- --- --- --- --- --- --- --- --- --- --g tca t-- --- --- --- --- --- --- --- --- --- --- --- --- --- --- --- -aa gtt agt taa gtc ata agt tag tta agt cat aag tta gtt aag tca taa gtt agt taa gtc ata agt tag tta agt cat aag tta gtt tat agt cta aca ctt cta att ttt ata cct tga tct ttt tct cta att atg aaa aag taa atc att atg aag atg gat g-a aat gga cga gat tgt gcg cat cgt taa cga tag tat gtg gta cgt acc taa cgc att tat gga cga cgg tga taa tga agg tca cat ttc tgt caa taa tgt ctg tca tat gta tct cgc att ctt tga tgt gga tat atc atc tta tct gtt taa att agt tat taa aca ctg cga tct gaa taa acg act aaa atg tgg taa ctc tcc att aca ttg cta tac gat gaa tac acg att taa tcc atc tgt att aaa gat att gtt acg cca cgg cat gcg taa ctt tga tag caa gga taa aaa agg aca tat tcc tct aca cca cta tct gat tca ttc act atc aat cga taa caa gat ctt tga tat act aac gga ccc cat tga tga ctt tag taa atc atc cga tct att gct gtg tta tct tag ata taa att caa tgg gag ctt aaa cta tta cgt tct gta caa att att gac taa agg atc tga ccc taa ttg cgt cga tga gga tgg act cac ttc tct tca tta cta ctg taa aca cat atc cgc gtt cca cga aag caa tta tta caa gtc aaa gag tca cac taa gat gcg agc tga gaa gcg att cat cta cgc gat aat aga tca tgg agc aaa cat taa cgc ggt tac gaa aat cgg aaa tac gcc gtt aca cac tta cct tca aca gta tac caa aca tag tcc tcg tgt ggt gta tgc tct ttt atc tcg agg agc cga tac gag gat acg taa taa tct taa ttg tac acc cat cat gga ata cat aaa gaa cga ttg tgc aac agg tca tat tct cat aat gtt act caa ttg gca cga aca aaa ata cgg gaa att aca aaa gga aga agg aca aca tct act tta tct att cat aaa aca taa tca agg ata tgg aag tcg ctc tct caa tat act acg gta tct act aga tag att cga cat tca gaa aga cga ata cta taa tac aat gac tcc tct tca tac cgc ctt cca gaa ttg caa taa caa tgt tgc ctc ata cct cgt ata cat tgg ata cga cat caa cct tcc gac taa aga cga taa gac agt att cga ctt ggt gtt tga aaa cag aaa cat tat ata caa ggc gga tgt cgt taa tga cat tat cca cca cag act gaa agt atc tct acc tat gat taa atc gtt gtt cta caa gat gtc gga gtt ctc tcc cta cga cga tca cta cgt aaa gaa gat aat agc cta ctg cct att aag gga cga gtc att tgc gga act aca tac taa att ctg ttt aaa cga gga cta taa aag tgt att tat gaa aaa tat atc att cga taa gat aga ttc cat cat cga aaa atg tag tcg tga cat aag tct cct caa aga gat tcg aat ctc aga cac cga ctt gta tac ggt att gag aac aga aga cat tcg gta tca cac ata tct cga agc cat aca ttc aga caa acg cat ttc att tcc cat gta cga cga tct cat aga aca gtg tca tct atc gat gga gca taa aag taa act cgt cga caa agc act caa taa att aga gtc tac cat cga tag tca atc tag act atc gta ttt gcc tcc gga aat tat gcg caa tat cat aac caa gct aag cga cta cca tct aaa cag tat gtt gta cgg aaa gaa cca tta caa ata tta tcc atg ata gaa aga aaa tat tta aaa aat aat cta tat gat tgg aga agt agg aaa caa aca gta aca aga cga cga tta cta c-a tta tta aat cat gag gtc cgt att ata ctc gta tat att gtt tct ctc atg tat aat aat aaa cgg aag aga tat agc acc aca tgc acc atc caa tgg aaa gtg taa aga caa cga ata cag aag ccg taa tct atg ttg tct atc gtg tcc tcc ggg aac tta cgc ttt cag att atg tga tag caa gac taa tac aca atg tac acc gtg tgg ttc gga tac ctt tac atc tca caa taa tca ttt aca ggc ttg tct aag ttg taa cgg aag atg tga tag taa tca ggt aga gac gcg atc gtg taa cac gac tca caa tag aat ctg tga atg ctc tcc agg ata tta ttg tct tct caa agg agc atc agg gtg tag aac atg tat ttc taa aac aaa gtg tgg aat agg ata cgg agt atc cgg ata cac gtc tac cgg aga cgt cat ctg ttc tcc gtg tgg tcc cgg aac ata ttc tca cac cgt ctc ttc cac aga taa atg cga acc cgt cgt aac cag caa tac att taa cta tat cga tgt gga aat taa cct gta tcc agt caa cga cac atc gtg tac tcg gac gac cac tac cgg tct cag cga atc cat ctc aac gtc gga act aac tat tac cat gaa tca taa aga ttg tga tcc agt ctt tcg tgc aga ata ctt ctc tgt cct taa taa tgt agc aac ttc agg att ctt tac agg aga aaa tag ata tca gaa tac ttc aaa gat atg tac tct gaa ttt cga gat taa atg taa caa caa aga ttc atc ttc caa aca gtt aac gaa aac aaa gaa tga tac tat cat gcc gca ttc aga gac ggt aac tct agt ggg cga ctg tct atc tag cgt cga cat cta cat act ata tag taa tac caa tac tca aga cta cga aaa tga tac aat ctc tta tca tat ggg taa tgt tct cga tgt caa tag cca tat gcc cgc tag ttg cga tat aca taa act gat cac taa ttc cca gaa tcc cac cca ctt ata gta ag- -tt ttt tta cct ata aat aat aaa tac aat aat taa ttt ctc gta aaa gta gaa aat ata ttc taa ttt att ata tgg taa gaa agt aga atc atc tag aac agt aat caa tca ata gca atc atg aaa caa tat att gtc ctg gca tgc atg tgc cta gtg gca gct gct atg cct act agt ctt caa caa tct tca tcc tcg tgt act gaa gaa gaa aac aaa cat cat atg gga atc gat gtt att atc aaa gtc aca aag caa gac caa aca ccg acc aat gat aag att tgt caa tcc gta acg gaa gtt aca gag a-- --- --- --- --- --- -cc gaa gat gat gag gta tcc gaa gaa gtt gta aaa gga gat ccc acc act tat tac act atc gtc ggt gcg ggt ctt aac atg aac ttt gga ttc acc aaa tgc cca aag att tta tcc atc tcc gaa tcc tct gat gga aac act gtg aat act aga ttg tcc agc gtg tca ccg gga caa ggt aag gac tct ccc gcg atc acg cgt gaa gaa gct ctg gct atg atc aaa gac tgt gag atg tct atc gac atc aga tgt agc gaa gaa gag aaa gac agt gac atc aag acc cat cca gta ctt ggg tct aac atc tca cat aag aaa gtg agt tac aaa gat atc atc ggt tca acg atc gtt gat aca aaa tgt gtc aag aac cta gag ttt agc gta cgt atc gga gac atg tgt gag gaa tca tct gaa ctt gaa gtc aag gat gga ttc aag tat gtc gac gga tcg gca tct gaa ggt gca acc gat gat act tca ctc atc gat tca aca aaa ctc aaa gca tgt gtc tga atc gat aac tct att cat ctg aaa atg gat gag ttg ggt taa tcg aac gat tca gac acc gca cca cga att aaa aaa g-- -ac cgg gca cta tat tcc ggt ttg caa aac aaa aat att taa cta cat tca caa aaa gtt acc tct cgt tac ttc ttc ttt ctg ttt caa tat gtg ata cga tat gat cac tat tcg tat tct ct- --- --- tgg tct cat aaa aaa gtt tta c-- --a aaa aaa aaa aaa a-t att ttt att ctc ttt ctc tct tcg at- ggt ctc aca aaa ata tta aac ctc ttt ctg atg tct caa cta ttt cgt aaa cga taa cgt cca aca ata tat tct cgt aga gct tat caa cat cct tat acc aat cta ggt tgt cag aca att gca tca taa aat aat gtt tat aat tta cac gtt aac atc ata taa taa acg tat ata gtt aat att ttt gga ata taa atg atc tgt aaa atc cat gta ggg gac act gct cac gtt ttt tct cta gta cat aat ttc aca caa gtt ttt ata cag aca aat taa ttc tcg tcc ata ta- ttt taa aac att gac ttt tgt act aag aaa aat atc ttg act aac cat ctc ttt ctc tct tcg atg ggt ctc aca aaa ata tta aac ctc ttt ctg atg g-- --- --- --- --- --- --- --- --- --- --- --- --- --- --- --- --- --- --- --- --- --- --- --- --- --- --- --- --- --- --- --- --- --- --- --- --- --- --- --- --- --- --- --- --- --- --- agt cgt aaa aag ttt tta tcc ttt ctc tct tcg ata ggt ctc --- --- --- --- --- --- --- --- --- --- --- --- --- --- --- --- --- --- --- --- --- --- --- --- --- --- --- --- --- --- --- --- --- --- --- --- --- --- --- --- --- --- --- --- --- --- --- --- --- --- --- --- --- --- --- --- --- --- --- --- --- --- --- --- --- --- --- --- --- --- aca aaa ata tta aac ctc ttt ctg atg gtc tct ata aa

>ON619836.1_Monkeypox_virus_isolate_MPXV_UK_2022_2_complete_genome

atc gag gac gac ttc aac tat gac agc tcg gtg gcg tct gcc agc gtg tac ata cga atg gca ttt cta aga aaa gtc tac ggt atc ctt -tc tac aat ttc ctt taa caa cgg caa cag ctg cag tat ttt tat act ttg aat gca -tc gga cat tta tac aag gga gtc ctg ttc taa tat tgg cat caa tgt tcg gat cta tag gct tga ttt tcg cat tga ctt tac aca gac atg aac atc ccc tga atc tgt aca tac ttt gtg gat tta cac tgt tag aat ctc taa cgc tgg cct ctg ttg tta ctt tct atg atg cac gta tcg tta tgc aag ctt tca tgt tga cta ctg cag tgt ttc ttg ctc tga cta cat ata ctc tac aat caa aga gag att tca gta aac ttg taa cag gat tgt ttg ctg ctt tct gga ttt taa ttt tgt cag gag tct tga gga taa agt tta aaa tag aat taa taa aga aca tat agg tca -tt ttt taa aca tgg ata gaa acc aag gtt gtt agt taa taa tat aca aga tat ttt ttc tca ctc tga tcc atg taa acc aag gac gag aga cac tct cat tcc tca ttc acg aca cca tta aaa atg gaa att aaa gcc ctc tat taa gca cag acg gct aca ggt cta cca tca ggt t-a cct tcg tct acc ttc aca atg gcc tct cct tgt gcc cag ttc agt ccc tgt cat tgc cac gct act aag gac tcc ctg aat acc gtg act gac gtc aga cat tgt ctg act gaa tac atc ctg tgg gtt tct cat aga tgg acc cat aga gaa agc gca ggg cct ctc tac agg ctt ctc atc tct ttc aga att gat gca atg gag cta ttt ggt agc gag ttg aag gag ttc tcg aat tca ctt ccg tgg gac aat atc gac aat tgc gtg gag atc att aaa tgt ttc atc aga aat gac tcc atg aaa acc gcc aaa gaa ctt tgt gca ata att gga ctt tgt act caa tca gct att gtc act gga aga gtc ttc aat gat aag tat atc gac ata cta ctt atg ctg cga aag att ctg aac gag aac gac tat ctc acc ctc ttg gat cat atc ctc act gct aaa tac taa atc tcc ttc atg ctc tct cac taa tac tct tac tca cta cac ttt tta tca tct tat gat gaa tga ttg cct tca tca ttt ttt cgt gga ata taa tat agg aat aat tag cac cag aat agc tat gga tat ctc gtt aag aat att ctc tat aag aga cat aat gta gac ata gtt att ata tcc ttc tta gat aag tgt tac gct act gga aag ttt cca tcg tta tta tta cat gaa gat gat ata att aaa cca aca ttg aga ttg gct ctt atg tta gct gga ttg aat tac tgt aat aaa tgc atc gag tat aga ggg ata tag caa ttc tcg ata ata gtc atg caa tat ttg aat gag act gat aat tta ggt aat aca gta ct- --- aca cac ata tct ttc tag att ata tat cgt taa aaa tct gta aga tgt ata ttt ctc ata agt atc cac tgt gta ata tta tta atg gat ata tag ata acg caa tag gga cta ata gta ttg taa aag ata taa tcg act att tac gta cat atc cag ata tct ata ttc cta cta gtt tgc tgc gta gtt gca tca ttg ata tgc atg -at tta tca gga ttc aga gat gaa tta cta agt aaa cta caa tcc cac aat aag taa gaa tca aat atc aaa aac tca ctt ttg att ttt cta gtc tta agt aat aca tat att tat taa tag acc tat gaa ata aaa aaa ggt aac aat gga ttc gcg tat agc tat tta cgt att agt atc ggc atc tct ttt gta tct tgt taa ttg tca caa act agt aca tta ctt caa tct gaa aat aaa tgg aag tga tat aac taa tac agc aga tat att gct gga caa tta tcc aat tat gac ctt tga tgg aaa gga tat tta tcc atc tat ctc gtt cat ggt cgg taa taa act ttt cct aga tct tta taa aaa tat ctt tga aga att ttt cag act att tcg agt atc tgt aag tag tca ata cga gga att aga ata tta tta ttc atg tga tta tac taa caa ccg tcc tac aat taa aca aca tta ctt tta taa cgg cga cga ata tac tga aat tga tag atc gaa aaa agc cac taa taa aaa cag ttg gtt aat tac ttc agg ctt tag act aca aaa atg gtt cga tag cga aga ttg tat aat tta tct cag atc ttt agt tag aag aat gga aga cag taa caa aaa cag taa aaa aac tta gta ctt aga tat cg- --a aaa aat ata ttt ttg tag act ctt gag aat aga agg aaa aca tgt aca taa tta taa aaa atg aaa atc aat ggc gaa taa gac agt gcg att cgc acc atg gag tcg gta gat ttc atg gct gtc gat gag cag ttt cac gac gac ctc gat ctt tgg tca tta tct ttg gta gat gat tat aaa aaa cat gga tta ggt gtt gac tgt tat gtt cta gaa cca gtt gtt gac agg aaa ata ttt gat aga ttt ctc ctt gaa cca att tgt gat cct gta gat gtt ctg tat gat tat ttt agg att cat aga gat aat att gat cag tat ata gta gat aga ctg ttt gca tat att aca tat aaa gat att ata tct gca tta gtg tca aag aat tat atg gaa gat att ttc tct ata att att aag aat tgt aat tct gtg caa gat ctc tta ctt tac tat cta tct aat gca tat gta gaa ata gac att gtt gat ctt atg gta gat cat ggg gct gta ata tat aaa ata gaa tgc ttg aat gcc tat ttt agg gga ata tgt aaa aag gaa agt agt gtt gtt gag ttt att ttg aat tgt ggt atc cca gat gaa aat gat gtt aaa tta gat cta tat aaa ata att cag tat act agg gga ttc ctt gta gat gaa ccc aca gta tta gaa att tat aag ctt tgt atc cca tat att gaa gat atc aat caa cta gat gct ggt gga agg acc ttg ctt tat cgc gct atc tat gca ggt tat ata gat tta gta tca tgg cta tta gaa aat gga gca aat gtc aac gca gta atg agt aat gga tat aca tgt ctt gac gtg gcc gtg gat agg gga tct gtc atc gcc cgt agg gaa gca cat ctt aaa ata tta gaa ata ttg ctt aga gaa cca ttg tct att gac tgt ata aaa tta gct ata ctt aat aat aca att gaa aac cat gat gtg ata aag ctc tgt atc aag tat ttt atg atg gta gat tat tca ctt tgt aat gtg tat gca tca tca ctc ttt gat tat ata att gat tgt aaa caa gaa ttg gag tac att agg cag atg aaa att cat aat aca acc atg tat gag tta atc tat aat aga gac aaa aac aag cat gct tcc cat att cta cat agg tat tct aaa cat cca gtt ttg aca cag tgt atc act aaa gga ttc aag att tac aca gaa gta acc gag cag gtc act aaa gct cta aac aga cgt gct cta ata gat gag ata ata aac aat gta tca act gat gac aat ctc cta tca aaa ctt cca tta gaa att agg gat cta att gtt tca caa gct gtc ata tag agt tct atc cac cca cct ttc ttg aaa tga gtt aat --- --- --- --- --- --- --- --- --- --- --- --- --- --- --- --- --- --- --- --- --- --- --- --- --- --- --- --- --- --- --- --- --- --- --- --- --- --- --- --- --- --- --- --- --- --- --- --- --- --- --- --- --- --- --- --- --- --- --- --- --- --- --- --- --- --- --- --- --- --- --- --- --- --- --- --- --- --- --- --- --- --- --- --- --- --- --- --- --- --- --- --- --- --- --- --- --- --- --- --- --- --- --- --- --- -ag tca taa gtt agt taa --- --- --- --- --- -gt cat aag tta gtt aag tca t-- --- --- --- --- --- --- --- --- --- --- --- --- --- --- --- -aa gtt agt taa gtc ata agt tag tta agt cat aag tta gtt aag tca taa gtt agt taa gtc ata agt tag tta agt cat aag tta gtt tat agt cta aca ctt cta att ttt ata cct tga tct ttt tct cta att atg aaa aag taa atc att atg aag atg gat g-a aat gga cga gat tgt gcg cat cgt taa cga tag tat gtg gta cgt acc taa cgc att tat gga cga cgg tga taa tga agg tca cat ttc tgt caa taa tgt ctg tca tat gta tct cgc att ctt tga tgt gga tat atc atc tca tct gtt taa att agt tat taa aca ctg cga tct gaa taa acg act aaa atg tgg taa ctc tcc att aca ttg cta tac gat gaa tac acg att taa tcc atc tgt att aaa gat att gtt acg cca cgg cat gcg taa ctt tga tag caa gga taa aaa agg aca tat tcc tct aca cca cta tct gat tca ttc act atc aat cga taa caa gat ctt tga tat act aac gga ccc cat tga tga ctt tag taa atc atc cga tct att gct gtg tta tct tag ata taa att caa tgg gag ctt aaa cta tta cgt tct gta caa att att gac taa agg atc tga ccc taa ttg cgt cga tga gga tgg act cac ttc tct tca tta cta ctg taa aca cat atc cgc gtt cca cga aag caa tta tta caa gtc aaa gag tca cac taa gat gcg agc tga gaa gcg att cat cta cgc gat aat aga tca tgg agc aaa cat taa cgc ggt tac gaa aat cgg aaa tac gcc gtt aca cac tta cct tca aca gta tac caa aca tag tcc tcg tgt ggt gta tgc tct ttt atc tcg agg agc cga tac gag gat acg taa taa tct taa ttg tac acc cat cat gga ata cat aaa gaa cga ttg tgc aac agg tca tat tct cat aat gtt act caa ttg gca cga aca aaa ata cgg gaa att aca aaa gga aga agg aca aca tct act tta tct att cat aaa aca taa tca agg ata tgg aag tcg ctc tct caa tat act acg gta tct act aga tag att cga cat tca gaa aga cga ata cta taa tac aat gac tcc tct tca tac cgc ctt cca gaa ttg caa taa caa tgt tgc ctc ata cct cgt ata cat tgg ata cga cat caa cct tcc gac taa aga cga taa gac agt att cga ctt ggt gtt tga aaa cag aaa cat tat ata caa ggc gga tgt cgt taa tga cat tat cca cca cag act gaa agt atc tct acc tat gat taa atc gtt gtt cta caa gat gtc gga gtt ctc tcc cta cga cga tca cta cgt aaa gaa gat aat agc cta ctg cct att aag gga cga gtc att tgc gga act aca tac taa att ctg ttt aaa cga gga cta taa aag tgt att tat gaa aaa tat atc att cga taa gat aga ttc cat cat cga aaa atg tag tcg tga cat aag tct cct caa aga gat tcg aat ctc aga cac cga ctt gta tac ggt att gag aac aga aga cat tcg gta tca cac ata tct cga agc cat aca ttc aga caa acg cat ttc att tcc cat gta cga cga tct cat aga aca gtg tca tct atc gat gga gca taa aag taa act cgt cga caa agc act caa taa att aga gtc tac cat cga tag tca atc tag act atc gta ttt gcc tcc gga aat tat gcg caa tat cat aac caa gct aag cga cta cca tct aaa cag tat gtt gta cgg aaa gaa cca tta caa ata tta tcc atg ata gaa aga aaa tat tta aaa aat aat cta tat gat tgg aga agt agg aaa caa aca gta aca aga cga cga tta cta c-a tta tta aat cat gag gtc cgt att ata ctc gta tat att gtt tct ctc atg tat aat aat aaa cgg aag aga tat agc acc aca tgc acc atc caa tgg aaa gtg taa aga caa cga ata cag aag ccg taa tct atg ttg tct atc gtg tcc tcc ggg aac tta cgc ttt cag att atg tga tag caa gac taa tac aca atg tac acc gtg tgg ttc gga tac ctt tac atc tca caa taa tca ttt aca ggc ttg tct aag ttg taa cgg aag atg tga tag taa tca ggt aga gac gcg atc gtg taa cac gac tca caa tag aat ctg tga atg ctc tcc agg ata tta ttg tct tct caa agg agc atc agg gtg tag aac atg tat ttc taa aac aaa gtg tgg aat agg ata cgg agt atc cgg ata cac gtc tac cgg aga cgt cat ctg ttc tcc gtg tgg tcc cgg aac ata ttc tca cac cgt ctc ttc cac aga taa atg cga acc cgt cgt aac cag caa tac att taa cta tat cga tgt gga aat taa cct gta tcc agt caa cga cac atc gtg tac tcg gac gac cac tac cgg tct cag cga atc cat ctc aac gtc gga act aac tat tac cat gaa tca taa aga ttg tga tcc agt ctt tcg tgc aga ata ctt ctc tgt cct taa taa tgt agc aac ttc agg att ctt tac agg aga aaa tag ata tca gaa tac ttc aaa gat atg tac tct gaa ttt cga gat taa atg taa caa caa aga ttc atc ttc caa aca gtt aac gaa aac aaa gaa tga tac tat cat gcc gca ttc aga gac ggt aac tct agt ggg cga ctg tct atc tag cgt cga cat cta cat act ata tag taa tac caa tac tca aga cta cga aaa tga tac aat ctc tta tca tat ggg taa tgt tct cga tgt caa tag cca tat gcc cgc tag ttg cga tat aca taa act gat cac taa ttc cca gaa tcc cac cca ctt ata gta ag- -tt ttt tta cct ata aat aat aaa tac aat aat taa ttt ctc gta aaa gta gaa aat ata ttc taa ttt att ata tgg taa gaa agt aga atc atc tag aac agt aat caa tca ata gca atc atg aaa caa tat att gtc ctg gca tgc atg tgc cta gtg gca gct gct atg cct act agt ctt caa caa tct tca tcc tcg tgt act gaa gaa gaa aac aaa cat cat atg gga atc gat gtt att atc aaa gtc aca aag caa gac caa aca ccg acc aat gat aag att tgt caa tcc gta acg gaa gtt aca gag a-- --- --- --- --- --- -cc gaa gat gat gag gta tcc gaa gaa gtt gta aaa gga gat ccc acc act tat tac act atc gtc ggt gcg ggt ctt aac atg aac ttt gga ttc acc aaa tgc cca aag att tta tcc atc tcc gaa tcc tct gat gga aac act gtg aat act aga ttg tcc agc gtg tca ccg gga caa ggt aag gac tct ccc gcg atc acg cgt gaa gaa gct ctg gct atg atc aaa gac tgt gag atg tct atc gac atc aga tgt agc gaa gaa gag aaa gac agt gac atc aag acc cat cca gta ctt ggg tct aac atc tca cat aag aaa gtg agt tac aaa gat atc atc ggt tca acg atc gtt gat aca aaa tgt gtc aag aac cta gag ttt agc gta cgt atc gga gac atg tgt gag gaa tca tct gaa ctt gaa gtc aag gat gga ttc aag tat gtc gac gga tcg gca tct gaa ggt gca acc gat gat act tca ctc atc gat tca aca aaa ctc aaa gca tgt gtc tga atc gat aac tct att cat ctg aaa atg gat gag ttg ggt taa tcg aac gat tca gac acc gca cca cga att aaa aaa g-- -ac cgg gca cta tat tcc ggt ttg caa aac aaa aat att taa cta cat tca caa aaa gtt acc tct cgt tac ttc ttc ttt ctg ttt caa tat gtg ata cga tat gat cac tat tcg tat tct ct- --- --- tgg tct cat aaa aaa gtt tta c-- --? aaa aaa aaa aaa a-t att ttt att ctc ttt ctc tct tcg at- ggt ctc aca aaa ata tta aac ctc ttt ctg atg tct caa cta ttt cgt aaa cga taa cgt cca aca ata tat tct cgt aga gct tat caa cat cct tat acc aat cta ggt tgt cag aca att gca tca taa aat aat gtt tat aat tta cac gtt aac atc ata taa taa acg tat ata gtt aat att ttt gga ata taa atg atc tgt aaa atc cat gta ggg gac act gct cac gtt ttt tct cta gta cat aat ttc aca caa gtt ttt ata cag aca aat taa ttc tcg tcc ata ta- ttt taa aac att gac ttt tgt act aag aaa aat atc ttg act aac cat ctc ttt ctc tct tcg atg ggt ctc aca aaa ata tta aac ctc ttt ctg atg g-- --- --- --- --- --- --- --- --- --- --- --- --- --- --- --- --- --- --- --- --- --- --- --- --- --- --- --- --- --- --- --- --- --- --- --- --- --- --- --- --- --- --- --- --- --- --- agt cgt aaa aag ttt tta tcc ttt ctc tct tcg ata ggt ctc --- --- --- --- --- --- --- --- --- --- --- --- --- --- --- --- --- --- --- --- --- --- --- --- --- --- --- --- --- --- --- --- --- --- --- --- --- --- --- --- --- --- --- --- --- --- --- --- --- --- --- --- --- --- --- --- --- --- --- --- --- --- --- --- --- --- --- --- --- --- aca aaa ata tta aac ctc ttt ctg atg gtc tct ata aa

>ON619838.1_Monkeypox_virus_isolate_MPXV_UK_2022_4_complete_genome

atc gag gac gac ttc aac tat gac agc tcg gtg gcg tct gcc agc gtg tac ata cga atg gca ttt cta aga aaa gtc tac ggt atc ctt -tc tac aat ttc ctt taa caa cgg caa cag ctg cag tat ttt tat act ttg aat gca -tc gga cat tta tac aag gga gtc ctg ttc taa tat tgg cat caa tgt tcg gat cta tag gct tga ttt tcg cat tga ctt tac aca gac atg aac atc ccc tga atc tgt aca tac ttt gtg gat tta cac tgt tag aat ctc taa cgc tgg cct ctg ttg tta ctt tct atg atg cac gta tcg tta tgc aag ctt tca tgt tga cta ctg cag tgt ttc ttg ctc tga cta cat ata ctc tac aat caa aga gag att tca gta aac ttg taa cag gat tgt ttg ctg ctt tct gga ttt taa ttt tgt cag gag tct tga gga taa agt tta aaa tag aat taa taa aga aca tat agg tca -tt ttt taa aca tgg ata gaa acc aag gtt gtt agt taa taa tat aca aga tat ttt ttc tca ctc tga tcc atg taa acc aag gac gag aga cac tct cat tcc tca ttc acg aca cca tta aaa atg gaa att aaa gcc ctc tat taa gca cag acg gct aca ggt cta cca tca ggt t-a cct tcg tct acc ttc aca atg gcc tct cct tgt gcc cag ttc agt ccc tgt cat tgc cac gct act aag gac tcc ctg aat acc gtg act gac gtc aga cat tgt ctg act gaa tac atc ctg tgg gtt tct cat aga tgg acc cat aga gaa agc gca ggg cct ctc tac agg ctt ctc atc tct ttc aga att gat gca atg gag cta ttt ggt agc gag ttg aag gag ttc tcg aat tca ctt ccg tgg gac aat atc gac aat tgc gtg gag atc att aaa tgt ttc atc aga aat gac tcc atg aaa acc gcc aaa gaa ctt tgt gca ata att gga ctt tgt act caa tca gct att gtc act gga aga gtc ttc aat gat aag tat atc gac ata cta ctt atg ctg cga aag att ctg aac gag aac gac tat ctc acc ctc ttg gat cat atc ctc act gct aaa tac taa atc tcc ttc atg ctc tct cac taa tac tct tac tca cta cac ttt tta tca tct tat gat gaa tga ttg cct tca tca ttt ttt cgt gga ata taa tat agg aat aat tag cac cag aat agc tat gga tat ctc gtt aag aat att ctc tat aag aga cat aat gta gac ata gtt att ata tcc ttc tta gat aag tgt tac gct act gga aag ttt cca tcg tta tta tta cat gaa gat gat ata att aaa cca aca ttg aga ttg gct ctt atg tta gct gga ttg aat tac tgt aat aaa tgc atc gag tat aga ggg ata tag caa ttc tcg ata ata gtc atg caa tat ttg aat gag act gat aat tta ggt aat aca gta ct- --- aca cac ata tct ttc tag att ata tat cgt taa aaa tct gta aga tgt ata ttt ctc ata agt atc cac tgt gta ata tta tta atg gat ata tag ata acg caa tag gga cta ata gta ttg taa aag ata taa tcg act att tac gta cat atc cag ata tct ata ttc cta cta gtt tgc tgc gta gtt gca tca ttg ata tgc atg -at tta tca gga ttc aga gat gaa tta cta agt aaa cta caa tcc cac aat aag taa gaa tca aat atc aaa aac tca ctt ttg att ttt cta gtc tta agt aat aca tat att tat taa tag acc tat gaa ata aaa aaa ggt aac aat gga ttc gcg tat agc tat tta cgt att agt atc ggc atc tct ttt gta tct tgt taa ttg tca caa act agt aca tta ctt caa tct gaa aat aaa tgg aag tga tat aac taa tac agc aga tat att gct gga caa tta tcc aat tat gac ctt tga tgg aaa gga tat tta tcc atc tat ctc gtt cat ggt cgg taa taa act ttt cct aga tct tta taa aaa tat ctt tga aga att ttt cag act att tcg agt atc tgt aag tag tca ata cga gga att aga ata tta tta ttc atg tga tta tac taa caa ccg tcc tac aat taa aca aca tta ctt tta taa cgg cga cga ata tac tga aat tga tag atc gaa aaa agc cac taa taa aaa cag ttg gtt aat tac ttc agg ctt tag act aca aaa atg gtt cga tag cga aga ttg tat aat tta tct cag atc ttt agt tag aag aat gga aga cag taa caa aaa cag taa aaa aac tta gta ctt aga tat cg- --a aaa aat ata ttt ttg tag act ctt gag aat aga agg aaa aca tgt aca taa tta taa aaa atg aaa atc aat ggc gaa taa gac agt gcg att cgc acc atg gag tcg gta gat ttc atg gct gtc gat gag cag ttt cac gac gac ctc gat ctt tgg tca tta tct ttg gta gat gat tat aaa aaa cat gga tta ggt gtt gac tgt tat gtt cta gaa cca gtt gtt gac agg aaa ata ttt gat aga ttt ctc ctt gaa cca att tgt gat cct gta gat gtt ctg tat gat tat ttt agg att cat aga gat aat att gat cag tat ata gta gat aga ctg ttt gca tat att aca tat aaa gat att ata tct gca tta gtg tca aag aat tat atg gaa gat att ttc tct ata att att aag aat tgt aat tct gtg caa gat ctc tta ctt tac tat cta tct aat gca tat gta gaa ata gac att gtt gat ctt atg gta gat cat ggg gct gta ata tat aaa ata gaa tgc ttg aat gcc tat ttt agg gga ata tgt aaa aag gaa agt agt gtt gtt gag ttt att ttg aat tgt ggt atc cca gat gaa aat gat gtt aaa tta gat cta tat aaa ata att cag tat act agg gga ttc ctt gta gat gaa ccc aca gta tta gaa att tat aag ctt tgt atc cca tat att gaa gat atc aat caa cta gat gct ggt gga agg acc ttg ctt tat cgc gct atc tat gca ggt tat ata gat tta gta tca tgg cta tta gaa aat gga gca aat gtc aac gca gta atg agt aat gga tat aca tgt ctt gac gtg gcc gtg gat agg gga tct gtc atc gcc cgt agg gaa gca cat ctt aaa ata tta gaa ata ttg ctt aga gaa cca ttg tct att gac tgt ata aaa tta gct ata ctt aat aat aca att gaa aac cat gat gtg ata aag ctc tgt atc aag tat ttt atg atg gta gat tat tca ctt tgt aat gtg tat gca tca tca ctc ttt gat tat ata att gat tgt aaa caa gaa ttg gag tac att agg cag atg aaa att cat aat aca acc atg tat gag tta atc tat aat aga gac aaa aac aag cat gct tcc cat att cta cat agg tat tct aaa cat cca gtt ttg aca cag tgt atc act aaa gga ttc aag att tac aca gaa gta acc gag cag gtc act aaa gct cta aac aga cgt gct cta ata gat gag ata ata aac aat gta tca act gat gac aat ctc cta tca aaa ctt cca tta gaa att agg gat cta att gtt tca caa gct gtc ata tag agt tct atc cac cca cct ttc ttg aaa tga gtt aat --- --- --- --- --- --- --- --- --- --- --- --- --- --- --- --- --- --- --- --- --- --- --- --- --- --- --- --- --- --- --- --- --- --- --- --- --- --- --- --- --- --- --- --- --- --- --- --- --- --- --- --- --- --- --- --- --- --- --- --- --- --- --- --- --- --- --- --- --- --- --- --- --- --- --- --- --- --- --- --- --- --- --- --- --- --- --- --- --- --- --- --- --- --- --- --- --- --- --- --- --- --- --- --- --- -ag tca taa gtt agt taa --- --- --- --- --- -gt cat aag tta gtt aag tca t-- --- --- --- --- --- --- --- --- --- --- --- --- --- --- --- -aa gtt agt taa gtc ata agt tag tta agt cat aag tta gtt aag tca taa gtt agt taa gtc ata agt tag tta agt cat aag tta gtt tat agt cta aca ctt cta att ttt ata cct tga tct ttt tct cta att atg aaa aag taa atc att atg aag atg gat g-a aat gga cga gat tgt gcg cat cgt taa cga tag tat gtg gta cgt acc taa cgc att tat gga cga cgg tga taa tga agg tca cat ttc tgt caa taa tgt ctg tca tat gta tct cgc att ctt tga tgt gga tat atc atc tca tct gtt taa att agt tat taa aca ctg cga tct gaa taa acg act aaa atg tgg taa ctc tcc att aca ttg cta tac gat gaa tac acg att taa tcc atc tgt att aaa gat att gtt acg cca cgg cat gcg taa ctt tga tag caa gga taa aaa agg aca tat tcc tct aca cca cta tct gat tca ttc act atc aat cga taa caa gat ctt tga tat act aac gga ccc cat tga tga ctt tag taa atc atc cga tct att gct gtg tta tct tag ata taa att caa tgg gag ctt aaa cta tta cgt tct gta caa att att gac taa agg atc tga ccc taa ttg cgt cga tga gga tgg act cac ttc tct tca tta cta ctg taa aca cat atc cgc gtt cca cga aag caa tta tta caa gtc aaa gag tca cac taa gat gcg agc tga gaa gcg att cat cta cgc gat aat aga tca tgg agc aaa cat taa cgc ggt tac gaa aat cgg aaa tac gcc gtt aca cac tta cct tca aca gta tac caa aca tag tcc tcg tgt ggt gta tgc tct ttt atc tcg agg agc cga tac gag gat acg taa taa tct taa ttg tac acc cat cat gga ata cat aaa gaa cga ttg tgc aac agg tca tat tct cat aat gtt act caa ttg gca cga aca aaa ata cgg gaa att aca aaa gga aga agg aca aca tct act tta tct att cat aaa aca taa tca agg ata tgg aag tcg ctc tct caa tat act acg gta tct act aga tag att cga cat tca gaa aga cga ata cta taa tac aat gac tcc tct tca tac cgc ctt cca gaa ttg caa taa caa tgt tgc ctc ata cct cgt ata cat tgg ata cga cat caa cct tcc gac taa aga cga taa gac agt att cga ctt ggt gtt tga aaa cag aaa cat tat ata caa ggc gga tgt cgt taa tga cat tat cca cca cag act gaa agt atc tct acc tat gat taa atc gtt gtt cta caa gat gtc gga gtt ctc tcc cta cga cga tca cta cgt aaa gaa gat aat agc cta ctg cct att aag gga cga gtc att tgc gga act aca tac taa att ctg ttt aaa cga gga cta taa aag tgt att tat gaa aaa tat atc att cga taa gat aga ttc cat cat cga aaa atg tag tcg tga cat aag tct cct caa aga gat tcg aat ctc aga cac cga ctt gta tac ggt att gag aac aga aga cat tcg gta tca cac ata tct cga agc cat aca ttc aga caa acg cat ttc att tcc cat gta cga cga tct cat aga aca gtg tca tct atc gat gga gca taa aag taa act cgt cga caa agc act caa taa att aga gtc tac cat cga tag tca atc tag act atc gta ttt gcc tcc gga aat tat gcg caa tat cat aac caa gct aag cga cta cca tct aaa cag tat gtt gta cgg aaa gaa cca tta caa ata tta tcc atg ata gaa aga aaa tat tta aaa aat aat cta tat gat tgg aga agt agg aaa caa aca gta aca aga cga cga tta cta c-a tta tta aat cat gag gtc cgt att ata ctc gta tat att gtt tct ctc atg tat aat aat aaa cgg aag aga tat agc acc aca tgc acc atc caa tgg aaa gtg taa aga caa cga ata cag aag ccg taa tct atg ttg tct atc gtg tcc tcc ggg aac tta cgc ttt cag att atg tga tag caa gac taa tac aca atg tac acc gtg tgg ttc gga tac ctt tac atc tca caa taa tca ttt aca ggc ttg tct aag ttg taa cgg aag atg tga tag taa tca ggt aga gac gcg atc gtg taa cac gac tca caa tag aat ctg tga atg ctc tcc agg ata tta ttg tct tct caa agg agc atc agg gtg tag aac atg tat ttc taa aac aaa gtg tgg aat agg ata cgg agt atc cgg ata cac gtc tac cgg aga cgt cat ctg ttc tcc gtg tgg tcc cgg aac ata ttc tca cac cgt ctc ttc cac aga taa atg cga acc cgt cgt aac cag caa tac att taa cta tat cga tgt gga aat taa cct gta tcc agt caa cga cac atc gtg tac tcg gac gac cac tac cgg tct cag cga atc cat ctc aac gtc gga act aac tat tac cat gaa tca taa aga ttg tga tcc agt ctt tcg tgc aga ata ctt ctc tgt cct taa taa tgt agc aac ttc agg att ctt tac agg aga aaa tag ata tca gaa tac ttc aaa gat atg tac tct gaa ttt cga gat taa atg taa caa caa aga ttc atc ttc caa aca gtt aac gaa aac aaa gaa tga tac tat cat gcc gca ttc aga gac ggt aac tct agt ggg cga ctg tct atc tag cgt cga cat cta cat act ata tag taa tac caa tac tca aga cta cga aaa tga tac aat ctc tta tca tat ggg taa tgt tct cga tgt caa tag cca tat gcc cgc tag ttg cga tat aca taa act gat cac taa ttc cca gaa tcc cac cca ctt ata gta ag- -tt ttt tta cct ata aat aat aaa tac aat aat taa ttt ctc gta aaa gta gaa aat ata ttc taa ttt att ata tgg taa gaa agt aga atc atc tag aac agt aat caa tca ata gca atc atg aaa caa tat att gtc ctg gca tgc atg tgc cta gtg gca gct gct atg cct act agt ctt caa caa tct tca tcc tcg tgt act gaa gaa gaa aac aaa cat cat atg gga atc gat gtt att atc aaa gtc aca aag caa gac caa aca ccg acc aat gat aag att tgt caa tcc gta acg gaa gtt aca gag a-- --- --- --- --- --- -cc gaa gat gat gag gta tcc gaa gaa gtt gta aaa gga gat ccc acc act tat tac act atc gtc ggt gcg ggt ctt aac atg aac ttt gga ttc acc aaa tgc cca aag att tta tcc atc tcc gaa tcc tct gat gga aac act gtg aat act aga ttg tcc agc gtg tca ccg gga caa ggt aag gac tct ccc gcg atc acg cgt gaa gaa gct ctg gct atg atc aaa gac tgt gag atg tct atc gac atc aga tgt agc gaa gaa gag aaa gac agt gac atc aag acc cat cca gta ctt ggg tct aac atc tca cat aag aaa gtg agt tac aaa gat atc atc ggt tca acg atc gtt gat aca aaa tgt gtc aag aac cta gag ttt agc gta cgt atc gga gac atg tgt gag gaa tca tct gaa ctt gaa gtc aag gat gga ttc aag tat gtc gac gga tcg gca tct gaa ggt gca acc gat gat act tca ctc atc gat tca aca aaa ctc aaa gca tgt gtc tga atc gat aac tct att cat ctg aaa atg gat gag ttg ggt taa tcg aac gat tca gac acc gca cca cga att aaa aaa g-- -ac cgg gca cta tat tcc ggt ttg caa aac aaa aat att taa cta cat tca caa aaa gtt acc tct cgt tac ttc ttc ttt ctg ttt caa tat gtg ata cga tat gat cac tat tcg tat tct ct- --- --- tgg tct cat aaa aaa gtt tta c-- --? aaa aaa aaa aaa a-t att ttt att ctc ttt ctc tct tcg at- ggt ctc aca aaa ata tta aac ctc ttt ctg atg tct caa cta ttt cgt aaa cga taa cgt cca aca ata tat tct cgt aga gct tat caa cat cct tat acc aat cta ggt tgt cag aca att gca tca taa aat aat gtt tat aat tta cac gtt aac atc ata taa taa acg tat ata gtt aat att ttt gga ata taa atg atc tgt aaa atc cat gta ggg gac act gct cac gtt ttt tct cta gta cat aat ttc aca caa gtt ttt ata cag aca aat taa ttc tcg tcc ata ta- ttt taa aac att gac ttt tgt act aag aaa aat atc ttg act aac cat ctc ttt ctc tct tcg atg ggt ctc aca aaa ata tta aac ctc ttt ctg atg g-- --- --- --- --- --- --- --- --- --- --- --- --- --- --- --- --- --- --- --- --- --- --- --- --- --- --- --- --- --- --- --- --- --- --- --- --- --- --- --- --- --- --- --- --- --- --- agt cgt aaa aag ttt tta tcc ttt ctc tct tcg ata ggt ctc --- --- --- --- --- --- --- --- --- --- --- --- --- --- --- --- --- --- --- --- --- --- --- --- --- --- --- --- --- --- --- --- --- --- --- --- --- --- --- --- --- --- --- --- --- --- --- --- --- --- --- --- --- --- --- --- --- --- --- --- --- --- --- --- --- --- --- --- --- --- aca aaa ata tta aac ctc ttt ctg atg gtc tct ata aa

>ON619835.1_Monkeypox_virus_isolate_MPXV_UK_2022_1_complete_genome

atc gag gac gac ttc aac tat gac agc tcg gtg gcg tct gcc agc gtg tac ata cga atg gca ttt cta aga aaa gtc tac ggt atc ctt -tc tac aat ttc ctt taa caa cgg caa cag ctg cag tat ttt tat act ttg aat gca -tc gga cat tta tac aag gga gtc ctg ttc taa tat tgg cat caa tgt tcg gat cta tag gct tga ttt tcg cat tga ctt tac aca gac atg aac atc ccc tga atc tgt aca tac ttt gtg gat tta cac tgt tag aat ctc taa cgc tgg cct ctg ttg tta ctt tct atg atg cac gta tcg tta tgc aag ctt tca tgt tga cta ctg cag tgt ttc ttg ctc tga cta cat ata ctc tac aat caa aga gag att tca gta aac ttg taa cag gat tgt ttg ctg ctt tct gga ttt taa ttt tgt cag gag tct tga gga taa agt tta aaa tag aat taa taa aga aca tat agg tca -tt ttt taa aca tgg ata gaa acc aag gtt gtt agt taa taa tat aca aga tat ttt ttc tca ctc tga tcc atg taa acc aag gac gag aga cac tct cat tcc tca ttc acg aca cca tta aaa atg gaa att aaa gcc ctc tat taa gca cag acg gct aca ggt cta cca tca ggt t-a cct tcg tct acc ttc aca atg gcc tct cct tgt gcc cag ttc agt ccc tgt cat tgc cac gct act aag gac tcc ctg aat acc gtg act gac gtc aga cat tgt ctg act gaa tac atc ctg tgg gtt tct cat aga tgg acc cat aga gaa agc gca ggg cct ctc tac agg ctt ctc atc tct ttc aga att gat gca atg gag cta ttt ggt agc gag ttg aag gag ttc tcg aat tca ctt ccg tgg gac aat atc gac aat tgc gtg gag atc att aaa tgt ttc atc aga aat gac tcc atg aaa acc gcc aaa gaa ctt tgt gca ata att gga ctt tgt act caa tca gct att gtc act gga aga gtc ttc aat gat aag tat atc gac ata cta ctt atg ctg cga aag att ctg aac gag aac gac tat ctc acc ctc ttg gat cat atc ctc act gct aaa tac taa atc tcc ttc atg ctc tct cac taa tac tct tac tca cta cac ttt tta tca tct tat gat gaa tga ttg cct tca tca ttt ttt cgt gga ata taa tat agg aat aat tag cac cag aat agc tat gga tat ctc gtt aag aat att ctc tat aag aga cat aat gta gac ata gtt att ata tcc ttc tta gat aag tgt tac gct act gga aag ttt cca tcg tta tta tta cat gaa gat gat ata att aaa cca aca ttg aga ttg gct ctt atg tta gct gga ttg aat tac tgt aat aaa tgc atc gag tat aga ggg ata tag caa ttc tcg ata ata gtc atg caa tat ttg aat gag act gat aat tta ggt aat aca gta ct- --- aca cac ata tct ttc tag att ata tat cgt taa aaa tct gta aga tgt ata ttt ctc ata agt atc cac tgt gta ata tta tta atg gat ata tag ata acg caa tag gga cta ata gta ttg taa aag ata taa tcg act att tac gta cat atc cag ata tct ata ttc cta cta gtt tgc tgc gta gtt gca tca ttg ata tgc atg -at tta tca gga ttc aga gat gaa tta cta agt aaa cta caa tcc cac aat aag taa gaa tca aat atc aaa aac tca ctt ttg att ttt cta gtc tta agt aat aca tat att tat taa tag acc tat gaa ata aaa aaa ggt aac aat gga ttc gcg tat agc tat tta cgt att agt atc ggc atc tct ttt gta tct tgt taa ttg tca caa act agt aca tta ctt caa tct gaa aat aaa tgg aag tga tat aac taa tac agc aga tat att gct gga caa tta tcc aat tat gac ctt tga tgg aaa gga tat tta tcc atc tat ctc gtt cat ggt cgg taa taa act ttt cct aga tct tta taa aaa tat ctt tga aga att ttt cag act att tcg agt atc tgt aag tag tca ata cga gga att aga ata tta tta ttc atg tga tta tac taa caa ccg tcc tac aat taa aca aca tta ctt tta taa cgg cga cga ata tac tga aat tga tag atc gaa aaa agc cac taa taa aaa cag ttg gtt aat tac ttc agg ctt tag act aca aaa atg gtt cga tag cga aga ttg tat aat tta tct cag atc ttt agt tag aag aat gga aga cag taa caa aaa cag taa aaa aac tta gta ctt aga tat cg- --a aaa aat ata ttt ttg tag act ctt gag aat aga agg aaa aca tgt aca taa tta taa aaa atg aaa atc aat ggc gaa taa gac agt gcg att cgc acc atg gag tcg gta gat ttc atg gct gtc gat gag cag ttt cac gac gac ctc gat ctt tgg tca tta tct ttg gta gat gat tat aaa aaa cat gga tta ggt gtt gac tgt tat gtt cta gaa cca gtt gtt gac agg aaa ata ttt gat aga ttt ctc ctt gaa cca att tgt gat cct gta gat gtt ctg tat gat tat ttt agg att cat aga gat aat att gat cag tat ata gta gat aga ctg ttt gca tat att aca tat aaa gat att ata tct gca tta gtg tca aag aat tat atg gaa gat att ttc tct ata att att aag aat tgt aat tct gtg caa gat ctc tta ctt tac tat cta tct aat gca tat gta gaa ata gac att gtt gat ctt atg gta gat cat ggg gct gta ata tat aaa ata gaa tgc ttg aat gcc tat ttt agg gga ata tgt aaa aag gaa agt agt gtt gtt gag ttt att ttg aat tgt ggt atc cca gat gaa aat gat gtt aaa tta gat cta tat aaa ata att cag tat act agg gga ttc ctt gta gat gaa ccc aca gta tta gaa att tat aag ctt tgt atc cca tat att gaa gat atc aat caa cta gat gct ggt gga agg acc ttg ctt tat cgc gct atc tat gca ggt tat ata gat tta gta tca tgg cta tta gaa aat gga gca aat gtc aac gca gta atg agt aat gga tat aca tgt ctt gac gtg gcc gtg gat agg gga tct gtc atc gcc cgt agg gaa gca cat ctt aaa ata tta gaa ata ttg ctt aga gaa cca ttg tct att gac tgt ata aaa tta gct ata ctt aat aat aca att gaa aac cat gat gtg ata aag ctc tgt atc aag tat ttt atg atg gta gat tat tca ctt tgt aat gtg tat gca tca tca ctc ttt gat tat ata att gat tgt aaa caa gaa ttg gag tac att agg cag atg aaa att cat aat aca acc atg tat gag tta atc tat aat aga gac aaa aac aag cat gct tcc cat att cta cat agg tat tct aaa cat cca gtt ttg aca cag tgt atc act aaa gga ttc aag att tac aca gaa gta acc gag cag gtc act aaa gct cta aac aga cgt gct cta ata gat gag ata ata aac aat gta tca act gat gac aat ctc cta tca aaa ctt cca tta gaa att agg gat cta att gtt tca caa gct gtc ata tag agt tct atc cac cca cct ttc ttg aaa tga gtt aat --- --- --- --- --- --- --- --- --- --- --- --- --- --- --- --- --- --- --- --- --- --- --- --- --- --- --- --- --- --- --- --- --- --- --- --- --- --- --- --- --- --- --- --- --- --- --- --- --- --- --- --- --- --- --- --- --- --- --- --- --- --- --- --- --- --- --- --- --- --- --- --- --- --- --- --- --- --- --- --- --- --- --- --- --- --- --- --- --- --- --- --- --- --- --- --- --- --- --- --- --- --- --- --- --- -ag tca taa gtt agt taa --- --- --- --- --- -gt cat aag tta gtt aag tca t-- --- --- --- --- --- --- --- --- --- --- --- --- --- --- --- -aa gtt agt taa gtc ata agt tag tta agt cat aag tta gtt aag tca taa gtt agt taa gtc ata agt tag tta agt cat aag tta gtt tat agt cta aca ctt cta att ttt ata cct tga tct ttt tct cta att atg aaa aag taa atc att atg aag atg gat g-a aat gga cga gat tgt gcg cat cgt taa cga tag tat gtg gta cgt acc taa cgc att tat gga cga cgg tga taa tga agg tca cat ttc tgt caa taa tgt ctg tca tat gta tct cgc att ctt tga tgt gga tat atc atc tca tct gtt taa att agt tat taa aca ctg cga tct gaa taa acg act aaa atg tgg taa ctc tcc att aca ttg cta tac gat gaa tac acg att taa tcc atc tgt att aaa gat att gtt acg cca cgg cat gcg taa ctt tga tag caa gga taa aaa agg aca tat tcc tct aca cca cta tct gat tca ttc act atc aat cga taa caa gat ctt tga tat act aac gga ccc cat tga tga ctt tag taa atc atc cga tct att gct gtg tta tct tag ata taa att caa tgg gag ctt aaa cta tta cgt tct gta caa att att gac taa agg atc tga ccc taa ttg cgt cga tga gga tgg act cac ttc tct tca tta cta ctg taa aca cat atc cgc gtt cca cga aag caa tta tta caa gtc aaa gag tca cac taa gat gcg agc tga gaa gcg att cat cta cgc gat aat aga tca tgg agc aaa cat taa cgc ggt tac gaa aat cgg aaa tac gcc gtt aca cac tta cct tca aca gta tac caa aca tag tcc tcg tgt ggt gta tgc tct ttt atc tcg agg agc cga tac gag gat acg taa taa tct taa ttg tac acc cat cat gga ata cat aaa gaa cga ttg tgc aac agg tca tat tct cat aat gtt act caa ttg gca cga aca aaa ata cgg gaa att aca aaa gga aga agg aca aca tct act tta tct att cat aaa aca taa tca agg ata tgg aag tcg ctc tct caa tat act acg gta tct act aga tag att cga cat tca gaa aga cga ata cta taa tac aat gac tcc tct tca tac cgc ctt cca gaa ttg caa taa caa tgt tgc ctc ata cct cgt ata cat tgg ata cga cat caa cct tcc gac taa aga cga taa gac agt att cga ctt ggt gtt tga aaa cag aaa cat tat ata caa ggc gga tgt cgt taa tga cat tat cca cca cag act gaa agt atc tct acc tat gat taa atc gtt gtt cta caa gat gtc gga gtt ctc tcc cta cga cga tca cta cgt aaa gaa gat aat agc cta ctg cct att aag gga cga gtc att tgc gga act aca tac taa att ctg ttt aaa cga gga cta taa aag tgt att tat gaa aaa tat atc att cga taa gat aga ttc cat cat cga aaa atg tag tcg tga cat aag tct cct caa aga gat tcg aat ctc aga cac cga ctt gta tac ggt att gag aac aga aga cat tcg gta tca cac ata tct cga agc cat aca ttc aga caa acg cat ttc att tcc cat gta cga cga tct cat aga aca gtg tca tct atc gat gga gca taa aag taa act cgt cga caa agc act caa taa att aga gtc tac cat cga tag tca atc tag act atc gta ttt gcc tcc gga aat tat gcg caa tat cat aac caa gct aag cga cta cca tct aaa cag tat gtt gta cgg aaa gaa cca tta caa ata tta tcc atg ata gaa aga aaa tat tta aaa aat aat cta tat gat tgg aga agt agg aaa caa aca gta aca aga cga cga tta cta c-a tta tta aat cat gag gtc cgt att ata ctc gta tat att gtt tct ctc atg tat aat aat aaa cgg aag aga tat agc acc aca tgc acc atc caa tgg aaa gtg taa aga caa cga ata cag aag ccg taa tct atg ttg tct atc gtg tcc tcc ggg aac tta cgc ttt cag att atg tga tag caa gac taa tac aca atg tac acc gtg tgg ttc gga tac ctt tac atc tca caa taa tca ttt aca ggc ttg tct aag ttg taa cgg aag atg tga tag taa tca ggt aga gac gcg atc gtg taa cac gac tca caa tag aat ctg tga atg ctc tcc agg ata tta ttg tct tct caa agg agc atc agg gtg tag aac atg tat ttc taa aac aaa gtg tgg aat agg ata cgg agt atc cgg ata cac gtc tac cgg aga cgt cat ctg ttc tcc gtg tgg tcc cgg aac ata ttc tca cac cgt ctc ttc cac aga taa atg cga acc cgt cgt aac cag caa tac att taa cta tat cga tgt gga aat taa cct gta tcc agt caa cga cac atc gtg tac tcg gac gac cac tac cgg tct cag cga atc cat ctc aac gtc gga act aac tat tac cat gaa tca taa aga ttg tga tcc agt ctt tcg tgc aga ata ctt ctc tgt cct taa taa tgt agc aac ttc agg att ctt tac agg aga aaa tag ata tca gaa tac ttc aaa gat atg tac tct gaa ttt cga gat taa atg taa caa caa aga ttc atc ttc caa aca gtt aac gaa aac aaa gaa tga tac tat cat gcc gca ttc aga gac ggt aac tct agt ggg cga ctg tct atc tag cgt cga cat cta cat act ata tag taa tac caa tac tca aga cta cga aaa tga tac aat ctc tta tca tat ggg taa tgt tct cga tgt caa tag cca tat gcc cgc tag ttg cga tat aca taa act gat cac taa ttc cca gaa tcc cac cca ctt ata gta ag- -tt ttt tta cct ata aat aat aaa tac aat aat taa ttt ctc gta aaa gta gaa aat ata ttc taa ttt att ata tgg taa gaa agt aga atc atc tag aac agt aat caa tca ata gca atc atg aaa caa tat att gtc ctg gca tgc atg tgc cta gtg gca gct gct atg cct act agt ctt caa caa tct tca tcc tcg tgt act gaa gaa gaa aac aaa cat cat atg gga atc gat gtt att atc aaa gtc aca aag caa gac caa aca ccg acc aat gat aag att tgt caa tcc gta acg gaa gtt aca gag a-- --- --- --- --- --- -cc gaa gat gat gag gta tcc gaa gaa gtt gta aaa gga gat ccc acc act tat tac act atc gtc ggt gcg ggt ctt aac atg aac ttt gga ttc acc aaa tgc cca aag att tta tcc atc tcc gaa tcc tct gat gga aac act gtg aat act aga ttg tcc agc gtg tca ccg gga caa ggt aag gac tct ccc gcg atc acg cgt gaa gaa gct ctg gct atg atc aaa gac tgt gag atg tct atc gac atc aga tgt agc gaa gaa gag aaa gac agt gac atc aag acc cat cca gta ctt ggg tct aac atc tca cat aag aaa gtg agt tac aaa gat atc atc ggt tca acg atc gtt gat aca aaa tgt gtc aag aac cta gag ttt agc gta cgt atc gga gac atg tgt gag gaa tca tct gaa ctt gaa gtc aag gat gga ttc aag tat gtc gac gga tcg gca tct gaa ggt gca acc gat gat act tca ctc atc gat tca aca aaa ctc aaa gca tgt gtc tga atc gat aac tct att cat ctg aaa atg gat gag ttg ggt taa tcg aac gat tca gac acc gca cca cga att aaa aaa g-- -ac cgg gca cta tat tcc ggt ttg caa aac aaa aat att taa cta cat tca caa aaa gtt acc tct cgt tac ttc ttc ttt ctg ttt caa tat gtg ata cga tat gat cac tat tcg tat tct ct- --- --- tgg tct cat aaa aaa gtt tta c-- --? aaa aaa aaa aaa a-t att ttt att ctc ttt ctc tct tcg at- ggt ctc aca aaa ata tta aac ctc ttt ctg atg tct caa cta ttt cgt aaa cga taa cgt cca aca ata tat tct cgt aga gct tat caa cat cct tat acc aat cta ggt tgt cag aca att gca tca taa aat aat gtt tat aat tta cac gtt aac atc ata taa taa acg tat ata gtt aat att ttt gga ata taa atg atc tgt aaa atc cat gta ggg gac act gct cac gtt ttt tct cta gta cat aat ttc aca caa gtt ttt ata cag aca aat taa ttc tcg tcc ata ta- ttt taa aac att gac ttt tgt act aag aaa aat atc ttg act aac cat ctc ttt ctc tct tcg atg ggt ctc aca aaa ata tta aac ctc ttt ctg atg g-- --- --- --- --- --- --- --- --- --- --- --- --- --- --- --- --- --- --- --- --- --- --- --- --- --- --- --- --- --- --- --- --- --- --- --- --- --- --- --- --- --- --- --- --- --- --- agt cgt aaa aag ttt tta tcc ttt ctc tct tcg ata ggt ctc --- --- --- --- --- --- --- --- --- --- --- --- --- --- --- --- --- --- --- --- --- --- --- --- --- --- --- --- --- --- --- --- --- --- --- --- --- --- --- --- --- --- --- --- --- --- --- --- --- --- --- --- --- --- --- --- --- --- --- --- --- --- --- --- --- --- --- --- --- --- aca aaa ata tta aac ctc ttt ctg atg gtc tct ata aa

>ON682269.2_Monkeypox_virus_isolate_MPXV/Germany/2022/RKI07_complete_genome

atc gag gac gac ttc aac tat gac agc tcg gtg gcg tct gcc agc gtg tac ata cga atg gca ttt cta aga aaa gtc tac ggt atc ctt -tc tac aat ttc ctt taa caa cgg caa cag ctg cag tat ttt tat act ttg aat gca -tc gga cat tta tac aag gga gtc ctg ttc taa tat tgg cat caa tgt tcg gat cta tag gct tga ttt tcg cat tga ctt tac aca gac atg aac atc ccc tga atc tgt aca tac ttt gtg gat tta cac tgt tag aat ctc taa cgc tgg cct ctg ttg tta ctt tct atg atg cac gta tcg tta tgc aag ctt tca tgt tga cta ctg cag tgt ttc ttg ctc tga cta cat ata ctc tac aat caa aga gag att tca gta aac ttg taa cag gat tgt ttg ctg ctt tct gga ttt taa ttt tgt cag gag tct tga gga taa agt tta aaa tag aat taa taa aga aca tat agg tca -tt ttt taa aca tgg ata gaa acc aag gtt gtt agt taa taa tat aca aga tat ttt ttc tca ctc tga tcc atg taa acc aag gac gag aga cac tct cat tcc tca ttc acg aca cca tta aaa atg gaa att aaa gcc ctc tat taa gca cag acg gct aca ggt cta cca tca ggt t-a cct tcg tct acc ttc aca atg gcc tct cct tgt gcc cag ttc agt ccc tgt cat tgc cac gct act aag gac tcc ctg aat acc gtg act gac gtc aga cat tgt ctg act gaa tac atc ctg tgg gtt tct cat aga tgg acc cat aga gaa agc gca ggg cct ctc tac agg ctt ctc atc tct ttc aga att gat gca atg gag cta ttt ggt agc gag ttg aag gag ttc tcg aat tca ctt ccg tgg gac aat atc gac aat tgc gtg gag atc att aaa tgt ttc atc aga aat gac tcc atg aaa acc gcc aaa gaa ctt tgt gca ata att gga ctt tgt act caa tca gct att gtc act gga aga gtc ttc aat gat aag tat atc gac ata cta ctt atg ctg cga aag att ctg aac gag aac gac tat ctc acc ctc ttg gat cat atc ctc act gct aaa tac taa atc tcc ttc atg ctc tct cac taa tac tct tac tca cta cac ttt tta tca tct tat gat gaa tga ttg cct tca tca ttt ttt cgt gga ata taa tat agg aat aat tag cac cag aat agc tat gga tat ctc gtt aag aat att ctc tat aag aga cat aat gta gac ata gtt att ata tcc ttc tta gat aag tgt tac gct act gga aag ttt cca tcg tta tta tta cat gaa gat gat ata att aaa cca aca ttg aga ttg gct ctt atg tta gct gga ttg aat tac tgt aat aaa tgc atc gag tat aga ggg ata tag caa ttc tcg ata ata gtc atg caa tat ttg aat gag act gat aat tta ggt aat aca gta ct- --- aca cac ata tct ttc tag att ata tat cgt taa aaa tct gta aga tgt ata ttt ctc ata agt atc cac tgt gta ata tta tta atg gat ata tag ata acg caa tag gga cta ata gta ttg taa aag ata taa tcg act att tac gta cat atc cag ata tct ata ttc cta cta gtt tgc tgc gta gtt gca tca ttg ata tgc atg -at tta tca gga ttc aga gat gaa tta cta agt aaa cta caa tcc cac aat aag taa gaa tca aat atc aaa aac tca ctt ttg att ttt cta gtc tta agt aat aca tat att tat taa tag acc tat gaa ata aaa aaa ggt aac aat gga ttc gcg tat agc tat tta cgt att agt atc ggc atc tct ttt gta tct tgt taa ttg tca caa act agt aca tta ctt caa tct gaa aat aaa tgg aag tga tat aac taa tac agc aga tat att gct gga caa tta tcc aat tat gac ctt tga tgg aaa gga tat tta tcc atc tat ctc gtt cat ggt cgg taa taa act ttt cct aga tct tta taa aaa tat ctt tga aga att ttt cag act att tcg agt atc tgt aag tag tca ata cga gga att aga ata tta tta ttc atg tga tta tac taa caa ccg tcc tac aat taa aca aca tta ctt tta taa cgg cga cga ata tac tga aat tga tag atc gaa aaa agc cac taa taa aaa cag ttg gtt aat tac ttc agg ctt tag act aca aaa atg gtt cga tag cga aga ttg tat aat tta tct cag atc ttt agt tag aag aat gga aga cag taa caa aaa cag taa aaa aac tta gta ctt aga tat cg- --a aaa aat ata ttt ttg tag act ctt gag aat aga agg aaa aca tgt aca taa tta taa aaa atg aaa atc aat ggc gaa taa gac agt gcg att cgc acc atg gag tcg gta gat ttc atg gct gtc gat gag cag ttt cac gac gac ctc gat ctt tgg tca tta tct ttg gta gat gat tat aaa aaa cat gga tta ggt gtt gac tgt tat gtt cta gaa cca gtt gtt gac agg aaa ata ttt gat aga ttt ctc ctt gaa cca att tgt gat cct gta gat gtt ctg tat gat tat ttt agg att cat aga gat aat att gat cag tat ata gta gat aga ctg ttt gca tat att aca tat aaa gat att ata tct gca tta gtg tca aag aat tat atg gaa gat att ttc tct ata att att aag aat tgt aat tct gtg caa gat ctc tta ctt tac tat cta tct aat gca tat gta gaa ata gac att gtt gat ctt atg gta gat cat ggg gct gta ata tat aaa ata gaa tgc ttg aat gcc tat ttt agg gga ata tgt aaa aag gaa agt agt gtt gtt gag ttt att ttg aat tgt ggt atc cca gat gaa aat gat gtt aaa tta gat cta tat aaa ata att cag tat act agg gga ttc ctt gta gat gaa ccc aca gta tta gaa att tat aag ctt tgt atc cca tat att gaa gat atc aat caa cta gat gct ggt gga agg acc ttg ctt tat cgc gct atc tat gca ggt tat ata gat tta gta tca tgg cta tta gaa aat gga gca aat gtc aac gca gta atg agt aat gga tat aca tgt ctt gac gtg gcc gtg gat agg gga tct gtc atc gcc cgt agg gaa gca cat ctt aaa ata tta gaa ata ttg ctt aga gaa cca ttg tct att gac tgt ata aaa tta gct ata ctt aat aat aca att gaa aac cat gat gtg ata aag ctc tgt atc aag tat ttt atg atg gta gat tat tca ctt tgt aat gtg tat gca tca tca ctc ttt gat tat ata att gat tgt aaa caa gaa ttg gag tac att agg cag atg aaa att cat aat aca acc atg tat gag tta atc tat aat aga gac aaa aac aag cat gct tcc cat att cta cat agg tat tct aaa cat cca gtt ttg aca cag tgt atc act aaa gga ttc aag att tac aca gaa gta acc gag cag gtc act aaa gct cta aac aga cgt gct cta ata gat gag ata ata aac aat gta tca act gat gac aat ctc cta tca aaa ctt cca tta gaa att agg gat cta att gtt tca caa gct gtc ata tag agt tct atc cac cca cct ttc ttg aaa tga gtt aat agt cat aag tt- --- --- --- --- --- --- --- --- --- --- --- --- --- --- --- --- --- --- --- --- --- --- --- --- --- --- --- --- --- --- --- --- --- --- --- --- --- --- --- --- --- --- --- --- --- --- --- --- --- --- --- --- --- --- --- --- --- --- --- --- --- --- --- --- --- --- --- --- --- --- --- --- --- --- --- --- --- --- --- --- --- --- --- --- --- --- --- --- --- agt taa gtc ata agt tag tta agt cat aag tta gtt aag tca taa gtt agt taa --- --- --- --- --- -gt cat aag tta gtt aag tca t-- --- --- --- --- --- --- --- --- --- --- --- --- --- --- --- -aa gtt agt taa gtc ata agt tag tta agt cat aag tta gtt aag tca taa gtt agt taa gtc ata agt tag tta agt cat aag tta gtt tat agt cta aca ctt cta att ttt ata cct tga tct ttt tct cta att atg aaa aag taa atc att atg aag atg gat g-a aat gga cga gat tgt gcg cat cgt taa cga tag tat gtg gta cgt acc taa cgc att tat gga cga cgg tga taa tga agg tca cat ttc tgt caa taa tgt ctg tca tat gta tct cgc att ctt tga tgt gga tat atc atc tca tct gtt taa att agt tat taa aca ctg cga tct gaa taa acg act aaa atg tgg taa ctc tcc att aca ttg cta tac gat gaa tac acg att taa tcc atc tgt att aaa gat att gtt acg cca cgg cat gcg taa ctt tga tag caa gga taa aaa agg aca tat tcc tct aca cca cta tct gat tca ttc act atc aat cga taa caa gat ctt tga tat act aac gga ccc cat tga tga ctt tag taa atc atc cga tct att gct gtg tta tct tag ata taa att caa tgg gag ctt aaa cta tta cgt tct gta caa att att gac taa agg atc tga ccc taa ttg cgt cga tga gga tgg act cac ttc tct tca tta cta ctg taa aca cat atc cgc gtt cca cga aag caa tta tta caa gtc aaa gag tca cac taa gat gcg agc tga gaa gcg att cat cta cgc gat aat aga tca tgg agc aaa cat taa cgc ggt tac gaa aat cgg aaa tac gcc gtt aca cac tta cct tca aca gta tac caa aca tag tcc tcg tgt ggt gta tgc tct ttt atc tcg agg agc cga tac gag gat acg taa taa tct taa ttg tac acc cat cat gga ata cat aaa gaa cga ttg tgc aac agg tca tat tct cat aat gtt act caa ttg gca cga aca aaa ata cgg gaa att aca aaa gga aga agg aca aca tct act tta tct att cat aaa aca taa tca agg ata tgg aag tcg ctc tct caa tat act acg gta tct act aga tag att cga cat tca gaa aga cga ata cta taa tac aat gac tcc tct tca tac cgc ctt cca gaa ttg caa taa caa tgt tgc ctc ata cct cgt ata cat tgg ata cga cat caa cct tcc gac taa aga cga taa gac agt att cga ctt ggt gtt tga aaa cag aaa cat tat ata caa ggc gga tgt cgt taa tga cat tat cca cca cag act gaa agt atc tct acc tat gat taa atc gtt gtt cta caa gat gtc gga gtt ctc tcc cta cga cga tca cta cgt aaa gaa gat aat agc cta ctg cct att aag gga cga gtc att tgc gga act aca tac taa att ctg ttt aaa cga gga cta taa aag tgt att tat gaa aaa tat atc att cga taa gat aga ttc cat cat cga aaa atg tag tcg tga cat aag tct cct caa aga gat tcg aat ctc aga cac cga ctt gta tac ggt att gag aac aga aga cat tcg gta tca cac ata tct cga agc cat aca ttc aga caa acg cat ttc att tcc cat gta cga cga tct cat aga aca gtg tca tct atc gat gga gca taa aag taa act cgt cga caa agc act caa taa att aga gtc tac cat cga tag tca atc tag act atc gta ttt gcc tcc gga aat tat gcg caa tat cat aac caa gct aag cga cta cca tct aaa cag tat gtt gta cgg aaa gaa cca tta caa ata tta tcc atg ata gaa aga aaa tat tta aaa aat aat cta tat gat tgg aga agt agg aaa caa aca gta aca aga cga cga tta cta c-a tta tta aat cat gag gtc cgt att ata ctc gta tat att gtt tct ctc atg tat aat aat aaa cgg aag aga tat agc acc aca tgc acc atc caa tgg aaa gtg taa aga caa cga ata cag aag ccg taa tct atg ttg tct atc gtg tcc tcc ggg aac tta cgc ttt cag att atg tga tag caa gac taa tac aca atg tac acc gtg tgg ttc gga tac ctt tac atc tca caa taa tca ttt aca ggc ttg tct aag ttg taa cgg aag atg tga tag taa tca ggt aga gac gcg atc gtg taa cac gac tca caa tag aat ctg tga atg ctc tcc agg ata tta ttg tct tct caa agg agc atc agg gtg tag aac atg tat ttc taa aac aaa gtg tgg aat agg ata cgg agt atc cgg ata cac gtc tac cgg aga cgt cat ctg ttc tcc gtg tgg tcc cgg aac ata ttc tca cac cgt ctc ttc cac aga taa atg cga acc cgt cgt aac cag caa tac att taa cta tat cga tgt gga aat taa cct gta tcc agt caa cga cac atc gtg tac tcg gac gac cac tac cgg tct cag cga atc cat ctc aac gtc gga act aac tat tac cat gaa tca taa aga ttg tga tcc agt ctt tcg tgc aga ata ctt ctc tgt cct taa taa tgt agc aac ttc agg att ctt tac agg aga aaa tag ata tca gaa tac ttc aaa gat atg tac tct gaa ttt cga gat taa atg taa caa caa aga ttc atc ttc caa aca gtt aac gaa aac aaa gaa tga tac tat cat gcc gca ttc aga gac ggt aac tct agt ggg cga ctg tct atc tag cgt cga cat cta cat act ata tag taa tac caa tac tca aga cta cga aaa tga tac aat ctc tta tca tat ggg taa tgt tct cga tgt caa tag cca tat gcc cgc tag ttg cga tat aca taa act gat cac taa ttc cca gaa tcc cac cca ctt ata gta ag- -tt ttt tta cct ata aat aat aaa tac aat aat taa ttt ctc gta aaa gta gaa aat ata ttc taa ttt att ata tgg taa gaa agt aga atc atc tag aac agt aat caa tca ata gca atc atg aaa caa tat att gtc ctg gca tgc atg tgc cta gtg gca gct gct atg cct act agt ctt caa caa tct tca tcc tcg tgt act gaa gaa gaa aac aaa cat cat atg gga atc gat gtt att atc aaa gtc aca aag caa gac caa aca ccg acc aat gat aag att tgt caa tcc gta acg gaa gtt aca gag a-- --- --- --- --- --- -cc gaa gat gat gag gta tcc gaa gaa gtt gta aaa gga gat ccc acc act tat tac act atc gtc ggt gcg ggt ctt aac atg aac ttt gga ttc acc aaa tgc cca aag att tta tcc atc tcc gaa tcc tct gat gga aac act gtg aat act aga ttg tcc agc gtg tca ccg gga caa ggt aag gac tct ccc gcg atc acg cgt gaa gaa gct ctg gct atg atc aaa gac tgt gag atg tct atc gac atc aga tgt agc gaa gaa gag aaa gac agt gac atc aag acc cat cca gta ctt ggg tct aac atc tca cat aag aaa gtg agt tac aaa gat atc atc ggt tca acg atc gtt gat aca aaa tgt gtc aag aac cta gag ttt agc gta cgt atc gga gac atg tgt gag gaa tca tct gaa ctt gaa gtc aag gat gga ttc aag tat gtc gac gga tcg gca tct gaa ggt gca acc gat gat act tca ctc atc gat tca aca aaa ctc aaa gca tgt gtc tga atc gat aac tct att cat ctg aaa atg gat gag ttg ggt taa tcg aac gat tca gac acc gca cca cga att aaa aaa g-- -ac cgg gca cta tat tcc ggt ttg caa aac aaa aat att taa cta cat tca caa aaa gtt acc tct cgt tac ttc ttc ttt ctg ttt caa tat gtg ata cga tat gat cac tat tcg tat tct ct- --- --- tgg tct cat aaa aaa gtt tta c-- --a aaa aaa aaa aaa a-t att ttt att ctc ttt ctc tct tcg at- ggt ctc aca aaa ata tta aac ctc ttt ctg atg tct caa cta ttt cgt aaa cga taa cgt cca aca ata tat tct cgt aga gct tat caa cat cct tat acc aat cta ggt tgt cag aca att gca tca taa aat aat gtt tat aat tta cac gtt aac atc ata taa taa acg tat ata gtt aat att ttt gga ata taa atg atc tgt aaa atc cat gta ggg gac act gct cac gtt ttt tct cta gta cat aat ttc aca caa gtt ttt ata cag aca aat taa ttc tcg tcc ata ta- ttt taa aac att gac ttt tgt act aag aaa aat atc ttg act aac cat ctc ttt ctc tct tcg atg ggt ctc aca aaa ata tta aac ctc ttt ctg atg g-- --- --- --- --- --- --- --- --- --- --- --- --- --- --- --- --- --- --- --- --- --- --- --- --- --- --- --- --- --- --- --- --- --- --- --- --- --- --- --- --- --- --- --- --- --- --- agt cgt aaa aag ttt tta tcc ttt ctc tct tcg ata ggt ctc --- --- --- --- --- --- --- --- --- --- --- --- --- --- --- --- --- --- --- --- --- --- --- --- --- --- --- --- --- --- --- --- --- --- --- --- --- --- --- --- --- --- --- --- --- --- --- --- --- --- --- --- --- --- --- --- --- --- --- --- --- --- --- --- --- --- --- --- --- --- aca aaa ata tta aac ctc ttt ctg atg gtc tct ata aa

>KJ642616.1_Monkeypox_virus_strain_PCH_complete_genome

atc gag gac gac ttc aac tat gac agc tcg gtg gcg tct gcc agc gtg tac ata cga atg gca ttt cta aga aaa gtc tac ggt atc ctt -tc tac aat ttc ctt taa caa cgg caa cag ctg cag tat ttt tat act ttg aat gca -tc gga cat tta tac aag gga gtc ctg ttc taa tat tgg cat caa tgt tcg gat cta tag gct tga ttt tcg cat tga ctt tac aca gac atg aac atc ccc tga atc tgt aca tac ttt gtg gat tta cac tgt tag aat ctc taa cgc tgg cct ctg ttg tta ctt tct atg atg cac gta tcg tta tgc aag ctt tca tgt tga cta ctg cag tgt ttc ttg ctc tga cta cat ata ctc tac aat caa aga gag att tca gta aac ttg taa cag gat tgt ttg ctg ctt tct gga ttt taa ttt tgt cag gag tct tga gga taa agt tta aaa tag aat taa taa aga aca tat agg tca -tt ttt taa aca tgg ata gaa acc aag gtt gtt agt taa taa tat aca aga tat ttt ttc tca ctc tga tcc atg taa acc aag gac gag aga cac tct cat tcc tca ttc acg aca cca tta aaa atg gaa att aaa gcc ctc tat taa gca cag acg gct aca ggt cta cca tca ggt t-a cct tcg tct acc ttc aca atg gcc tct cct tgt gcc cag ttc agt ccc tgt cat tgc cac gct act aag gac tcc ctg aat acc gtg act gat gtc aga cat tgt ctg act gaa tac atc ctg tgg gtt tct cat aga tgg acc cat aga gaa agc gca ggg cct ctc tac agg ctt ctc atc tct ttc aga att gat gca atg gag cta ttt ggt agc gag ttg aag gag ttc tcg gat tca ctt ccg tgg gac aat atc gac aat tgc gtg gag atc att aaa tgt ttc atc aga aat gac tcc atg aaa acc gcc aaa gaa ctt tgt gca ata att gga ctt tgt act caa tca gct att gtc act gga aga gtc ttc aat gat aag tat atc gac ata cta ctt atg ctg cga aag att ctg aac gag aac gac tat ctc acc ctc ttg gat cat atc ctc act gct aaa tac taa atc tcc ttc atg ctc tct cac taa tac tct tac tca cta cac ttt tta tca tct tat gat gaa tga ttg cct tca tca -tt ttt cgt gga ata taa tat agg aat aat tag cac cag aat agc tat gga tat ctc gtt aag aat att ctc gat aag aga cat aat gta gac ata gtt att ata tcc ttc tta gat aag tgt tac gct act gga aag ttt cca tcg tta tta tta cat gaa gat gat ata att aaa cca aca ttg aga ttg gct ctt atg tta gct gga ttg aat tac tgt aat aaa tgc atc gag tat aga ggg ata tag caa ttc tcg ata ata gtc atg caa tat ttg aat gag act gat aat tta ggt aat aca gta ct- --- aca cac ata tct ttc tag att ata tat cgt taa aaa tct gta aga ggt ata ttt ctc ata agt atc cac tgt gta ata tta tta atg gat ata tag ata acg caa tag gga cta ata gta ttg taa aag ata taa tcg act att tac gta cat atc cag ata tct ata ttc cta cta gtt tgc tgc gta gtt gca tca ttg ata tgc atg -at tta tca gga ttc aga gat gaa tta cta agt aaa cta caa tcc cac aat aag taa gaa tca aat atc aaa aac tca ctt ttg att ttt cta gtc tta agt aat aca tat att tat taa tag acc tat gaa ata aaa aaa ggt aac aat gga ttc gcg tat agc tat tta cgt att agt atc ggc atc tct ttt gta tct tgt taa ttg tca caa act agt aca tta ctt caa tct gaa aat aaa tgg aag tga tat aac taa tac agc aga tat att gct gga caa tta tcc aat tat gac ctt tga tgg aaa gga tat tta tcc atc tat ctc gtt cat ggt cgg taa taa act ttt cct aga tct tta taa aaa tat ctt tga aga att ttt cag act att tcg agt atc tgt aag tag tca ata cga gga att aga ata tta tta ttc atg tga tta tac taa caa ccg tcc tac aat taa aca aca tta ctt tta taa cgg cga aga ata tac tga aat tga tag atc gaa aaa agc cac taa taa aaa cag ttg gtt aat tac ttc agg ctt tag act aca aaa atg gtt cga tag cga aga ttg tat aat tta tct cag atc ttt agt tag aag aat gga aga cag taa caa aaa cag taa aaa aac tta gta ctt aga tat cg- -aa aaa aat ata ttt ttg tag act ctt gag aat aga agg aaa aca tgt aca taa tta taa aaa atg aaa atc aat ggc gaa taa gac agt gcg att cgc acc atg gag tcg gta gat ttc atg gct gtc gat gag cag ttt cac gac gac ctc gat ctt tgg tca tta tct ttg gta gat gat tat aaa aaa cat gga tta ggt gtt gac tgt tat gtt cta gaa cca gtt gtt gac agg aaa ata ttt gat aga ttt ctc ctt gaa cca att tgt gat cct gta gat gtt ctg tat gat tat ttt agg att cat aga gat aat att gat cag tat ata gta gat aga ctg ttt gca tat att aca tat aaa gat att ata tct gca tta gtg tca aag aat tat atg gaa gat att ttc tct ata att att aag aat tgt aat tct gta caa gat ctc tta ctt tac tat cta tct aat gca tat gta gaa ata gac att gtt gat ctt atg gta gat cat ggg gct gta ata tat aaa ata gaa tgc ttg aat gcc tat ttt agg gga ata tgt aaa aag gaa agt agt gtt gtt gag ttt att ttg aat tgt ggt atc cca gat gaa aat gat gtt aaa tta gat cta tat aaa ata att cag tat act agg gga ttc ctt gta gat gaa ccc aca gta tta gaa att tat aag ctt tgt atc cca tat att gaa gat atc aat caa cta gat gct ggt gga agg acc ttg ctt tat cgc gct atc tat gca ggt tat ata gat tta gta tca ttg cta tta gaa aat gga gca aat gtc aac gca gta atg agt aat gga tat aca tgt ctt gac gtg gcc gtg gat agg gga tct gtc atc gcc cgt agg gaa gca cat ctt aaa ata tta gaa ata ttg ctt aga gaa cca ttg tct att gac tgt ata aaa tta gct ata ctt aat aat aca att gaa aac cat gat gtg ata aag ctc tgt atc aag tat ttt atg atg gta gat tat tca ctt tgt aat gtg tat gca tca tca ctc ttt gat tat ata att gat tgt aaa caa gaa ttg gag tac att agg cag atg aaa att cat aat aca acc atg tat gag tta atc tat aat aga gac aaa aac aag cat gct tcc cat att cta cat agg tat tct aaa cat cca gtt ttg aca cag tgt atc act aaa gga ttc aag att tac aca gaa gta acc gag cag gtc act aaa gct cta aac aga cgt gct cta ata gat gag ata ata aac aat gta tca act gat gac aat ctc cta tca aaa ctt cca tta gaa att agg gat cta att gtt tca caa gct gtc ata tag agt tct atc cac cca cct ttc ttg aaa tga gtt aat --- --- --- --- --- --- --- --- --- --- --- --- --- --- --- --- --- --- --- --- --- --- --- --- --- --- --- --- --- --- --- --- --- --- --- --- --- --- --- --- --- --- --- --- --- --- --- --- --- --- --- --- --- --- --- --- --- --- --- --- --- --- --- --- --- --- --- --- --- --- --- --- --- --- --- --- --- --- --- --- --- --- --- --- --- --- --- --- --- --- --- --- --- --- --- --- --- --- --- --- --- --- --- --- --- -ag tca taa gtt agt taa --- --- --- --- --- -gt cat aag tta gtt aag tca taa gtt agt taa gtc ata agt tag tta agt cat aag tta gtt aag tca taa gtt agt taa gtc ata agt tag tta agt cat aag tta gtt aag tca taa gtt agt taa gtc ata agt tag tt- --t tat aag tta gtt tat agt cta aca ctt cta att ttt ata cct tga tct ttt tct cta att atg aaa aag taa atc att atg aag atg gat g-a aat gga cga gat tgt gcg cat cgt taa cga tag tat gtg gta cgt acc taa cgc att tat gga cga cgg tga taa tga agg tca cat ttc tgt caa taa tgt ctg tca tat gta tct cgc att ctt tga tgt gga tat atc atc tca tct gtt taa att agt tat taa aca ctg cga tct gaa taa acg act aaa atg tgg taa ctc tcc att aca ttg cta tac gat gaa tac acg att taa tcc atc tgt att aaa gat att gtt acg cca cgg cat gcg taa ctt tga tag caa gga taa aaa agg aca tat tcc tct aca cca cta tct gat tca ttc act atc aat cga taa caa gat ctt tga tat act aac gga ccc cat tga tga ctt tag taa atc atc cga tct att gct gtg tta tct tag ata taa att caa tgg gag ctt aaa cta tta cgt tct gta caa att att gac taa agg atc tga ccc taa ttg cgt cga tga gga tgg act cac ttc tct tca tta cta ctg taa aca cat atc cgc gtt cca cga aag caa tta tta caa gtc aaa gag tca cac taa gat gcg agc tga gaa gcg att cat cta cgc gat aat aga tca tgg agc aaa cat taa cgc ggt tac gaa aat cgg aaa tac gcc gtt aca cac tta cct tca aca gta tac caa aca tag tcc tcg tgt ggt gta tgc tct ttt atc tcg agg agc tga tac gag gat acg taa taa tct tga ttg tac acc cat cat gga ata cat aaa gaa cga ttg tgc aac agg tca tat tct cat aat gtt act caa ttg gca cga aca aaa ata cgg gaa att aca aaa gga aga agg aca aca tct act tta tct att cat aaa aca taa tca agg ata tgg aag tcg ctc tct caa tat act acg gta tct act aga tag att cga cat tca gaa aga cga ata cta taa tac aat gac tcc tct tca tac cgc ctt cca gaa ttg caa taa caa tgt tgc ctc ata cct cgt ata cat cgg ata cga cat caa cct tcc gac taa aga cga taa gac agt att cga ctt ggt gtt tga aaa cag aaa cat cat ata caa ggc gga tgt cgt taa tga cat cat cca cca cag act gaa agt atc tct acc tat gat taa atc gtt gtt cta caa gat gtc gga gtt ctc tcc cta cga cga tca cta cgt aaa gaa gat aat agc cta ctg cct att aag gga cga gtc att tgc gga act aca tac taa att ctg ttt aaa cga gga cta taa aag tgt att tat gaa aaa tat atc att cga taa gat aga ttc cat cat cga aaa atg tag tcg tga cat aag tct cct caa aga gat tcg aat ctc aga cac cga ctt gta tac ggt att gag aac aga aga cat ccg gta tca cac cta tct cga agc cat aca ttc aga caa acg cat ttc att tcc cat gta tga cga tct cat aga aca gtg tca tct atc gat gga gca taa aag taa act cgt cga caa agc act caa taa att aga gtc tac cat cga taa tca atc tag act atc gta ttt gcc tcc gga aat tat gcg caa tat cat aac caa gct aag cga cta cca tct aaa cag tat gtt gta cgg aaa gaa cca tta caa ata tta tcc atg ata gaa aga aaa tat tta aaa aat aat cta tat gat tgg aga agt agg aaa caa aca gta aca aga cga cga tta cta c-a --a tta aat cat gag gtc cgt att ata ctc gta tat att gtt tct ctc atg tat aat aat aaa cgg aag aga tat agc acc aca tgc acc atc caa tgg aaa gtg taa aga caa cga ata cag aag ccg taa tct atg ttg tct atc gtg tcc tcc ggg aac tta cgc ttc cag att atg tga tag caa gac taa tac aca atg tac acc gtg tgg ttc gga tac ctt tac atc tca caa taa tca ttt aca ggc ttg tct aag ttg taa cgg aag atg tga tag taa tca ggt aga gac gcg atc gtg taa cac gac tca caa tag aat ctg tga atg ctc tcc agg ata tta ttg tct tct caa agg agc atc agg gtg tag aac atg tat ttc taa aac aaa gtg tgg aat agg ata cgg agt atc cgg ata cac gtc tac cgg aga cgt cat ctg ttc tcc gtg tgg tcc cgg aac ata ttc tca cac cgt ctc ttc cac aga taa atg cga acc cgt cgt aac cag caa tac att taa cta tat cga tgt gga aat taa cct gta tcc agt caa cga cac atc gtg tac tcg gac gac cac tac cgg tct cag cga atc cat ctc aac gtc gga act aac tat tac cat gaa tca taa aga ttg tga tcc agt ctt tcg tgc aga ata ctt ctc tgt cct taa taa tgt agc aac ttc agg att ctt tac agg aga aaa tag ata tca gaa tac ttc aaa gat atg tac tct gaa ttt cga gat taa atg taa caa caa aga ttc atc ttc caa aca gtt aac gaa aac aaa gaa tga tac tat cat gcc gca ttc aga gac ggt aac tct agt ggg cga ctg tct atc tag cgt cga cat cta cat act ata tag taa tac caa tac tca aga cta cga aac gga tac aat ctc tta tca tat ggg taa tgt tct cga tgt caa tag cca tat gcc cgc tag ttg cga tat aca taa act gat cac taa ttc cca gaa tcc cac cca ctt ata gta ag- -tt ttt tta ccc ata aat aat aaa tac aat aat taa ttt ctc gta aaa gta gaa aat ata ttc taa ttt att ata tgg taa gaa agt aga atc atc tag aac agt aat caa tca ata gca atc atg aaa caa tat att gtc ctg gca tgc atg tgc cta gtg gca gct gct atg cct act agt ctt caa caa tct tca tcc tcg tgt act gaa gaa gaa aac aaa cat cat atg gga atc gat gtt att atc aaa gtc aca aag caa gac caa aca ccg acc aat gat aag att tgt caa tcc gta acg gaa gtt aca gag a-- --- --- --- --- --- -cc gaa gat gat gag gta tcc gaa gaa gtt gta aaa gga gat ccc acc act tat tac act atc gtc ggt gcg ggt ctt aac atg aac ttt gga ttc acc aaa tgc cca aag att tca tcc atc tcc gaa tcc tct gat gga aac act gtg aat act aga ttg tcc agc gtg tca ccg gga caa ggt aag gac tct ccc gcg atc acg cgt gaa gaa gct ctg gct atg atc aaa gac tgt gag atg tct atc gac atc aga tgt agc gaa gaa gag aaa gac agt gac atc aag acc cat cca gta ctt ggg tct aac atc tca cat aag aaa gtg agt tac aaa gat atc atc ggt tca acg atc gtt gat aca aaa tgt gtc aag aac cta gag ttt agc gta cgt atc gga gac atg tgt gag gaa tca tct gaa ctt gaa gtc aag gat gga ttc aag tat gtc gac gga tcg gca tct gaa ggt gca acc gat gat act tca ctc atc gat tca aca aaa ctc aaa gca tgt gtc tga atc gat aac tct att cat ctg aaa atg gat gag ttg ggt taa tcg aac gat tca gac acc gca cca cga att aaa aaa g-- -ac cgg gca cta tat tcc ggt ttg caa aac aaa aat att taa cta cat tta caa aaa gtt acc tct cgt tac ttc ttc ttt ctg ttt caa tat gtg ata cga tat gat cac tat tcg tat tct ct- --- --- tgg tct cat aaa aaa gtt tta c-- --a aaa aaa aaa aaa a-t att ttt att ctc ttt ctc tct tcg at- ggt ctc aca aaa ata tta aac ctc ttt ctg atg tct caa cta ttt cgt aaa cga taa cgt cca aca ata tat tct cgt aga gct tat caa cat cct tat acc aat cta ggt tgt cag aca att gca tca taa aat aat gtt tat aat tta cac gtt aac atc ata taa taa acg tat ata gtt aat att ttt gga ata taa atg atc tgt aaa atc cat gta ggg gac act gct cac gtt ttt tct cta gta cat aat ttc aca caa gtt ttt ata cag aca aat taa ttc tcg tcc ata ta- ttt taa aac att gac ttt tgt act aag aaa aat atc ttg act aac cat ctc ttt ctc tct tcg at- --- --- --- --- --- --- --- --- --- --- --- --- --- --- --- --- --- --- --- --- --- --- --- --- --- --- --- --- --- --- --- --- --- --- --- --- --- --- --- --- --- --- --- --- --- --- --- --- --- --- --- --- --- --- --- --- --- --- --- --- --- --- --- --- --- --- --- --- --- --- --- --- --- --- --- --- --- --- --- --- --- --- --- --- --- --- --- --- --- --- --- --- --- --- --- --- --- --- --- --- --- --- --- --- --- --- --- --- --- --- --- --- --- --- --- --- --- --- --- --- --- --- --- --- --- --- --- --- --- --- --- --- --- --- --- --- --- --- --- --- --- --- --- --- --- --- --- --- --- --- --- --- --- --- --

>AY753185.1_Monkeypox_virus_strain_COP-58_complete_genome

atc gag gac gac ttc aac tat gac agc tcg gtg gcg tct gcc agc gtg tac ata cga atg gca ttt cta aga aaa gtc tac ggt atc ctt -tc tac aat ttc ctt taa caa cgg caa cag ctg cag tat ttt tat act ttg aat gca -tc gga cat tta tac aag gga gtc ctg ttc taa tat tgg cat caa tgt tcg gat cta tag gct tga ttt tcg cat tga ctt tac aca gac atg aac atc ccc tga atc tgt aca tac ttt gtg gat tta cac tgt tag aat ctc taa cgc tgg cct ctg ttg tta ctt tct atg atg cac gta tcg tta tgc aag ctt tca tgt tga cta ctg cag tgt ttc ttg ctc tga cta cat ata ctc tac aat caa aga gag att tca gta aac ttg taa cag gat tgt ttg ctg ctt tct gga ttt taa ttt tgt cag gag tct tga gga taa agt tta aaa tag aat taa taa aga aca tat agg tca -tt ttt taa aca tgg ata gaa acc aag gtt gtt agt taa taa tat aca aga tat ttt ttc tca ctc tga tcc atg taa acc aag gac gag aga cac tct cat tcc tca ttc acg aca cca tta aaa atg gaa att aaa gcc ctc tat taa gca cag acg gct aca ggt cta cca tca ggt t-a cct tcg tct acc ttc aca atg gcc tct cct tgt gcc cag ttc agt ccc tgt cat tgc cac gct act aag gac tcc ctg aat acc gtg act gat gtc aga cat tgt ctg act gaa tac atc ctg tgg gtt tct cat aga tgg acc cat aga gaa agc gca ggg cct ctc tac agg ctt ctc atc tct ttc aga att gat gca atg gag cta ttt ggt agc gag ttg aag gag ttc tcg gat tca ctt ccg tgg gac aat atc gac aat tgc gtg gag atc att aaa tgt ttc atc aga aat gac tcc atg aaa acc gcc aaa gaa ctt tgt gca ata att gga ctt tgt act caa tca gct att gtc act gga aga gtc ttc aat gat aag tat atc gac ata cta ctt atg ctg cga aag att ctg aac gag aac gac tat ctc acc ctc ttg gat cat atc ctc act gct aaa tac taa atc tcc ttc atg ctc tct cac taa tac tct tac tca cta cac ttt tta tca tct tat gat gaa tga ttg cct tca tca -tt ttt cgt gga ata taa tat agg aat aat tag cac cag aat agc tat gga tat ctc gtt aag aat att ctc gat aag aga cat aat gta gac ata gtt att ata tcc ttc tta gat aag tgt tac gct act gga aag ttt cca tcg tta tta tta cat gaa gat gat ata att aaa cca aca ttg aga ttg gct ctt atg tta gct gga ttg aat tac tgt aat aaa tgc atc gag tat aga ggg ata tag caa ttc tcg ata ata gtc atg caa tat ttg aat gag act gat aat tta ggt aat aca gta ct- --- aca cac ata tct ttc tag att ata tat cgt taa aaa tct gta aga ggt ata ttt ctc ata agt atc cac tgt gta ata tta tta atg gat ata tag ata acg caa tag gga cta ata gta ttg taa aag ata taa tcg act att tac gta cat atc cag ata tct ata ttc cta cta gtt tgc tgc gta gtt gca tca ttg ata tgc atg -at tta tca gga ttc aga gat gaa tta cta agt aaa cta caa tcc cac aat aag taa gaa tca aat atc aaa aac tca ctt ttg att ttt cta gtc tta agt aat aca tat att tat taa tag acc tat gaa ata aaa aaa ggt aac aat gga ttc gcg tat agc tat tta cgt att agt atc ggc atc tct ttt gta tct tgt taa ttg tca caa act agt aca tta ctt caa tct gaa aat aaa tgg aag tga tat aac taa tac agc aga tat att gct gga caa tta tcc aat tat gac ctt tga tgg aaa gga tat tta tcc atc tat ctc gtt cat ggt cgg taa taa act ttt cct aga tct tta taa aaa tat ctt tga aga att ttt cag act att tcg agt atc tgt aag tag tca ata cga gga att aga ata tta tta ttc atg tga tta tac taa caa ccg tcc tac aat taa aca aca tta ctt tta taa cgg cga aga ata tac tga aat tga tag atc gaa aaa agc cac taa taa aaa cag ttg gtt aat tac ttc agg ctt tag act aca aaa atg gtt cga tag cga aga ttg tat aat tta tct cag atc ttt agt tag aag aat gga aga cag taa caa aaa cag taa aaa aac tta gta ctt aga tat cg- -aa aaa aat ata ttt ttg tag act ctt gag aat aga agg aaa aca tgt aca taa tta taa aaa atg aaa atc aat ggc gaa taa gac agt gcg att cgc acc atg gag tcg gta gat ttc atg gct gtc gat gag cag ttt cac gac gac ctc gat ctt tgg tca tta tct ttg gta gat gat tat aaa aaa cat gga tta ggt gtt gac tgt tat gtt cta gaa cca gtt gtt gac agg aaa ata ttt gat aga ttt ctc ctt gaa cca att tgt gat cct gta gat gtt ctg tat gat tat ttt agg att cat aga gat aat att gat cag tat ata gta gat aga ctg ttt gca tat att aca tat aaa gat att ata tct gca tta gtg tca aag aat tat atg gaa gat att ttc tct ata att att aag aat tgt aat tct gta caa gat ctc tta ctt tac tat cta tct aat gca tat gta gaa ata gac att gtt gat ctt atg gta gat cat ggg gct gta ata tat aaa ata gaa tgc ttg aat gcc tat ttt agg gga ata tgt aaa aag gaa agt agt gtt gtt gag ttt att ttg aat tgt ggt atc cca gat gaa aat gat gtt aaa tta gat cta tat aaa ata att cag tat act agg gga ttc ctt gta gat gaa ccc aca gta tta gaa att tat aag ctt tgt atc cca tat att gaa gat atc aat caa cta gat gct ggt gga agg acc ttg ctt tat cgc gct atc tat gca ggt tat ata gat tta gta tca ttg cta tta gaa aat gga gca aat gtc aac gca gta atg agt aat gga tat aca tgt ctt gac gtg gcc gtg gat agg gga tct gtc atc gcc cgt agg gaa gca cat ctt aaa ata tta gaa ata ttg ctt aga gaa cca ttg tct att gac tgt ata aaa tta gct ata ctt aat aat aca att gaa aac cat gat gtg ata aag ctc tgt atc aag tat ttt atg atg gta gat tat tca ctt tgt aat gtg tat gca tca tca ctc ttt gat tat ata att gat tgt aaa caa gaa ttg gag tac att agg cag atg aaa att cat aat aca acc atg tat gag tta atc tat aat aga gac aaa aac aag cat gct tcc cat att cta cat agg tat tct aaa cat cca gtt ttg aca cag tgt atc act aaa gga ttc aag att tac aca gaa gta acc gag cag gtc act aaa gct cta aac aga cgt gct cta ata gat gag ata ata aac aat gta tca act gat gac aat ctc cta tca aaa ctt cca tta gaa att agg gat cta att gtt tca caa gct gtc ata tag agt tct atc cac cca cct ttc ttg aaa tga gtt aat agt cat aag tta gtt aag tca taa gtt agt taa gtc ata agt tag tta agt cat aag tt- --- --- --- --- --- --- --- --- --- --- --- --- --- --- --- --- --- --- --- --- --- --- --- --- --- --- --- --- --- --- --- --- --- --- --- --- --- --- --- --- --- --- --- --- --- --- --- --- --- --- --- --- --- --- --- --- --- agt taa gtc ata agt tag tta agt cat aag tta gtt aag tca taa gtt agt taa gtc ata agt tag tta agt cat aag tta gtt aag tca taa gtt agt taa --- --- --- --- --- -gt cat aag tta gtt aag tca taa gtt agt taa gtc ata agt tag tta agt cat aag tta gtt aag tca taa gtt agt taa gtc ata agt tag tta agt cat aag tta gtt aag tca taa gtt agt taa gtc ata agt tag tt- --t tat aag tta gtt tat agt cta aca ctt cta att ttt ata cct tga tct ttt tct cta att atg aaa aag taa atc att atg aag atg gat g-a aat gga cga gat tgt gcg cat cgt taa cga tag tat gtg gta cgt acc taa cgc att tat gga cga cgg tga taa tga agg tca cat ttc tgt caa taa tgt ctg tca tat gta tct cgc att ctt tga tgt gga tat atc atc tca tct gtt taa att agt tat taa aca ctg cga tct gaa taa acg act aaa atg tgg taa ctc tcc att aca ttg cta tac gat gaa tac acg att taa tcc atc tgt att aaa gat att gtt acg cca cgg cat gcg taa ctt tga tag caa gga taa aaa agg aca tat tcc tct aca cca cta tct gat tca ttc act atc aat cga taa caa gat ctt tga tat act aac gga ccc cat tga tga ctt tag taa atc atc cga tct att gct gtg tta tct tag ata taa att caa tgg gag ctt aaa cta tta cgt tct gta caa att att gac taa agg atc tga ccc taa ttg cgt cga tga gga tgg act cac ttc tct tca tta cta ctg taa aca cat atc cgc gtt cca cga aag caa tta tta caa gtc aaa gag tca cac taa gat gcg agc tga gaa gcg att cat cta cgc gat aat aga tca tgg agc aaa cat taa cgc ggt tac gaa aat cgg aaa tac gcc gtt aca cac tta cct tca aca gta tac caa aca tag tcc tcg tgt ggt gta tgc tct ttt atc tcg agg agc tga tac gag gat acg taa taa tct tga ttg tac acc cat cat gga ata cat aaa gaa cga ttg tgc aac agg tca tat tct cat aat gtt act caa ttg gca cga aca aaa ata cgg gaa att aca aaa gga aga agg aca aca tct act tta tct att cat aaa aca taa tca agg ata tgg aag tcg ctc tct caa tat act acg gta tct act aga tag att cga cat tca gaa aga cga ata cta taa tac aat gac tcc tct tca tac cgc ctt cca gaa ttg caa taa caa tgt tgc ctc ata cct cgt ata cat cgg ata cga cat caa cct tcc gac taa aga cga taa gac agt att cga ctt ggt gtt tga aaa cag aaa cat cat ata caa ggc gga tgt cgt taa tga cat cat cca cca cag act gaa agt atc tct acc tat gat taa atc gtt gtt cta caa gat gtc gga gtt ctc tcc cta cga cga tca cta cgt aaa gaa gat aat agc cta ctg cct att aag gga cga gtc att tgc gga act aca tac taa att ctg ttt aaa cga gga cta taa aag tgt att tat gaa aaa tat atc att cga taa gat aga ttc cat cat cga aaa atg tag tcg tga cat aag tct cct caa aga gat tcg aat ctc aga cac cga ctt gta tac ggt att gag aac aga aga cat ccg gta tca cac cta tct cga agc cat aca ttc aga caa acg cat ttc att tcc cat gta tga cga tct cat aga aca gtg tca tct atc gat gga gca taa aag taa act cgt cga caa agc act caa taa att aga gtc tac cat cga taa tca atc tag act atc gta ttt gcc tcc gga aat tat gcg caa tat cat aac caa gct aag cga cta cca tct aaa cag tat gtt gta cgg aaa gaa cca tta caa ata tta tcc atg ata gaa aga aaa tat tta aaa aat aat cta tat gat tgg aga agt agg aaa caa aca gta aca aga cga cga tta cta c-a --a tta aat cat gag gtc cgt att ata ctc gta tat att gtt tct ctc atg tat aat aat aaa cgg aag aga tat agc acc aca tgc acc atc caa tgg aaa gtg taa aga caa cga ata cag aag ccg taa tct atg ttg tct atc gtg tcc tcc ggg aac tta cgc ttc cag att atg tga tag caa gac taa tac aca atg tac acc gtg tgg ttc gga tac ctt tac atc tca caa taa tca ttt aca ggc ttg tct aag ttg taa cgg aag atg tga tag taa tca ggt aga gac gcg atc gtg taa cac gac tca caa tag aat ctg tga atg ctc tcc agg ata tta ttg tct tct caa agg agc atc agg gtg tag aac atg tat ttc taa aac aaa gtg tgg aat agg ata cgg agt atc cgg ata cac gtc tac cgg aga cgt cat ctg ttc tcc gtg tgg tcc cgg aac ata ttc tca cac cgt ctc ttc cac aga taa atg cga acc cgt cgt aac cag caa tac att taa cta tat cga tgt gga aat taa cct gta tcc agt caa cga cac atc gtg tac tcg gac gac cac tac cgg tct cag cga atc cat ctc aac gtc gga act aac tat tac cat gaa tca taa aga ttg tga tcc agt ctt tcg tgc aga ata ctt ctc tgt cct taa taa tgt agc aac ttc agg att ctt tac agg aga aaa tag ata tca gaa tac ttc aaa gat atg tac tct gaa ttt cga gat taa atg taa caa caa aga ttc atc ttc caa aca gtt aac gaa aac aaa gaa tga tac tat cat gcc gca ttc aga gac ggt aac tct agt ggg cga ctg tct atc tag cgt cga cat cta cat act ata tag taa tac caa tac tca aga cta cga aac gga tac aat ctc tta tca tat ggg taa tgt tct cga tgt caa tag cca tat gcc cgc tag ttg cga tat aca taa act gat cac taa ttc cca gaa tcc cac cca ctt ata gta ag- -tt ttt tta ccc ata aat aat aaa tac aat aat taa ttt ctc gta aaa gta gaa aat ata ttc taa ttt att ata tgg taa gaa agt aga atc atc tag aac agt aat caa tca ata gca atc atg aaa caa tat att gtc ctg gca tgc atg tgc cta gtg gca gct gct atg cct act agt ctt caa caa tct tca tcc tcg tgt act gaa gaa gaa aac aaa cat cat atg gga atc gat gtt att atc aaa gtc aca aag caa gac caa aca ccg acc aat gat aag att tgt caa tcc gta acg gaa gtt aca gag a-- --- --- --- --- --- -cc gaa gat gat gag gta tcc gaa gaa gtt gta aaa gga gat ccc acc act tat tac act atc gtc ggt gcg ggt ctt aac atg aac ttt gga ttc acc aaa tgc cca aag att tca tcc atc tcc gaa tcc tct gat gga aac act gtg aat act aga ttg tcc agc gtg tca ccg gga caa ggt aag gac tct ccc gcg atc acg cgt gaa gaa gct ctg gct atg atc aaa gac tgt gag atg tct atc gac atc aga tgt agc gaa gaa gag aaa gac agt gac atc aag acc cat cca gta ctt ggg tct aac atc tca cat aag aaa gtg agt tac aaa gat atc atc ggt tca acg atc gtt gat aca aaa tgt gtc aag aac cta gag ttt agc gta cgt atc gga gac atg tgt gag gaa tca tct gaa ctt gaa gtc aag gat gga ttc aag tat gtc gac gga tcg gca tct gaa ggt gca acc gat gat act tca ctc atc gat tca aca aaa ctc aaa gca tgt gtc tga atc gat aac tct att cat ctg aaa atg gat gag ttg ggt taa tcg aac gat tca gac acc gca cca cga att aaa aaa g-- -ac cgg gca cta tat tcc ggt ttg caa aac aaa aat att taa cta cat tta caa aaa gtt acc tct cgt tac ttc ttc ttt ctg ttt caa tat gtg ata cga tat gat cac tat tcg tat tct ct- --- --- tgg tct cat aaa aaa gtt tta c-- -aa aaa aaa aaa aaa a-t att ttt att cta ttt ctc tct tcg at- ggt ctc aca aaa ata tta aac ctc ttt ctg atg tct caa cta ttt cgt aaa cga taa cgt cca aca ata tat tct cgt aga gct tat caa cat cct tat acc aat cta ggt tgt cag aca att gca tca taa aat aat gtt tat aat tta cac gtt aac atc ata taa taa acg tat ata gtt aat att ttt gga ata taa atg atc tgt aaa atc cat gta ggg gac act gct cac gtt ttt tct cta gta cat aat ttc aca caa gtt ttt ata cag aca aat taa ttc tcg tcc ata ta- ttt taa aac att gac ttt tgt act aag aaa aat atc ttg act aac cat ctc ttt ctc tct tcg atg ggt ctc aca aaa ata tta aac ctc ttt ctg atg g-- --- --- --- --- --- --- --- --- --- --- --- --- --- --- --- --- --- --- --- --- --- --- --- --- --- --- --- --- --- --- --- --- --- --- --- --- --- --- --- --- --- --- --- --- --- --- agt cgt aaa aag ttt tta tcc ttt ctc tct tcg atg ggt ctc --- --- --- --- --- --- --- --- --- --- --- --- --- --- --- --- --- --- --- --- --- --- --- --- --- --- --- --- --- --- --- --- --- --- --- --- --- --- --- --- --- --- --- --- --- --- --- --- --- --- --- --- --- --- --- --- --- --- --- --- --- --- --- --- --- --- --- --- --- --- aca aaa ata tta aac ctc ttt ctg atg gtc tct ata aa

>AY603973.1_Monkeypox_virus_strain_MPXV-WRAIR7-61_complete_genome

atc gag gac gac ttc aac tat gac agc tcg gtg gcg tct gcc agc gtg tac ata cga atg gca ttt cta aga aaa gtc tac ggt atc ctt -tc tac aat ttc ctt taa caa cgg caa cag ctg cag tat ttt tat act ttg aat gca -tc gga cat tta tac aag gga gtc ctg ttc taa tat tgg cat caa tgt tcg gat cta tag gct tga ttt tcg cat tga ctt tac aca gac atg aac atc ccc tga atc tgt aca tac ttt gtg gat tta cac tgt tag aat ctc taa cgc tgg cct ctg ttg tta ctt tct atg atg cac gta tcg tta tgc aag ctt tca tgt tga cta ctg cag tgt ttc ttg ctc tga cta cat ata ctc tac aat caa aga gag att tca gta aac ttg taa cag gat tgt ttg ctg ctt tct gga ttt taa ttt tgt cag gag tct tga gga taa agt tta aaa tag aat taa taa aga aca tat agg tca -tt ttt taa aca tgg ata gaa acc aag gtt gtt agt taa taa tat aca aga tat ttt ttc tca ctc tga tcc atg taa acc aag gac gag aga cac tct cat tcc tca ttc acg aca cca tta aaa atg gaa att aaa gcc ctc tat taa gca cag acg gct aca ggt cta cca tca ggt t-a cct tcg tct acc ttc aca atg gcc tct cct tgt gcc cag ttc agt ccc tgt cat tgc cac gct act aag gac tcc ctg aat acc gtg act gat gtc aga cat tgt ctg act gaa tac atc ctg tgg gtt tct cat aga tgg acc cat aga gaa agc gca ggg cct ctc tac agg ctt ctc atc tct ttc aga att gat gca atg gag cta ttt ggt agc gag ttg aag gag ttc tcg gat tca ctt ccg tgg gac aat atc gac aat tgc gtg gag atc att aaa tgt ttc atc aga aat gac tcc atg aaa acc gcc aaa gaa ctt tgt gca ata att gga ctt tgt act caa tca gct att gtc act gga aga gtc ttc aat gat aag tat atc gac ata cta ctt atg ctg cga aag att ctg aac gag aac gac tat ctc acc ctc ttg gat cat atc ctc act gct aaa tac taa atc tcc ttc atg ctc tct cac taa tac tct tac tca cta cac ttt tta tca tct tat gat gaa tga ttg cct tca tca -tt ttt cgt gga ata taa tat agg aat aat tag cac cag aat agc tat gga tat ctc gtt aag aat att ctc gat aag aga cat aat gta gac ata gtt att ata tcc ttc tta gat aag tgt tac gct act gga aag ttt cca tcg tta tta tta cat gaa gat gat ata att aaa cca aca ttg aga ttg gct ctt atg tta gct gga ttg aat tac tgt aat aaa tgc atc gag tat aga ggg ata tag caa ttc tcg ata ata gtc atg caa tat ttg aat gag act gat aat tta ggt aat aca gta ct- --- aca cac ata tct ttc tag att ata tat cgt taa aaa tct gta aga ggt ata ttt ctc ata agt atc cac tgt gta ata tta tta atg gat ata tag ata acg caa tag gga cta ata gta ttg taa aag ata taa tcg act att tac gta cat atc cag ata tct ata ttc cta cta gtt tgc tgc gta gtt gca tca ttg ata tgc atg -at tta tca gga ttc aga gat gaa tta cta agt aaa cta caa tcc cac aat aag taa gaa tca aat atc aaa aac tca ctt ttg att ttt cta gtc tta agt aat aca tat att tat taa tag acc tat gaa ata aaa aaa ggt aac aat gga ttc gcg tat agc tat tta cgt att agt atc ggc atc tct ttt gta tct tgt taa ttg tca caa act agt aca tta ctt caa tct gaa aat aaa tgg aag tga tat aac taa tac agc aga tat att gct gga caa tta tcc aat tat gac ctt tga tgg aaa gga tat tta tcc atc tat ctc gtt cat ggt cgg taa taa act ttt cct aga tct tta taa aaa tat ctt tga aga att ttt cag act att tcg agt atc tgt aag tag tca ata cga gga att aga ata tta tta ttc atg tga tta tac taa caa ccg tcc tac aat taa aca aca tta ctt tta taa cgg cga aga ata tac tga aat tga tag atc gaa aaa agc cac taa taa aaa cag ttg gtt aat tac ttc agg ctt tag act aca aaa atg gtt cga tag cga aga ttg tat aat tta tct cag atc ttt agt tag aag aat gga aga cag taa caa aaa cag taa aaa aac tta gta ctt aga tat cg- -aa aaa aat ata ttt ttg tag act ctt gag aat aga agg aaa aca tgt aca taa tta taa aaa atg aaa atc aat ggc gaa taa gac agt gcg att cgc acc atg gag tcg gta gat ttc atg gct gtc gat gag cag ttt cac gac gac ctc gat ctt tgg tca tta tct ttg gta gat gat tat aaa aaa cat gga tta ggt gtt gac tgt tat gtt cta gaa cca gtt gtt gac agg aaa ata ttt gat aga ttt ctc ctt gaa cca att tgt gat cct gta gat gtt ctg tat gat tat ttt agg att cat aga gat aat att gat cag tat ata gta gat aga ctg ttt gca tat att aca tat aaa gat att ata tct gca tta gtg tca aag aat tat atg gaa gat att ttc tct ata att att aag aat tgt aat tct gta caa gat ctc tta ctt tac tat cta tct aat gca tat gta gaa ata gac att gtt gat ctt atg gta gat cat ggg gct gta ata tat aaa ata gaa tgc ttg aat gcc tat ttt agg gga ata tgt aaa aag gaa agt agt gtt gtt gag ttt att ttg aat tgt ggt atc cca gat gaa aat gat gtt aaa tta gat cta tat aaa ata att cag tat act agg gga ttc ctt gta gat gaa ccc aca gta tta gaa att tat aag ctt tgt atc cca tat att gaa gat atc aat caa cta gat gct ggt gga agg acc ttg ctt tat cgc gct atc tat gca ggt tat ata gat tta gta tca ttg cta tta gaa aat gga gca aat gtc aac gca gta atg agt aat gga tat aca tgt ctt gac gtg gcc gtg gat agg gga tct gtc atc gcc cgt agg gaa gca cat ctt aaa ata tta gaa ata ttg ctt aga gaa cca ttg tct att gac tgt ata aaa tta gct ata ctt aat aat aca att gaa aac cat gat gtg ata aag ctc tgt atc aag tat ttt atg atg gta gat tat tca ctt tgt aat gtg tat gca tca tca ctc ttt gat tat ata att gat tgt aaa caa gaa ttg gag tac att agg cag atg aaa att cat aat aca acc atg tat gag tta atc tat aat aga gac aaa aac aag cat gct tcc cat att cta cat agg tat tct aaa cat cca gtt ttg aca cag tgt atc act aaa gga ttc aag att tac aca gaa gta acc gag cag gtc act aaa gct cta aac aga cgt gct cta ata gat gag ata ata aac aat gta tca act gat gac aat ctc cta tca aaa ctt cca tta gaa att agg gat cta att gtt tca caa gct gtc ata tag agt tct atc cac cca cct ttc ttg aaa tga gtt aat agt cat aag tt- --- --- --- --- --- --- --- --- --- --- --- --- --- --- --- --- --- --- --- --- --- --- --- --- --- --- --- --- --- --- --- --- --- --- --- --- --- --- --- --- --- --- --- --- --- --- --- --- --- --- --- --- --- --- --- --- --- --- --- --- --- --- --- --- --- --- --- --- --- --- --- --- --- --- --- --- --- --- --- --- --- --- --- --- --- --- --- --- --- agt taa gtc ata agt tag tta agt cat aag tta gtt aag tca taa gtt agt taa --- --- --- --- --- -gt cat aag tta gtt aag tca taa gtt agt taa gtc ata agt tag tta agt cat aag tta gtt aag tca taa gtt agt taa gtc ata agt tag tta agt cat aag tta gtt aag tca taa gtt agt taa gtc ata agt tag tt- --t tat aag tta gtt tat agt cta aca ctt cta att ttt ata cct tga tct ttt tct cta att atg aaa aag taa atc att atg aag atg gat g-a aat gga cga gat tgt gcg cat cgt taa cga tag tat gtg gta cgt acc taa cgc att tat gga cga cgg tga taa tga agg tca cat ttc tgt caa taa tgt ctg tca tat gta tct cgc att ctt tga tgt gga tat atc atc tca tct gtt taa att agt tat taa aca ctg cga tct gaa taa acg act aaa atg tgg taa ctc tcc att aca ttg cta tac gat gaa tac acg att taa tcc atc tgt att aaa gat att gtt acg cca cgg cat gcg taa ctt tga tag caa gga taa aaa agg aca tat tcc tct aca cca cta tct gat tca ttc act atc aat cga taa caa gat ctt tga tat act aac gga ccc cat tga tga ctt tag taa atc atc cga tct att gct gtg tta tct tag ata taa att caa tgg gag ctt aaa cta tta cgt tct gta caa att att gac taa agg atc tga ccc taa ttg cgt cga tga gga tgg act cac ttc tct tca tta cta ctg taa aca cat atc cgc gtt cca cga aag caa tta tta caa gtc aaa gag tca cac taa gat gcg agc tga gaa gcg att cat cta cgc gat aat aga tca tgg agc aaa cat taa cgc ggt tac gaa aat cgg aaa tac gcc gtt aca cac tta cct tca aca gta tac caa aca tag tcc tcg tgt ggt gta tgc tct ttt atc tcg agg agc tga tac gag gat acg taa taa tct tga ttg tac acc cat cat gga ata cat aaa gaa cga ttg tgc aac agg tca tat tct cat aat gtt act caa ttg gca cga aca aaa ata cgg gaa att aca aaa gga aga agg aca aca tct act tta tct att cat aaa aca taa tca agg ata tgg aag tcg ctc tct caa tat act acg gta tct act aga tag att cga cat tca gaa aga cga ata cta taa tac aat gac tcc tct tca tac cgc ctt cca gaa ttg caa taa caa tgt tgc ctc ata cct cgt ata cat cgg ata cga cat caa cct tcc gac taa aga cga taa gac agt att cga ctt ggt gtt tga aaa cag aaa cat cat ata caa ggc gga tgt cgt taa tga cat cat cca cca cag act gaa agt atc tct acc tat gat taa atc gtt gtt cta caa gat gtc gga gtt ctc tcc cta cga cga tca cta cgt aaa gaa gat aat agc cta ctg cct att aag gga cga gtc att tgc gga act aca tac taa att ctg ttt aaa cga gga cta taa aag tgt att tat gaa aaa tat atc att cga taa gat aga ttc cat cat cga aaa atg tag tcg tga cat aag tct cct caa aga gat tcg aat ctc aga cac cga ctt gta tac ggt att gag aac aga aga cat ccg gta tca cac cta tct cga agc cat aca ttc aga caa acg cat ttc att tcc cat gta tga cga tct cat aga aca gtg tca tct atc gat gga gca taa aag taa act cgt cga caa agc act caa taa att aga gtc tac cat cga taa tca atc tag act atc gta ttt gcc tcc gga aat tat gcg caa tat cat aac caa gct aag cga cta cca tct aaa cag tat gtt gta cgg aaa gaa cca tta caa ata tta tcc atg ata gaa aga aaa tat tta aaa aat aat cta tat gat tgg aga agt agg aaa caa aca gta aca aga cga cga tta cta c-a --a tta aat cat gag gtc cgt att ata ctc gta tat att gtt tct ctc atg tat aat aat aaa cgg aag aga tat agc acc aca tgc acc atc caa tgg aaa gtg taa aga caa cga ata cag aag ccg taa tct atg ttg tct atc gtg tcc tcc ggg aac tta cgc ttc cag att atg tga tag caa gac taa tac aca atg tac acc gtg tgg ttc gga tac ctt tac atc tca caa taa tca ttt aca ggc ttg tct aag ttg taa cgg aag atg tga tag taa tca ggt aga gac gcg atc gtg taa cac gac tca caa tag aat ctg tga atg ctc tcc agg ata tta ttg tct tct caa agg agc atc agg gtg tag aac atg tat ttc taa aac aaa gtg tgg aat agg ata cgg agt atc cgg ata cac gtc tac cgg aga cgt cat ctg ttc tcc gtg tgg tcc cgg aac ata ttc tca cac cgt ctc ttc cac aga taa atg cga acc cgt cgt aac cag caa tac att taa cta tat cga tgt gga aat taa cct gta tcc agt caa cga cac atc gtg tac tcg gac gac cac tac cgg tct cag cga atc cat ctc aac gtc gga act aac tat tac cat gaa tca taa aga ttg tga tcc agt ctt tcg tgc aga ata ctt ctc tgt cct taa taa tgt agc aac ttc agg att ctt tac agg aga aaa tag ata tca gaa tac ttc aaa gat atg tac tct gaa ttt cga gat taa atg taa caa caa aga ttc atc ttc caa aca gtt aac gaa aac aaa gaa tga tac tat cat gcc gca ttc aga gac ggt aac tct agt ggg cga ctg tct atc tag cgt cga cat cta cat act ata tag taa tac caa tac tca aga cta cga aac gga tac aat ctc tta tca tat ggg taa tgt tct cga tgt caa tag cca tat gcc cgc tag ttg cga tat aca taa act gat cac taa ttc cca gaa tcc cac cca ctt ata gta ag- -tt ttt tta ccc ata aat aat aaa tac aat aat taa ttt ctc gta aaa gta gaa aat ata ttc taa ttt att ata tgg taa gaa agt aga atc atc tag aac agt aat caa tca ata gca atc atg aaa caa tat att gtc ctg gca tgc atg tgc cta gtg gca gct gct atg cct act agt ctt caa caa tct tca tcc tcg tgt act gaa gaa gaa aac aaa cat cat atg gga atc gat gtt att atc aaa gtc aca aag caa gac caa aca ccg acc aat gat aag att tgt caa tcc gta acg gaa gtt aca gag a-- --- --- --- --- --- -cc gaa gat gat gag gta tcc gaa gaa gtt gta aaa gga gat ccc acc act tat tac act atc gtc ggt gcg ggt ctt aac atg aac ttt gga ttc acc aaa tgc cca aag att tca tcc atc tcc gaa tcc tct gat gga aac act gtg aat act aga ttg tcc agc gtg tca ccg gga caa ggt aag gac tct ccc gcg atc acg cgt gaa gaa gct ctg gct atg atc aaa gac tgt gag atg tct atc gac atc aga tgt agc gaa gaa gag aaa gac agt gac atc aag acc cat cca gta ctt ggg tct aac atc tca cat aag aaa gtg agt tac aaa gat atc atc ggt tca acg atc gtt gat aca aaa tgt gtc aag aac cta gag ttt agc gta cgt atc gga gac atg tgt gag gaa tca tct gaa ctt gaa gtc aag gat gga ttc aag tat gtc gac gga tcg gca tct gaa ggt gca acc gat gat act tca ctc atc gat tca aca aaa ctc aaa gca tgt gtc tga atc gat aac tct att cat ctg aaa atg gat gag ttg ggt taa tcg aac gat tca gac act gca cca cga att aaa aaa g-- -ac cgg gca cta tat tcc ggt ttg caa aac aaa aat att taa cta cat tta caa aaa gtt acc tct cgt tac ttc ttc ttt ctg ttt caa tat gtg ata cga tat gat cac tat tcg tat tct ct- --- --- tgg tct cat aaa aaa gtt tta c-- -aa aaa aaa aaa aaa a-t att ttt att cta ttt ctc tct tcg at- ggt ctc aca aaa ata tta aac ctc ttt ctg atg tct caa cta ttt cgt aaa cga taa cgt cca aca ata tat tct cgt aga gct tat caa cat cct tat acc aat cta ggt tgt cag aca att gca tca taa aat aat gtt tat aat tta cac gtt aac atc ata taa taa acg tat ata gtt aat att ttt gga ata taa atg atc tgt aaa atc cat gta ggg gac act gct cac gtt ttt tct cta gta cat aat ttc aca caa gtt ttt ata cag aca aat taa ttc tcg tcc ata ta- ttt taa aac att gac ttt tgt act aag aaa aat atc ttg act aac cat ctc ttt ctc tct tcg atg ggt ctc aca aaa ata tta aac ctc ttt ctg atg g-- --- --- --- --- --- --- --- --- --- --- --- --- --- --- --- --- --- --- --- --- --- --- --- --- --- --- --- --- --- --- --- --- --- --- --- --- --- --- --- --- --- --- --- --- --- --- agt cgt aaa aag ttt tta tcc ttt ctc tct tcg atg ggt ctc --- --- --- --- --- --- --- --- --- --- --- --- --- --- --- --- --- --- --- --- --- --- --- --- --- --- --- --- --- --- --- --- --- --- --- --- --- --- --- --- --- --- --- --- --- --- --- --- --- --- --- --- --- --- --- --- --- --- --- --- --- --- --- --- --- --- --- --- --- --- aca aaa ata tta aac ctc ttt ctg atg gtc tct ata aa

>ON619837.1_Monkeypox_virus_isolate_MPXV_UK_2022_3_complete_genome

atc gag gac gac ttc aac tat gac agc tcg gtg gcg tct gcc agc gtg tac ata cga atg gca ttt cta aga aaa gtc tac ggt atc ctt -tc tac aat ttc ctt taa caa cgg caa cag ctg cag tat ttt tat act ttg aat gca -tc gga cat tta tac aag gga gtc ctg ttc taa tat tgg cat caa tgt tcg gat cta tag gct tga ttt tcg cat tga ctt tac aca gac atg aac atc ccc tga atc tgt aca tac ttt gtg gat tta cac tgt tag aat ctc taa cgc tgg cct ctg ttg tta ctt tct atg atg cac gta tcg tta tgc aag ctt tca tgt tga cta ctg cag tgt ttc ttg ctc tga cta cat ata ctc tac aat caa aga gag att tca gta aac ttg taa cag gat tgt ttg ctg ctt tct gga ttt taa ttt tgt cag gag tct tga gga taa agt tta aaa tag aat taa taa aga aca tat agg tca -tt ttt taa aca tgg ata gaa acc aag gtt gtt agt taa taa tat aca aga tat ttt ttc tca ctc tga tcc atg taa acc aag gac gag aga cac tct cat tcc tca ttc acg aca cca tta aaa atg gaa att aaa gcc ctc tat taa gca cag acg gct aca ggt cta cca tca ggt t-a cct tcg tct acc ttc aca atg gcc tct cct tgt gcc cag ttc agt ccc tgt cat tgc cac gct act aag gac tcc ctg aat acc gtg act gac gtc aga cat tgt ctg act gaa tac atc ctg tgg gtt tct cat aga tgg acc cat aga gaa agc gca ggg cct ctc tac agg ctt ctc atc tct ttc aga att gat gca atg gag cta ttt ggt agc gag ttg aag gag ttc tcg aat tca ctt ccg tgg gac aat atc gac aat tgc gtg gag atc att aaa tgt ttc atc aga aat gac tcc atg aaa acc gcc aaa gaa ctt tgt gca ata att gga ctt tgt act caa tca gct att gtc act gga aga gtc ttc aat gat aag tat atc gac ata cta ctt atg ctg cga aag att ctg aac gag aac gac tat ctc acc ctc ttg gat cat atc ctc act gct aaa tac taa atc tcc ttc atg ctc tct cac taa tac tct tac tca cta cac ttt tta tca tct tat gat gaa tga ttg cct tca tca ttt ttt cgt gga ata taa tat agg aat aat tag cac cag aat agc tat gga tat ctc gtt aag aat att ctc tat aag aga cat aat gta gac ata gtt att ata tcc ttc tta gat aag tgt tac gct act gga aag ttt cca tcg tta tta tta cat gaa gat gat ata att aaa cca aca ttg aga ttg gct ctt atg tta gct gga ttg aat tac tgt aat aaa tgc atc gag tat aga ggg ata tag caa ttc tcg ata ata gtc atg caa tat ttg aat gag act gat aat tta ggt aat aca gta ct- --- aca cac ata tct ttc tag att ata tat cgt taa aaa tct gta aga tgt ata ttt ctc ata agt atc cac tgt gta ata tta tta atg gat ata tag ata acg caa tag gga cta ata gta ttg taa aag ata taa tcg act att tac gta cat atc cag ata tct ata ttc cta cta gtt tgc tgc gta gtt gca tca ttg ata tgc atg -at tta tca gga ttc aga gat gaa tta cta agt aaa cta caa tcc cac aat aag taa gaa tca aat atc aaa aac tca ctt ttg att ttt cta gtc tta agt aat aca tat att tat taa tag acc tat gaa ata aaa aaa ggt aac aat gga ttc gcg tat agc tat tta cgt att agt atc ggc atc tct ttt gta tct tgt taa ttg tca caa act agt aca tta ctt caa tct gaa aat aaa tgg aag tga tat aac taa tac agc aga tat att gct gga caa tta tcc aat tat gac ctt tga tgg aaa gga tat tta tcc atc tat ctc gtt cat ggt cgg taa taa act ttt cct aga tct tta taa aaa tat ctt tga aga att ttt cag act att tcg agt atc tgt aag tag tca ata cga gga att aga ata tta tta ttc atg tga tta tac taa caa ccg tcc tac aat taa aca aca tta ctt tta taa cgg cga cga ata tac tga aat tga tag atc gaa aaa agc cac taa taa aaa cag ttg gtt aat tac ttc agg ctt tag act aca aaa atg gtt cga tag cga aga ttg tat aat tta tct cag atc ttt agt tag aag aat gga aga cag taa caa aaa cag taa aaa aac tta gta ctt aga tat cg- --a aaa aat ata ttt ttg tag act ctt gag aat aga agg aaa aca tgt aca taa tta taa aaa atg aaa atc aat ggc gaa taa gac agt gcg att cgc acc atg gag tcg gta gat ttc atg gct gtc gat gag cag ttt cac gac gac ctc gat ctt tgg tca tta tct ttg gta gat gat tat aaa aaa cat gga tta ggt gtt gac tgt tat gtt cta gaa cca gtt gtt gac agg aaa ata ttt gat aga ttt ctc ctt gaa cca att tgt gat cct gta gat gtt ctg tat gat tat ttt agg att cat aga gat aat att gat cag tat ata gta gat aga ctg ttt gca tat att aca tat aaa gat att ata tct gca tta gtg tca aag aat tat atg gaa gat att ttc tct ata att att aag aat tgt aat tct gtg caa gat ctc tta ctt tac tat cta tct aat gca tat gta gaa ata gac att gtt gat ctt atg gta gat cat ggg gct gta ata tat aaa ata gaa tgc ttg aat gcc tat ttt agg gga ata tgt aaa aag gaa agt agt gtt gtt gag ttt att ttg aat tgt ggt atc cca gat gaa aat gat gtt aaa tta gat cta tat aaa ata att cag tat act agg gga ttc ctt gta gat gaa ccc aca gta tta gaa att tat aag ctt tgt atc cca tat att gaa gat atc aat caa cta gat gct ggt gga agg acc ttg ctt tat cgc gct atc tat gca ggt tat ata gat tta gta tca tgg cta tta gaa aat gga gca aat gtc aac gca gta atg agt aat gga tat aca tgt ctt gac gtg gcc gtg gat agg gga tct gtc atc gcc cgt agg gaa gca cat ctt aaa ata tta gaa ata ttg ctt aga gaa cca ttg tct att gac tgt ata aaa tta gct ata ctt aat aat aca att gaa aac cat gat gtg ata aag ctc tgt atc aag tat ttt atg atg gta gat tat tca ctt tgt aat gtg tat gca tca tca ctc ttt gat tat ata att gat tgt aaa caa gaa ttg gag tac att agg cag atg aaa att cat aat aca acc atg tat gag tta atc tat aat aga gac aaa aac aag cat gct tcc cat att cta cat agg tat tct aaa cat cca gtt ttg aca cag tgt atc act aaa gga ttc aag att tac aca gaa gta acc gag cag gtc act aaa gct cta aac aga cgt gct cta ata gat gag ata ata aac aat gta tca act gat gac aat ctc cta tca aaa ctt cca tta gaa att agg gat cta att gtt tca caa gct gtc ata tag agt tct atc cac cca cct ttc ttg aaa tga gtt aat --- --- --- --- --- --- --- --- --- --- --- --- --- --- --- --- --- --- --- --- --- --- --- --- --- --- --- --- --- --- --- --- --- --- --- --- --- --- --- --- --- --- --- --- --- --- --- --- --- --- --- --- --- --- --- --- --- --- --- --- --- --- --- --- --- --- --- --- --- --- --- --- --- --- --- --- --- --- --- --- --- --- --- --- --- --- --- --- --- --- --- --- --- --- --- --- --- --- --- --- --- --- --- --- --- -ag tca taa gtt agt taa --- --- --- --- --- -gt cat aag tta gtt aag tca t-- --- --- --- --- --- --- --- --- --- --- --- --- --- --- --- -aa gtt agt taa gtc ata agt tag tta agt cat aag tta gtt aag tca taa gtt agt taa gtc ata agt tag tta agt cat aag tta gtt tat agt cta aca ctt cta att ttt ata cct tga tct ttt tct cta att atg aaa aag taa atc att atg aag atg gat g-a aat gga cga gat tgt gcg cat cgt taa cga tag tat gtg gta cgt acc taa cgc att tat gga cga cgg tga taa tga agg tca cat ttc tgt caa taa tgt ctg tca tat gta tct cgc att ctt tga tgt gga tat atc atc tca tct gtt taa att agt tat taa aca ctg cga tct gaa taa acg act aaa atg tgg taa ctc tcc att aca ttg cta tac gat gaa tac acg att taa tcc atc tgt att aaa gat att gtt acg cca cgg cat gcg taa ctt tga tag caa gga taa aaa agg aca tat tcc tct aca cca cta tct gat tca ttc act atc aat cga taa caa gat ctt tga tat act aac gga ccc cat tga tga ctt tag taa atc atc cga tct att gct gtg tta tct tag ata taa att caa tgg gag ctt aaa cta tta cgt tct gta caa att att gac taa agg atc tga ccc taa ttg cgt cga tga gga tgg act cac ttc tct tca tta cta ctg taa aca cat atc cgc gtt cca cga aag caa tta tta caa gtc aaa gag tca cac taa gat gcg agc tga gaa gcg att cat cta cgc gat aat aga tca tgg agc aaa cat taa cgc ggt tac gaa aat cgg aaa tac gcc gtt aca cac tta cct tca aca gta tac caa aca tag tcc tcg tgt ggt gta tgc tct ttt atc tcg agg agc cga tac gag gat acg taa taa tct taa ttg tac acc cat cat gga ata cat aaa gaa cga ttg tgc aac agg tca tat tct cat aat gtt act caa ttg gca cga aca aaa ata cgg gaa att aca aaa gga aga agg aca aca tct act tta tct att cat aaa aca taa tca agg ata tgg aag tcg ctc tct caa tat act acg gta tct act aga tag att cga cat tca gaa aga cga ata cta taa tac aat gac tcc tct tca tac cgc ctt cca gaa ttg caa taa caa tgt tgc ctc ata cct cgt ata cat tgg ata cga cat caa cct tcc gac taa aga cga taa gac agt att cga ctt ggt gtt tga aaa cag aaa cat tat ata caa ggc gga tgt cgt taa tga cat tat cca cca cag act gaa agt atc tct acc tat gat taa atc gtt gtt cta caa gat gtc gga gtt ctc tcc cta cga cga tca cta cgt aaa gaa gat aat agc cta ctg cct att aag gga cga gtc att tgc gga act aca tac taa att ctg ttt aaa cga gga cta taa aag tgt att tat gaa aaa tat atc att cga taa gat aga ttc cat cat cga aaa atg tag tcg tga cat aag tct cct caa aga gat tcg aat ctc aga cac cga ctt gta tac ggt att gag aac aga aga cat tcg gta tca cac ata tct cga agc cat aca ttc aga caa acg cat ttc att tcc cat gta cga cga tct cat aga aca gtg tca tct atc gat gga gca taa aag taa act cgt cga caa agc act caa taa att aga gtc tac cat cga tag tca atc tag act atc gta ttt gcc tcc gga aat tat gcg caa tat cat aac caa gct aag cga cta cca tct aaa cag tat gtt gta cgg aaa gaa cca tta caa ata tta tcc atg ata gaa aga aaa tat tta aaa aat aat cta tat gat tgg aga agt agg aaa caa aca gta aca aga cga cga tta cta c-a tta tta aat cat gag gtc cgt att ata ctc gta tat att gtt tct ctc atg tat aat aat aaa cgg aag aga tat agc acc aca tgc acc atc caa tgg aaa gtg taa aga caa cga ata cag aag ccg taa tct atg ttg tct atc gtg tcc tcc ggg aac tta cgc ttt cag att atg tga tag caa gac taa tac aca atg tac acc gtg tgg ttc gga tac ctt tac atc tca caa taa tca ttt aca ggc ttg tct aag ttg taa cgg aag atg tga tag taa tca ggt aga gac gcg atc gtg taa cac gac tca caa tag aat ctg tga atg ctc tcc agg ata tta ttg tct tct caa agg agc atc agg gtg tag aac atg tat ttc taa aac aaa gtg tgg aat agg ata cgg agt atc cgg ata cac gtc tac cgg aga cgt cat ctg ttc tcc gtg tgg tcc cgg aac ata ttc tca cac cgt ctc ttc cac aga taa atg cga acc cgt cgt aac cag caa tac att taa cta tat cga tgt gga aat taa cct gta tcc agt caa cga cac atc gtg tac tcg gac gac cac tac cgg tct cag cga atc cat ctc aac gtc gga act aac tat tac cat gaa tca taa aga ttg tga tcc agt ctt tcg tgc aga ata ctt ctc tgt cct taa taa tgt agc aac ttc agg att ctt tac agg aga aaa tag ata tca gaa tac ttc aaa gat atg tac tct gaa ttt cga gat taa atg taa caa caa aga ttc atc ttc caa aca gtt aac gaa aac aaa gaa tga tac tat cat gcc gca ttc aga gac ggt aac tct agt ggg cga ctg tct atc tag cgt cga cat cta cat act ata tag taa tac caa tac tca aga cta cga aaa tga tac aat ctc tta tca tat ggg taa tgt tct cga tgt caa tag cca tat gcc cgc tag ttg cga tat aca taa act gat cac taa ttc cca gaa tcc cac cca ctt ata gta ag- -tt ttt tta cct ata aat aat aaa tac aat aat taa ttt ctc gta aaa gta gaa aat ata ttc taa ttt att ata tgg taa gaa agt aga atc atc tag aac agt aat caa tca ata gca atc atg aaa caa tat att gtc ctg gca tgc atg tgc cta gtg gca gct gct atg cct act agt ctt caa caa tct tca tcc tcg tgt act gaa gaa gaa aac aaa cat cat atg gga atc gat gtt att atc aaa gtc aca aag caa gac caa aca ccg acc aat gat aag att tgt caa tcc gta acg gaa gtt aca gag a-- --- --- --- --- --- -cc gaa gat gat gag gta tcc gaa gaa gtt gta aaa gga gat ccc acc act tat tac act atc gtc ggt gcg ggt ctt aac atg aac ttt gga ttc acc aaa tgc cca aag att tta tcc atc tcc gaa tcc tct gat gga aac act gtg aat act aga ttg tcc agc gtg tca ccg gga caa ggt aag gac tct ccc gcg atc acg cgt gaa gaa gct ctg gct atg atc aaa gac tgt gag atg tct atc gac atc aga tgt agc gaa gaa gag aaa gac agt gac atc aag acc cat cca gta ctt ggg tct aac atc tca cat aag aaa gtg agt tac aaa gat atc atc ggt tca acg atc gtt gat aca aaa tgt gtc aag aac cta gag ttt agc gta cgt atc gga gac atg tgt gag gaa tca tct gaa ctt gaa gtc aag gat gga ttc aag tat gtc gac gga tcg gca tct gaa ggt gca acc gat gat act tca ctc atc gat tca aca aaa ctc aaa gca tgt gtc tga atc gat aac tct att cat ctg aaa atg gat gag ttg ggt taa tcg aac gat tca gac acc gca cca cga att aaa aaa g-- -ac cgg gca cta tat tcc ggt ttg caa aac aaa aat att taa cta cat tca caa aaa gtt acc tct cgt tac ttc ttc ttt ctg ttt caa tat gtg ata cga tat gat cac tat tcg tat tct ct- --- --- tgg tct cat aaa aaa gtt tta c-- --? aaa aaa aaa aaa a-t att ttt att ctc ttt ctc tct tcg at- ggt ctc aca aaa ata tta aac ctc ttt ctg atg tct caa cta ttt cgt aaa cga taa cgt cca aca ata tat tct cgt aga gct tat caa cat cct tat acc aat cta ggt tgt cag aca att gca tca taa aat aat gtt tat aat tta cac gtt aac atc ata taa taa acg tat ata gtt aat att ttt gga ata taa atg atc tgt aaa atc cat gta ggg gac act gct cac gtt ttt tct cta gta cat aat ttc aca caa gtt ttt ata cag aca aat taa ttc tcg tcc ata ta- ttt taa aac att gac ttt tgt act aag aaa aat atc ttg act aac cat ctc ttt ctc tct tcg atg ggt ctc aca aaa ata tta aac ctc ttt ctg atg g-- --- --- --- --- --- --- --- --- --- --- --- --- --- --- --- --- --- --- --- --- --- --- --- --- --- --- --- --- --- --- --- --- --- --- --- --- --- --- --- --- --- --- --- --- --- --- agt cgt aaa aag ttt tta tcc ttt ctc tct tcg ata ggt ctc --- --- --- --- --- --- --- --- --- --- --- --- --- --- --- --- --- --- --- --- --- --- --- --- --- --- --- --- --- --- --- --- --- --- --- --- --- --- --- --- --- --- --- --- --- --- --- --- --- --- --- --- --- --- --- --- --- --- --- --- --- --- --- --- --- --- --- --- --- --- aca aaa ata tta aac ctc ttt ctg atg gtc tct ata aa

>ON585037.1_Monkeypox_virus_isolate_Monkeypox/PT0005/2022_complete_genome

atc gag gac gac ttc aac tat gac agc tcg gtg gcg tct gcc agc gtg tac ata cga atg gca ttt cta aga aaa gtc tac ggt atc ctt -tc tac aat ttc ctt taa caa cgg caa cag ctg cag tat ttt tat act ttg aat gca -tc gga cat tta tac aag gga gtc ctg ttc taa tat tgg cat caa tgt tcg gat cta tag gct tga ttt tcg cat tga ctt tac aca gac atg aac atc ccc tga atc tgt aca tac ttt gtg gat tta cac tgt tag aat ctc taa cgc tgg cct ctg ttg tta ctt tct atg atg cac gta tcg tta tgc aag ctt tca tgt tga cta ctg cag tgt ttc ttg ctc tga cta cat ata ctc tac aat caa aga gag att tca gta aac ttg taa cag gat tgt ttg ctg ctt tct gga ttt taa ttt tgt cag gag tct tga gga taa agt tta aaa tag aat taa taa aga aca tat agg tca -tt ttt taa aca tgg ata gaa acc aag gtt gtt agt taa taa tat aca aga tat ttt ttc tca ctc tga tcc atg taa acc aag gac gag aga cac tct cat tcc tca ttc acg aca cca tta aaa atg gaa att aaa gcc ctc tat taa gca cag acg gct aca ggt cta cca tca ggt t-a cct tcg tct acc ttc aca atg gcc tct cct tgt gcc cag ttc agt ccc tgt cat tgc cac gct act aag gac tcc ctg aat acc gtg act gac gtc aga cat tgt ctg act gaa tac atc ctg tgg gtt tct cat aga tgg acc cat aga gaa agc gca ggg cct ctc tac agg ctt ctc atc tct ttc aga att gat gca atg gag cta ttt ggt agc gag ttg aag gag ttc tcg aat tca ctt ccg tgg gac aat atc gac aat tgc gtg gag atc att aaa tgt ttc atc aga aat gac tcc atg aaa acc gcc aaa gaa ctt tgt gca ata att gga ctt tgt act caa tca gct att gtc act gga aga gtc ttc aat gat aag tat atc gac ata cta ctt atg ctg cga aag att ctg aac gag aac gac tat ctc acc ctc ttg gat cat atc ctc act gct aaa tac taa atc tcc ttc atg ctc tct cac taa tac tct tac tca cta cac ttt tta tca tct tat gat gaa tga ttg cct tca tca ttt ttt cgt gga ata taa tat agg aat aat tag cac cag aat agc tat gga tat ctc gtt aag aat att ctc tat aag aga cat aat gta gac ata gtt att ata tcc ttc tta gat aag tgt tac gct act gga aag ttt cca tcg tta tta tta cat gaa gat gat ata att aaa cca aca ttg aga ttg gct ctt atg tta gct gga ttg aat tac tgt aat aaa tgc atc gag tat aga ggg ata tag caa ttc tcg ata ata gtc atg caa tat ttg aat gag act gat aat tta ggt aat aca gta ct- --- aca cac ata tct ttc tag att ata tat cgt taa aaa tct gta aga tgt ata ttt ctc ata agt atc cac tgt gta ata tta tta atg gat ata tag ata acg caa tag gga cta ata gta ttg taa aag ata taa tcg act att tac gta cat atc cag ata tct ata ttc cta cta gtt tgc tgc gta gtt gca tca ttg ata tgc atg -at tta tca gga ttc aga gat gaa tta cta agt aaa cta caa tcc cac aat aag taa gaa tca aat atc aaa aac tca ctt ttg att ttt cta gtc tta agt aat aca tat att tat taa tag acc tat gaa ata aaa aaa ggt aac aat gga ttc gcg tat agc tat tta cgt att agt atc ggc atc tct ttt gta tct tgt taa ttg tca caa act agt aca tta ctt caa tct gaa aat aaa tgg aag tga tat aac taa tac agc aga tat att gct gga caa tta tcc aat tat gac ctt tga tgg aaa gga tat tta tcc atc tat ctc gtt cat ggt cgg taa taa act ttt cct aga tct tta taa aaa tat ctt tga aga att ttt cag act att tcg agt atc tgt aag tag tca ata cga gga att aga ata tta tta ttc atg tga tta tac taa caa ccg tcc tac aat taa aca aca tta ctt tta taa cgg cga cga ata tac tga aat tga tag atc gaa aaa agc cac taa taa aaa cag ttg gtt aat tac ttc agg ctt tag act aca aaa atg gtt cga tag cga aga ttg tat aat tta tct cag atc ttt agt tag aag aat gga aga cag taa caa aaa cag taa aaa aac tta gta ctt aga tat cg- --a aaa aat ata ttt ttg tag act ctt gag aat aga agg aaa aca tgt aca taa tta taa aaa atg aaa atc aat ggc gaa taa gac agt gcg att cgc acc atg gag tcg gta gat ttc atg gct gtc gat gag cag ttt cac gac gac ctc gat ctt tgg tca tta tct ttg gta gat gat tat aaa aaa cat gga tta ggt gtt gac tgt tat gtt cta gaa cca gtt gtt gac agg aaa ata ttt gat aga ttt ctc ctt gaa cca att tgt gat cct gta gat gtt ctg tat gat tat ttt agg att cat aga gat aat att gat cag tat ata gta gat aga ctg ttt gca tat att aca tat aaa gat att ata tct gca tta gtg tca aag aat tat atg gaa gat att ttc tct ata att att aag aat tgt aat tct gtg caa gat ctc tta ctt tac tat cta tct aat gca tat gta gaa ata gac att gtt gat ctt atg gta gat cat ggg gct gta ata tat aaa ata gaa tgc ttg aat gcc tat ttt agg gga ata tgt aaa aag gaa agt agt gtt gtt gag ttt att ttg aat tgt ggt atc cca gat gaa aat gat gtt aaa tta gat cta tat aaa ata att cag tat act agg gga ttc ctt gta gat gaa ccc aca gta tta gaa att tat aag ctt tgt atc cca tat att gaa gat atc aat caa cta gat gct ggt gga agg acc ttg ctt tat cgc gct atc tat gca ggt tat ata gat tta gta tca tgg cta tta gaa aat gga gca aat gtc aac gca gta atg agt aat gga tat aca tgt ctt gac gtg gcc gtg gat agg gga tct gtc atc gcc cgt agg gaa gca cat ctt aaa ata tta gaa ata ttg ctt aga gaa cca ttg tct att gac tgt ata aaa tta gct ata ctt aat aat aca att gaa aac cat gat gtg ata aag ctc tgt atc aag tat ttt atg atg gta gat tat tca ctt tgt aat gtg tat gca tca tca ctc ttt gat tat ata att gat tgt aaa caa gaa ttg gag tac att agg cag atg aaa att cat aat aca acc atg tat gag tta atc tat aat aga gac aaa aac aag cat gct tcc cat att cta cat agg tat tct aaa cat cca gtt ttg aca cag tgt atc act aaa gga ttc aag att tac aca gaa gta acc gag cag gtc act aaa gct cta aac aga cgt gct cta ata gat gag ata ata aac aat gta tca act gat gac aat ctc cta tca aaa ctt cca tta gaa att agg gat cta att gtt tca caa gct gtc ata tag agt tct atc cac cca cct ttc ttg aaa tga gtt aat --- --- --- --- --- --- --- --- --- --- --- --- --- --- --- --- --- --- --- --- --- --- --- --- --- --- --- --- --- --- --- --- --- --- --- --- --- --- --- --- --- --- --- --- --- --- --- --- --- --- --- --- --- --- --- --- --- --- --- --- --- --- --- --- --- --- --- --- --- --- --- --- --- --- --- --- --- --- --- --- --- --- --- --- --- --- --- --- --- --- --- --- --- --- --- --- --- --- --- --- --- --- --- --- --- -ag tca taa gtt agt taa --- --- --- --- --- -gt cat aag tta gtt aag tca t-- --- --- --- --- --- --- --- --- --- --- --- --- --- --- --- -aa gtt agt taa gtc ata agt tag tta agt cat aag tta gtt aag tca taa gtt agt taa gtc ata agt tag tta agt cat aag tta gtt tat agt cta aca ctt cta att ttt ata cct tga tct ttt tct cta att atg aaa aag taa atc att atg aag atg gat g-a aat gga cga gat tgt gcg cat cgt taa cga tag tat gtg gta cgt acc taa cgc att tat gga cga cgg tga taa tga agg tca cat ttc tgt caa taa tgt ctg tca tat gta tct cgc att ctt tga tgt gga tat atc atc tca tct gtt taa att agt tat taa aca ctg cga tct gaa taa acg act aaa atg tgg taa ctc tcc att aca ttg cta tac gat gaa tac acg att taa tcc atc tgt att aaa gat att gtt acg cca cgg cat gcg taa ctt tga tag caa gga taa aaa agg aca tat tcc tct aca cca cta tct gat tca ttc act atc aat cga taa caa gat ctt tga tat act aac gga ccc cat tga tga ctt tag taa atc atc cga tct att gct gtg tta tct tag ata taa att caa tgg gag ctt aaa cta tta cgt tct gta caa att att gac taa agg atc tga ccc taa ttg cgt cga tga gga tgg act cac ttc tct tca tta cta ctg taa aca cat atc cgc gtt cca cga aag caa tta tta caa gtc aaa gag tca cac taa gat gcg agc tga gaa gcg att cat cta cgc gat aat aga tca tgg agc aaa cat taa cgc ggt tac gaa aat cgg aaa tac gcc gtt aca cac tta cct tca aca gta tac caa aca tag tcc tcg tgt ggt gta tgc tct ttt atc tcg agg agc cga tac gag gat acg taa taa tct taa ttg tac acc cat cat gga ata cat aaa gaa cga ttg tgc aac agg tca tat tct cat aat gtt act caa ttg gca cga aca aaa ata cgg gaa att aca aaa gga aga agg aca aca tct act tta tct att cat aaa aca taa tca agg ata tgg aag tcg ctc tct caa tat act acg gta tct act aga tag att cga cat tca gaa aga cga ata cta taa tac aat gac tcc tct tca tac cgc ctt cca gaa ttg caa taa caa tgt tgc ctc ata cct cgt ata cat tgg ata cga cat caa cct tcc gac taa aga cga taa gac agt att cga ctt ggt gtt tga aaa cag aaa cat tat ata caa ggc gga tgt cgt taa tga cat tat cca cca cag act gaa agt atc tct acc tat gat taa atc gtt gtt cta caa gat gtc gga gtt ctc tcc cta cga cga tca cta cgt aaa gaa gat aat agc cta ctg cct att aag gga cga gtc att tgc gga act aca tac taa att ctg ttt aaa cga gga cta taa aag tgt att tat gaa aaa tat atc att cga taa gat aga ttc cat cat cga aaa atg tag tcg tga cat aag tct cct caa aga gat tcg aat ctc aga cac cga ctt gta tac ggt att gag aac aga aga cat tcg gta tca cac ata tct cga agc cat aca ttc aga caa acg cat ttc att tcc cat gta cga cga tct cat aga aca gtg tca tct atc gat gga gca taa aag taa act cgt cga caa agc act caa taa att aga gtc tac cat cga tag tca atc tag act atc gta ttt gcc tcc gga aat tat gcg caa tat cat aac caa gct aag cga cta cca tct aaa cag tat gtt gta cgg aaa gaa cca tta caa ata tta tcc atg ata gaa aga aaa tat tta aaa aat aat cta tat gat tgg aga agt agg aaa caa aca gta aca aga cga cga tta cta c-a tta tta aat cat gag gtc cgt att ata ctc gta tat att gtt tct ctc atg tat aat aat aaa cgg aag aga tat agc acc aca tgc acc atc caa tgg aaa gtg taa aga caa cga ata cag aag ccg taa tct atg ttg tct atc gtg tcc tcc ggg aac tta cgc ttt cag att atg tga tag caa gac taa tac aca atg tac acc gtg tgg ttc gga tac ctt tac atc tca caa taa tca ttt aca ggc ttg tct aag ttg taa cgg aag atg tga tag taa tca ggt aga gac gcg atc gtg taa cac gac tca caa tag aat ctg tga atg ctc tcc agg ata tta ttg tct tct caa agg agc atc agg gtg tag aac atg tat ttc taa aac aaa gtg tgg aat agg ata cgg agt atc cgg ata cac gtc tac cgg aga cgt cat ctg ttc tcc gtg tgg tcc cgg aac ata ttc tca cac cgt ctc ttc cac aga taa atg cga acc cgt cgt aac cag caa tac att taa cta tat cga tgt gga aat taa cct gta tcc agt caa cga cac atc gtg tac tcg gac gac cac tac cgg tct cag cga atc cat ctc aac gtc gga act aac tat tac cat gaa tca taa aga ttg tga tcc agt ctt tcg tgc aga ata ctt ctc tgt cct taa taa tgt agc aac ttc agg att ctt tac agg aga aaa tag ata tca gaa tac ttc aaa gat atg tac tct gaa ttt cga gat taa atg taa caa caa aga ttc atc ttc caa aca gtt aac gaa aac aaa gaa tga tac tat cat gcc gca ttc aga gac ggt aac tct agt ggg cga ctg tct atc tag cgt cga cat cta cat act ata tag taa tac caa tac tca aga cta cga aaa tga tac aat ctc tta tca tat ggg taa tgt tct cga tgt caa tag cca tat gcc cgc tag ttg cga tat aca taa act gat cac taa ttc cca gaa tcc cac cca ctt ata gta ag- -tt ttt tta cct ata aat aat aaa tac aat aat taa ttt ctc gta aaa gta gaa aat ata ttc taa ttt att ata tgg taa gaa agt aga atc atc tag aac agt aat caa tca ata gca atc atg aaa caa tat att gtc ctg gca tgc atg tgc cta gtg gca gct gct atg cct act agt ctt caa caa tct tca tcc tcg tgt act gaa gaa gaa aac aaa cat cat atg gga atc gat gtt att atc aaa gtc aca aag caa gac caa aca ccg acc aat gat aag att tgt caa tcc gta acg gaa gtt aca gag a-- --- --- --- --- --- -cc gaa gat gat gag gta tcc gaa gaa gtt gta aaa gga gat ccc acc act tat tac act atc gtc ggt gcg ggt ctt aac atg aac ttt gga ttc acc aaa tgc cca aag att tta tcc atc tcc gaa tcc tct gat gga aac act gtg aat act aga ttg tcc agc gtg tca ccg gga caa ggt aag gac tct ccc gcg atc acg cgt gaa gaa gct ctg gct atg atc aaa gac tgt gag atg tct atc gac atc aga tgt agc gaa gaa gag aaa gac agt gac atc aag acc cat cca gta ctt ggg tct aac atc tca cat aag aaa gtg agt tac aaa gat atc atc ggt tca acg atc gtt gat aca aaa tgt gtc aag aac cta gag ttt agc gta cgt atc gga gac atg tgt gag gaa tca tct gaa ctt gaa gtc aag gat gga ttc aag tat gtc gac gga tcg gca tct gaa ggt gca acc gat gat act tca ctc atc gat tca aca aaa ctc aaa gca tgt gtc tga atc gat aac tct att cat ctg aaa atg gat gag ttg ggt taa tcg aac gat tca gac acc gca cca cga att aaa aaa g-- -ac cgg gca cta tat tcc ggt ttg caa aac aaa aat att taa cta cat tca caa aaa gtt acc tct cgt tac ttc ttc ttt ctg ttt caa tat gtg ata cga tat gat cac tat tcg tat tct ct- --- --- tgg tct cat aaa aaa gtt tta c-- --a aaa aaa aaa aaa a-t att ttt att ctc ttt ctc tct tcg at- ggt ctc aca aaa ata tta aac ctc ttt ctg atg tct caa cta ttt cgt aaa cga taa cgt cca aca ata tat tct cgt aga gct tat caa cat cct tat acc aat cta ggt tgt cag aca att gca tca taa aat aat gtt tat aat tta cac gtt aac atc ata taa taa acg tat ata gtt aat att ttt gga ata taa atg atc tgt aaa atc cat gta ggg gac act gct cac gtt ttt tct cta gta cat aat ttc aca caa gtt ttt ata cag aca aat taa ttc tcg tcc ata ta- ttt taa aac att gac ttt tgt act aag aaa aat atc ttg act aac cat ctc ttt ctc tct tcg atg ggt ctc aca aaa ata tta aac ctc ttt ctg atg g-- --- --- --- --- --- --- --- --- --- --- --- --- --- --- --- --- --- --- --- --- --- --- --- --- --- --- --- --- --- --- --- --- --- --- --- --- --- --- --- --- --- --- --- --- --- --- agt cgt aaa aag ttt tta tcc ttt ctc tct tcg ata ggt ctc --- --- --- --- --- --- --- --- --- --- --- --- --- --- --- --- --- --- --- --- --- --- --- --- --- --- --- --- --- --- --- --- --- --- --- --- --- --- --- --- --- --- --- --- --- --- --- --- --- --- --- --- --- --- --- --- --- --- --- --- --- --- --- --- --- --- --- --- --- --- aca aaa ata tta aac ctc ttt ctg atg gtc tct ata aa

>AY741551.1_Monkeypox_virus_isolate_Sierra_Leone_complete_genome

atc gag gac gac ttc aac tat gac agc tcg gtg gcg tct gcc agc gtg tac ata cga atg gca ttt cta aga aaa gtc tac ggt atc ctt -tc tac aat ttc ctt taa caa cgg caa cag ctg cag tat ttt tat act ttg aat gca -tc gga cat tta tac aag gga gtc ctg ttc taa tat tgg cat caa tgt tcg gat cta tag gct tga ttt tcg cat tga ctt tac aca gac atg aac atc ccc tga atc tgt aca tac ttt gtg gat tta cac tgt tag aat ctc taa cgc tgg cct ctg ttg tta ctt tct atg atg cac gta tcg tta tgc aag ctt tca tgt tga cta ctg cag tgt ttc ttg ctc tga cta cat ata ctc tat aat caa aga gag att tca gta aac ttg taa cag gat tgt ttg ctg ctt tct gga ttt taa ttt tgt cag gag tct tga gga taa agt tta aaa tag aat taa taa aga aca tat agg tca -tt ttt taa aca tgg ata gaa acc aag gtt gtt agt taa taa tat aca aga tat ttt ttc tca ctc tga tcc atg taa acc aag gac gag aga cac tct cat tcc tca ttc acg aca cca tta aaa atg gaa att aaa gcc ctc tat taa gca cag acg gct aca ggt cta cca tca ggt t-a cct tcg tct acc ttc aca atg gcc tct cct tgt gcc cag ttc agt ccc tgt cat tgc cac gct act aag gac tcc ctg aat acc gtg act gat gtc aga cat tgt ctg act gaa tac atc ctg tgg gtt tct cat aga tgg acc cat aga gaa agc gca ggg cct ctc tac agg ctt ctc atc tct ttc aga att gat gca atg gag cta ttt ggt agc gag ttg aag gag ttc tcg gat tca ctt ccg tgg gac aat atc gac aat tgc gtg gag atc att aaa tgt ttc atc aga aat gac tcc atg aaa acc gcc aaa gaa ctt tgt gca ata att gga ctt tgt act caa tca gct att gtc act gga aga gtc ttc aat gat aag tat atc gac ata cta ctt atg ctg cga aag att ctg aac gag aac gac tat ctc acc ctc ttg gat cat atc ctc act gct aaa tac taa atc tcc ttc atg ctc tct cac taa tac tct tac tca cta cac ttt tta tca tct tat gat gaa tga ttg cct tca tca -tt ttt cgt gga ata taa tat agg aat aat tag cac cag aat agc tat gga tat ctc gtt aag aat att ctc gat aag aga cat aat gta gac ata gtt att ata tcc ttc tta gat aag tgt tac gct act gga aag ttt cca tcg tta tta ttg cat gaa gat gat ata att aaa cca aca ttg aga ttg gct ctt atg tta gct gga ttg aat tac tgt aat aaa tgc atc gag tat aga ggg ata tag caa ttc tcg ata ata gtc atg caa tat ttg aat gag act gat aat tta ggt aat aca gta ct- --- aca cac ata tct ttc tag att ata tat cgt taa aaa tct gta aga ggt ata ttt ctc ata agt atc cac tgt gta ata tta tta atg gat ata tag ata acg caa tag gga cta ata gta ttg taa aag ata taa tcg act att tac gta cat atc cag ata tct ata ttc cta cta gtt tgc tgc gta gtt gca tca ttg ata tgc atg -at tta tca gga ttc aga gat gaa tta cta agt aaa cta caa tcc cac aat aag taa gaa tca aat atc aaa aac tta ctt ttg att ttt cta gtc tta agt aat aca tat att tat taa tag acc tat gaa ata aaa aaa ggt aac aat gga ttc gcg tat agc tat tta cgt att agt atc ggc atc tct ttt gta tct tgt taa ttg tca caa act agt aca tta ctt caa tct gaa aat aaa tgg aag tga tat aac taa tac agc aga tat att gct gga caa tta tcc aat tat gac ctt tga tgg aaa gga tat tta tcc atc tat ctc gtt cat ggt cgg taa taa act ttt cct aga tct tta taa aaa tat ctt tga aga att ttt cag act att tcg agt atc tgt aag tag tca ata cga gga att aga ata tta tta ttc atg tga tta tac taa caa ccg tcc tac aat taa aca aca tta ctt tta taa cgg cga aga ata tac tga aat tga tag atc gaa aaa agc cac taa taa aaa cag ttg gtt aat tac ttc agg ctt tag act aca aaa atg gtt cga tag cga aga ttg tat aat tta tct cag atc ttt agt tag aag aat gga aga cag taa caa aaa cag taa aaa aac tta gta ctt aga tat cg- -aa aaa aat ata ttt ttg tag act ctt gag aat aga agg aaa aca tgt aca taa tta taa aaa atg aaa atc aat ggc gaa taa gac agt gcg att cgc acc atg gag tcg gta gat ttc atg gct gtc gat gag cag ttt cac gac gac ctc gat ctt tgg tca tta tct ttg gta gat gat tat aaa aaa cat gga tta ggt gtt gac tgt tat gtt cta gaa cca gtt gtt gac agg aaa ata ttt gat aga ttt ctc ctt gaa cca att tgt gat cct gta gat gtt ctg tat gat tat ttt agg att cat aga gat aat att gat cag tat ata gta gat aga ctg ttt gca tat att aca tat aaa gat att ata tct gca tta gtg tca aag aat tat atg gaa gat att ttc tct ata att att aag aat tgt aat tct gtg caa gat ctc tta ctt tac tat cta tct aat gaa tat gta gaa ata gac att gtt gat ctt atg gta gat cat ggg gct gta ata tat aaa ata gaa tgc ttg aat gcc tat ttt agg gga ata tgt aaa aag gaa agt agt gtt gtt gag ttt att ttg aat tgt ggt atc cca gat gaa aat gat gtt aaa tta gat cta tat aaa ata att cag tat act agg gga ttc ctt gta gat gaa ccc aca gta tta gaa att tat aag ctt tgt atc cca tat att gaa gat atc aat caa cta gat gct ggt gga agg acc ttg ctt tat cgc gct atc tat gca ggt tat ata gat tta gta tca ttg cta tta gaa aat gga gca aat gtc aac gca gta atg agt aat gga tat aca tgt ctt gac gtg gcc gtg gat agg gga tct gtc atc gcc cgt agg gaa gca cat ctt aaa ata tta gaa ata ttg ctt aga gaa cca ttg tct att gac tgt ata aaa tta gct ata ctt aat aat aca att gaa aac cat gat gtg ata aag ctc tgt atc aag tat ttt atg atg gta gat tat tca ctt tgt aat gtg tat gca tca tca ctc ttt gat tat ata att gat tgt aaa caa gaa ttg gag tac att agg cag atg aaa att cat aat aca acc atg tat gag tta atc tat aat aga gac aaa aac aag cat gct tcc cat att cta cat agg tat tct aaa cat cca gtt ttg aca cag tgt atc act aaa gga ttc aag att tac aca gaa gta acc gag cag gtc act aaa gct cta aac aga cgt gct cta ata gat gag ata ata aac aat gta tca act gat gac aat ctc cta tca aaa ctt cca tta gaa att agg gat cta atc gtt tca caa gct gtc ata tag agt tct atc cac cca cct ttc ttg aaa tga gtt aat --- --- --- --- --- --- --- --- --- --- --- --- --- --- --- --- --- --- --- --- --- --- --- --- --- --- --- --- --- --- --- --- --- --- --- --- --- --- --- --- --- --- --- --- --- --- --- --- --- --- --- --- --- --- --- --- --- --- --- --- --- --- --- --- --- --- --- --- --- --- --- --- --- --- --- --- --- --- --- --- --- --- --- --- --- --- --- --- --- --- --- --- --- --- --- --- --- --- --- --- --- --- --- --- --- --- --- --- --- --- --- --- --- --- --- --- --- --- --- --- --- --- --- --- --- --- --- --- --- --- --- --- --- --- --- --- --- --- --- --- --- --- --- --- --- --- --- --- --- --- --- --- --- -ag tca taa gtt agt taa gtc ata agt tag tt- --t tat aag tta gtt tat agt cta aca ctt cta att ttt ata cct tga tct ttt tct cta att atg aaa aag taa atc att atg aag atg gat g-a aat gga cga gat tgt gcg cat cgt taa cga tag tat gtg gta cgt acc taa cgc att tat gga cga cgg tga taa tga agg tca cat ttc tgt caa taa tgt ctg tca tat gta tct cgc att ctt tga tgt gga tat atc atc tca tct gtt taa att agt tat taa aca ctg cga tct gaa taa acg act aaa atg tgg taa ctc tcc att aca ttg cta tac gat gaa tac acg att taa tcc atc tgt att aaa gat att gtt acg cca cgg cat gcg taa ctt tga tag caa gga taa aaa agg aca tat tcc tct aca cca cta tct gat tca ttc act atc aat cga taa caa gat ctt tga tat act aac gga ccc cat tga tga ctt tag taa atc atc cga tct att gct gtg tta tct tag ata taa att caa tgg gag ctt aaa cta tta cgt tct gta caa att att gac taa agg atc tga ccc taa ttg cgt cga tgc gga tgg act cac ttc tct tca tta cta ctg taa aca cat atc cgc gtt cca cga aag caa tta tta caa gtc aaa gag tca cac taa gat gcg agc tga gaa gcg att cat cta cgc gat aat aga tca tgg agc aaa cat taa cgc ggt tac gaa aat cgg aaa tac gcc gtt aca cac tta cct tca aca gta tac caa aca tag tcc tcg tgt ggt gta tgc tct ttt atc tcg agg agc tga tac gag gat acg taa taa tct tga ttg tac acc cat cat gga ata cat aaa gaa cga ttg tgc aac agg tca tat tct cat aat gtt act caa ttg gca cga aca aaa ata cgg gaa att aca aaa gga aga agg aca aca tct act tta tct att cat aaa aca taa tca agg ata tgg aag tcg ctc tct caa tat act acg gta tct act aga tag att cga cat tca gaa aga cga ata cta taa tac aat gac tcc tct tca tac cgc ctt cca gaa ttg caa taa caa tgt tgc ctc ata cct cgt ata cat cgg ata cga cat caa cct tcc gac taa aga cga taa gac agt att cga ctt ggt gtt tga aaa cag aaa cat cat ata caa ggc gga tgt tgt taa tga cat cat cca cca cag act gaa agt atc tct acc tat gat taa atc gtt gtt cta caa gat gtc gga gtt ctc tcc cta cga cga tca cta cgt aaa gaa gat aat agc cta ctg cct att aag gga cga gtc att tgc gga act aca tac taa att ctg ttt aaa cga gga cta taa aag tgt att tat gaa aaa tat atc att cga taa gat aga ttc cat cat cga aaa atg tag tcg tga cat aag tct cct caa aga gat tcg aat ctc aga cac cga ctt gta tac ggt att gag aac aga aga cat ccg gta tca cac cta tct cga agc cat aca ttc aga caa acg cat ttc att tcc cat gta cga cga tct cat aga aca gtg tca tct atc gat gga gca taa aag taa act cgt cga caa agc act caa taa att aga gtc tac cat cga taa tca atc tag act atc gta ttt gcc tcc gga aat tat gcg caa tat cat aac caa gct aag cga cta cca tct aaa cag tat gtt gta cgg aaa gaa cca tta caa ata tta tcc atg ata gaa aga aaa tat tta aaa aat aat cta tat gat tgg aga agt agg aaa caa aca gta aca aga cga cga tta cta c-a tta tta aat cat gag gtc cgt att ata ctc gta tat att gtt tct ctc atg tat aat aat aaa cgg aag aga tat agc acc aca tgc acc atc caa tgg aaa gtg taa aga caa cga ata cag aag ccg taa tct atg ttg tct atc gtg tcc tcc ggg aac tta cgc ttc cag att atg tga tag caa gac taa tac aca atg tac acc gtg tgg ttc gga tac ctt tac atc tca caa taa tca ttt aca ggc ttg tct aag ttg taa cgg aag atg tga tag taa tca ggt aga gac gcg atc gtg taa cac gac tca caa tag aat ctg tga atg ctc tcc agg ata tta ttg tct tct caa agg agc att agg gtg tag aac atg tat ttc taa aac aaa gtg tgg aat agg ata cgg agt atc cgg ata cac gtc tac cgg aga cgt cat ctg ttc tcc gtg tgg tcc cgg aac ata ttc tca cac cgt ctc ttc cac aga taa atg cga acc cgt cgt aac cag caa tac att taa cta tat cga tgt gga aat taa cct gta tcc agt caa cga cac atc gtg tac tcg gac gac cac tac cgg tct cag cga atc cat ctc aac gtc gga act aac tat tac cat gaa tca taa aga ttg tga tcc agt ctt tcg tgc aga ata ctt ctc tgt cct taa taa tgt agc aac ttc agg att ctt tac agg aga aaa tag ata tca gaa tac ttc aaa gat atg tac tct gaa ttt cga gat taa atg taa caa caa aga ttc atc ttc caa aca gtt aac gaa aac aaa gaa tga tac tat cat gcc gca ttc aga gac ggt aac tct agt ggg cga ctg tct atc tag cgt cga cat cta cat act ata tag taa tac caa tac tca aga cta cga aac gga tac aat ctc tta tca tat ggg taa tgt tct cga tgt caa tag cca tat gcc cgc tag ttg cga tat aca taa act gat cac taa ttc cca gaa tcc cac cca ctt ata gta ag- -tt ttt tta ccc ata aat aat aaa tac aat aat taa ttt ctc gta aaa gta gaa aat ata ttc taa ttt att ata tgg taa gaa agt aga atc atc tag aac agt aat caa tca ata gca atc atg aaa caa tat att gtc ctg gca tgc atg tgc cta gtg gca gct gct atg cct act agt ctt caa caa tct tca tcc tcg tgt act gaa gaa gaa aac aaa cat cat atg gga atc gat gtt att atc aaa gtc aca aag caa gac caa aca ccg acc aat gat aag att tgt caa tcc gta acg gaa gtt aca gag a-- --- --- --- --- --- -cc gaa gat gat gag gta tcc gaa gaa gtt gta aaa gga gat ccc acc act tat tac act atc gtc ggt gcg ggt ctt aac atg aac ttt gga ttc acc aaa tgc cca aag att tca tcc atc tcc gaa tcc tct gat gga aac act gtg aat act aga ttg tcc agc gtg tca ccg gga caa ggt aag gac tct ccc gcg atc acg cgt gaa gaa gct ctg gct atg atc aaa gac tgt gag atg tct atc gac atc aga tgt agc gaa gaa gag aaa gac agt gac atc aag acc cat cca gta ctt ggg tct aac atc tca cat aag aaa gtg agt tac aaa gat atc atc ggt tca acg atc gtt gat aca aaa tgt gtc aag aac cta gag ttt agc gta cgt atc gga gac atg tgt gag gaa tca tct gaa ctt gaa gtc aag gat gga ttc aag tat gtc gac gga tcg gca tct gaa ggt gca acc gat gat act tca ctc atc gat tca aca aaa ctc aaa gca tgt gtc tga atc gat aac tct att cat ctg aaa atg gat gag ttg ggt taa tcg aac gat tca gac acc gca cca cga att aaa aaa g-- -ac cgg gca cta tat tcc ggt ttg caa aac aaa aat att taa cta cat tta caa aaa gtt acc tct cgt tac ttc ttc ttt ctg ttt caa tat gtg ata cga tat gat cac tat tcg tat tct ct- --- --- tgg tct cat aaa aaa gtt tta caa aaa aaa aaa aaa aaa a-t att ttt att cta ttt ctc tct tcg at- ggt ctc aca aaa ata tta aac ctc ttt ctg atg tct caa cta ttt cgt aaa cga taa cgt cca aca ata tat tct cgt aga gct tat caa cat cct tat acc aat cta ggt tgt cag aca att gca tca taa aat aat gtt tat aat tta cac gtt aac atc ata taa taa acg tat ata gtt aat att ttt gga ata taa atg atc tgt aaa atc cat gta ggg gac act gct cac gtt ttt tct cta gta cat aat ttc aca caa gtt ttt ata cag aca aat taa ttc tcg tcc ata ta- ttt taa aac att gac ttt tgt act aag aaa aat atc ttg act aac cat ctc ttt ctc tct tcg atg ggt ctc aca aaa ata tta aac ctc ttt ctg atg g-- --- --- --- --- --- --- --- --- --- --- --- --- --- --- --- --- --- --- --- --- --- --- --- --- --- --- --- --- --- --- --- --- --- --- --- --- --- --- --- --- --- --- --- --- --- --- agt cgt aaa aag ttt tta tcc ttt ctc tct tcg atg ggt ctc --- --- --- --- --- --- --- --- --- --- --- --- --- --- --- --- --- --- --- --- --- --- --- --- --- --- --- --- --- --- --- --- --- --- --- --- --- --- --- --- --- --- --- --- --- --- --- --- --- --- --- --- --- --- --- --- --- --- --- --- --- --- --- --- --- --- --- --- --- --- aca aaa ata tta aac ctc ttt ctg atg gtc tct ata aa

>ON622718.1_Monkeypox_virus_isolate_MPXV/ES0001/HUGTiP/2022_partial_genome

atc gag gac gac ttc aac tat gac agc tcg gtg gcg tct gcc agc gtg tac ata cga atg gca ttt cta aga aaa gtc tac ggt atc ctt -tc tac aat ttc ctt taa caa cgg caa cag ctg cag tat ttt tat act ttg aat gca -tc gga cat tta tac aag gga gtc ctg ttc taa tat tgg cat caa tgt tcg gat cta tag gct tga ttt tcg cat tga ctt tac aca gac atg aac atc ccc tga atc tgt aca tac ttt gtg gat tta cac tgt tag aat ctc taa cgc tgg cct ctg ttg tta ctt tct atg atg cac gta tcg tta tgc aag ctt tca tgt tga cta ctg cag tgt ttc ttg ctc tga cta cat ata ctc tac aat caa aga gag att tca gta aac ttg taa cag gat tgt ttg ctg ctt tct gga ttt taa ttt tgt cag gag tct tga gga taa agt tta aaa tag aat taa taa aga aca tat agg tca -tt ttt taa aca tgg ata gaa acc aag gtt gtt agt taa taa tat aca aga tat ttt ttc tca ctc tga tcc atg taa acc aag gac gag aga cac tct cat tcc tca ttc acg aca cca tta aaa atg gaa att aaa gcc ctc tat taa gca cag acg gct aca ggt cta cca tca ggt t-a cct tcg tct acc ttc aca atg gcc tct cct tgt gcc cag ttc agt ccc tgt cat tgc cac gct act aag gac tcc ctg aat acc gtg act gac gtc aga cat tgt ctg act gaa tac atc ctg tgg gtt tct cat aga tgg acc cat aga gaa agc gca ggg cct ctc tac agg ctt ctc atc tct ttc aga att gat gca atg gag cta ttt ggt agc gag ttg aag gag ttc tcg aat tca ctt ccg tgg gac aat atc gac aat tgc gtg gag atc att aaa tgt ttc atc aga aat gac tcc atg aaa acc gcc aaa gaa ctt tgt gca ata att gga ctt tgt act caa tca gct att gtc act gga aga gtc ttc aat gat aag tat atc gac ata cta ctt atg ctg cga aag att ctg aac gag aac gac tat ctc acc ctc ttg gat cat atc ctc act gct aaa tac taa atc tcc ttc atg ctc tct cac taa tac tct tac tca cta cac ttt tta tca tct tat gat gaa tga ttg cct tca tca ttt ttt cgt gga ata taa tat agg aat aat tag cac cag aat agc tat gga tat ctc gtt aag aat att ctc tat aag aga cat aat gta gac ata gtt att ata tcc ttc tta gat aag tgt tac gct act gga aag ttt cca tcg tta tta tta cat gaa gat gat ata att aaa cca aca ttg aga ttg gct ctt atg tta gct gga ttg aat tac tgt aat aaa tgc atc gag tat aga ggg ata tag caa ttc tcg ata ata gtc atg caa tat ttg aat gag act gat aat tta ggt aat aca gta ct- --- aca cac ata tct ttc tag att ata tat cgt taa aaa tct gta aga tgt ata ttt ctc ata agt atc cac tgt gta ata tta tta atg gat ata tag ata acg caa tag gga cta ata gta ttg taa aag ata taa tcg act att tac gta cat atc cag ata tct ata ttc cta cta gtt tgc tgc gta gtt gca tca ttg ata tgc atg -at tta tca gga ttc aga gat gaa tta cta agt aaa cta caa tcc cac aat aag taa gaa tca aat atc aaa aac tca ctt ttg att ttt cta gtc tta agt aat aca tat att tat taa tag acc tat gaa ata aaa aaa ggt aac aat gga ttc gcg tat agc tat tta cgt att agt atc ggc atc tct ttt gta tct tgt taa ttg tca caa act agt aca tta ctt caa tct gaa aat aaa tgg aag tga tat aac taa tac agc aga tat att gct gga caa tta tcc aat tat gac ctt tga tgg aaa gga tat tta tcc atc tat ctc gtt cat ggt cgg taa taa act ttt cct aga tct tta taa aaa tat ctt tga aga att ttt cag act att tcg agt atc tgt aag tag tca ata cga gga att aga ata tta tta ttc atg tga tta tac taa caa ccg tcc tac aat taa aca aca tta ctt tta taa cgg cga cga ata tac tga aat tga tag atc gaa aaa agc cac taa taa aaa cag ttg gtt aat tac ttc agg ctt tag act aca aaa atg gtt cga tag cga aga ttg tat aat tta tct cag atc ttt agt tag aag aat gga aga cag taa caa aaa cag taa aaa aac tta gta ctt aga tat cg- --a aaa aat ata ttt ttg tag act ctt gag aat aga agg aaa aca tgt aca taa tta taa aaa atg aaa atc aat ggc gaa taa gac agt gcg att cgc acc atg gag tcg gta gat ttc atg gct gtc gat gag cag ttt cac gac gac ctc gat ctt tgg tca tta tct ttg gta gat gat tat aaa aaa cat gga tta ggt gtt gac tgt tat gtt cta gaa cca gtt gtt gac agg aaa ata ttt gat aga ttt ctc ctt gaa cca att tgt gat cct gta gat gtt ctg tat gat tat ttt agg att cat aga gat aat att gat cag tat ata gta gat aga ctg ttt gca tat att aca tat aaa gat att ata tct gca tta gtg tca aag aat tat atg gaa gat att ttc tct ata att att aag aat tgt aat tct gtg caa gat ctc tta ctt tac tat cta tct aat gca tat gta gaa ata gac att gtt gat ctt atg gta gat cat ggg gct gta ata tat aaa ata gaa tgc ttg aat gcc tat ttt agg gga ata tgt aaa aag gaa agt agt gtt gtt gag ttt att ttg aat tgt ggt atc cca gat gaa aat gat gtt aaa tta gat cta tat aaa ata att tag tat act agg gga ttc ctt gta gat gaa ccc aca gta tta gaa att tat aag ctt tgt atc cca tat att gaa gat atc aat caa cta gat gct ggt gga agg acc ttg ctt tat cgc gct atc tat gca ggt tat ata gat tta gta tca tgg cta tta gaa aat gga gca aat gtc aac gca gta atg agt aat gga tat aca tgt ctt gac gtg gcc gtg gat agg gga tct gtc atc gcc cgt agg gaa gca cat ctt aaa ata tta gaa ata ttg ctt aga gaa cca ttg tct att gac tgt ata aaa tta gct ata ctt aat aat aca att gaa aac cat gat gtg ata aag ctc tgt atc aag tat ttt atg atg gta gat tat tca ctt tgt aat gtg tat gca tca tca ctc ttt gat tat ata att gat tgt aaa caa gaa ttg gag tac att agg cag atg aaa att cat aat aca acc atg tat gag tta atc tat aat aga gac aaa aac aag cat gct tcc cat att cta cat agg tat tct aaa cat cca gtt ttg aca cag tgt atc act aaa gga ttc aag att tac aca gaa gta acc gag cag gtc act aaa gct cta aac aga cgt gct cta ata gat gag ata ata aac aat gta tca act gat gac aat ctc cta tca aaa ctt cca tta gaa att agg gat cta att gtt tca caa gct gtc ata tag agt tct atc cac cca cct ttc ttg aaa tga gtt aat --- --- --- --- --- --- --- --- --- --- --- --- --- --- --- --- --- --- --- --- --- --- --- --- --- --- --- --- --- --- --- --- --- --- --- --- --- --- --- --- --- --- --- --- --- --- --- --- --- --- --- --- --- --- --- --- --- --- --- --- --- --- --- --- --- --- --- --- --- --- --- --- --- --- --- --- --- --- --- --- --- --- --- --- --- --- --- --- --- --- --- --- --- --- --- --- --- --- --- --- --- --- --- --- --- -ag tca taa gtt agt taa --- --- --- --- --- -gt cat aag tta gtt aag tca t-- --- --- --- --- --- --- --- --- --- --- --- --- --- --- --- -aa gtt agt taa gtc ata agt tag tta agt cat aag tta gtt aag tca taa gtt agt taa gtc ata agt tag tta agt cat aag tta gtt tat agt cta aca ctt cta att ttt ata cct tga tct ttt tct cta att atg aaa aag taa atc att atg aag atg gat g-a aat gga cga gat tgt gcg cat cgt taa cga tag tat gtg gta cgt acc taa cgc att tat gga cga cgg tga taa tga agg tca cat ttc tgt caa taa tgt ctg tca tat gta tct cgc att ctt tga tgt gga tat atc atc tca tct gtt taa att agt tat taa aca ctg cga tct gaa taa acg act aaa atg tgg taa ctc tcc att aca ttg cta tac gat gaa tac acg att taa tcc atc tgt att aaa gat att gtt acg cca cgg cat gcg taa ctt tga tag caa gga taa aaa agg aca tat tcc tct aca cca cta tct gat tca ttc act atc aat cga taa caa gat ctt tga tat act aac gga ccc cat tga tga ctt tag taa atc atc cga tct att gct gtg tta tct tag ata taa att caa tgg gag ctt aaa cta tta cgt tct gta caa att att gac taa agg atc tga ccc taa ttg cgt cga tga gga tgg act cac ttc tct tca tta cta ctg taa aca cat atc cgc gtt cca cga aag caa tta tta caa gtc aaa gag tca cac taa gat gcg agc tga gaa gcg att cat cta cgc gat aat aga tca tgg agc aaa cat taa cgc ggt tac gaa aat cgg aaa tac gcc gtt aca cac tta cct tca aca gta tac caa aca tag tcc tcg tgt ggt gta tgc tct ttt atc tcg agg agc cga tac gag gat acg taa taa tct taa ttg tac acc cat cat gga ata cat aaa gaa cga ttg tgc aac agg tca tat tct cat aat gtt act caa ttg gca cga aca aaa ata cgg gaa att aca aaa gga aga agg aca aca tct act tta tct att cat aaa aca taa tca agg ata tgg aag tcg ctc tct caa tat act acg gta tct act aga tag att cga cat tca gaa aga cga ata cta taa tac aat gac tcc tct tca tac cgc ctt cca gaa ttg caa taa caa tgt tgc ctc ata cct cgt ata cat tgg ata cga cat caa cct tcc gac taa aga cga taa gac agt att cga ctt ggt gtt tga aaa cag aaa cat tat ata caa ggc gga tgt cgt taa tga cat tat cca cca cag act gaa agt atc tct acc tat gat taa atc gtt gtt cta caa gat gtc gga gtt ctc tcc cta cga cga tca cta cgt aaa gaa gat aat agc cta ctg cct att aag gga cga gtc att tgc gga act aca tac taa att ctg ttt aaa cga gga cta taa aag tgt att tat gaa aaa tat atc att cga taa gat aga ttc cat cat cga aaa atg tag tcg tga cat aag tct cct caa aga gat tcg aat ctc aga cac cga ctt gta tac ggt att gag aac aga aga cat tcg gta tca cac ata tct cga agc cat aca ttc aga caa acg cat ttc att tcc cat gta cga cga tct cat aga aca gtg tca tct atc gat gga gca taa aag taa act cgt cga caa agc act caa taa att aga gtc tac cat cga tag tca atc tag act atc gta ttt gcc tcc gga aat tat gcg caa tat cat aac caa gct aag cga cta cca tct aaa cag tat gtt gta cgg aaa gaa cca tta caa ata tta tcc atg ata gaa aga aaa tat tta aaa aat aat cta tat gat tgg aga agt agg aaa caa aca gta aca aga cga cga tta cta c-a tta tta aat cat gag gtc cgt att ata ctc gta tat att gtt tct ctc atg tat aat aat aaa cgg aag aga tat agc acc aca tgc acc atc caa tgg aaa gtg taa aga caa cga ata cag aag ccg taa tct atg ttg tct atc gtg tcc tcc ggg aac tta cgc ttt cag att atg tga tag caa gac taa tac aca atg tac acc gtg tgg ttc gga tac ctt tac atc tca caa taa tca ttt aca ggc ttg tct aag ttg taa cgg aag atg tga tag taa tca ggt aga gac gcg atc gtg taa cac gac tca caa tag aat ctg tga atg ctc tcc agg ata tta ttg tct tct caa agg agc atc agg gtg tag aac atg tat ttc taa aac aaa gtg tgg aat agg ata cgg agt atc cgg ata cac gtc tac cgg aga cgt cat ctg ttc tcc gtg tgg tcc cgg aac ata ttc tca cac cgt ctc ttc cac aga taa atg cga acc cgt cgt aac cag caa tac att taa cta tat cga tgt gga aat taa cct gta tcc agt caa cga cac atc gtg tac tcg gac gac cac tac cgg tct cag cga atc cat ctc aac gtc gga act aac tat tac cat gaa tca taa aga ttg tga tcc agt ctt tcg tgc aga ata ctt ctc tgt cct taa taa tgt agc aac ttc agg att ctt tac agg aga aaa tag ata tca gaa tac ttc aaa gat atg tac tct gaa ttt cga gat taa atg taa caa caa aga ttc atc ttc caa aca gtt aac gaa aac aaa gaa tga tac tat cat gcc gca ttc aga gac ggt aac tct agt ggg cga ctg tct atc tag cgt cga cat cta cat act ata tag taa tac caa tac tca aga cta cga aaa tga tac aat ctc tta tca tat ggg taa tgt tct cga tgt caa tag cca tat gcc cgc tag ttg cga tat aca taa act gat cac taa ttc cca gaa tcc cac cca ctt ata gta ag- -tt ttt tta cct ata aat aat aaa tac aat aat taa ttt ctc gta aaa gta gaa aat ata ttc taa ttt att ata tgg taa gaa agt aga atc atc tag aac agt aat caa tca ata gca atc atg aaa caa tat att gtc ctg gca tgc atg tgc cta gtg gca gct gct atg cct act agt ctt caa caa tct tca tcc tcg tgt act gaa gaa gaa aac aaa cat cat atg gga atc gat gtt att atc aaa gtc aca aag caa gac caa aca ccg acc aat gat aag att tgt caa tcc gta acg gaa gtt aca gag a-- --- --- --- --- --- -cc gaa gat gat gag gta tcc gaa gaa gtt gta aaa gga gat ccc acc act tat tac act atc gtc ggt gcg ggt ctt aac atg aac ttt gga ttc acc aaa tgc cca aag att tta tcc atc tcc gaa tcc tct gat gga aac act gtg aat act aga ttg tcc agc gtg tca ccg gga caa ggt aag gac tct ccc gcg atc acg cgt gaa gaa gct ctg gct atg atc aaa gac tgt gag atg tct atc gac atc aga tgt agc gaa gaa gag aaa gac agt gac atc aag acc cat cca gta ctt ggg tct aac atc tca cat aag aaa gtg agt tac aaa gat atc atc ggt tca acg atc gtt gat aca aaa tgt gtc aag aac cta gag ttt agc gta cgt atc gga gac atg tgt gag gaa tca tct gaa ctt gaa gtc aag gat gga ttc aag tat gtc gac gga tcg gca tct gaa ggt gca acc gat gat act tca ctc atc gat tca aca aaa ctc aaa gca tgt gtc tga atc gat aac tct att cat ctg aaa atg gat gag ttg ggt taa tcg aac gat tca gac acc gca cca cga att aaa aaa g-- -ac cgg gca cta tat tcc ggt ttg caa aac aaa aat att taa cta cat tca caa aaa gtt acc tct cgt tac ttc ttc ttt ctg ttt caa tat gtg ata cga tat gat cac tat tcg tat tct ct- --- --- tgg tct cat aaa aaa gtt tta ca- -ma aaa aaa aaa aaa a-t att ttt att ctc ttt ctc tct tcg at- ggt ctc aca aaa ata tta aac ctc ttt ctg atg tct caa cta ttt cgt aaa cga taa cgt cca aca ata tat tct cgt aga gct tat caa cat cct tat acc aat cta ggt tgt cag aca att gca tca taa aat aat gtt tat aat tta cac gtt aac atc ata taa taa acg tat ata gtt aat att ttt gga ata taa atg atc tgt aaa atc cat gta ggg gac act gct cac gtt ttt tct cta gta cat aat ttc aca caa gtt ttt ata cag aca aat taa ttc tcg tcc ata tat ttt taa aac att gac ttt tgt act aag aaa aat atc ttg act aac cat ctc ttt ctc tct tcg atg ggt ctc aca aaa ata tta aac ctc tt- --- --- --- --- --- --- --- --- --- --- --- --- --- --- --- --- --- --- --- --- --- --- --- --- --- --- --- --- --- --- --- --- --- --- --- --- --- --- --- --- --- --- --- --- --- --- --- --- --- --- --- --- --- --- --- --- --- --- --- --- --- --- --- --- --- --- --- --- --- --- --- --- --- --- --- --- --- --- --- --- --- --- --- --- --- --- --- --- --- --- --- --- --- --- --- --- --- --- --- --- --- --- --- --- --- --- --- --- --- --- --- --- --- --- --- --- --- --- --- --- --- --- --- --- --- --- --- --- --- --- --- --- --- --- --- --- --- --- --- --- --- --- --- --- --- --

>MT903346.1_Monkeypox_virus_isolate_MPXV-USA2003_099_Gambian_Rat

atc gag gac gac ttc aac tat gac agc tcg gtg gcg tct gcc agc gtg tac ata cga atg gca ttt cta aga aaa gtc tac ggt att ctt -tc tac aat ttc ctt taa caa cgg caa cag ctg cag tat ttt tat act ttg aat gca -tc gga cat tta tac aag gga gtc ctg ttc taa tat tgg cat caa tgt tcg gat cta tag gct tga ttt tcg cat tga ctt tac aca gac atg aac atc ccc tga atc tgt aca tac ttt gtg gat tta cac tgt tag aat ctc taa cgc tgg cct ctg ttg tta ctt tct atg atg cac gta tcg tta tgc aag ctt tca tgt tga cta ctg cag tgt ttc ttg ctc tga cta cat ata ctc tac aat caa aga gag att tca gta aac ttg taa cag gat tgt ttg ctg ctt tct gga ttt taa ttt tgt cag gag tct tga gga taa agt tta aaa tag aat taa taa aga aca tat agg tca ttt ttt taa aca tgg ata gaa acc aag gtt gtt agt taa taa tat aca aga tat ttt ttc tca ctc tga tcc atg taa acc aag gac gag aga cac tct cat tcc tca ttc acg aca cca tta aaa atg gaa att aaa gcc ctc tat taa gca cag acg gct aca ggt cta cca tca ggt t-a cct tcg tct acc ttc aca atg gcc tct cct tgt gcc cag ttc agt ccc tgt cat tgc cac gct act aag gac tcc ctg aat acc gtg act gat gtc aga cat tgt ctg act gaa tac atc ctg tgg gtt tct cat aga tgg acc cat aga gaa agc gca ggg cct ctc tac agg ctt ctc atc tct ttc aga att gat gca atg gag cta ttt ggt agc gag ttg aag gag ttc tcg gat tca ctt ccg tgg gac aat atc gac aat tgc gtg gag atc att aaa tgt ttc atc aga aat gac tcc atg aaa acc gcc aaa gaa ctt tgt gca ata att gga ctt tgt act caa tca gct att gtc act gga aga gtc ttc aat gat aag tat atc gac ata cta ctt atg ctg cga aag att ctg aac gag aac gac tat ctc aca ctc ttg gat cat atc ctc act gct aaa tac taa atc tcc ttc atg ctc tct cac taa tac tct tac tca cta cac ttt tta tca tct tat gat gaa tga ttg cct tca tca -tt ttt cgt gga ata taa tat agg aat aat tag cac cag aat agc tat gga tat ctc gtt aag aat att ctc gat aag aga cat aat gta gac ata gtt att ata tcc ttc tta gat aag tgt tac gct act gga aag ttt cca tcg tta tta tta cat gaa gat gat ata att aaa cca aca ttg aga ttg gct ctt atg tta gct gga ttg aat tac tgt aat aaa tgc atc gag tat aga ggg ata tag caa ttc tcg ata ata gtc atg caa tat ttg aat gag act gat aat tta ggt aat aca gta ct- --- aca cac ata tct ttc tag att ata tat cgt taa aaa tct gta aga ggt ata ttt ctc ata agt atc cac tgt gta ata tta tta atg gat ata tag ata acg caa tag gga cta ata gta ttg taa aag ata taa tcg act att tac gta cat atc cag ata tct ata ttc cta cta gtt tgc tgc gta gtt gca tca ttg ata tgc atg -at tta tca gga ttc aga gat gaa tta cta agt aaa cta caa tcc cac aat aag taa gaa tca aat atc aaa aac tca ctt ttg att ttt cta gtc tta agt aat aca tat att tat taa tag acc tat gaa ata aaa aaa ggt aac aat gga ttc gcg tat agc tat tta cgt att agt atc ggc atc tct ttt gta tct tgt taa ttg tca caa act agt aca tta ctt caa tct gaa aat aaa tgg aag tga tat aac taa tac agc aga tat att gct gga caa tta tcc aat tat gac ctt tga tgg aaa gga tat tta tcc atc tat ctc gtt cat ggt cgg taa taa act ttt cct aga tct tta taa aaa tat ctt tga aga att ttt cag act att tcg agt atc tgt aag tag tca ata cga gga att aga ata tta tta ttc atg tga tta tac taa caa ccg tcc tac aat taa aca aca tta ctt tta taa cgg cga aga ata tac tga aat tga tag atc gaa aaa agc cac taa taa aaa cag ttg gtt aat tac ttc agg ctt tag act aca aaa atg gtt cga tag cga aga ttg tat aat tta tct cag atc ttt agt tag aag aat gga aga cag taa caa aaa cag taa aaa aac tta gta ctt aga tat cg- -aa aaa aat ata ttt ttg tag act ctt gag aat aga agg aaa aca tgt aca taa tta taa aaa atg aaa atc aat ggc gaa taa gac agt gcg att cgc acc atg gag tcg gta gat ttc atg gct gtc gat gag cag ttt cac gac gac ctc gat ctt tgg tca tta tct ttg gta gat gat tat aaa aaa cat gga tta ggt gtt gac tgt tat gtt cta gaa cca gtt gtt gac agg aaa ata ttt gat aga ttt ctc ctt gaa cca att tgt gat cct gta gat gtt ctg tat gat tat ttt agg att cat aga gat aat att gat cag tat ata gta gat aga ctg ttt gca tat att aca tat aaa gat att ata tct gca tta gtg tca aag aat tat atg gaa gat att ttc tct ata att att aag aat tgt aat tct gtg caa gat ctc tta ctt tac tat cta tct aat gca tat gta gaa ata gac att gtt gat ctt atg gta gat cat ggg gct gta ata tat aaa ata gaa tgc ttg aat gcc tat ttt agg gga ata tgt aaa aag gaa agt agt gtt gtt gag ttt att ttg aat tgt ggt atc cca gat gaa aat gat gtt aaa tta gat cta tat aaa ata att cag tat act agg gga ttc ctt gta gat gaa ccc aca gta tta gaa att tat aag ctt tgt atc cca tat att gaa gat atc aat caa cta gat gct ggt gga agg acc ttg ctt tat cgc gct atc tat gca ggt tat ata gat tta gta tca tgg cta tta gaa aat gga gca aat gtc aac gca gta atg agt aat gga tat aca tgt ctt gac gtg gcc gtg gat agg gga tct gtc atc gcc cgt agg gaa gca cat ctt aaa ata tta gaa ata ttg ctt aga gaa cca ttg tct att gac tgt ata aaa tta gct ata ctt aat aat aca att gaa aac cat gat gtg ata aag ctc tgt atc aag tat ttt atg atg gta gat tat tca ctt tgt aat gtg tat gca tca tca ctc ttt gat tat ata att gat tgt aaa caa gaa ttg gag tac att agg cag atg aaa att cat aat aca acc atg tat gag tta atc tat aat aga gac aaa aac aag cat gct tcc cat att cta cat agg tat tct aaa cat cca gtt ttg aca cag tgt atc act aaa gga ttc aag att tac aca gaa gta acc gag cag gtc act aaa gct cta aac aga cgt gct cta ata gat gag ata ata aac aat gta tca act gat gac aat ctc cta tca aaa ctt cca tta gaa att agg gat cta att gtt tca caa gct gtc ata tag agt tct atc cac cca cct ttc ttg aaa tga gtt aat --- --- --- --- --- --- --- --- --- --- --- --- --- --- --- --- --- --- --- --- --- --- --- --- --- --- --- --- --- --- --- --- --- --- --- --- --- --- --- --- --- --- --- --- --- --- --- --- --- --- --- --- --- --- --- --- --- --- --- --- --- --- --- --- --- --- --- --- --- --- --- --- --- --- --- --- --- --- --- --- --- --- --- --- --- --- --- --- --- --- --- --- --- --- --- --- --- --- --- --- --- --- --- --- --- -ag tca taa gtt agt taa --- --- --- --- --- -gt cat aag tta gtt aag tca taa gtt agt t-- --- --- --- --- --- --t tat aag --- --- --- --- taa gtt agt taa gtc ata agt tag tt- --t tat aag --- --- --- --- taa gtt agt taa gtc ata agt tag tt- --t tat aag tta gtt tat agt cta aca ctt cta att ttt ata cct tga tct ttt tct cta att atg aaa aag taa atc att atg aag atg gat g-a aat gga cga gat tgt gcg cat cgt taa cga tag tat gtg gta cgt acc taa cgc att tat gga cga cgg tga taa tga agg tca cat ttc tgt caa taa tgt ctg tca tat gta tct cgc att ctt tga tgt gga tat atc atc tca tct gtt taa att agt tat taa aca ctg cga tct gaa taa acg act aaa atg tgg taa ctc tcc att aca ttg cta tac gat gaa tac acg att taa tcc atc tgt att aaa gat att gtt acg cca cgg cat gcg taa ctt tga tag caa gga taa aaa agg aca tat tcc tct aca cca cta tct gat tca ttc act atc aat cga taa caa gat ctt tga tat act aac gga ccc cat tga tga ctt tag taa atc atc cga tct att gct gtg tta tct tag ata taa att caa tgg gag ctt aaa cta tta cgt tct gta caa att att gac taa agg atc tga ccc taa ttg cgt cga tga gga tgg act cac ttc tct tca tta cta ctg taa aca cat atc cgc gtt cca cga aag caa tta tta caa gtc aaa gag tca cac taa gat gcg agc tga gaa gcg att cat cta cgc gat aat aga tca tgg agc aaa cat taa cgc ggt tac gaa aat cgg aaa tac gcc gtt aca cac tta cct tca aca gta tac caa aca tag tcc tcg tgt ggt gta tgc tct ttt atc tcg agg agc cga tac gag gat acg taa taa tct tga ttg tac acc cat cat gga ata cat aaa gaa cga ttg tgc aac agg tca tat tct cat aat gtt act caa ttg gca cga aca aaa ata cgg gaa att aca aaa gga aga agg aca aca tct act tta tct att cat aaa aca taa tca agg ata tgg aag tcg ctc tct caa tat act acg gta tct act aga tag att cga cat tca gaa aga cga ata cta taa tac aat gac tcc tct tca tac cgc ctt cca gaa ttg caa taa caa tgt tgc ctc ata cct cgt ata cat cgg ata cga cat caa cct tcc gac taa aga cga taa gac agt att cga ctt ggt gtt tga aaa cag aaa cat cat ata caa ggc gga tgt cgt taa tga cat cat cca cca cag act gaa agt atc tct acc tat gat taa atc gtt gtt cta caa gat gtc gga gtt ctc tcc cta cga cga tca cta cgt aaa gaa gat aat agc cta ctg cct att aag gta cga gtc att tgc gga act aca tac taa att ctg ttt aaa cga gga cta taa aag tgt att tat gaa aaa tat atc att cga taa gat aga ttc cat cat cga aaa atg tag tcg tga cat aag tct cct caa aga gat tcg aat ctc aga cac cga ctt gta tac ggt att gag aac aga aga cat ccg gta tca cac cta tct cga agc cat aca ttc aga caa acg cat ttc att tcc cat gta cga cga tct cat aga aca gtg tca tct atc gat gga gca taa aag taa act cgt cga caa agc act caa taa att aga gtc tac cat cga tag tca atc tag act atc gta ttt gcc tcc gga aat tat gcg caa tat cat aac caa gct aag cga cta cca tct aaa cag tat gtt gta cgg aaa gaa cca tta caa ata tta tcc atg ata gaa aga aaa tat tta aaa aat aat cta tat gat tgg aga agt agg aaa caa aca gta aca aga cga cga tta cta c-a tta tta aat cat gag gtc cgt att ata ctc gta tat att gtt tct ctc atg tat aat aat aaa cgg aag aga tat agc acc aca tgc acc atc caa tgg aaa gtg taa aga caa cga ata cag aag ccg taa tct atg ttg tct atc gtg tcc tcc ggg aac tta cgc ttc cag att atg tga tag caa gac taa tac aca atg tac acc gtg tgg ttc gga tac ctt tac atc tca caa taa tca ttt aca ggc ttg tct aag ttg taa cgg aag atg tga tag taa tca ggt aga gac gcg atc gtg taa cac gac tca caa tag aat ctg tga atg ctc tcc agg ata tta ttg tct tct caa agg agc atc agg gtg tag aac atg tat ttc taa aac aaa gtg tgg aat agg ata cgg agt atc cgg ata cac gtc tac cgg aga cgt cat ctg ttc tcc gtg tgg tcc cgg aac ata ttc tca cac cgt ctc ttc cac aga taa atg cga acc cgt cgt aac cag caa tac att taa cta tat cga tgt gga aat taa cct gta tcc agt caa cga cac atc gtg tac tcg gac gac cac tac cgg tct cag cga atc cat ctc aac gtc gga act aac tat tac cat gaa tca taa aga ttg tga tcc agt ctt tcg tgc aga ata ctt ctc tgt cct taa taa tgt agc aac ttc agg att ctt tac agg aga aaa tag ata tca gaa tac ttc aaa gat atg tac tct gaa ttt cga gat taa atg taa caa caa aga ttc atc ttc caa aca gtt aac gaa aac aaa gaa tga tac tat cat gcc gca ttc aga gac ggt aac tct agt ggg cga ctg tct atc tag cgt cga cat cta cat act ata tag taa tac caa tac tca aga cta cga aac gga tac aat ctc tta tca tat ggg taa tgt tct cga tgt caa tag cca tat gcc cgc tag ttg cga tat aca taa act gat cac taa ttc cca gaa tcc cac cca ctt ata gta ag- -tt ttt tta ccc ata aat aat aaa tac aat aat taa ttt ctc gta aaa gta gaa aat ata ttc taa ttt att ata tgg taa gaa agt aga atc atc tag aac agt aat caa tca ata gca atc atg aaa caa tat att gtc ctg gca tgc atg tgc cta gtg gca gct gct atg cct act agt ctt caa caa tct tca tcc tcg tgt act gaa gaa gaa aac aaa cat cat atg gga atc gat gtt att atc aaa gtc aca aag caa gac caa aca ccg acc aat gat aag att tgt caa tcc gta acg gaa gtt aca gag a-- --- --- --- --- --- -cc gaa gat gat gag gta tcc gaa gaa gtt gta aaa gga gat ccc acc act tat tac act atc gtc ggt gcg ggt ctt aac atg aac ttt gga ttc acc aaa tgc cca aag att tca tcc atc tcc gaa tcc tct gat gga aac act gtg aat act aga ttg tcc agc gtg tca ccg gga caa ggt aag gac tct ccc gcg atc acg cgt gaa gaa gct ctg gct atg atc aaa gac tgt gag atg tct atc gac atc aga tgt agc gaa gaa gag aaa gac agt gac atc aag acc cat cca gta ctt ggg tct aac atc tca cat aag aaa gtg agt tac aaa gat atc atc ggt tca acg atc gtt gat aca aaa tgt gtc aag aac cta gag ttt agc gta cgt atc gga gac atg tgt gag gaa tca tct gaa ctt gaa gtc aag gat gga ttc aag tat gtc gac gga tcg gca tct gaa ggt gca acc gat gat act tca ctc atc gat tca aca aaa ctc aaa gca tgt gtc tga atc gat aac tct att cat ctg aaa atg gat gag ttg ggt taa tcg aac gat tca gac acc gca cca cga att aaa aaa g-- -ac cgg gca cta tat tcc ggt ttg caa aac aaa aat att taa cta cat tca caa aaa gtt acc tct cgt tac ttc ttc ttt ctg ttt caa tat gtg ata cga tat gat cac tat tcg tat tct ct- --- --- tgg tct cat aaa aaa gtt tta c-- --- --- -aa aaa aaa a-t att ttt att cta ttt ctc tct tcg at- ggt ctc aca aaa ata tta aac ctc ttt ctg atg tct caa cta ttt cgt aaa cga taa cgt cca aca ata tat tct cgt aga gct tat caa cat cct tat acc aat cta ggt tgt cag aca att gca tca taa aat aat gtt tat aat tta cac gtt aac atc ata taa taa acg tat ata gtt aat att ttt gga ata taa atg atc tgt aaa atc cat gta ggg gac act act cac gtt ttt tct cta gta cat aat ttc aca caa gtt ttt ata cag aca aat taa ttc tcg tcc ata ta- ttt taa aac att gac ttt tgt act aag aaa aat atc ttg act aac cat ctc ttt ctc tct tcg atg ggt ctc aca aaa ata tta aac ctc ttt ctg atg gag tcg taa aaa gtt ttt atc ctt tct ctc ttc gat ggg tct cac aaa aat att aaa cct ctt tct gat gga gtc gta aaa agt ttt tat cct ttc tct ctt cga tgg gtc tca caa aaa tat taa acc tct ttc tga tgg agt cgt aaa aag ttt tta tcc ttt ctc tct tcg atg ggt ctc --- --- --- --- --- --- --- --- --- --- --- --- --- --- --- --- --- --- --- --- --- --- --- --- --- --- --- --- --- --- --- --- --- --- --- --- --- --- --- --- --- --- --- --- --- --- --- --- --- --- --- --- --- --- --- --- --- --- --- --- --- --- --- --- --- --- --- --- --- --- aca aaa ata tta aac ctc ttt ctg atg gtc tct ata aa

>DQ011157.1_Monkeypox_virus_strain_USA_2003_039_complete_genome

atc gag gac gac ttc aac tat gac agc tcg gtg gcg tct gcc agc gtg tac ata cga atg gca ttt cta aga aaa gtc tac ggt att ctt -tc tac aat ttc ctt taa caa cgg caa cag ctg cag tat ttt tat act ttg aat gca -tc gga cat tta tac aag gga gtc ctg ttc taa tat tgg cat caa tgt tcg gat cta tag gct tga ttt tcg cat tga ctt tac aca gac atg aac atc ccc tga atc tgt aca tac ttt gtg gat tta cac tgt tag aat ctc taa cgc tgg cct ctg ttg tta ctt tct atg atg cac gta tcg tta tgc aag ctt tca tgt tga cta ctg cag tgt ttc ttg ctc tga cta cat ata ctc tac aat caa aga gag att tca gta aac ttg taa cag gat tgt ttg ctg ctt tct gga ttt taa ttt tgt cag gag tct tga gga taa agt tta aaa tag aat taa taa aga aca tat agg tca ttt ttt taa aca tgg ata gaa acc aag gtt gtt agt taa taa tat aca aga tat ttt ttc tca ctc tga tcc atg taa acc aag gac gag aga cac tct cat tcc tca ttc acg aca cca tta aaa atg gaa att aaa gcc ctc tat taa gca cag acg gct aca ggt cta cca tca ggt t-a cct tcg tct acc ttc aca atg gcc tct cct tgt gcc cag ttc agt ccc tgt cat tgc cac gct act aag gac tcc ctg aat acc gtg act gat gtc aga cat tgt ctg act gaa tac atc ctg tgg gtt tct cat aga tgg acc cat aga gaa agc gca ggg cct ctc tac agg ctt ctc atc tct ttc aga att gat gca atg gag cta ttt ggt agc gag ttg aag gag ttc tcg gat tca ctt ccg tgg gac aat atc gac aat tgc gtg gag atc att aaa tgt ttc atc aga aat gac tcc atg aaa acc gcc aaa gaa ctt tgt gca ata att gga ctt tgt act caa tca gct att gtc act gga aga gtc ttc aat gat aag tat atc gac ata cta ctt atg ctg cga aag att ctg aac gag aac gac tat ctc aca ctc ttg gat cat atc ctc act gct aaa tac taa atc tcc ttc atg ctc tct cac taa tac tct tac tca cta cac ttt tta tca tct tat gat gaa tga ttg cct tca tca -tt ttt cgt gga ata taa tat agg aat aat tag cac cag aat agc tat gga tat ctc gtt aag aat att ctc gat aag aga cat aat gta gac ata gtt att ata tcc ttc tta gat aag tgt tac gct act gga aag ttt cca tcg tta tta tta cat gaa gat gat ata att aaa cca aca ttg aga ttg gct ctt atg tta gct gga ttg aat tac tgt aat aaa tgc atc gag tat aga ggg ata tag caa ttc tcg ata ata gtc atg caa tat ttg aat gag act gat aat tta ggt aat aca gta ct- --- aca cac ata tct ttc tag att ata tat cgt taa aaa tct gta aga ggt ata ttt ctc ata agt atc cac tgt gta ata tta tta atg gat ata tag ata acg caa tag gga cta ata gta ttg taa aag ata taa tcg act att tac gta cat atc cag ata tct ata ttc cta cta gtt tgc tgc gta gtt gca tca ttg ata tgc atg -at tta tca gga ttc aga gat gaa tta cta agt aaa cta caa tcc cac aat aag taa gaa tca aat atc aaa aac tca ctt ttg att ttt cta gtc tta agt aat aca tat att tat taa tag acc tat gaa ata aaa aaa ggt aac aat gga ttc gcg tat agc tat tta cgt att agt atc ggc atc tct ttt gta tct tgt taa ttg tca caa act agt aca tta ctt caa tct gaa aat aaa tgg aag tga tat aac taa tac agc aga tat att gct gga caa tta tcc aat tat gac ctt tga tgg aaa gga tat tta tcc atc tat ctc gtt cat ggt cgg taa taa act ttt cct aga tct tta taa aaa tat ctt tga aga att ttt cag act att tcg agt atc tgt aag tag tca ata cga gga att aga ata tta tta ttc atg tga tta tac taa caa ccg tcc tac aat taa aca aca tta ctt tta taa cgg cga aga ata tac tga aat tga tag atc gaa aaa agc cac taa taa aaa cag ttg gtt aat tac ttc agg ctt tag act aca aaa atg gtt cga tag cga aga ttg tat aat tta tct cag atc ttt agt tag aag aat gga aga cag taa caa aaa cag taa aaa aac tta gta ctt aga tat cg- -aa aaa aat ata ttt ttg tag act ctt gag aat aga agg aaa aca tgt aca taa tta taa aaa atg aaa atc aat ggc gaa taa gac agt gcg att cgc acc atg gag tcg gta gat ttc atg gct gtc gat gag cag ttt cac gac gac ctc gat ctt tgg tca tta tct ttg gta gat gat tat aaa aaa cat gga tta ggt gtt gac tgt tat gtt cta gaa cca gtt gtt gac agg aaa ata ttt gat aga ttt ctc ctt gaa cca att tgt gat cct gta gat gtt ctg tat gat tat ttt agg att cat aga gat aat att gat cag tat ata gta gat aga ctg ttt gca tat att aca tat aaa gat att ata tct gca tta gtg tca aag aat tat atg gaa gat att ttc tct ata att att aag aat tgt aat tct gtg caa gat ctc tta ctt tac tat cta tct aat gca tat gta gaa ata gac att gtt gat ctt atg gta gat cat ggg gct gta ata tat aaa ata gaa tgc ttg aat gcc tat ttt agg gga ata tgt aaa aag gaa agt agt gtt gtt gag ttt att ttg aat tgt ggt atc cca gat gaa aat gat gtt aaa tta gat cta tat aaa ata att cag tat act agg gga ttc ctt gta gat gaa ccc aca gta tta gaa att tat aag ctt tgt atc cca tat att gaa gat atc aat caa cta gat gct ggt gga agg acc ttg ctt tat cgc gct atc tat gca ggt tat ata gat tta gta tca tgg cta tta gaa aat gga gca aat gtc aac gca gta atg agt aat gga tat aca tgt ctt gac gtg gcc gtg gat agg gga tct gtc atc gcc cgt agg gaa gca cat ctt aaa ata tta gaa ata ttg ctt aga gaa cca ttg tct att gac tgt ata aaa tta gct ata ctt aat aat aca att gaa aac cat gat gtg ata aag ctc tgt atc aag tat ttt atg atg gta gat tat tca ctt tgt aat gtg tat gca tca tca ctc ttt gat tat ata att gat tgt aaa caa gaa ttg gag tac att agg cag atg aaa att cat aat aca acc atg tat gag tta atc tat aat aga gac aaa aac aag cat gct tcc cat att cta cat agg tat tct aaa cat cca gtt ttg aca cag tgt atc act aaa gga ttc aag att tac aca gaa gta acc gag cag gtc act aaa gct cta aac aga cgt gct cta ata gat gag ata ata aac aat gta tca act gat gac aat ctc cta tca aaa ctt cca tta gaa att agg gat cta att gtt tca caa gct gtc ata tag agt tct atc cac cca cct ttc ttg aaa tga gtt aat --- --- --- --- --- --- --- --- --- --- --- --- --- --- --- --- --- --- --- --- --- --- --- --- --- --- --- --- --- --- --- --- --- --- --- --- --- --- --- --- --- --- --- --- --- --- --- --- --- --- --- --- --- --- --- --- --- --- --- --- --- --- --- --- --- --- --- --- --- --- --- --- --- --- --- --- --- --- --- --- --- --- --- --- --- --- --- --- --- --- --- --- --- --- --- --- --- --- --- --- --- --- --- --- --- -ag tca taa gtt agt taa --- --- --- --- --- -gt cat aag tta gtt aag tca taa gtt agt t-- --- --- --- --- --- --t tat aag --- --- --- --- taa gtt agt taa gtc ata agt tag tt- --t tat aag --- --- --- --- taa gtt agt taa gtc ata agt tag tt- --t tat aag tta gtt tat agt cta aca ctt cta att ttt ata cct tga tct ttt tct cta att atg aaa aag taa atc att atg aag atg gat g-a aat gga cga gat tgt gcg cat cgt taa cga tag tat gtg gta cgt acc taa cgc att tat gga cga cgg tga taa tga agg tca cat ttc tgt caa taa tgt ctg tca tat gta tct cgc att ctt tga tgt gga tat atc atc tca tct gtt taa att agt tat taa aca ctg cga tct gaa taa acg act aaa atg tgg taa ctc tcc att aca ttg cta tac gat gaa tac acg att taa tcc atc tgt att aaa gat att gtt acg cca cgg cat gcg taa ctt tga tag caa gga taa aaa agg aca tat tcc tct aca cca cta tct gat tca ttc act atc aat cga taa caa gat ctt tga tat act aac gga ccc cat tga tga ctt tag taa atc atc cga tct att gct gtg tta tct tag ata taa att caa tgg gag ctt aaa cta tta cgt tct gta caa att att gac taa agg atc tga ccc taa ttg cgt cga tga gga tgg act cac ttc tct tca tta cta ctg taa aca cat atc cgc gtt cca cga aag caa tta tta caa gtc aaa gag tca cac taa gat gcg agc tga gaa gcg att cat cta cgc gat aat aga tca tgg agc aaa cat taa cgc ggt tac gaa aat cgg aaa tac gcc gtt aca cac tta cct tca aca gta tac caa aca tag tcc tcg tgt ggt gta tgc tct ttt atc tcg agg agc cga tac gag gat acg taa taa tct tga ttg tac acc cat cat gga ata cat aaa gaa cga ttg tgc aac agg tca tat tct cat aat gtt act caa ttg gca cga aca aaa ata cgg gaa att aca aaa gga aga agg aca aca tct act tta tct att cat aaa aca taa tca agg ata tgg aag tcg ctc tct caa tat act acg gta tct act aga tag att cga cat tca gaa aga cga ata cta taa tac aat gac tcc tct tca tac cgc ctt cca gaa ttg caa taa caa tgt tgc ctc ata cct cgt ata cat cgg ata cga cat caa cct tcc gac taa aga cga taa gac agt att cga ctt ggt gtt tga aaa cag aaa cat cat ata caa ggc gga tgt cgt taa tga cat cat cca cca cag act gaa agt atc tct acc tat gat taa atc gtt gtt cta caa gat gtc gga gtt ctc tcc cta cga cga tca cta cgt aaa gaa gat aat agc cta ctg cct att aag gta cga gtc att tgc gga act aca tac taa att ctg ttt aaa cga gga cta taa aag tgt att tat gaa aaa tat atc att cga taa gat aga ttc cat cat cga aaa atg tag tcg tga cat aag tct cct caa aga gat tcg aat ctc aga cac cga ctt gta tac ggt att gag aac aga aga cat ccg gta tca cac cta tct cga agc cat aca ttc aga caa acg cat ttc att tcc cat gta cga cga tct cat aga aca gtg tca tct atc gat gga gca taa aag taa act cgt cga caa agc act caa taa att aga gtc tac cat cga tag tca atc tag act atc gta ttt gcc tcc gga aat tat gcg caa tat cat aac caa gct aag cga cta cca tct aaa cag tat gtt gta cgg aaa gaa cca tta caa ata tta tcc atg ata gaa aga aaa tat tta aaa aat aat cta tat gat tgg aga agt agg aaa caa aca gta aca aga cga cga tta cta c-a tta tta aat cat gag gtc cgt att ata ctc gta tat att gtt tct ctc atg tat aat aat aaa cgg aag aga tat agc acc aca tgc acc atc caa tgg aaa gtg taa aga caa cga ata cag aag ccg taa tct atg ttg tct atc gtg tcc tcc ggg aac tta cgc ttc cag att atg tga tag caa gac taa tac aca atg tac acc gtg tgg ttc gga tac ctt tac atc tca caa taa tca ttt aca ggc ttg tct aag ttg taa cgg aag atg tga tag taa tca ggt aga gac gcg atc gtg taa cac gac tca caa tag aat ctg tga atg ctc tcc agg ata tta ttg tct tct caa agg agc atc agg gtg tag aac atg tat ttc taa aac aaa gtg tgg aat agg ata cgg agt atc cgg ata cac gtc tac cgg aga cgt cat ctg ttc tcc gtg tgg tcc cgg aac ata ttc tca cac cgt ctc ttc cac aga taa atg cga acc cgt cgt aac cag caa tac att taa cta tat cga tgt gga aat taa cct gta tcc agt caa cga cac atc gtg tac tcg gac gac cac tac cgg tct cag cga atc cat ctc aac gtc gga act aac tat tac cat gaa tca taa aga ttg tga tcc agt ctt tcg tgc aga ata ctt ctc tgt cct taa taa tgt agc aac ttc agg att ctt tac agg aga aaa tag ata tca gaa tac ttc aaa gat atg tac tct gaa ttt cga gat taa atg taa caa caa aga ttc atc ttc caa aca gtt aac gaa aac aaa gaa tga tac tat cat gcc gca ttc aga gac ggt aac tct agt ggg cga ctg tct atc tag cgt cga cat cta cat act ata tag taa tac caa tac tca aga cta cga aac gga tac aat ctc tta tca tat ggg taa tgt tct cga tgt caa tag cca tat gcc cgc tag ttg cga tat aca taa act gat cac taa ttc cca gaa tcc cac cca ctt ata gta ag- -tt ttt tta ccc ata aat aat aaa tac aat aat taa ttt ctc gta aaa gta gaa aat ata ttc taa ttt att ata tgg taa gaa agt aga atc atc tag aac agt aat caa tca ata gca atc atg aaa caa tat att gtc ctg gca tgc atg tgc cta gtg gca gct gct atg cct act agt ctt caa caa tct tca tcc tcg tgt act gaa gaa gaa aac aaa cat cat atg gga atc gat gtt att atc aaa gtc aca aag caa gac caa aca ccg acc aat gat aag att tgt caa tcc gta acg gaa gtt aca gag a-- --- --- --- --- --- -cc gaa gat gat gag gta tcc gaa gaa gtt gta aaa gga gat ccc acc act tat tac act atc gtc ggt gcg ggt ctt aac atg aac ttt gga ttc acc aaa tgc cca aag att tca tcc atc tcc gaa tcc tct gat gga aac act gtg aat act aga ttg tcc agc gtg tca ccg gga caa ggt aag gac tct ccc gcg atc acg cgt gaa gaa gct ctg gct atg atc aaa gac tgt gag atg tct atc gac atc aga tgt agc gaa gaa gag aaa gac agt gac atc aag acc cat cca gta ctt ggg tct aac atc tca cat aag aaa gtg agt tac aaa gat atc atc ggt tca acg atc gtt gat aca aaa tgt gtc aag aac cta gag ttt agc gta cgt atc gga gac atg tgt gag gaa tca tct gaa ctt gaa gtc aag gat gga ttc aag tat gtc gac gga tcg gca tct gaa ggt gca acc gat gat act tca ctc atc gat tca aca aaa ctc aaa gca tgt gtc tga atc gat aac tct att cat ctg aaa atg gat gag ttg ggt taa tcg aac gat tca gac acc gca cca cga att aaa aaa g-- -ac cgg gca cta tat tcc ggt ttg caa aac aaa aat att taa cta cat tca caa aaa gtt acc tct cgt tac ttc ttc ttt ctg ttt caa tat gtg ata cga tat gat cac tat tcg tat tct ct- --- --- tgg tct cat aaa aaa gtt tta c-- --- --- aaa aaa aaa a-t att ttt att cta ttt ctc tct tcg at- ggt ctc aca aaa ata tta aac ctc ttt ctg atg tct caa cta ttt cgt aaa cga taa cgt cca aca ata tat tct cgt aga gct tat caa cat cct tat acc aat cta ggt tgt cag aca att gca tca taa aat aat gtt tat aat tta cac gtt aac atc ata taa taa acg tat ata gtt aat att ttt gga ata taa atg atc tgt aaa atc cat gta ggg gac act act cac gtt ttt tct cta gta cat aat ttc aca caa gtt ttt ata cag aca aat taa ttc tcg tcc ata ta- ttt taa aac att gac ttt tgt act aag aaa aat atc ttg act aac cat ctc ttt ctc tct tcg atg ggt ctc aca aaa ata tta aac ctc ttt ctg atg gag tcg taa aaa gtt ttt atc ctt tct ctc ttc gat ggg tct cac aaa aat att aaa cct ctt tct gat gga gtc gta aaa agt ttt tat cct ttc tct ctt cga tgg gtc tca caa aaa tat taa acc tct ttc tga tgg agt cgt aaa aag ttt tta tcc ttt ctc tct tcg atg ggt ctc --- --- --- --- --- --- --- --- --- --- --- --- --- --- --- --- --- --- --- --- --- --- --- --- --- --- --- --- --- --- --- --- --- --- --- --- --- --- --- --- --- --- --- --- --- --- --- --- --- --- --- --- --- --- --- --- --- --- --- --- --- --- --- --- --- --- --- --- --- --- aca aaa ata tta aac ctc ttt ctg atg gtc tct ata aa

>DQ011153.1_Monkeypox_virus_strain_USA_2003_044_complete_genome

atc gag gac gac ttc aac tat gac agc tcg gtg gcg tct gcc agc gtg tac ata cga atg gca ttt cta aga aaa gtc tac ggt att ctt -tc tac aat ttc ctt taa caa cgg caa cag ctg cag tat ttt tat act ttg aat gca -tc gga cat tta tac aag gga gtc ctg ttc taa tat tgg cat caa tgt tcg gat cta tag gct tga ttt tcg cat tga ctt tac aca gac atg aac atc ccc tga atc tgt aca tac ttt gtg gat tta cac tgt tag aat ctc taa cgc tgg cct ctg ttg tta ctt tct atg atg cac gta tcg tta tgc aag ctt tca tgt tga cta ctg cag tgt ttc ttg ctc tga cta cat ata ctc tac aat caa aga gag att tca gta aac ttg taa cag gat tgt ttg ctg ctt tct gga ttt taa ttt tgt cag gag tct tga gga taa agt tta aaa tag aat taa taa aga aca tat agg tca ttt ttt taa aca tgg ata gaa acc aag gtt gtt agt taa taa tat aca aga tat ttt ttc tca ctc tga tcc atg taa acc aag gac gag aga cac tct cat tcc tca ttc acg aca cca tta aaa atg gaa att aaa gcc ctc tat taa gca cag acg gct aca ggt cta cca tca ggt t-a cct tcg tct acc ttc aca atg gcc tct cct tgt gcc cag ttc agt ccc tgt cat tgc cac gct act aag gac tcc ctg aat acc gtg act gat gtc aga cat tgt ctg act gaa tac atc ctg tgg gtt tct cat aga tgg acc cat aga gaa agc gca ggg cct ctc tac agg ctt ctc atc tct ttc aga att gat gca atg gag cta ttt ggt agc gag ttg aag gag ttc tcg gat tca ctt ccg tgg gac aat atc gac aat tgc gtg gag atc att aaa tgt ttc atc aga aat gac tcc atg aaa acc gcc aaa gaa ctt tgt gca ata att gga ctt tgt act caa tca gct att gtc act gga aga gtc ttc aat gat aag tat atc gac ata cta ctt atg ctg cga aag att ctg aac gag aac gac tat ctc aca ctc ttg gat cat atc ctc act gct aaa tac taa atc tcc ttc atg ctc tct cac taa tac tct tac tca cta cac ttt tta tca tct tat gat gaa tga ttg cct tca tca -tt ttt cgt gga ata taa tat agg aat aat tag cac cag aat agc tat gga tat ctc gtt aag aat att ctc gat aag aga cat aat gta gac ata gtt att ata tcc ttc tta gat aag tgt tac gct act gga aag ttt cca tcg tta tta tta cat gaa gat gat ata att aaa cca aca ttg aga ttg gct ctt atg tta gct gga ttg aat tac tgt aat aaa tgc atc gag tat aga ggg ata tag caa ttc tcg ata ata gtc atg caa tat ttg aat gag act gat aat tta ggt aat aca gta ct- --- aca cac ata tct ttc tag att ata tat cgt taa aaa tct gta aga ggt ata ttt ctc ata agt atc cac tgt gta ata tta tta atg gat ata tag ata acg caa tag gga cta ata gta ttg taa aag ata taa tcg act att tac gta cat atc cag ata tct ata ttc cta cta gtt tgc tgc gta gtt gca tca ttg ata tgc atg -at tta tca gga ttc aga gat gaa tta cta agt aaa cta caa tcc cac aat aag taa gaa tca aat atc aaa aac tca ctt ttg att ttt cta gtc tta agt aat aca tat att tat taa tag acc tat gaa ata aaa aaa ggt aac aat gga ttc gcg tat agc tat tta cgt att agt atc ggc atc tct ttt gta tct tgt taa ttg tca caa act agt aca tta ctt caa tct gaa aat aaa tgg aag tga tat aac taa tac agc aga tat att gct gga caa tta tcc aat tat gac ctt tga tgg aaa gga tat tta tcc atc tat ctc gtt cat ggt cgg taa taa act ttt cct aga tct tta taa aaa tat ctt tga aga att ttt cag act att tcg agt atc tgt aag tag tca ata cga gga att aga ata tta tta ttc atg tga tta tac taa caa ccg tcc tac aat taa aca aca tta ctt tta taa cgg cga aga ata tac tga aat tga tag atc gaa aaa agc cac taa taa aaa cag ttg gtt aat tac ttc agg ctt tag act aca aaa atg gtt cga tag cga aga ttg tat aat tta tct cag atc ttt agt tag aag aat gga aga cag taa caa aaa cag taa aaa aac tta gta ctt aga tat cg- -aa aaa aat ata ttt ttg tag act ctt gag aat aga agg aaa aca tgt aca taa tta taa aaa atg aaa atc aat ggc gaa taa gac agt gcg att cgc acc atg gag tcg gta gat ttc atg gct gtc gat gag cag ttt cac gac gac ctc gat ctt tgg tca tta tct ttg gta gat gat tat aaa aaa cat gga tta ggt gtt gac tgt tat gtt cta gaa cca gtt gtt gac agg aaa ata ttt gat aga ttt ctc ctt gaa cca att tgt gat cct gta gat gtt ctg tat gat tat ttt agg att cat aga gat aat att gat cag tat ata gta gat aga ctg ttt gca tat att aca tat aaa gat att ata tct gca tta gtg tca aag aat tat atg gaa gat att ttc tct ata att att aag aat tgt aat tct gtg caa gat ctc tta ctt tac tat cta tct aat gca tat gta gaa ata gac att gtt gat ctt atg gta gat cat ggg gct gta ata tat aaa ata gaa tgc ttg aat gcc tat ttt agg gga ata tgt aaa aag gaa agt agt gtt gtt gag ttt att ttg aat tgt ggt atc cca gat gaa aat gat gtt aaa tta gat cta tat aaa ata att cag tat act agg gga ttc ctt gta gat gaa ccc aca gta tta gaa att tat aag ctt tgt atc cca tat att gaa gat atc aat caa cta gat gct ggt gga agg acc ttg ctt tat cgc gct atc tat gca ggt tat ata gat tta gta tca tgg cta tta gaa aat gga gca aat gtc aac gca gta atg agt aat gga tat aca tgt ctt gac gtg gcc gtg gat agg gga tct gtc atc gcc cgt agg gaa gca cat ctt aaa ata tta gaa ata ttg ctt aga gaa cca ttg tct att gac tgt ata aaa tta gct ata ctt aat aat aca att gaa aac cat gat gtg ata aag ctc tgt atc aag tat ttt atg atg gta gat tat tca ctt tgt aat gtg tat gca tca tca ctc ttt gat tat ata att gat tgt aaa caa gaa ttg gag tac att agg cag atg aaa att cat aat aca acc atg tat gag tta atc tat aat aga gac aaa aac aag cat gct tcc cat att cta cat agg tat tct aaa cat cca gtt ttg aca cag tgt atc act aaa gga ttc aag att tac aca gaa gta acc gag cag gtc act aaa gct cta aac aga cgt gct cta ata gat gag ata ata aac aat gta tca act gat gac aat ctc cta tca aaa ctt cca tta gaa att agg gat cta att gtt tca caa gct gtc ata tag agt tct atc cac cca cct ttc ttg aaa tga gtt aat --- --- --- --- --- --- --- --- --- --- --- --- --- --- --- --- --- --- --- --- --- --- --- --- --- --- --- --- --- --- --- --- --- --- --- --- --- --- --- --- --- --- --- --- --- --- --- --- --- --- --- --- --- --- --- --- --- --- --- --- --- --- --- --- --- --- --- --- --- --- --- --- --- --- --- --- --- --- --- --- --- --- --- --- --- --- --- --- --- --- --- --- --- --- --- --- --- --- --- --- --- --- --- --- --- -ag tca taa gtt agt taa --- --- --- --- --- -gt cat aag tta gtt aag tca taa gtt agt t-- --- --- --- --- --- --t tat aag --- --- --- --- taa gtt agt taa gtc ata agt tag tt- --t tat aag --- --- --- --- taa gtt agt taa gtc ata agt tag tt- --t tat aag tta gtt tat agt cta aca ctt cta att ttt ata cct tga tct ttt tct cta att atg aaa aag taa atc att atg aag atg gat g-a aat gga cga gat tgt gcg cat cgt taa cga tag tat gtg gta cgt acc taa cgc att tat gga cga cgg tga taa tga agg tca cat ttc tgt caa taa tgt ctg tca tat gta tct cgc att ctt tga tgt gga tat atc atc tca tct gtt taa att agt tat taa aca ctg cga tct gaa taa acg act aaa atg tgg taa ctc tcc att aca ttg cta tac gat gaa tac acg att taa tcc atc tgt att aaa gat att gtt acg cca cgg cat gcg taa ctt tga tag caa gga taa aaa agg aca tat tcc tct aca cca cta tct gat tca ttc act atc aat cga taa caa gat ctt tga tat act aac gga ccc cat tga tga ctt tag taa atc atc cga tct att gct gtg tta tct tag ata taa att caa tgg gag ctt aaa cta tta cgt tct gta caa att att gac taa agg atc tga ccc taa ttg cgt cga tga gga tgg act cac ttc tct tca tta cta ctg taa aca cat atc cgc gtt cca cga aag caa tta tta caa gtc aaa gag tca cac taa gat gcg agc tga gaa gcg att cat cta cgc gat aat aga tca tgg agc aaa cat taa cgc ggt tac gaa aat cgg aaa tac gcc gtt aca cac tta cct tca aca gta tac caa aca tag tcc tcg tgt ggt gta tgc tct ttt atc tcg agg agc cga tac gag gat acg taa taa tct tga ttg tac acc cat cat gga ata cat aaa gaa cga ttg tgc aac agg tca tat tct cat aat gtt act caa ttg gca cga aca aaa ata cgg gaa att aca aaa gga aga agg aca aca tct act tta tct att cat aaa aca taa tca agg ata tgg aag tcg ctc tct caa tat act acg gta tct act aga tag att cga cat tca gaa aga cga ata cta taa tac aat gac tcc tct tca tac cgc ctt cca gaa ttg caa taa caa tgt tgc ctc ata cct cgt ata cat cgg ata cga cat caa cct tcc gac taa aga cga taa gac agt att cga ctt ggt gtt tga aaa cag aaa cat cat ata caa ggc gga tgt cgt taa tga cat cat cca cca cag act gaa agt atc tct acc tat gat taa atc gtt gtt cta caa gat gtc gga gtt ctc tcc cta cga cga tca cta cgt aaa gaa gat aat agc cta ctg cct att aag gta cga gtc att tgc gga act aca tac taa att ctg ttt aaa cga gga cta taa aag tgt att tat gaa aaa tat atc att cga taa gat aga ttc cat cat cga aaa atg tag tcg tga cat aag tct cct caa aga gat tcg aat ctc aga cac cga ctt gta tac ggt att gag aac aga aga cat ccg gta tca cac cta tct cga agc cat aca ttc aga caa acg cat ttc att tcc cat gta cga cga tct cat aga aca gtg tca tct atc gat gga gca taa aag taa act cgt cga caa agc act caa taa att aga gtc tac cat cga tag tca atc tag act atc gta ttt gcc tcc gga aat tat gcg caa tat cat aac caa gct aag cga cta cca tct aaa cag tat gtt gta cgg aaa gaa cca tta caa ata tta tcc atg ata gaa aga aaa tat tta aaa aat aat cta tat gat tgg aga agt agg aaa caa aca gta aca aga cga cga tta cta c-a tta tta aat cat gag gtc cgt att ata ctc gta tat att gtt tct ctc atg tat aat aat aaa cgg aag aga tat agc acc aca tgc acc atc caa tgg aaa gtg taa aga caa cga ata cag aag ccg taa tct atg ttg tct atc gtg tcc tcc ggg aac tta cgc ttc cag att atg tga tag caa gac taa tac aca atg tac acc gtg tgg ttc gga tac ctt tac atc tca caa taa tca ttt aca ggc ttg tct aag ttg taa cgg aag atg tga tag taa tca ggt aga gac gcg atc gtg taa cac gac tca caa tag aat ctg tga atg ctc tcc agg ata tta ttg tct tct caa agg agc atc agg gtg tag aac atg tat ttc taa aac aaa gtg tgg aat agg ata cgg agt atc cgg ata cac gtc tac cgg aga cgt cat ctg ttc tcc gtg tgg tcc cgg aac ata ttc tca cac cgt ctc ttc cac aga taa atg cga acc cgt cgt aac cag caa tac att taa cta tat cga tgt gga aat taa cct gta tcc agt caa cga cac atc gtg tac tcg gac gac cac tac cgg tct cag cga atc cat ctc aac gtc gga act aac tat tac cat gaa tca taa aga ttg tga tcc agt ctt tcg tgc aga ata ctt ctc tgt cct taa taa tgt agc aac ttc agg att ctt tac agg aga aaa tag ata tca gaa tac ttc aaa gat atg tac tct gaa ttt cga gat taa atg taa caa caa aga ttc atc ttc caa aca gtt aac gaa aac aaa gaa tga tac tat cat gcc gca ttc aga gac ggt aac tct agt ggg cga ctg tct atc tag cgt cga cat cta cat act ata tag taa tac caa tac tca aga cta cga aac gga tac aat ctc tta tca tat ggg taa tgt tct cga tgt caa tag cca tat gcc cgc tag ttg cga tat aca taa act gat cac taa ttc cca gaa tcc cac cca ctt ata gta ag- -tt ttt tta ccc ata aat aat aaa tac aat aat taa ttt ctc gta aaa gta gaa aat ata ttc taa ttt att ata tgg taa gaa agt aga atc atc tag aac agt aat caa tca ata gca atc atg aaa caa tat att gtc ctg gca tgc atg tgc cta gtg gca gct gct atg cct act agt ctt caa caa tct tca tcc tcg tgt act gaa gaa gaa aac aaa cat cat atg gga atc gat gtt att atc aaa gtc aca aag caa gac caa aca ccg acc aat gat aag att tgt caa tcc gta acg gaa gtt aca gag a-- --- --- --- --- --- -cc gaa gat gat gag gta tcc gaa gaa gtt gta aaa gga gat ccc acc act tat tac act atc gtc ggt gcg ggt ctt aac atg aac ttt gga ttc acc aaa tgc cca aag att tca tcc atc tcc gaa tcc tct gat gga aac act gtg aat act aga ttg tcc agc gtg tca ccg gga caa ggt aag gac tct ccc gcg atc acg cgt gaa gaa gct ctg gct atg atc aaa gac tgt gag atg tct atc gac atc aga tgt agc gaa gaa gag aaa gac agt gac atc aag acc cat cca gta ctt ggg tct aac atc tca cat aag aaa gtg agt tac aaa gat atc atc ggt tca acg atc gtt gat aca aaa tgt gtc aag aac cta gag ttt agc gta cgt atc gga gac atg tgt gag gaa tca tct gaa ctt gaa gtc aag gat gga ttc aag tat gtc gac gga tcg gca tct gaa ggt gca acc gat gat act tca ctc atc gat tca aca aaa ctc aaa gca tgt gtc tga atc gat aac tct att cat ctg aaa atg gat gag ttg ggt taa tcg aac gat tca gac acc gca cca cga att aaa aaa g-- -ac cgg gca cta tat tcc ggt ttg caa aac aaa aat att taa cta cat tca caa aaa gtt acc tct cgt tac ttc ttc ttt ctg ttt caa tat gtg ata cga tat gat cac tat tcg tat tct ct- --- --- tgg tct cat aaa aaa gtt tta c-- --- --- aaa aaa aaa a-t att ttt att cta ttt ctc tct tcg at- ggt ctc aca aaa ata tta aac ctc ttt ctg atg tct caa cta ttt cgt aaa cga taa cgt cca aca ata tat tct cgt aga gct tat caa cat cct tat acc aat cta ggt tgt cag aca att gca tca taa aat aat gtt tat aat tta cac gtt aac atc ata taa taa acg tat ata gtt aat att ttt gga ata taa atg atc tgt aaa atc cat gta ggg gac act act cac gtt ttt tct cta gta cat aat ttc aca caa gtt ttt ata cag aca aat taa ttc tcg tcc ata ta- ttt taa aac att gac ttt tgt act aag aaa aat atc ttg act aac cat ctc ttt ctc tct tcg atg ggt ctc aca aaa ata tta aac ctc ttt ctg atg gag tcg taa aaa gtt ttt atc ctt tct ctc ttc gat ggg tct cac aaa aat att aaa cct ctt tct gat gga gtc gta aaa agt ttt tat cct ttc tct ctt cga tgg gtc tca caa aaa tat taa acc tct ttc tga tgg agt cgt aaa aag ttt tta tcc ttt ctc tct tcg atg ggt ctc --- --- --- --- --- --- --- --- --- --- --- --- --- --- --- --- --- --- --- --- --- --- --- --- --- --- --- --- --- --- --- --- --- --- --- --- --- --- --- --- --- --- --- --- --- --- --- --- --- --- --- --- --- --- --- --- --- --- --- --- --- --- --- --- --- --- --- --- --- --- aca aaa ata tta aac ctc ttt ctg atg gtc tct ata aa

>JX878422.1_Monkeypox_virus_isolate_DRC_07-0287_complete_genome

atc gag gac gac ttc aac tat gac agc tcg gtg gcg tct gcc agc gtg cac ata cga atg gca ttt cta aga aaa gtc tac ggt atc ctt gtc tac aat ttc ctt taa caa cgg caa cag ctg cag tat ttt tat act ttg aat gca ttc gga cat tta tac aag gga gtc ctg ttc taa tat tgg cat caa tgt tcg gat cta tag gct tga ttt tcg cat tga ctt tac aca gac ata aac atc ccc tga atc tgt acc tac ttt gtg gat tta cac tgt tag aat ctc taa cgc tgg cct ctg ttg tta ctt tct atg atg cgc gta tcg tta tac aag ctt tca tgt tga cta ctg cag tgt ttc ttg ctc tga cta cat gta ctc tac aat caa aga gag att tca gta aac ttg taa cag gat tgt ttg ctg ctt tct gga ttt taa ttt tgt cag gag tct tga gga taa agt tta aaa tag aat taa taa aga aca tat agg tca -tt ttt taa aca tgg ata gaa acc aag gtt gtt agt taa taa tat aca aga tat ttt ttc tca ctc tga tcc atg taa acc aag gac gag aga cac tct cat tcc tca ttc acg aca cca tta aaa atg gaa att aaa gcc ctc tat taa gca cag acg gct aca ggt cta cca tca ggt taa cct tcg tct acc ttc aca atg gcc ttt cct tgt gcc cag ttc agt ccc tgt cat tgc cac gct act aag gac tcc ctg aat acc gtg act gac gtc aga cat tgt ctg act gaa tac atc ctg tgg gtt tct cat aga tgg acc cat aga gaa agc gca ggg cct ctc tac agg ctt ctc atc tct ttc aga att gat gca atg gag cta ttt ggt agc gag ttg aag gag ttc tcg gat tca ctt ccg tgg gac aat atc gac aat tgc gtg gag atc att aaa tgt ttc atc aga aat gac tcc atg aaa acc gcc aaa gaa ctt tgt gca ata att gga ctt tgt act caa tca gct att gtc act gga aga gtc ttc aat gat aag tat atc gac ata cta ctt atg ctg cga aag att ctg aat gag aac gac tat ctc acc ctc ttg gat cat atc ctc act gct aaa tac taa atc tcc ttc atg ctc tct cac taa tac tct tac tca cta cac ttt tta tca tct tat gat gaa tga ttg cct tca tca -tt ttt cgt gga ata taa tat agg aat aat tag cac cag aat agc tat gga tat ctc gtt aag aat att ctc gat aag aga cat aat gta gac ata gtt att ata tcc ttc tta gat aag tgt tac gct act gga aag ttt cca tcg tta tta tta cgt gaa gat gat atc att aaa cca aca ttg aga ttg gct ctt atg tta gct gga ttg aat tac tgt aat aaa tgc atc gag tat aga ggg ata tag caa ttc tcg ata ata gtc atg caa tat ttg aat gag act gat aat tta ggt aat aca gta ct- -ac aca cac ata tct ttc tag att ata tat cgt taa aaa tct gta aga ggt ata ttt ctc ata agt atc cac tgt gta ata tta tta atg gat ata tag ata aca caa tag gga cta ata gta ttg taa aag ata taa tcg act att t-- gta cat atc cag ata tct ata ttc cta cta gtt tgc tgc gta gtt gca tca ttg ata tgc atg -at tta tca gga ttc aga gat gaa tta cta agt aaa cta caa tcc cac aat aag taa gaa tca aat atc aaa aac tca ctt ttg att ttt cta gtc tta agt aat aca tat att tat taa tag acc tat gaa at- aaa aaa gtt aac aat gga ttc gcg tat agc tat tta cgt att agt atc ggc atc tct ttt gta tct tgt taa ttg tca caa act agt aca tta ctt caa tct gaa aat aaa tgg aag tga tat aac taa tac agc aga tat att act gga caa tta tcc aat tat gac ctt tga tgg aaa gga tat tta tcc atc tat cgc gtt cat ggt cgg taa taa act ttt cct aga tct tta taa aaa tat ctt tgt aga att ttt cag act att tcg agt atc tgt aag tag tca ata cga gga att aga ata tta tta ttc atg tga tta tac taa caa ccg tcc tac aat taa aca aca tta ctt tta taa cgg cga aga ata tac tga aat tga tag atc gaa aaa agc cac taa taa aaa cag ttg gtt aat tac ttc agg ctt tag act aca aaa atg gtt cga tag cga aga ttg tat aat tta tct cag atc ttt agt tag aag aat gga aga cag taa caa aaa cag taa aaa aac tta gta ctt aga tat cg- aaa aaa aat ata ttt ttg tag act ctt gag aat aga agg aaa aca tgt aca taa tta taa aaa atg aaa atc aat ggc gaa taa gac agt gcg att cgc gcc atg gag tcg gca gat ttc atg gct gtc gat gag cag ttt cac gac gac ctc gat ctt tgg tca tta tct ttg gta gat gac tat aaa aaa cat gga tta ggt gtt gac tgt tat gtt cta gaa cca gtt gtt gac agg aaa ata ttt gat aga ttt ctc ctt gaa cca att tgt gat cct gta gat gtt ctg tat gat tat ttt agg att cat aga gat aat att gat cag tat ata gta gat aga ctg ttt gca tat att aca tat aaa gat att ata tct gca tta gtg tca aag aat tat atg gaa gat att ttc tct ata att att aag aat tgt aat tct gta caa gat ctc tta ctt tac tat cta tct aat gca tat gta gaa ata gac att gtt gat ttt atg gta gat cat ggg gct gta ata tat aaa ata gaa tgc ttg aat gcc tat ttt agg gga ata tgt aaa aag gaa agt agt gtt gtt gag ttt att ttg aat tgt ggt atc cca gat gaa aat gat gtt aaa tta gat cta tat aaa ata att cag tat act agg gga ttc ctt gta gat gaa ccc aca gta tta gaa att tat aag ctt tgt atc cca tat att gaa gat atc aat caa cta gat gct ggt gga agg acc ttg ctt tat cgc gct atc tat gca ggt tat ata gat tta gta tca tgg cta tta gaa aat gga gca aat gtc aac gca gta atg agt aat gga tat aca tgt ctt gac gtg gcc gtg gat agg gga tct gtc atc gcc cgt agg gaa gca cat ctt aaa ata tta gaa ata ttg ctt aga gaa cca ttg tct att gac tgt ata aaa tta gct ata ctt aat aat aca att gaa aac cat gat gtg ata aag ctc tgt atc aag tat ttt atg atg gta gat tat tca ctt tgt aat gtg tat gca tca tca ctc ttt gat tat ata att gat tgt aaa caa gaa ttg gag tac att agg cag atg aaa att cat aat aca acc atg tat gag tta atc tat aat aga gac aaa aac aag cat gct tcc cat att cta cat agg tat tct aaa cat cca gtt ttg aca cag tgt atc act aaa gga ttc aag att tac aca gaa gta acc gag cag gtc act aaa gct cta aac aga cgt gct cta ata gat gag ata ata aac aat gta tca act gat gac aat ctc cta tca aaa ctt cca tta gaa att agg gat cta att gtt tca caa gct gtc ata tag agt tct ctc cac cca cct ttc ttg aaa tga gtt aat --- --- --- --- --- --- --- --- --- --- --- --- --- --- --- --- --- --- --- --- --- --- --- --- --- --- --- --- --- --- --- --- --- --- --- --- --- --- --- --- --- --- --- --- --- --- --- --- --- --- --- --- --- --- --- --- --- --- --- --- --- --- --- --- --- --- --- --- --- --- --- --- --- --- --- --- --- --- --- --- --- --- --- --- --- --- --- --- --- --- --- --- --- --- --- --- --- --- --- --- --- --- --- --- --- --- --- --- --- --- --- --- --- --- --- --- --- --- --- --- --- --- --- --- --- --- --- --- --- --- --- --- --- --- --- --- --- --- --- --- --- --- --- --- --- --- --- --- --- --- --- --- --- -ag tca taa gtt agt taa gtc ata agt tag tt- --t tat aag tta gtt tat agt cta aca ctt cta att ttt ata cct tga tct ttt tct cta att atg aaa aag taa atc att atg aag atg gat gaa aat gga cga gat tgt gcg cat cgt taa cga tag tat gtg gta cgt acc taa cgc att tat gga cga cgg tga taa tga agg tca cat ttc tgt caa taa tgt ctg tca tat gta tct cgc att ctt tga tgt gga tat atc atc tca tct gtt taa att agt tat taa aca ctg cga tct gaa taa acg act aaa atg tgg taa ctc tcc att aca ttg cta tac gat gaa tac acg att taa tcc atc tgt att aaa gat att gtt acg cca cgg cat gcg taa ctt tga tag caa gga taa aaa agg aca tat tcc tct aca cca cta tct gat tca ttc act atc aat cga taa caa gat ctt tga tat act aac gga ccc cat tga tga ctt tag taa atc atc cga tct att gct gtg tta tct tag ata taa att caa tgg gag ctt aaa cta tta cgt tct gta caa att att gac taa agg atc tga ccc taa ttg cgt cga tga gga tgg act cac ttc tct tca tta cta ctg taa aca cat atc cgt gtt cca cga aag caa tta tta caa gtc aaa gag tca cac taa gat gcg agc tga gaa gcg att cat cta cgc gat aat aga tca tgg agc aaa cat taa cgc ggt tac gaa aat cgg aaa tac gcc gtt aca cac tta cct tca aca gta tac caa aca tag tcc tcg tgt ggt gta tgc tct ttt atc tcg agg agc cga tac gag gat acg taa taa tct tga ttg tac acc cat cat gga ata cat aaa gaa cga ttg tgc aac agg tca tat tct cat aat gtt act caa ttg gca cga aca aaa ata cgg gaa att aca aaa gga aga agg aca aca tct act tta tct att cat aaa aca taa tca agg ata tgg aag tcg ctc tct caa tat act acg gta tct act aga tag att cga cat tca gaa aga cga ata cta taa tac aat gac tcc tct tca tac cgc ctt cca gaa ttg caa taa caa tgt tgc ctc ata cct cgt ata cat cgg ata cga cat caa cct tcc gac taa aga cga taa gac agt att cga ctt ggt gtt tga aaa cag aaa cat cat ata caa ggc gga tgt cgt taa tga cat cat cca cca cag act gaa agt atc tct acc tat gat taa atc gtt gtt cta caa gat gtc gga gtt ctc tcc cta cga cga tca cta cgt aaa gaa gat aat agc cta ctg cct att aag gga cga gtc att tgc gga act aca tac taa att ctg ttt aaa cga gga cta taa aag tgt att tat gaa aaa tat atc att cga taa gat aga ttc cat cat cga aaa atg tag tcg tga cat aag tct cct caa aga gat tcg aat ctc aga cac caa ctt gta tac ggt att gag aac aga aga cat ccg gta tca cac cta tct cga agc cat aca ttc aga caa acg cat ttc att tcc cat gta cga cga tct cat aga aca gtg tca tct atc gat gga gca taa aag taa act cgt cga caa agc act caa taa att aga gtc tac cat cga tag tca atc tag act atc gta ttt gcc tcc gga aat tat gcg caa tat cat aac caa gct aag cga cta cca tct aaa cag tat gtt gta cgg aaa gaa cca tta caa ata tta tcc atg ata gaa aga aaa tat tta aaa aat aat cta tat gat tgg aga agt agg aaa caa aca gta aca aga cga cga tta cta c-a tta tta aat cat gag gtc cgt att ata ctc gta tat att gtt tct ctc atg tat aat aat aaa cgg aag aga tat agc acc aca tgc acc atc caa tgg aaa atg taa aga caa cga ata cag aag ccg taa tct atg ttg tct atc gtg tcc tcc ggg aac tta cgc ttc cag att atg tga tag caa gac taa tac aca atg tac gcc gtg tgg ttc gga tac ctt tac atc tca caa taa tca ttt aca ggc ttg tct aag ttg taa cgg aag atg tga tag taa tca ggt aga gac gcg atc gtg taa cac gac tca caa tag aat ctg tga atg ctc tcc agg ata tta ttg tct tct caa agg atc atc agg gtg tag aac atg tat ttc taa aac aaa gtg tgg aat agg ata cgg agt atc cgg ata cac gtc tac cgg aga cgt cat ctg ttc tcc gtg tgg tcc cgg aac ata ttc tca cac cgt ctc ttc cac aga taa atg cga acc --- cgt aac cag caa tac att taa cta tat cga tgt gga aat taa cct gta tcc agt caa cga cac atc gtg tac tcg gac gac cac tac cgg tct cag cga atc cat ctc aac gtc gga act aac tat tac cat gaa tca taa aga ttg tga tcc cgt ctt tcg tgc aga ata ctt ctc tgt cct taa taa tgt agc aac ttc agg att ctt tac agg aga aaa tag ata tca gaa tac ttc aaa gat atg tac tct gaa ttt cga gat taa atg taa caa caa aga ttc atc ttc caa aca gtt aac gaa aac aaa gaa tga tac tat cat gcc gca ttc aga gac ggt aac tct agt ggg cga ctg tct atc tag cgt cga cat cta cat act ata tag taa tac caa tac tca aga cta cga aac gga tac aat ctc tta tca tat ggg taa tgt tct cga tgt caa tag cca tat gcc cgc tag ttg tga tat aca taa act gat cac taa ttc cca gaa tcc cac cca ctt ata gta ag- ttt ttt tta ccc ata aat aat aaa tac aat aat taa ttt ctc gta aaa gta gaa aat ata ttc taa ttt att ata tgg taa gaa agt aga atc atc tag aac agt aat caa tca ata gca atc atg aaa caa tat att gtc ctg gca tgc atg tgc cta gtg gca gct gct atg cct act agt ctt caa caa tcc tca tcc tcg tgt act gaa gaa gaa aac aaa cat cat atg gga atc gat gtt att atc aaa gtc aca aag caa gac caa aca ccg acc aat gat aag att tgt caa tcc gta acg gaa gtt aca gag acc gaa gat gat gag gta tcc gaa gat gat gag gta tcc gaa gaa gtt gta aaa gga gat ccc acc act tat tac act atc gtc ggc gcg ggt ctt aac atg aac ttt gga ttc acc aaa tgc cca aag att tca tcc atc tcc gaa tcc tct gat gga aac act gtg aat act aga ttg tcc agc gtg tca cca gga caa ggt aag gac tct ccc gcg atc acg cgt gaa gaa gct ctg gct atg atc aaa gac tgt gag atg tct atc gac atc aga tgt agc gaa gaa gag aaa gac agc gac atc aag acc cat cca gta ctt ggg tct aac atc tca cat aag aaa gtg agt tac aaa gat atc atc ggt tca acg atc gtt gat aca aaa tgt gtt aag aac cta gag ttt agc gta cgt atc gga gac atg tgt gag gaa tca tct gaa ctt gaa gtc aag gat gga ttc aag tat gtc gac gga tcg gca tct gaa ggt gca acc gat gat act tca ctc atc gat tca aca aaa ctc aaa gca tgt gtc tga atc gat aac tct att cat ctg aaa atg gat gag ttg ggt taa tca aac gat tca gac acc gca cca cga att aaa aaa gtg tac cgg gta cta tat tcc ggt ttg caa aac aaa aat gtt taa cta cat tca caa aaa gtt acc tct cgt tac ttc ttc ttt ctg ttt caa tat gtg ata cga tat gat cac tat tcg tat tct ctt tct ctc ttg tct cat aaa aaa gtt tta c-- --- --- --- aaa aaa a-t att ttt att ctc ttt ctc tct tcg at- ggt ctc aca aaa ata tta aac ctc ttt ctg atg tct caa cta ttt cgt aaa cga taa cgt cca aca ata tat tct cgt aga gct tat caa tat cct tat gcc aat --- ggt tgt cag aca att gca tca taa aat aat gtt tat aat tta cac gtt aac atc ata taa taa acg tat ata gtt aat att ttt gga ata taa atg atc tgt aaa atc cat gta ggg gac act gct cac gtt ttt tct cta gta cat aat ttc aca caa gtt ttt ata cag aca aat taa ttc tcg tcc ata ta- ttt taa aac att gac ttt tgt act aag aaa aat atc ttg act aac cat ctc ttt ctc tcg tcg atg ggt ctc aca aaa ata tta aac ctc ttt ctg atg g-- --- --- --- --- --- --- --- --- --- --- --- --- --- --- --- --- --- --- --- --- --- --- --- --- --- --- --- --- --- --- --- --- --- --- --- --- --- --- --- --- --- --- --- --- --- --- agt cgt aaa aag ttt tta tcc ttt ctc tct tcg atg ggt ctc aca aaa ata tta aac ctc ttt ctg atg gag tcg taa aaa gtt ttt atc ctt tct ctc ttc gat ggg tct cac aaa aat att aaa cct ctt tct gat gga gtc gta aaa agt ttt tat cct ttc tct ctt cga tgg gtc tca caa aaa tat taa acc tct ttc tga tgg agt cgt aaa aag ttt tta tcc ttt ctc tcg tcg atg ggt ctc aca aaa ata tta aac ctc ttt ctg atg gtc tct ata aa

>KP849471.1_Monkeypox_virus_isolate_Yambuku_DRC_1985_complete_genome

atc gag gac gac ttc aac tat gac agc tcg gtg gcg tct gcc agc gtg cac ata cga atg gca ttt cta aga aaa gtc tac ggt atc ctt gtc tac aat ttc ctt taa caa cgg caa cag ctg cag tat ttt tat act ttg aat gca ttc gga cat tta tac aag gga gtc ctg ttc taa tat tgg cat caa tgt tcg gat cta tag gct tga ttt tcg cat tga ctt tac aca gac ata aac atc ccc tga atc tgt acc tac ttt gtg gat tta cac tgt tag aat ctc taa cgc tgg cct ctg ttg tta ctt tct atg atg cgc gta tcg tta tac aag ctt tca tgt tga cta ctg cag tgt ttc ttg ctc tga cta cat gta ctc tac aat caa aga gag att tca gta aac ttg taa cag gat tgt ttg ctg ctt tct gga ttt taa ttt tgt cag gag tct tga gga taa agt tta aaa tag aat taa taa aga aca tat agg tca -tt ttt taa aca tgg ata gaa acc aag gtt gtt agt taa taa tat aca aga tat ttt ttc tca ctc tga tcc atg taa acc aag gac gag aga cac tct cat tcc tca ttc acg aca cca tta aaa atg gaa att aaa gcc ctc tat taa gca cag acg gct aca ggt cta cca tca ggt taa cct tcg tct acc ttc aca atg gcc ttt cct tgt gcc cag ttc agt ccc tgt cat tgc cac gct act aag gac tcc ctg aat acc gtg act gac gtc aga cat tgt ctg act gaa tac atc ctg tgg gtt tct cat aga tgg acc cat aga gaa agc gca ggg cct ctc tac agg ctt ctc atc tct ttc aga att gat gca atg gag cta ttt ggt agc gag ttg aag gag ttc tcg gat tca ctt ccg tgg gac aat atc gac aat tgc gtg gag atc att aaa tgt ttc atc aga aat gac tcc atg aaa acc gcc aaa gaa ctt tgt gca ata att gga ctt tgt act caa tca gct att gtc act gga aga gtc ttc aat gat aag tat atc gac ata cta ctt atg ctg cga aag att ctg aat gag aac gac tat ctc acc ctc ttg gat cat atc ctc act gct aaa tac taa atc tcc ttc atg ctc tct cac taa tac tct tac tca cta cac ttt tta tca tct tat gat gaa tga ttg cct tca tca -tt ttt cgt gga ata taa tat agg aat aat tag cac cag aat agc tat gga tat ctc gtt aag aat att ctc gat aag aga cat aat gta gac ata gtt att ata tcc ttc tta gat aag tgt tac gct act gga aag ttt cca tcg tta tta tta cgt gaa gat gat atc att aaa cca aca ttg aga ttg gct ctt atg tta gct gga ttg aat tac tgt aat aaa tgc atc gag tat aga ggg ata tag caa ttc tcg ata ata gtc atg caa tat ttg aat gag act gat aat tta ggt aat aca gta ct- -ac aca cac ata tct ttc tag att ata tat cgt taa aaa tct gta aga ggt ata ttt ctc ata agt atc cac tgt gta ata tta tta atg gat ata tag ata aca caa tag gga cta ata gta ttg taa aag ata taa tcg act att t-- gta cat atc cag ata tct ata ttc cta cta gtt tgc tgc gta gtt gca tca ttg ata tgc atg -at tta tca gga ttc aga gat gaa tta cta agt aaa cta caa tcc cac aat aag taa gaa tca aat atc aaa aac tca ctt ttg att ttt cta gtc tta agt aat aca tat att tat taa tag acc tat gaa at- aaa aaa gtt aac aat gga ttc gcg tat agc tat tta cgt att agt atc ggc atc tct ttt gta tct tgt taa ttg tca caa act agt aca tta ctt caa tct gaa aat aaa tgg aag tga tat aac taa tac agc aga tat att gct gga caa tta tcc aat tat gac ctt tga tgg aaa gga tat tta tcc atc tat cgc gtt cat ggt cgg taa taa act ttt cct aga tct tta taa aaa tat ctt tgt aga att ttt cag act att tcg agt atc tgt aag tag tca ata cga gga att aga ata tta tta ttc atg tga tta tac taa caa ccg tcc tac aat taa aca aca tta ctt tta taa cgg cga aga ata tac tga aat tga tag atc gaa aaa agc cac taa taa aaa cag ttg gtt aat tac ttc agg ctt tag act aca aaa atg gtt cga tag cga aga ttg tat aat tta tct cag atc ttt agt tag aag aat gga aga cag taa caa aaa cag taa aaa aac tta gta ctt aga tat cg- aaa aaa aat ata ttt ttg tag act ctt gag aat aga agg aaa aca tgt aca taa tta taa aaa atg aaa atc aat ggc gaa taa gac agt gcg att cgc gcc atg gag tcg gca gat ttc atg gct gtc gat gag cag ttt cac gac gac ctc gat ctt tgg tca tta tct ttg gta gat gac tat aaa aaa cat gga tta ggt gtt gac tgt tat gtt cta gaa cca gtt gtt gac agg aaa ata ttt gat aga ttt ctc ctt gaa cca att tgt gat cct gta gat gtt ctg tat gat tat ttt agg att cat aga gat aat att gat cag tat ata gta gat aga ctg ttt gca tat att aca tat aaa gat att ata tct gca tta gtg tca aag aat tat atg gaa gat att ttc tct ata att att aag aat tgt aat tct gta caa gat ctc tta ctt tac tat cta tct aat gca tat gta gaa ata gac att gtt gat ttt atg gta gat cat ggg gct gta ata tat aaa ata gaa tgc ttg aat gcc tat ttt agg gga ata tgt aaa aag gaa agt agt gtt gtt gag ttt att ttg aat tgt ggt atc cca gat gaa aat gat gtt aaa tta gat cta tat aaa ata att cag tat act agg gga ttc ctt gta gat gaa ccc aca gta tta gaa att tat aag ctt tgt atc cca tat att gaa gat atc aat caa cta gat gct ggt gga agg acc ttg ctt tat cgc gct atc tat gca ggt tat ata gat tta gta tca tgg cta tta gaa aat gga gca aat gtc aac gca gta atg agt aat gga tat aca tgt ctt gac gtg gcc gtg gat agg gga tct gtc atc gcc cgt agg gaa gca cat ctt aaa ata tta gaa ata ttg ctt aga gaa cca ttg tct att gac tgt ata aaa tta gct ata ctt aat aat aca att gaa aac cat gat gtg ata aag ctc tgt atc aag tat ttt atg atg gta gat tat tca ctt tgt aat gtg tat gca tca tca ctc ttt gat tat ata att gat tgt aaa caa gaa ttg gag tac att agg cag atg aaa att cat aat aca acc atg tat gag tta atc tat aat aga gac aaa aac aag cat gct tcc cat att cta cat agg tat tct aaa cat cca gtt ttg aca cag tgt atc act aaa gga ttc aag att tac aca gaa gta acc gag cag gtc act aaa gct cta aac aga cgt gct cta ata gat gag ata ata aac aat gta tca act gat gac aat ctc cta tca aaa ctt cca tta gaa att agg gat cta att gtt tca caa gct gtc ata tag agt tct ctc cac cca cct ttc ttg aaa tga gtt aat --- --- --- --- --- --- --- --- --- --- --- --- --- --- --- --- --- --- --- --- --- --- --- --- --- --- --- --- --- --- --- --- --- --- --- --- --- --- --- --- --- --- --- --- --- --- --- --- --- --- --- --- --- --- --- --- --- --- --- --- --- --- --- --- --- --- --- --- --- --- --- --- --- --- --- --- --- --- --- --- --- --- --- --- --- --- --- --- --- --- --- --- --- --- --- --- --- --- --- --- --- --- --- --- --- --- --- --- --- --- --- --- --- --- --- --- --- --- --- --- --- --- --- --- --- --- --- --- --- --- --- --- --- --- --- --- --- --- --- --- --- --- --- --- --- --- --- --- --- --- --- --- --- -ag tca taa gtt agt taa gtc ata agt tag tt- --t tat aag tta gtt tat agt cta aca ctt cta att ttt ata cct tga tct ttt tct cta att atg aaa aag taa atc att atg aag atg gat gaa aat gga cga gat tgt gcg cat cgt taa cga tag tat gtg gta cgt acc taa cgc att tat gga cga cgg tga taa tga agg tca cat ttc tgt caa taa tgt ctg tca tat gta tct cgc att ctt tga tgt gga tat atc atc tca tct gtt taa att agt tat taa aca ctg cga tct gaa taa acg act aaa atg tgg taa ctc tcc att aca ttg cta tac gat gaa tac acg att taa tcc atc tgt att aaa gat att gtt acg cca cgg cat gcg taa ctt tga tag caa gga taa aaa agg aca tat tcc tct aca cca cta tct gat tca ttc act atc aat cga taa caa gat ctt tga tat act aac gga ccc cat tga tga ctt tag taa atc atc cga tct att gct gtg tta tct tag ata taa att caa tgg gag ctt aaa cta tta cgt tct gta caa att att gac taa agg atc tga ccc taa ttg cgt cga tga gga tgg act cac ttc tct tca tta cta ctg taa aca cat atc cgt gtt cca cga aag caa tta tta caa gtc aaa gag tca cac taa gat gcg agc tga gaa gcg att cat tta cgc gat aat aga tca tgg agc aaa cat taa cgc ggt tac gaa aat cgg aaa tac gcc gtt aca cac tta cct tca aca gta tac caa aca tag tcc tcg tgt ggt gta tgc tct ttt atc tcg agg agc cga tac gag gat acg taa taa tct tga ttg tac acc cat cat gga ata cat aaa gaa cga ttg tgc aac agg tca tat tct cat aat gtt act caa ttg gca cga aca aaa ata cgg gaa att aca aaa gga aga agg aca aca tct act tta tct att cat aaa aca taa tca agg ata tgg aag tcg ctc tct caa tat act acg gta tct act aga tag att cga cat tca gaa aga cga ata cta taa tac aat gac tcc tct tca tac cgc ctt cca gaa ttg caa taa caa tgt tgc ctc ata cct cgt ata cat cgg ata cga cat caa cct tcc gac taa aga cga taa gac agt att cga ctt ggt gtt tga aaa cag aaa cat cat ata caa ggc gga tgt cgt taa tga cat cat cca cca cag act gaa agt atc tct acc tat gat taa atc gtt gtt cta caa gat gtc gga gtt ctc tcc cta cga cga tca cta cgt aaa gaa gat aat agc cta ctg cct att aag gga cga gtc att tgc gga act aca tac taa att ctg ttt aaa cga gga cta taa aag tgt att tat gaa aaa tat atc att cga taa gat aga ttc cat cat cga aaa atg tag tcg tga cat aag tct cct caa aga gat tcg aat ctc aga cac caa ctt gta tac ggt att gag aac aga aga cat ccg gta tca cac cta tct cga agc cat aca ttc aga caa acg cat ttc att tcc cat gta cga cga tct cat aga aca gtg tca tct atc gat gga gca taa aag taa act cgt cga caa agc act caa taa att aga gtc tac cat cga tag tca atc tag act atc gta ttt gcc tcc gga aat tat gcg caa tat cat aac caa gct aag cga cta cca tct aaa cag tat gtt gta cgg aaa gaa cca tta caa ata tta tcc atg ata gaa aga aaa tat tta aaa aat aat cta tat gat tgg aga agt agg aaa caa aca gta aca aga cga cga tta cta c-a tta tta aat cat gag gtc cgt att ata ctc gta tat att gtt tct ctc atg tat aat aat aaa cgg aag aga tat agc acc aca tgc acc atc caa tgg aaa atg taa aga caa cga ata cag aag ccg taa tct atg ttg tct atc gtg tcc tcc ggg aac tta cgc ttc cag att atg tga tag caa gac taa tac aca atg tac gcc gtg tgg ttc gga tac ctt tac atc tca caa taa tca ttt aca ggc ttg tct aag ttg taa cgg aag atg tga tag taa tca ggt aga gac gcg atc gtg taa cac gac tca caa tag aat ctg tga atg ctc tcc agg ata tta ttg tct tct caa agg atc atc agg gtg tag aac atg tat ttc taa aac aaa gtg tgg aat agg ata cgg agt atc cgg ata cac gtc tac cgg aga cgt cat ctg ttc tcc gtg tgg tcc cgg aac ata ttc tca cac cgt ctc ttc cac aga taa atg cga acc --- cgt aac cag caa tac att taa cta tat cga tgt gga aat taa cct gta tcc agt caa cga cac atc gtg tac tcg gac gac cac tac cgg tct cag cga atc cat ctc aac gtc gga act aac tat tac cat gaa tca taa aga ttg tga tcc cgt ctt tcg tgc aga ata ctt ctc tgt cct taa taa tgt agc aac ttc agg att ctt tac agg aga aaa tag ata tca gaa tac ttc aaa gat atg tac tct gaa ttt cga gat taa atg taa caa caa aga ttc atc ttc caa aca gtt aac gaa aac aaa gaa tga tac tat cat gcc gca ttc aga gac ggt aac tct agt ggg cga ctg tct atc tag cgt cga cat cta cat act ata tag taa tac caa tac tca aga cta cga aac gga tac aat ctc tta tca tat ggg taa tgt tct cga tgt caa tag cca tat gcc cgc tag ttg cga tat aca taa act gat cac taa ttc cca gaa tcc cac cca ctt ata gta ag- ttt ttt tta ccc ata aat aat aaa tac aat aat taa ttt ctc gta aaa gta gaa aat ata ttc taa ttt att ata tgg taa gaa agt aga atc atc tag aac agt aat caa tca ata gca atc atg aaa caa tat att gtc ctg gca tgc atg tgc cta gtg gca gct gct atg cct act agt ctt caa caa tcc tca tcc tcg tgt act gaa gaa gaa aac aaa cat cat atg gga atc gat gtt att atc aaa gtc aca aag caa gac caa aca ccg acc aat gat aag att tgt caa tcc gta acg gaa gtt aca gag a-- --- --- --- --- --- -cc gaa gat gat gag gta tcc gaa gaa gtt gta aaa gga gat ccc acc act tat tac act atc gtc ggc gcg ggt ctt aac atg aac ttt gga ttc acc aaa tgc cca aag att tca tcc atc tcc gaa tcc tct gat gga aac act gtg aat act aga ttg tcc agc gtg tca cca gga caa ggt aag gac tct ccc gcg atc acg cgt gaa gaa gct ctg gct atg atc aaa gac tgt gag atg tct atc gac atc aga tgt agc gaa gaa gag aaa gac agc gac atc aag acc cat cca gta ctt ggg tct aac atc tca cat aag aaa gtg agt tac aaa gat atc atc ggt tca acg atc gtt gat aca aaa tgt gtt aag aac cta gag ttt agc gta cgt atc gga gac atg tgt gag gaa tca tct gaa ctt gaa gtc aag gat gga ttc aag tat gtc gac gga tcg gca tct gaa ggt gca acc gat gat act tca ctc atc gat tca aca aaa ctc aaa gca tgt gtc tga atc gat aac tct att cat ctg aaa atg gat gag ttg ggt taa tca aac gat tca gac acc gca cca cga att aaa aaa gtg tac cgg gta cta tat tcc ggt ttg caa aac aaa aat gtt taa cta cat tca caa aaa gtt acc tct cgt tac ttc ttc ttt ctg ttt caa tat gtg ata cga tat gat cac tat tcg tat tct ctt tct ctc ttg tct cat aaa aaa gtt tta c-- --- --- --- aaa aaa a-t att ttt att ctc ttt ctc tct tcg at- ggt ctc aca aaa ata tta aac ctc ttt ctg atg tct caa cta ttt cgt aaa cga taa cgt cca aca ata tat tct cgt aga gct tat caa tat cct tat gcc aat --- ggt tgt cag aca att gca tca taa aat aat gtt tat aat tta cac gtt aac atc ata taa taa acg tat ata gtt aat att ttt gga ata taa atg atc tgt aaa atc cat gta ggg gac act gct cac gtt ttt tct cta gta cat aat ttc aca caa gtt ttt ata cag aca aat taa ttc tcg tcc ata ta- ttt taa aac att gac ttt tgt act aag aaa aat atc ttg act aac cat ctc ttt ctc tcg tcg atg ggt ctc aca aaa ata tta aac ctc ttt ctg atg g-- --- --- --- --- --- --- --- --- --- --- --- --- --- --- --- --- --- --- --- --- --- --- --- --- --- --- --- --- --- --- --- --- --- --- --- --- --- --- --- --- --- --- --- --- --- --- agt cgt aaa aag ttt tta tcc ttt ctc tct tcg atg ggt ctc --- --- --- --- --- --- --- --- --- --- --- --- --- --- --- --- --- --- --- --- --- --- --- -ac aaa aat att aaa cct ctt tct gat gga gtc gta aaa agt ttt tat cct ttc tct ctt cga tgg gtc tca caa aaa tat taa acc tct ttc tga tgg agt cgt aaa aag ttt tta tcc ttt ctc tct tcg atg ggt ctc aca aaa ata tta aac ctc ttt ctg atg gtc tct ata aa

>DQ011155.1_Monkeypox_virus_strain_Zaire_1979-005_complete_genome

atc gag gac gac ttc aac tat gac agc tcg gtg gcg tct gcc agc gtg cac ata cga atg gca ttt cta aga aaa gtc tac ggt atc ctt gtc tac aat ttc ctt taa caa cgg caa cag ctg cag tat ttt tat act ttg aat gca ttc gga cat tta tac aag gga gtc ctg ttc taa tat tgg cat caa tgt tcg gat cta tag gct tga ttt tcg cat tga ctt tac aca gac ata aac atc ccc tga atc tgt acc tac ttt gtg gat tta cac tgt tag aat ctc taa cgc tgg cct ctg ttg tta ctt tct atg atg cgc gta tcg tta tac aag ctt tca tgt tga cta ctg cag tgt ttc ttg ctc tga cta cat gta ctc tac aat caa aga gag att tca gta aac ttg taa cag gat tgt ttg ctg ctt tct gga ttt taa ttt tgt cag gag tct tga gga taa agt tta aaa tag aat taa taa aga aca tat agg tca -tt ttt taa aca tgg ata gaa acc aag gtt gtt agt taa taa tat aca aga tat ttt ttc tca ctc tga tcc atg taa acc aag gac gag aga cac tct cat tcc tca ttc acg aca cca tta aaa atg gaa att aaa gcc ctc tat taa gca cag acg gct aca ggt cta cca tca ggt taa cct tcg tct acc ttc aca atg gcc ttt cct tgt gcc cag ttc agt ccc tgt cat tgc cac gct act aag gac tcc ctg aat acc gtg act gac gtc aga cat tgt ctg act gaa tac atc ctg tgg gtt tct cat aga tgg acc cat aga gaa agc gca ggg cct ctc tac agg ctt ctc atc tct ttc aga att gat gca atg gag cta ttt ggt agc gag ttg aag gag ttc tcg gat tca ctt ccg tgg gac aat atc gac aat tgc gtg gag atc att aaa tgt ttc atc aga aat gac tcc atg aaa acc gcc aaa gaa ctt tgt gca ata att gga ctt tgt act caa tca gct att gtc act gga aga gtc ttc aat gat aag tat atc gac ata cta ctt atg ctg cga aag att ctg aat gag aac gac tat ctc acc ctc ttg gat cat atc ctc act gct aaa tac taa atc tcc ttc atg ctc tct cac taa tac tct tac tca cta cac ttt tta tca tct tat gat gaa tga ttg cct tca tca -tt ttt cgt gga ata taa tat agg aat aat tag cac cag aat agc tat gga tat ctc gtt aag aat att ctc gat aag aga cat aat gta gac ata gtt att ata tcc ttc tta gat aag tgt tac gct act gga aag ttt cca tcg tta tta tta cgt gaa gat gat atc att aaa cca aca ttg aga ttg gct ctt atg tta gct gga ttg aat tac tgt aat aaa tgc atc gag tat aga ggg ata tag caa ttc tcg ata ata gtc atg caa tat ttg aat gag act gat aat tta ggt aat aca gta ct- -ac aca cac ata tct ttc tag att ata tat cgt taa aaa tct gta aga ggt ata ttt ctc ata agt atc cac tgt gta ata tta tta atg gat ata tag ata aca caa tag gga cta ata gta ttg taa aag ata taa tcg act att t-- gta cat atc cag ata tct ata ttc cta cta gtt tgc tgc gta gtt gca tca ttg ata tgc atg aat tta tca gga ttc aga gat gaa tta cta agt aaa cta caa tcc cac aat aag taa gaa tca aat atc aaa aac tca ctt ttg att ttt cta gtc tta agt aat aca tat att tat taa tag acc tat gaa at- aaa aaa gtt aac aat gga ttc gcg tat agc tat tta cgt att agt atc ggc atc tct ttt gta tct tgt taa ttg tca caa act agt aca tta ctt caa tct gaa aat aaa tgg aag tga tat aac taa tac agc aga tat att gct gga caa tta tcc aat tat gac ctt tga tgg aaa gga tat tta tcc atc tat cgc gtt cat ggt cgg taa taa act ttt cct aga tct tta taa aaa tat ctt tgt aga att ttt cag act att tcg agt atc tgt aag tag tca ata cga gga att aga ata tta tta ttc atg tga tta tac taa caa ccg tcc tac aat taa aca aca tta ctt tta taa cgg cga aga ata tac tga aat tga tag atc gaa aaa agc cac taa taa aaa cag ttg gtt aat tac ttc agg ctt tag act aca aaa atg gtt cga tag cga aga ttg tat aat tta tct cag atc ttt agt tag aag aat gga aga cag taa caa aaa cag taa aaa aac tta gta ctt aga tat cg- aaa aaa aat ata ttt ttg tag act ctt gag aat aga agg aaa aca tgt aca taa tta taa aaa atg aaa atc aat ggc gaa taa gac agt gcg att cgc gcc atg gag tcg gta gat ttc atg gct gtc gat gag cag ttt cac gac gac ctc gat ctt tgg tca tta tct ttg gta gat gac tat aaa aaa cat gga tta ggt gtt gac tgt tat gtt cta gaa cca gtt gtt gac agg aaa ata ttt gat aga ttt ctc ctt gaa cca att tgt gat cct gta gat gtt ctg tat gat tat ttt agg att cat aga gat aat att gat cag tat ata gta gat aga ctg ttt gca tat att aca tat aaa gat att ata tct gca tta gtg tca aag aat tat atg gaa gat att ttc tct ata att att aag aat tgt aat tct gta caa gat ctc tta ctt tac tat cta tct aat gca tat gta gaa ata gac att gtt gat ttt atg gta gat cat ggg gct gta ata tat aaa ata gaa tgc ttg aat gcc tat ttt agg gga ata tgt aaa aag gaa agt agt gtt gtt gag ttt att ttg aat tgt ggt atc cca gat gaa aat gat gtt aaa tta gat cta tat aaa ata att cag tat act agg gga ttc ctt gta gat gaa ccc aca gta tta gaa att tat aag ctt tgt atc cca tat att gaa gat atc aat caa cta gat gct ggt gga agg acc ttg ctt tat cgc gct atc tat gca ggt tat ata gat tta gta tca tgg cta tta gaa aat gga gca aat gtc aac gca gta atg agt aat gga tat aca tgt ctt gac gtg gcc gtg gat agg gga tct gtc atc gcc cgt agg gaa gca cat ctt aaa ata tta gaa ata ttg ctt aga gaa cca ttg tct att gac tgt ata aaa tta gct ata ctt aat aat aca att gaa aac cat gat gtg ata aag ctc tgt atc aag tat ttt atg atg gta gat tat tca ctt tgt aat gtg tat gca tca tca ctc ttt gat tat ata att gat tgt aaa caa gaa ttg gag tac att agg cag atg aaa att cat aat aca acc atg tat gag tta atc tat aat aga gac aaa aac aag cat gct tcc cat att cta cat agg tat tct aaa cat cca gtt ttg aca cag tgt atc act aaa gga ttc aag att tac aca gaa gta acc gag cag gtc act aaa gct cta aac aga cgt gct cta ata gat gag ata ata aac aat gta tca act gat gac aat ctc cta tca aaa ctt cca tta gaa att agg gat cta att gtt tca caa gct gtc ata tag agt tct ctc cac cca cct ttc ttg aaa tga gtt aat --- --- --- --- --- --- --- --- --- --- --- --- --- --- --- --- --- --- --- --- --- --- --- --- --- --- --- --- --- --- --- --- --- --- --- --- --- --- --- --- --- --- --- --- --- --- --- --- --- --- --- --- --- --- --- --- --- --- --- --- --- --- --- --- --- --- --- --- --- --- --- --- --- --- --- --- --- --- --- --- --- --- --- --- --- --- --- --- --- --- --- --- --- --- --- --- --- --- --- --- --- --- --- --- --- --- --- --- --- --- --- --- --- --- --- --- --- --- --- --- --- --- --- --- --- --- --- --- --- --- --- --- --- --- --- --- --- --- --- --- --- --- --- --- --- --- --- --- --- --- --- --- --- -ag tca taa gtt agt taa gtc ata agt tag tt- --t tat aag tta gtt tat agt cta aca ctt cta att ttt ata cct tga tct ttt tct cta att atg aaa aag taa atc att atg aag atg gat gaa aat gga cga gat tgt gcg cat cgt taa cga tag tat gtg gta cgt acc taa cgc att tat gga cga cgg tga taa tga agg tca cat ttc tgt caa taa tgt ctg tca tat gta tct cgc att ctt tga tgt gga tat atc atc tca tct gtt taa att agt tat taa aca ctg cga tct gaa taa acg act aaa atg tgg taa ctc tcc att aca ttg cta tac gat gaa tac acg att taa tcc atc tgt att aaa gat att gtt acg cca cgg cat gcg taa ctt tga tag caa gga taa aaa agg aca tat tcc tct aca cca cta tct gat tca ttc act atc aat cga taa caa gat ctt tga tat act aac gga ccc cat tga tga ctt tag taa atc atc cga tct att gct gtg tta tct tag ata taa att caa tgg gag ctt aaa cta tta cgt tct gta caa att att gac taa agg atc tga ccc taa ttg cgt cga tga gga tgg act cac ttc tct tca tta cta ctg taa aca cat atc cgt gtt cca cga aag caa tta tta caa gtc aaa gag tca cac taa gat gcg agc tga gaa gcg att cat cta cgc gat aat aga tca tgg agc aaa cat taa cgc ggt tac gaa aat cgg aaa tac gcc gtt aca cac tta cct tca aca gta tac caa aca tag tcc tcg tgt ggt gta tgc tct ttt atc tcg agg agc cga tac gag gat acg taa taa tct tga ttg tac acc cat cat gga ata cat aaa gaa cga ttg tgc aac agg tca tat tct cat aat gtt act caa ttg gca cga aca aaa ata cgg gaa att aca aaa gga aga agg aca aca tct act tta tct att cat aaa aca taa tca agg ata tgg aag tcg ctc tct caa tat act acg gta tct act aga tag att cga cat tca gaa aga cga ata cta taa tac aat gac tcc tct tca tac cgc ctt cca gaa ttg caa taa caa tgt tgc ctc ata cct cgt ata cat cgg ata cga cat caa cct tcc gac taa aga cga taa gac agt att cga ctt ggt gtt tga aaa cag aaa cat cat ata caa ggc gga tgt cgt taa tga cat cat cca cca cag act gaa agt atc tct acc tat gat taa atc gtt gtt cta caa gat gtc gga gtt ctc tcc cta cga cga tca cta cgt aaa gaa gat aat agc cta ctg cct att aag gga cga gtc att tgc gga act aca tac taa att ctg ttt aaa cga gga cta taa aag tgt att tat gaa aaa tat atc att cga taa gat aga ttc cat cat cga aaa atg tag tcg tga cat aag tct cct caa aga gat tcg aat ctc aga cac caa ctt gta tac ggt att gag aac aga aga cat ccg gta tca cac cta tct cga agc cat aca ttc aga caa acg cat ttc att tcc cat gta cga cga tct cat aga aca gtg tca tct atc gat gga gca taa aag taa act cgt cga caa agc act caa taa att aga gtc tac cat cga tag tca atc tag act atc gta ttt gcc tcc gga aat tat gcg caa tat cat aac caa gct aag cga cta cca tct aaa cag tat gtt gta cgg aaa gaa cca tta caa ata tta tcc atg ata gaa aga aaa tat tta aaa aat aat cta tat gat tgg aga agt agg aaa caa aca gta aca aga cga cga tta cta c-a tta tta aat cat gag gtc cgt att ata ctc gta tat att gtt tct ctc atg tat aat aat aaa cgg aag aga tat agc acc aca tgc acc atc caa tgg aaa atg taa aga caa cga ata cag aag ccg taa tct atg ttg tct atc gtg tcc tcc ggg aac tta cgc ttc cag att atg tga tag caa gac taa tac aca atg tac gcc gtg tgg ttc gga tac ctt tac atc tca caa taa tca ttt aca ggc ttg tct aag ttg taa cgg aag atg tga tag taa tca ggt aga gac gcg atc gtg taa cac gac tca caa tag aat ctg tga atg ctc tcc agg ata tta ttg tct tct caa agg atc atc agg gtg tag aac atg tat ttc taa aac aaa gtg tgg aat agg ata cgg agt atc cgg ata cac gtc tac cgg aga cgt cat ctg ttc tcc gtg tgg tcc cgg aac ata ttc tca cac cgt ctc ttc cac aga taa atg cga acc --- cgt aac cag caa tac att taa cta tat cga tgt gga aat taa cct gta tcc agt caa cga cac atc gtg tac tcg gac gac cac tac cgg tct cag cga atc cat ctc aac gtc gga act aac tat tac cat gaa tca taa aga ttg tga tcc cgt ctt tcg tgc aga ata ctt ctc tgt cct taa taa tgt agc aac ttc agg att ctt tac agg aga aaa tag ata tca gaa tac ttc aaa gat atg tac tct gaa ttt cga gat taa atg taa caa caa aga ttc atc ttc caa aca gtt aac gaa aac aaa gaa tga tac tat cat gcc gca ttc aga gac ggt aac tct agt ggg cga ctg tct atc tag cgt cga cat cta cat act ata tag taa tac caa tac tca aga cta cga aac gga tac aat ctc tta tca tat ggg taa tgt tct cga tgt caa tag cca tat gcc cgc tag ttg cga tat aca taa act gat cac taa ttc cca gaa tcc cac cca ctt ata gta ag- ttt ttt tta ccc ata aat aat aaa tac aat aat taa ttt ctc gta aaa gta gaa aat ata ttc taa ttt att ata tgg taa gaa agt aga atc atc tag aac agt aat caa tca ata gca atc atg aaa caa tat att gtc ctg gca tgc atg tgc cta gtg gca gct gct atg cct act agt ctt caa caa tcc tca tcc tcg tgt act gaa gaa gaa aac aaa cat cat atg gga atc gat gtt att atc aaa gtc aca aag caa gac caa aca ccg acc aat gat aag att tgt caa tcc gta acg gaa gtt aca gag a-- --- --- --- --- --- -cc gaa gat gat gag gta tcc gaa gaa gtt gta aaa gga gat ccc acc act tat tac aat atc gtc ggc gcg ggt ctt aac atg aac ttt gga ttc acc aaa tgc cca aag att tca tcc atc tcc gaa tcc tct gat gga aac act gtg aat act aga ttg tcc agc gtg tca cca gga caa ggt aag gac tct ccc gcg atc acg cgt gaa gaa gct ctg gct atg atc aaa gac tgt gag atg tct atc gac atc aga tgt agc gaa gaa gag aaa gac agc gac atc aag acc cat cca gta ctt ggg tct aac atc tca cat aag aaa gtg agt tac aaa gat atc atc ggt tca acg atc gtt gat aca aaa tgt gtt aag aac cta gag ttt agc gta cgt atc gga gac atg tgt gag gaa tca tct gaa ctt gaa gtc aag gat gga ttc aag tat gtc gac gga tcg gca tct gaa ggt gca acc gat gat act tca ctc atc gat tca aca aaa ctc aaa gca tgt gtc tga atc gat aac tct att cat ctg aaa atg gat gag ttg ggt taa tcg aac gat tca gac acc gca cca cga att aaa aaa gtg tac cgg gta cta tat tcc ggt ttg caa aac aaa aat gtt taa cta cat tca caa aaa gtt acc tct cgt tac ttc ttc ttt ctg ttt caa tat gtg ata cga tat gat cac tat tcg tat tct ctt tct ctc ttg tct cat aaa aaa gtt tta c-- --- --- --- aaa aaa a-t att ttt att ctc ttt ctc tct tcg at- ggt ctc aca aaa ata tta aac ctc ttt ctg atg tct caa cta ttt cgt aaa cga taa cgt cca aca ata tat tct cgt aga gct tat caa tat cct tat gcc aat --- ggt tgt cag aca att gca tca taa aat aat gtt tat aat tta cac gtt aac atc ata taa taa acg tat ata gtt aat att ttt gga ata taa atg atc tgt aaa atc cat gta ggg gac act gct cac gtt ttt tct cta gta cat aat ttc aca caa gtt ttt ata cag aca aat taa ttc tcg tcc ata ta- ttt taa aac att gac ttt tgt act aag aaa aat atc ttg act aac cat ctc ttt ctc tct tcg atg ggt ctc aca aaa ata tta aac ctc ttt ctg atg g-- --- --- --- --- --- --- --- --- --- --- --- --- --- --- --- --- --- --- --- --- --- --- --- --- --- --- --- --- --- --- --- --- --- --- --- --- --- --- --- --- --- --- --- --- --- --- agt cgt aaa aag ttt tta tcc ttt ctc tct tcg atg ggt ctc --- --- --- --- --- --- --- --- --- --- --- --- --- --- --- --- --- --- --- --- --- --- --- --- --- --- --- --- --- --- --- --- --- --- --- --- --- --- --- --- --- --- --- --- --- --- --a caa aaa tat taa acc tct ttc tga tgg agt cgt aaa aag ttt tta tcc ttt ctc tct tcg atg ggt ctc aca aaa ata tta aac ctc ttt ctg atg gtc tct ata aa

>KP849469.1_Monkeypox_virus_isolate_Boende_DRC_2008_complete_genome

atc gag gac gac ttc aac tat gac agc tcg gtg gcg tct gcc agc gtg cac ata cga atg gca ttt cta aga aaa gtc tac ggt atc ctt gtc tac aat ttc ctt taa caa cgg caa cag ctg cag tat ttt tat act ttg aat gca ttc gga cat tta tac aag gga gtc ctg ttc taa tat tgg cat caa tgt tcg gat cta tag gct tga ttt tcg cat tga ctt tac aca gac ata aac atc ccc tga atc tgt acc tac ttt gtg gat tta cac tgt tag aat ctc taa cgc tgg cct ctg ttg tta ctt tct atg atg cgc gta tcg tta tac aag ctt tca tgt tga cta ctg cag tgt ttc ttg ctc tga cta cat gta ctc tac aat caa aga gag att tca gta aac ttg taa cag gat tgt ttg ctg ctt tct gga ttt taa ttt tgt cag gag tct tga gga taa agt tta aaa tag aat taa taa aga aca tat agg tca -tt ttt taa aca tgg ata gaa acc aag gtt gtt agt taa t-a tat aca aga tat ttt ttc tca ctc tga tcc atg taa acc aag gac gag aga cac tct cat tcc tca ttc acg aca cca tta aaa atg gaa att aaa gcc ctc tat taa gca cag acg gct aca ggt cta cca tca ggt taa cct tcg tct acc ttc aca atg gcc ttt cct tgt gcc cag ttc agt ccc tgt cat tgc cac gct act aag gac tcc ctg aat acc gtg act gac gtc aga cat tgt ctg act gaa tac atc ctg tgg gtt tct cat aga tgg acc cat aga gaa agc gca ggg cct ctc tac agg ctt ctc atc tct ttc aga att gat gca atg gag cta ttt ggt agc gag ttg aag gag ttc tcg gat tca ctt ccg tgg gac aat atc gac aat tgc gtg gag atc att aaa tgt ttc atc aga aat gac tcc atg aaa acc gcc aaa gaa ctt tgt gca ata att gga ctt tgt act caa tca gct att gtc act gga aga gtc ttc aat gat aag tat atc gac ata cta ctt ata ctg cga aag att ctg aat gag aac gac tat ctc acc ctc ttg gat cat atc ctc act gct aaa tac taa atc tcc ttc atg ctc tct cac taa tac tct tac tca cta cac ttt tta tca tct tat gat gaa tga ttg cct tca tca -tt ttt cgt gga ata taa tat agg aat aat tag cac cag aat agc tat gga tat ctc gtt aag aat att ctc gat aag aga cat aat gta gac ata gtt att ata tcc ttc tta gat aag tgt tac gct act gga aag ttt cca tcg tta tta tta cgt gaa gat gat atc att aaa cca aca ttg aga ttg gct ctt atg tta gct gga ttg aat tac tgt aat aaa tgc atc gag tat aga ggg ata tag caa ttc tcg ata ata gtc atg caa tat ttg aat gag act gat aat tta ggt aat aca gta ct- -ac aca cac ata tct ttc tag att ata tat cgt taa aaa tct gta aga ggt ata ttt ctc ata agt atc cac tgt gta ata tta tta atg gat ata tag ata aca caa tag gga cta ata gta ttg taa aag ata taa tcg act att t-- gta cat atc cag ata tct ata ttc cta cta gtt tgc tgc gta gtt gca tca ttg ata tgc atg -at tta tca gga ttc aga gat gaa tta cta agt aaa cta caa tcc cac aat aag taa gaa tca aat atc aaa aac tca ctt ttg att ttt cta gtc tta agt aat aca tat att tat taa tag acc tat gaa at- aaa aaa gtt aac aat gga ttc gcg tat agc tat tta cgt att agt atc ggc atc tct ttt gta tct tgt taa ttg tca caa act agt aca tta ctt caa tct gaa aat aaa tgg aag tga tat aac taa tac agc aga tat att gct gga caa tta tcc aat tat gac ctt tga tgg aaa gga tat tta tcc atc tat cgc gtt cat ggt cgg taa taa act ttt cct aga tct tta taa aaa tat ctt tgt aga att ttt cag act att tcg agt atc tgt aag tag tca ata cga gga att aga ata tta tta ttc atg tga tta tac taa caa ccg tcc tac aat taa aca aca tta ctt tta taa cgg cga aga ata tac tga aat tga tag atc gaa aaa agc cac taa taa aaa cag ttg gtt aat tac ttc agg ctt tag act aca aaa atg gtt cga tag cga aga ttg tat aat tta tct cag atc ttt agt tag aag aat gga aga cag taa caa aaa cag taa aaa aac tta gta ctt aga tat cg- aaa aaa aat ata ttt ttg tag act ctt gag aat aga agg -aa aca tgt aca taa tta taa aaa atg aaa atc aat ggc gaa taa gac agt gcg att cgc gcc atg gag tcg gca gat ttc atg gct gtc gat gag cag ttt cac gac gac ctc gat ctt tgg tca tta tct ttg gta gat gac tat aaa aaa cat gga tta ggt gtt gac tgt tat gtt cta gaa cca gtt gtt gac agg aaa ata ttt gat aga ttt ctc ctt gaa cca att tgt gat cct gta gat gtt ctg tat gat tat ttt agg att cat aga gat aat att gat cag tat ata gta gat aga ctg ttt gca tat att aca tat aaa gat att ata tct gca tta gtg tca aag aat tat atg gaa gat att ttc tct ata att att aag aat tgt aat tct gta caa gat ctc tta ctt tac tat cta tct aat gca tat gta gaa ata gac att gtt gat ttt atg gta gat cat ggg gct gta ata tat aaa ata gaa tgc ttg aat gcc tat ttt agg gga ata tgt aaa aag gaa agt agt gtt gtt gag ttt att ttg aat cgt ggt atc cca gat gaa aat gat gtt aaa tta gat cta tat aaa ata att cag tat act agg gga ttc ctt gta gat gaa ccc aca gta tta gaa att tat aag ctt tgt atc cca tat att gaa gat atc aat caa cta gat gct ggt gga agg acc ttg ctt tat cgc gct atc tat gca ggt tat ata gat tta gta tca tgg cta tta gaa aat gga gca aat gtc aac gca gta atg agt aat gga tat aca tgt ctt gac gtg gcc gtg gat agg gga tct gtc atc gcc cgt agg gaa gca cat ctt aaa ata tta gaa ata ttg ctt aga gaa cca ttg tct att gac tgt ata aaa tta gct ata ctt aat aat aca att gaa aac cat gat gtg ata aag ctc tgt atc aag tat ttt atg atg gta gat tat tca ctt tgt aat gtg tat gca tca tca ctc ttt gat tat ata att gat tgt aaa caa gaa ttg gag tac att agg cag atg aaa att cat aat aca acc atg tat gag tta atc tat aat aga gac aaa aac aag cat gct tcc cat att cta cat agg tat tct aaa cat cca gtt ttg aca cag tgt atc act aaa gga ttc aag att tac aca gaa gta acc gag cag gtc act aaa gct cta aac aga cgt gct cta ata gat gag ata ata aac aat gta tca act gat gac aat ctc cta tca aaa ctt cca tta gaa att agg gat cta att gtt tca caa gct gtc ata tag agt tct ctc cac cca cct ttc ttg aaa tga gtt aat --- --- --- --- --- --- --- --- --- --- --- --- --- --- --- --- --- --- --- --- --- --- --- --- --- --- --- --- --- --- --- --- --- --- --- --- --- --- --- --- --- --- --- --- --- --- --- --- --- --- --- --- --- --- --- --- --- --- --- --- --- --- --- --- --- --- --- --- --- --- --- --- --- --- --- --- --- --- --- --- --- --- --- --- --- --- --- --- --- --- --- --- --- --- --- --- --- --- --- --- --- --- --- --- --- --- --- --- --- --- --- --- --- --- --- --- --- --- --- --- --- --- --- --- --- --- --- --- --- --- --- --- --- --- --- --- --- --- --- --- --- --- --- --- --- --- --- --- --- --- --- --- --- -ag tca taa gtt agt taa gtc ata agt tag tt- --t tat aag tta gtt tat agt cta aca ctt cta att ttt ata cct tga tct ttt tct cta att atg aaa aag taa atc att atg aag atg gat gaa aat gga cga gat tgt gcg cat cgt taa cga tag tat gtg gta cgt acc taa cgc att tat gga cga cgg tga taa tga agg tca cat ttc tgt caa taa tgt ctg tca tat gta tct cgc att ctt tga tgt gga tat atc atc tca tct gtt taa att agt tat taa aca ctg cga tct gaa taa acg act aaa atg tgg taa ctc tcc att aca ttg cta tac gat gaa tac acg att taa tcc atc tgt att aaa gat att gtt acg cca cgg cat gcg taa ctt tga tag caa gga taa aaa agg aca tat tcc tct aca cca cta tct gat tca ttc act atc aat cga taa caa gat ctt tga tat act aac gga ccc cat tga tga ctt tag taa atc atc cga tct att gct gtg tta tct tag ata taa att caa tgg gag ctt aaa cta tta cgt tct gta caa att att gac taa agg atc tga ccc taa ttg cgt cga tga gga tgg act cac ttc tct tca tta cta ctg taa aca cat atc cgt gtt cca cga aag caa tta tta caa gtc aaa gag tca cac taa gat gcg agc tga gaa gcg att cat cta cgc gat aat aga tca tgg agc aaa cat taa cgc ggt tac gaa aat cgg aaa tac gcc gtt aca cac tta cct tca aca gta tac caa aca tag tcc tcg tgt ggt gta tgc tct ttt atc tcg agg agc cga tac gag gat acg taa taa tct tga ttg tac acc cat cat gga ata cat aaa gaa cga ttg tgc aac agg tca tat tct cat aat gtt act caa ttg gca cga aca aaa ata cgg gaa att aca aaa gga aga agg aca aca tct act tta tct att cat aaa aca taa tca agg ata tgg aag tcg ctc tct caa tat act acg gta tct act aga tag att cga cat tca gaa aga cga ata cta taa tac aat gac tcc tct tca tac cgc ctt cca gaa ttg caa taa caa tgt tgc ctc ata cct cgt ata cat cgg ata cga cat caa cct tcc gac taa aga cga taa gac agt att cga ctt ggt gtt tga aaa cag aaa cat cat ata caa ggc gga tgt cgt taa tga cat cat cca cca cag act gaa agt atc tct acc tat gat taa atc gtt gtt cta caa gat gtc gga gtt ctc tcc cta cga cga tca cta cgt aaa gaa gat aat agc cta ctg cat att aag gga cga gtc att tgc gga act aca tac taa att ctg ttt aaa cga gga cta taa aag tgt att tat gaa aaa tat atc att cga taa gat aga ttc cat cat cga aaa atg tag tcg tga cat aag tct cct caa aga gat tcg aat ctc aga cac caa ctt gta tac ggt att gag aac aga aga cat ccg gta tca cac cta tct cga agc cat aca ttc aga caa acg cat ttc att tcc cat gta cga cga tct cat aga aca gtg tca tct atc gat gga gca taa aag taa act cgt cga caa agc act caa taa att aga gtc tac cat cga tag tca atc tag act atc gta ttt gcc tcc gga aat tat gcg caa tat cat aac caa gct aag cga cta cca tct aaa cag tat gtt gta cgg aaa gaa cca tta caa ata tta tcc atg ata gaa aga aaa tat tta aaa aat aat cta tat gat tgg aga agt agg aaa caa aca gta aca aga cga cga tta cta c-a tta tta aat cat gag gtc cgt att ata ctc gta tat att gtt tct ctc atg tat aat aat aaa cgg aag aga tat agc acc aca tgc acc atc caa tgg aaa atg taa aga caa cga ata cag aag ccg taa tct atg ttg tct atc gtg tcc tcc ggg aac tta cgc ttc cag att atg tga tag caa gac taa tac aca atg tac gcc gtg tgg ttc gga tac ctt tac atc tca caa taa tca ttt aca ggc ttg tct aag ttg taa cgg aag atg tga tag taa tca ggt aga gac gcg atc gtg taa cac gac tca caa tag aat ctg tga atg ctc tcc agg ata tta ttg tct tct caa agg atc atc agg gtg tag aac atg tat ttc taa aac aaa gtg tgg aat agg ata cgg agt atc cgg ata cac gtc tac cgg aga cgt cat ctg ttc tcc gtg tgg tcc cgg aac ata ttc tca cac cgt ctc ttc cac aga taa atg cga acc --- cgt aac cag caa tac att taa cta tat cga tgt gga aat taa cct gta tcc agt caa cga cac atc gtg tac tcg gac gac cac tac cgg tct cag cga atc cat ctc aac gtc gga act aac tat tac cat gaa tca taa aga ttg tga tcc cgt ctt tcg tgc aga ata ctt ctc tgt cct taa taa tgt agc aac ttc agg att ctt tac agg aga aaa tag ata tca gaa tac ttc aaa gat atg tac tct gaa ttt cga gat taa atg taa caa caa aga ttc atc ttc caa aca gtt aac gaa aac aaa gaa tga tac tat cat gcc gca ttc aga gac ggt aac tct agt ggg cga ctg tct atc tag cgt cga cat cta cat act ata tag taa tac caa tac tca aga cta cga aac gga tac aat ctc tta tca tat ggg taa tgt tct cga tgt caa tag cca tat gcc cgc tag ttg cga tat aca taa act gat cac taa ttc cca gaa tcc cac cca ctt ata gta ag- ttt ttt tta ccc ata aat aat aaa tac aat aat taa ttt ctc gta aaa gta gaa aat ata ttc taa ttt att ata tgg taa gaa agt aga atc atc tag aac agt aat caa tca ata gca atc atg aaa caa tat att gtc ctg gca tgc atg tgc cta gtg gca gct gct atg cct act agt ctt caa caa tcc tca tcc tcg tgt act gaa gaa gaa aac aaa cat cat atg gga atc gat gtt att atc aaa gtc aca aag caa gac caa aca ccg acc aat gat aag att tgt caa tcc gta acg gaa gtt aca gag acc gaa gat gat gag gta tcc gaa gat gat gag gta tcc gaa gaa gtt gta aaa gga gat ccc acc act tat tac aat atc gtc ggc gcg ggt ctt aac atg aac ttt gga ttc acc aaa tgc cca aag att tca tcc atc tcc gaa tcc tct gat gga aac act gtg aat act aga ttg tcc agc gtg tca cca gga caa ggt aag gac tct ccc gcg atc acg cgt gaa gaa gct ctg gct atg atc aaa gac tgt gag atg tct atc gac atc aga tgt agc gaa gaa gag aaa gac agc gac atc aag acc cat cca gta ctt ggg tct aac atc tca cat aag aaa gtg agt tac aaa gat atc atc ggt tca acg atc gtt gat aca aaa tgt gtt aag aac cta gag ttt agc gta cgt atc gga gac atg tgt gag gaa tca tct gaa ctt gaa gtc aag gat gga ttc aag tat gtc gat gga tcg gca tct gaa ggt gca acc gat gat act tca ctc atc gat tca aca aaa ctc aaa gca tgt gtc tga atc gat aac tct att cat ctg aaa atg gat gag ttg ggt taa tcg aac gat tca gac acc gca cca cga att aaa aaa gtg tac cgg gta cta tat tcc ggt ttg caa aac aaa aat gtt taa cta cat tca caa aaa gtt acc tct cgt tac ttc ttc ttt ctg ttt caa tat gtg ata cga tat gat cac tat tcg tat tct ctt tct ctc ttg tct cat aaa aaa gtt tta c-- --- --- --- aaa aaa a-t att ttt att ctc ttt ctc tct tcg at- ggt ctc aca aaa ata tta aac ctc ttt ctg atg tct caa cta ttt cgt aaa cga taa cgt cca aca ata tat tct cgt aga gct tat caa tat cct tat gcc aat --- ggt tgt cag aca att gca tca taa aat aat gtt tat aat tta cac gtt aac atc ata taa taa acg tat ata gtt aat att ttt gga ata taa atg atc tgt aaa atc cat gta ggg gac act gct cac gtt ttt tct cta gta cat aat ttc aca caa gtt ttt ata cag aca aat taa ttc tcg tcc ata ta- ttt taa aac att gac ttt tgt act aag aaa aat atc ttg act aac cat ctc ttt ctc tcg tcg atg ggt ctc aca aaa ata tta aac ctc ttt ctg atg g-- --- --- --- --- --- --- --- --- --- --- --- --- --- --- --- --- --- --- --- --- --- --- --- --- --- --- --- --- --- --- --- --- --- --- --- --- --- --- --- --- --- --- --- --- --- --- agt cgt aaa aag ttt tta tcc ttt ctc tcg tcg atg ggt ctc --- --- --- --- --- --- --- --- --- --- --- --- --- --- --- --- --- --- --- --- --- --- --- -ac aaa aat att aaa cct ctt tct gat gga gtc gta aaa agt ttt tat cct ttc tct ctt cga tgg gtc tca caa aaa tat taa acc tct ttc tga tgg agt cgt aaa aag ttt tta tcc ttt ctc tct tcg atg ggt ctc aca aaa ata tta aac ctc ttt ctg atg gtc tct ata aa

>KJ642619.1_Monkeypox_virus_strain_Gabon-1988_complete_genome

atc gag gac gac ttc aac tat ggc agc tcg gtg gcg tct gcc agc gtg cac ata cga atg gca ttt cta aga aaa gtc tac ggt atc ctt gtc tac aat ttc ctt taa caa cgg caa cag ctg cag tat ttt tat act ttg aat gca ttc gga cat tta tac aag gga gtc ctg ttc taa tat tgg cat caa tgt tcg gat cta tag gct tga ttt tcg cat tga ctt tac aca gac ata aac atc ccc tga atc tgt acc tac ttt gtg gat tta cac tgt tag aat ctc taa cgc tgg cct ctg ttg tta ctt tct atg atg cgc gta tcg tta tac aag ctt tca tgt tga cta ctg cag tgt ttc ttg ctc tga cta cat gta ctc tac aat caa aga gag att tca gta aac ttg taa cag gat tgt ttg ctg ctt tct gga ttt taa ttt tgt cag gag tct tga gga taa agt tta aaa tag aat taa taa aga aca tat agg tca -tt ttt taa aca tgg ata gaa acc aag gtt gtt agt taa taa tat aca aga tat ttt ttc tca ctc tga tcc atg taa acc aag gac gag aga cac tct cat tcc tca ttc acg aca cca tta aaa atg gaa att aaa gcc ctc tat taa gca cag acg gct aca ggt cta cca tca ggt taa cct tcg tct acc ttc aca atg gcc ttt cct tgt gcc cag ttc agt ccc tgt cat tgc cac gct act aag gac tcc ctg aat acc gtg act gac gtc aga cat tgt ctg act gaa tac atc ctg tgg gtt tct cat aga tgg acc cat aga gaa agc gca ggg cct ctc tac agg ctt ctc atc tct ttc aga att gat gca atg gag cta ttt ggt agc gag ttg aag gag ttc tcg gat tca ctt ccg tgg gac aat atc gac aat tgc gtg gag atc att aaa tgt ttc atc aga aat gac tcc atg aaa acc gcc aaa gaa ctt tgt gca ata att gga ctt tgt act caa tca gct att gtc act gga aga gtc ttc aat gat aag tat atc gac ata cta ctt atg ctg cga aag att ctg aat gag aac gac tat ctc acc ctc ttg gat cat atc ctc act gct aaa tac taa atc tcc ttc atg ctc tct cac taa tac tct tac tca cta cac ttt tta tca tct tat gat gaa tga ttg cct tca tca -tt ttt cgt gga ata taa tat agg aat aat tag cac cag aat agc tat gga tat ctc gtt aag aat att ctc gat aag aga cat aat gta gac ata gtt att ata tcc ttc tta gat aag tgt tac gct act gga aag ttt cca tcg tta tta tta cgt gaa gat gat atc att aaa cca aca ttg aga ttg gct ctt atg tta gct gga ttg aat tac tgt aat aaa tgc atc gag tat aga ggg ata tag caa ttc tcg ata ata gtc atg caa tat ttg aat gag act gat aat tta ggt aat aca gta cta cac aca cac ata tct ttc tag att ata tat cgt taa aaa tct gta aga ggt ata ttt ctc ata agt atc cac tgt gta ata tta tta atg gat ata tag ata aca caa tag gga cta ata gta ttg taa aag ata taa tcg act att t-- gta cat atc cag ata tct ata ttc cta cta gtt tgc tgc gta gtt gca tca ttg ata tgc atg -at tta tca gga ttc aga gat gaa tta cta agt aaa cta caa tcc cac aat aag taa gaa tca aat atc aaa aac tca ctt ttg att ttt cta gtc tta agt aat aca tat att tat taa tag acc tat gaa at- aaa aaa gtt aac aat gga ttc gcg tat agc tat tta cgt att agt atc ggt atc tct ttt gta tct tgt taa ttg tca caa act agt aca tta ctt caa tct gaa aat aaa tgg aag tga tat aac taa tac agc aga tat att gct gga caa tta tcc aat tat gac ctt tga tgg aaa gga tat tta tcc atc tat cgc gtt cat ggt cgg taa taa act ttt cct aga tct tta taa aaa tat ctt tgt aga att ttt cag act att tcg agt atc tgt aag tag tca ata cga gga att aga ata tta tta ttc atg tga tta tac taa caa ccg tcc tac aat taa aca aca tta ctt tta taa cgg cga aga ata tac tga aat tga tag atc gaa aaa agc cac taa taa aaa cag ttg gtt aat tac ttc agg ctt tag act aca aaa atg gtt cga tag cga aga ttg tat aat tta tct cag atc ttt agt tag aag aat gga aga cag taa caa aaa cag t-a aaa aac tta gta ctt aga tat cg- aaa aaa aat ata ttt ttg tag act ctt g-- --- aga agg aaa aca tgt aca taa tta taa aaa atg aaa atc aat ggc gaa taa gac agt gcg att cgc gcc atg gag tcg gca gat ttc atg gct gtc gat gag cag ttt cac gac gac ctc gat ctt tgg tca tta tct ttg gta gat gac tat aaa aaa cat gga tta ggt gtt gac tgt tat gtt cta gaa cca gtt gtt gac agg aaa ata ttt gat aga ttt ctc ctt gaa cca att tgt gat cct gta gat gtt ctg tat gat tat ttt agg att cat aga gat aat att gat cag tat ata gta gat aga ctg ttt gca tat att aca tat aaa gat att ata tct gca tta gtg tca aag aat tat atg gaa gat att ttc tct ata att att aag aat tgt aat tct gta caa gat ctc tta ctt tac tat cta tct aat gca tat gta gaa ata gac att gtt gat ttt atg gta gat cat ggg gct gta ata tat aaa ata gaa tgc ttg aat gcc tat ttt agg gga ata tgt aaa aag gaa agt agt gtt gtt gag ttt att ttg aat tgt ggt atc cca gat gaa aat gat gtt aaa tta gat cta tat aaa ata att cag tat act agg gga ttc ctt gta gat gaa ccc aca gta tta gaa att tat aag ctt tgt atc cca tat att gaa gat atc aat caa cta gat gct ggt gga agg acc ttg ctt tat cgc gct atc tat gca ggt tat ata gat tta gta tca tgg cta tta gaa aat gga gca aat gtc aac gca gta atg agt aat gga tat aca tgt ctt gac gtg gcc gtg gat agg gga tct gtc atc gcc cgt agg gaa gca cat ctt aaa ata tta gaa ata ttg ctt aga gaa cca ttg tct att gac tgt ata aaa tta gct ata ctt aat aat aca att gaa aac cat gat gtg ata aag ctc tgt atc aag tat ttt atg atg gta gat tat tca ctt tgt aat gtg tat gca tca tca ctc ttt gat tat ata att gat tgt aaa caa gaa ttg gag tac att agg cag atg aaa att cat aat aca acc atg tat gag tta atc tat aat aga gac aaa aac aag cat gct tcc cat att cta cat agg tat tct aaa cat cca gtt ttg aca cag tgt atc act aaa gga ttc aag att tac aca gaa gta acc gag cag gtc act aaa gct cta aac aga cgt gct cta ata gat gag ata ata aac aat gta tca act gat gac aat ctc cta tca aaa ctt cca tta gaa att agg gat cta att gtt tca caa gct gtc ata tag agt tct ctc cac cca cct ttc ttg aaa tga gtt aat --- --- --- --- --- --- --- --- --- --- --- --- --- --- --- --- --- --- --- --- --- --- --- --- --- --- --- --- --- --- --- --- --- --- --- --- --- --- --- --- --- --- --- --- --- --- --- --- --- --- --- --- --- --- --- --- --- --- --- --- --- --- --- --- --- --- --- --- --- --- --- --- --- --- --- --- --- --- --- --- --- --- --- --- --- --- --- --- --- --- --- --- --- --- --- --- --- --- --- --- --- --- --- --- --- --- --- --- --- --- --- --- --- --- --- --- --- --- --- --- --- --- --- --- --- --- --- --- --- --- --- --- --- --- --- --- --- --- --- --- --- --- --- --- --- --- --- --- --- --- --- --- --- --- --- --- --- --- --a gtc ata agt tag tt- --t tat aag tta gtt tat agt cta aca ctt cta att ttt ata cct tga tct ttt tct cta att atg aaa aag taa atc att atg aag atg gat gaa aat gga cga gat tgt gcg cat cgt taa cga tag tat gtg gta cgt acc taa cgc att tat gga cga cgg tga taa tga agg tca cat ttc tgt caa taa tgt ctg tca tat gta tct cgc att ctt tga tgt gga tat atc atc tca tct gtt taa att agt tat taa aca ctg cga tct gaa taa acg act aaa atg tgg taa ctc tcc att aca ttg cta tac gat gaa tac acg att taa tcc atc tgt att aaa gat att gtt acg cca cgg cat gcg taa ctt tga tag caa gga taa aaa agg aca tat tcc tct aca cca cta tct gat tca ttc act atc aat cga taa caa gat ctt tga tat act aac gga ccc cat tga tga ctt tag taa atc atc cga tct att gct gtg tta tct tag ata taa att caa tgg gag ctt aaa cta tta cgt tct gta caa att att gac taa agg atc tga ccc taa ttg cgt cga tga gga tgg act cac ttc tct tca tta cta ctg taa aca cat atc cgt gtt cca cga aag caa tta tta caa gtc aaa gag tca cac taa gat gcg agc tga gaa gcg att cat cta cgc gat aat aga tca tgg agc aaa cat taa cgc ggt tac gaa aat cgg aaa tac gcc gtt aca cac tta cct tca aca gta tac caa aca tag tcc tcg tgt ggt gta tgc tct ttt atc tcg agg agc cga tac gag gat acg taa taa tct tga ttg tac acc cat cat gga ata cat aaa gaa cga ttg tgc aac agg tca tat tct cat aat gtt act caa ttg gca cga aca aaa ata cgg gaa att aca aaa gga aga agg aca aca tct act tta tct att cat aaa aca taa tca agg ata tgg aag tcg ctc tct caa tat act acg gta tct act aga tag att cga cat tca gaa aga cga ata cta taa tac aat gac tcc tct tca tac cgc ctt cca gaa ttg caa taa caa tgt tgc ctc ata cct cgt ata cat cgg ata cga cat caa cct tcc gac taa aga cga taa gac agt att cga ctt ggt gtt tga aaa cag aaa cat cat ata caa ggc gga tgt cgt taa tga cat cat cca cca cag act gaa agt atc tct acc tat gat taa atc gtt gtt cta caa gat gtc gga gtt ctc tcc cta cga cga tca cta cgt aaa gaa gat aat agc cta ctg cct att aag gga cga gtc att tgc gga act aca tac taa att ctg ttt aaa cga gga cta taa aag tgt att tat gaa aaa tat atc att cga taa gat aga ttc cat cat cga aaa atg tag tcg tga cat aag tct cct caa aga gat tcg aat ctc aga cac caa ctt gta tac ggt att gag aac aga aga cat ccg gta tca cac cta tct cga agc cat aca ttc aga caa acg cat ttc att tcc cat gta cga cga tct cat aga aca gtg tca tct atc gat gga gca taa aag taa act cgt cga caa agc act caa taa att aga gtc tac cat cga tag tca atc tag act atc gta ttt gcc tcc gga aat tat gcg caa tat cat aac caa gct aag cga cta cca tct aaa cag tat gtt gta cgg aaa gaa cca tta caa ata tta tcc atg ata gaa aga aaa tat tta aaa aat aat cta tat gat tgg aga agt agg aaa caa aca gta aca aga cga cga tta cta c-a tta tta aat cat gag gtc cgt att ata ctc gta tat att gtt tct ctc atg tat aat aat aaa cgg aag aga tat agc acc aca tgc acc atc caa tgg aaa atg taa aga caa cga ata cag aag ccg taa tct atg ttg tct atc gtg tcc tcc ggg aac tta cgc ttc cag att atg tga tag caa gac taa tac aca atg tac gcc gtg tgg ttc gga tac ctt tac atc tca caa taa tca ttt aca ggc ttg tct aag ttg taa cgg aag atg tga tag taa tca ggt aga gac gcg atc gtg taa cac gac tca caa tag aat ctg tga atg ctc tcc agg ata tta ttg tct tct caa agg atc atc agg gtg tag aac atg tat ttc taa aac aaa gtg tgg aat agg ata cgg agt atc cgg ata cac gtc tac cgg aga cgt cat ctg ttc tcc gtg tgg tcc cgg aac ata ttc tca cac cgt ctc ttc cac aga taa atg cga acc --- cgt aac cag caa tac att taa cta tat cga tgt gga aat taa cct gta tcc agt caa cga cac atc gtg tac tcg gac gac cac tac cgg tct cag cga atc cat ctc aac gtc gga act aac tat tac cat gaa tca taa aga ttg tga tcc cgt ctt tcg tgc aga ata ctt ctc tgt cct taa taa tgt agc aac ttc agg att ctt tac agg aga aaa tag ata tca gaa tac ttc aaa gat atg tac tct gaa ttt cga gat taa atg taa caa caa aga ttc atc ttc caa aca gtt aac gaa aac aaa gaa tga tac tat cat gcc gca ttc aga gac ggt aac tct agt ggg cga ctg tct atc tag cgt tga cat cta cat act ata tag taa tac caa tac tca aga cta cga aac gga tac aat ctc tta tca tat ggg taa tgt tct cga tgt caa tag cca tat gcc cgc tag ttg cga tat aca taa act gat cac taa ttc cca gaa tcc cac cca ctt ata gta ag- ttt ttt tta ccc ata aat aat aaa tac aat aat taa ttt ctc gta aaa gta gaa aat ata ttc taa ttt att ata tgg taa gaa agt aga atc atc tag aac agt aat caa tca ata gca atc atg aaa caa tat att gtc ctg gca tgc atg tgc cta gtg gca gct gct atg cct act agt ctt caa caa tcc tca tcc tcg tgt act gaa gaa gaa aac aaa cat cat atg gga atc gat gtt att atc aaa gtc aca aag caa gac caa aca ccg acc aat gat aag att tgt caa tcc gta acg gaa gtt aca gag a-- --- --- --- --- --- -cc gaa gat gat gag gta tcc gaa gaa gtt gta aaa gga gat ccc acc act tat tac act atc gtc ggc gcg ggt ctt aac atg aac ttt gga ttc acc aaa tgc cca aag att tca tcc atc tcc gaa tcc tct gat gga aac act gtg aat act aga ttg tcc agc gtg tca cca gga caa ggt aag gac tct ccc gcg atc acg cgt gaa gaa gct ctg gct atg atc aaa gac tgt gag atg tct atc gac atc aga tgt agc gaa gaa gag aaa gac agc gac atc aag acc cat cca gta ctt ggg tct aac atc tca cat aag aaa gtg agt tac aaa gat atc atc ggt tca acg atc gtt gat aca aaa tgt gtt aag aac cta gag ttt agc gta cgt atc gga gac atg tgt gag gaa tca tct gaa ctt gaa gtc aag gat gga ttc aag tat gtc gac gga tcg gca tct gaa ggt gca acc gat gat act tca ctc atc gat tca aca aaa ctc aaa gca tgt gtc tga atc gat aac tct att cat ctg aaa atg gat gag ttg ggt taa tcg aac gat tca gac acc gca cca cga att aaa aaa gtg tac cgg gta cta tat tcc ggt ttg caa aac aaa aat gtt taa cta cat tca caa aaa gtt acc tct cgt tac ttc ttc ttt ctg ttt caa tat gtg ata cga tat gat cac tat tcg tat tct ctt tct ctc ttg tct cat aaa aaa gtt tta c-- --- --- --- aaa aaa a-t att ttt att ctc ttt ctc tct tcg at- ggt ctc aca aaa ata tta aac ctc ttt ctg atg tct caa cta ttt cgt aaa cga taa cgt cca aca ata tat tct cgt aga gct tat caa tat cct tat gcc aat cta ggt tgt cag aca att gca tca taa aat aat gtt tat aat tta cac gtt aac atc ata taa taa acg tat ata gtt aat att ttt gga ata taa atg atc tgt aaa atc cat gta ggg gac act gct cac gtt ttt tct cta gta cat aat ttc aca caa gtt ttt ata cag aca aat taa ttc tcg tcc ata ta- ttt taa aac att gac ttt tgt act aag aaa aat atc ttg act aac cat ctc ttt ctc tcg tcg atg ggt ctc aca aaa ata tta aac ctc ttt ctg atg g-- --- --- --- --- --- --- --- --- --- --- --- --- --- --- --- --- --- --- --- --- --- --- --- --- --- --- --- --- --- --- --- --- --- --- --- --- --- --- --- --- --- --- --- --- --- --- agt cgt aaa aag ttt tta tcc ttt ctc tct tcg atg ggt ctc --- --- --- --- --- --- --- --- --- --- --- --- --- --- --- --- --- --- --- --- --- --- --- --- --- --- --- --- --- --- --- --- --- --- --- --- --- --- --- --- --- --- --- --- --- --- --- --- --- --- --- --- --- --- --- --- --- --- --- --- --- --- --- --- --- --- --- --- --- --- aca aaa ata tta aac ctc ttt ctg atg gtc tct ata aa

>DQ011154.1_Monkeypox_virus_strain_Congo_2003_358_complete_genome

atc gag gac gac ttc aac tat gac agc tcg gtg gcg tct gcc agc gtg cac ata cga atg gca ttt cta aga aaa gtc tac ggt atc ctt gtc tac aat ttc ctt taa caa cgg caa cag ctg cag tat ttt tat act ttg aat gca ttc gga cat tta tac aag gga gtc ctg ttc taa tat tgg cat caa tgt tcg gat cta tag gct tga ttt tcg cat tga ctt tac aca gac ata aac atc ccc tga atc tgt acc tac ttt gtg gat tta cac tgt tag aat ctc taa cgc tgg cct ctg ttg tta ctt tct atg atg cgc gta tcg tta tac aag ctt tca tgt tga cta ctg cag tgt ttc ttg ctc tga cta cat gta ctc tac aat caa aga gag att tca gta aac ttg taa cag gat tgt ttg ctg att tct gga ttt taa ttt tgt cag gag tct tga gga taa agt tta aaa tag aat taa taa aga aca tat agg tca -tt ttt taa aca tgg ata gaa acc aag gtt gtt agt taa taa tat aca aga tat ttt ttc tca ctc tga tcc atg taa acc aag gac gag aga cac tct cat tcc tca ttc acg aca cca tta aaa atg gaa att aaa gcc ctc tat taa gca cag acg gct aca ggt cta cca tca ggt taa cct tcg tct acc ttc aca atg gcc ttt cct tgt gcc cag ttc agt ccc tgt cat tgc cac gct act aag gac tcc ctg aat acc gtg act gac gtc aga cat tgt ctg act gaa tac atc ctg tgg gtt tct cat aga tgg acc cat aga gaa agc gca ggg cct ctc tac agg ctt ctc atc tct ttc aga att gat gca atg gag cta ttt ggt agc gag ttg aag gag ttc tcg gat tca ctt ccg tgg gac aat atc gac aat tgc gtg gag atc att aaa tgt ttc atc aga aat gac tcc atg aaa acc gcc aaa gaa ctt tgt gca ata att gga ctt tgt act caa tca gct att gtc act gga aga gtc ttc aat gat aag tat atc gac ata cta ctt ata ctg cga aag att ctg aat gag aac gac tat ctc acc ctc ttg gat cat atc ctc act gct aaa tac taa atc tcc ttc atg ctc tct cac taa tac tct tac tca cta cac ttt tta tca tct tat gat gaa tga ttg cct tca tca -tt ttt cgt gga ata taa tat agg aat aat tag cac cag aat agc tat gga tat ctc gtt aag aat att ctc gat aag aga cat aat gta gac ata gtt att ata tcc ttc tta gat aag tgt tac gct act gga aag ttt cca tcg tta tta tta cgt gaa gat gat atc att aaa cca aca ttg aga ttg gct ctt atg tta gct gga ttg aat tac tgt aat aaa tgc atc gag tat aga ggg ata tag caa ttc tcg ata ata gtc atg caa tat ttg aat gag act gat aat tta ggt aat aca gta ct- -ac aca cac ata tct ttc tag att ata tat cgt taa aaa tct gta aga ggt ata ttt ctc ata agt atc cac tgt gta ata tta tta atg gat ata tag ata aca caa tag gga cta ata gta ttg taa aag ata taa tcg act att t-- gta cat atc cag ata tct ata ttc cta cta gtt tgc tgc gta gtt gca tca ttg ata tgc atg -at tta tca gga ttc aga gat gaa tta cta agt aaa cta caa tcc cac aat aag taa gaa tca aat atc aaa aac tca ctt ttg att ttt cta gtc tta agt aat aca tat att tat taa tag acc tat gaa at- aaa aaa gtt aac aat gga ttc gcg tat agc tat tta cgt att agt atc ggc atc tct ttt gta tct tgt taa ttg tca caa act agt aca tta ctt caa tct gaa aat aaa tgg aag tga tat aac taa tac agc aga tat att gct gga caa tta tcc aat tat gac ctt tga tgg aaa gga tat tta tcc atc tat cgc gtt cat ggt cgg taa taa act ttt cct aga tct tta taa aaa tat ctt tgt aga att ttt cag act att tcg agt atc tgt aag tag tca ata cga gga att aga ata tta tta ttc atg tga tta tac taa caa ccg tcc tac aat taa aca aca tta ctt tta taa cgg cga aga ata tac tga aat tga tag atc gaa aaa agc cac taa taa aaa cag ttg gtt aat tac ttc agg ctt tag act aca aaa atg gtt cga tag cga aga ttg tat aat tta tct cag atc ttt agt tag aag aat gga aga cag taa caa aaa cag taa aaa aac tta gta ctt aga tat cg- aaa aaa aat ata ttt ttg tag act ctt gag aat aga agg aaa aca tgt aca taa tta taa aaa atg aaa atc aat ggc gaa taa gac agt gcg att cgc gcc atg gag tcg gca gat ttc atg gct gtc gat gag cag ttt cac gac gac ctc gat ctt tgg tca tta tct ttg gta gat gac tat aaa aaa cat gga tta ggt gtt gac tgt tat gtt cta gaa cca gtt gtt gac agg aaa ata ttt gat aga ttt ctc ctt gaa cca att tgt gat cct gta gat gtt ctg tat gat tat ttt agg att cat aga gat aat att gat cag tat ata gta gat aga ctg ttt gca tat att aca tat aaa gat att ata tct gca tta gtg tca aag aat tat atg gaa gat att ttc tct ata att att aag aat tgt aat tct gta caa gat ctc tta ctt tac tat cta tct aat gca tat gta gaa ata gac att gtt gat ttt atg gta gat cat ggg gct gta ata tat aaa ata gaa tgc ttg aat gcc tat ttt agg gga ata tgt aaa aag gaa agt agt gtt gtt gag ttt att ttg aat tgt ggt atc cca gat gaa aat gat gtt aaa tta gat cta tat aaa ata att cag tat act agg gga ttc ctt gta gat gaa ccc aca gta tta gaa att tat aag ctt tgt atc cca tat att gaa gat atc aat caa cta gat gct ggt gga agg acc ttg ctt tat cgc gct atc tat gca ggt tat ata gat tta gta tca tgg cta tta gaa aat gga gca aat gtc aac gca gta atg agt aat gga tat aca tgt ctt gac gtg gcc gtg gat agg gga tct gtc atc gcc cgt agg gaa gca cat ctt aaa ata tta gaa ata ttg ctt aga gaa cca ttg tct att gac tgt ata aaa tta gct ata ctt aat aat aca att gaa aac cat gat gtg ata aag ctc tgt atc aag tat ttt atg atg gta gat tat tca ctt tgt aat gtg tat gca tca tca ctc ttt gat tat ata att gat tgt aaa caa gaa ttg gag tac att agg cag atg aaa att cat aat aca acc atg tat gag tta atc tat aat aga gac aaa aac aag cat gct tcc cat att cta cat agg tat tct aaa cat cca gtt ttg aca cag tgt atc act aaa gga ttc aag att tac aca gaa gta acc gag cag gtc act aaa gct cta aac aga cgt gct cta ata gat gag ata ata aac aat gta tca act gat gac aat ctc cta tca aaa ctt cca tta gaa att agg gat cta att gtt tca caa gct gtc ata tag agt tct ctc cac cca cct ttc ttg aaa tga gtt aat --- --- --- --- --- --- --- --- --- --- --- --- --- --- --- --- --- --- --- --- --- --- --- --- --- --- --- --- --- --- --- --- --- --- --- --- --- --- --- --- --- --- --- --- --- --- --- --- --- --- --- --- --- --- --- --- --- --- --- --- --- --- --- --- --- --- --- --- --- --- --- --- --- --- --- --- --- --- --- --- --- --- --- --- --- --- --- --- --- --- --- --- --- --- --- --- --- --- --- --- --- --- --- --- --- --- --- --- --- --- --- --- --- --- --- --- --- --- --- --- --- --- --- --- --- --- --- --- --- --- --- --- --- --- --- --- --- --- --- --- --- --- --- --- --- --- --- --- --- --- --- --- --- -ag tca taa gtt agt taa gtc ata agt tag tt- --t tat aag tta gtt tat agt cta aca ctt cta att ttt ata cct tga tct ttt tct cta att atg aaa aag taa atc att atg aag atg gat gaa aat gga cga gat tgt gcg cat cgt taa cga tag tat gtg gta cgt acc taa cgc att tat gga cga cgg tga taa tga agg tca cat ttc tgt caa taa tgt ctg tca tat gta tct cgc att ctt tga tgt gga tat atc atc tca tct gtt taa att agt tat taa aca ctg cga tct gaa taa acg act aaa atg tgg taa ctc tcc att aca ttg cta tac gat gaa tac acg att taa tcc atc tgt att aaa gat att gtt acg cca cgg cat gcg taa ctt tga tag caa gga taa aaa agg aca tat tcc tct aca cca cta tct gat tca ttc act atc aat cga taa caa gat ctt tga tat act aac gga ccc cat tga tga ctt tag taa atc atc cga tct att gct gtg tta tct tag ata taa att caa tgg gag ctt aaa cta tta cgt tct gta caa att att gac taa agg atc tga ccc taa ttg cgt cga tga gga tgg act cac ttc tct tca tta cta ctg taa aca cat atc cgt gtt cca cga aag caa tta tta caa gtc aaa gag tca cac taa gat gcg agc tga gaa gcg att cat cta cgc gat aat aga tca tgg agc aaa cat taa cgc ggt tac gaa aat cgg aaa tac gcc gtt aca cac tta cct tca aca gta tac caa aca tag tcc tcg tgt ggt gta tgc tct ttt atc tcg agg agc cga tac gag gat acg taa taa tct tga ttg tac acc cat cat gga ata cat aaa gaa cga ttg tgc aac agg tca tat tct cat aat gtt act caa ttg gca cga aca aaa ata cgg gaa att aca aaa gga aga agg aca aca tct act tta tct att cat aaa aca taa tca agg ata tgg aag tcg ctc tct caa tat act acg gta tct act aga tag att cga cat tca gaa aga cga ata cta taa tac aat gac tcc tct tca tac cgc ctt cca gaa ttg caa taa caa tgt tgc ctc ata cct cgt ata cat cgg ata cga cat caa cct tcc gac taa aga cga taa gac agt att cga ctt ggt gtt tga aaa cag aaa cat cat ata caa ggc gga tgt cgt taa tga cat cat cca cca cag act gaa agt atc tct acc tat gat taa atc gtt gtt cta caa gat gtc gga gtt ctc tcc cta cga cga tca cta cgt aaa gaa gat aat agc cta ctg cat att aag gga cga gtc att tgc gga act aca tac taa att ctg ttt aaa cga gga cta taa aag tgt att tat gaa aaa tat atc att cga taa gat aga ttc cat cat cga aaa atg tag tcg tga cat aag tct cct caa aga gat tcg aat ctc aga cac caa ctt gta tac ggt att gag aac aga aga cat ccg gta tca cac cta tct cga agc cat aca ttc aga caa acg cat ttc att tcc cat gta cga cga tct cat aga aca gtg tca tct atc gat gga gca taa aag taa act cgt cga caa agc act caa taa att aga gtc tac cat cga tag tca atc tag act atc gta ttt gcc tcc gga aat tat gcg caa tat cat aac caa gct aag cga cta cca tct aaa cag tat gtt gta cgg aaa gaa cca tta caa ata tta tcc atg ata gaa aga aaa tat tta aaa aat aat cta tat gat tgg aga agt agg aaa caa aca gta aca aga cga cga tta cta c-a tta tta aat cat gag gtc cgt att ata ctc gta tat att gtt tct ctc atg tat aat aat aaa cgg aag aga tat agc acc aca tgc acc atc caa tgg aaa atg taa aga caa cga ata cag aag ccg taa tct atg ttg tct atc gtg tcc tcc ggg aac tta cgc ttc cag att atg tga tag caa gac taa tac aca atg tac gcc gtg tgg ttc gga tac ctt tac atc tca caa taa tca ttt aca ggc ttg tct aag ttg taa cgg aag atg tga tag taa tca ggt aga gac gcg atc gtg taa cac gac tca caa tag aat ctg tga atg ctc tcc agg ata tta ttg tct tct caa agg atc atc agg gtg tag aac atg tat ttc taa aac aaa gtg tgg aat agg ata cgg agt atc cgg ata cac gtc tac cgg aga cgt cat ctg ttc tcc gtg tgg tcc cgg aac ata ttc tca cac cgt ctc ttc cac aga taa atg cga acc --- cgt aac cag caa tac att taa cta tat cga tgt gga aat taa cct gta tcc agt caa cga cac atc gtg tac tcg gac gac cac tac cgg tct cag cga atc cat ctc aac gtc gga act aac tat tac cat gaa tca taa aga ttg tga tcc cgt ctt tcg tgc aga ata ctt ctc tgt cct taa taa tgt agc aac ttc agg att ctt tac agg aga aaa tag ata tca gaa tac ttc aaa gat atg tac tct gaa ttt cga gat taa atg taa caa caa aga ttc atc ttc caa aca gtt aac gaa aac aaa gaa tga tac tat cat gcc gca ttc aga gac ggt aac tct agt ggg cga ctg tct atc tag cgt cga cat cta cat act ata tag taa tac caa tac tca aga cta cga aac gga tac aat ctc tta tca tat ggg taa tgt tct cga tgt caa tag cca tat gcc cgc tag ttg cga tat aca taa act gat cac taa ttc cca gaa tcc cac cca ctt ata gta ag- ttt ttt tta ccc ata aat aat aaa tac aat aat taa ttt ctc gta aaa gta gaa aat ata ttc taa ttt att ata tgg taa gaa agt aga atc atc tag aac agt aat caa tca ata gca atc atg aaa caa tat att gtc ctg gca tgc atg tgc cta gtg gca gct gct atg cct act agt ctt caa caa tcc tca tcc tcg tgt act gaa gaa gaa aac aaa cat cat atg gga atc gat gtt att atc aaa gtc aca aag caa gac caa aca ccg acc aat gat aag att tgt caa tcc gta acg gaa gtt aca gag acc gaa gat gat gag gta tcc gaa gat gat gag gta tcc gaa gaa gtt gta aaa gga gat ccc acc act tat tac aat atc gtc ggc gcg ggt ctt aac atg aac ttt gga ttc acc aaa tgc cca aag att tca tcc atc tcc gaa tcc tct gat gga aac act gtg aat act aga ttg tcc agc gtg tca cca gga caa ggt aag gac tct ccc gcg atc acg cgt gaa gaa gct ctg gct atg atc aaa gac tgt gag atg tct atc gac atc aga tgt agc gaa gaa gag aaa gac agc gac atc aag acc cat cca gta ctt ggg tct aac atc tca cat aag aaa gtg agt tac aaa gat atc atc ggt tca acg atc gtt gat aca aaa tgt gtt aag aac cta gag ttt agc gta cgt atc gga gac atg tgt gag gaa tca tct gaa ctt gaa gtc aag gat gga ttc aag tat gtc gat gga tcg gca tct gaa ggt gca acc gat gat act tca ctc atc gat tca aca aaa ctc aaa gca tgt gtc tga atc gat aac tct att cat ctg aaa atg gat gag ttg ggt taa tcg aac gat tca gac acc gca cca cga att aaa aaa gtg tac cgg gta cta tat tcc ggt ttg caa aac aaa aat gtt taa cta cat tca caa aaa gtt acc tct cgt tac ttc ttc ttt ctg ttt caa tat gtg ata cga tat gat cac tat tcg tat tct ctt tct ctc ttg tct cat aaa aaa gtt tta c-- --- --- --- aaa aaa a-t att ttt att ctc ttt ctc tct tcg at- ggt ctc aca aaa ata tta aac ctc ttt ctg atg tct caa cta ttt cgt aaa cga taa cgt cca aca ata tat tct cgt aga gct tat caa tat cct tat gcc aat --- ggt tgt cag aca att gca tca taa aat aat gtt tat aat tta cac gtt aac atc ata taa taa acg tat ata gtt aat att ttt gga ata taa atg atc tgt aaa atc cat gta ggg gac act gct cac gtt ttt tct cta gta cat aat ttc aca caa gtt ttt ata cag aca aat taa ttc tcg tcc ata ta- ttt taa aac att gac ttt tgt act aag aaa aat atc ttg act aac cat ctc ttt ctc tcg tcg atg ggt ctc aca aaa ata tta aac ctc ttt ctg atg g-- --- --- --- --- --- --- --- --- --- --- --- --- --- --- --- --- --- --- --- --- --- --- --- --- --- --- --- --- --- --- --- --- --- --- --- --- --- --- --- --- --- --- --- --- --- --- agt cgt aaa aag ttt tta tcc ttt ctc tct tcg atg ggt ctc --- --- --- --- --- --- --- --- --- --- --- --- --- --- --- --- --- --- --- --- --- --- --- -ac aaa aat att aaa cct ctt tct gat gga gtc gta aaa agt ttt tat cct ttc tct ctt cga tgg gtc tca caa aaa tat taa acc tct ttc tga tgg agt cgt aaa aag ttt tta tcc ttt ctc tct tcg atg ggt ctc aca aaa ata tta aac ctc ttt ctg atg gtc tct ata aa

>JX878424.1_Monkeypox_virus_isolate_DRC_07-0338_complete_genome

atc gag gac gac ttc aac tat gac agc tcg gtg gcg tct gcc agc gtg cac ata cga atg gca ttt cta aga aaa gtc tac ggt atc ctt gtc tac aat ttc ctt taa caa cgg caa cag ctg cag tat ttt tat act ttg aat gca ttc gga cat tta tac aag gga gtc ctg ttc taa tat tgg cat caa tgt tcg gat cta tag gct tga ttt tcg cat tga ctt tac aca gac ata aac atc ccc tga atc tgt acc tac ttt gtg gat tta cac tgt tag aat ctc taa cgc tgg cct ctg ttg tta ctt tct atg atg cgc gta tcg tta tac aag ctt tca tgt tga cta ctg cag tgt ttc ttg ctc tga cta cat gta ctc tac aat caa aga gag att tca gta aac ttg taa cag gat tgt ttg ctg ctt tct gga ttt taa ttt tgt cag gag tct tga gga taa agt tta aaa tag aat taa taa aga aca tat agg tca -tt ttt taa aca tgg ata gaa acc aag gtt gtt agt taa taa tat aca aga tat ttt ttc tca ctc tga tcc atg taa acc aag gac gag aga cac tct cat tcc tca ttc acg aca cca tta aaa atg gaa att aaa gcc ctc tat taa gca cag acg gct aca ggt cta cca tca ggt taa cct tcg tct acc ttc aca atg gcc ttt cct tgt gcc cag ttc agt ccc tgt cat tgc cac gct act aag gac tcc ctg aat acc gtg act gac gtc aga cat tgt ctg act gaa tac atc ctg tgg gtt tct cat aga tgg acc cat aga gaa agc gca ggg cct ctc tac agg ctt ctc atc tct ttc aga att gat gca atg gag cta ttt ggt agc gag ttg aag gag ttc tcg gat tca ctt ccg tgg gac aat atc gac aat tgc gtg gag atc att aaa tgt ttc atc aga aat gac tcc atg aaa acc gcc aaa gaa ctt tgt gca ata att gga ctt tgt act caa tca gct att gtc act gga aga gtc ttc aat gat aag tat atc gac ata cta ctt ata ctg cga aag att ctg aat gag aac gac tat ctc acc ctc ttg gat cat atc ctc act gct aaa tac taa atc tcc ttc atg ctc tct cac taa tac tct tac tca cta cac ttt tta tca tct tat gat gaa tga ttg cct tca tca -tt ttt cgt gga ata taa tat agg aat aat tag cac cag aat agc tat gga tat ctc gtt aag aat att ctc gat aag aga cat aat gta gac ata gtt att ata tcc ttc tta gat aag tgt tac gct act gga aag ttt cca tcg tta --- --- --- --- --- --- --- --- --- --- --- --- --- --- --- --- --- --- --- --- --- --- --- --- --- --- --- --- --- --- --- --- --- --- --- --- --- --- --- --- --- --- --- --- --- --- --- --- --- --- --- --- --- --- --- --- --- --- --- --- --- --- --- --- --- --- --- --- --- --- --- --- --- --- --- --- --- --- --- --- --- --- --- --- --- --- --- --- --- --- --- --- --- --- --- --- --- --- --- --- --- --- --- --- --- --- --- --- --- --- --- --- --- --- --- --- --- --- --- --- --- --- --- --- --- --- --- --- --- --- --- --- --- --- --- --- --- --- --- --- --- --- --- --- --- --- --- --- --- --- --- --- --- --- --- --- --- --- --- --- --- --- --- --- --- --- --- --- --- --- --- --- --- --- --- --- --- --- --- --- --- --- --- --- --- --- --- --- --- --- --- --- --- --- --- --- --- --- --- --- --- --- --- --- --- --- --- --- --- aag tga tat aac taa tac agc aga tat att gct gga caa tta tcc aat tat gac ctt tga tgg aaa gga tat tta tcc atc tat cgc gtt cat ggt cgg taa taa act ttt cct aga tct tta taa aaa tat ctt tgt aga att ttt cag act att tcg agt atc tgt aag tag tca ata cga gga att aga ata tta tta ttc atg tga tta tac taa caa ccg tcc tac aat taa aca aca tta ctt tta taa cgg cga aga ata tac tga aat tga tag atc gaa aaa agc cac taa taa aaa cag ttg gtt aat tac ttc agg ctt tag act aca aaa atg gtt cga tag cga aga ttg tat aat tta tct cag atc ttt agt tag aag aat gga aga cag taa caa aaa cag taa aaa aac tta gta ctt aga tat cg- aaa aaa aat ata ttt ttg tag act ctt gag aat aga agg aaa aca tgt aca taa tta taa aaa atg aaa atc aat ggc gaa taa gac agt gcg att cgc gcc atg gag tcg gca gat ttc atg gct gtc gat gag cag ttt cac gac gac ctc gat ctt tgg tca tta tct ttg gta gat gac tat aaa aaa cat gga tta ggt gtt gac tgt tat gtt cta gaa cca gtt gtt gac agg aaa ata ttt gat aga ttt ctc ctt gaa cca att tgt gat cct gta gat gtt ctg tat gat tat ttt agg att cat aga gat aat att gat cag tat ata gta gat aga ctg ttt gca tat att aca tat aaa gat att ata tct gca tta gtg tca aag aat tat atg gaa gat att ttc tct ata att att aag aat tgt aat tct gta caa gat ctc tta ctt tac tat cta tct aat gca tat gta gaa ata gac att gtt gat ttt atg gta gat cat ggg gct gta ata tat aaa ata gaa tgc ttg aat gcc tat ttt agg gga ata tgt aaa aag gaa agt agt gtt gtt gag ttt att ttg aat tgt ggt atc cca gat gaa aat gat gtt aaa tta gat cta tat aaa ata att cag tat act agg gga ttc ctt gta gat gaa ccc aca gta tta gaa att tat aag ctt tgt atc cca tat att gaa gat atc aat caa cta gat gct ggt gga agg acc ttg ctt tat cgc gct atc tat gca ggt tat ata gat tta gta tca tgg cta tta gaa aat gga gca aat gtc aac gca gta atg agt aat gga tat aca tgt ctt gac gtg gcc gtg gat agg gga tct gtc atc gcc cgt agg gaa gca cat ctt aaa ata tta gaa ata ttg ctt aga gaa cca ttg tct att gac tgt ata aaa tta gct ata ctt aat aat aca att gaa aac cat gat gtg ata aag ctc tgt atc aag tat ttt atg atg gta gat tat tca ctt tgt aat gtg tat gca tca tca ctc ttt gat tat ata att gat tgt aaa caa gaa ttg gag tac att agg cag atg aaa att cat aat aca acc atg tat gag tta atc tat aat aga gac aaa aac aag cat gct tcc cat att cta cat agg tat tct aaa cat cca gtt ttg aca cag tgt atc act aaa gga ttc aag att tac aca gaa gta acc gag cag gtc act aaa gct cta aac aga cgt gct cta ata gat gag ata ata aac aat gta tca act gat gac aat ctc cta tca aaa ctt cca tta gaa att agg gat cta att gtt tca caa gct gtc ata tag agt tct ctc cac cca cct ttc ttg aaa tga gtt aat --- --- --- --- --- --- --- --- --- --- --- --- --- --- --- --- --- --- --- --- --- --- --- --- --- --- --- --- --- --- --- --- --- --- --- --- --- --- --- --- --- --- --- --- --- --- --- --- --- --- --- --- --- --- --- --- --- --- --- --- --- --- --- --- --- --- --- --- --- --- --- --- --- --- --- --- --- --- --- --- --- --- --- --- --- --- --- --- --- --- --- --- --- --- --- --- --- --- --- --- --- --- --- --- --- --- --- --- --- --- --- --- --- --- --- --- --- --- --- --- --- --- --- --- --- --- --- --- --- --- --- --- --- --- --- --- --- --- --- --- --- --- --- --- --- --- --- --- --- --- --- --- --- -ag tca taa gtt agt taa gtc ata agt tag tt- --t tat aag tta gtt tat agt cta aca ctt cta att ttt ata cct tga tct ttt tct cta att atg aaa aag taa atc att atg aag atg gat gaa aat gga cga gat tgt gcg cat cgt taa cga tag tat gtg gta cgt acc taa cgc att tat gga cga cgg tga taa tga agg tca cat ttc tgt caa taa tgt ctg tca tat gta tct cgc att ctt tga tgt gga tat atc atc tca tct gtt taa att agt tat taa aca ctg cga tct gaa taa acg act aaa atg tgg taa ctc tcc att aca ttg cta tac gat gaa tac acg att taa tcc atc tgt att aaa gat att gtt acg cca cgg cat gcg taa ctt tga tag caa gga taa aaa agg aca tat tcc tct aca cca cta tct gat tca ttc act atc aat cga taa caa gat ctt tga tat act aac gga ccc cat tga tga ctt tag taa atc atc cga tct att gct gtg tta tct tag ata taa att caa tgg gag ctt aaa cta tta cgt tct gta caa att att gac taa agg atc tga ccc taa ttg cgt cga tga gga tgg act cac ttc tct tca tta cta ctg taa aca cat atc cgt gtt cca cga aag caa tta tta caa gtc aaa gag tca cac taa gat gcg agc tga gaa gcg att cat cta cgc gat aat aga tca tgg agc aaa cat taa cgc ggt tac gaa aat cgg aaa tac gcc gtt aca cac tta cct tca aca gta tac caa aca tag tcc tcg tgt ggt gta tgc tct ttt atc tcg agg agc cga tac gag gat acg taa taa tct tga ttg tac acc cat cat gga ata cat aaa gaa cga ttg tgc aac agg tca tat tct cat aat gtt act caa ttg gca cga aca aaa ata cgg gaa att aca aaa gga aga agg aca aca tct act tta tct att cat aaa aca taa tca agg ata tgg aag tcg ctc tct caa tat act acg gta tct act aga tag att cga cat tca gaa aga cga ata cta taa tac aat gac tcc tct tca tac cgc ctt cca gaa ttg caa taa caa tgt tgc ctc ata cct cgt ata cat cgg ata cga cat caa cct tcc gac taa aga cga taa gac agt att cga ctt ggt gtt tga aaa cag aaa cat cat ata caa ggc gga tgt cgt taa tga cat cat cca cca cag act gaa agt atc tct acc tat gat taa atc gtt gtt cta caa gat gtc gga gtt ctc tcc cta cga cga tca cta cgt aaa gaa gat aat agc cta ctg cat att aag gga cga gtc att tgc gga act aca tac taa att ctg ttt aaa cga gga cta taa aag tgt att tat gaa aaa tat atc att cga taa gat aga ttc cat cat cga aaa atg tag tcg tga cat aag tct cct caa aga gat tcg aat ctc aga cac caa ctt gta tac ggt att gag aac aga aga cat ccg gta tca cac cta tct cga agc cat aca ttc aga caa acg cat ttc att tcc cat gta cga cga tct cat aga aca gtg tca tct atc gat gga gca taa aag taa act cgt cga caa agc act caa taa att aga gtc tac cat cga tag tca atc tag act atc gta ttt gcc tcc gga aat tat gcg caa tat cat aac caa gct aag cga cta cca tct aaa cag tat gtt gta cgg aaa gaa cca tta caa ata tta tcc atg ata gaa aga aaa tat tta aaa aat aat cta tat gat tgg aga agt agg aaa caa aca gta aca aga cga cga tta cta c-a tta tta aat cat gag gtc cgt att ata ctc gta tat att gtt tct ctc atg tat aat aat aaa cgg aag aga tat agc acc aca tgc acc atc caa tgg aaa atg taa aga caa cga ata cag aag ccg taa tct atg ttg tct atc gtg tcc tcc ggg aac tta cgc ttc cag att atg tga tag caa gac taa tac aca atg tac gcc gtg tgg ttc gga tac ctt tac atc tca caa taa tca ttt aca ggc ttg tct aag ttg taa cgg aag atg tga tag taa tca ggt aga gac gcg atc gtg taa cac gac tca caa tag aat ctg tga atg ctc tcc agg ata tta ttg tct tct caa agg atc atc agg gtg tag aac atg tat ttc taa aac aaa gtg tgg aat agg ata cgg agt atc cgg ata cac gtc tac cgg aga cgt cat ctg ttc tcc gtg tgg tcc cgg aac ata ttc tca cac cgt ctc ttc cac aga taa atg cga acc --- cgt aac cag caa tac att taa cta tat cga tgt gga aat taa cct gta tcc agt caa cga cac atc gtg tac tcg gac gac cac tac cgg tct cag cga atc cat ctc aac gtc gga act aac tat tac cat gaa tca taa aga ttg tga tcc cgt ctt tcg tgc aga ata ctt ctc tgt cct taa taa tgt agc aac ttc agg att ctt tac agg aga aaa tag ata tca gaa tac ttc aaa gat atg tac tct gaa ttt cga gat taa atg taa caa caa aga ttc atc ttc caa aca gtt aac gaa aac aaa gaa tga tac tat cat gcc gca ttc aga gac ggt aac tct agt ggg cga ctg tct atc tag cgt cga cat cta cat act ata tag taa tac caa tac tca aga cta cga aac gga tac aat ctc tta tca tat ggg taa tgt tct cga tgt caa tag cca tat gcc cgc tag ttg cga tat aca taa act gat cac taa ttc cca gaa tcc cac cca ctt ata gta ag- ttt ttt tta ccc ata aat aat aaa tac aat aat taa ttt ctc gta aaa gta gaa aat ata ttc taa ttt att ata tgg taa gaa agt aga atc atc tag aac agt aat caa tca ata gca atc atg aaa caa tat att gtc ctg gca tgc atg tgc cta gtg gca gct gct atg cct act agt ctt caa caa tcc tca tcc tcg tgt act gaa gaa gaa aac aaa cat cat atg gga atc gat gtt att atc aaa gtc aca aag caa gac caa aca ccg acc aat gat aag att tgt caa tcc gta acg gaa gtt aca gag acc gaa gat gat gag gta tcc gaa gat gat gag gta tcc gaa gaa gtt gta aaa gga gat ccc acc act tat tac aat atc gtc ggc gcg ggt ctt aac atg aac ttt gga ttc acc aaa tgc cca aag att tca tcc atc tcc gaa tcc tct gat gga aac act gtg aat act aga ttg tcc agc gtg tca cca gga caa ggt aag gac tct ccc gcg atc acg cgt gaa gaa gct ctg gct atg atc aaa gac tgt gag atg tct atc gac atc aga tgt agc gaa gaa gag aaa gac agc gac atc aag acc cat cca gta ctt ggg tct aac atc tca cat aag aaa gtg agt tac aaa gat atc atc ggt tca acg atc gtt gat aca aaa tgt gtt aag aac cta gag ttt agc gta cgt atc gga gac atg tgt gag gaa tca tct gaa ctt gaa gtc aag gat gga ttc aag tat gtc gat gga tcg gca tct gaa ggt gca acc gat gat act tca ctc atc gat tca aca aaa ctc aaa gca tgt gtc tga atc gat aac tct att cat ctg aaa atg gat gag ttg ggt taa tcg aac gat tca gac acc gca cca cga att aaa aaa gtg tac cgg gta cta tat tcc ggt ttg caa aac aaa aat gtt taa cta cat tca caa aaa gtt acc tct cgt tac ttc ttc ttt ctg ttt caa tat gtg ata cga tat gat cac tat tcg tat tct ctt tct ctc ttg tct cat aaa aaa gtt tta c-- --- --- --- aaa aaa a-t att ttt att ctc ttt ctc tct tcg at- ggt ctc aca aaa ata tta aac ctc ttt ctg atg tct caa cta ttt cgt aaa cga taa cgt cca aca ata tat tct cgt aga gct tat caa tat cct tat gcc aat --- ggt tgt cag aca att gca tca taa aat aat gtt tat aat tta cac gtt aac atc ata taa taa acg tat ata gtt aat att ttt gga ata taa atg atc tgt aaa atc cat gta ggg gac act gct cac gtt ttt tct cta gta cat aat ttc aca caa gtt ttt ata cag aca aat taa ttc tcg tcc ata ta- ttt taa aac att gac ttt tgt act aag aaa aat atc ttg act aac cat ctc ttt ct- --- --- --- --- --- --- --- --- --- --- --- --- --- --- --- --- --- --- --- --- --- --- --- --- --- --- --- --- --- --- --- --- --- --- --- --- --- --- --- --- --- --- --- --- --- --- --- --- --- --- --- --- --- --- --- --- --- --- --- --- --- --- --- --- --- --- --- --- --- --c tcg tcg atg ggt ctc --- --- --- --- --- --- --- --- --- --- --- --- --- --- --- --- --- --- --- --- --- --- --- -ac aaa aat att aaa cct ctt tct gat gga gtc gta aaa agt ttt tat cct ttc tc- --t cga tgg gtc tca caa aaa tat taa acc tct ttc tga tgg agt cgt aaa aag ttt tta tcc ttt ctc tcg tcg atg ggt ctc aca aaa ata tta aac ctc ttt ctg atg gtc tct ata aa

>JX878407.1_Monkeypox_virus_isolate_DRC_06-0950_complete_genome

atc gag gac gac ttc aac tat gac agc tcg gtg gcg tct gcc agc gtg cac ata cga atg gca ttt cta aga aaa gtc tac ggt atc ctt gtc tac aat ttc ctt taa caa cgg caa cag ctg cag tat ttt tat act ttg aat gca ttc gga cat tta tac aag gga gtc ctg ttc taa tat tgg cat caa tgt tcg gat cta tag gct tga ttt tcg cat tga ctt tac aca gac ata aac atc ccc tga atc tgt acc tac ttt gtg gat tta cac tgt tag aat ctc taa cgc tgg cct ctg ttg tta ctt tct atg atg cgc gta tcg tta tac aag ctt tca tgt tga cta ctg cag tgt ttc ttg ctc tga cta cat gta ctc tac aat caa aga gag att tca gta aac ttg taa cag gat tgt ttg ctg ctt tct gga ttt taa ttt tgt cag gag tct tga gga taa agt tta aaa tag aat taa taa aga aca tat agg tca -tt ttt taa aca tgg ata gaa acc aag gtt gtt agt taa taa tat aca aga tat ttt ttc tca ctc tga tcc atg taa acc aag gac gag aga cac tct cat tcc tca ttc acg aca cca tta aaa atg gaa att aaa gcc ctc tat taa gca cag acg gct aca ggt cta cca tca ggt taa cct tcg tct acc ttc aca atg gcc ttt cct tgt gcc cag ttc agt ccc tgt cat tgc cac gct act aag gac tcc ctg aat acc gtg act gac gtc aga cat tgt ctg act gaa tac atc ctg tgg gtt tct cat aga tgg acc cat aga gaa agc gca ggg cct ctc tac agg ctt ctc atc tct ttc aga att gat gca atg gag cta ttt ggt agc gag ttg aag gag ttc tcg gat tca ctt ccg tgg gac aat atc gac aat tgc gtg gag atc att aaa tgt ttc atc aga aat gac tcc atg aaa acc gcc aaa gaa ctt tgt gca ata att gga ctt tgt act caa tca gct att gtc act gga aga gtc ttc aat gat aag tat atc gac ata cta ctt ata ctg cga aag att ctg aat gag aac gac tat ctc acc ctc ttg gat cat atc ctc act gct aaa tac taa atc tcc ttc atg ctc tct cac taa tac tct tac tca cta cac ttt tta tca tct tat gat gaa tga ttg cct tca tca -tt ttt cgt gga ata taa tat agg aat aat tag cac cag aat agc tat gga tat ctc gtt aag aat att ctc gat aag aga cat aat gta gac ata gtt att ata tcc ttc tta gat aag tgt tac gct act gga aag ttt cca tcg tta --- --- --- --- --- --- --- --- --- --- --- --- --- --- --- --- --- --- --- --- --- --- --- --- --- --- --- --- --- --- --- --- --- --- --- --- --- --- --- --- --- --- --- --- --- --- --- --- --- --- --- --- --- --- --- --- --- --- --- --- --- --- --- --- --- --- --- --- --- --- --- --- --- --- --- --- --- --- --- --- --- --- --- --- --- --- --- --- --- --- --- --- --- --- --- --- --- --- --- --- --- --- --- --- --- --- --- --- --- --- --- --- --- --- --- --- --- --- --- --- --- --- --- --- --- --- --- --- --- --- --- --- --- --- --- --- --- --- --- --- --- --- --- --- --- --- --- --- --- --- --- --- --- --- --- --- --- --- --- --- --- --- --- --- --- --- --- --- --- --- --- --- --- --- --- --- --- --- --- --- --- --- --- --- --- --- --- --- --- --- --- --- --- --- --- --- --- --- --- --- --- --- --- --- --- --- --- --- --- aag tga tat aac taa tac agc aga tat att gct gga caa tta tcc aat tat gac ctt tga tgg aaa gga tat tta tcc atc tat cgc gtt cat ggt cgg taa taa act ttt cct aga tct tta taa aaa tat ctt tgt aga att ttt cag act att tcg agt atc tgt aag tag tca ata cga gga att aga ata tta tta ttc atg tga tta tac taa caa ccg tcc tac aat taa aca aca tta ctt tta taa cgg cga aga ata tac tga aat tga tag atc gaa aaa agc cac taa taa aaa cag ttg gtt aat tac ttc agg ctt tag act aca aaa atg gtt cga tag cga aga ttg tat aat tta tct cag atc ttt agt tag aag aat gga aga cag taa caa aaa cag taa aaa aac tta gta ctt aga tat cg- aaa aaa aat ata ttt ttg tag act ctt gag aat aga agg aaa aca tgt aca taa tta taa aaa atg aaa atc aat ggc gaa taa gac agt gcg att cgc gcc atg gag tcg gca gat ttc atg gct gtc gat gag cag ttt cac gac gac ctc gat ctt tgg tca tta tct ttg gta gat gac tat aaa aaa cat gga tta ggt gtt gac tgt tat gtt cta gaa cca gtt gtt gac agg aaa ata ttt gat aga ttt ctc ctt gaa cca att tgt gat cct gta gat gtt ctg tat gat tat ttt agg att cat aga gat aat att gat cag tat ata gta gat aga ctg ttt gca tat att aca tat aaa gat att ata tct gca tta gtg tca aag aat tat atg gaa gat att ttc tct ata att att aag aat tgt aat tct gta caa gat ctc tta ctt tac tat cta tct aat gca tat gta gaa ata gac att gtt gat ttt atg gta gat cat ggg gct gta ata tat aaa ata gaa tgc ttg aat gcc tat ttt agg gga ata tgt aaa aag gaa agt agt gtt gtt gag ttt att ttg aat tgt ggt atc cca gat gaa aat gat gtt aaa tta gat cta tat aaa ata att cag tat act agg gga ttc ctt gta gat gaa ccc aca gta tta gaa att tat aag ctt tgt atc cca tat att gaa gat atc aat caa cta gat gct ggt gga agg acc ttg ctt tat cgc gct atc tat gca ggt tat ata gat tta gta tca tgg cta tta gaa aat gga gca aat gtc aac gca gta atg agt aat gga tat aca tgt ctt gac gtg gcc gtg gat agg gga tct gtc atc gcc cgt agg gaa gca cat ctt aaa ata tta gaa ata ttg ctt aga gaa cca ttg tct att gac tgt ata aaa tta gct ata ctt aat aat aca att gaa aac cat gat gtg ata aag ctc tgt atc aag tat ttt atg atg gta gat tat tca ctt tgt aat gtg tat gca tca tca ctc ttt gat tat ata att gat tgt aaa caa gaa ttg gag tac att agg cag atg aaa att cat aat aca acc atg tat gag tta atc tat aat aga gac aaa aac aag cat gct tcc cat att cta cat agg tat tct aaa cat cca gtt ttg aca cag tgt atc act aaa gga ttc aag att tac aca gaa gta acc gag cag gtc act aaa gct cta aac aga cgt gct cta ata gat gag ata ata aac aat gta tca act gat gac aat ctc cta tca aaa ctt cca tta gaa att agg gat cta att gtt tca caa gct gtc ata tag agt tct ctc cac cca cct ttc ttg aaa tga gtt aat --- --- --- --- --- --- --- --- --- --- --- --- --- --- --- --- --- --- --- --- --- --- --- --- --- --- --- --- --- --- --- --- --- --- --- --- --- --- --- --- --- --- --- --- --- --- --- --- --- --- --- --- --- --- --- --- --- --- --- --- --- --- --- --- --- --- --- --- --- --- --- --- --- --- --- --- --- --- --- --- --- --- --- --- --- --- --- --- --- --- --- --- --- --- --- --- --- --- --- --- --- --- --- --- --- --- --- --- --- --- --- --- --- --- --- --- --- --- --- --- --- --- --- --- --- --- --- --- --- --- --- --- --- --- --- --- --- --- --- --- --- --- --- --- --- --- --- --- --- --- --- --- --- -ag tca taa gtt agt taa gtc ata agt tag tt- --t tat aag tta gtt tat agt cta aca ctt cta att ttt ata cct tga tct ttt tct cta att atg aaa aag taa atc att atg aag atg gat gaa aat gga cga gat tgt gcg cat cgt taa cga tag tat gtg gta cgt acc taa cgc att tat gga cga cgg tga taa tga agg tca cat ttc tgt caa taa tgt ctg tca tat gta tct cgc att ctt tga tgt gga tat atc atc tca tct gtt taa att agt tat taa aca ctg cga tct gaa taa acg act aaa atg tgg taa ctc tcc att aca ttg cta tac gat gaa tac acg att taa tcc atc tgt att aaa gat att gtt acg cca cgg cat gcg taa ctt tga tag caa gga taa aaa agg aca tat tcc tct aca cca cta tct gat tca ttc act atc aat cga taa caa gat ctt tga tat act aac gga ccc cat tga tga ctt tag taa atc atc cga tct att gct gtg tta tct tag ata taa att caa tgg gag ctt aaa cta tta cgt tct gta caa att att gac taa agg atc tga ccc taa ttg cgt cga tga gga tgg act cac ttc tct tca tta cta ctg taa aca cat atc cgt gtt cca cga aag caa tta tta caa gtc aaa gag tca cac taa gat gcg agc tga gaa gcg att cat cta cgc gat aat aga tca tgg agc aaa cat taa cgc ggt tac gaa aat cgg aaa tac gcc gtt aca cac tta cct tca aca gta tac caa aca tag tcc tcg tgt ggt gta tgc tct ttt atc tcg agg agc cga tac gag gat acg taa taa tct tga ttg tac acc cat cat gga ata cat aaa gaa cga ttg tgc aac agg tca tat tct cat aat gtt act caa ttg gca cga aca aaa ata cgg gaa att aca aaa gga aga agg aca aca tct act tta tct att cat aaa aca taa tca agg ata tgg aag tcg ctc tct caa tat act acg gta tct act aga tag att cga cat tca gaa aga cga ata cta taa tac aat gac tcc tct tca tac cgc ctt cca gaa ttg caa taa caa tgt tgc ctc ata cct cgt ata cat cgg ata cga cat caa cct tcc gac taa aga cga taa gac agt att cga ctt ggt gtt tga aaa cag aaa cat cat ata caa ggc gga tgt cgt taa tga cat cat cca cca cag act gaa agt atc tct acc tat gat taa atc gtt gtt cta caa gat gtc gga gtt ctc tcc cta cga cga tca cta cgt aaa gaa gat aat agc cta ctg cat att aag gga cga gtc att tgc gga act aca tac taa att ctg ttt aaa cga gga cta taa aag tgt att tat gaa aaa tat atc att cga taa gat aga ttc cat cat cga aaa atg tag tcg tga cat aag tct cct caa aga gat tcg aat ctc aga cac caa ctt gta tac ggt att gag aac aga aga cat ccg gta tca cac cta tct cga agc cat aca ttc aga caa acg cat ttc att tcc cat gta cga cga tct cat aga aca gtg tca tct atc gat gga gca taa aag taa act cgt cga caa agc act caa taa att aga gtc tac cat cga tag tca atc tag act atc gta ttt gcc tcc gga aat tat gcg caa tat cat aac caa gct aag cga cta cca tct aaa cag tat gtt gta cgg aaa gaa cca tta caa ata tta tcc atg ata gaa aga aaa tat tta aaa aat aat cta tat gat tgg aga agt agg aaa caa aca gta aca aga cga cga tta cta c-a tta tta aat cat gag gtc cgt att ata ctc gta tat att gtt tct ctc atg tat aat aat aaa cgg aag aga tat agc acc aca tgc acc atc caa tgg aaa atg taa aga caa cga ata cag aag ccg taa tct atg ttg tct atc gtg tcc tcc ggg aac tta cgc ttc cag att atg tga tag caa gac taa tac aca atg tac gcc gtg tgg ttc gga tac ctt tac atc tca caa taa tca ttt aca ggc ttg tct aag ttg taa cgg aag atg tga tag taa tca ggt aga gac gcg atc gtg taa cac gac tca caa tag aat ctg tga atg ctc tcc agg ata tta ttg tct tct caa agg atc atc agg gtg tag aac atg tat ttc taa aac aaa gtg tgg aat agg ata cgg agt atc cgg ata cac gtc tac cgg aga cgt cat ctg ttc tcc gtg tgg tcc cgg aac ata ttc tca cac cgt ctc ttc cac aga taa atg cga acc --- cgt aac cag caa tac att taa cta tat cga tgt gga aat taa cct gta tcc agt caa cga cac atc gtg tac tcg gac gac cac tac cgg tct cag cga atc cat ctc aac gtc gga act aac tat tac cat gaa tca taa aga ttg tga tcc cgt ctt tcg tgc aga ata ctt ctc tgt cct taa taa tgt agc aac ttc agg att ctt tac agg aga aaa tag ata tca gaa tac ttc aaa gat atg tac tct gaa ttt cga gat taa atg taa caa caa aga ttc atc ttc caa aca gtt aac gaa aac aaa gaa tga tac tat cat gcc gca ttc aga gac ggt aac tct agt ggg cga ctg tct atc tag cgt cga cat cta cat act ata tag taa tac caa tac tca aga cta cga aac gga tac aat ctc tta tca tat ggg taa tgt tct cga tgt caa tag cca tat gcc cgc tag ttg cga tat aca taa act gat cac taa ttc cca gaa tcc cac cca ctt ata gta ag- ttt ttt tta ccc ata aat aat aaa tac aat aat taa ttt ctc gta aaa gta gaa aat ata ttc taa ttt att ata tgg taa gaa agt aga atc atc tag aac agt aat caa tca ata gca atc atg aaa caa tat att gtc ctg gca tgc atg tgc cta gtg gca gct gct atg cct act agt ctt caa caa tcc tca tcc tcg tgt act gaa gaa gaa aac aaa cat cat atg gga atc gat gtt att atc aaa gtc aca aag caa gac caa aca ccg acc aat gat aag att tgt caa tcc gta acg gaa gtt aca gag acc gaa gat gat gag gta tcc gaa gat gat gag gta tcc gaa gaa gtt gta aaa gga gat ccc acc act tat tac aat atc gtc ggc gcg ggt ctt aac atg aac ttt gga ttc acc aaa tgc cca aag att tca tcc atc tcc gaa tcc tct gat gga aac act gtg aat act aga ttg tcc agc gtg tca cca gga caa ggt aag gac tct ccc gcg atc acg cgt gaa gaa gct ctg gct atg atc aaa gac tgt gag atg tct atc gac atc aga tgt agc gaa gaa gag aaa gac agc gac atc aag acc cat cca gta ctt ggg tct aac atc tca cat aag aaa gtg agt tac aaa gat atc atc ggt tca acg atc gtt gat aca aaa tgt gtt aag aac cta gag ttt agc gta cgt atc gga gac atg tgt gag gaa tca tct gaa ctt gaa gtc aag gat gga ttc aag tat gtc gat gga tcg gca tct gaa ggt gca acc gat gat act tca ctc atc gat tca aca aaa ctc aaa gca tgt gtc tga atc gat aac tct att cat ctg aaa atg gat gag ttg ggt taa tcg aac gat tca gac acc gca cca cga att aaa aaa gtg tac cgg gta cta tat tcc ggt ttg caa aac aaa aat gtt taa cta cat tca caa aaa gtt acc tct cgt tac ttc ttc ttt ctg ttt caa tat gtg ata cga tat gat cac tat tcg tat tct ctt tct ctc ttg tct cat aaa aaa gtt tta c-- --- --- --- aaa aaa a-t att ttt att ctc ttt ctc tct tcg at- ggt ctc aca aaa ata tta aac ctc ttt ctg atg tct caa cta ttt cgt aaa cga taa cgt cca aca ata tat tct cgt aga gct tat caa tat cct tat gcc aat --- ggt tgt cag aca att gca tca taa aat aat gtt tat aat tta cac gtt aac atc ata taa taa acg tat ata gtt aat att ttt gga ata taa atg atc tgt aaa atc cat gta ggg gac act gct cac gtt ttt tct cta gta cat aat ttc aca caa gtt ttt ata cag aca aat taa ttc tcg tcc ata ta- ttt taa aac att gac ttt tgt act aag aaa aat atc ttg act aac cat ctc ttt ct- --- --- --- --- --- --- --- --- --- --- --- --- --- --- --- --- --- --- --- --- --- --- --- --- --- --- --- --- --- --- --- --- --- --- --- --- --- --- --- --- --- --- --- --- --- --- --- --- --- --- --- --- --- --- --- --- --- --- --- --- --- --- --- --- --- --- --- --- --- --c tcg tcg atg ggt ctc --- --- --- --- --- --- --- --- --- --- --- --- --- --- --- --- --- --- --- --- --- --- --- -ac aaa aat att aaa cct ctt tct gat gga gtc gta aaa agt ttt tat cct ttc tct cgt cga tgg gtc tca caa aaa tat taa acc tct ttc tga tgg agt cgt aaa aag ttt tta tcc ttt ctc tcg tcg atg ggt ctc aca aaa ata tta aac ctc ttt ctg atg gtc tct ata aa

>JX878423.1_Monkeypox_virus_isolate_DRC_07-0337_complete_genome

atc gag gac gac ttc aac tat gac agc tcg gtg gcg tct gcc agc gtg cac ata cga atg gca ttt cta aga aaa gtc tac ggt atc ctt gtc tac aat ttc ctt taa caa cgg caa cag ctg cag tat ttt tat act ttg aat gca ttc gga cat tta tac aag gga gtc ctg ttc taa tat tgg cat caa tgt tcg gat cta tag gct tga ttt tcg cat tga ctt tac aca gac ata aac atc ccc tga atc tgt acc tac ttt gtg gat tta cac tgt tag aat ctc taa cgc tgg cct ctg ttg tta ctt tct atg atg cgc gta tcg tta tac aag ctt tca tgt tga cta ctg cag tgt ttc ttg ctc tga cta cat gta ctc tac aat caa aga gag att tca gta aac ttg taa cag gat tgt ttg ctg ctt tct gga ttt taa ttt tgt cag gag tct tga gga taa agt tta aaa tag aat taa taa aga aca tat agg tca -tt ttt taa aca tgg ata gaa acc aag gtt gtt agt taa taa tat aca aga tat ttt ttc tca ctc tga tcc atg taa acc aag gac gag aga cac tct cat tcc tca ttc acg aca cca tta aaa atg gaa att aaa gcc ctc tat taa gca cag acg gct aca ggt cta cca tca ggt taa cct tcg tct acc ttc aca atg gcc ttt cct tgt gcc cag ttc agt ccc tgt cat tgc cac gct act aag gac tcc ctg aat acc gtg act gac gtc aga cat tgt ctg act gaa tac atc ctg tgg gtt tct cat aga tgg acc cat aga gaa agc gca ggg cct ctc tac agg ctt ctc atc tct ttc aga att gat gca atg gag cta ttt ggt agc gag ttg aag gag ttc tcg gat tca ctt ccg tgg gac aat atc gac aat tgc gtg gag atc att aaa tgt ttc atc aga aat gac tcc atg aaa acc gcc aaa gaa ctt tgt gca ata att gga ctt tgt act caa tca gct att gtc act gga aga gtc ttc aat gat aag tat atc gac ata cta ctt ata ctg cga aag att ctg aat gag aac gac tat ctc acc ctc ttg gat cat atc ctc act gct aaa tac taa atc tcc ttc atg ctc tct cac taa tac tct tac tca cta cac ttt tta tca tct tat gat gaa tga ttg cct tca tca -tt ttt cgt gga ata taa tat agg aat aat tag cac cag aat agc tat gga tat ctc gtt aag aat att ctc gat aag aga cat aat gta gac ata gtt att ata tcc ttc tta gat aag tgt tac gct act gga aag ttt cca tcg tta --- --- --- --- --- --- --- --- --- --- --- --- --- --- --- --- --- --- --- --- --- --- --- --- --- --- --- --- --- --- --- --- --- --- --- --- --- --- --- --- --- --- --- --- --- --- --- --- --- --- --- --- --- --- --- --- --- --- --- --- --- --- --- --- --- --- --- --- --- --- --- --- --- --- --- --- --- --- --- --- --- --- --- --- --- --- --- --- --- --- --- --- --- --- --- --- --- --- --- --- --- --- --- --- --- --- --- --- --- --- --- --- --- --- --- --- --- --- --- --- --- --- --- --- --- --- --- --- --- --- --- --- --- --- --- --- --- --- --- --- --- --- --- --- --- --- --- --- --- --- --- --- --- --- --- --- --- --- --- --- --- --- --- --- --- --- --- --- --- --- --- --- --- --- --- --- --- --- --- --- --- --- --- --- --- --- --- --- --- --- --- --- --- --- --- --- --- --- --- --- --- --- --- --- --- --- --- --- --- aag tga tat aac taa tac agc aga tat att gct gga caa tta tcc aat tat gac ctt tga tgg aaa gga tat tta tcc atc tat cgc gtt cat ggt cgg taa taa act ttt cct aga tct tta taa aaa tat ctt tgt aga att ttt cag act att tcg agt atc tgt aag tag tca ata cga gga att aga ata tta tta ttc atg tga tta tac taa caa ccg tcc tac aat taa aca aca tta ctt tta taa cgg cga aga ata tac tga aat tga tag atc gaa aaa agc cac taa taa aaa cag ttg gtt aat tac ttc agg ctt tag act aca aaa atg gtt cga tag cga aga ttg tat aat tta tct cag atc ttt agt tag aag aat gga aga cag taa caa aaa cag taa aaa aac tta gta ctt aga tat cg- aaa aaa aat ata ttt ttg tag act ctt gag aat aga agg aaa aca tgt aca taa tta taa aaa atg aaa atc aat ggc gaa taa gac agt gcg att cgc gcc atg gag tcg gca gat ttc atg gct gtc gat gag cag ttt cac gac gac ctc gat ctt tgg tca tta tct ttg gta gat gac tat aaa aaa cat gga tta ggt gtt gac tgt tat gtt cta gaa cca gtt gtt gac agg aaa ata ttt gat aga ttt ctc ctt gaa cca att tgt gat cct gta gat gtt ctg tat gat tat ttt agg att cat aga gat aat att gat cag tat ata gta gat aga ctg ttt gca tat att aca tat aaa gat att ata tct gca tta gtg tca aag aat tat atg gaa gat att ttc tct ata att att aag aat tgt aat tct gta caa gat ctc tta ctt tac tat cta tct aat gca tat gta gaa ata gac att gtt gat ttt atg gta gat cat ggg gct gta ata tat aaa ata gaa tgc ttg aat gcc tat ttt agg gga ata tgt aaa aag gaa agt agt gtt gtt gag ttt att ttg aat tgt ggt atc cca gat gaa aat gat gtt aaa tta gat cta tat aaa ata att cag tat act agg gga ttc ctt gta gat gaa ccc aca gta tta gaa att tat aag ctt tgt atc cca tat att gaa gat atc aat caa cta gat gct ggt gga agg acc ttg ctt tat cgc gct atc tat gca ggt tat ata gat tta gta tca tgg cta tta gaa aat gga gca aat gtc aac gca gta atg agt aat gga tat aca tgt ctt gac gtg gcc gtg gat agg gga tct gtc atc gcc cgt agg gaa gca cat ctt aaa ata tta gaa ata ttg ctt aga gaa cca ttg tct att gac tgt ata aaa tta gct ata ctt aat aat aca att gaa aac cat gat gtg ata aag ctc tgt atc aag tat ttt atg atg gta gat tat tca ctt tgt aat gtg tat gca tca tca ctc ttt gat tat ata att gat tgt aaa caa gaa ttg gag tac att agg cag atg aaa att cat aat aca acc atg tat gag tta atc tat aat aga gac aaa aac aag cat gct tcc cat att cta cat agg tat tct aaa cat cca gtt ttg aca cag tgt atc act aaa gga ttc aag att tac aca gaa gta acc gag cag gtc act aaa gct cta aac aga cgt gct cta ata gat gag ata ata aac aat gta tca act gat gac aat ctc cta tca aaa ctt cca tta gaa att agg gat cta att gtt tca caa gct gtc ata tag agt tct ctc cac cca cct ttc ttg aaa tga gtt aat --- --- --- --- --- --- --- --- --- --- --- --- --- --- --- --- --- --- --- --- --- --- --- --- --- --- --- --- --- --- --- --- --- --- --- --- --- --- --- --- --- --- --- --- --- --- --- --- --- --- --- --- --- --- --- --- --- --- --- --- --- --- --- --- --- --- --- --- --- --- --- --- --- --- --- --- --- --- --- --- --- --- --- --- --- --- --- --- --- --- --- --- --- --- --- --- --- --- --- --- --- --- --- --- --- --- --- --- --- --- --- --- --- --- --- --- --- --- --- --- --- --- --- --- --- --- --- --- --- --- --- --- --- --- --- --- --- --- --- --- --- --- --- --- --- --- --- --- --- --- --- --- --- -ag tca taa gtt agt taa gtc ata agt tag tt- --t tat aag tta gtt tat agt cta aca ctt cta att ttt ata cct tga tct ttt tct cta att atg aaa aag taa atc att atg aag atg gat gaa aat gga cga gat tgt gcg cat cgt taa cga tag tat gtg gta cgt acc taa cgc att tat gga cga cgg tga taa tga agg tca cat ttc tgt caa taa tgt ctg tca tat gta tct cgc att ctt tga tgt gga tat atc atc tca tct gtt taa att agt tat taa aca ctg cga tct gaa taa acg act aaa atg tgg taa ctc tcc att aca ttg cta tac gat gaa tac acg att taa tcc atc tgt att aaa gat att gtt acg cca cgg cat gcg taa ctt tga tag caa gga taa aaa agg aca tat tcc tct aca cca cta tct gat tca ttc act atc aat cga taa caa gat ctt tga tat act aac gga ccc cat tga tga ctt tag taa atc atc cga tct att gct gtg tta tct tag ata taa att caa tgg gag ctt aaa cta tta cgt tct gta caa att att gac taa agg atc tga ccc taa ttg cgt cga tga gga tgg act cac ttc tct tca tta cta ctg taa aca cat atc cgt gtt cca cga aag caa tta tta caa gtc aaa gag tca cac taa gat gcg agc tga gaa gcg att cat cta cgc gat aat aga tca tgg agc aaa cat taa cgc ggt tac gaa aat cgg aaa tac gcc gtt aca cac tta cct tca aca gta tac caa aca tag tcc tcg tgt ggt gta tgc tct ttt atc tcg agg agc cga tac gag gat acg taa taa tct tga ttg tac acc cat cat gga ata cat aaa gaa cga ttg tgc aac agg tca tat tct cat aat gtt act caa ttg gca cga aca aaa ata cgg gaa att aca aaa gga aga agg aca aca tct act tta tct att cat aaa aca taa tca agg ata tgg aag tcg ctc tct caa tat act acg gta tct act aga tag att cga cat tca gaa aga cga ata cta taa tac aat gac tcc tct tca tac cgc ctt cca gaa ttg caa taa caa tgt tgc ctc ata cct cgt ata cat cgg ata cga cat caa cct tcc gac taa aga cga taa gac agt att cga ctt ggt gtt tga aaa cag aaa cat cat ata caa ggc gga tgt cgt taa tga cat cat cca cca cag act gaa agt atc tct acc tat gat taa atc gtt gtt cta caa gat gtc gga gtt ctc tcc cta cga cga tca cta cgt aaa gaa gat aat agc cta ctg cat att aag gga cga gtc att tgc gga act aca tac taa att ctg ttt aaa cga gga cta taa aag tgt att tat gaa aaa tat atc att cga taa gat aga ttc cat cat cga aaa atg tag tcg tga cat aag tct cct caa aga gat tcg aat ctc aga cac caa ctt gta tac ggt att gag aac aga aga cat ccg gta tca cac cta tct cga agc cat aca ttc aga caa acg cat ttc att tcc cat gta cga cga tct cat aga aca gtg tca tct atc gat gga gca taa aag taa act cgt cga caa agc act caa taa att aga gtc tac cat cga tag tca atc tag act atc gta ttt gcc tcc gga aat tat gcg caa tat cat aac caa gct aag cga cta cca tct aaa cag tat gtt gta cgg aaa gaa cca tta caa ata tta tcc atg ata gaa aga aaa tat tta aaa aat aat cta tat gat tgg aga agt agg aaa caa aca gta aca aga cga cga tta cta c-a tta tta aat cat gag gtc cgt att ata ctc gta tat att gtt tct ctc atg tat aat aat aaa cgg aag aga tat agc acc aca tgc acc atc caa tgg aaa atg taa aga caa cga ata cag aag ccg taa tct atg ttg tct atc gtg tcc tcc ggg aac tta cgc ttc cag att atg tga tag caa gac taa tac aca atg tac gcc gtg tgg ttc gga tac ctt tac atc tca caa taa tca ttt aca ggc ttg tct aag ttg taa cgg aag atg tga tag taa tca ggt aga gac gcg atc gtg taa cac gac tca caa tag aat ctg tga atg ctc tcc agg ata tta ttg tct tct caa agg atc atc agg gtg tag aac atg tat ttc taa aac aaa gtg tgg aat agg ata cgg agt atc cgg ata cac gtc tac cgg aga cgt cat ctg ttc tcc gtg tgg tcc cgg aac ata ttc tca cac cgt ctc ttc cac aga taa atg cga acc --- cgt aac cag caa tac att taa cta tat cga tgt gga aat taa cct gta tcc agt caa cga cac atc gtg tac tcg gac gac cac tac cgg tct cag cga atc cat ctc aac gtc gga act aac tat tac cat gaa tca taa aga ttg tga tcc cgt ctt tcg tgc aga ata ctt ctc tgt cct taa taa tgt agc aac ttc agg att ctt tac agg aga aaa tag ata tca gaa tac ttc aaa gat atg tac tct gaa ttt cga gat taa atg taa caa caa aga ttc atc ttc caa aca gtt aac gaa aac aaa gaa tga tac tat cat gcc gca ttc aga gac ggt aac tct agt ggg cga ctg tct atc tag cgt cga cat cta cat act ata tag taa tac caa tac tca aga cta cga aac gga tac aat ctc tta tca tat ggg taa tgt tct cga tgt caa tag cca tat gcc cgc tag ttg cga tat aca taa act gat cac taa ttc cca gaa tcc cac cca ctt ata gta ag- ttt ttt tta ccc ata aat aat aaa tac aat aat taa ttt ctc gta aaa gta gaa aat ata ttc taa ttt att ata tgg taa gaa agt aga atc atc tag aac agt aat caa tca ata gca atc atg aaa caa tat att gtc ctg gca tgc atg tgc cta gtg gca gct gct atg cct act agt ctt caa caa tcc tca tcc tcg tgt act gaa gaa gaa aac aaa cat cat atg gga atc gat gtt att atc aaa gtc aca aag caa gac caa aca ccg acc aat gat aag att tgt caa tcc gta acg gaa gtt aca gag acc gaa gat gat gag gta tcc gaa gat gat gag gta tcc gaa gaa gtt gta aaa gga gat ccc acc act tat tac aat atc gtc ggc gcg ggt ctt aac atg aac ttt gga ttc acc aaa tgc cca aag att tca tcc atc tcc gaa tcc tct gat gga aac act gtg aat act aga ttg tcc agc gtg tca cca gga caa ggt aag gac tct ccc gcg atc acg cgt gaa gaa gct ctg gct atg atc aaa gac tgt gag atg tct atc gac atc aga tgt agc gaa gaa gag aaa gac agc gac atc aag acc cat cca gta ctt ggg tct aac atc tca cat aag aaa gtg agt tac aaa gat atc atc ggt tca acg atc gtt gat aca aaa tgt gtt aag aac cta gag ttt agc gta cgt atc gga gac atg tgt gag gaa tca tct gaa ctt gaa gtc aag gat gga ttc aag tat gtc gat gga tcg gca tct gaa ggt gca acc gat gat act tca ctc atc gat tca aca aaa ctc aaa gca tgt gtc tga atc gat aac tct att cat ctg aaa atg gat gag ttg ggt taa tcg aac gat tca gac acc gca cca cga att aaa aaa gtg tac cgg gta cta tat tcc ggt ttg caa aac aaa aat gtt taa cta cat tca caa aaa gtt acc tct cgt tac ttc ttc ttt ctg ttt caa tat gtg ata cga tat gat cac tat tcg tat tct ctt tct ctc ttg tct cat aaa aaa gtt tta c-- --- --- --- aaa aaa a-t att ttt att ctc ttt ctc tct tcg at- ggt ctc aca aaa ata tta aac ctc ttt ctg atg tct caa cta ttt cgt aaa cga taa cgt cca aca ata tat tct cgt aga gct tat caa tat cct tat gcc aat --- ggt tgt cag aca att gca tca taa aat aat gtt tat aat tta cac gtt aac atc ata taa taa acg tat ata gtt aat att ttt gga ata taa atg atc tgt aaa atc cat gta ggg gac act gct cac gtt ttt tct cta gta cat aat ttc aca caa gtt ttt ata cag aca aat taa ttc tcg tcc ata ta- ttt taa aac att gac ttt tgt act aag aaa aat atc ttg act aac cat ctc ttt ctc tcg tcg atg ggt ctc aca aaa ata tta aac ctc ttt ctg atg g-- --- --- --- --- --- --- --- --- --- --- --- --- --- --- --- --- --- --- --- --- --- --- --- --- --- --- --- --- --- --- --- --- --- --- --- --- --- --- --- --- --- --- --- --- --- --- agt cgt aaa aag ttt tta tcc ttt ctc tcg tcg atg ggt ctc --- --- --- --- --- --- --- --- --- --- --- --- --- --- --- --- --- --- --- --- --- --- --- -ac aaa aat att aaa cct ctt tct gat gga gtc gta aaa agt ttt tat cct ttc tct cgt cga tgg gtc tca caa aaa tat taa acc tct ttc tga tgg agt cgt aaa aag ttt tta tcc ttt ctc tcg tcg atg ggt ctc aca aaa ata tta aac ctc ttt ctg atg gtc tct ata aa

>JX878408.1_Monkeypox_virus_isolate_DRC_06-0970_complete_genome

atc gag gac gac ttc aac tat gac agc tcg gtg gcg tct gcc agc gtg cac ata cga atg gca ttt cta aga aaa gtc tac ggt atc ctt gtc tac aat ttc ctt taa caa cgg caa cag ctg cag tat ttt tat act ttg aat gca ttc gga cat tta tac aag gga gtc ctg ttc taa tat tgg cat caa tgt tcg gat cta tag gct tga ttt tcg cat tga ctt tac aca gac ata aac atc ccc tga atc tgt acc tac ttt gtg gat tta cac tgt tag aat ctc taa cgc tgg cct ctg ttg tta ctt tct atg atg cgc gta tcg tta tac aag ctt tca tgt tga cta ctg cag tgt ttc ttg ctc tga cta cat gta ctc tac aat caa aga gag att tca gta aac ttg taa cag gat tgt ttg ctg ctt tct gga ttt taa ttt tgt cag gag tct tga gga taa agt tta aaa tag aat taa taa aga aca tat agg tca -tt ttt taa aca tgg ata gaa acc aag gtt gtt agt taa taa tat aca aga tat ttt ttc tca ctc tga tcc atg taa acc aag gac gag aga cac tct cat tcc tca ttc acg aca cca tta aaa atg gaa att aaa gcc ctc tat taa gca cag acg gct aca ggt cta cca tca ggt taa cct tcg tct acc ttc aca atg gcc ttt cct tgt gcc cag ttc agt ccc tgt cat tgc cac gct act aag gac tcc ctg aat acc gtg act gac gtc aga cat tgt ctg act gaa tac atc ctg tgg gtt tct cat aga tgg acc cat aga gaa agc gca ggg cct ctc tac agg ctt ctc atc tct ttc aga att gat gca atg gag cta ttt ggt agc gag ttg aag gag ttc tcg gat tca ctt ccg tgg gac aat atc gac aat tgc gtg gag atc att aaa tgt ttc atc aga aat gac tcc atg aaa acc gcc aaa gaa ctt tgt gca ata att gga ctt tgt act caa tca gct att gtc act gga aga gtc ttc aat gat aag tat atc gac ata cta ctt atg ctg cga aag att ctg aat gag aac gac tat ctc acc ctc ttg gat cat atc ctc act gct aaa tac taa atc tcc ttc atg ctc tct cac taa tac tct tac tca cta cac ttt tta tca tct tat gat gaa tga ttg cct tca tca -tt ttt cgt gga ata taa tat agg aat aat tag cac cag aat agc tat gga tat ctc gtt aag aat att ctc gat aag aga cat aat gta gac ata gtt att ata tcc ttc tta gat aag tgt tac gct act gga aag ttt cca tcg tta tta tta cgt gaa gat gat atc att aaa cca aca ttg aga ttg gct ctt atg tta gct gga ttg aat tac tgt aat aaa tgc atc gag tat aga ggg ata tag caa ttc tcg ata ata gtc atg caa tat ttg aat gag act gat aat tta ggt aat aca gta ct- -ac aca cac ata tct ttc tag att ata tat cgt taa aaa tct gta aga ggt ata ttt ctc ata agt atc cac tgt gta ata tta tta atg gat ata tag ata aca caa tag gga cta ata gta ttg taa aag ata taa tcg act att t-- gta cat atc cag ata tct ata ttc cta cta gtt tgc tgc gta gtt gca tca ttg ata tgc atg -at tta tca gga ttc aga gat gaa tta cta agt aaa cta caa tcc cac aat aag taa gaa tca aat atc aaa aac tca ctt ttg att ttt cta gtc tta agt aat aca tat att tat taa tag acc tat gaa at- aaa aaa gtt aac aat gga ttc gcg tat agc tat tta cgt att agt atc ggc atc tct ttt gta tct tgt taa ttg tca caa act agt aca tta ctt caa tct gaa aat aaa tgg aag tga tat aac taa tac agc aga tat att gct gga caa tta tcc aat tat gac ctt tga tgg aaa gga tat tta tcc atc tat cgc gtt cat ggt cgg taa taa act ttt cct aga tct tta taa aaa tat ctt tgt aga att ttt cag act att tcg agt atc tgt aag tag tca ata cga gga att aga ata tta tta ttc atg tga tta tac taa caa ccg tcc tac aat taa aca aca tta ctt tta taa cgg cga aga ata tac tga aat tga tag atc gaa aaa agc cac taa taa aaa cag ttg gtt aat tac ttc agg ctt tag act aca aaa atg gtt cga tag cga aga ttg tat aat tta tct cag atc ttt agt tag aag aat gga aga cag taa caa aaa cag taa aaa aac tta gta ctt aga tat cg- aaa aaa aat ata ttt ttg tag act ctt gag aat aga agg aaa aca tgt aca taa tta taa aaa atg aaa atc aat ggc gaa taa gac agt gcg att cgc gcc atg gag tcg gca gat ttc atg gct gtc gat gag cag ttt cac gac gac ctc gat ctt tgg tca tta tct ttg gta gat gac tat aaa aaa cat gga tta ggt gtt gac tgt tat gtt cta gaa cca gtt gtt gac agg aaa ata ttt gat aga ttt ctc ctt gaa cca att tgt gat cct gta gat gtt ctg tat gat tat ttt agg att cat aga gat aat att gat cag tat ata gta gat aga ctg ttt gca tat att aca tat aaa gat att ata tct gca tta gtg tca aag aat tat atg gaa gat att ttc tct ata att att aag aat tgt aat tct gta caa gat ctc tta ctt tac tat cta tct aat gca tat gta gaa ata gac att gtt gat ttt atg gta gat cat ggg gct gta ata tat aaa ata gaa tgc ttg aat gcc tat ttt agg gga ata tgt aaa aag gaa agt agt gtt gtt gag ttt att ttg aat tgt ggt atc cca gat gaa aat gat gtt aaa tta gat cta tat aaa ata att cag tat act agg gga ttc ctt gta gat gaa ccc aca gta tta gaa att tat aag ctt tgt atc cca tat att gaa gat atc aat caa cta gat gct ggt gga agg acc ttg ctt tat cgc gct atc tat gca ggt tat ata gat tta gta tca tgg cta tta gaa aat gga gca aat gtc aac gca gta atg agt aat gga tat aca tgt ctt gac gtg gcc gtg gat agg gga tct gtc atc gcc cgt agg gaa aca cat ctt aaa ata tta gaa ata ttg ctt aga gaa cca ttg tct att gac tgt ata aaa tta gct ata ctt aat aat aca att gaa aac cat gat gtg ata aag ctc tgt atc aag tat ttt atg atg gta gat tat tca ctt tgt aat gtg tat gca tca tca ctc ttt gat tat ata att gat tgt aaa caa gaa ttg gag tac att agg cag atg aaa att cat aat aca acc atg tat gag tta atc tat aat aga gac aaa aac aag cat gct tcc cat att cta cat agg tat tct aaa cat cca gtt ttg aca cag tgt atc act aaa gga ttc aag att tac aca gaa gta acc gag cag gtc act aaa gct cta aac aga cgt gct cta ata gat gag ata ata aac aat gta tca act gat gac aat ctc cta tca aaa ctt cca tta gaa att agg gat cta att gtt tca caa gct gtc ata tag agt tct ctc cac cca cct ttc ttg aaa tga gtt aat --- --- --- --- --- --- --- --- --- --- --- --- --- --- --- --- --- --- --- --- --- --- --- --- --- --- --- --- --- --- --- --- --- --- --- --- --- --- --- --- --- --- --- --- --- --- --- --- --- --- --- --- --- --- --- --- --- --- --- --- --- --- --- --- --- --- --- --- --- --- --- --- --- --- --- --- --- --- --- --- --- --- --- --- --- --- --- --- --- --- --- --- --- --- --- --- --- --- --- --- --- --- --- --- --- --- --- --- --- --- --- --- --- --- --- --- --- --- --- --- --- --- --- --- --- --- --- --- --- --- --- --- --- --- --- --- --- --- --- --- --- --- --- --- --- --- --- --- --- --- --- --- --- -ag tca taa gtt agt taa gtc ata agt tag tt- --t tat aag tta gtt tat agt cta aca ctt cta att ttt ata cct tga tct ttt tct cta att atg aaa aag taa atc att atg aag atg gat gaa aat gga cga gat tgt gcg cat cgt taa cga tag tat gtg gta cgt acc taa cgc att tat gga cga cgg tga taa tga agg tca cat ttc tgt caa taa tgt ctg tca tat gta tct cgc att ctt tga tgt gga tat atc atc tca tct gtt taa att agt tat taa aca ctg cga tct gaa taa acg act aaa atg tgg taa ctc tcc att aca ttg cta tac gat gaa tac acg att taa tcc atc tgt att aaa gat att gtt acg cca cgg cat gcg taa ctt tga tag caa gga taa aaa agg aca tat tcc tct aca cca cta tct gat tca ttc act atc aat cga taa caa gat ctt tga tat act aac gga ccc cat tga tga ctt tag taa atc atc cga tct att gct gtg tta tct tag ata taa att caa tgg gag ctt aaa cta tta cgt tct gta caa att att gac taa agg atc tga ccc taa ttg cgt cga tga gga tgg act cac ttc tct tca tta cta ctg taa aca cat atc cgt gtt cca cga aag caa tta tta caa gtc aaa gag tca cac taa gat gcg agc tga gaa gcg att cat cta cgc gat aat aga tca tgg agc aaa cat taa cgc ggt tac gaa aat cgg aaa tac gcc gtt aca cac tta cct tca aca gta tac caa aca tag tcc tcg tgt ggt gta tgc tct ttt atc tcg agg agc cga tac gag gat acg taa taa tct tga ttg tac acc cat cat gga ata cat aaa gaa cga ttg tgc aac agg tca tat tct cat aat gtt act caa ttg gca cga aca aaa ata cgg gaa att aca aaa gga aga agg aca aca tct act tta tct att cat aaa aca taa tca agg ata tgg aag tcg ctc tct caa tat act acg gta tct act aga tag att cga cat tca gaa aga cga ata cta taa tac aat gac tcc tct tca tac cgc ctt cca gaa ttg caa taa caa tgt tgc ctc ata cct cgt ata cat cgg ata cga cat caa cct tcc gac taa aga cga taa gac agt att cga ctt ggt gtt tga aaa cag aaa cat cat ata caa ggc gga tgt cgt taa tga cat cat cca cca cag act gaa agt atc tct acc tat gat taa atc gtt gtt cta caa gat gtc gga gtt ctc tcc cta cga cga tca cta cgt aaa gaa gat aat agc cta ctg cct att aag gga cga gtc att tgc gga act aca tac taa att ctg ttt aaa cga gga cta taa aag tgt att tat gaa aaa tat atc att cga taa gat aga ttc cat cat cga aaa atg tag tcg tga cat aag tct cct caa aga gat tcg aat ctc aga cac caa ctt gta tac ggt att gag aac aga aga cat ccg gta tca cac cta tct cga agc cat aca ttc aga caa acg cat ttc att tcc cat gta cga cga tct cat aga aca gtg tca tct atc gat gga gca taa aag taa act cgt cga caa agc act caa taa att aga gtc tac cat cga tag tca atc tag act atc gta ttt gcc tcc gga aat tat gcg caa tat cat aac caa gct aag cga cta cca tct aaa cag tat gtt gta cgg aaa gaa cca tta caa ata tta tcc atg ata gaa aga aaa tat tta aaa aat aat cta tat gat tgg aga agt agg aaa caa aca gta aca aga cga cga tta cta c-a tta tta aat cat gag gtc cgt att ata ctc gta tat att gtt tct ctc atg tat aat aat aaa cgg aag aga tat agc acc aca tgc acc atc caa tgg aaa atg taa aga caa cga ata cag aag ccg taa tct atg ttg tct atc gtg tcc tcc ggg aac tta cgc ttc cag att atg tga tag caa gac taa tac aca atg tac gcc gtg tgg ttc gga tac ctt tac atc tca caa taa tca ttt aca ggc ttg tct aag ttg taa cgg aag atg tga tag taa tca ggt aga gac gcg atc gtg taa cac gac tca caa tag aat ctg tga atg ctc tcc agg ata tta ttg tct tct caa agg atc atc agg gtg tag aac atg tat ttc taa aac aaa gtg tgg aat agg ata cgg agt atc cgg ata cac gtc tac cgg aga cgt cat ctg ttc tcc gtg tgg tcc cgg aac ata ttc tca cac cgt ctc ttc cac aga taa atg cga acc --- cgt aac cag caa tac att taa cta tat cga tgt gga aat taa cct gta tcc agt caa cga cac atc gtg tac tcg gac gac cac tac cgg tct cag cga atc cat ctc aac gtc gga act aac tat tac cat gaa tca taa aga ttg tga tcc cgt ctt tcg tgc aga ata ctt ctc tgt cct taa taa tgt agc aac ttc agg att ctt tac agg aga aaa tag ata tca gaa tac ttc aaa gat atg tac tct gaa ttt cga gat taa atg taa caa caa aga ttc atc ttc caa aca gtt aac gaa aac aaa gaa tga tac tat cat gcc gca ttc aga gac ggt aac tct agt ggg cga ctg tct atc tag cgt cga cat cta cat act ata tag taa tac caa tac tca aga cta cga aac gga tac aat ctc tta tca tat ggg taa tgt tct cga tgt caa tag cca tat gcc cgc tag ttg cga tat aca taa act gat cac taa ttc cca gaa tcc cac cca ctt ata gta agt ttt ttt tta ccc ata aat aat aaa tac aat aat taa ttt ctc gta aaa gta gaa aat ata ttc taa ttt att ata tgg taa gaa agt aga atc atc tag aac agt aat caa tca ata gca atc atg aaa caa tat att gtc ctg gca tgc atg tgc cta gtg gca gct gct atg cct act agt ctt caa caa tcc tca tcc tcg tgt act gaa gaa gaa aac aaa cat cat atg gga atc gat gtt att atc aaa gtc aca aag caa gac caa aca ccg acc aat gat aag att tgt caa tcc gta acg gaa gtt aca gag a-- --- --- --- --- --- -cc gaa gat gat gag gta tcc gaa gaa gtt gta aaa gga gat ccc acc act tat tac act atc gtc ggc gcg ggt ctt aac atg aac ttt gga ttc acc aaa tgc cca aag att tca tcc atc tcc gaa tcc tct gat gga aac act gtg aat act aga ttg tcc agc gtg tca cca gga caa ggt aag gac tct ccc gcg atc acg cgt gaa gaa gct ctg gct atg atc aaa gac tgt gag atg tct atc gac atc aga tgt agc gaa gaa gag aaa gac agc gac atc aag acc cat cca gta ctt ggg tct aac atc tca cat aag aaa gtg agt tac aaa gat atc atc ggt tca acg atc gtt gat aca aaa tgt gtt aag aac cta gag ttt agc gta cgt atc gga gac atg tgt gag gaa tca tct gaa ctt gaa gtc aag gat gga ttc aag tat gtc gac gga tcg gca tct gaa ggt gca acc gat gat act tca ctc atc gat tca aca aaa ctc aaa gca tgt gtc tga atc gat aac tct att cat ctg aaa atg gat gag ttg ggt taa tcg aac gat tca gac acc gca cca cga att aaa aaa gtg tac cgg gta cta tat tcc ggt ttg caa aac aaa aat gtt taa cta cat tca caa aaa gtt acc tct cgt tac ttc ttc ttt ctg ttt caa tat gtg ata cga tat gat cac tat tcg tat tct ctt tct ctc ttg tct cat -aa aaa gtt tta c-- --- --- --- aaa aaa a-t att ttt att ctc ttt ctc tct tcg at- ggt ctc aca aaa ata tta aac ctc ttt ctg atg tct caa cta ttt cgt aaa cga taa cgt cca aca ata tat tct cgt aga gct tat caa tat cct tat gcc aat cta ggt tgt cag aca att gca tca taa aat aat gtt tat aat tta cac gtt aac atc ata taa taa acg tat ata gtt aat att ttt gga ata taa atg atc tgt aaa atc cat gta ggg gac act gct cac gtt ttt tct cta gta cat aat ttc aca caa gtt ttt ata cag aca aat taa ttc tcg tcc ata ta- ttt taa aac att gac ttt tgt act aag aaa aat atc ttg act aac cat ctc ttt ctc tcg tcg atg ggt ctc aca aaa ata tta aac ctc ttt ctg atg g-- --- --- --- --- --- --- --- --- --- --- --- --- --- --- --- --- --- --- --- --- --- --- --- --- --- --- --- --- --- --- --- --- --- --- --- --- --- --- --- --- --- --- --- --- --- --- agt cgt aaa aag ttt tta tcc ttt ctc tct tcg atg ggt ctc --- --- --- --- --- --- --- --- --- --- --- --- --- --- --- --- --- --- --- --- --- --- --- --- --- --- --- --- --- --- --- --- --- --- --- --- --- --- --- --- --- --- --- --- --- --- --- --- --- --- --- --- --- --- --- --- --- --- --- --- --- --- --- --- --- --- --- --- --- --- aca aaa ata tta aac ctc ttt ctg atg gtc tct ata aa

>JX878420.1_Monkeypox_virus_isolate_DRC_07-0283_complete_genome

atc gag gac gac ttc aac tat gac agc tcg gtg gcg tct gcc agc gtg cac ata cga atg gca ttt cta aga aaa gtc tac ggt atc ctt gtc tac aat ttc ctt taa caa cgg caa cag ctg cag tat ttt tat act ttg aat gca ttc gga cat tta tac aag gga gtc ctg ttc taa tat tgg cat caa tgt tcg gat cta tag gct tga ttt tcg cat tga ctt tac aca gac ata aac atc ccc tga atc tgt acc tac ttt gtg gat tta cac tgt tag aat ctc taa cgc tgg cct ctg ttg tta ctt tct atg atg cgc gta tcg tta tac aag ctt tca tgt tga cta ctg cag tgt ttc ttg ctc tga cta cat gta ctc tac aat caa aga gag att tca gta aac ttg taa cag gat tgt ttg ctg ctt tct gga ttt taa ttt tgt cag gag tct tga gga taa agt tta aaa tag aat taa taa aga aca tat agg tca -tt ttt taa aca tgg ata gaa acc aag gtt gtt agt taa taa tat aca aga tat ttt ttc tca ctc tga tcc atg taa acc aag gac gag aga cac tct cat tcc tca ttc acg aca cca tta aaa atg gaa att aaa gcc ctc tat taa gca cag acg gct aca ggt cta cca tca ggt taa cct tcg tct acc ttc aca atg gcc ttt cct tgt gcc cag ttc agt ccc tgt cat tgc cac gct act aag gac tcc ctg aat acc gtg act gac gtc aga cat tgt ctg act gaa tac atc ctg tgg gtt tct cat aga tgg acc cat aga gaa agc gca ggg cct ctc tac agg ctt ctc atc tct ttc aga att gat gca atg gag cta ttt ggt agc gag ttg aag gag ttc tcg gat tca ctt ccg tgg gac aat atc gac aat tgc gtg gag atc att aaa tgt ttc atc aga aat gac tcc atg aaa acc gcc aaa gaa ctt tgt gca ata att gga ctt tgt act caa tca gct att gtc act gga aga gtc ttc aat gat aag tat atc gac ata cta ctt atg ctg cga aag att ctg aat gag aac gac tat ctc acc ctc ttg gat cat atc ctc act gct aaa tac taa atc tcc ttc atg ctc tct cac taa tac tct tac tca cta cac ttt tta tca tct tat gat gaa tga ttg cct tca tca -tt ttt cgt gga ata taa tat agg aat aat tag cac cag aat agc tat gga tat ctc gtt aag aat att ctc gat aag aga cat aat gta gac ata gtt att ata tcc ttc tta gat aag tgt tac gct act gga aag ttt cca tcg tta tta tta cgt gaa gat gat atc att aaa cca aca ttg aga ttg gct ctt atg tta gct gga ttg aat tac tgt aat aaa tgc atc gag tat aga ggg ata tag caa ttc tcg ata ata gtc atg caa tat ttg aat gag act gat aat tta ggt aat aca gta ct- -ac aca cac ata tct ttc tag att ata tat cgt taa aaa tct gta aga ggt ata ttt ctc ata agt atc cac tgt gta ata tta tta atg gat ata tag ata aca caa tag gga cta ata gta ttg taa aag ata taa tcg act att t-- gta cat atc cag ata tct ata ttc cta cta gtt tgc tgc gta gtt gca tca ttg ata tgc atg -at tta tca gga ttc aga gat gaa tta cta agt aaa cta caa tcc cac aat aag taa gaa tca aat atc aaa aac tca ctt ttg att ttt cta gtc tta agt aat aca tat att tat taa tag acc tat gaa at- aaa aaa gtt aac aat gga ttc gcg tat agc tat tta cgt att agt atc ggc atc tct ttt gta tct tgt taa ttg tca caa act agt aca tta ctt caa tct gaa aat aaa tgg aag tga tat aac taa tac agc aga tat att gct gga caa tta tcc aat tat gac ctt tga tgg aaa gga tat tta tcc atc tat cgc gtt cat ggt cgg taa taa act ttt cct aga tct tta taa aaa tat ctt tgt aga att ttt cag act att tcg agt atc tgt aag tag tca ata cga gga att aga ata tta tta ttc atg tga tta tac taa caa ccg tcc tac aat taa aca aca tta ctt tta taa cgg cga aga ata tac tga aat tga tag atc gaa aaa agc cac taa taa aaa cag ttg gtt aat tac ttc agg ctt tag act aca aaa atg gtt cga tag cga aga ttg tat aat tta tct cag atc ttt agt tag aag aat gga aga cag taa caa aaa cag taa aaa aac tta gta ctt aga tat cg- aaa aaa aat ata ttt ttg tag act ctt gag aat aga agg aaa aca tgt aca taa tta taa aaa atg aaa atc aat ggc gaa taa gac agt gcg att cgc gcc atg gag tcg gca gat ttc atg gct gtc gat gag cag ttt cac gac gac ctc gat ctt tgg tca tta tct ttg gta gat gac tat aaa aaa cat gga tta ggt gtt gac tgt tat gtt cta gaa cca gtt gtt gac agg aaa ata ttt gat aga ttt ctc ctt gaa cca att tgt gat cct gta gat gtt ctg tat gat tat ttt agg att cat aga gat aat att gat cag tat ata gta gat aga ctg ttt gca tat att aca tat aaa gat att ata tct gca tta gtg tca aag aat tat atg gaa gat att ttc tct ata att att aag aat tgt aat tct gta caa gat ctc tta ctt tac tat cta tct aat gca tat gta gaa ata gac att gtt gat ttt atg gta gat cat ggg gct gta ata tat aaa ata gaa tgc ttg aat gcc tat ttt agg gga ata tgt aaa aag gaa agt agt gtt gtt gag ttt att ttg aat tgt ggt atc cca gat gaa aat gat gtt aaa tta gat cta tat aaa ata att cag tat act agg gga ttc ctt gta gat gaa ccc aca gta tta gaa att tat aag ctt tgt atc cca tat att gaa gat atc aat caa cta gat gct ggt gga agg acc ttg ctt tat cgc gct atc tat gca ggt tat ata gat tta gta tca tgg cta tta gaa aat gga gca aat gtc aac gca gta atg agt aat gga tat aca tgt ctt gac gtg gcc gtg gat agg gga tct gtc atc gcc cgt agg gaa aca cat ctt aaa ata tta gaa ata ttg ctt aga gaa cca ttg tct att gac tgt ata aaa tta gct ata ctt aat aat aca att gaa aac cat gat gtg ata aag ctc tgt atc aag tat ttt atg atg gta gat tat tca ctt tgt aat gtg tat gca tca tca ctc ttt gat tat ata att gat tgt aaa caa gaa ttg gag tac att agg cag atg aaa att cat aat aca acc atg tat gag tta atc tat aat aga gac aaa aac aag cat gct tcc cat att cta cat agg tat tct aaa cat cca gtt ttg aca cag tgt atc act aaa gga ttc aag att tac aca gaa gta acc gag cag gtc act aaa gct cta aac aga cgt gct cta ata gat gag ata ata aac aat gta tca act gat gac aat ctc cta tca aaa ctt cca tta gaa att agg gat cta att gtt tca caa gct gtc ata tag agt tct ctc cac cca cct ttc ttg aaa tga gtt aat --- --- --- --- --- --- --- --- --- --- --- --- --- --- --- --- --- --- --- --- --- --- --- --- --- --- --- --- --- --- --- --- --- --- --- --- --- --- --- --- --- --- --- --- --- --- --- --- --- --- --- --- --- --- --- --- --- --- --- --- --- --- --- --- --- --- --- --- --- --- --- --- --- --- --- --- --- --- --- --- --- --- --- --- --- --- --- --- --- --- --- --- --- --- --- --- --- --- --- --- --- --- --- --- --- --- --- --- --- --- --- --- --- --- --- --- --- --- --- --- --- --- --- --- --- --- --- --- --- --- --- --- --- --- --- --- --- --- --- --- --- --- --- --- --- --- --- --- --- --- --- --- --- -ag tca taa gtt agt taa gtc ata agt tag tt- --t tat aag tta gtt tat agt cta aca ctt cta att ttt ata cct tga tct ttt tct cta att atg aaa aag taa atc att atg aag atg gat gaa aat gga cga gat tgt gcg cat cgt taa cga tag tat gtg gta cgt acc taa cgc att tat gga cga cgg tga taa tga agg tca cat ttc tgt caa taa tgt ctg tca tat gta tct cgc att ctt tga tgt gga tat atc atc tca tct gtt taa att agt tat taa aca ctg cga tct gaa taa acg act aaa atg tgg taa ctc tcc att aca ttg cta tac gat gaa tac acg att taa tcc atc tgt att aaa gat att gtt acg cca cgg cat gcg taa ctt tga tag caa gga taa aaa agg aca tat tcc tct aca cca cta tct gat tca ttc act atc aat cga taa caa gat ctt tga tat act aac gga ccc cat tga tga ctt tag taa atc atc cga tct att gct gtg tta tct tag ata taa att caa tgg gag ctt aaa cta tta cgt tct gta caa att att gac taa agg atc tga ccc taa ttg cgt cga tga gga tgg act cac ttc tct tca tta cta ctg taa aca cat atc cgt gtt cca cga aag caa tta tta caa gtc aaa gag tca cac taa gat gcg agc tga gaa gcg att cat cta cgc gat aat aga tca tgg agc aaa cat taa cgc ggt tac gaa aat cgg aaa tac gcc gtt aca cac tta cct tca aca gta tac caa aca tag tcc tcg tgt ggt gta tgc tct ttt atc tcg agg agc cga tac gag gat acg taa taa tct tga ttg tac acc cat cat gga ata cat aaa gaa cga ttg tgc aac agg tca tat tct cat aat gtt act caa ttg gca cga aca aaa ata cgg gaa att aca aaa gga aga agg aca aca tct act tta tct att cat aaa aca taa tca agg ata tgg aag tcg ctc tct caa tat act acg gta tct act aga tag att cga cat tca gaa aga cga ata cta taa tac aat gac tcc tct tca tac cgc ctt cca gaa ttg caa taa caa tgt tgc ctc ata cct cgt ata cat cgg ata cga cat caa cct tcc gac taa aga cga taa gac agt att cga ctt ggt gtt tga aaa cag aaa cat cat ata caa ggc gga tgt cgt taa tga cat cat cca cca cag act gaa agt atc tct acc tat gat taa atc gtt gtt cta caa gat gtc gga gtt ctc tcc cta cga cga tca cta cgt aaa gaa gat aat agc cta ctg cct att aag gga cga gtc att tgc gga act aca tac taa att ctg ttt aaa cga gga cta taa aag tgt att tat gaa aaa tat atc att cga taa gat aga ttc cat cat cga aaa atg tag tcg tga cat aag tct cct caa aga gat tcg aat ctc aga cac caa ctt gta tac ggt att gag aac aga aga cat ccg gta tca cac cta tct cga agc cat aca ttc aga caa acg cat ttc att tcc cat gta cga cga tct cat aga aca gtg tca tct atc gat gga gca taa aag taa act cgt cga caa agc act caa taa att aga gtc tac cat cga tag tca atc tag act atc gta ttt gcc tcc gga aat tat gcg caa tat cat aac caa gct aag cga cta cca tct aaa cag tat gtt gta cgg aaa gaa cca tta caa ata tta tcc atg ata gaa aga aaa tat tta aaa aat aat cta tat gat tgg aga agt agg aaa caa aca gta aca aga cga cga tta cta c-a tta tta aat cat gag gtc cgt att ata ctc gta tat att gtt tct ctc atg tat aat aat aaa cgg aag aga tat agc acc aca tgc acc atc caa tgg aaa atg taa aga caa cga ata cag aag ccg taa tct atg ttg tct atc gtg tcc tcc ggg aac tta cgc ttc cag att atg tga tag caa gac taa tac aca atg tac gcc gtg tgg ttc gga tac ctt tac atc tca caa taa tca ttt aca ggc ttg tct aag ttg taa cgg aag atg tga tag taa tca ggt aga gac gcg atc gtg taa cac gac tca caa tag aat ctg tga atg ctc tcc agg ata tta ttg tct tct caa agg atc atc agg gtg tag aac atg tat ttc taa aac aaa gtg tgg aat agg ata cgg agt atc cgg ata cac gtc tac cgg aga cgt cat ctg ttc tcc gtg tgg tcc cgg aac ata ttc tca cac cgt ctc ttc cac aga taa atg cga acc --- cgt aac cag caa tac att taa cta tat cga tgt gga aat taa cct gta tcc agt caa cga cac atc gtg tac tcg gac gac cac tac cgg tct cag cga atc cat ctc aac gtc gga act aac tat tac cat gaa tca taa aga ttg tga tcc cgt ctt tcg tgc aga ata ctt ctc tgt cct taa taa tgt agc aac ttc agg att ctt tac agg aga aaa tag ata tca gaa tac ttc aaa gat atg tac tct gaa ttt cga gat taa atg taa caa caa aga ttc atc ttc caa aca gtt aac gaa aac aaa gaa tga tac tat cat gcc gca ttc aga gac ggt aac tct agt ggg cga ctg tct atc tag cgt cga cat cta cat act ata tag taa tac caa tac tca aga cta cga aac gga tac aat ctc tta tca tat ggg taa tgt tct cga tgt caa tag cca tat gcc cgc tag ttg cga tat aca taa act gat cac taa ttc cca gaa tcc cac cca ctt ata gta ag- ttt ttt tta ccc ata aat aat aaa tac aat aat taa ttt ctc gta aaa gta gaa aat ata ttc taa ttt att ata tgg taa gaa agt aga atc atc tag aac agt aat caa tca ata gca atc atg aaa caa tat att gtc ctg gca tgc atg tgc cta gtg gca gct gct atg cct act agt ctt caa caa tcc tca tcc tcg tgt act gaa gaa gaa aac aaa cat cat atg gga atc gat gtt att atc aaa gtc aca aag caa gac caa aca ccg acc aat gat aag att tgt caa tcc gta acg gaa gtt aca gag a-- --- --- --- --- --- -cc gaa gat gat gag gta tcc gaa gaa gtt gta aaa gga gat ccc acc act tat tac act atc gtc ggc gcg ggt ctt aac atg aac ttt gga ttc acc aaa tgc cca aag att tca tcc atc tcc gaa tcc tct gat gga aac act gtg aat act aga ttg tcc agc gtg tca cca gga caa ggt aag gac tct ccc gcg atc acg cgt gaa gaa gct ctg gct atg atc aaa gac tgt gag atg tct atc gac atc aga tgt agc gaa gaa gag aaa gac agc gac atc aag acc cat cca gta ctt ggg tct aac atc tca cat aag aaa gtg agt tac aaa gat atc atc ggt tca acg atc gtt gat aca aaa tgt gtt aag aac cta gag ttt agc gta cgt atc gga gac atg tgt gag gaa tca tct gaa ctt gaa gtc aag gat gga ttc aag tat gtc gac gga tcg gca tct gaa ggt gca acc gat gat act tca ctc atc gat tca aca aaa ctc aaa gca tgt gtc tga atc gat aac tct att cat ctg aaa atg gat gag ttg ggt taa tcg aac gat tca gac acc gca cca cga att aaa aaa gtg tac cgg gta cta tat tcc ggt ttg caa aac aaa aat gtt taa cta cat tca caa aaa gtt acc tct cgt tac ttc ttc ttt ctg ttt caa tat gtg ata cga tat gat cac tat tcg tat tct ctt tct ctc ttg tct cat -aa aaa gtt tta c-- --- --- --- aaa aaa a-t att ttt att ctc ttt ctc tct tcg at- ggt ctc aca aaa ata tta aac ctc ttt ctg atg tct caa cta ttt cgt aaa cga taa cgt cca aca ata tat tct cgt aga gct tat caa tat cct tat gcc aat cta ggt tgt cag aca att gca tca taa aat aat gtt tat aat tta cac gtt aac atc ata taa taa acg tat ata gtt aat att ttt gga ata taa atg atc tgt aaa atc cat gta ggg gac act gct cac gtt ttt tct cta gta cat aat ttc aca caa gtt ttt ata cag aca aat taa ttc tcg tcc ata ta- ttt taa aac att gac ttt tgt act aag aaa aat atc ttg act aac cat ctc ttt ctc tcg tcg atg ggt ctc aca aaa ata tta aac ctc ttt ctg atg g-- --- --- --- --- --- --- --- --- --- --- --- --- --- --- --- --- --- --- --- --- --- --- --- --- --- --- --- --- --- --- --- --- --- --- --- --- --- --- --- --- --- --- --- --- --- --- agt cgt aaa aag ttt tta tcc ttt ctc tct tcg atg ggt ctc --- --- --- --- --- --- --- --- --- --- --- --- --- --- --- --- --- --- --- --- --- --- --- --- --- --- --- --- --- --- --- --- --- --- --- --- --- --- --- --- --- --- --- --- --- --- --- --- --- --- --- --- --- --- --- --- --- --- --- --- --- --- --- --- --- --- --- --- --- --- aca aaa ata tta aac ctc ttt ctg atg gtc tct ata aa

>JX878419.1_Monkeypox_virus_isolate_DRC_07-0275_complete_genome

atc gag gac gac ttc aac tat gac agc tcg gtg gcg tct gcc agc gtg cac ata cga atg gca ttt cta aga aaa gtc tac ggt atc ctt gtc tac aat ttc ctt taa caa cgg caa cag ctg cag tat ttt tat act ttg aat gca ttc gga cat tta tac aag gga gtc ctg ttc taa tat tgg cat caa tgt tcg gat cta tag gct tga ttt tcg cat tga ctt tac aca gac ata aac atc ccc tga atc tgt acc tac ttt gtg gat tta cac tgt tag aat ctc taa cgc tgg cct ctg ttg tta ctt tct atg atg cgc gta tcg tta tac aag ctt tca tgt tga cta ctg cag tgt ttc ttg ctc tga cta cat gta ctc tac aat caa aga gag att tca gta aac ttg taa cag gat tgt ttg ctg ctt tct gga ttt taa ttt tgt cag gag tct tga gga taa agt tta aaa tag aat taa taa aga aca tat agg tca -tt ttt taa aca tgg ata gaa acc aag gtt gtt agt taa taa tat aca aga tat ttt ttc tca ctc tga tcc atg taa acc aag gac gag aga cac tct cat tcc tca ttc acg aca cca tta aaa atg gaa att aaa gcc ctc tat taa gca cag acg gct aca ggt cta cca tca ggt taa cct tcg tct acc ttc aca atg gcc ttt cct tgt gcc cag ttc agt ccc tgt cat tgc cac gct act aag gac tcc ctg aat acc gtg act gac gtc aga cat tgt ctg act gaa tac atc ctg tgg gtt tct cat aga tgg acc cat aga gaa agc gca ggg cct ctc tac agg ctt ctc atc tct ttc aga att gat gca atg gag cta ttt ggt agc gag ttg aag gag ttc tcg gat tca ctt ccg tgg gac aat atc gac aat tgc gtg gag atc att aaa tgt ttc atc aga aat gac tcc atg aaa acc gcc aaa gaa ctt tgt gca ata att gga ctt tgt act caa tca gct att gtc act gga aga gtc ttc aat gat aag tat atc gac ata cta ctt atg ctg cga aag att ctg aat gag aac gac tat ctc acc ctc ttg gat cat atc ctc act gct aaa tac taa atc tcc ttc atg ctc tct cac taa tac tct tac tca cta cac ttt tta tca tct tat gat gaa tga ttg cct tca tca -tt ttt cgt gga ata taa tat agg aat aat tag cac cag aat agc tat gga tat ctc gtt aag aat att ctc gat aag aga cat aat gta gac ata gtt att ata tcc ttc tta gat aag tgt tac gct act gga aag ttt cca tcg tta tta tta cgt gaa gat gat atc att aaa cca aca ttg aga ttg gct ctt atg tta gct gga ttg aat tac tgt aat aaa tgc atc gag tat aga ggg ata tag caa ttc tcg ata ata gtc atg caa tat ttg aat gag act gat aat tta ggt aat aca gta ct- -ac aca cac ata tct ttc tag att ata tat cgt taa aaa tct gta aga ggt ata ttt ctc ata agt atc cac tgt gta ata tta tta atg gat ata tag ata aca caa tag gga cta ata gta ttg taa aag ata taa tcg act att t-- gta cat atc cag ata tct ata ttc cta cta gtt tgc tgc gta gtt gca tca ttg ata tgc atg -at tta tca gga ttc aga gat gaa tta cta agt aaa cta caa tcc cac aat aag taa gaa tca aat atc aaa aac tca ctt ttg att ttt cta gtc tta agt aat aca tat att tat taa tag acc tat gaa at- aaa aaa gtt aac aat gga ttc gcg tat agc tat tta cgt att agt atc ggc atc tct ttt gta tct tgt taa ttg tca caa act agt aca tta ctt caa tct gaa aat aaa tgg aag tga tat aac taa tac agc aga tat att gct gga caa tta tcc aat tat gac ctt tga tgg aaa gga tat tta tcc atc tat cgc gtt cat ggt cgg taa taa act ttt cct aga tct tta taa aaa tat ctt tgt aga att ttt cag act att tcg agt atc tgt aag tag tca ata cga gga att aga ata tta tta ttc atg tga tta tac taa caa ccg tcc tac aat taa aca aca tta ctt tta taa cgg cga aga ata tac tga aat tga tag atc gaa aaa agc cac taa taa aaa cag ttg gtt aat tac ttc agg ctt tag act aca aaa atg gtt cga tag cga aga ttg tat aat tta tct cag atc ttt agt tag aag aat gga aga cag taa caa aaa cag taa aaa aac tta gta ctt aga tat cg- aaa aaa aat ata ttt ttg tag act ctt gag aat aga agg aaa aca tgt aca taa tta taa aaa atg aaa atc aat ggc gaa taa gac agt gcg att cgc gcc atg gag tcg gca gat ttc atg gct gtc gat gag cag ttt cac gac gac ctc gat ctt tgg tca tta tct ttg gta gat gac tat aaa aaa cat gga tta ggt gtt gac tgt tat gtt cta gaa cca gtt gtt gac agg aaa ata ttt gat aga ttt ctc ctt gaa cca att tgt gat cct gta gat gtt ctg tat gat tat ttt agg att cat aga gat aat att gat cag tat ata gta gat aga ctg ttt gca tat att aca tat aaa gat att ata tct gca tta gtg tca aag aat tat atg gaa gat att ttc tct ata att att aag aat tgt aat tct gta caa gat ctc tta ctt tac tat cta tct aat gca tat gta gaa ata gac att gtt gat ttt atg gta gat cat ggg gct gta ata tat aaa ata gaa tgc ttg aat gcc tat ttt agg gga ata tgt aaa aag gaa agt agt gtt gtt gag ttt att ttg aat tgt ggt atc cca gat gaa aat gat gtt aaa tta gat cta tat aaa ata att cag tat act agg gga ttc ctt gta gat gaa ccc aca gta tta gaa att tat aag ctt tgt atc cca tat att gaa gat atc aat caa cta gat gct ggt gga agg acc ttg ctt tat cgc gct atc tat gca ggt tat ata gat tta gta tca tgg cta tta gaa aat gga gca aat gtc aac gca gta atg agt aat gga tat aca tgt ctt gac gtg gcc gtg gat agg gga tct gtc atc gcc cgt agg gaa aca cat ctt aaa ata tta gaa ata ttg ctt aga gaa cca ttg tct att gac tgt ata aaa tta gct ata ctt aat aat aca att gaa aac cat gat gtg ata aag ctc tgt atc aag tat ttt atg atg gta gat tat tca ctt tgt aat gtg tat gca tca tca ctc ttt gat tat ata att gat tgt aaa caa gaa ttg gag tac att agg cag atg aaa att cat aat aca acc atg tat gag tta atc tat aat aga gac aaa aac aag cat gct tcc cat att cta cat agg tat tct aaa cat cca gtt ttg aca cag tgt atc act aaa gga ttc aag att tac aca gaa gta acc gag cag gtc act aaa gct cta aac aga cgt gct cta ata gat gag ata ata aac aat gta tca act gat gac aat ctc cta tca aaa ctt cca tta gaa att agg gat cta att gtt tca caa gct gtc ata tag agt tct ctc cac cca cct ttc ttg aaa tga gtt aat --- --- --- --- --- --- --- --- --- --- --- --- --- --- --- --- --- --- --- --- --- --- --- --- --- --- --- --- --- --- --- --- --- --- --- --- --- --- --- --- --- --- --- --- --- --- --- --- --- --- --- --- --- --- --- --- --- --- --- --- --- --- --- --- --- --- --- --- --- --- --- --- --- --- --- --- --- --- --- --- --- --- --- --- --- --- --- --- --- --- --- --- --- --- --- --- --- --- --- --- --- --- --- --- --- --- --- --- --- --- --- --- --- --- --- --- --- --- --- --- --- --- --- --- --- --- --- --- --- --- --- --- --- --- --- --- --- --- --- --- --- --- --- --- --- --- --- --- --- --- --- --- --- -ag tca taa gtt agt taa gtc ata agt tag tt- --t tat aag tta gtt tat agt cta aca ctt cta att ttt ata cct tga tct ttt tct cta att atg aaa aag taa atc att atg aag atg gat gaa aat gga cga gat tgt gcg cat cgt taa cga tag tat gtg gta cgt acc taa cgc att tat gga cga cgg tga taa tga agg tca cat ttc tgt caa taa tgt ctg tca tat gta tct cgc att ctt tga tgt gga tat atc atc tca tct gtt taa att agt tat taa aca ctg cga tct gaa taa acg act aaa atg tgg taa ctc tcc att aca ttg cta tac gat gaa tac acg att taa tcc atc tgt att aaa gat att gtt acg cca cgg cat gcg taa ctt tga tag caa gga taa aaa agg aca tat tcc tct aca cca cta tct gat tca ttc act atc aat cga taa caa gat ctt tga tat act aac gga ccc cat tga tga ctt tag taa atc atc cga tct att gct gtg tta tct tag ata taa att caa tgg gag ctt aaa cta tta cgt tct gta caa att att gac taa agg atc tga ccc taa ttg cgt cga tga gga tgg act cac ttc tct tca tta cta ctg taa aca cat atc cgt gtt cca cga aag caa tta tta caa gtc aaa gag tca cac taa gat gcg agc tga gaa gcg att cat cta cgc gat aat aga tca tgg agc aaa cat taa cgc ggt tac gaa aat cgg aaa tac gcc gtt aca cac tta cct tca aca gta tac caa aca tag tcc tcg tgt ggt gta tgc tct ttt atc tcg agg agc cga tac gag gat acg taa taa tct tga ttg tac acc cat cat gga ata cat aaa gaa cga ttg tgc aac agg tca tat tct cat aat gtt act caa ttg gca cga aca aaa ata cgg gaa att aca aaa gga aga agg aca aca tct act tta tct att cat aaa aca taa tca agg ata tgg aag tcg ctc tct caa tat act acg gta tct act aga tag att cga cat tca gaa aga cga ata cta taa tac aat gac tcc tct tca tac cgc ctt cca gaa ttg caa taa caa tgt tgc ctc ata cct cgt ata cat cgg ata cga cat caa cct tcc gac taa aga cga taa gac agt att cga ctt ggt gtt tga aaa cag aaa cat cat ata caa ggc gga tgt cgt taa tga cat cat cca cca cag act gaa agt atc tct acc tat gat taa atc gtt gtt cta caa gat gtc gga gtt ctc tcc cta cga cga tca cta cgt aaa gaa gat aat agc cta ctg cct att aag gga cga gtc att tgc gga act aca tac taa att ctg ttt aaa cga gga cta taa aag tgt att tat gaa aaa tat atc att cga taa gat aga ttc cat cat cga aaa atg tag tcg tga cat aag tct cct caa aga gat tcg aat ctc aga cac caa ctt gta tac ggt att gag aac aga aga cat ccg gta tca cac cta tct cga agc cat aca ttc aga caa acg cat ttc att tcc cat gta cga cga tct cat aga aca gtg tca tct atc gat gga gca taa aag taa act cgt cga caa agc act caa taa att aga gtc tac cat cga tag tca atc tag act atc gta ttt gcc tcc gga aat tat gcg caa tat cat aac caa gct aag cga cta cca tct aaa cag tat gtt gta cgg aaa gaa cca tta caa ata tta tcc atg ata gaa aga aaa tat tta aaa aat aat cta tat gat tgg aga agt agg aaa caa aca gta aca aga cga cga tta cta c-a tta tta aat cat gag gtc cgt att ata ctc gta tat att gtt tct ctc atg tat aat aat aaa cgg aag aga tat agc acc aca tgc acc atc caa tgg aaa atg taa aga caa cga ata cag aag ccg taa tct atg ttg tct atc gtg tcc tcc ggg aac tta cgc ttc cag att atg tga tag caa gac taa tac aca atg tac gcc gtg tgg ttc gga tac ctt tac atc tca caa taa tca ttt aca ggc ttg tct aag ttg taa cgg aag atg tga tag taa tca ggt aga gac gcg atc gtg taa cac gac tca caa tag aat ctg tga atg ctc tcc agg ata tta ttg tct tct caa agg atc atc agg gtg tag aac atg tat ttc taa aac aaa gtg tgg aat agg ata cgg agt atc cgg ata cac gtc tac cgg aga cgt cat ctg ttc tcc gtg tgg tcc cgg aac ata ttc tca cac cgt ctc ttc cac aga taa atg cga acc --- cgt aac cag caa tac att taa cta tat cga tgt gga aat taa cct gta tcc agt caa cga cac atc gtg tac tcg gac gac cac tac cgg tct cag cga atc cat ctc aac gtc gga act aac tat tac cat gaa tca taa aga ttg tga tcc cgt ctt tcg tgc aga ata ctt ctc tgt cct taa taa tgt agc aac ttc agg att ctt tac agg aga aaa tag ata tca gaa tac ttc aaa gat atg tac tct gaa ttt cga gat taa atg taa caa caa aga ttc atc ttc caa aca gtt aac gaa aac aaa gaa tga tac tat cat gcc gca ttc aga gac ggt aac tct agt ggg cga ctg tct atc tag cgt cga cat cta cat act ata tag taa tac caa tac tca aga cta cga aac gga tac aat ctc tta tca tat ggg taa tgt tct cga tgt caa tag cca tat gcc cgc tag ttg cga tat aca taa act gat cac taa ttc cca gaa tcc cac cca ctt ata gta agt ttt ttt tta ccc ata aat aat aaa tac aat aat taa ttt ctc gta aaa gta gaa aat ata ttc taa ttt att ata tgg taa gaa agt aga atc atc tag aac agt aat caa tca ata gca atc atg aaa caa tat att gtc ctg gca tgc atg tgc cta gtg gca gct gct atg cct act agt ctt caa caa tcc tca tcc tcg tgt act gaa gaa gaa aac aaa cat cat atg gga atc gat gtt att atc aaa gtc aca aag caa gac caa aca ccg acc aat gat aag att tgt caa tcc gta acg gaa gtt aca gag a-- --- --- --- --- --- -cc gaa gat gat gag gta tcc gaa gaa gtt gta aaa gga gat ccc acc act tat tac act atc gtc ggc gcg ggt ctt aac atg aac ttt gga ttc acc aaa tgc cca aag att tca tcc atc tcc gaa tcc tct gat gga aac act gtg aat act aga ttg tcc agc gtg tca cca gga caa ggt aag gac tct ccc gcg atc acg cgt gaa gaa gct ctg gct atg atc aaa gac tgt gag atg tct atc gac atc aga tgt agc gaa gaa gag aaa gac agc gac atc aag acc cat cca gta ctt ggg tct aac atc tca cat aag aaa gtg agt tac aaa gat atc atc ggt tca acg atc gtt gat aca aaa tgt gtt aag aac cta gag ttt agc gta cgt atc gga gac atg tgt gag gaa tca tct gaa ctt gaa gtc aag gat gga ttc aag tat gtc gac gga tcg gca tct gaa ggt gca acc gat gat act tca ctc atc gat tca aca aaa ctc aaa gca tgt gtc tga atc gat aac tct att cat ctg aaa atg gat gag ttg ggt taa tcg aac gat tca gac acc gca cca cga att aaa aaa gtg tac cgg gta cta tat tcc ggt ttg caa aac aaa aat gtt taa cta cat tca caa aaa gtt acc tct cgt tac ttc ttc ttt ctg ttt caa tat gtg ata cga tat gat cac tat tcg tat tct ctt tct ctc ttg tct cat -aa aaa gtt tta c-- --- --- --- aaa aaa a-t att ttt att ctc ttt ctc tct tcg at- ggt ctc aca aaa ata tta aac ctc ttt ctg atg tct caa cta ttt cgt aaa cga taa cgt cca aca ata tat tct cgt aga gct tat caa tat cct tat gcc aat cta ggt tgt cag aca att gca tca taa aat aat gtt tat aat tta cac gtt aac atc ata taa taa acg tat ata gtt aat att ttt gga ata taa atg atc tgt aaa atc cat gta ggg gac act gct cac gtt ttt tct cta gta cat aat ttc aca caa gtt ttt ata cag aca aat taa ttc tcg tcc ata ta- ttt taa aac att gac ttt tgt act aag aaa aat atc ttg act aac cat ctc ttt ctc tcg tcg atg ggt ctc aca aaa ata tta aac ctc ttt ctg atg g-- --- --- --- --- --- --- --- --- --- --- --- --- --- --- --- --- --- --- --- --- --- --- --- --- --- --- --- --- --- --- --- --- --- --- --- --- --- --- --- --- --- --- --- --- --- --- agt cgt aaa aag ttt tta tcc ttt ctc tct tcg atg ggt ctc --- --- --- --- --- --- --- --- --- --- --- --- --- --- --- --- --- --- --- --- --- --- --- --- --- --- --- --- --- --- --- --- --- --- --- --- --- --- --- --- --- --- --- --- --- --- --- --- --- --- --- --- --- --- --- --- --- --- --- --- --- --- --- --- --- --- --- --- --- --- aca aaa ata tta aac ctc ttt ctg atg gtc tct ata aa

>KJ642612.1_Monkeypox_virus_strain_Ikubi_complete_genome

atc gag gac gac ttc aac tat gac agc tcg gtg gcg tct gcc agc gtg cac ata cga atg gca ttt cta aga aaa gtc tac ggt atc ctt gtc tac aat ttc ctt taa caa cgg caa cag ctg cag tat ttt tat act ttg aat gca ttc gga cat tta tac aag gga gtc ctg ttc taa tat tgg cat caa tgt tcg gat cta tag gct tga ttt tcg cat tga ctt tac aca gac ata aac atc ccc tga atc tgt acc tac ttt gtg gat tta cac tgt tag aat ctc taa cgc tgg cct ctg ttg tta ctt tct atg atg cgc gta tcg tta tac aag ctt tca tgt tga cta ctg cag tgt ttc ttg ctc tga cta cat gta ctc tac aat caa aga gag att tca gta aac ttg taa cag gat tgt ttg ctg ctt tct gga ttt taa ttt tgt cag gag tct tga gga taa agt tta aaa tag aat taa taa aga aca tat agg tca -tt ttt taa aca tgg ata gaa acc aag gtt gtt agt taa taa tat aca aga tat ttt ttc tca ctc tga tcc atg taa acc aag gac gag aga cac tct cat tcc tca ttc acg aca cca tta aaa atg gaa att aaa gcc ctc tat taa gca cag acg gct aca ggt cta cca tca ggt taa cct tcg tct acc ttc aca atg gcc ttt cct tgt gcc cag ttc agt ccc tgt cat tgc cac gct act aag gac tcc ctg aat acc gtg act gac gtc aga cat tgt ctg act gaa tac atc ctg tgg gtt tct cat aga tgg acc cat aga gaa agc gca ggg cct ctc tac agg ctt ctc atc tct ttc aga att gat gca atg gag cta ttt ggt agc gag ttg aag gag ttc tcg gat tca ctt ccg tgg gac aat atc gac aat tgc gtg gag atc att aaa tgt ttc atc aga aat gac tcc atg aaa acc gcc aaa gaa ctt tgt gca ata att gga ctt tgt act caa tca gct att gtc act gga aga gtc ttc aat gat aag tat atc gac ata cta ctt atg ctg cga aag att ctg aat gag aac gac tat ctc acc ctc ttg gat cat atc ctc act gct aaa tac taa atc tcc ttc atg ctc tct cac taa tac tct tac tca cta cac ttt tta tca tct tat gat gaa tga ttg cct tca tca -tt ttt cgt gga ata taa tat agg aat aat tag cac cag aat agc tat gga tat ctc gtt aag aat att ctc gat aag aga cat aat gta gac ata gtt att ata tcc ttc tta gat aag tgt tac gct act gga aag ttt cca tcg tta tta tta cgt gaa gat gat atc att aaa cca aca ttg aga ttg gct ctt atg tta gct gga ttg aat tac tgt aat aaa tgc atc gag tat aga ggg ata tag caa ttc tcg ata ata gtc atg caa tat ttg aat gag act gat aat tta ggt aat aca gta ct- -ac aca cac ata tct ttc tag att ata tat cgt taa aaa tct gta aga ggt ata ttt ctc ata agt atc cac tgt gta ata tta tta atg gat ata tag ata aca caa tag gga cta ata gta ttg taa aag ata taa tcg act att t-- gta cat atc cag ata tct ata ttc cta cta gtt tgc tgc gta gtt gca tca ttg ata tgc atg -at tta tca gga ttc aga gat gaa tta cta agt aaa cta cga tcc cac aat aag taa gaa tca aat atc aaa aac tca ctt ttg att ttt cta gtc tta agt aat aca tat att tat taa tag acc tat gaa at- aaa aaa gtt aac aat gga ttc gcg tat agc tat tta cgt att agt atc ggc atc tct ttt gta tct tgt taa ttg tca caa act agt aca tta ctt caa tct gaa aat aaa tgg aag tga tat aac taa tac agc aga tat att gct gga caa tta tcc aat tat gac ctt tga tgg aaa gga tat tta tcc atc tat cgc gtt cat ggt cgg taa taa act ttt cct aga tct tta taa aaa tat ctt tgt aga att ttt cag act att tcg agt atc tgt aag tag tca ata cga gga att aga ata tta tta ttc atg tga tta tac taa caa ccg tcc tac aat taa aca aca tta ctt tta taa cgg cga aga ata tac tga aat tga tag atc gaa aaa agc cac taa taa aaa cag ttg gtt aat tac ttc agg ctt tag act aca aaa atg gtt cga tag cga aga ttg tat aat tta tct cag atc ttt agt tag aag aat gga aga cag taa caa aaa cag taa aaa aac tta gta ctt aga tat cg- aaa aaa aat ata ttt ttg tag act ctt gag aat aga agg aaa aca tgt aca taa tta taa aaa atg aaa atc aat ggc gaa taa gac agt gcg att cgc gcc atg gag tcg gca gat ttc atg gct gtc gat gag cag ttt cac gac gac ctc gat ctt tgg tca tta tct ttg gta gat gac tat aaa aaa cat gga tta ggt gtt gac tgt tat gtt cta gaa cca gtt gtt gac agg aaa ata ttt gat aga ttt ctc ctt gaa cca att tgt gat cct gta gat gtt ctg tat gat tat ttt agg att cat aga gat aat att gat cag tat ata gta gat aga ctg ttt gca tat att aca tat aaa gat att ata tct gca tta gtg tca aag aat tat atg gaa gat att ttc tct ata att att aag aat tgt aat tct gta caa gat ctc tta ctt tac tat cta tct aat gca tat gta gaa ata gac att gtt gat ttt atg gta gat cat ggg gct gta ata tat aaa ata gaa tgc ttg aat gcc tat ttt agg gga ata tgt aaa aag gaa agt agt gtt gtt gag ttt att ttg aat tgt ggt atc cca gat gaa aat gat gtt aaa tta gat cta tat aaa ata att cag tat act agg gga ttc ctt gta gat gaa ccc aca gta tta gaa att tat aag ctt tgt atc cca tat att gaa gat atc aat caa cta gat gct ggt gga agg acc ttg ctt tat cgc gct atc tat gca ggt tat ata gat tta gta tca tgg cta tta gaa aat gga gca aat gtc aac gca gta atg agt aat gga tat aca tgt ctt gac gtg gcc gtg gat agg gga tct gtc atc gcc cgt agg gaa gca cat ctt aaa ata tta gaa ata ttg ctt aga gaa cca ttg tct att gac tgt ata aaa tta gct ata ctt aat aat aca att gaa aac cat gat gtg ata aag ctc tgt atc aag tat ttt atg atg gta gat tat tca ctt tgt aat gtg tat gca tca tca ctc ttt gat tat ata att gat tgt aaa caa gaa ttg gag tac att agg cag atg aaa att cat aat aca acc atg tat gag tta atc tat aat aga gac aaa aac aag cat gct tcc cat att cta cat agg tat tct aaa cat cca gtt ttg aca cag tgt atc act aaa gga ttc aag att tac aca gaa gta acc gag cag gtc act aaa gct cta aac aga cgt gct cta ata gat gag ata ata aac aat gta tca act gat gac aat ctc cta tca aaa ctt cca tta gaa att agg gat cta att gtt tca caa gct gtc ata tag agt tct ctc cac cca cct ttc ttg aaa tga gtt aat --- --- --- --- --- --- --- --- --- --- --- --- --- --- --- --- --- --- --- --- --- --- --- --- --- --- --- --- --- --- --- --- --- --- --- --- --- --- --- --- --- --- --- --- --- --- --- --- --- --- --- --- --- --- --- --- --- --- --- --- --- --- --- --- --- --- --- --- --- --- --- --- --- --- --- --- --- --- --- --- --- --- --- --- --- --- --- --- --- --- --- --- --- --- --- --- --- --- --- --- --- --- --- --- --- --- --- --- --- --- --- --- --- --- --- --- --- --- --- --- --- --- --- --- --- --- --- --- --- --- --- --- --- --- --- --- --- --- --- --- --- --- --- --- --- --- --- --- --- --- --- --- --- -ag tca taa gtt agt taa gtc ata agt tag tt- --t tat aag tta gtt tat agt cta aca ctt cta att ttt ata cct tga tct ttt tct cta att atg aaa aag taa atc att atg aag atg gat gaa aat gga cga gat tgt gcg cat cgt taa cga tag tat gtg gta cgt acc taa cgc att tat gga cga cgg tga taa tga agg tca cat ttc tgt caa taa tgt ctg tca tat gta tct cgc att ctt tga tgt gga tat atc atc tca tct gtt taa att agt tat taa aca ctg cga tct gaa taa acg act aaa atg tgg taa ctc tcc att aca ttg cta tac gat gaa tac acg att taa tcc atc tgt att aaa gat att gtt acg cca cgg cat gcg taa ctt tga tag caa gga taa aaa agg aca tat tcc tct aca cca cta tct gat tca ttc act atc aat cga taa caa gat ctt tga tat act aac gga ccc cat tga tga ctt tag taa atc atc cga tct att gct gtg tta tct tag ata taa att caa tgg gag ctt aaa cta tta cgt tct gta caa att att gac taa agg atc tga ccc taa ttg cgt cga tga gga tgg act cac ttc tct tca tta cta ctg taa aca cat atc cgt gtt cca cga aag caa tta tta caa gtc aaa gag tca cac taa gat gcg agc tga gaa gcg att cat cta cgc gat aat aga tca tgg agc aaa cat taa cgc ggt tac gaa aat cgg aaa tac gcc gtt aca cac tta cct tca aca gta tac caa aca tag tcc tcg tgt ggt gta tgc tct ttt atc tcg agg agc cga tac gag gat acg taa taa tct tga ttg tac acc cat cat gga ata cat aaa gaa cga ttg tgc aac agg tca tat tct cat aat gtt act caa ttg gca cga aca aaa ata cgg gaa att aca aaa gga aga agg aca aca tct act tta tct att cat aaa aca taa tca agg ata tgg aag tcg ctc tct caa tat act acg gta tct act aga tag att cga cat tca gaa aga cga ata cta taa tac aat gac tcc tct tca tac cgc ctt cca gaa ttg caa taa caa tgt tgc ctc ata cct cgt ata cat cgg ata cga cat caa cct tcc gac taa aga cga taa gac agt att cga ctt ggt gtt tga aaa cag aaa cat cat ata caa ggc gga tgt cgt taa tga cat cat cca cca cag act gaa agt atc tct acc tat gat taa atc gtt gtt cta caa gat gtc gga gtt ctc tcc cta cga cga tca cta cgt aaa gaa gat aat agc cta ctg cct att aag gga cga gtc att tgc gga act aca tac taa att ctg ttt aaa cga gga cta taa aag tgt att tat gaa aaa tat atc att cga taa gat aga ttc cat cat cga aaa atg tag tcg tga cat aag tct cct caa aga gat tcg aat ctc aga cac caa ctt gta tac ggt att gag aac aga aga cat ccg gta tca cac cta tct cga agc cat aca ttc aga caa acg cat ttc att tcc cat gta cga cga tct cat aga aca gtg tca tct atc gat gga gca taa aag taa act cgt cga caa agc act caa taa att aga gtc tac cat cga tag tca atc tag act atc gta ttt gcc tcc gga aat tat gcg caa tat cat aac caa gct aag cga cta cca tct aaa cag tat gtt gta cgg aaa gaa cca tta caa ata tta tcc atg ata gaa aga aaa tat tta aaa aat aat cta tat gat tgg aga agt agg aaa caa aca gta aca aga cga cga tta cta c-a tta tta aat cat gag gtc cgt att ata ctc gta tat att gtt tct ctc atg tat aat aat aaa cgg aag aga tat agc acc aca tgc acc atc caa tgg aaa atg taa aga caa cga ata cag aag ccg taa tct atg ttg tct atc gtg tcc tcc ggg aac tta cgc ttc cag att atg tga tag caa gac taa tac aca atg tac gcc gtg tgg ttc gga tac ctt tac atc tca caa taa tca ttt aca ggc ttg tct aag ttg taa cgg aag atg tga tag taa tca ggt aga gac gcg atc gtg taa cac gac tca caa tag aat ctg tga atg ctc tcc agg ata tta ttg tct tct caa agg atc atc agg gtg tag aac atg tat ttc taa aac aaa gtg tgg aat agg ata cgg agt atc cgg ata cac gtc tac cgg aga cgt cat ctg ttc tcc gtg tgg tcc cgg aac ata ttc tca cac cgt ctc ttc cac aga taa atg cga acc --- cgt aac cag caa tac att taa cta tat cga tgt gga aat taa cct gta tcc agt caa cga cac atc gtg tac tcg gac gac cac tac cgg tct cag cga atc cat ctc aac gtc gga act aac tat tac cat gaa tca taa aga ttg tga tcc cgt ctt tcg tgc aga ata ctt ctc tgt cct taa taa tgt agc aac ttc agg att ctt tac agg aga aaa tag ata tca gaa tac ttc aaa gat atg tac tct gaa ttt cga gat taa atg taa caa caa aga ttc atc ttc caa aca gtt aac gaa aac aaa gaa tga tac tat cat gcc gca ttc aga gac ggt aac tct agt ggg cga ctg tct atc tag cgt cga cat cta cat act ata tag taa tac caa tac tca aga cta cga aac gga tac aat ctc tta tca tat ggg taa tgt tct cga tgt caa tag cca tat gcc cgc tag ttg cga tat aca taa act gat cac taa ttc cca gaa tcc cac cca ctt ata gta agt ttt ttt tta ccc ata aat aat aaa tac aat aat taa ttt ctc gta aaa gta gaa aat ata ttc taa ttt att ata tgg taa gaa agt aga atc atc tag aac agt aat caa tca ata gca atc atg aaa caa tat att gtc ctg gca tgc atg tgc cta gtg gca gct gct atg cct act agt ctt caa caa tcc tca tcc tcg tgt act gaa gaa gaa aac aaa cat cat atg gga atc gat gtt att atc aaa gtc aca aag caa gac caa aca ccg acc aat gat aag att tgt caa tcc gta acg gaa gtt aca gag a-- --- --- --- --- --- -cc gaa gat gat gag gta tcc gaa gaa gtt gta aaa gga gat ccc acc act tat tac act atc gtc ggc gcg ggt ctt aac atg aac ttt gga ttc acc aaa tgc cca aag att tca tcc atc tcc gaa tcc tct gat gga aac act gtg aat act aga ttg tcc agc gtg tca cca gga caa ggt aag gac tct ccc gcg atc acg cgt gaa gaa gct ctg gct atg atc aaa gac tgt gag atg tct atc gac atc aga tgt agc gaa gaa gag aaa gac agc gac atc aag acc cat cca gta ctt ggg tct aac atc tca cat aag aaa gtg agt tac aaa gat atc atc ggt tca acg atc gtt gat aca aaa tgt gtt aag aac cta gag ttt agc gta cgt atc gga gac atg tgt gag gaa tca tct gaa ctt gaa gtc aag gat gga ttc aag tat gtc gac gga tcg gca tct gaa ggt gca acc gat gat act tca ctc atc gat tca aca aaa ctc aaa gca tgt gtc tga atc gat aac tct att cat ctg aaa atg gat gag ttg ggt taa tcg aac gat tca gac acc gca cca cga att aaa aaa gtg tac cgg gta cta tat tcc ggt ttg caa aac aaa aat gtt taa cta cat tca caa aaa gtt acc tct cgt tac ttc ttc ttt ctg ttt caa tat gtg ata cga tat gat cac tat tcg tat tct ctt tct ctc ttg tct cat aaa aaa gtt tta c-- --- --- --- aaa aaa a-t att ttt att ctc ttt ctc tct tcg at- ggt ctc aca aaa ata tta aac ctc ttt ctg atg tct caa cta ttt cgt aaa cga taa cgt cca aca ata tat tct cgt aga gct tat caa tat cct tat gcc aat cta ggt tgt cag aca att gca tca taa aat aat gtt tat aat tta cac gtt aac atc ata taa taa acg tat ata gtt aat att ttt gga ata taa atg atc tgt aaa atc cat gta ggg gac act gct cac gtt ttt tct cta gta cat aat ttc aca caa gtt ttt ata cag aca aat taa ttc tcg tcc ata ta- ttt taa aac att gac ttt tgt act aag aaa aat atc ttg act aac cat ctc ttt ctc tct tcg atg ggt ctc aca aaa ata tta aac ctc ttt ctg atg g-- --- --- --- --- --- --- --- --- --- --- --- --- --- --- --- --- --- --- --- --- --- --- --- --- --- --- --- --- --- --- --- --- --- --- --- --- --- --- --- --- --- --- --- --- --- --- agt cgt aaa aag ttt tta tcc ttt ctc tct tcg atg ggt --- --- --- --- --- --- --- --- --- --- --- --- --- --- --- --- --- --- --- --- --- --- --- --- --- --- --- --- --- --- --- --- --- --- --- --- --- --- --- --- --- --- --- --- --- --- --- --- --- --- --- --- --- --- --- --- --- --- --- --- --- --- --- --- --- --- --- --- --- --- --- --- --- --- --- --- --- --- --- --- --- --- --- --

>JX878426.1_Monkeypox_virus_isolate_DRC_07-0450_complete_genome

atc gag gac gac ttc aac tat gac agc tcg gtg gcg tct gcc agc gtg cac ata cga atg gca ttt cta aga aaa gtc tac ggt atc ctt gtc tac aat ttc ctt taa caa cgg caa cag ctg cag tat ttt tat act ttg aat gca ttc gga cat tta tac aag gga gtc ctg ttc taa tat tgg cat caa tgt tcg gat cta tag gct tga ttt tcg cat tga ctt tac aca gac ata aac atc ccc tga atc tgt acc tac ttt gtg gat tta cac tgt tag aat ctc taa cgc tgg cct ctg ttg tta ctt tct atg atg cgc gta tcg tta tac aag ctt tca tgt tga cta ctg cag tgt ttc ttg ctc tga cta cat gta ctc tac aat caa aga gag att tca gta aac ttg taa cag gat tgt ttg ctg ctt tct gga ttt taa ttt tgt cag gag tct tga gga taa agt tta aaa tag aat taa taa aga aca tat agg tca -tt ttt taa aca tgg ata gaa acc aag gtt gtt agt taa taa tat aca aga tat ttt ttc tca ctc tga tcc atg taa acc aag gac gag aga cac tct cat tcc tca ttc acg aca cca tta aaa atg gaa att aaa gcc ctc tat taa gca cag acg gct aca ggt cta cca tca ggt taa cct tcg tct acc ttc aca atg gcc ttt cct tgt gcc cag ttc agt ccc tgt cat tgc cac gct act aag gac tcc ctg aat acc gtg act gac gtc aga cat tgt ctg act gaa tac atc ctg tgg gtt tct cat aga tgg acc cat aga gaa agc gca ggg cct ctc tac agg ctt ctc atc tct ttc aga att gat gca atg gag cta ttt ggt agc gag ttg aag gag ttc tcg gat tca ctt ccg tgg gac aat atc gac aat tgc gtg gag atc att aaa tgt ttc atc aga aat gac tcc atg aaa acc gcc aaa gaa ctt tgt gca ata att gga ctt tgt act caa tca gct att gtc act gga aga gtc ttc aat gat aag tat atc gac ata cta ctt ata ctg cga aag att ctg aat gag aac gac tat ctc acc ctc ttg gat cat atc ctc act gct aaa tac taa atc tcc ttc atg ctc tct cac taa tac tct tac tca cta cac ttt tta tca tct tat gat gaa tga ttg cct tca tca -tt ttt cgt gga ata taa tat agg aat aat tag cac cag aat agc tat gga tat ctc gtt aag aat att ctc gat aag aga cat aat gta gac ata gtt att ata tcc ttc tta gat aag tgt tac gct act gga aag ttt cca tcg tta tta tta cgt gaa gat gat atc att aaa cca aca ttg aga ttg gct ctt atg tta gct gga ttg aat tac tgt aat aaa tgc atc gag tat aga ggg ata tag caa ttc tcg ata ata gtc atg caa tat ttg aat gag act gat aat tta ggt aat aca gta ct- -ac aca cac ata tct ttc tag att ata tat cgt taa aaa tct gta aga ggt ata ttt ctc ata agt atc cac tgt gta ata tta tta atg gat ata tag ata aca caa tag gga cta ata gta ttg taa aag ata taa tcg act att t-- gta cat atc cag ata tct ata ttc cta cta gtt tgc tgc gta gtt gca tca ttg ata tgc atg -at tta tca gga ttc aga gat gaa tta cta agt aaa cta caa tcc cac aat aag taa gaa tca aat atc aaa aac tca ctt ttg att ttt cta gtc tta agt aat aca tat att tat taa tag acc tat gaa at- aaa aaa gtt aac aat gga ttc gcg tat agc tat tta cgt att agt atc ggc atc tct ttt gta tct tgt taa ttg tca caa act agt aca tta ctt caa tct gaa aat aaa tgg aag tga tat aac taa tac agc aga tat att gct gga caa tta tcc aat tat gac ctt tga tgg aaa gga tat tta tcc atc tat cgc gtt cat ggt cgg taa taa act ttt cct aga tct tta taa aaa tat ctt tgt aga att ttt cag act att tcg agt atc tgt aag tag tca ata cga gga att aga ata tta tta ttc atg tga tta tac taa caa ccg tcc tac aat taa aca aca tta ctt tta taa cgg cga aga ata tac tga aat tga tag atc gaa aaa agc cac taa taa aaa cag ttg gtt aat tac ttc agg ctt tag act aca aaa atg gtt cga tag cga aga ttg tat aat tta tct cag atc ttt agt tag aag aat gga aga cag taa caa aaa cag taa aaa aac tta gta ctt aga tat cg- aaa aaa aat ata ttt ttg tag act ctt gag aat aga agg aaa aca tgt aca taa tta taa aaa atg aaa atc aat ggc gaa taa gac agt gcg att cgc gcc atg gag tcg gca gat ttc atg gct gtc gat gag cag ttt cac gac gac ctc gat ctt tgg tca tta tct ttg gta gat gac tat aaa aaa cat gga tta ggt gtt gac tgt tat gtt cta gaa cca gtt gtt gac agg aaa ata ttt gat aga ttt ctc ctt gaa cca att tgt gat cct gta gat gtt ctg tat gat tat ttt agg att cat aga gat aat att gat cag tat ata gta gat aga ctg ttt gca tat att aca tat aaa gat att ata tct gca tta gtg tca aag aat tat atg gaa gat att ttc tct ata att att aag aat tgt aat tct gta caa gat ctc tta ctt tac tat cta tct aat gca tat gta gaa ata gac att gtt gat ttt atg gta gat cat ggg gct gta ata tat aaa ata gaa tgc ttg aat gcc tat ttt agg gga ata tgt aaa aag gaa agt agt gtt gtt gag ttt att ttg aat tgt ggt atc cca gat gaa aat gat gtt aaa tta gat cta tat aaa ata att cag tat act agg gga ttc ctt gta gat gaa ccc aca gta tta gaa att tat aag ctt tgt atc cca tat att gaa gat atc aat caa cta gat gct ggt gga agg acc ttg ctt tat cgc gct atc tat gca ggt tat ata gat tta gta tca tgg cta tta gaa aat gga gca aat gtc aac gca gta atg agt aat gga tat aca tgt ctt gac gtg gcc gtg gat agg gga tct gtc atc gcc cgt agg gaa aca cat ctt aaa ata tta gaa ata ttg ctt aga gaa cca ttg tct att gac tgt ata aaa tta gct ata ctt aat aat aca att gaa aac cat gat gtg ata aag ctc tgt atc aag tat ttt atg atg gta gat tat tca ctt tgt aat gtg tat gca tca tca ctc ttt gat tat ata att gat tgt aaa caa gaa ttg gag tac att agg cag atg aaa att cat aat aca acc atg tat gag tta atc tat aat aga gac aaa aac aag cat gct tcc cat att cta cat agg tat tct aaa cat cca gtt ttg aca cag tgt atc act aaa gga ttc aag att tac aca gaa gta acc gag cag gtc act aaa gct cta aac aga cgt gct cta ata gat gag ata ata aac aat gta tca act gat gac aat ctc cta tca aaa ctt cca tta gaa att agg gat cta att gtt tca caa gct gtc ata tag agt tct ctc cac cca cct ttc ttg aaa tga gtt aat --- --- --- --- --- --- --- --- --- --- --- --- --- --- --- --- --- --- --- --- --- --- --- --- --- --- --- --- --- --- --- --- --- --- --- --- --- --- --- --- --- --- --- --- --- --- --- --- --- --- --- --- --- --- --- --- --- --- --- --- --- --- --- --- --- --- --- --- --- --- --- --- --- --- --- --- --- --- --- --- --- --- --- --- --- --- --- --- --- --- --- --- --- --- --- --- --- --- --- --- --- --- --- --- --- --- --- --- --- --- --- --- --- --- --- --- --- --- --- --- --- --- --- --- --- --- --- --- --- --- --- --- --- --- --- --- --- --- --- --- --- --- --- --- --- --- --- --- --- --- --- --- --- -ag tca taa gtt agt taa gtc ata agt tag tt- --t tat aag tta gtt tat agt cta aca ctt cta att ttt ata cct tga tct ttt tct cta att atg aaa aag taa atc att atg aag atg gat gaa aat gga cga gat tgt gcg cat cgt taa cga tag tat gtg gta cgt acc taa cgc att tat gga cga cgg tga taa tga agg tca cat ttc tgt caa taa tgt ctg tca tat gta tct cgc att ctt tga tgt gga tat atc atc tca tct gtt taa att agt tat taa aca ctg cga tct gaa taa acg act aaa atg tgg taa ctc tcc att aca ttg cta tac gat gaa tac acg att taa tcc atc tgt att aaa gat att gtt acg cca cgg cat gcg taa ctt tga tag caa gga taa aaa agg aca tat tcc tct aca cca cta tct gat tca ttc act atc aat cga taa caa gat ctt tga tat act aac gga ccc cat tga tga ctt tag taa atc atc cga tct att gct gtg tta tct tag ata taa att caa tgg gag ctt aaa cta tta cgt tct gta caa att att gac taa agg atc tga ccc taa ttg cgt cga tga gga tgg act cac ttc tct tca tta cta ctg taa aca cat atc cgt gtt cca cga aag caa tta tta caa gtc aaa gag tca cac taa gat gcg agc tga gaa gcg att cat cta cgc gat aat aga tca tgg agc aaa cat taa cgc ggt tac gaa aat cgg aaa tac gcc gtt aca cac tta cct tca aca gta tac caa aca tag tcc tcg tgt ggt gta tgc tct ttt atc tcg agg agc cga tac gag gat acg taa taa tct tga ttg tac acc cat cat gga ata cat aaa gaa cga ttg tgc aac agg tca tat tct cat aat gtt act caa ttg gca cga aca aaa ata cgg gaa att aca aaa gga aga agg aca aca tct act tta tct att cat aaa aca taa tca agg ata tgg aag tcg ctc tct caa tat act acg gta tct act aga tag att cga cat tca gaa aga cga ata cta taa tac aat gac tcc tct tca tac cgc ctt cca gaa ttg caa taa caa tgt tgc ctc ata cct cgt ata cat cgg ata cga cat caa cct tcc gac taa aga cga taa gac agt att cga ctt ggt gtt tga aaa cag aaa cat cat ata caa ggc gga tgt cgt taa tga cat cat cca cca cag act gaa agt atc tct acc tat gat taa atc gtt gtt cta caa gat gtc gga gtt ctc tcc cta cga cga tca cta cgt aaa gaa gat aat agc cta ctg cct att aag gga cga gtc att tgc gga act aca tac taa att ctg ttt aaa cga gga cta taa aag tgt att tat gaa aaa tat atc att cga taa gat aga ttc cat cat cga aaa atg tag tcg tga cat aag tct cct caa aga gat tcg aat ctc aga cac caa ctt gta tac ggt att gag aac aga aga cat ccg gta tca cac cta tct cga agc cat aca ttc aga caa acg cat ttc att tcc cat gta cga cga tct cat aga aca gtg tca tct atc gat gga gca taa aag taa act cgt cga caa agc act caa taa att aga gtc tac cat cga tag tca atc tag act atc gta ttt gcc tcc gga aat tat gcg caa tat cat aac caa gct aag cga cta cca tct aaa cag tat gtt gta cgg aaa gaa cca tta caa ata tta tcc atg ata gaa aga aaa tat tta aaa aat aat cta tat gat tgg aga agt agg aaa caa aca gta aca aga cga cga tta cta c-a tta tta aat cat gag gtc cgt att ata ctc gta tat att gtt tct ctc atg tat aat aat aaa cgg aag aga tat agc acc aca tgc acc atc caa tgg aaa atg taa aga caa cga ata cag aag ccg taa tct atg ttg tct atc gtg tcc tcc ggg aac tta cgc ttc cag att atg tga tag caa gac taa tac aca atg tac gcc gtg tgg ttc gga tac ctt tac atc tca caa taa tca ttt aca ggc ttg tct aag ttg taa cgg aag atg tga tag taa tca ggt aga gac gcg atc gtg taa cac gac tca caa tag aat ctg tga atg ctc tcc agg ata tta ttg tct tct caa agg atc atc agg gtg tag aac atg tat ttc taa aac aaa gtg tgg aat agg ata cgg agt atc cgg ata cac gtc tac cgg aga cgt cat ctg ttc tcc gtg tgg tcc cgg aac ata ttc tca cac cgt ctc ttc cac aga taa atg cga acc --- cgt aac cag caa tac att taa cta tat cga tgt gga aat taa cct gta tcc agt caa cga cac atc gtg tac tcg gac gac cac tac cgg tct cag cga atc cat ctc aac gtc gga act aac tat tac cat gaa tca taa aga ttg tga tcc cgt ctt tcg tgc aga ata ctt ctc tgt cct taa taa tgt agc aac ttc agg att ctt tac agg aga aaa tag ata tca gaa tac ttc aaa gat atg tac tct gaa ttt cga gat taa atg taa caa caa aga ttc atc ttc caa aca gtt aac gaa aac aaa gaa tga tac tat cat gcc gca ttc aga gac ggt aac tct agt ggg cga ctg tct atc tag cgt cga cat cta cat act ata tag taa tac caa tac tca aga cta cga aac gga tac aat ctc tta tca tat ggg taa tgt tct cga tgt caa tag cca tat gcc cgc tag ttg cga tat aca taa act gat cac taa ttc cca gaa tcc cac cca ctt ata gta agt ttt ttt tta ccc ata aat aat aaa tac aat aat taa ttt ctc gta aaa gta gaa aat ata ttc taa ttt att ata tgg taa gaa agt aga atc atc tag aac agt aat caa tca ata gca atc atg aaa caa tat att gtc ctg gca tgc atg tgc cta gtg gca gct gct atg cct act agt ctt caa caa tcc tca tcc tcg tgt act gaa gaa gaa aac aaa cat cat atg gga atc gat gtt att atc aaa gtc aca aag caa gac caa aca ccg acc aat gat aag att tgt caa tcc gta acg gaa gtt aca gag a-- --- --- --- --- --- -cc gaa gat gat gag gta tcc gaa gaa gtt gta aaa gga gat ccc acc act tat tac act atc gtc ggc gcg ggt ctt aac atg aac ttt gga ttc acc aaa tgc cca aag att tca tcc atc tcc gaa tcc tct gat gga aac act gtg aat act aga ttg tcc agc gtg tca cca gga caa ggt aag gac tct ccc gcg atc acg cgt gaa gaa gct ctg gct atg atc aaa gac tgt gag atg tct atc gac atc aga tgt agc gaa gaa gag aaa gac agc gac atc aag acc cat cca gta ctt ggg tct aac atc tca cat aag aaa gtg agt tac aaa gat atc atc ggt tca acg atc gtt gat aca aaa tgt gtt aag aac cta gag ttt agc gta cgt atc gga gac atg tgt gag gaa tca tct gaa ctt gaa gtc aag gat gga ttc aag tat gtc gac gga tcg gca tct gaa ggt gca acc gat gat act tca ctc atc gat tca aca aaa ctc aaa gca tgt gtc tga atc gat aac tct att cat ctg aaa atg gat gag ttg ggt taa tcg aac gat tca gac acc gca cca cga att aaa aaa gtg tac cgg gta cta tat tcc ggt ttg caa aac aaa aat gtt taa cta cat tca caa aaa gtt acc tct cgt tac ttc ttc ttt ctg ttt caa tat gtg ata cga tat gat cac tat tcg tat tct ctt tct ctc ttg tct cat -aa aaa gtt tta c-- --- --- --- aaa aaa a-t att ttt att ctc ttt ctc tct tcg at- ggt ctc aca aaa ata tta aac ctc ttt ctg atg tct caa cta ttt cgt aaa cga taa cgt cca aca ata tat tct cgt aga gct tat caa tat cct tat gcc aat cta ggt tgt cag aca att gca tca taa aat aat gtt tat aat tta cac gtt aac atc ata taa taa acg tat ata gtt aat att ttt gga ata taa atg atc tgt aaa atc cat gta ggg gac act gct cac gtt ttt tct cta gta cat aat ttc aca caa gtt ttt ata cag aca aat taa ttc tcg tcc ata ta- ttt taa aac att gac ttt tgt act aag aaa aat atc ttg act aac cat ctc ttt ctc tcg tcg atg ggt ctc aca aaa ata tta aac ctc ttt ctg atg g-- --- --- --- --- --- --- --- --- --- --- --- --- --- --- --- --- --- --- --- --- --- --- --- --- --- --- --- --- --- --- --- --- --- --- --- --- --- --- --- --- --- --- --- --- --- --- agt cgt aaa aag ttt tta tcc ttt ctc tct tcg atg ggt ctc --- --- --- --- --- --- --- --- --- --- --- --- --- --- --- --- --- --- --- --- --- --- --- --- --- --- --- --- --- --- --- --- --- --- --- --- --- --- --- --- --- --- --- --- --- --- --- --- --- --- --- --- --- --- --- --- --- --- --- --- --- --- --- --- --- --- --- --- --- --- aca aaa ata tta aac ctc ttt ctg atg gtc tct ata aa

>MN702449.1_Monkeypox_virus_strain_B2_contig_SPADES

atc gag gac gac ttc aac tat gac agc tcg gtg gcg tct gcc agc gtg cac ata cga atg gca ttt cta aga aaa gtc tac ggt atc ctt gtc tac aat ttc ctt taa caa cgg caa cag ctg cag tat ttt tat act ttg aat gca ttc gga cat tta tac aag gga gtc ctg ttc taa tat tgg cat caa tgt tcg gat cta tag gct tga ttt tcg cat tga ctt tac aca gac ata aac atc ccc tga atc tgt acc tac ttt gtg gat tta cac tgt tag aat ctc taa cgc tgg cct ctg ttg tta ctt tct atg atg cgc gta tcg tta tac aag ctt tca tgt tga cta ctg cag tgt ttc ttg ctc tga cta cat gta ctc tac aat caa aga gag att tca gta aac ttg taa cag gat tgt ttg ctg ctt tct gga ttt taa ttt tgt cag gag tct tga gga taa agt tta aaa tag aat taa taa aga aca tat agg tca -tt ttt taa aca tgg ata gaa acc aag gtt gtt agt taa taa tat aca aga tat ttt ttc tca ctc tga tcc atg taa acc aag gac gag aga cac tct cat tcc tca ttc acg aca cca tta aaa atg gaa att aaa gcc ctc tat taa gca cag acg gct aca ggt cta cca tca ggt taa cct tcg tct acc ttc aca atg gcc ttt cct tgt gcc cag ttc agt ccc tgt cat tgc cac gct act aag gac tcc ctg aat acc gtg act gac gtc aga cat tgt ctg act gaa tac atc ctg tgg gtt tct cat aga tgg acc cat aga gaa agc gca ggg cct ctc tac agg ctt ctc atc tct ttc aga att gat gca atg gag cta ttt ggt agc gag ttg aag gag ttc tcg gat tca ctt ccg tgg gac aat atc gac aat tgc gtg gag atc att aaa tgt ttc atc aga aat gac tcc atg aaa acc gcc aaa gaa ctt tgt gca ata att gga ctt tgt act caa tca gct att gtc act gga aga gtc ttc aat gat aag tat atc gac ata cta ctt atg ctg cga aag att ctg aat gag aac gac tat ctc acc ctc ttg gat cat atc ctc act gct aaa tac taa atc tcc ttc atg ctc tct cac taa tac tct tac tca cta cac ttt tta tca tct tat gat gaa tga ttg cct tca tca -tt ttt cgt gga ata taa tat agg aat aat tag cac cag aat agc tat gga tat ctc gtt aag aat att ctc gat aag aga cat aat gta gac ata gtt att ata tcc ttc tta gat aag tgt tac gct act gga aag ttt cca tcg tta tta tta cgt gaa gat gat atc att aaa cca aca ttg aga ttg gct ctt atg tta gct gga ttg aat tac tgt aat aaa tgc atc gag tat aga ggg ata tag caa ttc tcg ata ata gtc atg caa tat ttg aat gag act gat aat tta ggt aat aca gta ct- -ac aca cac ata tct ttc tag att ata tat cgt taa aaa tct gta aga ggt ata ttt ctc ata agt atc cac tgt gta ata tta tta atg gat ata tag ata aca caa tag gga cta ata gta ttg taa aag ata taa tcg act att t-- gta cat atc cag ata tct ata ttc cta cta gtt tgc tgc gta gtt gca tca ttg ata tgc atg -at tta tca gga ttc aga gat gaa tta cta agt aaa cta caa tcc cac aat aag taa gaa tca aat atc aaa aac tca ctt ttg att ttt cta gtc tta agt aat aca tat att tat taa tag acc tat gaa at- aaa aaa gtt aac aat gga ttc gcg tat agc tat tta cgt att agt atc ggc atc tct ttt gta tct tgt taa ttg tca caa act agt aca tta ctt caa tct gaa aat aaa tgg aag tga tat aac taa tac agc aga tat att gct gga caa tta tcc aat tat gac ctt tga tgg aaa gga tat tta tcc atc tat cgc gtt cat ggt cgg taa taa act ttt cct aga tct tta taa aaa tat ctt tgt aga att ttt cag act att tcg agt atc tgt aag tag tca ata cga gga att aga ata tta tta ttc atg tga tta tac taa caa ccg tcc tac aat taa aca aca tta ctt tta taa cgg cga aga ata tac tga aat tga tag atc gaa aaa agc cac taa taa aaa cag ttg gtt aat tac ttc agg ctt tag act aca aaa atg gtt cga tag cga aga ttg tat aat tta tct cag atc ttt agt tag aag aat gga aga cag taa caa aaa cag taa aaa aac tta gta ctt aga tat cga aaa aaa aat ata ttt ttg tag act ctt gag aat aga agg aaa aca tgt aca taa tta taa aaa atg aaa atc aat ggc gaa taa gac agt gcg att cgc gcc atg gag tcg gca gat ttc atg gct gtc gat gag cag ttt cac gac gac ctc gat ctt tgg tca tta tct ttg gta gat gac tat aaa aaa cat gga tta ggt gtt gac tgt tat gtt cta gaa cca gtt gtt gac agg aaa ata ttt gat aga ttt ctc ctt gaa cca att tgt gat cct gta gat gtt ctg tat gat tat ttt agg att cat aga gat aat att gat cag tat ata gta gat aga ctg ttt gca tat att aca tat aaa gat att ata tct gca tta gtg tca aag aat tat atg gaa gat att ttc tct ata att att aag aat tgt aat tct gta caa gat ctc tta ctt tac tat cta tct aat gca tat gta gaa ata gac att gtt gat ttt atg gta gat cat ggg gct gta ata tat aaa ata gaa tgc ttg aat gcc tat ttt agg gga ata tgt aaa aag gaa agt agt gtt gtt gag ttt att ttg aat tgt ggt atc cca gat gaa aat gat gtt aaa tta gat cta tat aaa ata att cag tat act agg gga ttc ctt gta gat gaa ccc aca gta tta gaa att tat aag ctt tgt atc cca tat att gaa gat atc aat caa cta gat gct ggt gga agg acc ttg ctt tat cgc gct atc tat gca ggt tat ata gat tta gta tca tgg cta tta gaa aat gga gca aat gtc aac gca gta atg agt aat gga tat aca tgt ctt gac gtg gcc gtg gat agg gga tct gtc atc gcc cgt agg gaa gca cat ctt aaa ata tta gaa ata ttg ctt aga gaa cca ttg tct att gac tgt ata aaa tta gct ata ctt aat aat aca att gaa aac cat gat gtg ata aag ctc tgt atc aag tat ttt atg atg gta gat tat tca ctt tgt aat gtg tat gca tca tca ctc ttt gat tat ata att gat tgt aaa caa gaa ttg gag tac att agg cag atg aaa att cat aat aca acc atg tat gag tta atc tat aat aga gac aaa aac aag cat gct tcc cat att cta cat agg tat tct aaa cat cca gtt ttg aca cag tgt atc act aaa gga ttc aag att tac aca gaa gta acc gag cag gtc act aaa gct cta aac aga cgt gct cta ata gat gag ata ata aac aat gta tca act gat gac aat ctc cta tca aaa ctt cca tta gaa att agg gat cta att gtt tca caa gct gtc ata tag agt tct ctc cac cca cct ttc ttg aaa tga gtt aat --- --- --- --- --- --- --- --- --- --- --- --- --- --- --- --- --- --- --- --- --- --- --- --- --- --- --- --- --- --- --- --- --- --- --- --- --- --- --- --- --- --- --- --- --- --- --- --- --- --- --- --- --- --- --- --- --- --- --- --- --- --- --- --- --- --- --- --- --- --- --- --- --- --- --- --- --- --- --- --- --- --- --- --- --- --- --- --- --- --- --- --- --- --- --- --- --- --- --- --- --- --- --- --- --- --- --- --- --- --- --- --- --- --- --- --- --- --- --- --- --- --- --- --- --- --- --- --- --- --- --- --- --- --- --- --- --- --- --- --- --- --- --- --- --- --- --- --- --- --- --- --- --- -ag tca taa gtt agt taa gtc ata agt tag tt- --t tat aag tta gtt tat agt cta aca ctt cta att ttt ata cct tga tct ttt tct cta att atg aaa aag taa atc att atg aag atg gat gaa aat gga cga gat tgt gcg cat cgt taa cga tag tat gtg gta cgt acc taa cgc att tat gga cga cgg tga taa tga agg tca cat ttc tgt caa taa tgt ctg tca tat gta tct cgc att ctt tga tgt gga tat atc atc tca tct gtt taa att agt tat taa aca ctg cga tct gaa taa acg act aaa atg tgg taa ctc tcc att aca ttg cta tac gat gaa tac acg att taa tcc atc tgt att aaa gat att gtt acg cca cgg cat gcg taa ctt tga tag caa gga taa aaa agg aca tat tcc tct aca cca cta tct gat tca ttc act atc aat cga taa caa gat ctt tga tat act aac gga ccc cat tga tga ctt tag taa atc atc cga tct att gct gtg tta tct tag ata taa att caa tgg gag ctt aaa cta tta cgt tct gta caa att att gac taa agg atc tga ccc taa ttg cgt cga tga gga tgg act cac ttc tct tca tta cta ctg taa aca cat atc cgt gtt cca cga aag caa tta tta caa gtc aaa gag tca cac taa gat gcg agc tga gaa gcg att cat cta cgc gat aat aga tca tgg agc aaa cat taa cgc ggt tac gaa aat cgg aaa tac gcc gtt aca cac tta cct tca aca gta tac caa aca tag tcc tcg tgt ggt gta tgc tct ttt atc tcg agg agc cga tac gag gat acg taa taa tct tga ttg tac acc cat cat gga ata cat aaa gaa cga ttg tgc aac agg tca tat tct cat aat gtt act caa ttg gca cga aca aaa ata cgg gaa att aca aaa gga aga agg aca aca tct act tta tct att cat aaa aca taa tca agg ata tgg aag tcg ctc tct caa tat act acg gta tct act aga tag att cga cat tca gaa aga cga ata cta taa tac aat gac tcc tct tca tac cgc ctt cca gaa ttg caa taa caa tgt tgc ctc ata cct cgt ata cat cgg ata cga cat caa cct tcc gac taa aga cga taa gac agt att cga ctt ggt gtt tga aaa cag aaa cat cat ata caa ggc gga tgt cgt taa tga cat cat cca cca cag act gaa agt atc tct acc tat gat taa atc gtt gtt cta caa gat gtc gga gtt ctc tcc cta cga cga tca cta cgt aaa gaa gat aat agc cta ctg cct att aag gga cga gtc att tgc gga act aca tac taa att ctg ttt aaa cga gga cta taa aag tgt att tat gaa aaa tat atc att cga taa gat aga ttc cat cat cga aaa atg tag tcg tga cat aag tct cct caa aga gat tcg aat ctc aga cac caa ctt gta tac ggt att gag aac aga aga cat ccg gta tca cac cta tct cga agc cat aca ttc aga caa acg cat ttc att tcc cat gta cga cga tct cat aga aca gtg tca tct atc gat gga gca taa aag taa act cgt cga caa agc act caa taa att aga gtc tac cat cga tag tca atc tag act atc gta ttt gcc tcc gga aat tat gcg caa tat cat aac caa gct aag cga cta cca tct aaa cag tat gtt gta cgg aaa gaa cca tta caa ata tta tcc atg ata gaa aga aaa tat tta aaa aat aat cta tat gat tgg aga agt agg aaa caa aca gta aca aga cga cga tta cta c-a tta tta aat cat gag gtc cgt att ata ctc gta tat att gtt tct ctc atg tat aat aat aaa cgg aag aga tat agc acc aca tgc acc atc caa tgg aaa atg taa aga caa cga ata cag aag ccg taa tct atg ttg tct atc gtg tcc tcc ggg aac tta cgc ttc cag att atg tga tag caa gac taa tac aca atg tac gcc gtg tgg ttc gga tac ctt tac atc tca caa taa tca ttt aca ggc ttg tct aag ttg taa cgg aag atg tga tag taa tca ggt aga gac gcg atc gtg taa cac gac tca caa tag aat ctg tga atg ctc tcc agg ata tta ttg tct tct caa agg atc atc agg gtg tag aac atg tat ttc taa aac aaa gtg tgg aat agg ata cgg agt atc cgg ata cac gtc tac cgg aga cgt cat ctg ttc tcc gtg tgg tcc cgg aac ata ttc tca cac cgt ctc ttc cac aga taa atg cga acc --- cgt aac cag caa tac att taa cta tat cga tgt gga aat taa cct gta tcc agt caa cga cac atc gtg tac tcg gac gac cac tac cgg tct cag cga atc cat ctc aac gtc gga act aac tat tac cat gaa tca taa aga ttg tga tcc cgt ctt tcg tgc aga ata ctt ctc tgt cct taa taa tgt agc aac ttc agg att ctt tac agg aga aaa tag ata tca gaa tac ttc aaa gat atg tac tct gaa ttt cga gat taa atg taa caa caa aga ttc atc ttc caa aca gtt aac gaa aac aaa gaa tga tac tat cat gcc gca ttc aga gac ggt aac tct agt ggg cga ctg tct atc tag cgt cga cat cta cat act ata tag taa tac caa tac tca aga cta cga aac gga tac aat ctc tta tca tat ggg taa tgt tct cga tgt caa tag cca tat gcc cgc tag ttg cga tat aca taa act gat cac taa ttc cca gaa tcc cac cca ctt ata gta ag- -tt ttt tta ccc ata aat aat aaa tac aat aat taa ttt ctc gta aaa gta gaa aat ata ttc taa ttt att ata tgg taa gaa agt aga atc atc tag aac agt aat caa tca ata gca atc atg aaa caa tat att gtc ctg gca tgc atg tgc cta gtg gca gct gct atg cct act agt ctt caa caa tcc tca tcc tcg tgt act gaa gaa gaa aac aaa cat cat atg gga atc gat gtt att atc aaa gtc aca aag caa gac caa aca ccg acc aat gat aag att tgt caa tcc gta acg gaa gtt aca gag acc gaa gat gat gag gta tcc gaa gat gat gag gta tcc gaa gaa gtt gta aaa gga gat ccc acc act tat tac aat atc gtc ggc gcg ggt ctt aac atg aac ttt gga ttc acc aaa tgc cca aag att tca tcc atc tcc gaa tcc tct gat gga aac act gtg aat act aga ttg tcc agc gtg tca cca gga caa ggt aag gac tct ccc gcg atc acg cgt gaa gaa gct ctg gct atg atc aaa gac tgt gag atg tct atc gac atc aga tgt agc gaa gaa gag aaa gac agc gac atc aag acc cat cca gta ctt ggg tct aac atc tca cat aag aaa gtg agt tac aaa gat atc atc ggt tca acg atc gtt gat aca aaa tgt gtt aag aac cta gag ttt agc gta cgt atc gga gac atg tgt gag gaa tca tct gaa ctt gaa gtc aag gat gga ttc aag tat gtc gac gga tcg gca tct gaa ggt gca acc gat gat act tca ctc atc gat tca aca aaa ctc aaa gca tgt gtc tga atc gat aac tct att cat ctg aaa atg gat gag ttg ggt taa tcg aac gat tca gac acc gca cca cga att aaa aaa gtg tac cgg gta cta tat tcc ggt ttg caa aac aaa aat gtt taa cta cat tca caa aaa gtt acc tct cgt tac ttc ttc ttt ctg ttt caa tat gtg ata cga tat gat cac tat tcg tat tct ctt tct ctc ttg tct cat aaa aaa gtt tta c-- --- --- --- aaa aaa a-t att ttt att ctc ttt ctc tct tcg at- ggt ctc aca aaa ata tta aac ctc ttt ctg atg tct caa cta ttt cgt aaa cga taa cgt cca aca ata tat tct cgt aga gct tat caa tat cct tat gcc aat --- ggt tgt cag aca att gca tca taa aat aat gtt tat aat tta cac gtt aac atc ata taa taa acg tat ata gtt aat att ttt gga ata taa atg atc tgt aaa atc cat gta ggg gac act gct cac gtt ttt tct cta gta cat aat ttc aca caa gtt ttt ata cag aca aat taa ttc tcg tcc ata ta- ttt taa aac att gac ttt tgt act aag aaa aat atc ttg act aac cat ctc ttt ctc tcg tcg atg ggt ctc aca aaa ata tta aac ctc ttt ctg atg g-- --- --- --- --- --- --- --- --- --- --- --- --- --- --- --- --- --- --- --- --- --- --- --- --- --- --- --- --- --- --- --- --- --- --- --- --- --- --- --- --- --- --- --- --- --- --- agt cgt aaa aag ttt tta tcc ttt ctc tct tcg atg ggt ctc --- --- --- --- --- --- --- --- --- --- --- --- --- --- --- --- --- --- --- --- --- --- --- --- --- --- --- --- --- --- --- --- --- --- --- --- --- --- --- --- --- --- --- --- --- --- --a caa aaa tat taa acc tct ttc tga tgg agt cgt aaa aag ttt tta tcc ttt ctc tct tcg atg g-- --- --- --- --- --- --- --- --- --- --- --- --- --- --

>JX878418.1_Monkeypox_virus_isolate_DRC_07-0120_complete_genome

atc gag gac gac ttc aac tat gac agc tcg gtg gcg tct gcc agc gtg cac ata cga atg gca ttt cta aga aaa gtc tac ggt atc ctt gtc tac aat ttc ctt taa caa cgg caa cag ctg cag tat ttt tat act ttg aat gca ttc gga cat tta tac aag gga gtc ctg ttc taa tat tgg cat caa tgt tcg gat cta tag gct tga ttt tcg cat tga ctt tac aca gac ata aac atc ccc tga atc tgt acc tac ttt gtg gat tta cac tgt tag aat ctc taa cgc tgg cct ctg ttg tta ctt tct atg atg cgc gta tcg tta tac aag ctt tca tgt tga cta ctg cag tgt ttc ttg ctc tga cta cat gta ctc tac aat caa aga gag att tca gta aac ttg taa cag gat tgt ttg ctg ctt tct gga ttt taa ttt tgt cag gag tct tga gga taa agt tta aaa tag aat taa taa aga aca tat agg tca -tt ttt taa aca tgg ata gaa acc aag gtt gtt agt taa taa tat aca aga tat ttt ttc tca ctc tga tcc atg taa acc aag gac gag aga cac tct cat tcc tca ttc acg aca cca tta aaa atg gaa att aaa gcc ctc tat taa gca cag acg gct aca ggt cta cca tca ggt taa cct tcg tct acc ttc aca atg gcc ttt cct tgt gcc cag ttc agt ccc tgt cat tgc cac gct act aag gac tcc ctg aat acc gtg act gac gtc aga cat tgt ctg act gaa tac atc ctg tgg gtt tct cat aga tgg acc cat aga gaa agc gca ggg cct ctc tac agg ctt ctc atc tct ttc aga att gat gca atg gag cta ttt ggt agc gag ttg aag gag ttc tcg gat tca ctt ccg tgg gac aat atc gac aat tgc gtg gag atc att aaa tgt ttc atc aga aat gac tcc atg aaa acc gcc aaa gaa ctt tgt gca ata att gga ctt tgt act caa tca gct att gtc act gga aga gtc ttc aat gat aag tat atc gac ata cta ctt atg ctg cga aag att ctg aat gag aac gac tat ctc acc ctc ttg gat cat atc ctc act gct aaa tac taa atc tcc ttc atg ctc tct cac taa tac tct tac tca cta cac ttt tta tca tct tat gat gaa tga ttg cct tca tca -tt ttt cgt gga ata taa tat agg aat aat tag cac cag aat agc tat gga tat ctc gtt aag aat att ctc gat aag aga cat aat gta gac ata gtt att ata tcc ttc tta gat aag tgt tac gct act gga aag ttt cca tcg tta tta tta cgt gaa gat gat atc att aaa cca aca ttg aga ttg gct ctt atg tta gct gga ttg aat tac tgt aat aaa tgc atc gag tat aga ggg ata tag caa ttc tcg ata ata gtc atg caa tat ttg aat gag act gat aat tta ggt aat aca gta ct- -ac aca cac ata tct ttc tag att ata tat cgt taa aaa tct gta aga ggt ata ttt ctc ata agt atc cac tgt gta ata tta tta atg gat ata tag ata aca caa tag gga cta ata gta ttg taa aag ata taa tcg act att t-- gta cat atc cag ata tct ata ttc cta cta gtt tgc tgc gta gtt gca tca ttg ata tgc atg -at tta tca gga ttc aga gat gaa tta cta agt aaa cta caa tcc cac aat aag taa gaa tca aat atc aaa aac tca ctt ttg att ttt cta gtc tta agt aat aca tat att tat taa tag acc tat gaa at- aaa aaa gtt aac aat gga ttc gcg tat agc tat tta cgt att agt atc ggc atc tct ttt gta tct tgt taa ttg tca caa act agt aca tta ctt caa tct gaa aat aaa tgg aag tga tat aac taa tac agc aga tat att gct gga caa tta tcc aat tat gac ctt tga tgg aaa gga tat tta tcc atc tat cgc gtt cat ggt cgg taa taa act ttt cct aga tct tta taa aaa tat ctt tgt aga att ttt cag act att tcg agt atc tgt aag tag tca ata cga gga att aga ata tta tta ttc atg tga tta tac taa caa ccg tcc tac aat taa aca aca tta ctt tta taa cgg cga aga ata tac tga aat tga tag atc gaa aaa agc cac taa taa aaa cag ttg gtt aat tac ttc agg ctt tag act aca aaa atg gtt cga tag cga aga ttg tat aat tta tct cag atc ttt agt tag aag aat gga aga cag taa caa aaa cag taa aaa aac tta gta ctt aga tat cg- aaa aaa aat ata ttt ttg tag act ctt gag aat aga agg aaa aca tgt aca taa tta taa aaa atg aaa atc aat ggc gaa taa gac agt gcg att cgc gcc atg gag tcg gca gat ttc atg gct gtc gat gag cag ttt cac gac gac ctc gat ctt tgg tca tta tct ttg gta gat gac tat aaa aaa cat gga tta ggt gtt gac tgt tat gtt cta gaa cca gtt gtt gac agg aaa ata ttt gat aga ttt ctc ctt gaa cca att tgt gat cct gta gat gtt ctg tat gat tat ttt agg att cat aga gat aat att gat cag tat ata gta gat aga ctg ttt gca tat att aca tat aaa gat att ata tct gca tta gtg tca aag aat tat atg gaa gat att ttc tct ata att att aag aat tgt aat tct gta caa gat ctc tta ctt tac tat cta tct aat gca tat gta gaa ata gac att gtt gat ttt atg gta gat cat ggg gct gta ata tat aaa ata gaa tgc ttg aat gcc tat ttt agg gga ata tgt aaa aag gaa agt agt gtt gtt gag ttt att ttg aat tgt ggt atc cca gat gaa aat gat gtt aaa tta gat cta tat aaa ata att cag tat act agg gga ttc ctt gta gat gaa ccc aca gta tta gaa att tat aag ctt tgt atc cca tat att gaa gat atc aat caa cta gat gct ggt gga agg acc ttg ctt tat cgc gct atc tat gca ggt tat ata gat tta gta tca tgg cta tta gaa aat gga gca aat gtc aac gca gta atg agt aat gga tat aca tgt ctt gac gtg gcc gtg gat agg gga tct gtc atc gcc cgt agg gaa aca cat ctt aaa ata tta gaa ata ttg ctt aga gaa cca ttg tct att gac tgt ata aaa tta gct ata ctt aat aat aca att gaa aac cat gat gtg ata aag ctc tgt atc aag tat ttt atg atg gta gat tat tca ctt tgt aat gtg tat gca tca tca ctc ttt gat tat ata att gat tgt aaa caa gaa ttg gag tac att agg cag atg aaa att cat aat aca acc atg tat gag tta atc tat aat aga gac aaa aac aag cat gct tcc cat att cta cat agg tat tct aaa cat cca gtt ttg aca cag tgt atc act aaa gga ttc aag att tac aca gaa gta acc gag cag gtc act aaa gct cta aac aga cgt gct cta ata gat gag ata ata aac aat gta tca act gat gac aat ctc cta tca aaa ctt cca tta gaa att agg gat cta att gtt tca caa gct gtc ata tag agt tct ctc cac cca cct ttc ttg aaa tga gtt aat --- --- --- --- --- --- --- --- --- --- --- --- --- --- --- --- --- --- --- --- --- --- --- --- --- --- --- --- --- --- --- --- --- --- --- --- --- --- --- --- --- --- --- --- --- --- --- --- --- --- --- --- --- --- --- --- --- --- --- --- --- --- --- --- --- --- --- --- --- --- --- --- --- --- --- --- --- --- --- --- --- --- --- --- --- --- --- --- --- --- --- --- --- --- --- --- --- --- --- --- --- --- --- --- --- --- --- --- --- --- --- --- --- --- --- --- --- --- --- --- --- --- --- --- --- --- --- --- --- --- --- --- --- --- --- --- --- --- --- --- --- --- --- --- --- --- --- --- --- --- --- --- --- -ag tca taa gtt agt taa gtc ata agt tag tt- --t tat aag tta gtt tat agt cta aca ctt cta att ttt ata cct tga tct ttt tct cta att atg aaa aag taa atc att atg aag atg gat gaa aat gga cga gat tgt gcg cat cgt taa cga tag tat gtg gta cgt acc taa cgc att tat gga cga cgg tga taa tga agg tca cat ttc tgt caa taa tgt ctg tca tat gta tct cgc att ctt tga tgt gga tat atc atc tca tct gtt taa att agt tat taa aca ctg cga tct gaa taa acg act aaa atg tgg taa ctc tcc att aca ttg cta tac gat gaa tac acg att taa tcc atc tgt att aaa gat att gtt acg cca cgg cat gcg taa ctt tga tag caa gga taa aaa agg aca tat tcc tct aca cca cta tct gat tca ttc act atc aat cga taa caa gat ctt tga tat act aac gga ccc cat tga tga ctt tag taa atc atc cga tct att gct gtg tta tct tag ata taa att caa tgg gag ctt aaa cta tta cgt tct gta caa att att gac taa agg atc tga ccc taa ttg cgt cga tga gga tgg act cac ttc tct tca tta cta ctg taa aca cat atc cgt gtt cca cga aag caa tta tta caa gtc aaa gag tca cac taa gat gcg agc tga gaa gcg att cat cta cgc gat aat aga tca tgg agc aaa cat taa cgc ggt tac gaa aat cgg aaa tac gcc gtt aca cac tta cct tca aca gta tac caa aca tag tcc tcg tgt ggt gta tgc tct ttt atc tcg agg agc cga tac gag gat acg taa taa tct tga ttg tac acc cat cat gga ata cat aaa gaa cga ttg tgc aac agg tca tat tct cat aat gtt act caa ttg gca cga aca aaa ata cgg gaa att aca aaa gga aga agg aca aca tct act tta tct att cat aaa aca taa tca agg ata tgg aag tcg ctc tct caa tat act acg gta tct act aga tag att cga cat tca gaa aga cga ata cta taa tac aat gac tcc tct tca tac cgc ctt cca gaa ttg caa taa caa tgt tgc ctc ata cct cgt ata cat cgg ata cga cat caa cct tcc gac taa aga cga taa gac agt att cga ctt ggt gtt tga aaa cag aaa cat cat ata caa ggc gga tgt cgt taa tga cat cat cca cca cag act gaa agt atc tct acc tat gat taa atc gtt gtt cta caa gat gtc gga gtt ctc tcc cta cga cga tca cta cgt aaa gaa gat aat agc cta ctg cct att aag gga cga gtc att tgc gga act aca tac taa att ctg ttt aaa cga gga cta taa aag tgt att tat gaa aaa tat atc att cga taa gat aga ttc cat cat cga aaa atg tag tcg tga cat aag tct cct caa aga gat tcg aat ctc aga cac caa ctt gta tac ggt att gag aac aga aga cat ccg gta tca cac cta tct cga agc cat aca ttc aga caa acg cat ttc att tcc cat gta cga cga tct cat aga aca gtg tca tct atc gat gga gca taa aag taa act cgt cga caa agc act caa taa att aga gtc tac cat cga tag tca atc tag act atc gta ttt gcc tcc gga aat tat gcg caa tat cat aac caa gct aag cga cta cca tct aaa cag tat gtt gta cgg aaa gaa cca tta caa ata tta tcc atg ata gaa aga aaa tat tta aaa aat aat cta tat gat tgg aga agt agg aaa caa aca gta aca aga cga cga tta cta c-a tta tta aat cat gag gtc cgt att ata ctc gta tat att gtt tct ctc atg tat aat aat aaa cgg aag aga tat agc acc aca tgc acc atc caa tgg aaa atg taa aga caa cga ata cag aag ccg taa tct atg ttg tct atc gtg tcc tcc ggg aac tta cgc ttc cag att atg tga tag caa gac taa tac aca atg tac gcc gtg tgg ttc gga tac ctt tac atc tca caa taa tca ttt aca ggc ttg tct aag ttg taa cgg aag atg tga tag taa tca ggt aga gac gcg atc gtg taa cac gac tca caa tag aat ctg tga atg ctc tcc agg ata tta ttg tct tct caa agg atc atc agg gtg tag aac atg tat ttc taa aac aaa gtg tgg aat agg ata cgg agt atc cgg ata cac gtc tac cgg aga cgt cat ctg ttc tcc gtg tgg tcc cgg aac ata ttc tca cac cgt ctc ttc cac aga taa atg cga acc --- cgt aac cag caa tac att taa cta tat cga tgt gga aat taa cct gta tcc agt caa cga cac atc gtg tac tcg gac gac cac tac cgg tct cag cga atc cat ctc aac gtc gga act aac tat tac cat gaa tca taa aga ttg tga tcc cgt ctt tcg tgc aga ata ctt ctc tgt cct taa taa tgt agc aac ttc agg att ctt tac agg aga aaa tag ata tca gaa tac ttc aaa gat atg tac tct gaa ttt cga gat taa atg taa caa caa aga ttc atc ttc caa aca gtt aat gaa aac aaa gaa tga tac tat cat gcc gca ttc aga gac ggt aac tct agt ggg cga ctg tct atc tag cgt cga cat cta cat act ata tag taa tac caa tac tca aga cta cga aac gga tac aat ctc tta tca tat ggg taa tgt tct cga tgt caa tag cca tat gcc cgc tag ttg cga tat aca taa act gat cac taa ttc cca gaa tcc cac cca ctt ata gta agt ttt ttt tta ccc ata aat aat aaa tac aat aat taa ttt ctc gta aaa gta gaa aat ata ttc taa ttt att ata tgg taa gaa agt aga atc atc tag aac agt aat caa tca ata gca atc atg aaa caa tat att gtc ctg gca tgc atg tgc cta gtg gca gct gct atg cct act agt ctt caa caa tcc tca tcc tcg tgt act gaa gaa gaa aac aaa cat cat atg gga atc gat gtt att atc aaa gtc aca aag caa gac caa aca ccg acc aat gat aag att tgt caa tcc gta acg gaa gtt aca gag a-- --- --- --- --- --- -cc gaa gat gat gag gta tcc gaa gaa gtt gta aaa gga gat ccc acc act tat tac act atc gtc ggc gcg ggt ctt aac atg aac ttt gga ttc acc aaa tgc cca aag att tca tcc atc tcc gaa tcc tct gat gga aac act gtg aat act aga ttg tcc agc gtg tca cca gga caa ggt aag gac tct ccc gcg atc acg cgt gaa gaa gct ctg gct atg atc aaa gac tgt gag atg tct atc gac atc aga tgt agc gaa gaa gag aaa gac agc gac atc aag acc cat cca gta ctt ggg tct aac atc tca cat aag aaa gtg agt tac aaa gat atc atc ggt tca acg atc gtt gat aca aaa tgt gtt aag aac cta gag ttt agc gta cgt atc gga gac atg tgt gag gaa tca tct gaa ctt gaa gtc aag gat gga ttc aag tat gtc gac gga tcg gca tct gaa ggt gca acc gat gat act tca ctc atc gat tca aca aaa ctc aaa gca tgt gtc tga atc gat aac tct att cat ctg aaa atg gat gag ttg ggt taa tcg aac gat tca gac acc gca cca cga att aaa aaa gtg tac cgg gta cta tat tcc ggt ttg caa aac aaa aat gtt taa cta cat tca caa aaa gtt acc tct cgt tac ttc ttc ttt ctg ttt caa tat gtg ata cga tat gat cac tat tcg tat tct ctt tct ctc ttg tct cat -aa aaa gtt tta c-- --- --- --- aaa aaa a-t att ttt att ctc ttt ctc tct tcg at- ggt ctc aca aaa ata tta aac ctc ttt ctg atg tct caa cta ttt cgt aaa cga taa cgt cca aca ata tat tct cgt aga gct tat caa tat cct tat gcc aat cta ggt tgt cag aca att gca tca taa aat aat gtt tat aat tta cac gtt aac atc ata taa taa acg tat ata gtt aat att ttt gga ata taa atg atc tgt aaa atc cat gta ggg gac act gct cac gtt ttt tct cta gta cat aat ttc aca caa gtt ttt ata cag aca aat taa ttc tcg tcc ata ta- ttt taa aac att gac ttt tgt act aag aaa aat atc ttg act aac cat ctc ttt ctc tcg tcg atg ggt ctc aca aaa ata tta aac ctc ttt ctg atg g-- --- --- --- --- --- --- --- --- --- --- --- --- --- --- --- --- --- --- --- --- --- --- --- --- --- --- --- --- --- --- --- --- --- --- --- --- --- --- --- --- --- --- --- --- --- --- agt cgt aaa aag ttt tta tcc ttt ctc tct tcg atg ggt ctc --- --- --- --- --- --- --- --- --- --- --- --- --- --- --- --- --- --- --- --- --- --- --- --- --- --- --- --- --- --- --- --- --- --- --- --- --- --- --- --- --- --- --- --- --- --- --- --- --- --- --- --- --- --- --- --- --- --- --- --- --- --- --- --- --- --- --- --- --- --- aca aaa ata tta aac ctc ttt ctg atg gtc tct ata aa

>MN702445.1_Monkeypox_virus_strain_A4_contig_SPADES

atc gag gac gac ttc aac tat gac agc tcg gtg gcg tct gcc agc gtg cac ata cga atg gca ttt cta aga aaa gtc tac ggt atc ctt gtc tac aat ttc ctt taa caa cgg caa cag ctg cag tat ttt tat act ttg aat gca ttc gga cat tta tac aag gga gtc ctg ttc taa tat tgg cat caa tgt tcg gat cta tag gct tga ttt tcg cat tga ctt tac aca gac ata aac atc ccc tga atc tgt acc tac ttt gtg gat tta cac tgt tag aat ctc taa cgc tgg cct ctg ttg tta ctt tct atg atg cgc gta tcg tta tac aag ctt tca tgt tga cta ctg cag tgt ttc ttg ctc tga cta cat gta ctc tac aat caa aga gag att tca gta aac ttg taa cag gat tgt ttg ctg ctt tct gga ttt taa ttt tgt cag gag tct tga gga taa agt tta aaa tag aat taa taa aga aca tat agg tca -tt ttt taa aca tgg ata gaa acc aag gtt gtt agt taa taa tat aca aga tat ttt ttc tca ctc tga tcc atg taa acc aag gac gag aga cac tct cat tcc tca ttc acg aca cca tta aaa atg gaa att aaa gcc ctc tat taa gca cag acg gct aca ggt cta cca tca ggt taa cct tcg tct acc ttc aca atg gcc ttt cct tgt gcc cag ttc agt ccc tgt cat tgc cac gct act aag gac tcc ctg aat acc gtg act gac gtc aga cat tgt ctg act gaa tac atc ctg tgg gtt tct cat aga tgg acc cat aga gaa agc gca ggg cct ctc tac agg ctt ctc atc tct ttc aga att gat gca atg gag cta ttt ggt agc gag ttg aag gag ttc tcg gat tca ctt ccg tgg gac aat atc gac aat tgc gtg gag atc att aaa tgt ttc atc aga aat gac tcc atg aaa acc gcc aaa gaa ctt tgt gca ata att gga ctt tgt act caa tca gct att gtc act gga aga gtc ttc aat gat aag tat atc gac ata cta ctt atg ctg cga aag att ctg aat gag aac gac tat ctc acc ctc ttg gat cat atc ctc act gct aaa tac taa atc tcc ttc atg ctc tct cac taa tac tct tac tca cta cac ttt tta tca tct tat gat gaa tga ttg cct tca tca -tt ttt cgt gga ata taa tat agg aat aat tag cac cag aat agc tat gga tat ctc gtt aag aat att ctc gat aag aga cat aat gta gac ata gtt att ata tcc ttc tta gat aag tgt tac gct act gga aag ttt cca tcg tta tta tta cgt gaa gat gat atc att aaa cca aca ttg aga ttg gct ctt atg tta gct gga ttg aat tac tgt aat aaa tgc atc gag tat aga ggg ata tag caa ttc tcg ata ata gtc atg caa tat ttg aat gag act gat aat tta ggt aat aca gta ct- -ac aca cac ata tct ttc tag att ata tat cgt taa aaa tct gta aga ggt ata ttt ctc ata agt atc cac tgt gta ata tta tta atg gat ata tag ata aca caa tag gga cta ata gta ttg taa aag ata taa tcg act att t-- gta cat atc cag ata tct ata ttc cta cta gtt tgc tgc gta gtt gca tca ttg ata tgc atg -at tta tca gga ttc aga gat gaa tta cta agt aaa cta caa tcc cac aat aag taa gaa tca aat atc aaa aac tca ctt ttg att ttt cta gtc tta agt aat aca tat att tat taa tag acc tat gaa at- aaa aaa gtt aac aat gga ttc gcg tat agc tat tta cgt att agt atc ggc atc tct ttt gta tct tgt taa ttg tca caa act agt aca tta ctt caa tct gaa aat aaa tgg aag tga tat aac taa tac agc aga tat att gct gga caa tta tcc aat tat gac ctt tga tgg aaa gga tat tta tcc atc tat cgc gtt cat ggt cgg taa taa act ttt cct aga tct tta taa aaa tat ctt tgt aga att ttt cag act att tcg agt atc tgt aag tag tca ata cga gga att aga ata tta tta ttc atg tga tta tac taa caa ccg tcc tac aat taa aca aca tta ctt tta taa cgg cga aga ata tac tga aat tga tag atc gaa aaa agc cac taa taa aaa cag ttg gtt aat tac ttc agg ctt tag act aca aaa atg gtt cga tag cga aga ttg tat aat tta tct cag atc ttt agt tag aag aat gga aga cag taa caa aaa cag taa aaa aac tta gta ctt aga tat cga aaa aaa aat ata ttt ttg tag act ctt gag aat aga agg aaa aca tgt aca taa tta taa aaa atg aaa atc aat ggc gaa taa gac agt gcg att cgc gcc atg gag tcg gca gat ttc atg gct gtc gat gag cag ttt cac gac gac ctc gat ctt tgg tca tta tct ttg gta gat gac tat aaa aaa cat gga tta ggt gtt gac tgt tat gtt cta gaa cca gtt gtt gac agg aaa ata ttt gat aga ttt ctc ctt gaa cca att tgt gat cct gta gat gtt ctg tat gat tat ttt agg att cat aga gat aat att gat cag tat ata gta gat aga ctg ttt gca tat att aca tat aaa gat att ata tct gca tta gtg tca aag aat tat atg gaa gat att ttc tct ata att att aag aat tgt aat tct gta caa gat ctc tta ctt tac tat cta tct aat gca tat gta gaa ata gac att gtt gat ttt atg gta gat cat ggg gct gta ata tat aaa ata gaa tgc ttg aat gcc tat ttt agg gga ata tgt aaa aag gaa agt agt gtt gtt gag ttt att ttg aat tgt ggt atc cca gat gaa aat gat gtt aaa tta gat cta tat aaa ata att cag tat act agg gga ttc ctt gta gat gaa ccc aca gta tta gaa att tat aag ctt tgt atc cca tat att gaa gat atc aat caa cta gat gct ggt gga agg acc ttg ctt tat cgc gct atc tat gca ggt tat ata gat tta gta tca tgg cta tta gaa aat gga gca aat gtc aac gca gta atg agt aat gga tat aca tgt ctt gac gtg gcc gtg gat agg gga tct gtc atc gcc cgt agg gaa gca cat ctt aaa ata tta gaa ata ttg ctt aga gaa cca ttg tct att gac tgt ata aaa tta gct ata ctt aat aat aca att gaa aac cat gat gtg ata aag ctc tgt atc aag tat ttt atg atg gta gat tat tca ctt tgt aat gtg tat gca tca tca ctc ttt gat tat ata att gat tgt aaa caa gaa ttg gag tac att agg cag atg aaa att cat aat aca acc atg tat gag tta atc tat aat aga gac aaa aac aag cat gct tcc cat att cta cat agg tat tct aaa cat cca gtt ttg aca cag tgt atc act aaa gga ttc aag att tac aca gaa gta acc gag cag gtc act aaa gct cta aac aga cgt gct cta ata gat gag ata ata aac aat gta tca act gat gac aat ctc cta tca aaa ctt cca tta gaa att agg gat cta att gtt tca caa gct gtc ata tag agt tct ctc cac cca cct ttc ttg aaa tga gtt aat --- --- --- --- --- --- --- --- --- --- --- --- --- --- --- --- --- --- --- --- --- --- --- --- --- --- --- --- --- --- --- --- --- --- --- --- --- --- --- --- --- --- --- --- --- --- --- --- --- --- --- --- --- --- --- --- --- --- --- --- --- --- --- --- --- --- --- --- --- --- --- --- --- --- --- --- --- --- --- --- --- --- --- --- --- --- --- --- --- --- --- --- --- --- --- --- --- --- --- --- --- --- --- --- --- --- --- --- --- --- --- --- --- --- --- --- --- --- --- --- --- --- --- --- --- --- --- --- --- --- --- --- --- --- --- --- --- --- --- --- --- --- --- --- --- --- --- --- --- --- --- --- --- -ag tca taa gtt agt taa gtc ata agt tag tt- --t tat aag tta gtt tat agt cta aca ctt cta att ttt ata cct tga tct ttt tct cta att atg aaa aag taa atc att atg aag atg gat gaa aat gga cga gat tgt gcg cat cgt taa cga tag tat gtg gta cgt acc taa cgc att tat gga cga cgg tga taa tga agg tca cat ttc tgt caa taa tgt ctg tca tat gta tct cgc att ctt tga tgt gga tat atc atc tca tct gtt taa att agt tat taa aca ctg cga tct gaa taa acg act aaa atg tgg taa ctc tcc att aca ttg cta tac gat gaa tac acg att taa tcc atc tgt att aaa gat att gtt acg cca cgg cat gcg taa ctt tga tag caa gga taa aaa agg aca tat tcc tct aca cca cta tct gat tca ttc act atc aat cga taa caa gat ctt tga tat act aac gga ccc cat tga tga ctt tag taa atc atc cga tct att gct gtg tta tct tag ata taa att caa tgg gag ctt aaa cta tta cgt tct gta caa att att gac taa agg atc tga ccc taa ttg cgt cga tga gga tgg act cac ttc tct tca tta cta ctg taa aca cat atc cgt gtt cca cga aag caa tta tta caa gtc aaa gag tca cac taa gat gcg agc tga gaa gcg att cat cta cgc gat aat aga tca tgg agc aaa cat taa cgc ggt tac gaa aat cgg aaa tac gcc gtt aca cac tta cct tca aca gta tac caa aca tag tcc tcg tgt ggt gta tgc tct ttt atc tcg agg agc cga tac gag gat acg taa taa tct tga ttg tac acc cat cat gga ata cat aaa gaa cga ttg tgc aac agg tca tat tct cat aat gtt act caa ttg gca cga aca aaa ata cgg gaa att aca aaa gga aga agg aca aca tct act tta tct att cat aaa aca taa tca agg ata tgg aag tcg ctc tct caa tat act acg gta tct act aga tag att cga cat tca gaa aga cga ata cta taa tac aat gac tcc tct tca tac cgc ctt cca gaa ttg caa taa caa tgt tgc ctc ata cct cgt ata cat cgg ata cga cat caa cct tcc gac taa aga cga taa gac agt att cga ctt ggt gtt tga aaa cag aaa cat cat ata caa ggc gga tgt cgt taa tga cat cat cca cca cag act gaa agt atc tct acc tat gat taa atc gtt gtt cta caa gat gtc gga gtt ctc tcc cta cga cga tca cta cgt aaa gaa gat aat agc cta ctg cct att aag gga cga gtc att tgc gga act aca tac taa att ctg ttt aaa cga gga cta taa aag tgt att tat gaa aaa tat atc att cga taa gat aga ttc cat cat cga aaa atg tag tcg tga cat aag tct cct caa aga gat tcg aat ctc aga cac caa ctt gta tac ggt att gag aac aga aga cat ccg gta tca cac cta tct cga agc cat aca ttc aga caa acg cat ttc att tcc cat gta cga cga tct cat aga aca gtg tca tct atc gat gga gca taa aag taa act cgt cga caa agc act caa taa att aga gtc tac cat cga tag tca atc tag act atc gta ttt gcc tcc gga aat tat gcg caa tat cat aac caa gct aag cga cta cca tct aaa cag tat gtt gta cgg aaa gaa cca tta caa ata tta tcc atg ata gaa aga aaa tat tta aaa aat aat cta tat gat tgg aga agt agg aaa caa aca gta aca aga cga cga tta cta c-a tta tta aat cat gag gtc cgt att ata ctc gta tat att gtt tct ctc atg tat aat aat aaa cgg aag aga tat agc acc aca tgc acc atc caa tgg aaa atg taa aga caa cga ata cag aag ccg taa tct atg ttg tct atc gtg tcc tcc ggg aac tta cgc ttc cag att atg tga tag caa gac taa tac aca atg tac gcc gtg tgg ttc gga tac ctt tac atc tca caa taa tca ttt aca ggc ttg tct aag ttg taa cgg aag atg tga tag taa tca ggt aga gac gcg atc gtg taa cac gac tca caa tag aat ctg tga atg ctc tcc agg ata tta ttg tct tct caa agg atc atc agg gtg tag aac atg tat ttc taa aac aaa gtg tgg aat agg ata cgg agt atc cgg ata cac gtc tac cgg aga cgt cat ctg ttc tcc gtg tgg tcc cgg aac ata ttc tca cac cgt ctc ttc cac aga taa atg cga acc --- cgt aac cag caa tac att taa cta tat cga tgt gga aat taa cct gta tcc agt caa cga cac atc gtg tac tcg gac gac cac tac cgg tct cag cga atc cat ctc aac gtc gga act aac tat tac cat gaa tca taa aga ttg tga tcc cgt ctt tcg tgc aga ata ctt ctc tgt cct taa taa tgt agc aac ttc agg att ctt tac agg aga aaa tag ata tca gaa tac ttc aaa gat atg tac tct gaa ttt cga gat taa atg taa caa caa aga ttc atc ttc caa aca gtt aac gaa aac aaa gaa tga tac tat cat gcc gca ttc aga gac ggt aac tct agt ggg cga ctg tct atc tag cgt cga cat cta cat act ata tag taa tac caa tac tca aga cta cga aac gga tac aat ctc tta tca tat ggg taa tgt tct cga tgt caa tag cca tat gcc cgc tag ttg cga tat aca taa act gat cac taa ttc cca gaa tcc cac cca ctt ata gta ag- -tt ttt tta ccc ata aat aat aaa tac aat aat taa ttt ctc gta aaa gta gaa aat ata ttc taa ttt att ata tgg taa gaa agt aga atc atc tag aac agt aat caa tca ata gca atc atg aaa caa tat att gtc ctg gca tgc atg tgc cta gtg gca gct gct atg cct act agt ctt caa caa tcc tca tcc tcg tgt act gaa gaa gaa aac aaa cat cat atg gga atc gat gtt att atc aaa gtc aca aag caa gac caa aca ccg acc aat gat aag att tgt caa tcc gta acg gaa gtt aca gag acc gaa gat gat gag gta tcc gaa gat gat gag gta tcc gaa gaa gtt gta aaa gga gat ccc acc act tat tac aat atc gtc ggc gcg ggt ctt aac atg aac ttt gga ttc acc aaa tgc cca aag att tca tcc atc tcc gaa tcc tct gat gga aac act gtg aat act aga ttg tcc agc gtg tca cca gga caa ggt aag gac tct ccc gcg atc acg cgt gaa gaa gct ctg gct atg atc aaa gac tgt gag atg tct atc gac atc aga tgt agc gaa gaa gag aaa gac agc gac atc aag acc cat cca gta ctt ggg tct aac atc tca cat aag aaa gtg agt tac aaa gat atc atc ggt tca acg atc gtt gat aca aaa tgt gtt aag aac cta gag ttt agc gta cgt atc gga gac atg tgt gag gaa tca tct gaa ctt gaa gtc aag gat gga ttc aag tat gtc gac gga tcg gca tct gaa ggt gca acc gat gat act tca ctc atc gat tca aca aaa ctc aaa gca tgt gtc tga atc gat aac tct att cat ctg aaa atg gat gag ttg ggt taa tcg aac gat tca gac acc gca cca cga att aaa aaa gtg tac cgg gta cta tat tcc ggt ttg caa aac aaa aat gtt taa cta cat tca caa aaa gtt acc tct cgt tac ttc ttc ttt ctg ttt caa tat gtg ata cga tat gat cac tat tcg tat tct ctt tct ctc ttg tct cat aaa aaa gtt tta c-- --- --- --- aaa aaa a-t att ttt att ctc ttt ctc tct tcg at- ggt ctc aca aaa ata tta aac ctc ttt ctg atg tct caa cta ttt cgt aaa cga taa cgt cca aca ata tat tct cgt aga gct tat caa tat cct tat gcc aat --- ggt tgt cag aca att gca tca taa aat aat gtt tat aat tta cac gtt aac atc ata taa taa acg tat ata gtt aat att ttt gga ata taa atg atc tgt aaa atc cat gta ggg gac act gct cac gtt ttt tct cta gta cat aat ttc aca caa gtt ttt ata cag aca aat taa ttc tcg tcc ata ta- ttt taa aac att gac ttt tgt act aag aaa aat atc ttg act aac cat ctc ttt ctc tcg tcg atg ggt ctc aca aaa ata tta aac ctc ttt ctg atg g-- --- --- --- --- --- --- --- --- --- --- --- --- --- --- --- --- --- --- --- --- --- --- --- --- --- --- --- --- --- --- --- --- --- --- --- --- --- --- --- --- --- --- --- --- --- --- agt cgt aaa aag ttt tta tcc ttt ctc tct tcg atg ggt ctc --- --- --- --- --- --- --- --- --- --- --- --- --- --- --- --- --- --- --- --- --- --- --- --- --- --- --- --- --- --- --- --- --- --- --- --- --- --- --- --- --- --- --- --- --- --- --a caa aaa tat taa acc tct ttc tga tgg agt cgt aaa aag ttt tta tcc ttt ctc tct tcg atg ggt ctc aca aaa ata tta aac ctc ttt ctg atg gtc tct ata aa

>MN702444.1_Monkeypox_virus_strain_A5_contig_SPADES

atc gag gac gac ttc aac tat gac agc tcg gtg gcg tct gcc agc gtg cac ata cga atg gca ttt cta aga aaa gtc tac ggt atc ctt gtc tac aat ttc ctt taa caa cgg caa cag ctg cag tat ttt tat act ttg aat gca ttc gga cat tta tac aag gga gtc ctg ttc taa tat tgg cat caa tgt tcg gat cta tag gct tga ttt tcg cat tga ctt tac aca gac ata aac atc ccc tga atc tgt acc tac ttt gtg gat tta cac tgt tag aat ctc taa cgc tgg cct ctg ttg tta ctt tct atg atg cgc gta tcg tta tac aag ctt tca tgt tga cta ctg cag tgt ttc ttg ctc tga cta cat gta ctc tac aat caa aga gag att tca gta aac ttg taa cag gat tgt ttg ctg ctt tct gga ttt taa ttt tgt cag gag tct tga gga taa agt tta aaa tag aat taa taa aga aca tat agg tca -tt ttt taa aca tgg ata gaa acc aag gtt gtt agt taa taa tat aca aga tat ttt ttc tca ctc tga tcc atg taa acc aag gac gag aga cac tct cat tcc tca ttc acg aca cca tta aaa atg gaa att aaa gcc ctc tat taa gca cag acg gct aca ggt cta cca tca ggt taa cct tcg tct acc ttc aca atg gcc ttt cct tgt gcc cag ttc agt ccc tgt cat tgc cac gct act aag gac tcc ctg aat acc gtg act gac gtc aga cat tgt ctg act gaa tac atc ctg tgg gtt tct cat aga tgg acc cat aga gaa agc gca ggg cct ctc tac agg ctt ctc atc tct ttc aga att gat gca atg gag cta ttt ggt agc gag ttg aag gag ttc tcg gat tca ctt ccg tgg gac aat atc gac aat tgc gtg gag atc att aaa tgt ttc atc aga aat gac tcc atg aaa acc gcc aaa gaa ctt tgt gca ata att gga ctt tgt act caa tca gct att gtc act gga aga gtc ttc aat gat aag tat atc gac ata cta ctt atg ctg cga aag att ctg aat gag aac gac tat ctc acc ctc ttg gat cat atc ctc act gct aaa tac taa atc tcc ttc atg ctc tct cac taa tac tct tac tca cta cac ttt tta tca tct tat gat gaa tga ttg cct tca tca -tt ttt cgt gga ata taa tat agg aat aat tag cac cag aat agc tat gga tat ctc gtt aag aat att ctc gat aag aga cat aat gta gac ata gtt att ata tcc ttc tta gat aag tgt tac gct act gga aag ttt cca tcg tta tta tta cgt gaa gat gat atc att aaa cca aca ttg aga ttg gct ctt atg tta gct gga ttg aat tac tgt aat aaa tgc atc gag tat aga ggg ata tag caa ttc tcg ata ata gtc atg caa tat ttg aat gag act gat aat tta ggt aat aca gta ct- -ac aca cac ata tct ttc tag att ata tat cgt taa aaa tct gta aga ggt ata ttt ctc ata agt atc cac tgt gta ata tta tta atg gat ata tag ata aca caa tag gga cta ata gta ttg taa aag ata taa tcg act att t-- gta cat atc cag ata tct ata ttc cta cta gtt tgc tgc gta gtt gca tca ttg ata tgc atg -at tta tca gga ttc aga gat gaa tta cta agt aaa cta caa tcc cac aat aag taa gaa tca aat atc aaa aac tca ctt ttg att ttt cta gtc tta agt aat aca tat att tat taa tag acc tat gaa at- aaa aaa gtt aac aat gga ttc gcg tat agc tat tta cgt att agt atc ggc atc tct ttt gta tct tgt taa ttg tca caa act agt aca tta ctt caa tct gaa aat aaa tgg aag tga tat aac taa tac agc aga tat att gct gga caa tta tcc aat tat gac ctt tga tgg aaa gga tat tta tcc atc tat cgc gtt cat ggt cgg taa taa act ttt cct aga tct tta taa aaa tat ctt tgt aga att ttt cag act att tcg agt atc tgt aag tag tca ata cga gga att aga ata tta tta ttc atg tga tta tac taa caa ccg tcc tac aat taa aca aca tta ctt tta taa cgg cga aga ata tac tga aat tga tag atc gaa aaa agc cac taa taa aaa cag ttg gtt aat tac ttc agg ctt tag act aca aaa atg gtt cga tag cga aga ttg tat aat tta tct cag atc ttt agt tag aag aat gga aga cag taa caa aaa cag taa aaa aac tta gta ctt aga tat cga aaa aaa aat ata ttt ttg tag act ctt gag aat aga agg aaa aca tgt aca taa tta taa aaa atg aaa atc aat ggc gaa taa gac agt gcg att cgc gcc atg gag tcg gca gat ttc atg gct gtc gat gag cag ttt cac gac gac ctc gat ctt tgg tca tta tct ttg gta gat gac tat aaa aaa cat gga tta ggt gtt gac tgt tat gtt cta gaa cca gtt gtt gac agg aaa ata ttt gat aga ttt ctc ctt gaa cca att tgt gat cct gta gat gtt ctg tat gat tat ttt agg att cat aga gat aat att gat cag tat ata gta gat aga ctg ttt gca tat att aca tat aaa gat att ata tct gca tta gtg tca aag aat tat atg gaa gat att ttc tct ata att att aag aat tgt aat tct gta caa gat ctc tta ctt tac tat cta tct aat gca tat gta gaa ata gac att gtt gat ttt atg gta gat cat ggg gct gta ata tat aaa ata gaa tgc ttg aat gcc tat ttt agg gga ata tgt aaa aag gaa agt agt gtt gtt gag ttt att ttg aat tgt ggt atc cca gat gaa aat gat gtt aaa tta gat cta tat aaa ata att cag tat act agg gga ttc ctt gta gat gaa ccc aca gta tta gaa att tat aag ctt tgt atc cca tat att gaa gat atc aat caa cta gat gct ggt gga agg acc ttg ctt tat cgc gct atc tat gca ggt tat ata gat tta gta tca tgg cta tta gaa aat gga gca aat gtc aac gca gta atg agt aat gga tat aca tgt ctt gac gtg gcc gtg gat agg gga tct gtc atc gcc cgt agg gaa gca cat ctt aaa ata tta gaa ata ttg ctt aga gaa cca ttg tct att gac tgt ata aaa tta gct ata ctt aat aat aca att gaa aac cat gat gtg ata aag ctc tgt atc aag tat ttt atg atg gta gat tat tca ctt tgt aat gtg tat gca tca tca ctc ttt gat tat ata att gat tgt aaa caa gaa ttg gag tac att agg cag atg aaa att cat aat aca acc atg tat gag tta atc tat aat aga gac aaa aac aag cat gct tcc cat att cta cat agg tat tct aaa cat cca gtt ttg aca cag tgt atc act aaa gga ttc aag att tac aca gaa gta acc gag cag gtc act aaa gct cta aac aga cgt gct cta ata gat gag ata ata aac aat gta tca act gat gac aat ctc cta tca aaa ctt cca tta gaa att agg gat cta att gtt tca caa gct gtc ata tag agt tct ctc cac cca cct ttc ttg aaa tga gtt aat --- --- --- --- --- --- --- --- --- --- --- --- --- --- --- --- --- --- --- --- --- --- --- --- --- --- --- --- --- --- --- --- --- --- --- --- --- --- --- --- --- --- --- --- --- --- --- --- --- --- --- --- --- --- --- --- --- --- --- --- --- --- --- --- --- --- --- --- --- --- --- --- --- --- --- --- --- --- --- --- --- --- --- --- --- --- --- --- --- --- --- --- --- --- --- --- --- --- --- --- --- --- --- --- --- --- --- --- --- --- --- --- --- --- --- --- --- --- --- --- --- --- --- --- --- --- --- --- --- --- --- --- --- --- --- --- --- --- --- --- --- --- --- --- --- --- --- --- --- --- --- --- --- -ag tca taa gtt agt taa gtc ata agt tag tt- --t tat aag tta gtt tat agt cta aca ctt cta att ttt ata cct tga tct ttt tct cta att atg aaa aag taa atc att atg aag atg gat gaa aat gga cga gat tgt gcg cat cgt taa cga tag tat gtg gta cgt acc taa cgc att tat gga cga cgg tga taa tga agg tca cat ttc tgt caa taa tgt ctg tca tat gta tct cgc att ctt tga tgt gga tat atc atc tca tct gtt taa att agt tat taa aca ctg cga tct gaa taa acg act aaa atg tgg taa ctc tcc att aca ttg cta tac gat gaa tac acg att taa tcc atc tgt att aaa gat att gtt acg cca cgg cat gcg taa ctt tga tag caa gga taa aaa agg aca tat tcc tct aca cca cta tct gat tca ttc act atc aat cga taa caa gat ctt tga tat act aac gga ccc cat tga tga ctt tag taa atc atc cga tct att gct gtg tta tct tag ata taa att caa tgg gag ctt aaa cta tta cgt tct gta caa att att gac taa agg atc tga ccc taa ttg cgt cga tga gga tgg act cac ttc tct tca tta cta ctg taa aca cat atc cgt gtt cca cga aag caa tta tta caa gtc aaa gag tca cac taa gat gcg agc tga gaa gcg att cat cta cgc gat aat aga tca tgg agc aaa cat taa cgc ggt tac gaa aat cgg aaa tac gcc gtt aca cac tta cct tca aca gta tac caa aca tag tcc tcg tgt ggt gta tgc tct ttt atc tcg agg agc cga tac gag gat acg taa taa tct tga ttg tac acc cat cat gga ata cat aaa gaa cga ttg tgc aac agg tca tat tct cat aat gtt act caa ttg gca cga aca aaa ata cgg gaa att aca aaa gga aga agg aca aca tct act tta tct att cat aaa aca taa tca agg ata tgg aag tcg ctc tct caa tat act acg gta tct act aga tag att cga cat tca gaa aga cga ata cta taa tac aat gac tcc tct tca tac cgc ctt cca gaa ttg caa taa caa tgt tgc ctc ata cct cgt ata cat cgg ata cga cat caa cct tcc gac taa aga cga taa gac agt att cga ctt ggt gtt tga aaa cag aaa cat cat ata caa ggc gga tgt cgt taa tga cat cat cca cca cag act gaa agt atc tct acc tat gat taa atc gtt gtt cta caa gat gtc gga gtt ctc tcc cta cga cga tca cta cgt aaa gaa gat aat agc cta ctg cct att aag gga cga gtc att tgc gga act aca tac taa att ctg ttt aaa cga gga cta taa aag tgt att tat gaa aaa tat atc att cga taa gat aga ttc cat cat cga aaa atg tag tcg tga cat aag tct cct caa aga gat tcg aat ctc aga cac caa ctt gta tac ggt att gag aac aga aga cat ccg gta tca cac cta tct cga agc cat aca ttc aga caa acg cat ttc att tcc cat gta cga cga tct cat aga aca gtg tca tct atc gat gga gca taa aag taa act cgt cga caa agc act caa taa att aga gtc tac cat cga tag tca atc tag act atc gta ttt gcc tcc gga aat tat gcg caa tat cat aac caa gct aag cga cta cca tct aaa cag tat gtt gta cgg aaa gaa cca tta caa ata tta tcc atg ata gaa aga aaa tat tta aaa aat aat cta tat gat tgg aga agt agg aaa caa aca gta aca aga cga cga tta cta c-a tta tta aat cat gag gtc cgt att ata ctc gta tat att gtt tct ctc atg tat aat aat aaa cgg aag aga tat agc acc aca tgc acc atc caa tgg aaa atg taa aga caa cga ata cag aag ccg taa tct atg ttg tct atc gtg tcc tcc ggg aac tta cgc ttc cag att atg tga tag caa gac taa tac aca atg tac gcc gtg tgg ttc gga tac ctt tac atc tca caa taa tca ttt aca ggc ttg tct aag ttg taa cgg aag atg tga tag taa tca ggt aga gac gcg atc gtg taa cac gac tca caa tag aat ctg tga atg ctc tcc agg ata tta ttg tct tct caa agg atc atc agg gtg tag aac atg tat ttc taa aac aaa gtg tgg aat agg ata cgg agt atc cgg ata cac gtc tac cgg aga cgt cat ctg ttc tcc gtg tgg tcc cgg aac ata ttc tca cac cgt ctc ttc cac aga taa atg cga acc --- cgt aac cag caa tac att taa cta tat cga tgt gga aat taa cct gta tcc agt caa cga cac atc gtg tac tcg gac gac cac tac cgg tct cag cga atc cat ctc aac gtc gga act aac tat tac cat gaa tca taa aga ttg tga tcc cgt ctt tcg tgc aga ata ctt ctc tgt cct taa taa tgt agc aac ttc agg att ctt tac agg aga aaa tag ata tca gaa tac ttc aaa gat atg tac tct gaa ttt cga gat taa atg taa caa caa aga ttc atc ttc caa aca gtt aac gaa aac aaa gaa tga tac tat cat gcc gca ttc aga gac ggt aac tct agt ggg cga ctg tct atc tag cgt cga cat cta cat act ata tag taa tac caa tac tca aga cta cga aac gga tac aat ctc tta tca tat ggg taa tgt tct cga tgt caa tag cca tat gcc cgc tag ttg cga tat aca taa act gat cac taa ttc cca gaa tcc cac cca ctt ata gta ag- -tt ttt tta ccc ata aat aat aaa tac aat aat taa ttt ctc gta aaa gta gaa aat ata ttc taa ttt att ata tgg taa gaa agt aga atc atc tag aac agt aat caa tca ata gca atc atg aaa caa tat att gtc ctg gca tgc atg tgc cta gtg gca gct gct atg cct act agt ctt caa caa tcc tca tcc tcg tgt act gaa gaa gaa aac aaa cat cat atg gga atc gat gtt att atc aaa gtc aca aag caa gac caa aca ccg acc aat gat aag att tgt caa tcc gta acg gaa gtt aca gag acc gaa gat gat gag gta tcc gaa gat gat gag gta tcc gaa gaa gtt gta aaa gga gat ccc acc act tat tac aat atc gtc ggc gcg ggt ctt aac atg aac ttt gga ttc acc aaa tgc cca aag att tca tcc atc tcc gaa tcc tct gat gga aac act gtg aat act aga ttg tcc agc gtg tca cca gga caa ggt aag gac tct ccc gcg atc acg cgt gaa gaa gct ctg gct atg atc aaa gac tgt gag atg tct atc gac atc aga tgt agc gaa gaa gag aaa gac agc gac atc aag acc cat cca gta ctt ggg tct aac atc tca cat aag aaa gtg agt tac aaa gat atc atc ggt tca acg atc gtt gat aca aaa tgt gtt aag aac cta gag ttt agc gta cgt atc gga gac atg tgt gag gaa tca tct gaa ctt gaa gtc aag gat gga ttc aag tat gtc gac gga tcg gca tct gaa ggt gca acc gat gat act tca ctc atc gat tca aca aaa ctc aaa gca tgt gtc tga atc gat aac tct att cat ctg aaa atg gat gag ttg ggt taa tcg aac gat tca gac acc gca cca cga att aaa aaa gtg tac cgg gta cta tat tcc ggt ttg caa aac aaa aat gtt taa cta cat tca caa aaa gtt acc tct cgt tac ttc ttc ttt ctg ttt caa tat gtg ata cga tat gat cac tat tcg tat tct ctt tct ctc ttg tct cat aaa aaa gtt tta c-- --- --- --- aaa aaa a-t att ttt att ctc ttt ctc tct tcg at- ggt ctc aca aaa ata tta aac ctc ttt ctg atg tct caa cta ttt cgt aaa cga taa cgt cca aca ata tat tct cgt aga gct tat caa tat cct tat gcc aat --- ggt tgt cag aca att gca tca taa aat aat gtt tat aat tta cac gtt aac atc ata taa taa acg tat ata gtt aat att ttt gga ata taa atg atc tgt aaa atc cat gta ggg gac act gct cac gtt ttt tct cta gta cat aat ttc aca caa gtt ttt ata cag aca aat taa ttc tcg tcc ata ta- ttt taa aac att gac ttt tgt act aag aaa aat atc ttg act aac cat ctc ttt ctc tcg tcg atg ggt ctc aca aaa ata tta aac ctc ttt ctg atg g-- --- --- --- --- --- --- --- --- --- --- --- --- --- --- --- --- --- --- --- --- --- --- --- --- --- --- --- --- --- --- --- --- --- --- --- --- --- --- --- --- --- --- --- --- --- --- agt cgt aaa aag ttt tta tcc ttt ctc tct tcg atg g-- --- --- --- --- --- --- --- --- --- --- --- --- --- --- --- --- --- --- --- --- --- --- --- --- --- --- --- --- --- --- --- --- --- --- --- --- --- --- --- --- --- --- --- --- --- --- --- --- --- --- --- --- --- --- --- --- --- --- --- --- --- --- --- --- --- --- --- --- --- --- --- --- --- --- --- --- --- --- --- --- --- --- --- --

>KJ642613.1_Monkeypox_virus_strain_Congo_8_complete_genome

atc gag gac gac ttc aac tat gac agc tcg gtg gcg tct gcc agc gtg cac ata cga atg gca ttt cta aga aaa gtc tac ggt atc ctt gtc tac aat ttc ctt taa caa cgg caa cag ctg cag tat ttt tat act ttg aat gca ttc gga cat tta tac aag gga gtc ctg ttc taa tat tgg cat caa tgt tcg gat cta tag gct tga ttt tcg cat tga ctt tac aca gac ata aac atc ccc tga atc tgt acc tac ttt gtg gat tta cac tgt tag aat ctc taa cgc tgg cct ctg ttg tta ctt tct atg atg cgc gta tcg tta tac aag ctt tca tgt tga cta ctg cag tgt ttc ttg ctc tga cta cat gta ctc tac aat caa aga gag att tca gta aac ttg taa cag gat tgt ttg ctg ctt tct gga ttt taa ttt tgt cag gag tct tga gga taa agt tta aaa tag aat taa taa aga aca tat agg tca -tt ttt taa aca tgg ata gaa acc aag gtt gtt agt taa taa tat aca aga tat ttt ttc tca ctc tga tcc atg taa acc aag gac gag aga cac tct cat tcc tca ttc acg aca cca tta aaa atg gaa att aaa gcc ctc tat taa gca cag acg gct aca ggt cta cca tca ggt taa cct tcg tct acc ttc aca atg gcc ttt cct tgt gcc cag ttc agt ccc tgt cat tgc cac gct act aag gac tcc ctg aat acc gtg act gac gtc aga cat tgt ctg act gaa tac atc ctg tgg gtt tct cat aga tgg acc cat aga gaa agc gca ggg cct ctc tac agg ctt ctc atc tct ttc aga att gat gca atg gag cta ttt ggt agc gag ttg aag gag ttc tcg gat tca ctt ccg tgg gac aat atc gac aat tgc gtg gag atc att aaa tgt ttc atc aga aat gac tcc atg aaa acc gcc aaa gaa ctt tgt gca ata att gga ctt tgt act caa tca gct att gtc act gga aga gtc ttc aat gat aag tat atc gac ata cta ctt atg ctg cga aag att ctg aat gag aac gac tat ctc acc ctc ttg gat cat atc ctc act gct aaa tac taa atc tcc ttc atg ctc tct cac taa tac tct tac tca cta cac ttt tta tca tct tat gat gaa tga ttg cct tca tca -tt ttt cgt gga ata taa tat agg aat aat tag cac cag aat agc tat gga tat ctc gtt aag aat att ctc gat aag aga cat aat gta gac ata gtt att ata tcc ttc tta gat aag tgt tac gct act gga aag ttt cca tcg tta tta tta cgt gaa gat gat atc att aaa cca aca ttg aga ttg gct ctt atg tta gct gga ttg aat tac tgt aat aaa tgc atc gag tat aga ggg ata tag caa ttc tcg ata ata gtc atg caa tat ttg aat gag act gat aat tta ggt aat aca gta ct- -ac aca cac ata tct ttc tag att ata tat cgt taa aaa tct gta aga ggt ata ttt ctc ata agt atc cac tgt gta ata tta tta atg gat ata tag ata aca caa tag gga cta ata gta ttg taa aag ata taa tcg act att t-- gta cat atc cag ata tct ata ttc cta cta gtt tgc tgc gta gtt gca tca ttg ata tgc atg -at tta tca gga ttc aga gat gaa tta cta agt aaa cta caa tcc cac aat aag taa gaa tca aat atc aaa aac tca ctt ttg att ttt cta gtc tta agt aat aca tat att tat taa tag acc tat gaa at- aaa aaa gtt aac aat gga ttc gcg tat agc tat tta cgt att agt atc ggc atc tct ttt gta tct tgt taa ttg tca caa act agt aca tta ctt caa tct gaa aat aaa tgg aag tga tat aac taa tac agc aga tat att gct gga caa tta tcc aat tat gac ctt tga tgg aaa gga tat tta tcc atc tat cgc gtt cat ggt cgg taa taa act ttt cct aga tct tta taa aaa tat ctt tgt aga att ttt cag act att tcg agt atc tgt aag tag tca ata cga gga att aga ata tta tta ttc atg tga tta tac taa caa ccg tcc tac aat taa aca aca tta ctt tta taa cgg cga aga ata tac tga aat tga tag atc gaa aaa agc cac taa taa aaa cag ttg gtt aat tac ttc agg ctt tag act aca aaa atg gtt cga tag cga aga ttg tat aat tta tct cag atc ttt agt tag aag aat gga aga cag taa caa aaa cag taa aaa aac tta gta ctt aga tat cg- aaa aaa aat ata ttt ttg tag act ctt gag aat aga agg aaa aca tgt aca taa tta taa aaa atg aaa atc aat ggc gaa taa gac agt gcg att cgc gcc atg gag tcg gca gat ttc atg gct gtc gat gag cag ttt cac gac gac ctc gat ctt tgg tca tta tct ttg gta gat gac tat aaa aaa cat gga tta ggt gtt gac tgt tat gtt cta gaa cca gtt gtt gac agg aaa ata ttt gat aga ttt ctc ctt gaa cca att tgt gat cct gta gat gtt ctg tat gat tat ttt agg att cat aga gat aat att gat cag tat ata gta gat aga ctg ttt gca tat att aca tat aaa gat att ata tct gca tta gtg tca aag aat tat atg gaa gat att ttc tct ata att att aag aat tgt aat tct gta caa gat ctc tta ctt tac tat cta tct aat gca tat gta gaa ata gac att gtt gat ttt atg gta gat cat ggg gct gta ata tat aaa ata gaa tgc ttg aat gcc tat ttt agg gga ata tgt aaa aag gaa agt agt gtt gtt gag ttt att ttg aat tgt ggt atc cca gat gaa aat gat gtt aaa tta gat cta tat aaa ata att cag tat act agg gga ttc ctt gta gat gaa ccc aca gta tta gaa att tat aag ctt tgt atc cca tat att gaa gat atc aat caa cta gat gct ggt gga agg acc ttg ctt tat cgc gct atc tat gca ggt tat ata gat tta gta tca tgg cta tta gaa aat gga gca aat gtc aac gca gta atg agt aat gga tat aca tgt ctt gac gtg gcc gtg gat agg gga tct gtc atc gcc cgt agg gaa gca cat ctt aaa ata tta gaa ata ttg ctt aga gaa cca ttg tct att gac tgt ata aaa tta gct ata ctt aat aat aca att gaa aac cat gat gtg ata aag ctc tgt atc aag tat ttt atg atg gta gat tat tca ctt tgt aat gtg tat gca tca tca ctc ttt gat tat ata att gat tgt aaa caa gaa ttg gag tac att agg cag atg aaa att cat aat aca acc atg tat gag tta atc tat aat aga gac aaa aac aag cat gct tcc cat att cta cat agg tat tct aaa cat cca gtt ttg aca cag tgt atc act aaa gga ttc aag att tac aca gaa gta acc gag cag gtc act aaa gct cta aac aga cgt gct cta ata gat gag ata ata aac aat gta tca act gat gac aat ctc cta tca aaa ctt cca tta gaa att agg gat cta att gtt tca caa gct gtc ata tag agt tct ctc cac cca cct ttc ttg aaa tga gtt aat --- --- --- --- --- --- --- --- --- --- --- --- --- --- --- --- --- --- --- --- --- --- --- --- --- --- --- --- --- --- --- --- --- --- --- --- --- --- --- --- --- --- --- --- --- --- --- --- --- --- --- --- --- --- --- --- --- --- --- --- --- --- --- --- --- --- --- --- --- --- --- --- --- --- --- --- --- --- --- --- --- --- --- --- --- --- --- --- --- --- --- --- --- --- --- --- --- --- --- --- --- --- --- --- --- --- --- --- --- --- --- --- --- --- --- --- --- --- --- --- --- --- --- --- --- --- --- --- --- --- --- --- --- --- --- --- --- --- --- --- --- --- --- --- --- --- --- --- --- --- --- --- --- -ag tca taa gtt agt taa gtc ata agt tag tt- --t tat aag tta gtt tat agt cta aca ctt cta att ttt ata cct tga tct ttt tct cta att atg aaa aag taa atc att atg aag atg gat gaa aat gga cga gat tgt gcg cat cgt taa cga tag tat gtg gta cgt acc taa cgc att tat gga cga cgg tga taa tga agg tca cat ttc tgt caa taa tgt ctg tca tat gta tct cgc att ctt tga tgt gga tat atc atc tca tct gtt taa att agt tat taa aca ctg cga tct gaa taa acg act aaa atg tgg taa ctc tcc att aca ttg cta tac gat gaa tac acg att taa tcc atc tgt att aaa gat att gtt acg cca cgg cat gcg taa ctt tga tag caa gga taa aaa agg aca tat tcc tct aca cca cta tct gat tca ttc act atc aat cga taa caa gat ctt tga tat act aac gga ccc cat tga tga ctt tag taa atc atc cga tct att gct gtg tta tct tag ata taa att caa tgg gag ctt aaa cta tta cgt tct gta caa att att gac taa agg atc tga ccc taa ttg cgt cga tga gga tgg act cac ttc tct tca tta cta ctg taa aca cat atc cgt gtt cca cga aag caa tta tta caa gtc aaa gag tca cac taa gat gcg agc tga gaa gcg att cat cta cgc gat aat aga tca tgg agc aaa cat taa cgc ggt tac gaa aat cgg aaa tac gcc gtt aca cac tta cct tca aca gta tac caa aca tag tcc tcg tgt ggt gta tgc tct ttt atc tcg agg agc cga tac gag gat acg taa taa tct tga ttg tac acc cat cat gga ata cat aaa gaa cga ttg tgc aac agg tca tat tct cat aat gtt act caa ttg gca cga aca aaa ata cgg gaa att aca aaa gga aga agg aca aca tct act tta tct att cat aaa aca taa tca agg ata tgg aag tcg ctc tct caa tat act acg gta tct act aga tag att cga cat tca gaa aga cga ata cta taa tac aat gac tcc tct tca tac cgc ctt cca gaa ttg caa taa caa tgt tgc ctc ata cct cgt ata cat cgg ata cga cat caa cct tcc gac taa aga cga taa gac agt att cga ctt ggt gtt tga aaa cag aaa cat cat ata caa ggc gga tgt cgt taa tga cat cat cca cca cag act gaa agt atc tct acc tat gat taa atc gtt gtt cta caa gat gtc gga gtt ctc tcc cta cga cga tca cta cgt aaa gaa gat aat agc cta ctg cct att aag gga cga gtc att tgc gga act aca tac taa att ctg ttt aaa cga gga cta taa aag tgt att tat gaa aaa tat atc att cga taa gat aga ttc cat cat cga aaa atg tag tcg tga cat aag tct cct caa aga gat tcg aat ctc aga cac caa ctt gta tac ggt att gag aac aga aga cat ccg gta tca cac cta tct cga agc cat aca ttc aga caa acg cat ttc att tcc cat gta cga cga tct cat aga aca gtg tca tct atc gat gga gca taa aag taa act cgt cga caa agc act caa taa att aga gtc tac cat cga tag tca atc tag act atc gta ttt gcc tcc gga aat tat gcg caa tat cat aac caa gct aag cga cta cca tct aaa cag tat gtt gta cgg aaa gaa cca tta caa ata tta tcc atg ata gaa aga aaa tat tta aaa aat aat cta tat gat tgg aga agt agg aaa caa aca gta aca aga cga cga tta cta c-a tta tta aat cat gag gtc cgt att ata ctc gta tat att gtt tct ctc atg tat aat aat aaa cgg aag aga tat agc acc aca tgc acc atc caa tgg aaa atg taa aga caa cga ata cag aag ccg taa tct atg ttg tct atc gtg tcc tcc ggg aac tta cgc ttc cag att atg tga tag caa gac taa tac aca atg tac gcc gtg tgg ttc gga tac ctt tac atc tca caa taa tca ttt aca ggc ttg tct aag ttg taa cgg aag atg tga tag taa tca ggt aga gac gcg atc gtg taa cac gac tca caa tag aat ctg tga atg ctc tcc agg ata tta ttg tct tct caa agg atc atc agg gtg tag aac atg tat ttc taa aac aaa gtg tgg aat agg ata cgg agt atc cgg ata cac gtc tac cgg aga cgt cat ctg ttc tcc gtg tgg tcc cgg aac ata ttc tca cac cgt ctc ttc cac aga taa atg cga acc --- cgt aac cag caa tac att taa cta tat cga tgt gga aat taa cct gta tcc agt caa cga cac atc gtg tac tcg gac gac cac tac cgg tct cag cga atc cat ctc aac gtc gga act aac tat tac cat gaa tca taa aga ttg tga tcc cgt ctt tcg tgc aga ata ctt ctc tgt cct taa taa tgt agc aac ttc agg att ctt tac agg aga aaa tag ata tca gaa tac ttc aaa gat atg tac tct gaa ttt cga gat taa atg taa caa caa aga ttc atc ttc caa aca gtt aac gaa aac aaa gaa tga tac tat cat gcc gca ttc aga gac ggt aac tct agt ggg cga ctg tct atc tag cgt cga cat cta cat act ata tag taa tac caa tac tca aga cta cga aac gga tac aat ctc tta tca tat ggg taa tgt tct cga tgt caa tag cca tat gcc cgc tag ttg cga tat aca taa act gat cac taa ttc cca gaa tcc cac cca ctt ata gta ag- ttt ttt tta ccc ata aat aat aaa tac aat aat taa ttt ctc gta aaa gta gaa aat ata ttc taa ttt att ata tgg taa gaa agt aga atc atc tag aac agt aat caa tca ata gca atc atg aaa caa tat att gtc ctg gca tgc atg tgc cta gtg gca gct gct atg cct act agt ctt caa caa tcc tca tcc tcg tgt act gaa gaa gaa aac aaa cat cat atg gga atc gat gtt att atc aaa gtc aca aag caa gac caa aca ccg acc aat gat aag att tgt caa tcc gta acg gaa gtt aca gag acc gaa gat gat gag gta tcc gaa gat gat gag gta tcc gaa gaa gtt gta aaa gga gat ccc acc act tat tac aat atc gtc ggc gcg ggt ctt aac atg aac ttt gga ttc acc aaa tgc cca aag att tca tcc atc tcc gaa tcc tct gat gga aac act gtg aat act aga ttg tcc agc gtg tca cca gga caa ggt aag gac tct ccc gcg atc acg cgt gaa gaa gct ctg gct atg atc aaa gac tgt gag atg tct atc gac atc aga tgt agc gaa gaa gag aaa gac agc gac atc aag acc cat cca gta ctt ggg tct aac atc tca cat aag aaa gtg agt tac aaa gat atc atc ggt tca acg atc gtt gat aca aaa tgt gtt aag aac cta gag ttt agc gta cgt atc gga gac atg tgt gag gaa tca tct gaa ctt gaa gtc aag gat gga ttc aag tat gtc gac gga tcg gca tct gaa ggt gca acc gat gat act tca ctc atc gat tca aca aaa ctc aaa gca tgt gtc tga atc gat aac tct att cat ctg aaa atg gat gag ttg ggt taa tcg aac gat tca gac acc gca cca cga att aaa aaa gtg tac cgg gta cta tat tcc ggt ttg caa aac aaa aat gtt taa cta cat tca caa aaa gtt acc tct cgt tac ttc ttc ttt ctg ttt caa tat gtg ata cga tat gat cac tat tcg tat tct ctt tct ctc ttg tct cat aaa aaa gtt tta c-- --- --- --- aaa aaa a-t att ttt atc c-- ttt ctc tct tcg atg ggt ctc aca aaa ata tta aac ctc ttt ctg atg tct caa cta ttt cgt aaa cga taa cgt cca aca ata tat tct cgt aga gct tat caa tat cct tat gcc aat --- ggt tgt cag aca att gca tca taa aat aat gtt tat aat tta cac gtt aac atc ata taa taa acg tat ata gtt aat att ttt gga ata taa atg atc tgt aaa atc cat gta ggg gac act gct cac gtt ttt tct cta gta cat aat ttc aca caa gtt ttt ata cag aca aat taa ttc tcg tcc ata ta- ttt taa aac att gac ttt tgt act aag aaa aat atc ttg act aac cat ctc ttt ctc tcg tcg atg ggt --- --- --- --- --- --- --- --- --- --- --- --- --- --- --- --- --- --- --- --- --- --- --- --- --- --- --- --- --- --- --- --- --- --- --- --- --- --- --- --- --- --- --- --- --- --- --- --- --- --- --- --- --- --- --- --- --- --- --- --- --- --- --- --- --- --- --- --- --- --- --- --- --- --- --- --- --- --- --- --- --- --- --- --- --- --- --- --- --- --- --- --- --- --- --- --- --- --- --- --- --- --- --- --- --- --- --- --- --- --- --- --- --- --- --- --- --- --- --- --- --- --- --- --- --- --- --- --- --- --- --- --- --- --- --- --- --- --- --- --- --- --- --- --- --- --- --- --- --- --- --- --- --- --

>HM172544.1_Monkeypox_virus_strain_Zaire_1979-005_complete_genome

atc gag gac gac ttc aac tat gac agc tcg gtg gcg tct gcc agc gtg cac ata cga atg gca ttt cta aga aaa gtc tac ggt atc ctt gtc tac aat ttc ctt taa caa cgg caa cag ctg cag tat ttt tat act ttg aat gca ttc gga cat tta tac aag gga gtc ctg ttc taa tat tgg cat caa tgt tcg gat cta tag gct tga ttt tcg cat tga ctt tac aca gac ata aac atc ccc tga atc tgt acc tac ttt gtg gat tta cac tgt tag aat ctc taa cgc tgg cct ctg ttg tta ctt tct atg atg cgc gta tcg tta tac aag ctt tca tgt tga cta ctg cag tgt ttc ttg ctc tga cta cat gta ctc tac aat caa aga gag att tca gta aac ttg taa cag gat tgt ttg ctg ctt tct gga ttt taa ttt tgt cag gag tct tga gga taa agt tta aaa tag aat taa taa aga aca tat agg tca -tt ttt taa aca tgg ata gaa acc aag gtt gtt agt taa taa tat aca aga tat ttt ttc tca ctc tga tcc atg taa acc aag gac gag aga cac tct cat tcc tca ttc acg aca cca tta aaa atg gaa att aaa gcc ctc tat taa gca cag acg gct aca ggt cta cca tca ggt taa cct tcg tct acc ttc aca atg gcc ttt cct tgt gcc cag ttc agt ccc tgt cat tgc cac gct act aag gac tcc ctg aat acc gtg act gac gtc aga cat tgt ctg act gaa tac atc ctg tgg gtt tct cat aga tgg acc cat aga gaa agc gca ggg cct ctc tac agg ctt ctc atc tct ttc aga att gat gca atg gag cta ttt ggt agc gag ttg aag gag ttc tcg gat tca ctt ccg tgg gac aat atc gac aat tgc gtg gag atc att aaa tgt ttc atc aga aat gac tcc atg aaa acc gcc aaa gaa ctt tgt gca ata att gga ctt tgt act caa tca gct att gtc act gga aga gtc ttc aat gat aag tat atc gac ata cta ctt atg ctg cga aag att ctg aat gag aac gac tat ctc acc ctc ttg gat cat atc ctc act gct aaa tac taa atc tcc ttc atg ctc tct cac taa tac tct tac tca cta cac ttt tta tca tct tat gat gaa tga ttg cct tca tca -tt ttt cgt gga ata taa tat agg aat aat tag cac cag aat agc tat gga tat ctc gtt aag aat att ctc gat aag aga cat aat gta gac ata gtt atc ata tcc ttc tta gat aag tgt tac gct act gga aag ttt cca tcg tta tta tta cgt gaa gat gat atc att aaa cca aca ttg aga ttg gct ctt atg tta gct gga ttg aat tac tgt aat aaa tgc atc gag tat aga ggg ata tag caa ttc tcg ata ata gtc atg caa tat ttg aat gag act gat aat tta ggt aat aca gta ct- -ac aca cac ata tct ttc tag att ata tat cgt taa aaa tct gta aga ggt ata ttt ctc ata agt atc cac tgt gta ata tta tta atg gat ata tag ata aca caa tag gga cta ata gta ttg taa aag ata taa tcg act att t-- gta cat atc cag ata tct ata ttc cta cta gtt tgc tgc gta gtt gca tca ttg ata tgc atg -at tta tca gga ttc aga gat gaa tta cta agt aaa cta caa tcc cac aat aag taa gaa tca aat atc aaa aac tca ctt ttg att ttt cta gtc tta agt aat aca tat att tat taa tag acc tat gaa at- aaa aaa gtt aac aat gga ttc gcg tat agc tat tta cgt att agt atc ggc atc tct ttt gta tct tgt taa ttg tca caa act agt aca tta ctt caa tct gaa aat aaa tgg aag tga tat aac taa tac agc aga tat att gct gga caa tta tcc aat tat gac ctt tga tgg aaa gga tat tta tcc atc tat cgc gtt cat ggt cgg taa taa act ttt cct aga tct tta taa aaa tat ctt tgt aga att ttt cag act att tcg agt atc tgt aag tag tca ata cga gga att aga ata tta tta ttc atg tga tta tac taa caa ccg tcc tac aat taa aca aca tta ctt tta taa cgg cga aga ata tac tga aat tga tag atc gaa aaa agc cac taa taa aaa cag ttg gtt aat tac ttc agg ctt tag act aca aaa atg gtt cga tag cga aga ttg tat aat tta tct cag atc ttt agt tag aag aat gga aga cag taa caa aaa cag taa aaa aac tta gta ctt aga tat cg- aaa aaa aat ata ttt ttg tag act ctt gag aat aga agg aaa aca tgt aca taa tta taa aaa atg aaa atc aat ggc gaa taa gac agt gcg att cgc gcc atg gag tcg gta gat ttc atg gct gtc gat gag cag ttt cac gac gac ctc gat ctt tgg tca tta tct ttg gta gat gac tat aaa aaa cat gga tta ggt gtt gac tgt tat gtt cta gaa cca gtt gtt gac agg aaa ata ttt gat aga ttt ctc ctt gaa cca att tgt gat cct gta gat gtt ctg tat gat tat ttt agg att cat aga gat aat att gat cag tat ata gta gat aga ctg ttt gca tat att aca tat aaa gat att ata tct gca tta gtg tca aag aat tat atg gaa gat att ttc tct ata att att aag aat tgt aat tct gta caa gat ctc tta ctt tac tat cta tct aat gca tat gta gaa ata gac att gtt gat ttt atg gta gat cat ggg gct gta ata tat aaa ata gaa tgc ttg aat gcc tat ttt agg gga ata tgt aaa aag gaa agt agt gtt gtt gag ttt att ttg aat tgt ggt atc cca gat gaa aat gat gtt aaa tta gat cta tat aaa ata att cag tat act agg gga ttc ctt gta gat gaa ccc aca gta tta gaa att tat aag ctt tgt atc cca tat att gaa gat atc aat caa cta gat gct ggt gga agg acc ttg ctt tat cgc gct atc tat gca ggt tat ata gat tta gta tca tgg cta tta gaa aat gga gca aat gtc aac gca gta atg agt aat gga tat aca tgt ctt gac gtg gcc gtg gat agg gga tct gtc atc gcc cgt agg gaa gca cat ctt aaa ata tta gaa ata ttg ctt aga gaa cca ttg tct att gac tgt ata aaa tta gct ata ctt aat aat aca att gaa aac cat gat gtg ata aag ctc tgt atc aag tat ttt atg atg gta gat tat tca ctt tgt aat gtg tat gca tca tca ctc ttt gat tat ata att gat tgt aaa caa gaa ttg gag tac att agg cag atg aaa att cat aat aca acc atg tat gag tta atc tat aat aga gac aaa aac aag cat gct tcc cat att cta cat agg tat tct aaa cat cca gtt ttg aca cag tgt atc act aaa gga ttc aag att tac aca gaa gta acc gag cag gtc act aaa gct cta aac aga cgt gct cta ata gat gag ata ata aac aat gta tca act gat gac aat ctc cta tca aaa ctt cca tta gaa att agg gat cta att gtt tca caa gct gtc ata tag agt tct ctc cac cca cct ttc ttg aaa tga gtt aat --- --- --- --- --- --- --- --- --- --- --- --- --- --- --- --- --- --- --- --- --- --- --- --- --- --- --- --- --- --- --- --- --- --- --- --- --- --- --- --- --- --- --- --- --- --- --- --- --- --- --- --- --- --- --- --- --- --- --- --- --- --- --- --- --- --- --- --- --- --- --- --- --- --- --- --- --- --- --- --- --- --- --- --- --- --- --- --- --- --- --- --- --- --- --- --- --- --- --- --- --- --- --- --- --- --- --- --- --- --- --- --- --- --- --- --- --- --- --- --- --- --- --- --- --- --- --- --- --- --- --- --- --- --- --- --- --- --- --- --- --- --- --- --- --- --- --- --- --- --- --- --- --- -ag tca taa gtt agt taa gtc ata agt tag tt- --t tat aag tta gtt tat agt cta aca ctt cta att ttt ata cct tga tct ttt tct cta att atg aaa aag taa atc att atg aag atg gat gaa aat gga cga gat tgt gcg cat cgt taa cga tag tat gtg gta cgt acc taa cgc att tat gga cga cgg tga taa tga agg tca cat ttc tgt caa taa tgt ctg tca tat gta tct cgc att ctt tga tgt gga tat atc atc tca tct gtt taa att agt tat taa aca ctg cga tct gaa taa acg act aaa atg tgg taa ctc tcc att aca ttg cta tac gat gaa tac acg att taa tcc atc tgt att aaa gat att gtt acg cca cgg cat gcg taa ctt tga tag caa gga taa aaa agg aca tat tcc tct aca cca cta tct gat tca ttc act atc aat cga taa caa gat ctt tga tat act aac gga ccc cat tga tga ctt tag taa atc atc cga tct att gct gtg tta tct tag ata taa att caa tgg gag ctt aaa cta tta cgt tct gta caa att att gac taa agg atc tga ccc taa ttg cgt cga tga gga tgg act cac ttc tct tca tta cta ctg taa aca cat atc cgt gtt cca cga aag caa tta tta caa gtc aaa gag tca cac taa gat gcg agc tga gaa gcg att cat cta cgc gat aat aga tca tgg agc aaa cat taa cgc ggt tac gaa aat cgg aaa tac gcc gtt aca cac tta cct tca aca gta tac caa aca tag tcc tcg tgt ggt gta tgc tct ttt atc tcg agg agc cga tac gag gat acg taa taa tct tga ttg tac acc cat cat gga ata cat aaa gaa cga ttg tgc aac agg tca tat tct cat aat gtt act caa ttg gca cga aca aaa ata cgg gaa att aca aaa gga aga agg aca aca tct act tta tct att cat aaa aca taa tca agg ata tgg aag tcg ctc tct caa tat act acg gta tct act aga tag att cga cat tca gaa aga cga ata cta taa tac aat gac tcc tct tca tac cgc ctt cca gaa ttg caa taa caa tgt tgc ctc ata cct cgt ata cat cgg ata cga cat caa cct tcc gac taa aga cga taa gac agt att cga ctt ggt gtt tga aaa cag aaa cat cat ata caa ggc gga tgt cgt taa tga cat cat cca cca cag act gaa agt atc tct acc tat gat taa atc gtt gtt cta caa gat gtc gga gtt ctc tcc cta cga cga tca cta cgt aaa gaa gat aat agc cta ctg cct att aag gga cga gtc att tgc gga act aca tac taa att ctg ttt aaa cga gga cta taa aag tgt att tat gaa aaa tat atc att cga taa gat aga ttc cat cat cga aaa atg tag tcg tga cat aag tct cct caa aga gat tcg aat ctc aga cac caa ctt gta tac ggt att gag aac aga aga cat ccg gta tca cac cta tct cga agc cat aca ttc aga caa acg cat ttc att tcc cat gta cga cga tct cat aga aca gtg tca tct atc gat gga gca taa aag taa act cgt cga caa agc act caa taa att aga gtc tac cat cga tag tca atc tag act atc gta ttt gcc tcc gga aat tat gcg caa tat cat aac caa gct aag cga cta cca tct aaa cag tat gtt gta cgg aaa gaa cca tta caa ata tta tcc atg ata gaa aga aaa tat tta aaa aat aat cta tat gat tgg aga agt agg aaa caa aca gta aca aga cga cga tta cta c-a tta tta aat cat gag gtc cgt att ata ctc gta tat att gtt tct ctc atg tat aat aat aaa cgg aag aga tat agc acc aca tgc acc atc caa tgg aaa atg taa aga caa cga ata cag aag ccg taa tct atg ttg tct atc gtg tcc tcc ggg aac tta cgc ttc cag att atg tga tag caa gac taa tac aca atg tac gcc gtg tgg ttc gga tac ctt tac atc tca caa taa tca ttt aca ggc ttg tct aag ttg taa cgg aag atg tga tag taa tca ggt aga gac gcg atc gtg taa cac gac tca caa tag aat ctg tga atg ctc tcc agg ata tta ttg tct tct caa agg atc atc agg gtg tag aac atg tat ttc taa aac aaa gtg tgg aat agg ata cgg agt atc cgg ata cac gtc tac cgg aga cgt cat ctg ttc tcc gtg tgg tcc cgg aac ata ttc tca cac cgt ctc ttc cac aga taa atg cga acc --- cgt aac cag caa tac att taa cta tat cga tgt gga aat taa cct gta tcc agt caa cga cac atc gtg tac tcg gac gac cac tac cgg tct cag cga atc cat ctc aac gtc gga act aac tat tac cat gaa tca taa aga ttg tga tcc cgt ctt tcg tgc aga ata ctt ctc tgt cct taa taa tgt agc aac ttc agg att ctt tac agg aga aaa tag ata tca gaa tac ttc aaa gat atg tac tct gaa ttt cga gat taa atg taa caa caa aga ttc atc ttc caa aca gtt aac gaa aac aaa gaa tga tac tat cat gcc gca ttc aga gac ggt aac tct agt ggg cga ctg tct atc tag cgt cga cat cta cat act ata tag taa tac caa tac tca aga cta cga aac gga tac aat ctc tta tca tat ggg taa tgt tct cga tgt caa tag cca tat gcc cgc tag ttg cga tat aca taa act gat cac taa ttc cca gaa tcc cac cca ctt ata gta ag- ttt ttt tta ccc ata aat aat aaa tac aat aat taa ttt ctc gta aaa gta gaa aat ata ttc taa ttt att ata tgg taa gaa agt aga atc atc tag aac agt aat caa tca ata gca atc atg aaa caa tat att gtc ctg gca tgc atg tgc cta gtg gca gct gct atg cct act agt ctt caa caa tcc tca tcc tcg tgt act gaa gaa gaa aac aaa cat cat atg gga atc gat gtt att atc aaa gtc aca aag caa gac caa aca ccg acc aat gat aag att tgt caa tcc gta acg gaa gtt aca gag a-- --- --- --- --- --- -cc gaa gat gat gag gta tcc gaa gaa gtt gta aaa gga gat ccc acc act tat tac aat atc gtc ggc gcg ggt ctt aac atg aac ttt gga ttc acc aaa tgc cca aag att tca tcc atc tcc gaa tcc tct gat gga aac act gtg aat act aga ttg tcc agc gtg tca cca gga caa ggt aag gac tct ccc gcg atc acg cgt gaa gaa gct ctg gct atg atc aaa gac tgt gag atg tct atc gac atc aga tgt agc gaa gaa gag aaa gac agc gac atc aag acc cat cca gta ctt ggg tct aac atc tca cat aag aaa gtg agt tac aaa gat atc atc ggt tca acg atc gtt gat aca aaa tgt gtt aag aac cta gag ttt agc gta cgt atc gga gac atg tgt gag gaa tca tct gaa ctt gaa gtc aag gat gga ttc aag tat gtc gac gga tcg gca tct gaa ggt gca acc gat gat act tca ctc atc gat tca aca aaa ctc aaa gca tgt gtc tga atc gat aac tct att cat ctg aaa atg gat gag ttg ggt taa tcg aac gat tca gac acc gca cca cga att aaa aaa gtg tac cgg gta cta tat tcc ggt ttg caa aac aaa aat gtt taa cta cat tca caa aaa gtt acc tct cgt tac ttc ttc ttt ctg ttt caa tat gtg ata cga tat gat cac tat tcg tat tct ctt tct ctc ttg tct cat aaa aaa gtt tta c-- --- --- --- aaa aaa a-t att ttt att ctc ttt ctc tct tcg at- ggt ctc aca aaa ata tta aac ctc ttt ctg atg tct caa cta ttt cgt aaa cga taa cgt cca aca ata tat tct cgt aga gct tat caa tat cct tat gcc aat --- ggt tgt cag aca att gca tca taa aat aat gtt tat aat tta cac gtt aac atc ata taa taa acg tat ata gtt aat att ttt gga ata taa atg atc tgt aaa atc cat gta ggg gac act gct cac gtt ttt tct cta gta cat aat ttc aca caa gtt ttt ata cag aca aat taa ttc tcg tcc ata ta- ttt taa aac att gac ttt tgt act aag aaa aat atc ttg act aac cat ctc ttt ctc tct tcg atg ggt ctc aca aaa ata tta aac ctc ttt ctg atg g-- --- --- --- --- --- --- --- --- --- --- --- --- --- --- --- --- --- --- --- --- --- --- --- --- --- --- --- --- --- --- --- --- --- --- --- --- --- --- --- --- --- --- --- --- --- --- agt cgt aaa aag ttt tta tcc ttt ctc tct tcg atg ggt ctc --- --- --- --- --- --- --- --- --- --- --- --- --- --- --- --- --- --- --- --- --- --- --- --- --- --- --- --- --- --- --- --- --- --- --- --- --- --- --- --- --- --- --- --- --- --- --a caa aaa tat taa acc tct ttc tga tgg agt cgt aaa aag ttt tta tcc ttt ctc tct tcg atg ggt ctc aca aaa ata tta aac ctc ttt ctg atg gtc tct ata aa

>AF380138.1_Monkeypox_virus_strain_Zaire-96-I-16_complete_genome

atc gag gac gac ttc aac tat gac agc tcg gtg gcg tct gcc agc gtg cac ata cga atg gca ttt cta aga aaa gtc tac ggt atc ctt gtc tac aat ttc ctt taa caa cgg caa cag ctg cag tat ttt tat act ttg aat gca ttc gga cat tta tac aag gga gtc ctg ttc taa tat tgg cat caa tgt tcg gat cta tag gct tga ttt tcg cat tga ctt tac aca gac ata aac atc ccc tga atc tgt acc tac ttt gtg gat tta cac tgt tag aat ctc taa cgc tgg cct ctg ttg tta ctt tct atg atg cgc gta tcg tta tac aag ctt tca tgt tga cta ctg cag tgt ttc ttg ctc tga cta cat gta ctc tac aat caa aga gag att tca gta aac ttg taa cag gat tgt ttg ctg ctt tct gga ttt taa ttt tgt cag gag tct tga gga taa agt tta aaa tag aat taa taa aga aca tat agg tca -tt ttt taa aca tgg ata gaa acc aag gtt gtt agt taa taa tat aca aga tat ttt ttc tca ctc tga tcc atg taa acc aag gac gag aga cac tct cat tcc tca ttc acg aca cca tta aaa atg gaa att aaa gcc ctc tat taa gca cag acg gct aca ggt cta cca tca ggt taa cct tcg tct acc ttc aca atg gcc ttt cct tgt gcc cag ttc agt ccc tgt cat tgc cac gct act aag gac tcc ctg aat acc gtg act gac gtc aga cat tgt ctg act gaa tac atc ctg tgg gtt tct cat aga tgg acc cat aga gaa agc gca ggg cct ctc tac agg ctt ctc atc tct ttc aga att gat gca atg gag cta ttt ggt agc gag ttg aag gag ttc tcg gat tca ctt ccg tgg gac aat atc gac aat tgc gtg gag atc att aaa tgt ttc atc aga aat gac tcc atg aaa acc gcc aaa gaa ctt tgt gca ata att gga ctt tgt act caa tca gct att gtc act gga aga gtc ttc aat gat aag tat atc gac ata cta ctt atg ctg cga aag att ctg aat gag aac gac tat ctc acc ctc ttg gat cat atc ctc act gct aaa tac taa atc tcc ttc atg ctc tct cac taa tac tct tac tca cta cac ttt tta tca tct tat gat gaa tga ttg cct tca tca -tt ttt cgt gga ata taa tat agg aat aat tag cac cag aat agc tat gga tat ctc gtt aag aat att ctc gat aag aga cat aat gta gac ata gtt att ata tcc ttc tta gat aag tgt tac gct act gga aag ttt cca tcg tta tta tta cgt gaa gat gat atc att aaa cca aca ttg aga ttg gct ctt atg tta gct gga ttg aat tac tgt aat aaa tgc atc gag tat aga ggg ata tag caa ttc tcg ata ata gtc atg caa tat ttg aat gag act gat aat tta ggt aat aca gta ct- -ac aca cac ata tct ttc tag att ata tat cgt taa aaa tct gta aga ggt ata ttt ctc ata agt atc cac tgt gta ata tta tta atg gat ata tag ata aca caa tag gga cta ata gta ttg taa aag ata taa tcg act att t-- gta cat atc cag ata tct ata ttc cta cta gtt tgc tgc gta gtt gca tca ttg ata tgc atg -at tta tca gga ttc aga gat gaa tta cta agt aaa cta caa tcc cac aat aag taa gaa tca aat atc aaa aac tca ctt ttg att ttt cta gtc tta agt aat aca tat att tat taa tag acc tat gaa at- aaa aaa gtt aac aat gga ttc gcg tat agc tat tta cgt att agt atc ggc atc tct ttt gta tct tgt taa ttg tca caa act agt aca tta ctt caa tct gaa aat aaa tgg aag tga tat aac taa tac agc aga tat att gct gga caa tta tcc aat tat gac ctt tga tgg aaa gga tat tta tcc atc tat cgc gtt cat ggt cgg taa taa act ttt cct aga tct tta taa aaa tat ctt tgt aga att ttt cag act att tcg agt atc tgt aag tag tca ata cga gga att aga ata tta tta ttc atg tga tta tac taa caa ccg tcc tac aat taa aca aca tta ctt tta taa cgg cga aga ata tac tga aat tga tag atc gaa aaa agc cac taa taa aaa cag ttg gtt aat tac ttc agg ctt tag act aca aaa atg gtt cga tag cga aga ttg tat aat tta tct cag atc ttt agt tag aag aat gga aga cag taa caa aaa cag taa aaa aac tta gta ctt aga tat cg- aaa aaa aat ata ttt ttg tag act ctt gag aat aga agg aaa aca tgt aca taa tta taa aaa atg aaa atc aat ggc gaa taa gac agt gcg att cgc gcc atg gag tcg gca gat ttc atg gct gtc gat gag cag ttt cac gac gac ctc gat ctt tgg tca tta tct ttg gta gat gac tat aaa aaa cat gga tta ggt gtt gac tgt tat gtt cta gaa cca gtt gtt gac agg aaa ata ttt gat aga ttt ctc ctt gaa cca att tgt gat cct gta gat gtt ctg tat gat tat ttt agg att cat aga gat aat att gat cag tat ata gta gat aga ctg ttt gca tat att aca tat aaa gat att ata tct gca tta gtg tca aag aat tat atg gaa gat att ttc tct ata att att aag aat tgt aat tct gta caa gat ctc tta ctt tac tat cta tct aat gca tat gta gaa ata gac att gtt gat ttt atg gta gat cat ggg gct gta ata tat aaa ata gaa tgc ttg aat gcc tat ttt agg gga ata tgt aaa aag gaa agt agt gtt gtt gag ttt att ttg aat tgt ggt atc cca gat gaa aat gat gtt aaa tta gat cta tat aaa ata att cag tat act agg gga ttc ctt gta gat gaa ccc aca gta tta gaa att tat aag ctt tgt atc cca tat att gaa gat atc aat caa cta gat gct ggt gga agg acc ttg ctt tat cgc gct atc tat gca ggt tat ata gat tta gta tca tgg cta tta gaa aat gga gca aat gtc aac gca gta atg agt aat gga tat aca tgt ctt gac gtg gcc gtg gat agg gga tct gtc atc gcc cgt agg gaa aca cat ctt aaa ata tta gaa ata ttg ctt aga gaa cca ttg tct att gac tgt ata aaa tta gct ata ctt aat aat aca att gaa aac cat gat gtg ata aag ctc tgt atc aag tat ttt atg atg gta gat tat tca ctt tgt aat gtg tat gca tca tca ctc ttt gat tat ata att gat tgt aaa caa gaa ttg gag tac att agg cag atg aaa att cat aat aca acc atg tat gag tta atc tat aat aga gac aaa aac aag cat gct tcc cat att cta cat agg tat tct aaa cat cca gtt ttg aca cag tgt atc act aaa gga ttc aag att tac aca gaa gta acc gag cag gtc act aaa gct cta aac aga cgt gct cta ata gat gag ata ata aac aat gta tca act gat gac aat ctc cta tca aaa ctt cca tta gaa att agg gat cta att gtt tca caa gct gtc ata tag agt tct ctc cac cca cct ttc ttg aaa tga gtt aat --- --- --- --- --- --- --- --- --- --- --- --- --- --- --- --- --- --- --- --- --- --- --- --- --- --- --- --- --- --- --- --- --- --- --- --- --- --- --- --- --- --- --- --- --- --- --- --- --- --- --- --- --- --- --- --- --- --- --- --- --- --- --- --- --- --- --- --- --- --- --- --- --- --- --- --- --- --- --- --- --- --- --- --- --- --- --- --- --- --- --- --- --- --- --- --- --- --- --- --- --- --- --- --- --- --- --- --- --- --- --- --- --- --- --- --- --- --- --- --- --- --- --- --- --- --- --- --- --- --- --- --- --- --- --- --- --- --- --- --- --- --- --- --- --- --- --- --- --- --- --- --- --- -ag tca taa gtt agt taa gtc ata agt tag tt- --t tat aag tta gtt tat agt cta aca ctt cta att ttt ata cct tga tct ttt tct cta att atg aaa aag taa atc att atg aag atg gat gaa aat gga cga gat tgt gcg cat cgt taa cga tag tat gtg gta cgt acc taa cgc att tat gga cga cgg tga taa tga agg tca cat ttc tgt caa taa tgt ctg tca tat gta tct cgc att ctt tga tgt gga tat atc atc tca tct gtt taa att agt tat taa aca ctg cga tct gaa taa acg act aaa atg tgg taa ctc tcc att aca ttg cta tac gat gaa tac acg att taa tcc atc tgt att aaa gat att gtt acg cca cgg cat gcg taa ctt tga tag caa gga taa aaa agg aca tat tcc tct aca cca cta tct gat tca ttc act atc aat cga taa caa gat ctt tga tat act aac gga ccc cat tga tga ctt tag taa atc atc cga tct att gct gtg tta tct tag ata taa att caa tgg gag ctt aaa cta tta cgt tct gta caa att att gac taa agg atc tga ccc taa ttg cgt cga tga gga tgg act cac ttc tct tca tta cta ctg taa aca cat atc cgt gtt cca cga aag caa tta tta caa gtc aaa gag tca cac taa gat gcg agc tga gaa gcg att cat cta cgc gat aat aga tca tgg agc aaa cat taa cgc ggt tac gaa aat cgg aaa tac gcc gtt aca cac tta cct tca aca gta tac caa aca tag tcc tcg tgt ggt gta tgc tct ttt atc tcg agg agc cga tac gag gat acg taa taa tct tga ttg tac acc cat cat gga ata cat aaa gaa cga ttg tgc aac agg tca tat tct cat aat gtt act caa ttg gca cga aca aaa ata cgg gaa att aca aaa gga aga agg aca aca tct act tta tct att cat aaa aca taa tca agg ata tgg aag tcg ctc tct caa tat act acg gta tct act aga tag att cga cat tca gaa aga cga ata cta taa tac aat gac tcc tct tca tac cgc ctt cca gaa ttg caa taa caa tgt tgc ctc ata cct cgt ata cat cgg ata cga cat caa cct tcc gac taa aga cga taa gac agt att cga ctt ggt gtt tga aaa cag aaa cat cat ata caa ggc gga tgt cgt taa tga cat cat cca cca cag act gaa agt atc tct acc tat gat taa atc gtt gtt cta caa gat gtc gga gtt ctc tcc cta cga cga tca cta cgt aaa gaa gat aat agc cta ctg cct att aag gga cga gtc att tgc gga act aca tac taa att ctg ttt aaa cga gga cta taa aag tgt att tat gaa aaa tat atc att cga taa gat aga ttc cat cat cga aaa atg tag tcg tga cat aag tct cct caa aga gat tcg aat ctc aga cac caa ctt gta tac ggt att gag aac aga aga cat ccg gta tca cac cta tct cga agc cat aca ttc aga caa acg cat ttc att tcc cat gta cga cga tct cat aga aca gtg tca tct atc gat gga gca taa aag taa act cgt cga caa agc act caa taa att aga gtc tac cat cga tag tca atc tag act atc gta ttt gcc tcc gga aat tat gcg caa tat cat aac caa gct aag cga cta cca tct aaa cag tat gtt gta cgg aaa gaa cca tta caa ata tta tcc atg ata gaa aga aaa tat tta aaa aat aat cta tat gat tgg aga agt agg aaa caa aca gta aca aga cga cga tta cta cta tta tta aat cat gag gtc cgt att ata ctc gta tat att gtt tct ctc atg tat aat aat aaa cgg aag aga ttt agc acc aca tgc acc atc caa tgg aaa atg taa aga caa cga ata cag aag ccg taa tct atg ttg tct atc gtg tcc tcc ggg aac tta cgc ttc cag att atg tga tag caa gac taa tac aca atg tac gcc gtg tgg ttc gga tac ctt tac atc tca caa taa tca ttt aca ggc ttg tct aag ttg taa cgg aag atg tga tag taa tca ggt aga gac gcg atc gtg taa cac gac tca caa tag aat ctg tga atg ctc tcc agg ata tta ttg tct tct caa agg atc atc agg gtg tag aac atg tat ttc taa aac aaa gtg tgg aat agg ata cgg agt atc cgg ata cac gtc tac cgg aga cgt cat ctg ttc tcc gtg tgg tcc cgg aac ata ttc tca cac cgt ctc ttc cac aga taa atg cga acc --- cgt aac cag caa tac att taa cta tat cga tgt gga aat taa cct gta tcc agt caa cga cac atc gtg tac tcg gac gac cac tac cgg tct cag cga atc cat ctc aac gtc gga act aac tat tac cat gaa tca taa aga ttg tga tcc cgt ctt tcg tgc aga ata ctt ctc tgt cct taa taa tgt agc aac ttc agg att ctt tac agg aga aaa tag ata tca gaa tac ttc aaa gat atg tac tct gaa ttt cga gat taa atg taa caa caa aga ttc atc ttc caa aca gtt aac gaa aac aaa gaa tga tac tat cat gcc gca ttc aga gac ggt aac tct agt ggg cga ctg tct atc tag cgt cga cat cta cat act ata tag taa tac caa tac tca aga cta cga aac gga tac aat ctc tta tca tat ggg taa tgt tct cga tgt caa tag cca tat gcc cgc tag ttg cga tat aca taa act gat cac taa ttc cca gaa tcc cac cca ctt ata gta agt ttt ttt tta ccc ata aat aat aaa tac aat aat taa ttt ctc gta aaa gta gaa aat ata ttc taa ttt att ata tgg taa gaa agt aga atc atc tag aac agt aat caa tca ata gca atc atg aaa caa tat att gtc ctg gca tgc atg tgc cta gtg gca gct gct atg cct act agt ctt caa caa tcc tca tcc tcg tgt act gaa gaa gaa aac aaa cat cat atg gga atc gat gtt att atc aaa gtc aca aag caa gac caa aca ccg acc aat gat aag att tgt caa tcc gta acg gaa gtt aca gag a-- --- --- --- --- --- -cc gaa gat gat gag gta tcc gaa gaa gtt gta aaa gga gat ccc acc act tat tac act atc gtc ggc gcg ggt ctt aac atg aac ttt gga ttc acc aaa tgc cca aag att tca tcc atc tcc gaa tcc tct gat gga aac act gtg aat act aga ttg tcc agc gtg tca cca gga caa ggt aag gac tct ccc gcg atc acg cgt gaa gaa gct ctg gct atg atc aaa gac tgt gag atg tct atc gac atc aga tgt agc gaa gaa gag aaa gac agc gac atc aag acc cat cca gta ctt ggg tct aac atc tca cat aag aaa gtg agt tac aaa gat atc atc ggt tca acg atc gtt gat aca aaa tgt gtt aag aac cta gag ttt agc gta cgt atc gga gac atg tgt gag gaa tca tct gaa ctt gaa gtc aag gat gga ttc aag tat gtc gac gga tcg gca tct gaa ggt gca acc gat gat act tca ctc atc gat tca aca aaa ctc aaa gca tgt gtc tga atc gat aac tct att cat ctg aaa atg gat gag ttg ggt taa tcg aac gat tca gac acc gca cca cga att aaa aaa gtg tac cgg gta cta tat tcc ggt ttg caa aac aaa aat gtt taa cta cat tca caa aaa gtt acc tct cgt tac ttc ttc ttt ctg ttt caa tat gtg ata cga tat gat cac tat tcg tat tct ctt tct ctc ttg tct cat -aa aaa gtt tta c-- --- --- --- aaa aaa a-t att ttt att ctc ttt ctc tct tcg at- ggt ctc aca aaa ata tta aac ctc ttt ctg atg tct caa cta ttt cgt aaa cga taa cgt cca aca ata tat tct cgt aga gct tat caa tat cct tat gcc aat cta ggt tgt cag aca att gca tca taa aat aat gtt tat aat tta cac gtt aac atc ata taa taa acg tat ata gtt aat att ttt gga ata taa atg atc tgt aaa atc cat gta ggg gac act gct cac gtt ttt tct cta gta cat aat ttc aca caa gtt ttt ata cag aca aat taa ttc tcg tcc ata ta- ttt taa aac att gac ttt tgt act aag aaa aat atc ttg act aac cat ctc ttt ctc tcg tcg atg ggt ctc aca aaa ata tta aac ctc ttt ctg atg g-- --- --- --- --- --- --- --- --- --- --- --- --- --- --- --- --- --- --- --- --- --- --- --- --- --- --- --- --- --- --- --- --- --- --- --- --- --- --- --- --- --- --- --- --- --- --- agt cgt aaa aag ttt tta tcc ttt ctc tct tcg atg ggt ctc --- --- --- --- --- --- --- --- --- --- --- --- --- --- --- --- --- --- --- --- --- --- --- --- --- --- --- --- --- --- --- --- --- --- --- --- --- --- --- --- --- --- --- --- --- --- --- --- --- --- --- --- --- --- --- --- --- --- --- --- --- --- --- --- --- --- --- --- --- --- aca aaa ata tta aac ctc ttt ctg atg gtc tct ata aa

>HQ857562.1_Monkeypox_virus_strain_V79-I-005_complete_genome

atc gag gac gac ttc aac tat gac agc tcg gtg gcg tct gcc agc gtg cac ata cga atg gca ttt cta aga aaa gtc tac ggt atc ctt gtc tac aat ttc ctt taa caa cgg caa cag ctg cag tat ttt tat act ttg aat gca ttc gga cat tta tac aag gga gtc ctg ttc taa tat tgg cat caa tgt tcg gat cta tag gct tga ttt tcg cat tga ctt tac aca gac ata aac atc ccc tga atc tgt acc tac ttt gtg gat tta cac tgt tag aat ctc taa cgc tgg cct ctg ttg tta ctt tct atg atg cgc gta tcg tta tac aag ctt tca tgt tga cta ctg cag tgt ttc ttg ctc tga cta cat gta ctc tac aat caa aga gag att tca gta aac ttg taa cag gat tgt ttg ctg ctt tct gga ttt taa ttt tgt cag gag tct tga gga taa agt tta aaa tag aat taa taa aga aca tat agg tca -tt ttt taa aca tgg ata gaa acc aag gtt gtt agt taa taa tat aca aga tat ttt ttc tca ctc tga tcc atg taa acc aag gac gag aga cac tct cat tcc tca ttc acg aca cca tta aaa atg gaa att aaa gcc ctc tat taa gca cag acg gcy aca ggt cta cca tca ggt taa cct tcg tct acc ttc aca atg gcc ttt cct tgt gcc cag ttc agt ccc tgt cat tgc cac gct act aag gac tcc ctg aat acc gtg act gac gtc aga cat tkt ctg act gaa tac atc ctg tgg gtt tct cat aga tgg acc cat aga gaa agc gca ggg cct ctc tac agg ctt ctc atc tct ttc aga att gat gca atg gag cta ttt ggt agc gag ttg aag gag ttc tcg gat tca ctt ccg tgg gac aat atc gac aat tgc gtg gag atc att aaa tgt ttc atc aga aat gac tcc atg aaa acc gcc aaa gaa ctt tgt gca ata att gga ctt tgt act caa tca gct att gtc act gga aga gtc ttc aat gat aag tat atc gac ata cta ctt atg ctg cga aag att ctg aat gag aac gac tat ctc acc ctc ttg gat cat atc ctc act gct aaa tac taa atc tcc ttc atg ctc tct cac taa tac tct tac tca cta cac ttt tta tca tct tat gat gaa tga ttg cct tca tca -tt ttt cgt gga ata taa tat agg aat aat tag cac cag aat agc tat gga tat ctc gtt aag aat att ctc gat aag aga cat aat gta gac ata gtt att ata tcc ttc tta gat aag tgt tac gct act gga aag ttt cca tcg tta tta tta cgt gaa gat gat atc att aaa cca aca ttg aga ttg gct ctt atg tta gct gga ttg aat tac tgt aat aaa tgc atc gag tat aga ggg ata tag caa ttc tcg ata ata gtc atg caa tat ttg aat gag act gat aat tta ggt aat aca gta ct- -ac aca cac ata tct ttc tag att ata tat cgt taa aaa tct gta aga ggt ata ttt ctc ata agt atc cac tgt gta ata tta tta atg gat ata tag ata aca caa tag gga cta ata gta ttg taa aag ata taa tcg act att t-- gta cat atc cag ata tct ata ttc cta cta gtt tgc tgc gta gtt gca tca ttg ata tgc atg -at tta tca gga ttc aga gat gaa tta cta agt aaa cta caa tcc cac aat aag taa gaa tca aat atc aaa aac tca ctt ttg att ttt cta gtc tta agt aat aca tat att tat taa tag acc tat gaa at- aaa aaa gtt aac aat gga ttc gcg tat agc tat tta cgt att agt atc ggc atc tct ttt gta tct tgt taa ttg tca caa act agt aca tta ctt caa tct gaa aat aaa tgg aag tga tat aac taa tac agc aga tat att gct gga caa tta tcc aat tat gac ctt tga tgg aaa gga tat tta tcc atc tat cgc gtt cat ggt cgg taa taa act ttt cct aga tct tta taa aaa tat ctt tgt aga att ttt cag act att tcg agt atc tgt aag tag tca ata cga gga att aga ata tta tta ttc atg tga tta tac taa caa ccg tcc tac aat taa aca aca tta ctt tta taa cgg cga aga ata tac tga aat tga tag atc gaa aaa agc cac taa taa aaa cag ttg gtt aat tac ttc agg ctt tag act aca aaa atg gtt cga tag cga aga ttg tat aat tta tct cag atc ttt agt tag aag aat gga aga cag taa caa aaa cag taa aaa aac tta gta ctt aga tat cg- aaa aaa aat ata ttt ttg tag act ctt gag aat aga agg aaa aca tgt aca taa tta taa aaa atg aaa atc aat ggc gaa taa gac agt gcg att cgc gcc atg gag tcg gta gat ttc atg gct gtc gat gag cag ttt cac gac gac ctc gat ctt tgg tca tta tct ttg gta gat gac tat aaa aaa cat gga tta ggt gtt gac tgt tat gtt cta gaa cca gtt gtt gac agg aaa ata ttt gat aga ttt ctc ctt gaa cca att tgt gat cct gta gat gtt ctg tat gat tat ttt agg att cat aga gat aat att gat cag tat ata gta gat aga ctg ttt gca tat att aca tat aaa gat att ata tct gca tta gtg tca aag aat tat atg gaa gat att ttc tct ata att att aag aat tgt aat tct gta caa gat ctc tta ctt tac tat cta tct aat gca tat gta gaa ata gac att gtt gat ttt atg gta gat cat ggg gct gta ata tat aaa ata gaa tgc ttg aat gcc tat ttt agg gga ata tgt aaa aag gaa agt agt gtt gtt gag ttt att ttg aat tgt ggt atc cca gat gaa aat gat gtt aaa tta gat cta tat aaa ata att cag tat act agg gga ttc ctt gta gat gaa ccc aca gta tta gaa att tat aag ctt tgt atc cca tat att gaa gat atc aat caa cta gat gct ggt gga agg acc ttg ctt tat cgc gct atc tat gca ggt tat ata gat tta gta tca tgg cta tta gaa aat gga gca aat gtc aac gca gta atg agt aat gga tat aca tgt ctt gac gtg gcc gtg gat agg gga tct gtc atc gcc cgt agg gaa gca cat ctt aaa ata tta gaa ata ttg ctt aga gaa cca ttg tct att gac tgt ata aaa tta gct ata ctt aat aat aca att gaa aac cat gat gtg ata aag ctc tgt atc aag tat ttt atg atg gta gat tat tca ctt tgt aat gtg tat gca tca tca ctc ttt gat tat ata att gat tgt aaa caa gaa ttg gag tac att agg cag atg aaa att cat aat aca acc atg tat gag tta atc tat aat aga gac aaa aac aag cat gct tcc cat att cta cat agg tat tct aaa cat cca gtt ttg aca cag tgt atc act aaa gga ttc aag att tac aca gaa gta acc gag cag gtc act aaa gct cta aac aga cgt gct cta ata gat gag ata ata aac aat gta tca act gat gac aat ctc cta tca aaa ctt cca tta gaa att agg gat cta att gtt tca caa gct gtc ata tag agt tct ctc cac cca cct ttc ttg aaa tga gtt aat --- --- --- --- --- --- --- --- --- --- --- --- --- --- --- --- --- --- --- --- --- --- --- --- --- --- --- --- --- --- --- --- --- --- --- --- --- --- --- --- --- --- --- --- --- --- --- --- --- --- --- --- --- --- --- --- --- --- --- --- --- --- --- --- --- --- --- --- --- --- --- --- --- --- --- --- --- --- --- --- --- --- --- --- --- --- --- --- --- --- --- --- --- --- --- --- --- --- --- --- --- --- --- --- --- --- --- --- --- --- --- --- --- --- --- --- --- --- --- --- --- --- --- --- --- --- --- --- --- --- --- --- --- --- --- --- --- --- --- --- --- --- --- --- --- --- --- --- --- --- --- --- --- -ag tca taa gtt agt taa gtc ata agt tag tt- --t tat aag tta gtt tat agt cta aca ctt cta att ttt ata cct tga tct ttt tct cta att atg aaa aag taa atc att atg aag atg gat gaa aat gga cga gat tgt gcg cat cgt taa cga tag tat gtg gta cgt acc taa cgc att tat gga cga cgg tga taa tga agg tca cat ttc tgt caa taa tgt ctg tca tat gta tct cgc att ctt tga tgt gga tat atc atc tca tct gtt taa att agt tat taa aca ctg cga tct gaa taa acg act aaa atg tgg taa ctc tcc att aca ttg cta tac gat gaa tac acg att taa tcc atc tgt att aaa gat att gtt acg cca cgg cat gcg taa ctt tga tag caa gga taa aaa agg aca tat tcc tct aca cca cta tct gat tca ttc act atc aat cga taa caa gat ctt tga tat act aac gga ccc cat tga tga ctt tag taa atc atc cga tct att gct gtg tta tct tag ata taa att caa tgg gag ctt aaa cta tta cgt tct gta caa att att gac taa agg atc tga ccc taa ttg cgt cga tga gga tgg act cac ttc tct tca tta cta ctg taa aca cat atc cgt gtt cca cga aag caa tta tta caa gtc aaa gag tca cac taa gat gcg agc tga gaa gcg att cat cta cgc gat aat aga tca tgg agc aaa cat taa cgc ggt tac gaa aat cgg aaa tac gcc gtt aca cac tta cct tca aca gta tac caa aca tag tcc tcg tgt ggt gta tgc tct ttt atc tcg agg agc cga tac gag gat acg taa taa tct tga ttg tac acc cat cat gga ata cat aaa gaa cga ttg tgc aac agg tca tat tct cat aat gtt act caa ttg gca cga aca aaa ata cgg gaa att aca aaa gga aga agg aca aca tct act tta tct att cat aaa aca taa tca agg ata tgg aag tcg ctc tct caa tat act acg gta tct act aga tag att cga cat tca gaa aga cga ata cta taa tac aat gac tcc tct tca tac cgc ctt cca gaa ttg caa taa caa tgt tgc ctc ata cct cgt ata cat cgg ata cga cat caa cct tcc gac taa aga cga taa gac agt att cga ctt ggt gtt tga aaa cag aaa cat cat ata caa ggc gga tgt cgt taa tga cat cat cca cca cag act gaa agt atc tct acc tat gat taa atc gtt gtt cta caa gat gtc gga gtt ctc tcc cta cga cga tca cta cgt aaa gaa gat aat agc cta ctg cct att aag gga cga gtc att tgc gga act aca tac taa att ctg ttt aaa cga gga cta taa aag tgt att tat gaa aaa tat atc att cga taa gat aga ttc cat cat cga aaa atg tag tcg tga cat aag tct cct caa aga gat tcg aat ctc aga cac caa ctt gta tac ggt att gag aac aga aga cat ccg gta tca cac cta tct cga agc cat aca ttc aga caa acg cat ttc att tcc cat gta cga cga tct cat aga aca gtg tca tct atc gat gga gca taa aag taa act cgt cga caa agc act caa taa att aga gtc tac cat cga tag tca atc tag act atc gta ttt gcc tcc gga aat tat gcg caa tat cat aac caa gct aag cga cta cca tct aaa cag tat gtt gta cgg aaa gaa cca tta caa ata tta tcc atg ata gaa aga aaa tat tta aaa aat aat cta tat gat tgg aga agt agg aaa caa aca gta aca aga cga cga tta cta caa tta tta aat cat gag gtc cgt att ata ctc gta tat att gtt tct ctc atg tat aat aat aaa cgg aag aga tat agc acc aca tgc acc atc caa tgg aaa atg taa aga caa cga ata cag aag ccg taa tct atg ttg tct atc gtg tcc tcc ggg aac tta cgc ttc cag att atg tga tag caa gac taa tac aca atg tac gcc gtg tgg ttc gga tac ctt tac atc tca caa taa tca ttt aca ggc ttg tct aag ttg taa cgg aag atg tga tag taa tca ggt aga gac gcg atc gtg taa cac gac tca caa tag aat ctg tga atg ctc tcc agg ata tta ttg tct tct caa agg atc atc agg gtg tag aac atg tat ttc taa aac aaa gtg tgg aat agg ata cgg agt atc cgg ata cac gtc tac cgg aga cgt cat ctg ttc tcc gtg tgg tcc cgg aac ata ttc tca cac cgt ctc ttc cac aga taa atg cga acc --- cgt aac cag caa tac att taa cta tat cga tgt gga aat taa cct gta tcc agt caa cga cac atc gtg tac tcg gac gac cac tac cgg tct cag cga atc cat ctc aac gtc gga act aac tat tac cat gaa tca taa aga ttg tga tcc cgt ctt tcg tgc aga ata ctt ctc tgt cct taa taa tgt agc aac ttc agg att ctt tac agg aga aaa tag ata tca gaa tac ttc aaa gat atg tac tct gaa ttt cga gat taa atg taa caa caa aga ttc atc ttc caa aca gtt aac gaa aac aaa gaa tga tac tat cat gcc gca ttc aga gac ggt aac tct agt ggg cga ctg tct atc tag cgt cga cat cta cat act ata tag taa tac caa tac tca aga cta cga aac gga tac aat ctc tta tca tat ggg taa tgt tct cga tgt caa tag cca tat gcc cgc tag ttg cga tat aca taa act gat cac taa ttc cca gaa tcc cac cca ctt ata gta agt ttt ttt tta ccc ata aat aat aaa tac aat aat taa ttt ctc gta aaa gta gaa aat ata ttc taa ttt att ata tgg taa gaa agt aga atc atc tag aac agt aat caa tca ata gca atc atg aaa caa tat att gtc ctg gca tgc atg tgc cta gtg gca gct gct atg cct act agt ctt caa caa tcc tca tcc tcg tgt act gaa gaa gaa aac aaa cat cat atg gga atc gat gtt att atc aaa gtc aca aag caa gac caa aca ccg acc aat gat aag att tgt caa tcc gta acg gaa gtt aca gag a-- --- --- --- --- --- -cc gaa gat gat gag gta tcc gaa gaa gtt gta aaa gga gat ccc acc act tat tac aat atc gtc ggc gcg ggt ctt aac atg aac ttt gga ttc acc aaa tgc cca aag att tca tcc atc tcc gaa tcc tct gat gga aac act gtg aat act aga ttg tcc agc gtg tca cca gga caa ggt aag gac tct ccc gcg atc acg cgt gaa gaa gct ctg gct atg atc aaa gac tgt gag atg tct atc gac atc aga tgt agc gaa gaa gag aaa gac agc gac atc aag acc cat cca gta ctt ggg tct aac atc tca cat aag aaa gtg agt tac aaa gat atc atc ggt tca acg atc gtt gat aca aaa tgt gtt aag aac cta gag ttt agc gta cgt atc gga gac atg tgt gag gaa tca tct gaa ctt gaa gtc aag gat gga ttc aag tat gtc gac gga tcg gca tct gaa ggt gca acc gat gat act tca ctc atc gat tca aca aaa ctc aaa gca tgt gtc tga atc gat aac tct att cat ctg aaa atg gat gag ttg ggt taa tcg aac gat tca gac acc gca cca cga att aaa aaa gtg tac cgg gta cta tat tcc ggt ttg caa aac aaa aat gtt taa cta cat tca caa aaa gtt acc tct cgt tac ttc ttc ttt ctg ttt caa tat gtg ata cga tat gat cac tat tcg tat tct ctt tct ctc ttg tct cat -aa aaa gtt tta c-- --- --- --- aaa aaa a-t att ttt att ctc ttt ctc tct tcg at- ggt ctc aca aaa ata tta aac ctc ttt ctg atg tct caa cta ttt cgt aaa cga taa cgt cca aca ata tat tct cgt aga gct tat caa tat cct tat gcc aat ggt ggt tgt cag aca att gca tca taa aat aat gtt tat aat tta cac gtt aac atc ata taa taa acg tat ata gtt aat att ttt gga ata taa atg atc tgt aaa atc cat gta ggg gac act gct cac gtt ttt tct cta gta cat aat ttc aca caa gtt ttt ata cag aca aat taa ttc tcg tcc ata ta- ttt taa aac att gac ttt tgt act aag aaa aat atc ttg act aac cat ctc ttt ctc tct tcg atg ggt ctc aca aaa ata tta aac ctc ttt ctg atg g-- --- --- --- --- --- --- --- --- --- --- --- --- --- --- --- --- --- --- --- --- --- --- --- --- --- --- --- --- --- --- --- --- --- --- --- --- --- --- --- --- --- --- --- --- --- --- agt cgt aaa aag ttt tta tcc ttt ctc tct tcg atg ggt ctc --- --- --- --- --- --- --- --- --- --- --- --- --- --- --- --- --- --- --- --- --- --- --- --- --- --- --- --- --- --- --- --- --- --- --- --- --- --- --- --- --- --- --- --- --- --- --- --- --- --- --- --- --- --- --- --- --- --- --- --- --- --- --- --- --- --- --- --- --- --- aca aaa ata tta aac ctc ttt ctg atg gtc tct ata aa

>HQ857563.1_Monkeypox_virus_strain_D14L_knockout_complete_genome

atc gag gac gac ttc aac tat gac agc tcg gkg gcg tct gcc agc gtg cac ata cga atg gca ttt cta aga aaa gtc tac ggt atc ctt gtc tac aat ttc ctt taa caa cgg caa cag ctg cag tat ttt tat act ttg aat gca ttc gga cat tta tac aag gga gtc ctg ttc taa tat tgg cat caa tgt tcg gat cta tag gct tga ttt tcg cat tga ctt tac aca gac ata aac atc ccc tga atc tgt acc tac ttt gtg gat tta cac tgt tag aat ctc taa cgc tgg cct ctg ttg tta ctt tct atg atg cgc gta tcg tta tac aag ctt tca tgt tga cta ctg cag tgt ttc ttg ctc tga cta cat gta ctc tac aat caa aga gag att tca gta aac ttg taa cag gat tgt ttg ctg ctt tct gga ttt taa ttt tgt cag gag tct tga gga taa agt tta aaa tag aat taa taa aga aca tat agg tca -tt ttt taa aca tgg ata gaa acc aag gtt gtt agt taa taa tat aca aga tat ttt ttc tca ctc tga tcc atg taa acc aag gac gag aga cac tct cat tcc tca ttc acg aca cca tta aaa atg gaa att aaa gcc ctc tat taa gca cag acg gct aca ggt cta cca tca ggt taa cct tcg tct acc ttc aca atg gcc ttt cct tgt gcc cag ttc agt ccc tgt cat tgc cac gct act aag gac tcc ctg aat acc gtg act gac gtc aga cat tgt ctg act gaa tac atc ctg tgg gtt tct cat aga tgg acc cat aga gaa agc gca ggg cct ctc tac agg ctt ctc atc tct ttc aga att gat gca atg gag cta ttt ggt agc gag ttg aag gag ttc tcg gat tca ctt ccg tgg gac aat atc gac aat tgc gtg gag atc att aaa tgt ttc atc aga aat gac tcc atg aaa mcc gcc aaa gaa ctt tgt gca ata att gga ctt tgt act caa tca gct att gtc act gga aga gtc ttc aat gat aag tat atc gac ata cta ctt atg ctg cga aag att ctg aat gag aac gac tat ctc acc ctc ttg gat cat atc ctc act gct aaa tac taa atc tcc ttc atg ctc tct cac taa tac tct tac tca cta cac ttt tta tca tct tat gat gaa tga ttg cct tca tca -tt ttt cgt gga ata taa tat agg aat aat tag cac cag aat agc tat gga tat ctc gtt aag aat att ctc gat aag aga cat aat gta gac ata gtt att ata tcc ttc tta gat aag tgt tac gct act gga aag ttt cca tcg tta tta tta cgt gaa gat gat atc att aaa cca aca ttg aga ttg gct ctt atg tta gct gga ttg aat tac tgt aat aaa tgc atc gag tat aga ggg ata tag caa ttc tcg ata ata gtc atg caa tat ttg aat gag act gat aat tta ggt aat aca gta ct- -ac aca cac ata tct ttc tag att ata tat cgt taa aaa tct gta aga ggt ata ttt ctc ata agt atc cac tgt gta ata tta tta atg gat ata tag ata aca caa tag gga cta ata gta ttg taa aag ata taa tcg act att t-- gta cat atc cag ata tct ata ttc cta cta gtt tgc tgc gta gtt gca tca ttg ata tgc atg -at tta tca gga ttc aga gat gaa tta cta agt aaa cta caa tcc cac aat aag taa gaa tca aat atc aaa aac tca ctt ttg att ttt cta gtc tta agt aat aca tat att tat taa tag acc tat gaa at- aaa aaa gtt aac aat gga ttc gcg tat agc tat tta cgt att agt atc ggc atc tct ttt kta tct tgt taa ttg tca caa act agt aca tta ctt caa tct gaa aat aaa tgg aag tga tat aac taa tac agc aga tat att gct gga caa tta tcc aat tat gac ctt tga tgg aaa gga tat tta tcc atc tat cgc gtt cat ggt cgg taa taa act ttt cct aga tct tta taa aaa tat ctt tgt aga att ttt cag act att tcg agt atc tgt aag tag tca ata cga gga att aga ata tta tta ttc atg tga tta tac taa caa ccg tcc tac aat taa aca aca tta ctt tta taa cgg cga aga ata tac tga aat tga tag atc gaa aaa agc cac taa taa aaa cag ttg gtt aat tac ttc agg ctt tag act aca aaa atg gtt cga tag cga aga ttg tat aat tta tct cag atc ttt agt tag aag aat gga aga cag taa caa aaa cag taa aaa aac tta gta ctt aga tat cg- aaa aaa aat ata ttt ttg tag act ctt gag aat aga agg aaa aca tgt aca taa tta taa aaa atg aaa atc aat ggc gaa taa gac agt gcg att cgc gcc atg gag tcg gga gat ttc atg gct gtc gat gag cag ttt cac gac gac ctc gat ctt tgg tca tta tct ttg gta gat gac tat aaa aaa cat gga tta ggt gtt gac tgt tat gtt cta gaa cca gtt gtt gac agg aaa ata ttt gat aga ttt ctc ctt gaa cca att tgt gat cct gta gat gtt ctg tat gat tat ttt agg att cat aga gat aat att gat cag tat ata gta gat aga ctg ttt gca tat att aca tat aaa gat att ata tct gca tta gtg tca aag aat tat atg gaa gat att ttc tct ata att att aag aat tgt aat tct gta caa gat ctc tta ctt tac tat cta tct aat gca tat gta gaa ata gac att gtt gat ttt atg gta gat cat ggg gct gta ata tat aaa ata gaa tgc ttg aat gcc tat ttt agg gga ata tgt aaa aag gaa agt agt gtt gtt gag ttt att ttg aat tgt ggt atc cca gat gaa aat gat gtt aaa tta gat cta tat aaa ata att cag tat act agg gga ttc ctt gta gat gaa ccc aca gta tta gaa att tat aag ctt tgt atc cca tat att gaa gat atc aat caa cta gat gct ggt gga agg acc ttg ctt tat cgc gct atc tat gca ggt tat ata gat tta gta tca tgg cta tta gaa aat gga gca aat gtc aac gca gta atg agt aat gga tat aca tgt ctt gac gtg gcc gtg gat agg gga tct gtc atc gcc cgt agg gaa gca cat ctt aaa ata tta gaa ata ttg ctt aga gaa cca ttg tct att gac tgt ata aaa tta gct ata ctt aat aat aca att gaa aac cat gat gtg ata aag ctc tgt atc aag tat ttt atg atg gta gat tat tca ctt tgt aat gtg tat gca tca tca ctc ttt gat tat ata att gat tgt aaa caa gaa ttg gag tac att agg cag atg aaa att cat aat aca acc atg tat gag tta atc tat aat aga gac aaa aac aag cat gct tcc cat att cta cat agg tat tct aaa cat cca gtt ttg aca cag tgt atc act aaa gga ttc aag att tac aca gaa gta acc gag cag gtc act aaa gct cta aac aga cgt gct cta ata gat gag ata ata aac aat gta tca act gat gac aat ctc cta tca aaa ctt cca tta gaa att agg gat cta att gtt tca caa gct gtc ata tag agt tct ctc cac cca cct ttc ttg aaa tga gtt aat --- --- --- --- --- --- --- --- --- --- --- --- --- --- --- --- --- --- --- --- --- --- --- --- --- --- --- --- --- --- --- --- --- --- --- --- --- --- --- --- --- --- --- --- --- --- --- --- --- --- --- --- --- --- --- --- --- --- --- --- --- --- --- --- --- --- --- --- --- --- --- --- --- --- --- --- --- --- --- --- --- --- --- --- --- --- --- --- --- --- --- --- --- --- --- --- --- --- --- --- --- --- --- --- --- --- --- --- --- --- --- --- --- --- --- --- --- --- --- --- --- --- --- --- --- --- --- --- --- --- --- --- --- --- --- --- --- --- --- --- --- --- --- --- --- --- --- --- --- --- --- --- --- -ag tca taa gtt agt taa gtc ata agt tag tt- --t tat aag tta gtt tat agt cta aca ctt cta att ttt ata cct tga tct ttt tct cta att atg aaa aag taa atc att atg aag atg gat gaa aat gga cga gat tgt gcg cat cgt taa cga tag tat gtg gta cgt acc taa cgc att tat gga cga cgg tga taa tga agg tca cat ttc tgt caa taa tgt ctg tca tat gta tct cgc att ctt tga tgt gga tat atc atc tca tct gtt taa att agt tat taa aca ctg cga tct gaa taa acg act aaa atg tgg taa ctc tcc att aca ttg cta tac gat gaa tac acg att taa tcc atc tgt att aaa gat att gtt acg cca cgg cat gcg taa ctt tga tag caa gga taa aaa agg aca tat tcc tct aca cca cta tct gat tca ttc act atc aat cga taa caa gat ctt tga tat act aac gga ccc cat tga tga ctt tag taa atc atc cga tct att gct gtg tta tct tag ata taa att caa tgg gag ctt aaa cta tta cgt tct gta caa att att gac taa agg atc tga ccc taa ttg cgt cga tga gga tgg act cac ttc tct tca tta cta ctg taa aca cat atc cgt gtt cca cga aag caa tta tta caa gtc aaa gag tca cac taa gat gcg agc tga gaa gcg att cat cta cgc gat aat aga tca tgg agc aaa cat taa cgc ggt tac gaa aat cgg aaa tac gcc gtt aca cac tta cct tca aca gta tac caa aca tag tcc tcg tgt ggt gta tgc tct ttt atc tcg agg agc cga tac gag gat acg taa taa tct tga ttg tac acc cat cat gga ata cat aaa gaa cga ttg tgc aac agg tca tat tct cat aat gtt act caa ttg gca cga aca aaa ata cgg gaa att aca aaa gga aga agg aca aca tct act tta tct att cat aaa aca taa tca agg ata tgg aag tcg ctc tct caa tat act acg gta tct act aga tag att cga cat tca gaa aga cga ata cta taa tac aat gac tcc tct tca tac cgc ctt cca gaa ttg caa taa caa tgt tgc ctc ata cct cgt ata cat cgg ata cga cat caa cct tcc gac taa aga cga taa gac agt att cga ctt ggt gtt tga aaa cag aaa cat cat ata caa ggc gga tgt cgt taa tga cat cat cca cca cag act gaa agt atc tct acc tat gat taa atc gtt gtt cta caa gat gtc gga gtt ctc tcc cta cga cga tca cta cgt aaa gaa gat aat agc cta ctg cct att aag gga cga gtc att tgc gga act aca tac taa att ctg ttt aaa cga gga cta taa aag tgt att tat gaa aaa tat atc att cga taa gat aga ttc cat cat cga aaa atg tag tcg tga cat aag tct cct caa aga gat tcg aat ctc aga cac caa ctt gta tac ggt att gag aac aga aga cat ccg gta tca cac cta tct cga agc cat aca ttc aga caa acg cat ttc att tcc cat gta cga cga tct cat aga aca gtg tca tct atc gat gga gca taa aag taa act cgt cga caa agc act caa taa att aga gtc tac cat cga tag tca atc tag act atc gta ttt gcc tcc gga aat tat gcg caa tat cat aac caa gct aag cga cta cca tct aaa cag tat gtt gta cgg aaa gaa cca tta caa ata tta tcc atg ata gaa aga aaa tat tta aaa aat aat cta tat gat tgg aga agt agg aaa caa aca gta aca aga cga cga tta cta caa tta tta aat cat gag gtc cgt att ata ctc gta tat att gtt tct ctc atg tat aat aat aaa cgg aag aga tat agc acc aca tgc acc atc caa tgg aaa atg taa aga caa cga ata cag aag ccg taa tct atg ttg tct atc gtg tcc tcc ggg aac tta cgc ttc cag att atg tga tag caa gac taa tac aca atg tac gcc gtg tgg ttc gga tac ctt tac atc tca caa taa tca ttt aca ggc ttg tct aag ttg taa cgg aag atg tga tag taa tca ggt aga gac gcg atc gtg taa cac gac tca caa tag aat ctg tga atg ctc tcc agg ata tta ttg tct tct caa agg atc atc agg gtg tag aac atg tat ttc taa aac aaa gtg tgg aat agg ata cgg agt atc cgg ata cac gtc tac cgg aga cgt cat ctg ttc tcc gtg tgg tcc cgg aac ata ttc tca cac cgt ctc ttc cac aga taa atg cga acc --- cgt aac cag caa tac att taa cta tat cga tgt gga aat taa cct gta tcc agt caa cga cac atc gtg tac tcg gac gac cac tac cgg tct cag cga atc cat ctc aac gtc gga act aac tat tac cat gaa tca taa aga ttg tga tcc cgt ctt tcg tgc aga ata ctt ctc tgt cct taa taa tgt agc aac ttc agg att ctt tac agg aga aaa tag ata tca gaa tac ttc aaa gat atg tac tct gaa ttt cga gat taa atg taa caa caa aga ttc atc ttc caa aca gtt aac gaa aac aaa gaa tga tac tat cat gcc gca ttc aga gac ggt aac tct agt ggg cga ctg tct atc tag cgt cga cat cta cat act ata tag taa tac caa tac tca aga cta cga aac gga tac aat ctc tta tca trt ggg taa tgt tct cga tgt cra tag cca tat gcc cgc tag ttg cga tat aca taa act gat cac taa ttc cca gaa tcc cac cca ctt ata gta agt ttt ttt tta ccc ata aat aat aaa tac aat aat taa ttt ctc gta aaa gta gaa aat ata ttc taa ttt att ata tgg taa gaa agt aga atc atc tag aac agt aat caa tca ata gca atc atg aaa caa tat att gtc ctg gca tgc atg tgc cta gtg gca gct gct atg cct act agt ctt caa caa tcc tca tcc tcg tgt act gaa gaa gaa aac aaa cat cat atg gga atc gat gtt att atc aaa gtc aca aag caa gac caa aca ccg acc aat gat aag att tgt caa tcc gta acg gaa gtt aca gag a-- --- --- --- --- --- -cc gaa gat gat gag gta tcc gaa gaa gtt gta aaa gga gat ccc acc act tat tac aat atc gtc ggc gcg ggt ctt aac atg aac ttt gga ttc acc aaa tgc cca aag att tca tcc atc tcc gaa tcc tct gat gga aac act gtg aat act aga ttg tcc agc gtg tca cca gga caa ggt aag gac tct ccc gcg atc acg cgt gaa gaa gct ctg gct atg atc aaa gac tgt gag atg tct atc gac atc aga tgt agc gaa gaa gag aaa gac agc gac atc aag acc cat cca gta ctt ggg tct aac atc tca cat aag aaa gtg agt tac aaa gat atc atc ggt tca acg atc gtt gat aca aaa tgt gtt aag aac cta gag ttt agc gta cgt atc gga gac atg tgt gag gaa tca tct gaa ctt gaa gtc aag gat gga ttc aag tat gtc gac gga tcg gca tct gaa ggt gca acc gat gat act tca ctc atc gat tca aca aaa ctc aaa gca tgt gtc tga atc gat aac tct att cat ctg aaa atg gat gag ttg ggt taa tcg aac gat tca gac acc gca cca cga att aaa aaa gtg tac cgg gta cta tat tcc ggt ttg caa aac aaa aat gtt taa cta cat tca caa aaa gtt acc tct cgt tac ttc ttc ttt ctg ttt caa tat gtg ata cga tat gat cac tat tcg tat tct ctt tct ctc ttg tct cat -aa aaa gtt tta c-- --- --- --- aaa aaa a-t att ttt att ctc ttt ctc tct tcg at- ggt ctc aca aaa ata tta aac ctc ttt ctg atg tct caa cta ttt cgt aaa cga taa cgt cca aca ata tat tct cgt aga gct tat caa tat cct tat gcc aat ggt ggt tgt cag aca att gca tca taa aat aat gtt tat aat tta cac gtt aac atc ata taa taa acg tat ata gtt aat att ttt gga ata taa atg atc tgt aaa atc cat gta ggg gac act gct cac gtt ttt tct cta gta cat aat ttc aca caa gtt ttt ata cag aca aat taa ttc tcg tcc ata ta- ttt taa aac att gac ttt tgt act aag aaa aat atc ttg act aac cat ctc ttt ctc tct tcg atg ggt ctc aca aaa ata tta aac ctc ttt ctg atg g-- --- --- --- --- --- --- --- --- --- --- --- --- --- --- --- --- --- --- --- --- --- --- --- --- --- --- --- --- --- --- --- --- --- --- --- --- --- --- --- --- --- --- --- --- --- --- agt cgt aaa aag ttt tta tcc ttt ctc tct tcg atg ggt ctc --- --- --- --- --- --- --- --- --- --- --- --- --- --- --- --- --- --- --- --- --- --- --- --- --- --- --- --- --- --- --- --- --- --- --- --- --- --- --- --- --- --- --- --- --- --- --- --- --- --- --- --- --- --- --- --- --- --- --- --- --- --- --- --- --- --- --- --- --- --- aca aaa ata tta aac ctc ttt ctg atg gtc tct ata aa
