## Supplementary Data 4 for "Genomic regions insertion and deletion in Monkeypox virus causing multi-country outbreak-2022"

>ON674051.1_Monkeypox_virus_isolate_MPXV_USA_2022_FL001_complete_genome

ttt tat atc act acg gac ata aac cat tgt ata ttt ttt atg ttt att agt gta cac att ttg gaa gta agt tc- --- --- --- --- --- --- --- --- --- --- --- --- --- --- --- --- --- --- --- --- --- --- --- --- --- --- --- --- --- --- --- --- --- --- --- --- --- --- --- --- --- --- --- --- --- --- --- --- --- --- --- --- --- --- --- --- --- --- --- --- --- --- --- --- --- --- --- --- --- --- --- --- --- --- --- --- --- --- --- --- --- --- --- --- --- --- --- --- --- --- --- --- --- --- --- --- --- --- --- --- --- --- --- --- --- --- --- --- --- --- --- --- --- --- --- --- --- --- --- --- --- --- --- --- --- --- --- --- --- --- --- --- --- --- --- --- --- --- --- --- --- --- --- --- --- --- --- --- --- --- --- --- --- --- --- --- --- --- --- --- --- --- --- --- --- --- --- --- --- --- --- --- --- --- --- --- --- --- --- --- --- --- --- --- --- --- --- --- --- --- --- --- --- --- --- --- --- --- --- --- --- --- --- --- --- --- --- --- --- --- --- --- --- --- --- --- --- --- --- --- --- --- --- --- --- --- --- --- --- --- --- --- --- --- --- --- --- --- --- --- --- --- --- --- --- --- --- --- --- --- --- --- --- --- --- --- --- --- --- --- --- --- --- --- --- --- --- --- --- --- --- --- --- --- --- --- --- --- --- --- --- --- --- --- --- --- --- --- --- --- --- --- --- --- --- --- --- --- --- --- --- --- --- --- --- --- --- --- --- --- --- --- --- --- --- --- --- --- --- --- --- --- --- --- --- --- --- --- --- --- --- --- --- --- --- --- --- --- --- --- --- --- --- --- --- --- --- --- --- --- --- --- --- --- --- --- --- --- --- --- --- --- --- --- --- --- --- --- --- --- --- --- --- --- --- --- --- --- --- --- --- --- --- --- --- --- --- --- --- --- --- --- --- --- --- --- --- --- --- --- --- --- --- --- --- --- --- --- --- --- --- --- --- --- --- --- --- --- --- --- --- --- --- --- --- --- --- --- --- --- --- --- --- --- --- --- --- --- --- --- --- --- --- --- --- --- --- --- --- --- --- --- --- --- --- --- --- --- --- --- --- --- --- --- --- --- --- --- --- --- --- --- --- --- --- --- --- --- --- --- --- --- --- --- --- --- --- --- --- --- --- --- --- --- --- --- --- --- --- --- --- --- --- --- --- --- --- --- --- --- --- --- --- --- --- --- --- --- --- --- --- --- --- --- --- --- --- --- --- --- --- --- --- --- --- --- --- --- --- --- --- --- --- --- --- --- --- --- --- --- --- --- --- --- --- --- --- --- --- --- --- --- --- --- --- --- --- --- --- --- --- --- --- --- --- --- --- --- --- --- --- --- --- --- --- --- --- --- --- --- --- --- --- --- --- --- --- --- --- --- --- --- --- --- --- --- --- --- --- --- --- --- --- --- --- --- --- --- --- --- --- --- --- --- --- --- --- --- --- --- --- --- --- --- --- --- --- --- --- --- --- --- --- --- --- --- --- --- --- --- --- --- -ct gga tcg gat gtc acc gca gta ata ttg ttg att att tct gac atc gac gta tta tat agt ttt tta att cca tat ctt ttt t

>NC_063383.1_Monkeypox_virus_complete_genome

ttt tat atc act acg gac ata aac cat tgt ata ttt ttt atg ttt att agt gta cac att ttg gaa gta agt tc- --- --- --- --- --- --- --- --- --- --- --- --- --- --- --- --- --- --- --- --- --- --- --- --- --- --- --- --- --- --- --- --- --- --- --- --- --- --- --- --- --- --- --- --- --- --- --- --- --- --- --- --- --- --- --- --- --- --- --- --- --- --- --- --- --- --- --- --- --- --- --- --- --- --- --- --- --- --- --- --- --- --- --- --- --- --- --- --- --- --- --- --- --- --- --- --- --- --- --- --- --- --- --- --- --- --- --- --- --- --- --- --- --- --- --- --- --- --- --- --- --- --- --- --- --- --- --- --- --- --- --- --- --- --- --- --- --- --- --- --- --- --- --- --- --- --- --- --- --- --- --- --- --- --- --- --- --- --- --- --- --- --- --- --- --- --- --- --- --- --- --- --- --- --- --- --- --- --- --- --- --- --- --- --- --- --- --- --- --- --- --- --- --- --- --- --- --- --- --- --- --- --- --- --- --- --- --- --- --- --- --- --- --- --- --- --- --- --- --- --- --- --- --- --- --- --- --- --- --- --- --- --- --- --- --- --- --- --- --- --- --- --- --- --- --- --- --- --- --- --- --- --- --- --- --- --- --- --- --- --- --- --- --- --- --- --- --- --- --- --- --- --- --- --- --- --- --- --- --- --- --- --- --- --- --- --- --- --- --- --- --- --- --- --- --- --- --- --- --- --- --- --- --- --- --- --- --- --- --- --- --- --- --- --- --- --- --- --- --- --- --- --- --- --- --- --- --- --- --- --- --- --- --- --- --- --- --- --- --- --- --- --- --- --- --- --- --- --- --- --- --- --- --- --- --- --- --- --- --- --- --- --- --- --- --- --- --- --- --- --- --- --- --- --- --- --- --- --- --- --- --- --- --- --- --- --- --- --- --- --- --- --- --- --- --- --- --- --- --- --- --- --- --- --- --- --- --- --- --- --- --- --- --- --- --- --- --- --- --- --- --- --- --- --- --- --- --- --- --- --- --- --- --- --- --- --- --- --- --- --- --- --- --- --- --- --- --- --- --- --- --- --- --- --- --- --- --- --- --- --- --- --- --- --- --- --- --- --- --- --- --- --- --- --- --- --- --- --- --- --- --- --- --- --- --- --- --- --- --- --- --- --- --- --- --- --- --- --- --- --- --- --- --- --- --- --- --- --- --- --- --- --- --- --- --- --- --- --- --- --- --- --- --- --- --- --- --- --- --- --- --- --- --- --- --- --- --- --- --- --- --- --- --- --- --- --- --- --- --- --- --- --- --- --- --- --- --- --- --- --- --- --- --- --- --- --- --- --- --- --- --- --- --- --- --- --- --- --- --- --- --- --- --- --- --- --- --- --- --- --- --- --- --- --- --- --- --- --- --- --- --- --- --- --- --- --- --- --- --- --- --- --- --- --- --- --- --- --- --- --- --- --- --- --- --- --- --- --- --- --- --- --- --- --- --- --- --- --- --- --- --- --- --- --- --- --- --- --- --- --- --- --- -ct gga tcg gat gtc acc gca gta ata ttg ttg att att tct gac atc gac gta tta tat agt ttt tta att cca tat ctt ttt t

>ON676704.1_Monkeypox_virus_isolate_MPXV_USA_2022_FL002_complete_genome

ttt tat atc act acg gac ata aac cat tgt ata ttt ttt atg ttt att agt gta cac att ttg gaa gta agt tc- --- --- --- --- --- --- --- --- --- --- --- --- --- --- --- --- --- --- --- --- --- --- --- --- --- --- --- --- --- --- --- --- --- --- --- --- --- --- --- --- --- --- --- --- --- --- --- --- --- --- --- --- --- --- --- --- --- --- --- --- --- --- --- --- --- --- --- --- --- --- --- --- --- --- --- --- --- --- --- --- --- --- --- --- --- --- --- --- --- --- --- --- --- --- --- --- --- --- --- --- --- --- --- --- --- --- --- --- --- --- --- --- --- --- --- --- --- --- --- --- --- --- --- --- --- --- --- --- --- --- --- --- --- --- --- --- --- --- --- --- --- --- --- --- --- --- --- --- --- --- --- --- --- --- --- --- --- --- --- --- --- --- --- --- --- --- --- --- --- --- --- --- --- --- --- --- --- --- --- --- --- --- --- --- --- --- --- --- --- --- --- --- --- --- --- --- --- --- --- --- --- --- --- --- --- --- --- --- --- --- --- --- --- --- --- --- --- --- --- --- --- --- --- --- --- --- --- --- --- --- --- --- --- --- --- --- --- --- --- --- --- --- --- --- --- --- --- --- --- --- --- --- --- --- --- --- --- --- --- --- --- --- --- --- --- --- --- --- --- --- --- --- --- --- --- --- --- --- --- --- --- --- --- --- --- --- --- --- --- --- --- --- --- --- --- --- --- --- --- --- --- --- --- --- --- --- --- --- --- --- --- --- --- --- --- --- --- --- --- --- --- --- --- --- --- --- --- --- --- --- --- --- --- --- --- --- --- --- --- --- --- --- --- --- --- --- --- --- --- --- --- --- --- --- --- --- --- --- --- --- --- --- --- --- --- --- --- --- --- --- --- --- --- --- --- --- --- --- --- --- --- --- --- --- --- --- --- --- --- --- --- --- --- --- --- --- --- --- --- --- --- --- --- --- --- --- --- --- --- --- --- --- --- --- --- --- --- --- --- --- --- --- --- --- --- --- --- --- --- --- --- --- --- --- --- --- --- --- --- --- --- --- --- --- --- --- --- --- --- --- --- --- --- --- --- --- --- --- --- --- --- --- --- --- --- --- --- --- --- --- --- --- --- --- --- --- --- --- --- --- --- --- --- --- --- --- --- --- --- --- --- --- --- --- --- --- --- --- --- --- --- --- --- --- --- --- --- --- --- --- --- --- --- --- --- --- --- --- --- --- --- --- --- --- --- --- --- --- --- --- --- --- --- --- --- --- --- --- --- --- --- --- --- --- --- --- --- --- --- --- --- --- --- --- --- --- --- --- --- --- --- --- --- --- --- --- --- --- --- --- --- --- --- --- --- --- --- --- --- --- --- --- --- --- --- --- --- --- --- --- --- --- --- --- --- --- --- --- --- --- --- --- --- --- --- --- --- --- --- --- --- --- --- --- --- --- --- --- --- --- --- --- --- --- --- --- --- --- --- --- --- --- --- --- --- --- --- --- --- --- --- --- --- --- --- --- --- --- --- --- --- --- -ct gga tcg gat gtc acc gca gta ata ttg ttg att att tct gac atc gac gta tta tat agt ttt tta att cca tat ctt ttt t

>ON563414.3_Monkeypox_virus_isolate_MPXV_USA_2022_MA001_complete_genome

ttt tat atc act acg gac ata aac cat tgt ata ttt ttt atg ttt att agt gta cac att ttg gaa gta agt tc- --- --- --- --- --- --- --- --- --- --- --- --- --- --- --- --- --- --- --- --- --- --- --- --- --- --- --- --- --- --- --- --- --- --- --- --- --- --- --- --- --- --- --- --- --- --- --- --- --- --- --- --- --- --- --- --- --- --- --- --- --- --- --- --- --- --- --- --- --- --- --- --- --- --- --- --- --- --- --- --- --- --- --- --- --- --- --- --- --- --- --- --- --- --- --- --- --- --- --- --- --- --- --- --- --- --- --- --- --- --- --- --- --- --- --- --- --- --- --- --- --- --- --- --- --- --- --- --- --- --- --- --- --- --- --- --- --- --- --- --- --- --- --- --- --- --- --- --- --- --- --- --- --- --- --- --- --- --- --- --- --- --- --- --- --- --- --- --- --- --- --- --- --- --- --- --- --- --- --- --- --- --- --- --- --- --- --- --- --- --- --- --- --- --- --- --- --- --- --- --- --- --- --- --- --- --- --- --- --- --- --- --- --- --- --- --- --- --- --- --- --- --- --- --- --- --- --- --- --- --- --- --- --- --- --- --- --- --- --- --- --- --- --- --- --- --- --- --- --- --- --- --- --- --- --- --- --- --- --- --- --- --- --- --- --- --- --- --- --- --- --- --- --- --- --- --- --- --- --- --- --- --- --- --- --- --- --- --- --- --- --- --- --- --- --- --- --- --- --- --- --- --- --- --- --- --- --- --- --- --- --- --- --- --- --- --- --- --- --- --- --- --- --- --- --- --- --- --- --- --- --- --- --- --- --- --- --- --- --- --- --- --- --- --- --- --- --- --- --- --- --- --- --- --- --- --- --- --- --- --- --- --- --- --- --- --- --- --- --- --- --- --- --- --- --- --- --- --- --- --- --- --- --- --- --- --- --- --- --- --- --- --- --- --- --- --- --- --- --- --- --- --- --- --- --- --- --- --- --- --- --- --- --- --- --- --- --- --- --- --- --- --- --- --- --- --- --- --- --- --- --- --- --- --- --- --- --- --- --- --- --- --- --- --- --- --- --- --- --- --- --- --- --- --- --- --- --- --- --- --- --- --- --- --- --- --- --- --- --- --- --- --- --- --- --- --- --- --- --- --- --- --- --- --- --- --- --- --- --- --- --- --- --- --- --- --- --- --- --- --- --- --- --- --- --- --- --- --- --- --- --- --- --- --- --- --- --- --- --- --- --- --- --- --- --- --- --- --- --- --- --- --- --- --- --- --- --- --- --- --- --- --- --- --- --- --- --- --- --- --- --- --- --- --- --- --- --- --- --- --- --- --- --- --- --- --- --- --- --- --- --- --- --- --- --- --- --- --- --- --- --- --- --- --- --- --- --- --- --- --- --- --- --- --- --- --- --- --- --- --- --- --- --- --- --- --- --- --- --- --- --- --- --- --- --- --- --- --- --- --- --- --- --- --- --- --- --- --- --- --- --- --- --- --- --- --- --- --- --- --- --- --- --- --- --- --- --- --- --- --- --- --- -ct gga tcg gat gtc acc gca gta ata ttg ttg att att tct gac atc gac gta tta tat agt ttt tta att cca tat ctt ttt t

>ON676705.1_Monkeypox_virus_isolate_MPXV_USA_2022_UT001_complete_genome

ttt tat atc act acg gac ata aac cat tgt ata ttt ttt atg ttt att agt gta cac att ttg gaa gta agt tc- --- --- --- --- --- --- --- --- --- --- --- --- --- --- --- --- --- --- --- --- --- --- --- --- --- --- --- --- --- --- --- --- --- --- --- --- --- --- --- --- --- --- --- --- --- --- --- --- --- --- --- --- --- --- --- --- --- --- --- --- --- --- --- --- --- --- --- --- --- --- --- --- --- --- --- --- --- --- --- --- --- --- --- --- --- --- --- --- --- --- --- --- --- --- --- --- --- --- --- --- --- --- --- --- --- --- --- --- --- --- --- --- --- --- --- --- --- --- --- --- --- --- --- --- --- --- --- --- --- --- --- --- --- --- --- --- --- --- --- --- --- --- --- --- --- --- --- --- --- --- --- --- --- --- --- --- --- --- --- --- --- --- --- --- --- --- --- --- --- --- --- --- --- --- --- --- --- --- --- --- --- --- --- --- --- --- --- --- --- --- --- --- --- --- --- --- --- --- --- --- --- --- --- --- --- --- --- --- --- --- --- --- --- --- --- --- --- --- --- --- --- --- --- --- --- --- --- --- --- --- --- --- --- --- --- --- --- --- --- --- --- --- --- --- --- --- --- --- --- --- --- --- --- --- --- --- --- --- --- --- --- --- --- --- --- --- --- --- --- --- --- --- --- --- --- --- --- --- --- --- --- --- --- --- --- --- --- --- --- --- --- --- --- --- --- --- --- --- --- --- --- --- --- --- --- --- --- --- --- --- --- --- --- --- --- --- --- --- --- --- --- --- --- --- --- --- --- --- --- --- --- --- --- --- --- --- --- --- --- --- --- --- --- --- --- --- --- --- --- --- --- --- --- --- --- --- --- --- --- --- --- --- --- --- --- --- --- --- --- --- --- --- --- --- --- --- --- --- --- --- --- --- --- --- --- --- --- --- --- --- --- --- --- --- --- --- --- --- --- --- --- --- --- --- --- --- --- --- --- --- --- --- --- --- --- --- --- --- --- --- --- --- --- --- --- --- --- --- --- --- --- --- --- --- --- --- --- --- --- --- --- --- --- --- --- --- --- --- --- --- --- --- --- --- --- --- --- --- --- --- --- --- --- --- --- --- --- --- --- --- --- --- --- --- --- --- --- --- --- --- --- --- --- --- --- --- --- --- --- --- --- --- --- --- --- --- --- --- --- --- --- --- --- --- --- --- --- --- --- --- --- --- --- --- --- --- --- --- --- --- --- --- --- --- --- --- --- --- --- --- --- --- --- --- --- --- --- --- --- --- --- --- --- --- --- --- --- --- --- --- --- --- --- --- --- --- --- --- --- --- --- --- --- --- --- --- --- --- --- --- --- --- --- --- --- --- --- --- --- --- --- --- --- --- --- --- --- --- --- --- --- --- --- --- --- --- --- --- --- --- --- --- --- --- --- --- --- --- --- --- --- --- --- --- --- --- --- --- --- --- --- --- --- --- --- --- --- --- --- --- --- --- --- --- --- --- --- --- --- --- --- --- --- --- --- --- --- --- --- --- --- --- -ct gga tcg gat gtc acc gca gta ata ttg ttg att att tct gac atc gac gta tta tat agt ttt tta att cca tat ctt ttt t

>MN648051.1_Monkeypox_virus_strain_Israel_2018

ttt tat atc act acg gac ata aac cat tgt ata ttt ttt atg ttt att agt gta cac att ttg gaa gta agt tc- --- --- --- --- --- --- --- --- --- --- --- --- --- --- --- --- --- --- --- --- --- --- --- --- --- --- --- --- --- --- --- --- --- --- --- --- --- --- --- --- --- --- --- --- --- --- --- --- --- --- --- --- --- --- --- --- --- --- --- --- --- --- --- --- --- --- --- --- --- --- --- --- --- --- --- --- --- --- --- --- --- --- --- --- --- --- --- --- --- --- --- --- --- --- --- --- --- --- --- --- --- --- --- --- --- --- --- --- --- --- --- --- --- --- --- --- --- --- --- --- --- --- --- --- --- --- --- --- --- --- --- --- --- --- --- --- --- --- --- --- --- --- --- --- --- --- --- --- --- --- --- --- --- --- --- --- --- --- --- --- --- --- --- --- --- --- --- --- --- --- --- --- --- --- --- --- --- --- --- --- --- --- --- --- --- --- --- --- --- --- --- --- --- --- --- --- --- --- --- --- --- --- --- --- --- --- --- --- --- --- --- --- --- --- --- --- --- --- --- --- --- --- --- --- --- --- --- --- --- --- --- --- --- --- --- --- --- --- --- --- --- --- --- --- --- --- --- --- --- --- --- --- --- --- --- --- --- --- --- --- --- --- --- --- --- --- --- --- --- --- --- --- --- --- --- --- --- --- --- --- --- --- --- --- --- --- --- --- --- --- --- --- --- --- --- --- --- --- --- --- --- --- --- --- --- --- --- --- --- --- --- --- --- --- --- --- --- --- --- --- --- --- --- --- --- --- --- --- --- --- --- --- --- --- --- --- --- --- --- --- --- --- --- --- --- --- --- --- --- --- --- --- --- --- --- --- --- --- --- --- --- --- --- --- --- --- --- --- --- --- --- --- --- --- --- --- --- --- --- --- --- --- --- --- --- --- --- --- --- --- --- --- --- --- --- --- --- --- --- --- --- --- --- --- --- --- --- --- --- --- --- --- --- --- --- --- --- --- --- --- --- --- --- --- --- --- --- --- --- --- --- --- --- --- --- --- --- --- --- --- --- --- --- --- --- --- --- --- --- --- --- --- --- --- --- --- --- --- --- --- --- --- --- --- --- --- --- --- --- --- --- --- --- --- --- --- --- --- --- --- --- --- --- --- --- --- --- --- --- --- --- --- --- --- --- --- --- --- --- --- --- --- --- --- --- --- --- --- --- --- --- --- --- --- --- --- --- --- --- --- --- --- --- --- --- --- --- --- --- --- --- --- --- --- --- --- --- --- --- --- --- --- --- --- --- --- --- --- --- --- --- --- --- --- --- --- --- --- --- --- --- --- --- --- --- --- --- --- --- --- --- --- --- --- --- --- --- --- --- --- --- --- --- --- --- --- --- --- --- --- --- --- --- --- --- --- --- --- --- --- --- --- --- --- --- --- --- --- --- --- --- --- --- --- --- --- --- --- --- --- --- --- --- --- --- --- --- --- --- --- --- --- --- --- --- --- --- --- --- --- --- --- --- --- --- --- --- --- --- --- --- --- -ct gga tcg gat gtc acc gca gta ata ttg ttg att att tct gac atc gac gta tta tat agt ttt tta att cca tat ctt ttt t

>ON644344.1_Monkeypox_virus_isolate_MPXV_FVG-ITA_01_2022_complete_genome

ttt tat atc act acg gac ata aac cat tgt ata ttt ttt atg ttt att agt gta cac att ttg gaa gta agt tc- --- --- --- --- --- --- --- --- --- --- --- --- --- --- --- --- --- --- --- --- --- --- --- --- --- --- --- --- --- --- --- --- --- --- --- --- --- --- --- --- --- --- --- --- --- --- --- --- --- --- --- --- --- --- --- --- --- --- --- --- --- --- --- --- --- --- --- --- --- --- --- --- --- --- --- --- --- --- --- --- --- --- --- --- --- --- --- --- --- --- --- --- --- --- --- --- --- --- --- --- --- --- --- --- --- --- --- --- --- --- --- --- --- --- --- --- --- --- --- --- --- --- --- --- --- --- --- --- --- --- --- --- --- --- --- --- --- --- --- --- --- --- --- --- --- --- --- --- --- --- --- --- --- --- --- --- --- --- --- --- --- --- --- --- --- --- --- --- --- --- --- --- --- --- --- --- --- --- --- --- --- --- --- --- --- --- --- --- --- --- --- --- --- --- --- --- --- --- --- --- --- --- --- --- --- --- --- --- --- --- --- --- --- --- --- --- --- --- --- --- --- --- --- --- --- --- --- --- --- --- --- --- --- --- --- --- --- --- --- --- --- --- --- --- --- --- --- --- --- --- --- --- --- --- --- --- --- --- --- --- --- --- --- --- --- --- --- --- --- --- --- --- --- --- --- --- --- --- --- --- --- --- --- --- --- --- --- --- --- --- --- --- --- --- --- --- --- --- --- --- --- --- --- --- --- --- --- --- --- --- --- --- --- --- --- --- --- --- --- --- --- --- --- --- --- --- --- --- --- --- --- --- --- --- --- --- --- --- --- --- --- --- --- --- --- --- --- --- --- --- --- --- --- --- --- --- --- --- --- --- --- --- --- --- --- --- --- --- --- --- --- --- --- --- --- --- --- --- --- --- --- --- --- --- --- --- --- --- --- --- --- --- --- --- --- --- --- --- --- --- --- --- --- --- --- --- --- --- --- --- --- --- --- --- --- --- --- --- --- --- --- --- --- --- --- --- --- --- --- --- --- --- --- --- --- --- --- --- --- --- --- --- --- --- --- --- --- --- --- --- --- --- --- --- --- --- --- --- --- --- --- --- --- --- --- --- --- --- --- --- --- --- --- --- --- --- --- --- --- --- --- --- --- --- --- --- --- --- --- --- --- --- --- --- --- --- --- --- --- --- --- --- --- --- --- --- --- --- --- --- --- --- --- --- --- --- --- --- --- --- --- --- --- --- --- --- --- --- --- --- --- --- --- --- --- --- --- --- --- --- --- --- --- --- --- --- --- --- --- --- --- --- --- --- --- --- --- --- --- --- --- --- --- --- --- --- --- --- --- --- --- --- --- --- --- --- --- --- --- --- --- --- --- --- --- --- --- --- --- --- --- --- --- --- --- --- --- --- --- --- --- --- --- --- --- --- --- --- --- --- --- --- --- --- --- --- --- --- --- --- --- --- --- --- --- --- --- --- --- --- --- --- --- --- --- --- --- --- --- --- --- --- --- --- --- --- --- --- --- --- --- --- -ct gga tcg gat gtc acc gca gta ata ttg ttg att att tct gac atc gac gta tta tat agt ttt tta att cca tat ctt ttt t

>MT903343.1_Monkeypox_virus_isolate_MPXV-UK_P1

ttt tat atc act acg gac ata aac cat tgt ata ttt ttt atg ttt att agt gta cac att ttg gaa gta agt tc- --- --- --- --- --- --- --- --- --- --- --- --- --- --- --- --- --- --- --- --- --- --- --- --- --- --- --- --- --- --- --- --- --- --- --- --- --- --- --- --- --- --- --- --- --- --- --- --- --- --- --- --- --- --- --- --- --- --- --- --- --- --- --- --- --- --- --- --- --- --- --- --- --- --- --- --- --- --- --- --- --- --- --- --- --- --- --- --- --- --- --- --- --- --- --- --- --- --- --- --- --- --- --- --- --- --- --- --- --- --- --- --- --- --- --- --- --- --- --- --- --- --- --- --- --- --- --- --- --- --- --- --- --- --- --- --- --- --- --- --- --- --- --- --- --- --- --- --- --- --- --- --- --- --- --- --- --- --- --- --- --- --- --- --- --- --- --- --- --- --- --- --- --- --- --- --- --- --- --- --- --- --- --- --- --- --- --- --- --- --- --- --- --- --- --- --- --- --- --- --- --- --- --- --- --- --- --- --- --- --- --- --- --- --- --- --- --- --- --- --- --- --- --- --- --- --- --- --- --- --- --- --- --- --- --- --- --- --- --- --- --- --- --- --- --- --- --- --- --- --- --- --- --- --- --- --- --- --- --- --- --- --- --- --- --- --- --- --- --- --- --- --- --- --- --- --- --- --- --- --- --- --- --- --- --- --- --- --- --- --- --- --- --- --- --- --- --- --- --- --- --- --- --- --- --- --- --- --- --- --- --- --- --- --- --- --- --- --- --- --- --- --- --- --- --- --- --- --- --- --- --- --- --- --- --- --- --- --- --- --- --- --- --- --- --- --- --- --- --- --- --- --- --- --- --- --- --- --- --- --- --- --- --- --- --- --- --- --- --- --- --- --- --- --- --- --- --- --- --- --- --- --- --- --- --- --- --- --- --- --- --- --- --- --- --- --- --- --- --- --- --- --- --- --- --- --- --- --- --- --- --- --- --- --- --- --- --- --- --- --- --- --- --- --- --- --- --- --- --- --- --- --- --- --- --- --- --- --- --- --- --- --- --- --- --- --- --- --- --- --- --- --- --- --- --- --- --- --- --- --- --- --- --- --- --- --- --- --- --- --- --- --- --- --- --- --- --- --- --- --- --- --- --- --- --- --- --- --- --- --- --- --- --- --- --- --- --- --- --- --- --- --- --- --- --- --- --- --- --- --- --- --- --- --- --- --- --- --- --- --- --- --- --- --- --- --- --- --- --- --- --- --- --- --- --- --- --- --- --- --- --- --- --- --- --- --- --- --- --- --- --- --- --- --- --- --- --- --- --- --- --- --- --- --- --- --- --- --- --- --- --- --- --- --- --- --- --- --- --- --- --- --- --- --- --- --- --- --- --- --- --- --- --- --- --- --- --- --- --- --- --- --- --- --- --- --- --- --- --- --- --- --- --- --- --- --- --- --- --- --- --- --- --- --- --- --- --- --- --- --- --- --- --- --- --- --- --- --- --- --- --- --- --- --- --- --- --- --- --- --- --- --- -ct gga tcg gat gtc acc gca gta ata ttg ttg att att tct gac atc gac gta tta tat agt ttt tta att cca tat ctt ttt t

>ON631963.1_Monkeypox_virus_isolate_MPxV/VIDRL01/2022_complete_genome

ttt tat atc act acg gac ata aac cat tgt ata ttt ttt atg ttt att agt gta cac att ttg gaa gta agt tc- --- --- --- --- --- --- --- --- --- --- --- --- --- --- --- --- --- --- --- --- --- --- --- --- --- --- --- --- --- --- --- --- --- --- --- --- --- --- --- --- --- --- --- --- --- --- --- --- --- --- --- --- --- --- --- --- --- --- --- --- --- --- --- --- --- --- --- --- --- --- --- --- --- --- --- --- --- --- --- --- --- --- --- --- --- --- --- --- --- --- --- --- --- --- --- --- --- --- --- --- --- --- --- --- --- --- --- --- --- --- --- --- --- --- --- --- --- --- --- --- --- --- --- --- --- --- --- --- --- --- --- --- --- --- --- --- --- --- --- --- --- --- --- --- --- --- --- --- --- --- --- --- --- --- --- --- --- --- --- --- --- --- --- --- --- --- --- --- --- --- --- --- --- --- --- --- --- --- --- --- --- --- --- --- --- --- --- --- --- --- --- --- --- --- --- --- --- --- --- --- --- --- --- --- --- --- --- --- --- --- --- --- --- --- --- --- --- --- --- --- --- --- --- --- --- --- --- --- --- --- --- --- --- --- --- --- --- --- --- --- --- --- --- --- --- --- --- --- --- --- --- --- --- --- --- --- --- --- --- --- --- --- --- --- --- --- --- --- --- --- --- --- --- --- --- --- --- --- --- --- --- --- --- --- --- --- --- --- --- --- --- --- --- --- --- --- --- --- --- --- --- --- --- --- --- --- --- --- --- --- --- --- --- --- --- --- --- --- --- --- --- --- --- --- --- --- --- --- --- --- --- --- --- --- --- --- --- --- --- --- --- --- --- --- --- --- --- --- --- --- --- --- --- --- --- --- --- --- --- --- --- --- --- --- --- --- --- --- --- --- --- --- --- --- --- --- --- --- --- --- --- --- --- --- --- --- --- --- --- --- --- --- --- --- --- --- --- --- --- --- --- --- --- --- --- --- --- --- --- --- --- --- --- --- --- --- --- --- --- --- --- --- --- --- --- --- --- --- --- --- --- --- --- --- --- --- --- --- --- --- --- --- --- --- --- --- --- --- --- --- --- --- --- --- --- --- --- --- --- --- --- --- --- --- --- --- --- --- --- --- --- --- --- --- --- --- --- --- --- --- --- --- --- --- --- --- --- --- --- --- --- --- --- --- --- --- --- --- --- --- --- --- --- --- --- --- --- --- --- --- --- --- --- --- --- --- --- --- --- --- --- --- --- --- --- --- --- --- --- --- --- --- --- --- --- --- --- --- --- --- --- --- --- --- --- --- --- --- --- --- --- --- --- --- --- --- --- --- --- --- --- --- --- --- --- --- --- --- --- --- --- --- --- --- --- --- --- --- --- --- --- --- --- --- --- --- --- --- --- --- --- --- --- --- --- --- --- --- --- --- --- --- --- --- --- --- --- --- --- --- --- --- --- --- --- --- --- --- --- --- --- --- --- --- --- --- --- --- --- --- --- --- --- --- --- --- --- --- --- --- --- --- --- --- --- --- --- --- --- --- --- --- -ct gga tcg gat gtc acc gca gta ata ttg ttg att att tct gac atc gac gta tta tat agt ttt tta att cca tat ctt ttt t

>ON568298.1_Monkeypox_virus_isolate_MPXV-BY-IMB25241_complete_genome

ttt tat atc act acg gac ata aac cat tgt ata ttt ttt atg ttt att agt gta cac att ttg gaa gta agt tc- --- --- --- --- --- --- --- --- --- --- --- --- --- --- --- --- --- --- --- --- --- --- --- --- --- --- --- --- --- --- --- --- --- --- --- --- --- --- --- --- --- --- --- --- --- --- --- --- --- --- --- --- --- --- --- --- --- --- --- --- --- --- --- --- --- --- --- --- --- --- --- --- --- --- --- --- --- --- --- --- --- --- --- --- --- --- --- --- --- --- --- --- --- --- --- --- --- --- --- --- --- --- --- --- --- --- --- --- --- --- --- --- --- --- --- --- --- --- --- --- --- --- --- --- --- --- --- --- --- --- --- --- --- --- --- --- --- --- --- --- --- --- --- --- --- --- --- --- --- --- --- --- --- --- --- --- --- --- --- --- --- --- --- --- --- --- --- --- --- --- --- --- --- --- --- --- --- --- --- --- --- --- --- --- --- --- --- --- --- --- --- --- --- --- --- --- --- --- --- --- --- --- --- --- --- --- --- --- --- --- --- --- --- --- --- --- --- --- --- --- --- --- --- --- --- --- --- --- --- --- --- --- --- --- --- --- --- --- --- --- --- --- --- --- --- --- --- --- --- --- --- --- --- --- --- --- --- --- --- --- --- --- --- --- --- --- --- --- --- --- --- --- --- --- --- --- --- --- --- --- --- --- --- --- --- --- --- --- --- --- --- --- --- --- --- --- --- --- --- --- --- --- --- --- --- --- --- --- --- --- --- --- --- --- --- --- --- --- --- --- --- --- --- --- --- --- --- --- --- --- --- --- --- --- --- --- --- --- --- --- --- --- --- --- --- --- --- --- --- --- --- --- --- --- --- --- --- --- --- --- --- --- --- --- --- --- --- --- --- --- --- --- --- --- --- --- --- --- --- --- --- --- --- --- --- --- --- --- --- --- --- --- --- --- --- --- --- --- --- --- --- --- --- --- --- --- --- --- --- --- --- --- --- --- --- --- --- --- --- --- --- --- --- --- --- --- --- --- --- --- --- --- --- --- --- --- --- --- --- --- --- --- --- --- --- --- --- --- --- --- --- --- --- --- --- --- --- --- --- --- --- --- --- --- --- --- --- --- --- --- --- --- --- --- --- --- --- --- --- --- --- --- --- --- --- --- --- --- --- --- --- --- --- --- --- --- --- --- --- --- --- --- --- --- --- --- --- --- --- --- --- --- --- --- --- --- --- --- --- --- --- --- --- --- --- --- --- --- --- --- --- --- --- --- --- --- --- --- --- --- --- --- --- --- --- --- --- --- --- --- --- --- --- --- --- --- --- --- --- --- --- --- --- --- --- --- --- --- --- --- --- --- --- --- --- --- --- --- --- --- --- --- --- --- --- --- --- --- --- --- --- --- --- --- --- --- --- --- --- --- --- --- --- --- --- --- --- --- --- --- --- --- --- --- --- --- --- --- --- --- --- --- --- --- --- --- --- --- --- --- --- --- --- --- --- --- --- --- --- --- --- --- --- --- --- --- --- --- --- --- --- --- -ct gga tcg gat gtc acc gca gta ata ttg ttg att att tct gac atc gac gta tta tat agt ttt tta att cca tat ctt ttt t

>ON622722.2_Monkeypox_virus_isolate_MPXV_FR_HCL0001_2022_complete_genome

ttt tat atc act acg gac ata aac cat tgt ata ttt ttt atg ttt att agt gta cac att ttg gaa gta agt tc- --- --- --- --- --- --- --- --- --- --- --- --- --- --- --- --- --- --- --- --- --- --- --- --- --- --- --- --- --- --- --- --- --- --- --- --- --- --- --- --- --- --- --- --- --- --- --- --- --- --- --- --- --- --- --- --- --- --- --- --- --- --- --- --- --- --- --- --- --- --- --- --- --- --- --- --- --- --- --- --- --- --- --- --- --- --- --- --- --- --- --- --- --- --- --- --- --- --- --- --- --- --- --- --- --- --- --- --- --- --- --- --- --- --- --- --- --- --- --- --- --- --- --- --- --- --- --- --- --- --- --- --- --- --- --- --- --- --- --- --- --- --- --- --- --- --- --- --- --- --- --- --- --- --- --- --- --- --- --- --- --- --- --- --- --- --- --- --- --- --- --- --- --- --- --- --- --- --- --- --- --- --- --- --- --- --- --- --- --- --- --- --- --- --- --- --- --- --- --- --- --- --- --- --- --- --- --- --- --- --- --- --- --- --- --- --- --- --- --- --- --- --- --- --- --- --- --- --- --- --- --- --- --- --- --- --- --- --- --- --- --- --- --- --- --- --- --- --- --- --- --- --- --- --- --- --- --- --- --- --- --- --- --- --- --- --- --- --- --- --- --- --- --- --- --- --- --- --- --- --- --- --- --- --- --- --- --- --- --- --- --- --- --- --- --- --- --- --- --- --- --- --- --- --- --- --- --- --- --- --- --- --- --- --- --- --- --- --- --- --- --- --- --- --- --- --- --- --- --- --- --- --- --- --- --- --- --- --- --- --- --- --- --- --- --- --- --- --- --- --- --- --- --- --- --- --- --- --- --- --- --- --- --- --- --- --- --- --- --- --- --- --- --- --- --- --- --- --- --- --- --- --- --- --- --- --- --- --- --- --- --- --- --- --- --- --- --- --- --- --- --- --- --- --- --- --- --- --- --- --- --- --- --- --- --- --- --- --- --- --- --- --- --- --- --- --- --- --- --- --- --- --- --- --- --- --- --- --- --- --- --- --- --- --- --- --- --- --- --- --- --- --- --- --- --- --- --- --- --- --- --- --- --- --- --- --- --- --- --- --- --- --- --- --- --- --- --- --- --- --- --- --- --- --- --- --- --- --- --- --- --- --- --- --- --- --- --- --- --- --- --- --- --- --- --- --- --- --- --- --- --- --- --- --- --- --- --- --- --- --- --- --- --- --- --- --- --- --- --- --- --- --- --- --- --- --- --- --- --- --- --- --- --- --- --- --- --- --- --- --- --- --- --- --- --- --- --- --- --- --- --- --- --- --- --- --- --- --- --- --- --- --- --- --- --- --- --- --- --- --- --- --- --- --- --- --- --- --- --- --- --- --- --- --- --- --- --- --- --- --- --- --- --- --- --- --- --- --- --- --- --- --- --- --- --- --- --- --- --- --- --- --- --- --- --- --- --- --- --- --- --- --- --- --- --- --- --- --- --- --- --- --- --- --- --- --- --- --- --- --- --- --- -ct gga tcg gat gtc acc gca gta ata ttg ttg att att tct gac atc gac gta tta tat agt ttt tta att cca tat ctt ttt t

>ON627808.1_Monkeypox_virus_isolate_MPX/human/USA/UT-UPHL-82200022/2022_complete_genome

ttt tat atc act acg gac ata aac cat tgt ata ttt ttt atg ttt att agt gta cac att ttg gaa gta agt tc- --- --- --- --- --- --- --- --- --- --- --- --- --- --- --- --- --- --- --- --- --- --- --- --- --- --- --- --- --- --- --- --- --- --- --- --- --- --- --- --- --- --- --- --- --- --- --- --- --- --- --- --- --- --- --- --- --- --- --- --- --- --- --- --- --- --- --- --- --- --- --- --- --- --- --- --- --- --- --- --- --- --- --- --- --- --- --- --- --- --- --- --- --- --- --- --- --- --- --- --- --- --- --- --- --- --- --- --- --- --- --- --- --- --- --- --- --- --- --- --- --- --- --- --- --- --- --- --- --- --- --- --- --- --- --- --- --- --- --- --- --- --- --- --- --- --- --- --- --- --- --- --- --- --- --- --- --- --- --- --- --- --- --- --- --- --- --- --- --- --- --- --- --- --- --- --- --- --- --- --- --- --- --- --- --- --- --- --- --- --- --- --- --- --- --- --- --- --- --- --- --- --- --- --- --- --- --- --- --- --- --- --- --- --- --- --- --- --- --- --- --- --- --- --- --- --- --- --- --- --- --- --- --- --- --- --- --- --- --- --- --- --- --- --- --- --- --- --- --- --- --- --- --- --- --- --- --- --- --- --- --- --- --- --- --- --- --- --- --- --- --- --- --- --- --- --- --- --- --- --- --- --- --- --- --- --- --- --- --- --- --- --- --- --- --- --- --- --- --- --- --- --- --- --- --- --- --- --- --- --- --- --- --- --- --- --- --- --- --- --- --- --- --- --- --- --- --- --- --- --- --- --- --- --- --- --- --- --- --- --- --- --- --- --- --- --- --- --- --- --- --- --- --- --- --- --- --- --- --- --- --- --- --- --- --- --- --- --- --- --- --- --- --- --- --- --- --- --- --- --- --- --- --- --- --- --- --- --- --- --- --- --- --- --- --- --- --- --- --- --- --- --- --- --- --- --- --- --- --- --- --- --- --- --- --- --- --- --- --- --- --- --- --- --- --- --- --- --- --- --- --- --- --- --- --- --- --- --- --- --- --- --- --- --- --- --- --- --- --- --- --- --- --- --- --- --- --- --- --- --- --- --- --- --- --- --- --- --- --- --- --- --- --- --- --- --- --- --- --- --- --- --- --- --- --- --- --- --- --- --- --- --- --- --- --- --- --- --- --- --- --- --- --- --- --- --- --- --- --- --- --- --- --- --- --- --- --- --- --- --- --- --- --- --- --- --- --- --- --- --- --- --- --- --- --- --- --- --- --- --- --- --- --- --- --- --- --- --- --- --- --- --- --- --- --- --- --- --- --- --- --- --- --- --- --- --- --- --- --- --- --- --- --- --- --- --- --- --- --- --- --- --- --- --- --- --- --- --- --- --- --- --- --- --- --- --- --- --- --- --- --- --- --- --- --- --- --- --- --- --- --- --- --- --- --- --- --- --- --- --- --- --- --- --- --- --- --- --- --- --- --- --- --- --- --- --- --- --- --- --- --- --- --- --- --- --- --- --- --- --- --- --- -ct gga tcg gat gtc acc gca gta ata ttg ttg att att tct gac atc gac gta tta tat agt ttt tta att cca tat ctt ttt t

>ON676703.1_Monkeypox_virus_isolate_MPXV_USA_2022_CA001_complete_genome

ttt tat atc act acg gac ata aac cat tgt ata ttt ttt atg ttt att agt gta cac att ttg gaa gta agt tc- --- --- --- --- --- --- --- --- --- --- --- --- --- --- --- --- --- --- --- --- --- --- --- --- --- --- --- --- --- --- --- --- --- --- --- --- --- --- --- --- --- --- --- --- --- --- --- --- --- --- --- --- --- --- --- --- --- --- --- --- --- --- --- --- --- --- --- --- --- --- --- --- --- --- --- --- --- --- --- --- --- --- --- --- --- --- --- --- --- --- --- --- --- --- --- --- --- --- --- --- --- --- --- --- --- --- --- --- --- --- --- --- --- --- --- --- --- --- --- --- --- --- --- --- --- --- --- --- --- --- --- --- --- --- --- --- --- --- --- --- --- --- --- --- --- --- --- --- --- --- --- --- --- --- --- --- --- --- --- --- --- --- --- --- --- --- --- --- --- --- --- --- --- --- --- --- --- --- --- --- --- --- --- --- --- --- --- --- --- --- --- --- --- --- --- --- --- --- --- --- --- --- --- --- --- --- --- --- --- --- --- --- --- --- --- --- --- --- --- --- --- --- --- --- --- --- --- --- --- --- --- --- --- --- --- --- --- --- --- --- --- --- --- --- --- --- --- --- --- --- --- --- --- --- --- --- --- --- --- --- --- --- --- --- --- --- --- --- --- --- --- --- --- --- --- --- --- --- --- --- --- --- --- --- --- --- --- --- --- --- --- --- --- --- --- --- --- --- --- --- --- --- --- --- --- --- --- --- --- --- --- --- --- --- --- --- --- --- --- --- --- --- --- --- --- --- --- --- --- --- --- --- --- --- --- --- --- --- --- --- --- --- --- --- --- --- --- --- --- --- --- --- --- --- --- --- --- --- --- --- --- --- --- --- --- --- --- --- --- --- --- --- --- --- --- --- --- --- --- --- --- --- --- --- --- --- --- --- --- --- --- --- --- --- --- --- --- --- --- --- --- --- --- --- --- --- --- --- --- --- --- --- --- --- --- --- --- --- --- --- --- --- --- --- --- --- --- --- --- --- --- --- --- --- --- --- --- --- --- --- --- --- --- --- --- --- --- --- --- --- --- --- --- --- --- --- --- --- --- --- --- --- --- --- --- --- --- --- --- --- --- --- --- --- --- --- --- --- --- --- --- --- --- --- --- --- --- --- --- --- --- --- --- --- --- --- --- --- --- --- --- --- --- --- --- --- --- --- --- --- --- --- --- --- --- --- --- --- --- --- --- --- --- --- --- --- --- --- --- --- --- --- --- --- --- --- --- --- --- --- --- --- --- --- --- --- --- --- --- --- --- --- --- --- --- --- --- --- --- --- --- --- --- --- --- --- --- --- --- --- --- --- --- --- --- --- --- --- --- --- --- --- --- --- --- --- --- --- --- --- --- --- --- --- --- --- --- --- --- --- --- --- --- --- --- --- --- --- --- --- --- --- --- --- --- --- --- --- --- --- --- --- --- --- --- --- --- --- --- --- --- --- --- --- --- --- --- --- --- --- --- --- --- --- --- --- --- --- --- --- --- --- -ct gga tcg gat gtc acc gca gta ata ttg ttg att att tct gac atc gac gta tta tat agt ttt tta att cca tat ctt ttt t

>ON676706.1_Monkeypox_virus_isolate_MPXV_USA_2022_UT002_complete_genome

ttt tat atc act acg gac ata aac cat tgt ata ttt ttt atg ttt att agt gta cac att ttg gaa gta agt tc- --- --- --- --- --- --- --- --- --- --- --- --- --- --- --- --- --- --- --- --- --- --- --- --- --- --- --- --- --- --- --- --- --- --- --- --- --- --- --- --- --- --- --- --- --- --- --- --- --- --- --- --- --- --- --- --- --- --- --- --- --- --- --- --- --- --- --- --- --- --- --- --- --- --- --- --- --- --- --- --- --- --- --- --- --- --- --- --- --- --- --- --- --- --- --- --- --- --- --- --- --- --- --- --- --- --- --- --- --- --- --- --- --- --- --- --- --- --- --- --- --- --- --- --- --- --- --- --- --- --- --- --- --- --- --- --- --- --- --- --- --- --- --- --- --- --- --- --- --- --- --- --- --- --- --- --- --- --- --- --- --- --- --- --- --- --- --- --- --- --- --- --- --- --- --- --- --- --- --- --- --- --- --- --- --- --- --- --- --- --- --- --- --- --- --- --- --- --- --- --- --- --- --- --- --- --- --- --- --- --- --- --- --- --- --- --- --- --- --- --- --- --- --- --- --- --- --- --- --- --- --- --- --- --- --- --- --- --- --- --- --- --- --- --- --- --- --- --- --- --- --- --- --- --- --- --- --- --- --- --- --- --- --- --- --- --- --- --- --- --- --- --- --- --- --- --- --- --- --- --- --- --- --- --- --- --- --- --- --- --- --- --- --- --- --- --- --- --- --- --- --- --- --- --- --- --- --- --- --- --- --- --- --- --- --- --- --- --- --- --- --- --- --- --- --- --- --- --- --- --- --- --- --- --- --- --- --- --- --- --- --- --- --- --- --- --- --- --- --- --- --- --- --- --- --- --- --- --- --- --- --- --- --- --- --- --- --- --- --- --- --- --- --- --- --- --- --- --- --- --- --- --- --- --- --- --- --- --- --- --- --- --- --- --- --- --- --- --- --- --- --- --- --- --- --- --- --- --- --- --- --- --- --- --- --- --- --- --- --- --- --- --- --- --- --- --- --- --- --- --- --- --- --- --- --- --- --- --- --- --- --- --- --- --- --- --- --- --- --- --- --- --- --- --- --- --- --- --- --- --- --- --- --- --- --- --- --- --- --- --- --- --- --- --- --- --- --- --- --- --- --- --- --- --- --- --- --- --- --- --- --- --- --- --- --- --- --- --- --- --- --- --- --- --- --- --- --- --- --- --- --- --- --- --- --- --- --- --- --- --- --- --- --- --- --- --- --- --- --- --- --- --- --- --- --- --- --- --- --- --- --- --- --- --- --- --- --- --- --- --- --- --- --- --- --- --- --- --- --- --- --- --- --- --- --- --- --- --- --- --- --- --- --- --- --- --- --- --- --- --- --- --- --- --- --- --- --- --- --- --- --- --- --- --- --- --- --- --- --- --- --- --- --- --- --- --- --- --- --- --- --- --- --- --- --- --- --- --- --- --- --- --- --- --- --- --- --- --- --- --- --- --- --- --- --- --- --- --- --- --- --- --- --- --- --- --- --- --- --- --- --- --- -ct gga tcg gat gtc acc gca gta ata ttg ttg att att tct gac atc gac gta tta tat agt ttt tta att cca tat ctt ttt t

>ON649879.1_Monkeypox_virus_isolate_MPXV_ISR001_2022_complete_genome

ttt tat atc act acg gac ata aac cat tgt ata ttt ttt atg ttt att agt gta cac att ttg gaa gta agt tc- --- --- --- --- --- --- --- --- --- --- --- --- --- --- --- --- --- --- --- --- --- --- --- --- --- --- --- --- --- --- --- --- --- --- --- --- --- --- --- --- --- --- --- --- --- --- --- --- --- --- --- --- --- --- --- --- --- --- --- --- --- --- --- --- --- --- --- --- --- --- --- --- --- --- --- --- --- --- --- --- --- --- --- --- --- --- --- --- --- --- --- --- --- --- --- --- --- --- --- --- --- --- --- --- --- --- --- --- --- --- --- --- --- --- --- --- --- --- --- --- --- --- --- --- --- --- --- --- --- --- --- --- --- --- --- --- --- --- --- --- --- --- --- --- --- --- --- --- --- --- --- --- --- --- --- --- --- --- --- --- --- --- --- --- --- --- --- --- --- --- --- --- --- --- --- --- --- --- --- --- --- --- --- --- --- --- --- --- --- --- --- --- --- --- --- --- --- --- --- --- --- --- --- --- --- --- --- --- --- --- --- --- --- --- --- --- --- --- --- --- --- --- --- --- --- --- --- --- --- --- --- --- --- --- --- --- --- --- --- --- --- --- --- --- --- --- --- --- --- --- --- --- --- --- --- --- --- --- --- --- --- --- --- --- --- --- --- --- --- --- --- --- --- --- --- --- --- --- --- --- --- --- --- --- --- --- --- --- --- --- --- --- --- --- --- --- --- --- --- --- --- --- --- --- --- --- --- --- --- --- --- --- --- --- --- --- --- --- --- --- --- --- --- --- --- --- --- --- --- --- --- --- --- --- --- --- --- --- --- --- --- --- --- --- --- --- --- --- --- --- --- --- --- --- --- --- --- --- --- --- --- --- --- --- --- --- --- --- --- --- --- --- --- --- --- --- --- --- --- --- --- --- --- --- --- --- --- --- --- --- --- --- --- --- --- --- --- --- --- --- --- --- --- --- --- --- --- --- --- --- --- --- --- --- --- --- --- --- --- --- --- --- --- --- --- --- --- --- --- --- --- --- --- --- --- --- --- --- --- --- --- --- --- --- --- --- --- --- --- --- --- --- --- --- --- --- --- --- --- --- --- --- --- --- --- --- --- --- --- --- --- --- --- --- --- --- --- --- --- --- --- --- --- --- --- --- --- --- --- --- --- --- --- --- --- --- --- --- --- --- --- --- --- --- --- --- --- --- --- --- --- --- --- --- --- --- --- --- --- --- --- --- --- --- --- --- --- --- --- --- --- --- --- --- --- --- --- --- --- --- --- --- --- --- --- --- --- --- --- --- --- --- --- --- --- --- --- --- --- --- --- --- --- --- --- --- --- --- --- --- --- --- --- --- --- --- --- --- --- --- --- --- --- --- --- --- --- --- --- --- --- --- --- --- --- --- --- --- --- --- --- --- --- --- --- --- --- --- --- --- --- --- --- --- --- --- --- --- --- --- --- --- --- --- --- --- --- --- --- --- --- --- --- --- --- --- --- --- --- --- --- --- --- --- --- --- --- --- --- --- --- --- -ct gga tcg gat gtc acc gca gta ata ttg ttg att att tct gac atc gac gta tta tat agt ttt tta att cca tat ctt ttt t

>ON602722.1_Monkeypox_virus_isolate_MPXV_FRA_2022_TLS67_complete_genome

ttt tat atc act acg gac ata aac cat tgt ata ttt ttt atg ttt att agt gta cac att ttg gaa gta agt tc- --- --- --- --- --- --- --- --- --- --- --- --- --- --- --- --- --- --- --- --- --- --- --- --- --- --- --- --- --- --- --- --- --- --- --- --- --- --- --- --- --- --- --- --- --- --- --- --- --- --- --- --- --- --- --- --- --- --- --- --- --- --- --- --- --- --- --- --- --- --- --- --- --- --- --- --- --- --- --- --- --- --- --- --- --- --- --- --- --- --- --- --- --- --- --- --- --- --- --- --- --- --- --- --- --- --- --- --- --- --- --- --- --- --- --- --- --- --- --- --- --- --- --- --- --- --- --- --- --- --- --- --- --- --- --- --- --- --- --- --- --- --- --- --- --- --- --- --- --- --- --- --- --- --- --- --- --- --- --- --- --- --- --- --- --- --- --- --- --- --- --- --- --- --- --- --- --- --- --- --- --- --- --- --- --- --- --- --- --- --- --- --- --- --- --- --- --- --- --- --- --- --- --- --- --- --- --- --- --- --- --- --- --- --- --- --- --- --- --- --- --- --- --- --- --- --- --- --- --- --- --- --- --- --- --- --- --- --- --- --- --- --- --- --- --- --- --- --- --- --- --- --- --- --- --- --- --- --- --- --- --- --- --- --- --- --- --- --- --- --- --- --- --- --- --- --- --- --- --- --- --- --- --- --- --- --- --- --- --- --- --- --- --- --- --- --- --- --- --- --- --- --- --- --- --- --- --- --- --- --- --- --- --- --- --- --- --- --- --- --- --- --- --- --- --- --- --- --- --- --- --- --- --- --- --- --- --- --- --- --- --- --- --- --- --- --- --- --- --- --- --- --- --- --- --- --- --- --- --- --- --- --- --- --- --- --- --- --- --- --- --- --- --- --- --- --- --- --- --- --- --- --- --- --- --- --- --- --- --- --- --- --- --- --- --- --- --- --- --- --- --- --- --- --- --- --- --- --- --- --- --- --- --- --- --- --- --- --- --- --- --- --- --- --- --- --- --- --- --- --- --- --- --- --- --- --- --- --- --- --- --- --- --- --- --- --- --- --- --- --- --- --- --- --- --- --- --- --- --- --- --- --- --- --- --- --- --- --- --- --- --- --- --- --- --- --- --- --- --- --- --- --- --- --- --- --- --- --- --- --- --- --- --- --- --- --- --- --- --- --- --- --- --- --- --- --- --- --- --- --- --- --- --- --- --- --- --- --- --- --- --- --- --- --- --- --- --- --- --- --- --- --- --- --- --- --- --- --- --- --- --- --- --- --- --- --- --- --- --- --- --- --- --- --- --- --- --- --- --- --- --- --- --- --- --- --- --- --- --- --- --- --- --- --- --- --- --- --- --- --- --- --- --- --- --- --- --- --- --- --- --- --- --- --- --- --- --- --- --- --- --- --- --- --- --- --- --- --- --- --- --- --- --- --- --- --- --- --- --- --- --- --- --- --- --- --- --- --- --- --- --- --- --- --- --- --- --- --- --- --- --- --- --- --- --- --- --- --- --- --- --- --- -ct gga tcg gat gtc acc gca gta ata ttg ttg att att tct gac atc gac gta tta tat agt ttt tta att cca tat ctt ttt t

>ON675438.1_Monkeypox_virus_isolate_MPXV_USA_2022_VA001_complete_genome

ttt tat atc act acg gac ata aac cat tgt ata ttt ttt atg ttt att agt gta cac att ttg gaa gta agt tc- --- --- --- --- --- --- --- --- --- --- --- --- --- --- --- --- --- --- --- --- --- --- --- --- --- --- --- --- --- --- --- --- --- --- --- --- --- --- --- --- --- --- --- --- --- --- --- --- --- --- --- --- --- --- --- --- --- --- --- --- --- --- --- --- --- --- --- --- --- --- --- --- --- --- --- --- --- --- --- --- --- --- --- --- --- --- --- --- --- --- --- --- --- --- --- --- --- --- --- --- --- --- --- --- --- --- --- --- --- --- --- --- --- --- --- --- --- --- --- --- --- --- --- --- --- --- --- --- --- --- --- --- --- --- --- --- --- --- --- --- --- --- --- --- --- --- --- --- --- --- --- --- --- --- --- --- --- --- --- --- --- --- --- --- --- --- --- --- --- --- --- --- --- --- --- --- --- --- --- --- --- --- --- --- --- --- --- --- --- --- --- --- --- --- --- --- --- --- --- --- --- --- --- --- --- --- --- --- --- --- --- --- --- --- --- --- --- --- --- --- --- --- --- --- --- --- --- --- --- --- --- --- --- --- --- --- --- --- --- --- --- --- --- --- --- --- --- --- --- --- --- --- --- --- --- --- --- --- --- --- --- --- --- --- --- --- --- --- --- --- --- --- --- --- --- --- --- --- --- --- --- --- --- --- --- --- --- --- --- --- --- --- --- --- --- --- --- --- --- --- --- --- --- --- --- --- --- --- --- --- --- --- --- --- --- --- --- --- --- --- --- --- --- --- --- --- --- --- --- --- --- --- --- --- --- --- --- --- --- --- --- --- --- --- --- --- --- --- --- --- --- --- --- --- --- --- --- --- --- --- --- --- --- --- --- --- --- --- --- --- --- --- --- --- --- --- --- --- --- --- --- --- --- --- --- --- --- --- --- --- --- --- --- --- --- --- --- --- --- --- --- --- --- --- --- --- --- --- --- --- --- --- --- --- --- --- --- --- --- --- --- --- --- --- --- --- --- --- --- --- --- --- --- --- --- --- --- --- --- --- --- --- --- --- --- --- --- --- --- --- --- --- --- --- --- --- --- --- --- --- --- --- --- --- --- --- --- --- --- --- --- --- --- --- --- --- --- --- --- --- --- --- --- --- --- --- --- --- --- --- --- --- --- --- --- --- --- --- --- --- --- --- --- --- --- --- --- --- --- --- --- --- --- --- --- --- --- --- --- --- --- --- --- --- --- --- --- --- --- --- --- --- --- --- --- --- --- --- --- --- --- --- --- --- --- --- --- --- --- --- --- --- --- --- --- --- --- --- --- --- --- --- --- --- --- --- --- --- --- --- --- --- --- --- --- --- --- --- --- --- --- --- --- --- --- --- --- --- --- --- --- --- --- --- --- --- --- --- --- --- --- --- --- --- --- --- --- --- --- --- --- --- --- --- --- --- --- --- --- --- --- --- --- --- --- --- --- --- --- --- --- --- --- --- --- --- --- --- --- --- --- --- --- --- --- --- --- --- --- --- --- --- -ct gga tcg gat gtc acc gca gta ata ttg ttg att att tct gac atc gac gta tta tat agt ttt tta att cca tat ctt ttt t

>ON631241.1_Monkeypox_virus_isolate_2022/2_SLO_complete_genome

ttt tat atc act acg gac ata aac cat tgt ata ttt ttt atg ttt att agt gta cac att ttg gaa gta agt tc- --- --- --- --- --- --- --- --- --- --- --- --- --- --- --- --- --- --- --- --- --- --- --- --- --- --- --- --- --- --- --- --- --- --- --- --- --- --- --- --- --- --- --- --- --- --- --- --- --- --- --- --- --- --- --- --- --- --- --- --- --- --- --- --- --- --- --- --- --- --- --- --- --- --- --- --- --- --- --- --- --- --- --- --- --- --- --- --- --- --- --- --- --- --- --- --- --- --- --- --- --- --- --- --- --- --- --- --- --- --- --- --- --- --- --- --- --- --- --- --- --- --- --- --- --- --- --- --- --- --- --- --- --- --- --- --- --- --- --- --- --- --- --- --- --- --- --- --- --- --- --- --- --- --- --- --- --- --- --- --- --- --- --- --- --- --- --- --- --- --- --- --- --- --- --- --- --- --- --- --- --- --- --- --- --- --- --- --- --- --- --- --- --- --- --- --- --- --- --- --- --- --- --- --- --- --- --- --- --- --- --- --- --- --- --- --- --- --- --- --- --- --- --- --- --- --- --- --- --- --- --- --- --- --- --- --- --- --- --- --- --- --- --- --- --- --- --- --- --- --- --- --- --- --- --- --- --- --- --- --- --- --- --- --- --- --- --- --- --- --- --- --- --- --- --- --- --- --- --- --- --- --- --- --- --- --- --- --- --- --- --- --- --- --- --- --- --- --- --- --- --- --- --- --- --- --- --- --- --- --- --- --- --- --- --- --- --- --- --- --- --- --- --- --- --- --- --- --- --- --- --- --- --- --- --- --- --- --- --- --- --- --- --- --- --- --- --- --- --- --- --- --- --- --- --- --- --- --- --- --- --- --- --- --- --- --- --- --- --- --- --- --- --- --- --- --- --- --- --- --- --- --- --- --- --- --- --- --- --- --- --- --- --- --- --- --- --- --- --- --- --- --- --- --- --- --- --- --- --- --- --- --- --- --- --- --- --- --- --- --- --- --- --- --- --- --- --- --- --- --- --- --- --- --- --- --- --- --- --- --- --- --- --- --- --- --- --- --- --- --- --- --- --- --- --- --- --- --- --- --- --- --- --- --- --- --- --- --- --- --- --- --- --- --- --- --- --- --- --- --- --- --- --- --- --- --- --- --- --- --- --- --- --- --- --- --- --- --- --- --- --- --- --- --- --- --- --- --- --- --- --- --- --- --- --- --- --- --- --- --- --- --- --- --- --- --- --- --- --- --- --- --- --- --- --- --- --- --- --- --- --- --- --- --- --- --- --- --- --- --- --- --- --- --- --- --- --- --- --- --- --- --- --- --- --- --- --- --- --- --- --- --- --- --- --- --- --- --- --- --- --- --- --- --- --- --- --- --- --- --- --- --- --- --- --- --- --- --- --- --- --- --- --- --- --- --- --- --- --- --- --- --- --- --- --- --- --- --- --- --- --- --- --- --- --- --- --- --- --- --- --- --- --- --- --- --- --- --- --- --- --- --- --- --- --- --- --- --- --- --- --- --- -ct gga tcg gat gtc acc gca gta ata ttg ttg att att tct gac atc gac gta tta tat agt ttt tta att cca tat ctt ttt t

>ON622712.1_Monkeypox_virus_isolate_MPX/UZ_REGA_1/Belgium/2022_complete_genome

ttt tat atc act acg gac ata aac cat tgt ata ttt ttt atg ttt att agt gta cac att ttg gaa gta agt tc- --- --- --- --- --- --- --- --- --- --- --- --- --- --- --- --- --- --- --- --- --- --- --- --- --- --- --- --- --- --- --- --- --- --- --- --- --- --- --- --- --- --- --- --- --- --- --- --- --- --- --- --- --- --- --- --- --- --- --- --- --- --- --- --- --- --- --- --- --- --- --- --- --- --- --- --- --- --- --- --- --- --- --- --- --- --- --- --- --- --- --- --- --- --- --- --- --- --- --- --- --- --- --- --- --- --- --- --- --- --- --- --- --- --- --- --- --- --- --- --- --- --- --- --- --- --- --- --- --- --- --- --- --- --- --- --- --- --- --- --- --- --- --- --- --- --- --- --- --- --- --- --- --- --- --- --- --- --- --- --- --- --- --- --- --- --- --- --- --- --- --- --- --- --- --- --- --- --- --- --- --- --- --- --- --- --- --- --- --- --- --- --- --- --- --- --- --- --- --- --- --- --- --- --- --- --- --- --- --- --- --- --- --- --- --- --- --- --- --- --- --- --- --- --- --- --- --- --- --- --- --- --- --- --- --- --- --- --- --- --- --- --- --- --- --- --- --- --- --- --- --- --- --- --- --- --- --- --- --- --- --- --- --- --- --- --- --- --- --- --- --- --- --- --- --- --- --- --- --- --- --- --- --- --- --- --- --- --- --- --- --- --- --- --- --- --- --- --- --- --- --- --- --- --- --- --- --- --- --- --- --- --- --- --- --- --- --- --- --- --- --- --- --- --- --- --- --- --- --- --- --- --- --- --- --- --- --- --- --- --- --- --- --- --- --- --- --- --- --- --- --- --- --- --- --- --- --- --- --- --- --- --- --- --- --- --- --- --- --- --- --- --- --- --- --- --- --- --- --- --- --- --- --- --- --- --- --- --- --- --- --- --- --- --- --- --- --- --- --- --- --- --- --- --- --- --- --- --- --- --- --- --- --- --- --- --- --- --- --- --- --- --- --- --- --- --- --- --- --- --- --- --- --- --- --- --- --- --- --- --- --- --- --- --- --- --- --- --- --- --- --- --- --- --- --- --- --- --- --- --- --- --- --- --- --- --- --- --- --- --- --- --- --- --- --- --- --- --- --- --- --- --- --- --- --- --- --- --- --- --- --- --- --- --- --- --- --- --- --- --- --- --- --- --- --- --- --- --- --- --- --- --- --- --- --- --- --- --- --- --- --- --- --- --- --- --- --- --- --- --- --- --- --- --- --- --- --- --- --- --- --- --- --- --- --- --- --- --- --- --- --- --- --- --- --- --- --- --- --- --- --- --- --- --- --- --- --- --- --- --- --- --- --- --- --- --- --- --- --- --- --- --- --- --- --- --- --- --- --- --- --- --- --- --- --- --- --- --- --- --- --- --- --- --- --- --- --- --- --- --- --- --- --- --- --- --- --- --- --- --- --- --- --- --- --- --- --- --- --- --- --- --- --- --- --- --- --- --- --- --- --- --- --- --- --- --- --- --- --- --- --- --- -ct gga tcg gat gtc acc gca gta ata ttg ttg att att tct gac atc gac gta tta tat agt ttt tta att cca tat ctt ttt t

>ON609725.2_Monkeypox_virus_isolate_SLO_complete_genome

ttt tat atc act acg gac ata aac cat tgt ata ttt ttt atg ttt att agt gta cac att ttg gaa gta agt tc- --- --- --- --- --- --- --- --- --- --- --- --- --- --- --- --- --- --- --- --- --- --- --- --- --- --- --- --- --- --- --- --- --- --- --- --- --- --- --- --- --- --- --- --- --- --- --- --- --- --- --- --- --- --- --- --- --- --- --- --- --- --- --- --- --- --- --- --- --- --- --- --- --- --- --- --- --- --- --- --- --- --- --- --- --- --- --- --- --- --- --- --- --- --- --- --- --- --- --- --- --- --- --- --- --- --- --- --- --- --- --- --- --- --- --- --- --- --- --- --- --- --- --- --- --- --- --- --- --- --- --- --- --- --- --- --- --- --- --- --- --- --- --- --- --- --- --- --- --- --- --- --- --- --- --- --- --- --- --- --- --- --- --- --- --- --- --- --- --- --- --- --- --- --- --- --- --- --- --- --- --- --- --- --- --- --- --- --- --- --- --- --- --- --- --- --- --- --- --- --- --- --- --- --- --- --- --- --- --- --- --- --- --- --- --- --- --- --- --- --- --- --- --- --- --- --- --- --- --- --- --- --- --- --- --- --- --- --- --- --- --- --- --- --- --- --- --- --- --- --- --- --- --- --- --- --- --- --- --- --- --- --- --- --- --- --- --- --- --- --- --- --- --- --- --- --- --- --- --- --- --- --- --- --- --- --- --- --- --- --- --- --- --- --- --- --- --- --- --- --- --- --- --- --- --- --- --- --- --- --- --- --- --- --- --- --- --- --- --- --- --- --- --- --- --- --- --- --- --- --- --- --- --- --- --- --- --- --- --- --- --- --- --- --- --- --- --- --- --- --- --- --- --- --- --- --- --- --- --- --- --- --- --- --- --- --- --- --- --- --- --- --- --- --- --- --- --- --- --- --- --- --- --- --- --- --- --- --- --- --- --- --- --- --- --- --- --- --- --- --- --- --- --- --- --- --- --- --- --- --- --- --- --- --- --- --- --- --- --- --- --- --- --- --- --- --- --- --- --- --- --- --- --- --- --- --- --- --- --- --- --- --- --- --- --- --- --- --- --- --- --- --- --- --- --- --- --- --- --- --- --- --- --- --- --- --- --- --- --- --- --- --- --- --- --- --- --- --- --- --- --- --- --- --- --- --- --- --- --- --- --- --- --- --- --- --- --- --- --- --- --- --- --- --- --- --- --- --- --- --- --- --- --- --- --- --- --- --- --- --- --- --- --- --- --- --- --- --- --- --- --- --- --- --- --- --- --- --- --- --- --- --- --- --- --- --- --- --- --- --- --- --- --- --- --- --- --- --- --- --- --- --- --- --- --- --- --- --- --- --- --- --- --- --- --- --- --- --- --- --- --- --- --- --- --- --- --- --- --- --- --- --- --- --- --- --- --- --- --- --- --- --- --- --- --- --- --- --- --- --- --- --- --- --- --- --- --- --- --- --- --- --- --- --- --- --- --- --- --- --- --- --- --- --- --- --- --- --- --- --- --- --- --- --- --- --- --- --- --- --- --- --- -ct gga tcg gat gtc acc gca gta ata ttg ttg att att tct gac atc gac gta tta tat agt ttt tta att cca tat ctt ttt t

>ON622713.1_Monkeypox_virus_isolate_MPX/UZ_REGA_2/Belgium/2022_complete_genome

ttt tat atc act acg gac ata aac cat tgt ata ttt ttt atg ttt att agt gta cac att ttg gaa gta agt tc- --- --- --- --- --- --- --- --- --- --- --- --- --- --- --- --- --- --- --- --- --- --- --- --- --- --- --- --- --- --- --- --- --- --- --- --- --- --- --- --- --- --- --- --- --- --- --- --- --- --- --- --- --- --- --- --- --- --- --- --- --- --- --- --- --- --- --- --- --- --- --- --- --- --- --- --- --- --- --- --- --- --- --- --- --- --- --- --- --- --- --- --- --- --- --- --- --- --- --- --- --- --- --- --- --- --- --- --- --- --- --- --- --- --- --- --- --- --- --- --- --- --- --- --- --- --- --- --- --- --- --- --- --- --- --- --- --- --- --- --- --- --- --- --- --- --- --- --- --- --- --- --- --- --- --- --- --- --- --- --- --- --- --- --- --- --- --- --- --- --- --- --- --- --- --- --- --- --- --- --- --- --- --- --- --- --- --- --- --- --- --- --- --- --- --- --- --- --- --- --- --- --- --- --- --- --- --- --- --- --- --- --- --- --- --- --- --- --- --- --- --- --- --- --- --- --- --- --- --- --- --- --- --- --- --- --- --- --- --- --- --- --- --- --- --- --- --- --- --- --- --- --- --- --- --- --- --- --- --- --- --- --- --- --- --- --- --- --- --- --- --- --- --- --- --- --- --- --- --- --- --- --- --- --- --- --- --- --- --- --- --- --- --- --- --- --- --- --- --- --- --- --- --- --- --- --- --- --- --- --- --- --- --- --- --- --- --- --- --- --- --- --- --- --- --- --- --- --- --- --- --- --- --- --- --- --- --- --- --- --- --- --- --- --- --- --- --- --- --- --- --- --- --- --- --- --- --- --- --- --- --- --- --- --- --- --- --- --- --- --- --- --- --- --- --- --- --- --- --- --- --- --- --- --- --- --- --- --- --- --- --- --- --- --- --- --- --- --- --- --- --- --- --- --- --- --- --- --- --- --- --- --- --- --- --- --- --- --- --- --- --- --- --- --- --- --- --- --- --- --- --- --- --- --- --- --- --- --- --- --- --- --- --- --- --- --- --- --- --- --- --- --- --- --- --- --- --- --- --- --- --- --- --- --- --- --- --- --- --- --- --- --- --- --- --- --- --- --- --- --- --- --- --- --- --- --- --- --- --- --- --- --- --- --- --- --- --- --- --- --- --- --- --- --- --- --- --- --- --- --- --- --- --- --- --- --- --- --- --- --- --- --- --- --- --- --- --- --- --- --- --- --- --- --- --- --- --- --- --- --- --- --- --- --- --- --- --- --- --- --- --- --- --- --- --- --- --- --- --- --- --- --- --- --- --- --- --- --- --- --- --- --- --- --- --- --- --- --- --- --- --- --- --- --- --- --- --- --- --- --- --- --- --- --- --- --- --- --- --- --- --- --- --- --- --- --- --- --- --- --- --- --- --- --- --- --- --- --- --- --- --- --- --- --- --- --- --- --- --- --- --- --- --- --- --- --- --- --- --- --- --- --- --- --- --- --- --- --- --- --- --- --- -ct gga tcg gat gtc acc gca gta ata ttg ttg att att tct gac atc gac gta tta tat agt ttt tta att cca tat ctt ttt t

>ON676707.1_Monkeypox_virus_isolate_MPXV_USA_2021_TX_complete_genome

ttt tat atc act acg gac ata aac cat tgt ata ttt ttt atg ttt att agt gta cac att ttg gaa gta agt tc- --- --- --- --- --- --- --- --- --- --- --- --- --- --- --- --- --- --- --- --- --- --- --- --- --- --- --- --- --- --- --- --- --- --- --- --- --- --- --- --- --- --- --- --- --- --- --- --- --- --- --- --- --- --- --- --- --- --- --- --- --- --- --- --- --- --- --- --- --- --- --- --- --- --- --- --- --- --- --- --- --- --- --- --- --- --- --- --- --- --- --- --- --- --- --- --- --- --- --- --- --- --- --- --- --- --- --- --- --- --- --- --- --- --- --- --- --- --- --- --- --- --- --- --- --- --- --- --- --- --- --- --- --- --- --- --- --- --- --- --- --- --- --- --- --- --- --- --- --- --- --- --- --- --- --- --- --- --- --- --- --- --- --- --- --- --- --- --- --- --- --- --- --- --- --- --- --- --- --- --- --- --- --- --- --- --- --- --- --- --- --- --- --- --- --- --- --- --- --- --- --- --- --- --- --- --- --- --- --- --- --- --- --- --- --- --- --- --- --- --- --- --- --- --- --- --- --- --- --- --- --- --- --- --- --- --- --- --- --- --- --- --- --- --- --- --- --- --- --- --- --- --- --- --- --- --- --- --- --- --- --- --- --- --- --- --- --- --- --- --- --- --- --- --- --- --- --- --- --- --- --- --- --- --- --- --- --- --- --- --- --- --- --- --- --- --- --- --- --- --- --- --- --- --- --- --- --- --- --- --- --- --- --- --- --- --- --- --- --- --- --- --- --- --- --- --- --- --- --- --- --- --- --- --- --- --- --- --- --- --- --- --- --- --- --- --- --- --- --- --- --- --- --- --- --- --- --- --- --- --- --- --- --- --- --- --- --- --- --- --- --- --- --- --- --- --- --- --- --- --- --- --- --- --- --- --- --- --- --- --- --- --- --- --- --- --- --- --- --- --- --- --- --- --- --- --- --- --- --- --- --- --- --- --- --- --- --- --- --- --- --- --- --- --- --- --- --- --- --- --- --- --- --- --- --- --- --- --- --- --- --- --- --- --- --- --- --- --- --- --- --- --- --- --- --- --- --- --- --- --- --- --- --- --- --- --- --- --- --- --- --- --- --- --- --- --- --- --- --- --- --- --- --- --- --- --- --- --- --- --- --- --- --- --- --- --- --- --- --- --- --- --- --- --- --- --- --- --- --- --- --- --- --- --- --- --- --- --- --- --- --- --- --- --- --- --- --- --- --- --- --- --- --- --- --- --- --- --- --- --- --- --- --- --- --- --- --- --- --- --- --- --- --- --- --- --- --- --- --- --- --- --- --- --- --- --- --- --- --- --- --- --- --- --- --- --- --- --- --- --- --- --- --- --- --- --- --- --- --- --- --- --- --- --- --- --- --- --- --- --- --- --- --- --- --- --- --- --- --- --- --- --- --- --- --- --- --- --- --- --- --- --- --- --- --- --- --- --- --- --- --- --- --- --- --- --- --- --- --- --- --- --- --- --- --- --- --- --- --- --- --- --- -ct gga tcg gat gtc acc gca gta ata ttg ttg att att tct gac atc gac gta tta tat agt ttt tta att cca tat ctt ttt t

>MT903345.1_Monkeypox_virus_isolate_MPXV-UK_P3

ttt tat atc act acg gac ata aac cat tgt ata ttt ttt atg ttt att agt gta cac att ttg gaa gta agt tc- --- --- --- --- --- --- --- --- --- --- --- --- --- --- --- --- --- --- --- --- --- --- --- --- --- --- --- --- --- --- --- --- --- --- --- --- --- --- --- --- --- --- --- --- --- --- --- --- --- --- --- --- --- --- --- --- --- --- --- --- --- --- --- --- --- --- --- --- --- --- --- --- --- --- --- --- --- --- --- --- --- --- --- --- --- --- --- --- --- --- --- --- --- --- --- --- --- --- --- --- --- --- --- --- --- --- --- --- --- --- --- --- --- --- --- --- --- --- --- --- --- --- --- --- --- --- --- --- --- --- --- --- --- --- --- --- --- --- --- --- --- --- --- --- --- --- --- --- --- --- --- --- --- --- --- --- --- --- --- --- --- --- --- --- --- --- --- --- --- --- --- --- --- --- --- --- --- --- --- --- --- --- --- --- --- --- --- --- --- --- --- --- --- --- --- --- --- --- --- --- --- --- --- --- --- --- --- --- --- --- --- --- --- --- --- --- --- --- --- --- --- --- --- --- --- --- --- --- --- --- --- --- --- --- --- --- --- --- --- --- --- --- --- --- --- --- --- --- --- --- --- --- --- --- --- --- --- --- --- --- --- --- --- --- --- --- --- --- --- --- --- --- --- --- --- --- --- --- --- --- --- --- --- --- --- --- --- --- --- --- --- --- --- --- --- --- --- --- --- --- --- --- --- --- --- --- --- --- --- --- --- --- --- --- --- --- --- --- --- --- --- --- --- --- --- --- --- --- --- --- --- --- --- --- --- --- --- --- --- --- --- --- --- --- --- --- --- --- --- --- --- --- --- --- --- --- --- --- --- --- --- --- --- --- --- --- --- --- --- --- --- --- --- --- --- --- --- --- --- --- --- --- --- --- --- --- --- --- --- --- --- --- --- --- --- --- --- --- --- --- --- --- --- --- --- --- --- --- --- --- --- --- --- --- --- --- --- --- --- --- --- --- --- --- --- --- --- --- --- --- --- --- --- --- --- --- --- --- --- --- --- --- --- --- --- --- --- --- --- --- --- --- --- --- --- --- --- --- --- --- --- --- --- --- --- --- --- --- --- --- --- --- --- --- --- --- --- --- --- --- --- --- --- --- --- --- --- --- --- --- --- --- --- --- --- --- --- --- --- --- --- --- --- --- --- --- --- --- --- --- --- --- --- --- --- --- --- --- --- --- --- --- --- --- --- --- --- --- --- --- --- --- --- --- --- --- --- --- --- --- --- --- --- --- --- --- --- --- --- --- --- --- --- --- --- --- --- --- --- --- --- --- --- --- --- --- --- --- --- --- --- --- --- --- --- --- --- --- --- --- --- --- --- --- --- --- --- --- --- --- --- --- --- --- --- --- --- --- --- --- --- --- --- --- --- --- --- --- --- --- --- --- --- --- --- --- --- --- --- --- --- --- --- --- --- --- --- --- --- --- --- --- --- --- --- --- --- --- --- --- --- --- --- --- --- --- --- --- --- --- --- --- -ct gga tcg gat gtc acc gca gta ata ttg ttg att att tct gac atc gac gta tta tat agt ttt tta att cca tat ctt ttt t

>MT903344.1_Monkeypox_virus_isolate_MPXV-UK_P2

ttt tat atc act acg gac ata aac cat tgt ata ttt ttt atg ttt att agt gta cac att ttg gaa gta agt tc- --- --- --- --- --- --- --- --- --- --- --- --- --- --- --- --- --- --- --- --- --- --- --- --- --- --- --- --- --- --- --- --- --- --- --- --- --- --- --- --- --- --- --- --- --- --- --- --- --- --- --- --- --- --- --- --- --- --- --- --- --- --- --- --- --- --- --- --- --- --- --- --- --- --- --- --- --- --- --- --- --- --- --- --- --- --- --- --- --- --- --- --- --- --- --- --- --- --- --- --- --- --- --- --- --- --- --- --- --- --- --- --- --- --- --- --- --- --- --- --- --- --- --- --- --- --- --- --- --- --- --- --- --- --- --- --- --- --- --- --- --- --- --- --- --- --- --- --- --- --- --- --- --- --- --- --- --- --- --- --- --- --- --- --- --- --- --- --- --- --- --- --- --- --- --- --- --- --- --- --- --- --- --- --- --- --- --- --- --- --- --- --- --- --- --- --- --- --- --- --- --- --- --- --- --- --- --- --- --- --- --- --- --- --- --- --- --- --- --- --- --- --- --- --- --- --- --- --- --- --- --- --- --- --- --- --- --- --- --- --- --- --- --- --- --- --- --- --- --- --- --- --- --- --- --- --- --- --- --- --- --- --- --- --- --- --- --- --- --- --- --- --- --- --- --- --- --- --- --- --- --- --- --- --- --- --- --- --- --- --- --- --- --- --- --- --- --- --- --- --- --- --- --- --- --- --- --- --- --- --- --- --- --- --- --- --- --- --- --- --- --- --- --- --- --- --- --- --- --- --- --- --- --- --- --- --- --- --- --- --- --- --- --- --- --- --- --- --- --- --- --- --- --- --- --- --- --- --- --- --- --- --- --- --- --- --- --- --- --- --- --- --- --- --- --- --- --- --- --- --- --- --- --- --- --- --- --- --- --- --- --- --- --- --- --- --- --- --- --- --- --- --- --- --- --- --- --- --- --- --- --- --- --- --- --- --- --- --- --- --- --- --- --- --- --- --- --- --- --- --- --- --- --- --- --- --- --- --- --- --- --- --- --- --- --- --- --- --- --- --- --- --- --- --- --- --- --- --- --- --- --- --- --- --- --- --- --- --- --- --- --- --- --- --- --- --- --- --- --- --- --- --- --- --- --- --- --- --- --- --- --- --- --- --- --- --- --- --- --- --- --- --- --- --- --- --- --- --- --- --- --- --- --- --- --- --- --- --- --- --- --- --- --- --- --- --- --- --- --- --- --- --- --- --- --- --- --- --- --- --- --- --- --- --- --- --- --- --- --- --- --- --- --- --- --- --- --- --- --- --- --- --- --- --- --- --- --- --- --- --- --- --- --- --- --- --- --- --- --- --- --- --- --- --- --- --- --- --- --- --- --- --- --- --- --- --- --- --- --- --- --- --- --- --- --- --- --- --- --- --- --- --- --- --- --- --- --- --- --- --- --- --- --- --- --- --- --- --- --- --- --- --- --- --- --- --- --- --- --- --- --- --- --- --- --- --- --- --- --- --- --- --- -ct gga tcg gat gtc acc gca gta ata ttg ttg att att tct gac atc gac gta tta tat agt ttt tta att cca tat ctt ttt t

>MT903342.1_Monkeypox_virus_isolate_MPXV-Singapore

ttt tat atc act acg gac ata aac cat tgt ata ttt ttt atg ttt att agt gta cac att ttg gaa gta agt tc- --- --- --- --- --- --- --- --- --- --- --- --- --- --- --- --- --- --- --- --- --- --- --- --- --- --- --- --- --- --- --- --- --- --- --- --- --- --- --- --- --- --- --- --- --- --- --- --- --- --- --- --- --- --- --- --- --- --- --- --- --- --- --- --- --- --- --- --- --- --- --- --- --- --- --- --- --- --- --- --- --- --- --- --- --- --- --- --- --- --- --- --- --- --- --- --- --- --- --- --- --- --- --- --- --- --- --- --- --- --- --- --- --- --- --- --- --- --- --- --- --- --- --- --- --- --- --- --- --- --- --- --- --- --- --- --- --- --- --- --- --- --- --- --- --- --- --- --- --- --- --- --- --- --- --- --- --- --- --- --- --- --- --- --- --- --- --- --- --- --- --- --- --- --- --- --- --- --- --- --- --- --- --- --- --- --- --- --- --- --- --- --- --- --- --- --- --- --- --- --- --- --- --- --- --- --- --- --- --- --- --- --- --- --- --- --- --- --- --- --- --- --- --- --- --- --- --- --- --- --- --- --- --- --- --- --- --- --- --- --- --- --- --- --- --- --- --- --- --- --- --- --- --- --- --- --- --- --- --- --- --- --- --- --- --- --- --- --- --- --- --- --- --- --- --- --- --- --- --- --- --- --- --- --- --- --- --- --- --- --- --- --- --- --- --- --- --- --- --- --- --- --- --- --- --- --- --- --- --- --- --- --- --- --- --- --- --- --- --- --- --- --- --- --- --- --- --- --- --- --- --- --- --- --- --- --- --- --- --- --- --- --- --- --- --- --- --- --- --- --- --- --- --- --- --- --- --- --- --- --- --- --- --- --- --- --- --- --- --- --- --- --- --- --- --- --- --- --- --- --- --- --- --- --- --- --- --- --- --- --- --- --- --- --- --- --- --- --- --- --- --- --- --- --- --- --- --- --- --- --- --- --- --- --- --- --- --- --- --- --- --- --- --- --- --- --- --- --- --- --- --- --- --- --- --- --- --- --- --- --- --- --- --- --- --- --- --- --- --- --- --- --- --- --- --- --- --- --- --- --- --- --- --- --- --- --- --- --- --- --- --- --- --- --- --- --- --- --- --- --- --- --- --- --- --- --- --- --- --- --- --- --- --- --- --- --- --- --- --- --- --- --- --- --- --- --- --- --- --- --- --- --- --- --- --- --- --- --- --- --- --- --- --- --- --- --- --- --- --- --- --- --- --- --- --- --- --- --- --- --- --- --- --- --- --- --- --- --- --- --- --- --- --- --- --- --- --- --- --- --- --- --- --- --- --- --- --- --- --- --- --- --- --- --- --- --- --- --- --- --- --- --- --- --- --- --- --- --- --- --- --- --- --- --- --- --- --- --- --- --- --- --- --- --- --- --- --- --- --- --- --- --- --- --- --- --- --- --- --- --- --- --- --- --- --- --- --- --- --- --- --- --- --- --- --- --- --- --- --- --- --- --- --- --- --- --- --- --- --- --- --- --- -ct gga tcg gat gtc acc gca gta ata ttg ttg att att tct gac atc gac gta tta tat agt ttt tta att cca tat ctt ttt t

>ON676708.1_Monkeypox_virus_isolate_MPXV_USA_2021_MD_complete_genome

ttt tat atc act acg gac ata aac cat tgt ata ttt ttt atg ttt att agt gta cac att ttg gaa gta agt tc- --- --- --- --- --- --- --- --- --- --- --- --- --- --- --- --- --- --- --- --- --- --- --- --- --- --- --- --- --- --- --- --- --- --- --- --- --- --- --- --- --- --- --- --- --- --- --- --- --- --- --- --- --- --- --- --- --- --- --- --- --- --- --- --- --- --- --- --- --- --- --- --- --- --- --- --- --- --- --- --- --- --- --- --- --- --- --- --- --- --- --- --- --- --- --- --- --- --- --- --- --- --- --- --- --- --- --- --- --- --- --- --- --- --- --- --- --- --- --- --- --- --- --- --- --- --- --- --- --- --- --- --- --- --- --- --- --- --- --- --- --- --- --- --- --- --- --- --- --- --- --- --- --- --- --- --- --- --- --- --- --- --- --- --- --- --- --- --- --- --- --- --- --- --- --- --- --- --- --- --- --- --- --- --- --- --- --- --- --- --- --- --- --- --- --- --- --- --- --- --- --- --- --- --- --- --- --- --- --- --- --- --- --- --- --- --- --- --- --- --- --- --- --- --- --- --- --- --- --- --- --- --- --- --- --- --- --- --- --- --- --- --- --- --- --- --- --- --- --- --- --- --- --- --- --- --- --- --- --- --- --- --- --- --- --- --- --- --- --- --- --- --- --- --- --- --- --- --- --- --- --- --- --- --- --- --- --- --- --- --- --- --- --- --- --- --- --- --- --- --- --- --- --- --- --- --- --- --- --- --- --- --- --- --- --- --- --- --- --- --- --- --- --- --- --- --- --- --- --- --- --- --- --- --- --- --- --- --- --- --- --- --- --- --- --- --- --- --- --- --- --- --- --- --- --- --- --- --- --- --- --- --- --- --- --- --- --- --- --- --- --- --- --- --- --- --- --- --- --- --- --- --- --- --- --- --- --- --- --- --- --- --- --- --- --- --- --- --- --- --- --- --- --- --- --- --- --- --- --- --- --- --- --- --- --- --- --- --- --- --- --- --- --- --- --- --- --- --- --- --- --- --- --- --- --- --- --- --- --- --- --- --- --- --- --- --- --- --- --- --- --- --- --- --- --- --- --- --- --- --- --- --- --- --- --- --- --- --- --- --- --- --- --- --- --- --- --- --- --- --- --- --- --- --- --- --- --- --- --- --- --- --- --- --- --- --- --- --- --- --- --- --- --- --- --- --- --- --- --- --- --- --- --- --- --- --- --- --- --- --- --- --- --- --- --- --- --- --- --- --- --- --- --- --- --- --- --- --- --- --- --- --- --- --- --- --- --- --- --- --- --- --- --- --- --- --- --- --- --- --- --- --- --- --- --- --- --- --- --- --- --- --- --- --- --- --- --- --- --- --- --- --- --- --- --- --- --- --- --- --- --- --- --- --- --- --- --- --- --- --- --- --- --- --- --- --- --- --- --- --- --- --- --- --- --- --- --- --- --- --- --- --- --- --- --- --- --- --- --- --- --- --- --- --- --- --- --- --- --- --- --- --- --- --- --- --- --- --- --- --- --- --- -ct gga tcg gat gtc acc gca gta ata ttg ttg att att tct gac atc gac gta tta tat agt ttt tta att cca tat ctt ttt t

>ON585033.1_Monkeypox_virus_isolate_Monkeypox/PT0006/2022_complete_genome

ttt tat atc act acg gac ata aac cat tgt ata ttt ttt atg ttt att agt gta cac att ttg gaa gta agt tc- --- --- --- --- --- --- --- --- --- --- --- --- --- --- --- --- --- --- --- --- --- --- --- --- --- --- --- --- --- --- --- --- --- --- --- --- --- --- --- --- --- --- --- --- --- --- --- --- --- --- --- --- --- --- --- --- --- --- --- --- --- --- --- --- --- --- --- --- --- --- --- --- --- --- --- --- --- --- --- --- --- --- --- --- --- --- --- --- --- --- --- --- --- --- --- --- --- --- --- --- --- --- --- --- --- --- --- --- --- --- --- --- --- --- --- --- --- --- --- --- --- --- --- --- --- --- --- --- --- --- --- --- --- --- --- --- --- --- --- --- --- --- --- --- --- --- --- --- --- --- --- --- --- --- --- --- --- --- --- --- --- --- --- --- --- --- --- --- --- --- --- --- --- --- --- --- --- --- --- --- --- --- --- --- --- --- --- --- --- --- --- --- --- --- --- --- --- --- --- --- --- --- --- --- --- --- --- --- --- --- --- --- --- --- --- --- --- --- --- --- --- --- --- --- --- --- --- --- --- --- --- --- --- --- --- --- --- --- --- --- --- --- --- --- --- --- --- --- --- --- --- --- --- --- --- --- --- --- --- --- --- --- --- --- --- --- --- --- --- --- --- --- --- --- --- --- --- --- --- --- --- --- --- --- --- --- --- --- --- --- --- --- --- --- --- --- --- --- --- --- --- --- --- --- --- --- --- --- --- --- --- --- --- --- --- --- --- --- --- --- --- --- --- --- --- --- --- --- --- --- --- --- --- --- --- --- --- --- --- --- --- --- --- --- --- --- --- --- --- --- --- --- --- --- --- --- --- --- --- --- --- --- --- --- --- --- --- --- --- --- --- --- --- --- --- --- --- --- --- --- --- --- --- --- --- --- --- --- --- --- --- --- --- --- --- --- --- --- --- --- --- --- --- --- --- --- --- --- --- --- --- --- --- --- --- --- --- --- --- --- --- --- --- --- --- --- --- --- --- --- --- --- --- --- --- --- --- --- --- --- --- --- --- --- --- --- --- --- --- --- --- --- --- --- --- --- --- --- --- --- --- --- --- --- --- --- --- --- --- --- --- --- --- --- --- --- --- --- --- --- --- --- --- --- --- --- --- --- --- --- --- --- --- --- --- --- --- --- --- --- --- --- --- --- --- --- --- --- --- --- --- --- --- --- --- --- --- --- --- --- --- --- --- --- --- --- --- --- --- --- --- --- --- --- --- --- --- --- --- --- --- --- --- --- --- --- --- --- --- --- --- --- --- --- --- --- --- --- --- --- --- --- --- --- --- --- --- --- --- --- --- --- --- --- --- --- --- --- --- --- --- --- --- --- --- --- --- --- --- --- --- --- --- --- --- --- --- --- --- --- --- --- --- --- --- --- --- --- --- --- --- --- --- --- --- --- --- --- --- --- --- --- --- --- --- --- --- --- --- --- --- --- --- --- --- --- --- --- --- --- --- --- --- --- --- --- --- --- --- --- --- --- -ct gga tcg gat gtc acc gca gta ata ttg ttg att att tct gac atc gac gta tta tat agt ttt tta att cca tat ctt ttt t

>ON585035.1_Monkeypox_virus_isolate_Monkeypox/PT0009/2022_complete_genome

ttt tat atc act acg gac ata aac cat tgt ata ttt ttt atg ttt att agt gta cac att ttg gaa gta agt tc- --- --- --- --- --- --- --- --- --- --- --- --- --- --- --- --- --- --- --- --- --- --- --- --- --- --- --- --- --- --- --- --- --- --- --- --- --- --- --- --- --- --- --- --- --- --- --- --- --- --- --- --- --- --- --- --- --- --- --- --- --- --- --- --- --- --- --- --- --- --- --- --- --- --- --- --- --- --- --- --- --- --- --- --- --- --- --- --- --- --- --- --- --- --- --- --- --- --- --- --- --- --- --- --- --- --- --- --- --- --- --- --- --- --- --- --- --- --- --- --- --- --- --- --- --- --- --- --- --- --- --- --- --- --- --- --- --- --- --- --- --- --- --- --- --- --- --- --- --- --- --- --- --- --- --- --- --- --- --- --- --- --- --- --- --- --- --- --- --- --- --- --- --- --- --- --- --- --- --- --- --- --- --- --- --- --- --- --- --- --- --- --- --- --- --- --- --- --- --- --- --- --- --- --- --- --- --- --- --- --- --- --- --- --- --- --- --- --- --- --- --- --- --- --- --- --- --- --- --- --- --- --- --- --- --- --- --- --- --- --- --- --- --- --- --- --- --- --- --- --- --- --- --- --- --- --- --- --- --- --- --- --- --- --- --- --- --- --- --- --- --- --- --- --- --- --- --- --- --- --- --- --- --- --- --- --- --- --- --- --- --- --- --- --- --- --- --- --- --- --- --- --- --- --- --- --- --- --- --- --- --- --- --- --- --- --- --- --- --- --- --- --- --- --- --- --- --- --- --- --- --- --- --- --- --- --- --- --- --- --- --- --- --- --- --- --- --- --- --- --- --- --- --- --- --- --- --- --- --- --- --- --- --- --- --- --- --- --- --- --- --- --- --- --- --- --- --- --- --- --- --- --- --- --- --- --- --- --- --- --- --- --- --- --- --- --- --- --- --- --- --- --- --- --- --- --- --- --- --- --- --- --- --- --- --- --- --- --- --- --- --- --- --- --- --- --- --- --- --- --- --- --- --- --- --- --- --- --- --- --- --- --- --- --- --- --- --- --- --- --- --- --- --- --- --- --- --- --- --- --- --- --- --- --- --- --- --- --- --- --- --- --- --- --- --- --- --- --- --- --- --- --- --- --- --- --- --- --- --- --- --- --- --- --- --- --- --- --- --- --- --- --- --- --- --- --- --- --- --- --- --- --- --- --- --- --- --- --- --- --- --- --- --- --- --- --- --- --- --- --- --- --- --- --- --- --- --- --- --- --- --- --- --- --- --- --- --- --- --- --- --- --- --- --- --- --- --- --- --- --- --- --- --- --- --- --- --- --- --- --- --- --- --- --- --- --- --- --- --- --- --- --- --- --- --- --- --- --- --- --- --- --- --- --- --- --- --- --- --- --- --- --- --- --- --- --- --- --- --- --- --- --- --- --- --- --- --- --- --- --- --- --- --- --- --- --- --- --- --- --- --- --- --- --- --- --- --- --- --- --- --- --- --- --- --- --- --- --- --- --- --- --- -ct gga tcg gat gtc acc gca gta ata ttg ttg att att tct gac atc gac gta tta tat agt ttt tta att cca tat ctt ttt t

>ON649725.1_Monkeypox_virus_isolate_Monkeypox/PT0011/2022_complete_genome

ttt tat atc act acg gac ata aac cat tgt ata ttt ttt atg ttt att agt gta cac att ttg gaa gta agt tc- --- --- --- --- --- --- --- --- --- --- --- --- --- --- --- --- --- --- --- --- --- --- --- --- --- --- --- --- --- --- --- --- --- --- --- --- --- --- --- --- --- --- --- --- --- --- --- --- --- --- --- --- --- --- --- --- --- --- --- --- --- --- --- --- --- --- --- --- --- --- --- --- --- --- --- --- --- --- --- --- --- --- --- --- --- --- --- --- --- --- --- --- --- --- --- --- --- --- --- --- --- --- --- --- --- --- --- --- --- --- --- --- --- --- --- --- --- --- --- --- --- --- --- --- --- --- --- --- --- --- --- --- --- --- --- --- --- --- --- --- --- --- --- --- --- --- --- --- --- --- --- --- --- --- --- --- --- --- --- --- --- --- --- --- --- --- --- --- --- --- --- --- --- --- --- --- --- --- --- --- --- --- --- --- --- --- --- --- --- --- --- --- --- --- --- --- --- --- --- --- --- --- --- --- --- --- --- --- --- --- --- --- --- --- --- --- --- --- --- --- --- --- --- --- --- --- --- --- --- --- --- --- --- --- --- --- --- --- --- --- --- --- --- --- --- --- --- --- --- --- --- --- --- --- --- --- --- --- --- --- --- --- --- --- --- --- --- --- --- --- --- --- --- --- --- --- --- --- --- --- --- --- --- --- --- --- --- --- --- --- --- --- --- --- --- --- --- --- --- --- --- --- --- --- --- --- --- --- --- --- --- --- --- --- --- --- --- --- --- --- --- --- --- --- --- --- --- --- --- --- --- --- --- --- --- --- --- --- --- --- --- --- --- --- --- --- --- --- --- --- --- --- --- --- --- --- --- --- --- --- --- --- --- --- --- --- --- --- --- --- --- --- --- --- --- --- --- --- --- --- --- --- --- --- --- --- --- --- --- --- --- --- --- --- --- --- --- --- --- --- --- --- --- --- --- --- --- --- --- --- --- --- --- --- --- --- --- --- --- --- --- --- --- --- --- --- --- --- --- --- --- --- --- --- --- --- --- --- --- --- --- --- --- --- --- --- --- --- --- --- --- --- --- --- --- --- --- --- --- --- --- --- --- --- --- --- --- --- --- --- --- --- --- --- --- --- --- --- --- --- --- --- --- --- --- --- --- --- --- --- --- --- --- --- --- --- --- --- --- --- --- --- --- --- --- --- --- --- --- --- --- --- --- --- --- --- --- --- --- --- --- --- --- --- --- --- --- --- --- --- --- --- --- --- --- --- --- --- --- --- --- --- --- --- --- --- --- --- --- --- --- --- --- --- --- --- --- --- --- --- --- --- --- --- --- --- --- --- --- --- --- --- --- --- --- --- --- --- --- --- --- --- --- --- --- --- --- --- --- --- --- --- --- --- --- --- --- --- --- --- --- --- --- --- --- --- --- --- --- --- --- --- --- --- --- --- --- --- --- --- --- --- --- --- --- --- --- --- --- --- --- --- --- --- --- --- --- --- --- --- --- --- --- --- --- --- --- --- --- --- --- --- -ct gga tcg gat gtc acc gca gta ata ttg ttg att att tct gac atc gac gta tta tat agt ttt tta att cca tat ctt ttt t

>ON649724.1_Monkeypox_virus_isolate_Monkeypox/PT0014/2022_complete_genome

ttt tat atc act acg gac ata aac cat tgt ata ttt ttt atg ttt att agt gta cac att ttg gaa gta agt tc- --- --- --- --- --- --- --- --- --- --- --- --- --- --- --- --- --- --- --- --- --- --- --- --- --- --- --- --- --- --- --- --- --- --- --- --- --- --- --- --- --- --- --- --- --- --- --- --- --- --- --- --- --- --- --- --- --- --- --- --- --- --- --- --- --- --- --- --- --- --- --- --- --- --- --- --- --- --- --- --- --- --- --- --- --- --- --- --- --- --- --- --- --- --- --- --- --- --- --- --- --- --- --- --- --- --- --- --- --- --- --- --- --- --- --- --- --- --- --- --- --- --- --- --- --- --- --- --- --- --- --- --- --- --- --- --- --- --- --- --- --- --- --- --- --- --- --- --- --- --- --- --- --- --- --- --- --- --- --- --- --- --- --- --- --- --- --- --- --- --- --- --- --- --- --- --- --- --- --- --- --- --- --- --- --- --- --- --- --- --- --- --- --- --- --- --- --- --- --- --- --- --- --- --- --- --- --- --- --- --- --- --- --- --- --- --- --- --- --- --- --- --- --- --- --- --- --- --- --- --- --- --- --- --- --- --- --- --- --- --- --- --- --- --- --- --- --- --- --- --- --- --- --- --- --- --- --- --- --- --- --- --- --- --- --- --- --- --- --- --- --- --- --- --- --- --- --- --- --- --- --- --- --- --- --- --- --- --- --- --- --- --- --- --- --- --- --- --- --- --- --- --- --- --- --- --- --- --- --- --- --- --- --- --- --- --- --- --- --- --- --- --- --- --- --- --- --- --- --- --- --- --- --- --- --- --- --- --- --- --- --- --- --- --- --- --- --- --- --- --- --- --- --- --- --- --- --- --- --- --- --- --- --- --- --- --- --- --- --- --- --- --- --- --- --- --- --- --- --- --- --- --- --- --- --- --- --- --- --- --- --- --- --- --- --- --- --- --- --- --- --- --- --- --- --- --- --- --- --- --- --- --- --- --- --- --- --- --- --- --- --- --- --- --- --- --- --- --- --- --- --- --- --- --- --- --- --- --- --- --- --- --- --- --- --- --- --- --- --- --- --- --- --- --- --- --- --- --- --- --- --- --- --- --- --- --- --- --- --- --- --- --- --- --- --- --- --- --- --- --- --- --- --- --- --- --- --- --- --- --- --- --- --- --- --- --- --- --- --- --- --- --- --- --- --- --- --- --- --- --- --- --- --- --- --- --- --- --- --- --- --- --- --- --- --- --- --- --- --- --- --- --- --- --- --- --- --- --- --- --- --- --- --- --- --- --- --- --- --- --- --- --- --- --- --- --- --- --- --- --- --- --- --- --- --- --- --- --- --- --- --- --- --- --- --- --- --- --- --- --- --- --- --- --- --- --- --- --- --- --- --- --- --- --- --- --- --- --- --- --- --- --- --- --- --- --- --- --- --- --- --- --- --- --- --- --- --- --- --- --- --- --- --- --- --- --- --- --- --- --- --- --- --- --- --- --- --- --- --- --- --- --- --- --- --- --- --- --- --- --- --- --- -ct gga tcg gat gtc acc gca gta ata ttg ttg att att tct gac atc gac gta tta tat agt ttt tta att cca tat ctt ttt t

>ON649722.1_Monkeypox_virus_isolate_Monkeypox/PT0022/2022_complete_genome

ttt tat atc act acg gac ata aac cat tgt ata ttt ttt atg ttt att agt gta cac att ttg gaa gta agt tc- --- --- --- --- --- --- --- --- --- --- --- --- --- --- --- --- --- --- --- --- --- --- --- --- --- --- --- --- --- --- --- --- --- --- --- --- --- --- --- --- --- --- --- --- --- --- --- --- --- --- --- --- --- --- --- --- --- --- --- --- --- --- --- --- --- --- --- --- --- --- --- --- --- --- --- --- --- --- --- --- --- --- --- --- --- --- --- --- --- --- --- --- --- --- --- --- --- --- --- --- --- --- --- --- --- --- --- --- --- --- --- --- --- --- --- --- --- --- --- --- --- --- --- --- --- --- --- --- --- --- --- --- --- --- --- --- --- --- --- --- --- --- --- --- --- --- --- --- --- --- --- --- --- --- --- --- --- --- --- --- --- --- --- --- --- --- --- --- --- --- --- --- --- --- --- --- --- --- --- --- --- --- --- --- --- --- --- --- --- --- --- --- --- --- --- --- --- --- --- --- --- --- --- --- --- --- --- --- --- --- --- --- --- --- --- --- --- --- --- --- --- --- --- --- --- --- --- --- --- --- --- --- --- --- --- --- --- --- --- --- --- --- --- --- --- --- --- --- --- --- --- --- --- --- --- --- --- --- --- --- --- --- --- --- --- --- --- --- --- --- --- --- --- --- --- --- --- --- --- --- --- --- --- --- --- --- --- --- --- --- --- --- --- --- --- --- --- --- --- --- --- --- --- --- --- --- --- --- --- --- --- --- --- --- --- --- --- --- --- --- --- --- --- --- --- --- --- --- --- --- --- --- --- --- --- --- --- --- --- --- --- --- --- --- --- --- --- --- --- --- --- --- --- --- --- --- --- --- --- --- --- --- --- --- --- --- --- --- --- --- --- --- --- --- --- --- --- --- --- --- --- --- --- --- --- --- --- --- --- --- --- --- --- --- --- --- --- --- --- --- --- --- --- --- --- --- --- --- --- --- --- --- --- --- --- --- --- --- --- --- --- --- --- --- --- --- --- --- --- --- --- --- --- --- --- --- --- --- --- --- --- --- --- --- --- --- --- --- --- --- --- --- --- --- --- --- --- --- --- --- --- --- --- --- --- --- --- --- --- --- --- --- --- --- --- --- --- --- --- --- --- --- --- --- --- --- --- --- --- --- --- --- --- --- --- --- --- --- --- --- --- --- --- --- --- --- --- --- --- --- --- --- --- --- --- --- --- --- --- --- --- --- --- --- --- --- --- --- --- --- --- --- --- --- --- --- --- --- --- --- --- --- --- --- --- --- --- --- --- --- --- --- --- --- --- --- --- --- --- --- --- --- --- --- --- --- --- --- --- --- --- --- --- --- --- --- --- --- --- --- --- --- --- --- --- --- --- --- --- --- --- --- --- --- --- --- --- --- --- --- --- --- --- --- --- --- --- --- --- --- --- --- --- --- --- --- --- --- --- --- --- --- --- --- --- --- --- --- --- --- --- --- --- --- --- --- --- --- --- --- --- --- --- --- --- --- --- --- --- --- --- --- -ct gga tcg gat gtc acc gca gta ata ttg ttg att att tct gac atc gac gta tta tat agt ttt tta att cca tat ctt ttt t

>ON649721.1_Monkeypox_virus_isolate_Monkeypox/PT0021/2022_complete_genome

ttt tat atc act acg gac ata aac cat tgt ata ttt ttt atg ttt att agt gta cac att ttg gaa gta agt tc- --- --- --- --- --- --- --- --- --- --- --- --- --- --- --- --- --- --- --- --- --- --- --- --- --- --- --- --- --- --- --- --- --- --- --- --- --- --- --- --- --- --- --- --- --- --- --- --- --- --- --- --- --- --- --- --- --- --- --- --- --- --- --- --- --- --- --- --- --- --- --- --- --- --- --- --- --- --- --- --- --- --- --- --- --- --- --- --- --- --- --- --- --- --- --- --- --- --- --- --- --- --- --- --- --- --- --- --- --- --- --- --- --- --- --- --- --- --- --- --- --- --- --- --- --- --- --- --- --- --- --- --- --- --- --- --- --- --- --- --- --- --- --- --- --- --- --- --- --- --- --- --- --- --- --- --- --- --- --- --- --- --- --- --- --- --- --- --- --- --- --- --- --- --- --- --- --- --- --- --- --- --- --- --- --- --- --- --- --- --- --- --- --- --- --- --- --- --- --- --- --- --- --- --- --- --- --- --- --- --- --- --- --- --- --- --- --- --- --- --- --- --- --- --- --- --- --- --- --- --- --- --- --- --- --- --- --- --- --- --- --- --- --- --- --- --- --- --- --- --- --- --- --- --- --- --- --- --- --- --- --- --- --- --- --- --- --- --- --- --- --- --- --- --- --- --- --- --- --- --- --- --- --- --- --- --- --- --- --- --- --- --- --- --- --- --- --- --- --- --- --- --- --- --- --- --- --- --- --- --- --- --- --- --- --- --- --- --- --- --- --- --- --- --- --- --- --- --- --- --- --- --- --- --- --- --- --- --- --- --- --- --- --- --- --- --- --- --- --- --- --- --- --- --- --- --- --- --- --- --- --- --- --- --- --- --- --- --- --- --- --- --- --- --- --- --- --- --- --- --- --- --- --- --- --- --- --- --- --- --- --- --- --- --- --- --- --- --- --- --- --- --- --- --- --- --- --- --- --- --- --- --- --- --- --- --- --- --- --- --- --- --- --- --- --- --- --- --- --- --- --- --- --- --- --- --- --- --- --- --- --- --- --- --- --- --- --- --- --- --- --- --- --- --- --- --- --- --- --- --- --- --- --- --- --- --- --- --- --- --- --- --- --- --- --- --- --- --- --- --- --- --- --- --- --- --- --- --- --- --- --- --- --- --- --- --- --- --- --- --- --- --- --- --- --- --- --- --- --- --- --- --- --- --- --- --- --- --- --- --- --- --- --- --- --- --- --- --- --- --- --- --- --- --- --- --- --- --- --- --- --- --- --- --- --- --- --- --- --- --- --- --- --- --- --- --- --- --- --- --- --- --- --- --- --- --- --- --- --- --- --- --- --- --- --- --- --- --- --- --- --- --- --- --- --- --- --- --- --- --- --- --- --- --- --- --- --- --- --- --- --- --- --- --- --- --- --- --- --- --- --- --- --- --- --- --- --- --- --- --- --- --- --- --- --- --- --- --- --- --- --- --- --- --- --- --- --- --- --- --- --- --- --- --- --- --- --- --- --- --- --- --- -ct gga tcg gat gtc acc gca gta ata ttg ttg att att tct gac atc gac gta tta tat agt ttt tta att cca tat ctt ttt t

>ON649720.1_Monkeypox_virus_isolate_Monkeypox/PT0024/2022_complete_genome

ttt tat atc act acg gac ata aac cat tgt ata ttt ttt atg ttt att agt gta cac att ttg gaa gta agt tc- --- --- --- --- --- --- --- --- --- --- --- --- --- --- --- --- --- --- --- --- --- --- --- --- --- --- --- --- --- --- --- --- --- --- --- --- --- --- --- --- --- --- --- --- --- --- --- --- --- --- --- --- --- --- --- --- --- --- --- --- --- --- --- --- --- --- --- --- --- --- --- --- --- --- --- --- --- --- --- --- --- --- --- --- --- --- --- --- --- --- --- --- --- --- --- --- --- --- --- --- --- --- --- --- --- --- --- --- --- --- --- --- --- --- --- --- --- --- --- --- --- --- --- --- --- --- --- --- --- --- --- --- --- --- --- --- --- --- --- --- --- --- --- --- --- --- --- --- --- --- --- --- --- --- --- --- --- --- --- --- --- --- --- --- --- --- --- --- --- --- --- --- --- --- --- --- --- --- --- --- --- --- --- --- --- --- --- --- --- --- --- --- --- --- --- --- --- --- --- --- --- --- --- --- --- --- --- --- --- --- --- --- --- --- --- --- --- --- --- --- --- --- --- --- --- --- --- --- --- --- --- --- --- --- --- --- --- --- --- --- --- --- --- --- --- --- --- --- --- --- --- --- --- --- --- --- --- --- --- --- --- --- --- --- --- --- --- --- --- --- --- --- --- --- --- --- --- --- --- --- --- --- --- --- --- --- --- --- --- --- --- --- --- --- --- --- --- --- --- --- --- --- --- --- --- --- --- --- --- --- --- --- --- --- --- --- --- --- --- --- --- --- --- --- --- --- --- --- --- --- --- --- --- --- --- --- --- --- --- --- --- --- --- --- --- --- --- --- --- --- --- --- --- --- --- --- --- --- --- --- --- --- --- --- --- --- --- --- --- --- --- --- --- --- --- --- --- --- --- --- --- --- --- --- --- --- --- --- --- --- --- --- --- --- --- --- --- --- --- --- --- --- --- --- --- --- --- --- --- --- --- --- --- --- --- --- --- --- --- --- --- --- --- --- --- --- --- --- --- --- --- --- --- --- --- --- --- --- --- --- --- --- --- --- --- --- --- --- --- --- --- --- --- --- --- --- --- --- --- --- --- --- --- --- --- --- --- --- --- --- --- --- --- --- --- --- --- --- --- --- --- --- --- --- --- --- --- --- --- --- --- --- --- --- --- --- --- --- --- --- --- --- --- --- --- --- --- --- --- --- --- --- --- --- --- --- --- --- --- --- --- --- --- --- --- --- --- --- --- --- --- --- --- --- --- --- --- --- --- --- --- --- --- --- --- --- --- --- --- --- --- --- --- --- --- --- --- --- --- --- --- --- --- --- --- --- --- --- --- --- --- --- --- --- --- --- --- --- --- --- --- --- --- --- --- --- --- --- --- --- --- --- --- --- --- --- --- --- --- --- --- --- --- --- --- --- --- --- --- --- --- --- --- --- --- --- --- --- --- --- --- --- --- --- --- --- --- --- --- --- --- --- --- --- --- --- --- --- --- --- --- --- --- --- --- --- --- --- --- --- --- --- -ct gga tcg gat gtc acc gca gta ata ttg ttg att att tct gac atc gac gta tta tat agt ttt tta att cca tat ctt ttt t

>ON649723.1_Monkeypox_virus_isolate_Monkeypox/PT0013/2022_complete_genome

ttt tat atc act acg gac ata aac cat tgt ata ttt ttt atg ttt att agt gta cac att ttg gaa gta agt tc- --- --- --- --- --- --- --- --- --- --- --- --- --- --- --- --- --- --- --- --- --- --- --- --- --- --- --- --- --- --- --- --- --- --- --- --- --- --- --- --- --- --- --- --- --- --- --- --- --- --- --- --- --- --- --- --- --- --- --- --- --- --- --- --- --- --- --- --- --- --- --- --- --- --- --- --- --- --- --- --- --- --- --- --- --- --- --- --- --- --- --- --- --- --- --- --- --- --- --- --- --- --- --- --- --- --- --- --- --- --- --- --- --- --- --- --- --- --- --- --- --- --- --- --- --- --- --- --- --- --- --- --- --- --- --- --- --- --- --- --- --- --- --- --- --- --- --- --- --- --- --- --- --- --- --- --- --- --- --- --- --- --- --- --- --- --- --- --- --- --- --- --- --- --- --- --- --- --- --- --- --- --- --- --- --- --- --- --- --- --- --- --- --- --- --- --- --- --- --- --- --- --- --- --- --- --- --- --- --- --- --- --- --- --- --- --- --- --- --- --- --- --- --- --- --- --- --- --- --- --- --- --- --- --- --- --- --- --- --- --- --- --- --- --- --- --- --- --- --- --- --- --- --- --- --- --- --- --- --- --- --- --- --- --- --- --- --- --- --- --- --- --- --- --- --- --- --- --- --- --- --- --- --- --- --- --- --- --- --- --- --- --- --- --- --- --- --- --- --- --- --- --- --- --- --- --- --- --- --- --- --- --- --- --- --- --- --- --- --- --- --- --- --- --- --- --- --- --- --- --- --- --- --- --- --- --- --- --- --- --- --- --- --- --- --- --- --- --- --- --- --- --- --- --- --- --- --- --- --- --- --- --- --- --- --- --- --- --- --- --- --- --- --- --- --- --- --- --- --- --- --- --- --- --- --- --- --- --- --- --- --- --- --- --- --- --- --- --- --- --- --- --- --- --- --- --- --- --- --- --- --- --- --- --- --- --- --- --- --- --- --- --- --- --- --- --- --- --- --- --- --- --- --- --- --- --- --- --- --- --- --- --- --- --- --- --- --- --- --- --- --- --- --- --- --- --- --- --- --- --- --- --- --- --- --- --- --- --- --- --- --- --- --- --- --- --- --- --- --- --- --- --- --- --- --- --- --- --- --- --- --- --- --- --- --- --- --- --- --- --- --- --- --- --- --- --- --- --- --- --- --- --- --- --- --- --- --- --- --- --- --- --- --- --- --- --- --- --- --- --- --- --- --- --- --- --- --- --- --- --- --- --- --- --- --- --- --- --- --- --- --- --- --- --- --- --- --- --- --- --- --- --- --- --- --- --- --- --- --- --- --- --- --- --- --- --- --- --- --- --- --- --- --- --- --- --- --- --- --- --- --- --- --- --- --- --- --- --- --- --- --- --- --- --- --- --- --- --- --- --- --- --- --- --- --- --- --- --- --- --- --- --- --- --- --- --- --- --- --- --- --- --- --- --- --- --- --- --- --- --- --- --- --- --- --- --- --- --- --- --- --- --- -ct gga tcg gat gtc acc gca gta ata ttg ttg att att tct gac atc gac gta tta tat agt ttt tta att cca tat ctt ttt t

>ON649718.1_Monkeypox_virus_isolate_Monkeypox/PT0018/2022_complete_genome

ttt tat atc act acg gac ata aac cat tgt ata ttt ttt atg ttt att agt gta cac att ttg gaa gta agt tc- --- --- --- --- --- --- --- --- --- --- --- --- --- --- --- --- --- --- --- --- --- --- --- --- --- --- --- --- --- --- --- --- --- --- --- --- --- --- --- --- --- --- --- --- --- --- --- --- --- --- --- --- --- --- --- --- --- --- --- --- --- --- --- --- --- --- --- --- --- --- --- --- --- --- --- --- --- --- --- --- --- --- --- --- --- --- --- --- --- --- --- --- --- --- --- --- --- --- --- --- --- --- --- --- --- --- --- --- --- --- --- --- --- --- --- --- --- --- --- --- --- --- --- --- --- --- --- --- --- --- --- --- --- --- --- --- --- --- --- --- --- --- --- --- --- --- --- --- --- --- --- --- --- --- --- --- --- --- --- --- --- --- --- --- --- --- --- --- --- --- --- --- --- --- --- --- --- --- --- --- --- --- --- --- --- --- --- --- --- --- --- --- --- --- --- --- --- --- --- --- --- --- --- --- --- --- --- --- --- --- --- --- --- --- --- --- --- --- --- --- --- --- --- --- --- --- --- --- --- --- --- --- --- --- --- --- --- --- --- --- --- --- --- --- --- --- --- --- --- --- --- --- --- --- --- --- --- --- --- --- --- --- --- --- --- --- --- --- --- --- --- --- --- --- --- --- --- --- --- --- --- --- --- --- --- --- --- --- --- --- --- --- --- --- --- --- --- --- --- --- --- --- --- --- --- --- --- --- --- --- --- --- --- --- --- --- --- --- --- --- --- --- --- --- --- --- --- --- --- --- --- --- --- --- --- --- --- --- --- --- --- --- --- --- --- --- --- --- --- --- --- --- --- --- --- --- --- --- --- --- --- --- --- --- --- --- --- --- --- --- --- --- --- --- --- --- --- --- --- --- --- --- --- --- --- --- --- --- --- --- --- --- --- --- --- --- --- --- --- --- --- --- --- --- --- --- --- --- --- --- --- --- --- --- --- --- --- --- --- --- --- --- --- --- --- --- --- --- --- --- --- --- --- --- --- --- --- --- --- --- --- --- --- --- --- --- --- --- --- --- --- --- --- --- --- --- --- --- --- --- --- --- --- --- --- --- --- --- --- --- --- --- --- --- --- --- --- --- --- --- --- --- --- --- --- --- --- --- --- --- --- --- --- --- --- --- --- --- --- --- --- --- --- --- --- --- --- --- --- --- --- --- --- --- --- --- --- --- --- --- --- --- --- --- --- --- --- --- --- --- --- --- --- --- --- --- --- --- --- --- --- --- --- --- --- --- --- --- --- --- --- --- --- --- --- --- --- --- --- --- --- --- --- --- --- --- --- --- --- --- --- --- --- --- --- --- --- --- --- --- --- --- --- --- --- --- --- --- --- --- --- --- --- --- --- --- --- --- --- --- --- --- --- --- --- --- --- --- --- --- --- --- --- --- --- --- --- --- --- --- --- --- --- --- --- --- --- --- --- --- --- --- --- --- --- --- --- --- --- --- --- --- --- --- --- --- --- --- --- --- --- --- -ct gga tcg gat gtc acc gca gta ata ttg ttg att att tct gac atc gac gta tta tat agt ttt tta att cca tat ctt ttt t

>ON649719.1_Monkeypox_virus_isolate_Monkeypox/PT0012/2022_complete_genome

ttt tat atc act acg gac ata aac cat tgt ata ttt ttt atg ttt att agt gta cac att ttg gaa gta agt tc- --- --- --- --- --- --- --- --- --- --- --- --- --- --- --- --- --- --- --- --- --- --- --- --- --- --- --- --- --- --- --- --- --- --- --- --- --- --- --- --- --- --- --- --- --- --- --- --- --- --- --- --- --- --- --- --- --- --- --- --- --- --- --- --- --- --- --- --- --- --- --- --- --- --- --- --- --- --- --- --- --- --- --- --- --- --- --- --- --- --- --- --- --- --- --- --- --- --- --- --- --- --- --- --- --- --- --- --- --- --- --- --- --- --- --- --- --- --- --- --- --- --- --- --- --- --- --- --- --- --- --- --- --- --- --- --- --- --- --- --- --- --- --- --- --- --- --- --- --- --- --- --- --- --- --- --- --- --- --- --- --- --- --- --- --- --- --- --- --- --- --- --- --- --- --- --- --- --- --- --- --- --- --- --- --- --- --- --- --- --- --- --- --- --- --- --- --- --- --- --- --- --- --- --- --- --- --- --- --- --- --- --- --- --- --- --- --- --- --- --- --- --- --- --- --- --- --- --- --- --- --- --- --- --- --- --- --- --- --- --- --- --- --- --- --- --- --- --- --- --- --- --- --- --- --- --- --- --- --- --- --- --- --- --- --- --- --- --- --- --- --- --- --- --- --- --- --- --- --- --- --- --- --- --- --- --- --- --- --- --- --- --- --- --- --- --- --- --- --- --- --- --- --- --- --- --- --- --- --- --- --- --- --- --- --- --- --- --- --- --- --- --- --- --- --- --- --- --- --- --- --- --- --- --- --- --- --- --- --- --- --- --- --- --- --- --- --- --- --- --- --- --- --- --- --- --- --- --- --- --- --- --- --- --- --- --- --- --- --- --- --- --- --- --- --- --- --- --- --- --- --- --- --- --- --- --- --- --- --- --- --- --- --- --- --- --- --- --- --- --- --- --- --- --- --- --- --- --- --- --- --- --- --- --- --- --- --- --- --- --- --- --- --- --- --- --- --- --- --- --- --- --- --- --- --- --- --- --- --- --- --- --- --- --- --- --- --- --- --- --- --- --- --- --- --- --- --- --- --- --- --- --- --- --- --- --- --- --- --- --- --- --- --- --- --- --- --- --- --- --- --- --- --- --- --- --- --- --- --- --- --- --- --- --- --- --- --- --- --- --- --- --- --- --- --- --- --- --- --- --- --- --- --- --- --- --- --- --- --- --- --- --- --- --- --- --- --- --- --- --- --- --- --- --- --- --- --- --- --- --- --- --- --- --- --- --- --- --- --- --- --- --- --- --- --- --- --- --- --- --- --- --- --- --- --- --- --- --- --- --- --- --- --- --- --- --- --- --- --- --- --- --- --- --- --- --- --- --- --- --- --- --- --- --- --- --- --- --- --- --- --- --- --- --- --- --- --- --- --- --- --- --- --- --- --- --- --- --- --- --- --- --- --- --- --- --- --- --- --- --- --- --- --- --- --- --- --- --- --- --- --- --- --- --- --- --- --- --- --- --- --- --- -ct gga tcg gat gtc acc gca gta ata ttg ttg att att tct gac atc gac gta tta tat agt ttt tta att cca tat ctt ttt t

>ON682267.1_Monkeypox_virus_isolate_MPXV/Germany/2022/RKI010_complete_genome

ttt tat atc act acg gac ata aac cat tgt ata ttt ttt atg ttt att agt gta cac att ttg gaa gta agt tc- --- --- --- --- --- --- --- --- --- --- --- --- --- --- --- --- --- --- --- --- --- --- --- --- --- --- --- --- --- --- --- --- --- --- --- --- --- --- --- --- --- --- --- --- --- --- --- --- --- --- --- --- --- --- --- --- --- --- --- --- --- --- --- --- --- --- --- --- --- --- --- --- --- --- --- --- --- --- --- --- --- --- --- --- --- --- --- --- --- --- --- --- --- --- --- --- --- --- --- --- --- --- --- --- --- --- --- --- --- --- --- --- --- --- --- --- --- --- --- --- --- --- --- --- --- --- --- --- --- --- --- --- --- --- --- --- --- --- --- --- --- --- --- --- --- --- --- --- --- --- --- --- --- --- --- --- --- --- --- --- --- --- --- --- --- --- --- --- --- --- --- --- --- --- --- --- --- --- --- --- --- --- --- --- --- --- --- --- --- --- --- --- --- --- --- --- --- --- --- --- --- --- --- --- --- --- --- --- --- --- --- --- --- --- --- --- --- --- --- --- --- --- --- --- --- --- --- --- --- --- --- --- --- --- --- --- --- --- --- --- --- --- --- --- --- --- --- --- --- --- --- --- --- --- --- --- --- --- --- --- --- --- --- --- --- --- --- --- --- --- --- --- --- --- --- --- --- --- --- --- --- --- --- --- --- --- --- --- --- --- --- --- --- --- --- --- --- --- --- --- --- --- --- --- --- --- --- --- --- --- --- --- --- --- --- --- --- --- --- --- --- --- --- --- --- --- --- --- --- --- --- --- --- --- --- --- --- --- --- --- --- --- --- --- --- --- --- --- --- --- --- --- --- --- --- --- --- --- --- --- --- --- --- --- --- --- --- --- --- --- --- --- --- --- --- --- --- --- --- --- --- --- --- --- --- --- --- --- --- --- --- --- --- --- --- --- --- --- --- --- --- --- --- --- --- --- --- --- --- --- --- --- --- --- --- --- --- --- --- --- --- --- --- --- --- --- --- --- --- --- --- --- --- --- --- --- --- --- --- --- --- --- --- --- --- --- --- --- --- --- --- --- --- --- --- --- --- --- --- --- --- --- --- --- --- --- --- --- --- --- --- --- --- --- --- --- --- --- --- --- --- --- --- --- --- --- --- --- --- --- --- --- --- --- --- --- --- --- --- --- --- --- --- --- --- --- --- --- --- --- --- --- --- --- --- --- --- --- --- --- --- --- --- --- --- --- --- --- --- --- --- --- --- --- --- --- --- --- --- --- --- --- --- --- --- --- --- --- --- --- --- --- --- --- --- --- --- --- --- --- --- --- --- --- --- --- --- --- --- --- --- --- --- --- --- --- --- --- --- --- --- --- --- --- --- --- --- --- --- --- --- --- --- --- --- --- --- --- --- --- --- --- --- --- --- --- --- --- --- --- --- --- --- --- --- --- --- --- --- --- --- --- --- --- --- --- --- --- --- --- --- --- --- --- --- --- --- --- --- --- --- --- --- --- --- --- --- --- --- --- --- --- -ct gga tcg gat gtc acc gca gta ata ttg ttg att att tct gac atc gac gta tta tat agt ttt tta att cca tat ctt ttt t

>ON649717.1_Monkeypox_virus_isolate_Monkeypox/PT0019/2022_partial_genome

ttt tat atc act acg gac ata aac cat tgt ata ttt ttt atg ttt att agt gta cac att ttg gaa gta agt tc- --- --- --- --- --- --- --- --- --- --- --- --- --- --- --- --- --- --- --- --- --- --- --- --- --- --- --- --- --- --- --- --- --- --- --- --- --- --- --- --- --- --- --- --- --- --- --- --- --- --- --- --- --- --- --- --- --- --- --- --- --- --- --- --- --- --- --- --- --- --- --- --- --- --- --- --- --- --- --- --- --- --- --- --- --- --- --- --- --- --- --- --- --- --- --- --- --- --- --- --- --- --- --- --- --- --- --- --- --- --- --- --- --- --- --- --- --- --- --- --- --- --- --- --- --- --- --- --- --- --- --- --- --- --- --- --- --- --- --- --- --- --- --- --- --- --- --- --- --- --- --- --- --- --- --- --- --- --- --- --- --- --- --- --- --- --- --- --- --- --- --- --- --- --- --- --- --- --- --- --- --- --- --- --- --- --- --- --- --- --- --- --- --- --- --- --- --- --- --- --- --- --- --- --- --- --- --- --- --- --- --- --- --- --- --- --- --- --- --- --- --- --- --- --- --- --- --- --- --- --- --- --- --- --- --- --- --- --- --- --- --- --- --- --- --- --- --- --- --- --- --- --- --- --- --- --- --- --- --- --- --- --- --- --- --- --- --- --- --- --- --- --- --- --- --- --- --- --- --- --- --- --- --- --- --- --- --- --- --- --- --- --- --- --- --- --- --- --- --- --- --- --- --- --- --- --- --- --- --- --- --- --- --- --- --- --- --- --- --- --- --- --- --- --- --- --- --- --- --- --- --- --- --- --- --- --- --- --- --- --- --- --- --- --- --- --- --- --- --- --- --- --- --- --- --- --- --- --- --- --- --- --- --- --- --- --- --- --- --- --- --- --- --- --- --- --- --- --- --- --- --- --- --- --- --- --- --- --- --- --- --- --- --- --- --- --- --- --- --- --- --- --- --- --- --- --- --- --- --- --- --- --- --- --- --- --- --- --- --- --- --- --- --- --- --- --- --- --- --- --- --- --- --- --- --- --- --- --- --- --- --- --- --- --- --- --- --- --- --- --- --- --- --- --- --- --- --- --- --- --- --- --- --- --- --- --- --- --- --- --- --- --- --- --- --- --- --- --- --- --- --- --- --- --- --- --- --- --- --- --- --- --- --- --- --- --- --- --- --- --- --- --- --- --- --- --- --- --- --- --- --- --- --- --- --- --- --- --- --- --- --- --- --- --- --- --- --- --- --- --- --- --- --- --- --- --- --- --- --- --- --- --- --- --- --- --- --- --- --- --- --- --- --- --- --- --- --- --- --- --- --- --- --- --- --- --- --- --- --- --- --- --- --- --- --- --- --- --- --- --- --- --- --- --- --- --- --- --- --- --- --- --- --- --- --- --- --- --- --- --- --- --- --- --- --- --- --- --- --- --- --- --- --- --- --- --- --- --- --- --- --- --- --- --- --- --- --- --- --- --- --- --- --- --- --- --- --- --- --- --- --- --- --- --- --- --- --- --- --- --- --- --- -ct gga tcg gat gtc acc gca gta ata ttg ttg att att tct gac atc gac gta tta tat agt ttt tta att cca tat ctt ttt t

>ON649708.1_Monkeypox_virus_isolate_Monkeypox/PT0023/2022_partial_genome

ttt tat atc act acg gac ata aac cat tgt ata ttt ttt atg ttt att agt gta cac att ttg gaa gta agt tc- --- --- --- --- --- --- --- --- --- --- --- --- --- --- --- --- --- --- --- --- --- --- --- --- --- --- --- --- --- --- --- --- --- --- --- --- --- --- --- --- --- --- --- --- --- --- --- --- --- --- --- --- --- --- --- --- --- --- --- --- --- --- --- --- --- --- --- --- --- --- --- --- --- --- --- --- --- --- --- --- --- --- --- --- --- --- --- --- --- --- --- --- --- --- --- --- --- --- --- --- --- --- --- --- --- --- --- --- --- --- --- --- --- --- --- --- --- --- --- --- --- --- --- --- --- --- --- --- --- --- --- --- --- --- --- --- --- --- --- --- --- --- --- --- --- --- --- --- --- --- --- --- --- --- --- --- --- --- --- --- --- --- --- --- --- --- --- --- --- --- --- --- --- --- --- --- --- --- --- --- --- --- --- --- --- --- --- --- --- --- --- --- --- --- --- --- --- --- --- --- --- --- --- --- --- --- --- --- --- --- --- --- --- --- --- --- --- --- --- --- --- --- --- --- --- --- --- --- --- --- --- --- --- --- --- --- --- --- --- --- --- --- --- --- --- --- --- --- --- --- --- --- --- --- --- --- --- --- --- --- --- --- --- --- --- --- --- --- --- --- --- --- --- --- --- --- --- --- --- --- --- --- --- --- --- --- --- --- --- --- --- --- --- --- --- --- --- --- --- --- --- --- --- --- --- --- --- --- --- --- --- --- --- --- --- --- --- --- --- --- --- --- --- --- --- --- --- --- --- --- --- --- --- --- --- --- --- --- --- --- --- --- --- --- --- --- --- --- --- --- --- --- --- --- --- --- --- --- --- --- --- --- --- --- --- --- --- --- --- --- --- --- --- --- --- --- --- --- --- --- --- --- --- --- --- --- --- --- --- --- --- --- --- --- --- --- --- --- --- --- --- --- --- --- --- --- --- --- --- --- --- --- --- --- --- --- --- --- --- --- --- --- --- --- --- --- --- --- --- --- --- --- --- --- --- --- --- --- --- --- --- --- --- --- --- --- --- --- --- --- --- --- --- --- --- --- --- --- --- --- --- --- --- --- --- --- --- --- --- --- --- --- --- --- --- --- --- --- --- --- --- --- --- --- --- --- --- --- --- --- --- --- --- --- --- --- --- --- --- --- --- --- --- --- --- --- --- --- --- --- --- --- --- --- --- --- --- --- --- --- --- --- --- --- --- --- --- --- --- --- --- --- --- --- --- --- --- --- --- --- --- --- --- --- --- --- --- --- --- --- --- --- --- --- --- --- --- --- --- --- --- --- --- --- --- --- --- --- --- --- --- --- --- --- --- --- --- --- --- --- --- --- --- --- --- --- --- --- --- --- --- --- --- --- --- --- --- --- --- --- --- --- --- --- --- --- --- --- --- --- --- --- --- --- --- --- --- --- --- --- --- --- --- --- --- --- --- --- --- --- --- --- --- --- --- --- --- --- --- --- --- --- --- --- --- --- --- --- --- --- --- --- -ct gga tcg gat gtc acc gca gta ata ttg ttg att att tct gac atc gac gta tta tat agt ttt tta att cca tat ctt ttt t

>ON649709.1_Monkeypox_virus_isolate_Monkeypox/PT0028/2022_complete_genome

ttt tat atc act acg gac ata aac cat tgt ata ttt ttt atg ttt att agt gta cac att ttg gaa gta agt tc- --- --- --- --- --- --- --- --- --- --- --- --- --- --- --- --- --- --- --- --- --- --- --- --- --- --- --- --- --- --- --- --- --- --- --- --- --- --- --- --- --- --- --- --- --- --- --- --- --- --- --- --- --- --- --- --- --- --- --- --- --- --- --- --- --- --- --- --- --- --- --- --- --- --- --- --- --- --- --- --- --- --- --- --- --- --- --- --- --- --- --- --- --- --- --- --- --- --- --- --- --- --- --- --- --- --- --- --- --- --- --- --- --- --- --- --- --- --- --- --- --- --- --- --- --- --- --- --- --- --- --- --- --- --- --- --- --- --- --- --- --- --- --- --- --- --- --- --- --- --- --- --- --- --- --- --- --- --- --- --- --- --- --- --- --- --- --- --- --- --- --- --- --- --- --- --- --- --- --- --- --- --- --- --- --- --- --- --- --- --- --- --- --- --- --- --- --- --- --- --- --- --- --- --- --- --- --- --- --- --- --- --- --- --- --- --- --- --- --- --- --- --- --- --- --- --- --- --- --- --- --- --- --- --- --- --- --- --- --- --- --- --- --- --- --- --- --- --- --- --- --- --- --- --- --- --- --- --- --- --- --- --- --- --- --- --- --- --- --- --- --- --- --- --- --- --- --- --- --- --- --- --- --- --- --- --- --- --- --- --- --- --- --- --- --- --- --- --- --- --- --- --- --- --- --- --- --- --- --- --- --- --- --- --- --- --- --- --- --- --- --- --- --- --- --- --- --- --- --- --- --- --- --- --- --- --- --- --- --- --- --- --- --- --- --- --- --- --- --- --- --- --- --- --- --- --- --- --- --- --- --- --- --- --- --- --- --- --- --- --- --- --- --- --- --- --- --- --- --- --- --- --- --- --- --- --- --- --- --- --- --- --- --- --- --- --- --- --- --- --- --- --- --- --- --- --- --- --- --- --- --- --- --- --- --- --- --- --- --- --- --- --- --- --- --- --- --- --- --- --- --- --- --- --- --- --- --- --- --- --- --- --- --- --- --- --- --- --- --- --- --- --- --- --- --- --- --- --- --- --- --- --- --- --- --- --- --- --- --- --- --- --- --- --- --- --- --- --- --- --- --- --- --- --- --- --- --- --- --- --- --- --- --- --- --- --- --- --- --- --- --- --- --- --- --- --- --- --- --- --- --- --- --- --- --- --- --- --- --- --- --- --- --- --- --- --- --- --- --- --- --- --- --- --- --- --- --- --- --- --- --- --- --- --- --- --- --- --- --- --- --- --- --- --- --- --- --- --- --- --- --- --- --- --- --- --- --- --- --- --- --- --- --- --- --- --- --- --- --- --- --- --- --- --- --- --- --- --- --- --- --- --- --- --- --- --- --- --- --- --- --- --- --- --- --- --- --- --- --- --- --- --- --- --- --- --- --- --- --- --- --- --- --- --- --- --- --- --- --- --- --- --- --- --- --- --- --- --- --- --- --- --- --- --- --- --- --- --- --- --- --- --- -ct gga tcg gat gtc acc gca gta ata ttg ttg att att tct gac atc gac gta tta tat agt ttt tta att cca tat ctt ttt t

>ON614676.1_Monkeypox_virus_isolate_INMI-Pt1_partial_genome

ttt tat atc act acg gac ata aac cat tgt ata ttt ttt atg ttt att agt gta cac att ttg gaa gta agt tc- --- --- --- --- --- --- --- --- --- --- --- --- --- --- --- --- --- --- --- --- --- --- --- --- --- --- --- --- --- --- --- --- --- --- --- --- --- --- --- --- --- --- --- --- --- --- --- --- --- --- --- --- --- --- --- --- --- --- --- --- --- --- --- --- --- --- --- --- --- --- --- --- --- --- --- --- --- --- --- --- --- --- --- --- --- --- --- --- --- --- --- --- --- --- --- --- --- --- --- --- --- --- --- --- --- --- --- --- --- --- --- --- --- --- --- --- --- --- --- --- --- --- --- --- --- --- --- --- --- --- --- --- --- --- --- --- --- --- --- --- --- --- --- --- --- --- --- --- --- --- --- --- --- --- --- --- --- --- --- --- --- --- --- --- --- --- --- --- --- --- --- --- --- --- --- --- --- --- --- --- --- --- --- --- --- --- --- --- --- --- --- --- --- --- --- --- --- --- --- --- --- --- --- --- --- --- --- --- --- --- --- --- --- --- --- --- --- --- --- --- --- --- --- --- --- --- --- --- --- --- --- --- --- --- --- --- --- --- --- --- --- --- --- --- --- --- --- --- --- --- --- --- --- --- --- --- --- --- --- --- --- --- --- --- --- --- --- --- --- --- --- --- --- --- --- --- --- --- --- --- --- --- --- --- --- --- --- --- --- --- --- --- --- --- --- --- --- --- --- --- --- --- --- --- --- --- --- --- --- --- --- --- --- --- --- --- --- --- --- --- --- --- --- --- --- --- --- --- --- --- --- --- --- --- --- --- --- --- --- --- --- --- --- --- --- --- --- --- --- --- --- --- --- --- --- --- --- --- --- --- --- --- --- --- --- --- --- --- --- --- --- --- --- --- --- --- --- --- --- --- --- --- --- --- --- --- --- --- --- --- --- --- --- --- --- --- --- --- --- --- --- --- --- --- --- --- --- --- --- --- --- --- --- --- --- --- --- --- --- --- --- --- --- --- --- --- --- --- --- --- --- --- --- --- --- --- --- --- --- --- --- --- --- --- --- --- --- --- --- --- --- --- --- --- --- --- --- --- --- --- --- --- --- --- --- --- --- --- --- --- --- --- --- --- --- --- --- --- --- --- --- --- --- --- --- --- --- --- --- --- --- --- --- --- --- --- --- --- --- --- --- --- --- --- --- --- --- --- --- --- --- --- --- --- --- --- --- --- --- --- --- --- --- --- --- --- --- --- --- --- --- --- --- --- --- --- --- --- --- --- --- --- --- --- --- --- --- --- --- --- --- --- --- --- --- --- --- --- --- --- --- --- --- --- --- --- --- --- --- --- --- --- --- --- --- --- --- --- --- --- --- --- --- --- --- --- --- --- --- --- --- --- --- --- --- --- --- --- --- --- --- --- --- --- --- --- --- --- --- --- --- --- --- --- --- --- --- --- --- --- --- --- --- --- --- --- --- --- --- --- --- --- --- --- --- --- --- --- --- --- --- --- --- --- --- --- --- --- --- --- --- --- -ct gga tcg gat gtc acc gca gta ata ttg ttg att att tct gac atc gac gta tta tat agt ttt tta att cca tat ctt ttt t

>ON649712.1_Monkeypox_virus_isolate_Monkeypox/PT0025/2022_partial_genome

ttt tat atc act acg gac ata aac cat tgt ata ttt ttt atg ttt att agt gta cac att ttg gaa gta agt tc- --- --- --- --- --- --- --- --- --- --- --- --- --- --- --- --- --- --- --- --- --- --- --- --- --- --- --- --- --- --- --- --- --- --- --- --- --- --- --- --- --- --- --- --- --- --- --- --- --- --- --- --- --- --- --- --- --- --- --- --- --- --- --- --- --- --- --- --- --- --- --- --- --- --- --- --- --- --- --- --- --- --- --- --- --- --- --- --- --- --- --- --- --- --- --- --- --- --- --- --- --- --- --- --- --- --- --- --- --- --- --- --- --- --- --- --- --- --- --- --- --- --- --- --- --- --- --- --- --- --- --- --- --- --- --- --- --- --- --- --- --- --- --- --- --- --- --- --- --- --- --- --- --- --- --- --- --- --- --- --- --- --- --- --- --- --- --- --- --- --- --- --- --- --- --- --- --- --- --- --- --- --- --- --- --- --- --- --- --- --- --- --- --- --- --- --- --- --- --- --- --- --- --- --- --- --- --- --- --- --- --- --- --- --- --- --- --- --- --- --- --- --- --- --- --- --- --- --- --- --- --- --- --- --- --- --- --- --- --- --- --- --- --- --- --- --- --- --- --- --- --- --- --- --- --- --- --- --- --- --- --- --- --- --- --- --- --- --- --- --- --- --- --- --- --- --- --- --- --- --- --- --- --- --- --- --- --- --- --- --- --- --- --- --- --- --- --- --- --- --- --- --- --- --- --- --- --- --- --- --- --- --- --- --- --- --- --- --- --- --- --- --- --- --- --- --- --- --- --- --- --- --- --- --- --- --- --- --- --- --- --- --- --- --- --- --- --- --- --- --- --- --- --- --- --- --- --- --- --- --- --- --- --- --- --- --- --- --- --- --- --- --- --- --- --- --- --- --- --- --- --- --- --- --- --- --- --- --- --- --- --- --- --- --- --- --- --- --- --- --- --- --- --- --- --- --- --- --- --- --- --- --- --- --- --- --- --- --- --- --- --- --- --- --- --- --- --- --- --- --- --- --- --- --- --- --- --- --- --- --- --- --- --- --- --- --- --- --- --- --- --- --- --- --- --- --- --- --- --- --- --- --- --- --- --- --- --- --- --- --- --- --- --- --- --- --- --- --- --- --- --- --- --- --- --- --- --- --- --- --- --- --- --- --- --- --- --- --- --- --- --- --- --- --- --- --- --- --- --- --- --- --- --- --- --- --- --- --- --- --- --- --- --- --- --- --- --- --- --- --- --- --- --- --- --- --- --- --- --- --- --- --- --- --- --- --- --- --- --- --- --- --- --- --- --- --- --- --- --- --- --- --- --- --- --- --- --- --- --- --- --- --- --- --- --- --- --- --- --- --- --- --- --- --- --- --- --- --- --- --- --- --- --- --- --- --- --- --- --- --- --- --- --- --- --- --- --- --- --- --- --- --- --- --- --- --- --- --- --- --- --- --- --- --- --- --- --- --- --- --- --- --- --- --- --- --- --- --- --- --- --- --- --- --- --- --- --- --- --- --- --- --- -ct gga tcg gat gtc acc gca gta ata ttg ttg att att tct gac atc gac gta tta tat agt ttt tta att cca tat ctt ttt t

>ON595760.2_Monkeypox_virus_isolate_MPXV-CH-38134631/2022_partial_genome

ttt tat atc act acg gac ata aac cat tgt ata ttt ttt atg ttt att agt gta cac att ttg gaa gta agt tc- --- --- --- --- --- --- --- --- --- --- --- --- --- --- --- --- --- --- --- --- --- --- --- --- --- --- --- --- --- --- --- --- --- --- --- --- --- --- --- --- --- --- --- --- --- --- --- --- --- --- --- --- --- --- --- --- --- --- --- --- --- --- --- --- --- --- --- --- --- --- --- --- --- --- --- --- --- --- --- --- --- --- --- --- --- --- --- --- --- --- --- --- --- --- --- --- --- --- --- --- --- --- --- --- --- --- --- --- --- --- --- --- --- --- --- --- --- --- --- --- --- --- --- --- --- --- --- --- --- --- --- --- --- --- --- --- --- --- --- --- --- --- --- --- --- --- --- --- --- --- --- --- --- --- --- --- --- --- --- --- --- --- --- --- --- --- --- --- --- --- --- --- --- --- --- --- --- --- --- --- --- --- --- --- --- --- --- --- --- --- --- --- --- --- --- --- --- --- --- --- --- --- --- --- --- --- --- --- --- --- --- --- --- --- --- --- --- --- --- --- --- --- --- --- --- --- --- --- --- --- --- --- --- --- --- --- --- --- --- --- --- --- --- --- --- --- --- --- --- --- --- --- --- --- --- --- --- --- --- --- --- --- --- --- --- --- --- --- --- --- --- --- --- --- --- --- --- --- --- --- --- --- --- --- --- --- --- --- --- --- --- --- --- --- --- --- --- --- --- --- --- --- --- --- --- --- --- --- --- --- --- --- --- --- --- --- --- --- --- --- --- --- --- --- --- --- --- --- --- --- --- --- --- --- --- --- --- --- --- --- --- --- --- --- --- --- --- --- --- --- --- --- --- --- --- --- --- --- --- --- --- --- --- --- --- --- --- --- --- --- --- --- --- --- --- --- --- --- --- --- --- --- --- --- --- --- --- --- --- --- --- --- --- --- --- --- --- --- --- --- --- --- --- --- --- --- --- --- --- --- --- --- --- --- --- --- --- --- --- --- --- --- --- --- --- --- --- --- --- --- --- --- --- --- --- --- --- --- --- --- --- --- --- --- --- --- --- --- --- --- --- --- --- --- --- --- --- --- --- --- --- --- --- --- --- --- --- --- --- --- --- --- --- --- --- --- --- --- --- --- --- --- --- --- --- --- --- --- --- --- --- --- --- --- --- --- --- --- --- --- --- --- --- --- --- --- --- --- --- --- --- --- --- --- --- --- --- --- --- --- --- --- --- --- --- --- --- --- --- --- --- --- --- --- --- --- --- --- --- --- --- --- --- --- --- --- --- --- --- --- --- --- --- --- --- --- --- --- --- --- --- --- --- --- --- --- --- --- --- --- --- --- --- --- --- --- --- --- --- --- --- --- --- --- --- --- --- --- --- --- --- --- --- --- --- --- --- --- --- --- --- --- --- --- --- --- --- --- --- --- --- --- --- --- --- --- --- --- --- --- --- --- --- --- --- --- --- --- --- --- --- --- --- --- --- --- --- --- --- --- --- --- --- --- --- --- --- --- --- --- --- --- -ct gga tcg gat gtc acc gca gta ata ttg ttg att att tct rac atc gac gta tta tat agt ttt tta att cca tat ctt ttt t

>MT903337.1_Monkeypox_virus_isolate_MPXV-M2940_FCT

ttt tat atc act acg gac ata aac cat tgt ata ttt ttt atg ttt att agt gta cac att ttg gaa gta agt tc- --- --- --- --- --- --- --- --- --- --- --- --- --- --- --- --- --- --- --- --- --- --- --- --- --- --- --- --- --- --- --- --- --- --- --- --- --- --- --- --- --- --- --- --- --- --- --- --- --- --- --- --- --- --- --- --- --- --- --- --- --- --- --- --- --- --- --- --- --- --- --- --- --- --- --- --- --- --- --- --- --- --- --- --- --- --- --- --- --- --- --- --- --- --- --- --- --- --- --- --- --- --- --- --- --- --- --- --- --- --- --- --- --- --- --- --- --- --- --- --- --- --- --- --- --- --- --- --- --- --- --- --- --- --- --- --- --- --- --- --- --- --- --- --- --- --- --- --- --- --- --- --- --- --- --- --- --- --- --- --- --- --- --- --- --- --- --- --- --- --- --- --- --- --- --- --- --- --- --- --- --- --- --- --- --- --- --- --- --- --- --- --- --- --- --- --- --- --- --- --- --- --- --- --- --- --- --- --- --- --- --- --- --- --- --- --- --- --- --- --- --- --- --- --- --- --- --- --- --- --- --- --- --- --- --- --- --- --- --- --- --- --- --- --- --- --- --- --- --- --- --- --- --- --- --- --- --- --- --- --- --- --- --- --- --- --- --- --- --- --- --- --- --- --- --- --- --- --- --- --- --- --- --- --- --- --- --- --- --- --- --- --- --- --- --- --- --- --- --- --- --- --- --- --- --- --- --- --- --- --- --- --- --- --- --- --- --- --- --- --- --- --- --- --- --- --- --- --- --- --- --- --- --- --- --- --- --- --- --- --- --- --- --- --- --- --- --- --- --- --- --- --- --- --- --- --- --- --- --- --- --- --- --- --- --- --- --- --- --- --- --- --- --- --- --- --- --- --- --- --- --- --- --- --- --- --- --- --- --- --- --- --- --- --- --- --- --- --- --- --- --- --- --- --- --- --- --- --- --- --- --- --- --- --- --- --- --- --- --- --- --- --- --- --- --- --- --- --- --- --- --- --- --- --- --- --- --- --- --- --- --- --- --- --- --- --- --- --- --- --- --- --- --- --- --- --- --- --- --- --- --- --- --- --- --- --- --- --- --- --- --- --- --- --- --- --- --- --- --- --- --- --- --- --- --- --- --- --- --- --- --- --- --- --- --- --- --- --- --- --- --- --- --- --- --- --- --- --- --- --- --- --- --- --- --- --- --- --- --- --- --- --- --- --- --- --- --- --- --- --- --- --- --- --- --- --- --- --- --- --- --- --- --- --- --- --- --- --- --- --- --- --- --- --- --- --- --- --- --- --- --- --- --- --- --- --- --- --- --- --- --- --- --- --- --- --- --- --- --- --- --- --- --- --- --- --- --- --- --- --- --- --- --- --- --- --- --- --- --- --- --- --- --- --- --- --- --- --- --- --- --- --- --- --- --- --- --- --- --- --- --- --- --- --- --- --- --- --- --- --- --- --- --- --- --- --- --- --- --- --- --- --- --- --- --- --- --- --- --- --- --- --- -ct gga tcg gat gtc acc gca gta ata ttg ttg att att tct gac atc gac gta tta tat agt ttt tta att cca tat ctt ttt t

>MT903339.1_Monkeypox_virus_isolate_MPXV-M3021_Delta

ttt tat atc act acg gac ata aac cat tgt ata ttt ttt atg ttt att agt gta cac att ttg gaa gta agt tc- --- --- --- --- --- --- --- --- --- --- --- --- --- --- --- --- --- --- --- --- --- --- --- --- --- --- --- --- --- --- --- --- --- --- --- --- --- --- --- --- --- --- --- --- --- --- --- --- --- --- --- --- --- --- --- --- --- --- --- --- --- --- --- --- --- --- --- --- --- --- --- --- --- --- --- --- --- --- --- --- --- --- --- --- --- --- --- --- --- --- --- --- --- --- --- --- --- --- --- --- --- --- --- --- --- --- --- --- --- --- --- --- --- --- --- --- --- --- --- --- --- --- --- --- --- --- --- --- --- --- --- --- --- --- --- --- --- --- --- --- --- --- --- --- --- --- --- --- --- --- --- --- --- --- --- --- --- --- --- --- --- --- --- --- --- --- --- --- --- --- --- --- --- --- --- --- --- --- --- --- --- --- --- --- --- --- --- --- --- --- --- --- --- --- --- --- --- --- --- --- --- --- --- --- --- --- --- --- --- --- --- --- --- --- --- --- --- --- --- --- --- --- --- --- --- --- --- --- --- --- --- --- --- --- --- --- --- --- --- --- --- --- --- --- --- --- --- --- --- --- --- --- --- --- --- --- --- --- --- --- --- --- --- --- --- --- --- --- --- --- --- --- --- --- --- --- --- --- --- --- --- --- --- --- --- --- --- --- --- --- --- --- --- --- --- --- --- --- --- --- --- --- --- --- --- --- --- --- --- --- --- --- --- --- --- --- --- --- --- --- --- --- --- --- --- --- --- --- --- --- --- --- --- --- --- --- --- --- --- --- --- --- --- --- --- --- --- --- --- --- --- --- --- --- --- --- --- --- --- --- --- --- --- --- --- --- --- --- --- --- --- --- --- --- --- --- --- --- --- --- --- --- --- --- --- --- --- --- --- --- --- --- --- --- --- --- --- --- --- --- --- --- --- --- --- --- --- --- --- --- --- --- --- --- --- --- --- --- --- --- --- --- --- --- --- --- --- --- --- --- --- --- --- --- --- --- --- --- --- --- --- --- --- --- --- --- --- --- --- --- --- --- --- --- --- --- --- --- --- --- --- --- --- --- --- --- --- --- --- --- --- --- --- --- --- --- --- --- --- --- --- --- --- --- --- --- --- --- --- --- --- --- --- --- --- --- --- --- --- --- --- --- --- --- --- --- --- --- --- --- --- --- --- --- --- --- --- --- --- --- --- --- --- --- --- --- --- --- --- --- --- --- --- --- --- --- --- --- --- --- --- --- --- --- --- --- --- --- --- --- --- --- --- --- --- --- --- --- --- --- --- --- --- --- --- --- --- --- --- --- --- --- --- --- --- --- --- --- --- --- --- --- --- --- --- --- --- --- --- --- --- --- --- --- --- --- --- --- --- --- --- --- --- --- --- --- --- --- --- --- --- --- --- --- --- --- --- --- --- --- --- --- --- --- --- --- --- --- --- --- --- --- --- --- --- --- --- --- --- --- --- --- --- --- --- --- --- --- --- --- --- --- -ct gga tcg gat gtc acc gca gta ata ttg ttg att att tct gac atc gac gta tta tat agt ttt tta att cca tat ctt ttt t

>MT903338.1_Monkeypox_virus_isolate_MPXV-M2957_Lagos

ttt tat atc act acg gac ata aac cat tgt ata ttt ttt atg ttt att agt gta cac att ttg gaa gta agt tc- --- --- --- --- --- --- --- --- --- --- --- --- --- --- --- --- --- --- --- --- --- --- --- --- --- --- --- --- --- --- --- --- --- --- --- --- --- --- --- --- --- --- --- --- --- --- --- --- --- --- --- --- --- --- --- --- --- --- --- --- --- --- --- --- --- --- --- --- --- --- --- --- --- --- --- --- --- --- --- --- --- --- --- --- --- --- --- --- --- --- --- --- --- --- --- --- --- --- --- --- --- --- --- --- --- --- --- --- --- --- --- --- --- --- --- --- --- --- --- --- --- --- --- --- --- --- --- --- --- --- --- --- --- --- --- --- --- --- --- --- --- --- --- --- --- --- --- --- --- --- --- --- --- --- --- --- --- --- --- --- --- --- --- --- --- --- --- --- --- --- --- --- --- --- --- --- --- --- --- --- --- --- --- --- --- --- --- --- --- --- --- --- --- --- --- --- --- --- --- --- --- --- --- --- --- --- --- --- --- --- --- --- --- --- --- --- --- --- --- --- --- --- --- --- --- --- --- --- --- --- --- --- --- --- --- --- --- --- --- --- --- --- --- --- --- --- --- --- --- --- --- --- --- --- --- --- --- --- --- --- --- --- --- --- --- --- --- --- --- --- --- --- --- --- --- --- --- --- --- --- --- --- --- --- --- --- --- --- --- --- --- --- --- --- --- --- --- --- --- --- --- --- --- --- --- --- --- --- --- --- --- --- --- --- --- --- --- --- --- --- --- --- --- --- --- --- --- --- --- --- --- --- --- --- --- --- --- --- --- --- --- --- --- --- --- --- --- --- --- --- --- --- --- --- --- --- --- --- --- --- --- --- --- --- --- --- --- --- --- --- --- --- --- --- --- --- --- --- --- --- --- --- --- --- --- --- --- --- --- --- --- --- --- --- --- --- --- --- --- --- --- --- --- --- --- --- --- --- --- --- --- --- --- --- --- --- --- --- --- --- --- --- --- --- --- --- --- --- --- --- --- --- --- --- --- --- --- --- --- --- --- --- --- --- --- --- --- --- --- --- --- --- --- --- --- --- --- --- --- --- --- --- --- --- --- --- --- --- --- --- --- --- --- --- --- --- --- --- --- --- --- --- --- --- --- --- --- --- --- --- --- --- --- --- --- --- --- --- --- --- --- --- --- --- --- --- --- --- --- --- --- --- --- --- --- --- --- --- --- --- --- --- --- --- --- --- --- --- --- --- --- --- --- --- --- --- --- --- --- --- --- --- --- --- --- --- --- --- --- --- --- --- --- --- --- --- --- --- --- --- --- --- --- --- --- --- --- --- --- --- --- --- --- --- --- --- --- --- --- --- --- --- --- --- --- --- --- --- --- --- --- --- --- --- --- --- --- --- --- --- --- --- --- --- --- --- --- --- --- --- --- --- --- --- --- --- --- --- --- --- --- --- --- --- --- --- --- --- --- --- --- --- --- --- --- --- --- --- --- --- --- --- --- --- --- --- --- --- --- --- --- --- -ct gga tcg gat gtc acc gca gta ata ttg ttg att att tct gac atc gac gta tta tat agt ttt tta att cca tat ctt ttt t

>KJ642617.1_Monkeypox_virus_strain_Nigeria-SE-1971_complete_genome

ttt tat atc act acg gac ata aac cat tgt ata ttt ttt atg ttt att agt gta cac att ttg gaa gta agt tc- --- --- --- --- --- --- --- --- --- --- --- --- --- --- --- --- --- --- --- --- --- --- --- --- --- --- --- --- --- --- --- --- --- --- --- --- --- --- --- --- --- --- --- --- --- --- --- --- --- --- --- --- --- --- --- --- --- --- --- --- --- --- --- --- --- --- --- --- --- --- --- --- --- --- --- --- --- --- --- --- --- --- --- --- --- --- --- --- --- --- --- --- --- --- --- --- --- --- --- --- --- --- --- --- --- --- --- --- --- --- --- --- --- --- --- --- --- --- --- --- --- --- --- --- --- --- --- --- --- --- --- --- --- --- --- --- --- --- --- --- --- --- --- --- --- --- --- --- --- --- --- --- --- --- --- --- --- --- --- --- --- --- --- --- --- --- --- --- --- --- --- --- --- --- --- --- --- --- --- --- --- --- --- --- --- --- --- --- --- --- --- --- --- --- --- --- --- --- --- --- --- --- --- --- --- --- --- --- --- --- --- --- --- --- --- --- --- --- --- --- --- --- --- --- --- --- --- --- --- --- --- --- --- --- --- --- --- --- --- --- --- --- --- --- --- --- --- --- --- --- --- --- --- --- --- --- --- --- --- --- --- --- --- --- --- --- --- --- --- --- --- --- --- --- --- --- --- --- --- --- --- --- --- --- --- --- --- --- --- --- --- --- --- --- --- --- --- --- --- --- --- --- --- --- --- --- --- --- --- --- --- --- --- --- --- --- --- --- --- --- --- --- --- --- --- --- --- --- --- --- --- --- --- --- --- --- --- --- --- --- --- --- --- --- --- --- --- --- --- --- --- --- --- --- --- --- --- --- --- --- --- --- --- --- --- --- --- --- --- --- --- --- --- --- --- --- --- --- --- --- --- --- --- --- --- --- --- --- --- --- --- --- --- --- --- --- --- --- --- --- --- --- --- --- --- --- --- --- --- --- --- --- --- --- --- --- --- --- --- --- --- --- --- --- --- --- --- --- --- --- --- --- --- --- --- --- --- --- --- --- --- --- --- --- --- --- --- --- --- --- --- --- --- --- --- --- --- --- --- --- --- --- --- --- --- --- --- --- --- --- --- --- --- --- --- --- --- --- --- --- --- --- --- --- --- --- --- --- --- --- --- --- --- --- --- --- --- --- --- --- --- --- --- --- --- --- --- --- --- --- --- --- --- --- --- --- --- --- --- --- --- --- --- --- --- --- --- --- --- --- --- --- --- --- --- --- --- --- --- --- --- --- --- --- --- --- --- --- --- --- --- --- --- --- --- --- --- --- --- --- --- --- --- --- --- --- --- --- --- --- --- --- --- --- --- --- --- --- --- --- --- --- --- --- --- --- --- --- --- --- --- --- --- --- --- --- --- --- --- --- --- --- --- --- --- --- --- --- --- --- --- --- --- --- --- --- --- --- --- --- --- --- --- --- --- --- --- --- --- --- --- --- --- --- --- --- --- --- --- --- --- --- --- --- --- --- --- --- --- --- --- --- -ct gga tcg gat gtc acc gca gta ata ttg ttg att att tct gac atc gac gta tta tat agt ttt tta att cca tat ctt ttt t

>ON615424.1_Monkeypox_virus_isolate_MPXV_2022_NL001_partial_genome

ttt tat atc act acg gac ata aac cat tgt ata ttt ttt atg ttt att agt gta cac att ttg gaa gta agt tc- --- --- --- --- --- --- --- --- --- --- --- --- --- --- --- --- --- --- --- --- --- --- --- --- --- --- --- --- --- --- --- --- --- --- --- --- --- --- --- --- --- --- --- --- --- --- --- --- --- --- --- --- --- --- --- --- --- --- --- --- --- --- --- --- --- --- --- --- --- --- --- --- --- --- --- --- --- --- --- --- --- --- --- --- --- --- --- --- --- --- --- --- --- --- --- --- --- --- --- --- --- --- --- --- --- --- --- --- --- --- --- --- --- --- --- --- --- --- --- --- --- --- --- --- --- --- --- --- --- --- --- --- --- --- --- --- --- --- --- --- --- --- --- --- --- --- --- --- --- --- --- --- --- --- --- --- --- --- --- --- --- --- --- --- --- --- --- --- --- --- --- --- --- --- --- --- --- --- --- --- --- --- --- --- --- --- --- --- --- --- --- --- --- --- --- --- --- --- --- --- --- --- --- --- --- --- --- --- --- --- --- --- --- --- --- --- --- --- --- --- --- --- --- --- --- --- --- --- --- --- --- --- --- --- --- --- --- --- --- --- --- --- --- --- --- --- --- --- --- --- --- --- --- --- --- --- --- --- --- --- --- --- --- --- --- --- --- --- --- --- --- --- --- --- --- --- --- --- --- --- --- --- --- --- --- --- --- --- --- --- --- --- --- --- --- --- --- --- --- --- --- --- --- --- --- --- --- --- --- --- --- --- --- --- --- --- --- --- --- --- --- --- --- --- --- --- --- --- --- --- --- --- --- --- --- --- --- --- --- --- --- --- --- --- --- --- --- --- --- --- --- --- --- --- --- --- --- --- --- --- --- --- --- --- --- --- --- --- --- --- --- --- --- --- --- --- --- --- --- --- --- --- --- --- --- --- --- --- --- --- --- --- --- --- --- --- --- --- --- --- --- --- --- --- --- --- --- --- --- --- --- --- --- --- --- --- --- --- --- --- --- --- --- --- --- --- --- --- --- --- --- --- --- --- --- --- --- --- --- --- --- --- --- --- --- --- --- --- --- --- --- --- --- --- --- --- --- --- --- --- --- --- --- --- --- --- --- --- --- --- --- --- --- --- --- --- --- --- --- --- --- --- --- --- --- --- --- --- --- --- --- --- --- --- --- --- --- --- --- --- --- --- --- --- --- --- --- --- --- --- --- --- --- --- --- --- --- --- --- --- --- --- --- --- --- --- --- --- --- --- --- --- --- --- --- --- --- --- --- --- --- --- --- --- --- --- --- --- --- --- --- --- --- --- --- --- --- --- --- --- --- --- --- --- --- --- --- --- --- --- --- --- --- --- --- --- --- --- --- --- --- --- --- --- --- --- --- --- --- --- --- --- --- --- --- --- --- --- --- --- --- --- --- --- --- --- --- --- --- --- --- --- --- --- --- --- --- --- --- --- --- --- --- --- --- --- --- --- --- --- --- --- --- --- --- --- --- --- --- --- --- --- --- --- --- --- --- --- --- --- --- --- -ct gga tcg gat gtc acc gca gta ata ttg ttg att att tct gac atc gac gta tta tat agt ttt tta att cca tat ctt ttt t

>ON585034.1_Monkeypox_virus_isolate_Monkeypox/PT0007/2022_complete_genome

ttt tat atc act acg gac ata aac cat tgt ata ttt ttt atg ttt att agt gta cac att ttg gaa gta agt tc- --- --- --- --- --- --- --- --- --- --- --- --- --- --- --- --- --- --- --- --- --- --- --- --- --- --- --- --- --- --- --- --- --- --- --- --- --- --- --- --- --- --- --- --- --- --- --- --- --- --- --- --- --- --- --- --- --- --- --- --- --- --- --- --- --- --- --- --- --- --- --- --- --- --- --- --- --- --- --- --- --- --- --- --- --- --- --- --- --- --- --- --- --- --- --- --- --- --- --- --- --- --- --- --- --- --- --- --- --- --- --- --- --- --- --- --- --- --- --- --- --- --- --- --- --- --- --- --- --- --- --- --- --- --- --- --- --- --- --- --- --- --- --- --- --- --- --- --- --- --- --- --- --- --- --- --- --- --- --- --- --- --- --- --- --- --- --- --- --- --- --- --- --- --- --- --- --- --- --- --- --- --- --- --- --- --- --- --- --- --- --- --- --- --- --- --- --- --- --- --- --- --- --- --- --- --- --- --- --- --- --- --- --- --- --- --- --- --- --- --- --- --- --- --- --- --- --- --- --- --- --- --- --- --- --- --- --- --- --- --- --- --- --- --- --- --- --- --- --- --- --- --- --- --- --- --- --- --- --- --- --- --- --- --- --- --- --- --- --- --- --- --- --- --- --- --- --- --- --- --- --- --- --- --- --- --- --- --- --- --- --- --- --- --- --- --- --- --- --- --- --- --- --- --- --- --- --- --- --- --- --- --- --- --- --- --- --- --- --- --- --- --- --- --- --- --- --- --- --- --- --- --- --- --- --- --- --- --- --- --- --- --- --- --- --- --- --- --- --- --- --- --- --- --- --- --- --- --- --- --- --- --- --- --- --- --- --- --- --- --- --- --- --- --- --- --- --- --- --- --- --- --- --- --- --- --- --- --- --- --- --- --- --- --- --- --- --- --- --- --- --- --- --- --- --- --- --- --- --- --- --- --- --- --- --- --- --- --- --- --- --- --- --- --- --- --- --- --- --- --- --- --- --- --- --- --- --- --- --- --- --- --- --- --- --- --- --- --- --- --- --- --- --- --- --- --- --- --- --- --- --- --- --- --- --- --- --- --- --- --- --- --- --- --- --- --- --- --- --- --- --- --- --- --- --- --- --- --- --- --- --- --- --- --- --- --- --- --- --- --- --- --- --- --- --- --- --- --- --- --- --- --- --- --- --- --- --- --- --- --- --- --- --- --- --- --- --- --- --- --- --- --- --- --- --- --- --- --- --- --- --- --- --- --- --- --- --- --- --- --- --- --- --- --- --- --- --- --- --- --- --- --- --- --- --- --- --- --- --- --- --- --- --- --- --- --- --- --- --- --- --- --- --- --- --- --- --- --- --- --- --- --- --- --- --- --- --- --- --- --- --- --- --- --- --- --- --- --- --- --- --- --- --- --- --- --- --- --- --- --- --- --- --- --- --- --- --- --- --- --- --- --- --- --- --- --- --- --- --- --- --- --- --- --- --- --- --- --- --- --- --- --- -ct gga tcg gat gtc acc gca gta ata ttg ttg att att tct gac atc gac gta tta tat agt ttt tta att cca tat ctt ttt t

>ON649713.1_Monkeypox_virus_isolate_Monkeypox/PT0020/2022_complete_genome

ttt tat atc act acg gac ata aac cat tgt ata ttt ttt atg ttt att agt gta cac att ttg gaa gta agt tc- --- --- --- --- --- --- --- --- --- --- --- --- --- --- --- --- --- --- --- --- --- --- --- --- --- --- --- --- --- --- --- --- --- --- --- --- --- --- --- --- --- --- --- --- --- --- --- --- --- --- --- --- --- --- --- --- --- --- --- --- --- --- --- --- --- --- --- --- --- --- --- --- --- --- --- --- --- --- --- --- --- --- --- --- --- --- --- --- --- --- --- --- --- --- --- --- --- --- --- --- --- --- --- --- --- --- --- --- --- --- --- --- --- --- --- --- --- --- --- --- --- --- --- --- --- --- --- --- --- --- --- --- --- --- --- --- --- --- --- --- --- --- --- --- --- --- --- --- --- --- --- --- --- --- --- --- --- --- --- --- --- --- --- --- --- --- --- --- --- --- --- --- --- --- --- --- --- --- --- --- --- --- --- --- --- --- --- --- --- --- --- --- --- --- --- --- --- --- --- --- --- --- --- --- --- --- --- --- --- --- --- --- --- --- --- --- --- --- --- --- --- --- --- --- --- --- --- --- --- --- --- --- --- --- --- --- --- --- --- --- --- --- --- --- --- --- --- --- --- --- --- --- --- --- --- --- --- --- --- --- --- --- --- --- --- --- --- --- --- --- --- --- --- --- --- --- --- --- --- --- --- --- --- --- --- --- --- --- --- --- --- --- --- --- --- --- --- --- --- --- --- --- --- --- --- --- --- --- --- --- --- --- --- --- --- --- --- --- --- --- --- --- --- --- --- --- --- --- --- --- --- --- --- --- --- --- --- --- --- --- --- --- --- --- --- --- --- --- --- --- --- --- --- --- --- --- --- --- --- --- --- --- --- --- --- --- --- --- --- --- --- --- --- --- --- --- --- --- --- --- --- --- --- --- --- --- --- --- --- --- --- --- --- --- --- --- --- --- --- --- --- --- --- --- --- --- --- --- --- --- --- --- --- --- --- --- --- --- --- --- --- --- --- --- --- --- --- --- --- --- --- --- --- --- --- --- --- --- --- --- --- --- --- --- --- --- --- --- --- --- --- --- --- --- --- --- --- --- --- --- --- --- --- --- --- --- --- --- --- --- --- --- --- --- --- --- --- --- --- --- --- --- --- --- --- --- --- --- --- --- --- --- --- --- --- --- --- --- --- --- --- --- --- --- --- --- --- --- --- --- --- --- --- --- --- --- --- --- --- --- --- --- --- --- --- --- --- --- --- --- --- --- --- --- --- --- --- --- --- --- --- --- --- --- --- --- --- --- --- --- --- --- --- --- --- --- --- --- --- --- --- --- --- --- --- --- --- --- --- --- --- --- --- --- --- --- --- --- --- --- --- --- --- --- --- --- --- --- --- --- --- --- --- --- --- --- --- --- --- --- --- --- --- --- --- --- --- --- --- --- --- --- --- --- --- --- --- --- --- --- --- --- --- --- --- --- --- --- --- --- --- --- --- --- --- --- --- --- --- --- --- --- --- --- --- --- --- --- --- --- --- --- -ct gga tcg gat gtc acc gca gta ata ttg ttg att att tct gac atc gac gta tta tat agt ttt tta att cca tat ctt ttt t

>MG693723.1_Monkeypox_virus_isolate_MPXV_Nig_2017_297957_partial_genome

ttt tat atc act acg gac ata aac cat tgt ata ttt ttt atg ttt att agt gta cac att ttg gaa gta agt tc- --- --- --- --- --- --- --- --- --- --- --- --- --- --- --- --- --- --- --- --- --- --- --- --- --- --- --- --- --- --- --- --- --- --- --- --- --- --- --- --- --- --- --- --- --- --- --- --- --- --- --- --- --- --- --- --- --- --- --- --- --- --- --- --- --- --- --- --- --- --- --- --- --- --- --- --- --- --- --- --- --- --- --- --- --- --- --- --- --- --- --- --- --- --- --- --- --- --- --- --- --- --- --- --- --- --- --- --- --- --- --- --- --- --- --- --- --- --- --- --- --- --- --- --- --- --- --- --- --- --- --- --- --- --- --- --- --- --- --- --- --- --- --- --- --- --- --- --- --- --- --- --- --- --- --- --- --- --- --- --- --- --- --- --- --- --- --- --- --- --- --- --- --- --- --- --- --- --- --- --- --- --- --- --- --- --- --- --- --- --- --- --- --- --- --- --- --- --- --- --- --- --- --- --- --- --- --- --- --- --- --- --- --- --- --- --- --- --- --- --- --- --- --- --- --- --- --- --- --- --- --- --- --- --- --- --- --- --- --- --- --- --- --- --- --- --- --- --- --- --- --- --- --- --- --- --- --- --- --- --- --- --- --- --- --- --- --- --- --- --- --- --- --- --- --- --- --- --- --- --- --- --- --- --- --- --- --- --- --- --- --- --- --- --- --- --- --- --- --- --- --- --- --- --- --- --- --- --- --- --- --- --- --- --- --- --- --- --- --- --- --- --- --- --- --- --- --- --- --- --- --- --- --- --- --- --- --- --- --- --- --- --- --- --- --- --- --- --- --- --- --- --- --- --- --- --- --- --- --- --- --- --- --- --- --- --- --- --- --- --- --- --- --- --- --- --- --- --- --- --- --- --- --- --- --- --- --- --- --- --- --- --- --- --- --- --- --- --- --- --- --- --- --- --- --- --- --- --- --- --- --- --- --- --- --- --- --- --- --- --- --- --- --- --- --- --- --- --- --- --- --- --- --- --- --- --- --- --- --- --- --- --- --- --- --- --- --- --- --- --- --- --- --- --- --- --- --- --- --- --- --- --- --- --- --- --- --- --- --- --- --- --- --- --- --- --- --- --- --- --- --- --- --- --- --- --- --- --- --- --- --- --- --- --- --- --- --- --- --- --- --- --- --- --- --- --- --- --- --- --- --- --- --- --- --- --- --- --- --- --- --- --- --- --- --- --- --- --- --- --- --- --- --- --- --- --- --- --- --- --- --- --- --- --- --- --- --- --- --- --- --- --- --- --- --- --- --- --- --- --- --- --- --- --- --- --- --- --- --- --- --- --- --- --- --- --- --- --- --- --- --- --- --- --- --- --- --- --- --- --- --- --- --- --- --- --- --- --- --- --- --- --- --- --- --- --- --- --- --- --- --- --- --- --- --- --- --- --- --- --- --- --- --- --- --- --- --- --- --- --- --- --- --- --- --- --- --- --- --- --- --- --- --- --- --- --- --- --- --- --- --- --- -ct gga tcg gat gtc acc gca gta ata ttg ttg att att tct gac atc gac gta tta tat agt ttt tta att cca tat ctt ttt t

>ON585031.1_Monkeypox_virus_isolate_Monkeypox/PT0003/2022_complete_genome

ttt tat atc act acg gac ata aac cat tgt ata ttt ttt atg ttt att agt gta cac att ttg gaa gta agt tc- --- --- --- --- --- --- --- --- --- --- --- --- --- --- --- --- --- --- --- --- --- --- --- --- --- --- --- --- --- --- --- --- --- --- --- --- --- --- --- --- --- --- --- --- --- --- --- --- --- --- --- --- --- --- --- --- --- --- --- --- --- --- --- --- --- --- --- --- --- --- --- --- --- --- --- --- --- --- --- --- --- --- --- --- --- --- --- --- --- --- --- --- --- --- --- --- --- --- --- --- --- --- --- --- --- --- --- --- --- --- --- --- --- --- --- --- --- --- --- --- --- --- --- --- --- --- --- --- --- --- --- --- --- --- --- --- --- --- --- --- --- --- --- --- --- --- --- --- --- --- --- --- --- --- --- --- --- --- --- --- --- --- --- --- --- --- --- --- --- --- --- --- --- --- --- --- --- --- --- --- --- --- --- --- --- --- --- --- --- --- --- --- --- --- --- --- --- --- --- --- --- --- --- --- --- --- --- --- --- --- --- --- --- --- --- --- --- --- --- --- --- --- --- --- --- --- --- --- --- --- --- --- --- --- --- --- --- --- --- --- --- --- --- --- --- --- --- --- --- --- --- --- --- --- --- --- --- --- --- --- --- --- --- --- --- --- --- --- --- --- --- --- --- --- --- --- --- --- --- --- --- --- --- --- --- --- --- --- --- --- --- --- --- --- --- --- --- --- --- --- --- --- --- --- --- --- --- --- --- --- --- --- --- --- --- --- --- --- --- --- --- --- --- --- --- --- --- --- --- --- --- --- --- --- --- --- --- --- --- --- --- --- --- --- --- --- --- --- --- --- --- --- --- --- --- --- --- --- --- --- --- --- --- --- --- --- --- --- --- --- --- --- --- --- --- --- --- --- --- --- --- --- --- --- --- --- --- --- --- --- --- --- --- --- --- --- --- --- --- --- --- --- --- --- --- --- --- --- --- --- --- --- --- --- --- --- --- --- --- --- --- --- --- --- --- --- --- --- --- --- --- --- --- --- --- --- --- --- --- --- --- --- --- --- --- --- --- --- --- --- --- --- --- --- --- --- --- --- --- --- --- --- --- --- --- --- --- --- --- --- --- --- --- --- --- --- --- --- --- --- --- --- --- --- --- --- --- --- --- --- --- --- --- --- --- --- --- --- --- --- --- --- --- --- --- --- --- --- --- --- --- --- --- --- --- --- --- --- --- --- --- --- --- --- --- --- --- --- --- --- --- --- --- --- --- --- --- --- --- --- --- --- --- --- --- --- --- --- --- --- --- --- --- --- --- --- --- --- --- --- --- --- --- --- --- --- --- --- --- --- --- --- --- --- --- --- --- --- --- --- --- --- --- --- --- --- --- --- --- --- --- --- --- --- --- --- --- --- --- --- --- --- --- --- --- --- --- --- --- --- --- --- --- --- --- --- --- --- --- --- --- --- --- --- --- --- --- --- --- --- --- --- --- --- --- --- --- --- --- --- --- --- --- --- --- --- --- --- --- --- --- --- -ct gga tcg gat gtc acc gca gta ata ttg ttg att att tct gac atc gac gta tta tat agt ttt tta att cca tat ctt ttt t

>ON585038.1_Monkeypox_virus_isolate_Monkeypox/PT0008/2022_complete_genome

ttt tat atc act acg gac ata aac cat tgt ata ttt ttt atg ttt att agt gta cac att ttg gaa gta agt tc- --- --- --- --- --- --- --- --- --- --- --- --- --- --- --- --- --- --- --- --- --- --- --- --- --- --- --- --- --- --- --- --- --- --- --- --- --- --- --- --- --- --- --- --- --- --- --- --- --- --- --- --- --- --- --- --- --- --- --- --- --- --- --- --- --- --- --- --- --- --- --- --- --- --- --- --- --- --- --- --- --- --- --- --- --- --- --- --- --- --- --- --- --- --- --- --- --- --- --- --- --- --- --- --- --- --- --- --- --- --- --- --- --- --- --- --- --- --- --- --- --- --- --- --- --- --- --- --- --- --- --- --- --- --- --- --- --- --- --- --- --- --- --- --- --- --- --- --- --- --- --- --- --- --- --- --- --- --- --- --- --- --- --- --- --- --- --- --- --- --- --- --- --- --- --- --- --- --- --- --- --- --- --- --- --- --- --- --- --- --- --- --- --- --- --- --- --- --- --- --- --- --- --- --- --- --- --- --- --- --- --- --- --- --- --- --- --- --- --- --- --- --- --- --- --- --- --- --- --- --- --- --- --- --- --- --- --- --- --- --- --- --- --- --- --- --- --- --- --- --- --- --- --- --- --- --- --- --- --- --- --- --- --- --- --- --- --- --- --- --- --- --- --- --- --- --- --- --- --- --- --- --- --- --- --- --- --- --- --- --- --- --- --- --- --- --- --- --- --- --- --- --- --- --- --- --- --- --- --- --- --- --- --- --- --- --- --- --- --- --- --- --- --- --- --- --- --- --- --- --- --- --- --- --- --- --- --- --- --- --- --- --- --- --- --- --- --- --- --- --- --- --- --- --- --- --- --- --- --- --- --- --- --- --- --- --- --- --- --- --- --- --- --- --- --- --- --- --- --- --- --- --- --- --- --- --- --- --- --- --- --- --- --- --- --- --- --- --- --- --- --- --- --- --- --- --- --- --- --- --- --- --- --- --- --- --- --- --- --- --- --- --- --- --- --- --- --- --- --- --- --- --- --- --- --- --- --- --- --- --- --- --- --- --- --- --- --- --- --- --- --- --- --- --- --- --- --- --- --- --- --- --- --- --- --- --- --- --- --- --- --- --- --- --- --- --- --- --- --- --- --- --- --- --- --- --- --- --- --- --- --- --- --- --- --- --- --- --- --- --- --- --- --- --- --- --- --- --- --- --- --- --- --- --- --- --- --- --- --- --- --- --- --- --- --- --- --- --- --- --- --- --- --- --- --- --- --- --- --- --- --- --- --- --- --- --- --- --- --- --- --- --- --- --- --- --- --- --- --- --- --- --- --- --- --- --- --- --- --- --- --- --- --- --- --- --- --- --- --- --- --- --- --- --- --- --- --- --- --- --- --- --- --- --- --- --- --- --- --- --- --- --- --- --- --- --- --- --- --- --- --- --- --- --- --- --- --- --- --- --- --- --- --- --- --- --- --- --- --- --- --- --- --- --- --- --- --- --- --- --- --- --- --- --- --- --- --- --- --- --- --- --- -ct gga tcg gat gtc acc gca gta ata ttg ttg att att tct gac atc gac gta tta tat agt ttt tta att cca tat ctt ttt t

>DQ011156.1_Monkeypox_virus_strain_Liberia_1970_184_complete_genome

ttt tat atc act acg gac ata aac cat tgt ata att ttt atg ttt att agt gta cac att ttg gaa gta agt tc- --- --- --- --- --- --- --- --- --- --- --- --- --- --- --- --- --- --- --- --- --- --- --- --- --- --- --- --- --- --- --- --- --- --- --- --- --- --- --- --- --- --- --- --- --- --- --- --- --- --- --- --- --- --- --- --- --- --- --- --- --- --- --- --- --- --- --- --- --- --- --- --- --- --- --- --- --- --- --- --- --- --- --- --- --- --- --- --- --- --- --- --- --- --- --- --- --- --- --- --- --- --- --- --- --- --- --- --- --- --- --- --- --- --- --- --- --- --- --- --- --- --- --- --- --- --- --- --- --- --- --- --- --- --- --- --- --- --- --- --- --- --- --- --- --- --- --- --- --- --- --- --- --- --- --- --- --- --- --- --- --- --- --- --- --- --- --- --- --- --- --- --- --- --- --- --- --- --- --- --- --- --- --- --- --- --- --- --- --- --- --- --- --- --- --- --- --- --- --- --- --- --- --- --- --- --- --- --- --- --- --- --- --- --- --- --- --- --- --- --- --- --- --- --- --- --- --- --- --- --- --- --- --- --- --- --- --- --- --- --- --- --- --- --- --- --- --- --- --- --- --- --- --- --- --- --- --- --- --- --- --- --- --- --- --- --- --- --- --- --- --- --- --- --- --- --- --- --- --- --- --- --- --- --- --- --- --- --- --- --- --- --- --- --- --- --- --- --- --- --- --- --- --- --- --- --- --- --- --- --- --- --- --- --- --- --- --- --- --- --- --- --- --- --- --- --- --- --- --- --- --- --- --- --- --- --- --- --- --- --- --- --- --- --- --- --- --- --- --- --- --- --- --- --- --- --- --- --- --- --- --- --- --- --- --- --- --- --- --- --- --- --- --- --- --- --- --- --- --- --- --- --- --- --- --- --- --- --- --- --- --- --- --- --- --- --- --- --- --- --- --- --- --- --- --- --- --- --- --- --- --- --- --- --- --- --- --- --- --- --- --- --- --- --- --- --- --- --- --- --- --- --- --- --- --- --- --- --- --- --- --- --- --- --- --- --- --- --- --- --- --- --- --- --- --- --- --- --- --- --- --- --- --- --- --- --- --- --- --- --- --- --- --- --- --- --- --- --- --- --- --- --- --- --- --- --- --- --- --- --- --- --- --- --- --- --- --- --- --- --- --- --- --- --- --- --- --- --- --- --- --- --- --- --- --- --- --- --- --- --- --- --- --- --- --- --- --- --- --- --- --- --- --- --- --- --- --- --- --- --- --- --- --- --- --- --- --- --- --- --- --- --- --- --- --- --- --- --- --- --- --- --- --- --- --- --- --- --- --- --- --- --- --- --- --- --- --- --- --- --- --- --- --- --- --- --- --- --- --- --- --- --- --- --- --- --- --- --- --- --- --- --- --- --- --- --- --- --- --- --- --- --- --- --- --- --- --- --- --- --- --- --- --- --- --- --- --- --- --- --- --- --- --- --- --- --- --- --- --- --- --- --- --- --- --- --- --- --- --- --- --- --- -ct gga tcg gat gtc acc gca gta ata ttg ttg att att tct gac atc gac gta tta tat agt ttt tta att cca tat ctt ttt t

>KP849470.1_Monkeypox_virus_isolate_Cote_dIvoire_1971_complete_genome

ttt tat atc act acg gac ata aac cat tgt ata att ttt atg ttt att agt gta cac att ttg gaa gta agt tc- --- --- --- --- --- --- --- --- --- --- --- --- --- --- --- --- --- --- --- --- --- --- --- --- --- --- --- --- --- --- --- --- --- --- --- --- --- --- --- --- --- --- --- --- --- --- --- --- --- --- --- --- --- --- --- --- --- --- --- --- --- --- --- --- --- --- --- --- --- --- --- --- --- --- --- --- --- --- --- --- --- --- --- --- --- --- --- --- --- --- --- --- --- --- --- --- --- --- --- --- --- --- --- --- --- --- --- --- --- --- --- --- --- --- --- --- --- --- --- --- --- --- --- --- --- --- --- --- --- --- --- --- --- --- --- --- --- --- --- --- --- --- --- --- --- --- --- --- --- --- --- --- --- --- --- --- --- --- --- --- --- --- --- --- --- --- --- --- --- --- --- --- --- --- --- --- --- --- --- --- --- --- --- --- --- --- --- --- --- --- --- --- --- --- --- --- --- --- --- --- --- --- --- --- --- --- --- --- --- --- --- --- --- --- --- --- --- --- --- --- --- --- --- --- --- --- --- --- --- --- --- --- --- --- --- --- --- --- --- --- --- --- --- --- --- --- --- --- --- --- --- --- --- --- --- --- --- --- --- --- --- --- --- --- --- --- --- --- --- --- --- --- --- --- --- --- --- --- --- --- --- --- --- --- --- --- --- --- --- --- --- --- --- --- --- --- --- --- --- --- --- --- --- --- --- --- --- --- --- --- --- --- --- --- --- --- --- --- --- --- --- --- --- --- --- --- --- --- --- --- --- --- --- --- --- --- --- --- --- --- --- --- --- --- --- --- --- --- --- --- --- --- --- --- --- --- --- --- --- --- --- --- --- --- --- --- --- --- --- --- --- --- --- --- --- --- --- --- --- --- --- --- --- --- --- --- --- --- --- --- --- --- --- --- --- --- --- --- --- --- --- --- --- --- --- --- --- --- --- --- --- --- --- --- --- --- --- --- --- --- --- --- --- --- --- --- --- --- --- --- --- --- --- --- --- --- --- --- --- --- --- --- --- --- --- --- --- --- --- --- --- --- --- --- --- --- --- --- --- --- --- --- --- --- --- --- --- --- --- --- --- --- --- --- --- --- --- --- --- --- --- --- --- --- --- --- --- --- --- --- --- --- --- --- --- --- --- --- --- --- --- --- --- --- --- --- --- --- --- --- --- --- --- --- --- --- --- --- --- --- --- --- --- --- --- --- --- --- --- --- --- --- --- --- --- --- --- --- --- --- --- --- --- --- --- --- --- --- --- --- --- --- --- --- --- --- --- --- --- --- --- --- --- --- --- --- --- --- --- --- --- --- --- --- --- --- --- --- --- --- --- --- --- --- --- --- --- --- --- --- --- --- --- --- --- --- --- --- --- --- --- --- --- --- --- --- --- --- --- --- --- --- --- --- --- --- --- --- --- --- --- --- --- --- --- --- --- --- --- --- --- --- --- --- --- --- --- --- --- --- --- --- --- --- --- --- --- --- --- --- --- --- -ct gga tcg gat gtc acc gca gta ata ttg ttg att att tct gac atc gac gta tta tat agt ttt tta att cca tat ctt ttt t

>ON682264.2_Monkeypox_virus_isolate_MPXV/Germany/2022/RKI05_complete_genome

ttt tat atc act acg gac ata aac cat tgt ata ttt ttt atg ttt att agt gta cac att ttg gaa gta agt tc- --- --- --- --- --- --- --- --- --- --- --- --- --- --- --- --- --- --- --- --- --- --- --- --- --- --- --- --- --- --- --- --- --- --- --- --- --- --- --- --- --- --- --- --- --- --- --- --- --- --- --- --- --- --- --- --- --- --- --- --- --- --- --- --- --- --- --- --- --- --- --- --- --- --- --- --- --- --- --- --- --- --- --- --- --- --- --- --- --- --- --- --- --- --- --- --- --- --- --- --- --- --- --- --- --- --- --- --- --- --- --- --- --- --- --- --- --- --- --- --- --- --- --- --- --- --- --- --- --- --- --- --- --- --- --- --- --- --- --- --- --- --- --- --- --- --- --- --- --- --- --- --- --- --- --- --- --- --- --- --- --- --- --- --- --- --- --- --- --- --- --- --- --- --- --- --- --- --- --- --- --- --- --- --- --- --- --- --- --- --- --- --- --- --- --- --- --- --- --- --- --- --- --- --- --- --- --- --- --- --- --- --- --- --- --- --- --- --- --- --- --- --- --- --- --- --- --- --- --- --- --- --- --- --- --- --- --- --- --- --- --- --- --- --- --- --- --- --- --- --- --- --- --- --- --- --- --- --- --- --- --- --- --- --- --- --- --- --- --- --- --- --- --- --- --- --- --- --- --- --- --- --- --- --- --- --- --- --- --- --- --- --- --- --- --- --- --- --- --- --- --- --- --- --- --- --- --- --- --- --- --- --- --- --- --- --- --- --- --- --- --- --- --- --- --- --- --- --- --- --- --- --- --- --- --- --- --- --- --- --- --- --- --- --- --- --- --- --- --- --- --- --- --- --- --- --- --- --- --- --- --- --- --- --- --- --- --- --- --- --- --- --- --- --- --- --- --- --- --- --- --- --- --- --- --- --- --- --- --- --- --- --- --- --- --- --- --- --- --- --- --- --- --- --- --- --- --- --- --- --- --- --- --- --- --- --- --- --- --- --- --- --- --- --- --- --- --- --- --- --- --- --- --- --- --- --- --- --- --- --- --- --- --- --- --- --- --- --- --- --- --- --- --- --- --- --- --- --- --- --- --- --- --- --- --- --- --- --- --- --- --- --- --- --- --- --- --- --- --- --- --- --- --- --- --- --- --- --- --- --- --- --- --- --- --- --- --- --- --- --- --- --- --- --- --- --- --- --- --- --- --- --- --- --- --- --- --- --- --- --- --- --- --- --- --- --- --- --- --- --- --- --- --- --- --- --- --- --- --- --- --- --- --- --- --- --- --- --- --- --- --- --- --- --- --- --- --- --- --- --- --- --- --- --- --- --- --- --- --- --- --- --- --- --- --- --- --- --- --- --- --- --- --- --- --- --- --- --- --- --- --- --- --- --- --- --- --- --- --- --- --- --- --- --- --- --- --- --- --- --- --- --- --- --- --- --- --- --- --- --- --- --- --- --- --- --- --- --- --- --- --- --- --- --- --- --- --- --- --- --- --- --- --- --- --- --- --- --- --- --- --- --- -ct gga tcg gat gtc acc gca gta ata ttg ttg att att tct gac atc gac gta tta tat agt ttt tta att cca tat ctt ttt t

>ON682268.1_Monkeypox_virus_isolate_MPXV/Germany/2022/RKI08_complete_genome

ttt tat atc act acg gac ata aac cat tgt ata ttt ttt atg ttt att agt gta cac att ttg gaa gta agt tc- --- --- --- --- --- --- --- --- --- --- --- --- --- --- --- --- --- --- --- --- --- --- --- --- --- --- --- --- --- --- --- --- --- --- --- --- --- --- --- --- --- --- --- --- --- --- --- --- --- --- --- --- --- --- --- --- --- --- --- --- --- --- --- --- --- --- --- --- --- --- --- --- --- --- --- --- --- --- --- --- --- --- --- --- --- --- --- --- --- --- --- --- --- --- --- --- --- --- --- --- --- --- --- --- --- --- --- --- --- --- --- --- --- --- --- --- --- --- --- --- --- --- --- --- --- --- --- --- --- --- --- --- --- --- --- --- --- --- --- --- --- --- --- --- --- --- --- --- --- --- --- --- --- --- --- --- --- --- --- --- --- --- --- --- --- --- --- --- --- --- --- --- --- --- --- --- --- --- --- --- --- --- --- --- --- --- --- --- --- --- --- --- --- --- --- --- --- --- --- --- --- --- --- --- --- --- --- --- --- --- --- --- --- --- --- --- --- --- --- --- --- --- --- --- --- --- --- --- --- --- --- --- --- --- --- --- --- --- --- --- --- --- --- --- --- --- --- --- --- --- --- --- --- --- --- --- --- --- --- --- --- --- --- --- --- --- --- --- --- --- --- --- --- --- --- --- --- --- --- --- --- --- --- --- --- --- --- --- --- --- --- --- --- --- --- --- --- --- --- --- --- --- --- --- --- --- --- --- --- --- --- --- --- --- --- --- --- --- --- --- --- --- --- --- --- --- --- --- --- --- --- --- --- --- --- --- --- --- --- --- --- --- --- --- --- --- --- --- --- --- --- --- --- --- --- --- --- --- --- --- --- --- --- --- --- --- --- --- --- --- --- --- --- --- --- --- --- --- --- --- --- --- --- --- --- --- --- --- --- --- --- --- --- --- --- --- --- --- --- --- --- --- --- --- --- --- --- --- --- --- --- --- --- --- --- --- --- --- --- --- --- --- --- --- --- --- --- --- --- --- --- --- --- --- --- --- --- --- --- --- --- --- --- --- --- --- --- --- --- --- --- --- --- --- --- --- --- --- --- --- --- --- --- --- --- --- --- --- --- --- --- --- --- --- --- --- --- --- --- --- --- --- --- --- --- --- --- --- --- --- --- --- --- --- --- --- --- --- --- --- --- --- --- --- --- --- --- --- --- --- --- --- --- --- --- --- --- --- --- --- --- --- --- --- --- --- --- --- --- --- --- --- --- --- --- --- --- --- --- --- --- --- --- --- --- --- --- --- --- --- --- --- --- --- --- --- --- --- --- --- --- --- --- --- --- --- --- --- --- --- --- --- --- --- --- --- --- --- --- --- --- --- --- --- --- --- --- --- --- --- --- --- --- --- --- --- --- --- --- --- --- --- --- --- --- --- --- --- --- --- --- --- --- --- --- --- --- --- --- --- --- --- --- --- --- --- --- --- --- --- --- --- --- --- --- --- --- --- --- --- --- --- --- --- --- --- --- --- --- --- --- --- -ct gga tcg gat gtc acc gca gta ata ttg ttg att att tct gac atc gac gta tta tat agt ttt tta att cca tat ctt ttt t

>ON682270.1_Monkeypox_virus_isolate_MPXV/Germany/2022/RKI06_complete_genome

ttt tat atc act acg gac ata aac cat tgt ata ttt ttt atg ttt att agt gta cac att ttg gaa gta agt tc- --- --- --- --- --- --- --- --- --- --- --- --- --- --- --- --- --- --- --- --- --- --- --- --- --- --- --- --- --- --- --- --- --- --- --- --- --- --- --- --- --- --- --- --- --- --- --- --- --- --- --- --- --- --- --- --- --- --- --- --- --- --- --- --- --- --- --- --- --- --- --- --- --- --- --- --- --- --- --- --- --- --- --- --- --- --- --- --- --- --- --- --- --- --- --- --- --- --- --- --- --- --- --- --- --- --- --- --- --- --- --- --- --- --- --- --- --- --- --- --- --- --- --- --- --- --- --- --- --- --- --- --- --- --- --- --- --- --- --- --- --- --- --- --- --- --- --- --- --- --- --- --- --- --- --- --- --- --- --- --- --- --- --- --- --- --- --- --- --- --- --- --- --- --- --- --- --- --- --- --- --- --- --- --- --- --- --- --- --- --- --- --- --- --- --- --- --- --- --- --- --- --- --- --- --- --- --- --- --- --- --- --- --- --- --- --- --- --- --- --- --- --- --- --- --- --- --- --- --- --- --- --- --- --- --- --- --- --- --- --- --- --- --- --- --- --- --- --- --- --- --- --- --- --- --- --- --- --- --- --- --- --- --- --- --- --- --- --- --- --- --- --- --- --- --- --- --- --- --- --- --- --- --- --- --- --- --- --- --- --- --- --- --- --- --- --- --- --- --- --- --- --- --- --- --- --- --- --- --- --- --- --- --- --- --- --- --- --- --- --- --- --- --- --- --- --- --- --- --- --- --- --- --- --- --- --- --- --- --- --- --- --- --- --- --- --- --- --- --- --- --- --- --- --- --- --- --- --- --- --- --- --- --- --- --- --- --- --- --- --- --- --- --- --- --- --- --- --- --- --- --- --- --- --- --- --- --- --- --- --- --- --- --- --- --- --- --- --- --- --- --- --- --- --- --- --- --- --- --- --- --- --- --- --- --- --- --- --- --- --- --- --- --- --- --- --- --- --- --- --- --- --- --- --- --- --- --- --- --- --- --- --- --- --- --- --- --- --- --- --- --- --- --- --- --- --- --- --- --- --- --- --- --- --- --- --- --- --- --- --- --- --- --- --- --- --- --- --- --- --- --- --- --- --- --- --- --- --- --- --- --- --- --- --- --- --- --- --- --- --- --- --- --- --- --- --- --- --- --- --- --- --- --- --- --- --- --- --- --- --- --- --- --- --- --- --- --- --- --- --- --- --- --- --- --- --- --- --- --- --- --- --- --- --- --- --- --- --- --- --- --- --- --- --- --- --- --- --- --- --- --- --- --- --- --- --- --- --- --- --- --- --- --- --- --- --- --- --- --- --- --- --- --- --- --- --- --- --- --- --- --- --- --- --- --- --- --- --- --- --- --- --- --- --- --- --- --- --- --- --- --- --- --- --- --- --- --- --- --- --- --- --- --- --- --- --- --- --- --- --- --- --- --- --- --- --- --- --- --- --- --- --- --- --- --- --- --- --- --- --- --- --- -ct gga tcg gat gtc acc gca gta ata ttg ttg att att tct gac atc gac gta tta tat agt ttt tta att cca tat ctt ttt t

>ON637938.1_Monkeypox_virus_isolate_MPXV/Germany/2022/RKI01_complete_genome

ttt tat atc act acg gac ata aac cat tgt ata ttt ttt atg ttt att agt gta cac att ttg gaa gta agt tc- --- --- --- --- --- --- --- --- --- --- --- --- --- --- --- --- --- --- --- --- --- --- --- --- --- --- --- --- --- --- --- --- --- --- --- --- --- --- --- --- --- --- --- --- --- --- --- --- --- --- --- --- --- --- --- --- --- --- --- --- --- --- --- --- --- --- --- --- --- --- --- --- --- --- --- --- --- --- --- --- --- --- --- --- --- --- --- --- --- --- --- --- --- --- --- --- --- --- --- --- --- --- --- --- --- --- --- --- --- --- --- --- --- --- --- --- --- --- --- --- --- --- --- --- --- --- --- --- --- --- --- --- --- --- --- --- --- --- --- --- --- --- --- --- --- --- --- --- --- --- --- --- --- --- --- --- --- --- --- --- --- --- --- --- --- --- --- --- --- --- --- --- --- --- --- --- --- --- --- --- --- --- --- --- --- --- --- --- --- --- --- --- --- --- --- --- --- --- --- --- --- --- --- --- --- --- --- --- --- --- --- --- --- --- --- --- --- --- --- --- --- --- --- --- --- --- --- --- --- --- --- --- --- --- --- --- --- --- --- --- --- --- --- --- --- --- --- --- --- --- --- --- --- --- --- --- --- --- --- --- --- --- --- --- --- --- --- --- --- --- --- --- --- --- --- --- --- --- --- --- --- --- --- --- --- --- --- --- --- --- --- --- --- --- --- --- --- --- --- --- --- --- --- --- --- --- --- --- --- --- --- --- --- --- --- --- --- --- --- --- --- --- --- --- --- --- --- --- --- --- --- --- --- --- --- --- --- --- --- --- --- --- --- --- --- --- --- --- --- --- --- --- --- --- --- --- --- --- --- --- --- --- --- --- --- --- --- --- --- --- --- --- --- --- --- --- --- --- --- --- --- --- --- --- --- --- --- --- --- --- --- --- --- --- --- --- --- --- --- --- --- --- --- --- --- --- --- --- --- --- --- --- --- --- --- --- --- --- --- --- --- --- --- --- --- --- --- --- --- --- --- --- --- --- --- --- --- --- --- --- --- --- --- --- --- --- --- --- --- --- --- --- --- --- --- --- --- --- --- --- --- --- --- --- --- --- --- --- --- --- --- --- --- --- --- --- --- --- --- --- --- --- --- --- --- --- --- --- --- --- --- --- --- --- --- --- --- --- --- --- --- --- --- --- --- --- --- --- --- --- --- --- --- --- --- --- --- --- --- --- --- --- --- --- --- --- --- --- --- --- --- --- --- --- --- --- --- --- --- --- --- --- --- --- --- --- --- --- --- --- --- --- --- --- --- --- --- --- --- --- --- --- --- --- --- --- --- --- --- --- --- --- --- --- --- --- --- --- --- --- --- --- --- --- --- --- --- --- --- --- --- --- --- --- --- --- --- --- --- --- --- --- --- --- --- --- --- --- --- --- --- --- --- --- --- --- --- --- --- --- --- --- --- --- --- --- --- --- --- --- --- --- --- --- --- --- --- --- --- --- --- --- --- --- --- --- --- --- --- --- --- --- -ct gga tcg gat gtc acc gca gta ata ttg ttg att att tct gac atc gac gta tta tat agt ttt tta att cca tat ctt ttt t

>ON637939.1_Monkeypox_virus_isolate_MPXV/Germany/2022/RKI02_complete_genome

ttt tat atc act acg gac ata aac cat tgt ata ttt ttt atg ttt att agt gta cac att ttg gaa gta agt tc- --- --- --- --- --- --- --- --- --- --- --- --- --- --- --- --- --- --- --- --- --- --- --- --- --- --- --- --- --- --- --- --- --- --- --- --- --- --- --- --- --- --- --- --- --- --- --- --- --- --- --- --- --- --- --- --- --- --- --- --- --- --- --- --- --- --- --- --- --- --- --- --- --- --- --- --- --- --- --- --- --- --- --- --- --- --- --- --- --- --- --- --- --- --- --- --- --- --- --- --- --- --- --- --- --- --- --- --- --- --- --- --- --- --- --- --- --- --- --- --- --- --- --- --- --- --- --- --- --- --- --- --- --- --- --- --- --- --- --- --- --- --- --- --- --- --- --- --- --- --- --- --- --- --- --- --- --- --- --- --- --- --- --- --- --- --- --- --- --- --- --- --- --- --- --- --- --- --- --- --- --- --- --- --- --- --- --- --- --- --- --- --- --- --- --- --- --- --- --- --- --- --- --- --- --- --- --- --- --- --- --- --- --- --- --- --- --- --- --- --- --- --- --- --- --- --- --- --- --- --- --- --- --- --- --- --- --- --- --- --- --- --- --- --- --- --- --- --- --- --- --- --- --- --- --- --- --- --- --- --- --- --- --- --- --- --- --- --- --- --- --- --- --- --- --- --- --- --- --- --- --- --- --- --- --- --- --- --- --- --- --- --- --- --- --- --- --- --- --- --- --- --- --- --- --- --- --- --- --- --- --- --- --- --- --- --- --- --- --- --- --- --- --- --- --- --- --- --- --- --- --- --- --- --- --- --- --- --- --- --- --- --- --- --- --- --- --- --- --- --- --- --- --- --- --- --- --- --- --- --- --- --- --- --- --- --- --- --- --- --- --- --- --- --- --- --- --- --- --- --- --- --- --- --- --- --- --- --- --- --- --- --- --- --- --- --- --- --- --- --- --- --- --- --- --- --- --- --- --- --- --- --- --- --- --- --- --- --- --- --- --- --- --- --- --- --- --- --- --- --- --- --- --- --- --- --- --- --- --- --- --- --- --- --- --- --- --- --- --- --- --- --- --- --- --- --- --- --- --- --- --- --- --- --- --- --- --- --- --- --- --- --- --- --- --- --- --- --- --- --- --- --- --- --- --- --- --- --- --- --- --- --- --- --- --- --- --- --- --- --- --- --- --- --- --- --- --- --- --- --- --- --- --- --- --- --- --- --- --- --- --- --- --- --- --- --- --- --- --- --- --- --- --- --- --- --- --- --- --- --- --- --- --- --- --- --- --- --- --- --- --- --- --- --- --- --- --- --- --- --- --- --- --- --- --- --- --- --- --- --- --- --- --- --- --- --- --- --- --- --- --- --- --- --- --- --- --- --- --- --- --- --- --- --- --- --- --- --- --- --- --- --- --- --- --- --- --- --- --- --- --- --- --- --- --- --- --- --- --- --- --- --- --- --- --- --- --- --- --- --- --- --- --- --- --- --- --- --- --- --- --- --- --- --- --- --- --- --- --- --- --- --- -ct gga tcg gat gtc acc gca gta ata ttg ttg att att tct gac atc gac gta tta tat agt ttt tta att cca tat ctt ttt t

>ON682263.2_Monkeypox_virus_isolate_MPXV/Germany/2022/RKI03_complete_genome

ttt tat atc act acg gac ata aac cat tgt ata ttt ttt atg ttt att agt gta cac att ttg gaa gta agt tc- --- --- --- --- --- --- --- --- --- --- --- --- --- --- --- --- --- --- --- --- --- --- --- --- --- --- --- --- --- --- --- --- --- --- --- --- --- --- --- --- --- --- --- --- --- --- --- --- --- --- --- --- --- --- --- --- --- --- --- --- --- --- --- --- --- --- --- --- --- --- --- --- --- --- --- --- --- --- --- --- --- --- --- --- --- --- --- --- --- --- --- --- --- --- --- --- --- --- --- --- --- --- --- --- --- --- --- --- --- --- --- --- --- --- --- --- --- --- --- --- --- --- --- --- --- --- --- --- --- --- --- --- --- --- --- --- --- --- --- --- --- --- --- --- --- --- --- --- --- --- --- --- --- --- --- --- --- --- --- --- --- --- --- --- --- --- --- --- --- --- --- --- --- --- --- --- --- --- --- --- --- --- --- --- --- --- --- --- --- --- --- --- --- --- --- --- --- --- --- --- --- --- --- --- --- --- --- --- --- --- --- --- --- --- --- --- --- --- --- --- --- --- --- --- --- --- --- --- --- --- --- --- --- --- --- --- --- --- --- --- --- --- --- --- --- --- --- --- --- --- --- --- --- --- --- --- --- --- --- --- --- --- --- --- --- --- --- --- --- --- --- --- --- --- --- --- --- --- --- --- --- --- --- --- --- --- --- --- --- --- --- --- --- --- --- --- --- --- --- --- --- --- --- --- --- --- --- --- --- --- --- --- --- --- --- --- --- --- --- --- --- --- --- --- --- --- --- --- --- --- --- --- --- --- --- --- --- --- --- --- --- --- --- --- --- --- --- --- --- --- --- --- --- --- --- --- --- --- --- --- --- --- --- --- --- --- --- --- --- --- --- --- --- --- --- --- --- --- --- --- --- --- --- --- --- --- --- --- --- --- --- --- --- --- --- --- --- --- --- --- --- --- --- --- --- --- --- --- --- --- --- --- --- --- --- --- --- --- --- --- --- --- --- --- --- --- --- --- --- --- --- --- --- --- --- --- --- --- --- --- --- --- --- --- --- --- --- --- --- --- --- --- --- --- --- --- --- --- --- --- --- --- --- --- --- --- --- --- --- --- --- --- --- --- --- --- --- --- --- --- --- --- --- --- --- --- --- --- --- --- --- --- --- --- --- --- --- --- --- --- --- --- --- --- --- --- --- --- --- --- --- --- --- --- --- --- --- --- --- --- --- --- --- --- --- --- --- --- --- --- --- --- --- --- --- --- --- --- --- --- --- --- --- --- --- --- --- --- --- --- --- --- --- --- --- --- --- --- --- --- --- --- --- --- --- --- --- --- --- --- --- --- --- --- --- --- --- --- --- --- --- --- --- --- --- --- --- --- --- --- --- --- --- --- --- --- --- --- --- --- --- --- --- --- --- --- --- --- --- --- --- --- --- --- --- --- --- --- --- --- --- --- --- --- --- --- --- --- --- --- --- --- --- --- --- --- --- --- --- --- --- --- --- --- --- --- --- --- --- --- --- --- -ct gga tcg gat gtc acc gca gta ata ttg ttg att att tct gac atc gac gta tta tat agt ttt tta att cca tat ctt ttt t

>ON682266.1_Monkeypox_virus_isolate_MPXV/Germany/2022/RKI09_complete_genome

ttt tat atc act acg gac ata aac cat tgt ata ttt ttt atg ttt att agt gta cac att ttg gaa gta agt tc- --- --- --- --- --- --- --- --- --- --- --- --- --- --- --- --- --- --- --- --- --- --- --- --- --- --- --- --- --- --- --- --- --- --- --- --- --- --- --- --- --- --- --- --- --- --- --- --- --- --- --- --- --- --- --- --- --- --- --- --- --- --- --- --- --- --- --- --- --- --- --- --- --- --- --- --- --- --- --- --- --- --- --- --- --- --- --- --- --- --- --- --- --- --- --- --- --- --- --- --- --- --- --- --- --- --- --- --- --- --- --- --- --- --- --- --- --- --- --- --- --- --- --- --- --- --- --- --- --- --- --- --- --- --- --- --- --- --- --- --- --- --- --- --- --- --- --- --- --- --- --- --- --- --- --- --- --- --- --- --- --- --- --- --- --- --- --- --- --- --- --- --- --- --- --- --- --- --- --- --- --- --- --- --- --- --- --- --- --- --- --- --- --- --- --- --- --- --- --- --- --- --- --- --- --- --- --- --- --- --- --- --- --- --- --- --- --- --- --- --- --- --- --- --- --- --- --- --- --- --- --- --- --- --- --- --- --- --- --- --- --- --- --- --- --- --- --- --- --- --- --- --- --- --- --- --- --- --- --- --- --- --- --- --- --- --- --- --- --- --- --- --- --- --- --- --- --- --- --- --- --- --- --- --- --- --- --- --- --- --- --- --- --- --- --- --- --- --- --- --- --- --- --- --- --- --- --- --- --- --- --- --- --- --- --- --- --- --- --- --- --- --- --- --- --- --- --- --- --- --- --- --- --- --- --- --- --- --- --- --- --- --- --- --- --- --- --- --- --- --- --- --- --- --- --- --- --- --- --- --- --- --- --- --- --- --- --- --- --- --- --- --- --- --- --- --- --- --- --- --- --- --- --- --- --- --- --- --- --- --- --- --- --- --- --- --- --- --- --- --- --- --- --- --- --- --- --- --- --- --- --- --- --- --- --- --- --- --- --- --- --- --- --- --- --- --- --- --- --- --- --- --- --- --- --- --- --- --- --- --- --- --- --- --- --- --- --- --- --- --- --- --- --- --- --- --- --- --- --- --- --- --- --- --- --- --- --- --- --- --- --- --- --- --- --- --- --- --- --- --- --- --- --- --- --- --- --- --- --- --- --- --- --- --- --- --- --- --- --- --- --- --- --- --- --- --- --- --- --- --- --- --- --- --- --- --- --- --- --- --- --- --- --- --- --- --- --- --- --- --- --- --- --- --- --- --- --- --- --- --- --- --- --- --- --- --- --- --- --- --- --- --- --- --- --- --- --- --- --- --- --- --- --- --- --- --- --- --- --- --- --- --- --- --- --- --- --- --- --- --- --- --- --- --- --- --- --- --- --- --- --- --- --- --- --- --- --- --- --- --- --- --- --- --- --- --- --- --- --- --- --- --- --- --- --- --- --- --- --- --- --- --- --- --- --- --- --- --- --- --- --- --- --- --- --- --- --- --- --- --- --- --- --- --- --- --- --- --- --- --- --- --- -ct gga tcg gat gtc acc gca gta ata ttg ttg att att tct gac atc gac gta tta tat agt ttt tta att cca tat ctt ttt t

>ON619836.1_Monkeypox_virus_isolate_MPXV_UK_2022_2_complete_genome

ttt tat atc act acg gac ata aac cat tgt ata ttt ttt atg ttt att agt gta cac att ttg gaa gta agt tc- --- --- --- --- --- --- --- --- --- --- --- --- --- --- --- --- --- --- --- --- --- --- --- --- --- --- --- --- --- --- --- --- --- --- --- --- --- --- --- --- --- --- --- --- --- --- --- --- --- --- --- --- --- --- --- --- --- --- --- --- --- --- --- --- --- --- --- --- --- --- --- --- --- --- --- --- --- --- --- --- --- --- --- --- --- --- --- --- --- --- --- --- --- --- --- --- --- --- --- --- --- --- --- --- --- --- --- --- --- --- --- --- --- --- --- --- --- --- --- --- --- --- --- --- --- --- --- --- --- --- --- --- --- --- --- --- --- --- --- --- --- --- --- --- --- --- --- --- --- --- --- --- --- --- --- --- --- --- --- --- --- --- --- --- --- --- --- --- --- --- --- --- --- --- --- --- --- --- --- --- --- --- --- --- --- --- --- --- --- --- --- --- --- --- --- --- --- --- --- --- --- --- --- --- --- --- --- --- --- --- --- --- --- --- --- --- --- --- --- --- --- --- --- --- --- --- --- --- --- --- --- --- --- --- --- --- --- --- --- --- --- --- --- --- --- --- --- --- --- --- --- --- --- --- --- --- --- --- --- --- --- --- --- --- --- --- --- --- --- --- --- --- --- --- --- --- --- --- --- --- --- --- --- --- --- --- --- --- --- --- --- --- --- --- --- --- --- --- --- --- --- --- --- --- --- --- --- --- --- --- --- --- --- --- --- --- --- --- --- --- --- --- --- --- --- --- --- --- --- --- --- --- --- --- --- --- --- --- --- --- --- --- --- --- --- --- --- --- --- --- --- --- --- --- --- --- --- --- --- --- --- --- --- --- --- --- --- --- --- --- --- --- --- --- --- --- --- --- --- --- --- --- --- --- --- --- --- --- --- --- --- --- --- --- --- --- --- --- --- --- --- --- --- --- --- --- --- --- --- --- --- --- --- --- --- --- --- --- --- --- --- --- --- --- --- --- --- --- --- --- --- --- --- --- --- --- --- --- --- --- --- --- --- --- --- --- --- --- --- --- --- --- --- --- --- --- --- --- --- --- --- --- --- --- --- --- --- --- --- --- --- --- --- --- --- --- --- --- --- --- --- --- --- --- --- --- --- --- --- --- --- --- --- --- --- --- --- --- --- --- --- --- --- --- --- --- --- --- --- --- --- --- --- --- --- --- --- --- --- --- --- --- --- --- --- --- --- --- --- --- --- --- --- --- --- --- --- --- --- --- --- --- --- --- --- --- --- --- --- --- --- --- --- --- --- --- --- --- --- --- --- --- --- --- --- --- --- --- --- --- --- --- --- --- --- --- --- --- --- --- --- --- --- --- --- --- --- --- --- --- --- --- --- --- --- --- --- --- --- --- --- --- --- --- --- --- --- --- --- --- --- --- --- --- --- --- --- --- --- --- --- --- --- --- --- --- --- --- --- --- --- --- --- --- --- --- --- --- --- --- --- --- --- --- --- --- --- --- --- --- --- --- -ct gga tcg gat gtc acc gca gta ata ttg ttg att att tct gac atc gac gta tta tat agt ttt tta att cca tat ctt ttt t

>ON619838.1_Monkeypox_virus_isolate_MPXV_UK_2022_4_complete_genome

ttt tat atc act acg gac ata aac cat tgt ata ttt ttt atg ttt att agt gta cac att ttg gaa gta agt tc- --- --- --- --- --- --- --- --- --- --- --- --- --- --- --- --- --- --- --- --- --- --- --- --- --- --- --- --- --- --- --- --- --- --- --- --- --- --- --- --- --- --- --- --- --- --- --- --- --- --- --- --- --- --- --- --- --- --- --- --- --- --- --- --- --- --- --- --- --- --- --- --- --- --- --- --- --- --- --- --- --- --- --- --- --- --- --- --- --- --- --- --- --- --- --- --- --- --- --- --- --- --- --- --- --- --- --- --- --- --- --- --- --- --- --- --- --- --- --- --- --- --- --- --- --- --- --- --- --- --- --- --- --- --- --- --- --- --- --- --- --- --- --- --- --- --- --- --- --- --- --- --- --- --- --- --- --- --- --- --- --- --- --- --- --- --- --- --- --- --- --- --- --- --- --- --- --- --- --- --- --- --- --- --- --- --- --- --- --- --- --- --- --- --- --- --- --- --- --- --- --- --- --- --- --- --- --- --- --- --- --- --- --- --- --- --- --- --- --- --- --- --- --- --- --- --- --- --- --- --- --- --- --- --- --- --- --- --- --- --- --- --- --- --- --- --- --- --- --- --- --- --- --- --- --- --- --- --- --- --- --- --- --- --- --- --- --- --- --- --- --- --- --- --- --- --- --- --- --- --- --- --- --- --- --- --- --- --- --- --- --- --- --- --- --- --- --- --- --- --- --- --- --- --- --- --- --- --- --- --- --- --- --- --- --- --- --- --- --- --- --- --- --- --- --- --- --- --- --- --- --- --- --- --- --- --- --- --- --- --- --- --- --- --- --- --- --- --- --- --- --- --- --- --- --- --- --- --- --- --- --- --- --- --- --- --- --- --- --- --- --- --- --- --- --- --- --- --- --- --- --- --- --- --- --- --- --- --- --- --- --- --- --- --- --- --- --- --- --- --- --- --- --- --- --- --- --- --- --- --- --- --- --- --- --- --- --- --- --- --- --- --- --- --- --- --- --- --- --- --- --- --- --- --- --- --- --- --- --- --- --- --- --- --- --- --- --- --- --- --- --- --- --- --- --- --- --- --- --- --- --- --- --- --- --- --- --- --- --- --- --- --- --- --- --- --- --- --- --- --- --- --- --- --- --- --- --- --- --- --- --- --- --- --- --- --- --- --- --- --- --- --- --- --- --- --- --- --- --- --- --- --- --- --- --- --- --- --- --- --- --- --- --- --- --- --- --- --- --- --- --- --- --- --- --- --- --- --- --- --- --- --- --- --- --- --- --- --- --- --- --- --- --- --- --- --- --- --- --- --- --- --- --- --- --- --- --- --- --- --- --- --- --- --- --- --- --- --- --- --- --- --- --- --- --- --- --- --- --- --- --- --- --- --- --- --- --- --- --- --- --- --- --- --- --- --- --- --- --- --- --- --- --- --- --- --- --- --- --- --- --- --- --- --- --- --- --- --- --- --- --- --- --- --- --- --- --- --- --- --- --- --- --- --- --- --- --- --- --- --- --- --- -ct gga tcg gat gtc acc gca gta ata ttg ttg att att tct gac atc gac gta tta tat agt ttt tta att cca tat ctt ttt t

>ON619835.1_Monkeypox_virus_isolate_MPXV_UK_2022_1_complete_genome

ttt tat atc act acg gac ata aac cat tgt ata ttt ttt atg ttt att agt gta cac att ttg gaa gta agt tc- --- --- --- --- --- --- --- --- --- --- --- --- --- --- --- --- --- --- --- --- --- --- --- --- --- --- --- --- --- --- --- --- --- --- --- --- --- --- --- --- --- --- --- --- --- --- --- --- --- --- --- --- --- --- --- --- --- --- --- --- --- --- --- --- --- --- --- --- --- --- --- --- --- --- --- --- --- --- --- --- --- --- --- --- --- --- --- --- --- --- --- --- --- --- --- --- --- --- --- --- --- --- --- --- --- --- --- --- --- --- --- --- --- --- --- --- --- --- --- --- --- --- --- --- --- --- --- --- --- --- --- --- --- --- --- --- --- --- --- --- --- --- --- --- --- --- --- --- --- --- --- --- --- --- --- --- --- --- --- --- --- --- --- --- --- --- --- --- --- --- --- --- --- --- --- --- --- --- --- --- --- --- --- --- --- --- --- --- --- --- --- --- --- --- --- --- --- --- --- --- --- --- --- --- --- --- --- --- --- --- --- --- --- --- --- --- --- --- --- --- --- --- --- --- --- --- --- --- --- --- --- --- --- --- --- --- --- --- --- --- --- --- --- --- --- --- --- --- --- --- --- --- --- --- --- --- --- --- --- --- --- --- --- --- --- --- --- --- --- --- --- --- --- --- --- --- --- --- --- --- --- --- --- --- --- --- --- --- --- --- --- --- --- --- --- --- --- --- --- --- --- --- --- --- --- --- --- --- --- --- --- --- --- --- --- --- --- --- --- --- --- --- --- --- --- --- --- --- --- --- --- --- --- --- --- --- --- --- --- --- --- --- --- --- --- --- --- --- --- --- --- --- --- --- --- --- --- --- --- --- --- --- --- --- --- --- --- --- --- --- --- --- --- --- --- --- --- --- --- --- --- --- --- --- --- --- --- --- --- --- --- --- --- --- --- --- --- --- --- --- --- --- --- --- --- --- --- --- --- --- --- --- --- --- --- --- --- --- --- --- --- --- --- --- --- --- --- --- --- --- --- --- --- --- --- --- --- --- --- --- --- --- --- --- --- --- --- --- --- --- --- --- --- --- --- --- --- --- --- --- --- --- --- --- --- --- --- --- --- --- --- --- --- --- --- --- --- --- --- --- --- --- --- --- --- --- --- --- --- --- --- --- --- --- --- --- --- --- --- --- --- --- --- --- --- --- --- --- --- --- --- --- --- --- --- --- --- --- --- --- --- --- --- --- --- --- --- --- --- --- --- --- --- --- --- --- --- --- --- --- --- --- --- --- --- --- --- --- --- --- --- --- --- --- --- --- --- --- --- --- --- --- --- --- --- --- --- --- --- --- --- --- --- --- --- --- --- --- --- --- --- --- --- --- --- --- --- --- --- --- --- --- --- --- --- --- --- --- --- --- --- --- --- --- --- --- --- --- --- --- --- --- --- --- --- --- --- --- --- --- --- --- --- --- --- --- --- --- --- --- --- --- --- --- --- --- --- --- --- --- --- --- --- --- --- --- --- --- --- --- --- --- -ct gga tcg gat gtc acc gca gta ata ttg ttg att att tct gac atc gac gta tta tat agt ttt tta att cca tat ctt ttt t

>ON645312.1_Monkeypox_virus_isolate_MPXV_GSTT_Patient1_partial_genome

ttt tat atc act acg gac ata aac cat tgt ata ttt ttt atg ttt att agt gta cac att ttg gaa gta agt tc- --- --- --- --- --- --- --- --- --- --- --- --- --- --- --- --- --- --- --- --- --- --- --- --- --- --- --- --- --- --- --- --- --- --- --- --- --- --- --- --- --- --- --- --- --- --- --- --- --- --- --- --- --- --- --- --- --- --- --- --- --- --- --- --- --- --- --- --- --- --- --- --- --- --- --- --- --- --- --- --- --- --- --- --- --- --- --- --- --- --- --- --- --- --- --- --- --- --- --- --- --- --- --- --- --- --- --- --- --- --- --- --- --- --- --- --- --- --- --- --- --- --- --- --- --- --- --- --- --- --- --- --- --- --- --- --- --- --- --- --- --- --- --- --- --- --- --- --- --- --- --- --- --- --- --- --- --- --- --- --- --- --- --- --- --- --- --- --- --- --- --- --- --- --- --- --- --- --- --- --- --- --- --- --- --- --- --- --- --- --- --- --- --- --- --- --- --- --- --- --- --- --- --- --- --- --- --- --- --- --- --- --- --- --- --- --- --- --- --- --- --- --- --- --- --- --- --- --- --- --- --- --- --- --- --- --- --- --- --- --- --- --- --- --- --- --- --- --- --- --- --- --- --- --- --- --- --- --- --- --- --- --- --- --- --- --- --- --- --- --- --- --- --- --- --- --- --- --- --- --- --- --- --- --- --- --- --- --- --- --- --- --- --- --- --- --- --- --- --- --- --- --- --- --- --- --- --- --- --- --- --- --- --- --- --- --- --- --- --- --- --- --- --- --- --- --- --- --- --- --- --- --- --- --- --- --- --- --- --- --- --- --- --- --- --- --- --- --- --- --- --- --- --- --- --- --- --- --- --- --- --- --- --- --- --- --- --- --- --- --- --- --- --- --- --- --- --- --- --- --- --- --- --- --- --- --- --- --- --- --- --- --- --- --- --- --- --- --- --- --- --- --- --- --- --- --- --- --- --- --- --- --- --- --- --- --- --- --- --- --- --- --- --- --- --- --- --- --- --- --- --- --- --- --- --- --- --- --- --- --- --- --- --- --- --- --- --- --- --- --- --- --- --- --- --- --- --- --- --- --- --- --- --- --- --- --- --- --- --- --- --- --- --- --- --- --- --- --- --- --- --- --- --- --- --- --- --- --- --- --- --- --- --- --- --- --- --- --- --- --- --- --- --- --- --- --- --- --- --- --- --- --- --- --- --- --- --- --- --- --- --- --- --- --- --- --- --- --- --- --- --- --- --- --- --- --- --- --- --- --- --- --- --- --- --- --- --- --- --- --- --- --- --- --- --- --- --- --- --- --- --- --- --- --- --- --- --- --- --- --- --- --- --- --- --- --- --- --- --- --- --- --- --- --- --- --- --- --- --- --- --- --- --- --- --- --- --- --- --- --- --- --- --- --- --- --- --- --- --- --- --- --- --- --- --- --- --- --- --- --- --- --- --- --- --- --- --- --- --- --- --- --- --- --- --- --- --- --- --- --- --- --- --- --- --- --- --- --- --- --- --- --- -ct gga tcg gat gtc acc gca gta ata ttg ttg att att tct gac atc gac gta tta tat agt ttt tta att cca tat ctt ttt t

>ON682269.2_Monkeypox_virus_isolate_MPXV/Germany/2022/RKI07_complete_genome

ttt tat atc act acg gac ata aac cat tgt ata ttt ttt atg ttt att agt gta cac att ttg gaa gta agt tc- --- --- --- --- --- --- --- --- --- --- --- --- --- --- --- --- --- --- --- --- --- --- --- --- --- --- --- --- --- --- --- --- --- --- --- --- --- --- --- --- --- --- --- --- --- --- --- --- --- --- --- --- --- --- --- --- --- --- --- --- --- --- --- --- --- --- --- --- --- --- --- --- --- --- --- --- --- --- --- --- --- --- --- --- --- --- --- --- --- --- --- --- --- --- --- --- --- --- --- --- --- --- --- --- --- --- --- --- --- --- --- --- --- --- --- --- --- --- --- --- --- --- --- --- --- --- --- --- --- --- --- --- --- --- --- --- --- --- --- --- --- --- --- --- --- --- --- --- --- --- --- --- --- --- --- --- --- --- --- --- --- --- --- --- --- --- --- --- --- --- --- --- --- --- --- --- --- --- --- --- --- --- --- --- --- --- --- --- --- --- --- --- --- --- --- --- --- --- --- --- --- --- --- --- --- --- --- --- --- --- --- --- --- --- --- --- --- --- --- --- --- --- --- --- --- --- --- --- --- --- --- --- --- --- --- --- --- --- --- --- --- --- --- --- --- --- --- --- --- --- --- --- --- --- --- --- --- --- --- --- --- --- --- --- --- --- --- --- --- --- --- --- --- --- --- --- --- --- --- --- --- --- --- --- --- --- --- --- --- --- --- --- --- --- --- --- --- --- --- --- --- --- --- --- --- --- --- --- --- --- --- --- --- --- --- --- --- --- --- --- --- --- --- --- --- --- --- --- --- --- --- --- --- --- --- --- --- --- --- --- --- --- --- --- --- --- --- --- --- --- --- --- --- --- --- --- --- --- --- --- --- --- --- --- --- --- --- --- --- --- --- --- --- --- --- --- --- --- --- --- --- --- --- --- --- --- --- --- --- --- --- --- --- --- --- --- --- --- --- --- --- --- --- --- --- --- --- --- --- --- --- --- --- --- --- --- --- --- --- --- --- --- --- --- --- --- --- --- --- --- --- --- --- --- --- --- --- --- --- --- --- --- --- --- --- --- --- --- --- --- --- --- --- --- --- --- --- --- --- --- --- --- --- --- --- --- --- --- --- --- --- --- --- --- --- --- --- --- --- --- --- --- --- --- --- --- --- --- --- --- --- --- --- --- --- --- --- --- --- --- --- --- --- --- --- --- --- --- --- --- --- --- --- --- --- --- --- --- --- --- --- --- --- --- --- --- --- --- --- --- --- --- --- --- --- --- --- --- --- --- --- --- --- --- --- --- --- --- --- --- --- --- --- --- --- --- --- --- --- --- --- --- --- --- --- --- --- --- --- --- --- --- --- --- --- --- --- --- --- --- --- --- --- --- --- --- --- --- --- --- --- --- --- --- --- --- --- --- --- --- --- --- --- --- --- --- --- --- --- --- --- --- --- --- --- --- --- --- --- --- --- --- --- --- --- --- --- --- --- --- --- --- --- --- --- --- --- --- --- --- --- --- --- --- --- --- --- --- --- --- --- --- -ct gga tcg gat gtc acc gca gta ata ttg ttg att att tct gac atc gac gta tta tat agt ttt tta att cca tat ctt ttt t

>KJ642616.1_Monkeypox_virus_strain_PCH_complete_genome

ttt tat atc act acg gac ata aac cat tgt ata att ttt atg ttt att agt gta cac att ttg gaa gta agt tc- --- --- --- --- --- --- --- --- --- --- --- --- --- --- --- --- --- --- --- --- --- --- --- --- --- --- --- --- --- --- --- --- --- --- --- --- --- --- --- --- --- --- --- --- --- --- --- --- --- --- --- --- --- --- --- --- --- --- --- --- --- --- --- --- --- --- --- --- --- --- --- --- --- --- --- --- --- --- --- --- --- --- --- --- --- --- --- --- --- --- --- --- --- --- --- --- --- --- --- --- --- --- --- --- --- --- --- --- --- --- --- --- --- --- --- --- --- --- --- --- --- --- --- --- --- --- --- --- --- --- --- --- --- --- --- --- --- --- --- --- --- --- --- --- --- --- --- --- --- --- --- --- --- --- --- --- --- --- --- --- --- --- --- --- --- --- --- --- --- --- --- --- --- --- --- --- --- --- --- --- --- --- --- --- --- --- --- --- --- --- --- --- --- --- --- --- --- --- --- --- --- --- --- --- --- --- --- --- --- --- --- --- --- --- --- --- --- --- --- --- --- --- --- --- --- --- --- --- --- --- --- --- --- --- --- --- --- --- --- --- --- --- --- --- --- --- --- --- --- --- --- --- --- --- --- --- --- --- --- --- --- --- --- --- --- --- --- --- --- --- --- --- --- --- --- --- --- --- --- --- --- --- --- --- --- --- --- --- --- --- --- --- --- --- --- --- --- --- --- --- --- --- --- --- --- --- --- --- --- --- --- --- --- --- --- --- --- --- --- --- --- --- --- --- --- --- --- --- --- --- --- --- --- --- --- --- --- --- --- --- --- --- --- --- --- --- --- --- --- --- --- --- --- --- --- --- --- --- --- --- --- --- --- --- --- --- --- --- --- --- --- --- --- --- --- --- --- --- --- --- --- --- --- --- --- --- --- --- --- --- --- --- --- --- --- --- --- --- --- --- --- --- --- --- --- --- --- --- --- --- --- --- --- --- --- --- --- --- --- --- --- --- --- --- --- --- --- --- --- --- --- --- --- --- --- --- --- --- --- --- --- --- --- --- --- --- --- --- --- --- --- --- --- --- --- --- --- --- --- --- --- --- --- --- --- --- --- --- --- --- --- --- --- --- --- --- --- --- --- --- --- --- --- --- --- --- --- --- --- --- --- --- --- --- --- --- --- --- --- --- --- --- --- --- --- --- --- --- --- --- --- --- --- --- --- --- --- --- --- --- --- --- --- --- --- --- --- --- --- --- --- --- --- --- --- --- --- --- --- --- --- --- --- --- --- --- --- --- --- --- --- --- --- --- --- --- --- --- --- --- --- --- --- --- --- --- --- --- --- --- --- --- --- --- --- --- --- --- --- --- --- --- --- --- --- --- --- --- --- --- --- --- --- --- --- --- --- --- --- --- --- --- --- --- --- --- --- --- --- --- --- --- --- --- --- --- --- --- --- --- --- --- --- --- --- --- --- --- --- --- --- --- --- --- --- --- --- --- --- --- --- --- --- --- --- --- --- --- --- --- --- --- -ct gga tcg gat gtc acc gca gta ata ttg ttg att att tct gac atc gac gta tta tat agt ttt tta att cca tat ctt ttt t

>AY753185.1_Monkeypox_virus_strain_COP-58_complete_genome

ttt tat atc act acg gac ata aac cat tgt ata att ttt atg ttt att agt gta cac att ttg gaa gta agt tc- --- --- --- --- --- --- --- --- --- --- --- --- --- --- --- --- --- --- --- --- --- --- --- --- --- --- --- --- --- --- --- --- --- --- --- --- --- --- --- --- --- --- --- --- --- --- --- --- --- --- --- --- --- --- --- --- --- --- --- --- --- --- --- --- --- --- --- --- --- --- --- --- --- --- --- --- --- --- --- --- --- --- --- --- --- --- --- --- --- --- --- --- --- --- --- --- --- --- --- --- --- --- --- --- --- --- --- --- --- --- --- --- --- --- --- --- --- --- --- --- --- --- --- --- --- --- --- --- --- --- --- --- --- --- --- --- --- --- --- --- --- --- --- --- --- --- --- --- --- --- --- --- --- --- --- --- --- --- --- --- --- --- --- --- --- --- --- --- --- --- --- --- --- --- --- --- --- --- --- --- --- --- --- --- --- --- --- --- --- --- --- --- --- --- --- --- --- --- --- --- --- --- --- --- --- --- --- --- --- --- --- --- --- --- --- --- --- --- --- --- --- --- --- --- --- --- --- --- --- --- --- --- --- --- --- --- --- --- --- --- --- --- --- --- --- --- --- --- --- --- --- --- --- --- --- --- --- --- --- --- --- --- --- --- --- --- --- --- --- --- --- --- --- --- --- --- --- --- --- --- --- --- --- --- --- --- --- --- --- --- --- --- --- --- --- --- --- --- --- --- --- --- --- --- --- --- --- --- --- --- --- --- --- --- --- --- --- --- --- --- --- --- --- --- --- --- --- --- --- --- --- --- --- --- --- --- --- --- --- --- --- --- --- --- --- --- --- --- --- --- --- --- --- --- --- --- --- --- --- --- --- --- --- --- --- --- --- --- --- --- --- --- --- --- --- --- --- --- --- --- --- --- --- --- --- --- --- --- --- --- --- --- --- --- --- --- --- --- --- --- --- --- --- --- --- --- --- --- --- --- --- --- --- --- --- --- --- --- --- --- --- --- --- --- --- --- --- --- --- --- --- --- --- --- --- --- --- --- --- --- --- --- --- --- --- --- --- --- --- --- --- --- --- --- --- --- --- --- --- --- --- --- --- --- --- --- --- --- --- --- --- --- --- --- --- --- --- --- --- --- --- --- --- --- --- --- --- --- --- --- --- --- --- --- --- --- --- --- --- --- --- --- --- --- --- --- --- --- --- --- --- --- --- --- --- --- --- --- --- --- --- --- --- --- --- --- --- --- --- --- --- --- --- --- --- --- --- --- --- --- --- --- --- --- --- --- --- --- --- --- --- --- --- --- --- --- --- --- --- --- --- --- --- --- --- --- --- --- --- --- --- --- --- --- --- --- --- --- --- --- --- --- --- --- --- --- --- --- --- --- --- --- --- --- --- --- --- --- --- --- --- --- --- --- --- --- --- --- --- --- --- --- --- --- --- --- --- --- --- --- --- --- --- --- --- --- --- --- --- --- --- --- --- --- --- --- --- --- --- --- --- --- --- --- --- --- --- --- --- --- --- --- -ct gga tcg gat gtc acc gca gta ata ttg ttg att att tct gac atc gac gta tta tat agt ttt tta att cca tat ctt ttt t

>AY603973.1_Monkeypox_virus_strain_MPXV-WRAIR7-61_complete_genome

ttt tat atc act acg gac ata aac cat tgt ata att ttt atg ttt att agt gta cac att ttg gaa gta agt tc- --- --- --- --- --- --- --- --- --- --- --- --- --- --- --- --- --- --- --- --- --- --- --- --- --- --- --- --- --- --- --- --- --- --- --- --- --- --- --- --- --- --- --- --- --- --- --- --- --- --- --- --- --- --- --- --- --- --- --- --- --- --- --- --- --- --- --- --- --- --- --- --- --- --- --- --- --- --- --- --- --- --- --- --- --- --- --- --- --- --- --- --- --- --- --- --- --- --- --- --- --- --- --- --- --- --- --- --- --- --- --- --- --- --- --- --- --- --- --- --- --- --- --- --- --- --- --- --- --- --- --- --- --- --- --- --- --- --- --- --- --- --- --- --- --- --- --- --- --- --- --- --- --- --- --- --- --- --- --- --- --- --- --- --- --- --- --- --- --- --- --- --- --- --- --- --- --- --- --- --- --- --- --- --- --- --- --- --- --- --- --- --- --- --- --- --- --- --- --- --- --- --- --- --- --- --- --- --- --- --- --- --- --- --- --- --- --- --- --- --- --- --- --- --- --- --- --- --- --- --- --- --- --- --- --- --- --- --- --- --- --- --- --- --- --- --- --- --- --- --- --- --- --- --- --- --- --- --- --- --- --- --- --- --- --- --- --- --- --- --- --- --- --- --- --- --- --- --- --- --- --- --- --- --- --- --- --- --- --- --- --- --- --- --- --- --- --- --- --- --- --- --- --- --- --- --- --- --- --- --- --- --- --- --- --- --- --- --- --- --- --- --- --- --- --- --- --- --- --- --- --- --- --- --- --- --- --- --- --- --- --- --- --- --- --- --- --- --- --- --- --- --- --- --- --- --- --- --- --- --- --- --- --- --- --- --- --- --- --- --- --- --- --- --- --- --- --- --- --- --- --- --- --- --- --- --- --- --- --- --- --- --- --- --- --- --- --- --- --- --- --- --- --- --- --- --- --- --- --- --- --- --- --- --- --- --- --- --- --- --- --- --- --- --- --- --- --- --- --- --- --- --- --- --- --- --- --- --- --- --- --- --- --- --- --- --- --- --- --- --- --- --- --- --- --- --- --- --- --- --- --- --- --- --- --- --- --- --- --- --- --- --- --- --- --- --- --- --- --- --- --- --- --- --- --- --- --- --- --- --- --- --- --- --- --- --- --- --- --- --- --- --- --- --- --- --- --- --- --- --- --- --- --- --- --- --- --- --- --- --- --- --- --- --- --- --- --- --- --- --- --- --- --- --- --- --- --- --- --- --- --- --- --- --- --- --- --- --- --- --- --- --- --- --- --- --- --- --- --- --- --- --- --- --- --- --- --- --- --- --- --- --- --- --- --- --- --- --- --- --- --- --- --- --- --- --- --- --- --- --- --- --- --- --- --- --- --- --- --- --- --- --- --- --- --- --- --- --- --- --- --- --- --- --- --- --- --- --- --- --- --- --- --- --- --- --- --- --- --- --- --- --- --- --- --- --- --- --- --- --- --- --- --- --- --- --- --- --- --- --- --- --- -ct gga tcg gat gtc acc gca gta ata ttg ttg att att tct gac atc gac gta tta tat agt ttt tta att cca tat ctt ttt t

>ON619837.1_Monkeypox_virus_isolate_MPXV_UK_2022_3_complete_genome

ttt tat atc act acg gac ata aac cat tgt ata ttt ttt atg ttt att agt gta cac att ttg gaa gta agt tc- --- --- --- --- --- --- --- --- --- --- --- --- --- --- --- --- --- --- --- --- --- --- --- --- --- --- --- --- --- --- --- --- --- --- --- --- --- --- --- --- --- --- --- --- --- --- --- --- --- --- --- --- --- --- --- --- --- --- --- --- --- --- --- --- --- --- --- --- --- --- --- --- --- --- --- --- --- --- --- --- --- --- --- --- --- --- --- --- --- --- --- --- --- --- --- --- --- --- --- --- --- --- --- --- --- --- --- --- --- --- --- --- --- --- --- --- --- --- --- --- --- --- --- --- --- --- --- --- --- --- --- --- --- --- --- --- --- --- --- --- --- --- --- --- --- --- --- --- --- --- --- --- --- --- --- --- --- --- --- --- --- --- --- --- --- --- --- --- --- --- --- --- --- --- --- --- --- --- --- --- --- --- --- --- --- --- --- --- --- --- --- --- --- --- --- --- --- --- --- --- --- --- --- --- --- --- --- --- --- --- --- --- --- --- --- --- --- --- --- --- --- --- --- --- --- --- --- --- --- --- --- --- --- --- --- --- --- --- --- --- --- --- --- --- --- --- --- --- --- --- --- --- --- --- --- --- --- --- --- --- --- --- --- --- --- --- --- --- --- --- --- --- --- --- --- --- --- --- --- --- --- --- --- --- --- --- --- --- --- --- --- --- --- --- --- --- --- --- --- --- --- --- --- --- --- --- --- --- --- --- --- --- --- --- --- --- --- --- --- --- --- --- --- --- --- --- --- --- --- --- --- --- --- --- --- --- --- --- --- --- --- --- --- --- --- --- --- --- --- --- --- --- --- --- --- --- --- --- --- --- --- --- --- --- --- --- --- --- --- --- --- --- --- --- --- --- --- --- --- --- --- --- --- --- --- --- --- --- --- --- --- --- --- --- --- --- --- --- --- --- --- --- --- --- --- --- --- --- --- --- --- --- --- --- --- --- --- --- --- --- --- --- --- --- --- --- --- --- --- --- --- --- --- --- --- --- --- --- --- --- --- --- --- --- --- --- --- --- --- --- --- --- --- --- --- --- --- --- --- --- --- --- --- --- --- --- --- --- --- --- --- --- --- --- --- --- --- --- --- --- --- --- --- --- --- --- --- --- --- --- --- --- --- --- --- --- --- --- --- --- --- --- --- --- --- --- --- --- --- --- --- --- --- --- --- --- --- --- --- --- --- --- --- --- --- --- --- --- --- --- --- --- --- --- --- --- --- --- --- --- --- --- --- --- --- --- --- --- --- --- --- --- --- --- --- --- --- --- --- --- --- --- --- --- --- --- --- --- --- --- --- --- --- --- --- --- --- --- --- --- --- --- --- --- --- --- --- --- --- --- --- --- --- --- --- --- --- --- --- --- --- --- --- --- --- --- --- --- --- --- --- --- --- --- --- --- --- --- --- --- --- --- --- --- --- --- --- --- --- --- --- --- --- --- --- --- --- --- --- --- --- --- --- --- --- --- --- --- --- --- --- --- -ct gga tcg gat gtc acc gca gta ata ttg ttg att att tct gac atc gac gta tta tat agt ttt tta att cca tat ctt ttt t

>ON585037.1_Monkeypox_virus_isolate_Monkeypox/PT0005/2022_complete_genome

ttt tat atc act acg gac ata aac cat tgt ata ttt ttt atg ttt att agt gta cac att ttg gaa gta agt tc- --- --- --- --- --- --- --- --- --- --- --- --- --- --- --- --- --- --- --- --- --- --- --- --- --- --- --- --- --- --- --- --- --- --- --- --- --- --- --- --- --- --- --- --- --- --- --- --- --- --- --- --- --- --- --- --- --- --- --- --- --- --- --- --- --- --- --- --- --- --- --- --- --- --- --- --- --- --- --- --- --- --- --- --- --- --- --- --- --- --- --- --- --- --- --- --- --- --- --- --- --- --- --- --- --- --- --- --- --- --- --- --- --- --- --- --- --- --- --- --- --- --- --- --- --- --- --- --- --- --- --- --- --- --- --- --- --- --- --- --- --- --- --- --- --- --- --- --- --- --- --- --- --- --- --- --- --- --- --- --- --- --- --- --- --- --- --- --- --- --- --- --- --- --- --- --- --- --- --- --- --- --- --- --- --- --- --- --- --- --- --- --- --- --- --- --- --- --- --- --- --- --- --- --- --- --- --- --- --- --- --- --- --- --- --- --- --- --- --- --- --- --- --- --- --- --- --- --- --- --- --- --- --- --- --- --- --- --- --- --- --- --- --- --- --- --- --- --- --- --- --- --- --- --- --- --- --- --- --- --- --- --- --- --- --- --- --- --- --- --- --- --- --- --- --- --- --- --- --- --- --- --- --- --- --- --- --- --- --- --- --- --- --- --- --- --- --- --- --- --- --- --- --- --- --- --- --- --- --- --- --- --- --- --- --- --- --- --- --- --- --- --- --- --- --- --- --- --- --- --- --- --- --- --- --- --- --- --- --- --- --- --- --- --- --- --- --- --- --- --- --- --- --- --- --- --- --- --- --- --- --- --- --- --- --- --- --- --- --- --- --- --- --- --- --- --- --- --- --- --- --- --- --- --- --- --- --- --- --- --- --- --- --- --- --- --- --- --- --- --- --- --- --- --- --- --- --- --- --- --- --- --- --- --- --- --- --- --- --- --- --- --- --- --- --- --- --- --- --- --- --- --- --- --- --- --- --- --- --- --- --- --- --- --- --- --- --- --- --- --- --- --- --- --- --- --- --- --- --- --- --- --- --- --- --- --- --- --- --- --- --- --- --- --- --- --- --- --- --- --- --- --- --- --- --- --- --- --- --- --- --- --- --- --- --- --- --- --- --- --- --- --- --- --- --- --- --- --- --- --- --- --- --- --- --- --- --- --- --- --- --- --- --- --- --- --- --- --- --- --- --- --- --- --- --- --- --- --- --- --- --- --- --- --- --- --- --- --- --- --- --- --- --- --- --- --- --- --- --- --- --- --- --- --- --- --- --- --- --- --- --- --- --- --- --- --- --- --- --- --- --- --- --- --- --- --- --- --- --- --- --- --- --- --- --- --- --- --- --- --- --- --- --- --- --- --- --- --- --- --- --- --- --- --- --- --- --- --- --- --- --- --- --- --- --- --- --- --- --- --- --- --- --- --- --- --- --- --- --- --- --- --- --- --- --- --- --- --- --- --- --- --- -ct gga tcg gat gtc acc gca gta ata ttg ttg att att tct gac atc gac gta tta tat agt ttt tta att cca tat ctt ttt t

>ON585032.1_Monkeypox_virus_isolate_Monkeypox/PT0004/2022_complete_genome

ttt tat atc act acg gac ata aac cat tgt ata ttt ttt atg ttt att agt gta cac att ttg gaa gta agt tc- --- --- --- --- --- --- --- --- --- --- --- --- --- --- --- --- --- --- --- --- --- --- --- --- --- --- --- --- --- --- --- --- --- --- --- --- --- --- --- --- --- --- --- --- --- --- --- --- --- --- --- --- --- --- --- --- --- --- --- --- --- --- --- --- --- --- --- --- --- --- --- --- --- --- --- --- --- --- --- --- --- --- --- --- --- --- --- --- --- --- --- --- --- --- --- --- --- --- --- --- --- --- --- --- --- --- --- --- --- --- --- --- --- --- --- --- --- --- --- --- --- --- --- --- --- --- --- --- --- --- --- --- --- --- --- --- --- --- --- --- --- --- --- --- --- --- --- --- --- --- --- --- --- --- --- --- --- --- --- --- --- --- --- --- --- --- --- --- --- --- --- --- --- --- --- --- --- --- --- --- --- --- --- --- --- --- --- --- --- --- --- --- --- --- --- --- --- --- --- --- --- --- --- --- --- --- --- --- --- --- --- --- --- --- --- --- --- --- --- --- --- --- --- --- --- --- --- --- --- --- --- --- --- --- --- --- --- --- --- --- --- --- --- --- --- --- --- --- --- --- --- --- --- --- --- --- --- --- --- --- --- --- --- --- --- --- --- --- --- --- --- --- --- --- --- --- --- --- --- --- --- --- --- --- --- --- --- --- --- --- --- --- --- --- --- --- --- --- --- --- --- --- --- --- --- --- --- --- --- --- --- --- --- --- --- --- --- --- --- --- --- --- --- --- --- --- --- --- --- --- --- --- --- --- --- --- --- --- --- --- --- --- --- --- --- --- --- --- --- --- --- --- --- --- --- --- --- --- --- --- --- --- --- --- --- --- --- --- --- --- --- --- --- --- --- --- --- --- --- --- --- --- --- --- --- --- --- --- --- --- --- --- --- --- --- --- --- --- --- --- --- --- --- --- --- --- --- --- --- --- --- --- --- --- --- --- --- --- --- --- --- --- --- --- --- --- --- --- --- --- --- --- --- --- --- --- --- --- --- --- --- --- --- --- --- --- --- --- --- --- --- --- --- --- --- --- --- --- --- --- --- --- --- --- --- --- --- --- --- --- --- --- --- --- --- --- --- --- --- --- --- --- --- --- --- --- --- --- --- --- --- --- --- --- --- --- --- --- --- --- --- --- --- --- --- --- --- --- --- --- --- --- --- --- --- --- --- --- --- --- --- --- --- --- --- --- --- --- --- --- --- --- --- --- --- --- --- --- --- --- --- --- --- --- --- --- --- --- --- --- --- --- --- --- --- --- --- --- --- --- --- --- --- --- --- --- --- --- --- --- --- --- --- --- --- --- --- --- --- --- --- --- --- --- --- --- --- --- --- --- --- --- --- --- --- --- --- --- --- --- --- --- --- --- --- --- --- --- --- --- --- --- --- --- --- --- --- --- --- --- --- --- --- --- --- --- --- --- --- --- --- --- --- --- --- --- --- --- --- --- --- --- --- --- --- --- --- --- --- --- --- --- -ct gga tcg gat gtc acc gca gta ata ttg ttg att att tct gac atc gac gta tta tat agt ttt tta att cca tat ctt ttt t

>AY741551.1_Monkeypox_virus_isolate_Sierra_Leone_complete_genome

ttt tat atc act acg gac ata aac cat tgt ata att ttt atg ttt att agt gta cac att ttg gaa gta agt tc- --- --- --- --- --- --- --- --- --- --- --- --- --- --- --- --- --- --- --- --- --- --- --- --- --- --- --- --- --- --- --- --- --- --- --- --- --- --- --- --- --- --- --- --- --- --- --- --- --- --- --- --- --- --- --- --- --- --- --- --- --- --- --- --- --- --- --- --- --- --- --- --- --- --- --- --- --- --- --- --- --- --- --- --- --- --- --- --- --- --- --- --- --- --- --- --- --- --- --- --- --- --- --- --- --- --- --- --- --- --- --- --- --- --- --- --- --- --- --- --- --- --- --- --- --- --- --- --- --- --- --- --- --- --- --- --- --- --- --- --- --- --- --- --- --- --- --- --- --- --- --- --- --- --- --- --- --- --- --- --- --- --- --- --- --- --- --- --- --- --- --- --- --- --- --- --- --- --- --- --- --- --- --- --- --- --- --- --- --- --- --- --- --- --- --- --- --- --- --- --- --- --- --- --- --- --- --- --- --- --- --- --- --- --- --- --- --- --- --- --- --- --- --- --- --- --- --- --- --- --- --- --- --- --- --- --- --- --- --- --- --- --- --- --- --- --- --- --- --- --- --- --- --- --- --- --- --- --- --- --- --- --- --- --- --- --- --- --- --- --- --- --- --- --- --- --- --- --- --- --- --- --- --- --- --- --- --- --- --- --- --- --- --- --- --- --- --- --- --- --- --- --- --- --- --- --- --- --- --- --- --- --- --- --- --- --- --- --- --- --- --- --- --- --- --- --- --- --- --- --- --- --- --- --- --- --- --- --- --- --- --- --- --- --- --- --- --- --- --- --- --- --- --- --- --- --- --- --- --- --- --- --- --- --- --- --- --- --- --- --- --- --- --- --- --- --- --- --- --- --- --- --- --- --- --- --- --- --- --- --- --- --- --- --- --- --- --- --- --- --- --- --- --- --- --- --- --- --- --- --- --- --- --- --- --- --- --- --- --- --- --- --- --- --- --- --- --- --- --- --- --- --- --- --- --- --- --- --- --- --- --- --- --- --- --- --- --- --- --- --- --- --- --- --- --- --- --- --- --- --- --- --- --- --- --- --- --- --- --- --- --- --- --- --- --- --- --- --- --- --- --- --- --- --- --- --- --- --- --- --- --- --- --- --- --- --- --- --- --- --- --- --- --- --- --- --- --- --- --- --- --- --- --- --- --- --- --- --- --- --- --- --- --- --- --- --- --- --- --- --- --- --- --- --- --- --- --- --- --- --- --- --- --- --- --- --- --- --- --- --- --- --- --- --- --- --- --- --- --- --- --- --- --- --- --- --- --- --- --- --- --- --- --- --- --- --- --- --- --- --- --- --- --- --- --- --- --- --- --- --- --- --- --- --- --- --- --- --- --- --- --- --- --- --- --- --- --- --- --- --- --- --- --- --- --- --- --- --- --- --- --- --- --- --- --- --- --- --- --- --- --- --- --- --- --- --- --- --- --- --- --- --- --- --- --- --- --- --- --- --- --- --- -ct gga tcg gat gtc acc gca gta ata ttg ttg att att tct gac atc gac gta tta tat agt ttt tta att cca tat ctt ttt t

>ON622718.1_Monkeypox_virus_isolate_MPXV/ES0001/HUGTiP/2022_partial_genome

ttt tat atc act acg gac ata aac cat tgt ata ttt ttt atg ttt att agt gta cac att ttg gaa gta agt tc- --- --- --- --- --- --- --- --- --- --- --- --- --- --- --- --- --- --- --- --- --- --- --- --- --- --- --- --- --- --- --- --- --- --- --- --- --- --- --- --- --- --- --- --- --- --- --- --- --- --- --- --- --- --- --- --- --- --- --- --- --- --- --- --- --- --- --- --- --- --- --- --- --- --- --- --- --- --- --- --- --- --- --- --- --- --- --- --- --- --- --- --- --- --- --- --- --- --- --- --- --- --- --- --- --- --- --- --- --- --- --- --- --- --- --- --- --- --- --- --- --- --- --- --- --- --- --- --- --- --- --- --- --- --- --- --- --- --- --- --- --- --- --- --- --- --- --- --- --- --- --- --- --- --- --- --- --- --- --- --- --- --- --- --- --- --- --- --- --- --- --- --- --- --- --- --- --- --- --- --- --- --- --- --- --- --- --- --- --- --- --- --- --- --- --- --- --- --- --- --- --- --- --- --- --- --- --- --- --- --- --- --- --- --- --- --- --- --- --- --- --- --- --- --- --- --- --- --- --- --- --- --- --- --- --- --- --- --- --- --- --- --- --- --- --- --- --- --- --- --- --- --- --- --- --- --- --- --- --- --- --- --- --- --- --- --- --- --- --- --- --- --- --- --- --- --- --- --- --- --- --- --- --- --- --- --- --- --- --- --- --- --- --- --- --- --- --- --- --- --- --- --- --- --- --- --- --- --- --- --- --- --- --- --- --- --- --- --- --- --- --- --- --- --- --- --- --- --- --- --- --- --- --- --- --- --- --- --- --- --- --- --- --- --- --- --- --- --- --- --- --- --- --- --- --- --- --- --- --- --- --- --- --- --- --- --- --- --- --- --- --- --- --- --- --- --- --- --- --- --- --- --- --- --- --- --- --- --- --- --- --- --- --- --- --- --- --- --- --- --- --- --- --- --- --- --- --- --- --- --- --- --- --- --- --- --- --- --- --- --- --- --- --- --- --- --- --- --- --- --- --- --- --- --- --- --- --- --- --- --- --- --- --- --- --- --- --- --- --- --- --- --- --- --- --- --- --- --- --- --- --- --- --- --- --- --- --- --- --- --- --- --- --- --- --- --- --- --- --- --- --- --- --- --- --- --- --- --- --- --- --- --- --- --- --- --- --- --- --- --- --- --- --- --- --- --- --- --- --- --- --- --- --- --- --- --- --- --- --- --- --- --- --- --- --- --- --- --- --- --- --- --- --- --- --- --- --- --- --- --- --- --- --- --- --- --- --- --- --- --- --- --- --- --- --- --- --- --- --- --- --- --- --- --- --- --- --- --- --- --- --- --- --- --- --- --- --- --- --- --- --- --- --- --- --- --- --- --- --- --- --- --- --- --- --- --- --- --- --- --- --- --- --- --- --- --- --- --- --- --- --- --- --- --- --- --- --- --- --- --- --- --- --- --- --- --- --- --- --- --- --- --- --- --- --- --- --- --- --- --- --- --- --- --- --- --- --- --- --- --- --- --- -ct gga tcg gat gtc acc gca gta ata ttg ttg att att tct gac atc gac gta tta tat agt ttt tta att cca tat ctt ttt t

>MT903346.1_Monkeypox_virus_isolate_MPXV-USA2003_099_Gambian_Rat

ttt tat atc act acg gac ata aac cat tgt ata att ttt atg ttt att agt gta cac att ttg gaa gta agt tc- --- --- --- --- --- --- --- --- --- --- --- --- --- --- --- --- --- --- --- --- --- --- --- --- --- --- --- --- --- --- --- --- --- --- --- --- --- --- --- --- --- --- --- --- --- --- --- --- --- --- --- --- --- --- --- --- --- --- --- --- --- --- --- --- --- --- --- --- --- --- --- --- --- --- --- --- --- --- --- --- --- --- --- --- --- --- --- --- --- --- --- --- --- --- --- --- --- --- --- --- --- --- --- --- --- --- --- --- --- --- --- --- --- --- --- --- --- --- --- --- --- --- --- --- --- --- --- --- --- --- --- --- --- --- --- --- --- --- --- --- --- --- --- --- --- --- --- --- --- --- --- --- --- --- --- --- --- --- --- --- --- --- --- --- --- --- --- --- --- --- --- --- --- --- --- --- --- --- --- --- --- --- --- --- --- --- --- --- --- --- --- --- --- --- --- --- --- --- --- --- --- --- --- --- --- --- --- --- --- --- --- --- --- --- --- --- --- --- --- --- --- --- --- --- --- --- --- --- --- --- --- --- --- --- --- --- --- --- --- --- --- --- --- --- --- --- --- --- --- --- --- --- --- --- --- --- --- --- --- --- --- --- --- --- --- --- --- --- --- --- --- --- --- --- --- --- --- --- --- --- --- --- --- --- --- --- --- --- --- --- --- --- --- --- --- --- --- --- --- --- --- --- --- --- --- --- --- --- --- --- --- --- --- --- --- --- --- --- --- --- --- --- --- --- --- --- --- --- --- --- --- --- --- --- --- --- --- --- --- --- --- --- --- --- --- --- --- --- --- --- --- --- --- --- --- --- --- --- --- --- --- --- --- --- --- --- --- --- --- --- --- --- --- --- --- --- --- --- --- --- --- --- --- --- --- --- --- --- --- --- --- --- --- --- --- --- --- --- --- --- --- --- --- --- --- --- --- --- --- --- --- --- --- --- --- --- --- --- --- --- --- --- --- --- --- --- --- --- --- --- --- --- --- --- --- --- --- --- --- --- --- --- --- --- --- --- --- --- --- --- --- --- --- --- --- --- --- --- --- --- --- --- --- --- --- --- --- --- --- --- --- --- --- --- --- --- --- --- --- --- --- --- --- --- --- --- --- --- --- --- --- --- --- --- --- --- --- --- --- --- --- --- --- --- --- --- --- --- --- --- --- --- --- --- --- --- --- --- --- --- --- --- --- --- --- --- --- --- --- --- --- --- --- --- --- --- --- --- --- --- --- --- --- --- --- --- --- --- --- --- --- --- --- --- --- --- --- --- --- --- --- --- --- --- --- --- --- --- --- --- --- --- --- --- --- --- --- --- --- --- --- --- --- --- --- --- --- --- --- --- --- --- --- --- --- --- --- --- --- --- --- --- --- --- --- --- --- --- --- --- --- --- --- --- --- --- --- --- --- --- --- --- --- --- --- --- --- --- --- --- --- --- --- --- --- --- --- --- --- --- --- --- --- --- --- --- --- --- --- --- --- --- -ct gga tcg gat gtc acc gca gta ata ttg ttg att att tct gac atc gac gta tta tat agt ttt tta att cca tat ctt ttt t

>DQ011157.1_Monkeypox_virus_strain_USA_2003_039_complete_genome

ttt tat atc act acg gac ata aac cat tgt ata att ttt atg ttt att agt gta cac att ttg gaa gta agt tc- --- --- --- --- --- --- --- --- --- --- --- --- --- --- --- --- --- --- --- --- --- --- --- --- --- --- --- --- --- --- --- --- --- --- --- --- --- --- --- --- --- --- --- --- --- --- --- --- --- --- --- --- --- --- --- --- --- --- --- --- --- --- --- --- --- --- --- --- --- --- --- --- --- --- --- --- --- --- --- --- --- --- --- --- --- --- --- --- --- --- --- --- --- --- --- --- --- --- --- --- --- --- --- --- --- --- --- --- --- --- --- --- --- --- --- --- --- --- --- --- --- --- --- --- --- --- --- --- --- --- --- --- --- --- --- --- --- --- --- --- --- --- --- --- --- --- --- --- --- --- --- --- --- --- --- --- --- --- --- --- --- --- --- --- --- --- --- --- --- --- --- --- --- --- --- --- --- --- --- --- --- --- --- --- --- --- --- --- --- --- --- --- --- --- --- --- --- --- --- --- --- --- --- --- --- --- --- --- --- --- --- --- --- --- --- --- --- --- --- --- --- --- --- --- --- --- --- --- --- --- --- --- --- --- --- --- --- --- --- --- --- --- --- --- --- --- --- --- --- --- --- --- --- --- --- --- --- --- --- --- --- --- --- --- --- --- --- --- --- --- --- --- --- --- --- --- --- --- --- --- --- --- --- --- --- --- --- --- --- --- --- --- --- --- --- --- --- --- --- --- --- --- --- --- --- --- --- --- --- --- --- --- --- --- --- --- --- --- --- --- --- --- --- --- --- --- --- --- --- --- --- --- --- --- --- --- --- --- --- --- --- --- --- --- --- --- --- --- --- --- --- --- --- --- --- --- --- --- --- --- --- --- --- --- --- --- --- --- --- --- --- --- --- --- --- --- --- --- --- --- --- --- --- --- --- --- --- --- --- --- --- --- --- --- --- --- --- --- --- --- --- --- --- --- --- --- --- --- --- --- --- --- --- --- --- --- --- --- --- --- --- --- --- --- --- --- --- --- --- --- --- --- --- --- --- --- --- --- --- --- --- --- --- --- --- --- --- --- --- --- --- --- --- --- --- --- --- --- --- --- --- --- --- --- --- --- --- --- --- --- --- --- --- --- --- --- --- --- --- --- --- --- --- --- --- --- --- --- --- --- --- --- --- --- --- --- --- --- --- --- --- --- --- --- --- --- --- --- --- --- --- --- --- --- --- --- --- --- --- --- --- --- --- --- --- --- --- --- --- --- --- --- --- --- --- --- --- --- --- --- --- --- --- --- --- --- --- --- --- --- --- --- --- --- --- --- --- --- --- --- --- --- --- --- --- --- --- --- --- --- --- --- --- --- --- --- --- --- --- --- --- --- --- --- --- --- --- --- --- --- --- --- --- --- --- --- --- --- --- --- --- --- --- --- --- --- --- --- --- --- --- --- --- --- --- --- --- --- --- --- --- --- --- --- --- --- --- --- --- --- --- --- --- --- --- --- --- --- --- --- --- --- --- --- --- --- --- --- --- --- --- --- -ct gga tcg gat gtc acc gca gta ata ttg ttg att att tct gac atc gac gta tta tat agt ttt tta att cca tat ctt ttt t

>DQ011153.1_Monkeypox_virus_strain_USA_2003_044_complete_genome

ttt tat atc act acg gac ata aac cat tgt ata att ttt atg ttt att agt gta cac att ttg gaa gta agt tc- --- --- --- --- --- --- --- --- --- --- --- --- --- --- --- --- --- --- --- --- --- --- --- --- --- --- --- --- --- --- --- --- --- --- --- --- --- --- --- --- --- --- --- --- --- --- --- --- --- --- --- --- --- --- --- --- --- --- --- --- --- --- --- --- --- --- --- --- --- --- --- --- --- --- --- --- --- --- --- --- --- --- --- --- --- --- --- --- --- --- --- --- --- --- --- --- --- --- --- --- --- --- --- --- --- --- --- --- --- --- --- --- --- --- --- --- --- --- --- --- --- --- --- --- --- --- --- --- --- --- --- --- --- --- --- --- --- --- --- --- --- --- --- --- --- --- --- --- --- --- --- --- --- --- --- --- --- --- --- --- --- --- --- --- --- --- --- --- --- --- --- --- --- --- --- --- --- --- --- --- --- --- --- --- --- --- --- --- --- --- --- --- --- --- --- --- --- --- --- --- --- --- --- --- --- --- --- --- --- --- --- --- --- --- --- --- --- --- --- --- --- --- --- --- --- --- --- --- --- --- --- --- --- --- --- --- --- --- --- --- --- --- --- --- --- --- --- --- --- --- --- --- --- --- --- --- --- --- --- --- --- --- --- --- --- --- --- --- --- --- --- --- --- --- --- --- --- --- --- --- --- --- --- --- --- --- --- --- --- --- --- --- --- --- --- --- --- --- --- --- --- --- --- --- --- --- --- --- --- --- --- --- --- --- --- --- --- --- --- --- --- --- --- --- --- --- --- --- --- --- --- --- --- --- --- --- --- --- --- --- --- --- --- --- --- --- --- --- --- --- --- --- --- --- --- --- --- --- --- --- --- --- --- --- --- --- --- --- --- --- --- --- --- --- --- --- --- --- --- --- --- --- --- --- --- --- --- --- --- --- --- --- --- --- --- --- --- --- --- --- --- --- --- --- --- --- --- --- --- --- --- --- --- --- --- --- --- --- --- --- --- --- --- --- --- --- --- --- --- --- --- --- --- --- --- --- --- --- --- --- --- --- --- --- --- --- --- --- --- --- --- --- --- --- --- --- --- --- --- --- --- --- --- --- --- --- --- --- --- --- --- --- --- --- --- --- --- --- --- --- --- --- --- --- --- --- --- --- --- --- --- --- --- --- --- --- --- --- --- --- --- --- --- --- --- --- --- --- --- --- --- --- --- --- --- --- --- --- --- --- --- --- --- --- --- --- --- --- --- --- --- --- --- --- --- --- --- --- --- --- --- --- --- --- --- --- --- --- --- --- --- --- --- --- --- --- --- --- --- --- --- --- --- --- --- --- --- --- --- --- --- --- --- --- --- --- --- --- --- --- --- --- --- --- --- --- --- --- --- --- --- --- --- --- --- --- --- --- --- --- --- --- --- --- --- --- --- --- --- --- --- --- --- --- --- --- --- --- --- --- --- --- --- --- --- --- --- --- --- --- --- --- --- --- --- --- --- --- --- --- --- --- --- --- --- --- --- --- --- --- --- --- -ct gga tcg gat gtc acc gca gta ata ttg ttg att att tct gac atc gac gta tta tat agt ttt tta att cca tat ctt ttt t

>JX878410.1_Monkeypox_virus_isolate_DRC_06-1070_complete_genome

ttt tat atc act acg gac ata aac cat tgt ata att ttt atg ttt att agt gta cac att ttg gaa gta agt tcc ggc tgc cat gta ttt cct gga gag caa gta gat gat gaa gga acc aga tag ttt ata tcc ata ctt gca ctt aaa gtc tac att gta gtt gta tga gtg tat gat ctt tta agc cgc tag aag ttt tcc gtt tga tat agg atg tgg aca ttt aac aat ctg aca cgt ggg tgg att gga cca ttc tcc tcc tga aca cat gac acc aga gtt acc aat caa cga ata tcc act att gca act ata agt tac aat gct ccc atc gat ata aaa atc ctc gta tcc gtt atg tct tcc gtt gga tat aga tgg agg tga ttg gca ttt aac aga ttc gca aat agg tgc ctc agg att cca tac cat aga tcc agt aga tcc taa ttc aca ata cga ttt aga ttc acc gat caa atg ata tcc gct att aca aga gta cgt tat act aga gcc aaa gtc tac tcc gcc aat atc aag ttg gcc att atc gat atc tcg agg cga tgg gca tct ccg ttt aat aca ttg att aaa gag tgt cca tcc ggt acc ggt aca ttt agc ata tat ggg tcc cat ttt ttg ctt tct gta tcc agg tag aca tag ata ttc tat agt gtc tcc tat gtt gta att agc atc agt ctc tac act att ctt aaa ttt cat att aat ggg gcg tga cgg aat agt aca gta tga tag aac aca tcc tat tcc caa caa tgt cag gaa cgt cac gct ctc cac ctt cat att tat tta tcc gta aaa tgt tat cct gga cat cgt aca aat aat aaa aag ccc ata tat atg ttc gct att gta gaa att gtt ttt cac agt tgc tca aaa aca atg gca gtg act tat gag tta gtt aca ctt tgg agt ctc atc ttt agt aaa cat atc ata ata ttc gat att acg agt tga cat atc gaa caa att cca agt att tga ttt tgg ata ata ttc gta ttt tgc atc tgc tat aat taa gat ata atc acc aca aga aca cac gaa cgt ctt tcc tac atg gtt aaa gta cat gta caa ttc tat cca ttt gtc ttc ctt aac tat ata ttt gta tag ata att acg agt ctc atg agt aat tcc agt aat tgc ata gat gtc acc atc gta ttc tac agc ata aac tat act atg acg tct agg cat ggg aga ctt ttt tat cca acg att ttt agt gaa aca ttc cac atc gtt taa tac tac ata ttt ctc ata gtg gta taa act cca ccc att aca tat ata tca tcg ttt acg aat act gat gcg cct gaa tat cta gga gtg att aag ttt gga agt ctt ttc cat ttc gaa gtg ccg tgt ttc aaa tat tct gct ata ccc gtt gaa ata gaa aat tct aat cct cct att aca tat aac ttt cca tcg tta aca caa gta cta act tct gat ttt aac gac gac ata tta gta acc gtt ttc ca- --- ttt ttt ttg ttt taa gat cta ccc gcg ata cgg aat aaa cat gtc tat tgt taa tca tgc cgc caa taa tgt ata gac aat tat gta aaa cat ttg cat cat aga att gtc tat ctg tat tac cga cta tcg tcc aat att ctg ttc tag gag agt aat ggg tta ttg tgg ata tat aat cag agt ttt taa tga cta cta tat tat gtt tta tac cat ttc gtg tca cag ctt tgt aga ttt gga tat agt taa tcc caa caa tgc tat agc att gca tat agc att agt cat aaa ctt ggg atg taa aat gtt gat gat atc tac atc gtt tgg att ttt atg tat cca ctt taa taa tat tat agc gta aca tcc tca tga ttt acg tta acg ttt tcg tgt gat aag ata gtg gtc agt tca tcc ttt gat aat ttt cca aat tct gga tcg gat gtc acc gca gta ata ttg ttg att att tct gac atc gac gca tta tat agt ttt tta att cca tat ctt ttt t

>JX878428.1_Monkeypox_virus_isolate_DRC_07-0514_complete_genome

ttt tat atc act acg gac ata aac cat tgt ata att ttt atg ttt att agt gta cac att ttg gaa gta agt tcc ggc tgc cat gta ttt cct gga gag caa gta gat gat gaa gga acc aga tag ttt ata tcc ata ctt gca ctt aaa gtc tac att gta gtt gta tga gtg tat gat ctt tta agc cgc tag aag ttt tcc gtt tga tat agg atg tgg aca ttt aac aat ctg aca cgt ggg tgg att gga cca ttc tcc tcc tga aca cat gac acc aga gtt acc aat caa cga ata tcc act att gca act ata agt tac aat gct ccc atc gat ata aaa atc ctc gta tcc gtt atg tct tcc gtt gga tat aga tgg agg tga ttg gca ttt aac aga ttc gca aat agg tgc ctc agg att cca tac cat aga tcc agt aga tcc taa ttc aca ata cga ttt aga ttc acc gat caa atg ata tcc gct att aca aga gta cgt tat act aga gcc aaa gtc tac tcc gcc aat atc aag ttg gcc att atc gat atc tcg agg cga tgg gca tct ccg ttt aat aca ttg att aaa gag tgt cca tcc ggt acc ggt aca ttt agc ata tat ggg tcc cat ttt ttg ctt tct gta tcc agg tag aca tag ata ttc tat agt gtc tcc tat gtt gta att agc atc agt ctc tac act att ctt aaa ttt cat att aat ggg gcg tga cgg aat agt aca gta tga tag aac aca tcc tat tcc caa caa tgt cag gaa cgt cac gct ctc cac ctt cat att tat tta tcc gta aaa tgt tat cct gga cat cgt aca aat aat aaa aag ccc ata tat atg ttc gct att gta gaa att gtt ttt cac agt tgc tca aaa aca atg gca gtg act tat gag tta gtt aca ctt tgg agt ctc atc ttt agt aaa cat atc ata ata ttc gat att acg agt tga cat atc gaa caa att cca agt att tga ttt tgg ata ata ttc gta ttt tgc atc tgc tat aat taa gat ata atc acc aca aga aca cac gaa cgt ctt tcc tac atg gtt aaa gta cat gta caa ttc tat cca ttt gtc ttc ctt aac tat ata ttt gta tag ata att acg agt ctc atg agt aat tcc agt aat tgc ata gat gtc acc atc gta ttc tac agc ata aac tat act atg acg tct agg cat ggg aga ctt ttt tat cca acg att ttt agt gaa aca ttc cac atc gtt taa tac tac ata ttt ctc ata gtg gta taa act cca ccc att aca tat ata tca tcg ttt acg aat act gat gcg cct gaa tat cta gga gtg att aag ttt gga agt ctt ttc cat ttc gaa gtg ccg tgt ttc aaa tat tct gct ata ccc gtt gaa ata gaa aat tct aat cct cct att aca tat aac ttt cca tcg tta aca caa gta cta act tct gat ttt aac gac gac ata tta gta acc gtt ttc ca- --- ttt ttt ttg ttt taa gat cta ccc gcg ata cgg aat aaa cat gtc tat tgt taa tca tgc cgc caa taa tgt ata gac aat tat gta aaa cat ttg cat cat aga att gtc tat ctg tat tac cga cta tcg tcc aat att ctg ttc tag gag agt aat ggg tta ttg tgg ata tat aat cag agt ttt taa tga cta cta tat tat gtt tta tac cat ttc gtg tca cag ctt tgt aga ttt gga tat agt taa tcc caa caa tgc tat agc att gca tat agc att agt cat aaa ctt ggg atg taa aat gtt gat gat atc tac atc gtt tgg att ttt atg tat cca ctt taa taa tat tat agc gta aca tcc tca tga ttt acg tta acg ttt tcg tgt gat aag ata gtg gtc agt tca tcc ttt gat aat ttt cca aat tct gga tcg gat gtc acc gca gta ata ttg ttg att att tct gac atc gac gca tta tat agt ttt tta att cca tat ctt ttt t

>JX878427.1_Monkeypox_virus_isolate_DRC_07-0480_complete_genome

ttt tat atc act acg gac ata aac cat tgt ata att ttt atg ttt att agt gta cac att ttg gaa gta agt tcc ggc tgc cat gta ttt cct gga gag caa gta gat gat gaa gga acc aga tag ttt ata tcc ata ctt gca ctt aaa gtc tac att gta gtt gta tga gtg tat gat ctt tta agc cgc tag aag ttt tcc gtt tga tat agg atg tgg aca ttt aac aat ctg aca cgt ggg tgg att gga cca ttc tcc tcc tga aca cat gac acc aga gtt acc aat caa cga ata tcc act att gca act ata agt tac aat gct ccc atc gat ata aaa atc ctc gta tcc gtt atg tct tcc gtt gga tat aga tgg agg tga ttg gca ttt aac aga ttc gca aat agg tgc ctc agg att cca tac cat aga tcc agt aga tcc taa ttc aca ata cga ttt aga ttc acc gat caa atg ata tcc gct att aca aga gta cgt tat act aga gcc aaa gtc tac tcc gcc aat atc aag ttg gcc att atc gat atc tcg agg cga tgg gca tct ccg ttt aat aca ttg att aaa gag tgt cca tcc ggt acc ggt aca ttt agc ata tat ggg tcc cat ttt ttg ctt tct gta tcc agg tag aca tag ata ttc tat agt gtc tcc tat gtt gta att agc atc agt ctc tac act att ctt aaa ttt cat att aat ggg gcg tga cgg aat agt aca gta tga tag aac aca tcc tat tcc caa caa tgt cag gaa cgt cac gct ctc cac ctt cat att tat tta tcc gta aaa tgt tat cct gga cat cgt aca aat aat aaa aag ccc ata tat atg ttc gct att gta gaa att gtt ttt cac agt tgc tca aaa aca atg gca gtg act tat gag tta gtt aca ctt tgg agt ctc atc ttt agt aaa cat atc ata ata ttc gat att acg agt tga cat atc gaa caa att cca agt att tga ttt tgg ata ata ttc gta ttt tgc atc tgc tat aat taa gat ata atc acc aca aga aca cac gaa cgt ctt tcc tac atg gtt aaa gta cat gta caa ttc tat cca ttt gtc ttc ctt aac tat ata ttt gta tag ata att acg agt ctc atg agt aat tcc agt aat tgc ata gat gtc acc atc gta ttc tac agc ata aac tat act atg acg tct agg cat ggg aga ctt ttt tat cca acg att ttt agt gaa aca ttc cac atc gtt taa tac tac ata ttt ctc ata gtg gta taa act cca ccc att aca tat ata tca tcg ttt acg aat act gat gcg cct gaa tat cta gga gtg att aag ttt gga agt ctt ttc cat ttc gaa gtg ccg tgt ttc aaa tat tct gct ata ccc gtt gaa ata gaa aat tct aat cct cct att aca tat aac ttt cca tcg tta aca caa gta cta act tct gat ttt aac gac gac ata tta gta acc gtt ttc ca- --- ttt ttt ttg ttt taa gat cta ccc gcg ata cgg aat aaa cat gtc tat tgt taa tca tgc cgc caa taa tgt ata gac aat tat gta aaa cat ttg cat cat aga att gtc tat ctg tat tac cga cta tcg tcc aat att ctg ttc tag gag agt aat ggg tta ttg tgg ata tat aat cag agt ttt taa tga cta cta tat tat gtt tta tac cat ttc gtg tca cag ctt tgt aga ttt gga tat agt taa tcc caa caa tgc tat agc att gca tat agc att agt cat aaa ctt ggg atg taa aat gtt gat gat atc tac atc gtt tgg att ttt atg tat cca ctt taa taa tat tat agc gta aca tcc tca tga ttt acg tta acg ttt tcg tgt gat aag ata gtg gtc agt tca tcc ttt gat aat ttt cca aat tct gga tcg gat gtc acc gca gta ata ttg ttg att att tct gac atc gac gca tta tat agt ttt tta att cca tat ctt ttt t

>JX878421.1_Monkeypox_virus_isolate_DRC_07-0286_complete_genome

ttt tat atc act acg gac ata aac cat tgt ata att ttt atg ttt att agt gta cac att ttg gaa gta agt tcc ggc tgc cat gta ttt cct gga gag caa gta gat gat gaa gga acc aga tag ttt ata tcc ata ctt gca ctt aaa gtc tac att gta gtt gta tga gtg tat gat ctt tta agc cgc tag aag ttt tcc gtt tga tat agg atg tgg aca ttt aac aat ctg aca cgt ggg tgg att gga cca ttc tcc tcc tga aca cat gac acc aga gtt acc aat caa cga ata tcc act att gca act ata agt tac aat gct ccc atc gat ata aaa atc ctc gta tcc gtt atg tct tcc gtt gga tat aga tgg agg tga ttg gca ttt aac aga ttc gca aat agg tgc ctc agg att cca tac cat aga tcc agt aga tcc taa ttc aca ata cga ttt aga ttc acc gat caa atg ata tcc gct att aca aga gta cgt tat act aga gcc aaa gtc tac tcc gcc aat atc aag ttg gcc att atc gat atc tcg agg cga tgg gca tct ccg ttt aat aca ttg att aaa gag tgt cca tcc ggt acc ggt aca ttt agc ata tat ggg tcc cat ttt ttg ctt tct gta tcc agg tag aca tag ata ttc tat agt gtc tcc tat gtt gta att agc atc agt ctc tac act att ctt aaa ttt cat att aat ggg gcg tga cgg aat agt aca gta tga tag aac aca tcc tat tcc caa caa tgt cag gaa cgt cac gct ctc cac ctt cat att tat tta tcc gta aaa tgt tat cct gga cat cgt aca aat aat aaa aag ccc ata tat atg ttc gct att gta gaa att gtt ttt cac agt tgc tca aaa aca atg gca gtg act tat gag tta gtt aca ctt tgg agt ctc atc ttt agt aaa cat atc ata ata ttc gat att acg agt tga cat atc gaa caa att cca agt att tga ttt tgg ata ata ttc gta ttt tgc atc tgc tat aat taa gat ata atc acc aca aga aca cac gaa cgt ctt tcc tac atg gtt aaa gta cat gta caa ttc tat cca ttt gtc ttc ctt aac tat ata ttt gta tag ata att acg agt ctc atg agt aat tcc agt aat tgc ata gat gtc acc atc gta ttc tac agc ata aac tat act atg acg tct agg cat ggg aga ctt ttt tat cca acg att ttt agt gaa aca ttc cac atc gtt taa tac tac ata ttt ctc ata gtg gta taa act cca ccc att aca tat ata tca tcg ttt acg aat act gat gcg cct gaa tat cta gga gtg att aag ttt gga agt ctt ttc cat ttc gaa gtg ccg tgt ttc aaa tat tct gct ata ccc gtt gaa ata gaa aat tct aat cct cct att aca tat aac ttt cca tcg tta aca caa gta cta act tct gat ttt aac gac gac ata tta gta acc gtt ttc ca- --- ttt ttt ttg ttt taa gat cta ccc gcg ata cgg aat aaa cat gtc tat tgt taa tca tgc cgc caa taa tgt ata gac aat tat gta aaa cat ttg cat cat aga att gtc tat ctg tat tac cga cta tcg tcc aat att ctg ttc tag gag agt aat ggg tta ttg tgg ata tat aat cag agt ttt taa tga cta cta tat tat gtt tta tac cat ttc gtg tca cag ctt tgt aga ttt gga tat agt taa tcc caa caa tgc tat agc att gca tat agc att agt cat aaa ctt ggg atg taa aat gtt gat gat atc tac atc gtt tgg att ttt atg tat cca ctt taa taa tat tat agc gta aca tcc tca tga ttt acg tta acg ttt tcg tgt gat aag ata gtg gtc agt tca tcc ttt gat aat ttt cca aat tct gga tcg gat gtc acc gca gta ata ttg ttg att att tct gac atc gac gca tta tat agt ttt tta att cca tat ctt ttt t

>JX878415.1_Monkeypox_virus_isolate_DRC_07-0092_complete_genome

ttt tat atc act acg gac ata aac cat tgt ata att ttt atg ttt att agt gta cac att ttg gaa gta agt tcc ggc tgc cat gta ttt cct gga gag caa gta gat gat gaa gga acc aga tag ttt ata tcc ata ctt gca ctt aaa gtc tac att gta gtt gta tga gtg tat gat ctt tta agc cgc tag aag ttt tcc gtt tga tat agg atg tgg aca ttt aac aat ctg aca cgt ggg tgg att gga cca ttc tcc tcc tga aca cat gac acc aga gtt acc aat caa cga ata tcc act att gca act ata agt tac aat gct ccc atc gat ata aaa atc ctc gta tcc gtt atg tct tcc gtt gga tat aga tgg agg tga ttg gca ttt aac aga ttc gca aat agg tgc ctc agg att cca tac cat aga tcc agt aga tcc taa ttc aca ata cga ttt aga ttc acc gat caa atg ata tcc gct att aca aga gta cgt tat act aga gcc aaa gtc tac tcc gcc aat atc aag ttg gcc att atc gat atc tcg agg cga tgg gca tct ccg ttt aat aca ttg att aaa gag tgt cca tcc ggt acc ggt aca ttt agc ata tat ggg tcc cat ttt ttg ctt tct gta tcc agg tag aca tag ata ttc tat agt gtc tcc tat gtt gta att agc atc agt ctc tac act att ctt aaa ttt cat att aat ggg gcg tga cgg aat agt aca gta tga tag aac aca tcc tat tcc caa caa tgt cag gaa cgt cac gct ctc cac ctt cat att tat tta tcc gta aaa tgt tat cct gga cat cgt aca aat aat aaa aag ccc ata tat atg ttc gct att gta gaa att gtt ttt cac agt tgc tca aaa aca atg gca gtg act tat gag tta gtt aca ctt tgg agt ctc atc ttt agt aaa cat atc ata ata ttc gat att acg agt tga cat atc gaa caa att cca agt att tga ttt tgg ata ata ttc gta ttt tgc atc tgc tat aat taa gat ata atc acc aca aga aca cac gaa cgt ctt tcc tac atg gtt aaa gta cat gta caa ttc tat cca ttt gtc ttc ctt aac tat ata ttt gta tag ata att acg agt ctc atg agt aat tcc agt aat tgc ata gat gtc acc atc gta ttc tac agc ata aac tat act atg acg tct agg cat ggg aga ctt ttt tat cca acg att ttt agt gaa aca ttc cac atc gtt taa tac tac ata ttt ctc ata gtg gta taa act cca ccc att aca tat ata tca tcg ttt acg aat act gat gcg cct gaa tat cta gga gtg att aag ttt gga agt ctt ttc cat ttc gaa gtg ccg tgt ttc aaa tat tct gct ata ccc gtt gaa ata gaa aat tct aat cct cct att aca tat aac ttt cca tcg tta aca caa gta cta act tct gat ttt aac gac gac ata tta gta acc gtt ttc ca- --- ttt ttt ttg ttt taa gat cta ccc gcg ata cgg aat aaa cat gtc tat tgt taa tca tgc cgc caa taa tgt ata gac aat tat gta aaa cat ttg cat cat aga att gtc tat ctg tat tac cga cta tcg tcc aat att ctg ttc tag gag agt aat ggg tta ttg tgg ata tat aat cag agt ttt taa tga cta cta tat tat gtt tta tac cat ttc gtg tca cag ctt tgt aga ttt gga tat agt taa tcc caa caa tgc tat agc att gca tat agc att agt cat aaa ctt ggg atg taa aat gtt gat gat atc tac atc gtt tgg att ttt atg tat cca ctt taa taa tat tat agc gta aca tcc tca tga ttt acg tta acg ttt tcg tgt gat aag ata gtg gtc agt tca tcc ttt gat aat ttt cca aat tct gga tcg gat gtc acc gca gta ata ttg ttg att att tct gac atc gac gca tta tat agt ttt tta att cca tat ctt ttt t

>JX878414.1_Monkeypox_virus_isolate_DRC_07-0046_complete_genome

ttt tat atc act acg gac ata aac cat tgt ata att ttt atg ttt att agt gta cac att ttg gaa gta agt tcc ggc tgc cat gta ttt cct gga gag caa gta gat gat gaa gga acc aga tag ttt ata tcc ata ctt gca ctt aaa gtc tac att gta gtt gta tga gtg tat gat ctt tta agc cgc tag aag ttt tcc gtt tga tat agg atg tgg aca ttt aac aat ctg aca cgt ggg tgg att gga cca ttc tcc tcc tga aca cat gac acc aga gtt acc aat caa cga ata tcc act att gca act ata agt tac aat gct ccc atc gat ata aaa atc ctc gta tcc gtt atg tct tcc gtt gga tat aga tgg agg tga ttg gca ttt aac aga ttc gca aat agg tgc ctc agg att cca tac cat aga tcc agt aga tcc taa ttc aca ata cga ttt aga ttc acc gat caa atg ata tcc gct att aca aga gta cgt tat act aga gcc aaa gtc tac tcc gcc aat atc aag ttg gcc att atc gat atc tcg agg cga tgg gca tct ccg ttt aat aca ttg att aaa gag tgt cca tcc ggt acc ggt aca ttt agc ata tat ggg tcc cat ttt ttg ctt tct gta tcc agg tag aca tag ata ttc tat agt gtc tcc tat gtt gta att agc atc agt ctc tac act att ctt aaa ttt cat att aat ggg gcg tga cgg aat agt aca gta tga tag aac aca tcc tat tcc caa caa tgt cag gaa cgt cac gct ctc cac ctt cat att tat tta tcc gta aaa tgt tat cct gga cat cgt aca aat aat aaa aag ccc ata tat atg ttc gct att gta gaa att gtt ttt cac agt tgc tca aaa aca atg gca gtg act tat gag tta gtt aca ctt tgg agt ctc atc ttt agt aaa cat atc ata ata ttc gat att acg agt tga cat atc gaa caa att cca agt att tga ttt tgg ata ata ttc gta ttt tgc atc tgc tat aat taa gat ata atc acc aca aga aca cac gaa cgt ctt tcc tac atg gtt aaa gta cat gta caa ttc tat cca ttt gtc ttc ctt aac tat ata ttt gta tag ata att acg agt ctc atg agt aat tcc agt aat tgc ata gat gtc acc atc gta ttc tac agc ata aac tat act atg acg tct agg cat ggg aga ctt ttt tat cca acg att ttt agt gaa aca ttc cac atc gtt taa tac tac ata ttt ctc ata gtg gta taa act cca ccc att aca tat ata tca tcg ttt acg aat act gat gcg cct gaa tat cta gga gtg att aag ttt gga agt ctt ttc cat ttc gaa gtg ccg tgt ttc aaa tat tct gct ata ccc gtt gaa ata gaa aat tct aat cct cct att aca tat aac ttt cca tcg tta aca caa gta cta act tct gat ttt aac gac gac ata tta gta acc gtt ttc ca- --- ttt ttt ttg ttt taa gat cta ccc gcg ata cgg aat aaa cat gtc tat tgt taa tca tgc cgc caa taa tgt ata gac aat tat gta aaa cat ttg cat cat aga att gtc tat ctg tat tac cga cta tcg tcc aat att ctg ttc tag gag agt aat ggg tta ttg tgg ata tat aat cag agt ttt taa tga cta cta tat tat gtt tta tac cat ttc gtg tca cag ctt tgt aga ttt gga tat agt taa tcc caa caa tgc tat agc att gca tat agc att agt cat aaa ctt ggg atg taa aat gtt gat gat atc tac atc gtt tgg att ttt atg tat cca ctt taa taa tat tat agc gta aca tcc tca tga ttt acg tta acg ttt tcg tgt gat aag ata gtg gtc agt tca tcc ttt gat aat ttt cca aat tct gga tcg gat gtc acc gca gta ata ttg ttg att att tct gac atc gac gca tta tat agt ttt tta att cca tat ctt ttt t

>JX878413.1_Monkeypox_virus_isolate_DRC_07-0045_complete_genome

ttt tat atc act acg gac ata aac cat tgt ata att ttt atg ttt att agt gta cac att ttg gaa gta agt tcc ggc tgc cat gta ttt cct gga gag caa gta gat gat gaa gga acc aga tag ttt ata tcc ata ctt gca ctt aaa gtc tac att gta gtt gta tga gtg tat gat ctt tta agc cgc tag aag ttt tcc gtt tga tat agg atg tgg aca ttt aac aat ctg aca cgt ggg tgg att gga cca ttc tcc tcc tga aca cat gac acc aga gtt acc aat caa cga ata tcc act att gca act ata agt tac aat gct ccc atc gat ata aaa atc ctc gta tcc gtt atg tct tcc gtt gga tat aga tgg agg tga ttg gca ttt aac aga ttc gca aat agg tgc ctc agg att cca tac cat aga tcc agt aga tcc taa ttc aca ata cga ttt aga ttc acc gat caa atg ata tcc gct att aca aga gta cgt tat act aga gcc aaa gtc tac tcc gcc aat atc aag ttg gcc att atc gat atc tcg agg cga tgg gca tct ccg ttt aat aca ttg att aaa gag tgt cca tcc ggt acc ggt aca ttt agc ata tat ggg tcc cat ttt ttg ctt tct gta tcc agg tag aca tag ata ttc tat agt gtc tcc tat gtt gta att agc atc agt ctc tac act att ctt aaa ttt cat att aat ggg gcg tga cgg aat agt aca gta tga tag aac aca tcc tat tcc caa caa tgt cag gaa cgt cac gct ctc cac ctt cat att tat tta tcc gta aaa tgt tat cct gga cat cgt aca aat aat aaa aag ccc ata tat atg ttc gct att gta gaa att gtt ttt cac agt tgc tca aaa aca atg gca gtg act tat gag tta gtt aca ctt tgg agt ctc atc ttt agt aaa cat atc ata ata ttc gat att acg agt tga cat atc gaa caa att cca agt att tga ttt tgg ata ata ttc gta ttt tgc atc tgc tat aat taa gat ata atc acc aca aga aca cac gaa cgt ctt tcc tac atg gtt aaa gta cat gta caa ttc tat cca ttt gtc ttc ctt aac tat ata ttt gta tag ata att acg agt ctc atg agt aat tcc agt aat tgc ata gat gtc acc atc gta ttc tac agc ata aac tat act atg acg tct agg cat ggg aga ctt ttt tat cca acg att ttt agt gaa aca ttc cac atc gtt taa tac tac ata ttt ctc ata gtg gta taa act cca ccc att aca tat ata tca tcg ttt acg aat act gat gcg cct gaa tat cta gga gtg att aag ttt gga agt ctt ttc cat ttc gaa gtg ccg tgt ttc aaa tat tct gct ata ccc gtt gaa ata gaa aat tct aat cct cct att aca tat aac ttt cca tcg tta aca caa gta cta act tct gat ttt aac gac gac ata tta gta acc gtt ttc ca- --- ttt ttt ttg ttt taa gat cta ccc gcg ata cgg aat aaa cat gtc tat tgt taa tca tgc cgc caa taa tgt ata gac aat tat gta aaa cat ttg cat cat aga att gtc tat ctg tat tac cga cta tcg tcc aat att ctg ttc tag gag agt aat ggg tta ttg tgg ata tat aat cag agt ttt taa tga cta cta tat tat gtt tta tac cat ttc gtg tca cag ctt tgt aga ttt gga tat agt taa tcc caa caa tgc tat agc att gca tat agc att agt cat aaa ctt ggg atg taa aat gtt gat gat atc tac atc gtt tgg att ttt atg tat cca ctt taa taa tat tat agc gta aca tcc tca tga ttt acg tta acg ttt tcg tgt gat aag ata gtg gtc agt tca tcc ttt gat aat ttt cca aat tct gga tcg gat gtc acc gca gta ata ttg ttg att att tct gac atc gac gca tta tat agt ttt tta att cca tat ctt ttt t

>JX878422.1_Monkeypox_virus_isolate_DRC_07-0287_complete_genome

ttt tat atc act acg gac ata aac cat tgt ata att ttt atg ttt att agt gta cac att ttg gaa gta agt tcc ggc tgc cat gta ttt cct gga gag caa gta gat gat gaa gga acc aga tag ttt ata tcc ata ctt gca ctt aaa gtc tac att gta gtt gta tga gtg tat gat ctt tta agc cgc tag aag ttt tcc gtt tga tat agg atg tgg aca ttt aac aat ctg aca cgt ggg tgg att gga cca ttc tcc tcc tga aca cat gac acc aga gtt acc aat caa cga ata tcc act att gca act ata agt tac aat gct ccc atc gat ata aaa atc ctc gta tcc gtt atg tct tcc gtt gga tat aga tgg agg tga ttg gca ttt aac aga ttc gca aat agg tgc ctc agg att cca tac cat aga tcc agt aga tcc taa ttc aca ata cga ttt aga ttc acc gat caa atg ata tcc gct att aca aga gta cgt tat act aga gcc aaa gtc tac tcc gcc aat atc aag ttg gcc att atc gat atc tcg agg cga tgg gca tct ccg ttt aat aca ttg att aaa gag tgt cca tcc ggt acc ggt aca ttt agc ata tat ggg tcc cat ttt ttg ctt tct gta tcc agg tag aca tag ata ttc tat agt gtc tcc tat gtt gta att agc atc agt ctc tac act att ctt aaa ttt cat att aat ggg gcg tga cgg aat agt aca gta tga tag aac aca tcc tat tcc caa caa tgt cag gaa cgt cac gct ctc cac ctt cat att tat tta tcc gta aaa tgt tat cct gga cat cgt aca aat aat aaa aag ccc ata tat atg ttc gct att gta gaa att gtt ttt cac agt tgc tca aaa aca atg gca gtg act tat gag tta gtt aca ctt tgg agt ctc atc ttt agt aaa cat atc ata ata ttc gat att acg agt tga cat atc gaa caa att cca agt att tga ttt tgg ata ata ttc gta ttt tgc atc tgc tat aat taa gat ata atc acc aca aga aca cac gaa cgt ctt tcc tac atg gtt aaa gta cat gta caa ttc tat cca ttt gtc ttc ctt aac tat ata ttt gta tag ata att acg agt ctc atg agt aat tcc agt aat tgc ata gat gtc acc atc gta ttc tac agc ata aac tat act atg acg tct agg cat ggg aga ctt ttt tat cca acg att ttt agt gaa aca ttc cac atc gtt taa tac tac ata ttt ctc ata gtg gta taa act cca ccc att aca tat ata tca tcg ttt acg aat act gat gcg cct gaa tat cta gga gtg att aag ttt gga agt ctt ttc cat ttc gaa gtg ccg tgt ttc aaa tat tct gct ata ccc gtt gaa ata gaa aat tct aat cct cct att aca tat aac ttt cca tcg tta aca caa gta cta act tct gat ttt aac gac gac ata tta gta acc gtt ttc ca- --- ttt ttt ttg ttt taa gat cta ccc gcg ata cgg aat aaa cat gtc tat tgt taa tca tgc cgc caa taa tgt ata gac aat tat gta aaa cat ttg cat cat aga att gtc tat ctg tat tac cga cta tcg tcc aat att ctg ttc tag gag agt aat ggg tta ttg tgg ata tat aat cag agt ttt taa tga cta cta tat tat gtt tta tac cat ttc gtg tca cag ctt tgt aga ttt gga tat agt taa tcc caa caa tgc tat agc att gca tat agc att agt cat aaa ctt ggg atg taa aat gtt gat gat atc tac atc gtt tgg att ttt atg tat cca ctt taa taa tat tat agc gta aca tcc tca tga ttt acg tta acg ttt tcg tgt gat aag ata gtg gtc agt tca tcc ttt gat aat ttt cca aat tct gga tcg gat gtc acc gca gta ata ttg ttg att att tct gac atc gac gca tta tat agt ttt tta att cca tat ctt ttt t

>JX878416.1_Monkeypox_virus_isolate_DRC_07-0093_complete_genome

ttt tat atc act acg gac ata aac cat tgt ata att ttt atg ttt att agt gta cac att ttg gaa gta agt tcc ggc tgc cat gta ttt cct gga gag caa gta gat gat gaa gga acc aga tag ttt ata tcc ata ctt gca ctt aaa gtc tac att gta gtt gta tga gtg tat gat ctt tta agc cgc tag aag ttt tcc gtt tga tat agg atg tgg aca ttt aac aat ctg aca cgt ggg tgg att gga cca ttc tcc tcc tga aca cat gac acc aga gtt acc aat caa cga ata tcc act att gca act ata agt tac aat gct ccc atc gat ata aaa atc ctc gta tcc gtt atg tct tcc gtt gga tat aga tgg agg tga ttg gca ttt aac aga ttc gca aat agg tgc ctc agg att cca tac cat aga tcc agt aga tcc taa ttc aca ata cga ttt aga ttc acc gat caa atg ata tcc gct att aca aga gta cgt tat act aga gcc aaa gtc tac tcc gcc aat atc aag ttg gcc att atc gat atc tcg agg cga tgg gca tct ccg ttt aat aca ttg att aaa gag tgt cca tcc ggt acc ggt aca ttt agc ata tat ggg tcc cat ttt ttg ctt tct gta tcc agg tag aca tag ata ttc tat agt gtc tcc tat gtt gta att agc atc agt ctc tac act att ctt aaa ttt cat att aat ggg gcg tga cgg aat agt aca gta tga tag aac aca tcc tat tcc caa caa tgt cag gaa cgt cac gct ctc cac ctt cat att tat tta tcc gta aaa tgt tat cct gga cat cgt aca aat aat aaa aag ccc ata tat atg ttc gct att gta gaa att gtt ttt cac agt tgc tca aaa aca atg gca gtg act tat gag tta gtt aca ctt tgg agt ctc atc ttt agt aaa cat atc ata ata ttc gat att acg agt tga cat atc gaa caa att cca agt att tga ttt tgg ata ata ttc gta ttt tgc atc tgc tat aat taa gat ata atc acc aca aga aca cac gaa cgt ctt tcc tac atg gtt aaa gta cat gta caa ttc tat cca ttt gtc ttc ctt aac tat ata ttt gta tag ata att acg agt ctc atg agt aat tcc agt aat tgc ata gat gtc acc atc gta ttc tac agc ata aac tat act atg acg tct agg cat ggg aga ctt ttt tat cca acg att ttt agt gaa aca ttc cac atc gtt taa tac tac ata ttt ctc ata gtg gta taa act cca ccc att aca tat ata tca tcg ttt acg aat act gat gcg cct gaa tat cta gga gtg att aag ttt gga agt ctt ttc cat ttc gaa gtg ccg tgt ttc aaa tat tct gct ata ccc gtt gaa ata gaa aat tct aat cct cct att aca tat aac ttt cca tcg tta aca caa gta cta act tct gat ttt aac gac gac ata tta gta acc gtt ttc ca- --- ttt ttt ttg ttt taa gat cta ccc gcg ata cgg aat aaa cat gtc tat tgt taa tca tgc cgc caa taa tgt ata gac aat tat gta aaa cat ttg cat cat aga att gtc tat ctg tat tac cga cta tcg tcc aat att ctg ttc tag gag agt aat ggg tta ttg tgg ata tat aat cag agt ttt taa tga cta cta tat tat gtt tta tac cat ttc gtg tca cag ctt tgt aga ttt gga tat agt taa tcc caa caa tgc tat agc att gca tat agc att agt cat aaa ctt ggg atg taa aat gtt gat gat atc tac atc gtt tgg att ttt atg tat cca ctt taa taa tat tat agc gta aca tcc tca tga ttt acg tta acg ttt tcg tgt gat aag ata gtg gtc agt tca tcc ttt gat aat ttt cca aat tct gga tcg gat gtc acc gca gta ata ttg ttg att att tct gac atc gac gca tta tat agt ttt tta att cca tat ctt ttt t

>KP849471.1_Monkeypox_virus_isolate_Yambuku_DRC_1985_complete_genome

ttt tat atc act acg gac ata aac cat tgt ata att ttt atg ttt att agt gta cac att ttg gaa gta agt tcc ggc tgc cat gta ttt cct gga gag caa gta gat gat g-a gga acc aga tag ttt ata tcc ata ctt gca ctt aaa gtc tac att gta gtt gta tga gtg tat gat ctt tta agc cgc tag aag ttt tcc gtt tga tat agg atg tgg aca ttt aac aat ctg aca cgt ggg tgg att gga cca ttc tcc tcc tga aca cat gac acc aga gtt acc aat caa cga ata tcc act att gca act ata agt tac aat gct ccc atc gat ata aaa atc ctc gta tcc gtt atg tct tcc gtt gga tat aga tgg agg tga ttg gca ttt aac aga ttc gca aat agg tgc ctc agg att cca tac cat aga tcc agt aga tcc taa ttc aca ata cga ttt aga ttc acc gat caa atg ata tcc gct att aca aga gta cgt tat act aga gcc aaa gtc tac tcc gcc aat atc aag ttg gcc att atc gat atc tcg agg cga tgg gca tct ccg ttt aat aca ttg att aaa gag tgt cca tcc ggt acc ggt aca ttt agc ata tat ggg tcc cat ttt ttg ctt tct gta tcc agg tag aca tag ata ttc tat agt gtc tcc tat gtt gta att agc atc agt ctc tac act att ctt aaa ttt cat att aat ggg gcg tga cgg aat agt aca gta tga tag aac aca tcc tat tcc caa caa tgt cag gaa cgt cac gct ctc cac ctt cat att tat tta tcc gta aaa tgt tat cct gga cat cgt aca aat aat aaa aag ccc ata tat atg ttc gct att gta gaa att gtt ttt cac agt tgc tca aaa aca atg gca gtg act tat gag tta gtt aca ctt tgg agt ctc atc ttt agt aaa cat atc ata ata ttc gat att acg agt tga cat atc gaa caa att cca agt att tga ttt tgg ata ata ttc gta ttt tgc atc tgc tat aat taa gat ata atc acc aca aga aca cac gaa cgt ctt tcc tac atg gtt aaa gta cat gta caa ttc tat cca ttt gtc ttc ctt aac tat ata ttt gta tag ata att acg agt ctc atg agt aat tcc agt aat tgc ata gat gtc acc atc gta ttc tac agc ata aac tat act atg acg tct agg cat ggg aga ctt ttt tat cca acg att ttt agt gaa aca ttc cac atc gtt taa tac tac ata ttt ctc ata gtg gta taa act cca ccc att aca tat ata tca tcg ttt acg aat act gat gcg cct gaa tat cta gga gtg att aag ttt gga agt ctt ttc cat ttc gaa gtg ccg tgt ttc aaa tat tct gct ata ccc gtt gaa ata gaa aat tct aat cct cct att aca tat aac ttt cca tcg tta aca caa gta cta act tct gat ttt aac gac -ac ata tta gta acc gtt ttc ca- --- ttt ttt ttg ttt taa gat cta ccc gcg ata cgg aat aaa cat gtc tat tgt taa tca tgc cgc caa taa tgt ata gac aat tat gta aaa cat ttg cat cat aga att gtc tat ctg tat tac cga cta tcg tcc aat att ctg ttc tag gag agt aat ggg tta ttg tgg ata tat aat cag agt ttt taa tga cta cta tat tat gtt tta tac cat ttc gtg tca cag ctt tgt aga ttt gga tat agt taa tcc caa caa tgc tat agc att gca tat agc att agt cat aaa ctt ggg atg taa aat gtt gat gat atc tac atc gtt tgg att ttt atg tat cca ctt taa taa tat tat agc gta aca tcc tca tga ttt acg tta acg ttt tcg tgt gat aag ata gtg gtc agt tca tcc ttt gat aat ttt cca aat tct gga tcg gat gtc acc gca gta ata ttg ttg att att tct gac atc gac gca tta tat agt ttt tta att cca tat ctt ttt t

>DQ011155.1_Monkeypox_virus_strain_Zaire_1979-005_complete_genome

ttt tat atc act acg gac ata aac cat tgt ata att ttt atg ttt att agt gta cac att ttg gaa gta agt tcc ggc tgc cat gta ttt cct gga gag caa gta gat gat g-a gga acc aga tag ttt ata tcc ata ctt gca ctt aaa gtc tac att gta gtt gta tga gtg tat gat ctt tta agc cgc tag aag ttt tcc gtt tga tat agg atg tgg aca ttt aac aat ctg aca cgt ggg tgg att gga cca ttc tcc tcc tga aca cat gac acc aga gtt acc aat caa cga ata tcc act att gca act ata agt tac aat gct ccc atc gat ata aaa atc ctc gta tcc gtt atg tct tcc gtt gga tat aga tgg agg tga ttg gca ttt aac aga ttc gca aat agg tgc ctc agg att cca tac cat aga tcc agt aga tcc taa ttc aca ata cga ttt aga ttc acc gat caa atg ata tcc gct att aca aga gta cgt tat act aga gcc aaa gtc tac tcc gcc aat atc aag ttg gcc att atc gat atc tcg agg cga tgg gca tct ccg ttt aat aca ttg att aaa gag tgt cca tcc ggt acc ggt aca ttt agc ata tat ggg tcc cat ttt ttg ctt tct gta tcc agg tag aca tag ata ttc tat agt gtc tcc tat gtt gta att agc atc agt ctc tac act att ctt aaa ttt cat att aat ggg gcg tga cgg aat agt aca gta tga tag aac aca tcc tat tcc caa caa tgt cag gaa cgt cac gct ctc cac ctt cat att tat tta tcc gta aaa tgt tat cct gga cat cgt aca aat aat aaa aag ccc ata tat atg ttc gct att gta gaa att gtt ttt cac agt tgc tca aaa aca atg gca gtg act tat gag tta gtt aca ctt tgg agt ctc atc ttt agt aaa cat atc ata ata ttc gat att acg agt tga cat atc gaa caa att cca agt att tga ttt tgg ata ata ttc gta ttt tgc atc tgc tat aat taa gat ata atc acc aca aga aca cac gaa cgt ctt tcc tac atg gtt aaa gta cat gta caa ttc tat cca ttt gtc ttc ctt aac tat ata ttt gta tag ata att acg agt ctc atg agt aat tcc agt aat tgc ata gat gtc acc atc gta ttc tac agc ata aac tat act atg acg tct agg cat ggg aga ctt ttt tat cca acg att ttt agt gaa aca ttc cac atc gtt taa tac tac ata ttt ctc ata gtg gta taa act cca ccc att aca tat ata tca tcg ttt acg aat act gat gcg cct gaa tat cta gga gtg att aag ttt gga agt ctt ttc cat ttc gaa gtg ccg tgt ttc aaa tat tct gct ata ccc gtt gaa ata gaa aat tct aat cct cct att aca tat aac ttt cca tcg tta aca caa gta cta act tct gat ttt aac gac gac ata tta gta acc gtt ttc ca- --- -tt ttt ttg ttt taa gat cta ccc gcg ata cgg aat aaa cat gtc tat tgt taa tca tgc cgc caa taa tgt ata gac aat tat gta aaa cat ttg cat cat aga att gtc tat ctg tat tac cga cta tcg tcc aat att ctg ttc tag gag agt aat ggg tta ttg tga ata tat aat cag agt ttt taa tga cta cta tat tat gtt tta tac cat ttc gtg tca cag ctt tgt aga ttt gga tat agt taa tcc caa caa tgc tat agc att gca tat agc att agt cat aaa ctt ggg atg taa aat gtt gat gat atc tac atc gtt tgg att ttt atg tat cca ctt taa taa tat tat agc gta aca tcc tca tga ttt acg tta acg ttt tcg tgt gat aag ata gtg gtc agt tca tcc ttt gat aat ttt cca aat tct gga tcg gat gtc acc gca gta ata ttg ttg att att tct gac atc gac gca tta tat agt ttt tta att cca tat ctt ttt t

>KP849469.1_Monkeypox_virus_isolate_Boende_DRC_2008_complete_genome

ttt tat atc act acg gac ata aac cat tgt ata att ttt atg ttt att agt gta cac att ttg gaa gta agt tcc ggc tgc cat gta ttt cct gga gag caa gta gat gat g-a gga acc aga tag ttt ata tcc ata ctt gca ctt aaa gtc tac att gta gtt gta tga gtg tat gat ctt tta agc cgc tag aag ttt tcc gtt tga tat agg atg tgg aca ttt aac aat ctg aca cgt ggg tgg att gga cca ttc tcc tcc tga aca cat gac acc aga gtt acc aat caa cga ata tcc act att gca act ata agt tac aat gct ccc atc gat ata aaa atc ctc gta tcc gtt atg tct tcc gtt gga tat aga tgg agg tga ttg gca ttt aac aga ttc gca aat agg tgc ctc agg att cca tac cat aga tcc agt aga tcc taa ttc aca ata cga ttt aga ttc acc gat caa atg ata tcc gct att aca aga gta cgt tat act aga gcc aaa gtc tac tcc gcc aat atc aag ttg gcc att atc gat atc tcg agg cga tgg gca tct ccg ttt aat aca ttg att aaa gag tgt cca tcc ggt acc ggt aca ttt agc ata tat ggg tcc cat ttt ttg ctt tct gta tcc agg tag aca tag ata ttc tat agt gtc tcc tat gtt gta att agc atc agt ctc tac act att ctt aaa ttt cat att aat ggg gcg tga cgg aat agt aca gta tga tag aac aca tcc tat tcc caa caa tgt cag gaa cgt cac gct ctc cac ctt cat att tat tta tcc gta aaa tgt tat cct gga cat cgt aca aat aat aaa aag ccc ata tat atg ttc gct att gta gaa att gtt ttt cac agt tgc tca aaa aca atg gca gtg act tat gag tta gtt aca ctt tgg agt ctc atc ttt agt aaa cat atc ata ata ttc gat att acg agt tga cat atc gaa caa att cca agt att tga ttt tgg ata ata ttc gta ttt tgc atc tgc tat aat taa gat ata atc acc aca aga aca cac gaa cgt ctt tcc tac atg gtt aaa gta cat gta caa ttc tat cca ttt gtc ttc ctt aac tat ata ttt gta tag ata att acg agt ctc atg agt aat tcc agt aat tgc ata gat gtc acc atc gta ttc tac agc ata aac tat act atg acg tct agg cat ggg aga ctt ttt tat cca acg att ttt agt gaa aca ttc cac atc gtt taa tac tac ata ttt ctc ata gtg gta taa act cca ccc att aca tat ata tca tcg ttt acg aat act gat gcg cct gaa tat cta gga gtg att aag ttt gga agt ctt ttc cat ttc gaa gtg ccg tgt ttc aaa tat tct gct ata ccc gtt gaa ata gaa aat tct aat cct cct att aca tat aac ttt cca tcg tta aca caa gta cta act tct gat ttt aac gac gac ata tta gta acc gtt ttc ca- --- ttt ttt ttg ttt taa gat cta ccc gcg ata cgg aat aaa cat gtc tat tgt taa tca tgc cgc caa taa tgt ata gac aat tat gta aaa cat ttg cat cat aga att gtc tat ctg tat tac cga cta tcg tcc aat att ctg ttc tag gag agt aat ggg tta ttg tgg ata tat aat cag agt ttt taa tga cta cta tat tat gtt tta tac cat ttc gtg tca cag ctt tgt aga ttt gga tat agt taa tcc caa caa tgc tat agc att gca tat agc att agt cat aaa ctt ggg atg taa aat gtt gat gat atc tac atc gtt tgg att ttt atg tat cca ctt taa taa tat tat agc gta aca tcc tca tga ttt acg tta acg ttt tcg tgt gat aag ata gtg gtc agt tca tcc ttt gat aat ttt cca aat tct gga tcg gat gtc acc gca gta ata ttg ttg att att tct gac atc gac gca tta tat agt ttt tta att cca tat ctt ttt t

>KJ642619.1_Monkeypox_virus_strain_Gabon-1988_complete_genome

ttt tat atc act acg gac ata aac cat tgt ata att ttt atg ttt att agt gta cac att ttg gaa gta agt tcc ggc tgc cat gta ttt cct gga gag caa gta gat gat g-a gga acc aga tag ttt ata tcc ata ctt gca ctt aaa gtc tac att gta gtt gta tga gtg tat gat ctt tta agc cgc tag aag ttt tcc gtt tga tat agg atg tgg aca ttt aac aat ctg aca cgt ggg tgg att gga cca ttc tcc tcc tga aca cat gac acc aga gtt acc aat caa cga ata tcc act att gca act ata agt tac aat gct ccc atc gat ata aaa atc ctc gta tcc gtt atg tct tcc gtt gga tat aga tgg agg tga ttg gca ttt aac aga ttc gca aat agg tgc ctc agg att cca tac cat aga tcc agt aga tcc taa ttc aca ata cga ttt aga ttc acc gat caa atg ata tcc gct att aca aga gta cgt tat act aga gcc aaa gtc tac tcc gcc aat atc aag ttg gcc att atc gat atc tcg agg cga tgg gca tct ccg ttt aat aca ttg att aaa gag tgt cca tcc ggt acc ggt aca ttt agc ata tat ggg tcc cat ttt ttg ctt tct gta tcc agg tag aca tag ata ttc tat agt gtc tcc tat gtt gta att agc atc agt ctc tac act att ctt aaa ttt cat att aat ggg gcg tga cgg aat agt aca gta tga tag aac aca tcc tat tcc caa caa tgt cag gaa cgt cac gct ctc cac ctt cat att tat tta tcc gta aaa tgt tat cct gga cat cgt aca aat aat aaa aag ccc ata tat atg ttc gct att gta gaa att gtt ttt cac agt tgc tca aaa aca atg gca gtg act tat gag tta gtt aca ctt tgg agt ctc atc ttt agt aaa cat atc ata ata ttc gat att acg agt tga cat atc gaa caa att cca agt att tga ttt tgg ata ata ttc gta ttt tgc atc tgc tat aat taa gat ata atc acc aca aga aca cac gaa cgt ctt tcc tac atg gtt aaa gta cat gta caa ttc tat cca ttt gtc ttc ctt aac tat ata ttt gta tag ata att acg agt ctc atg agt aat tcc agt aat tgc ata gat gtc acc atc gta ttc tac agc ata aac tat act atg acg tct agg cat ggg aga ctt ttt tat cca acg att ttt agt gaa aca ttc cac atc gtt taa tac tac ata ttt ctc ata gtg gta taa act cca ccc att aca tat ata tca tcg ttt acg aat act gat gcg cct gaa tat cta gga gtg att aag ttt gga agt ctt ttc cat ttc gaa gtg ccg tgt ttc aaa tat tct gct ata ccc gtt gaa ata gaa aat tct aat cct cct att aca tat aac ttt cca tcg tta aca caa gta cta act tct gat ttt aac gac gac ata tta gta acc gtt ttc ca- --- ttt ttt ttg ttt taa gat cta ccc gcg ata cgg aat aaa cat gtc tat tgt taa tca tgc cgc caa taa tgt ata gac aat tat gta aaa cat ttg cat cat aga att gtc tat ctg tat tac cga cta tcg tcc aat att ctg ttc tag gag agt aat ggg tta ttg tgg ata tat aat cag agt ttt taa tga cta cta tat tat gtt tta tac cat ttc gtg tca cag ctt tgt aga ttt gga tat agt taa tcc caa caa tgc tat agc att gca tat agc att agt cat aaa ctt ggg atg taa aat gtt gat gat atc tac atc gtt tgg att ttt atg tat cca ctt taa taa tat tat agc gta aca tcc tca tga ttt acg tta acg ttt tcg tgt gat aag ata gtg gtc agt tca tcc ttt gat aat ttt cca aat tct gga tcg gat gtc acc gca gta ata ttg ttg att att tct gac atc gac gca tta tat agt ttt tta att cca tat ctt ttt t

>JX878411.1_Monkeypox_virus_isolate_DRC_06-1075_complete_genome

ttt tat atc act acg gac ata aac cat tgt ata att ttt atg ttt att agt gta cac att ttg gaa gta agt tcc ggc tgc cat gta ttt cct gga gag caa gta gat gat gaa gga acc aga tag ttt ata tcc ata ctt gca ctt aaa gtc tac att gta gtt gta tga gtg tat gat ctt tta agc cgc tag aag ttt tcc gtt tga tat agg atg tgg aca ttt aac aat ctg aca cgt ggg tgg att gga cca ttc tcc tcc tga aca cat gac acc aga gtt acc aat caa cga ata tcc act att gca act ata agt tac aat gct ccc atc gat ata aaa atc ctc gta tcc gtt atg tct tcc gtt gga tat aga tgg agg tga ttg gca ttt aac aga ttc gca aat agg tgc ctc agg att cca tac cat aga tcc agt aga tcc taa ttc aca ata cga ttt aga ttc acc gat caa atg ata tcc gct att aca aga gta cgt tat act aga gcc aaa gtc tac tcc gcc aat atc aag ttg gcc att atc gat atc tcg agg cga tgg gca tct ccg ttt aat aca ttg att aaa gag tgt cca tcc ggt acc ggt aca ttt agc ata tat ggg tcc cat ttt ttg ctt tct gta tcc agg tag aca tag ata ttc tat agt gtc tcc tat gtt gta att agc atc agt ctc tac act att ctt aaa ttt cat att aat ggg gcg tga cgg aat agt aca gta tga tag aac aca tcc tat tcc caa caa tgt cag gaa cgt cac gct ctc cac ctt cat att tat tta tcc gta aaa tgt tat cct gga cat cgt aca aat aat aaa aag ccc ata tat atg ttc gct att gta gaa att gtt ttt cac agt tgc tca aaa aca atg gca gtg act tat gag tta gtt aca ctt tgg agt ctc atc ttt agt aaa cat atc ata ata ttc gat att acg agt tga cat atc gaa caa att cca agt att tga ttt tgg ata ata ttc gta ttt tgc atc tgc tat aat taa gat ata atc acc aca aga aca cac gaa cgt ctt tcc tac atg gtt aaa gta cat gta caa ttc tat cca ttt gtc ttc ctt aac tat ata ttt gta tag ata att acg agt ctc atg agt aat tcc agt aat tgc ata gat gtc acc atc gta ttc tac agc ata aac tat act atg acg tct agg cat ggg aga ctt ttt tat cca acg att ttt agt gaa aca ttc cac atc gtt taa tac tac ata ttt ctc ata gtg gta taa act cca ccc att aca tat ata tca tcg ttt acg aat act gat gcg cct gaa tat cta gga gta att aag ttt gga agt ctt ttc cat ttc gaa gtg ccg tgt ttc aaa tat tct gct ata ccc gtt gaa ata gaa aat tct aat cct cct att aca tat aac ttt cca tcg tta aca caa gta cta act tct gat ttt aac gac gac ata tta gta acc gtt ttc ca- --- ttt ttt ttg ttt taa gat cta ccc gcg ata cgg aat aaa cat gtc tat tgt taa tca tgc cgc caa taa tgt ata gac aat tat gta aaa cat ttg cat cat aga att gtc tat ctg tat tac cga cta tcg tcc aat att ctg ttc tag gag agt aat ggg tta ttg tgg ata tat aat cag agt ttt taa tga cta cta tat tat gtt tta tac cat ttc gtg tca cag ctt tgt aga ttt gga tat agt taa tcc caa caa tgc tat agc att gca tat agc att agt cat aaa ctt ggg atg taa aat gtt gat gat atc tac atc gtt tgg att ttt atg tat cca ctt taa taa tat tat agc gta aca tcc tca tga ttt acg tta acg ttt tcg tgt gat aag ata gtg gtc agt tca tcc ttt gat aat ttt cca aat tct gga tcg gat gtc acc gca gta ata ttg ttg att att tct gac atc gac gca tta tat agt ttt tta att cca tat ctt ttt t

>JX878409.1_Monkeypox_virus_isolate_DRC_06-0999_complete_genome

ttt tat atc act acg gac ata aac cat tgt ata att ttt atg ttt att agt gta cac att ttg gaa gta agt tcc ggc tgc cat gta ttt cct gga gag caa gta gat gat gaa gga acc aga tag ttt ata tcc ata ctt gca ctt aaa gtc tac att gta gtt gta tga gtg tat gat ctt tta agc cgc tag aag ttt tcc gtt tga tat agg atg tgg aca ttt aac aat ctg aca cgt ggg tgg att gga cca ttc tcc tcc tga aca cat gac acc aga gtt acc aat caa cga ata tcc act att gca act ata agt tac aat gct ccc atc gat ata aaa atc ctc gta tcc gtt atg tct tcc gtt gga tat aga tgg agg tga ttg gca ttt aac aga ttc gca aat agg tgc ctc agg att cca tac cat aga tcc agt aga tcc taa ttc aca ata cga ttt aga ttc acc gat caa atg ata tcc gct att aca aga gta cgt tat act aga gcc aaa gtc tac tcc gcc aat atc aag ttg gcc att atc gat atc tcg agg cga tgg gca tct ccg ttt aat aca ttg att aaa gag tgt cca tcc ggt acc ggt aca ttt agc ata tat ggg tcc cat ttt ttg ctt tct gta tcc agg tag aca tag ata ttc tat agt gtc tcc tat gtt gta att agc atc agt ctc tac act att ctt aaa ttt cat att aat ggg gcg tga cgg aat agt aca gta tga tag aac aca tcc tat tcc caa caa tgt cag gaa cgt cac gct ctc cac ctt cat att tat tta tcc gta aaa tgt tat cct gga cat cgt aca aat aat aaa aag ccc ata tat atg ttc gct att gta gaa att gtt ttt cac agt tgc tca aaa aca atg gca gtg act tat gag tta gtt aca ctt tgg agt ctc atc ttt agt aaa cat atc ata ata ttc gat att acg agt tga cat atc gaa caa att cca agt att tga ttt tgg ata ata ttc gta ttt tgc atc tgc tat aat taa gat ata atc acc aca aga aca cac gaa cgt ctt tcc tac atg gtt aaa gta cat gta caa ttc tat cca ttt gtc ttc ctt aac tat ata ttt gta tag ata att acg agt ctc atg agt aat tcc agt aat tgc ata gat gtc acc atc gta ttc tac agc ata aac tat act atg acg tct agg cat ggg aga ctt ttt tat cca acg att ttt agt gaa aca ttc cac atc gtt taa tac tac ata ttt ctc ata gtg gta taa act cca ccc att aca tat ata tca tcg ttt acg aat act gat gcg cct gaa tat cta gga gta att aag ttt gga agt ctt ttc cat ttc gaa gtg ccg tgt ttc aaa tat tct gct ata ccc gtt gaa ata gaa aat tct aat cct cct att aca tat aac ttt cca tcg tta aca caa gta cta act tct gat ttt aac gac gac ata tta gta acc gtt ttc ca- --- ttt ttt ttg ttt taa gat cta ccc gcg ata cgg aat aaa cat gtc tat tgt taa tca tgc cgc caa taa tgt ata gac aat tat gta aaa cat ttg cat cat aga att gtc tat ctg tat tac cga cta tcg tcc aat att ctg ttc tag gag agt aat ggg tta ttg tgg ata tat aat cag agt ttt taa tga cta cta tat tat gtt tta tac cat ttc gtg tca cag ctt tgt aga ttt gga tat agt taa tcc caa caa tgc tat agc att gca tat agc att agt cat aaa ctt ggg atg taa aat gtt gat gat atc tac atc gtt tgg att ttt atg tat cca ctt taa taa tat tat agc gta aca tcc tca tga ttt acg tta acg ttt tcg tgt gat aag ata gtg gtc agt tca tcc ttt gat aat ttt cca aat tct gga tcg gat gtc acc gca gta ata ttg ttg att att tct gac atc gac gca tta tat agt ttt tta att cca tat ctt ttt t

>DQ011154.1_Monkeypox_virus_strain_Congo_2003_358_complete_genome

ttt tat atc act acg gac ata aac cat tgt ata att ttt atg ttt att agt gta cac att ttg gaa gta agt tcc ggc tgc cat gta ttt cct gga gag caa gta gat gat g-a gga acc aga tag ttt ata tcc ata ctt gca ctt aaa gtc tac att gta gtt gta tga gtg tat gat ctt tta agc cgc tag aag ttt tcc gtt tga tat agg atg tgg aca ttt aac aat ctg aca cgt ggg tgg att gga cca ttc tcc tcc tga aca cat gac acc aga gtt acc aat caa cga ata tcc act att gca act ata agt tac aat gct ccc atc gat ata aaa atc ctc gta tcc gtt atg tct tcc gtt gga tat aga tgg agg tga ttg gca ttt aac aga ttc gca aat agg tgc ctc agg att cca tac cat aga tcc agt aga tcc taa ttc aca ata cga ttt aga ttc acc gat caa atg ata tcc gct att aca aga gta cgt tat act aga gcc aaa gtc tac tcc gcc aat atc aag ttg gcc att atc gat atc tcg agg cga tgg gca tct ccg ttt aat aca ttg att aaa gag tgt cca tcc ggt acc ggt aca ttt agc ata tat ggg tcc cat ttt ttg ctt tct gta tcc agg tag aca tag ata ttc tat agt gtc tcc tat gtt gta att agc atc agt ctc tac act att ctt aaa ttt cat att aat ggg gcg tga cgg aat agt aca gta tga tag aac aca tcc tat tcc caa caa tgt cag gaa cgt cac gct ctc cac ctt cat att tat tta tcc gta aaa tgt tat cct gga cat cgt aca aat aat aaa aag ccc ata tat atg ttc gct att gta gaa att gtt ttt cac agt tgc tca aaa aca atg gca gtg act tat gag tta gtt aca ctt tgg agt ctc atc ttt agt aaa cat atc ata ata ttc gat att acg agt tga cat atc gaa caa att cca agt att tga ttt tgg ata ata ttc gta ttt tgc atc tgc tat aat taa gat ata atc acc aca aga aca cac gaa cgt ctt tcc tac atg gtt aaa gta cat gta caa ttc tat cca ttt gtc ttc ctt aac tat ata ttt gta tag ata att acg agt ctc atg agt aat tcc agt aat tgc ata gat gtc acc atc gta ttc tac agc ata aac tat act atg acg tct agg cat ggg aga ctt ttt tat cca acg att ttt agt gaa aca ttc cac atc gtt taa tac tac ata ttt ctc ata gtg gta taa act cca ccc att aca tat ata tca tcg ttt acg aat act gat gcg cct gaa tat cta gga gtg att aag ttt gga agt ctt ttc cat ttc gaa gtg ccg tgt ttc aaa tat tct gct ata ccc gtt gaa ata gaa aat tct aat cct cct att aca tat aac ttt cca tcg tta aca caa gta cta act tct gat ttt aac gac gac ata tta gta acc gtt ttc ca- --- ttt ttt ttg ttt taa gat cta ccc gcg ata cgg aat aaa cat gtc tat tgt taa tca tgc cgc caa taa tgt ata gac aat tat gta aaa cat ttg cat cat aga att gtc tat ctg tat tac cga cta tcg tcc aat att ctg ttc tag gag agt aat ggg tta ttg tgg ata tat aat cag agt ttt taa tga cta cta tat tat gtt tta tac cat ttc gtg tca cag ctt tgt aga ttt gga tat agt taa tcc caa caa tgc tat agc att gca tat agc att agt cat aaa ctt ggg atg taa aat gtt gat gat atc tac atc gtt tgg att ttt atg tat cca ctt taa taa tat tat agc gta aca tcc tca tga ttt acg tta acg ttt tcg tgt gat aag ata gtg gtc agt tca tcc ttt gat aat ttt cca aat tct gga tcg gat gtc acc gca gta ata ttg ttg att att tct gac atc gac gca tta tat agt ttt tta att cca tat ctt ttt t

>JX878412.1_Monkeypox_virus_isolate_DRC_06-1076_complete_genome

ttt tat atc act acg gac ata aac cat tgt ata att ttt atg ttt att agt gta cac att ttg gaa gta agt tcc ggc tgc cat gta ttt cct gga gag caa gta gat gat gaa gga acc aga tag ttt ata tcc ata ctt gca ctt aaa gtc tac att gta gtt gta tga gtg tat gat ctt tta agc cgc tag aag ttt tcc gtt tga tat agg atg tgg aca ttt aac aat ctg aca cgt ggg tgg att gga cca ttc tcc tcc tga aca cat gac acc aga gtt acc aat caa cga ata tcc act att gca act ata agt tac aat gct ccc atc gat ata aaa atc ctc gta tcc gtt atg tct tcc gtt gga tat aga tgg agg tga ttg gca ttt aac aga ttc gca aat agg tgc ctc agg att cca tac cat aga tcc agt aga tcc taa ttc aca ata cga ttt aga ttc acc gat caa atg ata tcc gct att aca aga gta cgt tat act aga gcc aaa gtc tac tcc gcc aat atc aag ttg gcc att atc gat atc tcg agg cga tgg gca tct ccg ttt aat aca ttg att aaa gag tgt cca tcc ggt acc ggt aca ttt agc ata tat ggg tcc cat ttt ttg ctt tct gta tcc agg tag aca tag ata ttc tat agt gtc tcc tat gtt gta att agc atc agt ctc tac act att ctt aaa ttt cat att aat ggg gcg tga cgg aat agt aca gta tga tag aac aca tcc tat tcc caa caa tgt cag gaa cgt cac gct ctc cac ctt cat att tat tta tcc gta aaa tgt tat cct gga cat cgt aca aat aat aaa aag ccc ata tat atg ttc gct att gta gaa att gtt ttt cac agt tgc tca aaa aca atg gca gtg act tat gag tta gtt aca ctt tgg agt ctc atc ttt agt aaa cat atc ata ata ttc gat att acg agt tga cat atc gaa caa att cca agt att tga ttt tgg ata ata ttc gta ttt tgc atc tgc tat aat taa gat ata atc acc aca aga aca cac gaa cgt ctt tcc tac atg gtt aaa gta cat gta caa ttc tat cca ttt gtc ttc ctt aac tat ata ttt gta tag ata att acg agt ctc atg agt aat tcc agt aat tgc ata gat gtc acc atc gta ttc tac agc ata aac tat act atg acg tct agg cat ggg aga ctt ttt tat cca acg att ttt agt gaa aca ttc cac atc gtt taa tac tac ata ttt ctc ata gtg gta taa act cca ccc att aca tat ata tca tcg ttt acg aat act gat gcg cct gaa tat cta gga gta att aag ttt gga agt ctt ttc cat ttc gaa gtg ccg tgt ttc aaa tat tct gct ata ccc gtt gaa ata gaa aat tct aat cct cct att aca tat aac ttt cca tcg tta aca caa gta cta act tct gat ttt aac gac gac ata tta gta acc gtt ttc ca- --- ttt ttt ttg ttt taa gat cta ccc gcg ata cgg aat aaa cat gtc tat tgt taa tca tgc cgc caa taa tgt ata gac aat tat gta aaa cat ttg cat cat aga att gtc tat ctg tat tac cga cta tcg tcc aat att ctg ttc tag gag agt aat ggg tta ttg tgg ata tat aat cag agt ttt taa tga cta cta tat tat gtt tta tac cat ttc gtg tca cag ctt tgt aga ttt gga tat agt taa tcc caa caa tgc tat agc att gca tat agc att agt cat aaa ctt ggg atg taa aat gtt gat gat atc tac atc gtt tgg att ttt atg tat cca ctt taa taa tat tat agc gta aca tcc tca tga ttt acg tta acg ttt tcg tgt gat aag ata gtg gtc agt tca tcc ttt gat aat ttt cca aat tct gga tcg gat gtc acc gca gta ata ttg ttg att att tct gac atc gac gca tta tat agt ttt tta att cca tat ctt ttt t

>MN702453.1_Monkeypox_virus_strain_A1_contig_SPADES

ttt tat atc act acg gac ata aac cat tgt ata att ttt atg ttt att agt gta cac att ttg gaa gta agt tcc ggc tgc cat gta ttt cct gga gag caa gta gat gat g-a gga acc aga tag ttt ata tcc ata ctt gca ctt aaa gtc tac att gta gtt gta tga gtg tat gat ctt tta agc cgc tag aag ttt tcc gtt tga tat agg atg tgg aca ttt aac aat ctg aca cgt ggg tgg att gga cca ttc tcc tcc tga aca cat gac acc aga gtt acc aat caa cga ata tcc act att gca act ata agt tac aat gct ccc atc gat ata aaa atc ctc gta tcc gtt atg tct tcc gtt gga tat aga tgg agg tga ttg gca ttt aac aga ttc gca aat agg tgc ctc agg att cca tac cat aga tcc agt aga tcc taa ttc aca ata cga ttt aga ttc acc gat caa atg ata tcc gct att aca aga gta cgt tat act aga gcc aaa gtc tac tcc gcc aat atc aag ttg gcc att atc gat atc tcg agg cga tgg gca tct ccg ttt aat aca ttg att aaa gag tgt cca tcc ggt acc ggt aca ttt agc ata tat ggg tcc cat ttt ttg ctt tct gta tcc agg tag aca tag ata ttc tat agt gtc tcc tat gtt gta att agc atc agt ctc tac act att ctt aaa ttt cat att aat ggg gcg tga cgg aat agt aca gta tga tag aac aca tcc tat tcc caa caa tgt cag gaa cgt cac gct ctc cac ctt cat att tat tta tcc gta aaa tgt tat cct gga cat cgt aca aat aat aaa aag ccc ata tat atg ttc gct att gta gaa att gtt ttt cac agt tgc tca aaa aca atg gca gtg act tat gag tta gtt aca ctt tgg agt ctc atc ttt agt aaa cat atc ata ata ttc gat att acg agt tga cat atc gaa caa att cca agt att tga ttt tgg ata ata ttc gta ttt tgc atc tgc tat aat taa gat ata atc acc aca aga aca cac gaa cgt ctt tcc tac atg gtt aaa gta cat gta caa ttc tat cca ttt gtc ttc ctt aac tat ata ttt gta tag ata att acg agt ctc atg agt aat tcc agt aat tgc ata gat gtc acc atc gta ttc tac agc ata aac tat act atg acg tct agg cat ggg aga ctt ttt tat cca acg att ttt agt gaa aca ttc cac atc gtt taa tac tac ata ttt ctc ata gtg gta taa act cca ccc att aca tat ata tca tcg ttt acg aat act gat gcg cct gaa tat cta gga gtg att aag ttt gga agt ctt ttc cat ttc gaa gtg ccg tgt ttc aaa tat tct gct ata ccc gtt gaa ata gaa aat tct aat cct cct att aca tat aac ttt cca tcg tta aca caa gta cta act tct gat ttt aac gac gac ata tta gta acc gtt ttc ca- --- ttt ttt ttg ttt taa gat cta ccc gcg ata cgg aat aaa cat gtc tat tgt taa tca tgc cgc caa taa tgt ata gac aat tat gta aaa cat ttg cat cat aga att gtc tat ctg tat tac cga cta tcg tcc aat att ctg ttc tag gag agt aat ggg tta ttg tgg ata tat aat cag agt ttt taa tga cta cta tat tat gtt tta tac cat ttc gtg tca cag ctt tgt aga ttt gga tat agt taa tcc caa caa tgc tat agc att gca tat agc att agt cat aaa ctt ggg atg taa aat gtt gat gat atc tac atc gtt tgg att ttt atg tat cca ctt taa taa tat tat agc gta aca tcc tca tga ttt acg tta acg ttt tcg tgt gat aag ata gtg gtc agt tca tcc ttt gat aat ttt cca aat tct gga tcg gat gtc acc gca gta ata ttg ttg att att tct gac atc gac gca tta tat agt ttt tta att cca tat ctt ttt t

>MN702452.1_Monkeypox_virus_strain_A2_contig_SPADES

ttt tat atc act acg gac ata aac cat tgt ata att ttt atg ttt att agt gta cac att ttg gaa gta agt tcc ggc tgc cat gta ttt cct gga gag caa gta gat gat g-a gga acc aga tag ttt ata tcc ata ctt gca ctt aaa gtc tac att gta gtt gta tga gtg tat gat ctt tta agc cgc tag aag ttt tcc gtt tga tat agg atg tgg aca ttt aac aat ctg aca cgt ggg tgg att gga cca ttc tcc tcc tga aca cat gac acc aga gtt acc aat caa cga ata tcc act att gca act ata agt tac aat gct ccc atc gat ata aaa atc ctc gta tcc gtt atg tct tcc gtt gga tat aga tgg agg tga ttg gca ttt aac aga ttc gca aat agg tgc ctc agg att cca tac cat aga tcc agt aga tcc taa ttc aca ata cga ttt aga ttc acc gat caa atg ata tcc gct att aca aga gta cgt tat act aga gcc aaa gtc tac tcc gcc aat atc aag ttg gcc att atc gat atc tcg agg cga tgg gca tct ccg ttt aat aca ttg att aaa gag tgt cca tcc ggt acc ggt aca ttt agc ata tat ggg tcc cat ttt ttg ctt tct gta tcc agg tag aca tag ata ttc tat agt gtc tcc tat gtt gta att agc atc agt ctc tac act att ctt aaa ttt cat att aat ggg gcg tga cgg aat agt aca gta tga tag aac aca tcc tat tcc caa caa tgt cag gaa cgt cac gct ctc cac ctt cat att tat tta tcc gta aaa tgt tat cct gga cat cgt aca aat aat aaa aag ccc ata tat atg ttc gct att gta gaa att gtt ttt cac agt tgc tca aaa aca atg gca gtg act tat gag tta gtt aca ctt tgg agt ctc atc ttt agt aaa cat atc ata ata ttc gat att acg agt tga cat atc gaa caa att cca agt att tga ttt tgg ata ata ttc gta ttt tgc atc tgc tat aat taa gat ata atc acc aca aga aca cac gaa cgt ctt tcc tac atg gtt aaa gta cat gta caa ttc tat cca ttt gtc ttc ctt aac tat ata ttt gta tag ata att acg agt ctc atg agt aat tcc agt aat tgc ata gat gtc acc atc gta ttc tac agc ata aac tat act atg acg tct agg cat ggg aga ctt ttt tat cca acg att ttt agt gaa aca ttc cac atc gtt taa tac tac ata ttt ctc ata gtg gta taa act cca ccc att aca tat ata tca tcg ttt acg aat act gat gcg cct gaa tat cta gga gtg att aag ttt gga agt ctt ttc cat ttc gaa gtg ccg tgt ttc aaa tat tct gct ata ccc gtt gaa ata gaa aat tct aat cct cct att aca tat aac ttt cca tcg tta aca caa gta cta act tct gat ttt aac gac gac ata tta gta acc gtt ttc ca- --- ttt ttt ttg ttt taa gat cta ccc gcg ata cgg aat aaa cat gtc tat tgt taa tca tgc cgc caa taa tgt ata gac aat tat gta aaa cat ttg cat cat aga att gtc tat ctg tat tac cga cta tcg tcc aat att ctg ttc tag gag agt aat ggg tta ttg tgg ata tat aat cag agt ttt taa tga cta cta tat tat gtt tta tac cat ttc gtg tca cag ctt tgt aga ttt gga tat agt taa tcc caa caa tgc tat agc att gca tat agc att agt cat aaa ctt ggg atg taa aat gtt gat gat atc tac atc gtt tgg att ttt atg tat cca ctt taa taa tat tat agc gta aca tcc tca tga ttt acg tta acg ttt tcg tgt gat aag ata gtg gtc agt tca tcc ttt gat aat ttt cca aat tct gga tcg gat gtc acc gca gta ata ttg ttg att att tct gac atc gac gca tta tat agt ttt tta att cca tat ctt ttt t

>MN702448.1_Monkeypox_virus_strain_015c_contig_SPADES

ttt tat atc act acg gac ata aac cat tgt ata att ttt atg ttt att agt gta cac att ttg gaa gta agt tcc ggc tgc cat gta ttt cct gga gag caa gta gat gat g-a gga acc aga tag ttt ata tcc ata ctt gca ctt aaa gtc tac att gta gtt gta tga gtg tat gat ctt tta agc cgc tag aag ttt tcc gtt tga tat agg atg tgg aca ttt aac aat ctg aca cgt ggg tgg att gga cca ttc tcc tcc tga aca cat gac acc aga gtt acc aat caa cga ata tcc act att gca act ata agt tac aat gct ccc atc gat ata aaa atc ctc gta tcc gtt atg tct tcc gtt gga tat aga tgg agg tga ttg gca ttt aac aga ttc gca aat agg tgc ctc agg att cca tac cat aga tcc agt aga tcc taa ttc aca ata cga ttt aga ttc acc gat caa atg ata tcc gct att aca aga gta cgt tat act aga gcc aaa gtc tac tcc gcc aat atc aag ttg gcc att atc gat atc tcg agg cga tgg gca tct ccg ttt aat aca ttg att aaa gag tgt cca tcc ggt acc ggt aca ttt agc ata tat ggg tcc cat ttt ttg ctt tct gta tcc agg tag aca tag ata ttc tat agt gtc tcc tat gtt gta att agc atc agt ctc tac act att ctt aaa ttt cat att aat ggg gcg tga cgg aat agt aca gta tga tag aac aca tcc tat tcc caa caa tgt cag gaa cgt cac gct ctc cac ctt cat att tat tta tcc gta aaa tgt tat cct gga cat cgt aca aat aat aaa aag ccc ata tat atg ttc gct att gta gaa att gtt ttt cac agt tgc tca aaa aca atg gca gtg act tat gag tta gtt aca ctt tgg agt ctc atc ttt agt aaa cat atc ata ata ttc gat att acg agt tga cat atc gaa caa att cca agt att tga ttt tgg ata ata ttc gta ttt tgc atc tgc tat aat taa gat ata atc acc aca aga aca cac gaa cgt ctt tcc tac atg gtt aaa gta cat gta caa ttc tat cca ttt gtc ttc ctt aac tat ata ttt gta tag ata att acg agt ctc atg agt aat tcc agt aat tgc ata gat gtc acc atc gta ttc tac agc ata aac tat act atg acg tct agg cat ggg aga ctt ttt tat cca acg att ttt agt gaa aca ttc cac atc gtt taa tac tac ata ttt ctc ata gtg gta taa act cca ccc att aca tat ata tca tcg ttt acg aat act gat gcg cct gaa tat cta gga gtg att aag ttt gga agt ctt ttc cat ttc gaa gtg ccg tgt ttc aaa tat tct gct ata ccc gtt gaa ata gaa aat tct aat cct cct att aca tat aac ttt cca tcg tta aca caa gta cta act tct gat ttt aac gac gac ata tta gta acc gtt ttc ca- --- ttt ttt ttg ttt taa gat cta ccc gcg ata cgg aat aaa cat gtc tat tgt taa tca tgc cgc caa taa tgt ata gac aat tat gta aaa cat ttg cat cat aga att gtc tat ctg tat tac cga cta tcg tcc aat att ctg ttc tag gag agt aat ggg tta ttg tgg ata tat aat cag agt ttt taa tga cta cta tat tat gtt tta tac cat ttc gtg tca cag ctt tgt aga ttt gga tat agt taa tcc caa caa tgc tat agc att gca tat agc att agt cat aaa ctt ggg atg taa aat gtt gat gat atc tac atc gtt tgg att ttt atg tat cca ctt taa taa tat tat agc gta aca tcc tca tga ttt acg tta acg ttt tcg tgt gat aag ata gtg gtc agt tca tcc ttt gat aat ttt cca aat tct gga tcg gat gtc acc gca gta ata ttg ttg att att tct gac atc gac gca tta tat agt ttt tta att cca tat ctt ttt t

>MN702447.1_Monkeypox_virus_strain_18_contig_SPADES

ttt tat atc act acg gac ata aac cat tgt ata att ttt atg ttt att agt gta cac att ttg gaa gta agt tcc ggc tgc cat gta ttt cct gga gag caa gta gat gat g-a gga acc aga tag ttt ata tcc ata ctt gca ctt aaa gtc tac att gta gtt gta tga gtg tat gat ctt tta agc cgc tag aag ttt tcc gtt tga tat agg atg tgg aca ttt aac aat ctg aca cgt ggg tgg att gga cca ttc tcc tcc tga aca cat gac acc aga gtt acc aat caa cga ata tcc act att gca act ata agt tac aat gct ccc atc gat ata aaa atc ctc gta tcc gtt atg tct tcc gtt gga tat aga tgg agg tga ttg gca ttt aac aga ttc gca aat agg tgc ctc agg att cca tac cat aga tcc agt aga tcc taa ttc aca ata cga ttt aga ttc acc gat caa atg ata tcc gct att aca aga gta cgt tat act aga gcc aaa gtc tac tcc gcc aat atc aag ttg gcc att atc gat atc tcg agg cga tgg gca tct ccg ttt aat aca ttg att aaa gag tgt cca tcc ggt acc ggt aca ttt agc ata tat ggg tcc cat ttt ttg ctt tct gta tcc agg tag aca tag ata ttc tat agt gtc tcc tat gtt gta att agc atc agt ctc tac act att ctt aaa ttt cat att aat ggg gcg tga cgg aat agt aca gta tga tag aac aca tcc tat tcc caa caa tgt cag gaa cgt cac gct ctc cac ctt cat att tat tta tcc gta aaa tgt tat cct gga cat cgt aca aat aat aaa aag ccc ata tat atg ttc gct att gta gaa att gtt ttt cac agt tgc tca aaa aca atg gca gtg act tat gag tta gtt aca ctt tgg agt ctc atc ttt agt aaa cat atc ata ata ttc gat att acg agt tga cat atc gaa caa att cca agt att tga ttt tgg ata ata ttc gta ttt tgc atc tgc tat aat taa gat ata atc acc aca aga aca cac gaa cgt ctt tcc tac atg gtt aaa gta cat gta caa ttc tat cca ttt gtc ttc ctt aac tat ata ttt gta tag ata att acg agt ctc atg agt aat tcc agt aat tgc ata gat gtc acc atc gta ttc tac agc ata aac tat act atg acg tct agg cat ggg aga ctt ttt tat cca acg att ttt agt gaa aca ttc cac atc gtt taa tac tac ata ttt ctc ata gtg gta taa act cca ccc att aca tat ata tca tcg ttt acg aat act gat gcg cct gaa tat cta gga gtg att aag ttt gga agt ctt ttc cat ttc gaa gtg ccg tgt ttc aaa tat tct gct ata ccc gtt gaa ata gaa aat tct aat cct cct att aca tat aac ttt cca tcg tta aca caa gta cta act tct gat ttt aac gac gac ata tta gta acc gtt ttc ca- --- ttt ttt ttg ttt taa gat cta ccc gcg ata cgg aat aaa cat gtc tat tgt taa tca tgc cgc caa taa tgt ata gac aat tat gta aaa cat ttg cat cat aga att gtc tat ctg tat tac cga cta tcg tcc aat att ctg ttc tag gag agt aat ggg tta ttg tgg ata tat aat cag agt ttt taa tga cta cta tat tat gtt tta tac cat ttc gtg tca cag ctt tgt aga ttt gga tat agt taa tcc caa caa tgc tat agc att gca tat agc att agt cat aaa ctt ggg atg taa aat gtt gat gat atc tac atc gtt tgg att ttt atg tat cca ctt taa taa tat tat agc gta aca tcc tca tga ttt acg tta acg ttt tcg tgt gat aag ata gtg gtc agt tca tcc ttt gat aat ttt cca aat tct gga tcg gat gtc acc gca gta ata ttg ttg att att tct gac atc gac gca tta tat agt ttt tta att cca tat ctt ttt t

>JX878424.1_Monkeypox_virus_isolate_DRC_07-0338_complete_genome

ttt tat atc act acg gac ata aac cat tgt ata att ttt atg ttt att agt gta cac att ttg gaa gta agt tcc ggc tgc cat gta ttt cct gga gag caa gta gat gat g-a gga acc aga tag ttt ata tcc ata ctt gca ctt aaa gtc tac att gta gtt gta tga gtg tat gat ctt tta agc cgc tag aag ttt tcc gtt tga tat agg atg tgg aca ttt aac aat ctg aca cgt ggg tgg att gga cca ttc tcc tcc tga aca cat gac acc aga gtt acc aat caa cga ata tcc act att gca act ata agt tac aat gct ccc atc gat ata aaa atc ctc gta tcc gtt atg tct tcc gtt gga tat aga tgg agg tga ttg gca ttt aac aga ttc gca aat agg tgc ctc agg att cca tac cat aga tcc agt aga tcc taa ttc aca ata cga ttt aga ttc acc gat caa atg ata tcc gct att aca aga gta cgt tat act aga gcc aaa gtc tac tcc gcc aat atc aag ttg gcc att atc gat atc tcg agg cga tgg gca tct ccg ttt aat aca ttg att aaa gag tgt cca tcc ggt acc ggt aca ttt agc ata tat ggg tcc cat ttt ttg ctt tct gta tcc agg tag aca tag ata ttc tat agt gtc tcc tat gtt gta att agc atc agt ctc tac act att ctt aaa ttt cat att aat ggg gcg tga cgg aat agt aca gta tga tag aac aca tcc tat tcc caa caa tgt cag gaa cgt cac gct ctc cac ctt cat att tat tta tcc gta aaa tgt tat cct gga cat cgt aca aat aat aaa aag ccc ata tat atg ttc gct att gta gaa att gtt ttt cac agt tgc tca aaa aca atg gca gtg act tat gag tta gtt aca ctt tgg agt ctc atc ttt agt aaa cat atc ata ata ttc gat att acg agt tga cat atc gaa caa att cca agt att tga ttt tgg ata ata ttc gta ttt tgc atc tgc tat aat taa gat ata atc acc aca aga aca cac gaa cgt ctt tcc tac atg gtt aaa gta cat gta caa ttc tat cca ttt gtc ttc ctt aac tat ata ttt gta tag ata att acg agt ctc atg agt aat tcc agt aat tgc ata gat gtc acc atc gta ttc tac agc ata aac tat act atg acg tct agg cat ggg aga ctt ttt tat cca acg att ttt agt gaa aca ttc cac atc gtt taa tac tac ata ttt ctc ata gtg gta taa act cca ccc att aca tat ata tca tcg ttt acg aat act gat gcg cct gaa tat cta gga gtg att aag ttt gga agt ctt ttc cat ttc gaa gtg ccg tgt ttc aaa tat tct gct ata ccc gtt gaa ata gaa aat tct aat cct cct att aca tat aac ttt cca tcg tta aca caa gta cta act tct gat ttt aac gac gac ata tta gta acc gtt ttc ca- --- ttt ttt ttg ttt taa gat cta ccc gcg ata cgg aat aaa cat gtc tat tgt taa tca tgc cgc caa taa tgt ata gac aat tat gta aaa cat ttg cat cat aga att gtc tat ctg tat tac cga cta tcg tcc aat att ctg ttc tag gag agt aat ggg tta ttg tgg ata tat aat cag agt ttt taa tga cta cta tat tat gtt tta tac cat ttc gtg tca cag ctt tgt aga ttt gga tat agt taa tcc caa caa tgc tat agc att gca tat agc att agt cat aaa ctt ggg atg taa aat gtt gat gat atc tac atc gtt tgg att ttt atg tat cca ctt taa taa tat tat agc gta aca tcc tca tga ttt acg tta acg ttt tcg tgt gat aag ata gtg gtc agt tca tcc ttt gat aat ttt cca aat tct gga tcg gat gtc acc gca gta ata ttg ttg att att tct gac atc gac gca tta tat agt ttt tta att cca tat ctt ttt t

>KC257460.1_Monkeypox_virus_strain_DRC_Yandongi_1985_complete_genome

ttt tat atc act acg gac ata aac cat tgt ata att ttt atg ttt att agt gta cac att ttg gaa gta agt tcc ggc tgc cat gta ttt cct gga gag caa gta gat gat g-a gga acc aga tag ttt ata tcc ata ctt gca ctt aaa gtc tac att gta gtt gta tga gtg tat gat ctt tta agc cgc tag aag ttt tcc gtt tga tat agg atg tgg aca ttt aac aat ctg aca cgt ggg tgg att gga cca ttc tcc tcc tga aca cat gac acc aga gtt acc aat caa cga ata tcc act att gca act ata agt tac aat gct ccc atc gat ata aaa atc ctc gta tcc gtt atg tct tcc gtt gga tat aga tgg agg tga ttg gca ttt aac aga ttc gca aat agg tgc ctc agg att cca tac cat aga tcc agt aga tcc taa ttc aca ata cga ttt aga ttc acc gat caa atg ata tcc gct att aca aga gta cgt tat act aga gcc aaa gtc tac tcc gcc aat atc aag ttg gcc att atc gat atc tcg agg cga tgg gca tct ccg ttt aat aca ttg att aaa gag tgt cca tcc ggt acc ggt aca ttt agc ata tat ggg tcc cat ttt ttg ctt tct gta tcc agg tag aca tag ata ttc tat agt gtc tcc tat gtt gta att agc atc agt ctc tac act att ctt aaa ttt cat att aat ggg gcg tga cgg aat agt aca gta tga tag aac aca tcc tat tcc caa caa tgt cag gaa cgt cac gct ctc cac ctt cat att tat tta tcc gta aaa tgt tat cct gga cat cgt aca aat aat aaa aag ccc ata tat atg ttc gct att gta gaa att gtt ttt cac agt tgc tca aaa aca atg gca gtg act tat gag tta gtt aca ctt tgg agt ctc atc ttt agt aaa cat atc ata ata ttc gat att acg agt tga cat atc gaa caa att cca agt att tga ttt tgg ata ata ttc gta ttt tgc atc tgc tat aat taa gat ata atc acc aca aga aca cac gaa cgt ctt tcc tac atg gtt aaa gta cat gta caa ttc tat cca ttt gtc ttc ctt aac tat ata ttt gta tag ata att acg agt ctc atg agt aat tcc agt aat tgc ata gat gtc acc atc gta ttc tac agc ata aac tat act atg acg tct agg cat ggg aga ctt ttt tat cca acg att ttt agt gaa aca ttc cac atc gtt taa tac tac ata ttt ctc ata gtg gta taa act cca ccc att aca tat ata tca tcg ttt acg aat act gat gcg cct gaa tat cta gga gtg att aag ttt gga agt ctt ttc cat ttc gaa gtg ccg tgt ttc aaa tat tct gct ata ccc gtt gaa ata gaa aat tct aat cct cct att aca tat aac ttt cca tcg tta aca caa gta cta act tct gat ttt aac gac gac ata tta gta acc gtt ttc ca- ttt ttt ttt ttg ttt taa gat cta ccc gcg ata cgg aat aaa cat gtc tat tgt taa tca tgc cgc caa taa tgt ata gac aat tat gta aaa cat ttg cat cat aga att gtc tat ctg tat tac cga cta tcg tcc aat att ctg ttc tag gag agt aat ggg tta ttg tga ata tat aat cag agt ttt taa tga cta cta tat tat gtt tta tac cat ttc gtg tca cag ctt tgt aga ttt gga tat agt taa tcc caa caa tgc tat agc att gca tat agc att agt cat aaa ctt ggg atg taa aat gtt gat gat atc tac atc gtt tgg att ttt atg tat cca ctt taa taa tat tat agc gta aca tcc tca tga ttt acg tta acg ttt tcg tgt gat aag ata gtg gtc agt tca tcc ttt gat aat ttt cca aat tct gga tcg gat gtc acc gca gta ata ttg ttg att att tct gac atc gac gca tta tat agt ttt tta att cca tat ctt ttt t

>JX878429.1_Monkeypox_virus_isolate_DRC_07-0662_complete_genome

ttt tat atc act acg gac ata aac cat tgt ata att ttt atg ttt att agt gta cac att ttg gaa gta agt tcc ggc tgc cat gta ttt cct gga gag caa gta gat gat g-a gga acc aga tag ttt ata tcc ata ctt gca ctt aaa gtc tac att gta gtt gta tga gtg tat gat ctt tta agc cgc tag aag ttt tcc gtt tga tat agg atg tgg aca ttt aac aat ctg aca cgt ggg tgg att gga cca ttc tcc tcc tga aca cat gac acc aga gtt acc aat caa cga ata tcc act att gca act ata agt tac aat gct ccc atc gat ata aaa atc ctc gta tcc gtt atg tct tcc gtt gga tat aga tgg agg tga ttg gca ttt aac aga ttc gca aat agg tgc ctc agg att cca tac cat aga tcc agt aga tcc taa ttc aca ata cga ttt aga ttc acc gat caa atg ata tcc gct att aca aga gta cgt tat act aga gcc aaa gtc tac tcc gcc aat atc aag ttg gcc att atc gat atc tcg agg cga tgg gca tct ccg ttt aat aca ttg att aaa gag tgt cca tcc ggt acc ggt aca ttt agc ata tat ggg tcc cat ttt ttg ctt tct gta tcc agg tag aca tag ata ttc tat agt gtc tcc tat gtt gta att agc atc agt ctc tac act att ctt aaa ttt cat att aat ggg gcg tga cgg aat agt aca gta tga tag aac aca tcc tat tcc caa caa tgt cag gaa cgt cac gct ctc cac ctt cat att tat tta tcc gta aaa tgt tat cct gga cat cgt aca aat aat aaa aag ccc ata tat atg ttc gct att gta gaa att gtt ttt cac agt tgc tca aaa aca atg gca gtg act tat gag tta gtt aca ctt tgg agt ctc atc ttt agt aaa cat atc ata ata ttc gat att acg agt tga cat atc gaa caa att cca agt att tga ttt tgg ata ata ttc gta ttt tgc atc tgc tat aat taa gat ata atc acc aca aga aca cac gaa cgt ctt tcc tac atg gtt aaa gta cat gta caa ttc tat cca ttt gtc ttc ctt aac tat ata ttt gta tag ata att acg agt ctc atg agt aat tcc agt aat tgc ata gat gtc acc atc gta ttc tac agc ata aac tat act atg acg tct agg cat ggg aga ctt ttt tat cca acg att ttt agt gaa aca ttc cac atc gtt taa tac tac ata ttt ctc ata gtg gta taa act cca ccc att aca tat ata tca tcg ttt acg aat act gat gcg cct gaa tat cta gga gtg att aag ttt gga agt ctt ttc cat ttc gaa gtg ccg tgt ttc aaa tat tct gct ata ccc gtt gaa ata gaa aat tct aat cct cct att aca tat aac ttt cca tcg tta aca caa gta cta act tct gat ttt aac gac gac ata tta gta acc gtt ttc ca- --- ttt ttt ttg ttt taa gat cta ccc gcg ata cgg aat aaa cat gtc tat tgt taa tca tgc cgc caa taa tgt ata gac aat tat gta aaa cat ttg cat cat aga att gtc tat ctg tat tac cga cta tcg tcc aat att ctg ttc tag gag agt aat ggg tta ttg tgg ata tat aat cag agt ttt taa tga cta cta tat tat gtt tta tac cat ttc gtg tca cag ctt tgt aga ttt gga tat agt taa tcc caa caa tgc tat agc att gca tat agc att agt cat aaa ctt ggg atg taa aat gtt gat gat atc tac atc gtt tgg att ttt atg tat cca ctt taa taa tat tat agc gta aca tcc tca tga ttt acg tta acg ttt tcg tgt gat aag ata gtg gtc agt tca tcc ttt gat aat ttt cca aat tct gga tcg gat gtc acc gca gta ata ttg ttg att att tct gac atc gac gca tta tat agt ttt tta att cca tat ctt ttt t

>JX878425.1_Monkeypox_virus_isolate_DRC_07-0354_complete_genome

ttt tat atc act acg gac ata aac cat tgt ata att ttt atg ttt att agt gta cac att ttg gaa gta agt tcc ggc tgc cat gta ttt cct gga gag caa gta gat gat g-a gga acc aga tag ttt ata tcc ata ctt gca ctt aaa gtc tac att gta gtt gta tga gtg tat gat ctt tta agc cgc tag aag ttt tcc gtt tga tat agg atg tgg aca ttt aac aat ctg aca cgt ggg tgg att gga cca ttc tcc tcc tga aca cat gac acc aga gtt acc aat caa cga ata tcc act att gca act ata agt tac aat gct ccc atc gat ata aaa atc ctc gta tcc gtt atg tct tcc gtt gga tat aga tgg agg tga ttg gca ttt aac aga ttc gca aat agg tgc ctc agg att cca tac cat aga tcc agt aga tcc taa ttc aca ata cga ttt aga ttc acc gat caa atg ata tcc gct att aca aga gta cgt tat act aga gcc aaa gtc tac tcc gcc aat atc aag ttg gcc att atc gat atc tcg agg cga tgg gca tct ccg ttt aat aca ttg att aaa gag tgt cca tcc ggt acc ggt aca ttt agc ata tat ggg tcc cat ttt ttg ctt tct gta tcc agg tag aca tag ata ttc tat agt gtc tcc tat gtt gta att agc atc agt ctc tac act att ctt aaa ttt cat att aat ggg gcg tga cgg aat agt aca gta tga tag aac aca tcc tat tcc caa caa tgt cag gaa cgt cac gct ctc cac ctt cat att tat tta tcc gta aaa tgt tat cct gga cat cgt aca aat aat aaa aag ccc ata tat atg ttc gct att gta gaa att gtt ttt cac agt tgc tca aaa aca atg gca gtg act tat gag tta gtt aca ctt tgg agt ctc atc ttt agt aaa cat atc ata ata ttc gat att acg agt tga cat atc gaa caa att cca agt att tga ttt tgg ata ata ttc gta ttt tgc atc tgc tat aat taa gat ata atc acc aca aga aca cac gaa cgt ctt tcc tac atg gtt aaa gta cat gta caa ttc tat cca ttt gtc ttc ctt aac tat ata ttt gta tag ata att acg agt ctc atg agt aat tcc agt aat tgc ata gat gtc acc atc gta ttc tac agc ata aac tat act atg acg tct agg cat ggg aga ctt ttt tat cca acg att ttt agt gaa aca ttc cac atc gtt taa tac tac ata ttt ctc ata gtg gta taa act cca ccc att aca tat ata tca tcg ttt acg aat act gat gcg cct gaa tat cta gga gtg att aag ttt gga agt ctt ttc cat ttc gaa gtg ccg tgt ttc aaa tat tct gct ata ccc gtt gaa ata gaa aat tct aat cct cct att aca tat aac ttt cca tcg tta aca caa gta cta act tct gat ttt aac gac gac ata tta gta acc gtt ttc ca- --- ttt ttt ttg ttt taa gat cta ccc gcg ata cgg aat aaa cat gtc tat tgt taa tca tgc cgc caa taa tgt ata gac aat tat gta aaa cat ttg cat cat aga att gtc tat ctg tat tac cga cta tcg tcc aat att ctg ttc tag gag agt aat ggg tta ttg tgg ata tat aat cag agt ttt taa tga cta cta tat tat gtt tta tac cat ttc gtg tca cag ctt tgt aga ttt gga tat agt taa tcc caa caa tgc tat agc att gca tat agc att agt cat aaa ctt ggg atg taa aat gtt gat gat atc tac atc gtt tgg att ttt atg tat cca ctt taa taa tat tat agc gta aca tcc tca tga ttt acg tta acg ttt tcg tgt gat aag ata gtg gtc agt tca tcc ttt gat aat ttt cca aat tct gga tcg gat gtc acc gca gta ata ttg ttg att att tct gac atc gac gca tta tat agt ttt tta att cca tat ctt ttt t

>JX878407.1_Monkeypox_virus_isolate_DRC_06-0950_complete_genome

ttt tat atc act acg gac ata aac cat tgt ata att ttt atg ttt att agt gta cac att ttg gaa gta agt tcc ggc tgc cat gta ttt cct gga gag caa gta gat gat g-a gga acc aga tag ttt ata tcc ata ctt gca ctt aaa gtc tac att gta gtt gta tga gtg tat gat ctt tta agc cgc tag aag ttt tcc gtt tga tat agg atg tgg aca ttt aac aat ctg aca cgt ggg tgg att gga cca ttc tcc tcc tga aca cat gac acc aga gtt acc aat caa cga ata tcc act att gca act ata agt tac aat gct ccc atc gat ata aaa atc ctc gta tcc gtt atg tct tcc gtt gga tat aga tgg agg tga ttg gca ttt aac aga ttc gca aat agg tgc ctc agg att cca tac cat aga tcc agt aga tcc taa ttc aca ata cga ttt aga ttc acc gat caa atg ata tcc gct att aca aga gta cgt tat act aga gcc aaa gtc tac tcc gcc aat atc aag ttg gcc att atc gat atc tcg agg cga tgg gca tct ccg ttt aat aca ttg att aaa gag tgt cca tcc ggt acc ggt aca ttt agc ata tat ggg tcc cat ttt ttg ctt tct gta tcc agg tag aca tag ata ttc tat agt gtc tcc tat gtt gta att agc atc agt ctc tac act att ctt aaa ttt cat att aat ggg gcg tga cgg aat agt aca gta tga tag aac aca tcc tat tcc caa caa tgt cag gaa cgt cac gct ctc cac ctt cat att tat tta tcc gta aaa tgt tat cct gga cat cgt aca aat aat aaa aag ccc ata tat atg ttc gct att gta gaa att gtt ttt cac agt tgc tca aaa aca atg gca gtg act tat gag tta gtt aca ctt tgg agt ctc atc ttt agt aaa cat atc ata ata ttc gat att acg agt tga cat atc gaa caa att cca agt att tga ttt tgg ata ata ttc gta ttt tgc atc tgc tat aat taa gat ata atc acc aca aga aca cac gaa cgt ctt tcc tac atg gtt aaa gta cat gta caa ttc tat cca ttt gtc ttc ctt aac tat ata ttt gta tag ata att acg agt ctc atg agt aat tcc agt aat tgc ata gat gtc acc atc gta ttc tac agc ata aac tat act atg acg tct agg cat ggg aga ctt ttt tat cca acg att ttt agt gaa aca ttc cac atc gtt taa tac tac ata ttt ctc ata gtg gta taa act cca ccc att aca tat ata tca tcg ttt acg aat act gat gcg cct gaa tat cta gga gtg att aag ttt gga agt ctt ttc cat ttc gaa gtg ccg tgt ttc aaa tat tct gct ata ccc gtt gaa ata gaa aat tct aat cct cct att aca tat aac ttt cca tcg tta aca caa gta cta act tct gat ttt aac gac gac ata tta gta acc gtt ttc ca- --- ttt ttt ttg ttt taa gat cta ccc gcg ata cgg aat aaa cat gtc tat tgt taa tca tgc cgc caa taa tgt ata gac aat tat gta aaa cat ttg cat cat aga att gtc tat ctg tat tac cga cta tcg tcc aat att ctg ttc tag gag agt aat ggg tta ttg tgg ata tat aat cag agt ttt taa tga cta cta tat tat gtt tta tac cat ttc gtg tca cag ctt tgt aga ttt gga tat agt taa tcc caa caa tgc tat agc att gca tat agc att agt cat aaa ctt ggg atg taa aat gtt gat gat atc tac atc gtt tgg att ttt atg tat cca ctt taa taa tat tat agc gta aca tcc tca tga ttt acg tta acg ttt tcg tgt gat aag ata gtg gtc agt tca tcc ttt gat aat ttt cca aat tct gga tcg gat gtc acc gca gta ata ttg ttg att att tct gac atc gac gca tta tat agt ttt tta att cca tat ctt ttt t

>MN702451.1_Monkeypox_virus_strain_A6_contig_SPADES

ttt tat atc act acg gac ata aac cat tgt ata att ttt atg ttt att agt gta cac att ttg gaa gta agt tcc ggc tgc cat gta ttt cct gga gag caa gta gat gat g-a gga acc aga tag ttt ata tcc ata ctt gca ctt aaa gtc tac att gta gtt gta tga gtg tat gat ctt tta agc cgc tag aag ttt tcc gtt tga tat agg atg tgg aca ttt aac aat ctg aca cgt ggg tgg att gga cca ttc tcc tcc tga aca cat gac acc aga gtt acc aat caa cga ata tcc act att gca act ata agt tac aat gct ccc atc gat ata aaa atc ctc gta tcc gtt atg tct tcc gtt gga tat aga tgg agg tga ttg gca ttt aac aga ttc gca aat agg tgc ctc agg att cca tac cat aga tcc agt aga tcc taa ttc aca ata cga ttt aga ttc acc gat caa atg ata tcc gct att aca aga gta cgt tat act aga gcc aaa gtc tac tcc gcc aat atc aag ttg gcc att atc gat atc tcg agg cga tgg gca tct ccg ttt aat aca ttg att aaa gag tgt cca tcc ggt acc ggt aca ttt agc ata tat ggg tcc cat ttt ttg ctt tct gta tcc agg tag aca tag ata ttc tat agt gtc tcc tat gtt gta att agc atc agt ctc tac act att ctt aaa ttt cat att aat ggg gcg tga cgg aat agt aca gta tga tag aac aca tcc tat tcc caa caa tgt cag gaa cgt cac gct ctc cac ctt cat att tat tta tcc gta aaa tgt tat cct gga cat cgt aca aat aat aaa aag ccc ata tat atg ttc gct att gta gaa att gtt ttt cac agt tgc tca aaa aca atg gca gtg act tat gag tta gtt aca ctt tgg agt ctc atc ttt agt aaa cat atc ata ata ttc gat att acg agt tga cat atc gaa caa att cca agt att tga ttt tgg ata ata ttc gta ttt tgc atc tgc tat aat taa gat ata atc acc aca aga aca cac gaa cgt ctt tcc tac atg gtt aaa gta cat gta caa ttc tat cca ttt gtc ttc ctt aac tat ata ttt gta tag ata att acg agt ctc atg agt aat tcc agt aat tgc ata gat gtc acc atc gta ttc tac agc ata aac tat act atg acg tct agg cat ggg aga ctt ttt tat cca acg att ttt agt gaa aca ttc cac atc gtt taa tac tac ata ttt ctc ata gtg gta taa act cca ccc att aca tat ata tca tcg ttt acg aat act gat gcg cct gaa tat cta gga gtg att aag ttt gga agt ctt ttc cat ttc gaa gtg ccg tgt ttc aaa tat tct gct ata ccc gtt gaa ata gaa aat tct aat cct cct att aca tat aac ttt cca tcg tta aca caa gta cta act tct gat ttt aac gac gac ata tta gta acc gtt ttc ca- --- ttt ttt ttg ttt taa gat cta ccc gcg ata cgg aat aaa cat gtc tat tgt taa tca tgc cgc caa taa tgt ata gac aat tat gta aaa cat ttg cat cat aga att gtc tat ctg tat tac cga cta tcg tcc aat att ctg ttc tag gag agt aat ggg tta ttg tgg ata tat aat cag agt ttt taa tga cta cta tat tat gtt tta tac cat ttc gtg tca cag ctt tgt aga ttt gga tat agt taa tcc caa caa tgc tat agc att gca tat agc att agt cat aaa ctt ggg atg taa aat gtt gat gat atc tac atc gtt tgg att ttt atg tat cca ctt taa taa tat tat agc gta aca tcc tca tga ttt acg tta acg ttt tcg tgt gat aag ata gtg gtc agt tca tcc ttt gat aat ttt cca aat tct gga tcg gat gtc acc gca gta ata ttg ttg att att tct gac atc gac gca tta tat agt ttt tta att cca tat ctt ttt t

>JX878423.1_Monkeypox_virus_isolate_DRC_07-0337_complete_genome

ttt tat atc act acg gac ata aac cat tgt ata att ttt atg ttt att agt gta cac att ttg gaa gta agt tcc ggc tgc cat gta ttt cct gga gag caa gta gat gat g-a gga acc aga tag ttt ata tcc ata ctt gca ctt aaa gtc tac att gta gtt gta tga gtg tat gat ctt tta agc cgc tag aag ttt tcc gtt tga tat agg atg tgg aca ttt aac aat ctg aca cgt ggg tgg att gga cca ttc tcc tcc tga aca cat gac acc aga gtt acc aat caa cga ata tcc act att gca act ata agt tac aat gct ccc atc gat ata aaa atc ctc gta tcc gtt atg tct tcc gtt gga tat aga tgg agg tga ttg gca ttt aac aga ttc gca aat agg tgc ctc agg att cca tac cat aga tcc agt aga tcc taa ttc aca ata cga ttt aga ttc acc gat caa atg ata tcc gct att aca aga gta cgt tat act aga gcc aaa gtc tac tcc gcc aat atc aag ttg gcc att atc gat atc tcg agg cga tgg gca tct ccg ttt aat aca ttg att aaa gag tgt cca tcc ggt acc ggt aca ttt agc ata tat ggg tcc cat ttt ttg ctt tct gta tcc agg tag aca tag ata ttc tat agt gtc tcc tat gtt gta att agc atc agt ctc tac act att ctt aaa ttt cat att aat ggg gcg tga cgg aat agt aca gta tga tag aac aca tcc tat tcc caa caa tgt cag gaa cgt cac gct ctc cac ctt cat att tat tta tcc gta aaa tgt tat cct gga cat cgt aca aat aat aaa aag ccc ata tat atg ttc gct att gta gaa att gtt ttt cac agt tgc tca aaa aca atg gca gtg act tat gag tta gtt aca ctt tgg agt ctc atc ttt agt aaa cat atc ata ata ttc gat att acg agt tga cat atc gaa caa att cca agt att tga ttt tgg ata ata ttc gta ttt tgc atc tgc tat aat taa gat ata atc acc aca aga aca cac gaa cgt ctt tcc tac atg gtt aaa gta cat gta caa ttc tat cca ttt gtc ttc ctt aac tat ata ttt gta tag ata att acg agt ctc atg agt aat tcc agt aat tgc ata gat gtc acc atc gta ttc tac agc ata aac tat act atg acg tct agg cat ggg aga ctt ttt tat cca acg att ttt agt gaa aca ttc cac atc gtt taa tac tac ata ttt ctc ata gtg gta taa act cca ccc att aca tat ata tca tcg ttt acg aat act gat gcg cct gaa tat cta gga gtg att aag ttt gga agt ctt ttc cat ttc gaa gtg ccg tgt ttc aaa tat tct gct ata ccc gtt gaa ata gaa aat tct aat cct cct att aca tat aac ttt cca tcg tta aca caa gta cta act tct gat ttt aac gac gac ata tta gta acc gtt ttc ca- --- ttt ttt ttg ttt taa gat cta ccc gcg ata cgg aat aaa cat gtc tat tgt taa tca tgc cgc caa taa tgt ata gac aat tat gta aaa cat ttg cat cat aga att gtc tat ctg tat tac cga cta tcg tcc aat att ctg ttc tag gag agt aat ggg tta ttg tgg ata tat aat cag agt ttt taa tga cta cta tat tat gtt tta tac cat ttc gtg tca cag ctt tgt aga ttt gga tat agt taa tcc caa caa tgc tat agc att gca tat agc att agt cat aaa ctt ggg atg taa aat gtt gat gat atc tac atc gtt tgg att ttt atg tat cca ctt taa taa tat tat agc gta aca tcc tca tga ttt acg tta acg ttt tcg tgt gat aag ata gtg gtc agt tca tcc ttt gat aat ttt cca aat tct gga tcg gat gtc acc gca gta ata ttg ttg att att tct gac atc gac gca tta tat agt ttt tta att cca tat ctt ttt t

>MN702450.1_Monkeypox_virus_strain_B1_contig_SPADES

ttt tat atc act acg gac ata aac cat tgt ata att ttt atg ttt att agt gta cac att ttg gaa gta agt tcc ggc tgc cat gta ttt cct gga gag caa gta gat gat g-a gga acc aga tag ttt ata tcc ata ctt gca ctt aaa gtc tac att gta gtt gta tga gtg tat gat ctt tta agc cgc tag aag ttt tcc gtt tga tat agg atg tgg aca ttt aac aat ctg aca cgt ggg tgg att gga cca ttc tcc tcc tga aca cat gac acc aga gtt acc aat caa cga ata tcc act att gca act ata agt tac aat gct ccc atc gat ata aaa atc ctc gta tcc gtt atg tct tcc gtt gga tat aga tgg agg tga ttg gca ttt aac aga ttc gca aat agg tgc ctc agg att cca tac cat aga tcc agt aga tcc taa ttc aca ata cga ttt aga ttc acc gat caa atg ata tcc gct att aca aga gta cgt tat act aga gcc aaa gtc tac tcc gcc aat atc aag ttg gcc att atc gat atc tcg agg cga tgg gca tct ccg ttt aat aca ttg att aaa gag tgt cca tcc ggt acc ggt aca ttt agc ata tat ggg tcc cat ttt ttg ctt tct gta tcc agg tag aca tag ata ttc tat agt gtc tcc tat gtt gta att agc atc agt ctc tac act att ctt aaa ttt cat att aat ggg gcg tga cgg aat agt aca gta tga tag aac aca tcc tat tcc caa caa tgt cag gaa cgt cac gct ctc cac ctt cat att tat tta tcc gta aaa tgt tat cct gga cat cgt aca aat aat aaa aag ccc ata tat atg ttc gct att gta gaa att gtt ttt cac agt tgc tca aaa aca atg gca gtg act tat gag tta gtt aca ctt tgg agt ctc atc ttt agt aaa cat atc ata ata ttc gat att acg agt tga cat atc gaa caa att cca agt att tga ttt tgg ata ata ttc gta ttt tgc atc tgc tat aat taa gat ata atc acc aca aga aca cac gaa cgt ctt tcc tac atg gtt aaa gta cat gta caa ttc tat cca ttt gtc ttc ctt aac tat ata ttt gta tag ata att acg agt ctc atg agt aat tcc agt aat tgc ata gat gtc acc atc gta ttc tac agc ata aac tat act atg acg tct agg cat ggg aga ctt ttt tat cca acg att ttt agt gaa aca ttc cac atc gtt taa tac tac ata ttt ctc ata gtg gta taa act cca ccc att aca tat ata tca tcg ttt acg aat act gat gcg cct gaa tat cta gga gtg att aag ttt gga agt ctt ttc cat ttc gaa gtg ccg tgt ttc aaa tat tct gct ata ccc gtt gaa ata gaa aat tct aat cct cct att aca tat aac ttt cca tcg tta aca caa gta cta act tct gat ttt aac gac gac ata tta gta acc gtt ttc ca- --- ttt ttt ttg ttt taa gat cta ccc gcg ata cgg aat aaa cat gtc tat tgt taa tca tgc cgc caa taa tgt ata gac aat tat gta aaa cat ttg cat cat aga att gtc tat ctg tat tac cga cta tcg tcc aat att ctg ttc tag gag agt aat ggg tta ttg tgg ata tat aat cag agt ttt taa tga cta cta tat tat gtt tta tac cat ttc gtg tca cag ctt tgt aga ttt gga tat agt taa tcc caa caa tgc tat agc att gca tat agc att agt cat aaa ctt ggg atg taa aat gtt gat gat atc tac atc gtt tgg att ttt atg tat cca ctt taa taa tat tat agc gta aca tcc tca tga ttt acg tta acg ttt tcg tgt gat aag ata gtg gtc agt tca tcc ttt gat aat ttt cca aat tct gga tcg gat gtc acc gca gta ata ttg ttg att att tct gac atc gac gca tta tat agt ttt tta att cca tat ctt ttt t

>JX878408.1_Monkeypox_virus_isolate_DRC_06-0970_complete_genome

ttt tat atc act acg gac ata aac cat tgt ata att ttt atg ttt att agt gta cac att ttg gaa gta agt tcc ggc tgc cat gta ttt cct gga gag caa gta gat gat g-a gga acc aga tag ttt ata tcc ata ctt gca ctt aaa gtc tac att gta gtt gta tga gag tat gat ctt tta agc cgc tag aag ttt tcc gtt tga tat agg atg tgg aca ttt aac aat ctg aca cgt ggg tgg att gga cca ttc tcc tcc tga aca cat gac acc aga gtt acc aat caa cga ata tcc act att gca act ata agt tac aat gct ccc atc gat ata aaa atc ctc gta tcc gtt atg tct tcc gtt gga tat aga tgg agg tga ttg gca ttt aac aga ttc gca aat agg tgc ctc agg att cca tac cat aga tcc agt aga tcc taa ttc aca ata cga ttt aga ttc acc gat caa atg ata tcc gct att aca aga gta cgt tat act aga gcc aaa gtc tac tcc gcc aat atc aag ttg gcc att atc gat atc tcg agg cga tgg gca tct ccg ttt aat aca ttg att aaa gag tgt cca tcc ggt acc ggt aca ttt agc ata tat ggg tcc cat ttt ttg ctt tct gta tcc agg tag aca tag ata ttc tat agt gtc tcc tat gtt gta att agc atc agt ctc tac act att ctt aaa ttt cat att aat ggg gcg tga cgg aat agt aca gta tga tag aac aca tcc tat tcc caa caa tgt cag gaa cgt cac gct ctc cac ctt cat att tat tta tcc gta aaa tgt tat cct gga cat cgt aca aat aat aaa aag ccc ata tat atg ttc gct att gta gaa att gtt ttt cac agt tgc tca aaa aca atg gca gtg act tat gag tta gtt aca ctt tgg agt ctc atc ttt agt aaa cat atc ata ata ttc gat att acg agt tga cat atc gaa caa att cca agt att tga ttt tgg ata ata ttc gta ttt tgc atc tgc tat aat taa gat ata atc acc aca aga aca cac gaa cgt ctt tcc tac atg gtt aaa gta cat gta caa ttc tat cca ttt gtc ttc ctt aac tat ata ttt gta tag ata att acg agt ctc atg agt aat tcc agt aat tgc ata gat gtc acc atc gta ttc tac agc ata aac tat act atg acg tct agg cat ggg aga ctt ttt tat cca acg att ttt agt gaa aca ttc cac atc gtt taa tac tac ata ttt ctc ata gtg gta taa act cca ccc att aca tat ata tca tcg ttt acg aat act gat gcg cct gaa tat cta gga gtg att aag ttt gga agt ctt ttc cat ttc gaa gtg ccg tgt ttc aaa tat tct gct ata ccc gtt gaa ata gaa aat tct aat cct cct att aca tat aac ttt cca tcg tta aca caa gta cta act tct gat ttt aac gac gac ata tta gta acc gtt ttc ca- --- ttt ttt ttg ttt taa gat cta ccc gcg ata cgg aat aaa cat gtc tat tgt taa tca tgc cgc caa taa tgt ata gac aat tat gta aaa cat ttg cat cat aga att gtc tat ctg tat tac cga cta tcg tcc aat att ctg ttc tag gag agt aat ggg tta ttg tgg ata tat aat cag agt ttt taa tga cta cta tat tat gtt tta tac cat ttc gtg tca cag ctt tgt aga ttt gga tat agt taa tcc caa caa tgc tat agc att gca tat agc att agt cat aaa ctt ggg atg taa aat gtt gat gat atc tac atc gtt tgg att ttt atg tat cca ctt taa taa tat tat agc gta aca tcc tca tga ttt acg tta acg ttt tcg tgt gat aag ata gtg gtc agt tca tcc ttt gat aat ttt cca aat tct gga tcg gat gtc acc gca gta ata ttg ttg att att tct gac atc gac gca tta tat agt ttt tta att cca tat ctt ttt t

>JX878420.1_Monkeypox_virus_isolate_DRC_07-0283_complete_genome

ttt tat atc act acg gac ata aac cat tgt ata att ttt atg ttt att agt gta cac att ttg gaa gta agt tcc ggc tgc cat gta ttt cct gga gag caa gta gat gat g-a gga acc aga tag ttt ata tcc ata ctt gca ctt aaa gtc tac att gta gtt gta tga gag tat gat ctt tta agc cgc tag aag ttt tcc gtt tga tat agg atg tgg aca ttt aac aat ctg aca cgt ggg tgg att gga cca ttc tcc tcc tga aca cat gac acc aga gtt acc aat caa cga ata tcc act att gca act ata agt tac aat gct ccc atc gat ata aaa atc ctc gta tcc gtt atg tct tcc gtt gga tat aga tgg agg tga ttg gca ttt aac aga ttc gca aat agg tgc ctc agg att cca tac cat aga tcc agt aga tcc taa ttc aca ata cga ttt aga ttc acc gat caa atg ata tcc gct att aca aga gta cgt tat act aga gcc aaa gtc tac tcc gcc aat atc aag ttg gcc att atc gat atc tcg agg cga tgg gca tct ccg ttt aat aca ttg att aaa gag tgt cca tcc ggt acc ggt aca ttt agc ata tat ggg tcc cat ttt ttg ctt tct gta tcc agg tag aca tag ata ttc tat agt gtc tcc tat gtt gta att agc atc agt ctc tac act att ctt aaa ttt cat att aat ggg gcg tga cgg aat agt aca gta tga tag aac aca tcc tat tcc caa caa tgt cag gaa cgt cac gct ctc cac ctt cat att tat tta tcc gta aaa tgt tat cct gga cat cgt aca aat aat aaa aag ccc ata tat atg ttc gct att gta gaa att gtt ttt cac agt tgc tca aaa aca atg gca gtg act tat gag tta gtt aca ctt tgg agt ctc atc ttt agt aaa cat atc ata ata ttc gat att acg agt tga cat atc gaa caa att cca agt att tga ttt tgg ata ata ttc gta ttt tgc atc tgc tat aat taa gat ata atc acc aca aga aca cac gaa cgt ctt tcc tac atg gtt aaa gta cat gta caa ttc tat cca ttt gtc ttc ctt aac tat ata ttt gta tag ata att acg agt ctc atg agt aat tcc agt aat tgc ata gat gtc acc atc gta ttc tac agc ata aac tat act atg acg tct agg cat ggg aga ctt ttt tat cca acg att ttt agt gaa aca ttc cac atc gtt taa tac tac ata ttt ctc ata gtg gta taa act cca ccc att aca tat ata tca tcg ttt acg aat act gat gcg cct gaa tat cta gga gtg att aag ttt gga agt ctt ttc cat ttc gaa gtg ccg tgt ttc aaa tat tct gct ata ccc gtt gaa ata gaa aat tct aat cct cct att aca tat aac ttt cca tcg tta aca caa gta cta act tct gat ttt aac gac gac ata tta gta acc gtt ttc ca- --- ttt ttt ttg ttt taa gat cta ccc gcg ata cgg aat aaa cat gtc tat tgt taa tca tgc cgc caa taa tgt ata gac aat tat gta aaa cat ttg cat cat aga att gtc tat ctg tat tac cga cta tcg tcc aat att ctg ttc tag gag agt aat ggg tta ttg tgg ata tat aat cag agt ttt taa tga cta cta tat tat gtt tta tac cat ttc gtg tca cag ctt tgt aga ttt gga tat agt taa tcc caa caa tgc tat agc att gca tat agc att agt cat aaa ctt ggg atg taa aat gtt gat gat atc tac atc gtt tgg att ttt atg tat cca ctt taa taa tat tat agc gta aca tcc tca tga ttt acg tta acg ttt tcg tgt gat aag ata gtg gtc agt tca tcc ttt gat aat ttt cca aat tct gga tcg gat gtc acc gca gta ata ttg ttg att att tct gac atc gac gca tta tat agt ttt tta att cca tat ctt ttt t

>JX878419.1_Monkeypox_virus_isolate_DRC_07-0275_complete_genome

ttt tat atc act acg gac ata aac cat tgt ata att ttt atg ttt att agt gta cac att ttg gaa gta agt tcc ggc tgc cat gta ttt cct gga gag caa gta gat gat g-a gga acc aga tag ttt ata tcc ata ctt gca ctt aaa gtc tac att gta gtt gta tga gag tat gat ctt tta agc cgc tag aag ttt tcc gtt tga tat agg atg tgg aca ttt aac aat ctg aca cgt ggg tgg att gga cca ttc tcc tcc tga aca cat gac acc aga gtt acc aat caa cga ata tcc act att gca act ata agt tac aat gct ccc atc gat ata aaa atc ctc gta tcc gtt atg tct tcc gtt gga tat aga tgg agg tga ttg gca ttt aac aga ttc gca aat agg tgc ctc agg att cca tac cat aga tcc agt aga tcc taa ttc aca ata cga ttt aga ttc acc gat caa atg ata tcc gct att aca aga gta cgt tat act aga gcc aaa gtc tac tcc gcc aat atc aag ttg gcc att atc gat atc tcg agg cga tgg gca tct ccg ttt aat aca ttg att aaa gag tgt cca tcc ggt acc ggt aca ttt agc ata tat ggg tcc cat ttt ttg ctt tct gta tcc agg tag aca tag ata ttc tat agt gtc tcc tat gtt gta att agc atc agt ctc tac act att ctt aaa ttt cat att aat ggg gcg tga cgg aat agt aca gta tga tag aac aca tcc tat tcc caa caa tgt cag gaa cgt cac gct ctc cac ctt cat att tat tta tcc gta aaa tgt tat cct gga cat cgt aca aat aat aaa aag ccc ata tat atg ttc gct att gta gaa att gtt ttt cac agt tgc tca aaa aca atg gca gtg act tat gag tta gtt aca ctt tgg agt ctc atc ttt agt aaa cat atc ata ata ttc gat att acg agt tga cat atc gaa caa att cca agt att tga ttt tgg ata ata ttc gta ttt tgc atc tgc tat aat taa gat ata atc acc aca aga aca cac gaa cgt ctt tcc tac atg gtt aaa gta cat gta caa ttc tat cca ttt gtc ttc ctt aac tat ata ttt gta tag ata att acg agt ctc atg agt aat tcc agt aat tgc ata gat gtc acc atc gta ttc tac agc ata aac tat act atg acg tct agg cat ggg aga ctt ttt tat cca acg att ttt agt gaa aca ttc cac atc gtt taa tac tac ata ttt ctc ata gtg gta taa act cca ccc att aca tat ata tca tcg ttt acg aat act gat gcg cct gaa tat cta gga gtg att aag ttt gga agt ctt ttc cat ttc gaa gtg ccg tgt ttc aaa tat tct gct ata ccc gtt gaa ata gaa aat tct aat cct cct att aca tat aac ttt cca tcg tta aca caa gta cta act tct gat ttt aac gac gac ata tta gta acc gtt ttc ca- --- ttt ttt ttg ttt taa gat cta ccc gcg ata cgg aat aaa cat gtc tat tgt taa tca tgc cgc caa taa tgt ata gac aat tat gta aaa cat ttg cat cat aga att gtc tat ctg tat tac cga cta tcg tcc aat att ctg ttc tag gag agt aat ggg tta ttg tgg ata tat aat cag agt ttt taa tga cta cta tat tat gtt tta tac cat ttc gtg tca cag ctt tgt aga ttt gga tat agt taa tcc caa caa tgc tat agc att gca tat agc att agt cat aaa ctt ggg atg taa aat gtt gat gat atc tac atc gtt tgg att ttt atg tat cca ctt taa taa tat tat agc gta aca tcc tca tga ttt acg tta acg ttt tcg tgt gat aag ata gtg gtc agt tca tcc ttt gat aat ttt cca aat tct gga tcg gat gtc acc gca gta ata ttg ttg att att tct gac atc gac gca tta tat agt ttt tta att cca tat ctt ttt t

>JX878417.1_Monkeypox_virus_isolate_DRC_07-0104_complete_genome

ttt tat atc act acg gac ata aac cat tgt ata att ttt atg ttt att agt gta cac att ttg gaa gta agt tcc ggc tgc cat gta ttt cct gga gag caa gta gat gat g-a gga acc aga tag ttt ata tcc ata ctt gca ctt aaa gtc tac att gta gtt gta tga gtg tat gat ctt tta agc cgc tag aag ttt tcc gtt tga tat agg atg tgg aca ttt aac aat ctg aca cgt ggg tgg att gga cca ttc tcc tcc tga aca cat gac acc aga gtt acc aat caa cga ata tcc act att gca act ata agt tac aat gct ccc atc gat ata aaa atc ctc gta tcc gtt atg tct tcc gtt gga tat aga tgg agg tga ttg gca ttt aac aga ttc gca aat agg tgc ctc agg att cca tac cat aga tcc agt aga tcc taa ttc aca ata cga ttt aga ttc acc gat caa atg ata tcc gct att aca aga gta cgt tat act aga gcc aaa gtc tac tcc gcc aat atc aag ttg acc att atc gat atc tcg agg cga tgg gca tct ccg ttt aat aca ttg att aaa gag tgt cca tcc ggt acc ggt aca ttt agc ata tat ggg tcc cat ttt ttg ctt tct gta tcc agg tag aca tag ata ttc tat agt gtc tcc tat gtt gta att agc atc agt ctc tac act att ctt aaa ttt cat att aat ggg gcg tga cgg aat agt aca gta tga tag aac aca tcc tat tcc caa caa tgt cag gaa cgt cac gct ctc cac ctt cat att tat tta tcc gta aaa tgt tat cct gga cat cgt aca aat aat aaa aag ccc ata tat atg ttc gct att gta gaa att gtt ttt cac agt tgc tca aaa aca atg gca gtg act tat gag ttc gtt aca ctt tgg agt ctc atc ttt agt aaa cat atc ata ata ttc gat att acg agt tga cat atc gaa caa att cca agt att tga ttt tgg ata ata ttc gta ttt tgc atc tgc tat aat taa gat ata atc acc aca aga aca cac gaa cgt ctt tcc tac atg gtt aaa gta cat gta caa ttc tat cca ttt gtc ttc ctt aac tat ata ttt gta tag ata att acg agt ctc atg agt aat tcc agt aat tgc ata gat gtc acc atc gta ttc tac agc ata aac tat act atg acg tct agg cat ggg aga ctt ttt tat cca acg att ttt agt gaa aca ttc cac atc gtt taa tac tac ata ttt ctc ata gtg gta taa act cca ccc att aca tat ata tca tcg ttt acg aat act gat gcg cct gaa tat cta gga gtg att aag ttt gga agt ctt ttc cat ttc gaa gtg ccg tgt ttc aaa tat tct gct ata ccc gtt gaa ata gaa aat tct aat cct cct att aca tat aac ttt cca tcg tta aca caa gta cta act tct gat ttt aac gac gac ata tta gta acc gtt ttc ca- --- ttt ttt ttg ttt taa gat cta ccc gcg ata cgg aat aaa cat gtc tat tgt taa tca tgc cgc caa taa tgt ata gac aat tat gta aaa cat ttg cat cat aga att gtc tat ctg tat tac cga cta tcg tcc aat att ctg ttc tag gag agt aat ggg tta ttg tgg ata tat aat cag agt ttt taa tga cta cta tat tat gtt tta tac cat ttc gtg tca cag ctt tgt aga ttt gga tat agt taa tcc caa caa tgc tat agc att gca tat agc att agt cat aaa ctt ggg atg taa aat gtt gat gat atc tac atc gtt tgg att ttt atg tat cca ctt taa taa tat tat agc gta aca tcc tca tga ttt acg tta acg ttt tcg tgt gat aag ata gtg gtc agt tca tcc ttt gat aat ttt cca aat tct gga tcg gat gtc acc gca gta ata ttg ttg att att tct gac atc gac gca tta tat agt ttt tta att cca tat ctt ttt t

>KJ642612.1_Monkeypox_virus_strain_Ikubi_complete_genome

ttt tat atc act acg gac ata aac cat tgt ata att ttt atg ttt att agt gta cac att ttg gaa gta agt tcc ggc tgc cat gta ttt cct gga gag caa gta gat gat g-a gga acc aga tag ttt ata tcc ata ctt gca ctt aaa gtc tac att gta gtt gta tga gag tat gat ctt tta agc cgc tag aag ttt tcc gtt tga tat agg atg tgg aca ttt aac aat ctg aca cgt ggg tgg att gga cca ttc tcc tcc tga aca cat gac acc aga gtt acc aat caa cga ata tcc act att gca act ata agt tac aat gct ccc atc gat ata aaa atc ctc gta tcc gtt atg tct tcc gtt gga tat aga tgg agg tga ttg gca ttt aac aga ttc gca aat agg tgc ctc agg att cca tac cat aga tcc agt aga tcc taa ttc aca ata cga ttt aga ttc acc gat caa atg ata tcc gct att aca aga gta cgt tat act aga gcc aaa gtc tac tcc gcc aat atc aag ttg gcc att atc gat atc tcg agg cga tgg gca tct ccg ttt aat aca ttg att aaa gag tgt cca tcc ggt acc ggt aca ttt agc ata tat ggg tcc cat ttt ttg ctt tct gta tcc agg tag aca tag ata ttc tat agt gtc tcc tat gtt gta att agc atc agt ctc tac act att ctt aaa ttt cat att aat ggg gcg tga cgg aat agt aca gta tga tag aac aca tcc tat tcc caa caa tgt cag gaa cgt cac gct ctc cac ctt cat att tat tta tcc gta aaa tgt tat cct gga cat cgt aca aat aat aaa aag ccc ata tat atg ttc gct att gta gaa att gtt ttt cac agt tgc tca aaa aca atg gca gtg act tat gag tta gtt aca ctt tgg agt ctc atc ttt agt aaa cat atc ata ata ttc gat att acg agt tga cat atc gaa caa att cca agt att tga ttt tgg ata ata ttc gta ttt tgc atc tgc tat aat taa gat ata atc acc aca aga aca cac gaa cgt ctt tcc tac atg gtt aaa gta cat gta caa ttc tat cca ttt gtc ttc ctt aac tat ata ttt gta tag ata att acg agt ctc atg agt aat tcc agt aat tgc ata gat gtc acc atc gta ttc tac agc ata aac tat act atg acg tct agg cat ggg aga ctt ttt tat cca acg att ttt agt gaa aca ttc cac atc gtt taa tac tac ata ttt ctc ata gtg gta taa act cca ccc att aca tat ata tca tcg ttt acg aat act gat gcg cct gaa tat cta gga gtg att aag ttt gga agt ctt ttc cat ttc gaa gtg ccg tgt ttc aaa tat tct gct ata ccc gtt gaa ata gaa aat tct aat cct cct att aca tat aac ttt cca tcg tta aca caa gta cta act tct gat ttt aac gac gac ata tta gta acc gtt ttc ca- --- ttt ttt ttg ttt taa gat cta ccc gcg ata cgg aat aaa cat gtc tat tgt taa tca tgc cgc caa taa tgt ata gac aat tat gta aaa cat ttg cat cat aga att gtc tat ctg tat tac cga cta tcg tcc aat att ctg ttc tag gag agt aat ggg tta ttg tgg ata tat aat cag agt ttt taa tga cta cta tat tat gtt tta tac cat ttc gtg tca cag ctt tgt aga ttt gga tat agt taa tcc caa caa tgc tat agc att gca tat agc att agt cat aaa ctt ggg atg taa aat gtt gat gat atc tac atc gtt tgg att ttt atg tat cca ctt taa taa tat tat agc gta aca tcc tca tga ttt acg tta acg ttt tcg tgt gat aag ata gtg gtc agt tca tcc ttt gat aat ttt cca aat tct gga tcg gat gtc acc gca gta ata ttg ttg att att tct gac atc gac gca tta tat agt ttt tta att cca tat ctt ttt t

>JX878426.1_Monkeypox_virus_isolate_DRC_07-0450_complete_genome

ttt tat atc act acg gac ata aac cat tgt ata att ttt atg ttt att agt gta cac att ttg gaa gta agt tcc ggc tgc cat gta ttt cct gga gag caa gta gat gat g-a gga acc aga tag ttt ata tcc ata ctt gca ctt aaa gtc tac att gta gtt gta tga gag tat gat ctt tta agc cgc tag aag ttt tcc gtt tga tat agg atg tgg aca ttt aac aat ctg aca cgt ggg tgg att gga cca ttc tcc tcc tga aca cat gac acc aga gtt acc aat caa cga ata tcc act att gca act ata agt tac aat gct ccc atc gat ata aaa atc ctc gta tcc gtt atg tct tcc gtt gga tat aga tgg agg tga ttg gca ttt aac aga ttc gca aat agg tgc ctc agg att cca tac cat aga tcc agt aga tcc taa ttc aca ata cga ttt aga ttc acc gat caa atg ata tcc gct att aca aga gta cgt tat act aga gcc aaa gtc tac tcc gcc aat atc aag ttg gcc att atc gat atc tcg agg cga tgg gca tct ccg ttt aat aca ttg att aaa gag tgt cca tcc ggt acc ggt aca ttt agc ata tat ggg tcc cat ttt ttg ctt tct gta tcc agg tag aca tag ata ttc tat agt gtc tcc tat gtt gta att agc atc agt ctc tac act att ctt aaa ttt cat att aat ggg gcg tga cgg aat agt aca gta tga tag aac aca tcc tat tcc caa caa tgt cag gaa cgt cac gct ctc cac ctt cat att tat tta tcc gta aaa tgt tat cct gga cat cgt aca aat aat aaa aag ccc ata tat atg ttc gct att gta gaa att gtt ttt cac agt tgc tca aaa aca atg gca gtg act tat gag tta gtt aca ctt tgg agt ctc atc ttt agt aaa cat atc ata ata ttc gat att acg agt tga cat atc gaa caa att cca agt att tga ttt tgg ata ata ttc gta ttt tgc atc tgc tat aat taa gat ata atc acc aca aga aca cac gaa cgt ctt tcc tac atg gtt aaa gta cat gta caa ttc tat cca ttt gtc ttc ctt aac tat ata ttt gta tag ata att acg agt ctc atg agt aat tcc agt aat tgc ata gat gtc acc atc gta ttc tac agc ata aac tat act atg acg tct agg cat ggg aga ctt ttt tat cca acg att ttt agt gaa aca ttc cac atc gtt taa tac tac ata ttt ctc ata gtg gta taa act cca ccc att aca tat ata tca tcg ttt acg aat act gat gcg cct gaa tat cta gga gtg att aag ttt gga agt ctt ttc cat ttc gaa gtg ccg tgt ttc aaa tat tct gct ata ccc gtt gaa ata gaa aat tct aat cct cct att aca tat aac ttt cca tcg tta aca caa gta cta act tct gat ttt aac gac gac ata tta gta acc gtt ttc ca- --- ttt ttt ttg ttt taa gat cta ccc gcg ata cgg aat aaa cat gtc tat tgt taa tca tgc cgc caa taa tgt ata gac aat tat gta aaa cat ttg cat cat aga att gtc tat ctg tat tac cga cta tcg tcc aat att ctg ttc tag gag agt aat ggg tta ttg tgg ata tat aat cag agt ttt taa tga cta cta tat tat gtt tta tac cat ttc gtg tca cag ctt tgt aga ttt gga tat agt taa tcc caa caa tgc tat agc att gca tat agc att agt cat aaa ctt ggg atg taa aat gtt gat gat atc tac atc gtt tgg att ttt atg tat cca ctt taa taa tat tat agc gta aca tcc tca tga ttt acg tta acg ttt tcg tgt gat aag ata gtg gtc agt tca tcc ttt gat aat ttt cca aat tct gga tcg gat gtc acc gca gta ata ttg ttg att att tct gac atc gac gca tta tat agt ttt tta att cca tat ctt ttt t

>KC257459.1_Monkeypox_virus_strain_Sudan_2005_01_complete_genome

ttt tat atc act acg gac ata aac cat tgt ata att ttt atg ttt att agt gta cac att ttg gaa gta agt tcc ggc tgc cat gta ttt cct gga gag caa gta gat gat g-a gga acc aga tag ttt ata tcc ata ctt gca ctt aaa gtc tac att gta gtt gta tga gtg tat gat ctt tta agc cgc tag aag ttt tcc gtt tga tat agg atg tgg aca ttt aac aat ctg aca cgt ggg tgg att gga cca ttc tcc tcc tga aca cat gac acc aga gtt acc aat caa cga ata tcc act att gca act ata agt tac aat gct ccc atc gat ata aaa atc ctc gta tcc gtt atg tct tcc gtt gga tat aga tgg agg tga ttg gca ttt aac aga ttc gca aat agg tgc ctc agg att cca tac cat aga tcc agt aga tcc taa ttc aca ata cga ttt aga ttc acc gat caa atg ata tcc gct att aca aga gta cgt tat act aga gcc aaa gtc tac tcc gcc aat atc aag ttg gcc att atc gat atc tcg agg cga tgg gca tct ccg ttt aat aca ttg att aaa gag tgt cca tcc ggt acc ggt aca ttt agc ata tat ggg tcc cat ttt ttg ctt tct gta tcc agg tag aca tag ata ttc tat agt gtc tcc tat gtt gta att agc atc agt ctc tac act att ctt aaa ttt cat att aat ggg gcg tga cgg aat agt aca gta tga tag aac aca tcc tat tcc caa caa tgt cag gaa cgt cac gct ctc cac ctt cat att tat tta tcc gta aaa tgt tat cct gga cat cgt aca aat aat aaa aag ccc ata tat atg ttc gct att gta gaa att gtt ttt cac agt tgc tca aaa aca atg gca gtg act tat gag tta gtt aca ctt tgg agt ctc atc ttt agt aaa cat atc ata ata ttc gat att acg agt tga cat atc gaa caa att cca agt att tga ttt tgg ata ata ttc gta ttt tgc atc tgc tat aat taa gat ata atc acc aca aga aca cac gaa cgt ctt tcc tac atg gtt aaa gta cat gta caa ttc tat cca ttt gtc ttc ctt aac tat ata ttt gta tag ata att acg agt ctc atg agt aat tcc agt aat tgc ata gat gtc acc atc gta ttc tac agc ata aac tat act atg acg tct agg cat ggg aga ctt ttt tat cca acg att ttt agt gaa aca ttc cac atc gtt taa tac tac ata ttt ctc ata gtg gta taa act cca ccc att aca tat ata tca tcg ttt acg aat act gat gcg cct gaa tat cta gga gtg att aag ttt gga agt ctt ttc cat ttc gaa gtg ccg tgt ttc aaa tat tct gct ata ccc gtt gaa ata gaa aat tct aat cct cct att aca tat aac ttt cca tcg tta aca caa gta cta act tct gat ttt aac gac gac ata tta gta acc gtt ttc ca- --t ttt ttt ttg ttt taa gat cta ccc gcg ata cgg aat aaa cat gtc tat tgt taa tca tgc cgc caa taa tgt ata gac aat tat gta aaa cat ttg cat cat aga att gtc tat ctg tat tac cga cta tcg tcc aat att ctg ttc tag gag agt aat ggg tta ttg tga ata tat aat cag agt ttt taa tga cta cta tat tat gtt tta tac cat ttc gtg tca cag ctt tgt aga ttt gga tat agt taa tcc caa caa tgc tat agc att gca tat agc att agt cat aaa ctt ggg atg taa aat gtt gat gat atc tac atc gtt tgg att ttt atg tat cca ctt taa taa tat tat agc gta aca tcc tca tga ttt acg tta acg ttt tcg tgt gat aag ata gtg gtc agt tca tcc ttt gat aat ttt cca aat tct gga tcg gat gtc acc gca gta ata ttg ttg att att tct gac atc gac gca tta tat agt ttt tta att cca tat ctt ttt t

>MN702449.1_Monkeypox_virus_strain_B2_contig_SPADES

ttt tat atc act acg gac ata aac cat tgt ata att ttt atg ttt att agt gta cac att ttg gaa gta agt tcc ggc tgc cat gta ttt cct gga gag caa gta gat gat g-a gga acc aga tag ttt ata tcc ata ctt gca ctt aaa gtc tac att gta gtt gta tga gtg tat gat ctt tta agc cgc tag aag ttt tcc gtt tga tat agg atg tgg aca ttt aac aat ctg aca cgt ggg tgg att gga cca ttc tcc tcc tga aca cat gac acc aga gtt acc aat caa cga ata tcc act att gca act ata agt tac aat gct ccc atc gat ata aaa atc ctc gta tcc gtt atg tct tcc gtt gga tat aga tgg agg tga ttg gca ttt aac aga ttc gca aat agg tgc ctc agg att cca tac cat aga tcc agt aga tcc taa ttc aca ata cga ttt aga ttc acc gat caa atg ata tcc gct att aca aga gta cgt tat act aga gcc aaa gtc tac tcc gcc aat atc aag ttg gcc att atc gat atc tcg agg cga tgg gca tct ccg ttt aat aca ttg att aaa gag tgt cca tcc ggt acc ggt aca ttt agc ata tat ggg tcc cat ttt ttg ctt tct gta tcc agg tag aca tag ata ttc tat agt gtc tcc tat gtt gta att agc atc agt ctc tac act att ctt aaa ttt cat att aat ggg gcg tga cgg aat agt aca gta tga tag aac aca tcc tat tcc caa caa tgt cag gaa cgt cac gct ctc cac ctt cat att tat tta tcc gta aaa tgt tat cct gga cat cgt aca aat aat aaa aag ccc ata tat atg ttc gct att gta gaa att gtt ttt cac agt tgc tca aaa aca atg gca gtg act tat gag tta gtt aca ctt tgg agt ctc atc ttt agt aaa cat atc ata ata ttc gat att acg agt tga cat atc gaa caa att cca agt att tga ttt tgg ata ata ttc gta ttt tgc atc tgc tat aat taa gat ata atc acc aca aga aca cac gaa cgt ctt tcc tac atg gtt aaa gta cat gta caa ttc tat cca ttt gtc ttc ctt aac tat ata ttt gta tag ata att acg agt ctc atg agt aat tcc agt aat tgc ata gat gtc acc atc gta ttc tac agc ata aac tat act atg acg tct agg cat ggg aga ctt ttt tat cca acg att ttt agt gaa aca ttc cac atc gtt taa tac tac ata ttt ctc ata gtg gta taa act cca ccc att aca tat ata tca tcg ttt acg aat act gat gcg cct gaa tat cta gga gtg att aag ttt gga agt ctt ttc cat ttc gaa gtg ccg tgt ttc aaa tat tct gct ata ccc gtt gaa ata gaa aat tct aat cct cct att aca tat aac ttt cca tcg tta aca caa gta cta act tct gat ttt aac gac gac ata tta gta acc gtt ttc cat ttt ttt ttt ttg ttt taa gat cta ccc gcg ata cgg aat aaa cat gtc tat tgt taa tca tgc cgc caa taa tgt ata gac aat tat gta aaa cat ttg cat cat aga att gtc tat ctg tat tac cga cta tcg tcc aat att ctg ttc tag gag agt aat ggg tta ttg tga ata tat aat cag agt ttt taa tga cta cta tat tat gtt tta tac cat ttc gtg tca cag ctt tgt aga ttt gga tat agt taa tcc caa caa tgc tat agc att gca tat agc att agt cat aaa ctt ggg atg taa aat gtt gat gat atc tac atc gtt tgg att ttt ata tat cca ctt taa taa tat tat agc gta aca tcc tca tga ttt acg tta acg ttt tcg tgt gat aag ata gtg gtc agt tca tcc ttt gat aat ttt cca aat tct gga tcg gat gtc acc gca gta ata ttg ttg att att tct gac atc gac gca tta tat agt ttt tta att cca tat ctt ttt t

>MN702446.1_Monkeypox_virus_strain_38c_contig_SPADES

ttt tat atc act acg gac ata aac cat tgt ata att ttt atg ttt att agt gta cac att ttg gaa gta agt tcc ggc tgc cat gta ttt cct gga gag caa gta gat gat g-a gga acc aga tag ttt ata tcc ata ctt gca ctt aaa gtc tac att gta gtt gta tga gtg tat gat ctt tta agc cgc tag aag ttt tcc gtt tga tat agg atg tgg aca ttt aac aat ctg aca cgt ggg tgg att gga cca ttc tcc tcc tga aca cat gac acc aga gtt acc aat caa cga ata tcc act att gca act ata agt tac aat gct ccc atc gat ata aaa atc ctc gta tcc gtt atg tct tcc gtt gga tat aga tgg agg tga ttg gca ttt aac aga ttc gca aat agg tgc ctc agg att cca tac cat aga tcc agt aga tcc taa ttc aca ata cga ttt aga ttc acc gat caa atg ata tcc gct att aca aga gta cgt tat act aga gcc aaa gtc tac tcc gcc aat atc aag ttg gcc att atc gat atc tcg agg cga tgg gca tct ccg ttt aat aca ttg att aaa gag tgt cca tcc ggt acc ggt aca ttt agc ata tat ggg tcc cat ttt ttg ctt tct gta tcc agg tag aca tag ata ttc tat agt gtc tcc tat gtt gta att agc atc agt ctc tac act att ctt aaa ttt cat att aat ggg gcg tga cgg aat agt aca gta tga tag aac aca tcc tat tcc caa caa tgt cag gaa cgt cac gct ctc cac ctt cat att tat tta tcc gta aaa tgt tat cct gga cat cgt aca aat aat aaa aag ccc ata tat atg ttc gct att gta gaa att gtt ttt cac agt tgc tca aaa aca atg gca gtg act tat gag tta gtt aca ctt tgg agt ctc atc ttt agt aaa cat atc ata ata ttc gat att acg agt tga cat atc gaa caa att cca agt att tga ttt tgg ata ata ttc gta ttt tgc atc tgc tat aat taa gat ata atc acc aca aga aca cac gaa cgt ctt tcc tac atg gtt aaa gta cat gta caa ttc tat cca ttt gtc ttc ctt aac tat ata ttt gta tag ata att acg agt ctc atg agt aat tcc agt aat tgc ata gat gtc acc atc gta ttc tac agc ata aac tat act atg acg tct agg cat ggg aga ctt ttt tat cca acg att ttt agt gaa aca ttc cac atc gtt taa tac tac ata ttt ctc ata gtg gta taa act cca ccc att aca tat ata tca tcg ttt acg aat act gat gcg cct gaa tat cta gga gtg att aag ttt gga agt ctt ttc cat ttc gaa gtg ccg tgt ttc aaa tat tct gct ata ccc gtt gaa ata gaa aat tct aat cct cct att aca tat aac ttt cca tcg tta aca caa gta cta act tct gat ttt aac gac gac ata tta gta acc gtt ttc ca- -tt ttt ttt ttg ttt taa gat cta ccc gcg ata cgg aat aaa cat gtc tat tgt taa tca tgc cgc caa taa tgt ata gac aat tat gta aaa cat ttg cat cat aga att gtc tat ctg tat tac cga cta tcg tcc aat att ctg ttc tag gag agt aat ggg tta ttg tga ata tat aat cag agt ttt taa tga cta cta tat tat gtt tta tac cat ttc gtg tca cag ctt tgt aga ttt gga tat agt taa tcc caa caa tgc tat agc att gca tat agc att agt cat aaa ctt ggg atg taa aat gtt gat gat atc tac atc gtt tgg att ttt ata tat cca ctt taa taa tat tat agc gta aca tcc tca tga ttt acg tta acg ttt tcg tgt gat aag ata gtg gtc agt tca tcc ttt gat aat ttt cca aat tct gga tcg gat gtc acc gca gta ata ttg ttg att att tct gac atc gac gca tta tat agt ttt tta att cca tat ctt ttt t

>JX878418.1_Monkeypox_virus_isolate_DRC_07-0120_complete_genome

ttt tat atc act acg gac ata aac cat tgt ata att ttt atg ttt att agt gta cac att ttg gaa gta agt tcc ggc tgc cat gta ttt cct gga gag caa gta gat gat g-a gga acc aga tag ttt ata tcc ata ctt gca ctt aaa gtc tac att gta gtt gta tga gag tat gat ctt tta agc cgc tag aag ttt tcc gtt tga tat agg atg tgg aca ttt aac aat ctg aca cgt ggg tgg att gga cca ttc tcc tcc tga aca cat gac acc aga gtt acc aat caa cga ata tcc act att gca act ata agt tac aat gct ccc atc gat ata aaa atc ctc gta tcc gtt atg tct tcc gtt gga tat aga tgg agg tga ttg gca ttt aac aga ttc gca aat agg tgc ctc agg att cca tac cat aga tcc agt aga tcc taa ttc aca ata cga ttt aga ttc acc gat caa atg ata tcc gct att aca aga gta cgt tat act aga gcc aaa gtc tac tcc gcc aat atc aag ttg gcc att atc gat atc tcg agg cga tgg gca tct ccg ttt aat aca ttg att aaa gag tgt cca tcc ggt acc ggt aca ttt agc ata tat ggg tcc cat ttt ttg ctt tct gta tcc agg tag aca tag ata ttc tat agt gtc tcc tat gtt gta att agc atc agt ctc tac act att ctt aaa ttt cat att aat ggg gcg tga cgg aat agt aca gta tga tag aac aca tcc tat tcc caa caa tgt cag gaa cgt cac gct ctc cac ctt cat att tat tta tcc gta aaa tgt tat cct gga cat cgt aca aat aat aaa aag ccc ata tat atg ttc gct att gta gaa att gtt ttt cac agt tgc tca aaa aca atg gca gtg act tat gag tta gtt aca ctt tgg agt ctc atc ttt agt aaa cat atc ata ata ttc gat att acg agt tga cat atc gaa caa att cca agt att tga ttt tgg ata ata ttc gta ttt tgc atc tgc tat aat taa gat ata atc acc aca aga aca cac gaa cgt ctt tcc tac atg gtt aaa gta cat gta caa ttc tat cca ttt gtc ttc ctt aac tat ata ttt gta tag ata att acg agt ctc atg agt aat tcc agt aat tgc ata gat gtc acc atc gta ttc tac agc ata aac tat act atg acg tct agg cat ggg aga ctt ttt tat cca acg att ttt agt gaa aca ttc cac atc gtt taa tac tac ata ttt ctc ata gtg gta taa act cca ccc att aca tat ata tca tcg ttt acg aat act gat gcg cct gaa tat cta gga gtg att aag ttt gga agt ctt ttc cat ttc gaa gtg ccg tgt ttc aaa tat tct gct ata ccc gtt gaa ata gaa aat tct aat cct cct att aca tat aac ttt cca tcg tta aca caa gta cta act tct gat ttt aac gac gac ata tta gta acc gtt ttc ca- --- ttt ttt ttg ttt taa gat cta ccc gcg ata cgg aat aaa cat gtc tat tgt taa tca tgc cgc caa taa tgt ata gac aat tat gta aaa cat ttg cat cat aga att gtc tat ctg tat tac cga cta tcg tcc aat att ctg ttc tag gag agt aat ggg tta ttg tgg ata tat aat cag agt ttt taa tga cta cta tat tat gtt tta tac cat ttc gtg tca cag ctt tgt aga ttt gga tat agt taa tcc caa caa tgc tat agc att gca tat agc att agt cat aaa ctt ggg atg taa aat gtt gat gat atc tac atc gtt tgg att ttt atg tat cca ctt taa taa tat tat agc gta aca tcc tca tga ttt acg tta acg ttt tcg tgt gat aag ata gtg gtc agt tca tcc ttt gat aat ttt cca aat tct gga tcg gat gtc acc gca gta ata ttg ttg att att tct gac atc gac gca tta tat agt ttt tta att cca tat ctt ttt t

>MN702445.1_Monkeypox_virus_strain_A4_contig_SPADES

ttt tat atc act acg gac ata aac cat tgt ata att ttt atg ttt att agt gta cac att ttg gaa gta agt tcc ggc tgc cat gta ttt cct gga gag caa gta gat gat g-a gga acc aga tag ttt ata tcc ata ctt gca ctt aaa gtc tac att gta gtt gta tga gtg tat gat ctt tta agc cgc tag aag ttt tcc gtt tga tat agg atg tgg aca ttt aac aat ctg aca cgt ggg tgg att gga cca ttc tcc tcc tga aca cat gac acc aga gtt acc aat caa cga ata tcc act att gca act ata agt tac aat gct ccc atc gat ata aaa atc ctc gta tcc gtt atg tct tcc gtt gga tat aga tgg agg tga ttg gca ttt aac aga ttc gca aat agg tgc ctc agg att cca tac cat aga tcc agt aga tcc taa ttc aca ata cga ttt aga ttc acc gat caa atg ata tcc gct att aca aga gta cgt tat act aga gcc aaa gtc tac tcc gcc aat atc aag ttg gcc att atc gat atc tcg agg cga tgg gca tct ccg ttt aat aca ttg att aaa gag tgt cca tcc ggt acc ggt aca ttt agc ata tat ggg tcc cat ttt ttg ctt tct gta tcc agg tag aca tag ata ttc tat agt gtc tcc tat gtt gta att agc atc agt ctc tac act att ctt aaa ttt cat att aat ggg gcg tga cgg aat agt aca gta tga tag aac aca tcc tat tcc caa caa tgt cag gaa cgt cac gct ctc cac ctt cat att tat tta tcc gta aaa tgt tat cct gga cat cgt aca aat aat aaa aag ccc ata tat atg ttc gct att gta gaa att gtt ttt cac agt tgc tca aaa aca atg gca gtg act tat gag tta gtt aca ctt tgg agt ctc atc ttt agt aaa cat atc ata ata ttc gat att acg agt tga cat atc gaa caa att cca agt att tga ttt tgg ata ata ttc gta ttt tgc atc tgc tat aat taa gat ata atc acc aca aga aca cac gaa cgt ctt tcc tac atg gtt aaa gta cat gta caa ttc tat cca ttt gtc ttc ctt aac tat ata ttt gta tag ata att acg agt ctc atg agt aat tcc agt aat tgc ata gat gtc acc atc gta ttc tac agc ata aac tat act atg acg tct agg cat ggg aga ctt ttt tat cca acg att ttt agt gaa aca ttc cac atc gtt taa tac tac ata ttt ctc ata gtg gta taa act cca ccc att aca tat ata tca tcg ttt acg aat act gat gcg cct gaa tat cta gga gtg att aag ttt gga agt ctt ttc cat ttc gaa gtg ccg tgt ttc aaa tat tct gct ata ccc gtt gaa ata gaa aat tct aat cct cct att aca tat aac ttt cca tcg tta aca caa gta cta act tct gat ttt aac gac gac ata tta gta acc gtt ttc ca- -tt ttt ttt ttg ttt taa gat cta ccc gcg ata cgg aat aaa cat gtc tat tgt taa tca tgc cgc caa taa tgt ata gac aat tat gta aaa cat ttg cat cat aga att gtc tat ctg tat tac cga cta tcg tcc aat att ctg ttc tag gag agt aat ggg tta ttg tga ata tat aat cag agt ttt taa tga cta cta tat tat gtt tta tac cat ttc gtg tca cag ctt tgt aga ttt gga tat agt taa tcc caa caa tgc tat agc att gca tat agc att agt cat aaa ctt ggg atg taa aat gtt gat gat atc tac atc gtt tgg att ttt ata tat cca ctt taa taa tat tat agc gta aca tcc tca tga ttt acg tta acg ttt tcg tgt gat aag ata gtg gtc agt tca tcc ttt gat aat ttt cca aat tct gga tcg gat gtc acc gca gta ata ttg ttg att att tct gac atc gac aca tta tat agt ttt tta att cca tat ctt ttt t

>MN702444.1_Monkeypox_virus_strain_A5_contig_SPADES

ttt tat atc act acg gac ata aac cat tgt ata att ttt atg ttt att agt gta cac att ttg gaa gta agt tcc ggc tgc cat gta ttt cct gga gag caa gta gat gat g-a gga acc aga tag ttt ata tcc ata ctt gca ctt aaa gtc tac att gta gtt gta tga gtg tat gat ctt tta agc cgc tag aag ttt tcc gtt tga tat agg atg tgg aca ttt aac aat ctg aca cgt ggg tgg att gga cca ttc tcc tcc tga aca cat gac acc aga gtt acc aat caa cga ata tcc act att gca act ata agt tac aat gct ccc atc gat ata aaa atc ctc gta tcc gtt atg tct tcc gtt gga tat aga tgg agg tga ttg gca ttt aac aga ttc gca aat agg tgc ctc agg att cca tac cat aga tcc agt aga tcc taa ttc aca ata cga ttt aga ttc acc gat caa atg ata tcc gct att aca aga gta cgt tat act aga gcc aaa gtc tac tcc gcc aat atc aag ttg gcc att atc gat atc tcg agg cga tgg gca tct ccg ttt aat aca ttg att aaa gag tgt cca tcc ggt acc ggt aca ttt agc ata tat ggg tcc cat ttt ttg ctt tct gta tcc agg tag aca tag ata ttc tat agt gtc tcc tat gtt gta att agc atc agt ctc tac act att ctt aaa ttt cat att aat ggg gcg tga cgg aat agt aca gta tga tag aac aca tcc tat tcc caa caa tgt cag gaa cgt cac gct ctc cac ctt cat att tat tta tcc gta aaa tgt tat cct gga cat cgt aca aat aat aaa aag ccc ata tat atg ttc gct att gta gaa att gtt ttt cac agt tgc tca aaa aca atg gca gtg act tat gag tta gtt aca ctt tgg agt ctc atc ttt agt aaa cat atc ata ata ttc gat att acg agt tga cat atc gaa caa att cca agt att tga ttt tgg ata ata ttc gta ttt tgc atc tgc tat aat taa gat ata atc acc aca aga aca cac gaa cgt ctt tcc tac atg gtt aaa gta cat gta caa ttc tat cca ttt gtc ttc ctt aac tat ata ttt gta tag ata att acg agt ctc atg agt aat tcc agt aat tgc ata gat gtc acc atc gta ttc tac agc ata aac tat act atg acg tct agg cat ggg aga ctt ttt tat cca acg att ttt agt gaa aca ttc cac atc gtt taa tac tac ata ttt ctc ata gtg gta taa act cca ccc att aca tat ata tca tcg ttt acg aat act gat gcg cct gaa tat cta gga gtg att aag ttt gga agt ctt ttc cat ttc gaa gtg ccg tgt ttc aaa tat tct gct ata ccc gtt gaa ata gaa aat tct aat cct cct att aca tat aac ttt cca tcg tta aca caa gta cta act tct gat ttt aac gac gac ata tta gta acc gtt ttc ca- -tt ttt ttt ttg ttt taa gat cta ccc gcg ata cgg aat aaa cat gtc tat tgt taa tca tgc cgc caa taa tgt ata gac aat tat gta aaa cat ttg cat cat aga att gtc tat ctg tat tac cga cta tcg tcc aat att ctg ttc tag gag agt aat ggg tta ttg tga ata tat aat cag agt ttt taa tga cta cta tat tat gtt tta tac cat ttc gtg tca cag ctt tgt aga ttt gga tat agt taa tcc caa caa tgc tat agc att gca tat agc att agt cat aaa ctt ggg atg taa aat gtt gat gat atc tac atc gtt tgg att ttt ata tat cca ctt taa taa tat tat agc gta aca tcc tca tga ttt acg tta acg ttt tcg tgt gat aag ata gtg gtc agt tca tcc ttt gat aat ttt cca aat tct gga tcg gat gtc acc gca gta ata ttg ttg att att tct gac atc gac aca tta tat agt ttt tta att cca tat ctt ttt t

>KJ642613.1_Monkeypox_virus_strain_Congo_8_complete_genome

ttt tat atc act acg gac ata aac cat tgt ata att ttt atg ttt att agt gta cac att ttg gaa gta agt tcc ggc tgc cat gta ttt cct gga gag caa gta gat gat g-a gga acc aga tag ttt ata tcc ata ctt gca ctt aaa gtc tac att gta gtt gta tga gtg tat gat ctt tta agc cgc tag aag ttt tcc gtt tga tat agg atg tgg aca ttt aac aat ctg aca cgt ggg tgg att gga cca ttc tcc tcc tga aca cat gac acc aga gtt acc aat caa cga ata tcc act att gca act ata agt tac aat gct ccc atc gat ata aaa atc ctc gta tcc gtt atg tct tcc gtt gga tat aga tgg agg tga ttg gca ttt aac aga ttc gca aat agg tgc ctc agg att cca tac cat aga tcc agt aga tcc taa ttc aca ata cga ttt aga ttc acc gat caa atg ata tcc gct att aca aga gta cgt tat act aga gcc aaa gtc tac tcc gcc aat atc aag ttg gcc att atc gat atc tcg agg cga tgg gca tct ccg ttt aat aca ttg att aaa gag tgt cca tcc ggt acc ggt aca ttt agc ata tat ggg tcc cat ttt ttg ctt tct gta tcc agg tag aca tag ata ttc tat agt gtc tcc tat gtt gta att agc atc agt ctc tac act att ctt aaa ttt cat att aat ggg gcg tga cgg aat agt aca gta tga tag aac aca tcc tat tcc caa caa tgt cag gaa cgt cac gct ctc cac ctt cat att tat tta tcc gta aaa tgt tat cct gga cat cgt aca aat aat aaa aag ccc ata tat atg ttc gct att gta gaa att gtt ttt cac agt tgc tca aaa aca atg gca gtg act tat gag tta gtt aca ctt tgg agt ctc atc ttt agt aaa cat atc ata ata ttc gat att acg agt tga cat atc gaa caa att cca agt att tga ttt tgg ata ata ttc gta ttt tgc atc tgc tat aat taa gat ata atc acc aca aga aca cac gaa cgt ctt tcc tac atg gtt aaa gta cat gta caa ttc tat cca ttt gtc ttc ctt aac tat ata ttt gta tag ata att acg agt ctc atg agt aat tcc agt aat tgc ata gat gtc acc atc gta ttc tac agc ata aac tat act atg acg tct agg cat ggg aga ctt ttt tat cca acg att ttt agt gaa aca ttc cac atc gtt taa tac tac ata ttt ctc ata gtg gta taa act cca ccc att aca tat ata tca tcg ttt acg aat act gat gcg cct gaa tat cta gga gtg att aag ttt gga agt ctt ttc cat ttc gaa gtg ccg tgt ttc aaa tat tct gct ata ccc gtt gaa ata gaa aat tct aat cct cct att aca tat aac ttt cca tcg tta aca caa gta cta act tct gat ttt aac gac gac ata tta gta acc gtt ttc ca- --- ttt ttt ttg ttt taa gat cta ccc gcg ata cgg aat aaa cat gtc tat tgt taa tca tgc cgc caa taa tgt ata gac aat tat gta aaa cat ttg cat cat aga att gtc tat ctg tat tac cga cta tcg tcc aat att ctg ttc tag gag agt aat ggg tta ttg tgg ata tat aat cag agt ttt taa tga cta cta tat tat gtt tta tac cat ttc gtg tca cag ctt tgt aga ttt gga tat agt taa tcc caa caa tgc tat agc att gca tat agc att agt cat aaa ctt ggg atg taa aat gtt gat gat atc tac atc gtt tgg att ttt atg tat cca ctt taa taa tat tat agc gta aca tcc tca tga ttt acg tta acg ttt tcg tgt gat aag ata gtg gtc agt tca tcc ttt gat aat ttt cca aat tct gga tcg gat gtc acc gca gta ata ttg ttg att att tct gac atc gac gca tta tat agt ttt tta att cca tat ctt ttt t

>HM172544.1_Monkeypox_virus_strain_Zaire_1979-005_complete_genome

ttt tat atc act acg gac ata aac cat tgt ata att ttt atg ttt att agt gta cac att ttg gaa gta agt tcc ggc tgc cat gta ttt cct gga gag caa gta gat gat g-a gga acc aga tag ttt ata tcc ata ctt gta ctt aaa gtc tac att gta gtt gta tga gtg tat gat ctt tta agc cgc tag aag ttt tcc gtt tga tat agg atg tgg aca ttt aac aat ctg aca cgt ggg tgg att gga cca ttc tcc tcc tga aca cat gac acc aga gtt acc aat caa cga ata tcc act att gca act ata agt tac aat gct ccc atc gat ata aaa atc ctc gta tcc gtt atg tct tcc gtt gga tat aga tgg agg tga ttg gca ttt aac aga ttc gca aat agg tgc ctc agg att cca tac cat aga tcc agt aga tcc taa ttc aca ata cga ttt aga ttc acc gat caa atg ata tcc gct att aca aga gta cgt tat act aga gcc aaa gtc tac tcc gcc aat atc aag ttg gcc att atc gat atc tcg agg cga tgg gca tct ccg ttt aat aca ttg att aaa gag tgt cca tcc ggt acc ggt aca ttt agc ata tat ggg tcc cat ttt ttg ctt tct gta tcc agg tag aca tag ata ttc tat agt gtc tcc tat gtt gta att agc atc agt ctc tac act att ctt aaa ttt cat att aat ggg gcg tga cgg aat agt aca gta tga tag aac aca tcc tat tcc caa caa tgt cag gaa cgt cac gct ctc cac ctt cat att tat tta tcc gta aaa tgt tat cct gga cat cgt aca aat aat aaa aag ccc ata tat atg ttc gct att gta gaa att gtt ttt cac agt tgc tca aaa aca atg gca gtg act tat gag tta gtt aca ctt tgg agt ctc atc ttt agt aaa cat atc ata ata ttc gat att acg agt tga cat atc gaa caa att cca agt att tga ttt tgg ata ata ttc gta ttt tgc atc tgc tat aat taa gat ata atc acc aca aga aca cac gaa cgt ctt tcc tac atg gtt aaa gta cat gta caa ttc tat cca ttt gtc ttc ctt aac tat ata ttt gta tag ata att acg agt ctc atg agt aat tcc agt aat tgc ata gat gtc acc atc gta ttc tac agc ata aac tat act atg acg tct agg cat ggg aga ctt ttt tat cca acg att ttt agt gaa aca ttc cac atc gtt taa tac tac ata ttt ctc ata gtg gta taa act cca ccc att aca tat ata tca tcg ttt acg aat act gat gcg cct gaa tat cta gga gtg att aag ttt gga agt ctt ttc cat ttc gaa gtg ccg tgt ttc aaa tat tct gct ata ccc gtt gaa ata gaa aat tct aat cct cct att aca tat aac ttt cca tcg tta aca caa gta cta act tct gat ttt aac gac gac ata tta gta acc gtt ttc ca- --- -tt ttt ttg ttt taa gat cta ccc gcg ata cgg aat aaa cat gtc tat tgt taa tca tgc cgc caa taa tgt ata gac aat tat gta aaa cat ttg cat cat aga att gtc tat ctg tat tac cga cta tcg tcc aat att ctg ttc tag gag agt aat ggg tta ttg tga ata tat aat cag agt ttt taa tga cta cta tat tat gtt tta tac cat ttc gtg tca cag ctt tgt aga ttt gga tat agt taa tcc caa caa tgc tat agc att gca tat agc att agt cat aaa ctt ggg atg taa aat gtt gat gat atc tac atc gtt tgg att ttt atg tat cca ctt taa taa tat tat agc gta aca tcc tca tga ttt acg tta acg ttt tcg tgt gat aag ata gtg gtc agt tca tcc ttt gat aat ttt cca aat tct gga tcg gat gtc acc gca gta ata ttg ttg att att tct gac atc gac gca tta tat agt ttt tta att cca tat ctt ttt t

>AF380138.1_Monkeypox_virus_strain_Zaire-96-I-16_complete_genome

ttt tat atc act acg gac ata aac cat tgt ata att ttt atg ttt att agt gta cac att ttg gaa gta agt tcc ggc tgc cat gta ttt cct gga gag caa gta gat gat g-a gga acc aga tag ttt ata tcc ata ctt gca ctt aaa gtc tac att gta gtt gta tga gag tat gat ctt tta agc cgc tag aag ttt tcc gtt tga tat agg atg tgg aca ttt aac aat ctg aca cgt ggg tgg att gga cca ttc tcc tcc tga aca cat gac acc aga gtt acc aat caa cga ata tcc act att gca act ata agt tac aat gct ccc atc gat ata aaa atc ctc gta tcc gtt atg tct tcc gtt gga tat aga tgg agg tga ttg gca ttt aac aga ttc gca aat agg tgc ctc agg att cca tac cat aga tcc agt aga tcc taa ttc aca ata cga ttt aga ttc acc gat caa atg ata tcc gct att aca aga gta cgt tat act aga gcc aaa gtc tac tcc gcc aat atc aag ttg gcc att atc gat atc tcg agg cga tgg gca tct ccg ttt aat aca ttg att aaa gag tgt cca tcc ggt acc ggt aca ttt agc ata tat ggg tcc cat ttt ttg ctt tct gta tcc agg tag aca tag ata ttc tat agt gtc tcc tat gtt gta att agc atc agt ctc tac act att ctt aaa ttt cat att aat ggg gcg tga cgg aat agt aca gta tga tag aac aca tcc tat tcc caa caa tgt cag gaa cgt cac gct ctc cac ctt cat att tat tta tcc gta aaa tgt tat cct gga cat cgt aca aat aat aaa aag ccc ata tat atg ttc gct att gta gaa att gtt ttt cac agt tgc tca aaa aca atg gca gtg act tat gag tta gtt aca ctt tgg agt ctc atc ttt agt aaa cat atc ata ata ttc gat att acg agt tga cat atc gaa caa att cca agt att tga ttt tgg ata ata ttc gta ttt tgc atc tgc tat aat taa gat ata atc acc aca aga aca cac gaa cgt ctt tcc tac atg gtt aaa gta cat gta caa ttc tat cca ttt gtc ttc ctt aac tat ata ttt gta tag ata att acg agt ctc atg agt aat tcc agt aat tgc ata gat gtc acc atc gta ttc tac agc ata aac tat act atg acg tct agg cat ggg aga ctt ttt tat cca acg att ttt agt gaa aca ttc cac atc gtt taa tac tac ata ttt ctc ata gtg gta taa act cca ccc att aca tat ata tca tcg ttt acg aat act gat gcg cct gaa tat cta gga gtg att aag ttt gga agt ctt ttc cat ttc gaa gtg ccg tgt ttc aaa tat tct gct ata ccc gtt gaa ata gaa aat tct aat cct cct att aca tat aac ttt cca tcg tta aca caa gta cta act tct gat ttt aac gac gac ata tta gta acc gtt ttc ca- --- ttt ttt ttg ttt taa gat cta ccc gcg ata cgg aat aaa cat gtc tat tgt taa tca tgc cgc caa taa tgt ata gac aat tat gta aaa cat ttg cat cat aga att gtc tat ctg tat tac cga cta tcg tcc aat att ctg ttc tag gag agt aat ggg tta ttg tgg ata tat aat cag agt ttt taa tga cta cta tat tat gtt tta tac cat ttc gtg tca cag ctt tgt aga ttt gga tat agt taa tcc caa caa tgc tat agc att gca tat agc att agt cat aaa ctt ggg atg taa aat gtt gat gat atc tac atc gtt tgg att ttt atg tat cca ctt taa taa tat tat agc gta aca tcc tca tga ttt acg tta acg ttt tcg tgt gat aag ata gtg gtc agt tca tcc ttt gat aat ttt cca aat tct gga tcg gat gtc acc gca gta ata ttg ttg att att tct gac atc gac gca tta tat agt ttt tta att cca tat ctt ttt t

>HQ857562.1_Monkeypox_virus_strain_V79-I-005_complete_genome

ttt tat atc act acg gac ata aac cat tgt ata att ttt atg ttt att agt gta cac att ttg gaa gta agt tcc ggc tgc cat gta ttt cct gga gag caa gta gat gat g-a gga acc aga tag ttt ata tcc ata ctt gca ctt aaa gtc tac att gta gtt gta tga gtg tat gat ctt tta agc cgc tag aag ttt tcc gtt tga tat agg atg tgg aca ttt aac aat ctg aca cgt ggg tgg att gga cca ttc tcc tcc tga aca cat gac acc aga gtt acc aat caa cga ata tcc act att gca act ata agt tac aat gct ccc atc gat ata aaa atc ctc gta tcc gtt atg tct tcc gtt ggr tat aga tgg rgg kga ttg gca ttt aac aga ttc gca aat agg tgc ctc agg att cca tac cat aga tcc agt aga tcc taa ttc aca ata cga ttt aga ttc acc gat caa atg ata tcc gct att aca aga gta cgt tat act aga gcc aaa gtc tmc ycc gcc aat atc aag ttg gcc att atc gat atc tcg agg cga tgg gca tct ccg ttt aat aca ttg att aaa gag tgt cca tcc ggt acc ggt aca ttt agc ata tat ggg tcc cat ttt ttg ctt tct gta tcc agg tag aca tag ata ttc tat agt gtc tcc tat gtt gta att agc atc agt ctc tac act att ctt aaa ttt cat att aat ggg gcg tga cgg aat agt aca gta tga tag aac aca tcc tat tcc caa caa tgt cag gaa cgt cac gct ctc cac ctt cat att tat tta tcc gta aaa tgt tat cct gga cat cgt aca aat aat aaa aag ccc ata tat atg ttc gct att gta gaa att gtt ttt cac agt tgc tca aaa aca atg gca gtg act tat gag tta gtt aca ctt tgg agt ctc atc ttt agt aaa cat atc ata ata ttc gat att acg agt tga cat atc gaa caa att cca agt att tga ttt tgg ata ata ttc gta ttt tgc atc tgc tat aat taa gat ata atc acc aca aga aca cac gaa cgt ctt tcc tac atg gtt aaa gta cat gta caa ttc tat cca ttt gtc ttc ctt aac tat ata ttt gta tag ata att acg agt ctc atg agt aat tcc agt aat tgc ata gat gtc acc atc gta ttc tac agc ata aac tat act atg acg tct agg cat ggg aga ctt ttt tat cca acg att ttt agt gaa aca ttc cac atc gtt taa tac tac ata ttt ctc ata gtg gta taa act cca ccc att aca tat ata tca tcg ttt acg aat act gat gcg cct gaa tat cta gga gtg att aag ttt gga agt ctt ttc cat ttc gaa gtg ccg tgt ttc aaa tat tct gct ata ccc gtt gaa ata gaa aat tct aat cct cct att aca tat aac ttt cca tcg tta aca caa gta cta act tct gat ttt aac gac gac ata tta gta acc gtt tty cm- --- wtt ttt tkk ttt taa gat cta ccc gcg ata cgg aat aaa cat gtc tat tgt taa tca tgc cgc caa taa tgt ata gac aat tat gta aaa cat ttg cat cat aga att gtc tat ctg tat tac cga cta tcg tcc aat att ctg ttc tag gag agt aat ggg tta ttg tga ata tat aat cag agt ttt taa tga cta cta tat tat gtt tta tac cat ttc gtg tca cag ctt tgt aga ttt gga tat agt taa tcc caa caa tgc tat agc att gca tat agc att agt cat aaa ctt ggg atg taa aat gtt gat gat atc tac atc gtt tgg att ttt atg tat cca ctt taa taa tat tat agc gta aca tcc tca tga ttt acg tta acg ttt tcg tgt gat aag ata gtg gtc agt tca tcc ttt gat aat ttt cca aat tct gga tcg gat gtc acc gca gta ata ttg ttg att att tct gac atc gac gca tta tat agt ttt tta att cca tat ctt ttt t
