## Supplementary Data 5 for "Genomic regions insertion and deletion in Monkeypox virus causing multi-country outbreak-2022"

>JX878428.1:19173-21126_Monkeypox_virus_isolate_DRC_07-0514_complete_genome

cgg ctg cca tgt att tcc tgg aga gca agt --- aga tga tga agg aac cag ata gtt tat atc cat act tgc act taa agt cta cat tgt agt tgt atg agt gta tga tct ttt aag -cc gct aga -ag ttt tcc gtt tga tat --- agg atg tgg aca ttt aac aat ctg aca cgt ggg tgg att gga cca ttc tcc tcc tga aca cat gac acc aga gtt acc aat caa cga ata tcc act att gca act ata agt tac aat gct ccc atc gat ata aaa atc ctc gta tcc gtt atg tct tcc gtt gga tat aga tgg agg tga ttg gca ttt aac aga ttc gca aat agg tgc ctc agg att cca tac cat aga tcc agt aga tcc taa ttc aca ata cga ttt aga ttc acc gat caa atg ata tcc gct att aca aga gta cgt tat act aga gcc aaa gtc tac tcc gcc aat atc aag ttg gcc att atc gat atc tcg agg cga tgg gca tct ccg ttt aat aca ttg att aaa gag tgt cca tcc ggt acc ggt aca ttt agc ata tat ggg tcc cat ttt ttg ctt tct gta tcc agg tag aca tag a-t att cta tag tgt ctc cta tgt tgt a-- --- --- --- --- --- -at tag cat cag tc- --t cta cac tat tct taa att tca tat taa tgg ggc gtg acg gaa tag tac agt atg ata gaa cac atc cta ttc cca aca atg tca gga acg tca cgc tct cca cct tca tat tta ttt atc cgt --a aaa tgt tat cct gga cat cgt aca aat aat a-a aaa gcc cat ata tat gtt cgc tat tgt aga aat tgt ttt tca cag ttg ctc aaa aac aat ggc agt gac tta tga gtt a-g tta cac ttt gga gtc tca tct tta gta aac ata tca taa tat tcg ata tta cga gtt gac ata tcg aac aaa ttc caa gta ttt gat ttt gga taa tat tcg tat ttt gca tct gct ata att aag ata taa tca cca caa gaa cac acg aac gtc ttt cct aca tgg tta aag tac atg tac aat tct atc cat ttg tct tcc tta act ata tat ttg tat aga taa tta cga gtc tca tga gta att cca gta att gca tag atg tca cca tcg tat tct aca gca taa act ata cta tga cgt cta ggc atg gga gac -tt ttt tat cca acg att ttt agt gaa aca ttc cac atc gtt taa tac tac ata ttt ctc ata -gt ggt ata aac tcc acc cat tac --a tat ata tca tcg ttt acg aat act gat gcg cct gaa tat cta gga gtg att aag ttt gga agt ctt ttc cat ttc gaa gtg ccg tgt ttc aaa tat tct gct ata ccc gtt gaa ata gaa aat tct aat cct cct att aca tat aac ttt cca tcg tta aca caa gta cta act tct gat ttt aac gac gac ata tta gta acc gtt ttc ca- --- --- --- --- --- --- -tt ttt ttt gtt tta aga tct acc cgc -ga tac gga ata aac atg tct att gtt aat cat gcc gcc aat aat gta tag aca att atg taa aac att tgc atc ata gaa ttg --- -tc tat ctg tat tac cga cta tcg tcc aat att ctg ttc tag gag agt aat ggg tta ttg tgg ata tat aat cag agt ttt taa tga cta cta tat tat gtt tta tac cat ttc gtg tca c-a gct ttg tag att tgg ata tag tta atc cca aca atg cta tag cat tgc ata tag cat tag tca taa act tgg gat gta aaa tgt tga tga tat cta cat cgt ttg gat ttt tat gta tcc act tta ata ata tta tag c-g taa cat cct cat ga- ttt acg tta acg ttt tcg tgt gat aag ata gtg gtc agt tca tcc ttt gat aat ttt cca aat t

>JX878427.1:19175-21128_Monkeypox_virus_isolate_DRC_07-0480_complete_genome

cgg ctg cca tgt att tcc tgg aga gca agt --- aga tga tga agg aac cag ata gtt tat atc cat act tgc act taa agt cta cat tgt agt tgt atg agt gta tga tct ttt aag -cc gct aga -ag ttt tcc gtt tga tat --- agg atg tgg aca ttt aac aat ctg aca cgt ggg tgg att gga cca ttc tcc tcc tga aca cat gac acc aga gtt acc aat caa cga ata tcc act att gca act ata agt tac aat gct ccc atc gat ata aaa atc ctc gta tcc gtt atg tct tcc gtt gga tat aga tgg agg tga ttg gca ttt aac aga ttc gca aat agg tgc ctc agg att cca tac cat aga tcc agt aga tcc taa ttc aca ata cga ttt aga ttc acc gat caa atg ata tcc gct att aca aga gta cgt tat act aga gcc aaa gtc tac tcc gcc aat atc aag ttg gcc att atc gat atc tcg agg cga tgg gca tct ccg ttt aat aca ttg att aaa gag tgt cca tcc ggt acc ggt aca ttt agc ata tat ggg tcc cat ttt ttg ctt tct gta tcc agg tag aca tag a-t att cta tag tgt ctc cta tgt tgt a-- --- --- --- --- --- -at tag cat cag tc- --t cta cac tat tct taa att tca tat taa tgg ggc gtg acg gaa tag tac agt atg ata gaa cac atc cta ttc cca aca atg tca gga acg tca cgc tct cca cct tca tat tta ttt atc cgt --a aaa tgt tat cct gga cat cgt aca aat aat a-a aaa gcc cat ata tat gtt cgc tat tgt aga aat tgt ttt tca cag ttg ctc aaa aac aat ggc agt gac tta tga gtt a-g tta cac ttt gga gtc tca tct tta gta aac ata tca taa tat tcg ata tta cga gtt gac ata tcg aac aaa ttc caa gta ttt gat ttt gga taa tat tcg tat ttt gca tct gct ata att aag ata taa tca cca caa gaa cac acg aac gtc ttt cct aca tgg tta aag tac atg tac aat tct atc cat ttg tct tcc tta act ata tat ttg tat aga taa tta cga gtc tca tga gta att cca gta att gca tag atg tca cca tcg tat tct aca gca taa act ata cta tga cgt cta ggc atg gga gac -tt ttt tat cca acg att ttt agt gaa aca ttc cac atc gtt taa tac tac ata ttt ctc ata -gt ggt ata aac tcc acc cat tac --a tat ata tca tcg ttt acg aat act gat gcg cct gaa tat cta gga gtg att aag ttt gga agt ctt ttc cat ttc gaa gtg ccg tgt ttc aaa tat tct gct ata ccc gtt gaa ata gaa aat tct aat cct cct att aca tat aac ttt cca tcg tta aca caa gta cta act tct gat ttt aac gac gac ata tta gta acc gtt ttc ca- --- --- --- --- --- --- -tt ttt ttt gtt tta aga tct acc cgc -ga tac gga ata aac atg tct att gtt aat cat gcc gcc aat aat gta tag aca att atg taa aac att tgc atc ata gaa ttg --- -tc tat ctg tat tac cga cta tcg tcc aat att ctg ttc tag gag agt aat ggg tta ttg tgg ata tat aat cag agt ttt taa tga cta cta tat tat gtt tta tac cat ttc gtg tca c-a gct ttg tag att tgg ata tag tta atc cca aca atg cta tag cat tgc ata tag cat tag tca taa act tgg gat gta aaa tgt tga tga tat cta cat cgt ttg gat ttt tat gta tcc act tta ata ata tta tag c-g taa cat cct cat ga- ttt acg tta acg ttt tcg tgt gat aag ata gtg gtc agt tca tcc ttt gat aat ttt cca aat t

>JX878422.1:19173-21126_Monkeypox_virus_isolate_DRC_07-0287_complete_genome

cgg ctg cca tgt att tcc tgg aga gca agt --- aga tga tga agg aac cag ata gtt tat atc cat act tgc act taa agt cta cat tgt agt tgt atg agt gta tga tct ttt aag -cc gct aga -ag ttt tcc gtt tga tat --- agg atg tgg aca ttt aac aat ctg aca cgt ggg tgg att gga cca ttc tcc tcc tga aca cat gac acc aga gtt acc aat caa cga ata tcc act att gca act ata agt tac aat gct ccc atc gat ata aaa atc ctc gta tcc gtt atg tct tcc gtt gga tat aga tgg agg tga ttg gca ttt aac aga ttc gca aat agg tgc ctc agg att cca tac cat aga tcc agt aga tcc taa ttc aca ata cga ttt aga ttc acc gat caa atg ata tcc gct att aca aga gta cgt tat act aga gcc aaa gtc tac tcc gcc aat atc aag ttg gcc att atc gat atc tcg agg cga tgg gca tct ccg ttt aat aca ttg att aaa gag tgt cca tcc ggt acc ggt aca ttt agc ata tat ggg tcc cat ttt ttg ctt tct gta tcc agg tag aca tag a-t att cta tag tgt ctc cta tgt tgt a-- --- --- --- --- --- -at tag cat cag tc- --t cta cac tat tct taa att tca tat taa tgg ggc gtg acg gaa tag tac agt atg ata gaa cac atc cta ttc cca aca atg tca gga acg tca cgc tct cca cct tca tat tta ttt atc cgt --a aaa tgt tat cct gga cat cgt aca aat aat a-a aaa gcc cat ata tat gtt cgc tat tgt aga aat tgt ttt tca cag ttg ctc aaa aac aat ggc agt gac tta tga gtt a-g tta cac ttt gga gtc tca tct tta gta aac ata tca taa tat tcg ata tta cga gtt gac ata tcg aac aaa ttc caa gta ttt gat ttt gga taa tat tcg tat ttt gca tct gct ata att aag ata taa tca cca caa gaa cac acg aac gtc ttt cct aca tgg tta aag tac atg tac aat tct atc cat ttg tct tcc tta act ata tat ttg tat aga taa tta cga gtc tca tga gta att cca gta att gca tag atg tca cca tcg tat tct aca gca taa act ata cta tga cgt cta ggc atg gga gac -tt ttt tat cca acg att ttt agt gaa aca ttc cac atc gtt taa tac tac ata ttt ctc ata -gt ggt ata aac tcc acc cat tac --a tat ata tca tcg ttt acg aat act gat gcg cct gaa tat cta gga gtg att aag ttt gga agt ctt ttc cat ttc gaa gtg ccg tgt ttc aaa tat tct gct ata ccc gtt gaa ata gaa aat tct aat cct cct att aca tat aac ttt cca tcg tta aca caa gta cta act tct gat ttt aac gac gac ata tta gta acc gtt ttc ca- --- --- --- --- --- --- -tt ttt ttt gtt tta aga tct acc cgc -ga tac gga ata aac atg tct att gtt aat cat gcc gcc aat aat gta tag aca att atg taa aac att tgc atc ata gaa ttg --- -tc tat ctg tat tac cga cta tcg tcc aat att ctg ttc tag gag agt aat ggg tta ttg tgg ata tat aat cag agt ttt taa tga cta cta tat tat gtt tta tac cat ttc gtg tca c-a gct ttg tag att tgg ata tag tta atc cca aca atg cta tag cat tgc ata tag cat tag tca taa act tgg gat gta aaa tgt tga tga tat cta cat cgt ttg gat ttt tat gta tcc act tta ata ata tta tag c-g taa cat cct cat ga- ttt acg tta acg ttt tcg tgt gat aag ata gtg gtc agt tca tcc ttt gat aat ttt cca aat t

>JX878421.1:19385-21338_Monkeypox_virus_isolate_DRC_07-0286_complete_genome

cgg ctg cca tgt att tcc tgg aga gca agt --- aga tga tga agg aac cag ata gtt tat atc cat act tgc act taa agt cta cat tgt agt tgt atg agt gta tga tct ttt aag -cc gct aga -ag ttt tcc gtt tga tat --- agg atg tgg aca ttt aac aat ctg aca cgt ggg tgg att gga cca ttc tcc tcc tga aca cat gac acc aga gtt acc aat caa cga ata tcc act att gca act ata agt tac aat gct ccc atc gat ata aaa atc ctc gta tcc gtt atg tct tcc gtt gga tat aga tgg agg tga ttg gca ttt aac aga ttc gca aat agg tgc ctc agg att cca tac cat aga tcc agt aga tcc taa ttc aca ata cga ttt aga ttc acc gat caa atg ata tcc gct att aca aga gta cgt tat act aga gcc aaa gtc tac tcc gcc aat atc aag ttg gcc att atc gat atc tcg agg cga tgg gca tct ccg ttt aat aca ttg att aaa gag tgt cca tcc ggt acc ggt aca ttt agc ata tat ggg tcc cat ttt ttg ctt tct gta tcc agg tag aca tag a-t att cta tag tgt ctc cta tgt tgt a-- --- --- --- --- --- -at tag cat cag tc- --t cta cac tat tct taa att tca tat taa tgg ggc gtg acg gaa tag tac agt atg ata gaa cac atc cta ttc cca aca atg tca gga acg tca cgc tct cca cct tca tat tta ttt atc cgt --a aaa tgt tat cct gga cat cgt aca aat aat a-a aaa gcc cat ata tat gtt cgc tat tgt aga aat tgt ttt tca cag ttg ctc aaa aac aat ggc agt gac tta tga gtt a-g tta cac ttt gga gtc tca tct tta gta aac ata tca taa tat tcg ata tta cga gtt gac ata tcg aac aaa ttc caa gta ttt gat ttt gga taa tat tcg tat ttt gca tct gct ata att aag ata taa tca cca caa gaa cac acg aac gtc ttt cct aca tgg tta aag tac atg tac aat tct atc cat ttg tct tcc tta act ata tat ttg tat aga taa tta cga gtc tca tga gta att cca gta att gca tag atg tca cca tcg tat tct aca gca taa act ata cta tga cgt cta ggc atg gga gac -tt ttt tat cca acg att ttt agt gaa aca ttc cac atc gtt taa tac tac ata ttt ctc ata -gt ggt ata aac tcc acc cat tac --a tat ata tca tcg ttt acg aat act gat gcg cct gaa tat cta gga gtg att aag ttt gga agt ctt ttc cat ttc gaa gtg ccg tgt ttc aaa tat tct gct ata ccc gtt gaa ata gaa aat tct aat cct cct att aca tat aac ttt cca tcg tta aca caa gta cta act tct gat ttt aac gac gac ata tta gta acc gtt ttc ca- --- --- --- --- --- --- -tt ttt ttt gtt tta aga tct acc cgc -ga tac gga ata aac atg tct att gtt aat cat gcc gcc aat aat gta tag aca att atg taa aac att tgc atc ata gaa ttg --- -tc tat ctg tat tac cga cta tcg tcc aat att ctg ttc tag gag agt aat ggg tta ttg tgg ata tat aat cag agt ttt taa tga cta cta tat tat gtt tta tac cat ttc gtg tca c-a gct ttg tag att tgg ata tag tta atc cca aca atg cta tag cat tgc ata tag cat tag tca taa act tgg gat gta aaa tgt tga tga tat cta cat cgt ttg gat ttt tat gta tcc act tta ata ata tta tag c-g taa cat cct cat ga- ttt acg tta acg ttt tcg tgt gat aag ata gtg gtc agt tca tcc ttt gat aat ttt cca aat t

>JX878416.1:19319-21272_Monkeypox_virus_isolate_DRC_07-0093_complete_genome

cgg ctg cca tgt att tcc tgg aga gca agt --- aga tga tga agg aac cag ata gtt tat atc cat act tgc act taa agt cta cat tgt agt tgt atg agt gta tga tct ttt aag -cc gct aga -ag ttt tcc gtt tga tat --- agg atg tgg aca ttt aac aat ctg aca cgt ggg tgg att gga cca ttc tcc tcc tga aca cat gac acc aga gtt acc aat caa cga ata tcc act att gca act ata agt tac aat gct ccc atc gat ata aaa atc ctc gta tcc gtt atg tct tcc gtt gga tat aga tgg agg tga ttg gca ttt aac aga ttc gca aat agg tgc ctc agg att cca tac cat aga tcc agt aga tcc taa ttc aca ata cga ttt aga ttc acc gat caa atg ata tcc gct att aca aga gta cgt tat act aga gcc aaa gtc tac tcc gcc aat atc aag ttg gcc att atc gat atc tcg agg cga tgg gca tct ccg ttt aat aca ttg att aaa gag tgt cca tcc ggt acc ggt aca ttt agc ata tat ggg tcc cat ttt ttg ctt tct gta tcc agg tag aca tag a-t att cta tag tgt ctc cta tgt tgt a-- --- --- --- --- --- -at tag cat cag tc- --t cta cac tat tct taa att tca tat taa tgg ggc gtg acg gaa tag tac agt atg ata gaa cac atc cta ttc cca aca atg tca gga acg tca cgc tct cca cct tca tat tta ttt atc cgt --a aaa tgt tat cct gga cat cgt aca aat aat a-a aaa gcc cat ata tat gtt cgc tat tgt aga aat tgt ttt tca cag ttg ctc aaa aac aat ggc agt gac tta tga gtt a-g tta cac ttt gga gtc tca tct tta gta aac ata tca taa tat tcg ata tta cga gtt gac ata tcg aac aaa ttc caa gta ttt gat ttt gga taa tat tcg tat ttt gca tct gct ata att aag ata taa tca cca caa gaa cac acg aac gtc ttt cct aca tgg tta aag tac atg tac aat tct atc cat ttg tct tcc tta act ata tat ttg tat aga taa tta cga gtc tca tga gta att cca gta att gca tag atg tca cca tcg tat tct aca gca taa act ata cta tga cgt cta ggc atg gga gac -tt ttt tat cca acg att ttt agt gaa aca ttc cac atc gtt taa tac tac ata ttt ctc ata -gt ggt ata aac tcc acc cat tac --a tat ata tca tcg ttt acg aat act gat gcg cct gaa tat cta gga gtg att aag ttt gga agt ctt ttc cat ttc gaa gtg ccg tgt ttc aaa tat tct gct ata ccc gtt gaa ata gaa aat tct aat cct cct att aca tat aac ttt cca tcg tta aca caa gta cta act tct gat ttt aac gac gac ata tta gta acc gtt ttc ca- --- --- --- --- --- --- -tt ttt ttt gtt tta aga tct acc cgc -ga tac gga ata aac atg tct att gtt aat cat gcc gcc aat aat gta tag aca att atg taa aac att tgc atc ata gaa ttg --- -tc tat ctg tat tac cga cta tcg tcc aat att ctg ttc tag gag agt aat ggg tta ttg tgg ata tat aat cag agt ttt taa tga cta cta tat tat gtt tta tac cat ttc gtg tca c-a gct ttg tag att tgg ata tag tta atc cca aca atg cta tag cat tgc ata tag cat tag tca taa act tgg gat gta aaa tgt tga tga tat cta cat cgt ttg gat ttt tat gta tcc act tta ata ata tta tag c-g taa cat cct cat ga- ttt acg tta acg ttt tcg tgt gat aag ata gtg gtc agt tca tcc ttt gat aat ttt cca aat t

>JX878415.1:19245-21198_Monkeypox_virus_isolate_DRC_07-0092_complete_genome

cgg ctg cca tgt att tcc tgg aga gca agt --- aga tga tga agg aac cag ata gtt tat atc cat act tgc act taa agt cta cat tgt agt tgt atg agt gta tga tct ttt aag -cc gct aga -ag ttt tcc gtt tga tat --- agg atg tgg aca ttt aac aat ctg aca cgt ggg tgg att gga cca ttc tcc tcc tga aca cat gac acc aga gtt acc aat caa cga ata tcc act att gca act ata agt tac aat gct ccc atc gat ata aaa atc ctc gta tcc gtt atg tct tcc gtt gga tat aga tgg agg tga ttg gca ttt aac aga ttc gca aat agg tgc ctc agg att cca tac cat aga tcc agt aga tcc taa ttc aca ata cga ttt aga ttc acc gat caa atg ata tcc gct att aca aga gta cgt tat act aga gcc aaa gtc tac tcc gcc aat atc aag ttg gcc att atc gat atc tcg agg cga tgg gca tct ccg ttt aat aca ttg att aaa gag tgt cca tcc ggt acc ggt aca ttt agc ata tat ggg tcc cat ttt ttg ctt tct gta tcc agg tag aca tag a-t att cta tag tgt ctc cta tgt tgt a-- --- --- --- --- --- -at tag cat cag tc- --t cta cac tat tct taa att tca tat taa tgg ggc gtg acg gaa tag tac agt atg ata gaa cac atc cta ttc cca aca atg tca gga acg tca cgc tct cca cct tca tat tta ttt atc cgt --a aaa tgt tat cct gga cat cgt aca aat aat a-a aaa gcc cat ata tat gtt cgc tat tgt aga aat tgt ttt tca cag ttg ctc aaa aac aat ggc agt gac tta tga gtt a-g tta cac ttt gga gtc tca tct tta gta aac ata tca taa tat tcg ata tta cga gtt gac ata tcg aac aaa ttc caa gta ttt gat ttt gga taa tat tcg tat ttt gca tct gct ata att aag ata taa tca cca caa gaa cac acg aac gtc ttt cct aca tgg tta aag tac atg tac aat tct atc cat ttg tct tcc tta act ata tat ttg tat aga taa tta cga gtc tca tga gta att cca gta att gca tag atg tca cca tcg tat tct aca gca taa act ata cta tga cgt cta ggc atg gga gac -tt ttt tat cca acg att ttt agt gaa aca ttc cac atc gtt taa tac tac ata ttt ctc ata -gt ggt ata aac tcc acc cat tac --a tat ata tca tcg ttt acg aat act gat gcg cct gaa tat cta gga gtg att aag ttt gga agt ctt ttc cat ttc gaa gtg ccg tgt ttc aaa tat tct gct ata ccc gtt gaa ata gaa aat tct aat cct cct att aca tat aac ttt cca tcg tta aca caa gta cta act tct gat ttt aac gac gac ata tta gta acc gtt ttc ca- --- --- --- --- --- --- -tt ttt ttt gtt tta aga tct acc cgc -ga tac gga ata aac atg tct att gtt aat cat gcc gcc aat aat gta tag aca att atg taa aac att tgc atc ata gaa ttg --- -tc tat ctg tat tac cga cta tcg tcc aat att ctg ttc tag gag agt aat ggg tta ttg tgg ata tat aat cag agt ttt taa tga cta cta tat tat gtt tta tac cat ttc gtg tca c-a gct ttg tag att tgg ata tag tta atc cca aca atg cta tag cat tgc ata tag cat tag tca taa act tgg gat gta aaa tgt tga tga tat cta cat cgt ttg gat ttt tat gta tcc act tta ata ata tta tag c-g taa cat cct cat ga- ttt acg tta acg ttt tcg tgt gat aag ata gtg gtc agt tca tcc ttt gat aat ttt cca aat t

>JX878414.1:19457-21410_Monkeypox_virus_isolate_DRC_07-0046_complete_genome

cgg ctg cca tgt att tcc tgg aga gca agt --- aga tga tga agg aac cag ata gtt tat atc cat act tgc act taa agt cta cat tgt agt tgt atg agt gta tga tct ttt aag -cc gct aga -ag ttt tcc gtt tga tat --- agg atg tgg aca ttt aac aat ctg aca cgt ggg tgg att gga cca ttc tcc tcc tga aca cat gac acc aga gtt acc aat caa cga ata tcc act att gca act ata agt tac aat gct ccc atc gat ata aaa atc ctc gta tcc gtt atg tct tcc gtt gga tat aga tgg agg tga ttg gca ttt aac aga ttc gca aat agg tgc ctc agg att cca tac cat aga tcc agt aga tcc taa ttc aca ata cga ttt aga ttc acc gat caa atg ata tcc gct att aca aga gta cgt tat act aga gcc aaa gtc tac tcc gcc aat atc aag ttg gcc att atc gat atc tcg agg cga tgg gca tct ccg ttt aat aca ttg att aaa gag tgt cca tcc ggt acc ggt aca ttt agc ata tat ggg tcc cat ttt ttg ctt tct gta tcc agg tag aca tag a-t att cta tag tgt ctc cta tgt tgt a-- --- --- --- --- --- -at tag cat cag tc- --t cta cac tat tct taa att tca tat taa tgg ggc gtg acg gaa tag tac agt atg ata gaa cac atc cta ttc cca aca atg tca gga acg tca cgc tct cca cct tca tat tta ttt atc cgt --a aaa tgt tat cct gga cat cgt aca aat aat a-a aaa gcc cat ata tat gtt cgc tat tgt aga aat tgt ttt tca cag ttg ctc aaa aac aat ggc agt gac tta tga gtt a-g tta cac ttt gga gtc tca tct tta gta aac ata tca taa tat tcg ata tta cga gtt gac ata tcg aac aaa ttc caa gta ttt gat ttt gga taa tat tcg tat ttt gca tct gct ata att aag ata taa tca cca caa gaa cac acg aac gtc ttt cct aca tgg tta aag tac atg tac aat tct atc cat ttg tct tcc tta act ata tat ttg tat aga taa tta cga gtc tca tga gta att cca gta att gca tag atg tca cca tcg tat tct aca gca taa act ata cta tga cgt cta ggc atg gga gac -tt ttt tat cca acg att ttt agt gaa aca ttc cac atc gtt taa tac tac ata ttt ctc ata -gt ggt ata aac tcc acc cat tac --a tat ata tca tcg ttt acg aat act gat gcg cct gaa tat cta gga gtg att aag ttt gga agt ctt ttc cat ttc gaa gtg ccg tgt ttc aaa tat tct gct ata ccc gtt gaa ata gaa aat tct aat cct cct att aca tat aac ttt cca tcg tta aca caa gta cta act tct gat ttt aac gac gac ata tta gta acc gtt ttc ca- --- --- --- --- --- --- -tt ttt ttt gtt tta aga tct acc cgc -ga tac gga ata aac atg tct att gtt aat cat gcc gcc aat aat gta tag aca att atg taa aac att tgc atc ata gaa ttg --- -tc tat ctg tat tac cga cta tcg tcc aat att ctg ttc tag gag agt aat ggg tta ttg tgg ata tat aat cag agt ttt taa tga cta cta tat tat gtt tta tac cat ttc gtg tca c-a gct ttg tag att tgg ata tag tta atc cca aca atg cta tag cat tgc ata tag cat tag tca taa act tgg gat gta aaa tgt tga tga tat cta cat cgt ttg gat ttt tat gta tcc act tta ata ata tta tag c-g taa cat cct cat ga- ttt acg tta acg ttt tcg tgt gat aag ata gtg gtc agt tca tcc ttt gat aat ttt cca aat t

>JX878413.1:19315-21268_Monkeypox_virus_isolate_DRC_07-0045_complete_genome

cgg ctg cca tgt att tcc tgg aga gca agt --- aga tga tga agg aac cag ata gtt tat atc cat act tgc act taa agt cta cat tgt agt tgt atg agt gta tga tct ttt aag -cc gct aga -ag ttt tcc gtt tga tat --- agg atg tgg aca ttt aac aat ctg aca cgt ggg tgg att gga cca ttc tcc tcc tga aca cat gac acc aga gtt acc aat caa cga ata tcc act att gca act ata agt tac aat gct ccc atc gat ata aaa atc ctc gta tcc gtt atg tct tcc gtt gga tat aga tgg agg tga ttg gca ttt aac aga ttc gca aat agg tgc ctc agg att cca tac cat aga tcc agt aga tcc taa ttc aca ata cga ttt aga ttc acc gat caa atg ata tcc gct att aca aga gta cgt tat act aga gcc aaa gtc tac tcc gcc aat atc aag ttg gcc att atc gat atc tcg agg cga tgg gca tct ccg ttt aat aca ttg att aaa gag tgt cca tcc ggt acc ggt aca ttt agc ata tat ggg tcc cat ttt ttg ctt tct gta tcc agg tag aca tag a-t att cta tag tgt ctc cta tgt tgt a-- --- --- --- --- --- -at tag cat cag tc- --t cta cac tat tct taa att tca tat taa tgg ggc gtg acg gaa tag tac agt atg ata gaa cac atc cta ttc cca aca atg tca gga acg tca cgc tct cca cct tca tat tta ttt atc cgt --a aaa tgt tat cct gga cat cgt aca aat aat a-a aaa gcc cat ata tat gtt cgc tat tgt aga aat tgt ttt tca cag ttg ctc aaa aac aat ggc agt gac tta tga gtt a-g tta cac ttt gga gtc tca tct tta gta aac ata tca taa tat tcg ata tta cga gtt gac ata tcg aac aaa ttc caa gta ttt gat ttt gga taa tat tcg tat ttt gca tct gct ata att aag ata taa tca cca caa gaa cac acg aac gtc ttt cct aca tgg tta aag tac atg tac aat tct atc cat ttg tct tcc tta act ata tat ttg tat aga taa tta cga gtc tca tga gta att cca gta att gca tag atg tca cca tcg tat tct aca gca taa act ata cta tga cgt cta ggc atg gga gac -tt ttt tat cca acg att ttt agt gaa aca ttc cac atc gtt taa tac tac ata ttt ctc ata -gt ggt ata aac tcc acc cat tac --a tat ata tca tcg ttt acg aat act gat gcg cct gaa tat cta gga gtg att aag ttt gga agt ctt ttc cat ttc gaa gtg ccg tgt ttc aaa tat tct gct ata ccc gtt gaa ata gaa aat tct aat cct cct att aca tat aac ttt cca tcg tta aca caa gta cta act tct gat ttt aac gac gac ata tta gta acc gtt ttc ca- --- --- --- --- --- --- -tt ttt ttt gtt tta aga tct acc cgc -ga tac gga ata aac atg tct att gtt aat cat gcc gcc aat aat gta tag aca att atg taa aac att tgc atc ata gaa ttg --- -tc tat ctg tat tac cga cta tcg tcc aat att ctg ttc tag gag agt aat ggg tta ttg tgg ata tat aat cag agt ttt taa tga cta cta tat tat gtt tta tac cat ttc gtg tca c-a gct ttg tag att tgg ata tag tta atc cca aca atg cta tag cat tgc ata tag cat tag tca taa act tgg gat gta aaa tgt tga tga tat cta cat cgt ttg gat ttt tat gta tcc act tta ata ata tta tag c-g taa cat cct cat ga- ttt acg tta acg ttt tcg tgt gat aag ata gtg gtc agt tca tcc ttt gat aat ttt cca aat t

>JX878410.1:19943-21896_Monkeypox_virus_isolate_DRC_06-1070_complete_genome

cgg ctg cca tgt att tcc tgg aga gca agt --- aga tga tga agg aac cag ata gtt tat atc cat act tgc act taa agt cta cat tgt agt tgt atg agt gta tga tct ttt aag -cc gct aga -ag ttt tcc gtt tga tat --- agg atg tgg aca ttt aac aat ctg aca cgt ggg tgg att gga cca ttc tcc tcc tga aca cat gac acc aga gtt acc aat caa cga ata tcc act att gca act ata agt tac aat gct ccc atc gat ata aaa atc ctc gta tcc gtt atg tct tcc gtt gga tat aga tgg agg tga ttg gca ttt aac aga ttc gca aat agg tgc ctc agg att cca tac cat aga tcc agt aga tcc taa ttc aca ata cga ttt aga ttc acc gat caa atg ata tcc gct att aca aga gta cgt tat act aga gcc aaa gtc tac tcc gcc aat atc aag ttg gcc att atc gat atc tcg agg cga tgg gca tct ccg ttt aat aca ttg att aaa gag tgt cca tcc ggt acc ggt aca ttt agc ata tat ggg tcc cat ttt ttg ctt tct gta tcc agg tag aca tag a-t att cta tag tgt ctc cta tgt tgt a-- --- --- --- --- --- -at tag cat cag tc- --t cta cac tat tct taa att tca tat taa tgg ggc gtg acg gaa tag tac agt atg ata gaa cac atc cta ttc cca aca atg tca gga acg tca cgc tct cca cct tca tat tta ttt atc cgt --a aaa tgt tat cct gga cat cgt aca aat aat a-a aaa gcc cat ata tat gtt cgc tat tgt aga aat tgt ttt tca cag ttg ctc aaa aac aat ggc agt gac tta tga gtt a-g tta cac ttt gga gtc tca tct tta gta aac ata tca taa tat tcg ata tta cga gtt gac ata tcg aac aaa ttc caa gta ttt gat ttt gga taa tat tcg tat ttt gca tct gct ata att aag ata taa tca cca caa gaa cac acg aac gtc ttt cct aca tgg tta aag tac atg tac aat tct atc cat ttg tct tcc tta act ata tat ttg tat aga taa tta cga gtc tca tga gta att cca gta att gca tag atg tca cca tcg tat tct aca gca taa act ata cta tga cgt cta ggc atg gga gac -tt ttt tat cca acg att ttt agt gaa aca ttc cac atc gtt taa tac tac ata ttt ctc ata -gt ggt ata aac tcc acc cat tac --a tat ata tca tcg ttt acg aat act gat gcg cct gaa tat cta gga gtg att aag ttt gga agt ctt ttc cat ttc gaa gtg ccg tgt ttc aaa tat tct gct ata ccc gtt gaa ata gaa aat tct aat cct cct att aca tat aac ttt cca tcg tta aca caa gta cta act tct gat ttt aac gac gac ata tta gta acc gtt ttc ca- --- --- --- --- --- --- -tt ttt ttt gtt tta aga tct acc cgc -ga tac gga ata aac atg tct att gtt aat cat gcc gcc aat aat gta tag aca att atg taa aac att tgc atc ata gaa ttg --- -tc tat ctg tat tac cga cta tcg tcc aat att ctg ttc tag gag agt aat ggg tta ttg tgg ata tat aat cag agt ttt taa tga cta cta tat tat gtt tta tac cat ttc gtg tca c-a gct ttg tag att tgg ata tag tta atc cca aca atg cta tag cat tgc ata tag cat tag tca taa act tgg gat gta aaa tgt tga tga tat cta cat cgt ttg gat ttt tat gta tcc act tta ata ata tta tag c-g taa cat cct cat ga- ttt acg tta acg ttt tcg tgt gat aag ata gtg gtc agt tca tcc ttt gat aat ttt cca aat t

>JX878412.1:19871-21824_Monkeypox_virus_isolate_DRC_06-1076_complete_genome

cgg ctg cca tgt att tcc tgg aga gca agt --- aga tga tga agg aac cag ata gtt tat atc cat act tgc act taa agt cta cat tgt agt tgt atg agt gta tga tct ttt aag -cc gct aga -ag ttt tcc gtt tga tat --- agg atg tgg aca ttt aac aat ctg aca cgt ggg tgg att gga cca ttc tcc tcc tga aca cat gac acc aga gtt acc aat caa cga ata tcc act att gca act ata agt tac aat gct ccc atc gat ata aaa atc ctc gta tcc gtt atg tct tcc gtt gga tat aga tgg agg tga ttg gca ttt aac aga ttc gca aat agg tgc ctc agg att cca tac cat aga tcc agt aga tcc taa ttc aca ata cga ttt aga ttc acc gat caa atg ata tcc gct att aca aga gta cgt tat act aga gcc aaa gtc tac tcc gcc aat atc aag ttg gcc att atc gat atc tcg agg cga tgg gca tct ccg ttt aat aca ttg att aaa gag tgt cca tcc ggt acc ggt aca ttt agc ata tat ggg tcc cat ttt ttg ctt tct gta tcc agg tag aca tag a-t att cta tag tgt ctc cta tgt tgt a-- --- --- --- --- --- -at tag cat cag tc- --t cta cac tat tct taa att tca tat taa tgg ggc gtg acg gaa tag tac agt atg ata gaa cac atc cta ttc cca aca atg tca gga acg tca cgc tct cca cct tca tat tta ttt atc cgt --a aaa tgt tat cct gga cat cgt aca aat aat a-a aaa gcc cat ata tat gtt cgc tat tgt aga aat tgt ttt tca cag ttg ctc aaa aac aat ggc agt gac tta tga gtt a-g tta cac ttt gga gtc tca tct tta gta aac ata tca taa tat tcg ata tta cga gtt gac ata tcg aac aaa ttc caa gta ttt gat ttt gga taa tat tcg tat ttt gca tct gct ata att aag ata taa tca cca caa gaa cac acg aac gtc ttt cct aca tgg tta aag tac atg tac aat tct atc cat ttg tct tcc tta act ata tat ttg tat aga taa tta cga gtc tca tga gta att cca gta att gca tag atg tca cca tcg tat tct aca gca taa act ata cta tga cgt cta ggc atg gga gac -tt ttt tat cca acg att ttt agt gaa aca ttc cac atc gtt taa tac tac ata ttt ctc ata -gt ggt ata aac tcc acc cat tac --a tat ata tca tcg ttt acg aat act gat gcg cct gaa tat cta gga gta att aag ttt gga agt ctt ttc cat ttc gaa gtg ccg tgt ttc aaa tat tct gct ata ccc gtt gaa ata gaa aat tct aat cct cct att aca tat aac ttt cca tcg tta aca caa gta cta act tct gat ttt aac gac gac ata tta gta acc gtt ttc ca- --- --- --- --- --- --- -tt ttt ttt gtt tta aga tct acc cgc -ga tac gga ata aac atg tct att gtt aat cat gcc gcc aat aat gta tag aca att atg taa aac att tgc atc ata gaa ttg --- -tc tat ctg tat tac cga cta tcg tcc aat att ctg ttc tag gag agt aat ggg tta ttg tgg ata tat aat cag agt ttt taa tga cta cta tat tat gtt tta tac cat ttc gtg tca c-a gct ttg tag att tgg ata tag tta atc cca aca atg cta tag cat tgc ata tag cat tag tca taa act tgg gat gta aaa tgt tga tga tat cta cat cgt ttg gat ttt tat gta tcc act tta ata ata tta tag c-g taa cat cct cat ga- ttt acg tta acg ttt tcg tgt gat aag ata gtg gtc agt tca tcc ttt gat aat ttt cca aat t

>JX878411.1:19941-21894_Monkeypox_virus_isolate_DRC_06-1075_complete_genome

cgg ctg cca tgt att tcc tgg aga gca agt --- aga tga tga agg aac cag ata gtt tat atc cat act tgc act taa agt cta cat tgt agt tgt atg agt gta tga tct ttt aag -cc gct aga -ag ttt tcc gtt tga tat --- agg atg tgg aca ttt aac aat ctg aca cgt ggg tgg att gga cca ttc tcc tcc tga aca cat gac acc aga gtt acc aat caa cga ata tcc act att gca act ata agt tac aat gct ccc atc gat ata aaa atc ctc gta tcc gtt atg tct tcc gtt gga tat aga tgg agg tga ttg gca ttt aac aga ttc gca aat agg tgc ctc agg att cca tac cat aga tcc agt aga tcc taa ttc aca ata cga ttt aga ttc acc gat caa atg ata tcc gct att aca aga gta cgt tat act aga gcc aaa gtc tac tcc gcc aat atc aag ttg gcc att atc gat atc tcg agg cga tgg gca tct ccg ttt aat aca ttg att aaa gag tgt cca tcc ggt acc ggt aca ttt agc ata tat ggg tcc cat ttt ttg ctt tct gta tcc agg tag aca tag a-t att cta tag tgt ctc cta tgt tgt a-- --- --- --- --- --- -at tag cat cag tc- --t cta cac tat tct taa att tca tat taa tgg ggc gtg acg gaa tag tac agt atg ata gaa cac atc cta ttc cca aca atg tca gga acg tca cgc tct cca cct tca tat tta ttt atc cgt --a aaa tgt tat cct gga cat cgt aca aat aat a-a aaa gcc cat ata tat gtt cgc tat tgt aga aat tgt ttt tca cag ttg ctc aaa aac aat ggc agt gac tta tga gtt a-g tta cac ttt gga gtc tca tct tta gta aac ata tca taa tat tcg ata tta cga gtt gac ata tcg aac aaa ttc caa gta ttt gat ttt gga taa tat tcg tat ttt gca tct gct ata att aag ata taa tca cca caa gaa cac acg aac gtc ttt cct aca tgg tta aag tac atg tac aat tct atc cat ttg tct tcc tta act ata tat ttg tat aga taa tta cga gtc tca tga gta att cca gta att gca tag atg tca cca tcg tat tct aca gca taa act ata cta tga cgt cta ggc atg gga gac -tt ttt tat cca acg att ttt agt gaa aca ttc cac atc gtt taa tac tac ata ttt ctc ata -gt ggt ata aac tcc acc cat tac --a tat ata tca tcg ttt acg aat act gat gcg cct gaa tat cta gga gta att aag ttt gga agt ctt ttc cat ttc gaa gtg ccg tgt ttc aaa tat tct gct ata ccc gtt gaa ata gaa aat tct aat cct cct att aca tat aac ttt cca tcg tta aca caa gta cta act tct gat ttt aac gac gac ata tta gta acc gtt ttc ca- --- --- --- --- --- --- -tt ttt ttt gtt tta aga tct acc cgc -ga tac gga ata aac atg tct att gtt aat cat gcc gcc aat aat gta tag aca att atg taa aac att tgc atc ata gaa ttg --- -tc tat ctg tat tac cga cta tcg tcc aat att ctg ttc tag gag agt aat ggg tta ttg tgg ata tat aat cag agt ttt taa tga cta cta tat tat gtt tta tac cat ttc gtg tca c-a gct ttg tag att tgg ata tag tta atc cca aca atg cta tag cat tgc ata tag cat tag tca taa act tgg gat gta aaa tgt tga tga tat cta cat cgt ttg gat ttt tat gta tcc act tta ata ata tta tag c-g taa cat cct cat ga- ttt acg tta acg ttt tcg tgt gat aag ata gtg gtc agt tca tcc ttt gat aat ttt cca aat t

>JX878409.1:19800-21753_Monkeypox_virus_isolate_DRC_06-0999_complete_genome

cgg ctg cca tgt att tcc tgg aga gca agt --- aga tga tga agg aac cag ata gtt tat atc cat act tgc act taa agt cta cat tgt agt tgt atg agt gta tga tct ttt aag -cc gct aga -ag ttt tcc gtt tga tat --- agg atg tgg aca ttt aac aat ctg aca cgt ggg tgg att gga cca ttc tcc tcc tga aca cat gac acc aga gtt acc aat caa cga ata tcc act att gca act ata agt tac aat gct ccc atc gat ata aaa atc ctc gta tcc gtt atg tct tcc gtt gga tat aga tgg agg tga ttg gca ttt aac aga ttc gca aat agg tgc ctc agg att cca tac cat aga tcc agt aga tcc taa ttc aca ata cga ttt aga ttc acc gat caa atg ata tcc gct att aca aga gta cgt tat act aga gcc aaa gtc tac tcc gcc aat atc aag ttg gcc att atc gat atc tcg agg cga tgg gca tct ccg ttt aat aca ttg att aaa gag tgt cca tcc ggt acc ggt aca ttt agc ata tat ggg tcc cat ttt ttg ctt tct gta tcc agg tag aca tag a-t att cta tag tgt ctc cta tgt tgt a-- --- --- --- --- --- -at tag cat cag tc- --t cta cac tat tct taa att tca tat taa tgg ggc gtg acg gaa tag tac agt atg ata gaa cac atc cta ttc cca aca atg tca gga acg tca cgc tct cca cct tca tat tta ttt atc cgt --a aaa tgt tat cct gga cat cgt aca aat aat a-a aaa gcc cat ata tat gtt cgc tat tgt aga aat tgt ttt tca cag ttg ctc aaa aac aat ggc agt gac tta tga gtt a-g tta cac ttt gga gtc tca tct tta gta aac ata tca taa tat tcg ata tta cga gtt gac ata tcg aac aaa ttc caa gta ttt gat ttt gga taa tat tcg tat ttt gca tct gct ata att aag ata taa tca cca caa gaa cac acg aac gtc ttt cct aca tgg tta aag tac atg tac aat tct atc cat ttg tct tcc tta act ata tat ttg tat aga taa tta cga gtc tca tga gta att cca gta att gca tag atg tca cca tcg tat tct aca gca taa act ata cta tga cgt cta ggc atg gga gac -tt ttt tat cca acg att ttt agt gaa aca ttc cac atc gtt taa tac tac ata ttt ctc ata -gt ggt ata aac tcc acc cat tac --a tat ata tca tcg ttt acg aat act gat gcg cct gaa tat cta gga gta att aag ttt gga agt ctt ttc cat ttc gaa gtg ccg tgt ttc aaa tat tct gct ata ccc gtt gaa ata gaa aat tct aat cct cct att aca tat aac ttt cca tcg tta aca caa gta cta act tct gat ttt aac gac gac ata tta gta acc gtt ttc ca- --- --- --- --- --- --- -tt ttt ttt gtt tta aga tct acc cgc -ga tac gga ata aac atg tct att gtt aat cat gcc gcc aat aat gta tag aca att atg taa aac att tgc atc ata gaa ttg --- -tc tat ctg tat tac cga cta tcg tcc aat att ctg ttc tag gag agt aat ggg tta ttg tgg ata tat aat cag agt ttt taa tga cta cta tat tat gtt tta tac cat ttc gtg tca c-a gct ttg tag att tgg ata tag tta atc cca aca atg cta tag cat tgc ata tag cat tag tca taa act tgg gat gta aaa tgt tga tga tat cta cat cgt ttg gat ttt tat gta tcc act tta ata ata tta tag c-g taa cat cct cat ga- ttt acg tta acg ttt tcg tgt gat aag ata gtg gtc agt tca tcc ttt gat aat ttt cca aat t

>KP849469.1:19265-21217_Monkeypox_virus_isolate_Boende_DRC_2008_complete_genome

cgg ctg cca tgt att tcc tgg aga gca agt --- aga tga tg- agg aac cag ata gtt tat atc cat act tgc act taa agt cta cat tgt agt tgt atg agt gta tga tct ttt aag -cc gct aga -ag ttt tcc gtt tga tat --- agg atg tgg aca ttt aac aat ctg aca cgt ggg tgg att gga cca ttc tcc tcc tga aca cat gac acc aga gtt acc aat caa cga ata tcc act att gca act ata agt tac aat gct ccc atc gat ata aaa atc ctc gta tcc gtt atg tct tcc gtt gga tat aga tgg agg tga ttg gca ttt aac aga ttc gca aat agg tgc ctc agg att cca tac cat aga tcc agt aga tcc taa ttc aca ata cga ttt aga ttc acc gat caa atg ata tcc gct att aca aga gta cgt tat act aga gcc aaa gtc tac tcc gcc aat atc aag ttg gcc att atc gat atc tcg agg cga tgg gca tct ccg ttt aat aca ttg att aaa gag tgt cca tcc ggt acc ggt aca ttt agc ata tat ggg tcc cat ttt ttg ctt tct gta tcc agg tag aca tag a-t att cta tag tgt ctc cta tgt tgt a-- --- --- --- --- --- -at tag cat cag tc- --t cta cac tat tct taa att tca tat taa tgg ggc gtg acg gaa tag tac agt atg ata gaa cac atc cta ttc cca aca atg tca gga acg tca cgc tct cca cct tca tat tta ttt atc cgt --a aaa tgt tat cct gga cat cgt aca aat aat a-a aaa gcc cat ata tat gtt cgc tat tgt aga aat tgt ttt tca cag ttg ctc aaa aac aat ggc agt gac tta tga gtt a-g tta cac ttt gga gtc tca tct tta gta aac ata tca taa tat tcg ata tta cga gtt gac ata tcg aac aaa ttc caa gta ttt gat ttt gga taa tat tcg tat ttt gca tct gct ata att aag ata taa tca cca caa gaa cac acg aac gtc ttt cct aca tgg tta aag tac atg tac aat tct atc cat ttg tct tcc tta act ata tat ttg tat aga taa tta cga gtc tca tga gta att cca gta att gca tag atg tca cca tcg tat tct aca gca taa act ata cta tga cgt cta ggc atg gga gac -tt ttt tat cca acg att ttt agt gaa aca ttc cac atc gtt taa tac tac ata ttt ctc ata -gt ggt ata aac tcc acc cat tac --a tat ata tca tcg ttt acg aat act gat gcg cct gaa tat cta gga gtg att aag ttt gga agt ctt ttc cat ttc gaa gtg ccg tgt ttc aaa tat tct gct ata ccc gtt gaa ata gaa aat tct aat cct cct att aca tat aac ttt cca tcg tta aca caa gta cta act tct gat ttt aac gac gac ata tta gta acc gtt ttc ca- --- --- --- --- --- --- -tt ttt ttt gtt tta aga tct acc cgc -ga tac gga ata aac atg tct att gtt aat cat gcc gcc aat aat gta tag aca att atg taa aac att tgc atc ata gaa ttg --- -tc tat ctg tat tac cga cta tcg tcc aat att ctg ttc tag gag agt aat ggg tta ttg tgg ata tat aat cag agt ttt taa tga cta cta tat tat gtt tta tac cat ttc gtg tca c-a gct ttg tag att tgg ata tag tta atc cca aca atg cta tag cat tgc ata tag cat tag tca taa act tgg gat gta aaa tgt tga tga tat cta cat cgt ttg gat ttt tat gta tcc act tta ata ata tta tag c-g taa cat cct cat ga- ttt acg tta acg ttt tcg tgt gat aag ata gtg gtc agt tca tcc ttt gat aat ttt cca aat t

>KJ642619.1:18900-20852_Monkeypox_virus_strain_Gabon-1988_complete_genome

cgg ctg cca tgt att tcc tgg aga gca agt --- aga tga tg- agg aac cag ata gtt tat atc cat act tgc act taa agt cta cat tgt agt tgt atg agt gta tga tct ttt aag -cc gct aga -ag ttt tcc gtt tga tat --- agg atg tgg aca ttt aac aat ctg aca cgt ggg tgg att gga cca ttc tcc tcc tga aca cat gac acc aga gtt acc aat caa cga ata tcc act att gca act ata agt tac aat gct ccc atc gat ata aaa atc ctc gta tcc gtt atg tct tcc gtt gga tat aga tgg agg tga ttg gca ttt aac aga ttc gca aat agg tgc ctc agg att cca tac cat aga tcc agt aga tcc taa ttc aca ata cga ttt aga ttc acc gat caa atg ata tcc gct att aca aga gta cgt tat act aga gcc aaa gtc tac tcc gcc aat atc aag ttg gcc att atc gat atc tcg agg cga tgg gca tct ccg ttt aat aca ttg att aaa gag tgt cca tcc ggt acc ggt aca ttt agc ata tat ggg tcc cat ttt ttg ctt tct gta tcc agg tag aca tag a-t att cta tag tgt ctc cta tgt tgt a-- --- --- --- --- --- -at tag cat cag tc- --t cta cac tat tct taa att tca tat taa tgg ggc gtg acg gaa tag tac agt atg ata gaa cac atc cta ttc cca aca atg tca gga acg tca cgc tct cca cct tca tat tta ttt atc cgt --a aaa tgt tat cct gga cat cgt aca aat aat a-a aaa gcc cat ata tat gtt cgc tat tgt aga aat tgt ttt tca cag ttg ctc aaa aac aat ggc agt gac tta tga gtt a-g tta cac ttt gga gtc tca tct tta gta aac ata tca taa tat tcg ata tta cga gtt gac ata tcg aac aaa ttc caa gta ttt gat ttt gga taa tat tcg tat ttt gca tct gct ata att aag ata taa tca cca caa gaa cac acg aac gtc ttt cct aca tgg tta aag tac atg tac aat tct atc cat ttg tct tcc tta act ata tat ttg tat aga taa tta cga gtc tca tga gta att cca gta att gca tag atg tca cca tcg tat tct aca gca taa act ata cta tga cgt cta ggc atg gga gac -tt ttt tat cca acg att ttt agt gaa aca ttc cac atc gtt taa tac tac ata ttt ctc ata -gt ggt ata aac tcc acc cat tac --a tat ata tca tcg ttt acg aat act gat gcg cct gaa tat cta gga gtg att aag ttt gga agt ctt ttc cat ttc gaa gtg ccg tgt ttc aaa tat tct gct ata ccc gtt gaa ata gaa aat tct aat cct cct att aca tat aac ttt cca tcg tta aca caa gta cta act tct gat ttt aac gac gac ata tta gta acc gtt ttc ca- --- --- --- --- --- --- -tt ttt ttt gtt tta aga tct acc cgc -ga tac gga ata aac atg tct att gtt aat cat gcc gcc aat aat gta tag aca att atg taa aac att tgc atc ata gaa ttg --- -tc tat ctg tat tac cga cta tcg tcc aat att ctg ttc tag gag agt aat ggg tta ttg tgg ata tat aat cag agt ttt taa tga cta cta tat tat gtt tta tac cat ttc gtg tca c-a gct ttg tag att tgg ata tag tta atc cca aca atg cta tag cat tgc ata tag cat tag tca taa act tgg gat gta aaa tgt tga tga tat cta cat cgt ttg gat ttt tat gta tcc act tta ata ata tta tag c-g taa cat cct cat ga- ttt acg tta acg ttt tcg tgt gat aag ata gtg gtc agt tca tcc ttt gat aat ttt cca aat t

>KJ642618.1:16354-18306_Monkeypox_virus_strain_Cameroon-1990_complete_genome

cgg ctg cca tgt att tcc tgg aga gca agt --- aga tga tg- agg aac cag ata gtt tat atc cat act tgc act taa agt cta cat tgt agt tgt atg agt gta tga tct ttt aag -cc gct aga -ag ttt tcc gtt tga tat --- agg atg tgg aca ttt aac aat ctg aca cgt ggg tgg att gga cca ttc tcc tcc tga aca cat gac acc aga gtt acc aat caa cga ata tcc act att gca act ata agt tac aat gct ccc atc gat ata aaa atc ctc gta tcc gtt atg tct tcc gtt gga tat aga tgg agg tga ttg gca ttt aac aga ttc gca aat agg tgc ctc agg att cca tac cat aga tcc agt aga tcc taa ttc aca ata cga ttt aga ttc acc gat caa atg ata tcc gct att aca aga gta cgt tat act aga gcc aaa gtc tac tcc gcc aat atc aag ttg gcc att atc gat atc tcg agg cga tgg gca tct ccg ttt aat aca ttg att aaa gag tgt cca tcc ggt acc ggt aca ttt agc ata tat ggg tcc cat ttt ttg ctt tct gta tcc agg tag aca tag a-t att cta tag tgt ctc cta tgt tgt a-- --- --- --- --- --- -at tag cat cag tc- --t cta cac tat tct taa att tca tat taa tgg ggc gtg acg gaa tag tac agt atg ata gaa cac atc cta ttc cca aca atg tca gga acg tca cgc tct cca cct tca tat tta ttt atc cgt --a aaa tgt tat cct gga cat cgt aca aat aat a-a aaa gcc cat ata tat gtt cgc tat tgt aga aat tgt ttt tca cag ttg ctc aaa aac aat ggc agt gac tta tga gtt a-g tta cac ttt gga gtc tca tct tta gta aac ata tca taa tat tcg ata tta cga gtt gac ata tcg aac aaa ttc caa gta ttt gat ttt gga taa tat tcg tat ttt gca tct gct ata att aag ata taa tca cca caa gaa cac acg aac gtc ttt cct aca tgg tta aag tac atg tac aat tct atc cat ttg tct tcc tta act ata tat ttg tat aga taa tta cga gtc tca tga gta att cca gta att gca tag atg tca cca tcg tat tct aca gca taa act ata cta tga cgt cta ggc atg gga gac -tt ttt tat cca acg att ttt agt gaa aca ttc cac atc gtt taa tac tac ata ttt ctc ata -gt ggt ata aac tcc acc cat tac --a tat ata tca tcg ttt acg aat act gat gcg cct gaa tat cta gga gtg att aag ttt gga agt ctt ttc cat ttc gaa gtg ccg tgt ttc aaa tat tct gct ata ccc gtt gaa ata gaa aat tct aat cct cct att aca tat aac ttt cca tcg tta aca caa gta cta act tct gat ttt aac gac gac ata tta gta acc gtt ttc ca- --- --- --- --- --- --- -tt ttt ttt gtt tta aga tct acc cgc -ga tac gga ata aac atg tct att gtt aat cat gcc gcc aat aat gta tag aca att atg taa aac att tgc atc ata gaa ttg --- -tc tat ctg tat tac cga cta tcg tcc aat att ctg ttc tag gag agt aat ggg tta ttg tgg ata tat aat cag agt ttt taa tga cta cta tat tat gtt tta tac cat ttc gtg tca c-a gct ttg tag att tgg ata tag tta atc cca aca atg cta tag cat tgc ata tag cat tag tca taa act tgg gat gta aaa tgt tga tga tat cta cat cgt ttg gat ttt tat gta tcc act tta ata ata tta tag c-g taa cat cct cat ga- ttt acg tta acg ttt tcg tgt gat aag ata gtg gtc agt tca tcc ttt gat aat ttt cca aat t

>KJ642613.1:18764-20716_Monkeypox_virus_strain_Congo_8_complete_genome

cgg ctg cca tgt att tcc tgg aga gca agt --- aga tga tg- agg aac cag ata gtt tat atc cat act tgc act taa agt cta cat tgt agt tgt atg agt gta tga tct ttt aag -cc gct aga -ag ttt tcc gtt tga tat --- agg atg tgg aca ttt aac aat ctg aca cgt ggg tgg att gga cca ttc tcc tcc tga aca cat gac acc aga gtt acc aat caa cga ata tcc act att gca act ata agt tac aat gct ccc atc gat ata aaa atc ctc gta tcc gtt atg tct tcc gtt gga tat aga tgg agg tga ttg gca ttt aac aga ttc gca aat agg tgc ctc agg att cca tac cat aga tcc agt aga tcc taa ttc aca ata cga ttt aga ttc acc gat caa atg ata tcc gct att aca aga gta cgt tat act aga gcc aaa gtc tac tcc gcc aat atc aag ttg gcc att atc gat atc tcg agg cga tgg gca tct ccg ttt aat aca ttg att aaa gag tgt cca tcc ggt acc ggt aca ttt agc ata tat ggg tcc cat ttt ttg ctt tct gta tcc agg tag aca tag a-t att cta tag tgt ctc cta tgt tgt a-- --- --- --- --- --- -at tag cat cag tc- --t cta cac tat tct taa att tca tat taa tgg ggc gtg acg gaa tag tac agt atg ata gaa cac atc cta ttc cca aca atg tca gga acg tca cgc tct cca cct tca tat tta ttt atc cgt --a aaa tgt tat cct gga cat cgt aca aat aat a-a aaa gcc cat ata tat gtt cgc tat tgt aga aat tgt ttt tca cag ttg ctc aaa aac aat ggc agt gac tta tga gtt a-g tta cac ttt gga gtc tca tct tta gta aac ata tca taa tat tcg ata tta cga gtt gac ata tcg aac aaa ttc caa gta ttt gat ttt gga taa tat tcg tat ttt gca tct gct ata att aag ata taa tca cca caa gaa cac acg aac gtc ttt cct aca tgg tta aag tac atg tac aat tct atc cat ttg tct tcc tta act ata tat ttg tat aga taa tta cga gtc tca tga gta att cca gta att gca tag atg tca cca tcg tat tct aca gca taa act ata cta tga cgt cta ggc atg gga gac -tt ttt tat cca acg att ttt agt gaa aca ttc cac atc gtt taa tac tac ata ttt ctc ata -gt ggt ata aac tcc acc cat tac --a tat ata tca tcg ttt acg aat act gat gcg cct gaa tat cta gga gtg att aag ttt gga agt ctt ttc cat ttc gaa gtg ccg tgt ttc aaa tat tct gct ata ccc gtt gaa ata gaa aat tct aat cct cct att aca tat aac ttt cca tcg tta aca caa gta cta act tct gat ttt aac gac gac ata tta gta acc gtt ttc ca- --- --- --- --- --- --- -tt ttt ttt gtt tta aga tct acc cgc -ga tac gga ata aac atg tct att gtt aat cat gcc gcc aat aat gta tag aca att atg taa aac att tgc atc ata gaa ttg --- -tc tat ctg tat tac cga cta tcg tcc aat att ctg ttc tag gag agt aat ggg tta ttg tgg ata tat aat cag agt ttt taa tga cta cta tat tat gtt tta tac cat ttc gtg tca c-a gct ttg tag att tgg ata tag tta atc cca aca atg cta tag cat tgc ata tag cat tag tca taa act tgg gat gta aaa tgt tga tga tat cta cat cgt ttg gat ttt tat gta tcc act tta ata ata tta tag c-g taa cat cct cat ga- ttt acg tta acg ttt tcg tgt gat aag ata gtg gtc agt tca tcc ttt gat aat ttt cca aat t

>JX878429.1:19315-21267_Monkeypox_virus_isolate_DRC_07-0662_complete_genome

cgg ctg cca tgt att tcc tgg aga gca agt --- aga tga tg- agg aac cag ata gtt tat atc cat act tgc act taa agt cta cat tgt agt tgt atg agt gta tga tct ttt aag -cc gct aga -ag ttt tcc gtt tga tat --- agg atg tgg aca ttt aac aat ctg aca cgt ggg tgg att gga cca ttc tcc tcc tga aca cat gac acc aga gtt acc aat caa cga ata tcc act att gca act ata agt tac aat gct ccc atc gat ata aaa atc ctc gta tcc gtt atg tct tcc gtt gga tat aga tgg agg tga ttg gca ttt aac aga ttc gca aat agg tgc ctc agg att cca tac cat aga tcc agt aga tcc taa ttc aca ata cga ttt aga ttc acc gat caa atg ata tcc gct att aca aga gta cgt tat act aga gcc aaa gtc tac tcc gcc aat atc aag ttg gcc att atc gat atc tcg agg cga tgg gca tct ccg ttt aat aca ttg att aaa gag tgt cca tcc ggt acc ggt aca ttt agc ata tat ggg tcc cat ttt ttg ctt tct gta tcc agg tag aca tag a-t att cta tag tgt ctc cta tgt tgt a-- --- --- --- --- --- -at tag cat cag tc- --t cta cac tat tct taa att tca tat taa tgg ggc gtg acg gaa tag tac agt atg ata gaa cac atc cta ttc cca aca atg tca gga acg tca cgc tct cca cct tca tat tta ttt atc cgt --a aaa tgt tat cct gga cat cgt aca aat aat a-a aaa gcc cat ata tat gtt cgc tat tgt aga aat tgt ttt tca cag ttg ctc aaa aac aat ggc agt gac tta tga gtt a-g tta cac ttt gga gtc tca tct tta gta aac ata tca taa tat tcg ata tta cga gtt gac ata tcg aac aaa ttc caa gta ttt gat ttt gga taa tat tcg tat ttt gca tct gct ata att aag ata taa tca cca caa gaa cac acg aac gtc ttt cct aca tgg tta aag tac atg tac aat tct atc cat ttg tct tcc tta act ata tat ttg tat aga taa tta cga gtc tca tga gta att cca gta att gca tag atg tca cca tcg tat tct aca gca taa act ata cta tga cgt cta ggc atg gga gac -tt ttt tat cca acg att ttt agt gaa aca ttc cac atc gtt taa tac tac ata ttt ctc ata -gt ggt ata aac tcc acc cat tac --a tat ata tca tcg ttt acg aat act gat gcg cct gaa tat cta gga gtg att aag ttt gga agt ctt ttc cat ttc gaa gtg ccg tgt ttc aaa tat tct gct ata ccc gtt gaa ata gaa aat tct aat cct cct att aca tat aac ttt cca tcg tta aca caa gta cta act tct gat ttt aac gac gac ata tta gta acc gtt ttc ca- --- --- --- --- --- --- -tt ttt ttt gtt tta aga tct acc cgc -ga tac gga ata aac atg tct att gtt aat cat gcc gcc aat aat gta tag aca att atg taa aac att tgc atc ata gaa ttg --- -tc tat ctg tat tac cga cta tcg tcc aat att ctg ttc tag gag agt aat ggg tta ttg tgg ata tat aat cag agt ttt taa tga cta cta tat tat gtt tta tac cat ttc gtg tca c-a gct ttg tag att tgg ata tag tta atc cca aca atg cta tag cat tgc ata tag cat tag tca taa act tgg gat gta aaa tgt tga tga tat cta cat cgt ttg gat ttt tat gta tcc act tta ata ata tta tag c-g taa cat cct cat ga- ttt acg tta acg ttt tcg tgt gat aag ata gtg gtc agt tca tcc ttt gat aat ttt cca aat t

>JX878425.1:19455-21407_Monkeypox_virus_isolate_DRC_07-0354_complete_genome

cgg ctg cca tgt att tcc tgg aga gca agt --- aga tga tg- agg aac cag ata gtt tat atc cat act tgc act taa agt cta cat tgt agt tgt atg agt gta tga tct ttt aag -cc gct aga -ag ttt tcc gtt tga tat --- agg atg tgg aca ttt aac aat ctg aca cgt ggg tgg att gga cca ttc tcc tcc tga aca cat gac acc aga gtt acc aat caa cga ata tcc act att gca act ata agt tac aat gct ccc atc gat ata aaa atc ctc gta tcc gtt atg tct tcc gtt gga tat aga tgg agg tga ttg gca ttt aac aga ttc gca aat agg tgc ctc agg att cca tac cat aga tcc agt aga tcc taa ttc aca ata cga ttt aga ttc acc gat caa atg ata tcc gct att aca aga gta cgt tat act aga gcc aaa gtc tac tcc gcc aat atc aag ttg gcc att atc gat atc tcg agg cga tgg gca tct ccg ttt aat aca ttg att aaa gag tgt cca tcc ggt acc ggt aca ttt agc ata tat ggg tcc cat ttt ttg ctt tct gta tcc agg tag aca tag a-t att cta tag tgt ctc cta tgt tgt a-- --- --- --- --- --- -at tag cat cag tc- --t cta cac tat tct taa att tca tat taa tgg ggc gtg acg gaa tag tac agt atg ata gaa cac atc cta ttc cca aca atg tca gga acg tca cgc tct cca cct tca tat tta ttt atc cgt --a aaa tgt tat cct gga cat cgt aca aat aat a-a aaa gcc cat ata tat gtt cgc tat tgt aga aat tgt ttt tca cag ttg ctc aaa aac aat ggc agt gac tta tga gtt a-g tta cac ttt gga gtc tca tct tta gta aac ata tca taa tat tcg ata tta cga gtt gac ata tcg aac aaa ttc caa gta ttt gat ttt gga taa tat tcg tat ttt gca tct gct ata att aag ata taa tca cca caa gaa cac acg aac gtc ttt cct aca tgg tta aag tac atg tac aat tct atc cat ttg tct tcc tta act ata tat ttg tat aga taa tta cga gtc tca tga gta att cca gta att gca tag atg tca cca tcg tat tct aca gca taa act ata cta tga cgt cta ggc atg gga gac -tt ttt tat cca acg att ttt agt gaa aca ttc cac atc gtt taa tac tac ata ttt ctc ata -gt ggt ata aac tcc acc cat tac --a tat ata tca tcg ttt acg aat act gat gcg cct gaa tat cta gga gtg att aag ttt gga agt ctt ttc cat ttc gaa gtg ccg tgt ttc aaa tat tct gct ata ccc gtt gaa ata gaa aat tct aat cct cct att aca tat aac ttt cca tcg tta aca caa gta cta act tct gat ttt aac gac gac ata tta gta acc gtt ttc ca- --- --- --- --- --- --- -tt ttt ttt gtt tta aga tct acc cgc -ga tac gga ata aac atg tct att gtt aat cat gcc gcc aat aat gta tag aca att atg taa aac att tgc atc ata gaa ttg --- -tc tat ctg tat tac cga cta tcg tcc aat att ctg ttc tag gag agt aat ggg tta ttg tgg ata tat aat cag agt ttt taa tga cta cta tat tat gtt tta tac cat ttc gtg tca c-a gct ttg tag att tgg ata tag tta atc cca aca atg cta tag cat tgc ata tag cat tag tca taa act tgg gat gta aaa tgt tga tga tat cta cat cgt ttg gat ttt tat gta tcc act tta ata ata tta tag c-g taa cat cct cat ga- ttt acg tta acg ttt tcg tgt gat aag ata gtg gtc agt tca tcc ttt gat aat ttt cca aat t

>JX878424.1:19100-21052_Monkeypox_virus_isolate_DRC_07-0338_complete_genome

cgg ctg cca tgt att tcc tgg aga gca agt --- aga tga tg- agg aac cag ata gtt tat atc cat act tgc act taa agt cta cat tgt agt tgt atg agt gta tga tct ttt aag -cc gct aga -ag ttt tcc gtt tga tat --- agg atg tgg aca ttt aac aat ctg aca cgt ggg tgg att gga cca ttc tcc tcc tga aca cat gac acc aga gtt acc aat caa cga ata tcc act att gca act ata agt tac aat gct ccc atc gat ata aaa atc ctc gta tcc gtt atg tct tcc gtt gga tat aga tgg agg tga ttg gca ttt aac aga ttc gca aat agg tgc ctc agg att cca tac cat aga tcc agt aga tcc taa ttc aca ata cga ttt aga ttc acc gat caa atg ata tcc gct att aca aga gta cgt tat act aga gcc aaa gtc tac tcc gcc aat atc aag ttg gcc att atc gat atc tcg agg cga tgg gca tct ccg ttt aat aca ttg att aaa gag tgt cca tcc ggt acc ggt aca ttt agc ata tat ggg tcc cat ttt ttg ctt tct gta tcc agg tag aca tag a-t att cta tag tgt ctc cta tgt tgt a-- --- --- --- --- --- -at tag cat cag tc- --t cta cac tat tct taa att tca tat taa tgg ggc gtg acg gaa tag tac agt atg ata gaa cac atc cta ttc cca aca atg tca gga acg tca cgc tct cca cct tca tat tta ttt atc cgt --a aaa tgt tat cct gga cat cgt aca aat aat a-a aaa gcc cat ata tat gtt cgc tat tgt aga aat tgt ttt tca cag ttg ctc aaa aac aat ggc agt gac tta tga gtt a-g tta cac ttt gga gtc tca tct tta gta aac ata tca taa tat tcg ata tta cga gtt gac ata tcg aac aaa ttc caa gta ttt gat ttt gga taa tat tcg tat ttt gca tct gct ata att aag ata taa tca cca caa gaa cac acg aac gtc ttt cct aca tgg tta aag tac atg tac aat tct atc cat ttg tct tcc tta act ata tat ttg tat aga taa tta cga gtc tca tga gta att cca gta att gca tag atg tca cca tcg tat tct aca gca taa act ata cta tga cgt cta ggc atg gga gac -tt ttt tat cca acg att ttt agt gaa aca ttc cac atc gtt taa tac tac ata ttt ctc ata -gt ggt ata aac tcc acc cat tac --a tat ata tca tcg ttt acg aat act gat gcg cct gaa tat cta gga gtg att aag ttt gga agt ctt ttc cat ttc gaa gtg ccg tgt ttc aaa tat tct gct ata ccc gtt gaa ata gaa aat tct aat cct cct att aca tat aac ttt cca tcg tta aca caa gta cta act tct gat ttt aac gac gac ata tta gta acc gtt ttc ca- --- --- --- --- --- --- -tt ttt ttt gtt tta aga tct acc cgc -ga tac gga ata aac atg tct att gtt aat cat gcc gcc aat aat gta tag aca att atg taa aac att tgc atc ata gaa ttg --- -tc tat ctg tat tac cga cta tcg tcc aat att ctg ttc tag gag agt aat ggg tta ttg tgg ata tat aat cag agt ttt taa tga cta cta tat tat gtt tta tac cat ttc gtg tca c-a gct ttg tag att tgg ata tag tta atc cca aca atg cta tag cat tgc ata tag cat tag tca taa act tgg gat gta aaa tgt tga tga tat cta cat cgt ttg gat ttt tat gta tcc act tta ata ata tta tag c-g taa cat cct cat ga- ttt acg tta acg ttt tcg tgt gat aag ata gtg gtc agt tca tcc ttt gat aat ttt cca aat t

>JX878423.1:19171-21123_Monkeypox_virus_isolate_DRC_07-0337_complete_genome

cgg ctg cca tgt att tcc tgg aga gca agt --- aga tga tg- agg aac cag ata gtt tat atc cat act tgc act taa agt cta cat tgt agt tgt atg agt gta tga tct ttt aag -cc gct aga -ag ttt tcc gtt tga tat --- agg atg tgg aca ttt aac aat ctg aca cgt ggg tgg att gga cca ttc tcc tcc tga aca cat gac acc aga gtt acc aat caa cga ata tcc act att gca act ata agt tac aat gct ccc atc gat ata aaa atc ctc gta tcc gtt atg tct tcc gtt gga tat aga tgg agg tga ttg gca ttt aac aga ttc gca aat agg tgc ctc agg att cca tac cat aga tcc agt aga tcc taa ttc aca ata cga ttt aga ttc acc gat caa atg ata tcc gct att aca aga gta cgt tat act aga gcc aaa gtc tac tcc gcc aat atc aag ttg gcc att atc gat atc tcg agg cga tgg gca tct ccg ttt aat aca ttg att aaa gag tgt cca tcc ggt acc ggt aca ttt agc ata tat ggg tcc cat ttt ttg ctt tct gta tcc agg tag aca tag a-t att cta tag tgt ctc cta tgt tgt a-- --- --- --- --- --- -at tag cat cag tc- --t cta cac tat tct taa att tca tat taa tgg ggc gtg acg gaa tag tac agt atg ata gaa cac atc cta ttc cca aca atg tca gga acg tca cgc tct cca cct tca tat tta ttt atc cgt --a aaa tgt tat cct gga cat cgt aca aat aat a-a aaa gcc cat ata tat gtt cgc tat tgt aga aat tgt ttt tca cag ttg ctc aaa aac aat ggc agt gac tta tga gtt a-g tta cac ttt gga gtc tca tct tta gta aac ata tca taa tat tcg ata tta cga gtt gac ata tcg aac aaa ttc caa gta ttt gat ttt gga taa tat tcg tat ttt gca tct gct ata att aag ata taa tca cca caa gaa cac acg aac gtc ttt cct aca tgg tta aag tac atg tac aat tct atc cat ttg tct tcc tta act ata tat ttg tat aga taa tta cga gtc tca tga gta att cca gta att gca tag atg tca cca tcg tat tct aca gca taa act ata cta tga cgt cta ggc atg gga gac -tt ttt tat cca acg att ttt agt gaa aca ttc cac atc gtt taa tac tac ata ttt ctc ata -gt ggt ata aac tcc acc cat tac --a tat ata tca tcg ttt acg aat act gat gcg cct gaa tat cta gga gtg att aag ttt gga agt ctt ttc cat ttc gaa gtg ccg tgt ttc aaa tat tct gct ata ccc gtt gaa ata gaa aat tct aat cct cct att aca tat aac ttt cca tcg tta aca caa gta cta act tct gat ttt aac gac gac ata tta gta acc gtt ttc ca- --- --- --- --- --- --- -tt ttt ttt gtt tta aga tct acc cgc -ga tac gga ata aac atg tct att gtt aat cat gcc gcc aat aat gta tag aca att atg taa aac att tgc atc ata gaa ttg --- -tc tat ctg tat tac cga cta tcg tcc aat att ctg ttc tag gag agt aat ggg tta ttg tgg ata tat aat cag agt ttt taa tga cta cta tat tat gtt tta tac cat ttc gtg tca c-a gct ttg tag att tgg ata tag tta atc cca aca atg cta tag cat tgc ata tag cat tag tca taa act tgg gat gta aaa tgt tga tga tat cta cat cgt ttg gat ttt tat gta tcc act tta ata ata tta tag c-g taa cat cct cat ga- ttt acg tta acg ttt tcg tgt gat aag ata gtg gtc agt tca tcc ttt gat aat ttt cca aat t

>JX878407.1:19100-21052_Monkeypox_virus_isolate_DRC_06-0950_complete_genome

cgg ctg cca tgt att tcc tgg aga gca agt --- aga tga tg- agg aac cag ata gtt tat atc cat act tgc act taa agt cta cat tgt agt tgt atg agt gta tga tct ttt aag -cc gct aga -ag ttt tcc gtt tga tat --- agg atg tgg aca ttt aac aat ctg aca cgt ggg tgg att gga cca ttc tcc tcc tga aca cat gac acc aga gtt acc aat caa cga ata tcc act att gca act ata agt tac aat gct ccc atc gat ata aaa atc ctc gta tcc gtt atg tct tcc gtt gga tat aga tgg agg tga ttg gca ttt aac aga ttc gca aat agg tgc ctc agg att cca tac cat aga tcc agt aga tcc taa ttc aca ata cga ttt aga ttc acc gat caa atg ata tcc gct att aca aga gta cgt tat act aga gcc aaa gtc tac tcc gcc aat atc aag ttg gcc att atc gat atc tcg agg cga tgg gca tct ccg ttt aat aca ttg att aaa gag tgt cca tcc ggt acc ggt aca ttt agc ata tat ggg tcc cat ttt ttg ctt tct gta tcc agg tag aca tag a-t att cta tag tgt ctc cta tgt tgt a-- --- --- --- --- --- -at tag cat cag tc- --t cta cac tat tct taa att tca tat taa tgg ggc gtg acg gaa tag tac agt atg ata gaa cac atc cta ttc cca aca atg tca gga acg tca cgc tct cca cct tca tat tta ttt atc cgt --a aaa tgt tat cct gga cat cgt aca aat aat a-a aaa gcc cat ata tat gtt cgc tat tgt aga aat tgt ttt tca cag ttg ctc aaa aac aat ggc agt gac tta tga gtt a-g tta cac ttt gga gtc tca tct tta gta aac ata tca taa tat tcg ata tta cga gtt gac ata tcg aac aaa ttc caa gta ttt gat ttt gga taa tat tcg tat ttt gca tct gct ata att aag ata taa tca cca caa gaa cac acg aac gtc ttt cct aca tgg tta aag tac atg tac aat tct atc cat ttg tct tcc tta act ata tat ttg tat aga taa tta cga gtc tca tga gta att cca gta att gca tag atg tca cca tcg tat tct aca gca taa act ata cta tga cgt cta ggc atg gga gac -tt ttt tat cca acg att ttt agt gaa aca ttc cac atc gtt taa tac tac ata ttt ctc ata -gt ggt ata aac tcc acc cat tac --a tat ata tca tcg ttt acg aat act gat gcg cct gaa tat cta gga gtg att aag ttt gga agt ctt ttc cat ttc gaa gtg ccg tgt ttc aaa tat tct gct ata ccc gtt gaa ata gaa aat tct aat cct cct att aca tat aac ttt cca tcg tta aca caa gta cta act tct gat ttt aac gac gac ata tta gta acc gtt ttc ca- --- --- --- --- --- --- -tt ttt ttt gtt tta aga tct acc cgc -ga tac gga ata aac atg tct att gtt aat cat gcc gcc aat aat gta tag aca att atg taa aac att tgc atc ata gaa ttg --- -tc tat ctg tat tac cga cta tcg tcc aat att ctg ttc tag gag agt aat ggg tta ttg tgg ata tat aat cag agt ttt taa tga cta cta tat tat gtt tta tac cat ttc gtg tca c-a gct ttg tag att tgg ata tag tta atc cca aca atg cta tag cat tgc ata tag cat tag tca taa act tgg gat gta aaa tgt tga tga tat cta cat cgt ttg gat ttt tat gta tcc act tta ata ata tta tag c-g taa cat cct cat ga- ttt acg tta acg ttt tcg tgt gat aag ata gtg gtc agt tca tcc ttt gat aat ttt cca aat t

>MN702453.1:18794-20746_Monkeypox_virus_strain_A1_contig_SPADES

cgg ctg cca tgt att tcc tgg aga gca agt --- aga tga tg- agg aac cag ata gtt tat atc cat act tgc act taa agt cta cat tgt agt tgt atg agt gta tga tct ttt aag -cc gct aga -ag ttt tcc gtt tga tat --- agg atg tgg aca ttt aac aat ctg aca cgt ggg tgg att gga cca ttc tcc tcc tga aca cat gac acc aga gtt acc aat caa cga ata tcc act att gca act ata agt tac aat gct ccc atc gat ata aaa atc ctc gta tcc gtt atg tct tcc gtt gga tat aga tgg agg tga ttg gca ttt aac aga ttc gca aat agg tgc ctc agg att cca tac cat aga tcc agt aga tcc taa ttc aca ata cga ttt aga ttc acc gat caa atg ata tcc gct att aca aga gta cgt tat act aga gcc aaa gtc tac tcc gcc aat atc aag ttg gcc att atc gat atc tcg agg cga tgg gca tct ccg ttt aat aca ttg att aaa gag tgt cca tcc ggt acc ggt aca ttt agc ata tat ggg tcc cat ttt ttg ctt tct gta tcc agg tag aca tag a-t att cta tag tgt ctc cta tgt tgt a-- --- --- --- --- --- -at tag cat cag tc- --t cta cac tat tct taa att tca tat taa tgg ggc gtg acg gaa tag tac agt atg ata gaa cac atc cta ttc cca aca atg tca gga acg tca cgc tct cca cct tca tat tta ttt atc cgt --a aaa tgt tat cct gga cat cgt aca aat aat a-a aaa gcc cat ata tat gtt cgc tat tgt aga aat tgt ttt tca cag ttg ctc aaa aac aat ggc agt gac tta tga gtt a-g tta cac ttt gga gtc tca tct tta gta aac ata tca taa tat tcg ata tta cga gtt gac ata tcg aac aaa ttc caa gta ttt gat ttt gga taa tat tcg tat ttt gca tct gct ata att aag ata taa tca cca caa gaa cac acg aac gtc ttt cct aca tgg tta aag tac atg tac aat tct atc cat ttg tct tcc tta act ata tat ttg tat aga taa tta cga gtc tca tga gta att cca gta att gca tag atg tca cca tcg tat tct aca gca taa act ata cta tga cgt cta ggc atg gga gac -tt ttt tat cca acg att ttt agt gaa aca ttc cac atc gtt taa tac tac ata ttt ctc ata -gt ggt ata aac tcc acc cat tac --a tat ata tca tcg ttt acg aat act gat gcg cct gaa tat cta gga gtg att aag ttt gga agt ctt ttc cat ttc gaa gtg ccg tgt ttc aaa tat tct gct ata ccc gtt gaa ata gaa aat tct aat cct cct att aca tat aac ttt cca tcg tta aca caa gta cta act tct gat ttt aac gac gac ata tta gta acc gtt ttc ca- --- --- --- --- --- --- -tt ttt ttt gtt tta aga tct acc cgc -ga tac gga ata aac atg tct att gtt aat cat gcc gcc aat aat gta tag aca att atg taa aac att tgc atc ata gaa ttg --- -tc tat ctg tat tac cga cta tcg tcc aat att ctg ttc tag gag agt aat ggg tta ttg tgg ata tat aat cag agt ttt taa tga cta cta tat tat gtt tta tac cat ttc gtg tca c-a gct ttg tag att tgg ata tag tta atc cca aca atg cta tag cat tgc ata tag cat tag tca taa act tgg gat gta aaa tgt tga tga tat cta cat cgt ttg gat ttt tat gta tcc act tta ata ata tta tag c-g taa cat cct cat ga- ttt acg tta acg ttt tcg tgt gat aag ata gtg gtc agt tca tcc ttt gat aat ttt cca aat t

>MN702452.1:18794-20746_Monkeypox_virus_strain_A2_contig_SPADES

cgg ctg cca tgt att tcc tgg aga gca agt --- aga tga tg- agg aac cag ata gtt tat atc cat act tgc act taa agt cta cat tgt agt tgt atg agt gta tga tct ttt aag -cc gct aga -ag ttt tcc gtt tga tat --- agg atg tgg aca ttt aac aat ctg aca cgt ggg tgg att gga cca ttc tcc tcc tga aca cat gac acc aga gtt acc aat caa cga ata tcc act att gca act ata agt tac aat gct ccc atc gat ata aaa atc ctc gta tcc gtt atg tct tcc gtt gga tat aga tgg agg tga ttg gca ttt aac aga ttc gca aat agg tgc ctc agg att cca tac cat aga tcc agt aga tcc taa ttc aca ata cga ttt aga ttc acc gat caa atg ata tcc gct att aca aga gta cgt tat act aga gcc aaa gtc tac tcc gcc aat atc aag ttg gcc att atc gat atc tcg agg cga tgg gca tct ccg ttt aat aca ttg att aaa gag tgt cca tcc ggt acc ggt aca ttt agc ata tat ggg tcc cat ttt ttg ctt tct gta tcc agg tag aca tag a-t att cta tag tgt ctc cta tgt tgt a-- --- --- --- --- --- -at tag cat cag tc- --t cta cac tat tct taa att tca tat taa tgg ggc gtg acg gaa tag tac agt atg ata gaa cac atc cta ttc cca aca atg tca gga acg tca cgc tct cca cct tca tat tta ttt atc cgt --a aaa tgt tat cct gga cat cgt aca aat aat a-a aaa gcc cat ata tat gtt cgc tat tgt aga aat tgt ttt tca cag ttg ctc aaa aac aat ggc agt gac tta tga gtt a-g tta cac ttt gga gtc tca tct tta gta aac ata tca taa tat tcg ata tta cga gtt gac ata tcg aac aaa ttc caa gta ttt gat ttt gga taa tat tcg tat ttt gca tct gct ata att aag ata taa tca cca caa gaa cac acg aac gtc ttt cct aca tgg tta aag tac atg tac aat tct atc cat ttg tct tcc tta act ata tat ttg tat aga taa tta cga gtc tca tga gta att cca gta att gca tag atg tca cca tcg tat tct aca gca taa act ata cta tga cgt cta ggc atg gga gac -tt ttt tat cca acg att ttt agt gaa aca ttc cac atc gtt taa tac tac ata ttt ctc ata -gt ggt ata aac tcc acc cat tac --a tat ata tca tcg ttt acg aat act gat gcg cct gaa tat cta gga gtg att aag ttt gga agt ctt ttc cat ttc gaa gtg ccg tgt ttc aaa tat tct gct ata ccc gtt gaa ata gaa aat tct aat cct cct att aca tat aac ttt cca tcg tta aca caa gta cta act tct gat ttt aac gac gac ata tta gta acc gtt ttc ca- --- --- --- --- --- --- -tt ttt ttt gtt tta aga tct acc cgc -ga tac gga ata aac atg tct att gtt aat cat gcc gcc aat aat gta tag aca att atg taa aac att tgc atc ata gaa ttg --- -tc tat ctg tat tac cga cta tcg tcc aat att ctg ttc tag gag agt aat ggg tta ttg tgg ata tat aat cag agt ttt taa tga cta cta tat tat gtt tta tac cat ttc gtg tca c-a gct ttg tag att tgg ata tag tta atc cca aca atg cta tag cat tgc ata tag cat tag tca taa act tgg gat gta aaa tgt tga tga tat cta cat cgt ttg gat ttt tat gta tcc act tta ata ata tta tag c-g taa cat cct cat ga- ttt acg tta acg ttt tcg tgt gat aag ata gtg gtc agt tca tcc ttt gat aat ttt cca aat t

>MN702451.1:12658-14610_Monkeypox_virus_strain_A6_contig_SPADES

cgg ctg cca tgt att tcc tgg aga gca agt --- aga tga tg- agg aac cag ata gtt tat atc cat act tgc act taa agt cta cat tgt agt tgt atg agt gta tga tct ttt aag -cc gct aga -ag ttt tcc gtt tga tat --- agg atg tgg aca ttt aac aat ctg aca cgt ggg tgg att gga cca ttc tcc tcc tga aca cat gac acc aga gtt acc aat caa cga ata tcc act att gca act ata agt tac aat gct ccc atc gat ata aaa atc ctc gta tcc gtt atg tct tcc gtt gga tat aga tgg agg tga ttg gca ttt aac aga ttc gca aat agg tgc ctc agg att cca tac cat aga tcc agt aga tcc taa ttc aca ata cga ttt aga ttc acc gat caa atg ata tcc gct att aca aga gta cgt tat act aga gcc aaa gtc tac tcc gcc aat atc aag ttg gcc att atc gat atc tcg agg cga tgg gca tct ccg ttt aat aca ttg att aaa gag tgt cca tcc ggt acc ggt aca ttt agc ata tat ggg tcc cat ttt ttg ctt tct gta tcc agg tag aca tag a-t att cta tag tgt ctc cta tgt tgt a-- --- --- --- --- --- -at tag cat cag tc- --t cta cac tat tct taa att tca tat taa tgg ggc gtg acg gaa tag tac agt atg ata gaa cac atc cta ttc cca aca atg tca gga acg tca cgc tct cca cct tca tat tta ttt atc cgt --a aaa tgt tat cct gga cat cgt aca aat aat a-a aaa gcc cat ata tat gtt cgc tat tgt aga aat tgt ttt tca cag ttg ctc aaa aac aat ggc agt gac tta tga gtt a-g tta cac ttt gga gtc tca tct tta gta aac ata tca taa tat tcg ata tta cga gtt gac ata tcg aac aaa ttc caa gta ttt gat ttt gga taa tat tcg tat ttt gca tct gct ata att aag ata taa tca cca caa gaa cac acg aac gtc ttt cct aca tgg tta aag tac atg tac aat tct atc cat ttg tct tcc tta act ata tat ttg tat aga taa tta cga gtc tca tga gta att cca gta att gca tag atg tca cca tcg tat tct aca gca taa act ata cta tga cgt cta ggc atg gga gac -tt ttt tat cca acg att ttt agt gaa aca ttc cac atc gtt taa tac tac ata ttt ctc ata -gt ggt ata aac tcc acc cat tac --a tat ata tca tcg ttt acg aat act gat gcg cct gaa tat cta gga gtg att aag ttt gga agt ctt ttc cat ttc gaa gtg ccg tgt ttc aaa tat tct gct ata ccc gtt gaa ata gaa aat tct aat cct cct att aca tat aac ttt cca tcg tta aca caa gta cta act tct gat ttt aac gac gac ata tta gta acc gtt ttc ca- --- --- --- --- --- --- -tt ttt ttt gtt tta aga tct acc cgc -ga tac gga ata aac atg tct att gtt aat cat gcc gcc aat aat gta tag aca att atg taa aac att tgc atc ata gaa ttg --- -tc tat ctg tat tac cga cta tcg tcc aat att ctg ttc tag gag agt aat ggg tta ttg tgg ata tat aat cag agt ttt taa tga cta cta tat tat gtt tta tac cat ttc gtg tca c-a gct ttg tag att tgg ata tag tta atc cca aca atg cta tag cat tgc ata tag cat tag tca taa act tgg gat gta aaa tgt tga tga tat cta cat cgt ttg gat ttt tat gta tcc act tta ata ata tta tag c-g taa cat cct cat ga- ttt acg tta acg ttt tcg tgt gat aag ata gtg gtc agt tca tcc ttt gat aat ttt cca aat t

>MN702450.1:12642-14594_Monkeypox_virus_strain_B1_contig_SPADES

cgg ctg cca tgt att tcc tgg aga gca agt --- aga tga tg- agg aac cag ata gtt tat atc cat act tgc act taa agt cta cat tgt agt tgt atg agt gta tga tct ttt aag -cc gct aga -ag ttt tcc gtt tga tat --- agg atg tgg aca ttt aac aat ctg aca cgt ggg tgg att gga cca ttc tcc tcc tga aca cat gac acc aga gtt acc aat caa cga ata tcc act att gca act ata agt tac aat gct ccc atc gat ata aaa atc ctc gta tcc gtt atg tct tcc gtt gga tat aga tgg agg tga ttg gca ttt aac aga ttc gca aat agg tgc ctc agg att cca tac cat aga tcc agt aga tcc taa ttc aca ata cga ttt aga ttc acc gat caa atg ata tcc gct att aca aga gta cgt tat act aga gcc aaa gtc tac tcc gcc aat atc aag ttg gcc att atc gat atc tcg agg cga tgg gca tct ccg ttt aat aca ttg att aaa gag tgt cca tcc ggt acc ggt aca ttt agc ata tat ggg tcc cat ttt ttg ctt tct gta tcc agg tag aca tag a-t att cta tag tgt ctc cta tgt tgt a-- --- --- --- --- --- -at tag cat cag tc- --t cta cac tat tct taa att tca tat taa tgg ggc gtg acg gaa tag tac agt atg ata gaa cac atc cta ttc cca aca atg tca gga acg tca cgc tct cca cct tca tat tta ttt atc cgt --a aaa tgt tat cct gga cat cgt aca aat aat a-a aaa gcc cat ata tat gtt cgc tat tgt aga aat tgt ttt tca cag ttg ctc aaa aac aat ggc agt gac tta tga gtt a-g tta cac ttt gga gtc tca tct tta gta aac ata tca taa tat tcg ata tta cga gtt gac ata tcg aac aaa ttc caa gta ttt gat ttt gga taa tat tcg tat ttt gca tct gct ata att aag ata taa tca cca caa gaa cac acg aac gtc ttt cct aca tgg tta aag tac atg tac aat tct atc cat ttg tct tcc tta act ata tat ttg tat aga taa tta cga gtc tca tga gta att cca gta att gca tag atg tca cca tcg tat tct aca gca taa act ata cta tga cgt cta ggc atg gga gac -tt ttt tat cca acg att ttt agt gaa aca ttc cac atc gtt taa tac tac ata ttt ctc ata -gt ggt ata aac tcc acc cat tac --a tat ata tca tcg ttt acg aat act gat gcg cct gaa tat cta gga gtg att aag ttt gga agt ctt ttc cat ttc gaa gtg ccg tgt ttc aaa tat tct gct ata ccc gtt gaa ata gaa aat tct aat cct cct att aca tat aac ttt cca tcg tta aca caa gta cta act tct gat ttt aac gac gac ata tta gta acc gtt ttc ca- --- --- --- --- --- --- -tt ttt ttt gtt tta aga tct acc cgc -ga tac gga ata aac atg tct att gtt aat cat gcc gcc aat aat gta tag aca att atg taa aac att tgc atc ata gaa ttg --- -tc tat ctg tat tac cga cta tcg tcc aat att ctg ttc tag gag agt aat ggg tta ttg tgg ata tat aat cag agt ttt taa tga cta cta tat tat gtt tta tac cat ttc gtg tca c-a gct ttg tag att tgg ata tag tta atc cca aca atg cta tag cat tgc ata tag cat tag tca taa act tgg gat gta aaa tgt tga tga tat cta cat cgt ttg gat ttt tat gta tcc act tta ata ata tta tag c-g taa cat cct cat ga- ttt acg tta acg ttt tcg tgt gat aag ata gtg gtc agt tca tcc ttt gat aat ttt cca aat t

>MN702448.1:12117-14069_Monkeypox_virus_strain_015c_contig_SPADES

cgg ctg cca tgt att tcc tgg aga gca agt --- aga tga tg- agg aac cag ata gtt tat atc cat act tgc act taa agt cta cat tgt agt tgt atg agt gta tga tct ttt aag -cc gct aga -ag ttt tcc gtt tga tat --- agg atg tgg aca ttt aac aat ctg aca cgt ggg tgg att gga cca ttc tcc tcc tga aca cat gac acc aga gtt acc aat caa cga ata tcc act att gca act ata agt tac aat gct ccc atc gat ata aaa atc ctc gta tcc gtt atg tct tcc gtt gga tat aga tgg agg tga ttg gca ttt aac aga ttc gca aat agg tgc ctc agg att cca tac cat aga tcc agt aga tcc taa ttc aca ata cga ttt aga ttc acc gat caa atg ata tcc gct att aca aga gta cgt tat act aga gcc aaa gtc tac tcc gcc aat atc aag ttg gcc att atc gat atc tcg agg cga tgg gca tct ccg ttt aat aca ttg att aaa gag tgt cca tcc ggt acc ggt aca ttt agc ata tat ggg tcc cat ttt ttg ctt tct gta tcc agg tag aca tag a-t att cta tag tgt ctc cta tgt tgt a-- --- --- --- --- --- -at tag cat cag tc- --t cta cac tat tct taa att tca tat taa tgg ggc gtg acg gaa tag tac agt atg ata gaa cac atc cta ttc cca aca atg tca gga acg tca cgc tct cca cct tca tat tta ttt atc cgt --a aaa tgt tat cct gga cat cgt aca aat aat a-a aaa gcc cat ata tat gtt cgc tat tgt aga aat tgt ttt tca cag ttg ctc aaa aac aat ggc agt gac tta tga gtt a-g tta cac ttt gga gtc tca tct tta gta aac ata tca taa tat tcg ata tta cga gtt gac ata tcg aac aaa ttc caa gta ttt gat ttt gga taa tat tcg tat ttt gca tct gct ata att aag ata taa tca cca caa gaa cac acg aac gtc ttt cct aca tgg tta aag tac atg tac aat tct atc cat ttg tct tcc tta act ata tat ttg tat aga taa tta cga gtc tca tga gta att cca gta att gca tag atg tca cca tcg tat tct aca gca taa act ata cta tga cgt cta ggc atg gga gac -tt ttt tat cca acg att ttt agt gaa aca ttc cac atc gtt taa tac tac ata ttt ctc ata -gt ggt ata aac tcc acc cat tac --a tat ata tca tcg ttt acg aat act gat gcg cct gaa tat cta gga gtg att aag ttt gga agt ctt ttc cat ttc gaa gtg ccg tgt ttc aaa tat tct gct ata ccc gtt gaa ata gaa aat tct aat cct cct att aca tat aac ttt cca tcg tta aca caa gta cta act tct gat ttt aac gac gac ata tta gta acc gtt ttc ca- --- --- --- --- --- --- -tt ttt ttt gtt tta aga tct acc cgc -ga tac gga ata aac atg tct att gtt aat cat gcc gcc aat aat gta tag aca att atg taa aac att tgc atc ata gaa ttg --- -tc tat ctg tat tac cga cta tcg tcc aat att ctg ttc tag gag agt aat ggg tta ttg tgg ata tat aat cag agt ttt taa tga cta cta tat tat gtt tta tac cat ttc gtg tca c-a gct ttg tag att tgg ata tag tta atc cca aca atg cta tag cat tgc ata tag cat tag tca taa act tgg gat gta aaa tgt tga tga tat cta cat cgt ttg gat ttt tat gta tcc act tta ata ata tta tag c-g taa cat cct cat ga- ttt acg tta acg ttt tcg tgt gat aag ata gtg gtc agt tca tcc ttt gat aat ttt cca aat t

>MN702447.1:12117-14069_Monkeypox_virus_strain_18_contig_SPADES

cgg ctg cca tgt att tcc tgg aga gca agt --- aga tga tg- agg aac cag ata gtt tat atc cat act tgc act taa agt cta cat tgt agt tgt atg agt gta tga tct ttt aag -cc gct aga -ag ttt tcc gtt tga tat --- agg atg tgg aca ttt aac aat ctg aca cgt ggg tgg att gga cca ttc tcc tcc tga aca cat gac acc aga gtt acc aat caa cga ata tcc act att gca act ata agt tac aat gct ccc atc gat ata aaa atc ctc gta tcc gtt atg tct tcc gtt gga tat aga tgg agg tga ttg gca ttt aac aga ttc gca aat agg tgc ctc agg att cca tac cat aga tcc agt aga tcc taa ttc aca ata cga ttt aga ttc acc gat caa atg ata tcc gct att aca aga gta cgt tat act aga gcc aaa gtc tac tcc gcc aat atc aag ttg gcc att atc gat atc tcg agg cga tgg gca tct ccg ttt aat aca ttg att aaa gag tgt cca tcc ggt acc ggt aca ttt agc ata tat ggg tcc cat ttt ttg ctt tct gta tcc agg tag aca tag a-t att cta tag tgt ctc cta tgt tgt a-- --- --- --- --- --- -at tag cat cag tc- --t cta cac tat tct taa att tca tat taa tgg ggc gtg acg gaa tag tac agt atg ata gaa cac atc cta ttc cca aca atg tca gga acg tca cgc tct cca cct tca tat tta ttt atc cgt --a aaa tgt tat cct gga cat cgt aca aat aat a-a aaa gcc cat ata tat gtt cgc tat tgt aga aat tgt ttt tca cag ttg ctc aaa aac aat ggc agt gac tta tga gtt a-g tta cac ttt gga gtc tca tct tta gta aac ata tca taa tat tcg ata tta cga gtt gac ata tcg aac aaa ttc caa gta ttt gat ttt gga taa tat tcg tat ttt gca tct gct ata att aag ata taa tca cca caa gaa cac acg aac gtc ttt cct aca tgg tta aag tac atg tac aat tct atc cat ttg tct tcc tta act ata tat ttg tat aga taa tta cga gtc tca tga gta att cca gta att gca tag atg tca cca tcg tat tct aca gca taa act ata cta tga cgt cta ggc atg gga gac -tt ttt tat cca acg att ttt agt gaa aca ttc cac atc gtt taa tac tac ata ttt ctc ata -gt ggt ata aac tcc acc cat tac --a tat ata tca tcg ttt acg aat act gat gcg cct gaa tat cta gga gtg att aag ttt gga agt ctt ttc cat ttc gaa gtg ccg tgt ttc aaa tat tct gct ata ccc gtt gaa ata gaa aat tct aat cct cct att aca tat aac ttt cca tcg tta aca caa gta cta act tct gat ttt aac gac gac ata tta gta acc gtt ttc ca- --- --- --- --- --- --- -tt ttt ttt gtt tta aga tct acc cgc -ga tac gga ata aac atg tct att gtt aat cat gcc gcc aat aat gta tag aca att atg taa aac att tgc atc ata gaa ttg --- -tc tat ctg tat tac cga cta tcg tcc aat att ctg ttc tag gag agt aat ggg tta ttg tgg ata tat aat cag agt ttt taa tga cta cta tat tat gtt tta tac cat ttc gtg tca c-a gct ttg tag att tgg ata tag tta atc cca aca atg cta tag cat tgc ata tag cat tag tca taa act tgg gat gta aaa tgt tga tga tat cta cat cgt ttg gat ttt tat gta tcc act tta ata ata tta tag c-g taa cat cct cat ga- ttt acg tta acg ttt tcg tgt gat aag ata gtg gtc agt tca tcc ttt gat aat ttt cca aat t

>AY743598.1:13452-15404_Monkeypox_virus_strain_Congo-8_D2L

cgg ctg cca tgt att tcc tgg aga gca agt --- aga tga tg- agg aac cag ata gtt tat atc cat act tgc act taa agt cta cat tgt agt tgt atg agt gta tga tct ttt aag -cc gct aga -ag ttt tcc gtt tga tat --- agg atg tgg aca ttt aac aat ctg aca cgt ggg tgg att gga cca ttc tcc tcc tga aca cat gac acc aga gtt acc aat caa cga ata tcc act att gca act ata agt tac aat gct ccc atc gat ata aaa atc ctc gta tcc gtt atg tct tcc gtt gga tat aga tgg agg tga ttg gca ttt aac aga ttc gca aat agg tgc ctc agg att cca tac cat aga tcc agt aga tcc taa ttc aca ata cga ttt aga ttc acc gat caa atg ata tcc gct att aca aga gta cgt tat act aga gcc aaa gtc tac tcc gcc aat atc aag ttg gcc att atc gat atc tcg agg cga tgg gca tct ccg ttt aat aca ttg att aaa gag tgt cca tcc ggt acc ggt aca ttt agc ata tat ggg tcc cat ttt ttg ctt tct gta tcc agg tag aca tag a-t att cta tag tgt ctc cta tgt tgt a-- --- --- --- --- --- -at tag cat cag tc- --t cta cac tat tct taa att tca tat taa tgg ggc gtg acg gaa tag tac agt atg ata gaa cac atc cta ttc cca aca atg tca gga acg tca cgc tct cca cct tca tat tta ttt atc cgt --a aaa tgt tat cct gga cat cgt aca aat aat a-a aaa gcc cat ata tat gtt cgc tat tgt aga aat tgt ttt tca cag ttg ctc aaa aac aat ggc agt gac tta tga gtt a-g tta cac ttt gga gtc tca tct tta gta aac ata tca taa tat tcg ata tta cga gtt gac ata tcg aac aaa ttc caa gta ttt gat ttt gga taa tat tcg tat ttt gca tct gct ata att aag ata taa tca cca caa gaa cac acg aac gtc ttt cct aca tgg tta aag tac atg tac aat tct atc cat ttg tct tcc tta act ata tat ttg tat aga taa tta cga gtc tca tga gta att cca gta att gca tag atg tca cca tcg tat tct aca gca taa act ata cta tga cgt cta ggc atg gga gac -tt ttt tat cca acg att ttt agt gaa aca ttc cac atc gtt taa tac tac ata ttt ctc ata -gt ggt ata aac tcc acc cat tac --a tat ata tca tcg ttt acg aat act gat gcg cct gaa tat cta gga gtg att aag ttt gga agt ctt ttc cat ttc gaa gtg ccg tgt ttc aaa tat tct gct ata ccc gtt gaa ata gaa aat tct aat cct cct att aca tat aac ttt cca tcg tta aca caa gta cta act tct gat ttt aac gac gac ata tta gta acc gtt ttc ca- --- --- --- --- --- --- -tt ttt ttt gtt tta aga tct acc cgc -ga tac gga ata aac atg tct att gtt aat cat gcc gcc aat aat gta tag aca att atg taa aac att tgc atc ata gaa ttg --- -tc tat ctg tat tac cga cta tcg tcc aat att ctg ttc tag gag agt aat ggg tta ttg tgg ata tat aat cag agt ttt taa tga cta cta tat tat gtt tta tac cat ttc gtg tca c-a gct ttg tag att tgg ata tag tta atc cca aca atg cta tag cat tgc ata tag cat tag tca taa act tgg gat gta aaa tgt tga tga tat cta cat cgt ttg gat ttt tat gta tcc act tta ata ata tta tag c-g taa cat cct cat ga- ttt acg tta acg ttt tcg tgt gat aag ata gtg gtc agt tca tcc ttt gat aat ttt cca aat t

>DQ011154.1:19095-21047_Monkeypox_virus_strain_Congo_2003_358_complete_genome

cgg ctg cca tgt att tcc tgg aga gca agt --- aga tga tg- agg aac cag ata gtt tat atc cat act tgc act taa agt cta cat tgt agt tgt atg agt gta tga tct ttt aag -cc gct aga -ag ttt tcc gtt tga tat --- agg atg tgg aca ttt aac aat ctg aca cgt ggg tgg att gga cca ttc tcc tcc tga aca cat gac acc aga gtt acc aat caa cga ata tcc act att gca act ata agt tac aat gct ccc atc gat ata aaa atc ctc gta tcc gtt atg tct tcc gtt gga tat aga tgg agg tga ttg gca ttt aac aga ttc gca aat agg tgc ctc agg att cca tac cat aga tcc agt aga tcc taa ttc aca ata cga ttt aga ttc acc gat caa atg ata tcc gct att aca aga gta cgt tat act aga gcc aaa gtc tac tcc gcc aat atc aag ttg gcc att atc gat atc tcg agg cga tgg gca tct ccg ttt aat aca ttg att aaa gag tgt cca tcc ggt acc ggt aca ttt agc ata tat ggg tcc cat ttt ttg ctt tct gta tcc agg tag aca tag a-t att cta tag tgt ctc cta tgt tgt a-- --- --- --- --- --- -at tag cat cag tc- --t cta cac tat tct taa att tca tat taa tgg ggc gtg acg gaa tag tac agt atg ata gaa cac atc cta ttc cca aca atg tca gga acg tca cgc tct cca cct tca tat tta ttt atc cgt --a aaa tgt tat cct gga cat cgt aca aat aat a-a aaa gcc cat ata tat gtt cgc tat tgt aga aat tgt ttt tca cag ttg ctc aaa aac aat ggc agt gac tta tga gtt a-g tta cac ttt gga gtc tca tct tta gta aac ata tca taa tat tcg ata tta cga gtt gac ata tcg aac aaa ttc caa gta ttt gat ttt gga taa tat tcg tat ttt gca tct gct ata att aag ata taa tca cca caa gaa cac acg aac gtc ttt cct aca tgg tta aag tac atg tac aat tct atc cat ttg tct tcc tta act ata tat ttg tat aga taa tta cga gtc tca tga gta att cca gta att gca tag atg tca cca tcg tat tct aca gca taa act ata cta tga cgt cta ggc atg gga gac -tt ttt tat cca acg att ttt agt gaa aca ttc cac atc gtt taa tac tac ata ttt ctc ata -gt ggt ata aac tcc acc cat tac --a tat ata tca tcg ttt acg aat act gat gcg cct gaa tat cta gga gtg att aag ttt gga agt ctt ttc cat ttc gaa gtg ccg tgt ttc aaa tat tct gct ata ccc gtt gaa ata gaa aat tct aat cct cct att aca tat aac ttt cca tcg tta aca caa gta cta act tct gat ttt aac gac gac ata tta gta acc gtt ttc ca- --- --- --- --- --- --- -tt ttt ttt gtt tta aga tct acc cgc -ga tac gga ata aac atg tct att gtt aat cat gcc gcc aat aat gta tag aca att atg taa aac att tgc atc ata gaa ttg --- -tc tat ctg tat tac cga cta tcg tcc aat att ctg ttc tag gag agt aat ggg tta ttg tgg ata tat aat cag agt ttt taa tga cta cta tat tat gtt tta tac cat ttc gtg tca c-a gct ttg tag att tgg ata tag tta atc cca aca atg cta tag cat tgc ata tag cat tag tca taa act tgg gat gta aaa tgt tga tga tat cta cat cgt ttg gat ttt tat gta tcc act tta ata ata tta tag c-g taa cat cct cat ga- ttt acg tta acg ttt tcg tgt gat aag ata gtg gtc agt tca tcc ttt gat aat ttt cca aat t

>KP849471.1:19186-21137_Monkeypox_virus_isolate_Yambuku_DRC_1985_complete_genome

cgg ctg cca tgt att tcc tgg aga gca agt --- aga tga tg- agg aac cag ata gtt tat atc cat act tgc act taa agt cta cat tgt agt tgt atg agt gta tga tct ttt aag -cc gct aga -ag ttt tcc gtt tga tat --- agg atg tgg aca ttt aac aat ctg aca cgt ggg tgg att gga cca ttc tcc tcc tga aca cat gac acc aga gtt acc aat caa cga ata tcc act att gca act ata agt tac aat gct ccc atc gat ata aaa atc ctc gta tcc gtt atg tct tcc gtt gga tat aga tgg agg tga ttg gca ttt aac aga ttc gca aat agg tgc ctc agg att cca tac cat aga tcc agt aga tcc taa ttc aca ata cga ttt aga ttc acc gat caa atg ata tcc gct att aca aga gta cgt tat act aga gcc aaa gtc tac tcc gcc aat atc aag ttg gcc att atc gat atc tcg agg cga tgg gca tct ccg ttt aat aca ttg att aaa gag tgt cca tcc ggt acc ggt aca ttt agc ata tat ggg tcc cat ttt ttg ctt tct gta tcc agg tag aca tag a-t att cta tag tgt ctc cta tgt tgt a-- --- --- --- --- --- -at tag cat cag tc- --t cta cac tat tct taa att tca tat taa tgg ggc gtg acg gaa tag tac agt atg ata gaa cac atc cta ttc cca aca atg tca gga acg tca cgc tct cca cct tca tat tta ttt atc cgt --a aaa tgt tat cct gga cat cgt aca aat aat a-a aaa gcc cat ata tat gtt cgc tat tgt aga aat tgt ttt tca cag ttg ctc aaa aac aat ggc agt gac tta tga gtt a-g tta cac ttt gga gtc tca tct tta gta aac ata tca taa tat tcg ata tta cga gtt gac ata tcg aac aaa ttc caa gta ttt gat ttt gga taa tat tcg tat ttt gca tct gct ata att aag ata taa tca cca caa gaa cac acg aac gtc ttt cct aca tgg tta aag tac atg tac aat tct atc cat ttg tct tcc tta act ata tat ttg tat aga taa tta cga gtc tca tga gta att cca gta att gca tag atg tca cca tcg tat tct aca gca taa act ata cta tga cgt cta ggc atg gga gac -tt ttt tat cca acg att ttt agt gaa aca ttc cac atc gtt taa tac tac ata ttt ctc ata -gt ggt ata aac tcc acc cat tac --a tat ata tca tcg ttt acg aat act gat gcg cct gaa tat cta gga gtg att aag ttt gga agt ctt ttc cat ttc gaa gtg ccg tgt ttc aaa tat tct gct ata ccc gtt gaa ata gaa aat tct aat cct cct att aca tat aac ttt cca tcg tta aca caa gta cta act tct gat ttt aac gac -ac ata tta gta acc gtt ttc ca- --- --- --- --- --- --- -tt ttt ttt gtt tta aga tct acc cgc -ga tac gga ata aac atg tct att gtt aat cat gcc gcc aat aat gta tag aca att atg taa aac att tgc atc ata gaa ttg --- -tc tat ctg tat tac cga cta tcg tcc aat att ctg ttc tag gag agt aat ggg tta ttg tgg ata tat aat cag agt ttt taa tga cta cta tat tat gtt tta tac cat ttc gtg tca c-a gct ttg tag att tgg ata tag tta atc cca aca atg cta tag cat tgc ata tag cat tag tca taa act tgg gat gta aaa tgt tga tga tat cta cat cgt ttg gat ttt tat gta tcc act tta ata ata tta tag c-g taa cat cct cat ga- ttt acg tta acg ttt tcg tgt gat aag ata gtg gtc agt tca tcc ttt gat aat ttt cca aat t

>KJ642612.1:17183-19135_Monkeypox_virus_strain_Ikubi_complete_genome

cgg ctg cca tgt att tcc tgg aga gca agt --- aga tga tg- agg aac cag ata gtt tat atc cat act tgc act taa agt cta cat tgt agt tgt atg aga gta tga tct ttt aag -cc gct aga -ag ttt tcc gtt tga tat --- agg atg tgg aca ttt aac aat ctg aca cgt ggg tgg att gga cca ttc tcc tcc tga aca cat gac acc aga gtt acc aat caa cga ata tcc act att gca act ata agt tac aat gct ccc atc gat ata aaa atc ctc gta tcc gtt atg tct tcc gtt gga tat aga tgg agg tga ttg gca ttt aac aga ttc gca aat agg tgc ctc agg att cca tac cat aga tcc agt aga tcc taa ttc aca ata cga ttt aga ttc acc gat caa atg ata tcc gct att aca aga gta cgt tat act aga gcc aaa gtc tac tcc gcc aat atc aag ttg gcc att atc gat atc tcg agg cga tgg gca tct ccg ttt aat aca ttg att aaa gag tgt cca tcc ggt acc ggt aca ttt agc ata tat ggg tcc cat ttt ttg ctt tct gta tcc agg tag aca tag a-t att cta tag tgt ctc cta tgt tgt a-- --- --- --- --- --- -at tag cat cag tc- --t cta cac tat tct taa att tca tat taa tgg ggc gtg acg gaa tag tac agt atg ata gaa cac atc cta ttc cca aca atg tca gga acg tca cgc tct cca cct tca tat tta ttt atc cgt --a aaa tgt tat cct gga cat cgt aca aat aat a-a aaa gcc cat ata tat gtt cgc tat tgt aga aat tgt ttt tca cag ttg ctc aaa aac aat ggc agt gac tta tga gtt a-g tta cac ttt gga gtc tca tct tta gta aac ata tca taa tat tcg ata tta cga gtt gac ata tcg aac aaa ttc caa gta ttt gat ttt gga taa tat tcg tat ttt gca tct gct ata att aag ata taa tca cca caa gaa cac acg aac gtc ttt cct aca tgg tta aag tac atg tac aat tct atc cat ttg tct tcc tta act ata tat ttg tat aga taa tta cga gtc tca tga gta att cca gta att gca tag atg tca cca tcg tat tct aca gca taa act ata cta tga cgt cta ggc atg gga gac -tt ttt tat cca acg att ttt agt gaa aca ttc cac atc gtt taa tac tac ata ttt ctc ata -gt ggt ata aac tcc acc cat tac --a tat ata tca tcg ttt acg aat act gat gcg cct gaa tat cta gga gtg att aag ttt gga agt ctt ttc cat ttc gaa gtg ccg tgt ttc aaa tat tct gct ata ccc gtt gaa ata gaa aat tct aat cct cct att aca tat aac ttt cca tcg tta aca caa gta cta act tct gat ttt aac gac gac ata tta gta acc gtt ttc ca- --- --- --- --- --- --- -tt ttt ttt gtt tta aga tct acc cgc -ga tac gga ata aac atg tct att gtt aat cat gcc gcc aat aat gta tag aca att atg taa aac att tgc atc ata gaa ttg --- -tc tat ctg tat tac cga cta tcg tcc aat att ctg ttc tag gag agt aat ggg tta ttg tgg ata tat aat cag agt ttt taa tga cta cta tat tat gtt tta tac cat ttc gtg tca c-a gct ttg tag att tgg ata tag tta atc cca aca atg cta tag cat tgc ata tag cat tag tca taa act tgg gat gta aaa tgt tga tga tat cta cat cgt ttg gat ttt tat gta tcc act tta ata ata tta tag c-g taa cat cct cat ga- ttt acg tta acg ttt tcg tgt gat aag ata gtg gtc agt tca tcc ttt gat aat ttt cca aat t

>JX878426.1:18898-20850_Monkeypox_virus_isolate_DRC_07-0450_complete_genome

cgg ctg cca tgt att tcc tgg aga gca agt --- aga tga tg- agg aac cag ata gtt tat atc cat act tgc act taa agt cta cat tgt agt tgt atg aga gta tga tct ttt aag -cc gct aga -ag ttt tcc gtt tga tat --- agg atg tgg aca ttt aac aat ctg aca cgt ggg tgg att gga cca ttc tcc tcc tga aca cat gac acc aga gtt acc aat caa cga ata tcc act att gca act ata agt tac aat gct ccc atc gat ata aaa atc ctc gta tcc gtt atg tct tcc gtt gga tat aga tgg agg tga ttg gca ttt aac aga ttc gca aat agg tgc ctc agg att cca tac cat aga tcc agt aga tcc taa ttc aca ata cga ttt aga ttc acc gat caa atg ata tcc gct att aca aga gta cgt tat act aga gcc aaa gtc tac tcc gcc aat atc aag ttg gcc att atc gat atc tcg agg cga tgg gca tct ccg ttt aat aca ttg att aaa gag tgt cca tcc ggt acc ggt aca ttt agc ata tat ggg tcc cat ttt ttg ctt tct gta tcc agg tag aca tag a-t att cta tag tgt ctc cta tgt tgt a-- --- --- --- --- --- -at tag cat cag tc- --t cta cac tat tct taa att tca tat taa tgg ggc gtg acg gaa tag tac agt atg ata gaa cac atc cta ttc cca aca atg tca gga acg tca cgc tct cca cct tca tat tta ttt atc cgt --a aaa tgt tat cct gga cat cgt aca aat aat a-a aaa gcc cat ata tat gtt cgc tat tgt aga aat tgt ttt tca cag ttg ctc aaa aac aat ggc agt gac tta tga gtt a-g tta cac ttt gga gtc tca tct tta gta aac ata tca taa tat tcg ata tta cga gtt gac ata tcg aac aaa ttc caa gta ttt gat ttt gga taa tat tcg tat ttt gca tct gct ata att aag ata taa tca cca caa gaa cac acg aac gtc ttt cct aca tgg tta aag tac atg tac aat tct atc cat ttg tct tcc tta act ata tat ttg tat aga taa tta cga gtc tca tga gta att cca gta att gca tag atg tca cca tcg tat tct aca gca taa act ata cta tga cgt cta ggc atg gga gac -tt ttt tat cca acg att ttt agt gaa aca ttc cac atc gtt taa tac tac ata ttt ctc ata -gt ggt ata aac tcc acc cat tac --a tat ata tca tcg ttt acg aat act gat gcg cct gaa tat cta gga gtg att aag ttt gga agt ctt ttc cat ttc gaa gtg ccg tgt ttc aaa tat tct gct ata ccc gtt gaa ata gaa aat tct aat cct cct att aca tat aac ttt cca tcg tta aca caa gta cta act tct gat ttt aac gac gac ata tta gta acc gtt ttc ca- --- --- --- --- --- --- -tt ttt ttt gtt tta aga tct acc cgc -ga tac gga ata aac atg tct att gtt aat cat gcc gcc aat aat gta tag aca att atg taa aac att tgc atc ata gaa ttg --- -tc tat ctg tat tac cga cta tcg tcc aat att ctg ttc tag gag agt aat ggg tta ttg tgg ata tat aat cag agt ttt taa tga cta cta tat tat gtt tta tac cat ttc gtg tca c-a gct ttg tag att tgg ata tag tta atc cca aca atg cta tag cat tgc ata tag cat tag tca taa act tgg gat gta aaa tgt tga tga tat cta cat cgt ttg gat ttt tat gta tcc act tta ata ata tta tag c-g taa cat cct cat ga- ttt acg tta acg ttt tcg tgt gat aag ata gtg gtc agt tca tcc ttt gat aat ttt cca aat t

>JX878420.1:18890-20842_Monkeypox_virus_isolate_DRC_07-0283_complete_genome

cgg ctg cca tgt att tcc tgg aga gca agt --- aga tga tg- agg aac cag ata gtt tat atc cat act tgc act taa agt cta cat tgt agt tgt atg aga gta tga tct ttt aag -cc gct aga -ag ttt tcc gtt tga tat --- agg atg tgg aca ttt aac aat ctg aca cgt ggg tgg att gga cca ttc tcc tcc tga aca cat gac acc aga gtt acc aat caa cga ata tcc act att gca act ata agt tac aat gct ccc atc gat ata aaa atc ctc gta tcc gtt atg tct tcc gtt gga tat aga tgg agg tga ttg gca ttt aac aga ttc gca aat agg tgc ctc agg att cca tac cat aga tcc agt aga tcc taa ttc aca ata cga ttt aga ttc acc gat caa atg ata tcc gct att aca aga gta cgt tat act aga gcc aaa gtc tac tcc gcc aat atc aag ttg gcc att atc gat atc tcg agg cga tgg gca tct ccg ttt aat aca ttg att aaa gag tgt cca tcc ggt acc ggt aca ttt agc ata tat ggg tcc cat ttt ttg ctt tct gta tcc agg tag aca tag a-t att cta tag tgt ctc cta tgt tgt a-- --- --- --- --- --- -at tag cat cag tc- --t cta cac tat tct taa att tca tat taa tgg ggc gtg acg gaa tag tac agt atg ata gaa cac atc cta ttc cca aca atg tca gga acg tca cgc tct cca cct tca tat tta ttt atc cgt --a aaa tgt tat cct gga cat cgt aca aat aat a-a aaa gcc cat ata tat gtt cgc tat tgt aga aat tgt ttt tca cag ttg ctc aaa aac aat ggc agt gac tta tga gtt a-g tta cac ttt gga gtc tca tct tta gta aac ata tca taa tat tcg ata tta cga gtt gac ata tcg aac aaa ttc caa gta ttt gat ttt gga taa tat tcg tat ttt gca tct gct ata att aag ata taa tca cca caa gaa cac acg aac gtc ttt cct aca tgg tta aag tac atg tac aat tct atc cat ttg tct tcc tta act ata tat ttg tat aga taa tta cga gtc tca tga gta att cca gta att gca tag atg tca cca tcg tat tct aca gca taa act ata cta tga cgt cta ggc atg gga gac -tt ttt tat cca acg att ttt agt gaa aca ttc cac atc gtt taa tac tac ata ttt ctc ata -gt ggt ata aac tcc acc cat tac --a tat ata tca tcg ttt acg aat act gat gcg cct gaa tat cta gga gtg att aag ttt gga agt ctt ttc cat ttc gaa gtg ccg tgt ttc aaa tat tct gct ata ccc gtt gaa ata gaa aat tct aat cct cct att aca tat aac ttt cca tcg tta aca caa gta cta act tct gat ttt aac gac gac ata tta gta acc gtt ttc ca- --- --- --- --- --- --- -tt ttt ttt gtt tta aga tct acc cgc -ga tac gga ata aac atg tct att gtt aat cat gcc gcc aat aat gta tag aca att atg taa aac att tgc atc ata gaa ttg --- -tc tat ctg tat tac cga cta tcg tcc aat att ctg ttc tag gag agt aat ggg tta ttg tgg ata tat aat cag agt ttt taa tga cta cta tat tat gtt tta tac cat ttc gtg tca c-a gct ttg tag att tgg ata tag tta atc cca aca atg cta tag cat tgc ata tag cat tag tca taa act tgg gat gta aaa tgt tga tga tat cta cat cgt ttg gat ttt tat gta tcc act tta ata ata tta tag c-g taa cat cct cat ga- ttt acg tta acg ttt tcg tgt gat aag ata gtg gtc agt tca tcc ttt gat aat ttt cca aat t

>JX878419.1:18891-20843_Monkeypox_virus_isolate_DRC_07-0275_complete_genome

cgg ctg cca tgt att tcc tgg aga gca agt --- aga tga tg- agg aac cag ata gtt tat atc cat act tgc act taa agt cta cat tgt agt tgt atg aga gta tga tct ttt aag -cc gct aga -ag ttt tcc gtt tga tat --- agg atg tgg aca ttt aac aat ctg aca cgt ggg tgg att gga cca ttc tcc tcc tga aca cat gac acc aga gtt acc aat caa cga ata tcc act att gca act ata agt tac aat gct ccc atc gat ata aaa atc ctc gta tcc gtt atg tct tcc gtt gga tat aga tgg agg tga ttg gca ttt aac aga ttc gca aat agg tgc ctc agg att cca tac cat aga tcc agt aga tcc taa ttc aca ata cga ttt aga ttc acc gat caa atg ata tcc gct att aca aga gta cgt tat act aga gcc aaa gtc tac tcc gcc aat atc aag ttg gcc att atc gat atc tcg agg cga tgg gca tct ccg ttt aat aca ttg att aaa gag tgt cca tcc ggt acc ggt aca ttt agc ata tat ggg tcc cat ttt ttg ctt tct gta tcc agg tag aca tag a-t att cta tag tgt ctc cta tgt tgt a-- --- --- --- --- --- -at tag cat cag tc- --t cta cac tat tct taa att tca tat taa tgg ggc gtg acg gaa tag tac agt atg ata gaa cac atc cta ttc cca aca atg tca gga acg tca cgc tct cca cct tca tat tta ttt atc cgt --a aaa tgt tat cct gga cat cgt aca aat aat a-a aaa gcc cat ata tat gtt cgc tat tgt aga aat tgt ttt tca cag ttg ctc aaa aac aat ggc agt gac tta tga gtt a-g tta cac ttt gga gtc tca tct tta gta aac ata tca taa tat tcg ata tta cga gtt gac ata tcg aac aaa ttc caa gta ttt gat ttt gga taa tat tcg tat ttt gca tct gct ata att aag ata taa tca cca caa gaa cac acg aac gtc ttt cct aca tgg tta aag tac atg tac aat tct atc cat ttg tct tcc tta act ata tat ttg tat aga taa tta cga gtc tca tga gta att cca gta att gca tag atg tca cca tcg tat tct aca gca taa act ata cta tga cgt cta ggc atg gga gac -tt ttt tat cca acg att ttt agt gaa aca ttc cac atc gtt taa tac tac ata ttt ctc ata -gt ggt ata aac tcc acc cat tac --a tat ata tca tcg ttt acg aat act gat gcg cct gaa tat cta gga gtg att aag ttt gga agt ctt ttc cat ttc gaa gtg ccg tgt ttc aaa tat tct gct ata ccc gtt gaa ata gaa aat tct aat cct cct att aca tat aac ttt cca tcg tta aca caa gta cta act tct gat ttt aac gac gac ata tta gta acc gtt ttc ca- --- --- --- --- --- --- -tt ttt ttt gtt tta aga tct acc cgc -ga tac gga ata aac atg tct att gtt aat cat gcc gcc aat aat gta tag aca att atg taa aac att tgc atc ata gaa ttg --- -tc tat ctg tat tac cga cta tcg tcc aat att ctg ttc tag gag agt aat ggg tta ttg tgg ata tat aat cag agt ttt taa tga cta cta tat tat gtt tta tac cat ttc gtg tca c-a gct ttg tag att tgg ata tag tta atc cca aca atg cta tag cat tgc ata tag cat tag tca taa act tgg gat gta aaa tgt tga tga tat cta cat cgt ttg gat ttt tat gta tcc act tta ata ata tta tag c-g taa cat cct cat ga- ttt acg tta acg ttt tcg tgt gat aag ata gtg gtc agt tca tcc ttt gat aat ttt cca aat t

>JX878418.1:18898-20850_Monkeypox_virus_isolate_DRC_07-0120_complete_genome

cgg ctg cca tgt att tcc tgg aga gca agt --- aga tga tg- agg aac cag ata gtt tat atc cat act tgc act taa agt cta cat tgt agt tgt atg aga gta tga tct ttt aag -cc gct aga -ag ttt tcc gtt tga tat --- agg atg tgg aca ttt aac aat ctg aca cgt ggg tgg att gga cca ttc tcc tcc tga aca cat gac acc aga gtt acc aat caa cga ata tcc act att gca act ata agt tac aat gct ccc atc gat ata aaa atc ctc gta tcc gtt atg tct tcc gtt gga tat aga tgg agg tga ttg gca ttt aac aga ttc gca aat agg tgc ctc agg att cca tac cat aga tcc agt aga tcc taa ttc aca ata cga ttt aga ttc acc gat caa atg ata tcc gct att aca aga gta cgt tat act aga gcc aaa gtc tac tcc gcc aat atc aag ttg gcc att atc gat atc tcg agg cga tgg gca tct ccg ttt aat aca ttg att aaa gag tgt cca tcc ggt acc ggt aca ttt agc ata tat ggg tcc cat ttt ttg ctt tct gta tcc agg tag aca tag a-t att cta tag tgt ctc cta tgt tgt a-- --- --- --- --- --- -at tag cat cag tc- --t cta cac tat tct taa att tca tat taa tgg ggc gtg acg gaa tag tac agt atg ata gaa cac atc cta ttc cca aca atg tca gga acg tca cgc tct cca cct tca tat tta ttt atc cgt --a aaa tgt tat cct gga cat cgt aca aat aat a-a aaa gcc cat ata tat gtt cgc tat tgt aga aat tgt ttt tca cag ttg ctc aaa aac aat ggc agt gac tta tga gtt a-g tta cac ttt gga gtc tca tct tta gta aac ata tca taa tat tcg ata tta cga gtt gac ata tcg aac aaa ttc caa gta ttt gat ttt gga taa tat tcg tat ttt gca tct gct ata att aag ata taa tca cca caa gaa cac acg aac gtc ttt cct aca tgg tta aag tac atg tac aat tct atc cat ttg tct tcc tta act ata tat ttg tat aga taa tta cga gtc tca tga gta att cca gta att gca tag atg tca cca tcg tat tct aca gca taa act ata cta tga cgt cta ggc atg gga gac -tt ttt tat cca acg att ttt agt gaa aca ttc cac atc gtt taa tac tac ata ttt ctc ata -gt ggt ata aac tcc acc cat tac --a tat ata tca tcg ttt acg aat act gat gcg cct gaa tat cta gga gtg att aag ttt gga agt ctt ttc cat ttc gaa gtg ccg tgt ttc aaa tat tct gct ata ccc gtt gaa ata gaa aat tct aat cct cct att aca tat aac ttt cca tcg tta aca caa gta cta act tct gat ttt aac gac gac ata tta gta acc gtt ttc ca- --- --- --- --- --- --- -tt ttt ttt gtt tta aga tct acc cgc -ga tac gga ata aac atg tct att gtt aat cat gcc gcc aat aat gta tag aca att atg taa aac att tgc atc ata gaa ttg --- -tc tat ctg tat tac cga cta tcg tcc aat att ctg ttc tag gag agt aat ggg tta ttg tgg ata tat aat cag agt ttt taa tga cta cta tat tat gtt tta tac cat ttc gtg tca c-a gct ttg tag att tgg ata tag tta atc cca aca atg cta tag cat tgc ata tag cat tag tca taa act tgg gat gta aaa tgt tga tga tat cta cat cgt ttg gat ttt tat gta tcc act tta ata ata tta tag c-g taa cat cct cat ga- ttt acg tta acg ttt tcg tgt gat aag ata gtg gtc agt tca tcc ttt gat aat ttt cca aat t

>JX878408.1:18898-20850_Monkeypox_virus_isolate_DRC_06-0970_complete_genome

cgg ctg cca tgt att tcc tgg aga gca agt --- aga tga tg- agg aac cag ata gtt tat atc cat act tgc act taa agt cta cat tgt agt tgt atg aga gta tga tct ttt aag -cc gct aga -ag ttt tcc gtt tga tat --- agg atg tgg aca ttt aac aat ctg aca cgt ggg tgg att gga cca ttc tcc tcc tga aca cat gac acc aga gtt acc aat caa cga ata tcc act att gca act ata agt tac aat gct ccc atc gat ata aaa atc ctc gta tcc gtt atg tct tcc gtt gga tat aga tgg agg tga ttg gca ttt aac aga ttc gca aat agg tgc ctc agg att cca tac cat aga tcc agt aga tcc taa ttc aca ata cga ttt aga ttc acc gat caa atg ata tcc gct att aca aga gta cgt tat act aga gcc aaa gtc tac tcc gcc aat atc aag ttg gcc att atc gat atc tcg agg cga tgg gca tct ccg ttt aat aca ttg att aaa gag tgt cca tcc ggt acc ggt aca ttt agc ata tat ggg tcc cat ttt ttg ctt tct gta tcc agg tag aca tag a-t att cta tag tgt ctc cta tgt tgt a-- --- --- --- --- --- -at tag cat cag tc- --t cta cac tat tct taa att tca tat taa tgg ggc gtg acg gaa tag tac agt atg ata gaa cac atc cta ttc cca aca atg tca gga acg tca cgc tct cca cct tca tat tta ttt atc cgt --a aaa tgt tat cct gga cat cgt aca aat aat a-a aaa gcc cat ata tat gtt cgc tat tgt aga aat tgt ttt tca cag ttg ctc aaa aac aat ggc agt gac tta tga gtt a-g tta cac ttt gga gtc tca tct tta gta aac ata tca taa tat tcg ata tta cga gtt gac ata tcg aac aaa ttc caa gta ttt gat ttt gga taa tat tcg tat ttt gca tct gct ata att aag ata taa tca cca caa gaa cac acg aac gtc ttt cct aca tgg tta aag tac atg tac aat tct atc cat ttg tct tcc tta act ata tat ttg tat aga taa tta cga gtc tca tga gta att cca gta att gca tag atg tca cca tcg tat tct aca gca taa act ata cta tga cgt cta ggc atg gga gac -tt ttt tat cca acg att ttt agt gaa aca ttc cac atc gtt taa tac tac ata ttt ctc ata -gt ggt ata aac tcc acc cat tac --a tat ata tca tcg ttt acg aat act gat gcg cct gaa tat cta gga gtg att aag ttt gga agt ctt ttc cat ttc gaa gtg ccg tgt ttc aaa tat tct gct ata ccc gtt gaa ata gaa aat tct aat cct cct att aca tat aac ttt cca tcg tta aca caa gta cta act tct gat ttt aac gac gac ata tta gta acc gtt ttc ca- --- --- --- --- --- --- -tt ttt ttt gtt tta aga tct acc cgc -ga tac gga ata aac atg tct att gtt aat cat gcc gcc aat aat gta tag aca att atg taa aac att tgc atc ata gaa ttg --- -tc tat ctg tat tac cga cta tcg tcc aat att ctg ttc tag gag agt aat ggg tta ttg tgg ata tat aat cag agt ttt taa tga cta cta tat tat gtt tta tac cat ttc gtg tca c-a gct ttg tag att tgg ata tag tta atc cca aca atg cta tag cat tgc ata tag cat tag tca taa act tgg gat gta aaa tgt tga tga tat cta cat cgt ttg gat ttt tat gta tcc act tta ata ata tta tag c-g taa cat cct cat ga- ttt acg tta acg ttt tcg tgt gat aag ata gtg gtc agt tca tcc ttt gat aat ttt cca aat t

>AF380138.1:18952-20904_Monkeypox_virus_strain_Zaire-96-I-16_complete_genome

cgg ctg cca tgt att tcc tgg aga gca agt --- aga tga tg- agg aac cag ata gtt tat atc cat act tgc act taa agt cta cat tgt agt tgt atg aga gta tga tct ttt aag -cc gct aga -ag ttt tcc gtt tga tat --- agg atg tgg aca ttt aac aat ctg aca cgt ggg tgg att gga cca ttc tcc tcc tga aca cat gac acc aga gtt acc aat caa cga ata tcc act att gca act ata agt tac aat gct ccc atc gat ata aaa atc ctc gta tcc gtt atg tct tcc gtt gga tat aga tgg agg tga ttg gca ttt aac aga ttc gca aat agg tgc ctc agg att cca tac cat aga tcc agt aga tcc taa ttc aca ata cga ttt aga ttc acc gat caa atg ata tcc gct att aca aga gta cgt tat act aga gcc aaa gtc tac tcc gcc aat atc aag ttg gcc att atc gat atc tcg agg cga tgg gca tct ccg ttt aat aca ttg att aaa gag tgt cca tcc ggt acc ggt aca ttt agc ata tat ggg tcc cat ttt ttg ctt tct gta tcc agg tag aca tag a-t att cta tag tgt ctc cta tgt tgt a-- --- --- --- --- --- -at tag cat cag tc- --t cta cac tat tct taa att tca tat taa tgg ggc gtg acg gaa tag tac agt atg ata gaa cac atc cta ttc cca aca atg tca gga acg tca cgc tct cca cct tca tat tta ttt atc cgt --a aaa tgt tat cct gga cat cgt aca aat aat a-a aaa gcc cat ata tat gtt cgc tat tgt aga aat tgt ttt tca cag ttg ctc aaa aac aat ggc agt gac tta tga gtt a-g tta cac ttt gga gtc tca tct tta gta aac ata tca taa tat tcg ata tta cga gtt gac ata tcg aac aaa ttc caa gta ttt gat ttt gga taa tat tcg tat ttt gca tct gct ata att aag ata taa tca cca caa gaa cac acg aac gtc ttt cct aca tgg tta aag tac atg tac aat tct atc cat ttg tct tcc tta act ata tat ttg tat aga taa tta cga gtc tca tga gta att cca gta att gca tag atg tca cca tcg tat tct aca gca taa act ata cta tga cgt cta ggc atg gga gac -tt ttt tat cca acg att ttt agt gaa aca ttc cac atc gtt taa tac tac ata ttt ctc ata -gt ggt ata aac tcc acc cat tac --a tat ata tca tcg ttt acg aat act gat gcg cct gaa tat cta gga gtg att aag ttt gga agt ctt ttc cat ttc gaa gtg ccg tgt ttc aaa tat tct gct ata ccc gtt gaa ata gaa aat tct aat cct cct att aca tat aac ttt cca tcg tta aca caa gta cta act tct gat ttt aac gac gac ata tta gta acc gtt ttc ca- --- --- --- --- --- --- -tt ttt ttt gtt tta aga tct acc cgc -ga tac gga ata aac atg tct att gtt aat cat gcc gcc aat aat gta tag aca att atg taa aac att tgc atc ata gaa ttg --- -tc tat ctg tat tac cga cta tcg tcc aat att ctg ttc tag gag agt aat ggg tta ttg tgg ata tat aat cag agt ttt taa tga cta cta tat tat gtt tta tac cat ttc gtg tca c-a gct ttg tag att tgg ata tag tta atc cca aca atg cta tag cat tgc ata tag cat tag tca taa act tgg gat gta aaa tgt tga tga tat cta cat cgt ttg gat ttt tat gta tcc act tta ata ata tta tag c-g taa cat cct cat ga- ttt acg tta acg ttt tcg tgt gat aag ata gtg gtc agt tca tcc ttt gat aat ttt cca aat t

>KC257459.1:19438-21391_Monkeypox_virus_strain_Sudan_2005_01_complete_genome

cgg ctg cca tgt att tcc tgg aga gca agt --- aga tga tg- agg aac cag ata gtt tat atc cat act tgc act taa agt cta cat tgt agt tgt atg agt gta tga tct ttt aag -cc gct aga -ag ttt tcc gtt tga tat --- agg atg tgg aca ttt aac aat ctg aca cgt ggg tgg att gga cca ttc tcc tcc tga aca cat gac acc aga gtt acc aat caa cga ata tcc act att gca act ata agt tac aat gct ccc atc gat ata aaa atc ctc gta tcc gtt atg tct tcc gtt gga tat aga tgg agg tga ttg gca ttt aac aga ttc gca aat agg tgc ctc agg att cca tac cat aga tcc agt aga tcc taa ttc aca ata cga ttt aga ttc acc gat caa atg ata tcc gct att aca aga gta cgt tat act aga gcc aaa gtc tac tcc gcc aat atc aag ttg gcc att atc gat atc tcg agg cga tgg gca tct ccg ttt aat aca ttg att aaa gag tgt cca tcc ggt acc ggt aca ttt agc ata tat ggg tcc cat ttt ttg ctt tct gta tcc agg tag aca tag a-t att cta tag tgt ctc cta tgt tgt a-- --- --- --- --- --- -at tag cat cag tc- --t cta cac tat tct taa att tca tat taa tgg ggc gtg acg gaa tag tac agt atg ata gaa cac atc cta ttc cca aca atg tca gga acg tca cgc tct cca cct tca tat tta ttt atc cgt --a aaa tgt tat cct gga cat cgt aca aat aat a-a aaa gcc cat ata tat gtt cgc tat tgt aga aat tgt ttt tca cag ttg ctc aaa aac aat ggc agt gac tta tga gtt a-g tta cac ttt gga gtc tca tct tta gta aac ata tca taa tat tcg ata tta cga gtt gac ata tcg aac aaa ttc caa gta ttt gat ttt gga taa tat tcg tat ttt gca tct gct ata att aag ata taa tca cca caa gaa cac acg aac gtc ttt cct aca tgg tta aag tac atg tac aat tct atc cat ttg tct tcc tta act ata tat ttg tat aga taa tta cga gtc tca tga gta att cca gta att gca tag atg tca cca tcg tat tct aca gca taa act ata cta tga cgt cta ggc atg gga gac -tt ttt tat cca acg att ttt agt gaa aca ttc cac atc gtt taa tac tac ata ttt ctc ata -gt ggt ata aac tcc acc cat tac --a tat ata tca tcg ttt acg aat act gat gcg cct gaa tat cta gga gtg att aag ttt gga agt ctt ttc cat ttc gaa gtg ccg tgt ttc aaa tat tct gct ata ccc gtt gaa ata gaa aat tct aat cct cct att aca tat aac ttt cca tcg tta aca caa gta cta act tct gat ttt aac gac gac ata tta gta acc gtt ttc ca- --- --- --- --- --- --- ttt ttt ttt gtt tta aga tct acc cgc -ga tac gga ata aac atg tct att gtt aat cat gcc gcc aat aat gta tag aca att atg taa aac att tgc atc ata gaa ttg --- -tc tat ctg tat tac cga cta tcg tcc aat att ctg ttc tag gag agt aat ggg tta ttg tga ata tat aat cag agt ttt taa tga cta cta tat tat gtt tta tac cat ttc gtg tca c-a gct ttg tag att tgg ata tag tta atc cca aca atg cta tag cat tgc ata tag cat tag tca taa act tgg gat gta aaa tgt tga tga tat cta cat cgt ttg gat ttt tat gta tcc act tta ata ata tta tag c-g taa cat cct cat ga- ttt acg tta acg ttt tcg tgt gat aag ata gtg gtc agt tca tcc ttt gat aat ttt cca aat t

>JX878417.1:19446-21398_Monkeypox_virus_isolate_DRC_07-0104_complete_genome

cgg ctg cca tgt att tcc tgg aga gca agt --- aga tga tg- agg aac cag ata gtt tat atc cat act tgc act taa agt cta cat tgt agt tgt atg agt gta tga tct ttt aag -cc gct aga -ag ttt tcc gtt tga tat --- agg atg tgg aca ttt aac aat ctg aca cgt ggg tgg att gga cca ttc tcc tcc tga aca cat gac acc aga gtt acc aat caa cga ata tcc act att gca act ata agt tac aat gct ccc atc gat ata aaa atc ctc gta tcc gtt atg tct tcc gtt gga tat aga tgg agg tga ttg gca ttt aac aga ttc gca aat agg tgc ctc agg att cca tac cat aga tcc agt aga tcc taa ttc aca ata cga ttt aga ttc acc gat caa atg ata tcc gct att aca aga gta cgt tat act aga gcc aaa gtc tac tcc gcc aat atc aag ttg acc att atc gat atc tcg agg cga tgg gca tct ccg ttt aat aca ttg att aaa gag tgt cca tcc ggt acc ggt aca ttt agc ata tat ggg tcc cat ttt ttg ctt tct gta tcc agg tag aca tag a-t att cta tag tgt ctc cta tgt tgt a-- --- --- --- --- --- -at tag cat cag tc- --t cta cac tat tct taa att tca tat taa tgg ggc gtg acg gaa tag tac agt atg ata gaa cac atc cta ttc cca aca atg tca gga acg tca cgc tct cca cct tca tat tta ttt atc cgt --a aaa tgt tat cct gga cat cgt aca aat aat a-a aaa gcc cat ata tat gtt cgc tat tgt aga aat tgt ttt tca cag ttg ctc aaa aac aat ggc agt gac tta tga gtt c-g tta cac ttt gga gtc tca tct tta gta aac ata tca taa tat tcg ata tta cga gtt gac ata tcg aac aaa ttc caa gta ttt gat ttt gga taa tat tcg tat ttt gca tct gct ata att aag ata taa tca cca caa gaa cac acg aac gtc ttt cct aca tgg tta aag tac atg tac aat tct atc cat ttg tct tcc tta act ata tat ttg tat aga taa tta cga gtc tca tga gta att cca gta att gca tag atg tca cca tcg tat tct aca gca taa act ata cta tga cgt cta ggc atg gga gac -tt ttt tat cca acg att ttt agt gaa aca ttc cac atc gtt taa tac tac ata ttt ctc ata -gt ggt ata aac tcc acc cat tac --a tat ata tca tcg ttt acg aat act gat gcg cct gaa tat cta gga gtg att aag ttt gga agt ctt ttc cat ttc gaa gtg ccg tgt ttc aaa tat tct gct ata ccc gtt gaa ata gaa aat tct aat cct cct att aca tat aac ttt cca tcg tta aca caa gta cta act tct gat ttt aac gac gac ata tta gta acc gtt ttc ca- --- --- --- --- --- --- -tt ttt ttt gtt tta aga tct acc cgc -ga tac gga ata aac atg tct att gtt aat cat gcc gcc aat aat gta tag aca att atg taa aac att tgc atc ata gaa ttg --- -tc tat ctg tat tac cga cta tcg tcc aat att ctg ttc tag gag agt aat ggg tta ttg tgg ata tat aat cag agt ttt taa tga cta cta tat tat gtt tta tac cat ttc gtg tca c-a gct ttg tag att tgg ata tag tta atc cca aca atg cta tag cat tgc ata tag cat tag tca taa act tgg gat gta aaa tgt tga tga tat cta cat cgt ttg gat ttt tat gta tcc act tta ata ata tta tag c-g taa cat cct cat ga- ttt acg tta acg ttt tcg tgt gat aag ata gtg gtc agt tca tcc ttt gat aat ttt cca aat t

>DQ011155.1:19076-21027_Monkeypox_virus_strain_Zaire_1979-005_complete_genome

cgg ctg cca tgt att tcc tgg aga gca agt --- aga tga tg- agg aac cag ata gtt tat atc cat act tgc act taa agt cta cat tgt agt tgt atg agt gta tga tct ttt aag -cc gct aga -ag ttt tcc gtt tga tat --- agg atg tgg aca ttt aac aat ctg aca cgt ggg tgg att gga cca ttc tcc tcc tga aca cat gac acc aga gtt acc aat caa cga ata tcc act att gca act ata agt tac aat gct ccc atc gat ata aaa atc ctc gta tcc gtt atg tct tcc gtt gga tat aga tgg agg tga ttg gca ttt aac aga ttc gca aat agg tgc ctc agg att cca tac cat aga tcc agt aga tcc taa ttc aca ata cga ttt aga ttc acc gat caa atg ata tcc gct att aca aga gta cgt tat act aga gcc aaa gtc tac tcc gcc aat atc aag ttg gcc att atc gat atc tcg agg cga tgg gca tct ccg ttt aat aca ttg att aaa gag tgt cca tcc ggt acc ggt aca ttt agc ata tat ggg tcc cat ttt ttg ctt tct gta tcc agg tag aca tag a-t att cta tag tgt ctc cta tgt tgt a-- --- --- --- --- --- -at tag cat cag tc- --t cta cac tat tct taa att tca tat taa tgg ggc gtg acg gaa tag tac agt atg ata gaa cac atc cta ttc cca aca atg tca gga acg tca cgc tct cca cct tca tat tta ttt atc cgt --a aaa tgt tat cct gga cat cgt aca aat aat a-a aaa gcc cat ata tat gtt cgc tat tgt aga aat tgt ttt tca cag ttg ctc aaa aac aat ggc agt gac tta tga gtt a-g tta cac ttt gga gtc tca tct tta gta aac ata tca taa tat tcg ata tta cga gtt gac ata tcg aac aaa ttc caa gta ttt gat ttt gga taa tat tcg tat ttt gca tct gct ata att aag ata taa tca cca caa gaa cac acg aac gtc ttt cct aca tgg tta aag tac atg tac aat tct atc cat ttg tct tcc tta act ata tat ttg tat aga taa tta cga gtc tca tga gta att cca gta att gca tag atg tca cca tcg tat tct aca gca taa act ata cta tga cgt cta ggc atg gga gac -tt ttt tat cca acg att ttt agt gaa aca ttc cac atc gtt taa tac tac ata ttt ctc ata -gt ggt ata aac tcc acc cat tac --a tat ata tca tcg ttt acg aat act gat gcg cct gaa tat cta gga gtg att aag ttt gga agt ctt ttc cat ttc gaa gtg ccg tgt ttc aaa tat tct gct ata ccc gtt gaa ata gaa aat tct aat cct cct att aca tat aac ttt cca tcg tta aca caa gta cta act tct gat ttt aac gac gac ata tta gta acc gtt ttc ca- --- --- --- --- --- --- --t ttt ttt gtt tta aga tct acc cgc -ga tac gga ata aac atg tct att gtt aat cat gcc gcc aat aat gta tag aca att atg taa aac att tgc atc ata gaa ttg --- -tc tat ctg tat tac cga cta tcg tcc aat att ctg ttc tag gag agt aat ggg tta ttg tga ata tat aat cag agt ttt taa tga cta cta tat tat gtt tta tac cat ttc gtg tca c-a gct ttg tag att tgg ata tag tta atc cca aca atg cta tag cat tgc ata tag cat tag tca taa act tgg gat gta aaa tgt tga tga tat cta cat cgt ttg gat ttt tat gta tcc act tta ata ata tta tag c-g taa cat cct cat ga- ttt acg tta acg ttt tcg tgt gat aag ata gtg gtc agt tca tcc ttt gat aat ttt cca aat t

>HM172544.1:19078-21029_Monkeypox_virus_strain_Zaire_1979-005_complete_genome

cgg ctg cca tgt att tcc tgg aga gca agt --- aga tga tg- agg aac cag ata gtt tat atc cat act tgt act taa agt cta cat tgt agt tgt atg agt gta tga tct ttt aag -cc gct aga -ag ttt tcc gtt tga tat --- agg atg tgg aca ttt aac aat ctg aca cgt ggg tgg att gga cca ttc tcc tcc tga aca cat gac acc aga gtt acc aat caa cga ata tcc act att gca act ata agt tac aat gct ccc atc gat ata aaa atc ctc gta tcc gtt atg tct tcc gtt gga tat aga tgg agg tga ttg gca ttt aac aga ttc gca aat agg tgc ctc agg att cca tac cat aga tcc agt aga tcc taa ttc aca ata cga ttt aga ttc acc gat caa atg ata tcc gct att aca aga gta cgt tat act aga gcc aaa gtc tac tcc gcc aat atc aag ttg gcc att atc gat atc tcg agg cga tgg gca tct ccg ttt aat aca ttg att aaa gag tgt cca tcc ggt acc ggt aca ttt agc ata tat ggg tcc cat ttt ttg ctt tct gta tcc agg tag aca tag a-t att cta tag tgt ctc cta tgt tgt a-- --- --- --- --- --- -at tag cat cag tc- --t cta cac tat tct taa att tca tat taa tgg ggc gtg acg gaa tag tac agt atg ata gaa cac atc cta ttc cca aca atg tca gga acg tca cgc tct cca cct tca tat tta ttt atc cgt --a aaa tgt tat cct gga cat cgt aca aat aat a-a aaa gcc cat ata tat gtt cgc tat tgt aga aat tgt ttt tca cag ttg ctc aaa aac aat ggc agt gac tta tga gtt a-g tta cac ttt gga gtc tca tct tta gta aac ata tca taa tat tcg ata tta cga gtt gac ata tcg aac aaa ttc caa gta ttt gat ttt gga taa tat tcg tat ttt gca tct gct ata att aag ata taa tca cca caa gaa cac acg aac gtc ttt cct aca tgg tta aag tac atg tac aat tct atc cat ttg tct tcc tta act ata tat ttg tat aga taa tta cga gtc tca tga gta att cca gta att gca tag atg tca cca tcg tat tct aca gca taa act ata cta tga cgt cta ggc atg gga gac -tt ttt tat cca acg att ttt agt gaa aca ttc cac atc gtt taa tac tac ata ttt ctc ata -gt ggt ata aac tcc acc cat tac --a tat ata tca tcg ttt acg aat act gat gcg cct gaa tat cta gga gtg att aag ttt gga agt ctt ttc cat ttc gaa gtg ccg tgt ttc aaa tat tct gct ata ccc gtt gaa ata gaa aat tct aat cct cct att aca tat aac ttt cca tcg tta aca caa gta cta act tct gat ttt aac gac gac ata tta gta acc gtt ttc ca- --- --- --- --- --- --- --t ttt ttt gtt tta aga tct acc cgc -ga tac gga ata aac atg tct att gtt aat cat gcc gcc aat aat gta tag aca att atg taa aac att tgc atc ata gaa ttg --- -tc tat ctg tat tac cga cta tcg tcc aat att ctg ttc tag gag agt aat ggg tta ttg tga ata tat aat cag agt ttt taa tga cta cta tat tat gtt tta tac cat ttc gtg tca c-a gct ttg tag att tgg ata tag tta atc cca aca atg cta tag cat tgc ata tag cat tag tca taa act tgg gat gta aaa tgt tga tga tat cta cat cgt ttg gat ttt tat gta tcc act tta ata ata tta tag c-g taa cat cct cat ga- ttt acg tta acg ttt tcg tgt gat aag ata gtg gtc agt tca tcc ttt gat aat ttt cca aat t

>KC257460.1:18875-20830_Monkeypox_virus_strain_DRC_Yandongi_1985_complete_genome

cgg ctg cca tgt att tcc tgg aga gca agt --- aga tga tg- agg aac cag ata gtt tat atc cat act tgc act taa agt cta cat tgt agt tgt atg agt gta tga tct ttt aag -cc gct aga -ag ttt tcc gtt tga tat --- agg atg tgg aca ttt aac aat ctg aca cgt ggg tgg att gga cca ttc tcc tcc tga aca cat gac acc aga gtt acc aat caa cga ata tcc act att gca act ata agt tac aat gct ccc atc gat ata aaa atc ctc gta tcc gtt atg tct tcc gtt gga tat aga tgg agg tga ttg gca ttt aac aga ttc gca aat agg tgc ctc agg att cca tac cat aga tcc agt aga tcc taa ttc aca ata cga ttt aga ttc acc gat caa atg ata tcc gct att aca aga gta cgt tat act aga gcc aaa gtc tac tcc gcc aat atc aag ttg gcc att atc gat atc tcg agg cga tgg gca tct ccg ttt aat aca ttg att aaa gag tgt cca tcc ggt acc ggt aca ttt agc ata tat ggg tcc cat ttt ttg ctt tct gta tcc agg tag aca tag a-t att cta tag tgt ctc cta tgt tgt a-- --- --- --- --- --- -at tag cat cag tc- --t cta cac tat tct taa att tca tat taa tgg ggc gtg acg gaa tag tac agt atg ata gaa cac atc cta ttc cca aca atg tca gga acg tca cgc tct cca cct tca tat tta ttt atc cgt --a aaa tgt tat cct gga cat cgt aca aat aat a-a aaa gcc cat ata tat gtt cgc tat tgt aga aat tgt ttt tca cag ttg ctc aaa aac aat ggc agt gac tta tga gtt a-g tta cac ttt gga gtc tca tct tta gta aac ata tca taa tat tcg ata tta cga gtt gac ata tcg aac aaa ttc caa gta ttt gat ttt gga taa tat tcg tat ttt gca tct gct ata att aag ata taa tca cca caa gaa cac acg aac gtc ttt cct aca tgg tta aag tac atg tac aat tct atc cat ttg tct tcc tta act ata tat ttg tat aga taa tta cga gtc tca tga gta att cca gta att gca tag atg tca cca tcg tat tct aca gca taa act ata cta tga cgt cta ggc atg gga gac -tt ttt tat cca acg att ttt agt gaa aca ttc cac atc gtt taa tac tac ata ttt ctc ata -gt ggt ata aac tcc acc cat tac --a tat ata tca tcg ttt acg aat act gat gcg cct gaa tat cta gga gtg att aag ttt gga agt ctt ttc cat ttc gaa gtg ccg tgt ttc aaa tat tct gct ata ccc gtt gaa ata gaa aat tct aat cct cct att aca tat aac ttt cca tcg tta aca caa gta cta act tct gat ttt aac gac gac ata tta gta acc gtt ttc ca- --- --- --- --- --- -tt ttt ttt ttt gtt tta aga tct acc cgc -ga tac gga ata aac atg tct att gtt aat cat gcc gcc aat aat gta tag aca att atg taa aac att tgc atc ata gaa ttg --- -tc tat ctg tat tac cga cta tcg tcc aat att ctg ttc tag gag agt aat ggg tta ttg tga ata tat aat cag agt ttt taa tga cta cta tat tat gtt tta tac cat ttc gtg tca c-a gct ttg tag att tgg ata tag tta atc cca aca atg cta tag cat tgc ata tag cat tag tca taa act tgg gat gta aaa tgt tga tga tat cta cat cgt ttg gat ttt tat gta tcc act tta ata ata tta tag c-g taa cat cct cat ga- ttt acg tta acg ttt tcg tgt gat aag ata gtg gtc agt tca tcc ttt gat aat ttt cca aat t

>MN702446.1:18752-20706_Monkeypox_virus_strain_38c_contig_SPADES

cgg ctg cca tgt att tcc tgg aga gca agt --- aga tga tg- agg aac cag ata gtt tat atc cat act tgc act taa agt cta cat tgt agt tgt atg agt gta tga tct ttt aag -cc gct aga -ag ttt tcc gtt tga tat --- agg atg tgg aca ttt aac aat ctg aca cgt ggg tgg att gga cca ttc tcc tcc tga aca cat gac acc aga gtt acc aat caa cga ata tcc act att gca act ata agt tac aat gct ccc atc gat ata aaa atc ctc gta tcc gtt atg tct tcc gtt gga tat aga tgg agg tga ttg gca ttt aac aga ttc gca aat agg tgc ctc agg att cca tac cat aga tcc agt aga tcc taa ttc aca ata cga ttt aga ttc acc gat caa atg ata tcc gct att aca aga gta cgt tat act aga gcc aaa gtc tac tcc gcc aat atc aag ttg gcc att atc gat atc tcg agg cga tgg gca tct ccg ttt aat aca ttg att aaa gag tgt cca tcc ggt acc ggt aca ttt agc ata tat ggg tcc cat ttt ttg ctt tct gta tcc agg tag aca tag a-t att cta tag tgt ctc cta tgt tgt a-- --- --- --- --- --- -at tag cat cag tc- --t cta cac tat tct taa att tca tat taa tgg ggc gtg acg gaa tag tac agt atg ata gaa cac atc cta ttc cca aca atg tca gga acg tca cgc tct cca cct tca tat tta ttt atc cgt --a aaa tgt tat cct gga cat cgt aca aat aat a-a aaa gcc cat ata tat gtt cgc tat tgt aga aat tgt ttt tca cag ttg ctc aaa aac aat ggc agt gac tta tga gtt a-g tta cac ttt gga gtc tca tct tta gta aac ata tca taa tat tcg ata tta cga gtt gac ata tcg aac aaa ttc caa gta ttt gat ttt gga taa tat tcg tat ttt gca tct gct ata att aag ata taa tca cca caa gaa cac acg aac gtc ttt cct aca tgg tta aag tac atg tac aat tct atc cat ttg tct tcc tta act ata tat ttg tat aga taa tta cga gtc tca tga gta att cca gta att gca tag atg tca cca tcg tat tct aca gca taa act ata cta tga cgt cta ggc atg gga gac -tt ttt tat cca acg att ttt agt gaa aca ttc cac atc gtt taa tac tac ata ttt ctc ata -gt ggt ata aac tcc acc cat tac --a tat ata tca tcg ttt acg aat act gat gcg cct gaa tat cta gga gtg att aag ttt gga agt ctt ttc cat ttc gaa gtg ccg tgt ttc aaa tat tct gct ata ccc gtt gaa ata gaa aat tct aat cct cct att aca tat aac ttt cca tcg tta aca caa gta cta act tct gat ttt aac gac gac ata tta gta acc gtt ttc ca- --- --- --- --- --- --t ttt ttt ttt gtt tta aga tct acc cgc -ga tac gga ata aac atg tct att gtt aat cat gcc gcc aat aat gta tag aca att atg taa aac att tgc atc ata gaa ttg --- -tc tat ctg tat tac cga cta tcg tcc aat att ctg ttc tag gag agt aat ggg tta ttg tga ata tat aat cag agt ttt taa tga cta cta tat tat gtt tta tac cat ttc gtg tca c-a gct ttg tag att tgg ata tag tta atc cca aca atg cta tag cat tgc ata tag cat tag tca taa act tgg gat gta aaa tgt tga tga tat cta cat cgt ttg gat ttt tat ata tcc act tta ata ata tta tag c-g taa cat cct cat ga- ttt acg tta acg ttt tcg tgt gat aag ata gtg gtc agt tca tcc ttt gat aat ttt cca aat t

>MN702445.1:12606-14560_Monkeypox_virus_strain_A4_contig_SPADES

cgg ctg cca tgt att tcc tgg aga gca agt --- aga tga tg- agg aac cag ata gtt tat atc cat act tgc act taa agt cta cat tgt agt tgt atg agt gta tga tct ttt aag -cc gct aga -ag ttt tcc gtt tga tat --- agg atg tgg aca ttt aac aat ctg aca cgt ggg tgg att gga cca ttc tcc tcc tga aca cat gac acc aga gtt acc aat caa cga ata tcc act att gca act ata agt tac aat gct ccc atc gat ata aaa atc ctc gta tcc gtt atg tct tcc gtt gga tat aga tgg agg tga ttg gca ttt aac aga ttc gca aat agg tgc ctc agg att cca tac cat aga tcc agt aga tcc taa ttc aca ata cga ttt aga ttc acc gat caa atg ata tcc gct att aca aga gta cgt tat act aga gcc aaa gtc tac tcc gcc aat atc aag ttg gcc att atc gat atc tcg agg cga tgg gca tct ccg ttt aat aca ttg att aaa gag tgt cca tcc ggt acc ggt aca ttt agc ata tat ggg tcc cat ttt ttg ctt tct gta tcc agg tag aca tag a-t att cta tag tgt ctc cta tgt tgt a-- --- --- --- --- --- -at tag cat cag tc- --t cta cac tat tct taa att tca tat taa tgg ggc gtg acg gaa tag tac agt atg ata gaa cac atc cta ttc cca aca atg tca gga acg tca cgc tct cca cct tca tat tta ttt atc cgt --a aaa tgt tat cct gga cat cgt aca aat aat a-a aaa gcc cat ata tat gtt cgc tat tgt aga aat tgt ttt tca cag ttg ctc aaa aac aat ggc agt gac tta tga gtt a-g tta cac ttt gga gtc tca tct tta gta aac ata tca taa tat tcg ata tta cga gtt gac ata tcg aac aaa ttc caa gta ttt gat ttt gga taa tat tcg tat ttt gca tct gct ata att aag ata taa tca cca caa gaa cac acg aac gtc ttt cct aca tgg tta aag tac atg tac aat tct atc cat ttg tct tcc tta act ata tat ttg tat aga taa tta cga gtc tca tga gta att cca gta att gca tag atg tca cca tcg tat tct aca gca taa act ata cta tga cgt cta ggc atg gga gac -tt ttt tat cca acg att ttt agt gaa aca ttc cac atc gtt taa tac tac ata ttt ctc ata -gt ggt ata aac tcc acc cat tac --a tat ata tca tcg ttt acg aat act gat gcg cct gaa tat cta gga gtg att aag ttt gga agt ctt ttc cat ttc gaa gtg ccg tgt ttc aaa tat tct gct ata ccc gtt gaa ata gaa aat tct aat cct cct att aca tat aac ttt cca tcg tta aca caa gta cta act tct gat ttt aac gac gac ata tta gta acc gtt ttc ca- --- --- --- --- --- --t ttt ttt ttt gtt tta aga tct acc cgc -ga tac gga ata aac atg tct att gtt aat cat gcc gcc aat aat gta tag aca att atg taa aac att tgc atc ata gaa ttg --- -tc tat ctg tat tac cga cta tcg tcc aat att ctg ttc tag gag agt aat ggg tta ttg tga ata tat aat cag agt ttt taa tga cta cta tat tat gtt tta tac cat ttc gtg tca c-a gct ttg tag att tgg ata tag tta atc cca aca atg cta tag cat tgc ata tag cat tag tca taa act tgg gat gta aaa tgt tga tga tat cta cat cgt ttg gat ttt tat ata tcc act tta ata ata tta tag c-g taa cat cct cat ga- ttt acg tta acg ttt tcg tgt gat aag ata gtg gtc agt tca tcc ttt gat aat ttt cca aat t

>MN702444.1:12611-14565_Monkeypox_virus_strain_A5_contig_SPADES

cgg ctg cca tgt att tcc tgg aga gca agt --- aga tga tg- agg aac cag ata gtt tat atc cat act tgc act taa agt cta cat tgt agt tgt atg agt gta tga tct ttt aag -cc gct aga -ag ttt tcc gtt tga tat --- agg atg tgg aca ttt aac aat ctg aca cgt ggg tgg att gga cca ttc tcc tcc tga aca cat gac acc aga gtt acc aat caa cga ata tcc act att gca act ata agt tac aat gct ccc atc gat ata aaa atc ctc gta tcc gtt atg tct tcc gtt gga tat aga tgg agg tga ttg gca ttt aac aga ttc gca aat agg tgc ctc agg att cca tac cat aga tcc agt aga tcc taa ttc aca ata cga ttt aga ttc acc gat caa atg ata tcc gct att aca aga gta cgt tat act aga gcc aaa gtc tac tcc gcc aat atc aag ttg gcc att atc gat atc tcg agg cga tgg gca tct ccg ttt aat aca ttg att aaa gag tgt cca tcc ggt acc ggt aca ttt agc ata tat ggg tcc cat ttt ttg ctt tct gta tcc agg tag aca tag a-t att cta tag tgt ctc cta tgt tgt a-- --- --- --- --- --- -at tag cat cag tc- --t cta cac tat tct taa att tca tat taa tgg ggc gtg acg gaa tag tac agt atg ata gaa cac atc cta ttc cca aca atg tca gga acg tca cgc tct cca cct tca tat tta ttt atc cgt --a aaa tgt tat cct gga cat cgt aca aat aat a-a aaa gcc cat ata tat gtt cgc tat tgt aga aat tgt ttt tca cag ttg ctc aaa aac aat ggc agt gac tta tga gtt a-g tta cac ttt gga gtc tca tct tta gta aac ata tca taa tat tcg ata tta cga gtt gac ata tcg aac aaa ttc caa gta ttt gat ttt gga taa tat tcg tat ttt gca tct gct ata att aag ata taa tca cca caa gaa cac acg aac gtc ttt cct aca tgg tta aag tac atg tac aat tct atc cat ttg tct tcc tta act ata tat ttg tat aga taa tta cga gtc tca tga gta att cca gta att gca tag atg tca cca tcg tat tct aca gca taa act ata cta tga cgt cta ggc atg gga gac -tt ttt tat cca acg att ttt agt gaa aca ttc cac atc gtt taa tac tac ata ttt ctc ata -gt ggt ata aac tcc acc cat tac --a tat ata tca tcg ttt acg aat act gat gcg cct gaa tat cta gga gtg att aag ttt gga agt ctt ttc cat ttc gaa gtg ccg tgt ttc aaa tat tct gct ata ccc gtt gaa ata gaa aat tct aat cct cct att aca tat aac ttt cca tcg tta aca caa gta cta act tct gat ttt aac gac gac ata tta gta acc gtt ttc ca- --- --- --- --- --- --t ttt ttt ttt gtt tta aga tct acc cgc -ga tac gga ata aac atg tct att gtt aat cat gcc gcc aat aat gta tag aca att atg taa aac att tgc atc ata gaa ttg --- -tc tat ctg tat tac cga cta tcg tcc aat att ctg ttc tag gag agt aat ggg tta ttg tga ata tat aat cag agt ttt taa tga cta cta tat tat gtt tta tac cat ttc gtg tca c-a gct ttg tag att tgg ata tag tta atc cca aca atg cta tag cat tgc ata tag cat tag tca taa act tgg gat gta aaa tgt tga tga tat cta cat cgt ttg gat ttt tat ata tcc act tta ata ata tta tag c-g taa cat cct cat ga- ttt acg tta acg ttt tcg tgt gat aag ata gtg gtc agt tca tcc ttt gat aat ttt cca aat t

>MN702449.1:12614-14570_Monkeypox_virus_strain_B2_contig_SPADES

cgg ctg cca tgt att tcc tgg aga gca agt --- aga tga tg- agg aac cag ata gtt tat atc cat act tgc act taa agt cta cat tgt agt tgt atg agt gta tga tct ttt aag -cc gct aga -ag ttt tcc gtt tga tat --- agg atg tgg aca ttt aac aat ctg aca cgt ggg tgg att gga cca ttc tcc tcc tga aca cat gac acc aga gtt acc aat caa cga ata tcc act att gca act ata agt tac aat gct ccc atc gat ata aaa atc ctc gta tcc gtt atg tct tcc gtt gga tat aga tgg agg tga ttg gca ttt aac aga ttc gca aat agg tgc ctc agg att cca tac cat aga tcc agt aga tcc taa ttc aca ata cga ttt aga ttc acc gat caa atg ata tcc gct att aca aga gta cgt tat act aga gcc aaa gtc tac tcc gcc aat atc aag ttg gcc att atc gat atc tcg agg cga tgg gca tct ccg ttt aat aca ttg att aaa gag tgt cca tcc ggt acc ggt aca ttt agc ata tat ggg tcc cat ttt ttg ctt tct gta tcc agg tag aca tag a-t att cta tag tgt ctc cta tgt tgt a-- --- --- --- --- --- -at tag cat cag tc- --t cta cac tat tct taa att tca tat taa tgg ggc gtg acg gaa tag tac agt atg ata gaa cac atc cta ttc cca aca atg tca gga acg tca cgc tct cca cct tca tat tta ttt atc cgt --a aaa tgt tat cct gga cat cgt aca aat aat a-a aaa gcc cat ata tat gtt cgc tat tgt aga aat tgt ttt tca cag ttg ctc aaa aac aat ggc agt gac tta tga gtt a-g tta cac ttt gga gtc tca tct tta gta aac ata tca taa tat tcg ata tta cga gtt gac ata tcg aac aaa ttc caa gta ttt gat ttt gga taa tat tcg tat ttt gca tct gct ata att aag ata taa tca cca caa gaa cac acg aac gtc ttt cct aca tgg tta aag tac atg tac aat tct atc cat ttg tct tcc tta act ata tat ttg tat aga taa tta cga gtc tca tga gta att cca gta att gca tag atg tca cca tcg tat tct aca gca taa act ata cta tga cgt cta ggc atg gga gac -tt ttt tat cca acg att ttt agt gaa aca ttc cac atc gtt taa tac tac ata ttt ctc ata -gt ggt ata aac tcc acc cat tac --a tat ata tca tcg ttt acg aat act gat gcg cct gaa tat cta gga gtg att aag ttt gga agt ctt ttc cat ttc gaa gtg ccg tgt ttc aaa tat tct gct ata ccc gtt gaa ata gaa aat tct aat cct cct att aca tat aac ttt cca tcg tta aca caa gta cta act tct gat ttt aac gac gac ata tta gta acc gtt ttc ca- --- --- --- --- --- ttt ttt ttt ttt gtt tta aga tct acc cgc -ga tac gga ata aac atg tct att gtt aat cat gcc gcc aat aat gta tag aca att atg taa aac att tgc atc ata gaa ttg --- -tc tat ctg tat tac cga cta tcg tcc aat att ctg ttc tag gag agt aat ggg tta ttg tga ata tat aat cag agt ttt taa tga cta cta tat tat gtt tta tac cat ttc gtg tca c-a gct ttg tag att tgg ata tag tta atc cca aca atg cta tag cat tgc ata tag cat tag tca taa act tgg gat gta aaa tgt tga tga tat cta cat cgt ttg gat ttt tat ata tcc act tta ata ata tta tag c-g taa cat cct cat ga- ttt acg tta acg ttt tcg tgt gat aag ata gtg gtc agt tca tcc ttt gat aat ttt cca aat t

>HQ857562.1:18944-20896_Monkeypox_virus_strain_V79-I-005_complete_genome

cgg ctg cca tgt att tcc tgg aga gca agt --- aga tga tg- agg aac cag ata gtt tat atc cat act tgc act taa agt cta cat tgt agt tgt atg agt gta tga tct ttt aag -cc gct aga -ag ttt tcc gtt tga tat --- agg atg tgg aca ttt aac aat ctg aca cgt ggg tgg att gga cca ttc tcc tcc tga aca cat gac acc aga gtt acc aat caa cga ata tcc act att gca act ata agt tac aat gct ccc atc gat ata aaa atc ctc gta tcc gtt atg tct tcc gtt ggr tat aga tgg rgg kga ttg gca ttt aac aga ttc gca aat agg tgc ctc agg att cca tac cat aga tcc agt aga tcc taa ttc aca ata cga ttt aga ttc acc gat caa atg ata tcc gct att aca aga gta cgt tat act aga gcc aaa gtc tmc ycc gcc aat atc aag ttg gcc att atc gat atc tcg agg cga tgg gca tct ccg ttt aat aca ttg att aaa gag tgt cca tcc ggt acc ggt aca ttt agc ata tat ggg tcc cat ttt ttg ctt tct gta tcc agg tag aca tag a-t att cta tag tgt ctc cta tgt tgt a-- --- --- --- --- --- -at tag cat cag tc- --t cta cac tat tct taa att tca tat taa tgg ggc gtg acg gaa tag tac agt atg ata gaa cac atc cta ttc cca aca atg tca gga acg tca cgc tct cca cct tca tat tta ttt atc cgt --a aaa tgt tat cct gga cat cgt aca aat aat a-a aaa gcc cat ata tat gtt cgc tat tgt aga aat tgt ttt tca cag ttg ctc aaa aac aat ggc agt gac tta tga gtt a-g tta cac ttt gga gtc tca tct tta gta aac ata tca taa tat tcg ata tta cga gtt gac ata tcg aac aaa ttc caa gta ttt gat ttt gga taa tat tcg tat ttt gca tct gct ata att aag ata taa tca cca caa gaa cac acg aac gtc ttt cct aca tgg tta aag tac atg tac aat tct atc cat ttg tct tcc tta act ata tat ttg tat aga taa tta cga gtc tca tga gta att cca gta att gca tag atg tca cca tcg tat tct aca gca taa act ata cta tga cgt cta ggc atg gga gac -tt ttt tat cca acg att ttt agt gaa aca ttc cac atc gtt taa tac tac ata ttt ctc ata -gt ggt ata aac tcc acc cat tac --a tat ata tca tcg ttt acg aat act gat gcg cct gaa tat cta gga gtg att aag ttt gga agt ctt ttc cat ttc gaa gtg ccg tgt ttc aaa tat tct gct ata ccc gtt gaa ata gaa aat tct aat cct cct att aca tat aac ttt cca tcg tta aca caa gta cta act tct gat ttt aac gac gac ata tta gta acc gtt tty cm- --- --- --- --- --- --- -wt ttt ttk ktt tta aga tct acc cgc -ga tac gga ata aac atg tct att gtt aat cat gcc gcc aat aat gta tag aca att atg taa aac att tgc atc ata gaa ttg --- -tc tat ctg tat tac cga cta tcg tcc aat att ctg ttc tag gag agt aat ggg tta ttg tga ata tat aat cag agt ttt taa tga cta cta tat tat gtt tta tac cat ttc gtg tca c-a gct ttg tag att tgg ata tag tta atc cca aca atg cta tag cat tgc ata tag cat tag tca taa act tgg gat gta aaa tgt tga tga tat cta cat cgt ttg gat ttt tat gta tcc act tta ata ata tta tag c-g taa cat cct cat ga- ttt acg tta acg ttt tcg tgt gat aag ata gtg gtc agt tca tcc ttt gat aat ttt cca aat t

>KC813493.1:26727-28684_Cowpox_virus_strain_HumLit08/1_complete_genome

cgg ctt cca tgt att tcc tgg aga gca agt --- aga tga tg- agg aac cag ata gtt tat atc cgt act tgc act taa agt cta cat tgt cgt tgt atg a-- gta tga tct ttt aaa ccc gct aga caa gta tcc gtt tga tat tgt agg atg tgg aca ttt aac aat ctg aca cgt ggg tgg atc gga cca ttc tcc tcc tga aca cag gac acc aga gtt acc aat caa cga ata tcc act att gca act ata cgt tac aac gct tcc atc ggt ata aaa atc ctc gta tcc gtt atg tct tcc gtt gga tat aga tgg agg tga ttg gca ttt aac aga ttc gca aat agg tgc ctc ggg att cca tac cat aga tcc agt aga tcc taa ttc aca ata cga ttt aga ttc acc gat caa atg ata tcc gct att aca aga gta cgt tat act aga gcc aaa gtc tac tcc gcc aat atc aag ttg gcc att atc gat atc tcg agg cga tgg gca tct ccg ttt aat aca ttg att aaa gag tgt cca tcc agt acc tgt aca ttt agc ata tat agg tcc cat ttt ttg ctt tct gta tcc agg tag aca tag a-t att cta tag tgt ctc cta tgt tgt a-- --- --- --- --- --- -at tag cat cag tc- --t cca cac tat tct taa att tca tat taa tgg gac gtg aag gaa tag gac agc atg ata gaa cgc atc cta ttc cca aca atg tca gga acg tca cgc tct cca cct tca tat tta ttt atc cgt -aa aaa tgt tat cct gga cat cgt aca aat aat a-a aaa gcc ca- -ta tat gtt cgc tat tgt aga aat tgt ttt tca cag ttg ctc aaa aac gat ggc aat gac tta tga gtt acg tta cac ttt gga gtc tca tct tta gta aac ata tca taa tat tcg ata tta cga gtt gac ata tcg aac aaa ttc caa gta ttt gat ttt gga taa tat tcg tat ttt gca tct gct ata att aag ata taa tcg ccg caa gaa cac acg aac atc ttt cct aca tgg tta aag tac atg tac aat tct atc cat ttg tct tcc tta act ata tat ttg tat aga taa tta cga gtc tca tga gta att cca gta att gca tag atg tca cca tcg tac tct aca gca taa act ata cta tga tgt cta ggc atg gga gac -tt ttt tat cca acg att ttt agt gaa aca ttc cac atc gtt taa tac tac ata ttt ctc ata cgt ggt ata aac tcc acc cat tac --a tat ata tca tcg ttt acg aat acc gat gcg cct gaa tat cta gga gta att aag ttt gga agt ctt atc cat ttc gaa gtg ccg tgt ttc aaa tat tct gcc aca ccc gtt gaa ata gaa aat tct aat cct cct att aca tat aac ttt cca tcg tta aca caa gta cta act tct gat ttt aac gac gac ata tta gta act gtt ttc ca- --- --- --- --- --- --- --t ttt ttc gtt tta aga tct acc cgc -ga tac gga ata aac atg tct att gtt aat cat gcc gcc aat aat gta tag aca att atg taa aac att tgc att ata gaa ttg --- -tc tat ctg tat tac cga cta tcg tcc aat att ctg ttc tag gag agt aat ggg tta ttg tgg ata tat aat cag agt ttt taa tga cta cta tat tat gtt tta tac cat ttc gtg tca ctg gct ttg tag att tgg ata tag tta atc cca aca atg ata tag cat tgc gca tag tat tag tca taa act tgg gat gta aaa tgt tga tga tat cta cat cgt ttg gat ttt tat gta tcc act tta ata ata tca tag ctg taa cat cct cat ga- ttt acg tta acg tct tca tgg gat aag ata gtt gtc agt tca tcc ttt gat aat ttt cca aat t

>HQ420893.1:33953-35916_Cowpox_virus_strain_Finland_2000_MAN_complete_genome

cgg ctt cca tgt att tcc tgg aga gca agt --- aga tga tg- aag aac cag ata ggt tat atc cgt act tgc act taa agt cta cat tgt cgt tgt atg a-- gta tga tct ttt aaa ccc gct aga caa gta tcc gtt tga tat tgt agg atg tgg aca ttt aac aat ctg aca cgt ggg tgg atc gga cca ttc tcc tcc tga aca cag gac acc aga gtt acc aat caa cga ata tcc act att gca act ata agt tac aac gct tcc atc ggt ata aaa atc ctc gta tcc gtt atg tct tcc gtt gga tat aga tgg agg tga ttg gca ttt aac aga ttc gca aat agg tgc ctc ggg att cca tac cat aga tcc agt aga tcc taa ttc aca ata cga ttt aga ttc acc gat caa atg ata tcc gct att aca aga gta cgt tat act aga gcc aaa gtc tac tcc gcc aat atc aag ttg gcc att atc gat atc tcg agg cga tgg gca tct ccg ttt aat aca ttg att aaa gag tgt cca tcc agt acc tat aca ttt agc ata tat agg tcc cat ttt ttg ctt tct gta tcc agg tag aca tag a-t att cta tag tgt ctc cta tgt tgt a-- --- --- --- -at tag cat tag cat cag tc- --t cca cac tat tct taa att tca tat taa tgg gac gtg acg gaa tag tac agc atg ata gaa cgc atc cta ttc cca aca atg tca gga acg tca cgc tct cca cct tca tat tta ttt atc cgt -aa aaa tgt tat cct gga cat cgt aca aat aat a-a aaa gcc ca- -ta tat gtt cgc tat tgt aga aat tgt ttt tca cag ttg ctc aaa aac gat ggc aat gac tta tga gtt acg tta cac ttt gga gtc tca tct tta gta aac ata tca taa tat tcg ata tta cga gtt gac ata tcg aac aaa ttc caa gta ttt gat ttt gga taa tat tcg tat ttt gca tct gct ata att aag ata taa tcg ccg caa gaa cac acg aac atc ttt cct aca tgg tta aag tac atg tac aat tct atc cat ttg tct tcc tta act ata tat ttg tat aga taa tta cga gtc tca tga gta atc cca gta att gca tag atg tca cca tcg tac tct aca gca taa act ata cta tga tgt cta ggc atg gga gac -tt ttt tat cca acg att ttt agt gaa aca ttc cac atc gtt taa tac tac ata ttt ctc ata cgt ggt ata aac tcc acc cat tac --a tat ata tca tcg ttt acg aat acc gat gcg cct gaa tat cta gga gta att aag ttt gga agt ctt atc cat ttc gaa gtg ccg tgt ttc aaa tat tct gcc aca ccc gtt gaa ata gaa aat tct aat cct cct att aca tat aac ttt cca tcg tta aca caa gta cta act tct gat ttt aac gac gac ata tta gta act gtt ttc ca- --- --- --- --- --- --- --t ttt ttc gtt tta aga tct acc cgc -ga tac gga ata aac atg tct att gtt aat cat gcc gcc aat aat gta tag aca att atg taa aac att tgc att ata gaa ttg --- -tc tat ctg tat tac cga cta tcg tcc aat att ctg ttc tag gag agt aat ggg tta ttg tgg ata tat aat cag agt ttt taa tga cta cta tat tat gtt tta tac cat ttc gtg tca ctg gct ttg tag att tgg ata tag tta atc cca aca atg ata tag cat tgc gca tag tat tag tca taa act tgg gat gta aaa tgt tga tga tat cta cat cgt ttg gat ttt tat gta tcc act tta ata ata tca tag ctg taa cat cct cat ga- ttt acg tta acg tct tcg tgg gat aag ata gtt gtc agt tca tcc ttt gat aat ttt cca aat t

>LT993232.1:33071-35035_Cowpox_virus_isolate_Ger/2015/Human2_genome_assembly_complete_genome:_monopartite

cgg ctt cca tgt att tcc tgg aga gca agt --- aga tga tg- agg aac cag ata gtt tat atc cgt act tgc act taa agt cta cat tgt cgt tgt atg a-- gta tga tct ttt aaa ccc gct aga caa gta tcc gtt tga tat tgt agg atg tgg aca ttt aac aat ctg aca cgt ggg tgg atc gga cca ttc tcc tcc tga aca cag gac acc aga gtt acc aat caa cga ata tcc act att gca act ata agt tac aac gct ccc atc ggt ata aaa atc ctc gta tcc gtt atg tct tcc gtt gga tat aga tgg agg gga ttg gca ttt aac aga ttc aca aat agg tgc ctc ggg att cca tac cat aga tcc agt aga tcc taa ttc aca ata cga ttt aga ttc acc gat caa atg ata tcc gct att aca aga gta cgt tat act aga gcc aaa gtc tac tcc acc aat atc aag ttg gcc att atc gat atc tcg agg cga tgg gca tct ccg ttt aat aca ttg att aaa gag tgt cca tcc agt acc tgt aca ttt agc ata tat agg tcc cat ttt ttg ctt tct gta tcc agg tag aca tag a-t att cta tag tgt ctc cta tgt tgt a-- --- --- --- -at tag cat tag cat cag tc- --t cca cac tat tct taa att tca tat taa tgg gtc gtg acg gaa tag tac agc atg ata gaa cgc atc cta ttc cca aca atg tca gga acg tca cgc tct cca cct tca tat tta ttt atc cgt -aa aaa tgt tat ccc gga cat tgt acc aat aat aaa aaa gcc ca- -ta tat gtt cgc tat tgt aga aat tgt ttt tca cag ttg ctc aaa aac gat ggc agt gac tta tga gtt acg tta cac ttt gga gtc tca tct tta gta aac ata tca taa tat tcg ata tta cga gtt gac ata tcg aac aaa ttc caa gta ttt gat ttt gga taa tat tcg tat ttt gca tct gct ata att aag ata taa tca ccg caa gaa cac acg aac atc ttt cct aca tgg tta aag tac atg tac aat tct atc cat ttg tct tcc tta act ata tat ttg tat aga taa tta cga gtc tca tga gta att cca gta att gca tag atg tca cca tcg tac tct aca gca taa act ata cta tga tgt cta ggc atg gga gac -tt ttt tat cca acg att ttt agt gaa aca ttc cat atc gtt taa tac tac ata ttt ctc ata agt agt ata aac tcc acc cat tac --a tat ata tca tcg ttt acg aat acc gat gcg cct gaa tat cta gga gta att aag ttt gga agt ctt atc cat ttc gaa gtg ccg tgt ttc aaa tat tct gct aca ccc gtt gaa ata gaa aat tct aat cct cct att aca tat aac ttt cca tcg tta aca caa gta cta act tct gat ttt aac gac gac ata tta gta acc gtt ttc ca- --- --- --- --- --- --- --t ttt ttc gtt tta aga tct acc cgc -ga tac gga ata aac atg tct att gtt aat cat gcc gcc aat aat gta tag aca att atg taa aac att tgc att ata gaa ttg --- -tc tat ctg tat tac cga cta tcg tcc aat att ctg ttc tag gag agt aat ggg tta ttg tgg ata tat aat cag agt ttt taa tga cta cta tat tat gtt tta tac cat ttc gtg tca ctg gct ttg tag att tgg ata tag tta atc cca aca atg ata tag cat tgc gca tag tat tag tca taa act tgg gat gta aaa tgt tga tga tat cta cat cgt ttg gat ttt tat gta tcc act tta ata ata tca tag ctg taa cat cct cat ga- ttt acg tta acg tct tcg tgg gat aag ata gtt gtc agt tca tcc ttt gat aat ttt cca aat t

>LT896730.1:34504-36468_Cowpox_virus_isolate_Ger/2010/Racoon_genome_assembly_complete_genome:_monopartite

cgg ctt cca tgt att tcc tgg aga gca agt --- aga tga tg- agg aac cag ata gtt tat atc cgt act tgc act taa agt cta cat tgt cgt tgt atg a-- gta tga tct ttt aaa ccc gct aga caa gta tcc gtt tga tat tgt agg atg tgg aca ttt aac aat ctg aca cgt ggg tgg atc gga cca ttc tcc tcc tga aca cag gac acc aga gtt acc aat caa cga ata tcc act att gca act ata agt tac aac gct ccc atc ggt ata aaa atc ctc gta tcc gtt atg tct tcc gtt gga tat aga tgg agg gga ttg gca ttt aac aga ttc aca aat agg tgc ctc ggg att cca tac cat aga tcc agt aga tcc taa ttc aca ata cga ttt aga ttc acc gat caa atg ata tcc gct att aca aga gta cgt tat act aga gcc aaa gtc tac tcc acc aat atc aag ttg gcc att atc gat atc tcg agg cga tgg gca tct ccg ttt aat aca ttg att aaa gag tgt cca tcc agt acc tgt aca ttt agc ata tat agg tcc cat ttt ttg ctt tct gta tcc agg tag aca tag a-t att cta tag tgt ctc cta tgt tgt a-- --- --- --- -at tag cat tag cat cag tc- --t cca cac tat tct taa att tca tat taa tgg gtc gtg acg gaa tag tac agc atg ata gaa cgc atc cta ttc cca aca atg tca gga acg tca cgc tct cca cct tca tat tta ttt atc cgt -aa aaa tgt tat ccc gga cat tgt acc aat aat aaa aaa gcc ca- -ta tat gtt cgc tat tgt aga aat tgt ttt tca cag ttg ctc aaa aac gat ggc agt gac tta tga gtt acg tta cac ttt gga gtc tca tct tta gta aac ata tca taa tat tcg ata tta cga gtt gac ata tcg aac aaa ttc caa gta ttt gat ttt gga taa tat tcg tat ttt gca tct gct ata att aag ata taa tca ccg caa gaa cac acg aac atc ttt cct aca tgg tta aag tac atg tac aat tct atc cat ttg tct tcc tta act ata tat ttg tat aga taa tta cga gtc tcg tga gta att cca gta att gca tag atg tca cca tcg tac tct aca gca taa act ata cta tga tgt cta ggc atg gga gac -tt ttt tat cca acg att ttt agt gaa aca ttc cac atc gtt taa tac tac ata ttt ctc ata cgt ggt ata aac tcc acc cat tac --a tat ata tca tcg ttt acg aat acc gat gcg cct gaa tat cta gga gta att aag ttt gga agt ctt atc cat ttc gaa gtg ccg tgt ttc aaa tat tct gcc aca ccc gtt gaa ata gaa aat tct aat cct cct att aca tat aac ttt cca tcg tta aca caa gta cta act tct gat ttt aac gac gac ata tta gta acc gtt ttc ca- --- --- --- --- --- --- --t ttt ttc gtt tta aga tct acc cgc -ga tac gga ata aac atg tct att gtt aat cat gcc gcc aat aat gta tag aca att atg taa aac att tgc att ata gaa ttg --- -tc tat ctg tat tac cga cta tcg tcc aat att ctg ttc tag gag agt aat ggg tta ttg tgg ata tat aat cag agt ttt taa tga cta cta tat tat gtt tta tac cat ttc gtg tca ctg gct ttg tag att tgg ata tag tta atc cca aca atg ata tag cat tgc gca tag tat tag tca taa act tgg gat gta aaa tgt tga tga tat cta cat cgt ttg gat ttt tat gta tcc act tta ata ata tca tag ctg taa cat cct cat ga- ttt acg tta acg tct tcg tgg gat aag ata gtt gtc agt tca tcc ttt gat aat ttt cca aat t

>LT896729.1:34430-36394_Cowpox_virus_isolate_Ger/2010/Cat_genome_assembly_complete_genome:_monopartite

cgg ctt cca tgt att tcc tgg aga gca agt --- aga tga tg- agg aac cag ata gtt tat atc cgt act tgc act taa agt cta cat tgt cgt tgt atg a-- gta tga tct ttt aaa ccc gct aga caa gta tcc gtt tga tat tgt agg atg tgg aca ttt aac aat ctg aca cgt ggg tgg atc gga cca ttc tcc tcc tga aca cag gac acc aga gtt acc aat caa cga ata tcc act att gca act ata agt tac aac gct ccc atc ggt ata aaa atc ctc gta tcc gtt atg tct tcc gtt gga tat aga tgg agg gga ttg gca ttt aac aga ttc aca aat agg tgc ctc ggg att cca tac cat aga tcc agt aga tcc taa ttc aca ata cga ttt aga ttc acc gat caa atg ata tcc gct att aca aga gta cgt tat act aga gcc aaa gtc tac tcc acc aat atc aag ttg gcc att atc gat atc tcg agg cga tgg gca tct ccg ttt aat aca ttg att aaa gag tgt cca tcc agt acc tgt aca ttt agc ata tat agg tcc cat ttt ttg ctt tct gta tcc agg tag aca tag a-t att cta tag tgt ctc cta tgt tgt a-- --- --- --- -at tag cat tag cat cag tc- --t cca cac tat tct taa att tca tat taa tgg gtc gtg acg gaa tag tac agc atg ata gaa cgc atc cta ttc cca aca atg tca gga acg tca cgc tct cca cct tca tat tta ttt atc cgt -aa aaa tgt tat ccc gga cat tgt acc aat aat aaa aaa gcc ca- -ta tat gtt cgc tat tgt aga aat tgt ttt tca cag ttg ctc aaa aac gat ggc agt gac tta tga gtt acg tta cac ttt gga gtc tca tct tta gta aac ata tca taa tat tcg ata tta cga gtt gac ata tcg aac aaa ttc caa gta ttt gat ttt gga taa tat tcg tat ttt gca tct gct ata att aag ata taa tca ccg caa gaa cac acg aac atc ttt cct aca tgg tta aag tac atg tac aat tct atc cat ttg tct tcc tta act ata tat ttg tat aga taa tta cga gtc tcg tga gta att cca gta att gca tag atg tca cca tcg tac tct aca gca taa act ata cta tga tgt cta ggc atg gga gac -tt ttt tat cca acg att ttt agt gaa aca ttc cac atc gtt taa tac tac ata ttt ctc ata cgt ggt ata aac tcc acc cat tac --a tat ata tca tcg ttt acg aat acc gat gcg cct gaa tat cta gga gta att aag ttt gga agt ctt atc cat ttc gaa gtg ccg tgt ttc aaa tat tct gcc aca ccc gtt gaa ata gaa aat tct aat cct cct att aca tat aac ttt cca tcg tta aca caa gta cta act tct gat ttt aac gac gac ata tta gta acc gtt ttc ca- --- --- --- --- --- --- --t ttt ttc gtt tta aga tct acc cgc -ga tac gga ata aac atg tct att gtt aat cat gcc gcc aat aat gta tag aca att atg taa aac att tgc att ata gaa ttg --- -tc tat ctg tat tac cga cta tcg tcc aat att ctg ttc tag gag agt aat ggg tta ttg tgg ata tat aat cag agt ttt taa tga cta cta tat tat gtt tta tac cat ttc gtg tca ctg gct ttg tag att tgg ata tag tta atc cca aca atg ata tag cat tgc gca tag tat tag tca taa act tgg gat gta aaa tgt tga tga tat cta cat cgt ttg gat ttt tat gta tcc act tta ata ata tca tag ctg taa cat cct cat ga- ttt acg tta acg tct tcg tgg gat aag ata gtt gtc agt tca tcc ttt gat aat ttt cca aat t

>LT896720.1:33041-35005_Cowpox_virus_isolate_Ger/2015/Human1_genome_assembly_complete_genome:_monopartite

cgg ctt cca tgt att tcc tgg aga gca agt --- aga tga tg- agg aac cag ata gtt tat atc cgt act tgc act taa agt cta cat tgt cgt tgt atg a-- gta tga tct ttt aaa ccc gct aga caa gta tcc gtt tga tat tgt agg atg tgg aca ttt aac aat ctg aca cgt ggg tgg atc gga cca ttc tcc tcc tga aca cag gac acc aga gtt acc aat caa cga ata tcc act att gca act ata agt tac aac gct ccc atc ggt ata aaa atc ctc gta tcc gtt atg tct tcc gtt gga tat aga tgg agg gga ttg gca ttt aac aga ttc aca aat agg tgc ctc ggg att cca tac cat aga tcc agt aga tcc taa ttc aca ata cga ttt aga ttc acc gat caa atg ata tcc gct att aca aga gta cgt tat act aga gcc aaa gtc tac tcc acc aat atc aag ttg gcc att atc gat atc tcg agg cga tgg gca tct ccg ttt aat aca ttg att aaa gag tgt cca tcc agt acc tgt aca ttt agc ata tat agg tcc cat ttt ttg ctt tct gta tcc agg tag aca tag a-t att cta tag tgt ctc cta tgt tgt a-- --- --- --- -at tag cat tag cat cag tc- --t cca cac tat tct taa att tca tat taa tgg gtc gtg acg gaa tag tac agc atg ata gaa cgc atc cta ttc cca aca atg tca gga acg tca cgc tct cca cct tca tat tta ttt atc cgt -aa aaa tgt tat ccc gga cat tgt acc aat aat aaa aaa gcc ca- -ta tat gtt cgc tat tgt aga aat tgt ttt tca cag ttg ctc aaa aac gat ggc agt gac tta tga gtt acg tta cac ttt gga gtc tca tct tta gta aac ata tca taa tat tcg ata tta cga gtt gac ata tcg aac aaa ttc caa gta ttt gat ttt gga taa tat tcg tat ttt gca tct gct ata att aag ata taa tca ccg caa gaa cac acg aac atc ttt cct aca tgg tta aag tac atg tac aat tct atc cat ttg tct tcc tta act ata tat ttg tat aga taa tta cga gtc tcg tga gta att cca gta att gca tag atg tca cca tcg tac tct aca gca taa act ata cta tga tgt cta ggc atg gga gac -tt ttt tat cca acg att ttt agt gaa aca ttc cac atc gtt taa tac tac ata ttt ctc ata cgt ggt ata aac tcc acc cat tac --a tat ata tca tcg ttt acg aat acc gat gcg cct gaa tat cta gga gta att aag ttt gga agt ctt atc cat ttc gaa gtg ccg tgt ttc aaa tat tct gcc aca ccc gtt gaa ata gaa aat tct aat cct cct att aca tat aac ttt cca tcg tta aca caa gta cta act tct gat ttt aac gac gac ata tta gta acc gtt ttc ca- --- --- --- --- --- --- --t ttt ttc gtt tta aga tct acc cgc -ga tac gga ata aac atg tct att gtt aat cat gcc gcc aat aat gta tag aca att atg taa aac att tgc att ata gaa ttg --- -tc tat ctg tat tac cga cta tcg tcc aat att ctg ttc tag gag agt aat ggg tta ttg tgg ata tat aat cag agt ttt taa tga cta cta tat tat gtt tta tac cat ttc gtg tca ctg gct ttg tag att tgg ata tag tta atc cca aca atg ata tag cat tgc gca tag tat tag tca taa act tgg gat gta aaa tgt tga tga tat cta cat cgt ttg gat ttt tat gta tcc act tta ata ata tca tag ctg taa cat cct cat ga- ttt acg tta acg tct tcg tgg gat aag ata gtt gtc agt tca tcc ttt gat aat ttt cca aat t

>DQ121394.1:21389-23352_Vaccinia_virus_strain_Lister_clone_VACV107_complete_genome

cgg ctt cca tgt att tcc tgg aga gca agt --- aga tga tg- agg aac cag ata gtt tat atc cgt act tgc act taa agt cta cat tgt cgt tgt atg a-- gta tga tct ttt aaa ccc gct aga caa gta tcc gtt tga tat tgt agg atg tgg aca ttt aac aat ctg aca cgt ggg tgg atc gga cca ttc tcc tcc tga aca cag gac acc aga gtt acc aat caa cga ata tcc act att gca act ata agt tac aac gct ccc atc ggt ata aaa atc ctc gta tcc gtt atg tct tcc gtt gga tat aga tgg agg gga ttg gca ttt aac aga ttc aca aat agg tgc ctc ggg att cca tac cat aga tcc agt aga tcc taa ttc aca ata cga ttt aga ttc acc gat caa atg ata tcc gct att aca aga gta cgt tat act aga gcc aaa gtc tac tcc acc aat atc aag ttg gcc att atc gat atc tcg agg cga tgg gca tct ccg ttt aat aca ttg att aaa gag tgt cca tcc agt acc tgt aca ttt agc ata tat agg tcc cat ttt ttg ctt tct gta tcc agg tag aca tag a-t att cta tag tgt ctc cta tgt tgt a-- --- --- --- -at tag cat tag cat cag tc- --t cca cac tat tct taa att tca tat taa tgg gtc gtg acg gaa tag tac agc atg ata gaa cgc atc cta ttc cca aca atg tca gga acg tca cgc tct cca cct tca tat tta ttt atc cgt -aa aaa tgt tat cct gga cat cgt aca aat aat a-a aaa gcc ca- -ta tat gtt cgc tat tgt aga aat tgt ttt tca cag ttg ctc aaa aac gat ggc agt gac tta tga gtt acg tta cac ttt gga gtc tca tct tta gta aac ata tca taa tat tcg ata tta cga gtt gac ata tcg aac aaa ttc caa gta ttt gat ttt gga taa tat tcg tat ttt gca tct gct ata att aag ata taa tca ccg caa gaa cac acg aac atc ttt cct aca tgg tta aag tac atg tac aat tct atc cat ttg tct tcc tta act ata tat ttg tat aga taa tta cga gtc tcg tga gta att cca gta att aca tag atg tcg ccg tcg tac tct aca gca taa act ata cta tga tgt cta ggc atg gga gac -tt ttt tat cca acg att ttt agt gaa aca ttc cac atc gtt taa tac tac ata ttt ctc ata cgt ggt ata aac tcc acc cat tac --a tat ata tca tcg ttt acg aat acc gac gcg cct gaa tat cta gga gta att aag ttt gga agt ctt atc cat ttc gaa gtg ccg tgt ttc aaa tat tct gcc aca ccc gtt gaa ata gaa aat tct aat cct cct att aca tat aac ttt cca tcg tta aca caa gta cta act tct gat ttt aac gac gac ata tta gta acc gtt ttc ca- --- --- --- --- --- --- --t ttt ttc gtt tta aga tct acc cgc -ga tac gga ata aac atg tct att gtt aat cat gcc gcc aat aat gta tag aca att atg taa aac att tgc att ata gaa ttg --- -tc tat ctg tat tac cga cta tcg tcc aat att ctg ttc tag gag agt aat ggg tta ttg tgg ata tat aat cag agt ttt taa tga cta cta tat tat gtt tta tac cat ttc gtg tca ctg gct ttg tag att tgg ata tag tta atc cca aca atg ata tag cat tgc gca tag tat tag tca taa act tgg gat gta aaa tgt tga tga tat cta cat cgt ttg gat ttt tat gta tcc act tta ata ata tca tag ctg taa cat cct cat ga- ttt acg tta acg tct tcg tgg gat aag ata gtt gtc agt tca tcc ttt gat aat ttt cca aat t

>KM044310.1:11170-13133_Vaccinia_virus_strain_Tashkent_clone_TKT4_complete_genome

cgg ctt cca tgt att tcc tgg aga gca agt --- aga tga tg- agg aac cag ata gtt tat atc cgt act tgc act taa agt cta cat tgt cgt tgt atg a-- gta tga tct ttt aaa ccc gct aga caa gta tcc gtt tga tat tgt agg atg tgg aca ttt aac aat ctg aca cgt ggg tgg atc gga cca ttc tcc tcc tga aca cag gac acc aga gtt acc aat caa cga ata tcc act att gca act ata agt tac aac gct ccc atc ggt ata aaa atc ctc gta tcc gtt atg tct tcc gtt gga tat aga tgg agg gga ttg gca ttt aac aga ttc aca aat agg tgc ctc ggg att cca tac cat aga tcc agt aga tcc taa ttc aca ata cga ttt aga ttc acc gat caa atg ata tcc gct att aca aga gta cgt tat act aga gcc aaa gtc tac tcc acc aat atc aag ttg gcc att atc gat atc tcg agg cga tgg gca tct ccg ttt aat aca ttg att aaa gag tgt cca tcc agt acc tgt aca ttt agc ata tat agg tcc cat ttt ttg ctt tct gta tcc agg tag aca tag a-t att cta tag tgt ctc cta tgt tgt a-- --- --- --- -at tag cat tag cat cag tc- --t cca cac tat tct taa att tca tat taa tgg gtc gtg acg gaa tag tac agc atg ata gaa cgc atc cta ttc cca aca atg tca gga acg tca cgc tct cca cct tca tat tta ttt atc cgt -aa aaa tgt tat cct gga cat cgt aca aat aat a-a aaa gcc ca- -ta tat gtt cgc tat tgt aga aat tgt ttt tca cag ttg ctc aaa aac gat ggc agt gac tta tga gtt acg tta cac ttt gga gtc tca tct tta gta aac ata tca taa tat tcg ata tta cga gtt gac ata tcg aac aaa ttc caa gta ttt gat ttt gga taa tat tcg tat ttt gca tct gct ata att aag ata taa tca ccg caa gaa cac acg aac atc ttt cct aca tgg tta aag tac atg tac aat tct atc cat ttg tct tcc tta act ata tat ttg tat aga taa tta cga gtc tcg tga gta att cca gta att aca tag atg tcg ccg tcg tac tct aca gca taa act ata cta tga tgt cta ggc atg gga gac -tt ttt tat cca acg att ttt agt gaa aca ttc cac atc gtt taa tac tac ata ttt ttc ata cgt ggt ata aac tcc acc cat tac --a tat ata tca tcg ttt acg aat acc gac gcg cct gaa tat cta gga gta att aag ttt gga agt ctt atc cat ttc gaa gtg ccg tgt ttc aaa tat tct gcc aca ccc gtt gaa ata gaa aat tct aat cct cct att aca tat aac ttt cca tcg tta aca caa gta cta act tct gat ttt aac gac gac ata tta gta acc gtt ttc ca- --- --- --- --- --- --- --t ttt ttc gtt tta aga tct acc cgc -ga tac gga ata aac atg tct att gtt aat cat gcc gcc aat aat gta tag aca att atg taa aac att tgc att ata gaa ttg --- -tc tat ctg tat tac cga cta tcg tcc aat att ctg ttc tag gag agt aat ggg tta ttg tgg ata tat aat cag agt ttt taa tga cta cta tat tat gtt tta tac cat ttc gtg tca ctg gct ttg tag att tgg ata tag tta atc cca aca atg ata tag cat tgc gca tag tat tag tca taa act tgg gat gta aaa tgt tga tga tat cta cat cgt ttg gat ttt tat gta tcc act tta ata ata tca tag ctg taa cat cct cat ga- ttt acg tta acg tct tcg tgg gat aag ata gtt gtc agt tca tcc ttt gat aat ttt cca aat t

>KM044309.1:11603-13566_Vaccinia_virus_strain_Tashkent_clone_TKT3_complete_genome

cgg ctt cca tgt att tcc tgg aga gca agt --- aga tga tg- agg aac cag ata gtt tat atc cgt act tgc act taa agt cta cat tgt cgt tgt atg a-- gta tga tct ttt aaa ccc gct aga caa gta tcc gtt tga tat tgt agg atg tgg aca ttt aac aat ctg aca cgt ggg tgg atc gga cca ttc tcc tcc tga aca cag gac acc aga gtt acc aat caa cga ata tcc act att gca act ata agt tac aac gct ccc atc ggt ata aaa atc ctc gta tcc gtt atg tct tcc gtt gga tat aga tgg agg gga ttg gca ttt aac aga ttc aca aat agg tgc ctc ggg att cca tac cat aga tcc agt aga tcc taa ttc aca ata cga ttt aga ttc acc gat caa atg ata tcc gct att aca aga gta cgt tat act aga gcc aaa gtc tac tcc acc aat atc aag ttg gcc att atc gat atc tcg agg cga tgg gca tct ccg ttt aat aca ttg att aaa gag tgt cca tcc agt acc tgt aca ttt agc ata tat agg tcc cat ttt ttg ctt tct gta tcc agg tag aca tag a-t att cta tag tgt ctc cta tgt tgt a-- --- --- --- -at tag cat tag cat cag tc- --t cca cac tat tct taa att tca tat taa tgg gtc gtg acg gaa tag tac agc atg ata gaa cgc atc cta ttc cca aca atg tca gga acg tca cgc tct cca cct tca tat tta ttt atc cgt -aa aaa tgt tat cct gga cat cgt aca aat aat a-a aaa gcc ca- -ta tat gtt cgc tat tgt aga aat tgt ttt tca cag ttg ctc aaa aac gat ggc agt gac tta tga gtt acg tta cac ttt gga gtc tca tct tta gta aac ata tca taa tat tcg ata tta cga gtt gac ata tcg aac aaa ttc caa gta ttt gat ttt gga taa tat tcg tat ttt gca tct gct ata att aag ata taa tca ccg caa gaa cac acg aac atc ttt cct aca tgg tta aag tac atg tac aat tct atc cat ttg tct tcc tta act ata tat ttg tat aga taa tta cga gtc tcg tga gta att cca gta att aca tag atg tcg ccg tcg tac tct aca gca taa act ata cta tga tgt cta ggc atg gga gac -tt ttt tat cca acg att ttt agt gaa aca ttc cac atc gtt taa tac tac ata ttt ttc ata cgt ggt ata aac tcc acc cat tac --a tat ata tca tcg ttt acg aat acc gac gcg cct gaa tat cta gga gta att aag ttt gga agt ctt atc cat ttc gaa gtg ccg tgt ttc aaa tat tct gcc aca ccc gtt gaa ata gaa aat tct aat cct cct att aca tat aac ttt cca tcg tta aca caa gta cta act tct gat ttt aac gac gac ata tta gta acc gtt ttc ca- --- --- --- --- --- --- --t ttt ttc gtt tta aga tct acc cgc -ga tac gga ata aac atg tct att gtt aat cat gcc gcc aat aat gta tag aca att atg taa aac att tgc att ata gaa ttg --- -tc tat ctg tat tac cga cta tcg tcc aat att ctg ttc tag gag agt aat ggg tta ttg tgg ata tat aat cag agt ttt taa tga cta cta tat tat gtt tta tac cat ttc gtg tca ctg gct ttg tag att tgg ata tag tta atc cca aca atg ata tag cat tgc gca tag tat tag tca taa act tgg gat gta aaa tgt tga tga tat cta cat cgt ttg gat ttt tat gta tcc act tta ata ata tca tag ctg taa cat cct cat ga- ttt acg tta acg tct tcg tgg gat aag ata gtt gtc agt tca tcc ttt gat aat ttt cca aat t

>LT896725.1:35493-37459_Cowpox_virus_isolate_Ger/2014/Cat2_genome_assembly_complete_genome:_monopartite

cgg ctt cca tgt att tcc tgg aga gca agt --- aga tga tg- agg aac cag ata gtt tat atc cgt act tgc act taa agt cta cat tgt cgt tgt atg a-- gta tga tct ttt aaa ccc gct aga caa gta tcc gtt tga tat tgt agg atg tgg aca ttt aac aat ctg aca cgt ggg tgg atc gga cca ttc tcc tcc tga aca cag gac acc aga gtt acc aat caa cga ata tcc act att gca act ata agt tac aac gct ccc atc ggt ata aaa atc ctc gta tcc gtt atg tct tcc gtt gga tat aga tgg agg gga ttg gca ttt aac aga ttc aca aat agg tgc ctc ggg att cca tac cat aga tcc agt aga tcc taa ttc aca ata cga ttt aga ttc acc gat caa atg ata tcc gct att aca aga gta cgt tat act aga gcc aaa gtc tac tcc acc aat atc aag ttg gcc att atc gat atc tcg agg cga tgg gca tct ccg ttt aat aca ttg att aaa gag tgt cca tcc agt acc tgt aca ttt agc ata tat agg tcc cat ttt ttg ctt tct gta tcc agg tag aca tag a-t att cta tag tgt ctc cta tgt tgt a-- --- --- --- -at tag cat tag cat cag tc- --t cca cac tat tct taa att tca tat taa tgg gtc gtg acg gaa tag tac agc atg ata gaa cgc atc cta ttc cca aca atg tca gga acg tca cgc tct cca cct tca tat tta ttt atc cgt -aa aaa tgt tat ccc gga cat tgt acc aat aat aaa aaa gcc ca- -ta tat gtt cgc tat tgt aga aat tgt ttt tca cag ttg ctc aaa aac gat ggc agt gac tta tga gtt acg tta cac ttt gga gtc tca tct tta gta aac ata tca taa tat tcg ata tta cga gtt gac ata tcg aac aaa ttc caa gta ttt gat ttt gga taa tat tcg tat ttt gca tct gct ata att aag ata taa tca ccg caa gaa cac acg aac atc ttt cct aca tgg tta aag tac atg tac aat tct atc cat ttg tct tcc tta act ata tat ttg tat aga taa tta cga gtc tcg tga gta att cca gta att gca tag atg tca cca tcg tac tct aca gca taa act ata cta tga tgt cta ggc atg gga gac -tt ttt tat cca acg att ttt agt gaa aca ttc cac atc gtt taa tac tac ata ttt ctc ata cgt ggt ata aac tcc acc cat tac ata tat ata tca tcg ttt acg aat acc gat gcg cct gaa tat cta gga gta att aag ttt gga agt ctt atc cat ttc gaa gtg ccg tgt ttc aaa tat tct gcc aca ccc gtt gaa ata gaa aat tct aat cct cct att aca tat aac ttt cca tcg tta aca caa gta cta act tct gat ttt aac gac gac ata tta gta acc gtt ttc ca- --- --- --- --- --- --- --t ttt ttc gtt tta aga tct acc cgc -ga tac gga ata aac atg tct att gtt aat cat gcc gcc aat aat gta tag aca att atg taa aac att tgc att ata gaa ttg --- -tc tat ctg tat tac cga cta tcg tcc aat att ctg ttc tag gag agt aat ggg tta ttg tgg ata tat aat cag agt ttt taa tga cta cta tat tat gtt tta tac cat ttc gtg tca ctg gct ttg tag att tgg ata tag tta atc cca aca atg ata tag cat tgc gca tag tat tag tca taa act tgg gat gta aaa tgt tga tga tat cta cat cgt ttg gat ttt tat gta tcc act tta ata ata tca tag ctg taa cat cct cat ga- ttt acg tta acg tct tcg tgg gat aag ata gtt gtc agt tca tcc ttt gat aat ttt cca aat t

>JX489137.1:18078-20036_Vaccinia_virus_strain_Tiantan_clone_TT10_complete_genome

cgg ctt cca tgt att tcc tgg aga gca agt --- aga tga tg- agg aac cag ata gtt tat atc cgt act tgc act taa agt tta cat tgt cgt tgt atg a-- gta tga tct ttt aaa ccc gct aga caa gta tcc gtt tga tat tgt agg atg tgg aca ttt aac aat ctg aca cgt ggg tgg atc gga cca ttc tcc tcc tga aca cag gac acc aga gtt acc aat caa cga ata tcc act att gca act ata agt tac aac gct ccc atc ggt ata aaa atc ctc gta tcc gtt atg tct tcc gtt gga tat aga tgg agg gga ttg gca ttt aac aga ttc aca aat agg tgc ctc ggg att cca tac cat aga tcc agt aga tcc taa ttc aca ata cga ttt aga ttc acc gat caa atg ata tcc gct att aca aga gta cgt tat act aga gcc aaa gtc tac tcc acc aat atc aag ttg gcc att atc gat atc tcg agg cga tgg gca tct ccg ttt aat aca ttg att aaa gag tgt cca tcc agt acc tgt aca ttt agc ata tat agg tcc cat ttt ttg ctt tct gta tcc agg tag aca tag a-t att cta tag tgt ctc cta tgt tgt a-- --- --- --- -at tag cat t-- --- -ag tt- --t cca cac tat tct taa att tca tat taa tgg gac gtg aag gaa tag gac agt atg ata gaa cgc atc cta ttc cca aca atg tca gga acg tca cgc tct cca cct tca tat tta ttt atc cgt -aa aaa tgt tat cct gga cat cgt aca aat aat aaa aaa gcc ca- -ta tat gtt cgc tat tgt aga aat tgt ttt tca cag ttg ctc aaa aac gat ggc agt gac tta tga gtt acg tta cac ttt gga gtc tca tct tta gta aac ata tca taa tat tcg ata tta cga gtt gac ata tcg aac aaa ttc caa gta ttt gat ttt gga taa tat tcg tat ttt gca tct gct ata att aag ata taa tca ccg caa gaa cac acg aac atc ttt cct aca tgg tta aag tac atg tac aat tct atc cat ttg tct tcc tta act ata tat ttg tat aga taa tta cga gtc tcg tga gta att cca gta att aca tag atg tcg ccg tcg tac tct aca gca taa act ata cta tga tgt cta ggc atg gga gac -tt ttt tat cca acg att ttt agt gaa aca ttc cac atc gtt taa tac tac ata ttt ttc ata cgt ggt ata aac tcc acc cat tac --a tat ata tca tcg ttt acg aat acc gac gcg cct gaa tat cta gga gta att aag ttt gga agt ctt atc cat ttc gaa gtg ccg tgt ttc aaa tat tct gcc aca ccc gtt gaa ata gaa aat tct aat cct cct att aca tat aac ttt cca tcg tta aca caa gta cta act tct gat ttt aac gac gac ata tta gta acc gtt ttc ca- --- --- --- --- --- --- --t ttt ttc gtt tca aga tct acc cgc -ga tac gga ata aac atg tct att gtt aat cat gcc gcc aat aat gta tag aca att atg taa aac att tgc att ata gaa ttg --- -tc tat ctg tat tac cga cta tcg tcc aat att ctg ttc tag gag agt aat ggg tta ttg tgg ata tat aat cag agt ttt taa tga cta cta tat tat gtt tta tac cat ttc gtg tca ctg gct ttg tag att tgg ata tag tta atc cca aca atg ata tag cat tgc gca tag tat tag tca taa act tgg gat gta aaa tgt tga tga tat cta cat cgt ttg gat ttt tat gta tcc act tta ata ata tca tag ctg taa cat cct cat ga- ttt acg tta acg tct tcg tgg gat aag ata gtt gtc agt tca tcc ttt gat aat ttt cca aat t

>EU410304.1:17942-19905_Synthetic_Vaccinia_virus_clone_GLV-1h68

cgg ctt cca tgt att tcc tgg aga gca agt --- aga tga tg- agg aac cag ata gtt tat atc cgt act tgc act taa agt cta cat tgt cgt tgt atg a-- gta tga tct ttt aaa ccc gct aga caa gta tcc gtt tga tat tgt agg atg tgg aca ttt aac aat ctg aca cgt ggg tgg atc gga cca ttc tcc tcc tga aca cag gac act aga gtt acc aat caa cga ata tcc act att gca act ata agt tac aac gct ccc atc ggt ata aaa atc ctc gta tcc gtt atg tct tcc gtt gga tat aga tgg agg gga ttg gca ttt aac aga ttc aca aat agg tgc ctc ggg att cca tac cat aga tcc agt aga tcc taa ttc aca ata cga ttt aga ttc acc gat caa ctg ata tcc gct att aca aga gta cgt tat act aga gcc aaa gtc tac tcc acc aat atc aag ttg gcc att atc gat atc tcg agg cga tgg gca tct ccg ttt aat aca ttg att aaa gag tgt cca tcc agt acc tgt aca ttt agc ata tat agg tcc cat ttt ttg ctt tct gta tcc agg tag aca tag a-t att cta tag tgt ctc cta tgt tgt a-- --- --- --- -at tag cat tag cat cag tc- --t cca cac tat tct taa att tca tat taa tgg gtc gtg acg gaa tag tac agc atg ata gaa cgc atc cta ttc cca aca atg tca gga acg tca cgc tct cca cct tca tat tta ttt atc cgt -aa aaa tgt tat cct gga cat cgt aca aat aat a-a aaa gcc ca- -ta tat gtt cgc tat tgt aga aat tgt ttt tca cag ttg ctc aaa aac gat ggc agt gac tta tga gtt acg tta cac ttt gga gtc tca tct tta gta aac ata tca taa tat tcg ata tta cga gtt gac ata tcg aac aaa ttc caa gta ttt gat ttt gga taa tat tcg tat ttt gca tct gct ata att aag ata taa tca ccg caa gaa cac acg aac atc ttt cct aca tgg tta aag tac atg tac aat tct atc cat ttg tct tcc tta act ata tat ttg tat aga taa tta cga gtc tcg tga gta att cca gta att aca tag atg tcg ccg tcg tac tct aca gca taa act ata cta tga tgt cta ggc atg gga gac -tt ttt tat cca acg att ttt agt gaa aca ttc cac atc gtt taa tac tac ata ttt ctc ata cgt ggt ata aac tcc acc cat tac --a tat ata tca tcg ttt acg aat acc gac gcg cct gaa tat cta gga gta att aag ttt gga agt ctt atc cat ttc gaa gtg ccg tgt ttc aaa tat tct gcc aca ccc gtt gaa ata gaa aat tct aat cct cct att aca tat aac ttt cca tcg tta aca caa gta cta act tct gat ttt aac gac gac ata tta gta acc gtt ttc ca- --- --- --- --- --- --- --t ttt ttc gtt tta aga tct acc cgc -ga tac gga ata aac atg tct att gtt aat cat gcc gcc aat aat gta tag aca att atg taa aac att tgc att ata gaa ttg --- -tc tat ctg tat tac cga cta tcg tcc aat att ctg ttc tag gag agt aat ggg tta ttg tgg ata tat aat cag agt ttt taa tga cta cta tat tat gtt tta tac cat ttc gtg tca ctg gct ttg tag att tgg ata tag tta atc cca aca atg ata tag cat tgc gca tag tat tag tca taa act tgg gat gta aaa tgt tga tga tat cta cat cgt ttg gat ttt tat gta tcc act tta ata ata tca tag ctg taa cat cct cat ga- ttt acg tta acg tct tcg tgg gat aag ata gtt gtc agt tca tcc ttt gat aat ttt cca aat t

>AY243312.1:18701-20664_Vaccinia_virus_WR_complete_genome

cgg ctt cca tgt att tcc tgg aga gca agt --- aga tga tg- agg aac cag ata gtt tat atc cgt act tgc act taa agt cta cat tgt cgt tgt atg a-- gta tga tct ttt aaa ccc gct aga caa gta tcc gtt tga tat tgt agg atg tgg aca ttt aac aat ctg aca cgt ggg tgg atc gga cca ttc tcc tcc tga aca cag gac acc aga gtt acc aat caa cga ata tcc act att gca act ata agt tac aac gct ccc atc ggt ata aaa atc ctc gta tcc gtt atg tct tcc gtt gga tat aga tgg agg gga ttg gca ttt aac aga ttc aca aat agg tgc ctc ggg att cca tac cat aga tcc agt aga tcc taa ttc aca ata cga ttt aga ttc acc gat caa atg ata tcc gct att aca aga gta cgt tat act aga gcc aaa gtc tac tcc acc aat atc aag ttg gcc att atc gat atc tcg agg cga tgg gca tct ccg ttt aat aca ttg att aaa gag tgt cca tcc agt acc tgt aca ttt agc ata tat agg tcc cat ttt ttg ctt tct gta tcc agg tag aca tag a-t att cta tag tgt ctc cta tgt tgt a-- --- --- --- -at tag cat tag cat cag tc- --t cca cac tat tct taa att tca tat taa tgg gtc gtg acg gaa tag tac agc atg ata gaa cgc atc cta ttc cca aca atg tca gga acg tca cgc tct cca cct tca tat tta ttt atc cgt -aa aaa tgt tat cct gga cat cgt aca aat aat a-a aaa gcc ca- -ta tat gtt cgc tat tgt aga aat tgt ttt tca cag ttg ctc aaa aac gat ggc agt gac tta tga gtt acg tta cac ttt gga gtc tca tct tta gta aac ata tca taa tat tcg ata tta cga gtt gac ata tcg aac aaa ttc caa gta ttt gat ttt gga taa tat tcg tat ttt gca tct gct ata att aag ata taa tca ccg caa gaa cac acg aac atc ttt cct aca tgg tta aag tac atg tat aat tct atc cat ttg tct tcc tta act ata tat ttg tat aga taa tta cga gtc tcg tga gta att cca gta att aca tag atg tcg ccg tcg tac tct aca gca taa act ata cta tga tgt cta ggc atg gga gac -tt ttt tat cca acg att ttt agt gaa aca ttc cac atc gtt taa tac tac ata ttt ttc ata cgt ggt ata aac tcc acc cat tac --a tat ata tca tcg ttt acg aat acc gac gcg cct gaa tat cta gga gta att aag ttt gga agt ctt atc cat ttc gaa gtg ccg tgt ttc aaa tat tct gcc aca ccc gtt gaa ata gaa aat tct aat cct cct att aca tat aac ttt cca tcg tta aca caa gta cta act tct gat ttt aac gac gac ata tta gta acc gtt ttc ca- --- --- --- --- --- --- --t ttt ttc gtt tca aga tct acc cgc -ga tac gga ata aac atg tct att gtt aat cat gcc gcc aat aat gta tag aca att atg taa aac att tgc att ata gaa ttg --- -tc tat ctg tat tac cga cta tcg tcc aat att ctg ttc tag gag agt aat ggg tta ttg tgg ata tat aat cag agt ttt taa tga cta cta tat tat gtt tta tac cat ttc gtg tca ctg gct ttg tag att tgg ata tag tta atc cca aca atg ata tag cat tgc gca tag tat tag tca taa act tgg gat gta aaa tgt tga tga tat cta cat cgt ttg gat ttt tat gta tcc act tta ata ata tca tag ctg taa cat cct cat ga- ttt acg tta acg ttt tcg tgg gat aag ata gtt gtc agt tca tcc ttt gat aat ttt cca aat t

>M22812.1:9562-11525_Vaccinia_virus_genome_left_end

cgg ctt cca tgt att tcc tgg aga gca agt --- aga tga tg- agg aac cag ata gtt tat atc cgt act tgc act taa agt cta cat tgt cgt tgt atg a-- gta tga tct ttt aaa ccc gct aga caa gta tcc gtt tga tat tgt agg atg tgg aca ttt aac aat ctg aca cgt ggg tgg atc gga cca ttc tcc tcc tga aca cag gac acc aga gtt acc aat caa cga ata tcc act att gca act ata agt tac aac gct ccc atc ggt ata aaa atc ctc gta tcc gtt atg tct tcc gtt gga tat aga tgg agg gga ttg gca ttt aac aga ttc aca aat agg tgc ctc ggg att cca tac cat aga tcc agt aga tcc taa ttc aca ata cga ttt aga ttc acc gat caa atg ata tcc gct att aca aga gta cgt tat act aga gcc aaa gtc tac tcc acc aat atc aag ttg gcc att atc gat atc tcg agg cga tgg gca tct ccg ttt aat aca ttg att aaa gag tgt cca tcc agt acc tgt aca ttt agc ata tat agg tcc cat ttt ttg ctt tct gta tcc agg tag aca tag a-t att cta tag tgt ctc cta tgt tgt a-- --- --- --- -at tag cat tag cat cag tc- --t cca cac tat tct taa att tca tat taa tgg gtc gtg acg gaa tag tac agc atg ata gaa cgc atc cta ttc cca aca atg tca gga acg tca cgc tct cca cct tca tat tta ttt atc cgt -aa aaa tgt tat cct gga cat cgt aca aat aat a-a aaa gcc ca- -ta tat gtt cgc tat tgt aga aat tgt ttt tca cag ttg ctc aaa aac gat ggc agt gac tta tga gtt acg tta cac ttt gga gtc tca tct tta gta aac ata tca taa tat tcg ata tta cga gtt gac ata tcg aac aaa ttc caa gta ttt gat ttt gga taa tat tcg tat ttt gca tct gct ata att aag ata taa tca ccg caa gaa cac acg aac atc ttt cct aca tgg tta aag tac atg tat aat tct atc cat ttg tct tcc tta act ata tat ttg tat aga taa tta cga gtc tcg tga gta att cca gta att aca tag atg tcg ccg tcg tac tct aca gca taa act ata cta tga tgt cta ggc atg gga gac -tt ttt tat cca acg att ttt agt gaa aca ttc cac atc gtt taa tac tac ata ttt ttc ata cgt ggt ata aac tcc acc cat tac --a tat ata tca tcg ttt acg aat acc gac gcg cct gaa tat cta gga gta att aag ttt gga agt ctt atc cat ttc gaa gtg ccg tgt ttc aaa tat tct gcc aca ccc gtt gaa ata gaa aat tct aat cct cct att aca tat aac ttt cca tcg tta aca caa gta cta act tct gat ttt aac gac gac ata tta gta acc gtt ttc ca- --- --- --- --- --- --- --t ttt ttc gtt tca aga tct acc cgc -ga tac gga ata aac atg tct att gtt aat cat gcc gcc aat aat gta tag aca att atg taa aac att tgc att ata gaa ttg --- -tc tat ctg tat tac cga cta tcg tcc aat att ctg ttc tag gag agt aat ggg tta ttg tgg ata tat aat cag agt ttt taa tga cta cta tat tat gtt tta tac cat ttc gtg tca ctg gct ttg tag att tgg ata tag tta atc cca aca atg ata tag cat tgc gca tag tat tag tca taa act tgg gat gta aaa tgt tga tga tat cta cat cgt ttg gat ttt tat gta tcc act tta ata ata tca tag ctg taa cat cct cat ga- ttt acg tta acg ttt tcg tgg gat aag ata gtt gtc agt tca tcc ttt gat aat ttt cca aat t

>MK035755.1:35164-37128_Cowpox_virus_strain_CPXV/75/01

cgg ctt cca tgt att tcc tgg aga gca agt --- aga tga tg- agg aac cag ata gtt tat atc cgt act tgc act taa agt cta cat tgt cgt tgt atg a-- gta tga tct ttt aaa ccc gct aga caa gta tcc gtt tga tat tgt agg atg tgg aca ttt aac aat ctg aca cgt ggg tgg atc gga cca ttc tcc tcc tga aca cag gac acc aga gtt acc aat caa cga ata tcc gct att aca act ata cgt tac aac gct tcc atc ggt ata aaa atc ctc gta tcc gtt atg tct tcc gtt gga tat aga tgg agg tga ttg gca ttt aac aga ttc gca aat agg tgc ctc ggg att cca tac cat aga tcc agt aga tcc taa ttc aca ata cga ttt aga ttc acc gat caa atg ata tcc gct att aca aga gta cgt tat act aga gcc aaa gtc tac tcc gcc aat atc aag ttg gcc att atc gat atc tcg agg cga tgg gca tct ccg ttt aat aca ttg att aaa gag tgt cca tcc agt acc tgt aca ttt agc ata tat agg tcc cat ttt ttg ctt tct gta tcc agg tag aca tag a-t att cta tag tgt ctc cta tgt tgt a-- --- --- --- -at tag cat tag cat cag tc- --t cca cac tat tct taa att tca tat taa tgg gtc gtg acg gaa tag tac agc atg ata gaa cgc atc cta ttc cca aca atg tca gga acg tca cgc tct cca cct tca tat tta ttt atc cgt -aa aaa tgt tat cct gga cat cgt acc aat aat aaa aaa gcc ca- -ta tat gtt cgc tat tgt aga aat tgt ttt tca cag ttg ctc aaa aac gat ggc agt gac tta tga gtt acg tta cac ttt gga gtc tca tct tta gta aac ata tca taa tat tcg ata tta cga gtt gac ata tcg aac aaa ttc caa gta ttt gat ttt gga taa tat tcg tat ttt gca tct gct ata att aag ata taa tcg ccg caa gaa cac acg aac atc ttt cct aca tgg tta aag tac atg tac aat tct atc cat ttg tct tcc tta act ata tat ttg tat aga taa tta cga gtc tca tga gta att cca gta att gca tag atg tca cca tcg tac tct aca gca taa act ata cta tga tgt cta ggc atg gga gac -tt ttt tat cca acg att ttt agt gaa aca ttc cac atc gtt taa tac tac ata ttt ttc ata cgt ggc ata aac tcc acc cat tac --a tat ata tca tcg ttt acg aat acc gat gcg cct gaa tat cta gga gta att aag ttt gga agt ctt atc cat ttc gaa gtg ccg tgt ttc aaa tat tct gcc aca ccc gtt gaa ata gaa aat tct aat cct cct att aca tat aac ttt cca tcg tta aca caa gta cta act tct gat ttt aac gac gac ata tta gta act gtt ttc ca- --- --- --- --- --- --- --t ttt ttc gtt tta aga tct acc cgc -ga tac gga ata aac atg tct att gtt aat cat gcc gcc aat aat gta tag aca att ata taa aac att tgc att ata gaa ttg --- -tc tat ctg tat tac cga cga ttg tcc aat att ctg ttc tag gag agt aat ggg tta ttg tgg ata tat aat cag agt ttt taa tga cta cta tat tat gtt tta tac cat ttc gtg tca cag gct ttg tag att tgg ata tag tta atc cca aca atg ata tag cat tgc aca tag tat tag tca taa act tgg gat gta aaa tgt tga tga tat cta cat cgt ttg gat ttt tat gta tcc act tta ata ata tca tag ctg taa cat cct cat ga- ttt acg tta acg tct tcg tgg gat aag ata gtt gtc agt tca tcc ttt gat aat ttt cca aat t

>LC315596.1:25860-27823_BAC_cloning_vector_pLC16m8.8S-BAC_DNA_complete_sequence

cgg ctt cca tgt att tcc tgg aga gca agt --- aga tga tg- agg aac cag ata gtt tat atc cgt act tgc act taa agt cta cat tgt cgt tgt atg a-- gta tga tct ttt aaa ccc gct aga caa gta tcc gtt tga tat tgt agg atg tgg aca ttt aac aat ctg aca cgt ggg tgg atc gga cca ttc tcc tcc tga aca cag gac act aga gtt acc aat caa cga ata tcc act att gca act ata agt tac aac gct ccc atc ggt ata aaa atc ctc gta tcc gtt atg tct tcc gtt gga tat aga tgg agg gga ttg gca ttt aac aga ttc aca aat agg tgc ctc ggg att cca tac cat aga tcc agt aga tcc taa ttc aca ata cga ttt aga ttc acc gat caa atg ata tcc gct att aca aga gta cgt tat act aga gcc aaa gtc tac tcc acc aat atc aag ttg gcc att atc gat atc tcg agg cga tgg gca tct ccg ttt aat aca ttg att aaa gag tgt cca tcc agt acc tgt aca ttt agc ata tat agg tcc cat ttt ttg ctt tct gta tcc agg tag aca tag a-t att cta tag tgt ctc cta tgt tgt a-- --- --- --- -at tag cat tag cat cag tc- --t cca cac tat tct taa att tca tat taa tgg gtc gtg acg gaa tag tac agc atg ata gaa cgc atc cta ttc cca aca atg tca gga acg tca cgc tct cca cct tca tat tta ttt atc cgt -aa aaa tgt tat cct gga cat cgt aca aat aat a-a aaa gcc ca- -ta tat gtt cgc tat tgt aga aat tgt ttt tca cag ttg ctc aaa aac gat ggc agt gac tta tga gtt acg tta cac ttt gga gtc tca tct tta gta aac ata tca taa tat tcg ata tta cga gtt gac ata tcg aac aaa ttc caa gta ttt gat ttt gga taa tat tcg tat ttt gca tct gct ata att aag ata taa tca ccg caa gaa cac acg aac atc ttt cct aca tgg tta aag tac atg tac aat tct atc cat ttg tct tcc tta act ata tat ttg tat aga taa tta cga gtc tcg tga gta att cca gta att aca tag atg tcg ccg tcg tac tct aca gca taa act ata cta tga tgt cta ggc atg gga gac -tt ttt tat cca acg att ttt agt gaa aca ttc cac atc gtt taa tac tac ata ttt ttc ata cgt ggt ata aac tcc acc cat tac --a tat ata tca tcg ttt acg aat acc gac gcg cct gaa tat cta gga gta att aag ttt gga agt ctt atc cat ttc gaa gtg ccg tgt ttc aaa tat tct gcc aca ccc gtt gaa ata gaa aat tct aat cct cct att aca tat aac ttt cca tcg tta aca caa gta cta act tct gat ttt aac gac gac ata tta gta acc gtt ttc ca- --- --- --- --- --- --- --t ttt ttc gtt tca aga tct acc cgc -ga tac gga ata aac atg tct att gtt aat cat gcc gcc aat aat gta tag aca att atg taa aac att tgc att ata gaa ttg --- -tc tat ctg tat tac cga cta tcg tcc aat att ctg ttc tag gag agt aat ggg tta ttg tgg ata tat aat cag agt ttt taa tga cta cta tat tat gtt tta tac cat ttc gtg tca ctg gct ttg tag att tgg ata tag tta atc cca aca atg ata tag cat tgc gca tag tat tag tca taa act tgg gat gta aaa tgt tga tga tat cta cat cgt ttg gat ttt tat gta tcc act tta ata ata tca tag ctg taa cat cct cat ga- ttt acg tta acg tct tcg tgg gat aag ata gtt gtc agt tca tcc ttt gat aat ttt cca aat t

>LT966077.1:18753-20716_Vaccinia_virus_WR_strain_WR_-_ATCC_VR-2035_genome_assembly_complete_genome:_monopartite

cgg ctt cca tgt att tcc tgg aga gca agt --- aga tga tg- agg aac cag ata gtt tat atc cgt act tgc act taa agt cta cat tgt cgt tgt atg a-- gta tga tct ttt aaa ccc gct aga caa gta tcc gtt taa tat tgt agg atg tgg aca ttt aac aat ctg aca cgt ggg tgg atc gga cca ttc tcc tcc tga aca cag gac acc aga gtt acc aat caa cga ata tcc act att gca act ata agt tac aac gct ccc atc ggt ata aaa atc ctc gta tcc gtt atg tct tcc gtt gga tat aga tgg agg gga ttg gca ttt aac aga ttc aca aat agg tgc ctc ggg att cca tac cat aga tcc agt aga tcc taa ttc aca ata cga ttt aga ttc acc gat caa atg ata tcc gct att aca aga gta cgt tat act aga gcc aaa gtc tac tcc acc aat atc aag ttg gcc att atc gat atc tcg agg cga tgg gca tct ccg ttt aat aca ttg att aaa gag tgt cca tcc agt acc tgt aca ttt agc ata tat agg tcc cat ttt ttg ctt tct gta tcc agg tag aca tag a-t att cta tag tgt ctc cta tgt tgt a-- --- --- --- -at tag cat tag cat cag tc- --t cca cac tat tct taa att tca tat taa tgg gtc gtg acg gaa tag tac agc atg ata gaa cgc atc cta ttc cca aca atg tca gga acg tca cgc tct cca cct tca tat tta ttt atc cgt -aa aaa tgt tat cct gga cat cgt aca aat aat a-a aaa gcc ca- -ta tat gtt cgc tat tgt aga aat tgt ttt tca cag ttg ctc aaa aac gat ggc agt gac tta tga gtt acg tta cac ttt gga gtc tca tct tta gta aac ata tca taa tat tcg ata tta cga gtt gac ata tcg aac aaa ttc caa gta ttt gat ttt gga taa tat tcg tat ttt gca tct gct ata att aag ata taa tca ccg caa gaa cac acg aac atc ttt cct aca tgg tta aag tac atg tat aat tct atc cat ttg tct tcc tta act ata tat ttg tat aga taa tta cga gtc tcg tga gta att cca gta att aca tag atg tcg ccg tcg tac tct aca gca taa act ata cta tga tgt cta ggc atg gga gac -tt ttt tat cca acg att ttt agt gaa aca ttc cac atc gtt taa tac tac ata ttt ttc ata cgt ggt ata aac tcc acc cat tac --a tat ata tca tcg ttt acg aat acc gac gcg cct gaa tat cta gga gta att aag ttt gga agt ctt atc cat ttc gaa gtg ccg tgt ttc aaa tat tct gcc aca ccc gtt gaa ata gaa aat tct aat cct cct att aca tat aac ttt cca tcg tta aca caa gta cta act tct gat ttt aac gac gac ata tta gta acc gtt ttc ca- --- --- --- --- --- --- --t ttt ttc gtt tca aga tct acc cgc -ga tac gga ata aac atg tct att gtt aat cat gcc gcc aat aat gta tag aca att atg taa aac att tgc att ata gaa ttg --- -tc tat ctg tat tac cga cta tcg tcc aat att ctg ttc tag gag agt aat ggg tta ttg tgg ata tat aat cag agt ttt taa tga cta cta tat tat gtt tta tac cat ttc gtg tca ctg gct ttg tag att tgg ata tag tta atc cca aca atg ata tag cat tgc gca tag tat tag tca taa act tgg gat gta aaa tgt tga tga tat cta cat cgt ttg gat ttt tat gta tcc act tta ata ata tca tag ctg taa cat cct cat ga- ttt acg tta acg ttt tcg tgg gat aag ata gtt gtc agt tca tcc ttt gat aat ttt cca aat t

>LT896719.1:34099-36062_Cowpox_virus_isolate_Ger/2013/Alpaca_genome_assembly_complete_genome:_monopartite

cgg ctt cca tgt att tcc tgg aga gca agt --- aga tga tg- agg aac cag ata gtt tat atc cgt act tgc act taa agt cta cat tgt cgt tgt atg a-- gta tga tct ttt aaa ccc gct aga caa gta tcc gtt tga tat tgt agg atg tgg aca ttt aac aat ctg aca cgt ggg tgg ata gga cca ttc tcc tcc tga aca cag gac acc aga gtt acc aat caa cga ata tcc act att gca act ata agt tac aac gct ccc atc ggt ata aaa atc ctc gta tcc gtt atg tct tcc gtt gga tat aga tgg agg gga ttg gca ttt aac aga ttc aca aat agg tgc ctc ggg att cca tac cat aga tcc agt aga tcc taa ttc aca ata cga ttt aga ttc acc gat caa atg ata tcc gct att aca aga gta cgt tat act aga gcc aaa gtc tac tcc acc aat atc aag ttg gcc att atc gat atc tcg agg cga tgg gca tct ccg ttt aat aca ttg att aaa gag tgt cca tcc agt acc tgt aca ttt agc ata tat agg tcc cat ttt ttg ctt tct gta tcc agg tag aca tag a-t att cta tag tgt ctc cta tgt tgt a-- --- --- --- -at tag cat tag cat cag tc- --t cca cac tat tct taa att tca tat taa tgg gtc gtg acg gaa tag tac agc atg ata gaa cgc atc cta ttc cca aca atg tca gga acg tca cgc tct cca cct tca tat tta ttt atc cgt -aa aaa tgt tat cct gga cat agt aca aat aat a-a aaa gcc ca- -ta tat gtt cgc tat tgt aga aat tgt ttt tca cag ttg ctc aaa aac gat ggc agt gac tta tga gtt acg tta cac ttt gga gtc tca tct tta gta aac ata tca taa tat tcg ata tta cga gtt gac ata tcg aac aaa ttc caa gta ttt gat ttt gga taa tat tcg tat ttt gca tct gct ata att aag ata taa tca ccg caa gaa cac acg aac atc ttt cct aca tgg tta aag tac atg tac aat tct atc cat ttg tct tcc tta act ata tat ttg tat aga taa tta cga gtc tcg tga gta att cca gta att gca tag atg tca cca tcg tac tct aca gca taa act ata cta tga tgt cta ggc atg gga gac -tt ttt tat cca acg att ttt agt gaa aca ttc tac atc gtt taa tac tac ata ttt ctc ata cgt ggt ata aac tcc acc cat tac --a tat ata tca tcg ttt acg aat acc gat gcg cct gaa tat cta gga gta att aag ttt gga agt ctt atc cat ttc gaa gtg ccg tgt ttc aaa tat tct gcc aca ccc gtt gaa ata gaa aat tct aat cct cct att aca tat aac ttt cca tcg tta aca caa gta cta act tct gat ttt aac gac gac ata tta gta act gtt ttc ca- --- --- --- --- --- --- --t ttt ttc gtt tta aga tct acc cgc -ga tac gga ata aac atg tct att gtt aat cat gcc gcc aat aat gta tag aca att ata taa aac att tgc att ata gaa ttg --- -tc tat ctg tat tac cga cga ttg tcc aat att ctg ttc tag gag agt aat ggg tta ttg tgg ata tat aat cag agt ttt taa tga cta cta tat tat gtt tta tac cat ttc gtg tca cag gct ttg tag att tgg ata tag tta atc cca aca atg ata tag cat tgc aca tag tat tag tca taa act tgg gat gta aaa tgt tga tga tat cta cat cgt ttg gat ttt tat gta tcc act tta ata ata tca tag ctg taa cat cct cgt ga- ttt acg ttg acg ctt tcg tgg gat aag ata gtt gtt agt tca tcc ttt gat aat ttt cca aat t

>KY549147.1:27110-29074_Cowpox_virus_isolate_CPXV_K780

cgg ctt cca tgt att tcc tgg aga gca agt --- aga tga tg- agg aac cag ata gtt tat atc cgt act tgc act taa agt cta cat tgt cgt tgt atg a-- gta tga tct ttt aaa ccc gct aga caa gta tcc gtt tga tat tgt agg atg tgg aca ttt aac aat ctg aca cgt ggg tgg atc gga cca ttc tcc tcc tga aca cag gac acc aga gtt acc aat caa cga ata tcc gct att aca act ata cgt tac aac gct tcc atc ggt ata aaa atc ctc gta tcc gtt atg tct tcc gtt gga tat aga tgg agg tga ttg gca ttt aac aga ttc gca aat agg tgc ctc ggg att cca tac cat aga tcc agt aga tcc taa ttc aca ata cga ttt aga ttc acc gat caa atg ata tcc gct att aca aga gta cgt tat act aga gcc aaa gtc tac tcc gcc aat atc aag ttg gcc att atc gat atc tcg agg cga tgg gca tct ccg ttt aat aca ttg att aaa gag tgt cca tcc agt acc tgt aca ttt agc ata tat agg tcc cat ttt ttg ctt tct gta tcc agg tag aca tag a-t att cta tag tgt ctc cta tgt tgt a-- --- --- --- -at tag cat tag cat cag tc- --t cca cac tat tct taa att tca tat taa tgg gtc gtg acg gaa tag tac agc atg ata gaa cgc atc cta ttc cca aca atg tca gga acg tca cgc tct cca cct tca tat tta ttt atc cgt -aa aaa tgt tat cct gga cat cgt acc aat aat aaa aaa gcc ca- -ta tat gtt cgc tat tgt aga aat tgt ttt tca cag ttg ctc aaa aac gat ggc agt gac tta tga gtt acg tta cac ttt gga gtc tca tct tta gta aac ata tca taa tat tcg ata tta cga gtt gac ata tcg aac aaa ttc caa gta ttt gat ttt gga taa tat tcg tat ttt gca tct gct ata att aag ata taa tcg ccg caa gaa cac acg aac atc ttt cct aca tgg tta aag tac atg tac aat tct atc cat ttg tct tcc tta act ata tat ttg tat aga taa tta cga gtc tca tga gta att cca gta att gca tag atg tca cca tcg tac tct aca gca taa act ata cta tga tgt cta ggc atg gga gac -tt ttt tat cca acg att ttt agt gaa aca ttc cac atc gtt taa tac tac ata ttt ttc ata cgt ggc ata aac tcc acc cat tac --a tat ata tca tcg ttt acg aat acc gat gcg cct gaa tat cta gga gta att aag ttt gga agt ctt atc cat ttc gaa gtg ccg tgt ttc aaa tat tct gcc aca ccc gtt gaa ata gaa aat tct aat cct cct att aca tat aac ttt cca tcg tta aca caa gta cta act tct gat ttt aac gac gac ata tta gta act gtt ttc ca- --- --- --- --- --- --- --t ttt ttc gtt tta aga tct acc cgc -ga tac gga ata aac atg tct att gtt aat cat gcc gcc aat aat gta tag aca att ata taa aac att tgc att ata gaa ttg --- -tc tat ctg tat tac cga cga ttg tcc aat att ctg ttc tag gag agt aat ggg tta ttg tgg ata tat aat cag agt ttt taa tga cta cta tat tat gtt tta tac cat ttc gtg tca cag gct ttg tag att tgg ata tag tta atc cca aca atg ata tag cat tgc aca tag tat tag tca taa act tgg gat gta aaa tgt tga tga tat cta cat cgt ttg gat ttt tat gta tcc act tta ata ata tca tag ctg taa cat cct cat ga- ttt acg tta acg tct tcg tgg gat aag ata gtt gtc agt tca tcc ttt gat aat ttt cca aat t

>KY549146.1:27302-29266_Cowpox_virus_isolate_CPXV_K779

cgg ctt cca tgt att tcc tgg aga gca agt --- aga tga tg- agg aac cag ata gtt tat atc cgt act tgc act taa agt cta cat tgt cgt tgt atg a-- gta tga tct ttt aaa ccc gct aga caa gta tcc gtt tga tat tgt agg atg tgg aca ttt aac aat ctg aca cgt ggg tgg atc gga cca ttc tcc tcc tga aca cag gac acc aga gtt acc aat caa cga ata tcc gct att aca act ata cgt tac aac gct tcc atc ggt ata aaa atc ctc gta tcc gtt atg tct tcc gtt gga tat aga tgg agg tga ttg gca ttt aac aga ttc gca aat agg tgc ctc ggg att cca tac cat aga tcc agt aga tcc taa ttc aca ata cga ttt aga ttc acc gat caa atg ata tcc gct att aca aga gta cgt tat act aga gcc aaa gtc tac tcc gcc aat atc aag ttg gcc att atc gat atc tcg agg cga tgg gca tct ccg ttt aat aca ttg att aaa gag tgt cca tcc agt acc tgt aca ttt agc ata tat agg tcc cat ttt ttg ctt tct gta tcc agg tag aca tag a-t att cta tag tgt ctc cta tgt tgt a-- --- --- --- -at tag cat tag cat cag tc- --t cca cac tat tct taa att tca tat taa tgg gtc gtg acg gaa tag tac agc atg ata gaa cgc atc cta ttc cca aca atg tca gga acg tca cgc tct cca cct tca tat tta ttt atc cgt -aa aaa tgt tat cct gga cat cgt acc aat aat aaa aaa gcc ca- -ta tat gtt cgc tat tgt aga aat tgt ttt tca cag ttg ctc aaa aac gat ggc agt gac tta tga gtt acg tta cac ttt gga gtc tca tct tta gta aac ata tca taa tat tcg ata tta cga gtt gac ata tcg aac aaa ttc caa gta ttt gat ttt gga taa tat tcg tat ttt gca tct gct ata att aag ata taa tcg ccg caa gaa cac acg aac atc ttt cct aca tgg tta aag tac atg tac aat tct atc cat ttg tct tcc tta act ata tat ttg tat aga taa tta cga gtc tca tga gta att cca gta att gca tag atg tca cca tcg tac tct aca gca taa act ata cta tga tgt cta ggc atg gga gac -tt ttt tat cca acg att ttt agt gaa aca ttc cac atc gtt taa tac tac ata ttt ttc ata cgt ggc ata aac tcc acc cat tac --a tat ata tca tcg ttt acg aat acc gat gcg cct gaa tat cta gga gta att aag ttt gga agt ctt atc cat ttc gaa gtg ccg tgt ttc aaa tat tct gcc aca ccc gtt gaa ata gaa aat tct aat cct cct att aca tat aac ttt cca tcg tta aca caa gta cta act tct gat ttt aac gac gac ata tta gta act gtt ttc ca- --- --- --- --- --- --- --t ttt ttc gtt tta aga tct acc cgc -ga tac gga ata aac atg tct att gtt aat cat gcc gcc aat aat gta tag aca att ata taa aac att tgc att ata gaa ttg --- -tc tat ctg tat tac cga cga ttg tcc aat att ctg ttc tag gag agt aat ggg tta ttg tgg ata tat aat cag agt ttt taa tga cta cta tat tat gtt tta tac cat ttc gtg tca cag gct ttg tag att tgg ata tag tta atc cca aca atg ata tag cat tgc aca tag tat tag tca taa act tgg gat gta aaa tgt tga tga tat cta cat cgt ttg gat ttt tat gta tcc act tta ata ata tca tag ctg taa cat cct cat ga- ttt acg tta acg tct tcg tgg gat aag ata gtt gtc agt tca tcc ttt gat aat ttt cca aat t

>KY463519.1:25662-27626_Cowpox_virus_strain_Germany_1971_EP1

cgg ctt cca tgt att tcc tgg aga gca agt --- aga tga tg- agg aac cag ata gtt tat atc cgt act tgc acg taa agt cta cat tgt cgt tgt atg a-- gta tga tct ttt aaa ccc gct aga caa gta tcc gtt tga tat tgt agg atg tgg aca ttt aac aat ctg aca cgt ggg tgg atc gga cca ttc tcc tcc tga aca cag gac acc aga gtt acc aat caa cga ata tcc act att aca act ata cgt tac aac gct tcc atc ggt ata aaa atc ctc gta tcc gtt atg tct tcc gtt gga tat aga tgg agg tga ttg gca ttt aac aga ttc gca aat agg tgc ctc ggg att cca tac cat aga tcc agt aga tcc taa ttc aca ata cga ttt aga ttc acc gat caa atg ata tcc gct att aca aga gta cgt tat act aga gcc aaa gtc tac tcc gcc aat atc aag ttg gcc att atc gat atc tcg agg cga tgg gca tct ccg ttt aat aca ttg att aaa gag tgt cca tcc agt acc tgt aca ttt agc ata tat agg tcc cat ttt ttg ctt tct gta tcc agg tag aca tag a-t att cta tag tgt ctc cta tgt tgt a-- --- --- --- -at tag cat tat cat cag ta- --t cca cac tat tct taa att tca tat taa tgg gtc gtg acg gaa tag tac agc atg ata gaa cgc atc cta ttc cca aca atg tca gga acg tca cgc tct cca cct tca tat tta ttt atc cgt -aa aaa tgt tat cct gga cat cgt acc aat aat aaa aaa gcc ca- -ta tat gtt cgc tat tgt aga aat tgt ttt tca cag ttg ctc aaa aac gat ggc agt gac tta tga gtt acg tta cac ttt gga gtc tca ttt tta gta aac ata tca taa tat tcg ata tta cga gtt gac ata tcg aac aaa ttc caa gta ttt gat ttt gga taa tat tcg tat ttt gca tct gct ata att aag ata taa tcg ccg caa gaa cac acg aac atc ttt cct aca tgg tta aag tac atg tac aat tct atc cat ttg tct tcc tta act ata tat ttg tat aga taa tta cga gtc tcg tga gta att cca gta att gca tag atg tca cca tcg tac tct aca gca taa act ata cta tga tgt cta ggc atg gga gac -tt ttt tat cca acg att ttt agt gaa aca ttc cac atc gtt taa tac tac ata ttt ctc ata cgt ggt ata aac tcc acc cat tac --a tat ata tca tcg ttt acg aat acc gat gcg cct gaa tat cta gga gta att aag ttt gga agt ctt atc cat ttc gaa gtg ccg tgt ttc aaa tat tct gcc aca ccc gtt gaa ata gaa aat tct aat cct cct att aca tat aac ttt cca tcg tta aca caa gta cta act tct gat ttt aac gac gac ata tta gta acc gtt ttc ca- --- --- --- --- --- --- --t ttt ttc gtt tta aga tct acc cgc -ga tac gga ata aac atg tct att gtt aat cat gcc gcc aat aat gta tag aca att ata taa aac att tgc att ata gaa ttg --- -tc tat ctg tat tac cga cga ttg tcc aat att ctg ttc tag gag agt aat ggg tta ttg tgg ata tat aat cag agt ttt taa tga cta cta tat tat gtt tta tac cat ttc gtg tca cag gct ttg tag att tgg ata tag tta atc cca aca atg ata tag cat tgc aca tag tat tag tca taa act tgg gat gta aaa tgt tga tga tat cta cat cgt ttg gat ttt tat gta tcc act tta ata ata tca tag ctg taa cat cct cat ga- ttt acg tta acg tct tcg tgg gat aag ata gtt gtc agt tca tcc ttt gat aat ttt cca aat t

>KC813509.1:22168-24131_Cowpox_virus_strain_HumBer07/1_complete_genome

cgg ctt cca tgt att tcc tgg aga gca agt --- aga tga tg- agg aac cag ata gtt tat atc cgt act tgc act taa agt cta cat tgt cgt tgt atg a-- gta tga tct ttt aaa ccc gct aga caa gta tcc gtt tga tat tgt agg atg tgg aca ttt aac aat ctg aca cgt ggg tgg atc gga cca ttc tcc tcc tga aca cag gac acc aga gtt acc aat caa cga ata tcc act att gca act ata agt tac aac gct ccc atc ggt ata aaa atc ctc gta tcc gtt atg tct tcc gtt gga tat aga tgg agg gga ttg gca ttt aac aga ttc aca aat agg tgc ctc ggg att cca tac cat aga tcc agt aga tcc taa ttc aca ata cga ttt aga ttc acc gat caa atg ata tcc gct att aca aga gta cgt tat act aga gcc aaa gtc tac tcc acc aat atc aag ttg gcc att atc gat atc tcg agg cga tgg gca tct ccg ttt aat aca ttg att aaa gag tgt cca tcc agt acc tgt aca ttt agc ata tat agg tcc cat ttt ttg ctt tct gta tcc agg tag aca tag a-t att cta tag tgt ctc cta tgt tgt a-- --- --- --- -at tag cat tag cat cag tc- --t cca cac tat tct taa att tca tat taa tgg gtc gtg acg gaa tag tac agc atg ata gaa cgc atc cta ttc cca aca atg tca gga acg tca cgc tct cca cct tca tat tta ttt atc cgt -aa aaa tgt tat cct gga cat agt aca aat aat a-a aaa gcc ca- -ta tat gtt cgc tat tgt aga aat tgt ttt tca cag ttg ctc aaa aac gat ggc agt gac tta tga gtt acg tta cac ttt gga gtc tca tct tta gta aac ata tca taa tat tcg ata tta cga gtt gac ata tcg aac aaa ttc caa gta ttt gat ttt gga taa tat tcg tat ttt gca tct gct ata att aag ata taa tca ccg caa gaa cac acg aac atc ttt cct aca tgg tta aag tac atg tac aat tct atc cat ttg tct tcc tta act ata tat ttg tat aga taa tta cga gtc tcg tga gta att cca gta att gca tag atg tca cca tcg tac tct aca gca taa act ata cta tga tgt cta ggc atg gga gac -tt ttt tat cca acg att ttt agt gaa aca ttc tac atc gtt taa tac tac ata ttt ctc ata cgt ggt ata aac tcc acc cat tac --a tat ata tca tcg ttt acg aat acc gat gcg cct gaa tat cta gga gta att aag ttt gga agt ctt atc cat ttc gaa gtg ccg tgt ttc aaa tat tct gcc aca ccc gtt gaa ata gaa aat tct aat cct cct att aca tat aac ttt cca tcg tta aca caa gta cta act tct gat ttt aac gac gac ata tta gta act gtt ttc ca- --- --- --- --- --- --- --t ttt ttc gtt tta aga tct acc cgc -ga tac gga ata aac atg tct att gtt aat cat gcc gcc aat aat gta tag aca att ata taa aac att tgc att ata gaa ttg --- -tc tat ctg tat tac cga cga ttg tcc aat att ctg ttc tag gag agt aat ggg tta ttg tgg ata tat aat cag agt ttt taa tga cta cta tat tat gtt tta tac cat ttc gtg tca cag gct ttg tag att tgg ata tag tta atc cca aca atg ata tag cat tgc aca tag tat tag tca taa act tgg gat gta aaa tgt tga tga tat cta cat cgt ttg gat ttt tat gta tcc act tta ata ata tca tag ctg taa cat cct cgt ga- ttt acg ttg acg ctt tcg tgg gat aag ata gtt gtt agt tca tcc ttt gat aat ttt cca aat t

>KC813506.1:35501-37464_Cowpox_virus_strain_CatPot07/1_complete_genome

cgg ctt cca tgt att tcc tgg aga gca agt --- aga tga tg- agg aac cag ata gtt tat atc cgt act tgc act taa agt cta cat tgt cgt tgt atg a-- gta tga tct ttt aaa ccc gct aga caa gta tcc gtt tga tat tgt agg atg tgg aca ttt aac aat ctg aca cgt ggg tgg atc gga cca ttc tcc tcc tga aca cag gac acc aga gtt acc aat caa cga ata tcc act att gca act ata agt tac aac gct ccc atc ggt ata aaa atc ctc gta tcc gtt atg tct tcc gtt gga tat aga tgg agg gga ttg gca ttt aac aga ttc aca aat agg tgc ctc ggg att cca tac cat aga tcc agt aga tcc taa ttc aca ata cga ttt aga ttc acc gat caa atg ata tcc gct att aca aga gta cgt tat act aga gcc aaa gtc tac tcc acc aat atc aag ttg gcc att atc gat atc tcg agg cga tgg gca tct ccg ttt aat aca ttg att aaa gag tgt cca tcc agt acc tgt aca ttt agc ata tat agg tcc cat ttt ttg ctt tct gta tcc agg tag aca tag a-t att cta tag tgt ctc cta tgt tgt a-- --- --- --- -at tag cat tag cat cag tc- --t cca cac tat tct taa att tca tat taa tgg gtc gtg acg gaa tag tac agc atg ata gaa cgc atc cta ttc cca aca atg tca gga acg tca cgc tct cca cct tca tat tta ttt atc cgt -aa aaa tgt tat cct gga cat agt aca aat aat a-a aaa gcc ca- -ta tat gtt cgc tat tgt aga aat tgt ttt tca cag ttg ctc aaa aac gat ggc agt gac tta tga gtt acg tta cac ttt gga gtc tca tct tta gta aac ata tca taa tat tcg ata tta cga gtt gac ata tcg aac aaa ttc caa gta ttt gat ttt gga taa tat tcg tat ttt gca tct gct ata att aag ata taa tca ccg caa gaa cac acg aac atc ttt cct aca tgg tta aag tac atg tac aat tct atc cat ttg tct tcc tta act ata tat ttg tat aga taa tta cga gtc tcg tga gta att cca gta att gca tag atg tca cca tcg tac tct aca gca taa act ata cta tga tgt cta ggc atg gga gac -tt ttt tat cca acg att ttt agt gaa aca ttc tac atc gtt taa tac tac ata ttt ctc ata cgt ggt ata aac tcc acc cat tac --a tat ata tca tcg ttt acg aat acc gat gcg cct gaa tat cta gga gta att aag ttt gga agt ctt atc cat ttc gaa gtg ccg tgt ttc aaa tat tct gcc aca ccc gtt gaa ata gaa aat tct aat cct cct att aca tat aac ttt cca tcg tta aca caa gta cta act tct gat ttt aac gac gac ata tta gta act gtt ttc ca- --- --- --- --- --- --- --t ttt ttc gtt tta aga tct acc cgc -ga tac gga ata aac atg tct att gtt aat cat gcc gcc aat aat gta tag aca att ata taa aac att tgc att ata gaa ttg --- -tc tat ctg tat tac cga cga ttg tcc aat att ctg ttc tag gag agt aat ggg tta ttg tgg ata tat aat cag agt ttt taa tga cta cta tat tat gtt tta tac cat ttc gtg tca cag gct ttg tag att tgg ata tag tta atc cca aca atg ata tag cat tgc aca tag tat tag tca taa act tgg gat gta aaa tgt tga tga tat cta cat cgt ttg gat ttt tat gta tcc act tta ata ata tca tag ctg taa cat cct cgt ga- ttt acg ttg acg ctt tcg tgg gat aag ata gtt gtt agt tca tcc ttt gat aat ttt cca aat t

>KC813502.1:27351-29314_Cowpox_virus_strain_CatBer07/1_complete_genome

cgg ctt cca tgt att tcc tgg aga gca agt --- aga tga tg- agg aac cag ata gtt tat atc cgt act tgc act taa agt cta cat tgt cgt tgt atg a-- gta tga tct ttt aaa ccc gct aga caa gta tcc gtt tga tat tgt agg atg tgg aca ttt aac aat ctg aca cgt ggg tgg atc gga cca ttc tcc tcc tga aca cag gac acc aga gtt acc aat caa cga ata tcc act att gca act ata agt tac aac gct ccc atc ggt ata aaa atc ctc gta tcc gtt atg tct tcc gtt gga tat aga tgg agg gga ttg gca ttt aac aga ttc aca aat agg tgc ctc ggg att cca tac cat aga tcc agt aga tcc taa ttc aca ata cga ttt aga ttc acc gat caa atg ata tcc gct att aca aga gta cgt tat act aga gcc aaa gtc tac tcc acc aat atc aag ttg gcc att atc gat atc tcg agg cga tgg gca tct ccg ttt aat aca ttg att aaa gag tgt cca tcc agt acc tgt aca ttt agc ata tat agg tcc cat ttt ttg ctt tct gta tcc agg tag aca tag a-t att cta tag tgt ctc cta tgt tgt a-- --- --- --- -at tag cat tag cat cag tc- --t cca cac tat tct taa att tca tat taa tgg gtc gtg acg gaa tag tac agc atg ata gaa cgc atc cta ttc cca aca atg tca gga acg tca cgc tct cca cct tca tat tta ttt atc cgt -aa aaa tgt tat cct gga cat agt aca aat aat a-a aaa gcc ca- -ta tat gtt cgc tat tgt aga aat tgt ttt tca cag ttg ctc aaa aac gat ggc agt gac tta tga gtt acg tta cac ttt gga gtc tca tct tta gta aac ata tca taa tat tcg ata tta cga gtt gac ata tcg aac aaa ttc caa gta ttt gat ttt gga taa tat tcg tat ttt gca tct gct ata att aag ata taa tca ccg caa gaa cac acg aac atc ttt cct aca tgg tta aag tac atg tac aat tct atc cat ttg tct tcc tta act ata tat ttg tat aga taa tta cga gtc tcg tga gta att cca gta att gca tag atg tca cca tcg tac tct aca gca taa act ata cta tga tgt cta ggc atg gga gac -tt ttt tat cca acg att ttt agt gaa aca ttc tac atc gtt taa tac tac ata ttt ctc ata cgt ggt ata aac tcc acc cat tac --a tat ata tca tcg ttt acg aat acc gat gcg cct gaa tat cta gga gta att aag ttt gga agt ctt atc cat ttc gaa gtg ccg tgt ttc aaa tat tct gcc aca ccc gtt gaa ata gaa aat tct aat cct cct att aca tat aac ttt cca tcg tta aca caa gta cta act tct gat ttt aac gac gac ata tta gta act gtt ttc ca- --- --- --- --- --- --- --t ttt ttc gtt tta aga tct acc cgc -ga tac gga ata aac atg tct att gtt aat cat gcc gcc aat aat gta tag aca att ata taa aac att tgc att ata gaa ttg --- -tc tat ctg tat tac cga cga ttg tcc aat att ctg ttc tag gag agt aat ggg tta ttg tgg ata tat aat cag agt ttt taa tga cta cta tat tat gtt tta tac cat ttc gtg tca cag gct ttg tag att tgg ata tag tta atc cca aca atg ata tag cat tgc aca tag tat tag tca taa act tgg gat gta aaa tgt tga tga tat cta cat cgt ttg gat ttt tat gta tcc act tta ata ata tca tag ctg taa cat cct cgt ga- ttt acg ttg acg ctt tcg tgg gat aag ata gtt gtt agt tca tcc ttt gat aat ttt cca aat t

>KC813495.1:28353-30316_Cowpox_virus_strain_HumMag07/1_complete_genome

cgg ctt cca tgt att tcc tgg aga gca agt --- aga tga tg- agg aac cag ata gtt tat atc cgt act tgc act taa agt cta cat tgt cgt tgt atg a-- gta tga tct ttt aaa ccc gct aga caa gta tcc gtt tga tat tgt agg atg tgg aca ttt aac aat ctg aca cgt ggg tgg atc gga cca ttc tcc tcc tga aca cag gac acc aga gtt acc aat caa cga ata tcc act att gca act ata agt tac aac gct ccc atc ggt ata aaa atc ctc gta tcc gtt atg tct tcc gtt gga tat aga tgg agg gga ttg gca ttt aac aga ttc aca aat agg tgc ctc ggg att cca tac cat aga tcc agt aga tcc taa ttc aca ata cga ttt aga ttc acc gat caa atg ata tcc gct att aca aga gta cgt tat act aga gcc aaa gtc tac tcc acc aat atc aag ttg gcc att atc gat atc tcg agg cga tgg gca tct ccg ttt aat aca ttg att aaa gag tgt cca tcc agt acc tgt aca ttt agc ata tat agg tcc cat ttt ttg ctt tct gta tcc agg tag aca tag a-t att cta tag tgt ctc cta tgt tgt a-- --- --- --- -at tag cat tag cat cag tc- --t cca cac tat tct taa att tca tat taa tgg gtc gtg acg gaa tag tac agc atg ata gaa cgc atc cta ttc cca aca atg tca gga acg tca cgc tct cca cct tca tat tta ttt atc cgt -aa aaa tgt tat cct gga cat agt aca aat aat a-a aaa gcc ca- -ta tat gtt cgc tat tgt aga aat tgt ttt tca cag ttg ctc aaa aac gat ggc agt gac tta tga gtt acg tta cac ttt gga gtc tca tct tta gta aac ata tca taa tat tcg ata tta cga gtt gac ata tcg aac aaa ttc caa gta ttt gat ttt gga taa tat tcg tat ttt gca tct gct ata att aag ata taa tca ccg caa gaa cac acg aac atc ttt cct aca tgg tta aag tac atg tac aat tct atc cat ttg tct tcc tta act ata tat ttg tat aga taa tta cga gtc tcg tga gta att cca gta att gca tag atg tca cca tcg tac tct aca gca taa act ata cta tga tgt cta ggc atg gga gac -tt ttt tat cca acg att ttt agt gaa aca ttc tac atc gtt taa tac tac ata ttt ctc ata cgt ggt ata aac tcc acc cat tac --a tat ata tca tcg ttt acg aat acc gat gcg cct gaa tat cta gga gta att aag ttt gga agt ctt atc cat ttc gaa gtg ccg tgt ttc aaa tat tct gcc aca ccc gtt gaa ata gaa aat tct aat cct cct att aca tat aac ttt cca tcg tta aca caa gta cta act tct gat ttt aac gac gac ata tta gta act gtt ttc ca- --- --- --- --- --- --- --t ttt ttc gtt tta aga tct acc cgc -ga tac gga ata aac atg tct att gtt aat cat gcc gcc aat aat gta tag aca att ata taa aac att tgc att ata gaa ttg --- -tc tat ctg tat tac cga cga ttg tcc aat att ctg ttc tag gag agt aat ggg tta ttg tgg ata tat aat cag agt ttt taa tga cta cta tat tat gtt tta tac cat ttc gtg tca cag gct ttg tag att tgg ata tag tta atc cca aca atg ata tag cat tgc aca tag tat tag tca taa act tgg gat gta aaa tgt tga tga tat cta cat cgt ttg gat ttt tat gta tcc act tta ata ata tca tag ctg taa cat cct cgt ga- ttt acg ttg acg ctt tcg tgg gat aag ata gtt gtt agt tca tcc ttt gat aat ttt cca aat t

>KC813491.1:27940-29903_Cowpox_virus_strain_BeaBer04/1_complete_genome

cgg ctt cca tgt att tcc tgg aga gca agt --- aga tga tg- agg aac cag ata gtt tat atc cgt act tgc act taa agt cta cat tgt cgt tgt atg a-- gta tga tct ttt aaa ccc gct aga caa gta tcc gtt tga tat tgt agg atg tgg aca ttt aac aat ctg aca cgt ggg tgg atc gga cca ttc tcc tcc tga aca cag gac acc aga gtt acc aat caa cga ata tcc act att gca act ata agt tac aac gct ccc atc ggt ata aaa atc ctc gta tcc gtt atg tct tcc gtt gga tat aga tgg agg gga ttg gca ttt aac aga ttc aca aat agg tgc ctc ggg att cca tac cat aga tcc agt aga tcc taa ttc aca ata cga ttt aga ttc acc gat caa atg ata tcc gct att aca aga gta cgt tat act aga gcc aaa gtc tac tcc acc aat atc aag ttg gcc att atc gat atc tcg agg cga tgg gca tct ccg ttt aat aca ttg att aaa gag tgt cca tcc agt acc tgt aca ttt agc ata tat agg tcc cat ttt ttg ctt tct gta tcc agg tag aca tag a-t att cta tag tgt ctc cta tgt tgt a-- --- --- --- -at tag cat tag cat cag tc- --t cca cac tat tct taa att tca tat taa tgg gtc gtg acg gaa tag tac agc atg ata gaa cgc atc cta ttc cca aca atg tca gga acg tca cgc tct cca cct tca tat tta ttt atc cgt -aa aaa tgt tat cct gga cat agt aca aat aat a-a aaa gcc ca- -ta tat gtt cgc tat tgt aga aat tgt ttt tca cag ttg ctc aaa aac gat ggc agt gac tta tga gtt acg tta cac ttt gga gtc tca tct tta gta aac ata tca taa tat tcg ata tta cga gtt gac ata tcg aac aaa ttc caa gta ttt gat ttt gga taa tat tcg tat ttt gca tct gct ata att aag ata taa tca ccg caa gaa cac acg aac atc ttt cct aca tgg tta aag tac atg tac aat tct atc cat ttg tct tcc tta act ata tat ttg tat aga taa tta cga gtc tcg tga gta att cca gta att gca tag atg tca cca tcg tac tct aca gca taa act ata cta tga tgt cta ggc atg gga gac -tt ttt tat cca acg att ttt agt gaa aca ttc tac atc gtt taa tac tac ata ttt ctc ata cgt ggt ata aac tcc acc cat tac --a tat ata tca tcg ttt acg aat acc gat gcg cct gaa tat cta gga gta att aag ttt gga agt ctt atc cat ttc gaa gtg ccg tgt ttc aaa tat tct gcc aca ccc gtt gaa ata gaa aat tct aat cct cct att aca tat aac ttt cca tcg tta aca caa gta cta act tct gat ttt aac gac gac ata tta gta act gtt ttc ca- --- --- --- --- --- --- --t ttt ttc gtt tta aga tct acc cgc -ga tac gga ata aac atg tct att gtt aat cat gcc gcc aat aat gta tag aca att ata taa aac att tgc att ata gaa ttg --- -tc tat ctg tat tac cga cga ttg tcc aat att ctg ttc tag gag agt aat ggg tta ttg tgg ata tat aat cag agt ttt taa tga cta cta tat tat gtt tta tac cat ttc gtg tca cag gct ttg tag att tgg ata tag tta atc cca aca atg ata tag cat tgc aca tag tat tag tca taa act tgg gat gta aaa tgt tga tga tat cta cat cgt ttg gat ttt tat gta tcc act tta ata ata tca tag ctg taa cat cct cgt ga- ttt acg ttg acg ctt tcg tgg gat aag ata gtt gtt agt tca tcc ttt gat aat ttt cca aat t

>JX489135.1:17591-19554_Vaccinia_virus_strain_Tiantan_clone_TT8_complete_genome

cgg ctg cca tgt att tcc tgg aga gca agt --- aga tga tg- agg aac cag ata ggt tat atc cgt act tgc act taa agt tta cat tgt cgt tgt atg a-- gta tga tct ttt aaa ccc gct aga caa gta tcc gtt tga tat tgt agg atg tgg aca ttt aac aat ctg aca cgt ggg tgg atc gga cca ttc tcc tcc tga aca cag gac acc aga gtt acc aat caa cga ata tcc act att gca act ata agt tac aac gct ccc atc ggt ata aaa atc ctc gta tcc gtt atg tct tcc gtt gga tat aga tgg agg gga ttg gca ttt aac aga ttc aca aat agg tgc ctc ggg att cca tac cat aga tcc agt aga tcc taa ttc aca ata cga ttt aga ttc acc gat caa atg ata tcc gct att aca aga gta cgt tat act aga gcc aaa gtc tac tcc acc aat atc aag ttg gcc att atc gat atc tcg agg cga tgg gca tct ccg ttt aat aca ttg att aaa gag tgt cca tcc agt acc tgt aca ttt agc ata tat agg tcc cat ttt ttg ctt tct gta tcc agg tag aca tag a-t att cta tag tgt ctc cta tgt tgt a-- --- --- --- -at tag cat tag cat cag tc- --t cca cac tat tct taa att tca tat taa tgg gtc gtg acg gaa tag tac agt atg ata gaa cgc atc cta ttc cca aca atg tca gga acg tca cgc tct cca cct tca tat tta ttt atc cgt -aa aaa tgt tat cct gga cat cgt aca aat aat a-a aaa gcc ca- -ta tat gtt cgc tat tgt aga aat tgt ttt tca cag ttg ctc aaa aac gat ggc agt gac tta tga gtt acg tta cac ttt gga gtc tca tct tta gta aac ata tca taa tat tcg ata tta cga gtt gac ata tcg aac aaa ttc caa gta ttt gat ttt gga taa tat tcg tat ttt gca tct gct ata att aag ata taa tca ccg caa gaa cac acg aac atc ttt cct aca tgg tta aag tac atg tac aat tct atc cat ttg tct tcc tta act ata tat ttg tat aga taa tta cga gtc tcg tga gta att cca gta att aca tag atg tcg ccg tcg tac tct aca gca taa act ata cta tga tgt cta ggc atg gga gac -tt ttt tat cca acg att ttt agt gaa aca ttc tac atc gtt taa tac tac ata ttt ctc ata cgt ggt ata aac tcc acc cat tac --a tat ata tca tcg ttt acg aat acc gat gcg cct gaa tat cta gga gta att aag ttt gga agt ctt atc cat ttt gaa gtg ccg tgt ttc aaa tat tct gcc aca ccc gtt gaa ata gaa aat tct aat cct cct att aca tat aac ttt cca ttg tta aca caa gta cta act tct gat ttt aac gac gac ata tta gta acc gtt ttc ca- --- --- --- --- --- --- --t ttt ttc gtt tca aga tct acc cgc -ga tac gga ata aac atg tct att gtt aat cat gcc gcc aat aat gta tag aca att atg taa aac att tgc att ata gaa ttg --- -tc tat ctg tat tac cga cta tcg tcc aat att ctg ttc tag gag agt aat ggg tta ttg tgg ata tat aat cag agt ttt taa tga cta cta tat tat gtt tta tac cat ttc gtg tca ctg gct ttg tag att tgg ata tag tta atc cca aca atg ata tag cat tgc gca tag tat tag tca taa act tgg gat gta aaa tgt tga tga tat cta cat cgt ttg gat ttt tat gta tcc act tta ata ata tca tag ctg taa cat cct cat ga- ttt acg tta acg tct tcg tgg gat aag ata gtt gtc agt tca tcc ttt gat aat ttt cca aat t

>HQ420898.1:34538-36502_Cowpox_virus_strain_Germany_2002_MKY_complete_genome

cgg ctt cca tgt att tcc tgg aga gca agt --- aga tga tg- agg aac cag ata gtt tat atc cgt act tgc act taa agt cta cat tgt cgt tgt atg a-- gta tga tct ttt aaa ccc gct aga caa gta tcc gtt tga tat tgt agg atg tgg aca ttt aac aat ctg aca cgt ggg tgg atc gga cca ttc tcc tcc tga aca cag gac acc aga gtt acc aat caa cga ata tcc gct att aca act ata cgt tac aac gct tcc atc ggt ata aaa atc ctc gta tcc gtt atg tct tcc gtt gga tat aga tgg agg tga ttg gca ttt aac aga ttc gca aat agg tgc ctc ggg att cca tac cat aga tcc agt aga tcc taa ttc aca ata cga ttt aga ttc acc gat caa atg ata tcc gct att aca aga gta cgt tat act aga gcc aaa gtc tac tcc gcc aat atc aag ttg gcc att atc gat atc tcg agg cga tgg gca tct ccg ttt aat aca ttg att aaa gag tgt cca tcc agt acc tgt aca ttt agc ata tat agg tcc cat ttt ttg ctt tct gta tcc agg tag aca tag a-t att cta tag tgt ctc cta tgt tgt a-- --- --- --- -at tag cat tag cat cag tc- --t cca cac tat tct taa att tca tat taa tgg gtc gtg acg gaa tag tac agc atg ata gaa cgc atc cta ttc cca aca atg tca gga acg tca cgc tct cca cct tca tat tta ttt atc cgt -aa aaa tgt tat cct gga cat cgt acc aat aat aaa aaa gcc ca- -ta tat gtt cgc tat tgt aga aat tgt ttt tca cag ttg ctc aaa aac gat ggc agt gac tta tga gtt acg tta cac ttt gga gtc tca tct tta gta aac ata tca taa tat tcg ata tta cga gtt gac ata tcg aac aaa ttc caa gta ttt gat ttt gga taa tat tcg tat ttt gca tct gct ata att aag ata taa tcg ccg caa gaa cac acg aac atc ttt cct aca tgg tta aag tac atg tac aat tct atc cat ttg tct tcc tta act ata tat ttg tat aga taa tta cga gtc tcg tga gta att cca gta att gca tag atg tca cca tcg tac tct aca gca tac act ata cta tga tgt cta ggc atg gga gac -tt ttt tat cca acg att ttt agt gaa aca ttc cac atc gtt taa tac tac ata ttt ctc ata cgt ggt ata aac tcc acc cat tac --a tat ata tca tcg ttt acg aat acc gat gcg cct gaa tat cta gga gta att aag ttt gga agt ctt atc cat ttc gaa gtg ccg tgt ttc aaa tat tct gcc aca ccc gtt gaa ata gaa aat tct aat cct cct att aca tat aac ttt cca tcg tta aca caa gta cta act tct gat ttt aac gac gac ata tta gta act gtt ttc ca- --- --- --- --- --- --- --t ttt ttc gtt tta aga tct acc cgc -ga tac gga ata aac atg tct att gtt aat cat gcc gcc aat aat gta tag aca att ata taa aac att tgc att ata gaa ttg --- -tc tat ctg tat tac cga cga ttg tcc aat att ctg ttc tag gag agt aat ggg tta ttg tgg ata tat aat cag agt ttt taa tga cta cta tat tat gtt tta tac cat ttc gtg tca cag gct ttg tag att tgg ata tag tta atc cca aca atg ata tag cat tgc aca tag tat tag tca taa act tgg gat gta aaa tgt tga tga tat cta cat cgt ttg gat ttt tat gta tcc act tta ata ata tca tag ctg taa cat cct cat ga- ttt acg tta acg tct tcg tgg gat aag ata gtt gtc agt tca tcc ttt gat aat ttt cca aat t

>HQ420895.1:35808-37772_Cowpox_virus_strain_Germany_1980_EP4_complete_genome

cgg ctt cca tgt att tcc tgg aga gca agt --- aga tga tg- agg aac cag ata gtt tat atc cgt act tgc act taa agt cta cat tgt cgt tgt atg a-- gta tga tct ttt aaa ccc gct aga caa gta tcc gtt tga tat tgt agg atg tgg aca ttt aac aat ctg aca cgt ggg tgg atc gga cca ttc tcc tcc tga aca cag gac acc aga gtt acc aat caa cga ata tcc gct att aca act ata cgt tac aac gct tcc atc ggt ata aaa atc ctc gta tcc gtt atg tct tcc gtt gga tat aga tgg agg tga ttg gca ttt aac aga ttc gca aat agg tgc ctc ggg att cca tac cat aga tcc agt aga tcc taa ttc aca ata cga ttt aga ttc acc gat caa atg ata tcc gct att aca aga gta cgt tat act aga gcc aaa gtc tac tcc gcc aat atc aag ttg gcc att atc gat atc tcg agg cga tgg gca tct ccg ttt aat aca ttg att aaa gag tgt cca tcc agt acc tgt aca ttt agc ata tat agg tcc cat ttt ttg ctt tct gta tcc agg tag aca tag a-t att cta tag tgt ctc cta tgt tgt a-- --- --- --- -at tag cat tag cat cag tc- --t cca cac tat tct taa att tca tat taa tgg gtc gtg acg gaa tag tac agc atg ata gaa cgc atc cta ttc cca aca atg tca gga acg tca cgc tct cca cct tca tat tta ttt atc cgt -aa aaa tgt tat cct gga cat cgt acc aat aat aaa aaa gcc ca- -ta tat gtt cgc tat tgt aga aat tgt ttt tca cag ttg ctc aaa aac gat ggc agt gac tta tga gtt acg tta cac ttt gga gtc tca tct tta gta aac ata tca taa tat tcg ata tta cga gtt gac ata tcg aac aaa ttc caa gta ttt gat ttt gga taa tat tcg tat ttt gca tct gct ata att aag ata taa tcg ccg caa gaa cac acg aac atc ttt cct aca tgg tta aag tac atg tac aat tct atc cat ttg tct tcc tta act ata tat ttg tat aga taa tta cga gtc tca tga gta att cca gta att gca tag atg tca cca tcg tac tct aca gca taa act ata cta tga tgt cta ggc atg gga gac -tt ttt tat cca acg att ttt agt gaa aca ttc cac atc gtt taa tac tac ata ttt ttc ata cgt ggc ata aac tcc acc cat tac --a tat ata tca tcg ttt acg aat acc gat gcg cct gaa tat cta gga gta att aag ttt gga agt ctt atc cat ttc gaa gtg ccg tgt ttc aaa tat tct gcc aca ccc gtt gaa ata gaa aat tct aat cct cct att aca tat aac ttt cca tcg tta aca caa gta cta act tct gat ttt aac gac gac ata tta gta act gtt ttc ca- --- --- --- --- --- --- --t ttt ttc gtt tta aga tct acc cgc -ga tac gga ata aac atg tct att gtt aat cat gcc gcc aat aat gta tag aca att ata taa aac att tgc att ata gaa ttg --- -tc tat ctg tat tac cga cga ttg tcc aat att ctg ttc tag gag agt aat ggg tta ttg tgg ata tat aat cag agt ttt taa tga cta cta tat tat gtt tta tac cat ttc gtg tca cag gct ttg tag att tgg ata tag tta atc cca aca atg ata tag cat tgc aca tag tat tag tca taa act tgg gat gta aaa tgt tga tga tat cta cat cgt ttg gat ttt tat gta tcc act tta ata ata tca tag ctg taa cat cct cat ga- ttt acg tta acg tct tcg tgg gat aag ata gtt gtc agt tca tcc ttt gat aat ttt cca aat t

>AY678275.1:21300-23263_Vaccinia_virus_strain_LC16m8_complete_genome

cgg ctt cca tgt att tcc tgg aga gca agt --- aga tga tg- agg aac cag ata gtt tat atc cgt act tgc act taa agt cta cat tgt cgt tgt atg a-- gta tga tct ttt aaa ccc gct aga caa gta tcc gtt tga tat tgt agg atg tgg aca ttt aac aat ctg aca cgt ggg tgg atc gga cca ttc tcc tcc tga aca cag gac act aga gtt acc aat caa cga ata tcc act att gca act ata agt tac aac gct ccc atc ggt ata aaa atc ctc gta tcc gtt atg tct tcc gtt gga tat aga tgg agg gga ttg gca ttt aac aga ttc aca aat agg tgc ctc ggg att cca tac cat aga tcc agt aga tcc taa ttc aca ata cga ttt aga ttc acc gat caa atg ata tcc gct att aca aga gta cgt tat act aga gcc aaa gtc tac tcc acc aat atc aag ttg gcc att atc gat atc tcg agg cga tgg gca tct ccg ttt aat aca ttg att aaa gag tgt cca tcc agt acc tgt aca ttt agc ata tat agg tcc cat ttt ttg ctt tct gta tcc agg tag aca tag a-t att cta tag tgt ctc cta tgt tgt a-- --- --- --- -at tag cat tag cat cag tc- --t cca cac tat tct taa att tca tat taa tgg gtc gtg acg gaa tag tac agc atg ata gaa cgc atc cta ttc cca aca atg tca gga acg tca cgc tct cca cct tca tat tta ttt atc cgt -aa aaa tgt tat cct gga cat cgt aca aat aat a-a aaa gcc ca- -ta tat gtt cgc tat tgt aga aat tgt ttt tca cag ttg ctc aaa aac gat ggc agt gac tta tga gtt acg tta cac ttt gga gtc tca tct tta gta aac ata tca taa tat tcg ata tta cga gtt gac ata tcg aac aaa ttc caa gta ttt gat ttt gga taa tat tcg tat ttt gca tct gct ata att aag ata taa tca ccg caa gaa cac acg aac atc ttt cct aca tgg tta aag tac atg tac aat tct atc cat ttg tct tcc tta act ata tat ttg tat aga taa tta cga gtc tcg tga gta att cca gta att aca tag atg tcg ccg tcg tac tct aca gca taa act ata cta tga tgt cta ggc atg gga gac -tt ttt tat cca acg att ttt agt gaa aca ttc cac atc gtt taa tac tac ata ttt ttc ata cgt ggt ata aac tcc acc cat tac --a tat ata tca tcg ttt acg aat acc gac gcg cct gaa tat cta gga gta att aag ttt gga agt ctt atc cat ttc gaa gtg ccg tgt ttc aaa tat tct gcc aca ccc gtt gaa ata gaa aat tct aat cct cct att aca tat aac ttt cca tcg tta aca caa gta cta act tct gat ttt aac gac gac ata tta gta acc gtt ttc ca- --- --- --- --- --- --- --t ttt ttc gtt tca aga tct acc cgc -ga tac gga ata aac atg tct att gtt aat cat gcc gcc aat aat gta tag aca att atg taa aac att tgc att ata gaa ttg --- -tc tat ctg tat tac cga cta tcg tcc aat att ctg ttc tag gag agt aat ggg tta ttg tgg ata tat aat cag agt ttt taa tga cta cta tat tat gtt tta tac cat ttc gtg tca ctg gct ttg tag att tgg ata tag tta atc cca aca atg ata tag cat tgc gca tag tat tag tca taa act tgg gat gta aaa tgt tga tga tat cta cat cgt ttg gat ttt tat gta tcc act tta ata ata tca tag ctg taa cat cct cat ga- ttt acg tta acg tct tcg tgg gat aag ata gtt gtc agt tca tcc ttt gat aat ttt cca aat t

>AY678277.1:21300-23263_Vaccinia_virus_strain_LC16mO_complete_genome

cgg ctt cca tgt att tcc tgg aga gca agt --- aga tga tg- agg aac cag ata gtt tat atc cgt act tgc act taa agt cta cat tgt cgt tgt atg a-- gta tga tct ttt aaa ccc gct aga caa gta tcc gtt tga tat tgt agg atg tgg aca ttt aac aat ctg aca cgt ggg tgg atc gga cca ttc tcc tcc tga aca cag gac act aga gtt acc aat caa cga ata tcc act att gca act ata agt tac aac gct ccc atc ggt ata aaa atc ctc gta tcc gtt atg tct tcc gtt gga tat aga tgg agg gga ttg gca ttt aac aga ttc aca aat agg tgc ctc ggg att cca tac cat aga tcc agt aga tcc taa ttc aca ata cga ttt aga ttc acc gat caa atg ata tcc gct att aca aga gta cgt tat act aga gcc aaa gtc tac tcc acc aat atc aag ttg gcc att atc gat atc tcg agg cga tgg gca tct ccg ttt aat aca ttg att aaa gag tgt cca tcc agt acc tgt aca ttt agc ata tat agg tcc cat ttt ttg ctt tct gta tcc agg tag aca tag a-t att cta tag tgt ctc cta tgt tgt a-- --- --- --- -at tag cat tag cat cag tc- --t cca cac tat tct taa att tca tat taa tgg gtc gtg acg gaa tag tac agc atg ata gaa cgc atc cta ttc cca aca atg tca gga acg tca cgc tct cca cct tca tat tta ttt atc cgt -aa aaa tgt tat cct gga cat cgt aca aat aat a-a aaa gcc ca- -ta tat gtt cgc tat tgt aga aat tgt ttt tca cag ttg ctc aaa aac gat ggc agt gac tta tga gtt acg tta cac ttt gga gtc tca tct tta gta aac ata tca taa tat tcg ata tta cga gtt gac ata tcg aac aaa ttc caa gta ttt gat ttt gga taa tat tcg tat ttt gca tct gct ata att aag ata taa tca ccg caa gaa cac acg aac atc ttt cct aca tgg tta aag tac atg tac aat tct atc cat ttg tct tcc tta act ata tat ttg tat aga taa tta cga gtc tcg tga gta att cca gta att aca tag atg tcg ccg tcg tac tct aca gca taa act ata cta tga tgt cta ggc atg gga gac -tt ttt tat cca acg att ttt agt gaa aca ttc cac atc gtt taa tac tac ata ttt ttc ata cgt ggt ata aac tcc acc cat tac --a tat ata tca tcg ttt acg aat acc gac gcg cct gaa tat cta gga gta att aag ttt gga agt ctt atc cat ttc gaa gtg ccg tgt ttc aaa tat tct gcc aca ccc gtt gaa ata gaa aat tct aat cct cct att aca tat aac ttt cca tcg tta aca caa gta cta act tct gat ttt aac gac gac ata tta gta acc gtt ttc ca- --- --- --- --- --- --- --t ttt ttc gtt tca aga tct acc cgc -ga tac gga ata aac atg tct att gtt aat cat gcc gcc aat aat gta tag aca att atg taa aac att tgc att ata gaa ttg --- -tc tat ctg tat tac cga cta tcg tcc aat att ctg ttc tag gag agt aat ggg tta ttg tgg ata tat aat cag agt ttt taa tga cta cta tat tat gtt tta tac cat ttc gtg tca ctg gct ttg tag att tgg ata tag tta atc cca aca atg ata tag cat tgc gca tag tat tag tca taa act tgg gat gta aaa tgt tga tga tat cta cat cgt ttg gat ttt tat gta tcc act tta ata ata tca tag ctg taa cat cct cat ga- ttt acg tta acg tct tcg tgg gat aag ata gtt gtc agt tca tcc ttt gat aat ttt cca aat t

>AY678276.1:21233-23196_Vaccinia_virus_strain_Lister_complete_genome

cgg ctt cca tgt att tcc tgg aga gca agt --- aga tga tg- agg aac cag ata gtt tat atc cgt act tgc act taa agt cta cat tgt cgt tgt atg a-- gta tga tct ttt aaa ccc gct aga caa gta tcc gtt tga tat tgt agg atg tgg aca ttt aac aat ctg aca cgt ggg tgg atc gga cca ttc tcc tcc tga aca cag gac acc aga gtt acc aat caa cga ata tcc act att gca act ata agt tac aac gct tcc atc ggt ata aaa atc ctc gta tcc gtt atg tct tcc gtt gga tat aga tgg agg gga ttg gca ttt aac aga ttc aca aat agg tgc ctc ggg att cca tac cat aga tcc agt aga tcc taa ttc aca ata cga ttt aga ttc acc gat caa atg ata tcc gct att aca aga gta cgt tat act aga gcc aaa gtc tac tcc acc aat atc aag ttg gcc att atc gat atc tcg agg cga tgg gca tct ccg ttt aat aca ttg att aaa gag tgt cca tcc agt acc tgt aca ttt agc ata tat agg tcc cat ttt ttg ctt tct gta tcc agg tag aca tag a-t att cta tag tgt ctc cta tgt tgt a-- --- --- --- -at tag cat tag cat cag tc- --t cca cac tat tct taa att tca tat taa tgg gtc gtg acg gaa tag tac agc atg ata gaa cgc atc cta ttc cca aca atg tca gga acg tca cgc tct cca cct tca tat tta ttt atc cgt -aa aaa tgt tat cct gga cat cgt aca aat aat a-a aaa gcc ca- -ta tat gtt cgc tat tgt aga aat tgt ttt tca cag ttg ctc aaa aac gat ggc agt gac tta tga gtt acg tta cac ttt gga gtc tca tct tta gta aac ata tca taa tat tcg ata tta cga gtt gac ata tcg aac aaa ttc caa gta ttt gat ttt gga taa tat tcg tat ttt gca tct gct ata att aag ata taa tca ccg caa gaa cac acg aac atc ttt cct aca tgg tta aag tac atg tac aat tct atc cat ttg tct tcc tta act ata tat ttg tat aga taa tta cga gtc tcg tga gta att cca gta att aca tag atg tcg ccg tcg tac tct aca gca taa act ata cta tga tgt cta ggc atg gga gac -tt ttt tat cca acg att ttt agt gaa aca ttc cac atc gtt taa tac tac ata ttt ttc ata cgt ggt ata aac tcc acc cat tac --a tat ata tca tcg ttt acg aat acc gac gcg cct gaa tat cta gga gta att aag ttt gga agt ctt atc cat ttc gaa gtg ccg tgt ttc aaa tat tct gcc aca ccc gtt gaa ata gaa aat tct aat cct cct att aca tat aac ttt cca tcg tta aca caa gta cta act tct gat ttt aac gac gac ata tta gta acc gtt ttc ca- --- --- --- --- --- --- --t ttt ttc gtt tca aga tct acc cgc -ga tac gga ata aac atg tct att gtt aat cat gcc gcc aat aat gta tag aca att atg taa aac att tgc att ata gaa ttg --- -tc tat ctg tat tac cga cta tcg tcc aat att ctg ttc tag gag agt aat ggg tta ttg tgg ata tat aat cag agt ttt taa tga cta cta tat tat gtt tta tac cat ttc gtg tca ctg gct ttg tag att tgg ata tag tta atc cca aca atg ata tag cat tgc gca tag tat tag tca taa act tgg gat gta aaa tgt tga tga tat cta cat cgt ttg gat ttt tat gta tcc act tta ata ata tca tag ctg taa cat cct cat ga- ttt acg tta acg tct tcg tgg gat aag ata gtt gtc agt tca tcc ttt gat aat ttt cca aat t

>MK035758.1:35000-36964_Cowpox_virus_strain_CPXV/Moritz_2015/3a

cgg ctt cca tgt att tcc tgg aga gca agt --- aga tga tg- agg aac cag ata gtt tat atc cgt act tgc act taa agt cta cat tgt cgt tgt atg a-- gta tga tct ttt aaa ccc gct aga caa gta tcc gtt tga tat tgt agg atg tgg aca ttt aac aat ctg aca cgt ggg tgg atc tga cca ttc tcc tcc tga aca cag gac acc aga gtt acc aat caa cga ata tcc gct att aca act ata cgt tac aac gct tcc atc ggt ata aaa atc ctc gta tcc gtt atg tct tcc gtt gga tat aga tgg agg tga ttg gca ttt aac aga ttc gca aat agg tgc ctc ggg att cca tac cat aga tcc agt aga tcc taa ttc aca ata cga ttt aga ttc acc gat caa atg ata tcc gct att aca aga gta cgt tat act aga gcc aaa gtc tac tcc gcc aat atc aag ttg gcc att atc gat atc tcg agg cga tgg gca tct ccg ttt aat aca ttg att aaa gag tgt cca tcc agt acc tgt aca ttt agc ata tat agg tcc cat ttt ttg ctt tct gta tcc agg tag aca tag a-t att cta tag tgt ctc cta tgt tgt a-- --- --- --- -at tag cat tag cat cag tc- --t cca cac tat tct taa att tca tat taa tgg gtc gtg acg gaa tag tac agc atg ata gaa cgc atc cta ttc cca aca atg tca gga acg tca cgc tct cca cct tca tat tta ttt atc cgt -aa aaa tgt tat cct gga cat cgt acc aat aat aaa aaa gcc ca- -ta tat gtt cgc tat tgt aga aat tgt ttt tca cag ttg ctc aaa aac gat ggc agt gac tta tga gtt acg tta cac ttt gga gtc tca tct tta gta aac ata tca taa tat tcg ata tta cga gtt gac ata tcg aac aaa ttc caa gta ttt gat ttt gga taa tat tcg tat ttt gca tct gct ata att aag ata taa tcg ccg caa gaa cac acg aac atc ttt cct aca tgg tta aag tac atg tac aat tct atc cat ttg tct tcc tta act ata tat ttg tat aga taa tta cga gtc tca tga gta att cca gta att gca tag atg tca cca tcg tac tct aca gca taa act ata cta tga tgt cta ggc atg gga gac -tt ttt tat cca acg att ttt agt gaa aca ttc cac atc gtt taa tac tac ata ttt ttc ata cgt ggc ata aac tcc acc cat tac --a tat ata tca tcg ttt acg aat acc gat gcg cct gaa tat cta gga gta att aag ttt gga agt ctt atc cat ttc gaa gtg ccg tgt ttc aaa tat tct gcc aca ccc gtt gaa ata gaa aat tct aat cct cct att aca tat aac ttt cca tcg tta aca caa gta cta act tct gat ttt aac gac gac ata tta gta act gtt ttc ca- --- --- --- --- --- --- --t ttt ttc gtt tta aga tct acc cgc -ga tac gga ata aac atg tct att gtt aat cat gcc gcc aat aat gta tag aca att ata taa aac att tgc att ata gaa ttg --- -tc tat ctg tat tac cga cga ttg tcc aat att ctg ttc tag gag agt aat ggg tta ttg tgg ata tat aat cag agt ttt taa tga cta cta tat tat gtt tta tac cat ttc gtg tca cag gct ttg tag att tgg ata tag tta atc cca aca atg ata tag cat tgc aca tag tat tag tca taa act tgg gat gta aaa tgt tga tga tat cta cat cgt ttg gat ttt tat gta tcc act tta ata ata tca tag ctg taa cat cct cat ga- ttt acg tta acg tct tcg tgg gat aag ata gtt gtc agt tca tcc ttt gat aat ttt cca aat t

>MK035756.1:34880-36844_Cowpox_virus_strain_CPXV/Leo_2015/5

cgg ctt cca tgt att tcc tgg aga gca agt --- aga tga tg- agg aac cag ata gtt tat atc cgt act tgc act taa agt cta cat tgt cgt tgt atg a-- gta tga tct ttt aaa ccc gct aga caa gta tcc gtt tga tat tgt agg atg tgg aca ttt aac aat ctg aca cgt ggg tgg atc tga cca ttc tcc tcc tga aca cag gac acc aga gtt acc aat caa cga ata tcc gct att aca act ata cgt tac aac gct tcc atc ggt ata aaa atc ctc gta tcc gtt atg tct tcc gtt gga tat aga tgg agg tga ttg gca ttt aac aga ttc gca aat agg tgc ctc ggg att cca tac cat aga tcc agt aga tcc taa ttc aca ata cga ttt aga ttc acc gat caa atg ata tcc gct att aca aga gta cgt tat act aga gcc aaa gtc tac tcc gcc aat atc aag ttg gcc att atc gat atc tcg agg cga tgg gca tct ccg ttt aat aca ttg att aaa gag tgt cca tcc agt acc tgt aca ttt agc ata tat agg tcc cat ttt ttg ctt tct gta tcc agg tag aca tag a-t att cta tag tgt ctc cta tgt tgt a-- --- --- --- -at tag cat tag cat cag tc- --t cca cac tat tct taa att tca tat taa tgg gtc gtg acg gaa tag tac agc atg ata gaa cgc atc cta ttc cca aca atg tca gga acg tca cgc tct cca cct tca tat tta ttt atc cgt -aa aaa tgt tat cct gga cat cgt acc aat aat aaa aaa gcc ca- -ta tat gtt cgc tat tgt aga aat tgt ttt tca cag ttg ctc aaa aac gat ggc agt gac tta tga gtt acg tta cac ttt gga gtc tca tct tta gta aac ata tca taa tat tcg ata tta cga gtt gac ata tcg aac aaa ttc caa gta ttt gat ttt gga taa tat tcg tat ttt gca tct gct ata att aag ata taa tcg ccg caa gaa cac acg aac atc ttt cct aca tgg tta aag tac atg tac aat tct atc cat ttg tct tcc tta act ata tat ttg tat aga taa tta cga gtc tca tga gta att cca gta att gca tag atg tca cca tcg tac tct aca gca taa act ata cta tga tgt cta ggc atg gga gac -tt ttt tat cca acg att ttt agt gaa aca ttc cac atc gtt taa tac tac ata ttt ttc ata cgt ggc ata aac tcc acc cat tac --a tat ata tca tcg ttt acg aat acc gat gcg cct gaa tat cta gga gta att aag ttt gga agt ctt atc cat ttc gaa gtg ccg tgt ttc aaa tat tct gcc aca ccc gtt gaa ata gaa aat tct aat cct cct att aca tat aac ttt cca tcg tta aca caa gta cta act tct gat ttt aac gac gac ata tta gta act gtt ttc ca- --- --- --- --- --- --- --t ttt ttc gtt tta aga tct acc cgc -ga tac gga ata aac atg tct att gtt aat cat gcc gcc aat aat gta tag aca att ata taa aac att tgc att ata gaa ttg --- -tc tat ctg tat tac cga cga ttg tcc aat att ctg ttc tag gag agt aat ggg tta ttg tgg ata tat aat cag agt ttt taa tga cta cta tat tat gtt tta tac cat ttc gtg tca cag gct ttg tag att tgg ata tag tta atc cca aca atg ata tag cat tgc aca tag tat tag tca taa act tgg gat gta aaa tgt tga tga tat cta cat cgt ttg gat ttt tat gta tcc act tta ata ata tca tag ctg taa cat cct cat ga- ttt acg tta acg tct tcg tgg gat aag ata gtt gtc agt tca tcc ttt gat aat ttt cca aat t

>MK035754.1:35336-37300_Cowpox_virus_strain_CPXV/Cookie_S963_15

cgg ctt cca tgt att tcc tgg aga gca agt --- aga tga tg- agg aac cag ata gtt tat atc cgt act tgc act taa agt cta cat tgt cgt tgt atg a-- gta tga tct ttt aaa ccc gct aga caa gta tcc gtt tga tat tgt agg atg tgg aca ttt aac aat ctg aca cgt ggg tgg atc tga cca ttc tcc tcc tga aca cag gac acc aga gtt acc aat caa cga ata tcc gct att aca act ata cgt tac aac gct tcc atc ggt ata aaa atc ctc gta tcc gtt atg tct tcc gtt gga tat aga tgg agg tga ttg gca ttt aac aga ttc gca aat agg tgc ctc ggg att cca tac cat aga tcc agt aga tcc taa ttc aca ata cga ttt aga ttc acc gat caa atg ata tcc gct att aca aga gta cgt tat act aga gcc aaa gtc tac tcc gcc aat atc aag ttg gcc att atc gat atc tcg agg cga tgg gca tct ccg ttt aat aca ttg att aaa gag tgt cca tcc agt acc tgt aca ttt agc ata tat agg tcc cat ttt ttg ctt tct gta tcc agg tag aca tag a-t att cta tag tgt ctc cta tgt tgt a-- --- --- --- -at tag cat tag cat cag tc- --t cca cac tat tct taa att tca tat taa tgg gtc gtg acg gaa tag tac agc atg ata gaa cgc atc cta ttc cca aca atg tca gga acg tca cgc tct cca cct tca tat tta ttt atc cgt -aa aaa tgt tat cct gga cat cgt acc aat aat aaa aaa gcc ca- -ta tat gtt cgc tat tgt aga aat tgt ttt tca cag ttg ctc aaa aac gat ggc agt gac tta tga gtt acg tta cac ttt gga gtc tca tct tta gta aac ata tca taa tat tcg ata tta cga gtt gac ata tcg aac aaa ttc caa gta ttt gat ttt gga taa tat tcg tat ttt gca tct gct ata att aag ata taa tcg ccg caa gaa cac acg aac atc ttt cct aca tgg tta aag tac atg tac aat tct atc cat ttg tct tcc tta act ata tat ttg tat aga taa tta cga gtc tca tga gta att cca gta att gca tag atg tca cca tcg tac tct aca gca taa act ata cta tga tgt cta ggc atg gga gac -tt ttt tat cca acg att ttt agt gaa aca ttc cac atc gtt taa tac tac ata ttt ttc ata cgt ggc ata aac tcc acc cat tac --a tat ata tca tcg ttt acg aat acc gat gcg cct gaa tat cta gga gta att aag ttt gga agt ctt atc cat ttc gaa gtg ccg tgt ttc aaa tat tct gcc aca ccc gtt gaa ata gaa aat tct aat cct cct att aca tat aac ttt cca tcg tta aca caa gta cta act tct gat ttt aac gac gac ata tta gta act gtt ttc ca- --- --- --- --- --- --- --t ttt ttc gtt tta aga tct acc cgc -ga tac gga ata aac atg tct att gtt aat cat gcc gcc aat aat gta tag aca att ata taa aac att tgc att ata gaa ttg --- -tc tat ctg tat tac cga cga ttg tcc aat att ctg ttc tag gag agt aat ggg tta ttg tgg ata tat aat cag agt ttt taa tga cta cta tat tat gtt tta tac cat ttc gtg tca cag gct ttg tag att tgg ata tag tta atc cca aca atg ata tag cat tgc aca tag tat tag tca taa act tgg gat gta aaa tgt tga tga tat cta cat cgt ttg gat ttt tat gta tcc act tta ata ata tca tag ctg taa cat cct cat ga- ttt acg tta acg tct tcg tgg gat aag ata gtt gtc agt tca tcc ttt gat aat ttt cca aat t

>MK035752.1:35103-37067_Cowpox_virus_strain_CPXV/Sammy_2015/4a

cgg ctt cca tgt att tcc tgg aga gca agt --- aga tga tg- agg aac cag ata gtt tat atc cgt act tgc act taa agt cta cat tgt cgt tgt atg a-- gta tga tct ttt aaa ccc gct aga caa gta tcc gtt tga tat tgt agg atg tgg aca ttt aac aat ctg aca cgt ggg tgg atc tga cca ttc tcc tcc tga aca cag gac acc aga gtt acc aat caa cga ata tcc gct att aca act ata cgt tac aac gct tcc atc ggt ata aaa atc ctc gta tcc gtt atg tct tcc gtt gga tat aga tgg agg tga ttg gca ttt aac aga ttc gca aat agg tgc ctc ggg att cca tac cat aga tcc agt aga tcc taa ttc aca ata cga ttt aga ttc acc gat caa atg ata tcc gct att aca aga gta cgt tat act aga gcc aaa gtc tac tcc gcc aat atc aag ttg gcc att atc gat atc tcg agg cga tgg gca tct ccg ttt aat aca ttg att aaa gag tgt cca tcc agt acc tgt aca ttt agc ata tat agg tcc cat ttt ttg ctt tct gta tcc agg tag aca tag a-t att cta tag tgt ctc cta tgt tgt a-- --- --- --- -at tag cat tag cat cag tc- --t cca cac tat tct taa att tca tat taa tgg gtc gtg acg gaa tag tac agc atg ata gaa cgc atc cta ttc cca aca atg tca gga acg tca cgc tct cca cct tca tat tta ttt atc cgt -aa aaa tgt tat cct gga cat cgt acc aat aat aaa aaa gcc ca- -ta tat gtt cgc tat tgt aga aat tgt ttt tca cag ttg ctc aaa aac gat ggc agt gac tta tga gtt acg tta cac ttt gga gtc tca tct tta gta aac ata tca taa tat tcg ata tta cga gtt gac ata tcg aac aaa ttc caa gta ttt gat ttt gga taa tat tcg tat ttt gca tct gct ata att aag ata taa tcg ccg caa gaa cac acg aac atc ttt cct aca tgg tta aag tac atg tac aat tct atc cat ttg tct tcc tta act ata tat ttg tat aga taa tta cga gtc tca tga gta att cca gta att gca tag atg tca cca tcg tac tct aca gca taa act ata cta tga tgt cta ggc atg gga gac -tt ttt tat cca acg att ttt agt gaa aca ttc cac atc gtt taa tac tac ata ttt ttc ata cgt ggc ata aac tcc acc cat tac --a tat ata tca tcg ttt acg aat acc gat gcg cct gaa tat cta gga gta att aag ttt gga agt ctt atc cat ttc gaa gtg ccg tgt ttc aaa tat tct gcc aca ccc gtt gaa ata gaa aat tct aat cct cct att aca tat aac ttt cca tcg tta aca caa gta cta act tct gat ttt aac gac gac ata tta gta act gtt ttc ca- --- --- --- --- --- --- --t ttt ttc gtt tta aga tct acc cgc -ga tac gga ata aac atg tct att gtt aat cat gcc gcc aat aat gta tag aca att ata taa aac att tgc att ata gaa ttg --- -tc tat ctg tat tac cga cga ttg tcc aat att ctg ttc tag gag agt aat ggg tta ttg tgg ata tat aat cag agt ttt taa tga cta cta tat tat gtt tta tac cat ttc gtg tca cag gct ttg tag att tgg ata tag tta atc cca aca atg ata tag cat tgc aca tag tat tag tca taa act tgg gat gta aaa tgt tga tga tat cta cat cgt ttg gat ttt tat gta tcc act tta ata ata tca tag ctg taa cat cct cat ga- ttt acg tta acg tct tcg tgg gat aag ata gtt gtc agt tca tcc ttt gat aat ttt cca aat t

>MK035751.1:35593-37557_Cowpox_virus_strain_CPXV/S2216/04

cgg ctt cca tgt att tcc tgg aga gca agt --- aga tga tg- agg aac cag ata gtt tat atc cgt act tgc act taa agt cta cat tgt cgt tgt atg a-- gta tga tct ttt aaa ccc gct aga caa gta tcc gtt tga tat tgt agg atg tgg aca ttt aac aat ctg aca cgt ggg tgg atc tga cca ttc tcc tcc tga aca cag gac acc aga gtt acc aat caa cga ata tcc gct att aca act ata cgt tac aac gct tcc atc ggt ata aaa atc ctc gta tcc gtt atg tct tcc gtt gga tat aga tgg agg tga ttg gca ttt aac aga ttc gca aat agg tgc ctc ggg att cca tac cat aga tcc agt aga tcc taa ttc aca ata cga ttt aga ttc acc gat caa atg ata tcc gct att aca aga gta cgt tat act aga gcc aaa gtc tac tcc gcc aat atc aag ttg gcc att atc gat atc tcg agg cga tgg gca tct ccg ttt aat aca ttg att aaa gag tgt cca tcc agt acc tgt aca ttt agc ata tat agg tcc cat ttt ttg ctt tct gta tcc agg tag aca tag a-t att cta tag tgt ctc cta tgt tgt a-- --- --- --- -at tag cat tag cat cag tc- --t cca cac tat tct taa att tca tat taa tgg gtc gtg acg gaa tag tac agc atg ata gaa cgc atc cta ttc cca aca atg tca gga acg tca cgc tct cca cct tca tat tta ttt atc cgt -aa aaa tgt tat cct gga cat cgt acc aat aat aaa aaa gcc ca- -ta tat gtt cgc tat tgt aga aat tgt ttt tca cag ttg ctc aaa aac gat ggc agt gac tta tga gtt acg tta cac ttt gga gtc tca tct tta gta aac ata tca taa tat tcg ata tta cga gtt gac ata tcg aac aaa ttc caa gta ttt gat ttt gga taa tat tcg tat ttt gca tct gct ata att aag ata taa tcg ccg caa gaa cac acg aac atc ttt cct aca tgg tta aag tac atg tac aat tct atc cat ttg tct tcc tta act ata tat ttg tat aga taa tta cga gtc tca tga gta att cca gta att gca tag atg tca cca tcg tac tct aca gca taa act ata cta tga tgt cta ggc atg gga gac -tt ttt tat cca acg att ttt agt gaa aca ttc cac atc gtt taa tac tac ata ttt ttc ata cgt ggc ata aac tcc acc cat tac --a tat ata tca tcg ttt acg aat acc gat gcg cct gaa tat cta gga gta att aag ttt gga agt ctt atc cat ttc gaa gtg ccg tgt ttc aaa tat tct gcc aca ccc gtt gaa ata gaa aat tct aat cct cct att aca tat aac ttt cca tcg tta aca caa gta cta act tct gat ttt aac gac gac ata tta gta act gtt ttc ca- --- --- --- --- --- --- --t ttt ttc gtt tta aga tct acc cgc -ga tac gga ata aac atg tct att gtt aat cat gcc gcc aat aat gta tag aca att ata taa aac att tgc att ata gaa ttg --- -tc tat ctg tat tac cga cga ttg tcc aat att ctg ttc tag gag agt aat ggg tta ttg tgg ata tat aat cag agt ttt taa tga cta cta tat tat gtt tta tac cat ttc gtg tca cag gct ttg tag att tgg ata tag tta atc cca aca atg ata tag cat tgc aca tag tat tag tca taa act tgg gat gta aaa tgt tga tga tat cta cat cgt ttg gat ttt tat gta tcc act tta ata ata tca tag ctg taa cat cct cat ga- ttt acg tta acg tct tcg tgg gat aag ata gtt gtc agt tca tcc ttt gat aat ttt cca aat t

>LT896731.1:34361-36331_Cowpox_virus_isolate_Ger/2015/Cat4_genome_assembly_complete_genome:_monopartite

cgg ctt cca tgt att tcc tgg aga gca agt --- aga tga tg- agg aac cag ata gtt tat atc cgt act tgc act taa agt cta cat tgt cgt tgt atg a-- gta tga tct ttt aaa ccc gct aga caa gta tcc gtt tga tat tgt agg atg tgg aca ttt aac aat ctg aca cgt ggg tgg atc gga cca ttc tcc tcc tga aca cag gac acc aga gtt acc aat caa cga ata tcc act att gca act ata agt tac aac gct ccc atc ggt ata aaa atc ctc gta tcc gtt atg tct tcc gtt gga tat aga tgg agg gga ttg gca ttt aac aga ttc aca aat agg tgc ctc ggg att cca tac cat aga tcc agt aga tcc taa ttc aca ata cga ttt aga ttc acc gat caa atg ata tcc gct att aca aga gta cgt tat act aga gcc aaa gtc tac tcc acc aat atc aag ttg gcc att atc gat atc tcg agg cga tgg gca tct ccg ttt aat aca ttg att aaa gag tgt cca tcc agt acc tgt aca ttt agc ata tat agg tcc cat ttt ttg ctt tct gta tcc agg tag aca tag a-t att cta tag tgt ctc cta tgt tgt aat tag c-- --- -at tag cat tag cat cag tc- --t cca cac tat tct taa att tca tat taa tgg gtc gtg acg gaa tag tac agc atg ata gaa cgc atc cta ttc cca aca atg tca gga acg tca cgc tct cca cct tca tat tta ttt atc cgt -aa aaa tgt tat ccc gga cat tgt acc aat aat aaa aaa gcc ca- -ta tat gtt cgc tat tgt aga aat tgt ttt tca cag ttg ctc aaa aac gat ggc agt gac tta tga gtt acg tta cac ttt gga gtc tca tct tta gta aac ata tca taa tat tcg ata tta cga gtt gac ata tcg aac aaa ttc caa gta ttt gat ttt gga taa tat tcg tat ttt gca tct gct ata att aag ata taa tca ccg caa gaa cac acg aac atc ttt cct aca tgg tta aag tac atg tac aat tct atc cat ttg tct tcc tta act ata tat ttg tat aga taa tta cga gtc tcg tga gta att cca gta att gca tag atg tca cca tcg tac tct aca gca taa act ata cta tga tgt cta ggc atg gga gac -tt ttt tat cca acg att ttt agt gaa aca ttc cac atc gtt taa tac tac ata ttt ctc ata cgt ggt ata aac tcc acc cat tac --a tat ata tca tcg ttt acg aat acc gat gcg cct gaa tat cta gga gta att aag ttt gga agt ctt atc cat ttc gaa gtg ccg tgt ttc aaa tat tct gcc aca ccc gtt gaa ata gaa aat tct aat cct cct att aca tat aac ttt cca tcg tta aca caa gta cta act tct gat ttt aac gac gac ata tta gta acc gtt ttc ca- --- --- --- --- --- --- --t ttt ttc gtt tta aga tct acc cgc -ga tac gga ata aac atg tct att gtt aat cat gcc gcc aat aat gta tag aca att atg taa aac att tgc att ata gaa ttg --- -tc tat ctg tat tac cga cta tcg tcc aat att ctg ttc tag gag agt aat ggg tta ttg tgg ata tat aat cag agt ttt taa tga cta cta tat tat gtt tta tac cat ttc gtg tca ctg gct ttg tag att tgg ata tag tta atc cca aca atg ata tag cat tgc gca tag tat tag tca taa act tgg gat gta aaa tgt tga tga tat cta cat cgt ttg gat ttt tat gta tcc act tta ata ata tca tag ctg taa cat cct cat ga- ttt acg tta acg tct tcg tgg gat aag ata gtt gtc agt tca tcc ttt gat aat ttt cca aat t

>LT896728.1:34811-36775_Cowpox_virus_isolate_Ger/2010/Rat_genome_assembly_complete_genome:_monopartite

cgg ctt cca tgt att tcc tgg aga gca agt --- aga tga tg- agg aac cag ata gtt tat atc cgt act tgc act taa agt cta cat tgt cgt tgt atg a-- gta tga tct ttt aaa ccc gct aga caa gta tcc gtt tga tat tgt agg atg tgg aca ttt aac aat ctg aca cgt ggg tgg atc gga cca ttc tcc tcc tga aca cag gac acc aga gtt acc aat caa cga ata tcc gct att aca act ata cgt tac aac gct tcc atc ggt ata aaa atc ctc gta tcc gtt atg tct tcc gtt gga tat aga tgg agg tga ttg gca ttt aac aga ttc gca aat agg tgc ctc ggg att cca tac cat aga tcc agt aga tcc taa ttc gca ata cga ttt aga ttc acc gat caa atg ata tcc gct att aca aga gta cgt tat act aga gcc aaa gtc tac tcc gcc aat atc aag ttg gcc att atc gat atc tcg agg cga tgg gca tct ccg ttt aat aca ttg att aaa gag tgt cca tcc agt acc tgt aca ttt agc ata tat agg tcc cat ttt ttg ctt tct gta tcc agg tag aca tag a-t att cta tag tgt ctc cta tgt tgt a-- --- --- --- -at tag cat tag cat cag tc- --t cca cac tat tct taa att tca tat taa tgg gtc gtg acg gaa tag tac agc atg ata gaa cgc atc cta ttc cca aca atg tca gga acg tca cgc tct cca cct tca tat tta ttt atc cgt -aa aaa tgt tat cct gga cat cgt acc aat aat aaa aaa gcc ca- -ta tat gtt cgc tat tgt aga aat tgt ttt tca cag ttg ctc aaa aac gat ggc agt gac tta tga gtt acg tta cac ttt gga gtc tca tct tta gta aac ata tca taa tat tcg ata tta cga gtt gac ata tcg aac aaa ttc caa gta ttt gat ttt gga taa tat tcg tat ttt gca tct gct ata att aag ata taa tcg ccg caa gaa cac acg aac atc ttt cct aca tgg tta aag tac atg tac aat tct atc cat ttg tct tcc tta act ata tat ttg tat aga taa tta cga gtc tca tga gta att cca gta att gca tag atg tca cca tcg tac tct aca gca taa act ata cta tga tgt cta ggc atg gga gac -tt ttt tat cca acg att ttt agt gaa aca ttc cac atc gtt taa tac tac ata ttt ttc ata cgt ggc ata aac tcc acc cat tac --a tat ata tca tcg ttt acg aat acc gat gcg cct gaa tat cta gga gta att aag ttt gga agt ctt atc cat ttc gaa gtg ccg tgt ttc aaa tat tct gcc aca ccc gtt gaa ata gaa aat tct aat cct cct att aca tat aac ttt cca tcg tta aca caa gta cta act tct gat ttt aac gac gac ata tta gta act gtt ttc ca- --- --- --- --- --- --- --t ttt ttc gtt tta aga tct acc cgc -ga tac gga ata aac atg tct att gtt aat cat gcc gcc aat aat gta tag aca att ata taa aac att tgc att ata gaa ttg --- -tc tat ctg tat tac cga cga ttg tcc aat att ctg ttc tag gag agt aat ggg tta ttg tgg ata tat aat cag agt ttt taa tga cta cta tat tat gtt tta tac cat ttc gtg tca cag gct ttg tag att tgg ata tag tta atc cca aca atg ata tag cat tgc aca tag tat tag tca taa act tgg gat gta aaa tgt tga tga tat cta cat cgt ttg gat ttt tat gta tcc act tta ata ata tca tag ctg taa cat cct cat ga- ttt acg tta acg tct tcg tgg gat aag ata gtt gtc agt tca tcc ttt gat aat ttt cca aat t

>LT896723.1:34866-36835_Cowpox_virus_isolate_Ger/2014/Cat1_genome_assembly_complete_genome:_monopartite

cgg ctt cca tgt att tcc tgg aga gca agt --- aga tga tg- agg aac cag ata gtt tat atc cgt act tgc act taa agt cta cat tgt cgt tgt atg a-- gta tga tct ttt aaa ccc gct aga caa gta tcc gtt tga tat tgt agg atg tgg aca ttt aac aat ctg aca cgt ggg tgg atc gga cca ttc tcc tcc tga aca cag gac acc aga gtt acc aat caa cga ata tcc act att gca act ata cgt tac aac gct tcc atc ggt ata aaa atc ctc gta tcc gtt atg tct tcc gtt gga tat aga tgg agg tga ttg gca ttt aac aga ttc gca aat agg tgc ctc ggg att cca tac cat aga tcc agt aga tcc taa ttc gca ata cga ttt aga ttc acc gat caa atg ata tcc gct att aca aga gta cgt tat act aga gcc aaa gtc tac tcc gcc aat atc aag ttg gcc att atc gat atc tcg agg cga tgg gca tct ccg ttt aat aca ttg att aaa gag tgt cca tcc agt acc tgt aca ttt agc ata tat agg tcc cat ttt ttg ctt tct gta tcc agg tag aca tag a-t att cta tag tgt ctc cta tgt tgt aat tag c-- --- -at tag cat tag cat cag tc- --t cca cac tat tct taa att tca tat taa tgg gtc gtg acg gaa tag tac agc atg ata gaa cgc atc cta ttc cca aca atg tca gga acg tca cgc tct cca cct tca tat tta ttt atc cgt -aa aaa tgt tat cct gga cat cgt aca aat aat a-a aaa gcc ca- -ta tat gtt cgc tat tgt aga aat tgt ttt tca cag ttg ctc aaa aac gat ggc agt gac tta tga gtt acg tta cac ttt gga gtc tca tct tta gta aac ata tca taa tat tcg ata tta cga gtt gac ata tcg aac aaa ttc caa gta ttt gat ttt gga taa tat tcg tat ttt gca tct gct ata att aag ata taa tca ccg caa gaa cac acg aac atc ttt cct aca tgg tta aag tac atg tac aat tct atc cat ttg tct tcc tta act ata tat ttg tat aga taa tta cga gtc tcg tga gta att cca gta att gca tag atg tca cca tcg tac tct aca gca taa act ata cta tga tgt cta ggc atg gga gac -tt ttt tat cca acg att ttt agt gaa aca ttc cac atc gtt taa tac tac ata ttt ctc ata cgt ggt ata aac tcc acc cat tac --a tat ata tca tcg ttt acg aat acc gat gcg cct gaa tat cta gga gta att aag ttt gga agt ctt atc cat ttc gaa gtg ccg tgt ttc aaa tat tct gcc aca ccc gtt gaa ata gaa aat tct aat cct cct att aca tat aac ttt cca tcg tta aca caa gta cta act tct gat ttt aac gac gac ata tta gta acc gtt ttc ca- --- --- --- --- --- --- --t ttt ttc gtt tta aga tct acc cgc -ga tac gga ata aac atg tct att gtt aat cat gcc gcc aat aat gta tag aca att ata taa aac att tgc att ata gaa ttg --- -tc tat ctg tat tac cga cga ttg tcc aat att ctg ttc tag gag agt aat ggg tta ttg tgg ata tat aat cag agt ttt taa tga cta cta tat tat gtt tta tac cat ttc gtg tca cag gct ttg tag att tgg ata tag tta atc cca aca atg ata tag cat tgc aca tag tat tag tca taa act tgg gat gta aaa tgt tga tga tat cta cat cgt ttg gat ttt tat gta tcc act tta ata ata tca tag ctg taa cat cct cgt ga- ttt acg ttg acg ctt tcg tgg gat aag ata gtt gtt agt tca tcc ttt gat aat ttt cca aat t

>KC813511.1:28219-30182_Cowpox_virus_strain_HumGri07/1_complete_genome

cgg ctt cca tgt att tcc tgg aga gca agt --- aga tga tg- agg aac cag ata gtt tat atc cgt act tgc act taa agt cta cat tgt cgt tgt atg a-- gta tga tct ttt aaa ccc gct aga caa gta tcc gtt tga tat tgt agg atg tgg aca ttt aac aat ctg aca cgt ggg tgg atc gga cca ttc tcc tcc tga aca cag gac acc aga gtt acc aat caa cga ata tcc act att gca act ata agt tac aac gct ccc atc ggt ata aaa atc ctc gta tcc gtt atg tct tcc gtt gga tat aga tgg agg gga ttg gca ttt aac aga ttc aca aat agg tgc ctc ggg att cca tac cat aga tcc agt aga tcc taa ttc gca ata cga ttt aga ttc acc gat caa atg ata tcc gct att aca aga gta cgt tat act aga gcc aaa gtc tac tcc acc aat atc aag ttg gcc att atc gat atc tcg agg cga tgg gca tct ccg ttt aat aca ttg att aaa gag tgt cca tcc agt acc tgt aca ttt agc ata tat agg tcc cat ttt ttg ctt tct gta tcc agg tag aca tag a-t att cta tag tgt ctc cta tgt tgt a-- --- --- --- -at tag cat tag cat cag tc- --t cca cac tat tct taa att tca tat taa tgg gtc gtg acg gaa tag tac agc atg ata gaa cgc atc cta ttc cca aca atg tca gga acg tca cgc tct cca cct tca tat tta ttt atc cgt -aa aaa tgt tat cct gga cat cgt aca aat aat a-a aaa gcc ca- -ta tat gtt cgc tat tgt aga aat tgt ttt tca cag ttg ctc aaa aac gat ggc agt gac tta tga gtt acg tta cac ttt gga gtc tca tct tta gta aac ata tca taa tat tcg ata tta cga gtt gac ata tcg aac aaa ttc caa gta ttt gat ttt gga taa tat tcg tat ttt gca tct gct ata att aag ata taa tca ccg caa gaa cac acg aac atc ttt cct aca tgg tta aag tac atg tac aat tct atc cat ttg tct tcc tta act ata tat ttg tat aga taa tta cga gtc tcg tga gta att cca gta att gca tag atg tca cca tcg tac tct aca gca taa act ata cta tga tgt cta ggc atg gga gac -tt ttt tat cca acg att ttt agt gaa aca ttc cac atc gtt taa tac tac ata ttt ctc ata cgt ggt ata aac tcc acc cat tac --a tat ata tca tcg ttt acg aat acc gat gcg cct gaa tat cta gga gta att aag ttt gga agt ctt atc cat ttc gaa gtg ccg tgt ttc aaa tat tct gcc aca ccc gtt gaa ata gaa aat tct aat cct cct att aca tat aac ttt cca tcg tta aca caa gta cta act tct gat ttt aac gac gac ata tta gta acc gtt ttc ca- --- --- --- --- --- --- --t ttt ttc gtt tta aga tct acc agc -ga cac gga ata aac atg tct att att aat cat gcc gcc aat aat gta tag aca att atg taa aac att tgc att ata gaa ttg --- -tc tat ctg tat tac cga cta tag tcc aat att ctg ttc tag gag agt aat ggg tta ttg tgg ata tat aat cag agt ttt taa tga cta cta tat tat gtt tta tac cat ttc gtg tca ccg gct ttg tag att tgg ata tag tta atc cca aca atg ata tag cat tgc gca tag tat tag tca taa act tgg gat gta aaa tgt tga tga tat cta cat cgt ttg gat ttt tat gta tcc act tta ata ata tca tag ctg taa cat cct cgt ga- ttt acg ttg acg ctt tcg tgg gat aag ata gtt gtt agt tca tcc ttt gat aat ttt cca aat t

>KC813507.1:16667-18630_Cowpox_virus_strain_EleGri07/1_complete_genome

cgg ctt cca tgt att tcc tgg aga gca agt --- aga tga tg- agg aac cag ata gtt tat atc cgt act tgc act taa agt cta cat tgt cgt tgt atg a-- gta tga tct ttt aaa ccc gct aga caa gta tcc gtt tga tat tgt agg atg tgg aca ttt aac aat ctg aca cgt ggg tgg atc gga cca ttc tcc tcc tga aca cag gac acc aga gtt acc aat caa cga ata tcc act att gca act ata agt tac aac gct ccc atc ggt ata aaa atc ctc gta tcc gtt atg tct tcc gtt gga tat aga tgg agg gga ttg gca ttt aac aga ttc aca aat agg tgc ctc ggg att cca tac cat aga tcc agt aga tcc taa ttc gca ata cga ttt aga ttc acc gat caa atg ata tcc gct att aca aga gta cgt tat act aga gcc aaa gtc tac tcc acc aat atc aag ttg gcc att atc gat atc tcg agg cga tgg gca tct ccg ttt aat aca ttg att aaa gag tgt cca tcc agt acc tgt aca ttt agc ata tat agg tcc cat ttt ttg ctt tct gta tcc agg tag aca tag a-t att cta tag tgt ctc cta tgt tgt a-- --- --- --- -at tag cat tag cat cag tc- --t cca cac tat tct taa att tca tat taa tgg gtc gtg acg gaa tag tac agc atg ata gaa cgc atc cta ttc cca aca atg tca gga acg tca cgc tct cca cct tca tat tta ttt atc cgt -aa aaa tgt tat cct gga cat cgt aca aat aat a-a aaa gcc ca- -ta tat gtt cgc tat tgt aga aat tgt ttt tca cag ttg ctc aaa aac gat ggc agt gac tta tga gtt acg tta cac ttt gga gtc tca tct tta gta aac ata tca taa tat tcg ata tta cga gtt gac ata tcg aac aaa ttc caa gta ttt gat ttt gga taa tat tcg tat ttt gca tct gct ata att aag ata taa tca ccg caa gaa cac acg aac atc ttt cct aca tgg tta aag tac atg tac aat tct atc cat ttg tct tcc tta act ata tat ttg tat aga taa tta cga gtc tcg tga gta att cca gta att gca tag atg tca cca tcg tac tct aca gca taa act ata cta tga tgt cta ggc atg gga gac -tt ttt tat cca acg att ttt agt gaa aca ttc cac atc gtt taa tac tac ata ttt ctc ata cgt ggt ata aac tcc acc cat tac --a tat ata tca tcg ttt acg aat acc gat gcg cct gaa tat cta gga gta att aag ttt gga agt ctt atc cat ttc gaa gtg ccg tgt ttc aaa tat tct gcc aca ccc gtt gaa ata gaa aat tct aat cct cct att aca tat aac ttt cca tcg tta aca caa gta cta act tct gat ttt aac gac gac ata tta gta acc gtt ttc ca- --- --- --- --- --- --- --t ttt ttc gtt tta aga tct acc agc -ga cac gga ata aac atg tct att att aat cat gcc gcc aat aat gta tag aca att atg taa aac att tgc att ata gaa ttg --- -tc tat ctg tat tac cga cta tag tcc aat att ctg ttc tag gag agt aat ggg tta ttg tgg ata tat aat cag agt ttt taa tga cta cta tat tat gtt tta tac cat ttc gtg tca ccg gct ttg tag att tgg ata tag tta atc cca aca atg ata tag cat tgc gca tag tat tag tca taa act tgg gat gta aaa tgt tga tga tat cta cat cgt ttg gat ttt tat gta tcc act tta ata ata tca tag ctg taa cat cct cgt ga- ttt acg ttg acg ctt tcg tgg gat aag ata gtt gtt agt tca tcc ttt gat aat ttt cca aat t

>AF095689.1:18243-20206_Vaccinia_virus_(strain_Tian_Tan)_complete_genome

cgg ctt cca tgt att tcc tgg aga gca agt --- aga tga tg- agg aac cag ata gtt tat atc cgt act tgc act taa agt cta cat tgt cgt tgt atg a-- gta tga tct ttt aaa ccc gct aga caa gta tcc gtt tga tat tgt agg atg tgg aca ttt aac aat ctg aca cgt ggg tgg atc gga cca ttc tcc tcc tga aca cag gac acc aga gtt acc aat caa cga ata tcc act att gca act ata agt tac aac gct tcc atc ggt ata aaa atc ctc gta tcc gtt atg tct tcc gtt gga tat aga tgg agg gga ttg gca ttt aac aga ttc aca aat agg tgc ctc ggg att cca tac cat aga tcc agt aga tcc taa ttc aca ata cga ttt aga ttc acc gat caa atg ata tcc gct att aca aga gta cgt tat act aga gcc aaa gtc tac tcc acc aat atc aag ttg gcc att atc gat atc tcg agg cga tgg gca tct ccg ttt aat aca ttg att aaa gag tgt cca tcc agt acc tgt aca ttt agc ata tat agg tcc cat ttt ttg ctt tct gta tcc agg tag aca tag a-t att cta tag tgt ctc cta tgt tgt a-- --- --- --- -at tag cat tag cat cag tc- --t cca cac tat tct taa att tca tat taa tgg gtc gtg acg gaa tag tac agc atg ata gaa cgc atc cta ttc cca aca atg tca gga acg tca cgc tct cca cct tca tat tta ttt atc cgt -aa aaa tgt tat cct gga cat cgt aca aat aat a-a aaa gcc ca- -ta tat gtt cgc tat tgt aga aat tgt ttt tca cag ttg ctc aaa aac gat ggc agt gac tta tga gtt acg tta cac ttt gga gtc tca tct tta gta aac ata tca taa tat tcg ata tta cga gtt gac ata tcg aac aaa ttc caa gta ttt gat ttt gga taa tat tcg tat ttt gca tct gct ata att aag ata taa tca ccg caa gaa cac acg aac atc ttt cct aca tgg tta aag tac atg tac aat tct atc cat ttg tct tcc tta act ata tat ttg tat aga taa tta cga gtc tcg tga gta att cca gta att aca tag atg tcg ccg tcg tac tct aca gca taa act ata cta tga tgt cta ggc atg gga gac -tt ttt tat cca acg att ttt agt gaa aca ttc cac atc gtt taa tac tac ata ttt ttc ata cgt ggt ata aac tcc acc cat tac --a tat ata tca tcg ttt acg aat acc gac gcg cct gaa tat cta gga gta att aag ttt gga agt ctt atc cat ttc gaa gtg ccg tgt ttc aaa tat tct gcc aca ccc gtt gaa ata gaa aat tct aat cct cct att aca tat aac ttt cca ttg tta aca caa gta cta act tct gat ttt aac gac gac ata tta gta acc gtt ttc ca- --- --- --- --- --- --- --t ttt ttc gtt tca aga tct acc cgc -ga tac gga ata aac atg tct att gtt aat cat gcc gcc aat aat gta tag aca att atg taa aac att tgc att ata gaa ttg --- -tc tat ctg tat tac cga cta tcg tcc aat att ctg ttc tag gag agt aat ggg tta ttg tgg ata tat aat cag agt ttt taa tga cta cta tat tat gtt tta tac cat ttc gtg tca ctg gct ttg tag att tgg ata tag tta atc cca aca atg ata tag cat tgc gca tag tat tag tca taa act tgg gat gta aaa tgt tga tga tat cta cat cgt ttg gat ttt tat gta tcc act tta ata ata tca tag ctg taa cat cct cat ga- ttt acg tta acg tct tcg tgg gat aag ata gtt gtc agt tca tcc ttt gat aat ttt cca aat t

>M35027.1:21784-23747_Vaccinia_virus_Copenhagen_complete_genome

cgg ctt cca tgt att tcc tgg aga gca agt --- aga tga tg- agg aac cag ata gtt tat atc cgt act tgc act taa agt cta cat tgt cgt tgt atg a-- gta tga tct ttt aaa ccc gct aga caa gta tcc gtt tga tat tgt agg atg tgg aca ttt aac aat ctg aca cgt ggg tgg atc gga cca ttc tcc tcc tga aca cag gac acc aga gtt acc aat caa cga ata tcc act att gca act ata agt tac aac gct tcc atc ggt ata aaa atc ctc gta tcc gtt atg tct tcc gtt gga tat aga tgg agg gga ttg gca ttt aac aga ttc aca aat agg tgc ctc ggg att cca tac cat aga tcc agt aga tcc taa ttc aca ata cga ttt aga ttc acc gat caa atg ata tcc gct att aca aga gta cgt tat act aga gcc aaa gtc tac tcc acc aat atc aag ttg gcc att atc gat atc tcg agg cga tgg gca tct ccg ttt aat aca ttg att aaa gag tgt cca tcc agt acc tgt aca ttt agc ata tat agg tcc cat ttt ttg ctt tct gta tcc agg tag aca tag a-t att cta tag tgt ctc cta tgt tgt a-- --- --- --- -at tag cat tag cat cag tc- --t cca cac tat tct taa att tca tat taa tgg gtc gtg acg gaa tag tac agc atg ata gaa cgc atc cta ttc cca aca atg tca gga acg tca cgc tct cca cct tca tat tta ttt atc cgt -aa aaa tgt tat cct gga cat cgt aca aat aat a-a aaa gcc ca- -ta tat gtt cgc tat tgt aga aat tgt ttt tca cag ttg ctc aaa aac gat ggc agt gac tta tga gtt acg tta cac ttt gga gtc tca tct tta gta aac ata tca taa tat tcg ata tta cga gtt gac ata tcg aac aaa ttc caa gta ttt gat ttt gga taa tat tcg tat ttt gca tct gct ata att aag ata taa tca ccg caa gaa cac acg aac atc ttt cct aca tgg tta aag tac atg tac aat tct atc cat ttg tct tcc tta act ata tat ttg tat aga taa tta cga gtc tcg tga gta att cca gta att aca tag atg tcg ccg tcg tac tct aca gca taa act ata cta tga tgt cta ggc atg gga gac -tt ttt tat cca acg att ttt agt gaa aca ttc cac atc gtt taa tac tac ata ttt ttc ata cgt ggt ata aac tcc acc cat tac --a tat ata tca tcg ttt acg aat acc gac gcg cct gaa tat cta gga gta att aag ttt gga agt ctt atc cat ttc gaa gtg ccg tgt ttc aaa tat tct gcc aca ccc gtt gaa ata gaa aat tct aat cct cct att aca tat aac ttt cca tcg tta aca caa gta cta act tct gat ttt aac gac gac ata tta gta acc gtt ttc ca- --- --- --- --- --- --- --t ttt ttc gtt tca aga tct acc cgc -ga tac gga ata aac atg tct att gtt aat cat gcc gcc aat aat gta tag aca att atg taa aac att tgc att ata gaa ttg --- -tc tat ctg tat tac cga cta tcg tcc aat att ctg tcc tag gag agt aat ggg tta ttg tgg ata tat aat cag agt ttt taa tga cta cta tat tat gtt tta tac cat ttc gtg tca ctg gct ttg tag att tgg ata tag tta atc cca aca atg ata tag cat tgc gca tag tat tag tca taa act tgg gat gta aaa tgt tga tga tat cta cat cgt ttg gat ttt tat gta tcc act tta ata ata tca tag ctg taa cat cct cat ga- ttt acg tta acg tct tcg tgg gat aag ata gtt gtc agt tca tcc ttt gat aat ttt cca aat t

>MK314711.1:15818-17779_Vaccinia_virus_strain_44/47.1_rMVA

cgg ctt cca tgt att tcc tgg aga gca agt --- aga tga tg- agg aac cag ata gtt tat atc cgt act tgc act taa agt cta cat tgt cgt tgt atg a-- gta tga tct ttt aaa ccc gct aga caa gta tcc gtt tga tat tgt agg atg tgg aca ttt aac aat ctg aca cgt ggg tgg atc gga cca ttc tcc tcc tga aca cag gac acc aga gtt acc aat caa cga ata tcc act att gca act ata agt tac aac gct ccc atc ggt ata aaa atc ctc gta tcc gtt atg tct tcc gtt gga tat aga tgg agg gga ttg gca ttt aac aga ttc aca aat agg tgc ctc ggg att cca tac cat aga tcc agt aga tcc taa ttc aca ata cga ttt aga ttc acc gat caa atg ata tcc gct att aca aga gta cgt tat act aga gcc aaa gtc tac tcc acc aat atc aag ttg gcc att atc gat atc tcg agg cga tgg gca tct ccg ttt aat aca ttg att aaa gag tgt cca tcc agt acc tgt aca ttt agc ata tat agg tcc cat ttt ttg ctt tct gta tcc agg tag aca tag a-t att cta tag tgt ctc cta tgt tgt a-- --- --- --- -at tag cat tag cat cag tc- --t cca cac tat tct taa att tca tat taa tgg gtc gtg acg gaa tag tac agc atg ata gaa cgc atc cta ttc cca aca atg tca gga acg tca cgc tct cca cct tca tat tta ttt atc cgt -aa aaa tgt tat cct gga cat cgt aca aat aat --- aaa gcc ca- -ta tat gtt cgc tat tgt aga aat tgt ttt tca cag ttg ctc aaa aac gat ggc agt gac tta tga gtt acg tta cac ttt gga gtc tca tct tta gta aac ata tca taa tat tcg ata tta cga gtt gac ata tcg aac aaa ttc caa gta ttt gat ttt gga taa tat tcg tat ttt gca tct gct ata att aag ata taa tca ccg caa gaa cac acg aac atc ttt cct aca tgg tta aag tac atg tac aat tct atc cat ttg tct tcc tta act ata tat ttg tat aga taa tta cga gtc tcg tga gta att cca gta att aca tag atg tcg ccg tcg tac tct aca gca taa act ata cta tga tgt cta ggc atg gga gac -tt ttt tat cca acg att ttt agt gaa aca ttc cac atc gtt taa tac tac ata ttt ttc ata cgt ggt ata aac tcc acc cat tac --a tat ata tca tcg ttt acg aat acc gac gcg cct gaa tat cta gga gta att aag ttt gga agt ctt atc cat ttc gaa gtg ccg tgt ttc aaa tat tct gcc aca ccc gtt gaa ata gaa aat tct aat cct cct att aca tat aac ttt cca tcg tta aca caa gta cta act tct gat ttt aac gac gac ata tta gta acc gtt ttc ca- --- --- --- --- --- --- --t ttt ttc gtt tca aga tct acc cgc -ga tac gga ata aac atg tct att gtt aat cat gcc gcc aat aat gta tag aca att atg taa aac att tgc att ata gaa ttg --- -tc tat ctg tat tac cga cta tcg tcc aat att ctg ttc tag gag agt aat ggg tta ttg tgg ata tat aat cag agt ttt taa tga cta cta tat tat gtt tta tac cat ttc gtg tca ctg gct ttg tag att tgg ata tag tta atc cca aca atg ata tag cat tgc gca tag tat tag tca taa act tgg gat gta aaa tgt tga tga tat cta cat cgt ttg gat ttt tat gta tcc act tta ata ata tca tag ctg taa cat cct cat ga- ttt acg tta acg tct tcg tgg gat aag ata gtt gtc agt tca tcc ttt gat aat ttt cca aat t

>MG663594.1:17429-19390_Vaccinia_virus_strain_51.1rMVA

cgg ctt cca tgt att tcc tgg aga gca agt --- aga tga tg- agg aac cag ata gtt tat atc cgt act tgc act taa agt cta cat tgt cgt tgt atg a-- gta tga tct ttt aaa ccc gct aga caa gta tcc gtt tga tat tgt agg atg tgg aca ttt aac aat ctg aca cgt ggg tgg atc gga cca ttc tcc tcc tga aca cag gac acc aga gtt acc aat caa cga ata tcc act att gca act ata agt tac aac gct ccc atc ggt ata aaa atc ctc gta tcc gtt atg tct tcc gtt gga tat aga tgg agg gga ttg gca ttt aac aga ttc aca aat agg tgc ctc ggg att cca tac cat aga tcc agt aga tcc taa ttc aca ata cga ttt aga ttc acc gat caa atg ata tcc gct att aca aga gta cgt tat act aga gcc aaa gtc tac tcc acc aat atc aag ttg gcc att atc gat atc tcg agg cga tgg gca tct ccg ttt aat aca ttg att aaa gag tgt cca tcc agt acc tgt aca ttt agc ata tat agg tcc cat ttt ttg ctt tct gta tcc agg tag aca tag a-t att cta tag tgt ctc cta tgt tgt a-- --- --- --- -at tag cat tag cat cag tc- --t cca cac tat tct taa att tca tat taa tgg gtc gtg acg gaa tag tac agc atg ata gaa cgc atc cta ttc cca aca atg tca gga acg tca cgc tct cca cct tca tat tta ttt atc cgt -aa aaa tgt tat cct gga cat cgt aca aat aat --- aaa gcc ca- -ta tat gtt cgc tat tgt aga aat tgt ttt tca cag ttg ctc aaa aac gat ggc agt gac tta tga gtt acg tta cac ttt gga gtc tca tct tta gta aac ata tca taa tat tcg ata tta cga gtt gac ata tcg aac aaa ttc caa gta ttt gat ttt gga taa tat tcg tat ttt gca tct gct ata att aag ata taa tca ccg caa gaa cac acg aac atc ttt cct aca tgg tta aag tac atg tac aat tct atc cat ttg tct tcc tta act ata tat ttg tat aga taa tta cga gtc tcg tga gta att cca gta att aca tag atg tcg ccg tcg tac tct aca gca taa act ata cta tga tgt cta ggc atg gga gac -tt ttt tat cca acg att ttt agt gaa aca ttc cac atc gtt taa tac tac ata ttt ttc ata cgt ggt ata aac tcc acc cat tac --a tat ata tca tcg ttt acg aat acc gac gcg cct gaa tat cta gga gta att aag ttt gga agt ctt atc cat ttc gaa gtg ccg tgt ttc aaa tat tct gcc aca ccc gtt gaa ata gaa aat tct aat cct cct att aca tat aac ttt cca tcg tta aca caa gta cta act tct gat ttt aac gac gac ata tta gta acc gtt ttc ca- --- --- --- --- --- --- --t ttt ttc gtt tca aga tct acc cgc -ga tac gga ata aac atg tct att gtt aat cat gcc gcc aat aat gta tag aca att atg taa aac att tgc att ata gaa ttg --- -tc tat ctg tat tac cga cta tcg tcc aat att ctg ttc tag gag agt aat ggg tta ttg tgg ata tat aat cag agt ttt taa tga cta cta tat tat gtt tta tac cat ttc gtg tca ctg gct ttg tag att tgg ata tag tta atc cca aca atg ata tag cat tgc gca tag tat tag tca taa act tgg gat gta aaa tgt tga tga tat cta cat cgt ttg gat ttt tat gta tcc act tta ata ata tca tag ctg taa cat cct cat ga- ttt acg tta acg tct tcg tgg gat aag ata gtt gtc agt tca tcc ttt gat aat ttt cca aat t

>KJ125439.1:21851-23814_Vaccinia_virus_strain_IHD-W_clone_IHDW1_complete_genome

cgg ctt cca tgt att tcc tgg aga gca agt --- aga tga tg- agg aac cag ata ggt tat atc cgt act tgc act taa agt tta cat tgt cgt tgt atg a-- gta tga tct ttt aaa ccc gct aga caa gta tcc gtt tga tat tgt agg atg tgg aca ttt aac aat ctg aca cgt ggg tgg atc gga cca ttc tcc tcc tga aca cag gac acc aga gtt acc aat caa cga ata tcc act att gca act ata agt tac aac gct tcc atc ggt ata aaa atc ctc gta tcc gtt atg tct tcc gtt gga tat aga tgg agg gga ttg gca ttt aac aga ttc aca aat agg tgc ctc ggg att cca tac cat aga tcc agt aga tcc taa ttc aca ata cga ttt aga ttc acc gat caa atg ata tcc gct att aca aga gta cgt tat act aga gcc aaa gtc tac tcc gcc aat atc aag ttg gcc att atc gat atc tcg agg cga tgg gca tct ccg ttt aat aca ttg att aaa gag tgt cca tcc agt acc tgt aca ttt agc ata tat agg tcc cat ttt ttg ctt tct gta tcc agg tag aca tag a-t att cta tag tgt ctc cta tgt tgt a-- --- --- --- -at tag cat tag cat cag tc- --t cca cac tat tct taa att tca tat taa tgg gtc gtg acg gaa tag tac agc atg ata gaa cgc atc cta ttc cca aca atg tca gga acg tca cgc tct cca cct tca tat tta ttt atc cgt -aa aaa tgt tat cct gga cat cgt aca aat aat a-a aaa gcc ca- -ta tat gtt cgc tat tgt aga aat tgt ttt tca cag ttg ctc aaa aac gat ggc agt gac tta tga gtt acg tta cac ttt gga gtc tca tct tta gta aac ata tca taa tat tcg ata tta cga gtt gac ata tcg aac aaa ttc caa gta ttt gat ttt gga taa tat tcg tat ttt gca tct gct ata att aag ata taa tca ccg caa gaa cac acg aac atc ttt cct aca tgg tta aag tac atg tac aat tct atc cat ttg tct tcc tta act ata tat ttg tat aga taa tta cga gtc tcg tga gta att cca gta att aca tag atg tcg ccg tcg tac tct aca gca taa act ata cta tga tgt cta ggc atg gga gac -tt ttt tat cca acg att ttt agt gaa aca ttc cac atc gtt taa tac tac ata ttt ttc ata cgt ggt ata aac tcc acc cat tac --a tat ata tca tcg ttt acg aat acc gac gcg cct gaa tat cta gga gta att aag ttt gga agt ctt atc cat ttc gaa gtg ccg tgt ttc aaa tat tct gcc aca ccc gtt gaa ata gaa aat tct aat cct cct att aca tat aac ttt cca ttg tta aca caa gta cta act tct gat ttt aac gac gac ata tta gta acc gtt ttc ca- --- --- --- --- --- --- --t ttt ttc gtt tca aga tct acc cgc -ga tac gga ata aac atg tct att gtt aat cat gcc gcc aat aat gta tag aca att atg taa aac att tgc att ata gaa ttg --- -tc tat ctg tat tac cga cta tcg tcc aat att ctg ttc tag gag agt aat ggg tta ttg tgg ata tat aat cag agt ttt taa tga cta cta tat tat gtt tta tac cat ttc gtg tca ctg gct ttg tag att tgg ata tag tta atc cca aca atg ata tag cat tgc gca tag tat tag tca taa act tgg gat gta aaa tgt tga tga tat cta cat cgt ttg gat ttt tat gta tcc act tta ata ata tca tag ctg taa cat cct cat ga- ttt acg tta acg tct tcg tgg gat aag ata gtt gtc agt tca tcc ttt gat aat ttt cca aat t

>JX489138.1:17970-19933_Vaccinia_virus_strain_Tiantan_clone_TT11_complete_genome

cgg ctt cca tgt att tcc tgg aga gca agt --- aga tga tg- agg aac cag ata gtt tat atc cgt act tgc act taa agt tta cat tgt cgt tgt atg a-- gta tga tct ttt aaa ccc gct aga caa gta tcc gtt tga tat tgt agg atg tgg aca ttt aac aat ttg aca cgt ggg tgg atc gga cca ttc tcc tcc tga aca cag gac acc aga gtt acc aat caa cga ata tcc act att gca act ata agt tac aac gct ccc atc ggt ata aaa atc ctc gta tcc gtt atg tct tcc gtt gga tat aga tgg agg gga ttg gca ttt aac aga ttc aca aat agg tgc ctc ggg att cca tac cat aga tcc agt aga tcc taa ttc aca ata cga ttt aga ttc acc gat caa atg ata tcc gct att aca aga gta cgt tat act aga gcc aaa gtc tac tcc acc aat atc aag ttg gcc att atc gat atc tcg agg cga tgg gca tct ccg ttt aat aca ttg att aaa gag tgt cca tcc agt acc tgt aca ttt agc ata tat agg tcc cat ttt ttg ctt tct gta tcc agg tag aca tag a-t att cta tag tgt ctc cta tgt tgt a-- --- --- --- -at tag cat tag cat cag tc- --t cca cac tat tct taa att tca tat taa tgg gtc gtg acg gaa tag tac agc atg ata gaa cgc atc cta ttc cca aca atg tca gga acg tca cgc tct cca cct tca tat tta ttt atc cgt -aa aaa tgt tat cct gga cat cgt aca aat aat a-a aaa gcc ca- -ta tat gtt cgc tat tgt aga aat tgt ttt tca cag ttg ctc aaa aac gat ggc agt gac tta tga gtt acg tta cac ttt gga gtt tca tct tta gta aac ata tca taa tat tcg ata tta cga gtt gac ata tcg aac aaa ttc caa gta ttt gat ttt gga taa tat tcg tat ttt gca tct gct ata att aag ata taa tca ccg caa gaa cac acg aac atc ttt cct aca tgg tta aag tac atg tac aat tct atc cat ttg tct tcc tta act ata tat ttg tat aga taa tta cga gtc tcg tga gta att cca gta att aca tag atg tcg ccg tcg tac tct aca gca taa act ata cta tga tgt cta ggc atg gga gac -tt ttt tat cca acg att ttt agt gaa aca ttc cac atc gtt taa tac tac ata ttt ttc ata cgt ggt ata aac tcc acc cat tac --a tat ata tca tcg ttt acg aat acc gac gcg cct gaa tat cta gga gta att aag ttt gga agt ctt atc cat ttc gaa gtg ccg tgt ttc aaa tat tct gcc aca ccc gtt gaa ata gaa aat tct aat cct cct att aca tat aac ttt cca tcg tta aca caa gta cta act tct gat ttt aac gac gac ata tta gta acc gtt ttc ca- --- --- --- --- --- --- --t ttt ttc gtt tca aga tct acc cgc -ga tac gga ata aac atg tct att gtt aat cat gcc gcc aat aat gta tag aca att atg taa aac att tgc att ata gaa ttg --- -tc tat ctg tat tac cga cta tcg tcc aat att ctg ttc tag gag agt aat ggg tta ttg tgg ata tat aat cag agt ttt taa tga cta cta tat tat gtt tta tac cat ttc gtg tca ctg gct ttg tag att tgg ata tag tta atc cca aca atg ata tag cat tgc gca tag tat tag tca taa act tgg gat gta aaa tgt tga tga tat cta cat cgt ttg gat ttt tat gta tcc act tta ata ata tca tag ctg taa cat cct cat ga- ttt acg tta acg tct tcg tgg gat aag ata gtt gtc agt tca tcc ttt gat aat ttt cca aat t

>JX489136.1:18056-20013_Vaccinia_virus_strain_Tiantan_clone_TT9_complete_genome

cgg ctg cca tgt att tcc tgg aga gca agt --- aga tga tg- agg aac cag ata gtt tat atc cgt act tgc act taa agt tta cat tgt cgt tgt atg a-- gta tga tct ttt aaa ccc gct aga caa gta tcc gtt tga tat tgt agg atg tgg aca ttt aac aat ctg aca cgt ggg tgg atc gga cca ttc tcc tcc tga aca cag gac acc aga gtt acc aat caa cga ata tcc act att gca act ata agt tac aac gct ccc atc ggt ata aaa atc ctc gta tcc gtt atg tct tcc gtt gga tat aga tgg agg gga ttg gca ttt aac aga ttc aca aat agg tgc ctc ggg att cca tac cat aga tcc agt aaa tcc taa ttc aca ata cga ttt aga ttc tcc tat caa ctg ata tcc gct att aca aga gta cgt tat act aga gcc aaa gtc tac tcc gcc aat atc aag ttg gcc att atc gat atc tcg agg cga tgg gca tct ccg ttt aat aca ttg att aaa gag tgt cca tcc agt acc tgt aca ttt agc ata tat agg tcc cat ttt ttg ctt tct gta tcc agg tag aca tag a-t att cta tag tgt ctc cta tgt tgt a-- --- --- --- -at tag cat t-- --- -ag tt- --t cca cac tat tct taa att tca tat taa tgg gac gtg aag gaa tag gac agt atg ata gaa cgc atc cta ttc cca aca atg tca gga acg tca cgc tct cca cct tca tat tta ttt atc cgt -aa aaa tgt tat cct gga cat cgt aca aat aat a-a aaa gcc ca- -ta tat gtt cgc tat tgt aga aat tgt ttt tca cag ttg ctc aaa aac gat ggc agt gac tta tga gtt acg tta cac ttt gga gtc tca tct tta gta aac ata tca taa tat tcg ata tta cga gtt gac ata tcg aac aaa ttc caa gta ttt gat ttt gga taa tat tcg tat ttt gca tct gct ata att aag ata taa tca ccg caa gaa cac acg aac atc ttt cct aca tgg tta aag tac atg tat aat tct atc cat ttg tct tcc tta act ata tat ttg tat aga taa tta cga gtc tcg tga gta att cca gta att aca tag atg tcg ccg tcg tac tct aca gca taa act ata cta tga tgt cta ggc atg gga gac -tt ttt tat cca acg att ttt agt gaa aca ttc cac atc gtt taa tac tac ata ttt ttc ata cgt ggt ata aac tcc acc cat tac --a tat ata tca tcg ttt acg aat acc gac gcg cct gaa tat cta gga gta att aag ttt gga agt ctt atc cat ttc gaa gtg ccg tgt ttc aaa tat tct gcc aca ccc gtt gaa ata gaa aat tct aat cct cct att aca tat aac ttt cca ttg tta aca caa gta cta act tct gat ttt aac gac gac ata tta gta acc gtt ttc ca- --- --- --- --- --- --- --t ttt ttc gtt tca aga tct acc cgc -ga tac gga ata aac atg tct att gtt aat cat gcc gcc aat aat gta tag aca att atg taa aac att tgc att ata gaa ttg --- -tc tat ctg tat tac cga cta tcg tcc aat att ctg ttc tag gag agt aat ggg tta ttg tgg ata tat aat cag agt ttt taa tga cta cta tat tat gtt tta tac cat ttc gtg tca ctg gct ttg tag att tgg ata tag tta atc cca aca atg ata tag cat tgc gca tag tat tag tca taa act tgg gat gta aaa tgt tga tga tat cta cat cgt ttg gat ttt tat gta tcc act tta ata ata tca tag ctg taa cat cct cat ga- ttt acg tta acg tct tcg tgg gat aag ata gtt gtc agt tca tcc ttt gat aat ttt cca aat t

>KC201194.1:22116-24079_Vaccinia_virus_strain_IHD-W_partial_sequence

cgg ctt cca tgt att tcc tgg aga gca agt --- aga tga tg- agg aac cag ata ggt tat atc cgt act tgc act taa agt tta cat tgt cgt tgt atg a-- gta tga tct ttt aaa ccc gct aga caa gta tcc gtt tga tat tgt agg atg tgg aca ttt aac aat ctg aca cgt ggg tgg atc gga cca ttc tcc tcc tga aca cag gac acc aga gtt acc aat caa cga ata tcc act att gca act ata agt tac aac gct tcc atc ggt ata aaa atc ctc gta tcc gtt atg tct tcc gtt gga tat aga tgg agg gga ttg gca ttt aac aga ttc aca aat agg tgc ctc ggg att cca tac cat aga tcc agt aga tcc taa ttc aca ata cga ttt aga ttc acc gat caa atg ata tcc gct att aca aga gta cgt tat act aga gcc aaa gtc tac tcc gcc aat atc aag ttg gcc att atc gat atc tcg agg cga tgg gca tct ccg ttt aat aca ttg att aaa gag tgt cca tcc agt acc tgt aca ttt agc ata tat agg tcc cat ttt ttg ctt tct gta tcc agg tag aca tag a-t att cta tag tgt ctc cta tgt tgt a-- --- --- --- -at tag cat tag cat cag tc- --t cca cac tat tct taa att tca tat taa tgg gtc gtg acg gaa tag tac agc atg ata gaa cgc atc cta ttc cca aca atg tca gga acg tca cgc tct cca cct tca tat tta ttt atc cgt -aa aaa tgt tat cct gga cat cgt aca aat aat a-a aaa gcc ca- -ta tat gtt cgc tat tgt aga aat tgt ttt tca cag ttg ctc aaa aac gat ggc agt gac tta tga gtt acg tta cac ttt gga gtc tca tct tta gta aac ata tca taa tat tcg ata tta cga gtt gac ata tcg aac aaa ttc caa gta ttt gat ttt gga taa tat tcg tat ttt gca tct gct ata att aag ata taa tca ccg caa gaa cac acg aac atc ttt cct aca tgg tta aag tac atg tac aat tct atc cat ttg tct tcc tta act ata tat ttg tat aga taa tta cga gtc tcg tga gta att cca gta att aca tag atg tcg ccg tcg tac tct aca gca taa act ata cta tga tgt cta ggc atg gga gac -tt ttt tat cca acg att ttt agt gaa aca ttc cac atc gtt taa tac tac ata ttt ttc ata cgt ggt ata aac tcc acc cat tac --a tat ata tca tcg ttt acg aat acc gac gcg cct gaa tat cta gga gta att aag ttt gga agt ctt atc cat ttc gaa gtg ccg tgt ttc aaa tat tct gcc aca ccc gtt gaa ata gaa aat tct aat cct cct att aca tat aac ttt cca ttg tta aca caa gta cta act tct gat ttt aac gac gac ata tta gta acc gtt ttc ca- --- --- --- --- --- --- --t ttt ttc gtt tca aga tct acc cgc -ga tac gga ata aac atg tct att gtt aat cat gcc gcc aat aat gta tag aca att atg taa aac att tgc att ata gaa ttg --- -tc tat ctg tat tac cga cta tcg tcc aat att ctg ttc tag gag agt aat ggg tta ttg tgg ata tat aat cag agt ttt taa tga cta cta tat tat gtt tta tac cat ttc gtg tca ctg gct ttg tag att tgg ata tag tta atc cca aca atg ata tag cat tgc gca tag tat tag tca taa act tgg gat gta aaa tgt tga tga tat cta cat cgt ttg gat ttt tat gta tcc act tta ata ata tca tag ctg taa cat cct cat ga- ttt acg tta acg tct tcg tgg gat aag ata gtt gtc agt tca tcc ttt gat aat ttt cca aat t

>MK239755.1:11916-13879_Buffalopox_virus_isolate_NIV116234

cgg ctt cca tgt att tcc tgg aga gca agt --- aga tga tg- agg aac cag ata gtt tat atc cgt act tgc act taa agt cta cat tgt cgt tgt atg a-- gta tga tct ttt aaa ccc gct aga caa gta tcc gtt tga tat tgt agg atg tgg aca ttt aac aat ctg aca cgt ggg tgg atc gga cca ttc tcc tcc tga aca cag gac act aga gtt acc aat caa cga ata tcc act att gca act ata agt tac aac gct tcc atc ggt ata aaa atc ctc gta tcc gtt atg tct tcc gtt gga tat aga tgg agg gga ttg gca ttt aac aga ttc aca aat agg tgc ctc ggg att cca tac cat aga tcc agt aga tcc taa ttc aca ata cga ttt aga ttc acc gat caa atg ata tcc gct att aca aga gta cgt tat act aga gcc aaa gtc tac tcc acc aat atc aag ttg gcc att atc gat atc tcg agg cga tgg gca tct ccg ttt aat aca ttg att aaa gag tgt cca tcc agt acc tgt aca ttt agc ata tat agg tcc cat ttt ttg ctt tct gta tcc agg tag aca tag a-t att cta tag tgt ctc cta tgt tgt a-- --- --- --- -at tag cat tag cat cag tc- --t cca cac tat tct taa att tca tat taa tgg gtc gtg acg gaa tag tac agc atg ata gaa cgc atc cta ttc cca aca atg tca gga acg tca cgc tct cca cct tca tat tta ttt atc cgt -aa aaa tgt tat cct gga cat cgt aca aat aat a-a aaa gcc ca- -ta tat gtt cgc tat tgt aga aat tgt ttt tca cag ttg ctc aaa aac gat ggc agt gac tta tga gtt acg tta cac ttt gga gtc tca tct tta gta aac ata tca taa tat tcg ata tta cga gtt gac ata tcg aac aaa ttc caa gta ttt gat ttt gga taa tat tcg tat ttt gca tct gct ata att aag ata taa tca ccg caa gaa cac acg aac atc ttt cct aca tgg tta aag tac atg tac aat tct atc cat ttg tct tcc tta act ata tat ttg tat aga taa tta cga gtc tcg tga gta att cca gta att aca tag atg tcg ccg tcg tac tct aca gca taa act ata cta tga tgt cta ggc atg gga gac -tt ttt tat cca acg att ttt agt gaa aca ttc cac atc gtt taa tac tac ata ttt ttc ata cgt ggt ata aac tcc acc cat tac --a tat ata tca tcg ttt acg aat acc gac gcg cct gaa tat cta gga gta att aag ttt gga agt att atc cat ttc gaa gtg ccg tgt ttc aaa tat tct gcc aca ccc gtt gaa ata gaa aat tct aat cct cct att aca tat aac ttt cca tcg tta aca caa gta cta act tct gat ttt aac gac gac ata tta gta acc gtt ttc ca- --- --- --- --- --- --- --t ttt ttc gtt tca aga tct acc cgc -ga tac gga ata aac atg tct att gtt aat cat gcc gcc aat aat gta tag aca att atg taa aac att tgc att ata gaa ttg --- -tc tat ctg tat tac cga cta tcg tcc aat att ctg ttc tag gag agt aat ggg tta ttg tgg ata tat aat cag agt ttt taa tga cta cta tat tat gtt tta tac cat ttc gtg tca ctg gct ttg tag att tgg ata tag tta atc cca aca atg ata tag cat tgc gca tag tat tag tca taa act tgg gat gta aaa tgt tga tga tat cta cat cgt ttg gat ttt tat gta tcc act tta ata ata tca tag ctg taa cat cct cat ga- ttt acg tta acg tct tcg tgg gat aag ata gtt gtc agt tca tcc ttt gat aat ttt cca aat t

>AY484669.1:23341-25304_Rabbitpox_virus_complete_genome

cgg ctg cca tgt att tcc tgg aga gca agt --- aga tga tg- agg aac cag ata ggt tat atc cgt act tgc act taa agt tta cat tgt cgt tgt atg a-- gta tga tct ttt aaa ccc gct aga caa gta tcc gtt tga tat tgt agg atg tgg aca ttt aac aat ctg aca cgt ggg tgg atc gga cca ttc tcc tcc tga aca cag gac acc aga gtt acc aat caa cga ata tcc act att gca act ata agt tac aac gct ccc atc ggt ata aaa atc ctc gta tcc gtt atg tct tcc gtt gga tat aga tgg agg gga ttg gca ttt aac aga ttc aca aat agg tgc ctc ggg att cca tac cat aga tcc agt aga tcc taa ttc aca ata cga ttt aga ttc acc gat caa atg ata tcc gct att aca aga gta cgt tat act aga gcc aaa gtc tac tcc gcc aat atc aag ttg gcc att atc gat atc tcg agg cga tgg gca tct ccg ttt aat aca ttg att aaa gag tgt cca tcc agt acc tgt aca ttt agc ata tat agg tcc cat ttt ttg ctt tct gta tcc agg tag aca tag a-t att cta tag tgt ctc cta tgt tgt a-- --- --- --- -at tag cat tag cat cag tc- --t cca cac tat tct taa att tca tat taa tgg gtc gtg acg gaa tag tac agc atg ata gaa cgc atc cta ttc cca aca atg tca gga acg tca cgc tct cca cct tca tat tta ttt atc cgt -aa aaa tgt tat cct gga cat cgt aca aat aat a-a aaa gcc ca- -ta tat gtt cgc tat tgt aga aat tgt ttt tca cag ttg ctc aaa aac gat ggc agt gac tta tga gtt acg tta cac ttt gga gtt tca tct tta gta aac ata tca taa tat tcg ata tta cga gtt gac ata tcg aac aaa ttc caa gta ttt gat ttt gga taa tat tcg tat ttt gca tct gct ata att aag ata taa tca ccg caa gaa cac acg aac atc ttt cct aca tgg tta aag tac atg tac aat tct atc cat ttg tct tcc tta act ata tat ttg tat aga taa tta cga gtc tcg tga gta att cca gta att aca tag atg tcg ccg tcg tac tct aca gca taa act ata cta tga tgt cta ggc atg gga gac -tt ttt tat cca acg att ttt agt gaa aca ttc cac atc gtt taa tac tac ata ttt ttc ata cgt ggt ata aac tcc acc cat tac --a tat ata tca tcg ttt acg aat acc gac gcg cct gaa tat cta gga gta att aag ttt gga agt ctt atc cat ttt gaa gtg ccg tgt ttc aaa tat tct gct aca ccc gtt gaa ata gaa aat tct aat cct cct att aca tat aac ttt cca ttg tta aca caa gta cta act tct gat ttt aac gac gac ata tta gta acc gtt ttc ca- --- --- --- --- --- --- --t ttt ttc gtt tca aga tct acc cgc -ga tac gga ata aac atg tct att gtt aat cat gcc gcc aat aat gta tag aca att atg taa aac att tgc att ata gaa ttg --- -tc tat ctg tat tac cga cta tcg tcc aat att ctg ttc tag gag agt aat ggg tta ttg tgg ata tat aat cag agt ttt taa tga cta cta tat tat gtt tta tac cat ttc gtg tca ctg gct ttg tag att tgg ata tag tta atc cca aca atg ata tag cat tgc gca tag tat tag tta taa act tgg gat gta aaa tgt tga tga tat cta cat cgt ttg gat ttt tat gta tcc act tta ata ata tca tag ctg taa cat cct cat ga- ttt acg tta acg tct tcg tgg gat aag ata gtt gtc agt tca tcc ttt gat aat ttt cca aat t

>MG599038.1:21951-23914_Buffalopox_virus_isolate_Karachi_2005

cgg ctt cca tgt att tcc tag aga gca agt --- aga tga tg- agg aac cag ata gtt tat atc cgt act tgc act taa agt cta cat tgt cgt tgt atg a-- gta tga tct ttt aaa ccc gct aga caa gta tcc gtt tga tat tgt agg atg tgg aca ttt aac aat ctg aca cgt ggg tgg atc gga cca ttc tcc tcc tga aca cag gac act aga gtt acc aat caa cga ata tcc act att gca act ata agt tac aac gct tcc atc ggt ata aaa atc ctc gta tcc gtt atg tct tcc gtt gga tat aga tgg agg gga ttg gca ttt aac aga ttc aca aat agg tgc ctc ggg att cca tac cat aga tcc agt aga tcc taa ttc aca ata cga ttt aga ttc acc gat caa atg ata tcc gct att aca aga gta cgt tat act aga gcc aaa gtc tac tcc acc aat atc aag ttg gcc att atc gat atc tcg agg cga tgg gca tct ccg ttt aat aca ttg att aaa gag tgt cca tcc agt acc tgt aca ttt agc ata tat agg tcc cat ttt ttg ctt tct gta tcc agg tag aca tag a-t att cta tag tgt ctc cta tgt tgt a-- --- --- --- -at tag cat tag cat cag tc- --t cca cac tat tct taa att tca tat taa tgg gtc gtg acg gaa tag tac agc atg ata gaa cgc atc cta ttc cca aca atg tca gga acg tca cgc tct cca cct tca tat tta ttt atc cgt -aa aaa tgt tat cct gga cat cgt aca aat aat a-a aaa gcc ca- -ta tat gtt cgc tat tgt aga aat tgt ttt tca cag ttg ctc aaa aac gat ggc agt gac tta tga gtt acg tta cac ttt gga gtc tca tct tta gta aac ata tca taa tat tcg ata tta cga gtt gac ata tcg aac aaa ttc caa gta ttt gat ttt gga taa tat tcg tat ttt gca tct gct ata att aag ata taa tca ccg caa gaa cac acg aac atc ttt cct aca tgg tta aag tac atg tac aat tct atc cat ttg tct tcc tta act ata tat ttg tat aga taa tta cga gtc tcg tga gta att cca gta att aca tag atg tcg ccg tcg tac tct aca gca taa act ata cta tga tgt cta ggc atg gga gac -tt ttt tat cca acg att ttt agt gaa aca ttc cac atc gtt taa tac tac ata ttt ttc ata cgt ggt ata aac tcc acc cat tac --a tat ata tca tcg ttt acg aat acc gac gcg cct gaa tat cta gga gta att aag ttt gga agt att atc cat ttc gaa gtg ccg tgt ttc aaa tat tct gcc aca ccc gtt gaa ata gaa aat tct aat cct cct att aca tat aac ttt cca tcg tta aca caa gta cta act tct gat ttt aac gac gac ata tta gta acc gtt ttc ca- --- --- --- --- --- --- --t ttt ttc gtt tca aga tct acc cgc -ga tac gga ata aac atg tct att gtt aat cat gcc gcc aat aat gta tag aca att atg taa aac att tgc att ata gaa ttg --- -tc tat ctg tat tac cga cta tcg tcc aat att ctg ttc tag gag agt aat ggg tta ttg tgg ata tat aat cag agt ttt taa tga cta cta tat tat gtt tta tac cat ttc gtg tca ctg gct ttg tag att tgg ata tag tta atc cca aca atg ata tag cat tgc gca tag tat tag tca taa act tgg gat gta aaa tgt tga tga tat cta cat cgt ttg gat ttt tat gta tcc act tta ata ata tca tag ctg taa cat cct cat ga- ttt acg tta acg tct tcg tgg gat aag ata gtt gtc agt tca tcc ttt gat aat ttt cca aat t

>LT896718.1:34869-36833_Cowpox_virus_isolate_Ger/2010/Alpaca_genome_assembly_complete_genome:_monopartite

cgg ctt cca tgt att tcc tgg aga gca agt --- aga tga tg- agg aac cag ata gtt tat atc cgt act tgc act taa agt cta cat tgt cgt tgt atg a-- gta tga tct ttt aaa ccc gct aga caa gta tcc gtt tga tat tgt agg atg tgg aca ttt aac aat ctg aca cgt ggg tgg atc gga cca ttc tcc tcc tga aca cag gac acc aga gtt acc aat caa cga ata tcc act att aca act ata cgt tac aac gct tcc atc ggt ata aaa atc ctc gta tcc gtt atg tct tcc gtt gga tat aga tgg agg tga ttg gca ttt aac aga ttc gca aat agg tgc ctc ggg att cca tac cat aga tcc agt aga tcc taa ttc aca ata cga ttt aga ttc acc gat caa atg ata tcc gct att aca aga gta cgt tat act aga gcc aaa gtc tac tcc gcc aat atc aag ttg gcc att atc gat atc tcg agg cga tgg gca tct ccg ttt aat aca ttg att aaa gag tgt cca tcc agt acc tgt aca ttt agc ata tat agg tcc cat ttt ttg ctt tct gta tcc agg tag aca tag a-t att cta tag tgt ctc cta tgt tgt a-- --- --- --- -at tag cat tag cat cag tc- --t cca cac tat tct taa att tca tat taa tgg gtc gtg acg gaa tag tac agc atg ata gaa cgc atc cta ttc cca aca atg tca gga atg tca cgc tct cca cct tca tat tta ttt atc cgt -aa aaa tgt tat cct gga cat cgt acc aat aat aaa aaa gcc ca- -ta tat gtt cgc tat tgt aga aat tgt ttt tca cag ttg ctc aaa aac gat ggc agt gac tta tga gtt acg tta cac ttt gga gtc tca tct tta gta aac ata tca taa tat tcg ata tta cga gtt gac ata tcg aac aaa ttc caa gta ttt gat ttt gga taa tat tcg tat ttt gca tct gct ata att aag ata taa tcg ccg caa gaa cac acg aac atc ttt cct aca tgg tta aag tac atg tac aat tct atc cat ttg tct tcc tta act ata tat ttg tat aga taa tta cga gtc tcg tgg gta att cca gta att gca tag atg tca cca tcg tac tct aca gca tac act ata cta tga tgt cta ggc atg gga gac -tt ttt tat cca acg att ttt agt gaa aca ttc cac atc gtt taa tac tac ata ttt ctc ata cgt ggt ata aac tcc acc cat tac --a tat ata tca tcg ttt acg aat acc gat gcg cct gaa tat cta gga gta att aag ttt gga agt ctt atc cat ttc gaa gtg ccg tgt ttc aaa tat tct gcc aca ccc gtt gaa ata gaa aat tct aat cct cct att aca tat aac ttt cca tcg tta aca caa gta cta act tct gat ttt aac gac gac ata tta gta acc gtt ttc ca- --- --- --- --- --- --- --t ttt ttc gtt tta aga tct acc agc -ga cac gga ata aac atg tct att att aat cat gcc gcc aat aat gta tag aca att atg taa aac att tgc att ata gaa ttg --- -tc tat ctg tat tac cga cta tag tcc aat att ctg ttc tag gag agt aat ggg tta ttg tgg ata tat aat cag agt ttt taa tga cta cta tat tat gtt tta tac cat ttc gtg tca ccg gct ttg tag att tgg ata tag tta atc cca aca atg ata tag cat tgc gca tag tat tag tca taa act tgg gat gta aaa tgt tga tga tat ctg cat cgt ttg gat ttt tat gta tcc act tta ata ata tta tag ctg taa cat cct cat ga- ttt acg tta acg tct tcg tgg gat aag ata gtt gtc agt tca tcc ttt gat aat ttt cca aat t

>KP233807.1:19061-21024_Vaccinia_virus_strain_L-IVP_complete_genome

cgg ctt cca tgt att tcc tgg aga gca agt --- aga tga tg- agg aac cag ata gtt tat atc cgt act tgc act taa agt cta cat tgt cgt tgt atg a-- gta tga tct ttt aaa ccc gct aga caa gta tcc gtt tga tat tgt agg atg tgg aca ttt aac aat ctg aca cgt ggg tgg atc gga cca ttc tcc tcc tga aca cag gac act aga gtt acc aat caa cga ata tcc act att gca act ata agt tac aac gct ccc atc ggt ata aaa atc ctc gta tcc gtt atg tct tcc gtt gga tat aga tgg agg gga ttg gca ttt aac aga ttc aca aat agg tgc ctc ggg att cca tac cat aga tcc agt aga tcc taa ttc aca ata cga ttt aga ttc tcc tat caa ctg ata tcc gct att aca aga gta cgt tat act aga gcc aaa gtc tac tcc acc aat atc aag ttg gcc att atc gat atc tcg agg cga tgg gca tct ccg ttt aat aca ttg att aaa gag tgt cca tcc agt acc tgt aca ttt agc ata tat agg tcc cat ttt ttg ctt tct gta tcc agg tag aca tag a-t att cta tag tgt ctc cta tgt tgt a-- --- --- --- -at tag cat tag cat cag tc- --t cca cac tat tct taa att tca tat taa tgg gtc gtg acg gaa tag tac agc atg ata gaa cgc atc cta ttc cca aca atg tca gga acg tca cgc tct cca cct tca tat tta ttt atc cgt -aa aaa tgt tat cct gga cat cgt aca aat aat a-a aaa gcc ca- -ta tat gtt cgc tat tgt aga aat tgt ttt tca cag ttg ctc aaa aac gat ggc agt gac tta tga gtt acg tta cac ttt gga gtc tca tct tta gta aac ata tca taa tat tcg ata tta cga gtt gac ata tcg aac aaa ttc caa gta ttt gat ttt gga taa tat tcg tat ttt gca tct gct ata att aag ata taa tca ccg caa gaa cac acg aac atc ttt cct aca tgg tta aag tac atg tac aat tct atc cat ttg tct tcc tta act ata tat ttg tat aga taa tta cga gtc tcg tga gta att cca gta att aca tag atg tcg ccg tcg tac tct aca gca taa act ata cta tga tgt cta ggc atg gga gac -tt ttt tat cca acg att ttt agt gaa aca ttc cac atc gtt taa tac tac ata ttt ttc ata cgt ggt ata aac tcc acc cat tac --a tat ata tca tcg ttt acg aat acc gac gcg cct gaa tat cta gga gta att aag ttt gga agt ctt atc cat ttc gaa gtg ccg tgt ttc aaa tat tct gcc aca ccc gtt gaa ata gaa aat tct aat cct cct att aca tat aac ttt cca tcg tta aca caa gta cta act tct gat ttt aac gac gac ata tta gta acc gtt ttc ca- --- --- --- --- --- --- --t ttt ttc gtt tca aga tct acc cgc -ga tac gga ata aac atg tct att gtt aat cat gcc gcc aat aat gta tag aca att atg taa aac att tgc att ata gaa ttg --- -tc tat ctg tat tac cga cta tcg tcc aat att ctg ttc tag gag agt aat ggg tta ttg tgg ata tat aat cag agt ttt taa tga cta cta tat tat gtt tta tac cat ttc gtg tca ctg gct ttg tag att tgg ata tag tta atc cca aca atg ata tag cat tgc gca tag tat tag tca taa act tgg gat gta aaa tgt tga tga tat cta cat cgt ttg gat ttt tat gta tcc act tta ata ata tca tag ctg taa cat cct cat ga- ttt acg tta acg tct tcg tgg gat aag ata gtt gtc agt tca tcc ttt gat aat ttt cca aat t

>OK422495.1:8001-9964_Buffalopox_virus_isolate_BPXV/buffalo/India/2011/Meerut-1/P50/A3_complete_genome

cgg ctt cca tgt att tcc tag aga gca agt --- aga tga tg- agg aac cag ata gtt tat atc cgt act tgc act taa agt cta cat tgt cgt tgt atg a-- gta tga tct ttt aaa ccc gct aga caa gta tcc gtt tga tat tgt agg atg tgg aca ttt aac aat ctg aca cgt ggg tgg atc gga cca ttc tcc tcc tga aca cag gac act aga gtt acc aat caa cga ata tcc act att gca act ata agt tac aac gct tcc atc ggt ata aaa atc ctc gta tcc gtt atg tct tcc gtt gga tat aga tgg agg gga ttg gca ttt aac aga ttc aca aat agg tgc ctc ggg att cca tac cat aga tcc agt aga tcc taa ttc aca ata cga ttt aga ttc acc gat caa atg ata tcc gct att aca aga gta cgt tat act aga gcc aaa gtc tac tcc acc aat atc aag ttg gcc att atc gat atc tcg agg cga tgg gca tct ccg ttt aat aca ttg att aaa gag tgt cca tcc agt acc tgt aca ttt agc ata tat agg tcc cat ttt ttg ctt tct gta tcc agg tag aca tag a-t att cta tag tgt ctc cta tgt tgt a-- --- --- --- -at tag cat tag cat cag tc- --t cca cac tat tct taa att tca tat taa tgg gtc gtg acg gaa tag tac agc atg ata gaa cgc atc cta ttc cca aca atg tca gga acg tca cgc tct cca cct tca tat tta ttt atc cgt -aa aaa tgt tat cct gga cat cgt aca aat aat a-a aaa gcc ca- -ta tat gtt cgc tat tgt aga aat tgt ttt tca cag ttg ctc aaa aac gat ggc agt gac tta tga gtt acg tta cac ttt gga gtc tca tct tta gta aac ata tca taa tat tcg ata tta cga gtt gac ata tcg aac aaa ttc caa gta ttt gat ttt gga taa tat tcg tat ttt gca tct gct ata att aag ata taa tca ccg caa gaa cac acg aac atc ttt cct aca tgg tta aag tac atg tac aat tct atc cat ttg tct tcc tta act ata tat ttg tat aga taa tta cga gtc tcg tga gta att cca gta att aca tag atg tcg ccg tcg tac tct aca gca taa act ata cta tga tgt cta ggc atg gga gac -tt ttt tat cca acg att ttt agt gaa aca ttc cac atc gtt taa tac tac ata ttt ttc ata cgt ggt ata aac tcc acc cat tac --a tat ata tca tcg ttt acg aat acc gac gcg cct gaa tat cta gga gta att aag ttt gga agt att atc cat ttc gaa gtg ccg tgt ttc aaa tat tct gcc aca ccc gtt gaa ata gaa aat tct aat cct cct att aca tat aac ttt cca tcg tta aca caa gta cta act tct gat ttt aac gac gac ata tta gta acc gtt ttc ca- --- --- --- --- --- --- --t ttt ttc gtt tca aga tct acc cgc -ga tac gga ata aac atg tct att gtt aat cat gcc gcc aat aat gta tag aca att atg taa aac att tgc att ata gaa ttg --- -tc tat ctg tat tac cga cta tcg tcc aat att ctg ttc tag gag agt aat ggg tta ttg tgg ata tat aat cag agt ttt taa tga cta cta tat tat gtt tta tac cat ttc gtg tca ctg gct ttg tag att tgg ata tag tta atc cca aca atg ata tag cat tgc gca tag tat tag tca taa act tgg gat gta aaa tgt tga tga tat cta cat cgt ttg gat ttt tat gta tcc act tta ata ata tca tag ctg taa cat cct cat ga- ttt acg tta acg tct tcg tgg gat aag ata gtt gtc agt tca tcc ttt gat aat ttt cca aat t

>KC813496.1:22501-24465_Cowpox_virus_strain_HumPad07/1_complete_genome

cgg ctt cca tgt att tcc tgg aga gca agt --- aga tga tg- agg aac cag ata gtt tat atc cgt act tgc act taa agt cta cat tgt cgt tgt atg a-- gta tga tct ttt aaa ccc gct aga caa gta tcc gtt tga tat tgt agg atg tgg aca ttt aac aat ctg aca cgt ggg tgg atc gga cca ttc tcc tcc tga aca cag gac acc aga gtt acc aat caa cga ata tcc gct att aca act ata cgt tac aac gct tcc atc ggt ata aaa atc ctc gta tcc gtt atg tct tcc gtt gga tat aga tgg agg tga ttg gca ttt aac aga ttc gca aat agg tgc ctc ggg att cca tac cat aga tcc agt aga tcc taa ttc aca ata cga ttt aga ttc acc gat caa atg ata tcc gct att aca aga gta cgt tat act aga gcc aaa gtc tac tcc gcc aat atc aag ttg gcc att atc gat atc tcg agg cga tgg gca tct ccg ttt aat aca ttg att aaa gag tgt cca tcc agt acc tgt aca ttt agc ata tat agg tcc cat ttt ttg ctt tct gta tcc agg tag aca tag a-t att cta tag tgt ctc cta tgt tgt a-- --- --- --- -at tag cat tag cat cag tc- --t cca cac tat tct taa att tca tat taa tgg gtc gtg acg gaa tag tac agc atg ata gaa cgc atc cta ttc cca aca atg tca gga acg tca cgc tct cca cct tca tat tta ttt atc cgt -aa aaa tgt tat cct gga cat cgt acc aat aat aaa aaa gcc ca- -ta tat gtt cgc tat tgt aga aat tgt ttt tca cag ttg ctc aaa aac gat ggc agt gac tta tga gtt acg tta cac ttt gga gtc tca tct tta gta aac ata tca taa tat tcg ata tta cga gtt gac ata tcg aac aaa ttc caa gta ttt gat ttt gga taa tat tcg tat ttt gca tct gct ata att aag ata taa tcg ccg caa gaa cac acg aac atc ttt cct aca tgg tta aag tac atg tac aat tct atc cat ttg tct tcc tta act ata tat ttg tat aga taa tta cga gtc tca tga gta att cca gta att gca tag atg tca cca tcg tac tct aca gca taa act ata cta tga tgt cta ggc atg gga gac -tt ttt tat cca acg att ttt agt gaa aca ttc cac atc gtt taa tac tac ata ttt ttc ata cgt ggc ata aac tcc acc cat tac --a tat ata tca tcg ttt acg aat acc gat gcg cct gaa tat cta gga gta att aag ttt gga agt ctt atc cat ttc gaa gtg ccg tgt ttc aaa tat tct gcc aca ccc gtt gaa ata gaa aat tct aat cct cct att aca tat aac ttt cca tcg tta aca caa gta cta act tct gat ttt aac gac gac ata tta gta act gtt ttc ca- --- --- --- --- --- --- --t ttt ttc gtt tta aga tct acc cgc -ga tac gga ata aac atg tct att gtt aat cat gcc gcc aat aat gta tag aca att ata taa aac att tgc att ata gaa ttg --- -tc tat ctg tat tac cga cga ttg tcc aat att ctg ttc tag gag agt aat ggg tta ttg tgg ata tat aat cag agt ttt taa tga cta cta tat tat gtt tta tac cat ttc gtg tca ctg gct ttg tag att tgg ata tag tta atc cca aca atg ata tag cat tgc gca tag tat tag tca taa act tgg gat gta aaa tgt tga taa tat cta cat cgt ttg gat ttt tat gta tcc act tta ata ata tca tag ctg taa cat cct cat ga- ttt acg tta acg tct tcg tgg gat aag ata gtt gtc agt tca tcc ttt gat aat ttt cca aat t

>MK239754.1:14218-16181_Buffalopox_virus_isolate_NIV116233

cgg ctt cca tgt att tcc tgg aga gca agt --- aga tga tg- agg aac cag ata gtt tat atc cgt act tgc act taa agt cta cat tgt cgt tgt atg a-- gta tga tct ttt aaa ccc gct aga caa gta tcc gtt tga tat tgt agg atg tgg aca ttt aac aat ctg aca cgt ggg tgg atc gga cca ttc tcc tcc tga aca cag gac act aga gtt acc aat caa cga ata tcc act att gca act ata agt tac aac gct tcc atc ggt ata aaa atc ctc gta tcc gtt atg tct tcc gtt gga tat aga tgg agg gga ttg gca ttt aac aga ttc aca aat agg tgc ctc ggg att cca tac cat aga tcc agt aga tcc taa ttc aca ata cga ttt aga ttc acc gat caa atg ata tcc gct att aca aga gta cgt tat act aga gcc aaa gtc tac tcc acc aat atc aag ttg gcc att atc gat atc tcg agg cga tgg gca tct ccg ttt aat aca ttg att aaa gag tgt cca tcc agt acc tgt aca ttt agc ata tat agg tcc cat ttt ttg ctt tct gta tcc agg tag aca tag a-t att cta tag tgt ctc cta tgt tgt a-- --- --- --- -at tag cat tag cat cag tc- --t cca cac tat tct taa att tca tat taa tgg gtc gtg acg gaa tag tac agc atg ata gaa cgc atc cta ttc cca aca atg tca gga acg tca cgc tct cca cct tca tat tta ttt atc cgt -aa aaa tgt tat cct gga cat cgt aca aat aat a-a aaa gcc ca- -ta tat gtt cgc tat tgt aga aat tgt ttt tca cag ttg ctc aaa aac gat ggc agt gac tta tga gtt acg tta cac ttt gga gtc tca tct tta gta aac ata tca taa tat tcg ata tta cga gtt gac ata tcg aac aaa ttc caa gta ttt gat ttt gga taa tat tcg tat ttt gca tct gct ata att aag ata taa tca ccg caa gaa cac acg aac atc ttt cct aca tgg tta aag tac atg tac aat tct atc cat ttg tct tcc tta act ata tat ttg tat aga taa tta cga gtc tcg tga gta att cca gta att aca tag atg tcg ccg tcg tac tct aca gca taa act ata cta tga tgt cta ggc atg gga gac -tt ttt tat cca acg att ttt agt gaa aca ttc cac atc gtt taa tac tac ata ttt ttc ata cgt ggt ata aac tcc acc cat tac --a tat ata tca tcg ttt acg aat acc gac gcg cct gaa tat cta gga gta att aag ttt gga agt att atc cat ttc gaa gtg ccg tgt ttc aaa tat tct gcc aca ccc gtt gaa ata gaa aat tct aat cct cct att aca tat aac ttt cca tcg tta aca caa gta cta act tct gat ttt aac gac gac ata tta gta acc gtt ttc ca- --- --- --- --- --- --- --t ttt ttc gtt tca aga tct acc cgc -ga tac gga ata aac atg tct att gtt aat cat gcc gcc aat aat gta tag aca att atg taa aac att tgc att ata gaa ttg --- -tc tat ctg tat tac cga cta ttg tcc aat att ctg ttc tag gag agt aat ggg tta ttg tgg ata tat aat cag agt ttt taa tga cta cta tat tat gtt tta tac cat ttc gtg tca ctg gct ttg tag att tgg ata tag tta atc cca aca atg ata tag cat tgc gca tag tat tag tca taa act tgg gat gta aaa tgt tga tga tat cta cat cgt ttg gat ttt tat gta tcc act tta ata ata tca tag ctg taa cat cct cat ga- ttt acg tta acg tct tcg tgg gat aag ata gtt gtc agt tca tcc ttt gat aat ttt cca aat t

>MK239753.1:14030-15993_Buffalopox_virus_isolate_NIV143262

cgg ctt cca tgt att tcc tgg aga gca agt --- aga tga tg- agg aac cag ata gtt tat atc cgt act tgc act taa agt cta cat tgt cgt tgt atg a-- gta tga tct ttt aaa ccc gct aga caa gta tcc gtt tga tat tgt agg atg tgg aca ttt aac aat ctg aca cgt ggg tgg atc gga cca ttc tcc tcc tga aca cag gac act aga gtt acc aat caa cga ata tcc act att gca act ata agt tac aac gct tcc atc ggt ata aaa atc ctc gta tcc gtt atg tct tcc gtt gga tat aga tgg agg gga ttg gca ttt aac aga ttc aca aat agg tgc ctc ggg att cca tac cat aga tcc agt aga tcc taa ttc aca ata cga ttt aga ttc acc gat caa atg ata tcc gct att aca aga gta cgt tat act aga gcc aaa gtc tac tcc acc aat atc aag ttg gcc att atc gat atc tcg agg cga tgg gca tct ccg ttt aat aca ttg att aaa gag tgt cca tcc agt acc tgt aca ttt agc ata tat agg tcc cat ttt ttg ctt tct gta tcc agg tag aca tag a-t att cta tag tgt ctc cta tgt tgt a-- --- --- --- -at tag cat tag cat cag tc- --t cca cac tat tct taa att tca tat taa tgg gtc gtg acg gaa tag tac agc atg ata gaa cgc atc cta ttc cca aca atg tca gga acg tca cgc tct cca cct tca tat tta ttt atc cgt -aa aaa tgt tat cct gga cat cgt aca aat aat a-a aaa gcc ca- -ta tat gtt cgc tat tgt aga aat tgt ttt tca cag ttg ctc aaa aac gat ggc agt gac tta tga gtt acg tta cac ttt gga gtc tca tct tta gta aac ata tca taa tat tcg ata tta cga gtt gac ata tcg aac aaa ttc caa gta ttt gat ttt gga taa tat tcg tat ttt gca tct gct ata att aag ata taa tca ccg caa gaa cac acg aac atc ttt cct aca tgg tta aag tac atg tac aat tct atc cat ttg tct tcc tta act ata tat ttg tat aga taa tta cga gtc tcg tga gta att cca gta att aca tag atg tcg ccg tcg tac tct aca gca taa act ata cta tga tgt cta ggc atg gga gac -tt ttt tat cca acg att ttt agt gaa aca ttc cac atc gtt taa tac tac ata ttt ttc ata cgt ggt ata aac tcc acc cat tac --a tat ata tca tcg ttt acg aat acc gac gcg cct gaa tat cta gga gta att aag ttt gga agt att atc cat ttc gaa gtg ccg tgt ttc aaa tat tct gcc aca ccc gtt gaa ata gaa aat tct aat cct cct att aca tat aac ttt cca tcg tta aca caa gta cta act tct gat ttt aac gac gac ata tta gta acc gtt ttc ca- --- --- --- --- --- --- --t ttt ttc gtt tca aga tct acc cgc -ga tac gga ata aac atg tct att gtt aat cat gcc gcc aat aat gta tag aca att atg taa aac att tgc att ata gaa ttg --- -tc tat ctg tat tac cga cta ttg tcc aat att ctg ttc tag gag agt aat ggg tta ttg tgg ata tat aat cag agt ttt taa tga cta cta tat tat gtt tta tac cat ttc gtg tca ctg gct ttg tag att tgg ata tag tta atc cca aca atg ata tag cat tgc gca tag tat tag tca taa act tgg gat gta aaa tgt tga tga tat cta cat cgt ttg gat ttt tat gta tcc act tta ata ata tca tag ctg taa cat cct cat ga- ttt acg tta acg tct tcg tgg gat aag ata gtt gtc agt tca tcc ttt gat aat ttt cca aat t

>KX781953.1:20766-22729_Vaccinia_virus_strain_L-IVP_clone_14

cgg ctt cca tgt att tcc tgg aga gca agt --- aga tga tg- agg aac cag ata gtt tat atc cgt act tgc act taa agt cta cat tgt cgt tgt atg a-- gta tga tct ttt aaa ccc gct aga caa gta tcc gtt tga tat tgt agg atg tgg aca ttt aac aat ctg aca cgt ggg tgg atc gga cca ttc tcc tcc tga aca cag gac acc aga gtt acc aat caa cga ata tcc act att gca act ata agt tac aac gct ccc atc ggt ata aaa atc ctc gta tcc gtt atg tct tcc gtt gga tat aga tgg agg gga ttg gca ttt aac aga ttc aca aat agg tgc ctc ggg att cca tac cat aga tcc agt aaa tcc taa ttc aca ata cga ttt aga ttc tcc tat caa ctg ata tcc gct att aca aga gta cgt tat act aga gcc aaa gtc tac tcc acc aat atc aag ttg gcc att atc gat atc tcg agg cga tgg gca tct ccg ttt aat aca ttg att aaa gag tgt cca tcc agt acc tgt aca ttt agc ata tat agg tcc cat ttt ttg ctt tct gta tcc agg tag aca tag a-t att cta tag tgt ctc cta tgt tgt a-- --- --- --- -at tag cat tag cat cag tc- --t cca cac tat tct taa att tca tat taa tgg gtc gtg acg gaa tag tac agc atg ata gaa cgc atc cta ttc cca aca atg tca gga acg tca cgc tct cca cct tca tat tta ttt atc cgt -aa aaa tgt tat cct gga cat cgt aca aat aat a-a aaa gcc ca- -ta tat gtt cgc tat tgt aga aat tgt ttt tca cag ttg ctc aaa aac gat ggc agt gac tta tga gtt acg tta cac ttt gga gtc tca tct tta gta aac ata tca taa tat tcg ata tta cga gtt gac ata tcg aac aaa ttc caa gta ttt gat ttt gga taa tat tcg tat ttt gca tct gct ata att aag ata taa tca ccg caa gaa cac acg aac atc ttt cct aca tgg tta aag tac atg tac aat tct atc cat ttg tct tcc tta act ata tat ttg tat aga taa tta cga gtc tcg tga gta att cca gta att aca tag atg tcg ccg tcg tac tct aca gca taa act ata cta tga tgt cta ggc atg gga gac -tt ttt tat cca acg att ttt agt gaa aca ttc cac atc gtt taa tac tac ata ttt ttc ata cgt ggt ata aac tcc acc tat tac --a tat ata tca tcg ttt acg aat acc gac gcg cct gaa tat cta gga gta att aag ttt gga agt ctt atc cat ttc gaa gtg ccg tgt ttc aaa tat tct gcc aca ccc gtt gaa ata gaa aat tct aat cct cct att aca tat aac ttt cca tcg tta aca caa gta cta act tct gat ttt aac gac gac ata tta gta acc gtt ttc ca- --- --- --- --- --- --- --t ttt ttc gtt tca aga tct acc cgc -ga tac gga ata aac atg tct att gtt aat cat gcc gcc aat aat gta tag aca att atg taa aac att tgc att ata gaa ttg --- -tc tat ctg tat tac cga cta tcg tcc aat att ctg ttc tag gag agt aat ggg tta ttg tgg ata tat aat cag agt ttt taa tga cta cta tat tat gtt tta tac cat ttc gtg tca ctg gct ttg tag att tgg ata tag tta atc cca aca atg ata tag cat tgc gca tag tat tag tca taa act tgg gat gta aaa tgt tga tga tat cta cat cgt ttg gat ttt tat gta tcc act tta ata ata tca tag ctg taa cat cct cat ga- ttt acg tta acg tct tcg tgg gat aag ata gtt gtc agt tca tcc ttt gat aat ttt cca aat t

>KX061501.1:20584-22547_Vaccinia_virus_strain_Lister_clone_Lister_Butantan

cgg ctt cca tgt att tcc tgg aga gca agt --- aga tga tg- agg aac cag ata gtt tat atc cgt act tgc act taa agt cta cat tgt cgt tgt atg a-- gta tga tct ttt aaa ccc gct aga caa gta tcc gtt tga tat tgt agg atg tgg aca ttt aac aat ctg aca cgt ggg tgg atc gga cca ttc tcc tcc tga aca cag gac act aga gtt acc aat caa cga ata tcc act att gca act ata agt tac aac gct ccc atc ggt ata aaa atc ctc gta tcc gtt atg tct tcc gtt gga tat aga tgg agg gga ttg gca ttt aac aga ttc aca aat agg tgc ctc ggg att cca tac cat aga tcc agt aga tcc taa ttc aca ata cga ttt aga ttc tcc tat caa ctg ata tcc gct att aca aga gta cgt tat act aga gcc aaa gtc tac tcc acc aat atc aag ttg gcc att atc gat atc tcg agg cga tgg gca tct ccg ttt aat aca ttg att aaa gag tgt cca tcc agt act tgt aca ttt agc ata tat agg tcc cat ttt ttg ctt tct gta tcc agg tag aca tag a-t att cta tag tgt ctc cta tgt tgt a-- --- --- --- -at tag cat tag cat cag tc- --t cca cac tat tct taa att tca tat taa tgg gtc gtg acg gaa tag tac agc atg ata gaa cgc atc cta ttc cca aca atg tca gga acg tca cgc tct cca cct tca tat tta ttt atc cgt -aa aaa tgt tat cct gga cat cgt aca aat aat a-a aaa gcc ca- -ta tat gtt cgc tat tgt aga aat tgt ttt tca cag ttg ctc aaa aac gat ggc agt gac tta tga gtt acg tta cac ttt gga gtc tca tct tta gta aac ata tca taa tat tcg ata tta cga gtt gac ata tcg aac aaa ttc caa gta ttt gat ttt gga taa tat tcg tat ttt gca tct gct ata att aag ata taa tca ccg caa gaa cac acg aac atc ttt cct aca tgg tta aag tac atg tac aat tct atc cat ttg tct tcc tta act ata tat ttg tat aga taa tta cga gtc tcg tga gta att cca gta att aca tag atg ttg ccg tcg tac tct aca gca taa act ata cta tga tgt cta ggc atg gga gac -tt ttt tat cca acg att ttt agt gaa aca ttc cac atc gtt taa tac tac ata ttt ctc ata cgt ggt ata aac tcc acc cat tac --a tat ata tca tcg ttt acg aat acc gac gcg cct gaa tat cta gga gta att aag ttt gga agt ctt atc cat ttc gaa gtg ccg tgt ttc aaa tat tct gcc aca ccc gtt gaa ata gaa aat tct aat cct cct att aca tat aac ttt cca tcg tta aca caa gta cta act tct gat ttt aac gac gac ata tta gta acc gtt ttc ca- --- --- --- --- --- --- --t ttt ttc gtt tca aga tct acc cgc -ga tac gga ata aac atg tct att gtt aat cat gcc gcc aat aat gta tag aca att atg taa aac att tgc att ata gaa ttg --- -tc tat ctg tat tac cga cta tcg tcc aat att ctg ttc tag gag agt aat ggg tta ttg tgg ata tat aat cag agt ttt taa tga cta cta tat tat gtt tta tac cat ttc gtg tca ctg gct ttg tag att tgg ata tag tta atc cca aca atg ata tag cat tgc gca tag tat tag tca taa act tgg gat gta aaa tgt tga tga tat cta cat cgt ttg gat ttt tat gta tcc act tta ata ata tca tag ctg taa cat cct cat ga- ttt acg tta acg tct tcg tgg gat aag ata gtt gtc agt tca tcc ttt gat aat ttt cca aat t

>KF866253.1:20550-22513_Vaccinia_virus_WAU86/88-1_complete_genome

cgg ctt cca tgt att tcc tgg aga gca agt --- aga tga tg- agg aac cag ata gtt tat atc cgt act tgc act taa agt cta cat tgt cgt tgt atg a-- gta tga tct ttt aaa ccc gct aga caa gta tcc gtt tga tat tgt agg atg tgg aca ttt aac aat ttg aca cgt ggg tgg atc gga cca ttc tcc tcc tga aca cag gac act aga gtt acc aat caa cga ata tcc act att gca act ata agt tac aac gct ccc atc ggt ata aaa atc ctc gta tcc gtt atg tct tcc gtt gga tat aga tgg agg gga ttg gca ttt aac aga ttc aca aat agg tgc ctc ggg att cca tac cat aga tcc agt aga tcc taa ttc aca ata cga ttt aga ttc tcc tat caa ctg ata tcc gct att aca aga gta cgt tat act aga gcc aaa gtc tac tcc acc aat atc aag ttg gcc att atc gat atc tcg agg cga tgg gca tct ccg ttt aat aca ttg att aaa gag tgt cca tcc agt acc tgt aca ttt agc ata tat agg tcc cat ttt ttg ctt tct gta tcc agg tag aca tag a-t att cta tag tgt ctc cta tgt tgt a-- --- --- --- -at tag cat tag cat cag tc- --t cca cac tat tct taa att tca tat taa tgg gtc gtg acg gaa tag tac agc atg ata gaa cgc atc cta ttc cca aca atg tca gga acg tca cgc tct cca cct tca tat tta ttt atc cgt -aa aaa tgt tat cct gga cat cgt aca aat aat a-a aaa gcc ca- -ta tat gtt cgc tat tgt aga aat tgt ttt tca cag ttg ctc aaa aac gat ggc agt gac tta tga gtt acg tta cac ttt gga gtc tca tct tta gta aac ata tca taa tat tcg ata tta cga gtt gac ata tcg aac aaa ttc caa gta ttt gat ttt gga taa tat tcg tat ttt gca tct gct ata att aag ata taa tca ccg caa gaa cac acg aac atc ttt cct aca tgg tta aag tac atg tac aat tct atc cat ttg tct tcc tta act ata tat ttg tat aga taa tta cga gtc tcg tga gta att cca gta att aca tag atg tcg ccg tcg tac tct aca gca taa act ata cta tga tgt cta ggc atg gga gac -tt ttt tat cca acg att ttt agt gaa aca ttc cac atc gtt taa tac tac ata ttt ttc ata cgt ggt ata aac tcc acc cat tac --a tat ata tca tcg ttt acg aat acc gac gcg cct gaa tat cta gga gta att aag ttt gga agt ctt atc cat ttc gaa gtg ccg tgt ttc aaa tat tct gcc aca ccc gtt gaa ata gaa aat tct aat cct cct att aca tat aac ttt cca tcg tta aca caa gta cta act tct gat ttt aac gac gac ata tta gta acc gtt ttc ca- --- --- --- --- --- --- --t ttt ttc gtt tca aga tct acc cgc -ga tac gga ata aac atg tct att gtt aat cat gcc gcc aat aat gta tag aca att atg taa aac att tgc att ata gaa ttg --- -tc tat ctg tat tac cga cta tcg tcc aat att ctg ttc tag gag agt aat ggg tta ttg tgg ata tat aat cag agt ttt taa tga cta cta tat tat gtt tta tac cat ttc gtg tca ctg gct ttg tag att tgg ata tag tta atc cca aca atg ata tag cat tgc gca tag tat tag tca taa act tgg gat gta aaa tgt tga tga tat cta cat cgt ttg gat ttt tat gta tcc act tta ata ata tca tag ctg taa cat cct cat ga- ttt acg tta acg tct tcg tgg gat aag ata gtt gtc agt tca tcc ttt gat aat ttt cca aat t

>KC813504.1:27673-29642_Cowpox_virus_strain_RatHei09/1_complete_genome

cgg ctt cca tgt att tcc tgg aga gca agt --- aga tga tg- agg aac cag ata gtt tat atc cgt act tgc act taa agt cta cat tgt cgt tgt atg a-- gta tga tct ttt aaa ccc gct aga caa gta tcc gtt tga tat tgt agg atg tgg aca ttt aac aat ctg aca cgt ggg tgg atc gga cca ttc tcc tcc tga aca cag gac acc aga gtt acc aat caa cga ata tcc act att gca act ata cgt tac aac gct tcc atc ggt ata aaa atc ctc gta tcc gtt atg tct tcc gtt gga tat aga tgg agg tga ttg gca ttt aac aga ttc gca aat agg tgc ctc ggg att cca tac cat aga tcc agt aga tcc taa ttc gca ata cga ttt aga ttc acc gat caa atg ata tcc gct att aca aga gta cgt tat act aga gcc aaa gtc tac tcc gcc aat atc aag ttg gcc att atc gat atc tcg agg cga tgg gca tct ccg ttt aat aca ttg att aaa gag tgt cca tcc agt acc tgt aca ttt agc ata tat agg tcc cat ttt ttg ctt tct gta tcc agg tag aca tag a-t att cta tag tgt ctc cta tgt tgt aat tag c-- --- -at tag cat tag cat cag tc- --t cca cac tat tct taa att tca tat taa tgg gtc gtg acg gaa tag tac agc atg ata gaa cgc atc cta ttc cca aca atg tca gga acg tca cgc tct cca cct tca tat tta ttt atc cgt -aa aaa tgt tat cct gga cat cgt aca aat aat a-a aaa gcc ca- -ta tat gtt cgc tat tgt aga aat tgt ttt tca cag ttg ctc aaa aac gat ggc agt gac tta tga gtt atg tta cac ttt gga gtc tca tct tta gta aac ata tca taa tat tcg ata tta cga gtt gac ata tcg aac aaa ttc caa gta ttt gat ttt gga taa tat tcg tat ttt gca tct gct ata att aag ata taa tca ccg caa gaa cac acg aac atc ttt cct aca tgg tta aag tac atg tac aat tct atc cat ttg tct tcc tta act ata tat ttg tat aga taa tta cga gtc tca tga gta att cca gta att gca tag atg tca cca tcg tac tct aca gca taa act ata cta tga tgt cta ggc atg gga gac -tt ttt tat cca acg att ttt agt gaa aca ttc cac atc gtt taa tac tac ata ttt ttc ata cgt ggt ata aac tcc acc cat tac --a tat ata tca tcg ttt acg aat acc gac gcg cct gaa tat cta gga gta att aag ttt gga agt ctt atc cat ttc gaa gtg ccg tgt ttc aaa tat tct gct acg cct gtt gaa ata gaa aat tct aat cct cct att aca tat aac ttt cca tca tta aca caa gta cta act tct gat ttt aac gac gac ata tta gta act gtt ttc ca- --- --- --- --- --- --- --t ttt ttc gtt tta aga tct acc cgc -ga tac gga ata aac atg tct att gtt aat cat gcc gcc aat aat gta tag aca att ata taa aac att tgc att ata gaa ttg --- -tc tat ctg tat tac cga cga ttg tcc aat att ctg ttc tag gag agt aat ggg tta ttg tgg ata tat aat cag agt ttt taa tga cta cta tat tat gtt tta tac cat ttc gtg tca ctg gct ttg tag att tgg ata tag tta atc cca aca atg ata tag cat tgc gca tag tat tag tca taa act tgg gat gta aaa tgt tga tga tat cta cat cgt ttg gat ttt tat gta tcc act tta ata ata tca tag ctg taa cat cct cat ga- ttt acg tta acg tct tcg tgg gat aag ata gtt gtc agt tca tcc ttt gat aat ttt cca aat t

>KC813500.1:16754-18723_Cowpox_virus_strain_MonKre08/4_complete_genome

cgg ctt cca tgt att tcc tgg aga gca agt --- aga tga tg- agg aac cag ata gtt tat atc cgt act tgc act taa agt cta cat tgt cgt tgt atg a-- gta tga tct ttt aaa ccc gct aga caa gta tcc gtt tga tat tgt agg atg tgg aca ttt aac aat ctg aca cgt ggg tgg atc gga cca ttc tcc tcc tga aca cag gac acc aga gtt acc aat caa cga ata tcc act att gca act ata cgt tac aac gct tcc atc ggt ata aaa atc ctc gta tcc gtt atg tct tcc gtt gga tat aga tgg agg tga ttg gca ttt aac aga ttc gca aat agg tgc ctc ggg att cca tac cat aga tcc agt aga tcc taa ttc gca ata cga ttt aga ttc acc gat caa atg ata tcc gct att aca aga gta cgt tat act aga gcc aaa gtc tac tcc gcc aat atc aag ttg gcc att atc gat atc tcg agg cga tgg gca tct ccg ttt aat aca ttg att aaa gag tgt cca tcc agt acc tgt aca ttt agc ata tat agg tcc cat ttt ttg ctt tct gta tcc agg tag aca tag a-t att cta tag tgt ctc cta tgt tgt aat tag c-- --- -at tag cat tag cat cag tc- --t cca cac tat tct taa att tca tat taa tgg gtc gtg acg gaa tag tac agc atg ata gaa cgc atc cta ttc cca aca atg tca gga acg tca cgc tct cca cct tca tat tta ttt atc cgt -aa aaa tgt tat cct gga cat cgt aca aat aat a-a aaa gcc ca- -ta tat gtt cgc tat tgt aga aat tgt ttt tca cag ttg ctc aaa aac gat ggc agt gac tta tga gtt atg tta cac ttt gga gtc tca tct tta gta aac ata tca taa tat tcg ata tta cga gtt gac ata tcg aac aaa ttc caa gta ttt gat ttt gga taa tat tcg tat ttt gca tct gct ata att aag ata taa tca ccg caa gaa cac acg aac atc ttt cct aca tgg tta aag tac atg tac aat tct atc cat ttg tct tcc tta act ata tat ttg tat aga taa tta cga gtc tca tga gta att cca gta att gca tag atg tca cca tcg tac tct aca gca taa act ata cta tga tgt cta ggc atg gga gac -tt ttt tat cca acg att ttt agt gaa aca ttc cac atc gtt taa tac tac ata ttt ttc ata cgt ggt ata aac tcc acc cat tac --a tat ata tca tcg ttt acg aat acc gac gcg cct gaa tat cta gga gta att aag ttt gga agt ctt atc cat ttc gaa gtg ccg tgt ttc aaa tat tct gct acg cct gtt gaa ata gaa aat tct aat cct cct att aca tat aac ttt cca tca tta aca caa gta cta act tct gat ttt aac gac gac ata tta gta act gtt ttc ca- --- --- --- --- --- --- --t ttt ttc gtt tta aga tct acc cgc -ga tac gga ata aac atg tct att gtt aat cat gcc gcc aat aat gta tag aca att ata taa aac att tgc att ata gaa ttg --- -tc tat ctg tat tac cga cga ttg tcc aat att ctg ttc tag gag agt aat ggg tta ttg tgg ata tat aat cag agt ttt taa tga cta cta tat tat gtt tta tac cat ttc gtg tca ctg gct ttg tag att tgg ata tag tta atc cca aca atg ata tag cat tgc gca tag tat tag tca taa act tgg gat gta aaa tgt tga tga tat cta cat cgt ttg gat ttt tat gta tcc act tta ata ata tca tag ctg taa cat cct cat ga- ttt acg tta acg tct tcg tgg gat aag ata gtt gtc agt tca tcc ttt gat aat ttt cca aat t

>KC813498.1:35726-37695_Cowpox_virus_strain_JagKre08/2_complete_genome

cgg ctt cca tgt att tcc tgg aga gca agt --- aga tga tg- agg aac cag ata gtt tat atc cgt act tgc act taa agt cta cat tgt cgt tgt atg a-- gta tga tct ttt aaa ccc gct aga caa gta tcc gtt tga tat tgt agg atg tgg aca ttt aac aat ctg aca cgt ggg tgg atc gga cca ttc tcc tcc tga aca cag gac acc aga gtt acc aat caa cga ata tcc act att gca act ata cgt tac aac gct tcc atc ggt ata aaa atc ctc gta tcc gtt atg tct tcc gtt gga tat aga tgg agg tga ttg gca ttt aac aga ttc gca aat agg tgc ctc ggg att cca tac cat aga tcc agt aga tcc taa ttc gca ata cga ttt aga ttc acc gat caa atg ata tcc gct att aca aga gta cgt tat act aga gcc aaa gtc tac tcc gcc aat atc aag ttg gcc att atc gat atc tcg agg cga tgg gca tct ccg ttt aat aca ttg att aaa gag tgt cca tcc agt acc tgt aca ttt agc ata tat agg tcc cat ttt ttg ctt tct gta tcc agg tag aca tag a-t att cta tag tgt ctc cta tgt tgt aat tag c-- --- -at tag cat tag cat cag tc- --t cca cac tat tct taa att tca tat taa tgg gtc gtg acg gaa tag tac agc atg ata gaa cgc atc cta ttc cca aca atg tca gga acg tca cgc tct cca cct tca tat tta ttt atc cgt -aa aaa tgt tat cct gga cat cgt aca aat aat a-a aaa gcc ca- -ta tat gtt cgc tat tgt aga aat tgt ttt tca cag ttg ctc aaa aac gat ggc agt gac tta tga gtt atg tta cac ttt gga gtc tca tct tta gta aac ata tca taa tat tcg ata tta cga gtt gac ata tcg aac aaa ttc caa gta ttt gat ttt gga taa tat tcg tat ttt gca tct gct ata att aag ata taa tca ccg caa gaa cac acg aac atc ttt cct aca tgg tta aag tac atg tac aat tct atc cat ttg tct tcc tta act ata tat ttg tat aga taa tta cga gtc tca tga gta att cca gta att gca tag atg tca cca tcg tac tct aca gca taa act ata cta tga tgt cta ggc atg gga gac -tt ttt tat cca acg att ttt agt gaa aca ttc cac atc gtt taa tac tac ata ttt ttc ata cgt ggt ata aac tcc acc cat tac --a tat ata tca tcg ttt acg aat acc gac gcg cct gaa tat cta gga gta att aag ttt gga agt ctt atc cat ttc gaa gtg ccg tgt ttc aaa tat tct gct acg cct gtt gaa ata gaa aat tct aat cct cct att aca tat aac ttt cca tca tta aca caa gta cta act tct gat ttt aac gac gac ata tta gta act gtt ttc ca- --- --- --- --- --- --- --t ttt ttc gtt tta aga tct acc cgc -ga tac gga ata aac atg tct att gtt aat cat gcc gcc aat aat gta tag aca att ata taa aac att tgc att ata gaa ttg --- -tc tat ctg tat tac cga cga ttg tcc aat att ctg ttc tag gag agt aat ggg tta ttg tgg ata tat aat cag agt ttt taa tga cta cta tat tat gtt tta tac cat ttc gtg tca ctg gct ttg tag att tgg ata tag tta atc cca aca atg ata tag cat tgc gca tag tat tag tca taa act tgg gat gta aaa tgt tga tga tat cta cat cgt ttg gat ttt tat gta tcc act tta ata ata tca tag ctg taa cat cct cat ga- ttt acg tta acg tct tcg tgg gat aag ata gtt gtc agt tca tcc ttt gat aat ttt cca aat t

>KC813497.1:35408-37377_Cowpox_virus_strain_JagKre08/1_complete_genome

cgg ctt cca tgt att tcc tgg aga gca agt --- aga tga tg- agg aac cag ata gtt tat atc cgt act tgc act taa agt cta cat tgt cgt tgt atg a-- gta tga tct ttt aaa ccc gct aga caa gta tcc gtt tga tat tgt agg atg tgg aca ttt aac aat ctg aca cgt ggg tgg atc gga cca ttc tcc tcc tga aca cag gac acc aga gtt acc aat caa cga ata tcc act att gca act ata cgt tac aac gct tcc atc ggt ata aaa atc ctc gta tcc gtt atg tct tcc gtt gga tat aga tgg agg tga ttg gca ttt aac aga ttc gca aat agg tgc ctc ggg att cca tac cat aga tcc agt aga tcc taa ttc gca ata cga ttt aga ttc acc gat caa atg ata tcc gct att aca aga gta cgt tat act aga gcc aaa gtc tac tcc gcc aat atc aag ttg gcc att atc gat atc tcg agg cga tgg gca tct ccg ttt aat aca ttg att aaa gag tgt cca tcc agt acc tgt aca ttt agc ata tat agg tcc cat ttt ttg ctt tct gta tcc agg tag aca tag a-t att cta tag tgt ctc cta tgt tgt aat tag c-- --- -at tag cat tag cat cag tc- --t cca cac tat tct taa att tca tat taa tgg gtc gtg acg gaa tag tac agc atg ata gaa cgc atc cta ttc cca aca atg tca gga acg tca cgc tct cca cct tca tat tta ttt atc cgt -aa aaa tgt tat cct gga cat cgt aca aat aat a-a aaa gcc ca- -ta tat gtt cgc tat tgt aga aat tgt ttt tca cag ttg ctc aaa aac gat ggc agt gac tta tga gtt atg tta cac ttt gga gtc tca tct tta gta aac ata tca taa tat tcg ata tta cga gtt gac ata tcg aac aaa ttc caa gta ttt gat ttt gga taa tat tcg tat ttt gca tct gct ata att aag ata taa tca ccg caa gaa cac acg aac atc ttt cct aca tgg tta aag tac atg tac aat tct atc cat ttg tct tcc tta act ata tat ttg tat aga taa tta cga gtc tca tga gta att cca gta att gca tag atg tca cca tcg tac tct aca gca taa act ata cta tga tgt cta ggc atg gga gac -tt ttt tat cca acg att ttt agt gaa aca ttc cac atc gtt taa tac tac ata ttt ttc ata cgt ggt ata aac tcc acc cat tac --a tat ata tca tcg ttt acg aat acc gac gcg cct gaa tat cta gga gta att aag ttt gga agt ctt atc cat ttc gaa gtg ccg tgt ttc aaa tat tct gct acg cct gtt gaa ata gaa aat tct aat cct cct att aca tat aac ttt cca tca tta aca caa gta cta act tct gat ttt aac gac gac ata tta gta act gtt ttc ca- --- --- --- --- --- --- --t ttt ttc gtt tta aga tct acc cgc -ga tac gga ata aac atg tct att gtt aat cat gcc gcc aat aat gta tag aca att ata taa aac att tgc att ata gaa ttg --- -tc tat ctg tat tac cga cga ttg tcc aat att ctg ttc tag gag agt aat ggg tta ttg tgg ata tat aat cag agt ttt taa tga cta cta tat tat gtt tta tac cat ttc gtg tca ctg gct ttg tag att tgg ata tag tta atc cca aca atg ata tag cat tgc gca tag tat tag tca taa act tgg gat gta aaa tgt tga tga tat cta cat cgt ttg gat ttt tat gta tcc act tta ata ata tca tag ctg taa cat cct cat ga- ttt acg tta acg tct tcg tgg gat aag ata gtt gtc agt tca tcc ttt gat aat ttt cca aat t

>KC813492.1:27529-29498_Cowpox_virus_strain_HumLan08/1_complete_genome

cgg ctt cca tgt att tcc tgg aga gca agt --- aga tga tg- agg aac cag ata gtt tat atc cgt act tgc act taa agt cta cat tgt cgt tgt atg a-- gta tga tct ttt aaa ccc gct aga caa gta tcc gtt tga tat tgt agg atg tgg aca ttt aac aat ctg aca cgt ggg tgg atc gga cca ttc tcc tcc tga aca cag gac acc aga gtt acc aat caa cga ata tcc act att gca act ata cgt tac aac gct tcc atc ggt ata aaa atc ctc gta tcc gtt atg tct tcc gtt gga tat aga tgg agg tga ttg gca ttt aac aga ttc gca aat agg tgc ctc ggg att cca tac cat aga tcc agt aga tcc taa ttc gca ata cga ttt aga ttc acc gat caa atg ata tcc gct att aca aga gta cgt tat act aga gcc aaa gtc tac tcc gcc aat atc aag ttg gcc att atc gat atc tcg agg cga tgg gca tct ccg ttt aat aca ttg att aaa gag tgt cca tcc agt acc tgt aca ttt agc ata tat agg tcc cat ttt ttg ctt tct gta tcc agg tag aca tag a-t att cta tag tgt ctc cta tgt tgt aat tag c-- --- -at tag cat tag cat cag tc- --t cca cac tat tct taa att tca tat taa tgg gtc gtg acg gaa tag tac agc atg ata gaa cgc atc cta ttc cca aca atg tca gga acg tca cgc tct cca cct tca tat tta ttt atc cgt -aa aaa tgt tat cct gga cat cgt aca aat aat a-a aaa gcc ca- -ta tat gtt cgc tat tgt aga aat tgt ttt tca cag ttg ctc aaa aac gat ggc agt gac tta tga gtt atg tta cac ttt gga gtc tca tct tta gta aac ata tca taa tat tcg ata tta cga gtt gac ata tcg aac aaa ttc caa gta ttt gat ttt gga taa tat tcg tat ttt gca tct gct ata att aag ata taa tca ccg caa gaa cac acg aac atc ttt cct aca tgg tta aag tac atg tac aat tct atc cat ttg tct tcc tta act ata tat ttg tat aga taa tta cga gtc tca tga gta att cca gta att gca tag atg tca cca tcg tac tct aca gca taa act ata cta tga tgt cta ggc atg gga gac -tt ttt tat cca acg att ttt agt gaa aca ttc cac atc gtt taa tac tac ata ttt ttc ata cgt ggt ata aac tcc acc cat tac --a tat ata tca tcg ttt acg aat acc gac gcg cct gaa tat cta gga gta att aag ttt gga agt ctt atc cat ttc gaa gtg ccg tgt ttc aaa tat tct gct acg cct gtt gaa ata gaa aat tct aat cct cct att aca tat aac ttt cca tca tta aca caa gta cta act tct gat ttt aac gac gac ata tta gta act gtt ttc ca- --- --- --- --- --- --- --t ttt ttc gtt tta aga tct acc cgc -ga tac gga ata aac atg tct att gtt aat cat gcc gcc aat aat gta tag aca att ata taa aac att tgc att ata gaa ttg --- -tc tat ctg tat tac cga cga ttg tcc aat att ctg ttc tag gag agt aat ggg tta ttg tgg ata tat aat cag agt ttt taa tga cta cta tat tat gtt tta tac cat ttc gtg tca ctg gct ttg tag att tgg ata tag tta atc cca aca atg ata tag cat tgc gca tag tat tag tca taa act tgg gat gta aaa tgt tga tga tat cta cat cgt ttg gat ttt tat gta tcc act tta ata ata tca tag ctg taa cat cct cat ga- ttt acg tta acg tct tcg tgg gat aag ata gtt gtc agt tca tcc ttt gat aat ttt cca aat t

>MH341447.1:20976-22939_Vaccinia_virus_strain_L-IVP_oncoQ

cgg ctt cca tgt att tcc tgg aga gca agt --- aga tga tg- agg aac cag ata gtt tat atc cgt act tgc act taa agt cta cat tgt cgt tgt atg a-- gta tga tct ttt aaa ccc gct aga caa gta tcc gtt tga tat tgt agg atg tgg aca ttt aac aat ctg aca cgt ggg tgg atc gga cca ttc tcc tcc tga aca cag gac act aga gtt acc aat caa cga ata tcc act att gca act ata agt tac aac gct ccc atc ggt ata aaa atc ctc gta tcc gtt atg tct tcc gtt gga tat aga tgg agg gga ttg gca ttt aac aga ttc aca aat agg tgc ctc ggg att cca tac cat aga tcc agt aaa tcc taa ttc aca ata cga ttt aga ttc tcc tat caa ctg ata tcc gct att aca aga gta cgt tat act aga gcc aaa gtc tac tcc acc aat atc aag ttg gcc att atc gat atc tcg agg cga tgg gca tct ccg ttt aat aca ttg att aaa gag tgt cca tcc agt acc tgt aca ttt agc ata tat agg tcc cat ttt ttg ctt tct gta tcc agg tag aca tag a-t att cta tag tgt ctc cta tgt tgt a-- --- --- --- -at tag cat tag cat cag tc- --t cca cac tat tct taa att tca tat taa tgg gtc gtg acg gaa tag tac agc atg ata gaa cgc atc cta ttc cca aca atg tca gga acg tca cgc tct cca cct tca tat tta ttt atc cgt -aa aaa tgt tat cct gga cat cgt aca aat aat a-a aaa gcc ca- -ta tat gtt cgc tat tgt aga aat tgt ttt tca cag ttg ctc aaa aac gat ggc agt gac tta tga gtt acg tta cac ttt gga gtc tca tct tta gta aac ata tca taa tat tcg ata tta cga gtt gac ata tcg aac aaa ttc caa gta ttt gat ttt gga taa tat tcg tat ttt gca tct gct ata att aag ata taa tca ccg caa gaa cac acg aac atc ttt cct aca tgg tta aag tac atg tac aat tct atc cat ttg tct tcc tta act ata tat ttg tat aga taa tta cga gtc tcg tga gta att cca gta att aca tag atg tcg ccg tcg tac tct aca gca taa act ata cta tga tgt cta ggc atg gga gac -tt ttt tat cca acg att ttt agt gaa aca ttc cac atc gtt taa tac tac ata ttt ttc ata cgt ggt ata aac tcc acc tat tac --a tat ata tca tcg ttt acg aat acc gac gcg cct gaa tat cta gga gta att aag ttt gga agt ctt atc cat ttc gaa gtg ccg tgt ttc aaa tat tct gcc aca ccc gtt gaa ata gaa aat tct aat cct cct att aca tat aac ttt cca tcg tta aca caa gta cta act tct gat ttt aac gac gac ata tta gta acc gtt ttc ca- --- --- --- --- --- --- --t ttt ttc gtt tca aga tct acc cgc -ga tac gga ata aac atg tct att gtt aat cat gcc gcc aat aat gta tag aca att atg taa aac att tgc att ata gaa ttg --- -tc tat ctg tat tac cga cta tcg tcc aat att ctg ttc tag gag agt aat ggg tta ttg tgg ata tat aat cag agt ttt taa tga cta cta tat tat gtt tta tac cat ttc gtg tca ctg gct ttg tag att tgg ata tag tta atc cca aca atg ata tag cat tgc gca tag tat tag tca taa act tgg gat gta aaa tgt tga tga tat cta cat cgt ttg gat ttt tat gta tcc act tta ata ata tca tag ctg taa cat cct cat ga- ttt acg tta acg tct tcg tgg gat aag ata gtt gtc agt tca tcc ttt gat aat ttt cca aat t

>MH341446.1:20842-22805_Vaccinia_virus_strain_L-IVP_oncoM

cgg ctt cca tgt att tcc tgg aga gca agt --- aga tga tg- agg aac cag ata gtt tat atc cgt act tgc act taa agt cta cat tgt cgt tgt atg a-- gta tga tct ttt aaa ccc gct aga caa gta tcc gtt tga tat tgt agg atg tgg aca ttt aac aat ctg aca cgt ggg tgg atc gga cca ttc tcc tcc tga aca cag gac act aga gtt acc aat caa cga ata tcc act att gca act ata agt tac aac gct ccc atc ggt ata aaa atc ctc gta tcc gtt atg tct tcc gtt gga tat aga tgg agg gga ttg gca ttt aac aga ttc aca aat agg tgc ctc ggg att cca tac cat aga tcc agt aaa tcc taa ttc aca ata cga ttt aga ttc tcc tat caa ctg ata tcc gct att aca aga gta cgt tat act aga gcc aaa gtc tac tcc acc aat atc aag ttg gcc att atc gat atc tcg agg cga tgg gca tct ccg ttt aat aca ttg att aaa gag tgt cca tcc agt acc tgt aca ttt agc ata tat agg tcc cat ttt ttg ctt tct gta tcc agg tag aca tag a-t att cta tag tgt ctc cta tgt tgt a-- --- --- --- -at tag cat tag cat cag tc- --t cca cac tat tct taa att tca tat taa tgg gtc gtg acg gaa tag tac agc atg ata gaa cgc atc cta ttc cca aca atg tca gga acg tca cgc tct cca cct tca tat tta ttt atc cgt -aa aaa tgt tat cct gga cat cgt aca aat aat a-a aaa gcc ca- -ta tat gtt cgc tat tgt aga aat tgt ttt tca cag ttg ctc aaa aac gat ggc agt gac tta tga gtt acg tta cac ttt gga gtc tca tct tta gta aac ata tca taa tat tcg ata tta cga gtt gac ata tcg aac aaa ttc caa gta ttt gat ttt gga taa tat tcg tat ttt gca tct gct ata att aag ata taa tca ccg caa gaa cac acg aac atc ttt cct aca tgg tta aag tac atg tac aat tct atc cat ttg tct tcc tta act ata tat ttg tat aga taa tta cga gtc tcg tga gta att cca gta att aca tag atg tcg ccg tcg tac tct aca gca taa act ata cta tga tgt cta ggc atg gga gac -tt ttt tat cca acg att ttt agt gaa aca ttc cac atc gtt taa tac tac ata ttt ttc ata cgt ggt ata aac tcc acc tat tac --a tat ata tca tcg ttt acg aat acc gac gcg cct gaa tat cta gga gta att aag ttt gga agt ctt atc cat ttc gaa gtg ccg tgt ttc aaa tat tct gcc aca ccc gtt gaa ata gaa aat tct aat cct cct att aca tat aac ttt cca tcg tta aca caa gta cta act tct gat ttt aac gac gac ata tta gta acc gtt ttc ca- --- --- --- --- --- --- --t ttt ttc gtt tca aga tct acc cgc -ga tac gga ata aac atg tct att gtt aat cat gcc gcc aat aat gta tag aca att atg taa aac att tgc att ata gaa ttg --- -tc tat ctg tat tac cga cta tcg tcc aat att ctg ttc tag gag agt aat ggg tta ttg tgg ata tat aat cag agt ttt taa tga cta cta tat tat gtt tta tac cat ttc gtg tca ctg gct ttg tag att tgg ata tag tta atc cca aca atg ata tag cat tgc gca tag tat tag tca taa act tgg gat gta aaa tgt tga tga tat cta cat cgt ttg gat ttt tat gta tcc act tta ata ata tca tag ctg taa cat cct cat ga- ttt acg tta acg tct tcg tgg gat aag ata gtt gtc agt tca tcc ttt gat aat ttt cca aat t

>MH341445.1:20782-22745_Vaccinia_virus_strain_L-IVP_oncoB

cgg ctt cca tgt att tcc tgg aga gca agt --- aga tga tg- agg aac cag ata gtt tat atc cgt act tgc act taa agt cta cat tgt cgt tgt atg a-- gta tga tct ttt aaa ccc gct aga caa gta tcc gtt tga tat tgt agg atg tgg aca ttt aac aat ctg aca cgt ggg tgg atc gga cca ttc tcc tcc tga aca cag gac act aga gtt acc aat caa cga ata tcc act att gca act ata agt tac aac gct ccc atc ggt ata aaa atc ctc gta tcc gtt atg tct tcc gtt gga tat aga tgg agg gga ttg gca ttt aac aga ttc aca aat agg tgc ctc ggg att cca tac cat aga tcc agt aaa tcc taa ttc aca ata cga ttt aga ttc tcc tat caa ctg ata tcc gct att aca aga gta cgt tat act aga gcc aaa gtc tac tcc acc aat atc aag ttg gcc att atc gat atc tcg agg cga tgg gca tct ccg ttt aat aca ttg att aaa gag tgt cca tcc agt acc tgt aca ttt agc ata tat agg tcc cat ttt ttg ctt tct gta tcc agg tag aca tag a-t att cta tag tgt ctc cta tgt tgt a-- --- --- --- -at tag cat tag cat cag tc- --t cca cac tat tct taa att tca tat taa tgg gtc gtg acg gaa tag tac agc atg ata gaa cgc atc cta ttc cca aca atg tca gga acg tca cgc tct cca cct tca tat tta ttt atc cgt -aa aaa tgt tat cct gga cat cgt aca aat aat a-a aaa gcc ca- -ta tat gtt cgc tat tgt aga aat tgt ttt tca cag ttg ctc aaa aac gat ggc agt gac tta tga gtt acg tta cac ttt gga gtc tca tct tta gta aac ata tca taa tat tcg ata tta cga gtt gac ata tcg aac aaa ttc caa gta ttt gat ttt gga taa tat tcg tat ttt gca tct gct ata att aag ata taa tca ccg caa gaa cac acg aac atc ttt cct aca tgg tta aag tac atg tac aat tct atc cat ttg tct tcc tta act ata tat ttg tat aga taa tta cga gtc tcg tga gta att cca gta att aca tag atg tcg ccg tcg tac tct aca gca taa act ata cta tga tgt cta ggc atg gga gac -tt ttt tat cca acg att ttt agt gaa aca ttc cac atc gtt taa tac tac ata ttt ttc ata cgt ggt ata aac tcc acc tat tac --a tat ata tca tcg ttt acg aat acc gac gcg cct gaa tat cta gga gta att aag ttt gga agt ctt atc cat ttc gaa gtg ccg tgt ttc aaa tat tct gcc aca ccc gtt gaa ata gaa aat tct aat cct cct att aca tat aac ttt cca tcg tta aca caa gta cta act tct gat ttt aac gac gac ata tta gta acc gtt ttc ca- --- --- --- --- --- --- --t ttt ttc gtt tca aga tct acc cgc -ga tac gga ata aac atg tct att gtt aat cat gcc gcc aat aat gta tag aca att atg taa aac att tgc att ata gaa ttg --- -tc tat ctg tat tac cga cta tcg tcc aat att ctg ttc tag gag agt aat ggg tta ttg tgg ata tat aat cag agt ttt taa tga cta cta tat tat gtt tta tac cat ttc gtg tca ctg gct ttg tag att tgg ata tag tta atc cca aca atg ata tag cat tgc gca tag tat tag tca taa act tgg gat gta aaa tgt tga tga tat cta cat cgt ttg gat ttt tat gta tcc act tta ata ata tca tag ctg taa cat cct cat ga- ttt acg tta acg tct tcg tgg gat aag ata gtt gtc agt tca tcc ttt gat aat ttt cca aat t

>KC207811.1:21018-22981_Vaccinia_virus_strain_TianTan_clone_TP5_complete_genome

cgg ctt cca tgt att tcc tgg aga gca agt --- aga tga tg- agg aac cag ata gtt tat atc cgt act tgc act taa agt tta cat tgt cgt tgt atg a-- gta tga tct ttt aaa ccc gct aga caa gta tcc gtt tga tat tgt agg atg tgg aca ttt aac aat ctg aca cgt ggg tgg atc gga cca ttc tcc tcc tga aca cag gac acc aga gtt acc aat caa cga ata tcc act att gca act ata agt tac aac gct ccc atc ggt ata aaa atc ctc gta tcc gtt atg tct tcc gtt gga tat aga tgg agg gga ttg gca ttt aac aga ttc aca aat agg tgc ctc ggg att cca tac cat aga tcc agt aaa tcc taa ttc aca ata cga ttt aga ttc tcc tat caa ctg ata tcc gct att aca aga gta cgt tat act aga gcc aaa gtc tac tcc acc aat atc aag ttg gcc att atc gat atc tcg agg cga tgg gca tct ccg ttt aat aca ttg att aaa gag tgt cca tcc agt acc tgt aca ttt agc ata tat agg tcc cat ttt ttg ctt tct gta tcc agg tag aca tag a-t att cta tag tgt ctc cta tgt tgt a-- --- --- --- -at tag cat tag cat cag tc- --t cca cac tat tct taa att tca tat taa tgg gac gtg aag gaa tag gac agt atg ata gaa cgc atc cta ttc cca aca atg tca gga acg tca cgc tct cca cct tca tat tta ttt atc cgt -aa aaa tgt tat cct gga cat cgt aca aat aat a-a aaa gcc ca- -ta tat gtt cgc tat tgt aga aat tgt ttt tca cag ttg ctc aaa aac gat ggc agt gac tta tga gtt acg tta cac ttt gga gtc tca tct tta gta aac ata tca taa tat tcg ata tta cga gtt gac ata tcg aac aaa ttc caa gta ttt gat ttt gga taa tat tcg tat ttt gca tct gct ata att aag ata taa tca ccg caa gaa cac acg aac atc ttt cct aca tgg tta aag tac atg tac aat tct atc cat ttg tct tcc tta act ata tat ttg tat aga taa tta cga gtc tcg tga gta att cca gta att aca tag atg tcg ccg tcg tac tct aca gca taa act ata cta tga tgt cta ggc atg gga gac -tt ttt tat cca acg att ttt agt gaa aca ttc cac atc gtt taa tac tac ata ttt ttc ata cgt ggt ata aac tcc acc cat tac --a tat ata tca tcg ttt acg aat acc gac gcg cct gaa tat cta gga gta att aag ttt gga agt ctt atc cat ttc gaa gtg ccg tgt ttc aaa tat tct gcc aca ccc gtt gaa ata gaa aat tct aat cct cct att aca tat aac ttt cca ttg tta aca caa gta cta act tct gat ttt aac gac gac ata tta gta acc gtt ttc ca- --- --- --- --- --- --- --t ttt ttc gtt tta aga tct acc cgc -ga tac gga ata aac atg tct att gtt aat cat gcc gcc aat aat gta tag aca att atg taa aac att tgc att ata gaa ttg --- -tc tat ctg tat tac cga cta tcg tcc aat att ctg ttc tag gag agt aat ggg tta ttg tgg ata tat aat cag agt ttt taa tga cta cta tat tat gtt tta tac cat ttc gtg tca ctg gct ttg tag att tgg ata tag tta atc cca aca atg ata tag cat tgc gca tag tat tag tca taa act tgg gat gta aaa tgt tga tga tat cta cat cgt ttg gat ttt tat gta tcc act tta ata ata tca tag ctg taa cat cct cat ga- ttt acg tta acg tct tcg tgg gat aag ata gtt gtc agt tca tcc ttt gat aat ttt cca aat t

>KC207810.1:15311-17274_Vaccinia_virus_strain_TianTan_clone_TP3_complete_genome

cgg ctt cca tgt att tcc tgg aga gca agt --- aga tga tg- agg aac cag ata gtt tat atc cgt act tgc act taa agt tta cat tgt cgt tgt atg a-- gta tga tct ttt aaa ccc gct aga caa gta tcc gtt tga tat tgt agg atg tgg aca ttt aac aat ctg aca cgt ggg tgg atc gga cca ttc tcc tcc tga aca cag gac acc aga gtt acc aat caa cga ata tcc act att gca act ata agt tac aac gct ccc atc ggt ata aaa atc ctc gta tcc gtt atg tct tcc gtt gga tat aga tgg agg gga ttg gca ttt aac aga ttc aca aat agg tgc ctc ggg att cca tac cat aga tcc agt aaa tcc taa ttc aca ata cga ttt aga ttc tcc tat caa ctg ata tcc gct att aca aga gta cgt tat act aga gcc aaa gtc tac tcc acc aat atc aag ttg gcc att atc gat atc tcg agg cga tgg gca tct ccg ttt aat aca ttg att aaa gag tgt cca tcc agt acc tgt aca ttt agc ata tat agg tcc cat ttt ttg ctt tct gta tcc agg tag aca tag a-t att cta tag tgt ctc cta tgt tgt a-- --- --- --- -at tag cat tag cat cag tc- --t cca cac tat tct taa att tca tat taa tgg gac gtg aag gaa tag gac agt atg ata gaa cgc atc cta ttc cca aca atg tca gga acg tca cgc tct cca cct tca tat tta ttt atc cgt -aa aaa tgt tat cct gga cat cgt aca aat aat a-a aaa gcc ca- -ta tat gtt cgc tat tgt aga aat tgt ttt tca cag ttg ctc aaa aac gat ggc agt gac tta tga gtt acg tta cac ttt gga gtc tca tct tta gta aac ata tca taa tat tcg ata tta cga gtt gac ata tcg aac aaa ttc caa gta ttt gat ttt gga taa tat tcg tat ttt gca tct gct ata att aag ata taa tca ccg caa gaa cac acg aac atc ttt cct aca tgg tta aag tac atg tac aat tct atc cat ttg tct tcc tta act ata tat ttg tat aga taa tta cga gtc tcg tga gta att cca gta att aca tag atg tcg ccg tcg tac tct aca gca taa act ata cta tga tgt cta ggc atg gga gac -tt ttt tat cca acg att ttt agt gaa aca ttc cac atc gtt taa tac tac ata ttt ttc ata cgt ggt ata aac tcc acc cat tac --a tat ata tca tcg ttt acg aat acc gac gcg cct gaa tat cta gga gta att aag ttt gga agt ctt atc cat ttc gaa gtg ccg tgt ttc aaa tat tct gcc aca ccc gtt gaa ata gaa aat tct aat cct cct att aca tat aac ttt cca ttg tta aca caa gta cta act tct gat ttt aac gac gac ata tta gta acc gtt ttc ca- --- --- --- --- --- --- --t ttt ttc gtt tta aga tct acc cgc -ga tac gga ata aac atg tct att gtt aat cat gcc gcc aat aat gta tag aca att atg taa aac att tgc att ata gaa ttg --- -tc tat ctg tat tac cga cta tcg tcc aat att ctg ttc tag gag agt aat ggg tta ttg tgg ata tat aat cag agt ttt taa tga cta cta tat tat gtt tta tac cat ttc gtg tca ctg gct ttg tag att tgg ata tag tta atc cca aca atg ata tag cat tgc gca tag tat tag tca taa act tgg gat gta aaa tgt tga tga tat cta cat cgt ttg gat ttt tat gta tcc act tta ata ata tca tag ctg taa cat cct cat ga- ttt acg tta acg tct tcg tgg gat aag ata gtt gtc agt tca tcc ttt gat aat ttt cca aat t

>JX489139.1:18154-20118_Vaccinia_virus_strain_Tiantan_clone_TT12_complete_genome

cgg ctt cca tgt att tcc tgg aga gca agt --- aga tga tg- agg aac cag ata gtt tat atc cgt act tgc act taa agt tta cat tgt cgt tgt atg a-- gta tga tct ttt aaa ccc gct aga caa gta tcc gtt tga tat tgt agg atg tgg aca ttt aac aat ctg aca cgt ggg tgg atc gga cca ttc tcc tcc tga aca cag gac acc aga gtt acc aat caa cga ata tcc act att gca act ata agt tac aac gct ccc atc ggt ata aaa atc ctc gta tcc gtt atg tct tcc gtt gga tat aga tgg agg gga ttg gca ttt aac aga ttc aca aat agg tgc ctc ggg att cca tac cat aga tcc agt aaa tcc taa ttc aca ata cga ttt aga ttc tcc tat caa ctg ata tcc gct att aca aga gta cgt tat act aga gcc aaa gtc tac tcc acc aat atc aag ttg gcc att atc gat atc tcg agg cga tgg gca tct ccg ttt aat aca ttg att aaa gag tgt cca tcc agt acc tgt aca ttt agc ata tat agg tcc cat ttt ttg ctt tct gta tcc agg tag aca tag a-t att cta tag tgt ctc cta tgt tgt a-- --- --- --- -at tag cat tag cat cag tc- --t cca cac tat tct taa att tca tat taa tgg gtc gtg acg gaa tag tac agc atg ata gaa cgc atc cta ttc cca aca atg tca gga acg tca cgc tct cca ctt tca tat tta ttt atc cgt -aa aaa tgt tat cct gga cat cgt aca aat aat aaa aaa gcc ca- -ta tat gtt cgc tat tgt aga aat tgt ttt tca cag ttg ctc aaa aac gat ggc agt gac tta tga gtt acg tta cac ttt gga gtc tca tct tta gta aac ata tca taa tat tcg ata tta cga gtt gac ata tcg aac aaa ttc caa gta ttt gat ttt gga taa tat tcg tat ttt gca tct gct ata att aag ata taa tca ccg caa gaa cac acg aac atc ttt cct aca tgg tta aag tac atg tac aat tct atc cat ttg tct tcc tta act ata tat ttg tat aga taa tta cga gtc tcg tga gta att cca gta att aca tag atg tcg ccg tcg tac tct aca gca taa act ata cta tga tgt cta ggc atg gga gac -tt ttt tat cca acg att ttt agt gaa aca ttc cac atc gtt taa tac tac ata ttt ttc ata cgt ggt ata aac tcc acc cat tac --a tat ata tca tcg ttt acg aat acc gac gcg cct gaa tat cta gga gta att aag ttt gga agt ctt atc cat ttc gaa gtg ccg tgt ttc aaa tat tct gcc aca ccc gtt gaa ata gaa aat tct aat cct cct att aca tat aac ttt cca tcg tta aca caa gta cta act tct gat ttt aac gac gac ata tta gta acc gtt ttc ca- --- --- --- --- --- --- --t ttt ttc gtt tta aga tct acc cgc -ga tac gga ata aac atg tct att gtt aat cat gcc gcc aat aat gta tag aca att atg taa aac att tgc att ata gaa ttg --- -tc tat ctg tat tac cga cta tcg tcc aat att ctg ttc tag gag agt aat ggg tta ttg tgg ata tat aat cag agt ttt taa tga cta cta tat tat gtt tta tac cat ttc gtg tca ctg gct ttg tag att tgg ata tag tta atc cca aca atg ata tag cat tgc gca tag tat tag tca taa act tgg gat gta aaa tgt tga tga tat cta cat cgt ttg gat ttt tat gta tcc act tta ata ata tca tag ctg taa cat cct cat ga- ttt acg tta acg tct tcg tgg gat aag ata gtt gtc agt tca tcc ttt gat aat ttt cca aat t

>BK013343.1:42576-44534_TPA_asm:_Vaccinia_virus_isolate_VK12

cgg ctg cca tgt att tcc tgg aga gca agt --- aga tga tg- agg aac cag ata ggt tat atc cgt act tgc act taa agt tta cat tgt cgt tgt atg a-- gta tga tct ttt aaa ccc gct aga caa gta tcc gtt tga tat tgt agg atg tgg aca ttt aac aat ttg aca cgt ggg tgg atc gga cca ttc tcc tcc tga aca cag gac acc aga gtt acc aat caa cga ata tcc act att gca act ata agt tac aac gct tcc atc ggt ata aaa atc ctc gta tcc gtt atg tct tcc gtt gga tat aga tgg agg gga ttg gca ttt aac aga ttc aca aat agg tgc ctc ggg att cca tac cat aga tcc agt aaa tcc taa ttc aca ata cga ttt aga ttc tcc tat caa ctg ata tcc gct att aca aga gta cgt tat act aga gcc aaa gtc tac tcc gcc aat atc aag ttg gcc att atc gat atc tcg agg cga tgg gca tct ccg ttt aat aca ttg att aaa gag tgt cca tcc agt acc tgt aca ttt agc ata tat agg tcc cat ttt ttg ctt tct gta tcc agg tag aca tag a-t att cta tag tgt ctc cta tgt tgt a-- --- --- --- --- --- -at tag cat tag tt- --t cca cac tat tct taa att tca tat taa tgg gac gtg aag gaa tag gac agt atg ata gaa cgc atc cta ttc cca aca atg tca gga acg tca cgc tct cca cct tca tat tta ttt atc cgt -aa aaa tgt tat cct gga cat cgt aca aat aat aaa aaa gcc ca- -ta tat gtt tgc tat tgt aga aat tgt ttt tca cag ttg ctc aaa aac gat ggc agt gac tta tga gtt acg tta cac ttt gga gtt tca tct tta gta aac ata tca taa tat tcg ata tta cga gtt gac ata tcg aac aaa ttc caa gta ttt gat ttt gga taa tat tcg tat ttt gca tct gct ata att aag ata taa tca ccg caa gaa cac acg aac atc ttt cct aca tgg tta aag tac atg tat aat tct atc cat ttg tct tcc tta act ata tat ttg tat aga taa tta cga gtc tca taa gta att cca gta att gca tag atg tca cca tcg tac tct aca gca taa act ata cta tga tgt cta ggc atg gga gac -tt ttt tat cca acg att ttt agt gaa aca ttc tac atc gtt taa tac tac ata ttt ctc ata cgt ggt ata aac tcc acc cat tac --a tat ata tca tcg ttt acg aat acc gat gcg cct gaa tat cta gga gta att aag ttt gga agt ctt atc cat ttt gaa gtg ccg tgt ttc aaa tat tct gct aca ccc gtt gaa ata gaa aat tct aat cct cct att aca tat aac ttt cca ttg tta aca caa gta cta act tct gat ttt aac gac gac ata tta gta acc gtt ttc ca- --- --- --- --- --- --- --t ttt ttc gtt tca aga tct acc cgc -ga tac gga ata aac atg tct att gtt aat cat gcc gcc aat aat gta tag aca att atg taa aac att tgc att ata gaa ttg --- -tc tat ctg tat tac cga cta tcg tcc aat att ctg ttc tag gag agt aat ggg tta ttg tgg ata tat aat cag agt ttt taa tga cta cta tat tat gtt tta tac cat ttc gtg tca ctg gct ttg tag att tgg ata tag tta atc cca aca atg ata tag cat tgc gca tag tat tag tta taa act tgg gat gta aaa tgt tga tga tat cta cat cgt ttg gat ttt tat gta tcc act tta ata ata tca tag ctg taa cat cct cat ga- ttt acg tta acg tct tcg tgg gat aag ata gtt gtc agt tca tcc ttt gat aat ttt cca aat t

>MF477237.2:21857-23815_Vaccinia_virus_strain_Mulford_1902_complete_genome

cgg ctg cca tgt att tcc tgg aga gca agt --- aga tga tg- agg aac cag ata ggt tat atc cgt act tgc act taa agt tta cat tgt cgt tgt atg a-- gta tga tct ttt aaa ccc gct aga caa gta tcc gtt tga tat tgt agg atg tgg aca ttt aac aat ttg aca cgt ggg tgg atc gga cca ttc tcc tcc tga aca cag gac act aga gtt acc aat caa cga ata tcc act att gca act ata agt tac aac gct tcc atc ggt ata aaa atc ctc gta tcc gtt atg tct tcc gtt gga tat aga tgg agg gga ttg gca ttt aac aga ttc aca aat agg tgc ctc ggg att cca tac cat aga tcc agt aaa tcc taa ttc aca ata cga ttt aga ttc tcc tat caa ctg ata tcc gct att aca aga gta cgt tat act aga gcc aaa gtc tac tcc gcc aat atc aag ttg gcc att atc gat atc tcg agg cga tgg gca tct ccg ttt aat aca ttg att aaa gag tgt cca tcc agt acc tgt aca ttt agc ata tat agg tcc cat ttt ttg ctt tct gta tcc agg tag aca tag a-t att cta tag tgt ctc cta tgt tgt a-- --- --- --- --- --- -at tag cat tag tt- --t cca cac tat tct taa att tta tat taa tgg gac gtg aag gaa tag gac agt atg ata gaa cgc atc cta ttc cca aca atg tca gga acg tca cgc tct cca cct tca tat tta ttt atc cgt -aa aaa tgt tat cct gga cat cgt aca aat aat aaa aaa gcc ca- -ta tat gtt tgc tat tgt aga aat tgt ttt tca cag ttg ctc aaa aac gat ggc agt gac tta tga gtt acg tta cac ttt gga gtt tca tct tta gta aac ata tca taa tat tcg ata tta cga gtt gac ata tcg aac aaa ttc caa gta ttt gat ttt gga taa tat tcg tat ttt gca tct gct ata att aag ata taa tca ccg caa gaa cac acg aac atc ttt cct aca tgg tta aag tac atg tat aat tct atc cat ttg tct tcc tta act ata tat ttg tat aga taa tta cga gtc tca taa gta att cca gta att gca tag atg tca cca tcg tac tct aca gca taa act ata cta tga tgt cta ggc atg gga gac -tt ttt tat cca acg att ttt agt gaa aca ttc tac atc gtt taa tac tac ata ttt ctc ata cgt ggt ata aac tcc acc cat tac --a tat ata tca tcg ttt acg aat acc gat gcg cct gaa tat cta gga gta att aag ttt gga agt ctt atc cat ttt gaa gtg ccg tgt ttc aaa tat tct gct aca ccc gtt gaa ata gaa aat tct aat cct cct att aca tat aac ttt cca ttg tta aca caa gta cta act tct gat ttt aac gac gac ata tta gta acc gtt ttc ca- --- --- --- --- --- --- --t ttt ttc gtt tca aga tct acc cgc -ga tac gga ata aac atg tct att gtt aat cat gcc gcc aat aat gta tag aca att atg taa aac att tgc att ata gaa ttg --- -tc tat ctg tat tac cga cta tcg tcc aat att ctg ttc tag gag agt aat ggg tta ttg tgg ata tat aat cag agt ttt taa tga cta cta tat tat gtt tta tac cat ttc gtg tca ctg gct ttg tag att tgg ata tag tta atc cca aca atg ata tag cat tgc gca tag tat tag tta taa act tgg gat gta aaa tgt tga tga tat cta cat cgt ttg gat ttt tat gta tcc act tta ata ata tca tag ctg taa cat cct cat ga- ttt acg tta acg tct tcg tgg gat aag ata gtt gtc agt tca tcc ttt gat aat ttt cca aat t

>KY369926.1:34732-36683_Cowpox_virus_strain_Kostroma_2015

cgg ctg cca tgt att tcc tgg aga gca agt --- aga tga tg- agg aac cag ata gtt tat atc cgt act tgc act taa agt cta cat tgt cgt tgt atg a-- gta tga tct ttt aaa ccc gct aga caa gta tcc gtt tga tat tgt agg atg tgg aca ttt aac aat ctg aca cgt ggg tgg atc gga cca ttc tcc tcc tga aca cag gac acc aga gtt acc aat caa cga ata tcc act att gca act ata cgt tac aac gct tcc atc ggt ata aaa atc ctc gta tcc gtt atg tct tcc gtt gga tat aga tgg agg gga ttg gca ttt aac aga ttc aca aat ggg tgc ctc ggg att cca tac cat aga tcc agt aga tcc taa ttc aca ata cga ttt aga ttc acc gat caa atg ata tcc gct att aca aga gta cgt tat act aga gcc aaa gtc tac tcc acc aat atc aag ttg gcc att atc gat atc tcg agg cga tgg gca tct ccg ttt aat aca ttg att aaa gag tgt cca tcc agt acc tgt aca ttt agc ata tat agg tcc cat ttt ttg ctt tct gta tcc agg tag aca tag a-t att cta tag tgt ctc cta tat tat a-- --- --- --- --- --- --- --- -at ggg aa- --t cca ctg tac cct taa att tca tag taa tag gac gtg aag gaa tag gac agc atg ata gaa cgc atc cta ttc cca aca atg tca gga acg tca cgc tct cca cct tca tat tta ttt atc cgt -aa aaa tgt tat cct gga cat cgt aca aat aat a-a aaa gcc ca- -ta tat gtt cgc tat tgt aga aat tgt ttt tca cag ttg ctc aaa aac gat ggc agt gac tta tga gtt acg tta cac ttt gga gtc tca tct tta gta aac ata tca taa tat tcg ata tta cga gtt gac ata tcg aac aaa ttc caa gta ttt gat ttt gga taa tat tcg tat ttt gca tct gct ata att aag ata taa tcg ccg caa gaa cat acg aac atc ttt cct aca tgg tta aag tac atg tac aat tct atc cat ttg tct tcc tta act ata tat ttg tat aga taa tta cga gtc tca tga gta att cca gta att gca tag atg tca cca tcg tac tct aca gca taa act ata cta tga tgt cta ggc atg gga gac -tt ttt tat cca acg att ttt agt gaa aca ttc cac atc gtt taa tac tac ata ttt ctc ata cgt ggt ata aac tcc acc cat tac --a tat ata tca tcg ttt acg aat acc gat gcg cct gaa tat cta gga gta att aag ttt gga agt ctt atc cat ttc gaa gtg ccg tgt ttc aaa tat tct gcc aca ccc gtt gaa ata gaa aat tct aat cct cct att aca tat aac ttt cca tcg tta aca caa gta cta act tct gat ttt aac gac gac ata tta gta act gtt ttc ca- --- --- --- --- --- --- --t ttt ttc gtt tta aga tct acc cgc -ga tac gga ata aac atg tct att gtt aat cat gcc gcc aat aat gta tag aca att atg taa aac att tgc att ata gaa ttg --- -tc tat ctg tat tac cga cta tcg tcc aat att ctg ttc tag gag agt aat ggg tta ttg tgg ata tat aat cag agt ttt taa tga cta cta tat tat gtt tta tac cat ttc gtg tca ctg gct ttg tag att tgg ata tag tta atc cca aca atg ata tag cat tgc aca tag tat tag tca taa act tgg gat gta aaa tgt tga tga tat cta cat cgt ttg gat ttt tat gta tcc act tta ata ata tca tag ctg taa cat cct cat ga- ttt acg tta acg tct tcg tgg gat aag ata gtt gtc agt tca tcc ttt gat aat ttt cca aat t

>KT013210.1:12702-14660_Vaccinia_virus_strain_Cantagalo_isolate_Cantagalo_virus_CM01

cgg ctg cca tgt att tcc tgg aga gca agt --- aga tga tg- agg aac cag ata ggt tat atc cgt act tgc act taa agt tta cat tgt cgt tgt atg a-- gta tga tct ttt aaa ccc gct aga caa gta tcc gtt tga tat tgt agg atg tgg aca ttt aac aat ttg aca cgt ggg tgg atc gga cca ttc tcc tcc tga aca cag gac act aga gtt acc aat caa cga ata tcc act att gca act ata agt tac aac gct tcc atc ggt ata aaa atc ctc gta tcc gtt atg tct tcc gtt gga tat aga tgg agg gga ttg gca ttt aac aga ttc aca aat agg tgc ctc ggg att cca tac cat aga tcc agt aaa tcc taa ttc aca ata cga ttt aga ttc tcc tat caa ctg ata tcc gct att aca aga gta cgt tat act aga gcc aaa gtc tac tcc gcc aat atc aag ttg gcc att atc gat atc tcg agg cga tgg gca tct ccg ttt aat aca ttg att aaa gag tgt cca tcc agt acc tgt aca ttt agc ata tat agg tcc cat ttt ttg ctt tct gta tcc agg tag aca tag a-t att cta tag tgt ctc cta tgt tgt a-- --- --- --- --- --- -at tag cat tag tt- --t cca cac tat tct taa att tca tat taa tgg gac gtg aag gaa tag gac agt atg ata gaa cgc atc cta ttc cca aca atg tca gga acg tca cgc tct cca cct tca tat tta ttt atc cgt -aa aaa tgt tat cct gga cat cgt aca aat aat aaa aaa gcc ca- -ta tat gtt tgc tat tgt aga aat tgt ttt tca cag ttg ctc aaa aac gat ggc agt gac tta tga gtt acg tta cac ttt gga gtt tca tct tta gta aac ata tca taa tat tcg ata tta cga gtt gac ata tcg aac aaa ttc caa gta ttt gat ttt gga taa tat tcg tat ttt gca tct gct ata att aag ata taa tca ccg caa gaa cac acg aac atc ttt cct aca tgg tta aag tac atg tac aat tct atc cat ttg tct tcc tta act ata tat ttg tat aga taa tta cga gtc tcg tga gta att cca gta att aca tag atg tcg ccg tcg tac tct aca gca taa act ata cta tga tgt cta ggc atg gga gac -tt ttt tat cca acg att ttt agt gaa aca ttc cac atc gtt taa tac tac ata ttt ttc ata cgt ggt ata aac tcc acc cat tac --a tat ata tca tcg ttt acg aat acc gac gcg cct gaa tat cta gga gta att aag ttt gga agt ctt atc cat ttt gaa gtg ccg tgt ttc aaa tat tct gct aca ccc gtt gaa ata gaa aat tct aat cct cct att aca tat aac ttt cca ttg tta aca caa gta cta act tct gat ttt aac gac gac ata tta gta acc gtt ttc ca- --- --- --- --- --- --- --t ttt ttc gtt tca aga tct acc cgc -ga tac gga ata aac atg tct att gtt aat cat gcc gcc aat aat gta tag aca att atg taa aac att tgc att ata gaa ttg --- -tc tat ctg tat tac cga cta tcg tcc aat att ctg ttc tag gag agt aat ggg tta ttg tgg ata tat aat cag agt ttt taa tga cta cta tat tat gtt tta tac cat ttc gtg tca ctg gct ttg tag att tgg ata tag tta atc cca aca atg ata tag cat tgc gca tag tat tag tta taa act tgg gat gta aaa tgt tga tga tat cta cat cgt ttg gat ttt tat gta tcc act tta ata ata tca tag ctg taa cat cct cat ga- ttt acg tta acg tct tcg tgg gat aag ata gtt gtc agt tca tcc ttt gat aat ttt cca aat t

>MW018155.1:12308-14266_Vaccinia_virus_strain_Cantagalo_isolate_Cantagalo_CG04

cgg ctg cca tgt att tcc tgg aga gca agt --- aga tga tg- agg aac cag ata ggt tat atc cgt act tgc act taa agt tta cat tgt cgt tgt atg a-- gta tga tct ttt aaa ccc gct aga caa gta tcc gtt tga tat tgt agg atg tgg aca ttt aac aat ttg aca cgt ggg tgg atc gga cca ttc tcc tcc tga aca cag gac act aga gtt acc aat caa cga ata tcc act att gca act ata agt tac aac gct tcc atc ggt ata aaa atc ctc gta tcc gtt atg tct tcc gtt gga tat aga tgg agg gga ttg gca ttt aac aga ttc aca aat agg tgc ctc ggg att cca tac cat aga tcc agt aaa tcc taa ttc aca ata cga ttt aga ttc tcc tat caa ctg ata tcc gct att aca aga gta cgt tat act aga gcc aaa gtc tac tcc gcc aat atc aag ttg gcc att atc gat atc tcg agg cga tgg gca tct ccg ttt aat aca ttg att aaa gag tgt cca tcc agt acc tgt aca ttt agc ata tat agg tcc cat ttt ttg ctt tct gta tcc agg tag aca tag a-t att cta tag tgt ctc cta tgt tgt a-- --- --- --- --- --- -at tag cat tag tt- --t cca cac tat tct taa att tca tat taa tgg gac gtg aag gaa tag gac agt atg ata gaa cgc atc cta ttc cca aca atg tca gga acg tca cgc tct cca cct tca tat tta ttt atc cgt -aa aaa tgt tat cct gga cat cgt aca aat aat aaa aaa gcc ca- -ta tat gtt tgc tat tgt aga aat tgt ttt tca cag ttg ctc aaa aac gat ggc agt gac tta tga gtt acg tta cac ttt gga gtt tca tct tta gta aac ata tca taa tat tcg ata tta cga gtt gac ata tcg aac aaa ttc caa gta ttt gat ttt gga taa tat tcg tat ttt gca tct gct ata att aag ata taa tca ccg caa gaa cac acg aac atc ttt cct aca tgg tta aag tac atg tac aat tct atc cat ttg tct tcc tta act ata tat ttg tat aga taa tta cga gtc tcg tga gta att cca gta att aca tag atg tcg ccg tcg tac tct aca gca taa act ata cta tga tgt cta ggc atg gga gac -tt ttt tat cca acg att ttt agt gaa aca ttc cac atc gtt taa tac tac ata ttt ttc ata cgt ggt ata aac tcc acc cat tac --a tat ata tca tcg ttt acg aat acc gac gcg cct gaa tat cta gga gta att aag ttt gga agt ctt atc cat ttt gaa gtg ccg tgt ttc aaa tat tct gct aca ccc gtt gaa ata gaa aat tct aat cct cct att aca tat aac ttt cca ttg tta aca caa gta cta act tct gat ttt aac gac gac ata tta gta acc gtt ttc ca- --- --- --- --- --- --- --t ttt ttc gtt tca aga tct acc cgc -ga tac gga ata aac atg tct att gtt aat cat gcc gcc aat aat gta tag aca att atg taa aac att tgc att ata gaa ttg --- -tc tat ctg tat tac cga cta tcg tcc aat att ctg ttc tag gag agt aat ggg tta ttg tgg ata tat aat cag agt ttt taa tga cta cta tat tat gtt tta tac cat ttc gtg tca ctg gct ttg tag att tgg ata tag tta atc cca aca atg ata tag cat tgc gca tag tat tag tta taa act tgg gat gta aaa tgt tga tga tat cta cat cgt ttg gat ttt tat gta tcc act tta ata ata tca tag ctg taa cat cct cat ga- ttt acg tta acg tct tcg tgg gat aag ata gtt gtc agt tca tcc ttt gat aat ttt cca aat t

>MW018154.1:12677-14635_Vaccinia_virus_strain_Cantagalo_isolate_Cantagalo_VI04

cgg ctg cca tgt att tcc tgg aga gca agt --- aga tga tg- agg aac cag ata ggt tat atc cgt act tgc act taa agt tta cat tgt cgt tgt atg a-- gta tga tct ttt aaa ccc gct aga caa gta tcc gtt tga tat tgt agg atg tgg aca ttt aac aat ttg aca cgt ggg tgg atc gga cca ttc tcc tcc tga aca cag gac act aga gtt acc aat caa cga ata tcc act att gca act ata agt tac aac gct tcc atc ggt ata aaa atc ctc gta tcc gtt atg tct tcc gtt gga tat aga tgg agg gga ttg gca ttt aac aga ttc aca aat agg tgc ctc ggg att cca tac cat aga tcc agt aaa tcc taa ttc aca ata cga ttt aga ttc tcc tat caa ctg ata tcc gct att aca aga gta cgt tat act aga gcc aaa gtc tac tcc gcc aat atc aag ttg gcc att atc gat atc tcg agg cga tgg gca tct ccg ttt aat aca ttg att aaa gag tgt cca tcc agt acc tgt aca ttt agc ata tat agg tcc cat ttt ttg ctt tct gta tcc agg tag aca tag a-t att cta tag tgt ctc cta tgt tgt a-- --- --- --- --- --- -at tag cat tag tt- --t cca cac tat tct taa att tca tat taa tgg gac gtg aag gaa tag gac agt atg ata gaa cgc atc cta ttc cca aca atg tca gga acg tca cgc tct cca cct tca tat tta ttt atc cgt -aa aaa tgt tat cct gga cat cgt aca aat aat aaa aaa gcc ca- -ta tat gtt tgc tat tgt aga aat tgt ttt tca cag ttg ctc aaa aac gat ggc agt gac tta tga gtt acg tta cac ttt gga gtt tca tct tta gta aac ata tca taa tat tcg ata tta cga gtt gac ata tcg aac aaa ttc caa gta ttt gat ttt gga taa tat tcg tat ttt gca tct gct ata att aag ata taa tca ccg caa gaa cac acg aac atc ttt cct aca tgg tta aag tac atg tac aat tct atc cat ttg tct tcc tta act ata tat ttg tat aga taa tta cga gtc tcg tga gta att cca gta att aca tag atg tcg ccg tcg tac tct aca gca taa act ata cta tga tgt cta ggc atg gga gac -tt ttt tat cca acg att ttt agt gaa aca ttc cac atc gtt taa tac tac ata ttt ttc ata cgt ggt ata aac tcc acc cat tac --a tat ata tca tcg ttt acg aat acc gac gcg cct gaa tat cta gga gta att aag ttt gga agt ctt atc cat ttt gaa gtg ccg tgt ttc aaa tat tct gct aca ccc gtt gaa ata gaa aat tct aat cct cct att aca tat aac ttt cca ttg tta aca caa gta cta act tct gat ttt aac gac gac ata tta gta acc gtt ttc ca- --- --- --- --- --- --- --t ttt ttc gtt tca aga tct acc cgc -ga tac gga ata aac atg tct att gtt aat cat gcc gcc aat aat gta tag aca att atg taa aac att tgc att ata gaa ttg --- -tc tat ctg tat tac cga cta tcg tcc aat att ctg ttc tag gag agt aat ggg tta ttg tgg ata tat aat cag agt ttt taa tga cta cta tat tat gtt tta tac cat ttc gtg tca ctg gct ttg tag att tgg ata tag tta atc cca aca atg ata tag cat tgc gca tag tat tag tta taa act tgg gat gta aaa tgt tga tga tat cta cat cgt ttg gat ttt tat gta tcc act tta ata ata tca tag ctg taa cat cct cat ga- ttt acg tta acg tct tcg tgg gat aag ata gtt gtc agt tca tcc ttt gat aat ttt cca aat t

>MW018153.1:12309-14267_Vaccinia_virus_strain_Cantagalo_isolate_Cantagalo_MI233

cgg ctg cca tgt att tcc tgg aga gca agt --- aga tga tg- agg aac cag ata ggt tat atc cgt act tgc act taa agt tta cat tgt cgt tgt atg a-- gta tga tct ttt aaa ccc gct aga caa gta tcc gtt tga tat tgt agg atg tgg aca ttt aac aat ttg aca cgt ggg tgg atc gga cca ttc tcc tcc tga aca cag gac act aga gtt acc aat caa cga ata tcc act att gca act ata agt tac aac gct tcc atc ggt ata aaa atc ctc gta tcc gtt atg tct tcc gtt gga tat aga tgg agg gga ttg gca ttt aac aga ttc aca aat agg tgc ctc ggg att cca tac cat aga tcc agt aaa tcc taa ttc aca ata cga ttt aga ttc tcc tat caa ctg ata tcc gct att aca aga gta cgt tat act aga gcc aaa gtc tac tcc gcc aat atc aag ttg gcc att atc gat atc tcg agg cga tgg gca tct ccg ttt aat aca ttg att aaa gag tgt cca tcc agt acc tgt aca ttt agc ata tat agg tcc cat ttt ttg ctt tct gta tcc agg tag aca tag a-t att cta tag tgt ctc cta tgt tgt a-- --- --- --- --- --- -at tag cat tag tt- --t cca cac tat tct taa att tca tat taa tgg gac gtg aag gaa tag gac agt atg ata gaa cgc atc cta ttc cca aca atg tca gga acg tca cgc tct cca cct tca tat tta ttt atc cgt -aa aaa tgt tat cct gga cat cgt aca aat aat aaa aaa gcc ca- -ta tat gtt tgc tat tgt aga aat tgt ttt tca cag ttg ctc aaa aac gat ggc agt gac tta tga gtt acg tta cac ttt gga gtt tca tct tta gta aac ata tca taa tat tcg ata tta cga gtt gac ata tcg aac aaa ttc caa gta ttt gat ttt gga taa tat tcg tat ttt gca tct gct ata att aag ata taa tca ccg caa gaa cac acg aac atc ttt cct aca tgg tta aag tac atg tac aat tct atc cat ttg tct tcc tta act ata tat ttg tat aga taa tta cga gtc tcg tga gta att cca gta att aca tag atg tcg ccg tcg tac tct aca gca taa act ata cta tga tgt cta ggc atg gga gac -tt ttt tat cca acg att ttt agt gaa aca ttc cac atc gtt taa tac tac ata ttt ttc ata cgt ggt ata aac tcc acc cat tac --a tat ata tca tcg ttt acg aat acc gac gcg cct gaa tat cta gga gta att aag ttt gga agt ctt atc cat ttt gaa gtg ccg tgt ttc aaa tat tct gct aca ccc gtt gaa ata gaa aat tct aat cct cct att aca tat aac ttt cca ttg tta aca caa gta cta act tct gat ttt aac gac gac ata tta gta acc gtt ttc ca- --- --- --- --- --- --- --t ttt ttc gtt tca aga tct acc cgc -ga tac gga ata aac atg tct att gtt aat cat gcc gcc aat aat gta tag aca att atg taa aac att tgc att ata gaa ttg --- -tc tat ctg tat tac cga cta tcg tcc aat att ctg ttc tag gag agt aat ggg tta ttg tgg ata tat aat cag agt ttt taa tga cta cta tat tat gtt tta tac cat ttc gtg tca ctg gct ttg tag att tgg ata tag tta atc cca aca atg ata tag cat tgc gca tag tat tag tta taa act tgg gat gta aaa tgt tga tga tat cta cat cgt ttg gat ttt tat gta tcc act tta ata ata tca tag ctg taa cat cct cat ga- ttt acg tta acg tct tcg tgg gat aag ata gtt gtc agt tca tcc ttt gat aat ttt cca aat t

>X94355.2:34812-36763_Cowpox_virus_strain_GRI-90_complete_genome

cgg ctg cca tgt att tcc tgg aga gca agt --- aga tga tg- agg aac cag ata gtt tat atc cgt act tgc act taa agt cta cat tgt cgt tgt atg a-- gta tga tct ttt aaa ccc gct aga caa gta tcc gtt tga tat tgt agg atg tgg aca ttt aac aat ctg aca cgt ggg tgg atc gga cca ttc tcc tcc tga aca cag gac acc aga gtt acc aat caa cga ata tcc act att gca act ata cgt tac aac gct tcc atc ggt ata aaa atc ctc gta tcc gtt atg tct tcc gtt gga tat aga tgg agg gga ttg gca ttt aac aga ttc aca aat ggg tgc ctc ggg att cca tac cat aga tcc agt aga tcc taa ttc aca ata cga ttt aga ttc acc gat caa atg ata tcc gct att aca aga gta cgt tat act aga gcc aaa gtc tac tcc gcc aat atc aag ttg gcc att atc gat atc tcg agg cga tgg gca tct ccg ttt aat aca ttg att aaa gag tgt cca tcc agt acc tgt aca ttt agc ata tat agg tcc cat ttt ttg ctt tct gta tcc agg tag aca tag a-t att cta tag tgt ctc cta tat tat a-- --- --- --- --- --- --- --- -at ggg aa- --t cca ctg tac cct taa att tca tag taa tag gac gtg aag gaa tag gac agc atg ata gaa cgc atc cta ttc cca aca atg tca gga acg tca cgc tct cca cct tca tat tta ttt atc cgt -aa aaa tgt tat cct gga cat cgt aca aat aat a-a aaa gcc ca- -ta tat gtt cgc tat tgt aga aat tgt ttt tca cag ttg ctc aaa aac gat ggc agt gac tta tga gtt acg tta cac ttt gga gtc tca tct tta gta aac ata tca taa tat tcg ata tta cga gtt gac ata tcg aac aaa ttc caa gta ttt gat ttt gga taa tat tcg tat ttt gca tct gct ata att aag ata taa tcg ccg caa gaa cat acg aac atc ttt cct aca tgg tta aag tac atg tac aat tct atc cat ttg tct tcc tta act ata tat ttg tat aga taa tta cga gtc tca tga gta att cca gta att gca tag atg tca cca tcg tac tct aca gca taa act ata cta tga tgt cta ggc atg gga gac -tt ttt tat cca acg att ttt agt gaa aca ttc cac atc gtt taa tac tac ata ttt ctc ata cgt ggt ata aac tcc acc cat tac --a tat ata tca tcg ttt acg aat acc gat gcg cct gaa tat cta gga gta att aag ttt gga agt ctt atc cat ttc gaa gtg ccg tgt ttc aaa tat tct gcc aca ccc gtt gaa ata gaa aat tct aat cct cct att aca tat aac ttt cca tcg tta aca caa gta cta act tct gat ttt aac gac gac ata tta gta act gtt ttc ca- --- --- --- --- --- --- --t ttt ttc gtt tta aga tct acc cgc -ga tac gga ata aac atg tct att gtt aat cat gcc gcc aat aat gta tag aca att atg taa aac att tgc att ata gaa ttg --- -tc tat ctg tat tac cga cta tcg tcc aat att ctg ttc tag gag agt aat ggg tta ttg tgg ata tat aat cag agt ttt taa tga cta cta tat tat gtt tta tac cat ttc gtg tca ctg gct ttg tag att tgg ata tag tta atc cca aca atg ata tag cat tgc gca tag tat tag tca taa act tgg gat gta aaa tgt tga tga tat cta cat cgt ttg gat ttt tat gta tcc act tta ata ata tca tag ctg taa cat cct cat ga- ttt acg tta acg tct tcg tgg gat aag ata gtt gtc agt tca tcc ttt gat aat ttt cca aat t

>MK035759.1:34204-36156_Cowpox_virus_strain_CPXV/Cepad_332

cgg ctt cca tgt att tcc tgg aga gca agt --- aga tga tg- agg aac cag ata gtt tat atc cgt act tgc act taa agt cta cat tgt cgt tgt atg a-- gta tga tct ttt aaa ccc gct aga caa gta tcc gtt tga tat tgt agg atg tgg aca ttt aac aat ctg aca cgt ggg tgg atc gga cca ttc tcc tcc tga aca cag gac agc aga gtt acc aat taa cga ata tcc act att gca act ata agt tac aac gct tcc atc ggt ata aaa atc ctc gta tcc gtt atg tct tcc gtt gga tat aga tgg agg tga ttg gca ttt aac aga ttc gca aat agg tgc ctc ggg att cca tac cat aga tcc agt aga tcc taa ttc aca ata cga ttt aga ttc tcc tat caa ctg ata tcc gct att aca aga gta cgt tat act aga gcc aaa gtc tac tcc gcc aat atc aag ttg gcc att atc gat atc tcg agg cga tgg gca tct ccg ttt aat aca ttg att aaa gag tgt cca tcc agt acc tgt aca ttt agc ata tat agg tcc cat ttt ttg ctt tct gta tcc agg tag aca tag a-t att cta tag tgt ctc cta tat tat a-- --- --- --- -at ggg aat --- --- --- --- --- cca ctg tac cct taa att tca tag taa tag gac gtg aag gaa tag gac agc atg ata gaa cgc atc cta ttc cca aca atg tca gga acg tca cgc tct cca cct tca tat tta ttt atc cgt -aa aaa tgt tat cct gga cat cgt aca aat aat aaa aaa gcc ca- -ta tat gtt cgc tat tgt aga aat tgt ttt tca cag ttg ctc aaa aac gat ggc agt gac tta tga gtt acg tta cac ttt gga gtc tca tct tta gta aac ata tca taa tat tcg ata tta cga gtt gac ata tcg aac aaa ttc caa gta ttt gat ttt gga taa tat tcg tat ttt gca tca gct ata att aag ata taa tca ccg caa gaa cac acg aac atc ttt cct aca tgg tta aag tac atg tac aat tct atc cat ttg tct tcc tta act ata tat ttg tat aga taa tta cga gtc tcg tga gta att cca gta att gca tag atg tca cca tcg tac tct aca gca taa act ata cta tga tgt cta ggc atg gga gac -tt ttt tat cca acg att ttt agt gaa aca ttc cac gtc gtt taa tac tac ata ttt ctc ata cgt ggt ata aac tcc acc cat tac --a tat ata tca tcg ttt acg aat acc gat gct cct gaa tat cta gga gta att aag ttt gga agt ctt atc cat ttc gaa gtg ccg tgt ttc aaa tat tct gcc acg ccc gtt gaa ata gaa aat tct aat cct cct att aca tat aac ttt cca tcg tta aca caa gta cta act tct gat ttt aac gac gac ata tta gta acc gtt ttc ca- --- --- --- --- --- --- --t ttt ttc gtt tta aga tct acc cgc -ga tac gga ata aac atg tct att gtt aat cat gcc gcc aat aat gta tag aca att atg taa aac att tgc att ata gaa ttg --- -tc tat ctg tat tac cga cta tcg tcc aat att ctg ttc tag gag agt aat ggg tta ttg tgg ata tat aat cag agt ttt taa tga cta cta tat tat gtt tta tac cat ttc gtg tca cag gct ttg tag att tgg ata tag tta atc cca aca atg cta tag cat tgc gca tag cat tag tca taa act tgg gat gta aaa tgt tga tga tat cta cat cgt ttg gat ttt tat gta tcc act tta ata ata tta tag ctg taa cat cct cat ga- ttt acg tta acg ttt tcg tgg gat aag ata gtt gtt agt tca tcc ttt gat aat ttt cca aat t

>MK035757.1:34502-36454_Cowpox_virus_strain_CPXV/Rat_Koelle

cgg ctt cca tgt att tcc tgg aga gca agt --- aga tga tg- agg aac cag ata gtt tat atc cgt act tgc act taa agt cta cat tgt cgt tgt atg a-- gta tga tct ttt aaa ccc gct aga caa gta tcc gtt tga tat tgt agg atg tgg aca ttt aac aat ctg aca cgt ggg tgg atc gga cca ttc tcc tcc tga aca cag gac agc aga gtt acc aat taa cga ata tcc act att gca act ata agt tac aac gct tcc atc ggt ata aaa atc ctc gta tcc gtt atg tct tcc gtt gga tat aga tgg agg tga ttg gca ttt aac aga ttc gca aat agg tgc ctc ggg att cca tac cat aga tcc agt aga tcc taa ttc aca ata cga ttt aga ttc tcc tat caa ctg ata tcc gct att aca aga gta cgt tat act aga gcc aaa gtc tac tcc gcc aat atc aag ttg gcc att atc gat atc tcg agg cga tgg gca tct ccg ttt aat aca ttg att aaa gag tgt cca tcc agt acc tgt aca ttt agc ata tat agg tcc cat ttt ttg ctt tct gta tcc agg tag aca tag a-t att cta tag tgt ctc cta tat tat a-- --- --- --- -at ggg aat --- --- --- --- --- cca ctg tac cct taa att tca tag taa tag gac gtg aag gaa tag gac agc atg ata gaa cgc atc cta ttc cca aca atg tca gga acg tca cgc tct cca cct tca tat tta ttt atc cgt -aa aaa tgt tat cct gga cat cgt aca aat aat aaa aaa gcc ca- -ta tat gtt cgc tat tgt aga aat tgt ttt tca cag ttg ctc aaa aac gat ggc agt gac tta tga gtt acg tta cac ttt gga gtc tca tct tta gta aac ata tca taa tat tcg ata tta cga gtt gac ata tcg aac aaa ttc caa gta ttt gat ttt gga taa tat tcg tat ttt gca tca gct ata att aag ata taa tca ccg caa gaa cac acg aac atc ttt cct aca tgg tta aag tac atg tac aat tct atc cat ttg tct tcc tta act ata tat ttg tat aga taa tta cga gtc tcg tga gta att cca gta att gca tag atg tca cca tcg tac tct aca gca taa act ata cta tga tgt cta ggc atg gga gac -tt ttt tat cca acg att ttt agt gaa aca ttc cac gtc gtt taa tac tac ata ttt ctc ata cgt ggt ata aac tcc acc cat tac --a tat ata tca tcg ttt acg aat acc gat gct cct gaa tat cta gga gta att aag ttt gga agt ctt atc cat ttc gaa gtg ccg tgt ttc aaa tat tct gcc acg ccc gtt gaa ata gaa aat tct aat cct cct att aca tat aac ttt cca tcg tta aca caa gta cta act tct gat ttt aac gac gac ata tta gta acc gtt ttc ca- --- --- --- --- --- --- --t ttt ttc gtt tta aga tct acc cgc -ga tac gga ata aac atg tct att gtt aat cat gcc gcc aat aat gta tag aca att atg taa aac att tgc att ata gaa ttg --- -tc tat ctg tat tac cga cta tcg tcc aat att ctg ttc tag gag agt aat ggg tta ttg tgg ata tat aat cag agt ttt taa tga cta cta tat tat gtt tta tac cat ttc gtg tca cag gct ttg tag att tgg ata tag tta atc cca aca atg cta tag cat tgc gca tag cat tag tca taa act tgg gat gta aaa tgt tga tga tat cta cat cgt ttg gat ttt tat gta tcc act tta ata ata tta tag ctg taa cat cct cat ga- ttt acg tta acg ttt tcg tgg gat aag ata gtt gtt agt tca tcc ttt gat aat ttt cca aat t

>MK035753.1:34206-36158_Cowpox_virus_strain_CPXV/Cepad_333

cgg ctt cca tgt att tcc tgg aga gca agt --- aga tga tg- agg aac cag ata gtt tat atc cgt act tgc act taa agt cta cat tgt cgt tgt atg a-- gta tga tct ttt aaa ccc gct aga caa gta tcc gtt tga tat tgt agg atg tgg aca ttt aac aat ctg aca cgt ggg tgg atc gga cca ttc tcc tcc tga aca cag gac agc aga gtt acc aat taa cga ata tcc act att gca act ata agt tac aac gct tcc atc ggt ata aaa atc ctc gta tcc gtt atg tct tcc gtt gga tat aga tgg agg tga ttg gca ttt aac aga ttc gca aat agg tgc ctc ggg att cca tac cat aga tcc agt aga tcc taa ttc aca ata cga ttt aga ttc tcc tat caa ctg ata tcc gct att aca aga gta cgt tat act aga gcc aaa gtc tac tcc gcc aat atc aag ttg gcc att atc gat atc tcg agg cga tgg gca tct ccg ttt aat aca ttg att aaa gag tgt cca tcc agt acc tgt aca ttt agc ata tat agg tcc cat ttt ttg ctt tct gta tcc agg tag aca tag a-t att cta tag tgt ctc cta tat tat a-- --- --- --- -at ggg aat --- --- --- --- --- cca ctg tac cct taa att tca tag taa tag gac gtg aag gaa tag gac agc atg ata gaa cgc atc cta ttc cca aca atg tca gga acg tca cgc tct cca cct tca tat tta ttt atc cgt -aa aaa tgt tat cct gga cat cgt aca aat aat aaa aaa gcc ca- -ta tat gtt cgc tat tgt aga aat tgt ttt tca cag ttg ctc aaa aac gat ggc agt gac tta tga gtt acg tta cac ttt gga gtc tca tct tta gta aac ata tca taa tat tcg ata tta cga gtt gac ata tcg aac aaa ttc caa gta ttt gat ttt gga taa tat tcg tat ttt gca tca gct ata att aag ata taa tca ccg caa gaa cac acg aac atc ttt cct aca tgg tta aag tac atg tac aat tct atc cat ttg tct tcc tta act ata tat ttg tat aga taa tta cga gtc tcg tga gta att cca gta att gca tag atg tca cca tcg tac tct aca gca taa act ata cta tga tgt cta ggc atg gga gac -tt ttt tat cca acg att ttt agt gaa aca ttc cac gtc gtt taa tac tac ata ttt ctc ata cgt ggt ata aac tcc acc cat tac --a tat ata tca tcg ttt acg aat acc gat gct cct gaa tat cta gga gta att aag ttt gga agt ctt atc cat ttc gaa gtg ccg tgt ttc aaa tat tct gcc acg ccc gtt gaa ata gaa aat tct aat cct cct att aca tat aac ttt cca tcg tta aca caa gta cta act tct gat ttt aac gac gac ata tta gta acc gtt ttc ca- --- --- --- --- --- --- --t ttt ttc gtt tta aga tct acc cgc -ga tac gga ata aac atg tct att gtt aat cat gcc gcc aat aat gta tag aca att atg taa aac att tgc att ata gaa ttg --- -tc tat ctg tat tac cga cta tcg tcc aat att ctg ttc tag gag agt aat ggg tta ttg tgg ata tat aat cag agt ttt taa tga cta cta tat tat gtt tta tac cat ttc gtg tca cag gct ttg tag att tgg ata tag tta atc cca aca atg cta tag cat tgc gca tag cat tag tca taa act tgg gat gta aaa tgt tga tga tat cta cat cgt ttg gat ttt tat gta tcc act tta ata ata tta tag ctg taa cat cct cat ga- ttt acg tta acg ttt tcg tgg gat aag ata gtt gtt agt tca tcc ttt gat aat ttt cca aat t

>MK035750.1:24015-25967_Cowpox_virus_strain_CPXV/Compiegne_2009

cgg ctt cca tgt att tcc tgg aga gca agt --- aga tga tg- agg aac cag ata gtt tat atc cgt act tgc act taa agt cta cat tgt cgt tgt atg a-- gta tga tct ttt aaa ccc gct aga caa gta tcc gtt tga tat tgt agg atg tgg aca ttt aac aat ctg aca cgt ggg tgg atc gga cca ttc tcc tcc tga aca cag gac agc aga gtt acc aat taa cga ata tcc act att gca act ata agt tac aac gct tcc atc ggt ata aaa atc ctc gta tcc gtt atg tct tcc gtt gga tat aga tgg agg tga ttg gca ttt aac aga ttc gca aat agg tgc ctc ggg att cca tac cat aga tcc agt aga tcc taa ttc aca ata cga ttt aga ttc tcc tat caa ctg ata tcc gct att aca aga gta cgt tat act aga gcc aaa gtc tac tcc gcc aat atc aag ttg gcc att atc gat atc tcg agg cga tgg gca tct ccg ttt aat aca ttg att aaa gag tgt cca tcc agt acc tgt aca ttt agc ata tat agg tcc cat ttt ttg ctt tct gta tcc agg tag aca tag a-t att cta tag tgt ctc cta tat tat a-- --- --- --- -at ggg aat --- --- --- --- --- cca ctg tac cct taa att tca tag taa tag gac gtg aag gaa tag gac agc atg ata gaa cgc atc cta ttc cca aca atg tca gga acg tca cgc tct cca cct tca tat tta ttt atc cgt -aa aaa tgt tat cct gga cat cgt aca aat aat aaa aaa gcc ca- -ta tat gtt cgc tat tgt aga aat tgt ttt tca cag ttg ctc aaa aac gat ggc agt gac tta tga gtt acg tta cac ttt gga gtc tca tct tta gta aac ata tca taa tat tcg ata tta cga gtt gac ata tcg aac aaa ttc caa gta ttt gat ttt gga taa tat tcg tat ttt gca tca gct ata att aag ata taa tca ccg caa gaa cac acg aac atc ttt cct aca tgg tta aag tac atg tac aat tct atc cat ttg tct tcc tta act ata tat ttg tat aga taa tta cga gtc tcg tga gta att cca gta att gca tag atg tca cca tcg tac tct aca gca taa act ata cta tga tgt cta ggc atg gga gac -tt ttt tat cca acg att ttt agt gaa aca ttc cac gtc gtt taa tac tac ata ttt ctc ata cgt ggt ata aac tcc acc cat tac --a tat ata tca tcg ttt acg aat acc gat gct cct gaa tat cta gga gta att aag ttt gga agt ctt atc cat ttc gaa gtg ccg tgt ttc aaa tat tct gcc acg ccc gtt gaa ata gaa aat tct aat cct cct att aca tat aac ttt cca tcg tta aca caa gta cta act tct gat ttt aac gac gac ata tta gta acc gtt ttc ca- --- --- --- --- --- --- --t ttt ttc gtt tta aga tct acc cgc -ga tac gga ata aac atg tct att gtt aat cat gcc gcc aat aat gta tag aca att atg taa aac att tgc att ata gaa ttg --- -tc tat ctg tat tac cga cta tcg tcc aat att ctg ttc tag gag agt aat ggg tta ttg tgg ata tat aat cag agt ttt taa tga cta cta tat tat gtt tta tac cat ttc gtg tca cag gct ttg tag att tgg ata tag tta atc cca aca atg cta tag cat tgc gca tag cat tag tca taa act tgg gat gta aaa tgt tga tga tat cta cat cgt ttg gat ttt tat gta tcc act tta ata ata tta tag ctg taa cat cct cat ga- ttt acg tta acg ttt tcg tgg gat aag ata gtt gtt agt tca tcc ttt gat aat ttt cca aat t

>MK035749.1:19711-21663_Cowpox_virus_strain_CPXV/Rat_Biederstein

cgg ctt cca tgt att tcc tgg aga gca agt --- aga tga tg- agg aac cag ata gtt tat atc cgt act tgc act taa agt cta cat tgt cgt tgt atg a-- gta tga tct ttt aaa ccc gct aga caa gta tcc gtt tga tat tgt agg atg tgg aca ttt aac aat ctg aca cgt ggg tgg atc gga cca ttc tcc tcc tga aca cag gac agc aga gtt acc aat taa cga ata tcc act att gca act ata agt tac aac gct tcc atc ggt ata aaa atc ctc gta tcc gtt atg tct tcc gtt gga tat aga tgg agg tga ttg gca ttt aac aga ttc gca aat agg tgc ctc ggg att cca tac cat aga tcc agt aga tcc taa ttc aca ata cga ttt aga ttc tcc tat caa ctg ata tcc gct att aca aga gta cgt tat act aga gcc aaa gtc tac tcc gcc aat atc aag ttg gcc att atc gat atc tcg agg cga tgg gca tct ccg ttt aat aca ttg att aaa gag tgt cca tcc agt acc tgt aca ttt agc ata tat agg tcc cat ttt ttg ctt tct gta tcc agg tag aca tag a-t att cta tag tgt ctc cta tat tat a-- --- --- --- -at ggg aat --- --- --- --- --- cca ctg tac cct taa att tca tag taa tag gac gtg aag gaa tag gac agc atg ata gaa cgc atc cta ttc cca aca atg tca gga acg tca cgc tct cca cct tca tat tta ttt atc cgt -aa aaa tgt tat cct gga cat cgt aca aat aat aaa aaa gcc ca- -ta tat gtt cgc tat tgt aga aat tgt ttt tca cag ttg ctc aaa aac gat ggc agt gac tta tga gtt acg tta cac ttt gga gtc tca tct tta gta aac ata tca taa tat tcg ata tta cga gtt gac ata tcg aac aaa ttc caa gta ttt gat ttt gga taa tat tcg tat ttt gca tca gct ata att aag ata taa tca ccg caa gaa cac acg aac atc ttt cct aca tgg tta aag tac atg tac aat tct atc cat ttg tct tcc tta act ata tat ttg tat aga taa tta cga gtc tcg tga gta att cca gta att gca tag atg tca cca tcg tac tct aca gca taa act ata cta tga tgt cta ggc atg gga gac -tt ttt tat cca acg att ttt agt gaa aca ttc cac gtc gtt taa tac tac ata ttt ctc ata cgt ggt ata aac tcc acc cat tac --a tat ata tca tcg ttt acg aat acc gat gct cct gaa tat cta gga gta att aag ttt gga agt ctt atc cat ttc gaa gtg ccg tgt ttc aaa tat tct gcc acg ccc gtt gaa ata gaa aat tct aat cct cct att aca tat aac ttt cca tcg tta aca caa gta cta act tct gat ttt aac gac gac ata tta gta acc gtt ttc ca- --- --- --- --- --- --- --t ttt ttc gtt tta aga tct acc cgc -ga tac gga ata aac atg tct att gtt aat cat gcc gcc aat aat gta tag aca att atg taa aac att tgc att ata gaa ttg --- -tc tat ctg tat tac cga cta tcg tcc aat att ctg ttc tag gag agt aat ggg tta ttg tgg ata tat aat cag agt ttt taa tga cta cta tat tat gtt tta tac cat ttc gtg tca cag gct ttg tag att tgg ata tag tta atc cca aca atg cta tag cat tgc gca tag cat tag tca taa act tgg gat gta aaa tgt tga tga tat cta cat cgt ttg gat ttt tat gta tcc act tta ata ata tta tag ctg taa cat cct cat ga- ttt acg tta acg ttt tcg tgg gat aag ata gtt gtt agt tca tcc ttt gat aat ttt cca aat t

>MK035748.1:34119-36071_Cowpox_virus_strain_CPXV/Boa_Marl

cgg ctt cca tgt att tcc tgg aga gca agt --- aga tga tg- agg aac cag ata gtt tat atc cgt act tgc act taa agt cta cat tgt cgt tgt atg a-- gta tga tct ttt aaa ccc gct aga caa gta tcc gtt tga tat tgt agg atg tgg aca ttt aac aat ctg aca cgt ggg tgg atc gga cca ttc tcc tcc tga aca cag gac agc aga gtt acc aat taa cga ata tcc act att gca act ata agt tac aac gct tcc atc ggt ata aaa atc ctc gta tcc gtt atg tct tcc gtt gga tat aga tgg agg tga ttg gca ttt aac aga ttc gca aat agg tgc ctc ggg att cca tac cat aga tcc agt aga tcc taa ttc aca ata cga ttt aga ttc tcc tat caa ctg ata tcc gct att aca aga gta cgt tat act aga gcc aaa gtc tac tcc gcc aat atc aag ttg gcc att atc gat atc tcg agg cga tgg gca tct ccg ttt aat aca ttg att aaa gag tgt cca tcc agt acc tgt aca ttt agc ata tat agg tcc cat ttt ttg ctt tct gta tcc agg tag aca tag a-t att cta tag tgt ctc cta tat tat a-- --- --- --- -at ggg aat --- --- --- --- --- cca ctg tac cct taa att tca tag taa tag gac gtg aag gaa tag gac agc atg ata gaa cgc atc cta ttc cca aca atg tca gga acg tca cgc tct cca cct tca tat tta ttt atc cgt -aa aaa tgt tat cct gga cat cgt aca aat aat aaa aaa gcc ca- -ta tat gtt cgc tat tgt aga aat tgt ttt tca cag ttg ctc aaa aac gat ggc agt gac tta tga gtt acg tta cac ttt gga gtc tca tct tta gta aac ata tca taa tat tcg ata tta cga gtt gac ata tcg aac aaa ttc caa gta ttt gat ttt gga taa tat tcg tat ttt gca tca gct ata att aag ata taa tca ccg caa gaa cac acg aac atc ttt cct aca tgg tta aag tac atg tac aat tct atc cat ttg tct tcc tta act ata tat ttg tat aga taa tta cga gtc tcg tga gta att cca gta att gca tag atg tca cca tcg tac tct aca gca taa act ata cta tga tgt cta ggc atg gga gac -tt ttt tat cca acg att ttt agt gaa aca ttc cac gtc gtt taa tac tac ata ttt ctc ata cgt ggt ata aac tcc acc cat tac --a tat ata tca tcg ttt acg aat acc gat gct cct gaa tat cta gga gta att aag ttt gga agt ctt atc cat ttc gaa gtg ccg tgt ttc aaa tat tct gcc acg ccc gtt gaa ata gaa aat tct aat cct cct att aca tat aac ttt cca tcg tta aca caa gta cta act tct gat ttt aac gac gac ata tta gta acc gtt ttc ca- --- --- --- --- --- --- --t ttt ttc gtt tta aga tct acc cgc -ga tac gga ata aac atg tct att gtt aat cat gcc gcc aat aat gta tag aca att atg taa aac att tgc att ata gaa ttg --- -tc tat ctg tat tac cga cta tcg tcc aat att ctg ttc tag gag agt aat ggg tta ttg tgg ata tat aat cag agt ttt taa tga cta cta tat tat gtt tta tac cat ttc gtg tca cag gct ttg tag att tgg ata tag tta atc cca aca atg cta tag cat tgc gca tag cat tag tca taa act tgg gat gta aaa tgt tga tga tat cta cat cgt ttg gat ttt tat gta tcc act tta ata ata tta tag ctg taa cat cct cat ga- ttt acg tta acg ttt tcg tgg gat aag ata gtt gtt agt tca tcc ttt gat aat ttt cca aat t

>MK035747.1:34121-36073_Cowpox_virus_strain_CPXV/Rat_Marl

cgg ctt cca tgt att tcc tgg aga gca agt --- aga tga tg- agg aac cag ata gtt tat atc cgt act tgc act taa agt cta cat tgt cgt tgt atg a-- gta tga tct ttt aaa ccc gct aga caa gta tcc gtt tga tat tgt agg atg tgg aca ttt aac aat ctg aca cgt ggg tgg atc gga cca ttc tcc tcc tga aca cag gac agc aga gtt acc aat taa cga ata tcc act att gca act ata agt tac aac gct tcc atc ggt ata aaa atc ctc gta tcc gtt atg tct tcc gtt gga tat aga tgg agg tga ttg gca ttt aac aga ttc gca aat agg tgc ctc ggg att cca tac cat aga tcc agt aga tcc taa ttc aca ata cga ttt aga ttc tcc tat caa ctg ata tcc gct att aca aga gta cgt tat act aga gcc aaa gtc tac tcc gcc aat atc aag ttg gcc att atc gat atc tcg agg cga tgg gca tct ccg ttt aat aca ttg att aaa gag tgt cca tcc agt acc tgt aca ttt agc ata tat agg tcc cat ttt ttg ctt tct gta tcc agg tag aca tag a-t att cta tag tgt ctc cta tat tat a-- --- --- --- -at ggg aat --- --- --- --- --- cca ctg tac cct taa att tca tag taa tag gac gtg aag gaa tag gac agc atg ata gaa cgc atc cta ttc cca aca atg tca gga acg tca cgc tct cca cct tca tat tta ttt atc cgt -aa aaa tgt tat cct gga cat cgt aca aat aat aaa aaa gcc ca- -ta tat gtt cgc tat tgt aga aat tgt ttt tca cag ttg ctc aaa aac gat ggc agt gac tta tga gtt acg tta cac ttt gga gtc tca tct tta gta aac ata tca taa tat tcg ata tta cga gtt gac ata tcg aac aaa ttc caa gta ttt gat ttt gga taa tat tcg tat ttt gca tca gct ata att aag ata taa tca ccg caa gaa cac acg aac atc ttt cct aca tgg tta aag tac atg tac aat tct atc cat ttg tct tcc tta act ata tat ttg tat aga taa tta cga gtc tcg tga gta att cca gta att gca tag atg tca cca tcg tac tct aca gca taa act ata cta tga tgt cta ggc atg gga gac -tt ttt tat cca acg att ttt agt gaa aca ttc cac gtc gtt taa tac tac ata ttt ctc ata cgt ggt ata aac tcc acc cat tac --a tat ata tca tcg ttt acg aat acc gat gct cct gaa tat cta gga gta att aag ttt gga agt ctt atc cat ttc gaa gtg ccg tgt ttc aaa tat tct gcc acg ccc gtt gaa ata gaa aat tct aat cct cct att aca tat aac ttt cca tcg tta aca caa gta cta act tct gat ttt aac gac gac ata tta gta acc gtt ttc ca- --- --- --- --- --- --- --t ttt ttc gtt tta aga tct acc cgc -ga tac gga ata aac atg tct att gtt aat cat gcc gcc aat aat gta tag aca att atg taa aac att tgc att ata gaa ttg --- -tc tat ctg tat tac cga cta tcg tcc aat att ctg ttc tag gag agt aat ggg tta ttg tgg ata tat aat cag agt ttt taa tga cta cta tat tat gtt tta tac cat ttc gtg tca cag gct ttg tag att tgg ata tag tta atc cca aca atg cta tag cat tgc gca tag cat tag tca taa act tgg gat gta aaa tgt tga tga tat cta cat cgt ttg gat ttt tat gta tcc act tta ata ata tta tag ctg taa cat cct cat ga- ttt acg tta acg ttt tcg tgg gat aag ata gtt gtt agt tca tcc ttt gat aat ttt cca aat t

>MK035746.1:20165-22117_Cowpox_virus_strain_CPXV/Boy_Biederstein

cgg ctt cca tgt att tcc tgg aga gca agt --- aga tga tg- agg aac cag ata gtt tat atc cgt act tgc act taa agt cta cat tgt cgt tgt atg a-- gta tga tct ttt aaa ccc gct aga caa gta tcc gtt tga tat tgt agg atg tgg aca ttt aac aat ctg aca cgt ggg tgg atc gga cca ttc tcc tcc tga aca cag gac agc aga gtt acc aat taa cga ata tcc act att gca act ata agt tac aac gct tcc atc ggt ata aaa atc ctc gta tcc gtt atg tct tcc gtt gga tat aga tgg agg tga ttg gca ttt aac aga ttc gca aat agg tgc ctc ggg att cca tac cat aga tcc agt aga tcc taa ttc aca ata cga ttt aga ttc tcc tat caa ctg ata tcc gct att aca aga gta cgt tat act aga gcc aaa gtc tac tcc gcc aat atc aag ttg gcc att atc gat atc tcg agg cga tgg gca tct ccg ttt aat aca ttg att aaa gag tgt cca tcc agt acc tgt aca ttt agc ata tat agg tcc cat ttt ttg ctt tct gta tcc agg tag aca tag a-t att cta tag tgt ctc cta tat tat a-- --- --- --- -at ggg aat --- --- --- --- --- cca ctg tac cct taa att tca tag taa tag gac gtg aag gaa tag gac agc atg ata gaa cgc atc cta ttc cca aca atg tca gga acg tca cgc tct cca cct tca tat tta ttt atc cgt -aa aaa tgt tat cct gga cat cgt aca aat aat aaa aaa gcc ca- -ta tat gtt cgc tat tgt aga aat tgt ttt tca cag ttg ctc aaa aac gat ggc agt gac tta tga gtt acg tta cac ttt gga gtc tca tct tta gta aac ata tca taa tat tcg ata tta cga gtt gac ata tcg aac aaa ttc caa gta ttt gat ttt gga taa tat tcg tat ttt gca tca gct ata att aag ata taa tca ccg caa gaa cac acg aac atc ttt cct aca tgg tta aag tac atg tac aat tct atc cat ttg tct tcc tta act ata tat ttg tat aga taa tta cga gtc tcg tga gta att cca gta att gca tag atg tca cca tcg tac tct aca gca taa act ata cta tga tgt cta ggc atg gga gac -tt ttt tat cca acg att ttt agt gaa aca ttc cac gtc gtt taa tac tac ata ttt ctc ata cgt ggt ata aac tcc acc cat tac --a tat ata tca tcg ttt acg aat acc gat gct cct gaa tat cta gga gta att aag ttt gga agt ctt atc cat ttc gaa gtg ccg tgt ttc aaa tat tct gcc acg ccc gtt gaa ata gaa aat tct aat cct cct att aca tat aac ttt cca tcg tta aca caa gta cta act tct gat ttt aac gac gac ata tta gta acc gtt ttc ca- --- --- --- --- --- --- --t ttt ttc gtt tta aga tct acc cgc -ga tac gga ata aac atg tct att gtt aat cat gcc gcc aat aat gta tag aca att atg taa aac att tgc att ata gaa ttg --- -tc tat ctg tat tac cga cta tcg tcc aat att ctg ttc tag gag agt aat ggg tta ttg tgg ata tat aat cag agt ttt taa tga cta cta tat tat gtt tta tac cat ttc gtg tca cag gct ttg tag att tgg ata tag tta atc cca aca atg cta tag cat tgc gca tag cat tag tca taa act tgg gat gta aaa tgt tga tga tat cta cat cgt ttg gat ttt tat gta tcc act tta ata ata tta tag ctg taa cat cct cat ga- ttt acg tta acg ttt tcg tgg gat aag ata gtt gtt agt tca tcc ttt gat aat ttt cca aat t

>LN864565.1:34024-35976_Cowpox_virus_complete_genome_strain_Ratpox09

cgg ctt cca tgt att tcc tgg aga gca agt --- aga tga tg- agg aac cag ata gtt tat atc cgt act tgc act taa agt cta cat tgt cgt tgt atg a-- gta tga tct ttt aaa ccc gct aga caa gta tcc gtt tga tat tgt agg atg tgg aca ttt aac aat ctg aca cgt ggg tgg atc gga cca ttc tcc tcc tga aca cag gac agc aga gtt acc aat taa cga ata tcc act att gca act ata agt tac aac gct tcc atc ggt ata aaa atc ctc gta tcc gtt atg tct tcc gtt gga tat aga tgg agg tga ttg gca ttt aac aga ttc gca aat agg tgc ctc ggg att cca tac cat aga tcc agt aga tcc taa ttc aca ata cga ttt aga ttc tcc tat caa ctg ata tcc gct att aca aga gta cgt tat act aga gcc aaa gtc tac tcc gcc aat atc aag ttg gcc att atc gat atc tcg agg cga tgg gca tct ccg ttt aat aca ttg att aaa gag tgt cca tcc agt acc tgt aca ttt agc ata tat agg tcc cat ttt ttg ctt tct gta tcc agg tag aca tag a-t att cta tag tgt ctc cta tat tat a-- --- --- --- -at ggg aat --- --- --- --- --- cca ctg tac cct taa att tca tag taa tag gac gtg aag gaa tag gac agc atg ata gaa cgc atc cta ttc cca aca atg tca gga acg tca cgc tct cca cct tca tat tta ttt atc cgt -aa aaa tgt tat cct gga cat cgt aca aat aat aaa aaa gcc ca- -ta tat gtt cgc tat tgt aga aat tgt ttt tca cag ttg ctc aaa aac gat ggc agt gac tta tga gtt acg tta cac ttt gga gtc tca tct tta gta aac ata tca taa tat tcg ata tta cga gtt gac ata tcg aac aaa ttc caa gta ttt gat ttt gga taa tat tcg tat ttt gca tca gct ata att aag ata taa tca ccg caa gaa cac acg aac atc ttt cct aca tgg tta aag tac atg tac aat tct atc cat ttg tct tcc tta act ata tat ttg tat aga taa tta cga gtc tcg tga gta att cca gta att gca tag atg tca cca tcg tac tct aca gca taa act ata cta tga tgt cta ggc atg gga gac -tt ttt tat cca acg att ttt agt gaa aca ttc cac gtc gtt taa tac tac ata ttt ctc ata cgt ggt ata aac tcc acc cat tac --a tat ata tca tcg ttt acg aat acc gat gct cct gaa tat cta gga gta att aag ttt gga agt ctt atc cat ttc gaa gtg ccg tgt ttc aaa tat tct gcc acg ccc gtt gaa ata gaa aat tct aat cct cct att aca tat aac ttt cca tcg tta aca caa gta cta act tct gat ttt aac gac gac ata tta gta acc gtt ttc ca- --- --- --- --- --- --- --t ttt ttc gtt tta aga tct acc cgc -ga tac gga ata aac atg tct att gtt aat cat gcc gcc aat aat gta tag aca att atg taa aac att tgc att ata gaa ttg --- -tc tat ctg tat tac cga cta tcg tcc aat att ctg ttc tag gag agt aat ggg tta ttg tgg ata tat aat cag agt ttt taa tga cta cta tat tat gtt tta tac cat ttc gtg tca cag gct ttg tag att tgg ata tag tta atc cca aca atg cta tag cat tgc gca tag cat tag tca taa act tgg gat gta aaa tgt tga tga tat cta cat cgt ttg gat ttt tat gta tcc act tta ata ata tta tag ctg taa cat cct cat ga- ttt acg tta acg ttt tcg tgg gat aag ata gtt gtt agt tca tcc ttt gat aat ttt cca aat t

>KF179385.1:15831-17789_Vaccinia_virus_complete_genome

cgg ctg cca tgt att tcc tgg aga gca agt --- aga tga tg- agg aac cag ata ggt tat atc cgt act tgc act taa agt tta cat tgt cgt tgt atg a-- gta tga tct ttt aaa ccc gct aga caa gta tcc gtt tga tat tgt agg atg tgg aca ttt aac aat ttg aca cgt ggg tgg atc gga cca ttc tcc tcc tga aca cag gac act aga gtt acc aat caa cga ata tcc act att gca act ata agt tac aac gct tcc atc ggt ata aaa atc ctc gta tcc gtt atg tct tcc gtt gga tat aga tgg agg gga ttg gca ttt aac aga ttc aca aat agg tgc ctc ggg att cca tac cat aga tcc agt aaa tcc taa ttc aca ata cga ttt aga ttc tcc tat caa ctg ata tcc gct att aca aga gta cgt tat act aga gcc aaa gtc tac tcc gcc aat atc aag ttg gcc att atc gat atc tcg agg cga tgg gca tct ccg ttt aat aca ttg att aaa gag tgt cca tcc agt acc tgt aca ttt agc ata tat agg tcc cat ttt ttg ctt tct gta tcc agg tag aca tag a-t att cta tag tgt ctc cta tgt tgt a-- --- --- --- --- --- -at tag cat tag tt- --t cca cac tat tct taa att tca tat taa tgg gac gtg aag gaa tag gac agt atg ata gaa cgc atc cta ttc cca aca atg tca gga acg tca cgc tct cca cct tca tat tta ttt atc cgt -aa aaa tgt tat cct gga cat cgt aca aat aat aaa aaa gcc ca- -ta tat gtt tgc tat tgt aga aat tgt ttt tca cag ttg ctc aaa aac gat ggc agt gac tta tga gtt acg tta cac ttt gga gtt tca tct tta gta aac ata tca taa tat tcg ata tta cga gtt gac ata tcg aac aaa ttc caa gta ttt gat ttt gga taa tat tcg tat ttt gca tct gct ata att aag ata taa tca ccg caa gaa cac acg aac atc ttt cct aca tgg tta aag tac atg tac aat tct atc cat ttg tct tcc tta act ata tat ttg tat aga taa tta cga gtc tcg tga gta att cca gta att aca tag atg tcg ccg tcg tac tct aca gca taa act ata cta tga tgt cta ggc atg gga gac -tt ttt tat cca acg att ttt agt gaa aca ttc cac atc gtt taa tac tac ata ttt ttc ata cgt ggt ata aac tcc acc cat tac --a tat ata tca tcg ttt acg aat acc gac gcg cct gaa tat cta gga gta att aag ttt gga agt ctt atc cat ttt gaa gtg ccg tgt ttc aaa tat tct gct aca ccc gtt gaa ata gaa aat tct aat cct cct att aca tat aac ttt cca ttg tta aca caa gta cta act tct gat ttt aac gac gac ata tta gta acc gtt ttc ca- --- --- --- --- --- --- --t ttt ttc gtt tca aga tct acc cgc -aa tac gga ata aac atg tct att gtt aat cat gcc gcc aat aat gta tag aca att atg taa aac att tgc att ata gaa ttg --- -tc tat ctg tat tac cga cta tcg tcc aat att ctg ttc tag gag agt aat ggg tta ttg tgg ata tat aat cag agt ttt taa tga cta cta tat tat gtt tta tac cat ttc gtg tca ctg gct ttg tag att tgg ata tag tta atc cca aca atg ata tag cat tgc gca tag tat tag tta taa act tgg gat gta aaa tgt tga tga tat cta cat cgt ttg gat ttt tat gta tcc act tta ata ata tca tag ctg taa cat cct cat ga- ttt acg tta acg tct tcg tgg gat aag ata gtt gtc agt tca tcc ttt gat aat ttt cca aat t

>KC813512.1:26341-28293_Cowpox_virus_strain_HumKre08/1_complete_genome

cgg ctt cca tgt att tcc tgg aga gca agt --- aga tga tg- agg aac cag ata gtt tat atc cgt act tgc act taa agt cta cat tgt cgt tgt atg a-- gta tga tct ttt aaa ccc gct aga caa gta tcc gtt tga tat tgt agg atg tgg aca ttt aac aat ctg aca cgt ggg tgg atc gga cca ttc tcc tcc tga aca cag gac agc aga gtt acc aat taa cga ata tcc act att gca act ata agt tac aac gct tcc atc ggt ata aaa atc ctc gta tcc gtt atg tct tcc gtt gga tat aga tgg agg tga ttg gca ttt aac aga ttc gca aat agg tgc ctc ggg att cca tac cat aga tcc agt aga tcc taa ttc aca ata cga ttt aga ttc tcc tat caa ctg ata tcc gct att aca aga gta cgt tat act aga gcc aaa gtc tac tcc gcc aat atc aag ttg gcc att atc gat atc tcg agg cga tgg gca tct ccg ttt aat aca ttg att aaa gag tgt cca tcc agt acc tgt aca ttt agc ata tat agg tcc cat ttt ttg ctt tct gta tcc agg tag aca tag a-t att cta tag tgt ctc cta tat tat a-- --- --- --- -at ggg aat --- --- --- --- --- cca ctg tac cct taa att tca tag taa tag gac gtg aag gaa tag gac agc atg ata gaa cgc atc cta ttc cca aca atg tca gga acg tca cgc tct cca cct tca tat tta ttt atc cgt -aa aaa tgt tat cct gga cat cgt aca aat aat aaa aaa gcc ca- -ta tat gtt cgc tat tgt aga aat tgt ttt tca cag ttg ctc aaa aac gat ggc agt gac tta tga gtt acg tta cac ttt gga gtc tca tct tta gta aac ata tca taa tat tcg ata tta cga gtt gac ata tcg aac aaa ttc caa gta ttt gat ttt gga taa tat tcg tat ttt gca tca gct ata att aag ata taa tca ccg caa gaa cac acg aac atc ttt cct aca tgg tta aag tac atg tac aat tct atc cat ttg tct tcc tta act ata tat ttg tat aga taa tta cga gtc tcg tga gta att cca gta att gca tag atg tca cca tcg tac tct aca gca taa act ata cta tga tgt cta ggc atg gga gac -tt ttt tat cca acg att ttt agt gaa aca ttc cac gtc gtt taa tac tac ata ttt ctc ata cgt ggt ata aac tcc acc cat tac --a tat ata tca tcg ttt acg aat acc gat gct cct gaa tat cta gga gta att aag ttt gga agt ctt atc cat ttc gaa gtg ccg tgt ttc aaa tat tct gcc acg ccc gtt gaa ata gaa aat tct aat cct cct att aca tat aac ttt cca tcg tta aca caa gta cta act tct gat ttt aac gac gac ata tta gta acc gtt ttc ca- --- --- --- --- --- --- --t ttt ttc gtt tta aga tct acc cgc -ga tac gga ata aac atg tct att gtt aat cat gcc gcc aat aat gta tag aca att atg taa aac att tgc att ata gaa ttg --- -tc tat ctg tat tac cga cta tcg tcc aat att ctg ttc tag gag agt aat ggg tta ttg tgg ata tat aat cag agt ttt taa tga cta cta tat tat gtt tta tac cat ttc gtg tca cag gct ttg tag att tgg ata tag tta atc cca aca atg cta tag cat tgc gca tag cat tag tca taa act tgg gat gta aaa tgt tga tga tat cta cat cgt ttg gat ttt tat gta tcc act tta ata ata tta tag ctg taa cat cct cat ga- ttt acg tta acg ttt tcg tgg gat aag ata gtt gtt agt tca tcc ttt gat aat ttt cca aat t

>KC813508.1:26666-28618_Cowpox_virus_strain_HumAac09/1_complete_genome

cgg ctt cca tgt att tcc tgg aga gca agt --- aga tga tg- agg aac cag ata gtt tat atc cgt act tgc act taa agt cta cat tgt cgt tgt atg a-- gta tga tct ttt aaa ccc gct aga caa gta tcc gtt tga tat tgt agg atg tgg aca ttt aac aat ctg aca cgt ggg tgg atc gga cca ttc tcc tcc tga aca cag gac agc aga gtt acc aat taa cga ata tcc act att gca act ata agt tac aac gct tcc atc ggt ata aaa atc ctc gta tcc gtt atg tct tcc gtt gga tat aga tgg agg tga ttg gca ttt aac aga ttc gca aat agg tgc ctc ggg att cca tac cat aga tcc agt aga tcc taa ttc aca ata cga ttt aga ttc tcc tat caa ctg ata tcc gct att aca aga gta cgt tat act aga gcc aaa gtc tac tcc gcc aat atc aag ttg gcc att atc gat atc tcg agg cga tgg gca tct ccg ttt aat aca ttg att aaa gag tgt cca tcc agt acc tgt aca ttt agc ata tat agg tcc cat ttt ttg ctt tct gta tcc agg tag aca tag a-t att cta tag tgt ctc cta tat tat a-- --- --- --- -at ggg aat --- --- --- --- --- cca ctg tac cct taa att tca tag taa tag gac gtg aag gaa tag gac agc atg ata gaa cgc atc cta ttc cca aca atg tca gga acg tca cgc tct cca cct tca tat tta ttt atc cgt -aa aaa tgt tat cct gga cat cgt aca aat aat aaa aaa gcc ca- -ta tat gtt cgc tat tgt aga aat tgt ttt tca cag ttg ctc aaa aac gat ggc agt gac tta tga gtt acg tta cac ttt gga gtc tca tct tta gta aac ata tca taa tat tcg ata tta cga gtt gac ata tcg aac aaa ttc caa gta ttt gat ttt gga taa tat tcg tat ttt gca tca gct ata att aag ata taa tca ccg caa gaa cac acg aac atc ttt cct aca tgg tta aag tac atg tac aat tct atc cat ttg tct tcc tta act ata tat ttg tat aga taa tta cga gtc tcg tga gta att cca gta att gca tag atg tca cca tcg tac tct aca gca taa act ata cta tga tgt cta ggc atg gga gac -tt ttt tat cca acg att ttt agt gaa aca ttc cac gtc gtt taa tac tac ata ttt ctc ata cgt ggt ata aac tcc acc cat tac --a tat ata tca tcg ttt acg aat acc gat gct cct gaa tat cta gga gta att aag ttt gga agt ctt atc cat ttc gaa gtg ccg tgt ttc aaa tat tct gcc acg ccc gtt gaa ata gaa aat tct aat cct cct att aca tat aac ttt cca tcg tta aca caa gta cta act tct gat ttt aac gac gac ata tta gta acc gtt ttc ca- --- --- --- --- --- --- --t ttt ttc gtt tta aga tct acc cgc -ga tac gga ata aac atg tct att gtt aat cat gcc gcc aat aat gta tag aca att atg taa aac att tgc att ata gaa ttg --- -tc tat ctg tat tac cga cta tcg tcc aat att ctg ttc tag gag agt aat ggg tta ttg tgg ata tat aat cag agt ttt taa tga cta cta tat tat gtt tta tac cat ttc gtg tca cag gct ttg tag att tgg ata tag tta atc cca aca atg cta tag cat tgc gca tag cat tag tca taa act tgg gat gta aaa tgt tga tga tat cta cat cgt ttg gat ttt tat gta tcc act tta ata ata tta tag ctg taa cat cct cat ga- ttt acg tta acg ttt tcg tgg gat aag ata gtt gtt agt tca tcc ttt gat aat ttt cca aat t

>KC813505.1:26587-28539_Cowpox_virus_strain_RatKre08/2_complete_genome

cgg ctt cca tgt att tcc tgg aga gca agt --- aga tga tg- agg aac cag ata gtt tat atc cgt act tgc act taa agt cta cat tgt cgt tgt atg a-- gta tga tct ttt aaa ccc gct aga caa gta tcc gtt tga tat tgt agg atg tgg aca ttt aac aat ctg aca cgt ggg tgg atc gga cca ttc tcc tcc tga aca cag gac agc aga gtt acc aat taa cga ata tcc act att gca act ata agt tac aac gct tcc atc ggt ata aaa atc ctc gta tcc gtt atg tct tcc gtt gga tat aga tgg agg tga ttg gca ttt aac aga ttc gca aat agg tgc ctc ggg att cca tac cat aga tcc agt aga tcc taa ttc aca ata cga ttt aga ttc tcc tat caa ctg ata tcc gct att aca aga gta cgt tat act aga gcc aaa gtc tac tcc gcc aat atc aag ttg gcc att atc gat atc tcg agg cga tgg gca tct ccg ttt aat aca ttg att aaa gag tgt cca tcc agt acc tgt aca ttt agc ata tat agg tcc cat ttt ttg ctt tct gta tcc agg tag aca tag a-t att cta tag tgt ctc cta tat tat a-- --- --- --- -at ggg aat --- --- --- --- --- cca ctg tac cct taa att tca tag taa tag gac gtg aag gaa tag gac agc atg ata gaa cgc atc cta ttc cca aca atg tca gga acg tca cgc tct cca cct tca tat tta ttt atc cgt -aa aaa tgt tat cct gga cat cgt aca aat aat aaa aaa gcc ca- -ta tat gtt cgc tat tgt aga aat tgt ttt tca cag ttg ctc aaa aac gat ggc agt gac tta tga gtt acg tta cac ttt gga gtc tca tct tta gta aac ata tca taa tat tcg ata tta cga gtt gac ata tcg aac aaa ttc caa gta ttt gat ttt gga taa tat tcg tat ttt gca tca gct ata att aag ata taa tca ccg caa gaa cac acg aac atc ttt cct aca tgg tta aag tac atg tac aat tct atc cat ttg tct tcc tta act ata tat ttg tat aga taa tta cga gtc tcg tga gta att cca gta att gca tag atg tca cca tcg tac tct aca gca taa act ata cta tga tgt cta ggc atg gga gac -tt ttt tat cca acg att ttt agt gaa aca ttc cac gtc gtt taa tac tac ata ttt ctc ata cgt ggt ata aac tcc acc cat tac --a tat ata tca tcg ttt acg aat acc gat gct cct gaa tat cta gga gta att aag ttt gga agt ctt atc cat ttc gaa gtg ccg tgt ttc aaa tat tct gcc acg ccc gtt gaa ata gaa aat tct aat cct cct att aca tat aac ttt cca tcg tta aca caa gta cta act tct gat ttt aac gac gac ata tta gta acc gtt ttc ca- --- --- --- --- --- --- --t ttt ttc gtt tta aga tct acc cgc -ga tac gga ata aac atg tct att gtt aat cat gcc gcc aat aat gta tag aca att atg taa aac att tgc att ata gaa ttg --- -tc tat ctg tat tac cga cta tcg tcc aat att ctg ttc tag gag agt aat ggg tta ttg tgg ata tat aat cag agt ttt taa tga cta cta tat tat gtt tta tac cat ttc gtg tca cag gct ttg tag att tgg ata tag tta atc cca aca atg cta tag cat tgc gca tag cat tag tca taa act tgg gat gta aaa tgt tga tga tat cta cat cgt ttg gat ttt tat gta tcc act tta ata ata tta tag ctg taa cat cct cat ga- ttt acg tta acg ttt tcg tgg gat aag ata gtt gtt agt tca tcc ttt gat aat ttt cca aat t

>KC813503.1:34371-36323_Cowpox_virus_strain_RatGer09/1_complete_genome

cgg ctt cca tgt att tcc tgg aga gca agt --- aga tga tg- agg aac cag ata gtt tat atc cgt act tgc act taa agt cta cat tgt cgt tgt atg a-- gta tga tct ttt aaa ccc gct aga caa gta tcc gtt tga tat tgt agg atg tgg aca ttt aac aat ctg aca cgt ggg tgg atc gga cca ttc tcc tcc tga aca cag gac agc aga gtt acc aat taa cga ata tcc act att gca act ata agt tac aac gct tcc atc ggt ata aaa atc ctc gta tcc gtt atg tct tcc gtt gga tat aga tgg agg tga ttg gca ttt aac aga ttc gca aat agg tgc ctc ggg att cca tac cat aga tcc agt aga tcc taa ttc aca ata cga ttt aga ttc tcc tat caa ctg ata tcc gct att aca aga gta cgt tat act aga gcc aaa gtc tac tcc gcc aat atc aag ttg gcc att atc gat atc tcg agg cga tgg gca tct ccg ttt aat aca ttg att aaa gag tgt cca tcc agt acc tgt aca ttt agc ata tat agg tcc cat ttt ttg ctt tct gta tcc agg tag aca tag a-t att cta tag tgt ctc cta tat tat a-- --- --- --- -at ggg aat --- --- --- --- --- cca ctg tac cct taa att tca tag taa tag gac gtg aag gaa tag gac agc atg ata gaa cgc atc cta ttc cca aca atg tca gga acg tca cgc tct cca cct tca tat tta ttt atc cgt -aa aaa tgt tat cct gga cat cgt aca aat aat aaa aaa gcc ca- -ta tat gtt cgc tat tgt aga aat tgt ttt tca cag ttg ctc aaa aac gat ggc agt gac tta tga gtt acg tta cac ttt gga gtc tca tct tta gta aac ata tca taa tat tcg ata tta cga gtt gac ata tcg aac aaa ttc caa gta ttt gat ttt gga taa tat tcg tat ttt gca tca gct ata att aag ata taa tca ccg caa gaa cac acg aac atc ttt cct aca tgg tta aag tac atg tac aat tct atc cat ttg tct tcc tta act ata tat ttg tat aga taa tta cga gtc tcg tga gta att cca gta att gca tag atg tca cca tcg tac tct aca gca taa act ata cta tga tgt cta ggc atg gga gac -tt ttt tat cca acg att ttt agt gaa aca ttc cac gtc gtt taa tac tac ata ttt ctc ata cgt ggt ata aac tcc acc cat tac --a tat ata tca tcg ttt acg aat acc gat gct cct gaa tat cta gga gta att aag ttt gga agt ctt atc cat ttc gaa gtg ccg tgt ttc aaa tat tct gcc acg ccc gtt gaa ata gaa aat tct aat cct cct att aca tat aac ttt cca tcg tta aca caa gta cta act tct gat ttt aac gac gac ata tta gta acc gtt ttc ca- --- --- --- --- --- --- --t ttt ttc gtt tta aga tct acc cgc -ga tac gga ata aac atg tct att gtt aat cat gcc gcc aat aat gta tag aca att atg taa aac att tgc att ata gaa ttg --- -tc tat ctg tat tac cga cta tcg tcc aat att ctg ttc tag gag agt aat ggg tta ttg tgg ata tat aat cag agt ttt taa tga cta cta tat tat gtt tta tac cat ttc gtg tca cag gct ttg tag att tgg ata tag tta atc cca aca atg cta tag cat tgc gca tag cat tag tca taa act tgg gat gta aaa tgt tga tga tat cta cat cgt ttg gat ttt tat gta tcc act tta ata ata tta tag ctg taa cat cct cat ga- ttt acg tta acg ttt tcg tgg gat aag ata gtt gtt agt tca tcc ttt gat aat ttt cca aat t

>KC813501.1:26342-28294_Cowpox_virus_strain_RatAac09/1_complete_genome

cgg ctt cca tgt att tcc tgg aga gca agt --- aga tga tg- agg aac cag ata gtt tat atc cgt act tgc act taa agt cta cat tgt cgt tgt atg a-- gta tga tct ttt aaa ccc gct aga caa gta tcc gtt tga tat tgt agg atg tgg aca ttt aac aat ctg aca cgt ggg tgg atc gga cca ttc tcc tcc tga aca cag gac agc aga gtt acc aat taa cga ata tcc act att gca act ata agt tac aac gct tcc atc ggt ata aaa atc ctc gta tcc gtt atg tct tcc gtt gga tat aga tgg agg tga ttg gca ttt aac aga ttc gca aat agg tgc ctc ggg att cca tac cat aga tcc agt aga tcc taa ttc aca ata cga ttt aga ttc tcc tat caa ctg ata tcc gct att aca aga gta cgt tat act aga gcc aaa gtc tac tcc gcc aat atc aag ttg gcc att atc gat atc tcg agg cga tgg gca tct ccg ttt aat aca ttg att aaa gag tgt cca tcc agt acc tgt aca ttt agc ata tat agg tcc cat ttt ttg ctt tct gta tcc agg tag aca tag a-t att cta tag tgt ctc cta tat tat a-- --- --- --- -at ggg aat --- --- --- --- --- cca ctg tac cct taa att tca tag taa tag gac gtg aag gaa tag gac agc atg ata gaa cgc atc cta ttc cca aca atg tca gga acg tca cgc tct cca cct tca tat tta ttt atc cgt -aa aaa tgt tat cct gga cat cgt aca aat aat aaa aaa gcc ca- -ta tat gtt cgc tat tgt aga aat tgt ttt tca cag ttg ctc aaa aac gat ggc agt gac tta tga gtt acg tta cac ttt gga gtc tca tct tta gta aac ata tca taa tat tcg ata tta cga gtt gac ata tcg aac aaa ttc caa gta ttt gat ttt gga taa tat tcg tat ttt gca tca gct ata att aag ata taa tca ccg caa gaa cac acg aac atc ttt cct aca tgg tta aag tac atg tac aat tct atc cat ttg tct tcc tta act ata tat ttg tat aga taa tta cga gtc tcg tga gta att cca gta att gca tag atg tca cca tcg tac tct aca gca taa act ata cta tga tgt cta ggc atg gga gac -tt ttt tat cca acg att ttt agt gaa aca ttc cac gtc gtt taa tac tac ata ttt ctc ata cgt ggt ata aac tcc acc cat tac --a tat ata tca tcg ttt acg aat acc gat gct cct gaa tat cta gga gta att aag ttt gga agt ctt atc cat ttc gaa gtg ccg tgt ttc aaa tat tct gcc acg ccc gtt gaa ata gaa aat tct aat cct cct att aca tat aac ttt cca tcg tta aca caa gta cta act tct gat ttt aac gac gac ata tta gta acc gtt ttc ca- --- --- --- --- --- --- --t ttt ttc gtt tta aga tct acc cgc -ga tac gga ata aac atg tct att gtt aat cat gcc gcc aat aat gta tag aca att atg taa aac att tgc att ata gaa ttg --- -tc tat ctg tat tac cga cta tcg tcc aat att ctg ttc tag gag agt aat ggg tta ttg tgg ata tat aat cag agt ttt taa tga cta cta tat tat gtt tta tac cat ttc gtg tca cag gct ttg tag att tgg ata tag tta atc cca aca atg cta tag cat tgc gca tag cat tag tca taa act tgg gat gta aaa tgt tga tga tat cta cat cgt ttg gat ttt tat gta tcc act tta ata ata tta tag ctg taa cat cct cat ga- ttt acg tta acg ttt tcg tgg gat aag ata gtt gtt agt tca tcc ttt gat aat ttt cca aat t

>MW018156.1:16060-18018_Vaccinia_virus_strain_Cantagalo_isolate_Cantagalo_ALEH2

cgg ctg cca tgt att tcc tgg aga gca agt --- aga tga tg- agg aac cag ata ggt tat atc cgt act tgc act taa agt tta cat tgt cgt tgt atg a-- gta tga tct ttt aaa ccc gct aga caa gta tcc gtt tga tat tgt agg atg tgg aca ttt aac aat ttg aca tgt ggg tgg atc gga cca ttc tcc tcc tga aca cag gac act aga gtt acc aat caa cga ata tcc act att gca act ata agt tac aac gct tcc atc ggt ata aaa atc ctc gta tcc gtt atg tct tcc gtt gga tat aga tgg agg gga ttg gca ttt aac aga ttc aca aat agg tgc ctc ggg att cca tac cat aga tcc agt aaa tcc taa ttc aca ata cga ttt aga ttc tcc tat caa ctg ata tcc gct att aca aga gta cgt tat act aga gcc aaa gtc tac tcc gcc aat atc aag ttg gcc att atc gat atc tcg agg cga tgg gca tct ccg ttt aat aca ttg att aaa gag tgt cca tcc agt acc tgt aca ttt agc ata tat agg tcc cat ttt ttg ctt tct gta tcc agg tag aca tag a-t att cta tag tgt ctc cta tgt tgt a-- --- --- --- --- --- -at tag cat tag tt- --t cca cac tat tct taa att tca tat taa tgg gac gtg aag gaa tag gac agt atg ata gaa cgc atc cta ttc cca aca atg tca gga acg tca cgc tct cca cct tca tat tta ttt atc cgt -aa aaa tgt tat cct gga cat cgt aca aat aat aaa aaa gcc ca- -ta tat gtt tgc tat tgt aga aat tgt ttt tca cag ttg ctc aaa aac gat ggc agt gac tta tga gtt acg tta cac ttt gga gtt tca tct tta gta aac ata tca taa tat tcg ata tta cga gtt gac ata tcg aac aaa ttc caa gta ttt gat ttt gga taa tat tcg tat ttt gca tct gct ata att aag ata taa tca ccg caa gaa cac acg aac atc ttt cct aca tgg tta aag tac atg tac aat tct atc cat ttg tct tcc tta act ata tat ttg tat aga taa tta cga gtc tcg tga gta att cca gta att aca tag atg tcg ccg tcg tac tct aca gca taa act ata cta tga tgt cta ggc atg gga gac -tt ttt tat cca acg att ttt agt gaa aca ttc cac atc gtt taa tac tac ata ttt ttc ata cgt ggt ata aac tcc acc cat tac --a tat ata tca tcg ttt acg aat acc gac gcg cct gaa tat cta gga gta att aag ttt gga agt ctt atc cat ttt gaa gtg ccg tgt ttc aaa tat tct gct aca ccc gtt gaa ata gaa aat tct aat cct cct att aca tat aac ttt cca ttg tta aca caa gta cta act tct gat ttt aac gac gac ata tta gta acc gtt ttc ca- --- --- --- --- --- --- --t ttt ttc gtt tca aga tct acc cgc -ga tac gga ata aac atg tct att gtt aat cat gcc gcc aat aat gta tag aca att atg taa aac att tgc att ata gaa ttg --- -tc tat ctg tat tac cga cta tcg tcc aat att ctg ttc tag gag agt aat ggg tta ttg tgg ata tat aat cag agt ttt taa tga cta cta tat tat gtt tta tac cat ttc gtg tca ctg gct ttg tag att tgg ata tag tta atc cca aca atg ata tag cat tgc gca tag tat tag tta taa act tgg gat gta aaa tgt tga tga tat cta cat cgt ttg gat ttt tat gta tcc act tta ata ata tca tag ctg taa cat cct cat ga- ttt acg tta acg tct tcg tgg gat aag ata gtt gtc agt tca tcc ttt gat aat ttt cca aat t

>KY349117.1:29869-31833_Horsepox_virus_strain_MNR_clone_2-1

cgg ctg cca tgt att tcc tgg aga gca agt --- aga tga tg- agg aac cag ata ggt tat atc cgt act tgc act taa agt tta cat tgt cgt tgt atg a-- gta tga tct ttt aaa ccc gct aga caa gta tcc gtt tga tat tgt agg atg tgg aca ttt aac aat ttg aca cgt ggg tgg atc gga cca ttc tcc tcc tga aca cag gac acc aga gtt acc aat caa cga ata tcc act att gca act ata agt tac aac gct tcc atc ggt ata aaa atc ctc gta tcc gtt atg tct tcc gtt gga tat aga tgg agg gaa ttg gca ttt aac aga ttc aca aat agg tgc ctc ggg att cca tac cat aga tcc agt aaa tcc taa ttc aca ata cga ttt aga ttc tcc tat caa ctg ata tcc gct att aca aga gta cgt tat act aga gcc aaa gtc tac tcc gcc aat atc aag ttg gcc att atc gat atc tcg agg cga tgg gca tct ccg ttt aat aca ttg att aaa gag tgt cca tcc agt acc tgt aca ttt agc ata tat agg tcc cat ttt ttg ctt tct gta tcc agg tag aca tag a-t att cta tag tgt ctc cta tgt tgt a-- --- --- --- -at tag cat tag cat tag tt- --t tca cac tat tct taa att tca tat taa tgg gac gtg aag gaa tag gac agt atg ata gaa cgc atc cta ttc cca aca atg tca gga acg tca cgc tct cca cct tca tat tta ttt atc cgt -aa aaa tgt tat cct gga cat cgt aca aat aat aaa aaa gcc ca- -ta tat gtt tgc tat tgt aga aat tgt ttt tca cag ttg ctc aaa aac gat ggc agt gac tta tga gtt acg tta cac ttt gga gtt tca tct tta gta aac ata tca taa tat tcg ata tta cga gtt gac ata tcg aac aaa ttc caa gta ttt gat ttt gga taa tat tcg tat ttt gca tct gct ata att aag ata taa tca ccg caa gaa cac acg aac atc ttt cct aca tgg tta aag tac atg tat aat tct atc cat ttg tct tcc tta act ata tat ttg tat aga taa tta cga gtc tca taa gta att cca gta att gca tag atg tca cca tcg tac tct aca gca taa act ata cta tga tgt cta ggc atg gga gac -tt ttt tat cca acg att ttt agt gaa aca ttc tac atc gtt taa tac tac ata ttt ctc ata cgt ggt ata aac tcc acc cat tac --a tat ata tca tcg ttt acg aat acc gat gcg cct gaa tat cta gga gta att aag ttt gga agt ctt atc cat ttt gaa gtg ccg tgt ttc aaa tat tct gct aca ccc gtt gaa ata gaa aat tct aat cct cct att aca tat aac ttt cca ttg tta aca caa gta cta act tct gat ttt aac gac gac ata tta gta acc gtt ttc ca- --- --- --- --- --- --- --t ttt ttc gtt tca aga tct acc cgc -ga tac gga ata aac atg tct att gtt aat cat gcc gcc aat aat gta tag aca att atg taa aac att tgc att ata gaa ttg --- -tc tat ctg tat tac cga cta tcg tcc aat att ctg ttc tag gag agt aat ggg tta ttg tgg ata tat aat cag agt ttt taa tga cta cta tat tat gtt tta tac cat ttc gtg tca ctg gct ttg tag att tgg ata tag tta atc cca aca atg ata tag cat tgc gca tag tat tag tta taa act tgg gat gta aaa tgt tga tga tat cta cat cgt ttg gat ttt tat gta tcc act tta ata ata tca tag ctg taa cat cct cat ga- ttt acg tta acg tct tcg tgg gat aag ata gtt gtc agt tca tcc ttt gat aat ttt cca aat t

>DQ792504.1:29780-31744_Horsepox_virus_isolate_MNR-76_complete_genome

cgg ctg cca tgt att tcc tgg aga gca agt --- aga tga tg- agg aac cag ata ggt tat atc cgt act tgc act taa agt tta cat tgt cgt tgt atg a-- gta tga tct ttt aaa ccc gct aga caa gta tcc gtt tga tat tgt agg atg tgg aca ttt aac aat ttg aca cgt ggg tgg atc gga cca ttc tcc tcc tga aca cag gac acc aga gtt acc aat caa cga ata tcc act att gca act ata agt tac aac gct tcc atc ggt ata aaa atc ctc gta tcc gtt atg tct tcc gtt gga tat aga tgg agg gaa ttg gca ttt aac aga ttc aca aat agg tgc ctc ggg att cca tac cat aga tcc agt aaa tcc taa ttc aca ata cga ttt aga ttc tcc tat caa ctg ata tcc gct att aca aga gta cgt tat act aga gcc aaa gtc tac tcc gcc aat atc aag ttg gcc att atc gat atc tcg agg cga tgg gca tct ccg ttt aat aca ttg att aaa gag tgt cca tcc agt acc tgt aca ttt agc ata tat agg tcc cat ttt ttg ctt tct gta tcc agg tag aca tag a-t att cta tag tgt ctc cta tgt tgt a-- --- --- --- -at tag cat tag cat tag tt- --t tca cac tat tct taa att tca tat taa tgg gac gtg aag gaa tag gac agt atg ata gaa cgc atc cta ttc cca aca atg tca gga acg tca cgc tct cca cct tca tat tta ttt atc cgt -aa aaa tgt tat cct gga cat cgt aca aat aat aaa aaa gcc ca- -ta tat gtt tgc tat tgt aga aat tgt ttt tca cag ttg ctc aaa aac gat ggc agt gac tta tga gtt acg tta cac ttt gga gtt tca tct tta gta aac ata tca taa tat tcg ata tta cga gtt gac ata tcg aac aaa ttc caa gta ttt gat ttt gga taa tat tcg tat ttt gca tct gct ata att aag ata taa tca ccg caa gaa cac acg aac atc ttt cct aca tgg tta aag tac atg tat aat tct atc cat ttg tct tcc tta act ata tat ttg tat aga taa tta cga gtc tca taa gta att cca gta att gca tag atg tca cca tcg tac tct aca gca taa act ata cta tga tgt cta ggc atg gga gac -tt ttt tat cca acg att ttt agt gaa aca ttc tac atc gtt taa tac tac ata ttt ctc ata cgt ggt ata aac tcc acc cat tac --a tat ata tca tcg ttt acg aat acc gat gcg cct gaa tat cta gga gta att aag ttt gga agt ctt atc cat ttt gaa gtg ccg tgt ttc aaa tat tct gct aca ccc gtt gaa ata gaa aat tct aat cct cct att aca tat aac ttt cca ttg tta aca caa gta cta act tct gat ttt aac gac gac ata tta gta acc gtt ttc ca- --- --- --- --- --- --- --t ttt ttc gtt tca aga tct acc cgc -ga tac gga ata aac atg tct att gtt aat cat gcc gcc aat aat gta tag aca att atg taa aac att tgc att ata gaa ttg --- -tc tat ctg tat tac cga cta tcg tcc aat att ctg ttc tag gag agt aat ggg tta ttg tgg ata tat aat cag agt ttt taa tga cta cta tat tat gtt tta tac cat ttc gtg tca ctg gct ttg tag att tgg ata tag tta atc cca aca atg ata tag cat tgc gca tag tat tag tta taa act tgg gat gta aaa tgt tga tga tat cta cat cgt ttg gat ttt tat gta tcc act tta ata ata tca tag ctg taa cat cct cat ga- ttt acg tta acg tct tcg tgg gat aag ata gtt gtc agt tca tcc ttt gat aat ttt cca aat t

>LT896733.1:34470-36429_Cowpox_virus_isolate_Ger/2015/Cat3_genome_assembly_complete_genome:_monopartite

cgg ctt cca tgt att tcc tgg aga gca agt --- aga tga tg- agg aac cag ata gtt tat atc cgt act tgc act taa agt cta cat tgt cgt tgt atg a-- gta tga tct ttt aaa ccc gct aga caa gta tcc gtt tga tat tgt agg atg tgg aca ttt aac aat ctg aca cgt ggg tgg atc gga cca ttc tcc tcc tga aca cag gac acc aga gtt acc aat caa cga ata tcc act att gca act ata agt tac aac gct ccc atc ggt ata aaa atc ctc gta tcc gtt atg tct tcc gtt gga tat aga tgg agg tga ttg gca ttt aac aga ttc gca aat agg tgc ctc ggg att cca tac cat aga tcc agt aga tcc taa ttc aca ata cga ttt aga ttc acc gat caa atg ata tcc gct att aca aga gta cgt tat act aga gcc aaa gtc tac tcc gcc aat atc aag ttg gcc att atc gat atc tcg agg cga tgg gca tct ccg ttt aat aca ttg att aaa gag tgt cca tcc agt acc tgt aca ttt agc ata tat agg tcc cat ttt ttg ctt tct gta tcc agg tag aca tag a-t att cta tag tgt ctc cta tgt tgt a-- --- --- --- -at tag aat tag cat cag tc- --t cca cac tat tct taa att tca tat taa tgg gac gtg acg gaa tag tac agc atg ata gaa cgc atc cta ttc cca aca atg tca gga acg tca cgc tct cca cct tca tat tta ttt atc cgt -aa aaa tgt tat ccc agg cca ttg tcc aat aat a-a aaa gtc ca- --- --t gtt ctc tat tgt agg aat tgt ttt tca cag ttg ctc aaa aac gat ggt agt gac tta cga gtt aca tta cac ttt ggt gtc tcg tct tta gta aac ata tca taa tat tcg ata tta cga gtt gac ata tcg aac aaa ttc caa gta ttt gat ttt gga taa tat tcg tat ttt gca tct gct ata att aag ata taa tca ccg caa gaa cac acg aac atc ttt cct aca tgg tta aag tac atg tac aat tct atc cat ttg tct tcc tta act ata tat ttg tat aga taa tta cga gtc tca tga gta att cca gta att gca tag atg tca cca tcg tac tct aca gca taa act ata cta tga tgt cta ggc atg gga gac -tt ttt tat cca acg att ttt agt gaa aca ttc cat atc gtt taa tac tac ata ttt ctc ata agt agt ata aac tcc acc cat tac --a tat ata tca tcg ttt acg aat acc gat gcg cct gaa tat cta gga gta att aag ttt gga agt ctt atc cat ttc gaa gtg ccg tgt ttc aaa tat tct gct aca ccc gtt gaa ata gaa aat tct aat cct cct att aca tat aac ttt cca tcg tta aca caa gta cta act tct gat ttt aac gac gac ata tta gta acc gtt ttc ca- --- --- --- --- --- --- --t ttt ttc gtt tta aga tct acc agc -ga cac gga ata aac atg tct att att aat cat gcc gcc aat aat gta tag aca att atg taa aac att tgc att ata gaa ttg --- -tc tat ctg tat tac cga cta tag tcc aat att ctg ttc tag gag agt aat ggg tta ttg tgg ata tat aat cag agt ttt taa tga cta cta tat tat gtt tta tac cat ttc gtg tca ccg gct ttg tag att tgg ata tag tta atc cca aca atg ata tag cat tgc gca tag tat tag tca taa act tgg gat gta aaa tgt tga tga tat ctg cat cgt ttg gat ttt tat gta tcc act tta ata ata tta tag ctg taa cat cct cat ga- ttt acg tta acg tct tcg tgg gat aag ata gtt gtc agt tca tcc ttt gat aat ttt cca aat t

>LT896727.1:35146-37105_Cowpox_virus_isolate_Ger/2015/Cat2_genome_assembly_complete_genome:_monopartite

cgg ctt cca tgt att tcc tgg aga gca agt --- aga tga tg- agg aac cag ata gtt tat atc cgt act tgc act taa agt cta cat tgt cgt tgt atg a-- gta tga tct ttt aaa ccc gct aga caa gta tcc gtt tga tat tgt agg atg tgg aca ttt aac aat ctg aca cgt ggg tgg atc gga cca ttc tcc tcc tga aca cag gac acc aga gtt acc aat caa cga ata tcc act att gca act ata agt tac aac gct ccc atc ggt ata aaa atc ctc gta tcc gtt atg tct tcc gtt gga tat aga tgg agg tga ttg gca ttt aac aga ttc gca aat agg tgc ctc ggg att cca tac cat aga tcc agt aga tcc taa ttc aca ata cga ttt aga ttc acc gat caa atg ata tcc gct att aca aga gta cgt tat act aga gcc aaa gtc tac tcc gcc aat atc aag ttg gcc att atc gat atc tcg agg cga tgg gca tct ccg ttt aat aca ttg att aaa gag tgt cca tcc agt acc tgt aca ttt agc ata tat agg tcc cat ttt ttg ctt tct gta tcc agg tag aca tag a-t att cta tag tgt ctc cta tgt tgt a-- --- --- --- -at tag aat tag cat cag tc- --t cca cac tat tct taa att tca tat taa tgg gac gtg acg gaa tag tac agc atg ata gaa cgc atc cta ttc cca aca atg tca gga acg tca cgc tct cca cct tca tat tta ttt atc cgt -aa aaa tgt tat ccc agg cca ttg tcc aat aat a-a aaa gtc ca- --- --t gtt ctc tat tgt agg aat tgt ttt tca cag ttg ctc aaa aac gat ggt agt gac tta cga gtt aca tta cac ttt ggt gtc tcg tct tta gta aac ata tca taa tat tcg ata tta cga gtt gac ata tcg aac aaa ttc caa gta ttt gat ttt gga taa tat tcg tat ttt gca tct gct ata att aag ata taa tca ccg caa gaa cac acg aac atc ttt cct aca tgg tta aag tac atg tac aat tct atc cat ttg tct tcc tta act ata tat ttg tat aga taa tta cga gtc tca tga gta att cca gta att gca tag atg tca cca tcg tac tct aca gca taa act ata cta tga tgt cta ggc atg gga gac -tt ttt tat cca acg att ttt agt gaa aca ttc cat atc gtt taa tac tac ata ttt ctc ata agt agt ata aac tcc acc cat tac --a tat ata tca tcg ttt acg aat acc gat gcg cct gaa tat cta gga gta att aag ttt gga agt ctt atc cat ttc gaa gtg ccg tgt ttc aaa tat tct gct aca ccc gtt gaa ata gaa aat tct aat cct cct att aca tat aac ttt cca tcg tta aca caa gta cta act tct gat ttt aac gac gac ata tta gta acc gtt ttc ca- --- --- --- --- --- --- --t ttt ttc gtt tta aga tct acc agc -ga cac gga ata aac atg tct att att aat cat gcc gcc aat aat gta tag aca att atg taa aac att tgc att ata gaa ttg --- -tc tat ctg tat tac cga cta tag tcc aat att ctg ttc tag gag agt aat ggg tta ttg tgg ata tat aat cag agt ttt taa tga cta cta tat tat gtt tta tac cat ttc gtg tca ccg gct ttg tag att tgg ata tag tta atc cca aca atg ata tag cat tgc gca tag tat tag tca taa act tgg gat gta aaa tgt tga tga tat ctg cat cgt ttg gat ttt tat gta tcc act tta ata ata tta tag ctg taa cat cct cat ga- ttt acg tta acg tct tcg tgg gat aag ata gtt gtc agt tca tcc ttt gat aat ttt cca aat t

>LT896726.1:34274-36233_Cowpox_virus_isolate_Ger/2012/Alpaca_genome_assembly_complete_genome:_monopartite

cgg ctt cca tgt att tcc tgg aga gca agt --- aga tga tg- agg aac cag ata gtt tat atc cgt act tgc act taa agt cta cat tgt cgt tgt atg a-- gta tga tct ttt aaa ccc gct aga caa gta tcc gtt tga tat tgt agg atg tgg aca ttt aac aat ctg aca cgt ggg tgg atc gga cca ttc tcc tcc tga aca cag gac acc aga gtt acc aat caa cga ata tcc act att gca act ata agt tac aac gct ccc atc ggt ata aaa atc ctc gta tcc gtt atg tct tcc gtt gga tat aga tgg agg tga ttg gca ttt aac aga ttc gca aat agg tgc ctc ggg att cca tac cat aga tcc agt aga tcc taa ttc aca ata cga ttt aga ttc acc gat caa atg ata tcc gct att aca aga gta cgt tat act aga gcc aaa gtc tac tcc gcc aat atc aag ttg gcc att atc gat atc tcg agg cga tgg gca tct ccg ttt aat aca ttg att aaa gag tgt cca tcc agt acc tgt aca ttt agc ata tat agg tcc cat ttt ttg ctt tct gta tcc agg tag aca tag a-t att cta tag tgt ctc cta tgt tgt a-- --- --- --- -at tag aat tag cat cag tc- --t cca cac tat tct taa att tca tat taa tgg gac gtg acg gaa tag tac agc atg ata gaa cgc atc cta ttc cca aca atg tca gga acg tca cgc tct cca cct tca tat tta ttt atc cgt -aa aaa tgt tat ccc agg cca ttg tcc aat aat a-a aaa gtc ca- --- --t gtt ctc tat tgt agg aat tgt ttt tca cag ttg ctc aaa aac gat ggt agt gac tta cga gtt aca tta cac ttt ggt gtc tcg tct tta gta aac ata tca taa tat tcg ata tta cga gtt gac ata tcg aac aaa ttc caa gta ttt gat ttt gga taa tat tcg tat ttt gca tct gct ata att aag ata taa tca ccg caa gaa cac acg aac atc ttt cct aca tgg tta aag tac atg tac aat tct atc cat ttg tct tcc tta act ata tat ttg tat aga taa tta cga gtc tca tga gta att cca gta att gca tag atg tca cca tcg tac tct aca gca taa act ata cta tga tgt cta ggc atg gga gac -tt ttt tat cca acg att ttt agt gaa aca ttc cat atc gtt taa tac tac ata ttt ctc ata agt agt ata aac tcc acc cat tac --a tat ata tca tcg ttt acg aat acc gat gcg cct gaa tat cta gga gta att aag ttt gga agt ctt atc cat ttc gaa gtg ccg tgt ttc aaa tat tct gct aca ccc gtt gaa ata gaa aat tct aat cct cct att aca tat aac ttt cca tcg tta aca caa gta cta act tct gat ttt aac gac gac ata tta gta acc gtt ttc ca- --- --- --- --- --- --- --t ttt ttc gtt tta aga tct acc agc -ga cac gga ata aac atg tct att att aat cat gcc gcc aat aat gta tag aca att atg taa aac att tgc att ata gaa ttg --- -tc tat ctg tat tac cga cta tag tcc aat att ctg ttc tag gag agt aat ggg tta ttg tgg ata tat aat cag agt ttt taa tga cta cta tat tat gtt tta tac cat ttc gtg tca ccg gct ttg tag att tgg ata tag tta atc cca aca atg ata tag cat tgc gca tag tat tag tca taa act tgg gat gta aaa tgt tga tga tat ctg cat cgt ttg gat ttt tat gta tcc act tta ata ata tta tag ctg taa cat cct cat ga- ttt acg tta acg tct tcg tgg gat aag ata gtt gtc agt tca tcc ttt gat aat ttt cca aat t

>BK013340.1:14888-16852_TPA_asm:_Vaccinia_virus_isolate_VK02

cgg ctg cca tgt att tcc tgg aaa gca agt --- aga tga tg- agg aac cag ata ggt tat atc cgt act tgc act taa agt tta cat tgt cgt tgt atg a-- gta tga tct ttt aaa ccc gct aga caa gta tcc gtt tga tat tgt agg atg tgg aca ttt aac aat ttg aca cgt ggg tgg atc gga cca ttc tcc tcc tga aca cag gac acc aga gtt acc aat caa cga ata tcc act att gca act ata agt tac aac gct tcc atc ggt ata aaa atc ctc gta tcc gtt atg tct tcc gtt gga tat aga tgg agg gaa ttg gca ttt aac aga ttc aca aat agg tgc ctc ggg att cca tac cat aga tcc agt aaa tcc taa ttc aca ata cga ttt aga ttc tcc tat caa ctg ata tcc gct att aca aga gta cgt tat act aga gcc aaa gtc tac tcc gcc aat atc aag ttg gcc att atc gat atc tcg agg cga tgg gca tct ccg ttt aat aca ttg att aaa gag tgt cca ttc agt acc tgt aca ttt agc ata tat agg tcc cat ttt ttg ctt tct gta tcc agg tag aca tag a-t att cta tag tgt ctc cta tgt tgt a-- --- --- --- -at tag cat tag cat tag tt- --t tca cac tat tct taa att tca tat taa tgg gac gtg aag gaa tag gac agt atg ata gaa cgc atc cta ttc cca aca atg tca gga acg tca cgc tct cca cct tca tat tta ttt atc cgt -aa aaa tgt tat cct gga cat cgt aca aat aat aaa aaa gcc ca- -ta tat gtt tgc tat tgt aga aat tgt ttt tca cag ttg ctc aaa aac gat ggc agt gac tta tga gtt acg tta cac ttt gga gtt tca tct tta gta aac ata tca taa tat tcg ata tta cga gtt gac ata tcg aac aaa ttc caa gta ttt gat ttt gga taa tat tcg tat ttt gca tct gct ata att aag ata taa tca ccg caa gaa cac acg aac atc ttt cct aca tgg tta aag tac atg tat aat tct atc cat ttg tct tcc tta act ata tat ttg tat aga taa tta cga gtc tca taa gta att cca gta att gca tag atg tca cca tcg tac tct aca gca taa act ata cta tga tgt cta ggc atg gga gac -tt ttt tat cca acg att ttt agt gaa aca ttc tac atc gtt taa tac tac ata ttt ctc ata cgt ggt ata aac tcc acc cat tac --a tat ata tca tcg ttt acg aat acc gat gcg cct gaa tat cta gga gta att aag ttt gga agt ctt atc cat ttt gaa gtg ccg tgt ttc aaa tat tct gct aca ccc gtt gaa ata gaa aat tct aat cct cct att aca tat aac ttt cca ttg tta aca caa gta cta act tct gat ttt aac gac gac ata tta gta acc gtt ttc ca- --- --- --- --- --- --- --t ttt ttc gtt tca aga tct acc cgc -ga tac gga ata aac atg tct att gtt aat cat gcc gcc aat aat gta tag aca att atg taa aac att tgc att ata gaa ttg --- -tc tat ctg tat tac cga cta tcg tcc aat att ctg ttc tag gag agt aat ggg tta ttg tgg ata tat aat cag agt ttt taa tga cta cta tat tat gtt tta tac cat ttc gtg tca ctg gct ttg tag att tgg ata tag tta atc cca aca atg ata tag cat tgc gca tag tat tag tta taa act tgg gat gta aaa tgt tga tga tat cta cat cgt ttg gat ttt tat gta tcc act tta ata ata tca tag ctg taa cat cct cat ga- ttt acg tta acg tct tcg tgg gat aag ata gtt gtc agt tca tcc ttt gat aat ttt cca aat t

>AY313848.1:21667-23606_Vaccinia_virus_strain_Acambis_clone_3_complete_genome

cgg ctg cca tgt att tcc tgg aga gca agt --- aga tga tg- agg aac cag ata gtt tat atc cgt act tgc act taa agt cta cat tgt cgt tgt atg a-- gta tga tct ttt aaa ccc gct aga caa gta tcc gtt tga tat tgt agg atg tgg aca ttt aac aat ctg aca cgt ggg tgg atc gga cca ttc tcc tcc tga aca cag gac acc aga gtt acc aat caa cga ata tcc act att gca act ata agt tac aac gct ccc atc ggt ata aaa atc ctc gta tcc gtt atg tct tcc gtt gga tat aga tgg agg gga ttg gca ttt aac aga ttc aca aat agg tgc ctc ggg att cca tac cat aga tcc agt aga tcc taa ttc aca ata cga ttt aga ttc acc gat caa atg ata tcc gct att aca aga gta cgt tat act aga gcc aaa gtc tac tcc acc aat atc aag ttg gcc att atc gat atc tcg agg cga tgg gca tct ccg ttt aat aca ttg att aaa gag tgt cca tcc agt acc tgt aca ttt agc ata tat agg tcc cat ttt ttg ctt tct gta tcc agg tag aca tag a-t att cta tag tgt ctc cta tgt tgt a-- --- --- --- -at tag cat t-- --- -ag tt- --t cca cac tat tct taa att tca tat taa tgg gtc gtg acg gaa tag tac agc atg ata gaa cgc atc cta ttc cca aca atg tca gga acg tca cgc tct cca cct tca tat tta ttt atc cgt -aa aaa tgt tat cct gga cat cgt aca aat aat a-a aaa gcc ca- -ta tat gtt cgc tat tgt aga aat tgt ttt tca cag ttg ctc aaa aac gat ggc agt gac tta tga gtt --- --- --- --- --- --- tca tct tta gta aac ata tca taa tat tcg ata tta cga gtt gac ata tcg aac aaa ttc caa gta ttt gat ttt gga taa tat tcg tat ttt gca tct gct ata att aag ata taa tca ccg caa gaa cac acg aac atc ttt cct aca tgg tta aag tac atg tac aat tct atc cat ttg tct tcc tta act ata tat ttg tat aga taa tta cga gtc tcg tga gta att cca gta att aca tag atg tcg ccg tcg tac tct aca gca taa act ata cta tga tgt cta ggc atg gga gac -tt ttt tat cca acg att ttt agt gaa aca ttc tac atc gtt taa tac tac ata ttt ttc ata cgt ggt ata aac tcc acc cat tac --a tat ata tca tcg ttt acg aat acc gat gcg cct gaa tat cta gga gta att aag ttt gga agt ctt atc cat ttt gaa gtg ccg tgt ttc aaa tat tct gct aca ccc gtt gaa ata gaa aat tct aat cct cct att aca tat aac ttt cca tcg tta aca caa gta cta act tct gat ttt aac gac gac ata tta gta acc gtt ttc ca- --- --- --- --- --- --- --t ttt ttc gtt tta aga tct acc cgc -ga tac gga ata aac atg tct att gtt aat cat gcc gcc aat aat gta tag aca att atg taa aac att tgc att ata gaa ttg --- -tc tat ctg tat tac cga cta tcg tcc aat att ctg ttc tag gag agt aat ggg tta ttg tgg ata tat aat cag agt ttt taa tga cta cta tat tat gtt tta tac cat ttc gtg tca ctg gct ttg tag att tgg ata tag tta atc cca aca atg ata tag cat tgc gca tag tat tag tca taa act tgg gat gta aaa tgt tga tga tat cta cat cgt ttg gat ttt tat gta tcc act tta ata ata tca tag ctg taa cat cct cat ga- ttt acg tta acg tct tcg tgg gat aag ata gtt gtc agt tca tcc ttt gat aat ttt cca aat t

>LT896732.2:35213-37172_Cowpox_virus_isolate_Ger/2017/Alpaca2_genome_assembly_complete_genome:_monopartite

cgg ctt cca tgt att tcc tgg aga gca agt --- aga tga tg- agg aac cag ata gtt tat atc cgt act tgc act taa agt cta cat tgt cgt tgt atg a-- gta tga tct ttt aaa ccc gct aga caa gta tcc gtt tga tat tgt agg atg tgg aca ttt aac aat ctg aca cgt ggg tgg atc gga cca ttc tcc tcc tga aca cag gac acc aga gtt acc aat caa cga ata tcc act att gca act ata agt tac aac gct ccc atc ggt ata aaa atc ctc gta tcc gtt atg tct tcc gtt gga tat aga tgg agg tga ttg gca ttt aac aga ttc gca aat agg tgc ctc ggg att cca tac cat aga tcc agt aga tcc taa ttc aca ata cga ttt aga ttc acc gat caa atg ata tcc gct att aca aga gta cgt tat act aga gcc aaa gtc tac tcc gcc aat atc aag ttg gcc att atc gat atc tcg agg cga tgg gca tct ccg ttt aat aca ttg att aaa gag tgt cca tcc agt acc tgt aca ttt agc ata tat agg tcc cat ttt ttg ctt tct gta tcc agg tag aca tag a-t att cta tag tgt ctc cta tgt tgt a-- --- --- --- -at tag aat tag cat cag tc- --t cca cac tat tct taa att tca tat taa tgg gac gtg acg gaa tag tac agc atg ata gaa cgc atc cta ttc cca aca atg tca gga acg tca cgc tct cca cct tca tat tta ttt atc cgt -aa aaa tgt tat ccc agg cca ttg tcc aat aat a-a aaa gtc ca- --- --t gtt ctc tat tgt agg aat tgt ttt tca cag ttg ctc aaa aac gat ggt agt gac tta cga gtt aca tta cac ttt ggt gtc tcg tct tta gta aac ata tca taa tat tcg ata tta cga gtt gac ata tcg aac aaa ttc caa gta ttt gat ttt gga taa tat tcg tat ttt gca tct gct ata att aag ata taa tca ccg caa gaa cac acg aac atc ttt cct aca tgg tta aag tac atg tac aat tct atc cat ttg tct tcc tta act ata tat ttg tat aga taa tta cga gtc tca tga gta att cca gta att gca tag atg tca cca tcg tac tct aca gca taa act ata cta tga tgt cta ggc atg gga gac -tt ttt tat cca acg att ttt agt gaa aca ttc cat atc gtt taa tac tac ata ttt ctc ata agt agt ata aac tcc acc cat tac --a tat ata tca tcg ttt acg aat acc gat gcg cct gaa tat cta gga gta att aag ttt gga agt ctt atc cat ttc gaa gtg ccg tgt ttc aaa tat tct gct aca ccc gtt gaa ata gaa aat tct aat cct cct att aca tat aac ttt cca tcg tta aca caa gta cta act tct gat ttt aac gac gac ata tta gta acc gtt ttc ca- --- --- --- --- --- --- --t ttt ttc gtt tta aga tct acc agc -ga cac gga ata aac atg tct att att aat cat gcc gcc aat aat gta tag aca att atg taa aac att tgc att ata gaa ttg --- -tc tat ctg tat tac cga cta tag tcc aat att ctg ttc tag gag agt aat ggg tta ttg tgg ata tat aat cag agt ttt taa tga cta cta tat tat gtt tta tac cat ttc gtg tca ccg gct ttg tag att tgg ata tag tta atc cca aca atg ata tag cat tgc gca tag tat tag tca taa act tgg gat gta aaa tgt tga tga tat ctg cat cgt ttg gat ttt tat gta tcc act tta ata ata tca tag ctg taa cat cct cat ga- ttt acg tta acg tct tcg tgg gat aag ata gtt gtc agt tca tcc ttt gat aat ttt cca aat t

>LT993231.1:35157-37116_Cowpox_virus_isolate_Ger/2015/Prairie-dog_genome_assembly_complete_genome:_monopartite

cgg ctt cca tgt att tcc tgg aga gca agt --- aga tga tg- agg aac cag ata gtt tat atc cgt act tgc act taa agt cta cat tgt cgt tgt atg a-- gta tga tct ttt aaa ccc gct aga caa gta tcc gtt tga tat tgt agg atg tgg aca ttt aac aat ctg aca cgt ggg tgg atc gga cca ttc tcc tcc tga aca cag gac acc aga gtt acc aat caa cga ata tcc act att gca act ata agt tac aac gct ccc atc ggt ata aaa atc ctc gta tcc gtt atg tct tcc gtt gga tat aga tgg agg tga ttg gca ttt aac aga ttc gca aat agg tgc ctc ggg att cca tac cat aga tcc agt aga tcc taa ttc aca ata cga ttt aga ttc acc gat caa atg ata tcc gct att aca aga gta cgt tat act aga gcc aaa gtc tac tcc gcc aat atc aag ttg gcc att atc gat atc tcg agg cga tgg gca tct ccg ttt aat aca ttg att aaa gag tgt cca tcc agt acc tgt aca ttt agc ata tat agg tcc cat ttt ttg ctt tct gta tcc agg tag aca tag a-t att cta tag tgt ctc cta tgt tgt a-- --- --- --- -at tag aat tag cat cag tc- --t cca cac tat tct taa att tca tat taa tgg gac gtg acg gaa tag tac agc atg ata gaa cgc atc cta ttc cca aca atg tca gga acg tca cgc tct cca cct tca tat tta ttt atc cgt -aa aaa tgt tat ccc agg cca ttg tcc aat aat a-a aaa gtc ca- --- --t gtt ctc tat tgt agg aat tgt ttt tca cag ttg ctc aaa aac gat ggt agt gac tta cga gtt aca tta cac ttt ggt gtc tcg tct tta gta aac ata tca taa tat tcg ata tta cga gtt gac ata tcg aac aaa ttc caa gta ttt gat ttt gga taa tat tcg tat ttt gca tct gct ata att aag ata taa tca ccg caa gaa cac acg aac atc ttt cct aca tgg tta aag tac atg tac aat tct atc cat ttg tct tcc tta act ata tat ttg tat aga taa tta cga gtc tca tga gta att cca gta att gca tag atg tca cca tcg tac tct aca gca taa act ata cta tga tgt cta ggc atg gga gac -tt ttt tat cca acg att ttt agt gaa aca ttc cat atc gtt taa tac tac ata ttt ctc ata agt agt ata aac tcc acc cat tac --a tat ata tca tcg ttt acg aat acc gat gcg cct gaa tat cta gga gta att aag ttt gga agt ctt atc cat ttc gaa gtg ccg tgt ttc aaa tat tct gct aca ccc gtt gaa ata gaa aat tct aat cct cct att aca tat aac ttt cca tcg tta aca caa gta cta act tct gat ttt aac gac gac ata tta gta acc gtt ttc ca- --- --- --- --- --- --- --t ttt ttc gtt tta aga tct acc agc -ga cac gga ata aac atg tct att att aat cat gcc gcc aat aat gta tag aca att atg taa aac att tgc att ata gaa ttg --- -tc tat ctg tat tac cga cta tag tcc aat att ctg ttc tag gag agt aat ggg tta ttg tgg ata tat aat cag agt ttt taa tga cta cta tat tat gtt tta tac cat ttc gtg tca ccg gct ttg tag att tgg ata tag tta atc cca aca ata ata tag cat tgc gca tag tat tag tca taa act tgg gat gta aaa tgt tga tga tat ctg cat cgt ttg gat ttt tat gta tcc act tta ata ata tta tag ctg taa cat cct cat ga- ttt acg tta acg tct tcg tgg gat aag ata gtt gtc agt tca tcc ttt gat aat ttt cca aat t

>LT993230.1:34255-36214_Cowpox_virus_isolate_Ger/2017/Alpaca1_genome_assembly_complete_genome:_monopartite

cgg ctt cca tgt att tcc tgg aga gca agt --- aga tga tg- agg aac cag ata gtt tat atc cgt act tgc act taa agt cta cat tgt cgt tgt atg a-- gta tga tct ttt aaa ccc gct aga caa gta tcc gtt tga tat tgt agg atg tgg aca ttt aac aat ctg aca cgt ggg tgg atc gga cca ttc tcc tcc tga aca cag gac acc aga gtt acc aat caa cga ata tcc act att gca act ata agt tac aac gct ccc atc ggt ata aaa atc ctc gta tcc gtt atg tct tcc gtt gga tat aga tgg agg tga ttg gca ttt aac aga ttc gca aat agg tgc ctc ggg att cca tac cat aga tcc agt aga tcc taa ttc aca ata cga ttt aga ttc acc gat caa atg ata tcc gct att aca aga gta cgt tat act aga gcc aaa gtc tac tcc gcc aat atc aag ttg gcc att atc gat atc tcg agg cga tgg gca tct ccg ttt aat aca ttg att aaa gag tgt cca tcc agt acc tgt aca ttt agc ata tat agg tcc cat ttt ttg ctt tct gta tcc agg tag aca tag a-t att cta tag tgt ctc cta tgt tgt a-- --- --- --- -at tag aat tag cat cag tc- --t cca cac tat tct taa att tca tat taa tgg gac gtg acg gaa tag tac agc atg ata gaa cgc atc cta ttc cca aca atg tca gga acg tca cgc tct cca cct tca tat tta ttt atc cgt -aa aaa tgt tat ccc agg cca ttg tcc aat aat a-a aaa gtc ca- --- --t gtt ctc tat tgt agg aat tgt ttt tca cag ttg ctc aaa aac gat ggt agt gac tta cga gtt aca tta cac ttt ggt gtc tcg tct tta gta aac ata tca taa tat tcg ata tta cga gtt gac ata tcg aac aaa ttc caa gta ttt gat ttt gga taa tat tcg tat ttt gca tct gct ata att aag ata taa tca ccg caa gaa cac acg aac atc ttt cct aca tgg tta aag tac atg tac aat tct atc cat ttg tct tcc tta act ata tat ttg tat aga taa tta cga gtc tca tga gta att cca gta att gca tag atg tca cca tcg tac tct aca gca taa act ata cta tga tgt cta ggc atg gga gac -tt ttt tat cca acg att ttt agt gaa aca ttc cat atc gtt taa tac tac ata ttt ctc ata agt agt ata aac tcc acc cat tac --a tat ata tca tcg ttt acg aat acc gat gcg cct gaa tat cta gga gta att aag ttt gga agt ctt atc cat ttc gaa gtg ccg tgt ttc aaa tat tct gct aca ccc gtt gaa ata gaa aat tct aat cct cct att aca tat aac ttt cca tcg tta aca caa gta cta act tct gat ttt aac gac gac ata tta gta acc gtt ttc ca- --- --- --- --- --- --- --t ttt ttc gtt tta aga tct acc agc -ga cac gga ata aac atg tct att att aat cat gcc gcc aat aat gta tag aca att atg taa aac att tgc att ata gaa ttg --- -tc tat ctg tat tac cga cta tag tcc aat att ctg ttc tag gag agt aat ggg tta ttg tgg ata tat aat cag agt ttt taa tga cta cta tat tat gtt tta tac cat ttc gtg tca ccg gct ttg tag att tgg ata tag tta atc cca aca ata ata tag cat tgc gca tag tat tag tca taa act tgg gat gta aaa tgt tga tga tat ctg cat cgt ttg gat ttt tat gta tcc act tta ata ata tta tag ctg taa cat cct cat ga- ttt acg tta acg tct tcg tgg gat aag ata gtt gtc agt tca tcc ttt gat aat ttt cca aat t

>LT993228.1:34392-36351_Cowpox_virus_isolate_Ger/2017/common_vole_FMEimka_genome_assembly_complete_genome:_monopartite

cgg ctt cca tgt att tcc tgg aga gca agt --- aga tga tg- agg aac cag ata gtt tat atc cgt act tgc act taa agt cta cat tgt cgt tgt atg a-- gta tga tct ttt aaa ccc gct aga caa gta tcc gtt tga tat tgt agg atg tgg aca ttt aac aat ctg aca cgt ggg tgg atc gga cca ttc tcc tcc tga aca cag gac acc aga gtt acc aat caa cga ata tcc act att gca act ata agt tac aac gct ccc atc ggt ata aaa atc ctc gta tcc gtt atg tct tcc gtt gga tat aga tgg agg tga ttg gca ttt aac aga ttc gca aat agg tgc ctc ggg att cca tac cat aga tcc agt aga tcc taa ttc aca ata cga ttt aga ttc acc gat caa atg ata tcc gct att aca aga gta cgt tat act aga gcc aaa gtc tac tcc gcc aat atc aag ttg gcc att atc gat atc tcg agg cga tgg gca tct ccg ttt aat aca ttg att aaa gag tgt cca tcc agt acc tgt aca ttt agc ata tat agg tcc cat ttt ttg ctt tct gta tcc agg tag aca tag a-t att cta tag tgt ctc cta tgt tgt a-- --- --- --- -at tag aat tag cat cag tc- --t cca cac tat tct taa att tca tat taa tgg gac gtg acg gaa tag tac agc atg ata gaa cgc atc cta ttc cca aca atg tca gga acg tca cgc tct cca cct tca tat tta ttt atc cgt -aa aaa tgt tat ccc agg cca ttg tcc aat aat a-a aaa gtc ca- --- --t gtt ctc tat tgt agg aat tgt ttt tca cag ttg ctc aaa aac gat ggt agt gac tta cga gtt aca tta cac ttt ggt gtc tcg tct tta gta aac ata tca taa tat tcg ata tta cga gtt gac ata tcg aac aaa ttc caa gta ttt gat ttt gga taa tat tcg tat ttt gca tct gct ata att aag ata taa tca ccg caa gaa cac acg aac atc ttt cct aca tgg tta aag tac atg tac aat tct atc cat ttg tct tcc tta act ata tat ttg tat aga taa tta cga gtc tca tga gta att cca gta att gca tag atg tca cca tcg tac tct aca gca taa act ata cta tga tgt cta ggc atg gga gac -tt ttt tat cca acg att ttt agt gaa aca ttc cat atc gtt taa tac tac ata ttt ctc ata agt agt ata aac tcc acc cat tac --a tat ata tca tcg ttt acg aat acc gat gcg cct gaa tat cta gga gta att aag ttt gga agt ctt atc cat ttc gaa gtg ccg tgt ttc aaa tat tct gct aca ccc gtt gaa ata gaa aat tct aat cct cct att aca tat aac ttt cca tcg tta aca caa gta cta act tct gat ttt aac gac gac ata tta gta acc gtt ttc ca- --- --- --- --- --- --- --t ttt ttc gtt tta aga tct acc agc -ga cac gga ata aac atg tct att att aat cat gcc gcc aat aat gta tag aca att atg taa aac att tgc att ata gaa ttg --- -tc tat ctg tat tac cga cta tag tcc aat att ctg ttc tag gag agt aat ggg tta ttg tgg ata tat aat cag agt ttt taa tga cta cta tat tat gtt tta tac cat ttc gtg tca ccg gct ttg tag att tgg ata tag tta atc cca aca ata ata tag cat tgc gca tag tat tag tca taa act tgg gat gta aaa tgt tga tga tat ctg cat cgt ttg gat ttt tat gta tcc act tta ata ata tta tag ctg taa cat cct cat ga- ttt acg tta acg tct tcg tgg gat aag ata gtt gtc agt tca tcc ttt gat aat ttt cca aat t

>JN654981.1:21419-23359_Vaccinia_virus_strain_Dryvax_clone_DPP15_complete_genome

cgg ctg cca tgt att tcc tgg aga gca agt --- aga tga tg- agg aac cag ata gtt tat atc cgt act tgc act taa agt cta cat tgt cgt tgt atg a-- gta tga tct ttt aaa ccc gct aga caa gta tcc gtt tga tat tgt agg atg tgg aca ttt aac aat ctg aca cgt ggg tgg atc gga cca ttc tcc tcc tga aca cag gac acc aga gtt acc aat caa cga ata tcc act att gca act ata agt tac aac gct ccc atc ggt ata aaa atc ctc gta tcc gtt atg tct tcc gtt gga tat aga tgg agg gga ttg gca ttt aac aga ttc aca aat agg tgc ctc ggg att cca tac cat aga tcc agt aga tcc taa ttc aca ata cga ttt aga ttc acc gat caa atg ata tcc gct att aca aga gta cgt tat act aga gcc aaa gtc tac tcc acc aat atc aag ttg gcc att atc gat atc tcg agg cga tgg gca tct ccg ttt aat aca ttg att aaa gag tgt cca tcc agt acc tgt aca ttt agc ata tat agg tcc cat ttt ttg ctt tct gta tcc agg tag aca tag a-t att cta tag tgt ctc cta tgt tgt a-- --- --- --- -at tag cat t-- --- -ag tt- --t cca cac tat tct taa att tca tat taa tgg gtc gtg acg gaa tag gac agt atg ata gaa cgc atc cta ttc cca aca atg tca gga acg tca cgc tct cca cct tca tat tta ttt atc cgt -aa aaa tgt tat cct gga cat cgt aca aat aat aaa aaa gcc ca- -ta tat gtt tgc tat tgt aga aat tgt ttt tca cag ttg ctc aaa aac gat ggc agt gac tta tga gtt --- --- --- --- --- --- tca tct tta gta aac ata tca taa tat tcg ata tta cga gtt gac ata tcg aac aaa ttc caa gta ttt gat ttt gga taa tat tcg tat ttt gca tct gct ata att aag ata taa tca ccg caa gaa cac acg aac atc ttt cct aca tgg tta aag tac atg tac aat tct atc cat ttg tct tcc tta act ata tat ttg tat aga taa tta cga gtc tcg tga gta att cca gta att aca tag atg tcg ccg tcg tac tct aca gca taa act ata cta tga tgt cta ggc atg gga gac -tt ttt tat cca acg att ttt agt gaa aca ttc cac atc gtt taa tac tac ata ttt ttc ata cgt ggt ata aac tcc acc cat tac --a tat ata tca tcg ttt acg aat acc gac gcg cct gaa tat cta gga gta att aag ttt gga agt ctt atc cat ttc gaa gtg ccg tgt ttc aaa tat tct gct aca ccc gtt gaa ata gaa aat tct aat cct cct att aca tat aac ttt cca tcg tta aca caa gta cta act tct gat ttt aac gac gac ata tta gta acc gtt ttc ca- --- --- --- --- --- --- --t ttt ttc gtt tta aga tct acc cgc -ga tac gga ata aac atg tct att gtt aat cat gcc gcc aat aat gta tag aca att atg taa aac att tgc att ata gaa ttg --- -tc tat ctg tat tac cga cta tcg tcc aat att ctg ttc tag gag agt aat ggg tta ttg tgg ata tat aat cag agt ttt taa tga cta cta tat tat gtt tta tac cat ttc gtg tca ctg gct ttg tag att tgg ata tag tta atc cca aca atg ata tag cat tgc gca tag tat tag tca taa act tgg gat gta aaa tgt tga tga tat cta cat cgt ttg gat ttt tat gta tcc act tta ata ata tca tag ctg taa cat cct cat ga- ttt acg tta acg tct tcg tgg gat aag ata gtt gtc agt tca tcc ttt gat aat ttt cca aat t

>LN864566.1:34116-36080_Cowpox_virus_complete_genome_strain_FM2292

cgg ctt cca tgt att tcc tgg aga gca agt --- aga tga tg- aag aac cag ata ggt tat atc cgt act tgc act taa agt cta cat tgt cgt tgt atg a-- gta tga tct ttt aaa ccc gct aga caa gta tcc gtt tgg tat tgt agg atg ttg aca ttt aac aat ctg aca cgt ggg cgg atc tgt cca ctc tcc tcc tga aca cag gac acc aga gtt acc aat caa cga ata tcc act att gca act ata agt tac aac gct tcc atc ggt ata aaa atc ctc gta tcc gtt atg tct tcc gtt gga tat aga tgg agg tga ttg gca ttt aac aga ttc gca aat agg tgc ctc ggg att cca tac cat aga tcc agt aga tcc taa ttc aca ata cga ttt aga ttc acc gat caa atg ata tcc gct att aca aga gta cgt tat act aga gcc aaa gtc tac tcc gcc aat atc aag ttg gcc att atc gat atc tcg agg cga tgg gca tct ccg ttt aat aca ttg att aaa gag tgt cca tcc agt acc tgt aca ttt agc ata tat agg tcc cat ttt ttg ctt tct gta tcc agg tag aca tag a-t att cta tag tgt ctc cta tgt tgt a-- --- --- --- -at tag cat tag cat cag tc- --t cca cac tat tct taa att tca tat taa tgg gtc gtg acg gaa tag tac agc atg ata gaa cgc atc cta ttc cca aca atg tca gga acg tca cgc tct cca cct tca tat tta ttt atc cgt -aa aaa tgt tat ccc gga cat tgt aca aat aat aaa aaa gcc ca- -ta tat gtt cgc tat tgt aga aat tgt ttt tca cag ttg ctc aaa aac gat ggc agt gac tta tga gtt acg tta cac ttt gga gtc tca tct tta gta aac ata tca taa tat tcg ata tta cga gtt gac ata tcg aac aaa ttc caa gta ttt gat ttt gga taa tat tcg tat ttt gca tct gct ata att aag ata taa tca ccg caa gaa cac acg aac atc ttt cct aca tgg tta aag tac atg tac aat tct atc cat ttg tct tcc tta act ata tat ttg tac aga taa tta cga gtc tcg tga gta att cca gta att aca tag atg tcg ccg tcg tac tca acc gca taa act ata cta tga tgt cta ggc atg gga gac -tt ttt tat cca acg att ttt agt gaa aca ttc cac atc gtt taa tac tac ata ttt ttc ata cgt ggt ata aac tcc acc cat tac --a tat ata tca tcg ttt acg aat acc gac gcg cct gaa tat cta gga gta att aag ttt gga agt ctt atc cat ttc gaa gtg ccg tgt ttc aaa tat tct act acg cct gtt gaa ata gaa aat tct aat cct cct att aca tat aac ttt cca tca tta aca caa gta cta act tct gat ttt aac gac gac ata tta gta act gtt ttc ca- --- --- --- --- --- --- --t ttt ttc gtt tta aga tct acc cgc -ga tac gga ata aac atg tct att gtt aat cat gcc gcc aat aat gta tag aca att atg taa aac att tgc att ata gaa ttg --- -tc tat ctg tat tac cga cga ttg tcc aat att ctg ttc tag gag agt aat ggg tta ttg tgg ata tat aat cag agt ttt taa tga cta cta tat tat gtt tta tac cat ttc gtg tca cag gct ttg tag att tgg ata tag tta atc cca aca atg ata tag cat tgc aca tag tat tag tca taa act tgg gat gta aaa tgt tga tga tat cta cat cgt ttg gat ttt tat gta tcc act tta ata ata tca tag ctg taa cat cct cgt ga- ttt acg ttg acg ttt tcg tgg gat aag ata gtt gtt agt tca tcc ttt gat aat ttt cca aat t

>JN654985.1:21446-23385_Vaccinia_virus_strain_Dryvax_clone_DPP20_complete_genome

cgg ctg cca tgt att tcc tgg aga gca agt --- aga tga tg- agg aac cag ata gtt tat atc cgt act tgc act taa agt cta cat tgt cgt tgt atg a-- gta tga tct ttt aaa ccc gct aga caa gta tcc gtt tga tat tgt agg atg tgg aca ttt aac aat ctg aca cgt ggg tgg atc gga cca ttc tcc tcc tga aca cag gac acc aga gtt acc aat caa cga ata tcc act att gca act ata agt tac aac gct ccc atc ggt ata aaa atc ctc gta tcc gtt atg tct tcc gtt gga tat aga tgg agg gga ttg gca ttt aac aga ttc aca aat agg tgc ctc ggg att cca tac cat aga tcc agt aga tcc taa ttc aca ata cga ttt aga ttc tcc tat caa ctg ata tcc gct att aca aga gta cgt tat act aga gcc aaa gtc tac tcc gcc aat atc aag ttg gcc att atc gat atc tcg agg cga tgg gca tct ccg ttt aat aca ttg att aaa gag tgt cca tcc agt acc tgt aca ttt agc ata tat agg tcc cat ttt ttg ctt tct gta tcc agg tag aca tag a-t att cta tag tgt ctc cta tgt tgt a-- --- --- --- -at tag cat t-- --- -ag tt- --t cca cac tat tct taa att tca tat taa tgg gac gtg aag gaa tag gac agt atg ata gaa cgc atc cta ttc cca aca atg tca gga acg tca cgc tct cca cct tca tat tta ttt atc cgt -aa aaa tgt tat cct gga cat cgt aca aat aat a-a aaa gcc ca- -ta tat gtt cgc tat tgt aga aat tgt ttt tca cag ttg ctc aaa aac gat ggc agt gac tta tga gtt --- --- --- --- --- --- tca tct tta gta aac ata tca taa tat tcg ata tta cga gtt gac ata tcg aac aaa ttc caa gta ttt gat ttt gga taa tat tcg tat ttt gca tct gct ata att aag ata taa tca ccg caa gaa cac acg aac atc ttt cct aca tgg tta aag tac atg tac aat tct atc cat ttg tct tcc tta act ata tat ttg tat aga taa tta cga gtc tcg tga gta att cca gta att gca tag atg tca ccg tcg tac tct aca gca taa act ata cta tga tgt cta ggc atg gga gac -tt ttt tat cca acg att ttt agt gaa aca ttc tac atc gtt taa tac tac ata ttt ctc ata cgt ggt ata aac tcc acc cat tac --a tat ata tca tcg ttt acg aat acc gat gcg cct gaa tat cta gga gta att aag ttt gga agt ctt atc cat ttc gaa gtg ccg tgt ttc aaa tat tct gcc aca ccc gtt gaa ata gaa aat tct aat cct cct att aca tat aac ttt cca ttg tta aca caa gta cta act tct gat ttt aac gac gac ata tta gta acc gtt ttc ca- --- --- --- --- --- --- --t ttt ttc gtt tca aga tct acc cgc -ga tac gga ata aac atg tct att gtt aat cat gcc gcc aat aat gta tag aca att atg taa aac att tgc att ata gaa ttg --- -tc tat ctg tat tac cga cta tcg tcc aat att ctg ttc tag gag agt aat ggg tta ttg tgg ata tat aat cag agt ttt taa tga cta cta tat tat gtt tta tac cat ttc gtg tca ctg gct ttg tag att tgg ata tag tta atc cca aca atg ata tag cat tgc gca tag tat tag tca taa act tgg gat gta aaa tgt tga tga tat cta cat cgt ttg gat ttt tat gta tcc act tta ata ata tca tag ctg taa cat cct cat ga- ttt acg tta acg tct tcg tgg gat aag ata gtt gtc agt tca tcc ttt gat aat ttt cca aat t

>DQ437593.1:34483-36447_Cowpox_virus_strain_Germany_91-3_complete_genome

cgg ctt cca tgt att tcc tgg aga gca agt --- aga tga tg- aag aac cag ata ggt tat atc cgt act tgc act taa agt cta cat tgt cgt tgt atg a-- gta tga tct ttt aaa ccc gct aga caa gta tcc gtt tgg tat tgt agg atg ttg aca ttt aac aat ctg aca cgt ggg cgg atc tgt cca ctc tcc tcc tga aca cag gac acc aga gtt acc aat caa cga ata tcc act att gca act ata agt tac aac gct tcc atc ggt ata aaa atc ctc gta tcc gtt atg tct tcc gtt gga tat aga tgg agg tga ttg gca ttt aac aga ttc gca aat agg tgc ctc ggg att cca tac cat aga tcc agt aga tcc taa ttc aca ata cga ttt aga ttc acc gat caa atg ata tcc gct att aca aga gta cgt tat act aga gcc aaa gtc tac tcc gcc aat atc aag ttg gcc att atc gat atc tcg agg cga tgg gca tct ccg ttt aat aca ttg att aaa gag tgt cca tcc agt acc tgt aca ttt agc ata tat agg tcc cat ttt ttg ctt tct gta tcc agg tag aca tag a-t att cta tag tgt ctc cta tgt tgt a-- --- --- --- -at tag cat tag cat cag tc- --t cca cac tat tct taa att tca tat taa tgg gtc gtg acg gaa tag tac agc atg ata gaa cgc atc cta ttc cca aca atg tca gga acg tca cgc tct cca cct tca tat tta ttt atc cgt -aa aaa tgt tat ccc gga cat tgt aca aat aat aaa aaa gcc ca- -ta tat gtt cgc tat tgt aga aat tgt ttt tca cag ttg ctc aaa aac gat ggc agt gac tta tga gtt acg tta cac ttt gga gtc tca tct tta gta aac ata tca taa tat tcg ata tta cga gtt gac ata tcg aac aaa ttc caa gta ttt gat ttt gga taa tat tcg tat ttt gca tct gct ata att aag ata taa tca ccg caa gaa cac acg aac atc ttt cct aca tgg tta aag tac atg tac aat tct atc cat ttg tct tcc tta act ata tat ttg tac aga taa tta cga gtc tcg tga gta att cca gta att aca tag atg tcg ccg tcg tac tca acc gca taa act ata cta tga tgt cta ggc atg gga gac -tt ttt tat cca acg att ttt agt gaa aca ttc cac atc gtt taa tac tac ata ttt ttc ata cgt ggt ata aac tcc acc cat tac --a tat ata tca tcg ttt acg aat acc gac gcg cct gaa tat cta gga gta att aag ttt gga agt ctt atc cat ttc gaa gtg ccg tgt ttc aaa tat tct act acg cct gtt gaa ata gaa aat tct aat cct cct att aca tat aac ttt cca tca tta aca caa gta cta act tct gat ttt aac gac gac ata tta gta act gtt ttc ca- --- --- --- --- --- --- --t ttt ttc gtt tta aga tct acc cgc -ga tac gga ata aac atg tct att gtt aat cat gcc gcc aat aat gta tag aca att atg taa aac att tgc att ata gaa ttg --- -tc tat ctg tat tac cga cga ttg tcc aat att ctg ttc tag gag agt aat ggg tta ttg tgg ata tat aat cag agt ttt taa tga cta cta tat tat gtt tta tac cat ttc gtg tca cag gct ttg tag att tgg ata tag tta atc cca aca atg ata tag cat tgc aca tag tat tag tca taa act tgg gat gta aaa tgt tga tga tat cta cat cgt ttg gat ttt tat gta tcc act tta ata ata tca tag ctg taa cat cct cgt ga- ttt acg ttg acg ttt tcg tgg gat aag ata gtt gtt agt tca tcc ttt gat aat ttt cca aat t

>HQ420897.1:35565-37530_Cowpox_virus_strain_Germany_1998_2_complete_genome

cgg ctt cca tgt att tcc tgg aga gca agt --- aga tga tg- aag aac cag ata ggt tat atc cgt act tgc act taa agt cta cat tgt cgt tgt atg a-- gta tga tct ttt aaa ccc gct aga caa gta tcc gtt tgg tat tgt agg atg ttg aca ttt aac aat ctg aca cgt ggg cgg atc tgt cca ctc tcc tcc tga aca cag gac acc aga gtt acc aat caa cga ata tcc act att gca act ata agt tac aac gct tcc atc ggt ata aaa atc ctc gta tcc gtt atg tct tcc gtt gga tat aga tgg agg tga ttg gca ttt aac aga ttc gca aat agg tgc ctc ggg att cca tac cat aga tcc agt aga tcc taa ttc aca ata cga ttt aga ttc acc gat caa atg ata tcc gct att aca aga gta cgt tat act aga gcc aaa gtc tac tcc gcc aat atc aag ttg gcc att atc gat atc tcg agg cga tgg gca tct ccg ttt aat aca ttg att aaa gag tgt cca tcc agt acc tgt aca ttt agc ata tat agg tcc cat ttt ttg ctt tct gta tcc agg tag aca tag a-t att cta tag tgt ctc cta tgt tgt a-- --- --- --- -at tag cat tag cat cag tc- --t cca cac tat tct taa att tca tat taa tgg gtc gtg acg gaa tag tac agc atg ata gaa cgc atc cta ttc cca aca atg tca gga acg tca cgc tct cca cct tca tat tta ttt atc cgt aaa aaa tgt tat ccc gga cat tgt aca aat aat aaa aaa gcc ca- -ta tat gtt cgc tat tgt aga aat tgt ttt tca cag ttg ctc aaa aac gat ggc agt gac tta tga gtt acg tta cac ttt gga gtc tca tct tta gta aac ata tca taa tat tcg ata tta cga gtt gac ata tcg aac aaa ttc caa gta ttt gat ttt gga taa tat tcg tat ttt gca tct gct ata att aag ata taa tca ccg caa gaa cac acg aac atc ttt cct aca tgg tta aag tac atg tac aat tct atc cat ttg tct tcc tta act ata tat ttg tac aga taa tta cga gtc tcg tga gta att cca gta att aca tag atg tcg ccg tcg tac tca acc gca taa act ata cta tga tgt cta ggc atg gga gac -tt ttt tat cca acg att ttt agt gaa aca ttc cac atc gtt taa tac tac ata ttt ttc ata cgt ggt ata aac tcc acc cat tac --a tat ata tca tcg ttt acg aat acc gac gcg cct gaa tat cta gga gta att aag ttt gga agt ctt atc cat ttc gaa gtg ccg tgt ttc aaa tat tct act acg cct gtt gaa ata gaa aat tct aat cct cct att aca tat aac ttt cca tca tta aca caa gta cta act tct gat ttt aac gac gac ata tta gta act gtt ttc ca- --- --- --- --- --- --- --t ttt ttc gtt tta aga tct acc cgc -ga tac gga ata aac atg tct att gtt aat cat gcc gcc aat aat gta tag aca att atg taa aac att tgc att ata gaa ttg --- -tc tat ctg tat tac cga cga ttg tcc aat att ctg ttc tag gag agt aat ggg tta ttg tgg ata tat aat cag agt ttt taa tga cta cta tat tat gtt tta tac cat ttc gtg tca cag gct ttg tag att tgg ata tag tta atc cca aca atg ata tag cat tgc aca tag tat tag tca taa act tgg gat gta aaa tgt tga tga tat cta cat cgt ttg gat ttt tat gta tcc act tta ata ata tca tag ctg taa cat cct cgt ga- ttt acg ttg acg ttt tcg tgg gat aag ata gtt gtt agt tca tcc ttt gat aat ttt cca aat t

>MN369532.1:5868-7838_Vaccinia_virus_isolate_VK01

cgg ctg cca tgt att tcc tgg aga gca agt --- aga tga tg- agg aac cag ata ggt tat atc cgt act tgc act taa agt tta cat tgt cgt tgt atg a-- gta tga tct ttt aaa ccc gct aga caa gta tcc gtt tga tat tgt agg atg tgg aca ttt aac aat ttg aca cgt ggg tgg atc gga cca ttc tcc tcc tga aca cag gac acc aga gtt acc aat caa cga ata tcc act att gca act ata agt tac aac gct tcc atc ggt ata aaa atc ctc gta tcc gtt atg tct tcc gtt gga tat aga tgg agg gga ttg gca ttt aac aga ttc aca aat agg tgc ctc ggg att cca tac cat aga tcc agt aaa tcc taa ttc aca ata cga ttt aga ttc tcc tat caa ctg ata tcc gct att aca aga gta cgt tat act aga gcc aaa gtc tac tcc gcc aat atc aag ttg gcc att atc gat atc tca agg cga tgg gca tct ccg ttt aat aca ttg att aaa gag tgt cca tcc agt acc tgt aca ttt agc ata tat agg tcc cat ttt ttg ctt tct gta tcc agg tag aca tag a-t att cta tag tgt ctc cta tgt tgt aat tag c-- --- -at tag cat tag cat tag tt- --t cca cac tat tct taa att tca tat taa tgg gac gtg aag gaa tag gac agt atg ata gaa cgc atc cta ttc cca aca atg tca gga acg tca cgc tct cca cct tca tat tta ttt atc cgt -aa aaa tgt tat cct gga cat cgt aca aat aat aaa aaa gcc ca- -ta tat gtt tgc tat tgt aga aat tgt ttt tca cag ttg ctc aaa aac gat ggc agt gac tta tga gtt acg tta cac ttt gga gtt tca tct tta gta aac ata tca taa tat tcg ata tta cga gtt gac ata tcg aac aaa ttc caa gta ttt gat ttt gga taa tat tcg tat ttt gca tct gct ata att aag ata taa tca ccg caa gaa cac acg aac atc ttt cct aca tgg tta aag tac atg tat aat tct atc cat ttg tct tcc tta act ata tat ttg tat aga taa tta cga gtc tca taa gta att cca gta att gca tag atg tca cca tcg tac tct aca gca taa act ata cta tga tgt cta ggc atg gga gac -tt ttt tat cca acg att ttt agt gaa aca ttc tac atc gtt taa tac tac ata ttt ctc ata cgt ggt ata aac tcc acc cat tac --a tat ata tca tcg ttt acg aat acc aat gcg cct gaa tat cta gga gta att aag ttt gga agt ctt atc cat ttt gaa gtg ccg tgt ttc aaa tat tct gct aca ccc gtt gaa ata gaa aat tct aat cct cct att aca tat aac ttt cca ttg tta aca caa gta cta act tct gat ttt aac gac gac ata tta gta acc gtt ttc ca- --- --- --- --- --- --- --t ttt ttc gtt tca aga tct acc cgc -ga tac gga ata aac atg tct att gtt aat cat gcc gcc aat aat gta tag aca att atg taa aac att tgc att ata gaa ttg --- -tc tat ctg tat tac cga cta tcg tcc aat att ctg ttc tag gag agt aat ggg tta ttg tgg ata tat aat cag agt ttt taa tga cta cta tat tat gtt tta tac cat ttc gtg tca ctg gct ttg tag att tgg ata tag tta atc cca aca atg ata tag cat tgc gca tag tat tag tta taa act tgg gat gta aaa tgt tga tga tat cta cat cgt ttg gat ttt tat gta tcc act tta ata ata tca tag ctg taa cat cct cat ga- ttt acg tta acg tct tcg tgg gat aag ata gtt gtc agt tca tcc ttt gat aat ttt cca aat t

>LT896722.1:34455-36419_Cowpox_virus_isolate_Ger/2007/Vole_genome_assembly_complete_genome:_monopartite

cgg ctt cca tgt att tcc tgg aga gca agt --- aga tga tg- aag aac cag ata ggt tat atc cgt act tgc act taa agt cta cat tgt cgt tgt atg a-- gta tga tct ttt aaa ccc gct aga caa gta tcc gtt tgg tat tgt agg atg ttg aca ttt aac aat ctg aca cgt ggg cgg atc tgt cca ctc tcc tcc tga aca cag gac acc aga gtt acc aat caa cga ata tcc act att gca act ata agt tac aac gct tcc atc ggt ata aaa atc ctc gta tcc gtt atg tct tcc gtt gga tat aga tgg agg tga ttg gca ttt aac aga ttc gca aat agg tgt ctc ggg att cca tac cat aga tcc agt aga tcc taa ttc aca ata cga ttt aga ttc acc gat caa atg ata tcc gct att aca aga gta cgt tat act aga gcc aaa gtc tac tcc gcc aat atc aag ttg gcc att atc gat atc tcg agg cga tgg gca tct ccg ttt aat aca ttg att aaa gag tgt cca tcc agt acc tgt aca ttt agc ata tat agg tcc cat ttt ttg ctt tct gta tcc agg tag aca tag a-t att cta tag tgt ctc cta tgt tgt a-- --- --- --- -at tag cat tag cat cag tc- --t cca cac tat tct taa att tca tat taa tgg gtc gtg acg gaa tag tac agc atg ata gaa cgc atc cta ttc cca aca atg tca gga acg tca cgc tct cca cct tca tat tta ttt atc cgt -aa aaa tgt tat ccc gga cat tgt aca aat aat aaa aaa gcc ca- -ta tat gtt cgc tat tgt aga aat tgt ttt tca cag ttg ctc aaa aac gat ggc agt gac tta tga gtt acg tta cac ttt gga gtc tca tct tta gta aac ata tca taa tat tcg ata tta cga gtt gac ata tcg aac aaa ttc caa gta ttt gat ttt gga taa tat tcg tat ttt gca tct gct ata att aag ata taa tca ccg caa gaa cac acg aac atc ttt cct aca tgg tta aag tac atg tac aat tct atc cat ttg tct tcc tta act ata tat ttg tac aga taa tta cga gtc tcg tga gta att cca gta att aca tag atg tcg ccg tcg tac tca acc gca taa act ata cta tga tgt cta ggc atg gga gac -tt ttt tat cca acg att ttt agt gaa aca ttc cac atc gtt taa tac tac ata ttt ttc ata cgt ggt ata aac tcc acc cat tac --a tat ata tca tcg ttt acg aat acc gac gcg cct gaa tat cta gga gta att aag ttt gga agt ctt atc cat ttc gaa gtg ccg tgt ttc aaa tat tct act acg cct gtt gaa ata gaa aat tct aat cct cct att aca tat aac ttt cca tca tta aca caa gta cta act tct gat ttt aac gac gac ata tta gta act gtt ttc ca- --- --- --- --- --- --- --t ttt ttc gtt tta aga tct acc cgc -ga tac gga ata aac atg tct att gtt aat cat gcc gcc aat aat gta tag aca att atg taa aac att tgc att ata gaa ttg --- -tc tat ctg tat tac cga cga ttg tcc aat att ctg ttc tag gag agt aat ggg tta ttg tgg ata tat aat cag agt ttt taa tga cta cta tat tat gtt tta tac cat ttc gtg tca cag gct ttg tag att tgg ata tag tta atc cca aca atg ata tag cat tgc aca tag tat tag tca taa act tgg gat gta aaa tgt tga tga tat cta cat cgt ttg gat ttt tat gta tcc act tta ata ata tca tag ctg taa cat cct cgt ga- ttt acg ttg acg ttt tcg tgg gat aag ata gtt gtt agt tca tcc ttt gat aat ttt cca aat t

>BK013342.1:25763-27733_TPA_asm:_Vaccinia_virus_isolate_VK08

cgg ctg cca tgt att tcc tgg aga gca agt --- aga tga tg- agg aac cag ata ggt tat atc cgt act tgc act taa agt tta cat tgt cgt tgt atg a-- gta tga tct ttt aaa ccc gct aga caa gta tcc gtt tga tat tgt agg atg tgg aca ttt aac aat ttg aca cgt ggg tgg atc gga cca ttc tcc tcc tga aca cag gac acc aga gtt acc aat caa cga ata tcc act att gca act ata agt tac aac gct tcc atc ggt ata aaa atc ctc gta tcc gtt atg tct tcc gtt gga tat aga tgg agg gga ttg gca ttt aac aga ttc aca aat agg tgc ctc ggg att cca tac cat aga tcc agt aaa tcc taa ttc aca ata cga ttt aga ttc tcc tat caa ctg ata tcc gct att aca aga gta cgt tat act aga gcc aaa gtc tac tcc gcc aat atc aag ttg gcc att atc gat atc tca agg cga tgg gca tct ccg ttt aat aca ttg att aaa gag tgt cca tcc agt acc tgt aca ttt agc ata tat agg tcc cat ttt ttg ctt tct gta tcc agg tag aca tag a-t att cta tag tgt ctc cta tgt tgt aat tag c-- --- -at tag cat tag cat tag tt- --t cca cac tat tct taa att tca tat taa tgg gac gtg aag gaa tag gac agt atg ata gaa cgc atc cta ttc cca aca atg tca gga acg tca cgc tct cca cct tca tat tta ttt atc cgt -aa aaa tgt tat cct gga cat cgt aca aat aat aaa aaa gcc ca- -ta tat gtt tgc tat tgt aga aat tgt ttt tca cag ttg ctc aaa aac gat ggc agt gac tta tga gtt acg tta cac ttt gga gtt tca tct tta gta aac ata tca taa tat tcg ata tta cga gtt gac ata tcg aac aaa ttc caa gta ttt gat ttt gga taa tat tcg tat ttt gca tct gct ata att aag ata taa tca ccg caa gaa cac acg aac atc ttt cct aca tgg tta aag tac atg tat aat tct atc cat ttg tct tcc tta act ata tat ttg tat aga taa tta cga gtc tca taa gta att cca gta att gca tag atg tca cca tcg tac tct aca gca taa act ata cta tga tgt cta ggc atg gga gac -tt ttt tat cca acg att ttt agt gaa aca ttc tac atc gtt taa tac tac ata ttt ctc ata cgt ggt ata aac tcc acc cat tac --a tat ata tca tcg ttt acg aat acc aat gcg cct gaa tat cta gga gta att aag ttt gga agt ctt atc cat ttt gaa gtg ccg tgt ttc aaa tat tct gct aca ccc gtt gaa ata gaa aat tct aat cct cct att aca tat aac ttt cca ttg tta aca caa gta cta act tct gat ttt aac gac gac ata tta gta acc gtt ttc ca- --- --- --- --- --- --- --t ttt ttc gtt tca aga tct acc cgc -ga tac gga ata aac atg tct att gtt aat cat gcc gcc aat aat gta tag aca att atg taa aac att tgc att ata gaa ttg --- -tc tat ctg tat tac cga cta tcg tcc aat att ctg ttc tag gag agt aat ggg tta ttg tgg ata tat aat cag agt ttt taa tga cta cta tat tat gtt tta tac cat ttc gtg tca ctg gct ttg tag att tgg ata tag tta atc cca aca atg ata tag cat tgc gca tag tat tag tta taa act tgg gat gta aaa tgt tga tga tat cta cat cgt ttg gat ttt tat gta tcc act tta ata ata tca tag ctg taa cat cct cat ga- ttt acg tta acg tct tcg tgg gat aag ata gtt gtc agt tca tcc ttt gat aat ttt cca aat t

>BK013341.1:29842-31812_TPA_asm:_Horsepox_virus_isolate_VK05

cgg ctg cca tgt att tcc tgg aga gca agt --- aga tga tg- agg aac cag ata ggt tat atc cgt act tgc act taa agt tta cat tgt cgt tgt atg a-- gta tga tct ttt aaa ccc gct aga caa gta tcc gtt tga tat tgt agg atg tgg aca ttt aac aat ttg aca cgt ggg tgg atc gga cca ttc tcc tcc tga aca cag gac acc aga gtt acc aat caa cga ata tcc act att gca act ata agt tac aac gct tcc atc ggt ata aaa atc ctc gta tcc gtt atg tct tcc gtt gga tat aga tgg agg gaa ttg gca ttt aac aga ttc aca aat agg tgc ctc ggg att cca tac cat aga tcc agt aaa tcc taa ttc aca ata cga ttt aga ttc tcc tat caa ctg ata tcc gct att aca aga gta cgt tat act aga gcc aaa gtc tac tcc gcc aat atc aag ttg gcc att atc gat atc tcg agg cga tgg gca tct ccg ttt aat aca ttg att aaa gag tgt cca tcc agt acc tgt aca ttt agc ata tat agg tcc cat ttt ttg ctt tct gta tcc agg tag aca tag a-t att cta tag tgt ctc cta tgt tgt aat tag c-- --- -at tag cat tag cat tag tt- --t tca cac tat tct taa att tca tat taa tgg gac gtg aag gaa tag gac agt atg ata gaa cgc atc cta ttc cca aca atg tca gga acg tca cgc tct cca cct tca tat tta ttt atc cgt -aa aaa tgt tat cct gga cat cgt aca aat aat aaa aaa gcc ca- -ta tat gtt tgc tat tgt aga aat tgt ttt tca cag ttg ctc aaa aac gat ggc agt gac tta tga gtt acg tta cac ttt gga gtt tca tct tta gta aac ata tca taa tat tcg ata tta cga gtt gac ata tcg aac aaa ttc caa gta ttt gat ttt gga taa tat tcg tat ttt gca tct gct ata att aag ata taa tca ccg caa gaa cac acg aac atc ttt cct aca tgg tta aag tac atg tat aat tct atc cat ttg tct tcc tta act ata tat ttg tat aga taa tta cga gtc tca taa gta att cca gta att gca tag atg tca cca tcg tac tct aca gca taa act ata cta tga tgt cta ggc atg gga gac -tt ttt tat cca acg att ttt agt gaa aca ttc tac atc gtt taa tac tac ata ttt ctc ata cgt ggt ata aac tcc acc cat tac --a tat ata tca tcg ttt acg aat acc gat gcg cct gaa tat cta gga gta att aag ttt gga agt ttt atc cat ttt gaa gtg ccg tgt ttc aaa tat tct gct aca ccc gtt gaa ata gaa aat tct aat cct cct att aca tat aac ttt cca ttg tta aca caa gta cta act tct gat ttt aac gac gac ata tta gta acc gtt ttc ca- --- --- --- --- --- --- --t ttt ttc gtt tca aga tct acc cgc -ga tac gga ata aac atg tct att gtt aat cat gcc gcc aat aat gta tag aca att atg taa aac att tgc att ata gaa ttg --- -tc tat ctg tat tac cga cta tcg tcc aat att ctg ttc tag gag agt aat ggg tta ttg tgg ata tat aat cag agt ttt taa tga cta cta tat tat gtt tta tac cat ttc gtg tca ctg gct ttg tag att tgg ata tag tta atc cca aca atg ata tag cat tgc gca tag tat tag tta taa act tgg gat gta aaa tgt tga tga tat cta cat cgt ttg gat ttt tat gta tcc act tta ata ata tca tag ctg taa cat cct cat ga- ttt acg tta acg tct tcg tgg gat aag ata gtt gtc agt tca tcc ttt gat aat ttt cca aat t

>BK013339.1:35624-37594_TPA_asm:_Vaccinia_virus_isolate_VK01

cgg ctg cca tgt att tcc tgg aga gca agt --- aga tga tg- agg aac cag ata ggt tat atc cgt act tgc act taa agt tta cat tgt cgt tgt atg a-- gta tga tct ttt aaa ccc gct aga caa gta tcc gtt tga tat tgt agg atg tgg aca ttt aac aat ttg aca cgt ggg tgg atc gga cca ttc tcc tcc tga aca cag gac acc aga gtt acc aat caa cga ata tcc act att gca act ata agt tac aac gct tcc atc ggt ata aaa atc ctc gta tcc gtt atg tct tcc gtt gga tat aga tgg agg gga ttg gca ttt aac aga ttc aca aat agg tgc ctc ggg att cca tac cat aga tcc agt aaa tcc taa ttc aca ata cga ttt aga ttc tcc tat caa ctg ata tcc gct att aca aga gta cgt tat act aga gcc aaa gtc tac tcc gcc aat atc aag ttg gcc att atc gat atc tca agg cga tgg gca tct ccg ttt aat aca ttg att aaa gag tgt cca tcc agt acc tgt aca ttt agc ata tat agg tcc cat ttt ttg ctt tct gta tcc agg tag aca tag a-t att cta tag tgt ctc cta tgt tgt aat tag c-- --- -at tag cat tag cat tag tt- --t cca cac tat tct taa att tca tat taa tgg gac gtg aag gaa tag gac agt atg ata gaa cgc atc cta ttc cca aca atg tca gga acg tca cgc tct cca cct tca tat tta ttt atc cgt -aa aaa tgt tat cct gga cat cgt aca aat aat aaa aaa gcc ca- -ta tat gtt tgc tat tgt aga aat tgt ttt tca cag ttg ctc aaa aac gat ggc agt gac tta tga gtt acg tta cac ttt gga gtt tca tct tta gta aac ata tca taa tat tcg ata tta cga gtt gac ata tcg aac aaa ttc caa gta ttt gat ttt gga taa tat tcg tat ttt gca tct gct ata att aag ata taa tca ccg caa gaa cac acg aac atc ttt cct aca tgg tta aag tac atg tat aat tct atc cat ttg tct tcc tta act ata tat ttg tat aga taa tta cga gtc tca taa gta att cca gta att gca tag atg tca cca tcg tac tct aca gca taa act ata cta tga tgt cta ggc atg gga gac -tt ttt tat cca acg att ttt agt gaa aca ttc tac atc gtt taa tac tac ata ttt ctc ata cgt ggt ata aac tcc acc cat tac --a tat ata tca tcg ttt acg aat acc aat gcg cct gaa tat cta gga gta att aag ttt gga agt ctt atc cat ttt gaa gtg ccg tgt ttc aaa tat tct gct aca ccc gtt gaa ata gaa aat tct aat cct cct att aca tat aac ttt cca ttg tta aca caa gta cta act tct gat ttt aac gac gac ata tta gta acc gtt ttc ca- --- --- --- --- --- --- --t ttt ttc gtt tca aga tct acc cgc -ga tac gga ata aac atg tct att gtt aat cat gcc gcc aat aat gta tag aca att atg taa aac att tgc att ata gaa ttg --- -tc tat ctg tat tac cga cta tcg tcc aat att ctg ttc tag gag agt aat ggg tta ttg tgg ata tat aat cag agt ttt taa tga cta cta tat tat gtt tta tac cat ttc gtg tca ctg gct ttg tag att tgg ata tag tta atc cca aca atg ata tag cat tgc gca tag tat tag tta taa act tgg gat gta aaa tgt tga tga tat cta cat cgt ttg gat ttt tat gta tcc act tta ata ata tca tag ctg taa cat cct cat ga- ttt acg tta acg tct tcg tgg gat aag ata gtt gtc agt tca tcc ttt gat aat ttt cca aat t

>JN654978.1:21423-23363_Vaccinia_virus_strain_Dryvax_clone_DPP11_complete_genome

cgg ctg cca tgt att tcc tgg aga gca agt --- aga tga tg- agg aac cag ata gtt tat atc cgt act tgc act taa agt cta cat tgt cgt tgt atg a-- gta tga tct ttt aaa ccc gct aga caa gta tcc gtt tga tat tgt agg atg tgg aca ttt aac aat ctg aca cgt ggg tgg atc gga cca ttc tcc tcc tga aca cag gac acc aga gtt acc aat caa cga ata tcc act att gca act ata agt tac aac gct ccc atc ggt ata aaa atc ctc gta tcc gtt atg tct tcc gtt gga tat aga tgg agg gga ttg gca ttt aac aga ttc aca aat agg tgc ctc ggg att cca tac cat aga tcc agt aga tcc taa ttc aca ata cga ttt aga ttc acc gat caa atg ata tcc gct att aca aga gta cgt tat act aga gcc aaa gtc tac tcc acc aat atc aag ttg gcc att atc gat atc tcg agg cga tgg gca tct ccg ttt aat aca ttg att aaa gag tgt cca tcc agt acc tgt aca ttt agc ata tat agg tcc cat ttt ttg ctt tct gta tcc agg tag aca tag a-t att cta tag tgt ctc cta tgt tgt a-- --- --- --- -at tag cat t-- --- -ag tt- --t cca cac tat tct taa att tta tat taa tgg gac gtg aag gaa tag gac agt atg ata gaa cgc atc cta ttc cca aca atg tca gga acg tca cgc tct cca cct tca tat tta ttt atc cgt -aa aaa tgt tat cct gga cat cgt aca aat aat aaa aaa gcc ca- -ta tat gtt tgc tat tgt aga aat tgt ttt tca cag ttg ctc aaa aac gat ggc agt gac tta tga gtt --- --- --- --- --- --- tca tct tta gta aac ata tca taa tat tcg ata tta cga gtt gac ata tcg aac aaa ttc caa gta ttt gat ttt gga taa tat tcg tat ttt gca tct gct ata att aag ata taa tca ccg caa gaa cac acg aac atc ttt cct aca tgg tta aag tac atg tac aat tct atc cat ttg tct tcc tta act ata tat ttg tat aga taa tta cga gtc tcg tga gta att cca gta att aca tag atg tcg ccg tcg tac tct aca gca taa act ata cta tga tgt cta ggc atg gga gac -tt ttt tat cca acg att ttt agt gaa aca ttc cac atc gtt taa tac tac ata ttt ttc ata cgt ggt ata aac tcc acc cat tac --a tat ata tca tcg ttt acg aat acc gac gcg cct gaa tat cta gga gta att aag ttt gga agt ctt atc cat ttc gaa gtg ccg tgt ttc aaa tat tct gct aca ccc gtt gaa ata gaa aat tct aat cct cct att aca tat aac ttt cca tcg tta aca caa gta cta act tct gat ttt aac gac gac ata tta gta acc gtt ttc ca- --- --- --- --- --- --- --t ttt ttc gtt tta aga tct acc cgc -ga tac gga ata aac atg tct att gtt aat cat gcc gcc aat aat gta tag aca att atg taa aac att tgc att ata gaa ttg --- -tc tat ctg tat tac cga cta tcg tcc aat att ctg ttc tag gag agt aat ggg tta ttg tgg ata tat aat cag agt ttt taa tga cta cta tat tat gtt tta tac cat ttc gtg tca ctg gct ttg tag att tgg ata tag tta atc cca aca atg ata tag cat tgc gca tag tat tag tca taa act tgg gat gta aaa tgt tga tga tat cta cat cgt ttg gat ttt tat gta tcc act tta ata ata tca tag ctg taa cat cct cat ga- ttt acg tta acg tct tcg tgg gat aag ata gtt gtc agt tca tcc ttt gat aat ttt cca aat t

>DQ377945.1:24195-26134_Vaccinia_virus_strain_3737_complete_genome

cgg ctg cca tgt att tcc tgg aga gca agt --- aga tga tg- agg aac cag ata gtt tat atc cgt act tgc act taa agt cta cat tgt cgt tgt atg a-- gta tga tct ttt aaa ccc gct aga caa gta tcc gtt tga tat tgt agg atg tgg aca ttt aac aat ctg aca cgt ggg tgg atc gga cca ttc tcc tcc tga aca cag gac acc aga gtt acc aat caa cga ata tcc act att gca act ata agt tac aac gct ccc atc ggt ata aaa atc ctc gta tcc gtt atg tct tcc gtt gga tat aga tgg agg gga ttg gca ttt aac aga ttc aca aat agg tgc ctc ggg att cca tac cat aga tcc agt aga tcc taa ttc aca ata cga ttt aga ttc acc gat caa atg ata tcc gct att aca aga gta cgt tat act aga gcc aaa gtc tac tcc acc aat atc aag ttg gcc att atc gat atc tcg agg cga tgg gca tct ccg ttt aat aca ttg att aaa gag tgt cca tcc agt acc tgt aca ttt agc ata tat agg tcc cat ttt ttg ctt tct gta tcc agg tag aca tag a-t att cta tag tgt ctc cta tgt tgt a-- --- --- --- -at tag cat t-- --- -ag tt- --t cca cac tat tct taa att tca tat taa tgg gac gtg aag gaa tag gac agt atg ata gaa cgc atc cta ttc cca aca atg tca gga acg tca cgc tct cca cct tca tat tta ttt atc cgt -aa aaa tgt tat cct gga cat cgt aca aat aat a-a aaa gcc ca- -ta tat gtt cgc tat tgt aga aat tgt ttt tca cag ttg ctc aaa aac gat ggc agt gac tta tga gtt --- --- --- --- --- --- tca tct tta gta aac ata tca taa tat tcg ata tta cga gtt gac ata tcg aac aaa ttc caa gta ttt gat ttt gga taa tat tcg tat ttt gca tct gct ata att aag ata taa tca ccg caa gaa cac acg aac atc ttt cct aca tgg tta aag tac atg tac aat tct atc cat ttg tct tcc tta act ata tat ttg tat aga taa tta cga gtc tcg tga gta att cca gta att aca tag atg tcg ccg tcg tac tct aca gca taa act ata cta tga tgt cta ggc atg gga gac -tt ttt tat cca acg att ttt agt gaa aca ttc cac atc gtt taa tac tac ata ttt ttc ata cgt ggt ata aac tcc acc cat tac --a tat ata tca tcg ttt acg aat acc gac gcg cct gaa tat cta gga gta att aag ttt gga agt ctt atc cat ttt gaa gtg ccg tgt ttc aaa tat tct gct aca ccc gtt gaa ata gaa aat tct aat cct cct att aca tat aac ttt cca ttg tta aca caa gta cta act tct gat ttt aac gac gac ata tta gta acc gtt ttc ca- --- --- --- --- --- --- --t ttt ttc gtt tta aga tct acc cgc -ga tac gga ata aac atg tct att gtt aat cat gcc gcc aat aat gta tag aca att atg taa aac att tgc att ata gaa ttg --- -tc tat ctg tat tac cga cta tcg tcc aat att ctg ttc tag gag agt aat ggg tta ttg tgg ata tat aat cag agt ttt taa tga cta cta tat tat gtt tta tac cat ttc gtg tca ctg gct ttg tag att tgg ata tag tta atc cca aca atg ata tag cat tgc gca tag tat tag tta taa act tgg gat gta aaa tgt tga tga tat cta cat cgt ttg gat ttt tat gta tcc act tta ata ata tca tag ctg taa cat cct cat ga- ttt acg tta acg tct tcg tgg gat aag ata gtt gtc agt tca tcc ttt gat aat ttt cca aat t

>JN654986.1:21401-23347_Vaccinia_virus_strain_Dryvax_clone_DPP21_complete_genome

cgg ctg cca tgt att tcc tgg aga gca agt --- aga tga tg- agg aac cag ata gtt tat atc cgt act tgc act taa agt cta cat tgt cgt tgt atg a-- gta tga tct ttt aaa ccc gct aga caa gta tcc gtt tga tat tgt agg atg tgg aca ttt aac aat ctg aca cgt ggg tgg atc gga cca ttc tcc tcc tga aca cag gac acc aga gtt acc aat caa cga ata tcc act att gca act ata agt tac aac gct ccc atc ggt ata aaa atc ctc gta tcc gtt atg tct tcc gtt gga tat aga tgg agg gga ttg gca ttt aac aga ttc aca aat agg tgc ctc ggg att cca tac cat aga tcc agt aga tcc taa ttc aca ata cga ttt aga ttc acc gat caa atg ata tcc gct att aca aga gta cgt tat act aga gcc aaa gtc tac tcc acc aat atc aag ttg gcc att atc gat atc tcg agg cga tgg gca tct ccg ttt aat aca ttg att aaa gag tgt cca tcc agt acc tgt aca ttt agc ata tat agg tcc cat ttt ttg ctt tct gta tcc agg tag aca tag a-t att cta tag tgt ctc cta tgt tgt a-- --- --- --- -at tag cat tag cat cag tc- --t cca cac tat tct taa att tca tat taa tgg gtc gtg acg gaa tag tac agc atg ata gaa cgc atc cta ttc cca aca atg tca gga acg tca cgc tct cca cct tca tat tta ttt atc cgt -aa aaa tgt tat cct gga cat cgt aca aat aat aaa aaa gcc ca- -ta tat gtt tgc tat tgt aga aat tgt ttt tca cag ttg ctc aaa aac gat ggc agt gac tta tga gtt --- --- --- --- --- --- tca tct tta gta aac ata tca taa tat tcg ata tta cga gtt gac ata tcg aac aaa ttc caa gta ttt gat ttt gga taa tat tcg tat ttt gca tct gct ata att aag ata taa tca ccg caa gaa cac acg aac atc ttt cct aca tgg tta aag tac atg tac aat tct atc cat ttg tct tcc tta act ata tat ttg tat aga taa tta cga gtc tcg tga gta att cca gta att aca tag atg tcg ccg tcg tac tct aca gca taa act ata cta tga tgt cta ggc atg gga gac -tt ttt tat cca acg att ttt agt gaa aca ttc cac atc gtt taa tac tac ata ttt ttc ata cgt ggt ata aac tcc acc cat tac --a tat ata tca tcg ttt acg aat acc gat gcg cct gaa tat cta gga gta att aag ttt gga agt ctt atc cat ttc gaa gtg ccg tgt ttc aaa tat tct gcc aca ccc gtt gaa ata gaa aat tct aat cct cct att aca tat aac ttt cca tcg tta aca caa gta cta act tct gat ttt aac gac gac ata tta gta acc gtt ttc ca- --- --- --- --- --- --- --t ttt ttc gtt tta aga tct acc cgc -ga tac gga ata aac atg tct att gtt aat cat gcc gcc aat aat gta tag aca att atg taa aac att tgc att ata gaa ttg --- -tc tat ctg tat tac cga cta tcg tcc aat att ctg ttc tag gag agt aat ggg tta ttg tgg ata tat aat cag agt ttt taa tga cta cta tat tat gtt tta tac cat ttc gtg tca ctg gct ttg tag att tgg ata tag tta atc cca aca atg ata tag cat tgc gca tag tat tag tca taa act tgg gat gta aaa tgt tga tga tat cta cat cgt ttg gat ttt tat gta tcc act tta ata ata tca tag ctg taa cat cct cat ga- ttt acg tta acg tct tcg tgg gat aag ata gtt gtc agt tca tcc ttt gat aat ttt cca aat t

>JN654984.1:21428-23367_Vaccinia_virus_strain_Dryvax_clone_DPP19_complete_genome

cgg ctg cca tgt att tcc tgg aga gca agt --- aga tga tg- agg aac cag ata ggt tat atc cgt act tgc act taa agt cta cat tgt cgt tgt atg a-- gta tga tct ttt aaa ccc gct aga caa gta tcc gtt tga tat tgt agg atg tgg aca ttt aac aat ttg aca cgt ggg tgg atc gga cca ttc tcc tcc tga aca cag gac acc aga gtt acc aat caa cga ata tcc act att gca act ata agt tac aac gct ccc atc ggt ata aaa atc ctc gta tcc gtt atg tct tcc gtt gga tat aga tgg agg gga ttg gca ttt aac aga ttc aca aat agg tgc ctc ggg att cca tac cat aga tcc agt aga tcc taa ttc aca ata cga ttt aga ttc acc gat caa atg ata tcc gct att aca aga gta cgt tat act aga gcc aaa gtc tac tcc acc aat atc aag ttg gcc att atc gat atc tcg agg cga tgg gca tct ccg ttt aat aca ttg att aaa gag tgt cca tcc agt acc tgt aca ttt agc ata tat agg tcc cat ttt ttg ctt tct gta tcc agg tag aca tag a-t att cta tag tgt ctc cta tgt tgt a-- --- --- --- -at tag cat t-- --- -ag tt- --t cca cac tat tct taa att tta tat taa tgg gac gtg aag gaa tag gac agt atg ata gaa cgc atc cta ttc cca aca atg tca gga acg tca cgc tct cca cct tca tat tta ttt atc cgt -aa aaa tgt tat cct gga cat cgt aca aat aat a-a aaa gcc ca- -ta tat gtt cgc tat tgt aga aat tgt ttt tca cag ttg ctc aaa aac gat ggc agt gac tta tga gtt --- --- --- --- --- --- tca tct tta gta aac ata tca taa tat tcg ata tta cga gtt gac ata tcg aac aaa ttc caa gta ttt gat ttt gga taa tat tcg tat ttt gca tct gct ata att aag ata taa tca ccg caa gaa cac acg aac atc ttt cct aca tgg tta aag tac atg tac aat tct atc cat ttg tct tcc tta act ata tat ttg tat aga taa tta cga gtc tcg tga gta att cca gta att aca tag atg tcg ccg tcg tac tct aca gca taa act ata cta tga tgt cta ggc atg gga gac -tt ttt tat cca acg att ttt agt gaa aca ttc cac atc gtt taa tac tac ata ttt ttc ata cgt ggt ata aac tcc acc cat tac --a tat ata tca tcg ttt acg aat acc gac gcg cct gaa tat cta gga gta att aag ttt gga agt ctt atc cat ttc gaa gtg ccg tgt ttc aaa tat tct gcc aca ccc gtt gaa ata gaa aat tct aat cct cct att aca tat aac ttt cca tcg tta aca caa gta cta act tct gat ttt aac gac gac ata tta gta acc gtt ttc ca- --- --- --- --- --- --- --t ttt ttc gtt tta aga tct acc cgc -ga tac gga ata aac atg tct att gtt aat cat gcc gcc aat aat gta tag aca att atg taa aac att tgc att ata gaa ttg --- -tc tat ctg tat tac cga cta tcg tcc aat att ctg ttc tag gag agt aat ggg tta ttg tgg ata tat aat cag agt ttt taa tga cta cta tat tat gtt tta tac cat ttc gtg tca ctg gct ttg tag att tgg ata tag tta atc cca aca atg ata tag cat tgc gca tag tat tag tca taa act tgg gat gta aaa tgt tga tga tat cta cat cgt ttg gat ttt tat gta tcc act tta ata ata tca tag ctg taa cat cct cat ga- ttt acg tta acg tct tcg tgg gat aag ata gtt gtc agt tca tcc ttt gat aat ttt cca aat t

>JN654979.1:21508-23447_Vaccinia_virus_strain_Dryvax_clone_DPP12_complete_genome

cgg ctg cca tgt att tcc tgg aga gca agt --- aga tga tg- agg aac cag ata gtt tat atc cgt act tgc act taa agt cta cat tgt cgt tgt atg a-- gta tga tct ttt aaa ccc gct aga caa gta tcc gtt tga tat tgt agg atg tgg aca ttt aac aat ctg aca cgt ggg tgg atc gga cca ttc tcc tcc tga aca cag gac acc aga gtt acc aat caa cga ata tcc act att gca act ata agt tac aac gct ccc atc ggt ata aaa atc ctc gta tcc gtt atg tct tcc gtt gga tat aga tgg agg gga ttg gca ttt aac aga ttc aca aat agg tgc ctc ggg att cca tac cat aga tcc agt aga tcc taa ttc aca ata cga ttt aga ttc acc gat caa atg ata tcc gct att aca aga gta cgt tat act aga gcc aaa gtc tac tcc gcc aat atc aag ttg gcc att atc gat atc tcg agg cga tgg gca tct ccg ttt aat aca ttg att aaa gag tgt cca tcc agt acc tgt aca ttt agc ata tat agg tcc cat ttt ttg ctt tct gta tcc agg tag aca tag a-t att cta tag tgt ctc cta tgt tgt a-- --- --- --- -at tag cat t-- --- -ag tt- --t cca cac tat tct taa att tta tat taa tgg gac gtg aag gaa tag gac agt atg ata gaa cgc atc cta ttc cca aca atg tca gga acg tca cgc tct cca cct tca tat tta ttt atc cgt -aa aaa tgt tat cct gga cat cgt aca aat aat a-a aaa gcc ca- -ta tat gtt cgc tat tgt aga aat tgt ttt tca cag ttg ctc aaa aac gat ggc agt gac tta tga gtt --- --- --- --- --- --- tca tct tta gta aac ata tca taa tat tcg ata tta cga gtt gac ata tcg aac aaa ttc caa gta ttt gat ttt gga taa tat tcg tat ttt gca tct gct ata att aag ata taa tca ccg caa gaa cac acg aac atc ttt cct aca tgg tta aag tac atg tat aat tct atc cat ttg tct tcc tta act ata tat ttg tat aga taa tta cga gtc tcg tga gta att cca gta att aca tag atg tcg ccg tcg tac tct aca gca taa act ata cta tga tgt cta ggc atg gga gac -tt ttt tat cca acg att ttt agt gaa aca ttc cac atc gtt taa tac tac ata ttt ttc ata cgt ggt ata aac tcc acc cat tac --a tat ata tca tcg ttt acg aat acc gac gcg cct gaa tat cta gga gta att aag ttt gga agt ctt atc cat ttt gaa gtg ccg tgt ttc aaa tat tct gct aca ccc gtt gaa ata gaa aat tct aat cct cct att aca tat aac ttt cca ttg tta aca caa gta cta act tct gat ttt aac gac gac ata tta gta acc gtt ttc ca- --- --- --- --- --- --- --t ttt ttc gtt tca aga tct acc cgc -ga tac gga ata aac atg tct att gtt aat cat gcc gcc aat aat gta tag aca att atg taa aac att tgc att ata gaa ttg --- -tc tat ctg tat tac cga cta tcg tcc aat att ctg ttc tag gag agt aat ggg tta ttg tgg ata tat aat cag agt ttt taa tga cta cta tat tat gtt tta tac cat ttc gtg tca ctg gct ttg tag att tgg ata tag tta atc cca aca atg ata tag cat tgc gca tag tat tag tca taa act tgg gat gta aaa tgt tga tga tat cta cat cgt ttg gat ttt tat gta tcc act tta ata ata tca tag ctg taa cat cct cat ga- ttt acg tta acg tct tcg tgg gat aag ata gtt gtc agt tca tcc ttt gat aat ttt cca aat t

>LN879483.1:34205-36164_Cowpox_virus_strain_CPXV_Amadeus_2015_complete_genome

cgg ctt cca tgt att tcc tgg aga gca agt --- aga tga tg- agg aac cag ata gtt tat atc cgt act tgc act taa agt cta cat tgt cgt tgt atg a-- gta tga tct ttt aaa ccc gct aga caa gta tcc gtt tga tat tgt agg atg tgg aca ttt aac aat ctg aca cgt ggg tgg atc gga cca ttc tcc tcc tga aca cag gac acc aga gtt acc aat caa cga ata tcc gct att aca act ata cgt tac aac gct tcc atc ggt ata aaa atc ctc gta tcc gtt atg tct tcc gtt gga tat aga tgg agg tga ttg gca ttt aac aga ttc gca aat agg tgc ctc ggg att cca tac cat aga tcc agt aga tcc taa ttc aca ata cga ttt aga ttc acc gat caa atg ata tcc gct att aca aga gta cgt tat act aga gcc aaa gtc tac tcc gcc aat atc aag ttg gcc att atc gat atc tcg agg cga tgg gca tct ccg ttt aat aca ttg att aaa gag tgt cca tcc agt acc tgt aca ttt agc ata tat agg tcc cat ttt ttg ctt tct ata tcc agg caa aca tac a-t att cta tag tgt ctc cta tgt tgt a-- --- --- --- -at tag aat tag cat cat tc- --c cca cac tat tct taa att tca tat taa tgg gac gtg acg gaa tag tac agc atg ata gaa cgc atc cta ttc cca aca atg tca gga acg tca cgc tct cca cct tca tat tta ttt atc cgt -aa aaa tgt tat ccc agg cca ttg tcc aat aat a-a aaa gtc ca- --- --t gtt ctc tat tgt agg aat tgt ttt tca cag ttg ctc aaa aac gat ggc agt gac tta tga gtt acg tta cac ttt gga gtc tca tct tta gta aac ata tca taa tat tcg ata tta cga gtt gac ata tcg aac aaa ttc caa gta ttt gat ttt gga taa tat tcg tat ttt gca tct gct ata att aag ata taa tca ccg caa gaa cac acg aac atc ttt cct aca tgg tta aag tac atg tac aat tct atc cat ttg tct tcc tta act ata tat ttg tat aga taa tta cga gtc tcg tga gta att cca gta att gca tag atg tca cca tcg tac tct aca gca taa act ata cta tga tgt cta ggc atg gga gac -tt ttt tat cca acg att ttt agt gaa aca ttc tac atc gtt taa tac tac ata ttt ctc ata cgt ggt ata aac tcc acc cat tac --a tat ata tca tcg ttt acg aat acc gat gcg cct gaa tat cta gga gta att aag ttt gga agt ctt atc cat ttc gaa gtg ccg tgt ttc aaa tat tct gcc aca ccc gtt gaa ata gaa aat tct aat cct cct att aca tat aac ttt cca tcg tta aca caa gta cta act tct gat ttt aac gac gac ata tta gta act gtt ttc ca- --- --- --- --- --- --- --t ttt ttc gtt tta aga tct acc cgc -ga tac gga ata aac atg tct att att aat cat gcc gcc aat aat gta tag aca att atg taa aac att tgc att ata gaa ttg --- -tc tat ctg tat tac cga cta tag tcc aat att ctg ttc tag gag agt aat ggg tta ttg tgg ata tat aat cag agt ttt taa tga cta cta tat tat gtt tta tac cat ttc gtg tca ccg gct ttg tag att tgg ata tag tta atc cca aca atg ata tag cat tgc gca tag tat tag tca taa act tgg gat gta aaa tgt tga tga tat ctg cat cgt ttg gat ttt tat gta tcc act tta ata ata tta tag ctg taa cat cct cat ga- ttt acg tta acg tct tcg tgg gat aag ata gtt gtc agt tca tcc ttt gat aat ttt cca aat t

>MT227314.1:21619-23565_Vaccinia_virus_strain_Acambis/2019

cgg ctg cca tgt att tcc tgg aga gca agt --- aga tga tg- agg aac cag ata gtt tat atc cgt act tgc act taa agt cta cat tgt cgt tgt atg a-- gta tga tct ttt aaa ccc gct aga caa gta tcc gtt tga tat tgt agg atg tgg aca ttt aac aat ctg aca cgt ggg tgg atc gga cca ttc tcc tcc tga aca cag gac acc aga gtt acc aat caa cga ata tcc act att gca act ata agt tac aac gct ccc atc ggt ata aaa atc ctc gta tcc gtt atg tct tcc gtt gga tat aga tgg agg gga ttg gca ttt aac aga ttc aca aat agg tgc ctc ggg att cca tac cat aga tcc agt aga tcc taa ttc aca ata cga ttt aga ttc acc gat caa atg ata tcc gct att aca aga gta cgt tat act aga gcc aaa gtc tac tcc acc aat atc aag ttg gcc att atc gat atc tcg agg cga tgg gca tct ccg ttt aat aca ttg att aaa gag tgt cca tcc agt acc tgt aca ttt agc ata tat agg tcc cat ttt ttg ctt tct gta tcc agg tag aca tag a-t att cta tag tgt ctc cta tgt tgt a-- --- --- --- -at tag cat tag cat cag tc- --t cca cac tat tct taa att tca tat taa tgg gtc gtg acg gaa tag tac agc atg ata gaa cgc atc cta ttc cca aca atg tca gga acg tca cgc tct cca cct tca tat tta ttt atc cgt -aa aaa tgt tat cct gga cat cgt aca aat aat aaa aaa gcc ca- -ta tat gtt tgc tat tgt aga aat tgt ttt tca cag ttg ctc aaa aac gat ggc agt gac tta tga gtt --- --- --- --- --- --- tca tct tta gta aac ata tca taa tat tcg ata tta cga gtt gac ata tcg aac aaa ttc caa gta ttt gat ttt gga taa tat tcg tat ttt gca tct gct ata att aag ata taa tca ccg caa gaa cac acg aac atc ttt cct aca tgg tta aag tac atg tac aat tct atc cat ttg tct tcc tta act ata tat ttg tat aga taa tta cga gtc tcg tga gta att cca gta att aca tag atg tcg ccg tcg tac tct aca gca taa act ata cta tga tgt cta ggc atg gga gac -tt ttt tat cca acg att ttt agt gaa aca ttc cac atc gtt taa tac tac ata ttt ttc ata cgt ggt ata aac tcc acc cat tac --a tat ata tca tcg ttt acg aat acc gac gcg cct gaa tat cta gga gta att aag ttt gga agt ctt atc cat ttc gaa gtg ccg tgt ttc aaa tat tct gcc aca ccc gtt gaa ata gaa aat tct aat cct cct att aca tat aac ttt cca tcg tta aca caa gta cta act tct gat ttt aac gac gac ata tta gta acc gtt ttc ca- --- --- --- --- --- --- --t ttt ttc gtt tta aga tct acc cgc -ga tac gga ata aac atg tct att gtt aat cat gcc gcc aat aat gta tag aca att atg taa aac att tgc att ata gaa ttg --- -tc tat ctg tat tac cga cta tcg tcc aat att ctg ttc tag gag agt aat ggg tta ttg tgg ata tat aat cag agt ttt taa tga cta cta tat tat gtt tta tac cat ttc gtg tca ctg gct ttg tag att tgg ata tag tta atc cca aca atg ata tag cat tgc gca tag tat tag tca taa act tgg gat gta aaa tgt tga tga tat cta cat cgt ttg gat ttt tat gta tcc act tta ata ata tca tag ctg taa cat cct cat ga- ttt acg tta acg tct tcg tgg gat aag ata gtt gtc agt tca tcc ttt gat aat ttt cca aat t

>MN974381.1:21910-23856_Vaccinia_virus_strain_TNX-1200

cgg ctg cca tgt att tcc tgg aga gca agt --- aga tga tg- agg aac cag ata gtt tat atc cgt act tgc act taa agt cta cat tgt cgt tgt atg a-- gta tga tct ttt aaa ccc gct aga caa gta tcc gtt tga tat tgt agg atg tgg aca ttt aac aat ctg aca cgt ggg tgg atc gga cca ttc tcc tcc tga aca cag gac acc aga gtt acc aat caa cga ata tcc act att gca act ata agt tac aac gct ccc atc ggt ata aaa atc ctc gta tcc gtt atg tct tcc gtt gga tat aga tgg agg gga ttg gca ttt aac aga ttc aca aat agg tgc ctc ggg att cca tac cat aga tcc agt aga tcc taa ttc aca ata cga ttt aga ttc acc gat caa atg ata tcc gct att aca aga gta cgt tat act aga gcc aaa gtc tac tcc acc aat atc aag ttg gcc att atc gat atc tcg agg cga tgg gca tct ccg ttt aat aca ttg att aaa gag tgt cca tcc agt acc tgt aca ttt agc ata tat agg tcc cat ttt ttg ctt tct gta tcc agg tag aca tag a-t att cta tag tgt ctc cta tgt tgt a-- --- --- --- -at tag cat tag cat cag tc- --t cca cac tat tct taa att tca tat taa tgg gtc gtg acg gaa tag tac agc atg ata gaa cgc atc cta ttc cca aca atg tca gga acg tca cgc tct cca cct tca tat tta ttt atc cgt -aa aaa tgt tat cct gga cat cgt aca aat aat aaa aaa gcc ca- -ta tat gtt tgc tat tgt aga aat tgt ttt tca cag ttg ctc aaa aac gat ggc agt gac tta tga gtt --- --- --- --- --- --- tca tct tta gta aac ata tca taa tat tcg ata tta cga gtt gac ata tcg aac aaa ttc caa gta ttt gat ttt gga taa tat tcg tat ttt gca tct gct ata att aag ata taa tca ccg caa gaa cac acg aac atc ttt cct aca tgg tta aag tac atg tac aat tct atc cat ttg tct tcc tta act ata tat ttg tat aga taa tta cga gtc tcg tga gta att cca gta att aca tag atg tcg ccg tcg tac tct aca gca taa act ata cta tga tgt cta ggc atg gga gac -tt ttt tat cca acg att ttt agt gaa aca ttc cac atc gtt taa tac tac ata ttt ttc ata cgt ggt ata aac tcc acc cat tac --a tat ata tca tcg ttt acg aat acc gac gcg cct gaa tat cta gga gta att aag ttt gga agt ctt atc cat ttc gaa gtg ccg tgt ttc aaa tat tct gcc aca ccc gtt gaa ata gaa aat tct aat cct cct att aca tat aac ttt cca tcg tta aca caa gta cta act tct gat ttt aac gac gac ata tta gta acc gtt ttc ca- --- --- --- --- --- --- --t ttt ttc gtt tta aga tct acc cgc -ga tac gga ata aac atg tct att gtt aat cat gcc gcc aat aat gta tag aca att atg taa aac att tgc att ata gaa ttg --- -tc tat ctg tat tac cga cta tcg tcc aat att ctg ttc tag gag agt aat ggg tta ttg tgg ata tat aat cag agt ttt taa tga cta cta tat tat gtt tta tac cat ttc gtg tca ctg gct ttg tag att tgg ata tag tta atc cca aca atg ata tag cat tgc gca tag tat tag tca taa act tgg gat gta aaa tgt tga tga tat cta cat cgt ttg gat ttt tat gta tcc act tta ata ata tca tag ctg taa cat cct cat ga- ttt acg tta acg tct tcg tgg gat aag ata gtt gtc agt tca tcc ttt gat aat ttt cca aat t

>MN974380.1:21856-23802_Vaccinia_virus_strain_VACV_li

cgg ctg cca tgt att tcc tgg aga gca agt --- aga tga tg- agg aac cag ata gtt tat atc cgt act tgc act taa agt cta cat tgt cgt tgt atg a-- gta tga tct ttt aaa ccc gct aga caa gta tcc gtt tga tat tgt agg atg tgg aca ttt aac aat ctg aca cgt ggg tgg atc gga cca ttc tcc tcc tga aca cag gac acc aga gtt acc aat caa cga ata tcc act att gca act ata agt tac aac gct ccc atc ggt ata aaa atc ctc gta tcc gtt atg tct tcc gtt gga tat aga tgg agg gga ttg gca ttt aac aga ttc aca aat agg tgc ctc ggg att cca tac cat aga tcc agt aga tcc taa ttc aca ata cga ttt aga ttc acc gat caa atg ata tcc gct att aca aga gta cgt tat act aga gcc aaa gtc tac tcc acc aat atc aag ttg gcc att atc gat atc tcg agg cga tgg gca tct ccg ttt aat aca ttg att aaa gag tgt cca tcc agt acc tgt aca ttt agc ata tat agg tcc cat ttt ttg ctt tct gta tcc agg tag aca tag a-t att cta tag tgt ctc cta tgt tgt a-- --- --- --- -at tag cat tag cat cag tc- --t cca cac tat tct taa att tca tat taa tgg gtc gtg acg gaa tag tac agc atg ata gaa cgc atc cta ttc cca aca atg tca gga acg tca cgc tct cca cct tca tat tta ttt atc cgt -aa aaa tgt tat cct gga cat cgt aca aat aat aaa aaa gcc ca- -ta tat gtt tgc tat tgt aga aat tgt ttt tca cag ttg ctc aaa aac gat ggc agt gac tta tga gtt --- --- --- --- --- --- tca tct tta gta aac ata tca taa tat tcg ata tta cga gtt gac ata tcg aac aaa ttc caa gta ttt gat ttt gga taa tat tcg tat ttt gca tct gct ata att aag ata taa tca ccg caa gaa cac acg aac atc ttt cct aca tgg tta aag tac atg tac aat tct atc cat ttg tct tcc tta act ata tat ttg tat aga taa tta cga gtc tcg tga gta att cca gta att aca tag atg tcg ccg tcg tac tct aca gca taa act ata cta tga tgt cta ggc atg gga gac -tt ttt tat cca acg att ttt agt gaa aca ttc cac atc gtt taa tac tac ata ttt ttc ata cgt ggt ata aac tcc acc cat tac --a tat ata tca tcg ttt acg aat acc gac gcg cct gaa tat cta gga gta att aag ttt gga agt ctt atc cat ttc gaa gtg ccg tgt ttc aaa tat tct gcc aca ccc gtt gaa ata gaa aat tct aat cct cct att aca tat aac ttt cca tcg tta aca caa gta cta act tct gat ttt aac gac gac ata tta gta acc gtt ttc ca- --- --- --- --- --- --- --t ttt ttc gtt tta aga tct acc cgc -ga tac gga ata aac atg tct att gtt aat cat gcc gcc aat aat gta tag aca att atg taa aac att tgc att ata gaa ttg --- -tc tat ctg tat tac cga cta tcg tcc aat att ctg ttc tag gag agt aat ggg tta ttg tgg ata tat aat cag agt ttt taa tga cta cta tat tat gtt tta tac cat ttc gtg tca ctg gct ttg tag att tgg ata tag tta atc cca aca atg ata tag cat tgc gca tag tat tag tca taa act tgg gat gta aaa tgt tga tga tat cta cat cgt ttg gat ttt tat gta tcc act tta ata ata tca tag ctg taa cat cct cat ga- ttt acg tta acg tct tcg tgg gat aag ata gtt gtc agt tca tcc ttt gat aat ttt cca aat t

>MK910851.1:19549-21513_Camelpox_virus_strain_Negev2016

cgg ctg cca tgt att tcc tgg aga gca agt --- aga tga tg- agg aac cag ata gtt tat atc cgt act tgc acg taa agt cta cat tgt cat tat acg a-- ata tga tct ttt aaa ccc gct aga caa gta tcc gtt tga tat tgt agg atg tgg aca ttt aac aat ctg aca cgt ggg tgg atc gga cca ttc tcc tcc tga aca cag gac acc aga gtt acc aat caa cga ata tcc act att gca act ata agt tac aac gct tcc atc ggt ata aaa att atc gta tcc gtt atg tct tcc gtt gga tat aga tgg agg tga ttg gca ttt aac aga ttc gca aat agg tgc ctc ggg att cca tac cat aga tcc agt aga tcc taa ttc aca ata cga ttt aga ttc acc gat caa atg ata tcc gct att aca aga gta cgt tat act aga acc aaa gtc tac tcc acc aat atc aag ttg gcc att atc gat atc tcg agg cga tgg gca tct ccg ttt aat aca ttg att aaa gag tgt cca tcc agt acc tgt aca ttt agc ata tat agg tcc cat ttt ttg ctt tct gta tcc agg tag aca tag a-t att cta tag tgt ctc cta tgt tgt a-- --- --- --- -at tag tat tag cat aag tc- --t cca cac tat tct taa att tca tat taa tgg gtc gtg acg gaa tag tac agc atg ata gaa cgc ata cta ttc cca aca atg tca gga acg tca cgc tct cca cct tca tat tta ttt atc cgt -aa aaa tgt tat cct gga cac tgt acc aat aat aaa aaa gcc ca- -ta tat gtt cgc tat tgt aga aat tgt ttt tca cag ttg ctc aaa aac gat ggc agt gac tta tga gtt acg tta cac ttt gga gtc tca tct tta gta aac ata tca taa tat tcg ata tta cga gtt gac ata tcg aac aaa atc caa gta ttt gat ttt gga taa tat tcg tat ttt gca tct gct ata att aag ata taa tcg ccg caa gaa cac acg aac atc ttt cct aca tgg tta aag tac atg tac aat tct atc cat ttg tct tcc tta act ata tat ttg tat aga taa tta cga gtc tcg tga gta att cca gta att gca tag atg tca cca tcg tac tct ata gca taa act ata cta tga tgt cta ggc atg gga gac -tt ttt tat cca acg att ttt agt gaa aca ttc cac atc gtt taa tac tac ata ttt ttc ata cgt ggt ata aac tcc acc cat tac --a tat ata tca tcg ttt acg aat acc gac gcg cct gaa tat cta gga gta att aag ttt gga agt ctt atc cat ttc gaa gtt ccg tgt ttc aaa tat tct gct acg cct gtt gaa ata gaa aat tct aat cct cct att aca tat aac ttt cca tca tta aca caa gta cta act tct gat ttt aac gac gac ata tta gta act gtt ttc ca- --- --- --- --- --- --- --t ttt ttc gtt tta aga tct acc cgc -ga tac gga ata aac atg tct att gtt aat cat gcc gcc aat aat gta tag aca att ata taa aac att tgc att ata gaa ttg --- -tc tat ctg tat tac cga cga ttg tcc aat att ctg ttc tag gag agt aat ggg tta ttg tgg ata tat aat cag agt ttt taa tga cta cta tat tat gtt tta tac cat ttc gtg tca cag gct ttg tag att tgg ata tag tta atc cca aca atg ata tag cat tgc aca tag tat tag tca taa act tgg gat gta aaa tgt tga taa tat cta cat cgt ttg gat ttt tat gta tcc act tta ata ata tca tag ctg taa cat cct cgt ga- ttt acg ttg acg ttt tcg tgg gat aag ata gtt gtt agt tca tcc ttt gat aat ttt cca aat t

>MW960424.1:21717-23663_Vector_synVACV-Delta1Delta3-6_complete_sequence

cgg ctg cca tgt att tcc tgg aga gca agt --- aga tga tg- agg aac cag ata gtt tat atc cgt act tgc act taa agt cta cat tgt cgt tgt atg a-- gta tga tct ttt aaa ccc gct aga caa gta tcc gtt tga tat tgt agg atg tgg aca ttt aac aat ctg aca cgt ggg tgg atc gga cca ttc tcc tcc tga aca cag gac acc aga gtt acc aat caa cga ata tcc act att gca act ata agt tac aac gct ccc atc ggt ata aaa atc ctc gta tcc gtt atg tct tcc gtt gga tat aga tgg agg gga ttg gca ttt aac aga ttc aca aat agg tgc ctc ggg att cca tac cat aga tcc agt aga tcc taa ttc aca ata cga ttt aga ttc acc gat caa atg ata tcc gct att aca aga gta cgt tat act aga gcc aaa gtc tac tcc acc aat atc aag ttg gcc att atc gat atc tcg agg cga tgg gca tct ccg ttt aat aca ttg att aaa gag tgt cca tcc agt acc tgt aca ttt agc ata tat agg tcc cat ttt ttg ctt tct gta tcc agg tag aca tag a-t att cta tag tgt ctc cta tgt tgt a-- --- --- --- -at tag cat tag cat cag tc- --t cca cac tat tct taa att tca tat taa tgg gtc gtg acg gaa tag tac agc atg ata gaa cgc atc cta ttc cca aca atg tca gga acg tca cgc tct cca cct tca tat tta ttt atc cgt -aa aaa tgt tat cct gga cat cgt aca aat aat aaa aaa gcc ca- -ta tat gtt tgc tat tgt aga aat tgt ttt tca cag ttg ctc aaa aac gat ggc agt gac tta tga gtt --- --- --- --- --- --- tca tct tta gta aac ata tca taa tat tcg ata tta cga gtt gac ata tcg aac aaa ttc caa gta ttt gat ttt gga taa tat tcg tat ttt gca tct gct ata att aag ata taa tca ccg caa gaa cac acg aac atc ttt cct aca tgg tta aag tac atg tac aat tct atc cat ttg tct tcc tta act ata tat ttg tat aga taa tta cga gtc tcg tga gta att cca gta att aca tag atg tcg ccg tcg tac tct aca gca taa act ata cta tga tgt cta ggc atg gga gac -tt ttt tat cca acg att ttt agt gaa aca ttc cac atc gtt taa tac tac ata ttt ttc ata cgt ggt ata aac tcc acc cat tac --a tat ata tca tcg ttt acg aat acc gac gcg cct gaa tat cta gga gta att aag ttt gga agt ctt atc cat ttc gaa gtg ccg tgt ttc aaa tat tct gcc aca ccc gtt gaa ata gaa aat tct aat cct cct att aca tat aac ttt cca tcg tta aca caa gta cta act tct gat ttt aac gac gac ata tta gta acc gtt ttc ca- --- --- --- --- --- --- --t ttt ttc gtt tta aga tct acc cgc -ga tac gga ata aac atg tct att gtt aat cat gcc gcc aat aat gta tag aca att atg taa aac att tgc att ata gaa ttg --- -tc tat ctg tat tac cga cta tcg tcc aat att ctg ttc tag gag agt aat ggg tta ttg tgg ata tat aat cag agt ttt taa tga cta cta tat tat gtt tta tac cat ttc gtg tca ctg gct ttg tag att tgg ata tag tta atc cca aca atg ata tag cat tgc gca tag tat tag tca taa act tgg gat gta aaa tgt tga tga tat cta cat cgt ttg gat ttt tat gta tcc act tta ata ata tca tag ctg taa cat cct cat ga- ttt acg tta acg tct tcg tgg gat aag ata gtt gtc agt tca tcc ttt gat aat ttt cca aat t

>MW960423.1:21718-23664_Vector_synVACV-Delta3-6_complete_sequence

cgg ctg cca tgt att tcc tgg aga gca agt --- aga tga tg- agg aac cag ata gtt tat atc cgt act tgc act taa agt cta cat tgt cgt tgt atg a-- gta tga tct ttt aaa ccc gct aga caa gta tcc gtt tga tat tgt agg atg tgg aca ttt aac aat ctg aca cgt ggg tgg atc gga cca ttc tcc tcc tga aca cag gac acc aga gtt acc aat caa cga ata tcc act att gca act ata agt tac aac gct ccc atc ggt ata aaa atc ctc gta tcc gtt atg tct tcc gtt gga tat aga tgg agg gga ttg gca ttt aac aga ttc aca aat agg tgc ctc ggg att cca tac cat aga tcc agt aga tcc taa ttc aca ata cga ttt aga ttc acc gat caa atg ata tcc gct att aca aga gta cgt tat act aga gcc aaa gtc tac tcc acc aat atc aag ttg gcc att atc gat atc tcg agg cga tgg gca tct ccg ttt aat aca ttg att aaa gag tgt cca tcc agt acc tgt aca ttt agc ata tat agg tcc cat ttt ttg ctt tct gta tcc agg tag aca tag a-t att cta tag tgt ctc cta tgt tgt a-- --- --- --- -at tag cat tag cat cag tc- --t cca cac tat tct taa att tca tat taa tgg gtc gtg acg gaa tag tac agc atg ata gaa cgc atc cta ttc cca aca atg tca gga acg tca cgc tct cca cct tca tat tta ttt atc cgt -aa aaa tgt tat cct gga cat cgt aca aat aat aaa aaa gcc ca- -ta tat gtt tgc tat tgt aga aat tgt ttt tca cag ttg ctc aaa aac gat ggc agt gac tta tga gtt --- --- --- --- --- --- tca tct tta gta aac ata tca taa tat tcg ata tta cga gtt gac ata tcg aac aaa ttc caa gta ttt gat ttt gga taa tat tcg tat ttt gca tct gct ata att aag ata taa tca ccg caa gaa cac acg aac atc ttt cct aca tgg tta aag tac atg tac aat tct atc cat ttg tct tcc tta act ata tat ttg tat aga taa tta cga gtc tcg tga gta att cca gta att aca tag atg tcg ccg tcg tac tct aca gca taa act ata cta tga tgt cta ggc atg gga gac -tt ttt tat cca acg att ttt agt gaa aca ttc cac atc gtt taa tac tac ata ttt ttc ata cgt ggt ata aac tcc acc cat tac --a tat ata tca tcg ttt acg aat acc gac gcg cct gaa tat cta gga gta att aag ttt gga agt ctt atc cat ttc gaa gtg ccg tgt ttc aaa tat tct gcc aca ccc gtt gaa ata gaa aat tct aat cct cct att aca tat aac ttt cca tcg tta aca caa gta cta act tct gat ttt aac gac gac ata tta gta acc gtt ttc ca- --- --- --- --- --- --- --t ttt ttc gtt tta aga tct acc cgc -ga tac gga ata aac atg tct att gtt aat cat gcc gcc aat aat gta tag aca att atg taa aac att tgc att ata gaa ttg --- -tc tat ctg tat tac cga cta tcg tcc aat att ctg ttc tag gag agt aat ggg tta ttg tgg ata tat aat cag agt ttt taa tga cta cta tat tat gtt tta tac cat ttc gtg tca ctg gct ttg tag att tgg ata tag tta atc cca aca atg ata tag cat tgc gca tag tat tag tca taa act tgg gat gta aaa tgt tga tga tat cta cat cgt ttg gat ttt tat gta tcc act tta ata ata tca tag ctg taa cat cct cat ga- ttt acg tta acg tct tcg tgg gat aag ata gtt gtc agt tca tcc ttt gat aat ttt cca aat t

>MW960422.1:21733-23679_Vector_synVACV-Delta1-3_complete_sequence

cgg ctg cca tgt att tcc tgg aga gca agt --- aga tga tg- agg aac cag ata gtt tat atc cgt act tgc act taa agt cta cat tgt cgt tgt atg a-- gta tga tct ttt aaa ccc gct aga caa gta tcc gtt tga tat tgt agg atg tgg aca ttt aac aat ctg aca cgt ggg tgg atc gga cca ttc tcc tcc tga aca cag gac acc aga gtt acc aat caa cga ata tcc act att gca act ata agt tac aac gct ccc atc ggt ata aaa atc ctc gta tcc gtt atg tct tcc gtt gga tat aga tgg agg gga ttg gca ttt aac aga ttc aca aat agg tgc ctc ggg att cca tac cat aga tcc agt aga tcc taa ttc aca ata cga ttt aga ttc acc gat caa atg ata tcc gct att aca aga gta cgt tat act aga gcc aaa gtc tac tcc acc aat atc aag ttg gcc att atc gat atc tcg agg cga tgg gca tct ccg ttt aat aca ttg att aaa gag tgt cca tcc agt acc tgt aca ttt agc ata tat agg tcc cat ttt ttg ctt tct gta tcc agg tag aca tag a-t att cta tag tgt ctc cta tgt tgt a-- --- --- --- -at tag cat tag cat cag tc- --t cca cac tat tct taa att tca tat taa tgg gtc gtg acg gaa tag tac agc atg ata gaa cgc atc cta ttc cca aca atg tca gga acg tca cgc tct cca cct tca tat tta ttt atc cgt -aa aaa tgt tat cct gga cat cgt aca aat aat aaa aaa gcc ca- -ta tat gtt tgc tat tgt aga aat tgt ttt tca cag ttg ctc aaa aac gat ggc agt gac tta tga gtt --- --- --- --- --- --- tca tct tta gta aac ata tca taa tat tcg ata tta cga gtt gac ata tcg aac aaa ttc caa gta ttt gat ttt gga taa tat tcg tat ttt gca tct gct ata att aag ata taa tca ccg caa gaa cac acg aac atc ttt cct aca tgg tta aag tac atg tac aat tct atc cat ttg tct tcc tta act ata tat ttg tat aga taa tta cga gtc tcg tga gta att cca gta att aca tag atg tcg ccg tcg tac tct aca gca taa act ata cta tga tgt cta ggc atg gga gac -tt ttt tat cca acg att ttt agt gaa aca ttc cac atc gtt taa tac tac ata ttt ttc ata cgt ggt ata aac tcc acc cat tac --a tat ata tca tcg ttt acg aat acc gac gcg cct gaa tat cta gga gta att aag ttt gga agt ctt atc cat ttc gaa gtg ccg tgt ttc aaa tat tct gcc aca ccc gtt gaa ata gaa aat tct aat cct cct att aca tat aac ttt cca tcg tta aca caa gta cta act tct gat ttt aac gac gac ata tta gta acc gtt ttc ca- --- --- --- --- --- --- --t ttt ttc gtt tta aga tct acc cgc -ga tac gga ata aac atg tct att gtt aat cat gcc gcc aat aat gta tag aca att atg taa aac att tgc att ata gaa ttg --- -tc tat ctg tat tac cga cta tcg tcc aat att ctg ttc tag gag agt aat ggg tta ttg tgg ata tat aat cag agt ttt taa tga cta cta tat tat gtt tta tac cat ttc gtg tca ctg gct ttg tag att tgg ata tag tta atc cca aca atg ata tag cat tgc gca tag tat tag tca taa act tgg gat gta aaa tgt tga tga tat cta cat cgt ttg gat ttt tat gta tcc act tta ata ata tca tag ctg taa cat cct cat ga- ttt acg tta acg tct tcg tgg gat aag ata gtt gtc agt tca tcc ttt gat aat ttt cca aat t

>MW960421.1:21725-23671_Vector_synVACV-Delta5-6_complete_sequence

cgg ctg cca tgt att tcc tgg aga gca agt --- aga tga tg- agg aac cag ata gtt tat atc cgt act tgc act taa agt cta cat tgt cgt tgt atg a-- gta tga tct ttt aaa ccc gct aga caa gta tcc gtt tga tat tgt agg atg tgg aca ttt aac aat ctg aca cgt ggg tgg atc gga cca ttc tcc tcc tga aca cag gac acc aga gtt acc aat caa cga ata tcc act att gca act ata agt tac aac gct ccc atc ggt ata aaa atc ctc gta tcc gtt atg tct tcc gtt gga tat aga tgg agg gga ttg gca ttt aac aga ttc aca aat agg tgc ctc ggg att cca tac cat aga tcc agt aga tcc taa ttc aca ata cga ttt aga ttc acc gat caa atg ata tcc gct att aca aga gta cgt tat act aga gcc aaa gtc tac tcc acc aat atc aag ttg gcc att atc gat atc tcg agg cga tgg gca tct ccg ttt aat aca ttg att aaa gag tgt cca tcc agt acc tgt aca ttt agc ata tat agg tcc cat ttt ttg ctt tct gta tcc agg tag aca tag a-t att cta tag tgt ctc cta tgt tgt a-- --- --- --- -at tag cat tag cat cag tc- --t cca cac tat tct taa att tca tat taa tgg gtc gtg acg gaa tag tac agc atg ata gaa cgc atc cta ttc cca aca atg tca gga acg tca cgc tct cca cct tca tat tta ttt atc cgt -aa aaa tgt tat cct gga cat cgt aca aat aat aaa aaa gcc ca- -ta tat gtt tgc tat tgt aga aat tgt ttt tca cag ttg ctc aaa aac gat ggc agt gac tta tga gtt --- --- --- --- --- --- tca tct tta gta aac ata tca taa tat tcg ata tta cga gtt gac ata tcg aac aaa ttc caa gta ttt gat ttt gga taa tat tcg tat ttt gca tct gct ata att aag ata taa tca ccg caa gaa cac acg aac atc ttt cct aca tgg tta aag tac atg tac aat tct atc cat ttg tct tcc tta act ata tat ttg tat aga taa tta cga gtc tcg tga gta att cca gta att aca tag atg tcg ccg tcg tac tct aca gca taa act ata cta tga tgt cta ggc atg gga gac -tt ttt tat cca acg att ttt agt gaa aca ttc cac atc gtt taa tac tac ata ttt ttc ata cgt ggt ata aac tcc acc cat tac --a tat ata tca tcg ttt acg aat acc gac gcg cct gaa tat cta gga gta att aag ttt gga agt ctt atc cat ttc gaa gtg ccg tgt ttc aaa tat tct gcc aca ccc gtt gaa ata gaa aat tct aat cct cct att aca tat aac ttt cca tcg tta aca caa gta cta act tct gat ttt aac gac gac ata tta gta acc gtt ttc ca- --- --- --- --- --- --- --t ttt ttc gtt tta aga tct acc cgc -ga tac gga ata aac atg tct att gtt aat cat gcc gcc aat aat gta tag aca att atg taa aac att tgc att ata gaa ttg --- -tc tat ctg tat tac cga cta tcg tcc aat att ctg ttc tag gag agt aat ggg tta ttg tgg ata tat aat cag agt ttt taa tga cta cta tat tat gtt tta tac cat ttc gtg tca ctg gct ttg tag att tgg ata tag tta atc cca aca atg ata tag cat tgc gca tag tat tag tca taa act tgg gat gta aaa tgt tga tga tat cta cat cgt ttg gat ttt tat gta tcc act tta ata ata tca tag ctg taa cat cct cat ga- ttt acg tta acg tct tcg tgg gat aag ata gtt gtc agt tca tcc ttt gat aat ttt cca aat t

>MW960420.1:21733-23679_Vector_synVACV-Delta6_complete_sequence

cgg ctg cca tgt att tcc tgg aga gca agt --- aga tga tg- agg aac cag ata gtt tat atc cgt act tgc act taa agt cta cat tgt cgt tgt atg a-- gta tga tct ttt aaa ccc gct aga caa gta tcc gtt tga tat tgt agg atg tgg aca ttt aac aat ctg aca cgt ggg tgg atc gga cca ttc tcc tcc tga aca cag gac acc aga gtt acc aat caa cga ata tcc act att gca act ata agt tac aac gct ccc atc ggt ata aaa atc ctc gta tcc gtt atg tct tcc gtt gga tat aga tgg agg gga ttg gca ttt aac aga ttc aca aat agg tgc ctc ggg att cca tac cat aga tcc agt aga tcc taa ttc aca ata cga ttt aga ttc acc gat caa atg ata tcc gct att aca aga gta cgt tat act aga gcc aaa gtc tac tcc acc aat atc aag ttg gcc att atc gat atc tcg agg cga tgg gca tct ccg ttt aat aca ttg att aaa gag tgt cca tcc agt acc tgt aca ttt agc ata tat agg tcc cat ttt ttg ctt tct gta tcc agg tag aca tag a-t att cta tag tgt ctc cta tgt tgt a-- --- --- --- -at tag cat tag cat cag tc- --t cca cac tat tct taa att tca tat taa tgg gtc gtg acg gaa tag tac agc atg ata gaa cgc atc cta ttc cca aca atg tca gga acg tca cgc tct cca cct tca tat tta ttt atc cgt -aa aaa tgt tat cct gga cat cgt aca aat aat aaa aaa gcc ca- -ta tat gtt tgc tat tgt aga aat tgt ttt tca cag ttg ctc aaa aac gat ggc agt gac tta tga gtt --- --- --- --- --- --- tca tct tta gta aac ata tca taa tat tcg ata tta cga gtt gac ata tcg aac aaa ttc caa gta ttt gat ttt gga taa tat tcg tat ttt gca tct gct ata att aag ata taa tca ccg caa gaa cac acg aac atc ttt cct aca tgg tta aag tac atg tac aat tct atc cat ttg tct tcc tta act ata tat ttg tat aga taa tta cga gtc tcg tga gta att cca gta att aca tag atg tcg ccg tcg tac tct aca gca taa act ata cta tga tgt cta ggc atg gga gac -tt ttt tat cca acg att ttt agt gaa aca ttc cac atc gtt taa tac tac ata ttt ttc ata cgt ggt ata aac tcc acc cat tac --a tat ata tca tcg ttt acg aat acc gac gcg cct gaa tat cta gga gta att aag ttt gga agt ctt atc cat ttc gaa gtg ccg tgt ttc aaa tat tct gcc aca ccc gtt gaa ata gaa aat tct aat cct cct att aca tat aac ttt cca tcg tta aca caa gta cta act tct gat ttt aac gac gac ata tta gta acc gtt ttc ca- --- --- --- --- --- --- --t ttt ttc gtt tta aga tct acc cgc -ga tac gga ata aac atg tct att gtt aat cat gcc gcc aat aat gta tag aca att atg taa aac att tgc att ata gaa ttg --- -tc tat ctg tat tac cga cta tcg tcc aat att ctg ttc tag gag agt aat ggg tta ttg tgg ata tat aat cag agt ttt taa tga cta cta tat tat gtt tta tac cat ttc gtg tca ctg gct ttg tag att tgg ata tag tta atc cca aca atg ata tag cat tgc gca tag tat tag tca taa act tgg gat gta aaa tgt tga tga tat cta cat cgt ttg gat ttt tat gta tcc act tta ata ata tca tag ctg taa cat cct cat ga- ttt acg tta acg tct tcg tgg gat aag ata gtt gtc agt tca tcc ttt gat aat ttt cca aat t

>MW960419.1:21721-23667_Vector_synVACV-SFV_complete_sequence

cgg ctg cca tgt att tcc tgg aga gca agt --- aga tga tg- agg aac cag ata gtt tat atc cgt act tgc act taa agt cta cat tgt cgt tgt atg a-- gta tga tct ttt aaa ccc gct aga caa gta tcc gtt tga tat tgt agg atg tgg aca ttt aac aat ctg aca cgt ggg tgg atc gga cca ttc tcc tcc tga aca cag gac acc aga gtt acc aat caa cga ata tcc act att gca act ata agt tac aac gct ccc atc ggt ata aaa atc ctc gta tcc gtt atg tct tcc gtt gga tat aga tgg agg gga ttg gca ttt aac aga ttc aca aat agg tgc ctc ggg att cca tac cat aga tcc agt aga tcc taa ttc aca ata cga ttt aga ttc acc gat caa atg ata tcc gct att aca aga gta cgt tat act aga gcc aaa gtc tac tcc acc aat atc aag ttg gcc att atc gat atc tcg agg cga tgg gca tct ccg ttt aat aca ttg att aaa gag tgt cca tcc agt acc tgt aca ttt agc ata tat agg tcc cat ttt ttg ctt tct gta tcc agg tag aca tag a-t att cta tag tgt ctc cta tgt tgt a-- --- --- --- -at tag cat tag cat cag tc- --t cca cac tat tct taa att tca tat taa tgg gtc gtg acg gaa tag tac agc atg ata gaa cgc atc cta ttc cca aca atg tca gga acg tca cgc tct cca cct tca tat tta ttt atc cgt -aa aaa tgt tat cct gga cat cgt aca aat aat aaa aaa gcc ca- -ta tat gtt tgc tat tgt aga aat tgt ttt tca cag ttg ctc aaa aac gat ggc agt gac tta tga gtt --- --- --- --- --- --- tca tct tta gta aac ata tca taa tat tcg ata tta cga gtt gac ata tcg aac aaa ttc caa gta ttt gat ttt gga taa tat tcg tat ttt gca tct gct ata att aag ata taa tca ccg caa gaa cac acg aac atc ttt cct aca tgg tta aag tac atg tac aat tct atc cat ttg tct tcc tta act ata tat ttg tat aga taa tta cga gtc tcg tga gta att cca gta att aca tag atg tcg ccg tcg tac tct aca gca taa act ata cta tga tgt cta ggc atg gga gac -tt ttt tat cca acg att ttt agt gaa aca ttc cac atc gtt taa tac tac ata ttt ttc ata cgt ggt ata aac tcc acc cat tac --a tat ata tca tcg ttt acg aat acc gac gcg cct gaa tat cta gga gta att aag ttt gga agt ctt atc cat ttc gaa gtg ccg tgt ttc aaa tat tct gcc aca ccc gtt gaa ata gaa aat tct aat cct cct att aca tat aac ttt cca tcg tta aca caa gta cta act tct gat ttt aac gac gac ata tta gta acc gtt ttc ca- --- --- --- --- --- --- --t ttt ttc gtt tta aga tct acc cgc -ga tac gga ata aac atg tct att gtt aat cat gcc gcc aat aat gta tag aca att atg taa aac att tgc att ata gaa ttg --- -tc tat ctg tat tac cga cta tcg tcc aat att ctg ttc tag gag agt aat ggg tta ttg tgg ata tat aat cag agt ttt taa tga cta cta tat tat gtt tta tac cat ttc gtg tca ctg gct ttg tag att tgg ata tag tta atc cca aca atg ata tag cat tgc gca tag tat tag tca taa act tgg gat gta aaa tgt tga tga tat cta cat cgt ttg gat ttt tat gta tcc act tta ata ata tca tag ctg taa cat cct cat ga- ttt acg tta acg tct tcg tgg gat aag ata gtt gtc agt tca tcc ttt gat aat ttt cca aat t

>MW960418.1:21739-23685_Vector_synVACV-wt_complete_sequence

cgg ctg cca tgt att tcc tgg aga gca agt --- aga tga tg- agg aac cag ata gtt tat atc cgt act tgc act taa agt cta cat tgt cgt tgt atg a-- gta tga tct ttt aaa ccc gct aga caa gta tcc gtt tga tat tgt agg atg tgg aca ttt aac aat ctg aca cgt ggg tgg atc gga cca ttc tcc tcc tga aca cag gac acc aga gtt acc aat caa cga ata tcc act att gca act ata agt tac aac gct ccc atc ggt ata aaa atc ctc gta tcc gtt atg tct tcc gtt gga tat aga tgg agg gga ttg gca ttt aac aga ttc aca aat agg tgc ctc ggg att cca tac cat aga tcc agt aga tcc taa ttc aca ata cga ttt aga ttc acc gat caa atg ata tcc gct att aca aga gta cgt tat act aga gcc aaa gtc tac tcc acc aat atc aag ttg gcc att atc gat atc tcg agg cga tgg gca tct ccg ttt aat aca ttg att aaa gag tgt cca tcc agt acc tgt aca ttt agc ata tat agg tcc cat ttt ttg ctt tct gta tcc agg tag aca tag a-t att cta tag tgt ctc cta tgt tgt a-- --- --- --- -at tag cat tag cat cag tc- --t cca cac tat tct taa att tca tat taa tgg gtc gtg acg gaa tag tac agc atg ata gaa cgc atc cta ttc cca aca atg tca gga acg tca cgc tct cca cct tca tat tta ttt atc cgt -aa aaa tgt tat cct gga cat cgt aca aat aat aaa aaa gcc ca- -ta tat gtt tgc tat tgt aga aat tgt ttt tca cag ttg ctc aaa aac gat ggc agt gac tta tga gtt --- --- --- --- --- --- tca tct tta gta aac ata tca taa tat tcg ata tta cga gtt gac ata tcg aac aaa ttc caa gta ttt gat ttt gga taa tat tcg tat ttt gca tct gct ata att aag ata taa tca ccg caa gaa cac acg aac atc ttt cct aca tgg tta aag tac atg tac aat tct atc cat ttg tct tcc tta act ata tat ttg tat aga taa tta cga gtc tcg tga gta att cca gta att aca tag atg tcg ccg tcg tac tct aca gca taa act ata cta tga tgt cta ggc atg gga gac -tt ttt tat cca acg att ttt agt gaa aca ttc cac atc gtt taa tac tac ata ttt ttc ata cgt ggt ata aac tcc acc cat tac --a tat ata tca tcg ttt acg aat acc gac gcg cct gaa tat cta gga gta att aag ttt gga agt ctt atc cat ttc gaa gtg ccg tgt ttc aaa tat tct gcc aca ccc gtt gaa ata gaa aat tct aat cct cct att aca tat aac ttt cca tcg tta aca caa gta cta act tct gat ttt aac gac gac ata tta gta acc gtt ttc ca- --- --- --- --- --- --- --t ttt ttc gtt tta aga tct acc cgc -ga tac gga ata aac atg tct att gtt aat cat gcc gcc aat aat gta tag aca att atg taa aac att tgc att ata gaa ttg --- -tc tat ctg tat tac cga cta tcg tcc aat att ctg ttc tag gag agt aat ggg tta ttg tgg ata tat aat cag agt ttt taa tga cta cta tat tat gtt tta tac cat ttc gtg tca ctg gct ttg tag att tgg ata tag tta atc cca aca atg ata tag cat tgc gca tag tat tag tca taa act tgg gat gta aaa tgt tga tga tat cta cat cgt ttg gat ttt tat gta tcc act tta ata ata tca tag ctg taa cat cct cat ga- ttt acg tta acg tct tcg tgg gat aag ata gtt gtc agt tca tcc ttt gat aat ttt cca aat t

>MZ300860.1:19458-21422_Camelpox_virus_strain_D1621/20_complete_genome

cgg ctg cca tgt att tcc tgg aga gca agt --- aga tga tg- agg aac cag ata gtt tat atc cgt act tgc acg taa agt cta cat tgt cat tat acg a-- ata tga tct ttt aaa ccc gct aga caa gta tcc gtt tga tat tgt agg atg tgg aca ttt aac aat ctg aca cgt ggg tgg atc gga cca ttc tcc tcc tga aca cag gac acc aga gtt acc aat caa cga ata tcc act att gca act ata agt tac aac gct tcc atc ggt ata aaa att atc gta tcc gtt atg tct tcc gtt gga tat aga tgg agg tga ttg gca ttt aac aga ttc gca aat agg tgc ctc ggg att cca tac cat aga tcc agt aga tcc taa ttc aca ata cga ttt aga ttc acc gat caa atg ata tcc gct att aca aga gta cgt tat act aga acc aaa gtc tac tcc acc aat atc aag ttg gcc att atc gat atc tcg agg cga tgg gca tct ccg ttt aat aca ttg att aaa gag tgt cca tcc agt acc tgt aca ttt agc ata tat agg tcc cat ttt ttg ctt tct gta tcc agg tag aca tag a-t att cta tag tgt ctc cta tgt tgt a-- --- --- --- -at tag tat tag cat aag tc- --t cca cac tat tct taa att tca tat taa tgg gtc gtg acg gaa tag tac agc atg ata gaa cgc ata cta ttc cca aca atg tca gga acg tca cgc tct cca cct tca tat tta ttt atc cgt -aa aaa tgt tat cct gga cac tgt acc aat aat aaa aaa gcc ca- -ta tat gtt cgc tat tgt aga aat tgt ttt tca cag ttg ctc aaa aac gat ggc agt gac tta tga gtt acg tta cac ttt gga gtc tca tct tta gta aac ata tca taa tat tcg ata tta cga gtt gac ata tcg aac aaa atc caa gta ttt gat ttt gga taa tat tcg tat ttt gca tct gct ata att aag ata taa tcg ccg caa gaa cac acg aac atc ttt cct aca tgg tta aag tac atg tac aat tct atc cat ttg tct tcc tta act ata tat ttg tat aga taa tta cga gtc tcg tga gta att cca gta att gca tag atg tca cca tcg tac tct ata gca taa act ata cta tga tgt cta ggc atg gga gac -tt ttt tat cca acg att ttt agt gaa aca ttc cac atc gtt taa tac tac ata ttt ttc ata cgt ggt ata aac tcc acc cat tac --a tat ata tca tcg ttt acg aat acc gac gcg cct gaa tat cta gga gta att aag ttt gga agt ctt atc cat ttc gaa gtt ccg tgt ttc aaa tat tct gct acg cct gtt gaa ata gaa aat tct aat cct cct att aca tat aac ttt cca tca tta aca caa gta cta act tct gat ttt aac gac gac ata tta gta act gtt ttc ca- --- --- --- --- --- --- --t ttt ttc gtt tta aga tct acc cgc -ga tac gga ata aac atg tct att gtt aat cat gcc gcc aat aat gta tag aca att ata taa aac att tgc att ata gaa ttg --- -tc tat ctg tat tac cga cga ttg tcc aat att ctg ttc tag gag agt aat ggg tta ttg tgg ata tat aat cag agt ttt taa tga cta cta tat tat gtt tta tac cat ttc gtg tca cag gct ttg tag att tgg ata tag tta atc cca aca atg ata tag cat tgc aca tag tat tag tca taa act tgg gat gta aaa tgt tga taa tat cta cat cgt ttg gat ttt tat gta tcc act tta ata ata tca tag ctg taa cat cct cgt ga- ttt acg ttg acg ttt tcg tgg gat aag ata gtt gtt agt tca tcc ttt gat aat ttt cca aat t

>MZ300859.1:19459-21423_Camelpox_virus_strain_D1734/20_complete_genome

cgg ctg cca tgt att tcc tgg aga gca agt --- aga tga tg- agg aac cag ata gtt tat atc cgt act tgc acg taa agt cta cat tgt cat tat acg a-- ata tga tct ttt aaa ccc gct aga caa gta tcc gtt tga tat tgt agg atg tgg aca ttt aac aat ctg aca cgt ggg tgg atc gga cca ttc tcc tcc tga aca cag gac acc aga gtt acc aat caa cga ata tcc act att gca act ata agt tac aac gct tcc atc ggt ata aaa att atc gta tcc gtt atg tct tcc gtt gga tat aga tgg agg tga ttg gca ttt aac aga ttc gca aat agg tgc ctc ggg att cca tac cat aga tcc agt aga tcc taa ttc aca ata cga ttt aga ttc acc gat caa atg ata tcc gct att aca aga gta cgt tat act aga acc aaa gtc tac tcc acc aat atc aag ttg gcc att atc gat atc tcg agg cga tgg gca tct ccg ttt aat aca ttg att aaa gag tgt cca tcc agt acc tgt aca ttt agc ata tat agg tcc cat ttt ttg ctt tct gta tcc agg tag aca tag a-t att cta tag tgt ctc cta tgt tgt a-- --- --- --- -at tag tat tag cat aag tc- --t cca cac tat tct taa att tca tat taa tgg gtc gtg acg gaa tag tac agc atg ata gaa cgc ata cta ttc cca aca atg tca gga acg tca cgc tct cca cct tca tat tta ttt atc cgt -aa aaa tgt tat cct gga cac tgt acc aat aat aaa aaa gcc ca- -ta tat gtt cgc tat tgt aga aat tgt ttt tca cag ttg ctc aaa aac gat ggc agt gac tta tga gtt acg tta cac ttt gga gtc tca tct tta gta aac ata tca taa tat tcg ata tta cga gtt gac ata tcg aac aaa atc caa gta ttt gat ttt gga taa tat tcg tat ttt gca tct gct ata att aag ata taa tcg ccg caa gaa cac acg aac atc ttt cct aca tgg tta aag tac atg tac aat tct atc cat ttg tct tcc tta act ata tat ttg tat aga taa tta cga gtc tcg tga gta att cca gta att gca tag atg tca cca tcg tac tct ata gca taa act ata cta tga tgt cta ggc atg gga gac -tt ttt tat cca acg att ttt agt gaa aca ttc cac atc gtt taa tac tac ata ttt ttc ata cgt ggt ata aac tcc acc cat tac --a tat ata tca tcg ttt acg aat acc gac gcg cct gaa tat cta gga gta att aag ttt gga agt ctt atc cat ttc gaa gtt ccg tgt ttc aaa tat tct gct acg cct gtt gaa ata gaa aat tct aat cct cct att aca tat aac ttt cca tca tta aca caa gta cta act tct gat ttt aac gac gac ata tta gta act gtt ttc ca- --- --- --- --- --- --- --t ttt ttc gtt tta aga tct acc cgc -ga tac gga ata aac atg tct att gtt aat cat gcc gcc aat aat gta tag aca att ata taa aac att tgc att ata gaa ttg --- -tc tat ctg tat tac cga cga ttg tcc aat att ctg ttc tag gag agt aat ggg tta ttg tgg ata tat aat cag agt ttt taa tga cta cta tat tat gtt tta tac cat ttc gtg tca cag gct ttg tag att tgg ata tag tta atc cca aca atg ata tag cat tgc aca tag tat tag tca taa act tgg gat gta aaa tgt tga taa tat cta cat cgt ttg gat ttt tat gta tcc act tta ata ata tca tag ctg taa cat cct cgt ga- ttt acg ttg acg ttt tcg tgg gat aag ata gtt gtt agt tca tcc ttt gat aat ttt cca aat t

>MZ300858.1:19459-21423_Camelpox_virus_strain_D1795/20_complete_genome

cgg ctg cca tgt att tcc tgg aga gca agt --- aga tga tg- agg aac cag ata gtt tat atc cgt act tgc acg taa agt cta cat tgt cat tat acg a-- ata tga tct ttt aaa ccc gct aga caa gta tcc gtt tga tat tgt agg atg tgg aca ttt aac aat ctg aca cgt ggg tgg atc gga cca ttc tcc tcc tga aca cag gac acc aga gtt acc aat caa cga ata tcc act att gca act ata agt tac aac gct tcc atc ggt ata aaa att atc gta tcc gtt atg tct tcc gtt gga tat aga tgg agg tga ttg gca ttt aac aga ttc gca aat agg tgc ctc ggg att cca tac cat aga tcc agt aga tcc taa ttc aca ata cga ttt aga ttc acc gat caa atg ata tcc gct att aca aga gta cgt tat act aga acc aaa gtc tac tcc acc aat atc aag ttg gcc att atc gat atc tcg agg cga tgg gca tct ccg ttt aat aca ttg att aaa gag tgt cca tcc agt acc tgt aca ttt agc ata tat agg tcc cat ttt ttg ctt tct gta tcc agg tag aca tag a-t att cta tag tgt ctc cta tgt tgt a-- --- --- --- -at tag tat tag cat aag tc- --t cca cac tat tct taa att tca tat taa tgg gtc gtg acg gaa tag tac agc atg ata gaa cgc ata cta ttc cca aca atg tca gga acg tca cgc tct cca cct tca tat tta ttt atc cgt -aa aaa tgt tat cct gga cac tgt acc aat aat aaa aaa gcc ca- -ta tat gtt cgc tat tgt aga aat tgt ttt tca cag ttg ctc aaa aac gat ggc agt gac tta tga gtt acg tta cac ttt gga gtc tca tct tta gta aac ata tca taa tat tcg ata tta cga gtt gac ata tcg aac aaa atc caa gta ttt gat ttt gga taa tat tcg tat ttt gca tct gct ata att aag ata taa tcg ccg caa gaa cac acg aac atc ttt cct aca tgg tta aag tac atg tac aat tct atc cat ttg tct tcc tta act ata tat ttg tat aga taa tta cga gtc tcg tga gta att cca gta att gca tag atg tca cca tcg tac tct ata gca taa act ata cta tga tgt cta ggc atg gga gac -tt ttt tat cca acg att ttt agt gaa aca ttc cac atc gtt taa tac tac ata ttt ttc ata cgt ggt ata aac tcc acc cat tac --a tat ata tca tcg ttt acg aat acc gac gcg cct gaa tat cta gga gta att aag ttt gga agt ctt atc cat ttc gaa gtt ccg tgt ttc aaa tat tct gct acg cct gtt gaa ata gaa aat tct aat cct cct att aca tat aac ttt cca tca tta aca caa gta cta act tct gat ttt aac gac gac ata tta gta act gtt ttc ca- --- --- --- --- --- --- --t ttt ttc gtt tta aga tct acc cgc -ga tac gga ata aac atg tct att gtt aat cat gcc gcc aat aat gta tag aca att ata taa aac att tgc att ata gaa ttg --- -tc tat ctg tat tac cga cga ttg tcc aat att ctg ttc tag gag agt aat ggg tta ttg tgg ata tat aat cag agt ttt taa tga cta cta tat tat gtt tta tac cat ttc gtg tca cag gct ttg tag att tgg ata tag tta atc cca aca atg ata tag cat tgc aca tag tat tag tca taa act tgg gat gta aaa tgt tga taa tat cta cat cgt ttg gat ttt tat gta tcc act tta ata ata tca tag ctg taa cat cct cgt ga- ttt acg ttg acg ttt tcg tgg gat aag ata gtt gtt agt tca tcc ttt gat aat ttt cca aat t

>MZ300857.1:19458-21422_Camelpox_virus_strain_D1804/20_complete_genome

cgg ctg cca tgt att tcc tgg aga gca agt --- aga tga tg- agg aac cag ata gtt tat atc cgt act tgc acg taa agt cta cat tgt cat tat acg a-- ata tga tct ttt aaa ccc gct aga caa gta tcc gtt tga tat tgt agg atg tgg aca ttt aac aat ctg aca cgt ggg tgg atc gga cca ttc tcc tcc tga aca cag gac acc aga gtt acc aat caa cga ata tcc act att gca act ata agt tac aac gct tcc atc ggt ata aaa att atc gta tcc gtt atg tct tcc gtt gga tat aga tgg agg tga ttg gca ttt aac aga ttc gca aat agg tgc ctc ggg att cca tac cat aga tcc agt aga tcc taa ttc aca ata cga ttt aga ttc acc gat caa atg ata tcc gct att aca aga gta cgt tat act aga acc aaa gtc tac tcc acc aat atc aag ttg gcc att atc gat atc tcg agg cga tgg gca tct ccg ttt aat aca ttg att aaa gag tgt cca tcc agt acc tgt aca ttt agc ata tat agg tcc cat ttt ttg ctt tct gta tcc agg tag aca tag a-t att cta tag tgt ctc cta tgt tgt a-- --- --- --- -at tag tat tag cat aag tc- --t cca cac tat tct taa att tca tat taa tgg gtc gtg acg gaa tag tac agc atg ata gaa cgc ata cta ttc cca aca atg tca gga acg tca cgc tct cca cct tca tat tta ttt atc cgt -aa aaa tgt tat cct gga cac tgt acc aat aat aaa aaa gcc ca- -ta tat gtt cgc tat tgt aga aat tgt ttt tca cag ttg ctc aaa aac gat ggc agt gac tta tga gtt acg tta cac ttt gga gtc tca tct tta gta aac ata tca taa tat tcg ata tta cga gtt gac ata tcg aac aaa atc caa gta ttt gat ttt gga taa tat tcg tat ttt gca tct gct ata att aag ata taa tcg ccg caa gaa cac acg aac atc ttt cct aca tgg tta aag tac atg tac aat tct atc cat ttg tct tcc tta act ata tat ttg tat aga taa tta cga gtc tcg tga gta att cca gta att gca tag atg tca cca tcg tac tct ata gca taa act ata cta tga tgt cta ggc atg gga gac -tt ttt tat cca acg att ttt agt gaa aca ttc cac atc gtt taa tac tac ata ttt ttc ata cgt ggt ata aac tcc acc cat tac --a tat ata tca tcg ttt acg aat acc gac gcg cct gaa tat cta gga gta att aag ttt gga agt ctt atc cat ttc gaa gtt ccg tgt ttc aaa tat tct gct acg cct gtt gaa ata gaa aat tct aat cct cct att aca tat aac ttt cca tca tta aca caa gta cta act tct gat ttt aac gac gac ata tta gta act gtt ttc ca- --- --- --- --- --- --- --t ttt ttc gtt tta aga tct acc cgc -ga tac gga ata aac atg tct att gtt aat cat gcc gcc aat aat gta tag aca att ata taa aac att tgc att ata gaa ttg --- -tc tat ctg tat tac cga cga ttg tcc aat att ctg ttc tag gag agt aat ggg tta ttg tgg ata tat aat cag agt ttt taa tga cta cta tat tat gtt tta tac cat ttc gtg tca cag gct ttg tag att tgg ata tag tta atc cca aca atg ata tag cat tgc aca tag tat tag tca taa act tgg gat gta aaa tgt tga taa tat cta cat cgt ttg gat ttt tat gta tcc act tta ata ata tca tag ctg taa cat cct cgt ga- ttt acg ttg acg ttt tcg tgg gat aag ata gtt gtt agt tca tcc ttt gat aat ttt cca aat t

>MZ300856.1:19395-21359_Camelpox_virus_strain_D1865/20_complete_genome

cgg ctg cca tgt att tcc tgg aga gca agt --- aga tga tg- agg aac cag ata gtt tat atc cgt act tgc acg taa agt cta cat tgt cat tat acg a-- ata tga tct ttt aaa ccc gct aga caa gta tcc gtt tga tat tgt agg atg tgg aca ttt aac aat ctg aca cgt ggg tgg atc gga cca ttc tcc tcc tga aca cag gac acc aga gtt acc aat caa cga ata tcc act att gca act ata agt tac aac gct tcc atc ggt ata aaa att atc gta tcc gtt atg tct tcc gtt gga tat aga tgg agg tga ttg gca ttt aac aga ttc gca aat agg tgc ctc ggg att cca tac cat aga tcc agt aga tcc taa ttc aca ata cga ttt aga ttc acc gat caa atg ata tcc gct att aca aga gta cgt tat act aga acc aaa gtc tac tcc acc aat atc aag ttg gcc att atc gat atc tcg agg cga tgg gca tct ccg ttt aat aca ttg att aaa gag tgt cca tcc agt acc tgt aca ttt agc ata tat agg tcc cat ttt ttg ctt tct gta tcc agg tag aca tag a-t att cta tag tgt ctc cta tgt tgt a-- --- --- --- -at tag tat tag cat aag tc- --t cca cac tat tct taa att tca tat taa tgg gtc gtg acg gaa tag tac agc atg ata gaa cgc ata cta ttc cca aca atg tca gga acg tca cgc tct cca cct tca tat tta ttt atc cgt -aa aaa tgt tat cct gga cac tgt acc aat aat aaa aaa gcc ca- -ta tat gtt cgc tat tgt aga aat tgt ttt tca cag ttg ctc aaa aac gat ggc agt gac tta tga gtt acg tta cac ttt gga gtc tca tct tta gta aac ata tca taa tat tcg ata tta cga gtt gac ata tcg aac aaa atc caa gta ttt gat ttt gga taa tat tcg tat ttt gca tct gct ata att aag ata taa tcg ccg caa gaa cac acg aac atc ttt cct aca tgg tta aag tac atg tac aat tct atc cat ttg tct tcc tta act ata tat ttg tat aga taa tta cga gtc tcg tga gta att cca gta att gca tag atg tca cca tcg tac tct ata gca taa act ata cta tga tgt cta ggc atg gga gac -tt ttt tat cca acg att ttt agt gaa aca ttc cac atc gtt taa tac tac ata ttt ttc ata cgt ggt ata aac tcc acc cat tac --a tat ata tca tcg ttt acg aat acc gac gcg cct gaa tat cta gga gta att aag ttt gga agt ctt atc cat ttc gaa gtt ccg tgt ttc aaa tat tct gct acg cct gtt gaa ata gaa aat tct aat cct cct att aca tat aac ttt cca tca tta aca caa gta cta act tct gat ttt aac gac gac ata tta gta act gtt ttc ca- --- --- --- --- --- --- --t ttt ttc gtt tta aga tct acc cgc -ga tac gga ata aac atg tct att gtt aat cat gcc gcc aat aat gta tag aca att ata taa aac att tgc att ata gaa ttg --- -tc tat ctg tat tac cga cga ttg tcc aat att ctg ttc tag gag agt aat ggg tta ttg tgg ata tat aat cag agt ttt taa tga cta cta tat tat gtt tta tac cat ttc gtg tca cag gct ttg tag att tgg ata tag tta atc cca aca atg ata tag cat tgc aca tag tat tag tca taa act tgg gat gta aaa tgt tga taa tat cta cat cgt ttg gat ttt tat gta tcc act tta ata ata tca tag ctg taa cat cct cgt ga- ttt acg ttg acg ttt tcg tgg gat aag ata gtt gtt agt tca tcc ttt gat aat ttt cca aat t

>JN654976.1:21422-23362_Vaccinia_virus_strain_Dryvax_clone_DPP9_complete_genome

cgg ctg cca tgt att tcc tgg aga gca agt --- aga tga tg- agg aac cag ata gtt tat atc cgt act tgc act taa agt cta cat tgt cgt tgt atg a-- gta tga tct ttt aaa ccc gct aga caa gta tcc gtt tga tat tgt agg atg tgg aca ttt aac aat ctg aca cgt ggg tgg atc gga cca ttc tcc tcc tga aca cag gac acc aga gtt acc aat caa cga ata tcc act att gca act ata agt tac aac gct tcc atc ggt ata aaa atc ctc gta tcc gtt atg tct tcc gtt gga tat aga tgg agg gga ttg gca ttt aac aga ttc aca aat agg tgc ctc ggg att cca tac cat aga tcc agt aga tcc taa ttc aca ata cga ttt aga ttc acc gat caa atg ata tcc gct att aca aga gta cgt tat act aga gcc aaa gtc tac tcc acc aat atc aag ttg gcc att atc gat atc tcg agg cga tgg gca tct ccg ttt aat aca ttg att aaa gag tgt cca tcc agt acc tgt aca ttt agc ata tat agg tcc cat ttt ttg ctt tct gta tcc agg tag aca tag a-t att cta tag tgt ctc cta tgt tgt a-- --- --- --- -at tag cat t-- --- -ag tt- --t cca cac tat tct taa att tca tat taa tgg gac gtg aag gaa tag gac agc atg ata gaa cgc atc cta ttc cca aca atg tca gga acg tca cgc tct cca cct tca tat tta ttt atc cgt -aa aaa tgt tat cct gga cat cgt aca aat aat aaa aaa gcc ca- -ta tat gtt tgc tat tgt aga aat tgt ttt tca cag ttg ctc aaa aac gat ggc agt gac tta tga gtt --- --- --- --- --- --- tca tct tta gta aac ata tca taa tat tcg ata tta cga gtt gac ata tcg aac aaa ttc caa gta ttt gat ttt gga taa tat tcg tat ttt gca tct gct ata att aag ata taa tca ccg caa gaa cac acg aac atc ttt cct aca tgg tta aag tac atg tac aat tct atc cat ttg tct tcc tta act ata tat ttg tat aga taa tta cga gtc tcg tga gta att cca gta att aca tag atg tcg ccg tcg tac tct aca gca taa act ata cta tga tgt cta ggc atg gga gac -tt ttt tat cca acg att ttt agt gaa aca ttc cac atc gtt taa tac tac ata ttt ttc ata cgt ggt ata aac tcc acc cat tac --a tat ata tca tcg ttt acg aat acc gac gcg cct gaa tat cta gga gta att aag ttt gga agt ctt atc cat ttc gaa gtg ccg tgt ttc aaa tat tct gcc aca ccc gtt gaa ata gaa aat tct aat cct cct att aca tat aac ttt cca tcg tta aca caa gta cta act tct gat ttt aac gac gac ata tta gta acc gtt ttc ca- --- --- --- --- --- --- --t ttt ttc gtt tta aga tct acc cgc -ga tac gga ata aac atg tct att gtt aat cat gcc gcc aat aat gta tag aca att atg taa aac att tgc att ata gaa ttg --- -tc tat ctg tat tac cga cta tcg tcc aat att ctg ttc tag gag agt aat ggg tta ttg tgg ata tat aat cag agt ttt taa tga cta cta tat tat gtt tta tac cat ttc gtg tca ctg gct ttg tag att tgg ata tag tta atc cca aca atg ata tag cat tgc gca tag tat tag tca taa act tgg gat gta aaa tgt tga tga tat cta cat cgt ttg gat ttt tat gta tcc act tta ata ata tca tag ctg taa cat cct cat ga- ttt acg tta acg tct tcg tgg gat aag ata gtt gtc agt tca tcc ttt gat aat ttt cca aat t

>AY313847.1:21638-23584_Vaccinia_virus_strain_Acambis_clone_2000_complete_genome

cgg ctg cca tgt att tcc tgg aga gca agt --- aga tga tg- agg aac cag ata gtt tat atc cgt act tgc act taa agt cta cat tgt cgt tgt atg a-- gta tga tct ttt aaa ccc gct aga caa gta tcc gtt tga tat tgt agg atg tgg aca ttt aac aat ctg aca cgt ggg tgg atc gga cca ttc tcc tcc tga aca cag gac acc aga gtt acc aat caa cga ata tcc act att gca act ata agt tac aac gct ccc atc ggt ata aaa atc ctc gta tcc gtt atg tct tcc gtt gga tat aga tgg agg gga ttg gca ttt aac aga ttc aca aat agg tgc ctc ggg att cca tac cat aga tcc agt aga tcc taa ttc aca ata cga ttt aga ttc acc gat caa atg ata tcc gct att aca aga gta cgt tat act aga gcc aaa gtc tac tcc acc aat atc aag ttg gcc att atc gat atc tcg agg cga tgg gca tct ccg ttt aat aca ttg att aaa gag tgt cca tcc agt acc tgt aca ttt agc ata tat agg tcc cat ttt ttg ctt tct gta tcc agg tag aca tag a-t att cta tag tgt ctc cta tgt tgt a-- --- --- --- -at tag cat tag cat cag tc- --t cca cac tat tct taa att tca tat taa tgg gtc gtg acg gaa tag tac agc atg ata gaa cgc atc cta ttc cca aca atg tca gga acg tca cgc tct cca cct tca tat tta ttt atc cgt -aa aaa tgt tat cct gga cat cgt aca aat aat aaa aaa gcc ca- -ta tat gtt tgc tat tgt aga aat tgt ttt tca cag ttg ctc aaa aac gat ggc agt gac tta tga gtt --- --- --- --- --- --- tca tct tta gta aac ata tca taa tat tcg ata tta cga gtt gac ata tcg aac aaa ttc caa gta ttt gat ttt gga taa tat tcg tat ttt gca tct gct ata att aag ata taa tca ccg caa gaa cac acg aac atc ttt cct aca tgg tta aag tac atg tac aat tct atc cat ttg tct tcc tta act ata tat ttg tat aga taa tta cga gtc tcg tga gta att cca gta att aca tag atg tcg ccg tcg tac tct aca gca taa act ata cta tga tgt cta ggc atg gga gac -tt ttt tat cca acg att ttt agt gaa aca ttc cac atc gtt taa tac tac ata ttt ttc ata cgt ggt ata aac tcc acc cat tac --a tat ata tca tcg ttt acg aat acc gac gcg cct gaa tat cta gga gta att aag ttt gga agt ctt atc cat ttc gaa gtg ccg tgt ttc aaa tat tct gcc aca ccc gtt gaa ata gaa aat tct aat cct cct att aca tat aac ttt cca tcg tta aca caa gta cta act tct gat ttt aac gac gac ata tta gta acc gtt ttc ca- --- --- --- --- --- --- --t ttt ttc gtt tta aga tct acc cgc -ga tac gga ata aac atg tct att gtt aat cat gcc gcc aat aat gta tag aca att atg taa aac att tgc att ata gaa ttg --- -tc tat ctg tat tac cga cta tcg tcc aat att ctg ttc tag gag agt aat ggg tta ttg tgg ata tat aat cag agt ttt taa tga cta cta tat tat gtt tta tac cat ttc gtg tca ctg gct ttg tag att tgg ata tag tta atc cca aca atg ata tag cat tgc gca tag tat tag tca taa act tgg gat gta aaa tgt tga tga tat cta cat cgt ttg gat ttt tat gta tcc act tta ata ata tca tag ctg taa cat cct cat ga- ttt acg tta acg tct tcg tgg gat aag ata gtt gtc agt tca tcc ttt gat aat ttt cca aat t

>KJ125438.1:21456-23396_Vaccinia_virus_strain_Dryvax_clone_DPP25_complete_genome

cgg ctg cca tgt att tcc tgg aga gca agt --- aga tga tg- agg aac cag ata ggt tat atc cgt act tgc act taa agt tta cat tgt cgt tgt atg a-- gta tga tct ttt aaa ccc gct aga caa gta tcc gtt tga tat tgt agg atg tgg aca ttt aac aat ctg aca cgt ggg tgg atc gga cca ttc tcc tcc tga aca cag gac acc aga gtt acc aat caa cga ata tcc act att gca act ata agt tac aac gct ccc atc ggt ata aaa atc ctc gta tcc gtt atg tct tcc gtt gga tat aga tgg agg gga ttg gca ttt aac aga ttc aca aat agg tgc ctc ggg att cca tac cat aga tcc agt aga tcc taa ttc aca ata cga ttt aga ttc acc gat caa atg ata tcc gct att aca aga gta cgt tat act aga gcc aaa gtc tac tcc acc aat atc aag ttg gcc att atc gat atc tcg agg cga tgg gca tct ccg ttt aat aca ttg att aaa gag tgt cca tcc agt acc tgt aca ttt agc ata tat agg tcc cat ttt ttg ctt tct gta tcc agg tag aca tag a-t att cta tag tgt ctc cta tgt tgt a-- --- --- --- -at tag cat t-- --- -ag tt- --t cca cac tat tct taa att tta tat taa tgg gac gtg aag gaa tag gac agt atg ata gaa cgc atc cta ttc cca aca atg tca gga acg tca cgc tct cca cct tca tat tta ttt atc cgt -aa aaa tgt tat cct gga cat cgt aca aat aat aaa aaa gcc ca- -ta tat gtt tgc tat tgt aga aat tgt ttt tca cag ttg ctc aaa aac gat ggc agt gac tta tga gtt --- --- --- --- --- --- tca tct tta gta aac ata tca taa tat tcg ata tta cga gtt gac ata tcg aac aaa ttc caa gta ttt gat ttt gga taa tat tcg tat ttt gca tct gct ata att aag ata taa tca ccg caa gaa cac acg aac atc ttt cct aca tgg tta aag tac atg tac aat tct atc cat ttg tct tcc tta act ata tat ttg tat aga taa tta cga gtc tcg tga gta att cca gta att aca tag atg tcg ccg tcg tac tct aca gca taa act ata cta tga tgt cta ggc atg gga gac -tt ttt tat cca acg att ttt agt gaa aca ttc cac atc gtt taa tac tac ata ttt ttc ata cgt ggt ata aac tcc acc cat tac --a tat ata tca tcg ttt acg aat acc gac gcg cct gaa tat cta gga gta att aag ttt gga agt ctt atc cat ttc gaa gtg ccg tgt ttc aaa tat tct gcc aca ccc gtt gaa ata gaa aat tct aat cct cct att aca tat aac ttt cca tcg tta aca caa gta cta act tct gat ttt aac gac gac ata tta gta acc gtt ttc ca- --- --- --- --- --- --- --t ttt ttc gtt tta aga tct acc cgc -ga tac gga ata aac atg tct att gtt aat cat gcc gcc aat aat gta tag aca att atg taa aac att tgc att ata gaa ttg --- -tc tat ctg tat tac cga cta tcg tcc aat att ctg ttc tag gag agt aat ggg tta ttg tgg ata tat aat cag agt ttt taa tga cta cta tat tat gtt tta tac cat ttc gtg tca ctg gct ttg tag att tgg ata tag tta atc cca aca atg ata tag cat tgc gca tag tat tag tca taa act tgg gat gta aaa tgt tga tga tat cta cat cgt ttg gat ttt tat gta tcc act tta ata ata tca tag ctg taa cat cct cat ga- ttt acg tta acg tct tcg tgg gat aag ata gtt gtc agt tca tcc ttt gat aat ttt cca aat t

>JN654982.1:21525-23471_Vaccinia_virus_strain_Dryvax_clone_DPP16_complete_genome

cgg ctg cca tgt att tcc tgg aga gca agt --- aga tga tg- agg aac cag ata gtt tat atc cgt act tgc act taa agt cta cat tgt cgt tgt atg a-- gta tga tct ttt aaa ccc gct aga caa gta tcc gtt tga tat tgt agg atg tgg aca ttt aac aat ctg aca cgt ggg tgg atc gga cca ttc tcc tcc tga aca cag gac acc aga gtt acc aat caa cga ata tcc act att gca act ata agt tac aac gct ccc atc ggt ata aaa atc ctc gta tcc gtt atg tct tcc gtt gga tat aga tgg agg gga ttg gca ttt aac aga ttc aca aat agg tgc ctc ggg att cca tac cat aga tcc agt aga tcc taa ttc aca ata cga ttt aga ttc acc gat caa atg ata tcc gct att aca aga gta cgt tat act aga gcc aaa gtc tac tcc acc aat atc aag ttg gcc att atc gat atc tcg agg cga tgg gca tct ccg ttt aat aca ttg att aaa gag tgt cca tcc agt acc tgt aca ttt agc ata tat agg tcc cat ttt ttg ctt tct gta tcc agg tag aca tag a-t att cta tag tgt ctc cta tgt tgt a-- --- --- --- -at tag cat tag cat cag tc- --t cca cac tat tct taa att tca tat taa tgg gtc gtg acg gaa tag tac agc atg ata gaa cgc atc cta ttc cca aca atg tca gga acg tca cgc tct cca cct tca tat tta ttt atc cgt -aa aaa tgt tat cct gga cat cgt aca aat aat aaa aaa gcc ca- -ta tat gtt tgc tat tgt aga aat tgt ttt tca cag ttg ctc aaa aac gat ggc agt gac tta tga gtt --- --- --- --- --- --- tca tct tta gta aac ata tca taa tat tcg ata tta cga gtt gac ata tcg aac aaa ttc caa gta ttt gat ttt gga taa tat tcg tat ttt gca tct gct ata att aag ata taa tca ccg caa gaa cac acg aac atc ttt cct aca tgg tta aag tac atg tac aat tct atc cat ttg tct tcc tta act ata tat ttg tat aga taa tta cga gtc tcg tga gta att cca gta att aca tag atg tcg ccg tcg tac tct aca gca taa act ata cta tga tgt cta ggc atg gga gac -tt ttt tat cca acg att ttt agt gaa aca ttc cac atc gtt taa tac tac ata ttt ttc ata cgt ggt ata aac tcc acc cat tac --a tat ata tca tcg ttt acg aat acc gac gcg cct gaa tat cta gga gta att aag ttt gga agt ctt atc cat ttc gaa gtg ccg tgt ttc aaa tat tct gcc aca ccc gtt gaa ata gaa aat tct aat cct cct att aca tat aac ttt cca ttg tta aca caa gta cta act tct gat ttt aac gac gac ata tta gta acc gtt ttc ca- --- --- --- --- --- --- --t ttt ttc gtt tta aga tct acc cgc -ga tac gga ata aac atg tct att gtt aat cat gcc gcc aat aat gta tag aca att atg taa aac att tgc att ata gaa ttg --- -tc tat ctg tat tac cga cta tcg tcc aat att ctg ttc tag gag agt aat ggg tta ttg tgg ata tat aat cag agt ttt taa tga cta cta tat tat gtt tta tac cat ttc gtg tca ctg gct ttg tag att tgg ata tag tta atc cca aca atg ata tag cat tgc gca tag tat tag tca taa act tgg gat gta aaa tgt tga tga tat cta cat cgt ttg gat ttt tat gta tcc act tta ata ata tca tag ctg taa cat cct cat ga- ttt acg tta acg tct tcg tgg gat aag ata gtt gtc agt tca tcc ttt gat aat ttt cca aat t

>DQ439815.1:23932-25877_Vaccinia_virus_strain_DUKE_complete_genome

cgg ctg cca tgt att tcc tgg aga gca agt --- aga tga tg- agg aac cag ata gtt tat atc cgt act tgc act taa agt cta cat tgt cgt tgt atg a-- gta tga tct ttt aaa ccc gct aga caa gta tcc gtt tga tat tgt agg atg tgg aca ttt aac aat ctg aca cgt ggg tgg atc gga cca ttc tcc tcc tga aca cag gac acc aga gtt acc aat caa cga ata tcc act att gca act ata agt tac aac gct ccc atc ggt ata aaa atc ctc gta tcc gtt atg tct tcc gtt gga tat aga tgg agg gga ttg gca ttt aac aga ttc aca aat agg tgc ctc ggg att cca tac cat aga tcc agt aaa tcc taa ttc aca ata cga ttt aga ttc tcc tat caa ctg ata tcc gct att aca aga gta cgt tat act aga gcc aaa gtc tac tcc gcc aat atc aag ttg gcc att atc gat atc tcg agg cga tgg gca tct ccg ttt aat aca ttg att aaa gag tgt cca tcc agt acc tgt aca ttt agc ata tat agg tcc cat ttt ttg ctt tct gta tcc agg tag aca tag a-t att cta tag tgt ctc cta tgt tgt a-- --- --- --- -at tag cat tag cat cag tc- --t cca cac tat tct taa att tca tat taa tgg gtc gtg acg gaa tag tac agc atg ata gaa cgc atc cta ttc cca aca atg tca gga acg tca cgc tct cca cct tca tat tta ttt atc cgt -aa aaa tgt tat cct gga cat cgt aca aat aat a-a aaa gcc ca- -ta tat gtt cgc tat tgt aga aat tgt ttt tca cag ttg ctc aaa aac gat ggc agt gac tta tga gtt --- --- --- --- --- --- tca tct tta gta aac ata tca taa tat tcg ata tta cga gtt gac ata tcg aac aaa ttc caa gta ttt gat ttt gga taa tat tcg tat ttt gca tct gct ata att aag ata taa tca ccg caa gaa cac acg aac atc ttt cct aca tgg tta aag tac atg tac aat tct atc cat ttg tct tcc tta act ata tat ttg tat aga taa tta cga gtc tcg tga gta att cca gta att aca tag atg tcg ccg tcg tac tct aca gca taa act ata cta tga tgt cta ggc atg gga gac -tt ttt tat cca acg att ttt agt gaa aca ttc tac atc gtt taa tac tac ata ttt ctc ata cgt ggt ata aac tcc acc cat tac --a tat ata tca tcg ttt acg aat acc gat gcg cct gaa tat cta gga gta att aag ttt gga agt ctt atc cat ttt gaa gtg ccg tgt ttc aaa tat tct gct aca ccc gtt gaa ata gaa aat tct aat cct cct att aca tat aac ttt cca ttg tta aca caa gta cta act tct gat ttt aac gac gac ata tta gta acc gtt ttc ca- --- --- --- --- --- --- --t ttt ttc gtt tta aga tct acc cgc -ga tac gga ata aac atg tct att gtt aat cat gcc gcc aat aat gta tag aca att atg taa aac att tgc att ata gaa ttg --- -tc tat ctg tat tac cga cta tcg tcc aat att ctg ttc tag gag agt aat ggg tta ttg tgg ata tat aat cag agt ttt taa tga cta cta tat tat gtt tta tac cat ttc gtg tca ctg gct ttg tag att tgg ata tag tta atc cca aca atg ata tag cat tgc gca tag tat tag tca taa act tgg gat gta aaa tgt tga tga tat cta cat cgt ttg gat ttt tat gta tcc act tta ata ata tca tag ctg taa cat cct cat ga- ttt acg tta acg tct tcg tgg gat aag ata gtt gtc agt tca tcc ttt gat aat ttt cca aat t

>JN654980.1:21383-23322_Vaccinia_virus_strain_Dryvax_clone_DPP13_complete_genome

cgg ctg cca tgt att tcc tgg aga gca agt --- aga tga tg- agg aac cag ata ggt tat atc cgt act tgc act taa agt cta cat tgt cgt tgt atg a-- gta tga tct ttt aaa ccc gct aga caa gta tcc gtt tga tat tgt agg atg tgg aca ttt aac aat ctg aca cgt ggg tgg atc gga cca ttc tcc tcc tga aca cag gac acc aga gtt acc aat caa cga ata tcc act att gca act ata agt tac aac gct tcc atc ggt ata aaa atc ctc gta tcc gtt atg tct tcc gtt gga tat aga tgg agg gga ttg gca ttt aac aga ttc aca aat agg tgc ctc ggg att cca tac cat aga tcc agt aga tcc taa ttc aca ata cga ttt aga ttc tcc tat caa ctg ata tcc gct att aca aga gta cgt tat act aga gcc aaa gtc tac tcc gcc aat atc aag ttg gcc att atc gat atc tcg agg cga tgg gca tct ccg ttt aat aca ttg att aaa gag tgt cca tcc agt acc tgt aca ttt agc ata tat agg tcc cat ttt ttg ctt tct gta tcc agg tag aca tag a-t att cta tag tgt ctc cta tgt tgt a-- --- --- --- -at tag cat t-- --- -ag tt- --t cca cac tat tct taa att tta tat taa tgg gac gtg aag gaa tag gac agt atg ata gaa cgc atc cta ttc cca aca atg tca gga acg tca cgc tct cca cct tca tat tta ttt atc cgt -aa aaa tgt tat cct gga cat cgt aca aat aat a-a aaa gcc ca- -ta tat gtt cgc tat tgt aga aat tgt ttt tca cag ttg ctc aaa aac gat ggc agt gac tta tga gtt --- --- --- --- --- --- tca tct tta gta aac ata tca taa tat tcg ata tta cga gtt gac ata tcg aac aaa ttc caa gta ttt gat ttt gga taa tat tcg tat ttt gca tct gct ata att aag ata taa tca ccg caa gaa cac acg aac atc ttt cct aca tgg tta aag tac atg tac aat tct atc cat ttg tct tcc tta act ata tat ttg tat aga taa tta cga gtc tcg tga gta att cca gta att aca tag atg tcg ccg tcg tac tct aca gca taa act ata cta tga tgt cta ggc atg gga gac -tt ttt tat cca acg att ttt agt gaa aca ttc cac atc gtt taa tac tac ata ttt ttc ata cgt ggt ata aac tcc acc cat tac --a tat ata tca tcg ttt acg aat acc gac gcg cct gaa tat cta gga gta att aag ttt gga agt ctt atc cat ttc gaa gtg ccg tgt ttc aaa tat tct gcc aca ccc gtt gaa ata gaa aat tct aat cct cct att aca tat aac ttt cca tcg tta aca caa gta cta act tct gat ttt aac gac gac ata tta gta acc gtt ttc ca- --- --- --- --- --- --- --t ttt ttc gtt tca aga tct acc cgc -ga tac gga ata aac atg tct att gtt aat cat gcc gcc aat aat gta tag aca att atg taa aac att tgc att ata gaa ttg --- -tc tat ctg tat tac cga cta tcg tcc aat att ctg ttc tag gag agt aat ggg tta ttg tgg ata tat aat cag agt ttt taa tga cta cta tat tat gtt tta tac cat ttc gtg tca ctg gct ttg tag att tgg ata tag tta atc cca aca atg ata tag cat tgc gca tag tat tag tca taa act tgg gat gta aaa tgt tga tga tat cta cat cgt ttg gat ttt tat gta tcc act tta ata ata tca tag ctg taa cat cct cat ga- ttt acg tta acg tct tcg tgg gat aag ata gtt gtc agt tca tcc ttt gat aat ttt cca aat t

>JN654977.1:21377-23323_Vaccinia_virus_strain_Dryvax_clone_DPP10_complete_genome

cgg ctg cca tgt att tcc tgg aga gca agt --- aga tga tg- agg aac cag ata ggt tat atc cgt act tgc act taa agt tta cat tgt cgt tgt atg a-- gta tga tct ttt aaa ccc gct aga caa gta tcc gtt tga tat tgt agg atg tgg aca ttt aac aat ctg aca cgt ggg tgg atc gga cca ttc tcc tcc tga aca cag gac acc aga gtt acc aat caa cga ata tcc act att gca act ata agt tac aac gct ccc atc ggt ata aaa atc ctc gta tcc gtt atg tct tcc gtt gga tat aga tgg agg gga ttg gca ttt aac aga ttc aca aat agg tgc ctc ggg att cca tac cat aga tcc agt aga tcc taa ttc aca ata cga ttt aga ttc acc gat caa atg ata tcc gct att aca aga gta cgt tat act aga gcc aaa gtc tac tcc acc aat atc aag ttg gcc att atc gat atc tcg agg cga tgg gca tct ccg ttt aat aca ttg att aaa gag tgt cca tcc agt acc tgt aca ttt agc ata tat agg tcc cat ttt ttg ctt tct gta tcc agg tag aca tag a-t att cta tag tgt ctc cta tgt tgt a-- --- --- --- -at tag cat tag cat cag tc- --t cca cac tat tct taa att tca tat taa tgg gac gtg aag gaa tag tac agc atg ata gaa cgc atc cta ttc cca aca atg tca gga acg tca cgc tct cca cct tca tat tta ttt atc cgt -aa aaa tgt tat cct gga cat cgt aca aat aat aaa aaa gcc ca- -ta tat gtt tgc tat tgt aga aat tgt ttt tca cag ttg ctc aaa aac gat ggc agt gac tta tga gtt --- --- --- --- --- --- tca tct tta gta aac ata tca taa tat tcg ata tta cga gtt gac ata tcg aac aaa ttc caa gta ttt gat ttt gga taa tat tcg tat ttt gca tct gct ata att aag ata taa tca ccg caa gaa cac acg aac atc ttt cct aca tgg tta aag tac atg tat aat tct atc cat ttg tct tcc tta act ata tat ttg tat aga taa tta cga gtc tcg tga gta att cca gta att aca tag atg tcg ccg tcg tac tct aca gca taa act ata cta tga tgt cta ggc atg gga gac -tt ttt tat cca acg att ttt agt gaa aca ttc tac atc gtt taa tac tac ata ttt ctc ata cgt ggt ata aac tcc acc cat tac --a tat ata tca tcg ttt acg aat acc gat gcg cct gaa tat cta gga gta att aag ttt gga agt ctt atc cat ttc gaa gtg ccg tgt ttc aaa tat tct gct aca ccc gtt gaa ata gaa aat tct aat cct cct att aca tat aac ttt cca tcg tta aca caa gta cta act tct gat ttt aac gac gac ata tta gta acc gtt ttc ca- --- --- --- --- --- --- --t ttt ttc gtt tta aga tct acc cgc -ga tac gga ata aac atg tct att gtt aat cat gcc gcc aat aat gta tag aca att atg taa aac att tgc att ata gaa ttg --- -tc tat ctg tat tac cga cta tcg tcc aat att ctg ttc tag gag agt aat ggg tta ttg tgg ata tat aat cag agt ttt taa tga cta cta tat tat gtt tta tac cat ttc gtg tca ctg gct ttg tag att tgg ata tag tta atc cca aca atg ata tag cat tgc gca tag tat tag tca taa act tgg gat gta aaa tgt tga tga tat cta cat cgt ttg gat ttt tat gta tcc act tta ata ata tca tag ctg taa cat cct cat ga- ttt acg tta acg tct tcg tgg gat aag ata gtt gtc agt tca tcc ttt gat aat ttt cca aat t

>JN654983.1:21521-23467_Vaccinia_virus_strain_Dryvax_clone_DPP17_complete_genome

cgg ctg cca tgt att tcc tgg aga gca agt --- aga tga tg- agg aac cag ata ggt tat atc cgt act tgc act taa agt tta cat tgt cgt tgt atg a-- gta tga tct ttt aaa ccc gct aga caa gta tcc gtt tga tat tgt agg atg tgg aca ttt aac aat ctg aca cgt ggg tgg atc gga cca ttc tcc tcc tga aca cag gac acc aga gtt acc aat caa cga ata tcc act att gca act ata agt tac aac gct ccc atc ggt ata aaa atc ctc gta tcc gtt atg tct tcc gtt gga tat aga tgg agg gga ttg gca ttt aac aga ttc aca aat agg tgc ctc ggg att cca tac cat aga tcc agt aga tcc taa ttc aca ata cga ttt aga ttc acc gat caa atg ata tcc gct att aca aga gta cgt tat act aga gcc aaa gtc tac tcc acc aat atc aag ttg gcc att atc gat atc tcg agg cga tgg gca tct ccg ttt aat aca ttg att aaa gag tgt cca tcc agt acc tgt aca ttt agc ata tat agg tcc cat ttt ttg ctt tct gta tcc agg tag aca tag a-t att cta tag tgt ctc cta tgt tgt a-- --- --- --- -at tag cat tag cat cag tc- --t cca cac tat tct taa att tca tat taa tgg gac gtg aag gaa tag gac agt atg ata gaa cgc atc cta ttc cca aca atg tca gga acg tca cgc tct cca cct tca tat tta ttt atc cgt -aa aaa tgt tat cct gga cat cgt aca aat aat aaa aaa gcc ca- -ta tat gtt tgc tat tgt aga aat tgt ttt tca cag ttg ctc aaa aac gat ggc agt gac tta tga gtt --- --- --- --- --- --- tca tct tta gta aac ata tca taa tat tcg ata tta cga gtt gac ata tcg aac aaa ttc caa gta ttt gat ttt gga taa tat tcg tat ttt gca tct gct ata att aag ata taa tca ccg caa gaa cac acg aac atc ttt cct aca tgg tta aag tac atg tac aat tct atc cat ttg tct tcc tta act ata tat ttg tat aga taa tta cga gtc tcg tga gta att cca gta att aca tag atg tcg ccg tcg tac tct aca gca taa act ata cta tga tgt cta ggc atg gga gac -tt ttt tat cca acg att ttt agt gaa aca ttc tac atc gtt taa tac tac ata ttt ctc ata cgt ggt ata aac tcc acc cat tac --a tat ata tca tcg ttt acg aat acc gat gcg cct gaa tat cta gga gta att aag ttt gga agt ctt atc cat ttt gaa gtg ccg tgt ttc aaa tat tct gct aca ccc gtt gaa ata gaa aat tct aat cct cct att aca tat aac ttt cca tcg tta aca caa gta cta act tct gat ttt aac gac gac ata tta gta acc gtt ttc ca- --- --- --- --- --- --- --t ttt ttc gtt tca aga tct acc cgc -ga tac gga ata aac atg tct att gtt aat cat gcc gcc aat aat gta tag aca att atg taa aac att tgc att ata gaa ttg --- -tc tat ctg tat tac cga cta tcg tcc aat att ctg ttc tag gag agt aat ggg tta ttg tgg ata tat aat cag agt ttt taa tga cta cta tat tat gtt tta tac cat ttc gtg tca ctg gct ttg tag att tgg ata tag tta atc cca aca atg ata tag cat tgc gca tag tat tag tca taa act tgg gat gta aaa tgt tga tga tat cta cat cgt ttg gat ttt tat gta tcc act tta ata ata tca tag ctg taa cat cct cat ga- ttt acg tta acg tct tcg tgg gat aag ata gtt gtc agt tca tcc ttt gat aat ttt cca aat t

>KY549143.2:30321-32274_Cowpox_virus_isolate_CPXV_Catpox3L97

cgg ctt cca tgt att tcc tgg aga gca agt --- aga tga tg- agg aac cag ata gtt tat atc cgt gtc tgc act taa agt cta cat tgt cgt tgt atg a-- gta tga tct ttt aaa ccc gct aga caa gta tcc gtt tga tat tgt agg atg tgg aca ttt aac aat ctg aca cgt ggg tgg atc gga cca ttc tcc tcc tga aca cag gat gcc aga gtt acc aat taa cga ata tcc act att gca act ata agt tac aac gct tcc atc ggt ata aaa atc ctc gta tcc gtt atg tct tcc gtt gga tat aga tgg agg tga ttg gca ttt aac aga ttc gca aat agg tgc ctc ggg att cca tac cat aga tcc agt aga tcc taa ttc aca ata cga ttt aga ttc tcc tat caa ctg ata tcc gct att aca aga gta cgt tat act aga gcc aaa gtc tac tcc gcc aat atc aag ttg gcc att atc gat atc tcg agg cga tgg gca tct ccg ttt aat aca ttg att aaa gag tgt cca tcc agt acc tgt aca ttt agc ata tat agg tcc cat ttt ttg ctt tct gta tcc agg tag aca tag a-t att cta tag tgt ctc cta tat tat a-- --- --- --- -at tag cat tag cat cag tc- --c cca cac tat tct taa att tca tag taa tag gac gtg aag gaa tag gac agc atg ata gaa cgc atc cta ttc cca aca atg tca gga acg tca cgc tct cca cct tca tat tta ttt atc cgt -aa aaa tgt tat cct atg cat cgt ac- aat aat a-a aaa gtc at- --- --- --- -gc tat tgt aga aat tgt ttt tca cag ttg ctc aaa aac gat ggc agt gac tta tga gtt acg tta cac ttt gga gtc tca tct tta gta aac ata tca taa tat tcg ata tta cga gtt gac ata tcg aac aaa ttc caa gta ttt gat ttt gga taa tat tcg tat ttt gca tct gct ata att aag ata taa tca ccg caa gaa cac acg aac atc ttt cct aca tgg tta aag tac atg tac aat tct atc cat ttg tct tcc tta act ata tat ttg tat aga taa tta cga gtc tcg tga gta att cca gta att gca tag atg tca cca tcg tac tct aca gca taa act ata cta tga tgt cta ggc atg gga gac -tt ttt tat cca acg att ttt agt gaa aca ttc cac gtc gtt taa tac tac ata ttt ctc ata cgt ggt ata aac tcc acc cat tac --a tat ata tca tcg ttt acg aat acc gat gct cct gaa tat cta gga gta att aag ttt gga agt ctt atc cat ttc gaa gtg ccg tgt ttc aaa tat tct gcc acg ccc gtt gaa ata gaa aat tct aat cct cct att aca tat aac ttt cca tcg tta aca caa gta cta act tct gat ttt aac gac gac ata tta gta acc gtt ttc ca- --- --- --- --- --- --- --t ttt ttc gtt tta aga tct acc cgc -ga tac gga ata aac atg tct att gtt aat cat gcc gcc aat aat gta tag aca att atg taa aac att tgc att ata gaa ttg --- -tc tat ctg tat tac cga cta tcg tcc aat att ctg ttc tag gag agt aat ggg tta ttg tgg ata tat aat cag agt ttt taa tga cta cta tat tat gtt tta tac cat ttc gtg tca cag gct ttg tag att tgg ata tag tta atc cca aca atg cta tag cat tgc gca tag cat tag tca taa act tgg gat gta aaa tgt tga tga tat cta cat cgt ttg gat ttt tat gta tcc act tta ata ata tta tag ctg taa cat cct cat ga- ttt acg tta acg ttt tcg tgg gat aag ata gtt gtt agt tca tcc ttt gat aat ttt cca aat t

>KP768318.1:19852-21822_Camelpox_virus_strain_0408151v_complete_genome

cgg ctg cca tgt att tcc tgg aga gca agt --- aga tga tg- agg aac cag ata gtt tat atc cgt act tgc acg taa agt cta cat tgt cat tat acg a-- ata tga tct ttt aaa ccc gct aga caa gta tcc gtt tga tat tgt agg atg tgg aca ttt aac aat ctg aca cgt ggg tgg atc gga cca ttc tcc tcc tga aca cag gac acc aga gtt acc aat caa cga ata tcc act att gca act ata agt tac aac gct tcc atc ggt ata aaa att atc gta tcc gtt atg tct tcc gtt gga tat aga tgg agg tga ttg gca ttt aac aga ttc gca aat agg tgc ctc ggg att cca tac cat aga tcc agt aga tcc taa ttc aca ata cga ttt aga ttc acc gat caa atg ata tcc gct att aca aga gta cgt tat act aga gcc aaa gtc tac tcc acc aat atc aag ttg gcc att atc gat atc tcg agg cga tgg gca tct ccg ttt aat aca ttg att aaa gag tgt cca tcc agt acc tgt aca ttt agc ata tat agg tcc cat ttt ttg ctt tct gta tcc agg tag aca tag a-t att cta tag tgt ctc cta tgt tgt aat tag t-- --- -at tag tat tag cat aag tc- --t cca cac tat tct taa att tca tat taa tgg gtc gtg acg gaa tag tac agc atg ata gaa cgc ata cta ttc cca aca atg tca gga acg tca cgc tct cca cct tca tat tta ttt atc cgt -aa aaa tgt tat cct gga cac tgt acc aat aat aaa aaa tcc ca- -ta tat gtt cgc tat tgt aga aat tgt ttt tca cag ttg ctc aaa aac gat ggc agt gac tta tga gtt acg tta cac ttt gga gtc tca tct tta gta aac ata tca taa tat tcg ata tta cga gtt gac ata tcg aac aaa atc caa gta ttt gat ttt gga taa tat tcg tat ttt gca tct gct ata att aag ata taa tcg ccg caa gaa cac acg aac atc ttt cct aca tgg tta aag tac atg tac aat tct atc cat ttg tct tcc tta act ata tat ttg tat aga taa tta cga gtc tcg tga gta att cca gta att gca tag atg tca cca tcg tac tct ata gca taa act ata cta tga tgt cta ggc atg gga gac -tt ttt tat cca acg att ttt agt gaa aca ttc cac atc gtt taa tac tac ata ttt ttc ata cgt ggt ata aac tcc acc cat tac --a tat ata tca tcg ttt acg aat acc gac gcg cct gaa tat cta gga gta att aag ttt gga agt ctt atc cat ttc gaa gtt ccg tgt ttc aaa tat tct gct acg cct gtt gaa ata gaa aat tct aat cct cct att aca tat aac ttt cca tca tta aca caa gta cta act tct gat ttt aac gac gac ata tta gta act gtt ttc ca- --- --- --- --- --- --- --t ttt ttc gtt tta aga tct acc cgc -ga tac gga ata aac atg tct att gtt aat cat gcc gcc aat aat gta tag aca att ata taa aac att tgc att ata gaa ttg --- -tc tat ctg tat tac cga cga ttg tcc aat att ctg ttc tag gag agt aat ggg tta ttg tgg ata tat aat cag agt ttt taa tga cta cta tat tat gtt tta tac cat ttc gtg tca cag gct ttg tag att tgg ata tag tta atc cca aca atg ata tag cat tgc aca tag tat tag tca taa act tgg gat gta aaa tgt tga taa tat cta cat cgt ttg gat ttt tat gta tcc act tta ata ata tca tag ctg taa cat cct cgt ga- ttt acg ttg acg ttt tcg tgg gat aag ata gtt gtt agt tca tcc ttt gat aat ttt cca aat t

>AF438165.1:21498-23468_Camelpox_virus_M-96_from_Kazakhstan_complete_genome

cgg ctg cca tgt att tcc tgg aga gca agt --- aga tga tg- agg aac cag ata gtt tat atc cgt act tgc acg taa agt cta cat tgt cat tat acg a-- ata tga tct ttt aaa ccc gct aga caa gta tcc gtt tga tat tgt agg atg tgg aca ttt aac aat ctg aca cgt ggg tgg atc gga cca ttc tcc tcc tga aca cag gac acc aga gtt acc aat caa cga ata tcc act att gca act ata agt tac aac gct tcc atc ggt ata aaa att atc gta tcc gtt atg tct tcc gtt gga tat aga tgg agg tga ttg gca ttt aac aga ttc gca aat agg tgc ctc ggg att cca tac cat aga tcc agt aga tcc taa ttc aca ata cga ttt aga ttc acc gat caa atg ata tcc gct att aca aga gta cgt tat act aga gcc aaa gtc tac tcc acc aat atc aag ttg gcc att atc gat atc tcg agg cga tgg gca tct ccg ttt aat aca ttg att aaa gag tgt cca tcc agt acc tgt aca ttt agc ata tat agg tcc cat ttt ttg ctt tct gta tcc agg tag aca tag a-t att cta tag tgt ctc cta tgt tgt aat tag t-- --- -at tag tat tag cat aag tc- --t cca cac tat tct taa att tca tat taa tgg gtc gtg acg gaa tag tac agc atg ata gaa cgc ata cta ttc cca aca atg tca gga acg tca cgc tct cca cct tca tat tta ttt atc cgt -aa aaa tgt tat cct gga cac tgt acc aat aat aaa aaa tcc ca- -ta tat gtt cgc tat tgt aga aat tgt ttt tca cag ttg ctc aaa aac gat ggc agt gac tta tga gtt acg tta cac ttt gga gtc tca tct tta gta aac ata tca taa tat tcg ata tta cga gtt gac ata tcg aac aaa atc caa gta ttt gat ttt gga taa tat tcg tat ttt gca tct gct ata att aag ata taa tcg ccg caa gaa cac acg aac atc ttt cct aca tgg tta aag tac atg tac aat tct atc cat ttg tct tcc tta act ata tat ttg tat aga taa tta cga gtc tcg tga gta att cca gta att gca tag atg tca cca tcg tac tct ata gca taa act ata cta tga tgt cta ggc atg gga gac -tt ttt tat cca acg att ttt agt gaa aca ttc cac atc gtt taa tac tac ata ttt ttc ata cgt ggt ata aac tcc acc cat tac --a tat ata tca tcg ttt acg aat acc gac gcg cct gaa tat cta gga gta att aag ttt gga agt ctt atc cat ttc gaa gtt ccg tgt ttc aaa tat tct gct acg cct gtt gaa ata gaa aat tct aat cct cct att aca tat aac ttt cca tca tta aca caa gta cta act tct gat ttt aac gac gac ata tta gta act gtt ttc ca- --- --- --- --- --- --- --t ttt ttc gtt tta aga tct acc cgc -ga tac gga ata aac atg tct att gtt aat cat gcc gcc aat aat gta tag aca att ata taa aac att tgc att ata gaa ttg --- -tc tat ctg tat tac cga cga ttg tcc aat att ctg ttc tag gag agt aat ggg tta ttg tgg ata tat aat cag agt ttt taa tga cta cta tat tat gtt tta tac cat ttc gtg tca cag gct ttg tag att tgg ata tag tta atc cca aca atg ata tag cat tgc aca tag tat tag tca taa act tgg gat gta aaa tgt tga taa tat cta cat cgt ttg gat ttt tat gta tcc act tta ata ata tca tag ctg taa cat cct cgt ga- ttt acg ttg acg ttt tcg tgg gat aag ata gtt gtt agt tca tcc ttt gat aat ttt cca aat t

>AY009089.1:19810-21780_Camelpox_virus_CMS_complete_genome

cgg ctg cca tgt att tcc tgg aga gca agt --- aga tga tg- agg aac cag ata gtt tat atc cgt act tgc acg taa agt cta cat tgt cat tat acg a-- ata tga tct ttt aaa ccc gct aga caa gta tcc gtt tga tat tgt agg atg tgg aca ttt aac aat ctg aca cgt ggg tgg atc gga cca ttc tcc tcc tga aca cag gac acc aga gtt acc aat caa cga ata tcc act att gca act ata agt tac aac gct tcc atc ggt ata aaa att atc gta tcc gtt atg tct tcc gtt gga tat aga tgg agg tga ttg gca ttt aac aga ttc gca aat agg tgc ctc ggg att cca tac cat aga tcc agt aga tcc taa ttc aca ata cga ttt aga ttc acc gat caa atg ata tcc gct att aca aga gta cgt tat act aga gcc aaa gtc tac tcc acc aat atc aag ttg gcc att atc gat atc tcg agg cga tgg gca tct ccg ttt aat aca ttg att aaa gag tgt cca tcc agt acc tgt aca ttt agc ata tat agg tcc cat ttt ttg ctt tct gta tcc agg tag aca tag a-t att cta tag tgt ctc cta tgt tgt aat tag t-- --- -at tag tat tag cat aag tc- --t cca cac tat tct taa att tca tat taa tgg gtc gtg acg gaa tag tac agc atg ata gaa cgc ata cta ttc cca aca atg tca gga acg tca cgc tct cca cct tca tat tta ttt atc cgt -aa aaa tgt tat cct gga cac tgt acc aat aat aaa aaa tcc ca- -ta tat gtt cgc tat tgt aga aat tgt ttt tca cag ttg ctc aaa aac gat ggc agt gac tta tga gtt acg tta cac ttt gga gtc tca tct tta gta aac ata tca taa tat tcg ata tta cga gtt gac ata tcg aac aaa atc caa gta ttt gat ttt gga taa tat tcg tat ttt gca tct gct ata att aag ata taa tcg ccg caa gaa cac acg aac atc ttt cct aca tgg tta aag tac atg tac aat tct atc cat ttg tct tcc tta act ata tat ttg tat aga taa tta cga gtc tcg tga gta att cca gta att gca tag atg tca cca tcg tac tct ata gca taa act ata cta tga tgt cta ggc atg gga gac -tt ttt tat cca acg att ttt agt gaa aca ttc cac atc gtt taa tac tac ata ttt ttc ata cgt ggt ata aac tcc acc cat tac --a tat ata tca tcg ttt acg aat acc gac gcg cct gaa tat cta gga gta att aag ttt gga agt ctt atc cat ttc gaa gtt ccg tgt ttc aaa tat tct gct acg cct gtt gaa ata gaa aat tct aat cct cct att aca tat aac ttt cca tca tta aca caa gta cta act tct gat ttt aac gac gac ata tta gta act gtt ttc ca- --- --- --- --- --- --- --t ttt ttc gtt tta aga tct acc cgc -ga tac gga ata aac atg tct att gtt aat cat gcc gcc aat aat gta tag aca att ata taa aac att tgc att ata gaa ttg --- -tc tat ctg tat tac cga cga ttg tcc aat att ctg ttc tag gag agt aat ggg tta ttg tgg ata tat aat cag agt ttt taa tga cta cta tat tat gtt tta tac cat ttc gtg tca cag gct ttg tag att tgg ata tag tta atc cca aca atg ata tag cat tgc aca tag tat tag tca taa act tgg gat gta aaa tgt tga taa tat cta cat cgt ttg gat ttt tat gta tcc act tta ata ata tca tag ctg taa cat cct cgt ga- ttt acg ttg acg ttt tcg tgg gat aag ata gtt gtt agt tca tcc ttt gat aat ttt cca aat t

>LR800245.1:15680-17650_Variola_virus_genome_assembly_complete_genome:_monopartite

cgg ttg cca tgt att tcc tgg aga gca agt --- aga tga tg- agg aac cag ata gtt tat atc cgt act tgc acg taa agt cta cat tgt cat tat acg a-- ata tga tct ttt aaa ccc gct aga caa gta tcc gtt taa tat tgt agg atg tgg aca ttt aac aat ctg aca cat ggg tgg atc gga cca ttc tcc tcc tga aca cag gac act aga gtt acc aat caa cga ata tcc act att gca act ata agt tac aac gct tcc atc ggt ata aaa atc ctc gta tcc gtt atg tct tcc gtt gga tat aga tgg agg taa ttg gca ttt aac aga ttc gca aat agg tgc ctt ggg att cca tac cat aga tcc agt aga tcc taa ttc aca ata cga ttt ata ttc acc gat caa atg ata tcc gct att aca aga gta cgt tat act aga gcc aaa gtc tac tcc gcc aat atc aag atg gcc att atc gat atc tcg agg cga tgg gca tct ccg ttt aat aca ttg att aaa gag tgt cca tcc agt acc tgt aca ttt agc ata tat agg tcc cat ttt ttg ctt tct gta tcc agg tag aca tag a-t att cta tag tgt ctc cta tgt tgt aat tag c-- --- -at tag cat tag cat cag tc- --t cca cac tat tct taa att tca tat taa tgg gtc gtg acg gaa tag tac agc atg ata aaa cgc atc cta ttc cca aca atg tca gga acg tca cgc tct cca cct tca tat tta ttt atc cgt -aa aaa tgt tat cct gga cat tgt gcc aat aat aaa aaa gcc ca- -ta tat gtt cgc tat tgt aga aat tgt ttt tca tag ttg ctc aaa aac gat ggc ggt gac tta tga gtt acg tta cac ttt gga gtc tca tct tta gta aac ata tca taa tat tcg ata tta cga gtt gac ata tcg aac aaa ttc caa gta ttt gat ttt gga taa tat tcg tat ttt gca tct gct ata att aag ata taa tcg ccg caa gaa cac acg aac atc ttt cct aca tgg tta aag tac atg tac aat tct atc cat ttg tct tcc tta act ata tat ttg tat aga taa tta cga gtc tcg tga gta att cca gta att gta tag atg tca cca tcg tac tct aca gca taa act ata cta tga tgt cta ggc atg gga gac -tt ttt tat cca acg att ttt agt gaa aca ttc cac atc gtt taa tac tac ata ttt ttc ata cgt ggt ata aac tcc acc cat tac --a tat ata tca tcg ttt acg aat acc gac gcg cct gaa tat cta gga gta att aag ttt gga agt ctt atc cat ttc gaa gtg ccg tgt ttc aaa tat tct gct aca cct gtt gaa ata gaa aat ttt aat cct cct att aca tat aac ttt cca tca tta aca caa gta cta act tct gat ttt aac gac gac ata tta gta act gtt ttc ca- --- --- --- --- --- --- --t ttt ttc gtt tta aga tct acc cgc -ga tac gga ata aac atg tct att gtt aat cat gcc gcc aat aat gta tag aca att ata taa aac att tgc att ata gaa ttg --- -tc tat ctg tat tac cga cga ttg tcc aat att ctg ttc tag gag agt aat ggg tta ttg tgg ata tat aat cag agt ttt taa tga cta cta tat tat gtt tta tac cat ttc gtg tca cag gct ttg tag att tgg ata tag tta atc cca aca atg ata tag cat tgc aca tag tat tag tca taa act tgg gat gta aaa tgt tga tga tat cta cat cgt ttg gat ttt tat gta tcc act tta ata ata tca tag ctg taa cat cct cgt ga- ttt acg ttg acg ttt tcg tgg gat aag ata gtt gtt agt tca tcc ttt gat aat ttt cca aat t

>LR800244.1:15673-17643_Variola_virus_genome_assembly_complete_genome:_monopartite

cgg ttg cca tgt att tcc tgg aga gca agt --- aga tga tg- agg aac cag ata gtt tat atc cgt act tgc acg taa agt cta cat tgt cat tat acg a-- ata tga tct ttt aaa ccc gct aga caa gta tcc gtt taa tat tgt agg atg tgg aca ttt aac aat ctg aca cat ggg tgg atc gga cca ttc tcc tcc tga aca cag gac act aga gtt acc aat caa cga ata tcc act att gca act ata agt tac aac gct tcc atc ggt ata aaa atc ctc gta tcc gtt atg tct tcc gtt gga tat aga tgg agg taa ttg gca ttt aac aga ttc gca aat agg tgc ctt ggg att cca tac cat aga tcc agt aga tcc taa ttc aca ata cga ttt ata ttc acc gat caa atg ata tcc gct att aca aga gta cgt tat act aga gcc aaa gtc tac tcc gcc aat atc aag atg gcc att atc gat atc tcg agg cga tgg gca tct ccg ttt aat aca ttg att aaa gag tgt cca tcc agt acc tgt aca ttt agc ata tat agg tcc cat ttt ttg ctt tct gta tcc agg tag aca tag a-t att cta tag tgt ctc cta tgt tgt aat tag c-- --- -at tag cgt tag cat cag tc- --t cca cac tat tct taa att tca tat taa tgg gtc gtg acg gaa tag tac agc atg ata aaa cgc atc cta ttc cca aca atg tca gga acg tca cgc tct cca cct tca tat tta ttt atc cgt -aa aaa tgt tat cct gga cat tgt gcc aat aat aaa aaa gcc ca- -ta tat gtt cgc tat tgt aga aat tgt ttt tca tag ttg ctc aaa aac gat ggc ggt gac tta tga gtt acg tta cac ttt gga gtc tca tct tta gta aac ata tca taa tat tcg ata tta cga gtt gac ata tcg aac aaa ttc caa gta ttt gat ttt gga taa tat tcg tat ttt gca tct gct ata att aag ata taa tcg ccg caa gaa cac acg aac atc ttt cct aca tgg tta aag tac atg tac aat tct atc cat ttg tct tcc tta act ata tat ttg tat aga taa tta cga gtc tcg tga gta att cca gta att gta tag atg tca cca tcg tac tct aca gca taa act ata cta tga tgt cta ggc atg gga gac -tt ttt tat cca acg att ttt agt gaa aca ttc cac atc gtt taa tac tac ata ttt ttc ata cgt ggt ata aac tcc acc cat tac --a tat ata tca tcg ttt acg aat acc gac gcg cct gaa tat cta gga gta att aag ttt gga agt ctt atc cat ttc gaa gtg ccg tgt ttc aaa tat tct gct aca cct gtt gaa ata gaa aat ttt aat cct cct att aca tat aac ttt cca tca tta aca caa gta cta act tct gat ttt aac gac gac ata tta gta act gtt ttc ca- --- --- --- --- --- --- --t ttt ttc gtt tta aga tct acc cgc -ga tac gga ata aac atg tct att gtt aat cat gcc gcc aat aat gta tag aca att ata taa aac att tgc att ata gaa ttg --- -tc tat ctg tat tac cga cga ttg tcc aat att ctg ttc tag gag agt aat ggg tta ttg tgg ata tat aat cag agt ttt taa tga cta cta tat tat gtt tta tac cat ttc gtg tca cag gct ttg tag att tgg ata tag tta atc cca aca atg ata tag cat tgc aca tag tat tag tca taa act tgg gat gta aaa tgt tga tga tat cta cat cgt ttg gat ttt tat gta tcc act tta ata ata tca tag ctg taa cat cct cgt ga- ttt acg ttg acg ttt tcg tgg gat aag ata gtt gtt agt tca tcc ttt gat aat ttt cca aat t

>AM501482.1:19025-20958_Vaccinia_virus_Ankara_strain_chorioallantois_vaccinia_virus_Ankara_(CVA)_complete_coding_genome

cgg ctt cca tgt att tcc tgg aga gca agt --- aga tga tg- agg aac cag ata gtt tat atc cgt act tgc act taa agt cta cat tgt cgt tgt atg a-- gta tga tct ttt aaa ccc gct aga caa gta tcc gtt tga tat tgt agg atg tgg aca ttt aac aat ctg aca cgt ggg tgg atc gga cca ttc tcc tcc tga aca cag gac acc aga gtt acc aat caa cga ata tcc act att gca act ata agt tac aac gct ccc atc ggt ata aaa atc ctc gta tcc gtt atg tct tcc gtt gga tat aga tgg agg gga ttg gca ttt aac aga ttc aca aat agg tgc ctc ggg att cca tac cat aga tcc agt aga tcc taa ttc aca ata cga ttt aga ttc acc gat caa atg ata tcc gct att aca aga gta cgt tat act aga gcc aaa gtc tac tcc acc aat atc aag ttg gcc att atc gat atc tcg agg cga tgg gca tct ccg ttt aat aca ttg att aaa gag tgt cca tcc agt acc tgt aca ttt agc ata tat agg tcc cat ttt ttg ctt tct gta tcc agg tag aca tag a-t att cta tag tgt ctc cta tgt tgt a-- --- --- --- -at tag cat tag cat cag tc- --t cca cac tat tct taa att tca tat taa tgg gtc gtg acg gaa tag tac agc atg ata gaa cgc atc cta ttc cca aca atg tca gga acg tca cgc tct cca ctt tca tat tta ttt atc cgt -aa aaa tgt tat cct gga cat cgt aca aat aat a-a aaa gcc ca- -ta tat gtt cgc tat tgt aga aat tgt ttt tca cag ttg ctc aaa aac gat ggc agt gac tta tga gtt --- --- --- --- --- --- tca tct tta gta aac ata tca taa tat tcg ata tta cga gtt gac ata tcg aac aaa ttc caa gta ttt gat ttt gga taa tat tcg tat ttt gca tct gct ata att aag ata taa tca ccg caa gaa cac acg aac atc ttt cct aca tgg tta aag tac atg tac aat tct atc cat ttg tct tcc tta act ata tat ttg tat aga taa tta cga gtc tcg tga gta att cca gta att gca tag atg tcg ccg tcg tac tct aca gca taa act ata cta tga tgt cta ggc atg gga gac -tt ttt tat cca acg att ttt agt gaa aca ttc tac atc gtt taa tac tac ata ttt ctc ata cgt ggt ata aac tcc acc cat tac --a tat ata tca tcg ttt acg aat acc gac gcg cct gaa tat cta gga gta att aag ttt gga agt ctt atc cat ttc gaa gtg ccg tgt ttc aaa tat tct gct aca ccc gtt gaa ata gaa aat tct aat cct cct att aca tat aac ttt cca tcg tta aca caa gta cta act tct gat ttt aac gac gac ata tta gta acc gtt ttc ca- --- --- --- --- --- --- --t ttt ttc gtt tca aga tct acc cgc -ga tac gga ata aac atg tct att gtt aat cat gcc gcc aat aat gta tag aca att atg taa aac att tgc att ata gaa ttg --- -tc tat ctg tat tac cga cta tcg tcc aat att ctg ttc tag gag agt aat ggg tta ttg tgg ata tat aat cag agt ttt taa tga cta cta tat tat gtt tta tac cat ttc gtg tca ctg gct ttg tag att tgg ata tag tta atc cca aca atg ata tag c-- --- --- --- -at tag tca taa act tgg gat gta aaa tgt tga tga tat cta cat cgt ttg gat ttt tat gta tcc act tta ata ata tca tag ctg taa cat cct cat ga- ttt acg tta acg tct tcg tgg gat aag ata gtt gtc agt tca tcc ttt gat aat ttt cca aat t

>LR800247.1:15603-17575_Variola_virus_genome_assembly_complete_genome:_monopartite

cgg ttg cca tgt att tcc tgg aga gca agt --- aga tga tg- agg aac cag ata gtt tat atc cgt acg tgc acg taa agt cta cat tgt cat tat acg a-- ata tga tct ttt aaa ccc gct aga caa gta tcc gtt taa tat tgt agg atg tgg aca ttt aac aat ctg aca cgt ggg tgg atc gga cca ttc tcc tcc tga aca cag gac act aaa gtt acc aat caa cga ata tcc act att gca act ata agt tac gac gct tcc atc ggt ata aaa atc ctc gta tcc gtt atg tct tcc gtt gga tat aga tgg agg taa ttg gca ttt aac aga ttc gca aat agg tgc ctt ggg att cca tac cat aga tcc agt aga tcc taa ttc aca ata cga ttt ata ttc acc gat caa atg ata tcc gct att aca aga gta cgt tat act aga gcc aaa gtc tac tcc gcc aat atc gag atg gcc att atc gat atc tcg agg cga tgg gca tct ccg ttt aat aca ttg att aaa gag tgt cca tcc agt acc tgt aca ttt agc ata tat agg tcc cat ttt ttg ctt tct gta tcc agg tag aca tag a-t att cta tag tgt ctc cta tgt tgt aat tag c-- --- -at tag cat tag cat cag tc- --t cca cac tat tct taa att tca tat taa tgg gtc gtg acg gaa tag tac agc atg ata gaa cgc atc cta ttc cca aca atg tca gga acg tca cgc tct cca cct tca tat tta ttt atc cgt -aa aaa tgt tat cct gga cat tgt gcc aat aat aaa aaa gcc ca- -ta tat gtt cgy tat tgt aga aat tgt ttt tca tag ttg ctc aaa aac gat ggc ggt gac tta tga gtt acg tta cac ttt gga gtc tca tct tta gta aac ata tca taa tat ttg ata tta cga gtt gac ata tcg aac aaa ttc caa gta ttt gat ttt gga taa tat tcg tat ttt gca tct gct ata att aag ata taa tcg ccg caa gaa cac acg aac atc ttt cct aca tgg tta aag tac atg tac aat tct atc cat ttg tct tcc tta act ata tat ttg tat aga taa tta cga gtc tcg tga gta att cca gta att gca tag atg tca cca tca tac tct aca gca taa act ata cta tga tgt cta ggc atg gga gac ttt ttt tat cca acg att ttt agt gaa aca ttc cac atc gtt taa tac tac ata ttt ttc ata cgt ggt ata aac tcc acc cat tac --a tat ata tca tcg ttt acr aat acc gac gcg cct gaa tat cta gga gta att aag ttt gga agt ctt atc cat ttc gaa gtg ccg tgt ttc aaa tat tct gct aca cct gtt gaa ata gaa aat ttt aat cct tct att aca tat aac ttt cca tca tta aca caa gta cta act tct gat ttt aac gac gac ata tta gta act gtt ttc ca- --- --- --- --- --- --- -tt ttt ttc gtt tta aga tct acc cgc -ga tac gga ata aac atg tct att gtt aat cat gcc gcc aat aat gta tag aca att ata taa aac att tgc att ata gaa ttg --- -tc tat ctg tat tac cga cga ttg tcc aat att ctg ttc tag gag agt aat ggg tta ttg tgg ata tat aat cag agt ttt taa tga cta cta tat tat gtt tta tac cat ttc gtg tca cag gct ttg tag att tgg ata tag tta atc cca aca atg ata tag cat tgc aca tag tat tag tca taa act tgg gat gta aaa tgt tga tga tat cta cat cgt ttg gat ttt tat gta tcc act tta ata ata tca tag ctg taa cat cct cgt ga- ttt acg ttg acg ttt tcg tgg gat aag ata gtt gtt agt tca tcc ttt gat aat ttt cca aat t

>LR800246.1:15629-17601_Variola_virus_genome_assembly_complete_genome:_monopartite

cgg ttg cca tgt att tcc tgg aga gca agt --- aga tga tg- agg aac cag ata gtt tat atc cgt acg tgc acg taa agt cta cat tgt cat tat acg a-- ata tga tct ttt aaa ccc gct aga caa gta tcc gtt taa tat tgt agg atg tgg aca ttt aac aat ctg aca cgt ggg tgg atc gga cca ttc tcc tcc tga aca cag gac act aaa gtt acc aat caa cga ata tcc act att gca act ata agt tac gac gct tcc atc ggt ata aaa atc ctc gta tcc gtt atg tct tcc gtt gga tat aga tgg agg taa ttg gca ttt aac aga ttc gca aat agg tgc ctt ggg att cca tac cat aga tcc agt aga tcc taa ttc aca ata cga ttt ata ttc acc gat caa atg ata tcc gct att aca aga gta cgt tat act aga gcc aaa gtc tac tcc gcc aat atc gag atg gcc att atc gat atc tcg agg cga tgg gca tct ccg ttt aat aca ttg att aaa gag tgt cca tcc agt acc tgt aca ttt agc ata tat agg tcc cat ttt ttg ctt tct gta tcc agg tag aca tag a-t att cta tag tgt ctc cta tgt tgt aat tag c-- --- -at tag cat tag cat cag tc- --t cca cac tat tct taa att tca tat taa tgg gtc gtg acg gaa tag tac agc atg ata gaa cgc atc cta ttc cca aca atg tca gga acg tca cgc tct cca cct tca tat tta ttt atc cgt -aa aaa tgt tat cct gga cat tgt gcc aat aat aaa aaa gcc ca- -ta tat gtt cgc tat tgt aga aat tgt ttt tca tag ttg ctc aaa aac gat ggc ggt gac tta tga gtt acg tta cac ttt gga gtc tca tct tta gta aac ata tca taa tat ttg ata tta cra gtt gac ata tcg aac aaa ttc caa rta ttt gat ttt gga taa tat tcg tat ttt gca tct gct ata att aag ata taa tcg ctg caa gaa cac acg aac atc ttt cct aca tgg tta aag tac atg tac aat tct atc cat ttg tct tcc tta act ata tat ttg tat aga taa tta cga gtc tcg tga gta att cca gta att gca tag atg tca cca tca tac tct aca gca taa act ata cta tga tgt cta ggc atg gga gac ttt ttt tat cca acg att ttt agt gaa aca ttc cac atc gtt taa tac tac ata ttt ttc ata cgt ggt ata aac tcc acc cat tac --a tat ata tca tcg ttt acg aat acc gac gcg cct gaa tat cta gga gta att aag ttt gga agt ctt atc cat ttc gaa gtg ccg tgt ttc aaa tat tct gct aca cct gtt gaa ata gaa aat ttt aat cct tct att aca tat aac ttt cca tca tta aca caa gta cta act tct gat ttt aac gac gac ata tta gta act gtt ttc ca- --- --- --- --- --- --- -tt ttt ttc gtt tta aga tct acc cgc -ga tac gga ata aac atg tct att gtt aat cat gcc gcc aat aat gta tag aca att ata taa aac att tgc att ata gaa ttg --- -tc tat ctg tat tac cga cga ttg tcc aat att ctg ttc tag gag agt aat ggg tta ttg tgg ata tat aat cag agt ttt taa tga cta cta tat tat gtt tta tac cat ttc gtg tca cag gct ttg tag att tgg ata tag tta atc cca aca atg ata tag cat tgc aca tag tat tag tca taa act tgg gat gta aaa tgt tga tga tat cta cat cgt ttg gat ttt tat gta tcc act tta ata ata tca tag ctg taa cat cct cgt ga- ttt acg ttg acg ttt tcg tgg gat aag ata gtt gtt agt tca tcc ttt gat aat ttt cca aat t

>KY549148.1:35214-37179_Cowpox_virus_isolate_CPXV_1639

cgg ctg cca tgt att tcc tgg aga gca agt --- aga tga tg- agg aac cag ata gtt tat atc cgt gcc tgc acg taa agt gta cat tgt cgt tgt gtg a-- gta tga tct ttt aaa ccc gct aga caa gta tcc gtt tgt tat tga agg atg tgg gca ttt aac gat ctg aca cgt ggg tgg atc gga cca ttc tcc tcc tga aca cag gac acc aga gtt acc aat caa cga ata tcc act att gca act ata cgt tac aac gct tcc ttc ggt ata aaa atc ctc gta tcc gtt atg tct tcc gtt gga tat aga tgg agg gga ttg gca ttt aac aga ttc aca aat ggg tgc ctc ggg att cca cac cat aga tcc agt ata tcc tag ttc aca ata cga ttt aga ttc tcc tat caa ctg ata tcc gct att gca aga gta cgt tat act aga gcc aaa ctc tac tcc atc aat atc aat ttg acc att atc aat atc tcg agg tga tgg aca ttt ccg ttt aat aca ttg att aaa gag tgt cca tcc agt acc tgt aca ttt agc ata tat agg tcc cat ttt ttg ctt tct gta tcc agg tag aca tag a-t att cta tag tgt ctc cta tgt tgt a-- --- --- --- -at tag cat tag cat cag tc- --c cca cac tat tct taa att tca tat taa tgg gac gtg atg gaa tag tac agc atg ata gca cgc atc cta ttc cca aca atg tca gga acg tca cgc tct cca cct tca tat tta ttt atc cgt -aa aaa tgt tat ccc agg cca ttg tcc aat aat a-a aaa gct ctc ata tat gtt cgc tat tgt aaa aat tgt ttt tca cag ttg ctc aaa aac gat ggt agt gac tta cga gtt acg tta cac ttt gga gtc tca tct tta gtg aac ata tca taa tat tcg ata tta cga gtt gac ata tcg aac aaa ttc caa gta ttt gat ttt gga taa tat tcg tat ttt gca tct gct ata att aag ata taa tca ccg caa gaa cac acg aac atc ttt cct aca tgg tta aag tac atg tac aat tct atc cat ttg tct tcc tta act ata tat ttg tat aga taa tta cga gtc tca tga gta att cca gta att aca tag atg tca ccg tcg tac tct aca gca taa act ata cta tga tgt cta ggc atg gga gac -tt ttt tat cca acg att ttt agt gaa aca ttc cac atc gtt taa tac tac ata ttt ctc ata cgt ggt ata aac tcc acc cat tac --a tat ata tca tcg ttt acg aat acc gat gcg cct gaa tat cta gga gta att aag ttt gga agt ctt atc cat ttc gaa gtg tcg tgt ttc aaa tat tct gct aca ccc gtt gaa ata gaa aat tct aat cct cct att aca tat aac ttt cca tcg tta aca caa gta cta act tct gat ttt aac gac gac ata tta gta acc gtt ttc ca- --- --- --- --- --- --- --t ttt ttc gtt tta aga tct aca cgc -ga tac gga ata aac atg tct att gtt aat cat gcc gcc aat aat gta tag aca att atg taa aac att tgc att ata gaa ttg --- -tc tat ctg tat tac cga cta tcg tcc aat att ctg ttc tag gag agt aat ggg tta ttg tgg ata tat aat cag agt ttt taa tga cta cta tat tat gtt tta tac cat ttc gtg tca ccg gct tta tag att tgg ata tag tta atc cca aca atg ata tag cat tgc gca tag cat tag tca taa act tgg gat gta aaa tgt tga tga tat cta cat cgt ttg gat ttt tat gta tcc act tta ata ata tca tag ctg taa cat cct cat ga- ttt acg tta acg tct tcg tgt gat aag ata gtt gtc agt tca tcc ttt gat aat ttt cca aat t

>HQ420900.1:34051-36016_Cowpox_virus_strain_UK2000_K2984_complete_genome

cgg ctg cca tgt att tcc tgg aga gca agt --- aga tga tg- agg aac cag ata gtt tat atc cgt gcc tgc acg taa agt gta cat tgt cgt tgt gtg a-- gta tga tct ttt aaa ccc gct aga caa gta tcc gtt tgt tat tga agg atg tgg gca ttt aac gat ctg aca cgt ggg tgg atc gga cca ttc tcc tcc tga aca cag gac acc aga gtt acc aat caa cga ata tcc act att gca act ata cgt tac aac gct tcc ttc ggt ata aaa atc ctc gta tcc gtt atg tct tcc gtt gga tat aga tgg agg gga ttg gca ttt aac aga ttc aca aat ggg tgc ctc ggg att cca cac cat aga tcc agt ata tcc tag ttc aca ata cga ttt aga ttc tcc tat caa ctg ata tcc gct att gca aga gta cgt tat act aga gcc aaa ctc tac tcc atc aat atc aat ttg acc att atc aat atc tcg agg tga tgg aca ttt ccg ttt aat aca ttg att aaa gag tgt cca tcc agt acc tgt aca ttt agc ata tat agg tcc cat ttt ttg ctt tct gta tcc agg tag aca tag a-t att cta tag tgt ctc cta tgt tgt a-- --- --- --- -at tag cat tag cat cag tc- --c cca cac tat tct taa att tca tat taa tgg gac gtg atg gaa tag tac agc atg ata gca cgc atc cta ttc cca aca atg tca gga acg tca cgc tct cca cct tca tat tta ttt atc cgt -aa aaa tgt tat ccc agg cca ttg tcc aat aat a-a aaa gct ctc ata tat gtt cgc tat tgt aaa aat tgt ttt tca cag ttg ctc aaa aac gat ggt agt gac tta cga gtt acg tta cac ttt gga gtc tca tct tta gta aac ata tca taa tat tcg ata tta cga gtt gac ata tcg aac aaa ttc caa gta ttt gat ttt gga taa tat tcg tat ttt gca tct gct ata att aag ata taa tca ccg caa gaa cac acg aac atc ttt cct aca tgg tta aag tac atg tac aat tct atc cat ttg tct tcc tta act ata tat ttg tat aga taa tta cga gtc tca tga gta att cca gta att aca tag atg tca ccg tcg tac tct aca gca taa act ata cta tga tgt cta ggc atg gga gac -tt ttt tat cca acg att ttt agt gaa aca ttc cac atc gtt taa tac tac ata ttt ctc ata cgt ggt ata aac tcc acc cat tac --a tat ata tca tcg ttt acg aat acc gat gcg cct gaa tat cta gga gta att aag ttt gga agt ctt atc cat ttc gaa gtg tcg tgt ttc aaa tat tct gct aca ccc gtt gaa ata gaa aat tct aat cct cct att aca tat aac ttt cca tcg tta aca caa gta cta act tct gat ttt aac gac gac ata tta gta acc gtt ttc ca- --- --- --- --- --- --- --t ttt ttc gtt tta aga tct acc cgc -ga tac gga ata aac atg tct att gtt aat cat gcc gcc aat aat gta tag aca att atg taa aac att tgc att ata gaa ttg --- -tc tat ctg tat tac cga cga tcg tcc aat att ctg ttc tag gag agt aat ggg tta ttg tgg ata tat aat cag agt ttt taa tga cta cta tat tat gtt tta tac cat ttc gtg tca ccg gct tta tag att tgg ata tag tta atc cca aca atg ata tag cat tgc gca tag cat tag tca taa act tgg gat gta aaa tgt tga tga tat cta cat cgt ttg gat ttt tat gta tcc act tta ata ata tca tag ctg taa cat cct cat ga- ttt acg tta acg tct tcg tgg gat aag ata gtt gtc aat tca tcc ttt gat aat ttt cca aat t

>KY549149.1:35813-37778_Cowpox_virus_isolate_CPXV_K2739

cgg ctg cca tgt att tcc tgg aga gca agt --- aga tga tg- agg aac cag ata gtt tat atc cgt gcc tgc acg taa agt gta cat tgt cgt tgt gtg a-- gta tga tct ttt aaa ccc gct aga caa gta tcc gtt tgt tat tga agg atg tgg gca ttt aac gat ctg aca cgt ggg tgg atc gga cca ttc tcc tcc tga aca cag gac acc aga gtt acc aat caa cga ata tcc act att gca act ata cgt tac aac gct tcc ttc ggt ata aaa atc ctc gta tcc gtt atg tct tcc gtt gga tat aga tgg agg gga tgg gca ttt aac aga ttc aca aat ggg tgc ctc ggg att cca cac cat aga tcc agt ata tcc tag ttc aca ata cga ttt aga ttc tcc tat caa ctg ata tcc gct att gca aga gta cgt tat act aga gcc aaa ctc tac tcc atc aat atc aat ttg acc att atc aat atc tcg agg tga tgg aca ttt ccg ttt aat aca ttg att aaa gag tgt cca tcc agt acc tgt aca ttt agc ata tat agg tcc cat ttt ttg ctt tct gta tcc agg tag aca tag a-t att cta tag tgt ctc cta tgt tgt a-- --- --- --- -at tag cat tag cat cag tc- --c cca cac tat tct taa att tca tat taa tgg gac gtg acg gaa tag tac agc atg ata gca cgc atc cta ttc cca aca atg tca gga acg tca cgc tct cca cct tca tat tta ttt atc cgt -aa aaa tgt tat ccc agg cca ttg tcc aat aat a-a aaa gct ctc ata tat gtt cgc tat tgt aaa aat tgt ttt tca cag ttg ctc aaa aac gat ggt agt gac tta cga gtt acg tta cac ttt gga gtc tca tct tta gta aac ata tca taa tat tcg ata tta cga gtt gac ata tcg aac aaa ttc caa gta ttt gat ttt gga taa tat tcg tat ttt gca tct gct ata att aag ata taa tca ccg caa gaa cac acg aac atc ttt cct aca tgg tta aag tac atg tac aat tct atc cat ttg tct tcc tta act ata tat ttg tat aga taa tta cga gtc tca tga gta att cca gta aat aca tag atg tca ccg tcg tac tct aca gca taa act ata cta tga tgt cta ggc atg gga gac -tt ttt tat cca acg att ttt agt gaa aca ttc cac atc gtt taa tac tac ata ttt ctc ata cgt ggt ata aac tcc acc cat tac --a tat ata tca tcg ttt acg aat acc gat gcg cct gaa tat cta gga gta att aag ttt gga agt ctt atc cat ttc gaa gtg tcg tgt ttc aaa tat tct gct aca ccc gtt gaa ata gaa aat tct aat cct cct att aca tat aac ttt cca tcg tta aca caa gta cta act tct gat ttt aac gac gac ata tta gta acc gtt ttc ca- --- --- --- --- --- --- --t ttt ttc gtt tta aga tct acc cgc -ga cac gga ata aac atg tct att gtt aat cat gcc gcc aat aat gta tag aca att atg taa aac att tgc att ata gaa ttg --- -tc tat ctg tat tac caa cga tcg tcc aat att ctg ttc tag gag agt aat ggg tta ttg tgg ata tat aat cag agt ttt taa tga cta cta tat tat gtt tta tac cat ttc gtg tca ccg gct tta tag att tgg ata tag tta atc cca aca atg ata tag cat tgc gca tag cat tag tca taa act tgg gat gta aaa tgt tga tga tat cta cat cgt ttg gat ttt tat gta tcc act tta ata ata tca tag ctg taa cat cct cat ga- ttt acg tta acg tct tca tgg gat aag ata gtt gtc aat tca tcc ttt gat aat ttt cca aat t

>KY549151.1:30706-32665_Cowpox_virus_isolate_Norwayfeline

cgg ctg cca tgt att tcc tgg aga gca agt --- aga tga tg- agg aac cag ata gtt tat atc cgt gcc tgc atg taa agt cta cat tgt cgt tgt gtg a-- gta tga tct ttt aaa ccc gct aga caa gta tcc gtt tgt tat tga agg atg tgg gca ttt aac gat ctg aca cgt ggg tgg atc gga cca ttc tcc tcc tga aca cac gat gcc aga gtt acc aat caa cga ata tcc ttt att gca act ata agt tac aac gct ccc atg ggt ata aaa atc ctc gta tcc gtt atg tct tcc gtt gga tat aga tgg agg gga ttg gca ttt aac aga ttc aca aat ggg tgc ctc ggg att cca tac cat aga tcc agt ata tcc tag ttc aca ata cga ttt aga ttc tcc tat caa ctg ata tcc gct att gca aga gta cgt tat act aga gcc aaa ctc tac tcc acc aat atc aat ttg acc att atc aat atc tcg agg aga tgg aca ttt acg ttt aat aca ttg att aaa gag tgt cca tcc agt acc tgt aca ttt agc ata tat agg tcc cat ttt ttg ctt tct gta tcc agg tag aca tag a-t att cta tag tgt ctc cta tgt tgt a-- --- --- --- -at tag aat tag cat tag tc- --t ctg cac tat tct taa att tca tat taa tgg gac gtg acg gaa tag tac agc atg ata gaa cgc atc cta ttc cca aca atg tca gga acg tca cgc tct cca cct tca tat tta ttt atc cgt -aa aaa tgt tat ccc agg cca ttg tcc aat aat a-a aaa gtc ca- --- --t gtt ctc tat tgt agg aat tgt ttt tca cag ttg ctc aaa aac gat ggt agt gac tta cga gtt aca tta cac ttt gga gtc tca tct tta gta aac ata tca taa tat tcg ata tta cga gtt gac ata tcg aac aaa ttc caa gta ttt gat ttt gga taa tat tcg tat ttt gca tct gct ata att aag ata taa tca ccg caa gaa cac acg aac atc ttt cct aca tgg tta aag tac atg tac aat tct atc cat ttg tct tcc tta act ata tat ttg tat aga taa tta cga gtc tca tga gta att cca gta att aca tag atg tcg ccg tcg tac tct aca gca taa act ata cta tga tgt cta ggc ata gga gac -tt ttt tat cca acg att ttt agt gaa aca ttc cac atc gtt taa tac tac ata ttt ctc ata cgt ggt ata aac tcc acc cat tac --a tat ata tca tcg ttt acg aat acc gat gcg cct gaa tat cta gga gta att aag ttt gga agt ctt atc cat ctc gaa gtg ccg tgt ttc aaa tat tct gct aca ccc gtt gaa ata gaa aat tct aat cct cct att aca tat aac ttt cca tcg tta aca caa gta cta act tct gat ttt aac gac gac ata tta gta acc gtt ttc ca- --- --- --- --- --- --- --t ttt tta gtt tta aga tct acc cgc -ga tac gga ata aac atg tct att gtt aat cat gcc gcc aat aat gta tag aca att atg taa aac att tgc att ata gaa ttg --- -tc tat ctg tat tac caa cga tcg tcc aat att ctg ttc tag gag agt aat ggg tta ttg tgg ata tat aat cag agt ttt taa tga tca cta tat tat gtt tta tac cat ttc gtg tca cag gct ttg tag att tgg ata tag tta atc cca aca atg cta tag cat tgc gca tag cat tag tca taa act tag gat gta aaa tgt tga tga tat cta cat cgt ttg gat ttt tat gta tcc act tta ata ata tta tag ctg taa cat cct cat ga- ttt acg tta acg tct tcg tgg gat aag ata gtt gtc agt tca tcc ttt gat aat ttt cca aat t

>KY549144.1:35276-37241_Cowpox_virus_isolate_CPXV_Catpox5wv1

cgg ctg cca tgt att tcc tgg aga gca agt --- aga tga tg- agg aac cag ata gtt tat atc cgt gcc tgc acg taa agt gta cat tgt cgt tgt gtg a-- gta tga tct ttt aaa ccc gct aga caa gta tcc gtt tgt tat tga agg atg tgg gca ttt aac gat ctg aca cgt ggg tgg atc gga cca ttc tcc tcc tga aca cac gat gcc aga gtt acc aat caa cga ata tcc act att gca act ata agt tac aac gct tcc atc ggt ata aaa atc ctc gta tcc gtt atg cct tcc att ggt tac aga tgg agg gga tgg gca ttt aac aga ttc aca aat ggg tgc ctc ggg att cca cac cat aga tcc agt ata tcc tag ttc aca ata cga ttt aga ttc tcc tat caa ctg ata tcc gct att gca aga gta cgt tat act aga gcc aaa ctc tac tcc atc aat atc aat ttg acc att atc aat atc tcg agg tga tgg aca ttt ccg ttt aat aca ttg att aaa gag tgt cca tcc agt acc tgt aca ttt agc ata tat agg tcc cat ttt ttg ctt tct gta tcc agg tag aca tag a-t att cta tag tgt ctc cta tgt tgt a-- --- --- --- -at tag tat tag cat cag tc- --c cca cac tat tct taa att tca tat taa tgg gac gtg acg gaa tag tac agc atg ata gca cgc atc cta ttc cca aca atg tca gga acg tca cgc tct cca cct tca tat tta ttt atc cgt -aa aaa tgt tat ccc agg cca ttg tcc aat aat a-a aaa gct ctc ata tat gtt cgc tat tgt aaa aat tgt ttt tca cag ttg ctc aaa aac gat ggt agt gac tta cga gtt acg tta cac ttt gga gtc tca tct tta gta aac ata tca taa tat tcg ata tta cga gtt gac ata tcg aac aaa ttc caa gta ttt gat ttt gga taa tat tcg tat ttt gca tct gct ata att aag ata taa tca ccg caa gaa cac acg aac atc ttt cct aca tgg tta aag tac atg tac aat tct atc cat ttg tct tcc tta act ata tat ttg tat aga taa tta cga gtc tca tga gta att cca gta att aca tag atg tca ccg tcg tac tct aca gca taa act ata cta tga tgt cta ggc atg gga gac -tt ttt tat cca acg att ttt agt gaa aca ttc cac atc gtt taa tac tac ata ttt ctc ata cgt ggt ata aac tcc acc cat tac --a tat ata tca tcg ttt acg aat acc gac gcg cct gaa tat cta gga gta att aag ttt gga agt ctt atc cat ttt gaa gtg tcg tgt ttc aaa tat tct gct aca ccc gtt gaa ata gaa aat tct aat cct cct att aca tat aac ttt cca tcg tta aca caa gta cta act tct gat ttt aac gac gac ata tta gta acc gtt ttc ca- --- --- --- --- --- --- --t ttt ttc gtt tta aga tct aca cgc -ga tac gga ata aac atg tct att gtt aat cat gcc gcc aat aat gta tag aga att atg taa aac att tgc att ata gaa ttg --- -tc tat ctg tat tac cga cga tcg tcc aat att ctg ttc tag gag agt aat ggg tta ttg tgg ata tat aat cag agt ttt taa tga tca cta tat tat gtt tta tac cat ttc gtg tca ccg gct tta tag att tgg ata tag tta atc cca aca atg ata tag cat tgc gca tag cat tag tca taa act tgg gat gta aaa tgt tga tga tat cta cat cgt ttg gat ttt tat gta tcc act tta ata ata tca tag ctg taa cat cct cat ga- ttt acg tta acg tct tcg tgt gat aag ata gtt gtc agt tca tcc ttt gat aat ttt cca aat t

>HQ420894.1:33092-35051_Cowpox_virus_strain_France_2001_Nancy_complete_genome

cgg ctt cca tgt att tcc tgg aga gca agt --- aga tga tg- aag aac cag ata ggt tat atc cgt act tgc act taa agt cta cat tgt cgt tgt atg a-- gta tga tct ttt aaa ccc gct aga caa gta tcc gtt tga tat tgt agg atg tgg aca ttt aac aat ctg aca cgt ggg tgg atc gga cca ttc tcc tcc tga aca cag gac acc aga gtt acc aat caa cga ata tcc act att gca act ata agt tac aac gct tcc atc ggt ata aaa atc ctc gta tcc gtt atg cct tcc att ggt tac aga tgg agg gga tgg aca ttt aac aga ttc aca aat ggg tgc ctc ggg att cca cac cat aga tcc agt ata tcc tag ttc aca ata cga ttt aga ttc tcc gat caa ctg ata tcc gct att gca aga gta cgt tat act aga gcc aaa ctc tac tcc acc aat atc aat ttg acc att ttc gat atc tcg agg tga tgg aca ttt ccg ttt aat aca ttg att aaa gag tgt cca tcc agt acc tgt aca ttt agc ata tat agg tcc cat ttt ttg ctt tct gta tcc agg tag aca tag a-t att cta tag tgt ctc cta tgt tgt a-- --- --- --- -at tag cat tag cat cag tc- --c cca cac tat tct taa att tca tat taa tgg gac gtg acg gaa tag tac agc atg ata gca cgc atc cta ttc cca aca atg tca gga acg tca cgc tct cca cct tca tat tta ttt atc cgt -aa aaa tgt tat ccc agg cca ttg tcc aat aat a-a aaa gtc ca- --- --t gtt ctc tat tgt agg aat tgt ttt tca cag ttg ctc aaa aac gat ggt agt gac tta cga gtt acg tta cac ttt gga gtc tca tct tta gta aac ata tca taa tat tcg ata tta cga gtt gac ata tcg aac aaa ttc caa gta ttt gat ttt gga taa tat tcg tat ttt gca tct gct ata att aag ata taa tca ccg caa gaa cat acg aac atc ttt cct aca tgg tta aag tac atg tac aat tct atc cat ttg tct tcc tta act ata tat ttg tat aga taa tta cga gtc tca tga gta att cca gta att aca tag atg tcg ccg tcg tac tct aca gca taa act ata cta tga tgt cta ggc atg gga gac -tt ttt tat cca acg att ttt agt gaa aca ttc cac atc gtt taa tac tac ata ttt ctc ata cgt ggt ata aac tcc acc cat tac --a tat ata tca tcg ttt acg aat acc gat gcg cct gaa tat cta gga gta att aag ttt gga agt ctt atc cat ctc gaa gtg ccg tgt ttc aaa tat tct gct aca cct gtt gaa ata gaa aat tct aat cct cct att aca tat aac ttt cca tcg tta aca caa gta cta act tct gat ttt aac gac gac ata tta gta acc gtt ttc ca- --- --- --- --- --- --- --t ttt ttc gtt tta aga tct acc agc -ga cac gga ata aac atg tct att att aat cat gcc gcc aat aat gta tag aca att atg taa aac att tgc att ata gaa ttg --- -tc tat ctg tat tac caa cga tcg tcc aat att ctg ttc tag gag agt aat ggg tta ttg tgg ata tat aat cag agt ttt taa tga tca cta tat tat gtt tta tac cat ttc gtg tca ccg gct ttg tag att tgg ata tag tta atc cca aca atg ata tag cat tgc gca tag tat tag tca taa act tgg gat gta aaa tgt tga tga tat cta cat cgt ttg gat ttt tat gta tcc act tta ata ata tta tag ctg taa cat cct cat ga- ttt acg tta acg tct tcg tgg gat aag ata gtt gtc agt tca tcc ttt gat aat ttt cca aat t

>KY554976.1:25426-27372_Ectromelia_virus_isolate_Hampstead

cgg ctg cca tgt att ttc cgg aga gca agt --- aga tga tg- agg aac cag ata gtt tat atc cgt act tgc act taa agt cta cat tgt cgt tgt atg a-- gta tga tct ttt aaa ccc gct aga caa gta tcc gtt tga tat tgt agg atg tgg aca ttt aac aat ctg aca cgt ggg tgg atc gga cca ttc tcc tcc tga aca cag gac acc aga gtt acc aac caa tga ata tcc act att gca act ata cgt tac aac gct tcc ttc ggt ata aaa atc ctc gta tcc gtt atg tct tcc gtt gga tat aga tgg agg gga ttg gca ttt aac aga ttc gca aat ggg tgc ctc ggg att cca tac cat aga tcc agt aga tcc tag ttc aca ata cga ttt aga ttc tcc gat caa ctg ata tcc gct att gca aga gta cgt tat act aga gcc aaa gtc tgt tcc acc aat atc aag ttg acc att atc gat atc tcg agg tga tgg aca tct ccg ttt aat aca ttg att aaa gag tgt cca tcc att gcc tgt gca ttt aac ata tat gga tcc cat ttt ttg ctt tct gta tcc agg tag aca tac a-t att cta tag tgt ctc cta ctt tgt a-- --- --- --- --- --- -ag tat cat cag tc- --c cca cat tat tct taa att tca tat taa tgg gac gtg acg gaa tag tac agc atg ata gca cgc atc cta ttc cca aca atg tca gaa acg tca cgc tct cca cct tca tat tta ttt atc cgt -aa aaa tgt tat cct atg cat tgt -cc aat aat --a aaa gtc at- --- --- --- -gc tat tgt agg aat ttt ttt tca cag ttg ctc aaa aac gat ggt agt gac tta tga gtt aca tta cat ttt gga gtc tca tct tta gta aac ata tca taa tat tcg ata tta cga gtt gac gta tct aac aaa ttc caa gta ttt gat ttt gga taa tat tcg tat ttt gca tct gct ata att aag ata taa tca cca caa gaa cac acg aac atc ttt cct aca tgg tta aag tac atg tat aat tct atc cat ttg tct tca tta act ata tat ttg tat aga taa tta cga gtc tca tga gta att cca gta att gca tag atg tca cca tcg tac tct aca gca taa act ata cta tga tgt cta ggc atg gga gac -tt ttt tat cca acg att ttt agt gaa aca ttc cac atc att taa tac tac ata ttt ctc ata cgt ggt ata aac tcc acc cat tac --a tat ata tca tcg ttt acg aat acc gat gct cct gaa tat cta gga gta att aag ttt gga agt ctt atc cat ttc gaa gcg cca tgt ttc aaa tat tct gct aca gcc gtt gaa ata gaa aat tct aat cct cct att aca tat aac ttt cca ttg tta aca caa gta cta act tct gat ttt aac gac gac ata tta gtg acc gtt ttc ca- --- --- --- --- --- --- --t ttt ttc gtt tta aga tct acc cgc -ga tac gga ata aac atg tct att gtt aat cat gcc gcc aat aat gta tag aca att atg taa aac att tgc att ata gaa ttg --- -tc tat ctg tat tac cga cta tcg tcc aat att ctg ttc tag gag agt aat ggg tta ttg tgg ata tat aat cag agt ttt taa tga cta cta tat tat gtt tta tac cat ttc gtg tca cag gct ttg tag att tgg ata tag tta atc cca aca atg cta tag cat tgc gca tag cat tag tca taa act tgg gat gta aaa tgt tga tga tat cta cat cgt ttg gat ttt tat gta tcc act tta ata ata tta tag ctg taa cat cct cat ga- ttt acg tta acg ttt tcg tgg gat aag ata gtt gtt agt tca tcc ttt gat aat ttt cca aat t

>OU343159.1:26318-28264_Ectromelia_virus_strain_MouKre_genome_assembly_chromosome:_1

cgg ctg cca tgt att ttc cgg aga gca agt --- aga tga tg- agg aac cag ata gtt tat atc cgt act tgc act taa agt cta cat tgt cgt tgt atg a-- gta tga tct ttt aaa ccc gct aga caa gta tcc gtt tga tat tgt agg atg tgg aca ttt aac aat ctg aca cgt ggg tgg atc gga cca ttc tcc tcc tga aca cag gac acc aga gtt acc aac caa tga ata tcc act att gca act ata cgt tac aac gct tcc ttc ggt ata aaa atc ctc gta tcc gtt atg tct tcc gtt gga tat aga tgg agg gga ttg gca ttt aac aga ttc gca aat ggg tgc ctc ggg att cca tac cat aga tcc agt aga tcc tag ttc aca ata cga ttt aga ttc tcc gat caa ctg ata tcc gct att gca aga gta cgt tat act aga gcc aaa gtc tgt tcc acc aat atc aag ttg acc att atc gat atc tcg agg tga tgg aca tct ccg ttt aat aca ttg att aaa gag tgt cca tcc att gcc tgt gca ttt aac ata tat gga tcc cat ttt ttg ctt tct gta tcc agg tag aca tac a-t att cta tag tgt ctc cta ctt tgt a-- --- --- --- --- --- -ag tat cat cag tc- --c cca cat tat tct taa att tca tat taa tgg gac gtg acg gaa tag tac agc atg ata gca cgc atc cta ttc cca aca atg tca gaa acg tca cgc tct cca cct tca tat tta ttt atc cgt -aa aaa tgt tat cct atg cat tgt -cc aat aat --a aaa gtc at- --- --- --- -gc tat tgt agg aat ttt ttt tca cag ttg ctc aaa aac gat ggt agt gac tta tga gtt aca tta cat ttt gga gtc tca tct tta gta aac ata tca taa tat tcg ata tta cga gtt gac gta tct aac aaa ttc caa gta ttt gat ttt gga taa tat tcg tat ttt gca tct gct ata att aag ata taa tca cca caa gaa cac acg aac atc ttt cct aca tgg tta aag tac atg tat aat tct atc cat ttg tct tca tta act ata tat ttg tat aga taa tta cga gtc tca tga gta att cca gta att gca tag atg tca cca tcg tac tct aca gca taa act ata cta tga tgt cta ggc atg gga gac -tt ttt tat cca acg att ttt agt gaa aca ttc cac atc att taa tac tac ata ttt ctc ata cgt ggt ata aac tcc acc cat tac --a tat ata tca tcg ttt acg aat acc gat gct cct gaa tat cta gga gta att aag ttt gga agt ctt atc cat ttc gaa gcg cca tgt ttc aaa tat tct gct aca gcc gtt gaa ata gaa aat tct aat cct cct att aca tat aac ttt cca ttg tta aca caa gta cta act tct gat ttt aac gac gac ata tta gtg acc gtt ttc ca- --- --- --- --- --- --- --t ttt ttc gtt tta aga tct acc cgc -ga tac gga ata aac atg tct att gtt aat cat gcc gcc aat aat gta tag aca att atg taa aac att tgc att ata gaa ttg --- -tc tat ctg tat tac cga cta tcg tcc aat att ctg ttc tag gag agt aat ggg tta ttg tgg ata tat aat cag agt ttt taa tga cta cta tat tat gtt tta tac cat ttc gtg tca cag gct ttg tag att tgg ata tag tta atc cca aca atg cta tag cat tgc gca tag cat tag tca taa act tgg gat gta aaa tgt tga tga tat cta cat cgt ttg gat ttt tat gta tcc act tta ata ata tta tag ctg taa cat cct cat ga- ttt acg tta acg ttt tcg tgg gat aag ata gtt gtt agt tca tcc ttt gat aat ttt cca aat t

>OU343158.1:25459-27405_Ectromelia_virus_strain_Mp1_genome_assembly_chromosome:_1

cgg ctg cca tgt att ttc cgg aga gca agt --- aga tga tg- agg aac cag ata gtt tat atc cgt act tgc act taa agt cta cat tgt cgt tgt atg a-- gta tga tct ttt aaa ccc gct aga caa gta tcc gtt tga tat tgt agg atg tgg aca ttt aac aat ctg aca cgt ggg tgg atc gga cca ttc tcc tcc tga aca cag gac acc aga gtt acc aac caa tga ata tcc act att gca act ata cgt tac aac gct tcc ttc ggt ata aaa atc ctc gta tcc gtt atg tct tcc gtt gga tat aga tgg agg gga ttg gca ttt aac aga ttc gca aat ggg tgc ctc ggg att cca tac cat aga tcc agt aga tcc tag ttc aca ata cga ttt aga ttc tcc gat caa ctg ata tcc gct att gca aga gta cgt tat act aga gcc aaa gtc tgt tcc acc aat atc aag ttg acc att atc gat atc tcg agg tga tgg aca tct ccg ttt aat aca ttg att aaa gag tgt cca tcc att gcc tgt gca ttt aac ata tat gga tcc cat ttt ttg ctt tct gta tcc agg tag aca tac a-t att cta tag tgt ctc cta ctt tgt a-- --- --- --- --- --- -ag tat cat cag tc- --c cca cat tat tct taa att tca tat taa tgg gac gtg acg gaa tag tac agc atg ata gca cgc atc cta ttc cca aca atg tca gaa acg tca cgc tct cca cct tca tat tta ttt atc cgt -aa aaa tgt tat cct atg cat tgt -cc aat aat --a aaa gtc at- --- --- --- -gc tat tgt agg aat ttt ttt tca cag ttg ctc aaa aac gat ggt agt gac tta tga gtt aca tta cat ttt gga gtc tca tct tta gta aac ata tca taa tat tcg ata tta cga gtt gac gta tct aac aaa ttc caa gta ttt gat ttt gga taa tat tcg tat ttt gca tct gct ata att aag ata taa tca cca caa gaa cac acg aac atc ttt cct aca tgg tta aag tac atg tat aat tct atc cat ttg tct tca tta act ata tat ttg tat aga taa tta cga gtc tca tga gta att cca gta att gca tag atg tca cca tcg tac tct aca gca taa act ata cta tga tgt cta ggc atg gga gac -tt ttt tat cca acg att ttt agt gaa aca ttc cac atc att taa tac tac ata ttt ctc ata cgt ggt ata aac tcc acc cat tac --a tat ata tca tcg ttt acg aat acc gat gct cct gaa tat cta gga gta att aag ttt gga agt ctt atc cat ttc gaa gcg cca tgt ttc aaa tat tct gct aca gcc gtt gaa ata gaa aat tct aat cct cct att aca tat aac ttt cca ttg tta aca caa gta cta act tct gat ttt aac gac gac ata tta gtg acc gtt ttc ca- --- --- --- --- --- --- --t ttt ttc gtt tta aga tct acc cgc -ga tac gga ata aac atg tct att gtt aat cat gcc gcc aat aat gta tag aca att atg taa aac att tgc att ata gaa ttg --- -tc tat ctg tat tac cga cta tcg tcc aat att ctg ttc tag gag agt aat ggg tta ttg tgg ata tat aat cag agt ttt taa tga cta cta tat tat gtt tta tac cat ttc gtg tca cag gct ttg tag att tgg ata tag tta atc cca aca atg cta tag cat tgc gca tag cat tag tca taa act tgg gat gta aaa tgt tga tga tat cta cat cgt ttg gat ttt tat gta tcc act tta ata ata tta tag ctg taa cat cct cat ga- ttt acg tta acg ttt tcg tgg gat aag ata gtt gtt agt tca tcc ttt gat aat ttt cca aat t

>OU343157.1:25662-27608_Ectromelia_virus_strain_Mp4_genome_assembly_chromosome:_1

cgg ctg cca tgt att ttc cgg aga gca agt --- aga tga tg- agg aac cag ata gtt tat atc cgt act tgc act taa agt cta cat tgt cgt tgt atg a-- gta tga tct ttt aaa ccc gct aga caa gta tcc gtt tga tat tgt agg atg tgg aca ttt aac aat ctg aca cgt ggg tgg atc gga cca ttc tcc tcc tga aca cag gac acc aga gtt acc aac caa tga ata tcc act att gca act ata cgt tac aac gct tcc ttc ggt ata aaa atc ctc gta tcc gtt atg tct tcc gtt gga tat aga tgg agg gga ttg gca ttt aac aga ttc gca aat ggg tgc ctc ggg att cca tac cat aga tcc agt aga tcc tag ttc aca ata cga ttt aga ttc tcc gat caa ctg ata tcc gct att gca aga gta cgt tat act aga gcc aaa gtc tgt tcc acc aat atc aag ttg acc att atc gat atc tcg agg tga tgg aca tct ccg ttt aat aca ttg att aaa gag tgt cca tcc att gcc tgt gca ttt aac ata tat gga tcc cat ttt ttg ctt tct gta tcc agg tag aca tac a-t att cta tag tgt ctc cta ctt tgt a-- --- --- --- --- --- -ag tat cat cag tc- --c cca cat tat tct taa att tca tat taa tgg gac gtg acg gaa tag tac agc atg ata gca cgc atc cta ttc cca aca atg tca gaa acg tca cgc tct cca cct tca tat tta ttt atc cgt -aa aaa tgt tat cct atg cat tgt -cc aat aat --a aaa gtc at- --- --- --- -gc tat tgt agg aat ttt ttt tca cag ttg ctc aaa aac gat ggt agt gac tta tga gtt aca tta cat ttt gga gtc tca tct tta gta aac ata tca taa tat tcg ata tta cga gtt gac gta tct aac aaa ttc caa gta ttt gat ttt gga taa tat tcg tat ttt gca tct gct ata att aag ata taa tca cca caa gaa cac acg aac atc ttt cct aca tgg tta aag tac atg tat aat tct atc cat ttg tct tca tta act ata tat ttg tat aga taa tta cga gtc tca tga gta att cca gta att gca tag atg tca cca tcg tac tct aca gca taa act ata cta tga tgt cta ggc atg gga gac -tt ttt tat cca acg att ttt agt gaa aca ttc cac atc att taa tac tac ata ttt ctc ata cgt ggt ata aac tcc acc cat tac --a tat ata tca tcg ttt acg aat acc gat gct cct gaa tat cta gga gta att aag ttt gga agt ctt atc cat ttc gaa gcg cca tgt ttc aaa tat tct gct aca gcc gtt gaa ata gaa aat tct aat cct cct att aca tat aac ttt cca ttg tta aca caa gta cta act tct gat ttt aac gac gac ata tta gtg acc gtt ttc ca- --- --- --- --- --- --- --t ttt ttc gtt tta aga tct acc cgc -ga tac gga ata aac atg tct att gtt aat cat gcc gcc aat aat gta tag aca att atg taa aac att tgc att ata gaa ttg --- -tc tat ctg tat tac cga cta tcg tcc aat att ctg ttc tag gag agt aat ggg tta ttg tgg ata tat aat cag agt ttt taa tga cta cta tat tat gtt tta tac cat ttc gtg tca cag gct ttg tag att tgg ata tag tta atc cca aca atg cta tag cat tgc gca tag cat tag tca taa act tgg gat gta aaa tgt tga tga tat cta cat cgt ttg gat ttt tat gta tcc act tta ata ata tta tag ctg taa cat cct cat ga- ttt acg tta acg ttt tcg tgg gat aag ata gtt gtt agt tca tcc ttt gat aat ttt cca aat t

>OU343156.1:25451-27397_Ectromelia_virus_strain_Mill-Hill_genome_assembly_chromosome:_1

cgg ctg cca tgt att ttc cgg aga gca agt --- aga tga tg- agg aac cag ata gtt tat atc cgt act tgc act taa agt cta cat tgt cgt tgt atg a-- gta tga tct ttt aaa ccc gct aga caa gta tcc gtt tga tat tgt agg atg tgg aca ttt aac aat ctg aca cgt ggg tgg atc gga cca ttc tcc tcc tga aca cag gac acc aga gtt acc aac caa tga ata tcc act att gca act ata cgt tac aac gct tcc ttc ggt ata aaa atc ctc gta tcc gtt atg tct tcc gtt gga tat aga tgg agg gga ttg gca ttt aac aga ttc gca aat ggg tgc ctc ggg att cca tac cat aga tcc agt aga tcc tag ttc aca ata cga ttt aga ttc tcc gat caa ctg ata tcc gct att gca aga gta cgt tat act aga gcc aaa gtc tgt tcc acc aat atc aag ttg acc att atc gat atc tcg agg tga tgg aca tct ccg ttt aat aca ttg att aaa gag tgt cca tcc att gcc tgt gca ttt aac ata tat gga tcc cat ttt ttg ctt tct gta tcc agg tag aca tac a-t att cta tag tgt ctc cta ctt tgt a-- --- --- --- --- --- -ag tat cat cag tc- --c cca cat tat tct taa att tca tat taa tgg gac gtg acg gaa tag tac agc atg ata gca cgc atc cta ttc cca aca atg tca gaa acg tca cgc tct cca cct tca tat tta ttt atc cgt -aa aaa tgt tat cct atg cat tgt -cc aat aat --a aaa gtc at- --- --- --- -gc tat tgt agg aat ttt ttt tca cag ttg ctc aaa aac gat ggt agt gac tta tga gtt aca tta cat ttt gga gtc tca tct tta gta aac ata tca taa tat tcg ata tta cga gtt gac gta tct aac aaa ttc caa gta ttt gat ttt gga taa tat tcg tat ttt gca tct gct ata att aag ata taa tca cca caa gaa cac acg aac atc ttt cct aca tgg tta aag tac atg tat aat tct atc cat ttg tct tca tta act ata tat ttg tat aga taa tta cga gtc tca tga gta att cca gta att gca tag atg tca cca tcg tac tct aca gca taa act ata cta tga tgt cta ggc atg gga gac -tt ttt tat cca acg att ttt agt gaa aca ttc cac atc att taa tac tac ata ttt ctc ata cgt ggt ata aac tcc acc cat tac --a tat ata tca tcg ttt acg aat acc gat gct cct gaa tat cta gga gta att aag ttt gga agt ctt atc cat ttc gaa gcg cca tgt ttc aaa tat tct gct aca gcc gtt gaa ata gaa aat tct aat cct cct att aca tat aac ttt cca ttg tta aca caa gta cta act tct gat ttt aac gac gac ata tta gtg acc gtt ttc ca- --- --- --- --- --- --- --t ttt ttc gtt tta aga tct acc cgc -ga tac gga ata aac atg tct att gtt aat cat gcc gcc aat aat gta tag aca att atg taa aac att tgc att ata gaa ttg --- -tc tat ctg tat tac cga cta tcg tcc aat att ctg ttc tag gag agt aat ggg tta ttg tgg ata tat aat cag agt ttt taa tga cta cta tat tat gtt tta tac cat ttc gtg tca cag gct ttg tag att tgg ata tag tta atc cca aca atg cta tag cat tgc gca tag cat tag tca taa act tgg gat gta aaa tgt tga tga tat cta cat cgt ttg gat ttt tat gta tcc act tta ata ata tta tag ctg taa cat cct cat ga- ttt acg tta acg ttt tcg tgg gat aag ata gtt gtt agt tca tcc ttt gat aat ttt cca aat t

>OU343155.1:25556-27502_Ectromelia_virus_strain_Mp5_genome_assembly_chromosome:_1

cgg ctg cca tgt att ttc cgg aga gca agt --- aga tga tg- agg aac cag ata gtt tat atc cgt act tgc act taa agt cta cat tgt cgt tgt atg a-- gta tga tct ttt aaa ccc gct aga caa gta tcc gtt tga tat tgt agg atg tgg aca ttt aac aat ctg aca cgt ggg tgg atc gga cca ttc tcc tcc tga aca cag gac acc aga gtt acc aac caa tga ata tcc act att gca act ata cgt tac aac gct tcc ttc ggt ata aaa atc ctc gta tcc gtt atg tct tcc gtt gga tat aga tgg agg gga ttg gca ttt aac aga ttc gca aat ggg tgc ctc ggg att cca tac cat aga tcc agt aga tcc tag ttc aca ata cga ttt aga ttc tcc gat caa ctg ata tcc gct att gca aga gta cgt tat act aga gcc aaa gtc tgt tcc acc aat atc aag ttg acc att atc gat atc tcg agg tga tgg aca tct ccg ttt aat aca ttg att aaa gag tgt cca tcc att gcc tgt gca ttt aac ata tat gga tcc cat ttt ttg ctt tct gta tcc agg tag aca tac a-t att cta tag tgt ctc cta ctt tgt a-- --- --- --- --- --- -ag tat cat cag tc- --c cca cat tat tct taa att tca tat taa tgg gac gtg acg gaa tag tac agc atg ata gca cgc atc cta ttc cca aca atg tca gaa acg tca cgc tct cca cct tca tat tta ttt atc cgt -aa aaa tgt tat cct atg cat tgt -cc aat aat --a aaa gtc at- --- --- --- -gc tat tgt agg aat ttt ttt tca cag ttg ctc aaa aac gat ggt agt gac tta tga gtt aca tta cat ttt gga gtc tca tct tta gta aac ata tca taa tat tcg ata tta cga gtt gac gta tct aac aaa ttc caa gta ttt gat ttt gga taa tat tcg tat ttt gca tct gct ata att aag ata taa tca cca caa gaa cac acg aac atc ttt cct aca tgg tta aag tac atg tat aat tct atc cat ttg tct tca tta act ata tat ttg tat aga taa tta cga gtc tca tga gta att cca gta att gca tag atg tca cca tcg tac tct aca gca taa act ata cta tga tgt cta ggc atg gga gac -tt ttt tat cca acg att ttt agt gaa aca ttc cac atc att taa tac tac ata ttt ctc ata cgt ggt ata aac tcc acc cat tac --a tat ata tca tcg ttt acg aat acc gat gct cct gaa tat cta gga gta att aag ttt gga agt ctt atc cat ttc gaa gcg cca tgt ttc aaa tat tct gct aca gcc gtt gaa ata gaa aat tct aat cct cct att aca tat aac ttt cca ttg tta aca caa gta cta act tct gat ttt aac gac gac ata tta gtg acc gtt ttc ca- --- --- --- --- --- --- --t ttt ttc gtt tta aga tct acc cgc -ga tac gga ata aac atg tct att gtt aat cat gcc gcc aat aat gta tag aca att atg taa aac att tgc att ata gaa ttg --- -tc tat ctg tat tac cga cta tcg tcc aat att ctg ttc tag gag agt aat ggg tta ttg tgg ata tat aat cag agt ttt taa tga cta cta tat tat gtt tta tac cat ttc gtg tca cag gct ttg tag att tgg ata tag tta atc cca aca atg cta tag cat tgc gca tag cat tag tca taa act tgg gat gta aaa tgt tga tga tat cta cat cgt ttg gat ttt tat gta tcc act tta ata ata tta tag ctg taa cat cct cat ga- ttt acg tta acg ttt tcg tgg gat aag ata gtt gtt agt tca tcc ttt gat aat ttt cca aat t

>OU343113.1:25556-27502_Ectromelia_virus_strain_Ishibashi_genome_assembly_chromosome:_1

cgg ctg cca tgt att ttc cgg aga gca agt --- aga tga tg- agg aac cag ata gtt tat atc cgt act tgc act taa agt cta cat tgt cgt tgt atg a-- gta tga tct ttt aaa ccc gct aga caa gta tcc gtt tga tat tgt agg atg tgg aca ttt aac aat ctg aca cgt ggg tgg atc gga cca ttc tcc tcc tga aca cag gac acc aga gtt acc aac caa tga ata tcc act att gca act ata cgt tac aac gct tcc ttc ggt ata aaa atc ctc gta tcc gtt atg tct tcc gtt gga tat aga tgg agg gga ttg gca ttt aac aga ttc gca aat ggg tgc ctc ggg att cca tac cat aga tcc agt aga tcc tag ttc aca ata cga ttt aga ttc tcc gat caa ctg ata tcc gct att gca aga gta cgt tat act aga gcc aaa gtc tgt tcc acc aat atc aag ttg acc att atc gat atc tcg agg tga tgg aca tct ccg ttt aat aca ttg att aaa gag tgt cca tcc att gcc tgt gca ttt aac ata tat gga tcc cat ttt ttg ctt tct gta tcc agg tag aca tac a-t att cta tag tgt ctc cta ctt tgt a-- --- --- --- --- --- -ag tat cat cag tc- --c cca cat tat tct taa att tca tat taa tgg gac gtg acg gaa tag tac agc atg ata gca cgc atc cta ttc cca aca atg tca gaa acg tca cgc tct cca cct tca tat tta ttt atc cgt -aa aaa tgt tat cct atg cat tgt -cc aat aat --a aaa gtc at- --- --- --- -gc tat tgt agg aat ttt ttt tca cag ttg ctc aaa aac gat ggt agt gac tta tga gtt aca tta cat ttt gga gtc tca tct tta gta aac ata tca taa tat tcg ata tta cga gtt gac gta tct aac aaa ttc caa gta ttt gat ttt gga taa tat tcg tat ttt gca tct gct ata att aag ata taa tca cca caa gaa cac acg aac atc ttt cct aca tgg tta aag tac atg tat aat tct atc cat ttg tct tca tta act ata tat ttg tat aga taa tta cga gtc tca tga gta att cca gta att gca tag atg tca cca tcg tac tct aca gca taa act ata cta tga tgt cta ggc atg gga gac -tt ttt tat cca acg att ttt agt gaa aca ttc cac atc att taa tac tac ata ttt ctc ata cgt ggt ata aac tcc acc cat tac --a tat ata tca tcg ttt acg aat acc gat gct cct gaa tat cta gga gta att aag ttt gga agt ctt atc cat ttc gaa gcg cca tgt ttc aaa tat tct gct aca gcc gtt gaa ata gaa aat tct aat cct cct att aca tat aac ttt cca ttg tta aca caa gta cta act tct gat ttt aac gac gac ata tta gtg acc gtt ttc ca- --- --- --- --- --- --- --t ttt ttc gtt tta aga tct acc cgc -ga tac gga ata aac atg tct att gtt aat cat gcc gcc aat aat gta tag aca att atg taa aac att tgc att ata gaa ttg --- -tc tat ctg tat tac cga cta tcg tcc aat att ctg ttc tag gag agt aat ggg tta ttg tgg ata tat aat cag agt ttt taa tga cta cta tat tat gtt tta tac cat ttc gtg tca cag gct ttg tag att tgg ata tag tta atc cca aca atg cta tag cat tgc gca tag cat tag tca taa act tgg gat gta aaa tgt tga tga tat cta cat cgt ttg gat ttt tat gta tcc act tta ata ata tta tag ctg taa cat cct cat ga- ttt acg tta acg ttt tcg tgg gat aag ata gtt gtt agt tca tcc ttt gat aat ttt cca aat t

>OU343112.1:25463-27409_Ectromelia_virus_strain_Hampstead_Egg_genome_assembly_chromosome:_1

cgg ctg cca tgt att ttc cgg aga gca agt --- aga tga tg- agg aac cag ata gtt tat atc cgt act tgc act taa agt cta cat tgt cgt tgt atg a-- gta tga tct ttt aaa ccc gct aga caa gta tcc gtt tga tat tgt agg atg tgg aca ttt aac aat ctg aca cgt ggg tgg atc gga cca ttc tcc tcc tga aca cag gac acc aga gtt acc aac caa tga ata tcc act att gca act ata cgt tac aac gct tcc ttc ggt ata aaa atc ctc gta tcc gtt atg tct tcc gtt gga tat aga tgg agg gga ttg gca ttt aac aga ttc gca aat ggg tgc ctc ggg att cca tac cat aga tcc agt aga tcc tag ttc aca ata cga ttt aga ttc tcc gat caa ctg ata tcc gct att gca aga gta cgt tat act aga gcc aaa gtc tgt tcc acc aat atc aag ttg acc att atc gat atc tcg agg tga tgg aca tct ccg ttt aat aca ttg att aaa gag tgt cca tcc att gcc tgt gca ttt aac ata tat gga tcc cat ttt ttg ctt tct gta tcc agg tag aca tac a-t att cta tag tgt ctc cta ctt tgt a-- --- --- --- --- --- -ag tat cat cag tc- --c cca cat tat tct taa att tca tat taa tgg gac gtg acg gaa tag tac agc atg ata gca cgc atc cta ttc cca aca atg tca gaa acg tca cgc tct cca cct tca tat tta ttt atc cgt -aa aaa tgt tat cct atg cat tgt -cc aat aat --a aaa gtc at- --- --- --- -gc tat tgt agg aat ttt ttt tca cag ttg ctc aaa aac gat ggt agt gac tta tga gtt aca tta cat ttt gga gtc tca tct tta gta aac ata tca taa tat tcg ata tta cga gtt gac gta tct aac aaa ttc caa gta ttt gat ttt gga taa tat tcg tat ttt gca tct gct ata att aag ata taa tca cca caa gaa cac acg aac atc ttt cct aca tgg tta aag tac atg tat aat tct atc cat ttg tct tca tta act ata tat ttg tat aga taa tta cga gtc tca tga gta att cca gta att gca tag atg tca cca tcg tac tct aca gca taa act ata cta tga tgt cta ggc atg gga gac -tt ttt tat cca acg att ttt agt gaa aca ttc cac atc att taa tac tac ata ttt ctc ata cgt ggt ata aac tcc acc cat tac --a tat ata tca tcg ttt acg aat acc gat gct cct gaa tat cta gga gta att aag ttt gga agt ctt atc cat ttc gaa gcg cca tgt ttc aaa tat tct gct aca gcc gtt gaa ata gaa aat tct aat cct cct att aca tat aac ttt cca ttg tta aca caa gta cta act tct gat ttt aac gac gac ata tta gtg acc gtt ttc ca- --- --- --- --- --- --- --t ttt ttc gtt tta aga tct acc cgc -ga tac gga ata aac atg tct att gtt aat cat gcc gcc aat aat gta tag aca att atg taa aac att tgc att ata gaa ttg --- -tc tat ctg tat tac cga cta tcg tcc aat att ctg ttc tag gag agt aat ggg tta ttg tgg ata tat aat cag agt ttt taa tga cta cta tat tat gtt tta tac cat ttc gtg tca cag gct ttg tag att tgg ata tag tta atc cca aca atg cta tag cat tgc gca tag cat tag tca taa act tgg gat gta aaa tgt tga tga tat cta cat cgt ttg gat ttt tat gta tcc act tta ata ata tta tag ctg taa cat cct cat ga- ttt acg tta acg ttt tcg tgg gat aag ata gtt gtt agt tca tcc ttt gat aat ttt cca aat t

>AF012825.2:26803-28749_Ectromelia_virus_strain_Moscow_complete_genome

cgg ctg cca tgt att ttc cgg aga gca agt --- aga tga tg- agg aac cag ata gtt tat atc cgt act tgc act taa agt cta cat tgt cgt tgt atg a-- gta tga tct ttt aaa ccc gct aga caa gta tcc gtt tga tat tgt agg atg tgg aca ttt aac aat ctg aca cgt ggg tgg atc gga cca ttc tcc tcc tga aca cag gac acc aga gtt acc aac caa tga ata tcc act att gca act ata cgt tac aac gct tcc ttc ggt ata aaa atc ctc gta tcc gtt atg tct tcc gtt gga tat aga tgg agg gga ttg gca ttt aac aga ttc gca aat ggg tgc ctc ggg att cca tac cat aga tcc agt aga tcc tag ttc aca ata cga ttt aga ttc tcc gat caa ctg ata tcc gct att gca aga gta cgt tat act aga gcc aaa gtc tgt tcc acc aat atc aag ttg acc att atc gat atc tcg agg tga tgg aca tct ccg ttt aat aca ttg att aaa gag tgt cca tcc att gcc tgt gca ttt aac ata tat gga tcc cat ttt ttg ctt tct gta tcc agg tag aca tac a-t att cta tag tgt ctc cta ctt tgt a-- --- --- --- --- --- -ag tat cat cag tc- --c cca cat tat tct taa att tca tat taa tgg gac gtg acg gaa tag tac agc atg ata gca cgc atc cta ttc cca aca atg tca gaa acg tca cgc tct cca cct tca tat tta ttt atc cgt -aa aaa tgt tat cct atg cat tgt -cc aat aat --a aaa gtc at- --- --- --- -gc tat tgt agg aat ttt ttt tca cag ttg ctc aaa aac gat ggt agt gac tta tga gtt aca tta cat ttt gga gtc tca tct tta gta aac ata tca taa tat tcg ata tta cga gtt gac gta tct aac aaa ttc caa gta ttt gat ttt gga taa tat tcg tat ttt gca tct gct ata att aag ata taa tca cca caa gaa cac acg aac atc ttt cct aca tgg tta aag tac atg tat aat tct atc cat ttg tct tca tta act ata tat ttg tat aga taa tta cga gtc tca tga gta att cca gta att gca tag atg tca cca tcg tac tct aca gca taa act ata cta tga tgt cta ggc atg gga gac -tt ttt tat cca acg att ttt agt gaa aca ttc cac atc att taa tac tac ata ttt ctc ata cgt ggt ata aac tcc acc cat tac --a tat ata tca tcg ttt acg aat acc gat gct cct gaa tat cta gga gta att aag ttt gga agt ctt atc cat ttc gaa gcg cca tgt ttc aaa tat tct gct aca gcc gtt gaa ata gaa aat tct aat cct cct att aca tat aac ttt cca ttg tta aca caa gta cta act tct gat ttt aac gac gac ata tta gtg acc gtt ttc ca- --- --- --- --- --- --- --t ttt ttc gtt tta aga tct acc cgc -ga tac gga ata aac atg tct att gtt aat cat gcc gcc aat aat gta tag aca att atg taa aac att tgc att ata gaa ttg --- -tc tat ctg tat tac cga cta tcg tcc aat att ctg ttc tag gag agt aat ggg tta ttg tgg ata tat aat cag agt ttt taa tga cta cta tat tat gtt tta tac cat ttc gtg tca cag gct ttg tag att tgg ata tag tta atc cca aca atg cta tag cat tgc gca tag cat tag tca taa act tgg gat gta aaa tgt tga tga tat cta cat cgt ttg gat ttt tat gta tcc act tta ata ata tta tag ctg taa cat cct cat ga- ttt acg tta acg ttt tcg tgg gat aag ata gtt gtt agt tca tcc ttt gat aat ttt cca aat t

>KY549145.1:35829-37782_Cowpox_virus_isolate_CPXV_K428

cgg ctg cca tgt att tcc tgg aga gca agt --- aga tga tg- agg aac cag ata gtt tat atc cgt gcc tgc acg taa agt gta cat tgt cgt tgt gtg a-- gta tga tct ttt aaa ccc gct aga caa gta tcc gtt tgt tat tga agg atg tgg gca ttt aac gat ctg aca cgt ggg tgg atc gga cca ttc tcc tcc tga aca cag gac acc aga gtt acc aat caa cga ata tcc act att gca act ata cgt tac aac gct tcc ttc ggt ata aaa atc ctc gta tcc gtt atg tct tcc gtt gga tat aga tgg agg gga ttg gca ttt aac aga ttc aca aat ggg tgc ctc ggg att cca cac cat aga tcc agt ata tcc tag ttc aca ata cga ttt aga ttc tcc tat caa ctg ata tcc gct att gca aga gta cgt tat act aga gcc aaa ctc tac tcc atc aat atc aat ttg acc att atc aat atc tcg agg tga tgg aca ttt ccg ttt aat aca ttg att aaa gag tgt cca tcc agt acc tgt aca ttt agc ata tat agg tcc cat ttt ttg ctt tct gta tcc agg tag aca tag a-t att cta tag tgt ctc cta tgt tgt a-- --- --- --- -at tag cat tag cat cag tc- --c cca cac tat tct taa att tca tat taa tgg gac gtg atg gaa tag tac agc atg ata gca cgc atc cta ttc cca aca atg tca gga acg tca cgc tct cca cct tca tat tta ttt atc cgt -aa aaa tgt tat cct atg -ca ttg tcc aat aat a-a aaa gtc at- --- --- --- -gc tat tgt agg aat tgt ttt tca cag ttg ctc aaa aac gat ggt agt gac tta cga gtt acg tta cac ttt gga gtc tca tct tta gtg aac ata tca taa tat tcg ata tta cga gtt gac ata tcg aac aaa ttc caa gta ttt gat ttt gga taa tat tcg tat ttt gca tct gct ata att aag ata taa tca ccg caa gaa cac acg aac atc ttt cct aca tgg tta aag tac atg tac aat tct atc cat ttg tct tcc tta act ata tat ttg tat aga taa tta cga gtc tca tga gta att cca gta att aca tag atg tca ccg tcg tac tct aca gca taa act ata cta tga tgt cta ggc atg gga gac -tt ttt tat cca acg att ttt agt gaa aca ttc cac atc gtt taa tac tac ata ttt ctc ata cgt ggt ata aac tcc acc cat tac --a tat ata tca tcg ttt acg aat acc gat gcg cct gaa tat cta gga gta att aag ttt gga agt ctt atc cat ttc gaa gtg tcg tgt ttc aaa tat tct gct aca ccc gtt gaa ata gaa aat tct aat cct cct att aca tat aac ttt cca tcg tta aca caa gta cta act tct gat ttt aac gac gac ata tta gta acc gtt ttc ca- --- --- --- --- --- --- --t ttt ttc gtt tta aga tct acc cgc -ga cac gga ata aac atg tct att gtt aat cat tcc gcc aat aat gta tag aca att atg taa aac att tgc att ata gaa ttg --- -tc tat ctg tat tac cga cta tcg tcc aat att ctg ttc tag gag agt aat ggg tta ttg tgg ata tat aat cag agt ttt taa tga cta cta tat tat gtt tta tac cat ttc gtg tca ccg gct tta tag att tgg ata tag tta atc cca aca atg ata tag cat tgc gca tag cat tag tca taa act tgg gat gta aaa tgt tga tga tat cta cat cgt ttg gat ttt tat gta tcc act tta ata ata tca tag ctg taa cat cct cat ga- ttt acg tta acg tct tcg tgt gat aag ata gtt gtc agt tca tcc ttt gat aat ttt cca aat t

>KJ563295.1:24686-26632_Ectromelia_virus_Naval_complete_genome

cgg ctg cca tgt att ttc cgg aga gca agt --- aga tga tg- agg aac cag ata gtt tat atc cgt act tgc act taa agt ata cat tgt cgt tgt atg a-- gta tga tct ttt aaa ccc gct aga caa gta tcc gtt tga tat tgt agg atg tgg aca ttt aac aat ctg aca cgt ggg tgg atc gga cca ttc tcc tcc tga aca cag gac acc aga gtt acc aac caa tga ata tcc act att gca act ata cgt tac aac gct tcc ttc ggt ata aaa atc ctc gta tcc gtt atg tct tcc gtt gga tat aga tgg agg gga ttg gca ttt aac aga ttc gca aat ggg tgc ctc ggg att cca tac cat aga tcc agt aga tcc tag ttc aca ata cga ttt aga ttc tcc gat caa ctg ata tcc gct att gca aga gta cgt tat act aga gcc aaa gtc tgt tcc acc aat atc aag ttg acc att atc gat atc tcg agg tga tgg aca tct ccg ttt aat aca ttg att aaa gag tgt cca tcc att gcc tgt gca ttt aac ata tat gga tcc cat ttt ttg ctt tct gta tcc agg tag aca tac a-t att cta tag tat ctc cta ctt tgt a-- --- --- --- --- --- -ag tat cat cag tt- --c cca cat tat tct taa att tca tat taa tgg gac gtg acg gaa tag tac agc atg ata gca cac atc cta ttc cca aca atg tca gga acg tca cgc tct cca cct tca tat tta ttt atc cgt -aa aaa tgt tat cct atg cat tgt -cc aat aat --a aaa gtc ac- --- --- --- -gc tat tgt agg aat ttt ttt tca cag ttg ctc aaa aac gat ggt agt gac tta tga gtt aca tta cat ttt gga gtc tca tct tta gta aac ata tca taa tat tcg ata tta cga gtt gac gca tct aac aaa ttc caa gta ttt gat ttt gga taa tat tcg tat ttt gca tct gct ata att aag ata taa tca cca caa gaa cac acg aac atc ttt cct aca tgg tta aag tac atg tat aat tct atc cat ttg tct tca tta act ata tat ttg tat aga taa tta cga gtc tca tga gta att cca gta att gca tag atg tca cca tcg tac tct aca gca taa act ata cta tga tgt cta ggc atg gga gac -tt ttt tat cca acg att ttt agt gaa aca ttc cac atc att taa tac tac ata ttt ctc ata cgt ggt ata aac tcc acc cat tac --a tat ata tca tcg ttt acg aat acc gat gct cct gaa tat cta gga gta att aag ttt gga agt ctt atc cat ttc gaa gcg cca tgt ttc aaa tat tct gct aca gcc gtt gaa ata gaa aat tct aat cct cct att aca tat aac ttt cca ttg tta aca caa gta cta act tct gat ttt aac gac gac ata tta gtg acc gtt ttc ca- --- --- --- --- --- --- --t ttt ttc gtt tta aga tct acc cgc -ga tac gga ata aac atg tct att gtt aat cat gcc gcc aat aat gta tag aca att atg taa aac att tgc att ata gaa ttg --- -tc tat ctg tat tac cga cta tcg tcc aat att ctg ttc tag gag agt aat ggg tta ttg tgg ata tat aat cag agt ttt taa tga cta cta tat tat gtt tta tac cat ttc gtg tca cag gct ttg tag att tgg ata tag tta atc cca aca atg cta tag cat tgc gca tag cat tag tca taa act tgg gat gta aaa tgt tga tga tat cta cat cgt ttg gat ttt tat gta tcc act tta ata ata tta tag ctg taa cat cct cat ga- ttt acg tta acg ttt tcg tgg gat aag ata gtt gtt agt tca tcc ttt gat aat ttt cca aat t

>MN912466.1:24030-25976_Ectromelia_virus_WH

cgg ctg cca tgt att ttc cgg aga gca agt --- aga tga tg- agg aac cag ata gtt tat atc cgt act tgc act taa agt ata cat tgt cgt tgt atg a-- gta tga tct ttt aaa ccc gct aga caa gta tcc gtt tga tat tgt agg atg tgg aca ttt aac aat ctg aca cgt ggg tgg atc gga cca ttc tcc tcc tga aca cag gac acc aga gtt acc aac caa tga ata tcc act att gca act ata cgt tac aac gct tcc ttc ggt ata aaa atc ctc gta tcc gtt atg tct tcc gtt gga tat aga tgg agg gga ttg gca ttt aac aga ttc gca aat ggg tgc ctc ggg att cca tac cat aga tcc agt aga tcc tag ttc aca ata cga ttt aga ttc tcc gat caa ctg ata tcc gct att gca aga gta cgt tat act aga gcc aaa gtc tgt tcc acc aat atc aag ttg acc att atc gat atc tcg agg tga tgg aca tct ccg ttt aat aca ttg att aaa gag tgt cca tcc att gcc tgt gca ttt aac ata tat gga tcc cat ttt ttg ctt tct gta tcc agg tag aca tac a-t att cta tag tat ctc cta ctt tgt a-- --- --- --- --- --- -ag tat cat cag tt- --c cca cat tat tct taa att tca tat taa tgg gac gtg acg gaa tag tac agc atg ata gca cac atc cta ttc cca aca atg tca gga acg tca cgc tct cca cct tca tat tta ttt atc cgt -aa aaa tgt tat cct atg cat tgt -cc aat aat --a aaa gtc ac- --- --- --- -gc tat tgt agg aat ttt ttt tca cag ttg ctc aaa aac gat ggt agt gac tta tga gtt aca tta cat ttt gga gtc tca tct tta gta aac ata tca taa tat tcg ata tta cga gtt gac gca tct aac aaa ttc caa gta ttt gat ttt gga taa tat tcg tat ttt gca tct gct ata att aag ata taa tca cca caa gaa cac acg aac atc ttt cct aca tgg tta aag tac atg tat aat tct atc cat ttg tct tca tta act ata tat ttg tat aga taa tta cga gtc tca tga gta att cca gta att gca tag atg tca cca tcg tac tct aca gca taa act ata cta tga tgt cta ggc atg gga gac -tt ttt tat cca acg att ttt agt gaa aca ttc cac atc att taa tac tac ata ttt ctc ata cgt ggt ata aac tcc acc cat tac --a tat ata tca tcg ttt acg aat acc gat gct cct gaa tat cta gga gta att aag ttt gga agt ctt atc cat ttc gaa gcg cca tgt ttc aaa tat tct gct aca gcc gtt gaa ata gaa aat tct aat cct cct att aca tat aac ttt cca ttg tta aca caa gta cta act tct gat ttt aac gac gac ata tta gtg acc gtt ttc ca- --- --- --- --- --- --- --t ttt ttc gtt tta aga tct acc cgc -ga tac gga ata aac atg tct att gtt aat cat gcc gcc aat aat gta tag aca att atg taa aac att tgc att ata gaa ttg --- -tc tat ctg tat tac cga cta tcg tcc aat att ctg ttc tag gag agt aat ggg tta ttg tgg ata tat aat cag agt ttt taa tga cta cta tat tat gtt tta tac cat ttc gtg tca cag gct ttg tag att tgg ata tag tta atc cca aca atg cta tag cat tgc gca tag cat tag tca taa act tgg gat gta aaa tgt tga tga tat cta cat cgt ttg gat ttt tat gta tcc act tta ata ata tta tag ctg taa cat cct cat ga- ttt acg tta acg ttt tcg tgg gat aag ata gtt gtt agt tca tcc ttt gat aat ttt cca aat t

>JQ410350.1:24341-26287_Ectromelia_virus_ERPV_culture-collection_ATCC:VR-1431_complete_genome

cgg ctg cca tgt att ttc cgg aga gca agt --- aga tga tg- agg aac cag ata gtt tat atc cgt act tgc act taa agt ata cat tgt cgt tgt atg a-- gta tga tct ttt aaa ccc gct aga caa gta tcc gtt tga tat tgt agg atg tgg aca ttt aac aat ctg aca cgt ggg tgg atc gga cca ttc tcc tcc tga aca cag gac acc aga gtt acc aac caa tga ata tcc act att gca act ata cgt tac aac gct tcc ttc ggt ata aaa atc ctc gta tcc gtt atg tct tcc gtt gga tat aga tgg agg gga ttg gca ttt aac aga ttc gca aat ggg tgc ctc ggg att cca tac cat aga tcc agt aga tcc tag ttc aca ata cga ttt aga ttc tcc gat caa ctg ata tcc gct att gca aga gta cgt tat act aga gcc aaa gtc tgt tcc acc aat atc aag ttg acc att atc gat atc tcg agg tga tgg aca tct ccg ttt aat aca ttg att aaa gag tgt cca tcc att gcc tgt gca ttt aac ata tat gga tcc cat ttt ttg ctt tct gta tcc agg tag aca tac a-t att cta tag tat ctc cta ctt tgt a-- --- --- --- --- --- -ag tat cat cag tt- --c cca cat tat tct taa att tca tat taa tgg gac gtg acg gaa tag tac agc atg ata gca cac atc cta ttc cca aca atg tca gga acg tca cgc tct cca cct tca tat tta ttt atc cgt -aa aaa tgt tat cct atg cat tgt -cc aat aat --a aaa gtc ac- --- --- --- -gc tat tgt agg aat ttt ttt tca cag ttg ctc aaa aac gat ggt agt gac tta tga gtt aca tta cat ttt gga gtc tca tct tta gta aac ata tca taa tat tcg ata tta cga gtt gac gca tct aac aaa ttc caa gta ttt gat ttt gga taa tat tcg tat ttt gca tct gct ata att aag ata taa tca cca caa gaa cac acg aac atc ttt cct aca tgg tta aag tac atg tat aat tct atc cat ttg tct tca tta act ata tat ttg tat aga taa tta cga gtc tca tga gta att cca gta att gca tag atg tca cca tcg tac tct aca gca taa act ata cta tga tgt cta ggc atg gga gac -tt ttt tat cca acg att ttt agt gaa aca ttc cac atc att taa tac tac ata ttt ctc ata cgt ggt ata aac tcc acc cat tac --a tat ata tca tcg ttt acg aat acc gat gct cct gaa tat cta gga gta att aag ttt gga agt ctt atc cat ttc gaa gcg cca tgt ttc aaa tat tct gct aca gcc gtt gaa ata gaa aat tct aat cct cct att aca tat aac ttt cca ttg tta aca caa gta cta act tct gat ttt aac gac gac ata tta gtg acc gtt ttc ca- --- --- --- --- --- --- --t ttt ttc gtt tta aga tct acc cgc -ga tac gga ata aac atg tct att gtt aat cat gcc gcc aat aat gta tag aca att atg taa aac att tgc att ata gaa ttg --- -tc tat ctg tat tac cga cta tcg tcc aat att ctg ttc tag gag agt aat ggg tta ttg tgg ata tat aat cag agt ttt taa tga cta cta tat tat gtt tta tac cat ttc gtg tca cag gct ttg tag att tgg ata tag tta atc cca aca atg cta tag cat tgc gca tag cat tag tca taa act tgg gat gta aaa tgt tga tga tat cta cat cgt ttg gat ttt tat gta tcc act tta ata ata tta tag ctg taa cat cct cat ga- ttt acg tta acg ttt tcg tgg gat aag ata gtt gtt agt tca tcc ttt gat aat ttt cca aat t

>KY549150.2:35064-37017_Cowpox_virus_isolate_CPXV_K4207

cgg ctg cca tgt att tcc tgg aga gca agt --- aga tga tg- agg aac cag ata gtt tat atc cgt gcc tgc acg taa agt gta cat tgt cgt tgt gtg a-- gta tga tct ttt aaa ccc gct aga caa gta tcc gtt tgt tat tga agg atg tgg gca ttt aac gat ctg aca cgt ggg tgg atc gga cca ttc tcc tcc tga aca cag gac acc aga gtt acc aat caa cga ata tcc act att gca act ata cgt tac aac gct tcc ttc ggt ata aaa atc ctc gta tcc gtt atg tct tcc gtt gga tat aga tgg agg gga tgg gca ttt aac aga ttc aca aat ggg tgc ctc ggg att cca cac cat aga tcc agt ata tcc tag ttc aca ata cga ttt aga ttc tcc tat caa ctg ata tcc gct att gca aga gta cgt tat act aga gcc aaa ctc tac tcc atc aat atc aat ttg acc att atc aat atc tcg agg cga tgg aca ttt ccg ttt aat aca ttg att aaa gag tgt cca tcc agt acc tgt aca ttt agc ata tat agg tcc cat ttt ttg ctt tct gta tcc agg tag aca tag a-t att cta tag tgt ctc cta tgt tgt a-- --- --- --- -at tag cat tag cat cag tc- --c cca cac tat tct taa att tca tat taa tgg gac gtg acg gaa tag tac agc atg ata gca cgc atc cta ttc cca aca atg tca gga acg tca cgc tct cca cct tca tat tta ttt atc cgt -aa aaa tgt tat ccc agg -ca ttg tcc aat aat a-a aaa gtc at- --- --- --- -gc tat tgt aaa aat tgt ttt tca cag ttg ctc aaa aac gat ggt agt gac tta cga gtt acg tta cac ttt gga gtc tca tct tta gta aac ata tca taa tat tcg ata tta cga gtt gac ata tcg aac aaa ttc caa gta ttt gat ttt gga taa tat tcg tat ttt gca tct gct ata att aag ata taa tca ccg caa gaa cac acg aac atc ttt cct aca tgg tta aag tac atg tac aat tct atc cat ttg tct tcc tta act ata tat ttg tat aga taa tta cga gtc tca tga gta att cca gta att aca tag atg tca ccg tcg tac tct aca gca taa act ata cta tga tgt cta ggc atg gga gac -tt ttt tat cca acg att ttt agt gaa aca ttc cac atc gtt taa tac tac ata ttt ctc ata cgt ggt ata aac tcc acc cat tac --a tat ata tca tcg ttt acg aat acc gat gcg cct gaa tat cta gga gta att aag ttt gga agt ctt atc cat ttc gaa gtg tcg tgt ttc aaa tat tct gct aca ccc gtt gaa ata gaa aat tct aat cct cct att aca tat aac ttt cca tcg tta aca caa gta cta act tct gat ttt aac gac gac ata tta gta acc gtt ttc ca- --- --- --- --- --- --- --t ttt ttc gtt tta aga tct acc cgc -ga cac gga ata aac atg tct att gtt aat cat gcc gcc aat aat gta tag aca att atg taa aac att tgc att ata gaa ttg --- -tc tat ctg tat tac caa cga tcg tcc aat att ctg ttc tag gag agt aat ggg tta ttg tgg ata tat aat cag agt ttt taa tga tca cta tat tat gtt tta tac cat ttc gtg tca ccg gct tta tag att tgg ata tag tta atc cca aca atg ata tag cat tgc gca tag cat tag tca taa act tgg gat gta aaa tgt tga tga tat cta cat cgt ttg gat ttt tat gta tcc act tta ata ata tca tag ctg taa cat cct cat ga- ttt acg tta acg tct tcg tgg gat aag ata gtt gtc aat tca tcc ttt gat aat ttt cca aat t

>KC813499.1:33422-35381_Cowpox_virus_strain_MarLei07/1_complete_genome

cgg ctt cca tgt att tcc tgg aga gca agt --- aga tga tg- aag aac cag ata ggt tat atc cgt act tgc act taa agt cta cat tgt cgt tgt atg a-- gta tga tct ttt aaa ccc gct aga caa gta tcc gtt tgt tat tga agg atg tgg gca ttt aac gat ctg aca cgt ggg tgg atc ggt cca ttc tcc tcc tga aca cac gat gcc aga gtt acc aat caa cga ata tcc ttt att gca act ata agt tac aac gct tcc atc ggt ata aaa atc ctc gta tcc gtt atg tct tcc att ggt tac aga tgg agg gga tgg aca ttt aac aga ttc aca aat ggg tgc ctc ggg att cca cac cat aga tcc agt ata tcc tag ttc aca ata cga ttt aga ttc tcc gat caa ctg ata tcc gct att gca aga gta cgt tat act aga gcc aaa ctc tac tcc acc aat atc aat ttg acc att ttc gat atc tcg agg tga tgg aca ttt ccg ttt aat aca ttg att aaa gag tgt cca tcc agt acc tgt aca ttt agc ata tat agg tcc cat ttt ttg ctt tct gta tcc agg tag aca tag a-t att cta tag tgt ctc cta tgt tgt a-- --- --- --- -at tag cat tag cat cag tc- --c cca cac tat tct taa att tca tat taa tgg gac gtg acg gaa tag tac agc atg ata gca cgc atc cta ttc cca aca atg tca gga acg tca cgc tct cca cct tca tat tta ttt atc cgt -aa aaa tgt tat ccc agg cca ttg tcc aat aat a-a aaa gtc ca- --- --t gtt ctc tat tgt agg aat tgt ttt tca cag ttg ctc aaa aac gat ggt agt gac tta cga gtt acg tta cac ttt gga gtc tca tct tta gta aac ata tca taa tat tcg ata tta cga gtt gac ata tcg aac aaa ttc caa gta ttt gat ttt gga taa tat tcg tat ttt gca tct gct ata att aag ata taa tca ccg caa gaa cac acg aac atc ttt cct aca tgg tta aag tac atg tac aat tct atc cat ttg tct tcc tta act ata tat ttg tat aga taa tta cga gtc tca tga gta att cca gta att aca tag atg tcg ccg tcg tac tct aca gca taa act ata cta tga tgt cta ggc atg gga gac -tt ttt tat cca acg att ttt agt gaa aca ttc cac atc gtt taa tac tac ata ttt ctc ata cgt ggt ata aac tcc acc cat tac --a tat ata tca tcg ttt acg aat acc gat gcg cct gaa tat cta gga gta att aag ttt gga agt ctt atc cat ctc gaa gtg ccg tgt ttc aaa tat tct gct aca cct gtt gaa ata gaa aat tct aat cct cct att aca tat aac ttt cca tcg tta aca caa gta cta act tct gat ttt aac gac gac ata tta gta acc gtt ttc ca- --- --- --- --- --- --- --t ttt ttc gtt tta aga tct act cgc -ga cac gga ata aac atg tct att gtt aat cat gcc gcc aat aag gta tag aca att atg taa aac att tgc att ata gaa ttg --- -tc tat ctg tat tac caa cga tcg tcc aat att ctg ttc tag gag agt aat ggg tta ttg tgg ata tat aat cag agt ttt taa tga cta cta tat tat gtt tta tac cat ttc gtg tca ccg gct ttg tag att tgg ata tag tta atc cca aca atg ata tag cat tgc gca tag tat tag tca taa act tgg gat gta aaa tgt tga tga tat ctg cat cgt ttg gat ttt tat gta tcc act tta ata ata tta tag ctg taa cat cct cat ga- ttt acg tta acg tct tcg ttg gat aag ata gtt gtc agt tca tcc ttt gat aat ttt cca aat t

>HQ420899.1:33482-35441_Cowpox_virus_strain_Norway_1994_MAN_complete_genome

cgg ctg cca tgt att tcc tgg aga gca agt --- aga tga tg- atg aac cag ata gtt tat atc cgt gcc tgc atg taa agt cta cat tgt cgt tgt gtg a-- gta tga tct ttt aaa ccc gct aga caa gta tcc gtt tgt tat tga agg atg tgg gca ttt aac gat ctg aca cgt ggg tgg atc gga cca ttc tcc tcc tga aca cac gat gcc aga gtt acc aat caa cga ata tcc ttt att gca act ata agt tac aac gct tcc atc ggt ata aaa atc ctc gta tcc gtt atg cct tcc att ggt tac aga tgg agg gga tgg aca ttt aac aga ttc aca aat ggg tgc ctc ggg att cca cac cat aga tcc agt ata tcc tag ttc aca ata cga ttt aga ttc tcc tat caa ctg ata tcc gct att gca aga gta cgt tat act aga gcc aaa atc tac tcc acc aat atc aat ttg acc att atc aat atc tcg agg aga tgg aca ttt acg ttt aat aca ttg att aaa gag tgt cca tcc agt acc tgt aca ttt agc ata tat agg tcc cat ttt ttg ctt tct gta tcc agg tag aca tag a-t att cta tag tgt ctc cta tgt tgt a-- --- --- --- -at tag aat tag cat tag tc- --t ctg cac tat tct taa att tca tat taa tgg gac gtg acg gaa tag tac agc atg ata gaa cgc atc cta ttc cca aca atg tca gga acg tca cgc tct cca cct tca tat tta ttt atc cgt -aa aaa tgt tat ccc agg cca ttg tcc aat aat a-a aaa gtc ca- --- --t gtt ctc tat tgt agg aat tgt ttt tca cag ttg ctc aaa aac gat ggt agt gac tta cga gtt aca tta cac ttt gga gtc tca tct tta gta aac ata tca taa tat tcg ata tta cga gtt gac ata tcg aac aaa ttc caa gta ttt gat ttt gga taa tat tcg tat ttt gca tct gct ata att aag ata taa tca ccg caa gaa cac acg aac atc ttt cct aca tgg tta aag tac atg tac aat tct atc cat ttg tct tcc tta act ata tat ttg tat aga taa tta cga gtc tca tga gta att cca gta att aca tag atg tcg ccg tcg tac tct aca gca taa act ata cta tga tgt cta ggc ata gga gac -tt ttt tat cca acg att ttt agt gaa aca ttc cac atc gtt taa tac tac ata ttt ctc ata cgt ggt ata aac tcc acc cat tac --a tat ata tca tcg ttt acg aat acc gat gcg cct gaa tat cta gga gta att aag ttt gga agt ctt atc cat ttc gaa gtg tcg tgt ttc aaa tat tct gct aca ccc gtt gaa ata gaa aat tct aat cct cct att aca tat aac ttt cca tcg tta aca caa gta cta act tct gat ttt aac gac gac ata tta gta acc gtt ttc ca- --- --- --- --- --- --- --t ttt tta gtt tta aga tct acc cgc -ga tac gga ata aac atg tct att gtt aat cat gcc gcc aat aat gta tag aca att atg taa aac att tgc att ata gaa ttg --- -tc tat ctg tat tac caa cga tcg tcc aat att ctg ttc tag gag agt aat ggg tta ttg tgg ata tat aat cag agt ttt taa tga tca cta tat tat gtt tta tac cat ttc gtg tca cag gct ttg tag att tgg ata tag tta atc cca aca atg cta tag cat tgc gca tag cat tag tca taa act tag gat gta aaa tgt tga tga tat cta cat cgt ttg gat ttt tat gta tcc act tta ata ata tta tag ctg taa cat cct cat ga- ttt acg tta acg tct tcg tgg gat aag ata gtt gtc agt tca tcc ttt gat aat ttt cca aat t

>AF482758.2:35149-37102_Cowpox_virus_strain_Brighton_Red_complete_genome

cgg ctg cca tgt att tcc tgg aga gca agt --- aga tga tg- agg aac cag ata gtt tat atc cgt gtc tgc act taa agt cta cat tgt cgt tgt gtg a-- gta tga tct ttt aaa ccc gct aga caa gta tcc gtt tgt tat tga agg atg tgg gca ttt aac gat ctg aca cgt ggg tgg atc gga cca ttc tcc tcc tga aca cac gat gcc aga gtt acc aat caa cga ata tcc act att gca act ata cgt tac aac gct tcc atc ggt ata aaa atc ctc gta tcc gtt atg cct tcc att ggt tac aga tgg agg gga tgg gca ttt aac aga ttc aca aat ggg tgc ctc ggg att cca cac cat aga tcc agt ata tcc tag ttc aca ata cga ttt aga ttc tcc tat caa ctg ata tcc gct att gca aga gta cgt tat act aga gcc aaa ctc tac tcc acc aat atc aat ttg acc att atc aat atc tcg agg tga tgg aca ttt ccg ttt aat aca ttg att aaa gag tgt cca tcc agt acc tgt aca ttt agc ata tat agg tcc cat ttt ttg ctt tct gta tcc agg tag aca tag a-t att cta tag tgt ctc cta tgt tgt a-- --- --- --- -at tag cat tag cat cag tc- --c cca cac tat tct taa att tca tat taa tgg gac gtg acg gaa tag tac agc atg ata gca cgc atc cta ttc cca aca atg tca gga acg tca cgc tct cca cct tca tat tta ttt atc cgt -aa aaa tgt tat cct atg -ca ttg tcc aat aat a-a aaa gtc at- --- --- --- -gc tat tgt agg aat tgt ttt tca cag ttg ctc aaa aac gat ggt agt gac tta cga gtt acg tta cac ttt gga gtc tca tct tta gta aac ata tca taa tat tcg ata tta cga gtt gac ata tcg aac aaa ttc caa gta ttt gat ttt gga taa tat tcg tat ttt gca tct gct ata att aag ata taa tca ccg caa gaa cac acg aac atc ttt cct aca tgg tta aag tac atg tac aat tct atc cat ttg tct tcc tta act ata tat ttg tat aga taa tta cga gtc tca tga gta att cca gta att aca tag atg tca ccg tcg tac tct aca gca taa act ata cta tga tgt cta ggc atg gga gac -tt ttt tat cca acg att ttt agt gaa aca ttc cac atc gtt taa tac tac ata ttt ctc ata cgt ggt ata aac tcc acc cat tac --a tat ata tca tcg ttt acg aat acc gat gcg cct gaa tat cta gga gta att aag ttt gga agt ctt atc cat ttc gaa gtg tcg tgt ttc aaa tat tct gct aca ccc gtt gaa ata gaa aat tct aat cct cct att aca tat aac ttt cca tcg tta aca caa gta cta act tct gat ttt aac gac gac ata tta gta acc gtt ttc ca- --- --- --- --- --- --- --t ttt ttc gtt tta aga tct acc cgc -ga cac gga ata aac atg tct att gtt aat cat tcc gcc aat aat gta tag aca att atg taa aac att tgc att ata gaa ttg --- -tc tat ctg tat tac cga cta tcg tcc aat att ctg ttc tag gag agt aat ggg tta ttg tgg ata tat aat cag agt ttt taa tga tca cta tat tat gtt tta tac cat ttc gtg tca ccg gct tta tag att tgg ata tag tta atc cca aca atg ata tag cat tgc gca tag cat tag tca taa act tgg gat gta aaa tgt tga tga tat cta cat cgt ttg gat ttt tat gta tcc act tta ata ata tca tag ctg taa cat cct cat ga- ttt acg tta acg tct tcg tgt gat aag ata gtt gtc agt tca tcc ttt gat aat ttt cca aat t

>HQ420896.1:33444-35397_Cowpox_virus_strain_Germany_1990_2_complete_genome

cgg ctg cca tgt att tcc tgg aga gca agt --- aga tga tg- agg aac cag ata gtt tat atc cgt gcc tgc atg taa agt cta cat tgt cgt tgt gtg a-- gta tga tct ttt aaa ccc gct aga caa gta tcc gtt tgt tat tga agg atg tgg gca ttt aac gat ctg aca cgt ggg tgg atc ggt cca ttc tcc tcc tga aca cac gat gcc aga gtt acc aat caa cga ata tcc ttt att gca act ata agt tac aac gct tcc atc ggt ata aaa atc ctc gta tcc gtt atg cct tcc att ggt tac aga tgg agg gga tgg aca ttt aac aga ttc aca aat ggg tgc ctc agg att cca cac cat aga tcc agt ata tcc tag ttc aca ata cga ttt aga ttc tcc gat caa ctg ata tcc gct att gca aga gta cgt tat act aga gcc aaa ctc tac tcc acc aat atc aat ttg acc att ttc gat atc tcg agg tga tgg aca ttt ccg ttt aat aca ttg att aaa gag tgt cca tcc agt acc tgt aca ttt agc ata tat agg tcc cat ttt ttg ctt tct gta tcc agg tag aca tag a-t att cta tag tgt ctc cta ctt tgt a-- --- --- --- --- --- -ag tat cat cag tc- --c cca cac tat tct taa att tca tat taa tgg gac gtg acg gaa tag tac agc atg ata gca cgc atc cta ttc cca aca atg tca gga acg tca cgc tct cca cct tca tat tta ttt atc cgt -aa aaa tgt tat ccc agg cca ttg tcc aat aat a-a aaa gtc ca- --- --t gtt ctc tat tgt agg aat tgt ttt tca cag ttg ctc aaa aac gat ggt agt gac tta cga gtt aca tta cac ttt ggt gtc tcg tct tta gta aac ata tca taa tat tcg ata tta cga gtt gac ata tcg aac aaa ttc caa gta ttt gat ttt gga taa tat tcg tat ttt gca tct gct ata att aag ata taa tca ccg caa gaa cac acg aac atc ttt cct aca tgg tta aag tac atg tac aat tct atc cat ttg tct tcc tta act ata tat ttg tat aga taa tta cga gtc tca tga gta att cca gta att aca tag atg tca ccg tca tac tct acg gca taa act ata cta tga tgt cta ggc atg gga gac -tt ttt tat cca acg att ttt agt gaa aca ttc cat atc gtt taa tac tac ata ttt ctc ata cgt agt ata aac tcc acc cat tac --a tat ata tca tcg ttt acg aat acc gat gcg cct gaa tat cta gga gta att aag ttt gga agc ctt atc cat ctc gaa gtg ccg tgt ttc aaa tat tct gct aca cct gtt gaa ata gaa aat tct aat cct cct att aca tat aac ttt cca tcg tta aca caa gta cta act tct gat ttt aac gac gac ata tta gta acc gtt ttc ca- --- --- --- --- --- --- --t ttt ttc gtt tta aga tct acc cgc -ga cac gga ata aac atg tct att gtt aat cat gcc gcc aat aat gta tag aca att atg taa aac att tgc att ata gaa ttg --- -tc tat ctg tat tac cga cta tcg tcc aat att ctg ttc tag gag agt aat ggg tta ttg tgg ata tat aat cag agt ttt taa tga cta cta tat tat gtt tta tac cat ttc gtg tca ccg gct ttg tag att tgg ata tag tta atc cca aca atg ata tag cat cgc gca tag cat tag tca taa act tgg gat gta aaa tgt tga tga tat cta cat cgt ttg gat ttt tat gta tcc act tta ata ata tta tag ctg taa cat cct cat ga- ttt acg tta acg tct tcg tgg gat aag ata gtt gtc agt tca tcc ttt gat aat ttt cca aat t

>LT883663.1:32519-34484_Cowpox_virus_strain_France_Amiens_2016_genome_assembly_complete_genome:_monopartite

cgg ctg cca tgt att tcc tgg aga gca agt --- aga tga tg- agg aac cag ata gtt tat atc cat act tgc acg taa agt gta cat tgt cgt tgt gtg a-- gta tga tct ttt aaa ccc gct aga caa gta tcc gtt tgt tat tga agg atg tgg gca ttt aac gat ctg aca cgt ggg cgg atc gga cca ttc tcc tcc tga aca cac gat gcc aga gtt acc aat caa cga ata tcc ttt att gca act ata agt tac aac gct tcc atc gat ata aaa atc ctc gta tcc gtt atg cct tcc att ggt tac aga tgg agg gga tgg aca ttt aac aga ttc aca aat ggg tgc ctc ggg att cca cac cat aga tcc agt ata tcc tag ttc aca ata cga ttt aga ttc tcc gat caa ctg ata tcc gct att gca aga gta cgt tat act aga gcc aaa ctc tac tcc acc aat atc aat ttg acc att ttc gat atc tcg agg tga tgg aca ttt ccg ttt aat aca ttg att aaa gag tgt cca tcc agt acc tgt aca ttt agc ata tat agg tcc cat ttt ttg ctt tct gta tcc agg tag aca tag a-t att cta tag tgt ctc cta tgt tgt aat tag c-- --- -at tag cat tag cat cag tc- --c cca cac tat tct taa att tca tat taa tgg gac gtg acg gaa tag tac agc atg ata gaa cgc atc cta ttc cca aca atg tca gga acg tca cgc tct cca cct tca tat tta ttt atc cgt -aa aaa tgt tat ccc agg cca ttg tcc aat aat a-a aaa gtc ca- --- --t gtt ctc tat tgt agg aat tgt ttt tca cag ttg ctc aaa aac gat ggt agt gac tta cga gtt aca tta cac ttt ggt gtc tcg tct tta gta aac ata tca taa tat tcg ata tta cga gtt gac ata tcg aac aaa ttc caa gta ttt gat ttt gga taa tat tcg tat ttt gca tct gct ata att aag ata taa tca ccg caa gaa cac acg aac atc ttt cct aca tgg tta aag tac atg tac aat tct atc cat ttg tct tcc tta act ata tat ttg tat aga taa tta cga gtc tca tga gta att cca gta att aca tag atg tcg ccg tcg tac tct aca gca taa act ata cta tga tgt cta ggc atg gga gac -tt ttt tat cca acg att ttt agt aaa aca ttc cat atc gtt taa tac tac ata ttt ctc ata cgt ggt ata aac tcc acc cat tac --a tat ata tca tcg ttt acg aat acc gat gcg cct gaa tat cta gga gta att aag ttt gga agt ctt atc cat ctc gaa gtg ccg tgt ttc aaa tat tct gct aca cct gtt gaa ata gaa aat tct aat cct cct att aca tat aac ttt cca tcg tta aca caa gta cta act tct gat ttt aac gac gac ata tta gta acc gtt ttc ca- --- --- --- --- --- --- --t ttt ttc gtt tta aga tct act cgc -ga cac gga ata aac atg tct att gtt aat cat gcc gcc aat aag gta tag aca att atg taa aac att tgc att ata gaa ttg --- -tc tat ctg tat tac cga cta tcg tcc aat att ctg ttc tag gag agt aat ggg tta ttg tgg ata tat aat cag agt ttt taa tga cta cta tat tat gtt tta tac cat ttc gtg tca ctg gct ttg tag att tgg ata tag tta atc cca aca atg ata tag cat tgc gca tag tat tag tca taa act tgg gat gta aaa tgt tga tga tat tta cat cgt ttg gat ttt tat gta tcc act tta ata ata tta tag ctg taa cat cct cat ga- ttt acg tta acg tct tcg ttg gat aag ata gtt gtc agt tca tcc ttt gat aat ttt cca aat t

>NC_055231.1:27166-29113_Orthopoxvirus_Abatino_complete_genome

cgg ctg cca tgt att tcc tgg aga gca agt --- aga tga tg- agg aac cag ata gtt tat atc cat act tgc acg taa agt gta cat agt cat tat acg a-- ata tga tct ttt aaa ccc gct aga caa gta tcc gtt tgc tat tgt agg atg ttg aca ttt aac aat ctg aca cgt ggg cgg atc tgt cca ctc tcc tcc tga aca cag gac acc cga gtt acc aac caa cga ata tcc act att gca act ata cgt tac aac gct tcc ttc ggt ata aaa atc ctc gta tcc gtt atg tct tcc gtt gga tat aga tgg agg gga ttg gca ttt aac aga ttc aca aat ggg tgc ctc ggg att cca tac cat aga tcc agt aga tcc tag ttc aca ata cga ttt aga ttc tcc gat caa ttt ata tcc act att aca aga gta cgt tat act gga tcc aaa gtc tgt tcc acc aat atc aag ttg acc att atc gat atc tcg agg tga tgg aca tct ccg ttt aag aca ttg att aaa gag tgt cca tcc att gcc tgt gca ttt agc ata tat gga tcc cat ttt ttg ctt tct gta ccc agg tag aca tac a-t att cta tag tgt ctc cta ctt tgt a-- --- --- --- --- --- -ag tat cat cag tc- --c cca cac tat tct taa att tca tat taa tgg gac gtg acg gaa tag tac agc atg ata gca cgc atc cta ttc cca aca atg tca gga acg tca cgc tct cca cct tca tat tta ttt atc cgt -aa aaa tgt tat cct atg caa tgt -cc aat aat a-a aaa ggc at- --- --- --- -gc tat tgt agg aat tgt ttt tca cag ttg ctc aaa aac gat ggt agt gac tta tga gtt aca tta cac ttt gga gtc tca tct tta gta aac ata tca taa tat tcg ata tta cga gtt gac ata tct aac aaa ttc caa gta ttt gat ttt gga taa tat tcg tat ttt gca tct gct ata att aag ata taa tcg ccg caa gaa cac acg aac atc ttt cct aca tgg tta aag tac atg tac aat tct atc cat ttg tct tcc tta act ata tat ttg tat aga taa tta cga gtc tca tga gta att cca gta att gca tag atg tca ccg tcg tac tct aca gca taa act ata cta tga tgt cta ggc atg gga gac -tt ttt tat cca acg att ttt agt gaa aca ttc cat atc gtt taa tac tac ata ttt ctc ata cgt ggt ata aac tcc acc cat tac --a tat ata tca tcg ttt acg aat acc gat gcg cct gaa tat cta gga gta att aag ttt gga agt ctt atc cat ttc gaa gtg ccg tgt ttc aaa tat tct gct aca cct gtt gaa ata gaa aat tct aat cct cct att aca tat aac ttt cca tcg tta aca caa gta cta act tct gat ttt aac gac gac ata tta gta acc gtt ttc ca- --- --- --- --- --- --- --t ttt ttc gtt tta aga tct acc cgt -ga cac gga ata aac atg cct att gtt aat cat gcc gcc aat aat gta tag aca att atg taa aac att tgc att ata gaa ttg --- -tc tat ctg tat tac cga cta tcg tcc aac att ctg ttc tag gag agt aat ggg tta ttg tgg ata tat aat cag agt ttt taa tga cta cta tat tat gtt tta tac cat ttc gtg tca cag gct ttg tag att cgg ata tag tta atc cca aca atg cta tag cat tgc gca tag cat tag tca taa act cgg gat gta aaa tgt tga tga tat cta cat cgt ttg gat ttt tat gta tcc act tta ata ata tta tag ctg taa cat cct cat ga- ttt acg tta acg ttt tcg tgg gat aag ata gtt gtc agt tca tcc ttt gat aat ttt cca aat t

>LT896724.1:33566-35525_Cowpox_virus_isolate_Ger/2015/Cat1_genome_assembly_complete_genome:_monopartite

cgg ctg cca tgt att tcc tgg aga gca agt --- aga tga tg- agg aac cag ata gtt tat atc cgt gcc tgc atg taa agt cta cat tgt cgt tgt gtg a-- gta tga tct ttt aaa ccc gct aga caa gta tcc gtt tgt tat tga agg atg tgg aca ttt aac gat ctg aca cgt ggg tgg atc ggt cca ttc tcc tcc tga aca cac gat gcc aga gtt acc aat caa cga ata tcc ttt att gca act ata agt tac aac gct tcc atc ggt ata aaa atc ctc gta tcc gtt atg cct tcc att ggt tac aga tgg agg gga tgg aca ttt aac aga ttc aca aat ggg tgc ctc ggg att cca cac cat aga tcc agt ata tcc tag ttc aca ata cga ttt aga ttc tcc gat caa ctg ata tcc gct att gca aga gta cgt tat act aga gcc aaa ctc tac tcc acc gat atc aat ttg acc att ttc gat atc tcg agg tga tgg aca ttt ccg ttt aat aca ttg att aaa gag tgt cca tcc agt acc tgt aca ttt agc ata tat agg tcc cat ttt ttg ctt tct gta tcc ggg tag aca tag a-t att cta tag tgt ctc cta tgt tgt a-- --- --- --- -at tag aat tag cat tag tc- --t ctg cac tat tct taa att tca tat taa tgg gac gtg acg gaa tag tac agc atg ata gca cgc atc cta ttc cca aca atg tca gga acg tca cgc tct cca cct tca tat tta ttt atc cgt -aa aaa tgt tat ccc agg cca ttg tcc aat aat a-a aaa gtc ca- --- --t gtt ctc tat tgt agg aat tgt ttt tca cag ttg ctc aaa aac gat ggt agt gac tta cga gtt aca tta cac ttt ggt gtc tcg tct tta gta aac ata tca taa tat tcg ata tta cga gtt gac ata tcg aac aaa ttc caa gta ttt gat ttt gga taa tat tcg tat ttt gca tct gct ata att aag ata taa tca ccg caa gaa cac acg aac atc ttt cct aca tgg tta aag tac atg tac aat tct atc cat ttg tct tcc tta act ata tat ttg tat aga taa tta cga gtc tca tga gta att cca gta att aca tag atg tca ccg tca tac tct acg gca taa act ata cta tga tgt cta ggc atg gga gac -tt ttt tat cca acg att ttt agt gaa aca ttc cat atc gtt taa tac tac ata ttt ctc ata cgt agt ata aac tcc acc cat tac --a tat ata tca tcg ttt acg aat acc gat gcg cct gaa tat cta gga gta att aag ttt gga agt ctt atc cat ttc gaa gtg ccg tgt ttc aaa tat tct gct aca ccc gtt gaa ata gaa aat tct aat cct cct att aca tat aac ttt cca tcg tta aca caa gta cta act tct gat ttt aac gac gac ata tta gta acc gtt ttc ca- --- --- --- --- --- --- --t ttt ttc gtt tta aga tct acc agc -ga cac gga ata aac atg tct att att aat cat gcc gcc aat aat gta tag aca att atg taa aac att tgc att ata gaa ttg --- -tc tat ctg tat tac cga cta tag tcc aat att ctg ttc tag gag agt aat ggg tta ttg tgg ata tat aat cag agt ttt taa tga cta cta tat tat gtt tta tac cat ttc gtg tca ccg gct ttg tag att tgg ata tag tta atc cca aca atg ata tag cat tgc gca tag tat tag tca taa act tgg gat gta aaa tgt tga tga tat ctg cat cgt ttg gat ttt tat gta tcc act tta ata ata tta tag ctg taa cat cct cat ga- ttt acg tta acg tct tcg tgg gat aag ata gtt gtc agt tca tcc ttt gat aat ttt cca aat t

>MG012795.1:7081-9037_Vaccinia_virus_isolate_CEyV1

cgg ctg cca tgt att tcc tgg aga gca agt --- aga tga tg- agg aac cag ata ggt tat atc cgt act tgc act caa agt tta cat tgt cga tgt atg a-- gta tga tct ttt aaa ccc gct aga caa gta tgc gtt tga tat tgt agg atg tgg aca ttt aac aat ttg aca cgt ggg tga atc gga cca ttc tcc tcc tga aca cag gac act aga gtt acc aat caa cga ata tct act act gca act ata cgt tac aac gct tcc atc ggt aga aaa atc ctc gta tcc gtt atg tct tcc gtt gga tat aga tgg atg gga ttg tca ttt aac aga ttc aca aat agg tgc ctc ggg att cca tac cat aga tcc agt aaa tcc taa ttc aca ata caa ttt aga ttc tcc tat caa ctg ata tcc gct att aca aga gta cgt tat act aga gcg aaa gtc tac tcc gcc aat atc aag ttg gcc att atc gat atc tcg agg cga tgg gca tct ccg ttt aat aca ttg att aaa gag tgt cca tcc agt act tgt aca ttt agc ata tat cgg tcc cat ttt ttg ctt cct gta tcc agg tag aca taa a-t att cta tag tgt ctc cta tgt tgt a-- --- --- --- --- --- -at tag cat tag tt- --t cca cac tat tct taa att tcc tat taa tgg gac gtg aag gaa tag gac agt atg aga gaa cgc att cta ttc cca aca atg tca gga acg tca cgc tct cca cct tca tat ttt ttt ttc cgt -aa aaa tgt tat cct gga cat cgt aca aat aat aaa aaa gcc ca- -ta tat gtt tgc tat tgt aga aat tgt ttt tca cac ttg ctc aaa aac gat ggc agt gat tta tga gtt acg taa cac ttt gga gtt tca tct tta gta aac ata tca taa tat ctg ata tta cga gtt gac ata tcg aac aaa ctc caa gta ttt gat ttt gga taa tat tcg tat ttt gca tct gct ata att aag ata taa tca ccg gaa gaa cac acg aac atc ttt cct aca tgg tta aag tac gtg tac tat tct atc cat ttg tct tcc tta act ata tat ttg gat aga taa tta cga gtc tcg tga gta att cca ata att aca tag atg tcg ccg tcg tac tct aca gca taa act ata cta tga tgt cta ggc atg gga gac -tt ttt tat cca ccg att ttt agt gaa aca ttc aac atc gtt taa tac tac ata ttt ttc ata cgt ggt ata aac tcc acc cat tac --a gat ata tca tcg ttt acg aat acc gac gcg cct gaa tat cta gga gta att aag ttt gga agt ctt atc cat ttt gaa gtg ccg tgt ttc aaa tat tct gct aca ccc gtt gaa ata gaa att tct aat cct cct att aca tat aac ttt cca ttg tta aca caa gta cga act tct gat ttt aac gac gac ata tta gta acc gtt ttc ca- --- --- --- --- --- --- --t ttt ttc gtt tca aga tct acc cgc -aa aac gga ata aac atg tct att gtt aat cat gcc gcc aag aat gta tag aca att atg taa aac att tgc att ata gaa ttg --- -tc tat cag tac tgc cga cta tcg tcc aat att ctg ttc tag gag agt aat ggg tta ttg tgg ata tat aat cag agt ttt taa tga cta cta tat tat gtt tta tac cat ttc gtg tca ctg gct ttg tag att tcg ata tag tta atc cta aca atg ata tag cat tgc gca tcg tat tag tta tta act tgg gat gta aaa tat tga tga tat cta gat cgt ttg gat ttt tat gta tcc act tta ata ata tct tag ctg taa cat cct cat ga- ttt acg tta acg tct tcg tgg gat aag ata gtt gtc agt tca tcc ttt gat aat ttt cca aa- -

>LT896721.1:34303-36250_Cowpox_virus_isolate_Ger_2010_MKY_genome_assembly_complete_genome:_monopartite

cgg ctg cca tgt att tcc tgg aga gca agt --- aga tga tg- agg aac cat ata gtt tat atc cat act tgc acg taa agt gta cat agt cat tat acg a-- ata tga tct ttt aaa ccc gct aga caa gta tcc gtt tgg tat tgt agg atg ttg aca ttt aac aat ctg aca cgt ggg cgg atc tgt cca ttc tcc tcc tga aca caa gat acc cga gtt acc aac caa cga ata tcc act att gca act ata agt tac aac gct tcc atc tgt ata aaa atc ctc gta tcc gtt atg tct tcc att ggt tac aga tgg agg gga ttg gca ttt aag aga ttc gca aat ggg tgc ccc ggg att cca cac cat aga tcc agt aga tcc taa ttc aca ata cga ttt aga ttc tcc gat caa ctg ata tcc gcc att gca aga gta cgt tat act aga gcc aaa gtc tac tcc act aat atc aat ttg acc att ttc gat atc tcg agg tga tgg aca tct ccg ttt aat aca ttg att aaa gag tga cca tcc att gcc tgt gca ttt agc ata tat gga tcc cat ttt ttg ctt tct gta tcc agg tag aca ttc a-t att cta tag tgt ctc cta ctt tgt a-- --- --- --- --- --- -ag tac cat cag tc- --c cca cac tat tct taa att tca tat taa tgg gac gtg acg gaa tag tac agc atg ata gca cgc atc cta ttc cca aca atg tca gga acg tca cgc tct cca cct tca tat tta ttt atc cgt -aa aaa tgt tat cct atg cat tga -cc aat aat a-a aaa gtc at- --- --- --- -gc tat tgt agg aat tgt ttt tca cag ttg ctc aaa aac gat gat agc gac tta tga gtt aca tta cac ttt gga gtc tca tct tta gta aac ata tca taa tat tcg ata tta cga gtt gac ata tct aac aaa ttc caa gta ttt gat ttt gga taa tat tcg tat ttt gca tct gct ata att aag ata taa tcg ccg caa gaa cac acg aac atc ttt cct aca tgg tta aag tac atg tac aat tct atc cat ttg tct tcc tta act ata tat ttg tat aga taa tta cga gtc tca tga gta att cca gta att aca tag atg tca ccg tcg tac tct acg gca taa act ata cta tga tgt cta ggc atg gga gac -tt ttt tat cca acg att ttt agt gaa aca ttc cat atc gtt taa tac tac ata ttt ctc ata cgt ggt ata aac tcc acc cat tac --a tat ata tca tcg ttt acg aat acc gat gcg cct gaa tat cta gga gta att aag ttt gga agc ctt atc cat ttc gaa gtg ccg tgt ttc aaa tat tct gcc aca cct gtt gaa ata gaa aat tct aat cct cct att aca tat aac ttt cca tcg tta aca caa gta cta act tct gat ttt aac gac gac ata tta gta acc gtt ttc ca- --- --- --- --- --- --- --t ttt ttc gtt tta aga tct acc cgc -ga tac gga ata aac atg tct att gtt aat cat gcc gcc aat aat gta tag aca att atg taa aac att tgc att ata gaa ttg --- -tc tat ctg tat tac cga cta tcg tcc aat att ctg ttc tag gag agt aat ggg tta ttg tgg ata tat aat cag agt ttt taa tga cta cta tat tat gtt tta tac cat ttc gtg tca ctg gct ttg tag att tgg ata tag tta atc cca aca atg ata tag cat tgc gca tag tat tag tca taa act tgg gat gta aaa tgt tga tga tat cta cat cgt ttg gat ttt tat gta tcc act tta ata ata tca tag ctg taa cat cct cat ga- ttt acg tta acg tct tcg tgg gat aag ata gtt gtc agt tca tcc ttt gat aat ttt cca aat t

>KC813510.1:34211-36158_Cowpox_virus_strain_HumGra07/1_complete_genome

cgg ctg cca tgt att tcc tgg aga gca agt --- aga tga tg- aag aac cag ata ggt tat atc cgt act tgc act taa agt gta cat tgt cgt tgt atg a-- gta tga tct ttt aaa ccc gct aga caa gta tcc gtt tag tat tgt agg atg tgg aca ttt aac aat ctg aca cgt ggg tgg atc tgt cca ctc tcc tcc tga aca cag gac acc cga gtt acc aac caa cga ata tcc act att gca act ata agt tac aac gct tcc atc tgt ata aaa atc ctc gta tcc gtt atg cct tcc att ggt tac aga tgg agg gga tgg gca ttt aag aga ttc gca aat ggg tgc ccc ggg att cca cac cat aga tcc agt ata tcc tag ttc aca ata cga ttt aga ttc tcc gat caa ctg ata tcc gct att gca aga gta cgt tat gct aga gcc aaa gtc tac tcc act aat atc aat ttg acc att ttc gat atc tcg agg cga tgg aca tct ccg ttt aat aca ttg att aaa gag tgt cca tcc att gcc tgt gca ttt agc ata tat gga tcc cat ttt ttg ctt tct gta tcc agg tag aca ttc a-t att cta tag tgt ctc cta ctt tgt a-- --- --- --- --- --- -ag tac cat cag tc- --c cca cac tat tct taa att tca tat taa tgg gac gtg acg gaa tag tac agc atg ata gca cgc atc cta ttc cca aca atg tca gga acg tca cgc tct cca cct tca tat tta ttt atc cgt -aa aaa cgt tat ctt ata cat tgt -cc aat aat a-a aaa gtc at- --- --- --- -gc tat tgt agg aat tgt ttt tca cag ttg ctc aaa aac gat gat agc gac tta tga gtt aca tta cac ttt gga gtc tca tct tta gta aac ata tca taa tat tcg ata tta cga gtt gac ata tct aac aaa ttc caa gta ttt gat ttt gga taa tat tcg tat ttt gca tct gct ata att aag ata taa tcg ccg caa gaa cac acg aac atc ttt cct aca tgg tta aag tac atg tac aat tct atc cat ttg tct tcc tta act ata tat ttg tat aga taa tta cga gtc tca tga gta att cca gta att aca tag atg tca cca tcg tac tct acg gca tac act ata cta tga tgt cta ggc atg gga gac -tt ttt tat cca acg att ttt agt gaa aca ttc cat atc gtt taa tac tac ata ttt ctc ata cgt ggt ata aac tcc acc cat tac --a tat ata tca tcg ttt acg aat acc gat gcg cct gaa tat cta gga gta att aag ttt gga agc ctt atc cat ttc gaa gtg ccg tgt ttc aaa tat tct gct acg cct gtt gaa ata gaa aat tct aat cct cct att aca tat aac ttt cca tca tta aca caa gta cta act tct gat ttt aac gac gac ata tta gta act gtt ttc ca- --- --- --- --- --- --- --t ttt ttc gtt tta aga tct acc cgc -ga tac gga ata aac atg tct att gtt aat cat gcc gcc aat aat gta tag aca att ata taa aac att tgc att ata gaa ttg --- -tc tat ctg tat tac cga cga ttg tcc aat att ctg ttc tag gag agt aat ggg tta ttg tgg ata tat aat cag agt ttt taa tga cta cta tat tat gtt tta tac cat ttc gtg tca cag gct ttg tag att tgg ata tag tta atc cca aca atg ata tag cat tgc aca tag tat tag tca taa act tgg gat gta aaa tgt tga tga tat cta cat cgt ttg gat ttt tat gta tcc act tta ata ata tca tag ctg taa cat cct cgt ga- ttt acg ttg acg ttt tcg tgg gat aag ata gtt gtt agt tca tcc ttt gat aat ttt cca aat t

>KY569020.1:24580-26544_Cowpox_virus_strain_CPXV_CheNuru_DK_2012

cgg ctg cca tgt att tcc tgg aga gca agt --- aga tga tg- agg aac cag ata gtt tat atc cgt gcc tgc atg taa agt gta cat tgt cgt tgt gtg a-- ata tga tct ttt aaa ccc gct aga caa gta tcc gtt tgt tat tga agg atg tgg gca ttt aac gat ctg aca cgt ggg cgg atc gga cca ttc tcc tcc tga aca cac gat gcc aga gtt acc aat caa cga ata tcc ttt att gca act ata agt tac aac gct tcc atc gat ata aaa atc ctc gta gcc gtt atg cct tcc att ggt tac aga tgg agg gga tgg aca ttt aac aga ttc aca aat ggg tgc ctc ggg att cca cac cat aga tcc agt ata tcc tag ttc aca ata cga ttt aga ttc tcc tat caa ctg ata tcc gct att gca aga gta cgt tat act aga gcc aaa ctc tac tcc acc aat atc aat ttg acc att ttc gat atc tcg agg tga tgg aca ttt acg ttt aat aca ttg att aaa gag tgt cca tcc agt acc tgt aca ttt agc ata tat agg tcc cat ttt ttg ctt tct gta tcc agg tag aca tag a-t att cta tag tgt ctc cta tgt tgt aat tag --- --- -at tag aat tag cat tag tc- --t ctg cac tat tct taa att tca tat taa tgg gac gtg acg gaa tag tac agc atg ata gaa cgc atc cta ttc cca aca atg tca gga acg tca cgc tct cca cct tca tat tta ttt atc cgt -aa aaa tgt tat ccc agg cca ttg tcc aat aat a-a aaa gtc ca- --- --t gtt atc tat tgt agg aat tgt ttt tca cag ttg ctc aaa aac gat ggt agt gac tta cga gtt aca tta cac ttt ggt gtc tcg tct tta gta aac ata tca taa tat tcg ata tta cga gtt gac ata tcg aac aaa ttc caa gta ttt gat ttt gga taa tat tcg tat ttt gca tct gct ata att aag ata taa tca ccg caa gaa cac acg aac atc ttt cct aca tgg tta aag tac atg tac aat tct atc cat ttg tct tcc tta act ata tat ttg tat aga taa tta cga gtc tca tga gta att cca gta att aca tag atg tca ccg tca tac tct acg gca taa act ata cta tga tgt cta ggc atg gga gac -tt ttt tat cca act att ttt agt gaa aca ttc cac atc gtt taa tac tac ata ttt ctc ata cgt ggt ata aac tcc acc cat tac --a tat ata tca tcg ttt acg aat acc gat gca cct gaa tat cta gga gta att aag ttt gga agt ctt atc cat ctc gaa gtg ccg tgt ttc aaa tat tct gct aca cct gtt gaa ata gaa aat tct aat cct cct att aca tat aac ttt cca tcg tta aca caa gta cta act tct gat ttt aac gac gac ata tta gta acc gtt ttc ca- --- --- --- --- --- --- --t ttt ttc gtt tta aga tct acc cgc -ga cac gga ata aac atg tct att gtt aat cat gcc gcc aat aat gta tag aca att atg taa aac att tgc att ata gaa ttg --- -tc tat ctg tat tac caa cga tcg tcc aat att ctg ttc tag gag agt aat ggg tta ttg tgg ata tat aat cag agt ttt taa tga tca cta tat tat gtt tta tac cat ttc gtg tca ccg gct ttg tag att tgg ata tag tta atc cca aca atg cta tag cat tgc gca tag cat tag tca taa act tgg gat gta aaa tgt tga tga tat ctg cat cgt ttg gat ttt tat gta tcc act tta ata ata tta tag ctg taa cat cct cat ga- ttt acg tta acg tct tcg tgg gat aag ata gtt gtc agt tca tcc ttt gat aat ttt cca aat t

>LT993226.1:33502-35468_Cowpox_virus_isolate_Ger/2014/Human_genome_assembly_complete_genome:_monopartite

cgg ctg cca tgt att tcc tgg aga gca agt --- aga tga tg- agg aac cag ata gtt tat atc cgt gcc tgc atg taa agt cta cat tgt cgt tgt gtg a-- gta tga tct ttt aaa ccc gct aga caa gta tcc gtt tgt tat tga agg atg tgg aca ttt aac gat ctg aca cgt ggg tgg atc ggt cca ttc tcc tcc tga aca cac gat gcc aga gtt acc aat caa cga ata tcc ttt att gca act ata agt tac aac gct tcc atc ggt ata aaa atc ctc gta tcc gtt atg cct tcc att ggt tac aga tgg agg gga tgg aca ttt aac aga ttc aca aat ggg tgc ctc ggg att cca cac cat aga tcc agt ata tcc tag ttc aca ata cga tgt aga ttc tcc tat caa ctg ata tcc gct att gca aga gta cgt tat act aga gcc aaa ctc tac tcc acc aat atc aat ttg acc att ttc gat atc tcg agg tga tgg aca ttt ccg ttt aat aca ttg att aaa gag tgt cca tcc agt acc tgt aca ttt agc ata tat agg tcc cat ttt ttg ctt tct gta tcc agg tag aca tag att att cta tag tgt ctc cta tgt tgt aat tag a-- --- -at tag aat tag cat tag tc- --t ctg cac tat tct taa att tca tat taa tgg gac gtg acg gaa tag tac agc atg ata gca cgc atc cta ttc cca aca atg tca gga acg tca cgc tct cca cct tca tat tta ttt atc cgt -aa aaa tgt tat ccc agg cca ttg tcc aat aat a-a aaa gtc ca- --- --t gtt ctc tat tgt agg aat tgt ttt tca cag ttg ctc aaa aac gat ggt agt gac tta cga gtt aca tta cac ttt gga gtc tca tct tta gta aac ata tca taa tat tcg ata tta cga gtt gac ata tcg aac aaa ttc caa gta ttt gat ttt gga taa tat tcg tat ttt gca tct gct ata att aag ata taa tca ccg caa gaa cac acg aac atc ttt cct aca tgg tta aag tac atg tac aat tct atc cat ttg tct tcc tta act ata tat ttg tat aga tag tta cga gtc tca tga gta att cca gta att aca tag atg tcg ccg tcg tac tct aca gca taa act ata cta tga tgt cta ggc atg gga gac -tt ttt tat cca acg att ttt agt gaa aca ttc cat atc gtt taa tac tac ata ttt ctc ata cgt ggt ata aac tcc acc cat tac --a tat ata tca tcg ttt acg aat acc gat gcg cct gaa tat cta gga gta att aag ttt gga agt ctt atc cat ctc gaa gtg ccg tgt ttc aaa tat tct gcc aca cct gtt gaa ata gaa aat tct aat cct cct att aca tat aac ttt cca tcg tta aca caa gta cta act tct gat ttt aac gac gac ata tta gta acc gtt ttc ca- --- --- --- --- --- --- --t ttt ttc gtt tta aga tct acc agc -ga cac gga ata aac atg tct att att aat cat gcc gcc aat aat gta tag aca att atg taa aac att tgc att ata gaa ttg --- -tc tat ctg tat tac cga cta tcg tcc aat att ctg ttc tag gag agt aat ggg tta ttg tgg ata tat aat cag agt ttt taa tga tca cta tat tat gtt tta tac cat ttc gtg tca ccg gct ttg tag att tgg ata tag tta atc cca aca atg ata tag cat tgc gca tag tat tag tca taa act tga gat gta aaa tgt tga tga tat ctg cat cgt ttg gat ttt tat gta tcc act tta ata ata tta tag ctg taa cat cct cat ga- ttt acg tta acg tct tcg tgg gat aag ata gtt gtc agt tca tcc ttt gat aat ttt cca aat t

>KY569022.1:24558-26523_Cowpox_virus_strain_CPXV_CheTopCut_DK_2011

cgg ctg cca tgt att tcc tgg aga gca agt --- aga tga tg- agg aac cag ata gtt tat atc cgt gcc tgc atg taa agt gta cat tgt cgt tgt gtg a-- ata tga tct ttt aaa ccc gct aga caa gta tcc gtt tgt tat tga agg atg tgg gca ttt aac gat ctg aca cgt ggg cgg atc gga cca ttc tcc tcc tga aca cac gat gcc aga gtt acc aat caa cga ata tcc ttt att gca act ata agt tac aac gct tcc atc gat ata aaa atc ctc gta gcc gtt atg cct tcc att ggt tac aga tgg agg gga tgg aca ttt aac aga ttc aca aat ggg tgc ctc ggg att cca cac cat aga tcc agt ata tcc tag ttc aca ata cga ttt aga ttc tcc tat caa ctg ata tcc gct att gca aga gta cgt tat act aga gcc aaa ctc tac tcc acc aat atc aat ttg acc att ttc gat atc tcg agg tga tgg aca ttt acg ttt aat aca ttg att aaa gag tgt cca tcc agt acc tgt aca ttt agc ata tat agg tcc cat ttt ttg ctt tct gta tcc agg tag aca tag a-t att cta tag tgt ctc cta tgt tgt aat tag a-- --- -tt tag aat tag cat tag tc- --t ctg cac tat tct taa att tca tat taa tgg gac gtg acg gaa tag tac agc atg ata gaa cgc atc cta ttc cca aca atg tca gga acg tca cgc tct cca cct tca tat tta ttt atc cgt -aa aaa tgt tat ccc agg cca ttg tcc aat aat a-a aaa gtc ca- --- --t gtt atc tat tgt agg aat tgt ttt tca cag ttg ctc aaa aac gat ggt agt gac tta cga gtt aca tta cac ttt ggt gtc tcg tct tta gta aac ata tca taa tat tcg ata tta cga gtt gac ata tcg aac aaa ttc caa gta ttt gat ttt gga taa tat tcg tat ttt gca tct gct ata att aag ata taa tca ccg caa gaa cac acg aac atc ttt cct aca tgg tta aag tac atg tac aat tct atc cat ttg tct tcc tta act ata tat ttg tat aga taa tta cga gtc tca tga gta att cca gta att aca tag atg tca ccg tca tac tct acg gca taa act ata cta tga tgt cta ggc atg gga gac -tt ttt tat cca act att ttt agt gaa aca ttc cac atc gtt taa tac tac ata ttt ctc ata cgt ggt ata aac tcc acc cat tac --a tat ata tca tcg ttt acg aat acc gat gca cct gaa tat cta gga gta att aag ttt gga agt ctt atc cat ctc gaa gtg ccg tgt ttc aaa tat tct gct aca cct gtt gaa ata gaa aat tct aat cct cct att aca tat aac ttt cca tcg tta aca caa gta cta act tct gat ttt aac gac gac ata tta gta acc gtt ttc ca- --- --- --- --- --- --- --t ttt ttc gtt tta aga tct acc cgc -ga cac gga ata aac atg tct att gtt aat cat gcc gcc aat aat gta tag aca att atg taa aac att tgc att ata gaa ttg --- -tc tat ctg tat tac caa cga tcg tcc aat att ctg ttc tag gag agt aat ggg tta ttg tgg ata tat aat cag agt ttt taa tga tca cta tat tat gtt tta tac cat ttc gtg tca ccg gct ttg tag att tgg ata tag tta atc cca aca atg cta tag cat tgc gca tag cat tag tca taa act tgg gat gta aaa tgt tga tga tat ctg cat cgt ttg gat ttt tat gta tcc act tta ata ata tta tag ctg taa cat cct cat ga- ttt acg tta acg tct tcg tgg gat aag ata gtt gtc agt tca tcc ttt gat aat ttt cca aat t

>KY569018.1:24598-26563_Cowpox_virus_strain_CPXV_CheHurley_DK_2012

cgg ctg cca tgt att tcc tgg aga gca agt --- aga tga tg- agg aac cag ata gtt tat atc cgt gcc tgc atg taa agt gta cat tgt cgt tgt gtg a-- ata tga tct ttt aaa ccc gct aga caa gta tcc gtt tgt tat tga agg atg tgg gca ttt aac gat ctg aca cgt ggg cgg atc gga cca ttc tcc tcc tga aca cac aat gcc aga gtt acc aat caa cga ata tcc ttt att gca act ata agt tac aac gct tcc atc gat ata aaa atc ctc gta gcc gtt atg cct tcc att ggt tac aga tgg agg gga tgg aca ttt aac aga ttc aca aat ggg tgc ctc ggg att cca cac cat aga tcc agt ata tcc tag ttc aca ata cga ttt aga ttc tcc tat caa ctg ata tcc gct att gca aga gta cgt tat act aga gcc aaa ctc tac tcc acc aat atc aat ttg acc att ttc gat atc tcg agg tga tgg aca ttt acg ttt aat aca ttg att aaa gag tgt cca tcc agt acc tgt aca ttt agc ata tat agg tcc cat ttt ttg ctt tct gta tcc agg tag aca tag a-t att cta tag tgt ctc cta tgt tgt aat tag a-- --- -at tag aat tag cat tag tc- --t ctg cac tat tct taa att tca tat taa tgg gac gtg acg gaa tag tac agc atg ata gaa cgc atc cta ttc cca aca atg tca gga acg tca cgc tct cca cct tca tat tta ttt atc cgt -aa aaa tgt tat ccc agg cca ttg tcc aat aat a-a aaa gtc ca- --- --t gtt atc tat tgt agg aat tgt ttt tca cag ttg ctc aaa aac gat ggt agt gac tta cga gtt aca tta cac ttt ggt gtc tcg tct tta gta aac ata tca taa tat tcg ata tta cga gtt gac ata tcg aac aaa ttc caa gta ttt gat ttt gga taa tat tcg tat ttt gca tct gct ata att aag ata taa tca ccg caa gaa cac acg aac atc ttt cct aca tgg tta aag tac atg tac aat tct atc cat ttg tct tcc tta act ata tat ttg tat aga taa tta cga gtc tca tga gta att cca gta att aca tag atg tca ccg tca tac tct acg gca taa act ata cta tga tgt cta ggc atg gga gac -tt ttt tat cca acg att ttt agt gaa aca ttc cac atc gtt taa tac tac ata ttt ctc ata cgt ggt ata aac tcc acc cat tac --a tat ata tca tcg ttt acg aat acc gat gca cct gaa tat cta gga gta att aag ttt gga agt ctt atc cat ctc gaa gtg ccg tgt ttc aaa tat tct gct aca cct gtt gaa ata gaa aat tct aat cct cct att aca tat aac ttt cca tcg tta aca caa gta cta act tct gat ttt aac gac gac ata tta gta acc gtt ttc ca- --- --- --- --- --- --- --t ttt ttc gtt tta aga tct acc cgc -ga cac gga ata aac atg tct att gtt aat cat gcc gcc aat aat gta tag aca att atg taa aac att tgc att ata gaa ttg --- -tc tat ctg tat tac caa cga tcg tcc aat att ctg ttc tag gag agt aat ggg tta ttg tgg ata tat aat cag agt ttt taa tga tca cta tat tat gtt tta tac cat ttc gtg tca ccg gct ttg tag att tgg ata tag tta atc cca aca atg cta tag cat tgc gca tag cat tag tca taa act tgg gat gta aaa tgt tga tga tat ctg cat cgt ttg gat ttt tat gta tcc act tta ata ata tta tag ctg taa cat cct cat ga- ttt acg tta acg tct tcg tgg gat aag ata gtt gtc agt tca tcc ttt gat aat ttt cca aat t

>LR812035.1:34140-36087_Cowpox_virus_isolate_GerMygEK_938/17_genome_assembly_complete_genome:_monopartite

cgg ctg cca tgt att tcc tgg aga gca agt --- aga tga tg- agg aac cat ata gtt tat atc cat act tgc acg taa agt gta cat agt cat tat acg a-- ata tga tct ttt aaa ccc gct aga caa gta tcc gtt tgg tat tgt agg atg ttg aca ttt aat aat ctg aca cgt ggg cgg atc tgt cca ttc tcc tcc tga aca caa gat acc cga gtt acc aac caa cga ata tcc act att gca act ata agt tac aac gct tcc atc tgt ata aaa atc ctc gta tcc gtt atg tct tcc att ggt tac aga tgg agg gga ttg gca ttt aag aga ttc gca aat ggg tgc ccc ggg att cca cac cat aga tcc agt aga tcc taa ttc aca ata cga ttt aga ttc tcc gat caa ctg ata tcc gcc att gca aga gta cgt tat act aga gcc aaa gtc tac tcc act aat atc aat ttg acc att ttc gat atc tcg agg tga tgg aca tct ccg ttt aat aca ttg att aaa gag tga cca tcc att gcc tgt gca ttt agc ata tat gga tcc cat ttt ttg ctt tct gta tcc agg tag aca ttc a-t att cta tag tgt ctc cta ctt tgt a-- --- --- --- --- --- -ag tac cat cag tc- --c cca cac tat tct taa att tca tat taa tgg gac gtg acg gaa tag tac agc atg ata gca cgc atc cta ttc cca aca atg tca gga acg tca cgc tct cca cct tca tat tta ttt atc cgt -aa aaa tgt tat cct atg cat tga -cc aat aat a-a aaa gtc at- --- --- --- -gc tat tgt agg aat tgt ttt tca cag ttg ctc aaa aac gat gat agc gac tta tga gtt aca tta cac ttt gga gtc tca tct tta gta aac ata tca taa tat tcg ata tta cga gtt gac ata tct aac aaa ttc caa gta ttt gat ttt gga taa tat tcg tat ttt gca tct gct ata att aag ata taa tcg ccg caa gaa cac acg aac atc ttt cct aca tgg tta aag tac atg tac aat tct atc cat ttg tct tcc tta act ata tat ttg tat aga taa tta cga gtc tca tga gta att cca gta att aca tag atg tca ccg tcg tac tct acg gca taa act ata cta tga tgt cta ggc atg gga gac -tt ttt tat cca acg att ttt agt gaa aca ttc cat atc gtt taa tac tac ata ttt ctc ata cgt ggt ata aac tcc acc cat tac --a tat ata tca tcg ttt acg aat acc gat gcg cct gaa tat cta gga gta att aag ttt gga agc ctt atc cat ttc gaa gtg ccg tgt ttt aaa tat tct gcc aca cct gtt gaa ata gaa aat tct aat cct cct att aca tat aac ttt cca tcg tta aca caa gta cta act tct gat ttt aac gac gac ata tta gta acc gtt ttc ca- --- --- --- --- --- --- --t ttt ttc gtt tta aga tct acc cgc -ga tac gga ata aac atg tct att gtt aat cat gcc gcc aat aat gta tag aca att atg taa aac att tgc att ata gaa ttg --- -tc tat ctg tat tac cga cta tcg tcc aat att ctg ttc tag gag agt aat ggg tta ttg tgg ata tat aat cag agt ttt taa tga cta cta tat tat gtt tta tac cat ttc gtg tca ctg gct ttg tag att tgg ata tag tta atc cca aca atg ata tag cat tgc gca tag tat tag tca taa act tgg gat gta aaa tgt tga tga tat cta cat cgt ttg gat ttt tat gta tcc act tta ata ata tca tag ctg taa cat cct cat ga- ttt acg tta acg tct tcg tgg gat aag ata gtt gtc agt tca tcc ttt gat aat ttt cca aat t

>KY569021.1:24607-26572_Cowpox_virus_strain_CPXV_CheGrey_DK_2010

cgg ctg cca tgt att tcc tgg aga gca agt --- aga tga tg- agg aac cag ata gtt tat atc cgt gcc tgc atg taa agt gta cat tgt cgt tgt gtg a-- ata tga tct ttt aaa ccc gct aga caa gta tcc gtt tgt tat tga agg atg tgg gca ttt aac gat ctg aca cgt ggg cgg atc gga cca ttc tcc tcc tga aca cac aat gcc aga gtt acc aat caa cga ata tcc ttt att gca act ata agt tac aac gct tcc atc gat ata aaa atc ctc gta gcc gtt atg cct tcc att ggt tac aga tgg agg gga tgg aca ttt aac aga ttc aca aat ggg tgc ctc ggg att cca cac cat aga tcc agt ata tcc tag ttc aca ata cga ttt aga ttc tcc tat caa ctg ata tcc gct att gca aga gta cgt tat act aga gcc aaa ctc tac tcc acc aat atc aat ttg acc att ttc gat atc tcg agg tga tgg aca ttt acg ttt aat aca ttg att aaa gag tgt cca tcc agt acc tgt aca ttt agc ata tat agg tcc cat ttt ttg ctt tct gta tcc agg tag aca tag a-t att cta tag tgt ctc cta tgt tgt aat tag a-- --- -at tag aat tag cat tag tc- --t ctg cac tat tct taa att tca tat taa tgg gac gtg acg gaa tag tac agc atg ata gaa cgc atc cta ttc cca aca atg tca gga acg tca cgc tct cca cct tca tat tta ttt atc cgt -aa aaa tgt tat ccc agg cca ttg tcc aat aat a-a aaa gtc ca- --- --t gtt atc tat tgt agg aat tgt ttt tca cag ttg ctc aaa aac gat ggt agt gac tta cga gtt aca tta cac ttt ggt gtc tcg tct tta gta aac ata tca taa tat tcg ata tta cga gtt gac ata tcg aac aaa ttc caa gta ttt gat ttt gga taa tat tcg tat ttt gca tct gct ata att aag ata taa tca ccg caa gaa cac acg aac atc ttt cct aca tgg tta aag tac atg tac aat tct atc cat ttg tct tcc tta act ata tat ttg tat aga taa tta cga gtc tca tga gta att cca gta att aca tag atg tca ccg tca tac tct acg gca taa act ata cta tga tgt cta ggc atg gga gac -tt ttt tat cca act att ttt agt gaa aca ttc cac atc gtt taa tac tac ata ttt ctc ata cgt ggt ata aac tcc acc cat tac --a tat ata tca tcg ttt acg aat acc gat gca cct gaa tat cta gga gta att aag ttt gga agt ctt atc cat ctc gaa gtg ccg tgt ttc aaa tat tct gct aca cct gtt gaa ata gaa aat tct aat cct cct att aca tat aac ttt cca tcg tta aca caa gta cta act tct gat ttt aac gac gac ata tta gta acc gtt ttc ca- --- --- --- --- --- --- --t ttt ttc gtt tta aga tct acc cgc -ga cac gga ata aac atg tct att gtt aat cat gcc gcc aat aat gta tag aca att atg taa aac att tgc att ata gaa ttg --- -tc tat ctg tat tac caa cga tcg tcc aat att ctg ttc tag gag agt aat ggg tta ttg tgg ata tat aat cag agt ttt taa tga tca cta tat tat gtt tta tac cat ttc gtg tca ccg gct ttg tag att tgg ata tag tta atc cca aca atg cta tag cat tgc gca tag cat tag tca taa act tgg gat gta aaa tgt tga tga tat ctg cat cgt ttg gat ttt tat gta tcc act tta ata ata tta tag ctg taa cat cct cat ga- ttt acg tta acg tct tcg tgg gat aag ata gtt gtc agt tca tcc ttt gat aat ttt cca aat t

>KY569019.1:33153-35118_Cowpox_virus_strain_CPXV_CheNova_DK_2014

cgg ctg cca tgt att tcc tgg aga gca agt --- aga tga tg- agg aac cag ata gtt tat atc cgt gcc tgc atg taa agt gta cat tgt cgt tgt gtg a-- ata tga tct ttt aaa ccc gct aga caa gta tcc gtt tgt tat tga agg atg tgg gca ttt aac gat ctg aca cgt ggg cgg atc gga cca ttc tcc tcc tga aca cac aat gcc aga gtt acc aat caa cga ata tcc ttt att gca act ata agt tac aac gct tcc atc gat ata aaa atc ctc gta gcc gtt atg cct tcc att ggt tac aga tgg agg gga tgg aca ttt aac aga ttc aca aat ggg tgc ctc ggg att cca cac cat aga tcc agt ata tcc tag ttc aca ata cga ttt aga ttc tcc tat caa ctg ata tcc gct att gca aga gta cgt tat act aga gcc aaa ctc tac tcc acc aat atc aat ttg acc att ttc gat atc tcg agg tga tgg aca ttt acg ttt aat aca ttg att aaa gag tgt cca tcc agt acc tgt aca ttt agc ata tat agg tcc cat ttt ttg ctt tct gta tcc agg tag aca tag a-t att cta tag tgt ctc cta tgt tgt aat tag a-- --- -at tag aat tag cat tag tc- --t ctg cac tat tct taa att tca tat taa tgg gac gtg acg gaa tag tac agc atg ata gaa cgc atc cta ttc cca aca atg tca gga acg tca cgc tct cca cct tca tat tta ttt atc cgt -aa aaa tgt tat ccc agg cca ttg tcc aat aat a-a aaa gtc ca- --- --t gtt atc tat tgt agg aat tgt ttt tca cag ttg ctc aaa aac gat ggt agt gac tta cga gtt aca tta cac ttt ggt gtc tcg tct tta gta aac ata tca taa tat tcg ata tta cga gtt gac ata tcg aac aaa ttc caa gta ttt gat ttt gga taa tat tcg tat ttt gca tct gct ata att aag ata taa tca ccg caa gaa cac acg aac atc ttt cct aca tgg tta aag tac atg tac aat tct atc cat ttg tct tcc tta act ata tat ttg tat aga taa tta cga gtc tca tga gta att cca gta att aca tag atg tca ccg tca tac tct acg gca taa act ata cta tga tgt cta ggc atg gga gac -tt ttt tat cca act att ttt agt gaa aca ttc cac atc gtt taa tac tac ata ttt ctc ata cgt ggt ata aac tcc acc cat tac --a tat ata tca tcg ttt acg aat acc gat gca cct gaa tat cta gga gta att aag ttt gga agt ctt atc cat ctc gaa gtg ccg tgt ttc aaa tat tct gct aca cct gtt gaa ata gaa aat tct aat cct cct att aca tat aac ttt cca tcg tta aca caa gta cta act tct gat ttt aac gac gac ata tta gta acc gtt ttc ca- --- --- --- --- --- --- --t ttt ttc gtt tta aga tct acc cgc -ga cac gga ata aac atg tct att gtt aat cat gcc gcc aat aat gta tag aca att atg taa aac att tgc att ata gaa ttg --- -tc tat ctg tat tac caa cga tcg tcc aat att ctg ttc tag gag agt aat ggg tta ttg tgg ata tat aat cag agt ttt taa tga tca cta tat tat gtt tta tac cat ttc gtg tca ccg gct ttg tag att tgg ata tag tta atc cca aca atg cta tag cat tgc gca tag cat tag tca taa act tgg gat gta aaa tgt tga tga tat ctg cat cgt ttg gat ttt tat gta tcc act tta ata ata tta tag ctg taa cat cct cat ga- ttt acg tta acg tct tcg tgg gat aag ata gtt gtc agt tca tcc ttt gat aat ttt cca aat t

>MG012796.1:18701-20664_Vaccinia_virus_isolate_CEyV2

cgg ctt cca tgt att tcc tgg aga gca ggt --- aga tga tg- agg aac cag ata gtt tat atc cgt att tac act taa agt tta cat tgt cgt ggt atg a-- gtg tga tct ttt aaa ccc gct aga caa gtg tcc ttt tgt ttt tgt tgc atg tgg aca ttt aac aat ctg aca cgt ggg tgg atc gga cca ttc tcc tcc tga aca cag gac acc aga gtt acc aat caa cga ata tcc act att gca act ata agt tac aac gct ccc atc ggt ata ata atc ctc gta tcc gtt atg tct tcc gtt gga tat aga tgg agg gga ttg gca ttt aac aga ttc aca aac agg tgc ctc ggg att cca tac cat aga tcc agt aga tcc taa ttc ata ata cga tta aga ttc acc gat caa atg ata tcc gct att aca agt gta cgt tat act aga gac aaa gtc tac tcc acc aat atc aag ttg gcc att atc gat atc tcg agg cga tgg gca tct ccg ttt aat aca ttg att aaa gag tgt cca tcc agt acc tgt aca ttt agc ata ttt agg tcc cat ttt ttg ctt tct gta ttc agg tag aca tac a-t ata cta tag tgt ctc cta tgt tgt a-- --- --- --- -aa tag cat tag cat cag tc- --t cca cgc tat tct taa att tcg tat taa tgg gtc gtg acg gaa cag tac agc atg ata gaa cgc atc cta ttt cca aca atg tca gga acg tca cgc tct cca cct tca tat tta ttt atc cgt -aa aga tgt tat ctt gga cat cgt aca aat aat a-a aaa gcc ca- -ta tat gtt cgc tat tgt aga aat tgt ttt tca cac ttg cgc aaa aac gat gtc agt gac tta tgt gtt acg tta cac ttt gga gtc tta tct tta gta aac att tca taa tat tct ata tta cga gtt gac ata tcg aac aaa ttc caa gta ttt gat ttt gga taa tat tcg tat tat gca tct gct ata att aag ata tta tca ccg caa gat cac acg aac atc ttt cct aca tgg tta gag tac atg tat aat tct atc cat ttg tct tcc tta act ata tat ctg tat aga taa tta cga gtc tcg tga gta att cca gta att aca tag agg tcg ccg tcg tac tct aca cca taa act ata cga tga tgt cta ggc aag gga gaa -tt ttt tat cca acg att ttt agt gaa aca ttc aac atc ggt taa tac tac aaa ttt ttc ata cgt ggt ata aac tcc agc cat tac --a tat tta tca acg ttt acg aat tcc gac gcg cct gaa tat cta gga gtt att aag ttt gga agt ctt atc cat ttc gaa gta ccg tgt ttc aaa tat tct acc aca ccc gtt gaa ata gaa aat act aat cct cct atg aca tat aac ttt cca tcg tta aca caa gta cta act tct gat ctt aac gac gac aca tta gta acc gtt ttc ca- --- --- --- --- --- --- --t ttt ttc gtt tcg aga tct acc cgc -ga tac gga ata aac atg act att gtt aat cat gcc gcc aat aat gta tag aca att atg taa aac att tgc att ata gaa ttg --- -tc tat ctg tat tac cga cta tcg tcc aat att ctg ttc tag gag agt aaa ggg tta ttg tgg ata tat aat cag agt ttt taa tga cta cta tat tat gtt tta tac cat ttc gtg tca ctg gct ttg tag att tgg atc tag tta atc cca aca atg ata tga cat tgc gca tag tat tag tca taa act tgg gat gta aaa tct tga tga tat cta cat cgt ttg aat ttt tat cga tcc act tta ata ata tca tag atg taa cat cct cat ga- ttt acg tta acg ttt tcg tgg gat aag ata gtt gtc agt tca ccc ttt gat aat ttt cca aat t

>BK010317.1:13031-14980_TPA_asm:_Variola_virus_isolate_VD21

cgg ttg cca tgt att tcc tgg aga gca agt --- aga tga tg- agg aac cag ata gtt tat atc cgt act tgc acg taa agt cta cat tgt cat tat acg a-- ata tga tct ttt aaa ccc gct aga caa gta tcc gtt taa tat tgt agg atg tgg aca ttt aac aat ctg aca cgt ggg tgg att aga cca ttc tcc tcc tga aca cag gac acc aga gtt acc aat caa cga ata tcc act att gca act ata agt tac aac gct tcc atc ggt ata aaa atc att ata tcc gtt atg tct tcc gtt gga tat aga tgg agg taa ttg gca ttt aac aga ttc gca aat agg tgc ctt ggg att cca tac cat aga tcc agt aga tcc taa ttc aca ata cga ttt ata ttc acc gat caa ata ata tcc gct att aca aga gta cgt tat act aga gcc aaa gtc tac tcc gcc aat atc aag atg gcc att atc gat atc tcg agg cga cgg gca tct ccg ttt aat aca ttg att aaa gag tgt cca tcc agt acc tgt aca ttt agc ata tat agg tcc cat ttt ttg ctt tct gta tcc agg tag aca tag a-t att cta tag tgt ctc cta tgt tgt a-- --- --- --- -at tag cat tag cat cag tc- --t cca cac tat tct taa att tca tat taa tgg gtc gtg acg gaa tag tac agc atg ata gaa cgc atc cta ttc cca aca atg tca gga acg tca ctc tct cca cct tca tat tta ttt atc cgt -aa aaa tgt tat cct gga cat tgt gcc aat aat aaa aaa gcc ca- -ta tat gtt cgc tat tgt aga aat tgt ttt tca tag ttg ctc aaa aac gat ggc ggt gac tta tga gtc acg tta cac ttt gga gtc tca tct tta gta aac ata tca taa tat tcg ata tta cga gtt gac aca ttg aac aaa ttc caa gta ttt gat ttt gga taa tat tcg tat ttt gca tct gct ata att aag ata taa tcg cca caa gaa cac gcg aac atc ttt cct aca tgg tta aag tac atg tac aat tct atc cat ttg tct tcc tta act ata tat ttg tat aga taa tta cga gtc tcg tgt gta att cca gta att gca tag atg tta cca tcg tac tct aca gca taa act ata cta taa tgt cta ggc atg gga gac -tt ttt tat cca aca att ttt agt gaa aca ttc cac atc gtt taa tat tac ata ttt ttc ata cgt ggt ata aac tcc gcc cat tac --a tat ata tca tca ttt acg aat acc gac gcg tct gaa tat cta gga gta att aag ttt gga tgt ctt atc cat ttc aaa gtg tag cgt ttt aaa tat tct gct aca cct gtt gaa ata gaa aat ttt aat cct cct att aca tat aac ttt cca tca tta aca cac gta cta act ttt gat ttt aac gac gac ata tta gta act gtt ttc ca- --- --- --- --- --- --- -tt ttt ttc gtt tta aga tct acc cac gga tac gaa ata aac atg tct att gtt aat cat gct gcc aat aat gta tag aca att ata taa aac att tgc att ata gaa ttg tct atc tat ctg tat tac cga cga ttg tcc aat att ctg ttc tag gag agt aat ggg tta ttg tgg ata tat aat cag agt ttt taa tga cta cta tat tat gtt tta tac cat ttc gtg tca cag gct ttg tag att tgg a-- --- --- --- --- --- --- -ta tag cat tgc aca tag tat tag tca taa act tgg gat gta aaa tgt tga tga tat cta cat cgt ttg gat ttt tat gta tcc act tta ata ata tca tag ctg taa cat cct cgt ga- ttt acg ttg acg ttt tcg tgg gat aag ata gtt att agt tca tcc ttt gat aat ttt cca aat t

>DQ441437.1:12933-14878_Variola_virus_strain_Sierra_Leone_1969_(V68-258)_complete_genome

cgg ttg cca tgt att tcc tgg aga gca agt --- aga tga tg- agg aac cag ata gtt tat atc cgt act tgc acg taa agt cta cat tgt cat tat acg a-- ata tga tct ttt aaa ccc gct aga caa gta tcc gtt taa tat tgt agg atg tgg aca ttt aac aat ctg aca cgt ggg tgg att aga cca ttc tcc tcc tga aca cag gac acc aga gtt acc aat caa cga ata tcc act att gca act ata agt tac aac gct tcc atc ggt ata aaa atc att ata tcc gtt atg tct tcc gtt gga tat aga tgg agg taa ttg gca ttt aac aga ttc gca aat agg tgc ctt ggg att cca tac cat aga tcc agt aga tcc taa ttt aca ata cga ttt ata ttc acc gat caa ata ata tcc gct att aca aga gta cgt tat act aga gcc aaa gtc tac tcc gcc aat atc aag atg gcc att atc gat atc tcg agg cga cgg gca tct ccg ttt aat aca ttg att aaa gag tgt cca tcc agt acc tgt aca ttt agc ata tat agg tcc cat ttt ttg ctt tct gta tcc agg tag aca tag a-t att cta tag tgt ctc cta tgt tgt a-- --- --- --- -at tag cat tag cat cag tc- --t cca cac tat tct taa att tca tat taa tgg gtc gtg acg gaa tag tac agc atg ata gaa cgc atc cta ttc cca aca atg tca gga acg tca ctc tct cca cct tca tat tta ttt atc cgt -aa aaa tgt tat cct gga cat tgt gtc aat aat aaa aaa gct ca- -ta tat gtt tgc tat tgt aga aat tgt ttt tca tag ttg ctc aaa aac gat ggc ggt gac tta tga gtc acg tta cac ttt gga gtc tca tct tta gta aac ata tca taa tat tcg ata tta cga gtt gac ata ttg aac aaa ttc caa gta ttt gat ttt gga taa tat tcg tat ttt gca tct gct ata att aag ata taa tcg cca caa gaa cac acg aac atc ttt cct aca tgg tta aag tac atg tac aat tct atc cat ttg tct tcc tta act ata tat ttg tat aga taa tta cga gtc tcg tgt gta att cca gta att gca tag atg tta cca tcg tac tct aca gca taa act ata cta tga tgt cta ggc atg gga gac -tt ttt tat cca aca att ttt agt gaa aca ttc cac atc gtt taa tat tac ata ttt ttc ata cgt ggt ata aac tcc acc cat tac --a tat ata tca tca ttt acg aat acc gac gcg tct gaa tat cta gga gta att aag ttt gga tgt ctt atc cat ttc gaa gtg tag cgt ttt aaa tat tct gct aca cct gtt gaa ata gaa aat ttt aat cct cct att aca tat aac ttt cca tca tta aca cac gta ct- act ttt gat ttt aac gac gac ata tta gta act gtt ttc ca- --- --- --- --- --- --- -tt ttt ttc gtt tta aga tct acc cac gga tac gaa ata aac atg tct att gtt aat cat gcc gcc aat aat gta tag aca att ata taa aac att tgc att ata gaa ttg tct atc tat ctg tat tac cga caa ttg tcc aat att ctg ttc tag gag agt aat ggg tta ttg tgg ata tat aat cag agt ttt taa tga cta cta tat tat gtt tta tac cat ttc gtg tca cag gct ttg tag att tgg a-- --- --- --- --- --- --- -ta tag cat tgc aca tag tat tag tca taa act tgg gat gta aaa tgt tga tga tat cta cat cgt ttg gat ttt tat gta tcc act tta ata ata tca tag ctg taa cat cct tgt gat ttt acg ttg acg ttt tcg tgg gat aag ata gtt --- -gt tca tcc ttt gat aat ttt cca aat t

>DQ441416.1:13002-14947_Variola_virus_strain_Benin_Dahomey_1968_(v68-59)_complete_genome

cgg ttg cca tgt att tcc tgg aga gca agt --- aga tga tg- agg aac cag ata gtt tat atc cgt act tgc acg taa agt cta cat tgt cat tat acg a-- ata tga tct ttt aaa ccc gct aga caa gta tcc gtt taa tat tgt agg atg tgg aca ttt aac aat ctg aca cgt ggg tgg att aga cca ttc tcc tcc tga aca cag gac acc aga gtt acc aat caa cga ata tcc act att gca act ata agt tac aac gct tcc atc ggt ata aaa atc att ata tcc gtt atg tct tcc gtt gga tat aga tgg agg taa ttg gca ttt aac aga ttc gca aat agg tgc ctt ggg att cca tac cat aga tcc agt aga tcc taa ttt aca ata cga ttt ata ttc acc gat caa ata ata tcc gct att aca aga gta cgt tat act aga gcc aaa gtc tac tcc gcc aat atc aag atg gcc att atc gat atc tcg agg cga cgg gca tct ccg ttt aat aca ttg att aaa gag tgt cca tcc agt acc tgt aca ttt agc ata tat agg tcc cat ttt ttg ctt tct gta tcc agg tag aca tag a-t att cta tag tgt ctc cta tgt tgt a-- --- --- --- -at tag cat tag cat cag tc- --t cca cac tat tct taa att tca tat taa tgg gtc gtg acg gaa tag tac agc atg ata gaa cgc atc cta ttc cca aca atg tca gga acg tca ctc tct cca cct tca tat tta ttt atc cgt -aa aaa tgt tat cct gga cat tgt gtc aat aat aaa aaa gct ca- -ta tat gtt tgc tat tgt aga aat tgt ttt tca tag ttg ctc aaa aac gat ggc ggt gac tta tga gtc acg tta cac ttt gga gtc tca tct tta gta aac ata tca taa tat tcg ata tta cga gtt gac ata ttg aac aaa ttc caa gta ttt gat ttt gga taa tat tcg tat ttt gca tct gct ata att aag ata taa tcg cca caa gaa cac acg aac atc ttt cct aca tgg tta aag tac atg tac aat tct atc cat ttg tct tcc tta act ata tat ttg tat aga taa tta cga gtc tcg tgt gta att cca gta att gca tag atg tta cca tcg tac tct aca gca taa act ata cta tga tgt cta ggc atg gga gac -tt ttt tat cca aca att ttt agt gaa aca ttc cac atc gtt taa tat tac ata ttt ttc ata cgt ggt ata aac tcc acc cat tac --a tat ata tca tca ttt acg aat acc gac gcg tct gaa tat cta gga gta att aag ttt gga tgt ctt atc cat ttc gaa gtg tag cgt ttt aaa tat tct gct aca cct gtt gaa ata gaa aat ttt aat cct cct att aca tat aac ttt cca tca tta aca cac gta ct- act ttt gat ttt aac gac gac ata tta gta act gtt ttc ca- --- --- --- --- --- --- -tt ttt ttc gtt tta aga tct acc cac gga tac gaa ata aac atg tct att gtt aat cat gcc gcc aat aat gta tag aca att ata taa aac att tgc att ata gaa ttg tct atc tat ctg tat tac cga caa ttg tcc aat att ctg ttc tag gag agt aat ggg tta ttg tgg ata tat aat cag agt ttt taa tga cta cta tat tat gtt tta tac cat ttc gtg tca cag gct ttg tag att tgg a-- --- --- --- --- --- --- -ta tag cat tgc aca tag tat tag tca taa act tgg gat gta aaa tgt tga tga tat cta cat cgt ttg gat ttt tat gta tcc act tta ata ata tca tag ctg taa cat cct tgt gat ttt acg ttg acg ttt tcg tgg gat aag ata gtt --- -gt tca tcc ttt gat aat ttt cca aat t

>KC813494.1:25876-27847_Cowpox_virus_strain_HumLue09/1_complete_genome

cgg ctg cca tgt att tcc tgg aga gca agt --- aga tga tg- agg aac cag ata gtt tat atc cgt gcc tgc atg taa agt gta cat tgt cgt tgt gtg a-- ata tga tct ttt aaa ccc gct aga caa gta tcc gtt tgt tat tga agg atg tgg gca ttt aac gat ctg aca cgt ggg tgg atc ggt cca ttc tcc tcc tga aca cac gat gcc aga gtt acc aat caa cga ata tcc ttt att gca act ata agt tac aac gct tcc atc ggt ata aaa atc ctc gta tcc gtt atg cct tcc att ggt tac aga tgg agg gga tgg aca ttt aac aga ttc aca aat ggg tgc ctc ggg att cca cac cat aga tcc agt ata tcc tag ttc aca ata cga ttt aga ttc tcc tat caa ctg ata tcc gct att gca aga gta cgt tat act aga gcc aaa ctc tac tcc acc aat atc aat ttg acc att atc aat atc tcg agg aga tgg aca ttt ccg ttt aat aca ttg att aaa gag tgt cca tcc agt acc tgt aca ttt agc ata tat agg tcc cat ttt ttg ctt tct gta tcc agg tag aca tag a-t att cta tag tgt ctc cta tgt tgt aat tag aat tag aat tag aat tag cat tag tc- --t ctg cac tat tct taa att tca tat taa tgg gac gtg acg gaa tag tac agc atg ata gaa cgc atc cta ttc cca aca atg tca gga acg tca cgc tct cca cct tca tat tta ttt atc cgt -aa aaa tgt tat ccc agg cca ttg tcc aat aat a-a aaa gtc ca- --- --t gtt ctc tat tgt agg aat tgt ttt tca cag ttg ctc aaa aac gat ggt agt gac tta cga gtt aca tta cac ttt ggt gtc tca tct tta gta aac ata tca taa tat tcg ata tta cga gtt gac ata tct aac aaa ttc caa gta ttt gat ttt gga taa tat tcg tat ttt gca tct gct ata att aag ata taa tca ccg caa gaa cac acg aac atc ttt cct aca tgg tta aag tac atg tac aat tct atc cat ttg tct tcc tta act ata tat ttg tat aga taa tta cga gtc tca tga gta att cca gta att aca tag ata tca ccg tca tac tct acg gca taa act ata cta tga tgt cta ggc atg gga gac -tt ttt tat cca acg att ttt agt gaa aca ttc cat atc gtt taa tac tac ata ttt ctc ata cgt agt ata aac tcc acc cat tac --a tat ata tca tcg ttt acg aat aca gat gcg cct gaa tat cta gga gta att aag ttt gga agc ctt atc cat ctc gaa gtg ccg tgt ttc aaa tat tct gct aca cct gtt gaa ata gaa aat tct aat cct cct att aca tat aac ttt cca tcg tta aca caa gta cta act tct gat ttt aac gac gac ata tta gta acc gtt ttc ca- --- --- --- --- --- --- --t ttt ttc gtt tta aga tct acc agc -ga cac gga ata aac atg tct att gtt aat cat gcc acc aat aag gta tag aca att atg taa aac att tgc att ata gaa ttg --- -tc tat ctg tat tac caa cga tcg tcc aat att ctg ttc tag gag agt aat ggg tta ttg tgg ata tat aat cag agt ttt taa tga tca cta tat tat gtt tta tac cat ttc gtg tca ccg gct ttg tag att tgg ata tag tta atc cca aca atg ata tag cat tgc gca tag tat tag tca taa act tgg gat gta aaa tgt tga tga tat ctg cat cgt ttg gat ttt tat gta tcc act tta ata ata tta tag ctg taa cat cct cat ga- ttt acg tta acg tct tcg tgt gat aag ata gtt gtc agt tca tcc ttt gat aat ttt cca aat t

>DQ441434.1:12865-14810_Variola_virus_strain_Niger_1969_(001_importation_from_Nigeria)_complete_genome

cgg ttg cca tgt att tcc tgg aga gca agt --- aga tga tg- agg aac cag ata gtt tat atc cgt act tgc acg taa agt cta cat tgt cat tat acg a-- ata tga tct ttt aaa ccc gct aga caa gta tcc gtt taa tat tgt agg atg tgg aca ttt aac aat ctg aca cgt ggg tgg att aga cca ttc tcc tcc tga aca cag gac acc aga gtt acc aat caa cga ata tcc act att gca act ata agt tac aac gct tcc atc ggt ata aaa atc att ata tcc gtt atg tct tcc gtt gga tat aga tgg agg taa ttg gca ttt aac aga ttc gca aat agg tgc ctt ggg att cca tac cat aga tcc agt aga tcc taa ttt aca ata cga ttt ata ttc acc gat caa ata ata tcc gct att aca aga gta cgt tat act aga gcc aaa gtc tac tcc gcc aat atc aag atg gcc att atc gat atc tcg agg cga cgg gca tct ccg ttt aat aca ttg att aaa gag tgt cca tcc agt acc tgt aca ttt agc ata tat agg tcc cat ttt ttg ctt tct gta tcc agg tag aca tag a-t att cta tag tgt ctc cta tgt tgt a-- --- --- --- -at tag cat tag cat cag tc- --t cca cac tat tct taa att tca tat taa tgg gtc gtg acg gaa tag tac agc atg ata gaa cgc atc cta ttc cca aca atg tca gga acg tca ctc tct cca cct tca tat tta ttt atc cgt -aa aaa tgt tat cct gga cat tgt gtc aat aat aaa aaa gct ca- -ta tat gtt tgc tat tgt aga aat tgt ttt tca tag ttg ctc aaa aac gat ggc ggt gac tta tga gtc acg tta cac ttt gga gtc tca tct tta gta aac ata tca taa tat tcg ata tta cga gtt gac ata ttg aac aaa ttc caa gta ttt gat ttt gga taa tat tcg tat ttt gca tct gct ata att aag ata taa tcg cca caa gaa cac acg aac atc ttt cct aca tgg tta aag tac atg tac aat tct atc cat ttg tct tcc tta act ata tat ttg tat aga taa tta cga gtc tcg tgt gta att cca gta att gca tag atg tta cca tcg tac tct aca gca taa act ata cta tga tgt cta ggc atg gga gac -tt ttt tat cca aca att ttt agt gaa aca ttc cac atc gtt taa tat tac ata ttt ttc ata cgt ggt ata aac tcc acc cat tac --a tat ata tca tca ttt acg aat acc gac gcg tct gaa tat cta gga gta att aag ttt gga tgt ctt atc cat ttc gaa gtg tag cgt ttt aaa tat tct gct aca cct gtt gaa ata gaa aat ttt aat cct cct att aca tat aac ttt cca tca tta aca cac gta ct- act ttt gat ttt aac gac gac ata tta gta act gtt ttc ca- --- --- --- --- --- --- -tt ttt ttc gtt tta aga tct acc cac gga tac gaa ata aac atg tct att gtt aat cat gcc gcc aat aat gta tag aca att ata taa aac att tgc att ata gaa ttg tct atc tat ctg tat tac cga caa ttg tcc aat att ctg ttc tag gag agt aat ggg tta ttg tgg ata tat aat cag agt ttt taa tga cta cta tat tat gtt tta tat cat ttc gtg tca cag gct ttg tag att tgg a-- --- --- --- --- --- --- -ta tag cat tgc aca tag tat tag tca taa act tgg gat gta aaa tgt tga tga tat cta cat cgt ttg gat ttt tat gta tcc act tta ata ata tca tag ctg taa cat cct tgt gat ttt acg ttg acg ttt tcg tgg gat aag ata gtt --- -gt tca tcc ttt gat aat ttt cca aat t

>DQ441432.1:12236-14181_Variola_virus_strain_Korea_1947_(Lee_Masterseed)_complete_genome

cgg ttg cca tgt att tcc tgg aga gca agt --- aga tga tg- agg aac cag ata gtt tat atc cgt act tgc acg taa agt cta cat tgt cat tat acg a-- ata tga tct ttt aaa ccc gct aga caa gta tcc gtt taa tat tgt agg atg tgg aca ttt aac aat ctg aca cgt ggg tgg att aga cca ttc tcc tcc tga aca cag gac acc aga gtt acc aat caa cga ata tcc act att gca act ata agt tac aac gct tcc atc ggt ata aaa atc att ata tcc gtt atg tct tcc gtt gga tat aga tgg agg taa ttg gca ttt aac aga ttc gca aat agg tgc ctt ggg att cca tac cat aga tcc agt aga tcc taa ttt aca ata cga ttt ata ttc acc gat caa ata ata tcc gct att aca aga gta cgt tat act aga gcc aaa gtc tac tcc gcc aat atc aag atg gcc att atc gat atc tcg agg cga cgg gca tct ccg ttt aat aca ttg att aaa gag tgt cca tcc agt acc tgt aca ttt agc ata tat agg tcc cat ttt ttg ctt tct gta tcc agg tag aca tag a-t att cta tag tgt ctc cta tgt tgt a-- --- --- --- -at tag cat tag cat cag tc- --t cca cac tat tct taa att tca tat taa tgg gtc gtg acg gaa tag tac agc atg ata gaa cgc atc cta ttc cca aca atg tca gga acg tca ctc tct cca cct tca tat tta ttt atc cgt -aa aaa tgt tat cct gga cat tgt gcc aat aat aaa aaa gcc ca- -ta tat gtt cgc tat tgt aga aat tgt ttt tca tag ttg ctc aaa aac gat ggc ggt gac tta tga gtc acg tta cac ttt gga gtc tca tct tta gta aac ata tca taa tat tcg ata tta cga gtt gac aca ttg aac aaa -tc caa gta ttt gat ttt gga taa tat tcg tat ttt gca tct gct ata att aag ata taa tcg cca caa aaa cac gcg aac atc ttt cct aca tgg tta aag tac atg tac aat tct atc cat ttg tct tcc tta act ata tat ttg tat aga taa tta cga gtc tcg tgt gta att cca gta att gca tag atg tta cca tcg tac tct aca gca taa act ata cta tga tgt cta ggc atg gga gac -tt ttt tat cca aca att ttt agt gaa aca ttc cac atc gtt taa tat tac ata ttt ttc ata cgt ggt ata aac tcc acc cat tac --a tat ata tca tca ttt acg aat acc gac gcg tct gaa tat cta gga gta att aag ttt gga tgt ctt atc cat ttc aaa gtg tag cgt ttt aaa tat tct gct aca cct gtt gaa ata gaa aat ttt aat cct cct att aca tat aac ttt cca tca tta aca cac gta cta act ttt gat ttt aac gac gac ata tta gta act gtt ttc ca- --- --- --- --- --- --- ttt ttt ttc gtt tta aga tct acc cac gga tac gaa ata aac atg tct att gtt aat cat gcc acc aat aat gta tag aca att ata taa aac att tgc att ata gaa ttg tct atc tat ctg tat tac cga cga ttg tcc aat att ctg ttc tag gag agt aat ggg tta ttg tgg ata tat aat cag agt ttt taa tga cta cta tat tat gtt tta tac cat ttc gtg tca cag gct ttg tag att tgg a-- --- --- --- --- --- --- -ta tag cat tgc aca tag tat tag tca taa act tgg gat gta aac tgt tga tga tat cta cat cgt ttg gat ttt tat gta tcc act tta ata ata tca tag ctg taa cat cct cgt ga- ttt acg ttg acg ttt tcg tgg gat aag ata gtt --- -gt tca tcc ttt gat aat ttt cca aat t

>DQ441431.1:12090-14036_Variola_virus_strain_Japan_1951_(Stillwell_Masterseed)_complete_genome

cgg ttg cca tgt att tcc tgg aga gca agt --- aga tga tg- agg aac cag ata gtt tat atc cgt act tgc acg taa agt cta cat tgt cat tat acg a-- ata tga tct ttt aaa ccc gct aga caa gta tcc gtt taa tat tgt agg atg tgg aca ttt aac aat ctg aca cgt ggg tgg att aga cca ttc tcc tcc tga aca cag gac acc aga gtt acc aat caa cga ata tcc act att gca act ata agt tac aac gct tcc atc ggt ata aaa atc att ata tcc gtt atg tct tcc gtt gga tat aga tgg agg taa ttg gca ttt aac aga ttc gca aat agg tgc ctt ggg att cca tac cat aga tcc agt aga tcc taa ttt aca ata cga ttt ata ttc acc gat caa ata ata tcc gct att aca aga gta cgt tat act aga gcc aaa gtc tac tcc gcc aat atc aag atg gcc att atc gat atc tcg agg cga cgg gca tct ccg ttt aat aca ttg att aaa gag tgt cca tcc agt acc tgt aca ttt agc ata tat agg tcc cat ttt ttg ctt tct gta tcc agg tag aca tag a-t att cta tag tgt ctc cta tgt tgt a-- --- --- --- -at tag cat tag cat cag tc- --t cca cac tat tct taa att tca tat taa tgg gtc gtg acg gaa tag tac agc atg ata gaa cgc atc cta ttc cca aca atg tca gga acg tca ctc tct cca cct tca tat tta ttt atc cgt -aa aaa tgt tat cct gga cat tgt gcc aat aat aaa aaa gcc ca- -ta tat gtt cgc tat tgt aga aat tgt ttt tca tag ttg ctc aaa aac gat ggc ggt gac tta tga gtc acg tta cac ttt gga gtc tca tct tta gta aac ata tca taa tat tcg ata tta cga gtt gac aca ttg aac aaa -tc caa gta ttt gat ttt gga taa tat tcg tat ttt gca tct gct ata att aag ata taa tcg cca caa aaa cac gcg aac atc ttt cct aca tgg tta aag tac atg tac aat tct atc cat ttg tct tcc tta act ata tat ttg tat aga taa tta cga gtc tcg tgt gta att cca gta att gca tag atg tta cca tcg tac tct aca gca taa act ata cta tga tgt cta ggc atg gga gac -tt ttt tat cca aca att ttt agt gaa aca ttc cac atc gtt taa tat tac ata ttt ttc ata cgt ggt ata aac tcc acc cat tac --a tat ata tca tca ttt acg aat acc gac gcg gct gaa tat cta gga gta att aag ttt gga tgt ctt atc cat ttc gaa gtg tag cgt ttt aaa tat tct gct aca cct gtt gaa ata gaa aat ttt aat cct cct att aca tat aac ttt cca tca tta aca cac gta cta act ttt gat ttt aac gac gac ata tta gta act gtt ttc ca- --- --- --- --- --- --t ttt ttt ttc gtt tta aga tct acc cac gga tac gaa ata aac atg tct att gtt aat cat gcc acc aat aat gta tag aca att ata taa aac att tgc att ata gaa ttg tct atc tat ctg tat tac cga cga ttg tcc aat att ctg ttc tag gag agt aat ggg tta ttg tgg ata tat aat cag agt ttt taa tga cta cta tat tat gtt tta tac cat ttc gtg tca cag gct ttg tag att tgg a-- --- --- --- --- --- --- -ta tag cat tgc aca tag tat tag tca taa act tgg gat gta aac tgt tga tga tat cta cat cgt ttg gat ttt tat gta tcc act tta ata ata tca tag ctg taa cat cct cgt ga- ttt acg ttg acg ttt tcg tgg gat aag ata gtt --- -gt tca tcc ttt gat aat ttt cca aat t

>DQ441430.1:12091-14037_Variola_virus_strain_Japan_1951_(Harper_Masterseed)_complete_genome

cgg ttg cca tgt att tcc tgg aga gca agt --- aga tga tg- agg aac cag ata gtt tat atc cgt act tgc acg taa agt cta cat tgt cat tat acg a-- ata tga tct ttt aaa ccc gct aga caa gta tcc gtt taa tat tgt agg atg tgg aca ttt aac aat ctg aca cgt ggg tgg att aga cca ttc tcc tcc tga aca cag gac acc aga gtt acc aat caa cga ata tcc act att gca act ata agt tac aac gct tcc atc ggt ata aaa atc att ata tcc gtt atg tct tcc gtt gga tat aga tgg agg taa ttg gca ttt aac aga ttc gca aat agg tgc ctt ggg att cca tac cat aga tcc agt aga tcc taa ttt aca ata cga ttt ata ttc acc gat caa ata ata tcc gct att aca aga gta cgt tat act aga gcc aaa gtc tac tcc gcc aat atc aag atg gcc att atc gat atc tcg agg cga cgg gca tct ccg ttt aat aca ttg att aaa gag tgt cca tcc agt acc tgt aca ttt agc ata tat agg tcc cat ttt ttg ctt tct gta tcc agg tag aca tag a-t att cta tag tgt ctc cta tgt tgt a-- --- --- --- -at tag cat tag cat cag tc- --t cca cac tat tct taa att tca tat taa tgg gtc gtg acg gaa tag tac agc atg ata gaa cgc atc cta ttc cca aca atg tca gga acg tca ctc tct cca cct tca tat tta ttt atc cgt -aa aaa tgt tat cct gga cat tgt gcc aat aat aaa aaa gcc ca- -ta tat gtt cgc tat tgt aga aat tgt ttt tca tag ttg ctc aaa aac gat ggc ggt gac tta tga gtc acg tta cac ttt gga gtc tca tct tta gta aac ata tca taa tat tcg ata tta cga gtt gac aca ttg aac aaa -tc caa gta ttt gat ttt gga taa tat tcg tat ttt gca tct gct ata att aag ata taa tcg cca caa aaa cac gcg aac atc ttt cct aca tgg tta aag tac atg tac aat tct atc cat ttg tct tcc tta act ata tat ttg tat aga taa tta cga gtc tcg tgt gta att cca gta att gca tag atg tta cca tcg tac tct aca gca taa act ata cta tga tgt cta ggc atg gga gac -tt ttt tat cca aca att ttt agt gaa aca ttc cac atc gtt taa tat tac ata ttt ttc ata cgt ggt ata aac tcc acc cat tac --a tat ata tca tca ttt acg aat acc gac gcg gct gaa tat cta gga gta att aag ttt gga tgt ctt atc cat ttc gaa gtg tag cgt ttt aaa tat tct gct aca cct gtt gaa ata gaa aat ttt aat cct cct att aca tat aac ttt cca tca tta aca cac gta cta act ttt gat ttt aac gac gac ata tta gta act gtt ttc ca- --- --- --- --- --- --t ttt ttt ttc gtt tta aga tct acc cac gga tac gaa ata aac atg tct att gtt aat cat gcc acc aat aat gta tag aca att ata taa aac att tgc att ata gaa ttg tct atc tat ctg tat tac cga cga ttg tcc aat att ctg ttc tag gag agt aat ggg tta ttg tgg ata tat aat cag agt ttt taa tga cta cta tat tat gtt tta tac cat ttc gtg tca cag gct ttg tag att tgg a-- --- --- --- --- --- --- -ta tag cat tgc aca tag tat tag tca taa act tgg gat gta aac tgt tga tga tat cta cat cgt ttg gat ttt tat gta tcc act tta ata ata tca tag ctg taa cat cct cgt ga- ttt acg ttg acg ttt tcg tgg gat aag ata gtt --- -gt tca tcc ttt gat aat ttt cca aat t

>DQ441429.1:12102-14048_Variola_virus_strain_Japan_1946_(Yamada_MS-2(A)_Tokyo)_complete_genome

cgg ttg cca tgt att tcc tgg aga gca agt --- aga tga tg- agg aac cag ata gtt tat atc cgt act tgc acg taa agt cta cat tgt cat tat acg a-- ata tga tct ttt aaa ccc gct aga caa gta tcc gtt taa tat tgt agg atg tgg aca ttt aac aat ctg aca cgt ggg tgg att aga cca ttc tcc tcc tga aca cag gac acc aga gtt acc aat caa cga ata tcc act att gca act ata agt tac aac gct tcc atc ggt ata aaa atc att ata tcc gtt atg tct tcc gtt gga tat aga tgg agg taa ttg gca ttt aac aga ttc gca aat agg tgc ctt ggg att cca tac cat aga tcc agt aga tcc taa ttt aca ata cga ttt ata ttc acc gat caa ata ata tcc gct att aca aga gta cgt tat act aga gcc aaa gtc tac tcc gcc aat atc aag atg gcc att atc gat atc tcg agg cga cgg gca tct ccg ttt aat aca ttg att aaa gag tgt cca tcc agt acc tgt aca ttt agc ata tat agg tcc cat ttt ttg ctt tct gta tcc agg tag aca tag a-t att cta tag tgt ctc cta tgt tgt a-- --- --- --- -at tag cat tag cat cag tc- --t cca cac tat tct taa att tca tat taa tgg gtc gtg acg gaa tag tac agc atg ata gaa cgc atc cta ttc cca aca atg tca gga acg tca ctc tct cca cct tca tat tta ttt atc cgt -aa aaa tgt tat cct gga cat tgt gcc aat aat aaa aaa gcc ca- -ta tat gtt cgc tat tgt aga aat tgt ttt tca tag ttg ctc aaa aac gat ggc ggt gac tta tga gtc acg tta cac ttt gga gtc tca tct tta gta aac ata tca taa tat tcg ata tta cga gtt gac aca ttg aac aaa -tc caa gta ttt gat ttt gga taa tat tcg tat ttt gca tct gct ata att aag ata taa tcg cca caa aaa cac gcg aac atc ttt cct aca tgg tta aag tac atg tac aat tct atc cat ttg tct tcc tta act ata tat ttg tat aga taa tta cga gtc tcg tgt gta att cca gta att gca tag atg tta cca tcg tac tct aca gca taa act ata cta tga tgt cta ggc atg gga gac -tt ttt tat cca aca att ttt agt gaa aca ttc cac atc gtt taa tat tac ata ttt ttc ata cgt ggt ata aac tcc acc cat tac --a tat ata tca tca ttt acg aat acc gac gcg gct gaa tat cta gga gta att aag ttt gga tgt ctt atc cat ttc gaa gtg tag cgt ttt aaa tat tct gct aca cct gtt gaa ata gaa aat ttt aat cct cct att aca tat aac ttt cca tca tta aca cac gta cta act ttt gat ttt aac gac gac ata tta gta act gtt ttc ca- --- --- --- --- --- --t ttt ttt ttc gtt tta aga tct acc cac gga tac gaa ata aac atg tct att gtt aat cat gcc acc aat aat gta tag aca att ata taa aac att tgc att ata gaa ttg tct atc tat ctg tat tac cga cga ttg tcc aat att ctg ttc tag gag agt aat ggg tta ttg tgg ata tat aat cag agt ttt taa tga cta cta tat tat gtt tta tac cat ttc gtg tca cag gct ttg tag att tgg a-- --- --- --- --- --- --- -ta tag cat tgc aca tag tat tag tca taa act tgg gat gta aac tgt tga tga tat cta cat cgt ttg gat ttt tat gta tcc act tta ata ata tca tag ctg taa cat cct cgt ga- ttt acg ttg acg ttt tcg tgg gat aag ata gtt --- -gt tca tcc ttt gat aat ttt cca aat t

>DQ441428.1:12102-14048_Variola_virus_strain_India_1953_(New_Delhi)_complete_genome

cgg ttg cca tgt att tcc tgg aga gca agt --- aga tga tg- agg aac cag ata gtt tat atc cgt act tgc acg taa agt cta cat tgt cat tat acg a-- ata tga tct ttt aaa ccc gct aga caa gta tcc gtt taa tat tgt agg atg tgg aca ttt aac aat ctg aca cgt ggg tgg att aga cca ttc tcc tcc tga aca cag gac acc aga gtt acc aat caa cga ata tcc act att gca act ata agt tac aac gct tcc atc ggt ata aaa atc att ata tcc gtt atg tct tcc gtt gga tat aga tgg agg taa ttg gca ttt aac aga ttc gca aat agg tgc ctt ggg att cca tac cat aga tcc agt aga tcc taa ttt aca ata cga ttt ata ttc acc gat caa ata ata tcc gct att aca aga gta cgt tat act aga gcc aaa gtc tac tcc gcc aat atc aag atg gcc att atc gat atc tcg agg cga cgg gca tct ccg ttt aat aca ttg att aaa gag tgt cca tcc agt acc tgt aca ttt agc ata tat agg tcc cat ttt ttg ctt tct gta tcc agg tag aca tag a-t att cta tag tgt ctc cta tgt tgt a-- --- --- --- -at tag cat tag cat cag tc- --t cca cac tat tct taa att tca tat taa tgg gtc gtg acg gaa tag tac agc atg ata gaa cgc atc cta ttc cca aca atg tca gga acg tca ctc tct cca cct tca tat tta ttt atc cgt -aa aaa tgt tat cct gga cat tgt gcc aat aat aaa aaa gcc ca- -ta tat gtt cgc tat tgt aga aat tgt ttt tca tag ttg ctc aaa aac gat ggc ggt gac tta tga gtc acg tta cac ttt gga gtc tca tct tta gta aac ata tca taa tat tcg ata tta cga gtt gac aca ttg aac aaa -tc caa gta ttt gat ttt gga taa tat tcg tat ttt gca tct gct ata att aag ata taa tcg cca caa aaa cac gcg aac atc ttt cct aca tgg tta aag tac atg tac aat tct atc cat ttg tct tcc tta act ata tat ttg tat aga taa tta cga gtc tcg tgt gta att cca gta att gca tag atg tta cca tcg tac tct aca gca taa act ata cta tga tgt cta ggc atg gga gac -tt ttt tat cca aca att ttt agt gaa aca ttc cac atc gtt taa tat tac ata ttt ttc ata cgt ggt ata aac tcc acc cat tac --a tat ata tca tca ttt acg aat acc gac gcg gct gaa tat cta gga gta att aag ttt gga tgt ctt atc cat ttc gaa gtg tag cgt ttt aaa tat tct gct aca cct gtt gaa ata gaa aat ttt aat cct cct att aca tat aac ttt cca tca tta aca cac gta cta act ttt gat ttt aac gac gac ata tta gta act gtt ttc ca- --- --- --- --- --- --t ttt ttt ttc gtt tta aga tct acc cac gga tac gaa ata aac atg tct att gtt aat cat gcc acc aat aat gta tag aca att ata taa aac att tgc att ata gaa ttg tct atc tat ctg tat tac cga cga ttg tcc aat att ctg ttc tag gag agt aat ggg tta ttg tgg ata tat aat cag agt ttt taa tga cta cta tat tat gtt tta tac cat ttc gtg tca cag gct ttg tag att tgg a-- --- --- --- --- --- --- -ta tag cat tgc aca tag tat tag tca taa act tgg gat gta aac tgt tga tga tat cta cat cgt ttg gat ttt tat gta tcc act tta ata ata tca tag ctg taa cat cct cgt ga- ttt acg ttg acg ttt tcg tgg gat aag ata gtt --- -gt tca tcc ttt gat aat ttt cca aat t

>DQ441426.1:12933-14878_Variola_virus_strain_Guinea_1969_(005)_complete_genome

cgg ttg cca tgt att tcc tgg aga gca agt --- aga tga tg- agg aac cag ata gtt tat atc cgt act tgc acg taa agt cta cat tgt cat tat acg a-- ata tga tct ttt aaa ccc gct aga caa gta tcc gtt taa tat tgt agg atg tgg aca ttt aac aat ctg aca cgt ggg tgg att aga cca ttc tcc tcc tga aca cag gac acc aga gtt acc aat caa cga ata tcc act att gca act ata agt tac aac gct tcc atc ggt ata aaa atc att ata tcc gtt atg tct tcc gtt gga tat aga tgg agg taa ttg gca ttt aac aga ttc gca aat agg tgc ctt ggg att cca tac cat aga tcc agt aga tcc taa ttt aca ata cga ttt ata ttc acc gat caa ata ata tcc gct att aca aga gta cgt tat act aga gcc aaa gtc tac tcc gcc aat atc aag atg gcc att atc gat atc tcg agg cga cgg gca tct ccg ttt aat aca ttg att aaa gag tgt cca tcc agt acc tgt aca ttt agc ata tat agg tcc cat ttt ttg ctt tct gta tcc agg tag aca tag a-t att cta tag tgt ctc cta tgt tgt a-- --- --- --- -at tag cat tag cat cag tc- --t cca cac tat tct taa att tca tat taa tgg gtc gtg acg gaa tag tac agc atg ata gaa cgc atc cta ttc cca aca atg tca gga acg tca ctc tct cca cct tca tat tta ttt atc cgt -aa aaa tgt tat cct gga cat tgt gtc aat aat aaa aaa gct ca- -ta tat gtt tgc tat tgt aga aat tgt ttt tca tag ttg ctc aaa aac gat ggc ggt gac tta tga gtc acg tta cac ttt gga gtc tca tct tta gta aac ata tca taa tat tcg ata tta cga gtt gac ata ttg aac aaa ttc caa gta ttt gat ttt gga taa tat tcg tat ttt gca tct gct ata att aag ata taa tcg cca caa gaa cac acg aac atc ttt cct aca tgg tta aag tac atg tac aat tct atc cat ttg tct tcc tta act ata tat ttg tat aga taa tta cga gtc tcg tgt gta att cca gta att gca tag atg tta cca tcg tac tct aca gca taa act ata cta tga tgt cta ggc atg gga gac -tt ttt tat cca aca att ttt agt gaa aca ttc cac atc gtt taa tat tac ata ttt ttc ata cgt ggt ata aac tcc acc cat tac --a tat ata tca tca ttt acg aat acc gac gcg tct gaa tat cta gga gta att aag ttt gga tgt ctt atc cat ttc gaa gtg tag cgt ttt aaa tat tct gct aca cct gtt gaa ata gaa aat ttt aat cct cct att aca tat aac ttt cca tca tta aca cac gta ct- act ttt gat ttt aac gac gac ata tta gta act gtt ttc ca- --- --- --- --- --- --- -tt ttt ttc gtt tta aga tct acc cac gga tac gaa ata aac atg tct att gtt aat cat gcc gcc aat aat gta tag aca att ata taa aac att tgc att ata gaa ttg tct atc tat ctg tat tac cga caa ttg tcc aat att ctg ttc tag gag agt aat ggg tta ttg tgg ata tat aat cag agt ttt taa tga cta cta tat tat gtt tta tac cat ttc gtg tca cag gct ttg tag att tgg a-- --- --- --- --- --- --- -ta tag cat tgc aca tag tat tag tca taa act tgg gat gta aaa tgt tga tga tat cta cat cgt ttg gat ttt tat gta tcc act tta ata ata tca tag ctg taa cat cct tgt gat ttt acg ttg atg ttt tcg tgg gat aag ata gtt --- -gt tca tcc ttt gat aat ttt cca aat t

>LT706528.1:11645-13592_Variola_virus_isolate_VARV_V563_genome_assembly_chromosome:_I

cgg ttg cca tgt att tcc tgg aga gca agt --- aga tga tg- agg aac cag ata gtt tat atc cgt act tgc acg taa agt cta cat tgt cat tat acg a-- ata tga tct ttt aaa ccc gct aga caa gta tcc gtt taa tat tgt agg atg tgg aca ttt aac aat ctg aca cgt ggg tgg att aga cca ttc tcc tcc tga aca cag gac acc aga gtt acc aat caa cga ata tcc act att gca act ata agt tac aac gct tcc atc ggt ata aaa atc att ata tcc gtt atg tct tcc gtt gga tat aga tgg agg taa ttg gca ttt aac aga ttc gca aat agg tgc ctt ggg att cca tac cat aga tcc agt aga tcc taa ttt aca ata cga ttt ata ttc acc gat caa ata ata tcc gct att aca aga gta cgt tat act aga gcc aaa gtc tac tcc gcc aat atc aag atg gcc att atc gat atc tcg agg cga cgg gca tct ccg ttt aat aca ttg att aaa gag tgt cca tcc agt acc tgt aca ttt agc ata tat agg tcc cat ttt ttg ctt tct gta tcc agg tag aca tag a-t att cta tag tgt ctc cta tgt tgt a-- --- --- --- -at tag cat tag cat cag tc- --t cca cac tat tct taa att tca tat taa tgg gtc gtg acg gaa tag tac agc atg ata gaa cgc atc cta ttc cca aca atg tca gga acg tca ctc tct cca cct tca tat tta ttt atc cgt -aa aaa tgt tat cct gga cat tgt gcc aat aat aaa aaa gcc ca- -ta tat gtt cgc tat tgt aga aat tgt ttt tca tag ttg ctc aaa aac gat ggc ggt gac tta tga gtc acg tta cac ttt gga gtc tca tct tta gta aac ata tca taa tat tcg ata tta cga gtt gac aca ttg aac aaa -tc caa gta ttt gat ttt gga taa tat tcg tat ttt gca tct gct ata att aag ata taa tcg cca caa aaa cac gcg aac atc ttt cct aca tgg tta aag tac atg tac aat tct atc cat ttg tct tcc tta act ata tat ttg tat aga taa tta cga gtc tcg tgt gta att cca gta att gca tag atg tta cca tcg tac tct aca gca taa act ata cta tga tgt cta ggc atg gga gac -tt ttt tat cca aca att ttt agt gaa aca ttc cac atc gtt taa tat tac ata ttt ttc ata cgt ggt ata aac tcc acc cat tac --a tat ata tca tca ttt acg aat acc gac gcg tct gaa tat cta gga gta att aag ttt gga tgt ctt atc cat ttc gaa gtg tag cgt ttt aaa tat tct gct aca cct gtt gaa ata gaa aat ttt aat cct cct att aca tat aac ttt cca tca tta aca cac gta cta act ttt gat ttt aac gac gac ata tta gta act gtt ttc ca- --- --- --- --- --- -tt ttt ttt ttc gtt tta aga tct acc cac gga tac gaa ata aac atg tct att gtt aat cat gcc acc aat aat gta tag aca att ata taa aac att tgc att ata gaa ttg tct atc tat ctg tat tac cga cga ttg tcc aat att ctg ttc tag gag agt aat ggg tta ttg tgg ata tat aat cag agt ttt taa tga cta cta tat tat gtt tta tac cat ttc gtg tca cag gct ttg tag att tgg a-- --- --- --- --- --- --- -ta tag cat tgc aca tag tat tag tca taa act tgg gat gta aac tgt tga tga tat cta cat cgt ttg gat ttt tat gta tcc act tta ata ata tca tag ctg taa cat cct cgt ga- ttt acg ttg acg ttt tcg tgg gat aag ata gtt --- -gt tca tcc ttt gat aat ttt cca aat t

>DQ437582.1:12291-14238_Variola_virus_strain_China_Horn_1948_complete_genome

cgg ttg cca tgt att tcc tgg aga gca agt --- aga tga tg- agg aac cag ata gtt tat atc cgt act tgc acg taa agt cta cat tgt cat tat acg a-- ata tga tct ttt aaa ccc gct aga caa gta tcc gtt taa tat tgt agg atg tgg aca ttt aac aat ctg aca cgt ggg tgg att aga cca ttc tcc tcc tga aca cag gac acc aga gtt acc aat caa cga ata tcc act att gca act ata agt tac aac gct tcc atc ggt ata aaa atc att ata tcc gtt atg tct tcc gtt gga tat aga tgg agg taa ttg gca ttt aac aga ttc gca aat agg tgc ctt ggg att cca tac cat aga tcc agt aga tcc taa ttt aca ata cga ttt ata ttc acc gat caa ata ata tcc gct att aca aga gta cgt tat act aga gcc aaa gtc tac tcc gcc aat atc aag atg gcc att atc gat atc tcg agg cga cgg gca tct ccg ttt aat aca ttg att aaa gag tgt cca tcc agt acc tgt aca ttt agc ata tat agg tcc cat ttt ttg ctt tct gta tcc agg tag aca tag a-t att cta tag tgt ctc cta tgt tgt a-- --- --- --- -at tag cat tag cat cag tc- --t cca cac tat tct taa att tca tat taa tgg gtc gtg acg gaa tag tac agc atg ata gaa cgc atc cta ttc cca aca atg tca gga acg tca ctc tct cca cct tca tat tta ttt atc cgt -aa aaa tgt tat cct gga cat tgt gcc aat aat aaa aaa gcc ca- -ta tat gtt cgc tat tgt aga aat tgt ttt tca tag ttg ctc aaa aac gat ggc ggt gac tta tga gtc acg tta cac ttt gga gtc tca tct tta gta aac ata tca taa tat tcg ata tta cga gtt gac aca ttg aac aaa -tc caa gta ttt gat ttt gga taa tat tcg tat ttt gca tct gct ata att aag ata taa tcg cca caa aaa cac gcg aac atc ttt cct aca tgg tta aag tac atg tac aat tct atc cat ttg tct tcc tta act ata tat ttg tat aga taa tta cga gtc tcg tgt gta att cca gta att gca tag atg tta cca tcg tac tct aca gca taa act ata cta tga tgt cta ggc atg gga gac -tt ttt tat cca aca att ttt agt gaa aca ttc cac atc gtt taa tat tac ata ttt ttc ata cgt ggt ata aac tcc acc cat tac --a tat ata tca tca ttt acg aat acc gac gcg tct gaa tat cta gga gta att aag ttt gga tgt ctt atc cat ttc gaa gtg tag cgt ttt aaa tat tct gct aca cct gtt gaa ata gaa aat ttt aat cct cct att aca tat aac ttt cca tca tta aca cac gta cta act ttt gat ttt aac gac gac ata tta gta act gtt ttc ca- --- --- --- --- --- -tt ttt ttt ttc gtt tta aga tct acc cac gga tac gaa ata aac atg tct att gtt aat cat gcc acc aat aat gta tag aca att ata taa aac att tgc att ata gaa ttg tct atc tat ctg tat tac cga cga ttg tcc aat att ctg ttc tag gag agt aat ggg tta ttg tgg ata tat aat cag agt ttt taa tga cta cta tat tat gtt tta tac cat ttc gtg tca cag gct ttg tag att tgg a-- --- --- --- --- --- --- -ta tag cat tgc aca tag tat tag tca taa act tgg gat gta aac tgt tga tga tat cta cat cgt ttg gat ttt tat gta tcc act tta ata ata tca tag ctg taa cat cct cgt ga- ttt acg ttg acg ttt tcg tgg gat aag ata gtt --- -gt tca tcc ttt gat aat ttt cca aat t

>EF611198.1:79-2023_Variola_virus_strain_Khateen_left_variable_genomic_sequence

cgg ttg cca tgt att tcc tgg aga gca agt --- aga tga tg- agg aac cag ata gtt tat atc cgt act tgc acg taa agt cta cat tgt cat tat acg a-- ata tga tct ttt aaa ccc gct aga caa gta tcc gtt taa tat tgt agg atg tgg aca ttt aac aat ctg aca cgt ggg tgg att aga cca ttc tcc tcc tga aca cag gac acc aga gtt acc aat caa cga ata tcc act att gca act ata agt tac aac gct tcc atc ggt ata aaa atc att ata tcc gtt atg tct tcc gtt gga tat aga tgg agg taa ttg gca ttt aac aga ttc gca aat agg tgc ctt ggg att cca tac cat aga tcc agt aga tcc taa ttt aca ata cga ttt ata ttc acc aat caa ata ata tcc gct att aca aga gta cgt tat act aga gcc aaa gtc tac tcc gcc aat atc aag atg gcc att atc gat atc tcg agg cga cgg gca tct ccg ttt aat aca ttg att aaa gag tgt cca tcc agt acc tgt aca ttt agc ata tat agg tcc cat ttt ttg ctt tct gta tcc agg tag aca tag a-t att cta tag tgt ctc cta tgt tgt a-- --- --- --- -at tag cat tag cat cag tc- --t cca cac tat tct taa att tca tat taa tgg gtc gtg acg gaa tag tac agc atg ata gaa cgc atc cta ttc cca aca atg tca gga acg tca ctc tct cca cct tca tat tta ttt atc cgt -aa aaa tgt tat cct gga cat tgt gcc aat aat aaa aaa gcc ca- -ta tat gtt cgc tat tgt aga aat tgt ttt tca tag ttg ctc aaa aac gat ggc ggt gac tta tga gtc acg tta cac ttt gga gtc tca tct tta gta aac ata tca taa tat tcg ata tta cga gtt gac aca ttg aac aaa -tc caa gta ttt gat ttt gga taa tat tcg tat ttt gca tct gct ata att aag ata taa tcg cca caa aaa cac gcg aac atc ttt cct aca tgg tta aag tac atg tac aat tct atc cat ttg tct tcc tta act ata tat ttg tat aga taa tta cga gtc tcg tgt gta att cca gta att gca tag atg tta cca tcg tac tct aca gca taa act ata cta tga tgt cta ggc atg gga gac -tt ttt tat cca aca att ttt agt gaa aca ttc cac atc gtt taa tat tac ata ttt ttc ata cgt ggt ata aac tcc acc cat tac --a tat ata tca tca ttt acg aat acc gac gcg tct gaa tat cta gga gta att aag ttt gga tgt ctt atc cat ttc gaa gtg tag cgt ttt aaa tat ttt gct aca cct gtt gaa ata gaa aat ttt aat cct cct att aca tat aac ttt cca tca tta aca cac gta cta act ttt gat ttt aac gac aac ata tta gta act gtt ttc ca- --- --- --- --- --- --- -tt ttt ttt ttt tta aga tct acc cac gga tac gaa ata aac atg tct att gtt aat cat gcc acc aat aat gta tag aca att ata taa aac att tgc att ata gaa ttg tct atc tat ctg tat tac cga cga ttg tcc aat att ctg ttc tag gag agt aat ggg tta ttg tgg ata tat aat cag agt ttt taa tga cta cta tat tat gtt tta tac cat ttc gtg tca cag gct ttg tag att tgg a-- --- --- --- --- --- --- -ta tag cat tgc aca tag tat tag tca taa act tgg gat gta aac tgt tga tga tat cta cat cgt ttg gat ttt tat gta tcc act tta ata ata tca tag ctg taa cat cct cgt ga- ttt acg ttg acg ttt tcg tgg gat aag ata gtt --- -gt tca tcc ttt gat aat ttt cca aat t

>EF611196.1:79-2022_Variola_virus_strain_Ind-4a_left_variable_genomic_sequence

cgg ttg cca tgt att tcc tgg aga gca agt --- aga tga tg- agg aac cag ata gtt tat atc cgt act tgc acg taa agt cta cat tgt cat tat acg a-- ata tga tct ttt aaa ccc gct aga caa gta tcc gtt taa tat tgt agg atg tgg aca ttt aac aat ctg aca cgt ggg tgg att aga cca ttc tcc tcc tga aca cag gac acc aga gtt acc aat caa cga ata tcc act att gca act ata agt tac aac gct tcc atc ggt ata aaa atc att ata tcc gtt atg tct tcc gtt gga tat aga tgg agg taa ttg gca ttt aac aga ttc gca aat agg tgc ctt ggg att cca tac cat aga tcc agt aga tcc taa ttt aca ata cga ttt ata ttc acc aat caa ata ata tcc gct att aca aga gta cgt tat act aga gcc aaa gtc tac tcc gcc aat atc aag atg gcc att atc gat atc tcg agg cga cgg gca tct ccg ttt aat aca ttg att aaa gag tgt cca tcc agt acc tgt aca ttt agc ata tat agg tcc cat ttt ttg ctt tct gta tcc agg tag aca tag a-t att cta tag tgt ctc cta tgt tgt a-- --- --- --- -at tag cat tag cat cag tc- --t cca cac tat tct taa att tca tat taa tgg gtc gtg acg gaa tag tac agc atg ata gaa cgc atc cta ttc cca aca atg tca gga acg tca ctc tct cca cct tca tat tta ttt atc cgt -aa aaa tgt tat cct gga cat tgt gcc aat aat aaa aaa gcc ca- -ta tat gtt cgc tat tgt aga aat tgt ttt tca tag ttg ctc aaa aac gat ggc ggt gac tta tga gtc acg tta cac ttt gga gtc tca tct tta gta aac ata tca taa tat tcg ata tta cga gtt gac aca ttg aac aaa -tc caa gta ttt gat ttt gga taa tat tcg tat ttt gca tct gct ata att aag ata taa tcg cca caa aaa cac gcg aac atc ttt cct aca tgg tta aag tac atg tac aat tct atc cat ttg tct tcc tta act ata tat ttg tat aga taa tta cga gtc tcg tgt gta att cca gta att gca tag atg tta cca tcg tac tct aca gca taa act ata cta tga tgt cta ggc atg gga gac -tt ttt tat cca aca att ttt agt gaa aca ttc cac atc gtt taa tat tac ata ttt ttc ata cgt ggt ata aac tcc acc cat tac --a tat ata tca tca ttt acg aat acc gac gcg tct gaa tat cta gga gta att aag ttt gga tgt ctt atc cat ttc gaa gtg tag cgt ttt aaa tat ttt gct aca cct gtt gaa ata gaa aat ttt aat cct cct att aca tat aac ttt cca tca tta aca cac gta cta act ttt gat ttt aac gac aac ata tta gta act gtt ttc ca- --- --- --- --- --- --- --t ttt ttt ttt tta aga tct acc cac gga tac gaa ata aac atg tct att gtt aat cat gcc acc aat aat gta tag aca att ata taa aac att tgc att ata gaa ttg tct atc tat ctg tat tac cga cga ttg tcc aat att ctg ttc tag gag agt aat ggg tta ttg tgg ata tat aat cag agt ttt taa tga cta cta tat tat gtt tta tac cat ttc gtg tca cag gct ttg tag att tgg a-- --- --- --- --- --- --- -ta tag cat tgc aca tag tat tag tca taa act tgg gat gta aac tgt tga tga tat cta cat cgt ttg gat ttt tat gta tcc act tta ata ata tca tag ctg taa cat cct cgt ga- ttt acg ttg acg ttt tcg tgg gat aag ata gtt --- -gt tca tcc ttt gat aat ttt cca aat t

>EF611195.1:79-2023_Variola_virus_strain_Ind-3a_left_variable_genomic_sequence

cgg ttg cca tgt att tcc tgg aga gca agt --- aga tga tg- agg aac cag ata gtt tat atc cgt act tgc acg taa agt cta cat tgt cat tat acg a-- ata tga tct ttt aaa ccc gct aga caa gta tcc gtt taa tat tgt agg atg tgg aca ttt aac aat ctg aca cgt ggg tgg att aga cca ttc tcc tcc tga aca cag gac acc aga gtt acc aat caa cga ata tcc act att gca act ata agt tac aac gct tcc atc ggt ata aaa atc att ata tcc gtt atg tct tcc gtt gga tat aga tgg agg taa ttg gca ttt aac aga ttc gca aat agg tgc ctt ggg att cca tac cat aga tcc agt aga tcc taa ttt aca ata cga ttt ata ttc acc aat caa ata ata tcc gct att aca aga gta cgt tat act aga gcc aaa gtc tac tcc gcc aat atc aag atg gcc att atc gat atc tcg agg cga cgg gca tct ccg ttt aat aca ttg att aaa gag tgt cca tcc agt acc tgt aca ttt agc ata tat agg tcc cat ttt ttg ctt tct gta tcc agg tag aca tag a-t att cta tag tgt ctc cta tgt tgt a-- --- --- --- -at tag cat tag cat cag tc- --t cca cac tat tct taa att tca tat taa tgg gtc gtg acg gaa tag tac agc atg ata gaa cgc atc cta ttc cca aca atg tca gga acg tca ctc tct cca cct tca tat tta ttt atc cgt -aa aaa tgt tat cct gga cat tgt gcc aat aat aaa aaa gcc ca- -ta tat gtt cgc tat tgt aga aat tgt ttt tca tag ttg ctc aaa aac gat ggc ggt gac tta tga gtc acg tta cac ttt gga gtc tca tct tta gta aac ata tca taa tat tcg ata tta cga gtt gac aca ttg aac aaa -tc caa gta ttt gat ttt gga taa tat tcg tat ttt gca tct gct ata att aag ata taa tcg cca caa aaa cac gcg aac atc ttt cct aca tgg tta aag tac atg tac aat tct atc cat ttg tct tcc tta act ata tat ttg tat aga taa tta cga gtc tcg tgt gta att cca gta att gca tag atg tta cca tcg tac tct aca gca taa act ata cta tga tgt cta ggc atg gga gac -tt ttt tat cca aca att ttt agt gaa aca ttc cac atc gtt taa tat tac ata ttt ttc ata cgt ggt ata aac tcc acc cat tac --a tat ata tca tca ttt acg aat acc gac gcg tct gaa tat cta gga gta att aag ttt gga tgt ctt atc cat ttc gaa gtg tag cgt ttt aaa tat ttt gct aca cct gtt gaa ata gaa aat ttt aat cct cct att aca tat aac ttt cca tca tta aca cac gta cta act ttt gat ttt aac gac aac ata tta gta act gtt ttc ca- --- --- --- --- --- --- -tt ttt ttt ttt tta aga tct acc cac gga tac gaa ata aac atg tct att gtt aat cat gcc acc aat aat gta tag aca att ata taa aac att tgc att ata gaa ttg tct atc tat ctg tat tac cga cga ttg tcc aat att ctg ttc tag gag agt aat ggg tta ttg tgg ata tat aat cag agt ttt taa tga cta cta tat tat gtt tta tac cat ttc gtg tca cag gct ttg tag att tgg a-- --- --- --- --- --- --- -ta tag cat tgc aca tag tat tag tca taa act tgg gat gta aac tgt tga tga tat cta cat cgt ttg gat ttt tat gta tcc act tta ata ata tca tag ctg taa cat cct cgt ga- ttt acg ttg acg ttt tcg tgg gat aag ata gtt --- -gt tca tcc ttt gat aat ttt cca aat t

>DQ437581.1:12236-14179_Variola_virus_strain_Bangladesh_1975_v75-550_Banu_complete_genome

cgg ttg cca tgt att tcc tgg aga gca agt --- aga tga tg- agg aac cag ata gtt tat atc cgt act tgc acg taa agt cta cat tgt cat tat acg a-- ata tga tct ttt aaa ccc gct aga caa gta tcc gtt taa tat tgt agg atg tgg aca ttt aac aat ctg aca cgt ggg tgg att aga cca ttc tcc tcc tga aca cag gac acc aga gtt acc aat caa cga ata tcc act att gca act ata agt tac aac gct tcc atc ggt ata aaa atc att ata tcc gtt atg tct tcc gtt gga tat aga tgg agg taa ttg gca ttt aac aga ttc gca aat agg tgc ctt ggg att cca tac cat aga tcc agt aga tcc taa ttt aca ata cga ttt ata ttc acc aat caa ata ata tcc gct att aca aga gta cgt tat act aga gcc aaa gtc tac tcc gcc aat atc aag atg gcc att atc gat atc tcg agg cga cgg gca tct ccg ttt aat aca ttg att aaa gag tgt cca tcc agt acc tgt aca ttt agc ata tat agg tcc cat ttt ttg ctt tct gta tcc agg tag aca tag a-t att cta tag tgt ctc cta tgt tgt a-- --- --- --- -at tag cat tag cat cag tc- --t cca cac tat tct taa att tca tat taa tgg gtc gtg acg gaa tag tac agc atg ata gaa cgc atc cta ttc cca aca atg tca gga acg tca ctc tct cca cct tca tat tta ttt atc cgt -aa aaa tgt tat cct gga cat tgt gcc aat aat aaa aaa gcc ca- -ta tat gtt cgc tat tgt aga aat tgt ttt tca tag ttg ctc aaa aac gat ggc ggt gac tta tga gtc acg tta cac ttt gga gtc tca tct tta gta aac ata tca taa tat tcg ata tta cga gtt gac aca ttg aac aaa -tc caa gta ttt gat ttt gga taa tat tcg tat ttt gca tct gct ata att aag ata taa tcg cca caa aaa cac gcg aac atc ttt cct aca tgg tta aag tac atg tac aat tct atc cat ttg tct tcc tta act ata tat ttg tat aga taa tta cga gtc tcg tgt gta att cca gta att gca tag atg tta cca tcg tac tct aca gca taa act ata cta tga tgt cta ggc atg gga gac -tt ttt tat cca aca att ttt agt gaa aca ttc cac atc gtt taa tat tac ata ttt ttc ata cgt ggt ata aac tcc acc cat tac --a tat ata tca tca ttt acg aat acc gac gcg tct gaa tat cta gga gta att aag ttt gga tgt ctt atc cat ttc gaa gtg tag cgt ttt aaa tat ttt gct aca cct gtt gaa ata gaa aat ttt aat cct cct att aca tat aac ttt cca tca tta aca cac gta cta act ttt gat ttt aac gac aac ata tta gta act gtt ttc ca- --- --- --- --- --- --- --t ttt ttt ttt tta aga tct acc cac gga tac gaa ata aac atg tct att gtt aat cat gcc acc aat aat gta tag aca att ata taa aac att tgc att ata gaa ttg tct atc tat ctg tat tac cga cga ttg tcc aat att ctg ttc tag gag agt aat ggg tta ttg tgg ata tat aat cag agt ttt taa tga cta cta tat tat gtt tta tac cat ttc gtg tca cag gct ttg tag att tgg a-- --- --- --- --- --- --- -ta tag cat tgc aca tag tat tag tca taa act tgg gat gta aac tgt tga tga tat cta cat cgt ttg gat ttt tat gta tcc act tta ata ata tca tag ctg taa cat cct cgt ga- ttt acg ttg acg ttt tcg tgg gat aag ata gtt --- -gt tca tcc ttt gat aat ttt cca aat t

>DQ441445.1:12101-14045_Variola_virus_strain_United_Kingdom_1946_Hinden_(Middlesex)_complete_genome

cgg ttg cca tgt att tcc tgg aga gca agt --- aga tga tg- agg aac cag ata gtt tat atc cgt act tgc acg taa agt cta cat tgt cat tat acg a-- ata tga tct ttt aaa ccc gct aga caa gta tcc gtt taa tat tgt agg atg tgg aca ttt aac aat ctg aca cgt ggg tgg att aga cca ttc tcc tcc tga aca cag gac acc aga gtt acc aat caa cga ata tcc act att gca act ata agt tac aac gct tcc atc ggt ata aaa atc att ata tcc gtt atg tct tcc gtt gga tat aga tgg agg taa ttg gca ttt aac aga ttc gca aat agg tgc ctt ggg att cca tac cat aga tcc agt aga tcc taa ttt aca ata cga ttt ata ttc acc aat caa ata ata tcc gct att aca aga gta cgt tat act aga gcc aaa gtc tac tcc gcc aat atc aag atg gcc att atc gat atc tcg agg cga cgg gca tct ccg ttt aat aca ttg att aaa gag tgt cca tcc agt acc tgt aca ttt agc ata tat agg tcc cat ttt ttg ctt tct gta tcc agg tag aca tag a-t att cta tag tgt ctc cta tgt tgt a-- --- --- --- -at tag cat tag cat cag tc- --t cca cac tat tct taa att tca tat taa tgg gtc gtg acg gaa tag tac agc atg ata gaa cgc atc cta ttc cca aca atg tca gga acg tca ctc tct cca cct tca tat tta ttt atc cgt -aa aaa tgt tat cct gga cat tgt gcc aat aat aaa aaa gcc ca- -ta tat gtt cgc tat tgt aga aat tgt ttt tca tag ttg ctc aaa aac gat ggc ggt gac tta tga gtc acg tta cac ttt gga gtc tca tct tta gta aac ata tca taa tat tcg ata tta cga gtt gac aca ttg aac aaa -tc caa gta ttt gat ttt gga taa tat tcg tat ttt gca tct gct ata att aag ata taa tcg cca caa aaa cac gcg aac atc ttt cct aca tgg tta aag tac atg tac aat tct atc cat ttg tct tcc tta act ata tat ttg tat aga taa tta cga gtc tcg tgt gta att cca gta att gca tag atg tta tca tcg tac tct aca gca taa act ata cta tga tgt cta ggc atg gga gac -tt ttt tat cca aca att ttt agt gaa aca ttc cac atc gtt taa tat tac ata ttt ttc ata cgt ggt ata aac tcc acc cat tac --a tat ata tca tca ttt acg aat acc gac gcg tct gaa tat cta gga gta att aag ttt gga tgt ctt atc cat ttc gaa gtg tag cgt ttt aaa tat ttt gct aca cct gtt gaa ata gaa aat ttt aat cct cct att aca tat aac ttt cca tca tta aca cac gta cta act ttt gat ttt aac gac gac ata tta gta act gtt ttc ca- --- --- --- --- --- --- -tt ttt ttt ttt tta aga tct acc cac gga tac gaa ata aac atg tct att gtt aat cat gcc acc aat aat gta tag aca att ata taa aac att tgc att ata gaa ttg tct atc tat ctg tat tac cga caa ttg tcc aat att ctg ttc tag gag agt aat ggg tta ttg tgg ata tat aat cag agt ttt taa tga cta cta tat tat gtt tta tac cat ttc gtg tca cag gct ttg tag att tgg a-- --- --- --- --- --- --- -ta tag cat tgc aca tag tat tag tca taa act tgg gat gta aac tgt tga tga tat cta cat cgt ttg gat ttt tat gta tcc act tta ata ata tca tag ctg taa cat cct cgt ga- ttt acg ttg acg ttt tcg tgg gat aag ata gtt --- -gt tca tcc ttt gat aat ttt cca aat t

>DQ441421.1:12305-14248_Variola_virus_strain_Bangladesh_1974_(Shahzaman)_complete_genome

cgg ttg cca tgt att tcc tgg aga gca agt --- aga tga tg- agg aac cag ata gtt tat atc cgt act tgc acg taa agt cta cat tgt cat tat acg a-- ata tga tct ttt aaa ccc gct aga caa gta tcc gtt taa tat tgt agg atg tgg aca ttt aac aat ctg aca cgt ggg tgg att aga cca ttc tcc tcc tga aca cag gac acc aga gtt acc aat caa cga ata tcc act att gca act ata agt tac aac gct tcc atc ggt ata aaa atc att ata tcc gtt atg tct tcc gtt gga tat aga tgg agg taa ttg gca ttt aac aga ttc gca aat agg tgc ctt ggg att cca tac cat aga tcc agt aga tcc taa ttt aca ata cga ttt ata ttc acc aat caa ata ata tcc gct att aca aga gta cgt tat act aga gcc aaa gtc tac tcc gcc aat atc aag atg gcc att atc gat atc tcg agg cga cgg gca tct ccg ttt aat aca ttg att aaa gag tgt cca tcc agt acc tgt aca ttt agc ata tat agg tcc cat ttt ttg ctt tct gta tcc agg tag aca tag a-t att cta tag tgt ctc cta tgt tgt a-- --- --- --- -at tag cat tag cat cag tc- --t cca cac tat tct taa att tca tat taa tgg gtc gtg acg gaa tag tac agc atg ata gaa cgc atc cta ttc cca aca atg tca gga acg tca ctc tct cca cct tca tat tta ttt atc cgt -aa aaa tgt tat cct gga cat tgt gcc aat aat aaa aaa gcc ca- -ta tat gtt cgc tat tgt aga aat tgt ttt tca tag ttg ctc aaa aac gat ggc ggt gac tta tga gtc acg tta cac ttt gga gtc tca tct tta gta aac ata tca taa tat tcg ata tta cga gtt gac aca ttg aac aaa -tc caa gta ttt gat ttt gga taa tat tcg tat ttt gca tct gct ata att aag ata taa tcg cca caa aaa cac gcg aac atc ttt cct aca tgg tta aag tac atg tac aat tct atc cat ttg tct tcc tta act ata tat ttg tat aga taa tta cga gtc tcg tgt gta att cca gta att gca tag atg tta cca tcg tac tct aca gca taa act ata cta tga tgt cta ggc atg gga gac -tt ttt tat cca aca att ttt agt gaa aca ttc cac atc gtt taa tat tac ata ttt ttc ata cgt ggt ata aac tcc acc cat tac --a tat ata tca tca ttt acg aat acc gac gcg tct gaa tat cta gga gta att aag ttt gga tgt ctt atc cat ttc gaa gtg tag cgt ttt aaa tat ttt gct aca cct gtt gaa ata gaa aat ttt aat cct cct att aca tat aac ttt cca tca tta aca cac gta cta act ttt gat ttt aac gac aac ata tta gta act gtt ttc ca- --- --- --- --- --- --- --t ttt ttt ttt tta aga tct acc cac gga tac gaa ata aac atg tct att gtt aat cat gcc acc aat aat gta tag aca att ata taa aac att tgc att ata gaa ttg tct atc tat ctg tat tac cga cga ttg tcc aat att ctg ttc tag gag agt aat ggg tta ttg tgg ata tat aat cag agt ttt taa tga cta cta tat tat gtt tta tac cat ttc gtg tca cag gct ttg tag att tgg a-- --- --- --- --- --- --- -ta tag cat tgc aca tag tat tag tca taa act tgg gat gta aac tgt tga tga tat cta cat cgt ttg gat ttt tat gta tcc act tta ata ata tca tag ctg taa cat cct cgt ga- ttt acg ttg acg ttt tcg tgg gat aag ata gtt --- -gt tca tcc ttt gat aat ttt cca aat t

>DQ441420.1:12305-14248_Variola_virus_strain_Bangladesh_1974_(nur_islam)_complete_genome

cgg ttg cca tgt att tcc tgg aga gca agt --- aga tga tg- agg aac cag ata gtt tat atc cgt act tgc acg taa agt cta cat tgt cat tat acg a-- ata tga tct ttt aaa ccc gct aga caa gta tcc gtt taa tat tgt agg atg tgg aca ttt aac aat ctg aca cgt ggg tgg att aga cca ttc tcc tcc tga aca cag gac acc aga gtt acc aat caa cga ata tcc act att gca act ata agt tac aac gct tcc atc ggt ata aaa atc att ata tcc gtt atg tct tcc gtt gga tat aga tgg agg taa ttg gca ttt aac aga ttc gca aat agg tgc ctt ggg att cca tac cat aga tcc agt aga tcc taa ttt aca ata cga ttt ata ttc acc aat caa ata ata tcc gct att aca aga gta cgt tat act aga gcc aaa gtc tac tcc gcc aat atc aag atg gcc att atc gat atc tcg agg cga cgg gca tct ccg ttt aat aca ttg att aaa gag tgt cca tcc agt acc tgt aca ttt agc ata tat agg tcc cat ttt ttg ctt tct gta tcc agg tag aca tag a-t att cta tag tgt ctc cta tgt tgt a-- --- --- --- -at tag cat tag cat cag tc- --t cca cac tat tct taa att tca tat taa tgg gtc gtg acg gaa tag tac agc atg ata gaa cgc atc cta ttc cca aca atg tca gga acg tca ctc tct cca cct tca tat tta ttt atc cgt -aa aaa tgt tat cct gga cat tgt gcc aat aat aaa aaa gcc ca- -ta tat gtt cgc tat tgt aga aat tgt ttt tca tag ttg ctc aaa aac gat ggc ggt gac tta tga gtc acg tta cac ttt gga gtc tca tct tta gta aac ata tca taa tat tcg ata tta cga gtt gac aca ttg aac aaa -tc caa gta ttt gat ttt gga taa tat tcg tat ttt gca tct gct ata att aag ata taa tcg cca caa aaa cac gcg aac atc ttt cct aca tgg tta aag tac atg tac aat tct atc cat ttg tct tcc tta act ata tat ttg tat aga taa tta cga gtc tcg tgt gta att cca gta att gca tag atg tta cca tcg tac tct aca gca taa act ata cta tga tgt cta ggc atg gga gac -tt ttt tat cca aca att ttt agt gaa aca ttc cac atc gtt taa tat tac ata ttt ttc ata cgt ggt ata aac tcc acc cat tac --a tat ata tca tca ttt acg aat acc gac gcg tct gaa tat cta gga gta att aag ttt gga tgt ctt atc cat ttc gaa gtg tag cgt ttt aaa tat ttt gct aca cct gtt gaa ata gaa aat ttt aat cct cct att aca tat aac ttt cca tca tta aca cac gta cta act ttt gat ttt aac gac aac ata tta gta act gtt ttc ca- --- --- --- --- --- --- --t ttt ttt ttt tta aga tct acc cac gga tac gaa ata aac atg tct att gtt aat cat gcc acc aat aat gta tag aca att ata taa aac att tgc att ata gaa ttg tct atc tat ctg tat tac cga cga ttg tcc aat att ctg ttc tag gag agt aat ggg tta ttg tgg ata tat aat cag agt ttt taa tga cta cta tat tat gtt tta tac cat ttc gtg tca cag gct ttg tag att tgg a-- --- --- --- --- --- --- -ta tag cat tgc aca tag tat tag tca taa act tgg gat gta aac tgt tga tga tat cta cat cgt ttg gat ttt tat gta tcc act tta ata ata tca tag ctg taa cat cct cgt ga- ttt acg ttg acg ttt tcg tgg gat aag ata gtt --- -gt tca tcc ttt gat aat ttt cca aat t

>L22579.1:12321-14264_Variola_major_virus_(strain_Bangladesh-1975)_complete_genome

cgg ttg cca tgt att tcc tgg aga gca agt --- aga tga tg- agg aac cag ata gtt tat atc cgt act tgc acg taa agt cta cat tgt cat tat acg a-- ata tga tct ttt aaa ccc gct aga caa gta tcc gtt taa tat tgt agg atg tgg aca ttt aac aat ctg aca cgt ggg tgg att aga cca ttc tcc tcc tga aca cag gac acc aga gtt acc aat caa cga ata tcc act att gca act ata agt tac aac gct tcc atc ggt ata aaa atc att ata tcc gtt atg tct tcc gtt gga tat aga tgg agg taa ttg gca ttt aac aga ttc gca aat agg tgc ctt ggg att cca tac cat aga tcc agt aga tcc taa ttt aca ata cga ttt ata ttc acc aat caa ata ata tcc gct att aca aga gta cgt tat act aga gcc aaa gtc tac tcc gcc aat atc aag atg gcc att atc gat atc tcg agg cga cgg gca tct ccg ttt aat aca ttg att aaa gag tgt cca tcc agt acc tgt aca ttt agc ata tat agg tcc cat ttt ttg ctt tct gta tcc agg tag aca tag a-t att cta tag tgt ctc cta tgt tgt a-- --- --- --- -at tag cat tag cat cag tc- --t cca cac tat tct taa att tca tat taa tgg gtc gtg acg gaa tag tac agc atg ata gaa cgc atc cta ttc cca aca atg tca gga acg tca ctc tct cca cct tca tat tta ttt atc cgt -aa aaa tgt tat cct gga cat tgt gcc aat aat aaa aaa gcc ca- -ta tat gtt cgc tat tgt aga aat tgt ttt tca tag ttg ctc aaa aac gat ggc ggt gac tta tga gtc acg tta cac ttt gga gtc tca tct tta gta aac ata tca taa tat tcg ata tta cga gtt gac aca ttg aac aaa -tc caa gta ttt gat ttt gga taa tat tcg tat ttt gca tct gct ata att aag ata taa tcg cca caa aaa cac gcg aac atc ttt cct aca tgg tta aag tac atg tac aat tct atc cat ttg tct tcc tta act ata tat ttg tat aga taa tta cga gtc tcg tgt gta att cca gta att gca tag atg tta cca tcg tac tct aca gca taa act ata cta tga tgt cta ggc atg gga gac -tt ttt tat cca aca att ttt agt gaa aca ttc cac atc gtt taa tat tac ata ttt ttc ata cgt ggt ata aac tcc acc cat tac --a tat ata tca tca ttt acg aat acc gac gcg tct gaa tat cta gga gta att aag ttt gga tgt ctt atc cat ttc gaa gtg tag cgt ttt aaa tat ttt gct aca cct gtt gaa ata gaa aat ttt aat cct cct att aca tat aac ttt cca tca tta aca cac gta cta act ttt gat ttt aac gac aac ata tta gta act gtt ttc ca- --- --- --- --- --- --- --t ttt ttt ttt tta aga tct acc cac gga tac gaa ata aac atg tct att gtt aat cat gcc acc aat aat gta tag aca att ata taa aac att tgc att ata gaa ttg tct atc tat ctg tat tac cga cga ttg tcc aat att ctg ttc tag gag agt aat ggg tta ttg tgg ata tat aat cag agt ttt taa tga cta cta tat tat gtt tta tac cat ttc gtg tca cag gct ttg tag att tgg a-- --- --- --- --- --- --- -ta tag cat tgc aca tag tat tag tca taa act tgg gat gta aac tgt tga tga tat cta cat cgt ttg gat ttt tat gta tcc act tta ata ata tca tag ctg taa cat cct cgt ga- ttt acg ttg acg ttt tcg tgg gat aag ata gtt --- -gt tca tcc ttt gat aat ttt cca aat t

>EF611202.1:79-2023_Variola_virus_strain_Ngami_left_variable_genomic_sequence

cgg ttg cca tgt att tcc tgg aga gca agt --- aga tga tg- agg aac cag ata gtt tat atc cgt act tgc acg taa agt cta cat tgt cat tat acg a-- ata tga tct ttt aaa ccc gct aga caa gta tcc gtt taa tat tgt agg atg tgg aca ttt aac aat ctg aca cgt ggg tgg att aga cca ttc tcc tcc tga aca cag gac acc aga gtt acc aat caa cga ata tcc act att gca act ata agt tac aac gct tcc atc ggt ata aaa atc att ata tcc gtt atg tct tcc gtt gga tat aga tgg agg taa ttg gca ttt aac aga ttc gca aat agg tgc ctt ggg att cca tac cat aga tcc agt aga tcc taa ttt aca ata cga ttt ata ttc acc aat caa ata ata tcc gct att aca aga gta cgt tat act aga gcc aaa gtc tac tcc gcc aat atc aag atg gcc att atc gat atc tcg agg cga cgg gca tct ccg ttt aat aca ttg att aaa gag tgt cca tcc agt acc tgt aca ttt agc ata tat agg tcc cat ttt ttg ctt tct gta tcc agg tag aca tag a-t att cta tag tgt ctc cta tgt tgt a-- --- --- --- -at tag cat tag cat cag tc- --t cca cac tat tct taa att tca tat taa tgg gtc gtg acg gaa tag tac agc atg ata gaa cgc atc cta ttc cca aca atg tca gga acg tca ctc tct cca cct tca tat tta ttt atc cgt -aa aaa tgt tat cct gga cat tgt gcc aat aat aaa aaa gcc ca- -ta tat gtt cgc tat tgt aga act tgt ttt tca tag ttg ctc aaa aac gat ggc ggt gac tta tga gtc acg tta cac ttt gga gtc tca tct tta gta aac ata tca taa tat tcg ata tta cga gtt gac aca ttg aac aaa -tc caa gta ttt gat ttt gga taa tat tcg tat ttt gca tct gct ata att aag ata taa tcg cca caa aaa cac gcg aac atc ttt cct aca tgg tta aag tac atg tac aat tct atc cat ttg tct tcc tta act ata tat ttg tat aga taa tta cga gtc tcg tgt gta att cca gta att gca tag atg tta cca tcg tac tct aca gca taa act ata cta tga tgt cta ggc atg gga gac -tt ttt tat cca aca att ttt agt gaa aca ttc cac atc gtt taa tat tac ata ttt ttc ata cgt ggt ata aac tcc acc cat tac --a tat ata tca tca ttt acg aat acc gac gcg tct gaa tat cta gga gta att aag ttt gga tgt ctt atc cat ttc gaa gtg tag cgt ttt aaa tat ttt gct aca cct gtt gaa ata gaa aat ttt aat cct cct att aca tat aac ttt cca tca tta aca cac gta cta act ttt gat ttt aac gac gac ata tta gta act gtt ttc ca- --- --- --- --- --- --- ttt ttt ttt gtt tta aga tct acc cac gga tac gaa ata aac atg tct att gtt aat cat gcc acc aat aat gta tag aca att ata taa aa- att tgc att ata gaa ttg tct atc tat ctg tat tac cga cga ttg tcc aat att ctg ttc tag gag agt aat ggg tta ttg tgg ata tat aat cag agt ttt taa tga cta cta tat tat gtt tta tac cat ttc gtg tca cag gct ttg tag att tgg a-- --- --- --- --- --- --- -ta tag cat tgc aca tag tat tag tca taa act tgg gat gta aac tgt tga tga tat cta cat cgt ttg gat ttt tat gta tcc act tta ata ata tca tag ctg taa cat cct cgt ga- ttt acg ttg acg ttt tcg tgg gat aag ata gtt --- -gt tca tcc ttt gat aat ttt cca aat t

>EF611200.1:79-2023_Variola_virus_strain_Mary_left_variable_genomic_sequence

cgg ttg cca tgt att tcc tgg aga gca agt --- aga tga tg- agg aac cag ata gtt tat atc cgt act tgc acg taa agt cta cat tgt cat tat acg a-- ata tga tct ttt aaa ccc gct aga caa gta tcc gtt taa tat tgt agg atg tgg aca ttt aac aat ctg aca cgt ggg tgg att aga cca ttc tcc tcc tga aca cag gac acc aga gtt acc aat caa cga ata tcc act att gca act ata agt tac aac gct tcc atc ggt ata aaa atc att ata tcc gtt atg tct tcc gtt gga tat aga tgg agg taa ttg gca ttt aac aga ttc gca aat agg tgc ctt ggg att cca tac cat aga tcc agt aga tcc taa ttt aca ata cga ttt ata ttc acc aat caa ata ata tcc gct att aca aga gta cgt tat act aga gcc aaa gtc tac tcc gcc aat atc aag atg gcc att atc gat atc tcg agg cga cgg gca tct ccg ttt aat aca ttg att aaa gag tgt cca tcc agt acc tgt aca ttt agc ata tat agg tcc cat ttt ttg ctt tct gta tcc agg tag aca tag a-t att cta tag tgt ctc cta tgt tgt a-- --- --- --- -at tag cat tag cat cag tc- --t cca cac tat tct taa att tca tat taa tgg gtc gtg acg gaa tag tac agc atg ata gaa cgc atc cta ttc cca aca atg tca gga acg tca ctc tct cca cct tca tat tta ttt atc cgt -aa aaa tgt tat cct gga cat tgt gcc aat aat aaa aaa gcc ca- -ta tat gtt cgc tat tgt aga act tgt ttt tca tag ttg ctc aaa aac gat ggc ggt gac tta tga gtc acg tta cac ttt gga gtc tca tct tta gta aac ata tca taa tat tcg ata tta cga gtt gac aca ttg aac aaa -tc caa gta ttt gat ttt gga taa tat tcg tat ttt gca tct gct ata att aag ata taa tcg cca caa aaa cac gcg aac atc ttt cct aca tgg tta aag tac atg tac aat tct atc cat ttg tct tcc tta act ata tat ttg tat aga taa tta cga gtc tcg tgt gta att cca gta att gca tag atg tta cca tcg tac tct aca gca taa act ata cta tga tgt cta ggc atg gga gac -tt ttt tat cca aca att ttt agt gaa aca ttc cac atc gtt taa tat tac ata ttt ttc ata cgt ggt ata aac tcc acc cat tac --a tat ata tca tca ttt acg aat acc gac gcg tct gaa tat cta gga gta att aag ttt gga tgt ctt atc cat ttc gaa gtg tag cgt ttt aaa tat ttt gct aca cct gtt gaa ata gaa aat ttt aat cct cct att aca tat aac ttt cca tca tta aca cac gta cta act ttt gat ttt aac gac gac ata tta gta act gtt ttc ca- --- --- --- --- --- --- ttt ttt ttt gtt tta aga tct acc cac gga tac gaa ata aac atg tct att gtt aat cat gcc acc aat aat gta tag aca att ata taa aa- att tgc att ata gaa ttg tct atc tat ctg tat tac cga cga ttg tcc aat att ctg ttc tag gag agt aat ggg tta ttg tgg ata tat aat cag agt ttt taa tga cta cta tat tat gtt tta tac cat ttc gtg tca cag gct ttg tag att tgg a-- --- --- --- --- --- --- -ta tag cat tgc aca tag tat tag tca taa act tgg gat gta aac tgt tga tga tat cta cat cgt ttg gat ttt tat gta tcc act tta ata ata tca tag ctg taa cat cct cgt ga- ttt acg ttg acg ttt tcg tgg gat aag ata gtt --- -gt tca tcc ttt gat aat ttt cca aat t

>EF611194.1:79-2023_Variola_virus_strain_Helder_left_variable_genomic_sequence

cgg ttg cca tgt att tcc tgg aga gca agt --- aga tga tg- agg aac cag ata gtt tat atc cgt act tgc acg taa agt cta cat tgt cat tat acg a-- ata tga tct ttt aaa ccc gct aga caa gta tcc gtt taa tat tgt agg atg tgg aca ttt aac aat ctg aca cgt ggg tgg att aga cca ttc tcc tcc tga aca cag gac acc aga gtt acc aat caa cga ata tcc act att gca act ata agt tac aac gct tcc atc ggt ata aaa atc att ata tcc gtt atg tct tcc gtt gga tat aga tgg agg taa ttg gca ttt aac aga ttc gca aat agg tgc ctt ggg att cca tac cat aga tcc agt aga tcc taa ttt aca ata cga ttt ata ttc acc aat caa ata ata tcc gct att aca aga gta cgt tat act aga gcc aaa gtc tac tcc gcc aat atc aag atg gcc att atc gat atc tcg agg cga cgg gca tct ccg ttt aat aca ttg att aaa gag tgt cca tcc agt acc tgt aca ttt agc ata tat agg tcc cat ttt ttg ctt tct gta tcc agg tag aca tag a-t att cta tag tgt ctc cta tgt tgt a-- --- --- --- -at tag cat tag cat cag tc- --t cca cac tat tct taa att tca tat taa tgg gtc gtg acg gaa tag tac agc atg ata gaa cgc atc cta ttc cca aca atg tca gga acg tca ctc tct cca cct tca tat tta ttt atc cgt -aa aaa tgt tat cct gga cat tgt gcc aat aat aaa aaa gcc ca- -ta tat gtt cgc tat tgt aga act tgt ttt tca tag ttg ctc aaa aac gat ggc ggt gac tta tga gtc acg tta cac ttt gga gtc tca tct tta gta aac ata tca taa tat tcg ata tta cga gtt gac aca ttg aac aaa -tc caa gta ttt gat ttt gga taa tat tcg tat ttt gca tct gct ata att aag ata taa tcg cca caa aaa cac gcg aac atc ttt cct aca tgg tta aag tac atg tac aat tct atc cat ttg tct tcc tta act ata tat ttg tat aga taa tta cga gtc tcg tgt gta att cca gta att gca tag atg tta cca tcg tac tct aca gca taa act ata cta tga tgt cta ggc atg gga gac -tt ttt tat cca aca att ttt agt gaa aca ttc cac atc gtt taa tat tac ata ttt ttc ata cgt ggt ata aac tcc acc cat tac --a tat ata tca tca ttt acg aat acc gac gcg tct gaa tat cta gga gta att aag ttt gga tgt ctt atc cat ttc gaa gtg tag cgt ttt aaa tat ttt gct aca cct gtt gaa ata gaa aat ttt aat cct cct att aca tat aac ttt cca tca tta aca cac gta cta act ttt gat ttt aac gac gac ata tta gta act gtt ttc ca- --- --- --- --- --- --- ttt ttt ttt gtt tta aga tct acc cac gga tac gaa ata aac atg tct att gtt aat cat gcc acc aat aat gta tag aca att ata taa aa- att tgc att ata gaa ttg tct atc tat ctg tat tac cga cga ttg tcc aat att ctg ttc tag gag agt aat ggg tta ttg tgg ata tat aat cag agt ttt taa tga cta cta tat tat gtt tta tac cat ttc gtg tca cag gct ttg tag att tgg a-- --- --- --- --- --- --- -ta tag cat tgc aca tag tat tag tca taa act tgg gat gta aac tgt tga tga tat cta cat cgt ttg gat ttt tat gta tcc act tta ata ata tca tag ctg taa cat cct cgt ga- ttt acg ttg acg ttt tcg tgg gat aag ata gtt --- -gt tca tcc ttt gat aat ttt cca aat t

>DQ437592.1:12101-14044_Variola_virus_strain_Syria_1972_V72-199_complete_genome

cgg ttg cca tgt att tcc tgg aga gca agt --- aga tga tg- agg aac cag ata gtt tat atc cgt act tgc acg taa agt cta cat tgt cat tat acg a-- ata tga tct ttt aaa ccc gct aga caa gta tcc gtt taa tat tgt agg atg tgg aca ttt aac aat ctg aca cgt ggg tgg att aga cca ttc tcc tcc tga aca cag gac acc aga gtt acc aat caa cga ata tcc act att gca act ata agt tac aac gct tcc atc ggt ata aaa atc att ata tcc gtt atg tct tcc gtt gga tat aga tgg agg taa ttg gca ttt aac aga ttc gca aat agg tgc ctt ggg att cca tac cat aga tcc agt aga tcc taa ttt aca ata cga ttt ata ttc acc aat caa ata ata tcc gct att aca aga gta cgt tat act aga gcc aaa gtc tac tcc gcc aat atc aag atg gcc att atc gat atc tcg agg cga cgg gca tct ccg ttt aat aca ttg att aaa gag tgt cca tcc agt acc tgt aca ttt agc ata tat agg tcc cat ttt ttg ctt tct gta tcc agg tag aca tag a-t att cta tag tgt ctc cta tgt tgt a-- --- --- --- -at tag cat tag cat cag tc- --t cca cac tat tct taa att tca tat taa tgg gtc gtg acg gaa tag tac agc atg ata gaa cgc atc cta ttc cca aca atg tca gga acg tca ctc tct cca cct tca tat tta ttt atc cgt -aa aaa tgt tat cct gga cat tgt gcc aat aat aaa aaa gcc ca- -ta tat gtt cgc tat tgt aga aat tgt ttt tca tag ttg ctc aaa aac gat ggc ggt gac tta tga gtc acg tta cac ttt gga gtc tca tct tta gta aac ata tca taa tat tcg ata tta cga gtt gac aca ttg aac aaa -tc caa gta ttt gat ttt gga taa tat tcg tat ttt gca tct gct ata att aag ata taa tcg cca caa aaa cac gcg aac atc ttt cct aca tgg tta aag tac atg tac aat tct atc cat ttg tct tcc tta act ata tat ttg tat aga taa tta cga gtc tcg tgt gta att cca gta att gca tag atg tta cca tcg tac tct aca gca taa act ata cta tga tgt cta ggc atg gga gac -tt ttt tat cca aca att ttt agt gaa aca ttc cac att gtt taa tat tac ata ttt ttc ata cgt ggt ata aac tcc acc cat tac --a tat ata tca tca ttt acg aat acc gac gcg tct gaa tat cta gga gta att aag ttt gga tgt ctt atc cat ttc gaa gtg tag cgt ttt aaa tat ttt gct aca cct gtt gaa ata gaa aat ttt aat cct cct att aca tat aac ttt cca tca tta aca cac gta cta act ttt gat ttt gac gac gac ata tta gta act gtt ttc ca- --- --- --- --- --- --- --t ttt ttt ttt tta aga tct acc cac gga tac gaa ata aac atg tct att gtt aat cat gcc acc aat aat gta tag aca att ata taa aac att tgc att ata gaa ttg tct atc tat ctg tat tac cga cga ttg tcc aat att ctg ttc tag gag agt aat ggg tta ttg tgg ata tat aat cag agt ttt taa tga cta cta tat tat gtt tta tac cat ttc gtg tca cag gct ttg tag att tgg a-- --- --- --- --- --- --- -ta tag cat tgc aca tag tat tag tca taa act tgg gat gta aac tgt tga tga tat cta cat cgt ttg gat ttt tat gta tcc act tta ata ata tca tag ctg taa cat cct cgt ga- ttt acg ttg acg ttt tcg tgg gat aag ata gtt --- -gt tca tcc ttt gat aat ttt cca aat t

>DQ441448.1:12101-14044_Variola_virus_strain_Yugoslavia_1972_V72-164_complete_genome

cgg ttg cca tgt att tcc tgg aga gca agt --- aga tga tg- agg aac cag ata gtt tat atc cgt act tgc acg taa agt cta cat tgt cat tat acg a-- ata tga tct ttt aaa ccc gct aga caa gta tcc gtt taa tat tgt agg atg tgg aca ttt aac aat ctg aca cgt ggg tgg att aga cca ttc tcc tcc tga aca cag gac acc aga gtt acc aat caa cga ata tcc act att gca act ata agt tac aac gct tcc atc ggt ata aaa atc att ata tcc gtt atg tct tcc gtt gga tat aga tgg agg taa ttg gca ttt aac aga ttc gca aat agg tgc ctt ggg att cca tac cat aga tcc agt aga tcc taa ttt aca ata cga ttt ata ttc acc aat caa ata ata tcc gct att aca aga gta cgt tat act aga gcc aaa gtc tac tcc gcc aat atc aag atg gcc att atc gat atc tcg agg cga cgg gca tct ccg ttt aat aca ttg att aaa gag tgt cca tcc agt acc tgt aca ttt agc ata tat agg tcc cat ttt ttg ctt tct gta tcc agg tag aca tag a-t att cta tag tgt ctc cta tgt tgt a-- --- --- --- -at tag cat tag cat cag tc- --t cca cac tat tct taa att tca tat taa tgg gtc gtg acg gaa tag tac agc atg ata gaa cgc atc cta ttc cca aca atg tca gga acg tca ctc tct cca cct tca tat tta ttt atc cgt -aa aaa tgt tat cct gga cat tgt gcc aat aat aaa aaa gcc ca- -ta tat gtt cgc tat tgt aga aat tgt ttt tca tag ttg ctc aaa aac gat ggc ggt gac tta tga gtc acg tta cac ttt gga gtc tca tct tta gta aac ata tca taa tat tcg ata tta cga gtt gac aca ttg aac aaa -tc caa gta ttt gat ttt gga taa tat tcg tat ttt gca tct gct ata att aag ata taa tcg cca caa aaa cac gcg aac atc ttt cct aca tgg tta aag tac atg tac aat tct atc cat ttg tct tcc tta act ata tat ttg tat aga taa tta cga gtc tcg tgt gta att cca gta att gca tag atg tta cca tcg tac tct aca gca taa act ata cta tga tgt cta ggc atg gga gac -tt ttt tat cca aca att ttt agt gaa aca ttc cac att gtt taa tat tac ata ttt ttc ata cgt ggt ata aac tcc acc cat tac --a tat ata tca tca ttt acg aat acc gac gcg tct gaa tat cta gga gta att aag ttt gga tgt ctt atc cat ttc gaa gtg tag cgt ttt aaa tat ttt gct aca cct gtt gaa ata gaa aat ttt aat cct cct att aca tat aac ttt cca tca tta aca cac gta cta act ttt gat ttt gac gac gac ata tta gta act gtt ttc ca- --- --- --- --- --- --- --t ttt ttt ttt tta aga tct acc cac gga tac gaa ata aac atg tct att gtt aat cat gcc acc aat aat gta tag aca att ata taa aac att tgc att ata gaa ttg tct atc tat ctg tat tac cga cga ttg tcc aat att ctg ttc tag gag agt aat ggg tta ttg tgg ata tat aat cag agt ttt taa tga cta cta tat tat gtt tta tac cat ttc gtg tca cag gct ttg tag att tgg a-- --- --- --- --- --- --- -ta tag cat tgc aca tag tat tag tca taa act tgg gat gta aac tgt tga tga tat cta cat cgt ttg gat ttt tat gta tcc act tta ata ata tca tag ctg taa cat cct cgt ga- ttt acg ttg acg ttt tcg tgg gat aag ata gtt --- -gt tca tcc ttt gat aat ttt cca aat t

>DQ437588.1:12103-14045_Variola_virus_strain_Nepal_1973_complete_genome

cgg ttg cca tgt att tcc tgg aga gca agt --- aga tga tg- agg aac cag ata gtt tat atc cgt act tgc acg taa agt cta cat tgt cat tat acg a-- ata tga tct ttt aaa ccc gct aga caa gta tcc gtt taa tat tgt agg atg tgg aca ttt aac aat ctg aca cgt ggg tgg att aga cca ttc tcc tcc tga aca cag gac acc aga gtt acc aat caa cga ata tcc act att gca act ata agt tac aac gct tcc atc ggt ata aaa atc att ata tcc gtt atg tct tcc gtt gga tat aga tgg agg taa ttg gca ttt aac aga ttc gca aat agg tgc ctt ggg att cca tac cat aga tcc agt aga tcc taa ttt aca ata cga ttt ata ttc acc aat caa ata ata tcc gct att aca aga gta cgt tat act aga gcc aaa gtc tac tcc gcc aat atc aag atg gcc att atc gat atc tcg agg cga cgg gca tct ccg ttt aat aca ttg att aaa gag tgt cca tcc agt acc tgt aca ttt agc ata tat agg tcc cat ttt ttg ctt tct gta tcc agg tag aca tag a-t att cta tag tgt ctc cta tgt tgt a-- --- --- --- -at tag cat tag cat cag tc- --t cca cac tat tct taa att tca tat taa tgg gtc gtg acg gaa tag tac agc atg ata gaa cgc atc cta ttc cca aca atg tca gga acg tca ctc tct cca cct tca tat tta ttt atc cgt -aa aaa tgt tat cct gga cat tgt gcc aat aat aaa aaa gcc ca- -ta tat gtt cgc tat tgt aga aat tgt ttt tca tag ttg ctc aaa aac gat ggc ggt gac tta tga gtc acg tta cac ttt gga gtc tca tct tta gta aac ata tca taa tat tcg ata tta cga gtt gac aca ttg aac aaa -tc caa gta ttt gat ttt gga taa tat tcg tat ttt gca tct gct ata att aag ata taa tcg cca caa aaa cac gcg aac atc ttt cct aca tgg tta aag tac atg tac aat tct atc cat ttg tct tcc tta act ata tat ttg tat aga taa tta cga gtc tcg tgt gta att cca gta att gca tag atg tta cca tcg tac tct aca gca taa act ata cta tga tgt cta ggc atg gga gac -tt ttt tat cca aca att ttt agt gaa aca ttc cac atc gtt taa tat tac ata ttt ttc ata cgt ggt ata aac tcc acc cat tac --a tat ata tca tca ttt acg aat acc gac gcg tct gaa tat cta gga gta att aag ttt gga tgt ctt atc cat ttc gaa gtg tag cgt ttt aaa tat ttt gct aca cct gtt gaa ata gaa aat ttt aat cct cct att aca tat aac ttt cca tca tta aca cac gta cta act ttt gat ttt aac gac aac ata tta gta act gtt ttc ca- --- --- --- --- --- --- --- ttt ttt ttt tta aga tct acc cac gga tac gaa ata aac atg tct att gtt aat cat gcc acc aat aat gta tag aca att ata taa aac att tgc att ata gaa ttg tct atc tat ctg tat tac cga cga ttg tcc aat att ctg ttc tag gag agt aat ggg tta ttg tgg ata tat aat cag agt ttt taa tga cta cta tat tat gtt tta tac cat ttc gtg tca cag gct ttg tag att tgg a-- --- --- --- --- --- --- -ta tag cat tgc aca tag tat tag tca taa act tgg gat gta aac tgt tga tga tat cta cat cgt ttg gat ttt tat gta tcc act tta ata ata tca tag ctg taa cat cct cgt ga- ttt acg ttg acg ttt tcg tgg gat aag ata gtt --- -gt tca tcc ttt gat aat ttt cca aat t

>DQ441447.1:12902-14844_Variola_virus_strain_United_Kingdom_1952_Butler_complete_genome

cgg ttg cca tgt att tcc tgg aga gca agt --- aga tga tg- agg aac cag ata gtt tat atc cgt act tgc acg taa agt cta cat tgt cat tat acg a-- ata tga tct ttt aaa ccc gct aga caa gta tcc gtt taa tat tgt agg ata tgg aca ttt aac aat ctg aca cgt ggg tgg att aga cca ttc tcc tcc tga aca cag gac acc aga gtt acc aat caa cga ata tcc act att gca act ata agt tac aac gct tcc atc ggt ata aaa atc att ata tcc gtt atg tct tcc gtt gga tat aga tgg agg taa ttg gca ttt aac aga ttc gca aat agg tgc ctt ggg att cca tac cat aga tcc agt aga tcc taa ttt aca ata cga ttt ata ttc acc gat caa ata ata tcc gct att aca aga gta cgt tat act aga gcc aaa gtc tac tcc gcc aat atc aag atg gcc att atc gat atc tcg agg cga cgg gca tct ccg ttt aat aca ttg att aaa gag tgt cca tcc agt acc tgt aca ttt agc ata tat agg tcc cat ttt ttg ctt tct gta tcc agg tag aca tag a-t att cta tag tgt ctc cta tgt tgt a-- --- --- --- -at tag cat tag cat cag tc- --t cca cac tat tct taa att tca tat taa tgg gtc gtg acg gaa tag tac agc atg ata gaa cgc atc cta ttc cca aca atg tca gga acg tca ctc tct cca cct tca tat tta ttt atc cgt -aa aaa tgt tat cct gga cat tgt gtc aat aat aaa aaa gct ca- -ta tat gtt tgc tat tgt aga aat tgt ttt tca tag ttg ctc aaa aac gat ggc ggt gac tta tga gtc acg tta cac ttt gga gtc tca tct tta gta aac ata tca taa tat tcg ata tta cga gtt gac ata ttg aac aaa ttc caa gta ttt gat ttt gga taa tat tcg tat ttt gca tct gct ata att aag ata taa tcg cca caa gaa cac acg aac atc ttt cct aca tgg tta aag tac atg tac aat tct atc cat ttg tct tcc tta act ata tat ttg tat aga taa tta cga gtc tcg tgt gta att cca gta att gca tag atg tta cca tcg tac tct aca gca taa act ata cta tga tgt cta ggc atg gga gac -tt ttt tat cca aca att ttt agt gaa aca ttc cac atc gtt taa tat tac ata ttt ttc ata cgt ggt ata aac tcc acc cat tac --a tat ata tca tca ttt acg aat acc gac gcg tct gaa tat cta gga gta att aag ttt gga tgt ctt atc cat ttc gaa gtg tag cgt ttt aaa tat tct gct aca cct gtt aaa ata gaa aat ttt aat cct cct att aca tat aac ttt cca tca tta aca cac gta ct- act ttt gat ttt aac gac gac ata tta gta act gtt ttc ca- --- --- --- --- --- --- -tt ttt ttc gtt tta aga ttt acc cac gga tac gaa ata aac atg tct att gtt aat cat gcc gcc aat aat gta tag aca att ata taa aac att tgc att ata gaa ttg tct atc tat ctg tat tac cga cga ttg tcc aat att ctg ttc tag gag agt aat ggg tta ttg tgg --a tat aat cag agt ttt taa tga cta cta tat tat gtt tta tac cat ttc gtg tca cag gct ttg tag att tgg a-- --- --- --- --- --- --- -ta tag cat tgc aca tag tat tag tca taa act tgg gat gta aaa tgt tga tga tat cta cat cgt ttg gat ttt tat gta tcc act tta ata ata tca tag ctg taa cat cct tgt ga- ttt acg ttg acg ttt tcg tgg gat aag ata gtt --- -gt tca tcc ttt gat aat ttt cca aat t

>DQ441419.1:12902-14844_Variola_virus_strain_Brazil_1966_(v66-39_Sao_Paulo)_complete_genome

cgg ttg cca tgt att tcc tgg aga gca agt --- aga tga tg- agg aac cag ata gtt tat atc cgt act tgc acg taa agt cta cat tgt cat tat acg a-- ata tga tct ttt aaa ccc gct aga caa gta tcc gtt taa tat tgt agg ata tgg aca ttt aac aat ctg aca cgt ggg tgg att aga cca ttc tcc tcc tga aca cag gac acc aga gtt acc aat caa cga ata tcc act att gca act ata agt tac aac gct tcc atc ggt ata aaa atc att ata tcc gtt atg tct tcc gtt gga tat aga tgg agg taa ttg gca ttt aac aga ttc gca aat agg tgc ctt ggg att cca tac cat aga tcc agt aga tcc taa ttt aca ata cga ttt ata ttc acc gat caa ata ata tcc gct att aca aga gta cgt tat act aga gcc aaa gtc tac tcc gcc aat atc aag atg gcc att atc gat atc tcg agg cga cgg gca tct ccg ttt aat aca ttg att aaa gag tgt cca tcc agt acc tgt aca ttt agc ata tat agg tcc cat ttt ttg ctt tct gta tcc agg tag aca tag a-t att cta tag tgt ctc cta tgt tgt a-- --- --- --- -at tag cat tag cat cag tc- --t cca cac tat tct taa att tca tat taa tgg gtc gtg acg gaa tag tac agc atg ata gaa cgc atc cta ttc cca aca atg tca gga acg tca ctc tct cca cct tca tat tta ttt atc cgt -aa aaa tgt tat cct gga cat tgt gtc aat aat aaa aaa gct ca- -ta tat gtt tgc tat tgt aga aat tgt ttt tca tag ttg ctc aaa aac gat ggc ggt gac tta tga gtc acg tta cac ttt gga gtc tca tct tta gta aac ata tca taa tat tcg ata tta cga gtt gac ata ttg aac aaa ttc caa gta ttt gat ttt gga taa tat tcg tat ttt gca tct gct ata att aag ata taa tcg cca caa gaa cac acg aac atc ttt cct aca tgg tta aag tac atg tac aat tct atc cat ttg tct tcc tta act ata tat ttg tat aga taa tta cga gtc tcg tgt gta att cca gta att gca tag atg tta cca tcg tac tct aca gca taa act ata cta tga tgt cta ggc atg gga gac -tt ttt tat cca aca att ttt agt gaa aca ttc cac atc gtt taa tat tac ata ttt ttc ata cgt ggt ata aac tcc acc cat tac --a tat ata tca tca ttt acg aat acc gac gcg tct gaa tat cta gga gta att aag ttt gga tgt ctt atc cat ttc gaa gtg tag cgt ttt aaa tat tct gct aca cct gtt aaa ata gaa aat ttt aat cct cct att aca tat aac ttt cca tca tta aca cac gta ct- act ttt gat ttt aac gac gac ata tta gta act gtt ttc ca- --- --- --- --- --- --- -tt ttt ttc gtt tta aga ttt acc cac gga tac gaa ata aac atg tct att gtt aat cat gcc gcc aat aat gta tag aca att ata taa aac att tgc att ata gaa ttg tct atc tat ctg tat tac cga cga ttg tcc aat att ctg ttc tag gag agt aat ggg tta ttg tgg --a tat aat cag agt ttt taa tga cta cta tat tat gtt tta tac cat ttc gtg tca cag gct ttg tag att tgg a-- --- --- --- --- --- --- -ta tag cat tgc aca tag tat tag tca taa act tgg gat gta aaa tgt tga tga tat cta cat cgt ttg gat ttt tat gta tcc act tta ata ata tca tag ctg taa cat cct tgt ga- ttt acg ttg acg ttt tcg tgg gat aag ata gtt --- -gt tca tcc ttt gat aat ttt cca aat t

>KT184691.1:19549-21441_Vaccinia_virus_strain_IOC_clone_B388

cgg ctg cca tgt att tcc tgg aga gca agt --- aga tga tg- agg aac cag ata ggt tat atc cgt act tgc act taa agt tta cat tgt cgt tgt atg a-- gta tga tct ttt aaa ccc gct aga caa gta tcc gtt tga tat tgt agg atg tgg aca ttt aac aat ttg aca cgt ggg tgg atc gga cca ttc tcc tcc tga aca cag gac act aga gtt acc aat caa cga ata tcc act att gca act ata agt tac aac gct tcc atc ggt ata aaa atc ctc gta tcc gtt atg tct tcc gtt gga tat aga tgg agg gga ttg gca ttt aac aga ttc aca aat agg tgc ctc ggg att cca tac cat aga tcc agt aaa tcc taa ttc aca ata cga ttt aga ttc tcc tat caa ctg ata tcc gct att aca aga gta cgt tat act aga gcc aaa gtc tac tcc gcc aat atc aag ttg gcc att atc gat atc tcg agg cga tgg gca tct ccg ttt aat aca ttg att aaa gag tgt cca tcc agt acc tgt aca ttt agc ata tat agg tcc cat ttt ttg ctt tct gta tcc agg tag aca tag a-t att cta tag tgt ctc cta tgt tgt a-- --- --- --- --- --- -at tag cat tag tt- --t cca cac tat tct taa att tca tat taa tgg gac gtg aag gaa tag gac agt atg ata gaa cgc atc cta ttc cca aca atg tca gga acg tca cgc tct cca cct tca tat tta ttt atc cgt -aa aaa tgt tat cct gga cat cgt aca aat aat aaa aaa gcc ca- -ta tat gtt tgc tat tgt aga aat tgt ttt tca cag ttg ctc aaa aac gat ggc agt gac tta tga gtt --- --- --- --- --- --- tca tct tta gta aac ata tca taa tat tcg ata tta cga gtt gac ata tcg aac aaa ttc caa gta ttt gat ttt gga taa tat tcg tat ttt gca tct gc- --- --- --- -ta taa tca ccg caa gaa cac acg aac atc ttt cct aca tgg tta aag tac atg tac aat tct atc cat ttg tct tcc tta act ata tat ttg tat aga taa tta cga gtc tcg tg- --- --- --a gta att aca tag atg tcg ccg tcg tac tct aca gca taa act ata cta tga tgt cta ggc atg gga gac -tt ttt tat cca acg att ttt agt gaa aca ttc cac atc gtt taa tac tac ata ttt ttc ata cgt ggt ata aac tcc acc cat tac --a tat ata tca tcg ttt acg aat acc gac gcg cct gaa tat cta gga gta att aag ttt gga agt ctt atc cat ttt gaa gtg ccg tgt ttc aaa tat tct gct aca ccc gtt gaa ata gaa aat tct aat cct cct att aca tat aac ttt cca ttg tta aca caa gta cta act tct gat ttt aac gac gac ata tta gta acc gtt ttc ca- --- --- --- --- --- --- --t ttt ttc gtt tca aga tct acc cgc -ga tac gga ata aac atg --- --- --- --- --- -cc gcc aat aat gta tag aca att atg taa aac att tgc att ata gaa ttg --- -tc tat ctg tat tac cga cta tcg tcc aat att ctg ttc tag gag agt aat ggg tta ttg tgg ata tat aat cag agt ttt taa tga cta cta tat tat gtt tta tac cat ttc gtg tca ctg gct ttg tag att tgg ata tag tta atc cca aca atg ata tag c-- --- --- --- -at tag tta taa act tgg gat gta aaa tgt tga tga tat cta cat cgt ttg gat ttt tat gta tcc act tta ata ata tca tag ctg taa cat cct cat ga- ttt acg tta acg tct tcg tgg gat aag ata gtt gtc agt tca tcc ttt gat aat ttt cca aat t

>EF611191.1:79-2021_Variola_virus_strain_Brazil_131_left_variable_genomic_sequence

cgg ttg cca tgt att tcc tgg aga gca agt --- aga tga tg- agg aac cag ata gtt tat atc cgt act tgc acg taa agt cta cat tgt cat tat acg a-- ata tga tct ttt aaa ccc gct aga caa gta tcc gtt taa tat tgt agg ata tgg aca ttt aac aat ctg aca cgt ggg tgg att aga cca ttc tcc tcc tga aca cag gac acc aga gtt acc aat caa cga ata tcc act att gca act ata agt tac aac gct tcc atc tgt ata aaa atc att ata tcc gtt atg tct tcc gtt gga tat aga tgg agg taa ttg gca ttt aac aga ttc gca aat agg tgc ctt ggg att cca tac cat aga tcc agt aga tcc taa ttt aca ata cga ttt ata ttc acc gat caa ata ata tcc gct att aca aga gta cgt tat act aga gcc aaa gtc tac tcc gcc aat atc aag atg gcc att atc gat atc tcg agg cga cgg gca tct ccg ttt aat aca ttg att aaa gag tgt cca tcc agt acc tgt aca ttt agc ata tat agg tcc cat ttt ttg ctt tct gta tcc agg tag aca tag a-t att cta tag tgt ctc cta tgt tgt a-- --- --- --- -at tag cat tag cat cag tc- --t cca cac tat tct taa att tca tat taa tgg gtc gtg acg gaa tag tac agc atg ata gaa cgc atc cta ttc cca aca atg tca gga acg tca ctc tct cca cct tca tat tta ttt atc cgt -aa aaa tgt tat cct gga cat tgt gtc aat aat aaa aaa gct ca- -ta tat gtt tgc tat tgt aga aat tgt ttt tca tag ttg ctc aaa aac gat ggc ggt gac tta tga gtc acg tta cac ttt gga gtc tca tct tta gta aac ata tca taa tat tcg ata tta cga gtt gac ata ttg aac aaa ttc caa gta ttt gat ttt gga taa tat tcg tat ttt gca tct gct ata att aag ata taa tcg cca caa gaa cac acg aac atc ttt cct aca tgg tta aag tac atg tac aat tct atc cat ttg tct tcc tta act ata tat ttg tat aga taa tta cga gtc tcg tgt gta att cca gta att gca tag atg tta cca tcg tac tct aca gca taa act ata cta tga tgt cta ggc atg gga gac -tt ttt tat cca aca att ttt agt gaa aca ttc cac atc gtt taa tat tac ata ttt ttc ata cgt ggt ata aac tcc acc cat tac --a tat ata tca tca ttt acg aat acc gac gcg tct gaa tat cta gga gta att aag ttt gga tgt ctt atc cat ttc gaa gtg tag cgt ttt aaa tat tct gct aca cct gtt aaa ata gaa aat ttt aat cct cct att aca tat aac ttt cca tca tta aca cac gta ct- act ttt gat ttt aac gac gac ata tta gta act gtt ttc ca- --- --- --- --- --- --- -tt ttt ttc gtt tta aga ttt acc cac gga tac gaa ata aac atg tct att gtt aat cat gcc gcc aat aat gta tag aca att ata taa aac att tgc att ata gaa ttg tct atc tat ctg tat tac cga cga ttg tcc aat att ctg ttc tag gag agt aat ggg tta ttg tgg --a tat aat cag agt ttt taa tga cta cta tat tat gtt tta tac cat ttc gtg tca cag gct ttg tag att tgg a-- --- --- --- --- --- --- -ta tag cat tgc aca tag tat tag tca taa act tgg gat gta aaa tgt tga tga tat cta cat cgt ttg gat ttt tat gta tcc act tta ata ata tca tag ctg taa cat cct tgt ga- ttt acg ttg acg ttt tcg tgg gat aag ata gtt --- -gt tca tcc ttt gat aat ttt cca aat t

>Y16780.1:12476-14418_variola_minor_virus_complete_genome

cgg ttg cca tgt att tcc tgg aga gca agt --- aga tga tg- agg aac cag ata gtt tat atc cgt act tgc acg taa agt cta cat tgt cat tat acg a-- ata tga tct ttt aaa ccc gct aga caa gta tcc gtt taa tat tgt agg ata tgg aca ttt aac aat ctg aca cgt ggg tgg att aga cca ttc tcc tcc tga aca cag gac acc aga gtt acc aat caa cga ata tcc act att gca act ata agt tac aac gct tcc atc tgt ata aaa atc att ata tcc gtt atg tct tcc gtt gga tat aga tgg agg taa ttg gca ttt aac aga ttc gca aat agg tgc ctt ggg att cca tac cat aga tcc agt aga tcc taa ttt aca ata cga ttt ata ttc acc gat caa ata ata tcc gct att aca aga gta cgt tat act aga gcc aaa gtc tac tcc gcc aat atc aag atg gcc att atc gat atc tcg agg cga cgg gca tct ccg ttt aat aca ttg att aaa gag tgt cca tcc agt acc tgt aca ttt agc ata tat agg tcc cat ttt ttg ctt tct gta tcc agg tag aca tag a-t att cta tag tgt ctc cta tgt tgt a-- --- --- --- -at tag cat tag cat cag tc- --t cca cac tat tct taa att tca tat taa tgg gtc gtg acg gaa tag tac agc atg ata gaa cgc atc cta ttc cca aca atg tca gga acg tca ctc tct cca cct tca tat tta ttt atc cgt -aa aaa tgt tat cct gga cat tgt gtc aat aat aaa aaa gct ca- -ta tat gtt tgc tat tgt aga aat tgt ttt tca tag ttg ctc aaa aac gat ggc ggt gac tta tga gtc acg tta cac ttt gga gtc tca tct tta gta aac ata tca taa tat tcg ata tta cga gtt gac ata ttg aac aaa ttc caa gta ttt gat ttt gga taa tat tcg tat ttt gca tct gct ata att aag ata taa tcg cca caa gaa cac acg aac atc ttt cct aca tgg tta aag tac atg tac aat tct atc cat ttg tct tcc tta act ata tat ttg tat aga taa tta cga gtc tcg tgt gta att cca gta att gca tag atg tta cca tcg tac tct aca gca taa act ata cta tga tgt cta ggc atg gga gac -tt ttt tat cca aca att ttt agt gaa aca ttc cac atc gtt taa tat tac ata ttt ttc ata cgt ggt ata aac tcc acc cat tac --a tat ata tca tca ttt acg aat acc gac gcg tct gaa tat cta gga gta att aag ttt gga tgt ctt atc cat ttc gaa gtg tag cgt ttt aaa tat tct gct aca cct gtt aaa ata gaa aat ttt aat cct cct att aca tat aac ttt cca tca tta aca cac gta ct- act ttt gat ttt aac gac gac ata tta gta act gtt ttc ca- --- --- --- --- --- --- -tt ttt ttc gtt tta aga ttt acc cac gga tac gaa ata aac atg tct att gtt aat cat gcc gcc aat aat gta tag aca att ata taa aac att tgc att ata gaa ttg tct atc tat ctg tat tac cga cga ttg tcc aat att ctg ttc tag gag agt aat ggg tta ttg tgg --a tat aat cag agt ttt taa tga cta cta tat tat gtt tta tac cat ttc gtg tca cag gct ttg tag att tgg a-- --- --- --- --- --- --- -ta tag cat tgc aca tag tat tag tca taa act tgg gat gta aaa tgt tga tga tat cta cat cgt ttg gat ttt tat gta tcc act tta ata ata tca tag ctg taa cat cct tgt ga- ttt acg ttg acg ttt tcg tgg gat aag ata gtt --- -gt tca tcc ttt gat aat ttt cca aat t

>U18338.1:12476-14418_Variola_virus_Garcia-1966_left_near-terminal_region

cgg ttg cca tgt att tcc tgg aga gca agt --- aga tga tg- agg aac cag ata gtt tat atc cgt act tgc acg taa agt cta cat tgt cat tat acg a-- ata tga tct ttt aaa ccc gct aga caa gta tcc gtt taa tat tgt agg ata tgg aca ttt aac aat ctg aca cgt ggg tgg att aga cca ttc tcc tcc tga aca cag gac acc aga gtt acc aat caa cga ata tcc act att gca act ata agt tac aac gct tcc atc tgt ata aaa atc att ata tcc gtt atg tct tcc gtt gga tat aga tgg agg taa ttg gca ttt aac aga ttc gca aat agg tgc ctt ggg att cca tac cat aga tcc agt aga tcc taa ttt aca ata cga ttt ata ttc acc gat caa ata ata tcc gct att aca aga gta cgt tat act aga gcc aaa gtc tac tcc gcc aat atc aag atg gcc att atc gat atc tcg agg cga cgg gca tct ccg ttt aat aca ttg att aaa gag tgt cca tcc agt acc tgt aca ttt agc ata tat agg tcc cat ttt ttg ctt tct gta tcc agg tag aca tag a-t att cta tag tgt ctc cta tgt tgt a-- --- --- --- -at tag cat tag cat cag tc- --t cca cac tat tct taa att tca tat taa tgg gtc gtg acg gaa tag tac agc atg ata gaa cgc atc cta ttc cca aca atg tca gga acg tca ctc tct cca cct tca tat tta ttt atc cgt -aa aaa tgt tat cct gga cat tgt gtc aat aat aaa aaa gct ca- -ta tat gtt tgc tat tgt aga aat tgt ttt tca tag ttg ctc aaa aac gat ggc ggt gac tta tga gtc acg tta cac ttt gga gtc tca tct tta gta aac ata tca taa tat tcg ata tta cga gtt gac ata ttg aac aaa ttc caa gta ttt gat ttt gga taa tat tcg tat ttt gca tct gct ata att aag ata taa tcg cca caa gaa cac acg aac atc ttt cct aca tgg tta aag tac atg tac aat tct atc cat ttg tct tcc tta act ata tat ttg tat aga taa tta cga gtc tcg tgt gta att cca gta att gca tag atg tta cca tcg tac tct aca gca taa act ata cta tga tgt cta ggc atg gga gac -tt ttt tat cca aca att ttt agt gaa aca ttc cac atc gtt taa tat tac ata ttt ttc ata cgt ggt ata aac tcc acc cat tac --a tat ata tca tca ttt acg aat acc gac gcg tct gaa tat cta gga gta att aag ttt gga tgt ctt atc cat ttc gaa gtg tag cgt ttt aaa tat tct gct aca cct gtt aaa ata gaa aat ttt aat cct cct att aca tat aac ttt cca tca tta aca cac gta ct- act ttt gat ttt aac gac gac ata tta gta act gtt ttc ca- --- --- --- --- --- --- -tt ttt ttc gtt tta aga ttt acc cac gga tac gaa ata aac atg tct att gtt aat cat gcc gcc aat aat gta tag aca att ata taa aac att tgc att ata gaa ttg tct atc tat ctg tat tac cga cga ttg tcc aat att ctg ttc tag gag agt aat ggg tta ttg tgg --a tat aat cag agt ttt taa tga cta cta tat tat gtt tta tac cat ttc gtg tca cag gct ttg tag att tgg a-- --- --- --- --- --- --- -ta tag cat tgc aca tag tat tag tca taa act tgg gat gta aaa tgt tga tga tat cta cat cgt ttg gat ttt tat gta tcc act tta ata ata tca tag ctg taa cat cct tgt ga- ttt acg ttg acg ttt tcg tgg gat aag ata gtt --- -gt tca tcc ttt gat aat ttt cca aat t

>DQ437587.1:12101-14047_Variola_virus_strain_Iran_1972_2602_Tabriz_complete_genome

cgg ttg cca tgt att tcc tgg aga gca agt --- aga tga tg- agg aac cag ata gtt tat atc cgt act tgc acg taa agt cta cat tgt cat tat acg a-- ata tga tct ttt aaa ccc gct aga caa gta tcc gtt taa tat tgt agg atg tgg aca ttt aac aat ctg aca cgt ggg tgg att aga cca ttc tcc tcc tga aca cag gac acc aga gtt acc aat caa cga ata tcc act att gca act ata agt tac aac gct tcc atc ggt ata aaa atc att ata tcc gtt atg tct tcc gtt gga tat aga tgg agg taa ttg gca ttt aac aga ttc gca aat agg tgc ctt ggg att cca tac cat aga tcc agt aga tcc taa ttt aca ata cga ttt ata ttc acc aat caa ata ata tcc gct att aca aga gta cgt tat act aga gcc aaa gtc tac tcc gcc aat atc aag atg gcc att atc gat atc tcg agg cga cgg gca tct ccg ttt aat aca ttg att aaa gag tgt cca tcc agt acc tgt aca ttt agc ata tat agg tcc cat ttt ttg ctt tct gta tcc agg tag aca tag a-t att cta tag tgt ctc cta tgt tgt a-- --- --- --- -at tag cat tag cat cag tc- --t cca cac tat tct taa att tca tat taa tgg gtc gtg acg gaa tag tac agc atg ata gaa cgc atc cta ttc cca aca atg tca gga acg tca ctc tct cca cct tca tat tta ttt atc cgt -aa aaa tgt tat cct gga cat tgt gcc aat aat aaa aaa gcc ca- -ta tat gtt cgc tat tgt aga aat tgt ttt tca tag ttg ctc aaa aac gat ggc ggt gac tta tga gtc acg tta cac ttt gga gtc tca tct tta gta aac ata tca taa tat tcg ata tta cga gtt gac aca ttg aac aaa -tc caa gta ttt gat ttt gga taa tat tcg tat ttt gca tct gct ata att aag ata taa tcg cca caa aaa cac gcg aac atc ttt cct aca tgg tta aag tac atg tac aat tct atc cat ttg tct tcc tta act ata tat ttg tat aga taa tta cga gtc tcg tgt gta att cca gta att gca tag atg tta cca tcg tac tct aca gca taa act ata cta tga tgt cta ggc atg gga gac -tt ttt tat cca aca att ttt agt gaa aca ttc cac att gtt taa tat tac ata ttt ttc ata cgt ggt ata aac tcc acc cat tac --a tat ata tca tca ttt acg aat acc gac gcg tct gaa tat cta gga gta att aag ttt gga tgt ctt atc cat ttc gaa gtg tag cgt ttt aaa tat ttt gct aca cct gtt gaa ata gaa aat ttt aat cct cct att aca tat aac ttt cca tca tta aca cac gta cta act ttt gat ttt gac gac gac ata tta gta act gtt ttc ca- --- --- --- --- --- --t ttt ttt ttt ttt tta aga tct acc cac gga tac gaa ata aac atg tct att gtt aat cat gcc acc aat aat gta tag aca att ata taa aac att tgc att ata gaa ttg tct atc tat ctg tat tac cga cga ttg tcc aat att ctg ttc tag gag agt aat ggg tta ttg tgg ata tat aat cag agt ttt taa tga cta cta tat tat gtt tta tac cat ttc gtg tca cag gct ttg tag att tgg a-- --- --- --- --- --- --- -ta tag cat tgc aca tag tat tag tca taa act tgg gat gta aac tgt tga tga tat cta cat cgt ttg gat ttt tat gta tcc act tta ata ata tca tag ctg taa cat cct cgt ga- ttt acg ttg acg ttt tcg tgg gat aag ata gtt --- -gt tca tcc ttt gat aat ttt cca aat t

>EF611203.1:79-2026_Variola_virus_strain_Rw-18_left_variable_genomic_sequence

cgg ttg cca tgt att tcc tgg aga gca agt --- aga tga tg- agg aac cag ata gtt tat atc cgt act tgc acg taa agt cta cat tgt cat tat acg a-- ata tga tct ttt aaa ccc gct aga caa gta tcc gtt taa tat tgt agg atg tgg aca ttt aac aat ctg aca cgt ggg tgg att aga cca ttc tcc tcc tga aca cag gac acc aga gtt acc aat caa cga ata tcc act att gca act ata agt tac aac gct tcc atc ggt ata aaa atc att ata tcc gtt atg tct tcc gtt gga tat aga tgg agg taa ttg gca ttt aac aga ttc gca aat agg tgc ctt ggg att cca tac cat aga tcc agt aga tcc taa ttt aca ata cga ttt ata ttc acc aat caa ata ata tcc gct att aca aga gta cgt tat act aga gcc aaa gtc tac tcc gcc aat atc aag atg gcc att atc gat atc tcg agg cga cgg gca tct ccg ttt aat aca ttg att aaa gag tgt cca tcc agt acc tgt aca ttt agc ata tat agg tcc cat ttt ttg ctt tct gta tcc agg tag aca tag a-t att cta tag tgt ctc cta tgt tgt a-- --- --- --- -at tag cat tag cat cag tc- --t cca cac tat tct taa att tca tat taa tgg gtc gtg acg gaa tag tac agc atg ata gaa cgc atc cta ttc cca aca atg tca gga acg tca ctc tct cca cct tca tat tta ttt atc cgt -aa aaa tgt tat cct gga cat tgt gcc aat aat aaa aaa gcc ca- -ta tat gtt cgc tat tgt aga act tgt ttt tca tag ttg ctc aaa aac gat ggc ggt gac tta tga gtc acg tta cac ttt gga gtc tca tct tta gta aac ata tca taa tat tcg ata tta cga gtt gac aca ttg aac aaa -tc caa gta ttt gat ttt gga taa tat tcg tat ttt gca tct gct ata att aag ata taa tcg cca caa aaa cac gcg aac atc ttt cct aca tgg tta aag tac atg tac aat tct atc cat ttg tct tcc tta act ata tat ttg tat aga taa tta cga gtc tcg tgt gta att cca gta att gca tag atg tta cca tcg tac tct aca gca taa act ata cta tga tgt cta ggc atg gga gac -tt ttt tat cca aca att ttt agt gaa aca ttc cac atc gtt taa tat tac ata ttt ttc ata cgt ggt ata aac tcc acc cat tac --a tat ata tca tca ttt acg aat acc gac gcg tct gaa tat cta gga gta att aag ttt gga tgt ctt atc cat ttc gaa gtg tag cgt ttt aaa tat ttt gct aca cct gtt gaa ata gaa aat ttt aat cct cct att aca tat aac ttt cca tca tta aca cac gta cta act ttt gat ttt aac gac gac ata tta gta act gtt ttc ca- --- --- --- --- --- ttt ttt ttt ttt gtt tta aga tct acc cac gga tac gaa ata aac atg tct att gtt aat cat gcc acc aat aat gta tag aca att ata taa aa- att tgc att ata gaa ttg tct atc tat ctg tat tac cga cga ttg tcc aat att ctg ttc tag gag agt aat ggg tta ttg tgg ata tat aat cag agt ttt taa tga cta cta tat tat gtt tta tac cat ttc gtg tca cag gct ttg tag att tgg a-- --- --- --- --- --- --- -ta tag cat tgc aca tag tat tag tca taa act tgg gat gta aac tgt tga tga tat cta cat cgt ttg gat ttt tat gta tcc act tta ata ata tca tag ctg taa cat cct cgt ga- ttt acg ttg acg ttt tcg tgg gat aag ata gtt --- -gt tca tcc ttt gat aat ttt cca aat t

>EF611201.1:79-2019_Variola_virus_strain_M-Sur-60_left_variable_genomic_sequence

cgg ttg cca tgt att tcc tgg aga gca agt --- aga tga tg- agg aac cag ata gtt tat atc cgt act tgc acg taa agt cta cat tgt cat tat acg a-- ata tga tct ttt aaa ccc gct aga caa gta tcc gtt taa tat tgt agg atg cgg aca ttt aac aat ctg aca cgt ggg tgg att aga cca ttc tcc tcc tga aca cag gac acc aga gtt acc aat caa cga ata tcc act att gca act ata agt tac aac gct tcc atc ggt ata aaa atc att ata tcc gtt atg tct tcc gtt gga tat aga tgg agg taa ttg gca ttt aac aga ttc gca aat agg tgc ctt ggg att cca tac cat aga tcc agt aga tcc taa ttt aca ata cga ttt ata ttc acc aat caa ata ata tcc gct att aca aga gta cgt tat act aga gcc aaa gtc tac tcc gcc aat atc aag atg gcc att atc gat atc tcg agg cga cgg gca tct ccg ttt aat aca ttg att aaa gag tgt cca tcc agt acc tgt aca ttt agc ata tat agg tcc cat ttt ttg ctt tct gta tcc agg tag aca tag a-t att cta tag tgt ctc cta tgt tgt a-- --- --- --- -at tag cat tag cat cag tc- --t cca cac tat tct taa att tca tat taa tgg gtc gtg acg gaa tag tac agc atg ata gaa cgc atc cta ttc cca aca atg tca gga acg tca ctc tct cca cct tca tat tta ttt atc cgt -aa aaa tgt tat cct gga cat tgt gcc aat aat aaa aaa gcc ca- -ta tat gtt cgc tat tgt aga aat tgt ttt tca tag ttg ctc aaa aac gat ggc ggt gac tta tga gtc acg tta cac ttt gga gtc tca tct tta gta aac ata tca taa tat tcg ata tta cga gtt gac aca ttg aac aaa -tc caa gta ttt gat ttt gga taa tat tcg tat ttt gca tct gct ata att aag ata taa tcg cca caa aaa cac gcg aac atc ttt cct aca tgg tta aag tac atg tac aat tct atc cat ttg tct tcc tta act ata tat ttg tat aga taa tta cga gtc tcg tgt gta att cca gta att gca tag atg tta cca tcg tac tct aca gca taa act ata cta tga tgt cta ggc atg gga gac -tt ttt tat cca aca att ttt agt gaa aca ttc cac atc gtt taa tat tac ata ttt ttc ata cgt ggt ata aac tcc acc cat tac --a tat ata tca tca ttt acg aat acc gac gcg tct gaa tat cta gga gta att aag ttt gga tgt ctt atc cat ttc gaa gtg tag cgt ttt aaa tat ttt gct aca cct gtt gaa ata gaa aat ttt aat cct cct att aca tat aac ttt cca tca tta aca cac gta cta act ttt gat ttt aac gac aac ata tta gta act gtt ttc ca- --- --- --- --- --- --- --- --t ttt ttt tta aga tct acc cac gga tac gaa ata aac atg tct att gtt aat cat gcc acc aat aat gta tag aca att ata taa aac att tgc att ata gaa ttg tct atc tat ctg tat tac cga cga ttg tcc aat att ctg ttc tag gag agt aat ggg tta ttg tgg ata tat aat cag agt ttt taa tga cta cta tat tat gtt tta tac cat ttc gtg tca cag gct ttg tag att tgg a-- --- --- --- --- --- --- -ta tag cat tgc aca tag tat tag tca taa act tgg gat gta aac tgt tga tga tat cta cat cgt ttg gat ttt tat gta tcc act tta ata ata tca tag ctg taa cat cct cgt ga- ttt acg ttg acg ttt tcg tgg gat aag ata gtt --- -gt tca tcc ttt gat aat ttt cca aat t

>EF611187.1:79-2019_Variola_virus_strain_M-Gavr-60_left_variable_genomic_sequence

cgg ttg cca tgt att tcc tgg aga gca agt --- aga tga tg- agg aac cag ata gtt tat atc cgt act tgc acg taa agt cta cat tgt cat tat acg a-- ata tga tct ttt aaa ccc gct aga caa gta tcc gtt taa tat tgt agg atg cgg aca ttt aac aat ctg aca cgt ggg tgg att aga cca ttc tcc tcc tga aca cag gac acc aga gtt acc aat caa cga ata tcc act att gca act ata agt tac aac gct tcc atc ggt ata aaa atc att ata tcc gtt atg tct tcc gtt gga tat aga tgg agg taa ttg gca ttt aac aga ttc gca aat agg tgc ctt ggg att cca tac cat aga tcc agt aga tcc taa ttt aca ata cga ttt ata ttc acc aat caa ata ata tcc gct att aca aga gta cgt tat act aga gcc aaa gtc tac tcc gcc aat atc aag atg gcc att atc gat atc tcg agg cga cgg gca tct ccg ttt aat aca ttg att aaa gag tgt cca tcc agt acc tgt aca ttt agc ata tat agg tcc cat ttt ttg ctt tct gta tcc agg tag aca tag a-t att cta tag tgt ctc cta tgt tgt a-- --- --- --- -at tag cat tag cat cag tc- --t cca cac tat tct taa att tca tat taa tgg gtc gtg acg gaa tag tac agc atg ata gaa cgc atc cta ttc cca aca atg tca gga acg tca ctc tct cca cct tca tat tta ttt atc cgt -aa aaa tgt tat cct gga cat tgt gcc aat aat aaa aaa gcc ca- -ta tat gtt cgc tat tgt aga aat tgt ttt tca tag ttg ctc aaa aac gat ggc ggt gac tta tga gtc acg tta cac ttt gga gtc tca tct tta gta aac ata tca taa tat tcg ata tta cga gtt gac aca ttg aac aaa -tc caa gta ttt gat ttt gga taa tat tcg tat ttt gca tct gct ata att aag ata taa tcg cca caa aaa cac gcg aac atc ttt cct aca tgg tta aag tac atg tac aat tct atc cat ttg tct tcc tta act ata tat ttg tat aga taa tta cga gtc tcg tgt gta att cca gta att gca tag atg tta cca tcg tac tct aca gca taa act ata cta tga tgt cta ggc atg gga gac -tt ttt tat cca aca att ttt agt gaa aca ttc cac atc gtt taa tat tac ata ttt ttc ata cgt ggt ata aac tcc acc cat tac --a tat ata tca tca ttt acg aat acc gac gcg tct gaa tat cta gga gta att aag ttt gga tgt ctt atc cat ttc gaa gtg tag cgt ttt aaa tat ttt gct aca cct gtt gaa ata gaa aat ttt aat cct cct att aca tat aac ttt cca tca tta aca cac gta cta act ttt gat ttt aac gac aac ata tta gta act gtt ttc ca- --- --- --- --- --- --- --- --t ttt ttt tta aga tct acc cac gga tac gaa ata aac atg tct att gtt aat cat gcc acc aat aat gta tag aca att ata taa aac att tgc att ata gaa ttg tct atc tat ctg tat tac cga cga ttg tcc aat att ctg ttc tag gag agt aat ggg tta ttg tgg ata tat aat cag agt ttt taa tga cta cta tat tat gtt tta tac cat ttc gtg tca cag gct ttg tag att tgg a-- --- --- --- --- --- --- -ta tag cat tgc aca tag tat tag tca taa act tgg gat gta aac tgt tga tga tat cta cat cgt ttg gat ttt tat gta tcc act tta ata ata tca tag ctg taa cat cct cgt ga- ttt acg ttg acg ttt tcg tgg gat aag ata gtt --- -gt tca tcc ttt gat aat ttt cca aat t

>EF611193.1:79-2031_Variola_virus_strain_Congo-9_left_variable_genomic_sequence

cgg ttg cca tgt att tcc tgg aga gca agt --- aga tga ta- agg aac cag ata gtt tat atc cgt act tgc acg taa agt cta cat tgt cat tat acg a-- ata tga tct ttt aaa ccc gct aga caa gta tcc gtt taa tat tgt agg atg tgg aca ttt aac aat ctg aca cgt ggg tgg att aga cca ttc tcc tcc tga aca cag gac acc aga gtt acc aat caa cga ata tcc act att gca act ata agt tac aac gct tcc atc ggt ata aaa atc att ata tcc gtt atg tct tcc gtt gga tat aga tgg agg taa ttg gca ttt aac aga ttc gca aat agg tgc ctt ggg att cca tac cat aga tcc agt aga tcc taa ttt aca ata cga ttt ata ttc acc aat caa ata ata tcc gct att aca aga gta cgt tat act aga gcc aaa gtc tac tcc gcc aat atc aag atg gcc att atc gat atc tcg agg cga cgg gca tct ccg ttt aat aca ttg att aaa gag tgt cca tcc agt acc tgt aca ttt agc ata tat agg tcc cat ttt ttg ctt tct gta tcc agg tag aca tag a-t att cta tag tgt ctc cta tgt tgt a-- --- --- --- -at tag cat tag cat cag tc- --t cca cac tat tct taa att tca tat taa tgg gtc gtg acg gaa tag tac agc atg ata gaa cgc atc cta ttc cca aca atg tca gga acg tca ctc tct cca cct tca tat tta ttt atc cgt -aa aaa tgt tat cct gga cat tgt gcc aat aat aaa aaa gcc ca- -ta tat gtt cgc tat tgt aga act tgt ttt tca tag ttg ctc aaa aac gat ggc ggt gac tta tga gtc acg tta cac ttt gga gtc tca tct tta gta aac ata tca taa tat tcg ata tta cga gtt gac aca ttg aac aaa -tc caa gta ttt gat ttt gga taa tat tcg tat ttt gca tct gct ata att aag ata taa tcg cca caa aaa cac gcg aac atc ttt cct aca tgg tta aag tac atg tac aat tct atc cat ttg tct tcc tta act ata tat ttg tat aga taa tta cga gtc tcg tgt gta att cca gta att gca tag atg tta cca tcg tac tct aca gca taa act ata cta tga tgt cta ggc atg gga gac -tt ttt tat cca aca att ttt agt gaa aca ttc cac atc gtt taa tat tac ata ttt ttc ata cgt ggt ata aac tcc acc cat tac --a tat ata tca tca ttt acg aat acc gac gcg tct gaa tat cta gga gta att aag ttt gga tgt ctt atc cat ttc gaa gtg tag cgt ttt aaa tat ttt gct aca cct gtt gaa ata gaa aat ttt aat cct cct att aca tat aac ttt cca tca tta aca cac gta cta act ttt gat ttt aac gac gac ata tta gta act gtt ttc ca- --- --- --- -tt ttt ttt ttt gtt ttt gtt tta aga tct acc cac gga tac gaa ata aac atg tct att gtt aat cat gcc acc aat aat gta tag aca att ata taa aa- att tgc att ata gaa ttg tct atc tat ctg tat tac cga cga ttg tcc aat att ctg ttc tag gag agt aat ggg tta ttg tgg ata tat aat cag agt ttt taa tga cta cta tat tat gtt tta tac cat ttc gtg tca cag gct ttg tag att tgg a-- --- --- --- --- --- --- -ta tag cat tgc aca tag tat tag tca taa act tgg gat gta aac tgt tga tga tat cta cat cgt ttg gat ttt tat gta tcc act tta ata ata tca tag ctg taa cat cct cgt ga- ttt acg ttg acg ttt tcg tgg gat aag ata gtt --- -gt tca tcc ttt gat aat ttt cca aat t

>DQ441436.1:12107-14059_Variola_virus_strain_South_Africa_1965_(103_Tvaal_Nelspruit)_complete_genome

cgg ttg cca tgt att tcc tgg aga gca agt --- aga tga tg- agg aac cag ata gtt tat atc cgt act tgc acg taa agt cta cat tgt cat tat acg a-- ata tga tct ttt aaa ccc gct aga caa gta tcc gtt taa tat tgt agg atg tgg aca ttt aac aat ctg aca cgt ggg tgg att aga cca ttc tcc tcc tga aca cag gac acc aga gtt acc aat caa cga ata tcc act att gca act ata agt tac aac gct tcc atc ggt ata aaa atc att ata tcc gtt atg tct tcc gtt gga tat aga tgg agg taa ttg gca ttt aac aga ttc gca aat agg tgc ctt ggg att cca tac cat aga tcc agt aga tcc taa ttt aca ata cga ttt ata ttc acc aat caa ata ata tcc gct att aca aga gta cgt tat act aga gcc aaa gtc tac tcc gcc aat atc aag atg gcc att atc gat atc tcg agg cga cgg gca tct ccg ttt aat aca ttg att aaa gag tgt cca tcc agt acc tgt aca ttt agc ata tat agg tcc cat ttt ttg ctt tct gta tcc agg tag aca tag a-t att cta tag tgt ctc cta tgt tgt a-- --- --- --- -at tag cat tag cat cag tc- --t cca cac tat tct taa att tca tat taa tgg gtc gtg acg gaa tag tac agc atg ata gaa cgc atc cta ttc cca aca atg tca gga acg tca ctc tct cca cct tca tat tta ttt atc cgt -aa aaa tgt tat cct gga cat tgt gcc aat aat aaa aaa gcc ca- -ta tat gtt cgc tat tgt aga act tgt ttt tca tag ttg ctc aaa aac gat ggc ggt gac tta tga gtc acg tta cac ttt gga gtc tca tct tta gta aac ata tca taa tat tcg ata tta cga gtt gac aca ttg aac aaa -tc caa gta ttt gat ttt gga taa tat tcg tat ttt gca tct gct ata att aag ata taa tcg cca caa aaa cac gcg aac atc ttt cct aca tgg tta aag tac atg tac aat tct atc cat ttg tct tcc tta act ata tat ttg tat aga taa tta cga gtc tcg tgt gta att cca gta att gca tag atg tta cca tcg tac tct aca gca taa act ata cta tga tgt cta ggc atg gga gac -tt ttt tat cca aca att ttt agt gaa aca ttc cac atc gtt taa tat tac ata ttt ttc ata cgt ggt ata aac tcc acc cat tac --a tat ata tca tca ttt acg aat acc gac gcg tct gaa tat cta gga gta att aag ttt gga tgt ctt atc cat ttc gaa gtg tag cgt ttt aaa tat ttt gct aca cct gtt gaa ata gaa aat ttt aat cct cct att aca tat aac ttt cca tca tta aca cac gta cta act ttt gat ttt aac gac gac ata tta gta act gtt ttc ca- --- --- --- -tt ttt ttt ttt ttt ttt gtt tta aga tct acc cac gga tac gaa ata aac atg tct att gtt aat cat gcc acc aat aat gta tag aca att ata taa aa- att tgc att ata gaa ttg tct atc tat ctg tat tac cga cga ttg tcc aat att ctg ttc tag gag agt aat ggg tta ttg tgg ata tat aat cag agt ttt taa tga cta cta tat tat gtt tta tac cat ttc gtg tca cag gct ttg tag att tgg a-- --- --- --- --- --- --- -ta tag cat tgc aca tag tat tag tca taa act tgg gat gta aac tgt tga tga tat cta cat cgt ttg gat ttt tat gta tcc act tta ata ata tca tag ctg taa cat cct cgt ga- ttt acg ttg acg ttt tcg tgg gat aag ata gtt --- -gt tca tcc ttt gat aat ttt cca aat t

>DQ441435.1:12106-14058_Variola_virus_strain_South_Africa_1965_(102_Natal_Ingwavuma)_complete_genome

cgg ttg cca tgt att tcc tgg aga gca agt --- aga tga tg- agg aac cag ata gtt tat atc cgt act tgc acg taa agt cta cat tgt cat tat acg a-- ata tga tct ttt aaa ccc gct aga caa gta tcc gtt taa tat tgt agg atg tgg aca ttt aac aat ctg aca cgt ggg tgg att aga cca ttc tcc tcc tga aca cag gac acc aga gtt acc aat caa cga ata tcc act att gca act ata agt tac aac gct tcc atc ggt ata aaa atc att ata tcc gtt atg tct tcc gtt gga tat aga tgg agg taa ttg gca ttt aac aga ttc gca aat agg tgc ctt ggg att cca tac cat aga tcc agt aga tcc taa ttt aca ata cga ttt ata ttc acc aat caa ata ata tcc gct att aca aga gta cgt tat act aga gcc aaa gtc tac tcc gcc aat atc aag atg gcc att atc gat atc tcg agg cga cgg gca tct ccg ttt aat aca ttg att aaa gag tgt cca tcc agt acc tgt aca ttt agc ata tat agg tcc cat ttt ttg ctt tct gta tcc agg tag aca tag a-t att cta tag tgt ctc cta tgt tgt a-- --- --- --- -at tag cat tag cat cag tc- --t cca cac tat tct taa att tca tat taa tgg gtc gtg acg gaa tag tac agc atg ata gaa cgc atc cta ttc cca aca atg tca gga acg tca ctc tct cca cct tca tat tta ttt atc cgt -aa aaa tgt tat cct gga cat tgt gcc aat aat aaa aaa gcc ca- -ta tat gtt cgc tat tgt aga act tgt ttt tca tag ttg ctc aaa aac gat ggc ggt gac tta tga gtc acg tta cac ttt gga gtc tca tct tta gta aac ata tca taa tat tcg ata tta cga gtt gac aca ttg aac aaa -tc caa gta ttt gat ttt gga taa tat tcg tat ttt gca tct gct ata att aag ata taa tcg cca caa aaa cac gcg aac atc ttt cct aca tgg tta aag tac atg tac aat tct atc cat ttg tct tcc tta act ata tat ttg tat aga taa tta cga gtc tcg tgt gta att cca gta att gca tag atg tta cca tcg tac tct aca gca taa act ata cta tga tgt cta ggc atg gga gac -tt ttt tat cca aca att ttt agt gaa aca ttc cac atc gtt taa tat tac ata ttt ttc ata cgt ggt ata aac tcc acc cat tac --a tat ata tca tca ttt acg aat acc gac gcg tct gaa tat cta gga gta att aag ttt gga tgt ctt atc cat ttc gaa gtg tag cgt ttt aaa tat ttt gct aca cct gtt gaa ata gaa aat ttt aat cct cct att aca tat aac ttt cca tca tta aca cac gta cta act ttt gat ttt aac gac gac ata tta gta act gtt ttc ca- --- --- --- -tt ttt ttt ttt ttt ttt gtt tta aga tct acc cac gga tac gaa ata aac atg tct att gtt aat cat gcc acc aat aat gta tag aca att ata taa aa- att tgc att ata gaa ttg tct atc tat ctg tat tac cga cga ttg tcc aat att ctg ttc tag gag agt aat ggg tta ttg tgg ata tat aat cag agt ttt taa tga cta cta tat tat gtt tta tac cat ttc gtg tca cag gct ttg tag att tgg a-- --- --- --- --- --- --- -ta tag cat tgc aca tag tat tag tca taa act tgg gat gta aac tgt tga tga tat cta cat cgt ttg gat ttt tat gta tcc act tta ata ata tca tag ctg taa cat cct cgt ga- ttt acg ttg acg ttt tcg tgg gat aag ata gtt --- -gt tca tcc ttt gat aat ttt cca aat t

>DQ441423.1:12302-14254_Variola_virus_strain_Congo_9_1970_(v74-227_Gispen)_complete_genome

cgg ttg cca tgt att tcc tgg aga gca agt --- aga tga ta- agg aac cag ata gtt tat atc cgt act tgc acg taa agt cta cat tgt cat tat acg a-- ata tga tct ttt aaa ccc gct aga caa gta tcc gtt taa tat tgt agg atg tgg aca ttt aac aat ctg aca cgt ggg tgg att aga cca ttc tcc tcc tga aca cag gac acc aga gtt acc aat caa cga ata tcc act att gca act ata agt tac aac gct tcc atc ggt ata aaa atc att ata tcc gtt atg tct tcc gtt gga tat aga tgg agg taa ttg gca ttt aac aga ttc gca aat agg tgc ctt ggg att cca tac cat aga tcc agt aga tcc taa ttt aca ata cga ttt ata ttc acc aat caa ata ata tcc gct att aca aga gta cgt tat act aga gcc aaa gtc tac tcc gcc aat atc aag atg gcc att atc gat atc tcg agg cga cgg gca tct ccg ttt aat aca ttg att aaa gag tgt cca tcc agt acc tgt aca ttt agc ata tat agg tcc cat ttt ttg ctt tct gta tcc agg tag aca tag a-t att cta tag tgt ctc cta tgt tgt a-- --- --- --- -at tag cat tag cat cag tc- --t cca cac tat tct taa att tca tat taa tgg gtc gtg acg gaa tag tac agc atg ata gaa cgc atc cta ttc cca aca atg tca gga acg tca ctc tct cca cct tca tat tta ttt atc cgt -aa aaa tgt tat cct gga cat tgt gcc aat aat aaa aaa gcc ca- -ta tat gtt cgc tat tgt aga act tgt ttt tca tag ttg ctc aaa aac gat ggc ggt gac tta tga gtc acg tta cac ttt gga gtc tca tct tta gta aac ata tca taa tat tcg ata tta cga gtt gac aca ttg aac aaa -tc caa gta ttt gat ttt gga taa tat tcg tat ttt gca tct gct ata att aag ata taa tcg cca caa aaa cac gcg aac atc ttt cct aca tgg tta aag tac atg tac aat tct atc cat ttg tct tcc tta act ata tat ttg tat aga taa tta cga gtc tcg tgt gta att cca gta att gca tag atg tta cca tcg tac tct aca gca taa act ata cta tga tgt cta ggc atg gga gac -tt ttt tat cca aca att ttt agt gaa aca ttc cac atc gtt taa tat tac ata ttt ttc ata cgt ggt ata aac tcc acc cat tac --a tat ata tca tca ttt acg aat acc gac gcg tct gaa tat cta gga gta att aag ttt gga tgt ctt atc cat ttc gaa gtg tag cgt ttt aaa tat ttt gct aca cct gtt gaa ata gaa aat ttt aat cct cct att aca tat aac ttt cca tca tta aca cac gta cta act ttt gat ttt aac gac gac ata tta gta act gtt ttc ca- --- --- --- -tt ttt ttt ttt gtt ttt gtt tta aga tct acc cac gga tac gaa ata aac atg tct att gtt aat cat gcc acc aat aat gta tag aca att ata taa aa- att tgc att ata gaa ttg tct atc tat ctg tat tac cga cga ttg tcc aat att ctg ttc tag gag agt aat ggg tta ttg tgg ata tat aat cag agt ttt taa tga cta cta tat tat gtt tta tac cat ttc gtg tca cag gct ttg tag att tgg a-- --- --- --- --- --- --- -ta tag cat tgc aca tag tat tag tca taa act tgg gat gta aac tgt tga tga tat cta cat cgt ttg gat ttt tat gta tcc act tta ata ata tca tag ctg taa cat cct cgt ga- ttt acg ttg acg ttt tcg tgg gat aag ata gtt --- -gt tca tcc ttt gat aat ttt cca aat t

>EF611188.1:79-2034_Variola_virus_strain_6-58_left_variable_genomic_sequence

cgg ttg cca tgt att tcc tgg aga gca agt --- aga tga tg- agg aac cag ata gtt tat atc cgt act tgc acg taa agt cta cat tgt cat tat acg a-- ata tga tct ttt aaa ccc gct aga caa gta tcc gtt taa tat tgt agg atg tgg aca ttt aac aat ctg aca cgt ggg tgg att aga cca ttc tcc tcc tga aca cag gac acc aga gtt acc aat caa cga ata tcc act att gca act ata agt tac aac gct tcc atc ggt ata aaa atc att ata tcc gtt atg tct tcc gtt gga tat aga tgg agg taa ttg gca ttt aac aga ttc gca aat agg tgc ctt ggg att cca tac cat aga tcc agt aga tcc taa ttt aca ata cga ttt ata ttc acc aat caa ata ata tcc gct att aca aga gta cgt tat act aga gcc aaa gtc tac tcc gcc aat atc aag atg gcc att atc gat atc tcg agg cga cgg gca tct ccg ttt aat aca ttg att aaa gag tgt cca tcc agt acc tgt aca ttt agc ata tat agg tcc cat ttt ttg ctt tct gta tcc agg tag aca tag a-t att cta tag tgt ctc cta tgt tgt a-- --- --- --- -at tag cat tag cat cag tc- --t cca cac tat tct taa att tca tat taa tgg gtc gtg acg gaa tag tac agc atg ata gaa cgc atc cta ttc cca aca atg tca gga acg tca ctc tct cca cct tca tat tta ttt atc cgt -aa aaa tgt tat cct gga cat tgt gcc aat aat aaa aaa gcc ca- -ta tat gtt cgc tat tgt aga aat tgt ttt tca tag ttg ctc aaa aac gat ggc ggt gac tta tga gtc acg tta cac ttt gga gtc tca tct tta gta aac ata tca taa tat tcg ata tta cga gtt gac aca ttg aac aaa -tc caa gta ttt gat ttt gga taa tat tcg tat ttt gca tct gct ata att aag ata taa tcg cca caa aaa cac gcg aac atc ttt cct aca tgg tta aag tac atg tac aat tct atc cat ttg tct tcc tta act ata tat ttg tat aga taa tta cga gtc tcg tgt gta att cca gta att gca tag atg tta cca tcg tac tct aca gca taa act ata cta tga tgt cta ggc atg gga gac -tt ttt tat cca aca att ttt agt gaa aca ttc cac atc gtt taa tat tac ata ttt ttc ata cgt ggt ata aac tcc acc cat tac --a tat ata tca tca ttt acg aat acc gac gcg tct gaa tat cta gga gta att aag ttt gga tgt ctt atc cat ttc gaa gtg tag cgt ttt aaa tat ttt gct aca cct gtt gaa ata gaa aat ttt aat tct cct att aca tat aac ttt cca tca tta aca cac gta cta act ttt gat ttt aac gac gac ata tta gta act gtt ttc ca- --- --- --t ttt ttt ttt ttt ttt ttt gtt tta aga tct acc cac gga tac gaa ata aac atg tct att gtt aat cat gcc acc aat aat gta tag aca att ata taa aac att tgc att ata gaa ttg tct atc tat ctg tat tac cga cga ttg tcc aat att ctg ttc tag gag agt aat ggg tta ttg tgg ata tat aat cag agt ttt taa tga cta cta tat tat gtt tta tac cat ttc gtg tca cag gct ttg tag att tgg a-- --- --- --- --- --- --- -ta tag cat tgc aca tag tat tag tca taa act tgg gat gta aac tgt tga tga tat cta cat cgt ttg gat ttt tat gta tcc act tta ata ata tca tag ctg taa cat cct cgt ga- ttt acg ttg acg ttt tcg tgg gat aag ata gtt --- -gt tca tcc ttt gat aat ttt cca aat t

>DQ437580.1:12101-14054_Variola_virus_strain_Afghanistan_1970_Variolator_4_complete_genome

cgg ttg cca tgt att tcc tgg aga gca agt --- aga tga tg- agg aac cag ata gtt tat atc cgt act tgc acg taa agt cta cat tgt cat tat acg a-- ata tga tct ttt aaa ccc gct aga caa gta tcc gtt taa tat tgt agg atg tgg aca ttt aac aat ctg aca cgt ggg tgg att aga cca ttc tcc tcc tga aca cag gac acc aga gtt acc aat caa cga ata tcc act att gca act ata agt tac aac gct tcc atc ggt ata aaa atc att ata tcc gtt atg tct tcc gtt gga tat aga tgg agg taa ttg gca ttt aac aga ttc gca aat agg tgc ctt ggg att cca tac cat aga tcc agt aga tcc taa ttt aca ata cga ttt ata ttc acc aat caa ata ata tcc gct att aca aga gta cgt tat act aga gcc aaa gtc tac tcc gcc aat atc aag atg gcc att atc gat atc tcg agg cga cgg gca tct ccg ttt aat aca ttg att aaa gag tgt cca tcc agt acc tgt aca ttt agc ata tat agg tcc cat ttt ttg ctt tct gta tcc agg tag aca tag a-t att cta tag tgt ctc cta tgt tgt a-- --- --- --- -at tag cat tag cat cag tc- --t cca cac tat tct taa att tca tat taa tgg gtc gtg acg gaa tag tac agc atg ata gaa cgc atc cta ttc cca aca atg tca gga acg tca ctc tct cca cct tca tat tta ttt atc cgt -aa aaa tgt tat cct gga cat tgt gcc aat aat aaa aaa gcc ca- -ta tat gtt cgc tat tgt aga aat tgt ttt tca tag ttg ctc aaa aac gat ggc ggt gac tta tga gtc acg tta cac ttt gga gtc tca tct tta gta aac ata tca taa tat tcg ata tta cga gtt gac aca ttg aac aaa -tc caa gta ttt gat ttt gga taa tat tcg tat ttt gca tct gct ata att aag ata taa tcg cca caa aaa cac gcg aac atc ttt cct aca tgg tta aag tac atg tac aat tct atc cat ttg tct tcc tta act ata tat ttg tat aga taa tta cga gtc tcg tgt gta att cca gta att gca tag atg tta cca tcg tac tct aca gca taa act ata cta tga tgt cta ggc atg gga gac -tt ttt tat cca aca att ttt agt gaa aca ttc cac att gtt taa tat tac ata ttt ttc ata cgt ggt ata aac tcc acc cat tac --a tat ata tca tca ttt acg aat acc gac gcg tct gaa tat cta gga gta att aag ttt gga tgt ctt atc cat ttc gaa gtg tag cgt ttt aaa tat ttt gct aca cct gtt gaa ata gaa aat ttt aat cct cct att aca tat aac ttt cca tca tta aca cac gta cta act ttt gat ttt gac gac gac ata tta gta act gtt ttc ca- --- --- --- -tt ttt ttt ttt ttt ttt ttt tta aga tct acc cac gga tac gaa ata aac atg tct att gtt aat cat gcc acc aat aat gta tag aca att ata taa aac att tgc att ata gaa ttg tct atc tat ctg tat tac cga cga ttg tcc aat att ctg ttc tag gag agt aat ggg tta ttg tgg ata tat aat cag agt ttt taa tga cta cta tat tat gtt tta tac cat ttc gtg tca cag gct ttg tag att tgg a-- --- --- --- --- --- --- -ta tag cat tgc aca tag tat tag tca taa act tgg gat gta aac tgt tga tga tat cta cat cgt ttg gat ttt tat gta tcc act tta ata ata tca tag ctg taa cat cct cgt ga- ttt acg ttg acg ttt tcg tgg gat aag ata gtt --- -gt tca tcc ttt gat aat ttt cca aat t

>DQ441433.1:12102-14057_Variola_virus_strain_Kuwait_1967_(K1629)_complete_genome

cgg ttg cca tgt att tcc tgg aga gca agt --- aga tga tg- agg aac cag ata gtt tat atc cgt act tgc acg taa agt cta cat tgt cat tat acg a-- ata tga tct ttt aaa ccc gct aga caa gta tcc gtt taa tat tgt agg atg tgg aca ttt aac aat ctg aca cgt ggg tgg att aga cca ttc tcc tcc tga aca cag gac acc aga gtt acc aat caa cga ata tcc act att gca act ata agt tac aac gct tcc atc ggt ata aaa atc att ata tcc gtt atg tct tcc gtt gga tat aga tgg agg taa ttg gca ttt aac aga ttc gca aat agg tgc ctt ggg att cca tac cat aga tcc agt aga tcc taa ttt aca ata cga ttt ata ttc acc aat caa ata ata tcc gct att aca aga gta cgt tat act aga gcc aaa gtc tac tcc gcc aat atc aag atg gcc att atc gat atc tcg agg cga cgg gca tct ccg ttt aat aca ttg att aaa gag tgt cca tcc agt acc tgt aca ttt agc ata tat agg tcc cat ttt ttg ctt tct gta tcc agg tag aca tag a-t att cta tag tgt ctc cta tgt tgt a-- --- --- --- -at tag cat tag cat cag tc- --t cca cac tat tct taa att tca tat taa tgg gtc gtg acg gaa tag tac agc atg ata gaa cgc atc cta ttc cca aca atg tca gga acg tca ctc tct cca cct tca tat tta ttt atc cgt -aa aaa tgt tat cct gga cat tgt gcc aat aat aaa aaa gcc ca- -ta tat gtt cgc tat tgt aga aat tgt ttt tca tag ttg ctc aaa aac gat ggc ggt gac tta tga gtc acg tta cac ttt gga gtc tca tct tta gta aac ata tca taa tat tcg ata tta cga gtt gac aca ttg aac aaa -tc caa gta ttt gat ttt gga taa tat tcg tat ttt gca tct gct ata att aag ata taa tcg cca caa aaa cac gcg aac atc ttt cct aca tgg tta aag tac atg tac aat tct atc cat ttg tct tcc tta act ata tat ttg tat aga taa tta cga gtc tcg tgt gta att cca gta att gca tag atg tta cca tcg tac tct aca gca taa act ata cta tga tgt cta ggc atg gga gac -tt ttt tat cca aca att ttt agt gaa aca ttc cac att gtt taa tat tac ata ttt ttc ata cgt ggt ata aac tcc acc cat tac --a tat ata tca tca ttt acg aat acc gac gcg tct gaa tat cta gga gta att aag ttt gga tgt ctt atc cat ttc gaa gtg tag cgt ttt aaa tat ttt gct aca cct gtt gaa ata gaa aat ttt aat cct cct att aca tat aac ttt cca tca tta aca cac gta cta act ttt gat ttt aac gac gac ata tta gta act gtt ttc ca- --- --- --t ttt ttt ttt ttt ttt ttt ttt tta aga tct acc cac gga tac gaa ata aac atg tct att gtt aat cat gcc acc aat aat gta tag aca att ata taa aac att tgc att ata gaa ttg tct atc tat ctg tat tac cga cga ttg tcc aat att ctg ttc tag gag agt aat ggg tta ttg tgg ata tat aat cag agt ttt taa tga cta cta tat tat gtt tta tac cat ttc gtg tca cag gct ttg tag att tgg a-- --- --- --- --- --- --- -ta tag cat tgc aca tag tat tag tca taa act tgg gat gta aac tgt tga tga tat cta cat cgt ttg gat ttt tat gta tcc act tta ata ata tca tag ctg taa cat cct cgt ga- ttt acg ttg acg ttt tcg tgg gat aag ata gtt --- -gt tca tcc ttt gat aat ttt cca aat t

>X69198.1:11624-13580_Variola_virus_DNA_complete_genome

cgg ttg cca tgt att tcc tgg aga gca agt --- aga tga tg- agg aac cag ata gtt tat atc cgt act tgc acg taa agt cta cat tgt cat tat acg a-- ata tga tct ttt aaa ccc gct aga caa gta tcc gtt taa tat tgt agg atg tgg aca ttt aac aat ctg aca cgt ggg tgg att aga cca ttc tcc tcc tga aca cag gac acc aga gtt acc aat caa cga ata tcc act att gca act ata agt tac aac gct tcc atc ggt ata aaa atc att ata tcc gtt atg tct tcc gtt gga tat aga tgg agg taa ttg gca ttt aac aga ttc gca aat agg tgc ctt ggg att cca tac cat aga tcc agt aga tcc taa ttt aca ata cga ttt ata ttc acc aat caa ata ata tcc gct att aca aga gta cgt tat act aga gcc aaa gtc tac tcc gcc aat atc aag atg gcc att atc gat atc tcg agg cga cgg gca tct ccg ttt aat aca ttg att aaa gag tgt cca tcc agt acc tgt aca ttt agc ata tat agg tcc cat ttt ttg ctt tct gta tcc agg tag aca tag a-t att cta tag tgt ctc cta tgt tgt a-- --- --- --- -at tag cat tag cat cag tc- --t cca cac tat tct taa atg tca tat taa tgg gtc gtg acg gaa tag tac agc atg ata gaa cgc atc cta ttc cca aca atg tca gga acg tca ctc tct cca cct tca tat tta ttt atc cgt -aa aaa tgt tat cct gga cat tgt gcc aat aat a-a aaa gcc ca- -ta tat gtt cgc tat tgt aga aat tgt ttt tca tag ttg ctc aaa aac gat ggc ggt gac tta tga gtc acg tta cac ttt gga gtc tca tct tta gta aac ata tca taa tat tcg ata tta cga gtt gac aca ttg aac aaa -tc caa gta ttt gat ttt gga taa tat tcg tat ttt gca tct gct ata att aag ata taa tcg cca caa aaa cac gcg aac atc ttt cct aca tgg tta aag tac atg tac aat tct atc cat ttg tct tcc tta act ata tat ttg tat aga taa tta cga gtc tcg tgt gta att cca gta att gca tag atg tta cca tcg tac tct aca gca taa act ata cta tga tgt cta ggc atg gga gac -tt ttt tat cca aca att ttt agt gaa aca ttc cac atc gtt taa tat tac ata ttt ttc ata cgt ggt ata aac tcc acc cat tac --a tat ata tca tca ttt acg aat acc gac gcg tct gaa tat cta gga gta att aag ttt gga tgt ctt atc cat ttc gaa gtg tag cgt ttt aaa tat ttt gct aca cct gtt gaa ata gaa aat ttt aat tct cct att aca tat aac ttt cca tca tta aca cac gta cta act ttt gat ttt aac gac gac ata tta gta act gtt ttc ca- --- --- ttt ttt ttt ttt ttt ttt ttt gtt tta aga tct acc cac gga tac gaa ata aac atg tct att gtt aat cat gcc acc aat aat gta tag aca att ata taa aac att tgc att ata gaa ttg tct atc tat ctg tat tac cga cga ttg tcc aat att ctg ttc tag gag agt aat ggg tta ttg tgg ata tat aat cag agt ttt taa tga cta cta tat tat gtt tta tac cat ttc gtg tca cag gct ttg tag att tgg a-- --- --- --- --- --- --- -ta tag cat tgc aca tag tat tag tca taa act tgg gat gta aac tgt tga tga tat cta cat cgt ttg gat ttt tat gta tcc act tta ata ata tca tag ctg taa cat cct cgt ga- ttt acg ttg acg ttt tcg tgg gat aag ata gtt --- -gt tca tcc ttt gat aat ttt cca aat t

>EF611205.1:79-2036_Variola_virus_strain_Kuw-5_left_variable_genomic_sequence

cgg ttg cca tgt att tcc tgg aga gca agt --- aga tga tg- agg aac cag ata gtt tat atc cgt act tgc acg taa agt cta cat tgt cat tat acg a-- ata tga tct ttt aaa ccc gct aga caa gta tcc gtt taa tat tgt agg atg tgg aca ttt aac aat ctg aca cgt ggg tgg att aga cca ttc tcc tcc tga aca cag gac acc aga gtt acc aat caa cga ata tcc act att gca act ata agt tac aac gct tcc atc ggt ata aaa atc att ata tcc gtt atg tct tcc gtt gga tat aga tgg agg taa ttg gca ttt aac aga ttc gca aat agg tgc ctt ggg att cca tac cat aga tcc agt aga tcc taa ttt aca ata cga ttt ata ttc acc aat caa ata ata tcc gct att aca aga gta cgt tat act aga gcc aaa gtc tac tcc gcc aat atc aag atg gcc att atc gat atc tcg agg cga cgg gca tct ccg ttt aat aca ttg att aaa gag tgt cca tcc agt acc tgt aca ttt agc ata tat agg tcc cat ttt ttg ctt tct gta tcc agg tag aca tag a-t att cta tag tgt ctc cta tgt tgt a-- --- --- --- -at tag cat tag cat cag tc- --t cca cac tat tct taa att tca tat taa tgg gtc gtg acg gaa tag tac agc atg ata gaa cgc atc cta ttc cca aca atg tca gga acg tca ctc tct cca cct tca tat tta ttt atc cgt -aa aaa tgt tat cct gga cat tgt gcc aat aat aaa aaa gcc ca- -ta tat gtt cgc tat tgt aga aat tgt ttt tca tag ttg ctc aaa aac gat ggc ggt gac tta tga gtc acg tta cac ttt gga gtc tca tct tta gta aac ata tca taa tat tcg ata tta cga gtt gac aca ttg aac aaa -tc caa gta ttt gat ttt gga taa tat tcg tat ttt gca tct gct ata att aag ata taa tcg cca caa aaa cac gcg aac atc ttt cct aca tgg tta aag tac atg tac aat tct atc cat ttg tct tcc tta act ata tat ttg tat aga taa tta cga gtc tcg tgt gta att cca gta att gca tag atg tta cca tcg tac tct aca gca taa act ata cta tga tgt cta ggc atg gga gac -tt ttt tat cca aca att ttt agt gaa aca ttc cac att gtt taa tat tac ata ttt ttc ata cgt ggt ata aac tcc acc cat tac --a tat ata tca tca ttt acg aat acc gac gcg tct gaa tat cta gga gta att aag ttt gga tgt ctt atc cat ttc gaa gtg tag cgt ttt aaa tat ttt gct aca cct gtt gaa ata gaa aat ttt aat cct cct att aca tat aac ttt cca tca tta aca cac gta cta act ttt gat ttt aac gac gac ata tta gta act gtt ttc ca- --- --- ttt ttt ttt ttt ttt ttt ttt ttt tta aga tct acc cac gga tac gaa ata aac atg tct att gtt aat cat gcc acc aat aat gta tag aca att ata taa aac att tgc att ata gaa ttg tct atc tat ctg tat tac cga cga ttg tcc aat att ctg ttc tag gag agt aat ggg tta ttg tgg ata tat aat cag agt ttt taa tga cta cta tat tat gtt tta tac cat ttc gtg tca cag gct ttg tag att tgg a-- --- --- --- --- --- --- -ta tag cat tgc aca tag tat tag tca taa act tgg gat gta aac tgt tga tga tat cta cat cgt ttg gat ttt tat gta tcc act tta ata ata tca tag ctg taa cat cct cgt ga- ttt acg ttg acg ttt tcg tgg gat aag ata gtt --- -gt tca tcc ttt gat aat ttt cca aat t

>EF611190.1:79-2037_Variola_virus_strain_Aziz_left_variable_genomic_sequence

cgg ttg cca tgt att tcc tgg aga gca agt --- aga tga tg- agg aac cag ata gtt tat atc cgt act tgc acg taa agt cta cat tgt cat tat acg a-- ata tga tct ttt aaa ccc gct aga caa gta tcc gtt taa tat tgt agg atg tgg aca ttt aac aat ctg aca cgt ggg tgg att aga cca ttc tcc tcc tga aca cag gac acc aga gtt acc aat caa cga ata tcc act att gca act ata agt tac aac gct tcc atc ggt ata aaa atc att ata tcc gtt atg tct tcc gtt gga tat aga tgg agg taa ttg gca ttt aac aga ttc gca aat agg tgc ctt ggg att cca tac cat aga tcc agt aga tcc taa ttt aca ata cga ttt ata ttc acc aat caa ata ata tcc gct att aca aga gta cgt tat act aga gcc aaa gtc tac tcc gcc aat atc aag atg gcc att atc gat atc tcg agg cga cgg gca tct ccg ttt aat aca ttg att aaa gag tgt cca tcc agt acc tgt aca ttt agc ata tat agg tcc cat ttt ttg ctt tct gta tcc agg tag aca tag a-t att cta tag tgt ctc cta tgt tgt a-- --- --- --- -at tag cat tag cat cag tc- --t cca cac tat tct taa att tca tat taa tgg gtc gtg acg gaa tag tac agc atg ata gaa cgc atc cta ttc cca aca atg tca gga acg tca ctc tct cca cct tca tat tta ttt atc cgt -aa aaa tgt tat cct gga cat tgt gcc aat aat aaa aaa gcc ca- -ta tat gtt cgc tat tgt aga aat tgt ttt tca tag ttg ctc aaa aac gat ggc ggt gac tta tga gtc acg tta cac ttt gga gtc tca tct tta gta aac ata tca taa tat tcg ata tta cga gtt gac aca ttg aac aaa -tc caa gta ttt gat ttt gga taa tat tcg tat ttt gca tct gct ata att aag ata taa tcg cca caa aaa cac gcg aac atc ttt cct aca tgg tta aag tac atg tac aat tct atc cat ttg tct tcc tta act ata tat ttg tat aga taa tta cga gtc tcg tgt gta att cca gta att gca tag atg tta cca tcg tac tct aca gca taa act ata cta tga tgt cta ggc atg gga gac -tt ttt tat cca aca att ttt agt gaa aca ttc cac att gtt taa tat tac ata ttt ttc ata cgt ggt ata aac tcc acc cat tac --a tat ata tca tca ttt acg aat acc gac gcg tct gaa tat cta gga gta att aag ttt gga tgt ctt atc cat ttc gaa gtg tag cgt ttt aaa tat ttt gct aca cct gtt gaa ata gaa aat ttt aat cct cct att aca tat aac ttt cca tca tta aca cac gta cta act ttt gat ttt aac gac gac ata tta gta act gtt ttc ca- --- --t ttt ttt ttt ttt ttt ttt ttt gtt tta aga tct acc cac gga tac gaa ata aac atg tct att gtt aat cat gcc acc aat aat gta tag aca att ata taa aac att tgc att ata gaa ttg tct atc tat ctg tat tac cga cga ttg tcc aat att ctg ttc tag gag agt aat ggg tta ttg tgg ata tat aat cag agt ttt taa tga cta cta tat tat gtt tta tac cat ttc gtg tca cag gct ttg tag att tgg a-- --- --- --- --- --- --- -ta tag cat tgc aca tag tat tag tca taa act tgg gat gta aac tgt tga tga tat cta cat cgt ttg gat ttt tat gta tcc act tta ata ata tca tag ctg taa cat cct cgt ga- ttt acg ttg acg ttt tcg tgg gat aag ata gtt --- -gt tca tcc ttt gat aat ttt cca aat t

>DQ441427.1:12101-14059_Variola_virus_strain_India_1953_(Kali-Muthu-M50_Madras)_complete_genome

cgg ttg cca tgt att tcc tgg aga gca agt --- aga tga tg- agg aac cag ata gtt tat atc cgt act tgc acg taa agt cta cat tgt cat tat acg a-- ata tga tct ttt aaa ccc gct aga caa gta tcc gtt taa tat tgt agg atg tgg aca ttt aac aat ctg gca cgt ggg tgg att aga cca ttc tcc tcc tga aca cag gac acc aga gtt acc aat caa cga ata tcc act att gca act ata agt tac aac gct tcc atc ggt ata aaa atc att ata tcc gtt atg tct tcc gtt gga tat aga tgg agg taa ttg gca ttt aac aga ttc gca aat agg tgc ctt ggg att cca tac cat aga tcc agt aga tcc taa ttt aca ata cga ttt ata ttc acc aat caa ata ata tcc gct att aca aga gta cgt tat act aga gcc aaa gtc tac tcc gcc aat atc aag atg gcc att atc gat atc tcg agg cga cgg gca tct ccg ttt aat aca ttg att aaa gag tgt cca tcc agt acc tgt aca ttt agc ata tat agg tcc cat ttt ttg ctt tct gta tcc agg tag aca tag a-t att cta tag tgt ctc cta tgt tgt a-- --- --- --- -at tag cat tag cat cag tc- --t cca cac tat tct taa att tca tat taa tgg gtc gtg acg gaa tag tac agc atg ata gaa cgc atc cta ttc cca aca atg tca gga acg tca ctc tct cca cct tca tat tta ttt atc cgt -aa aaa tgt tat cct gga cat tgt gcc aat aat aaa aaa gcc ca- -ta tat gtt cgc tat tgt aga aat tgt ttt tca tag ttg ctc aaa aac gat ggc ggt gac tta tga gtc acg tta cac ttt gga gtc tca tct tta gta aac ata tca taa tat tcg ata tta cga gtt gac aca ttg aac aaa -tc caa gta ttt gat ttt gga taa tat tcg tat ttt gca tct gct ata att aag ata taa tcg cca caa aaa cac gcg aac atc ttt cct aca tgg tta aag tac atg tac aat tct atc cat ttg tct tcc tta act ata tat ttg tat aga taa tta cga gtc tcg tgt gta att cca gta att gca tag atg tta cca tcg tac tct aca gca taa act ata cta tga tgt cta ggc atg gga gac -tt ttt tat cca aca att ttt agt gaa aca ttc cac atc gtt taa tat tac ata ttt ttc ata cgt ggt ata aac tcc acc cat tac --a tat ata tca tca ttt acg aat acc gac gcg tct gaa tat cta gga gta att aag ttt gga tgt ctt atc cat ttc gaa gtg tag cgt ttt aaa tat ttt gct aca cct gtt gaa ata gaa aat ttt aat cct cct att aca tat aac ttt cca tca tta aca cac gta cta act ttt gat ttt aac gac gac ata tta gta act gtt ttc ca- --- --t ttt ttt ttt ttt ttt ttt ttt gtt tta aga tct acc cac gga tac gaa ata aac atg tct att gtt aat cat gcc acc aat aat gta tag aca att ata taa aac att tgc att ata gaa ttg tct atc tat ctg tat tac cga cga ttg tcc aat att ctg ttc tag gag agt aat ggg tta ttg tgg ata tat aat cag agt ttt taa tga cta cta tat tat gtt tta tac cat ttc gtg tca cag gct ttg tag att tgg a-- --- --- --- --- --- --- -ta tag cat tgc aca tag tat tag tca taa act tgg gat gta aac tgt tga tga tat cta cat cgt ttg gat ttt tat gta tcc act tta ata ata tca tag ctg taa cat cct cgt ga- ttt acg ttg acg ttt tcg tgg gat aag ata gtt --- -gt tca tcc ttt gat aat ttt cca aat t

>DQ441441.1:12382-14338_Variola_virus_strain_Sudan_1947_(Rumbec)_complete_genome

cgg ttg cca tgt att tcc tgg aga gca agt --- aga tga tg- agg aac cag ata gtt tat atc cgt act tgc acg taa agt cta cat tgt cat tat acg a-- ata tga tct ttt aaa ccc gct aga caa gta tcc gtt taa tat tgt agg atg tgg aca ttt aac aat ctg aca cgt ggg tgg att aga cca ttc tcc tcc tga aca cag gac acc aga gtt acc aat caa cga ata tcc act att gca act ata agt tac aac gct tcc atc ggt ata aaa atc att ata tcc gtt atg tct tcc gtt gga tat aga tgg agg taa ttg gca ttt aac aga ttc gca aat agg tgc ctt ggg att cca tac cat aga tcc agt aga tcc taa ttt aca ata cga ttt ata ttc acc aat caa ata ata tcc gct att aca aga gta cgt tat act aga gcc aaa gtc tac tcc gcc aat atc aag atg gcc att atc gat atc tcg agg cga cgg gca tct ccg ttt aat aca ttg att aaa gag tgt cca tcc agt acc tgt aca ttt agc ata tat agg tcc cat ttt ttg ctt tct gta tcc agg tag aca tag a-t att cta tag tgt ctc cta tgt tgt a-- --- --- --- -at tag cat tag cat cag tc- --t cca cac tat tct taa att tca tat taa tgg gtc gtg acg gaa tag tac agc atg ata gaa cgc atc cta ttc cca aca atg tca gga acg tca ctc tct cca cct tca tat tta ttt atc cgt -aa aaa tgt tat cct gga cat tgt gcc aat aat aaa aaa gcc ca- -ta tat gtt cgc tat tgt aga act tgt ttt tca tag ttg ctc aaa aac gat ggc ggt gac tta tga gtc acg tta cac ttt gga gtc tca tct tta gta aac ata tca taa tat tcg ata tta cga gtt gac aca ttg aac aaa -tc caa gta ttt gat ttt gga taa tat tcg tat ttt gca tct gct ata att aag ata taa tcg cca caa aaa cac gcg aac atc ttt cct aca tgg tta aag tac atg tac aat tct atc cat ttg tct tcc tta act ata tat ttg tat aga taa tta cga gtc tcg tgt gta att cca gta att gca tag atg tta cca tcg tac tct aca gca taa act ata cta tga tgt cta ggc atg gga gac -tt ttt tat cca aca att ttt agt gaa aca ttc cac atc gtt taa tat tac ata ttt ttc ata cgt ggt ata aac tcc acc cat tac --a tat ata tca tca ttt acg aat acc gac gcg tct gaa tat cta gga gta att aag ttt gga tgt ctt atc cat ttc gaa gtg tag cgt ttt aaa tat ttt gct aca cct gtt gaa ata gaa aat ttt aat cct cct att aca tat aac ttt cca tca tta aca cac gta cta act ttt gat ttt aac gac gac ata tta gta act gtt ttc ca- --- --- ttt ttt ttt ttt ttt ttt ttt gtt tta aga tct acc cac gga tac gaa ata aac atg tct att gtt aat cat gcc acc aat aat gta tag aca att ata taa aa- att tgc att ata gaa ttg tct atc tat ctg tat tac cga cga ttg tcc aat att ctg ttc tag gag agt aat ggg tta ttg tgg ata tat aat cag agt ttt taa tga cta cta tat tat gtt tta tac cat ttc gtg tca cag gct ttg tag att tgg a-- --- --- --- --- --- --- -ta tag cat tgc aca tag tat tag tca taa act tgg gat gta aac tgt tga tga tat cta cat cgt ttg gat ttt tat gta tcc act tta ata ata tca tag ctg taa cat cct cgt ga- ttt acg ttg acg ttt tcg tgg gat aag ata gtt --- -gt tca tcc ttt gat aat ttt cca aat t

>DQ441440.1:12313-14268_Variola_virus_strain_Sudan_1947_(Juba)_complete_genome

cgg ttg cca tgt att tcc tgg aga gca agt --- aga tga tg- agg aac cag ata gtt tat atc cgt act tgc acg taa agt cta cat tgt cat tat acg a-- ata tga tct ttt aaa ccc gct aga caa gta tcc gtt taa tat tgt agg atg tgg aca ttt aac aat ctg aca cgt ggg tgg att aga cca ttc tcc tcc tga aca cag gac acc aga gtt acc aat caa cga ata tcc act att gca act ata agt tac aac gct tcc atc ggt ata aaa atc att ata tcc gtt atg tct tcc gtt gga tat aga tgg agg taa ttg gca ttt aac aga ttc gca aat agg tgc ctt ggg att cca tac cat aga tcc agt aga tcc taa ttt aca ata cga ttt ata ttc acc aat caa ata ata tcc gct att aca aga gta cgt tat act aga gcc aaa gtc tac tcc gcc aat atc aag atg gcc att atc gat atc tcg agg cga cgg gca tct ccg ttt aat aca ttg att aaa gag tgt cca tcc agt acc tgt aca ttt agc ata tat agg tcc cat ttt ttg ctt tct gta tcc agg tag aca tag a-t att cta tag tgt ctc cta tgt tgt a-- --- --- --- -at tag cat tag cat cag tc- --t cca cac tat tct taa att tca tat taa tgg gtc gtg acg gaa tag tac agc atg ata gaa cgc atc cta ttc cca aca atg tca gga acg tca ctc tct cca cct tca tat tta ttt atc cgt -aa aaa tgt tat cct gga cat tgt gcc aat aat aaa aaa gcc ca- -ta tat gtt cgc tat tgt aga act tgt ttt tca tag ttg ctc aaa aac gat ggc ggt gac tta tga gtc acg tta cac ttt gga gtc tca tct tta gta aac ata tca taa tat tcg ata tta cga gtt gac aca ttg aac aaa -tc caa gta ttt gat ttt gga taa tat tcg tat ttt gca tct gct ata att aag ata taa tcg cca caa aaa cac gcg aac atc ttt cct aca tgg tta aag tac atg tac aat tct atc cat ttg tct tcc tta act ata tat ttg tat aga taa tta cga gtc tcg tgt gta att cca gta att gca tag atg tta cca tcg tac tct aca gca taa act ata cta tga tgt cta ggc atg gga gac -tt ttt tat cca aca att ttt agt gaa aca ttc cac atc gtt taa tat tac ata ttt ttc ata cgt ggt ata aac tcc acc cat tac --a tat ata tca tca ttt acg aat acc gac gcg tct gaa tat cta gga gta att aag ttt gga tgt ctt atc cat ttc gaa gtg tag cgt ttt aaa tat ttt gct aca cct gtt gaa ata gaa aat ttt aat cct cct att aca tat aac ttt cca tca tta aca cac gta cta act ttt gat ttt aac gac gac ata tta gta act gtt ttc ca- --- --- -tt ttt ttt ttt ttt ttt ttt ttt tta aga tct acc cac gga tac gaa ata aac atg tct att gtt aat cat gcc acc aat aat gta tag aca att ata taa aa- att tgc att ata gaa ttg tct atc tat ctg tat tac cga cga ttg tcc aat att ctg ttc tag gag agt aat ggg tta ttg tgg ata tat aat cag agt ttt taa tga cta cta tat tat gtt tta tac cat ttc gtg tca cag gct ttg tag att tgg a-- --- --- --- --- --- --- -ta tag cat tgc aca tag tat tag tca taa act tgg gat gta aac tgt tga tga tat cta cat cgt ttg gat ttt tat gta tcc act tta ata ata tca tag ctg taa cat cct cgt ga- ttt acg ttg acg ttt tcg tgg gat aag ata gtt --- -gt tca tcc ttt gat aat ttt cca aat t

>EF611204.1:79-2038_Variola_virus_strain_Wzim_Ahmed_left_variable_genomic_sequence

cgg ttg cca tgt att tcc tgg aga gca agt --- aga tga tg- agg aac cag ata gtt tat atc cgt act tgc acg taa agt cta cat tgt cat tat acg a-- ata tga tct ttt aaa ccc gct aga caa gta tcc gtt taa tat tgt agg atg tgg aca ttt aac aat ctg aca cgt ggg tgg att aga cca ttc tcc tcc tga aca cag gac acc aga gtt acc aat caa cga ata tcc act att gca act ata agt tac aac gct tcc atc ggt ata aaa atc att ata tcc gtt atg tct tcc gtt gga tat aga tgg agg taa ttg gca ttt aac aga ttc gca aat agg tgc ctt ggg att cca tac cat aga tcc agt aga tcc taa ttt aca ata cga ttt ata ttc acc aat caa ata ata tcc gct att aca aga gta cgt tat act aga gcc aaa gtc tac tcc gcc aat atc aag atg gcc att atc gat atc tcg agg cga cgg gca tct ccg ttt aat aca ttg att aaa gag tgt cca tcc agt acc tgt aca ttt agc ata tat agg tcc cat ttt ttg ctt tct gta tcc agg tag aca tag a-t att cta tag tgt ctc cta tgt tgt a-- --- --- --- -at tag cat tag cat cag tc- --t cca cac tat tct taa att tca tat taa tgg gtc gtg acg gaa tag tac agc atg ata gaa cgc atc cta ttc cca aca atg tca gga acg tca ctc tct cca cct tca tat tta ttt atc cgt -aa aaa tgt tat cct gga cat tgt gcc aat aat aaa aaa gcc ca- -ta tat gtt cgc tat tgt aga aat tgt ttt tca tag ttg ctc aaa aac gat ggc ggt gac tta tga gtc acg tta cac ttt gga gtc tca tct tta gta aac ata tca taa tat tcg ata tta cga gtt gac aca ttg aac aaa -tc caa gta ttt gat ttt gga taa tat tcg tat ttt gca tct gct ata att aag ata taa tcg cca caa aaa cac gcg aac atc ttt cct aca tgg tta aag tac atg tac aat tct atc cat ttg tct tcc tta act ata tat ttg tat aga taa tta cga gtc tcg tgt gta att cca gta att gca tag atg tta cca tcg tac tct aca gca taa act ata cta tga tgt cta ggc atg gga gac -tt ttt tat cca aca att ttt agt gaa aca ttc cac att gtt taa tat tac ata ttt ttc ata cgt ggt ata aac tcc acc cat tac --a tat ata tca tca ttt acg aat acc gac gcg tct gaa tat cta gga gta att aag ttt gga tgt ctt atc cat ttc gaa gtg tag cgt ttt aaa tat ttt gct aca cct gtt gaa ata gaa aat ttt aat cct cct att aca tat aac ttt cca tca tta aca cac gta cta act ttt gat ttt aac gac gac ata tta gta act gtt ttc ca- --- -tt ttt ttt ttt ttt ttt ttt ttt gtt tta aga tct acc cac gga tac gaa ata aac atg tct att gtt aat cat gcc acc aat aat gta tag aca att ata taa aac att tgc att ata gaa ttg tct atc tat ctg tat tac cga cga ttg tcc aat att ctg ttc tag gag agt aat ggg tta ttg tgg ata tat aat cag agt ttt taa tga cta cta tat tat gtt tta tac cat ttc gtg tca cag gct ttg tag att tgg a-- --- --- --- --- --- --- -ta tag cat tgc aca tag tat tag tca taa act tgg gat gta aac tgt tga tga tat cta cat cgt ttg gat ttt tat gta tcc act tta ata ata tca tag ctg taa cat cct cgt ga- ttt acg ttg acg ttt tcg tgg gat aag ata gtt --- -gt tca tcc ttt gat aat ttt cca aat t

>DQ441446.1:12101-14062_Variola_virus_strain_United_Kingdom_1947_Higgins_(Staffordshire)_complete_genome

cgg ttg cca tgt att tcc tgg aga gca agt --- aga tga tg- agg aac cag ata gtt tat atc cgt act tgc acg taa agt cta cat tgt cat tat acg a-- ata tga tct ttt aaa ccc gct aga caa gta tcc gtt taa tat tgt agg atg tgg aca ttt aac aat ctg aca cgt ggg tgg att aga cca ttc tcc tcc tga aca cag gac acc aga gtt acc aat caa cga ata tcc act att gca act ata agt tac aac gct tcc atc ggt ata aaa atc att ata tcc gtt atg tct tcc gtt gga tat aga tgg agg taa ttg gca ttt aac aga ttc gca aat agg tgc ctt ggg att cca tac cat aga tcc agt aga tcc taa ttt aca ata cga ttt ata ttc acc aat caa ata ata tcc gct att aca aga gta cgt tat act aga gcc aaa gtc tac tcc gcc aat atc aag atg gcc att atc gat atc tcg agg cga cgg gca tct ccg ttt aat aca ttg att aaa gag tgt cca tcc agt acc tgt aca ttt agc ata tat agg tcc cat ttt ttg ctt tct gta tcc agg tag aca tag a-t att cta tag tgt ctc cta tgt tgt a-- --- --- --- -at tag cat tag cat cag tc- --t cca cac tat tct taa att tca tat taa tgg gtc gtg acg gaa tag tac agc atg ata gaa cgc atc cta ttc cca aca atg tca gga acg tca ctc tct cca cct tca tat tta ttt atc cgt -aa aaa tgt tat cct gga cat tgt gcc aat aat aaa aaa gcc ca- -ta tat gtt cgc tat tgt aga aat tgt ttt tca tag ttg ctc aaa aac gat ggc ggt gac tta tga gtc acg tta cac ttt gga gtc tca tct tta gta aac ata tca taa tat tcg ata tta cga gtt gac aca ttg aac aaa -tc caa gta ttt gat ttt gga taa tat tcg tat ttt gca tct gct ata att aag ata taa tcg cca caa aaa cac gcg aac atc ttt cct aca tgg tta aag tac atg tac aat tct atc cat ttg tct tcc tta act ata tat ttg tat aga taa tta cga gtc tcg tgt gta att cca gta att gca tag atg tta cca tcg tac tct aca gca taa act ata cta tga tgt cta ggc atg gga gac -tt ttt tat cca aca att ttt agt gaa aca ttc cac atc gtt taa tat tac ata ttt ttc ata cgt ggt ata aac tcc acc cat tac --a tat ata tca tca ttt acg aat acc gac gcg tct gaa tat cta gga gta att aag ttt gga tgt ctt atc cat ttc gaa gtg tag cgt ttt aaa tat ttt gct aca cct gtt gaa ata gaa aat ttt aat cct cct att aca tat aac ttt cca tca tta aca cac gta cta act ttt gat ttt aac gac gac ata tta gta act gtt ttc cat t-- -tt ttt ttt ttt ttt ttt ttt ttt gtt tta aga tct acc cac gga tac gaa ata aac atg tct att gtt aat cat gcc acc aat aat gta tag aca att ata taa aac att tgc att ata gaa ttg tct atc tat ctg tat tac cga cga ttg tcc aat att ctg ttc tag gag agt aat ggg tta ttg tgg ata tat aat cag agt ttt taa tga cta cta tat tat gtt tta tac cat ttc gtg tca cag gct ttg tag att tgg a-- --- --- --- --- --- --- -ta tag cat tgc aca tag tat tag tca taa act tgg gat gta aac tgt tga tga tat cta cat cgt ttg gat ttt tat gta tcc act tta ata ata tca tag ctg taa cat cct cgt ga- ttt acg ttg acg ttt tcg tgg gat aag ata gtt --- -gt tca tcc ttt gat aat ttt cca aat t

>KT184690.1:19897-21778_Vaccinia_virus_strain_IOC_clone_B141

cgg ctg cca tgt att tcc tgg aga gca agt --- aga tga tg- agg aac cag ata ggt tat atc cgt act tgc act taa agt tta cat tgt cgt tgt atg a-- gta tga tct ttt aaa ccc gct aga caa gta tcc gtt tga tat tgt agg atg tgg aca ttt aac aat ttg aca cgt ggg tgg atc gga cca ttc tcc tcc tga aca cag gac acc aga gtt acc aat caa cga ata tcc act att gca act ata agt tac aac gct tcc atc ggt ata aaa atc ctc gta tcc gtt atg tct tcc gtt gga tat aga tgg agg gga ttg gca ttt aac aga ttc aca aat agg tgc ctc ggg att cca tac cat aga tcc agt aaa tcc taa ttc aca ata cga ttt aga ttc tcc tat caa ctg ata tcc gct att aca aga gta cgt tat act aga gcc aaa gtc tac tcc gcc aat atc aag ttg gcc att atc gat atc tcg agg cga tgg gca tct ccg ttt aat aca ttg att aaa gag tgt cca tcc agt acc tgt aca ttt agc ata tat agg t-- --- --- --- ctt tct gta tcc agg tag aca tag a-t att cta tag tgt ctc cta tgt tgt a-- --- --- --- --- --- -at tag cat tag tt- --t cca cac tat tct taa att tca tat taa tgg gac gtg aag gaa tag gac agt atg ata gaa cgc atc cta ttc cca aca atg tca gga acg tca cgc tct cca cct tca tat tta ttt atc cgt -aa aaa tgt tat cct gga cat cgt aca aat aat aaa aaa gcc ca- -ta tat gtt tgc tat tgt aga aat tgt ttt tca cag ttg ctc aaa aac gat ggc agt gac tta tga gtt --- --- --- --- --- --- tca tct tta gta aac ata tca taa tat tcg ata tta cga gtt gac ata tcg aac aaa ttc caa gta ttt gat ttt gga taa tat tcg tat ttt gca tct gc- --- --- --- -ta taa tca ccg caa gaa cac acg aac atc ttt cct aca tgg tta aag tac atg tac aat tct atc cat ttg tct tcc tta act ata tat ttg tat aga taa tta cga gtc tcg tg- --- --- --a gta att aca tag atg tcg ccg tcg tac tct aca gca taa act ata cta tga tgt cta ggc atg gga gac -tt ttt tat cca acg att ttt agt gaa aca ttc tac atc gtt taa tac tac ata ttt ttc ata cgt ggt ata aac tcc acc cat tac --a tat ata tca tcg ttt acg aat acc gac gcg cct gaa tat cta gga gta att aag ttt gga agt ctt atc cat ttt gaa gtg ccg tgt ttc aaa tat tct gct aca ccc gtt gaa ata gaa aat tct aat cct cct att aca tat aac ttt cca ttg tta aca caa gta cta act tct gat ttt aac gac gac ata tta gta acc gtt ttc ca- --- --- --- --- --- --- --t ttt ttc gtt tta aga tct acc cgc -ga tac gga ata atc atg --- --- --- --- --- -cc gcc aat aat gta tag aca att atg taa aac att tgc att ata gaa ttg --- -tc tat ctg tat tac cga cta tcg tcc aat att ctg ttc tag gag agt aat ggg tta ttg tgg ata tat aat cag agt ttt taa tga cta cta tat tat gtt tta tac cat ttc gtg tca ctg gct ttg tag att tgg ata tag tta atc cca aca atg ata tag c-- --- --- --- -at tag tca taa act tgg gat gta aaa tgt tga tga tat cta cat cgt ttg gat ttt tat gta tcc act tta ata ata tca tag ctg taa cat cct cat ga- ttt acg tta acg tct tcg tgg gat aag ata gtt gtc agt tca tcc ttt gat aat ttt cca aat t

>EF611189.1:79-2037_Variola_virus_strain_13/62_left_variable_genomic_sequence

cgg ttg cca tgt att tcc tgg aga gca agt --- aga tga tg- agg aac cag ata gtt tat atc cgt act tgc acg taa agt cta cat tgt cat tat acg a-- ata tga tct ttt aaa ccc gct aga caa gta tcc gtt taa tat tgt agg atg tgg aca ttt aac aat ctg aca cgt ggg tgg att aga cca ttc tcc tcc tga aca cag gac acc aga gtt acc aat caa cga ata tcc act att gca act ata agt tac aac gct tcc atc ggt ata aaa atc att ata tcc gtt atg tct tcc gtt gga tat aga tgg agg taa ttg gca ttt aac aga ttc gca aat agg tgc ctt ggg att cca tac cat aga tcc agt aga tcc taa ttt aca ata cga ttt ata ttc acc aat caa ata ata tcc gct att aca aga gta cgt tat act aga gcc aaa gtc tac tcc gcc aat atc aag atg gcc att atc gat atc tcg agg cga cgg gca tct ccg ttt aat aca ttg att aaa gag tgt cca tcc agt acc tgt aca ttt agc ata tat agg tcc cat ttt ttg ctt tct gta tcc agg tag aca tag a-t att cta tag tgt ctc cta tgt tgt a-- --- --- --- -at tag cat tag cat cag tc- --t cca cac tat tct taa att tca tat taa tgg gtc gtg acg gaa tag tac agc atg ata gaa cgc atc cta ttc cca aca atg tca gga acg tca ctc tct cca cct tca tat tta ttt atc cgt -aa aaa tgt tat cct gga cat tgt gcc aat aat aaa aaa gcc ca- -ta tat gtt cgc tat tgt aga act tgt ttt tca tag ttg ctc aaa aac gat ggc ggt gac tta tga gtc acg tta cac ttt gga gtc tca tct tta gta aac ata tca taa tat tcg ata tta cga gtt gac aca ttg aac aaa -tc caa gta ttt gat ttt gga taa tat tcg tat ttt gca tct gct ata att aag ata taa tcg cca caa aaa cac gcg aac atc ttt cct aca tgg tta aag tac atg tac aat tct atc cat ttg tct tcc tta act ata tat ttg tat aga taa tta cga gtc tcg tgt gta att cca gta att gca tag atg tta cca tcg tac tct aca gca taa act ata cta tga tgt cta ggc atg gga gac -tt ttt tat cca aca att ttt agt gaa aca ttc cac atc gtt taa tat tac ata ttt ttc ata cgt ggt ata aac tcc acc cat tac --a tat ata tca tca ttt acg aat acc gac gcg tct gaa tat cta gga gta att aag ttt gga tgt ctt atc cat ttc gaa gtg tag cgt ttt aaa tat ttt gct aca cct gtt gaa ata gaa aat ttt aat cct cct att aca tat aac ttt cca tca tta aca cac gta cta act ttt gat ttt aac gac gac ata tta gta act gtt ttc ca- --- -tt ttt ttt ttt ttt ttt ttt ttt gtt tta aga tct acc cac gga tac gaa ata aac atg tct att gtt aat cat gcc acc aat aat gta tag aca att ata taa aa- att tgc att ata gaa ttg tct atc tat ctg tat tac cga cga ttg tcc aat att ctg ttc tag gag agt aat ggg tta ttg tgg ata tat aat cag agt ttt taa tga cta cta tat tat gtt tta tac cat ttc gtg tca cag gct ttg tag att tgg a-- --- --- --- --- --- --- -ta tag cat tgc aca tag tat tag tca taa act tgg gat gta aac tgt tga tga tat cta cat cgt ttg gat ttt tat gta tcc act tta ata ata tca tag ctg taa cat cct cgt ga- ttt acg ttg acg ttt tcg tgg gat aag ata gtt --- -gt tca tcc ttt gat aat ttt cca aat t

>DQ437586.1:12171-14132_Variola_virus_strain_India_1964_7125_Vellore_complete_genome

cgg ttg cca tgt att tcc tgg aga gca agt --- aga tga tg- agg aac cag ata gtt tat atc cgt act tgc acg taa agt cta cat tgt cat tat acg a-- ata tga tct ttt aaa ccc gct aga caa gta tcc gtt taa tat tgt agg atg tgg aca ttt aac aat ctg aca cgt ggg tgg att aga cca ttc tcc tcc tga aca cag gac acc aga gtt acc aat caa cga ata tcc act att gca act ata agt tac aac gct tcc atc ggt ata aaa atc att ata tcc gtt atg tct tcc gtt gga tat aga tgg agg taa ttg gca ttt aac aga ttc gca aat agg tgc ctt ggg att cca tac cat aga tcc agt aga tcc taa ttt aca ata cga ttt ata ttc acc aat caa ata ata tcc gct att aca aga gta cgt tat act aga gcc aaa gtc tac tcc gcc aat atc aag atg gcc att atc gat atc tcg agg cga cgg gca tct ccg ttt aat aca ttg att aaa gag tgt cca tcc agt acc tgt aca ttt agc ata tat agg tcc cat ttt ttg ctt tct gta tcc agg tag aca tag a-t att cta tag tgt ctc cta tgt tgt a-- --- --- --- -at tag cat tag cat cag tc- --t cca cac tat tct taa att tca tat taa tgg gtc gtg acg gaa tag tac agc atg ata gaa cgc atc cta ttc cca aca atg tca gga acg tca ctc tct cca cct tca tat tta ttt atc cgt -aa aaa tgt tat cct gga cat tgt gcc aat aat aaa aaa gcc ca- -ta tat gtt cgc tat tgt aga aat tgt ttt tca tag ttg ctc aaa aac gat ggc ggt gac tta tga gtc acg tta cac ttt gga gtc tca tct tta gta aac ata tca taa tat tcg ata tta cga gtt gac aca ttg aac aaa -tc caa gta ttt gat ttt gga taa tat tcg tat ttt gca tct gct ata att aag ata taa tcg cca caa aaa cac gcg aac atc ttt cct aca tgg tta aag tac atg tac aat tct atc cat ttg tct tcc tta act ata tat ttg tat aga taa tta cga gtc tcg tgt gta att cca gta att gca tag atg tta cca tcg tac tct aca gca taa act ata cta tga tgt cta ggc atg gga gac -tt ttt tat cca aca att ttt agt gaa aca ttc cac att gtt taa tat tac ata ttt ttc ata cgt ggt ata aac tcc acc cat tac --a tat ata tca tca ttt acg aat acc gac gcg tct gaa tat cta gga gta att aag ttt gga tgt ctt atc cat ttc gaa gtg tag cgt ttt aaa tat ttt gct aca cct gtt gaa ata gaa aat ttt aat cct cct att aca tat aac ttt cca tca tta aca cac gta cta act ttt gat ttt aac gac gac ata tta gta act gtt ttc cat t-- -tt ttt ttt ttt ttt ttt ttt ttt gtt tta aga tct acc cac gga tac gaa ata aac atg tct att gtt aat cat gcc acc aat aat gta tag aca att ata taa aac att tgc att ata gaa ttg tct atc tat ctg tat tac cga cga ttg tcc aat att ctg ttc tag gag agt aat ggg tta ttg tgg ata tat aat cag agt ttt taa tga cta cta tat tat gtt tta tac cat ttc gtg tca cag gct ttg tag att tgg a-- --- --- --- --- --- --- -ta tag cat tgc aca tag tat tag tca taa act tgg gat gta aac tgt tga tga tat cta cat cgt ttg gat ttt tat gta tcc act tta ata ata tca tag ctg taa cat cct cgt ga- ttt acg ttg acg ttt tcg tgg gat aag ata gtt --- -gt tca tcc ttt gat aat ttt cca aat t

>DQ437585.1:12310-14271_Variola_virus_strain_India_1964_7124_Vellore_complete_genome

cgg ttg cca tgt att tcc tgg aga gca agt --- aga tga tg- agg aac cag ata gtt tat atc cgt act tgc acg taa agt cta cat tgt cat tat acg a-- ata tga tct ttt aaa ccc gct aga caa gta tcc gtt taa tat tgt agg atg tgg aca ttt aac aat ctg aca cgt ggg tgg att aga cca ttc tcc tcc tga aca cag gac acc aga gtt acc aat caa cga ata tcc act att gca act ata agt tac aac gct tcc atc ggt ata aaa atc att ata tcc gtt atg tct tcc gtt gga tat aga tgg agg taa ttg gca ttt aac aga ttc gca aat agg tgc ctt ggg att cca tac cat aga tcc agt aga tcc taa ttt aca ata cga ttt ata ttc acc aat caa ata ata tcc gct att aca aga gta cgt tat act aga gcc aaa gtc tac tcc gcc aat atc aag atg gcc att atc gat atc tcg agg cga cgg gca tct ccg ttt aat aca ttg att aaa gag tgt cca tcc agt acc tgt aca ttt agc ata tat agg tcc cat ttt ttg ctt tct gta tcc agg tag aca tag a-t att cta tag tgt ctc cta tgt tgt a-- --- --- --- -at tag cat tag cat cag tc- --t cca cac tat tct taa att tca tat taa tgg gtc gtg acg gaa tag tac agc atg ata gaa cgc atc cta ttc cca aca atg tca gga acg tca ctc tct cca cct tca tat tta ttt atc cgt -aa aaa tgt tat cct gga cat tgt gcc aat aat aaa aaa gcc ca- -ta tat gtt cgc tat tgt aga aat tgt ttt tca tag ttg ctc aaa aac gat ggc ggt gac tta tga gtc acg tta cac ttt gga gtc tca tct tta gta aac ata tca taa tat tcg ata tta cga gtt gac aca ttg aac aaa -tc caa gta ttt gat ttt gga taa tat tcg tat ttt gca tct gct ata att aag ata taa tcg cca caa aaa cac gcg aac atc ttt cct aca tgg tta aag tac atg tac aat tct atc cat ttg tct tcc tta act ata tat ttg tat aga taa tta cga gtc tcg tgt gta att cca gta att gca tag atg tta cca tcg tac tct aca gca taa act ata cta tga tgt cta ggc atg gga gac -tt ttt tat cca aca att ttt agt gaa aca ttc cac att gtt taa tat tac ata ttt ttc ata cgt ggt ata aac tcc acc cat tac --a tat ata tca tca ttt acg aat acc gac gcg tct gaa tat cta gga gta att aag ttt gga tgt ctt atc cat ttc gaa gtg tag cgt ttt aaa tat ttt gct aca cct gtt gaa ata gaa aat ttt aat cct cct att aca tat aac ttt cca tca tta aca cac gta cta act ttt gat ttt aac gac gac ata tta gta act gtt ttc cat t-- -tt ttt ttt ttt ttt ttt ttt ttt gtt tta aga tct acc cac gga tac gaa ata aac atg tct att gtt aat cat gcc acc aat aat gta tag aca att ata taa aac att tgc att ata gaa ttg tct atc tat ctg tat tac cga cga ttg tcc aat att ctg ttc tag gag agt aat ggg tta ttg tgg ata tat aat cag agt ttt taa tga cta cta tat tat gtt tta tac cat ttc gtg tca cag gct ttg tag att tgg a-- --- --- --- --- --- --- -ta tag cat tgc aca tag tat tag tca taa act tgg gat gta aac tgt tga tga tat cta cat cgt ttg gat ttt tat gta tcc act tta ata ata tca tag ctg taa cat cct cgt ga- ttt acg ttg acg ttt tcg tgg gat aag ata gtt --- -gt tca tcc ttt gat aat ttt cca aat t

>DQ437589.1:12101-14062_Variola_virus_strain_Pakistan_1969_(Rafig_Lahore)_complete_sequence

cgg ttg cca tgt att tcc tgg aga gca agt --- aga tga tg- agg aac cag ata gtt tat atc cgt act tgc acg taa agt cta cat tgt cat tat acg a-- ata tga tct ttt aaa ccc gct aga caa gta tcc gtt taa tat tgt agg atg tgg aca ttt aac aat ctg aca cgt ggg tgg att aga cca ttc tcc tcc tga aca cag gac acc aga gtt acc aat caa cga ata tcc act att gca act ata agt tac aac gct tcc atc ggt ata aaa atc att ata tcc gtt atg tct tcc gtt gga tat aga tgg agg taa ttg gca ttt aac aga ttc gca aat agg tgc ctt ggg att cca tac cat aga tcc agt aga tcc taa ttt aca ata cga ttt ata ttc acc aat caa ata ata tcc gct att aca aga gta cgt tat act aga gcc aaa gtc tac tcc gcc aat atc aag atg gcc att atc gat atc tcg agg cga cgg gca tct ccg ttt aat aca ttg att aaa gag tgt cca tcc agt acc tgt aca ttt agc ata tat agg tcc cat ttt ttg ctt tct gta tcc agg tag aca tag a-t att cta tag tgt ctc cta tgt tgt a-- --- --- --- -at tag cat tag cat cag tc- --t cca cac tat tct taa att tca tat taa tgg gtc gtg acg gaa tag tac agc atg ata gaa cgc atc cta ttc cca aca atg tca gga acg tca ctc tct cca cct tca tat tta ttt atc cgt -aa aaa tgt tat cct gga cat tgt gcc aat aat aaa aaa gcc ca- -ta tat gtt cgc tat tgt aga aat tgt ttt tca tag ttg ctc aaa aac gat ggc ggt gac tta tga gtc acg tta cac ttt gga gtc tca tct tta gta aac ata tca taa tat tcg ata tta cga gtt gac aca ttg aac aaa -tc caa gta ttt gat ttt gga taa tat tcg tat ttt gca tct gct ata att aag ata taa tcg cca caa aaa cac gcg aac atc ttt cct aca tgg tta aag tac atg tac aat tct atc cat ttg tct tcc tta act ata tat ttg tat aga taa tta cga gtc tcg tgt gta att cca gta att gca tag atg tta cca tcg tac tct aca gca taa act ata cta tga tgt cta ggc atg gga gac -tt ttt tat cca aca att ttt agt gaa aca ttc cac att gtt taa tat tac ata ttt ttc ata cgt ggt ata aac tcc acc cat tac --a tat ata tca tca ttt acg aat acc gac gcg tct gaa tat cta gga gta att aag ttt gga tgt ctt atc cat ttc gaa gtg tag cgt ttt aaa tat ttt gct aca cct gtt gaa ata gaa aat ttt aat cct cct att aca tat aac ttt cca tca tta aca cac gta cta act ttt gat ttt gac gac gac ata tta gta act gtt ttc cat t-- -tt ttt ttt ttt ttt ttt ttt ttt gtt tta aga tct acc cac gga tac gaa ata aac atg tct att gtt aat cat gcc acc aat aat gta tag aca att ata taa aac att tgc att ata gaa ttg tct atc tat ctg tat tac cga cga ttg tcc aat att ctg ttc tag gag agt aat ggg tta ttg tgg ata tat aat cag agt ttt taa tga cta cta tat tat gtt tta tac cat ttc gtg tca cag gct ttg tag att tgg a-- --- --- --- --- --- --- -ta tag cat tgc aca tag tat tag tca taa act tgg gat gta aac tgt tga tga tat cta cat cgt ttg gat ttt tat gta tcc act tta ata ata tca tag ctg taa cat cct cgt ga- ttt acg ttg acg ttt tcg tgg gat aag ata gtt --- -gt tca tcc ttt gat aat ttt cca aat t

>EF611197.1:79-2042_Variola_virus_strain_India_71_left_variable_genomic_sequence

cgg ttg cca tgt att tcc tgg aga gca agt --- aga tga tg- agg aac cag ata gtt tat atc cgt act tgc acg taa agt cta cat tgt cat tat acg a-- ata tga tct ttt aaa ccc gct aga caa gta tcc gtt taa tat tgt agg atg tgg aca ttt aac aat ctg aca cgt ggg tgg att aga cca ttc tcc tcc tga aca cag gac acc aga gtt acc aat caa cga ata tcc act att gca act ata agt tac aac gct tcc atc ggt ata aaa atc att ata tcc gtt atg tct tcc gtt gga tat aga tgg agg taa ttg gca ttt aac aga ttc gca aat agg tgc ctt ggg att cca tac cat aga tcc agt aga tcc taa ttt aca ata cga ttt ata ttc acc aat caa ata ata tcc gct att aca aga gta cgt tat act aga gcc aaa gtc tac tcc gcc aat atc aag atg gcc att atc gat atc tag agg cga cgg gca tct ccg ttt aat aca ttg att aaa gag tgt cca tcc agt acc tgt aca ttt agc ata tat agg tcc cat ttt ttg ctt tct gta tcc agg tag aca tag a-t att cta tag tgt ctc cta tgt tgt a-- --- --- --- -at tag cat tag cat cag tc- --t cca cac tat tct taa att tca tat taa tgg gtc gtg acg gaa tag tac agc atg ata gaa cgc atc cta ttc cca aca atg tca gga acg tca ctc tct cca cct tca tat tta ttt atc cgt -aa aaa tgt tat cct gga cat tgt gcc aat aat aaa aaa gcc ca- -ta tat gtt cgc tat tgt aga aat tgt ttt tca tag ttg ctc aaa aac gat ggc ggt gac tta tga gtc acg tta cac ttt gga gtc tca tct tta gta aac ata tca taa tat tcg ata tta cga gtt gac aca ttg aac aaa -tc caa gta ttt gat ttt gga taa tat tcg tat ttt gca tct gct ata att aag ata taa tcg cca caa aaa cac gcg aac atc ttt cct aca tgg tta aag tac atg tac aat tct atc cat ttg tct tcc tta act ata tat ttg tat aga taa tta cga gtc tcg tgt gta att cca gta att gca tag atg tta cca tcg tac tct aca gca taa act ata cta tga tgt cta ggc atg gga gac -tt ttt tat cca aca att ttt agt gaa aca ttc cac atc gtt taa tat tac ata ttt ttc ata cgt ggt ata aac tcc acc cat tac --a tat ata tca tca ttt acg aat acc gac gcg tct gaa tat cta gga gta att aag ttt gga tgt ctt atc cat ttc gaa gtg tag cgt ttt aaa tat ttt gct aca cct gtt gaa ata gaa aat ttt aat cct cct att aca tat aac ttt cca tca tta aca cac gta cta act ttt gat ttt aac gac gac ata tta gta act gtt ttc cat tt- ttt ttt ttt ttt ttt ttt ttt ttt gtt tta aga tct acc cac gga tac gaa ata aac atg tct att gtt aat cat gcc acc aat aat gta tag aca att ata taa aac att tgc att ata gaa ttg tct atc tat ctg tat tac cga cga ttg tcc aat att ctg ttc tag gag agt aat ggg tta ttg tgg ata tat aat cag agt ttt taa tga cta cta tat tat gtt tta tac cat ttc gtg tca cag gct ttg tag att tgg a-- --- --- --- --- --- --- -ta tag cat tgc aca tag tat tag tca taa act tgg gat gta aac tgt tga tga tat cta cat cgt ttg gat ttt tat gta tcc act tta ata ata tca tag ctg taa cat cct cgt ga- ttt acg ttg acg ttt tcg tgg gat aag ata gtt --- -gt tca tcc ttt gat aat ttt cca aat t

>DQ437590.1:12312-14275_Variola_virus_strain_Somalia_1977_complete_genome

cgg ttg cca tgt att tcc tgg aga gca agt --- aga tga tg- agg aac cag ata gtt tat atc cgt act tgc acg taa agt cta cat tgt cat tat acg a-- ata tga tct ttt aaa ccc gct aga caa gta tcc gtt taa tat tgt agg atg tgg aca ttt aac aat ctg aca cgt ggg tgg att aga cca ttc tcc tcc tga aca cag gac acc aga gtt acc aat caa cga ata tcc act att gca act ata agt tac aac gct tcc atc ggt ata aaa atc att ata tcc gtt atg tct tcc gtt gga tat aga tgg agg taa ttg gca ttt aac aga ttc gca aat agg tgc ctt ggg att cca tac cat aga tcc agt aga tcc taa ttt aca ata cga ttt ata ttc acc aat caa ata ata tcc gct att aca aga gta cgt tat act aga gcc aaa gtc tac tcc gcc aat atc aag atg gcc att atc gat atc tcg agg cga cgg gca tct ccg ttt aat aca ttg att aaa gag tgt cca tcc agt acc tgt aca ttt agc ata tat agg tcc cat ttt ttg ctt tct gta tcc agg tag aca tag a-t att cta tag tgt ctc cta tgt tgt a-- --- --- --- -at tag cat tag cat cag tc- --t cca cac tat tct taa att tca tat taa tgg gtc gtg acg gaa tag tac agc atg ata gaa cgc atc cta ttc cca aca atg tca gga acg tca ctc tct cca cct tca tat tta ttt atc cgt -aa aaa tgt tat cct gga cat tgt gcc aat aat aaa aaa gcc ca- -ta tat gtt cgc tat tgt aga act tgt ttt tca tag ttg ctc aaa aac gat ggc ggt gac tta tga gtc acg tta cac ttt gga gtc tca tct tta gta aac ata tca taa tat tcg ata tta cga gtt gac aca ttg aac aaa -tc caa gta ttt gat ttt gga taa tat tcg tat ttt gca tct gct ata att aag ata taa tcg cca caa aaa cac gcg aac atc ttt cct aca tgg tta aag tac atg tac aat tct atc cat ttg tct tcc tta act ata tat ttg tat aga taa tta cga gtc tcg tgt gta att cca gta att gca tag atg tta cca tcg tac tct aca gca taa act ata cta tga tgt cta ggc atg gga gac -tt ttt tat cca aca att ttt agt gaa aca ttc cac atc gtt taa tat tac ata ttt ttc ata cgt ggt ata aac tcc acc cat tac --a tat ata tca tca ttt acg aat acc gac gcg tct gaa tat cta gga gta att aag ttt gga tgt ctt atc cat ttc gaa gtg tag cgt ttt aaa tat ttt gct aca cct gtt gaa ata gaa aat ttt aat cct cct att aca tat aac ttt cca tca tta aca cac gta cta act ttt gat ttt aac gac gac ata tta gta act gtt ttc cat ttt ttt ttt ttt ttt ttt ttt ttt ttt gtt tta aga tct acc cac gga tac gaa ata aac atg tct att gtt aat cat gcc acc aat aat gta tag aca att ata taa aa- att tgc att ata gaa ttg tct atc tat ctg tat tac cga cga ttg tcc aat att ctg ttc tag gag agt aat ggg tta ttg tgg ata tat aat cag agt ttt taa tga cta cta tat tat gtt tta tac cat ttc gtg tca cag gct ttg tag att tgg a-- --- --- --- --- --- --- -ta tag cat tgc aca tag tat tag tca taa act tgg gat gta aac tgt tga tga tat cta cat cgt ttg gat ttt tat gta tcc act tta ata ata tca tag ctg taa cat cct cgt ga- ttt acg ttg acg ttt tcg tgg gat aag ata gtt --- -gt tca tcc ttt gat aat ttt cca aat t

>DQ441443.1:12241-14204_Variola_virus_strain_Tanzania_1965_kembula_complete_genome

cgg ttg cca tgt att tcc tgg aga gca agt --- aga tga tg- agg aac cag ata gtt tat atc cgt act tgc acg taa agt cta cat tgt cat tat acg a-- ata tga tct ttt aaa ccc gct aga caa gta tcc gtt taa tat tgt agg atg tgg aca ttt aac aat ctg aca cgt ggg tgg att aga cca ttc tcc tcc tga aca cag gac acc aga gtt acc aat caa cga ata tcc act att gca act ata agt tac aac gct tcc atc ggt ata aaa atc att ata tcc gtt atg tct tcc gtt gga tat aga tgg agg taa ttg gca ttt aac aga ttc gca aat agg tgc ctt ggg att cca tac cat aga tcc agt aga tcc taa ttt aca ata cga ttt ata ttc acc aat caa ata ata tcc gct att aca aga gta cgt tat act aga gcc aaa gtc tac tcc gcc aat atc aag atg gcc att atc gat atc tcg agg cga cgg gca tct ccg ttt aat aca ttg att aaa gag tgt cca tcc agt acc tgt aca ttt agc ata tat agg tcc cat ttt ttg ctt tct gta tcc agg tag aca tag a-t att cta tag tgt ctc cta tgt tgt a-- --- --- --- -at tag cat tag cat cag tc- --t cca cac tat tct taa att tca tat taa tgg gtc gtg acg gaa tag tac agc atg ata gaa cgc atc cta ttc cca aca atg tca gga acg tca ctc tct cca cct tca tat tta ttt atc cgt -aa aaa tgt tat cct gga cat tgt gcc aat aat aaa aaa gcc ca- -ta tat gtt cgc tat tgt aga act tgt ttt tca tag ttg ctc aaa aac gat ggc ggt gac tta tga gtc acg tta cac ttt gga gtc tca tct tta gta aac ata tca taa tat tcg ata tta cga gtt gac aca ttg aac aaa -tc caa gta ttt gat ttt gga taa tat tcg tat ttt gca tct gct ata att aag ata taa tcg cca caa aaa cac gcg aac atc ttt cct aca tgg tta aag tac atg tac aat tct atc cat ttg tct tcc tta act ata tat ttg tat aga taa tta cga gtc tcg tgt gta att cca gta att gca tag atg tta cca tcg tac tct aca gca taa act ata cta tga tgt cta ggc atg gga gac -tt ttt tat cca aca att ttt agt gaa aca ttc cac atc gtt taa tat tac ata ttt ttc ata cgt ggt ata aac tcc acc cat tac --a tat ata tca tca ttt acg aat acc gac gcg tct gaa tat cta gga gta att aag ttt gga tgt ctt atc cat ttc gaa gtg tag cgt ttt aaa tat ttt gct aca cct gtt gaa ata gaa aat ttt aat cct cct att aca tat aac ttt cca tca tta aca cac gta cta act ttt gat ttt aac gac gac ata tta gta act gtt ttc cat ttt ttt ttt ttt ttt ttt ttt gtt ttt gtt tta aga tct acc cac gga tac gaa ata aac atg tct att gtt aat cat gcc acc aat aat gta tag aca att ata taa aa- att tgc att ata gaa ttg tct atc tat ctg tat tac cga cga ttg tcc aat att ctg ttc tag gag agt aat ggg tta ttg tgg ata tat aat cag agt ttt taa tga cta cta tat tat gtt tta tac cat ttc gtg tca cag gct ttg tag att tgg a-- --- --- --- --- --- --- -ta tag cat tgc aca tag tat tag tca taa act tgg gat gta aac tgt tga tga tat cta cat cgt ttg gat ttt tat gta tcc act tta ata ata tca tag ctg taa cat cct cgt ga- ttt acg ttg acg ttt tcg tgg gat aag ata gtt --- -gt tca tcc ttt gat aat ttt cca aat t

>DQ441439.1:11326-13289_Variola_virus_strain_Somalia_1977_(V77-1605)_complete_genome

cgg ttg cca tgt att tcc tgg aga gca agt --- aga tga tg- agg aac cag ata gtt tat atc cgt act tgc acg taa agt cta cat tgt cat tat acg a-- ata tga tct ttt aaa ccc gct aga caa gta tcc gtt taa tat tgt agg atg tgg aca ttt aac aat ctg aca cgt ggg tgg att aga cca ttc tcc tcc tga aca cag gac acc aga gtt acc aat caa cga ata tcc act att gca act ata agt tac aac gct tcc atc ggt ata aaa atc att ata tcc gtt atg tct tcc gtt gga tat aga tgg agg taa ttg gca ttt aac aga ttc gca aat agg tgc ctt ggg att cca tac cat aga tcc agt aga tcc taa ttt aca ata cga ttt ata ttc acc aat caa ata ata tcc gct att aca aga gta cgt tat act aga gcc aaa gtc tac tcc gcc aat atc aag atg gcc att atc gat atc tcg agg cga cgg gca tct ccg ttt aat aca ttg att aaa gag tgt cca tcc agt acc tgt aca ttt agc ata tat agg tcc cat ttt ttg ctt tct gta tcc agg tag aca tag a-t att cta tag tgt ctc cta tgt tgt a-- --- --- --- -at tag cat tag cat cag tc- --t cca cac tat tct taa att tca tat taa tgg gtc gtg acg gaa tag tac agc atg ata gaa cgc atc cta ttc cca aca atg tca gga acg tca ctc tct cca cct tca tat tta ttt atc cgt -aa aaa tgt tat cct gga cat tgt gcc aat aat aaa aaa gcc ca- -ta tat gtt cgc tat tgt aga act tgt ttt tca tag ttg ctc aaa aac gat ggc ggt gac tta tga gtc acg tta cac ttt gga gtc tca tct tta gta aac ata tca taa tat tcg ata tta cga gtt gac aca ttg aac aaa -tc caa gta ttt gat ttt gga taa tat tcg tat ttt gca tct gct ata att aag ata taa tcg cca caa aaa cac gcg aac atc ttt cct aca tgg tta aag tac atg tac aat tct atc cat ttg tct tcc tta act ata tat ttg tat aga taa tta cga gtc tcg tgt gta att cca gta att gca tag atg tta cca tcg tac tct aca gca taa act ata cta tga tgt cta ggc atg gga gac -tt ttt tat cca aca att ttt agt gaa aca ttc cac atc gtt taa tat tac ata ttt ttc ata cgt ggt ata aac tcc acc cat tac --a tat ata tca tca ttt acg aat acc gac gcg tct gaa tat cta gga gta att aag ttt gga tgt ctt atc cat ttc gaa gtg tag cgt ttt aaa tat ttt gct aca cct gtt gaa ata gaa aat ttt aat cct cct att aca tat aac ttt cca tca tta aca cac gta cta act ttt gat ttt aac gac gac ata tta gta act gtt ttc cat ttt ttt ttt ttt ttt ttt ttt ttt ttt gtt tta aga tct acc cac gga tac gaa ata aac atg tct att gtt aat cat gcc acc aat aat gta tag aca att ata taa aa- att tgc att ata gaa ttg tct atc tat ctg tat tac cga cga ttg tcc aat att ctg ttc tag gag agt aat ggg tta ttg tgg ata tat aat cag agt ttt taa tga cta cta tat tat gtt tta tac cat ttc gtg tca cag gct ttg tag att tgg a-- --- --- --- --- --- --- -ta tag cat tgc aca tag tat tag tca taa act tgg gat gta aac tgt tga tga tat cta cat cgt ttg gat ttt tat gta tcc act tta ata ata tca tag ctg taa cat cct cgt ga- ttt acg ttg acg ttt tcg tgg gat aag ata gtt --- -gt tca tcc ttt gat aat ttt cca aat t

>DQ441438.1:11326-13289_Variola_virus_strain_Somalia_1977_(V77-1252)_complete_genome

cgg ttg cca tgt att tcc tgg aga gca agt --- aga tga tg- agg aac cag ata gtt tat atc cgt act tgc acg taa agt cta cat tgt cat tat acg a-- ata tga tct ttt aaa ccc gct aga caa gta tcc gtt taa tat tgt agg atg tgg aca ttt aac aat ctg aca cgt ggg tgg att aga cca ttc tcc tcc tga aca cag gac acc aga gtt acc aat caa cga ata tcc act att gca act ata agt tac aac gct tcc atc ggt ata aaa atc att ata tcc gtt atg tct tcc gtt gga tat aga tgg agg taa ttg gca ttt aac aga ttc gca aat agg tgc ctt ggg att cca tac cat aga tcc agt aga tcc taa ttt aca ata cga ttt ata ttc acc aat caa ata ata tcc gct att aca aga gta cgt tat act aga gcc aaa gtc tac tcc gcc aat atc aag atg gcc att atc gat atc tcg agg cga cgg gca tct ccg ttt aat aca ttg att aaa gag tgt cca tcc agt acc tgt aca ttt agc ata tat agg tcc cat ttt ttg ctt tct gta tcc agg tag aca tag a-t att cta tag tgt ctc cta tgt tgt a-- --- --- --- -at tag cat tag cat cag tc- --t cca cac tat tct taa att tca tat taa tgg gtc gtg acg gaa tag tac agc atg ata gaa cgc atc cta ttc cca aca atg tca gga acg tca ctc tct cca cct tca tat tta ttt atc cgt -aa aaa tgt tat cct gga cat tgt gcc aat aat aaa aaa gcc ca- -ta tat gtt cgc tat tgt aga act tgt ttt tca tag ttg ctc aaa aac gat ggc ggt gac tta tga gtc acg tta cac ttt gga gtc tca tct tta gta aac ata tca taa tat tcg ata tta cga gtt gac aca ttg aac aaa -tc caa gta ttt gat ttt gga taa tat tcg tat ttt gca tct gct ata att aag ata taa tcg cca caa aaa cac gcg aac atc ttt cct aca tgg tta aag tac atg tac aat tct atc cat ttg tct tcc tta act ata tat ttg tat aga taa tta cga gtc tcg tgt gta att cca gta att gca tag atg tta cca tcg tac tct aca gca taa act ata cta tga tgt cta ggc atg gga gac -tt ttt tat cca aca att ttt agt gaa aca ttc cac atc gtt taa tat tac ata ttt ttc ata cgt ggt ata aac tcc acc cat tac --a tat ata tca tca ttt acg aat acc gac gcg tct gaa tat cta gga gta att aag ttt gga tgt ctt atc cat ttc gaa gtg tag cgt ttt aaa tat ttt gct aca cct gtt gaa ata gaa aat ttt aat cct cct att aca tat aac ttt cca tca tta aca cac gta cta act ttt gat ttt aac gac gac ata tta gta act gtt ttc cat ttt ttt ttt ttt ttt ttt ttt ttt ttt gtt tta aga tct acc cac gga tac gaa ata aac atg tct att gtt aat cat gcc acc aat aat gta tag aca att ata taa aa- att tgc att ata gaa ttg tct atc tat ctg tat tac cga cga ttg tcc aat att ctg ttc tag gag agt aat ggg tta ttg tgg ata tat aat cag agt ttt taa tga cta cta tat tat gtt tta tac cat ttc gtg tca cag gct ttg tag att tgg a-- --- --- --- --- --- --- -ta tag cat tgc aca tag tat tag tca taa act tgg gat gta aac tgt tga tga tat cta cat cgt ttg gat ttt tat gta tcc act tta ata ata tca tag ctg taa cat cct cgt ga- ttt acg ttg acg ttt tcg tgg gat aag ata gtt --- -gt tca tcc ttt gat aat ttt cca aat t

>LT706529.1:11524-13444_Variola_virus_isolate_VARV_V1588_genome_assembly_chromosome:_I

cgg ttg cca tgt att tcc tgg aga gca agt --- aga tga tg- agg aac cag ata gtt tat atc cgt act tgc acg taa agt cta cat tgt cat tat acg a-- ata tga tct ttt aaa ccc gct aga caa gta tcc gtt taa tat tgt agg atg tgg aca ttt aac aat ctg aca cgt ggg tgg att aga cca ttc tcc tcc tga aca cag gac acc aga gtt acc aat caa cga ata tcc act att gca act ata agt tac aac gct tcc atc ggt ata aaa atc att ata tcc gtt atg tct tcc gtt gga tat aga tgg agg taa ttg gca ttt aac aga ttc gca aat agg tgc ctt ggg att cca tac cat aga tcc agt aga tcc taa ttt aca ata cga ttt ata ttc acc gat caa ata ata tcc gct att aca aga gta cgt tat act aga gcc aaa gtc tac tcc gcc aat atc aag atg gcc att atc gat atc tcg agg cga cgg gca tct ccg ttt aat aca ttg att aaa gag tgt cca tcc agt acc tgt aca ttt agc ata tat agg tcc cat ttt ttg ctt tct gta tcc agg tag aca tag a-t att cta tag tgt ctc cta tgt tgt a-- --- --- --- -at tag cat tag cat cag tc- --t cca cac tat tct taa att tca tat taa tgg gtc gtg acg gaa tag tac agc atg ata gaa cgc atc cta ttc cca aca atg tca gga acg tca ctc tct cca cct tca tat tta ttt atc cgt -aa aaa tgt tat cct gga cat tgt gtc aat aat aaa aaa gct ca- -ta tat gtt tgc tat tgt aga aat tgt ttt tca tag ttg ctc aaa aac gat ggc ggt gac tta tga gtc acg tta cac ttt gga gtc tca tct tta gta aac ata tca taa tat tcg ata tta cga gtt gac ata ttg aac aaa ttc caa gta ttt gat ttt gga taa tat tcg tat ttt gca tct gct ata att aag ata taa tcg cca caa gaa cac acg aac atc ttt cct aca tgg tta aag tac atg tac aat tct atc cat ttg tct tcc tta act ata tat ttg tat aga taa tta cga gtc tcg tgt gta att cca gta att gca tag atg tta cca tcg tac tct aca gca taa act ata cta tga tgt cta ggc atg gga gac -tt ttt tat cca aca att ttt agt gaa aca ttc cac atc gtt taa tat tac ata ttt ttc ata cgt ggt ata aac tcc acc cat tac --a tat ata tca tca ttt acg aat act gac gcg tct gaa tat cta gga gta att aag ttt gga tgt ctt atc cat ttc gaa gtg tag cgt ttt aaa tat tct gct aca cct gtt aaa ata gaa aat ttt aat cct cct att aca tat aac ttt cca tca tta --- --- --- --- --- --- --- --- -ac gac gac ata tta gta act gtt ttc ca- --- --- --- --- --- --- -tt ttt ttc gtt tta aga tct acc cac gga tac gaa ata aac atg tct att gtt aat cat gcc gcc aat aat gta tag aca att ata taa aac att tgc att ata gaa ttg tct atc tat ctg tat tac cga cga ttg tcc aat att ctg ttc tag gag agt aat ggg tta ttg tgg ata tat aat cag agt ttt taa tga cta cta tat tat gtt tta tac cat ttc gtg tca cag gct ttg tag att tgg a-- --- --- --- --- --- --- -ta tag cat tgc aca tag tat tag tca taa act tgg gat gta aaa tgt tga tga tat cta cat cgt ttg gat ttt tat gta tcc act tta ata ata tca tag ctg taa cat cct tgt ga- ttt acg ttg acg ttt tcg tgg gat aag ata gtt --- -gt tca tcc ttt gat aat ttt cca aat t

>DQ441442.1:12103-14026_Variola_virus_strain_Sumatra_1970_V70-228_complete_genome

cgg ttg cca tgt att tcc tgg aga gca agt --- aga tga tg- agg aac cag ata gtt tat atc cgt act tgc acg taa agt cta cat tgt cat tat acg a-- ata tga tct ttt aaa ccc gct aga caa gta tcc gtt taa tat tgt agg atg tgg aca ttt aac aat ttg aca cgt ggg tgg att aga cca ttc tcc tcc tga aca cag gac acc aga gtt acc aat caa cga ata tcc act att gca act ata agt tac aac gct tcc atc ggt ata aaa atc att ata tcc gtt atg tct tcc gtt gga tat aga tgg agg taa ttg gca ttt aac aga ttc gca aat agg tgc ctt ggg att cca tac cat aga tcc agt aga tcc taa ttt aca ata cga ttt ata ttc acc aat caa ata ata tcc gct att aca aga gta cgt tat act aga gcc aaa gtc tac tcc gcc aat atc aag atg gcc att atc gat atc tcg agg cga cgg gca tct ccg ttt aat aca ttg att aaa gag tgt cca tcc agt acc tgt aca ttt agc ata tat agg tcc cat ttt ttg ctt tct gta tcc agg tag aca tag a-t att cta tag tgt ctc cta tgt tgt a-- --- --- --- -at tag cat tag cat cag tc- --t cca cac tat tct taa att tca tat taa tgg gtc gtg atg gaa tag tac agc atg ata gaa cgc atc cta ttc cca aca atg tca gga acg tca ctc tct cca cct tca tat tta ttt atc cgt -aa aaa tgt tat cct gga cat tgt gcc aat aat aaa aaa gcc ca- -ta tat gtt cgc tat tgt aga aat tgt ttt tca tag ttg ctc aaa aac gat ggc ggt gac tta tga gt- --- --- --- --- --- --c tca tct tta gta aac ata tca taa tat tcg ata tta cga gtt gac aca ttg aac aaa -tc caa gta ttt gat ttt gga taa tat tcg tat ttt gca tct gct ata att aag ata taa tcg cca caa aaa cac gcg aac atc ttt cct aca tgg tta aag tac atg tac aat tct atc cat ttg tct tcc tta act ata tat ttg tat aga taa tta cga gtc tcg tgt gta att cca gta att gca tag atg tta cca ttg tac tct aca gca taa act ata cta tga tgt cta ggc atg gga gac -tt ttt tat cca aca att ttt agt gaa aca ttc cac atc gtt taa tat tac ata ttt ttc ata cgt ggt ata aac tcc acc cat tac --a tat ata tca tca ttt acg aat acc gac gcg tct gaa tat cta gga gta att aag ttt gga tgt ctt atc cat ttc gaa gtg tag cgt ttt aaa tat ttt gct aca cct gtt gaa ata gaa aat ttt aat cct cct att aca tat aac ttt cca tca tta aca cac gta cta act ttt gat ttt aac gac gac ata tta gta act gtt ttc ca- --- --- --- --- --- --- --- -tt ttt ttt tta aga tct acc cac gga tac gaa ata aac atg tct att gtt aat cat gcc acc aat aat gta tag aca att ata taa aac att tgc att ata gaa ttg tct atc tat ctg tat tac cga cga ttg tcc aat att ctg ttc tag gag agt aat ggg tta ttg tgg ata tat aat cag agt ttt taa tga cta cta tat tat gtt tta tac cat ttc gtg tca cag gct ttg tag att tgg a-- --- --- --- --- --- --- -ta tag cat tgc aca tag tat tag tca taa act tgg gat gta aac tgt tga tga tat cta cat cgt ttg gat ttt tat gta tcc act tta ata ata tca tag ctg taa cat cct cgt ga- ttt acg ttg acg ttt tcg tgg gat aag ata gtt --- -gt tca tcc ttt gat aat ttt cca aat t

>DQ437591.1:12103-14025_Variola_virus_strain_Sumatra_1970_V70-222_complete_genome

cgg ttg cca tgt att tcc tgg aga gca agt --- aga tga tg- agg aac cag ata gtt tat atc cgt act tgc acg taa agt cta cat tgt cat tat acg a-- ata tga tct ttt aaa ccc gct aga caa gta tcc gtt taa tat tgt agg atg tgg aca ttt aac aat ttg aca cgt ggg tgg att aga cca ttc tcc tcc tga aca cag gac acc aga gtt acc aat caa cga ata tcc act att gca act ata agt tac aac gct tcc atc ggt ata aaa atc att ata tcc gtt atg tct tcc gtt gga tat aga tgg agg taa ttg gca ttt aac aga ttc gca aat agg tgc ctt ggg att cca tac cat aga tcc agt aga tcc taa ttt aca ata cga ttt ata ttc acc aat caa ata ata tcc gct att aca aga gta cgt tat act aga gcc aaa gtc tac tcc gcc aat atc aag atg gcc att atc gat atc tcg agg cga cgg gca tct ccg ttt aat aca ttg att aaa gag tgt cca tcc agt acc tgt aca ttt agc ata tat agg tcc cat ttt ttg ctt tct gta tcc agg tag aca tag a-t att cta tag tgt ctc cta tgt tgt a-- --- --- --- -at tag cat tag cat cag tc- --t cca cac tat tct taa att tca tat taa tgg gtc gtg atg gaa tag tac agc atg ata gaa cgc atc cta ttc cca aca atg tca gga acg tca ctc tct cca cct tca tat tta ttt atc cgt -aa aaa tgt tat cct gga cat tgt gcc aat aat aaa aaa gcc ca- -ta tat gtt cgc tat tgt aga aat tgt ttt tca tag ttg ctc aaa aac gat ggc ggt gac tta tga gt- --- --- --- --- --- --c tca tct tta gta aac ata tca taa tat tcg ata tta cga gtt gac aca ttg aac aaa -tc caa gta ttt gat ttt gga taa tat tcg tat ttt gca tct gct ata att aag ata taa tcg cca caa aaa cac gcg aac atc ttt cct aca tgg tta aag tac atg tac aat tct atc cat ttg tct tcc tta act ata tat ttg tat aga taa tta cga gtc tcg tgt gta att cca gta att gca tag atg tta cca ttg tac tct aca gca taa act ata cta tga tgt cta ggc atg gga gac -tt ttt tat cca aca att ttt agt gaa aca ttc cac atc gtt taa tat tac ata ttt ttc ata cgt ggt ata aac tcc acc cat tac --a tat ata tca tca ttt acg aat acc gac gcg tct gaa tat cta gga gta att aag ttt gga tgt ctt atc cat ttc gaa gtg tag cgt ttt aaa tat ttt gct aca cct gtt gaa ata gaa aat ttt aat cct cct att aca tat aac ttt cca tca tta aca cac gta cta act ttt gat ttt aac gac gac ata tta gta act gtt ttc ca- --- --- --- --- --- --- --- --t ttt ttt tta aga tct acc cac gga tac gaa ata aac atg tct att gtt aat cat gcc acc aat aat gta tag aca att ata taa aac att tgc att ata gaa ttg tct atc tat ctg tat tac cga cga ttg tcc aat att ctg ttc tag gag agt aat ggg tta ttg tgg ata tat aat cag agt ttt taa tga cta cta tat tat gtt tta tac cat ttc gtg tca cag gct ttg tag att tgg a-- --- --- --- --- --- --- -ta tag cat tgc aca tag tat tag tca taa act tgg gat gta aac tgt tga tga tat cta cat cgt ttg gat ttt tat gta tcc act tta ata ata tca tag ctg taa cat cct cgt ga- ttt acg ttg acg ttt tcg tgg gat aag ata gtt --- -gt tca tcc ttt gat aat ttt cca aat t

>DQ437584.1:12031-13969_Variola_virus_strain_Germany_1958_Heidelberg_complete_genome

cgg ttg cca tgt att tcc tgg aga gca agt --- aga tga tg- agg aac cag ata gtt tat atc cgt act tgc acg taa agt cta cat tgt cat tat acg a-- ata tga tct ttt aaa ccc gct aga caa gta tcc gtt taa tat tgt agg atg tgg aca ttt aac aat ctg aca cgt ggg tgg att aga cca ttc tcc tcc tga aca cag gac acc aga gtt acc aat caa cga ata tcc act att gca act ata agt tac aac gct tcc atc ggt ata aaa atc att ata tcc gtt atg tct tcc gtt gga tat aga tgg agg taa ttg gca ttt aac aga ttc gca aat agg tgc ctt ggg att cca tac cat aga tcc agt aga tcc taa ttt aca ata cga ttt ata ttc acc aat caa ata ata tcc gct att aca aga gta cgt tat act aga gcc aaa gtc tac tcc gcc aat atc aag atg gcc att atc gat atc tcg agg cga cgg gca tct ccg ttt aat aca ttg att aaa gag tgt cca tcc agt acc tgt aca ttt agc ata tat agg tcc cat ttt ttg ctt tct gta tcc agg tag aca tag a-t att cta tag tgt ctc cta tgt tgt a-- --- --- --- -at tag cat tag cat cag tc- --t cca cac tat tct taa att tca tat taa tgg gtc gtg acg gaa tag tac agc atg ata gaa cgc atc cta ttc cca aca atg tca gga acg tca ctc tct cca cct tca tat tta ttt atc cgt -aa aaa tgt tat cct gga cat tgt gcc aat aat aaa aaa gcc ca- -ta tat gtt cgc tat tgt aga aat tgt ttt tca tag ttg ctc aaa aac gat ggc ggt gac tta tga gt- --- --- --- --- --- --c tca tct tta gta aac ata tca taa tat tcg ata tta cga gtt gac aca ttg aac aaa -tc caa gta ttt gat ttt gga taa tat tcg tat ttt gca tct gct ata att aag ata taa tcg cca caa aaa cac gcg aac atc ttt cct aca tgg tta aag tac atg tac aat tct atc cat ttg tct tcc tta act ata tat ttg tat aga taa tta cga gtc tcg tgt gta att cca gta att gca tag atg tta cca tcg tac tct aca gca taa act ata cta tga tgt cta ggc atg gga gac -tt ttt tat cca aca att ttt agt gaa aca ttc cac atc gtt taa tat tac ata ttt ttc ata cgt ggt ata aac tcc acc cat tac --a tat ata tca tca ttt acg aat acc gac gcg tct gaa tat cta gga gta att aag ttt gga tgt ctt atc cat ttc gaa gtg tag cgt ttt aaa tat ttt gct aca cct gtt gaa ata gaa aat ttt aat cct cct att aca tat aac ttt cca tca tta aca cac gta cta act ttt gat ttt aac gac gac ata tta gta act gtt ttc ca- --- --- -tt ttt ttt ttt ttt ttt ttt gtt tta aga tct acc cac gga tac gaa ata aac atg tct att gtt aat cat gcc acc aat aat gta tag aca att ata taa aac att tgc att ata gaa ttg tct atc tat ctg tat tac cga cga ttg tcc aat att ctg ttc tag gag agt aat ggg tta ttg tgg ata tat aat cag agt ttt taa tga cta cta tat tat gtt tta tac cat ttc gtg tca cag gct ttg tag att tgg a-- --- --- --- --- --- --- -ta tag cat tgc aca tag tat tag tca taa act tgg gat gta aac tgt tga tga tat cta cat cgt ttg gat ttt tat gta tcc act tta ata ata tca tag ctg taa cat cct cgt ga- ttt acg ttg acg ttt tcg tgg gat aag ata gtt --- -gt tca tcc ttt gat aat ttt cca aat t

>MN244298.1:32803-34752_Akhmeta_virus_strain_A40_complete_genome

-gg ctg cca tgt att tcc tgg aga gca agt --- aga tga tg- agg aac cag ata gtt tat atc cgt att tgc acg taa agt aca cat tgt cgt tgt acg a-- gta tga tct ttt aaa ccc gct aga caa gta tcc gtt tgg tat tgt agg atg tgg aca ttt aac aat ctg aca cat ggg tgt atc tgt cca ctc tcc gcc tga aca cag gac acc aga gtt acc aat caa cga ata tcc act att gca act ata cgt tac gac gct tcc atc ggt ata aaa atc ctc gta tcc gtt atg tct tcc gtt gga tat aga tgg agg tga ttg gca ttt aac aga ttc aca agt agg tgc ctc ggg att cca cac cat aga tcc agt aga tcc taa ttc aca ata caa ttt aga ttc acc gat caa atg ata tcc gct att aca aga gta tgt tat act gga tcc aaa gtc cgt tcc act aat atc aag ttg acc att atc gat atc tcg agg cga tgg aca tct ccg ttt aat aca ttg att aac gag taa cca tcc att gcc tgt aca ttt agc ata tat ggg tcc cat ttt ttg ctt tct gta tcc tgg tag aca taa a-t att cta tag tgt ctc cta ctt tgt a-- --- --- --- --- --- -gt tct cat tgg tcg aat cca ctg tat tct taa att tca tat taa tgg gac gtg agg gaa tat tac agc atg ata gta cgc atc cta ttc cca cca atg tca gga atg tca cac act ccg act tca tat tta ttt atc cgt -aa aaa tgt tgt ccc atg tat cat agc gat ata a-a aag gtc tc- --- --- --- --t tat tgc agg aat tgt ttt tca cag ttg ttc aaa aac gat ggt agt gac tta tgc gtt aca tta cac ttt gga gtt tcg tct cta gta aac ata tca taa tat tcg ata tta cgc gtt gac ata ttg tac gaa ttc caa gta ttt gat ttt gga taa tat tcg tat ttt gca tct gct ata att aag ata taa tca ccg caa gaa cac acg aac atc ttt cct aca tgg tta aag tac atg tac aat tct atc cat ttg tct tcc tta act ata tat ttg tat aga taa tta cga gtc tcg tgc gta att cca gta att aca tag atg tcg ccg tcg tac tca act gca taa act ata cta tga tgt cta ggc atg gga gac -tt ttt tat cca acg att ttt agt gaa aca ttc cat atc gtt taa tac tac ata ttt ctc ata cgt ggt ata aac tcc acc cat tac --a tat ata tca tcg ttt acg aat acc gac gcg cct gaa tat cta gga gta att aag ttt gga agt ctt atc cat ttc gaa gtg ccg tgt ttc aaa tat tct gct aca cct gtt gaa ata gaa aat tct aat cct cct att aca tat aac ttt cca tca tta aca caa gta cta act tct gat ttt aac gac gac ata tta gta acc gtt tta ca- --- --- --- --- --- --- --t ttt ttc gtt tta aga tct acc cgc -ga tac gga ata aac atg tct att gtt aat cat gcc acc aat aat gta tag aca att atg taa cac att tgc gtt ata gaa ttg --- -tc tat ctg tat tac cga cga ttg tcc aat att ctg ttc tag gag agt aat gtg tta ttg tag ata tat aat cag agt ttt taa tgg cta tta tgt tag gtt tta tac cat ttc gtg tta cgg gtt tta cag att ggg ata tag tta atc ccg aca acg ata tag cct tac gca tat tat tag aca taa act tgg gat gta aaa tgt tga tga tat ccc tat cgt ttg gat ttt taa gca tcc act tta ata ata tca tag ctg taa cat cct cat ga- ttt atg tta acg tct tcg tgg gat aag ata gtt gtc agt tca tcc ttt gat aat ttt cca aat t

>MN244297.1:32676-34625_Akhmeta_virus_strain_A39_complete_genome

-gg ctg cca tgt att tcc tgg aga gca agt --- aga tga tg- agg aac cag ata gtt tat atc cgt att tgc acg taa agt aca cat tgt cgt tgt acg a-- gta tga tct ttt aaa ccc gct aga caa gta tcc gtt tgg tat tgt agg atg tgg aca ttt aac aat ctg aca cat ggg tgt atc tgt cca ctc tcc gcc tga aca cag gac acc aga gtt acc aat caa cga ata tcc act att gca act ata cgt tac gac gct tcc atc ggt ata aaa atc ctc gta tcc gtt atg tct tcc gtt gga tat aga tgg agg tga ttg gca ttt aac aga ttc aca agt agg tgc ctc ggg att cca cac cat aga tcc agt aga tcc taa ttc aca ata caa ttt aga ttc acc gat caa atg ata tcc gct att aca aga gta tgt tat act gga tcc aaa gtc cgt tcc act aat atc aag ttg acc att atc gat atc tcg agg cga tgg aca tct ccg ttt aat aca ttg att aac gag taa cca tcc att gcc tgt aca ttt agc ata tat ggg tcc cat ttt ttg ctt tct gta tcc tgg tag aca taa a-t att cta tag tgt ctc cta ctt tgt a-- --- --- --- --- --- -gt tct cat tgg tcg aat cca ctg tat tct taa att tca tat taa tgg gac gtg agg gaa tat tac agc atg ata gta cgc atc cta ttc cca cca atg tca gga atg tca cac act ccg act tca tat tta ttt atc cgt -aa aaa tgt tgt ccc atg tat cat agc gat ata a-a aag gtc tc- --- --- --- --t tat tgc agg aat tgt ttt tca cag ttg ttc aaa aac gat ggt agt gac tta tgc gtt aca tta cac ttt gga gtt tcg tct cta gta aac ata tca taa tat tcg ata tta cgc gtt gac ata ttg tac gaa ttc caa gta ttt gat ttt gga taa tat tcg tat ttt gca tct gct ata att aag ata taa tca ccg caa gaa cac acg aac atc ttt cct aca tgg tta aag tac atg tac aat tct atc cat ttg tct tcc tta act ata tat ttg tat aga taa tta cga gtc tcg tgc gta att cca gta att aca tag atg tcg ccg tcg tac tca act gca taa act ata cta tga tgt cta ggc atg gga gac -tt ttt tat cca acg att ttt agt gaa aca ttc cat atc gtt taa tac tac ata ttt ctc ata cgt ggt ata aac tcc acc cat tac --a tat ata tca tcg ttt acg aat acc gac gcg cct gaa tat cta gga gta att aag ttt gga agt ctt atc cat ttc gaa gtg ccg tgt ttc aaa tat tct gct aca cct gtt gaa ata gaa aat tct aat cct cct att aca tat aac ttt cca tca tta aca caa gta cta act tct gat ttt aac gac gac ata tta gta acc gtt tta ca- --- --- --- --- --- --- --t ttt ttc gtt tta aga tct acc cgc -ga tac gga ata aac atg tct att gtt aat cat gcc acc aat aat gta tag aca att atg taa cac att tgc gtt ata gaa ttg --- -tc tat ctg tat tac cga cga ttg tcc aat att ctg ttc tag gag agt aat gtg tta ttg tag ata tat aat cag agt ttt taa tgg cta tta tgt tag gtt tta tac cat ttc gtg tta cgg gtt tta cag att ggg ata tag tta atc ccg aca acg ata tag cct tac gca tat tat tag aca taa act tgg gat gta aaa tgt tga tga tat ccc tat cgt ttg gat ttt taa gca tcc act tta ata ata tca tag ctg taa cat cct cat ga- ttt atg tta acg tct tcg tgg gat aag ata gtt gtc agt tca tcc ttt gat aat ttt cca aat t

>MN244296.1:33378-35327_Akhmeta_virus_strain_G66_complete_genome

-gg ctg cca tgt att tcc tgg aga gca agt --- aga tga tg- agg aac cag ata gtt tat atc cgt att tgc acg taa agt aca cat tgt cgt tgt acg a-- gta tga tct ttt aaa ccc gct aga caa gta tcc gtt tgg tat tgt agg atg tgg aca ttt aac aat ctg aca cgt ggg tgt atc tgt cca ctc tcc gcc tga aca cag gac acc aga gtt acc aat caa cga ata tcc act att gca act ata cgt tac aac gct tcc atc ggt ata aaa atc ctc gta tcc gtt atg tct tcc gtt gga tat aga tgg agg tga ttg gca ttt aac aga ttc aca agt agg tgc ctc ggg att cca cac cat aga tcc agt aga tcc taa ttc aca ata caa ttt aga ttc acc gat caa atg ata tcc gct att aca aga gta tgt tat act gga tcc aaa gtc cgt tcc act aat atc aag ttg acc att atc gat atc tcg agg cga tgg aca tct ccg ttt aat aca ttg att aac gag taa cca tcc att gcc tgt aca ttt agc ata tat ggg tcc cat ttt ttg ctt tct gta tcc tgg tag aca taa a-t agt cta tag tgt ctc cta ctt tgt a-- --- --- --- --- --- -gc tct cat tgg tcg aat cca ctg tat tct taa att tca tat taa tgg gac gtg agg gaa tat tac agc atg ata gta cgc atc cta ttc cca tca atg tca gga atg tca cac act ccg act tca tat tta ttt atc cgt -aa aaa tgt tgt ccc atg tat cgt agc ggt ata a-a aag gtc tc- --- --- --- --t tat tgc agg aat tgt ttt tca cag ttg ttc aaa aac gat ggt agt gac tta tgc gtt aca tta cac ttt gga gtt tcg tct cta gta aac ata tca taa tat tcg ata tta cgc gtt gac ata ttg tac gaa ttc caa gta ttt gat ttt gga taa tat tcg tat ttt gca tct gct ata att aag ata taa tca ccg caa gaa cac acg aac atc ttt cct aca tgg tta aag tac atg tac aat tct atc cat ttg tct tcc tta act ata tat ttg tat aga taa tta cga gtc tcg tga gta att cca gta att aca tag atg tcg ccg tcg tac tca act gca taa act ata cta tga tgt cta ggc atg gga gac -tt ttt tat cca acg att ttt agt gaa aca ttc cac atc gtt taa tac tac ata ttt ctc ata cgt ggt ata aac tcc acc cat tac --a tat ata tca tcg ttt acg aat acc gac gcg cct gaa tat cta gga gta att aag ttt gga agt ctt atc cat ttc gaa gtg ccg tgt ttc aaa tat tct gct aca cct gtt gaa ata gaa aat tct aat cct cct att aca tat aac ttt cca tca tta aca caa gta cta act tct gat ttt aac gac gac ata tta gta acc gtt tta ca- --- --- --- --- --- --- --t ttt ttc gtt tta aga tct acc cgc -ga tac gga ata aac atg tct att gtt aat cat gcc acc aat aat ata tag aca att atg taa cac att tgc gtt ata gaa ttg --- -tc tat ctg tat tac cga cga ttg tcc aat att ctg ttc tag gag agt aat gtg tta ttg tag ata tat aat cag agt ttt taa tgg cta tta tgt tag gtt tta tac cat ttc gtg tta cgg gtt tta cag att ggg ata tag tta atc ccg aca gcg ata tag cct tac gca tat tat tag aca taa act tgg gat gta aaa tgt tga tga tat ccc tat cgt ttg gat ttt taa gca tcc act tta ata ata tca tag ctg taa cat cct cat ga- ttt atg tta acg tct tcg tgg gat aag ata gtt gtc agt tca tcc ttt gat aat ttt cca aat t

>MH607142.1:33365-35314_Akhmeta_virus_isolate_Akhmeta_2013-85_complete_genome

-gg ctg cca tgt att tcc tgg aga gca agt --- aga tga tg- agg aac cag ata gtt tat atc cgt att tgc acg taa agt aca cat tgt cgt tgt acg a-- gta tga tct ttt aaa ccc gct aga caa gta tcc gtt tgg tat tgt agg atg tgg aca ttt aac aat ctg aca cgt ggg tgt atc tgt cca ctc tcc gcc tga aca cag gac acc aga gtt acc aat caa cga ata tcc act att gca act ata cgt tac aac gct tcc atc ggt ata aaa atc ctc gta tcc gtt atg tct tcc gtt gga tat aga tgg agg tga ttg gca ttt aac aga ttc aca agt agg tgc ctc ggg att cca cac cat aga tcc agt aga tcc taa ttc aca ata caa ttt aga ttc acc gat caa atg ata tcc gct att aca aga gta tgt tat act gga tcc aaa gtc cgt tcc act aat atc aag ttg acc att atc gat atc tcg agg cga tgg aca tct ccg ttt aat aca ttg att aac gag taa cca tcc att gcc tgt aca ttt agc ata tat ggg tcc cat ttt ttg ctt tct gta tcc tgg tag aca taa a-t agt cta tag tgt ctc cta ctt tgt a-- --- --- --- --- --- -gc tct cat tgg tcg aat cca ctg tat tct taa att tca tat taa tgg gac gtg agg gaa tat tac agc atg ata gta cgc atc cta ttc cca tca atg tca gga atg tca cac act ccg act tca tat tta ttt atc cgt -aa aaa tgt tgt ccc atg tat cgt agc ggt ata a-a aag gtc tc- --- --- --- --t tat tgc agg aat tgt ttt tca cag ttg ttc aaa aac gat ggt agt gac tta tgc gtt aca tta cac ttt gga gtt tcg tct cta gta aac ata tca taa tat tcg ata tta cgc gtt gac ata ttg tac gaa ttc caa gta ttt gat ttt gga taa tat tcg tat ttt gca tct gct ata att aag ata taa tca ccg caa gaa cac acg aac atc ttt cct aca tgg tta aag tac atg tac aat tct atc cat ttg tct tcc tta act ata tat ttg tat aga taa tta cga gtc tcg tga gta att cca gta att aca tag atg tcg ccg tcg tac tca act gca taa act ata cta tga tgt cta ggc atg gga gac -tt ttt tat cca acg att ttt agt gaa aca ttc cac atc gtt taa tac tac ata ttt ctc ata cgt ggt ata aac tcc acc cat tac --a tat ata tca tcg ttt acg aat acc gac gcg cct gaa tat cta gga gta att aag ttt gga agt ctt atc cat ttc gaa gtg ccg tgt ttc aaa tat tct gct aca cct gtt gaa ata gaa aat tct aat cct cct att aca tat aac ttt cca tca tta aca caa gta cta act tct gat ttt aac gac gac ata tta gta acc gtt tta ca- --- --- --- --- --- --- --t ttt ttc gtt tta aga tct acc cgc -ga tac gga ata aac atg tct att gtt aat cat gcc acc aat aat ata tag aca att atg taa cac att tgc gtt ata gaa ttg --- -tc tat ctg tat tac cga cga ttg tcc aat att ctg ttc tag gag agt aat gtg tta ttg tag ata tat aat cag agt ttt taa tgg cta tta tgt tag gtt tta tac cat ttc gtg tta cgg gtt tta cag att ggg ata tag tta atc ccg aca gcg ata tag cct tac gca tat tat tag aca taa act tgg gat gta aaa tgt tga tga tat ccc tat cgt ttg gat ttt taa gca tcc act tta ata ata tca tag ctg taa cat cct cat ga- ttt atg tta acg tct tcg tgg gat aag ata gtt gtc agt tca tcc ttt gat aat ttt cca aat t

>NC_055230.1:33365-35314_Akhmeta_virus_isolate_Akhmeta_2013-88_complete_genome

-gg ctg cca tgt att tcc tgg aga gca agt --- aga tga tg- agg aac cag ata gtt tat atc cgt att tgc acg taa agt aca cat tgt cgt tgt acg a-- gta tga tct ttt aaa ccc gct aga caa gta tcc gtt tgg tat tgt agg atg tgg aca ttt aac aat ctg aca cgt ggg tgt atc tgt cca ctc tcc gcc tga aca cag gac acc aga gtt acc aat caa cga ata tcc act att gca act ata cgt tac aac gct tcc atc ggt ata aaa atc ctc gta tcc gtt atg tct tcc gtt gga tat aga tgg agg tga ttg gca ttt aac aga ttc aca agt agg tgc ctc ggg att cca cac cat aga tcc agt aga tcc taa ttc aca ata caa ttt aga ttc acc gat caa atg ata tcc gct att aca aga gta tgt tat act gga tcc aaa gtc cgt tcc act aat atc aag ttg acc att atc gat atc tcg agg cga tgg aca tct ccg ttt aat aca ttg att aac gag taa cca tcc att gcc tgt aca ttt agc ata tat ggg tcc cat ttt ttg ctt tct gta tcc tgg tag aca taa a-t agt cta tag tgt ctc cta ctt tgt a-- --- --- --- --- --- -gc tct cat tgg tcg aat cca ctg tat tct taa att tca tat taa tgg gac gtg agg gaa tat tac agc atg ata gta cgc atc cta ttc cca tca atg tca gga atg tca cac act ccg act tca tat tta ttt atc cgt -aa aaa tgt tgt ccc atg tat cgt agc ggt ata a-a aag gtc tc- --- --- --- --t tat tgc agg aat tgt ttt tca cag ttg ttc aaa aac gat ggt agt gac tta tgc gtt aca tta cac ttt gga gtt tcg tct cta gta aac ata tca taa tat tcg ata tta cgc gtt gac ata ttg tac gaa ttc caa gta ttt gat ttt gga taa tat tcg tat ttt gca tct gct ata att aag ata taa tca ccg caa gaa cac acg aac atc ttt cct aca tgg tta aag tac atg tac aat tct atc cat ttg tct tcc tta act ata tat ttg tat aga taa tta cga gtc tcg tga gta att cca gta att aca tag atg tcg ccg tcg tac tca act gca taa act ata cta tga tgt cta ggc atg gga gac -tt ttt tat cca acg att ttt agt gaa aca ttc cac atc gtt taa tac tac ata ttt ctc ata cgt ggt ata aac tcc acc cat tac --a tat ata tca tcg ttt acg aat acc gac gcg cct gaa tat cta gga gta att aag ttt gga agt ctt atc cat ttc gaa gtg ccg tgt ttc aaa tat tct gct aca cct gtt gaa ata gaa aat tct aat cct cct att aca tat aac ttt cca tca tta aca caa gta cta act tct gat ttt aac gac gac ata tta gta acc gtt tta ca- --- --- --- --- --- --- --t ttt ttc gtt tta aga tct acc cgc -ga tac gga ata aac atg tct att gtt aat cat gcc acc aat aat ata tag aca att atg taa cac att tgc gtt ata gaa ttg --- -tc tat ctg tat tac cga cga ttg tcc aat att ctg ttc tag gag agt aat gtg tta ttg tag ata tat aat cag agt ttt taa tgg cta tta tgt tag gtt tta tac cat ttc gtg tta cgg gtt tta cag att ggg ata tag tta atc ccg aca gcg ata tag cct tac gca tat tat tag aca taa act tgg gat gta aaa tgt tga tga tat ccc tat cgt ttg gat ttt taa gca tcc act tta ata ata tca tag ctg taa cat cct cat ga- ttt atg tta acg tct tcg tgg gat aag ata gtt gtc agt tca tcc ttt gat aat ttt cca aat t

>MH607143.1:30978-32927_Akhmeta_virus_isolate_Vani_2010_complete_genome

-gg ctg cca tgt att tcc tgg aga gca agt --- aga tga tg- agg aac cag ata gtt tat atc cgt att tgc acg taa agt aca cat tgt cgt tgt acg a-- gta tga tct ttt aaa ccc gct aga caa gta tcc gtt tgg tat tgt agg atg tgg aca ttt aac aat ctg aca cat ggg tgt atc tgt cca ctc tcc gcc tga aca cag gac acc aga gtt acc aat caa cga ata tcc act att gca act ata cgt tac gac gct tcc atc ggt ata aaa atc ctc gta tcc gtt atg tct tcc gtt gga tat aga tgg agg tga tcg gca ttt aac aga ttc aca agt agg tgc ctc ggg att cca cac cat aga tcc agt aga tcc taa ttc aca ata caa ttt aga ttc acc gat caa atg ata tcc gct att aca aga gta tgt tat act gga tcc aaa gtc cgt tcc act aat atc aag ttg acc att atc gat atc tcg agg cga tgg aca tct ccg ttt aat aca ttg att aac gag taa cca tcc att gcc tgt aca ttt agt ata tat ggg tcc cat ttt ttg ctt tct gta tcc tgg tag aca taa a-t att cta tag tgt ctc cta ctt tgt a-- --- --- --- --- --- -gt tct cat tgg tcg aat cca ctg tat tct taa att tca tat taa tgg gac gtg agg gaa tat tac agc atg ata gta cgc atc cta ttc cca cca atg tca gga atg tcg cac act ccg act tca tat tta ttt atc cgt -aa aaa tgt tgt ccc atg tat cat agc gat ata a-a aag gtc tc- --- --- --- --t tat tgc agg aat tgt ttt tca cag ttg ttc aaa aac gat ggt agt gac tta tgc gtt aca tta cac ttt gga gtt tcg tct cta gta aac ata tca taa tat tcg ata tta cgc gtt gac ata ttg tac gaa ttc caa gta ttt gat ttt gga taa tat tcg tat ttt gca tct gct ata att aag ata taa tca ccg caa gaa cac acg aac atc ttt cct aca tgg tta aag tac atg tac aat tct atc cat ttg tct tcc tta act ata tat ttg tat aga taa tta cga gtc tcg tgc gta att cca gta att aca tag atg tcg ccg tcg tac tca act gca taa act ata cta tga tgt cta ggc atg gga gac -tt ttt tat cca acg att ttt agt gaa aca ttc cat atc gtt taa tac tac ata ttt ctc ata cgt ggt ata aac tcc acc cat tac --a tat ata tca tcg ttt acg aat acc gac gcg cct gaa tat cta gga gta att aag ttt gga agt ctt atc cat ttc gaa gtg ccg tgt ttc aaa tat tct gct aca cct gtt gaa ata gaa aat tct aat cct cct att aca tat aac ttt cca tca tta aca caa gta cta act tct gat ttt aac gac gac ata tta gta acc gtt tta ca- --- --- --- --- --- --- --t ttt ttc gtt tta aga tct acc cgc -ga tac gga ata aac atg tct att gtt aat cat gcc acc aat aat gta tag aca att atg taa cac att tgc gtt ata gaa ttg --- -tc tat ctg tat tac cga cga ttg tcc aat att ctg ttc tag gag agt aat gtg tta ttg tag ata tat aat cag agt ttt taa tgg cta tta tgt tag gtt tta tac cat ttc gtg tta cgg gtt tta cag att ggg ata tag tta atc ccg aca gcg ata tag cct tac gca tat tat tag aca taa act tgg gat gta aaa tgt tga tga tat ccc tat cgt ttg gat ttt taa gca tcc act tta ata ata tca tag ctg taa cat cct cat ga- ttt atg tta acg tct tcg tgg gat aag ata gtt gtc agt tca tcc ttt gat aat ttt cca aat t

>DQ441444.1:12032-13559_Variola_virus_strain_United_Kingdom_1946_Harvey_complete_genome

cgg ttg cca tgt att tcc tgg aga gca agt --- aga tga tg- agg aac cag ata gtt tat atc cgt act tgc acg taa agt cta cat tgt cat tat acg a-- ata tga tct ttt aaa ccc gct aga caa gta tcc gtt taa tat tgt agg atg tgg aca ttt aac aat ctg aca cgt ggg tgg att aga cca ttc tcc tcc tga aca cag gac acc aga gtt acc aat caa cga ata tcc act att gca act ata agt tac aac gct tcc atc ggt ata aaa atc att ata tcc gtt atg tct tcc gtt gga tat aga tgg agg taa ttg gca ttt aac aga ttc gca aat agg tgc ctt ggg att cca tac cat aga tcc agt aga tcc taa ttt aca ata cga ttt ata ttc acc aat caa ata ata tcc gct att aca aga gta cgt tat act aga gcc aaa gtc tac tcc gcc aat atc aag atg gcc att atc gat atc tcg agg cga cgg gca tct ccg ttt aat aca ttg att aaa gag tgt cca tcc agt acc tgt aca ttt agc ata tat agg tcc cat ttt ttg ctt tct gta tcc agg tag aca tag a-t att cta tag tgt ctc cta tgt tgt a-- --- --- --- -at tag cat tag cat cag tc- --t cca cac tat tct taa att tca tat taa tgg gtc gtg acg gaa tag tac agc atg ata gaa cgc atc cta ttc cca aca atg tca gga acg tca ctc tct cca cct tca tat tta ttt atc cgt -aa aaa tgt tat cct gga cat tgt gcc aat aat aaa aaa gcc ca- -ta tat gtt cgc tat tgt aga aat tgt ttt tca tag ttg ctc aaa aac gat ggc ggt gac tta tga gtc acg tta cac ttt gga gtc tca tct tta gta aac ata tca taa tat tcg ata tta cga gtt gac aca ttg aac aaa -tc caa gta ttt gat ttt gga taa tat tcg tat ttt gca tct gct ata att aag ata taa tcg cca caa aaa cac gcg aac atc ttt cct aca tgg tta aag tac atg tac aat tct atc cat ttg tct tcc tta act ata tat ttg tat aga taa tta cga gtc tcg tgt gta att cca gta att gca tag atg tta cca tcg tac tct aca gca taa act ata cta tga tgt cta ggc atg gga gac -tt ttt tat cca aca att ttt agt gaa aca ttc cac atc gtt taa tat tac ata ttt ttc ata cgt ggt ata aac tcc acc cat tac --a tat ata tca tca ttt acg aat acc gac gcg tct gaa tat cta gga gta att aag ttt gga tgt ctt atc cat ttc gaa gtg tag cgt ttt aaa tat tct gct aca cct gtt gaa ata gaa aat ttt aat cct cct att aca tat aac ttt cca tca tta aca cac gta cta act ttt gat ttt aac gac gac ata tta gta act gtt ttc ca- --- --- --- --- --- --- -tt ttt ttt ttt tt- --- --- --- --- --- --- --- --- --- --- --- --- --- --- --- --- --- --- --- --- --- --- --- --- --- --- --- --- --- --- --- --- --- --- --- --- --- --- --- --- --- --- --- --- --- --- --- --- --- --- --- --- --- --- --- --- --- --- --- --- --- --- --- --- --- --- --- --- --- --- --- --- --- --- --- --- --- --- --- --- --- --- --- --- --- --- --- --- --- --- --- --- --- --- --- --- --- --- --- --- --- --- --- --- --- --- --- --- --- --- --- --- --- --- --- --- --- --- --- --- --- --- --- --- --- --- --- --- --- --- --- --- --- --- --- --- --- --- --- --- --- --- --- --- --- --- --- -

>KY358055.1:11556-13072_Variola_virus_strain_VD21_17th_century

cgg ttg cca tgt att tcc tgg aga gca agt --- aga tga tg- agg aac cag ata gtt tat atc cgt act tgc acg taa agt cta cat tgt cat tat acg a-- ata tga tct ttt aaa ccc gct aga caa gta tcc gtt taa tat tgt agg atg tgg aca ttt aac aat ctg aca cgt ggg tgg att aga cca ttc tcc tcc tga aca cag gac acc aga gtt acc aat caa cga ata tcc act att gca act ata agt tac aac gct tcc atc ggt ata aaa atc att ata tcc gtt atg tct tcc gtt gga tat aga tgg agg taa ttg gca ttt aac aga ttc gca aat agg tgc ctt ggg att cca tac cat aga tcc agt aga tcc taa ttc aca ata cga ttt ata ttc acc gat caa ata ata tcc gct att aca aga gta cgt tat act aga gcc aaa gtc tac tcc gcc aat atc aag atg gcc att atc gat atc tcg agg cga cgg gca tct ccg ttt aat aca ttg att aaa gag tgt cca tcc agt acc tgt aca ttt agc ata tat agg tcc cat ttt ttg ctt tct gta tcc agg tag aca tag a-t att cta tag tgt ctc cta tgt tgt a-- --- --- --- -at tag cat tag cat cag tc- --t cca cac tat tct taa att tca tat taa tgg gtc gtg acg gaa tag tac agc atg ata gaa cgc atc cta ttc cca aca atg tca gga acg tca ctc tct cca cct tca tat tta ttt atc cgt -aa aaa tgt tat cct gga cat tgt gcc aat aat aaa aaa gcc ca- -ta tat gtt cgc tat tgt aga aat tgt ttt tca tag ttg ctc aaa aac gat ggc ggt gac tta tga gtc acg tta cac ttt gga gtc tca tct tta gta aac ata tca taa tat tcg ata tta cga gtt gac aca ttg aac aaa ttc caa gta ttt gat ttt gga taa tat tcg tat ttt gca tct gct ata att aag ata taa tcg cca caa gaa cac gcg aac atc ttt cct aca tgg tta aag tac atg tac aat tct atc cat ttg tct tcc tta act ata tat ttg tat aga taa tta cga gtc tcg tgt gta att cca gta att gca tag atg tta cca tcg tac tct aca gca taa act ata cta taa tgt cta ggc atg gga gac -tt ttt tat cca aca att ttt agt gaa aca ttc cac atc gtt taa tat tac ata ttt ttc ata cgt ggt ata aac tcc rcc cat tac --a tat ata tca tca ttt acg aat acc gac gcg tct gaa tat cta gga gta att aag ttt gga tgt ctt atc cat ttc aaa gtg tag cgt ttt aaa tat tct gct aca cct gtt gaa ata gaa aat ttt aat cct cct att aca tat aac ttt cca tca tta aca cac gta cta act ttt gat ttt aac gac gac ata tta gta act gtt ttc ca- --- --- --- --- --- --- --- --- --- --- --- --- --- --- --- --- --- --- --- --- --- --- --- --- --- --- --- --- --- --- --- --- -t- --- --- --- --- --- --- --- --- --- --- --- --- --- --- --- --- --- --- --- --- --- --- --- --- --- --- --- --- --- --- --- --- --- --- --- --- --- --- --- --- --- --- --- --- --- --- --- --- --- --- --- --- --- --- --- --- --- --- --- --- --- --- --- --- --- --- --- --- --- --- --- --- --- --- --- --- --- --- --- --- --- --- --- --- --- --- --- --- --- --- --- --- --- --- --- --- --- --- --- --- --- --- --- --- --- --- --- --- --- --- --- --- --- --- --- --- --- --- --- --- --- --- --- --- --- -

>EF611192.1:79-1606_Variola_virus_strain_Congo-2_left_variable_genomic_sequence

cgg ttg cca tgt att tcc tgg aga gca agt --- aga tga ta- agg aac cag ata gtt tat atc cgt act tgc acg taa agt cta cat tgt cat tat acg a-- ata tga tct ttt aaa ccc gct aga caa gta tcc gtt taa tat tgt agg atg tgg aca ttt aac aat ctg aca cgt ggg tgg att aga cca ttc tcc tcc tga aca cag gac acc aga gtt acc aat caa cga ata tcc act att gca act ata agt tac aac gct tcc atc ggt ata aaa atc att ata tcc gtt atg tct tcc gtt gga tat aga tgg agg taa ttg gca ttt aac aga ttc gca aat agg tgc ctt ggg att cca tac cat aga tcc agt aga tcc taa ttt aca ata cga ttt ata ttc acc aat caa ata ata tcc gct att aca aga gta cgt tat act aga gcc aaa gtc tac tcc gcc aat atc aag atg gcc att atc gat atc tcg agg cga cgg gca tct ccg ttt aat aca ttg att aaa gag tgt cca tcc agt acc tgt aca ttt agc ata tat agg tcc cat ttt ttg ctt tct gta tcc agg tag aca tag a-t att cta tag tgt ctc cta tgt tgt a-- --- --- --- -at tag cat tag cat cag tc- --t cca cac tat tct taa att tca tat taa tgg gtc gtg acg gaa tag tac agc atg ata gaa cgc atc cta ttc cca aca atg tca gga acg tca ctc tct cca cct tca tat tta ttt atc cgt -aa aaa tgt tat cct gga cat tgt gcc aat aat aaa aaa gcc ca- -ta tat gtt cgc tat tgt aga act tgt ttt tca tag ttg ctc aaa aac gat ggc ggt gac tta tga gtc acg tta cac ttt gga gtc tca tct tta gta aac ata tca taa tat tcg ata tta cga gtt gac aca ttg aac aaa -tc caa gta ttt gat ttt gga taa tat tcg tat ttt gca tct gct ata att aag ata taa tcg cca caa aaa cac gcg aac atc ttt cct aca tgg tta aag tac atg tac aat tct atc cat ttg tct tcc tta act ata tat ttg tat aga taa tta cga gtc tcg tgt gta att cca gta att gca tag atg tta cca tcg tac tct aca gca taa act ata cta tga tgt cta ggc atg gga gac -tt ttt tat cca aca att ttt agt gaa aca ttc cac atc gtt taa tat tac ata ttt ttc ata cgt ggt ata aac tcc acc cat tac --a tat ata tca tca ttt acg aat acc gac gcg tct gaa tat cta gga gta att aag ttt gga tgt ctt atc cat ttc gaa gtg tag cgt ttt aaa tat ttt gct aca cct gtt gaa ata gaa aat ttt aat cct cct att aca tat aac ttt cca tca tta aca cac gta cta act ttt gat ttt aac gac gac ata tta gta act gtt ttc ca- --- --- --- --- --- --- -tt ttt ttt ttt tt- --- --- --- --- --- --- --- --- --- --- --- --- --- --- --- --- --- --- --- --- --- --- --- --- --- --- --- --- --- --- --- --- --- --- --- --- --- --- --- --- --- --- --- --- --- --- --- --- --- --- --- --- --- --- --- --- --- --- --- --- --- --- --- --- --- --- --- --- --- --- --- --- --- --- --- --- --- --- --- --- --- --- --- --- --- --- --- --- --- --- --- --- --- --- --- --- --- --- --- --- --- --- --- --- --- --- --- --- --- --- --- --- --- --- --- --- --- --- --- --- --- --- --- --- --- --- --- --- --- --- --- --- --- --- --- --- --- --- --- --- --- --- --- --- --- --- --- -

>DQ437583.1:12445-13972_Variola_virus_strain_Congo_1970_complete_genome

cgg ttg cca tgt att tcc tgg aga gca agt --- aga tga ta- agg aac cag ata gtt tat atc cgt act tgc acg taa agt cta cat tgt cat tat acg a-- ata tga tct ttt aaa ccc gct aga caa gta tcc gtt taa tat tgt agg atg tgg aca ttt aac aat ctg aca cgt ggg tgg att aga cca ttc tcc tcc tga aca cag gac acc aga gtt acc aat caa cga ata tcc act att gca act ata agt tac aac gct tcc atc ggt ata aaa atc att ata tcc gtt atg tct tcc gtt gga tat aga tgg agg taa ttg gca ttt aac aga ttc gca aat agg tgc ctt ggg att cca tac cat aga tcc agt aga tcc taa ttt aca ata cga ttt ata ttc acc aat caa ata ata tcc gct att aca aga gta cgt tat act aga gcc aaa gtc tac tcc gcc aat atc aag atg gcc att atc gat atc tcg agg cga cgg gca tct ccg ttt aat aca ttg att aaa gag tgt cca tcc agt acc tgt aca ttt agc ata tat agg tcc cat ttt ttg ctt tct gta tcc agg tag aca tag a-t att cta tag tgt ctc cta tgt tgt a-- --- --- --- -at tag cat tag cat cag tc- --t cca cac tat tct taa att tca tat taa tgg gtc gtg acg gaa tag tac agc atg ata gaa cgc atc cta ttc cca aca atg tca gga acg tca ctc tct cca cct tca tat tta ttt atc cgt -aa aaa tgt tat cct gga cat tgt gcc aat aat aaa aaa gcc ca- -ta tat gtt cgc tat tgt aga act tgt ttt tca tag ttg ctc aaa aac gat ggc ggt gac tta tga gtc acg tta cac ttt gga gtc tca tct tta gta aac ata tca taa tat tcg ata tta cga gtt gac aca ttg aac aaa -tc caa gta ttt gat ttt gga taa tat tcg tat ttt gca tct gct ata att aag ata taa tcg cca caa aaa cac gcg aac atc ttt cct aca tgg tta aag tac atg tac aat tct atc cat ttg tct tcc tta act ata tat ttg tat aga taa tta cga gtc tcg tgt gta att cca gta att gca tag atg tta cca tcg tac tct aca gca taa act ata cta tga tgt cta ggc atg gga gac -tt ttt tat cca aca att ttt agt gaa aca ttc cac atc gtt taa tat tac ata ttt ttc ata cgt ggt ata aac tcc acc cat tac --a tat ata tca tca ttt acg aat acc gac gcg tct gaa tat cta gga gta att aag ttt gga tgt ctt atc cat ttc gaa gtg tag cgt ttt aaa tat ttt gct aca cct gtt gaa ata gaa aat ttt aat cct cct att aca tat aac ttt cca tca tta aca cac gta cta act ttt gat ttt aac gac gac ata tta gta act gtt ttc ca- --- --- --- --- --- --- -tt ttt ttt ttt tt- --- --- --- --- --- --- --- --- --- --- --- --- --- --- --- --- --- --- --- --- --- --- --- --- --- --- --- --- --- --- --- --- --- --- --- --- --- --- --- --- --- --- --- --- --- --- --- --- --- --- --- --- --- --- --- --- --- --- --- --- --- --- --- --- --- --- --- --- --- --- --- --- --- --- --- --- --- --- --- --- --- --- --- --- --- --- --- --- --- --- --- --- --- --- --- --- --- --- --- --- --- --- --- --- --- --- --- --- --- --- --- --- --- --- --- --- --- --- --- --- --- --- --- --- --- --- --- --- --- --- --- --- --- --- --- --- --- --- --- --- --- --- --- --- --- --- --- -

>U18337.1:11672-13199_Variola_virus_Congo-1965_left_near-terminal_region

cgg ttg cca tgt att tcc tgg aga gca agt --- aga tga ta- agg aac cag ata gtt tat atc cgt act tgc acg taa agt cta cat tgt cat tat acg a-- ata tga tct ttt aaa ccc gct aga caa gta tcc gtt taa tat tgt agg atg tgg aca ttt aac aat ctg aca cgt ggg tgg att aga cca ttc tcc tcc tga aca cag gac acc aga gtt acc aat caa cga ata tcc act att gca act ata agt tac aac gct tcc atc ggt ata aaa atc att ata tcc gtt atg tct tcc gtt gga tat aga tgg agg taa ttg gca ttt aac aga ttc gca aat agg tgc ctt ggg att cca tac cat aga tcc agt aga tcc taa ttt aca ata cga ttt ata ttc acc aat caa ata ata tcc gct att aca aga gta cgt tat act aga gcc aaa gtc tac tcc gcc aat atc aag atg gcc att atc gat atc tcg agg cga cgg gca tct ccg ttt aat aca ttg att aaa gag tgt cca tcc agt acc tgt aca ttt agc ata tat agg tcc cat ttt ttg ctt tct gta tcc agg tag aca tag a-t att cta tag tgt ctc cta tgt tgt a-- --- --- --- -at tag cat tag cat cag tc- --t cca cac tat tct taa att tca tat taa tgg gtc gtg acg gaa tag tac agc atg ata gaa cgc atc cta ttc cca aca atg tca gga acg tca ctc tct cca cct tca tat tta ttt atc cgt -aa aaa tgt tat cct gga cat tgt gcc aat aat aaa aaa gcc ca- -ta tat gtt cgc tat tgt aga act tgt ttt tca tag ttg ctc aaa aac gat ggc ggt gac tta tga gtc acg tta cac ttt gga gtc tca tct tta gta aac ata tca taa tat tcg ata tta cga gtt gac aca ttg aac aaa -tc caa gta ttt gat ttt gga taa tat tcg tat ttt gca tct gct ata att aag ata taa tcg cca caa aaa cac gcg aac atc ttt cct aca tgg tta aag tac atg tac aat tct atc cat ttg tct tcc tta act ata tat ttg tat aga taa tta cga gtc tcg tgt gta att cca gta att gca tag atg tta cca tcg tac tct aca gca taa act ata cta tga tgt cta ggc atg gga gac -tt ttt tat cca aca att ttt agt gaa aca ttc cac atc gtt taa tat tac ata ttt ttc ata cgt ggt ata aac tcc acc cat tac --a tat ata tca tca ttt acg aat acc gac gcg tct gaa tat cta gga gta att aag ttt gga tgt ctt atc cat ttc gaa gtg tag cgt ttt aaa tat ttt gct aca cct gtt gaa ata gaa aat ttt aat cct cct att aca tat aac ttt cca tca tta aca cac gta cta act ttt gat ttt aac gac gac ata tta gta act gtt ttc ca- --- --- --- --- --- --- -tt ttt ttt ttt tt- --- --- --- --- --- --- --- --- --- --- --- --- --- --- --- --- --- --- --- --- --- --- --- --- --- --- --- --- --- --- --- --- --- --- --- --- --- --- --- --- --- --- --- --- --- --- --- --- --- --- --- --- --- --- --- --- --- --- --- --- --- --- --- --- --- --- --- --- --- --- --- --- --- --- --- --- --- --- --- --- --- --- --- --- --- --- --- --- --- --- --- --- --- --- --- --- --- --- --- --- --- --- --- --- --- --- --- --- --- --- --- --- --- --- --- --- --- --- --- --- --- --- --- --- --- --- --- --- --- --- --- --- --- --- --- --- --- --- --- --- --- --- --- --- --- --- --- -

>DQ441425.1:12523-14050_Variola_virus_strain_Ethiopia_1972_(Eth17_R14-1X-72_Addis)_complete_genome

cgg ttg cca tgt att tcc tgg aga gca agt --- aga tga tg- agg aac cag ata gtt tat atc cgt act tgc acg taa agt cta cat tgt cat tat acg a-- ata tga tct ttt aaa ccc gct aga caa gta tcc gtt taa tat tgt agg atg tgg aca ttt aac aat ctg aca cgt ggg tgg att aga cca ttc tcc tcc tga aca cag gac acc aga gtt acc aat caa cga ata tcc act att gca act ata agt tac aac gct tcc atc ggt ata aaa atc att ata tcc gtt atg tct tcc gtt gga tat aga tgg agg taa ttg gca ttt aac aga ttc gca aat agg tgc ctt ggg att cca tac cat aga tcc agt aga tcc taa ttt aca ata cga ttt ata ttc acc aat caa ata ata tcc gct att aca aga gta cgt tat act aga gcc aaa gtc tac tcc gcc aat atc aag atg gcc att atc gat atc tcg agg cga cgg gca tct ccg ttt aat aca ttg att aaa gag tgt cca tcc agt acc tgt aca ttt agc ata tat agg tcc cat ttt ttg ctt tct gta tcc agg tag aca tag a-t att cta tag tgt ctc cta tgt tgt a-- --- --- --- -at tag cat tag cat cag tc- --t cca cac tat tct taa att tca tat taa tgg gtc gtg acg gaa tag tac agc atg ata gaa cgc atc cta ttc cca aca atg tca gga acg tca ctc tct cca cct tca tat tta ttt atc cgt -aa aaa tgt tat cct gga cat tgt gcc aat aat aaa aaa gcc ca- -ta tat gtt cgc tat tgt aga act tgt ttt tca tag ttg ctc aaa aac gat ggc ggt gac tta tga gtc acg tta cac ttt gga gtc tca tct tta gta aac ata tca taa tat tcg ata tta cga gtt gac aca ttg aac aaa -tc caa gta ttt gat ttt gga taa tat tcg tat ttt gca tct gct ata att aag ata taa tcg cca caa aaa cac gcg aac atc ttt cct aca tgg tta aag tac atg tac aat tct atc cat ttg tct tcc tta act ata tat ttg tat aga taa tta cga gtc tcg tgt gta att cca gta att gca tag atg tta cca tcg tac tct aca gca taa act ata cta tga tgt cta ggc atg gga gac -tt ttt tat cca aca att ttt agt gaa aca ttc cac atc gtt taa tat tac ata ttt ttc ata cgt ggt ata aac tcc acc cat tac --a tat ata tca tca ttt acg aat acc gac gcg tct gaa tat cta gga gta att aag ttt gga tgt ctt atc cat ttc gaa gtg tag cgt ttt aaa tat ttt gct aca cct gtt gaa ata gaa aat ttt aat cct cct att aca tat aac ttt cca tca tta aca cac gta cta act ttt gat ttt aac gac gac ata tta gta act gtt ttc ct- --- --- --- --- --- --- -tt ttt ttt ttt tt- --- --- --- --- --- --- --- --- --- --- --- --- --- --- --- --- --- --- --- --- --- --- --- --- --- --- --- --- --- --- --- --- --- --- --- --- --- --- --- --- --- --- --- --- --- --- --- --- --- --- --- --- --- --- --- --- --- --- --- --- --- --- --- --- --- --- --- --- --- --- --- --- --- --- --- --- --- --- --- --- --- --- --- --- --- --- --- --- --- --- --- --- --- --- --- --- --- --- --- --- --- --- --- --- --- --- --- --- --- --- --- --- --- --- --- --- --- --- --- --- --- --- --- --- --- --- --- --- --- --- --- --- --- --- --- --- --- --- --- --- --- --- --- --- --- --- --- -

>DQ441424.1:12523-14050_Variola_virus_strain_Ethiopia_1972_(Eth16_R14-1X-72_Addis)_complete_genome

cgg ttg cca tgt att tcc tgg aga gca agt --- aga tga tg- agg aac cag ata gtt tat atc cgt act tgc acg taa agt cta cat tgt cat tat acg a-- ata tga tct ttt aaa ccc gct aga caa gta tcc gtt taa tat tgt agg atg tgg aca ttt aac aat ctg aca cgt ggg tgg att aga cca ttc tcc tcc tga aca cag gac acc aga gtt acc aat caa cga ata tcc act att gca act ata agt tac aac gct tcc atc ggt ata aaa atc att ata tcc gtt atg tct tcc gtt gga tat aga tgg agg taa ttg gca ttt aac aga ttc gca aat agg tgc ctt ggg att cca tac cat aga tcc agt aga tcc taa ttt aca ata cga ttt ata ttc acc aat caa ata ata tcc gct att aca aga gta cgt tat act aga gcc aaa gtc tac tcc gcc aat atc aag atg gcc att atc gat atc tcg agg cga cgg gca tct ccg ttt aat aca ttg att aaa gag tgt cca tcc agt acc tgt aca ttt agc ata tat agg tcc cat ttt ttg ctt tct gta tcc agg tag aca tag a-t att cta tag tgt ctc cta tgt tgt a-- --- --- --- -at tag cat tag cat cag tc- --t cca cac tat tct taa att tca tat taa tgg gtc gtg acg gaa tag tac agc atg ata gaa cgc atc cta ttc cca aca atg tca gga acg tca ctc tct cca cct tca tat tta ttt atc cgt -aa aaa tgt tat cct gga cat tgt gcc aat aat aaa aaa gcc ca- -ta tat gtt cgc tat tgt aga act tgt ttt tca tag ttg ctc aaa aac gat ggc ggt gac tta tga gtc acg tta cac ttt gga gtc tca tct tta gta aac ata tca taa tat tcg ata tta cga gtt gac aca ttg aac aaa -tc caa gta ttt gat ttt gga taa tat tcg tat ttt gca tct gct ata att aag ata taa tcg cca caa aaa cac gcg aac atc ttt cct aca tgg tta aag tac atg tac aat tct atc cat ttg tct tcc tta act ata tat ttg tat aga taa tta cga gtc tcg tgt gta att cca gta att gca tag atg tta cca tcg tac tct aca gca taa act ata cta tga tgt cta ggc atg gga gac -tt ttt tat cca aca att ttt agt gaa aca ttc cac atc gtt taa tat tac ata ttt ttc ata cgt ggt ata aac tcc acc cat tac --a tat ata tca tca ttt acg aat acc gac gcg tct gaa tat cta gga gta att aag ttt gga tgt ctt atc cat ttc gaa gtg tag cgt ttt aaa tat ttt gct aca cct gtt gaa ata gaa aat ttt aat cct cct att aca tat aac ttt cca tca tta aca cac gta cta act ttt gat ttt aac gac gac ata tta gta act gtt ttc ct- --- --- --- --- --- --- -tt ttt ttt ttt tt- --- --- --- --- --- --- --- --- --- --- --- --- --- --- --- --- --- --- --- --- --- --- --- --- --- --- --- --- --- --- --- --- --- --- --- --- --- --- --- --- --- --- --- --- --- --- --- --- --- --- --- --- --- --- --- --- --- --- --- --- --- --- --- --- --- --- --- --- --- --- --- --- --- --- --- --- --- --- --- --- --- --- --- --- --- --- --- --- --- --- --- --- --- --- --- --- --- --- --- --- --- --- --- --- --- --- --- --- --- --- --- --- --- --- --- --- --- --- --- --- --- --- --- --- --- --- --- --- --- --- --- --- --- --- --- --- --- --- --- --- --- --- --- --- --- --- --- -

>U18340.1:11883-13410_Variola_virus_Somalia-1977_left_variable_region

cgg ttg cca tgt att tcc tgg aga gca agt --- aga tga tg- agg aac cag ata gtt tat atc cgt act tgc acg taa agt cta cat tgt cat tat acg a-- ata tga tct ttt aaa ccc gct aga caa gta tcc gtt taa tat tgt agg atg tgg aca ttt aac aat ctg aca cgt ggg tgg att aga cca ttc tcc tcc tga aca cag gac acc aga gtt acc aat caa cga ata tcc act att gca act ata agt tac aac gct tcc atc ggt ata aaa atc att ata tcc gtt atg tct tcc gtt gga tat aga tgg agg taa ttg gca ttt aac aga ttc gca aat agg tgc ctt ggg att cca tac cat aga tcc agt aga tcc taa ttt aca ata cga ttt ata ttc acc aat caa ata ata tcc gct att aca aga gta cgt tat act aga gcc aaa gtc tac tcc gcc aat atc aag atg gcc att atc gat atc tcg agg cga cgg gca tct ccg ttt aat aca ttg att aaa gag tgt cca tcc agt acc tgt aca ttt agc ata tat agg tcc cat ttt ttg ctt tct gta tcc agg tag aca tag a-t att cta tag tgt ctc cta tgt tgt a-- --- --- --- -at tag cat tag cat cag tc- --t cca cac tat tct taa att tca tat taa tgg gtc gtg acg gaa tag tac agc atg ata gaa cgc atc cta ttc cca aca atg tca gga acg tca ctc tct cca cct tca tat tta ttt atc cgt -aa aaa tgt tat cct gga cat tgt gcc aat aat aaa aaa gcc ca- -ta tat gtt cgc tat tgt aga act tgt ttt tca tag ttg ctc aaa aac gat ggc ggt gac tta tga gtc acg tta cac ttt gga gtc tca tct tta gta aac ata tca taa tat tcg ata tta cga gtt gac aca ttg aac aaa -tc caa gta ttt gat ttt gga taa tat tcg tat ttt gca tct gct ata att aag ata taa tcg cca caa aaa cac gcg aac atc ttt cct aca tgg tta aag tac atg tac aat tct atc cat ttg tct tcc tta act ata tat ttg tat aga taa tta cga gtc tcg tgt gta att cca gta att gca tag atg tta cca tcg tac tct aca gca taa act ata cta tga tgt cta ggc atg gga gac -tt ttt tat cca aca att ttt agt gaa aca ttc cac atc gtt taa tat tac ata ttt ttc ata cgt ggt ata aac tcc acc cat tac --a tat ata tca tca ttt acg aat acc gac gcg tct gaa tat cta gga gta att aag ttt gga tgt ctt atc cat ttc gaa gtg tag cgt ttt aaa tat ttt gct aca cct gtt gaa ata gaa aat ttt aat cct cct att aca tat aac ttt cca tca tta aca cac gta cta act ttt ggt ttt aac gac gac ata tta gta act gtt ttc ca- --- --- --- --- --- --- -tt ttt ttt ttt tt- --- --- --- --- --- --- --- --- --- --- --- --- --- --- --- --- --- --- --- --- --- --- --- --- --- --- --- --- --- --- --- --- --- --- --- --- --- --- --- --- --- --- --- --- --- --- --- --- --- --- --- --- --- --- --- --- --- --- --- --- --- --- --- --- --- --- --- --- --- --- --- --- --- --- --- --- --- --- --- --- --- --- --- --- --- --- --- --- --- --- --- --- --- --- --- --- --- --- --- --- --- --- --- --- --- --- --- --- --- --- --- --- --- --- --- --- --- --- --- --- --- --- --- --- --- --- --- --- --- --- --- --- --- --- --- --- --- --- --- --- --- --- --- --- --- --- --- -

>DQ441418.1:12113-13640_Variola_virus_strain_Botswana_1973_(v73-225)_complete_genome

cgg ttg cca tgt att tcc tgg aga gca agt --- aga tga tg- agg aac cag ata gtt tat atc cgt act tgc acg taa agt cta cat tgt cat tat acg a-- ata tga tct ttt aaa ccc gct aga caa gta tcc gtt taa tat tgt agg atg tgg aca ttt aac aat ctg aca cgt ggg tgg att aga cca ttc tcc tcc tga aca cag gac acc aga gtt acc aat caa cga ata tcc act att gca act ata agt tac aac gct tcc atc ggt ata aaa atc att ata tcc gtt atg tct tcc gtt gga tat aga tgg agg taa ttg gca ttt aac aga ttc gca aat agg tgc ctt ggg att cca tac cat aga tcc agt aga tcc taa ttt aca ata cga ttt ata ttc acc aat caa ata ata tcc gct att aca aga gta cgt tat act aga gcc aaa gtc tac tcc gcc aat atc aag atg gcc att atc gat atc tcg agg cga cgg gca tct ccg ttt aat aca ttg att aaa gag tgt cca tcc agt acc tgt aca ttt agc ata tat agg tcc cat ttt ttg ctt tct gta tcc agg tag aca tag a-t att cta tag tgt ctc cta tgt tgt a-- --- --- --- -at tag cat tag cat cag tc- --t cca cac tat tct taa att tca tat taa tgg gtc gtg acg gaa tag tac agc atg ata gaa cgc atc cta ttc cca aca atg tca gga acg tca ctc tct cca cct tca tat tta ttt atc cgt -aa aaa tgt tat cct gga cat tgt gcc aat aat aaa aaa gcc ca- -ta tat gtt cgc tat tgt aga act tgt ttt tca tag ttg ctc aaa aac gat ggc ggt gac tta tga gtc acg tta cac ttt gga gtc tca tct tta gta aac ata tca taa tat tcg ata tta caa gtt gac aca ttg aac aaa -tc caa gta ttt gat ttt gga taa tat tcg tat ttt gca tct gct ata att aag ata taa tcg cca caa aaa cac gcg aac atc ttt cct aca tgg tta aag tac atg tac aat tct atc cat ttg tct tcc tta act ata tat ttg tat aga taa tta cga gtc tcg tgt gta att cca gta att gca tag atg tta cca tcg tac tct aca gca taa act ata cta tga tgt cta ggc atg gga gac -tt ttt tat cca aca att ttt agt gaa aca ttc cac atc gtt taa tat tac ata ttt ttc ata cgt ggt ata aac tcc acc cat tac --a tat ata tca tca ttt acg aat acc gac gcg tct gaa tat cta gga gta att aag ttt gga tgt ctt atc cat ttc gaa gtg tag cgt ttt aaa tat ttt gct aca cct gtt gaa ata gaa aat ttt aat cct cct att aca tat aac ttt cca tca tta aca cac gta cta act ttt gat ttt aac gac gat ata tta gta act gtt ttc ca- --- --- --- --- --- --- -tt ttt ttt ttt tt- --- --- --- --- --- --- --- --- --- --- --- --- --- --- --- --- --- --- --- --- --- --- --- --- --- --- --- --- --- --- --- --- --- --- --- --- --- --- --- --- --- --- --- --- --- --- --- --- --- --- --- --- --- --- --- --- --- --- --- --- --- --- --- --- --- --- --- --- --- --- --- --- --- --- --- --- --- --- --- --- --- --- --- --- --- --- --- --- --- --- --- --- --- --- --- --- --- --- --- --- --- --- --- --- --- --- --- --- --- --- --- --- --- --- --- --- --- --- --- --- --- --- --- --- --- --- --- --- --- --- --- --- --- --- --- --- --- --- --- --- --- --- --- --- --- --- --- -

>DQ441417.1:12113-13640_Variola_virus_strain_Botswana_1972_(v72-143)_complete_genome

cgg ttg cca tgt att tcc tgg aga gca agt --- aga tga tg- agg aac cag ata gtt tat atc cgt act tgc acg taa agt cta cat tgt cat tat acg a-- ata tga tct ttt aaa ccc gct aga caa gta tcc gtt taa tat tgt agg atg tgg aca ttt aac aat ctg aca cgt ggg tgg att aga cca ttc tcc tcc tga aca cag gac acc aga gtt acc aat caa cga ata tcc act att gca act ata agt tac aac gct tcc atc ggt ata aaa atc att ata tcc gtt atg tct tcc gtt gga tat aga tgg agg taa ttg gca ttt aac aga ttc gca aat agg tgc ctt ggg att cca tac cat aga tcc agt aga tcc taa ttt aca ata cga ttt ata ttc acc aat caa ata ata tcc gct att aca aga gta cgt tat act aga gcc aaa gtc tac tcc gcc aat atc aag atg gcc att atc gat atc tcg agg cga cgg gca tct ccg ttt aat aca ttg att aaa gag tgt cca tcc agt acc tgt aca ttt agc ata tat agg tcc cat ttt ttg ctt tct gta tcc agg tag aca tag a-t att cta tag tgt ctc cta tgt tgt a-- --- --- --- -at tag cat tag cat cag tc- --t cca cac tat tct taa att tca tat taa tgg gtc gtg acg gaa tag tac agc atg ata gaa cgc atc cta ttc cca aca atg tca gga acg tca ctc tct cca cct tca tat tta ttt atc cgt -aa aaa tgt tat cct gga cat tgt gcc aat aat aaa aaa gcc ca- -ta tat gtt cgc tat tgt aga act tgt ttt tca tag ttg ctc aaa aac gat ggc ggt gac tta tga gtc acg tta cac ttt gga gtc tca tct tta gta aac ata tca taa tat tcg ata tta caa gtt gac aca ttg aac aaa -tc caa gta ttt gat ttt gga taa tat tcg tat ttt gca tct gct ata att aag ata taa tcg cca caa aaa cac gcg aac atc ttt cct aca tgg tta aag tac atg tac aat tct atc cat ttg tct tcc tta act ata tat ttg tat aga taa tta cga gtc tcg tgt gta att cca gta att gca tag atg tta cca tcg tac tct aca gca taa act ata cta tga tgt cta ggc atg gga gac -tt ttt tat cca aca att ttt agt gaa aca ttc cac atc gtt taa tat tac ata ttt ttc ata cgt ggt ata aac tcc acc cat tac --a tat ata tca tca ttt acg aat acc gac gcg tct gaa tat cta gga gta att aag ttt gga tgt ctt atc cat ttc gaa gtg tag cgt ttt aaa tat ttt gct aca cct gtt gaa ata gaa aat ttt aat cct cct att aca tat aac ttt cca tca tta aca cac gta cta act ttt gat ttt aac gac gat ata tta gta act gtt ttc ca- --- --- --- --- --- --- -tt ttt ttt ttt tt- --- --- --- --- --- --- --- --- --- --- --- --- --- --- --- --- --- --- --- --- --- --- --- --- --- --- --- --- --- --- --- --- --- --- --- --- --- --- --- --- --- --- --- --- --- --- --- --- --- --- --- --- --- --- --- --- --- --- --- --- --- --- --- --- --- --- --- --- --- --- --- --- --- --- --- --- --- --- --- --- --- --- --- --- --- --- --- --- --- --- --- --- --- --- --- --- --- --- --- --- --- --- --- --- --- --- --- --- --- --- --- --- --- --- --- --- --- --- --- --- --- --- --- --- --- --- --- --- --- --- --- --- --- --- --- --- --- --- --- --- --- --- --- --- --- --- --- -

>MN240300.1:30811-32757_Alaskapox_virus_isolate_Alaska_2015

-gg ctg cca tgt atc tcc tag aga gca agt --- aga tga cg- agg aac cag ata gtt cat atc cgt gtt tgc acg taa agt gta cct tat cat tat acg a-- gta tga tct ttt aaa ccc gtt --- cac gta tcc gtt tga tat tgt agg ata ttg aca ttt aac aat ctg gca cgt tgg cgg atc tgt cca ttc tcc tcc cga aca caa gat acc aga gtt acc aat cag tga ata tcc ttt att gca act ata agt cac gac gct ccc atc ggt ata aaa atc ctc ata tcc gtt atg att tcc gtt ggt tac aga tgg agg tga ttg gca ttt aac cga ttc aca aat agg tgc ctc ggg att cca tac cat ata tcc ggc aga tcc tag ctc aca ata tga tct aga ttc tcc gat caa ttt ata tcc gct att aca aga gta agt tat act gga tcc aaa gtc cgt tcc act aat atc aag ttg acc att atc gat atc tcg agg cga tga aca ttt ccg ttt aat aca ttg att aag gag tga cca tcc att gct tgt aca ttt agc ata tat ggg tcc cat ttt ttg ctt tct gta tcc tgg tag aca taa a-t att cta tag tgt ctc cta tat tat a-- --- --- --- --- --- -at tct cat tgg ta- --g cca ctg tat tct taa att tca tat taa tgg gac gta agg gac tat tac agc atg ata gta cac atc cta ttc caa aca atg tca gga atg tga cgc att ccg act tca tat tta ttt atc cgt aaa aaa tgt tat acc gga cat tgt --- -at aat aaa aag gtc ta- --- --- aat tac tat tgt agg aat tgt ttt tca cag ttg ttc aaa aac gat gtc agt gac tta tgc gtt aca tta cac gtt gga gtc tcg tct cta gta aac ata tca taa tat tcg ata tta cgc gtt gac ata ttg tac gaa ttc caa gta ttt gat ttt gga taa tat tcg tat ttt gca tcc gct ata atc aaa ata taa tcg cca caa gaa cac atg aac atc ttt ccc aca tgg tta aag tac ata tac aat tct atc cat ttg tct tcg tta act ata tat ttg tac aga taa tta cgt gtt tca tga gtg atc cca gaa atg aca tag atg tcc ccg tcg tac tct act gca taa act ata cta tga tgt cta ggc ata gga gac -tt ttt tat cca acg att ttt agt aaa gca ttc cac atc gtt taa tat cat gta tct ctc ata cgt ggt ata cac tcc acc cat cac --a tac ata tcg tcg tct acg aat acc gac gcg cct gaa tat cta gga gtg att aag ttt gga agt ctt atc cat ttt gag gta tca tgt ttc aaa tat tca act act cct gtc gat ata gaa aat tcc aat cct cct att acg tat agc ttt cca tca tta aca caa gta cta act ttt gat ttt aac gac gac atg tta gta acc gtt ttc ca- --- --- --- --- --- --- --t ttc ttt gtt tta aga tct aaa cga -ga cac gga ata aat atg tct atg gtt aat cat tcc tcc agt aac ata tag aca att atg cat cac att cgc gct aaa gaa ttg --- -ta cat cag tat tac cga cta tag tcc aat atc ctg ttc tag ggg agt aat gag tta ttg tag ata tat aat cag agt ttt taa tgg cta tta tgt tac gtt tta tac cat ttc gtg tta cag gtt tta tag att ggg ata tat tta atc cca aca acg ata tgg cct tac gca tat tat tag aca taa act tgg gat gta aaa tgt tga tga tat cat ggt cat ttg gat ttt tac gca tcc act tta ata ata tta tag cca cga cat cct cat ta- ttt aca ttg ata tct tcg tgg gat aag ata gtt att agt tca tcc ttt gat aat tct cca aat t
