## Supplementary Data 6 for "Genomic regions insertion and deletion in Monkeypox virus causing multi-country outbreak-2022"

>ON674051.1_Monkeypox_virus_isolate_MPXV_USA_2022_FL001_complete_genome

tta cca cgt ctg tac tcg acg agc tca cgt tta aga gat tca att tcc agt ttg tat cgg tcc atg tct cca ttg cta cac cac cat tag att tac agg ctg cta gtt gtc gtt cga gat cag aaa tac gtg ttt tct tgg aat gga ttt cgt cga tgt act tgt cat gat tgg cat cga aac act tat taa gtt ctt ttt ttc aat tct acg att tta ttt ctt tcg cga gtc aat tcc ctc ctg tag taa cta tca gtt ttg tca gat tca cgc tct cta cgt aga ctt tct tgt aag tta cta att tgt tcc ctg gca tta ccg agt tca gtt tta tat gcc gaa tag agt tct gat tca tcc ttt gag aag atc tct agc gat cgt tca aga tcc ctg att cta gtc ttt agc cta ttt acc tcc tca gaa gat gct ccg tta ccg ttt tta caa tcg tta aga tgt cta tca aga tcc atg att cta tct ctt ttc cat atc agc att gat ttc att att acg ttc gca gtc gtt caa ctg tat t

>NC_063383.1_Monkeypox_virus_complete_genome

tta cca cgt ctg tac tcg acg agc tca cgt tta aga gat tca att tcc agt ttg tat cgg tcc atg tct cca ttg cta cac cac cat tag att tac agg ctg cta gtt gtc gtt cga gat cag aaa tac gtg ttt tct tgg aat gga ttt cgt cga tgt act tgt cat gat tgg cat cga aac act tat taa gtt ctt ttt ttc aat tct acg att tta ttt ctt tcg cga gtc aat tcc ctc ctg tag taa cta tca gtt ttg tca gat tca cgc tct cta cgt aga ctt tct tgt aag tta cta att tgt tcc ctg gca tta ccg agt tca gtt tta tat gcc gaa tag agt tct gat tca tcc ttt gag aag atc tct agc gat cgt tca aga tcc ctg att cta gtc ttt agc cta ttt acc tcc tca gaa gat gct ccg tta ccg ttt tta caa tcg tta aga tgt cta tca aga tcc atg att cta tct ctt ttc cat atc agc att gat ttc att att acg ttc gca gtc gtt caa ctg tat t

>ON676704.1_Monkeypox_virus_isolate_MPXV_USA_2022_FL002_complete_genome

tta cca cgt ctg tac tcg acg agc tca cgt tta aga gat tca att tcc agt ttg tat cgg tcc atg tct cca ttg cta cac cac cat tag att tac agg ctg cta gtt gtc gtt cga gat cag aaa tac gtg ttt tct tgg aat gga ttt cgt cga tgt act tgt cat gat tgg cat cga aac act tat taa gtt ctt ttt ttc aat tct acg att tta ttt ctt tcg cga gtc aat tcc ctc ctg tag taa cta tca gtt ttg tca gat tca cgc tct cta cgt aga ctt tct tgt aag tta cta att tgt tcc ctg gca tta ccg agt tca gtt tta tat gcc gaa tag agt tct gat tca tcc ttt gag aag atc tct agc gat cgt tca aga tcc ctg att cta gtc ttt agc cta ttt acc ttc tca gaa gat gct ccg tta ccg ttt tta caa tcg tta aga tgt cta tca aga tcc atg att cta tct ctt ttc cat atc agc att gat ttc att att acg ttc gca gtc gtt caa ctg tat t

>ON563414.3_Monkeypox_virus_isolate_MPXV_USA_2022_MA001_complete_genome

tta cca cgt ctg tac tcg acg agc tca cgt tta aga gat tca att tcc agt ttg tat cgg tcc atg tct cca ttg cta cac cac cat tag att tac agg ctg cta gtt gtc gtt cga gat cag aaa tac gtg ttt tct tgg aat gga ttt cgt cga tgt act tgt cat gat tgg cat cga aac act tat taa gtt ctt ttt ttc aat tct acg att tta ttt ctt tcg cga gtc aat tcc ctc ctg tag taa cta tca gtt ttg tca gat tca cgc tct cta cgt aga ctt tct tgt aag tta cta att tgt tcc ctg gca tta ccg agt tca gtt tta tat gcc gaa tag agt tct gat tca tcc ttt gag aag atc tct agc gat cgt tca aga tcc ctg att cta gtc ttt agc cta ttt acc tcc tca gaa gat gct ccg tta ccg ttt tta caa tcg tta aga tgt cta tca aga tcc atg att cta tct ctt ttc cat atc agc att gat ttc att att acg ttc gca gtc gtt caa ctg tat t

>ON676705.1_Monkeypox_virus_isolate_MPXV_USA_2022_UT001_complete_genome

tta cca cgt ctg tac tcg acg agc tca cgt tta aga gat tca att tcc agt ttg tat cgg tcc atg tct cca ttg cta cac cac cat tag att tac agg ctg cta gtt gtc gtt cga gat cag aaa tac gtg ttt tct tgg aat gga ttt cgt cga tgt act tgt cat gat tgg cat cga aac act tat taa gtt ctt ttt ttc aat tct acg att tta ttt ctt tcg cga gtc aat tcc ctc ctg tag taa cta tca gtt ttg tca gat tca cgc tct cta cgt aga ctt tct tgt aag tta cta att tgt tcc ctg gca tta ccg agt tca gtt tta tat gcc gaa tag agt tct gat tca tcc ttt gag aag atc tct agc gat cgt tca aga tcc ctg att cta gtc ttt agc cta ttt acc tcc tca gaa gat gct ccg tta ccg ttt tta caa tcg tta aga tgt cta tca aga tcc atg att cta tct ctt ttc cat atc agc att gat ttc att att acg ttc gca gtc gtt caa ctg tat t

>MN648051.1_Monkeypox_virus_strain_Israel_2018

tta cca cgt ctg tac tcg acg agc tca cgt tta aga gat tca att tcc agt ttg tat cgg tcc atg tct cca ttg cta cac cac cat tag att tac agg ctg cta gtt gtc gtt cga gat cag aaa tac gtg ttt tct tgg aat gga ttt cgt cga tgt act tgt cat gat tgg cat cga aac act tat taa gtt ctt ttt ttc aat tct acg att tta ttt ctt tcg cga gtc aat tcc ctc ctg tag taa cta tca gtt ttg tca gat tca cgc tct cta cgt aga ctt tct tgt aag tta cta att tgt tcc ctg gca tta ccg agt tca gtt tta tat gcc gaa tag agt tct gat tca tcc ttt gag aag atc tct agc gat cgt tca aga tcc ctg att cta gtc ttt agc cta ttt acc tcc tca gaa gat gct ccg tta ccg ttt tta caa tcg tta aga tgt cta tca aga tcc atg att cta tct ctt ttc cat atc agc att gat ttc att att acg ttc gca gtc gtt caa ctg tat t

>ON644344.1_Monkeypox_virus_isolate_MPXV_FVG-ITA_01_2022_complete_genome

tta cca cgt ctg tac tcg acg agc tca cgt tta aga gat tca att tcc agt ttg tat cgg tcc atg tct cca ttg cta cac cac cat tag att tac agg ctg cta gtt gtc gtt cga gat cag aaa tac gtg ttt tct tgg aat gga ttt cgt cga tgt act tgt cat gat tgg cat cga aac act tat taa gtt ctt ttt ttc aat tct acg att tta ttt ctt tcg cga gtc aat tcc ctc ctg tag taa cta tca gtt ttg tca gat tca cgc tct cta cgt aga ctt tct tgt aag tta cta att tgt tcc ctg gca tta ccg agt tca gtt tta tat gcc gaa tag agt tct gat tca tcc ttt gag aag atc tct agc gat cgt tca aga tcc ctg att cta gtc ttt agc cta ttt acc tcc tca gaa gat gct ccg tta ccg ttt tta caa tcg tta aga tgt cta tca aga tcc atg att cta tct ctt ttc cat atc agc att gat ttc att att acg ttc gca gtc gtt caa ctg tat t

>MT903343.1_Monkeypox_virus_isolate_MPXV-UK_P1

tta cca cgt ctg tac tcg acg agc tca cgt tta aga gat tca att tcc agt ttg tat cgg tcc atg tct cca ttg cta cac cac cat tag att tac agg ctg cta gtt gtc gtt cga gat cag aaa tac gtg ttt tct tgg aat gga ttt cgt cga tgt act tgt cat gat tgg cat cga aac act tat taa gtt ctt ttt ttc aat tct acg att tta ttt ctt tcg cga gtc aat tcc ctc ctg tag taa cta tca gtt ttg tca gat tca cgc tct cta cgt aga ctt tct tgt aag tta cta att tgt tcc ctg gca tta ccg agt tca gtt tta tat gcc gaa tag agt tct gat tca tcc ttt gag aag atc tct agc gat cgt tca aga tcc ctg att cta gtc ttt agc cta ttt acc tcc tca gaa gat gct ccg tta ccg ttt tta caa tcg tta aga tgt cta tca aga tcc atg att cta tct ctt ttc cat atc agc att gat ttc att att acg ttc gca gtc gtt caa ctg tat t

>ON631963.1_Monkeypox_virus_isolate_MPxV/VIDRL01/2022_complete_genome

tta cca cgt ctg tac tcg acg agc tca cgt tta aga gat tca att tcc agt ttg tat cgg tcc atg tct cca ttg cta cac cac cat tag att tac agg ctg cta gtt gtc gtt cga gat cag aaa tac gtg ttt tct tgg aat gga ttt cgt cga tgt act tgt cat gat tgg cat cga aac act tat taa gtt ctt ttt ttc aat tct acg att tta ttt ctt tcg cga gtc aat tcc ctc ctg tag taa cta tca gtt ttg tca gat tca cgc tct cta cgt aga ctt tct tgt aag tta cta att tgt tcc ctg gca tta ccg agt tca gtt tta tat gcc gaa tag agt tct gat tca tcc ttt gag aag atc tct agc gat cgt tca aga tcc ctg att cta gtc ttt agc cta ttt acc tcc tca gaa gat gct ccg tta ccg ttt tta caa tcg tta aga tgt cta tca aga tcc atg att cta tct ctt ttc cat atc agc att gat ttc att att acg ttc gca gtc gtt caa ctg tat t

>ON568298.1_Monkeypox_virus_isolate_MPXV-BY-IMB25241_complete_genome

tta cca cgt ctg tac tcg acg agc tca cgt tta aga gat tca att tcc agt ttg tat cgg tcc atg tct cca ttg cta cac cac cat tag att tac agg ctg cta gtt gtc gtt cga gat cag aaa tac gtg ttt tct tgg aat gga ttt cgt cga tgt act tgt cat gat tgg cat cga aac act tat taa gtt ctt ttt ttc aat tct acg att tta ttt ctt tcg cga gtc aat tcc ctc ctg tag taa cta tca gtt ttg tca gat tca cgc tct cta cgt aga ctt tct tgt aag tta cta att tgt tcc ctg gca tta ccg agt tca gtt tta tat gcc gaa tag agt tct gat tca tcc ttt gag aag atc tct agc gat cgt tca aga tcc ctg att cta gtc ttt agc cta ttt acc tcc tca gaa gat gct ccg tta ccg ttt tta caa tcg tta aga tgt cta tca aga tcc atg att cta tct ctt ttc cat atc agc att gat ttc att att acg ttc gca gtc gtt caa ctg tat t

>ON622722.2_Monkeypox_virus_isolate_MPXV_FR_HCL0001_2022_complete_genome

tta cca cgt ctg tac tcg acg agc tca cgt tta aga gat tca att tcc agt ttg tat cgg tcc atg tct cca ttg cta cac cac cat tag att tac agg ctg cta gtt gtc gtt cga gat cag aaa tac gtg ttt tct tgg aat gga ttt cgt cga tgt act tgt cat gat tgg cat cga aac act tat taa gtt ctt ttt ttc aat tct acg att tta ttt ctt tcg cga gtc aat tcc ctc ctg tag taa cta tca gtt ttg tca gat tca cgc tct cta cgt aga ctt tct tgt aag tta cta att tgt tcc ctg gca tta ccg agt tca gtt tta tat gcc gaa tag agt tct gat tca tcc ttt gag aag atc tct agc gat cgt tca aga tcc ctg att cta gtc ttt agc cta ttt acc tcc tca gaa gat gct ccg tta ccg ttt tta caa tcg tta aga tgt cta tca aga tcc atg att cta tct ctt ttc cat atc agc att gat ttc att att acg ttc gca gtc gtt caa ctg tat t

>ON627808.1_Monkeypox_virus_isolate_MPX/human/USA/UT-UPHL-82200022/2022_complete_genome

tta cca cgt ctg tac tcg acg agc tca cgt tta aga gat tca att tcc agt ttg tat cgg tcc atg tct cca ttg cta cac cac cat tag att tac agg ctg cta gtt gtc gtt cga gat cag aaa tac gtg ttt tct tgg aat gga ttt cgt cga tgt act tgt cat gat tgg cat cga aac act tat taa gtt ctt ttt ttc aat tct acg att tta ttt ctt tcg cga gtc aat tcc ctc ctg tag taa cta tca gtt ttg tca gat tca cgc tct cta cgt aga ctt tct tgt aag tta cta att tgt tcc ctg gca tta ccg agt tca gtt tta tat gcc gaa tag agt tct gat tca tcc ttt gag aag atc tct agc gat cgt tca aga tcc ctg att cta gtc ttt agc cta ttt acc tcc tca gaa gat gct ccg tta ccg ttt tta caa tcg tta aga tgt cta tca aga tcc atg att cta tct ctt ttc cat atc agc att gat ttc att att acg ttc gca gtc gtt caa ctg tat t

>ON676703.1_Monkeypox_virus_isolate_MPXV_USA_2022_CA001_complete_genome

tta cca cgt ctg tac tcg acg agc tca cgt tta aga gat tca att tcc agt ttg tat cgg tcc atg tct cca ttg cta cac cac cat tag att tac agg ctg cta gtt gtc gtt cga gat cag aaa tac gtg ttt tct tgg aat gga ttt cgt cga tgt act tgt cat gat tgg cat cga aac act tat taa gtt ctt ttt ttc aat tct acg att tta ttt ctt tcg cga gtc aat tcc ctc ctg tag taa cta tca gtt ttg tca gat tca cgc tct cta cgt aga ctt tct tgt aag tta cta att tgt tcc ctg gca tta ccg agt tca gtt tta tat gcc gaa tag agt tct gat tca tcc ttt gag aag atc tct agc gat cgt tca aga tcc ctg att cta gtc ttt agc cta ttt acc tcc tca gaa gat gct ccg tta ccg ttt tta caa tcg tta aga tgt cta tca aga tcc atg att cta tct ctt ttc cat atc agc att gat ttc att att acg ttc gca gtc gtt caa ctg tat t

>ON676706.1_Monkeypox_virus_isolate_MPXV_USA_2022_UT002_complete_genome

tta cca cgt ctg tac tcg acg agc tca cgt tta aga gat tca att tcc agt ttg tat cgg tcc atg tct cca ttg cta cac cac cat tag att tac agg ctg cta gtt gtc gtt cga gat cag aaa tac gtg ttt tct tgg aat gga ttt cgt cga tgt act tgt cat gat tgg cat cga aac act tat taa gtt ctt ttt ttc aat tct acg att tta ttt ctt tcg cga gtc aat tcc ctc ctg tag taa cta tca gtt ttg tca gat tca cgc tct cta cgt aga ctt tct tgt aag tta cta att tgt tcc ctg gca tta ccg agt tca gtt tta tat gcc gaa tag agt tct gat tca tcc ttt gag aag atc tct agc gat cgt tca aga tcc ctg att cta gtc ttt agc cta ttt acc tcc tca gaa gat gct ccg tta ccg ttt tta caa tcg tta aga tgt cta tca aga tcc atg att cta tct ctt ttc cat atc agc att gat ttc att att acg ttc gca gtc gtt caa ctg tat t

>ON649879.1_Monkeypox_virus_isolate_MPXV_ISR001_2022_complete_genome

tta cca cgt ctg tac tcg acg agc tca cgt tta aga gat tca att tcc agt ttg tat cgg tcc atg tct cca ttg cta cac cac cat tag att tac agg ctg cta gtt gtc gtt cga gat cag aaa tac gtg ttt tct tgg aat gga ttt cgt cga tgt act tgt cat gat tgg cat cga aac act tat taa gtt ctt ttt ttc aat tct acg att tta ttt ctt tcg cga gtc aat tcc ctc ctg tag taa cta tca gtt ttg tca gat tca cgc tct cta cgt aga ctt tct tgt aag tta cta att tgt tcc ctg gca tta ccg agt tca gtt tta tat gcc gaa tag agt tct gat tca tcc ttt gag aag atc tct agc gat cgt tca aga tcc ctg att cta gtc ttt agc cta ttt acc tcc tca gaa gat gct ccg tta ccg ttt tta caa tcg tta aga tgt cta tca aga tcc atg att cta tct ctt ttc cat atc agc att gat ttc att att acg ttc gca gtc gtt caa ctg tat t

>ON602722.1_Monkeypox_virus_isolate_MPXV_FRA_2022_TLS67_complete_genome

tta cca cgt ctg tac tcg acg agc tca cgt tta aga gat tca att tcc agt ttg tat cgg tcc atg tct cca ttg cta cac cac cat tag att tac agg ctg cta gtt gtc gtt cga gat cag aaa tac gtg ttt tct tgg aat gga ttt cgt cga tgt act tgt cat gat tgg cat cga aac act tat taa gtt c-t ttt ttc aat tct acg att tta ttt ctt tcg cga gtc aat tcc ctc ctg tag taa cta tca gtt ttg tca gat tca cgc tct cta cgt aga ctt tct tgt aag tta cta att tgt tcc ctg gca tta ccg agt tca gtt tta tat gcc gaa tag agt tct gat tca tcc ttt gag aag atc tct agc gat cgt tca aga tcc ctg att cta gtc ttt agc cta ttt acc tcc tca gaa gat gct ccg tta ccg ttt tta caa tcg tta aga tgt cta tca aga tcc atg att cta tct ctt ttc cat atc agc att gat ttc att att acg ttc gca gtc gtt caa ctg tat t

>ON675438.1_Monkeypox_virus_isolate_MPXV_USA_2022_VA001_complete_genome

tta cca cgt ctg tac tcg acg agc tca cgt tta aga gat tca att tcc agt ttg tat cgg tcc atg tct cca ttg cta cac cac cat tag att tac agg ctg cta gtt gtc gtt cga gat cag aaa tac gtg ttt tct tgg aat gga ttt cgt cga tgt act tgt cat gat tgg cat cga aac act tat taa gtt ctt ttt ttc aat tct acg att tta ttt ctt tcg cga gtc aat tcc ctc ctg tag taa cta tca gtt ttg tca gat tca cgc tct cta cgt aga ctt tct tgt aag tta cta att tgt tcc ctg gca tta ccg agt tca gtt tta tat gcc gaa tag agt tct gat tca tcc ttt gag aag atc tct agc gat cgt tca aga tcc ctg att cta gtc ttt agc cta ttt acc tcc tca gaa gat gct ccg tta ccg ttt tta caa tcg tta aga tgt cta tca aga tcc atg att cta tct ctt ttc cat atc agc att gat ttc att att acg ttc gca gtc gtt caa ctg tat t

>ON631241.1_Monkeypox_virus_isolate_2022/2_SLO_complete_genome

tta cca cgt ctg tac tcg acg agc tca cgt tta aga gat tca att tcc agt ttg tat cgg tcc atg tct cca ttg cta cac cac cat tag att tac agg ctg cta gtt gtc gtt cga gat cag aaa tac gtg ttt tct tgg aat gga ttt cgt cga tgt act tgt cat gat tgg cat cga aac act tat taa gtt ctt ttt ttc aat tct acg att tta ttt ctt tcg cga gtc aat tcc ctc ctg tag taa cta tca gtt ttg tca gat tca cgc tct cta cgt aga ctt tct tgt aag tta cta att tgt tcc ctg gca tta ccg agt tca gtt tta tat gcc gaa tag agt tct gat tca tcc ttt gag aag atc tct agc gat cgt tca aga tcc ctg att cta gtc ttt agc cta ttt acc tcc tca gaa gat gct ccg tta ccg ttt tta caa tcg tta aga tgt cta tca aga tcc atg att cta tct ctt ttc cat atc agc att gat ttc att att acg ttc gca gtc gtt caa ctg tat t

>ON622712.1_Monkeypox_virus_isolate_MPX/UZ_REGA_1/Belgium/2022_complete_genome

tta cca cgt ctg tac tcg acg agc tca cgt tta aga gat tca att tcc agt ttg tat cgg tcc atg tct cca ttg cta cac cac cat tag att tac agg ctg cta gtt gtc gtt cga gat cag aaa tac gtg ttt tct tgg aat gga ttt cgt cga tgt act tgt cat gat tgg cat cga aac act tat taa gtt ctt ttt ttc aat tct acg att tta ttt ctt tcg cga gtc aat tcc ctc ctg tag taa cta tca gtt ttg tca gat tca cgc tct cta cgt aga ctt tct tgt aag tta cta att tgt tcc ctg gca tta ccg agt tca gtt tta tat gcc gaa tag agt tct gat tca tcc ttt gag aag atc tct agc gat cgt tca aga tcc ctg att cta gtc ttt agc cta ttt acc tcc tca gaa gat gct ccg tta ccg ttt tta caa tcg tta aga tgt cta tca aga tcc atg att cta tct ctt ttc cat atc agc att gat ttc att att acg ttc gca gtc gtt caa ctg tat t

>ON609725.2_Monkeypox_virus_isolate_SLO_complete_genome

tta cca cgt ctg tac tcg acg agc tca cgt tta aga gat tca att tcc agt ttg tat cgg tcc atg tct cca ttg cta cac cac cat tag att tac agg ctg cta gtt gtc gtt cga gat cag aaa tac gtg ttt tct tgg aat gga ttt cgt cga tgt act tgt cat gat tgg cat cga aac act tat taa gtt ctt ttt ttc aat tct acg att tta ttt ctt tcg cga gtc aat tcc ctc ctg tag taa cta tca gtt ttg tca gat tca cgc tct cta cgt aga ctt tct tgt aag tta cta att tgt tcc ctg gca tta ccg agt tca gtt tta tat gcc gaa tag agt tct gat tca tcc ttt gag aag atc tct agc gat cgt tca aga tcc ctg att cta gtc ttt agc cta ttt acc tcc tca gaa gat gct ccg tta ccg ttt tta caa tcg tta aga tgt cta tca aga tcc atg att cta tct ctt ttc cat atc agc att gat ttc att att acg ttc gca gtc gtt caa ctg tat t

>ON622713.1_Monkeypox_virus_isolate_MPX/UZ_REGA_2/Belgium/2022_complete_genome

tta cca cgt ctg tac tcg acg agc tca cgt tta aga gat tca att tcc agt ttg tat cgg tcc atg tct cca ttg cta cac cac cat tag att tac agg ctg cta gtt gtc gtt cga gat cag aaa tac gtg ttt tct tgg aat gga ttt cgt cga tgt act tgt cat gat tgg cat cga aac act tat taa gtt ctt ttt ttc aat tct acg att tta ttt ctt tcg cga gtc aat tcc ctc ctg tag taa cta tca gtt ttg tca gat tca cgc tct cta cgt aga ctt tct tgt aag tta cta att tgt tcc ctg gca tta ccg agt tca gtt tta tat gcc gaa tag agt tct gat tca tcc ttt gag aag atc tct agc gat cgt tca aga tcc ctg att cta gtc ttt agc cta ttt acc tcc tca gaa gat gct ccg tta ccg ttt tta caa tcg tta aga tgt cta tca aga tcc atg att cta tct ctt ttc cat atc agc att gat ttc att att acg ttc gca gtc gtt caa ctg tat t

>ON676707.1_Monkeypox_virus_isolate_MPXV_USA_2021_TX_complete_genome

tta cca cgt ctg tac tcg acg agc tca cgt tta aga gat tca att tcc agt ttg tat cgg tcc atg tct cca ttg cta cac cac cat tag att tac agg ctg cta gtt gtc gtt cga gat cag aaa tac gtg ttt tct tgg aat gga ttt cgt cga tgt act tgt cat gat tgg cat cga aac act tat taa gtt ctt ttt ttc aat tct acg att tta ttt ctt tcg cga gtc aat tcc ctc ctg tag taa cta tca gtt ttg tca gat tca cgc tct cta cgt aga ctt tct tgt aag tta cta att tgt tcc ctg gca tta ccg agt tca gtt tta tat gcc gaa tag agt tct gat tca tcc ttt gag aag atc tct agc gat cgt tca aga tcc ctg att cta gtc ttt agc cta ttt acc tcc tca gaa gat gct ccg tta ccg ttt tta caa tcg tta aga tgt cta tca aga tcc atg att cta tct ctt ttc cat atc agc att gat ttc att att acg ttc gca gtc gtt caa ctg tat t

>MT903345.1_Monkeypox_virus_isolate_MPXV-UK_P3

tta cca cgt ctg tac tcg acg agc tca cgt tta aga gat tca att tcc agt ttg tat cgg tcc atg tct cca ttg cta cac cac cat tag att tac agg ctg cta gtt gtc gtt cga gat cag aaa tac gtg ttt tct tgg aat gga ttt cgt cga tgt act tgt cat gat tgg cat cga aac act tat taa gtt ctt ttt ttc aat tct acg att tta ttt ctt tcg cga gtc aat tcc ctc ctg tag taa cta tca gtt ttg tca gat tca cgc tct cta cgt aga ctt tct tgt aag tta cta att tgt tcc ctg gca tta ccg agt tca gtt tta tat gcc gaa tag agt tct gat tca tcc ttt gag aag atc tct agc gat cgt tca aga tcc ctg att cta gtc ttt agc cta ttt acc tcc tca gaa gat gct ccg tta ccg ttt tta caa tcg tta aga tgt cta tca aga tcc atg att cta tct ctt ttc cat atc agc att gat ttc att att acg ttc gca gtc gtt caa ctg tat t

>MT903344.1_Monkeypox_virus_isolate_MPXV-UK_P2

tta cca cgt ctg tac tcg acg agc tca cgt tta aga gat tca att tcc agt ttg tat cgg tcc atg tct cca ttg cta cac cac cat tag att tac agg ctg cta gtt gtc gtt cga gat cag aaa tac gtg ttt tct tgg aat gga ttt cgt cga tgt act tgt cat gat tgg cat cga aac act tat taa gtt ctt ttt ttc aat tct acg att tta ttt ctt tcg cga gtc aat tcc ctc ctg tag taa cta tca gtt ttg tca gat tca cgc tct cta cgt aga ctt tct tgt aag tta cta att tgt tcc ctg gca tta ccg agt tca gtt tta tat gcc gaa tag agt tct gat tca tcc ttt gag aag atc tct agc gat cgt tca aga tcc ctg att cta gtc ttt agc cta ttt acc tcc tca gaa gat gct ccg tta ccg ttt tta caa tcg tta aga tgt cta tca aga tcc atg att cta tct ctt ttc cat atc agc att gat ttc att att acg ttc gca gtc gtt caa ctg tat t

>MT903342.1_Monkeypox_virus_isolate_MPXV-Singapore

tta cca cgt ctg tac tcg acg agc tca cgt tta aga gat tca att tcc agt ttg tat cgg tcc atg tct cca ttg cta cac cac cat tag att tac agg ctg cta gtt gtc gtt cga gat cag aaa tac gtg ttt tct tgg aat gga ttt cgt cga tgt act tgt cat gat tgg cat cga aac act tat taa gtt ctt ttt ttc aat tct acg att tta ttt ctt tcg cga gtc aat tcc ctc ctg tag taa cta tca gtt ttg tca gat tca cgc tct cta cgt aga ctt tct tgt aag tta cta att tgt tcc ctg gca tta ccg agt tca gtt tta tat gcc gaa tag agt tct gat tca tcc ttt gag aag atc tct agc gat cgt tca aga tcc ctg att cta gtc ttt agc cta ttt acc tcc tca gaa gat gct ccg tta ccg ttt tta caa tcg tta aga tgt cta tca aga tcc atg att cta tct ctt ttc cat atc agc att gat ttc att att acg ttc gca gtc gtt caa ctg tat t

>ON676708.1_Monkeypox_virus_isolate_MPXV_USA_2021_MD_complete_genome

tta cca cgt ctg tac tcg acg agc tca cgt tta aga gat tca att tcc agt ttg tat cgg tcc atg tct cca ttg cta cac cac cat tag att tac agg ctg cta gtt gtc gtt cga gat cag aaa tac gtg ttt tct tgg aat gga ttt cgt cga tgt act tgt cat gat tgg cat cga aac act tat taa gtt ctt ttt ttc aat tct acg att tta ttt ctt tcg cga gtc aat tcc ctc ctg tag taa cta tca gtt ttg tca gat tca cgc ttt cta cgt aga ctt tct tgt aag tta cta att tgt tcc ctg gca tta ccg agt tca gtt tta tat gcc gaa tag agt tct gat tca tcc ttt gag aag atc tct agc gat cgt tca aga tcc ctg att cta gtc ttt agc cta ttt acc tcc tca gaa gat gct ccg tta ccg ttt tta caa tcg tta aga tgt cta tca aga tcc atg att cta tct ctt ttc cat atc agc att gat ttc att att acg ttc gca gtc gtt caa ctg tat t

>ON585033.1_Monkeypox_virus_isolate_Monkeypox/PT0006/2022_complete_genome

tta cca cgt ctg tac tcg acg agc tca cgt tta aga gat tca att tcc agt ttg tat cgg tcc atg tct cca ttg cta cac cac cat tag att tac agg ctg cta gtt gtc gtt cga gat cag aaa tac gtg ttt tct tgg aat gga ttt cgt cga tgt act tgt cat gat tgg cat cga aac act tat taa gtt ctt ttt ttc aat tct acg att tta ttt ctt tcg cga gtc aat tcc ctc ctg tag taa cta tca gtt ttg tca gat tca cgc tct cta cgt aga ctt tct tgt aag tta cta att tgt tcc ctg gca tta ccg agt tca gtt tta tat gcc gaa tag agt tct gat tca tcc ttt gag aag atc tct agc gat cgt tca aga tcc ctg att cta gtc ttt agc cta ttt acc tcc tca gaa gat gct ccg tta ccg ttt tta caa tcg tta aga tgt cta tca aga tcc atg att cta tct ctt ttc cat atc agc att gat ttc att att acg ttc gca gtc gtt caa ctg tat t

>ON585035.1_Monkeypox_virus_isolate_Monkeypox/PT0009/2022_complete_genome

tta cca cgt ctg tac tcg acg agc tca cgt tta aga gat tca att tcc agt ttg tat cgg tcc atg tct cca ttg cta cac cac cat tag att tac agg ctg cta gtt gtc gtt cga gat cag aaa tac gtg ttt tct tgg aat gga ttt cgt cga tgt act tgt cat gat tgg cat cga aac act tat taa gtt ctt ttt ttc aat tct acg att tta ttt ctt tcg cga gtc aat tcc ctc ctg tag taa cta tca gtt ttg tca gat tca cgc tct cta cgt aga ctt tct tgt aag tta cta att tgt tcc ctg gca tta ccg agt tca gtt tta tat gcc gaa tag agt tct gat tca tcc ttt gag aag atc tct agc gat cgt tca aga tcc ctg att cta gtc ttt agc cta ttt acc tcc tca gaa gat gct ccg tta ccg ttt tta caa tcg tta aga tgt cta tca aga tcc atg att cta tct ctt ttc cat atc agc att gat ttc att att acg ttc gca gtc gtt caa ctg tat t

>ON649725.1_Monkeypox_virus_isolate_Monkeypox/PT0011/2022_complete_genome

tta cca cgt ctg tac tcg acg agc tca cgt tta aga gat tca att tcc agt ttg tat cgg tcc atg tct cca ttg cta cac cac cat tag att tac agg ctg cta gtt gtc gtt cga gat cag aaa tac gtg ttt tct tgg aat gga ttt cgt cga tgt act tgt cat gat tgg cat cga aac act tat taa gtt ctt ttt ttc aat tct acg att tta ttt ctt tcg cga gtc aat tcc ctc ctg tag taa cta tca gtt ttg tca gat tca cgc tct cta cgt aga ctt tct tgt aag tta cta att tgt tcc ctg gca tta ccg agt tca gtt tta tat gcc gaa tag agt tct gat tca tcc ttt gag aag atc tct agc gat cgt tca aga tcc ctg att cta gtc ttt agc cta ttt acc tcc tca gaa gat gct ccg tta ccg ttt tta caa tcg tta aga tgt cta tca aga tcc atg att cta tct ctt ttc cat atc agc att gat ttc att att acg ttc gca gtc gtt caa ctg tat t

>ON649724.1_Monkeypox_virus_isolate_Monkeypox/PT0014/2022_complete_genome

tta cca cgt ctg tac tcg acg agc tca cgt tta aga gat tca att tcc agt ttg tat cgg tcc atg tct cca ttg cta cac cac cat tag att tac agg ctg cta gtt gtc gtt cga gat cag aaa tac gtg ttt tct tgg aat gga ttt cgt cga tgt act tgt cat gat tgg cat cga aac act tat taa gtt ctt ttt ttc aat tct acg att tta ttt ctt tcg cga gtc aat tcc ctc ctg tag taa cta tca gtt ttg tca gat tca cgc tct cta cgt aga ctt tct tgt aag tta cta att tgt tcc ctg gca tta ccg agt tca gtt tta tat gcc gaa tag agt tct gat tca tcc ttt gag aag atc tct agc gat cgt tca aga tcc ctg att cta gtc ttt agc cta ttt acc tcc tca gaa gat gct ccg tta ccg ttt tta caa tcg tta aga tgt cta tca aga tcc atg att cta tct ctt ttc cat atc agc att gat ttc att att acg ttc gca gtc gtt caa ctg tat t

>ON649722.1_Monkeypox_virus_isolate_Monkeypox/PT0022/2022_complete_genome

tta cca cgt ctg tac tcg acg agc tca cgt tta aga gat tca att tcc agt ttg tat cgg tcc atg tct cca ttg cta cac cac cat tag att tac agg ctg cta gtt gtc gtt cga gat cag aaa tac gtg ttt tct tgg aat gga ttt cgt cga tgt act tgt cat gat tgg cat cga aac act tat taa gtt ctt ttt ttc aat tct acg att tta ttt ctt tcg cga gtc aat tcc ctc ctg tag taa cta tca gtt ttg tca gat tca cgc tct cta cgt aga ctt tct tgt aag tta cta att tgt tcc ctg gca tta ccg agt tca gtt tta tat gcc gaa tag agt tct gat tca tcc ttt gag aag atc tct agc gat cgt tca aga tcc ctg att cta gtc ttt agc cta ttt acc tcc tca gaa gat gct ccg tta ccg ttt tta caa tcg tta aga tgt cta tca aga tcc atg att cta tct ctt ttc cat atc agc att gat ttc att att acg ttc gca gtc gtt caa ctg tat t

>ON649721.1_Monkeypox_virus_isolate_Monkeypox/PT0021/2022_complete_genome

tta cca cgt ctg tac tcg acg agc tca cgt tta aga gat tca att tcc agt ttg tat cgg tcc atg tct cca ttg cta cac cac cat tag att tac agg ctg cta gtt gtc gtt cga gat cag aaa tac gtg ttt tct tgg aat gga ttt cgt cga tgt act tgt cat gat tgg cat cga aac act tat taa gtt ctt ttt ttc aat tct acg att tta ttt ctt tcg cga gtc aat tcc ctc ctg tag taa cta tca gtt ttg tca gat tca cgc tct cta cgt aga ctt tct tgt aag tta cta att tgt tcc ctg gca tta ccg agt tca gtt tta tat gcc gaa tag agt tct gat tca tcc ttt gag aag atc tct agc gat cgt tca aga tcc ctg att cta gtc ttt agc cta ttt acc tcc tca gaa gat gct ccg tta ccg ttt tta caa tcg tta aga tgt cta tca aga tcc atg att cta tct ctt ttc cat atc agc att gat ttc att att acg ttc gca gtc gtt caa ctg tat t

>ON649720.1_Monkeypox_virus_isolate_Monkeypox/PT0024/2022_complete_genome

tta cca cgt ctg tac tcg acg agc tca cgt tta aga gat tca att tcc agt ttg tat cgg tcc atg tct cca ttg cta cac cac cat tag att tac agg ctg cta gtt gtc gtt cga gat cag aaa tac gtg ttt tct tgg aat gga ttt cgt cga tgt act tgt cat gat tgg cat cga aac act tat taa gtt ctt ttt ttc aat tct acg att tta ttt ctt tcg cga gtc aat tcc ctc ctg tag taa cta tca gtt ttg tca gat tca cgc tct cta cgt aga ctt tct tgt aag tta cta att tgt tcc ctg gca tta ccg agt tca gtt tta tat gcc gaa tag agt tct gat tca tcc ttt gag aag atc tct agc gat cgt tca aga tcc ctg att cta gtc ttt agc cta ttt acc tcc tca gaa gat gct ccg tta ccg ttt tta caa tcg tta aga tgt cta tca aga tcc atg att cta tct ctt ttc cat atc agc att gat ttc att att acg ttc gca gtc gtt caa ctg tat t

>ON649723.1_Monkeypox_virus_isolate_Monkeypox/PT0013/2022_complete_genome

tta cca cgt ctg tac tcg acg agc tca cgt tta aga gat tca att tcc agt ttg tat cgg tcc atg tct cca ttg cta cac cac cat tag att tac agg ctg cta gtt gtc gtt cga gat cag aaa tac gtg ttt tct tgg aat gga ttt cgt cga tgt act tgt cat gat tgg cat cga aac act tat taa gtt ctt ttt ttc aat tct acg att tta ttt ctt tcg cga gtc aat tcc ctc ctg tag taa cta tca gtt ttg tca gat tca cgc tct cta cgt aga ctt tct tgt aag tta cta att tgt tcc ctg gca tta ccg agt tca gtt tta tat gcc gaa tag agt tct gat tca tcc ttt gag aag atc tct agc gat cgt tca aga tcc ctg att cta gtc ttt agc cta ttt acc tcc tca gaa gat gct ccg tta ccg ttt tta caa tcg tta aga tgt cta tca aga tcc atg att cta tct ctt ttc cat atc agc att gat ttc att att acg ttc gca gtc gtt caa ctg tat t

>ON649718.1_Monkeypox_virus_isolate_Monkeypox/PT0018/2022_complete_genome

tta cca cgt ctg tac tcg acg agc tca cgt tta aga gat tca att tcc agt ttg tat cgg tcc atg tct cca ttg cta cac cac cat tag att tac agg ctg cta gtt gtc gtt cga gat cag aaa tac gtg ttt tct tgg aat gga ttt cgt cga tgt act tgt cat gat tgg cat cga aac act tat taa gtt ctt ttt ttc aat tct acg att tta ttt ctt tcg cga gtc aat tcc ctc ctg tag taa cta tca gtt ttg tca gat tca cgc tct cta cgt aga ctt tct tgt aag tta cta att tgt tcc ctg gca tta ccg agt tca gtt tta tat gcc gaa tag agt tct gat tca tcc ttt gag aag atc tct agc gat cgt tca aga tcc ctg att cta gtc ttt agc cta ttt acc tcc tca gaa gat gct ccg tta ccg ttt tta caa tcg tta aga tgt cta tca aga tcc atg att cta tct ctt ttc cat atc agc att gat ttc att att acg ttc gca gtc gtt caa ctg tat t

>ON649719.1_Monkeypox_virus_isolate_Monkeypox/PT0012/2022_complete_genome

tta cca cgt ctg tac tcg acg agc tca cgt tta aga gat tca att tcc agt ttg tat cgg tcc atg tct cca ttg cta cac cac cat tag att tac agg ctg cta gtt gtc gtt cga gat cag aaa tac gtg ttt tct tgg aat gga ttt cgt cga tgt act tgt cat gat tgg cat cga aac act tat taa gtt ctt ttt ttc aat tct acg att tta ttt ctt tcg cga gtc aat tcc ctc ctg tag taa cta tca gtt ttg tca gat tca cgc tct cta cgt aga ctt tct tgt aag tta cta att tgt tcc ctg gca tta ccg agt tca gtt tta tat gcc gaa tag agt tct gat tca tcc ttt gag aag atc tct agc gat cgt tca aga tcc ctg att cta gtc ttt agc cta ttt acc tcc tca gaa gat gct ccg tta ccg ttt tta caa tcg tta aga tgt cta tca aga tcc atg att cta tct ctt ttc cat atc agc att gat ttc att att acg ttc gca gtc gtt caa ctg tat t

>ON682267.1_Monkeypox_virus_isolate_MPXV/Germany/2022/RKI010_complete_genome

tta cca cgt ctg tac tcg acg agc tca cgt tta aga gat tca att tcc agt ttg tat cgg tcc atg tct cca ttg cta cac cac cat tag att tac agg ctg cta gtt gtc gtt cga gat cag aaa tac gtg ttt tct tgg aat gga ttt cgt cga tgt act tgt cat gat tgg cat cga aac act tat taa gtt ctt ttt ttc aat tct acg att tta ttt ctt tcg cga gtc aat tcc ctc ctg tag taa cta tca gtt ttg tca gat tca cgc tct cta cgt aga ctt tct tgt aag tta cta att tgt tcc ctg gca tta ccg agt tca gtt tta tat gcc gaa tag agt tct gat tca tcc ttt gag aag atc tct agc gat cgt tca aga tcc ctg att cta gtc ttt agc cta ttt acc tcc tca gaa gat gct ccg tta ccg ttt tta caa tcg tta aga tgt cta tca aga tcc atg att cta tct ctt ttc cat atc agc att gat ttc att att acg ttc gca gtc gtt caa ctg tat t

>ON649717.1_Monkeypox_virus_isolate_Monkeypox/PT0019/2022_partial_genome

tta cca cgt ctg tac tcg acg agc tca cgt tta aga gat tca att tcc agt ttg tat cgg tcc atg tct cca ttg cta cac cac cat tag att tac agg ctg cta gtt gtc gtt cga gat cag aaa tac gtg ttt tct tgg aat gga ttt cgt cga tgt act tgt cat gat tgg cat cga aac act tat taa gtt ctt ttt ttc aat tct acg att tta ttt ctt tcg cga gtc aat tcc ctc ctg tag taa cta tca gtt ttg tca gat tca cgc tct cta cgt aga ctt tct tgt aag tta cta att tgt tcc ctg gca tta ccg agt tca gtt tta tat gcc gaa tag agt tct gat tca tcc ttt gag aag atc tct agc gat cgt tca aga tcc ctg att cta gtc ttt agc cta ttt acc tcc tca gaa gat gct ccg tta ccg ttt tta caa tcg tta aga tgt cta tca aga tcc atg att cta tct ctt ttc cat atc agc att gat ttc att att acg ttc gca gtc gtt caa ctg tat t

>ON649708.1_Monkeypox_virus_isolate_Monkeypox/PT0023/2022_partial_genome

tta cca cgt ctg tac tcg acg agc tca cgt tta aga gat tca att tcc agt ttg tat cgg tcc atg tct cca ttg cta cac cac cat tag att tac agg ctg cta gtt gtc gtt cga gat cag aaa tac gtg ttt tct tgg aat gga ttt cgt cga tgt act tgt cat gat tgg cat cga aac act tat taa gtt ctt ttt ttc aat tct acg att tta ttt ctt tcg cga gtc aat tcc ctc ctg tag taa cta tca gtt ttg tca gat tca cgc tct cta cgt aga ctt tct tgt aag tta cta att tgt tcc ctg gca tta ccg agt tca gtt tta tat gcc gaa tag agt tct gat tca tcc ttt gag aag atc tct agc gat cgt tca aga tcc ctg att cta gtc ttt agc cta ttt acc tcc tca gaa gat gct ccg tta ccg ttt tta caa tcg tta aga tgt cta tca aga tcc atg att cta tct ctt ttc cat atc agc att gat ttc att att acg ttc gca gtc gtt caa ctg tat t

>ON649709.1_Monkeypox_virus_isolate_Monkeypox/PT0028/2022_complete_genome

tta cca cgt ctg tac tcg acg agc tca cgt tta aga gat tca att tcc agt ttg tat cgg tcc atg tct cca ttg cta cac cac cat tag att tac agg ctg cta gtt gtc gtt cga gat cag aaa tac gtg ttt tct tgg aat gga ttt cgt cga tgt act tgt cat gat tgg cat cga aac act tat taa gtt ctt ttt ttc aat tct acg att tta ttt ctt tcg cga gtc aat tcc ctc ctg tag taa cta tca gtt ttg tca gat tca cgc tct cta cgt aga ctt tct tgt aag tta cta att tgt tcc ctg gca tta ccg agt tca gtt tta tat gcc gaa tag agt tct gat tca tcc ttt gag aag atc tct agc gat cgt tca aga tcc ctg att cta gtc ttt agc cta ttt acc tcc tca gaa gat gct ccg tta ccg ttt tta caa tcg tta aga tgt cta tca aga tcc atg att cta tct ctt ttc cat atc agc att gat ttc att att acg ttc gca gtc gtt caa ctg tat t

>ON614676.1_Monkeypox_virus_isolate_INMI-Pt1_partial_genome

tta cca cgt ctg tac tcg acg agc tca cgt tta aga gat tca att tcc agt ttg tat cgg tcc atg tct cca ttg cta cac cac cat tag att tac agg ctg cta gtt gtc gtt cga gat cag aaa tac gtg ttt tct tgg aat gga ttt cgt cga tgt act tgt cat gat tgg cat cga aac act tat taa gtt ctt ttt ttc aat tct acg att tta ttt ctt tcg cga gtc aat tcc ctc ctg tag taa cta tca gtt ttg tca gat tca cgc tct cta cgt aga ctt tct tgt aag tta cta att tgt tcc ctg gca tta ccg agt tca gtt tta tat gcc gaa tag agt tct gat tca tcc ttt gag aag atc tct agc gat cgt tca aga tcc ctg att cta gtc ttt agc cta ttt acc tcc tca gaa gat gct ccg tta ccg ttt tta caa tcg tta aga tgt cta tca aga tcc atg att cta tct ctt ttc cat atc agc att gat ttc att att acg ttc gca gtc gtt caa ctg tat t

>ON649712.1_Monkeypox_virus_isolate_Monkeypox/PT0025/2022_partial_genome

tta cca cgt ctg tac tcg acg agc tca cgt tta aga gat tca att tcc agt ttg tat cgg tcc atg tct cca ttg cta cac cac cat tag att tac agg ctg cta gtt gtc gtt cga gat cag aaa tac gtg ttt tct tgg aat gga ttt cgt cga tgt act tgt cat gat tgg cat cga aac act tat taa gtt ctt ttt ttc aat tct acg att tta ttt ctt tcg cga gtc aat tcc ctc ctg tag taa cta tca gtt ttg tca gat tca cgc tct cta cgt aga ctt tct tgt aag tta cta att tgt tcc ctg gca tta ccg agt tca gtt tta tat gcc gaa tag agt tct gat tca tcc ttt gag aag atc tct agc gat cgt tca aga tcc ctg att cta gtc ttt agc cta ttt acc tcc tca gaa gat gct ccg tta ccg ttt tta caa tcg tta aga tgt cta tca aga tcc atg att cta tct ctt ttc cat atc agc att gat ttc att att acg ttc gca gtc gtt caa ctg tat t

>ON595760.2_Monkeypox_virus_isolate_MPXV-CH-38134631/2022_partial_genome

tta cca cgt ctg tac tcg acg agc tca cgt tta aga gat tca att tcc agt ttg tat cgg tcc atg tct cca ttg cta cac cac cat tag att tac agg ctg cta gtt gtc gtt cga gat cag aaa tac gtg ttt tct tgg aat gga ttt cgt cga tgt act tgt cat gat tgg cat cga aac act tat taa gtt ctt ttt ttc aat tct acg att tta ttt ctt tcg cga gtc aat tcc ctc ctg tag taa cta tca gtt ttg tca gat tca cgc tct cta cgt aga ctt tct tgt aag tta cta att tgt tcc ctg gca tta ccg agt tca gtt tta tat gcc gaa tag agt tct gat tca tcc ttt gag aag atc tct agc gat cgt tca aga tcc ctg att cta gtc ttt agc cta ttt acc tcc tca gaa gat gct ccg tta ccg ttt tta caa tcg tta aga tgt cta tca aga tcc atg att cta tct ctt ttc cat atc agc att gat ttc att att acg ttc gca gtc gtt caa ctg tat t

>MT903337.1_Monkeypox_virus_isolate_MPXV-M2940_FCT

tta cca cgt ctg tac tcg acg agc tca cgt tta aga gat tca att tcc agt ttg tat cgg tcc atg tct cca ttg cta cac cac cat tag att tac agg ctg cta gtt gtc gtt cga gat cag aaa tac gtg ttt tct tgg aat gga ttt cgt cga tgt act tgt cat gat tgg cat cga aac act tat taa gtt ctt ttt ttc aat tct acg att tta ttt ctt tcg cga gtc aat tcc ctc ctg tag taa cta tca gtt ttg tca gat tca cgc tct cta cgt aga ctt tct tgt aag tta cta att tgt tcc ctg gca tta ccg agt tca gtt tta tat gcc gaa tag agt tct gat tca tcc ttt gag aag atc tct agc gat cgt tca aga tcc ctg att cta gtc ttt agc cta ttt acc tcc tca gaa gat gct ccg tta ccg ttt tta caa tcg tta aga tgt cta tca aga tcc atg att cta tct ctt ttc cat atc agc att gat ttc att att acg ttc gca gtc gtt caa ctg tat t

>MT903339.1_Monkeypox_virus_isolate_MPXV-M3021_Delta

tta cca cgt ctg tac tcg acg agc tca cgt tta aga gat tca att tcc agt ttg tat cgg tcc atg tct cca ttg cta cac cac cat tag att tac agg ctg cta gtt gtc gtt cga gat cag aaa tac gtg ttt tct tgg aat gga ttt cgt cga tgt act tgt cat gat tgg cat cga aac act tat taa gtt ctt ttt ttc aat tct acg att tta ttt ctt tcg cga gtc aat tcc ctc ctg tag taa cta tca gtt ttg tca gat tca cgc tct cta cgt aga ctt tct tgt aag tta cta att tgt tcc ctg gca tta ccg agt tca gtt tta tat gcc gaa tag agt tct gat tca tcc ttt gag aag atc tct agc gat cgt tca aga tcc ctg att cta gtc ttt agc cta ttt acc tcc tca gaa gat gct ccg tta ccg ttt tta caa tcg tta aga tgt cta tca aga tcc atg att cta tct ctt ttc cat atc agc att gat ttc att att acg ttc gca gtc gtt caa ctg tat t

>MT903338.1_Monkeypox_virus_isolate_MPXV-M2957_Lagos

tta cca cgt ctg tac tcg acg agc tca cgt tta aga gat tca att tcc agt ttg tat cgg tcc atg tct cca ttg cta cac cac cat tag att tac agg ctg cta gtt gtc gtt cga gat cag aaa tac gtg ttt tct tgg aat gga ttt cgt cga tgt act tgt cat gat tgg cat cga aac act tat taa gtt ctt ttt ttc aat tct acg att tta ttt ctt tcg cga gtc aat tcc ctc ctg tag taa cta tca gtt ttg tca gat tca cgc tct cta cgt aga ctt tct tgt aag tta cta att tgt tcc ctg gca tta ccg agt tca gtt tta tat gcc gaa tag agt tct gat tca tcc ttt gag aag atc tct agc gat cgt tca aga tcc ctg att cta gtc ttt agc cta ttt acc tcc tca gaa gat gct ccg tta ccg ttt tta caa tcg tta aga tgt cta tca aga tcc atg att cta tct ctt ttc cat atc agc att gat ttc att att acg ttc gca gtc gtt caa ctg tat t

>KJ642617.1_Monkeypox_virus_strain_Nigeria-SE-1971_complete_genome

tta cca cgt ctg tac tcg acg agc tca cgt tta aga gat tca att tcc agt ttg tat cgg tcc atg tct cca ttg cta cac cac cat tag att tac agg ctg cta gtt gtc gtt cga gat cag aaa tac gtg ttt tct tgg aat gga ttt cgt cga tgt act tgt cat gat tgg cat cga aac act tat taa gtt ctt ttt ttc aat tct acg att tta ttt ctt tcg cga gtc aat tcc ctc ctg tag taa cta tca gtt ttg tca gat tca cgc tct cta cgt aga ctt tct tgt aag tta cta att tgt tcc ctg gca tta ccg agt tca gtt tta tat gcc gaa tag agt tct gat tca tcc ttt gag aag atc tct agc gat cgt tca aga tcc ctg att cta gtc ttt agc cta ttt acc tcc tca gaa gat gct ccg tta ccg ttt tta caa tcg tta aga tgt cta tca aga tcc atg att cta tct ctt ttc cat atc agc att gat ttc att att acg ttc gca gtc gtt caa ctg tat t

>ON615424.1_Monkeypox_virus_isolate_MPXV_2022_NL001_partial_genome

tta cca cgt ctg tac tcg acg agc tca cgt tta aga gat tca att tcc agt ttg tat cgg tcc atg tct cca ttg cta cac cac cat tag att tac agg ctg cta gtt gtc gtt cga gat cag aaa tac gtg ttt tct tgg aat gga ttt cgt cga tgt act tgt cat gat tgg cat cga aac act tat taa gtt ctt ttt ttc aat tct acg att tta ttt ctt tcg cga gtc aat tcc ctc ctg tag taa cta tca gtt ttg tca gat tca cgc tct cta cgt aga ctt tct tgt aag tta cta att tgt tcc ctg gca tta ccg agt tca gtt tta tat gcc gaa tag agt tct gat tca tcc ttt gag aag atc tct agc gat cgt tca aga tcc ctg att cta gtc ttt agc cta ttt acc tcc tca gaa gat gct ccg tta ccg ttt tta caa tcg tta aga tgt cta tca aga tcc atg att cta tct ctt ttc cat atc agc att gat ttc att att acg ttc gca gtc gtt caa ctg tat t

>ON585034.1_Monkeypox_virus_isolate_Monkeypox/PT0007/2022_complete_genome

tta cca cgt ctg tac tcg acg agc tca cgt tta aga gat tca att tcc agt ttg tat cgg tcc atg tct cca ttg cta cac cac cat tag att tac agg ctg cta gtt gtc gtt cga gat cag aaa tac gtg ttt tct tgg aat gga ttt cgt cga tgt act tgt cat gat tgg cat cga aac act tat taa gtt ctt ttt ttc aat tct acg att tta ttt ctt tcg cga gtc aat tcc ctc ctg tag taa cta tca gtt ttg tca gat tca cgc tct cta cgt aga ctt tct tgt aag tta cta att tgt tcc ctg gca tta ccg agt tca gtt tta tat gcc gaa tag agt tct gat tca tcc ttt gag aag atc tct agc gat cgt tca aga tcc ctg att cta gtc ttt agc cta ttt acc tcc tca gaa gat gct ccg tta ccg ttt tta caa tcg tta aga tgt cta tca aga tcc atg att cta tct ctt ttc cat atc agc att gat ttc att att acg ttc gca gtc gtt caa ctg tat t

>ON649713.1_Monkeypox_virus_isolate_Monkeypox/PT0020/2022_complete_genome

tta cca cgt ctg tac tcg acg agc tca cgt tta aga gat tca att tcc agt ttg tat cgg tcc atg tct cca ttg cta cac cac cat tag att tac agg ctg cta gtt gtc gtt cga gat cag aaa tac gtg ttt tct tgg aat gga ttt cgt cga tgt act tgt cat gat tgg cat cga aac act tat taa gtt ctt ttt ttc aat tct acg att tta ttt ctt tcg cga gtc aat tcc ctc ctg tag taa cta tca gtt ttg tca gat tca cgc tct cta cgt aga ctt tct tgt aag tta cta att tgt tcc ctg gca tta ccg agt tca gtt tta tat gcc gaa tag agt tct gat tca tcc ttt gag aag atc tct agc gat cgt tca aga tcc ctg att cta gtc ttt agc cta ttt acc tcc tca gaa gat gct ccg tta ccg ttt tta caa tcg tta aga tgt cta tca aga tcc atg att cta tct ctt ttc cat atc agc att gat ttc att att acg ttc gca gtc gtt caa ctg tat t

>MG693723.1_Monkeypox_virus_isolate_MPXV_Nig_2017_297957_partial_genome

tta cca cgt ctg tac tcg acg agc tca cgt tta aga gat tca att tcc agt ttg tat cgg tcc atg tct cca ttg cta cac cac cat tag att tac agg ctg cta gtt gtc gtt cga gat cag aaa tac gtg ttt tct tgg aat gga ttt cgt cga tgt act tgt cat gat tgg cat cga aac act tat taa gtt ctt ttt ttc aat tct acg att tta ttt ctt tcg cga gtc aat tcc ctc ctg tag taa cta tca gtt ttg tca gat tca cgc tct cta cgt aga ctt tct tgt aag tta cta att tgt tcc ctg gca tta ccg agt tca gtt tta tat gcc gaa tag agt tct gat tca tcc ttt gag aag atc tct agc gat cgt tca aga tcc ctg att cta gtc ttt agc cta ttt acc tcc tca gaa gat gct ccg tta ccg ttt tta caa tcg tta aga tgt cta tca aga tcc atg att cta tct ctt ttc cat atc agc att gat ttc att att acg ttc gca gtc gtt caa ctg tat t

>ON585031.1_Monkeypox_virus_isolate_Monkeypox/PT0003/2022_complete_genome

tta cca cgt ctg tac tcg acg agc tca cgt tta aga gat tca att tcc agt ttg tat cgg tcc atg tct cca ttg cta cac cac cat tag att tac agg ctg cta gtt gtc gtt cga gat cag aaa tac gtg ttt tct tgg aat gga ttt cgt cga tgt act tgt cat gat tgg cat cga aac act tat taa gtt ctt ttt ttc aat tct acg att tta ttt ctt tcg cga gtc aat tcc ctc ctg tag taa cta tca gtt ttg tca gat tca cgc tct cta cgt aga ctt tct tgt aag tta cta att tgt tcc ctg gca tta ccg agt tca gtt tta tat gcc gaa tag agt tct gat tca tcc ttt gag aag atc tct agc gat cgt tca aga tcc ctg att cta gtc ttt agc cta ttt acc tcc tca gaa gat gct ccg tta ccg ttt tta caa tcg tta aga tgt cta tca aga tcc atg att cta tct ctt ttc cat atc agc att gat ttc att att acg ttc gca gtc gtt caa ctg tat t

>ON585038.1_Monkeypox_virus_isolate_Monkeypox/PT0008/2022_complete_genome

tta cca cgt ctg tac tcg acg agc tca cgt tta aga gat tca att tcc agt ttg tat cgg tcc atg tct cca ttg cta cac cac cat tag att tac agg ctg cta gtt gtc gtt cga gat cag aaa tac gtg ttt tct tgg aat gga ttt cgt cga tgt act tgt cat gat tgg cat cga aac act tat taa gtt ctt ttt ttc aat tct acg att tta ttt ctt tcg cga gtc aat tcc ctc ctg tag taa cta tca gtt ttg tca gat tca cgc tct cta cgt aga ctt tct tgt aag tta cta att tgt tcc ctg gca tta ccg agt tca gtt tta tat gcc gaa tag agt tct gat tca tcc ttt gag aag atc tct agc gat cgt tca aga tcc ctg att cta gtc ttt agc cta ttt acc tcc tca gaa gat gct ccg tta ccg ttt tta caa tcg tta aga tgt cta tca aga tcc atg att cta tct ctt ttc cat atc agc att gat ttc att att acg ttc gca gtc gtt caa ctg tat t

>DQ011156.1_Monkeypox_virus_strain_Liberia_1970_184_complete_genome

tta cca cgt ctg tac tcg acg agc tca cgt tta aga gat tca att tcc agt ttg tat cgg tcc atg tct cca ttg cta cac cat cat tag att tac agg ctg tta gtt gtc gtt cga gat cag aaa tac gtg ttt tct tgg aat gga ttt cgt cga tgt act tgt cat gat tgg cat cga aac act tat taa gtt ctt ttt ttc aat tct acg att tta ttt ctt tcg cga gtc aat tcc ctc cta tag taa cta tca gtt ttg tca gat tca cgc tct cta cgt aga ctt tct tgc aag tta cta att tgt tcc ctg gca tta ccg agt tca gtt tta tat gcc gaa tag agt tct gat tca tcc ttt gag aag atc tct agc gat cgt tca aga tcc ctg att cta gtc ttt agc cta ttt acc tcc tca gaa gat gct ccg tta ccg ttt tta caa tcg tta aga tgt cta tca aga tcc atg att cta tct ctt ttc cat atc agc att gat ttc att att acg tcc gca gtc gtt caa ctg tat t

>KP849470.1_Monkeypox_virus_isolate_Cote_dIvoire_1971_complete_genome

tta cca cgt ctg tac tcg acg agc tca cgt tta aga gat tca att tcc agt ttg tat cgg tcc atg tct cca ttg cta cac cat cat tag att tac agg ctg tta gtt gtc gtt cga gat cag aaa tac gtg ttt tct tgg aat gga ttt cgt cga tgt act tgt cat gat tgg cat cga aac act tat taa gtt ctt ttt ttc aat tct acg att tta ttt ctt tcg cga gtc aat tcc ctc cta tag taa cta tca gtt ttg tca gat tca cgc tct cta cgt aga ctt tct tgc aag tta cta att tgt tcc ctg gca tta ccg agt tca gtt tta tat gcc gaa tag agt tct gat tca tcc ttt gag aag atc tct agc gat cgt tca aga tcc ctg att cta gtc ttt agc cta ttt acc tcc tca gaa gat gct ccg tta ccg ttt tta caa tcg tta aga tgt cta tca aga tcc atg att cta tct ctt ttc cat atc agc att gat ttc att att acg tcc gca gtc gtt caa ctg tat t

>ON682264.2_Monkeypox_virus_isolate_MPXV/Germany/2022/RKI05_complete_genome

tta cca cgt ctg tac tcg acg agc tca cgt tta aga gat tca att tcc agt ttg tat cgg tcc atg tct cca ttg cta cac cac cat tag att tac agg ctg cta gtt gtc gtt cga gat cag aaa tac gtg ttt tct tgg aat gga ttt cgt cga tgt act tgt cat gat tgg cat cga aac act tat taa gtt ctt ttt ttc aat tct acg att tta ttt ctt tcg cga gtc aat tcc ctc ctg tag taa cta tca gtt ttg tca gat tca cgc tct cta cgt aga ctt tct tgt aag tta cta att tgt tcc ctg gca tta ccg agt tca gtt tta tat gcc gaa tag agt tct gat tca tcc ttt gag aag atc tct agc gat cgt tca aga tcc ctg att cta gtc ttt agc cta ttt acc tcc tca gaa gat gct ccg tta ccg ttt tta caa tcg tta aga tgt cta tca aga tcc atg att cta tct ctt ttc cat atc agc att gat ttc att att acg ttc gca gtc gtt caa ctg tat t

>ON682268.1_Monkeypox_virus_isolate_MPXV/Germany/2022/RKI08_complete_genome

tta cca cgt ctg tac tcg acg agc tca cgt tta aga gat tca att tcc agt ttg tat cgg tcc atg tct cca ttg cta cac cac cat tag att tac agg ctg cta gtt gtc gtt cga gat cag aaa tac gtg ttt tct tgg aat gga ttt cgt cga tgt act tgt cat gat tgg cat cga aac act tat taa gtt ctt ttt ttc aat tct acg att tta ttt ctt tcg cga gtc aat tcc ctc ctg tag taa cta tca gtt ttg tca gat tca cgc tct cta cgt aga ctt tct tgt aag tta cta att tgt tcc ctg gca tta ccg agt tca gtt tta tat gcc gaa tag agt tct gat tca tcc ttt gag aag atc tct agc gat cgt tca aga tcc ctg att cta gtc ttt agc cta ttt acc tcc tca gaa gat gct ccg tta ccg ttt tta caa tcg tta aga tgt cta tca aga tcc atg att cta tct ctt ttc cat atc agc att gat ttc att att acg ttc gca gtc gtt caa ctg tat t

>ON682270.1_Monkeypox_virus_isolate_MPXV/Germany/2022/RKI06_complete_genome

tta cca cgt ctg tac tcg acg agc tca cgt tta aga gat tca att tcc agt ttg tat cgg tcc atg tct cca ttg cta cac cac cat tag att tac agg ctg cta gtt gtc gtt cga gat cag aaa tac gtg ttt tct tgg aat gga ttt cgt cga tgt act tgt cat gat tgg cat cga aac act tat taa gtt ctt ttt ttc aat tct acg att tta ttt ctt tcg cga gtc aat tcc ctc ctg tag taa cta tca gtt ttg tca gat tca cgc tct cta cgt aga ctt tct tgt aag tta cta att tgt tcc ctg gca tta ccg agt tca gtt tta tat gcc gaa tag agt tct gat tca tcc ttt gag aag atc tct agc gat cgt tca aga tcc ctg att cta gtc ttt agc cta ttt acc tcc tca gaa gat gct ccg tta ccg ttt tta caa tcg tta aga tgt cta tca aga tcc atg att cta tct ctt ttc cat atc agc att gat ttc att att acg ttc gca gtc gtt caa ctg tat t

>ON637938.1_Monkeypox_virus_isolate_MPXV/Germany/2022/RKI01_complete_genome

tta cca cgt ctg tac tcg acg agc tca cgt tta aga gat tca att tcc agt ttg tat cgg tcc atg tct cca ttg cta cac cac cat tag att tac agg ctg cta gtt gtc gtt cga gat cag aaa tac gtg ttt tct tgg aat gga ttt cgt cga tgt act tgt cat gat tgg cat cga aac act tat taa gtt ctt ttt ttc aat tct acg att tta ttt ctt tcg cga gtc aat tcc ctc ctg tag taa cta tca gtt ttg tca gat tca cgc tct cta cgt aga ctt tct tgt aag tta cta att tgt tcc ctg gca tta ccg agt tca gtt tta tat gcc gaa tag agt tct gat tca tcc ttt gag aag atc tct agc gat cgt tca aga tcc ctg att cta gtc ttt agc cta ttt acc tcc tca gaa gat gct ccg tta ccg ttt tta caa tcg tta aga tgt cta tca aga tcc atg att cta tct ctt ttc cat atc agc att gat ttc att att acg ttc gca gtc gtt caa ctg tat t

>ON637939.1_Monkeypox_virus_isolate_MPXV/Germany/2022/RKI02_complete_genome

tta cca cgt ctg tac tcg acg agc tca cgt tta aga gat tca att tcc agt ttg tat cgg tcc atg tct cca ttg cta cac cac cat tag att tac agg ctg cta gtt gtc gtt cga gat cag aaa tac gtg ttt tct tgg aat gga ttt cgt cga tgt act tgt cat gat tgg cat cga aac act tat taa gtt ctt ttt ttc aat tct acg att tta ttt ctt tcg cga gtc aat tcc ctc ctg tag taa cta tca gtt ttg tca gat tca cgc tct cta cgt aga ctt tct tgt aag tta cta att tgt tcc ctg gca tta ccg agt tca gtt tta tat gcc gaa tag agt tct gat tca tcc ttt gag aag atc tct agc gat cgt tca aga tcc ctg att cta gtc ttt agc cta ttt acc tcc tca gaa gat gct ccg tta ccg ttt tta caa tcg tta aga tgt cta tca aga tcc atg att cta tct ctt ttc cat atc agc att gat ttc att att acg ttc gca gtc gtt caa ctg tat t

>ON682263.2_Monkeypox_virus_isolate_MPXV/Germany/2022/RKI03_complete_genome

tta cca cgt ctg tac tcg acg agc tca cgt tta aga gat tca att tcc agt ttg tat cgg tcc atg tct cca ttg cta cac cac cat tag att tac agg ctg cta gtt gtc gtt cga gat cag aaa tac gtg ttt tct tgg aat gga ttt cgt cga tgt act tgt cat gat tgg cat cga aac act tat taa gtt ctt ttt ttc aat tct acg att tta ttt ctt tcg cga gtc aat tcc ctc ctg tag taa cta tca gtt ttg tca gat tca cgc tct cta cgt aga ctt tct tgt aag tta cta att tgt tcc ctg gca tta ccg agt tca gtt tta tat gcc gaa tag agt tct gat tca tcc ttt gag aag atc tct agc gat cgt tca aga tcc ctg att cta gtc ttt agc cta ttt acc tcc tca gaa gat gct ccg tta ccg ttt tta caa tcg tta aga tgt cta tca aga tcc atg att cta tct ctt ttc cat atc agc att gat ttc att att acg ttc gca gtc gtt caa ctg tat t

>ON682266.1_Monkeypox_virus_isolate_MPXV/Germany/2022/RKI09_complete_genome

tta cca cgt ctg tac tcg acg agc tca cgt tta aga gat tca att tcc agt ttg tat cgg tcc atg tct cca ttg cta cac cac cat tag att tac agg ctg cta gtt gtc gtt cga gat cag aaa tac gtg ttt tct tgg aat gga ttt cgt cga tgt act tgt cat gat tgg cat cga aac act tat taa gtt ctt ttt ttc aat tct acg att tta ttt ctt tcg cga gtc aat tcc ctc ctg tag taa cta tca gtt ttg tca gat tca cgc tct cta cgt aga ctt tct tgt aag tta cta att tgt tcc ctg gca tta ccg agt tca gtt tta tat gcc gaa tag agt tct gat tca tcc ttt gag aag atc tct agc gat cgt tca aga tcc ctg att cta gtc ttt agc cta ttt acc tcc tca gaa gat gct ccg tta ccg ttt tta caa tcg tta aga tgt cta tca aga tcc atg att cta tct ctt ttc cat atc agc att gat ttc att att acg ttc gca gtc gtt caa ctg tat t

>ON619836.1_Monkeypox_virus_isolate_MPXV_UK_2022_2_complete_genome

tta cca cgt ctg tac tcg acg agc tca cgt tta aga gat tca att tcc agt ttg tat cgg tcc atg tct cca ttg cta cac cac cat tag att tac agg ctg cta gtt gtc gtt cga gat cag aaa tac gtg ttt tct tgg aat gga ttt cgt cga tgt act tgt cat gat tgg cat cga aac act tat taa gtt ctt ttt ttc aat tct acg att tta ttt ctt tcg cga gtc aat tcc ctc ctg tag taa cta tca gtt ttg tca gat tca cgc tct cta cgt aga ctt tct tgt aag tta cta att tgt tcc ctg gca tta ccg agt tca gtt tta tat gcc gaa tag agt tct gat tca tcc ttt gag aag atc tct agc gat cgt tca aga tcc ctg att cta gtc ttt agc cta ttt acc tcc tca gaa gat gct ccg tta ccg ttt tta caa tcg tta aga tgt cta tca aga tcc atg att cta tct ctt ttc cat atc agc att gat ttc att att acg ttc gca gtc gtt caa ctg tat t

>ON619838.1_Monkeypox_virus_isolate_MPXV_UK_2022_4_complete_genome

tta cca cgt ctg tac tcg acg agc tca cgt tta aga gat tca att tcc agt ttg tat cgg tcc atg tct cca ttg cta cac cac cat tag att tac agg ctg cta gtt gtc gtt cga gat cag aaa tac gtg ttt tct tgg aat gga ttt cgt cga tgt act tgt cat gat tgg cat cga aac act tat taa gtt ctt ttt ttc aat tct acg att tta ttt ctt tcg cga gtc aat tcc ctc ctg tag taa cta tca gtt ttg tca gat tca cgc tct cta cgt aga ctt tct tgt aag tta cta att tgt tcc ctg gca tta ccg agt tca gtt tta tat gcc gaa tag agt tct gat tca tcc ttt gag aag atc tct agc gat cgt tca aga tcc ctg att cta gtc ttt agc cta ttt acc tcc tca gaa gat gct ccg tta ccg ttt tta caa tcg tta aga tgt cta tca aga tcc atg att cta tct ctt ttc cat atc agc att gat ttc att att acg ttc gca gtc gtt caa ctg tat t

>ON619835.1_Monkeypox_virus_isolate_MPXV_UK_2022_1_complete_genome

tta cca cgt ctg tac tcg acg agc tca cgt tta aga gat tca att tcc agt ttg tat cgg tcc atg tct cca ttg cta cac cac cat tag att tac agg ctg cta gtt gtc gtt cga gat cag aaa tac gtg ttt tct tgg aat gga ttt cgt cga tgt act tgt cat gat tgg cat cga aac act tat taa gtt ctt ttt ttc aat tct acg att tta ttt ctt tcg cga gtc aat tcc ctc ctg tag taa cta tca gtt ttg tca gat tca cgc tct cta cgt aga ctt tct tgt aag tta cta att tgt tcc ctg gca tta ccg agt tca gtt tta tat gcc gaa tag agt tct gat tca tcc ttt gag aag atc tct agc gat cgt tca aga tcc ctg att cta gtc ttt agc cta ttt acc tcc tca gaa gat gct ccg tta ccg ttt tta caa tcg tta aga tgt cta tca aga tcc atg att cta tct ctt ttc cat atc agc att gat ttc att att acg ttc gca gtc gtt caa ctg tat t

>ON682269.2_Monkeypox_virus_isolate_MPXV/Germany/2022/RKI07_complete_genome

tta cca cgt ctg tac tcg acg agc tca cgt tta aga gat tca att tcc agt ttg tat cgg tcc atg tct cca ttg cta cac cac cat tag att tac agg ctg cta gtt gtc gtt cga gat cag aaa tac gtg ttt tct tgg aat gga ttt cgt cga tgt act tgt cat gat tgg cat cga aac act tat taa gtt ctt ttt ttc aat tct acg att tta ttt ctt tcg cga gtc aat tcc ctc ctg tag taa cta tca gtt ttg tca gat tca cgc tct cta cgt aga ctt tct tgt aag tta cta att tgt tcc ctg gca tta ccg agt tca gtt tta tat gcc gaa tag agt tct gat tca tcc ttt gag aag atc tct agc gat cgt tca aga tcc ctg att cta gtc ttt agc cta ttt acc tcc tca gaa gat gct ccg tta ccg ttt tta caa tcg tta aga tgt cta tca aga tcc atg att cta tct ctt ttc cat atc agc att gat ttc att att acg ttc gca gtc gtt caa ctg tat t

>KJ642616.1_Monkeypox_virus_strain_PCH_complete_genome

tta cca cgt ctg tac tca acg agc tca cgt tta aga gat tca att tcc agt ttg tat cgg tcc atg tct cca ttg cta cac cac cat tag att tac agg ctg tta gtt gtc gtt cga gat cag aaa tac gtg ttt tct tgg aat gga ttt cgt cga tgt act tgt cat gat tgg cat cga aac act tat taa gtt ctt ttt ttc aat tct acg att tta ttt ctt tcg cga gtc aat tcc ctc cta tag taa cta tca gtt ttg tca gat tca cgc tct cta cgt aga ctt tct tgc aag tta cta att tgt tcc ctg gca tta ccg agt tca gtt tta tat gcc gaa tag agt tct gat tca tcc ttt gag aag atc tct agc gat cgt tca aga tcc ctg att cta gtc ttt agc cta ttt acc tcc tca gaa gat gct ccg tta ccg ttt tta caa tcg tta aga tgt cta tca aga tcc atg att cta tct ctt ttc cat atc agc att gat ttc att att acg tcc gca gtc gtt caa ctg tat t

>AY753185.1_Monkeypox_virus_strain_COP-58_complete_genome

tta cca cgt ctg tac tcg acg agc tca cgt tta aga gat tca att tcc agt ttg tat cgg tcc atg tct cca ttg cta cac ca- cat tag att tac agg ctg tta gtt gtc gtt cga gat cag aaa tac gtg ttt tct tgg aat gga ttt cgt cga tgt act tgt cat gat tgg cat cga aac act tat taa gtt ctt ttt ttc aat tct acg att tta ttt ctt tcg cga gtc aat tcc ctc cta tag taa cta tca gtt ttg tca gat tca cgc tct cta cgt aga ctt tct tgc aag tta cta att tgt tcc ctg gca tta ccg agt tca gtt tta tat gcc gaa tag agt tct gat tca tcc ttt gag aag atc tct agc gat cgt tca aga tcc ctg att cta gtc ttt agc cta ttt acc tcc tca gaa gat gct ccg tta ccg ttt tta caa tcg tta aga tgt cta tca aga tcc atg att cta tct ctt ttc cat atc agc att gat ttc att att acg tcc gca gtc gtt caa ctg tat t

>AY603973.1_Monkeypox_virus_strain_MPXV-WRAIR7-61_complete_genome

tta cca cgt ctg tac tcg acg agc tca cgt tta aga gat tca att tcc agt ttg tat cgg tcc atg tct cca ttg cta cac ca- cat tag att tac agg ctg tta gtt gtc gtt cga gat cag aaa tac gtg ttt tct tgg aat gga ttt cgt cga tgt act tgt cat gat tgg cat cga aac act tat taa gtt ctt ttt ttc aat tct acg att tta ttt ctt tcg cga gtc aat tcc ctc cta tag taa cta tca gtt ttg tca gat tca cgc tct cta cgt aga ctt tct tgc aag tta cta att tgt tcc ctg gca tta ccg agt tca gtt tta tat gcc gaa tag agt tct gat tca tcc ttt gag aag atc tct agc gat cgt tca aga tcc ctg att cta gtc ttt agc cta ttt acc tcc tca gaa gat gct ccg tta ccg ttt tta caa tcg tta aga tgt cta tca aga tcc atg att cta tct ctt ttc cat atc agc att gat ttc att att acg tcc gca gtc gtt caa ctg tat t

>ON619837.1_Monkeypox_virus_isolate_MPXV_UK_2022_3_complete_genome

tta cca cgt ctg tac tcg acg agc tca cgt tta aga gat tca att tcc agt ttg tat cgg tcc atg tct cca ttg cta cac cac cat tag att tac agg ctg cta gtt gtc gtt cga gat cag aaa tac gtg ttt tct tgg aat gga ttt cgt cga tgt act tgt cat gat tgg cat cga aac act tat taa gtt c?t ttt ttc aat tct acg att tta ttt ctt tcg cga gtc aat tcc ctc ctg tag taa cta tca gtt ttg tca gat tca cgc tct cta cgt aga ctt tct tgt aag tta cta att tgt tcc ctg gca tta ccg agt tca gtt tta tat gcc gaa tag agt tct gat tca tcc ttt gag aag atc tct agc gat cgt tca aga tcc ctg att cta gtc ttt agc cta ttt acc tcc tca gaa gat gct ccg tta ccg ttt tta caa tcg tta aga tgt cta tca aga tcc atg att cta tct ctt ttc cat atc agc att gat ttc att att acg ttc gca gtc gtt caa ctg tat t

>ON585037.1_Monkeypox_virus_isolate_Monkeypox/PT0005/2022_complete_genome

tta cca cgt ctg tac tcg acg agc tca cgt tta aga gat tca att tcc agt ttg tat cgg tcc atg tct cca ttg cta cac cac cat tag att tac agg ctg cta gtt gtc gtt cga gat cag aaa tac gtg ttt tct tgg aat gga ttt cgt cga tgt act tgt cat gat tgg cat cga aac act tat taa gtt ctt ttt ttc aat tct acg att tta ttt ctt tcg cga gtc aat tcc ctc ctg tag taa cta tca gtt ttg tca gat tca cgc tct cta cgt aga ctt tct tgt aag tta cta att tgt tcc ctg gca tta ccg agt tca gtt tta tat gcc gaa tag agt tct gat tca tcc ttt gag aag atc tct agc gat cgt tca aga tcc ctg att cta gtc ttt agc cta ttt acc tcc tca gaa gat gct ccg tta ccg ttt tta caa tcg tta aga tgt cta tca aga tcc atg att cta tct ctt ttc cat atc agc att gat ttc att att acg ttc gca gtc gtt caa ctg tat t

>ON585032.1_Monkeypox_virus_isolate_Monkeypox/PT0004/2022_complete_genome

tta cca cgt ctg tac tcg acg agc tca cgt tta aga gat tca att tcc agt ttg tat cgg tcc atg tct cca ttg cta cac cac cat tag att tac agg ctg cta gtt gtc gtt cga gat cag aaa tac gtg ttt tct tgg aat gga ttt cgt cga tgt act tgt cat gat tgg cat cga aac act tat taa gtt ctt ttt ttc aat tct acg att tta ttt ctt tcg cga gtc aat tcc ctc ctg tag taa cta tca gtt ttg tca gat tca cgc tct cta cgt aga ctt tct tgt aag tta cta att tgt tcc ctg gca tta ccg agt tca gtt tta tat gcc gaa tag agt tct gat tca tcc ttt gag aag atc tct agc gat cgt tca aga tcc ctg att cta gtc ttt agc cta ttt acc tcc tca gaa gat gct ccg tta ccg ttt tta caa tcg tta aga tgt cta tca aga tcc atg att cta tct ctt ttc cat atc agc att gat ttc att att acg ttc gca gtc gtt caa ctg tat t

>AY741551.1_Monkeypox_virus_isolate_Sierra_Leone_complete_genome

tta cca cgt ctg tac tcg acg agc tca cgt tta aga gat tca att tcc agt ttg tat cgg tcc atg tct cca ttg cta cac cat cat tag att tac agg ctg tta gtt atc gtt cga gat cag aaa tac gtg ttt tct tgg aat gga ttt cgt cga tgt act tgt cat gat tgg cat cga aac act tat taa gtt ctt ttt ttc aat tct acg att tta ttt ctt tcg cga gtc aat tcc ctc cta tag taa cta tca gtt ttg tca gat tca cgc tct cta cgt aga ctt tct tgc aag tta cta att tgt tcc ctg gca tta ccg agt tca gtt tta tat gcc gaa tag agt tct gat tca tcc ttt gag aag atc tct agc gat cgt tca aga tcc ctg att cta gtc ttt agc cta ttt acc tcc tca gaa gat gct ccg tta ccg ttt tta caa tcg tta aga tgt cta tca aga tcc atg att cta tct ctt ttc cat atc agc att gat ttc att att acg tcc gca gtc gtt caa ctg tat t

>ON622718.1_Monkeypox_virus_isolate_MPXV/ES0001/HUGTiP/2022_partial_genome

tta cca cgt ctg tac tcg acg agc tca cgt tta aga gat tca att tcc agt ttg tat cgg tcc atg tct cca ttg cta cac cac cat tag att tac agg ctg cta gtt gtc gtt cga gat cag aaa tac gtg ttt tct tgg aat gga ttt cgt cga tgt act tgt cat gat tgg cat cga aac act tat taa gtt ctt ttt ttc aat tct acg att tta ttt ctt tcg cga gtc aat tcc ctc ctg tag taa cta tca gtt ttg tca gat tca cgc tct cta cgt aga ctt tct tgt aag tta cta att tgt tcc ctg gca tta ccg agt tca gtt tta tat gcc gaa tag agt tct gat tca tcc ttt gag aag atc tct agc gat cgt tca aga tcc ctg att cta gtc ttt agc cta ttt acc tcc tca gaa gat gct ccg tta ccg ttt tta caa tcg tta aga tgt cta tca aga tcc atg att cta tct ctt ttc cat atc agc att gat ttc att att acg ttc gca gtc gtt caa ctg tat t

>MT903346.1_Monkeypox_virus_isolate_MPXV-USA2003_099_Gambian_Rat

tta ccg cgt ctg tac tcg acg agc tca cgt tta aga gat tca att tcc agt ttg tat cgg tcc atg tct cca ttg cta cac cac cat tag att tac agg ctg tta gtt gtc gtt cga gat cag aaa tac gtg ttt tct tgg aat gga ttt cgt cga tgt act tgt cat gat tgg cat cga aac act tat taa gtt ctt ttt ttc aat tct acg att tta ttt ctt tcg cga gtc aat tcc ctc cta tag taa cta tca gtt ttg tca gat tca cgc tct cta cgt aga ctt tct tgc aag tta cta att tgt tcc ctg gca tta ccg agt tca gtt tta tat gcc gaa tag agt tct gat tca tcc ttt gag aag atc tct agc gat cgt tca aga tcc ctg att cta gtc ttt agc cta ttt acc tcc tca gaa gat gct ccg tta ccg ttt tta caa tcg tta aga tgt cta tca aga tcc atg att cta tct ctt ttc cat atc agc att gat ttc att att acg tcc gca gtc gtt caa ctg tat t

>DQ011157.1_Monkeypox_virus_strain_USA_2003_039_complete_genome

tta ccg cgt ctg tac tcg acg agc tca cgt tta aga gat tca att tcc agt ttg tat cgg tcc atg tct cca ttg cta cac cac cat tag att tac agg ctg tta gtt gtc gtt cga gat cag aaa tac gtg ttt tct tgg aat gga ttt cgt cga tgt act tgt cat gat tgg cat cga aac act tat taa gtt ctt ttt ttc aat tct acg att tta ttt ctt tcg cga gtc aat tcc ctc cta tag taa cta tca gtt ttg tca gat tca cgc tct cta cgt aga ctt tct tgc aag tta cta att tgt tcc ctg gca tta ccg agt tca gtt tta tat gcc gaa tag agt tct gat tca tcc ttt gag aag atc tct agc gat cgt tca aga tcc ctg att cta gtc ttt agc cta ttt acc tcc tca gaa gat gct ccg tta ccg ttt tta caa tcg tta aga tgt cta tca aga tcc atg att cta tct ctt ttc cat atc agc att gat ttc att att acg tcc gca gtc gtt caa ctg tat t

>DQ011153.1_Monkeypox_virus_strain_USA_2003_044_complete_genome

tta ccg cgt ctg tac tcg acg agc tca cgt tta aga gat tca att tcc agt ttg tat cgg tcc atg tct cca ttg cta cac cac cat tag att tac agg ctg tta gtt gtc gtt cga gat cag aaa tac gtg ttt tct tgg aat gga ttt cgt cga tgt act tgt cat gat tgg cat cga aac act tat taa gtt ctt ttt ttc aat tct acg att tta ttt ctt tcg cga gtc aat tcc ctc cta tag taa cta tca gtt ttg tca gat tca cgc tct cta cgt aga ctt tct tgc aag tta cta att tgt tcc ctg gca tta ccg agt tca gtt tta tat gcc gaa tag agt tct gat tca tcc ttt gag aag atc tct agc gat cgt tca aga tcc ctg att cta gtc ttt agc cta ttt acc tcc tca gaa gat gct ccg tta ccg ttt tta caa tcg tta aga tgt cta tca aga tcc atg att cta tct ctt ttc cat atc agc att gat ttc att att acg tcc gca gtc gtt caa ctg tat t

>JX878410.1_Monkeypox_virus_isolate_DRC_06-1070_complete_genome

tta cca cgt ctg tac tcg gcg agc tca cgt tta aga gat tca att tcc agt ttg ta- --- --- --- --- --- --- --- --- --- --- --- --- --- --- --- --- --- --- --- --- --- --- --- --- --- --- --- --- --- --- --- --- --- --- --- --- --- --- --- --- --- --- --- --- --- --- --- --- --- --- --- --- --- --- --- --- --- --- --- --- --- --- --- --- --- --- --- --- --- --- --- --- --- --- --- --- --- --- --- --- --- --- --- --- --- --- --- --- --- --- --- --- --- --- --- --- --- --- --- --- --- --- --- --- --- --- --- --- --- --- --- --- --- --- --- --- --- --- --- --- --- --- --- --- --- --- --- --- --- --- --- --- --- --- --- --- --- --- --- --- --- --- --- --- --- --- --- --- --- --- --c att gat ttc att att acg tcc gca gtc gtt caa ctg tat t

>ON645312.1_Monkeypox_virus_isolate_MPXV_GSTT_Patient1_partial_genome

tta cca cgt ctg tac tcg acg agc tca cgt tta aga gat tca att tcc agt ttg ta- --- --- --- --- --- --- --- --- --- --- --- --- --- --- --- --- --- --- --- --- --- --- --- --- --- --- --- --- --- --- --- --- --- --- --- --- --- --- --- --- --- --- --- --- --- --- --- --- --- --- --- --- --- --- --- --- --- --- --- --- --- --- --- --- --- --- --- --- --- --- --- --- --- --- --- --- --- --- --- --- --- --- --- --- --- --- --- --- --- --- --- --- --- --- --- --- --- --- --- --- --- --- --- --- --- --- --- --- --- --- --- --- --- --- --- --- --- --- --- --- --- --- --- --- --- --- --- --- --- --- --- --- --- --- --- --- --- --- --- --- --- --- --- --- --- --- --- --- --- --- --c att gat ttc att att acg ttc gca gtc gtt caa ctg tat t

>JX878428.1_Monkeypox_virus_isolate_DRC_07-0514_complete_genome

tta cca cgt ctg tac tcg gcg agc tca cgt tta aga gat tca att tcc agt ttg ta- --- --- --- --- --- --- --- --- --- --- --- --- --- --- --- --- --- --- --- --- --- --- --- --- --- --- --- --- --- --- --- --- --- --- --- --- --- --- --- --- --- --- --- --- --- --- --- --- --- --- --- --- --- --- --- --- --- --- --- --- --- --- --- --- --- --- --- --- --- --- --- --- --- --- --- --- --- --- --- --- --- --- --- --- --- --- --- --- --- --- --- --- --- --- --- --- --- --- --- --- --- --- --- --- --- --- --- --- --- --- --- --- --- --- --- --- --- --- --- --- --- --- --- --- --- --- --- --- --- --- --- --- --- --- --- --- --- --- --- --- --- --- --- --- --- --- --- --- --- --- --c att gat ttc att att acg tcc gca gtc gtt caa ctg tat t

>JX878427.1_Monkeypox_virus_isolate_DRC_07-0480_complete_genome

tta cca cgt ctg tac tcg gcg agc tca cgt tta aga gat tca att tcc agt ttg ta- --- --- --- --- --- --- --- --- --- --- --- --- --- --- --- --- --- --- --- --- --- --- --- --- --- --- --- --- --- --- --- --- --- --- --- --- --- --- --- --- --- --- --- --- --- --- --- --- --- --- --- --- --- --- --- --- --- --- --- --- --- --- --- --- --- --- --- --- --- --- --- --- --- --- --- --- --- --- --- --- --- --- --- --- --- --- --- --- --- --- --- --- --- --- --- --- --- --- --- --- --- --- --- --- --- --- --- --- --- --- --- --- --- --- --- --- --- --- --- --- --- --- --- --- --- --- --- --- --- --- --- --- --- --- --- --- --- --- --- --- --- --- --- --- --- --- --- --- --- --- --c att gat ttc att att acg tcc gca gtc gtt caa ctg tat t

>JX878421.1_Monkeypox_virus_isolate_DRC_07-0286_complete_genome

tta cca cgt ctg tac tcg gcg agc tca cgt tta aga gat tca att tcc agt ttg ta- --- --- --- --- --- --- --- --- --- --- --- --- --- --- --- --- --- --- --- --- --- --- --- --- --- --- --- --- --- --- --- --- --- --- --- --- --- --- --- --- --- --- --- --- --- --- --- --- --- --- --- --- --- --- --- --- --- --- --- --- --- --- --- --- --- --- --- --- --- --- --- --- --- --- --- --- --- --- --- --- --- --- --- --- --- --- --- --- --- --- --- --- --- --- --- --- --- --- --- --- --- --- --- --- --- --- --- --- --- --- --- --- --- --- --- --- --- --- --- --- --- --- --- --- --- --- --- --- --- --- --- --- --- --- --- --- --- --- --- --- --- --- --- --- --- --- --- --- --- --- --c att gat ttc att att acg tcc gca gtc gtt caa ctg tat t

>JX878415.1_Monkeypox_virus_isolate_DRC_07-0092_complete_genome

tta cca cgt ctg tac tcg gcg agc tca cgt tta aga gat tca att tcc agt ttg ta- --- --- --- --- --- --- --- --- --- --- --- --- --- --- --- --- --- --- --- --- --- --- --- --- --- --- --- --- --- --- --- --- --- --- --- --- --- --- --- --- --- --- --- --- --- --- --- --- --- --- --- --- --- --- --- --- --- --- --- --- --- --- --- --- --- --- --- --- --- --- --- --- --- --- --- --- --- --- --- --- --- --- --- --- --- --- --- --- --- --- --- --- --- --- --- --- --- --- --- --- --- --- --- --- --- --- --- --- --- --- --- --- --- --- --- --- --- --- --- --- --- --- --- --- --- --- --- --- --- --- --- --- --- --- --- --- --- --- --- --- --- --- --- --- --- --- --- --- --- --- --c att gat ttc att att acg tcc gca gtc gtt caa ctg tat t

>JX878414.1_Monkeypox_virus_isolate_DRC_07-0046_complete_genome

tta cca cgt ctg tac tcg gcg agc tca cgt tta aga gat tca att tcc agt ttg ta- --- --- --- --- --- --- --- --- --- --- --- --- --- --- --- --- --- --- --- --- --- --- --- --- --- --- --- --- --- --- --- --- --- --- --- --- --- --- --- --- --- --- --- --- --- --- --- --- --- --- --- --- --- --- --- --- --- --- --- --- --- --- --- --- --- --- --- --- --- --- --- --- --- --- --- --- --- --- --- --- --- --- --- --- --- --- --- --- --- --- --- --- --- --- --- --- --- --- --- --- --- --- --- --- --- --- --- --- --- --- --- --- --- --- --- --- --- --- --- --- --- --- --- --- --- --- --- --- --- --- --- --- --- --- --- --- --- --- --- --- --- --- --- --- --- --- --- --- --- --- --c att gat ttc att att acg tcc gca gtc gtt caa ctg tat t

>JX878413.1_Monkeypox_virus_isolate_DRC_07-0045_complete_genome

tta cca cgt ctg tac tcg gcg agc tca cgt tta aga gat tca att tcc agt ttg ta- --- --- --- --- --- --- --- --- --- --- --- --- --- --- --- --- --- --- --- --- --- --- --- --- --- --- --- --- --- --- --- --- --- --- --- --- --- --- --- --- --- --- --- --- --- --- --- --- --- --- --- --- --- --- --- --- --- --- --- --- --- --- --- --- --- --- --- --- --- --- --- --- --- --- --- --- --- --- --- --- --- --- --- --- --- --- --- --- --- --- --- --- --- --- --- --- --- --- --- --- --- --- --- --- --- --- --- --- --- --- --- --- --- --- --- --- --- --- --- --- --- --- --- --- --- --- --- --- --- --- --- --- --- --- --- --- --- --- --- --- --- --- --- --- --- --- --- --- --- --- --c att gat ttc att att acg tcc gca gtc gtt caa ctg tat t

>JX878422.1_Monkeypox_virus_isolate_DRC_07-0287_complete_genome

tta cca cgt ctg tac tcg gcg agc tca cgt tta aga gat tca att tcc agt ttg ta- --- --- --- --- --- --- --- --- --- --- --- --- --- --- --- --- --- --- --- --- --- --- --- --- --- --- --- --- --- --- --- --- --- --- --- --- --- --- --- --- --- --- --- --- --- --- --- --- --- --- --- --- --- --- --- --- --- --- --- --- --- --- --- --- --- --- --- --- --- --- --- --- --- --- --- --- --- --- --- --- --- --- --- --- --- --- --- --- --- --- --- --- --- --- --- --- --- --- --- --- --- --- --- --- --- --- --- --- --- --- --- --- --- --- --- --- --- --- --- --- --- --- --- --- --- --- --- --- --- --- --- --- --- --- --- --- --- --- --- --- --- --- --- --- --- --- --- --- --- --- --c att gat ttc att att acg tcc gca gtc gtt caa ctg tat t

>JX878416.1_Monkeypox_virus_isolate_DRC_07-0093_complete_genome

tta cca cgt ctg tac tcg gcg agc tca cgt tta aga gat tca att tcc agt ttg ta- --- --- --- --- --- --- --- --- --- --- --- --- --- --- --- --- --- --- --- --- --- --- --- --- --- --- --- --- --- --- --- --- --- --- --- --- --- --- --- --- --- --- --- --- --- --- --- --- --- --- --- --- --- --- --- --- --- --- --- --- --- --- --- --- --- --- --- --- --- --- --- --- --- --- --- --- --- --- --- --- --- --- --- --- --- --- --- --- --- --- --- --- --- --- --- --- --- --- --- --- --- --- --- --- --- --- --- --- --- --- --- --- --- --- --- --- --- --- --- --- --- --- --- --- --- --- --- --- --- --- --- --- --- --- --- --- --- --- --- --- --- --- --- --- --- --- --- --- --- --- --c att gat ttc att att acg tcc gca gtc gtt caa ctg tat t

>KP849471.1_Monkeypox_virus_isolate_Yambuku_DRC_1985_complete_genome

tta cca cgt ctg tac tca gcg agc tca cgt tta aga gat tca att tcc agt ttg ta- --- --- --- --- --- --- --- --- --- --- --- --- --- --- --- --- --- --- --- --- --- --- --- --- --- --- --- --- --- --- --- --- --- --- --- --- --- --- --- --- --- --- --- --- --- --- --- --- --- --- --- --- --- --- --- --- --- --- --- --- --- --- --- --- --- --- --- --- --- --- --- --- --- --- --- --- --- --- --- --- --- --- --- --- --- --- --- --- --- --- --- --- --- --- --- --- --- --- --- --- --- --- --- --- --- --- --- --- --- --- --- --- --- --- --- --- --- --- --- --- --- --- --- --- --- --- --- --- --- --- --- --- --- --- --- --- --- --- --- --- --- --- --- --- --- --- --- --- --- --- --c att gat ttc att att acg tcc gca gtc gtt caa ctg tat t

>DQ011155.1_Monkeypox_virus_strain_Zaire_1979-005_complete_genome

tta cca cgt ctg tac tcg gcg agc tca cgt tta aga gat tca att tcc agt ttg ta- --- --- --- --- --- --- --- --- --- --- --- --- --- --- --- --- --- --- --- --- --- --- --- --- --- --- --- --- --- --- --- --- --- --- --- --- --- --- --- --- --- --- --- --- --- --- --- --- --- --- --- --- --- --- --- --- --- --- --- --- --- --- --- --- --- --- --- --- --- --- --- --- --- --- --- --- --- --- --- --- --- --- --- --- --- --- --- --- --- --- --- --- --- --- --- --- --- --- --- --- --- --- --- --- --- --- --- --- --- --- --- --- --- --- --- --- --- --- --- --- --- --- --- --- --- --- --- --- --- --- --- --- --- --- --- --- --- --- --- --- --- --- --- --- --- --- --- --- --- --- --c att gat ttc att att acg tcc gca gtc gtt --- ctg tat t

>KP849469.1_Monkeypox_virus_isolate_Boende_DRC_2008_complete_genome

tta cca cgt ctg tac tcg gcg agc tca cgt tta aga gat tca att tcc agt ttg ta- --- --- --- --- --- --- --- --- --- --- --- --- --- --- --- --- --- --- --- --- --- --- --- --- --- --- --- --- --- --- --- --- --- --- --- --- --- --- --- --- --- --- --- --- --- --- --- --- --- --- --- --- --- --- --- --- --- --- --- --- --- --- --- --- --- --- --- --- --- --- --- --- --- --- --- --- --- --- --- --- --- --- --- --- --- --- --- --- --- --- --- --- --- --- --- --- --- --- --- --- --- --- --- --- --- --- --- --- --- --- --- --- --- --- --- --- --- --- --- --- --- --- --- --- --- --- --- --- --- --- --- --- --- --- --- --- --- --- --- --- --- --- --- --- --- --- --- --- --- --- --c att gat ttc att att acg tcc gca gtc gtt caa ctg tat t

>KJ642619.1_Monkeypox_virus_strain_Gabon-1988_complete_genome

tta cca cgt ctg tac tcg gcg agc tca cgt tta aga gat tca att tcc agt ttg ta- --- --- --- --- --- --- --- --- --- --- --- --- --- --- --- --- --- --- --- --- --- --- --- --- --- --- --- --- --- --- --- --- --- --- --- --- --- --- --- --- --- --- --- --- --- --- --- --- --- --- --- --- --- --- --- --- --- --- --- --- --- --- --- --- --- --- --- --- --- --- --- --- --- --- --- --- --- --- --- --- --- --- --- --- --- --- --- --- --- --- --- --- --- --- --- --- --- --- --- --- --- --- --- --- --- --- --- --- --- --- --- --- --- --- --- --- --- --- --- --- --- --- --- --- --- --- --- --- --- --- --- --- --- --- --- --- --- --- --- --- --- --- --- --- --- --- --- --- --- --- --c att gat ttc att att acg tcc gca gtc gtt caa ctg tat t

>JX878411.1_Monkeypox_virus_isolate_DRC_06-1075_complete_genome

tta cca cgt ctg tac tcg gcg agc tca cgt tta aga gat tca att tcc agt ttg ta- --- --- --- --- --- --- --- --- --- --- --- --- --- --- --- --- --- --- --- --- --- --- --- --- --- --- --- --- --- --- --- --- --- --- --- --- --- --- --- --- --- --- --- --- --- --- --- --- --- --- --- --- --- --- --- --- --- --- --- --- --- --- --- --- --- --- --- --- --- --- --- --- --- --- --- --- --- --- --- --- --- --- --- --- --- --- --- --- --- --- --- --- --- --- --- --- --- --- --- --- --- --- --- --- --- --- --- --- --- --- --- --- --- --- --- --- --- --- --- --- --- --- --- --- --- --- --- --- --- --- --- --- --- --- --- --- --- --- --- --- --- --- --- --- --- --- --- --- --- --- --c att gat ttc att att acg tcc gca gtc gtt caa ctg tat t

>JX878409.1_Monkeypox_virus_isolate_DRC_06-0999_complete_genome

tta cca cgt ctg tac tcg gcg agc tca cgt tta aga gat tca att tcc agt ttg ta- --- --- --- --- --- --- --- --- --- --- --- --- --- --- --- --- --- --- --- --- --- --- --- --- --- --- --- --- --- --- --- --- --- --- --- --- --- --- --- --- --- --- --- --- --- --- --- --- --- --- --- --- --- --- --- --- --- --- --- --- --- --- --- --- --- --- --- --- --- --- --- --- --- --- --- --- --- --- --- --- --- --- --- --- --- --- --- --- --- --- --- --- --- --- --- --- --- --- --- --- --- --- --- --- --- --- --- --- --- --- --- --- --- --- --- --- --- --- --- --- --- --- --- --- --- --- --- --- --- --- --- --- --- --- --- --- --- --- --- --- --- --- --- --- --- --- --- --- --- --- --c att gat ttc att att acg tcc gca gtc gtt caa ctg tat t

>DQ011154.1_Monkeypox_virus_strain_Congo_2003_358_complete_genome

tta cca cgt ctg tac tcg gcg agc tca cgt tta aga gat tca att tcc agt ttg ta- --- --- --- --- --- --- --- --- --- --- --- --- --- --- --- --- --- --- --- --- --- --- --- --- --- --- --- --- --- --- --- --- --- --- --- --- --- --- --- --- --- --- --- --- --- --- --- --- --- --- --- --- --- --- --- --- --- --- --- --- --- --- --- --- --- --- --- --- --- --- --- --- --- --- --- --- --- --- --- --- --- --- --- --- --- --- --- --- --- --- --- --- --- --- --- --- --- --- --- --- --- --- --- --- --- --- --- --- --- --- --- --- --- --- --- --- --- --- --- --- --- --- --- --- --- --- --- --- --- --- --- --- --- --- --- --- --- --- --- --- --- --- --- --- --- --- --- --- --- --- --c att gat ttc att att acg tcc gca gtc gtt caa ctg tat t

>JX878412.1_Monkeypox_virus_isolate_DRC_06-1076_complete_genome

tta cca cgt ctg tac tcg gcg agc tca cgt tta aga gat tca att tcc agt ttg ta- --- --- --- --- --- --- --- --- --- --- --- --- --- --- --- --- --- --- --- --- --- --- --- --- --- --- --- --- --- --- --- --- --- --- --- --- --- --- --- --- --- --- --- --- --- --- --- --- --- --- --- --- --- --- --- --- --- --- --- --- --- --- --- --- --- --- --- --- --- --- --- --- --- --- --- --- --- --- --- --- --- --- --- --- --- --- --- --- --- --- --- --- --- --- --- --- --- --- --- --- --- --- --- --- --- --- --- --- --- --- --- --- --- --- --- --- --- --- --- --- --- --- --- --- --- --- --- --- --- --- --- --- --- --- --- --- --- --- --- --- --- --- --- --- --- --- --- --- --- --- --c att gat ttc att att acg tcc gca gtc gtt caa ctg tat t

>MN702453.1_Monkeypox_virus_strain_A1_contig_SPADES

tta cca cgt ctg tac tcg gcg agc tca cgt tta aga gat tca att tcc agt ttg ta- --- --- --- --- --- --- --- --- --- --- --- --- --- --- --- --- --- --- --- --- --- --- --- --- --- --- --- --- --- --- --- --- --- --- --- --- --- --- --- --- --- --- --- --- --- --- --- --- --- --- --- --- --- --- --- --- --- --- --- --- --- --- --- --- --- --- --- --- --- --- --- --- --- --- --- --- --- --- --- --- --- --- --- --- --- --- --- --- --- --- --- --- --- --- --- --- --- --- --- --- --- --- --- --- --- --- --- --- --- --- --- --- --- --- --- --- --- --- --- --- --- --- --- --- --- --- --- --- --- --- --- --- --- --- --- --- --- --- --- --- --- --- --- --- --- --- --- --- --- --- --c att gat ttc att att acg tcc gca gtc gtt caa ctg tat t

>MN702452.1_Monkeypox_virus_strain_A2_contig_SPADES

tta cca cgt ctg tac tcg gcg agc tca cgt tta aga gat tca att tcc agt ttg ta- --- --- --- --- --- --- --- --- --- --- --- --- --- --- --- --- --- --- --- --- --- --- --- --- --- --- --- --- --- --- --- --- --- --- --- --- --- --- --- --- --- --- --- --- --- --- --- --- --- --- --- --- --- --- --- --- --- --- --- --- --- --- --- --- --- --- --- --- --- --- --- --- --- --- --- --- --- --- --- --- --- --- --- --- --- --- --- --- --- --- --- --- --- --- --- --- --- --- --- --- --- --- --- --- --- --- --- --- --- --- --- --- --- --- --- --- --- --- --- --- --- --- --- --- --- --- --- --- --- --- --- --- --- --- --- --- --- --- --- --- --- --- --- --- --- --- --- --- --- --- --c att gat ttc att att acg tcc gca gtc gtt caa ctg tat t

>MN702448.1_Monkeypox_virus_strain_015c_contig_SPADES

tta cca cgt ctg tac tcg gcg agc tca cgt tta aga gat tca att tcc agt ttg ta- --- --- --- --- --- --- --- --- --- --- --- --- --- --- --- --- --- --- --- --- --- --- --- --- --- --- --- --- --- --- --- --- --- --- --- --- --- --- --- --- --- --- --- --- --- --- --- --- --- --- --- --- --- --- --- --- --- --- --- --- --- --- --- --- --- --- --- --- --- --- --- --- --- --- --- --- --- --- --- --- --- --- --- --- --- --- --- --- --- --- --- --- --- --- --- --- --- --- --- --- --- --- --- --- --- --- --- --- --- --- --- --- --- --- --- --- --- --- --- --- --- --- --- --- --- --- --- --- --- --- --- --- --- --- --- --- --- --- --- --- --- --- --- --- --- --- --- --- --- --- --c att gat ttc att att acg tcc gca gtc gtt caa ctg tat t

>MN702447.1_Monkeypox_virus_strain_18_contig_SPADES

tta cca cgt ctg tac tcg gcg agc tca cgt tta aga gat tca att tcc agt ttg ta- --- --- --- --- --- --- --- --- --- --- --- --- --- --- --- --- --- --- --- --- --- --- --- --- --- --- --- --- --- --- --- --- --- --- --- --- --- --- --- --- --- --- --- --- --- --- --- --- --- --- --- --- --- --- --- --- --- --- --- --- --- --- --- --- --- --- --- --- --- --- --- --- --- --- --- --- --- --- --- --- --- --- --- --- --- --- --- --- --- --- --- --- --- --- --- --- --- --- --- --- --- --- --- --- --- --- --- --- --- --- --- --- --- --- --- --- --- --- --- --- --- --- --- --- --- --- --- --- --- --- --- --- --- --- --- --- --- --- --- --- --- --- --- --- --- --- --- --- --- --- --c att gat ttc att att acg tcc gca gtc gtt caa ctg tat t

>JX878424.1_Monkeypox_virus_isolate_DRC_07-0338_complete_genome

tta cca cgt ctg tac tcg gcg agc tca cgt tta aga gat tca att tcc agt ttg ta- --- --- --- --- --- --- --- --- --- --- --- --- --- --- --- --- --- --- --- --- --- --- --- --- --- --- --- --- --- --- --- --- --- --- --- --- --- --- --- --- --- --- --- --- --- --- --- --- --- --- --- --- --- --- --- --- --- --- --- --- --- --- --- --- --- --- --- --- --- --- --- --- --- --- --- --- --- --- --- --- --- --- --- --- --- --- --- --- --- --- --- --- --- --- --- --- --- --- --- --- --- --- --- --- --- --- --- --- --- --- --- --- --- --- --- --- --- --- --- --- --- --- --- --- --- --- --- --- --- --- --- --- --- --- --- --- --- --- --- --- --- --- --- --- --- --- --- --- --- --- --c att gat ttc att att acg tcc gca gtc gtt caa ctg tat t

>KC257460.1_Monkeypox_virus_strain_DRC_Yandongi_1985_complete_genome

tta cca cgt ctg tac tcg gcg agc tca cgt tta aga gat tca att tcc agt ttg ta- --- --- --- --- --- --- --- --- --- --- --- --- --- --- --- --- --- --- --- --- --- --- --- --- --- --- --- --- --- --- --- --- --- --- --- --- --- --- --- --- --- --- --- --- --- --- --- --- --- --- --- --- --- --- --- --- --- --- --- --- --- --- --- --- --- --- --- --- --- --- --- --- --- --- --- --- --- --- --- --- --- --- --- --- --- --- --- --- --- --- --- --- --- --- --- --- --- --- --- --- --- --- --- --- --- --- --- --- --- --- --- --- --- --- --- --- --- --- --- --- --- --- --- --- --- --- --- --- --- --- --- --- --- --- --- --- --- --- --- --- --- --- --- --- --- --- --- --- --- --- --c att gat ttc att att acg tcc gca gtc gtt --- ctg tat t

>JX878429.1_Monkeypox_virus_isolate_DRC_07-0662_complete_genome

tta cca cgt ctg tac tcg gcg agc tca cgt tta aga gat tca att tcc agt ttg ta- --- --- --- --- --- --- --- --- --- --- --- --- --- --- --- --- --- --- --- --- --- --- --- --- --- --- --- --- --- --- --- --- --- --- --- --- --- --- --- --- --- --- --- --- --- --- --- --- --- --- --- --- --- --- --- --- --- --- --- --- --- --- --- --- --- --- --- --- --- --- --- --- --- --- --- --- --- --- --- --- --- --- --- --- --- --- --- --- --- --- --- --- --- --- --- --- --- --- --- --- --- --- --- --- --- --- --- --- --- --- --- --- --- --- --- --- --- --- --- --- --- --- --- --- --- --- --- --- --- --- --- --- --- --- --- --- --- --- --- --- --- --- --- --- --- --- --- --- --- --- --c att gat ttc att att acg tcc gca gtc gtt caa ctg tat t

>JX878425.1_Monkeypox_virus_isolate_DRC_07-0354_complete_genome

tta cca cgt ctg tac tcg gcg agc tca cgt tta aga gat tca att tcc agt ttg ta- --- --- --- --- --- --- --- --- --- --- --- --- --- --- --- --- --- --- --- --- --- --- --- --- --- --- --- --- --- --- --- --- --- --- --- --- --- --- --- --- --- --- --- --- --- --- --- --- --- --- --- --- --- --- --- --- --- --- --- --- --- --- --- --- --- --- --- --- --- --- --- --- --- --- --- --- --- --- --- --- --- --- --- --- --- --- --- --- --- --- --- --- --- --- --- --- --- --- --- --- --- --- --- --- --- --- --- --- --- --- --- --- --- --- --- --- --- --- --- --- --- --- --- --- --- --- --- --- --- --- --- --- --- --- --- --- --- --- --- --- --- --- --- --- --- --- --- --- --- --- --c att gat ttc att att acg tcc gca gtc gtt caa ctg tat t

>JX878407.1_Monkeypox_virus_isolate_DRC_06-0950_complete_genome

tta cca cgt ctg tac tcg gcg agc tca cgt tta aga gat tca att tcc agt ttg ta- --- --- --- --- --- --- --- --- --- --- --- --- --- --- --- --- --- --- --- --- --- --- --- --- --- --- --- --- --- --- --- --- --- --- --- --- --- --- --- --- --- --- --- --- --- --- --- --- --- --- --- --- --- --- --- --- --- --- --- --- --- --- --- --- --- --- --- --- --- --- --- --- --- --- --- --- --- --- --- --- --- --- --- --- --- --- --- --- --- --- --- --- --- --- --- --- --- --- --- --- --- --- --- --- --- --- --- --- --- --- --- --- --- --- --- --- --- --- --- --- --- --- --- --- --- --- --- --- --- --- --- --- --- --- --- --- --- --- --- --- --- --- --- --- --- --- --- --- --- --- --c att gat ttc att att acg tcc gca gtc gtt caa ctg tat t

>MN702451.1_Monkeypox_virus_strain_A6_contig_SPADES

tta cca cgt ctg tac tcg gcg agc tca cgt tta aga gat tca att tcc agt ttg ta- --- --- --- --- --- --- --- --- --- --- --- --- --- --- --- --- --- --- --- --- --- --- --- --- --- --- --- --- --- --- --- --- --- --- --- --- --- --- --- --- --- --- --- --- --- --- --- --- --- --- --- --- --- --- --- --- --- --- --- --- --- --- --- --- --- --- --- --- --- --- --- --- --- --- --- --- --- --- --- --- --- --- --- --- --- --- --- --- --- --- --- --- --- --- --- --- --- --- --- --- --- --- --- --- --- --- --- --- --- --- --- --- --- --- --- --- --- --- --- --- --- --- --- --- --- --- --- --- --- --- --- --- --- --- --- --- --- --- --- --- --- --- --- --- --- --- --- --- --- --- --c att gat ttc att att acg tcc gca gtc gtt caa ctg tat t

>JX878423.1_Monkeypox_virus_isolate_DRC_07-0337_complete_genome

tta cca cgt ctg tac tcg gcg agc tca cgt tta aga gat tca att tcc agt ttg ta- --- --- --- --- --- --- --- --- --- --- --- --- --- --- --- --- --- --- --- --- --- --- --- --- --- --- --- --- --- --- --- --- --- --- --- --- --- --- --- --- --- --- --- --- --- --- --- --- --- --- --- --- --- --- --- --- --- --- --- --- --- --- --- --- --- --- --- --- --- --- --- --- --- --- --- --- --- --- --- --- --- --- --- --- --- --- --- --- --- --- --- --- --- --- --- --- --- --- --- --- --- --- --- --- --- --- --- --- --- --- --- --- --- --- --- --- --- --- --- --- --- --- --- --- --- --- --- --- --- --- --- --- --- --- --- --- --- --- --- --- --- --- --- --- --- --- --- --- --- --- --c att gat ttc att att acg tcc gca gtc gtt caa ctg tat t

>MN702450.1_Monkeypox_virus_strain_B1_contig_SPADES

tta cca cgt ctg tac tcg gcg agc tca cgt tta aga gat tca att tcc agt ttg ta- --- --- --- --- --- --- --- --- --- --- --- --- --- --- --- --- --- --- --- --- --- --- --- --- --- --- --- --- --- --- --- --- --- --- --- --- --- --- --- --- --- --- --- --- --- --- --- --- --- --- --- --- --- --- --- --- --- --- --- --- --- --- --- --- --- --- --- --- --- --- --- --- --- --- --- --- --- --- --- --- --- --- --- --- --- --- --- --- --- --- --- --- --- --- --- --- --- --- --- --- --- --- --- --- --- --- --- --- --- --- --- --- --- --- --- --- --- --- --- --- --- --- --- --- --- --- --- --- --- --- --- --- --- --- --- --- --- --- --- --- --- --- --- --- --- --- --- --- --- --- --c att gat ttc att att acg tcc gca gtc gtt caa ctg tat t

>JX878408.1_Monkeypox_virus_isolate_DRC_06-0970_complete_genome

tta cca cgt ctg tac tcg gcg agc tca cgt tta aga gat tca att tcc agt ttg ta- --- --- --- --- --- --- --- --- --- --- --- --- --- --- --- --- --- --- --- --- --- --- --- --- --- --- --- --- --- --- --- --- --- --- --- --- --- --- --- --- --- --- --- --- --- --- --- --- --- --- --- --- --- --- --- --- --- --- --- --- --- --- --- --- --- --- --- --- --- --- --- --- --- --- --- --- --- --- --- --- --- --- --- --- --- --- --- --- --- --- --- --- --- --- --- --- --- --- --- --- --- --- --- --- --- --- --- --- --- --- --- --- --- --- --- --- --- --- --- --- --- --- --- --- --- --- --- --- --- --- --- --- --- --- --- --- --- --- --- --- --- --- --- --- --- --- --- --- --- --- --c att gat ttc att att acg tcc gca gtc gtt caa ctg tat t

>JX878420.1_Monkeypox_virus_isolate_DRC_07-0283_complete_genome

tta cca cgt ctg tac tcg gcg agc tca cgt tta aga gat tca att tcc agt ttg ta- --- --- --- --- --- --- --- --- --- --- --- --- --- --- --- --- --- --- --- --- --- --- --- --- --- --- --- --- --- --- --- --- --- --- --- --- --- --- --- --- --- --- --- --- --- --- --- --- --- --- --- --- --- --- --- --- --- --- --- --- --- --- --- --- --- --- --- --- --- --- --- --- --- --- --- --- --- --- --- --- --- --- --- --- --- --- --- --- --- --- --- --- --- --- --- --- --- --- --- --- --- --- --- --- --- --- --- --- --- --- --- --- --- --- --- --- --- --- --- --- --- --- --- --- --- --- --- --- --- --- --- --- --- --- --- --- --- --- --- --- --- --- --- --- --- --- --- --- --- --- --c att gat ttc att att acg tcc gca gtc gtt caa ctg tat t

>JX878419.1_Monkeypox_virus_isolate_DRC_07-0275_complete_genome

tta cca cgt ctg tac tcg gcg agc tca cgt tta aga gat tca att tcc agt ttg ta- --- --- --- --- --- --- --- --- --- --- --- --- --- --- --- --- --- --- --- --- --- --- --- --- --- --- --- --- --- --- --- --- --- --- --- --- --- --- --- --- --- --- --- --- --- --- --- --- --- --- --- --- --- --- --- --- --- --- --- --- --- --- --- --- --- --- --- --- --- --- --- --- --- --- --- --- --- --- --- --- --- --- --- --- --- --- --- --- --- --- --- --- --- --- --- --- --- --- --- --- --- --- --- --- --- --- --- --- --- --- --- --- --- --- --- --- --- --- --- --- --- --- --- --- --- --- --- --- --- --- --- --- --- --- --- --- --- --- --- --- --- --- --- --- --- --- --- --- --- --- --c att gat ttc att att acg tcc gca gtc gtt caa ctg tat t

>JX878417.1_Monkeypox_virus_isolate_DRC_07-0104_complete_genome

tta cca cgt ctg tac tcg gcg agc tca cgt tta aga gat tca att tcc agt ttg ta- --- --- --- --- --- --- --- --- --- --- --- --- --- --- --- --- --- --- --- --- --- --- --- --- --- --- --- --- --- --- --- --- --- --- --- --- --- --- --- --- --- --- --- --- --- --- --- --- --- --- --- --- --- --- --- --- --- --- --- --- --- --- --- --- --- --- --- --- --- --- --- --- --- --- --- --- --- --- --- --- --- --- --- --- --- --- --- --- --- --- --- --- --- --- --- --- --- --- --- --- --- --- --- --- --- --- --- --- --- --- --- --- --- --- --- --- --- --- --- --- --- --- --- --- --- --- --- --- --- --- --- --- --- --- --- --- --- --- --- --- --- --- --- --- --- --- --- --- --- --- --c att gat ttc att att acg tcc gca gtc gtt caa ctg tat t

>KJ642612.1_Monkeypox_virus_strain_Ikubi_complete_genome

tta cca cgt ctg tac tcg gcg agc tca cgt tta aga gat tca att tcc agt ttg ta- --- --- --- --- --- --- --- --- --- --- --- --- --- --- --- --- --- --- --- --- --- --- --- --- --- --- --- --- --- --- --- --- --- --- --- --- --- --- --- --- --- --- --- --- --- --- --- --- --- --- --- --- --- --- --- --- --- --- --- --- --- --- --- --- --- --- --- --- --- --- --- --- --- --- --- --- --- --- --- --- --- --- --- --- --- --- --- --- --- --- --- --- --- --- --- --- --- --- --- --- --- --- --- --- --- --- --- --- --- --- --- --- --- --- --- --- --- --- --- --- --- --- --- --- --- --- --- --- --- --- --- --- --- --- --- --- --- --- --- --- --- --- --- --- --- --- --- --- --- --- --c att gat ttc att att acg tcc gca gtc gtt caa ctg tat t

>JX878426.1_Monkeypox_virus_isolate_DRC_07-0450_complete_genome

tta cca cgt ctg tac tcg gcg agc tca cgt tta aga gat tca att tcc agt ttg ta- --- --- --- --- --- --- --- --- --- --- --- --- --- --- --- --- --- --- --- --- --- --- --- --- --- --- --- --- --- --- --- --- --- --- --- --- --- --- --- --- --- --- --- --- --- --- --- --- --- --- --- --- --- --- --- --- --- --- --- --- --- --- --- --- --- --- --- --- --- --- --- --- --- --- --- --- --- --- --- --- --- --- --- --- --- --- --- --- --- --- --- --- --- --- --- --- --- --- --- --- --- --- --- --- --- --- --- --- --- --- --- --- --- --- --- --- --- --- --- --- --- --- --- --- --- --- --- --- --- --- --- --- --- --- --- --- --- --- --- --- --- --- --- --- --- --- --- --- --- --- --c att gat ttc att att acg tcc gca gtc gtt caa ctg tat t

>KC257459.1_Monkeypox_virus_strain_Sudan_2005_01_complete_genome

tta cca cgt ctg tac tcg gcg agc tca cgt tta aga gat tca att tcc agt ttg ta- --- --- --- --- --- --- --- --- --- --- --- --- --- --- --- --- --- --- --- --- --- --- --- --- --- --- --- --- --- --- --- --- --- --- --- --- --- --- --- --- --- --- --- --- --- --- --- --- --- --- --- --- --- --- --- --- --- --- --- --- --- --- --- --- --- --- --- --- --- --- --- --- --- --- --- --- --- --- --- --- --- --- --- --- --- --- --- --- --- --- --- --- --- --- --- --- --- --- --- --- --- --- --- --- --- --- --- --- --- --- --- --- --- --- --- --- --- --- --- --- --- --- --- --- --- --- --- --- --- --- --- --- --- --- --- --- --- --- --- --- --- --- --- --- --- --- --- --- --- --- --c att gat ttc att att acg tcc aca gtc gtt --- ctg tat t

>MN702449.1_Monkeypox_virus_strain_B2_contig_SPADES

tta cca cgt ctg tac tcg gcg agc tca cgt tta aga tat tca att tcc agt ttg ta- --- --- --- --- --- --- --- --- --- --- --- --- --- --- --- --- --- --- --- --- --- --- --- --- --- --- --- --- --- --- --- --- --- --- --- --- --- --- --- --- --- --- --- --- --- --- --- --- --- --- --- --- --- --- --- --- --- --- --- --- --- --- --- --- --- --- --- --- --- --- --- --- --- --- --- --- --- --- --- --- --- --- --- --- --- --- --- --- --- --- --- --- --- --- --- --- --- --- --- --- --- --- --- --- --- --- --- --- --- --- --- --- --- --- --- --- --- --- --- --- --- --- --- --- --- --- --- --- --- --- --- --- --- --- --- --- --- --- --- --- --- --- --- --- --- --- --- --- --- --- --c att gat ttc att att acg tcc gca gtc gtt caa ctg tat t

>MN702446.1_Monkeypox_virus_strain_38c_contig_SPADES

tta cca cgt ctg tac tcg gcg agc tca cgt tta aga tat tca att tcc agt ttg ta- --- --- --- --- --- --- --- --- --- --- --- --- --- --- --- --- --- --- --- --- --- --- --- --- --- --- --- --- --- --- --- --- --- --- --- --- --- --- --- --- --- --- --- --- --- --- --- --- --- --- --- --- --- --- --- --- --- --- --- --- --- --- --- --- --- --- --- --- --- --- --- --- --- --- --- --- --- --- --- --- --- --- --- --- --- --- --- --- --- --- --- --- --- --- --- --- --- --- --- --- --- --- --- --- --- --- --- --- --- --- --- --- --- --- --- --- --- --- --- --- --- --- --- --- --- --- --- --- --- --- --- --- --- --- --- --- --- --- --- --- --- --- --- --- --- --- --- --- --- --- --c att gat ttc att att acg tcc gca gtc gtt caa ctg tat t

>JX878418.1_Monkeypox_virus_isolate_DRC_07-0120_complete_genome

tta cca cgt ctg tac tcg gcg agc tca cgt tta aga gat tca att tcc agt ttg ta- --- --- --- --- --- --- --- --- --- --- --- --- --- --- --- --- --- --- --- --- --- --- --- --- --- --- --- --- --- --- --- --- --- --- --- --- --- --- --- --- --- --- --- --- --- --- --- --- --- --- --- --- --- --- --- --- --- --- --- --- --- --- --- --- --- --- --- --- --- --- --- --- --- --- --- --- --- --- --- --- --- --- --- --- --- --- --- --- --- --- --- --- --- --- --- --- --- --- --- --- --- --- --- --- --- --- --- --- --- --- --- --- --- --- --- --- --- --- --- --- --- --- --- --- --- --- --- --- --- --- --- --- --- --- --- --- --- --- --- --- --- --- --- --- --- --- --- --- --- --- --c att gat ttc att att acg tcc gca gtc gtt caa ctg tat t

>MN702445.1_Monkeypox_virus_strain_A4_contig_SPADES

tta cca cgt ctg tac tcg gcg agc tca cgt tta aga tat tca att tcc agt ttg ta- --- --- --- --- --- --- --- --- --- --- --- --- --- --- --- --- --- --- --- --- --- --- --- --- --- --- --- --- --- --- --- --- --- --- --- --- --- --- --- --- --- --- --- --- --- --- --- --- --- --- --- --- --- --- --- --- --- --- --- --- --- --- --- --- --- --- --- --- --- --- --- --- --- --- --- --- --- --- --- --- --- --- --- --- --- --- --- --- --- --- --- --- --- --- --- --- --- --- --- --- --- --- --- --- --- --- --- --- --- --- --- --- --- --- --- --- --- --- --- --- --- --- --- --- --- --- --- --- --- --- --- --- --- --- --- --- --- --- --- --- --- --- --- --- --- --- --- --- --- --- --c att gat ttc att att acg tcc gca gtc gtt caa ctg tat t

>MN702444.1_Monkeypox_virus_strain_A5_contig_SPADES

tta cca cgt ctg tac tcg gcg agc tca cgt tta aga tat tca att tcc agt ttg ta- --- --- --- --- --- --- --- --- --- --- --- --- --- --- --- --- --- --- --- --- --- --- --- --- --- --- --- --- --- --- --- --- --- --- --- --- --- --- --- --- --- --- --- --- --- --- --- --- --- --- --- --- --- --- --- --- --- --- --- --- --- --- --- --- --- --- --- --- --- --- --- --- --- --- --- --- --- --- --- --- --- --- --- --- --- --- --- --- --- --- --- --- --- --- --- --- --- --- --- --- --- --- --- --- --- --- --- --- --- --- --- --- --- --- --- --- --- --- --- --- --- --- --- --- --- --- --- --- --- --- --- --- --- --- --- --- --- --- --- --- --- --- --- --- --- --- --- --- --- --- --c att gat ttc att att acg tcc gca gtc gtt caa ctg tat t

>KJ642613.1_Monkeypox_virus_strain_Congo_8_complete_genome

tta cca cgt ctg tac tcg gcg agc tca cgt tta aga gat tca att tcc agt ttg ta- --- --- --- --- --- --- --- --- --- --- --- --- --- --- --- --- --- --- --- --- --- --- --- --- --- --- --- --- --- --- --- --- --- --- --- --- --- --- --- --- --- --- --- --- --- --- --- --- --- --- --- --- --- --- --- --- --- --- --- --- --- --- --- --- --- --- --- --- --- --- --- --- --- --- --- --- --- --- --- --- --- --- --- --- --- --- --- --- --- --- --- --- --- --- --- --- --- --- --- --- --- --- --- --- --- --- --- --- --- --- --- --- --- --- --- --- --- --- --- --- --- --- --- --- --- --- --- --- --- --- --- --- --- --- --- --- --- --- --- --- --- --- --- --- --- --- --- --- --- --- --c att gat ttc att att acg tcc aca gtc gtt caa ctg tat t

>HM172544.1_Monkeypox_virus_strain_Zaire_1979-005_complete_genome

tta cca cgt ctg tac tcg gcg agc tca cgt tta aga gat tca att tcc agt ttg ta- --- --- --- --- --- --- --- --- --- --- --- --- --- --- --- --- --- --- --- --- --- --- --- --- --- --- --- --- --- --- --- --- --- --- --- --- --- --- --- --- --- --- --- --- --- --- --- --- --- --- --- --- --- --- --- --- --- --- --- --- --- --- --- --- --- --- --- --- --- --- --- --- --- --- --- --- --- --- --- --- --- --- --- --- --- --- --- --- --- --- --- --- --- --- --- --- --- --- --- --- --- --- --- --- --- --- --- --- --- --- --- --- --- --- --- --- --- --- --- --- --- --- --- --- --- --- --- --- --- --- --- --- --- --- --- --- --- --- --- --- --- --- --- --- --- --- --- --- --- --- --c att gat ttc att att acg tcc gca gtc gtt --- ctg tat t

>AF380138.1_Monkeypox_virus_strain_Zaire-96-I-16_complete_genome

tta cca cgt ctg tac tcg gcg agc tca cgt tta aga gat tca att tcc agt ttg ta- --- --- --- --- --- --- --- --- --- --- --- --- --- --- --- --- --- --- --- --- --- --- --- --- --- --- --- --- --- --- --- --- --- --- --- --- --- --- --- --- --- --- --- --- --- --- --- --- --- --- --- --- --- --- --- --- --- --- --- --- --- --- --- --- --- --- --- --- --- --- --- --- --- --- --- --- --- --- --- --- --- --- --- --- --- --- --- --- --- --- --- --- --- --- --- --- --- --- --- --- --- --- --- --- --- --- --- --- --- --- --- --- --- --- --- --- --- --- --- --- --- --- --- --- --- --- --- --- --- --- --- --- --- --- --- --- --- --- --- --- --- --- --- --- --- --- --- --- --- --- --c att gat ttc att att acg tcc gca gtc gtt caa ctg tat t

>HQ857562.1_Monkeypox_virus_strain_V79-I-005_complete_genome

tta cca cgt ctg tac tcg gcg agc tca cgt tta aga gat tca att tcc agt ttg ta- --- --- --- --- --- --- --- --- --- --- --- --- --- --- --- --- --- --- --- --- --- --- --- --- --- --- --- --- --- --- --- --- --- --- --- --- --- --- --- --- --- --- --- --- --- --- --- --- --- --- --- --- --- --- --- --- --- --- --- --- --- --- --- --- --- --- --- --- --- --- --- --- --- --- --- --- --- --- --- --- --- --- --- --- --- --- --- --- --- --- --- --- --- --- --- --- --- --- --- --- --- --- --- --- --- --- --- --- --- --- --- --- --- --- --- --- --- --- --- --- --- --- --- --- --- --- --- --- --- --- --- --- --- --- --- --- --- --- --- --- --- --- --- --- --- --- --- --- --- --- --c att gat ttc att att acg tcc gca gtc gtt ctk ctg tat t

>HQ857563.1_Monkeypox_virus_strain_D14L_knockout_complete_genome

tta cca cgt ctg tac tcg gcg agc tca cgt tta aga gat tca att tcc agt ttg ta- --- --- --- --- --- --- --- --- --- --- --- --- --- --- --- --- --- --- --- --- --- --- --- --- --- --- --- --- --- --- --- --- --- --- --- --- --- --- --- --- --- --- --- --- --- --- --- --- --- --- --- --- --- --- --- --- --- --- --- --- --- --- --- --- --- --- --- --- --- --- --- --- --- --- --- --- --- --- --- --- --- --- --- --- --- --- --- --- --- --- --- --- --- --- --- --- --- --- --- --- --- --- --- --- --- --- --- --- --- --- --- --- --- --- --- --- --- --- --- --- --- --- --- --- --- --- --- --- --- --- --- --- --- --- --- --- --- --- --- --- --- --- --- --- --- --- --- --- --- --- --c att gat ttc att att acg tcc gca gtc gtt ctk ctg tat t
