## Supplementary Data 7 for "Genomic regions insertion and deletion in Monkeypox virus causing multi-country outbreak-2022"

>MT250197.1:133708-134155_Monkeypox_virus_strain_Singapore_2019

gct gct agt tgt cgt tcg aga tca gaa ata cgt gtt ttc ttg gaa tgg att tcg tcg atg tac ttg tca tga ttg gca tcg aaa cac tta tta agt tct ttt ttt caa ttc tac gat ttt att tct ttc gcg agt caa ttc cct cct gta gta act atc agt ttt gtc aga ttc acg ctc tct acg tag act ttc ttg taa gtt act aat ttg ttc cct ggc att acc gag ttc agt ttt ata tgc cga ata gag ttc tga ttc atc ctt tga gaa gat ctc tag cga tcg ttc aag atc cct gat tct agt ctt tag cct att tac ctc ctc aga aga tgc tcc gtt acc gtt --- ttt aca atc gtt aag atg tct atc aag atc cat gat tct atc tct ttt cca tat cag cat tga ttt cat tat tac gtt cgc agt cgt tca act g

>MN648051.1:133749-134196_Monkeypox_virus_strain_Israel_2018

gct gct agt tgt cgt tcg aga tca gaa ata cgt gtt ttc ttg gaa tgg att tcg tcg atg tac ttg tca tga ttg gca tcg aaa cac tta tta agt tct ttt ttt caa ttc tac gat ttt att tct ttc gcg agt caa ttc cct cct gta gta act atc agt ttt gtc aga ttc acg ctc tct acg tag act ttc ttg taa gtt act aat ttg ttc cct ggc att acc gag ttc agt ttt ata tgc cga ata gag ttc tga ttc atc ctt tga gaa gat ctc tag cga tcg ttc aag atc cct gat tct agt ctt tag cct att tac ctc ctc aga aga tgc tcc gtt acc gtt --- ttt aca atc gtt aag atg tct atc aag atc cat gat tct atc tct ttt cca tat cag cat tga ttt cat tat tac gtt cgc agt cgt tca act g

>MG693724.1:126550-126997_Monkeypox_virus_isolate_MPXV_Nig_2017_298464_partial_genome

gct gct agt tgt cgt tcg aga tca gaa ata cgt gtt ttc ttg gaa tgg att tcg tcg atg tac ttg tca tga ttg gca tcg aaa cac tta tta agt tct ttt ttt caa ttc tac gat ttt att tct ttc gcg agt caa ttc cct cct gta gta act atc agt ttt gtc aga ttc acg ctc tct acg tag act ttc ttg taa gtt act aat ttg ttc cct ggc att acc gag ttc agt ttt ata tgc cga ata gag ttc tga ttc atc ctt tga gaa gat ctc tag cga tcg ttc aag atc cct gat tct agt ctt tag cct att tac ctc ctc aga aga tgc tcc gtt acc gtt --- ttt aca atc gtt aag atg tct atc aag atc cat gat tct atc tct ttt cca tat cag cat tga ttt cat tat tac gtt cgc agt cgt tca act g

>MG693723.1:127206-127653_Monkeypox_virus_isolate_MPXV_Nig_2017_297957_partial_genome

gct gct agt tgt cgt tcg aga tca gaa ata cgt gtt ttc ttg gaa tgg att tcg tcg atg tac ttg tca tga ttg gca tcg aaa cac tta tta agt tct ttt ttt caa ttc tac gat ttt att tct ttc gcg agt caa ttc cct cct gta gta act atc agt ttt gtc aga ttc acg ctc tct acg tag act ttc ttg taa gtt act aat ttg ttc cct ggc att acc gag ttc agt ttt ata tgc cga ata gag ttc tga ttc atc ctt tga gaa gat ctc tag cga tcg ttc aag atc cct gat tct agt ctt tag cct att tac ctc ctc aga aga tgc tcc gtt acc gtt --- ttt aca atc gtt aag atg tct atc aag atc cat gat tct atc tct ttt cca tat cag cat tga ttt cat tat tac gtt cgc agt cgt tca act g

>KJ642617.1:133708-134155_Monkeypox_virus_strain_Nigeria-SE-1971_complete_genome

gct gct agt tgt cgt tcg aga tca gaa ata cgt gtt ttc ttg gaa tgg att tcg tcg atg tac ttg tca tga ttg gca tcg aaa cac tta tta agt tct ttt ttt caa ttc tac gat ttt att tct ttc gcg agt caa ttc cct cct gta gta act atc agt ttt gtc aga ttc acg ctc tct acg tag act ttc ttg taa gtt act aat ttg ttc cct ggc att acc gag ttc agt ttt ata tgc cga ata gag ttc tga ttc atc ctt tga gaa gat ctc tag cga tcg ttc aag atc cct gat tct agt ctt tag cct att tac ctc ctc aga aga tgc tcc gtt acc gtt --- ttt aca atc gtt aag atg tct atc aag atc cat gat tct atc tct ttt cca tat cag cat tga ttt cat tat tac gtt cgc agt cgt tca act g

>ON682269.2:133524-133971_Monkeypox_virus_isolate_MPXV/Germany/2022/RKI07_complete_genome

gct gct agt tgt cgt tcg aga tca gaa ata cgt gtt ttc ttg gaa tgg att tcg tcg atg tac ttg tca tga ttg gca tcg aaa cac tta tta agt tct ttt ttt caa ttc tac gat ttt att tct ttc gcg agt caa ttc cct cct gta gta act atc agt ttt gtc aga ttc acg ctc tct acg tag act ttc ttg taa gtt act aat ttg ttc cct ggc att acc gag ttc agt ttt ata tgc cga ata gag ttc tga ttc atc ctt tga gaa gat ctc tag cga tcg ttc aag atc cct gat tct agt ctt tag cct att tac ctc ctc aga aga tgc tcc gtt acc gtt --- ttt aca atc gtt aag atg tct atc aag atc cat gat tct atc tct ttt cca tat cag cat tga ttt cat tat tac gtt cgc agt cgt tca act g

>ON682265.2:133524-133971_Monkeypox_virus_isolate_MPXV/Germany/2022/RKI04_partial_genome

gct gct agt tgt cgt tcg aga tca gaa ata cgt gtt ttc ttg gaa tgg att tcg tcg atg tac ttg tca tga ttg gca tcg aaa cac tta tta agt tct ttt ttt caa ttc tac gat ttt att tct ttc gcg agt caa ttc cct cct gta gta act atc agt ttt gtc aga ttc acg ctc tct acg tag act ttc ttg taa gtt act aat ttg ttc cct ggc att acc gag ttc agt ttt ata tgc cga ata gag ttc tga ttc atc ctt tga gaa gat ctc tag cga tcg ttc aag atc cct gat tct agt ctt tag cct att tac ctc ctc aga aga tgc tcc gtt acc gtt --- ttt aca atc gtt aag atg tct atc aag atc cat gat tct atc tct ttt cca tat cag cat tga ttt cat tat tac gtt cgc agt cgt tca act g

>ON682264.2:133524-133971_Monkeypox_virus_isolate_MPXV/Germany/2022/RKI05_complete_genome

gct gct agt tgt cgt tcg aga tca gaa ata cgt gtt ttc ttg gaa tgg att tcg tcg atg tac ttg tca tga ttg gca tcg aaa cac tta tta agt tct ttt ttt caa ttc tac gat ttt att tct ttc gcg agt caa ttc cct cct gta gta act atc agt ttt gtc aga ttc acg ctc tct acg tag act ttc ttg taa gtt act aat ttg ttc cct ggc att acc gag ttc agt ttt ata tgc cga ata gag ttc tga ttc atc ctt tga gaa gat ctc tag cga tcg ttc aag atc cct gat tct agt ctt tag cct att tac ctc ctc aga aga tgc tcc gtt acc gtt --- ttt aca atc gtt aag atg tct atc aag atc cat gat tct atc tct ttt cca tat cag cat tga ttt cat tat tac gtt cgc agt cgt tca act g

>ON682263.2:133556-134003_Monkeypox_virus_isolate_MPXV/Germany/2022/RKI03_complete_genome

gct gct agt tgt cgt tcg aga tca gaa ata cgt gtt ttc ttg gaa tgg att tcg tcg atg tac ttg tca tga ttg gca tcg aaa cac tta tta agt tct ttt ttt caa ttc tac gat ttt att tct ttc gcg agt caa ttc cct cct gta gta act atc agt ttt gtc aga ttc acg ctc tct acg tag act ttc ttg taa gtt act aat ttg ttc cct ggc att acc gag ttc agt ttt ata tgc cga ata gag ttc tga ttc atc ctt tga gaa gat ctc tag cga tcg ttc aag atc cct gat tct agt ctt tag cct att tac ctc ctc aga aga tgc tcc gtt acc gtt --- ttt aca atc gtt aag atg tct atc aag atc cat gat tct atc tct ttt cca tat cag cat tga ttt cat tat tac gtt cgc agt cgt tca act g

>ON682270.1:133524-133971_Monkeypox_virus_isolate_MPXV/Germany/2022/RKI06_complete_genome

gct gct agt tgt cgt tcg aga tca gaa ata cgt gtt ttc ttg gaa tgg att tcg tcg atg tac ttg tca tga ttg gca tcg aaa cac tta tta agt tct ttt ttt caa ttc tac gat ttt att tct ttc gcg agt caa ttc cct cct gta gta act atc agt ttt gtc aga ttc acg ctc tct acg tag act ttc ttg taa gtt act aat ttg ttc cct ggc att acc gag ttc agt ttt ata tgc cga ata gag ttc tga ttc atc ctt tga gaa gat ctc tag cga tcg ttc aag atc cct gat tct agt ctt tag cct att tac ctc ctc aga aga tgc tcc gtt acc gtt --- ttt aca atc gtt aag atg tct atc aag atc cat gat tct atc tct ttt cca tat cag cat tga ttt cat tat tac gtt cgc agt cgt tca act g

>ON682268.1:133524-133971_Monkeypox_virus_isolate_MPXV/Germany/2022/RKI08_complete_genome

gct gct agt tgt cgt tcg aga tca gaa ata cgt gtt ttc ttg gaa tgg att tcg tcg atg tac ttg tca tga ttg gca tcg aaa cac tta tta agt tct ttt ttt caa ttc tac gat ttt att tct ttc gcg agt caa ttc cct cct gta gta act atc agt ttt gtc aga ttc acg ctc tct acg tag act ttc ttg taa gtt act aat ttg ttc cct ggc att acc gag ttc agt ttt ata tgc cga ata gag ttc tga ttc atc ctt tga gaa gat ctc tag cga tcg ttc aag atc cct gat tct agt ctt tag cct att tac ctc ctc aga aga tgc tcc gtt acc gtt --- ttt aca atc gtt aag atg tct atc aag atc cat gat tct atc tct ttt cca tat cag cat tga ttt cat tat tac gtt cgc agt cgt tca act g

>ON682267.1:133524-133971_Monkeypox_virus_isolate_MPXV/Germany/2022/RKI010_complete_genome

gct gct agt tgt cgt tcg aga tca gaa ata cgt gtt ttc ttg gaa tgg att tcg tcg atg tac ttg tca tga ttg gca tcg aaa cac tta tta agt tct ttt ttt caa ttc tac gat ttt att tct ttc gcg agt caa ttc cct cct gta gta act atc agt ttt gtc aga ttc acg ctc tct acg tag act ttc ttg taa gtt act aat ttg ttc cct ggc att acc gag ttc agt ttt ata tgc cga ata gag ttc tga ttc atc ctt tga gaa gat ctc tag cga tcg ttc aag atc cct gat tct agt ctt tag cct att tac ctc ctc aga aga tgc tcc gtt acc gtt --- ttt aca atc gtt aag atg tct atc aag atc cat gat tct atc tct ttt cca tat cag cat tga ttt cat tat tac gtt cgc agt cgt tca act g

>ON682266.1:133524-133971_Monkeypox_virus_isolate_MPXV/Germany/2022/RKI09_complete_genome

gct gct agt tgt cgt tcg aga tca gaa ata cgt gtt ttc ttg gaa tgg att tcg tcg atg tac ttg tca tga ttg gca tcg aaa cac tta tta agt tct ttt ttt caa ttc tac gat ttt att tct ttc gcg agt caa ttc cct cct gta gta act atc agt ttt gtc aga ttc acg ctc tct acg tag act ttc ttg taa gtt act aat ttg ttc cct ggc att acc gag ttc agt ttt ata tgc cga ata gag ttc tga ttc atc ctt tga gaa gat ctc tag cga tcg ttc aag atc cct gat tct agt ctt tag cct att tac ctc ctc aga aga tgc tcc gtt acc gtt --- ttt aca atc gtt aag atg tct atc aag atc cat gat tct atc tct ttt cca tat cag cat tga ttt cat tat tac gtt cgc agt cgt tca act g

>ON676707.1:133590-134037_Monkeypox_virus_isolate_MPXV_USA_2021_TX_complete_genome

gct gct agt tgt cgt tcg aga tca gaa ata cgt gtt ttc ttg gaa tgg att tcg tcg atg tac ttg tca tga ttg gca tcg aaa cac tta tta agt tct ttt ttt caa ttc tac gat ttt att tct ttc gcg agt caa ttc cct cct gta gta act atc agt ttt gtc aga ttc acg ctc tct acg tag act ttc ttg taa gtt act aat ttg ttc cct ggc att acc gag ttc agt ttt ata tgc cga ata gag ttc tga ttc atc ctt tga gaa gat ctc tag cga tcg ttc aag atc cct gat tct agt ctt tag cct att tac ctc ctc aga aga tgc tcc gtt acc gtt --- ttt aca atc gtt aag atg tct atc aag atc cat gat tct atc tct ttt cca tat cag cat tga ttt cat tat tac gtt cgc agt cgt tca act g

>ON676706.1:133477-133924_Monkeypox_virus_isolate_MPXV_USA_2022_UT002_complete_genome

gct gct agt tgt cgt tcg aga tca gaa ata cgt gtt ttc ttg gaa tgg att tcg tcg atg tac ttg tca tga ttg gca tcg aaa cac tta tta agt tct ttt ttt caa ttc tac gat ttt att tct ttc gcg agt caa ttc cct cct gta gta act atc agt ttt gtc aga ttc acg ctc tct acg tag act ttc ttg taa gtt act aat ttg ttc cct ggc att acc gag ttc agt ttt ata tgc cga ata gag ttc tga ttc atc ctt tga gaa gat ctc tag cga tcg ttc aag atc cct gat tct agt ctt tag cct att tac ctc ctc aga aga tgc tcc gtt acc gtt --- ttt aca atc gtt aag atg tct atc aag atc cat gat tct atc tct ttt cca tat cag cat tga ttt cat tat tac gtt cgc agt cgt tca act g

>ON676705.1:133523-133970_Monkeypox_virus_isolate_MPXV_USA_2022_UT001_complete_genome

gct gct agt tgt cgt tcg aga tca gaa ata cgt gtt ttc ttg gaa tgg att tcg tcg atg tac ttg tca tga ttg gca tcg aaa cac tta tta agt tct ttt ttt caa ttc tac gat ttt att tct ttc gcg agt caa ttc cct cct gta gta act atc agt ttt gtc aga ttc acg ctc tct acg tag act ttc ttg taa gtt act aat ttg ttc cct ggc att acc gag ttc agt ttt ata tgc cga ata gag ttc tga ttc atc ctt tga gaa gat ctc tag cga tcg ttc aag atc cct gat tct agt ctt tag cct att tac ctc ctc aga aga tgc tcc gtt acc gtt --- ttt aca atc gtt aag atg tct atc aag atc cat gat tct atc tct ttt cca tat cag cat tga ttt cat tat tac gtt cgc agt cgt tca act g

>ON676703.1:133529-133976_Monkeypox_virus_isolate_MPXV_USA_2022_CA001_complete_genome

gct gct agt tgt cgt tcg aga tca gaa ata cgt gtt ttc ttg gaa tgg att tcg tcg atg tac ttg tca tga ttg gca tcg aaa cac tta tta agt tct ttt ttt caa ttc tac gat ttt att tct ttc gcg agt caa ttc cct cct gta gta act atc agt ttt gtc aga ttc acg ctc tct acg tag act ttc ttg taa gtt act aat ttg ttc cct ggc att acc gag ttc agt ttt ata tgc cga ata gag ttc tga ttc atc ctt tga gaa gat ctc tag cga tcg ttc aag atc cct gat tct agt ctt tag cct att tac ctc ctc aga aga tgc tcc gtt acc gtt --- ttt aca atc gtt aag atg tct atc aag atc cat gat tct atc tct ttt cca tat cag cat tga ttt cat tat tac gtt cgc agt cgt tca act g

>ON675438.1:133533-133980_Monkeypox_virus_isolate_MPXV_USA_2022_VA001_complete_genome

gct gct agt tgt cgt tcg aga tca gaa ata cgt gtt ttc ttg gaa tgg att tcg tcg atg tac ttg tca tga ttg gca tcg aaa cac tta tta agt tct ttt ttt caa ttc tac gat ttt att tct ttc gcg agt caa ttc cct cct gta gta act atc agt ttt gtc aga ttc acg ctc tct acg tag act ttc ttg taa gtt act aat ttg ttc cct ggc att acc gag ttc agt ttt ata tgc cga ata gag ttc tga ttc atc ctt tga gaa gat ctc tag cga tcg ttc aag atc cct gat tct agt ctt tag cct att tac ctc ctc aga aga tgc tcc gtt acc gtt --- ttt aca atc gtt aag atg tct atc aag atc cat gat tct atc tct ttt cca tat cag cat tga ttt cat tat tac gtt cgc agt cgt tca act g

>ON674051.1:133539-133986_Monkeypox_virus_isolate_MPXV_USA_2022_FL001_complete_genome

gct gct agt tgt cgt tcg aga tca gaa ata cgt gtt ttc ttg gaa tgg att tcg tcg atg tac ttg tca tga ttg gca tcg aaa cac tta tta agt tct ttt ttt caa ttc tac gat ttt att tct ttc gcg agt caa ttc cct cct gta gta act atc agt ttt gtc aga ttc acg ctc tct acg tag act ttc ttg taa gtt act aat ttg ttc cct ggc att acc gag ttc agt ttt ata tgc cga ata gag ttc tga ttc atc ctt tga gaa gat ctc tag cga tcg ttc aag atc cct gat tct agt ctt tag cct att tac ctc ctc aga aga tgc tcc gtt acc gtt --- ttt aca atc gtt aag atg tct atc aag atc cat gat tct atc tct ttt cca tat cag cat tga ttt cat tat tac gtt cgc agt cgt tca act g

>ON649879.1:133360-133807_Monkeypox_virus_isolate_MPXV_ISR001_2022_complete_genome

gct gct agt tgt cgt tcg aga tca gaa ata cgt gtt ttc ttg gaa tgg att tcg tcg atg tac ttg tca tga ttg gca tcg aaa cac tta tta agt tct ttt ttt caa ttc tac gat ttt att tct ttc gcg agt caa ttc cct cct gta gta act atc agt ttt gtc aga ttc acg ctc tct acg tag act ttc ttg taa gtt act aat ttg ttc cct ggc att acc gag ttc agt ttt ata tgc cga ata gag ttc tga ttc atc ctt tga gaa gat ctc tag cga tcg ttc aag atc cct gat tct agt ctt tag cct att tac ctc ctc aga aga tgc tcc gtt acc gtt --- ttt aca atc gtt aag atg tct atc aag atc cat gat tct atc tct ttt cca tat cag cat tga ttt cat tat tac gtt cgc agt cgt tca act g

>ON649725.1:133535-133982_Monkeypox_virus_isolate_Monkeypox/PT0011/2022_complete_genome

gct gct agt tgt cgt tcg aga tca gaa ata cgt gtt ttc ttg gaa tgg att tcg tcg atg tac ttg tca tga ttg gca tcg aaa cac tta tta agt tct ttt ttt caa ttc tac gat ttt att tct ttc gcg agt caa ttc cct cct gta gta act atc agt ttt gtc aga ttc acg ctc tct acg tag act ttc ttg taa gtt act aat ttg ttc cct ggc att acc gag ttc agt ttt ata tgc cga ata gag ttc tga ttc atc ctt tga gaa gat ctc tag cga tcg ttc aag atc cct gat tct agt ctt tag cct att tac ctc ctc aga aga tgc tcc gtt acc gtt --- ttt aca atc gtt aag atg tct atc aag atc cat gat tct atc tct ttt cca tat cag cat tga ttt cat tat tac gtt cgc agt cgt tca act g

>ON649724.1:133535-133982_Monkeypox_virus_isolate_Monkeypox/PT0014/2022_complete_genome

gct gct agt tgt cgt tcg aga tca gaa ata cgt gtt ttc ttg gaa tgg att tcg tcg atg tac ttg tca tga ttg gca tcg aaa cac tta tta agt tct ttt ttt caa ttc tac gat ttt att tct ttc gcg agt caa ttc cct cct gta gta act atc agt ttt gtc aga ttc acg ctc tct acg tag act ttc ttg taa gtt act aat ttg ttc cct ggc att acc gag ttc agt ttt ata tgc cga ata gag ttc tga ttc atc ctt tga gaa gat ctc tag cga tcg ttc aag atc cct gat tct agt ctt tag cct att tac ctc ctc aga aga tgc tcc gtt acc gtt --- ttt aca atc gtt aag atg tct atc aag atc cat gat tct atc tct ttt cca tat cag cat tga ttt cat tat tac gtt cgc agt cgt tca act g

>ON649723.1:133535-133982_Monkeypox_virus_isolate_Monkeypox/PT0013/2022_complete_genome

gct gct agt tgt cgt tcg aga tca gaa ata cgt gtt ttc ttg gaa tgg att tcg tcg atg tac ttg tca tga ttg gca tcg aaa cac tta tta agt tct ttt ttt caa ttc tac gat ttt att tct ttc gcg agt caa ttc cct cct gta gta act atc agt ttt gtc aga ttc acg ctc tct acg tag act ttc ttg taa gtt act aat ttg ttc cct ggc att acc gag ttc agt ttt ata tgc cga ata gag ttc tga ttc atc ctt tga gaa gat ctc tag cga tcg ttc aag atc cct gat tct agt ctt tag cct att tac ctc ctc aga aga tgc tcc gtt acc gtt --- ttt aca atc gtt aag atg tct atc aag atc cat gat tct atc tct ttt cca tat cag cat tga ttt cat tat tac gtt cgc agt cgt tca act g

>ON649722.1:133535-133982_Monkeypox_virus_isolate_Monkeypox/PT0022/2022_complete_genome

gct gct agt tgt cgt tcg aga tca gaa ata cgt gtt ttc ttg gaa tgg att tcg tcg atg tac ttg tca tga ttg gca tcg aaa cac tta tta agt tct ttt ttt caa ttc tac gat ttt att tct ttc gcg agt caa ttc cct cct gta gta act atc agt ttt gtc aga ttc acg ctc tct acg tag act ttc ttg taa gtt act aat ttg ttc cct ggc att acc gag ttc agt ttt ata tgc cga ata gag ttc tga ttc atc ctt tga gaa gat ctc tag cga tcg ttc aag atc cct gat tct agt ctt tag cct att tac ctc ctc aga aga tgc tcc gtt acc gtt --- ttt aca atc gtt aag atg tct atc aag atc cat gat tct atc tct ttt cca tat cag cat tga ttt cat tat tac gtt cgc agt cgt tca act g

>ON649721.1:133535-133982_Monkeypox_virus_isolate_Monkeypox/PT0021/2022_complete_genome

gct gct agt tgt cgt tcg aga tca gaa ata cgt gtt ttc ttg gaa tgg att tcg tcg atg tac ttg tca tga ttg gca tcg aaa cac tta tta agt tct ttt ttt caa ttc tac gat ttt att tct ttc gcg agt caa ttc cct cct gta gta act atc agt ttt gtc aga ttc acg ctc tct acg tag act ttc ttg taa gtt act aat ttg ttc cct ggc att acc gag ttc agt ttt ata tgc cga ata gag ttc tga ttc atc ctt tga gaa gat ctc tag cga tcg ttc aag atc cct gat tct agt ctt tag cct att tac ctc ctc aga aga tgc tcc gtt acc gtt --- ttt aca atc gtt aag atg tct atc aag atc cat gat tct atc tct ttt cca tat cag cat tga ttt cat tat tac gtt cgc agt cgt tca act g

>ON649720.1:133535-133982_Monkeypox_virus_isolate_Monkeypox/PT0024/2022_complete_genome

gct gct agt tgt cgt tcg aga tca gaa ata cgt gtt ttc ttg gaa tgg att tcg tcg atg tac ttg tca tga ttg gca tcg aaa cac tta tta agt tct ttt ttt caa ttc tac gat ttt att tct ttc gcg agt caa ttc cct cct gta gta act atc agt ttt gtc aga ttc acg ctc tct acg tag act ttc ttg taa gtt act aat ttg ttc cct ggc att acc gag ttc agt ttt ata tgc cga ata gag ttc tga ttc atc ctt tga gaa gat ctc tag cga tcg ttc aag atc cct gat tct agt ctt tag cct att tac ctc ctc aga aga tgc tcc gtt acc gtt --- ttt aca atc gtt aag atg tct atc aag atc cat gat tct atc tct ttt cca tat cag cat tga ttt cat tat tac gtt cgc agt cgt tca act g

>ON649719.1:133535-133982_Monkeypox_virus_isolate_Monkeypox/PT0012/2022_complete_genome

gct gct agt tgt cgt tcg aga tca gaa ata cgt gtt ttc ttg gaa tgg att tcg tcg atg tac ttg tca tga ttg gca tcg aaa cac tta tta agt tct ttt ttt caa ttc tac gat ttt att tct ttc gcg agt caa ttc cct cct gta gta act atc agt ttt gtc aga ttc acg ctc tct acg tag act ttc ttg taa gtt act aat ttg ttc cct ggc att acc gag ttc agt ttt ata tgc cga ata gag ttc tga ttc atc ctt tga gaa gat ctc tag cga tcg ttc aag atc cct gat tct agt ctt tag cct att tac ctc ctc aga aga tgc tcc gtt acc gtt --- ttt aca atc gtt aag atg tct atc aag atc cat gat tct atc tct ttt cca tat cag cat tga ttt cat tat tac gtt cgc agt cgt tca act g

>ON649718.1:133535-133982_Monkeypox_virus_isolate_Monkeypox/PT0018/2022_complete_genome

gct gct agt tgt cgt tcg aga tca gaa ata cgt gtt ttc ttg gaa tgg att tcg tcg atg tac ttg tca tga ttg gca tcg aaa cac tta tta agt tct ttt ttt caa ttc tac gat ttt att tct ttc gcg agt caa ttc cct cct gta gta act atc agt ttt gtc aga ttc acg ctc tct acg tag act ttc ttg taa gtt act aat ttg ttc cct ggc att acc gag ttc agt ttt ata tgc cga ata gag ttc tga ttc atc ctt tga gaa gat ctc tag cga tcg ttc aag atc cct gat tct agt ctt tag cct att tac ctc ctc aga aga tgc tcc gtt acc gtt --- ttt aca atc gtt aag atg tct atc aag atc cat gat tct atc tct ttt cca tat cag cat tga ttt cat tat tac gtt cgc agt cgt tca act g

>ON649717.1:133529-133976_Monkeypox_virus_isolate_Monkeypox/PT0019/2022_partial_genome

gct gct agt tgt cgt tcg aga tca gaa ata cgt gtt ttc ttg gaa tgg att tcg tcg atg tac ttg tca tga ttg gca tcg aaa cac tta tta agt tct ttt ttt caa ttc tac gat ttt att tct ttc gcg agt caa ttc cct cct gta gta act atc agt ttt gtc aga ttc acg ctc tct acg tag act ttc ttg taa gtt act aat ttg ttc cct ggc att acc gag ttc agt ttt ata tgc cga ata gag ttc tga ttc atc ctt tga gaa gat ctc tag cga tcg ttc aag atc cct gat tct agt ctt tag cct att tac ctc ctc aga aga tgc tcc gtt acc gtt --- ttt aca atc gtt aag atg tct atc aag atc cat gat tct atc tct ttt cca tat cag cat tga ttt cat tat tac gtt cgc agt cgt tca act g

>ON649716.1:133524-133971_Monkeypox_virus_isolate_Monkeypox/PT0016/2022_partial_genome

gct gct agt tgt cgt tcg aga tca gaa ata cgt gtt ttc ttg gaa tgg att tcg tcg atg tac ttg tca tga ttg gca tcg aaa cac tta tta agt tct ttt ttt caa ttc tac gat ttt att tct ttc gcg agt caa ttc cct cct gta gta act atc agt ttt gtc aga ttc acg ctc tct acg tag act ttc ttg taa gtt act aat ttg ttc cct ggc att acc gag ttc agt ttt ata tgc cga ata gag ttc tga ttc atc ctt tga gaa gat ctc tag cga tcg ttc aag atc cct gat tct agt ctt tag cct att tac ctc ctc aga aga tgc tcc gtt acc gtt --- ttt aca atc gtt aag atg tct atc aag atc cat gat tct atc tct ttt cca tat cag cat tga ttt cat tat tac gtt cgc agt cgt tca act g

>ON649715.1:133523-133970_Monkeypox_virus_isolate_Monkeypox/PT0017/2022_partial_genome

gct gct agt tgt cgt tcg aga tca gaa ata cgt gtt ttc ttg gaa tgg att tcg tcg atg tac ttg tca tga ttg gca tcg aaa cac tta tta agt tct ttt ttt caa ttc tac gat ttt att tct ttc gcg agt caa ttc cct cct gta gta act atc agt ttt gtc aga ttc acg ctc tct acg tag act ttc ttg taa gtt act aat ttg ttc cct ggc att acc gag ttc agt ttt ata tgc cga ata gag ttc tga ttc atc ctt tga gaa gat ctc tag cga tcg ttc aag atc cct gat tct agt ctt tag cct att tac ctc ctc aga aga tgc tcc gtt acc gtt --- ttt aca atc gtt aag atg tct atc aag atc cat gat tct atc tct ttt cca tat cag cat tga ttt cat tat tac gtt cgc agt cgt tca act g

>ON649714.1:133524-133971_Monkeypox_virus_isolate_Monkeypox/PT0015/2022_partial_genome

gct gct agt tgt cgt tcg aga tca gaa ata cgt gtt ttc ttg gaa tgg att tcg tcg atg tac ttg tca tga ttg gca tcg aaa cac tta tta agt tct ttt ttt caa ttc tac gat ttt att tct ttc gcg agt caa ttc cct cct gta gta act atc agt ttt gtc aga ttc acg ctc tct acg tag act ttc ttg taa gtt act aat ttg ttc cct ggc att acc gag ttc agt ttt ata tgc cga ata gag ttc tga ttc atc ctt tga gaa gat ctc tag cga tcg ttc aag atc cct gat tct agt ctt tag cct att tac ctc ctc aga aga tgc tcc gtt acc gtt --- ttt aca atc gtt aag atg tct atc aag atc cat gat tct atc tct ttt cca tat cag cat tga ttt cat tat tac gtt cgc agt cgt tca act g

>ON649713.1:132622-133069_Monkeypox_virus_isolate_Monkeypox/PT0020/2022_complete_genome

gct gct agt tgt cgt tcg aga tca gaa ata cgt gtt ttc ttg gaa tgg att tcg tcg atg tac ttg tca tga ttg gca tcg aaa cac tta tta agt tct ttt ttt caa ttc tac gat ttt att tct ttc gcg agt caa ttc cct cct gta gta act atc agt ttt gtc aga ttc acg ctc tct acg tag act ttc ttg taa gtt act aat ttg ttc cct ggc att acc gag ttc agt ttt ata tgc cga ata gag ttc tga ttc atc ctt tga gaa gat ctc tag cga tcg ttc aag atc cct gat tct agt ctt tag cct att tac ctc ctc aga aga tgc tcc gtt acc gtt --- ttt aca atc gtt aag atg tct atc aag atc cat gat tct atc tct ttt cca tat cag cat tga ttt cat tat tac gtt cgc agt cgt tca act g

>ON649712.1:133529-133976_Monkeypox_virus_isolate_Monkeypox/PT0025/2022_partial_genome

gct gct agt tgt cgt tcg aga tca gaa ata cgt gtt ttc ttg gaa tgg att tcg tcg atg tac ttg tca tga ttg gca tcg aaa cac tta tta agt tct ttt ttt caa ttc tac gat ttt att tct ttc gcg agt caa ttc cct cct gta gta act atc agt ttt gtc aga ttc acg ctc tct acg tag act ttc ttg taa gtt act aat ttg ttc cct ggc att acc gag ttc agt ttt ata tgc cga ata gag ttc tga ttc atc ctt tga gaa gat ctc tag cga tcg ttc aag atc cct gat tct agt ctt tag cct att tac ctc ctc aga aga tgc tcc gtt acc gtt --- ttt aca atc gtt aag atg tct atc aag atc cat gat tct atc tct ttt cca tat cag cat tga ttt cat tat tac gtt cgc agt cgt tca act g

>ON649711.1:133516-133963_Monkeypox_virus_isolate_Monkeypox/PT0026/2022_partial_genome

gct gct agt tgt cgt tcg aga tca gaa ata cgt gtt ttc ttg gaa tgg att tcg tcg atg tac ttg tca tga ttg gca tcg aaa cac tta tta agt tct ttt ttt caa ttc tac gat ttt att tct ttc gcg agt caa ttc cct cct gta gta act atc agt ttt gtc aga ttc acg ctc tct acg tag act ttc ttg taa gtt act aat ttg ttc cct ggc att acc gag ttc agt ttt ata tgc cga ata gag ttc tga ttc atc ctt tga gaa gat ctc tag cga tcg ttc aag atc cct gat tct agt ctt tag cct att tac ctc ctc aga aga tgc tcc gtt acc gtt --- ttt aca atc gtt aag atg tct atc aag atc cat gat tct atc tct ttt cca tat cag cat tga ttt cat tat tac gtt cgc agt cgt tca act g

>ON649710.1:133529-133976_Monkeypox_virus_isolate_Monkeypox/PT0027/2022_partial_genome

gct gct agt tgt cgt tcg aga tca gaa ata cgt gtt ttc ttg gaa tgg att tcg tcg atg tac ttg tca tga ttg gca tcg aaa cac tta tta agt tct ttt ttt caa ttc tac gat ttt att tct ttc gcg agt caa ttc cct cct gta gta act atc agt ttt gtc aga ttc acg ctc tct acg tag act ttc ttg taa gtt act aat ttg ttc cct ggc att acc gag ttc agt ttt ata tgc cga ata gag ttc tga ttc atc ctt tga gaa gat ctc tag cga tcg ttc aag atc cct gat tct agt ctt tag cct att tac ctc ctc aga aga tgc tcc gtt acc gtt --- ttt aca atc gtt aag atg tct atc aag atc cat gat tct atc tct ttt cca tat cag cat tga ttt cat tat tac gtt cgc agt cgt tca act g

>ON649709.1:133535-133982_Monkeypox_virus_isolate_Monkeypox/PT0028/2022_complete_genome

gct gct agt tgt cgt tcg aga tca gaa ata cgt gtt ttc ttg gaa tgg att tcg tcg atg tac ttg tca tga ttg gca tcg aaa cac tta tta agt tct ttt ttt caa ttc tac gat ttt att tct ttc gcg agt caa ttc cct cct gta gta act atc agt ttt gtc aga ttc acg ctc tct acg tag act ttc ttg taa gtt act aat ttg ttc cct ggc att acc gag ttc agt ttt ata tgc cga ata gag ttc tga ttc atc ctt tga gaa gat ctc tag cga tcg ttc aag atc cct gat tct agt ctt tag cct att tac ctc ctc aga aga tgc tcc gtt acc gtt --- ttt aca atc gtt aag atg tct atc aag atc cat gat tct atc tct ttt cca tat cag cat tga ttt cat tat tac gtt cgc agt cgt tca act g

>ON649708.1:133529-133976_Monkeypox_virus_isolate_Monkeypox/PT0023/2022_partial_genome

gct gct agt tgt cgt tcg aga tca gaa ata cgt gtt ttc ttg gaa tgg att tcg tcg atg tac ttg tca tga ttg gca tcg aaa cac tta tta agt tct ttt ttt caa ttc tac gat ttt att tct ttc gcg agt caa ttc cct cct gta gta act atc agt ttt gtc aga ttc acg ctc tct acg tag act ttc ttg taa gtt act aat ttg ttc cct ggc att acc gag ttc agt ttt ata tgc cga ata gag ttc tga ttc atc ctt tga gaa gat ctc tag cga tcg ttc aag atc cct gat tct agt ctt tag cct att tac ctc ctc aga aga tgc tcc gtt acc gtt --- ttt aca atc gtt aag atg tct atc aag atc cat gat tct atc tct ttt cca tat cag cat tga ttt cat tat tac gtt cgc agt cgt tca act g

>ON622722.2:133528-133975_Monkeypox_virus_isolate_MPXV_FR_HCL0001_2022_complete_genome

gct gct agt tgt cgt tcg aga tca gaa ata cgt gtt ttc ttg gaa tgg att tcg tcg atg tac ttg tca tga ttg gca tcg aaa cac tta tta agt tct ttt ttt caa ttc tac gat ttt att tct ttc gcg agt caa ttc cct cct gta gta act atc agt ttt gtc aga ttc acg ctc tct acg tag act ttc ttg taa gtt act aat ttg ttc cct ggc att acc gag ttc agt ttt ata tgc cga ata gag ttc tga ttc atc ctt tga gaa gat ctc tag cga tcg ttc aag atc cct gat tct agt ctt tag cct att tac ctc ctc aga aga tgc tcc gtt acc gtt --- ttt aca atc gtt aag atg tct atc aag atc cat gat tct atc tct ttt cca tat cag cat tga ttt cat tat tac gtt cgc agt cgt tca act g

>ON644344.1:133749-134196_Monkeypox_virus_isolate_MPXV_FVG-ITA_01_2022_complete_genome

gct gct agt tgt cgt tcg aga tca gaa ata cgt gtt ttc ttg gaa tgg att tcg tcg atg tac ttg tca tga ttg gca tcg aaa cac tta tta agt tct ttt ttt caa ttc tac gat ttt att tct ttc gcg agt caa ttc cct cct gta gta act atc agt ttt gtc aga ttc acg ctc tct acg tag act ttc ttg taa gtt act aat ttg ttc cct ggc att acc gag ttc agt ttt ata tgc cga ata gag ttc tga ttc atc ctt tga gaa gat ctc tag cga tcg ttc aag atc cct gat tct agt ctt tag cct att tac ctc ctc aga aga tgc tcc gtt acc gtt --- ttt aca atc gtt aag atg tct atc aag atc cat gat tct atc tct ttt cca tat cag cat tga ttt cat tat tac gtt cgc agt cgt tca act g

>ON595760.2:133435-133882_Monkeypox_virus_isolate_MPXV-CH-38134631/2022_partial_genome

gct gct agt tgt cgt tcg aga tca gaa ata cgt gtt ttc ttg gaa tgg att tcg tcg atg tac ttg tca tga ttg gca tcg aaa cac tta tta agt tct ttt ttt caa ttc tac gat ttt att tct ttc gcg agt caa ttc cct cct gta gta act atc agt ttt gtc aga ttc acg ctc tct acg tag act ttc ttg taa gtt act aat ttg ttc cct ggc att acc gag ttc agt ttt ata tgc cga ata gag ttc tga ttc atc ctt tga gaa gat ctc tag cga tcg ttc aag atc cct gat tct agt ctt tag cct att tac ctc ctc aga aga tgc tcc gtt acc gtt --- ttt aca atc gtt aag atg tct atc aag atc cat gat tct atc tct ttt cca tat cag cat tga ttt cat tat tac gtt cgc agt cgt tca act g

>ON637939.1:133592-134039_Monkeypox_virus_isolate_MPXV/Germany/2022/RKI02_complete_genome

gct gct agt tgt cgt tcg aga tca gaa ata cgt gtt ttc ttg gaa tgg att tcg tcg atg tac ttg tca tga ttg gca tcg aaa cac tta tta agt tct ttt ttt caa ttc tac gat ttt att tct ttc gcg agt caa ttc cct cct gta gta act atc agt ttt gtc aga ttc acg ctc tct acg tag act ttc ttg taa gtt act aat ttg ttc cct ggc att acc gag ttc agt ttt ata tgc cga ata gag ttc tga ttc atc ctt tga gaa gat ctc tag cga tcg ttc aag atc cct gat tct agt ctt tag cct att tac ctc ctc aga aga tgc tcc gtt acc gtt --- ttt aca atc gtt aag atg tct atc aag atc cat gat tct atc tct ttt cca tat cag cat tga ttt cat tat tac gtt cgc agt cgt tca act g

>ON637938.1:133592-134039_Monkeypox_virus_isolate_MPXV/Germany/2022/RKI01_complete_genome

gct gct agt tgt cgt tcg aga tca gaa ata cgt gtt ttc ttg gaa tgg att tcg tcg atg tac ttg tca tga ttg gca tcg aaa cac tta tta agt tct ttt ttt caa ttc tac gat ttt att tct ttc gcg agt caa ttc cct cct gta gta act atc agt ttt gtc aga ttc acg ctc tct acg tag act ttc ttg taa gtt act aat ttg ttc cct ggc att acc gag ttc agt ttt ata tgc cga ata gag ttc tga ttc atc ctt tga gaa gat ctc tag cga tcg ttc aag atc cct gat tct agt ctt tag cct att tac ctc ctc aga aga tgc tcc gtt acc gtt --- ttt aca atc gtt aag atg tct atc aag atc cat gat tct atc tct ttt cca tat cag cat tga ttt cat tat tac gtt cgc agt cgt tca act g

>ON631963.1:133554-134001_Monkeypox_virus_isolate_MPxV/VIDRL01/2022_complete_genome

gct gct agt tgt cgt tcg aga tca gaa ata cgt gtt ttc ttg gaa tgg att tcg tcg atg tac ttg tca tga ttg gca tcg aaa cac tta tta agt tct ttt ttt caa ttc tac gat ttt att tct ttc gcg agt caa ttc cct cct gta gta act atc agt ttt gtc aga ttc acg ctc tct acg tag act ttc ttg taa gtt act aat ttg ttc cct ggc att acc gag ttc agt ttt ata tgc cga ata gag ttc tga ttc atc ctt tga gaa gat ctc tag cga tcg ttc aag atc cct gat tct agt ctt tag cct att tac ctc ctc aga aga tgc tcc gtt acc gtt --- ttt aca atc gtt aag atg tct atc aag atc cat gat tct atc tct ttt cca tat cag cat tga ttt cat tat tac gtt cgc agt cgt tca act g

>ON609725.2:133580-134027_Monkeypox_virus_isolate_SLO_complete_genome

gct gct agt tgt cgt tcg aga tca gaa ata cgt gtt ttc ttg gaa tgg att tcg tcg atg tac ttg tca tga ttg gca tcg aaa cac tta tta agt tct ttt ttt caa ttc tac gat ttt att tct ttc gcg agt caa ttc cct cct gta gta act atc agt ttt gtc aga ttc acg ctc tct acg tag act ttc ttg taa gtt act aat ttg ttc cct ggc att acc gag ttc agt ttt ata tgc cga ata gag ttc tga ttc atc ctt tga gaa gat ctc tag cga tcg ttc aag atc cct gat tct agt ctt tag cct att tac ctc ctc aga aga tgc tcc gtt acc gtt --- ttt aca atc gtt aag atg tct atc aag atc cat gat tct atc tct ttt cca tat cag cat tga ttt cat tat tac gtt cgc agt cgt tca act g

>ON631241.1:133580-134027_Monkeypox_virus_isolate_2022/2_SLO_complete_genome

gct gct agt tgt cgt tcg aga tca gaa ata cgt gtt ttc ttg gaa tgg att tcg tcg atg tac ttg tca tga ttg gca tcg aaa cac tta tta agt tct ttt ttt caa ttc tac gat ttt att tct ttc gcg agt caa ttc cct cct gta gta act atc agt ttt gtc aga ttc acg ctc tct acg tag act ttc ttg taa gtt act aat ttg ttc cct ggc att acc gag ttc agt ttt ata tgc cga ata gag ttc tga ttc atc ctt tga gaa gat ctc tag cga tcg ttc aag atc cct gat tct agt ctt tag cct att tac ctc ctc aga aga tgc tcc gtt acc gtt --- ttt aca atc gtt aag atg tct atc aag atc cat gat tct atc tct ttt cca tat cag cat tga ttt cat tat tac gtt cgc agt cgt tca act g

>ON563414.3:133545-133992_Monkeypox_virus_isolate_MPXV_USA_2022_MA001_complete_genome

gct gct agt tgt cgt tcg aga tca gaa ata cgt gtt ttc ttg gaa tgg att tcg tcg atg tac ttg tca tga ttg gca tcg aaa cac tta tta agt tct ttt ttt caa ttc tac gat ttt att tct ttc gcg agt caa ttc cct cct gta gta act atc agt ttt gtc aga ttc acg ctc tct acg tag act ttc ttg taa gtt act aat ttg ttc cct ggc att acc gag ttc agt ttt ata tgc cga ata gag ttc tga ttc atc ctt tga gaa gat ctc tag cga tcg ttc aag atc cct gat tct agt ctt tag cct att tac ctc ctc aga aga tgc tcc gtt acc gtt --- ttt aca atc gtt aag atg tct atc aag atc cat gat tct atc tct ttt cca tat cag cat tga ttt cat tat tac gtt cgc agt cgt tca act g

>ON627808.1:133532-133979_Monkeypox_virus_isolate_MPX/human/USA/UT-UPHL-82200022/2022_complete_genome

gct gct agt tgt cgt tcg aga tca gaa ata cgt gtt ttc ttg gaa tgg att tcg tcg atg tac ttg tca tga ttg gca tcg aaa cac tta tta agt tct ttt ttt caa ttc tac gat ttt att tct ttc gcg agt caa ttc cct cct gta gta act atc agt ttt gtc aga ttc acg ctc tct acg tag act ttc ttg taa gtt act aat ttg ttc cct ggc att acc gag ttc agt ttt ata tgc cga ata gag ttc tga ttc atc ctt tga gaa gat ctc tag cga tcg ttc aag atc cct gat tct agt ctt tag cct att tac ctc ctc aga aga tgc tcc gtt acc gtt --- ttt aca atc gtt aag atg tct atc aag atc cat gat tct atc tct ttt cca tat cag cat tga ttt cat tat tac gtt cgc agt cgt tca act g

>ON622718.1:133016-133463_Monkeypox_virus_isolate_MPXV/ES0001/HUGTiP/2022_partial_genome

gct gct agt tgt cgt tcg aga tca gaa ata cgt gtt ttc ttg gaa tgg att tcg tcg atg tac ttg tca tga ttg gca tcg aaa cac tta tta agt tct ttt ttt caa ttc tac gat ttt att tct ttc gcg agt caa ttc cct cct gta gta act atc agt ttt gtc aga ttc acg ctc tct acg tag act ttc ttg taa gtt act aat ttg ttc cct ggc att acc gag ttc agt ttt ata tgc cga ata gag ttc tga ttc atc ctt tga gaa gat ctc tag cga tcg ttc aag atc cct gat tct agt ctt tag cct att tac ctc ctc aga aga tgc tcc gtt acc gtt --- ttt aca atc gtt aag atg tct atc aag atc cat gat tct atc tct ttt cca tat cag cat tga ttt cat tat tac gtt cgc agt cgt tca act g

>ON622713.1:133757-134204_Monkeypox_virus_isolate_MPX/UZ_REGA_2/Belgium/2022_complete_genome

gct gct agt tgt cgt tcg aga tca gaa ata cgt gtt ttc ttg gaa tgg att tcg tcg atg tac ttg tca tga ttg gca tcg aaa cac tta tta agt tct ttt ttt caa ttc tac gat ttt att tct ttc gcg agt caa ttc cct cct gta gta act atc agt ttt gtc aga ttc acg ctc tct acg tag act ttc ttg taa gtt act aat ttg ttc cct ggc att acc gag ttc agt ttt ata tgc cga ata gag ttc tga ttc atc ctt tga gaa gat ctc tag cga tcg ttc aag atc cct gat tct agt ctt tag cct att tac ctc ctc aga aga tgc tcc gtt acc gtt --- ttt aca atc gtt aag atg tct atc aag atc cat gat tct atc tct ttt cca tat cag cat tga ttt cat tat tac gtt cgc agt cgt tca act g

>ON622712.1:133757-134204_Monkeypox_virus_isolate_MPX/UZ_REGA_1/Belgium/2022_complete_genome

gct gct agt tgt cgt tcg aga tca gaa ata cgt gtt ttc ttg gaa tgg att tcg tcg atg tac ttg tca tga ttg gca tcg aaa cac tta tta agt tct ttt ttt caa ttc tac gat ttt att tct ttc gcg agt caa ttc cct cct gta gta act atc agt ttt gtc aga ttc acg ctc tct acg tag act ttc ttg taa gtt act aat ttg ttc cct ggc att acc gag ttc agt ttt ata tgc cga ata gag ttc tga ttc atc ctt tga gaa gat ctc tag cga tcg ttc aag atc cct gat tct agt ctt tag cct att tac ctc ctc aga aga tgc tcc gtt acc gtt --- ttt aca atc gtt aag atg tct atc aag atc cat gat tct atc tct ttt cca tat cag cat tga ttt cat tat tac gtt cgc agt cgt tca act g

>ON619838.1:133537-133984_Monkeypox_virus_isolate_MPXV_UK_2022_4_complete_genome

gct gct agt tgt cgt tcg aga tca gaa ata cgt gtt ttc ttg gaa tgg att tcg tcg atg tac ttg tca tga ttg gca tcg aaa cac tta tta agt tct ttt ttt caa ttc tac gat ttt att tct ttc gcg agt caa ttc cct cct gta gta act atc agt ttt gtc aga ttc acg ctc tct acg tag act ttc ttg taa gtt act aat ttg ttc cct ggc att acc gag ttc agt ttt ata tgc cga ata gag ttc tga ttc atc ctt tga gaa gat ctc tag cga tcg ttc aag atc cct gat tct agt ctt tag cct att tac ctc ctc aga aga tgc tcc gtt acc gtt --- ttt aca atc gtt aag atg tct atc aag atc cat gat tct atc tct ttt cca tat cag cat tga ttt cat tat tac gtt cgc agt cgt tca act g

>ON619836.1:133543-133990_Monkeypox_virus_isolate_MPXV_UK_2022_2_complete_genome

gct gct agt tgt cgt tcg aga tca gaa ata cgt gtt ttc ttg gaa tgg att tcg tcg atg tac ttg tca tga ttg gca tcg aaa cac tta tta agt tct ttt ttt caa ttc tac gat ttt att tct ttc gcg agt caa ttc cct cct gta gta act atc agt ttt gtc aga ttc acg ctc tct acg tag act ttc ttg taa gtt act aat ttg ttc cct ggc att acc gag ttc agt ttt ata tgc cga ata gag ttc tga ttc atc ctt tga gaa gat ctc tag cga tcg ttc aag atc cct gat tct agt ctt tag cct att tac ctc ctc aga aga tgc tcc gtt acc gtt --- ttt aca atc gtt aag atg tct atc aag atc cat gat tct atc tct ttt cca tat cag cat tga ttt cat tat tac gtt cgc agt cgt tca act g

>ON619835.1:133537-133984_Monkeypox_virus_isolate_MPXV_UK_2022_1_complete_genome

gct gct agt tgt cgt tcg aga tca gaa ata cgt gtt ttc ttg gaa tgg att tcg tcg atg tac ttg tca tga ttg gca tcg aaa cac tta tta agt tct ttt ttt caa ttc tac gat ttt att tct ttc gcg agt caa ttc cct cct gta gta act atc agt ttt gtc aga ttc acg ctc tct acg tag act ttc ttg taa gtt act aat ttg ttc cct ggc att acc gag ttc agt ttt ata tgc cga ata gag ttc tga ttc atc ctt tga gaa gat ctc tag cga tcg ttc aag atc cct gat tct agt ctt tag cct att tac ctc ctc aga aga tgc tcc gtt acc gtt --- ttt aca atc gtt aag atg tct atc aag atc cat gat tct atc tct ttt cca tat cag cat tga ttt cat tat tac gtt cgc agt cgt tca act g

>ON615424.1:133521-133968_Monkeypox_virus_isolate_MPXV_2022_NL001_partial_genome

gct gct agt tgt cgt tcg aga tca gaa ata cgt gtt ttc ttg gaa tgg att tcg tcg atg tac ttg tca tga ttg gca tcg aaa cac tta tta agt tct ttt ttt caa ttc tac gat ttt att tct ttc gcg agt caa ttc cct cct gta gta act atc agt ttt gtc aga ttc acg ctc tct acg tag act ttc ttg taa gtt act aat ttg ttc cct ggc att acc gag ttc agt ttt ata tgc cga ata gag ttc tga ttc atc ctt tga gaa gat ctc tag cga tcg ttc aag atc cct gat tct agt ctt tag cct att tac ctc ctc aga aga tgc tcc gtt acc gtt --- ttt aca atc gtt aag atg tct atc aag atc cat gat tct atc tct ttt cca tat cag cat tga ttt cat tat tac gtt cgc agt cgt tca act g

>ON614676.1:133527-133974_Monkeypox_virus_isolate_INMI-Pt1_partial_genome

gct gct agt tgt cgt tcg aga tca gaa ata cgt gtt ttc ttg gaa tgg att tcg tcg atg tac ttg tca tga ttg gca tcg aaa cac tta tta agt tct ttt ttt caa ttc tac gat ttt att tct ttc gcg agt caa ttc cct cct gta gta act atc agt ttt gtc aga ttc acg ctc tct acg tag act ttc ttg taa gtt act aat ttg ttc cct ggc att acc gag ttc agt ttt ata tgc cga ata gag ttc tga ttc atc ctt tga gaa gat ctc tag cga tcg ttc aag atc cct gat tct agt ctt tag cct att tac ctc ctc aga aga tgc tcc gtt acc gtt --- ttt aca atc gtt aag atg tct atc aag atc cat gat tct atc tct ttt cca tat cag cat tga ttt cat tat tac gtt cgc agt cgt tca act g

>ON585038.1:132621-133068_Monkeypox_virus_isolate_Monkeypox/PT0008/2022_complete_genome

gct gct agt tgt cgt tcg aga tca gaa ata cgt gtt ttc ttg gaa tgg att tcg tcg atg tac ttg tca tga ttg gca tcg aaa cac tta tta agt tct ttt ttt caa ttc tac gat ttt att tct ttc gcg agt caa ttc cct cct gta gta act atc agt ttt gtc aga ttc acg ctc tct acg tag act ttc ttg taa gtt act aat ttg ttc cct ggc att acc gag ttc agt ttt ata tgc cga ata gag ttc tga ttc atc ctt tga gaa gat ctc tag cga tcg ttc aag atc cct gat tct agt ctt tag cct att tac ctc ctc aga aga tgc tcc gtt acc gtt --- ttt aca atc gtt aag atg tct atc aag atc cat gat tct atc tct ttt cca tat cag cat tga ttt cat tat tac gtt cgc agt cgt tca act g

>ON585037.1:132616-133063_Monkeypox_virus_isolate_Monkeypox/PT0005/2022_complete_genome

gct gct agt tgt cgt tcg aga tca gaa ata cgt gtt ttc ttg gaa tgg att tcg tcg atg tac ttg tca tga ttg gca tcg aaa cac tta tta agt tct ttt ttt caa ttc tac gat ttt att tct ttc gcg agt caa ttc cct cct gta gta act atc agt ttt gtc aga ttc acg ctc tct acg tag act ttc ttg taa gtt act aat ttg ttc cct ggc att acc gag ttc agt ttt ata tgc cga ata gag ttc tga ttc atc ctt tga gaa gat ctc tag cga tcg ttc aag atc cct gat tct agt ctt tag cct att tac ctc ctc aga aga tgc tcc gtt acc gtt --- ttt aca atc gtt aag atg tct atc aag atc cat gat tct atc tct ttt cca tat cag cat tga ttt cat tat tac gtt cgc agt cgt tca act g

>ON585036.1:133524-133971_Monkeypox_virus_isolate_Monkeypox/PT0010/2022_partial_genome

gct gct agt tgt cgt tcg aga tca gaa ata cgt gtt ttc ttg gaa tgg att tcg tcg atg tac ttg tca tga ttg gca tcg aaa cac tta tta agt tct ttt ttt caa ttc tac gat ttt att tct ttc gcg agt caa ttc cct cct gta gta act atc agt ttt gtc aga ttc acg ctc tct acg tag act ttc ttg taa gtt act aat ttg ttc cct ggc att acc gag ttc agt ttt ata tgc cga ata gag ttc tga ttc atc ctt tga gaa gat ctc tag cga tcg ttc aag atc cct gat tct agt ctt tag cct att tac ctc ctc aga aga tgc tcc gtt acc gtt --- ttt aca atc gtt aag atg tct atc aag atc cat gat tct atc tct ttt cca tat cag cat tga ttt cat tat tac gtt cgc agt cgt tca act g

>ON585035.1:133535-133982_Monkeypox_virus_isolate_Monkeypox/PT0009/2022_complete_genome

gct gct agt tgt cgt tcg aga tca gaa ata cgt gtt ttc ttg gaa tgg att tcg tcg atg tac ttg tca tga ttg gca tcg aaa cac tta tta agt tct ttt ttt caa ttc tac gat ttt att tct ttc gcg agt caa ttc cct cct gta gta act atc agt ttt gtc aga ttc acg ctc tct acg tag act ttc ttg taa gtt act aat ttg ttc cct ggc att acc gag ttc agt ttt ata tgc cga ata gag ttc tga ttc atc ctt tga gaa gat ctc tag cga tcg ttc aag atc cct gat tct agt ctt tag cct att tac ctc ctc aga aga tgc tcc gtt acc gtt --- ttt aca atc gtt aag atg tct atc aag atc cat gat tct atc tct ttt cca tat cag cat tga ttt cat tat tac gtt cgc agt cgt tca act g

>ON585034.1:133529-133976_Monkeypox_virus_isolate_Monkeypox/PT0007/2022_complete_genome

gct gct agt tgt cgt tcg aga tca gaa ata cgt gtt ttc ttg gaa tgg att tcg tcg atg tac ttg tca tga ttg gca tcg aaa cac tta tta agt tct ttt ttt caa ttc tac gat ttt att tct ttc gcg agt caa ttc cct cct gta gta act atc agt ttt gtc aga ttc acg ctc tct acg tag act ttc ttg taa gtt act aat ttg ttc cct ggc att acc gag ttc agt ttt ata tgc cga ata gag ttc tga ttc atc ctt tga gaa gat ctc tag cga tcg ttc aag atc cct gat tct agt ctt tag cct att tac ctc ctc aga aga tgc tcc gtt acc gtt --- ttt aca atc gtt aag atg tct atc aag atc cat gat tct atc tct ttt cca tat cag cat tga ttt cat tat tac gtt cgc agt cgt tca act g

>ON585033.1:133535-133982_Monkeypox_virus_isolate_Monkeypox/PT0006/2022_complete_genome

gct gct agt tgt cgt tcg aga tca gaa ata cgt gtt ttc ttg gaa tgg att tcg tcg atg tac ttg tca tga ttg gca tcg aaa cac tta tta agt tct ttt ttt caa ttc tac gat ttt att tct ttc gcg agt caa ttc cct cct gta gta act atc agt ttt gtc aga ttc acg ctc tct acg tag act ttc ttg taa gtt act aat ttg ttc cct ggc att acc gag ttc agt ttt ata tgc cga ata gag ttc tga ttc atc ctt tga gaa gat ctc tag cga tcg ttc aag atc cct gat tct agt ctt tag cct att tac ctc ctc aga aga tgc tcc gtt acc gtt --- ttt aca atc gtt aag atg tct atc aag atc cat gat tct atc tct ttt cca tat cag cat tga ttt cat tat tac gtt cgc agt cgt tca act g

>ON585032.1:133529-133976_Monkeypox_virus_isolate_Monkeypox/PT0004/2022_complete_genome

gct gct agt tgt cgt tcg aga tca gaa ata cgt gtt ttc ttg gaa tgg att tcg tcg atg tac ttg tca tga ttg gca tcg aaa cac tta tta agt tct ttt ttt caa ttc tac gat ttt att tct ttc gcg agt caa ttc cct cct gta gta act atc agt ttt gtc aga ttc acg ctc tct acg tag act ttc ttg taa gtt act aat ttg ttc cct ggc att acc gag ttc agt ttt ata tgc cga ata gag ttc tga ttc atc ctt tga gaa gat ctc tag cga tcg ttc aag atc cct gat tct agt ctt tag cct att tac ctc ctc aga aga tgc tcc gtt acc gtt --- ttt aca atc gtt aag atg tct atc aag atc cat gat tct atc tct ttt cca tat cag cat tga ttt cat tat tac gtt cgc agt cgt tca act g

>ON585031.1:133529-133976_Monkeypox_virus_isolate_Monkeypox/PT0003/2022_complete_genome

gct gct agt tgt cgt tcg aga tca gaa ata cgt gtt ttc ttg gaa tgg att tcg tcg atg tac ttg tca tga ttg gca tcg aaa cac tta tta agt tct ttt ttt caa ttc tac gat ttt att tct ttc gcg agt caa ttc cct cct gta gta act atc agt ttt gtc aga ttc acg ctc tct acg tag act ttc ttg taa gtt act aat ttg ttc cct ggc att acc gag ttc agt ttt ata tgc cga ata gag ttc tga ttc atc ctt tga gaa gat ctc tag cga tcg ttc aag atc cct gat tct agt ctt tag cct att tac ctc ctc aga aga tgc tcc gtt acc gtt --- ttt aca atc gtt aag atg tct atc aag atc cat gat tct atc tct ttt cca tat cag cat tga ttt cat tat tac gtt cgc agt cgt tca act g

>ON585030.1:133529-133976_Monkeypox_virus_isolate_Monkeypox/PT0002/2022_partial_genome

gct gct agt tgt cgt tcg aga tca gaa ata cgt gtt ttc ttg gaa tgg att tcg tcg atg tac ttg tca tga ttg gca tcg aaa cac tta tta agt tct ttt ttt caa ttc tac gat ttt att tct ttc gcg agt caa ttc cct cct gta gta act atc agt ttt gtc aga ttc acg ctc tct acg tag act ttc ttg taa gtt act aat ttg ttc cct ggc att acc gag ttc agt ttt ata tgc cga ata gag ttc tga ttc atc ctt tga gaa gat ctc tag cga tcg ttc aag atc cct gat tct agt ctt tag cct att tac ctc ctc aga aga tgc tcc gtt acc gtt --- ttt aca atc gtt aag atg tct atc aag atc cat gat tct atc tct ttt cca tat cag cat tga ttt cat tat tac gtt cgc agt cgt tca act g

>ON568298.1:133676-134123_Monkeypox_virus_isolate_MPXV-BY-IMB25241_complete_genome

gct gct agt tgt cgt tcg aga tca gaa ata cgt gtt ttc ttg gaa tgg att tcg tcg atg tac ttg tca tga ttg gca tcg aaa cac tta tta agt tct ttt ttt caa ttc tac gat ttt att tct ttc gcg agt caa ttc cct cct gta gta act atc agt ttt gtc aga ttc acg ctc tct acg tag act ttc ttg taa gtt act aat ttg ttc cct ggc att acc gag ttc agt ttt ata tgc cga ata gag ttc tga ttc atc ctt tga gaa gat ctc tag cga tcg ttc aag atc cct gat tct agt ctt tag cct att tac ctc ctc aga aga tgc tcc gtt acc gtt --- ttt aca atc gtt aag atg tct atc aag atc cat gat tct atc tct ttt cca tat cag cat tga ttt cat tat tac gtt cgc agt cgt tca act g

>MT903345.1:133560-134007_Monkeypox_virus_isolate_MPXV-UK_P3

gct gct agt tgt cgt tcg aga tca gaa ata cgt gtt ttc ttg gaa tgg att tcg tcg atg tac ttg tca tga ttg gca tcg aaa cac tta tta agt tct ttt ttt caa ttc tac gat ttt att tct ttc gcg agt caa ttc cct cct gta gta act atc agt ttt gtc aga ttc acg ctc tct acg tag act ttc ttg taa gtt act aat ttg ttc cct ggc att acc gag ttc agt ttt ata tgc cga ata gag ttc tga ttc atc ctt tga gaa gat ctc tag cga tcg ttc aag atc cct gat tct agt ctt tag cct att tac ctc ctc aga aga tgc tcc gtt acc gtt --- ttt aca atc gtt aag atg tct atc aag atc cat gat tct atc tct ttt cca tat cag cat tga ttt cat tat tac gtt cgc agt cgt tca act g

>MT903344.1:133546-133993_Monkeypox_virus_isolate_MPXV-UK_P2

gct gct agt tgt cgt tcg aga tca gaa ata cgt gtt ttc ttg gaa tgg att tcg tcg atg tac ttg tca tga ttg gca tcg aaa cac tta tta agt tct ttt ttt caa ttc tac gat ttt att tct ttc gcg agt caa ttc cct cct gta gta act atc agt ttt gtc aga ttc acg ctc tct acg tag act ttc ttg taa gtt act aat ttg ttc cct ggc att acc gag ttc agt ttt ata tgc cga ata gag ttc tga ttc atc ctt tga gaa gat ctc tag cga tcg ttc aag atc cct gat tct agt ctt tag cct att tac ctc ctc aga aga tgc tcc gtt acc gtt --- ttt aca atc gtt aag atg tct atc aag atc cat gat tct atc tct ttt cca tat cag cat tga ttt cat tat tac gtt cgc agt cgt tca act g

>MT903343.1:133540-133987_Monkeypox_virus_isolate_MPXV-UK_P1

gct gct agt tgt cgt tcg aga tca gaa ata cgt gtt ttc ttg gaa tgg att tcg tcg atg tac ttg tca tga ttg gca tcg aaa cac tta tta agt tct ttt ttt caa ttc tac gat ttt att tct ttc gcg agt caa ttc cct cct gta gta act atc agt ttt gtc aga ttc acg ctc tct acg tag act ttc ttg taa gtt act aat ttg ttc cct ggc att acc gag ttc agt ttt ata tgc cga ata gag ttc tga ttc atc ctt tga gaa gat ctc tag cga tcg ttc aag atc cct gat tct agt ctt tag cct att tac ctc ctc aga aga tgc tcc gtt acc gtt --- ttt aca atc gtt aag atg tct atc aag atc cat gat tct atc tct ttt cca tat cag cat tga ttt cat tat tac gtt cgc agt cgt tca act g

>MT903342.1:133553-134000_Monkeypox_virus_isolate_MPXV-Singapore

gct gct agt tgt cgt tcg aga tca gaa ata cgt gtt ttc ttg gaa tgg att tcg tcg atg tac ttg tca tga ttg gca tcg aaa cac tta tta agt tct ttt ttt caa ttc tac gat ttt att tct ttc gcg agt caa ttc cct cct gta gta act atc agt ttt gtc aga ttc acg ctc tct acg tag act ttc ttg taa gtt act aat ttg ttc cct ggc att acc gag ttc agt ttt ata tgc cga ata gag ttc tga ttc atc ctt tga gaa gat ctc tag cga tcg ttc aag atc cct gat tct agt ctt tag cct att tac ctc ctc aga aga tgc tcc gtt acc gtt --- ttt aca atc gtt aag atg tct atc aag atc cat gat tct atc tct ttt cca tat cag cat tga ttt cat tat tac gtt cgc agt cgt tca act g

>MT903341.1:125069-125516_Monkeypox_virus_isolate_MPXV-M5320_M15_Bayelsa

gct gct agt tgt cgt tcg aga tca gaa ata cgt gtt ttc ttg gaa tgg att tcg tcg atg tac ttg tca tga ttg gca tcg aaa cac tta tta agt tct ttt ttt caa ttc tac gat ttt att tct ttc gcg agt caa ttc cct cct gta gta act atc agt ttt gtc aga ttc acg ctc tct acg tag act ttc ttg taa gtt act aat ttg ttc cct ggc att acc gag ttc agt ttt ata tgc cga ata gag ttc tga ttc atc ctt tga gaa gat ctc tag cga tcg ttc aag atc cct gat tct agt ctt tag cct att tac ctc ctc aga aga tgc tcc gtt acc gtt --- ttt aca atc gtt aag atg tct atc aag atc cat gat tct atc tct ttt cca tat cag cat tga ttt cat tat tac gtt cgc agt cgt tca act g

>NC_063383.1:133535-133982_Monkeypox_virus_complete_genome

gct gct agt tgt cgt tcg aga tca gaa ata cgt gtt ttc ttg gaa tgg att tcg tcg atg tac ttg tca tga ttg gca tcg aaa cac tta tta agt tct ttt ttt caa ttc tac gat ttt att tct ttc gcg agt caa ttc cct cct gta gta act atc agt ttt gtc aga ttc acg ctc tct acg tag act ttc ttg taa gtt act aat ttg ttc cct ggc att acc gag ttc agt ttt ata tgc cga ata gag ttc tga ttc atc ctt tga gaa gat ctc tag cga tcg ttc aag atc cct gat tct agt ctt tag cct att tac ctc ctc aga aga tgc tcc gtt acc gtt --- ttt aca atc gtt aag atg tct atc aag atc cat gat tct atc tct ttt cca tat cag cat tga ttt cat tat tac gtt cgc agt cgt tca act g

>MT903339.1:133702-134149_Monkeypox_virus_isolate_MPXV-M3021_Delta

gct gct agt tgt cgt tcg aga tca gaa ata cgt gtt ttc ttg gaa tgg att tcg tcg atg tac ttg tca tga ttg gca tcg aaa cac tta tta agt tct ttt ttt caa ttc tac gat ttt att tct ttc gcg agt caa ttc cct cct gta gta act atc agt ttt gtc aga ttc acg ctc tct acg tag act ttc ttg taa gtt act aat ttg ttc cct ggc att acc gag ttc agt ttt ata tgc cga ata gag ttc tga ttc atc ctt tga gaa gat ctc tag cga tcg ttc aag atc cct gat tct agt ctt tag cct att tac ctc ctc aga aga tgc tcc gtt acc gtt --- ttt aca atc gtt aag atg tct atc aag atc cat gat tct atc tct ttt cca tat cag cat tga ttt cat tat tac gtt cgc agt cgt tca act g

>MT903338.1:133701-134148_Monkeypox_virus_isolate_MPXV-M2957_Lagos

gct gct agt tgt cgt tcg aga tca gaa ata cgt gtt ttc ttg gaa tgg att tcg tcg atg tac ttg tca tga ttg gca tcg aaa cac tta tta agt tct ttt ttt caa ttc tac gat ttt att tct ttc gcg agt caa ttc cct cct gta gta act atc agt ttt gtc aga ttc acg ctc tct acg tag act ttc ttg taa gtt act aat ttg ttc cct ggc att acc gag ttc agt ttt ata tgc cga ata gag ttc tga ttc atc ctt tga gaa gat ctc tag cga tcg ttc aag atc cct gat tct agt ctt tag cct att tac ctc ctc aga aga tgc tcc gtt acc gtt --- ttt aca atc gtt aag atg tct atc aag atc cat gat tct atc tct ttt cca tat cag cat tga ttt cat tat tac gtt cgc agt cgt tca act g

>MT903337.1:133700-134147_Monkeypox_virus_isolate_MPXV-M2940_FCT

gct gct agt tgt cgt tcg aga tca gaa ata cgt gtt ttc ttg gaa tgg att tcg tcg atg tac ttg tca tga ttg gca tcg aaa cac tta tta agt tct ttt ttt caa ttc tac gat ttt att tct ttc gcg agt caa ttc cct cct gta gta act atc agt ttt gtc aga ttc acg ctc tct acg tag act ttc ttg taa gtt act aat ttg ttc cct ggc att acc gag ttc agt ttt ata tgc cga ata gag ttc tga ttc atc ctt tga gaa gat ctc tag cga tcg ttc aag atc cct gat tct agt ctt tag cct att tac ctc ctc aga aga tgc tcc gtt acc gtt --- ttt aca atc gtt aag atg tct atc aag atc cat gat tct atc tct ttt cca tat cag cat tga ttt cat tat tac gtt cgc agt cgt tca act g

>ON619837.1:133536-133983_Monkeypox_virus_isolate_MPXV_UK_2022_3_complete_genome

gct gct agt tgt cgt tcg aga tca gaa ata cgt gtt ttc ttg gaa tgg att tcg tcg atg tac ttg tca tga ttg gca tcg aaa cac tta tta agt tc? ttt ttt caa ttc tac gat ttt att tct ttc gcg agt caa ttc cct cct gta gta act atc agt ttt gtc aga ttc acg ctc tct acg tag act ttc ttg taa gtt act aat ttg ttc cct ggc att acc gag ttc agt ttt ata tgc cga ata gag ttc tga ttc atc ctt tga gaa gat ctc tag cga tcg ttc aag atc cct gat tct agt ctt tag cct att tac ctc ctc aga aga tgc tcc gtt acc gtt --- ttt aca atc gtt aag atg tct atc aag atc cat gat tct atc tct ttt cca tat cag cat tga ttt cat tat tac gtt cgc agt cgt tca act g

>ON585029.1:133739-134186_Monkeypox_virus_isolate_Monkeypox/PT0001/2022_partial_genome

gct gct agt tgt cgt tcg aga tca gaa ata cgt gtt ttc ttg gaa tgg att tcg tcg atg tac ttg tca tga ttg gca tcg aaa cac tta tta agt tc? ttt ttt caa ttc tac gat ttt att tct ttc gcg agt caa ttc cct cct gta gta act atc agt ttt gtc aga ttc acg ctc tct acg tag act ttc ttg taa gtt act aat ttg ttc cct ggc att acc gag ttc agt ttt ata tgc cga ata gag ttc tga ttc atc ctt tga gaa gat ctc tag cga tcg ttc aag atc cct gat tct agt ctt tag cct att tac ctc ctc aga aga tgc tcc gtt acc gtt --- ttt aca atc gtt aag atg tct atc aag atc cat gat tct atc tct ttt cca tat cag cat tga ttt cat tat tac gtt cgc agt cgt tca act g

>ON676708.1:133530-133977_Monkeypox_virus_isolate_MPXV_USA_2021_MD_complete_genome

gct gct agt tgt cgt tcg aga tca gaa ata cgt gtt ttc ttg gaa tgg att tcg tcg atg tac ttg tca tga ttg gca tcg aaa cac tta tta agt tct ttt ttt caa ttc tac gat ttt att tct ttc gcg agt caa ttc cct cct gta gta act atc agt ttt gtc aga ttc acg ctt tct acg tag act ttc ttg taa gtt act aat ttg ttc cct ggc att acc gag ttc agt ttt ata tgc cga ata gag ttc tga ttc atc ctt tga gaa gat ctc tag cga tcg ttc aag atc cct gat tct agt ctt tag cct att tac ctc ctc aga aga tgc tcc gtt acc gtt --- ttt aca atc gtt aag atg tct atc aag atc cat gat tct atc tct ttt cca tat cag cat tga ttt cat tat tac gtt cgc agt cgt tca act g

>ON676704.1:133529-133976_Monkeypox_virus_isolate_MPXV_USA_2022_FL002_complete_genome

gct gct agt tgt cgt tcg aga tca gaa ata cgt gtt ttc ttg gaa tgg att tcg tcg atg tac ttg tca tga ttg gca tcg aaa cac tta tta agt tct ttt ttt caa ttc tac gat ttt att tct ttc gcg agt caa ttc cct cct gta gta act atc agt ttt gtc aga ttc acg ctc tct acg tag act ttc ttg taa gtt act aat ttg ttc cct ggc att acc gag ttc agt ttt ata tgc cga ata gag ttc tga ttc atc ctt tga gaa gat ctc tag cga tcg ttc aag atc cct gat tct agt ctt tag cct att tac ctt ctc aga aga tgc tcc gtt acc gtt --- ttt aca atc gtt aag atg tct atc aag atc cat gat tct atc tct ttt cca tat cag cat tga ttt cat tat tac gtt cgc agt cgt tca act g

>ON622720.1:133524-133971_Monkeypox_virus_isolate_MPXV-CH-38156923/2022_partial_genome

gct gct agt tgt cgt tcg aga tca gaa ata cgt gtt ttc ttg gaa tgg att tcg tcg atg tac ttg tca tga ttg gca tcg aaa cac tta tta agt tc? ttt ttt caa ttc tac gat ttt att tct ttc gcg agt caa tt? cct cct gta gta act atc agt ttt gtc aga ttc acg ctc tct acg tag act ttc ttg taa gtt act aat ttg ttc cct ggc att acc gag ttc agt ttt ata tgc cga ata gag ttc tga ttc atc ctt tga gaa gat ctc tag cga tcg ttc aag atc cct gat tct agt ctt tag cct att tac ctc ctc aga aga tgc tcc gtt acc gtt --- ttt aca atc gtt aag atg tct atc aag atc cat gat tct atc tct ttt cca tat cag cat tga ttt cat tat tac gtt cgc agt cgt tca act g

>ON602722.1:133413-133859_Monkeypox_virus_isolate_MPXV_FRA_2022_TLS67_complete_genome

gct gct agt tgt cgt tcg aga tca gaa ata cgt gtt ttc ttg gaa tgg att tcg tcg atg tac ttg tca tga ttg gca tcg aaa cac tta tta agt tc- ttt ttt caa ttc tac gat ttt att tct ttc gcg agt caa ttc cct cct gta gta act atc agt ttt gtc aga ttc acg ctc tct acg tag act ttc ttg taa gtt act aat ttg ttc cct ggc att acc gag ttc agt ttt ata tgc cga ata gag ttc tga ttc atc ctt tga gaa gat ctc tag cga tcg ttc aag atc cct gat tct agt ctt tag cct att tac ctc ctc aga aga tgc tcc gtt acc gtt --- ttt aca atc gtt aag atg tct atc aag atc cat gat tct atc tct ttt cca tat cag cat tga ttt cat tat tac gtt cgc agt cgt tca act g

>KP849470.1:136123-136570_Monkeypox_virus_isolate_Cote_dIvoire_1971_complete_genome

gct gtt agt tgt cgt tcg aga tca gaa ata cgt gtt ttc ttg gaa tgg att tcg tcg atg tac ttg tca tga ttg gca tcg aaa cac tta tta agt tct ttt ttt caa ttc tac gat ttt att tct ttc gcg agt caa ttc cct cct ata gta act atc agt ttt gtc aga ttc acg ctc tct acg tag act ttc ttg caa gtt act aat ttg ttc cct ggc att acc gag ttc agt ttt ata tgc cga ata gag ttc tga ttc atc ctt tga gaa gat ctc tag cga tcg ttc aag atc cct gat tct agt ctt tag cct att tac ctc ctc aga aga tgc tcc gtt acc gtt --- ttt aca atc gtt aag atg tct atc aag atc cat gat tct atc tct ttt cca tat cag cat tga ttt cat tat tac gtc cgc agt cgt tca act g

>KJ642616.1:135180-135627_Monkeypox_virus_strain_PCH_complete_genome

gct gtt agt tgt cgt tcg aga tca gaa ata cgt gtt ttc ttg gaa tgg att tcg tcg atg tac ttg tca tga ttg gca tcg aaa cac tta tta agt tct ttt ttt caa ttc tac gat ttt att tct ttc gcg agt caa ttc cct cct ata gta act atc agt ttt gtc aga ttc acg ctc tct acg tag act ttc ttg caa gtt act aat ttg ttc cct ggc att acc gag ttc agt ttt ata tgc cga ata gag ttc tga ttc atc ctt tga gaa gat ctc tag cga tcg ttc aag atc cct gat tct agt ctt tag cct att tac ctc ctc aga aga tgc tcc gtt acc gtt --- ttt aca atc gtt aag atg tct atc aag atc cat gat tct atc tct ttt cca tat cag cat tga ttt cat tat tac gtc cgc agt cgt tca act g

>KJ642614.1:126741-127188_Monkeypox_virus_strain_UTC_complete_genome

gct gtt agt tgt cgt tcg aga tca gaa ata cgt gtt ttc ttg gaa tgg att tcg tcg atg tac ttg tca tga ttg gca tcg aaa cac tta tta agt tct ttt ttt caa ttc tac gat ttt att tct ttc gcg agt caa ttc cct cct ata gta act atc agt ttt gtc aga ttc acg ctc tct acg tag act ttc ttg caa gtt act aat ttg ttc cct ggc att acc gag ttc agt ttt ata tgc cga ata gag ttc tga ttc atc ctt tga gaa gat ctc tag cga tcg ttc aag atc cct gat tct agt ctt tag cct att tac ctc ctc aga aga tgc tcc gtt acc gtt --- ttt aca atc gtt aag atg tct atc aag atc cat gat tct atc tct ttt cca tat cag cat tga ttt cat tat tac gtc cgc agt cgt tca act g

>MT903346.1:135508-135955_Monkeypox_virus_isolate_MPXV-USA2003_099_Gambian_Rat

gct gtt agt tgt cgt tcg aga tca gaa ata cgt gtt ttc ttg gaa tgg att tcg tcg atg tac ttg tca tga ttg gca tcg aaa cac tta tta agt tct ttt ttt caa ttc tac gat ttt att tct ttc gcg agt caa ttc cct cct ata gta act atc agt ttt gtc aga ttc acg ctc tct acg tag act ttc ttg caa gtt act aat ttg ttc cct ggc att acc gag ttc agt ttt ata tgc cga ata gag ttc tga ttc atc ctt tga gaa gat ctc tag cga tcg ttc aag atc cct gat tct agt ctt tag cct att tac ctc ctc aga aga tgc tcc gtt acc gtt --- ttt aca atc gtt aag atg tct atc aag atc cat gat tct atc tct ttt cca tat cag cat tga ttt cat tat tac gtc cgc agt cgt tca act g

>DQ011157.1:135509-135956_Monkeypox_virus_strain_USA_2003_039_complete_genome

gct gtt agt tgt cgt tcg aga tca gaa ata cgt gtt ttc ttg gaa tgg att tcg tcg atg tac ttg tca tga ttg gca tcg aaa cac tta tta agt tct ttt ttt caa ttc tac gat ttt att tct ttc gcg agt caa ttc cct cct ata gta act atc agt ttt gtc aga ttc acg ctc tct acg tag act ttc ttg caa gtt act aat ttg ttc cct ggc att acc gag ttc agt ttt ata tgc cga ata gag ttc tga ttc atc ctt tga gaa gat ctc tag cga tcg ttc aag atc cct gat tct agt ctt tag cct att tac ctc ctc aga aga tgc tcc gtt acc gtt --- ttt aca atc gtt aag atg tct atc aag atc cat gat tct atc tct ttt cca tat cag cat tga ttt cat tat tac gtc cgc agt cgt tca act g

>DQ011156.1:135956-136403_Monkeypox_virus_strain_Liberia_1970_184_complete_genome

gct gtt agt tgt cgt tcg aga tca gaa ata cgt gtt ttc ttg gaa tgg att tcg tcg atg tac ttg tca tga ttg gca tcg aaa cac tta tta agt tct ttt ttt caa ttc tac gat ttt att tct ttc gcg agt caa ttc cct cct ata gta act atc agt ttt gtc aga ttc acg ctc tct acg tag act ttc ttg caa gtt act aat ttg ttc cct ggc att acc gag ttc agt ttt ata tgc cga ata gag ttc tga ttc atc ctt tga gaa gat ctc tag cga tcg ttc aag atc cct gat tct agt ctt tag cct att tac ctc ctc aga aga tgc tcc gtt acc gtt --- ttt aca atc gtt aag atg tct atc aag atc cat gat tct atc tct ttt cca tat cag cat tga ttt cat tat tac gtc cgc agt cgt tca act g

>DQ011153.1:135509-135956_Monkeypox_virus_strain_USA_2003_044_complete_genome

gct gtt agt tgt cgt tcg aga tca gaa ata cgt gtt ttc ttg gaa tgg att tcg tcg atg tac ttg tca tga ttg gca tcg aaa cac tta tta agt tct ttt ttt caa ttc tac gat ttt att tct ttc gcg agt caa ttc cct cct ata gta act atc agt ttt gtc aga ttc acg ctc tct acg tag act ttc ttg caa gtt act aat ttg ttc cct ggc att acc gag ttc agt ttt ata tgc cga ata gag ttc tga ttc atc ctt tga gaa gat ctc tag cga tcg ttc aag atc cct gat tct agt ctt tag cct att tac ctc ctc aga aga tgc tcc gtt acc gtt --- ttt aca atc gtt aag atg tct atc aag atc cat gat tct atc tct ttt cca tat cag cat tga ttt cat tat tac gtc cgc agt cgt tca act g

>AY753185.1:135722-136169_Monkeypox_virus_strain_COP-58_complete_genome

gct gtt agt tgt cgt tcg aga tca gaa ata cgt gtt ttc ttg gaa tgg att tcg tcg atg tac ttg tca tga ttg gca tcg aaa cac tta tta agt tct ttt ttt caa ttc tac gat ttt att tct ttc gcg agt caa ttc cct cct ata gta act atc agt ttt gtc aga ttc acg ctc tct acg tag act ttc ttg caa gtt act aat ttg ttc cct ggc att acc gag ttc agt ttt ata tgc cga ata gag ttc tga ttc atc ctt tga gaa gat ctc tag cga tcg ttc aag atc cct gat tct agt ctt tag cct att tac ctc ctc aga aga tgc tcc gtt acc gtt --- ttt aca atc gtt aag atg tct atc aag atc cat gat tct atc tct ttt cca tat cag cat tga ttt cat tat tac gtc cgc agt cgt tca act g

>AY603973.1:135479-135926_Monkeypox_virus_strain_MPXV-WRAIR7-61_complete_genome

gct gtt agt tgt cgt tcg aga tca gaa ata cgt gtt ttc ttg gaa tgg att tcg tcg atg tac ttg tca tga ttg gca tcg aaa cac tta tta agt tct ttt ttt caa ttc tac gat ttt att tct ttc gcg agt caa ttc cct cct ata gta act atc agt ttt gtc aga ttc acg ctc tct acg tag act ttc ttg caa gtt act aat ttg ttc cct ggc att acc gag ttc agt ttt ata tgc cga ata gag ttc tga ttc atc ctt tga gaa gat ctc tag cga tcg ttc aag atc cct gat tct agt ctt tag cct att tac ctc ctc aga aga tgc tcc gtt acc gtt --- ttt aca atc gtt aag atg tct atc aag atc cat gat tct atc tct ttt cca tat cag cat tga ttt cat tat tac gtc cgc agt cgt tca act g

>KJ642615.1:133827-134275_Monkeypox_virus_strain_W-Nigeria_complete_genome

gct gtt agt tgt cgt tcg aga tca gaa ata cgt gtt ttc ttg gaa tgg att tcg tcg atg tac ttg tca tga ttg gca tcg aaa cac tta tta agt tct ttt ttt caa ttc tac gat ttt att tct ttc gcg agt caa ttc cct cct ata gta act atc agt ttt gtc aga ttc acg ctc tct acg tag act ttc ttg caa gtt act aat ttg ttc cct ggc att acc gag ttc agt ttt ata tgc cga ata gag ttc tga ttc atc ctt tga gaa gat ctc tag cga tcg ttc aag atc cct gat tct agt ctt tag cct att tac ctc ctc aga aga tgc tcc gtt acc gtt --t ttt aca atc gtt aag atg tct atc aag atc cat gat tct atc tct ttt cca tat cag cat tga ttt cat tat tac gtc cgc agt cgt tca act g

>AY741551.1:135503-135950_Monkeypox_virus_isolate_Sierra_Leone_complete_genome

gct gtt agt tat cgt tcg aga tca gaa ata cgt gtt ttc ttg gaa tgg att tcg tcg atg tac ttg tca tga ttg gca tcg aaa cac tta tta agt tct ttt ttt caa ttc tac gat ttt att tct ttc gcg agt caa ttc cct cct ata gta act atc agt ttt gtc aga ttc acg ctc tct acg tag act ttc ttg caa gtt act aat ttg ttc cct ggc att acc gag ttc agt ttt ata tgc cga ata gag ttc tga ttc atc ctt tga gaa gat ctc tag cga tcg ttc aag atc cct gat tct agt ctt tag cct att tac ctc ctc aga aga tgc tcc gtt acc gtt --- ttt aca atc gtt aag atg tct atc aag atc cat gat tct atc tct ttt cca tat cag cat tga ttt cat tat tac gtc cgc agt cgt tca act g

>MG693725.1:125266-125713_Monkeypox_virus_isolate_MPX_Nig_2017_298481_partial_genome

gct gct agt tgt cgt tcg aga tca gaa ata cgt gtt ttc ttg gaa tgg att tcg tcg atg tac ttg tca tga ttg gca tcg aaa cac tta tta agt tct ttt ttt caa ttc tac gat ttt att tct ttc gcg agt caa ttc cct cct gta gta act atc agt ttt gtc aga ttc acg ctc tct acg tag act ttc ttg taa gtt act ??? ??? ??? cct ggc att acc gag ttc agt ttt ata tgc cga ata gag ttc tga ttc atc ctt tga gaa gat ctc tag cga tcg ttc aag atc cct gat tct agt ctt tag cct att tac ctc ctc aga aga tgc tcc gtt acc gtt --- ttt aca atc gtt aag atg tct atc aag atc cat gat tct atc tct ttt cca tat cag cat tga ttt cat tat tac gtt cgc agt cgt tca act g

>KC813510.1:151579-151979_Cowpox_virus_strain_HumGra07/1_complete_genome

gct gct agt tgt cgt tcg aga tca gaa ata cgg gtt ttc ttg gaa ttg att tcg tcg atg tac ttg gca tga ttg gca tcg aaa cac tta tta agt tc- ttt ttc caa ttc tac gat ttt att tct ttc gcg agt caa ttc cct cct gta gta act atc tgt ttt gtc aga ttc acg ctc tct acg tag act ttc ttg caa gtt act aat ttg ttc cct ggc acg tcc gag ttc agt ttt ata tgc cga ata gag ttc tga ttc atc ctt tga gca gat ctc tag cga tcg ttc aag atc cct gat tct agt ctt tag cct att tac ctc ctc aga aga tgt tcc gtt acc gtt gcg ttt aca ctc gtt aag ctg tct atc aag atc cat gat tct atc tct --- --- --- --- --- --- --- --- --- --- --- --- --- --- --- --- -

>MK035759.1:151597-151997_Cowpox_virus_strain_CPXV/Cepad_332

gct gct agt tgt cgt tcg aga tca gaa ata cgg gtt ttc ttg gaa ttg att tcg tcg atg tac ttg gca tga ttg gca tcg aaa cac tta tta agt tc- ttt ttc caa ttc tac gat ttt att tct ttc gcg agt caa ttc cct cct gta gta act atc tgt ttt gtc aga ttc acg ctc tct acg tag act ttc ttg caa gtt act aat ttg ttc cct ggc acg tcc gag ttc agt ttt ata tgc cga ata gag ttc tga ttc atc ctt tga gca gat ctc tag cga tcg ttc aag atc cct gat tct agt ctt tag cct att tac ctc ctc aga aga tgt tcc gtt acc gtt gcg ttt aca ctc gtt gag ctg tct atc aag atc cat gat tct atc tct --- --- --- --- --- --- --- --- --- --- --- --- --- --- --- --- -

>MK035757.1:151894-152294_Cowpox_virus_strain_CPXV/Rat_Koelle

gct gct agt tgt cgt tcg aga tca gaa ata cgg gtt ttc ttg gaa ttg att tcg tcg atg tac ttg gca tga ttg gca tcg aaa cac tta tta agt tc- ttt ttc caa ttc tac gat ttt att tct ttc gcg agt caa ttc cct cct gta gta act atc tgt ttt gtc aga ttc acg ctc tct acg tag act ttc ttg caa gtt act aat ttg ttc cct ggc acg tcc gag ttc agt ttt ata tgc cga ata gag ttc tga ttc atc ctt tga gca gat ctc tag cga tcg ttc aag atc cct gat tct agt ctt tag cct att tac ctc ctc aga aga tgt tcc gtt acc gtt gcg ttt aca ctc gtt gag ctg tct atc aag atc cat gat tct atc tct --- --- --- --- --- --- --- --- --- --- --- --- --- --- --- --- -

>MK035753.1:151599-151999_Cowpox_virus_strain_CPXV/Cepad_333

gct gct agt tgt cgt tcg aga tca gaa ata cgg gtt ttc ttg gaa ttg att tcg tcg atg tac ttg gca tga ttg gca tcg aaa cac tta tta agt tc- ttt ttc caa ttc tac gat ttt att tct ttc gcg agt caa ttc cct cct gta gta act atc tgt ttt gtc aga ttc acg ctc tct acg tag act ttc ttg caa gtt act aat ttg ttc cct ggc acg tcc gag ttc agt ttt ata tgc cga ata gag ttc tga ttc atc ctt tga gca gat ctc tag cga tcg ttc aag atc cct gat tct agt ctt tag cct att tac ctc ctc aga aga tgt tcc gtt acc gtt gcg ttt aca ctc gtt gag ctg tct atc aag atc cat gat tct atc tct --- --- --- --- --- --- --- --- --- --- --- --- --- --- --- --- -

>MK035750.1:141394-141794_Cowpox_virus_strain_CPXV/Compiegne_2009

gct gct agt tgt cgt tcg aga tca gaa ata cgg gtt ttc ttg gaa ttg att tcg tcg atg tac ttg gca tga ttg gca tcg aaa cac tta tta agt tc- ttt ttc caa ttc tac gat ttt att tct ttc gcg agt caa ttc cct cct gta gta act atc tgt ttt gtc aga ttc acg ctc tct acg tag act ttc ttg caa gtt act aat ttg ttc cct ggc acg tcc gag ttc agt ttt ata tgc cga ata gag ttc tga ttc atc ctt tga gca gat ctc tag cga tcg ttc aag atc cct gat tct agt ctt tag cct att tac ctc ctc aga aga tgt tcc gtt acc gtt gcg ttt aca ctc gtt gag ctg tct atc aag atc cat gat tct atc tct --- --- --- --- --- --- --- --- --- --- --- --- --- --- --- --- -

>MK035749.1:137103-137503_Cowpox_virus_strain_CPXV/Rat_Biederstein

gct gct agt tgt cgt tcg aga tca gaa ata cgg gtt ttc ttg gaa ttg att tcg tcg atg tac ttg gca tga ttg gca tcg aaa cac tta tta agt tc- ttt ttc caa ttc tac gat ttt att tct ttc gcg agt caa ttc cct cct gta gta act atc tgt ttt gtc aga ttc acg ctc tct acg tag act ttc ttg caa gtt act aat ttg ttc cct ggc acg tcc gag ttc agt ttt ata tgc cga ata gag ttc tga ttc atc ctt tga gca gat ctc tag cga tcg ttc aag atc cct gat tct agt ctt tag cct att tac ctc ctc aga aga tgt tcc gtt acc gtt gcg ttt aca ctc gtt gag ctg tct atc aag atc cat gat tct atc tct --- --- --- --- --- --- --- --- --- --- --- --- --- --- --- --- -

>MK035748.1:151511-151911_Cowpox_virus_strain_CPXV/Boa_Marl

gct gct agt tgt cgt tcg aga tca gaa ata cgg gtt ttc ttg gaa ttg att tcg tcg atg tac ttg gca tga ttg gca tcg aaa cac tta tta agt tc- ttt ttc caa ttc tac gat ttt att tct ttc gcg agt caa ttc cct cct gta gta act atc tgt ttt gtc aga ttc acg ctc tct acg tag act ttc ttg caa gtt act aat ttg ttc cct ggc acg tcc gag ttc agt ttt ata tgc cga ata gag ttc tga ttc atc ctt tga gca gat ctc tag cga tcg ttc aag atc cct gat tct agt ctt tag cct att tac ctc ctc aga aga tgt tcc gtt acc gtt gcg ttt aca ctc gtt gag ctg tct atc aag atc cat gat tct atc tct --- --- --- --- --- --- --- --- --- --- --- --- --- --- --- --- -

>MK035747.1:151513-151913_Cowpox_virus_strain_CPXV/Rat_Marl

gct gct agt tgt cgt tcg aga tca gaa ata cgg gtt ttc ttg gaa ttg att tcg tcg atg tac ttg gca tga ttg gca tcg aaa cac tta tta agt tc- ttt ttc caa ttc tac gat ttt att tct ttc gcg agt caa ttc cct cct gta gta act atc tgt ttt gtc aga ttc acg ctc tct acg tag act ttc ttg caa gtt act aat ttg ttc cct ggc acg tcc gag ttc agt ttt ata tgc cga ata gag ttc tga ttc atc ctt tga gca gat ctc tag cga tcg ttc aag atc cct gat tct agt ctt tag cct att tac ctc ctc aga aga tgt tcc gtt acc gtt gcg ttt aca ctc gtt gag ctg tct atc aag atc cat gat tct atc tct --- --- --- --- --- --- --- --- --- --- --- --- --- --- --- --- -

>MK035746.1:137557-137957_Cowpox_virus_strain_CPXV/Boy_Biederstein

gct gct agt tgt cgt tcg aga tca gaa ata cgg gtt ttc ttg gaa ttg att tcg tcg atg tac ttg gca tga ttg gca tcg aaa cac tta tta agt tc- ttt ttc caa ttc tac gat ttt att tct ttc gcg agt caa ttc cct cct gta gta act atc tgt ttt gtc aga ttc acg ctc tct acg tag act ttc ttg caa gtt act aat ttg ttc cct ggc acg tcc gag ttc agt ttt ata tgc cga ata gag ttc tga ttc atc ctt tga gca gat ctc tag cga tcg ttc aag atc cct gat tct agt ctt tag cct att tac ctc ctc aga aga tgt tcc gtt acc gtt gcg ttt aca ctc gtt gag ctg tct atc aag atc cat gat tct atc tct --- --- --- --- --- --- --- --- --- --- --- --- --- --- --- --- -

>LN864565.1:151415-151815_Cowpox_virus_complete_genome_strain_Ratpox09

gct gct agt tgt cgt tcg aga tca gaa ata cgg gtt ttc ttg gaa ttg att tcg tcg atg tac ttg gca tga ttg gca tcg aaa cac tta tta agt tc- ttt ttc caa ttc tac gat ttt att tct ttc gcg agt caa ttc cct cct gta gta act atc tgt ttt gtc aga ttc acg ctc tct acg tag act ttc ttg caa gtt act aat ttg ttc cct ggc acg tcc gag ttc agt ttt ata tgc cga ata gag ttc tga ttc atc ctt tga gca gat ctc tag cga tcg ttc aag atc cct gat tct agt ctt tag cct att tac ctc ctc aga aga tgt tcc gtt acc gtt gcg ttt aca ctc gtt gag ctg tct atc aag atc cat gat tct atc tct --- --- --- --- --- --- --- --- --- --- --- --- --- --- --- --- -

>KC813512.1:143733-144133_Cowpox_virus_strain_HumKre08/1_complete_genome

gct gct agt tgt cgt tcg aga tca gaa ata cgg gtt ttc ttg gaa ttg att tcg tcg atg tac ttg gca tga ttg gca tcg aaa cac tta tta agt tc- ttt ttc caa ttc tac gat ttt att tct ttc gcg agt caa ttc cct cct gta gta act atc tgt ttt gtc aga ttc acg ctc tct acg tag act ttc ttg caa gtt act aat ttg ttc cct ggc acg tcc gag ttc agt ttt ata tgc cga ata gag ttc tga ttc atc ctt tga gca gat ctc tag cga tcg ttc aag atc cct gat tct agt ctt tag cct att tac ctc ctc aga aga tgt tcc gtt acc gtt gcg ttt aca ctc gtt gag ctg tct atc aag atc cat gat tct atc tct --- --- --- --- --- --- --- --- --- --- --- --- --- --- --- --- -

>KC813508.1:144056-144456_Cowpox_virus_strain_HumAac09/1_complete_genome

gct gct agt tgt cgt tcg aga tca gaa ata cgg gtt ttc ttg gaa ttg att tcg tcg atg tac ttg gca tga ttg gca tcg aaa cac tta tta agt tc- ttt ttc caa ttc tac gat ttt att tct ttc gcg agt caa ttc cct cct gta gta act atc tgt ttt gtc aga ttc acg ctc tct acg tag act ttc ttg caa gtt act aat ttg ttc cct ggc acg tcc gag ttc agt ttt ata tgc cga ata gag ttc tga ttc atc ctt tga gca gat ctc tag cga tcg ttc aag atc cct gat tct agt ctt tag cct att tac ctc ctc aga aga tgt tcc gtt acc gtt gcg ttt aca ctc gtt gag ctg tct atc aag atc cat gat tct atc tct --- --- --- --- --- --- --- --- --- --- --- --- --- --- --- --- -

>KC813505.1:143979-144379_Cowpox_virus_strain_RatKre08/2_complete_genome

gct gct agt tgt cgt tcg aga tca gaa ata cgg gtt ttc ttg gaa ttg att tcg tcg atg tac ttg gca tga ttg gca tcg aaa cac tta tta agt tc- ttt ttc caa ttc tac gat ttt att tct ttc gcg agt caa ttc cct cct gta gta act atc tgt ttt gtc aga ttc acg ctc tct acg tag act ttc ttg caa gtt act aat ttg ttc cct ggc acg tcc gag ttc agt ttt ata tgc cga ata gag ttc tga ttc atc ctt tga gca gat ctc tag cga tcg ttc aag atc cct gat tct agt ctt tag cct att tac ctc ctc aga aga tgt tcc gtt acc gtt gcg ttt aca ctc gtt gag ctg tct atc aag atc cat gat tct atc tct --- --- --- --- --- --- --- --- --- --- --- --- --- --- --- --- -

>KC813503.1:151763-152163_Cowpox_virus_strain_RatGer09/1_complete_genome

gct gct agt tgt cgt tcg aga tca gaa ata cgg gtt ttc ttg gaa ttg att tcg tcg atg tac ttg gca tga ttg gca tcg aaa cac tta tta agt tc- ttt ttc caa ttc tac gat ttt att tct ttc gcg agt caa ttc cct cct gta gta act atc tgt ttt gtc aga ttc acg ctc tct acg tag act ttc ttg caa gtt act aat ttg ttc cct ggc acg tcc gag ttc agt ttt ata tgc cga ata gag ttc tga ttc atc ctt tga gca gat ctc tag cga tcg ttc aag atc cct gat tct agt ctt tag cct att tac ctc ctc aga aga tgt tcc gtt acc gtt gcg ttt aca ctc gtt gag ctg tct atc aag atc cat gat tct atc tct --- --- --- --- --- --- --- --- --- --- --- --- --- --- --- --- -

>KC813501.1:143734-144134_Cowpox_virus_strain_RatAac09/1_complete_genome

gct gct agt tgt cgt tcg aga tca gaa ata cgg gtt ttc ttg gaa ttg att tcg tcg atg tac ttg gca tga ttg gca tcg aaa cac tta tta agt tc- ttt ttc caa ttc tac gat ttt att tct ttc gcg agt caa ttc cct cct gta gta act atc tgt ttt gtc aga ttc acg ctc tct acg tag act ttc ttg caa gtt act aat ttg ttc cct ggc acg tcc gag ttc agt ttt ata tgc cga ata gag ttc tga ttc atc ctt tga gca gat ctc tag cga tcg ttc aag atc cct gat tct agt ctt tag cct att tac ctc ctc aga aga tgt tcc gtt acc gtt gcg ttt aca ctc gtt gag ctg tct atc aag atc cat gat tct atc tct --- --- --- --- --- --- --- --- --- --- --- --- --- --- --- --- -

>NC_055231.1:144625-145025_Orthopoxvirus_Abatino_complete_genome

gct gct agc tgt cgt tcg aga tca gaa ata cgg gtt ttc ttg gaa ttg att tcg tcg atg tac ttg gca tga ttg gca tcg aaa cac tta tta agt tc- ttt ttc caa ttc tac gat ttt att tct ttc gcg agt caa ttc cct cct gta gta act atc ggt ttt gtc aga ttc acg ctc tct acg tag act ttc ttg caa gtt act aat ttg ttc cct ggc acg tcc gag ttc agt ttt ata tgc cga ata gag ttc tga ttc atc ctt tga gca gat ctc tag aga tcg ttc aag atc cct gat tct agt ctt tag cct att tac ctc ctc aga aga tga tcc gtt acc gtt gcg ttt aca ctc gtt aag ctg tct atc aag atc cat gat tct atc tct --- --- --- --- --- --- --- --- --- --- --- --- --- --- --- --- -

>DQ437594.1:135696-136096_Taterapox_virus_strain_Dahomey_1968_complete_genome

gct gct agc tgt cgt tcg aga tca gaa ata cgg gtt ttc ttg gaa ttg att tcg tcg atg tac ttg gca tga ttg gca tcg aaa cac tta tta agt tc- ttt ttc caa ttc tac gat ttt att tct ttc gcg agt caa ttc cct cct gta gta gct atc ggt ttt gtc aga ttc acg ctc tct acg tag act ttc ttg caa gtt act aat ttg ttc cct ggc acg tcc gag ttc agt ttt ata tgc cga ata gag ttc tga ttc atc ctt tga gca gat ctc tag cga tcg ttc aag atc cct gat tct agt ctt tag cct att tac ctc ctc aga aga tgt tcc gtt acc gtt gcg ttt aca ctc gtt aag ctg tct atc aag atc cat gat tct atc tct --- --- --- --- --- --- --- --- --- --- --- --- --- --- --- --- -

>MN369532.1:123226-123626_Vaccinia_virus_isolate_VK01

gct gct agt tgt cgt tcg aga tca gaa ata cgg gtt ttc ttg gaa ttg att tcg tcg atg tac ttg gca tga ttg gca tcg aaa cac tta tta agt tc- ttt ttc caa ttc tac gat ttt att tct ttc gcg agt caa ttc cct cct gta gta act atc tgt ttt gtc aga ttc acg ctc tct acg tag act ttc ttg caa gtt act aat ttg ttc cct agc acg tcc gag ttc agt ttt ata tgc cga ata gag ttc tga ttc atc ctt tga gca gat ctc tag cga tcg ttt aag att cct gat tct agt ctt tag cct att tac ctc ctc aga aga tgt tcc gtt acc gtt gcg ttt aca ctc gtt aag ctg tct atc aag atc cat gat tct atc tct --- --- --- --- --- --- --- --- --- --- --- --- --- --- --- --- -

>LT896724.1:151288-151688_Cowpox_virus_isolate_Ger/2015/Cat1_genome_assembly_complete_genome:_monopartite

gct gct agt tgt cgt tcg aga tca gaa ata cgg gtt ttc ttg gaa ttg att tcg tcg atg tac ttg gca tga ttg gta tcg aaa cac tta tta agt tc- ttt ttc caa ttc tac gat ttt att tct ttc gcg agt caa ttc cct cct gta gta act atc tgt ttt gtc aga ttc acg ctc tct acg tag act ttc ttg caa gtt act aat ttg ttc cct ggc acg tcc gag ttc agt ttt ata tgc cga ata gag ttc tga ttc atc ctt tga gca gat ctc tag aga tcg ttc aag atc cct gat tct agt ttt tag cct att tac ctc ctc aga aga tgt tcc gtt acc gtt gcg ttt aca ctc gtt aag ctg tct atc aag atc cat gat tct atc tct --- --- --- --- --- --- --- --- --- --- --- --- --- --- --- --- -

>BK013342.1:143109-143509_TPA_asm:_Vaccinia_virus_isolate_VK08

gct gct agt tgt cgt tcg aga tca gaa ata cgg gtt ttc ttg gaa ttg att tcg tcg atg tac ttg gca tga ttg gca tcg aaa cac tta tta agt tc- ttt ttc caa ttc tac gat ttt att tct ttc gcg agt caa ttc cct cct gta gta act atc tgt ttt gtc aga ttc acg ctc tct acg tag act ttc ttg caa gtt act aat ttg ttc cct agc acg tcc gag ttc agt ttt ata tgc cga ata gag ttc tga ttc atc ctt tga gca gat ctc tag cga tcg ttt aag att cct gat tct agt ctt tag cct att tac ctc ctc aga aga tgt tcc gtt acc gtt gcg ttt aca ctc gtt aag ctg tct atc aag atc cat gat tct atc tct --- --- --- --- --- --- --- --- --- --- --- --- --- --- --- --- -

>BK013341.1:147188-147588_TPA_asm:_Horsepox_virus_isolate_VK05

gct gct agt tgt cgt tcg aga tca gaa ata cgg gtt ttc ttg gaa ttg att tcg tcg atg tac ttg gca tga ttg gca tcg aaa cac tta tta agt tc- ttt ttc caa ttc tac gat ttt att tct ttc gcg agt caa ttc cct cct gta gta act atc tgt ttt gtc aga ttc acg ctc tct acg tag act ttc ttg caa gtt act aat ttg ttc cct agc acg tcc gag ttc agt ttt ata tgc cga ata gag ttc tga ttc atc ctt tga gca gat ctc tag cga tcg ttt aag att cct gat tct agt ctt tag cct att tac ctc ctc aga aga tgt tcc gtt acc gtt gcg ttt aca ctc gtt aag ctg tct atc aag atc cat gat tct atc tct --- --- --- --- --- --- --- --- --- --- --- --- --- --- --- --- -

>BK013339.1:152982-153382_TPA_asm:_Vaccinia_virus_isolate_VK01

gct gct agt tgt cgt tcg aga tca gaa ata cgg gtt ttc ttg gaa ttg att tcg tcg atg tac ttg gca tga ttg gca tcg aaa cac tta tta agt tc- ttt ttc caa ttc tac gat ttt att tct ttc gcg agt caa ttc cct cct gta gta act atc tgt ttt gtc aga ttc acg ctc tct acg tag act ttc ttg caa gtt act aat ttg ttc cct agc acg tcc gag ttc agt ttt ata tgc cga ata gag ttc tga ttc atc ctt tga gca gat ctc tag cga tcg ttt aag att cct gat tct agt ctt tag cct att tac ctc ctc aga aga tgt tcc gtt acc gtt gcg ttt aca ctc gtt aag ctg tct atc aag atc cat gat tct atc tct --- --- --- --- --- --- --- --- --- --- --- --- --- --- --- --- -

>KY369926.1:151997-152397_Cowpox_virus_strain_Kostroma_2015

gct gct agc tgt cgt tcg aga tca gaa ata cgg gtt ttc ttg gaa ttg att tcg tcg atg tac ttg gca tga ttg gta tcg aaa cac tta tta agt tc- ttt ttc caa ttc tac gat ttt att tct ttc gcg agt caa ttc cct cct gta gta gct atc ggt ttt gtc aga ttc acg ctc tct acg tag act ttc ttg caa att act aat ttg ttc cct ggc acg tcc gag ttc agt ttt ata tgc cga ata gag ttc tga ttc atc ctt tga gca gat ctc tag aga tcg ttc aag atc cct gat tct agt ctt tag cct att tac ctc ctc aga aga tgc tcc gtt acc gtt gcg ttt aca ctc gtt aag ctg tct atc aag atc cat gat tct atc tct --- --- --- --- --- --- --- --- --- --- --- --- --- --- --- --- -

>BK013340.1:132242-132642_TPA_asm:_Vaccinia_virus_isolate_VK02

gct gct agt tgt cgt tcg aga tca gaa ata cgg gtt ttc ttg gaa ttg att tcg tcg atg tac ttg gca tga ttg gca tcg aaa cac tta tta agt tc- ttt ttc caa ttc tac gat ttt att tct ttc gcg agt caa ttc cct cct gta gta act atc tgt ttt gtc aga ttc acg ctc tct acg tag act ttc ttg caa gtt act aat ttg ttc cct agc acg tcc gag ttt agt ttt ata tgc cga ata gag ttc tga ttc atc ctt tga gca gat ctc tag cga tcg ttt aag att cct gat tct agt ctt tag cct att tac ctc ctc aga aga tgt tcc gtt acc gtt gcg ttt aca ctc gtt aag ctg tct atc aag atc cat gat tct atc tct --- --- --- --- --- --- --- --- --- --- --- --- --- --- --- --- -

>HQ420893.1:151277-151677_Cowpox_virus_strain_Finland_2000_MAN_complete_genome

gct gct agc tgt cgt tcg aga tca gaa ata cgg gtt ttc ttg gaa ttg att tcg tcg atg tac ttg gca tga ttg gta tcg aaa cac tta tta agt tc- ttt ttc caa ttc tac gat ttt att tct ttc gcg agt caa ttc cct cct gta gta gct atc ggt ttt gtc aga ttc acg ctc tct acg tag act ttc ttg caa gtt act aat ttg ttc cct ggc acg tcc gag ttc agt ttt ata tgc cga ata gag ttc tga ttc atc ctt tga gca gat ctc tag aga tcg ttc aag atc cct gat tct agt ctt tag cct att tac ctc ctc aga aga tgc tcc gtt acc gtt gcg ttt aca ctc gtt gag ctg tct atc aag atc cat gat tct atc tct --- --- --- --- --- --- --- --- --- --- --- --- --- --- --- --- -

>AF095689.1:135345-135745_Vaccinia_virus_(strain_Tian_Tan)_complete_genome

gct gct agt tgt cgt tcg aga tca gaa ata cgg gtt ttc ttg gaa ttg att tcg tcg atg tac ttg gca tga ttg gca tcg aaa cac tta tta agt tc- ttt ttc caa ttc tac gat ttt att tct ttc gcg agt caa ttc cct cct gta gta act atc tgt ttt gtc aga ttc acg ctc tct acg tag act ttc ttg caa gtt act aat ttg ttc cct agc acg tcc gag ttt agt ttt ata tgc cga ata gag ttc tga ttc atc ctt tga gca gat ctc tag cga tcg ttt aag att cct gat tct agt ctt tag cct att tac ctc ctc aga aga tgt tcc gtt acc gtt gcg ttt aca ctc gtt aag ctg tct atc aag atc cat gat tct atc tct --- --- --- --- --- --- --- --- --- --- --- --- --- --- --- --- -

>KY549143.2:147529-147929_Cowpox_virus_isolate_CPXV_Catpox3L97

gct gat agt tgt cgt tcg aga tca gaa ata cgg gtt ttc ttg gaa ttg att tcg tcg atg tac ttg gca tga ttg gca tcg aaa cac tta tta agt tc- ttt ttc caa ttc tac gat ttt att tct ttc gcg agt caa ttc cct cct gta gta act atc tgt ttt gtc aga ttc acg ctc tct acg tag act ttc ttg caa gtt act aat ttg ttc tct agc acg tcc gag ttc agt ttt ata tgc cga ata gag ttc tga ttc atc ctt tga gca gat ctc tag aga tcg ttc aag atc cct gat tct agt ctt tag cct att tac ctc ctc aga aga tgt tcc gtt acc gtt gcg ttt aca ctc gtt gag ctg tct atc aag atc cat gat tct atc tct --- --- --- --- --- --- --- --- --- --- --- --- --- --- --- --- -

>KC813493.1:144099-144499_Cowpox_virus_strain_HumLit08/1_complete_genome

gct gct agt tgt cgt tcg aga tca gaa ata cgg gtt ttc ttg gaa ttg att tcg tcg atg tac ttg gca tga ttg gca tcg aaa cac tta tta agt tc- ttt ttc caa ttc tac gat ttt att tct ttc gcg agt caa ttc cct cct gta gta act atc ggt ttt gtc aga ttc acg ctc tct ccg tag act ttc ttg caa gtt act aat ttg ttc cct ggc acg tcc gag ttc agt ttt ata tgc cga ata gag ttc tga ttc atc ctt tga gca gat ctc tag aga tcg ttc aag atc cct aat tct agt ttt tag cct att tac ctc ctc aga aga tgt tcc gtt acc gtt gcg ttt aca ctc gtt gag ctg tct atc aag atc cat gat tct atc tct --- --- --- --- --- --- --- --- --- --- --- --- --- --- --- --- -

>JX489139.1:135276-135676_Vaccinia_virus_strain_Tiantan_clone_TT12_complete_genome

gct gct agt tgt cgt tcg aga tca gaa ata cgg gtt ttc ttg gaa ttg att tcg tcg atg tac ttg gca tga ttg gca tcg aaa cac tta tta agt tc- ttt ttc caa ttc tac gat ttt att tct ttc gcg agt caa ttc cct cct gta gta act atc tgt ttt gtc aga ttc acg ctc tct acg tag act ttc ttg caa gtt act aat ttg ttc cct agc acg tcc gag ttt agt ttt ata tgc tga ata gag ttc tga ttc atc ctt tga gca gat ctc tag cga tcg ttt aag att cct gat tct agt ctt tag cct att tac ctc ctc aga aga tgt tcc gtt acc gtt gcg ttt aca ctc gtt aag ctg tct atc aag atc cat gat tct atc tct --- --- --- --- --- --- --- --- --- --- --- --- --- --- --- --- -

>JX489136.1:135126-135526_Vaccinia_virus_strain_Tiantan_clone_TT9_complete_genome

gct gct agt tgt cgt tcg aga tca gaa ata cgg gtt ttc ttg gaa ttg att tcg tcg atg tac ttg gca tga ttg gca tcg aaa cac tta tta agt tc- ttt ttc caa ttc tac gat ttt att tct ttc gcg agt caa ttc cct cct gta gta act atc tgt ttt gtc aga ttc acg ctc tct acg tag act ttc ttg caa gtt act aat ttg ttc cct agc acg tcc gag ttt agt ttt ata tgc tga ata gag ttc tga ttc atc ctt tga gca gat ctc tag cga tcg ttt aag att cct gat tct agt ctt tag cct att tac ctc ctc aga aga tgt tcc gtt acc gtt gcg ttt aca ctc gtt aag ctg tct atc aag atc cat gat tct atc tct --- --- --- --- --- --- --- --- --- --- --- --- --- --- --- --- -

>JX489135.1:134716-135116_Vaccinia_virus_strain_Tiantan_clone_TT8_complete_genome

gct gct agt tgt cgt tcg aga tca gaa ata cgg gtt ttc ttg gaa ttg att tcg tcg atg tac ttg gca tga ttg gca tcg aaa cac tta tta agt tc- ttt ttc caa ttc tac gat ttt att tct ttc gcg agt caa ttc cct cct gta gta act atc tgt ttt gtc aga ttc acg ctc tct acg tag act ttc ttg caa gtt act aat ttg ttc cct agc acg tcc gag ttt agt ttt ata tgc tga ata gag ttc tga ttc atc ctt tga gca gat ctc tag cga tcg ttt aag att cct gat tct agt ctt tag cct att tac ctc ctc aga aga tgt tcc gtt acc gtt gcg ttt aca ctc gtt aag ctg tct atc aag atc cat gat tct atc tct --- --- --- --- --- --- --- --- --- --- --- --- --- --- --- --- -

>BK013343.1:159950-160350_TPA_asm:_Vaccinia_virus_isolate_VK12

gct gct agt tgt cgt tcg aga tca gaa ata cgg gtt ttc ttg gaa ttg att tcg tcg atg tac ttg gca tga ttg gca tcg aaa cac tta tta agt tc- ttt ttc caa ttc tac gat ttt att tct ttc gcg agt caa ttc cct cct gta gta act atc tgt ttt gtc aga ttc acg ctc tct acg tag act ttc ttg caa gtt act aat ttg ttc cct agc acg tcc gag ttt agt ttt ata tgc tga ata gag ttc tga ttc atc ctt tga gca gat ctc tag cga tcg ttt aag att cct gat tct agt ctt tag cct att tac ctc ctc aga aga tgt tcc gtt acc gtt gcg ttt aca ctc gtt aag ctg tct atc aag atc cat gat tct atc tct --- --- --- --- --- --- --- --- --- --- --- --- --- --- --- --- -

>HQ420899.1:151266-151666_Cowpox_virus_strain_Norway_1994_MAN_complete_genome

gct gct agt tgt cgt tcg aga tca gaa ata cgg gtt ttc ttg gaa ttg att tcg tcg atg tac ttg gca tga ttg gca tcg aaa cac tta tta agt tc- ttt ttc caa ttc tat gat ttt att tct ttc gcg agt caa ttc cct cct gta gta act atc ggt ttt gtc aga ttc acg ctc ttt acg tag act ttc ttg caa gtt act aat ttg ttc cct tgc acg tcc gag ttc agt ttt ata tgc cga ata gag ttc tga ttc atc ctt tga gca gat ctc tag aga tcg ttc aag atc cct aat tct agt ttt tag cct att tac ctc ctc aga aga tgc tcc gtt acc gtt gcg ttt aca ctc gtt aag ctg tct atc aag atc cat gat tct atc tct --- --- --- --- --- --- --- --- --- --- --- --- --- --- --- --- -

>HQ407377.1:150409-150809_Cowpox_virus_strain_Austria_1999_complete_genome

gct gat agt tgt cgt tcg aga tca gaa ata cgg gtt ttc ttg gaa ttg att tcg tcg atg tac ttg gca tga ttg gca tcg aaa cac tta tta agt tc- ttt ttc caa ttc tac gat ttt att tct ttc gcg agt caa ttc cct cct gta gta act atc tgt ttt gtc aga ttc acg ctc tct acg tag act ttc ttg caa gtt act aat ttg ttc tct agc acg tcc gag ttc agt ttt ata tgc cga ata gag ttc tga ttc atc ctt tga gca gat ctc tag aga tcg ttc aag atc cct gat tct agt ctt tag cct att tac ctc ctc aga aga tgt tcc gtt acc gtt gcg ttt aca ctc gtt gag ctg tct atc aag atc cat gat tct atc tct --- --- --- --- --- --- --- --- --- --- --- --- --- --- --- --- -

>LR800245.1:130805-131205_Variola_virus_genome_assembly_complete_genome:_monopartite

gct gct agc tgt cgt tcg aga tca gaa ata cgg gtt ttc ttg gaa ttg att tcg tca atg tac ttg gca tga ttg aca tcg aaa cac tta tta agt tc- ttt ttc caa ttc tac gat ttt att tct ttc gcg agt caa ttc cct cct gta gta gct atc ggt ttt gtc aga ttc acg ctc tct acg tag act ttc ttg caa gtt act aat ttg ttc cct ggc acg tcc gag ttt agt ttt ata tgc cga ata gag ttc tga ttc atc ctt tga gca gat ctc tag cga tcg ttc aag atc cct gat tct agt ctt tag cct att tac ctc ctc aga aga tgt tcc gtt acc gtt gcg ttt aca ctc gtt aag ctg tct atc aag atc cat gat tat atc tct --- --- --- --- --- --- --- --- --- --- --- --- --- --- --- --- -

>LR800244.1:130798-131198_Variola_virus_genome_assembly_complete_genome:_monopartite

gct gct agc tgt cgt tcg aga tca gaa ata cgg gtt ttc ttg gaa ttg att tcg tca atg tac ttg gca tga ttg aca tcg aaa cac tta tta agt tc- ttt ttc caa ttc tac gat ttt att tct ttc gcg agt caa ttc cct cct gta gta gct atc ggt ttt gtc aga ttc acg ctc tct acg tag act ttc ttg caa gtt act aat ttg ttc cct ggc acg tcc gag ttt agt ttt ata tgc cga ata gag ttc tga ttc atc ctt tga gca gat ctc tag cga tcg ttc aag atc cct gat tct agt ctt tag cct att tac ctc ctc aga aga tgt tcc gtt acc gtt gcg ttt aca ctc gtt aag ctg tct atc aag atc cat gat tat atc tct --- --- --- --- --- --- --- --- --- --- --- --- --- --- --- --- -

>LT896723.1:152360-152760_Cowpox_virus_isolate_Ger/2014/Cat1_genome_assembly_complete_genome:_monopartite

gct gct agt tgt cgt tcg aga tca gaa ata cgg gtt ttc ttg gaa tcg att tcg tcg atg tac ttg gca tga ttg gta tcg aaa cac tta tta agt tc- ttt ttc caa ttc tac gat ttt att tct ttc gcg agt caa ttc cct cct gta gta act atc tgt ttt gtc aga ttc acg ctc tct acg tag act ttc ttg caa gtt act aat ttg ttc tct agc acg tcc gag ttc agt ttt ata tgc cga ata gag ttc tga ttc atc ctt tga gca gat ctc tag aga tcg ttc aag atc cct aat tct agt ttt tag cct att tac ctc ctc aga aga tgt tcc gtt acc gtt gcg ttt aca ctc gtt aag ctg tct atc aag atc cat gat tct atc tct --- --- --- --- --- --- --- --- --- --- --- --- --- --- --- --- -

>KY549151.1:148056-148456_Cowpox_virus_isolate_Norwayfeline

gct gct agt tgt cgt tcg aga tca gaa ata cgg gtt ttc ttg gaa ttg att tcg tcg atg tac ttg gca tga ttg gca tcg aaa cac tta tta agt tc- ttt ttc caa ttc tat gat ttt att tct ttc gcg agt caa ttc cct cct gta gta act atc ggt ttt gtc aga ttc acg ctc ttt acg tag act ttc ttg caa gtt act aat ttg ttc cct tgc acg tct gag ttc agt ttt ata tgc cga ata gag ttc tga ttc atc ctt tga gca gat ctc tag aga tcg ttc aag atc cct aat tct agt ttt tag cct att tac ctc ctc aga aga tgc tcc gtt acc gtt gcg ttt aca ctc gtt aag ctg tct atc aag atc cat gat tct atc tct --- --- --- --- --- --- --- --- --- --- --- --- --- --- --- --- -

>KY463519.1:143066-143466_Cowpox_virus_strain_Germany_1971_EP1

gct gct agt tgt cgt tcg aga tca gaa ata cgg gtt ttc ttg gaa tcg att tcg tcg atg tac ttg gca tga ttg gta tcg aaa cac tta tta agt tc- ttt ttc caa ttc tac gat ttt att tct ttc gcg agt caa ttc cct cct gta gta act atc tgt ttt gtc aga ttc acg ctc tct acg tag act ttc ttg caa gtt act aat ttg ttc tct agc acg tcc gag ttc agt ttt ata tgc cga ata gag ttc tga ttc atc ctt tga gca gat ctc tag aga tcg ttc aag atc cct aat tct agt ttt tag cct att tac ctc ctc aga aga tgt tcc gtt acc gtt gcg ttt aca ctc gtt aag ctg tct atc aag atc cat gat tct atc tct --- --- --- --- --- --- --- --- --- --- --- --- --- --- --- --- -

>KY349117.1:147249-147648_Horsepox_virus_strain_MNR_clone_2-1

gct gct agt tgt cgt tcg aga tca gaa ata cgg gtt ttc ttg -aa ttg att tcg tcg atg tac ttg gca tga ttg gca tcg aaa cac tta tta agt tc- ttt ttc caa ttc tac gat ttt att tct ttc gca agt caa ttc cct cct gta gta act atc tgt ttt gtt aga ttc acg ctc tct acg tag act ttc ttg caa gtt act aat ttg ttc cct agc acg tcc gag ttc agt ttt ata tgc cga ata gag ttc tga ttc atc ctt tga gca gat ctc tag cga tcg ttt aag att cct gat tct agt ctt tag cct att tac ctc ctc aga aga tgt tcc gtt acc gtt gcg ttt aca ctc gtt aag ctg tct atc aag atc cat gat tct atc tct --- --- --- --- --- --- --- --- --- --- --- --- --- --- --- --- -

>DQ792504.1:147160-147559_Horsepox_virus_isolate_MNR-76_complete_genome

gct gct agt tgt cgt tcg aga tca gaa ata cgg gtt ttc ttg -aa ttg att tcg tcg atg tac ttg gca tga ttg gca tcg aaa cac tta tta agt tc- ttt ttc caa ttc tac gat ttt att tct ttc gca agt caa ttc cct cct gta gta act atc tgt ttt gtt aga ttc acg ctc tct acg tag act ttc ttg caa gtt act aat ttg ttc cct agc acg tcc gag ttc agt ttt ata tgc cga ata gag ttc tga ttc atc ctt tga gca gat ctc tag cga tcg ttt aag att cct gat tct agt ctt tag cct att tac ctc ctc aga aga tgt tcc gtt acc gtt gcg ttt aca ctc gtt aag ctg tct atc aag atc cat gat tct atc tct --- --- --- --- --- --- --- --- --- --- --- --- --- --- --- --- -

>LR800247.1:130762-131162_Variola_virus_genome_assembly_complete_genome:_monopartite

gct act agc tgt cgt tcg aga tca gaa ata cgg gtt ttc ttg gaa ttg att tcg tcg atg tac ttg gca tga ttg aca tcg aaa cac tta tta agt tc- ttt ttc caa ttc tac gat ttt att tct ttc gcg agt caa ttc cct cct gta gta gct atc ggt ttt gtc aga ttc acg ctc tct acg tag act ttc ttg caa gtt act aat ttg ttc cct ggt acg tcc gag ttc agt ttt ata tac cga ata gag ttc tga ttc atc ctt tga gca gat ctc tag cga tcg ttc aag atc cct gat tct agt ctt tag cct att tac ctc ctc aaa aga tgt tcc gtt acc gtt gcg ttt aca ctc gtt aag ctg tct atc aag atc cat gat tct atc tct --- --- --- --- --- --- --- --- --- --- --- --- --- --- --- --- -

>LT993226.1:151330-151730_Cowpox_virus_isolate_Ger/2014/Human_genome_assembly_complete_genome:_monopartite

gct gca agt tgt cgt tcg aga tca gaa ata cgg gtt ttc ttg gaa ttg att tcg tcg atg tac ttg gca tga ttg gta tcg aaa cac tta tta agt tc- ttt ttc caa ttc tac gat ttt att tct ttc gcg agt caa ttc cct cct gta gta act atc tgt ttt gtc aga ttc acg ctc tct acg tag act ttc ttg caa gtt act aat ttg ttc tct agc acg tcc gag ttc agt ttt ata tgc cga ata gag ttc tga ttc atc ctt tga gca gat ctc tag aga tcg ttc aag atc cct aat tct agt ttt tag cct att tac ctc ctc aga aga tgt tcc gtt acc gtt gcg ttt aca ttc gtt aag ctg tct atc aag atc cat gat tct atc tct --- --- --- --- --- --- --- --- --- --- --- --- --- --- --- --- -

>LT896722.1:151777-152177_Cowpox_virus_isolate_Ger/2007/Vole_genome_assembly_complete_genome:_monopartite

gct gca agt tgt cgt tcg aga tca gaa ata cgg gtt ttc ttg gaa ttg att tcg tcg atg tac ttg gca tga ttg gta tcg aaa cac tta tta agt tc- ttt ttc caa ttc tac gat ttt att tct ttc gcg agt caa ttc cct cct gta gta act atc tgt ttt gtc aga ttc acg ctc tct acg tag act ttc ttg caa gtt act aat ttg ttc tct agc acg tcc gag ttc agt ttt ata tgc cga ata gag ttc tga ttc atc ctt tga gca gat ctc tag aga tcg ttc aag atc cct aat tct agt ttt tag cct att tac ctc ctc aga aga tgt tcc gtt acc gtt gcg ttt aca ttc gtt aag ctg tct atc aag atc cat gat tct atc tct --- --- --- --- --- --- --- --- --- --- --- --- --- --- --- --- -

>LT896721.1:151846-152246_Cowpox_virus_isolate_Ger_2010_MKY_genome_assembly_complete_genome:_monopartite

gct gct agt tgt cgt tcg aga tca gaa ata cgg gtt ttc ttg gaa ttg att tcg tcg atg tac ttg gca tga ttg gta tcg aaa cac tta tta agt tc- ttt ttc caa ttc tac gat ttt att tct ttc gcg agt caa ttc cct cct gta gta act atc tgt ttt gtc aga ttc acg ctc tct acg tag act ttc ttg caa gtt act aat ttg ttc tct agc acg tcc gag ttc agt ttt ata tgc aga ata gag ttc tga ttc atc ctt tga gca gat ctc tag aga tcg ttc aag atc cct aat tct agt ttt tag cct att tac ctc ctc aga aga tgt tcc gtt acc gtt gcg ttt aca ttc gtt aag ctg tct atc aag atc cat gat tct atc tct --- --- --- --- --- --- --- --- --- --- --- --- --- --- --- --- -

>KY549145.1:153378-153778_Cowpox_virus_isolate_CPXV_K428

gct gct agt tgt cgt tcg aga tca gaa ata cgg gtt ttc ttg gaa ttg att tcg tcg atg tac ttg gca tga ttg gta tcg aaa cac tta tta agt tc- ttt ttc caa ttc tac gat ttt att tct ttc acg agt caa ttc cct cct gta gta act atc ggt ttt gtc aga ttc acg ctc tct acg tag act ttc ttg caa gtt act aat ttg ttc tct ggc acg tcc gag ttc act ctt ata tgc tga ata gag ttc tga ttc atc ctt tga gca gat ctc tag aga tcg ttc aag atc cct gat tct agt ttt tag cct att tac ctc ctc aga aga tgc tcc gtt acc gtt gcg ttt aca ttc gtt aag ctg tct atc aag atc cat gat tct atc tct --- --- --- --- --- --- --- --- --- --- --- --- --- --- --- --- -

>LR812035.1:151721-152121_Cowpox_virus_isolate_GerMygEK_938/17_genome_assembly_complete_genome:_monopartite

gct gct agt tgt cgt tcg aga tca gaa ata cgg gtt ttc ttg gaa ttg att tcg tcg atg tac ttg gca tga ttg gta tcg aaa cac tta tta agt tc- ttt ttc caa ttc tac gat ttt att tct ttc gcg agt caa ttc cct cct gta gta act atc tgt ttt gtc aga ttc acg ctc tct acg tag act ttc ttg caa gtt act aat ttg ttc tct agc acg tcc gag ttc agt ttt ata tgc aga ata gag ttc tga ttc atc ctt tga gca gat ctc tag aga tcg ttc aag atc cct aat tct agt ttt tag cct att tac ctc ctc aga aga tgt tcc gtt acc gtt gcg ttt aca ttc gtt aag ctg tct atc aag atc cat gat tct atc tct --- --- --- --- --- --- --- --- --- --- --- --- --- --- --- --- -

>HQ420896.1:150864-151264_Cowpox_virus_strain_Germany_1990_2_complete_genome

gct gct agt tgt cgt tcg aga tca gaa ata cgg gtt ttc ttg gaa ttg att tcg tcg atg tac ttg gca tga ttg gta tcg aaa cac tta tta agt tc- ttt ttc caa ttc tac gat ttt att tct ttc gcg agt caa ttc cct cct gta gta act atc tgt ttt gtc aga ttc acg ctc tct acg tag act ttc ttg caa gtt act aat ttg ttc tct ggc acg tcc gag ttc act ctt ata tgc tga ata gag ttc tga ttc atc ctt tga gca gat ctc tag aga tcg ttc aag atc cct gat tct agt ttt tag cct att tac ctc ctc aga aga tgt tcc gtt acc gtt gcg ttt aca ttc gtt aag ctg tct atc aag atc cat gat tct atc tct --- --- --- --- --- --- --- --- --- --- --- --- --- --- --- --- -

>KY549150.2:152582-152982_Cowpox_virus_isolate_CPXV_K4207

gct gct agt tgt cgt tcg aga tca gaa ata cgg gtt ttc ttg gaa ttg att tcg tcg atg tac ttg gca tga ttg gta tcg aaa cac tta tta agt tc- ttt ttc caa ttc tac gat ttt att tct ttc acg agt caa ttc cct cct gta gta act atc ggt ttt gtc aga ttc acg ctc tct acg tag act ttc ttg caa gtt act aat ttg ttc tct ggc acg tcc gag ttc act ctt ata tgc tga ata gag ttc tga ttc atc ctt tga gca gat ctc tag aga tcg ttc aag atc cct gat tct agt ttt tag cct att tac ctc ctc aga aga tgt tcc gtt acc gtt gcg ttt aca ttc gtt aag ctg tct atc aag atc cat gat tct atc tct --- --- --- --- --- --- --- --- --- --- --- --- --- --- --- --- -

>LT883663.1:150116-150516_Cowpox_virus_strain_France_Amiens_2016_genome_assembly_complete_genome:_monopartite

gct gct agt tgt cgt tcg aga tca gaa ata cgg gtt ttc ttg gaa ttg att tcg tcg atg tac ttg gca tga ttg gta tcg aaa cac tta tta agt tc- ttt ttc caa ttc tac gat ttt att tct ttc acg agt caa ttc cct cct gta gta act atc ggt ttt gtc aga ttc acg ctc tct acg tag act ttc ttg caa gtt act aat ttg gtc tct ggc acg tcc gag ttc act ctt ata tgc tga ata gag ttc tga ttc atc ctt tga gca gat ctc tag aga tcg ttc aag atc cct gat tct agt ctt tag cct att tac ctc ctc aga aga tgt tcc gtt acc gtt gcg ttt aca ttc gtt aag ctg tct atc aag atc cat gat tct atc tct --- --- --- --- --- --- --- --- --- --- --- --- --- --- --- --- -

>KY569022.1:142056-142456_Cowpox_virus_strain_CPXV_CheTopCut_DK_2011

gct gct agt tgt cgt tcg aga tca gaa ata cgg gtt ttc ttg gaa tcg att tcg tcg atg tac ttg gca tga ttg gaa tcg aaa cac tta tta agt tc- ttt ttc caa ttc tac gat ttt att tct ttc acg agt caa ttc cct cct gta gta act atc ggt ttt gtc aga ttc acg ctc tct acg tag act ttc ttg caa gtt act aat ttg ttc tct ggc acg tcc gag ttc act ctt ata tgc tga ata gag ttc tga ttc atc ctt tga gca gat ctc tag aga tcg ttc aag atc cct gat tct agt ttt tag cct att tac ctc ctc gga aga tgc tcc gtt acc gtt gcg ttt aca ttc gtt aag ctg tct atc aag atc cat gat tct atc tct --- --- --- --- --- --- --- --- --- --- --- --- --- --- --- --- -

>KY569021.1:142095-142495_Cowpox_virus_strain_CPXV_CheGrey_DK_2010

gct gct agt tgt cgt tcg aga tca gaa ata cgg gtt ttc ttg gaa tcg att tcg tcg atg tac ttg gca tga ttg gaa tcg aaa cac tta tta agt tc- ttt ttc caa ttc tac gat ttt att tct ttc acg agt caa ttc cct cct gta gta act atc ggt ttt gtc aga ttc acg ctc tct acg tag act ttc ttg caa gtt act aat ttg ttc tct ggc acg tcc gag ttc act ctt ata tgc tga ata gag ttc tga ttc atc ctt tga gca gat ctc tag aga tcg ttc aag atc cct gat tct agt ttt tag cct att tac ctc ctc gga aga tgc tcc gtt acc gtt gcg ttt aca ttc gtt aag ctg tct atc aag atc cat gat tct atc tct --- --- --- --- --- --- --- --- --- --- --- --- --- --- --- --- -

>KY569020.1:141974-142374_Cowpox_virus_strain_CPXV_CheNuru_DK_2012

gct gct agt tgt cgt tcg aga tca gaa ata cgg gtt ttc ttg gaa tcg att tcg tcg atg tac ttg gca tga ttg gaa tcg aaa cac tta tta agt tc- ttt ttc caa ttc tac gat ttt att tct ttc acg agt caa ttc cct cct gta gta act atc ggt ttt gtc aga ttc acg ctc tct acg tag act ttc ttg caa gtt act aat ttg ttc tct ggc acg tcc gag ttc act ctt ata tgc tga ata gag ttc tga ttc atc ctt tga gca gat ctc tag aga tcg ttc aag atc cct gat tct agt ttt tag cct att tac ctc ctc gga aga tgc tcc gtt acc gtt gcg ttt aca ttc gtt aag ctg tct atc aag atc cat gat tct atc tct --- --- --- --- --- --- --- --- --- --- --- --- --- --- --- --- -

>KY569019.1:150654-151054_Cowpox_virus_strain_CPXV_CheNova_DK_2014

gct gct agt tgt cgt tcg aga tca gaa ata cgg gtt ttc ttg gaa tcg att tcg tcg atg tac ttg gca tga ttg gaa tcg aaa cac tta tta agt tc- ttt ttc caa ttc tac gat ttt att tct ttc acg agt caa ttc cct cct gta gta act atc ggt ttt gtc aga ttc acg ctc tct acg tag act ttc ttg caa gtt act aat ttg ttc tct ggc acg tcc gag ttc act ctt ata tgc tga ata gag ttc tga ttc atc ctt tga gca gat ctc tag aga tcg ttc aag atc cct gat tct agt ttt tag cct att tac ctc ctc gga aga tgc tcc gtt acc gtt gcg ttt aca ttc gtt aag ctg tct atc aag atc cat gat tct atc tct --- --- --- --- --- --- --- --- --- --- --- --- --- --- --- --- -

>KY569018.1:142007-142407_Cowpox_virus_strain_CPXV_CheHurley_DK_2012

gct gct agt tgt cgt tcg aga tca gaa ata cgg gtt ttc ttg gaa tcg att tcg tcg atg tac ttg gca tga ttg gaa tcg aaa cac tta tta agt tc- ttt ttc caa ttc tac gat ttt att tct ttc acg agt caa ttc cct cct gta gta act atc ggt ttt gtc aga ttc acg ctc tct acg tag act ttc ttg caa gtt act aat ttg ttc tct ggc acg tcc gag ttc act ctt ata tgc tga ata gag ttc tga ttc atc ctt tga gca gat ctc tag aga tcg ttc aag atc cct gat tct agt ttt tag cct att tac ctc ctc gga aga tgc tcc gtt acc gtt gcg ttt aca ttc gtt aag ctg tct atc aag atc cat gat tct atc tct --- --- --- --- --- --- --- --- --- --- --- --- --- --- --- --- -

>KY549148.1:152896-153296_Cowpox_virus_isolate_CPXV_1639

gct gct agt tgt cgt tcg aga tca gaa ata cgg gtt ttc ttg gaa ttg att tcg tcg atg tac ttg gca tga ttg gta tcg aaa cac tta tta agt tc- ttt ttc caa ttc tac gat ttt att tct ttc acg agt caa ttc cct cct gta gta act atc ggt ttt gtc aga ttc acg ctc tct acg tag act ttc ttg caa gtt act aat ttg ttc cct tgc acg tcc gag ttc act ctt ata tgc tga ata gag ttc tga ttc atc ctt tga gca gat ctc tag aga tcg ttc aag atc cct gat tct agt ctt tag cct att tac ctc ctc aga aga tgt tcc gtt acc gtc gcg ttt aca ttc gtt aag ctg tct atc aag atc cat gat tct atc tct --- --- --- --- --- --- --- --- --- --- --- --- --- --- --- --- -

>KY549147.1:144478-144878_Cowpox_virus_isolate_CPXV_K780

gct gct agt tgt cgt tcg aga tca gaa ata cgg gtt ttc ttg gaa tcg att tcg tcg atg tac ttg gca tga ttg gta tcg aaa cac tta tta agt tc- ttt ttc caa ttc tac gat ttt att tct ttc acg agt caa ttc cct cct gta gta act atc ggt ttt gtc aga ttc acg ctc tct acg tag act ttc ttg caa gtt act aat ttg ttc tct ggc acg tcc gag ttc act ctt ata tgc tga ata gag ttc tga ttc atc ctt tga gca gat ctc tag aga tcg ttc aag atc cct gat tct agt ttt tag cct att tac ctc ctc aga aga tgt tcc gtt acc gtt gcg ttt aca ttc gtt aag ctg tct atc aag atc cat gat tct atc tct --- --- --- --- --- --- --- --- --- --- --- --- --- --- --- --- -

>KY549146.1:144712-145112_Cowpox_virus_isolate_CPXV_K779

gct gct agt tgt cgt tcg aga tca gaa ata cgg gtt ttc ttg gaa tcg att tcg tcg atg tac ttg gca tga ttg gta tcg aaa cac tta tta agt tc- ttt ttc caa ttc tac gat ttt att tct ttc acg agt caa ttc cct cct gta gta act atc ggt ttt gtc aga ttc acg ctc tct acg tag act ttc ttg caa gtt act aat ttg ttc tct ggc acg tcc gag ttc act ctt ata tgc tga ata gag ttc tga ttc atc ctt tga gca gat ctc tag aga tcg ttc aag atc cct gat tct agt ttt tag cct att tac ctc ctc aga aga tgt tcc gtt acc gtt gcg ttt aca ttc gtt aag ctg tct atc aag atc cat gat tct atc tct --- --- --- --- --- --- --- --- --- --- --- --- --- --- --- --- -

>LN864566.1:151605-152005_Cowpox_virus_complete_genome_strain_FM2292

gct gca agt tgt cgt tcg aga tca gaa ata cgg gtt ttc ttg gaa ttg att tcg tcg atg tac ttg gca tga ttg gta tcg aaa cac tta tta agt tc- ttt ttc caa ttc tac gat ttt att tct ttc gcg agt caa ttc cct cct gta gta act atc tgt ttt gtc aga ttc acg ctc tct acg tag act ttc ttg caa gtt act aat ttg ttc tct ggc acg tcc gag ttc act ctt ata tgc tga ata gag ttc tga ttc atc ctt tga gca gat ctc tag aga tcg ttc aag atc cct gat tct agt ttt tag cct att tac ctc ctc aga aga tgt tcc gtt acc gtt gcg ttt aca ttc gtt aag ctg tct atc aag atc cat gat tct atc tct --- --- --- --- --- --- --- --- --- --- --- --- --- --- --- --- -

>KC813509.1:139673-140073_Cowpox_virus_strain_HumBer07/1_complete_genome

gct gct agt tgt cgt tcg aga tca gaa ata cgg gtt ttc ttg gaa tcg att tcg tcg atg tac ttg gca tga ttg gca tcg aaa cac tta tta agt tc- ttt ttc caa ttc tac gat ttt att tct ttc acg cgt caa ttc cct cct gta gta act atc ggt ttt gtc aga ttc acg ctc tct acg tag act ttc ttg caa gtt act aat ttg ttc tct ggc acg tcc gag ttc act ctt ata tgc tga ata gag ttc tga ttc atc ctt tga gca gat ctc tag aga tcg ttc aag atc cct gat tct agt ttt tag cct att tac ctc ctc aga aga tgt tcc gtt acc gtt gcg ttt aca ttc gtt aag ctg tct atc aag atc cat gat tct atc tct --- --- --- --- --- --- --- --- --- --- --- --- --- --- --- --- -

>KC813506.1:152900-153300_Cowpox_virus_strain_CatPot07/1_complete_genome

gct gct agt tgt cgt tcg aga tca gaa ata cgg gtt ttc ttg gaa tcg att tcg tcg atg tac ttg gca tga ttg gca tcg aaa cac tta tta agt tc- ttt ttc caa ttc tac gat ttt att tct ttc acg cgt caa ttc cct cct gta gta act atc ggt ttt gtc aga ttc acg ctc tct acg tag act ttc ttg caa gtt act aat ttg ttc tct ggc acg tcc gag ttc act ctt ata tgc tga ata gag ttc tga ttc atc ctt tga gca gat ctc tag aga tcg ttc aag atc cct gat tct agt ttt tag cct att tac ctc ctc aga aga tgt tcc gtt acc gtt gcg ttt aca ttc gtt aag ctg tct atc aag atc cat gat tct atc tct --- --- --- --- --- --- --- --- --- --- --- --- --- --- --- --- -

>KC813504.1:145156-145556_Cowpox_virus_strain_RatHei09/1_complete_genome

gct gct agt tgt cgt tcg aga tca gaa ata cgg gtt ttc ttg gaa tcg att tcg tcg atg tac ttg gca tga ttg gta tcg aaa cac tta tta agt tc- ttt ttc caa ttc tac gat ttt att tct ttc acg cgt caa ttc cct cct gta gta act atc ggt ttt gtc aga ttc acg ctc tct acg tag act ttc ttg caa gtt act aat ttg ttc cct ggc acg tcc gag ttc agt ttt ata tgc cga ata gag ttc tga ttc atc ctt tga gca gat ctc taa aga tcg ttc aag atc cct aat tct agt ttt tag cct att tac ctc ctc aga aga tgt tcc gtt acc gtt gcg ttt aca ctc gtt gag ctg tct atc aag atc cat gat tct atc tct --- --- --- --- --- --- --- --- --- --- --- --- --- --- --- --- -

>KC813502.1:144856-145256_Cowpox_virus_strain_CatBer07/1_complete_genome

gct gct agt tgt cgt tcg aga tca gaa ata cgg gtt ttc ttg gaa tcg att tcg tcg atg tac ttg gca tga ttg gca tcg aaa cac tta tta agt tc- ttt ttc caa ttc tac gat ttt att tct ttc acg cgt caa ttc cct cct gta gta act atc ggt ttt gtc aga ttc acg ctc tct acg tag act ttc ttg caa gtt act aat ttg ttc tct ggc acg tcc gag ttc act ctt ata tgc tga ata gag ttc tga ttc atc ctt tga gca gat ctc tag aga tcg ttc aag atc cct gat tct agt ttt tag cct att tac ctc ctc aga aga tgt tcc gtt acc gtt gcg ttt aca ttc gtt aag ctg tct atc aag atc cat gat tct atc tct --- --- --- --- --- --- --- --- --- --- --- --- --- --- --- --- -

>KC813500.1:134234-134634_Cowpox_virus_strain_MonKre08/4_complete_genome

gct gct agt tgt cgt tcg aga tca gaa ata cgg gtt ttc ttg gaa tcg att tcg tcg atg tac ttg gca tga ttg gta tcg aaa cac tta tta agt tc- ttt ttc caa ttc tac gat ttt att tct ttc acg cgt caa ttc cct cct gta gta act atc ggt ttt gtc aga ttc acg ctc tct acg tag act ttc ttg caa gtt act aat ttg ttc cct ggc acg tcc gag ttc agt ttt ata tgc cga ata gag ttc tga ttc atc ctt tga gca gat ctc taa aga tcg ttc aag atc cct aat tct agt ttt tag cct att tac ctc ctc aga aga tgt tcc gtt acc gtt gcg ttt aca ctc gtt gag ctg tct atc aag atc cat gat tct atc tct --- --- --- --- --- --- --- --- --- --- --- --- --- --- --- --- -

>KC813499.1:151055-151455_Cowpox_virus_strain_MarLei07/1_complete_genome

gct gct agt tgt cgt tcg aga tca gaa ata cgg gtt ttc ttg gaa ttg att tcg tcg atg tac ttg gca tga ttg gta tcg aaa cac tta tta agt tc- ttt ttc caa ttc tac gat ttt att tct ttc acg agt caa ttc cct cct gta gta act atc ggt ttt gtc aga ttc acg ctc tct acg tag act ttc ttg caa gtt act aat ttg gtc tct ggc acg tcc gag ttc act ctt ata tgc tga ata gag ttc tga ttc atc ctt tga gca gat ctc tag aga tcg ttc aag atc cct gat tct agt ctt tag cct att tac ctc ctc aga aga tgt tcc gtt acc gtt gcg ttt aca ttc gtt aag ctg tct atc aag atc cat gat tct atc tct --- --- --- --- --- --- --- --- --- --- --- --- --- --- --- --- -

>KC813498.1:153210-153610_Cowpox_virus_strain_JagKre08/2_complete_genome

gct gct agt tgt cgt tcg aga tca gaa ata cgg gtt ttc ttg gaa tcg att tcg tcg atg tac ttg gca tga ttg gta tcg aaa cac tta tta agt tc- ttt ttc caa ttc tac gat ttt att tct ttc acg cgt caa ttc cct cct gta gta act atc ggt ttt gtc aga ttc acg ctc tct acg tag act ttc ttg caa gtt act aat ttg ttc cct ggc acg tcc gag ttc agt ttt ata tgc cga ata gag ttc tga ttc atc ctt tga gca gat ctc taa aga tcg ttc aag atc cct aat tct agt ttt tag cct att tac ctc ctc aga aga tgt tcc gtt acc gtt gcg ttt aca ctc gtt gag ctg tct atc aag atc cat gat tct atc tct --- --- --- --- --- --- --- --- --- --- --- --- --- --- --- --- -

>KC813497.1:152892-153292_Cowpox_virus_strain_JagKre08/1_complete_genome

gct gct agt tgt cgt tcg aga tca gaa ata cgg gtt ttc ttg gaa tcg att tcg tcg atg tac ttg gca tga ttg gta tcg aaa cac tta tta agt tc- ttt ttc caa ttc tac gat ttt att tct ttc acg cgt caa ttc cct cct gta gta act atc ggt ttt gtc aga ttc acg ctc tct acg tag act ttc ttg caa gtt act aat ttg ttc cct ggc acg tcc gag ttc agt ttt ata tgc cga ata gag ttc tga ttc atc ctt tga gca gat ctc taa aga tcg ttc aag atc cct aat tct agt ttt tag cct att tac ctc ctc aga aga tgt tcc gtt acc gtt gcg ttt aca ctc gtt gag ctg tct atc aag atc cat gat tct atc tct --- --- --- --- --- --- --- --- --- --- --- --- --- --- --- --- -

>KC813495.1:145791-146191_Cowpox_virus_strain_HumMag07/1_complete_genome

gct gct agt tgt cgt tcg aga tca gaa ata cgg gtt ttc ttg gaa tcg att tcg tcg atg tac ttg gca tga ttg gca tcg aaa cac tta tta agt tc- ttt ttc caa ttc tac gat ttt att tct ttc acg cgt caa ttc cct cct gta gta act atc ggt ttt gtc aga ttc acg ctc tct acg tag act ttc ttg caa gtt act aat ttg ttc tct ggc acg tcc gag ttc act ctt ata tgc tga ata gag ttc tga ttc atc ctt tga gca gat ctc tag aga tcg ttc aag atc cct gat tct agt ttt tag cct att tac ctc ctc aga aga tgt tcc gtt acc gtt gcg ttt aca ttc gtt aag ctg tct atc aag atc cat gat tct atc tct --- --- --- --- --- --- --- --- --- --- --- --- --- --- --- --- -

>KC813492.1:145014-145414_Cowpox_virus_strain_HumLan08/1_complete_genome

gct gct agt tgt cgt tcg aga tca gaa ata cgg gtt ttc ttg gaa tcg att tcg tcg atg tac ttg gca tga ttg gta tcg aaa cac tta tta agt tc- ttt ttc caa ttc tac gat ttt att tct ttc acg cgt caa ttc cct cct gta gta act atc ggt ttt gtc aga ttc acg ctc tct acg tag act ttc ttg caa gtt act aat ttg ttc cct ggc acg tcc gag ttc agt ttt ata tgc cga ata gag ttc tga ttc atc ctt tga gca gat ctc taa aga tcg ttc aag atc cct aat tct agt ttt tag cct att tac ctc ctc aga aga tgt tcc gtt acc gtt gcg ttt aca ctc gtt gag ctg tct atc aag atc cat gat tct atc tct --- --- --- --- --- --- --- --- --- --- --- --- --- --- --- --- -

>KC813491.1:145310-145710_Cowpox_virus_strain_BeaBer04/1_complete_genome

gct gct agt tgt cgt tcg aga tca gaa ata cgg gtt ttc ttg gaa tcg att tcg tcg atg tac ttg gca tga ttg gca tcg aaa cac tta tta agt tc- ttt ttc caa ttc tac gat ttt att tct ttc acg cgt caa ttc cct cct gta gta act atc ggt ttt gtc aga ttc acg ctc tct acg tag act ttc ttg caa gtt act aat ttg ttc tct ggc acg tcc gag ttc act ctt ata tgc tga ata gag ttc tga ttc atc ctt tga gca gat ctc tag aga tcg ttc aag atc cct gat tct agt ttt tag cct att tac ctc ctc aga aga tgt tcc gtt acc gtt gcg ttt aca ttc gtt aag ctg tct atc aag atc cat gat tct atc tct --- --- --- --- --- --- --- --- --- --- --- --- --- --- --- --- -

>HQ420900.1:151600-152000_Cowpox_virus_strain_UK2000_K2984_complete_genome

gct gct agt tgt cgt tcg aga tca gaa ata cgg gtt ttc ttg gaa ttg att tcg tcg atg tac ttg gca tga ttg gta tcg aaa cac tta tta agt tc- ttt ttc caa ttc tac gat ttt att tct ttc acg agt caa ttc cct cct gta gta act atc ggt ttt gtc aga ttc acg ctc tct acg tag act ttc ttg caa gtt act aat ttg ttc cct tgc acg tcc gag ttc act ctt ata tgc tga ata gag ttc tga ttc atc ctt tga gca gat ctc tag aga tcg ttc aag atc cct gat tct agt ctt tag cct att tac ctc ctc aga aga tgt tcc gtt acc gtc gcg ttt aca ttc gtt aag ctg tct atc aag atc cat gat tct atc tct --- --- --- --- --- --- --- --- --- --- --- --- --- --- --- --- -

>HQ420897.1:152883-153283_Cowpox_virus_strain_Germany_1998_2_complete_genome

gct gct agt tgt cgt tcg aga tca gaa ata cgg gtt ttc ttg gaa ttg att tcg tcg atg tac ttg gca tga ttg gca tcg aaa cac tta tta agt tc- ttt ttc caa ttc tac gat ttt att tct ttc acg cgt caa ttc cct cct gta gta act atc ggt ttt gtc aga ttc acg ctc tct acg tag act ttc ttg caa gtt act aat ttg ttc tct ggc acg tcc gag ttc act ctt ata tgc tga ata gag ttc tga ttc atc ctt tga gca gat ctc tag aga tcg ttc aag atc cct gat tct agt ttt tag cct att tac ctc ctc aga aga tgt tcc gtt acc gtt gcg ttt aca ttc gtt aag ctg tct atc aag atc cat gat tct atc tct --- --- --- --- --- --- --- --- --- --- --- --- --- --- --- --- -

>HQ420894.1:150868-151268_Cowpox_virus_strain_France_2001_Nancy_complete_genome

gct gct agt tgt cgt tcg aga tca gaa ata cgg gtt ttc ttg gaa ttg att tcg tcg atg tac ttg gca tga ttg gta tcg aaa cac tta tta agt tc- ttt ttc caa ttc tac gat ttt att tct ttc acg agt caa ttc cct cct gta gta act atc ggt ttt gtc aga ttc acg ctc tct acg tag act ttc ttg caa gtt act aat ttg gtc tct ggc acg tcc gag ttc act ctt ata tgc tga ata gag ttc tga ttc atc ctt tga gca gat ctc tag aga tcg ttc aag atc cct gat tct agt ctt tag cct att tac ctc ctc aga aga tgt tcc gtt acc gtt gcg ttt aca ttc gtt aag ctg tct atc aag atc cat gat tct atc tct --- --- --- --- --- --- --- --- --- --- --- --- --- --- --- --- -

>DQ437593.1:151844-152244_Cowpox_virus_strain_Germany_91-3_complete_genome

gct gct agt tgt cgt tcg aga tca gaa ata cgg gtt ttc ttg gaa ttg att tcg tcg atg tac ttg gca tga ttg gta tcg aaa cac tta tta agt tc- ttt ttc caa ttc tac gat ttt att tct ttc gcg agt caa ttc cct cct gta gta act atc tgt ttt gtc aga ttc acg ctc tct acg tag act ttc ttg caa gtt act aat ttg ttc tct ggc acg tcc gag ttc act ctt ata tgc tga ata gag ttc tga ttc atc ctt tga gca gat ctc tag aga tcg ttc aag atc cct aat tct agt ttt tag cct att tac ctc ctc aga aga tgt tcc gtt acc gtt gcg ttt aca ttc gtt aag ctg tct atc aag atc cat gat tct atc tct --- --- --- --- --- --- --- --- --- --- --- --- --- --- --- --- -

>X94355.2:152095-152495_Cowpox_virus_strain_GRI-90_complete_genome

gct gct agc tgt cgt tcg aga tca gaa ata cgg gtt ttc ttg gaa ttg att tcg tcg atg tac ttg gca tga ttg gca tcg aaa cac tta tta agt tc- ttt ttc caa ttc tac gat ttt att tct ttc gcg agt caa ttc cct cct gta gta gct atc ggt ttt gtc aga ttc acg ctc tct acg tag act ttc ttg caa att act aat ttg ttc cct ggc acg tcc gag ttc agt ttt ata tgc cga ata gag ttc tga ttc atc ctt tga gca gat ctc tag aga tcg ttc aag atc cct aat tct agt ttt tag cct att tac ctc ctc aga aga tgt tcc gtt gcc gtt gcg ttt aca ctc gtt gag ctg tct atc aag atc cat gat tct atc tct --- --- --- --- --- --- --- --- --- --- --- --- --- --- --- --- -

>MK035758.1:152337-152737_Cowpox_virus_strain_CPXV/Moritz_2015/3a

gct gct agt tgt cgt tcg aga tca gaa ata cgg gtt ttc ttg gaa tcg att tcg tcg atg tac ttg gca tga ttg gta tcg aaa cac tta tta agt tc- ttt ttc caa ttc tac gat ttt att tct ttc acg cgt caa ttc cct cct gta gta act atc ggt ttt gtc aga ttc acg ctc tct acg tag act ttc ttg caa gtt act aat ttg ttc tct ggc acg tcc gag ttc act ctt ata tgc tga ata gag ttc tga ttc atc ctt tga gca gat ctc tag aga tcg ttc aag atc cct gat tct agt ttt tag cct att tac ctc ctc aga aga tgt tcc gtt acc gtt gcg ttt aca ttc gtt aag ctg tct atc aag atc cat gat tct atc tct --- --- --- --- --- --- --- --- --- --- --- --- --- --- --- --- -

>MK035756.1:152209-152609_Cowpox_virus_strain_CPXV/Leo_2015/5

gct gct agt tgt cgt tcg aga tca gaa ata cgg gtt ttc ttg gaa tcg att tcg tcg atg tac ttg gca tga ttg gta tcg aaa cac tta tta agt tc- ttt ttc caa ttc tac gat ttt att tct ttc acg cgt caa ttc cct cct gta gta act atc ggt ttt gtc aga ttc acg ctc tct acg tag act ttc ttg caa gtt act aat ttg ttc tct ggc acg tcc gag ttc act ctt ata tgc tga ata gag ttc tga ttc atc ctt tga gca gat ctc tag aga tcg ttc aag atc cct gat tct agt ttt tag cct att tac ctc ctc aga aga tgt tcc gtt acc gtt gcg ttt aca ttc gtt aag ctg tct atc aag atc cat gat tct atc tct --- --- --- --- --- --- --- --- --- --- --- --- --- --- --- --- -

>MK035755.1:152566-152966_Cowpox_virus_strain_CPXV/75/01

gct gct agt tgt cgt tcg aga tca gaa ata cgg gtt ttc ttg gaa tcg att tcg tcg atg tac ttg gca tga ttg gta tcg aaa cac tta tta agt tc- ttt ttc caa ttc tac gat ttt att tct ttc acg cgt caa ttc cct cct gta gta act atc ggt ttt gtc aga ttc acg ctc tct acg tag act ttc ttg caa gtt act aat ttg ttc tct ggc acg tcc gag ttc act ctt ata tgc tga ata gag ttc tga ttc atc ctt tga gca gat ctc tag aga tcg ttc aag atc cct gat tct agt ttt tag cct att tac ctc ctc aga aga tgt tcc gtt acc gtt gcg ttt aca ttc gtt aag ctg tct atc aag atc cat gat tct atc tct --- --- --- --- --- --- --- --- --- --- --- --- --- --- --- --- -

>MK035754.1:152665-153065_Cowpox_virus_strain_CPXV/Cookie_S963_15

gct gct agt tgt cgt tcg aga tca gaa ata cgg gtt ttc ttg gaa tcg att tcg tcg atg tac ttg gca tga ttg gta tcg aaa cac tta tta agt tc- ttt ttc caa ttc tac gat ttt att tct ttc acg cgt caa ttc cct cct gta gta act atc ggt ttt gtc aga ttc acg ctc tct acg tag act ttc ttg caa gtt act aat ttg ttc tct ggc acg tcc gag ttc act ctt ata tgc tga ata gag ttc tga ttc atc ctt tga gca gat ctc tag aga tcg ttc aag atc cct gat tct agt ttt tag cct att tac ctc ctc aga aga tgt tcc gtt acc gtt gcg ttt aca ttc gtt aag ctg tct atc aag atc cat gat tct atc tct --- --- --- --- --- --- --- --- --- --- --- --- --- --- --- --- -

>MK035752.1:152432-152832_Cowpox_virus_strain_CPXV/Sammy_2015/4a

gct gct agt tgt cgt tcg aga tca gaa ata cgg gtt ttc ttg gaa tcg att tcg tcg atg tac ttg gca tga ttg gta tcg aaa cac tta tta agt tc- ttt ttc caa ttc tac gat ttt att tct ttc acg cgt caa ttc cct cct gta gta act atc ggt ttt gtc aga ttc acg ctc tct acg tag act ttc ttg caa gtt act aat ttg ttc tct ggc acg tcc gag ttc act ctt ata tgc tga ata gag ttc tga ttc atc ctt tga gca gat ctc tag aga tcg ttc aag atc cct gat tct agt ttt tag cct att tac ctc ctc aga aga tgt tcc gtt acc gtt gcg ttt aca ttc gtt aag ctg tct atc aag atc cat gat tct atc tct --- --- --- --- --- --- --- --- --- --- --- --- --- --- --- --- -

>LT896732.2:152692-153092_Cowpox_virus_isolate_Ger/2017/Alpaca2_genome_assembly_complete_genome:_monopartite

gct gct agt tgt cgt tcg aga tca gaa ata cgg gtt ttc ttg gaa tcg att tcg tcg atg tac ttg gca tga ttg gta tcg aaa cac tta tta agt tc- ttt ttc caa ttc tac gat ttt att tct ttc acg cgt caa ttc cct cct gta gta act atc ggt ttt gtc aga ttc acg ctc tct acg tag act ttc ttg caa gtt act aat ttg ttc tct ggc acg tcc gag ttc act ctt ata tgc tga ata gag ttc tga ttc atc ctt tga gca gat ctc tag aga tcg ttc aag atc cct gat tct agt ttt tag cct att tac ctc ctc aga aga tgt tcc gtt acc gtt gcg ttt aca ttc gtt aag ctg tct atc aag atc cat gat tct atc tct --- --- --- --- --- --- --- --- --- --- --- --- --- --- --- --- -

>LT993232.1:150445-150845_Cowpox_virus_isolate_Ger/2015/Human2_genome_assembly_complete_genome:_monopartite

gct gct agt tgt cgt tcg aga tca gaa ata cgg gtt ttc ttg gaa tcg att tcg tcg atg tac ttg gca tga ttg gta tcg aaa cac tta tta agt tc- ttt ttc caa ttc tac gat ttt att tct ttc acg cgt caa ttc cct cct gta gta act atc ggt ttt gtc aga ttc acg ctc tct acg tag act ttc ttg caa gtt act aat ttg ttc tct ggc acg tcc gag ttc act ctt ata tgc tga ata gag ttc tga ttc atc ctt tga gca gat ctc tag aga tcg ttc aag atc cct gat tct agt ttt tag cct att tac ctc ctc aga aga tgt tcc gtt acc gtt gcg ttt aca ttc gtt aag ctg tct atc aag atc cat gat tct atc tct --- --- --- --- --- --- --- --- --- --- --- --- --- --- --- --- -

>LT993231.1:152542-152942_Cowpox_virus_isolate_Ger/2015/Prairie-dog_genome_assembly_complete_genome:_monopartite

gct gct agt tgt cgt tcg aga tca gaa ata cgg gtt ttc ttg gaa tcg att tcg tcg atg tac ttg gca tga ttg gta tcg aaa cac tta tta agt tc- ttt ttc caa ttc tac gat ttt att tct ttc acg cgt caa ttc cct cct gta gta act atc ggt ttt gtc aga ttc acg ctc tct acg tag act ttc ttg caa gtt act aat ttg ttc tct ggc acg tcc gag ttc act ctt ata tgc tga ata gag ttc tga ttc atc ctt tga gca gat ctc tag aga tcg ttc aag atc cct gat tct agt ttt tag cct att tac ctc ctc aga aga tgt tcc gtt acc gtt gcg ttt aca ttc gtt aag ctg tct atc aag atc cat gat tct atc tct --- --- --- --- --- --- --- --- --- --- --- --- --- --- --- --- -

>LT993230.1:151701-152101_Cowpox_virus_isolate_Ger/2017/Alpaca1_genome_assembly_complete_genome:_monopartite

gct gct agt tgt cgt tcg aga tca gaa ata cgg gtt ttc ttg gaa tcg att tcg tcg atg tac ttg gca tga ttg gta tcg aaa cac tta tta agt tc- ttt ttc caa ttc tac gat ttt att tct ttc acg cgt caa ttc cct cct gta gta act atc ggt ttt gtc aga ttc acg ctc tct acg tag act ttc ttg caa gtt act aat ttg ttc tct ggc acg tcc gag ttc act ctt ata tgc tga ata gag ttc tga ttc atc ctt tga gca gat ctc tag aga tcg ttc aag atc cct gat tct agt ttt tag cct att tac ctc ctc aga aga tgt tcc gtt acc gtt gcg ttt aca ttc gtt aag ctg tct atc aag atc cat gat tct atc tct --- --- --- --- --- --- --- --- --- --- --- --- --- --- --- --- -

>LT993228.1:151838-152238_Cowpox_virus_isolate_Ger/2017/common_vole_FMEimka_genome_assembly_complete_genome:_monopartite

gct gct agt tgt cgt tcg aga tca gaa ata cgg gtt ttc ttg gaa tcg att tcg tcg atg tac ttg gca tga ttg gta tcg aaa cac tta tta agt tc- ttt ttc caa ttc tac gat ttt att tct ttc acg cgt caa ttc cct cct gta gta act atc ggt ttt gtc aga ttc acg ctc tct acg tag act ttc ttg caa gtt act aat ttg ttc tct ggc acg tcc gag ttc act ctt ata tgc tga ata gag ttc tga ttc atc ctt tga gca gat ctc tag aga tcg ttc aag atc cct gat tct agt ttt tag cct att tac ctc ctc aga aga tgt tcc gtt acc gtt gcg ttt aca ttc gtt aag ctg tct atc aag atc cat gat tct atc tct --- --- --- --- --- --- --- --- --- --- --- --- --- --- --- --- -

>LT896733.1:151890-152290_Cowpox_virus_isolate_Ger/2015/Cat3_genome_assembly_complete_genome:_monopartite

gct gct agt tgt cgt tcg aga tca gaa ata cgg gtt ttc ttg gaa tcg att tcg tcg atg tac ttg gca tga ttg gta tcg aaa cac tta tta agt tc- ttt ttc caa ttc tac gat ttt att tct ttc acg cgt caa ttc cct cct gta gta act atc ggt ttt gtc aga ttc acg ctc tct acg tag act ttc ttg caa gtt act aat ttg ttc tct ggc acg tcc gag ttc act ctt ata tgc tga ata gag ttc tga ttc atc ctt tga gca gat ctc tag aga tcg ttc aag atc cct gat tct agt ttt tag cct att tac ctc ctc aga aga tgt tcc gtt acc gtt gcg ttt aca ttc gtt aag ctg tct atc aag atc cat gat tct atc tct --- --- --- --- --- --- --- --- --- --- --- --- --- --- --- --- -

>LT896731.1:151787-152187_Cowpox_virus_isolate_Ger/2015/Cat4_genome_assembly_complete_genome:_monopartite

gct gct agt tgt cgt tcg aga tca gaa ata cgg gtt ttc ttg gaa tcg att tcg tcg atg tac ttg gca tga ttg gta tcg aaa cac tta tta agt tc- ttt ttc caa ttc tac gat ttt att tct ttc acg cgt caa ttc cct cct gta gta act atc ggt ttt gtc aga ttc acg ctc tct acg tag act ttc ttg caa gtt act aat ttg ttc tct ggc acg tcc gag ttc act ctt ata tgc tga ata gag ttc tga ttc atc ctt tga gca gat ctc tag aga tcg ttc aag atc cct gat tct agt ttt tag cct att tac ctc ctc aga aga tgt tcc gtt acc gtt gcg ttt aca ttc gtt aag ctg tct atc aag atc cat gat tct atc tct --- --- --- --- --- --- --- --- --- --- --- --- --- --- --- --- -

>LT896730.1:151778-152178_Cowpox_virus_isolate_Ger/2010/Racoon_genome_assembly_complete_genome:_monopartite

gct gct agt tgt cgt tcg aga tca gaa ata cgg gtt ttc ttg gaa tcg att tcg tcg atg tac ttg gca tga ttg gta tcg aaa cac tta tta agt tc- ttt ttc caa ttc tac gat ttt att tct ttc acg cgt caa ttc cct cct gta gta act atc ggt ttt gtc aga ttc acg ctc tct acg tag act ttc ttg caa gtt act aat ttg ttc tct ggc acg tcc gag ttc act ctt ata tgc tga ata gag ttc tga ttc atc ctt tga gca gat ctc tag aga tcg ttc aag atc cct gat tct agt ttt tag cct att tac ctc ctc aga aga tgt tcc gtt acc gtt gcg ttt aca ttc gtt aag ctg tct atc aag atc cat gat tct atc tct --- --- --- --- --- --- --- --- --- --- --- --- --- --- --- --- -

>LT896729.1:151747-152147_Cowpox_virus_isolate_Ger/2010/Cat_genome_assembly_complete_genome:_monopartite

gct gct agt tgt cgt tcg aga tca gaa ata cgg gtt ttc ttg gaa tcg att tcg tcg atg tac ttg gca tga ttg gta tcg aaa cac tta tta agt tc- ttt ttc caa ttc tac gat ttt att tct ttc acg cgt caa ttc cct cct gta gta act atc ggt ttt gtc aga ttc acg ctc tct acg tag act ttc ttg caa gtt act aat ttg ttc tct ggc acg tcc gag ttc act ctt ata tgc tga ata gag ttc tga ttc atc ctt tga gca gat ctc tag aga tcg ttc aag atc cct gat tct agt ttt tag cct att tac ctc ctc aga aga tgt tcc gtt acc gtt gcg ttt aca ttc gtt aag ctg tct atc aag atc cat gat tct atc tct --- --- --- --- --- --- --- --- --- --- --- --- --- --- --- --- -

>LT896728.1:152177-152577_Cowpox_virus_isolate_Ger/2010/Rat_genome_assembly_complete_genome:_monopartite

gct gct agt tgt cgt tcg aga tca gaa ata cgg gtt ttc ttg gaa tcg att tcg tcg atg tac ttg gca tga ttg gta tcg aaa cac tta tta agt tc- ttt ttc caa ttc tac gat ttt att tct ttc acg cgt caa ttc cct cct gta gta act atc ggt ttt gtc aga ttc acg ctc tct acg tag act ttc ttg caa gtt act aat ttg ttc tct ggc acg tcc gag ttc act ctt ata tgc tga ata gag ttc tga ttc atc ctt tga gca gat ctc tag aga tcg ttc aag atc cct gat tct agt ttt tag cct att tac ctc ctc aga aga tgt tcc gtt acc gtt gcg ttt aca ttc gtt aag ctg tct atc aag atc cat gat tct atc tct --- --- --- --- --- --- --- --- --- --- --- --- --- --- --- --- -

>LT896727.1:152576-152976_Cowpox_virus_isolate_Ger/2015/Cat2_genome_assembly_complete_genome:_monopartite

gct gct agt tgt cgt tcg aga tca gaa ata cgg gtt ttc ttg gaa tcg att tcg tcg atg tac ttg gca tga ttg gta tcg aaa cac tta tta agt tc- ttt ttc caa ttc tac gat ttt att tct ttc acg cgt caa ttc cct cct gta gta act atc ggt ttt gtc aga ttc acg ctc tct acg tag act ttc ttg caa gtt act aat ttg ttc tct ggc acg tcc gag ttc act ctt ata tgc tga ata gag ttc tga ttc atc ctt tga gca gat ctc tag aga tcg ttc aag atc cct gat tct agt ttt tag cct att tac ctc ctc aga aga tgt tcc gtt acc gtt gcg ttt aca ttc gtt aag ctg tct atc aag atc cat gat tct atc tct --- --- --- --- --- --- --- --- --- --- --- --- --- --- --- --- -

>LT896726.1:151650-152050_Cowpox_virus_isolate_Ger/2012/Alpaca_genome_assembly_complete_genome:_monopartite

gct gct agt tgt cgt tcg aga tca gaa ata cgg gtt ttc ttg gaa tcg att tcg tcg atg tac ttg gca tga ttg gta tcg aaa cac tta tta agt tc- ttt ttc caa ttc tac gat ttt att tct ttc acg cgt caa ttc cct cct gta gta act atc ggt ttt gtc aga ttc acg ctc tct acg tag act ttc ttg caa gtt act aat ttg ttc tct ggc acg tcc gag ttc act ctt ata tgc tga ata gag ttc tga ttc atc ctt tga gca gat ctc tag aga tcg ttc aag atc cct gat tct agt ttt tag cct att tac ctc ctc aga aga tgt tcc gtt acc gtt gcg ttt aca ttc gtt aag ctg tct atc aag atc cat gat tct atc tct --- --- --- --- --- --- --- --- --- --- --- --- --- --- --- --- -

>LT896725.1:152942-153342_Cowpox_virus_isolate_Ger/2014/Cat2_genome_assembly_complete_genome:_monopartite

gct gct agt tgt cgt tcg aga tca gaa ata cgg gtt ttc ttg gaa ttg att tcg tcg atg tac ttg gca tga ttg gta tcg aaa cac tta tta agt tc- ttt ttc caa ttc tac gat ttt att tct ttc acg cgt caa ttc cct cct gta gta act atc ggt ttt gtc aga ttc acg ctc tct acg tag act ttc ttg caa gtt act aat ttg ttc tct ggc acg tcc gag ttc act ctt ata tgc tga ata gag ttc tga ttc atc ctt tga gca gat ctc tag aga tcg ttc aag atc cct gat tct agt ttt tag cct att tac ctc ctc aga aga tgt tcc gtt acc gtt gcg ttt aca ttc gtt aag ctg tct atc aag atc cat gat tct atc tct --- --- --- --- --- --- --- --- --- --- --- --- --- --- --- --- -

>LT896720.1:150453-150853_Cowpox_virus_isolate_Ger/2015/Human1_genome_assembly_complete_genome:_monopartite

gct gct agt tgt cgt tcg aga tca gaa ata cgg gtt ttc ttg gaa tcg att tcg tcg atg tac ttg gca tga ttg gta tcg aaa cac tta tta agt tc- ttt ttc caa ttc tac gat ttt att tct ttc acg cgt caa ttc cct cct gta gta act atc ggt ttt gtc aga ttc acg ctc tct acg tag act ttc ttg caa gtt act aat ttg ttc tct ggc acg tcc gag ttc act ctt ata tgc tga ata gag ttc tga ttc atc ctt tga gca gat ctc tag aga tcg ttc aag atc cct gat tct agt ttt tag cct att tac ctc ctc aga aga tgt tcc gtt acc gtt gcg ttt aca ttc gtt aag ctg tct atc aag atc cat gat tct atc tct --- --- --- --- --- --- --- --- --- --- --- --- --- --- --- --- -

>LT896719.1:151513-151913_Cowpox_virus_isolate_Ger/2013/Alpaca_genome_assembly_complete_genome:_monopartite

gct gct agt tgt cgt tcg aga tca gaa ata cgg gtt ttc ttg gaa tcg att tcg tcg atg tac ttg gca tga ttg gta tcg aaa cac tta tta agt tc- ttt ttc caa ttc tac gat ttt att tct ttc acg agt caa ttc cct cct gta gta act atc ggt ttt gtc aga ttc acg ctc tct acg tag act ttc ttg caa gtt act aat ttg ttc tct ggc acg tcc gag ttc act ctt ata tgc tga ata gag ttc tga ttc atc ctt tga gca gat ctc tag aga tcg ttc aag atc cct gat tct agt ttt tag cct att tac ctc ctc agt aga tgt tcc gtt acc gtt gcg ttt aca ttc gtt aag ctg tct atc aag atc cat gat tct atc tct --- --- --- --- --- --- --- --- --- --- --- --- --- --- --- --- -

>LT896718.1:152324-152724_Cowpox_virus_isolate_Ger/2010/Alpaca_genome_assembly_complete_genome:_monopartite

gct gct agt tgt cgt tcg aga tca gaa ata cgg gtt ttc ttg gaa tcg att tcg tcg atg tac ttg gca tga ttg gta tcg aaa cac tta tta agt tc- ttt ttc caa ttc tac gat ttt att tct ttc acg cgt caa ttc cct cct gta gta act atc ggt ttt gtc aga ttc acg ctc tct acg tag act ttc ttg caa gtt act aat ttg ttc tct ggc acg tcc gag ttc act ctt ata tgc tga ata gag ttc tga ttc atc ctt tga gca gat ctc tag aga tcg ttc aag atc cct gat tct agt ttt tag cct att tac ctc ctc aga aga tgt tcc gtt acc gtt gcg ttt aca ttc gtt aag ctg tct atc aag atc cat gat tct atc tct --- --- --- --- --- --- --- --- --- --- --- --- --- --- --- --- -

>KY549149.1:153449-153849_Cowpox_virus_isolate_CPXV_K2739

gct gct agt tgt cgt tcg aga tca gaa ata cgg gtt ttc ttg gaa ttg att tcg tcg atg tac ttg gca tga ttg gta tcg aaa cac tta tta agt tc- ttt ttc caa ttc tac gat ttt att tct ttc acg agt caa ttc cct cct gta gta act atc ggt ttt gtc aga ttc acg ctc tct acg tag act ttc ttg caa gtt act aat ttg ttc tct ggc acg tcc gag ttc act ctt ata tgc tga ata gag ttc tga ttc atc ctt tga gca gat ctc tag aga tcg ttc aag atc cct gat tct agt ttt tag cct att tac ctc ctc aga aga tgt tcc gtt acc gtc gcg ttt aca ttc gtt aag ctg tct atc aag atc cat gat tct atc tct --- --- --- --- --- --- --- --- --- --- --- --- --- --- --- --- -

>KY549144.1:153046-153446_Cowpox_virus_isolate_CPXV_Catpox5wv1

gct gct agt tgt cgt tcg aga tca gaa ata cgg gtt ttc ttg gaa ttg att tcg tcg atg tac ttg gca tga ttg gta tcg aaa cac tta tta agt tc- ttt ttc caa ttc tac gat ttt att tct ttc acg agt caa ttc cct cct gta gta act atc ggt ttt gtc aga ttc acg ctc tct acg tag act ttc ttg caa gtt act aat ttg ttc tct ggc acg tcc gag ttc act ctt ata tgc tga ata gag ttc tga ttc atc ctt tga gca gat ctc tag aga tcg ttc aag atc cct gat tct agt ttt tag cct att tac ctc ctc aga aga tgt tcc gtt acc gtc gcg ttt aca ttc gtt aag ctg tct atc aag atc cat gat tct atc tct --- --- --- --- --- --- --- --- --- --- --- --- --- --- --- --- -

>LN879483.1:151658-152058_Cowpox_virus_strain_CPXV_Amadeus_2015_complete_genome

gct gct agt tgt cgt tcg aga tca gaa ata cgg gtt ttc ttg gaa tcg att tcg tcg atg tac ttg gca tga ttg gta tcg aaa cac tta tta agt tc- ttt ttc caa ttc tac gat ttt att tct ttc acg agt caa ttc cct cct gta gta act atc ggt ttt gtc aga ttc acg ctc tct acg tag act ttc ttg caa gtt act aat ttg ttc tct ggc acg tcc gag ttc act ctt ata tgc tga ata gag ttc tga ttc atc ctt tga gca gat ctc tag aga tcg ttc aag atc cct gat tct agt ttt tag cct att tac ctc ctc agt aga tgt tcc gtt acc gtt gcg ttt aca ttc gtt aag ctg tct atc aag atc cat gat tct atc tct --- --- --- --- --- --- --- --- --- --- --- --- --- --- --- --- -

>KC813511.1:145670-146070_Cowpox_virus_strain_HumGri07/1_complete_genome

gct gct agt tgt cgt tcg aga tca gaa ata cgg gtt ttc ttg gaa tcg att tcg tcg atg tac ttg gca tga ttg gta tcg aaa cac tta tta agt tc- ttt ttc caa ttc tac gat ttt att tct ttc acg cgt caa ttc cct cct gta gta act atc ggt ttt gtc aga ttc acg ctc tct acg tag act ttc ttg caa gtt act aat ttg ttc tct ggc acg tcc gag ttc act ctt ata tgc tga ata gag ttc tga ttc atc ctt tga gca gat ctc tag aga tcg ttc aag atc cct gat tct agt ttt tag cct att tac ctc ctc aga aga tgt tcc gtt acc gtt gcg ttt aca ttc gtt aag ctg tct atc aag atc cat gat tct atc tct --- --- --- --- --- --- --- --- --- --- --- --- --- --- --- --- -

>KC813507.1:134142-134542_Cowpox_virus_strain_EleGri07/1_complete_genome

gct gct agt tgt cgt tcg aga tca gaa ata cgg gtt ttc ttg gaa tcg att tcg tcg atg tac ttg gca tga ttg gta tcg aaa cac tta tta agt tc- ttt ttc caa ttc tac gat ttt att tct ttc acg cgt caa ttc cct cct gta gta act atc ggt ttt gtc aga ttc acg ctc tct acg tag act ttc ttg caa gtt act aat ttg ttc tct ggc acg tcc gag ttc act ctt ata tgc tga ata gag ttc tga ttc atc ctt tga gca gat ctc tag aga tcg ttc aag atc cct gat tct agt ttt tag cct att tac ctc ctc aga aga tgt tcc gtt acc gtt gcg ttt aca ttc gtt aag ctg tct atc aag atc cat gat tct atc tct --- --- --- --- --- --- --- --- --- --- --- --- --- --- --- --- -

>KC813496.1:139804-140204_Cowpox_virus_strain_HumPad07/1_complete_genome

gct gct agt tgt cgt tcg aga tca gaa ata cgg gtt ttc ttg gaa tcg att tcg tcg atg tac ttg gca tga ttg gta tcg aaa cac tta tta agt tc- ttt ttc caa ttc tac gat ttt att tct ttc acg cgt caa ttc cct cct gta gta act atc ggt ttt gtc aga ttc acg ctc tct acg tag act ttc ttg caa gtt act aat ttg ttc tct ggc acg tcc gag ttc act ctt ata tgc tga ata gag ttc tga ttc atc ctt tga gca gat ctc tag aga tcg ttc aag atc cct gat tct agt ttt tag cct att tac ctc ctc aga aga tgt tcc gtt acc gtt gcg ttt aca ttc gtt aag ctg tct atc aag atc cat gat tct atc tct --- --- --- --- --- --- --- --- --- --- --- --- --- --- --- --- -

>KC813494.1:143293-143693_Cowpox_virus_strain_HumLue09/1_complete_genome

gct gct agt tgt cgt tcg aga tca gaa ata cgg gtt ttc ttg gaa tcg att tcg tcg atg tac ttg gca tga ttg gta tcg aaa cac tta tta agt tc- ttt ttc caa ttc tac gat ttt att tct ttc acg agt caa ttc cct cct gta gta act atc ggt ttt gtc aga ttc acg ctc tct acg tag act ttc ttg caa gtt act aat ttg ttc tct ggc acg tcc gag ttc act ctt ata tgc tga ata gag ttc tga ttc atc ctt tga gca gat ctc tag aga tcg ttc aag atc cct aat tct agt ttt tag cct att tac ctc ctc aga aga tgt tcc gtt acc gtt gcg ttt aca ttc gtt aag ctg tct atc aag atc cat gat tct atc tct --- --- --- --- --- --- --- --- --- --- --- --- --- --- --- --- -

>HQ420898.1:151944-152344_Cowpox_virus_strain_Germany_2002_MKY_complete_genome

gct gct agt tgt cgt tcg aga tca gaa ata cgg gtt ttc ttg gaa tcg att tcg tcg atg tac ttg gca tga ttg gta tcg aaa cac tta tta agt tc- ttt ttc caa ttc tac gat ttt att tct ttc acg agt caa ttc cct cct gta gta act atc ggt ttt gtc aga ttc acg ctc tct acg tag act ttc ttg caa gtt act aat ttg ttc tct ggc acg tcc gag ttc act ctt ata tgc tga ata gag ttc tga ttc atc ctt tga gca gat ctc tag aga tcg ttc aag atc cct gat tct agt ttt tag cct att tac ctc ctc agt aga tgt tcc gtt acc gtt gcg ttt aca ttc gtt aag ctg tct atc aag atc cat gat tct atc tct --- --- --- --- --- --- --- --- --- --- --- --- --- --- --- --- -

>HQ420895.1:153180-153580_Cowpox_virus_strain_Germany_1980_EP4_complete_genome

gct gct agt tgt cgt tcg aga tca gaa ata cgg gtt ttc ttg gaa tcg att tcg tcg atg tac ttg gca tga ttg gta tcg aaa cac tta tta agt tc- ttt ttc caa ttc tac gat ttt att tct ttc acg cgt caa ttc cct cct gta gta act atc ggt ttt gtc aga ttc acg ctc tct acg tag act ttc ttg caa gtt act aat ttg ttc tct ggc acg tcc gag ttc act ctt ata tgc tga ata gag ttc tga ttc atc ctt tga gca gat ctc tag aga tcg ttc aag atc cct gat tct agt ttt tag cct att tac ctc ctc aga aga tgt tcc gtt acc gtt gcg ttt aca ttc gtt aag ctg tct atc aag atc cat gat tct atc tct --- --- --- --- --- --- --- --- --- --- --- --- --- --- --- --- -

>AF482758.2:152667-153067_Cowpox_virus_strain_Brighton_Red_complete_genome

gct gct agt tgt cgt tcg aga tca gaa ata cgg gtt ttc ttg gaa ttg att tcg tcg atg tac ttg gca tga ttg gta tcg aaa cac tta tta agt tc- ttt ttc caa ttc tac gat ttt att tct ttc acg agt caa ttc cct cct gta gta act atc ggt ttt gtc aga ttc acg ctc tct acg tag act ttc ttg cag gtt act aat ttg ttc cct tgc acg tcc gag ttc act ctt ata tgc tga ata gag ttc tga ttc atc ctt tga gca gat ctc tag aga tcg ttc aag atc cct gat tct agt ctt tag cct att tac ctc ctc aga aga tgt tcc gtt acc gtc gcg ttt aca ttc gtt aag ctg tct atc aag atc cat gat tct atc tct --- --- --- --- --- --- --- --- --- --- --- --- --- --- --- --- -

>MK035751.1:152961-153361_Cowpox_virus_strain_CPXV/S2216/04

gct gct agt tgt cgt tcg aga tca gaa ata cgg gtt ttc ttg gaa tcg att tcg tcg atg tac ttg gca tga ttg gta tcg aaa cac tta tta agt tc- ttt ttc caa ttc tac gat ttt att tct ttc acg cgt caa ttc cct cct gta gta act atc ggt ttt gtc aga ttc acg ctc tct acg tag act ttc ttg caa gtt act aat ttg ttc tct ggt acg tcc gag ttc act ctt ata tgc tga ata gag ttc tga ttc atc ctt tga gca gat ctc tag aga tcg ttc aag atc cct gat tct agt ttt tag cct att tac ctc ctc aga aga tgt tcc gtt acc gtt gcg ttt aca ttc gtt aag ctg tct atc aag atc cat gat tct atc tct --- --- --- --- --- --- --- --- --- --- --- --- --- --- --- --- -

>MT227314.1:138568-138959_Vaccinia_virus_strain_Acambis/2019

gct gct agt tgt cgt tcg aga tca gaa ata cgg gtt ttc ttg gaa ttg att tcg tcg atg tac ttg gca --- --- --- tcg aaa cac tta tta agt tc- ttt ttc caa ttc tac gat ttt att tct ttc gcg agt caa ttc cct cct gta gta act atc tgt ttt gtc aga ttc acg ctc tct acg tag act ttc ttg caa gtt act aat ttg ttc cct agc acg tcc gag ttt agt ttt ata tgc cga ata gag ttc tga ttc atc ctt tga gca gat ctc tag aga tcg ttc aag atc cct gat tct agt ctt tag cct att tac ctc ctc gga aga tgc tcc gtt acc gtt gcg ttt aca ctc gtt aag ctg tct atc aag atc cat gat tct atc tct --- --- --- --- --- --- --- --- --- --- --- --- --- --- --- --- -

>MN974381.1:138859-139250_Vaccinia_virus_strain_TNX-1200

gct gct agt tgt cgt tcg aga tca gaa ata cgg gtt ttc ttg gaa ttg att tcg tcg atg tac ttg gca --- --- --- tcg aaa cac tta tta agt tc- ttt ttc caa ttc tac gat ttt att tct ttc gcg agt caa ttc cct cct gta gta act atc tgt ttt gtc aga ttc acg ctc tct acg tag act ttc ttg caa gtt act aat ttg ttc cct agc acg tcc gag ttt agt ttt ata tgc cga ata gag ttc tga ttc atc ctt tga gca gat ctc tag aga tcg ttc aag atc cct gat tct agt ctt tag cct att tac ctc ctc gga aga tgc tcc gtt acc gtt gcg ttt aca ctc gtt aag ctg tct atc aag atc cat gat tct atc tct --- --- --- --- --- --- --- --- --- --- --- --- --- --- --- --- -

>MN974380.1:138805-139196_Vaccinia_virus_strain_VACV_li

gct gct agt tgt cgt tcg aga tca gaa ata cgg gtt ttc ttg gaa ttg att tcg tcg atg tac ttg gca --- --- --- tcg aaa cac tta tta agt tc- ttt ttc caa ttc tac gat ttt att tct ttc gcg agt caa ttc cct cct gta gta act atc tgt ttt gtc aga ttc acg ctc tct acg tag act ttc ttg caa gtt act aat ttg ttc cct agc acg tcc gag ttt agt ttt ata tgc cga ata gag ttc tga ttc atc ctt tga gca gat ctc tag aga tcg ttc aag atc cct gat tct agt ctt tag cct att tac ctc ctc gga aga tgc tcc gtt acc gtt gcg ttt aca ctc gtt aag ctg tct atc aag atc cat gat tct atc tct --- --- --- --- --- --- --- --- --- --- --- --- --- --- --- --- -

>KJ125438.1:138307-138698_Vaccinia_virus_strain_Dryvax_clone_DPP25_complete_genome

gct gct agt tgt cgt tcg aga tca gaa ata cgg gtt ttc ttg gaa ttg att tcg tcg atg tac ttg gca --- --- --- tcg aaa cac tta tta agt tc- ttt ttc caa ttc tac gat ttt att tct ttc gcg agt caa ttc cct cct gta gta act atc tgt ttt gtc aga ttc acg ctc tct acg tag act ttc ttg caa gtt act aat ttg ttc cct agc acg tcc gag ttt agt ttt ata tgc cga ata gag ttc tga ttc atc ctt tga gca gat ctc tag aga tcg ttc aag atc cct gat tct agt ctt tag cct att tac ctc ctc gga aga tgc tcc gtt acc gtt gcg ttt aca ctc gtt aag ctg tct atc aag atc cat gat tct atc tct --- --- --- --- --- --- --- --- --- --- --- --- --- --- --- --- -

>MW960424.1:139417-139808_Vector_synVACV-Delta1Delta3-6_complete_sequence

gct gct agt tgt cgt tcg aga tca gaa ata cgg gtt ttc ttg gaa ttg att tcg tcg atg tac ttg gca --- --- --- tcg aaa cac tta tta agt tc- ttt ttc caa ttc tac gat ttt att tct ttc gcg agt caa ttc cct cct gta gta act atc tgt ttt gtc aga ttc acg ctc tct acg tag act ttc ttg caa gtt act aat ttg ttc cct agc acg tcc gag ttt agt ttt ata tgc cga ata gag ttc tga ttc atc ctt tga gca gat ctc tag aga tcg ttc aag atc cct gat tct agt ctt tag cct att tac ctc ctc gga aga tgc tcc gtt acc gtt gcg ttt aca ctc gtt aag ctg tct atc aag atc cat gat tct atc tct --- --- --- --- --- --- --- --- --- --- --- --- --- --- --- --- -

>MW960423.1:139418-139809_Vector_synVACV-Delta3-6_complete_sequence

gct gct agt tgt cgt tcg aga tca gaa ata cgg gtt ttc ttg gaa ttg att tcg tcg atg tac ttg gca --- --- --- tcg aaa cac tta tta agt tc- ttt ttc caa ttc tac gat ttt att tct ttc gcg agt caa ttc cct cct gta gta act atc tgt ttt gtc aga ttc acg ctc tct acg tag act ttc ttg caa gtt act aat ttg ttc cct agc acg tcc gag ttt agt ttt ata tgc cga ata gag ttc tga ttc atc ctt tga gca gat ctc tag aga tcg ttc aag atc cct gat tct agt ctt tag cct att tac ctc ctc gga aga tgc tcc gtt acc gtt gcg ttt aca ctc gtt aag ctg tct atc aag atc cat gat tct atc tct --- --- --- --- --- --- --- --- --- --- --- --- --- --- --- --- -

>MW960422.1:139432-139823_Vector_synVACV-Delta1-3_complete_sequence

gct gct agt tgt cgt tcg aga tca gaa ata cgg gtt ttc ttg gaa ttg att tcg tcg atg tac ttg gca --- --- --- tcg aaa cac tta tta agt tc- ttt ttc caa ttc tac gat ttt att tct ttc gcg agt caa ttc cct cct gta gta act atc tgt ttt gtc aga ttc acg ctc tct acg tag act ttc ttg caa gtt act aat ttg ttc cct agc acg tcc gag ttt agt ttt ata tgc cga ata gag ttc tga ttc atc ctt tga gca gat ctc tag aga tcg ttc aag atc cct gat tct agt ctt tag cct att tac ctc ctc gga aga tgc tcc gtt acc gtt gcg ttt aca ctc gtt aag ctg tct atc aag atc cat gat tct atc tct --- --- --- --- --- --- --- --- --- --- --- --- --- --- --- --- -

>MW960421.1:139425-139816_Vector_synVACV-Delta5-6_complete_sequence

gct gct agt tgt cgt tcg aga tca gaa ata cgg gtt ttc ttg gaa ttg att tcg tcg atg tac ttg gca --- --- --- tcg aaa cac tta tta agt tc- ttt ttc caa ttc tac gat ttt att tct ttc gcg agt caa ttc cct cct gta gta act atc tgt ttt gtc aga ttc acg ctc tct acg tag act ttc ttg caa gtt act aat ttg ttc cct agc acg tcc gag ttt agt ttt ata tgc cga ata gag ttc tga ttc atc ctt tga gca gat ctc tag aga tcg ttc aag atc cct gat tct agt ctt tag cct att tac ctc ctc gga aga tgc tcc gtt acc gtt gcg ttt aca ctc gtt aag ctg tct atc aag atc cat gat tct atc tct --- --- --- --- --- --- --- --- --- --- --- --- --- --- --- --- -

>MW960420.1:139433-139824_Vector_synVACV-Delta6_complete_sequence

gct gct agt tgt cgt tcg aga tca gaa ata cgg gtt ttc ttg gaa ttg att tcg tcg atg tac ttg gca --- --- --- tcg aaa cac tta tta agt tc- ttt ttc caa ttc tac gat ttt att tct ttc gcg agt caa ttc cct cct gta gta act atc tgt ttt gtc aga ttc acg ctc tct acg tag act ttc ttg caa gtt act aat ttg ttc cct agc acg tcc gag ttt agt ttt ata tgc cga ata gag ttc tga ttc atc ctt tga gca gat ctc tag aga tcg ttc aag atc cct gat tct agt ctt tag cct att tac ctc ctc gga aga tgc tcc gtt acc gtt gcg ttt aca ctc gtt aag ctg tct atc aag atc cat gat tct atc tct --- --- --- --- --- --- --- --- --- --- --- --- --- --- --- --- -

>MW960419.1:139421-139812_Vector_synVACV-SFV_complete_sequence

gct gct agt tgt cgt tcg aga tca gaa ata cgg gtt ttc ttg gaa ttg att tcg tcg atg tac ttg gca --- --- --- tcg aaa cac tta tta agt tc- ttt ttc caa ttc tac gat ttt att tct ttc gcg agt caa ttc cct cct gta gta act atc tgt ttt gtc aga ttc acg ctc tct acg tag act ttc ttg caa gtt act aat ttg ttc cct agc acg tcc gag ttt agt ttt ata tgc cga ata gag ttc tga ttc atc ctt tga gca gat ctc tag aga tcg ttc aag atc cct gat tct agt ctt tag cct att tac ctc ctc gga aga tgc tcc gtt acc gtt gcg ttt aca ctc gtt aag ctg tct atc aag atc cat gat tct atc tct --- --- --- --- --- --- --- --- --- --- --- --- --- --- --- --- -

>MW960418.1:139439-139830_Vector_synVACV-wt_complete_sequence

gct gct agt tgt cgt tcg aga tca gaa ata cgg gtt ttc ttg gaa ttg att tcg tcg atg tac ttg gca --- --- --- tcg aaa cac tta tta agt tc- ttt ttc caa ttc tac gat ttt att tct ttc gcg agt caa ttc cct cct gta gta act atc tgt ttt gtc aga ttc acg ctc tct acg tag act ttc ttg caa gtt act aat ttg ttc cct agc acg tcc gag ttt agt ttt ata tgc cga ata gag ttc tga ttc atc ctt tga gca gat ctc tag aga tcg ttc aag atc cct gat tct agt ctt tag cct att tac ctc ctc gga aga tgc tcc gtt acc gtt gcg ttt aca ctc gtt aag ctg tct atc aag atc cat gat tct atc tct --- --- --- --- --- --- --- --- --- --- --- --- --- --- --- --- -

>JN654984.1:138327-138718_Vaccinia_virus_strain_Dryvax_clone_DPP19_complete_genome

gct gct agt tgt cgt tcg aga tca gaa ata cgg gtt ttc ttg gaa ttg att tcg tcg atg tac ttg gca --- --- --- tcg aaa cac tta tta agt tc- ttt ttc caa ttc tac gat ttt att tct ttc gcg agt caa ttc cct cct gta gta act atc tgt ttt gtc aga ttc acg ctc tct acg tag act ttc ttg caa gtt act aat ttg ttc cct agc acg tcc gag ttt agt ttt ata tgc cga ata gag ttc tga ttc atc ctt tga gca gat ctc tag aga tcg ttc aag atc cct gat tct agt ctt tag cct att tac ctc ctc aga aga tgt tcc gtt acc gtt gcg ttt aca ctc gtt aag ctg tct atc aag atc cat gat tct atc tct --- --- --- --- --- --- --- --- --- --- --- --- --- --- --- --- -

>JN654983.1:138446-138837_Vaccinia_virus_strain_Dryvax_clone_DPP17_complete_genome

gct gct agt tgt cgt tcg aga tca gaa ata cgg gtt ttc ttg gaa ttg att tcg tcg atg tac ttg gca --- --- --- tcg aaa cac tta tta agt tc- ttt ttc caa ttc tac gat ttt att tct ttc gcg agt caa ttc cct cct gta gta act atc tgt ttt gtc aga ttc acg ctc tct acg tag act ttc ttg caa gtt act aat ttg ttc cct agc acg tcc gag ttt agt ttt ata tgc cga ata gag ttc tga ttc atc ctt tga gca gat ctc tag aga tcg ttc aag atc cct gat tct agt ctt tag cct att tac ctc ctc gga aga tgc tcc gtt acc gtt gcg ttt aca ctc gtt aag ctg tct atc aag atc cat gat tct atc tct --- --- --- --- --- --- --- --- --- --- --- --- --- --- --- --- -

>JN654980.1:138265-138656_Vaccinia_virus_strain_Dryvax_clone_DPP13_complete_genome

gct gct agt tgt cgt tcg aga tca gaa ata cgg gtt ttc ttg gaa ttg att tcg tcg atg tac ttg gca --- --- --- tcg aaa cac tta tta agt tc- ttt ttc caa ttc tac gat ttt att tct ttc gcg agt caa ttc cct cct gta gta act atc tgt ttt gtc aga ttc acg ctc tct acg tag act ttc ttg caa gtt act aat ttg ttc cct agc acg tcc gag ttt agt ttt ata tgc cga ata gag ttc tga ttc atc ctt tga gca gat ctc tag aga tcg ttc aag atc cct gat tct agt ctt tag cct att tac ctc ctc gga aga tgc tcc gtt acc gtt gcg ttt aca ctc gtt aag ctg tct atc aag atc cat gat tct atc tct --- --- --- --- --- --- --- --- --- --- --- --- --- --- --- --- -

>JN654976.1:138280-138671_Vaccinia_virus_strain_Dryvax_clone_DPP9_complete_genome

gct gct agt tgt cgt tcg aga tca gaa ata cgg gtt ttc ttg gaa ttg att tcg tcg atg tac ttg gca --- --- --- tcg aaa cac tta tta agt tc- ttt ttc caa ttc tac gat ttt att tct ttc gcg agt caa ttc cct cct gta gta act atc tgt ttt gtc aga ttc acg ctc tct acg tag act ttc ttg caa gtt act aat ttg ttc cct agc acg tcc gag ttt agt ttt ata tgc cga ata gag ttc tga ttc atc ctt tga gca gat ctc tag aga tcg ttc aag atc cct gat tct agt ctt tag cct att tac ctc ctc gga aga tgc tcc gtt acc gtt gcg ttt aca ctc gtt aag ctg tct atc aag atc cat gat tct atc tct --- --- --- --- --- --- --- --- --- --- --- --- --- --- --- --- -

>DQ439815.1:140880-141271_Vaccinia_virus_strain_DUKE_complete_genome

gct gct agt tgt cgt tcg aga tca gaa ata cgg gtt ttc ttg gaa ttg att tcg tcg atg tac ttg gca --- --- --- tcg aaa cac tta tta agt tc- ttt ttc caa ttc tac gat ttt att tct ttc gcg agt caa ttc cct cct gta gta act atc tgt ttt gtc aga ttc acg ctc tct acg tag act ttc ttg caa gtt act aat ttg ttc cct agc acg tcc gag ttt agt ttt ata tgc cga ata gag ttc tga ttc atc ctt tga gca gat ctc tag aga tcg ttc aag atc cct gat tct agt ctt tag cct att tac ctc ctc gga aga tgc tcc gtt acc gtt gcg ttt aca ctc gtt aag ctg tct atc aag atc cat gat tct atc tct --- --- --- --- --- --- --- --- --- --- --- --- --- --- --- --- -

>AY313847.1:138587-138978_Vaccinia_virus_strain_Acambis_clone_2000_complete_genome

gct gct agt tgt cgt tcg aga tca gaa ata cgg gtt ttc ttg gaa ttg att tcg tcg atg tac ttg gca --- --- --- tcg aaa cac tta tta agt tc- ttt ttc caa ttc tac gat ttt att tct ttc gcg agt caa ttc cct cct gta gta act atc tgt ttt gtc aga ttc acg ctc tct acg tag act ttc ttg caa gtt act aat ttg ttc cct agc acg tcc gag ttt agt ttt ata tgc cga ata gag ttc tga ttc atc ctt tga gca gat ctc tag aga tcg ttc aag atc cct gat tct agt ctt tag cct att tac ctc ctc gga aga tgc tcc gtt acc gtt gcg ttt aca ctc gtt aag ctg tct atc aag atc cat gat tct atc tct --- --- --- --- --- --- --- --- --- --- --- --- --- --- --- --- -

>AY313848.1:138629-139020_Vaccinia_virus_strain_Acambis_clone_3_complete_genome

gct gct agt tgt cgt tcg aga tca gaa ata cgg gtt ttc ttg gaa ttg att tcg tcg atg tac ttg gca --- --- --- tcg aaa cac tta tta agt tc- ttt ttc caa ttc tac gat ttt att tct ttc gcg agt caa ttc cct cct gta gta act atc tgt ttt gtc aga ttc acg ctc tct acg tag act ttc ttg caa gtt act aat ttg ttc cct agc acg tcc gag ttt agt ttt ata tgc cga ata gag ttc tga ttc atc ctt tga gca gat ctc tag aga tcg ttc aag atc cct gat tct agt ctt tag cct att tac ctc ctc gga aga tgc tcc gtt acc gtt gcg ttt aca ctc gtt aag ctg tct atc aag atc cat gat tct atc tct --- --- --- --- --- --- --- --- --- --- --- --- --- --- --- --- -

>DQ377945.1:141137-141528_Vaccinia_virus_strain_3737_complete_genome

gct gct agt tgt cgt tcg aga tca gaa ata cgg gtt ttc ttg gaa ttg att tcg tcg atg tac ttg gca --- --- --- tcg aaa cac tta tta agt tc- ttt ttc caa ttc tac gat ttt att tct ttc gcg agt caa ttc cct cct gta gta act atc tgt ttt gtc aga ttc acg ctc tct acg tag act ttc ttg caa gtt act aat ttg ttc cct agc acg tcc gag ttt agt ttt ata tgc cga ata gag ttc tga ttc atc ctt tga gca gat ctc tag aga tcg ttc aag atc cct gat tct agt ctt tag cct att tac ctc ctc gga aga tgc tcc gtt acc gtt gcg ttt aca ctc gtt aag ctg tct atc aag atc cat gat tct atc tct --- --- --- --- --- --- --- --- --- --- --- --- --- --- --- --- -

>BK010317.1:127954-128354_TPA_asm:_Variola_virus_isolate_VD21

gct gtt agc tgt cgt tca aga tca gaa ata cgg ttt ttc ttg gaa ttg att tcg tca atg tac tta aca tga tta aca tcg aaa cac tta tta agt tc- ttt ttc caa ttc tac gat ttt att tct ttc gcg agt caa ttc cct cct gta gta gct atc ggt ttt gtc aga ttc acg ctc tct acg tag act ttc ttg caa gtt act aat ttg ttc cct gac acg tcc gag ttc agt ttt ata tgc cga ata gag ttc tga ttc atc ctt tga gca gat ctc tag cga ttg ttt aag atc cct aat tct agt ctt tag cct att tac ctc ctc aga aga tgt tcc gtt acc gtt gcg ttt aca ctc gtt aag ttg tct atc aag atc cat gat tct atc tct --- --- --- --- --- --- --- --- --- --- --- --- --- --- --- --- -

>KY358055.1:126518-126918_Variola_virus_strain_VD21_17th_century

gct gtt agc tgt cgt tca aga tca gaa ata cgg ttt ttc ttg gaa ttg att tcg tca atg tac tta aca tga tta aca tcg aaa cac tta tta agt tc- ttt ttc caa ttc tac gat ttt att tct ttc gcg agt caa ttc cct cct gta gta gct atc ggt ttt gtc aga ttc acg ctc tct acg tag act ttc ttg caa gtt act aat ttg ttc cct gac acg tcc gag ttc agt ttt ata tgc cga ata gag ttc tga ttc atc ctt tga gca gat ctc tag cga ttg ttt aag atc cct aat tct agt ctt tag cct att tac ctc ctc aga aga tgt tcc gtt acc gtt gcg ttt aca ctc gtt aag ttg tct atc aag atc cat gat tct atc tct --- --- --- --- --- --- --- --- --- --- --- --- --- --- --- --- -

>MF477237.2:138889-139280_Vaccinia_virus_strain_Mulford_1902_complete_genome

gct gct agt tgt cgt tcg aga tca gaa ata cgg gtt ttc ttg gaa ttg att tcg tcg atg tac ttg gca --- --- --- tcg aaa cac tta tta agt tc- ttt ttc caa ttc tac gat ttt att tct ttc gcg agt caa ttc cct cct gta gta act atc tgt ttt gtc aga ttc acg ctc tct acg tag act ttc ttg caa gtt act aat ttg ttc cct agc acg tcc gag ttc agt ttt ata tgc tga ata gag ttc tga ttc atc ctt tga gca gat ctc tag cga tcg ttt aag att cct gat tct agt ctt tag cct att tac ctc ctc aga aga tgt tcc gtt acc gtt gcg ttt aca ctc gtt aag ctg tct atc aag atc cat gat tct atc tct --- --- --- --- --- --- --- --- --- --- --- --- --- --- --- --- -

>KC207811.1:138151-138542_Vaccinia_virus_strain_TianTan_clone_TP5_complete_genome

gct gct agt tgt cgt tcg aga tca gaa ata cgg gtt ttc ttg gaa ttg att tcg tcg atg tac ttg gca --- --- --- tcg aaa cac tta tta agt tc- ttt ttc caa ttc tac gat ttt att tct ttc gcg agt caa ttc cct cct gta gta act atc tgt ttt gtc aga ttc acg ctc tct acg tag act ttc ttg caa gtt act aat ttg ttc cct agc acg tcc gag ttc agt ttt ata tgc tga ata gag ttc tga ttc atc ctt tga gca gat ctc tag cga tcg ttt aag att cct gat tct agt ctt tag cct att tac ctc ctc aga aga tgt tcc gtt acc gtt gcg ttt aca ctc gtt aag ctg tct atc aag atc cat gat tct atc tct --- --- --- --- --- --- --- --- --- --- --- --- --- --- --- --- -

>KC207810.1:132443-132834_Vaccinia_virus_strain_TianTan_clone_TP3_complete_genome

gct gct agt tgt cgt tcg aga tca gaa ata cgg gtt ttc ttg gaa ttg att tcg tcg atg tac ttg gca --- --- --- tcg aaa cac tta tta agt tc- ttt ttc caa ttc tac gat ttt att tct ttc gcg agt caa ttc cct cct gta gta act atc tgt ttt gtc aga ttc acg ctc tct acg tag act ttc ttg caa gtt act aat ttg ttc cct agc acg tcc gag ttc agt ttt ata tgc tga ata gag ttc tga ttc atc ctt tga gca gat ctc tag cga tcg ttt aag att cct gat tct agt ctt tag cct att tac ctc ctc aga aga tgt tcc gtt acc gtt gcg ttt aca ctc gtt aag ctg tct atc aag atc cat gat tct atc tct --- --- --- --- --- --- --- --- --- --- --- --- --- --- --- --- -

>JX489137.1:135160-135551_Vaccinia_virus_strain_Tiantan_clone_TT10_complete_genome

gct gct agt tgt cgt tcg aga tca gaa ata cgg gtt ttc ttg gaa ttg att tcg tcg atg tac ttg gca --- --- --- tcg aaa cac tta tta agt tc- ttt ttc caa ttc tac gat ttt att tct ttc gcg agt caa ttc cct cct gta gta act atc tgt ttt gtc aga ttc acg ctc tct acg tag act ttc ttg caa gtt act aat ttg ttc cct agc acg tcc gag ttc agt ttt ata tgc tga ata gag ttc tga ttc atc ctt tga gca gat ctc tag cga tcg ttt aag att cct gat tct agt ctt tag cct att tac ctc ctc aga aga tgt tcc gtt acc gtt gcg ttt aca ctc gtt aag ctg tct atc aag atc cat gat tct atc tct --- --- --- --- --- --- --- --- --- --- --- --- --- --- --- --- -

>JN654986.1:138287-138678_Vaccinia_virus_strain_Dryvax_clone_DPP21_complete_genome

gct gct agt tgt cgt tcg aga tca gaa ata cgg gtt ttc ttg gaa ttg att tcg tcg atg tac ttg gca --- --- --- tcg aaa cac tta tta agt tc- ttt ttc caa ttc tac gat ttt att tct ttc gcg agt caa ttc cct cct gta gta act atc tgt ttt gtc aga ttc acg ctc tct acg tag act ttc ttg caa gtt act aat ttg ttc cct agc acg tcc gag ttt agt ttt ata tgc cga ata gag ttc tga ttc atc ctt tga gca gat ctc tag aga tcg ttc aag att cct gat tct agt ctt tag cct att tac ctc ctc gga aga tgc tcc gtt acc gtt gcg ttt aca ctc gtt aag ctg tct atc aag atc cat gat tct atc tct --- --- --- --- --- --- --- --- --- --- --- --- --- --- --- --- -

>JN654985.1:138370-138761_Vaccinia_virus_strain_Dryvax_clone_DPP20_complete_genome

gct gct agt tgt cgt tcg aga tca gaa ata cgg gtt ttc ttg gaa ttg att tcg tcg atg tac ttg gca --- --- --- tcg aaa cac tta tta agt tc- ttt ttc caa ttc tac aat ttt att tct ttc gcg agt caa ttc cct cct gta gta act atc tgt ttt gtc aga ttc acg ctc tct acg tag act ttc ttg caa gtt act aat ttg ttc cct agc acg tcc gag ttt agt ttt ata tgc cga ata gag ttc tga ttc atc ctt tga gca gat ctc tag aga tcg ttc aag atc cct gat tct agt ctt tag cct att tac ctc ctc gga aga tgc tcc gtt acc gtt gcg ttt aca ctc gtt aag ctg tct atc aag atc cat gat tct atc tct --- --- --- --- --- --- --- --- --- --- --- --- --- --- --- --- -

>JN654981.1:138282-138673_Vaccinia_virus_strain_Dryvax_clone_DPP15_complete_genome

gct gct agt tgt cgt tcg aga tca gaa ata cgg gtt ttc ttg gaa ttg att tcg tcg atg tac ttg gca --- --- --- tcg aaa cac tta tta agt tc- ttt ttc caa ttc tac gat ttt att tct ttc gcg agt caa ttc cct cct gta gta act atc tgt ttt gtc aga ttc acg ctc tct acg tag act ttc ttg caa gtt act aat ttg ttc tct agc acg tcc gag ttt agt ttt ata tgc cga ata gag ttc tga ttc atc ctt tga gca gat ctc tag aga tcg ttc aag atc cct gat tct agt ctt tag cct att tac ctc ctc gga aga tgc tcc gtt acc gtt gcg ttt aca ctc gtt aag ctg tct atc aag atc cat gat tct atc tct --- --- --- --- --- --- --- --- --- --- --- --- --- --- --- --- -

>JN654979.1:138470-138861_Vaccinia_virus_strain_Dryvax_clone_DPP12_complete_genome

gct gct agt tgt cgt tcg aga tca gaa ata cgg gtt ttc ttg gaa ttg att tcg tcg atg tac ttg gca --- --- --- tcg aaa cac tta tta agt tc- ttt ttc caa ttc tac aat ttt att tct ttc gcg agt caa ttc cct cct gta gta act atc tgt ttt gtc aga ttc acg ctc tct acg tag act ttc ttg caa gtt act aat ttg ttc cct agc acg tcc gag ttt agt ttt ata tgc cga ata gag ttc tga ttc atc ctt tga gca gat ctc tag aga tcg ttc aag atc cct gat tct agt ctt tag cct att tac ctc ctc gga aga tgc tcc gtt acc gtt gcg ttt aca ctc gtt aag ctg tct atc aag atc cat gat tct atc tct --- --- --- --- --- --- --- --- --- --- --- --- --- --- --- --- -

>JN654978.1:138276-138667_Vaccinia_virus_strain_Dryvax_clone_DPP11_complete_genome

gct gct agt tgt cgt tcg aga tca gaa ata cgg gtt ttc ttg gaa ttg att tcg tcg atg tac ttg gca --- --- --- tcg aaa cac tta tta agt tc- ttt ttc caa ttc tac gat ttt att tct ttc gcg agt caa ttc cct cct gta gta act atc tgt ttt gtc aga ttc acg ctc tct acg tag act ttc ttg caa gtt act aat ttg ttc tct agc acg tcc gag ttt agt ttt ata tgc cga ata gag ttc tga ttc atc ctt tga gca gat ctc tag aga tcg ttc aag atc cct gat tct agt ctt tag cct att tac ctc ctc gga aga tgc tcc gtt acc gtt gcg ttt aca ctc gtt aag ctg tct atc aag atc cat gat tct atc tct --- --- --- --- --- --- --- --- --- --- --- --- --- --- --- --- -

>JN654977.1:138234-138625_Vaccinia_virus_strain_Dryvax_clone_DPP10_complete_genome

gct gct agt tgt cgt tcg aga tca gaa ata cgg gtt ttc ttg gaa ttg att tcg tcg atg tac ttg gca --- --- --- tcg aaa cac tta tta agt tc- ttt ttc caa ttc tac aat ttt att tct ttc gcg agt caa ttc cct cct gta gta act atc tgt ttt gtc aga ttc acg ctc tct acg tag act ttc ttg caa gtt act aat ttg ttc cct agc acg tcc gag ttt agt ttt ata tgc cga ata gag ttc tga ttc atc ctt tga gca gat ctc tag aga tcg ttc aag atc cct gat tct agt ctt tag cct att tac ctc ctc gga aga tgc tcc gtt acc gtt gcg ttt aca ctc gtt aag ctg tct atc aag atc cat gat tct atc tct --- --- --- --- --- --- --- --- --- --- --- --- --- --- --- --- -

>AY484669.1:140566-140957_Rabbitpox_virus_complete_genome

gct gct agt tgt cgt tcg aga tca gaa ata cgg gtt ttc ttg gaa ttg att tcg tcg atg tac ttg gca --- --- --- tcg aaa cac tta tta agt tc- ttt ttc caa ttc tac gat ttt att tct ttc gcg agt caa ttc cct cct gta gta act atc tgt ttt gtc aga ttc acg ctc tct acg tag act ttc ttg caa gtt act aat ttg ttc cct agc acg tcc gag ttt agt ttt ata tgc tga ata gag ttc tga ttc atc ctt tga gca gat ctc tag aga tcg ttc aag atc cct gat tct agt ctt tag cct att tac ctc ctc gga aga tgc tcc gtt acc gtt gcg ttt aca ctc gtt aag ctg tct atc aag atc cat gat tct atc tct --- --- --- --- --- --- --- --- --- --- --- --- --- --- --- --- -

>DQ441437.1:128046-128446_Variola_virus_strain_Sierra_Leone_1969_(V68-258)_complete_genome

gct gtt agc tgt cgt tca aga tca gaa ata cgg ttt ttc ttg gaa ttg att tcg tca atg tac tta aca tga tta aca tcg aaa cac tta tta agt tc- ttt ttc caa ttc tac gat ttt att tct ttc gcg agt caa ttc cct cct gta gta gct atc ggt ttt gtc aga ttc acg ctc tct acg tag act ttc ttg caa gtt act aat ttg ttc cct gac acg tcc gag ttc agt ttt ata tgc cga ata gag ttc tga ttt atc ctt tga gca gat ctc tag cga ttg ttt aag atc cct aat tct agt ctt tag cct att tac ctc ctc aga aga tgt tcc gtt acc gtt gcg ttt aca ctc gtt aag ctg tct atc aag atc cat gat tct atc tct --- --- --- --- --- --- --- --- --- --- --- --- --- --- --- --- -

>DQ441434.1:127938-128338_Variola_virus_strain_Niger_1969_(001_importation_from_Nigeria)_complete_genome

gct gtt agc tgt cgt tca aga tca gaa ata cgg ttt ttc ttg gaa ttg att tcg tca atg tac tta aca tga tta aca tcg aaa cac tta tta agt tc- ttt ttc caa ttc tac gat ttt att tct ttc gcg agt caa ttc cct cct gta gta gct atc ggt ttt gtc aga ttc acg ctc tct acg tag act ttc ttg caa gtt act aat ttg ttc cct gac acg tcc gag ttc agt ttt ata tgc cga ata gag ttc tga ttt atc ctt tga gca gat ctc tag cga ttg ttt aag atc cct aat tct agt ctt tag cct att tac ctc ctc aga aga tgt tcc gtt acc gtt gcg ttt aca ctc gtt aag ctg tct atc aag atc cat gat tct atc tct --- --- --- --- --- --- --- --- --- --- --- --- --- --- --- --- -

>DQ441426.1:127875-128275_Variola_virus_strain_Guinea_1969_(005)_complete_genome

gct gtt agc tgt cgt tca aga tca gaa ata cgg ttt ttc ttg gaa ttg att tcg tca atg tac tta aca tga tta aca tcg aaa cac tta tta agt tc- ttt ttc caa ttc tac gat ttt att tct ttc gcg agt caa ttc cct cct gta gta gct atc ggt ttt gtc aga ttc acg ctc tct acg tag act ttc ttg caa gtt act aat ttg ttc cct gac acg tcc gag ttc agt ttt ata tgc cga ata gag ttc tga ttt atc ctt tga gca gat ctc tag cga ttg ttt aag atc cct aat tct agt ctt tag cct att tac ctc ctc aga aga tgt tcc gtt acc gtt gcg ttt aca ctc gtt aag ctg tct atc aag atc cat gat tct atc tct --- --- --- --- --- --- --- --- --- --- --- --- --- --- --- --- -

>LR800246.1:130755-131155_Variola_virus_genome_assembly_complete_genome:_monopartite

gct act agc tgt cgt tcg aga tca gaa ata cgg gtt ttc ttg gaa ttg att tcg tcg atg tac ttg gca ??? ??? ??? ??g aaa cac tta tta agt tc- ttt ttc caa ttc tac gat ttt att tct ttc gcg agt caa ttc cct cct gta gta gct atc ggt ttt gtc aga ttc acg ctc tct acg tag act ttc ttg caa gtt act aat ttg ttc cct ggt acg tcc gag ttt agt ttt ata tac cga ata gag ttc tga ttc atc ctt tga gca gat ctc tag cga tcg ttc aag atc cct gat tct agt ctt tag cct att tac ctc ctc aaa aga tgt tcc gtt acc gtt gcg ttt aca ctc gtt aag ctg tct atc aag atc cat gat tct atc tct --- --- --- --- --- --- --- --- --- --- --- --- --- --- --- --- -

>MK336431.1:118827-119218_Vaccinia_virus_isolate_KVAC103

gct gct agt tgt cgt tcg aga tca gaa ata cgg gtt ttc ttg gaa ttg att tcg tcg atg tac ttg gca --- --- --- tcg aaa cac tta tta agt tc- ttt ttc caa ttc tac gat ttt att tct ttc gcg agt caa ttc cct cct gta gta act atc tgt ttt gtc aga ttc acg ctc tct acg tag act ttc ttg caa gtt act aat ttg ttc cct agc acg tcc gag ttt agt ttt ata tgc tga ata gag ttc tga ttc atc ctt tga gca gat ctc tag cga tcg ttt aag att cct gat tct agt ctt tag cct att tac ctc ctc aga aga tgt tcc gtt acc gtt gcg ttt aca ctc gtt aag ctg tct atc aag atc cat gat tct atc tct --- --- --- --- --- --- --- --- --- --- --- --- --- --- --- --- -

>MH341447.1:138500-138891_Vaccinia_virus_strain_L-IVP_oncoQ

gct gct agt tgt cgt tcg aga tca gaa ata cgg gtt ttc ttg gaa ttg att tcg tcg atg tac ttg gca --- --- --- tcg aaa cac tta tta agt tc- ttt ttc caa ttc tac gat ttt att tct ttc gcg agt caa ttc cct cct gta gta act atc tgt ttt gtc aga ttc acg ctc tct acg tag act ttc ttg caa gtt act aat ttg ttc cct agc acg tcc gag ttt agt ttt ata tgc tga ata gag ttc tga ttc atc ctt tga gca gat ctc tag cga tcg ttt aag att cct gat tct agt ctt tag cct att tac ctc ctc aga aga tgt tcc gtt acc gtt gcg ttt aca ctc gtt aag ctg tct atc aag atc cat gat tct atc tct --- --- --- --- --- --- --- --- --- --- --- --- --- --- --- --- -

>MH341446.1:138366-138757_Vaccinia_virus_strain_L-IVP_oncoM

gct gct agt tgt cgt tcg aga tca gaa ata cgg gtt ttc ttg gaa ttg att tcg tcg atg tac ttg gca --- --- --- tcg aaa cac tta tta agt tc- ttt ttc caa ttc tac gat ttt att tct ttc gcg agt caa ttc cct cct gta gta act atc tgt ttt gtc aga ttc acg ctc tct acg tag act ttc ttg caa gtt act aat ttg ttc cct agc acg tcc gag ttt agt ttt ata tgc tga ata gag ttc tga ttc atc ctt tga gca gat ctc tag cga tcg ttt aag att cct gat tct agt ctt tag cct att tac ctc ctc aga aga tgt tcc gtt acc gtt gcg ttt aca ctc gtt aag ctg tct atc aag atc cat gat tct atc tct --- --- --- --- --- --- --- --- --- --- --- --- --- --- --- --- -

>MH341445.1:138306-138697_Vaccinia_virus_strain_L-IVP_oncoB

gct gct agt tgt cgt tcg aga tca gaa ata cgg gtt ttc ttg gaa ttg att tcg tcg atg tac ttg gca --- --- --- tcg aaa cac tta tta agt tc- ttt ttc caa ttc tac gat ttt att tct ttc gcg agt caa ttc cct cct gta gta act atc tgt ttt gtc aga ttc acg ctc tct acg tag act ttc ttg caa gtt act aat ttg ttc cct agc acg tcc gag ttt agt ttt ata tgc tga ata gag ttc tga ttc atc ctt tga gca gat ctc tag cga tcg ttt aag att cct gat tct agt ctt tag cct att tac ctc ctc aga aga tgt tcc gtt acc gtt gcg ttt aca ctc gtt aag ctg tct atc aag atc cat gat tct atc tct --- --- --- --- --- --- --- --- --- --- --- --- --- --- --- --- -

>LC315596.1:142955-143346_BAC_cloning_vector_pLC16m8.8S-BAC_DNA_complete_sequence

gct gct agt tgt cgt tcg aga tca gaa ata cgg gtt ttc ttg gaa ttg att tcg tcg atg tac ttg gca --- --- --- tcg aaa cac tta tta agt tc- ttt ttc caa ttc tac gat ttt att tct ttc gcg agt caa ttc cct cct gta gta act atc tgt ttt gtc aga ttc acg ctc tct acg tag act ttc ttg caa gtt act aat ttg ttc cct agc acg tcc gag ttt agt ttt ata tgc tga ata gag ttc tga ttc atc ctt tga gca gat ctc tag cga tcg ttt aag att cct gat tct agt ctt tag cct att tac ctc ctc aga aga tgt tcc gtt acc gtt gcg ttt aca ctc gtt aag ctg tct atc aag atc cat gat tct atc tct --- --- --- --- --- --- --- --- --- --- --- --- --- --- --- --- -

>LT966077.1:135861-136252_Vaccinia_virus_WR_strain_WR_-_ATCC_VR-2035_genome_assembly_complete_genome:_monopartite

gct gct agt tgt cgt tcg aga tca gaa ata cgg gtt ttc ttg gaa ttg att tcg tcg atg tac ttg gca --- --- --- tcg aaa cac tta tta agt tc- ttt ttc caa ttc tac gat ttt att tct ttc gcg agt caa ttc cct cct gta gta act atc tgt ttt gtc aga ttc acg ctc tct acg tag act ttc ttg caa gtt act aat ttg ttc cct agc acg tcc gag ttt agt ttt ata tgc tga ata gag ttc tga ttc atc ctt tga gca gat ctc tag cga tcg ttt aag att cct gat tct agt ctt tag cct att tac ctc ctc aga aga tgt tcc gtt acc gtt gcg ttt aca ctc gtt aag ctg tct atc aag atc cat gat tct atc tct --- --- --- --- --- --- --- --- --- --- --- --- --- --- --- --- -

>KX781953.1:137773-138164_Vaccinia_virus_strain_L-IVP_clone_14

gct gct agt tgt cgt tcg aga tca gaa ata cgg gtt ttc ttg gaa ttg att tcg tcg atg tac ttg gca --- --- --- tcg aaa cac tta tta agt tc- ttt ttc caa ttc tac gat ttt att tct ttc gcg agt caa ttc cct cct gta gta act atc tgt ttt gtc aga ttc acg ctc tct acg tag act ttc ttg caa gtt act aat ttg ttc cct agc acg tcc gag ttt agt ttt ata tgc tga ata gag ttc tga ttc atc ctt tga gca gat ctc tag cga tcg ttt aag att cct gat tct agt ctt tag cct att tac ctc ctc aga aga tgt tcc gtt acc gtt gcg ttt aca ctc gtt aag ctg tct atc aag atc cat gat tct atc tct --- --- --- --- --- --- --- --- --- --- --- --- --- --- --- --- -

>KT184691.1:136157-136548_Vaccinia_virus_strain_IOC_clone_B388

gct gct agt tgt cgt tcg aga tca gaa ata cgg gtt ttc ttg gaa ttg att tcg tcg atg tac ttg gca --- --- --- tcg aaa cac tta tta agt tc- ttt ttc caa ttc tac gat ttt att tct ttc gcg agt caa ttc cct cct gta gta act atc tgt ttt gtc aga ttc acg ctc tct acg tag act ttc ttg caa gtt act aat ttg ttc cct agc acg tcc gag ttt agt ttt ata tgc tga ata gag ttc tga ttc atc ctt tga gca gat ctc tag cga tcg ttt aag att cct gat tct agt ctt tag cct att tac ctc ctc aga aga tgt tcc gtt acc gtt gcg ttt aca ctc gtt aag ctg tct atc aag atc cat gat tct atc tct --- --- --- --- --- --- --- --- --- --- --- --- --- --- --- --- -

>KT184690.1:136537-136928_Vaccinia_virus_strain_IOC_clone_B141

gct gct agt tgt cgt tcg aga tca gaa ata cgg gtt ttc ttg gaa ttg att tcg tcg atg tac ttg gca --- --- --- tcg aaa cac tta tta agt tc- ttt ttc caa ttc tac gat ttt att tct ttc gcg agt caa ttc cct cct gta gta act atc tgt ttt gtc aga ttc acg ctc tct acg tag act ttc ttg caa gtt act aat ttg ttc cct agc acg tcc gag ttt agt ttt ata tgc tga ata gag ttc tga ttc atc ctt tga gca gat ctc tag cga tcg ttt aag att cct gat tct agt ctt tag cct att tac ctc ctc aga aga tgt tcc gtt acc gtt gcg ttt aca ctc gtt aag ctg tct atc aag atc cat gat tct atc tct --- --- --- --- --- --- --- --- --- --- --- --- --- --- --- --- -

>KP233807.1:136097-136488_Vaccinia_virus_strain_L-IVP_complete_genome

gct gct agt tgt cgt tcg aga tca gaa ata cgg gtt ttc ttg gaa ttg att tcg tcg atg tac ttg gca --- --- --- tcg aaa cac tta tta agt tc- ttt ttc caa ttc tac gat ttt att tct ttc gcg agt caa ttc cct cct gta gta act atc tgt ttt gtc aga ttc acg ctc tct acg tag act ttc ttg caa gtt act aat ttg ttc cct agc acg tcc gag ttt agt ttt ata tgc tga ata gag ttc tga ttc atc ctt tga gca gat ctc tag cga tcg ttt aag att cct gat tct agt ctt tag cct att tac ctc ctc aga aga tgt tcc gtt acc gtt gcg ttt aca ctc gtt aag ctg tct atc aag atc cat gat tct atc tct --- --- --- --- --- --- --- --- --- --- --- --- --- --- --- --- -

>KJ125439.1:138943-139334_Vaccinia_virus_strain_IHD-W_clone_IHDW1_complete_genome

gct gct agt tgt cgt tcg aga tca gaa ata cgg gtt ttc ttg gaa ttg att tcg tcg atg tac ttg gca --- --- --- tcg aaa cac tta tta agt tc- ttt ttc caa ttc tac gat ttt att tct ttc gcg agt caa ttc cct cct gta gta act atc tgt ttt gtc aga ttc acg ctc tct acg tag act ttc ttg caa gtt act aat ttg ttc cct agc acg tcc gag ttt agt ttt ata tgc tga ata gag ttc tga ttc atc ctt tga gca gat ctc tag cga tcg ttt aag att cct gat tct agt ctt tag cct att tac ctc ctc aga aga tgt tcc gtt acc gtt gcg ttt aca ctc gtt aag ctg tct atc aag atc cat gat tct atc tct --- --- --- --- --- --- --- --- --- --- --- --- --- --- --- --- -

>KJ399295.1:17-408_Vaccinia_virus_isolate_SFI1_acidophilic_type_inclusion_protein_gene_partial_cds

gct gct agt tgt cgt tcg aga tca gaa ata cgg gtt ttc ttg gaa ttg att tcg tcg atg tac ttg gca --- --- --- tcg aaa cac tta tta agt tc- ttt ttc caa ttc tac gat ttt att tct ttc gcg agt caa ttc cct cct gta gta act atc tgt ttt gtc aga ttc acg ctc tct acg tag act ttc ttg caa gtt act aat ttg ttc cct agc acg tcc gag ttt agt ttt ata tgc tga ata gag ttc tga ttc atc ctt tga gca gat ctc tag cga tcg ttt aag att cct gat tct agt ctt tag cct att tac ctc ctc aga aga tgt tcc gtt acc gtt gcg ttt aca ctc gtt aag ctg tct atc aag atc cat gat tct atc tct --- --- --- --- --- --- --- --- --- --- --- --- --- --- --- --- -

>JX489138.1:135102-135493_Vaccinia_virus_strain_Tiantan_clone_TT11_complete_genome

gct gct agt tgt cgt tcg aga tca gaa ata cgg gtt ttc ttg gaa ttg att tcg tcg atg tac ttg gca --- --- --- tcg aaa cac tta tta agt tc- ttt ttc caa ttc tac gat ttt att tct ttc gcg agt caa ttc cct cct gta gta act atc tgt ttt gtc aga ttc acg ctc tct acg tag act ttc ttg caa gtt act aat ttg ttc cct agc acg tcc gag ttt agt ttt ata tgc tga ata gag ttc tga ttc atc ctt tga gca gat ctc tag cga tcg ttt aag att cct gat tct agt ctt tag cct att tac ctc ctc aga aga tgt tcc gtt acc gtt gcg ttt aca ctc gtt aag ctg tct atc aag atc cat gat tct atc tct --- --- --- --- --- --- --- --- --- --- --- --- --- --- --- --- -

>JN654982.1:138481-138872_Vaccinia_virus_strain_Dryvax_clone_DPP16_complete_genome

gct gct agt tgt cgt tcg aga tca gaa ata cgg gtt ttc ttg gaa ttg att tcg tcg atg tac ttg gca --- --- --- tcg aaa cac tta tta agt tc- ttt ttc caa ttc tac aat ttt att tct ttc gcg agt caa ttc cct cct gta gta act atc tgt ttt gtc aga ttc acg ctc tct acg tag act ttc ttg caa gtt act aat ttg ttc cct agc acg tcc gag ttt agt ttt ata tgc cga ata gag ttc tga ttc atc ctt tga gca gat ctc tag cga tcg ttt aag att cct gat tct agt ctt tag cct att tac ctc ctc aga aga tgt tcc gtt acc gtt gcg ttt aca ctc gtt aag ctg tct atc aag atc cat gat tct atc tct --- --- --- --- --- --- --- --- --- --- --- --- --- --- --- --- -

>AM501482.1:136000-136391_Vaccinia_virus_Ankara_strain_chorioallantois_vaccinia_virus_Ankara_(CVA)_complete_coding_genome

gct gct agt tgt cgt tcg aga tca gaa ata cgg gtt ttc ttg gaa ttg att tcg tcg atg tac ttg gca --- --- --- tcg aaa cac tta tta agt tc- ttt ttc caa ttc tac gat ttt att tct ttc gcg agt caa ttc cct cct gta gta act atc tgt ttt gtc aga ttc acg ctc tct acg tag act ttc ttg caa gtt act aat ttg ttc cct agc acg tcc gag ttt agt ttt ata tgc tga ata gag ttc tga ttc atc ctt tga gca gat ctc tag cga tcg ttt aag att cct gat tct agt ctt tag cct att tac ctc ctc aga aga tgt tcc gtt acc gtt gcg ttt aca ctc gtt aag ctg tct atc aag atc cat gat tct atc tct --- --- --- --- --- --- --- --- --- --- --- --- --- --- --- --- -

>DQ121394.1:138438-138829_Vaccinia_virus_strain_Lister_clone_VACV107_complete_genome

gct gct agt tgt cgt tcg aga tca gaa ata cgg gtt ttc ttg gaa ttg att tcg tcg atg tac ttg gca --- --- --- tcg aaa cac tta tta agt tc- ttt ttc caa ttc tac gat ttt att tct ttc gcg agt caa ttc cct cct gta gta act atc tgt ttt gtc aga ttc acg ctc tct acg tag act ttc ttg caa gtt act aat ttg ttc cct agc acg tcc gag ttt agt ttt ata tgc tga ata gag ttc tga ttc atc ctt tga gca gat ctc tag cga tcg ttt aag att cct gat tct agt ctt tag cct att tac ctc ctc aga aga tgt tcc gtt acc gtt gcg ttt aca ctc gtt aag ctg tct atc aag atc cat gat tct atc tct --- --- --- --- --- --- --- --- --- --- --- --- --- --- --- --- -

>AY678275.1:138395-138786_Vaccinia_virus_strain_LC16m8_complete_genome

gct gct agt tgt cgt tcg aga tca gaa ata cgg gtt ttc ttg gaa ttg att tcg tcg atg tac ttg gca --- --- --- tcg aaa cac tta tta agt tc- ttt ttc caa ttc tac gat ttt att tct ttc gcg agt caa ttc cct cct gta gta act atc tgt ttt gtc aga ttc acg ctc tct acg tag act ttc ttg caa gtt act aat ttg ttc cct agc acg tcc gag ttt agt ttt ata tgc tga ata gag ttc tga ttc atc ctt tga gca gat ctc tag cga tcg ttt aag att cct gat tct agt ctt tag cct att tac ctc ctc aga aga tgt tcc gtt acc gtt gcg ttt aca ctc gtt aag ctg tct atc aag atc cat gat tct atc tct --- --- --- --- --- --- --- --- --- --- --- --- --- --- --- --- -

>AY678277.1:138394-138785_Vaccinia_virus_strain_LC16mO_complete_genome

gct gct agt tgt cgt tcg aga tca gaa ata cgg gtt ttc ttg gaa ttg att tcg tcg atg tac ttg gca --- --- --- tcg aaa cac tta tta agt tc- ttt ttc caa ttc tac gat ttt att tct ttc gcg agt caa ttc cct cct gta gta act atc tgt ttt gtc aga ttc acg ctc tct acg tag act ttc ttg caa gtt act aat ttg ttc cct agc acg tcc gag ttt agt ttt ata tgc tga ata gag ttc tga ttc atc ctt tga gca gat ctc tag cga tcg ttt aag att cct gat tct agt ctt tag cct att tac ctc ctc aga aga tgt tcc gtt acc gtt gcg ttt aca ctc gtt aag ctg tct atc aag atc cat gat tct atc tct --- --- --- --- --- --- --- --- --- --- --- --- --- --- --- --- -

>AY678276.1:138280-138671_Vaccinia_virus_strain_Lister_complete_genome

gct gct agt tgt cgt tcg aga tca gaa ata cgg gtt ttc ttg gaa ttg att tcg tcg atg tac ttg gca --- --- --- tcg aaa cac tta tta agt tc- ttt ttc caa ttc tac gat ttt att tct ttc gcg agt caa ttc cct cct gta gta act atc tgt ttt gtc aga ttc acg ctc tct acg tag act ttc ttg caa gtt act aat ttg ttc cct agc acg tcc gag ttt agt ttt ata tgc tga ata gag ttc tga ttc atc ctt tga gca gat ctc tag cga tcg ttt aag att cct gat tct agt ctt tag cct att tac ctc ctc aga aga tgt tcc gtt acc gtt gcg ttt aca ctc gtt aag ctg tct atc aag atc cat gat tct atc tct --- --- --- --- --- --- --- --- --- --- --- --- --- --- --- --- -

>AY243312.1:135756-136147_Vaccinia_virus_WR_complete_genome

gct gct agt tgt cgt tcg aga tca gaa ata cgg gtt ttc ttg gaa ttg att tcg tcg atg tac ttg gca --- --- --- tcg aaa cac tta tta agt tc- ttt ttc caa ttc tac gat ttt att tct ttc gcg agt caa ttc cct cct gta gta act atc tgt ttt gtc aga ttc acg ctc tct acg tag act ttc ttg caa gtt act aat ttg ttc cct agc acg tcc gag ttt agt ttt ata tgc tga ata gag ttc tga ttc atc ctt tga gca gat ctc tag cga tcg ttt aag att cct gat tct agt ctt tag cct att tac ctc ctc aga aga tgt tcc gtt acc gtt gcg ttt aca ctc gtt aag ctg tct atc aag atc cat gat tct atc tct --- --- --- --- --- --- --- --- --- --- --- --- --- --- --- --- -

>X57318.1:107-498_Vaccina_virus_genomic_DNA

gct gct agt tgt cgt tcg aga tca gaa ata cgg gtt ttc ttg gaa ttg att tcg tcg atg tac ttg gca --- --- --- tcg aaa cac tta tta agt tc- ttt ttc caa ttc tac gat ttt att tct ttc gcg agt caa ttc cct cct gta gta act atc tgt ttt gtc aga ttc acg ctc tct acg tag act ttc ttg caa gtt act aat ttg ttc cct agc acg tcc gag ttt agt ttt ata tgc tga ata gag ttc tga ttc atc ctt tga gca gat ctc tag cga tcg ttt aag att cct gat tct agt ctt tag cct att tac ctc ctc aga aga tgt tcc gtt acc gtt gcg ttt aca ctc gtt aag ctg tct atc aag atc cat gat tct atc tct --- --- --- --- --- --- --- --- --- --- --- --- --- --- --- --- -

>LT706529.1:126357-126757_Variola_virus_isolate_VARV_V1588_genome_assembly_chromosome:_I

gct gtt agc tgt cgt tca aga tca gaa ata cgg ttt ttc ttg gaa ttg att tcg tca atg tac tta aca tga tta aca tcg aaa cac tta tta agt tc- ttt ttc caa ttc tac gat ttt att tct ttc gcg agt caa ttc cct cct gta gta gct atc ggt ttt gtc aaa ttc acg ctc tct acg tag act ttc ttg caa gtt act aat ttg ttc cct gac acg tcc gag ttc agt ttt ata tgc cga ata gag ttc tga ttt atc ctt tga gca gat ctc tag cga ttg ttt aag atc cct aat tct agt ctt tag cct att tac ctc ctc aga aga tgt tcc gtt acc gtt gcg ttt aca ctc gtt aag ctg tct atc aag atc cat gat tct atc tct --- --- --- --- --- --- --- --- --- --- --- --- --- --- --- --- -

>DQ441447.1:128038-128438_Variola_virus_strain_United_Kingdom_1952_Butler_complete_genome

gct gtt agc tgt cgt tca aga tca gaa ata cgg ttt ttc ttg aaa ttg att tcg tca atg tac tta aca tga tta aca tcg aaa cac tta tta agt tc- ttt ttc caa ttc tac gat ttt att tct ttc gcg agt caa ttc cct cct gta gta gct atc ggt ttt gtc aga ttc acg ctc tct acg tag act ttc ttg caa gtt act aat ttg ttc cct gac acg tcc gag ttc agt ttt ata tgc cga ata gag ttc tga ttt atc ctt tga gca gat ctc tag cga ttg ttt aag atc cct aat tct agt ctt tag cct att tac ctc ctc aga aga tgt tcc gtt acc gtt gcg ttt aca ctc gtt aag ctg tct atc aag atc cat gat tct atc tct --- --- --- --- --- --- --- --- --- --- --- --- --- --- --- --- -

>DQ441419.1:127910-128310_Variola_virus_strain_Brazil_1966_(v66-39_Sao_Paulo)_complete_genome

gct gtt agc tgt cgt tca aga tca gaa ata cgg ttt ttc ttg aaa ttg att tcg tca atg tac tta aca tga tta aca tcg aaa cac tta tta agt tc- ttt ttc caa ttc tac gat ttt att tct ttc gcg agt caa ttc cct cct gta gta gct atc ggt ttt gtc aga ttc acg ctc tct acg tag act ttc ttg caa gtt act aat ttg ttc cct gac acg tcc gag ttc agt ttt ata tgc cga ata gag ttc tga ttt atc ctt tga gca gat ctc tag cga ttg ttt aag atc cct aat tct agt ctt tag cct att tac ctc ctc aga aga tgt tcc gtt acc gtt gcg ttt aca ctc gtt aag ctg tct atc aag atc cat gat tct atc tct --- --- --- --- --- --- --- --- --- --- --- --- --- --- --- --- -

>DQ441416.1:127953-128353_Variola_virus_strain_Benin_Dahomey_1968_(v68-59)_complete_genome

gct gtt agc tgt cgt tca aga tta gaa ata cgg ttt ttc ttg gaa ttg att tcg tca atg tac tta aca tga tta aca tcg aaa cac tta tta agt tc- ttt ttc caa ttc tac gat ttt att tct ttc gcg agt caa ttc cct cct gta gta gct atc ggt ttt gtc aga ttc acg ctc tct acg tag act ttc ttg caa gtt act aat ttg ttc cct gac acg tcc gag ttc agt ttt ata tgc cga ata gag ttc tga ttt atc ctt tga gca gat ctc tag cga ttg ttt aag atc cct aat tct agt ctt tag cct att tac ctc ctc aga aga tgt tcc gtt acc gtt gcg ttt aca ctc gtt aag ctg tct atc aag atc cat gat tct atc tct --- --- --- --- --- --- --- --- --- --- --- --- --- --- --- --- -

>Y16780.1:127584-127984_variola_minor_virus_complete_genome

gct gtt agc tgt cgt tca aga tca gaa ata cgg ttt ttc ttg aaa ttg att tcg tca atg tac tta aca tga tta aca tcg aaa cac tta tta agt tc- ttt ttc caa ttc tac gat ttt att tct ttc gcg agt caa ttc cct cct gta gta gct atc ggt ttt gtc aga ttc acg ctc tct acg tag act ttc ttg caa gtt act aat ttg ttc cct gac acg tcc gag ttc agt ttt ata tgc cga ata gag ttc tga ttt atc ctt tga gca gat ctc tag cga ttg ttt aag atc cct aat tct agt ctt tag cct att tac ctc ctc aga aga tgt tcc gtt acc gtt gcg ttt aca ctc gtt aag ctg tct atc aag atc cat gat tct atc tct --- --- --- --- --- --- --- --- --- --- --- --- --- --- --- --- -

>X76266.1:304-704_Variola_virus_(Garcia-1966)_A27LA28LA29LA30LA31LA32LA33LA34RA35LA36RA37RA38RA39RA40R_genes

gct gtt agc tgt cgt tca aga tca gaa ata cgg ttt ttc ttg aaa ttg att tcg tca atg tac tta aca tga tta aca tcg aaa cac tta tta agt tc- ttt ttc caa ttc tac gat ttt att tct ttc gcg agt caa ttc cct cct gta gta gct atc ggt ttt gtc aga ttc acg ctc tct acg tag act ttc ttg caa gtt act aat ttg ttc cct gac acg tcc gag ttc agt ttt ata tgc cga ata gag ttc tga ttt atc ctt tga gca gat ctc tag cga ttg ttt aag atc cct aat tct agt ctt tag cct att tac ctc ctc aga aga tgt tcc gtt acc gtt gcg ttt aca ctc gtt aag ctg tct atc aag atc cat gat tct atc tct --- --- --- --- --- --- --- --- --- --- --- --- --- --- --- --- -

>KX061501.1:137640-138031_Vaccinia_virus_strain_Lister_clone_Lister_Butantan

gct gct agt tgt cgt tcg aga tca gaa ata cgg gtt ttc ttg gaa ttg att tcg tcg atg tac ttg gca --- --- --- tcg aaa cac tta tta agt tc- ttt ttc caa ttc tac gat ttt att tct ttc gcg agt caa ttc cct cct gta gta act atc tgt ttt gtc aga ttc acg ctt tct acg tag act ttc ttg caa gtt act aat ttg ttc cct agc acg tcc gag ttt agt ttt ata tgc tga ata gag ttc tga ttc atc ctt tga gca gat ctc tag cga tcg ttt aag att cct gat tct agt ctt tag cct att tac ctc ctc aga aga tgt tcc gtt acc gtt gcg ttt aca ctc gtt aag ctg tct atc aag atc cat gat tct atc tct --- --- --- --- --- --- --- --- --- --- --- --- --- --- --- --- -

>KF866253.1:137617-138008_Vaccinia_virus_WAU86/88-1_complete_genome

gct gct agt tgt cgt tcg aga tca gaa ata cgg gtt ttc ttg gaa ttg att tcg tcg atg tac ttg gca --- --- --- tcg aaa cac tta tta agt tc- ttt ttc caa ttc tac gat ttt att tct ttc gcg agt caa ttc cct cct gta gta act atc tgt ttt gtc aaa ttc acg ctc tct acg tag act ttc ttg caa gtt act aat ttg ttc cct agc acg tcc gag ttt agt ttt ata tgc tga ata gag ttc tga ttc atc ctt tga gca gat ctc tag cga tcg ttt aag att cct gat tct agt ctt tag cct att tac ctc ctc aga aga tgt tcc gtt acc gtt gcg ttt aca ctc gtt aag ctg tct atc aag atc cat gat tct atc tct --- --- --- --- --- --- --- --- --- --- --- --- --- --- --- --- -

>EU410304.1:145550-145941_Synthetic_Vaccinia_virus_clone_GLV-1h68

gct gct agt tgt cgt tcg aga tca gaa ata cgg gtt ttc ttg gaa ttg att tcg tcg atg tac ttg gca --- --- --- tcg aaa cac tta tta agt tc- ttt ttc caa ttc tac gat ttt att tct ttc gcg agt caa ttc cct cct gta gta act atc tgt ttt gtc aga ttc acg ctc tct acg tag act ttc ttg caa gtt act aat ttg ttc cct agc acg tcc gag ttt agt ttt ata tgc tga ata gag ttc tga ttc atc ctt tga gca gat ctc tag cga tcg ttt aag att cct aat tct agt ctt tag cct att tac ctc ctc aga aga tgt tcc gtt acc gtt gcg ttt aca ctc gtt aag ctg tct atc aag atc cat gat tct atc tct --- --- --- --- --- --- --- --- --- --- --- --- --- --- --- --- -

>LT706528.1:126606-127006_Variola_virus_isolate_VARV_V563_genome_assembly_chromosome:_I

gct gtt agc tgt cgt tca aga tca gaa ata cgg ttt ttc ttg gaa ttg att tcg tca atg tac tta aca tga tta aca tcg aaa cac tta tta agt tc- ttt ttc caa ttc tac gat ttt att tct ttc gcg agt caa ttc cct cct gta gta gct atc ggt ttt gtc aga ttc acg ctc tct acg tag act ttc ttg caa gtt act aat ttg ttc cct gac acg tcc aag ttc agt ttt ata tgc cga ata gag ttc tga ttt atc ctt tga gca gat ctc tag cga ttg ttt aag atc act aat tct agt ctt tag cct att tac ctc ctc aga aga tgt tcc gtt acc gtt gcg ttt aca ctc gtt aag ctg tct atc aag atc cat gat tct atc tct --- --- --- --- --- --- --- --- --- --- --- --- --- --- --- --- -

>DQ437592.1:126961-127361_Variola_virus_strain_Syria_1972_V72-199_complete_genome

gct gtt agc tgt cgt tca aga tca gaa ata cgg ttt ttc ttg gaa ttg att tcg tca atg tac tta aca tga tta aca tcg aaa cac tta tta agt tc- ttt ttc caa ttc tac gat ttt att tct ttc gcg agt caa ttc cct cct gta gta gct atc ggt ttt gtc aga ttc acg ctc tct acg tag act ttc ttg caa gtt act aat ttg ttc cct gac acg tcc aag ttc agt ttt ata tgc cga ata gag ttc tga ttt atc ctt tga gca gat ctc tag cga ttg ttt aag atc act aat tct agt ctt tag cct att tac ctc ctc aga aga tgt tcc gtt acc gtt gcg ttt aca ctc gtt aag ctg tct atc aag atc cat gat tct atc tct --- --- --- --- --- --- --- --- --- --- --- --- --- --- --- --- -

>DQ437589.1:126977-127377_Variola_virus_strain_Pakistan_1969_(Rafig_Lahore)_complete_sequence

gct gtt agc tgt cgt tca aga tca gaa ata cgg ttt ttc ttg gaa ttg att tcg tca atg tac tta aca tga tta aca tcg aaa cac tta tta agt tc- ttt ttc caa ttc tac gat ttt att tct ttc gcg agt caa ttc cct cct gta gta gct atc ggt ttt gtc aga ttc acg ctc tct acg tag act ttc ttg caa gtt act aat ttg ttc cct gac acg tcc aag ttc agt ttt ata tgc cga ata gag ttc tga ttt atc ctt tga gca gat ctc tag cga ttg ttt aag atc act aat tct agt ctt tag cct att tac ctc ctc aga aga tgt tcc gtt acc gtt gcg ttt aca ctc gtt aag ctg tct atc aag atc cat gat tct atc tct --- --- --- --- --- --- --- --- --- --- --- --- --- --- --- --- -

>DQ437587.1:126964-127364_Variola_virus_strain_Iran_1972_2602_Tabriz_complete_genome

gct gtt agc tgt cgt tca aga tca gaa ata cgg ttt ttc ttg gaa ttg att tcg tca atg tac tta aca tga tta aca tcg aaa cac tta tta agt tc- ttt ttc caa ttc tac gat ttt att tct ttc gcg agt caa ttc cct cct gta gta gct atc ggt ttt gtc aga ttc acg ctc tct acg tag act ttc ttg caa gtt act aat ttg ttc cct gac acg tcc aag ttc agt ttt ata tgc cga ata gag ttc tga ttt atc ctt tga gca gat ctc tag cga ttg ttt aag atc act aat tct agt ctt tag cct att tac ctc ctc aga aga tgt tcc gtt acc gtt gcg ttt aca ctc gtt aag ctg tct atc aag atc cat gat tct atc tct --- --- --- --- --- --- --- --- --- --- --- --- --- --- --- --- -

>DQ437586.1:127040-127440_Variola_virus_strain_India_1964_7125_Vellore_complete_genome

gct gtt agc tgt cgt tca aga tca gaa ata cgg ttt ttc ttg gaa ttg att tcg tca atg tac tta aca tga tta aca tcg aaa cac tta tta agt tc- ttt ttc caa ttc tac gat ttt att tct ttc gcg agt caa ttc cct cct gta gta gct atc ggt ttt gtc aga ttc acg ctc tct acg tag act ttc ttg caa gtt act aat ttg ttc cct gac acg tcc aag ttc agt ttt ata tgc cga ata gag ttc tga ttt atc ctt tga gca gat ctc tag cga ttg ttt aag atc act aat tct agt ctt tag cct att tac ctc ctc aga aga tgt tcc gtt acc gtt gcg ttt aca ctc gtt aag ctg tct atc aag atc cat gat tct atc tct --- --- --- --- --- --- --- --- --- --- --- --- --- --- --- --- -

>DQ437585.1:127179-127579_Variola_virus_strain_India_1964_7124_Vellore_complete_genome

gct gtt agc tgt cgt tca aga tca gaa ata cgg ttt ttc ttg gaa ttg att tcg tca atg tac tta aca tga tta aca tcg aaa cac tta tta agt tc- ttt ttc caa ttc tac gat ttt att tct ttc gcg agt caa ttc cct cct gta gta gct atc ggt ttt gtc aga ttc acg ctc tct acg tag act ttc ttg caa gtt act aat ttg ttc cct gac acg tcc aag ttc agt ttt ata tgc cga ata gag ttc tga ttt atc ctt tga gca gat ctc tag cga ttg ttt aag atc act aat tct agt ctt tag cct att tac ctc ctc aga aga tgt tcc gtt acc gtt gcg ttt aca ctc gtt aag ctg tct atc aag atc cat gat tct atc tct --- --- --- --- --- --- --- --- --- --- --- --- --- --- --- --- -

>DQ437582.1:127278-127678_Variola_virus_strain_China_Horn_1948_complete_genome

gct gtt agc tgt cgt tca aga tca gaa ata cgg ttt ttc ttg gaa ttg att tcg tca atg tac tta aca tga tta aca tcg aaa cac tta tta agt tc- ttt ttc caa ttc tac gat ttt att tct ttc gcg agt caa ttc cct cct gta gta gct atc ggt ttt gtc aga ttc acg ctc tct acg tag act ttc ttg caa gtt act aat ttg ttc cct gac acg tcc aag ttc agt ttt ata tgc cga ata gag ttc tga ttt atc ctt tga gca gat ctc tag cga ttg ttt aag atc act aat tct agt ctt tag cct att tac ctc ctc aga aga tgt tcc gtt acc gtt gcg ttt aca ctc gtt aag ctg tct atc aag atc cat gat tct atc tct --- --- --- --- --- --- --- --- --- --- --- --- --- --- --- --- -

>DQ437580.1:126971-127371_Variola_virus_strain_Afghanistan_1970_Variolator_4_complete_genome

gct gtt agc tgt cgt tca aga tca gaa ata cgg ttt ttc ttg gaa ttg att tcg tca atg tac tta aca tga tta aca tcg aaa cac tta tta agt tc- ttt ttc caa ttc tac gat ttt att tct ttc gcg agt caa ttc cct cct gta gta gct atc ggt ttt gtc aga ttc acg ctc tct acg tag act ttc ttg caa gtt act aat ttg ttc cct gac acg tcc aag ttc agt ttt ata tgc cga ata gag ttc tga ttt atc ctt tga gca gat ctc tag cga ttg ttt aag atc act aat tct agt ctt tag cct att tac ctc ctc aga aga tgt tcc gtt acc gtt gcg ttt aca ctc gtt aag ctg tct atc aag atc cat gat tct atc tct --- --- --- --- --- --- --- --- --- --- --- --- --- --- --- --- -

>DQ441448.1:126961-127361_Variola_virus_strain_Yugoslavia_1972_V72-164_complete_genome

gct gtt agc tgt cgt tca aga tca gaa ata cgg ttt ttc ttg gaa ttg att tcg tca atg tac tta aca tga tta aca tcg aaa cac tta tta agt tc- ttt ttc caa ttc tac gat ttt att tct ttc gcg agt caa ttc cct cct gta gta gct atc ggt ttt gtc aga ttc acg ctc tct acg tag act ttc ttg caa gtt act aat ttg ttc cct gac acg tcc aag ttc agt ttt ata tgc cga ata gag ttc tga ttt atc ctt tga gca gat ctc tag cga ttg ttt aag atc act aat tct agt ctt tag cct att tac ctc ctc aga aga tgt tcc gtt acc gtt gcg ttt aca ctc gtt aag ctg tct atc aag atc cat gat tct atc tct --- --- --- --- --- --- --- --- --- --- --- --- --- --- --- --- -

>DQ441445.1:127026-127426_Variola_virus_strain_United_Kingdom_1946_Hinden_(Middlesex)_complete_genome

gct gtt agc tgt cgt tca aga tca gaa ata cgg ttt ttc ttg gaa ttg att tcg tca atg tac tta aca tga tta aca tcg aaa cac tta tta agt tc- ttt ttc caa ttc tac gat ttt att tct ttc gcg agt caa ttc cct cct gta gta gct atc ggt ttt gtc aga ttc acg ctc tct acg tag act ttc ttg caa gtt act aat ttg ttc cct gac acg tcc aag ttc agt ttt ata tgc cga ata gag ttc tga ttt atc ctt tga gca gat ctc tag cga ttg ttt aag atc act aat tct agt ctt tag cct att tac ctc ctc aga aga tgt tcc gtt acc gtt gcg ttt aca ctc gtt aag ctg tct atc aag atc cat gat tct atc tct --- --- --- --- --- --- --- --- --- --- --- --- --- --- --- --- -

>DQ441444.1:127007-127407_Variola_virus_strain_United_Kingdom_1946_Harvey_complete_genome

gct gtt agc tgt cgt tca aga tca gaa ata cgg ttt ttc ttg gaa ttg att tcg tca atg tac tta aca tga tta aca tcg aaa cac tta tta agt tc- ttt ttc caa ttc tac gat ttt att tct ttc gcg agt caa ttc cct cct gta gta gct atc ggt ttt gtc aga ttc acg ctc tct acg tag act ttc ttg caa gtt act aat ttg ttc cct gac acg tcc aag ttc agt ttt ata tgc cga ata gag ttc tga ttt atc ctt tga gca gat ctc tag cga ttg ttt aag atc act aat tct agt ctt tag cct att tac ctc ctc aga aga tgt tcc gtt acc gtt gcg ttt aca ctc gtt aag ctg tct atc aag atc cat gat tct atc tct --- --- --- --- --- --- --- --- --- --- --- --- --- --- --- --- -

>DQ441433.1:126972-127372_Variola_virus_strain_Kuwait_1967_(K1629)_complete_genome

gct gtt agc tgt cgt tca aga tca gaa ata cgg ttt ttc ttg gaa ttg att tcg tca atg tac tta aca tga tta aca tcg aaa cac tta tta agt tc- ttt ttc caa ttc tac gat ttt att tct ttc gcg agt caa ttc cct cct gta gta gct atc ggt ttt gtc aga ttc acg ctc tct acg tag act ttc ttg caa gtt act aat ttg ttc cct gac acg tcc aag ttc agt ttt ata tgc cga ata gag ttc tga ttt atc ctt tga gca gat ctc tag cga ttg ttt aag atc act aat tct agt ctt tag cct att tac ctc ctc aga aga tgt tcc gtt acc gtt gcg ttt aca ctc gtt aag ctg tct atc aag atc cat gat tct atc tct --- --- --- --- --- --- --- --- --- --- --- --- --- --- --- --- -

>DQ441432.1:127159-127559_Variola_virus_strain_Korea_1947_(Lee_Masterseed)_complete_genome

gct gtt agc tgt cgt tca aga tca gaa ata cgg ttt ttc ttg gaa ttg att tcg tca atg tac tta aca tga tta aca tcg aaa cac tta tta agt tc- ttt ttc caa ttc tac gat ttt att tct ttc gcg agt caa ttc cct cct gta gta gct atc ggt ttt gtc aga ttc acg ctc tct acg tag act ttc ttg caa gtt act aat ttg ttc cct gac acg tcc aag ttc agt ttt ata tgc cga ata gag ttc tga ttt atc ctt tga gca gat ctc tag cga ttg ttt aag atc act aat tct agt ctt tag cct att tac ctc ctc aga aga tgt tcc gtt acc gtt gcg ttt aca ctc gtt aag ctg tct atc aag atc cat gat tct atc tct --- --- --- --- --- --- --- --- --- --- --- --- --- --- --- --- -

>DQ441430.1:127042-127442_Variola_virus_strain_Japan_1951_(Harper_Masterseed)_complete_genome

gct gtt agc tgt cgt tca aga tca gaa ata cgg ttt ttc ttg gaa ttg att tcg tca atg tac tta aca tga tta aca tcg aaa cac tta tta agt tc- ttt ttc caa ttc tac gat ttt att tct ttc gcg agt caa ttc cct cct gta gta gct atc ggt ttt gtc aga ttc acg ctc tct acg tag act ttc ttg caa gtt act aat ttg ttc cct gac acg tcc aag ttc agt ttt ata tgc cga ata gag ttc tga ttt atc ctt tga gca gat ctc tag cga ttg ttt aag atc act aat tct agt ctt tag cct att tac ctc ctc aga aga tgt tcc gtt acc gtt gcg ttt aca ctc gtt aag ctg tct atc aag atc cat gat tct atc tct --- --- --- --- --- --- --- --- --- --- --- --- --- --- --- --- -

>DQ516056.1:654-1045_Vaccinia_virus_strain_SPAn232_nonfunctional_ATI_gene_partial_sequence

gct gct agt tgt cgt tcg aga tca gaa ata cgg gtt ttc ttg gaa ttg att tcg tcg atg tac ttg gca --- --- --- tcg aaa cac tta tta agt tc- ttt ttc caa ttc tac gat ttt att tct ttc gcg agt caa ttc cct cct gta gta act atc tgt ttt gtc aga ttc acg ctc tct acg tag act ttc ttg caa gtt act aat ctg ttc cct agc acg tcc gag ttt agt ttt ata tgc tga ata gag ttc tga ttc atc ctt tga gca gat ctc tag cga tcg ttt aag gtt cct gat tct agt ctt tag cct att tac ctc ctc aga aga tgt tcc gtt acc gtt gcg ttt aca ctc gtt aag ctg tct atc aag atc cat gat tct atc tct --- --- --- --- --- --- --- --- --- --- --- --- --- --- --- --- -

>DQ437590.1:127248-127648_Variola_virus_strain_Somalia_1977_complete_genome

gct gtt agc tgt cgt tca aga tca gaa ata cgg ttt ttc ttg gaa ttg att tcg tca atg tac tta aca tga tta aca tcg aaa cac tta tta agt tc- ttt ttc caa ttc tac gat ttt att tct ttc gcg agt caa ttc cct cct gta gta gct atc ggt ttt gtc aga ttc acg ctc tct acg tag act ttc ttg caa gtt act aat ttg ttc cct gac acg tcc aag ttc agt ttt ata tgc cga ata gag ttc tga ttt atc ctt taa gca gat ctc tag cga ttg ttt aag atc act aat tct agt ctt tag cct att tac ctc ctc aga aga tgt tcc gtt acc gtt gcg ttt aca ctc gtt aag ctg tct atc aag atc cat gat tct atc tct --- --- --- --- --- --- --- --- --- --- --- --- --- --- --- --- -

>DQ437588.1:126953-127353_Variola_virus_strain_Nepal_1973_complete_genome

gct gtt agc tgt cgt tca aga tca gaa ata cgg ttt ttc ttg gaa ttg att tcg tca atg tac tta aca tga tta aca tcg aaa cac tta tta agt tc- ttt ttc caa ttc tac gat ttt att tct ttc gcg agt caa ttc cct cct gta gta gct atc ggt ttt gtc aga ttc acg ctc tct acg tag act ttc ttg caa gtt act aat ttg ttc cct gac acg tcc aag ttc agt ttt ata tgc cga ata gag ttc tga ttt atc ctt taa gca gat ctc tag cga ttg ttt aag atc act aat tct agt ctt tag cct att tac ctc ctc aga aga tgt tcc gtt acc gtt gcg ttt aca ctc gtt aag ctg tct atc aag atc cat gat tct atc tct --- --- --- --- --- --- --- --- --- --- --- --- --- --- --- --- -

>DQ437583.1:127349-127749_Variola_virus_strain_Congo_1970_complete_genome

gct gtt agc tgt cgt tca aga tca gaa ata cgg ttt ttc ttg gaa ttg att tcg tca atg tac tta aca tga tta aca tcg aaa cac tta tta agt tc- ttt ttc caa ttc tac gat ttt att tct ttc gcg agt caa ttc cct cct gta gta gct atc ggt ttt gtc aga ttc acg ctc tct acg tag act ttc ttg caa gtt act aat ttg ttc cct gac acg tcc aag ttc agt ttt ata tgc cga ata gag ttc tga ttt atc ctt taa gca gat ctc tag cga ttg ttt aag atc act aat tct agt ctt tag cct att tac ctc ctc aga aga tgt tcc gtt acc gtt gcg ttt aca ctc gtt aag ctg tct atc aag atc cat gat tct atc tct --- --- --- --- --- --- --- --- --- --- --- --- --- --- --- --- -

>DQ441443.1:127118-127518_Variola_virus_strain_Tanzania_1965_kembula_complete_genome

gct gtt agc tgt cgt tca aga tca gaa ata cgg ttt ttc ttg gaa ttg att tcg tca atg tac tta aca tga tta aca tcg aaa cac tta tta agt tc- ttt ttc caa ttc tac gat ttt att tct ttc gcg agt caa ttc cct cct gta gta gct atc ggt ttt gtc aga ttc acg ctc tct acg tag act ttc ttg caa gtt act aat ttg ttc cct gac acg tcc aag ttc agt ttt ata tgc cga ata gag ttc tga ttt atc ctt taa gca gat ctc tag cga ttg ttt aag atc act aat tct agt ctt tag cct att tac ctc ctc aga aga tgt tcc gtt acc gtt gcg ttt aca ctc gtt aag ctg tct atc aag atc cat gat tct atc tct --- --- --- --- --- --- --- --- --- --- --- --- --- --- --- --- -

>DQ441441.1:127341-127741_Variola_virus_strain_Sudan_1947_(Rumbec)_complete_genome

gct gtt agc tgt cgt tca aga tca gaa ata cgg ttt ttc ttg gaa ttg att tcg tca atg tac tta aca tga tta aca tcg aaa cac tta tta agt tc- ttt ttc caa ttc tac gat ttt att tct ttc gcg agt caa ttc cct cct gta gta gct atc ggt ttt gtc aga ttc acg ctc tct acg tag act ttc ttg caa gtt act aat ttg ttc cct gac acg tcc aag ttc agt ttt ata tgc cga ata gag ttc tga ttt atc ctt taa gca gat ctc tag cga ttg ttt aag atc act aat tct agt ctt tag cct att tac ctc ctc aga aga tgt tcc gtt acc gtt gcg ttt aca ctc gtt aag ctg tct atc aag atc cat gat tct atc tct --- --- --- --- --- --- --- --- --- --- --- --- --- --- --- --- -

>DQ441440.1:127280-127680_Variola_virus_strain_Sudan_1947_(Juba)_complete_genome

gct gtt agc tgt cgt tca aga tca gaa ata cgg ttt ttc ttg gaa ttg att tcg tca atg tac tta aca tga tta aca tcg aaa cac tta tta agt tc- ttt ttc caa ttc tac gat ttt att tct ttc gcg agt caa ttc cct cct gta gta gct atc ggt ttt gtc aga ttc acg ctc tct acg tag act ttc ttg caa gtt act aat ttg ttc cct gac acg tcc aag ttc agt ttt ata tgc cga ata gag ttc tga ttt atc ctt taa gca gat ctc tag cga ttg ttt aag atc act aat tct agt ctt tag cct att tac ctc ctc aga aga tgt tcc gtt acc gtt gcg ttt aca ctc gtt aag ctg tct atc aag atc cat gat tct atc tct --- --- --- --- --- --- --- --- --- --- --- --- --- --- --- --- -

>DQ441439.1:126253-126653_Variola_virus_strain_Somalia_1977_(V77-1605)_complete_genome

gct gtt agc tgt cgt tca aga tca gaa ata cgg ttt ttc ttg gaa ttg att tcg tca atg tac tta aca tga tta aca tcg aaa cac tta tta agt tc- ttt ttc caa ttc tac gat ttt att tct ttc gcg agt caa ttc cct cct gta gta gct atc ggt ttt gtc aga ttc acg ctc tct acg tag act ttc ttg caa gtt act aat ttg ttc cct gac acg tcc aag ttc agt ttt ata tgc cga ata gag ttc tga ttt atc ctt taa gca gat ctc tag cga ttg ttt aag atc act aat tct agt ctt tag cct att tac ctc ctc aga aga tgt tcc gtt acc gtt gcg ttt aca ctc gtt aag ctg tct atc aag atc cat gat tct atc tct --- --- --- --- --- --- --- --- --- --- --- --- --- --- --- --- -

>DQ441438.1:126262-126662_Variola_virus_strain_Somalia_1977_(V77-1252)_complete_genome

gct gtt agc tgt cgt tca aga tca gaa ata cgg ttt ttc ttg gaa ttg att tcg tca atg tac tta aca tga tta aca tcg aaa cac tta tta agt tc- ttt ttc caa ttc tac gat ttt att tct ttc gcg agt caa ttc cct cct gta gta gct atc ggt ttt gtc aga ttc acg ctc tct acg tag act ttc ttg caa gtt act aat ttg ttc cct gac acg tcc aag ttc agt ttt ata tgc cga ata gag ttc tga ttt atc ctt taa gca gat ctc tag cga ttg ttt aag atc act aat tct agt ctt tag cct att tac ctc ctc aga aga tgt tcc gtt acc gtt gcg ttt aca ctc gtt aag ctg tct atc aag atc cat gat tct atc tct --- --- --- --- --- --- --- --- --- --- --- --- --- --- --- --- -

>DQ441436.1:127078-127478_Variola_virus_strain_South_Africa_1965_(103_Tvaal_Nelspruit)_complete_genome

gct gtt agc tgt cgt tca aga tca gaa ata cgg ttt ttc ttg gaa ttg att tcg tca atg tac tta aca tga tta aca tcg aaa cac tta tta agt tc- ttt ttc caa ttc tac gat ttt att tct ttc gcg agt caa ttc cct cct gta gta gct atc ggt ttt gtc aga ttc acg ctc tct acg tag act ttc ttg caa gtt act aat ttg ttc cct gac acg tcc aag ttc agt ttt ata tgc cga ata gag ttc tga ttt atc ctt taa gca gat ctc tag cga ttg ttt aag atc act aat tct agt ctt tag cct att tac ctc ctc aga aga tgt tcc gtt acc gtt gcg ttt aca ctc gtt aag ctg tct atc aag atc cat gat tct atc tct --- --- --- --- --- --- --- --- --- --- --- --- --- --- --- --- -

>DQ441435.1:127245-127645_Variola_virus_strain_South_Africa_1965_(102_Natal_Ingwavuma)_complete_genome

gct gtt agc tgt cgt tca aga tca gaa ata cgg ttt ttc ttg gaa ttg att tcg tca atg tac tta aca tga tta aca tcg aaa cac tta tta agt tc- ttt ttc caa ttc tac gat ttt att tct ttc gcg agt caa ttc cct cct gta gta gct atc ggt ttt gtc aga ttc acg ctc tct acg tag act ttc ttg caa gtt act aat ttg ttc cct gac acg tcc aag ttc agt ttt ata tgc cga ata gag ttc tga ttt atc ctt taa gca gat ctc tag cga ttg ttt aag atc act aat tct agt ctt tag cct att tac ctc ctc aga aga tgt tcc gtt acc gtt gcg ttt aca ctc gtt aag ctg tct atc aag atc cat gat tct atc tct --- --- --- --- --- --- --- --- --- --- --- --- --- --- --- --- -

>DQ441431.1:127050-127450_Variola_virus_strain_Japan_1951_(Stillwell_Masterseed)_complete_genome

gct gtt agc tgt cgt tca aga tca gaa ata cgg ttt ttc ttg gaa ttg att tcg tca atg tac tta aca tga tta aca tcg aaa cac tta tta agt tc- ttt ttc caa ttc tac gat ttt att tct ttc gcg agt caa ttc cct cct gta gta gct atc ggt ttt gtc aga ttc acg ctc tct acg tag act ttc ttg caa gtt act aat ttg ttc cct gac acg tcc aag ttc agt ttt ata tgc cga ata gag ttc tga ttt atc ctt tga gca gat ctc tag cga ttg ttt aag atc act aat tct agt ctt tag cct att tac ctc ctc aga aga tgt tcc gtt acc att gcg ttt aca ctc gtt aag ctg tct atc aag atc cat gat tct atc tct --- --- --- --- --- --- --- --- --- --- --- --- --- --- --- --- -

>DQ441429.1:127053-127453_Variola_virus_strain_Japan_1946_(Yamada_MS-2(A)_Tokyo)_complete_genome

gct gtt agc tgt cgt tca aga tca gaa ata cgg ttt ttc ttg gaa ttg att tcg tca atg tac tta aca tga tta aca tcg aaa cac tta tta agt tc- ttt ttc caa ttt tac gat ttt att tct ttc gcg agt caa ttc cct cct gta gta gct atc ggt ttt gtc aga ttc acg ctc tct acg tag act ttc ttg caa gtt act aat ttg ttc cct gac acg tcc aag ttc agt ttt ata tgc cga ata gag ttc tga ttt atc ctt tga gca gat ctc tag cga ttg ttt aag atc act aat tct agt ctt tag cct att tac ctc ctc aga aga tgt tcc gtt acc gtt gcg ttt aca ctc gtt aag ctg tct atc aag atc cat gat tct atc tct --- --- --- --- --- --- --- --- --- --- --- --- --- --- --- --- -

>DQ441428.1:127053-127453_Variola_virus_strain_India_1953_(New_Delhi)_complete_genome

gct gtt agc tgt cgt tca aga tca gaa ata cgg ttt ttc ttg gaa ttg att tcg tca atg tac tta aca tga tta aca tcg aaa cac tta tta agt tc- ttt ttc caa ttt tac gat ttt att tct ttc gcg agt caa ttc cct cct gta gta gct atc ggt ttt gtc aga ttc acg ctc tct acg tag act ttc ttg caa gtt act aat ttg ttc cct gac acg tcc aag ttc agt ttt ata tgc cga ata gag ttc tga ttt atc ctt tga gca gat ctc tag cga ttg ttt aag atc act aat tct agt ctt tag cct att tac ctc ctc aga aga tgt tcc gtt acc gtt gcg ttt aca ctc gtt aag ctg tct atc aag atc cat gat tct atc tct --- --- --- --- --- --- --- --- --- --- --- --- --- --- --- --- -

>DQ441427.1:127052-127452_Variola_virus_strain_India_1953_(Kali-Muthu-M50_Madras)_complete_genome

gct gtt agc tgt cgt tca aga tca gaa ata cgg ttt ttc ttg gaa ttg att tcg tca atg tac tta aca tga tta aca tcg aaa cac tta tta agt tc- ttt ttc caa ttc tac gat ttt att tct ttc gcg agt caa ttc cct cct gta gta gct atc ggt ttt gtc aga ttc acg ctc tct acg tag act ttc ttg caa gtt act aat ttg ttc cct gac acg tcc aag ttc agt ttt ata tgc cga ata gag ttc tga ttt atc ctt taa gca gat ctc tag cga ttg ttt aag atc act aat tct agt ctt tag cct att tac ctc ctc aga aga tgt tcc gtt acc gtt gcg ttt aca ctc gtt aag ctg tct atc aag atc cat gat tct atc tct --- --- --- --- --- --- --- --- --- --- --- --- --- --- --- --- -

>DQ441425.1:127443-127843_Variola_virus_strain_Ethiopia_1972_(Eth17_R14-1X-72_Addis)_complete_genome

gct gtt agc tgt cgt tca aga tca gaa ata cgg ttt ttc ttg gaa ttg att tcg tca atg tac tta aca tga tta aca tcg aaa cac tta tta agt tc- ttt ttc caa ttc tac gat ttt att tct ttc gcg agt caa ttc cct cct gta gta gct atc ggt ttt gtc aga ttc acg ctc tct acg tag act ttc ttg caa gtt act aat ttg ttc cct gac acg tcc aag ttc agt ttt ata tgc cga ata gag ttc tga ttt atc ctt taa gca gat ctc tag cga ttg ttt aag atc act aat tct agt ctt tag cct att tac ctc ctc aga aga tgt tcc gtt acc gtt gcg ttt aca ctc gtt aag ctg tct atc aag atc cat gat tct atc tct --- --- --- --- --- --- --- --- --- --- --- --- --- --- --- --- -

>DQ441424.1:127443-127843_Variola_virus_strain_Ethiopia_1972_(Eth16_R14-1X-72_Addis)_complete_genome

gct gtt agc tgt cgt tca aga tca gaa ata cgg ttt ttc ttg gaa ttg att tcg tca atg tac tta aca tga tta aca tcg aaa cac tta tta agt tc- ttt ttc caa ttc tac gat ttt att tct ttc gcg agt caa ttc cct cct gta gta gct atc ggt ttt gtc aga ttc acg ctc tct acg tag act ttc ttg caa gtt act aat ttg ttc cct gac acg tcc aag ttc agt ttt ata tgc cga ata gag ttc tga ttt atc ctt taa gca gat ctc tag cga ttg ttt aag atc act aat tct agt ctt tag cct att tac ctc ctc aga aga tgt tcc gtt acc gtt gcg ttt aca ctc gtt aag ctg tct atc aag atc cat gat tct atc tct --- --- --- --- --- --- --- --- --- --- --- --- --- --- --- --- -

>DQ441423.1:127178-127578_Variola_virus_strain_Congo_9_1970_(v74-227_Gispen)_complete_genome

gct gtt agc tgt cgt tca aga tca gaa ata cgg ttt ttc ttg gaa ttg att tcg tca atg tac tta aca tga tta aca tcg aaa cac tta tta agt tc- ttt ttc caa ttc tac gat ttt att tct ttc gcg agt caa ttc cct cct gta gta gct atc ggt ttt gtc aga ttc acg ctc tct acg tag act ttc ttg caa gtt act aat ttg ttc cct gac acg tcc aag ttc agt ttt ata tgc cga ata gag ttc tga ttt atc ctt taa gca gat ctc tag cga ttg ttt aag atc act aat tct agt ctt tag cct att tac ctc ctc aga aga tgt tcc gtt acc gtt gcg ttt aca ctc gtt aag ctg tct atc aag atc cat gat tct atc tct --- --- --- --- --- --- --- --- --- --- --- --- --- --- --- --- -

>DQ441418.1:127071-127471_Variola_virus_strain_Botswana_1973_(v73-225)_complete_genome

gct gtt agc tgt cgt tca aga tca gaa ata cgg ttt ttc ttg gaa ttg att tcg tca atg tac tta aca tga tta aca tcg aaa cac tta tta agt tc- ttt ttc caa ttc tac gat ttt att tct ttc gcg agt caa ttc cct cct gta gta gct atc ggt ttt gtc aga ttc acg ctc tct acg tag act ttc ttg caa gtt act aat ttg ttc cct gac acg tcc aag ttc agt ttt ata tgc cga ata gag ttc tga ttt atc ctt taa gca gat ctc tag cga ttg ttt aag atc act aat tct agt ctt tag cct att tac ctc ctc aga aga tgt tcc gtt acc gtt gcg ttt aca ctc gtt aag ctg tct atc aag atc cat gat tct atc tct --- --- --- --- --- --- --- --- --- --- --- --- --- --- --- --- -

>DQ441417.1:127081-127481_Variola_virus_strain_Botswana_1972_(v72-143)_complete_genome

gct gtt agc tgt cgt tca aga tca gaa ata cgg ttt ttc ttg gaa ttg att tcg tca atg tac tta aca tga tta aca tcg aaa cac tta tta agt tc- ttt ttc caa ttc tac gat ttt att tct ttc gcg agt caa ttc cct cct gta gta gct atc ggt ttt gtc aga ttc acg ctc tct acg tag act ttc ttg caa gtt act aat ttg ttc cct gac acg tcc aag ttc agt ttt ata tgc cga ata gag ttc tga ttt atc ctt taa gca gat ctc tag cga ttg ttt aag atc act aat tct agt ctt tag cct att tac ctc ctc aga aga tgt tcc gtt acc gtt gcg ttt aca ctc gtt aag ctg tct atc aag atc cat gat tct atc tct --- --- --- --- --- --- --- --- --- --- --- --- --- --- --- --- -

>X69198.1:126578-126978_Variola_virus_DNA_complete_genome

gct gtt agc tgt cgt tca aga tca gaa ata cgg ttt ttc ttg gaa ttg att tcg tca atg tac tta aca tga tta aca tcg aaa cac tta tta agt tc- ttt ttc caa ttc tac gat ttt att tct ttc gcg agt caa ttc cct cct gta gta gct atc ggt ttt gtc aga ttc acg ctc tct acg tag act ttc ttg caa gtt act aat ttg ttc cct gac acg tcc aag ttc agt ttt ata tgc cga ata gag ttc tga ttt atc ctt taa gca gat ctc tag cga ttg ttt aag atc act aat tct agt ctt tag cct att tac ctc ctc aga aga tgt tcc gtt acc gtt gcg ttt aca ctc gtt aag ctg tct atc aag atc cat gat tct atc tct --- --- --- --- --- --- --- --- --- --- --- --- --- --- --- --- -

>X67115.1:303-703_Variola_virus_(XhoI-_E_genome_fragment)_genes_for_fusion_protein_RNA_polymerase_subunit_RNA_binding_ribonucleoprotein_ATP/GTP_binding_protein

gct gtt agc tgt cgt tca aga tca gaa ata cgg ttt ttc ttg gaa ttg att tcg tca atg tac tta aca tga tta aca tcg aaa cac tta tta agt tc- ttt ttc caa ttc tac gat ttt att tct ttc gcg agt caa ttc cct cct gta gta gct atc ggt ttt gtc aga ttc acg ctc tct acg tag act ttc ttg caa gtt act aat ttg ttc cct gac acg tcc aag ttc agt ttt ata tgc cga ata gag ttc tga ttt atc ctt taa gca gat ctc tag cga ttg ttt aag atc act aat tct agt ctt tag cct att tac ctc ctc aga aga tgt tcc gtt acc gtt gcg ttt aca ctc gtt aag ctg tct atc aag atc cat gat tct atc tct --- --- --- --- --- --- --- --- --- --- --- --- --- --- --- --- -

>JQ361140.1:645-1036_Vaccinia_virus_isolate_SH2V_acidophilic_type_inclusion_gene_complete_cds_and_acidophilic_type_inclusion_gene_partial_cds

gct gct agt tgt cgt tcg aga tca gaa ata cgg gtt ttc ttg gaa ttg att tcg tcg atg tac ttg gca --- --- --- tcg aaa cac tta tta agt tc- ttt ttc caa ttc tac gat ttt att tct ttc gcg agt caa ttc cct cct gta gta act atc tgt ttt gtc aga tag acg cac tct acg tag act ttc ttg caa gtt act aat ttg ttc cct agc acg tcc gag ttt agt ttt ata tgc tga ata gag ttc tga ttc atc ctt tga gca gat ctc tag cga tcg ttt aag att cct gat tct agt ctt tag cct att tac ctc ctc aga aga tgt tcc gtt acc gtt gcg ttt aca ctc gtt aag ctg tct atc aag atc cat gat tct atc tct --- --- --- --- --- --- --- --- --- --- --- --- --- --- --- --- -

>DQ437591.1:127083-127483_Variola_virus_strain_Sumatra_1970_V70-222_complete_genome

gct gtt agc tgt cgt tca aga tca gaa ata cgg ttt ttc ttg gaa ttg att tcg tca atg tac tta aca tga tta aca tcg aaa cac tta tta agt tc- ttt ttc caa ttc tac gat ttt att tct ttc gcg agt caa ttc cct cct gta gta gct atc ggt ttt gtc aga ttc acg ctc tct acg tag act ttc ttg caa gtt act aat ttg ttc cct gac acg tcc aag ttc agt ttt ata tgc cga ata tag ttc tga ttt atc ctt taa gca gat ctc tag cga ttg ttt aag atc act aat tct agt ctt tag cct att tac ctc ctc aga aga tgt tcc gtt acc gtt gcg ttt aca ctc gtt aag ctg tct atc aag atc cat gat tct atc tct --- --- --- --- --- --- --- --- --- --- --- --- --- --- --- --- -

>DQ437584.1:127003-127403_Variola_virus_strain_Germany_1958_Heidelberg_complete_genome

gct gtt agc tgt cgt tca aga tca gaa ata cgg ttt ttc ttg gaa ttg att tcg tca atg tac tta aca tga tta aca tcg aaa cac tta tta agt tc- ttt ttc caa ttc tac gat ttt att tct ttc gcg agt caa ttc cct cct gta gta gct atc ggt ttt gtc aga ttt acg ctc tct acg tag act ttc ttg caa gtt act aat ttg ttc cct gac acg tcc aag ttc agt ttt ata tgc cga ata gag ttc tga ttt atc ctt taa gca gat ctc tag cga ttg ttt aag atc act aat tct agt ctt tag cct att tac ctc ctc aga aga tgt tcc gtt acc gtt gcg ttt aca ctc gtt aag ctg tct atc aag atc cat gat tct atc tct --- --- --- --- --- --- --- --- --- --- --- --- --- --- --- --- -

>DQ441446.1:127109-127509_Variola_virus_strain_United_Kingdom_1947_Higgins_(Staffordshire)_complete_genome

gct gtt agc tgt cgt tca aga tca gaa ata cgg ttt ttc ttg gaa ttg att tcg tca atg tac tta aca tga tta aca tcg aaa cac tta tta agt tc- ttt ttc caa ttc tac gat ttt att tct ttc gcg agt caa ttc cct cct gta gta gct atc ggt ttt gtc aga ttt acg ctc tct acg tag act ttc ttg caa gtt act aat ttg ttc cct gac acg tcc aag ttc agt ttt ata tgc cga ata gag ttc tga ttt atc ctt taa gca gat ctc tag cga ttg ttt aag atc act aat tct agt ctt tag cct att tac ctc ctc aga aga tgt tcc gtt acc gtt gcg ttt aca ctc gtt aag ctg tct atc aag atc cat gat tct atc tct --- --- --- --- --- --- --- --- --- --- --- --- --- --- --- --- -

>DQ441442.1:127048-127448_Variola_virus_strain_Sumatra_1970_V70-228_complete_genome

gct gtt agc tgt cgt tca aga tca gaa ata cgg ttt ttc ttg gaa ttg att tcg tca atg tac tta aca tga tta aca tcg aaa cac tta tta agt tc- ttt ttc caa ttc tac gat ttt att tct ttc gcg agt caa ttc cct cct gta gta gct atc ggt ttt gtc aga ttc acg ctc tct acg tag act ttc ttg caa gtt act aat ttg ttc cct gac acg tcc aag ttc agt ttt ata tgc cga ata tag ttc tga ttt atc ctt taa gca gat ctc tag cga ttg ttt aag atc act aat tct agt ctt tag cct att tac ctc ctc aga aga tgt tcc gtt acc gtt gcg ttt aca ctc gtt aag ctg tct atc aag atc cat gat tct atc tct --- --- --- --- --- --- --- --- --- --- --- --- --- --- --- --- -

>KM044310.1:128244-128632_Vaccinia_virus_strain_Tashkent_clone_TKT4_complete_genome

gct gct agt t-- -gt tcg aga tca gaa ata cgg gtt ttc ttg gaa ttg att tcg tcg atg tac ttg gca --- --- --- tcg aaa cac tta tta agt tc- ttt ttc caa ttc tac gat ttt att tct ttc gcg agt caa ttc cct cct gta gta act atc tgt ttt gtc aga ttc acg ctc ttt acg tag act ttc ttg caa gtt act aat ttg ttc cct agc acg tcc gag ttt agt ttt ata tgc tga ata gag ttc tga ttc atc ctt tga gca gat ctc tag cga tcg ttt aag att cct gat tct agt ctt tag cct att tac ctc ctc aga aga tgt tcc gtt acc gtt gcg ttt aca ctc gtt aag ctg tct atc aag atc cat gat tct atc tct --- --- --- --- --- --- --- --- --- --- --- --- --- --- --- --- -

>KM044309.1:128677-129065_Vaccinia_virus_strain_Tashkent_clone_TKT3_complete_genome

gct gct agt t-- -gt tcg aga tca gaa ata cgg gtt ttc ttg gaa ttg att tcg tcg atg tac ttg gca --- --- --- tcg aaa cac tta tta agt tc- ttt ttc caa ttc tac gat ttt att tct ttc gcg agt caa ttc cct cct gta gta act atc tgt ttt gtc aga ttc acg ctc ttt acg tag act ttc ttg caa gtt act aat ttg ttc cct agc acg tcc gag ttt agt ttt ata tgc tga ata gag ttc tga ttc atc ctt tga gca gat ctc tag cga tcg ttt aag att cct gat tct agt ctt tag cct att tac ctc ctc aga aga tgt tcc gtt acc gtt gcg ttt aca ctc gtt aag ctg tct atc aag atc cat gat tct atc tct --- --- --- --- --- --- --- --- --- --- --- --- --- --- --- --- -

>DQ437581.1:127122-127516_Variola_virus_strain_Bangladesh_1975_v75-550_Banu_complete_genome

gct gtt agc tgt cgt tca aga tca gaa ata cgg ttt ttc ttg gaa ttg att tcg tca atg tac tta aca tga tta aca tcg aaa cac tta tta agt tc- ttt ttc caa ttc tac gat ttt att tct ttc gcg agt caa ttc cct cct gta gta gct atc ggt ttt gtc aga ttc acg ctc tct acg tag act ttc ttg caa gtt act aat ttg ttc cct gac acg tcc aag ttc agt ttt ata tgc cga ata gag ttc tga ttt atc ctt taa gca gat ctc tag cga ttg ttt aag atc act aat tct agt ctt tag cct att tac ctc ctc aga aga tgt tcc gtt --- --- gcg ttt acg ctc gtt aag ctg tct atc aag atc cat gat tct atc tct --- --- --- --- --- --- --- --- --- --- --- --- --- --- --- --- -

>DQ441421.1:127199-127593_Variola_virus_strain_Bangladesh_1974_(Shahzaman)_complete_genome

gct gtt agc tgt cgt tca aga tca gaa ata cgg ttt ttc ttg gaa ttg att tcg tca atg tac tta aca tga tta aca tcg aaa cac tta tta agt tc- ttt ttc caa ttc tac gat ttt att tct ttc gcg agt caa ttc cct cct gta gta gct atc ggt ttt gtc aga ttc acg ctc tct acg tag act ttc ttg caa gtt act aat ttg ttc cct gac acg tcc aag ttc agt ttt ata tgc cga ata gag ttc tga ttt atc ctt taa gca gat ctc tag cga ttg ttt aag atc act aat tct agt ctt tag cct att tac ctc ctc aga aga tgt tcc gtt --- --- gcg ttt acg ctc gtt aag ctg tct atc aag atc cat gat tct atc tct --- --- --- --- --- --- --- --- --- --- --- --- --- --- --- --- -

>DQ441420.1:127199-127593_Variola_virus_strain_Bangladesh_1974_(nur_islam)_complete_genome

gct gtt agc tgt cgt tca aga tca gaa ata cgg ttt ttc ttg gaa ttg att tcg tca atg tac tta aca tga tta aca tcg aaa cac tta tta agt tc- ttt ttc caa ttc tac gat ttt att tct ttc gcg agt caa ttc cct cct gta gta gct atc ggt ttt gtc aga ttc acg ctc tct acg tag act ttc ttg caa gtt act aat ttg ttc cct gac acg tcc aag ttc agt ttt ata tgc cga ata gag ttc tga ttt atc ctt taa gca gat ctc tag cga ttg ttt aag atc act aat tct agt ctt tag cct att tac ctc ctc aga aga tgt tcc gtt --- --- gcg ttt acg ctc gtt aag ctg tct atc aag atc cat gat tct atc tct --- --- --- --- --- --- --- --- --- --- --- --- --- --- --- --- -

>L22579.1:127207-127601_Variola_major_virus_(strain_Bangladesh-1975)_complete_genome

gct gtt agc tgt cgt tca aga tca gaa ata cgg ttt ttc ttg gaa ttg att tcg tca atg tac tta aca tga tta aca tcg aaa cac tta tta agt tc- ttt ttc caa ttc tac gat ttt att tct ttc gcg agt caa ttc cct cct gta gta gct atc ggt ttt gtc aga ttc acg ctc tct acg tag act ttc ttg caa gtt act aat ttg ttc cct gac acg tcc aag ttc agt ttt ata tgc cga ata gag ttc tga ttt atc ctt taa gca gat ctc tag cga ttg ttt aag atc act aat tct agt ctt tag cct att tac ctc ctc aga aga tgt tcc gtt --- --- gcg ttt acg ctc gtt aag ctg tct atc aag atc cat gat tct atc tct --- --- --- --- --- --- --- --- --- --- --- --- --- --- --- --- -

>MG599038.1:139002-139386_Buffalopox_virus_isolate_Karachi_2005

gct gct agt tgt cgt tcg aga tca gaa ata cgg gtt ttc ttg gaa ttg att tcg tcg atg tac ttg gca --- --- --- tcg aaa cac tta tta agt tc- ttt ttc caa ttc tac gat ttt att tct ttc gcg agt caa ttc cct cct gta gta act atc tgt ttt gtc aga ttc acg ctc tct acg tag act ttc ttg caa gtt act aat ttg ttc cct agc acg tcc gag ttt agt ttt ata tgc tga ata gag ttc tga ttc atc ctt tga gca gat ctc tag cga tcg ttt aag att cct gat tct agt ctt tag cct att tac ctc ctc ag- --- --- tcc gtt acc gtt gcg ttt aca ctc gtt aag ctg tct atc aag atc cat gat tct atc tct --- --- --- --- --- --- --- --- --- --- --- --- --- --- --- --- -

>OK422495.1:125041-125425_Buffalopox_virus_isolate_BPXV/buffalo/India/2011/Meerut-1/P50/A3_complete_genome

gct gct agt tgt cgt tcg aga tca gaa ata cgg gtt ttc ttg gaa ttg att tcg tcg atg tac ttg gca --- --- --- tcg aaa cac tta tta agt tc- ttt ttc caa ttc tac gat ttt att tct ttc gcg agt caa ttc cct cct gta gta act atc tgt ttt gtc aga ttc acg ctc tct acg tag act ttc ttg caa gtt act aat ttg ttc cct agc acg tcc gag ttt agt ttt ata tgc tga ata gag ttc tga ttc atc ctt tga gca gat ctc tag cga tcg ttt aag att cct gat tct agt ctt tag cct att tac ctc ctc ag- --- --- tcc gtt acc gtt gcg ttt aca ctc gtt aag ctg tct atc aag atc cat gat tct atc tct --- --- --- --- --- --- --- --- --- --- --- --- --- --- --- --- -

>MK239755.1:128968-129352_Buffalopox_virus_isolate_NIV116234

gct gct agt tgt cgt tcg aga tca gaa ata cgg gtt ttc ttg gaa ttg att tcg tca atg tac ttg gca --- --- --- tcg aaa cac tta tta agt tc- ttt ttc caa ttc tac gat ttt att tct ttc gcg agt caa ttc cct cct gta gta act atc tgt ttt gtc aga ttc acg ctc tct acg tag act ttc ttg caa gtt act aat ttg ttc cct agc acg tcc gag ttt agt ttt ata tgc tga ata gag ttc tga ttc atc ctt tga gca gat ctc tag cga tcg ttt aag att cct gat tct agt ctt tag cct att tac ctc ctc ag- --- --- tcc gtt acc gtt gcg ttt aca ctc gtt aag ctg tct atc aag atc cat gat tct atc tct --- --- --- --- --- --- --- --- --- --- --- --- --- --- --- --- -

>MK239754.1:131278-131662_Buffalopox_virus_isolate_NIV116233

gct gct agt tgt cgt tcg aga tca gaa ata cgg gtt ttc ttg gaa ttg att tcg tca atg tac ttg gca --- --- --- tcg aaa cac tta tta agt tc- ttt ttc caa ttc tac gat ttt att tct ttc gcg agt caa ttc cct cct gta gta act atc tgt ttt gtc aga ttc acg ctc tct acg tag act ttc ttg caa gtt act aat ttg ttc cct agc acg tcc gag ttt agt ttt ata tgc tga ata gag ttc tga ttc atc ctt tga gca gat ctc tag cga tcg ttt aag att cct gat tct agt ctt tag cct att tac ctc ctc ag- --- --- tcc gtt acc gtt gcg ttt aca ctc gtt aag ctg tct atc aag atc cat gat tct atc tct --- --- --- --- --- --- --- --- --- --- --- --- --- --- --- --- -

>MK239753.1:131090-131474_Buffalopox_virus_isolate_NIV143262

gct gct agt tgt cgt tcg aga tca gaa ata cgg gtt ttc ttg gaa ttg att tcg tca atg tac ttg gca --- --- --- tcg aaa cac tta tta agt tc- ttt ttc caa ttc tac gat ttt att tct ttc gcg agt caa ttc cct cct gta gta act atc tgt ttt gtc aga ttc acg ctc tct acg tag act ttc ttg caa gtt act aat ttg ttc cct agc acg tcc gag ttt agt ttt ata tgc tga ata gag ttc tga ttc atc ctt tga gca gat ctc tag cga tcg ttt aag att cct gat tct agt ctt tag cct att tac ctc ctc ag- --- --- tcc gtt acc gtt gcg ttt aca ctc gtt aag ctg tct atc aag atc cat gat tct atc tct --- --- --- --- --- --- --- --- --- --- --- --- --- --- --- --- -

>MG012796.1:135756-136147_Vaccinia_virus_isolate_CEyV2

gct ggt agt tgt cgt tcg aga tca gaa ata cgg gtt ttc ttg gaa ttg att tcg tcg atg tcc ttt gca --- --- --- tcg aaa cac tta tta agt tc- ttt ttc caa ttg tac gat ttt att tct ttc gcg agt caa ttc cct cct gta gta act atc tgt ttg gtc aga ttc acg ccc tct acg tag att ctc ttg caa gtt act aat ttg ttc cct agc acg tcc gag ttt agt ttt ata tgc tga ata gag ttc tga ttc atc ctt tga gca gat ctc tgg cga tcg ttt aag att cct gat tct agt ctt tag act aat tac ctc ctc aga aga tgt tcc gtt acc ctt ggg ttt aca ctc gtt aag ctg tct atc aag atc cat gat tct atc tct --- --- --- --- --- --- --- --- --- --- --- --- --- --- --- --- -

>KT013210.1:129658-130037_Vaccinia_virus_strain_Cantagalo_isolate_Cantagalo_virus_CM01

gct gct agt tgt cgt tcg aga tca gaa ata cgg gtt ttc ttg gaa ttg att tcg tca atg tac ttg gca --- --- --- tcg aaa cac tta tta agt tc- ttt ttc caa ttc tac gat ttt att tct ttc gcg agt caa ttc cct cct gta gta act atc tgt ttt g-- --- --- --- -tc tct acg tag act ttc ttg caa gtt act aat ttg ttc cct agc acg tcc gag ttc agt ttt ata tgc tga ata gag ttc tga ttc atc ctt tga gca gat ctc tag aga tcg ttc aag atc cct gat tct agt ctt tag cct att tac ctc ctc aga aga tgt tcc gtt acc gtt gcg ttt aca ctc gtt aag ctg tct atc aag atc cat gat tct atc tct --- --- --- --- --- --- --- --- --- --- --- --- --- --- --- --- -

>KF179385.1:132776-133155_Vaccinia_virus_complete_genome

gct gct agt tgt cgt tcg aga tca gaa ata cgg gtt ttc ttg gaa ttg att tcg tca atg tac ttg gca --- --- --- tcg aaa cac tta tta agt tc- ttt ttc caa ttc tac gat ttt att tct ttc gcg agt caa ttc cct cct gta gta act atc tgt ttt g-- --- --- --- -tc tct acg tag act ttc ttg caa gtt act aat ttg ttc cct agc acg tcc gag ttc agt ttt ata tgc tga ata gag ttc tga ttc atc ctt tga gca gat ctc tag aga tcg ttc aag atc cct gat tct agt ctt tag cct att tac ctc ctc aga aga tgt tcc gtt acc gtt gcg ttt aca ctc gtt aag ctg tct atc aag atc cat gat tct atc tct --- --- --- --- --- --- --- --- --- --- --- --- --- --- --- --- -

>MW018156.1:133005-133384_Vaccinia_virus_strain_Cantagalo_isolate_Cantagalo_ALEH2

gct gct agt tgt cgt tcg aga tca gaa ata cgg gtt ttc ttg gaa ttg att tcg tca atg tac ttg gca --- --- --- tcg aaa cac tta tta agt tc- ttt ttc caa ttc tac gat ttt att tct ttc gcg agt caa ttc cct cct gta gta act atc tgt ttt g-- --- --- --- -tc tct acg tag act ttc ttg caa gtt act aat ttg ttc cct agc acg tcc gag ttc agt ttt ata tgc tga ata gag ttc tga ttc atc ctt tga gca gat ctc tag aga tcg ttc aag atc cct gat tct agt ctt tag cct att tac ctc ctc aga aga tgt tcc gtt acc gtt gcg ttt aca ctc gtt aag ctg tct atc aag atc cat gat tct atc tct --- --- --- --- --- --- --- --- --- --- --- --- --- --- --- --- -

>MW018155.1:129257-129636_Vaccinia_virus_strain_Cantagalo_isolate_Cantagalo_CG04

gct gct agt tgt cgt tcg aga tca gaa ata cgg gtt ttc ttg gaa ttg att tcg tca atg tac ttg gca --- --- --- tcg aaa cac tta tta agt tc- ttt ttc caa ttc tac gat ttt att tct ttc gcg agt caa ttc cct cct gta gta act atc tgt ttt g-- --- --- --- -tc tct acg tag act ttc ttg caa gtt act aat ttg ttc cct agc acg tcc gag ttc agt ttt ata tgc tga ata gag ttc tga ttc atc ctt tga gca gat ctc tag aga tcg ttc aag atc cct gat tct agt ctt tag cct att tac ctc ctc aga aga tgt tcc gtt acc gtt gcg ttt aca ctc gtt aag ctg tct atc aag atc cat gat tct atc tct --- --- --- --- --- --- --- --- --- --- --- --- --- --- --- --- -

>MW018154.1:129619-129998_Vaccinia_virus_strain_Cantagalo_isolate_Cantagalo_VI04

gct gct agt tgt cgt tcg aga tca gaa ata cgg gtt ttc ttg gaa ttg att tcg tca atg tac ttg gca --- --- --- tcg aaa cac tta tta agt tc- ttt ttc caa ttc tac gat ttt att tct ttc gcg agt caa ttc cct cct gta gta act atc tgt ttt g-- --- --- --- -tc tct acg tag act ttc ttg caa gtt act aat ttg ttc cct agc acg tcc gag ttc agt ttt ata tgc tga ata gag ttc tga ttc atc ctt tga gca gat ctc tag aga tcg ttc aag atc cct gat tct agt ctt tag cct att tac ctc ctc aga aga tgt tcc gtt acc gtt gcg ttt aca ctc gtt aag ctg tct atc aag atc cat gat tct atc tct --- --- --- --- --- --- --- --- --- --- --- --- --- --- --- --- -

>MW018153.1:129251-129630_Vaccinia_virus_strain_Cantagalo_isolate_Cantagalo_MI233

gct gct agt tgt cgt tcg aga tca gaa ata cgg gtt ttc ttg gaa ttg att tcg tca atg tac ttg gca --- --- --- tcg aaa cac tta tta agt tc- ttt ttc caa ttc tac gat ttt att tct ttc gcg agt caa ttc cct cct gta gta act atc tgt ttt g-- --- --- --- -tc tct acg tag act ttc ttg caa gtt act aat ttg ttc cct agc acg tcc gag ttc agt ttt ata tgc tga ata gag ttc tga ttc atc ctt tga gca gat ctc tag aga tcg ttc aag atc cct gat tct agt ctt tag cct att tac ctc ctc aga aga tgt tcc gtt acc gtt gcg ttt aca ctc gtt aag ctg tct atc aag atc cat gat tct atc tct --- --- --- --- --- --- --- --- --- --- --- --- --- --- --- --- -

>DQ516060.1:633-1012_Vaccinia_virus_strain_Passatempo_nonfunctional_ATI_gene_partial_sequence

gct gct agt tgt cgt tcg aga tca gaa ata cgg gtt ttc ttg gaa ttg att tcg tca atg tac ttg gca --- --- --- tcg aaa cac tta tta agt tc- ttt ttc caa ttc tac gat ttt att tct ttc gcg agt caa ttc cct cct gta gta act atc tgt ttt g-- --- --- --- -tc tct acg tag act ttc ttg caa gtt act aat ttg ttc cct agc acg tcc gag ttc agt ttt ata tgc tga ata gag ttc tga ttc atc ctt tga gca gat ctc tag aga tcg ttc aag atc cct gat tct agt ctt tag cct att tac ctc ctc aga aga tgt tcc gtt acc gtt gcg ttt aca ctc gtt aag ctg tct atc aag atc cat gat tct atc tct --- --- --- --- --- --- --- --- --- --- --- --- --- --- --- --- -

>DQ516057.1:633-1012_Vaccinia_virus_strain_Aracatuba_nonfunctional_ATI_gene_partial_sequence

gct gct agt tgt cgt tcg aga tca gaa ata cgg gtt ttc ttg gaa ttg att tcg tca atg tac ttg gca --- --- --- tcg aaa cac tta tta agt tc- ttt ttc caa ttc tac gat ttt att tct ttc gcg agt caa ttc cct cct gta gta act atc tgt ttt g-- --- --- --- -tc tct acg tag act ttc ttg caa gtt act aat ttg ttc cct agc acg tcc gag ttc agt ttt ata tgc tga ata gag ttc tga ttc atc ctt tga gca gat ctc tag aga tcg ttc aag atc cct gat tct agt ctt tag cct att tac ctc ctc aga aga tgt tcc gtt acc gtt gcg ttt aca ctc gtt aag ctg tct atc aag atc cat gat tct atc tct --- --- --- --- --- --- --- --- --- --- --- --- --- --- --- --- -

>JQ361142.1:624-1003_Vaccinia_virus_isolate_CARV_acidophilic_type_inclusion_gene_complete_cds_and_acidophilic_type_inclusion_gene_partial_cds

gct gct agt tga cgt tcg aga tca gaa ata cgg gtt ttc ttg gaa ttg att tcg tcg atg tac ttg gca --- --- --- tcg aaa cac tta tta agt tc- ttt ttc caa ttc tac gat ttt att tct ttc gcg agt caa ttc cct cct gta gta act atc tgt ttt g-- --- --- --- -tc tct acg tag act ttc ttg caa gtt act aat ttg ttc cct agc acg tcc gag ttc agt ttt ata tgc tga ata gag ttc tga ttc atc ctt tga gca gat ctc tag aga tcg ttc aag atc cct gat tct agt ctt tag cct att tac ctc ctc aga aga tgt tcc gtt acc gtt gcg ttt aca ctc gtt aag ctg tct atc aag acc cat gat tct atc tct --- --- --- --- --- --- --- --- --- --- --- --- --- --- --- --- -

>JQ361147.1:624-1003_Vaccinia_virus_isolate_PARV2_nonfunctional_acidophilic_type_inclusion_gene_complete_sequence_and_acidophilic_type_inclusion_gene_partial_cds

gct gct agt tga cgt tcg aga tca gaa ata cgg gtt ttc ttg gaa ttg att tcg tca atg tac ttg gca --- --- --- tcg aaa cac tta tta agt tc- ttt ttc caa ttc tac gat ttt att tct ttc gcg agt caa ttc cct cct gta gta act atc tgt ttt g-- --- --- --- -tc tct acg tag act ttc ttg caa gtt act aat ttg ttc cct agc acg tcc gag ttc agt ttt ata tgc tga ata gag ttc tga ttc atc ctt tga gca gat ctc tag aga tcg ttc aag atc cct gat tct agt ctt tag cct att tac ctc ctc aga aga tgt tcc gtt acc gtt gcg ttt aca ctc gtt aag ctg tct atc aag acc cat gat tct atc tct --- --- --- --- --- --- --- --- --- --- --- --- --- --- --- --- -

>JQ361146.1:624-1003_Vaccinia_virus_isolate_SB2005V_acidophilic_type_inclusion_gene_complete_cds_and_acidophilic_type_inclusion_gene_partial_cds

gct gct agt tga cgt tcg aga tca gaa ata cgg gtt ttc ttg gaa ttg att tcg tca gtg tac ttg gca --- --- --- tcg aaa cac tta tta agt tc- ttt ttc caa ttc tac gat ttt att tct ttc gcg agt caa ttc cct cct gta gta act atc tgt ttt g-- --- --- --- -tc tct acg tag act ttc ttg caa gtt act aat ttg ttc cct agc acg tcc gag ttc agt ttt ata tgc tga ata gag ttc tga ttc atc ctt tga gca gat ctc tag aga tcg ttc aag atc cct gat tct agt ctt tag cct att tac ctc ctc aga aga tgt tcc gtt acc gtt gcg ttt aca ctc gtt aag ctg tct atc aag acc cat gat tct atc tct --- --- --- --- --- --- --- --- --- --- --- --- --- --- --- --- -

>JQ361145.1:624-1003_Vaccinia_virus_isolate_BGV1_acidophilic_type_inclusion_gene_complete_cds_and_acidophilic_type_inclusion_gene_partial_cds

gct gct agt tga cgt tcg aga tca gaa ata cgg gtt ttc ttg gaa ttg att tcg tca atg tac ttg gca --- --- --- tcg aaa cac tta tta agt tc- ttt ttc caa ttc tac gat ttt att tct ttc gcg agt caa ttc cct cct gta gta act atc tgt ttt g-- --- --- --- -tc tct acg tag act ttc ttg caa gtt act aat ttg ttc cct agc acg tcc gag ttc agt ttt ata tgc tga ata gag ttc tga ttc atc ctt tga gca gat ctc tag aga tcg ttc aag atc cct gat tct agt ctt tag cct att tac ctc ctc aga aga tgt tcc gtt acc gtt gcg ttt aca ctc gtt aag ctg tct atc aag acc cat gat tct atc tct --- --- --- --- --- --- --- --- --- --- --- --- --- --- --- --- -

>JQ361144.1:624-1003_Vaccinia_virus_isolate_MUNV_acidophilic_type_inclusion_gene_complete_cds_and_acidophilic_type_inclusion_gene_partial_cds

gct gct agt tga cgt tcg aga tca gaa ata cgg gtt ttc ttg gaa ttg att tcg tca atg tac ttg gca --- --- --- tcg aaa cac tta tta agt tc- ttt ttc caa ttc tac gat ttt att tct ttc gcg agt caa ttc cct cct gta gta act atc tgt ttt g-- --- --- --- -tc tct acg tag act ttc ttg caa gtt act aat ttg ttc cct agc acg tcc gag ttc agt ttt ata tgc tga ata gag ttc tga ttc atc ctt tga gca gat ctc tag aga tcg ttc aag atc cct gat tct agt ctt tag cct att tac ctc ctc aga aga tgt tcc gtt acc gtt gcg ttt aca ctc gtt aag ctg tct atc aag acc cat gat tct atc tct --- --- --- --- --- --- --- --- --- --- --- --- --- --- --- --- -

>JQ361143.1:624-1003_Vaccinia_virus_isolate_ESV1_acidophilic_type_inclusion_gene_complete_cds_and_acidophilic_type_inclusion_gene_partial_cds

gct gct agt tga cgt tcg aga tca gaa ata cgg gtt ttc ttg gaa ttg att tcg tca atg tac ttg gca --- --- --- tcg aaa cac tta tta agt tc- ttt ttc caa ttc tac gat ttt att tct ttc gcg agt caa ttc cct cct gta gta act atc tgt ttt g-- --- --- --- -tc tct acg tag act ttc ttg caa gtt act aat ttg ttc cct agc acg tcc gag ttc agt ttt ata tgc tga ata gag ttc tga ttc atc ctt tga gca gat ctc tag aga tcg ttc aag atc cct gat tct agt ctt tag cct att tac ctc ctc aga aga tgt tcc gtt acc gtt gcg ttt aca ctc gtt aag ctg tct atc aag acc cat gat tct atc tct --- --- --- --- --- --- --- --- --- --- --- --- --- --- --- --- -

>JQ361141.1:624-1003_Vaccinia_virus_isolate_GOV1_acidophilic_type_inclusion_gene_complete_cds_and_acidophilic_type_inclusion_gene_partial_cds

gct gct agt tga cgt tcg aga tca gaa ata cgg gtt ttc ttg gaa ttg att tcg tca atg tac ttg gca --- --- --- tcg aaa cac tta tta agt tc- ttt ttc caa ttc tac gat ttt att tct ttc gcg agt caa ttc cct cct gta gta act atc tgt ttt g-- --- --- --- -tc tct acg tag act ttc ttg caa gtt act aat ttg ttc cct agc acg tcc gag ttc agt ttt ata tgc tga ata gag ttc tga ttc atc ctt tga gca gat ctc tag aga tcg ttc aag atc cct gat tct agt ctt tag cct att tac ctc ctc aga aga tgt tcc gtt acc gtt gcg ttt aca ctc gtt aag ctg tct atc aag acc cat gat tct atc tct --- --- --- --- --- --- --- --- --- --- --- --- --- --- --- --- -

>DQ516059.1:633-1012_Vaccinia_virus_strain_Muriae_nonfunctional_ATI_gene_partial_sequence

gct gct agt tgt cgt tcg aaa tca gaa ata cgg gtt ttc ttg gaa ttg att tcg tca atg tac ttg gca --- --- --- tcg aaa cac tta tta agt tc- ttt ttc caa ttc tac gat ttt att tct ttc gcg agt caa ttc cct cct gta gta act atc tgt ttt g-- --- --- --- -tc tct acg tag act ttc ttg caa gtt act aat ttg ttc cct agc acg tcc ggg ttc agt ctt ata tgc cga ata gag ttc tga ttc atc ctt tga gca gat ctc tag aga tcg ttc aag atc cct gat tct agt ctt tag cct att tac ctc ctc aga aga tgt tcc gtt acc gtt gcg ttt aca ctc gtt aag ctg tct atc aag atc cat gat tct atc tct --- --- --- --- --- --- --- --- --- --- --- --- --- --- --- --- -

>DQ516058.1:633-1012_Vaccinia_virus_strain_Guarani_P2_nonfunctional_ATI_gene_partial_sequence

gct gct agt tga cgt tcg aga tca gaa ata cgg gtt ttc ttg gaa ttg att tcg tca atg tac ttg gca --- --- --- tcg aaa cac tta tta agt tc- ttt ttc caa ttc tac gat ttt att tct ttc gcg agt caa ttc cct cct gta gta act atc tgt ttt g-- --- --- --- -tc tct acg tag act ttc ttg caa gtt act aat ttg ttc cct agc acg tcc gag ttc agt ttt ata tgc tga ata gag ttc tga ttc atc ctt tga gca gat ctc tag aga tcg ttc aag atc cct gat tct agt ctt tag cct att tac ctc ctc aga aga tgt tcc gtt acc gtt gcg ttt aca ctc gtt aag ctg tct atc aag acc cat gat tct atc tct --- --- --- --- --- --- --- --- --- --- --- --- --- --- --- --- -

>MG012795.1:124026-124405_Vaccinia_virus_isolate_CEyV1

gct gct agt agt cgt tcg aga tca gaa ata cgg gtt ttc ttg gaa ttg att tcg tca atg tac ttg gaa --- --- --- tag aaa cac tta tta agt tc- ttt ttc caa ttc tac gat ttt att tct ttc acg agt caa ttc cct cct gta gta act atc tgt ttt g-- --- --- --- -tc tct ccg tag act ttc ttg caa gtt act aat tgg ttc cct agc acg tcc gag ttc agt ttt ata tgc tgc ata gag ttc tga ttc atc ctt tga gca gat cac tag aga tcg ttc aag atc cct gat tct agt ctt tag cct att tac ctc ctc aga aga tgt tcc gtt acc gtt gcg ttt aca ctc gtt aag ttg tct atc aag ttc cat gat tct atc tct --- --- --- --- --- --- --- --- --- --- --- --- --- --- --- --- -
