## Supplementary Data 8 for "Genomic regions insertion and deletion in Monkeypox virus causing multi-country outbreak-2022"

>ON674051.1_Monkeypox_virus_isolate_MPXV_USA_2022_FL001_complete_genome

tgt tat tgt agt atg ata ata tca aaa aga tgg ata taa aga att tac tga cta cat gta cta ttt tac att act aca ttg gct acg gca tat ata cct att tcg tca ctt cca cac gct ccg gta aac ggg tgt cat gtg acg agg gag aat ctt gat aag agg cat aat caa tgt tgt aat ccg atg tcc acc tgg aga att tgc caa ggt cag atg tag agt tgg tag tga taa cac aaa atg tga aca ctg ccc acc tca tac ata tac cgc aat ccc caa tta ttc taa tag atg tca tca atg tag aaa atg ccc aac agg atc att tga taa ggt aaa gtg tac cgg aac aca gaa caa at- gtt cgt gtc atc ctg gtt ggt ata cgc tac tga ttc ttc aca gac tga aga ttg tcg aga ttt gtg tac caa aaa agg aga tgt cca tgc gga tac ttt ggt gga ata gat gaa gga aat cct att tgt aaa tcg tgt tgt gtt ggt gaa tat tgc gac tac cta cgt aat tat aga ctt gat cca ttt cct cca tgc aaa cta tct atc taa atg taa tta att atg att ttg atg ata atg tta cca tac att ata tca cta ctt ggt tag tgt gta tta ttt agt atg gaa gac cta tta ata att act tat ctt ttg acg atc ttg tta taa tta taa tat aaa aat act tat gac ata gta a-- --- -ct cat aat tgc tga cgc gat aaa ttc gta ata atc tgt ttt gtt caa att ttt ata agg aat cta cag gca taa aaa taa aaa tat aat cta taa tat act ctt aca acg cca tca tga ata gca gtg aat taa ttg ctg tta tta tgg att tag aaa tag tgg acg att ttg tga tat taa tat agt tat taa tga tga aag gat aaa cgc gca tag act cat cct atc tgg agc ctc cga ata ttt ttt cca ttc tgt ttt cca ata att tta tcg att cta atg aat acg aag tta atc taa gtc att tag att atc aaa gtg tta act att tga tcg att aca ttt atg gga tac ctt tga gcc taa cta aca ata acg tga aat ata ttc ttt caa ccg ctg att ttt tta caa att gga tct gtc att act gag tgc gaa aaa tac ata ctt aaa aat ctt tgt ttt aga aac tgt atc gat ttc --- --- tac ata tac gct gat aaa tat aat aac aag aaa ata gaa tta gca tcg ttt aac aca ata tta cga aat att ttg aga ctc atc aac aat gaa aac ttt aaa tac tta aca gag gaa tca atg ata aaa att tta agc gat gat atg tta tat ata aaa aat gag gat ttc acc cca ctg att ctt att aaa tgg tta gag agt aca cca acc atg tac cgt cga gtt act tag atg cct cag aat atc a-t ttc ttt ccc cac aag tta taa aat cac ttt ata gtc atc gac tgg tta gtt caa tct acg aat gta taa cat tct taa aca ata tag cat tct tgg ata aat cat ttc cta gat acc ata tca tcg agt tga tat cta tcg gta taa gta att cac atg ata aga ttt cca taa act gct aca atc ata aaa aaa ttc atg gga aat gat atc ttc acg tag ata tag gtg tag ttt cgc agt gac cgt cct gga taa tat tat cta tat gat ggg tgg ata tga tca gtc cct gta tag aag ttc aaa ggt tat agc gta caa tac atg tac taa ttc ttg gat ata tga tat acc aga gct aaa aat atc atc gtt cta att gcg gag gag ttg cca atg acg aat aca ttt att gta tag gcg gta tac gcg atc agg agt cat cgt tga tat cta gta tcg ata gat gga agc cat caa aac cat att ggc aga agt atg cta aaa tgt gcg aac caa aat gtg ata tgg ggt tgc gat ttt aaa cgg att aat ata tgt cat agg tgg agt cgt taa agg tga cac aca tat acc aac gca cta gag agt tta tca gaa gat gga tgg atg aat cat caa cgt ctt cca ata aaa atg tcc aat atg tcg acg att gtt cat gct gga aag att tat ata tct aga ggt tac aac aat agt agt gta gtt aat gta ata tcg aat ct- -ag tcc tta gct ata atc cga tat atg atg aat gga cca aat tat cat cat taa ata ttc cta gaa tta atc ctg ctc tat ggt cag tgt ata ata aat tat atg tag gag gag taa tat ctg atg atg ttc aaa cta ata cat ctg aaa cat acg ata aag aaa aa

>NC_063383.1_Monkeypox_virus_complete_genome

tgt tat tgt agt atg ata ata tca aaa aga tgg ata taa aga att tac tga cta cat gta cta ttt tac att act aca ttg gct acg gca tat ata cct att tcg tca ctt cca cac gct ccg gta aac ggg tgt cat gtg acg agg gag aat ctt gat aag agg cat aat caa tgt tgt aat ccg atg tcc acc tgg aga att tgc caa ggt cag atg tag agt tgg tag tga taa cac aaa atg tga aca ctg ccc acc tca tac ata tac cgc aat ccc caa tta ttc taa tag atg tca tca atg tag aaa atg ccc aac agg atc att tga taa ggt aaa gtg tac cgg aac aca gaa caa at- gtt cgt gtc atc ctg gtt ggt ata cgc tac tga ttc ttc aca gac tga aga ttg tcg aga ttt gtg tac caa aaa agg aga tgt cca tgc gga tac ttt ggt gga ata gat gaa gga aat cct att tgt aaa tcg tgt tgt gtt ggt gaa tat tgc gac tac cta cgt aat tat aga ctt gat cca ttt cct cca tgc aaa cta tct atc taa atg taa tta att atg att ttg atg ata atg tta cca tac att ata tca cta ctt ggt tag tgt gta tta ttt agt atg gaa gac cta tta ata att act tat ctt ttg acg atc ttg tta taa tta taa tat aaa aat act tat gac ata gta a-- --- -ct cat aat tgc tga cgc gat aaa ttc gta ata atc tgt ttt gtt caa att ttt ata agg aat cta cag gca taa aaa taa aaa tat aat cta taa tat act ctt aca acg cca tca tga ata gca gtg aat taa ttg ctg tta tta tgg att tag aaa tag tgg acg att ttg tga tat taa tat agt tat taa tga tga aag gat aaa cgc gca tag act cat cct atc tgg agc ctc cga ata ttt ttt cca ttc tgt ttt cca ata att tta tcg att cta atg aat acg aag tta atc taa gtc att tag att atc aaa gtg tta act att tga tcg att aca ttt atg gga tac ctt tga gcc taa cta aca ata acg tga aat ata ttc ttt caa ccg ctg att ttt tta caa att gga tct gtc att act gag tgc gaa aaa tac ata ctt aaa aat ctt tgt ttt aga aac tgt atc gat ttc --- --- tac ata tac gct gat aaa tat aat aac aag aaa ata gaa tta gca tcg ttt aac aca ata tta cga aat att ttg aga ctc atc aac aat gaa aac ttt aaa tac tta aca gag gaa tca atg ata aaa att tta agc gat gat atg tta tat ata aaa aat gag gat ttc acc cca ctg att ctc att aaa tgg tta gag agt aca cca acc atg tac cgt cga gtt act tag atg cct cag aat atc a-t ttc ttt ccc cac aag tta taa aat cac ttt ata gtc atc gac tgg tta gtt caa tct acg aat gta taa cat tct taa aca ata tag cat tct tgg ata aat cat ttc cta gat acc ata tca tcg agt tga tat cta tcg gta taa gta att cac atg ata aga ttt cca taa act gct aca atc ata aaa aaa ttc atg gga aat gat atc ttc acg tag ata tag gtg tag ttt cgc agt gac cgt cct gga taa tat tat cta tat gat ggg tgg ata tga tca gtc cct gta tag aag ttc aaa ggt tat agc gta caa tac atg tac taa ttc ttg gat ata tga tat acc aga gct aaa aat atc atc gtt cta att gcg gag gag ttg cca atg acg aat aca ttt att gta tag gcg gta tac gcg atc agg agt cat cgt tga tat cta gta tcg ata gat gga agc cat caa aac cat att ggc aga agt atg cta aaa tgt gcg aac caa aat gtg ata tgg ggt tgc gat ttt aaa cgg att aat ata tgt cat agg tgg agt cgt taa agg tga cac aca tat acc aac gca cta gag agt tta tca gaa gat gga tgg atg aat cat caa cgt ctt cca ata aaa atg tcc aat atg tcg acg att gtt cat gct gga aag att tat ata tct aga ggt tac aac aat agt agt gta gtt aat gta ata tcg aat ct- -ag tcc tta gct ata atc cga tat atg atg aat gga cca aat tat cat cat taa ata ttc cta gaa tta atc ctg ctc tat ggt cag tgt ata ata aat tat atg tag gag gag taa tat ctg atg atg ttc aaa cta ata cat ctg aaa cat acg ata aag aaa aa

>ON676704.1_Monkeypox_virus_isolate_MPXV_USA_2022_FL002_complete_genome

tgt tat tgt agt atg ata ata tca aaa aga tgg ata taa aga att tac tga cta cat gta cta ttt tac att act aca ttg gct acg gca tat ata cct att tcg tca ctt cca cac gct ccg gta aac ggg tgt cat gtg acg agg gag aat ctt gat aag agg cat aat caa tgt tgt aat ccg atg tcc acc tgg aga att tgc caa ggt cag atg tag agt tgg tag tga taa cac aaa atg tga aca ctg ccc acc tca tac ata tac cgc aat ccc caa tta ttc taa tag atg tca tca atg tag aaa atg ccc aac agg atc att tga taa ggt aaa gtg tac cgg aac aca gaa caa at- gtt cgt gtc atc ctg gtt ggt ata cgc tac tga ttc ttc aca gac tga aga ttg tcg aga ttt gtg tac caa aaa agg aga tgt cca tgc gga tac ttt ggt gga ata gat gaa gga aat cct att tgt aaa tcg tgt tgt gtt ggt gaa tat tgc gac tac cta cgt aat tat aga ctt gat cca ttt cct cca tgc aaa cta tct atc taa atg taa tta att atg att ttg atg ata atg tta cca tac att ata tca cta ctt ggt tag tgt gta tta ttt agt atg gaa gac cta tta ata att act tat ctt ttg acg atc ttg tta taa tta taa tat aaa aat act tat gac ata gta a-- --- -ct cat aat tgc tga cgc gat aaa ttc gta ata atc tgt ttt gtt caa att ttt ata agg aat cta cag gca taa aaa taa aaa tat aat cta taa tat act ctt aca acg cca tca tga ata gca gtg aat taa ttg ctg tta tta tgg att tag aaa tag tgg acg att ttg tga tat taa tat agt tat taa tga tga aag gat aaa cgc gca tag act cat cct atc tgg agc ctc cga ata ttt ttt cca ttc tgt ttt cca ata att tta tcg att cta atg aat acg aag tta atc taa gtc att tag att atc aaa gtg tta act att tga tcg att aca ttt atg gga tac ctt tga gcc taa cta aca ata acg tga aat ata ttc ttt caa ccg ctg att ttt tta caa att gga tct gtc att act gag tgc gaa aaa tac ata ctt aaa aat ctt tgt ttt aga aac tgt atc gat ttc --- --- tac ata tac gct gat aaa tat aat aac aag aaa ata gaa tta gca tcg ttt aac aca ata tta cga aat att ttg aga ctc atc aac aat gaa aac ttt aaa tac tta aca gag gaa tca atg ata aaa att tta agc gat gat atg tta tat ata aaa aat gag gat ttc acc cca ctg att ctc att aaa tgg tta gag agt aca cca acc atg tac cgt cga gtt act tag atg cct cag aat atc a-t ttc ttt ccc cac aag tta taa aat cac ttt ata gtc atc gac tgg tta gtt caa tct acg aat gta taa cat tct taa aca ata tag cat tct tgg ata aat cat ttc cta gat acc ata tca tcg agt tga tat cta tcg gta taa gta att cac atg ata aga ttt cca taa act gct aca atc ata aaa aaa ttc atg gga aat gat atc ttc acg tag ata tag gtg tag ttt cgc agt gac cgt cct gga taa tat tat cta tat gat ggg tgg ata tga tca gtc cct gta tag aag ttc aaa ggt tat agc gta caa tac atg tac taa ttc ttg gat ata tga tat acc aga gct aaa aat atc atc gtt cta att gcg gag gag ttg cca atg acg aat aca ttt att gta tag gcg gta tac gcg atc agg agt cat cgt tga tat cta gta tcg ata gat gga agc cat caa aac cat att ggc aga agt atg cta aaa tgt gcg aac caa aat gtg ata tgg ggt tgc gat ttt aaa cgg att aat ata tgt cat agg tgg agt cgt taa agg tga cac aca tat acc aac gca cta gag agt tta tca gaa gat gga tgg atg aat cat caa cgt ctt cca ata aaa atg tcc aat atg tcg acg att gtt cat gct gga aag att tat ata tct aga ggt tac aac aat agt agt gta gtt aat gta ata tcg aat ct- -ag tcc tta gct ata atc cga tat atg atg aat gga cca aat tat cat cat taa ata ttc cta gaa tta atc ctg ctc tat ggt cag tgt ata ata aat tat atg tag gag gag taa tat ctg atg atg ttc aaa cta ata cat ctg aaa cat acg ata aag aaa aa

>ON563414.3_Monkeypox_virus_isolate_MPXV_USA_2022_MA001_complete_genome

tgt tat tgt agt atg ata ata tca aaa aga tgg ata taa aga att tac tga cta cat gta cta ttt tac att act aca ttg gct acg gca tat ata cct att tcg tca ctt cca cac gct ccg gta aac ggg tgt cat gtg acg agg gag aat ctt gat aag agg cat aat caa tgt tgt aat ccg atg tcc acc tgg aga att tgc caa ggt cag atg tag agt tgg tag tga taa cac aaa atg tga aca ctg ccc acc tca tac ata tac cgc aat ccc caa tta ttc taa tag atg tca tca atg tag aaa atg ccc aac agg atc att tga taa ggt aaa gtg tac cgg aac aca gaa caa at- gtt cgt gtc atc ctg gtt ggt ata cgc tac tga ttc ttc aca gac tga aga ttg tcg aga ttt gtg tac caa aaa agg aga tgt cca tgc gga tac ttt ggt gga ata gat gaa gga aat cct att tgt aaa tcg tgt tgt gtt ggt gaa tat tgc gac tac cta cgt aat tat aga ctt gat cca ttt cct cca tgc aaa cta tct atc taa atg taa tta att atg att ttg atg ata atg tta cca tac att ata tca cta ctt ggt tag tgt gta tta ttt agt atg gaa gac cta tta ata att act tat ctt ttg acg atc ttg tta taa tta taa tat aaa aat act tat gac ata gta a-- --- -ct cat aat tgc tga cgc gat aaa ttc gta ata atc tgt ttt gtt caa att ttt ata agg aat cta cag gca taa aaa taa aaa tat aat cta taa tat act ctt aca acg cca tca tga ata gca gtg aat taa ttg ctg tta tta tgg att tag aaa tag tgg acg att ttg tga tat taa tat agt tat taa tga tga aag gat aaa cgc gca tag act cat cct atc tgg agc ctc cga ata ttt ttt cca ttc tgt ttt cca ata att tta tcg att cta atg aat acg aag tta atc taa gtc att tag att atc aaa gtg tta act att tga tcg att aca ttt atg gga tac ctt tga gcc taa cta aca ata acg tga aat ata ttc ttt caa ccg ctg att ttt tta caa att gga tct gtc att act gag tgc gaa aaa tac ata ctt aaa aat ctt tgt ttt aga aac tgt atc gat ttc --- --- tac ata tac gct gat aaa tat aat aac aag aaa ata gaa tta gca tcg ttt aac aca ata tta cga aat att ttg aga ctc atc aac aat gaa aac ttt aaa tac tta aca gag gaa tca atg ata aaa att tta agc gat gat atg tta tat ata aaa aat gag gat ttc acc cca ctg att ctc att aaa tgg tta gag agt aca cca acc atg tac cgt cga gtt act tag atg cct cag aat atc a-t ttc ttt ccc cac aag tta taa aat cac ttt ata gtc atc gac tgg tta gtt caa tct acg aat gta taa cat tct taa aca ata tag cat tct tgg ata aat cat ttc cta gat acc ata tca tcg agt tga tat cta tcg gta taa gta att cac atg ata aga ttt cca taa act gct aca atc ata aaa aaa ttc atg gga aat gat atc ttc acg tag ata tag gtg tag ttt cgc agt gac cgt cct gga taa tat tat cta tat gat ggg tgg ata tga tca gtc cct gta tag aag ttc aaa ggt tat agc gta caa tac atg tac taa ttc ttg gat ata tga tat acc aga gct aaa aat atc atc gtt cta att gcg gag gag ttg cca atg acg aat aca ttt att gta tag gcg gta tac gcg atc agg agt cat cgt tga tat cta gta tcg ata gat gga agc cat caa aac cat att ggc aga agt atg cta aaa tgt gcg aac caa aat gtg ata tgg ggt tgc gat ttt aaa cgg att aat ata tgt cat agg tgg agt cgt taa agg tga cac aca tat acc aac gca cta gag agt tta tca gaa gat gga tgg atg aat cat caa cgt ctt cca ata aaa atg tcc aat atg tcg acg att gtt cat gct gga aag att tat ata tct aga ggt tac aac aat agt agt gta gtt aat gta ata tcg aat ct- -ag tcc tta gct ata atc cga tat atg atg aat gga cca aat tat cat cat taa ata ttc cta gaa tta atc ctg ctc tat ggt cag tgt ata ata aat tat atg tag gag gag taa tat ctg atg atg ttc aaa cta ata cat ctg aaa cat acg ata aag aaa aa

>ON676705.1_Monkeypox_virus_isolate_MPXV_USA_2022_UT001_complete_genome

tgt tat tgt agt atg ata ata tca aaa aga tgg ata taa aga att tac tga cta cat gta cta ttt tac att act aca ttg gct acg gca tat ata cct att tcg tca ctt cca cac gct ccg gta aac ggg tgt cat gtg acg agg gag aat ctt gat aag agg cat aat caa tgt tgt aat ccg atg tcc acc tgg aga att tgc caa ggt cag atg tag agt tgg tag tga taa cac aaa atg tga aca ctg ccc acc tca tac ata tac cgc aat ccc caa tta ttc taa tag atg tca tca atg tag aaa atg ccc aac agg atc att tga taa ggt aaa gtg tac cgg aac aca gaa caa at- gtt cgt gtc atc ctg gtt ggt ata cgc tac tga ttc ttc aca gac tga aga ttg tcg aga ttt gtg tac caa aaa agg aga tgt cca tgc gga tac ttt ggt gga ata gat gaa gga aat cct att tgt aaa tcg tgt tgt gtt ggt gaa tat tgc gac tac cta cgt aat tat aga ctt gat cca ttt cct cca tgc aaa cta tct atc taa atg taa tta att atg att ttg atg ata atg tta cca tac att ata tca cta ctt ggt tag tgt gta tta ttt agt atg gaa gac cta tta ata att act tat ctt ttg acg atc ttg tta taa tta taa tat aaa aat act tat gac ata gta a-- --- -ct cat aat tgc tga cgc gat aaa ttc gta ata atc tgt ttt gtt caa att ttt ata agg aat cta cag gca taa aaa taa aaa tat aat cta taa tat act ctt aca acg cca tca tga ata gca gtg aat taa ttg ctg tta tta tgg att tag aaa tag tgg acg att ttg tga tat taa tat agt tat taa tga tga aag gat aaa cgc gca tag act cat cct atc tgg agc ctc cga ata ttt ttt cca ttc tgt ttt cca ata att tta tcg att cta atg aat acg aag tta atc taa gtc att tag att atc aaa gtg tta act att tga tcg att aca ttt atg gga tac ctt tga gcc taa cta aca ata acg tga aat ata ttc ttt caa ccg ctg att ttt tta caa att gga tct gtc att act gag tgc gaa aaa tac ata ctt aaa aat ctt tgt ttt aga aac tgt atc gat ttc --- --- tac ata tac gct gat aaa tat aat aac aag aaa ata gaa tta gca tcg ttt aac aca ata tta cga aat att ttg aga ctc atc aac aat gaa aac ttt aaa tac tta aca gag gaa tca atg ata aaa att tta agc gat gat atg tta tat ata aaa aat gag gat ttc acc cca ctg att ctc att aaa tgg tta gag agt aca cca acc atg tac cgt cga gtt act tag atg cct cag aat atc a-t ttc ttt ccc cac aag tta taa aat cac ttt ata gtc atc gac tgg tta gtt caa tct acg aat gta taa cat tct taa aca ata tag cat tct tgg ata aat cat ttc cta gat acc ata tca tcg agt tga tat cta tcg gta taa gta att cac atg ata aga ttt cca taa act gct aca atc ata aaa aaa ttc atg gga aat gat atc ttc acg tag ata tag gtg tag ttt cgc agt gac cgt cct gga taa tat tat cta tat gat ggg tgg ata tga tca gtc cct gta tag aag ttc aaa ggt tat agc gta caa tac atg tac taa ttc ttg gat ata tga tat acc aga gct aaa aat atc atc gtt cta att gcg gag gag ttg cca atg acg aat aca ttt att gta tag gcg gta tac gcg atc agg agt cat cgt tga tat cta gta tcg ata gat gga agc cat caa aac cat att ggc aga agt atg cta aaa tgt gcg aac caa aat gtg ata tgg ggt tgc gat ttt aaa cgg att aat ata tgt cat agg tgg agt cgt taa agg tga cac aca tat acc aac gca cta gag agt tta tca gaa gat gga tgg atg aat cat caa cgt ctt cca ata aaa atg tcc aat atg tcg acg att gtt cat gct gga aag att tat ata tct aga ggt tac aac aat agt agt gta gtt aat gta ata tcg aat ct- -ag tcc tta gct ata atc cga tat atg atg aat gga cca aat tat cat cat taa ata ttc cta gaa tta atc ctg ctc tat ggt cag tgt ata ata aat tat atg tag gag gag taa tat ctg atg atg ttc aaa cta ata cat ctg aaa cat acg ata aag aaa aa

>MN648051.1_Monkeypox_virus_strain_Israel_2018

tgt tat tgt agt atg ata ata tca aaa aga tgg ata taa aga att tac tga cta cat gta cta ttt tac att act aca ttg gct acg gca tat ata cct att tcg tca ctt cca cac gct ccg gta aac ggg tgt cat gtg acg agg gag aat ctt gat aag agg cat aat caa tgt tgt aat ccg atg tcc acc tgg aga att tgc caa ggt cag atg tag agt tgg tag tga taa cac aaa atg tga aca ctg ccc acc tca tac ata tac cgc aat ccc caa tta ttc taa tag atg tca tca atg tag aaa atg ccc aac agg atc att tga taa ggt aaa gtg tac cgg aac aca gaa caa at- gtt cgt gtc atc ctg gtt ggt ata cgc tac tga ttc ttc aca gac tga aga ttg tcg aga ttt gtg tac caa aaa agg aga tgt cca tgc gga tac ttt ggt gga ata gat gaa gga aat cct att tgt aaa tcg tgt tgt gtt ggt gaa tat tgc gac tac cta cgt aat tat aga ctt gat cca ttt cct cca tgc aaa cta tct atc taa atg taa tta att atg att ttg atg ata atg tta cca tac att ata tca cta ctt ggt tag tgt gta tta ttt agt atg gaa gac cta tta ata att act tat ctt ttg acg atc ttg tta taa tta taa tat aaa aat act tat gac ata gta a-- --- -ct cat aat tgc tga cgc gat aaa ttc gta ata atc tgt ttt gtt caa att ttt ata agg aat cta cag gca taa aaa taa aaa tat aat cta taa tat act ctt aca acg cca tca tga ata gca gtg aat taa ttg ctg tta tta tgg att tag aaa tag tgg acg att ttg tga tat taa tat agt tat taa tga tga aag gat aaa cgc gca tag act cat cct atc tgg agc ctc cga ata ttt ttt cca ttc tgt ttt cca ata att tta tcg att cta atg aat acg aag tta atc taa gtc att tag att atc aaa gtg tta act att tga tcg att aca ttt atg gga tac ctt tga gcc taa cta aca ata acg tga aat ata ttc ttt caa ccg ctg att ttt tta caa att gga tct gtc att act gag tgc gaa aaa tac ata ctt aaa aat ctt tgt ttt aga aac tgt atc gat ttc --- --- tac ata tac gct gat aaa tat aat aac aag aaa ata gaa tta gca tcg ttt aac aca ata tta cga aat att ttg aga ctc atc aac aat gaa aac ttt aaa tac tta aca gag gaa tca atg ata aaa att tta agc gat gat atg tta tat ata aaa aat gag gat ttc acc cca ctg att ctc att aaa tgg tta gag agt aca cca acc atg tac cgt cga gtt act tag atg cct cag aat atc a-t ttc ttt ccc cac aag tta taa aat cac ttt ata gtc atc gac tgg tta gtt caa tct acg aat gta taa cat tct taa aca ata tag cat tct tgg ata aat cat ttc cta gat acc ata tca tcg agt tga tat cta tcg gta taa gta att cac atg ata aga ttt cca taa act gct aca atc ata aaa aaa ttc atg gga aat gat atc ttc acg tag ata tag gtg tag ttt cgc agt gac cgt cct gga taa tat tat cta tat gat ggg tgg ata tga tca gtc cct gta tag aag ttc aaa ggt tat agc gta caa tac atg tac taa ttc ttg gat ata tga tat acc aga gct aaa aat atc atc gtt cta att gcg gag gag ttg cca atg acg aat aca ttt att gta tag gcg gta tac gcg atc agg agt cat cgt tga tat cta gta tcg ata gat gga agc cat caa aac cat att ggc aga agt atg cta aaa tgt gcg aac caa aat gtg ata tgg ggt tgc gat ttt aaa cgg att aat ata tgt cat agg tgg agt cgt taa agg tga cac aca tat acc aac gca cta gag agt tta tca gaa gat gga tgg atg aat cat caa cgt ctt cca ata aaa atg tcc aat atg tcg acg att gtt cat gct gga aag att tat ata tct aga ggt tac aac aat agt agt gta gtt aat gta ata tcg aat ct- -ag tcc tta gct ata atc cga tat atg atg aat gga cca aat tat cat cat taa ata ttc cta gaa tta atc ctg ctc tat ggt cag tgt ata ata aat tat atg tag gag gag taa tat ctg atg atg ttc aaa cta ata cat ctg aaa cat acg ata aag aaa aa

>ON644344.1_Monkeypox_virus_isolate_MPXV_FVG-ITA_01_2022_complete_genome

tgt tat tgt agt atg ata ata tca aaa aga tgg ata taa aga att tac tga cta cat gta cta ttt tac att act aca ttg gct acg gca tat ata cct att tcg tca ctt cca cac gct ccg gta aac ggg tgt cat gtg acg agg gag aat ctt gat aag agg cat aat caa tgt tgt aat ccg atg tcc acc tgg aga att tgc caa ggt cag atg tag agt tgg tag tga taa cac aaa atg tga aca ctg ccc acc tca tac ata tac cgc aat ccc caa tta ttc taa tag atg tca tca atg tag aaa atg ccc aac agg atc att tga taa ggt aaa gtg tac cgg aac aca gaa caa at- gtt cgt gtc atc ctg gtt ggt ata cgc tac tga ttc ttc aca gac tga aga ttg tcg aga ttt gtg tac caa aaa agg aga tgt cca tgc gga tac ttt ggt gga ata gat gaa gga aat cct att tgt aaa tcg tgt tgt gtt ggt gaa tat tgc gac tac cta cgt aat tat aga ctt gat cca ttt cct cca tgc aaa cta tct atc taa atg taa tta att atg att ttg atg ata atg tta cca tac att ata tca cta ctt ggt tag tgt gta tta ttt agt atg gaa gac cta tta ata att act tat ctt ttg acg atc ttg tta taa tta taa tat aaa aat act tat gac ata gta a-- --- -ct cat aat tgc tga cgc gat aaa ttc gta ata atc tgt ttt gtt caa att ttt ata agg aat cta cag gca taa aaa taa aaa tat aat cta taa tat act ctt aca acg cca tca tga ata gca gtg aat taa ttg ctg tta tta tgg att tag aaa tag tgg acg att ttg tga tat taa tat agt tat taa tga tga aag gat aaa cgc gca tag act cat cct atc tgg agc ctc cga ata ttt ttt cca ttc tgt ttt cca ata att tta tcg att cta atg aat acg aag tta atc taa gtc att tag att atc aaa gtg tta act att tga tcg att aca ttt atg gga tac ctt tga gcc taa cta aca ata acg tga aat ata ttc ttt caa ccg ctg att ttt tta caa att gga tct gtc att act gag tgc gaa aaa tac ata ctt aaa aat ctt tgt ttt aga aac tgt atc gat ttc --- --- tac ata tac gct gat aaa tat aat aac aag aaa ata gaa tta gca tcg ttt aac aca ata tta cga aat att ttg aga ctc atc aac aat gaa aac ttt aaa tac tta aca gag gaa tca atg ata aaa att tta agc gat gat atg tta tat ata aaa aat gag gat ttc acc cca ctg att ctc att aaa tgg tta gag agt aca cca acc atg tac cgt cga gtt act tag atg cct cag aat atc a-t ttc ttt ccc cac aag tta taa aat cac ttt ata gtc atc gac tgg tta gtt caa tct acg aat gta taa cat tct taa aca ata tag cat tct tgg ata aat cat ttc cta gat acc ata tca tcg agt tga tat cta tcg gta taa gta att cac atg ata aga ttt cca taa act gct aca atc ata aaa aaa ttc atg gga aat gat atc ttc acg tag ata tag gtg tag ttt cgc agt gac cgt cct gga taa tat tat cta tat gat ggg tgg ata tga tca gtc cct gta tag aag ttc aaa ggt tat agc gta caa tac atg tac taa ttc ttg gat ata tga tat acc aga gct aaa aat atc atc gtt cta att gcg gag gag ttg cca atg acg aat aca ttt att gta tag gcg gta tac gcg atc agg agt cat cgt tga tat cta gta tcg ata gat gga agc cat caa aac cat att ggc aga agt atg cta aaa tgt gcg aac caa aat gtg ata tgg ggt tgc gat ttt aaa cgg att aat ata tgt cat agg tgg agt cgt taa agg tga cac aca tat acc aac gca cta gag agt tta tca gaa gat gga tgg atg aat cat caa cgt ctt cca ata aaa atg tcc aat atg tcg acg att gtt cat gct gga aag att tat ata tct aga ggt tac aac aat agt agt gta gtt aat gta ata tcg aat ct- -ag tcc tta gct ata atc cga tat atg atg aat gga cca aat tat cat cat taa ata ttc cta gaa tta atc ctg ctc tat ggt cag tgt ata ata aat tat atg tag gag gag taa tat ctg atg atg ttc aaa cta ata cat ctg aaa cat acg ata aag aaa aa

>MT903343.1_Monkeypox_virus_isolate_MPXV-UK_P1

tgt tat tgt agt atg ata ata tca aaa aga tgg ata taa aga att tac tga cta cat gta cta ttt tac att act aca ttg gct acg gca tat ata cct att tcg tca ctt cca cac gct ccg gta aac ggg tgt cat gtg acg agg gag aat ctt gat aag agg cat aat caa tgt tgt aat ccg atg tcc acc tgg aga att tgc caa ggt cag atg tag agt tgg tag tga taa cac aaa atg tga aca ctg ccc acc tca tac ata tac cgc aat ccc caa tta ttc taa tag atg tca tca atg tag aaa atg ccc aac agg atc att tga taa ggt aaa gtg tac cgg aac aca gaa caa at- gtt cgt gtc atc ctg gtt ggt ata cgc tac tga ttc ttc aca gac tga aga ttg tcg aga ttt gtg tac caa aaa agg aga tgt cca tgc gga tac ttt ggt gga ata gat gaa gga aat cct att tgt aaa tcg tgt tgt gtt ggt gaa tat tgc gac tac cta cgt aat tat aga ctt gat cca ttt cct cca tgc aaa cta tct atc taa atg taa tta att atg att ttg atg ata atg tta cca tac att ata tca cta ctt ggt tag tgt gta tta ttt agt atg gaa gac cta tta ata att act tat ctt ttg acg atc ttg tta taa tta taa tat aaa aat act tat gac ata gta a-- --- -ct cat aat tgc tga cgc gat aaa ttc gta ata atc tgt ttt gtt caa att ttt ata agg aat cta cag gca taa aaa taa aaa tat aat cta taa tat act ctt aca acg cca tca tga ata gca gtg aat taa ttg ctg tta tta tgg att tag aaa tag tgg acg att ttg tga tat taa tat agt tat taa tga tga aag gat aaa cgc gca tag act cat cct atc tgg agc ctc cga ata ttt ttt cca ttc tgt ttt cca ata att tta tcg att cta atg aat acg aag tta atc taa gtc att tag att atc aaa gtg tta act att tga tcg att aca ttt atg gga tac ctt tga gcc taa cta aca ata acg tga aat ata ttc ttt caa ccg ctg att ttt tta caa att gga tct gtc att act gag tgc gaa aaa tac ata ctt aaa aat ctt tgt ttt aga aac tgt atc gat ttc --- --- tac ata tac gct gat aaa tat aat aac aag aaa ata gaa tta gca tcg ttt aac aca ata tta cga aat att ttg aga ctc atc aac aat gaa aac ttt aaa tac tta aca gag gaa tca atg ata aaa att tta agc gat gat atg tta tat ata aaa aat gag gat ttc acc cca ctg att ctc att aaa tgg tta gag agt aca cca acc atg tac cgt cga gtt act tag atg cct cag aat atc a-t ttc ttt ccc cac aag tta taa aat cac ttt ata gtc atc gac tgg tta gtt caa tct acg aat gta taa cat tct taa aca ata tag cat tct tgg ata aat cat ttc cta gat acc ata tca tcg agt tga tat cta tcg gta taa gta att cac atg ata aga ttt cca taa act gct aca atc ata aaa aaa ttc atg gga aat gat atc ttc acg tag ata tag gtg tag ttt cgc agt gac cgt cct gga taa tat tat cta tat gat ggg tgg ata tga tca gtc cct gta tag aag ttc aaa ggt tat agc gta caa tac atg tac taa ttc ttg gat ata tga tat acc aga gct aaa aat atc atc gtt cta att gcg gag gag ttg cca atg acg aat aca ttt att gta tag gcg gta tac gcg atc agg agt cat cgt tga tat cta gta tcg ata gat gga agc cat caa aac cat att ggc aga agt atg cta aaa tgt gcg aac caa aat gtg ata tgg ggt tgc gat ttt aaa cgg att aat ata tgt cat agg tgg agt cgt taa agg tga cac aca tat acc aac gca cta gag agt tta tca gaa gat gga tgg atg aat cat caa cgt ctt cca ata aaa atg tcc aat atg tcg acg att gtt cat gct gga aag att tat ata tct aga ggt tac aac aat agt agt gta gtt aat gta ata tcg aat ct- -ag tcc tta gct ata atc cga tat atg atg aat gga cca aat tat cat cat taa ata ttc cta gaa tta atc ctg ctc tat ggt cag tgt ata ata aat tat atg tag gag gag taa tat ctg atg atg ttc aaa cta ata cat ctg aaa cat acg ata aag aaa aa

>ON631963.1_Monkeypox_virus_isolate_MPxV/VIDRL01/2022_complete_genome

tgt tat tgt agt atg ata ata tca aaa aga tgg ata taa aga att tac tga cta cat gta cta ttt tac att act aca ttg gct acg gca tat ata cct att tcg tca ctt cca cac gct ccg gta aac ggg tgt cat gtg acg agg gag aat ctt gat aag agg cat aat caa tgt tgt aat ccg atg tcc acc tgg aga att tgc caa ggt cag atg tag agt tgg tag tga taa cac aaa atg tga aca ctg ccc acc tca tac ata tac cgc aat ccc caa tta ttc taa tag atg tca tca atg tag aaa atg ccc aac agg atc att tga taa ggt aaa gtg tac cgg aac aca gaa caa at- gtt cgt gtc atc ctg gtt ggt ata cgc tac tga ttc ttc aca gac tga aga ttg tcg aga ttt gtg tac caa aaa agg aga tgt cca tgc gga tac ttt ggt gga ata gat gaa gga aat cct att tgt aaa tcg tgt tgt gtt ggt gaa tat tgc gac tac cta cgt aat tat aga ctt gat cca ttt cct cca tgc aaa cta tct atc taa atg taa tta att atg att ttg atg ata atg tta cca tac att ata tca cta ctt ggt tag tgt gta tta ttt agt atg gaa gac cta tta ata att act tat ctt ttg acg atc ttg tta taa tta taa tat aaa aat act tat gac ata gta a-- --- -ct cat aat tgc tga cgc gat aaa ttc gta ata atc tgt ttt gtt caa att ttt ata agg aat cta cag gca taa aaa taa aaa tat aat cta taa tat act ctt aca acg cca tca tga ata gca gtg aat taa ttg ctg tta tta tgg att tag aaa tag tgg acg att ttg tga tat taa tat agt tat taa tga tga aag gat aaa cgc gca tag act cat cct atc tgg agc ctc cga ata ttt ttt cca ttc tgt ttt cca ata att tta tcg att cta atg aat acg aag tta atc taa gtc att tag att atc aaa gtg tta act att tga tcg att aca ttt atg gga tac ctt tga gcc taa cta aca ata acg tga aat ata ttc ttt caa ccg ctg att ttt tta caa att gga tct gtc att act gag tgc gaa aaa tac ata ctt aaa aat ctt tgt ttt aga aac tgt atc gat ttc --- --- tac ata tac gct gat aaa tat aat aac aag aaa ata gaa tta gca tcg ttt aac aca ata tta cga aat att ttg aga ctc atc aac aat gaa aac ttt aaa tac tta aca gag gaa tca atg ata aaa att tta agc gat gat atg tta tat ata aaa aat gag gat ttc acc cca ctg att ctc att aaa tgg tta gag agt aca cca acc atg tac cgt cga gtt act tag atg cct cag aat atc a-t ttc ttt ccc cac aag tta taa aat cac ttt ata gtc atc gac tgg tta gtt caa tct acg aat gta taa cat tct taa aca ata tag cat tct tgg ata aat cat ttc cta gat acc ata tca tcg agt tga tat cta tcg gta taa gta att cac atg ata aga ttt cca taa act gct aca atc ata aaa aaa ttc atg gga aat gat atc ttc acg tag ata tag gtg tag ttt cgc agt gac cgt cct gga taa tat tat cta tat gat ggg tgg ata tga tca gtc cct gta tag aag ttc aaa ggt tat agc gta caa tac atg tac taa ttc ttg gat ata tga tat acc aga gct aaa aat atc atc gtt cta att gcg gag gag ttg cca atg acg aat aca ttt att gta tag gcg gta tac gcg atc agg agt cat cgt tga tat cta gta tcg ata gat gga agc cat caa aac cat att ggc aga agt atg cta aaa tgt gcg aac caa aat gtg ata tgg ggt tgc gat ttt aaa cgg att aat ata tgt cat agg tgg agt cgt taa agg tga cac aca tat acc aac gca cta gag agt tta tca gaa gat gga tgg atg aat cat caa cgt ctt cca ata aaa atg tcc aat atg tcg acg att gtt cat gct gga aag att tat ata tct aga ggt tac aac aat agt agt gta gtt aat gta ata tcg aat ct- -ag tcc tta gct ata atc cga tat atg atg aat gga cca aat tat cat cat taa ata ttc cta gaa tta atc ctg ctc tat ggt cag tgt ata ata aat tat atg tag gag gag taa tat ctg atg atg ttc aaa cta ata cat ctg aaa cat acg ata aag aaa aa

>ON568298.1_Monkeypox_virus_isolate_MPXV-BY-IMB25241_complete_genome

tgt tat tgt agt atg ata ata tca aaa aga tgg ata taa aga att tac tga cta cat gta cta ttt tac att act aca ttg gct acg gca tat ata cct att tcg tca ctt cca cac gct ccg gta aac ggg tgt cat gtg acg agg gag aat ctt gat aag agg cat aat caa tgt tgt aat ccg atg tcc acc tgg aga att tgc caa ggt cag atg tag agt tgg tag tga taa cac aaa atg tga aca ctg ccc acc tca tac ata tac cgc aat ccc caa tta ttc taa tag atg tca tca atg tag aaa atg ccc aac agg atc att tga taa ggt aaa gtg tac cgg aac aca gaa caa at- gtt cgt gtc atc ctg gtt ggt ata cgc tac tga ttc ttc aca gac tga aga ttg tcg aga ttt gtg tac caa aaa agg aga tgt cca tgc gga tac ttt ggt gga ata gat gaa gga aat cct att tgt aaa tcg tgt tgt gtt ggt gaa tat tgc gac tac cta cgt aat tat aga ctt gat cca ttt cct cca tgc aaa cta tct atc taa atg taa tta att atg att ttg atg ata atg tta cca tac att ata tca cta ctt ggt tag tgt gta tta ttt agt atg gaa gac cta tta ata att act tat ctt ttg acg atc ttg tta taa tta taa tat aaa aat act tat gac ata gta a-- --- -ct cat aat tgc tga cgc gat aaa ttc gta ata atc tgt ttt gtt caa att ttt ata agg aat cta cag gca taa aaa taa aaa tat aat cta taa tat act ctt aca acg cca tca tga ata gca gtg aat taa ttg ctg tta tta tgg att tag aaa tag tgg acg att ttg tga tat taa tat agt tat taa tga tga aag gat aaa cgc gca tag act cat cct atc tgg agc ctc cga ata ttt ttt cca ttc tgt ttt cca ata att tta tcg att cta atg aat acg aag tta atc taa gtc att tag att atc aaa gtg tta act att tga tcg att aca ttt atg gga tac ctt tga gcc taa cta aca ata acg tga aat ata ttc ttt caa ccg ctg att ttt tta caa att gga tct gtc att act gag tgc gaa aaa tac ata ctt aaa aat ctt tgt ttt aga aac tgt atc gat ttc --- --- tac ata tac gct gat aaa tat aat aac aag aaa ata gaa tta gca tcg ttt aac aca ata tta cga aat att ttg aga ctc atc aac aat gaa aac ttt aaa tac tta aca gag gaa tca atg ata aaa att tta agc gat gat atg tta tat ata aaa aat gag gat ttc acc cca ctg att ctc att aaa tgg tta gag agt aca cca acc atg tac cgt cga gtt act tag atg cct cag aat atc a-t ttc ttt ccc cac aag tta taa aat cac ttt ata gtc atc gac tgg tta gtt caa tct acg aat gta taa cat tct taa aca ata tag cat tct tgg ata aat cat ttc cta gat acc ata tca tcg agt tga tat cta tcg gta taa gta att cac atg ata aga ttt cca taa act gct aca atc ata aaa aaa ttc atg gga aat gat atc ttc acg tag ata tag gtg tag ttt cgc agt gac cgt cct gga taa tat tat cta tat gat ggg tgg ata tga tca gtc cct gta tag aag ttc aaa ggt tat agc gta caa tac atg tac taa ttc ttg gat ata tga tat acc aga gct aaa aat atc atc gtt cta att gcg gag gag ttg cca atg acg aat aca ttt att gta tag gcg gta tac gcg atc agg agt cat cgt tga tat cta gta tcg ata gat gga agc cat caa aac cat att ggc aga agt atg cta aaa tgt gcg aac caa aat gtg ata tgg ggt tgc gat ttt aaa cgg att aat ata tgt cat agg tgg agt cgt taa agg tga cac aca tat acc aac gca cta gag agt tta tca gaa gat gga tgg atg aat cat caa cgt ctt cca ata aaa atg tcc aat atg tcg acg att gtt cat gct gga aag att tat ata tct aga ggt tac aac aat agt agt gta gtt aat gta ata tcg aat ct- -ag tcc tta gct ata atc cga tat atg atg aat gga cca aat tat cat cat taa ata ttc cta gaa tta atc ctg ctc tat ggt cag tgt ata ata aat tat atg tag gag gag taa tat ctg atg atg ttc aaa cta ata cat ctg aaa cat acg ata aag aaa aa

>ON622722.2_Monkeypox_virus_isolate_MPXV_FR_HCL0001_2022_complete_genome

tgt tat tgt agt atg ata ata tca aaa aga tgg ata taa aga att tac tga cta cat gta cta ttt tac att act aca ttg gct acg gca tat ata cct att tcg tca ctt cca cac gct ccg gta aac ggg tgt cat gtg acg agg gag aat ctt gat aag agg cat aat caa tgt tgt aat ccg atg tcc acc tgg aga att tgc caa ggt cag atg tag agt tgg tag tga taa cac aaa atg tga aca ctg ccc acc tca tac ata tac cgc aat ccc caa tta ttc taa tag atg tca tca atg tag aaa atg ccc aac agg atc att tga taa ggt aaa gtg tac cgg aac aca gaa caa at- gtt cgt gtc atc ctg gtt ggt ata cgc tac tga ttc ttc aca gac tga aga ttg tcg aga ttt gtg tac caa aaa agg aga tgt cca tgc gga tac ttt ggt gga ata gat gaa gga aat cct att tgt aaa tcg tgt tgt gtt ggt gaa tat tgc gac tac cta cgt aat tat aga ctt gat cca ttt cct cca tgc aaa cta tct atc taa atg taa tta att atg att ttg atg ata atg tta cca tac att ata tca cta ctt ggt tag tgt gta tta ttt agt atg gaa gac cta tta ata att act tat ctt ttg acg atc ttg tta taa tta taa tat aaa aat act tat gac ata gta a-- --- -ct cat aat tgc tga cgc gat aaa ttc gta ata atc tgt ttt gtt caa att ttt ata agg aat cta cag gca taa aaa taa aaa tat aat cta taa tat act ctt aca acg cca tca tga ata gca gtg aat taa ttg ctg tta tta tgg att tag aaa tag tgg acg att ttg tga tat taa tat agt tat taa tga tga aag gat aaa cgc gca tag act cat cct atc tgg agc ctc cga ata ttt ttt cca ttc tgt ttt cca ata att tta tcg att cta atg aat acg aag tta atc taa gtc att tag att atc aaa gtg tta act att tga tcg att aca ttt atg gga tac ctt tga gcc taa cta aca ata acg tga aat ata ttc ttt caa ccg ctg att ttt tta caa att gga tct gtc att act gag tgc gaa aaa tac ata ctt aaa aat ctt tgt ttt aga aac tgt atc gat ttc --- --- tac ata tac gct gat aaa tat aat aac aag aaa ata gaa tta gca tcg ttt aac aca ata tta cga aat att ttg aga ctc atc aac aat gaa aac ttt aaa tac tta aca gag gaa tca atg ata aaa att tta agc gat gat atg tta tat ata aaa aat gag gat ttc acc cca ctg att ctc att aaa tgg tta gag agt aca cca acc atg tac cgt cga gtt act tag atg cct cag aat atc a-t ttc ttt ccc cac aag tta taa aat cac ttt ata gtc atc gac tgg tta gtt caa tct acg aat gta taa cat tct taa aca ata tag cat tct tgg ata aat cat ttc cta gat acc ata tca tcg agt tga tat cta tcg gta taa gta att cac atg ata aga ttt cca taa act gct aca atc ata aaa aaa ttc atg gga aat gat atc ttc acg tag ata tag gtg tag ttt cgc agt gac cgt cct gga taa tat tat cta tat gat ggg tgg ata tga tca gtc cct gta tag aag ttc aaa ggt tat agc gta caa tac atg tac taa ttc ttg gat ata tga tat acc aga gct aaa aat atc atc gtt cta att gcg gag gag ttg cca atg acg aat aca ttt att gta tag gcg gta tac gcg atc agg agt cat cgt tga tat cta gta tcg ata gat gga agc cat caa aac cat att ggc aga agt atg cta aaa tgt gcg aac caa aat gtg ata tgg ggt tgc gat ttt aaa cgg att aat ata tgt cat agg tgg agt cgt taa agg tga cac aca tat acc aac gca cta gag agt tta tca gaa gat gga tgg atg aat cat caa cgt ctt cca ata aaa atg tcc aat atg tcg acg att gtt cat gct gga aag att tat ata tct aga ggt tac aac aat agt agt gta gtt aat gta ata tcg aat ct- -ag tcc tta gct ata atc cga tat atg atg aat gga cca aat tat cat cat taa ata ttc cta gaa tta atc ctg ctc tat ggt cag tgt ata ata aat tat atg tag gag gag taa tat ctg atg atg ttc aaa cta ata cat ctg aaa cat acg ata aag aaa aa

>ON627808.1_Monkeypox_virus_isolate_MPX/human/USA/UT-UPHL-82200022/2022_complete_genome

tgt tat tgt agt atg ata ata tca aaa aga tgg ata taa aga att tac tga cta cat gta cta ttt tac att act aca ttg gct acg gca tat ata cct att tcg tca ctt cca cac gct ccg gta aac ggg tgt cat gtg acg agg gag aat ctt gat aag agg cat aat caa tgt tgt aat ccg atg tcc acc tgg aga att tgc caa ggt cag atg tag agt tgg tag tga taa cac aaa atg tga aca ctg ccc acc tca tac ata tac cgc aat ccc caa tta ttc taa tag atg tca tca atg tag aaa atg ccc aac agg atc att tga taa ggt aaa gtg tac cgg aac aca gaa caa at- gtt cgt gtc atc ctg gtt ggt ata cgc tac tga ttc ttc aca gac tga aga ttg tcg aga ttt gtg tac caa aaa agg aga tgt cca tgc gga tac ttt ggt gga ata gat gaa gga aat cct att tgt aaa tcg tgt tgt gtt ggt gaa tat tgc gac tac cta cgt aat tat aga ctt gat cca ttt cct cca tgc aaa cta tct atc taa atg taa tta att atg att ttg atg ata atg tta cca tac att ata tca cta ctt ggt tag tgt gta tta ttt agt atg gaa gac cta tta ata att act tat ctt ttg acg atc ttg tta taa tta taa tat aaa aat act tat gac ata gta a-- --- -ct cat aat tgc tga cgc gat aaa ttc gta ata atc tgt ttt gtt caa att ttt ata agg aat cta cag gca taa aaa taa aaa tat aat cta taa tat act ctt aca acg cca tca tga ata gca gtg aat taa ttg ctg tta tta tgg att tag aaa tag tgg acg att ttg tga tat taa tat agt tat taa tga tga aag gat aaa cgc gca tag act cat cct atc tgg agc ctc cga ata ttt ttt cca ttc tgt ttt cca ata att tta tcg att cta atg aat acg aag tta atc taa gtc att tag att atc aaa gtg tta act att tga tcg att aca ttt atg gga tac ctt tga gcc taa cta aca ata acg tga aat ata ttc ttt caa ccg ctg att ttt tta caa att gga tct gtc att act gag tgc gaa aaa tac ata ctt aaa aat ctt tgt ttt aga aac tgt atc gat ttc --- --- tac ata tac gct gat aaa tat aat aac aag aaa ata gaa tta gca tcg ttt aac aca ata tta cga aat att ttg aga ctc atc aac aat gaa aac ttt aaa tac tta aca gag gaa tca atg ata aaa att tta agc gat gat atg tta tat ata aaa aat gag gat ttc acc cca ctg att ctc att aaa tgg tta gag agt aca cca acc atg tac cgt cga gtt act tag atg cct cag aat atc a-t ttc ttt ccc cac aag tta taa aat cac ttt ata gtc atc gac tgg tta gtt caa tct acg aat gta taa cat tct taa aca ata tag cat tct tgg ata aat cat ttc cta gat acc ata tca tcg agt tga tat cta tcg gta taa gta att cac atg ata aga ttt cca taa act gct aca atc ata aaa aaa ttc atg gga aat gat atc ttc acg tag ata tag gtg tag ttt cgc agt gac cgt cct gga taa tat tat cta tat gat ggg tgg ata tga tca gtc cct gta tag aag ttc aaa ggt tat agc gta caa tac atg tac taa ttc ttg gat ata tga tat acc aga gct aaa aat atc atc gtt cta att gcg gag gag ttg cca atg acg aat aca ttt att gta tag gcg gta tac gcg atc agg agt cat cgt tga tat cta gta tcg ata gat gga agc cat caa aac cat att ggc aga agt atg cta aaa tgt gcg aac caa aat gtg ata tgg ggt tgc gat ttt aaa cgg att aat ata tgt cat agg tgg agt cgt taa agg tga cac aca tat acc aac gca cta gag agt tta tca gaa gat gga tgg atg aat cat caa cgt ctt cca ata aaa atg tcc aat atg tcg acg att gtt cat gct gga aag att tat ata tct aga ggt tac aac aat agt agt gta gtt aat gta ata tcg aat ct- -ag tcc tta gct ata atc cga tat atg atg aat gga cca aat tat cat cat taa ata ttc cta gaa tta atc ctg ctc tat ggt cag tgt ata ata aat tat atg tag gag gag taa tat ctg atg atg ttc aaa cta ata cat ctg aaa cat acg ata aag aaa aa

>ON676703.1_Monkeypox_virus_isolate_MPXV_USA_2022_CA001_complete_genome

tgt tat tgt agt atg ata ata tca aaa aga tgg ata taa aga att tac tga cta cat gta cta ttt tac att act aca ttg gct acg gca tat ata cct att tcg tca ctt cca cac gct ccg gta aac ggg tgt cat gtg acg agg gag aat ctt gat aag agg cat aat caa tgt tgt aat ccg atg tcc acc tgg aga att tgc caa ggt cag atg tag agt tgg tag tga taa cac aaa atg tga aca ctg ccc acc tca tac ata tac cgc aat ccc caa tta ttc taa tag atg tca tca atg tag aaa atg ccc aac agg atc att tga taa ggt aaa gtg tac cgg aac aca gaa caa at- gtt cgt gtc atc ctg gtt ggt ata cgc tac tga ttc ttc aca gac tga aga ttg tcg aga ttt gtg tac caa aaa agg aga tgt cca tgc gga tac ttt ggt gga ata gat gaa gga aat cct att tgt aaa tcg tgt tgt gtt ggt gaa tat tgc gac tac cta cgt aat tat aga ctt gat cca ttt cct cca tgc aaa cta tct atc taa atg taa tta att atg att ttg atg ata atg tta cca tac att ata tca cta ctt ggt tag tgt gta tta ttt agt atg gaa gac cta tta ata att act tat ctt ttg acg atc ttg tta taa tta taa tat aaa aat act tat gac ata gta a-- --- -ct cat aat tgc tga cgc gat aaa ttc gta ata atc tgt ttt gtt caa att ttt ata agg aat cta cag gca taa aaa taa aaa tat aat cta taa tat act ctt aca acg cca tca tga ata gca gtg aat taa ttg ctg tta tta tgg att tag aaa tag tgg acg att ttg tga tat taa tat agt tat taa tga tga aag gat aaa cgc gca tag act cat cct atc tgg agc ctc cga ata ttt ttt cca ttc tgt ttt cca ata att tta tcg att cta atg aat acg aag tta atc taa gtc att tag att atc aaa gtg tta act att tga tcg att aca ttt atg gga tac ctt tga gcc taa cta aca ata acg tga aat ata ttc ttt caa ccg ctg att ttt tta caa att gga tct gtc att act gag tgc gaa aaa tac ata ctt aaa aat ctt tgt ttt aga aac tgt atc gat ttc --- --- tac ata tac gct gat aaa tat aat aac aag aaa ata gaa tta gca tcg ttt aac aca ata tta cga aat att ttg aga ctc atc aac aat gaa aac ttt aaa tac tta aca gag gaa tca atg ata aaa att tta agc gat gat atg tta tat ata aaa aat gag gat ttc acc cca ctg att ctc att aaa tgg tta gag agt aca cca acc atg tac cgt cga gtt act tag atg cct cag aat atc a-t ttc ttt ccc cac aag tta taa aat cac ttt ata gtc atc gac tgg tta gtt caa tct acg aat gta taa cat tct taa aca ata tag cat tct tgg ata aat cat ttc cta gat acc ata tca tcg agt tga tat cta tcg gta taa gta att cac atg ata aga ttt cca taa act gct aca atc ata aaa aaa ttc atg gga aat gat atc ttc acg tag ata tag gtg tag ttt cgc agt gac cgt cct gga taa tat tat cta tat gat ggg tgg ata tga tca gtc cct gta tag aag ttc aaa ggt tat agc gta caa tac atg tac taa ttc ttg gat ata tga tat acc aga gct aaa aat atc atc gtt cta att gcg gag gag ttg cca atg acg aat aca ttt att gta tag gcg gta tac gcg atc agg agt cat cgt tga tat cta gta tcg ata gat gga agc cat caa aac cat att ggc aga agt atg cta aaa tgt gcg aac caa aat gtg ata tgg ggt tgc gat ttt aaa cgg att aat ata tgt cat agg tgg agt cgt taa agg tga cac aca tat acc aac gca cta gag agt tta tca gaa gat gga tgg atg aat cat caa cgt ctt cca ata aaa atg tcc aat atg tcg acg att gtt cat gct gga aag att tat ata tct aga ggt tac aac aat agt agt gta gtt aat gta ata tcg aat ct- -ag tcc tta gct ata atc cga tat atg atg aat gga cca aat tat cat cat taa ata ttc cta gaa tta atc ctg ctc tat ggt cag tgt ata ata aat tat atg tag gag gag taa tat ctg atg atg ttc aaa cta ata cat ctg aaa cat acg ata aag aaa aa

>ON676706.1_Monkeypox_virus_isolate_MPXV_USA_2022_UT002_complete_genome

tgt tat tgt agt atg ata ata tca aaa aga tgg ata taa aga att tac tga cta cat gta cta ttt tac att act aca ttg gct acg gca tat ata cct att tcg tca ctt cca cac gct ccg gta aac ggg tgt cat gtg acg agg gag aat ctt gat aag agg cat aat caa tgt tgt aat ccg atg tcc acc tgg aga att tgc caa ggt cag atg tag agt tgg tag tga taa cac aaa atg tga aca ctg ccc acc tca tac ata tac cgc aat ccc caa tta ttc taa tag atg tca tca atg tag aaa atg ccc aac agg atc att tga taa ggt aaa gtg tac cgg aac aca gaa caa at- gtt cgt gtc atc ctg gtt ggt ata cgc tac tga ttc ttc aca gac tga aga ttg tcg aga ttt gtg tac caa aaa agg aga tgt cca tgc gga tac ttt ggt gga ata gat gaa gga aat cct att tgt aaa tcg tgt tgt gtt ggt gaa tat tgc gac tac cta cgt aat tat aga ctt gat cca ttt cct cca tgc aaa cta tct atc taa atg taa tta att atg att ttg atg ata atg tta cca tac att ata tca cta ctt ggt tag tgt gta tta ttt agt atg gaa gac cta tta ata att act tat ctt ttg acg atc ttg tta taa tta taa tat aaa aat act tat gac ata gta a-- --- -ct cat aat tgc tga cgc gat aaa ttc gta ata atc tgt ttt gtt caa att ttt ata agg aat cta cag gca taa aaa taa aaa tat aat cta taa tat act ctt aca acg cca tca tga ata gca gtg aat taa ttg ctg tta tta tgg att tag aaa tag tgg acg att ttg tga tat taa tat agt tat taa tga tga aag gat aaa cgc gca tag act cat cct atc tgg agc ctc cga ata ttt ttt cca ttc tgt ttt cca ata att tta tcg att cta atg aat acg aag tta atc taa gtc att tag att atc aaa gtg tta act att tga tcg att aca ttt atg gga tac ctt tga gcc taa cta aca ata acg tga aat ata ttc ttt caa ccg ctg att ttt tta caa att gga tct gtc att act gag tgc gaa aaa tac ata ctt aaa aat ctt tgt ttt aga aac tgt atc gat ttc --- --- tac ata tac gct gat aaa tat aat aac aag aaa ata gaa tta gca tcg ttt aac aca ata tta cga aat att ttg aga ctc atc aac aat gaa aac ttt aaa tac tta aca gag gaa tca atg ata aaa att tta agc gat gat atg tta tat ata aaa aat gag gat ttc acc cca ctg att ctc att aaa tgg tta gag agt aca cca acc atg tac cgt cga gtt act tag atg cct cag aat atc a-t ttc ttt ccc cac aag tta taa aat cac ttt ata gtc atc gac tgg tta gtt caa tct acg aat gta taa cat tct taa aca ata tag cat tct tgg ata aat cat ttc cta gat acc ata tca tcg agt tga tat cta tcg gta taa gta att cac atg ata aga ttt cca taa act gct aca atc ata aaa aaa ttc atg gga aat gat atc ttc acg tag ata tag gtg tag ttt cgc agt gac cgt cct gga taa tat tat cta tat gat ggg tgg ata tga tca gtc cct gta tag aag ttc aaa ggt tat agc gta caa tac atg tac taa ttc ttg gat ata tga tat acc aga gct aaa aat atc atc gtt cta att gcg gag gag ttg cca atg acg aat aca ttt att gta tag gcg gta tac gcg atc agg agt cat cgt tga tat cta gta tcg ata gat gga agc cat caa aac cat att ggc aga agt atg cta aaa tgt gcg aac caa aat gtg ata tgg ggt tgc gat ttt aaa cgg att aat ata tgt cat agg tgg agt cgt taa agg tga cac aca tat acc aac gca cta gag agt tta tca gaa gat gga tgg atg aat cat caa cgt ctt cca ata aaa atg tcc aat atg tcg acg att gtt cat gct gga aag att tat ata tct aga ggt tac aac aat agt agt gta gtt aat gta ata tcg aat ct- -ag tcc tta gct ata atc cga tat atg atg aat gga cca aat tat cat cat taa ata ttc cta gaa tta atc ctg ctc tat ggt cag tgt ata ata aat tat atg tag gag gag taa tat ctg atg atg ttc aaa cta ata cat ctg aaa cat acg ata aag aaa aa

>ON649879.1_Monkeypox_virus_isolate_MPXV_ISR001_2022_complete_genome

tgt tat tgt agt atg ata ata tca aaa aga tgg ata taa aga att tac tga cta cat gta cta ttt tac att act aca ttg gct acg gca tat ata cct att tcg tca ctt cca cac gct ccg gta aac ggg tgt cat gtg acg agg gag aat ctt gat aag agg cat aat caa tgt tgt aat ccg atg tcc acc tgg aga att tgc caa ggt cag atg tag agt tgg tag tga taa cac aaa atg tga aca ctg ccc acc tca tac ata tac cgc aat ccc caa tta ttc taa tag atg tca tca atg tag aaa atg ccc aac agg atc att tga taa ggt aaa gtg tac cgg aac aca gaa caa at- gtt cgt gtc atc ctg gtt ggt ata cgc tac tga ttc ttc aca gac tga aga ttg tcg aga ttt gtg tac caa aaa agg aga tgt cca tgc gga tac ttt ggt gga ata gat gaa gga aat cct att tgt aaa tcg tgt tgt gtt ggt gaa tat tgc gac tac cta cgt aat tat aga ctt gat cca ttt cct cca tgc aaa cta tct atc taa atg taa tta att atg att ttg atg ata atg tta cca tac att ata tca cta ctt ggt tag tgt gta tta ttt agt atg gaa gac cta tta ata att act tat ctt ttg acg atc ttg tta taa tta taa tat aaa aat act tat gac ata gta a-- --- -ct cat aat tgc tga cgc gat aaa ttc gta ata atc tgt ttt gtt caa att ttt ata agg aat cta cag gca taa aaa taa aaa tat aat cta taa tat act ctt aca acg cca tca tga ata gca gtg aat taa ttg ctg tta tta tgg att tag aaa tag tgg acg att ttg tga tat taa tat agt tat taa tga tga aag gat aaa cgc gca tag act cat cct atc tgg agc ctc cga ata ttt ttt cca ttc tgt ttt cca ata att tta tcg att cta atg aat acg aag tta atc taa gtc att tag att atc aaa gtg tta act att tga tcg att aca ttt atg gga tac ctt tga gcc taa cta aca ata acg tga aat ata ttc ttt caa ccg ctg att ttt tta caa att gga tct gtc att act gag tgc gaa aaa tac ata ctt aaa aat ctt tgt ttt aga aac tgt atc gat ttc --- --- tac ata tac gct gat aaa tat aat aac aag aaa ata gaa tta gca tcg ttt aac aca ata tta cga aat att ttg aga ctc atc aac aat gaa aac ttt aaa tac tta aca gag gaa tca atg ata aaa att tta agc gat gat atg tta tat ata aaa aat gag gat ttc acc cca ctg att ctc att aaa tgg tta gag agt aca cca acc atg tac cgt cga gtt act tag atg cct cag aat atc a-t ttc ttt ccc cac aag tta taa aat cac ttt ata gtc atc gac tgg tta gtt caa tct acg aat gta taa cat tct taa aca ata tag cat tct tgg ata aat cat ttc cta gat acc ata tca tcg agt tga tat cta tcg gta taa gta att cac atg ata aga ttt cca taa act gct aca atc ata aaa aaa ttc atg gga aat gat atc ttc acg tag ata tag gtg tag ttt cgc agt gac cgt cct gga taa tat tat cta tat gat ggg tgg ata tga tca gtc cct gta tag aag ttc aaa ggt tat agc gta caa tac atg tac taa ttc ttg gat ata tga tat acc aga gct aaa aat atc atc gtt cta att gcg gag gag ttg cca atg acg aat aca ttt att gta tag gcg gta tac gcg atc agg agt cat cgt tga tat cta gta tcg ata gat gga agc cat caa aac cat att ggc aga agt atg cta aaa tgt gcg aac caa aat gtg ata tgg ggt tgc gat ttt aaa cgg att aat ata tgt cat agg tgg agt cgt taa agg tga cac aca tat acc aac gca cta gag agt tta tca gaa gat gga tgg atg aat cat caa cgt ctt cca ata aaa atg tcc aat atg tcg acg att gtt cat gct gga aag att tat ata tct aga ggt tac aac aat agt agt gta gtt aat gta ata tcg aat ct- -ag tcc tta gct ata atc cga tat atg atg aat gga cca aat tat cat cat taa ata ttc cta gaa tta atc ctg ctc tat ggt cag tgt ata ata aat tat atg tag gag gag taa tat ctg atg atg ttc aaa cta ata cat ctg aaa cat acg ata aag aaa aa

>ON602722.1_Monkeypox_virus_isolate_MPXV_FRA_2022_TLS67_complete_genome

tgt tat tgt agt atg ata ata tc- aaa aga tgg ata taa aga att tac tga cta cat gta cta ttt tac att act aca ttg gct acg gca tat ata cct att tcg tca ctt cca cac gct ccg gta aac ggg tgt cat gtg acg agg gag aat ctt gat aag agg cat aat caa tgt tgt aat ccg atg tcc acc tgg aga att tgc caa ggt cag atg tag agt tgg tag tga taa cac aaa atg tga aca ctg ccc acc tca tac ata tac cgc aat ccc caa tta ttc taa tag atg tca tca atg tag aaa atg ccc aac agg atc att tga taa ggt aaa gtg tac cgg aac aca gaa caa at- gtt cgt gtc atc ctg gtt ggt ata cgc tac tga ttc ttc aca gac tga aga ttg tcg aga ttt gtg tac c-a aaa agg aga tgt cca tgc gga tac ttt ggt gga ata gat gaa gga aat cct att tgt aaa tcg tgt tgt gtt ggt gaa tat tgc gac tac cta cgt aat tat aga ctt gat cca ttt cct cca tgc aaa cta tct atc taa atg taa tta att atg att ttg atg ata atg tta cca tac att ata tca cta ctt ggt tag tgt gta tta ttt agt atg gaa gac cta tta ata att act tat ctt ttg acg atc ttg tta taa tta taa tat aaa aat act tat gac ata gta a-- --- -ct cat aat tgc tga cgc gat aaa ttc gta ata atc tgt ttt gtt caa att ttt ata agg aat cta cag gca taa aaa taa aaa tat aat cta taa tat act ctt aca acg cca tca tga ata gca gtg aat taa ttg ctg tta tta tgg att tag aaa tag tgg acg att ttg tga tat taa tat agt tat taa tga tga aag gat aaa cgc gca tag act cat cct atc tgg agc ctc cga ata ttt ttt cca ttc tgt ttt cca ata att tta tcg att cta atg aat acg aag tta atc taa gtc att tag att atc aaa gtg tta act att tga tcg att aca ttt atg gga tac ctt tga gcc taa cta aca ata acg tga aat ata ttc ttt caa ccg ctg att ttt tta caa att gga tct gtc att act gag tgc gaa aaa tac ata ctt aaa aat ctt tgt ttt aga aac tgt atc gat ttc --- --- tac ata tac gct gat aaa tat aat aac aag aaa ata gaa tta gca tcg ttt aac aca ata tta cga aat att ttg aga ctc atc aac aat gaa aac ttt aaa tac tta aca gag gaa tca atg ata aaa att tta agc gat gat atg tta tat ata aaa aat gag gat ttc acc cca ctg att ctc att aaa tgg tta gag agt aca cca acc atg tac cgt cga gtt act tag atg cct cag aat atc a-t ttc ttt ccc cac aag tta taa aat cac ttt ata gtc atc gac tgg tta gtt caa tct acg aat gta taa cat tct taa aca ata tag cat tct tgg ata aat cat ttc cta gat acc ata tca tcg agt tga tat cta tcg gta taa gta att cac atg ata aga ttt cca taa act gct aca atc at- aaa aaa ttc atg gga aat gat atc ttc acg tag ata tag gtg tag ttt cgc agt gac cgt cct gga taa tat tat cta tat gat ggg tgg ata tga tca gtc cct gta tag aag ttc aaa ggt tat agc gta caa tac atg tac taa ttc ttg gat ata tga tat acc aga gct aaa aat atc atc gtt cta att gcg gag gag ttg cca atg acg aat aca ttt att gta tag gcg gta tac gcg atc a?g agt cat cgt tga tat cta gta tcg ata gat gga agc cat caa aac cat att ggc aga agt atg cta aaa tgt gcg aac caa aat gtg ata tgg ggt tgc gat ttt aaa cgg att aat ata tgt cat agg tgg agt cgt taa agg tga cac aca tat acc aac gca cta gag agt tta tca gaa gat gga tgg atg aat cat caa cgt ctt cca ata aaa atg tcc aat atg tcg acg att gtt cat gct gga aag att tat ata tct aga ggt tac aac aat agt agt gta gtt aat gta ata tcg aat ct- -ag tcc tta gct ata atc cga tat atg atg aat gga cca aat tat cat cat taa ata ttc cta gaa tta atc ctg ctc tat ggt cag tgt ata ata aat tat atg tag gag gag taa tat ctg atg atg ttc aaa cta ata cat ctg aaa cat acg ata aag aaa aa

>ON675438.1_Monkeypox_virus_isolate_MPXV_USA_2022_VA001_complete_genome

tgt tat tgt agt atg ata ata tca aaa aga tgg ata taa aga att tac tga cta cat gta cta ttt tac att act aca ttg gct acg gca tat ata cct att ttg tca ctt cca cac gct ccg gta aac ggg tgt cat gtg acg agg gag aat ctt gat aag agg cat aat caa tgt tgt aat ccg atg tcc acc tgg aga att tgc caa ggt cag atg tag agt tgg tag tga taa cac aaa atg tga aca ctg ccc acc tca tac ata tac cgc aat ccc caa tta ttc taa tag atg tca tca atg tag aaa atg ccc aac agg atc att tga taa ggt aaa gtg tac cgg aac aca gaa caa at- gtt cgt gtc atc ctg gtt ggt ata cgc tac tga ttc ttc aca gac tga aga ttg tcg aga ttt gtg tac caa aaa agg aga tgt cca tgc gga tac ttt ggt gga ata gat gaa gga aat cct att tgt aaa tcg tgt tgt gtt ggt gaa tat tgc gac tac cta cgt aat tat aga ctt gat cca ttt cct cca tgc aaa cta tct atc taa atg taa tta att atg att ttg atg ata atg tta cca tac att ata tca cta ctt ggt tag tgt gta tta ttt agt atg gaa gac cta tta ata att act tat ctt ttg acg atc ttg tta taa tta taa tat aaa aat act tat gac ata gta a-- --- -ct cat aat tgc tga cgc gat aaa ttc gta ata atc tgt ttt gtt caa att ttt ata agg aat cta cag gca taa aaa taa aaa tat aat cta taa tat act ctt aca acg cca tca tga ata gca gtg aat taa ttg ctg tta tta tgg att tag aaa tag tgg acg att ttg tga tat taa tat agt tat taa tga tga aag gat aaa cgc gca tag act cat cct atc tgg agc ctc cga ata ttt ttt cca ttc tgt ttt cca ata att tta tcg att cta atg aat acg aag tta atc taa gtc att tag att atc aaa gtg tta act att tga tcg att aca ttt atg gga tac ctt tga gcc taa cta aca ata acg tga aat ata ttc ttt caa ccg ctg att ttt tta caa att gga tct gtc att act gag tgc gaa aaa tac ata ctt aaa aat ctt tgt ttt aga aac tgt atc gat ttc --- --- tac ata tac gct gat aaa tat aat aac aag aaa ata gaa tta gca tcg ttt aac aca ata tta cga aat att ttg aga ctc atc aac aat gaa aac ttt aaa tac tta aca gag gaa tca atg ata aaa att tta agc gat gat atg tta tat ata aaa aat gag gat ttc acc cca ctg att ctc att aaa tgg tta gag agt aca cca acc atg tac cgt cga gtt act tag atg cct cag aat atc a-t ttc ttt ccc cac aag tta taa aat cac ttt ata gtc atc gac tgg tta gtt caa tct acg aat gta taa cat tct taa aca ata tag cat tct tgg ata aat cat ttc cta gat acc ata tca tcg agt tga tat cta tcg gta taa gta att cac atg ata aga ttt cca taa act gct aca atc ata aaa aaa ttc atg gga aat gat atc ttc acg tag ata tag gtg tag ttt cgc agt gac cgt cct gga taa tat tat cta tat gat ggg tgg ata tga tca gtc cct gta tag aag ttc aaa ggt tat agc gta caa tac atg tac taa ttc ttg gat ata tga tat acc aga gct aaa aat atc atc gtt cta att gcg gag gag ttg cca atg acg aat aca ttt att gta tag gcg gta tac gcg atc agg agt cat cgt tga tat cta gta tcg ata gat gga agc cat caa aac cat att ggc aga agt atg cta aaa tgt gcg aac caa aat gtg ata tgg ggt tgc gat ttt aaa cgg att aat ata tgt cat agg tgg agt cgt taa agg tga cac aca tat acc aac gca cta gag agt tta tca gaa gat gga tgg atg aat cat caa cgt ctt cca ata aaa atg tcc aat atg tcg acg att gtt cat gct gga aag att tat ata tct aga ggt tac aac aat agt agt gta gtt aat gta ata tcg aat ct- -ag tcc tta gct ata atc cga tat atg atg aat gga cca aat tat cat cat taa ata ttc cta gaa tta atc ctg ctc tat ggt cag tgt ata ata aat tat atg tag gag gag taa tat ctg atg atg ttc aaa cta ata cat ctg aaa cat acg ata aag aaa aa

>ON631241.1_Monkeypox_virus_isolate_2022/2_SLO_complete_genome

tgt tat tgt agt atg ata ata tca aaa aga tgg ata taa aga att tac tga cta cat gta cta ttt tac att act aca ttg gct acg gca tat ata cct att tcg tca ctt cca cac gct ccg gta aac ggg tgt cat gtg acg agg gag aat ctt gat aag agg cat aat caa tgt tgt aat ccg atg tcc acc tgg aga att tgc caa ggt cag atg tag agt tgg tag tga taa cac aaa atg tga aca ctg ccc acc tca tac ata tac cgc aat ccc caa tta ttc taa tag atg tca tca atg tag aaa atg ccc aac agg atc att tga taa ggt aaa gtg tac cgg aac aca gaa caa at- gtt cgt gtc atc ctg gtt ggt ata cgc tac tga ttc ttc aca gac tga aga ttg tcg aga ttt gtg tac caa aaa agg aga tgt cca tgc gga tac ttt ggt gga ata gat gaa gga aat cct att tgt aaa tcg tgt tgt gtt ggt gaa tat tgc gac tac cta cgt aat tat aga ctt gat cca ttt cct cca tgc aaa cta tct atc taa atg taa tta att atg att ttg atg ata atg tta cca tac att ata tca cta ctt ggt tag tgt gta tta ttt agt atg gaa gac cta tta ata att act tat ctt ttg acg atc ttg tta taa tta taa tat aaa aat act tat gac ata gta a-- --- -ct cat aat tgc tga cgc gat aaa ttc gta ata atc tgt ttt gtt caa att ttt ata agg aat cta cag gca taa aaa taa aaa tat aat cta taa tat act ctt aca acg cca tca tga ata gca gtg aat taa ttg ctg tta tta tgg att tag aaa tag tgg acg att ttg tga tat taa tat agt tat taa tga tga aag gat aaa cgc gca tag act cat cct atc tgg agc ctc cga ata ttt ttt cca ttc tgt ttt cca ata att tta tcg att cta atg aat acg aag tta atc taa gtc att tag att atc aaa gtg tta act att tga tcg att aca ttt atg gga tac ctt tga gcc taa cta aca ata acg tga aat ata ttc ttt caa ccg ctg att ttt tta caa att gga tct gtc att act gag tgc gaa aaa tac ata ctt aaa aat ctt tgt ttt aga aac tgt atc gat ttc --- --- tac ata tac gct gat aaa tat aat aac aag aaa ata gaa tta gca tcg ttt aac aca ata tta cga aat att ttg aga ctc atc aac aat gaa aac ttt aaa tac tta aca gag gaa tca atg ata aaa att tta agc gat gat atg tta tat ata aaa aat gag gat ttc acc cca ctg att ctc att aaa tgg tta gag agt aca cca acc atg tac cgt cga gtt act tag atg cct cag aat atc a-t ttc ttt ccc cac aag tta taa aat cac ttt ata gtc atc gac tgg tta gtt caa tct acg aat gta taa cat tct taa aca ata tag cat tct tgg ata aat cat ttc cta gat acc ata tca tcg agt tga tat cta tcg gta taa gta att cac atg ata aga ttt cca taa act gct aca atc ata aaa aaa ttc atg gga aat gat atc ttc acg tag ata tag gtg tag ttt cgc agt gac cgt cct gga taa tat tat cta tat gat ggg tgg ata tga tca gtc cct gta tag aag ttc aaa ggt tat agc gta caa tac atg tac taa ttc ttg gat ata tga tat acc aga gct aaa aat atc atc gtt cta att gcg gag gag ttg cca atg acg aat aca ttt att gta tag gcg gta tac gcg atc agg agt cat cgt tga tat cta gta tcg ata gat gga agc cat caa aac cat att ggc aga agt atg cta aaa tgt gcg aac caa aat gtg ata tgg ggt tgc gat ttt aaa cgg att aat ata tgt cat agg tgg agt cgt taa agg tga cac aca tat acc aac gca cta gag agt tta tca gaa gat gga tgg atg aat cat caa cgt ctt cca ata aaa atg tcc aat atg tcg acg att gtt cat gct gga aag att tat ata tct aga ggt tac aac aat agt agt gta gtt aat gta ata tcg aat ct- -ag tcc tta gct ata atc cga tat atg atg aat gga cca aat tat cat cat taa ata ttc cta gaa tta atc ctg ctc tat ggt cag tgt ata ata aat tat atg tag gag gag taa tat ctg atg atg ttc aaa cta ata cat ctg aaa cat acg ata aag aaa aa

>ON622712.1_Monkeypox_virus_isolate_MPX/UZ_REGA_1/Belgium/2022_complete_genome

tgt tat tgt agt atg ata ata tca aaa aga tgg ata taa aga att tac tga cta cat gta cta ttt tac att act aca ttg gct acg gca tat ata cct att tcg tca ctt cca cac gct ccg gta aac ggg tgt cat gtg acg agg gag aat ctt gat aag agg cat aat caa tgt tgt aat ccg atg tcc acc tgg aga att tgc caa ggt cag atg tag agt tgg tag tga taa cac aaa atg tga aca ctg ccc acc tca tac ata tac cgc aat ccc caa tta ttc taa tag atg tca tca atg tag aaa atg ccc aac agg atc att tga taa ggt aaa gtg tac cgg aac aca gaa caa at- gtt cgt gtc atc ctg gtt ggt ata cgc tac tga ttc ttc aca gac tga aga ttg tcg aga ttt gtg tac caa aaa agg aga tgt cca tgc gga tac ttt ggt gga ata gat gaa gga aat cct att tgt aaa tcg tgt tgt gtt ggt gaa tat tgc gac tac cta cgt aat tat aga ctt gat cca ttt cct cca tgc aaa cta tct atc taa atg taa tta att atg att ttg atg ata atg tta cca tac att ata tca cta ctt ggt tag tgt gta tta ttt agt atg gaa gac cta tta ata att act tat ctt ttg acg atc ttg tta taa tta taa tat aaa aat act tat gac ata gta a-- --- -ct cat aat tgc tga cgc gat aaa ttc gta ata atc tgt ttt gtt caa att ttt ata agg aat cta cag gca taa aaa taa aaa tat aat cta taa tat act ctt aca acg cca tca tga ata gca gtg aat taa ttg ctg tta tta tgg att tag aaa tag tgg acg att ttg tga tat taa tat agt tat taa tga tga aag gat aaa cgc gca tag act cat cct atc tgg agc ctc cga ata ttt ttt cca ttc tgt ttt cca ata att tta tcg att cta atg aat acg aag tta atc taa gtc att tag att atc aaa gtg tta act att tga tcg att aca ttt atg gga tac ctt tga gcc taa cta aca ata acg tga aat ata ttc ttt caa ccg ctg att ttt tta caa att gga tct gtc att act gag tgc gaa aaa tac ata ctt aaa aat ctt tgt ttt aga aac tgt atc gat ttc --- --- tac ata tac gct gat aaa tat aat aac aag aaa ata gaa tta gca tcg ttt aac aca ata tta cga aat att ttg aga ctc atc aac aat gaa aac ttt aaa tac tta aca gag gaa tca atg ata aaa att tta agc gat gat atg tta tat ata aaa aat gag gat ttc acc cca ctg att ctc att aaa tgg tta gag agt aca cca acc atg tac cgt cga gtt act tag atg cct cag aat atc a-t ttc ttt ccc cac aag tta taa aat cac ttt ata gtc atc gac tgg tta gtt caa tct acg aat gta taa cat tct taa aca ata tag cat tct tgg ata aat cat ttc cta gat acc ata tca tcg agt tga tat cta tcg gta taa gta att cac atg ata aga ttt cca taa act gct aca atc ata aaa aaa ttc atg gga aat gat atc ttc acg tag ata tag gtg tag ttt cgc agt gac cgt cct gga taa tat tat cta tat gat ggg tgg ata tga tca gtc cct gta tag aag ttc aaa ggt tat agc gta caa tac atg tac taa ttc ttg gat ata tga tat acc aga gct aaa aat atc atc gtt cta att gcg gag gag ttg cca atg acg aat aca ttt att gta tag gcg gta tac gcg atc agg agt cat cgt tga tat cta gta tcg ata gat gga agc cat caa aac cat att ggc aga agt atg cta aaa tgt gcg aac caa aat gtg ata tgg ggt tgc gat ttt aaa cgg att aat ata tgt cat agg tgg agt cgt taa agg tga cac aca tat acc aac gca cta gag agt tta tca gaa gat gga tgg atg aat cat caa cgt ctt cca ata aaa atg tcc aat atg tcg acg att gtt cat gct gga aag att tat ata tct aga ggt tac aac aat agt agt gta gtt aat gta ata tcg aat ct- -ag tcc tta gct ata atc cga tat atg atg aat gga cca aat tat cat cat taa ata ttc cta gaa tta atc ctg ctc tat ggt cag tgt ata ata aat tat atg tag gag gag taa tat ctg atg atg ttc aaa cta ata cat ctg aaa cat acg ata aag aaa aa

>ON609725.2_Monkeypox_virus_isolate_SLO_complete_genome

tgt tat tgt agt atg ata ata tca aaa aga tgg ata taa aga att tac tga cta cat gta cta ttt tac att act aca ttg gct acg gca tat ata cct att tcg tca ctt cca cac gct ccg gta aac ggg tgt cat gtg acg agg gag aat ctt gat aag agg cat aat caa tgt tgt aat ccg atg tcc acc tgg aga att tgc caa ggt cag atg tag agt tgg tag tga taa cac aaa atg tga aca ctg ccc acc tca tac ata tac cgc aat ccc caa tta ttc taa tag atg tca tca atg tag aaa atg ccc aac agg atc att tga taa ggt aaa gtg tac cgg aac aca gaa caa at- gtt cgt gtc atc ctg gtt ggt ata cgc tac tga ttc ttc aca gac tga aga ttg tcg aga ttt gtg tac caa aaa agg aga tgt cca tgc gga tac ttt ggt gga ata gat gaa gga aat cct att tgt aaa tcg tgt tgt gtt ggt gaa tat tgc gac tac cta cgt aat tat aga ctt gat cca ttt cct cca tgc aaa cta tct atc taa atg taa tta att atg att ttg atg ata atg tta cca tac att ata tca cta ctt ggt tag tgt gta tta ttt agt atg gaa gac cta tta ata att act tat ctt ttg acg atc ttg tta taa tta taa tat aaa aat act tat gac ata gta a-- --- -ct cat aat tgc tga cgc gat aaa ttc gta ata atc tgt ttt gtt caa att ttt ata agg aat cta cag gca taa aaa taa aaa tat aat cta taa tat act ctt aca acg cca tca tga ata gca gtg aat taa ttg ctg tta tta tgg att tag aaa tag tgg acg att ttg tga tat taa tat agt tat taa tga tga aag gat aaa cgc gca tag act cat cct atc tgg agc ctc cga ata ttt ttt cca ttc tgt ttt cca ata att tta tcg att cta atg aat acg aag tta atc taa gtc att tag att atc aaa gtg tta act att tga tcg att aca ttt atg gga tac ctt tga gcc taa cta aca ata acg tga aat ata ttc ttt caa ccg ctg att ttt tta caa att gga tct gtc att act gag tgc gaa aaa tac ata ctt aaa aat ctt tgt ttt aga aac tgt atc gat ttc --- --- tac ata tac gct gat aaa tat aat aac aag aaa ata gaa tta gca tcg ttt aac aca ata tta cga aat att ttg aga ctc atc aac aat gaa aac ttt aaa tac tta aca gag gaa tca atg ata aaa att tta agc gat gat atg tta tat ata aaa aat gag gat ttc acc cca ctg att ctc att aaa tgg tta gag agt aca cca acc atg tac cgt cga gtt act tag atg cct cag aat atc a-t ttc ttt ccc cac aag tta taa aat cac ttt ata gtc atc gac tgg tta gtt caa tct acg aat gta taa cat tct taa aca ata tag cat tct tgg ata aat cat ttc cta gat acc ata tca tcg agt tga tat cta tcg gta taa gta att cac atg ata aga ttt cca taa act gct aca atc ata aaa aaa ttc atg gga aat gat atc ttc acg tag ata tag gtg tag ttt cgc agt gac cgt cct gga taa tat tat cta tat gat ggg tgg ata tga tca gtc cct gta tag aag ttc aaa ggt tat agc gta caa tac atg tac taa ttc ttg gat ata tga tat acc aga gct aaa aat atc atc gtt cta att gcg gag gag ttg cca atg acg aat aca ttt att gta tag gcg gta tac gcg atc agg agt cat cgt tga tat cta gta tcg ata gat gga agc cat caa aac cat att ggc aga agt atg cta aaa tgt gcg aac caa aat gtg ata tgg ggt tgc gat ttt aaa cgg att aat ata tgt cat agg tgg agt cgt taa agg tga cac aca tat acc aac gca cta gag agt tta tca gaa gat gga tgg atg aat cat caa cgt ctt cca ata aaa atg tcc aat atg tcg acg att gtt cat gct gga aag att tat ata tct aga ggt tac aac aat agt agt gta gtt aat gta ata tcg aat ct- -ag tcc tta gct ata atc cga tat atg atg aat gga cca aat tat cat cat taa ata ttc cta gaa tta atc ctg ctc tat ggt cag tgt ata ata aat tat atg tag gag gag taa tat ctg atg atg ttc aaa cta ata cat ctg aaa cat acg ata aag aaa aa

>ON622713.1_Monkeypox_virus_isolate_MPX/UZ_REGA_2/Belgium/2022_complete_genome

tgt tat tgt agt atg ata ata tca aaa aga tgg ata taa aga att tac tga cta cat gta cta ttt tac att act aca ttg gct acg gca tat ata cct att tcg tca ctt cca cac gct ccg gta aac ggg tgt cat gtg acg agg gag aat ctt gat aag agg cat aat caa tgt tgt aat ccg atg tcc acc tgg aga att tgc caa ggt cag atg tag agt tgg tag tga taa cac aaa atg tga aca ctg ccc acc tca tac ata tac cgc aat ccc caa tta ttc taa tag atg tca tca atg tag aaa atg ccc aac agg atc att tga taa ggt aaa gtg tac cgg aac aca gaa caa at- gtt cgt gtc atc ctg gtt ggt ata cgc tac tga ttc ttc aca gac tga aga ttg tcg aga ttt gtg tac caa aaa agg aga tgt cca tgc gga tac ttt ggt gga ata gat gaa gga aat cct att tgt aaa tcg tgt tgt gtt ggt gaa tat tgc gac tac cta cgt aat tat aga ctt gat cca ttt cct cca tgc aaa cta tct atc taa atg taa tta att atg att ttg atg ata atg tta cca tac att ata tca cta ctt ggt tag tgt gta tta ttt agt atg gaa gac cta tta ata att act tat ctt ttg acg atc ttg tta taa tta taa tat aaa aat act tat gac ata gta a-- --- -ct cat aat tgc tga cgc gat aaa ttc gta ata atc tgt ttt gtt caa att ttt ata agg aat cta cag gca taa aaa taa aaa tat aat cta taa tat act ctt aca acg cca tca tga ata gca gtg aat taa ttg ctg tta tta tgg att tag aaa tag tgg acg att ttg tga tat taa tat agt tat taa tga tga aag gat aaa cgc gca tag act cat cct atc tgg agc ctc cga ata ttt ttt cca ttc tgt ttt cca ata att tta tcg att cta atg aat acg aag tta atc taa gtc att tag att atc aaa gtg tta act att tga tcg att aca ttt atg gga tac ctt tga gcc taa cta aca ata acg tga aat ata ttc ttt caa ccg ctg att ttt tta caa att gga tct gtc att act gag tgc gaa aaa tac ata ctt aaa aat ctt tgt ttt aga aac tgt atc gat ttc --- --- tac ata tac gct gat aaa tat aat aac aag aaa ata gaa tta gca tcg ttt aac aca ata tta cga aat att ttg aga ctc atc aac aat gaa aac ttt aaa tac tta aca gag gaa tca atg ata aaa att tta agc gat gat atg tta tat ata aaa aat gag gat ttc acc cca ctg att ctc att aaa tgg tta gag agt aca cca acc atg tac cgt cga gtt act tag atg cct cag aat atc a-t ttc ttt ccc cac aag tta taa aat cac ttt ata gtc atc gac tgg tta gtt caa tct acg aat gta taa cat tct taa aca ata tag cat tct tgg ata aat cat ttc cta gat acc ata tca tcg agt tga tat cta tcg gta taa gta att cac atg ata aga ttt cca taa act gct aca atc ata aaa aaa ttc atg gga aat gat atc ttc acg tag ata tag gtg tag ttt cgc agt gac cgt cct gga taa tat tat cta tat gat ggg tgg ata tga tca gtc cct gta tag aag ttc aaa ggt tat agc gta caa tac atg tac taa ttc ttg gat ata tga tat acc aga gct aaa aat atc atc gtt cta att gcg gag gag ttg cca atg acg aat aca ttt att gta tag gcg gta tac gcg atc agg agt cat cgt tga tat cta gta tcg ata gat gga agc cat caa aac cat att ggc aga agt atg cta aaa tgt gcg aac caa aat gtg ata tgg ggt tgc gat ttt aaa cgg att aat ata tgt cat agg tgg agt cgt taa agg tga cac aca tat acc aac gca cta gag agt tta tca gaa gat gga tgg atg aat cat caa cgt ctt cca ata aaa atg tcc aat atg tcg acg att gtt cat gct gga aag att tat ata tct aga ggt tac aac aat agt agt gta gtt aat gta ata tcg aat ct- -ag tcc tta gct ata atc cga tat atg atg aat gga cca aat tat cat cat taa ata ttc cta gaa tta atc ctg ctc tat ggt cag tgt ata ata aat tat atg tag gag gag taa tat ctg atg atg ttc aaa cta ata cat ctg aaa cat acg ata aag aaa aa

>ON676707.1_Monkeypox_virus_isolate_MPXV_USA_2021_TX_complete_genome

tgt tat tgt agt atg ata ata tca aaa aga tgg ata taa aga att tac tga cta cat gta cta ttt tac att act aca ttg gct acg gca tat ata cct att tcg tca ctt cca cac gct ccg gta aac ggg tgt cat gtg acg agg gag aat ctt gat aag agg cat aat caa tgt tgt aat ccg atg tcc acc tgg aga att tgc caa ggt cag atg tag agt tgg tag tga taa cac aaa atg tga aca ctg ccc acc tca tac ata tac cgc aat ccc caa tta ttc taa tag atg tca tca atg tag aaa atg ccc aac agg atc att tga taa ggt aaa gtg tac cgg aac aca gaa caa at- gtt cgt gtc atc ctg gtt ggt ata cgc tac tga ttc ttc aca gac tga aga ttg tcg aga ttt gtg tac caa aaa agg aga tgt cca tgc gga tac ttt ggt gga ata gat gaa gga aat cct att tgt aaa tcg tgt tgt gtt ggt gaa tat tgc gac tac cta cgt aat tat aga ctt gat cca ttt cct cca tgc aaa cta tct atc taa atg taa tta att atg att ttg atg ata atg tta cca tac att ata tca cta ctt ggt tag tgt gta tta ttt agt atg gaa gac cta tta ata att act tat ctt ttg acg atc ttg tta taa tta taa tat aaa aat act tat gac ata gta a-- --- -ct cat aat tgc tga cgc gat aaa ttc gta ata atc tgt ttt gtt caa att ttt ata agg aat cta cag gca taa aaa taa aaa tat aat cta taa tat act ctt aca acg cca tca tga ata gca gtg aat taa ttg ctg tta tta tgg att tag aaa tag tgg acg att ttg tga tat taa tat agt tat taa tga tga aag gat aaa cgc gca tag act cat cct atc tgg agc ctc cga ata ttt ttt cca ttc tgt ttt cca ata att tta tcg att cta atg aat acg aag tta atc taa gtc att tag att atc aaa gtg tta act att tga tcg att aca ttt atg gga tac ctt tga gcc taa cta aca ata acg tga aat ata ttc ttt caa ccg ctg att ttt tta caa att gga tct gtc att act gag tgc gaa aaa tac ata ctt aaa aat ctt tgt ttt aga aac tgt atc gat ttc --- --- tac ata tac gct gat aaa tat aat aac aag aaa ata gaa tta gca tcg ttt aac aca ata tta cga aat att ttg aga ctc atc aac aat gaa aac ttt aaa tac tta aca gag gaa tca atg ata aaa att tta agc gat gat atg tta tat ata aaa aat gag gat ttc acc cca ctg att ctc att aaa tgg tta gag agt aca cca acc atg tac cgt cga gtt act tag atg cct cag aat atc a-t ttc ttt ccc cac aag tta taa aat cac ttt ata gtc atc gac tgg tta gtt caa tct acg aat gta taa cat tct taa aca ata tag cat tct tgg ata aat cat ttc cta gat acc ata tca tcg agt tga tat cta tcg gta taa gta att cac atg ata aga ttt cca taa act gct aca atc ata aaa aaa ttc atg gga aat gat atc ttc acg tag ata tag gtg tag ttt cgc agt gac cgt cct gga taa tat tat cta tat gat ggg tgg ata tga tca gtc cct gta tag aag ttc aaa ggt tat agc gta caa tac atg tac taa ttc ttg gat ata tga tat acc aga gct aaa aat atc atc gtt cta att gcg gag gag ttg cca atg acg aat aca ttt att gta tag gcg gta tac gcg atc agg agt cat cgt tga tat cta gta tcg ata gat gga agc cat caa aac cat att ggc aga agt atg cta aaa tgt gcg aac caa aat gtg ata tgg ggt tgc gat ttt aaa cgg att aat ata tgt cat agg tgg agt cgt taa agg tga cac aca tat acc aac gca cta gag agt tta tca gaa gat gga tgg atg aat cat caa cgt ctt cca ata aaa atg tcc aat atg tcg acg att gtt cat gct gga aag att tat ata tct aga ggt tac aac aat agt agt gta gtt aat gta ata tcg aat ct- -ag tcc tta gct ata atc cga tat atg atg aat gga cca aat tat cat cat taa ata ttc cta gaa tta atc ctg ctc tat ggt cag tgt ata ata aat tat atg tag gag gag taa tat ctg atg atg ttc aaa cta ata cat ctg aaa cat acg ata aag aaa aa

>MT903345.1_Monkeypox_virus_isolate_MPXV-UK_P3

tgt tat tgt agt atg ata ata tca aaa aga tgg ata taa aga att tac tga cta cat gta cta ttt tac att act aca ttg gct acg gca tat ata cct att tcg tca ctt cca cac gct ccg gta aac ggg tgt cat gtg acg agg gag aat ctt gat aag agg cat aat caa tgt tgt aat ccg atg tcc acc tgg aga att tgc caa ggt cag atg tag agt tgg tag tga taa cac aaa atg tga aca ctg ccc acc tca tac ata tac cgc aat ccc caa tta ttc taa tag atg tca tca atg tag aaa atg ccc aac agg atc att tga taa ggt aaa gtg tac cgg aac aca gaa caa at- gtt cgt gtc atc ctg gtt ggt ata cgc tac tga ttc ttc aca gac tga aga ttg tcg aga ttt gtg tac caa aaa agg aga tgt cca tgc gga tac ttt ggt gga ata gat gaa gga aat cct att tgt aaa tcg tgt tgt gtt ggt gaa tat tgc gac tac cta cgt aat tat aga ctt gat cca ttt cct cca tgc aaa cta tct atc taa atg taa tta att atg att ttg atg ata atg tta cca tac att ata tca cta ctt ggt tag tgt gta tta ttt agt atg gaa gac cta tta ata att act tat ctt ttg acg atc ttg tta taa tta taa tat aaa aat act tat gac ata gta a-- --- -ct cat aat tgc tga cgc gat aaa ttc gta ata atc tgt ttt gtt caa att ttt ata agg aat cta cag gca taa aaa taa aaa tat aat cta taa tat act ctt aca acg cca tca tga ata gca gtg aat taa ttg ctg tta tta tgg att tag aaa tag tgg acg att ttg tga tat taa tat agt tat taa tga tga aag gat aaa cgc gca tag act cat cct atc tgg agc ctc cga ata ttt ttt cca ttc tgt ttt cca ata att tta tcg att cta atg aat acg aag tta atc taa gtc att tag att atc aaa gtg tta act att tga tcg att aca ttt atg gga tac ctt tga gcc taa cta aca ata acg tga aat ata ttc ttt caa ccg ctg att ttt tta caa att gga tct gtc att act gag tgc gaa aaa tac ata ctt aaa aat ctt tgt ttt aga aac tgt atc gat ttc --- --- tac ata tac gct gat aaa tat aat aac aag aaa ata gaa tta gca tcg ttt aac aca ata tta cga aat att ttg aga ctc atc aac aat gaa aac ttt aaa tac tta aca gag gaa tca atg ata aaa att tta agc gat gat atg tta tat ata aaa aat gag gat ttc acc cca ctg att ctc att aaa tgg tta gag agt aca cca acc atg tac cgt cga gtt act tag atg cct cag aat atc a-t ttc ttt ccc cac aag tta taa aat cac ttt ata gtc atc gac tgg tta gtt caa tct acg aat gta taa cat tct taa aca ata tag cat tct tgg ata aat cat ttc cta gat acc ata tca tcg agt tga tat cta tcg gta taa gta att cac atg ata aga ttt cca taa act gct aca atc ata aaa aaa ttc atg gga aat gat atc ttc acg tag ata tag gtg tag ttt cgc agt gac cgt cct gga taa tat tat cta tat gat ggg tgg ata tga tca gtc cct gta tag aag ttc aaa ggt tat agc gta caa tac atg tac taa ttc ttg gat ata tga tat acc aga gct aaa aat atc atc gtt cta att gcg gag gag ttg cca atg acg aat aca ttt att gta tag gcg gta tac gcg atc agg agt cat cgt tga tat cta gta tcg ata gat gga agc cat caa aac cat att ggc aga agt atg cta aaa tgt gcg aac caa aat gtg ata tgg ggt tgc gat ttt aaa cgg att aat ata tgt cat agg tgg agt cgt taa agg tga cac aca tat acc aac gca cta gag agt tta tca gaa gat gga tgg atg aat cat caa cgt ctt cca ata aaa atg tcc aat atg tcg acg att gtt cat gct gga aag att tat ata tct aga ggt tac aac aat agt agt gta gtt aat gta ata tcg aat ct- -ag tcc tta gct ata atc cga tat atg atg aat gga cca aat tat cat cat taa ata ttc cta gaa tta atc ctg ctc tat ggt cag tgt ata ata aat tat atg tag gag gag taa tat ctg atg atg ttc aaa cta ata cat ctg aaa cat acg ata aag aaa aa

>MT903344.1_Monkeypox_virus_isolate_MPXV-UK_P2

tgt tat tgt agt atg ata ata tca aaa aga tgg ata taa aga att tac tga cta cat gta cta ttt tac att act aca ttg gct acg gca tat ata cct att tcg tca ctt cca cac gct ccg gta aac ggg tgt cat gtg acg agg gag aat ctt gat aag agg cat aat caa tgt tgt aat ccg atg tcc acc tgg aga att tgc caa ggt cag atg tag agt tgg tag tga taa cac aaa atg tga aca ctg ccc acc tca tac ata tac cgc aat ccc caa tta ttc taa tag atg tca tca atg tag aaa atg ccc aac agg atc att tga taa ggt aaa gtg tac cgg aac aca gaa caa at- gtt cgt gtc atc ctg gtt ggt ata cgc tac tga ttc ttc aca gac tga aga ttg tcg aga ttt gtg tac caa aaa agg aga tgt cca tgc gga tac ttt ggt gga ata gat gaa gga aat cct att tgt aaa tcg tgt tgt gtt ggt gaa tat tgc gac tac cta cgt aat tat aga ctt gat cca ttt cct cca tgc aaa cta tct atc taa atg taa tta att atg att ttg atg ata atg tta cca tac att ata tca cta ctt ggt tag tgt gta tta ttt agt atg gaa gac cta tta ata att act tat ctt ttg acg atc ttg tta taa tta taa tat aaa aat act tat gac ata gta a-- --- -ct cat aat tgc tga cgc gat aaa ttc gta ata atc tgt ttt gtt caa att ttt ata agg aat cta cag gca taa aaa taa aaa tat aat cta taa tat act ctt aca acg cca tca tga ata gca gtg aat taa ttg ctg tta tta tgg att tag aaa tag tgg acg att ttg tga tat taa tat agt tat taa tga tga aag gat aaa cgc gca tag act cat cct atc tgg agc ctc cga ata ttt ttt cca ttc tgt ttt cca ata att tta tcg att cta atg aat acg aag tta atc taa gtc att tag att atc aaa gtg tta act att tga tcg att aca ttt atg gga tac ctt tga gcc taa cta aca ata acg tga aat ata ttc ttt caa ccg ctg att ttt tta caa att gga tct gtc att act gag tgc gaa aaa tac ata ctt aaa aat ctt tgt ttt aga aac tgt atc gat ttc --- --- tac ata tac gct gat aaa tat aat aac aag aaa ata gaa tta gca tcg ttt aac aca ata tta cga aat att ttg aga ctc atc aac aat gaa aac ttt aaa tac tta aca gag gaa tca atg ata aaa att tta agc gat gat atg tta tat ata aaa aat gag gat ttc acc cca ctg att ctc att aaa tgg tta gag agt aca cca acc atg tac cgt cga gtt act tag atg cct cag aat atc a-t ttc ttt ccc cac aag tta taa aat cac ttt ata gtc atc gac tgg tta gtt caa tct acg aat gta taa cat tct taa aca ata tag cat tct tgg ata aat cat ttc cta gat acc ata tca tcg agt tga tat cta tcg gta taa gta att cac atg ata aga ttt cca taa act gct aca atc ata aaa aaa ttc atg gga aat gat atc ttc acg tag ata tag gtg tag ttt cgc agt gac cgt cct gga taa tat tat cta tat gat ggg tgg ata tga tca gtc cct gta tag aag ttc aaa ggt tat agc gta caa tac atg tac taa ttc ttg gat ata tga tat acc aga gct aaa aat atc atc gtt cta att gcg gag gag ttg cca atg acg aat aca ttt att gta tag gcg gta tac gcg atc agg agt cat cgt tga tat cta gta tcg ata gat gga agc cat caa aac cat att ggc aga agt atg cta aaa tgt gcg aac caa aat gtg ata tgg ggt tgc gat ttt aaa cgg att aat ata tgt cat agg tgg agt cgt taa agg tga cac aca tat acc aac gca cta gag agt tta tca gaa gat gga tgg atg aat cat caa cgt ctt cca ata aaa atg tcc aat atg tcg acg att gtt cat gct gga aag att tat ata tct aga ggt tac aac aat agt agt gta gtt aat gta ata tcg aat ct- -ag tcc tta gct ata atc cga tat atg atg aat gga cca aat tat cat cat taa ata ttc cta gaa tta atc ctg ctc tat ggt cag tgt ata ata aat tat atg tag gag gag taa tat ctg atg atg ttc aaa cta ata cat ctg aaa cat acg ata aag aaa aa

>MT903342.1_Monkeypox_virus_isolate_MPXV-Singapore

tgt tat tgt agt atg ata ata tca aaa aga tgg ata taa aga att tac tga cta cat gta cta ttt tac att act aca ttg gct acg gca tat ata cct att tcg tca ctt cca cac gct ccg gta aac ggg tgt cat gtg acg agg gag aat ctt gat aag agg cat aat caa tgt tgt aat ccg atg tcc acc tgg aga att tgc caa ggt cag atg tag agt tgg tag tga taa cac aaa atg tga aca ctg ccc acc tca tac ata tac cgc aat ccc caa tta ttc taa tag atg tca tca atg tag aaa atg ccc aac agg atc att tga taa ggt aaa gtg tac cgg aac aca gaa caa at- gtt cgt gtc atc ctg gtt ggt ata cgc tac tga ttc ttc aca gac tga aga ttg tcg aga ttt gtg tac caa aaa agg aga tgt cca tgc gga tac ttt ggt gga ata gat gaa gga aat cct att tgt aaa tcg tgt tgt gtt ggt gaa tat tgc gac tac cta cgt aat tat aga ctt gat cca ttt cct cca tgc aaa cta tct atc taa atg taa tta att atg att ttg atg ata atg tta cca tac att ata tca cta ctt ggt tag tgt gta tta ttt agt atg gaa gac cta tta ata att act tat ctt ttg acg atc ttg tta taa tta taa tat aaa aat act tat gac ata gta a-- --- -ct cat aat tgc tga cgc gat aaa ttc gta ata atc tgt ttt gtt caa att ttt ata agg aat cta cag gca taa aaa taa aaa tat aat cta taa tat act ctt aca acg cca tca tga ata gca gtg aat taa ttg ctg tta tta tgg att tag aaa tag tgg acg att ttg tga tat taa tat agt tat taa tga tga aag gat aaa cgc gca tag act cat cct atc tgg agc ctc cga ata ttt ttt cca ttc tgt ttt cca ata att tta tcg att cta atg aat acg aag tta atc taa gtc att tag att atc aaa gtg tta act att tga tcg att aca ttt atg gga tac ctt tga gcc taa cta aca ata acg tga aat ata ttc ttt caa ccg ctg att ttt tta caa att gga tct gtc att act gag tgc gaa aaa tac ata ctt aaa aat ctt tgt ttt aga aac tgt atc gat ttc --- --- tac ata tac gct gat aaa tat aat aac aag aaa ata gaa tta gca tcg ttt aac aca ata tta cga aat att ttg aga ctc atc aac aat gaa aac ttt aaa tac tta aca gag gaa tca atg ata aaa att tta agc gat gat atg tta tat ata aaa aat gag gat ttc acc cca ctg att ctc att aaa tgg tta gag agt aca cca acc atg tac cgt cga gtt act tag atg cct cag aat atc a-t ttc ttt ccc cac aag tta taa aat cac ttt ata gtc atc gac tgg tta gtt caa tct acg aat gta taa cat tct taa aca ata tag cat tct tgg ata aat cat ttc cta gat acc ata tca tcg agt tga tat cta tcg gta taa gta att cac atg ata aga ttt cca taa act gct aca atc ata aaa aaa ttc atg gga aat gat atc ttc acg tag ata tag gtg tag ttt cgc agt gac cgt cct gga taa tat tat cta tat gat ggg tgg ata tga tca gtc cct gta tag aag ttc aaa ggt tat agc gta caa tac atg tac taa ttc ttg gat ata tga tat acc aga gct aaa aat atc atc gtt cta att gcg gag gag ttg cca atg acg aat aca ttt att gta tag gcg gta tac gcg atc agg agt cat cgt tga tat cta gta tcg ata gat gga agc cat caa aac cat att ggc aga agt atg cta aaa tgt gcg aac caa aat gtg ata tgg ggt tgc gat ttt aaa cgg att aat ata tgt cat agg tgg agt cgt taa agg tga cac aca tat acc aac gca cta gag agt tta tca gaa gat gga tgg atg aat cat caa cgt ctt cca ata aaa atg tcc aat atg tcg acg att gtt cat gct gga aag att tat ata tct aga ggt tac aac aat agt agt gta gtt aat gta ata tcg aat ct- -ag tcc tta gct ata atc cga tat atg atg aat gga cca aat tat cat cat taa ata ttc cta gaa tta atc ctg ctc tat ggt cag tgt ata ata aat tat atg tag gag gag taa tat ctg atg atg ttc aaa cta ata cat ctg aaa cat acg ata aag aaa aa

>ON676708.1_Monkeypox_virus_isolate_MPXV_USA_2021_MD_complete_genome

tgt tat tgt agt atg ata ata tca aaa aga tgg ata taa aga att tac tga cta cat gta cta ttt tac att act aca ttg gct acg gca tat ata cct att tcg tca ctt cca cac gct ccg gta aac ggg tgt cat gtg acg agg gag aat ctt gat aag agg cat aat caa tgt tgt aat ccg atg tcc acc tgg aga att tgc caa ggt cag atg tag agt tgg tag tga taa cac aaa atg tga aca ctg ccc acc tca tac ata tac cgc aat ccc caa tta ttc taa tag atg tca tca atg tag aaa atg ccc aac agg atc att tga taa ggt aaa gtg tac cgg aac aca gaa caa at- gtt cgt gtc atc ctg gtt ggt ata cgc tac tga ttc ttc aca gac tga aga ttg tcg aga ttt gtg tac caa aaa agg aga tgt cca tgc gga tac ttt ggt gga ata gat gaa gga aat cct att tgt aaa tcg tgt tgt gtt ggt gaa tat tgc gac tac cta cgt aat tat aga ctt gat cca ttt cct cca tgc aaa cta tct atc taa atg taa tta att atg att ttg atg ata atg tta cca tac att ata tca cta ctt ggt tag tgt gta tta ttt agt atg gaa gac cta tta ata att act tat ctt ttg acg atc ttg tta taa tta taa tat aaa aat act tat gac ata gta a-- --- -ct cat aat tgc tga cgc gat aaa ttc gta ata atc tgt ttt gtt caa att ttt ata agg aat cta cag gca taa aaa taa aaa tat aat cta taa tat act ctt aca acg cca tca tga ata gca gtg aat taa ttg ctg tta tta tgg att tag aaa tag tgg acg att ttg tga tat taa tat agt tat taa tga tga aag gat aaa cgc gca tag act cat cct atc tgg agc ctc cga ata ttt ttt cca ttc tgt ttt cca ata att tta tcg att cta atg aat acg aag tta atc taa gtc att tag att atc aaa gtg tta act att tga tcg att aca ttt atg gga tac ctt tga gcc taa cta aca ata acg tga aat ata ttc ttt caa ccg ctg att ttt tta caa att gga tct gtc att act gag tgc gaa aaa tac ata ctt aaa aat ctt tgt ttt aga aac tgt atc gat ttc --- --- tac ata tac gct gat aaa tat aat aac aag aaa ata gaa tta gca tcg ttt aac aca ata tta cga aat att ttg aga ctc atc aac aat gaa aac ttt aaa tac tta aca gag gaa tca atg ata aaa att tta agc gat gat atg tta tat ata aaa aat gag gat ttc acc cca ctg att ctc att aaa tgg tta gag agt aca cca acc atg tac cgt cga gtt act tag atg cct cag aat atc a-t ttc ttt ccc cac aag tta taa aat cac ttt ata gtc atc gac tgg tta gtt caa tct acg aat gta taa cat tct taa aca ata tag cat tct tgg ata aat cat ttc cta gat acc ata tca tcg agt tga tat cta tcg gta taa gta att cac atg ata aga ttt cca taa act gct aca atc ata aaa aaa ttc atg gga aat gat atc ttc acg tag ata tag gtg tag ttt cgc agt gac cgt cct gga taa tat tat cta tat gat ggg tgg ata tga tca gtc cct gta tag aag ttc aaa ggt tat agc gta caa tac atg tac taa ttc ttg gat ata tga tat acc aga gct aaa aat atc atc gtt cta att gcg gag gag ttg cca atg acg aat aca ttt att gta tag gcg gta tac gcg atc agg agt cat cgt tga tat cta gta tcg ata gat gga agc cat caa aac cat att ggc aga agt atg cta aaa tgt gcg aac caa aat gtg ata tgg ggt tgc gat ttt aaa cgg att aat ata tgt cat agg tgg agt cgt taa agg tga cac aca tat acc aac gca cta gag agt tta tca gaa gat gga tgg atg aat cat caa cgt ctt cca ata aaa atg tcc aat atg tcg acg att gtt cat gct gga aag att tat ata tct aga ggt tac aac aat agt agt gta gtt aat gta ata tcg aat ct- -ag tcc tta gct ata atc cga tat atg atg aat gga cca aat tat cat cat taa ata ttc cta gaa tta atc ctg ctc tat ggt cag tgt ata ata aat tat atg tag gag gag taa tat ctg atg atg ttc aaa cta ata cat ctg aaa cat acg ata aag aaa aa

>ON585033.1_Monkeypox_virus_isolate_Monkeypox/PT0006/2022_complete_genome

tgt tat tgt agt atg ata ata tca aaa aga tgg ata taa aga att tac tga cta cat gta cta ttt tac att act aca ttg gct acg gca tat ata cct att tcg tca ctt cca cac gct ccg gta aac ggg tgt cat gtg acg agg gag aat ctt gat aag agg cat aat caa tgt tgt aat ccg atg tcc acc tgg aga att tgc caa ggt cag atg tag agt tgg tag tga taa cac aaa atg tga aca ctg ccc acc tca tac ata tac cgc aat ccc caa tta ttc taa tag atg tca tca atg tag aaa atg ccc aac agg atc att tga taa ggt aaa gtg tac cgg aac aca gaa caa at- gtt cgt gtc atc ctg gtt ggt ata cgc tac tga ttc ttc aca gac tga aga ttg tcg aga ttt gtg tac caa aaa agg aga tgt cca tgc gga tac ttt ggt gga ata gat gaa gga aat cct att tgt aaa tcg tgt tgt gtt ggt gaa tat tgc gac tac cta cgt aat tat aga ctt gat cca ttt cct cca tgc aaa cta tct atc taa atg taa tta att atg att ttg atg ata atg tta cca tac att ata tca cta ctt ggt tag tgt gta tta ttt agt atg gaa gac cta tta ata att act tat ctt ttg acg atc ttg tta taa tta taa tat aaa aat act tat gac ata gta a-- --- -ct cat aat tgc tga cgc gat aaa ttc gta ata atc tgt ttt gtt caa att ttt ata agg aat cta cag gca taa aaa taa aaa tat aat cta taa tat act ctt aca acg cca tca tga ata gca gtg aat taa ttg ctg tta tta tgg att tag aaa tag tgg acg att ttg tga tat taa tat agt tat taa tga tga aag gat aaa cgc gca tag act cat cct atc tgg agc ctc cga ata ttt ttt cca ttc tgt ttt cca ata att tta tcg att cta atg aat acg aag tta atc taa gtc att tag att atc aaa gtg tta act att tga tcg att aca ttt atg gga tac ctt tga gcc taa cta aca ata acg tga aat ata ttc ttt caa ccg ctg att ttt tta caa att gga tct gtc att act gag tgc gaa aaa tac ata ctt aaa aat ctt tgt ttt aga aac tgt atc gat ttc --- --- tac ata tac gct gat aaa tat aat aac aag aaa ata gaa tta gca tcg ttt aac aca ata tta cga aat att ttg aga ctc atc aac aat gaa aac ttt aaa tac tta aca gag gaa tca atg ata aaa att tta agc gat gat atg tta tat ata aaa aat gag gat ttc acc cca ctg att ctc att aaa tgg tta gag agt aca cca acc atg tac cgt cga gtt act tag atg cct cag aat atc a-t ttc ttt ccc cac aag tta taa aat cac ttt ata gtc atc gac tgg tta gtt caa tct acg aat gta taa cat tct taa aca ata tag cat tct tgg ata aat cat ttc cta gat acc ata tca tcg agt tga tat cta tcg gta taa gta att cac atg ata aga ttt cca taa act gct aca atc ata aaa aaa ttc atg gga aat gat atc ttc acg tag ata tag gtg tag ttt cgc agt gac cgt cct gga taa tat tat cta tat gat ggg tgg ata tga tca gtc cct gta tag aag ttc aaa ggt tat agc gta caa tac atg tac taa ttc ttg gat ata tga tat acc aga gct aaa aat atc atc gtt cta att gcg gag gag ttg cca atg acg aat aca ttt att gta tag gcg gta tac gcg atc agg agt cat cgt tga tat cta gta tcg ata gat gga agc cat caa aac cat att ggc aga agt atg cta aaa tgt gcg aac caa aat gtg ata tgg ggt tgc gat ttt aaa cgg att aat ata tgt cat agg tgg agt cgt taa agg tga cac aca tat acc aac gca cta gag agt tta tca gaa gat gga tgg atg aat cat caa cgt ctt cca ata aaa atg tcc aat atg tcg acg att gtt cat gct gga aag att tat ata tct aga ggt tac aac aat agt agt gta gtt aat gta ata tcg aat ct- -ag tcc tta gct ata atc cga tat atg atg aat gga cca aat tat cat cat taa ata ttc cta gaa tta atc ctg ctc tat ggt cag tgt ata ata aat tat atg tag gag gag taa tat ctg atg atg ttc aaa cta ata cat ctg aaa cat acg ata aag aaa aa

>ON585035.1_Monkeypox_virus_isolate_Monkeypox/PT0009/2022_complete_genome

tgt tat tgt agt atg ata ata tca aaa aga tgg ata taa aga att tac tga cta cat gta cta ttt tac att act aca ttg gct acg gca tat ata cct att tcg tca ctt cca cac gct ccg gta aac ggg tgt cat gtg acg agg gag aat ctt gat aag agg cat aat caa tgt tgt aat ccg atg tcc acc tgg aga att tgc caa ggt cag atg tag agt tgg tag tga taa cac aaa atg tga aca ctg ccc acc tca tac ata tac cgc aat ccc caa tta ttc taa tag atg tca tca atg tag aaa atg ccc aac agg atc att tga taa ggt aaa gtg tac cgg aac aca gaa caa at- gtt cgt gtc atc ctg gtt ggt ata cgc tac tga ttc ttc aca gac tga aga ttg tcg aga ttt gtg tac caa aaa agg aga tgt cca tgc gga tac ttt ggt gga ata gat gaa gga aat cct att tgt aaa tcg tgt tgt gtt ggt gaa tat tgc gac tac cta cgt aat tat aga ctt gat cca ttt cct cca tgc aaa cta tct atc taa atg taa tta att atg att ttg atg ata atg tta cca tac att ata tca cta ctt ggt tag tgt gta tta ttt agt atg gaa gac cta tta ata att act tat ctt ttg acg atc ttg tta taa tta taa tat aaa aat act tat gac ata gta a-- --- -ct cat aat tgc tga cgc gat aaa ttc gta ata atc tgt ttt gtt caa att ttt ata agg aat cta cag gca taa aaa taa aaa tat aat cta taa tat act ctt aca acg cca tca tga ata gca gtg aat taa ttg ctg tta tta tgg att tag aaa tag tgg acg att ttg tga tat taa tat agt tat taa tga tga aag gat aaa cgc gca tag act cat cct atc tgg agc ctc cga ata ttt ttt cca ttc tgt ttt cca ata att tta tcg att cta atg aat acg aag tta atc taa gtc att tag att atc aaa gtg tta act att tga tcg att aca ttt atg gga tac ctt tga gcc taa cta aca ata acg tga aat ata ttc ttt caa ccg ctg att ttt tta caa att gga tct gtc att act gag tgc gaa aaa tac ata ctt aaa aat ctt tgt ttt aga aac tgt atc gat ttc --- --- tac ata tac gct gat aaa tat aat aac aag aaa ata gaa tta gca tcg ttt aac aca ata tta cga aat att ttg aga ctc atc aac aat gaa aac ttt aaa tac tta aca gag gaa tca atg ata aaa att tta agc gat gat atg tta tat ata aaa aat gag gat ttc acc cca ctg att ctc att aaa tgg tta gag agt aca cca acc atg tac cgt cga gtt act tag atg cct cag aat atc a-t ttc ttt ccc cac aag tta taa aat cac ttt ata gtc atc gac tgg tta gtt caa tct acg aat gta taa cat tct taa aca ata tag cat tct tgg ata aat cat ttc cta gat acc ata tca tcg agt tga tat cta tcg gta taa gta att cac atg ata aga ttt cca taa act gct aca atc ata aaa aaa ttc atg gga aat gat atc ttc acg tag ata tag gtg tag ttt cgc agt gac cgt cct gga taa tat tat cta tat gat ggg tgg ata tga tca gtc cct gta tag aag ttc aaa ggt tat agc gta caa tac atg tac taa ttc ttg gat ata tga tat acc aga gct aaa aat atc atc gtt cta att gcg gag gag ttg cca atg acg aat aca ttt att gta tag gcg gta tac gcg atc agg agt cat cgt tga tat cta gta tcg ata gat gga agc cat caa aac cat att ggc aga agt atg cta aaa tgt gcg aac caa aat gtg ata tgg ggt tgc gat ttt aaa cgg att aat ata tgt cat agg tgg agt cgt taa agg tga cac aca tat acc aac gca cta gag agt tta tca gaa gat gga tgg atg aat cat caa cgt ctt cca ata aaa atg tcc aat atg tcg acg att gtt cat gct gga aag att tat ata tct aga ggt tac aac aat agt agt gta gtt aat gta ata tcg aat ct- -ag tcc tta gct ata atc cga tat atg atg aat gga cca aat tat cat cat taa ata ttc cta gaa tta atc ctg ctc tat ggt cag tgt ata ata aat tat atg tag gag gag taa tat ctg atg atg ttc aaa cta ata cat ctg aaa cat acg ata aag aaa aa

>ON649725.1_Monkeypox_virus_isolate_Monkeypox/PT0011/2022_complete_genome

tgt tat tgt agt atg ata ata tca aaa aga tgg ata taa aga att tac tga cta cat gta cta ttt tac att act aca ttg gct acg gca tat ata cct att tcg tca ctt cca cac gct ccg gta aac ggg tgt cat gtg acg agg gag aat ctt gat aag agg cat aat caa tgt tgt aat ccg atg tcc acc tgg aga att tgc caa ggt cag atg tag agt tgg tag tga taa cac aaa atg tga aca ctg ccc acc tca tac ata tac cgc aat ccc caa tta ttc taa tag atg tca tca atg tag aaa atg ccc aac agg atc att tga taa ggt aaa gtg tac cgg aac aca gaa caa at- gtt cgt gtc atc ctg gtt ggt ata cgc tac tga ttc ttc aca gac tga aga ttg tcg aga ttt gtg tac caa aaa agg aga tgt cca tgc gga tac ttt ggt gga ata gat gaa gga aat cct att tgt aaa tcg tgt tgt gtt ggt gaa tat tgc gac tac cta cgt aat tat aga ctt gat cca ttt cct cca tgc aaa cta tct atc taa atg taa tta att atg att ttg atg ata atg tta cca tac att ata tca cta ctt ggt tag tgt gta tta ttt agt atg gaa gac cta tta ata att act tat ctt ttg acg atc ttg tta taa tta taa tat aaa aat act tat gac ata gta a-- --- -ct cat aat tgc tga cgc gat aaa ttc gta ata atc tgt ttt gtt caa att ttt ata agg aat cta cag gca taa aaa taa aaa tat aat cta taa tat act ctt aca acg cca tca tga ata gca gtg aat taa ttg ctg tta tta tgg att tag aaa tag tgg acg att ttg tga tat taa tat agt tat taa tga tga aag gat aaa cgc gca tag act cat cct atc tgg agc ctc cga ata ttt ttt cca ttc tgt ttt cca ata att tta tcg att cta atg aat acg aag tta atc taa gtc att tag att atc aaa gtg tta act att tga tcg att aca ttt atg gga tac ctt tga gcc taa cta aca ata acg tga aat ata ttc ttt caa ccg ctg att ttt tta caa att gga tct gtc att act gag tgc gaa aaa tac ata ctt aaa aat ctt tgt ttt aga aac tgt atc gat ttc --- --- tac ata tac gct gat aaa tat aat aac aag aaa ata gaa tta gca tcg ttt aac aca ata tta cga aat att ttg aga ctc atc aac aat gaa aac ttt aaa tac tta aca gag gaa tca atg ata aaa att tta agc gat gat atg tta tat ata aaa aat gag gat ttc acc cca ctg att ctc att aaa tgg tta gag agt aca cca acc atg tac cgt cga gtt act tag atg cct cag aat atc a-t ttc ttt ccc cac aag tta taa aat cac ttt ata gtc atc gac tgg tta gtt caa tct acg aat gta taa cat tct taa aca ata tag cat tct tgg ata aat cat ttc cta gat acc ata tca tcg agt tga tat cta tcg gta taa gta att cac atg ata aga ttt cca taa act gct aca atc ata aaa aaa ttc atg gga aat gat atc ttc acg tag ata tag gtg tag ttt cgc agt gac cgt cct gga taa tat tat cta tat gat ggg tgg ata tga tca gtc cct gta tag aag ttc aaa ggt tat agc gta caa tac atg tac taa ttc ttg gat ata tga tat acc aga gct aaa aat atc atc gtt cta att gcg gag gag ttg cca atg acg aat aca ttt att gta tag gcg gta tac gcg atc agg agt cat cgt tga tat cta gta tcg ata gat gga agc cat caa aac cat att ggc aga agt atg cta aaa tgt gcg aac caa aat gtg ata tgg ggt tgc gat ttt aaa cgg att aat ata tgt cat agg tgg agt cgt taa agg tga cac aca tat acc aac gca cta gag agt tta tca gaa gat gga tgg atg aat cat caa cgt ctt cca ata aaa atg tcc aat atg tcg acg att gtt cat gct gga aag att tat ata tct aga ggt tac aac aat agt agt gta gtt aat gta ata tcg aat ct- -ag tcc tta gct ata atc cga tat atg atg aat gga cca aat tat cat cat taa ata ttc cta gaa tta atc ctg ctc tat ggt cag tgt ata ata aat tat atg tag gag gag taa tat ctg atg atg ttc aaa cta ata cat ctg aaa cat acg ata aag aaa aa

>ON649724.1_Monkeypox_virus_isolate_Monkeypox/PT0014/2022_complete_genome

tgt tat tgt agt atg ata ata tca aaa aga tgg ata taa aga att tac tga cta cat gta cta ttt tac att act aca ttg gct acg gca tat ata cct att tcg tca ctt cca cac gct ccg gta aac ggg tgt cat gtg acg agg gag aat ctt gat aag agg cat aat caa tgt tgt aat ccg atg tcc acc tgg aga att tgc caa ggt cag atg tag agt tgg tag tga taa cac aaa atg tga aca ctg ccc acc tca tac ata tac cgc aat ccc caa tta ttc taa tag atg tca tca atg tag aaa atg ccc aac agg atc att tga taa ggt aaa gtg tac cgg aac aca gaa caa at- gtt cgt gtc atc ctg gtt ggt ata cgc tac tga ttc ttc aca gac tga aga ttg tcg aga ttt gtg tac caa aaa agg aga tgt cca tgc gga tac ttt ggt gga ata gat gaa gga aat cct att tgt aaa tcg tgt tgt gtt ggt gaa tat tgc gac tac cta cgt aat tat aga ctt gat cca ttt cct cca tgc aaa cta tct atc taa atg taa tta att atg att ttg atg ata atg tta cca tac att ata tca cta ctt ggt tag tgt gta tta ttt agt atg gaa gac cta tta ata att act tat ctt ttg acg atc ttg tta taa tta taa tat aaa aat act tat gac ata gta a-- --- -ct cat aat tgc tga cgc gat aaa ttc gta ata atc tgt ttt gtt caa att ttt ata agg aat cta cag gca taa aaa taa aaa tat aat cta taa tat act ctt aca acg cca tca tga ata gca gtg aat taa ttg ctg tta tta tgg att tag aaa tag tgg acg att ttg tga tat taa tat agt tat taa tga tga aag gat aaa cgc gca tag act cat cct atc tgg agc ctc cga ata ttt ttt cca ttc tgt ttt cca ata att tta tcg att cta atg aat acg aag tta atc taa gtc att tag att atc aaa gtg tta act att tga tcg att aca ttt atg gga tac ctt tga gcc taa cta aca ata acg tga aat ata ttc ttt caa ccg ctg att ttt tta caa att gga tct gtc att act gag tgc gaa aaa tac ata ctt aaa aat ctt tgt ttt aga aac tgt atc gat ttc --- --- tac ata tac gct gat aaa tat aat aac aag aaa ata gaa tta gca tcg ttt aac aca ata tta cga aat att ttg aga ctc atc aac aat gaa aac ttt aaa tac tta aca gag gaa tca atg ata aaa att tta agc gat gat atg tta tat ata aaa aat gag gat ttc acc cca ctg att ctc att aaa tgg tta gag agt aca cca acc atg tac cgt cga gtt act tag atg cct cag aat atc a-t ttc ttt ccc cac aag tta taa aat cac ttt ata gtc atc gac tgg tta gtt caa tct acg aat gta taa cat tct taa aca ata tag cat tct tgg ata aat cat ttc cta gat acc ata tca tcg agt tga tat cta tcg gta taa gta att cac atg ata aga ttt cca taa act gct aca atc ata aaa aaa ttc atg gga aat gat atc ttc acg tag ata tag gtg tag ttt cgc agt gac cgt cct gga taa tat tat cta tat gat ggg tgg ata tga tca gtc cct gta tag aag ttc aaa ggt tat agc gta caa tac atg tac taa ttc ttg gat ata tga tat acc aga gct aaa aat atc atc gtt cta att gcg gag gag ttg cca atg acg aat aca ttt att gta tag gcg gta tac gcg atc agg agt cat cgt tga tat cta gta tcg ata gat gga agc cat caa aac cat att ggc aga agt atg cta aaa tgt gcg aac caa aat gtg ata tgg ggt tgc gat ttt aaa cgg att aat ata tgt cat agg tgg agt cgt taa agg tga cac aca tat acc aac gca cta gag agt tta tca gaa gat gga tgg atg aat cat caa cgt ctt cca ata aaa atg tcc aat atg tcg acg att gtt cat gct gga aag att tat ata tct aga ggt tac aac aat agt agt gta gtt aat gta ata tcg aat ct- -ag tcc tta gct ata atc cga tat atg atg aat gga cca aat tat cat cat taa ata ttc cta gaa tta atc ctg ctc tat ggt cag tgt ata ata aat tat atg tag gag gag taa tat ctg atg atg ttc aaa cta ata cat ctg aaa cat acg ata aag aaa aa

>ON649722.1_Monkeypox_virus_isolate_Monkeypox/PT0022/2022_complete_genome

tgt tat tgt agt atg ata ata tca aaa aga tgg ata taa aga att tac tga cta cat gta cta ttt tac att act aca ttg gct acg gca tat ata cct att tcg tca ctt cca cac gct ccg gta aac ggg tgt cat gtg acg agg gag aat ctt gat aag agg cat aat caa tgt tgt aat ccg atg tcc acc tgg aga att tgc caa ggt cag atg tag agt tgg tag tga taa cac aaa atg tga aca ctg ccc acc tca tac ata tac cgc aat ccc caa tta ttc taa tag atg tca tca atg tag aaa atg ccc aac agg atc att tga taa ggt aaa gtg tac cgg aac aca gaa caa at- gtt cgt gtc atc ctg gtt ggt ata cgc tac tga ttc ttc aca gac tga aga ttg tcg aga ttt gtg tac caa aaa agg aga tgt cca tgc gga tac ttt ggt gga ata gat gaa gga aat cct att tgt aaa tcg tgt tgt gtt ggt gaa tat tgc gac tac cta cgt aat tat aga ctt gat cca ttt cct cca tgc aaa cta tct atc taa atg taa tta att atg att ttg atg ata atg tta cca tac att ata tca cta ctt ggt tag tgt gta tta ttt agt atg gaa gac cta tta ata att act tat ctt ttg acg atc ttg tta taa tta taa tat aaa aat act tat gac ata gta a-- --- -ct cat aat tgc tga cgc gat aaa ttc gta ata atc tgt ttt gtt caa att ttt ata agg aat cta cag gca taa aaa taa aaa tat aat cta taa tat act ctt aca acg cca tca tga ata gca gtg aat taa ttg ctg tta tta tgg att tag aaa tag tgg acg att ttg tga tat taa tat agt tat taa tga tga aag gat aaa cgc gca tag act cat cct atc tgg agc ctc cga ata ttt ttt cca ttc tgt ttt cca ata att tta tcg att cta atg aat acg aag tta atc taa gtc att tag att atc aaa gtg tta act att tga tcg att aca ttt atg gga tac ctt tga gcc taa cta aca ata acg tga aat ata ttc ttt caa ccg ctg att ttt tta caa att gga tct gtc att act gag tgc gaa aaa tac ata ctt aaa aat ctt tgt ttt aga aac tgt atc gat ttc --- --- tac ata tac gct gat aaa tat aat aac aag aaa ata gaa tta gca tcg ttt aac aca ata tta cga aat att ttg aga ctc atc aac aat gaa aac ttt aaa tac tta aca gag gaa tca atg ata aaa att tta agc gat gat atg tta tat ata aaa aat gag gat ttc acc cca ctg att ctc att aaa tgg tta gag agt aca cca acc atg tac cgt cga gtt act tag atg cct cag aat atc a-t ttc ttt ccc cac aag tta taa aat cac ttt ata gtc atc gac tgg tta gtt caa tct acg aat gta taa cat tct taa aca ata tag cat tct tgg ata aat cat ttc cta gat acc ata tca tcg agt tga tat cta tcg gta taa gta att cac atg ata aga ttt cca taa act gct aca atc ata aaa aaa ttc atg gga aat gat atc ttc acg tag ata tag gtg tag ttt cgc agt gac cgt cct gga taa tat tat cta tat gat ggg tgg ata tga tca gtc cct gta tag aag ttc aaa ggt tat agc gta caa tac atg tac taa ttc ttg gat ata tga tat acc aga gct aaa aat atc atc gtt cta att gcg gag gag ttg cca atg acg aat aca ttt att gta tag gcg gta tac gcg atc agg agt cat cgt tga tat cta gta tcg ata gat gga agc cat caa aac cat att ggc aga agt atg cta aaa tgt gcg aac caa aat gtg ata tgg ggt tgc gat ttt aaa cgg att aat ata tgt cat agg tgg agt cgt taa agg tga cac aca tat acc aac gca cta gag agt tta tca gaa gat gga tgg atg aat cat caa cgt ctt cca ata aaa atg tcc aat atg tcg acg att gtt cat gct gga aag att tat ata tct aga ggt tac aac aat agt agt gta gtt aat gta ata tcg aat ct- -ag tcc tta gct ata atc cga tat atg atg aat gga cca aat tat cat cat taa ata ttc cta gaa tta atc ctg ctc tat ggt cag tgt ata ata aat tat atg tag gag gag taa tat ctg atg atg ttc aaa cta ata cat ctg aaa cat acg ata aag aaa aa

>ON649721.1_Monkeypox_virus_isolate_Monkeypox/PT0021/2022_complete_genome

tgt tat tgt agt atg ata ata tca aaa aga tgg ata taa aga att tac tga cta cat gta cta ttt tac att act aca ttg gct acg gca tat ata cct att tcg tca ctt cca cac gct ccg gta aac ggg tgt cat gtg acg agg gag aat ctt gat aag agg cat aat caa tgt tgt aat ccg atg tcc acc tgg aga att tgc caa ggt cag atg tag agt tgg tag tga taa cac aaa atg tga aca ctg ccc acc tca tac ata tac cgc aat ccc caa tta ttc taa tag atg tca tca atg tag aaa atg ccc aac agg atc att tga taa ggt aaa gtg tac cgg aac aca gaa caa at- gtt cgt gtc atc ctg gtt ggt ata cgc tac tga ttc ttc aca gac tga aga ttg tcg aga ttt gtg tac caa aaa agg aga tgt cca tgc gga tac ttt ggt gga ata gat gaa gga aat cct att tgt aaa tcg tgt tgt gtt ggt gaa tat tgc gac tac cta cgt aat tat aga ctt gat cca ttt cct cca tgc aaa cta tct atc taa atg taa tta att atg att ttg atg ata atg tta cca tac att ata tca cta ctt ggt tag tgt gta tta ttt agt atg gaa gac cta tta ata att act tat ctt ttg acg atc ttg tta taa tta taa tat aaa aat act tat gac ata gta a-- --- -ct cat aat tgc tga cgc gat aaa ttc gta ata atc tgt ttt gtt caa att ttt ata agg aat cta cag gca taa aaa taa aaa tat aat cta taa tat act ctt aca acg cca tca tga ata gca gtg aat taa ttg ctg tta tta tgg att tag aaa tag tgg acg att ttg tga tat taa tat agt tat taa tga tga aag gat aaa cgc gca tag act cat cct atc tgg agc ctc cga ata ttt ttt cca ttc tgt ttt cca ata att tta tcg att cta atg aat acg aag tta atc taa gtc att tag att atc aaa gtg tta act att tga tcg att aca ttt atg gga tac ctt tga gcc taa cta aca ata acg tga aat ata ttc ttt caa ccg ctg att ttt tta caa att gga tct gtc att act gag tgc gaa aaa tac ata ctt aaa aat ctt tgt ttt aga aac tgt atc gat ttc --- --- tac ata tac gct gat aaa tat aat aac aag aaa ata gaa tta gca tcg ttt aac aca ata tta cga aat att ttg aga ctc atc aac aat gaa aac ttt aaa tac tta aca gag gaa tca atg ata aaa att tta agc gat gat atg tta tat ata aaa aat gag gat ttc acc cca ctg att ctc att aaa tgg tta gag agt aca cca acc atg tac cgt cga gtt act tag atg cct cag aat atc a-t ttc ttt ccc cac aag tta taa aat cac ttt ata gtc atc gac tgg tta gtt caa tct acg aat gta taa cat tct taa aca ata tag cat tct tgg ata aat cat ttc cta gat acc ata tca tcg agt tga tat cta tcg gta taa gta att cac atg ata aga ttt cca taa act gct aca atc ata aaa aaa ttc atg gga aat gat atc ttc acg tag ata tag gtg tag ttt cgc agt gac cgt cct gga taa tat tat cta tat gat ggg tgg ata tga tca gtc cct gta tag aag ttc aaa ggt tat agc gta caa tac atg tac taa ttc ttg gat ata tga tat acc aga gct aaa aat atc atc gtt cta att gcg gag gag ttg cca atg acg aat aca ttt att gta tag gcg gta tac gcg atc agg agt cat cgt tga tat cta gta tcg ata gat gga agc cat caa aac cat att ggc aga agt atg cta aaa tgt gcg aac caa aat gtg ata tgg ggt tgc gat ttt aaa cgg att aat ata tgt cat agg tgg agt cgt taa agg tga cac aca tat acc aac gca cta gag agt tta tca gaa gat gga tgg atg aat cat caa cgt ctt cca ata aaa atg tcc aat atg tcg acg att gtt cat gct gga aag att tat ata tct aga ggt tac aac aat agt agt gta gtt aat gta ata tcg aat ct- -ag tcc tta gct ata atc cga tat atg atg aat gga cca aat tat cat cat taa ata ttc cta gaa tta atc ctg ctc tat ggt cag tgt ata ata aat tat atg tag gag gag taa tat ctg atg atg ttc aaa cta ata cat ctg aaa cat acg ata aag aaa aa

>ON649720.1_Monkeypox_virus_isolate_Monkeypox/PT0024/2022_complete_genome

tgt tat tgt agt atg ata ata tca aaa aga tgg ata taa aga att tac tga cta cat gta cta ttt tac att act aca ttg gct acg gca tat ata cct att tcg tca ctt cca cac gct ccg gta aac ggg tgt cat gtg acg agg gag aat ctt gat aag agg cat aat caa tgt tgt aat ccg atg tcc acc tgg aga att tgc caa ggt cag atg tag agt tgg tag tga taa cac aaa atg tga aca ctg ccc acc tca tac ata tac cgc aat ccc caa tta ttc taa tag atg tca tca atg tag aaa atg ccc aac agg atc att tga taa ggt aaa gtg tac cgg aac aca gaa caa at- gtt cgt gtc atc ctg gtt ggt ata cgc tac tga ttc ttc aca gac tga aga ttg tcg aga ttt gtg tac caa aaa agg aga tgt cca tgc gga tac ttt ggt gga ata gat gaa gga aat cct att tgt aaa tcg tgt tgt gtt ggt gaa tat tgc gac tac cta cgt aat tat aga ctt gat cca ttt cct cca tgc aaa cta tct atc taa atg taa tta att atg att ttg atg ata atg tta cca tac att ata tca cta ctt ggt tag tgt gta tta ttt agt atg gaa gac cta tta ata att act tat ctt ttg acg atc ttg tta taa tta taa tat aaa aat act tat gac ata gta a-- --- -ct cat aat tgc tga cgc gat aaa ttc gta ata atc tgt ttt gtt caa att ttt ata agg aat cta cag gca taa aaa taa aaa tat aat cta taa tat act ctt aca acg cca tca tga ata gca gtg aat taa ttg ctg tta tta tgg att tag aaa tag tgg acg att ttg tga tat taa tat agt tat taa tga tga aag gat aaa cgc gca tag act cat cct atc tgg agc ctc cga ata ttt ttt cca ttc tgt ttt cca ata att tta tcg att cta atg aat acg aag tta atc taa gtc att tag att atc aaa gtg tta act att tga tcg att aca ttt atg gga tac ctt tga gcc taa cta aca ata acg tga aat ata ttc ttt caa ccg ctg att ttt tta caa att gga tct gtc att act gag tgc gaa aaa tac ata ctt aaa aat ctt tgt ttt aga aac tgt atc gat ttc --- --- tac ata tac gct gat aaa tat aat aac aag aaa ata gaa tta gca tcg ttt aac aca ata tta cga aat att ttg aga ctc atc aac aat gaa aac ttt aaa tac tta aca gag gaa tca atg ata aaa att tta agc gat gat atg tta tat ata aaa aat gag gat ttc acc cca ctg att ctc att aaa tgg tta gag agt aca cca acc atg tac cgt cga gtt act tag atg cct cag aat atc a-t ttc ttt ccc cac aag tta taa aat cac ttt ata gtc atc gac tgg tta gtt caa tct acg aat gta taa cat tct taa aca ata tag cat tct tgg ata aat cat ttc cta gat acc ata tca tcg agt tga tat cta tcg gta taa gta att cac atg ata aga ttt cca taa act gct aca atc ata aaa aaa ttc atg gga aat gat atc ttc acg tag ata tag gtg tag ttt cgc agt gac cgt cct gga taa tat tat cta tat gat ggg tgg ata tga tca gtc cct gta tag aag ttc aaa ggt tat agc gta caa tac atg tac taa ttc ttg gat ata tga tat acc aga gct aaa aat atc atc gtt cta att gcg gag gag ttg cca atg acg aat aca ttt att gta tag gcg gta tac gcg atc agg agt cat cgt tga tat cta gta tcg ata gat gga agc cat caa aac cat att ggc aga agt atg cta aaa tgt gcg aac caa aat gtg ata tgg ggt tgc gat ttt aaa cgg att aat ata tgt cat agg tgg agt cgt taa agg tga cac aca tat acc aac gca cta gag agt tta tca gaa gat gga tgg atg aat cat caa cgt ctt cca ata aaa atg tcc aat atg tcg acg att gtt cat gct gga aag att tat ata tct aga ggt tac aac aat agt agt gta gtt aat gta ata tcg aat ct- -ag tcc tta gct ata atc cga tat atg atg aat gga cca aat tat cat cat taa ata ttc cta gaa tta atc ctg ctc tat ggt cag tgt ata ata aat tat atg tag gag gag taa tat ctg atg atg ttc aaa cta ata cat ctg aaa cat acg ata aag aaa aa

>ON649723.1_Monkeypox_virus_isolate_Monkeypox/PT0013/2022_complete_genome

tgt tat tgt agt atg ata ata tca aaa aga tgg ata taa aga att tac tga cta cat gta cta ttt tac att act aca ttg gct acg gca tat ata cct att tcg tca ctt cca cac gct ccg gta aac ggg tgt cat gtg acg agg gag aat ctt gat aag agg cat aat caa tgt tgt aat ccg atg tcc acc tgg aga att tgc caa ggt cag atg tag agt tgg tag tga taa cac aaa atg tga aca ctg ccc acc tca tac ata tac cgc aat ccc caa tta ttc taa tag atg tca tca atg tag aaa atg ccc aac agg atc att tga taa ggt aaa gtg tac cgg aac aca gaa caa at- gtt cgt gtc atc ctg gtt ggt ata cgc tac tga ttc ttc aca gac tga aga ttg tcg aga ttt gtg tac caa aaa agg aga tgt cca tgc gga tac ttt ggt gga ata gat gaa gga aat cct att tgt aaa tcg tgt tgt gtt ggt gaa tat tgc gac tac cta cgt aat tat aga ctt gat cca ttt cct cca tgc aaa cta tct atc taa atg taa tta att atg att ttg atg ata atg tta cca tac att ata tca cta ctt ggt tag tgt gta tta ttt agt atg gaa gac cta tta ata att act tat ctt ttg acg atc ttg tta taa tta taa tat aaa aat act tat gac ata gta a-- --- -ct cat aat tgc tga cgc gat aaa ttc gta ata atc tgt ttt gtt caa att ttt ata agg aat cta cag gca taa aaa taa aaa tat aat cta taa tat act ctt aca acg cca tca tga ata gca gtg aat taa ttg ctg tta tta tgg att tag aaa tag tgg acg att ttg tga tat taa tat agt tat taa tga tga aag gat aaa cgc gca tag act cat cct atc tgg agc ctc cga ata ttt ttt cca ttc tgt ttt cca ata att tta tcg att cta atg aat acg aag tta atc taa gtc att tag att atc aaa gtg tta act att tga tcg att aca ttt atg gga tac ctt tga gcc taa cta aca ata acg tga aat ata ttc ttt caa ccg ctg att ttt tta caa att gga tct gtc att act gag tgc gaa aaa tac ata ctt aaa aat ctt tgt ttt aga aac tgt atc gat ttc --- --- tac ata tac gct gat aaa tat aat aac aag aaa ata gaa tta gca tcg ttt aac aca ata tta cga aat att ttg aga ctc atc aac aat gaa aac ttt aaa tac tta aca gag gaa tca atg ata aaa att tta agc gat gat atg tta tat ata aaa aat gag gat ttc acc cca ctg att ctc att aaa tgg tta gag agt aca cca acc atg tac cgt cga gtt act tag atg cct cag aat atc a-t ttc ttt ccc cac aag tta taa aat cac ttt ata gtc atc gac tgg tta gtt caa tct acg aat gta taa cat tct taa aca ata tag cat tct tgg ata aat cat ttc cta gat acc ata tca tcg agt tga tat cta tcg gta taa gta att cac atg ata aga ttt cca taa act gct aca atc ata aaa aaa ttc atg gga aat gat atc ttc acg tag ata tag gtg tag ttt cgc agt gac cgt cct gga taa tat tat cta tat gat ggg tgg ata tga tca gtc cct gta tag aag ttc aaa ggt tat agc gta caa tac atg tac taa ttc ttg gat ata tga tat acc aga gct aaa aat atc atc gtt cta att gcg gag gag ttg cca atg acg aat aca ttt att gta tag gcg gta tac gcg atc agg agt cat cgt tga tat cta gta tcg ata gat gga agc cat caa aac cat att ggc aga agt atg cta aaa tgt gcg aac caa aat gtg ata tgg ggt tgc gat ttt aaa cgg att aat ata tgt cat agg tgg agt cgt taa agg tga cac aca tat acc aac gca cta gag agt tta tca gaa gat gga tgg atg aat cat caa cgt ctt cca ata aaa atg tcc aat atg tcg acg att gtt cat gct gga aag att tat ata tct aga ggt tac aac aat agt agt gta gtt aat gta ata tcg aat ct- -ag tcc tta gct ata atc cga tat atg atg aat gga cca aat tat cat cat taa ata ttc cta gaa tta atc ctg ctc tat ggt cag tgt ata ata aat tat atg tag gag gag taa tat ctg atg atg ttc aaa cta ata cat ctg aaa cat acg ata aag aaa aa

>ON649718.1_Monkeypox_virus_isolate_Monkeypox/PT0018/2022_complete_genome

tgt tat tgt agt atg ata ata tca aaa aga tgg ata taa aga att tac tga cta cat gta cta ttt tac att act aca ttg gct acg gca tat ata cct att tcg tca ctt cca cac gct ccg gta aac ggg tgt cat gtg acg agg gag aat ctt gat aag agg cat aat caa tgt tgt aat ccg atg tcc acc tgg aga att tgc caa ggt cag atg tag agt tgg tag tga taa cac aaa atg tga aca ctg ccc acc tca tac ata tac cgc aat ccc caa tta ttc taa tag atg tca tca atg tag aaa atg ccc aac agg atc att tga taa ggt aaa gtg tac cgg aac aca gaa caa at- gtt cgt gtc atc ctg gtt ggt ata cgc tac tga ttc ttc aca gac tga aga ttg tcg aga ttt gtg tac caa aaa agg aga tgt cca tgc gga tac ttt ggt gga ata gat gaa gga aat cct att tgt aaa tcg tgt tgt gtt ggt gaa tat tgc gac tac cta cgt aat tat aga ctt gat cca ttt cct cca tgc aaa cta tct atc taa atg taa tta att atg att ttg atg ata atg tta cca tac att ata tca cta ctt ggt tag tgt gta tta ttt agt atg gaa gac cta tta ata att act tat ctt ttg acg atc ttg tta taa tta taa tat aaa aat act tat gac ata gta a-- --- -ct cat aat tgc tga cgc gat aaa ttc gta ata atc tgt ttt gtt caa att ttt ata agg aat cta cag gca taa aaa taa aaa tat aat cta taa tat act ctt aca acg cca tca tga ata gca gtg aat taa ttg ctg tta tta tgg att tag aaa tag tgg acg att ttg tga tat taa tat agt tat taa tga tga aag gat aaa cgc gca tag act cat cct atc tgg agc ctc cga ata ttt ttt cca ttc tgt ttt cca ata att tta tcg att cta atg aat acg aag tta atc taa gtc att tag att atc aaa gtg tta act att tga tcg att aca ttt atg gga tac ctt tga gcc taa cta aca ata acg tga aat ata ttc ttt caa ccg ctg att ttt tta caa att gga tct gtc att act gag tgc gaa aaa tac ata ctt aaa aat ctt tgt ttt aga aac tgt atc gat ttc --- --- tac ata tac gct gat aaa tat aat aac aag aaa ata gaa tta gca tcg ttt aac aca ata tta cga aat att ttg aga ctc atc aac aat gaa aac ttt aaa tac tta aca gag gaa tca atg ata aaa att tta agc gat gat atg tta tat ata aaa aat gag gat ttc acc cca ctg att ctc att aaa tgg tta gag agt aca cca acc atg tac cgt cga gtt act tag atg cct cag aat atc a-t ttc ttt ccc cac aag tta taa aat cac ttt ata gtc atc gac tgg tta gtt caa tct acg aat gta taa cat tct taa aca ata tag cat tct tgg ata aat cat ttc cta gat acc ata tca tcg agt tga tat cta tcg gta taa gta att cac atg ata aga ttt cca taa act gct aca atc ata aaa aaa ttc atg gga aat gat atc ttc acg tag ata tag gtg tag ttt cgc agt gac cgt cct gga taa tat tat cta tat gat ggg tgg ata tga tca gtc cct gta tag aag ttc aaa ggt tat agc gta caa tac atg tac taa ttc ttg gat ata tga tat acc aga gct aaa aat atc atc gtt cta att gcg gag gag ttg cca atg acg aat aca ttt att gta tag gcg gta tac gcg atc agg agt cat cgt tga tat cta gta tcg ata gat gga agc cat caa aac cat att ggc aga agt atg cta aaa tgt gcg aac caa aat gtg ata tgg ggt tgc gat ttt aaa cgg att aat ata tgt cat agg tgg agt cgt taa agg tga cac aca tat acc aac gca cta gag agt tta tca gaa gat gga tgg atg aat cat caa cgt ctt cca ata aaa atg tcc aat atg tcg acg att gtt cat gct gga aag att tat ata tct aga ggt tac aac aat agt agt gta gtt aat gta ata tcg aat ct- -ag tcc tta gct ata atc cga tat atg atg aat gga cca aat tat cat cat taa ata ttc cta gaa tta atc ctg ctc tat ggt cag tgt ata ata aat tat atg tag gag gag taa tat ctg atg atg ttc aaa cta ata cat ctg aaa cat acg ata aag aaa aa

>ON649719.1_Monkeypox_virus_isolate_Monkeypox/PT0012/2022_complete_genome

tgt tat tgt agt atg ata ata tca aaa aga tgg ata taa aga att tac tga cta cat gta cta ttt tac att act aca ttg gct acg gca tat ata cct att tcg tca ctt cca cac gct ccg gta aac ggg tgt cat gtg acg agg gag aat ctt gat aag agg cat aat caa tgt tgt aat ccg atg tcc acc tgg aga att tgc caa ggt cag atg tag agt tgg tag tga taa cac aaa atg tga aca ctg ccc acc tca tac ata tac cgc aat ccc caa tta ttc taa tag atg tca tca atg tag aaa atg ccc aac agg atc att tga taa ggt aaa gtg tac cgg aac aca gaa caa at- gtt cgt gtc atc ctg gtt ggt ata cgc tac tga ttc ttc aca gac tga aga ttg tcg aga ttt gtg tac caa aaa agg aga tgt cca tgc gga tac ttt ggt gga ata gat gaa gga aat cct att tgt aaa tcg tgt tgt gtt ggt gaa tat tgc gac tac cta cgt aat tat aga ctt gat cca ttt cct cca tgc aaa cta tct atc taa atg taa tta att atg att ttg atg ata atg tta cca tac att ata tca cta ctt ggt tag tgt gta tta ttt agt atg gaa gac cta tta ata att act tat ctt ttg acg atc ttg tta taa tta taa tat aaa aat act tat gac ata gta a-- --- -ct cat aat tgc tga cgc gat aaa ttc gta ata atc tgt ttt gtt caa att ttt ata agg aat cta cag gca taa aaa taa aaa tat aat cta taa tat act ctt aca acg cca tca tga ata gca gtg aat taa ttg ctg tta tta tgg att tag aaa tag tgg acg att ttg tga tat taa tat agt tat taa tga tga aag gat aaa cgc gca tag act cat cct atc tgg agc ctc cga ata ttt ttt cca ttc tgt ttt cca ata att tta tcg att cta atg aat acg aag tta atc taa gtc att tag att atc aaa gtg tta act att tga tcg att aca ttt atg gga tac ctt tga gcc taa cta aca ata acg tga aat ata ttc ttt caa ccg ctg att ttt tta caa att gga tct gtc att act gag tgc gaa aaa tac ata ctt aaa aat ctt tgt ttt aga aac tgt atc gat ttc --- --- tac ata tac gct gat aaa tat aat aac aag aaa ata gaa tta gca tcg ttt aac aca ata tta cga aat att ttg aga ctc atc aac aat gaa aac ttt aaa tac tta aca gag gaa tca atg ata aaa att tta agc gat gat atg tta tat ata aaa aat gag gat ttc acc cca ctg att ctc att aaa tgg tta gag agt aca cca acc atg tac cgt cga gtt act tag atg cct cag aat atc a-t ttc ttt ccc cac aag tta taa aat cac ttt ata gtc atc gac tgg tta gtt caa tct acg aat gta taa cat tct taa aca ata tag cat tct tgg ata aat cat ttc cta gat acc ata tca tcg agt tga tat cta tcg gta taa gta att cac atg ata aga ttt cca taa act gct aca atc ata aaa aaa ttc atg gga aat gat atc ttc acg tag ata tag gtg tag ttt cgc agt gac cgt cct gga taa tat tat cta tat gat ggg tgg ata tga tca gtc cct gta tag aag ttc aaa ggt tat agc gta caa tac atg tac taa ttc ttg gat ata tga tat acc aga gct aaa aat atc atc gtt cta att gcg gag gag ttg cca atg acg aat aca ttt att gta tag gcg gta tac gcg atc agg agt cat cgt tga tat cta gta tcg ata gat gga agc cat caa aac cat att ggc aga agt atg cta aaa tgt gcg aac caa aat gtg ata tgg ggt tgc gat ttt aaa cgg att aat ata tgt cat agg tgg agt cgt taa agg tga cac aca tat acc aac gca cta gag agt tta tca gaa gat gga tgg atg aat cat caa cgt ctt cca ata aaa atg tcc aat atg tcg acg att gtt cat gct gga aag att tat ata tct aga ggt tac aac aat agt agt gta gtt aat gta ata tcg aat ct- -ag tcc tta gct ata atc cga tat atg atg aat gga cca aat tat cat cat taa ata ttc cta gaa tta atc ctg ctc tat ggt cag tgt ata ata aat tat atg tag gag gag taa tat ctg atg atg ttc aaa cta ata cat ctg aaa cat acg ata aag aaa aa

>ON682267.1_Monkeypox_virus_isolate_MPXV/Germany/2022/RKI010_complete_genome

tgt tat tgt agt atg ata ata tca aaa aga tgg ata taa aga att tac tga cta cat gta cta ttt tac att act aca ttg gct acg gca tat ata cct att tcg tca ctt cca cac gct ccg gta aac ggg tgt cat gtg acg agg gag aat ctt gat aag agg cat aat caa tgt tgt aat ccg atg tcc acc tgg aga att tgc caa ggt cag atg tag agt tgg tag tga taa cac aaa atg tga aca ctg ccc acc tca tac ata tac cgc aat ccc caa tta ttc taa tag atg tca tca atg tag aaa atg ccc aac agg atc att tga taa ggt aaa gtg tac cgg aac aca gaa caa at- gtt cgt gtc atc ctg gtt ggt ata cgc tac tga ttc ttc aca gac tga aga ttg tcg aga ttt gtg tac caa aaa agg aga tgt cca tgc gga tac ttt ggt gga ata gat gaa gga aat cct att tgt aaa tcg tgt tgt gtt ggt gaa tat tgc gac tac cta cgt aat tat aga ctt gat cca ttt cct cca tgc aaa cta tct atc taa atg taa tta att atg att ttg atg ata atg tta cca tac att ata tca cta ctt ggt tag tgt gta tta ttt agt atg gaa gac cta tta ata att act tat ctt ttg acg atc ttg tta taa tta taa tat aaa aat act tat gac ata gta a-- --- -ct cat aat tgc tga cgc gat aaa ttc gta ata atc tgt ttt gtt caa att ttt ata agg aat cta cag gca taa aaa taa aaa tat aat cta taa tat act ctt aca acg cca tca tga ata gca gtg aat taa ttg ctg tta tta tgg att tag aaa tag tgg acg att ttg tga tat taa tat agt tat taa tga tga aag gat aaa cgc gca tag act cat cct atc tgg agc ctc cga ata ttt ttt cca ttc tgt ttt cca ata att tta tcg att cta atg aat acg aag tta atc taa gtc att tag att atc aaa gtg tta act att tga tcg att aca ttt atg gga tac ctt tga gcc taa cta aca ata acg tga aat ata ttc ttt caa ccg ctg att ttt tta caa att gga tct gtc att act gag tgc gaa aaa tac ata ctt aaa aat ctt tgt ttt aga aac tgt atc gat ttc --- --- tac ata tac gct gat aaa tat aat aac aag aaa ata gaa tta gca tcg ttt aac aca ata tta cga aat att ttg aga ctc atc aac aat gaa aac ttt aaa tac tta aca gag gaa tca atg ata aaa att tta agc gat gat atg tta tat ata aaa aat gag gat ttc acc cca ctg att ctc att aaa tgg tta gag agt aca cca acc atg tac cgt cga gtt act tag atg cct cag aat atc a-t ttc ttt ccc cac aag tta taa aat cac ttt ata gtc atc gac tgg tta gtt caa tct acg aat gta taa cat tct taa aca ata tag cat tct tgg ata aat cat ttc cta gat acc ata tca tcg agt tga tat cta tcg gta taa gta att cac atg ata aga ttt cca taa act gct aca atc ata aaa aaa ttc atg gga aat gat atc ttc acg tag ata tag gtg tag ttt cgc agt gac cgt cct gga taa tat tat cta tat gat ggg tgg ata tga tca gtc cct gta tag aag ttc aaa ggt tat agc gta caa tac atg tac taa ttc ttg gat ata tga tat acc aga gct aaa aat atc atc gtt cta att gcg gag gag ttg cca atg acg aat aca ttt att gta tag gcg gta tac gcg atc agg agt cat cgt tga tat cta gta tcg ata gat gga agc cat caa aac cat att ggc aga agt atg cta aaa tgt gcg aac caa aat gtg ata tgg ggt tgc gat ttt aaa cgg att aat ata tgt cat agg tgg agt cgt taa agg tga cac aca tat acc aac gca cta gag agt tta tca gaa gat gga tgg atg aat cat caa cgt ctt cca ata aaa atg tcc aat atg tcg acg att gtt cat gct gga aag att tat ata tct aga ggt tac aac aat agt agt gta gtt aat gta ata tcg aat ct- -ag tcc tta gct ata atc cga tat atg atg aat gga cca aat tat cat cat taa ata ttc cta gaa tta atc ctg ctc tat ggt cag tgt ata ata aat tat atg tag gag gag taa tat ctg atg atg ttc aaa cta ata cat ctg aaa cat acg ata aag aaa aa

>ON649717.1_Monkeypox_virus_isolate_Monkeypox/PT0019/2022_partial_genome

tgt tat tgt agt atg ata ata tca aaa aga tgg ata taa aga att tac tga cta cat gta cta ttt tac att act aca ttg gct acg gca tat ata cct att tcg tca ctt cca cac gct ccg gta aac ggg tgt cat gtg acg agg gag aat ctt gat aag agg cat aat caa tgt tgt aat ccg atg tcc acc tgg aga att tgc caa ggt cag atg tag agt tgg tag tga taa cac aaa atg tga aca ctg ccc acc tca tac ata tac cgc aat ccc caa tta ttc taa tag atg tca tca atg tag aaa atg ccc aac agg atc att tga taa ggt aaa gtg tac cgg aac aca gaa caa at- gtt cgt gtc atc ctg gtt ggt ata cgc tac tga ttc ttc aca gac tga aga ttg tcg aga ttt gtg tac caa aaa agg aga tgt cca tgc gga tac ttt ggt gga ata gat gaa gga aat cct att tgt aaa tcg tgt tgt gtt ggt gaa tat tgc gac tac cta cgt aat tat aga ctt gat cca ttt cct cca tgc aaa cta tct atc taa atg taa tta att atg att ttg atg ata atg tta cca tac att ata tca cta ctt ggt tag tgt gta tta ttt agt atg gaa gac cta tta ata att act tat ctt ttg acg atc ttg tta taa tta taa tat aaa aat act tat gac ata gta a-- --- -ct cat aat tgc tga cgc gat aaa ttc gta ata atc tgt ttt gtt caa att ttt ata agg aat cta cag gca taa aaa taa aaa tat aat cta taa tat act ctt aca acg cca tca tga ata gca gtg aat taa ttg ctg tta tta tgg att tag aaa tag tgg acg att ttg tga tat taa tat agt tat taa tga tga aag gat aaa cgc gca tag act cat cct atc tgg agc ctc cga ata ttt ttt cca ttc tgt ttt cca ata att tta tcg att cta atg aat acg aag tta atc taa gtc att tag att atc aaa gtg tta act att tga tcg att aca ttt atg gga tac ctt tga gcc taa cta aca ata acg tga aat ata ttc ttt caa ccg ctg att ttt tta caa att gga tct gtc att act gag tgc gaa aaa tac ata ctt aaa aat ctt tgt ttt aga aac tgt atc gat ttc --- --- tac ata tac gct gat aaa tat aat aac aag aaa ata gaa tta gca tcg ttt aac aca ata tta cga aat att ttg aga ctc atc aac aat gaa aac ttt aaa tac tta aca gag gaa tca atg ata aaa att tta agc gat gat atg tta tat ata aaa aat gag gat ttc acc cca ctg att ctc att aaa tgg tta gag agt aca cca acc atg tac cgt cga gtt act tag atg cct cag aat atc a-t ttc ttt ccc cac aag tta taa aat cac ttt ata gtc atc gac tgg tta gtt caa tct acg aat gta taa cat tct taa aca ata tag cat tct tgg ata aat cat ttc cta gat acc ata tca tcg agt tga tat cta tcg gta taa gta att cac atg ata aga ttt cca taa act gct aca atc ata aaa aaa ttc atg gga aat gat atc ttc acg tag ata tag gtg tag ttt cgc agt gac cgt cct gga taa tat tat cta tat gat ggg tgg ata tga tca gtc cct gta tag aag ttc aaa ggt tat agc gta caa tac atg tac taa ttc ttg gat ata tga tat acc aga gct aaa aat atc atc gtt cta att gcg gag gag ttg cca atg acg aat aca ttt att gta tag gcg gta tac gcg atc agg agt cat cgt tga tat cta gta tcg ata gat gga agc cat caa aac cat att ggc aga agt atg cta aaa tgt gcg aac caa aat gtg ata tgg ggt tgc gat ttt aaa cgg att aat ata tgt cat agg tgg agt cgt taa agg tga cac aca tat acc aac gca cta gag agt tta tca gaa gat gga tgg atg aat cat caa cgt ctt cca ata aaa atg tcc aat atg tcg acg att gtt cat gct gga aag att tat ata tct aga ggt tac aac aat agt agt gta gtt aat gta ata tcg aat ct- -ag tcc tta gct ata atc cga tat atg atg aat gga cca aat tat cat cat taa ata ttc cta gaa tta atc ctg ctc tat ggt cag tgt ata ata aat tat atg tag gag gag taa tat ctg atg atg ttc aaa cta ata cat ctg aaa cat acg ata aag aaa aa

>ON649708.1_Monkeypox_virus_isolate_Monkeypox/PT0023/2022_partial_genome

tgt tat tgt agt atg ata ata tca aaa aga tgg ata taa aga att tac tga cta cat gta cta ttt tac att act aca ttg gct acg gca tat ata cct att tcg tca ctt cca cac gct ccg gta aac ggg tgt cat gtg acg agg gag aat ctt gat aag agg cat aat caa tgt tgt aat ccg atg tcc acc tgg aga att tgc caa ggt cag atg tag agt tgg tag tga taa cac aaa atg tga aca ctg ccc acc tca tac ata tac cgc aat ccc caa tta ttc taa tag atg tca tca atg tag aaa atg ccc aac agg atc att tga taa ggt aaa gtg tac cgg aac aca gaa caa at- gtt cgt gtc atc ctg gtt ggt ata cgc tac tga ttc ttc aca gac tga aga ttg tcg aga ttt gtg tac caa aaa agg aga tgt cca tgc gga tac ttt ggt gga ata gat gaa gga aat cct att tgt aaa tcg tgt tgt gtt ggt gaa tat tgc gac tac cta cgt aat tat aga ctt gat cca ttt cct cca tgc aaa cta tct atc taa atg taa tta att atg att ttg atg ata atg tta cca tac att ata tca cta ctt ggt tag tgt gta tta ttt agt atg gaa gac cta tta ata att act tat ctt ttg acg atc ttg tta taa tta taa tat aaa aat act tat gac ata gta a-- --- -ct cat aat tgc tga cgc gat aaa ttc gta ata atc tgt ttt gtt caa att ttt ata agg aat cta cag gca taa aaa taa aaa tat aat cta taa tat act ctt aca acg cca tca tga ata gca gtg aat taa ttg ctg tta tta tgg att tag aaa tag tgg acg att ttg tga tat taa tat agt tat taa tga tga aag gat aaa cgc gca tag act cat cct atc tgg agc ctc cga ata ttt ttt cca ttc tgt ttt cca ata att tta tcg att cta atg aat acg aag tta atc taa gtc att tag att atc aaa gtg tta act att tga tcg att aca ttt atg gga tac ctt tga gcc taa cta aca ata acg tga aat ata ttc ttt caa ccg ctg att ttt tta caa att gga tct gtc att act gag tgc gaa aaa tac ata ctt aaa aat ctt tgt ttt aga aac tgt atc gat ttc --- --- tac ata tac gct gat aaa tat aat aac aag aaa ata gaa tta gca tcg ttt aac aca ata tta cga aat att ttg aga ctc atc aac aat gaa aac ttt aaa tac tta aca gag gaa tca atg ata aaa att tta agc gat gat atg tta tat ata aaa aat gag gat ttc acc cca ctg att ctc att aaa tgg tta gag agt aca cca acc atg tac cgt cga gtt act tag atg cct cag aat atc a-t ttc ttt ccc cac aag tta taa aat cac ttt ata gtc atc gac tgg tta gtt caa tct acg aat gta taa cat tct taa aca ata tag cat tct tgg ata aat cat ttc cta gat acc ata tca tcg agt tga tat cta tcg gta taa gta att cac atg ata aga ttt cca taa act gct aca atc ata aaa aaa ttc atg gga aat gat atc ttc acg tag ata tag gtg tag ttt cgc agt gac cgt cct gga taa tat tat cta tat gat ggg tgg ata tga tca gtc cct gta tag aag ttc aaa ggt tat agc gta caa tac atg tac taa ttc ttg gat ata tga tat acc aga gct aaa aat atc atc gtt cta att gcg gag gag ttg cca atg acg aat aca ttt att gta tag gcg gta tac gcg atc agg agt cat cgt tga tat cta gta tcg ata gat gga agc cat caa aac cat att ggc aga agt atg cta aaa tgt gcg aac caa aat gtg ata tgg ggt tgc gat ttt aaa cgg att aat ata tgt cat agg tgg agt cgt taa agg tga cac aca tat acc aac gca cta gag agt tta tca gaa gat gga tgg atg aat cat caa cgt ctt cca ata aaa atg tcc aat atg tcg acg att gtt cat gct gga aag att tat ata tct aga ggt tac aac aat agt agt gta gtt aat gta ata tcg aat ct- -ag tcc tta gct ata atc cga tat atg atg aat gga cca aat tat cat cat taa ata ttc cta gaa tta atc ctg ctc tat ggt cag tgt ata ata aat tat atg tag gag gag taa tat ctg atg atg ttc aaa cta ata cat ctg aaa cat acg ata aag aaa aa

>ON649709.1_Monkeypox_virus_isolate_Monkeypox/PT0028/2022_complete_genome

tgt tat tgt agt atg ata ata tca aaa aga tgg ata taa aga att tac tga cta cat gta cta ttt tac att act aca ttg gct acg gca tat ata cct att tcg tca ctt cca cac gct ccg gta aac ggg tgt cat gtg acg agg gag aat ctt gat aag agg cat aat caa tgt tgt aat ccg atg tcc acc tgg aga att tgc caa ggt cag atg tag agt tgg tag tga taa cac aaa atg tga aca ctg ccc acc tca tac ata tac cgc aat ccc caa tta ttc taa tag atg tca tca atg tag aaa atg ccc aac agg atc att tga taa ggt aaa gtg tac cgg aac aca gaa caa at- gtt cgt gtc atc ctg gtt ggt ata cgc tac tga ttc ttc aca gac tga aga ttg tcg aga ttt gtg tac caa aaa agg aga tgt cca tgc gga tac ttt ggt gga ata gat gaa gga aat cct att tgt aaa tcg tgt tgt gtt ggt gaa tat tgc gac tac cta cgt aat tat aga ctt gat cca ttt cct cca tgc aaa cta tct atc taa atg taa tta att atg att ttg atg ata atg tta cca tac att ata tca cta ctt ggt tag tgt gta tta ttt agt atg gaa gac cta tta ata att act tat ctt ttg acg atc ttg tta taa tta taa tat aaa aat act tat gac ata gta a-- --- -ct cat aat tgc tga cgc gat aaa ttc gta ata atc tgt ttt gtt caa att ttt ata agg aat cta cag gca taa aaa taa aaa tat aat cta taa tat act ctt aca acg cca tca tga ata gca gtg aat taa ttg ctg tta tta tgg att tag aaa tag tgg acg att ttg tga tat taa tat agt tat taa tga tga aag gat aaa cgc gca tag act cat cct atc tgg agc ctc cga ata ttt ttt cca ttc tgt ttt cca ata att tta tcg att cta atg aat acg aag tta atc taa gtc att tag att atc aaa gtg tta act att tga tcg att aca ttt atg gga tac ctt tga gcc taa cta aca ata acg tga aat ata ttc ttt caa ccg ctg att ttt tta caa att gga tct gtc att act gag tgc gaa aaa tac ata ctt aaa aat ctt tgt ttt aga aac tgt atc gat ttc --- --- tac ata tac gct gat aaa tat aat aac aag aaa ata gaa tta gca tcg ttt aac aca ata tta cga aat att ttg aga ctc atc aac aat gaa aac ttt aaa tac tta aca gag gaa tca atg ata aaa att tta agc gat gat atg tta tat ata aaa aat gag gat ttc acc cca ctg att ctc att aaa tgg tta gag agt aca cca acc atg tac cgt cga gtt act tag atg cct cag aat atc a-t ttc ttt ccc cac aag tta taa aat cac ttt ata gtc atc gac tgg tta gtt caa tct acg aat gta taa cat tct taa aca ata tag cat tct tgg ata aat cat ttc cta gat acc ata tca tcg agt tga tat cta tcg gta taa gta att cac atg ata aga ttt cca taa act gct aca atc ata aaa aaa ttc atg gga aat gat atc ttc acg tag ata tag gtg tag ttt cgc agt gac cgt cct gga taa tat tat cta tat gat ggg tgg ata tga tca gtc cct gta tag aag ttc aaa ggt tat agc gta caa tac atg tac taa ttc ttg gat ata tga tat acc aga gct aaa aat atc atc gtt cta att gcg gag gag ttg cca atg acg aat aca ttt att gta tag gcg gta tac gcg atc agg agt cat cgt tga tat cta gta tcg ata gat gga agc cat caa aac cat att ggc aga agt atg cta aaa tgt gcg aac caa aat gtg ata tgg ggt tgc gat ttt aaa cgg att aat ata tgt cat agg tgg agt cgt taa agg tga cac aca tat acc aac gca cta gag agt tta tca gaa gat gga tgg atg aat cat caa cgt ctt cca ata aaa atg tcc aat atg tcg acg att gtt cat gct gga aag att tat ata tct aga ggt tac aac aat agt agt gta gtt aat gta ata tcg aat ct- -ag tcc tta gct ata atc cga tat atg atg aat gga cca aat tat cat cat taa ata ttc cta gaa tta atc ctg ctc tat ggt cag tgt ata ata aat tat atg tag gag gag taa tat ctg atg atg ttc aaa cta ata cat ctg aaa cat acg ata aag aaa aa

>ON614676.1_Monkeypox_virus_isolate_INMI-Pt1_partial_genome

tgt tat tgt agt atg ata ata tca aaa aga tgg ata taa aga att tac tga cta cat gta cta ttt tac att act aca ttg gct acg gca tat ata cct att tcg tca ctt cca cac gct ccg gta aac ggg tgt cat gtg acg agg gag aat ctt gat aag agg cat aat caa tgt tgt aat ccg atg tcc acc tgg aga att tgc caa ggt cag atg tag agt tgg tag tga taa cac aaa atg tga aca ctg ccc acc tca tac ata tac cgc aat ccc caa tta ttc taa tag atg tca tca atg tag aaa atg ccc aac agg atc att tga taa ggt aaa gtg tac cgg aac aca gaa caa at- gtt cgt gtc atc ctg gtt ggt ata cgc tac tga ttc ttc aca gac tga aga ttg tcg aga ttt gtg tac caa aaa agg aga tgt cca tgc gga tac ttt ggt gga ata gat gaa gga aat cct att tgt aaa tcg tgt tgt gtt ggt gaa tat tgc gac tac cta cgt aat tat aga ctt gat cca ttt cct cca tgc aaa cta tct atc taa atg taa tta att atg att ttg atg ata atg tta cca tac att ata tca cta ctt ggt tag tgt gta tta ttt agt atg gaa gac cta tta ata att act tat ctt ttg acg atc ttg tta taa tta taa tat aaa aat act tat gac ata gta a-- --- -ct cat aat tgc tga cgc gat aaa ttc gta ata atc tgt ttt gtt caa att ttt ata agg aat cta cag gca taa aaa taa aaa tat aat cta taa tat act ctt aca acg cca tca tga ata gca gtg aat taa ttg ctg tta tta tgg att tag aaa tag tgg acg att ttg tga tat taa tat agt tat taa tga tga aag gat aaa cgc gca tag act cat cct atc tgg agc ctc cga ata ttt ttt cca ttc tgt ttt cca ata att tta tcg att cta atg aat acg aag tta atc taa gtc att tag att atc aaa gtg tta act att tga tcg att aca ttt atg gga tac ctt tga gcc taa cta aca ata acg tga aat ata ttc ttt caa ccg ctg att ttt tta caa att gga tct gtc att act gag tgc gaa aaa tac ata ctt aaa aat ctt tgt ttt aga aac tgt atc gat ttc --- --- tac ata tac gct gat aaa tat aat aac aag aaa ata gaa tta gca tcg ttt aac aca ata tta cga aat att ttg aga ctc atc aac aat gaa aac ttt aaa tac tta aca gag gaa tca atg ata aaa att tta agc gat gat atg tta tat ata aaa aat gag gat ttc acc cca ctg att ctc att aaa tgg tta gag agt aca cca acc atg tac cgt cga gtt act tag atg cct cag aat atc a-t ttc ttt ccc cac aag tta taa aat cac ttt ata gtc atc gac tgg tta gtt caa tct acg aat gta taa cat tct taa aca ata tag cat tct tgg ata aat cat ttc cta gat acc ata tca tcg agt tga tat cta tcg gta taa gta att cac atg ata aga ttt cca taa act gct aca atc ata aaa aaa ttc atg gga aat gat atc ttc acg tag ata tag gtg tag ttt cgc agt gac cgt cct gga taa tat tat cta tat gat ggg tgg ata tga tca gtc cct gta tag aag ttc aaa ggt tat agc gta caa tac atg tac taa ttc ttg gat ata tga tat acc aga gct aaa aat atc atc gtt cta att gcg gag gag ttg cca atg acg aat aca ttt att gta tag gcg gta tac gcg atc agg agt cat cgt tga tat cta gta tcg ata gat gga agc cat caa aac cat att ggc aga agt atg cta aaa tgt gcg aac caa aat gtg ata tgg ggt tgc gat ttt aaa cgg att aat ata tgt cat agg tgg agt cgt taa agg tga cac aca tat acc aac gca cta gag agt tta tca gaa gat gga tgg atg aat cat caa cgt ctt cca ata aaa atg tcc aat atg tcg acg att gtt cat gct gga aag att tat ata tct aga ggt tac aac aat agt agt gta gtt aat gta ata tcg aat ct- -ag tcc tta gct ata atc cga tat atg atg aat gga cca aat tat cat cat taa ata ttc cta gaa tta atc ctg ctc tat ggt cag tgt ata ata aat tat atg tag gag gag taa tat ctg atg atg ttc aaa cta ata cat ctg aaa cat acg ata aag aaa aa

>ON649712.1_Monkeypox_virus_isolate_Monkeypox/PT0025/2022_partial_genome

tgt tat tgt agt atg ata ata tca aaa aga tgg ata taa aga att tac tga cta cat gta cta ttt tac att act aca ttg gct acg gca tat ata cct att tcg tca ctt cca cac gct ccg gta aac ggg tgt cat gtg acg agg gag aat ctt gat aag agg cat aat caa tgt tgt aat ccg atg tcc acc tgg aga att tgc caa ggt cag atg tag agt tgg tag tga taa cac aaa atg tga aca ctg ccc acc tca tac ata tac cgc aat ccc caa tta ttc taa tag atg tca tca atg tag aaa atg ccc aac agg atc att tga taa ggt aaa gtg tac cgg aac aca gaa caa at- gtt cgt gtc atc ctg gtt ggt ata cgc tac tga ttc ttc aca gac tga aga ttg tcg aga ttt gtg tac caa aaa agg aga tgt cca tgc gga tac ttt ggt gga ata gat gaa gga aat cct att tgt aaa tcg tgt tgt gtt ggt gaa tat tgc gac tac cta cgt aat tat aga ctt gat cca ttt cct cca tgc aaa cta tct atc taa atg taa tta att atg att ttg atg ata atg tta cca tac att ata tca cta ctt ggt tag tgt gta tta ttt agt atg gaa gac cta tta ata att act tat ctt ttg acg atc ttg tta taa tta taa tat aaa aat act tat gac ata gta a-- --- -ct cat aat tgc tga cgc gat aaa ttc gta ata atc tgt ttt gtt caa att ttt ata agg aat cta cag gca taa aaa taa aaa tat aat cta taa tat act ctt aca acg cca tca tga ata gca gtg aat taa ttg ctg tta tta tgg att tag aaa tag tgg acg att ttg tga tat taa tat agt tat taa tga tga aag gat aaa cgc gca tag act cat cct atc tgg agc ctc cga ata ttt ttt cca ttc tgt ttt cca ata att tta tcg att cta atg aat acg aag tta atc taa gtc att tag att atc aaa gtg tta act att tga tcg att aca ttt atg gga tac ctt tga gcc taa cta aca ata acg tga aat ata ttc ttt caa ccg ctg att ttt tta caa att gga tct gtc att act gag tgc gaa aaa tac ata ctt aaa aat ctt tgt ttt aga aac tgt atc gat ttc --- --- tac ata tac gct gat aaa tat aat aac aag aaa ata gaa tta gca tcg ttt aac aca ata tta cga aat att ttg aga ctc atc aac aat gaa aac ttt aaa tac tta aca gag gaa tca atg ata aaa att tta agc gat gat atg tta tat ata aaa aat gag gat ttc acc cca ctg att ctc att aaa tgg tta gag agt aca cca acc atg tac cgt cga gtt act tag atg cct cag aat atc a-t ttc ttt ccc cac aag tta taa aat cac ttt ata gtc atc gac tgg tta gtt caa tct acg aat gta taa cat tct taa aca ata tag cat tct tgg ata aat cat ttc cta gat acc ata tca tcg agt tga tat cta tcg gta taa gta att cac atg ata aga ttt cca taa act gct aca atc ata aaa aaa ttc atg gga aat gat atc ttc acg tag ata tag gtg tag ttt cgc agt gac cgt cct gga taa tat tat cta tat gat ggg tgg ata tga tca gtc cct gta tag aag ttc aaa ggt tat agc gta caa tac atg tac taa ttc ttg gat ata tga tat acc aga gct aaa aat atc atc gtt cta att gcg gag gag ttg cca atg acg aat aca ttt att gta tag gcg gta tac gcg atc agg agt cat cgt tga tat cta gta tcg ata gat gga agc cat caa aac cat att ggc aga agt atg cta aaa tgt gcg aac caa aat gtg ata tgg ggt tgc gat ttt aaa cgg att aat ata tgt cat agg tgg agt cgt taa agg tga cac aca tat acc aac gca cta gag agt tta tca gaa gat gga tgg atg aat cat caa cgt ctt cca ata aaa atg tcc aat atg tcg acg att gtt cat gct gga aag att tat ata tct aga ggt tac aac aat agt agt gta gtt aat gta ata tcg aat ct- -ag tcc tta gct ata atc cga tat atg atg aat gga cca aat tat cat cat taa ata ttc cta gaa tta atc ctg ctc tat ggt cag tgt ata ata aat tat atg tag gag gag taa tat ctg atg atg ttc aaa cta ata cat ctg aaa cat acg ata aag aaa aa

>ON595760.2_Monkeypox_virus_isolate_MPXV-CH-38134631/2022_partial_genome

tgt tat tgt agt atg ata ata tca aaa aga tgg ata taa aga att tac tga cta cat gta cta ttt tac att act aca ttg gct acg gca tat ata cct att tcg tca ctt cca cac gct ccg gta aac ggg tgt cat gtg acg agg gag aat ctt gat aag agg cat aat caa tgt tgt aat ccg atg tcc acc tgg aga att tgc caa ggt cag atg tag agt tgg tag tga taa cac aaa atg tga aca ctg ccc acc tca tac ata tac cgc aat ccc caa tta ttc taa tag atg tca tca atg tag aaa atg ccc aac agg atc att tga taa ggt aaa gtg tac cgg aac aca gaa caa at- gtt cgt gtc atc ctg gtt ggt ata cgc tac tga ttc ttc aca gac tga aga ttg tcg aga ttt gtg tac caa aaa agg aga tgt cca tgc gga tac ttt ggt gga ata gat gaa gga aat cct att tgt aaa tcg tgt tgt gtt ggt gaa tat tgc gac tac cta cgt aat tat aga ctt gat cca ttt cct cca tgc aaa cta tct atc taa atg taa tta att atg att ttg atg ata atg tta cca tac att ata tca cta ctt ggt tag tgt gta tta ttt agt atg gaa gac cta tta ata att act tat ctt ttg acg atc ttg tta taa tta taa tat aaa aat act tat gac ata gta a-- --- -ct cat aat tgc tga cgc gat aaa ttc gta ata atc tgt ttt gtt caa att ttt ata agg aat cta cag gca taa aaa taa aaa tat aat cta taa tat act ctt aca acg cca tca tga ata gca gtg aat taa ttg ctg tta tta tgg att tag aaa tag tgg acg att ttg tga tat taa tat agt tat taa tga tga aag gat aaa cgc gca tag act cat cct atc tgg agc ctc cga ata ttt ttt cca ttc tgt ttt cca ata att tta tcg att cta atg aat acg aag tta atc taa gtc att tag att atc aaa gtg tta act att tga tcg att aca ttt atg gga tac ctt tga gcc taa cta aca ata acg tga aat ata ttc ttt caa ccg ctg att ttt tta caa att gga tct gtc att act gag tgc gaa aaa tac ata ctt aaa aat ctt tgt ttt aga aac tgt atc gat ttc --- --- tac ata tac gct gat aaa tat aat aac aag aaa ata gaa tta gca tcg ttt aac aca ata tta cga aat att ttg aga ctc atc aac aat gaa aac ttt aaa tac tta aca gag gaa tca atg ata aaa att tta agc gat gat atg tta tat ata aaa aat gag gat ttc acc cca ctg att ctc att aaa tgg tta gag agt aca cca acc atg tac cgt cga gtt act tag atg cct cag aat atc a-t ttc ttt ccc cac aag tta taa aat cac ttt ata gtc atc gac tgg tta gtt caa tct acg aat gta taa cat tct taa aca ata tag cat tct tgg ata aat cat ttc cta gat acc ata tca tcg agt tga tat cta tcg gta taa gta att cac atg ata aga ttt cca taa act gct aca atc ata aaa aaa ttc atg gga aat gat atc ttc acg tag ata tag gtg tag ttt cgc agt gac cgt cct gga taa tat tat cta tat gat ggg tgg ata tga tca gtc cct gta tag aag ttc aaa ggt tat agc gta caa tac atg tac taa ttc ttg gat ata tga tat acc aga gct aaa aat atc atc gtt cta att gcg gag gag ttg cca atg acg aat aca ttt att gta tag gcg gta tac gcg atc agg agt cat cgt tga tat cta gta tcg ata gat gga agc cat caa aac cat att ggc aga agt atg cta aaa tgt gcg aac caa aat gtg ata tgg ggt tgc gat ttt aaa cgg att aat ata tgt cat agg tgg agt cgt taa agg tga cac aca tat acc aac gca cta gag agt tta tca gaa gat gga tgg atg aat cat caa cgt ctt cca ata aaa atg tcc aat atg tcg acg att gtt cat gct gga aag att tat ata tct aga ggt tac aac aat agt agt gta gtt aat gta ata tcg aat ct- -ag tcc tta gct ata atc cga tat atg atg aat gga cca aat tat cat cat taa ata ttc cta gaa tta atc ctg ctc tat ggt cag tgt ata ata aat tat atg tag gag gag taa tat ctg atg atg ttc aaa cta ata cat ctg aaa cat acg ata aag aaa aa

>MT903337.1_Monkeypox_virus_isolate_MPXV-M2940_FCT

tgt tat tgt agt atg ata ata tca aaa aga tgg ata taa aga att tac tga cta cat gta cta ttt tac att act aca ttg gct acg gca tat ata cct att tcg tca ctt cca cac gct ccg gta aac ggg tgt cat gtg acg agg gag aat ctt gat aag agg cat aat caa tgt tgt aat ccg atg tcc acc tgg aga att tgc caa ggt cag atg tag agt tgg tag tga taa cac aaa atg tga aca ctg ccc acc tca tac ata tac cgc aat ccc caa tta ttc taa tag atg tca tca atg tag aaa atg ccc aac agg atc att tga taa ggt aaa gtg tac cgg aac aca gaa caa at- gtt cgt gtc atc ctg gtt ggt ata cgc tac tga ttc ttc aca gac tga aga ttg tcg aga ttt gtg tac caa aaa agg aga tgt cca tgc gga tac ttt ggt gga ata gat gaa gga aat cct att tgt aaa tcg tgt tgt gtt ggt gaa tat tgc gac tac cta cgt aat tat aga ctt gat cca ttt cct cca tgc aaa cta tct atc taa atg taa tta att atg att ttg atg ata atg tta cca tac att ata tca cta ctt ggt tag tgt gta tta ttt agt atg gaa gac cta tta ata att act tat ctt ttg acg atc ttg tta taa tta taa tat aaa aat act tat gac ata gta a-- --- -ct cat aat tgc tga cgc gat aaa ttc gta ata atc tgt ttt gtt caa att ttt ata agg aat cta cag gca taa aaa taa aaa tat aat cta taa tat act ctt aca acg cca tca tga ata gca gtg aat taa ttg ctg tta tta tgg att tag aaa tag tgg acg att ttg tga tat taa tat agt tat taa tga tga aag gat aaa cgc gca tag act cat cct atc tgg agc ctc cga ata ttt ttt cca ttc tgt ttt cca ata att tta tcg att cta atg aat acg aag tta atc taa gtc att tag att atc aaa gtg tta act att tga tcg att aca ttt atg gga tac ctt tga gcc taa cta aca ata acg tga aat ata ttc ttt caa ccg ctg att ttt tta caa att gga tct gtc att act gag tgc gaa aaa tac ata ctt aaa aat ctt tgt ttt aga aac tgt atc gat ttc --- --- tac ata tac gct gat aaa tat aat aac aag aaa ata gaa tta gca tcg ttt aac aca ata tta cga aat att ttg aga ctc atc aac aat gaa aac ttt aaa tac tta aca gag gaa tca atg ata aaa att tta agc gat gat atg tta tat ata aaa aat gag gat ttc acc cca ctg att ctc att aaa tgg tta gag agt aca cca acc atg tac cgt cga gtt act tag atg cct cag aat atc a-t ttc ttt ccc cac aag tta taa aat cac ttt ata gtc atc gac tgg tta gtt caa tct acg aat gta taa cat tct taa aca ata tag cat tct tgg ata aat cat ttc cta gat acc ata tca tcg agt tga tat cta tcg gta taa gta att cac atg ata aga ttt cca taa act gct aca atc ata aaa aaa ttc atg gga aat gat atc ttc acg tag ata tag gtg tag ttt cgc agt gac cgt cct gga taa tat tat cta tat gat ggg tgg ata tga tca gtc cct gta tag aag ttc aaa ggt tat agc gta caa tac atg tac taa ttc ttg gat ata tga tat acc aga gct aaa aat atc atc gtt cta att gcg gag gag ttg cca atg acg aat aca ttt att gta tag gcg gta tac gcg atc agg agt cat cgt tga tat cta gta tcg ata gat gga agc cat caa aac cat att ggc aga agt atg cta aaa tgt gcg aac caa aat gtg ata tgg ggt tgc gat ttt aaa cgg att aat ata tgt cat agg tgg agt cgt taa agg tga cac aca tat acc aac gca cta gag agt tta tca gaa gat gga tgg atg aat cat caa cgt ctt cca ata aaa atg tcc aat atg tcg acg att gtt cat gct gga aag att tat ata tct aga ggt tac aac aat agt agt gta gtt aat gta ata tcg aat ct- -ag tcc tta gct ata atc cga tat atg atg aat gga cca aat tat cat cat taa ata ttc cta gaa tta atc ctg ctc tat ggt cag tgt ata ata aat tat atg tag gag gag taa tat ctg atg atg ttc aaa cta ata cat ctg aaa cat acg ata aag aaa aa

>MT903339.1_Monkeypox_virus_isolate_MPXV-M3021_Delta

tgt tat tgt agt atg ata ata tca aaa aga tgg ata taa aga att tac tga cta cat gta cta ttt tac att act aca ttg gct acg gca tat ata cct att tcg tca ctt cca cac gct ccg gta aac ggg tgt cat gtg acg agg gag aat ctt gat aag agg cat aat caa tgt tgt aat ccg atg tcc acc tgg aga att tgc caa ggt cag atg tag agt tgg tag tga taa cac aaa atg tga aca ctg ccc acc tca tac ata tac cgc aat ccc caa tta ttc taa tag atg tca tca atg tag aaa atg ccc aac agg atc att tga taa ggt aaa gtg tac cgg aac aca gaa caa at- gtt cgt gtc atc ctg gtt ggt ata cgc tac tga ttc ttc aca gac tga aga ttg tcg aga ttt gtg tac caa aaa agg aga tgt cca tgc gga tac ttt ggt gga ata gat gaa gga aat cct att tgt aaa tcg tgt tgt gtt ggt gaa tat tgc gac tac cta cgt aat tat aga ctt gat cca ttt cct cca tgc aaa cta tct atc taa atg taa tta att atg att ttg atg ata atg tta cca tac att ata tca cta ctt ggt tag tgt gta tta ttt agt atg gaa gac cta tta ata att act tat ctt ttg acg atc ttg tta taa tta taa tat aaa aat act tat gac ata gta a-- --- -ct cat aat tgc tga cgc gat aaa ttc gta ata atc tgt ttt gtt caa att ttt ata agg aat cta cag gca taa aaa taa aaa tat aat cta taa tat act ctt aca acg cca tca tga ata gca gtg aat taa ttg ctg tta tta tgg att tag aaa tag tgg acg att ttg tga tat taa tat agt tat taa tga tga aag gat aaa cgc gca tag act cat cct atc tgg agc ctc cga ata ttt ttt cca ttc tgt ttt cca ata att tta tcg att cta atg aat acg aag tta atc taa gtc att tag att atc aaa gtg tta act att tga tcg att aca ttt atg gga tac ctt tga gcc taa cta aca ata acg tga aat ata ttc ttt caa ccg ctg att ttt tta caa att gga tct gtc att act gag tgc gaa aaa tac ata ctt aaa aat ctt tgt ttt aga aac tgt atc gat ttc --- --- tac ata tac gct gat aaa tat aat aac aag aaa ata gaa tta gca tcg ttt aac aca ata tta cga aat att ttg aga ctc atc aac aat gaa aac ttt aaa tac tta aca gag gaa tca atg ata aaa att tta agc gat gat atg tta tat ata aaa aat gag gat ttc acc cca ctg att ctc att aaa tgg tta gag agt aca cca acc atg tac cgt cga gtt act tag atg cct cag aat atc a-t ttc ttt ccc cac aag tta taa aat cac ttt ata gtc atc gac tgg tta gtt caa tct acg aat gta taa cat tct taa aca ata tag cat tct tgg ata aat cat ttc cta gat acc ata tca tcg agt tga tat cta tcg gta taa gta att cac atg ata aga ttt cca taa act gct aca atc ata aaa aaa ttc atg gga aat gat atc ttc acg tag ata tag gtg tag ttt cgc agt gac cgt cct gga taa tat tat cta tat gat ggg tgg ata tga tca gtc cct gta tag aag ttc aaa ggt tat agc gta caa tac atg tac taa ttc ttg gat ata tga tat acc aga gct aaa aat atc atc gtt cta att gcg gag gag ttg cca atg acg aat aca ttt att gta tag gcg gta tac gcg atc agg agt cat cgt tga tat cta gta tcg ata gat gga agc cat caa aac cat att ggc aga agt atg cta aaa tgt gcg aac caa aat gtg ata tgg ggt tgc gat ttt aaa cgg att aat ata tgt cat agg tgg agt cgt taa agg tga cac aca tat acc aac gca cta gag agt tta tca gaa gat gga tgg atg aat cat caa cgt ctt cca ata aaa atg tcc aat atg tcg acg att gtt cat gct gga aag att tat ata tct aga ggt tac aac aat agt agt gta gtt aat gta ata tcg aat ct- -ag tcc tta gct ata atc cga tat atg atg aat gga cca aat tat cat cat taa ata ttc cta gaa tta atc ctg ctc tat ggt cag tgt ata ata aat tat atg tag gag gag taa tat ctg atg atg ttc aaa cta ata cat ctg aaa cat acg ata aag aaa aa

>MT903338.1_Monkeypox_virus_isolate_MPXV-M2957_Lagos

tgt tat tgt agt atg ata ata tca aaa aga tgg ata taa aga att tac tga cta cat gta cta ttt tac att act aca ttg gct acg gca tat ata cct att tcg tca ctt cca cac gct ccg gta aac ggg tgt cat gtg acg agg gag aat ctt gat aag agg cat aat caa tgt tgt aat ccg atg tcc acc tgg aga att tgc caa ggt cag atg tag agt tgg tag tga taa cac aaa atg tga aca ctg ccc acc tca tac ata tac cgc aat ccc caa tta ttc taa tag atg tca tca atg tag aaa atg ccc aac agg atc att tga taa ggt aaa gtg tac cgg aac aca gaa caa at- gtt cgt gtc atc ctg gtt ggt ata cgc tac tga ttc ttc aca gac tga aga ttg tcg aga ttt gtg tac caa aaa agg aga tgt cca tgc gga tac ttt ggt gga ata gat gaa gga aat cct att tgt aaa tcg tgt tgt gtt ggt gaa tat tgc gac tac cta cgt aat tat aga ctt gat cca ttt cct cca tgc aaa cta tct atc taa atg taa tta att atg att ttg atg ata atg tta cca tac att ata tca cta ctt ggt tag tgt gta tta ttt agt atg gaa gac cta tta ata att act tat ctt ttg acg atc ttg tta taa tta taa tat aaa aat act tat gac ata gta a-- --- -ct cat aat tgc tga cgc gat aaa ttc gta ata atc tgt ttt gtt caa att ttt ata agg aat cta cag gca taa aaa taa aaa tat aat cta taa tat act ctt aca acg cca tca tga ata gca gtg aat taa ttg ctg tta tta tgg att tag aaa tag tgg acg att ttg tga tat taa tat agt tat taa tga tga aag gat aaa cgc gca tag act cat cct atc tgg agc ctc cga ata ttt ttt cca ttc tgt ttt cca ata att tta tcg att cta atg aat acg aag tta atc taa gtc att tag att atc aaa gtg tta act att tga tcg att aca ttt atg gga tac ctt tga gcc taa cta aca ata acg tga aat ata ttc ttt caa ccg ctg att ttt tta caa att gga tct gtc att act gag tgc gaa aaa tac ata ctt aaa aat ctt tgt ttt aga aac tgt atc gat ttc --- --- tac ata tac gct gat aaa tat aat aac aag aaa ata gaa tta gca tcg ttt aac aca ata tta cga aat att ttg aga ctc atc aac aat gaa aac ttt aaa tac tta aca gag gaa tca atg ata aaa att tta agc gat gat atg tta tat ata aaa aat gag gat ttc acc cca ctg att ctc att aaa tgg tta gag agt aca cca acc atg tac cgt cga gtt act tag atg cct cag aat atc a-t ttc ttt ccc cac aag tta taa aat cac ttt ata gtc atc gac tgg tta gtt caa tct acg aat gta taa cat tct taa aca ata tag cat tct tgg ata aat cat ttc cta gat acc ata tca tcg agt tga tat cta tcg gta taa gta att cac atg ata aga ttt cca taa act gct aca atc ata aaa aaa ttc atg gga aat gat atc ttc acg tag ata tag gtg tag ttt cgc agt gac cgt cct gga taa tat tat cta tat gat ggg tgg ata tga tca gtc cct gta tag aag ttc aaa ggt tat agc gta caa tac atg tac taa ttc ttg gat ata tga tat acc aga gct aaa aat atc atc gtt cta att gcg gag gag ttg cca atg acg aat aca ttt att gta tag gcg gta tac gcg atc agg agt cat cgt tga tat cta gta tcg ata gat gga agc cat caa aac cat att ggc aga agt atg cta aaa tgt gcg aac caa aat gtg ata tgg ggt tgc gat ttt aaa cgg att aat ata tgt cat agg tgg agt cgt taa agg tga cac aca tat acc aac gca cta gag agt tta tca gaa gat gga tgg atg aat cat caa cgt ctt cca ata aaa atg tcc aat atg tcg acg att gtt cat gct gga aag att tat ata tct aga ggt tac aac aat agt agt gta gtt aat gta ata tcg aat ct- -ag tcc tta gct ata atc cga tat atg atg aat gga cca aat tat cat cat taa ata ttc cta gaa tta atc ctg ctc tat ggt cag tgt ata ata aat tat atg tag gag gag taa tat ctg atg atg ttc aaa cta ata cat ctg aaa cat acg ata aag aaa aa

>KJ642617.1_Monkeypox_virus_strain_Nigeria-SE-1971_complete_genome

tgt tat tgt agt atg ata ata tca aaa aga tgg ata taa aga att tac tga cta cat gta cta ttt tac att act aca ttg gct acg gca tat ata cct att tcg tca ctt cca cac gct ccg gta aac ggg tgt cat gtg acg agg gag aat ctt gat aag agg cat aat caa tgt tgt aat ccg atg tcc acc tgg aga att tgc caa ggt cag atg tag agt tgg tag tga taa cac aaa atg tga aca ctg ccc acc tca tac ata tac cgc aat ccc caa tta ttc taa tag atg tca tca atg tag aaa atg ccc aac agg atc att tga taa ggt aaa gtg tac cgg aac aca gaa caa at- gtt cgt gtc atc ctg gtt ggt ata cgc tac tga ttc ttc aca gac tga aga ttg tcg aga ttt gtg tac caa aaa agg aga tgt cca tgc gga tac ttt ggt gga ata gat gaa gga aat cct att tgt aaa tcg tgt tgt gtt ggt gaa tat tgc gac tac cta cgt aat tat aga ctt gat cca ttt cct cca tgc aaa cta tct atc taa atg taa tta att atg att ttg atg ata atg tta cca tac att ata tca cta ctt ggt tag tgt gta tta ttt agt atg gaa gac cta tta ata att act tat ctt ttg acg atc ttg tta taa tta taa tat aaa aat act tat gac ata gta a-- --- -ct cat aat tgc tga cgc gat aaa ttc gta ata atc tgt ttt gtt caa att ttt ata agg aat cta cag gca taa aaa taa aaa tat aat cta taa tat act ctt aca acg cca tca tga ata gca gtg aat taa ttg ctg tta tta tgg att tag aaa tag tgg acg att ttg tga tat taa tat agt tat taa tga tga aag gat aaa cgc gca tag act cat cct atc tgg agc ctc cga ata ttt ttt cca ttc tgt ttt cca ata att tta tcg att cta atg aat acg aag tta atc taa gtc att tag att atc aaa gtg tta act att tga tcg att aca ttt atg gga tac ctt tga gcc taa cta aca ata acg tga aat ata ttc ttt caa ccg ctg att ttt tta caa att gga tct gtc att act gag tgc gaa aaa tac ata ctt aaa aat ctt tgt ttt aga aac tgt atc gat ttc --- --- tac ata tac gct gat aaa tat aat aac aag aaa ata gaa tta gca tcg ttt aac aca ata tta cga aat att ttg aga ctc atc aac aat gaa aac ttt aaa tac tta aca gag gaa tca atg ata aaa att tta agc gat gat atg tta tat ata aaa aat gag gat ttc acc cca ctg att ctc att aaa tgg tta gag agt aca cca acc atg tac cgt cga gtt act tag atg cct cag aat atc a-t ttc ttt ccc cac aag tta taa aat cac ttt ata gtc atc gac tgg tta gtt caa tct acg aat gta taa cat tct taa aca ata tag cat tct tgg ata aat cat ttc cta gat acc ata tca tcg agt tga tat cta tcg gta taa gta att cac atg ata aga ttt cca taa act gct aca atc ata aaa aaa ttc atg gga aat gat atc ttc acg tag ata tag gtg tag ttt cgc agt gac cgt cct gga taa tat tat cta tat gat ggg tgg ata tga tca gtc cct gta tag aag ttc aaa ggt tat agc gta caa tac atg tac taa ttc ttg gat ata tga tat acc aga gct -aa aat atc atc gtt cta att gcg gag gag ttg cca atg acg aat aca ttt att gta tag gcg gta tac gcg atc agg agt cat cgt tga tat cta gta tcg ata gat gga agc cat caa aac cat att ggc aga agt atg cta aaa tgt gcg aac caa aat gtg ata tgg ggt tgc gat ttt aaa cgg att aat ata tgt cat agg tgg agt cgt taa agg tga cac aca tat acc aac gca cta gag agt tta tca gaa gat gga tgg atg aat cat caa cgt ctt cca ata aaa atg tcc aat atg tcg acg att gtt cat gct gga aag att tat ata tct aga ggt tac aac aat agt agt gta gtt aat gta ata tcg aat ct- -ag tcc tta gct ata atc cga tat atg atg aat gga cca aat tat cat cat taa ata ttc cta gaa tta atc ctg ctc tat ggt cag tgt ata ata aat tat atg tag gag gag taa tat ctg atg atg ttc aaa cta ata cat ctg aaa cat acg ata aag aaa aa

>ON615424.1_Monkeypox_virus_isolate_MPXV_2022_NL001_partial_genome

tgt tat tgt agt atg ata ata tca aaa aga tgg ata taa aga att tac tga cta cat gta cta ttt tac att act aca ttg gct acg gca tat ata cct att tcg tca ctt cca cac gct ccg gta aac ggg tgt cat gtg acg agg gag aat ctt gat aag agg cat aat caa tgt tgt aat ccg atg tcc acc tgg aga att tgc caa ggt cag atg tag agt tgg tag tga taa cac aaa atg tga aca ctg ccc acc tca tac ata tac cgc aat ccc caa tta ttc taa tag atg tca tca atg tag aaa atg ccc aac agg atc att tga taa ggt aaa gtg tac cgg aac aca gaa caa at- gtt cgt gtc atc ctg gtt ggt ata cgc tac tga ttc ttc aca gac tga aga ttg tcg aga ttt gtg tac caa aaa agg aga tgt cca tgc gga tac ttt ggt gga ata gat gaa gga aat cct att tgt aaa tcg tgt tgt gtt ggt gaa tat tgc gac tac cta cgt aat tat aga ctt gat cca ttt cct cca tgc aaa cta tct atc taa atg taa tta att atg att ttg atg ata atg tta cca tac att ata tca cta ctt ggt tag tgt gta tta ttt agt atg gaa gac cta tta ata att act tat ctt ttg acg atc ttg tta taa tta taa tat aaa aat act tat gac ata gta a-- --- -ct cat aat tgc tga cgc gat aaa ttc gta ata atc tgt ttt gtt caa att ttt ata agg aat cta cag gca taa aaa taa aaa tat aat cta taa tat act ctt aca acg cca tca tga ata gca gtg aat taa ttg ctg tta tta tgg att tag aaa tag tgg acg att ttg tga tat taa tat agt tat taa tga tga aag gat aaa cgc gca tag act cat cct atc tgg agc ctc cga ata ttt ttt cca ttc tgt ttt cca ata att tta tcg att cta atg aat acg aag tta atc taa gtc att tag att atc aaa gtg tta act att tga tcg att aca ttt atg gga tac ctt tga gcc taa cta aca ata acg tga aat ata ttc ttt caa ccg ctg att ttt tta caa att gga tct gtc att act gag tgc gaa aaa tac ata ctt aaa aat ctt tgt ttt aga aac tgt atc gat ttc --- --- tac ata tac gct gat aaa tat aat aac aag aaa ata gaa tta gca tcg ttt aac aca ata tta cga aat att ttg aga ctc atc aac aat gaa aac ttt aaa tac tta aca gag gaa tca atg ata aaa att tta agc gat gat atg tta tat ata aaa aat gag gat ttc acc cca ctg att ctc att aaa tgg tta gag agt aca cca acc atg tac cgt cga gtt act tag atg cct cag aat atc a-t ttc ttt ccc cac aag tta taa aat cac ttt ata gtc atc gac tgg tta gtt caa tct acg aat gta taa cat tct taa aca ata tag cat tct tgg ata aat cat ttc cta gat acc ata tca tcg agt tga tat cta tcg gta taa gta att cac atg ata aga ttt cca taa act gct aca atc ata aaa aaa ttc atg gga aat gat atc ttc acg tag ata tag gtg tag ttt cgc agt gac cgt cct gga taa tat tat cta tat gat ggg tgg ata tga tca gtc cct gta tag aag ttc aaa ggt tat agc gta caa tac atg tac taa ttc ttg gat ata tga tat acc aga gct aaa aat atc atc gtt cta att gcg gag gag ttg cca atg acg aat aca ttt att gta tag gcg gta tac gcg atc agg agt cat cgt tga tat cta gta tcg ata gat gga agc cat caa aac cat att ggc aga agt atg cta aaa tgt gcg aac caa aat gtg ata tgg ggt tgc gat ttt aaa cgg att aat ata tgt cat agg tgg agt cgt taa agg tga cac aca tat acc aac gca cta gag agt tta tca gaa gat gga tgg atg aat cat caa cgt ctt cca ata aaa atg tcc aat atg tcg acg att gtt cat gct gga aag att tat ata tct aga ggt tac aac aat agt agt gta gtt aat gta ata tcg aat ct- -ag tcc tta gct ata atc cga tat atg atg aat gga cca aat tat cat cat taa ata ttc cta gaa tta atc ctg ctc tat ggt cag tgt ata ata aat tat atg tag gag gag taa tat ctg atg atg ttc aaa cta ata cat ctg aaa cat acg ata aag aaa aa

>ON585034.1_Monkeypox_virus_isolate_Monkeypox/PT0007/2022_complete_genome

tgt tat tgt agt atg ata ata tca aaa aga tgg ata taa aga att tac tga cta cat gta cta ttt tac att act aca ttg gct acg gca tat ata cct att tcg tca ctt cca cac gct ccg gta aac ggg tgt cat gtg acg agg gag aat ctt gat aag agg cat aat caa tgt tgt aat ccg atg tcc acc tgg aga att tgc caa ggt cag atg tag agt tgg tag tga taa cac aaa atg tga aca ctg ccc acc tca tac ata tac cgc aat ccc caa tta ttc taa tag atg tca tca atg tag aaa atg ccc aac agg atc att tga taa ggt aaa gtg tac cgg aac aca gaa caa at- gtt cgt gtc atc ctg gtt ggt ata cgc tac tga ttc ttc aca gac tga aga ttg tcg aga ttt gtg tac caa aaa agg aga tgt cca tgc gga tac ttt ggt gga ata gat gaa gga aat cct att tgt aaa tcg tgt tgt gtt ggt gaa tat tgc gac tac cta cgt aat tat aga ctt gat cca ttt cct cca tgc aaa cta tct atc taa atg taa tta att atg att ttg atg ata atg tta cca tac att ata tca cta ctt ggt tag tgt gta tta ttt agt atg gaa gac cta tta ata att act tat ctt ttg acg atc ttg tta taa tta taa tat aaa aat act tat gac ata gta a-- --- -ct cat aat tgc tga cgc gat aaa ttc gta ata atc tgt ttt gtt caa att ttt ata agg aat cta cag gca taa aaa taa aaa tat aat cta taa tat act ctt aca acg cca tca tga ata gca gtg aat taa ttg ctg tta tta tgg att tag aaa tag tgg acg att ttg tga tat taa tat agt tat taa tga tga aag gat aaa cgc gca tag act cat cct atc tgg agc ctc cga ata ttt ttt cca ttc tgt ttt cca ata att tta tcg att cta atg aat acg aag tta atc taa gtc att tag att atc aaa gtg tta act att tga tcg att aca ttt atg gga tac ctt tga gcc taa cta aca ata acg tga aat ata ttc ttt caa ccg ctg att ttt tta caa att gga tct gtc att act gag tgc gaa aaa tac ata ctt aaa aat ctt tgt ttt aga aac tgt atc gat ttc --- --- tac ata tac gct gat aaa tat aat aac aag aaa ata gaa tta gca tcg ttt aac aca ata tta cga aat att ttg aga ctc atc aac aat gaa aac ttt aaa tac tta aca gag gaa tca atg ata aaa att tta agc gat gat atg tta tat ata aaa aat gag gat ttc acc cca ctg att ctc att aaa tgg tta gag agt aca cca acc atg tac cgt cga gtt act tag atg cct cag aat atc a-t ttc ttt ccc cac aag tta taa aat cac ttt ata gtc atc gac tgg tta gtt caa tct acg aat gta taa cat tct taa aca ata tag cat tct tgg ata aat cat ttc cta gat acc ata tca tcg agt tga tat cta tcg gta taa gta att cac atg ata aga ttt cca taa act gct aca atc ata aaa aaa ttc atg gga aat gat atc ttc acg tag ata tag gtg tag ttt cgc agt gac cgt cct gga taa tat tat cta tat gat ggg tgg ata tga tca gtc cct gta tag aag ttc aaa ggt tat agc gta caa tac atg tac taa ttc ttg gat ata tga tat acc aga gct aaa aat atc atc gtt cta att gcg gag gag ttg cca atg acg aat aca ttt att gta tag gcg gta tac gcg atc agg agt cat cgt tga tat cta gta tcg ata gat gga agc cat caa aac cat att ggc aga agt atg cta aaa tgt gcg aac caa aat gtg ata tgg ggt tgc gat ttt aaa cgg att aat ata tgt cat agg tgg agt cgt taa agg tga cac aca tat acc aac gca cta gag agt tta tca gaa gat gga tgg atg aat cat caa cgt ctt cca ata aaa atg tcc aat atg tcg acg att gtt cat gct gga aag att tat ata tct aga ggt tac aac aat agt agt gta gtt aat gta ata tcg aat ct- -ag tcc tta gct ata atc cga tat atg atg aat gga cca aat tat cat cat taa ata ttc cta gaa tta atc ctg ctc tat ggt cag tgt ata ata aat tat atg tag gag gag taa tat ctg atg atg ttc aaa cta ata cat ctg aaa cat acg ata aag aaa aa

>ON649713.1_Monkeypox_virus_isolate_Monkeypox/PT0020/2022_complete_genome

tgt tat tgt agt atg ata ata tca aaa aga tgg ata taa aga att tac tga cta cat gta cta ttt tac att act aca ttg gct acg gca tat ata cct att tcg tca ctt cca cac gct ccg gta aac ggg tgt cat gtg acg agg gag aat ctt gat aag agg cat aat caa tgt tgt aat ccg atg tcc acc tgg aga att tgc caa ggt cag atg tag agt tgg tag tga taa cac aaa atg tga aca ctg ccc acc tca tac ata tac cgc aat ccc caa tta ttc taa tag atg tca tca atg tag aaa atg ccc aac agg atc att tga taa ggt aaa gtg tac cgg aac aca gaa caa at- gtt cgt gtc atc ctg gtt ggt ata cgc tac tga ttc ttc aca gac tga aga ttg tcg aga ttt gtg tac caa aaa agg aga tgt cca tgc gga tac ttt ggt gga ata gat gaa gga aat cct att tgt aaa tcg tgt tgt gtt ggt gaa tat tgc gac tac cta cgt aat tat aga ctt gat cca ttt cct cca tgc aaa cta tct atc taa atg taa tta att atg att ttg atg ata atg tta cca tac att ata tca cta ctt ggt tag tgt gta tta ttt agt atg gaa gac cta tta ata att act tat ctt ttg acg atc ttg tta taa tta taa tat aaa aat act tat gac ata gta a-- --- -ct cat aat tgc tga cgc gat aaa ttc gta ata atc tgt ttt gtt caa att ttt ata agg aat cta cag gca taa aaa taa aaa tat aat cta taa tat act ctt aca acg cca tca tga ata gca gtg aat taa ttg ctg tta tta tgg att tag aaa tag tgg acg att ttg tga tat taa tat agt tat taa tga tga aag gat aaa cgc gca tag act cat cct atc tgg agc ctc cga ata ttt ttt cca ttc tgt ttt cca ata att tta tcg att cta atg aat acg aag tta atc taa gtc att tag att atc aaa gtg tta act att tga tcg att aca ttt atg gga tac ctt tga gcc taa cta aca ata acg tga aat ata ttc ttt caa ccg ctg att ttt tta caa att gga tct gtc att act gag tgc gaa aaa tac ata ctt aaa aat ctt tgt ttt aga aac tgt atc gat ttc --- --- tac ata tac gct gat aaa tat aat aac aag aaa ata gaa tta gca tcg ttt aac aca ata tta cga aat att ttg aga ctc atc aac aat gaa aac ttt aaa tac tta aca gag gaa tca atg ata aaa att tta agc gat gat atg tta tat ata aaa aat gag gat ttc acc cca ctg att ctc att aaa tgg tta gag agt aca cca acc atg tac cgt cga gtt act tag atg cct cag aat atc a-t ttc ttt ccc cac aag tta taa aat cac ttt ata gtc atc gac tgg tta gtt caa tct acg aat gta taa cat tct taa aca ata tag cat tct tgg ata aat cat ttc cta gat acc ata tca tcg agt tga tat cta tcg gta taa gta att cac atg ata aga ttt cca taa act gct aca atc ata aaa aaa ttc atg gga aat gat atc ttc acg tag ata tag gtg tag ttt cgc agt gac cgt cct gga taa tat tat cta tat gat ggg tgg ata tga tca gtc cct gta tag aag ttc aaa ggt tat agc gta caa tac atg tac taa ttc ttg gat ata tga tat acc aga gct aaa aat atc atc gtt cta att gcg gag gag ttg cca atg acg aat aca ttt att gta tag gcg gta tac gcg atc agg agt cat cgt tga tat cta gta tcg ata gat gga agc cat caa aac cat att ggc aga agt atg cta aaa tgt gcg aac caa aat gtg ata tgg ggt tgc gat ttt aaa cgg att aat ata tgt cat agg tgg agt cgt taa agg tga cac aca tat acc aac gca cta gag agt tta tca gaa gat gga tgg atg aat cat caa cgt ctt cca ata aaa atg tcc aat atg tcg acg att gtt cat gct gga aag att tat ata tct aga ggt tac aac aat agt agt gta gtt aat gta ata tcg aat ct- -ag tcc tta gct ata atc cga tat atg atg aat gga cca aat tat cat cat taa ata ttc cta gaa tta atc ctg ctc tat ggt cag tgt ata ata aat tat atg tag gag gag taa tat ctg atg atg ttc aaa cta ata cat ctg aaa cat acg ata aag aaa aa

>MG693723.1_Monkeypox_virus_isolate_MPXV_Nig_2017_297957_partial_genome

tgt tat tgt agt atg ata ata tca aaa aga tgg ata taa aga att tac tga cta cat gta cta ttt tac att act aca ttg gct acg gca tat ata cct att tcg tca ctt cca cac gct ccg gta aac ggg tgt cat gtg acg agg gag aat ctt gat aag agg cat aat caa tgt tgt aat ccg atg tcc acc tgg aga att tgc caa ggt cag atg tag agt tgg tag tga taa cac aaa atg tga aca ctg ccc acc tca tac ata tac cgc aat ccc caa tta ttc taa tag atg tca tca atg tag aaa atg ccc aac agg atc att tga taa ggt aaa gtg tac cgg aac aca gaa caa at- gtt cgt gtc atc ctg gtt ggt ata cgc tac tga ttc ttc aca gac tga aga ttg tcg aga ttt gtg tac caa aaa agg aga tgt cca tgc gga tac ttt ggt gga ata gat gaa gga aat cct att tgt aaa tcg tgt tgt gtt ggt gaa tat tgc gac tac cta cgt aat tat aga ctt gat cca ttt cct cca tgc aaa cta tct atc taa atg taa tta att atg att ttg atg ata atg tta cca tac att ata tca cta ctt ggt tag tgt gta tta ttt agt atg gaa gac cta tta ata att act tat ctt ttg acg atc ttg tta taa tta taa tat aaa aat act tat gac ata gta a-- --- -ct cat aat tgc tga cgc gat aaa ttc gta ata atc tgt ttt gtt caa att ttt ata agg aat cta cag gca taa aaa taa aaa tat aat cta taa tat act ctt aca acg cca tca tga ata gca gtg aat taa ttg ctg tta tta tgg att tag aaa tag tgg acg att ttg tga tat taa tat agt tat taa tga tga aag gat aaa cgc gca tag act cat cct atc tgg agc ctc cga ata ttt ttt cca ttc tgt ttt cca ata att tta tcg att cta atg aat acg aag tta atc taa gtc att tag att atc aaa gtg tta act att tga tcg att aca ttt atg gga tac ctt tga gcc taa cta aca ata acg tga aat ata ttc ttt caa ccg ctg att ttt tta caa att gga tct gtc att act gag tgc gaa aaa tac ata ctt aaa aat ctt tgt ttt aga aac tgt atc gat ttc --- --- tac ata tac gct gat aaa tat aat aac aag aaa ata gaa tta gca tcg ttt aac aca ata tta cga aat att ttg aga ctc atc aac aat gaa aac ttt aaa tac tta aca gag gaa tca atg ata aaa att tta agc gat gat atg tta tat ata aaa aat gag gat ttc acc cca ctg att ctc att aaa tgg tta gag agt aca cca acc atg tac cgt cga gtt act tag atg cct cag aat atc a-t ttc ttt ccc cac aag tta taa aat cac ttt ata gtc atc gac tgg tta gtt caa tct acg aat gta taa cat tct taa aca ata tag cat tct tgg ata aat cat ttc cta gat acc ata tca tcg agt tga tat cta tcg gta taa gta att cac atg ata aga ttt cca taa act gct aca atc ata aaa aaa ttc atg gga aat gat atc ttc acg tag ata tag gtg tag ttt cgc agt gac cgt cct gga taa tat tat cta tat gat ggg tgg ata tga tca gtc cct gta tag aag ttc aaa ggt tat agc gta caa tac atg tac taa ttc ttg gat ata tga tat acc aga gct aaa aat atc atc gtt cta att gcg gag gag ttg cca atg acg aat aca ttt att gta tag gcg gta tac gcg atc agg agt cat cgt tga tat cta gta tcg ata gat gga agc cat caa aac cat att ggc aga agt atg cta aaa tgt gcg aac caa aat gtg ata tgg ggt tgc gat ttt aaa cgg att aat ata tgt cat agg tgg agt cgt taa agg tga cac aca tat acc aac gca cta gag agt tta tca gaa gat gga tgg atg aat cat caa cgt ctt cca ata aaa atg tcc aat atg tcg acg att gtt cat gct gga aag att tat ata tct aga ggt tac aac aat agt agt gta gtt aat gta ata tcg aat ct- -ag tcc tta gct ata atc cga tat atg atg aat gga cca aat tat cat cat taa ata ttc cta gaa tta atc ctg ctc tat ggt cag tgt ata ata aat tat atg tag gag gag taa tat ctg atg atg ttc aaa cta ata cat ctg aaa cat acg ata aag aaa aa

>ON585031.1_Monkeypox_virus_isolate_Monkeypox/PT0003/2022_complete_genome

tgt tat tgt agt atg ata ata tca aaa aga tgg ata taa aga att tac tga cta cat gta cta ttt tac att act aca ttg gct acg gca tat ata cct att tcg tca ctt cca cac gct ccg gta aac ggg tgt cat gtg acg agg gag aat ctt gat aag agg cat aat caa tgt tgt aat ccg atg tcc acc tgg aga att tgc caa ggt cag atg tag agt tgg tag tga taa cac aaa atg tga aca ctg ccc acc tca tac ata tac cgc aat ccc caa tta ttc taa tag atg tca tca atg tag aaa atg ccc aac agg atc att tga taa ggt aaa gtg tac cgg aac aca gaa caa at- gtt cgt gtc atc ctg gtt ggt ata cgc tac tga ttc ttc aca gac tga aga ttg tcg aga ttt gtg tac caa aaa agg aga tgt cca tgc gga tac ttt ggt gga ata gat gaa gga aat cct att tgt aaa tcg tgt tgt gtt ggt gaa tat tgc gac tac cta cgt aat tat aga ctt gat cca ttt cct cca tgc aaa cta tct atc taa atg taa tta att atg att ttg atg ata atg tta cca tac att ata tca cta ctt ggt tag tgt gta tta ttt agt atg gaa gac cta tta ata att act tat ctt ttg acg atc ttg tta taa tta taa tat aaa aat act tat gac ata gta a-- --- -ct cat aat tgc tga cgc gat aaa ttc gta ata atc tgt ttt gtt caa att ttt ata agg aat cta cag gca taa aaa taa aaa tat aat cta taa tat act ctt aca acg cca tca tga ata gca gtg aat taa ttg ctg tta tta tgg att tag aaa tag tgg acg att ttg tga tat taa tat agt tat taa tga tga aag gat aaa cgc gca tag act cat cct atc tgg agc ctc cga ata ttt ttt cca ttc tgt ttt cca ata att tta tcg att cta atg aat acg aag tta atc taa gtc att tag att atc aaa gtg tta act att tga tcg att aca ttt atg gga tac ctt tga gcc taa cta aca ata acg tga aat ata ttc ttt caa ccg ctg att ttt tta caa att gga tct gtc att act gag tgc gaa aaa tac ata ctt aaa aat ctt tgt ttt aga aac tgt atc gat ttc --- --- tac ata tac gct gat aaa tat aat aac aag aaa ata gaa tta gca tcg ttt aac aca ata tta cga aat att ttg aga ctc atc aac aat gaa aac ttt aaa tac tta aca gag gaa tca atg ata aaa att tta agc gat gat atg tta tat ata aaa aat gag gat ttc acc cca ctg att ctc att aaa tgg tta gag agt aca cca acc atg tac cgt cga gtt act tag atg cct cag aat atc a-t ttc ttt ccc cac aag tta taa aat cac ttt ata gtc atc gac tgg tta gtt caa tct acg aat gta taa cat tct taa aca ata tag cat tct tgg ata aat cat ttc cta gat acc ata tca tcg agt tga tat cta tcg gta taa gta att cac atg ata aga ttt cca taa act gct aca atc ata aaa aaa ttc atg gga aat gat atc ttc acg tag ata tag gtg tag ttt cgc agt gac cgt cct gga taa tat tat cta tat gat ggg tgg ata tga tca gtc cct gta tag aag ttc aaa ggt tat agc gta caa tac atg tac taa ttc ttg gat ata tga tat acc aga gct aaa aat atc atc gtt cta att gcg gag gag ttg cca atg acg aat aca ttt att gta tag gcg gta tac gcg atc agg agt cat cgt tga tat cta gta tcg ata gat gga agc cat caa aac cat att ggc aga agt atg cta aaa tgt gcg aac caa aat gtg ata tgg ggt tgc gat ttt aaa cgg att aat ata tgt cat agg tgg agt cgt taa agg tga cac aca tat acc aac gca cta gag agt tta tca gaa gat gga tgg atg aat cat caa cgt ctt cca ata aaa atg tcc aat atg tcg acg att gtt cat gct gga aag att tat ata tct aga ggt tac aac aat agt agt gta gtt aat gta ata tcg aat ct- -ag tcc tta gct ata atc cga tat atg atg aat gga cca aat tat cat cat taa ata ttc cta gaa tta atc ctg ctc tat ggt cag tgt ata ata aat tat atg tag gag gag taa tat ctg atg atg ttc aaa cta ata cat ctg aaa cat acg ata aag aaa aa

>ON585038.1_Monkeypox_virus_isolate_Monkeypox/PT0008/2022_complete_genome

tgt tat tgt agt atg ata ata tca aaa aga tgg ata taa aga att tac tga cta cat gta cta ttt tac att act aca ttg gct acg gca tat ata cct att tcg tca ctt cca cac gct ccg gta aac ggg tgt cat gtg acg agg gag aat ctt gat aag agg cat aat caa tgt tgt aat ccg atg tcc acc tgg aga att tgc caa ggt cag atg tag agt tgg tag tga taa cac aaa atg tga aca ctg ccc acc tca tac ata tac cgc aat ccc caa tta ttc taa tag atg tca tca atg tag aaa atg ccc aac agg atc att tga taa ggt aaa gtg tac cgg aac aca gaa caa at- gtt cgt gtc atc ctg gtt ggt ata cgc tac tga ttc ttc aca gac tga aga ttg tcg aga ttt gtg tac caa aaa agg aga tgt cca tgc gga tac ttt ggt gga ata gat gaa gga aat cct att tgt aaa tcg tgt tgt gtt ggt gaa tat tgc gac tac cta cgt aat tat aga ctt gat cca ttt cct cca tgc aaa cta tct atc taa atg taa tta att atg att ttg atg ata atg tta cca tac att ata tca cta ctt ggt tag tgt gta tta ttt agt atg gaa gac cta tta ata att act tat ctt ttg acg atc ttg tta taa tta taa tat aaa aat act tat gac ata gta a-- --- -ct cat aat tgc tga cgc gat aaa ttc gta ata atc tgt ttt gtt caa att ttt ata agg aat cta cag gca taa aaa taa aaa tat aat cta taa tat act ctt aca acg cca tca tga ata gca gtg aat taa ttg ctg tta tta tgg att tag aaa tag tgg acg att ttg tga tat taa tat agt tat taa tga tga aag gat aaa cgc gca tag act cat cct atc tgg agc ctc cga ata ttt ttt cca ttc tgt ttt cca ata att tta tcg att cta atg aat acg aag tta atc taa gtc att tag att atc aaa gtg tta act att tga tcg att aca ttt atg gga tac ctt tga gcc taa cta aca ata acg tga aat ata ttc ttt caa ccg ctg att ttt tta caa att gga tct gtc att act gag tgc gaa aaa tac ata ctt aaa aat ctt tgt ttt aga aac tgt atc gat ttc --- --- tac ata tac gct gat aaa tat aat aac aag aaa ata gaa tta gca tcg ttt aac aca ata tta cga aat att ttg aga ctc atc aac aat gaa aac ttt aaa tac tta aca gag gaa tca atg ata aaa att tta agc gat gat atg tta tat ata aaa aat gag gat ttc acc cca ctg att ctc att aaa tgg tta gag agt aca cca acc atg tac cgt cga gtt act tag atg cct cag aat atc a-t ttc ttt ccc cac aag tta taa aat cac ttt ata gtc atc gac tgg tta gtt caa tct acg aat gta taa cat tct taa aca ata tag cat tct tgg ata aat cat ttc cta gat acc ata tca tcg agt tga tat cta tcg gta taa gta att cac atg ata aga ttt cca taa act gct aca atc ata aaa aaa ttc atg gga aat gat atc ttc acg tag ata tag gtg tag ttt cgc agt gac cgt cct gga taa tat tat cta tat gat ggg tgg ata tga tca gtc cct gta tag aag ttc aaa ggt tat agc gta caa tac atg tac taa ttc ttg gat ata tga tat acc aga gct aaa aat atc atc gtt cta att gcg gag gag ttg cca atg acg aat aca ttt att gta tag gcg gta tac gcg atc agg agt cat cgt tga tat cta gta tcg ata gat gga agc cat caa aac cat att ggc aga agt atg cta aaa tgt gcg aac caa aat gtg ata tgg ggt tgc gat ttt aaa cgg att aat ata tgt cat agg tgg agt cgt taa agg tga cac aca tat acc aac gca cta gag agt tta tca gaa gat gga tgg atg aat cat caa cgt ctt cca ata aaa atg tcc aat atg tcg acg att gtt cat gct gga aag att tat ata tct aga ggt tac aac aat agt agt gta gtt aat gta ata tcg aat ct- -ag tcc tta gct ata atc cga tat atg atg aat gga cca aat tat cat cat taa ata ttc cta gaa tta atc ctg ctc tat ggt cag tgt ata ata aat tat atg tag gag gag taa tat ctg atg atg ttc aaa cta ata cat ctg aaa cat acg ata aag aaa aa

>DQ011156.1_Monkeypox_virus_strain_Liberia_1970_184_complete_genome

tgt tat tgt agt atg ata ata tca aaa aga tgg ata taa aga att tac tga cta cat gta cta ttt tac att act aca ttg gct acg g-- tat ata cct act tcg tca ctt cca cac gct cca gaa aac ggg tgt cat gtg acg agg gag aat ctt gat aag agg cat aat caa tgt tgt aat ccg gtg tcc acc tgg aga att tgc caa ggt cag atg t-- agt tgg tag tga taa cac aaa atg tga aca ctg ccc acc tca tac ata tac cgc aat ccc caa tta ttc taa tag atg tca tca atg tat aaa atg ccc aac agg atc att tga taa ggt aaa gtg tac cgg aac aca gaa caa at- gtt cgt gtc atc ctg gtt ggt ata cgc tac tga ttc ttc aca gac tga aga ttg tcg aga ttt gtg tac caa aaa agg aga tgt cca tgc gga tac ttt ggt gga ata gat gaa gga aat cct att tgt aaa tcg tgt tgt gtt ggt gaa tat tgc gac tac cta cgt aat tat aga ctt gat cca ttt cct cca tgc aaa cta tct atc taa atg taa tta att atg att ttg atg ata atg tta cca tac att ata tcg cta ctt ggt tag tgt gta tta ttc agt atg gaa gac cta tta ata att act tat ctt ttg acg atc ttg tta taa tta taa tat aaa aat act tat gac ata gta aca gta act cat aat tgc tga cgc gat aaa ttc gta ata atc tgt ttt gtt caa att ttt ata agg aat cta cag gca taa aaa taa aaa tat aat cta taa tat act ctt aca acg cca tca tga ata gca gtg aat taa ttg ctg tta tta tgg att tag aaa tag tgg acg att ttg tga tat taa tat agt tat taa tga tga aag gat aaa cgc tca tag att cat cct atc tgg agc ctc cga ata ttt ttt cca ttc tgt ttt cca ata att tta tcg att cta atg aat acg aag tta atc taa gtc att tag att atc aaa gtg tta act att tga tcg att aca ttt atg gga tac ctt tga gcc taa cta acg ata acg tga aat ata ttc ttt caa ccg ctg att ttt tta caa att gga tct gcc att act gag tgc gaa aaa tac ata ctt aaa aat ctt tgt ttt aga aac tgt atc gat ttc --- --- tac ata tac gct gat aaa tat aat aac aag aaa ata gaa tta gca tcg ttt aac aca ata tta cga aat att ttg aga ctc atc aac aat gaa aac ttt aaa tac tta aca gag gaa tca --- --- --- --- --- --- -at gat atg tta tat ata aaa aat gag gat ttc acc cca ctg att ctc att aaa tgg tta gag agt aca cca acc atg tac cgt cga gtt act tag atg cct cag aat atc a-t ttc ttt ccc cac aag tta taa aat cac ttt ata gtc atc gac tgg tta gtt caa tct acg aat gta taa cat tct taa aca ata tag cat tct tgg ata aat cat ttc cta gat acc ata tca tcg agt tga tat cta tcg gta taa gta att cac atg ata aga ttt cca taa act gct aca atc ata aaa aaa ttc atg gga aat gat atc ttc acg tag at- --g gtg tag ttt cgc agt gac cgt cct gga taa tat tat cta tat gat ggg tgg ata tga tca gtc cct gta tag aag ttc aaa ggt tat agc gta caa tac atg tac taa ttc ttg gat ata tga tat acc aga gct -aa aat atc ctc gtt cta att gcg gag gag ttg cca atg acg aat aca ttt att gta tag gcg gta tac gcg atc agg agt cat cgt tga tat cta gta tcg ata gat gga agc cat caa aac cat att ggc aga agt atg cta aaa tgt gcg aac caa aat gtg ata tgg ggt tgc gat ttt aaa cgg att aat ata tgt cat agg tgg agt cgt taa agg tga cac aca tgt acc gac gca cta gag agt tta tca gaa gat gga tgg atg aat cat caa cgt ctt cca ata aaa atg tcc aat atg tcg acg att gtt cat gct gga aag att tat ata tct gga ggt tac aac aat agt agt gta gtt aat gta ata tcg aat cta gag tcc tta gct ata atc cga tat atg atg aat gga cca aat tat cat cat taa ata ttc cta gaa tta atc ctg ctc tat ggt cag tgt ata ata aat tat atg tag gag gag gaa tat ctg atg atg ttc aaa cta ata cat ctg aaa cat acg ata aag aaa aa

>KP849470.1_Monkeypox_virus_isolate_Cote_dIvoire_1971_complete_genome

tgt tat tgt agt atg ata ata tca aaa aga tgg ata taa aga att tac tga cta cat gta cta ttt tac att act aca ttg gct acg g-- tat ata cct act tcg tca ctt cca cac gct cca gaa aac ggg tgt cat gtg acg agg gag aat ctt gat aag agg cat aat caa tgt tgt aat ccg gtg tcc acc tgg aga att tgc caa ggt cag atg t-- agt tgg tag tga taa cac aaa atg tga aca ctg ccc acc tca tac ata tac cgc aat ccc caa tta ttc taa tag atg tca tca atg tat aaa atg ccc aac agg atc att tga taa ggt aaa gtg tac cgg aac aca gaa caa at- gtt cgt gtc atc ctg gtt ggt ata cgc tac tga ttc ttc aca gac tga aga ttg tcg aga ttt gtg tac caa aaa agg aga tgt cca tgc gga tac ttt ggt gga ata gat gaa gga aat cct att tgt aaa tcg tgt tgt gtt ggt gaa tat tgc gac tac cta cgt aat tat aga ctt gat cca ttt cct cca tgc aaa cta tct atc taa atg taa tta att atg att ttg atg ata atg tta cca tac att ata tcg cta ctt ggt tag tgt gta tta ttc agt atg gaa gac cta tta ata att act tat ctt ttg acg atc ttg tta taa tta taa tat aaa aat act tat gac ata gta aca gta act cat aat tgc tga cgc gat aaa ttc gta ata atc tgt ttt gtt caa att ttt ata agg aat cta cag gca taa aaa taa aaa tat aat cta taa tat act ctt aca acg cca tca tga ata gca gtg aat taa ttg ctg tta tta tgg att tag aaa tag tgg acg att ttg tga tat taa tat agt tat taa tga tga aag gat aaa cgc tca tag att cat cct atc tgg agc ctc cga ata ttt ttt cca ttc tgt ttt cca ata att tta tcg att cta atg aat acg aag tta atc taa gtc att tag att atc aaa gtg tta act att tga tcg att aca ttt atg gga tac ctt tga gcc taa cta acg ata acg tga aat ata ttc ttt caa ccg ctg att ttt tta caa att gga tct gcc att act gag tgc gaa aaa tac ata ctt aaa aat ctt tgt ttt aga aac tgt atc gat ttc --- --- tac ata tac gct gat aaa tat aat aac aag aaa ata gaa tta gca tcg ttt aac aca ata tta cga aat att ttg aga ctc atc aac aat gaa aac ttt aaa tac tta aca gag gaa tca --- --- --- --- --- --- -at gat atg tta tat ata aaa aat gag gat ttc acc cca ctg att ctc att aaa tgg tta gag agt aca cca acc atg tac cgt cga gtt act tag atg cct cag aat atc a-t ttc ttt ccc cac aag tta taa aat cac ttt ata gtc atc gac tgg tta gtt caa tct acg aat gta taa cat tct taa aca ata tag cat tct tgg ata aat cat ttc cta gat acc ata tca tcg agt tga tat cta tcg gta taa gta att cac atg ata aga ttt cca taa act gct aca atc ata aaa aaa ttc atg gga aat gat atc ttc acg tag at- --g gtg tag ttt cgc agt gac cgt cct gga taa tat tat cta tat gat ggg tgg ata tga tca gtc cct gta tag aag ttc aaa ggt tat agc gta caa tac atg tac taa ttc ttg gat ata tga tat acc aga gct -aa aat atc ctc gtt cta att gcg gag gag ttg cca atg acg aat aca ttt att gta tag gcg gta tac gcg atc agg agt cat cgt tga tat cta gta tcg ata gat gga agc cat caa aac cat att ggc aga agt atg cta aaa tgt gcg aac caa aat gtg ata tgg ggt tgc gat ttt aaa cgg att aat ata tgt cat agg tgg agt cgt taa agg tga cac aca tgt acc gac gca cta gag agt tta tca gaa gat gga tgg atg aat cat caa cgt ctt cca ata aaa atg tcc aat atg tcg acg att gtt cat gct gga aag att tat ata tct gga ggt tac aac aat agt agt gta gtt aat gta ata tcg aat cta gag tcc tta gct ata atc cga tat atg atg aat gga cca aat tat cat cat taa ata ttc cta gaa tta atc ctg ctc tat ggt cag tgt ata ata aat tat atg tag gag gag gaa tat ctg atg atg ttc aaa cta ata cat ctg aaa cat acg ata aag aaa aa

>ON682264.2_Monkeypox_virus_isolate_MPXV/Germany/2022/RKI05_complete_genome

tgt tat tgt agt atg ata ata tca aaa aga tgg ata taa aga att tac tga cta cat gta cta ttt tac att act aca ttg gct acg gca tat ata cct att tcg tca ctt cca cac gct ccg gta aac ggg tgt cat gtg acg agg gag aat ctt gat aag agg cat aat caa tgt tgt aat ccg atg tcc acc tgg aga att tgc caa ggt cag atg tag agt tgg tag tga taa cac aaa atg tga aca ctg ccc acc tca tac ata tac cgc aat ccc caa tta ttc taa tag atg tca tca atg tag aaa atg ccc aac agg atc att tga taa ggt aaa gtg tac cgg aac aca gaa caa at- gtt cgt gtc atc ctg gtt ggt ata cgc tac tga ttc ttc aca gac tga aga ttg tcg aga ttt gtg tac caa aaa agg aga tgt cca tgc gga tac ttt ggt gga ata gat gaa gga aat cct att tgt aaa tcg tgt tgt gtt ggt gaa tat tgc gac tac cta cgt aat tat aga ctt gat cca ttt cct cca tgc aaa cta tct atc taa atg taa tta att atg att ttg atg ata atg tta cca tac att ata tca cta ctt ggt tag tgt gta tta ttt agt atg gaa gac cta tta ata att act tat ctt ttg acg atc ttg tta taa tta taa tat aaa aat act tat gac ata gta a-- --- -ct cat aat tgc tga cgc gat aaa ttc gta ata atc tgt ttt gtt caa att ttt ata agg aat cta cag gca taa aaa taa aaa tat aat cta taa tat act ctt aca acg cca tca tga ata gca gtg aat taa ttg ctg tta tta tgg att tag aaa tag tgg acg att ttg tga tat taa tat agt tat taa tga tga aag gat aaa cgc gca tag act cat cct atc tgg agc ctc cga ata ttt ttt cca ttc tgt ttt cca ata att tta tcg att cta atg aat acg aag tta atc taa gtc att tag att atc aaa gtg tta act att tga tcg att aca ttt atg gga tac ctt tga gcc taa cta aca ata acg tga aat ata ttc ttt caa ccg ctg att ttt tta caa att gga tct gtc att act gag tgc gaa aaa tac ata ctt aaa aat ctt tgt ttt aga aac tgt atc gat ttc --- --- tac ata tac gct gat aaa tat aat aac aag aaa ata gaa tta gca tcg ttt aac aca ata tta cga aat att ttg aga ctc atc aac aat gaa aac ttt aaa tac tta aca gag gaa tca atg ata aaa att tta agc gat gat atg tta tat ata aaa aat gag gat ttc acc cca ctg att ctc att aaa tgg tta gag agt aca cca acc atg tac cgt cga gtt act tag atg cct cag aat atc a-t ttc ttt ccc cac aag tta taa aat cac ttt ata gtc atc gac tgg tta gtt caa tct acg aat gta taa cat tct taa aca ata tag cat tct tgg ata aat cat ttc cta gat acc ata tca tcg agt tga tat cta tcg gta taa gta att cac atg ata aga ttt cca taa act gct aca atc ata aaa aaa ttc atg gga aat gat atc ttc acg tag ata tag gtg tag ttt cgc agt gac cgt cct gga taa tat tat cta tat gat ggg tgg ata tga tca gtc cct gta tag aag ttc aaa ggt tat agc gta caa tac atg tac taa ttc ttg gat ata tga tat acc aga gct aaa aat atc atc gtt cta att gcg gag gag ttg cca atg acg aat aca ttt att gta tag gcg gta tac gcg atc agg agt cat cgt tga tat cta gta tcg ata gat gga agc cat caa aac cat att ggc aga agt atg cta aaa tgt gcg aac caa aat gtg ata tgg ggt tgc gat ttt aaa cgg att aat ata tgt cat agg tgg agt cgt taa agg tga cac aca tat acc aac gca cta gag agt tta tca gaa gat gga tgg atg aat cat caa cgt ctt cca ata aaa atg tcc aat atg tcg acg att gtt cat gct gga aag att tat ata tct aga ggt tac aac aat agt agt gta gtt aat gta ata tcg aat ct- -ag tcc tta gct ata atc cga tat atg atg aat gga cca aat tat cat cat taa ata ttc cta gaa tta atc ctg ctc tat ggt cag tgt ata ata aat tat atg tag gag gag taa tat ctg atg atg ttc aaa cta ata cat ctg aaa cat acg ata aag aaa aa

>ON682268.1_Monkeypox_virus_isolate_MPXV/Germany/2022/RKI08_complete_genome

tgt tat tgt agt atg ata ata tca aaa aga tgg ata taa aga att tac tga cta cat gta cta ttt tac att act aca ttg gct acg gca tat ata cct att tcg tca ctt cca cac gct ccg gta aac ggg tgt cat gtg acg agg gag aat ctt gat aag agg cat aat caa tgt tgt aat ccg atg tcc acc tgg aga att tgc caa ggt cag atg tag agt tgg tag tga taa cac aaa atg tga aca ctg ccc acc tca tac ata tac cgc aat ccc caa tta ttc taa tag atg tca tca atg tag aaa atg ccc aac agg atc att tga taa ggt aaa gtg tac cgg aac aca gaa caa at- gtt cgt gtc atc ctg gtt ggt ata cgc tac tga ttc ttc aca gac tga aga ttg tcg aga ttt gtg tac caa aaa agg aga tgt cca tgc gga tac ttt ggt gga ata gat gaa gga aat cct att tgt aaa tcg tgt tgt gtt ggt gaa tat tgc gac tac cta cgt aat tat aga ctt gat cca ttt cct cca tgc aaa cta tct atc taa atg taa tta att atg att ttg atg ata atg tta cca tac att ata tca cta ctt ggt tag tgt gta tta ttt agt atg gaa gac cta tta ata att act tat ctt ttg acg atc ttg tta taa tta taa tat aaa aat act tat gac ata gta a-- --- -ct cat aat tgc tga cgc gat aaa ttc gta ata atc tgt ttt gtt caa att ttt ata agg aat cta cag gca taa aaa taa aaa tat aat cta taa tat act ctt aca acg cca tca tga ata gca gtg aat taa ttg ctg tta tta tgg att tag aaa tag tgg acg att ttg tga tat taa tat agt tat taa tga tga aag gat aaa cgc gca tag act cat cct atc tgg agc ctc cga ata ttt ttt cca ttc tgt ttt cca ata att tta tcg att cta atg aat acg aag tta atc taa gtc att tag att atc aaa gtg tta act att tga tcg att aca ttt atg gga tac ctt tga gcc taa cta aca ata acg tga aat ata ttc ttt caa ccg ctg att ttt tta caa att gga tct gtc att act gag tgc gaa aaa tac ata ctt aaa aat ctt tgt ttt aga aac tgt atc gat ttc --- --- tac ata tac gct gat aaa tat aat aac aag aaa ata gaa tta gca tcg ttt aac aca ata tta cga aat att ttg aga ctc atc aac aat gaa aac ttt aaa tac tta aca gag gaa tca atg ata aaa att tta agc gat gat atg tta tat ata aaa aat gag gat ttc acc cca ctg att ctc att aaa tgg tta gag agt aca cca acc atg tac cgt cga gtt act tag atg cct cag aat atc a-t ttc ttt ccc cac aag tta taa aat cac ttt ata gtc atc gac tgg tta gtt caa tct acg aat gta taa cat tct taa aca ata tag cat tct tgg ata aat cat ttc cta gat acc ata tca tcg agt tga tat cta tcg gta taa gta att cac atg ata aga ttt cca taa act gct aca atc ata aaa aaa ttc atg gga aat gat atc ttc acg tag ata tag gtg tag ttt cgc agt gac cgt cct gga taa tat tat cta tat gat ggg tgg ata tga tca gtc cct gta tag aag ttc aaa ggt tat agc gta caa tac atg tac taa ttc ttg gat ata tga tat acc aga gct aaa aat atc atc gtt cta att gcg gag gag ttg cca atg acg aat aca ttt att gta tag gcg gta tac gcg atc agg agt cat cgt tga tat cta gta tcg ata gat gga agc cat caa aac cat att ggc aga agt atg cta aaa tgt gcg aac caa aat gtg ata tgg ggt tgc gat ttt aaa cgg att aat ata tgt cat agg tgg agt cgt taa agg tga cac aca tat acc aac gca cta gag agt tta tca gaa gat gga tgg atg aat cat caa cgt ctt cca ata aaa atg tcc aat atg tcg acg att gtt cat gct gga aag att tat ata tct aga ggt tac aac aat agt agt gta gtt aat gta ata tcg aat ct- -ag tcc tta gct ata atc cga tat atg atg aat gga cca aat tat cat cat taa ata ttc cta gaa tta atc ctg ctc tat ggt cag tgt ata ata aat tat atg tag gag gag taa tat ctg atg atg ttc aaa cta ata cat ctg aaa cat acg ata aag aaa aa

>ON682270.1_Monkeypox_virus_isolate_MPXV/Germany/2022/RKI06_complete_genome

tgt tat tgt agt atg ata ata tca aaa aga tgg ata taa aga att tac tga cta cat gta cta ttt tac att act aca ttg gct acg gca tat ata cct att tcg tca ctt cca cac gct ccg gta aac ggg tgt cat gtg acg agg gag aat ctt gat aag agg cat aat caa tgt tgt aat ccg atg tcc acc tgg aga att tgc caa ggt cag atg tag agt tgg tag tga taa cac aaa atg tga aca ctg ccc acc tca tac ata tac cgc aat ccc caa tta ttc taa tag atg tca tca atg tag aaa atg ccc aac agg atc att tga taa ggt aaa gtg tac cgg aac aca gaa caa at- gtt cgt gtc atc ctg gtt ggt ata cgc tac tga ttc ttc aca gac tga aga ttg tcg aga ttt gtg tac caa aaa agg aga tgt cca tgc gga tac ttt ggt gga ata gat gaa gga aat cct att tgt aaa tcg tgt tgt gtt ggt gaa tat tgc gac tac cta cgt aat tat aga ctt gat cca ttt cct cca tgc aaa cta tct atc taa atg taa tta att atg att ttg atg ata atg tta cca tac att ata tca cta ctt ggt tag tgt gta tta ttt agt atg gaa gac cta tta ata att act tat ctt ttg acg atc ttg tta taa tta taa tat aaa aat act tat gac ata gta a-- --- -ct cat aat tgc tga cgc gat aaa ttc gta ata atc tgt ttt gtt caa att ttt ata agg aat cta cag gca taa aaa taa aaa tat aat cta taa tat act ctt aca acg cca tca tga ata gca gtg aat taa ttg ctg tta tta tgg att tag aaa tag tgg acg att ttg tga tat taa tat agt tat taa tga tga aag gat aaa cgc gca tag act cat cct atc tgg agc ctc cga ata ttt ttt cca ttc tgt ttt cca ata att tta tcg att cta atg aat acg aag tta atc taa gtc att tag att atc aaa gtg tta act att tga tcg att aca ttt atg gga tac ctt tga gcc taa cta aca ata acg tga aat ata ttc ttt caa ccg ctg att ttt tta caa att gga tct gtc att act gag tgc gaa aaa tac ata ctt aaa aat ctt tgt ttt aga aac tgt atc gat ttc --- --- tac ata tac gct gat aaa tat aat aac aag aaa ata gaa tta gca tcg ttt aac aca ata tta cga aat att ttg aga ctc atc aac aat gaa aac ttt aaa tac tta aca gag gaa tca atg ata aaa att tta agc gat gat atg tta tat ata aaa aat gag gat ttc acc cca ctg att ctc att aaa tgg tta gag agt aca cca acc atg tac cgt cga gtt act tag atg cct cag aat atc a-t ttc ttt ccc cac aag tta taa aat cac ttt ata gtc atc gac tgg tta gtt caa tct acg aat gta taa cat tct taa aca ata tag cat tct tgg ata aat cat ttc cta gat acc ata tca tcg agt tga tat cta tcg gta taa gta att cac atg ata aga ttt cca taa act gct aca atc ata aaa aaa ttc atg gga aat gat atc ttc acg tag ata tag gtg tag ttt cgc agt gac cgt cct gga taa tat tat cta tat gat ggg tgg ata tga tca gtc cct gta tag aag ttc aaa ggt tat agc gta caa tac atg tac taa ttc ttg gat ata tga tat acc aga gct aaa aat atc atc gtt cta att gcg gag gag ttg cca atg acg aat aca ttt att gta tag gcg gta tac gcg atc agg agt cat cgt tga tat cta gta tcg ata gat gga agc cat caa aac cat att ggc aga agt atg cta aaa tgt gcg aac caa aat gtg ata tgg ggt tgc gat ttt aaa cgg att aat ata tgt cat agg tgg agt cgt taa agg tga cac aca tat acc aac gca cta gag agt tta tca gaa gat gga tgg atg aat cat caa cgt ctt cca ata aaa atg tcc aat atg tcg acg att gtt cat gct gga aag att tat ata tct aga ggt tac aac aat agt agt gta gtt aat gta ata tcg aat ct- -ag tcc tta gct ata atc cga tat atg atg aat gga cca aat tat cat cat taa ata ttc cta gaa tta atc ctg ctc tat ggt cag tgt ata ata aat tat atg tag gag gag taa tat ctg atg atg ttc aaa cta ata cat ctg aaa cat acg ata aag aaa aa

>ON637938.1_Monkeypox_virus_isolate_MPXV/Germany/2022/RKI01_complete_genome

tgt tat tgt agt atg ata ata tca aaa aga tgg ata taa aga att tac tga cta cat gta cta ttt tac att act aca ttg gct acg gca tat ata cct att tcg tca ctt cca cac gct ccg gta aac ggg tgt cat gtg acg agg gag aat ctt gat aag agg cat aat caa tgt tgt aat ccg atg tcc acc tgg aga att tgc caa ggt cag atg tag agt tgg tag tga taa cac aaa atg tga aca ctg ccc acc tca tac ata tac cgc aat ccc caa tta ttc taa tag atg tca tca atg tag aaa atg ccc aac agg atc att tga taa ggt aaa gtg tac cgg aac aca gaa caa at- gtt cgt gtc atc ctg gtt ggt ata cgc tac tga ttc ttc aca gac tga aga ttg tcg aga ttt gtg tac caa aaa agg aga tgt cca tgc gga tac ttt ggt gga ata gat gaa gga aat cct att tgt aaa tcg tgt tgt gtt ggt gaa tat tgc gac tac cta cgt aat tat aga ctt gat cca ttt cct cca tgc aaa cta tct atc taa atg taa tta att atg att ttg atg ata atg tta cca tac att ata tca cta ctt ggt tag tgt gta tta ttt agt atg gaa gac cta tta ata att act tat ctt ttg acg atc ttg tta taa tta taa tat aaa aat act tat gac ata gta a-- --- -ct cat aat tgc tga cgc gat aaa ttc gta ata atc tgt ttt gtt caa att ttt ata agg aat cta cag gca taa aaa taa aaa tat aat cta taa tat act ctt aca acg cca tca tga ata gca gtg aat taa ttg ctg tta tta tgg att tag aaa tag tgg acg att ttg tga tat taa tat agt tat taa tga tga aag gat aaa cgc gca tag act cat cct atc tgg agc ctc cga ata ttt ttt cca ttc tgt ttt cca ata att tta tcg att cta atg aat acg aag tta atc taa gtc att tag att atc aaa gtg tta act att tga tcg att aca ttt atg gga tac ctt tga gcc taa cta aca ata acg tga aat ata ttc ttt caa ccg ctg att ttt tta caa att gga tct gtc att act gag tgc gaa aaa tac ata ctt aaa aat ctt tgt ttt aga aac tgt atc gat ttc --- --- tac ata tac gct gat aaa tat aat aac aag aaa ata gaa tta gca tcg ttt aac aca ata tta cga aat att ttg aga ctc atc aac aat gaa aac ttt aaa tac tta aca gag gaa tca atg ata aaa att tta agc gat gat atg tta tat ata aaa aat gag gat ttc acc cca ctg att ctc att aaa tgg tta gag agt aca cca acc atg tac cgt cga gtt act tag atg cct cag aat atc a-t ttc ttt ccc cac aag tta taa aat cac ttt ata gtc atc gac tgg tta gtt caa tct acg aat gta taa cat tct taa aca ata tag cat tct tgg ata aat cat ttc cta gat acc ata tca tcg agt tga tat cta tcg gta taa gta att cac atg ata aga ttt cca taa act gct aca atc ata aaa aaa ttc atg gga aat gat atc ttc acg tag ata tag gtg tag ttt cgc agt gac cgt cct gga taa tat tat cta tat gat ggg tgg ata tga tca gtc cct gta tag aag ttc aaa ggt tat agc gta caa tac atg tac taa ttc ttg gat ata tga tat acc aga gct aaa aat atc atc gtt cta att gcg gag gag ttg cca atg acg aat aca ttt att gta tag gcg gta tac gcg atc agg agt cat cgt tga tat cta gta tcg ata gat gga agc cat caa aac cat att ggc aga agt atg cta aaa tgt gcg aac caa aat gtg ata tgg ggt tgc gat ttt aaa cgg att aat ata tgt cat agg tgg agt cgt taa agg tga cac aca tat acc aac gca cta gag agt tta tca gaa gat gga tgg atg aat cat caa cgt ctt cca ata aaa atg tcc aat atg tcg acg att gtt cat gct gga aag att tat ata tct aga ggt tac aac aat agt agt gta gtt aat gta ata tcg aat ct- -ag tcc tta gct ata atc cga tat atg atg aat gga cca aat tat cat cat taa ata ttc cta gaa tta atc ctg ctc tat ggt cag tgt ata ata aat tat atg tag gag gag taa tat ctg atg atg ttc aaa cta ata cat ctg aaa cat acg ata aag aaa aa

>ON637939.1_Monkeypox_virus_isolate_MPXV/Germany/2022/RKI02_complete_genome

tgt tat tgt agt atg ata ata tca aaa aga tgg ata taa aga att tac tga cta cat gta cta ttt tac att act aca ttg gct acg gca tat ata cct att tcg tca ctt cca cac gct ccg gta aac ggg tgt cat gtg acg agg gag aat ctt gat aag agg cat aat caa tgt tgt aat ccg atg tcc acc tgg aga att tgc caa ggt cag atg tag agt tgg tag tga taa cac aaa atg tga aca ctg ccc acc tca tac ata tac cgc aat ccc caa tta ttc taa tag atg tca tca atg tag aaa atg ccc aac agg atc att tga taa ggt aaa gtg tac cgg aac aca gaa caa at- gtt cgt gtc atc ctg gtt ggt ata cgc tac tga ttc ttc aca gac tga aga ttg tcg aga ttt gtg tac caa aaa agg aga tgt cca tgc gga tac ttt ggt gga ata gat gaa gga aat cct att tgt aaa tcg tgt tgt gtt ggt gaa tat tgc gac tac cta cgt aat tat aga ctt gat cca ttt cct cca tgc aaa cta tct atc taa atg taa tta att atg att ttg atg ata atg tta cca tac att ata tca cta ctt ggt tag tgt gta tta ttt agt atg gaa gac cta tta ata att act tat ctt ttg acg atc ttg tta taa tta taa tat aaa aat act tat gac ata gta a-- --- -ct cat aat tgc tga cgc gat aaa ttc gta ata atc tgt ttt gtt caa att ttt ata agg aat cta cag gca taa aaa taa aaa tat aat cta taa tat act ctt aca acg cca tca tga ata gca gtg aat taa ttg ctg tta tta tgg att tag aaa tag tgg acg att ttg tga tat taa tat agt tat taa tga tga aag gat aaa cgc gca tag act cat cct atc tgg agc ctc cga ata ttt ttt cca ttc tgt ttt cca ata att tta tcg att cta atg aat acg aag tta atc taa gtc att tag att atc aaa gtg tta act att tga tcg att aca ttt atg gga tac ctt tga gcc taa cta aca ata acg tga aat ata ttc ttt caa ccg ctg att ttt tta caa att gga tct gtc att act gag tgc gaa aaa tac ata ctt aaa aat ctt tgt ttt aga aac tgt atc gat ttc --- --- tac ata tac gct gat aaa tat aat aac aag aaa ata gaa tta gca tcg ttt aac aca ata tta cga aat att ttg aga ctc atc aac aat gaa aac ttt aaa tac tta aca gag gaa tca atg ata aaa att tta agc gat gat atg tta tat ata aaa aat gag gat ttc acc cca ctg att ctc att aaa tgg tta gag agt aca cca acc atg tac cgt cga gtt act tag atg cct cag aat atc a-t ttc ttt ccc cac aag tta taa aat cac ttt ata gtc atc gac tgg tta gtt caa tct acg aat gta taa cat tct taa aca ata tag cat tct tgg ata aat cat ttc cta gat acc ata tca tcg agt tga tat cta tcg gta taa gta att cac atg ata aga ttt cca taa act gct aca atc ata aaa aaa ttc atg gga aat gat atc ttc acg tag ata tag gtg tag ttt cgc agt gac cgt cct gga taa tat tat cta tat gat ggg tgg ata tga tca gtc cct gta tag aag ttc aaa ggt tat agc gta caa tac atg tac taa ttc ttg gat ata tga tat acc aga gct aaa aat atc atc gtt cta att gcg gag gag ttg cca atg acg aat aca ttt att gta tag gcg gta tac gcg atc agg agt cat cgt tga tat cta gta tcg ata gat gga agc cat caa aac cat att ggc aga agt atg cta aaa tgt gcg aac caa aat gtg ata tgg ggt tgc gat ttt aaa cgg att aat ata tgt cat agg tgg agt cgt taa agg tga cac aca tat acc aac gca cta gag agt tta tca gaa gat gga tgg atg aat cat caa cgt ctt cca ata aaa atg tcc aat atg tcg acg att gtt cat gct gga aag att tat ata tct aga ggt tac aac aat agt agt gta gtt aat gta ata tcg aat ct- -ag tcc tta gct ata atc cga tat atg atg aat gga cca aat tat cat cat taa ata ttc cta gaa tta atc ctg ctc tat ggt cag tgt ata ata aat tat atg tag gag gag taa tat ctg atg atg ttc aaa cta ata cat ctg aaa cat acg ata aag aaa aa

>ON682263.2_Monkeypox_virus_isolate_MPXV/Germany/2022/RKI03_complete_genome

tgt tat tgt agt atg ata ata tca aaa aga tgg ata taa aga att tac tga cta cat gta cta ttt tac att act aca ttg gct acg gca tat ata cct att tcg tca ctt cca cac gct ccg gta aac ggg tgt cat gtg acg agg gag aat ctt gat aag agg cat aat caa tgt tgt aat ccg atg tcc acc tgg aga att tgc caa ggt cag atg tag agt tgg tag tga taa cac aaa atg tga aca ctg ccc acc tca tac ata tac cgc aat ccc caa tta ttc taa tag atg tca tca atg tag aaa atg ccc aac agg atc att tga taa ggt aaa gtg tac cgg aac aca gaa caa at- gtt cgt gtc atc ctg gtt ggt ata cgc tac tga ttc ttc aca gac tga aga ttg tcg aga ttt gtg tac caa aaa agg aga tgt cca tgc gga tac ttt ggt gga ata gat gaa gga aat cct att tgt aaa tcg tgt tgt gtt ggt gaa tat tgc gac tac cta cgt aat tat aga ctt gat cca ttt cct cca tgc aaa cta tct atc taa atg taa tta att atg att ttg atg ata atg tta cca tac att ata tca cta ctt ggt tag tgt gta tta ttt agt atg gaa gac cta tta ata att act tat ctt ttg acg atc ttg tta taa tta taa tat aaa aat act tat gac ata gta a-- --- -ct cat aat tgc tga cgc gat aaa ttc gta ata atc tgt ttt gtt caa att ttt ata agg aat cta cag gca taa aaa taa aaa tat aat cta taa tat act ctt aca acg cca tca tga ata gca gtg aat taa ttg ctg tta tta tgg att tag aaa tag tgg acg att ttg tga tat taa tat agt tat taa tga tga aag gat aaa cgc gca tag act cat cct atc tgg agc ctc cga ata ttt ttt cca ttc tgt ttt cca ata att tta tcg att cta atg aat acg aag tta atc taa gtc att tag att atc aaa gtg tta act att tga tcg att aca ttt atg gga tac ctt tga gcc taa cta aca ata acg tga aat ata ttc ttt caa ccg ctg att ttt tta caa att gga tct gtc att act gag tgc gaa aaa tac ata ctt aaa aat ctt tgt ttt aga aac tgt atc gat ttc --- --- tac ata tac gct gat aaa tat aat aac aag aaa ata gaa tta gca tcg ttt aac aca ata tta cga aat att ttg aga ctc atc aac aat gaa aac ttt aaa tac tta aca gag gaa tca atg ata aaa att tta agc gat gat atg tta tat ata aaa aat gag gat ttc acc cca ctg att ctc att aaa tgg tta gag agt aca cca acc atg tac cgt cga gtt act tag atg cct cag aat atc a-t ttc ttt ccc cac aag tta taa aat cac ttt ata gtc atc gac tgg tta gtt caa tct acg aat gta taa cat tct taa aca ata tag cat tct tgg ata aat cat ttc cta gat acc ata tca tcg agt tga tat cta tcg gta taa gta att cac atg ata aga ttt cca taa act gct aca atc ata aaa aaa ttc atg gga aat gat atc ttc acg tag ata tag gtg tag ttt cgc agt gac cgt cct gga taa tat tat cta tat gat ggg tgg ata tga tca gtc cct gta tag aag ttc aaa ggt tat agc gta caa tac atg tac taa ttc ttg gat ata tga tat acc aga gct aaa aat atc atc gtt cta att gcg gag gag ttg cca atg acg aat aca ttt att gta tag gcg gta tac gcg atc agg agt cat cgt tga tat cta gta tcg ata gat gga agc cat caa aac cat att ggc aga agt atg cta aaa tgt gcg aac caa aat gtg ata tgg ggt tgc gat ttt aaa cgg att aat ata tgt cat agg tgg agt cgt taa agg tga cac aca tat acc aac gca cta gag agt tta tca gaa gat gga tgg atg aat cat caa cgt ctt cca ata aaa atg tcc aat atg tcg acg att gtt cat gct gga aag att tat ata tct aga ggt tac aac aat agt agt gta gtt aat gta ata tcg aat ct- -ag tcc tta gct ata atc cga tat atg atg aat gga cca aat tat cat cat taa ata ttc cta gaa tta atc ctg ctc tat ggt cag tgt ata ata aat tat atg tag gag gag taa tat ctg atg atg ttc aaa cta ata cat ctg aaa cat acg ata aag aaa aa

>ON682266.1_Monkeypox_virus_isolate_MPXV/Germany/2022/RKI09_complete_genome

tgt tat tgt agt atg ata ata tca aaa aga tgg ata taa aga att tac tga cta cat gta cta ttt tac att act aca ttg gct acg gca tat ata cct att tcg tca ctt cca cac gct ccg gta aac ggg tgt cat gtg acg agg gag aat ctt gat aag agg cat aat caa tgt tgt aat ccg atg tcc acc tgg aga att tgc caa ggt cag atg tag agt tgg tag tga taa cac aaa atg tga aca ctg ccc acc tca tac ata tac cgc aat ccc caa tta ttc taa tag atg tca tca atg tag aaa atg ccc aac agg atc att tga taa ggt aaa gtg tac cgg aac aca gaa caa at- gtt cgt gtc atc ctg gtt ggt ata cgc tac tga ttc ttc aca gac tga aga ttg tcg aga ttt gtg tac caa aaa agg aga tgt cca tgc gga tac ttt ggt gga ata gat gaa gga aat cct att tgt aaa tcg tgt tgt gtt ggt gaa tat tgc gac tac cta cgt aat tat aga ctt gat cca ttt cct cca tgc aaa cta tct atc taa atg taa tta att atg att ttg atg ata atg tta cca tac att ata tca cta ctt ggt tag tgt gta tta ttt agt atg gaa gac cta tta ata att act tat ctt ttg acg atc ttg tta taa tta taa tat aaa aat act tat gac ata gta a-- --- -ct cat aat tgc tga cgc gat aaa ttc gta ata atc tgt ttt gtt caa att ttt ata agg aat cta cag gca taa aaa taa aaa tat aat cta taa tat act ctt aca acg cca tca tga ata gca gtg aat taa ttg ctg tta tta tgg att tag aaa tag tgg acg att ttg tga tat taa tat agt tat taa tga tga aag gat aaa cgc gca tag act cat cct atc tgg agc ctc cga ata ttt ttt cca ttc tgt ttt cca ata att tta tcg att cta atg aat acg aag tta atc taa gtc att tag att atc aaa gtg tta act att tga tcg att aca ttt atg gga tac ctt tga gcc taa cta aca ata acg tga aat ata ttc ttt caa ccg ctg att ttt tta caa att gga tct gtc att act gag tgc gaa aaa tac ata ctt aaa aat ctt tgt ttt aga aac tgt atc gat ttc --- --- tac ata tac gct gat aaa tat aat aac aag aaa ata gaa tta gca tcg ttt aac aca ata tta cga aat att ttg aga ctc atc aac aat gaa aac ttt aaa tac tta aca gag gaa tca atg ata aaa att tta agc gat gat atg tta tat ata aaa aat gag gat ttc acc cca ctg att ctc att aaa tgg tta gag agt aca cca acc atg tac cgt cga gtt act tag atg cct cag aat atc a-t ttc ttt ccc cac aag tta taa aat cac ttt ata gtc atc gac tgg tta gtt caa tct acg aat gta taa cat tct taa aca ata tag cat tct tgg ata aat cat ttc cta gat acc ata tca tcg agt tga tat cta tcg gta taa gta att cac atg ata aga ttt cca taa act gct aca atc ata aaa aaa ttc atg gga aat gat atc ttc acg tag ata tag gtg tag ttt cgc agt gac cgt cct gga taa tat tat cta tat gat ggg tgg ata tga tca gtc cct gta tag aag ttc aaa ggt tat agc gta caa tac atg tac taa ttc ttg gat ata tga tat acc aga gct aaa aat atc atc gtt cta att gcg gag gag ttg cca atg acg aat aca ttt att gta tag gcg gta tac gcg atc agg agt cat cgt tga tat cta gta tcg ata gat gga agc cat caa aac cat att ggc aga agt atg cta aaa tgt gcg aac caa aat gtg ata tgg ggt tgc gat ttt aaa cgg att aat ata tgt cat agg tgg agt cgt taa agg tga cac aca tat acc aac gca cta gag agt tta tca gaa gat gga tgg atg aat cat caa cgt ctt cca ata aaa atg tcc aat atg tcg acg att gtt cat gct gga aag att tat ata tct aga ggt tac aac aat agt agt gta gtt aat gta ata tcg aat ct- -ag tcc tta gct ata atc cga tat atg atg aat gga cca aat tat cat cat taa ata ttc cta gaa tta atc ctg ctc tat ggt cag tgt ata ata aat tat atg tag gag gag taa tat ctg atg atg ttc aaa cta ata cat ctg aaa cat acg ata aag aaa aa

>ON619836.1_Monkeypox_virus_isolate_MPXV_UK_2022_2_complete_genome

tgt tat tgt agt atg ata ata tca aaa aga tgg ata taa aga att tac tga cta cat gta cta ttt tac att act aca ttg gct acg gca tat ata cct att tcg tca ctt cca cac gct ccg gta aac ggg tgt cat gtg acg agg gag aat ctt gat aag agg cat aat caa tgt tgt aat ccg atg tcc acc tgg aga att tgc caa ggt cag atg tag agt tgg tag tga taa cac aaa atg tga aca ctg ccc acc tca tac ata tac cgc aat ccc caa tta ttc taa tag atg tca tca atg tag aaa atg ccc aac agg atc att tga taa ggt aaa gtg tac cgg aac aca gaa caa at- gtt cgt gtc atc ctg gtt ggt ata cgc tac tga ttc ttc aca gac tga aga ttg tcg aga ttt gtg tac caa aaa agg aga tgt cca tgc gga tac ttt ggt gga ata gat gaa gga aat cct att tgt aaa tcg tgt tgt gtt ggt gaa tat tgc gac tac cta cgt aat tat aga ctt gat cca ttt cct cca tgc aaa cta tct atc taa atg taa tta att atg att ttg atg ata atg tta cca tac att ata tca cta ctt ggt tag tgt gta tta ttt agt atg gaa gac cta tta ata att act tat ctt ttg acg atc ttg tta taa tta taa tat aaa aat act tat gac ata gta a-- --- -ct cat aat tgc tga cgc gat aaa ttc gta ata atc tgt ttt gtt caa att ttt ata agg aat cta cag gca taa aaa taa aaa tat aat cta taa tat act ctt aca acg cca tca tga ata gca gtg aat taa ttg ctg tta tta tgg att tag aaa tag tgg acg att ttg tga tat taa tat agt tat taa tga tga aag gat aaa cgc gca tag act cat cct atc tgg agc ctc cga ata ttt ttt cca ttc tgt ttt cca ata att tta tcg att cta atg aat acg aag tta atc taa gtc att tag att atc aaa gtg tta act att tga tcg att aca ttt atg gga tac ctt tga gcc taa cta aca ata acg tga aat ata ttc ttt caa ccg ctg att ttt tta caa att gga tct gtc att act gag tgc gaa aaa tac ata ctt aaa aat ctt tgt ttt aga aac tgt atc gat ttc --- --- tac ata tac gct gat aaa tat aat aac aag aaa ata gaa tta gca tcg ttt aac aca ata tta cga aat att ttg aga ctc atc aac aat gaa aac ttt aaa tac tta aca gag gaa tca atg ata aaa att tta agc gat gat atg tta tat ata aaa aat gag gat ttc acc cca ctg att ctc att aaa tgg tta gag agt aca cca acc atg tac cgt cga gtt act tag atg cct cag aat atc a-t ttc ttt ccc cac aag tta taa aat cac ttt ata gtc atc gac tgg tta gtt caa tct acg aat gta taa cat tct taa aca ata tag cat tct tgg ata aat cat ttc cta gat acc ata tca tcg agt tga tat cta tcg gta taa gta att cac atg ata aga ttt cca taa act gct aca atc ata aaa aaa ttc atg gga aat gat atc ttc acg tag ata tag gtg tag ttt cgc agt gac cgt cct gga taa tat tat cta tat gat ggg tgg ata tga tca gtc cct gta tag aag ttc aaa ggt tat agc gta caa tac atg tac taa ttc ttg gat ata tga tat acc aga gct aaa aat atc atc gtt cta att gcg gag gag ttg cca atg acg aat aca ttt att gta tag gcg gta tac gcg atc agg agt cat cgt tga tat cta gta tcg ata gat gga agc cat caa aac cat att ggc aga agt atg cta aaa tgt gcg aac caa aat gtg ata tgg ggt tgc gat ttt aaa cgg att aat ata tgt cat agg tgg agt cgt taa agg tga cac aca tat acc aac gca cta gag agt tta tca gaa gat gga tgg atg aat cat caa cgt ctt cca ata aaa atg tcc aat atg tcg acg att gtt cat gct gga aag att tat ata tct aga ggt tac aac aat agt agt gta gtt aat gta ata tcg aat ct- -ag tcc tta gct ata atc cga tat atg atg aat gga cca aat tat cat cat taa ata ttc cta gaa tta atc ctg ctc tat ggt cag tgt ata ata aat tat atg tag gag gag taa tat ctg atg atg ttc aaa cta ata cat ctg aaa cat acg ata aag aaa aa

>ON619838.1_Monkeypox_virus_isolate_MPXV_UK_2022_4_complete_genome

tgt tat tgt agt atg ata ata tca aaa aga tgg ata taa aga att tac tga cta cat gta cta ttt tac att act aca ttg gct acg gca tat ata cct att tcg tca ctt cca cac gct ccg gta aac ggg tgt cat gtg acg agg gag aat ctt gat aag agg cat aat caa tgt tgt aat ccg atg tcc acc tgg aga att tgc caa ggt cag atg tag agt tgg tag tga taa cac aaa atg tga aca ctg ccc acc tca tac ata tac cgc aat ccc caa tta ttc taa tag atg tca tca atg tag aaa atg ccc aac agg atc att tga taa ggt aaa gtg tac cgg aac aca gaa caa at- gtt cgt gtc atc ctg gtt ggt ata cgc tac tga ttc ttc aca gac tga aga ttg tcg aga ttt gtg tac caa aaa agg aga tgt cca tgc gga tac ttt ggt gga ata gat gaa gga aat cct att tgt aaa tcg tgt tgt gtt ggt gaa tat tgc gac tac cta cgt aat tat aga ctt gat cca ttt cct cca tgc aaa cta tct atc taa atg taa tta att atg att ttg atg ata atg tta cca tac att ata tca cta ctt ggt tag tgt gta tta ttt agt atg gaa gac cta tta ata att act tat ctt ttg acg atc ttg tta taa tta taa tat aaa aat act tat gac ata gta a-- --- -ct cat aat tgc tga cgc gat aaa ttc gta ata atc tgt ttt gtt caa att ttt ata agg aat cta cag gca taa aaa taa aaa tat aat cta taa tat act ctt aca acg cca tca tga ata gca gtg aat taa ttg ctg tta tta tgg att tag aaa tag tgg acg att ttg tga tat taa tat agt tat taa tga tga aag gat aaa cgc gca tag act cat cct atc tgg agc ctc cga ata ttt ttt cca ttc tgt ttt cca ata att tta tcg att cta atg aat acg aag tta atc taa gtc att tag att atc aaa gtg tta act att tga tcg att aca ttt atg gga tac ctt tga gcc taa cta aca ata acg tga aat ata ttc ttt caa ccg ctg att ttt tta caa att gga tct gtc att act gag tgc gaa aaa tac ata ctt aaa aat ctt tgt ttt aga aac tgt atc gat ttc --- --- tac ata tac gct gat aaa tat aat aac aag aaa ata gaa tta gca tcg ttt aac aca ata tta cga aat att ttg aga ctc atc aac aat gaa aac ttt aaa tac tta aca gag gaa tca atg ata aaa att tta agc gat gat atg tta tat ata aaa aat gag gat ttc acc cca ctg att ctc att aaa tgg tta gag agt aca cca acc atg tac cgt cga gtt act tag atg cct cag aat atc a-t ttc ttt ccc cac aag tta taa aat cac ttt ata gtc atc gac tgg tta gtt caa tct acg aat gta taa cat tct taa aca ata tag cat tct tgg ata aat cat ttc cta gat acc ata tca tcg agt tga tat cta tcg gta taa gta att cac atg ata aga ttt cca taa act gct aca atc ata aaa aaa ttc atg gga aat gat atc ttc acg tag ata tag gtg tag ttt cgc agt gac cgt cct gga taa tat tat cta tat gat ggg tgg ata tga tca gtc cct gta tag aag ttc aaa ggt tat agc gta caa tac atg tac taa ttc ttg gat ata tga tat acc aga gct aaa aat atc atc gtt cta att gcg gag gag ttg cca atg acg aat aca ttt att gta tag gcg gta tac gcg atc agg agt cat cgt tga tat cta gta tcg ata gat gga agc cat caa aac cat att ggc aga agt atg cta aaa tgt gcg aac caa aat gtg ata tgg ggt tgc gat ttt aaa cgg att aat ata tgt cat agg tgg agt cgt taa agg tga cac aca tat acc aac gca cta gag agt tta tca gaa gat gga tgg atg aat cat caa cgt ctt cca ata aaa atg tcc aat atg tcg acg att gtt cat gct gga aag att tat ata tct aga ggt tac aac aat agt agt gta gtt aat gta ata tcg aat ct- -ag tcc tta gct ata atc cga tat atg atg aat gga cca aat tat cat cat taa ata ttc cta gaa tta atc ctg ctc tat ggt cag tgt ata ata aat tat atg tag gag gag taa tat ctg atg atg ttc aaa cta ata cat ctg aaa cat acg ata aag aaa aa

>ON619835.1_Monkeypox_virus_isolate_MPXV_UK_2022_1_complete_genome

tgt tat tgt agt atg ata ata tca aaa aga tgg ata taa aga att tac tga cta cat gta cta ttt tac att act aca ttg gct acg gca tat ata cct att tcg tca ctt cca cac gct ccg gta aac ggg tgt cat gtg acg agg gag aat ctt gat aag agg cat aat caa tgt tgt aat ccg atg tcc acc tgg aga att tgc caa ggt cag atg tag agt tgg tag tga taa cac aaa atg tga aca ctg ccc acc tca tac ata tac cgc aat ccc caa tta ttc taa tag atg tca tca atg tag aaa atg ccc aac agg atc att tga taa ggt aaa gtg tac cgg aac aca gaa caa at- gtt cgt gtc atc ctg gtt ggt ata cgc tac tga ttc ttc aca gac tga aga ttg tcg aga ttt gtg tac caa aaa agg aga tgt cca tgc gga tac ttt ggt gga ata gat gaa gga aat cct att tgt aaa tcg tgt tgt gtt ggt gaa tat tgc gac tac cta cgt aat tat aga ctt gat cca ttt cct cca tgc aaa cta tct atc taa atg taa tta att atg att ttg atg ata atg tta cca tac att ata tca cta ctt ggt tag tgt gta tta ttt agt atg gaa gac cta tta ata att act tat ctt ttg acg atc ttg tta taa tta taa tat aaa aat act tat gac ata gta a-- --- -ct cat aat tgc tga cgc gat aaa ttc gta ata atc tgt ttt gtt caa att ttt ata agg aat cta cag gca taa aaa taa aaa tat aat cta taa tat act ctt aca acg cca tca tga ata gca gtg aat taa ttg ctg tta tta tgg att tag aaa tag tgg acg att ttg tga tat taa tat agt tat taa tga tga aag gat aaa cgc gca tag act cat cct atc tgg agc ctc cga ata ttt ttt cca ttc tgt ttt cca ata att tta tcg att cta atg aat acg aag tta atc taa gtc att tag att atc aaa gtg tta act att tga tcg att aca ttt atg gga tac ctt tga gcc taa cta aca ata acg tga aat ata ttc ttt caa ccg ctg att ttt tta caa att gga tct gtc att act gag tgc gaa aaa tac ata ctt aaa aat ctt tgt ttt aga aac tgt atc gat ttc --- --- tac ata tac gct gat aaa tat aat aac aag aaa ata gaa tta gca tcg ttt aac aca ata tta cga aat att ttg aga ctc atc aac aat gaa aac ttt aaa tac tta aca gag gaa tca atg ata aaa att tta agc gat gat atg tta tat ata aaa aat gag gat ttc acc cca ctg att ctc att aaa tgg tta gag agt aca cca acc atg tac cgt cga gtt act tag atg cct cag aat atc a-t ttc ttt ccc cac aag tta taa aat cac ttt ata gtc atc gac tgg tta gtt caa tct acg aat gta taa cat tct taa aca ata tag cat tct tgg ata aat cat ttc cta gat acc ata tca tcg agt tga tat cta tcg gta taa gta att cac atg ata aga ttt cca taa act gct aca atc ata aaa aaa ttc atg gga aat gat atc ttc acg tag ata tag gtg tag ttt cgc agt gac cgt cct gga taa tat tat cta tat gat ggg tgg ata tga tca gtc cct gta tag aag ttc aaa ggt tat agc gta caa tac atg tac taa ttc ttg gat ata tga tat acc aga gct aaa aat atc atc gtt cta att gcg gag gag ttg cca atg acg aat aca ttt att gta tag gcg gta tac gcg atc agg agt cat cgt tga tat cta gta tcg ata gat gga agc cat caa aac cat att ggc aga agt atg cta aaa tgt gcg aac caa aat gtg ata tgg ggt tgc gat ttt aaa cgg att aat ata tgt cat agg tgg agt cgt taa agg tga cac aca tat acc aac gca cta gag agt tta tca gaa gat gga tgg atg aat cat caa cgt ctt cca ata aaa atg tcc aat atg tcg acg att gtt cat gct gga aag att tat ata tct aga ggt tac aac aat agt agt gta gtt aat gta ata tcg aat ct- -ag tcc tta gct ata atc cga tat atg atg aat gga cca aat tat cat cat taa ata ttc cta gaa tta atc ctg ctc tat ggt cag tgt ata ata aat tat atg tag gag gag taa tat ctg atg atg ttc aaa cta ata cat ctg aaa cat acg ata aag aaa aa

>ON682269.2_Monkeypox_virus_isolate_MPXV/Germany/2022/RKI07_complete_genome

tgt tat tgt agt atg ata ata tca aaa aga tgg ata taa aga att tac tga cta cat gta cta ttt tac att act aca ttg gct acg gca tat ata cct att tcg tca ctt cca cac gct ccg gta aac ggg tgt cat gtg acg agg gag aat ctt gat aag agg cat aat caa tgt tgt aat ccg atg tcc acc tgg aga att tgc caa ggt cag atg tag agt tgg tag tga taa cac aaa atg tga aca ctg ccc acc tca tac ata tac cgc aat ccc caa tta ttc taa tag atg tca tca atg tag aaa atg ccc aac agg atc att tga taa ggt aaa gtg tac cgg aac aca gaa caa at- gtt cgt gtc atc ctg gtt ggt ata cgc tac tga ttc ttc aca gac tga aga ttg tcg aga ttt gtg tac caa aaa agg aga tgt cca tgc gga tac ttt ggt gga ata gat gaa gga aat cct att tgt aaa tcg tgt tgt gtt ggt gaa tat tgc gac tac cta cgt aat tat aga ctt gat cca ttt cct cca tgc aaa cta tct atc taa atg taa tta att atg att ttg atg ata atg tta cca tac att ata tca cta ctt ggt tag tgt gta tta ttt agt atg gaa gac cta tta ata att act tat ctt ttg acg atc ttg tta taa tta taa tat aaa aat act tat gac ata gta a-- --- -ct cat aat tgc tga cgc gat aaa ttc gta ata atc tgt ttt gtt caa att ttt ata agg aat cta cag gca taa aaa taa aaa tat aat cta taa tat act ctt aca acg cca tca tga ata gca gtg aat taa ttg ctg tta tta tgg att tag aaa tag tgg acg att ttg tga tat taa tat agt tat taa tga tga aag gat aaa cgc gca tag act cat cct atc tgg agc ctc cga ata ttt ttt cca ttc tgt ttt cca ata att tta tcg att cta atg aat acg aag tta atc taa gtc att tag att atc aaa gtg tta act att tga tcg att aca ttt atg gga tac ctt tga gcc taa cta aca ata acg tga aat ata ttc ttt caa ccg ctg att ttt tta caa att gga tct gtc att act gag tgc gaa aaa tac ata ctt aaa aat ctt tgt ttt aga aac tgt atc gat ttc --- --- tac ata tac gct gat aaa tat aat aac aag aaa ata gaa tta gca tcg ttt aac aca ata tta cga aat att ttg aga ctc atc aac aat gaa aac ttt aaa tac tta aca gag gaa tca atg ata aaa att tta agc gat gat atg tta tat ata aaa aat gag gat ttc acc cca ctg att ctc att aaa tgg tta gag agt aca cca acc atg tac cgt cga gtt act tag atg cct cag aat atc a-t ttc ttt ccc cac aag tta taa aat cac ttt ata gtc atc gac tgg tta gtt caa tct acg aat gta taa cat tct taa aca ata tag cat tct tgg ata aat cat ttc cta gat acc ata tca tcg agt tga tat cta tcg gta taa gta att cac atg ata aga ttt cca taa act gct aca atc ata aaa aaa ttc atg gga aat gat atc ttc acg tag ata tag gtg tag ttt cgc agt gac cgt cct gga taa tat tat cta tat gat ggg tgg ata tga tca gtc cct gta tag aag ttc aaa ggt tat agc gta caa tac atg tac taa ttc ttg gat ata tga tat acc aga gct aaa aat atc atc gtt cta att gcg gag gag ttg cca atg acg aat aca ttt att gta tag gcg gta tac gcg atc agg agt cat cgt tga tat cta gta tcg ata gat gga agc cat caa aac cat att ggc aga agt atg cta aaa tgt gcg aac caa aat gtg ata tgg ggt tgc gat ttt aaa cgg att aat ata tgt cat agg tgg agt cgt taa agg tga cac aca tat acc aac gca cta gag agt tta tca gaa gat gga tgg atg aat cat caa cgt ctt cca ata aaa atg tcc aat atg tcg acg att gtt cat gct gga aag att tat ata tct aga ggt tac aac aat agt agt gta gtt aat gta ata tcg aat ct- -ag tcc tta gct ata atc cga tat atg atg aat gga cca aat tat cat cat taa ata ttc cta gaa tta atc ctg ctc tat ggt cag tgt ata ata aat tat atg tag gag gag taa tat ctg atg atg ttc aaa cta ata cat ctg aaa cat acg ata aag aaa aa

>KJ642616.1_Monkeypox_virus_strain_PCH_complete_genome

tgt tat tgt agt atg ata ata tca aaa aga tgg ata taa aga att tac tga cta cat gta cta ttt tac att act aca ttg gct acg g-- tat ata cct act tcg tca ctt caa cac gct cca gta aac ggg tgt cat gtg acg agg gag aat ctt gat aag agg cat aat caa tgt tgt aat ccg gtg tcc acc tgg aga att tgc caa ggt cag atg t-- agt tgg tag tga taa cac aaa atg tga aca ctg ccc acc tca tac ata tac cgc aat ccc caa tta ttc taa tag atg tca tca atg tag aaa atg ccc aac agg atc att tga taa ggt aaa gtg tac cgg aac aca gaa caa at- gtt cgt gtc atc ctg gtt ggt ata cgc tac tga ttc ttc aca gac tga aga ttg tcg aga ttt gtg tac caa aaa agg aga tgt cca tgc gga tac ttt ggt gga ata gat gaa gga aat cct att tgt aaa tcg tgt tgt gtt ggt gaa tat tgc gac tac cta cgt aat tat aga ctt gat cca ttt cct cca tgc aac cta tct atc taa atg taa tta att atg att ttg atg ata atg tta cca tac att ata tcg cta ctt ggt tag tgt gta tta ttc agt atg gaa gac cta tta ata att act tat ctt ttg acg atc ttg tta taa tta taa tat aaa aat act tat gac ata gta aca gta act cat aat tgc tga cgc gat aaa ttc gta ata atc tgt ttt gtt caa att ttt ata agg aat cta cag gca taa aaa taa aaa tat aat cta taa tat act ctt aca acg cca tca tga ata gca gtg aat taa ttg ctg tta tta tgg att tag aaa tag tgg acg att ttg tga tat taa tat agt tat taa tga tga aag gat aaa cgc tca tag att cat cct atc tgg agc ctc cga ata ttt ttt cca ttc tgt ttt cca ata att tta tcg att cta atg aat acg aag tta atc taa gtc att tag att atc aaa gtg tta act att tga tcg att aca ttt atg gga tac ctt tga gcc taa cta acg ata acg tga aat ata ttc ttt caa ccg ctg att ttt tta caa att gga tct gcc att act gag tgc gaa aaa tac ata ctt aaa aat ctt tgt ttt aga aac tgt atc gat ttc --- --- tac ata tac gct gat aaa tat aat aac aag aaa ata gaa tta gca tcg ttt aac aca cta tta cga aat att ttg aga ctc atc aac aat gaa aac ttt aaa tac tta aca gag gaa tca --- --- --- --- --- --- -at gat atg tta tat ata aaa aat gag gat ttc acc cca ctg att ctc att aaa tgg tta gag agt aca cca acc atg tac cgt cga gtt act tag atg cct cag aat atc a-t ttc ttt ccc cac aag tta taa aat cac ttt ata gtc atc gac tgg tta gtt caa tct acg aat gta taa cat tct taa aca ata tag cat tct tgg ata aat cat ttc cta gat acc ata tca tcg agt tga tat cta tcg gta taa gta att cac atg ata aga ttt cca taa act gct aca atc ata aaa aaa ttc atg gga aat gat atc ttc acg tag at- --g gtg tag ttt cgc agt gac cgt cct gga taa tat tat cta tat gat ggg tgg ata tga tca gtc cct gta tag aag ttc aaa ggt tat agc gta caa tac atg tac taa ttc ttg gat ata tga tat acc aga gct -aa aat atc ctc gtt cta att gcg gag gag ttg cca atg acg aat aca ttt att gta tag gcg gta tac gcg atc agg agt cat cgt tga tat cta gta tcg ata gat gga agc cat caa aac cat att ggc aga agt atg cta aaa tgt gcg aac caa aat gtg ata tgg ggt tgc gat ttt aaa cgg att aat ata tgt cat agg tgg agt tgt taa agg tga cac aca tgt acc gac gca cta gag agt tta tca gaa gat gga tgg atg aat cat caa cgt ctt cca ata aaa atg tcc aat atg tcg acg att gtt cat gct gga aag att tat ata tct gga ggt tac aac aat agt agt gta gtt aat gta ata tcg aat cta gag tcc tta gct ata atc cga tat atg atg aat gga cca aat tat cat cat taa ata ttc cta gaa tta atc ctg ctc tat ggt cag tgt ata ata aat tat atg tag gag gag gaa tat ctg atg atg ttc aaa cta ata cat ctg aaa cat acg ata aag aaa aa

>AY753185.1_Monkeypox_virus_strain_COP-58_complete_genome

tgt tat tgt agt atg ata ata tca aaa aga tgg ata taa aga att tac tga cta cat gta cta ttt tac att act aca ttg gct acg g-- tat ata cct act tcg tca ctt cca cac gct cca gta aac ggg tgt cat gtg acg agg gag aat ctt gat aag agg cat aat caa tgt tgt aat ccg gtg tcc acc tgg aga att tgc caa ggt cag atg t-- agt tgg tag tga taa cac aaa atg tga aca ctg ccc acc tca tac ata tac cgc aat ccc caa tta ttc taa tag atg tca tca atg tag aaa atg ccc aac agg atc att tga taa ggt aaa gtg tac cgg aac aca gaa caa at- gtt cgt gtc atc ctg gtt ggt ata cgc tac tga ttc ttc aca gac tga aga ttg tcg aga ttt gtg tac caa aaa agg aga tgt cca tgc gga tac ttt ggt gga ata gat gaa gga aat cct att tgt aaa tcg tgt tgt gtt ggt gaa tat tgc gac tac cta cgt aat tat aga ctt gat cca ttt cct cca tgc aac cta tct atc taa atg taa tta att atg att ttg atg ata atg tta cca tac att ata tcg cta ctt ggt tag tgt gta tta ttc agt atg gaa gac cta tta ata att act tat ctt ttg acg atc ttg tta taa tta taa tat aaa aat act tat gac ata gta aca gta act cat aat tgc tga cgc gat aaa ttc gta ata atc tgt ttt gtt caa att ttt ata agg aat cta cag gca taa aaa taa aaa tat aat cta taa tat act ctt aca acg cca tca tga ata gca gtg aat taa ttg ctg tta tta tgg att tag aaa tag tgg acg att ttg tga tat taa tat agt tat taa tga tga aag gat aaa cgc tca tag att cat cct atc tgg agc ctc cga ata ttt ttt cca ttc tgt ttt cca ata att tta tcg att cta atg aat acg aag tta atc taa gtc att tag att atc aaa gtg tta act att tga tcg att aca ttt atg gga tac ctt tga gcc taa cta acg ata acg tga aat ata ttc ttt caa ccg ctg att ttt tta caa att gga tct gcc att act gag tgc gaa aaa tac ata ctt aaa aat ctt tgt ttt aga aac tgt atc gat ttc --- --- tac ata tac gct gat aaa tat aat aac aag aaa ata gaa tta gca tcg ttt aac aca ata tta cga aat att ttg aga ctc atc aac aat gaa aac ttt aaa tac tta aca gag gaa tca --- --- --- --- --- --- -at gat atg tta tat ata aaa aat gag gat ttc acc cca ctg att ctc att aaa tgg tta gag agt aca cca acc atg tac cgt cga gtt act tag atg cct cag aat atc a-t ttc ttt ccc cac aag tta taa aat cac ttt ata gtc atc gac tgg tta gtt caa tct acg aat gta taa cat tct taa aca ata tag cat tct tgg ata aat cat ttc cta gat acc ata tca tcg agt tga tat cta tcg gta taa gta att cac atg ata aga ttt cca taa act gct aca atc ata aaa aaa ttc atg gga aat gat atc ttc acg tag at- --g gtg tag ttt cgc agt gac cgt cct gga taa tat tat cta tat gat ggg tgg ata tga tca gtc cct gta tag aag ttc aaa ggt tat agc gta caa tac atg tac taa ttc ttg gat ata tga tat acc aga gct -aa aat atc ctc gtt cta att gcg gag gag ttg cca atg acg aat aca ttt att gta tag gcg gta tac gcg atc agg agt cat cgt tga tat cta gta tcg ata gat gga agc cat caa aac cat att ggc aga agt atg cta aaa tgt gcg aac caa aat gtg ata tgg ggt tgc gat ttt aaa cgg att aat ata tgt cat agg tgg agt cgt taa agg tga cac aca tgt acc gac gca cta gag agt tta tca gaa gat gga tgg atg aat cat caa cgt ctt cca ata aaa atg tcc aat atg tcg acg att gtt cat gct gga aag att tat ata tct gga ggt tac aac aat agt agt gta gtt aat gta ata tcg aat cta gag tcc tta gct ata atc cga tat atg atg aat gga cca aat tat cat cat taa ata ttc cta gaa tta atc ctg ctc tat ggt cag tgt ata ata aat tat atg tag gag gag gaa tat ctg atg atg ttc aaa cta ata cat ctg aaa cat acg ata aag aaa aa

>AY603973.1_Monkeypox_virus_strain_MPXV-WRAIR7-61_complete_genome

tgt tat tgt agt atg ata ata tca aaa aga tgg ata taa aga att tac tga cta cat gta cta ttt tac att act aca ttg gct acg g-- tat ata cct act tcg tca ctt cca cac gct cca gta aac ggg tgt cat gtg acg agg gag aat ctt gat aag agg cat aat caa tgt tgt aat ccg gtg tcc acc tgg aga att tgc caa ggt cag atg t-- agt tgg tag tga taa cac aaa atg tga aca ctg ccc acc tca tac ata tac cgc aat ccc caa tta ttc taa tag atg tca tca atg tag aaa atg ccc aac agg atc att tga taa ggt aaa gtg tac cgg aac aca gaa caa at- gtt cgt gtc atc ctg gtt ggt ata cgc tac tga ttc ttc aca gac tga aga ttg tcg aga ttt gtg tac caa aaa agg aga tgt cca tgc gga tac ttt ggt gga ata gat gaa gga aat cct att tgt aaa tcg tgt tgt gtt ggt gaa tat tgc gac tac cta cgt aat tat aga ctt gat cca ttt cct cca tgc aac cta tct atc taa atg taa tta att atg att ttg atg ata atg tta cca tac att ata tcg cta ctt ggt tag tgt gta tta ttc agt atg gaa gac cta tta ata att act tat ctt ttg acg atc ttg tta taa tta taa tat aaa aat act tat gac ata gta aca gta act cat aat tgc tga cgc gat aaa ttc gta ata atc tgt ttt gtt caa att ttt ata agg aat cta cag gca taa aaa taa aaa tat aat cta taa tat act ctt aca acg cca tca tga ata gca gtg aat taa ttg ctg tta tta tgg att tag aaa tag tgg acg att ttg tga tat taa tat agt tat taa tga tga aag gat aaa cgc tca tag att cat cct atc tgg agc ctc cga ata ttt ttt cca ttc tgt ttt cca ata att tta tcg att cta atg aat acg aag tta atc taa gtc att tag att atc aaa gtg tta act att tga tcg att aca ttt atg gga tac ctt tga gcc taa cta acg ata acg tga aat ata ttc ttt caa ccg ctg att ttt tta caa att gga tct gcc att act gag tgc gaa aaa tac ata ctt aaa aat ctt tgt ttt aga aac tgt atc gat ttc --- --- tac ata tac gct gat aaa tat aat aac aag aaa ata gaa tta gca tcg ttt aac aca ata tta cga aat att ttg aga ctc atc aac aat gaa aac ttt aaa tac tta aca gag gaa tca --- --- --- --- --- --- -at gat atg tta tat ata aaa aat gag gat ttc acc cca ctg att ctc att aaa tgg tta gag agt aca cca acc atg tac cgt cga gtt act tag atg cct cag aat atc a-t ttc ttt ccc cac aag tta taa aat cac ttt ata gtc atc gac tgg tta gtt caa tct acg aat gta taa cat tct taa aca ata tag cat tct tgg ata aat cat ttc cta gat acc ata tca tcg agt tga tat cta tcg gta taa gta att cac atg ata aga ttt cca taa act gct aca atc ata aaa aaa ttc atg gga aat gat atc ttc acg tag at- --g gtg tag ttt cgc agt gac cgt cct gga taa tat tat cta tat gat ggg tgg ata tga tca gtc cct gta tag aag ttc aaa ggt tat agc gta caa tac atg tac taa ttc ttg gat ata tga tat acc aga gct -aa aat atc ctc gtt cta att gcg gag gag ttg cca atg acg aat aca ttt att gta tag gcg gta tac gcg atc agg agt cat cgt tga tat cta gta tcg ata gat gga agc cat caa aac cat att ggc aga agt atg cta aaa tgt gcg aac caa aat gtg ata tgg ggt tgc gat ttt aaa cgg att aat ata tgt cat agg tgg agt cgt taa agg tga cac aca tgt acc gac gca cta gag agt tta tca gaa gat gga tgg atg aat cat caa cgt ctt cca ata aaa atg tcc aat atg tcg acg att gtt cat gct gga aag att tat ata tct gga ggt tac aac aat agt agt gta gtt aat gta ata tcg aat cta gag tcc tta gct ata atc cga tat atg atg aat gga cca aat tat cat cat taa ata ttc cta gaa tta atc ctg ctc tat ggt cag tgt ata ata aat tat atg tag gag gag gaa tat ctg atg atg ttc aaa cta ata cat ctg aaa cat acg ata aag aaa aa

>ON619837.1_Monkeypox_virus_isolate_MPXV_UK_2022_3_complete_genome

tgt tat tgt agt atg ata ata tca aaa aga tgg ata taa aga att tac tga cta cat gta cta ttt tac att act aca ttg gct acg gca tat ata cct att tcg tca ctt cca cac gct ccg gta aac ggg tgt cat gtg acg agg gag aat ctt gat aag agg cat aat caa tgt tgt aat ccg atg tcc acc tgg aga att tgc caa ggt cag atg tag agt tgg tag tga taa cac aaa atg tga aca ctg ccc acc tca tac ata tac cgc aat ccc caa tta ttc taa tag atg tca tca atg tag aaa atg ccc aac agg atc att tga taa ggt aaa gtg tac cgg aac aca gaa caa at- gtt cgt gtc atc ctg gtt ggt ata cgc tac tga ttc ttc aca gac tga aga ttg tcg aga ttt gtg tac caa aaa agg aga tgt cca tgc gga tac ttt ggt gga ata gat gaa gga aat cct att tgt aaa tcg tgt tgt gtt ggt gaa tat tgc gac tac cta cgt aat tat aga ctt gat cca ttt cct cca tgc aaa cta tct atc taa atg taa tta att atg att ttg atg ata atg tta cca tac att ata tca cta ctt ggt tag tgt gta tta ttt agt atg gaa gac cta tta ata att act tat ctt ttg acg atc ttg tta taa tta taa tat aaa aat act tat gac ata gta a-- --- -ct cat aat tgc tga cgc gat aaa ttc gta ata atc tgt ttt gtt caa att ttt ata agg aat cta cag gca taa aaa taa aaa tat aat cta taa tat act ctt aca acg cca tca tga ata gca gtg aat taa ttg ctg tta tta tgg att tag aaa tag tgg acg att ttg tga tat taa tat agt tat taa tga tga aag gat aaa cgc gca tag act cat cct atc tgg agc ctc cga ata ttt ttt cca ttc tgt ttt cca ata att tta tcg att cta atg aat acg aag tta atc taa gtc att tag att atc aaa gtg tta act att tga tcg att aca ttt atg gga tac ctt tga gcc taa cta aca ata acg tga aat ata ttc ttt caa ccg ctg att ttt tta caa att gga tct gtc att act gag tgc gaa aaa tac ata ctt aaa aat ctt tgt ttt aga aac tgt atc gat ttc --- --- tac ata tac gct gat aaa tat aat aac aag aaa ata gaa tta gca tcg ttt aac aca ata tta cga aat att ttg aga ctc atc aac aat gaa aac ttt aaa tac tta aca gag gaa tca atg ata aaa att tta agc gat gat atg tta tat ata aaa aat gag gat ttc acc cca ctg att ctc att aaa tgg tta gag agt aca cca acc atg tac cgt cga gtt act tag atg cct cag aat atc a-t ttc ttt ccc cac aag tta taa aat cac ttt ata gtc atc gac tgg tta gtt caa tct acg aat gta taa cat tct taa aca ata tag cat tct tgg ata aat cat ttc cta gat acc ata tca tcg agt tga tat cta tcg gta taa gta att cac atg ata aga ttt cca taa act gct aca atc ata aaa aaa ttc atg gga aat gat atc ttc acg tag ata tag gtg tag ttt cgc agt gac cgt cct gga taa tat tat cta tat gat ggg tgg ata tga tca gtc cct gta tag aag ttc aaa ggt tat agc gta caa tac atg tac taa ttc ttg gat ata tga tat acc aga gct aaa aat atc atc gtt cta att gcg gag gag ttg cca atg acg aat aca ttt att gta tag gcg gta tac gcg atc agg agt cat cgt tga tat cta gta tcg ata gat gga agc cat caa aac cat att ggc aga agt atg cta aaa tgt gcg aac caa aat gtg ata tgg ggt tgc gat ttt aaa cgg att aat ata tgt cat agg tgg agt cgt taa agg tga cac aca tat acc aac gca cta gag agt tta tca gaa gat gga tgg atg aat cat caa cgt ctt cca ata aaa atg tcc aat atg tcg acg att gtt cat gct gga aag att tat ata tct aga ggt tac aac aat agt agt gta gtt aat gta ata tcg aat ct- -ag tcc tta gct ata atc cga tat atg atg aat gga cca aat tat cat cat taa ata ttc cta gaa tta atc ctg ctc tat ggt cag tgt ata ata aat tat atg tag gag gag taa tat ctg atg atg ttc aaa cta ata cat ctg aaa cat acg ata aag aaa aa

>ON585037.1_Monkeypox_virus_isolate_Monkeypox/PT0005/2022_complete_genome

tgt tat tgt agt atg ata ata tca aaa aga tgg ata taa aga att tac tga cta cat gta cta ttt tac att act aca ttg gct acg gca tat ata cct att tcg tca ctt cca cac gct ccg gta aac ggg tgt cat gtg acg agg gag aat ctt gat aag agg cat aat caa tgt tgt aat ccg atg tcc acc tgg aga att tgc caa ggt cag atg tag agt tgg tag tga taa cac aaa atg tga aca ctg ccc acc tca tac ata tac cgc aat ccc caa tta ttc taa tag atg tca tca atg tag aaa atg ccc aac agg atc att tga taa ggt aaa gtg tac cgg aac aca gaa caa at- gtt cgt gtc atc ctg gtt ggt ata cgc tac tga ttc ttc aca gac tga aga ttg tcg aga ttt gtg tac caa aaa agg aga tgt cca tgc gga tac ttt ggt gga ata gat gaa gga aat cct att tgt aaa tcg tgt tgt gtt ggt gaa tat tgc gac tac cta cgt aat tat aga ctt gat cca ttt cct cca tgc aaa cta tct atc taa atg taa tta att atg att ttg atg ata atg tta cca tac att ata tca cta ctt ggt tag tgt gta tta ttt agt atg gaa gac cta tta ata att act tat ctt ttg acg atc ttg tta taa tta taa tat aaa aat act tat gac ata gta a-- --- -ct cat aat tgc tga cgc gat aaa ttc gta ata atc tgt ttt gtt caa att ttt ata agg aat cta cag gca taa aaa taa aaa tat aat cta taa tat act ctt aca acg cca tca tga ata gca gtg aat taa ttg ctg tta tta tgg att tag aaa tag tgg acg att ttg tga tat taa tat agt tat taa tga tga aag gat aaa cgc gca tag act cat cct atc tgg agc ctc cga ata ttt ttt cca ttc tgt ttt cca ata att tta tcg att cta atg aat acg aag tta atc taa gtc att tag att atc aaa gtg tta act att tga tcg att aca ttt atg gga tac ctt tga gcc taa cta aca ata acg tga aat ata ttc ttt caa ccg ctg att ttt tta caa att gga tct gtc att act gag tgc gaa aaa tac ata ctt aaa aat ctt tgt ttt aga aac tgt atc gat ttc --- --- tac ata tac gct gat aaa tat aat aac aag aaa ata gaa tta gca tcg ttt aac aca ata tta cga aat att ttg aga ctc atc aac aat gaa aac ttt aaa tac tta aca gag gaa tca atg ata aaa att tta agc gat gat atg tta tat ata aaa aat gag gat ttc acc cca ctg att ctc att aaa tgg tta gag agt aca cca acc atg tac cgt cga gtt act tag atg cct cag aat atc a-t ttc ttt ccc cac aag tta taa aat cac ttt ata gtc atc gac tgg tta gtt caa tct acg aat gta taa cat tct taa aca ata tag cat tct tgg ata aat cat ttc cta gat acc ata tca tcg agt tga tat cta tcg gta taa gta att cac atg ata aga ttt cca taa act gct aca atc ata aaa aaa ttc atg gga aat gat atc ttc acg tag ata tag gtg tag ttt cgc agt gac cgt cct gga taa tat tat cta tat gat ggg tgg ata tga tca gtc cct gta tag aag ttc aaa ggt tat agc gta caa tac atg tac taa ttc ttg gat ata tga tat acc aga gct aaa aat atc atc gtt cta att gcg gag gag ttg cca atg acg aat aca ttt att gta tag gcg gta tac gcg atc agg agt cat cgt tga tat cta gta tcg ata gat gga agc cat caa aac cat att ggc aga agt atg cta aaa tgt gcg aac caa aat gtg ata tgg ggt tgc gat ttt aaa cgg att aat ata tgt cat agg tgg agt cgt taa agg tga cac aca tat acc aac gca cta gag agt tta tca gaa gat gga tgg atg aat cat caa cgt ctt cca ata aaa atg tcc aat atg tcg acg att gtt cat gct gga aag att tat ata tct aga ggt tac aac aat agt agt gta gtt aat gta ata tcg aat ct- -ag tcc tta gct ata atc cga tat atg atg aat gga cca aat tat cat cat taa ata ttc cta gaa tta atc ctg ctc tat ggt cag tgt ata ata aat tat atg tag gag gag taa tat ctg atg atg ttc aaa cta ata cat ctg aaa cat acg ata aag aaa aa

>ON585032.1_Monkeypox_virus_isolate_Monkeypox/PT0004/2022_complete_genome

tgt tat tgt agt atg ata ata tca aaa aga tgg ata taa aga att tac tga cta cat gta cta ttt tac att act aca ttg gct acg gca tat ata cct att tcg tca ctt cca cac gct ccg gta aac ggg tgt cat gtg acg agg gag aat ctt gat aag agg cat aat caa tgt tgt aat ccg atg tcc acc tgg aga att tgc caa ggt cag atg tag agt tgg tag tga taa cac aaa atg tga aca ctg ccc acc tca tac ata tac cgc aat ccc caa tta ttc taa tag atg tca tca atg tag aaa atg ccc aac agg atc att tga taa ggt aaa gtg tac cgg aac aca gaa caa at- gtt cgt gtc atc ctg gtt ggt ata cgc tac tga ttc ttc aca gac tga aga ttg tcg aga ttt gtg tac caa aaa agg aga tgt cca tgc gga tac ttt ggt gga ata gat gaa gga aat cct att tgt aaa tcg tgt tgt gtt ggt gaa tat tgc gac tac cta cgt aat tat aga ctt gat cca ttt cct cca tgc aaa cta tct atc taa atg taa tta att atg att ttg atg ata atg tta cca tac att ata tca cta ctt ggt tag tgt gta tta ttt agt atg gaa gac cta tta ata att act tat ctt ttg acg atc ttg tta taa tta taa tat aaa aat act tat gac ata gta a-- --- -ct cat aat tgc tga cgc gat aaa ttc gta ata atc tgt ttt gtt caa att ttt ata agg aat cta cag gca taa aaa taa aaa tat aat cta taa tat act ctt aca acg cca tca tga ata gca gtg aat taa ttg ctg tta tta tgg att tag aaa tag tgg acg att ttg tga tat taa tat agt tat taa tga tga aag gat aaa cgc gca tag act cat cct atc tgg agc ctc cga ata ttt ttt cca ttc tgt ttt cca ata att tta tcg att cta atg aat acg aag tta atc taa gtc att tag att atc aaa gtg tta act att tga tcg att aca ttt atg gga tac ctt tga gcc taa cta aca ata acg tga aat ata ttc ttt caa ccg ctg att ttt tta caa att gga tct gtc att act gag tgc gaa aaa tac ata ctt aaa aat ctt tgt ttt aga aac tgt atc gat ttc --- --- tac ata tac gct gat aaa tat aat aac aag aaa ata gaa tta gca tcg ttt aac aca ata tta cga aat att ttg aga ctc atc aac aat gaa aac ttt aaa tac tta aca gag gaa tca atg ata aaa att tta agc gat gat atg tta tat ata aaa aat gag gat ttc acc cca ctg att ctc att aaa tgg tta gag agt aca cca acc atg tac cgt cga gtt act tag atg cct cag aat atc a-t ttc ttt ccc cac aag tta taa aat cac ttt ata gtc atc gac tgg tta gtt caa tct acg aat gta taa cat tct taa aca ata tag cat tct tgg ata aat cat ttc cta gat acc ata tca tcg agt tga tat cta tcg gta taa gta att cac atg ata aga ttt cca taa act gct aca atc ata aaa aaa ttc atg gga aat gat atc ttc acg tag ata tag gtg tag ttt cgc agt gac cgt cct gga taa tat tat cta tat gat ggg tgg ata tga tca gtc cct gta tag aag ttc aaa ggt tat agc gta caa tac atg tac taa ttc ttg gat ata tga tat acc aga gct aaa aat atc atc gtt cta att gcg gag gag ttg cca atg acg aat aca ttt att gta tag gcg gta tac gcg atc agg agt cat cgt tga tat cta gta tcg ata gat gga agc cat caa aac cat att ggc aga agt atg cta aaa tgt gcg aac caa aat gtg ata tgg ggt tgc gat ttt aaa cgg att aat ata tgt cat agg tgg agt cgt taa agg tga cac aca tat acc aac gca cta gag agt tta tca gaa gat gga tgg atg aat cat caa cgt ctt cca ata aaa atg tcc aat atg tcg acg att gtt cat gct gga aag att tat ata tct aga ggt tac aac aat agt agt gta gtt aat gta ata tcg aat ct- -ag tcc tta gct ata atc cga tat atg atg aat gga cca aat tat cat cat taa ata ttc cta gaa tta atc ctg ctc tat ggt cag tgt ata ata aat tat atg tag gag gag taa tat ctg atg atg ttc aaa cta ata cat ctg aaa cat acg ata aag aaa aa

>AY741551.1_Monkeypox_virus_isolate_Sierra_Leone_complete_genome

tgt tat tgt agt atg ata ata tca aaa aga tgg ata taa aga att tac tga cta cat gta cta ttt tac att act aca ttg gct acg g-- tat ata cct act tcg tca ctt cca cac gct cca gta aac ggg tgt cat gtg acg agg gag aat ctt gat aag agg cat aat caa tgt tgt aat ccg gtg tcc acc tgg aga att tgc caa ggt cag atg t-- agt tgg tag tga taa cac aaa atg tga aca ctg ccc acc tca tac ata tac cgc aat ccc caa tta ttc taa tag atg tca tca atg tag aaa atg ccc aac agg atc att tga taa ggt aaa gtg tac cgg aac aca gaa caa atg gtt cgt gtc atc ctg gtt ggt ata cgc tac tga ttc ttc aca gac tga aga ttg tcg aga ttt gtg tac caa aaa agg aga tgt cca tgc gga tac ttt ggt gga ata gat gaa gga aat cct att tgt aaa tcg tgt tgt gtt ggt gaa tat tgc gac tac cta cgt aat tat aga ctt gat cca ttt cct cca tgc aaa cta tct atc taa atg taa tta att atg att ttg atg ata atg tta cca tac att ata tcg cta ctt ggt tag tgt gta tta ttc agt atg gaa gac cta tta ata att act tat ctt ttg acg atc ttg tta taa tta taa tat aaa aat act tat gac ata gta aca gta act cat aat tgc tga cgc gat aaa ttc gta ata atc tgt ttt gtt caa att ttt ata agg aat cta cag gca taa aaa taa aaa tat aat cta taa tat act ctt aca acg cca tca tga ata gca gtg aat taa ttg ctg tta tta tgg att tag aaa tag tgg acg att ttg tga tat taa tat agt tat taa tga tga aag gat aaa cgc tca tag att cat cct atc tgg agc ctc cga ata ttt ttt cca ttc tgt ttt cca ata att tta tcg att cta atg aat acg aag tta atc taa gtc att tag att atc aaa gtg tta act att tga tcg att aca ttt atg gga tac ctt tga gcc taa cta acg ata acg tga aat ata ttc ttt caa ccg ctg att ttt tta caa att gga tct gcc att act gag tgc gaa aaa tac ata ctt aaa aat ctt tgt ttt aga aac tgt atc gat ttc --- --- tac ata tac gct gat aaa tat aat aac aag aaa ata gaa tta gca tcg ttt aac aca ata tta cga aat att ttg aga ctc atc aac aat gaa aac ttt aaa tac tta aca gag gaa tca --- --- --- --- --- --- -at gat atg tta tat ata aaa aat gag gat ttc acc cca ctg att ctc att aaa tgg tta gag agt aca cca acc atg tac cgt cga gtt act tag atg cct cag aat atc att ttc ttt ccc cac aag tta taa aat cac ttt ata gtc atc gac tgg tta gtt caa tct acg aat gta taa cat tct taa aca ata tag cat tct tgg ata aat cat ttc cta gat acc ata tca tcg agt tga tat cta tcg gta taa gta att cac atg ata aga ttt cca taa act gct aca atc ata aaa aaa ttc atg gga aat gat atc ttc acg tag at- --g gtg tag ttt cgc agt gac cgt cct gga taa tat tat cta tat gat ggg tgg ata tga tca gtc cct gta tag aag ttc aaa ggt tat agc gta caa tac atg tac taa ttc ttg gat ata tga tat acc aga gct -aa aat atc ctc gtt cta att gcg gag gag ttg cca atg acg aat aca ttt att gta tag gcg gta tac gcg atc agg agt cat cgt tga tat cta gta tcg ata gat gga agc cat caa aac cat att ggc aga agt atg cta aaa tgt gcg aac caa aat gtg ata tgg ggt tgc gat ttt aaa cgg att aat ata tgt cat agg tgg agt cgt taa agg tga cac aca tgt acc gac gca cta gag agt tta tca gaa gat gga tgg atg aat cat caa cgt ctt cca ata aaa atg tcc aat atg tcg acg att gtt cat gct gga aag att tat ata tct gga ggt tac aac aat agt agt gta gtt aat gta ata tcg aat cta gag tcc tta gct ata atc cga tat atg atg aat gga cca aat tat cat cat taa ata ttc cta gaa tta atc ctg ctc tat ggt cag tgt ata ata aat tat atg tag gag gag gaa tat ctg atg atg ttc aaa cta ata cat ctg aaa cat acg ata aag aaa aa

>ON622718.1_Monkeypox_virus_isolate_MPXV/ES0001/HUGTiP/2022_partial_genome

tgt tat tgt agt atg ata ata tca aaa aga tgg ata taa aga att tac tga cta cat gta cta ttt tac att act aca ttg gct acg gca tat ata cct att tcg tca ctt cca cac gct ccg gta aac ggg tgt cat gtg acg agg gag aat ctt gat aag agg cat aat caa tgt tgt aat ccg atg tcc acc tgg aga att tgc caa ggt cag atg tag agt tgg tag tga taa cac aaa atg tga aca ctg ccc acc tca tac ata tac cgc aat ccc caa tta ttc taa tag atg tca tca atg tag aaa atg ccc aac agg atc att tga taa ggt aaa gtg tac cgg aac aca gaa caa at- gtt cgt gtc atc ctg gtt ggt ata cgc tac tga ttc ttc aca gac tga aga ttg tcg aga ttt gtg tac caa aaa agg aga tgt cca tgc gga tac ttt ggt gga ata gat gaa gga aat cct att tgt aaa tcg tgt tgt gtt ggt gaa tat tgc gac tac cta cgt aat tat aga ctt gat cca ttt cct cca tgc aaa cta tct atc taa atg taa tta att atg att ttg atg ata atg tta cca tac att ata tca cta ctt ggt tag tgt gta tta ttt agt atg gaa gac cta tta ata att act tat ctt ttg acg atc ttg tta taa tta taa tat aaa aat act tat gac ata gta a-- --- -ct cat aat tgc tga cgc gat aaa ttc gta ata atc tgt ttt g?? ??? ??? ??? ??? ??? ?at cta cag gca taa aaa taa aaa tat aat cta taa tat act ctt aca acg cca tca tga ata gca gtg aat taa ttg ctg tta tta tgg att tag aaa tag tgg acg att ttg tga tat taa tat agt tat taa tga tga aag gat aaa cgc gca tag act cat cct atc tgg agc ctc cga ata ttt ttt cca ttc tgt ttt cca ata att tta tcg att cta atg aat acg aag tta atc taa gtc att tag att atc aaa gtg tta act att tga tcg att aca ttt atg gga tac ctt tga gcc taa cta aca ata acg tga aat ata ttc ttt caa ccg ctg att ttt tta caa att gga tct gtc att act gag tgc gaa aaa tac ata ctt aaa aat ctt tgt ttt aga aac tgt atc gat ttc --- --- tac ata tac gct gat aaa tat aat aac aag aaa ata gaa tta gca tcg ttt aac aca ata tta cga aat att ttg aga ctc atc aac aat gaa aac ttt aaa tac tta aca gag gaa tca atg ata aaa att tta agc gat gat atg tta tat ata aaa aat gag gat ttc acc cca ctg att ctc att aaa tgg tta gag agt aca cca acc atg tac cgt cga gtt act tag atg cct cag aat atc a-t ttc ttt ccc cac aag tta taa aat cac ttt ata gtc atc gac tgg tta gtt caa tct acg aat gta taa cat tct taa aca ata tag cat tct tgg ata aat cat ttc cta gat acc ata tca tcg agt tga tat cta tcg gta taa gta att cac atg ata aga ttt cca taa act gct aca atc ata aaa aaa ttc atg gga aat gat atc ttc acg tag ata tag gtg tag ttt cgc agt gac cgt cct gga taa tat tat cta tat gat ggg tgg ata tga tca gtc cct gta tag aag ttc aaa ggt tat agc gta caa tac atg tac taa ttc ttg gat ata tga tat acc aga gct aaa aat atc atc gtt cta att gcg gag gag ttg cca atg acg aat aca ttt att gta tag gcg gta tac gcg atc agg agt cat cgt tga tat cta gta tcg ata gat gga agc cat caa aac cat att ggc aga agt atg cta aaa tgt gcg aac caa aat gtg ata tgg ggt tgc gat ttt aaa cgg att aat ata tgt cat agg tgg agt cgt taa agg tga cac aca tat acc aac gca cta gag agt tta tca gaa gat gga tgg atg aat cat caa cgt ctt cca ata aaa atg tcc aat atg tcg acg att gtt cat gct gga aag att tat ata tct aga ggt tac aac aat agt agt gta gtt aat gta ata tcg aat ct- -ag tcc tta gct ata atc cga tat atg atg aat gga cca aat tat cat cat taa ata ttc cta gaa tta atc ctg ctc tat ggt cag tgt ata ata aat tat atg tag gag gag taa tat ctg atg atg ttc aaa cta ata cat ctg aaa cat acg ata aag aaa aa

>MT903346.1_Monkeypox_virus_isolate_MPXV-USA2003_099_Gambian_Rat

tgt tat tgt agt atg ata ata tca aaa aga tgg ata taa aga att tac tga cta cat gta cta ttt tac att act aca ttg gct acg g-- tat ata cct act tcg tca ctt cca cac gct cca gta aac ggg tgt cat gtg acg agg gag aat ctt gat aag agg cat aat caa tgt tgt aat ccg gtg tcc acc tgg aga att tgc caa ggt cag atg t-- agt tgg tag tga taa cac aaa atg tga aca ctg ccc acc tca tac ata tac cgc aat ccc caa tta ttc taa tag atg tca tca atg tag aaa atg ccc aac agg atc att tga taa ggt aaa gtg tac cgg aac aca gaa caa at- gtt cgt gtc atc ctg gtt ggt ata cgc tac -ga ttc ttc aca gac tga aga ttg tcg aga ttt gtg tac caa aaa agg aga tgt cca tgc gga tac ttt ggt gga ata gat gaa gga aat cct att tgt aaa tcg tgt tgt gtt ggt gaa tat tgc gac tac cta cgt aat tat aga ctt gat cca ttt cct cca tgc aaa cta tct atc taa atg taa tta att atg att ttg atg ata atg tta cca tac att ata tcg cta ctt ggt tag tgt gta tta ttc agt atg gaa gac cta tta ata att act tat ctt ttg acg atc ttg tta taa tta taa tat aaa aat act tat gac ata gta aca gta act cat aat tgc tga cgc gat aaa ttc gta ata atc tgt ttt gtt caa att ttt ata agg aat cta cag gca taa aaa taa aaa tat aat cta taa tat act ctt aca acg cca tca tga ata gca gtg aat taa ttg ctg tta tta tgg att tag aaa tag tgg acg att ttg tga tat taa tat agt tat taa tga tga aag gat aaa cgc tca tag att cat cct atc tgg agc ctc cga ata ttt ttt cca ttc tgt ttt cca ata att tta tcg att cta atg aat acg aag tta atc taa gtc att tag att atc aaa gtg tta act att tga tcg att aca ttt atg gga tac ctt tga gcc taa cta acg ata acg tga aat ata ttc ttt caa ccg ctg att ttt tta caa att gga tct gcc att act gag tgc gaa aaa tac ata ctt aaa aat ctt tgt ttt aga aac tgt atc gat ttc tac ata tac ata tac gct gat aaa tat aat aac aag aaa ata gaa tta gca tcg ttt aac aca ata tta cga aat att ttg aga ctc atc aac aat gaa aac ttt aaa tac tta aca gag gaa tca --- --- --- --- --- --- -at gat atg tta tat ata aaa aat gag gat ttc acc cca ctg att ctc att aaa tgg tta gag agt aca cca acc atg tac cgt cga gtt act tag atg cct cag aat atc a-t ttc ttt ccc cac aag tta taa aat cac ttt ata gtc atc gac tgg tta gtt caa tct acg aat gta taa cat tct taa aca ata tag cat tct tgg ata aat cat ttc cta gat acc ata tca tcg agt tga tat cta tcg gta taa gta att cac atg ata aga ttt cca taa act gct aca atc ata aaa aaa ttc atg gga aat gat atc ttc acg tag at- --g gtg tag ttt cgc agt gac cgt cct gga taa tat tat cta tat gat ggg tgg ata tga tca gtc cct gta tag aag ttc aaa ggt tat agc gta caa tac atg tac taa ttc ttg gat ata tga tat acc aga gct -aa aat atc ctc gtt cta att gcg gag gag ttg cca atg acg aat aca ttt att gta tag gcg gta tac gcg atc agg agt cat cgt tga tat cta gta tcg ata gat gga agc cat caa aac cat att ggc aga agt atg cta aaa tgt gcg aac caa aat gtg ata tgg ggt tgc gat ttt aaa cgg att aat ata tgt cat agg tgg agt cgt taa agg tga cac aca tgt acc gac gca cta gag agt tta tca gaa gat gga tgg atg aat cat caa cgt ctt cca ata aaa atg tcc aat atg tcg acg att gtt cat gct gga aag att tat ata tct gga ggt tac aac aat agt agt gta gtt aat gta ata tcg aat cta gag tcc tta gct ata atc cga tat atg atg aat gga cca aat tat cat cat taa ata ttc cta gaa tta atc ctg ctc tat ggt cag tgt ata ata aat tat atg tag gag gag gaa tat ctg atg atg ttc aaa cta ata cat ctg aaa cat acg ata aag aaa aa

>DQ011157.1_Monkeypox_virus_strain_USA_2003_039_complete_genome

tgt tat tgt agt atg ata ata tca aaa aga tgg ata taa aga att tac tga cta cat gta cta ttt tac att act aca ttg gct acg g-- tat ata cct act tcg tca ctt cca cac gct cca gta aac ggg tgt cat gtg acg agg gag aat ctt gat aag agg cat aat caa tgt tgt aat ccg gtg tcc acc tgg aga att tgc caa ggt cag atg t-- agt tgg tag tga taa cac aaa atg tga aca ctg ccc acc tca tac ata tac cgc aat ccc caa tta ttc taa tag atg tca tca atg tag aaa atg ccc aac agg atc att tga taa ggt aaa gtg tac cgg aac aca gaa caa at- gtt cgt gtc atc ctg gtt ggt ata cgc tac -ga ttc ttc aca gac tga aga ttg tcg aga ttt gtg tac caa aaa agg aga tgt cca tgc gga tac ttt ggt gga ata gat gaa gga aat cct att tgt aaa tcg tgt tgt gtt ggt gaa tat tgc gac tac cta cgt aat tat aga ctt gat cca ttt cct cca tgc aaa cta tct atc taa atg taa tta att atg att ttg atg ata atg tta cca tac att ata tcg cta ctt ggt tag tgt gta tta ttc agt atg gaa gac cta tta ata att act tat ctt ttg acg atc ttg tta taa tta taa tat aaa aat act tat gac ata gta aca gta act cat aat tgc tga cgc gat aaa ttc gta ata atc tgt ttt gtt caa att ttt ata agg aat cta cag gca taa aaa taa aaa tat aat cta taa tat act ctt aca acg cca tca tga ata gca gtg aat taa ttg ctg tta tta tgg att tag aaa tag tgg acg att ttg tga tat taa tat agt tat taa tga tga aag gat aaa cgc tca tag att cat cct atc tgg agc ctc cga ata ttt ttt cca ttc tgt ttt cca ata att tta tcg att cta atg aat acg aag tta atc taa gtc att tag att atc aaa gtg tta act att tga tcg att aca ttt atg gga tac ctt tga gcc taa cta acg ata acg tga aat ata ttc ttt caa ccg ctg att ttt tta caa att gga tct gcc att act gag tgc gaa aaa tac ata ctt aaa aat ctt tgt ttt aga aac tgt atc gat ttc tac ata tac ata tac gct gat aaa tat aat aac aag aaa ata gaa tta gca tcg ttt aac aca ata tta cga aat att ttg aga ctc atc aac aat gaa aac ttt aaa tac tta aca gag gaa tca --- --- --- --- --- --- -at gat atg tta tat ata aaa aat gag gat ttc acc cca ctg att ctc att aaa tgg tta gag agt aca cca acc atg tac cgt cga gtt act tag atg cct cag aat atc a-t ttc ttt ccc cac aag tta taa aat cac ttt ata gtc atc gac tgg tta gtt caa tct acg aat gta taa cat tct taa aca ata tag cat tct tgg ata aat cat ttc cta gat acc ata tca tcg agt tga tat cta tcg gta taa gta att cac atg ata aga ttt cca taa act gct aca atc ata aaa aaa ttc atg gga aat gat atc ttc acg tag at- --g gtg tag ttt cgc agt gac cgt cct gga taa tat tat cta tat gat ggg tgg ata tga tca gtc cct gta tag aag ttc aaa ggt tat agc gta caa tac atg tac taa ttc ttg gat ata tga tat acc aga gct -aa aat atc ctc gtt cta att gcg gag gag ttg cca atg acg aat aca ttt att gta tag gcg gta tac gcg atc agg agt cat cgt tga tat cta gta tcg ata gat gga agc cat caa aac cat att ggc aga agt atg cta aaa tgt gcg aac caa aat gtg ata tgg ggt tgc gat ttt aaa cgg att aat ata tgt cat agg tgg agt cgt taa agg tga cac aca tgt acc gac gca cta gag agt tta tca gaa gat gga tgg atg aat cat caa cgt ctt cca ata aaa atg tcc aat atg tcg acg att gtt cat gct gga aag att tat ata tct gga ggt tac aac aat agt agt gta gtt aat gta ata tcg aat cta gag tcc tta gct ata atc cga tat atg atg aat gga cca aat tat cat cat taa ata ttc cta gaa tta atc ctg ctc tat ggt cag tgt ata ata aat tat atg tag gag gag gaa tat ctg atg atg ttc aaa cta ata cat ctg aaa cat acg ata aag aaa aa

>DQ011153.1_Monkeypox_virus_strain_USA_2003_044_complete_genome

tgt tat tgt agt atg ata ata tca aaa aga tgg ata taa aga att tac tga cta cat gta cta ttt tac att act aca ttg gct acg g-- tat ata cct act tcg tca ctt cca cac gct cca gta aac ggg tgt cat gtg acg agg gag aat ctt gat aag agg cat aat caa tgt tgt aat ccg gtg tcc acc tgg aga att tgc caa ggt cag atg t-- agt tgg tag tga taa cac aaa atg tga aca ctg ccc acc tca tac ata tac cgc aat ccc caa tta ttc taa tag atg tca tca atg tag aaa atg ccc aac agg atc att tga taa ggt aaa gtg tac cgg aac aca gaa caa at- gtt cgt gtc atc ctg gtt ggt ata cgc tac -ga ttc ttc aca gac tga aga ttg tcg aga ttt gtg tac caa aaa agg aga tgt cca tgc gga tac ttt ggt gga ata gat gaa gga aat cct att tgt aaa tcg tgt tgt gtt ggt gaa tat tgc gac tac cta cgt aat tat aga ctt gat cca ttt cct cca tgc aaa cta tct atc taa atg taa tta att atg att ttg atg ata atg tta cca tac att ata tcg cta ctt ggt tag tgt gta tta ttc agt atg gaa gac cta tta ata att act tat ctt ttg acg atc ttg tta taa tta taa tat aaa aat act tat gac ata gta aca gta act cat aat tgc tga cgc gat aaa ttc gta ata atc tgt ttt gtt caa att ttt ata agg aat cta cag gca taa aaa taa aaa tat aat cta taa tat act ctt aca acg cca tca tga ata gca gtg aat taa ttg ctg tta tta tgg att tag aaa tag tgg acg att ttg tga tat taa tat agt tat taa tga tga aag gat aaa cgc tca tag att cat cct atc tgg agc ctc cga ata ttt ttt cca ttc tgt ttt cca ata att tta tcg att cta atg aat acg aag tta atc taa gtc att tag att atc aaa gtg tta act att tga tcg att aca ttt atg gga tac ctt tga gcc taa cta acg ata acg tga aat ata ttc ttt caa ccg ctg att ttt tta caa att gga tct gcc att act gag tgc gaa aaa tac ata ctt aaa aat ctt tgt ttt aga aac tgt atc gat ttc tac ata tac ata tac gct gat aaa tat aat aac aag aaa ata gaa tta gca tcg ttt aac aca ata tta cga aat att ttg aga ctc atc aac aat gaa aac ttt aaa tac tta aca gag gaa tca --- --- --- --- --- --- -at gat atg tta tat ata aaa aat gag gat ttc acc cca ctg att ctc att aaa tgg tta gag agt aca cca acc atg tac cgt cga gtt act tag atg cct cag aat atc a-t ttc ttt ccc cac aag tta taa aat cac ttt ata gtc atc gac tgg tta gtt caa tct acg aat gta taa cat tct taa aca ata tag cat tct tgg ata aat cat ttc cta gat acc ata tca tcg agt tga tat cta tcg gta taa gta att cac atg ata aga ttt cca taa act gct aca atc ata aaa aaa ttc atg gga aat gat atc ttc acg tag at- --g gtg tag ttt cgc agt gac cgt cct gga taa tat tat cta tat gat ggg tgg ata tga tca gtc cct gta tag aag ttc aaa ggt tat agc gta caa tac atg tac taa ttc ttg gat ata tga tat acc aga gct -aa aat atc ctc gtt cta att gcg gag gag ttg cca atg acg aat aca ttt att gta tag gcg gta tac gcg atc agg agt cat cgt tga tat cta gta tcg ata gat gga agc cat caa aac cat att ggc aga agt atg cta aaa tgt gcg aac caa aat gtg ata tgg ggt tgc gat ttt aaa cgg att aat ata tgt cat agg tgg agt cgt taa agg tga cac aca tgt acc gac gca cta gag agt tta tca gaa gat gga tgg atg aat cat caa cgt ctt cca ata aaa atg tcc aat atg tcg acg att gtt cat gct gga aag att tat ata tct gga ggt tac aac aat agt agt gta gtt aat gta ata tcg aat cta gag tcc tta gct ata atc cga tat atg atg aat gga cca aat tat cat cat taa ata ttc cta gaa tta atc ctg ctc tat ggt cag tgt ata ata aat tat atg tag gag gag gaa tat ctg atg atg ttc aaa cta ata cat ctg aaa cat acg ata aag aaa aa

>ON645312.1_Monkeypox_virus_isolate_MPXV_GSTT_Patient1_partial_genome

tgt tat tgt agt atg ata ata tca aaa aga tgg ata taa aga at- --- --- --- --- --- --- --- --- --- --- --- --- --- --- --- --- --- --- --- --- --- --- --- --- --- --- --- --- --- --- --- --- --- --- --- --- --- --- --- --- --- --- --- --- --- --- --- --- --- --- --- --- --- --- --- --- --- --- --- --- --- --- --- --- --- --- --- --- --- --- --- --- --- --- --- --- --- --- --- --- --- --- --- --- --- --- --- --- --- --- --- --- --- --- --- --- --- --- --- --- --- --- --- --- --- --- --- --- --- --- --- --- --- --- --- --- --- --- --- --- --- --- --- --- --- --- --- --- --- --- --- --- --- --- --- --- --- --- --- --- --- --- --- --- --- --- --- --- --- --- --- --- --- --- --- --- --- --- --- --- --- --- --- --- --- --- --- --- --- --- --- --- --- --- --- --- --- --- --- --- --- --- --- --- --- --- --- --- --- --- --- --- --- --- --- --- --- --- --- --- --- --- --- --- --- --- --- --- --- --- --- --- --- --- --- --- --- --- --- --- --- --- --- --- --- --- --- --- --- --- --- --- --- --- --- --- --- --- --- --- --- --- --- --- --- --- --- --- --- --- --- --- --- --- --- --- --- --- --- --- --- --- --- --- --- --- --- --- --- --- --- --- --- --- --- --- --- --- --- --- --- --- --- --- --- --- --- --- --- --- --- --- --- --- --- --- --- --- --- --- --- --- --- --- --- --- --- --- --- --- --- --- --- --- --- --- --- --- --- --- --- --- --- --- --- --- --- --- --- --- --- --- --- --- --- --- --- --- --- --- --- --- --- --- --- --- --- --- --- --- --- --- --- --- --- --- --- --- --- --- --- --- --- --- --- --- --- --- --- --- --- --- --- --- --- --- --- --- --- --- --- --- --- --- --- --- --- --- --- --- --- --- --- --- --- --- --- --- --- --- --- --- --- --- --- --- --- --- --- --- --- --- --- --- --- --- --- --- --- --- --- --- --- --- --- --- --- --- --- --- --- --- --- --- --- --- --- --- --- --- --- --- --- --- --- --- --- --- --- --- --- --- --- --- --- --- --- --- --- --- --- --- --- --- --- --- --- --- --- --- --- --- --- --- --- --- --- --- --- --- --- --- --- --- --- --- --- --- --- --- --- --- --- --- --- --- --- --- --- --- --- --- --- --- --- --- --- --- --- --- --- --- --- --- --- --- --- --- --- --- --- --- --- --- --- --- --- --- --- --- --- --- --- --- --- --- --- --- --- --- --- --- --- --- --- --- --- --- --- --- --- --- --- --- --- --- --- --- --- --- --- --- --- --- --- --- --- --- --- --- --- --- --- --- --- --- --- --- --- --- --- --- --- --- --- --- --- --- --- --- --- --- --- --- --- --- --- --- --- --- --- --- --- --- --- --- --- --- --- --- --- --- --- --- --- --- --- --- --- --- --- --- --- --- --- --- --- --- --- --- --- --- --- --- --- --- --- --- --- --- --- --- --- --- --- --- --- --- --- --- --- --- --- --- --- --- --- --- --- --- --- --- --- --- --- --- --- --- --- --- --- --- --- --- --- --- --- --- --- --- --- --- --- --- --- --- --- --- --- --- --- --- --- --- --- --- --- --- --- --- --- --- --- --- --- --- --- --- --- --- --- --- --- --- --- --- --- --- --- --- --- --- --- --- --- --- --- --- --- --- --- --- --- --- --- --- --- --- --- --- --- --- --- --- --- --- --- --- --- --- --- --- --- --- --- --- --- --- --- --t ggt cag tgt ata ata aat tat atg tag gag gag taa tat ctg atg atg ttc aaa cta ata cat ctg aaa cat acg ata aag aaa aa

>JX878410.1_Monkeypox_virus_isolate_DRC_06-1070_complete_genome

tgt tat tgt agt atg ata ata tca aaa aga tgg ata taa aga at- --- --- --- --- --- --- --- --- --- --- --- --- --- --- --- --- --- --- --- --- --- --- --- --- --- --- --- --- --- --- --- --- --- --- --- --- --- --- --- --- --- --- --- --- --- --- --- --- --- --- --- --- --- --- --- --- --- --- --- --- --- --- --- --- --- --- --- --- --- --- --- --- --- --- --- --- --- --- --- --- --- --- --- --- --- --- --- --- --- --- --- --- --- --- --- --- --- --- --- --- --- --- --- --- --- --- --- --- --- --- --- --- --- --- --- --- --- --- --- --- --- --- --- --- --- --- --- --- --- --- --- --- --- --- --- --- --- --- --- --- --- --- --- --- --- --- --- --- --- --- --- --- --- --- --- --- --- --- --- --- --- --- --- --- --- --- --- --- --- --- --- --- --- --- --- --- --- --- --- --- --- --- --- --- --- --- --- --- --- --- --- --- --- --- --- --- --- --- --- --- --- --- --- --- --- --- --- --- --- --- --- --- --- --- --- --- --- --- --- --- --- --- --- --- --- --- --- --- --- --- --- --- --- --- --- --- --- --- --- --- --- --- --- --- --- --- --- --- --- --- --- --- --- --- --- --- --- --- --- --- --- --- --- --- --- --- --- --- --- --- --- --- --- --- --- --- --- --- --- --- --- --- --- --- --- --- --- --- --- --- --- --- --- --- --- --- --- --- --- --- --- --- --- --- --- --- --- --- --- --- --- --- --- --- --- --- --- --- --- --- --- --- --- --- --- --- --- --- --- --- --- --- --- --- --- --- --- --- --- --- --- --- --- --- --- --- --- --- --- --- --- --- --- --- --- --- --- --- --- --- --- --- --- --- --- --- --- --- --- --- --- --- --- --- --- --- --- --- --- --- --- --- --- --- --- --- --- --- --- --- --- --- --- --- --- --- --- --- --- --- --- --- --- --- --- --- --- --- --- --- --- --- --- --- --- --- --- --- --- --- --- --- --- --- --- --- --- --- --- --- --- --- --- --- --- --- --- --- --- --- --- --- --- --- --- --- --- --- --- --- --- --- --- --- --- --- --- --- --- --- --- --- --- --- --- --- --- --- --- --- --- --- --- --- --- --- --- --- --- --- --- --- --- --- --- --- --- --- --- --- --- --- --- --- --- --- --- --- --- --- --- --- --- --- --- --- --- --- --- --- --- --- --- --- --- --- --- --- --- --- --- --- --- --- --- --- --- --- --- --- --- --- --- --- --- --- --- --- --- --- --- --- --- --- --- --- --- --- --- --- --- --- --- --- --- --- --- --- --- --- --- --- --- --- --- --- --- --- --- --- --- --- --- --- --- --- --- --- --- --- --- --- --- --- --- --- --- --- --- --- --- --- --- --- --- --- --- --- --- --- --- --- --- --- --- --- --- --- --- --- --- --- --- --- --- --- --- --- --- --- --- --- --- --- --- --- --- --- --- --- --- --- --- --- --- --- --- --- --- --- --- --- --- --- --- --- --- --- --- --- --- --- --- --- --- --- --- --- --- --- --- --- --- --- --- --- --- --- --- --- --- --- --- --- --- --- --- --- --- --- --- --- --- --- --- --- --- --- --- --- --- --- --- --- --- --- --- --- --- --- --- --- --- --- --- --- --- --- --- --- --- --- --- --- --- --- --- --- --- --- --- --- --- --- --- --- --- --- --- --- --- --- --- --- --- --- --- --- --- --- --- --- --- --- --- --- --- --- --- --- --- --- --- --- --- --- --- --- --- --t ggt cag tgt ata ata aat tat atg tag gag gag gaa tat ctg atg atg ttc aaa cta ata cat ctg aaa cat acg ata aag aaa aa

>JX878428.1_Monkeypox_virus_isolate_DRC_07-0514_complete_genome

tgt tat tgt agt atg ata ata tca aaa aga tgg ata taa aga at- --- --- --- --- --- --- --- --- --- --- --- --- --- --- --- --- --- --- --- --- --- --- --- --- --- --- --- --- --- --- --- --- --- --- --- --- --- --- --- --- --- --- --- --- --- --- --- --- --- --- --- --- --- --- --- --- --- --- --- --- --- --- --- --- --- --- --- --- --- --- --- --- --- --- --- --- --- --- --- --- --- --- --- --- --- --- --- --- --- --- --- --- --- --- --- --- --- --- --- --- --- --- --- --- --- --- --- --- --- --- --- --- --- --- --- --- --- --- --- --- --- --- --- --- --- --- --- --- --- --- --- --- --- --- --- --- --- --- --- --- --- --- --- --- --- --- --- --- --- --- --- --- --- --- --- --- --- --- --- --- --- --- --- --- --- --- --- --- --- --- --- --- --- --- --- --- --- --- --- --- --- --- --- --- --- --- --- --- --- --- --- --- --- --- --- --- --- --- --- --- --- --- --- --- --- --- --- --- --- --- --- --- --- --- --- --- --- --- --- --- --- --- --- --- --- --- --- --- --- --- --- --- --- --- --- --- --- --- --- --- --- --- --- --- --- --- --- --- --- --- --- --- --- --- --- --- --- --- --- --- --- --- --- --- --- --- --- --- --- --- --- --- --- --- --- --- --- --- --- --- --- --- --- --- --- --- --- --- --- --- --- --- --- --- --- --- --- --- --- --- --- --- --- --- --- --- --- --- --- --- --- --- --- --- --- --- --- --- --- --- --- --- --- --- --- --- --- --- --- --- --- --- --- --- --- --- --- --- --- --- --- --- --- --- --- --- --- --- --- --- --- --- --- --- --- --- --- --- --- --- --- --- --- --- --- --- --- --- --- --- --- --- --- --- --- --- --- --- --- --- --- --- --- --- --- --- --- --- --- --- --- --- --- --- --- --- --- --- --- --- --- --- --- --- --- --- --- --- --- --- --- --- --- --- --- --- --- --- --- --- --- --- --- --- --- --- --- --- --- --- --- --- --- --- --- --- --- --- --- --- --- --- --- --- --- --- --- --- --- --- --- --- --- --- --- --- --- --- --- --- --- --- --- --- --- --- --- --- --- --- --- --- --- --- --- --- --- --- --- --- --- --- --- --- --- --- --- --- --- --- --- --- --- --- --- --- --- --- --- --- --- --- --- --- --- --- --- --- --- --- --- --- --- --- --- --- --- --- --- --- --- --- --- --- --- --- --- --- --- --- --- --- --- --- --- --- --- --- --- --- --- --- --- --- --- --- --- --- --- --- --- --- --- --- --- --- --- --- --- --- --- --- --- --- --- --- --- --- --- --- --- --- --- --- --- --- --- --- --- --- --- --- --- --- --- --- --- --- --- --- --- --- --- --- --- --- --- --- --- --- --- --- --- --- --- --- --- --- --- --- --- --- --- --- --- --- --- --- --- --- --- --- --- --- --- --- --- --- --- --- --- --- --- --- --- --- --- --- --- --- --- --- --- --- --- --- --- --- --- --- --- --- --- --- --- --- --- --- --- --- --- --- --- --- --- --- --- --- --- --- --- --- --- --- --- --- --- --- --- --- --- --- --- --- --- --- --- --- --- --- --- --- --- --- --- --- --- --- --- --- --- --- --- --- --- --- --- --- --- --- --- --- --- --- --- --- --- --- --- --- --- --- --- --- --- --- --- --- --- --- --- --- --- --- --- --- --- --- --- --- --- --- --- --- --- --- --- --- --- --- --- --- --- --- --- --- --- --- --- --t ggt cag tgt ata ata aat tat atg tag gag gag gaa tat ctg atg atg ttc aaa cta ata cat ctg aaa cat acg ata aag aaa aa

>JX878427.1_Monkeypox_virus_isolate_DRC_07-0480_complete_genome

tgt tat tgt agt atg ata ata tca aaa aga tgg ata taa aga at- --- --- --- --- --- --- --- --- --- --- --- --- --- --- --- --- --- --- --- --- --- --- --- --- --- --- --- --- --- --- --- --- --- --- --- --- --- --- --- --- --- --- --- --- --- --- --- --- --- --- --- --- --- --- --- --- --- --- --- --- --- --- --- --- --- --- --- --- --- --- --- --- --- --- --- --- --- --- --- --- --- --- --- --- --- --- --- --- --- --- --- --- --- --- --- --- --- --- --- --- --- --- --- --- --- --- --- --- --- --- --- --- --- --- --- --- --- --- --- --- --- --- --- --- --- --- --- --- --- --- --- --- --- --- --- --- --- --- --- --- --- --- --- --- --- --- --- --- --- --- --- --- --- --- --- --- --- --- --- --- --- --- --- --- --- --- --- --- --- --- --- --- --- --- --- --- --- --- --- --- --- --- --- --- --- --- --- --- --- --- --- --- --- --- --- --- --- --- --- --- --- --- --- --- --- --- --- --- --- --- --- --- --- --- --- --- --- --- --- --- --- --- --- --- --- --- --- --- --- --- --- --- --- --- --- --- --- --- --- --- --- --- --- --- --- --- --- --- --- --- --- --- --- --- --- --- --- --- --- --- --- --- --- --- --- --- --- --- --- --- --- --- --- --- --- --- --- --- --- --- --- --- --- --- --- --- --- --- --- --- --- --- --- --- --- --- --- --- --- --- --- --- --- --- --- --- --- --- --- --- --- --- --- --- --- --- --- --- --- --- --- --- --- --- --- --- --- --- --- --- --- --- --- --- --- --- --- --- --- --- --- --- --- --- --- --- --- --- --- --- --- --- --- --- --- --- --- --- --- --- --- --- --- --- --- --- --- --- --- --- --- --- --- --- --- --- --- --- --- --- --- --- --- --- --- --- --- --- --- --- --- --- --- --- --- --- --- --- --- --- --- --- --- --- --- --- --- --- --- --- --- --- --- --- --- --- --- --- --- --- --- --- --- --- --- --- --- --- --- --- --- --- --- --- --- --- --- --- --- --- --- --- --- --- --- --- --- --- --- --- --- --- --- --- --- --- --- --- --- --- --- --- --- --- --- --- --- --- --- --- --- --- --- --- --- --- --- --- --- --- --- --- --- --- --- --- --- --- --- --- --- --- --- --- --- --- --- --- --- --- --- --- --- --- --- --- --- --- --- --- --- --- --- --- --- --- --- --- --- --- --- --- --- --- --- --- --- --- --- --- --- --- --- --- --- --- --- --- --- --- --- --- --- --- --- --- --- --- --- --- --- --- --- --- --- --- --- --- --- --- --- --- --- --- --- --- --- --- --- --- --- --- --- --- --- --- --- --- --- --- --- --- --- --- --- --- --- --- --- --- --- --- --- --- --- --- --- --- --- --- --- --- --- --- --- --- --- --- --- --- --- --- --- --- --- --- --- --- --- --- --- --- --- --- --- --- --- --- --- --- --- --- --- --- --- --- --- --- --- --- --- --- --- --- --- --- --- --- --- --- --- --- --- --- --- --- --- --- --- --- --- --- --- --- --- --- --- --- --- --- --- --- --- --- --- --- --- --- --- --- --- --- --- --- --- --- --- --- --- --- --- --- --- --- --- --- --- --- --- --- --- --- --- --- --- --- --- --- --- --- --- --- --- --- --- --- --- --- --- --- --- --- --- --- --- --- --- --- --- --- --- --- --- --- --- --- --- --- --- --- --- --- --- --- --- --- --- --- --- --- --- --- --- --- --- --- --- --- --- --t ggt cag tgt ata ata aat tat atg tag gag gag gaa tat ctg atg atg ttc aaa cta ata cat ctg aaa cat acg ata aag aaa aa

>JX878421.1_Monkeypox_virus_isolate_DRC_07-0286_complete_genome

tgt tat tgt agt atg ata ata tca aaa aga tgg ata taa aga at- --- --- --- --- --- --- --- --- --- --- --- --- --- --- --- --- --- --- --- --- --- --- --- --- --- --- --- --- --- --- --- --- --- --- --- --- --- --- --- --- --- --- --- --- --- --- --- --- --- --- --- --- --- --- --- --- --- --- --- --- --- --- --- --- --- --- --- --- --- --- --- --- --- --- --- --- --- --- --- --- --- --- --- --- --- --- --- --- --- --- --- --- --- --- --- --- --- --- --- --- --- --- --- --- --- --- --- --- --- --- --- --- --- --- --- --- --- --- --- --- --- --- --- --- --- --- --- --- --- --- --- --- --- --- --- --- --- --- --- --- --- --- --- --- --- --- --- --- --- --- --- --- --- --- --- --- --- --- --- --- --- --- --- --- --- --- --- --- --- --- --- --- --- --- --- --- --- --- --- --- --- --- --- --- --- --- --- --- --- --- --- --- --- --- --- --- --- --- --- --- --- --- --- --- --- --- --- --- --- --- --- --- --- --- --- --- --- --- --- --- --- --- --- --- --- --- --- --- --- --- --- --- --- --- --- --- --- --- --- --- --- --- --- --- --- --- --- --- --- --- --- --- --- --- --- --- --- --- --- --- --- --- --- --- --- --- --- --- --- --- --- --- --- --- --- --- --- --- --- --- --- --- --- --- --- --- --- --- --- --- --- --- --- --- --- --- --- --- --- --- --- --- --- --- --- --- --- --- --- --- --- --- --- --- --- --- --- --- --- --- --- --- --- --- --- --- --- --- --- --- --- --- --- --- --- --- --- --- --- --- --- --- --- --- --- --- --- --- --- --- --- --- --- --- --- --- --- --- --- --- --- --- --- --- --- --- --- --- --- --- --- --- --- --- --- --- --- --- --- --- --- --- --- --- --- --- --- --- --- --- --- --- --- --- --- --- --- --- --- --- --- --- --- --- --- --- --- --- --- --- --- --- --- --- --- --- --- --- --- --- --- --- --- --- --- --- --- --- --- --- --- --- --- --- --- --- --- --- --- --- --- --- --- --- --- --- --- --- --- --- --- --- --- --- --- --- --- --- --- --- --- --- --- --- --- --- --- --- --- --- --- --- --- --- --- --- --- --- --- --- --- --- --- --- --- --- --- --- --- --- --- --- --- --- --- --- --- --- --- --- --- --- --- --- --- --- --- --- --- --- --- --- --- --- --- --- --- --- --- --- --- --- --- --- --- --- --- --- --- --- --- --- --- --- --- --- --- --- --- --- --- --- --- --- --- --- --- --- --- --- --- --- --- --- --- --- --- --- --- --- --- --- --- --- --- --- --- --- --- --- --- --- --- --- --- --- --- --- --- --- --- --- --- --- --- --- --- --- --- --- --- --- --- --- --- --- --- --- --- --- --- --- --- --- --- --- --- --- --- --- --- --- --- --- --- --- --- --- --- --- --- --- --- --- --- --- --- --- --- --- --- --- --- --- --- --- --- --- --- --- --- --- --- --- --- --- --- --- --- --- --- --- --- --- --- --- --- --- --- --- --- --- --- --- --- --- --- --- --- --- --- --- --- --- --- --- --- --- --- --- --- --- --- --- --- --- --- --- --- --- --- --- --- --- --- --- --- --- --- --- --- --- --- --- --- --- --- --- --- --- --- --- --- --- --- --- --- --- --- --- --- --- --- --- --- --- --- --- --- --- --- --- --- --- --- --- --- --- --- --- --- --- --- --- --- --- --- --- --- --- --- --- --- --- --- --- --- --- --- --t ggt cag tgt ata ata aat tat atg tag gag gag gaa tat ctg atg atg ttc aaa cta ata cat ctg aaa cat acg ata aag aaa aa

>JX878415.1_Monkeypox_virus_isolate_DRC_07-0092_complete_genome

tgt tat tgt agt atg ata ata tca aaa aga tgg ata taa aga at- --- --- --- --- --- --- --- --- --- --- --- --- --- --- --- --- --- --- --- --- --- --- --- --- --- --- --- --- --- --- --- --- --- --- --- --- --- --- --- --- --- --- --- --- --- --- --- --- --- --- --- --- --- --- --- --- --- --- --- --- --- --- --- --- --- --- --- --- --- --- --- --- --- --- --- --- --- --- --- --- --- --- --- --- --- --- --- --- --- --- --- --- --- --- --- --- --- --- --- --- --- --- --- --- --- --- --- --- --- --- --- --- --- --- --- --- --- --- --- --- --- --- --- --- --- --- --- --- --- --- --- --- --- --- --- --- --- --- --- --- --- --- --- --- --- --- --- --- --- --- --- --- --- --- --- --- --- --- --- --- --- --- --- --- --- --- --- --- --- --- --- --- --- --- --- --- --- --- --- --- --- --- --- --- --- --- --- --- --- --- --- --- --- --- --- --- --- --- --- --- --- --- --- --- --- --- --- --- --- --- --- --- --- --- --- --- --- --- --- --- --- --- --- --- --- --- --- --- --- --- --- --- --- --- --- --- --- --- --- --- --- --- --- --- --- --- --- --- --- --- --- --- --- --- --- --- --- --- --- --- --- --- --- --- --- --- --- --- --- --- --- --- --- --- --- --- --- --- --- --- --- --- --- --- --- --- --- --- --- --- --- --- --- --- --- --- --- --- --- --- --- --- --- --- --- --- --- --- --- --- --- --- --- --- --- --- --- --- --- --- --- --- --- --- --- --- --- --- --- --- --- --- --- --- --- --- --- --- --- --- --- --- --- --- --- --- --- --- --- --- --- --- --- --- --- --- --- --- --- --- --- --- --- --- --- --- --- --- --- --- --- --- --- --- --- --- --- --- --- --- --- --- --- --- --- --- --- --- --- --- --- --- --- --- --- --- --- --- --- --- --- --- --- --- --- --- --- --- --- --- --- --- --- --- --- --- --- --- --- --- --- --- --- --- --- --- --- --- --- --- --- --- --- --- --- --- --- --- --- --- --- --- --- --- --- --- --- --- --- --- --- --- --- --- --- --- --- --- --- --- --- --- --- --- --- --- --- --- --- --- --- --- --- --- --- --- --- --- --- --- --- --- --- --- --- --- --- --- --- --- --- --- --- --- --- --- --- --- --- --- --- --- --- --- --- --- --- --- --- --- --- --- --- --- --- --- --- --- --- --- --- --- --- --- --- --- --- --- --- --- --- --- --- --- --- --- --- --- --- --- --- --- --- --- --- --- --- --- --- --- --- --- --- --- --- --- --- --- --- --- --- --- --- --- --- --- --- --- --- --- --- --- --- --- --- --- --- --- --- --- --- --- --- --- --- --- --- --- --- --- --- --- --- --- --- --- --- --- --- --- --- --- --- --- --- --- --- --- --- --- --- --- --- --- --- --- --- --- --- --- --- --- --- --- --- --- --- --- --- --- --- --- --- --- --- --- --- --- --- --- --- --- --- --- --- --- --- --- --- --- --- --- --- --- --- --- --- --- --- --- --- --- --- --- --- --- --- --- --- --- --- --- --- --- --- --- --- --- --- --- --- --- --- --- --- --- --- --- --- --- --- --- --- --- --- --- --- --- --- --- --- --- --- --- --- --- --- --- --- --- --- --- --- --- --- --- --- --- --- --- --- --- --- --- --- --- --- --- --- --- --- --- --- --- --- --- --- --- --- --- --- --- --- --- --- --- --- --- --- --- --- --- --- --- --- --- --- --- --- --t ggt cag tgt ata ata aat tat atg tag gag gag gaa tat ctg atg atg ttc aaa cta ata cat ctg aaa cat acg ata aag aaa aa

>JX878414.1_Monkeypox_virus_isolate_DRC_07-0046_complete_genome

tgt tat tgt agt atg ata ata tca aaa aga tgg ata taa aga at- --- --- --- --- --- --- --- --- --- --- --- --- --- --- --- --- --- --- --- --- --- --- --- --- --- --- --- --- --- --- --- --- --- --- --- --- --- --- --- --- --- --- --- --- --- --- --- --- --- --- --- --- --- --- --- --- --- --- --- --- --- --- --- --- --- --- --- --- --- --- --- --- --- --- --- --- --- --- --- --- --- --- --- --- --- --- --- --- --- --- --- --- --- --- --- --- --- --- --- --- --- --- --- --- --- --- --- --- --- --- --- --- --- --- --- --- --- --- --- --- --- --- --- --- --- --- --- --- --- --- --- --- --- --- --- --- --- --- --- --- --- --- --- --- --- --- --- --- --- --- --- --- --- --- --- --- --- --- --- --- --- --- --- --- --- --- --- --- --- --- --- --- --- --- --- --- --- --- --- --- --- --- --- --- --- --- --- --- --- --- --- --- --- --- --- --- --- --- --- --- --- --- --- --- --- --- --- --- --- --- --- --- --- --- --- --- --- --- --- --- --- --- --- --- --- --- --- --- --- --- --- --- --- --- --- --- --- --- --- --- --- --- --- --- --- --- --- --- --- --- --- --- --- --- --- --- --- --- --- --- --- --- --- --- --- --- --- --- --- --- --- --- --- --- --- --- --- --- --- --- --- --- --- --- --- --- --- --- --- --- --- --- --- --- --- --- --- --- --- --- --- --- --- --- --- --- --- --- --- --- --- --- --- --- --- --- --- --- --- --- --- --- --- --- --- --- --- --- --- --- --- --- --- --- --- --- --- --- --- --- --- --- --- --- --- --- --- --- --- --- --- --- --- --- --- --- --- --- --- --- --- --- --- --- --- --- --- --- --- --- --- --- --- --- --- --- --- --- --- --- --- --- --- --- --- --- --- --- --- --- --- --- --- --- --- --- --- --- --- --- --- --- --- --- --- --- --- --- --- --- --- --- --- --- --- --- --- --- --- --- --- --- --- --- --- --- --- --- --- --- --- --- --- --- --- --- --- --- --- --- --- --- --- --- --- --- --- --- --- --- --- --- --- --- --- --- --- --- --- --- --- --- --- --- --- --- --- --- --- --- --- --- --- --- --- --- --- --- --- --- --- --- --- --- --- --- --- --- --- --- --- --- --- --- --- --- --- --- --- --- --- --- --- --- --- --- --- --- --- --- --- --- --- --- --- --- --- --- --- --- --- --- --- --- --- --- --- --- --- --- --- --- --- --- --- --- --- --- --- --- --- --- --- --- --- --- --- --- --- --- --- --- --- --- --- --- --- --- --- --- --- --- --- --- --- --- --- --- --- --- --- --- --- --- --- --- --- --- --- --- --- --- --- --- --- --- --- --- --- --- --- --- --- --- --- --- --- --- --- --- --- --- --- --- --- --- --- --- --- --- --- --- --- --- --- --- --- --- --- --- --- --- --- --- --- --- --- --- --- --- --- --- --- --- --- --- --- --- --- --- --- --- --- --- --- --- --- --- --- --- --- --- --- --- --- --- --- --- --- --- --- --- --- --- --- --- --- --- --- --- --- --- --- --- --- --- --- --- --- --- --- --- --- --- --- --- --- --- --- --- --- --- --- --- --- --- --- --- --- --- --- --- --- --- --- --- --- --- --- --- --- --- --- --- --- --- --- --- --- --- --- --- --- --- --- --- --- --- --- --- --- --- --- --- --- --- --- --- --- --- --- --- --- --- --- --- --- --- --- --- --- --- --- --- --- --- --- --- --- --t ggt cag tgt ata ata aat tat atg tag gag gag gaa tat ctg atg atg ttc aaa cta ata cat ctg aaa cat acg ata aag aaa aa

>JX878413.1_Monkeypox_virus_isolate_DRC_07-0045_complete_genome

tgt tat tgt agt atg ata ata tca aaa aga tgg ata taa aga at- --- --- --- --- --- --- --- --- --- --- --- --- --- --- --- --- --- --- --- --- --- --- --- --- --- --- --- --- --- --- --- --- --- --- --- --- --- --- --- --- --- --- --- --- --- --- --- --- --- --- --- --- --- --- --- --- --- --- --- --- --- --- --- --- --- --- --- --- --- --- --- --- --- --- --- --- --- --- --- --- --- --- --- --- --- --- --- --- --- --- --- --- --- --- --- --- --- --- --- --- --- --- --- --- --- --- --- --- --- --- --- --- --- --- --- --- --- --- --- --- --- --- --- --- --- --- --- --- --- --- --- --- --- --- --- --- --- --- --- --- --- --- --- --- --- --- --- --- --- --- --- --- --- --- --- --- --- --- --- --- --- --- --- --- --- --- --- --- --- --- --- --- --- --- --- --- --- --- --- --- --- --- --- --- --- --- --- --- --- --- --- --- --- --- --- --- --- --- --- --- --- --- --- --- --- --- --- --- --- --- --- --- --- --- --- --- --- --- --- --- --- --- --- --- --- --- --- --- --- --- --- --- --- --- --- --- --- --- --- --- --- --- --- --- --- --- --- --- --- --- --- --- --- --- --- --- --- --- --- --- --- --- --- --- --- --- --- --- --- --- --- --- --- --- --- --- --- --- --- --- --- --- --- --- --- --- --- --- --- --- --- --- --- --- --- --- --- --- --- --- --- --- --- --- --- --- --- --- --- --- --- --- --- --- --- --- --- --- --- --- --- --- --- --- --- --- --- --- --- --- --- --- --- --- --- --- --- --- --- --- --- --- --- --- --- --- --- --- --- --- --- --- --- --- --- --- --- --- --- --- --- --- --- --- --- --- --- --- --- --- --- --- --- --- --- --- --- --- --- --- --- --- --- --- --- --- --- --- --- --- --- --- --- --- --- --- --- --- --- --- --- --- --- --- --- --- --- --- --- --- --- --- --- --- --- --- --- --- --- --- --- --- --- --- --- --- --- --- --- --- --- --- --- --- --- --- --- --- --- --- --- --- --- --- --- --- --- --- --- --- --- --- --- --- --- --- --- --- --- --- --- --- --- --- --- --- --- --- --- --- --- --- --- --- --- --- --- --- --- --- --- --- --- --- --- --- --- --- --- --- --- --- --- --- --- --- --- --- --- --- --- --- --- --- --- --- --- --- --- --- --- --- --- --- --- --- --- --- --- --- --- --- --- --- --- --- --- --- --- --- --- --- --- --- --- --- --- --- --- --- --- --- --- --- --- --- --- --- --- --- --- --- --- --- --- --- --- --- --- --- --- --- --- --- --- --- --- --- --- --- --- --- --- --- --- --- --- --- --- --- --- --- --- --- --- --- --- --- --- --- --- --- --- --- --- --- --- --- --- --- --- --- --- --- --- --- --- --- --- --- --- --- --- --- --- --- --- --- --- --- --- --- --- --- --- --- --- --- --- --- --- --- --- --- --- --- --- --- --- --- --- --- --- --- --- --- --- --- --- --- --- --- --- --- --- --- --- --- --- --- --- --- --- --- --- --- --- --- --- --- --- --- --- --- --- --- --- --- --- --- --- --- --- --- --- --- --- --- --- --- --- --- --- --- --- --- --- --- --- --- --- --- --- --- --- --- --- --- --- --- --- --- --- --- --- --- --- --- --- --- --- --- --- --- --- --- --- --- --- --- --- --- --- --- --- --- --- --- --- --- --- --- --- --- --- --- --- --- --- --- --- --- --- --- --- --- --- --- --- --t ggt cag tgt ata ata aat tat atg tag gag gag gaa tat ctg atg atg ttc aaa cta ata cat ctg aaa cat acg ata aag aaa aa

>JX878422.1_Monkeypox_virus_isolate_DRC_07-0287_complete_genome

tgt tat tgt agt atg ata ata tca aaa aga tgg ata taa aga at- --- --- --- --- --- --- --- --- --- --- --- --- --- --- --- --- --- --- --- --- --- --- --- --- --- --- --- --- --- --- --- --- --- --- --- --- --- --- --- --- --- --- --- --- --- --- --- --- --- --- --- --- --- --- --- --- --- --- --- --- --- --- --- --- --- --- --- --- --- --- --- --- --- --- --- --- --- --- --- --- --- --- --- --- --- --- --- --- --- --- --- --- --- --- --- --- --- --- --- --- --- --- --- --- --- --- --- --- --- --- --- --- --- --- --- --- --- --- --- --- --- --- --- --- --- --- --- --- --- --- --- --- --- --- --- --- --- --- --- --- --- --- --- --- --- --- --- --- --- --- --- --- --- --- --- --- --- --- --- --- --- --- --- --- --- --- --- --- --- --- --- --- --- --- --- --- --- --- --- --- --- --- --- --- --- --- --- --- --- --- --- --- --- --- --- --- --- --- --- --- --- --- --- --- --- --- --- --- --- --- --- --- --- --- --- --- --- --- --- --- --- --- --- --- --- --- --- --- --- --- --- --- --- --- --- --- --- --- --- --- --- --- --- --- --- --- --- --- --- --- --- --- --- --- --- --- --- --- --- --- --- --- --- --- --- --- --- --- --- --- --- --- --- --- --- --- --- --- --- --- --- --- --- --- --- --- --- --- --- --- --- --- --- --- --- --- --- --- --- --- --- --- --- --- --- --- --- --- --- --- --- --- --- --- --- --- --- --- --- --- --- --- --- --- --- --- --- --- --- --- --- --- --- --- --- --- --- --- --- --- --- --- --- --- --- --- --- --- --- --- --- --- --- --- --- --- --- --- --- --- --- --- --- --- --- --- --- --- --- --- --- --- --- --- --- --- --- --- --- --- --- --- --- --- --- --- --- --- --- --- --- --- --- --- --- --- --- --- --- --- --- --- --- --- --- --- --- --- --- --- --- --- --- --- --- --- --- --- --- --- --- --- --- --- --- --- --- --- --- --- --- --- --- --- --- --- --- --- --- --- --- --- --- --- --- --- --- --- --- --- --- --- --- --- --- --- --- --- --- --- --- --- --- --- --- --- --- --- --- --- --- --- --- --- --- --- --- --- --- --- --- --- --- --- --- --- --- --- --- --- --- --- --- --- --- --- --- --- --- --- --- --- --- --- --- --- --- --- --- --- --- --- --- --- --- --- --- --- --- --- --- --- --- --- --- --- --- --- --- --- --- --- --- --- --- --- --- --- --- --- --- --- --- --- --- --- --- --- --- --- --- --- --- --- --- --- --- --- --- --- --- --- --- --- --- --- --- --- --- --- --- --- --- --- --- --- --- --- --- --- --- --- --- --- --- --- --- --- --- --- --- --- --- --- --- --- --- --- --- --- --- --- --- --- --- --- --- --- --- --- --- --- --- --- --- --- --- --- --- --- --- --- --- --- --- --- --- --- --- --- --- --- --- --- --- --- --- --- --- --- --- --- --- --- --- --- --- --- --- --- --- --- --- --- --- --- --- --- --- --- --- --- --- --- --- --- --- --- --- --- --- --- --- --- --- --- --- --- --- --- --- --- --- --- --- --- --- --- --- --- --- --- --- --- --- --- --- --- --- --- --- --- --- --- --- --- --- --- --- --- --- --- --- --- --- --- --- --- --- --- --- --- --- --- --- --- --- --- --- --- --- --- --- --- --- --- --- --- --- --- --- --- --- --- --- --- --- --- --- --- --- --- --- --- --- --- --- --- --- --t ggt cag tgt ata ata aat tat atg tag gag gag gaa tat ctg atg atg ttc aaa cta ata cat ctg aaa cat acg ata aag aaa aa

>JX878416.1_Monkeypox_virus_isolate_DRC_07-0093_complete_genome

tgt tat tgt agt atg ata ata tca aaa aga tgg ata taa aga at- --- --- --- --- --- --- --- --- --- --- --- --- --- --- --- --- --- --- --- --- --- --- --- --- --- --- --- --- --- --- --- --- --- --- --- --- --- --- --- --- --- --- --- --- --- --- --- --- --- --- --- --- --- --- --- --- --- --- --- --- --- --- --- --- --- --- --- --- --- --- --- --- --- --- --- --- --- --- --- --- --- --- --- --- --- --- --- --- --- --- --- --- --- --- --- --- --- --- --- --- --- --- --- --- --- --- --- --- --- --- --- --- --- --- --- --- --- --- --- --- --- --- --- --- --- --- --- --- --- --- --- --- --- --- --- --- --- --- --- --- --- --- --- --- --- --- --- --- --- --- --- --- --- --- --- --- --- --- --- --- --- --- --- --- --- --- --- --- --- --- --- --- --- --- --- --- --- --- --- --- --- --- --- --- --- --- --- --- --- --- --- --- --- --- --- --- --- --- --- --- --- --- --- --- --- --- --- --- --- --- --- --- --- --- --- --- --- --- --- --- --- --- --- --- --- --- --- --- --- --- --- --- --- --- --- --- --- --- --- --- --- --- --- --- --- --- --- --- --- --- --- --- --- --- --- --- --- --- --- --- --- --- --- --- --- --- --- --- --- --- --- --- --- --- --- --- --- --- --- --- --- --- --- --- --- --- --- --- --- --- --- --- --- --- --- --- --- --- --- --- --- --- --- --- --- --- --- --- --- --- --- --- --- --- --- --- --- --- --- --- --- --- --- --- --- --- --- --- --- --- --- --- --- --- --- --- --- --- --- --- --- --- --- --- --- --- --- --- --- --- --- --- --- --- --- --- --- --- --- --- --- --- --- --- --- --- --- --- --- --- --- --- --- --- --- --- --- --- --- --- --- --- --- --- --- --- --- --- --- --- --- --- --- --- --- --- --- --- --- --- --- --- --- --- --- --- --- --- --- --- --- --- --- --- --- --- --- --- --- --- --- --- --- --- --- --- --- --- --- --- --- --- --- --- --- --- --- --- --- --- --- --- --- --- --- --- --- --- --- --- --- --- --- --- --- --- --- --- --- --- --- --- --- --- --- --- --- --- --- --- --- --- --- --- --- --- --- --- --- --- --- --- --- --- --- --- --- --- --- --- --- --- --- --- --- --- --- --- --- --- --- --- --- --- --- --- --- --- --- --- --- --- --- --- --- --- --- --- --- --- --- --- --- --- --- --- --- --- --- --- --- --- --- --- --- --- --- --- --- --- --- --- --- --- --- --- --- --- --- --- --- --- --- --- --- --- --- --- --- --- --- --- --- --- --- --- --- --- --- --- --- --- --- --- --- --- --- --- --- --- --- --- --- --- --- --- --- --- --- --- --- --- --- --- --- --- --- --- --- --- --- --- --- --- --- --- --- --- --- --- --- --- --- --- --- --- --- --- --- --- --- --- --- --- --- --- --- --- --- --- --- --- --- --- --- --- --- --- --- --- --- --- --- --- --- --- --- --- --- --- --- --- --- --- --- --- --- --- --- --- --- --- --- --- --- --- --- --- --- --- --- --- --- --- --- --- --- --- --- --- --- --- --- --- --- --- --- --- --- --- --- --- --- --- --- --- --- --- --- --- --- --- --- --- --- --- --- --- --- --- --- --- --- --- --- --- --- --- --- --- --- --- --- --- --- --- --- --- --- --- --- --- --- --- --- --- --- --- --- --- --- --- --- --- --- --- --- --- --- --- --- --- --- --- --- --- --- --- --- --t ggt cag tgt ata ata aat tat atg tag gag gag gaa tat ctg atg atg ttc aaa cta ata cat ctg aaa cat acg ata aag aaa aa

>KP849471.1_Monkeypox_virus_isolate_Yambuku_DRC_1985_complete_genome

tgt tat tgt agt atg ata ata tca aaa aga tgg ata taa aga at- --- --- --- --- --- --- --- --- --- --- --- --- --- --- --- --- --- --- --- --- --- --- --- --- --- --- --- --- --- --- --- --- --- --- --- --- --- --- --- --- --- --- --- --- --- --- --- --- --- --- --- --- --- --- --- --- --- --- --- --- --- --- --- --- --- --- --- --- --- --- --- --- --- --- --- --- --- --- --- --- --- --- --- --- --- --- --- --- --- --- --- --- --- --- --- --- --- --- --- --- --- --- --- --- --- --- --- --- --- --- --- --- --- --- --- --- --- --- --- --- --- --- --- --- --- --- --- --- --- --- --- --- --- --- --- --- --- --- --- --- --- --- --- --- --- --- --- --- --- --- --- --- --- --- --- --- --- --- --- --- --- --- --- --- --- --- --- --- --- --- --- --- --- --- --- --- --- --- --- --- --- --- --- --- --- --- --- --- --- --- --- --- --- --- --- --- --- --- --- --- --- --- --- --- --- --- --- --- --- --- --- --- --- --- --- --- --- --- --- --- --- --- --- --- --- --- --- --- --- --- --- --- --- --- --- --- --- --- --- --- --- --- --- --- --- --- --- --- --- --- --- --- --- --- --- --- --- --- --- --- --- --- --- --- --- --- --- --- --- --- --- --- --- --- --- --- --- --- --- --- --- --- --- --- --- --- --- --- --- --- --- --- --- --- --- --- --- --- --- --- --- --- --- --- --- --- --- --- --- --- --- --- --- --- --- --- --- --- --- --- --- --- --- --- --- --- --- --- --- --- --- --- --- --- --- --- --- --- --- --- --- --- --- --- --- --- --- --- --- --- --- --- --- --- --- --- --- --- --- --- --- --- --- --- --- --- --- --- --- --- --- --- --- --- --- --- --- --- --- --- --- --- --- --- --- --- --- --- --- --- --- --- --- --- --- --- --- --- --- --- --- --- --- --- --- --- --- --- --- --- --- --- --- --- --- --- --- --- --- --- --- --- --- --- --- --- --- --- --- --- --- --- --- --- --- --- --- --- --- --- --- --- --- --- --- --- --- --- --- --- --- --- --- --- --- --- --- --- --- --- --- --- --- --- --- --- --- --- --- --- --- --- --- --- --- --- --- --- --- --- --- --- --- --- --- --- --- --- --- --- --- --- --- --- --- --- --- --- --- --- --- --- --- --- --- --- --- --- --- --- --- --- --- --- --- --- --- --- --- --- --- --- --- --- --- --- --- --- --- --- --- --- --- --- --- --- --- --- --- --- --- --- --- --- --- --- --- --- --- --- --- --- --- --- --- --- --- --- --- --- --- --- --- --- --- --- --- --- --- --- --- --- --- --- --- --- --- --- --- --- --- --- --- --- --- --- --- --- --- --- --- --- --- --- --- --- --- --- --- --- --- --- --- --- --- --- --- --- --- --- --- --- --- --- --- --- --- --- --- --- --- --- --- --- --- --- --- --- --- --- --- --- --- --- --- --- --- --- --- --- --- --- --- --- --- --- --- --- --- --- --- --- --- --- --- --- --- --- --- --- --- --- --- --- --- --- --- --- --- --- --- --- --- --- --- --- --- --- --- --- --- --- --- --- --- --- --- --- --- --- --- --- --- --- --- --- --- --- --- --- --- --- --- --- --- --- --- --- --- --- --- --- --- --- --- --- --- --- --- --- --- --- --- --- --- --- --- --- --- --- --- --- --- --- --- --- --- --- --- --- --- --- --- --- --- --- --- --- --- --- --- --- --- --- --- --- --- --- --- --t ggt cag tgt ata ata aat tat atg tag gag gag gaa tat ctg atg atg ttc aaa cta ata cat ctg aaa cat acg ata aag aaa aa

>DQ011155.1_Monkeypox_virus_strain_Zaire_1979-005_complete_genome

tgt tat tgt agt atg ata ata tca aaa aga tgg ata taa aga at- --- --- --- --- --- --- --- --- --- --- --- --- --- --- --- --- --- --- --- --- --- --- --- --- --- --- --- --- --- --- --- --- --- --- --- --- --- --- --- --- --- --- --- --- --- --- --- --- --- --- --- --- --- --- --- --- --- --- --- --- --- --- --- --- --- --- --- --- --- --- --- --- --- --- --- --- --- --- --- --- --- --- --- --- --- --- --- --- --- --- --- --- --- --- --- --- --- --- --- --- --- --- --- --- --- --- --- --- --- --- --- --- --- --- --- --- --- --- --- --- --- --- --- --- --- --- --- --- --- --- --- --- --- --- --- --- --- --- --- --- --- --- --- --- --- --- --- --- --- --- --- --- --- --- --- --- --- --- --- --- --- --- --- --- --- --- --- --- --- --- --- --- --- --- --- --- --- --- --- --- --- --- --- --- --- --- --- --- --- --- --- --- --- --- --- --- --- --- --- --- --- --- --- --- --- --- --- --- --- --- --- --- --- --- --- --- --- --- --- --- --- --- --- --- --- --- --- --- --- --- --- --- --- --- --- --- --- --- --- --- --- --- --- --- --- --- --- --- --- --- --- --- --- --- --- --- --- --- --- --- --- --- --- --- --- --- --- --- --- --- --- --- --- --- --- --- --- --- --- --- --- --- --- --- --- --- --- --- --- --- --- --- --- --- --- --- --- --- --- --- --- --- --- --- --- --- --- --- --- --- --- --- --- --- --- --- --- --- --- --- --- --- --- --- --- --- --- --- --- --- --- --- --- --- --- --- --- --- --- --- --- --- --- --- --- --- --- --- --- --- --- --- --- --- --- --- --- --- --- --- --- --- --- --- --- --- --- --- --- --- --- --- --- --- --- --- --- --- --- --- --- --- --- --- --- --- --- --- --- --- --- --- --- --- --- --- --- --- --- --- --- --- --- --- --- --- --- --- --- --- --- --- --- --- --- --- --- --- --- --- --- --- --- --- --- --- --- --- --- --- --- --- --- --- --- --- --- --- --- --- --- --- --- --- --- --- --- --- --- --- --- --- --- --- --- --- --- --- --- --- --- --- --- --- --- --- --- --- --- --- --- --- --- --- --- --- --- --- --- --- --- --- --- --- --- --- --- --- --- --- --- --- --- --- --- --- --- --- --- --- --- --- --- --- --- --- --- --- --- --- --- --- --- --- --- --- --- --- --- --- --- --- --- --- --- --- --- --- --- --- --- --- --- --- --- --- --- --- --- --- --- --- --- --- --- --- --- --- --- --- --- --- --- --- --- --- --- --- --- --- --- --- --- --- --- --- --- --- --- --- --- --- --- --- --- --- --- --- --- --- --- --- --- --- --- --- --- --- --- --- --- --- --- --- --- --- --- --- --- --- --- --- --- --- --- --- --- --- --- --- --- --- --- --- --- --- --- --- --- --- --- --- --- --- --- --- --- --- --- --- --- --- --- --- --- --- --- --- --- --- --- --- --- --- --- --- --- --- --- --- --- --- --- --- --- --- --- --- --- --- --- --- --- --- --- --- --- --- --- --- --- --- --- --- --- --- --- --- --- --- --- --- --- --- --- --- --- --- --- --- --- --- --- --- --- --- --- --- --- --- --- --- --- --- --- --- --- --- --- --- --- --- --- --- --- --- --- --- --- --- --- --- --- --- --- --- --- --- --- --- --- --- --- --- --- --- --- --- --- --- --- --- --- --- --- --- --- --- --- --- --- --- --- --- --- --- --- --- --- --t ggt cag tgt ata ata aat tat atg tag gag gag gaa tat ctg atg atg ttc aaa cta ata cat ctg aaa cat acg ata aag aaa aa

>KP849469.1_Monkeypox_virus_isolate_Boende_DRC_2008_complete_genome

tgt tat tgt agt atg ata ata tca aaa aga tgg ata taa aga at- --- --- --- --- --- --- --- --- --- --- --- --- --- --- --- --- --- --- --- --- --- --- --- --- --- --- --- --- --- --- --- --- --- --- --- --- --- --- --- --- --- --- --- --- --- --- --- --- --- --- --- --- --- --- --- --- --- --- --- --- --- --- --- --- --- --- --- --- --- --- --- --- --- --- --- --- --- --- --- --- --- --- --- --- --- --- --- --- --- --- --- --- --- --- --- --- --- --- --- --- --- --- --- --- --- --- --- --- --- --- --- --- --- --- --- --- --- --- --- --- --- --- --- --- --- --- --- --- --- --- --- --- --- --- --- --- --- --- --- --- --- --- --- --- --- --- --- --- --- --- --- --- --- --- --- --- --- --- --- --- --- --- --- --- --- --- --- --- --- --- --- --- --- --- --- --- --- --- --- --- --- --- --- --- --- --- --- --- --- --- --- --- --- --- --- --- --- --- --- --- --- --- --- --- --- --- --- --- --- --- --- --- --- --- --- --- --- --- --- --- --- --- --- --- --- --- --- --- --- --- --- --- --- --- --- --- --- --- --- --- --- --- --- --- --- --- --- --- --- --- --- --- --- --- --- --- --- --- --- --- --- --- --- --- --- --- --- --- --- --- --- --- --- --- --- --- --- --- --- --- --- --- --- --- --- --- --- --- --- --- --- --- --- --- --- --- --- --- --- --- --- --- --- --- --- --- --- --- --- --- --- --- --- --- --- --- --- --- --- --- --- --- --- --- --- --- --- --- --- --- --- --- --- --- --- --- --- --- --- --- --- --- --- --- --- --- --- --- --- --- --- --- --- --- --- --- --- --- --- --- --- --- --- --- --- --- --- --- --- --- --- --- --- --- --- --- --- --- --- --- --- --- --- --- --- --- --- --- --- --- --- --- --- --- --- --- --- --- --- --- --- --- --- --- --- --- --- --- --- --- --- --- --- --- --- --- --- --- --- --- --- --- --- --- --- --- --- --- --- --- --- --- --- --- --- --- --- --- --- --- --- --- --- --- --- --- --- --- --- --- --- --- --- --- --- --- --- --- --- --- --- --- --- --- --- --- --- --- --- --- --- --- --- --- --- --- --- --- --- --- --- --- --- --- --- --- --- --- --- --- --- --- --- --- --- --- --- --- --- --- --- --- --- --- --- --- --- --- --- --- --- --- --- --- --- --- --- --- --- --- --- --- --- --- --- --- --- --- --- --- --- --- --- --- --- --- --- --- --- --- --- --- --- --- --- --- --- --- --- --- --- --- --- --- --- --- --- --- --- --- --- --- --- --- --- --- --- --- --- --- --- --- --- --- --- --- --- --- --- --- --- --- --- --- --- --- --- --- --- --- --- --- --- --- --- --- --- --- --- --- --- --- --- --- --- --- --- --- --- --- --- --- --- --- --- --- --- --- --- --- --- --- --- --- --- --- --- --- --- --- --- --- --- --- --- --- --- --- --- --- --- --- --- --- --- --- --- --- --- --- --- --- --- --- --- --- --- --- --- --- --- --- --- --- --- --- --- --- --- --- --- --- --- --- --- --- --- --- --- --- --- --- --- --- --- --- --- --- --- --- --- --- --- --- --- --- --- --- --- --- --- --- --- --- --- --- --- --- --- --- --- --- --- --- --- --- --- --- --- --- --- --- --- --- --- --- --- --- --- --- --- --- --- --- --- --- --- --- --- --- --- --- --- --- --- --- --- --- --- --- --- --- --- --- --- --- --- --- --- --t ggt cag tgt ata ata aat tat atg tag gag gag gaa tat ctg atg atg ttc aaa cta ata cat ctg aaa cat acg ata aag aaa aa

>KJ642619.1_Monkeypox_virus_strain_Gabon-1988_complete_genome

tgt tat tgt agt atg ata ata tca aaa aga tgg ata taa aga at- --- --- --- --- --- --- --- --- --- --- --- --- --- --- --- --- --- --- --- --- --- --- --- --- --- --- --- --- --- --- --- --- --- --- --- --- --- --- --- --- --- --- --- --- --- --- --- --- --- --- --- --- --- --- --- --- --- --- --- --- --- --- --- --- --- --- --- --- --- --- --- --- --- --- --- --- --- --- --- --- --- --- --- --- --- --- --- --- --- --- --- --- --- --- --- --- --- --- --- --- --- --- --- --- --- --- --- --- --- --- --- --- --- --- --- --- --- --- --- --- --- --- --- --- --- --- --- --- --- --- --- --- --- --- --- --- --- --- --- --- --- --- --- --- --- --- --- --- --- --- --- --- --- --- --- --- --- --- --- --- --- --- --- --- --- --- --- --- --- --- --- --- --- --- --- --- --- --- --- --- --- --- --- --- --- --- --- --- --- --- --- --- --- --- --- --- --- --- --- --- --- --- --- --- --- --- --- --- --- --- --- --- --- --- --- --- --- --- --- --- --- --- --- --- --- --- --- --- --- --- --- --- --- --- --- --- --- --- --- --- --- --- --- --- --- --- --- --- --- --- --- --- --- --- --- --- --- --- --- --- --- --- --- --- --- --- --- --- --- --- --- --- --- --- --- --- --- --- --- --- --- --- --- --- --- --- --- --- --- --- --- --- --- --- --- --- --- --- --- --- --- --- --- --- --- --- --- --- --- --- --- --- --- --- --- --- --- --- --- --- --- --- --- --- --- --- --- --- --- --- --- --- --- --- --- --- --- --- --- --- --- --- --- --- --- --- --- --- --- --- --- --- --- --- --- --- --- --- --- --- --- --- --- --- --- --- --- --- --- --- --- --- --- --- --- --- --- --- --- --- --- --- --- --- --- --- --- --- --- --- --- --- --- --- --- --- --- --- --- --- --- --- --- --- --- --- --- --- --- --- --- --- --- --- --- --- --- --- --- --- --- --- --- --- --- --- --- --- --- --- --- --- --- --- --- --- --- --- --- --- --- --- --- --- --- --- --- --- --- --- --- --- --- --- --- --- --- --- --- --- --- --- --- --- --- --- --- --- --- --- --- --- --- --- --- --- --- --- --- --- --- --- --- --- --- --- --- --- --- --- --- --- --- --- --- --- --- --- --- --- --- --- --- --- --- --- --- --- --- --- --- --- --- --- --- --- --- --- --- --- --- --- --- --- --- --- --- --- --- --- --- --- --- --- --- --- --- --- --- --- --- --- --- --- --- --- --- --- --- --- --- --- --- --- --- --- --- --- --- --- --- --- --- --- --- --- --- --- --- --- --- --- --- --- --- --- --- --- --- --- --- --- --- --- --- --- --- --- --- --- --- --- --- --- --- --- --- --- --- --- --- --- --- --- --- --- --- --- --- --- --- --- --- --- --- --- --- --- --- --- --- --- --- --- --- --- --- --- --- --- --- --- --- --- --- --- --- --- --- --- --- --- --- --- --- --- --- --- --- --- --- --- --- --- --- --- --- --- --- --- --- --- --- --- --- --- --- --- --- --- --- --- --- --- --- --- --- --- --- --- --- --- --- --- --- --- --- --- --- --- --- --- --- --- --- --- --- --- --- --- --- --- --- --- --- --- --- --- --- --- --- --- --- --- --- --- --- --- --- --- --- --- --- --- --- --- --- --- --- --- --- --- --- --- --- --- --- --- --- --- --- --- --- --- --- --- --- --- --- --- --- --- --- --- --- --- --- --- --- --t ggt cag tgt ata ata aat tat atg tag gag gag gaa tat ctg atg atg ttc aaa cta ata cat ctg aaa cat acg ata aag aaa aa

>JX878411.1_Monkeypox_virus_isolate_DRC_06-1075_complete_genome

tgt tat tgt agt atg ata ata tca aaa aga tgg ata taa aga at- --- --- --- --- --- --- --- --- --- --- --- --- --- --- --- --- --- --- --- --- --- --- --- --- --- --- --- --- --- --- --- --- --- --- --- --- --- --- --- --- --- --- --- --- --- --- --- --- --- --- --- --- --- --- --- --- --- --- --- --- --- --- --- --- --- --- --- --- --- --- --- --- --- --- --- --- --- --- --- --- --- --- --- --- --- --- --- --- --- --- --- --- --- --- --- --- --- --- --- --- --- --- --- --- --- --- --- --- --- --- --- --- --- --- --- --- --- --- --- --- --- --- --- --- --- --- --- --- --- --- --- --- --- --- --- --- --- --- --- --- --- --- --- --- --- --- --- --- --- --- --- --- --- --- --- --- --- --- --- --- --- --- --- --- --- --- --- --- --- --- --- --- --- --- --- --- --- --- --- --- --- --- --- --- --- --- --- --- --- --- --- --- --- --- --- --- --- --- --- --- --- --- --- --- --- --- --- --- --- --- --- --- --- --- --- --- --- --- --- --- --- --- --- --- --- --- --- --- --- --- --- --- --- --- --- --- --- --- --- --- --- --- --- --- --- --- --- --- --- --- --- --- --- --- --- --- --- --- --- --- --- --- --- --- --- --- --- --- --- --- --- --- --- --- --- --- --- --- --- --- --- --- --- --- --- --- --- --- --- --- --- --- --- --- --- --- --- --- --- --- --- --- --- --- --- --- --- --- --- --- --- --- --- --- --- --- --- --- --- --- --- --- --- --- --- --- --- --- --- --- --- --- --- --- --- --- --- --- --- --- --- --- --- --- --- --- --- --- --- --- --- --- --- --- --- --- --- --- --- --- --- --- --- --- --- --- --- --- --- --- --- --- --- --- --- --- --- --- --- --- --- --- --- --- --- --- --- --- --- --- --- --- --- --- --- --- --- --- --- --- --- --- --- --- --- --- --- --- --- --- --- --- --- --- --- --- --- --- --- --- --- --- --- --- --- --- --- --- --- --- --- --- --- --- --- --- --- --- --- --- --- --- --- --- --- --- --- --- --- --- --- --- --- --- --- --- --- --- --- --- --- --- --- --- --- --- --- --- --- --- --- --- --- --- --- --- --- --- --- --- --- --- --- --- --- --- --- --- --- --- --- --- --- --- --- --- --- --- --- --- --- --- --- --- --- --- --- --- --- --- --- --- --- --- --- --- --- --- --- --- --- --- --- --- --- --- --- --- --- --- --- --- --- --- --- --- --- --- --- --- --- --- --- --- --- --- --- --- --- --- --- --- --- --- --- --- --- --- --- --- --- --- --- --- --- --- --- --- --- --- --- --- --- --- --- --- --- --- --- --- --- --- --- --- --- --- --- --- --- --- --- --- --- --- --- --- --- --- --- --- --- --- --- --- --- --- --- --- --- --- --- --- --- --- --- --- --- --- --- --- --- --- --- --- --- --- --- --- --- --- --- --- --- --- --- --- --- --- --- --- --- --- --- --- --- --- --- --- --- --- --- --- --- --- --- --- --- --- --- --- --- --- --- --- --- --- --- --- --- --- --- --- --- --- --- --- --- --- --- --- --- --- --- --- --- --- --- --- --- --- --- --- --- --- --- --- --- --- --- --- --- --- --- --- --- --- --- --- --- --- --- --- --- --- --- --- --- --- --- --- --- --- --- --- --- --- --- --- --- --- --- --- --- --- --- --- --- --- --- --- --- --- --- --- --- --- --- --- --- --- --- --- --- --- --- --- --- --- --- --t ggt cag tgt ata ata aat tat atg tag gag gag gaa tat ctg atg atg ttc aaa cta ata cat ctg aaa cat acg ata aag aaa aa

>JX878409.1_Monkeypox_virus_isolate_DRC_06-0999_complete_genome

tgt tat tgt agt atg ata ata tca aaa aga tgg ata taa aga at- --- --- --- --- --- --- --- --- --- --- --- --- --- --- --- --- --- --- --- --- --- --- --- --- --- --- --- --- --- --- --- --- --- --- --- --- --- --- --- --- --- --- --- --- --- --- --- --- --- --- --- --- --- --- --- --- --- --- --- --- --- --- --- --- --- --- --- --- --- --- --- --- --- --- --- --- --- --- --- --- --- --- --- --- --- --- --- --- --- --- --- --- --- --- --- --- --- --- --- --- --- --- --- --- --- --- --- --- --- --- --- --- --- --- --- --- --- --- --- --- --- --- --- --- --- --- --- --- --- --- --- --- --- --- --- --- --- --- --- --- --- --- --- --- --- --- --- --- --- --- --- --- --- --- --- --- --- --- --- --- --- --- --- --- --- --- --- --- --- --- --- --- --- --- --- --- --- --- --- --- --- --- --- --- --- --- --- --- --- --- --- --- --- --- --- --- --- --- --- --- --- --- --- --- --- --- --- --- --- --- --- --- --- --- --- --- --- --- --- --- --- --- --- --- --- --- --- --- --- --- --- --- --- --- --- --- --- --- --- --- --- --- --- --- --- --- --- --- --- --- --- --- --- --- --- --- --- --- --- --- --- --- --- --- --- --- --- --- --- --- --- --- --- --- --- --- --- --- --- --- --- --- --- --- --- --- --- --- --- --- --- --- --- --- --- --- --- --- --- --- --- --- --- --- --- --- --- --- --- --- --- --- --- --- --- --- --- --- --- --- --- --- --- --- --- --- --- --- --- --- --- --- --- --- --- --- --- --- --- --- --- --- --- --- --- --- --- --- --- --- --- --- --- --- --- --- --- --- --- --- --- --- --- --- --- --- --- --- --- --- --- --- --- --- --- --- --- --- --- --- --- --- --- --- --- --- --- --- --- --- --- --- --- --- --- --- --- --- --- --- --- --- --- --- --- --- --- --- --- --- --- --- --- --- --- --- --- --- --- --- --- --- --- --- --- --- --- --- --- --- --- --- --- --- --- --- --- --- --- --- --- --- --- --- --- --- --- --- --- --- --- --- --- --- --- --- --- --- --- --- --- --- --- --- --- --- --- --- --- --- --- --- --- --- --- --- --- --- --- --- --- --- --- --- --- --- --- --- --- --- --- --- --- --- --- --- --- --- --- --- --- --- --- --- --- --- --- --- --- --- --- --- --- --- --- --- --- --- --- --- --- --- --- --- --- --- --- --- --- --- --- --- --- --- --- --- --- --- --- --- --- --- --- --- --- --- --- --- --- --- --- --- --- --- --- --- --- --- --- --- --- --- --- --- --- --- --- --- --- --- --- --- --- --- --- --- --- --- --- --- --- --- --- --- --- --- --- --- --- --- --- --- --- --- --- --- --- --- --- --- --- --- --- --- --- --- --- --- --- --- --- --- --- --- --- --- --- --- --- --- --- --- --- --- --- --- --- --- --- --- --- --- --- --- --- --- --- --- --- --- --- --- --- --- --- --- --- --- --- --- --- --- --- --- --- --- --- --- --- --- --- --- --- --- --- --- --- --- --- --- --- --- --- --- --- --- --- --- --- --- --- --- --- --- --- --- --- --- --- --- --- --- --- --- --- --- --- --- --- --- --- --- --- --- --- --- --- --- --- --- --- --- --- --- --- --- --- --- --- --- --- --- --- --- --- --- --- --- --- --- --- --- --- --- --- --- --- --- --- --- --- --- --- --- --- --- --- --- --- --- --- --- --- --- --- --- --- --- --- --t ggt cag tgt ata ata aat tat atg tag gag gag gaa tat ctg atg atg ttc aaa cta ata cat ctg aaa cat acg ata aag aaa aa

>DQ011154.1_Monkeypox_virus_strain_Congo_2003_358_complete_genome

tgt tat tgt agt atg ata ata tca aaa aga tgg ata taa aga at- --- --- --- --- --- --- --- --- --- --- --- --- --- --- --- --- --- --- --- --- --- --- --- --- --- --- --- --- --- --- --- --- --- --- --- --- --- --- --- --- --- --- --- --- --- --- --- --- --- --- --- --- --- --- --- --- --- --- --- --- --- --- --- --- --- --- --- --- --- --- --- --- --- --- --- --- --- --- --- --- --- --- --- --- --- --- --- --- --- --- --- --- --- --- --- --- --- --- --- --- --- --- --- --- --- --- --- --- --- --- --- --- --- --- --- --- --- --- --- --- --- --- --- --- --- --- --- --- --- --- --- --- --- --- --- --- --- --- --- --- --- --- --- --- --- --- --- --- --- --- --- --- --- --- --- --- --- --- --- --- --- --- --- --- --- --- --- --- --- --- --- --- --- --- --- --- --- --- --- --- --- --- --- --- --- --- --- --- --- --- --- --- --- --- --- --- --- --- --- --- --- --- --- --- --- --- --- --- --- --- --- --- --- --- --- --- --- --- --- --- --- --- --- --- --- --- --- --- --- --- --- --- --- --- --- --- --- --- --- --- --- --- --- --- --- --- --- --- --- --- --- --- --- --- --- --- --- --- --- --- --- --- --- --- --- --- --- --- --- --- --- --- --- --- --- --- --- --- --- --- --- --- --- --- --- --- --- --- --- --- --- --- --- --- --- --- --- --- --- --- --- --- --- --- --- --- --- --- --- --- --- --- --- --- --- --- --- --- --- --- --- --- --- --- --- --- --- --- --- --- --- --- --- --- --- --- --- --- --- --- --- --- --- --- --- --- --- --- --- --- --- --- --- --- --- --- --- --- --- --- --- --- --- --- --- --- --- --- --- --- --- --- --- --- --- --- --- --- --- --- --- --- --- --- --- --- --- --- --- --- --- --- --- --- --- --- --- --- --- --- --- --- --- --- --- --- --- --- --- --- --- --- --- --- --- --- --- --- --- --- --- --- --- --- --- --- --- --- --- --- --- --- --- --- --- --- --- --- --- --- --- --- --- --- --- --- --- --- --- --- --- --- --- --- --- --- --- --- --- --- --- --- --- --- --- --- --- --- --- --- --- --- --- --- --- --- --- --- --- --- --- --- --- --- --- --- --- --- --- --- --- --- --- --- --- --- --- --- --- --- --- --- --- --- --- --- --- --- --- --- --- --- --- --- --- --- --- --- --- --- --- --- --- --- --- --- --- --- --- --- --- --- --- --- --- --- --- --- --- --- --- --- --- --- --- --- --- --- --- --- --- --- --- --- --- --- --- --- --- --- --- --- --- --- --- --- --- --- --- --- --- --- --- --- --- --- --- --- --- --- --- --- --- --- --- --- --- --- --- --- --- --- --- --- --- --- --- --- --- --- --- --- --- --- --- --- --- --- --- --- --- --- --- --- --- --- --- --- --- --- --- --- --- --- --- --- --- --- --- --- --- --- --- --- --- --- --- --- --- --- --- --- --- --- --- --- --- --- --- --- --- --- --- --- --- --- --- --- --- --- --- --- --- --- --- --- --- --- --- --- --- --- --- --- --- --- --- --- --- --- --- --- --- --- --- --- --- --- --- --- --- --- --- --- --- --- --- --- --- --- --- --- --- --- --- --- --- --- --- --- --- --- --- --- --- --- --- --- --- --- --- --- --- --- --- --- --- --- --- --- --- --- --- --- --- --- --- --- --- --- --- --- --- --- --- --- --- --- --- --- --- --- --- --- --- --- --- --- --- --t ggt cag tgt ata ata aat tat atg tag gag gag gaa tat ctg atg atg ttc aaa cta ata cat ctg aaa cat acg ata aag aaa aa

>JX878412.1_Monkeypox_virus_isolate_DRC_06-1076_complete_genome

tgt tat tgt agt atg ata ata tca aaa aga tgg ata taa aga at- --- --- --- --- --- --- --- --- --- --- --- --- --- --- --- --- --- --- --- --- --- --- --- --- --- --- --- --- --- --- --- --- --- --- --- --- --- --- --- --- --- --- --- --- --- --- --- --- --- --- --- --- --- --- --- --- --- --- --- --- --- --- --- --- --- --- --- --- --- --- --- --- --- --- --- --- --- --- --- --- --- --- --- --- --- --- --- --- --- --- --- --- --- --- --- --- --- --- --- --- --- --- --- --- --- --- --- --- --- --- --- --- --- --- --- --- --- --- --- --- --- --- --- --- --- --- --- --- --- --- --- --- --- --- --- --- --- --- --- --- --- --- --- --- --- --- --- --- --- --- --- --- --- --- --- --- --- --- --- --- --- --- --- --- --- --- --- --- --- --- --- --- --- --- --- --- --- --- --- --- --- --- --- --- --- --- --- --- --- --- --- --- --- --- --- --- --- --- --- --- --- --- --- --- --- --- --- --- --- --- --- --- --- --- --- --- --- --- --- --- --- --- --- --- --- --- --- --- --- --- --- --- --- --- --- --- --- --- --- --- --- --- --- --- --- --- --- --- --- --- --- --- --- --- --- --- --- --- --- --- --- --- --- --- --- --- --- --- --- --- --- --- --- --- --- --- --- --- --- --- --- --- --- --- --- --- --- --- --- --- --- --- --- --- --- --- --- --- --- --- --- --- --- --- --- --- --- --- --- --- --- --- --- --- --- --- --- --- --- --- --- --- --- --- --- --- --- --- --- --- --- --- --- --- --- --- --- --- --- --- --- --- --- --- --- --- --- --- --- --- --- --- --- --- --- --- --- --- --- --- --- --- --- --- --- --- --- --- --- --- --- --- --- --- --- --- --- --- --- --- --- --- --- --- --- --- --- --- --- --- --- --- --- --- --- --- --- --- --- --- --- --- --- --- --- --- --- --- --- --- --- --- --- --- --- --- --- --- --- --- --- --- --- --- --- --- --- --- --- --- --- --- --- --- --- --- --- --- --- --- --- --- --- --- --- --- --- --- --- --- --- --- --- --- --- --- --- --- --- --- --- --- --- --- --- --- --- --- --- --- --- --- --- --- --- --- --- --- --- --- --- --- --- --- --- --- --- --- --- --- --- --- --- --- --- --- --- --- --- --- --- --- --- --- --- --- --- --- --- --- --- --- --- --- --- --- --- --- --- --- --- --- --- --- --- --- --- --- --- --- --- --- --- --- --- --- --- --- --- --- --- --- --- --- --- --- --- --- --- --- --- --- --- --- --- --- --- --- --- --- --- --- --- --- --- --- --- --- --- --- --- --- --- --- --- --- --- --- --- --- --- --- --- --- --- --- --- --- --- --- --- --- --- --- --- --- --- --- --- --- --- --- --- --- --- --- --- --- --- --- --- --- --- --- --- --- --- --- --- --- --- --- --- --- --- --- --- --- --- --- --- --- --- --- --- --- --- --- --- --- --- --- --- --- --- --- --- --- --- --- --- --- --- --- --- --- --- --- --- --- --- --- --- --- --- --- --- --- --- --- --- --- --- --- --- --- --- --- --- --- --- --- --- --- --- --- --- --- --- --- --- --- --- --- --- --- --- --- --- --- --- --- --- --- --- --- --- --- --- --- --- --- --- --- --- --- --- --- --- --- --- --- --- --- --- --- --- --- --- --- --- --- --- --- --- --- --- --- --- --- --- --- --- --- --- --- --- --- --- --- --- --- --- --- --- --- --- --- --- --t ggt cag tgt ata ata aat tat atg tag gag gag gaa tat ctg atg atg ttc aaa cta ata cat ctg aaa cat acg ata aag aaa aa

>MN702453.1_Monkeypox_virus_strain_A1_contig_SPADES

tgt tat tgt agt atg ata ata tca aaa aga tgg ata taa aga at- --- --- --- --- --- --- --- --- --- --- --- --- --- --- --- --- --- --- --- --- --- --- --- --- --- --- --- --- --- --- --- --- --- --- --- --- --- --- --- --- --- --- --- --- --- --- --- --- --- --- --- --- --- --- --- --- --- --- --- --- --- --- --- --- --- --- --- --- --- --- --- --- --- --- --- --- --- --- --- --- --- --- --- --- --- --- --- --- --- --- --- --- --- --- --- --- --- --- --- --- --- --- --- --- --- --- --- --- --- --- --- --- --- --- --- --- --- --- --- --- --- --- --- --- --- --- --- --- --- --- --- --- --- --- --- --- --- --- --- --- --- --- --- --- --- --- --- --- --- --- --- --- --- --- --- --- --- --- --- --- --- --- --- --- --- --- --- --- --- --- --- --- --- --- --- --- --- --- --- --- --- --- --- --- --- --- --- --- --- --- --- --- --- --- --- --- --- --- --- --- --- --- --- --- --- --- --- --- --- --- --- --- --- --- --- --- --- --- --- --- --- --- --- --- --- --- --- --- --- --- --- --- --- --- --- --- --- --- --- --- --- --- --- --- --- --- --- --- --- --- --- --- --- --- --- --- --- --- --- --- --- --- --- --- --- --- --- --- --- --- --- --- --- --- --- --- --- --- --- --- --- --- --- --- --- --- --- --- --- --- --- --- --- --- --- --- --- --- --- --- --- --- --- --- --- --- --- --- --- --- --- --- --- --- --- --- --- --- --- --- --- --- --- --- --- --- --- --- --- --- --- --- --- --- --- --- --- --- --- --- --- --- --- --- --- --- --- --- --- --- --- --- --- --- --- --- --- --- --- --- --- --- --- --- --- --- --- --- --- --- --- --- --- --- --- --- --- --- --- --- --- --- --- --- --- --- --- --- --- --- --- --- --- --- --- --- --- --- --- --- --- --- --- --- --- --- --- --- --- --- --- --- --- --- --- --- --- --- --- --- --- --- --- --- --- --- --- --- --- --- --- --- --- --- --- --- --- --- --- --- --- --- --- --- --- --- --- --- --- --- --- --- --- --- --- --- --- --- --- --- --- --- --- --- --- --- --- --- --- --- --- --- --- --- --- --- --- --- --- --- --- --- --- --- --- --- --- --- --- --- --- --- --- --- --- --- --- --- --- --- --- --- --- --- --- --- --- --- --- --- --- --- --- --- --- --- --- --- --- --- --- --- --- --- --- --- --- --- --- --- --- --- --- --- --- --- --- --- --- --- --- --- --- --- --- --- --- --- --- --- --- --- --- --- --- --- --- --- --- --- --- --- --- --- --- --- --- --- --- --- --- --- --- --- --- --- --- --- --- --- --- --- --- --- --- --- --- --- --- --- --- --- --- --- --- --- --- --- --- --- --- --- --- --- --- --- --- --- --- --- --- --- --- --- --- --- --- --- --- --- --- --- --- --- --- --- --- --- --- --- --- --- --- --- --- --- --- --- --- --- --- --- --- --- --- --- --- --- --- --- --- --- --- --- --- --- --- --- --- --- --- --- --- --- --- --- --- --- --- --- --- --- --- --- --- --- --- --- --- --- --- --- --- --- --- --- --- --- --- --- --- --- --- --- --- --- --- --- --- --- --- --- --- --- --- --- --- --- --- --- --- --- --- --- --- --- --- --- --- --- --- --- --- --- --- --- --- --- --- --- --- --- --- --- --- --- --- --- --- --- --- --- --- --- --- --- --- --- --- --- --- --- --- --- --- --- --- --- --- --t ggt cag tgt ata ata aat tat atg tag gag gag gaa tat ctg atg atg ttc aaa cta ata cat ctg aaa cat acg ata aag aaa aa

>MN702452.1_Monkeypox_virus_strain_A2_contig_SPADES

tgt tat tgt agt atg ata ata tca aaa aga tgg ata taa aga at- --- --- --- --- --- --- --- --- --- --- --- --- --- --- --- --- --- --- --- --- --- --- --- --- --- --- --- --- --- --- --- --- --- --- --- --- --- --- --- --- --- --- --- --- --- --- --- --- --- --- --- --- --- --- --- --- --- --- --- --- --- --- --- --- --- --- --- --- --- --- --- --- --- --- --- --- --- --- --- --- --- --- --- --- --- --- --- --- --- --- --- --- --- --- --- --- --- --- --- --- --- --- --- --- --- --- --- --- --- --- --- --- --- --- --- --- --- --- --- --- --- --- --- --- --- --- --- --- --- --- --- --- --- --- --- --- --- --- --- --- --- --- --- --- --- --- --- --- --- --- --- --- --- --- --- --- --- --- --- --- --- --- --- --- --- --- --- --- --- --- --- --- --- --- --- --- --- --- --- --- --- --- --- --- --- --- --- --- --- --- --- --- --- --- --- --- --- --- --- --- --- --- --- --- --- --- --- --- --- --- --- --- --- --- --- --- --- --- --- --- --- --- --- --- --- --- --- --- --- --- --- --- --- --- --- --- --- --- --- --- --- --- --- --- --- --- --- --- --- --- --- --- --- --- --- --- --- --- --- --- --- --- --- --- --- --- --- --- --- --- --- --- --- --- --- --- --- --- --- --- --- --- --- --- --- --- --- --- --- --- --- --- --- --- --- --- --- --- --- --- --- --- --- --- --- --- --- --- --- --- --- --- --- --- --- --- --- --- --- --- --- --- --- --- --- --- --- --- --- --- --- --- --- --- --- --- --- --- --- --- --- --- --- --- --- --- --- --- --- --- --- --- --- --- --- --- --- --- --- --- --- --- --- --- --- --- --- --- --- --- --- --- --- --- --- --- --- --- --- --- --- --- --- --- --- --- --- --- --- --- --- --- --- --- --- --- --- --- --- --- --- --- --- --- --- --- --- --- --- --- --- --- --- --- --- --- --- --- --- --- --- --- --- --- --- --- --- --- --- --- --- --- --- --- --- --- --- --- --- --- --- --- --- --- --- --- --- --- --- --- --- --- --- --- --- --- --- --- --- --- --- --- --- --- --- --- --- --- --- --- --- --- --- --- --- --- --- --- --- --- --- --- --- --- --- --- --- --- --- --- --- --- --- --- --- --- --- --- --- --- --- --- --- --- --- --- --- --- --- --- --- --- --- --- --- --- --- --- --- --- --- --- --- --- --- --- --- --- --- --- --- --- --- --- --- --- --- --- --- --- --- --- --- --- --- --- --- --- --- --- --- --- --- --- --- --- --- --- --- --- --- --- --- --- --- --- --- --- --- --- --- --- --- --- --- --- --- --- --- --- --- --- --- --- --- --- --- --- --- --- --- --- --- --- --- --- --- --- --- --- --- --- --- --- --- --- --- --- --- --- --- --- --- --- --- --- --- --- --- --- --- --- --- --- --- --- --- --- --- --- --- --- --- --- --- --- --- --- --- --- --- --- --- --- --- --- --- --- --- --- --- --- --- --- --- --- --- --- --- --- --- --- --- --- --- --- --- --- --- --- --- --- --- --- --- --- --- --- --- --- --- --- --- --- --- --- --- --- --- --- --- --- --- --- --- --- --- --- --- --- --- --- --- --- --- --- --- --- --- --- --- --- --- --- --- --- --- --- --- --- --- --- --- --- --- --- --- --- --- --- --- --- --- --- --- --- --- --- --- --- --- --- --- --- --- --- --- --- --- --- --- --- --- --- --- --- --- --- --- --t ggt cag tgt ata ata aat tat atg tag gag gag gaa tat ctg atg atg ttc aaa cta ata cat ctg aaa cat acg ata aag aaa aa

>MN702448.1_Monkeypox_virus_strain_015c_contig_SPADES

tgt tat tgt agt atg ata ata tca aaa aga tgg ata taa aga at- --- --- --- --- --- --- --- --- --- --- --- --- --- --- --- --- --- --- --- --- --- --- --- --- --- --- --- --- --- --- --- --- --- --- --- --- --- --- --- --- --- --- --- --- --- --- --- --- --- --- --- --- --- --- --- --- --- --- --- --- --- --- --- --- --- --- --- --- --- --- --- --- --- --- --- --- --- --- --- --- --- --- --- --- --- --- --- --- --- --- --- --- --- --- --- --- --- --- --- --- --- --- --- --- --- --- --- --- --- --- --- --- --- --- --- --- --- --- --- --- --- --- --- --- --- --- --- --- --- --- --- --- --- --- --- --- --- --- --- --- --- --- --- --- --- --- --- --- --- --- --- --- --- --- --- --- --- --- --- --- --- --- --- --- --- --- --- --- --- --- --- --- --- --- --- --- --- --- --- --- --- --- --- --- --- --- --- --- --- --- --- --- --- --- --- --- --- --- --- --- --- --- --- --- --- --- --- --- --- --- --- --- --- --- --- --- --- --- --- --- --- --- --- --- --- --- --- --- --- --- --- --- --- --- --- --- --- --- --- --- --- --- --- --- --- --- --- --- --- --- --- --- --- --- --- --- --- --- --- --- --- --- --- --- --- --- --- --- --- --- --- --- --- --- --- --- --- --- --- --- --- --- --- --- --- --- --- --- --- --- --- --- --- --- --- --- --- --- --- --- --- --- --- --- --- --- --- --- --- --- --- --- --- --- --- --- --- --- --- --- --- --- --- --- --- --- --- --- --- --- --- --- --- --- --- --- --- --- --- --- --- --- --- --- --- --- --- --- --- --- --- --- --- --- --- --- --- --- --- --- --- --- --- --- --- --- --- --- --- --- --- --- --- --- --- --- --- --- --- --- --- --- --- --- --- --- --- --- --- --- --- --- --- --- --- --- --- --- --- --- --- --- --- --- --- --- --- --- --- --- --- --- --- --- --- --- --- --- --- --- --- --- --- --- --- --- --- --- --- --- --- --- --- --- --- --- --- --- --- --- --- --- --- --- --- --- --- --- --- --- --- --- --- --- --- --- --- --- --- --- --- --- --- --- --- --- --- --- --- --- --- --- --- --- --- --- --- --- --- --- --- --- --- --- --- --- --- --- --- --- --- --- --- --- --- --- --- --- --- --- --- --- --- --- --- --- --- --- --- --- --- --- --- --- --- --- --- --- --- --- --- --- --- --- --- --- --- --- --- --- --- --- --- --- --- --- --- --- --- --- --- --- --- --- --- --- --- --- --- --- --- --- --- --- --- --- --- --- --- --- --- --- --- --- --- --- --- --- --- --- --- --- --- --- --- --- --- --- --- --- --- --- --- --- --- --- --- --- --- --- --- --- --- --- --- --- --- --- --- --- --- --- --- --- --- --- --- --- --- --- --- --- --- --- --- --- --- --- --- --- --- --- --- --- --- --- --- --- --- --- --- --- --- --- --- --- --- --- --- --- --- --- --- --- --- --- --- --- --- --- --- --- --- --- --- --- --- --- --- --- --- --- --- --- --- --- --- --- --- --- --- --- --- --- --- --- --- --- --- --- --- --- --- --- --- --- --- --- --- --- --- --- --- --- --- --- --- --- --- --- --- --- --- --- --- --- --- --- --- --- --- --- --- --- --- --- --- --- --- --- --- --- --- --- --- --- --- --- --- --- --- --- --- --- --- --- --- --- --- --- --- --- --- --- --- --- --- --- --- --- --- --- --- --- --- --- --- --- --- --t ggt cag tgt ata ata aat tat atg tag gag gag gaa tat ctg atg atg ttc aaa cta ata cat ctg aaa cat acg ata aag aaa aa

>MN702447.1_Monkeypox_virus_strain_18_contig_SPADES

tgt tat tgt agt atg ata ata tca aaa aga tgg ata taa aga at- --- --- --- --- --- --- --- --- --- --- --- --- --- --- --- --- --- --- --- --- --- --- --- --- --- --- --- --- --- --- --- --- --- --- --- --- --- --- --- --- --- --- --- --- --- --- --- --- --- --- --- --- --- --- --- --- --- --- --- --- --- --- --- --- --- --- --- --- --- --- --- --- --- --- --- --- --- --- --- --- --- --- --- --- --- --- --- --- --- --- --- --- --- --- --- --- --- --- --- --- --- --- --- --- --- --- --- --- --- --- --- --- --- --- --- --- --- --- --- --- --- --- --- --- --- --- --- --- --- --- --- --- --- --- --- --- --- --- --- --- --- --- --- --- --- --- --- --- --- --- --- --- --- --- --- --- --- --- --- --- --- --- --- --- --- --- --- --- --- --- --- --- --- --- --- --- --- --- --- --- --- --- --- --- --- --- --- --- --- --- --- --- --- --- --- --- --- --- --- --- --- --- --- --- --- --- --- --- --- --- --- --- --- --- --- --- --- --- --- --- --- --- --- --- --- --- --- --- --- --- --- --- --- --- --- --- --- --- --- --- --- --- --- --- --- --- --- --- --- --- --- --- --- --- --- --- --- --- --- --- --- --- --- --- --- --- --- --- --- --- --- --- --- --- --- --- --- --- --- --- --- --- --- --- --- --- --- --- --- --- --- --- --- --- --- --- --- --- --- --- --- --- --- --- --- --- --- --- --- --- --- --- --- --- --- --- --- --- --- --- --- --- --- --- --- --- --- --- --- --- --- --- --- --- --- --- --- --- --- --- --- --- --- --- --- --- --- --- --- --- --- --- --- --- --- --- --- --- --- --- --- --- --- --- --- --- --- --- --- --- --- --- --- --- --- --- --- --- --- --- --- --- --- --- --- --- --- --- --- --- --- --- --- --- --- --- --- --- --- --- --- --- --- --- --- --- --- --- --- --- --- --- --- --- --- --- --- --- --- --- --- --- --- --- --- --- --- --- --- --- --- --- --- --- --- --- --- --- --- --- --- --- --- --- --- --- --- --- --- --- --- --- --- --- --- --- --- --- --- --- --- --- --- --- --- --- --- --- --- --- --- --- --- --- --- --- --- --- --- --- --- --- --- --- --- --- --- --- --- --- --- --- --- --- --- --- --- --- --- --- --- --- --- --- --- --- --- --- --- --- --- --- --- --- --- --- --- --- --- --- --- --- --- --- --- --- --- --- --- --- --- --- --- --- --- --- --- --- --- --- --- --- --- --- --- --- --- --- --- --- --- --- --- --- --- --- --- --- --- --- --- --- --- --- --- --- --- --- --- --- --- --- --- --- --- --- --- --- --- --- --- --- --- --- --- --- --- --- --- --- --- --- --- --- --- --- --- --- --- --- --- --- --- --- --- --- --- --- --- --- --- --- --- --- --- --- --- --- --- --- --- --- --- --- --- --- --- --- --- --- --- --- --- --- --- --- --- --- --- --- --- --- --- --- --- --- --- --- --- --- --- --- --- --- --- --- --- --- --- --- --- --- --- --- --- --- --- --- --- --- --- --- --- --- --- --- --- --- --- --- --- --- --- --- --- --- --- --- --- --- --- --- --- --- --- --- --- --- --- --- --- --- --- --- --- --- --- --- --- --- --- --- --- --- --- --- --- --- --- --- --- --- --- --- --- --- --- --- --- --- --- --- --- --- --- --- --- --- --- --- --- --- --- --- --- --- --- --- --- --- --- --- --- --- --- --- --- --- --- --t ggt cag tgt ata ata aat tat atg tag gag gag gaa tat ctg atg atg ttc aaa cta ata cat ctg aaa cat acg ata aag aaa aa

>JX878424.1_Monkeypox_virus_isolate_DRC_07-0338_complete_genome

tgt tat tgt agt atg ata ata tca aaa aga tgg ata taa aga at- --- --- --- --- --- --- --- --- --- --- --- --- --- --- --- --- --- --- --- --- --- --- --- --- --- --- --- --- --- --- --- --- --- --- --- --- --- --- --- --- --- --- --- --- --- --- --- --- --- --- --- --- --- --- --- --- --- --- --- --- --- --- --- --- --- --- --- --- --- --- --- --- --- --- --- --- --- --- --- --- --- --- --- --- --- --- --- --- --- --- --- --- --- --- --- --- --- --- --- --- --- --- --- --- --- --- --- --- --- --- --- --- --- --- --- --- --- --- --- --- --- --- --- --- --- --- --- --- --- --- --- --- --- --- --- --- --- --- --- --- --- --- --- --- --- --- --- --- --- --- --- --- --- --- --- --- --- --- --- --- --- --- --- --- --- --- --- --- --- --- --- --- --- --- --- --- --- --- --- --- --- --- --- --- --- --- --- --- --- --- --- --- --- --- --- --- --- --- --- --- --- --- --- --- --- --- --- --- --- --- --- --- --- --- --- --- --- --- --- --- --- --- --- --- --- --- --- --- --- --- --- --- --- --- --- --- --- --- --- --- --- --- --- --- --- --- --- --- --- --- --- --- --- --- --- --- --- --- --- --- --- --- --- --- --- --- --- --- --- --- --- --- --- --- --- --- --- --- --- --- --- --- --- --- --- --- --- --- --- --- --- --- --- --- --- --- --- --- --- --- --- --- --- --- --- --- --- --- --- --- --- --- --- --- --- --- --- --- --- --- --- --- --- --- --- --- --- --- --- --- --- --- --- --- --- --- --- --- --- --- --- --- --- --- --- --- --- --- --- --- --- --- --- --- --- --- --- --- --- --- --- --- --- --- --- --- --- --- --- --- --- --- --- --- --- --- --- --- --- --- --- --- --- --- --- --- --- --- --- --- --- --- --- --- --- --- --- --- --- --- --- --- --- --- --- --- --- --- --- --- --- --- --- --- --- --- --- --- --- --- --- --- --- --- --- --- --- --- --- --- --- --- --- --- --- --- --- --- --- --- --- --- --- --- --- --- --- --- --- --- --- --- --- --- --- --- --- --- --- --- --- --- --- --- --- --- --- --- --- --- --- --- --- --- --- --- --- --- --- --- --- --- --- --- --- --- --- --- --- --- --- --- --- --- --- --- --- --- --- --- --- --- --- --- --- --- --- --- --- --- --- --- --- --- --- --- --- --- --- --- --- --- --- --- --- --- --- --- --- --- --- --- --- --- --- --- --- --- --- --- --- --- --- --- --- --- --- --- --- --- --- --- --- --- --- --- --- --- --- --- --- --- --- --- --- --- --- --- --- --- --- --- --- --- --- --- --- --- --- --- --- --- --- --- --- --- --- --- --- --- --- --- --- --- --- --- --- --- --- --- --- --- --- --- --- --- --- --- --- --- --- --- --- --- --- --- --- --- --- --- --- --- --- --- --- --- --- --- --- --- --- --- --- --- --- --- --- --- --- --- --- --- --- --- --- --- --- --- --- --- --- --- --- --- --- --- --- --- --- --- --- --- --- --- --- --- --- --- --- --- --- --- --- --- --- --- --- --- --- --- --- --- --- --- --- --- --- --- --- --- --- --- --- --- --- --- --- --- --- --- --- --- --- --- --- --- --- --- --- --- --- --- --- --- --- --- --- --- --- --- --- --- --- --- --- --- --- --- --- --- --- --- --- --- --- --- --- --- --- --- --- --- --- --- --- --- --- --- --- --- --- --- --- --- --- --- --- --- --- --t ggt cag tgt ata ata aat tat atg tag gag gag gaa tat ctg atg atg ttc aaa cta ata cat ctg aaa cat acg ata aag aaa aa

>KC257460.1_Monkeypox_virus_strain_DRC_Yandongi_1985_complete_genome

tgt tat tgt agt atg ata ata tca aaa aga tgg ata taa aga at- --- --- --- --- --- --- --- --- --- --- --- --- --- --- --- --- --- --- --- --- --- --- --- --- --- --- --- --- --- --- --- --- --- --- --- --- --- --- --- --- --- --- --- --- --- --- --- --- --- --- --- --- --- --- --- --- --- --- --- --- --- --- --- --- --- --- --- --- --- --- --- --- --- --- --- --- --- --- --- --- --- --- --- --- --- --- --- --- --- --- --- --- --- --- --- --- --- --- --- --- --- --- --- --- --- --- --- --- --- --- --- --- --- --- --- --- --- --- --- --- --- --- --- --- --- --- --- --- --- --- --- --- --- --- --- --- --- --- --- --- --- --- --- --- --- --- --- --- --- --- --- --- --- --- --- --- --- --- --- --- --- --- --- --- --- --- --- --- --- --- --- --- --- --- --- --- --- --- --- --- --- --- --- --- --- --- --- --- --- --- --- --- --- --- --- --- --- --- --- --- --- --- --- --- --- --- --- --- --- --- --- --- --- --- --- --- --- --- --- --- --- --- --- --- --- --- --- --- --- --- --- --- --- --- --- --- --- --- --- --- --- --- --- --- --- --- --- --- --- --- --- --- --- --- --- --- --- --- --- --- --- --- --- --- --- --- --- --- --- --- --- --- --- --- --- --- --- --- --- --- --- --- --- --- --- --- --- --- --- --- --- --- --- --- --- --- --- --- --- --- --- --- --- --- --- --- --- --- --- --- --- --- --- --- --- --- --- --- --- --- --- --- --- --- --- --- --- --- --- --- --- --- --- --- --- --- --- --- --- --- --- --- --- --- --- --- --- --- --- --- --- --- --- --- --- --- --- --- --- --- --- --- --- --- --- --- --- --- --- --- --- --- --- --- --- --- --- --- --- --- --- --- --- --- --- --- --- --- --- --- --- --- --- --- --- --- --- --- --- --- --- --- --- --- --- --- --- --- --- --- --- --- --- --- --- --- --- --- --- --- --- --- --- --- --- --- --- --- --- --- --- --- --- --- --- --- --- --- --- --- --- --- --- --- --- --- --- --- --- --- --- --- --- --- --- --- --- --- --- --- --- --- --- --- --- --- --- --- --- --- --- --- --- --- --- --- --- --- --- --- --- --- --- --- --- --- --- --- --- --- --- --- --- --- --- --- --- --- --- --- --- --- --- --- --- --- --- --- --- --- --- --- --- --- --- --- --- --- --- --- --- --- --- --- --- --- --- --- --- --- --- --- --- --- --- --- --- --- --- --- --- --- --- --- --- --- --- --- --- --- --- --- --- --- --- --- --- --- --- --- --- --- --- --- --- --- --- --- --- --- --- --- --- --- --- --- --- --- --- --- --- --- --- --- --- --- --- --- --- --- --- --- --- --- --- --- --- --- --- --- --- --- --- --- --- --- --- --- --- --- --- --- --- --- --- --- --- --- --- --- --- --- --- --- --- --- --- --- --- --- --- --- --- --- --- --- --- --- --- --- --- --- --- --- --- --- --- --- --- --- --- --- --- --- --- --- --- --- --- --- --- --- --- --- --- --- --- --- --- --- --- --- --- --- --- --- --- --- --- --- --- --- --- --- --- --- --- --- --- --- --- --- --- --- --- --- --- --- --- --- --- --- --- --- --- --- --- --- --- --- --- --- --- --- --- --- --- --- --- --- --- --- --- --- --- --- --- --- --- --- --- --- --- --- --- --- --- --- --- --- --- --- --- --- --- --- --- --- --- --- --- --- --- --- --- --- --- --- --- --t ggt cag tgt ata ata aat tat atg tag gag gag gaa tat ctg atg atg ttc aaa cta ata cat ctg aaa cat acg ata aag aaa aa

>JX878429.1_Monkeypox_virus_isolate_DRC_07-0662_complete_genome

tgt tat tgt agt atg ata ata tca aaa aga tgg ata taa aga at- --- --- --- --- --- --- --- --- --- --- --- --- --- --- --- --- --- --- --- --- --- --- --- --- --- --- --- --- --- --- --- --- --- --- --- --- --- --- --- --- --- --- --- --- --- --- --- --- --- --- --- --- --- --- --- --- --- --- --- --- --- --- --- --- --- --- --- --- --- --- --- --- --- --- --- --- --- --- --- --- --- --- --- --- --- --- --- --- --- --- --- --- --- --- --- --- --- --- --- --- --- --- --- --- --- --- --- --- --- --- --- --- --- --- --- --- --- --- --- --- --- --- --- --- --- --- --- --- --- --- --- --- --- --- --- --- --- --- --- --- --- --- --- --- --- --- --- --- --- --- --- --- --- --- --- --- --- --- --- --- --- --- --- --- --- --- --- --- --- --- --- --- --- --- --- --- --- --- --- --- --- --- --- --- --- --- --- --- --- --- --- --- --- --- --- --- --- --- --- --- --- --- --- --- --- --- --- --- --- --- --- --- --- --- --- --- --- --- --- --- --- --- --- --- --- --- --- --- --- --- --- --- --- --- --- --- --- --- --- --- --- --- --- --- --- --- --- --- --- --- --- --- --- --- --- --- --- --- --- --- --- --- --- --- --- --- --- --- --- --- --- --- --- --- --- --- --- --- --- --- --- --- --- --- --- --- --- --- --- --- --- --- --- --- --- --- --- --- --- --- --- --- --- --- --- --- --- --- --- --- --- --- --- --- --- --- --- --- --- --- --- --- --- --- --- --- --- --- --- --- --- --- --- --- --- --- --- --- --- --- --- --- --- --- --- --- --- --- --- --- --- --- --- --- --- --- --- --- --- --- --- --- --- --- --- --- --- --- --- --- --- --- --- --- --- --- --- --- --- --- --- --- --- --- --- --- --- --- --- --- --- --- --- --- --- --- --- --- --- --- --- --- --- --- --- --- --- --- --- --- --- --- --- --- --- --- --- --- --- --- --- --- --- --- --- --- --- --- --- --- --- --- --- --- --- --- --- --- --- --- --- --- --- --- --- --- --- --- --- --- --- --- --- --- --- --- --- --- --- --- --- --- --- --- --- --- --- --- --- --- --- --- --- --- --- --- --- --- --- --- --- --- --- --- --- --- --- --- --- --- --- --- --- --- --- --- --- --- --- --- --- --- --- --- --- --- --- --- --- --- --- --- --- --- --- --- --- --- --- --- --- --- --- --- --- --- --- --- --- --- --- --- --- --- --- --- --- --- --- --- --- --- --- --- --- --- --- --- --- --- --- --- --- --- --- --- --- --- --- --- --- --- --- --- --- --- --- --- --- --- --- --- --- --- --- --- --- --- --- --- --- --- --- --- --- --- --- --- --- --- --- --- --- --- --- --- --- --- --- --- --- --- --- --- --- --- --- --- --- --- --- --- --- --- --- --- --- --- --- --- --- --- --- --- --- --- --- --- --- --- --- --- --- --- --- --- --- --- --- --- --- --- --- --- --- --- --- --- --- --- --- --- --- --- --- --- --- --- --- --- --- --- --- --- --- --- --- --- --- --- --- --- --- --- --- --- --- --- --- --- --- --- --- --- --- --- --- --- --- --- --- --- --- --- --- --- --- --- --- --- --- --- --- --- --- --- --- --- --- --- --- --- --- --- --- --- --- --- --- --- --- --- --- --- --- --- --- --- --- --- --- --- --- --- --- --- --- --- --- --- --- --- --- --- --- --- --- --- --- --- --- --- --- --- --- --- --- --- --- --t ggt cag tgt ata ata aat tat atg tag gag gag gaa tat ctg atg atg ttc aaa cta ata cat ctg aaa cat acg ata aag aaa aa

>JX878425.1_Monkeypox_virus_isolate_DRC_07-0354_complete_genome

tgt tat tgt agt atg ata ata tca aaa aga tgg ata taa aga at- --- --- --- --- --- --- --- --- --- --- --- --- --- --- --- --- --- --- --- --- --- --- --- --- --- --- --- --- --- --- --- --- --- --- --- --- --- --- --- --- --- --- --- --- --- --- --- --- --- --- --- --- --- --- --- --- --- --- --- --- --- --- --- --- --- --- --- --- --- --- --- --- --- --- --- --- --- --- --- --- --- --- --- --- --- --- --- --- --- --- --- --- --- --- --- --- --- --- --- --- --- --- --- --- --- --- --- --- --- --- --- --- --- --- --- --- --- --- --- --- --- --- --- --- --- --- --- --- --- --- --- --- --- --- --- --- --- --- --- --- --- --- --- --- --- --- --- --- --- --- --- --- --- --- --- --- --- --- --- --- --- --- --- --- --- --- --- --- --- --- --- --- --- --- --- --- --- --- --- --- --- --- --- --- --- --- --- --- --- --- --- --- --- --- --- --- --- --- --- --- --- --- --- --- --- --- --- --- --- --- --- --- --- --- --- --- --- --- --- --- --- --- --- --- --- --- --- --- --- --- --- --- --- --- --- --- --- --- --- --- --- --- --- --- --- --- --- --- --- --- --- --- --- --- --- --- --- --- --- --- --- --- --- --- --- --- --- --- --- --- --- --- --- --- --- --- --- --- --- --- --- --- --- --- --- --- --- --- --- --- --- --- --- --- --- --- --- --- --- --- --- --- --- --- --- --- --- --- --- --- --- --- --- --- --- --- --- --- --- --- --- --- --- --- --- --- --- --- --- --- --- --- --- --- --- --- --- --- --- --- --- --- --- --- --- --- --- --- --- --- --- --- --- --- --- --- --- --- --- --- --- --- --- --- --- --- --- --- --- --- --- --- --- --- --- --- --- --- --- --- --- --- --- --- --- --- --- --- --- --- --- --- --- --- --- --- --- --- --- --- --- --- --- --- --- --- --- --- --- --- --- --- --- --- --- --- --- --- --- --- --- --- --- --- --- --- --- --- --- --- --- --- --- --- --- --- --- --- --- --- --- --- --- --- --- --- --- --- --- --- --- --- --- --- --- --- --- --- --- --- --- --- --- --- --- --- --- --- --- --- --- --- --- --- --- --- --- --- --- --- --- --- --- --- --- --- --- --- --- --- --- --- --- --- --- --- --- --- --- --- --- --- --- --- --- --- --- --- --- --- --- --- --- --- --- --- --- --- --- --- --- --- --- --- --- --- --- --- --- --- --- --- --- --- --- --- --- --- --- --- --- --- --- --- --- --- --- --- --- --- --- --- --- --- --- --- --- --- --- --- --- --- --- --- --- --- --- --- --- --- --- --- --- --- --- --- --- --- --- --- --- --- --- --- --- --- --- --- --- --- --- --- --- --- --- --- --- --- --- --- --- --- --- --- --- --- --- --- --- --- --- --- --- --- --- --- --- --- --- --- --- --- --- --- --- --- --- --- --- --- --- --- --- --- --- --- --- --- --- --- --- --- --- --- --- --- --- --- --- --- --- --- --- --- --- --- --- --- --- --- --- --- --- --- --- --- --- --- --- --- --- --- --- --- --- --- --- --- --- --- --- --- --- --- --- --- --- --- --- --- --- --- --- --- --- --- --- --- --- --- --- --- --- --- --- --- --- --- --- --- --- --- --- --- --- --- --- --- --- --- --- --- --- --- --- --- --- --- --- --- --- --- --- --- --- --- --- --- --- --- --- --- --- --- --- --- --- --- --- --- --- --- --- --- --- --- --- --- --- --t ggt cag tgt ata ata aat tat atg tag gag gag gaa tat ctg atg atg ttc aaa cta ata cat ctg aaa cat acg ata aag aaa aa

>JX878407.1_Monkeypox_virus_isolate_DRC_06-0950_complete_genome

tgt tat tgt agt atg ata ata tca aaa aga tgg ata taa aga at- --- --- --- --- --- --- --- --- --- --- --- --- --- --- --- --- --- --- --- --- --- --- --- --- --- --- --- --- --- --- --- --- --- --- --- --- --- --- --- --- --- --- --- --- --- --- --- --- --- --- --- --- --- --- --- --- --- --- --- --- --- --- --- --- --- --- --- --- --- --- --- --- --- --- --- --- --- --- --- --- --- --- --- --- --- --- --- --- --- --- --- --- --- --- --- --- --- --- --- --- --- --- --- --- --- --- --- --- --- --- --- --- --- --- --- --- --- --- --- --- --- --- --- --- --- --- --- --- --- --- --- --- --- --- --- --- --- --- --- --- --- --- --- --- --- --- --- --- --- --- --- --- --- --- --- --- --- --- --- --- --- --- --- --- --- --- --- --- --- --- --- --- --- --- --- --- --- --- --- --- --- --- --- --- --- --- --- --- --- --- --- --- --- --- --- --- --- --- --- --- --- --- --- --- --- --- --- --- --- --- --- --- --- --- --- --- --- --- --- --- --- --- --- --- --- --- --- --- --- --- --- --- --- --- --- --- --- --- --- --- --- --- --- --- --- --- --- --- --- --- --- --- --- --- --- --- --- --- --- --- --- --- --- --- --- --- --- --- --- --- --- --- --- --- --- --- --- --- --- --- --- --- --- --- --- --- --- --- --- --- --- --- --- --- --- --- --- --- --- --- --- --- --- --- --- --- --- --- --- --- --- --- --- --- --- --- --- --- --- --- --- --- --- --- --- --- --- --- --- --- --- --- --- --- --- --- --- --- --- --- --- --- --- --- --- --- --- --- --- --- --- --- --- --- --- --- --- --- --- --- --- --- --- --- --- --- --- --- --- --- --- --- --- --- --- --- --- --- --- --- --- --- --- --- --- --- --- --- --- --- --- --- --- --- --- --- --- --- --- --- --- --- --- --- --- --- --- --- --- --- --- --- --- --- --- --- --- --- --- --- --- --- --- --- --- --- --- --- --- --- --- --- --- --- --- --- --- --- --- --- --- --- --- --- --- --- --- --- --- --- --- --- --- --- --- --- --- --- --- --- --- --- --- --- --- --- --- --- --- --- --- --- --- --- --- --- --- --- --- --- --- --- --- --- --- --- --- --- --- --- --- --- --- --- --- --- --- --- --- --- --- --- --- --- --- --- --- --- --- --- --- --- --- --- --- --- --- --- --- --- --- --- --- --- --- --- --- --- --- --- --- --- --- --- --- --- --- --- --- --- --- --- --- --- --- --- --- --- --- --- --- --- --- --- --- --- --- --- --- --- --- --- --- --- --- --- --- --- --- --- --- --- --- --- --- --- --- --- --- --- --- --- --- --- --- --- --- --- --- --- --- --- --- --- --- --- --- --- --- --- --- --- --- --- --- --- --- --- --- --- --- --- --- --- --- --- --- --- --- --- --- --- --- --- --- --- --- --- --- --- --- --- --- --- --- --- --- --- --- --- --- --- --- --- --- --- --- --- --- --- --- --- --- --- --- --- --- --- --- --- --- --- --- --- --- --- --- --- --- --- --- --- --- --- --- --- --- --- --- --- --- --- --- --- --- --- --- --- --- --- --- --- --- --- --- --- --- --- --- --- --- --- --- --- --- --- --- --- --- --- --- --- --- --- --- --- --- --- --- --- --- --- --- --- --- --- --- --- --- --- --- --- --- --- --- --- --- --- --- --- --- --- --- --- --- --- --- --- --- --- --- --- --- --- --- --- --- --- --- --t ggt cag tgt ata ata aat tat atg tag gag gag gaa tat ctg atg atg ttc aaa cta ata cat ctg aaa cat acg ata aag aaa aa

>MN702451.1_Monkeypox_virus_strain_A6_contig_SPADES

tgt tat tgt agt atg ata ata tca aaa aga tgg ata taa aga at- --- --- --- --- --- --- --- --- --- --- --- --- --- --- --- --- --- --- --- --- --- --- --- --- --- --- --- --- --- --- --- --- --- --- --- --- --- --- --- --- --- --- --- --- --- --- --- --- --- --- --- --- --- --- --- --- --- --- --- --- --- --- --- --- --- --- --- --- --- --- --- --- --- --- --- --- --- --- --- --- --- --- --- --- --- --- --- --- --- --- --- --- --- --- --- --- --- --- --- --- --- --- --- --- --- --- --- --- --- --- --- --- --- --- --- --- --- --- --- --- --- --- --- --- --- --- --- --- --- --- --- --- --- --- --- --- --- --- --- --- --- --- --- --- --- --- --- --- --- --- --- --- --- --- --- --- --- --- --- --- --- --- --- --- --- --- --- --- --- --- --- --- --- --- --- --- --- --- --- --- --- --- --- --- --- --- --- --- --- --- --- --- --- --- --- --- --- --- --- --- --- --- --- --- --- --- --- --- --- --- --- --- --- --- --- --- --- --- --- --- --- --- --- --- --- --- --- --- --- --- --- --- --- --- --- --- --- --- --- --- --- --- --- --- --- --- --- --- --- --- --- --- --- --- --- --- --- --- --- --- --- --- --- --- --- --- --- --- --- --- --- --- --- --- --- --- --- --- --- --- --- --- --- --- --- --- --- --- --- --- --- --- --- --- --- --- --- --- --- --- --- --- --- --- --- --- --- --- --- --- --- --- --- --- --- --- --- --- --- --- --- --- --- --- --- --- --- --- --- --- --- --- --- --- --- --- --- --- --- --- --- --- --- --- --- --- --- --- --- --- --- --- --- --- --- --- --- --- --- --- --- --- --- --- --- --- --- --- --- --- --- --- --- --- --- --- --- --- --- --- --- --- --- --- --- --- --- --- --- --- --- --- --- --- --- --- --- --- --- --- --- --- --- --- --- --- --- --- --- --- --- --- --- --- --- --- --- --- --- --- --- --- --- --- --- --- --- --- --- --- --- --- --- --- --- --- --- --- --- --- --- --- --- --- --- --- --- --- --- --- --- --- --- --- --- --- --- --- --- --- --- --- --- --- --- --- --- --- --- --- --- --- --- --- --- --- --- --- --- --- --- --- --- --- --- --- --- --- --- --- --- --- --- --- --- --- --- --- --- --- --- --- --- --- --- --- --- --- --- --- --- --- --- --- --- --- --- --- --- --- --- --- --- --- --- --- --- --- --- --- --- --- --- --- --- --- --- --- --- --- --- --- --- --- --- --- --- --- --- --- --- --- --- --- --- --- --- --- --- --- --- --- --- --- --- --- --- --- --- --- --- --- --- --- --- --- --- --- --- --- --- --- --- --- --- --- --- --- --- --- --- --- --- --- --- --- --- --- --- --- --- --- --- --- --- --- --- --- --- --- --- --- --- --- --- --- --- --- --- --- --- --- --- --- --- --- --- --- --- --- --- --- --- --- --- --- --- --- --- --- --- --- --- --- --- --- --- --- --- --- --- --- --- --- --- --- --- --- --- --- --- --- --- --- --- --- --- --- --- --- --- --- --- --- --- --- --- --- --- --- --- --- --- --- --- --- --- --- --- --- --- --- --- --- --- --- --- --- --- --- --- --- --- --- --- --- --- --- --- --- --- --- --- --- --- --- --- --- --- --- --- --- --- --- --- --- --- --- --- --- --- --- --- --- --- --- --- --- --- --- --- --- --- --- --- --- --- --- --- --- --- --- --- --- --- --- --- --- --- --t ggt cag tgt ata ata aat tat atg tag gag gag gaa tat ctg atg atg ttc aaa cta ata cat ctg aaa cat acg ata aag aaa aa

>JX878423.1_Monkeypox_virus_isolate_DRC_07-0337_complete_genome

tgt tat tgt agt atg ata ata tca aaa aga tgg ata taa aga at- --- --- --- --- --- --- --- --- --- --- --- --- --- --- --- --- --- --- --- --- --- --- --- --- --- --- --- --- --- --- --- --- --- --- --- --- --- --- --- --- --- --- --- --- --- --- --- --- --- --- --- --- --- --- --- --- --- --- --- --- --- --- --- --- --- --- --- --- --- --- --- --- --- --- --- --- --- --- --- --- --- --- --- --- --- --- --- --- --- --- --- --- --- --- --- --- --- --- --- --- --- --- --- --- --- --- --- --- --- --- --- --- --- --- --- --- --- --- --- --- --- --- --- --- --- --- --- --- --- --- --- --- --- --- --- --- --- --- --- --- --- --- --- --- --- --- --- --- --- --- --- --- --- --- --- --- --- --- --- --- --- --- --- --- --- --- --- --- --- --- --- --- --- --- --- --- --- --- --- --- --- --- --- --- --- --- --- --- --- --- --- --- --- --- --- --- --- --- --- --- --- --- --- --- --- --- --- --- --- --- --- --- --- --- --- --- --- --- --- --- --- --- --- --- --- --- --- --- --- --- --- --- --- --- --- --- --- --- --- --- --- --- --- --- --- --- --- --- --- --- --- --- --- --- --- --- --- --- --- --- --- --- --- --- --- --- --- --- --- --- --- --- --- --- --- --- --- --- --- --- --- --- --- --- --- --- --- --- --- --- --- --- --- --- --- --- --- --- --- --- --- --- --- --- --- --- --- --- --- --- --- --- --- --- --- --- --- --- --- --- --- --- --- --- --- --- --- --- --- --- --- --- --- --- --- --- --- --- --- --- --- --- --- --- --- --- --- --- --- --- --- --- --- --- --- --- --- --- --- --- --- --- --- --- --- --- --- --- --- --- --- --- --- --- --- --- --- --- --- --- --- --- --- --- --- --- --- --- --- --- --- --- --- --- --- --- --- --- --- --- --- --- --- --- --- --- --- --- --- --- --- --- --- --- --- --- --- --- --- --- --- --- --- --- --- --- --- --- --- --- --- --- --- --- --- --- --- --- --- --- --- --- --- --- --- --- --- --- --- --- --- --- --- --- --- --- --- --- --- --- --- --- --- --- --- --- --- --- --- --- --- --- --- --- --- --- --- --- --- --- --- --- --- --- --- --- --- --- --- --- --- --- --- --- --- --- --- --- --- --- --- --- --- --- --- --- --- --- --- --- --- --- --- --- --- --- --- --- --- --- --- --- --- --- --- --- --- --- --- --- --- --- --- --- --- --- --- --- --- --- --- --- --- --- --- --- --- --- --- --- --- --- --- --- --- --- --- --- --- --- --- --- --- --- --- --- --- --- --- --- --- --- --- --- --- --- --- --- --- --- --- --- --- --- --- --- --- --- --- --- --- --- --- --- --- --- --- --- --- --- --- --- --- --- --- --- --- --- --- --- --- --- --- --- --- --- --- --- --- --- --- --- --- --- --- --- --- --- --- --- --- --- --- --- --- --- --- --- --- --- --- --- --- --- --- --- --- --- --- --- --- --- --- --- --- --- --- --- --- --- --- --- --- --- --- --- --- --- --- --- --- --- --- --- --- --- --- --- --- --- --- --- --- --- --- --- --- --- --- --- --- --- --- --- --- --- --- --- --- --- --- --- --- --- --- --- --- --- --- --- --- --- --- --- --- --- --- --- --- --- --- --- --- --- --- --- --- --- --- --- --- --- --- --- --- --- --- --- --- --- --- --- --- --- --- --- --- --- --- --- --- --- --- --- --- --- --- --- --- --t ggt cag tgt ata ata aat tat atg tag gag gag gaa tat ctg atg atg ttc aaa cta ata cat ctg aaa cat acg ata aag aaa aa

>MN702450.1_Monkeypox_virus_strain_B1_contig_SPADES

tgt tat tgt agt atg ata ata tca aaa aga tgg ata taa aga at- --- --- --- --- --- --- --- --- --- --- --- --- --- --- --- --- --- --- --- --- --- --- --- --- --- --- --- --- --- --- --- --- --- --- --- --- --- --- --- --- --- --- --- --- --- --- --- --- --- --- --- --- --- --- --- --- --- --- --- --- --- --- --- --- --- --- --- --- --- --- --- --- --- --- --- --- --- --- --- --- --- --- --- --- --- --- --- --- --- --- --- --- --- --- --- --- --- --- --- --- --- --- --- --- --- --- --- --- --- --- --- --- --- --- --- --- --- --- --- --- --- --- --- --- --- --- --- --- --- --- --- --- --- --- --- --- --- --- --- --- --- --- --- --- --- --- --- --- --- --- --- --- --- --- --- --- --- --- --- --- --- --- --- --- --- --- --- --- --- --- --- --- --- --- --- --- --- --- --- --- --- --- --- --- --- --- --- --- --- --- --- --- --- --- --- --- --- --- --- --- --- --- --- --- --- --- --- --- --- --- --- --- --- --- --- --- --- --- --- --- --- --- --- --- --- --- --- --- --- --- --- --- --- --- --- --- --- --- --- --- --- --- --- --- --- --- --- --- --- --- --- --- --- --- --- --- --- --- --- --- --- --- --- --- --- --- --- --- --- --- --- --- --- --- --- --- --- --- --- --- --- --- --- --- --- --- --- --- --- --- --- --- --- --- --- --- --- --- --- --- --- --- --- --- --- --- --- --- --- --- --- --- --- --- --- --- --- --- --- --- --- --- --- --- --- --- --- --- --- --- --- --- --- --- --- --- --- --- --- --- --- --- --- --- --- --- --- --- --- --- --- --- --- --- --- --- --- --- --- --- --- --- --- --- --- --- --- --- --- --- --- --- --- --- --- --- --- --- --- --- --- --- --- --- --- --- --- --- --- --- --- --- --- --- --- --- --- --- --- --- --- --- --- --- --- --- --- --- --- --- --- --- --- --- --- --- --- --- --- --- --- --- --- --- --- --- --- --- --- --- --- --- --- --- --- --- --- --- --- --- --- --- --- --- --- --- --- --- --- --- --- --- --- --- --- --- --- --- --- --- --- --- --- --- --- --- --- --- --- --- --- --- --- --- --- --- --- --- --- --- --- --- --- --- --- --- --- --- --- --- --- --- --- --- --- --- --- --- --- --- --- --- --- --- --- --- --- --- --- --- --- --- --- --- --- --- --- --- --- --- --- --- --- --- --- --- --- --- --- --- --- --- --- --- --- --- --- --- --- --- --- --- --- --- --- --- --- --- --- --- --- --- --- --- --- --- --- --- --- --- --- --- --- --- --- --- --- --- --- --- --- --- --- --- --- --- --- --- --- --- --- --- --- --- --- --- --- --- --- --- --- --- --- --- --- --- --- --- --- --- --- --- --- --- --- --- --- --- --- --- --- --- --- --- --- --- --- --- --- --- --- --- --- --- --- --- --- --- --- --- --- --- --- --- --- --- --- --- --- --- --- --- --- --- --- --- --- --- --- --- --- --- --- --- --- --- --- --- --- --- --- --- --- --- --- --- --- --- --- --- --- --- --- --- --- --- --- --- --- --- --- --- --- --- --- --- --- --- --- --- --- --- --- --- --- --- --- --- --- --- --- --- --- --- --- --- --- --- --- --- --- --- --- --- --- --- --- --- --- --- --- --- --- --- --- --- --- --- --- --- --- --- --- --- --- --- --- --- --- --- --- --- --- --- --- --- --- --- --- --- --- --- --- --- --- --- --- --- --- --t ggt cag tgt ata ata aat tat atg tag gag gag gaa tat ctg atg atg ttc aaa cta ata cat ctg aaa cat acg ata aag aaa aa

>JX878408.1_Monkeypox_virus_isolate_DRC_06-0970_complete_genome

tgt tat tgt agt atg ata ata tca aaa aga tgg ata taa aga at- --- --- --- --- --- --- --- --- --- --- --- --- --- --- --- --- --- --- --- --- --- --- --- --- --- --- --- --- --- --- --- --- --- --- --- --- --- --- --- --- --- --- --- --- --- --- --- --- --- --- --- --- --- --- --- --- --- --- --- --- --- --- --- --- --- --- --- --- --- --- --- --- --- --- --- --- --- --- --- --- --- --- --- --- --- --- --- --- --- --- --- --- --- --- --- --- --- --- --- --- --- --- --- --- --- --- --- --- --- --- --- --- --- --- --- --- --- --- --- --- --- --- --- --- --- --- --- --- --- --- --- --- --- --- --- --- --- --- --- --- --- --- --- --- --- --- --- --- --- --- --- --- --- --- --- --- --- --- --- --- --- --- --- --- --- --- --- --- --- --- --- --- --- --- --- --- --- --- --- --- --- --- --- --- --- --- --- --- --- --- --- --- --- --- --- --- --- --- --- --- --- --- --- --- --- --- --- --- --- --- --- --- --- --- --- --- --- --- --- --- --- --- --- --- --- --- --- --- --- --- --- --- --- --- --- --- --- --- --- --- --- --- --- --- --- --- --- --- --- --- --- --- --- --- --- --- --- --- --- --- --- --- --- --- --- --- --- --- --- --- --- --- --- --- --- --- --- --- --- --- --- --- --- --- --- --- --- --- --- --- --- --- --- --- --- --- --- --- --- --- --- --- --- --- --- --- --- --- --- --- --- --- --- --- --- --- --- --- --- --- --- --- --- --- --- --- --- --- --- --- --- --- --- --- --- --- --- --- --- --- --- --- --- --- --- --- --- --- --- --- --- --- --- --- --- --- --- --- --- --- --- --- --- --- --- --- --- --- --- --- --- --- --- --- --- --- --- --- --- --- --- --- --- --- --- --- --- --- --- --- --- --- --- --- --- --- --- --- --- --- --- --- --- --- --- --- --- --- --- --- --- --- --- --- --- --- --- --- --- --- --- --- --- --- --- --- --- --- --- --- --- --- --- --- --- --- --- --- --- --- --- --- --- --- --- --- --- --- --- --- --- --- --- --- --- --- --- --- --- --- --- --- --- --- --- --- --- --- --- --- --- --- --- --- --- --- --- --- --- --- --- --- --- --- --- --- --- --- --- --- --- --- --- --- --- --- --- --- --- --- --- --- --- --- --- --- --- --- --- --- --- --- --- --- --- --- --- --- --- --- --- --- --- --- --- --- --- --- --- --- --- --- --- --- --- --- --- --- --- --- --- --- --- --- --- --- --- --- --- --- --- --- --- --- --- --- --- --- --- --- --- --- --- --- --- --- --- --- --- --- --- --- --- --- --- --- --- --- --- --- --- --- --- --- --- --- --- --- --- --- --- --- --- --- --- --- --- --- --- --- --- --- --- --- --- --- --- --- --- --- --- --- --- --- --- --- --- --- --- --- --- --- --- --- --- --- --- --- --- --- --- --- --- --- --- --- --- --- --- --- --- --- --- --- --- --- --- --- --- --- --- --- --- --- --- --- --- --- --- --- --- --- --- --- --- --- --- --- --- --- --- --- --- --- --- --- --- --- --- --- --- --- --- --- --- --- --- --- --- --- --- --- --- --- --- --- --- --- --- --- --- --- --- --- --- --- --- --- --- --- --- --- --- --- --- --- --- --- --- --- --- --- --- --- --- --- --- --- --- --- --- --- --- --- --- --- --- --- --- --- --- --- --- --- --- --- --- --- --- --- --- --- --- --- --- --- --- --- --- --t ggt cag tgt ata ata aat tat atg tag gag gag gaa tat ctg atg atg ttc aaa cta ata cat ctg aaa cat acg ata aag aaa aa

>JX878420.1_Monkeypox_virus_isolate_DRC_07-0283_complete_genome

tgt tat tgt agt atg ata ata tca aaa aga tgg ata taa aga at- --- --- --- --- --- --- --- --- --- --- --- --- --- --- --- --- --- --- --- --- --- --- --- --- --- --- --- --- --- --- --- --- --- --- --- --- --- --- --- --- --- --- --- --- --- --- --- --- --- --- --- --- --- --- --- --- --- --- --- --- --- --- --- --- --- --- --- --- --- --- --- --- --- --- --- --- --- --- --- --- --- --- --- --- --- --- --- --- --- --- --- --- --- --- --- --- --- --- --- --- --- --- --- --- --- --- --- --- --- --- --- --- --- --- --- --- --- --- --- --- --- --- --- --- --- --- --- --- --- --- --- --- --- --- --- --- --- --- --- --- --- --- --- --- --- --- --- --- --- --- --- --- --- --- --- --- --- --- --- --- --- --- --- --- --- --- --- --- --- --- --- --- --- --- --- --- --- --- --- --- --- --- --- --- --- --- --- --- --- --- --- --- --- --- --- --- --- --- --- --- --- --- --- --- --- --- --- --- --- --- --- --- --- --- --- --- --- --- --- --- --- --- --- --- --- --- --- --- --- --- --- --- --- --- --- --- --- --- --- --- --- --- --- --- --- --- --- --- --- --- --- --- --- --- --- --- --- --- --- --- --- --- --- --- --- --- --- --- --- --- --- --- --- --- --- --- --- --- --- --- --- --- --- --- --- --- --- --- --- --- --- --- --- --- --- --- --- --- --- --- --- --- --- --- --- --- --- --- --- --- --- --- --- --- --- --- --- --- --- --- --- --- --- --- --- --- --- --- --- --- --- --- --- --- --- --- --- --- --- --- --- --- --- --- --- --- --- --- --- --- --- --- --- --- --- --- --- --- --- --- --- --- --- --- --- --- --- --- --- --- --- --- --- --- --- --- --- --- --- --- --- --- --- --- --- --- --- --- --- --- --- --- --- --- --- --- --- --- --- --- --- --- --- --- --- --- --- --- --- --- --- --- --- --- --- --- --- --- --- --- --- --- --- --- --- --- --- --- --- --- --- --- --- --- --- --- --- --- --- --- --- --- --- --- --- --- --- --- --- --- --- --- --- --- --- --- --- --- --- --- --- --- --- --- --- --- --- --- --- --- --- --- --- --- --- --- --- --- --- --- --- --- --- --- --- --- --- --- --- --- --- --- --- --- --- --- --- --- --- --- --- --- --- --- --- --- --- --- --- --- --- --- --- --- --- --- --- --- --- --- --- --- --- --- --- --- --- --- --- --- --- --- --- --- --- --- --- --- --- --- --- --- --- --- --- --- --- --- --- --- --- --- --- --- --- --- --- --- --- --- --- --- --- --- --- --- --- --- --- --- --- --- --- --- --- --- --- --- --- --- --- --- --- --- --- --- --- --- --- --- --- --- --- --- --- --- --- --- --- --- --- --- --- --- --- --- --- --- --- --- --- --- --- --- --- --- --- --- --- --- --- --- --- --- --- --- --- --- --- --- --- --- --- --- --- --- --- --- --- --- --- --- --- --- --- --- --- --- --- --- --- --- --- --- --- --- --- --- --- --- --- --- --- --- --- --- --- --- --- --- --- --- --- --- --- --- --- --- --- --- --- --- --- --- --- --- --- --- --- --- --- --- --- --- --- --- --- --- --- --- --- --- --- --- --- --- --- --- --- --- --- --- --- --- --- --- --- --- --- --- --- --- --- --- --- --- --- --- --- --- --- --- --- --- --- --- --- --- --- --- --- --- --- --- --- --- --- --- --- --- --- --- --- --- --- --- --- --- --- --t ggt cag tgt ata ata aat tat atg tag gag gag gaa tat ctg atg atg ttc aaa cta ata cat ctg aaa cat acg ata aag aaa aa

>JX878419.1_Monkeypox_virus_isolate_DRC_07-0275_complete_genome

tgt tat tgt agt atg ata ata tca aaa aga tgg ata taa aga at- --- --- --- --- --- --- --- --- --- --- --- --- --- --- --- --- --- --- --- --- --- --- --- --- --- --- --- --- --- --- --- --- --- --- --- --- --- --- --- --- --- --- --- --- --- --- --- --- --- --- --- --- --- --- --- --- --- --- --- --- --- --- --- --- --- --- --- --- --- --- --- --- --- --- --- --- --- --- --- --- --- --- --- --- --- --- --- --- --- --- --- --- --- --- --- --- --- --- --- --- --- --- --- --- --- --- --- --- --- --- --- --- --- --- --- --- --- --- --- --- --- --- --- --- --- --- --- --- --- --- --- --- --- --- --- --- --- --- --- --- --- --- --- --- --- --- --- --- --- --- --- --- --- --- --- --- --- --- --- --- --- --- --- --- --- --- --- --- --- --- --- --- --- --- --- --- --- --- --- --- --- --- --- --- --- --- --- --- --- --- --- --- --- --- --- --- --- --- --- --- --- --- --- --- --- --- --- --- --- --- --- --- --- --- --- --- --- --- --- --- --- --- --- --- --- --- --- --- --- --- --- --- --- --- --- --- --- --- --- --- --- --- --- --- --- --- --- --- --- --- --- --- --- --- --- --- --- --- --- --- --- --- --- --- --- --- --- --- --- --- --- --- --- --- --- --- --- --- --- --- --- --- --- --- --- --- --- --- --- --- --- --- --- --- --- --- --- --- --- --- --- --- --- --- --- --- --- --- --- --- --- --- --- --- --- --- --- --- --- --- --- --- --- --- --- --- --- --- --- --- --- --- --- --- --- --- --- --- --- --- --- --- --- --- --- --- --- --- --- --- --- --- --- --- --- --- --- --- --- --- --- --- --- --- --- --- --- --- --- --- --- --- --- --- --- --- --- --- --- --- --- --- --- --- --- --- --- --- --- --- --- --- --- --- --- --- --- --- --- --- --- --- --- --- --- --- --- --- --- --- --- --- --- --- --- --- --- --- --- --- --- --- --- --- --- --- --- --- --- --- --- --- --- --- --- --- --- --- --- --- --- --- --- --- --- --- --- --- --- --- --- --- --- --- --- --- --- --- --- --- --- --- --- --- --- --- --- --- --- --- --- --- --- --- --- --- --- --- --- --- --- --- --- --- --- --- --- --- --- --- --- --- --- --- --- --- --- --- --- --- --- --- --- --- --- --- --- --- --- --- --- --- --- --- --- --- --- --- --- --- --- --- --- --- --- --- --- --- --- --- --- --- --- --- --- --- --- --- --- --- --- --- --- --- --- --- --- --- --- --- --- --- --- --- --- --- --- --- --- --- --- --- --- --- --- --- --- --- --- --- --- --- --- --- --- --- --- --- --- --- --- --- --- --- --- --- --- --- --- --- --- --- --- --- --- --- --- --- --- --- --- --- --- --- --- --- --- --- --- --- --- --- --- --- --- --- --- --- --- --- --- --- --- --- --- --- --- --- --- --- --- --- --- --- --- --- --- --- --- --- --- --- --- --- --- --- --- --- --- --- --- --- --- --- --- --- --- --- --- --- --- --- --- --- --- --- --- --- --- --- --- --- --- --- --- --- --- --- --- --- --- --- --- --- --- --- --- --- --- --- --- --- --- --- --- --- --- --- --- --- --- --- --- --- --- --- --- --- --- --- --- --- --- --- --- --- --- --- --- --- --- --- --- --- --- --- --- --- --- --- --- --- --- --- --- --- --- --- --- --- --- --- --- --- --- --- --- --- --- --- --- --- --- --- --- --- --- --- --- --t ggt cag tgt ata ata aat tat atg tag gag gag gaa tat ctg atg atg ttc aaa cta ata cat ctg aaa cat acg ata aag aaa aa

>JX878417.1_Monkeypox_virus_isolate_DRC_07-0104_complete_genome

tgt tat tgt agt atg ata ata tca aaa aga tgg ata taa aga at- --- --- --- --- --- --- --- --- --- --- --- --- --- --- --- --- --- --- --- --- --- --- --- --- --- --- --- --- --- --- --- --- --- --- --- --- --- --- --- --- --- --- --- --- --- --- --- --- --- --- --- --- --- --- --- --- --- --- --- --- --- --- --- --- --- --- --- --- --- --- --- --- --- --- --- --- --- --- --- --- --- --- --- --- --- --- --- --- --- --- --- --- --- --- --- --- --- --- --- --- --- --- --- --- --- --- --- --- --- --- --- --- --- --- --- --- --- --- --- --- --- --- --- --- --- --- --- --- --- --- --- --- --- --- --- --- --- --- --- --- --- --- --- --- --- --- --- --- --- --- --- --- --- --- --- --- --- --- --- --- --- --- --- --- --- --- --- --- --- --- --- --- --- --- --- --- --- --- --- --- --- --- --- --- --- --- --- --- --- --- --- --- --- --- --- --- --- --- --- --- --- --- --- --- --- --- --- --- --- --- --- --- --- --- --- --- --- --- --- --- --- --- --- --- --- --- --- --- --- --- --- --- --- --- --- --- --- --- --- --- --- --- --- --- --- --- --- --- --- --- --- --- --- --- --- --- --- --- --- --- --- --- --- --- --- --- --- --- --- --- --- --- --- --- --- --- --- --- --- --- --- --- --- --- --- --- --- --- --- --- --- --- --- --- --- --- --- --- --- --- --- --- --- --- --- --- --- --- --- --- --- --- --- --- --- --- --- --- --- --- --- --- --- --- --- --- --- --- --- --- --- --- --- --- --- --- --- --- --- --- --- --- --- --- --- --- --- --- --- --- --- --- --- --- --- --- --- --- --- --- --- --- --- --- --- --- --- --- --- --- --- --- --- --- --- --- --- --- --- --- --- --- --- --- --- --- --- --- --- --- --- --- --- --- --- --- --- --- --- --- --- --- --- --- --- --- --- --- --- --- --- --- --- --- --- --- --- --- --- --- --- --- --- --- --- --- --- --- --- --- --- --- --- --- --- --- --- --- --- --- --- --- --- --- --- --- --- --- --- --- --- --- --- --- --- --- --- --- --- --- --- --- --- --- --- --- --- --- --- --- --- --- --- --- --- --- --- --- --- --- --- --- --- --- --- --- --- --- --- --- --- --- --- --- --- --- --- --- --- --- --- --- --- --- --- --- --- --- --- --- --- --- --- --- --- --- --- --- --- --- --- --- --- --- --- --- --- --- --- --- --- --- --- --- --- --- --- --- --- --- --- --- --- --- --- --- --- --- --- --- --- --- --- --- --- --- --- --- --- --- --- --- --- --- --- --- --- --- --- --- --- --- --- --- --- --- --- --- --- --- --- --- --- --- --- --- --- --- --- --- --- --- --- --- --- --- --- --- --- --- --- --- --- --- --- --- --- --- --- --- --- --- --- --- --- --- --- --- --- --- --- --- --- --- --- --- --- --- --- --- --- --- --- --- --- --- --- --- --- --- --- --- --- --- --- --- --- --- --- --- --- --- --- --- --- --- --- --- --- --- --- --- --- --- --- --- --- --- --- --- --- --- --- --- --- --- --- --- --- --- --- --- --- --- --- --- --- --- --- --- --- --- --- --- --- --- --- --- --- --- --- --- --- --- --- --- --- --- --- --- --- --- --- --- --- --- --- --- --- --- --- --- --- --- --- --- --- --- --- --- --- --- --- --- --- --- --- --- --- --- --- --- --- --- --- --- --- --- --- --- --- --- --- --- --- --- --- --- --- --t ggt cag tgt ata ata aat tat atg tag gag gag gaa tat ctg atg atg ttc aaa cta ata cat ctg aaa cat acg ata aag aaa aa

>KJ642612.1_Monkeypox_virus_strain_Ikubi_complete_genome

tgt tat tgt agt atg ata ata tca aaa aga tgg ata taa aga at- --- --- --- --- --- --- --- --- --- --- --- --- --- --- --- --- --- --- --- --- --- --- --- --- --- --- --- --- --- --- --- --- --- --- --- --- --- --- --- --- --- --- --- --- --- --- --- --- --- --- --- --- --- --- --- --- --- --- --- --- --- --- --- --- --- --- --- --- --- --- --- --- --- --- --- --- --- --- --- --- --- --- --- --- --- --- --- --- --- --- --- --- --- --- --- --- --- --- --- --- --- --- --- --- --- --- --- --- --- --- --- --- --- --- --- --- --- --- --- --- --- --- --- --- --- --- --- --- --- --- --- --- --- --- --- --- --- --- --- --- --- --- --- --- --- --- --- --- --- --- --- --- --- --- --- --- --- --- --- --- --- --- --- --- --- --- --- --- --- --- --- --- --- --- --- --- --- --- --- --- --- --- --- --- --- --- --- --- --- --- --- --- --- --- --- --- --- --- --- --- --- --- --- --- --- --- --- --- --- --- --- --- --- --- --- --- --- --- --- --- --- --- --- --- --- --- --- --- --- --- --- --- --- --- --- --- --- --- --- --- --- --- --- --- --- --- --- --- --- --- --- --- --- --- --- --- --- --- --- --- --- --- --- --- --- --- --- --- --- --- --- --- --- --- --- --- --- --- --- --- --- --- --- --- --- --- --- --- --- --- --- --- --- --- --- --- --- --- --- --- --- --- --- --- --- --- --- --- --- --- --- --- --- --- --- --- --- --- --- --- --- --- --- --- --- --- --- --- --- --- --- --- --- --- --- --- --- --- --- --- --- --- --- --- --- --- --- --- --- --- --- --- --- --- --- --- --- --- --- --- --- --- --- --- --- --- --- --- --- --- --- --- --- --- --- --- --- --- --- --- --- --- --- --- --- --- --- --- --- --- --- --- --- --- --- --- --- --- --- --- --- --- --- --- --- --- --- --- --- --- --- --- --- --- --- --- --- --- --- --- --- --- --- --- --- --- --- --- --- --- --- --- --- --- --- --- --- --- --- --- --- --- --- --- --- --- --- --- --- --- --- --- --- --- --- --- --- --- --- --- --- --- --- --- --- --- --- --- --- --- --- --- --- --- --- --- --- --- --- --- --- --- --- --- --- --- --- --- --- --- --- --- --- --- --- --- --- --- --- --- --- --- --- --- --- --- --- --- --- --- --- --- --- --- --- --- --- --- --- --- --- --- --- --- --- --- --- --- --- --- --- --- --- --- --- --- --- --- --- --- --- --- --- --- --- --- --- --- --- --- --- --- --- --- --- --- --- --- --- --- --- --- --- --- --- --- --- --- --- --- --- --- --- --- --- --- --- --- --- --- --- --- --- --- --- --- --- --- --- --- --- --- --- --- --- --- --- --- --- --- --- --- --- --- --- --- --- --- --- --- --- --- --- --- --- --- --- --- --- --- --- --- --- --- --- --- --- --- --- --- --- --- --- --- --- --- --- --- --- --- --- --- --- --- --- --- --- --- --- --- --- --- --- --- --- --- --- --- --- --- --- --- --- --- --- --- --- --- --- --- --- --- --- --- --- --- --- --- --- --- --- --- --- --- --- --- --- --- --- --- --- --- --- --- --- --- --- --- --- --- --- --- --- --- --- --- --- --- --- --- --- --- --- --- --- --- --- --- --- --- --- --- --- --- --- --- --- --- --- --- --- --- --- --- --- --- --- --- --- --- --- --- --- --- --- --- --- --- --- --- --- --- --- --- --- --- --- --- --- --t ggt cag tgt ata ata aat tat atg tag gag gag gaa tat ctg atg atg ttc aaa cta ata cat ctg aaa cat acg ata aag aaa aa

>JX878426.1_Monkeypox_virus_isolate_DRC_07-0450_complete_genome

tgt tat tgt agt atg ata ata tca aaa aga tgg ata taa aga at- --- --- --- --- --- --- --- --- --- --- --- --- --- --- --- --- --- --- --- --- --- --- --- --- --- --- --- --- --- --- --- --- --- --- --- --- --- --- --- --- --- --- --- --- --- --- --- --- --- --- --- --- --- --- --- --- --- --- --- --- --- --- --- --- --- --- --- --- --- --- --- --- --- --- --- --- --- --- --- --- --- --- --- --- --- --- --- --- --- --- --- --- --- --- --- --- --- --- --- --- --- --- --- --- --- --- --- --- --- --- --- --- --- --- --- --- --- --- --- --- --- --- --- --- --- --- --- --- --- --- --- --- --- --- --- --- --- --- --- --- --- --- --- --- --- --- --- --- --- --- --- --- --- --- --- --- --- --- --- --- --- --- --- --- --- --- --- --- --- --- --- --- --- --- --- --- --- --- --- --- --- --- --- --- --- --- --- --- --- --- --- --- --- --- --- --- --- --- --- --- --- --- --- --- --- --- --- --- --- --- --- --- --- --- --- --- --- --- --- --- --- --- --- --- --- --- --- --- --- --- --- --- --- --- --- --- --- --- --- --- --- --- --- --- --- --- --- --- --- --- --- --- --- --- --- --- --- --- --- --- --- --- --- --- --- --- --- --- --- --- --- --- --- --- --- --- --- --- --- --- --- --- --- --- --- --- --- --- --- --- --- --- --- --- --- --- --- --- --- --- --- --- --- --- --- --- --- --- --- --- --- --- --- --- --- --- --- --- --- --- --- --- --- --- --- --- --- --- --- --- --- --- --- --- --- --- --- --- --- --- --- --- --- --- --- --- --- --- --- --- --- --- --- --- --- --- --- --- --- --- --- --- --- --- --- --- --- --- --- --- --- --- --- --- --- --- --- --- --- --- --- --- --- --- --- --- --- --- --- --- --- --- --- --- --- --- --- --- --- --- --- --- --- --- --- --- --- --- --- --- --- --- --- --- --- --- --- --- --- --- --- --- --- --- --- --- --- --- --- --- --- --- --- --- --- --- --- --- --- --- --- --- --- --- --- --- --- --- --- --- --- --- --- --- --- --- --- --- --- --- --- --- --- --- --- --- --- --- --- --- --- --- --- --- --- --- --- --- --- --- --- --- --- --- --- --- --- --- --- --- --- --- --- --- --- --- --- --- --- --- --- --- --- --- --- --- --- --- --- --- --- --- --- --- --- --- --- --- --- --- --- --- --- --- --- --- --- --- --- --- --- --- --- --- --- --- --- --- --- --- --- --- --- --- --- --- --- --- --- --- --- --- --- --- --- --- --- --- --- --- --- --- --- --- --- --- --- --- --- --- --- --- --- --- --- --- --- --- --- --- --- --- --- --- --- --- --- --- --- --- --- --- --- --- --- --- --- --- --- --- --- --- --- --- --- --- --- --- --- --- --- --- --- --- --- --- --- --- --- --- --- --- --- --- --- --- --- --- --- --- --- --- --- --- --- --- --- --- --- --- --- --- --- --- --- --- --- --- --- --- --- --- --- --- --- --- --- --- --- --- --- --- --- --- --- --- --- --- --- --- --- --- --- --- --- --- --- --- --- --- --- --- --- --- --- --- --- --- --- --- --- --- --- --- --- --- --- --- --- --- --- --- --- --- --- --- --- --- --- --- --- --- --- --- --- --- --- --- --- --- --- --- --- --- --- --- --- --- --- --- --- --- --- --- --- --- --- --- --- --- --- --- --- --- --- --- --- --- --- --- --- --- --- --- --- --- --- --- --- --t ggt cag tgt ata ata aat tat atg tag gag gag gaa tat ctg atg atg ttc aaa cta ata cat ctg aaa cat acg ata aag aaa aa

>KC257459.1_Monkeypox_virus_strain_Sudan_2005_01_complete_genome

tgt tat tgt agt atg ata ata tca aaa aga tgg ata taa aga at- --- --- --- --- --- --- --- --- --- --- --- --- --- --- --- --- --- --- --- --- --- --- --- --- --- --- --- --- --- --- --- --- --- --- --- --- --- --- --- --- --- --- --- --- --- --- --- --- --- --- --- --- --- --- --- --- --- --- --- --- --- --- --- --- --- --- --- --- --- --- --- --- --- --- --- --- --- --- --- --- --- --- --- --- --- --- --- --- --- --- --- --- --- --- --- --- --- --- --- --- --- --- --- --- --- --- --- --- --- --- --- --- --- --- --- --- --- --- --- --- --- --- --- --- --- --- --- --- --- --- --- --- --- --- --- --- --- --- --- --- --- --- --- --- --- --- --- --- --- --- --- --- --- --- --- --- --- --- --- --- --- --- --- --- --- --- --- --- --- --- --- --- --- --- --- --- --- --- --- --- --- --- --- --- --- --- --- --- --- --- --- --- --- --- --- --- --- --- --- --- --- --- --- --- --- --- --- --- --- --- --- --- --- --- --- --- --- --- --- --- --- --- --- --- --- --- --- --- --- --- --- --- --- --- --- --- --- --- --- --- --- --- --- --- --- --- --- --- --- --- --- --- --- --- --- --- --- --- --- --- --- --- --- --- --- --- --- --- --- --- --- --- --- --- --- --- --- --- --- --- --- --- --- --- --- --- --- --- --- --- --- --- --- --- --- --- --- --- --- --- --- --- --- --- --- --- --- --- --- --- --- --- --- --- --- --- --- --- --- --- --- --- --- --- --- --- --- --- --- --- --- --- --- --- --- --- --- --- --- --- --- --- --- --- --- --- --- --- --- --- --- --- --- --- --- --- --- --- --- --- --- --- --- --- --- --- --- --- --- --- --- --- --- --- --- --- --- --- --- --- --- --- --- --- --- --- --- --- --- --- --- --- --- --- --- --- --- --- --- --- --- --- --- --- --- --- --- --- --- --- --- --- --- --- --- --- --- --- --- --- --- --- --- --- --- --- --- --- --- --- --- --- --- --- --- --- --- --- --- --- --- --- --- --- --- --- --- --- --- --- --- --- --- --- --- --- --- --- --- --- --- --- --- --- --- --- --- --- --- --- --- --- --- --- --- --- --- --- --- --- --- --- --- --- --- --- --- --- --- --- --- --- --- --- --- --- --- --- --- --- --- --- --- --- --- --- --- --- --- --- --- --- --- --- --- --- --- --- --- --- --- --- --- --- --- --- --- --- --- --- --- --- --- --- --- --- --- --- --- --- --- --- --- --- --- --- --- --- --- --- --- --- --- --- --- --- --- --- --- --- --- --- --- --- --- --- --- --- --- --- --- --- --- --- --- --- --- --- --- --- --- --- --- --- --- --- --- --- --- --- --- --- --- --- --- --- --- --- --- --- --- --- --- --- --- --- --- --- --- --- --- --- --- --- --- --- --- --- --- --- --- --- --- --- --- --- --- --- --- --- --- --- --- --- --- --- --- --- --- --- --- --- --- --- --- --- --- --- --- --- --- --- --- --- --- --- --- --- --- --- --- --- --- --- --- --- --- --- --- --- --- --- --- --- --- --- --- --- --- --- --- --- --- --- --- --- --- --- --- --- --- --- --- --- --- --- --- --- --- --- --- --- --- --- --- --- --- --- --- --- --- --- --- --- --- --- --- --- --- --- --- --- --- --- --- --- --- --- --- --- --- --- --- --- --- --- --- --- --- --- --- --- --- --- --- --- --- --- --- --- --- --- --- --- --- --- --- --- --- --t ggt cag tgt ata ata aat tat atg tag gag gag gaa tat atg atg atg ttc aaa cta ata cat ctg aaa cat acg ata aag aaa aa

>MN702449.1_Monkeypox_virus_strain_B2_contig_SPADES

tgt tat tgt agt atg ata ata tca aaa aga tgg ata taa aga at- --- --- --- --- --- --- --- --- --- --- --- --- --- --- --- --- --- --- --- --- --- --- --- --- --- --- --- --- --- --- --- --- --- --- --- --- --- --- --- --- --- --- --- --- --- --- --- --- --- --- --- --- --- --- --- --- --- --- --- --- --- --- --- --- --- --- --- --- --- --- --- --- --- --- --- --- --- --- --- --- --- --- --- --- --- --- --- --- --- --- --- --- --- --- --- --- --- --- --- --- --- --- --- --- --- --- --- --- --- --- --- --- --- --- --- --- --- --- --- --- --- --- --- --- --- --- --- --- --- --- --- --- --- --- --- --- --- --- --- --- --- --- --- --- --- --- --- --- --- --- --- --- --- --- --- --- --- --- --- --- --- --- --- --- --- --- --- --- --- --- --- --- --- --- --- --- --- --- --- --- --- --- --- --- --- --- --- --- --- --- --- --- --- --- --- --- --- --- --- --- --- --- --- --- --- --- --- --- --- --- --- --- --- --- --- --- --- --- --- --- --- --- --- --- --- --- --- --- --- --- --- --- --- --- --- --- --- --- --- --- --- --- --- --- --- --- --- --- --- --- --- --- --- --- --- --- --- --- --- --- --- --- --- --- --- --- --- --- --- --- --- --- --- --- --- --- --- --- --- --- --- --- --- --- --- --- --- --- --- --- --- --- --- --- --- --- --- --- --- --- --- --- --- --- --- --- --- --- --- --- --- --- --- --- --- --- --- --- --- --- --- --- --- --- --- --- --- --- --- --- --- --- --- --- --- --- --- --- --- --- --- --- --- --- --- --- --- --- --- --- --- --- --- --- --- --- --- --- --- --- --- --- --- --- --- --- --- --- --- --- --- --- --- --- --- --- --- --- --- --- --- --- --- --- --- --- --- --- --- --- --- --- --- --- --- --- --- --- --- --- --- --- --- --- --- --- --- --- --- --- --- --- --- --- --- --- --- --- --- --- --- --- --- --- --- --- --- --- --- --- --- --- --- --- --- --- --- --- --- --- --- --- --- --- --- --- --- --- --- --- --- --- --- --- --- --- --- --- --- --- --- --- --- --- --- --- --- --- --- --- --- --- --- --- --- --- --- --- --- --- --- --- --- --- --- --- --- --- --- --- --- --- --- --- --- --- --- --- --- --- --- --- --- --- --- --- --- --- --- --- --- --- --- --- --- --- --- --- --- --- --- --- --- --- --- --- --- --- --- --- --- --- --- --- --- --- --- --- --- --- --- --- --- --- --- --- --- --- --- --- --- --- --- --- --- --- --- --- --- --- --- --- --- --- --- --- --- --- --- --- --- --- --- --- --- --- --- --- --- --- --- --- --- --- --- --- --- --- --- --- --- --- --- --- --- --- --- --- --- --- --- --- --- --- --- --- --- --- --- --- --- --- --- --- --- --- --- --- --- --- --- --- --- --- --- --- --- --- --- --- --- --- --- --- --- --- --- --- --- --- --- --- --- --- --- --- --- --- --- --- --- --- --- --- --- --- --- --- --- --- --- --- --- --- --- --- --- --- --- --- --- --- --- --- --- --- --- --- --- --- --- --- --- --- --- --- --- --- --- --- --- --- --- --- --- --- --- --- --- --- --- --- --- --- --- --- --- --- --- --- --- --- --- --- --- --- --- --- --- --- --- --- --- --- --- --- --- --- --- --- --- --- --- --- --- --- --- --- --- --- --- --- --- --- --- --- --- --- --- --- --- --- --- --- --- --- --- --- --- --t ggt cag tgt ata ata aat tat atg tag gag gag gaa tat ctg atg atg ttc aaa cta ata cat ctg aaa cat acg ata aag aaa aa

>MN702446.1_Monkeypox_virus_strain_38c_contig_SPADES

tgt tat tgt agt atg ata ata tca aaa aga tgg ata taa aga at- --- --- --- --- --- --- --- --- --- --- --- --- --- --- --- --- --- --- --- --- --- --- --- --- --- --- --- --- --- --- --- --- --- --- --- --- --- --- --- --- --- --- --- --- --- --- --- --- --- --- --- --- --- --- --- --- --- --- --- --- --- --- --- --- --- --- --- --- --- --- --- --- --- --- --- --- --- --- --- --- --- --- --- --- --- --- --- --- --- --- --- --- --- --- --- --- --- --- --- --- --- --- --- --- --- --- --- --- --- --- --- --- --- --- --- --- --- --- --- --- --- --- --- --- --- --- --- --- --- --- --- --- --- --- --- --- --- --- --- --- --- --- --- --- --- --- --- --- --- --- --- --- --- --- --- --- --- --- --- --- --- --- --- --- --- --- --- --- --- --- --- --- --- --- --- --- --- --- --- --- --- --- --- --- --- --- --- --- --- --- --- --- --- --- --- --- --- --- --- --- --- --- --- --- --- --- --- --- --- --- --- --- --- --- --- --- --- --- --- --- --- --- --- --- --- --- --- --- --- --- --- --- --- --- --- --- --- --- --- --- --- --- --- --- --- --- --- --- --- --- --- --- --- --- --- --- --- --- --- --- --- --- --- --- --- --- --- --- --- --- --- --- --- --- --- --- --- --- --- --- --- --- --- --- --- --- --- --- --- --- --- --- --- --- --- --- --- --- --- --- --- --- --- --- --- --- --- --- --- --- --- --- --- --- --- --- --- --- --- --- --- --- --- --- --- --- --- --- --- --- --- --- --- --- --- --- --- --- --- --- --- --- --- --- --- --- --- --- --- --- --- --- --- --- --- --- --- --- --- --- --- --- --- --- --- --- --- --- --- --- --- --- --- --- --- --- --- --- --- --- --- --- --- --- --- --- --- --- --- --- --- --- --- --- --- --- --- --- --- --- --- --- --- --- --- --- --- --- --- --- --- --- --- --- --- --- --- --- --- --- --- --- --- --- --- --- --- --- --- --- --- --- --- --- --- --- --- --- --- --- --- --- --- --- --- --- --- --- --- --- --- --- --- --- --- --- --- --- --- --- --- --- --- --- --- --- --- --- --- --- --- --- --- --- --- --- --- --- --- --- --- --- --- --- --- --- --- --- --- --- --- --- --- --- --- --- --- --- --- --- --- --- --- --- --- --- --- --- --- --- --- --- --- --- --- --- --- --- --- --- --- --- --- --- --- --- --- --- --- --- --- --- --- --- --- --- --- --- --- --- --- --- --- --- --- --- --- --- --- --- --- --- --- --- --- --- --- --- --- --- --- --- --- --- --- --- --- --- --- --- --- --- --- --- --- --- --- --- --- --- --- --- --- --- --- --- --- --- --- --- --- --- --- --- --- --- --- --- --- --- --- --- --- --- --- --- --- --- --- --- --- --- --- --- --- --- --- --- --- --- --- --- --- --- --- --- --- --- --- --- --- --- --- --- --- --- --- --- --- --- --- --- --- --- --- --- --- --- --- --- --- --- --- --- --- --- --- --- --- --- --- --- --- --- --- --- --- --- --- --- --- --- --- --- --- --- --- --- --- --- --- --- --- --- --- --- --- --- --- --- --- --- --- --- --- --- --- --- --- --- --- --- --- --- --- --- --- --- --- --- --- --- --- --- --- --- --- --- --- --- --- --- --- --- --- --- --- --- --- --- --- --- --- --- --- --- --- --- --- --- --- --- --- --- --- --- --- --- --- --- --- --- --- --- --- --- --- --- --- --t ggt cag tgt ata ata aat tat atg tag gag gag gaa tat ctg atg atg ttc aaa cta ata cat ctg aaa cat acg ata aag aaa aa

>JX878418.1_Monkeypox_virus_isolate_DRC_07-0120_complete_genome

tgt tat tgt agt atg ata ata tca aaa aga tgg ata taa aga at- --- --- --- --- --- --- --- --- --- --- --- --- --- --- --- --- --- --- --- --- --- --- --- --- --- --- --- --- --- --- --- --- --- --- --- --- --- --- --- --- --- --- --- --- --- --- --- --- --- --- --- --- --- --- --- --- --- --- --- --- --- --- --- --- --- --- --- --- --- --- --- --- --- --- --- --- --- --- --- --- --- --- --- --- --- --- --- --- --- --- --- --- --- --- --- --- --- --- --- --- --- --- --- --- --- --- --- --- --- --- --- --- --- --- --- --- --- --- --- --- --- --- --- --- --- --- --- --- --- --- --- --- --- --- --- --- --- --- --- --- --- --- --- --- --- --- --- --- --- --- --- --- --- --- --- --- --- --- --- --- --- --- --- --- --- --- --- --- --- --- --- --- --- --- --- --- --- --- --- --- --- --- --- --- --- --- --- --- --- --- --- --- --- --- --- --- --- --- --- --- --- --- --- --- --- --- --- --- --- --- --- --- --- --- --- --- --- --- --- --- --- --- --- --- --- --- --- --- --- --- --- --- --- --- --- --- --- --- --- --- --- --- --- --- --- --- --- --- --- --- --- --- --- --- --- --- --- --- --- --- --- --- --- --- --- --- --- --- --- --- --- --- --- --- --- --- --- --- --- --- --- --- --- --- --- --- --- --- --- --- --- --- --- --- --- --- --- --- --- --- --- --- --- --- --- --- --- --- --- --- --- --- --- --- --- --- --- --- --- --- --- --- --- --- --- --- --- --- --- --- --- --- --- --- --- --- --- --- --- --- --- --- --- --- --- --- --- --- --- --- --- --- --- --- --- --- --- --- --- --- --- --- --- --- --- --- --- --- --- --- --- --- --- --- --- --- --- --- --- --- --- --- --- --- --- --- --- --- --- --- --- --- --- --- --- --- --- --- --- --- --- --- --- --- --- --- --- --- --- --- --- --- --- --- --- --- --- --- --- --- --- --- --- --- --- --- --- --- --- --- --- --- --- --- --- --- --- --- --- --- --- --- --- --- --- --- --- --- --- --- --- --- --- --- --- --- --- --- --- --- --- --- --- --- --- --- --- --- --- --- --- --- --- --- --- --- --- --- --- --- --- --- --- --- --- --- --- --- --- --- --- --- --- --- --- --- --- --- --- --- --- --- --- --- --- --- --- --- --- --- --- --- --- --- --- --- --- --- --- --- --- --- --- --- --- --- --- --- --- --- --- --- --- --- --- --- --- --- --- --- --- --- --- --- --- --- --- --- --- --- --- --- --- --- --- --- --- --- --- --- --- --- --- --- --- --- --- --- --- --- --- --- --- --- --- --- --- --- --- --- --- --- --- --- --- --- --- --- --- --- --- --- --- --- --- --- --- --- --- --- --- --- --- --- --- --- --- --- --- --- --- --- --- --- --- --- --- --- --- --- --- --- --- --- --- --- --- --- --- --- --- --- --- --- --- --- --- --- --- --- --- --- --- --- --- --- --- --- --- --- --- --- --- --- --- --- --- --- --- --- --- --- --- --- --- --- --- --- --- --- --- --- --- --- --- --- --- --- --- --- --- --- --- --- --- --- --- --- --- --- --- --- --- --- --- --- --- --- --- --- --- --- --- --- --- --- --- --- --- --- --- --- --- --- --- --- --- --- --- --- --- --- --- --- --- --- --- --- --- --- --- --- --- --- --- --- --- --- --- --- --- --- --- --- --- --- --- --- --- --- --- --- --- --- --- --- --- --- --- --t ggt cag tgt ata ata aat tat atg tag gag gag gaa tat ctg atg atg ttc aaa cta ata cat ctg aaa cat acg ata aag aaa aa

>MN702445.1_Monkeypox_virus_strain_A4_contig_SPADES

tgt tat tgt agt atg ata ata tca aaa aga tgg ata taa aga at- --- --- --- --- --- --- --- --- --- --- --- --- --- --- --- --- --- --- --- --- --- --- --- --- --- --- --- --- --- --- --- --- --- --- --- --- --- --- --- --- --- --- --- --- --- --- --- --- --- --- --- --- --- --- --- --- --- --- --- --- --- --- --- --- --- --- --- --- --- --- --- --- --- --- --- --- --- --- --- --- --- --- --- --- --- --- --- --- --- --- --- --- --- --- --- --- --- --- --- --- --- --- --- --- --- --- --- --- --- --- --- --- --- --- --- --- --- --- --- --- --- --- --- --- --- --- --- --- --- --- --- --- --- --- --- --- --- --- --- --- --- --- --- --- --- --- --- --- --- --- --- --- --- --- --- --- --- --- --- --- --- --- --- --- --- --- --- --- --- --- --- --- --- --- --- --- --- --- --- --- --- --- --- --- --- --- --- --- --- --- --- --- --- --- --- --- --- --- --- --- --- --- --- --- --- --- --- --- --- --- --- --- --- --- --- --- --- --- --- --- --- --- --- --- --- --- --- --- --- --- --- --- --- --- --- --- --- --- --- --- --- --- --- --- --- --- --- --- --- --- --- --- --- --- --- --- --- --- --- --- --- --- --- --- --- --- --- --- --- --- --- --- --- --- --- --- --- --- --- --- --- --- --- --- --- --- --- --- --- --- --- --- --- --- --- --- --- --- --- --- --- --- --- --- --- --- --- --- --- --- --- --- --- --- --- --- --- --- --- --- --- --- --- --- --- --- --- --- --- --- --- --- --- --- --- --- --- --- --- --- --- --- --- --- --- --- --- --- --- --- --- --- --- --- --- --- --- --- --- --- --- --- --- --- --- --- --- --- --- --- --- --- --- --- --- --- --- --- --- --- --- --- --- --- --- --- --- --- --- --- --- --- --- --- --- --- --- --- --- --- --- --- --- --- --- --- --- --- --- --- --- --- --- --- --- --- --- --- --- --- --- --- --- --- --- --- --- --- --- --- --- --- --- --- --- --- --- --- --- --- --- --- --- --- --- --- --- --- --- --- --- --- --- --- --- --- --- --- --- --- --- --- --- --- --- --- --- --- --- --- --- --- --- --- --- --- --- --- --- --- --- --- --- --- --- --- --- --- --- --- --- --- --- --- --- --- --- --- --- --- --- --- --- --- --- --- --- --- --- --- --- --- --- --- --- --- --- --- --- --- --- --- --- --- --- --- --- --- --- --- --- --- --- --- --- --- --- --- --- --- --- --- --- --- --- --- --- --- --- --- --- --- --- --- --- --- --- --- --- --- --- --- --- --- --- --- --- --- --- --- --- --- --- --- --- --- --- --- --- --- --- --- --- --- --- --- --- --- --- --- --- --- --- --- --- --- --- --- --- --- --- --- --- --- --- --- --- --- --- --- --- --- --- --- --- --- --- --- --- --- --- --- --- --- --- --- --- --- --- --- --- --- --- --- --- --- --- --- --- --- --- --- --- --- --- --- --- --- --- --- --- --- --- --- --- --- --- --- --- --- --- --- --- --- --- --- --- --- --- --- --- --- --- --- --- --- --- --- --- --- --- --- --- --- --- --- --- --- --- --- --- --- --- --- --- --- --- --- --- --- --- --- --- --- --- --- --- --- --- --- --- --- --- --- --- --- --- --- --- --- --- --- --- --- --- --- --- --- --- --- --- --- --- --- --- --- --- --- --- --- --- --- --- --- --- --- --- --- --- --- --- --- --- --- --- --- --- --- --- --t ggt cag tgt ata ata aat tat atg tag gag gag gaa tat ctg atg atg ttc aaa cta ata cat ctg aaa cat acg ata aag aaa aa

>MN702444.1_Monkeypox_virus_strain_A5_contig_SPADES

tgt tat tgt agt atg ata ata tca aaa aga tgg ata taa aga at- --- --- --- --- --- --- --- --- --- --- --- --- --- --- --- --- --- --- --- --- --- --- --- --- --- --- --- --- --- --- --- --- --- --- --- --- --- --- --- --- --- --- --- --- --- --- --- --- --- --- --- --- --- --- --- --- --- --- --- --- --- --- --- --- --- --- --- --- --- --- --- --- --- --- --- --- --- --- --- --- --- --- --- --- --- --- --- --- --- --- --- --- --- --- --- --- --- --- --- --- --- --- --- --- --- --- --- --- --- --- --- --- --- --- --- --- --- --- --- --- --- --- --- --- --- --- --- --- --- --- --- --- --- --- --- --- --- --- --- --- --- --- --- --- --- --- --- --- --- --- --- --- --- --- --- --- --- --- --- --- --- --- --- --- --- --- --- --- --- --- --- --- --- --- --- --- --- --- --- --- --- --- --- --- --- --- --- --- --- --- --- --- --- --- --- --- --- --- --- --- --- --- --- --- --- --- --- --- --- --- --- --- --- --- --- --- --- --- --- --- --- --- --- --- --- --- --- --- --- --- --- --- --- --- --- --- --- --- --- --- --- --- --- --- --- --- --- --- --- --- --- --- --- --- --- --- --- --- --- --- --- --- --- --- --- --- --- --- --- --- --- --- --- --- --- --- --- --- --- --- --- --- --- --- --- --- --- --- --- --- --- --- --- --- --- --- --- --- --- --- --- --- --- --- --- --- --- --- --- --- --- --- --- --- --- --- --- --- --- --- --- --- --- --- --- --- --- --- --- --- --- --- --- --- --- --- --- --- --- --- --- --- --- --- --- --- --- --- --- --- --- --- --- --- --- --- --- --- --- --- --- --- --- --- --- --- --- --- --- --- --- --- --- --- --- --- --- --- --- --- --- --- --- --- --- --- --- --- --- --- --- --- --- --- --- --- --- --- --- --- --- --- --- --- --- --- --- --- --- --- --- --- --- --- --- --- --- --- --- --- --- --- --- --- --- --- --- --- --- --- --- --- --- --- --- --- --- --- --- --- --- --- --- --- --- --- --- --- --- --- --- --- --- --- --- --- --- --- --- --- --- --- --- --- --- --- --- --- --- --- --- --- --- --- --- --- --- --- --- --- --- --- --- --- --- --- --- --- --- --- --- --- --- --- --- --- --- --- --- --- --- --- --- --- --- --- --- --- --- --- --- --- --- --- --- --- --- --- --- --- --- --- --- --- --- --- --- --- --- --- --- --- --- --- --- --- --- --- --- --- --- --- --- --- --- --- --- --- --- --- --- --- --- --- --- --- --- --- --- --- --- --- --- --- --- --- --- --- --- --- --- --- --- --- --- --- --- --- --- --- --- --- --- --- --- --- --- --- --- --- --- --- --- --- --- --- --- --- --- --- --- --- --- --- --- --- --- --- --- --- --- --- --- --- --- --- --- --- --- --- --- --- --- --- --- --- --- --- --- --- --- --- --- --- --- --- --- --- --- --- --- --- --- --- --- --- --- --- --- --- --- --- --- --- --- --- --- --- --- --- --- --- --- --- --- --- --- --- --- --- --- --- --- --- --- --- --- --- --- --- --- --- --- --- --- --- --- --- --- --- --- --- --- --- --- --- --- --- --- --- --- --- --- --- --- --- --- --- --- --- --- --- --- --- --- --- --- --- --- --- --- --- --- --- --- --- --- --- --- --- --- --- --- --- --- --- --- --- --- --- --- --- --- --- --- --- --- --- --- --- --- --- --- --- --- --- --- --- --- --t ggt cag tgt ata ata aat tat atg tag gag gag gaa tat ctg atg atg ttc aaa cta ata cat ctg aaa cat acg ata aag aaa aa

>KJ642613.1_Monkeypox_virus_strain_Congo_8_complete_genome

tgt tat tgt agt atg ata ata tca aaa aga tgg ata taa aga at- --- --- --- --- --- --- --- --- --- --- --- --- --- --- --- --- --- --- --- --- --- --- --- --- --- --- --- --- --- --- --- --- --- --- --- --- --- --- --- --- --- --- --- --- --- --- --- --- --- --- --- --- --- --- --- --- --- --- --- --- --- --- --- --- --- --- --- --- --- --- --- --- --- --- --- --- --- --- --- --- --- --- --- --- --- --- --- --- --- --- --- --- --- --- --- --- --- --- --- --- --- --- --- --- --- --- --- --- --- --- --- --- --- --- --- --- --- --- --- --- --- --- --- --- --- --- --- --- --- --- --- --- --- --- --- --- --- --- --- --- --- --- --- --- --- --- --- --- --- --- --- --- --- --- --- --- --- --- --- --- --- --- --- --- --- --- --- --- --- --- --- --- --- --- --- --- --- --- --- --- --- --- --- --- --- --- --- --- --- --- --- --- --- --- --- --- --- --- --- --- --- --- --- --- --- --- --- --- --- --- --- --- --- --- --- --- --- --- --- --- --- --- --- --- --- --- --- --- --- --- --- --- --- --- --- --- --- --- --- --- --- --- --- --- --- --- --- --- --- --- --- --- --- --- --- --- --- --- --- --- --- --- --- --- --- --- --- --- --- --- --- --- --- --- --- --- --- --- --- --- --- --- --- --- --- --- --- --- --- --- --- --- --- --- --- --- --- --- --- --- --- --- --- --- --- --- --- --- --- --- --- --- --- --- --- --- --- --- --- --- --- --- --- --- --- --- --- --- --- --- --- --- --- --- --- --- --- --- --- --- --- --- --- --- --- --- --- --- --- --- --- --- --- --- --- --- --- --- --- --- --- --- --- --- --- --- --- --- --- --- --- --- --- --- --- --- --- --- --- --- --- --- --- --- --- --- --- --- --- --- --- --- --- --- --- --- --- --- --- --- --- --- --- --- --- --- --- --- --- --- --- --- --- --- --- --- --- --- --- --- --- --- --- --- --- --- --- --- --- --- --- --- --- --- --- --- --- --- --- --- --- --- --- --- --- --- --- --- --- --- --- --- --- --- --- --- --- --- --- --- --- --- --- --- --- --- --- --- --- --- --- --- --- --- --- --- --- --- --- --- --- --- --- --- --- --- --- --- --- --- --- --- --- --- --- --- --- --- --- --- --- --- --- --- --- --- --- --- --- --- --- --- --- --- --- --- --- --- --- --- --- --- --- --- --- --- --- --- --- --- --- --- --- --- --- --- --- --- --- --- --- --- --- --- --- --- --- --- --- --- --- --- --- --- --- --- --- --- --- --- --- --- --- --- --- --- --- --- --- --- --- --- --- --- --- --- --- --- --- --- --- --- --- --- --- --- --- --- --- --- --- --- --- --- --- --- --- --- --- --- --- --- --- --- --- --- --- --- --- --- --- --- --- --- --- --- --- --- --- --- --- --- --- --- --- --- --- --- --- --- --- --- --- --- --- --- --- --- --- --- --- --- --- --- --- --- --- --- --- --- --- --- --- --- --- --- --- --- --- --- --- --- --- --- --- --- --- --- --- --- --- --- --- --- --- --- --- --- --- --- --- --- --- --- --- --- --- --- --- --- --- --- --- --- --- --- --- --- --- --- --- --- --- --- --- --- --- --- --- --- --- --- --- --- --- --- --- --- --- --- --- --- --- --- --- --- --- --- --- --- --- --- --- --- --- --- --- --- --- --- --- --- --- --- --- --- --- --- --- --- --- --- --- --- --- --- --- --- --- --t ggt cag tgt ata ata aat tat atg tag gag gag gaa tat ctg atg atg ttc aaa cta ata cat ctg aaa cat acg ata aag aaa aa

>HM172544.1_Monkeypox_virus_strain_Zaire_1979-005_complete_genome

tgt tat tgt agt atg ata ata tca aaa aga tgg ata taa aga at- --- --- --- --- --- --- --- --- --- --- --- --- --- --- --- --- --- --- --- --- --- --- --- --- --- --- --- --- --- --- --- --- --- --- --- --- --- --- --- --- --- --- --- --- --- --- --- --- --- --- --- --- --- --- --- --- --- --- --- --- --- --- --- --- --- --- --- --- --- --- --- --- --- --- --- --- --- --- --- --- --- --- --- --- --- --- --- --- --- --- --- --- --- --- --- --- --- --- --- --- --- --- --- --- --- --- --- --- --- --- --- --- --- --- --- --- --- --- --- --- --- --- --- --- --- --- --- --- --- --- --- --- --- --- --- --- --- --- --- --- --- --- --- --- --- --- --- --- --- --- --- --- --- --- --- --- --- --- --- --- --- --- --- --- --- --- --- --- --- --- --- --- --- --- --- --- --- --- --- --- --- --- --- --- --- --- --- --- --- --- --- --- --- --- --- --- --- --- --- --- --- --- --- --- --- --- --- --- --- --- --- --- --- --- --- --- --- --- --- --- --- --- --- --- --- --- --- --- --- --- --- --- --- --- --- --- --- --- --- --- --- --- --- --- --- --- --- --- --- --- --- --- --- --- --- --- --- --- --- --- --- --- --- --- --- --- --- --- --- --- --- --- --- --- --- --- --- --- --- --- --- --- --- --- --- --- --- --- --- --- --- --- --- --- --- --- --- --- --- --- --- --- --- --- --- --- --- --- --- --- --- --- --- --- --- --- --- --- --- --- --- --- --- --- --- --- --- --- --- --- --- --- --- --- --- --- --- --- --- --- --- --- --- --- --- --- --- --- --- --- --- --- --- --- --- --- --- --- --- --- --- --- --- --- --- --- --- --- --- --- --- --- --- --- --- --- --- --- --- --- --- --- --- --- --- --- --- --- --- --- --- --- --- --- --- --- --- --- --- --- --- --- --- --- --- --- --- --- --- --- --- --- --- --- --- --- --- --- --- --- --- --- --- --- --- --- --- --- --- --- --- --- --- --- --- --- --- --- --- --- --- --- --- --- --- --- --- --- --- --- --- --- --- --- --- --- --- --- --- --- --- --- --- --- --- --- --- --- --- --- --- --- --- --- --- --- --- --- --- --- --- --- --- --- --- --- --- --- --- --- --- --- --- --- --- --- --- --- --- --- --- --- --- --- --- --- --- --- --- --- --- --- --- --- --- --- --- --- --- --- --- --- --- --- --- --- --- --- --- --- --- --- --- --- --- --- --- --- --- --- --- --- --- --- --- --- --- --- --- --- --- --- --- --- --- --- --- --- --- --- --- --- --- --- --- --- --- --- --- --- --- --- --- --- --- --- --- --- --- --- --- --- --- --- --- --- --- --- --- --- --- --- --- --- --- --- --- --- --- --- --- --- --- --- --- --- --- --- --- --- --- --- --- --- --- --- --- --- --- --- --- --- --- --- --- --- --- --- --- --- --- --- --- --- --- --- --- --- --- --- --- --- --- --- --- --- --- --- --- --- --- --- --- --- --- --- --- --- --- --- --- --- --- --- --- --- --- --- --- --- --- --- --- --- --- --- --- --- --- --- --- --- --- --- --- --- --- --- --- --- --- --- --- --- --- --- --- --- --- --- --- --- --- --- --- --- --- --- --- --- --- --- --- --- --- --- --- --- --- --- --- --- --- --- --- --- --- --- --- --- --- --- --- --- --- --- --- --- --- --- --- --- --- --- --- --- --- --- --- --- --- --- --- --- --- --- --- --- --- --t ggt cag tgt ata ata aat tat atg tag gag gag gaa tat ctg atg atg ttc aaa cta ata cat ctg aaa cat acg ata aag aaa aa

>AF380138.1_Monkeypox_virus_strain_Zaire-96-I-16_complete_genome

tgt tat tgt agt atg ata ata tca aaa aga tgg ata taa aga at- --- --- --- --- --- --- --- --- --- --- --- --- --- --- --- --- --- --- --- --- --- --- --- --- --- --- --- --- --- --- --- --- --- --- --- --- --- --- --- --- --- --- --- --- --- --- --- --- --- --- --- --- --- --- --- --- --- --- --- --- --- --- --- --- --- --- --- --- --- --- --- --- --- --- --- --- --- --- --- --- --- --- --- --- --- --- --- --- --- --- --- --- --- --- --- --- --- --- --- --- --- --- --- --- --- --- --- --- --- --- --- --- --- --- --- --- --- --- --- --- --- --- --- --- --- --- --- --- --- --- --- --- --- --- --- --- --- --- --- --- --- --- --- --- --- --- --- --- --- --- --- --- --- --- --- --- --- --- --- --- --- --- --- --- --- --- --- --- --- --- --- --- --- --- --- --- --- --- --- --- --- --- --- --- --- --- --- --- --- --- --- --- --- --- --- --- --- --- --- --- --- --- --- --- --- --- --- --- --- --- --- --- --- --- --- --- --- --- --- --- --- --- --- --- --- --- --- --- --- --- --- --- --- --- --- --- --- --- --- --- --- --- --- --- --- --- --- --- --- --- --- --- --- --- --- --- --- --- --- --- --- --- --- --- --- --- --- --- --- --- --- --- --- --- --- --- --- --- --- --- --- --- --- --- --- --- --- --- --- --- --- --- --- --- --- --- --- --- --- --- --- --- --- --- --- --- --- --- --- --- --- --- --- --- --- --- --- --- --- --- --- --- --- --- --- --- --- --- --- --- --- --- --- --- --- --- --- --- --- --- --- --- --- --- --- --- --- --- --- --- --- --- --- --- --- --- --- --- --- --- --- --- --- --- --- --- --- --- --- --- --- --- --- --- --- --- --- --- --- --- --- --- --- --- --- --- --- --- --- --- --- --- --- --- --- --- --- --- --- --- --- --- --- --- --- --- --- --- --- --- --- --- --- --- --- --- --- --- --- --- --- --- --- --- --- --- --- --- --- --- --- --- --- --- --- --- --- --- --- --- --- --- --- --- --- --- --- --- --- --- --- --- --- --- --- --- --- --- --- --- --- --- --- --- --- --- --- --- --- --- --- --- --- --- --- --- --- --- --- --- --- --- --- --- --- --- --- --- --- --- --- --- --- --- --- --- --- --- --- --- --- --- --- --- --- --- --- --- --- --- --- --- --- --- --- --- --- --- --- --- --- --- --- --- --- --- --- --- --- --- --- --- --- --- --- --- --- --- --- --- --- --- --- --- --- --- --- --- --- --- --- --- --- --- --- --- --- --- --- --- --- --- --- --- --- --- --- --- --- --- --- --- --- --- --- --- --- --- --- --- --- --- --- --- --- --- --- --- --- --- --- --- --- --- --- --- --- --- --- --- --- --- --- --- --- --- --- --- --- --- --- --- --- --- --- --- --- --- --- --- --- --- --- --- --- --- --- --- --- --- --- --- --- --- --- --- --- --- --- --- --- --- --- --- --- --- --- --- --- --- --- --- --- --- --- --- --- --- --- --- --- --- --- --- --- --- --- --- --- --- --- --- --- --- --- --- --- --- --- --- --- --- --- --- --- --- --- --- --- --- --- --- --- --- --- --- --- --- --- --- --- --- --- --- --- --- --- --- --- --- --- --- --- --- --- --- --- --- --- --- --- --- --- --- --- --- --- --- --- --- --- --- --- --- --- --- --- --- --- --- --- --- --- --- --- --- --- --- --- --- --- --- --- --- --- --- --- --- --- --t ggt cag tgt ata ata aat tat atg tag gag gag gaa tat ctg atg atg ttc aaa cta ata cat ctg aaa cat acg ata aag aaa aa

>HQ857562.1_Monkeypox_virus_strain_V79-I-005_complete_genome

tgt tat tgt agt atg ata ata tca aaa aga tgg ata taa aga at- --- --- --- --- --- --- --- --- --- --- --- --- --- --- --- --- --- --- --- --- --- --- --- --- --- --- --- --- --- --- --- --- --- --- --- --- --- --- --- --- --- --- --- --- --- --- --- --- --- --- --- --- --- --- --- --- --- --- --- --- --- --- --- --- --- --- --- --- --- --- --- --- --- --- --- --- --- --- --- --- --- --- --- --- --- --- --- --- --- --- --- --- --- --- --- --- --- --- --- --- --- --- --- --- --- --- --- --- --- --- --- --- --- --- --- --- --- --- --- --- --- --- --- --- --- --- --- --- --- --- --- --- --- --- --- --- --- --- --- --- --- --- --- --- --- --- --- --- --- --- --- --- --- --- --- --- --- --- --- --- --- --- --- --- --- --- --- --- --- --- --- --- --- --- --- --- --- --- --- --- --- --- --- --- --- --- --- --- --- --- --- --- --- --- --- --- --- --- --- --- --- --- --- --- --- --- --- --- --- --- --- --- --- --- --- --- --- --- --- --- --- --- --- --- --- --- --- --- --- --- --- --- --- --- --- --- --- --- --- --- --- --- --- --- --- --- --- --- --- --- --- --- --- --- --- --- --- --- --- --- --- --- --- --- --- --- --- --- --- --- --- --- --- --- --- --- --- --- --- --- --- --- --- --- --- --- --- --- --- --- --- --- --- --- --- --- --- --- --- --- --- --- --- --- --- --- --- --- --- --- --- --- --- --- --- --- --- --- --- --- --- --- --- --- --- --- --- --- --- --- --- --- --- --- --- --- --- --- --- --- --- --- --- --- --- --- --- --- --- --- --- --- --- --- --- --- --- --- --- --- --- --- --- --- --- --- --- --- --- --- --- --- --- --- --- --- --- --- --- --- --- --- --- --- --- --- --- --- --- --- --- --- --- --- --- --- --- --- --- --- --- --- --- --- --- --- --- --- --- --- --- --- --- --- --- --- --- --- --- --- --- --- --- --- --- --- --- --- --- --- --- --- --- --- --- --- --- --- --- --- --- --- --- --- --- --- --- --- --- --- --- --- --- --- --- --- --- --- --- --- --- --- --- --- --- --- --- --- --- --- --- --- --- --- --- --- --- --- --- --- --- --- --- --- --- --- --- --- --- --- --- --- --- --- --- --- --- --- --- --- --- --- --- --- --- --- --- --- --- --- --- --- --- --- --- --- --- --- --- --- --- --- --- --- --- --- --- --- --- --- --- --- --- --- --- --- --- --- --- --- --- --- --- --- --- --- --- --- --- --- --- --- --- --- --- --- --- --- --- --- --- --- --- --- --- --- --- --- --- --- --- --- --- --- --- --- --- --- --- --- --- --- --- --- --- --- --- --- --- --- --- --- --- --- --- --- --- --- --- --- --- --- --- --- --- --- --- --- --- --- --- --- --- --- --- --- --- --- --- --- --- --- --- --- --- --- --- --- --- --- --- --- --- --- --- --- --- --- --- --- --- --- --- --- --- --- --- --- --- --- --- --- --- --- --- --- --- --- --- --- --- --- --- --- --- --- --- --- --- --- --- --- --- --- --- --- --- --- --- --- --- --- --- --- --- --- --- --- --- --- --- --- --- --- --- --- --- --- --- --- --- --- --- --- --- --- --- --- --- --- --- --- --- --- --- --- --- --- --- --- --- --- --- --- --- --- --- --- --- --- --- --- --- --- --- --- --- --- --- --- --- --- --- --- --- --- --- --- --- --- --- --- --- --- --- --- --- --- --- --t ggt cag tgt ata ata aat tat atg tag gag gag gaa tat ctg atg atg ttc aaa cta ata cat ctg aaa cat acg ata aag aaa aa

>HQ857563.1_Monkeypox_virus_strain_D14L_knockout_complete_genome

tgt tat tgt agt atg ata ata tca aaa aga tgg ata taa aga at- --- --- --- --- --- --- --- --- --- --- --- --- --- --- --- --- --- --- --- --- --- --- --- --- --- --- --- --- --- --- --- --- --- --- --- --- --- --- --- --- --- --- --- --- --- --- --- --- --- --- --- --- --- --- --- --- --- --- --- --- --- --- --- --- --- --- --- --- --- --- --- --- --- --- --- --- --- --- --- --- --- --- --- --- --- --- --- --- --- --- --- --- --- --- --- --- --- --- --- --- --- --- --- --- --- --- --- --- --- --- --- --- --- --- --- --- --- --- --- --- --- --- --- --- --- --- --- --- --- --- --- --- --- --- --- --- --- --- --- --- --- --- --- --- --- --- --- --- --- --- --- --- --- --- --- --- --- --- --- --- --- --- --- --- --- --- --- --- --- --- --- --- --- --- --- --- --- --- --- --- --- --- --- --- --- --- --- --- --- --- --- --- --- --- --- --- --- --- --- --- --- --- --- --- --- --- --- --- --- --- --- --- --- --- --- --- --- --- --- --- --- --- --- --- --- --- --- --- --- --- --- --- --- --- --- --- --- --- --- --- --- --- --- --- --- --- --- --- --- --- --- --- --- --- --- --- --- --- --- --- --- --- --- --- --- --- --- --- --- --- --- --- --- --- --- --- --- --- --- --- --- --- --- --- --- --- --- --- --- --- --- --- --- --- --- --- --- --- --- --- --- --- --- --- --- --- --- --- --- --- --- --- --- --- --- --- --- --- --- --- --- --- --- --- --- --- --- --- --- --- --- --- --- --- --- --- --- --- --- --- --- --- --- --- --- --- --- --- --- --- --- --- --- --- --- --- --- --- --- --- --- --- --- --- --- --- --- --- --- --- --- --- --- --- --- --- --- --- --- --- --- --- --- --- --- --- --- --- --- --- --- --- --- --- --- --- --- --- --- --- --- --- --- --- --- --- --- --- --- --- --- --- --- --- --- --- --- --- --- --- --- --- --- --- --- --- --- --- --- --- --- --- --- --- --- --- --- --- --- --- --- --- --- --- --- --- --- --- --- --- --- --- --- --- --- --- --- --- --- --- --- --- --- --- --- --- --- --- --- --- --- --- --- --- --- --- --- --- --- --- --- --- --- --- --- --- --- --- --- --- --- --- --- --- --- --- --- --- --- --- --- --- --- --- --- --- --- --- --- --- --- --- --- --- --- --- --- --- --- --- --- --- --- --- --- --- --- --- --- --- --- --- --- --- --- --- --- --- --- --- --- --- --- --- --- --- --- --- --- --- --- --- --- --- --- --- --- --- --- --- --- --- --- --- --- --- --- --- --- --- --- --- --- --- --- --- --- --- --- --- --- --- --- --- --- --- --- --- --- --- --- --- --- --- --- --- --- --- --- --- --- --- --- --- --- --- --- --- --- --- --- --- --- --- --- --- --- --- --- --- --- --- --- --- --- --- --- --- --- --- --- --- --- --- --- --- --- --- --- --- --- --- --- --- --- --- --- --- --- --- --- --- --- --- --- --- --- --- --- --- --- --- --- --- --- --- --- --- --- --- --- --- --- --- --- --- --- --- --- --- --- --- --- --- --- --- --- --- --- --- --- --- --- --- --- --- --- --- --- --- --- --- --- --- --- --- --- --- --- --- --- --- --- --- --- --- --- --- --- --- --- --- --- --- --- --- --- --- --- --- --- --- --- --- --- --- --- --- --- --- --- --- --- --- --- --- --- --- --- --- --- --- --- --- --- --- --- --- --- --t ggt cag tgt ata ata aat tat atg tag gag gag gaa tat ctg atg atg ttc aaa cta ata cat ctg aaa cat acg ata aag aaa aa
