## Supplementary Data 9 for "Genomic regions insertion and deletion in Monkeypox virus causing multi-country outbreak-2022"

>ON674051.1:156355-158618_Monkeypox_virus_isolate_MPXV_USA_2022_FL001_complete_genome

tta ctg act aca --- tgt a-c tat tt- -ta cat tac tac att ggc tac ggc ata tat acc tat ttc gtc act tcc aca cgc tcc ggt aaa cgg gtg tca tgt gac gag gga ga- -at ctt gat aag agg cat aat caa tgt tgt aat ccg atg tcc acc tgg aga att tgc caa ggt cag atg tag agt tgg tag tga taa cac aaa atg tga aca ctg ccc acc tca tac ata tac cgc aat ccc caa tta ttc t-a ata gat gtc atc aat gta gaa aat gcc caa cag gat cat ttg ata agg taa agt gta ccg gaa cac aga ac- --a aat -gt tcg tgt cat cct ggt tgg ta- tac gct act gat tct tca cag act gaa gat tgt cga gat ttg tgt acc aaa aaa gga gat gtc cat gcg gat act ttg gtg gaa tag atg aa- --g gaa atc cta ttt gta aat cgt gtt g-- -tg ttg gtg aat att gcg act acc tac gta att ata gac ttg atc cat ttc ctc cat gca aac tat cta tct aaa tgt aat taa tta --- --- --- --- --- --- --- --- tga ttt --- --- --- -tg atg ata atg tta -cc ata c-a tt- --- -at atc act act tgg tta gtg tgt att att tag tat gga aga cct att aat aa- --- --- -tt act tat ctt ttg acg atc ttg tt- -at aat tat --- --- --- --- aat ata aaa --a tac tta tga --- --- cat agt aac tca t-- --- --- --- aat tgc tg- acg cga taa att cgt aat aat ctg ttt tgt tca aat ttt tat aag ga- --- --a tct aca ggc ata aaa at- aaa aat ata atc tat aat ata c-- tct tac a-- acg cca tca tga at- --a gca gtg aat taa ttg ctg tta tt- atg gat tta gaa ata gtg gac gat ttt gtg ata tta ata tag tta tta atg atg aaa gga taa acg cgc ata gac tca tcc tat ctg gag cct ccg aat att ttt tcc att ctg ttt tcc aat aat ttt atc gat tct aat gaa tac gaa gtt aat cta agt cat tta gat tat caa agt gtt aac tat ttg atc gat tac att tat ggg ata cct ttg agc cta act aac aat aac gtg aaa tat att ctt tca acc gct ga- -tt ttt tta caa att gga tct gtc att act gag tgc gaa aaa tac ata ctt aaa aat ctt tgt ttt aga aac tgt atc gat ttc --- --- tac ata tac gct gat aaa tat aat aac aag aaa ata gaa tta gca tcg ttt aac aca ata tta cga aat att ttg aga ctc atc aac aat gaa aac ttt aaa tac tta aca gag gaa tca atg ata aaa att tta agc gat gat atg tta tat ata aaa aat gag gat ttc acc cca ctg att ctt att aaa tgg tta gag agt aca c-- caa cca tgt acc gtc gag tta ctt aga tgc ctc aga ata tca -tt tct ttc ccc aca agt tat aaa atc act tta tag tca tcg act ggt tag ttc aat cta cga atg tat aac att ctt aaa caa tat agc att ctt gga taa atc att tcc tag ata cca tat cat cga gtt gat atc tat cgg tat aag taa ttc aca tga taa gat ttc cat aaa ctg cta caa tca t-- aaa aaa att cat ggg aaa tga tat ctt cac gta gat ata ggt gta gtt tcg cag tga ccg tcc tgg ata ata tta tct ata tga tgg gtg gat atg atc agt ccc tgt ata gaa gtt caa agg tta tag cgt aca ata cat gta cta att ctt gga tat atg ata tac cag agc taa aaa tat cat cgt tct aat tgc gga gga gtt gcc aat gac gaa tac att tat tgt ata ggc ggt ata cgc gat cag gag tca tcg ttg ata tct agt atc gat aga tgg aag cca tca aaa cca tat tgg cag aag tat gct aaa atg tgc gaa cca aaa tgt gat atg gg- gtt gcg att tta aac gga tta ata tat gtc ata ggt gga gtc gtt aaa ggt gac aca cat ata cca acg cac tag aga gtt tat cag aag atg gat gga tga atc atc aac gtc ttc caa taa aaa tgt cca ata tgt cga cga ttg ttc atg ctg gaa aga ttt ata tat cta gag gtt aca aca ata gta gtg tag tta atg taa tat cga atc t-- agt cct tag cta taa tcc gat ata tga tga atg gac caa att atc atc att aaa tat tcc tag aat taa tcc tgc tct a

>MN648051.1:156584-158847_Monkeypox_virus_strain_Israel_2018

tta ctg act aca --- tgt a-c tat tt- -ta cat tac tac att ggc tac ggc ata tat acc tat ttc gtc act tcc aca cgc tcc ggt aaa cgg gtg tca tgt gac gag gga ga- -at ctt gat aag agg cat aat caa tgt tgt aat ccg atg tcc acc tgg aga att tgc caa ggt cag atg tag agt tgg tag tga taa cac aaa atg tga aca ctg ccc acc tca tac ata tac cgc aat ccc caa tta ttc t-a ata gat gtc atc aat gta gaa aat gcc caa cag gat cat ttg ata agg taa agt gta ccg gaa cac aga ac- --a aat -gt tcg tgt cat cct ggt tgg ta- tac gct act gat tct tca cag act gaa gat tgt cga gat ttg tgt acc aaa aaa gga gat gtc cat gcg gat act ttg gtg gaa tag atg aa- --g gaa atc cta ttt gta aat cgt gtt g-- -tg ttg gtg aat att gcg act acc tac gta att ata gac ttg atc cat ttc ctc cat gca aac tat cta tct aaa tgt aat taa tta --- --- --- --- --- --- --- --- tga ttt --- --- --- -tg atg ata atg tta -cc ata c-a tt- --- -at atc act act tgg tta gtg tgt att att tag tat gga aga cct att aat aa- --- --- -tt act tat ctt ttg acg atc ttg tt- -at aat tat --- --- --- --- aat ata aaa --a tac tta tga --- --- cat agt aac tca t-- --- --- --- aat tgc tg- acg cga taa att cgt aat aat ctg ttt tgt tca aat ttt tat aag ga- --- --a tct aca ggc ata aaa at- aaa aat ata atc tat aat ata c-- tct tac a-- acg cca tca tga at- --a gca gtg aat taa ttg ctg tta tt- atg gat tta gaa ata gtg gac gat ttt gtg ata tta ata tag tta tta atg atg aaa gga taa acg cgc ata gac tca tcc tat ctg gag cct ccg aat att ttt tcc att ctg ttt tcc aat aat ttt atc gat tct aat gaa tac gaa gtt aat cta agt cat tta gat tat caa agt gtt aac tat ttg atc gat tac att tat ggg ata cct ttg agc cta act aac aat aac gtg aaa tat att ctt tca acc gct ga- -tt ttt tta caa att gga tct gtc att act gag tgc gaa aaa tac ata ctt aaa aat ctt tgt ttt aga aac tgt atc gat ttc --- --- tac ata tac gct gat aaa tat aat aac aag aaa ata gaa tta gca tcg ttt aac aca ata tta cga aat att ttg aga ctc atc aac aat gaa aac ttt aaa tac tta aca gag gaa tca atg ata aaa att tta agc gat gat atg tta tat ata aaa aat gag gat ttc acc cca ctg att ctc att aaa tgg tta gag agt aca c-- caa cca tgt acc gtc gag tta ctt aga tgc ctc aga ata tca -tt tct ttc ccc aca agt tat aaa atc act tta tag tca tcg act ggt tag ttc aat cta cga atg tat aac att ctt aaa caa tat agc att ctt gga taa atc att tcc tag ata cca tat cat cga gtt gat atc tat cgg tat aag taa ttc aca tga taa gat ttc cat aaa ctg cta caa tca t-- aaa aaa att cat ggg aaa tga tat ctt cac gta gat ata ggt gta gtt tcg cag tga ccg tcc tgg ata ata tta tct ata tga tgg gtg gat atg atc agt ccc tgt ata gaa gtt caa agg tta tag cgt aca ata cat gta cta att ctt gga tat atg ata tac cag agc taa aaa tat cat cgt tct aat tgc gga gga gtt gcc aat gac gaa tac att tat tgt ata ggc ggt ata cgc gat cag gag tca tcg ttg ata tct agt atc gat aga tgg aag cca tca aaa cca tat tgg cag aag tat gct aaa atg tgc gaa cca aaa tgt gat atg gg- gtt gcg att tta aac gga tta ata tat gtc ata ggt gga gtc gtt aaa ggt gac aca cat ata cca acg cac tag aga gtt tat cag aag atg gat gga tga atc atc aac gtc ttc caa taa aaa tgt cca ata tgt cga cga ttg ttc atg ctg gaa aga ttt ata tat cta gag gtt aca aca ata gta gtg tag tta atg taa tat cga atc t-- agt cct tag cta taa tcc gat ata tga tga atg gac caa att atc atc att aaa tat tcc tag aat taa tcc tgc tct a

>MG693723.1:150044-152307_Monkeypox_virus_isolate_MPXV_Nig_2017_297957_partial_genome

tta ctg act aca --- tgt a-c tat tt- -ta cat tac tac att ggc tac ggc ata tat acc tat ttc gtc act tcc aca cgc tcc ggt aaa cgg gtg tca tgt gac gag gga ga- -at ctt gat aag agg cat aat caa tgt tgt aat ccg atg tcc acc tgg aga att tgc caa ggt cag atg tag agt tgg tag tga taa cac aaa atg tga aca ctg ccc acc tca tac ata tac cgc aat ccc caa tta ttc t-a ata gat gtc atc aat gta gaa aat gcc caa cag gat cat ttg ata agg taa agt gta ccg gaa cac aga ac- --a aat -gt tcg tgt cat cct ggt tgg ta- tac gct act gat tct tca cag act gaa gat tgt cga gat ttg tgt acc aaa aaa gga gat gtc cat gcg gat act ttg gtg gaa tag atg aa- --g gaa atc cta ttt gta aat cgt gtt g-- -tg ttg gtg aat att gcg act acc tac gta att ata gac ttg atc cat ttc ctc cat gca aac tat cta tct aaa tgt aat taa tta --- --- --- --- --- --- --- --- tga ttt --- --- --- -tg atg ata atg tta -cc ata c-a tt- --- -at atc act act tgg tta gtg tgt att att tag tat gga aga cct att aat aa- --- --- -tt act tat ctt ttg acg atc ttg tt- -at aat tat --- --- --- --- aat ata aaa --a tac tta tga --- --- cat agt aac tca t-- --- --- --- aat tgc tg- acg cga taa att cgt aat aat ctg ttt tgt tca aat ttt tat aag ga- --- --a tct aca ggc ata aaa at- aaa aat ata atc tat aat ata c-- tct tac a-- acg cca tca tga at- --a gca gtg aat taa ttg ctg tta tt- atg gat tta gaa ata gtg gac gat ttt gtg ata tta ata tag tta tta atg atg aaa gga taa acg cgc ata gac tca tcc tat ctg gag cct ccg aat att ttt tcc att ctg ttt tcc aat aat ttt atc gat tct aat gaa tac gaa gtt aat cta agt cat tta gat tat caa agt gtt aac tat ttg atc gat tac att tat ggg ata cct ttg agc cta act aac aat aac gtg aaa tat att ctt tca acc gct ga- -tt ttt tta caa att gga tct gtc att act gag tgc gaa aaa tac ata ctt aaa aat ctt tgt ttt aga aac tgt atc gat ttc --- --- tac ata tac gct gat aaa tat aat aac aag aaa ata gaa tta gca tcg ttt aac aca ata tta cga aat att ttg aga ctc atc aac aat gaa aac ttt aaa tac tta aca gag gaa tca atg ata aaa att tta agc gat gat atg tta tat ata aaa aat gag gat ttc acc cca ctg att ctc att aaa tgg tta gag agt aca c-- caa cca tgt acc gtc gag tta ctt aga tgc ctc aga ata tca -tt tct ttc ccc aca agt tat aaa atc act tta tag tca tcg act ggt tag ttc aat cta cga atg tat aac att ctt aaa caa tat agc att ctt gga taa atc att tcc tag ata cca tat cat cga gtt gat atc tat cgg tat aag taa ttc aca tga taa gat ttc cat aaa ctg cta caa tca t-- aaa aaa att cat ggg aaa tga tat ctt cac gta gat ata ggt gta gtt tcg cag tga ccg tcc tgg ata ata tta tct ata tga tgg gtg gat atg atc agt ccc tgt ata gaa gtt caa agg tta tag cgt aca ata cat gta cta att ctt gga tat atg ata tac cag agc taa aaa tat cat cgt tct aat tgc gga gga gtt gcc aat gac gaa tac att tat tgt ata ggc ggt ata cgc gat cag gag tca tcg ttg ata tct agt atc gat aga tgg aag cca tca aaa cca tat tgg cag aag tat gct aaa atg tgc gaa cca aaa tgt gat atg gg- gtt gcg att tta aac gga tta ata tat gtc ata ggt gga gtc gtt aaa ggt gac aca cat ata cca acg cac tag aga gtt tat cag aag atg gat gga tga atc atc aac gtc ttc caa taa aaa tgt cca ata tgt cga cga ttg ttc atg ctg gaa aga ttt ata tat cta gag gtt aca aca ata gta gtg tag tta atg taa tat cga atc t-- agt cct tag cta taa tcc gat ata tga tga atg gac caa att atc atc att aaa tat tcc tag aat taa tcc tgc tct a

>ON682269.2:156360-158623_Monkeypox_virus_isolate_MPXV/Germany/2022/RKI07_complete_genome

tta ctg act aca --- tgt a-c tat tt- -ta cat tac tac att ggc tac ggc ata tat acc tat ttc gtc act tcc aca cgc tcc ggt aaa cgg gtg tca tgt gac gag gga ga- -at ctt gat aag agg cat aat caa tgt tgt aat ccg atg tcc acc tgg aga att tgc caa ggt cag atg tag agt tgg tag tga taa cac aaa atg tga aca ctg ccc acc tca tac ata tac cgc aat ccc caa tta ttc t-a ata gat gtc atc aat gta gaa aat gcc caa cag gat cat ttg ata agg taa agt gta ccg gaa cac aga ac- --a aat -gt tcg tgt cat cct ggt tgg ta- tac gct act gat tct tca cag act gaa gat tgt cga gat ttg tgt acc aaa aaa gga gat gtc cat gcg gat act ttg gtg gaa tag atg aa- --g gaa atc cta ttt gta aat cgt gtt g-- -tg ttg gtg aat att gcg act acc tac gta att ata gac ttg atc cat ttc ctc cat gca aac tat cta tct aaa tgt aat taa tta --- --- --- --- --- --- --- --- tga ttt --- --- --- -tg atg ata atg tta -cc ata c-a tt- --- -at atc act act tgg tta gtg tgt att att tag tat gga aga cct att aat aa- --- --- -tt act tat ctt ttg acg atc ttg tt- -at aat tat --- --- --- --- aat ata aaa --a tac tta tga --- --- cat agt aac tca t-- --- --- --- aat tgc tg- acg cga taa att cgt aat aat ctg ttt tgt tca aat ttt tat aag ga- --- --a tct aca ggc ata aaa at- aaa aat ata atc tat aat ata c-- tct tac a-- acg cca tca tga at- --a gca gtg aat taa ttg ctg tta tt- atg gat tta gaa ata gtg gac gat ttt gtg ata tta ata tag tta tta atg atg aaa gga taa acg cgc ata gac tca tcc tat ctg gag cct ccg aat att ttt tcc att ctg ttt tcc aat aat ttt atc gat tct aat gaa tac gaa gtt aat cta agt cat tta gat tat caa agt gtt aac tat ttg atc gat tac att tat ggg ata cct ttg agc cta act aac aat aac gtg aaa tat att ctt tca acc gct ga- -tt ttt tta caa att gga tct gtc att act gag tgc gaa aaa tac ata ctt aaa aat ctt tgt ttt aga aac tgt atc gat ttc --- --- tac ata tac gct gat aaa tat aat aac aag aaa ata gaa tta gca tcg ttt aac aca ata tta cga aat att ttg aga ctc atc aac aat gaa aac ttt aaa tac tta aca gag gaa tca atg ata aaa att tta agc gat gat atg tta tat ata aaa aat gag gat ttc acc cca ctg att ctc att aaa tgg tta gag agt aca c-- caa cca tgt acc gtc gag tta ctt aga tgc ctc aga ata tca -tt tct ttc ccc aca agt tat aaa atc act tta tag tca tcg act ggt tag ttc aat cta cga atg tat aac att ctt aaa caa tat agc att ctt gga taa atc att tcc tag ata cca tat cat cga gtt gat atc tat cgg tat aag taa ttc aca tga taa gat ttc cat aaa ctg cta caa tca t-- aaa aaa att cat ggg aaa tga tat ctt cac gta gat ata ggt gta gtt tcg cag tga ccg tcc tgg ata ata tta tct ata tga tgg gtg gat atg atc agt ccc tgt ata gaa gtt caa agg tta tag cgt aca ata cat gta cta att ctt gga tat atg ata tac cag agc taa aaa tat cat cgt tct aat tgc gga gga gtt gcc aat gac gaa tac att tat tgt ata ggc ggt ata cgc gat cag gag tca tcg ttg ata tct agt atc gat aga tgg aag cca tca aaa cca tat tgg cag aag tat gct aaa atg tgc gaa cca aaa tgt gat atg gg- gtt gcg att tta aac gga tta ata tat gtc ata ggt gga gtc gtt aaa ggt gac aca cat ata cca acg cac tag aga gtt tat cag aag atg gat gga tga atc atc aac gtc ttc caa taa aaa tgt cca ata tgt cga cga ttg ttc atg ctg gaa aga ttt ata tat cta gag gtt aca aca ata gta gtg tag tta atg taa tat cga atc t-- agt cct tag cta taa tcc gat ata tga tga atg gac caa att atc atc att aaa tat tcc tag aat taa tcc tgc tct a

>ON682265.2:156360-158623_Monkeypox_virus_isolate_MPXV/Germany/2022/RKI04_partial_genome

tta ctg act aca --- tgt a-c tat tt- -ta cat tac tac att ggc tac ggc ata tat acc tat ttc gtc act tcc aca cgc tcc ggt aaa cgg gtg tca tgt gac gag gga ga- -at ctt gat aag agg cat aat caa tgt tgt aat ccg atg tcc acc tgg aga att tgc caa ggt cag atg tag agt tgg tag tga taa cac aaa atg tga aca ctg ccc acc tca tac ata tac cgc aat ccc caa tta ttc t-a ata gat gtc atc aat gta gaa aat gcc caa cag gat cat ttg ata agg taa agt gta ccg gaa cac aga ac- --a aat -gt tcg tgt cat cct ggt tgg ta- tac gct act gat tct tca cag act gaa gat tgt cga gat ttg tgt acc aaa aaa gga gat gtc cat gcg gat act ttg gtg gaa tag atg aa- --g gaa atc cta ttt gta aat cgt gtt g-- -tg ttg gtg aat att gcg act acc tac gta att ata gac ttg atc cat ttc ctc cat gca aac tat cta tct aaa tgt aat taa tta --- --- --- --- --- --- --- --- tga ttt --- --- --- -tg atg ata atg tta -cc ata c-a tt- --- -at atc act act tgg tta gtg tgt att att tag tat gga aga cct att aat aa- --- --- -tt act tat ctt ttg acg atc ttg tt- -at aat tat --- --- --- --- aat ata aaa --a tac tta tga --- --- cat agt aac tca t-- --- --- --- aat tgc tg- acg cga taa att cgt aat aat ctg ttt tgt tca aat ttt tat aag ga- --- --a tct aca ggc ata aaa at- aaa aat ata atc tat aat ata c-- tct tac a-- acg cca tca tga at- --a gca gtg aat taa ttg ctg tta tt- atg gat tta gaa ata gtg gac gat ttt gtg ata tta ata tag tta tta atg atg aaa gga taa acg cgc ata gac tca tcc tat ctg gag cct ccg aat att ttt tcc att ctg ttt tcc aat aat ttt atc gat tct aat gaa tac gaa gtt aat cta agt cat tta gat tat caa agt gtt aac tat ttg atc gat tac att tat ggg ata cct ttg agc cta act aac aat aac gtg aaa tat att ctt tca acc gct ga- -tt ttt tta caa att gga tct gtc att act gag tgc gaa aaa tac ata ctt aaa aat ctt tgt ttt aga aac tgt atc gat ttc --- --- tac ata tac gct gat aaa tat aat aac aag aaa ata gaa tta gca tcg ttt aac aca ata tta cga aat att ttg aga ctc atc aac aat gaa aac ttt aaa tac tta aca gag gaa tca atg ata aaa att tta agc gat gat atg tta tat ata aaa aat gag gat ttc acc cca ctg att ctc att aaa tgg tta gag agt aca c-- caa cca tgt acc gtc gag tta ctt aga tgc ctc aga ata tca -tt tct ttc ccc aca agt tat aaa atc act tta tag tca tcg act ggt tag ttc aat cta cga atg tat aac att ctt aaa caa tat agc att ctt gga taa atc att tcc tag ata cca tat cat cga gtt gat atc tat cgg tat aag taa ttc aca tga taa gat ttc cat aaa ctg cta caa tca t-- aaa aaa att cat ggg aaa tga tat ctt cac gta gat ata ggt gta gtt tcg cag tga ccg tcc tgg ata ata tta tct ata tga tgg gtg gat atg atc agt ccc tgt ata gaa gtt caa agg tta tag cgt aca ata cat gta cta att ctt gga tat atg ata tac cag agc taa aaa tat cat cgt tct aat tgc gga gga gtt gcc aat gac gaa tac att tat tgt ata ggc ggt ata cgc gat cag gag tca tcg ttg ata tct agt atc gat aga tgg aag cca tca aaa cca tat tgg cag aag tat gct aaa atg tgc gaa cca aaa tgt gat atg gg- gtt gcg att tta aac gga tta ata tat gtc ata ggt gga gtc gtt aaa ggt gac aca cat ata cca acg cac tag aga gtt tat cag aag atg gat gga tga atc atc aac gtc ttc caa taa aaa tgt cca ata tgt cga cga ttg ttc atg ctg gaa aga ttt ata tat cta gag gtt aca aca ata gta gtg tag tta atg taa tat cga atc t-- agt cct tag cta taa tcc gat ata tga tga atg gac caa att atc atc att aaa tat tcc tag aat taa tcc tgc tct a

>ON682264.2:156359-158622_Monkeypox_virus_isolate_MPXV/Germany/2022/RKI05_complete_genome

tta ctg act aca --- tgt a-c tat tt- -ta cat tac tac att ggc tac ggc ata tat acc tat ttc gtc act tcc aca cgc tcc ggt aaa cgg gtg tca tgt gac gag gga ga- -at ctt gat aag agg cat aat caa tgt tgt aat ccg atg tcc acc tgg aga att tgc caa ggt cag atg tag agt tgg tag tga taa cac aaa atg tga aca ctg ccc acc tca tac ata tac cgc aat ccc caa tta ttc t-a ata gat gtc atc aat gta gaa aat gcc caa cag gat cat ttg ata agg taa agt gta ccg gaa cac aga ac- --a aat -gt tcg tgt cat cct ggt tgg ta- tac gct act gat tct tca cag act gaa gat tgt cga gat ttg tgt acc aaa aaa gga gat gtc cat gcg gat act ttg gtg gaa tag atg aa- --g gaa atc cta ttt gta aat cgt gtt g-- -tg ttg gtg aat att gcg act acc tac gta att ata gac ttg atc cat ttc ctc cat gca aac tat cta tct aaa tgt aat taa tta --- --- --- --- --- --- --- --- tga ttt --- --- --- -tg atg ata atg tta -cc ata c-a tt- --- -at atc act act tgg tta gtg tgt att att tag tat gga aga cct att aat aa- --- --- -tt act tat ctt ttg acg atc ttg tt- -at aat tat --- --- --- --- aat ata aaa --a tac tta tga --- --- cat agt aac tca t-- --- --- --- aat tgc tg- acg cga taa att cgt aat aat ctg ttt tgt tca aat ttt tat aag ga- --- --a tct aca ggc ata aaa at- aaa aat ata atc tat aat ata c-- tct tac a-- acg cca tca tga at- --a gca gtg aat taa ttg ctg tta tt- atg gat tta gaa ata gtg gac gat ttt gtg ata tta ata tag tta tta atg atg aaa gga taa acg cgc ata gac tca tcc tat ctg gag cct ccg aat att ttt tcc att ctg ttt tcc aat aat ttt atc gat tct aat gaa tac gaa gtt aat cta agt cat tta gat tat caa agt gtt aac tat ttg atc gat tac att tat ggg ata cct ttg agc cta act aac aat aac gtg aaa tat att ctt tca acc gct ga- -tt ttt tta caa att gga tct gtc att act gag tgc gaa aaa tac ata ctt aaa aat ctt tgt ttt aga aac tgt atc gat ttc --- --- tac ata tac gct gat aaa tat aat aac aag aaa ata gaa tta gca tcg ttt aac aca ata tta cga aat att ttg aga ctc atc aac aat gaa aac ttt aaa tac tta aca gag gaa tca atg ata aaa att tta agc gat gat atg tta tat ata aaa aat gag gat ttc acc cca ctg att ctc att aaa tgg tta gag agt aca c-- caa cca tgt acc gtc gag tta ctt aga tgc ctc aga ata tca -tt tct ttc ccc aca agt tat aaa atc act tta tag tca tcg act ggt tag ttc aat cta cga atg tat aac att ctt aaa caa tat agc att ctt gga taa atc att tcc tag ata cca tat cat cga gtt gat atc tat cgg tat aag taa ttc aca tga taa gat ttc cat aaa ctg cta caa tca t-- aaa aaa att cat ggg aaa tga tat ctt cac gta gat ata ggt gta gtt tcg cag tga ccg tcc tgg ata ata tta tct ata tga tgg gtg gat atg atc agt ccc tgt ata gaa gtt caa agg tta tag cgt aca ata cat gta cta att ctt gga tat atg ata tac cag agc taa aaa tat cat cgt tct aat tgc gga gga gtt gcc aat gac gaa tac att tat tgt ata ggc ggt ata cgc gat cag gag tca tcg ttg ata tct agt atc gat aga tgg aag cca tca aaa cca tat tgg cag aag tat gct aaa atg tgc gaa cca aaa tgt gat atg gg- gtt gcg att tta aac gga tta ata tat gtc ata ggt gga gtc gtt aaa ggt gac aca cat ata cca acg cac tag aga gtt tat cag aag atg gat gga tga atc atc aac gtc ttc caa taa aaa tgt cca ata tgt cga cga ttg ttc atg ctg gaa aga ttt ata tat cta gag gtt aca aca ata gta gtg tag tta atg taa tat cga atc t-- agt cct tag cta taa tcc gat ata tga tga atg gac caa att atc atc att aaa tat tcc tag aat taa tcc tgc tct a

>ON682263.2:156392-158655_Monkeypox_virus_isolate_MPXV/Germany/2022/RKI03_complete_genome

tta ctg act aca --- tgt a-c tat tt- -ta cat tac tac att ggc tac ggc ata tat acc tat ttc gtc act tcc aca cgc tcc ggt aaa cgg gtg tca tgt gac gag gga ga- -at ctt gat aag agg cat aat caa tgt tgt aat ccg atg tcc acc tgg aga att tgc caa ggt cag atg tag agt tgg tag tga taa cac aaa atg tga aca ctg ccc acc tca tac ata tac cgc aat ccc caa tta ttc t-a ata gat gtc atc aat gta gaa aat gcc caa cag gat cat ttg ata agg taa agt gta ccg gaa cac aga ac- --a aat -gt tcg tgt cat cct ggt tgg ta- tac gct act gat tct tca cag act gaa gat tgt cga gat ttg tgt acc aaa aaa gga gat gtc cat gcg gat act ttg gtg gaa tag atg aa- --g gaa atc cta ttt gta aat cgt gtt g-- -tg ttg gtg aat att gcg act acc tac gta att ata gac ttg atc cat ttc ctc cat gca aac tat cta tct aaa tgt aat taa tta --- --- --- --- --- --- --- --- tga ttt --- --- --- -tg atg ata atg tta -cc ata c-a tt- --- -at atc act act tgg tta gtg tgt att att tag tat gga aga cct att aat aa- --- --- -tt act tat ctt ttg acg atc ttg tt- -at aat tat --- --- --- --- aat ata aaa --a tac tta tga --- --- cat agt aac tca t-- --- --- --- aat tgc tg- acg cga taa att cgt aat aat ctg ttt tgt tca aat ttt tat aag ga- --- --a tct aca ggc ata aaa at- aaa aat ata atc tat aat ata c-- tct tac a-- acg cca tca tga at- --a gca gtg aat taa ttg ctg tta tt- atg gat tta gaa ata gtg gac gat ttt gtg ata tta ata tag tta tta atg atg aaa gga taa acg cgc ata gac tca tcc tat ctg gag cct ccg aat att ttt tcc att ctg ttt tcc aat aat ttt atc gat tct aat gaa tac gaa gtt aat cta agt cat tta gat tat caa agt gtt aac tat ttg atc gat tac att tat ggg ata cct ttg agc cta act aac aat aac gtg aaa tat att ctt tca acc gct ga- -tt ttt tta caa att gga tct gtc att act gag tgc gaa aaa tac ata ctt aaa aat ctt tgt ttt aga aac tgt atc gat ttc --- --- tac ata tac gct gat aaa tat aat aac aag aaa ata gaa tta gca tcg ttt aac aca ata tta cga aat att ttg aga ctc atc aac aat gaa aac ttt aaa tac tta aca gag gaa tca atg ata aaa att tta agc gat gat atg tta tat ata aaa aat gag gat ttc acc cca ctg att ctc att aaa tgg tta gag agt aca c-- caa cca tgt acc gtc gag tta ctt aga tgc ctc aga ata tca -tt tct ttc ccc aca agt tat aaa atc act tta tag tca tcg act ggt tag ttc aat cta cga atg tat aac att ctt aaa caa tat agc att ctt gga taa atc att tcc tag ata cca tat cat cga gtt gat atc tat cgg tat aag taa ttc aca tga taa gat ttc cat aaa ctg cta caa tca t-- aaa aaa att cat ggg aaa tga tat ctt cac gta gat ata ggt gta gtt tcg cag tga ccg tcc tgg ata ata tta tct ata tga tgg gtg gat atg atc agt ccc tgt ata gaa gtt caa agg tta tag cgt aca ata cat gta cta att ctt gga tat atg ata tac cag agc taa aaa tat cat cgt tct aat tgc gga gga gtt gcc aat gac gaa tac att tat tgt ata ggc ggt ata cgc gat cag gag tca tcg ttg ata tct agt atc gat aga tgg aag cca tca aaa cca tat tgg cag aag tat gct aaa atg tgc gaa cca aaa tgt gat atg gg- gtt gcg att tta aac gga tta ata tat gtc ata ggt gga gtc gtt aaa ggt gac aca cat ata cca acg cac tag aga gtt tat cag aag atg gat gga tga atc atc aac gtc ttc caa taa aaa tgt cca ata tgt cga cga ttg ttc atg ctg gaa aga ttt ata tat cta gag gtt aca aca ata gta gtg tag tta atg taa tat cga atc t-- agt cct tag cta taa tcc gat ata tga tga atg gac caa att atc atc att aaa tat tcc tag aat taa tcc tgc tct a

>ON682270.1:156360-158623_Monkeypox_virus_isolate_MPXV/Germany/2022/RKI06_complete_genome

tta ctg act aca --- tgt a-c tat tt- -ta cat tac tac att ggc tac ggc ata tat acc tat ttc gtc act tcc aca cgc tcc ggt aaa cgg gtg tca tgt gac gag gga ga- -at ctt gat aag agg cat aat caa tgt tgt aat ccg atg tcc acc tgg aga att tgc caa ggt cag atg tag agt tgg tag tga taa cac aaa atg tga aca ctg ccc acc tca tac ata tac cgc aat ccc caa tta ttc t-a ata gat gtc atc aat gta gaa aat gcc caa cag gat cat ttg ata agg taa agt gta ccg gaa cac aga ac- --a aat -gt tcg tgt cat cct ggt tgg ta- tac gct act gat tct tca cag act gaa gat tgt cga gat ttg tgt acc aaa aaa gga gat gtc cat gcg gat act ttg gtg gaa tag atg aa- --g gaa atc cta ttt gta aat cgt gtt g-- -tg ttg gtg aat att gcg act acc tac gta att ata gac ttg atc cat ttc ctc cat gca aac tat cta tct aaa tgt aat taa tta --- --- --- --- --- --- --- --- tga ttt --- --- --- -tg atg ata atg tta -cc ata c-a tt- --- -at atc act act tgg tta gtg tgt att att tag tat gga aga cct att aat aa- --- --- -tt act tat ctt ttg acg atc ttg tt- -at aat tat --- --- --- --- aat ata aaa --a tac tta tga --- --- cat agt aac tca t-- --- --- --- aat tgc tg- acg cga taa att cgt aat aat ctg ttt tgt tca aat ttt tat aag ga- --- --a tct aca ggc ata aaa at- aaa aat ata atc tat aat ata c-- tct tac a-- acg cca tca tga at- --a gca gtg aat taa ttg ctg tta tt- atg gat tta gaa ata gtg gac gat ttt gtg ata tta ata tag tta tta atg atg aaa gga taa acg cgc ata gac tca tcc tat ctg gag cct ccg aat att ttt tcc att ctg ttt tcc aat aat ttt atc gat tct aat gaa tac gaa gtt aat cta agt cat tta gat tat caa agt gtt aac tat ttg atc gat tac att tat ggg ata cct ttg agc cta act aac aat aac gtg aaa tat att ctt tca acc gct ga- -tt ttt tta caa att gga tct gtc att act gag tgc gaa aaa tac ata ctt aaa aat ctt tgt ttt aga aac tgt atc gat ttc --- --- tac ata tac gct gat aaa tat aat aac aag aaa ata gaa tta gca tcg ttt aac aca ata tta cga aat att ttg aga ctc atc aac aat gaa aac ttt aaa tac tta aca gag gaa tca atg ata aaa att tta agc gat gat atg tta tat ata aaa aat gag gat ttc acc cca ctg att ctc att aaa tgg tta gag agt aca c-- caa cca tgt acc gtc gag tta ctt aga tgc ctc aga ata tca -tt tct ttc ccc aca agt tat aaa atc act tta tag tca tcg act ggt tag ttc aat cta cga atg tat aac att ctt aaa caa tat agc att ctt gga taa atc att tcc tag ata cca tat cat cga gtt gat atc tat cgg tat aag taa ttc aca tga taa gat ttc cat aaa ctg cta caa tca t-- aaa aaa att cat ggg aaa tga tat ctt cac gta gat ata ggt gta gtt tcg cag tga ccg tcc tgg ata ata tta tct ata tga tgg gtg gat atg atc agt ccc tgt ata gaa gtt caa agg tta tag cgt aca ata cat gta cta att ctt gga tat atg ata tac cag agc taa aaa tat cat cgt tct aat tgc gga gga gtt gcc aat gac gaa tac att tat tgt ata ggc ggt ata cgc gat cag gag tca tcg ttg ata tct agt atc gat aga tgg aag cca tca aaa cca tat tgg cag aag tat gct aaa atg tgc gaa cca aaa tgt gat atg gg- gtt gcg att tta aac gga tta ata tat gtc ata ggt gga gtc gtt aaa ggt gac aca cat ata cca acg cac tag aga gtt tat cag aag atg gat gga tga atc atc aac gtc ttc caa taa aaa tgt cca ata tgt cga cga ttg ttc atg ctg gaa aga ttt ata tat cta gag gtt aca aca ata gta gtg tag tta atg taa tat cga atc t-- agt cct tag cta taa tcc gat ata tga tga atg gac caa att atc atc att aaa tat tcc tag aat taa tcc tgc tct a

>ON682268.1:156359-158622_Monkeypox_virus_isolate_MPXV/Germany/2022/RKI08_complete_genome

tta ctg act aca --- tgt a-c tat tt- -ta cat tac tac att ggc tac ggc ata tat acc tat ttc gtc act tcc aca cgc tcc ggt aaa cgg gtg tca tgt gac gag gga ga- -at ctt gat aag agg cat aat caa tgt tgt aat ccg atg tcc acc tgg aga att tgc caa ggt cag atg tag agt tgg tag tga taa cac aaa atg tga aca ctg ccc acc tca tac ata tac cgc aat ccc caa tta ttc t-a ata gat gtc atc aat gta gaa aat gcc caa cag gat cat ttg ata agg taa agt gta ccg gaa cac aga ac- --a aat -gt tcg tgt cat cct ggt tgg ta- tac gct act gat tct tca cag act gaa gat tgt cga gat ttg tgt acc aaa aaa gga gat gtc cat gcg gat act ttg gtg gaa tag atg aa- --g gaa atc cta ttt gta aat cgt gtt g-- -tg ttg gtg aat att gcg act acc tac gta att ata gac ttg atc cat ttc ctc cat gca aac tat cta tct aaa tgt aat taa tta --- --- --- --- --- --- --- --- tga ttt --- --- --- -tg atg ata atg tta -cc ata c-a tt- --- -at atc act act tgg tta gtg tgt att att tag tat gga aga cct att aat aa- --- --- -tt act tat ctt ttg acg atc ttg tt- -at aat tat --- --- --- --- aat ata aaa --a tac tta tga --- --- cat agt aac tca t-- --- --- --- aat tgc tg- acg cga taa att cgt aat aat ctg ttt tgt tca aat ttt tat aag ga- --- --a tct aca ggc ata aaa at- aaa aat ata atc tat aat ata c-- tct tac a-- acg cca tca tga at- --a gca gtg aat taa ttg ctg tta tt- atg gat tta gaa ata gtg gac gat ttt gtg ata tta ata tag tta tta atg atg aaa gga taa acg cgc ata gac tca tcc tat ctg gag cct ccg aat att ttt tcc att ctg ttt tcc aat aat ttt atc gat tct aat gaa tac gaa gtt aat cta agt cat tta gat tat caa agt gtt aac tat ttg atc gat tac att tat ggg ata cct ttg agc cta act aac aat aac gtg aaa tat att ctt tca acc gct ga- -tt ttt tta caa att gga tct gtc att act gag tgc gaa aaa tac ata ctt aaa aat ctt tgt ttt aga aac tgt atc gat ttc --- --- tac ata tac gct gat aaa tat aat aac aag aaa ata gaa tta gca tcg ttt aac aca ata tta cga aat att ttg aga ctc atc aac aat gaa aac ttt aaa tac tta aca gag gaa tca atg ata aaa att tta agc gat gat atg tta tat ata aaa aat gag gat ttc acc cca ctg att ctc att aaa tgg tta gag agt aca c-- caa cca tgt acc gtc gag tta ctt aga tgc ctc aga ata tca -tt tct ttc ccc aca agt tat aaa atc act tta tag tca tcg act ggt tag ttc aat cta cga atg tat aac att ctt aaa caa tat agc att ctt gga taa atc att tcc tag ata cca tat cat cga gtt gat atc tat cgg tat aag taa ttc aca tga taa gat ttc cat aaa ctg cta caa tca t-- aaa aaa att cat ggg aaa tga tat ctt cac gta gat ata ggt gta gtt tcg cag tga ccg tcc tgg ata ata tta tct ata tga tgg gtg gat atg atc agt ccc tgt ata gaa gtt caa agg tta tag cgt aca ata cat gta cta att ctt gga tat atg ata tac cag agc taa aaa tat cat cgt tct aat tgc gga gga gtt gcc aat gac gaa tac att tat tgt ata ggc ggt ata cgc gat cag gag tca tcg ttg ata tct agt atc gat aga tgg aag cca tca aaa cca tat tgg cag aag tat gct aaa atg tgc gaa cca aaa tgt gat atg gg- gtt gcg att tta aac gga tta ata tat gtc ata ggt gga gtc gtt aaa ggt gac aca cat ata cca acg cac tag aga gtt tat cag aag atg gat gga tga atc atc aac gtc ttc caa taa aaa tgt cca ata tgt cga cga ttg ttc atg ctg gaa aga ttt ata tat cta gag gtt aca aca ata gta gtg tag tta atg taa tat cga atc t-- agt cct tag cta taa tcc gat ata tga tga atg gac caa att atc atc att aaa tat tcc tag aat taa tcc tgc tct a

>ON682267.1:156359-158622_Monkeypox_virus_isolate_MPXV/Germany/2022/RKI010_complete_genome

tta ctg act aca --- tgt a-c tat tt- -ta cat tac tac att ggc tac ggc ata tat acc tat ttc gtc act tcc aca cgc tcc ggt aaa cgg gtg tca tgt gac gag gga ga- -at ctt gat aag agg cat aat caa tgt tgt aat ccg atg tcc acc tgg aga att tgc caa ggt cag atg tag agt tgg tag tga taa cac aaa atg tga aca ctg ccc acc tca tac ata tac cgc aat ccc caa tta ttc t-a ata gat gtc atc aat gta gaa aat gcc caa cag gat cat ttg ata agg taa agt gta ccg gaa cac aga ac- --a aat -gt tcg tgt cat cct ggt tgg ta- tac gct act gat tct tca cag act gaa gat tgt cga gat ttg tgt acc aaa aaa gga gat gtc cat gcg gat act ttg gtg gaa tag atg aa- --g gaa atc cta ttt gta aat cgt gtt g-- -tg ttg gtg aat att gcg act acc tac gta att ata gac ttg atc cat ttc ctc cat gca aac tat cta tct aaa tgt aat taa tta --- --- --- --- --- --- --- --- tga ttt --- --- --- -tg atg ata atg tta -cc ata c-a tt- --- -at atc act act tgg tta gtg tgt att att tag tat gga aga cct att aat aa- --- --- -tt act tat ctt ttg acg atc ttg tt- -at aat tat --- --- --- --- aat ata aaa --a tac tta tga --- --- cat agt aac tca t-- --- --- --- aat tgc tg- acg cga taa att cgt aat aat ctg ttt tgt tca aat ttt tat aag ga- --- --a tct aca ggc ata aaa at- aaa aat ata atc tat aat ata c-- tct tac a-- acg cca tca tga at- --a gca gtg aat taa ttg ctg tta tt- atg gat tta gaa ata gtg gac gat ttt gtg ata tta ata tag tta tta atg atg aaa gga taa acg cgc ata gac tca tcc tat ctg gag cct ccg aat att ttt tcc att ctg ttt tcc aat aat ttt atc gat tct aat gaa tac gaa gtt aat cta agt cat tta gat tat caa agt gtt aac tat ttg atc gat tac att tat ggg ata cct ttg agc cta act aac aat aac gtg aaa tat att ctt tca acc gct ga- -tt ttt tta caa att gga tct gtc att act gag tgc gaa aaa tac ata ctt aaa aat ctt tgt ttt aga aac tgt atc gat ttc --- --- tac ata tac gct gat aaa tat aat aac aag aaa ata gaa tta gca tcg ttt aac aca ata tta cga aat att ttg aga ctc atc aac aat gaa aac ttt aaa tac tta aca gag gaa tca atg ata aaa att tta agc gat gat atg tta tat ata aaa aat gag gat ttc acc cca ctg att ctc att aaa tgg tta gag agt aca c-- caa cca tgt acc gtc gag tta ctt aga tgc ctc aga ata tca -tt tct ttc ccc aca agt tat aaa atc act tta tag tca tcg act ggt tag ttc aat cta cga atg tat aac att ctt aaa caa tat agc att ctt gga taa atc att tcc tag ata cca tat cat cga gtt gat atc tat cgg tat aag taa ttc aca tga taa gat ttc cat aaa ctg cta caa tca t-- aaa aaa att cat ggg aaa tga tat ctt cac gta gat ata ggt gta gtt tcg cag tga ccg tcc tgg ata ata tta tct ata tga tgg gtg gat atg atc agt ccc tgt ata gaa gtt caa agg tta tag cgt aca ata cat gta cta att ctt gga tat atg ata tac cag agc taa aaa tat cat cgt tct aat tgc gga gga gtt gcc aat gac gaa tac att tat tgt ata ggc ggt ata cgc gat cag gag tca tcg ttg ata tct agt atc gat aga tgg aag cca tca aaa cca tat tgg cag aag tat gct aaa atg tgc gaa cca aaa tgt gat atg gg- gtt gcg att tta aac gga tta ata tat gtc ata ggt gga gtc gtt aaa ggt gac aca cat ata cca acg cac tag aga gtt tat cag aag atg gat gga tga atc atc aac gtc ttc caa taa aaa tgt cca ata tgt cga cga ttg ttc atg ctg gaa aga ttt ata tat cta gag gtt aca aca ata gta gtg tag tta atg taa tat cga atc t-- agt cct tag cta taa tcc gat ata tga tga atg gac caa att atc atc att aaa tat tcc tag aat taa tcc tgc tct a

>ON682266.1:156359-158622_Monkeypox_virus_isolate_MPXV/Germany/2022/RKI09_complete_genome

tta ctg act aca --- tgt a-c tat tt- -ta cat tac tac att ggc tac ggc ata tat acc tat ttc gtc act tcc aca cgc tcc ggt aaa cgg gtg tca tgt gac gag gga ga- -at ctt gat aag agg cat aat caa tgt tgt aat ccg atg tcc acc tgg aga att tgc caa ggt cag atg tag agt tgg tag tga taa cac aaa atg tga aca ctg ccc acc tca tac ata tac cgc aat ccc caa tta ttc t-a ata gat gtc atc aat gta gaa aat gcc caa cag gat cat ttg ata agg taa agt gta ccg gaa cac aga ac- --a aat -gt tcg tgt cat cct ggt tgg ta- tac gct act gat tct tca cag act gaa gat tgt cga gat ttg tgt acc aaa aaa gga gat gtc cat gcg gat act ttg gtg gaa tag atg aa- --g gaa atc cta ttt gta aat cgt gtt g-- -tg ttg gtg aat att gcg act acc tac gta att ata gac ttg atc cat ttc ctc cat gca aac tat cta tct aaa tgt aat taa tta --- --- --- --- --- --- --- --- tga ttt --- --- --- -tg atg ata atg tta -cc ata c-a tt- --- -at atc act act tgg tta gtg tgt att att tag tat gga aga cct att aat aa- --- --- -tt act tat ctt ttg acg atc ttg tt- -at aat tat --- --- --- --- aat ata aaa --a tac tta tga --- --- cat agt aac tca t-- --- --- --- aat tgc tg- acg cga taa att cgt aat aat ctg ttt tgt tca aat ttt tat aag ga- --- --a tct aca ggc ata aaa at- aaa aat ata atc tat aat ata c-- tct tac a-- acg cca tca tga at- --a gca gtg aat taa ttg ctg tta tt- atg gat tta gaa ata gtg gac gat ttt gtg ata tta ata tag tta tta atg atg aaa gga taa acg cgc ata gac tca tcc tat ctg gag cct ccg aat att ttt tcc att ctg ttt tcc aat aat ttt atc gat tct aat gaa tac gaa gtt aat cta agt cat tta gat tat caa agt gtt aac tat ttg atc gat tac att tat ggg ata cct ttg agc cta act aac aat aac gtg aaa tat att ctt tca acc gct ga- -tt ttt tta caa att gga tct gtc att act gag tgc gaa aaa tac ata ctt aaa aat ctt tgt ttt aga aac tgt atc gat ttc --- --- tac ata tac gct gat aaa tat aat aac aag aaa ata gaa tta gca tcg ttt aac aca ata tta cga aat att ttg aga ctc atc aac aat gaa aac ttt aaa tac tta aca gag gaa tca atg ata aaa att tta agc gat gat atg tta tat ata aaa aat gag gat ttc acc cca ctg att ctc att aaa tgg tta gag agt aca c-- caa cca tgt acc gtc gag tta ctt aga tgc ctc aga ata tca -tt tct ttc ccc aca agt tat aaa atc act tta tag tca tcg act ggt tag ttc aat cta cga atg tat aac att ctt aaa caa tat agc att ctt gga taa atc att tcc tag ata cca tat cat cga gtt gat atc tat cgg tat aag taa ttc aca tga taa gat ttc cat aaa ctg cta caa tca t-- aaa aaa att cat ggg aaa tga tat ctt cac gta gat ata ggt gta gtt tcg cag tga ccg tcc tgg ata ata tta tct ata tga tgg gtg gat atg atc agt ccc tgt ata gaa gtt caa agg tta tag cgt aca ata cat gta cta att ctt gga tat atg ata tac cag agc taa aaa tat cat cgt tct aat tgc gga gga gtt gcc aat gac gaa tac att tat tgt ata ggc ggt ata cgc gat cag gag tca tcg ttg ata tct agt atc gat aga tgg aag cca tca aaa cca tat tgg cag aag tat gct aaa atg tgc gaa cca aaa tgt gat atg gg- gtt gcg att tta aac gga tta ata tat gtc ata ggt gga gtc gtt aaa ggt gac aca cat ata cca acg cac tag aga gtt tat cag aag atg gat gga tga atc atc aac gtc ttc caa taa aaa tgt cca ata tgt cga cga ttg ttc atg ctg gaa aga ttt ata tat cta gag gtt aca aca ata gta gtg tag tta atg taa tat cga atc t-- agt cct tag cta taa tcc gat ata tga tga atg gac caa att atc atc att aaa tat tcc tag aat taa tcc tgc tct a

>ON676708.1:156365-158628_Monkeypox_virus_isolate_MPXV_USA_2021_MD_complete_genome

tta ctg act aca --- tgt a-c tat tt- -ta cat tac tac att ggc tac ggc ata tat acc tat ttc gtc act tcc aca cgc tcc ggt aaa cgg gtg tca tgt gac gag gga ga- -at ctt gat aag agg cat aat caa tgt tgt aat ccg atg tcc acc tgg aga att tgc caa ggt cag atg tag agt tgg tag tga taa cac aaa atg tga aca ctg ccc acc tca tac ata tac cgc aat ccc caa tta ttc t-a ata gat gtc atc aat gta gaa aat gcc caa cag gat cat ttg ata agg taa agt gta ccg gaa cac aga ac- --a aat -gt tcg tgt cat cct ggt tgg ta- tac gct act gat tct tca cag act gaa gat tgt cga gat ttg tgt acc aaa aaa gga gat gtc cat gcg gat act ttg gtg gaa tag atg aa- --g gaa atc cta ttt gta aat cgt gtt g-- -tg ttg gtg aat att gcg act acc tac gta att ata gac ttg atc cat ttc ctc cat gca aac tat cta tct aaa tgt aat taa tta --- --- --- --- --- --- --- --- tga ttt --- --- --- -tg atg ata atg tta -cc ata c-a tt- --- -at atc act act tgg tta gtg tgt att att tag tat gga aga cct att aat aa- --- --- -tt act tat ctt ttg acg atc ttg tt- -at aat tat --- --- --- --- aat ata aaa --a tac tta tga --- --- cat agt aac tca t-- --- --- --- aat tgc tg- acg cga taa att cgt aat aat ctg ttt tgt tca aat ttt tat aag ga- --- --a tct aca ggc ata aaa at- aaa aat ata atc tat aat ata c-- tct tac a-- acg cca tca tga at- --a gca gtg aat taa ttg ctg tta tt- atg gat tta gaa ata gtg gac gat ttt gtg ata tta ata tag tta tta atg atg aaa gga taa acg cgc ata gac tca tcc tat ctg gag cct ccg aat att ttt tcc att ctg ttt tcc aat aat ttt atc gat tct aat gaa tac gaa gtt aat cta agt cat tta gat tat caa agt gtt aac tat ttg atc gat tac att tat ggg ata cct ttg agc cta act aac aat aac gtg aaa tat att ctt tca acc gct ga- -tt ttt tta caa att gga tct gtc att act gag tgc gaa aaa tac ata ctt aaa aat ctt tgt ttt aga aac tgt atc gat ttc --- --- tac ata tac gct gat aaa tat aat aac aag aaa ata gaa tta gca tcg ttt aac aca ata tta cga aat att ttg aga ctc atc aac aat gaa aac ttt aaa tac tta aca gag gaa tca atg ata aaa att tta agc gat gat atg tta tat ata aaa aat gag gat ttc acc cca ctg att ctc att aaa tgg tta gag agt aca c-- caa cca tgt acc gtc gag tta ctt aga tgc ctc aga ata tca -tt tct ttc ccc aca agt tat aaa atc act tta tag tca tcg act ggt tag ttc aat cta cga atg tat aac att ctt aaa caa tat agc att ctt gga taa atc att tcc tag ata cca tat cat cga gtt gat atc tat cgg tat aag taa ttc aca tga taa gat ttc cat aaa ctg cta caa tca t-- aaa aaa att cat ggg aaa tga tat ctt cac gta gat ata ggt gta gtt tcg cag tga ccg tcc tgg ata ata tta tct ata tga tgg gtg gat atg atc agt ccc tgt ata gaa gtt caa agg tta tag cgt aca ata cat gta cta att ctt gga tat atg ata tac cag agc taa aaa tat cat cgt tct aat tgc gga gga gtt gcc aat gac gaa tac att tat tgt ata ggc ggt ata cgc gat cag gag tca tcg ttg ata tct agt atc gat aga tgg aag cca tca aaa cca tat tgg cag aag tat gct aaa atg tgc gaa cca aaa tgt gat atg gg- gtt gcg att tta aac gga tta ata tat gtc ata ggt gga gtc gtt aaa ggt gac aca cat ata cca acg cac tag aga gtt tat cag aag atg gat gga tga atc atc aac gtc ttc caa taa aaa tgt cca ata tgt cga cga ttg ttc atg ctg gaa aga ttt ata tat cta gag gtt aca aca ata gta gtg tag tta atg taa tat cga atc t-- agt cct tag cta taa tcc gat ata tga tga atg gac caa att atc atc att aaa tat tcc tag aat taa tcc tgc tct a

>ON676707.1:156409-158672_Monkeypox_virus_isolate_MPXV_USA_2021_TX_complete_genome

tta ctg act aca --- tgt a-c tat tt- -ta cat tac tac att ggc tac ggc ata tat acc tat ttc gtc act tcc aca cgc tcc ggt aaa cgg gtg tca tgt gac gag gga ga- -at ctt gat aag agg cat aat caa tgt tgt aat ccg atg tcc acc tgg aga att tgc caa ggt cag atg tag agt tgg tag tga taa cac aaa atg tga aca ctg ccc acc tca tac ata tac cgc aat ccc caa tta ttc t-a ata gat gtc atc aat gta gaa aat gcc caa cag gat cat ttg ata agg taa agt gta ccg gaa cac aga ac- --a aat -gt tcg tgt cat cct ggt tgg ta- tac gct act gat tct tca cag act gaa gat tgt cga gat ttg tgt acc aaa aaa gga gat gtc cat gcg gat act ttg gtg gaa tag atg aa- --g gaa atc cta ttt gta aat cgt gtt g-- -tg ttg gtg aat att gcg act acc tac gta att ata gac ttg atc cat ttc ctc cat gca aac tat cta tct aaa tgt aat taa tta --- --- --- --- --- --- --- --- tga ttt --- --- --- -tg atg ata atg tta -cc ata c-a tt- --- -at atc act act tgg tta gtg tgt att att tag tat gga aga cct att aat aa- --- --- -tt act tat ctt ttg acg atc ttg tt- -at aat tat --- --- --- --- aat ata aaa --a tac tta tga --- --- cat agt aac tca t-- --- --- --- aat tgc tg- acg cga taa att cgt aat aat ctg ttt tgt tca aat ttt tat aag ga- --- --a tct aca ggc ata aaa at- aaa aat ata atc tat aat ata c-- tct tac a-- acg cca tca tga at- --a gca gtg aat taa ttg ctg tta tt- atg gat tta gaa ata gtg gac gat ttt gtg ata tta ata tag tta tta atg atg aaa gga taa acg cgc ata gac tca tcc tat ctg gag cct ccg aat att ttt tcc att ctg ttt tcc aat aat ttt atc gat tct aat gaa tac gaa gtt aat cta agt cat tta gat tat caa agt gtt aac tat ttg atc gat tac att tat ggg ata cct ttg agc cta act aac aat aac gtg aaa tat att ctt tca acc gct ga- -tt ttt tta caa att gga tct gtc att act gag tgc gaa aaa tac ata ctt aaa aat ctt tgt ttt aga aac tgt atc gat ttc --- --- tac ata tac gct gat aaa tat aat aac aag aaa ata gaa tta gca tcg ttt aac aca ata tta cga aat att ttg aga ctc atc aac aat gaa aac ttt aaa tac tta aca gag gaa tca atg ata aaa att tta agc gat gat atg tta tat ata aaa aat gag gat ttc acc cca ctg att ctc att aaa tgg tta gag agt aca c-- caa cca tgt acc gtc gag tta ctt aga tgc ctc aga ata tca -tt tct ttc ccc aca agt tat aaa atc act tta tag tca tcg act ggt tag ttc aat cta cga atg tat aac att ctt aaa caa tat agc att ctt gga taa atc att tcc tag ata cca tat cat cga gtt gat atc tat cgg tat aag taa ttc aca tga taa gat ttc cat aaa ctg cta caa tca t-- aaa aaa att cat ggg aaa tga tat ctt cac gta gat ata ggt gta gtt tcg cag tga ccg tcc tgg ata ata tta tct ata tga tgg gtg gat atg atc agt ccc tgt ata gaa gtt caa agg tta tag cgt aca ata cat gta cta att ctt gga tat atg ata tac cag agc taa aaa tat cat cgt tct aat tgc gga gga gtt gcc aat gac gaa tac att tat tgt ata ggc ggt ata cgc gat cag gag tca tcg ttg ata tct agt atc gat aga tgg aag cca tca aaa cca tat tgg cag aag tat gct aaa atg tgc gaa cca aaa tgt gat atg gg- gtt gcg att tta aac gga tta ata tat gtc ata ggt gga gtc gtt aaa ggt gac aca cat ata cca acg cac tag aga gtt tat cag aag atg gat gga tga atc atc aac gtc ttc caa taa aaa tgt cca ata tgt cga cga ttg ttc atg ctg gaa aga ttt ata tat cta gag gtt aca aca ata gta gtg tag tta atg taa tat cga atc t-- agt cct tag cta taa tcc gat ata tga tga atg gac caa att atc atc att aaa tat tcc tag aat taa tcc tgc tct a

>ON676706.1:156295-158558_Monkeypox_virus_isolate_MPXV_USA_2022_UT002_complete_genome

tta ctg act aca --- tgt a-c tat tt- -ta cat tac tac att ggc tac ggc ata tat acc tat ttc gtc act tcc aca cgc tcc ggt aaa cgg gtg tca tgt gac gag gga ga- -at ctt gat aag agg cat aat caa tgt tgt aat ccg atg tcc acc tgg aga att tgc caa ggt cag atg tag agt tgg tag tga taa cac aaa atg tga aca ctg ccc acc tca tac ata tac cgc aat ccc caa tta ttc t-a ata gat gtc atc aat gta gaa aat gcc caa cag gat cat ttg ata agg taa agt gta ccg gaa cac aga ac- --a aat -gt tcg tgt cat cct ggt tgg ta- tac gct act gat tct tca cag act gaa gat tgt cga gat ttg tgt acc aaa aaa gga gat gtc cat gcg gat act ttg gtg gaa tag atg aa- --g gaa atc cta ttt gta aat cgt gtt g-- -tg ttg gtg aat att gcg act acc tac gta att ata gac ttg atc cat ttc ctc cat gca aac tat cta tct aaa tgt aat taa tta --- --- --- --- --- --- --- --- tga ttt --- --- --- -tg atg ata atg tta -cc ata c-a tt- --- -at atc act act tgg tta gtg tgt att att tag tat gga aga cct att aat aa- --- --- -tt act tat ctt ttg acg atc ttg tt- -at aat tat --- --- --- --- aat ata aaa --a tac tta tga --- --- cat agt aac tca t-- --- --- --- aat tgc tg- acg cga taa att cgt aat aat ctg ttt tgt tca aat ttt tat aag ga- --- --a tct aca ggc ata aaa at- aaa aat ata atc tat aat ata c-- tct tac a-- acg cca tca tga at- --a gca gtg aat taa ttg ctg tta tt- atg gat tta gaa ata gtg gac gat ttt gtg ata tta ata tag tta tta atg atg aaa gga taa acg cgc ata gac tca tcc tat ctg gag cct ccg aat att ttt tcc att ctg ttt tcc aat aat ttt atc gat tct aat gaa tac gaa gtt aat cta agt cat tta gat tat caa agt gtt aac tat ttg atc gat tac att tat ggg ata cct ttg agc cta act aac aat aac gtg aaa tat att ctt tca acc gct ga- -tt ttt tta caa att gga tct gtc att act gag tgc gaa aaa tac ata ctt aaa aat ctt tgt ttt aga aac tgt atc gat ttc --- --- tac ata tac gct gat aaa tat aat aac aag aaa ata gaa tta gca tcg ttt aac aca ata tta cga aat att ttg aga ctc atc aac aat gaa aac ttt aaa tac tta aca gag gaa tca atg ata aaa att tta agc gat gat atg tta tat ata aaa aat gag gat ttc acc cca ctg att ctc att aaa tgg tta gag agt aca c-- caa cca tgt acc gtc gag tta ctt aga tgc ctc aga ata tca -tt tct ttc ccc aca agt tat aaa atc act tta tag tca tcg act ggt tag ttc aat cta cga atg tat aac att ctt aaa caa tat agc att ctt gga taa atc att tcc tag ata cca tat cat cga gtt gat atc tat cgg tat aag taa ttc aca tga taa gat ttc cat aaa ctg cta caa tca t-- aaa aaa att cat ggg aaa tga tat ctt cac gta gat ata ggt gta gtt tcg cag tga ccg tcc tgg ata ata tta tct ata tga tgg gtg gat atg atc agt ccc tgt ata gaa gtt caa agg tta tag cgt aca ata cat gta cta att ctt gga tat atg ata tac cag agc taa aaa tat cat cgt tct aat tgc gga gga gtt gcc aat gac gaa tac att tat tgt ata ggc ggt ata cgc gat cag gag tca tcg ttg ata tct agt atc gat aga tgg aag cca tca aaa cca tat tgg cag aag tat gct aaa atg tgc gaa cca aaa tgt gat atg gg- gtt gcg att tta aac gga tta ata tat gtc ata ggt gga gtc gtt aaa ggt gac aca cat ata cca acg cac tag aga gtt tat cag aag atg gat gga tga atc atc aac gtc ttc caa taa aaa tgt cca ata tgt cga cga ttg ttc atg ctg gaa aga ttt ata tat cta gag gtt aca aca ata gta gtg tag tta atg taa tat cga atc t-- agt cct tag cta taa tcc gat ata tga tga atg gac caa att atc atc att aaa tat tcc tag aat taa tcc tgc tct a

>ON676705.1:156341-158604_Monkeypox_virus_isolate_MPXV_USA_2022_UT001_complete_genome

tta ctg act aca --- tgt a-c tat tt- -ta cat tac tac att ggc tac ggc ata tat acc tat ttc gtc act tcc aca cgc tcc ggt aaa cgg gtg tca tgt gac gag gga ga- -at ctt gat aag agg cat aat caa tgt tgt aat ccg atg tcc acc tgg aga att tgc caa ggt cag atg tag agt tgg tag tga taa cac aaa atg tga aca ctg ccc acc tca tac ata tac cgc aat ccc caa tta ttc t-a ata gat gtc atc aat gta gaa aat gcc caa cag gat cat ttg ata agg taa agt gta ccg gaa cac aga ac- --a aat -gt tcg tgt cat cct ggt tgg ta- tac gct act gat tct tca cag act gaa gat tgt cga gat ttg tgt acc aaa aaa gga gat gtc cat gcg gat act ttg gtg gaa tag atg aa- --g gaa atc cta ttt gta aat cgt gtt g-- -tg ttg gtg aat att gcg act acc tac gta att ata gac ttg atc cat ttc ctc cat gca aac tat cta tct aaa tgt aat taa tta --- --- --- --- --- --- --- --- tga ttt --- --- --- -tg atg ata atg tta -cc ata c-a tt- --- -at atc act act tgg tta gtg tgt att att tag tat gga aga cct att aat aa- --- --- -tt act tat ctt ttg acg atc ttg tt- -at aat tat --- --- --- --- aat ata aaa --a tac tta tga --- --- cat agt aac tca t-- --- --- --- aat tgc tg- acg cga taa att cgt aat aat ctg ttt tgt tca aat ttt tat aag ga- --- --a tct aca ggc ata aaa at- aaa aat ata atc tat aat ata c-- tct tac a-- acg cca tca tga at- --a gca gtg aat taa ttg ctg tta tt- atg gat tta gaa ata gtg gac gat ttt gtg ata tta ata tag tta tta atg atg aaa gga taa acg cgc ata gac tca tcc tat ctg gag cct ccg aat att ttt tcc att ctg ttt tcc aat aat ttt atc gat tct aat gaa tac gaa gtt aat cta agt cat tta gat tat caa agt gtt aac tat ttg atc gat tac att tat ggg ata cct ttg agc cta act aac aat aac gtg aaa tat att ctt tca acc gct ga- -tt ttt tta caa att gga tct gtc att act gag tgc gaa aaa tac ata ctt aaa aat ctt tgt ttt aga aac tgt atc gat ttc --- --- tac ata tac gct gat aaa tat aat aac aag aaa ata gaa tta gca tcg ttt aac aca ata tta cga aat att ttg aga ctc atc aac aat gaa aac ttt aaa tac tta aca gag gaa tca atg ata aaa att tta agc gat gat atg tta tat ata aaa aat gag gat ttc acc cca ctg att ctc att aaa tgg tta gag agt aca c-- caa cca tgt acc gtc gag tta ctt aga tgc ctc aga ata tca -tt tct ttc ccc aca agt tat aaa atc act tta tag tca tcg act ggt tag ttc aat cta cga atg tat aac att ctt aaa caa tat agc att ctt gga taa atc att tcc tag ata cca tat cat cga gtt gat atc tat cgg tat aag taa ttc aca tga taa gat ttc cat aaa ctg cta caa tca t-- aaa aaa att cat ggg aaa tga tat ctt cac gta gat ata ggt gta gtt tcg cag tga ccg tcc tgg ata ata tta tct ata tga tgg gtg gat atg atc agt ccc tgt ata gaa gtt caa agg tta tag cgt aca ata cat gta cta att ctt gga tat atg ata tac cag agc taa aaa tat cat cgt tct aat tgc gga gga gtt gcc aat gac gaa tac att tat tgt ata ggc ggt ata cgc gat cag gag tca tcg ttg ata tct agt atc gat aga tgg aag cca tca aaa cca tat tgg cag aag tat gct aaa atg tgc gaa cca aaa tgt gat atg gg- gtt gcg att tta aac gga tta ata tat gtc ata ggt gga gtc gtt aaa ggt gac aca cat ata cca acg cac tag aga gtt tat cag aag atg gat gga tga atc atc aac gtc ttc caa taa aaa tgt cca ata tgt cga cga ttg ttc atg ctg gaa aga ttt ata tat cta gag gtt aca aca ata gta gtg tag tta atg taa tat cga atc t-- agt cct tag cta taa tcc gat ata tga tga atg gac caa att atc atc att aaa tat tcc tag aat taa tcc tgc tct a

>ON676704.1:156347-158610_Monkeypox_virus_isolate_MPXV_USA_2022_FL002_complete_genome

tta ctg act aca --- tgt a-c tat tt- -ta cat tac tac att ggc tac ggc ata tat acc tat ttc gtc act tcc aca cgc tcc ggt aaa cgg gtg tca tgt gac gag gga ga- -at ctt gat aag agg cat aat caa tgt tgt aat ccg atg tcc acc tgg aga att tgc caa ggt cag atg tag agt tgg tag tga taa cac aaa atg tga aca ctg ccc acc tca tac ata tac cgc aat ccc caa tta ttc t-a ata gat gtc atc aat gta gaa aat gcc caa cag gat cat ttg ata agg taa agt gta ccg gaa cac aga ac- --a aat -gt tcg tgt cat cct ggt tgg ta- tac gct act gat tct tca cag act gaa gat tgt cga gat ttg tgt acc aaa aaa gga gat gtc cat gcg gat act ttg gtg gaa tag atg aa- --g gaa atc cta ttt gta aat cgt gtt g-- -tg ttg gtg aat att gcg act acc tac gta att ata gac ttg atc cat ttc ctc cat gca aac tat cta tct aaa tgt aat taa tta --- --- --- --- --- --- --- --- tga ttt --- --- --- -tg atg ata atg tta -cc ata c-a tt- --- -at atc act act tgg tta gtg tgt att att tag tat gga aga cct att aat aa- --- --- -tt act tat ctt ttg acg atc ttg tt- -at aat tat --- --- --- --- aat ata aaa --a tac tta tga --- --- cat agt aac tca t-- --- --- --- aat tgc tg- acg cga taa att cgt aat aat ctg ttt tgt tca aat ttt tat aag ga- --- --a tct aca ggc ata aaa at- aaa aat ata atc tat aat ata c-- tct tac a-- acg cca tca tga at- --a gca gtg aat taa ttg ctg tta tt- atg gat tta gaa ata gtg gac gat ttt gtg ata tta ata tag tta tta atg atg aaa gga taa acg cgc ata gac tca tcc tat ctg gag cct ccg aat att ttt tcc att ctg ttt tcc aat aat ttt atc gat tct aat gaa tac gaa gtt aat cta agt cat tta gat tat caa agt gtt aac tat ttg atc gat tac att tat ggg ata cct ttg agc cta act aac aat aac gtg aaa tat att ctt tca acc gct ga- -tt ttt tta caa att gga tct gtc att act gag tgc gaa aaa tac ata ctt aaa aat ctt tgt ttt aga aac tgt atc gat ttc --- --- tac ata tac gct gat aaa tat aat aac aag aaa ata gaa tta gca tcg ttt aac aca ata tta cga aat att ttg aga ctc atc aac aat gaa aac ttt aaa tac tta aca gag gaa tca atg ata aaa att tta agc gat gat atg tta tat ata aaa aat gag gat ttc acc cca ctg att ctc att aaa tgg tta gag agt aca c-- caa cca tgt acc gtc gag tta ctt aga tgc ctc aga ata tca -tt tct ttc ccc aca agt tat aaa atc act tta tag tca tcg act ggt tag ttc aat cta cga atg tat aac att ctt aaa caa tat agc att ctt gga taa atc att tcc tag ata cca tat cat cga gtt gat atc tat cgg tat aag taa ttc aca tga taa gat ttc cat aaa ctg cta caa tca t-- aaa aaa att cat ggg aaa tga tat ctt cac gta gat ata ggt gta gtt tcg cag tga ccg tcc tgg ata ata tta tct ata tga tgg gtg gat atg atc agt ccc tgt ata gaa gtt caa agg tta tag cgt aca ata cat gta cta att ctt gga tat atg ata tac cag agc taa aaa tat cat cgt tct aat tgc gga gga gtt gcc aat gac gaa tac att tat tgt ata ggc ggt ata cgc gat cag gag tca tcg ttg ata tct agt atc gat aga tgg aag cca tca aaa cca tat tgg cag aag tat gct aaa atg tgc gaa cca aaa tgt gat atg gg- gtt gcg att tta aac gga tta ata tat gtc ata ggt gga gtc gtt aaa ggt gac aca cat ata cca acg cac tag aga gtt tat cag aag atg gat gga tga atc atc aac gtc ttc caa taa aaa tgt cca ata tgt cga cga ttg ttc atg ctg gaa aga ttt ata tat cta gag gtt aca aca ata gta gtg tag tta atg taa tat cga atc t-- agt cct tag cta taa tcc gat ata tga tga atg gac caa att atc atc att aaa tat tcc tag aat taa tcc tgc tct a

>ON676703.1:156347-158610_Monkeypox_virus_isolate_MPXV_USA_2022_CA001_complete_genome

tta ctg act aca --- tgt a-c tat tt- -ta cat tac tac att ggc tac ggc ata tat acc tat ttc gtc act tcc aca cgc tcc ggt aaa cgg gtg tca tgt gac gag gga ga- -at ctt gat aag agg cat aat caa tgt tgt aat ccg atg tcc acc tgg aga att tgc caa ggt cag atg tag agt tgg tag tga taa cac aaa atg tga aca ctg ccc acc tca tac ata tac cgc aat ccc caa tta ttc t-a ata gat gtc atc aat gta gaa aat gcc caa cag gat cat ttg ata agg taa agt gta ccg gaa cac aga ac- --a aat -gt tcg tgt cat cct ggt tgg ta- tac gct act gat tct tca cag act gaa gat tgt cga gat ttg tgt acc aaa aaa gga gat gtc cat gcg gat act ttg gtg gaa tag atg aa- --g gaa atc cta ttt gta aat cgt gtt g-- -tg ttg gtg aat att gcg act acc tac gta att ata gac ttg atc cat ttc ctc cat gca aac tat cta tct aaa tgt aat taa tta --- --- --- --- --- --- --- --- tga ttt --- --- --- -tg atg ata atg tta -cc ata c-a tt- --- -at atc act act tgg tta gtg tgt att att tag tat gga aga cct att aat aa- --- --- -tt act tat ctt ttg acg atc ttg tt- -at aat tat --- --- --- --- aat ata aaa --a tac tta tga --- --- cat agt aac tca t-- --- --- --- aat tgc tg- acg cga taa att cgt aat aat ctg ttt tgt tca aat ttt tat aag ga- --- --a tct aca ggc ata aaa at- aaa aat ata atc tat aat ata c-- tct tac a-- acg cca tca tga at- --a gca gtg aat taa ttg ctg tta tt- atg gat tta gaa ata gtg gac gat ttt gtg ata tta ata tag tta tta atg atg aaa gga taa acg cgc ata gac tca tcc tat ctg gag cct ccg aat att ttt tcc att ctg ttt tcc aat aat ttt atc gat tct aat gaa tac gaa gtt aat cta agt cat tta gat tat caa agt gtt aac tat ttg atc gat tac att tat ggg ata cct ttg agc cta act aac aat aac gtg aaa tat att ctt tca acc gct ga- -tt ttt tta caa att gga tct gtc att act gag tgc gaa aaa tac ata ctt aaa aat ctt tgt ttt aga aac tgt atc gat ttc --- --- tac ata tac gct gat aaa tat aat aac aag aaa ata gaa tta gca tcg ttt aac aca ata tta cga aat att ttg aga ctc atc aac aat gaa aac ttt aaa tac tta aca gag gaa tca atg ata aaa att tta agc gat gat atg tta tat ata aaa aat gag gat ttc acc cca ctg att ctc att aaa tgg tta gag agt aca c-- caa cca tgt acc gtc gag tta ctt aga tgc ctc aga ata tca -tt tct ttc ccc aca agt tat aaa atc act tta tag tca tcg act ggt tag ttc aat cta cga atg tat aac att ctt aaa caa tat agc att ctt gga taa atc att tcc tag ata cca tat cat cga gtt gat atc tat cgg tat aag taa ttc aca tga taa gat ttc cat aaa ctg cta caa tca t-- aaa aaa att cat ggg aaa tga tat ctt cac gta gat ata ggt gta gtt tcg cag tga ccg tcc tgg ata ata tta tct ata tga tgg gtg gat atg atc agt ccc tgt ata gaa gtt caa agg tta tag cgt aca ata cat gta cta att ctt gga tat atg ata tac cag agc taa aaa tat cat cgt tct aat tgc gga gga gtt gcc aat gac gaa tac att tat tgt ata ggc ggt ata cgc gat cag gag tca tcg ttg ata tct agt atc gat aga tgg aag cca tca aaa cca tat tgg cag aag tat gct aaa atg tgc gaa cca aaa tgt gat atg gg- gtt gcg att tta aac gga tta ata tat gtc ata ggt gga gtc gtt aaa ggt gac aca cat ata cca acg cac tag aga gtt tat cag aag atg gat gga tga atc atc aac gtc ttc caa taa aaa tgt cca ata tgt cga cga ttg ttc atg ctg gaa aga ttt ata tat cta gag gtt aca aca ata gta gtg tag tta atg taa tat cga atc t-- agt cct tag cta taa tcc gat ata tga tga atg gac caa att atc atc att aaa tat tcc tag aat taa tcc tgc tct a

>ON649879.1:156178-158441_Monkeypox_virus_isolate_MPXV_ISR001_2022_complete_genome

tta ctg act aca --- tgt a-c tat tt- -ta cat tac tac att ggc tac ggc ata tat acc tat ttc gtc act tcc aca cgc tcc ggt aaa cgg gtg tca tgt gac gag gga ga- -at ctt gat aag agg cat aat caa tgt tgt aat ccg atg tcc acc tgg aga att tgc caa ggt cag atg tag agt tgg tag tga taa cac aaa atg tga aca ctg ccc acc tca tac ata tac cgc aat ccc caa tta ttc t-a ata gat gtc atc aat gta gaa aat gcc caa cag gat cat ttg ata agg taa agt gta ccg gaa cac aga ac- --a aat -gt tcg tgt cat cct ggt tgg ta- tac gct act gat tct tca cag act gaa gat tgt cga gat ttg tgt acc aaa aaa gga gat gtc cat gcg gat act ttg gtg gaa tag atg aa- --g gaa atc cta ttt gta aat cgt gtt g-- -tg ttg gtg aat att gcg act acc tac gta att ata gac ttg atc cat ttc ctc cat gca aac tat cta tct aaa tgt aat taa tta --- --- --- --- --- --- --- --- tga ttt --- --- --- -tg atg ata atg tta -cc ata c-a tt- --- -at atc act act tgg tta gtg tgt att att tag tat gga aga cct att aat aa- --- --- -tt act tat ctt ttg acg atc ttg tt- -at aat tat --- --- --- --- aat ata aaa --a tac tta tga --- --- cat agt aac tca t-- --- --- --- aat tgc tg- acg cga taa att cgt aat aat ctg ttt tgt tca aat ttt tat aag ga- --- --a tct aca ggc ata aaa at- aaa aat ata atc tat aat ata c-- tct tac a-- acg cca tca tga at- --a gca gtg aat taa ttg ctg tta tt- atg gat tta gaa ata gtg gac gat ttt gtg ata tta ata tag tta tta atg atg aaa gga taa acg cgc ata gac tca tcc tat ctg gag cct ccg aat att ttt tcc att ctg ttt tcc aat aat ttt atc gat tct aat gaa tac gaa gtt aat cta agt cat tta gat tat caa agt gtt aac tat ttg atc gat tac att tat ggg ata cct ttg agc cta act aac aat aac gtg aaa tat att ctt tca acc gct ga- -tt ttt tta caa att gga tct gtc att act gag tgc gaa aaa tac ata ctt aaa aat ctt tgt ttt aga aac tgt atc gat ttc --- --- tac ata tac gct gat aaa tat aat aac aag aaa ata gaa tta gca tcg ttt aac aca ata tta cga aat att ttg aga ctc atc aac aat gaa aac ttt aaa tac tta aca gag gaa tca atg ata aaa att tta agc gat gat atg tta tat ata aaa aat gag gat ttc acc cca ctg att ctc att aaa tgg tta gag agt aca c-- caa cca tgt acc gtc gag tta ctt aga tgc ctc aga ata tca -tt tct ttc ccc aca agt tat aaa atc act tta tag tca tcg act ggt tag ttc aat cta cga atg tat aac att ctt aaa caa tat agc att ctt gga taa atc att tcc tag ata cca tat cat cga gtt gat atc tat cgg tat aag taa ttc aca tga taa gat ttc cat aaa ctg cta caa tca t-- aaa aaa att cat ggg aaa tga tat ctt cac gta gat ata ggt gta gtt tcg cag tga ccg tcc tgg ata ata tta tct ata tga tgg gtg gat atg atc agt ccc tgt ata gaa gtt caa agg tta tag cgt aca ata cat gta cta att ctt gga tat atg ata tac cag agc taa aaa tat cat cgt tct aat tgc gga gga gtt gcc aat gac gaa tac att tat tgt ata ggc ggt ata cgc gat cag gag tca tcg ttg ata tct agt atc gat aga tgg aag cca tca aaa cca tat tgg cag aag tat gct aaa atg tgc gaa cca aaa tgt gat atg gg- gtt gcg att tta aac gga tta ata tat gtc ata ggt gga gtc gtt aaa ggt gac aca cat ata cca acg cac tag aga gtt tat cag aag atg gat gga tga atc atc aac gtc ttc caa taa aaa tgt cca ata tgt cga cga ttg ttc atg ctg gaa aga ttt ata tat cta gag gtt aca aca ata gta gtg tag tta atg taa tat cga atc t-- agt cct tag cta taa tcc gat ata tga tga atg gac caa att atc atc att aaa tat tcc tag aat taa tcc tgc tct a

>ON649725.1:156353-158616_Monkeypox_virus_isolate_Monkeypox/PT0011/2022_complete_genome

tta ctg act aca --- tgt a-c tat tt- -ta cat tac tac att ggc tac ggc ata tat acc tat ttc gtc act tcc aca cgc tcc ggt aaa cgg gtg tca tgt gac gag gga ga- -at ctt gat aag agg cat aat caa tgt tgt aat ccg atg tcc acc tgg aga att tgc caa ggt cag atg tag agt tgg tag tga taa cac aaa atg tga aca ctg ccc acc tca tac ata tac cgc aat ccc caa tta ttc t-a ata gat gtc atc aat gta gaa aat gcc caa cag gat cat ttg ata agg taa agt gta ccg gaa cac aga ac- --a aat -gt tcg tgt cat cct ggt tgg ta- tac gct act gat tct tca cag act gaa gat tgt cga gat ttg tgt acc aaa aaa gga gat gtc cat gcg gat act ttg gtg gaa tag atg aa- --g gaa atc cta ttt gta aat cgt gtt g-- -tg ttg gtg aat att gcg act acc tac gta att ata gac ttg atc cat ttc ctc cat gca aac tat cta tct aaa tgt aat taa tta --- --- --- --- --- --- --- --- tga ttt --- --- --- -tg atg ata atg tta -cc ata c-a tt- --- -at atc act act tgg tta gtg tgt att att tag tat gga aga cct att aat aa- --- --- -tt act tat ctt ttg acg atc ttg tt- -at aat tat --- --- --- --- aat ata aaa --a tac tta tga --- --- cat agt aac tca t-- --- --- --- aat tgc tg- acg cga taa att cgt aat aat ctg ttt tgt tca aat ttt tat aag ga- --- --a tct aca ggc ata aaa at- aaa aat ata atc tat aat ata c-- tct tac a-- acg cca tca tga at- --a gca gtg aat taa ttg ctg tta tt- atg gat tta gaa ata gtg gac gat ttt gtg ata tta ata tag tta tta atg atg aaa gga taa acg cgc ata gac tca tcc tat ctg gag cct ccg aat att ttt tcc att ctg ttt tcc aat aat ttt atc gat tct aat gaa tac gaa gtt aat cta agt cat tta gat tat caa agt gtt aac tat ttg atc gat tac att tat ggg ata cct ttg agc cta act aac aat aac gtg aaa tat att ctt tca acc gct ga- -tt ttt tta caa att gga tct gtc att act gag tgc gaa aaa tac ata ctt aaa aat ctt tgt ttt aga aac tgt atc gat ttc --- --- tac ata tac gct gat aaa tat aat aac aag aaa ata gaa tta gca tcg ttt aac aca ata tta cga aat att ttg aga ctc atc aac aat gaa aac ttt aaa tac tta aca gag gaa tca atg ata aaa att tta agc gat gat atg tta tat ata aaa aat gag gat ttc acc cca ctg att ctc att aaa tgg tta gag agt aca c-- caa cca tgt acc gtc gag tta ctt aga tgc ctc aga ata tca -tt tct ttc ccc aca agt tat aaa atc act tta tag tca tcg act ggt tag ttc aat cta cga atg tat aac att ctt aaa caa tat agc att ctt gga taa atc att tcc tag ata cca tat cat cga gtt gat atc tat cgg tat aag taa ttc aca tga taa gat ttc cat aaa ctg cta caa tca t-- aaa aaa att cat ggg aaa tga tat ctt cac gta gat ata ggt gta gtt tcg cag tga ccg tcc tgg ata ata tta tct ata tga tgg gtg gat atg atc agt ccc tgt ata gaa gtt caa agg tta tag cgt aca ata cat gta cta att ctt gga tat atg ata tac cag agc taa aaa tat cat cgt tct aat tgc gga gga gtt gcc aat gac gaa tac att tat tgt ata ggc ggt ata cgc gat cag gag tca tcg ttg ata tct agt atc gat aga tgg aag cca tca aaa cca tat tgg cag aag tat gct aaa atg tgc gaa cca aaa tgt gat atg gg- gtt gcg att tta aac gga tta ata tat gtc ata ggt gga gtc gtt aaa ggt gac aca cat ata cca acg cac tag aga gtt tat cag aag atg gat gga tga atc atc aac gtc ttc caa taa aaa tgt cca ata tgt cga cga ttg ttc atg ctg gaa aga ttt ata tat cta gag gtt aca aca ata gta gtg tag tta atg taa tat cga atc t-- agt cct tag cta taa tcc gat ata tga tga atg gac caa att atc atc att aaa tat tcc tag aat taa tcc tgc tct a

>ON649724.1:156353-158616_Monkeypox_virus_isolate_Monkeypox/PT0014/2022_complete_genome

tta ctg act aca --- tgt a-c tat tt- -ta cat tac tac att ggc tac ggc ata tat acc tat ttc gtc act tcc aca cgc tcc ggt aaa cgg gtg tca tgt gac gag gga ga- -at ctt gat aag agg cat aat caa tgt tgt aat ccg atg tcc acc tgg aga att tgc caa ggt cag atg tag agt tgg tag tga taa cac aaa atg tga aca ctg ccc acc tca tac ata tac cgc aat ccc caa tta ttc t-a ata gat gtc atc aat gta gaa aat gcc caa cag gat cat ttg ata agg taa agt gta ccg gaa cac aga ac- --a aat -gt tcg tgt cat cct ggt tgg ta- tac gct act gat tct tca cag act gaa gat tgt cga gat ttg tgt acc aaa aaa gga gat gtc cat gcg gat act ttg gtg gaa tag atg aa- --g gaa atc cta ttt gta aat cgt gtt g-- -tg ttg gtg aat att gcg act acc tac gta att ata gac ttg atc cat ttc ctc cat gca aac tat cta tct aaa tgt aat taa tta --- --- --- --- --- --- --- --- tga ttt --- --- --- -tg atg ata atg tta -cc ata c-a tt- --- -at atc act act tgg tta gtg tgt att att tag tat gga aga cct att aat aa- --- --- -tt act tat ctt ttg acg atc ttg tt- -at aat tat --- --- --- --- aat ata aaa --a tac tta tga --- --- cat agt aac tca t-- --- --- --- aat tgc tg- acg cga taa att cgt aat aat ctg ttt tgt tca aat ttt tat aag ga- --- --a tct aca ggc ata aaa at- aaa aat ata atc tat aat ata c-- tct tac a-- acg cca tca tga at- --a gca gtg aat taa ttg ctg tta tt- atg gat tta gaa ata gtg gac gat ttt gtg ata tta ata tag tta tta atg atg aaa gga taa acg cgc ata gac tca tcc tat ctg gag cct ccg aat att ttt tcc att ctg ttt tcc aat aat ttt atc gat tct aat gaa tac gaa gtt aat cta agt cat tta gat tat caa agt gtt aac tat ttg atc gat tac att tat ggg ata cct ttg agc cta act aac aat aac gtg aaa tat att ctt tca acc gct ga- -tt ttt tta caa att gga tct gtc att act gag tgc gaa aaa tac ata ctt aaa aat ctt tgt ttt aga aac tgt atc gat ttc --- --- tac ata tac gct gat aaa tat aat aac aag aaa ata gaa tta gca tcg ttt aac aca ata tta cga aat att ttg aga ctc atc aac aat gaa aac ttt aaa tac tta aca gag gaa tca atg ata aaa att tta agc gat gat atg tta tat ata aaa aat gag gat ttc acc cca ctg att ctc att aaa tgg tta gag agt aca c-- caa cca tgt acc gtc gag tta ctt aga tgc ctc aga ata tca -tt tct ttc ccc aca agt tat aaa atc act tta tag tca tcg act ggt tag ttc aat cta cga atg tat aac att ctt aaa caa tat agc att ctt gga taa atc att tcc tag ata cca tat cat cga gtt gat atc tat cgg tat aag taa ttc aca tga taa gat ttc cat aaa ctg cta caa tca t-- aaa aaa att cat ggg aaa tga tat ctt cac gta gat ata ggt gta gtt tcg cag tga ccg tcc tgg ata ata tta tct ata tga tgg gtg gat atg atc agt ccc tgt ata gaa gtt caa agg tta tag cgt aca ata cat gta cta att ctt gga tat atg ata tac cag agc taa aaa tat cat cgt tct aat tgc gga gga gtt gcc aat gac gaa tac att tat tgt ata ggc ggt ata cgc gat cag gag tca tcg ttg ata tct agt atc gat aga tgg aag cca tca aaa cca tat tgg cag aag tat gct aaa atg tgc gaa cca aaa tgt gat atg gg- gtt gcg att tta aac gga tta ata tat gtc ata ggt gga gtc gtt aaa ggt gac aca cat ata cca acg cac tag aga gtt tat cag aag atg gat gga tga atc atc aac gtc ttc caa taa aaa tgt cca ata tgt cga cga ttg ttc atg ctg gaa aga ttt ata tat cta gag gtt aca aca ata gta gtg tag tta atg taa tat cga atc t-- agt cct tag cta taa tcc gat ata tga tga atg gac caa att atc atc att aaa tat tcc tag aat taa tcc tgc tct a

>ON649723.1:156353-158616_Monkeypox_virus_isolate_Monkeypox/PT0013/2022_complete_genome

tta ctg act aca --- tgt a-c tat tt- -ta cat tac tac att ggc tac ggc ata tat acc tat ttc gtc act tcc aca cgc tcc ggt aaa cgg gtg tca tgt gac gag gga ga- -at ctt gat aag agg cat aat caa tgt tgt aat ccg atg tcc acc tgg aga att tgc caa ggt cag atg tag agt tgg tag tga taa cac aaa atg tga aca ctg ccc acc tca tac ata tac cgc aat ccc caa tta ttc t-a ata gat gtc atc aat gta gaa aat gcc caa cag gat cat ttg ata agg taa agt gta ccg gaa cac aga ac- --a aat -gt tcg tgt cat cct ggt tgg ta- tac gct act gat tct tca cag act gaa gat tgt cga gat ttg tgt acc aaa aaa gga gat gtc cat gcg gat act ttg gtg gaa tag atg aa- --g gaa atc cta ttt gta aat cgt gtt g-- -tg ttg gtg aat att gcg act acc tac gta att ata gac ttg atc cat ttc ctc cat gca aac tat cta tct aaa tgt aat taa tta --- --- --- --- --- --- --- --- tga ttt --- --- --- -tg atg ata atg tta -cc ata c-a tt- --- -at atc act act tgg tta gtg tgt att att tag tat gga aga cct att aat aa- --- --- -tt act tat ctt ttg acg atc ttg tt- -at aat tat --- --- --- --- aat ata aaa --a tac tta tga --- --- cat agt aac tca t-- --- --- --- aat tgc tg- acg cga taa att cgt aat aat ctg ttt tgt tca aat ttt tat aag ga- --- --a tct aca ggc ata aaa at- aaa aat ata atc tat aat ata c-- tct tac a-- acg cca tca tga at- --a gca gtg aat taa ttg ctg tta tt- atg gat tta gaa ata gtg gac gat ttt gtg ata tta ata tag tta tta atg atg aaa gga taa acg cgc ata gac tca tcc tat ctg gag cct ccg aat att ttt tcc att ctg ttt tcc aat aat ttt atc gat tct aat gaa tac gaa gtt aat cta agt cat tta gat tat caa agt gtt aac tat ttg atc gat tac att tat ggg ata cct ttg agc cta act aac aat aac gtg aaa tat att ctt tca acc gct ga- -tt ttt tta caa att gga tct gtc att act gag tgc gaa aaa tac ata ctt aaa aat ctt tgt ttt aga aac tgt atc gat ttc --- --- tac ata tac gct gat aaa tat aat aac aag aaa ata gaa tta gca tcg ttt aac aca ata tta cga aat att ttg aga ctc atc aac aat gaa aac ttt aaa tac tta aca gag gaa tca atg ata aaa att tta agc gat gat atg tta tat ata aaa aat gag gat ttc acc cca ctg att ctc att aaa tgg tta gag agt aca c-- caa cca tgt acc gtc gag tta ctt aga tgc ctc aga ata tca -tt tct ttc ccc aca agt tat aaa atc act tta tag tca tcg act ggt tag ttc aat cta cga atg tat aac att ctt aaa caa tat agc att ctt gga taa atc att tcc tag ata cca tat cat cga gtt gat atc tat cgg tat aag taa ttc aca tga taa gat ttc cat aaa ctg cta caa tca t-- aaa aaa att cat ggg aaa tga tat ctt cac gta gat ata ggt gta gtt tcg cag tga ccg tcc tgg ata ata tta tct ata tga tgg gtg gat atg atc agt ccc tgt ata gaa gtt caa agg tta tag cgt aca ata cat gta cta att ctt gga tat atg ata tac cag agc taa aaa tat cat cgt tct aat tgc gga gga gtt gcc aat gac gaa tac att tat tgt ata ggc ggt ata cgc gat cag gag tca tcg ttg ata tct agt atc gat aga tgg aag cca tca aaa cca tat tgg cag aag tat gct aaa atg tgc gaa cca aaa tgt gat atg gg- gtt gcg att tta aac gga tta ata tat gtc ata ggt gga gtc gtt aaa ggt gac aca cat ata cca acg cac tag aga gtt tat cag aag atg gat gga tga atc atc aac gtc ttc caa taa aaa tgt cca ata tgt cga cga ttg ttc atg ctg gaa aga ttt ata tat cta gag gtt aca aca ata gta gtg tag tta atg taa tat cga atc t-- agt cct tag cta taa tcc gat ata tga tga atg gac caa att atc atc att aaa tat tcc tag aat taa tcc tgc tct a

>ON649722.1:156353-158616_Monkeypox_virus_isolate_Monkeypox/PT0022/2022_complete_genome

tta ctg act aca --- tgt a-c tat tt- -ta cat tac tac att ggc tac ggc ata tat acc tat ttc gtc act tcc aca cgc tcc ggt aaa cgg gtg tca tgt gac gag gga ga- -at ctt gat aag agg cat aat caa tgt tgt aat ccg atg tcc acc tgg aga att tgc caa ggt cag atg tag agt tgg tag tga taa cac aaa atg tga aca ctg ccc acc tca tac ata tac cgc aat ccc caa tta ttc t-a ata gat gtc atc aat gta gaa aat gcc caa cag gat cat ttg ata agg taa agt gta ccg gaa cac aga ac- --a aat -gt tcg tgt cat cct ggt tgg ta- tac gct act gat tct tca cag act gaa gat tgt cga gat ttg tgt acc aaa aaa gga gat gtc cat gcg gat act ttg gtg gaa tag atg aa- --g gaa atc cta ttt gta aat cgt gtt g-- -tg ttg gtg aat att gcg act acc tac gta att ata gac ttg atc cat ttc ctc cat gca aac tat cta tct aaa tgt aat taa tta --- --- --- --- --- --- --- --- tga ttt --- --- --- -tg atg ata atg tta -cc ata c-a tt- --- -at atc act act tgg tta gtg tgt att att tag tat gga aga cct att aat aa- --- --- -tt act tat ctt ttg acg atc ttg tt- -at aat tat --- --- --- --- aat ata aaa --a tac tta tga --- --- cat agt aac tca t-- --- --- --- aat tgc tg- acg cga taa att cgt aat aat ctg ttt tgt tca aat ttt tat aag ga- --- --a tct aca ggc ata aaa at- aaa aat ata atc tat aat ata c-- tct tac a-- acg cca tca tga at- --a gca gtg aat taa ttg ctg tta tt- atg gat tta gaa ata gtg gac gat ttt gtg ata tta ata tag tta tta atg atg aaa gga taa acg cgc ata gac tca tcc tat ctg gag cct ccg aat att ttt tcc att ctg ttt tcc aat aat ttt atc gat tct aat gaa tac gaa gtt aat cta agt cat tta gat tat caa agt gtt aac tat ttg atc gat tac att tat ggg ata cct ttg agc cta act aac aat aac gtg aaa tat att ctt tca acc gct ga- -tt ttt tta caa att gga tct gtc att act gag tgc gaa aaa tac ata ctt aaa aat ctt tgt ttt aga aac tgt atc gat ttc --- --- tac ata tac gct gat aaa tat aat aac aag aaa ata gaa tta gca tcg ttt aac aca ata tta cga aat att ttg aga ctc atc aac aat gaa aac ttt aaa tac tta aca gag gaa tca atg ata aaa att tta agc gat gat atg tta tat ata aaa aat gag gat ttc acc cca ctg att ctc att aaa tgg tta gag agt aca c-- caa cca tgt acc gtc gag tta ctt aga tgc ctc aga ata tca -tt tct ttc ccc aca agt tat aaa atc act tta tag tca tcg act ggt tag ttc aat cta cga atg tat aac att ctt aaa caa tat agc att ctt gga taa atc att tcc tag ata cca tat cat cga gtt gat atc tat cgg tat aag taa ttc aca tga taa gat ttc cat aaa ctg cta caa tca t-- aaa aaa att cat ggg aaa tga tat ctt cac gta gat ata ggt gta gtt tcg cag tga ccg tcc tgg ata ata tta tct ata tga tgg gtg gat atg atc agt ccc tgt ata gaa gtt caa agg tta tag cgt aca ata cat gta cta att ctt gga tat atg ata tac cag agc taa aaa tat cat cgt tct aat tgc gga gga gtt gcc aat gac gaa tac att tat tgt ata ggc ggt ata cgc gat cag gag tca tcg ttg ata tct agt atc gat aga tgg aag cca tca aaa cca tat tgg cag aag tat gct aaa atg tgc gaa cca aaa tgt gat atg gg- gtt gcg att tta aac gga tta ata tat gtc ata ggt gga gtc gtt aaa ggt gac aca cat ata cca acg cac tag aga gtt tat cag aag atg gat gga tga atc atc aac gtc ttc caa taa aaa tgt cca ata tgt cga cga ttg ttc atg ctg gaa aga ttt ata tat cta gag gtt aca aca ata gta gtg tag tta atg taa tat cga atc t-- agt cct tag cta taa tcc gat ata tga tga atg gac caa att atc atc att aaa tat tcc tag aat taa tcc tgc tct a

>ON649721.1:156353-158616_Monkeypox_virus_isolate_Monkeypox/PT0021/2022_complete_genome

tta ctg act aca --- tgt a-c tat tt- -ta cat tac tac att ggc tac ggc ata tat acc tat ttc gtc act tcc aca cgc tcc ggt aaa cgg gtg tca tgt gac gag gga ga- -at ctt gat aag agg cat aat caa tgt tgt aat ccg atg tcc acc tgg aga att tgc caa ggt cag atg tag agt tgg tag tga taa cac aaa atg tga aca ctg ccc acc tca tac ata tac cgc aat ccc caa tta ttc t-a ata gat gtc atc aat gta gaa aat gcc caa cag gat cat ttg ata agg taa agt gta ccg gaa cac aga ac- --a aat -gt tcg tgt cat cct ggt tgg ta- tac gct act gat tct tca cag act gaa gat tgt cga gat ttg tgt acc aaa aaa gga gat gtc cat gcg gat act ttg gtg gaa tag atg aa- --g gaa atc cta ttt gta aat cgt gtt g-- -tg ttg gtg aat att gcg act acc tac gta att ata gac ttg atc cat ttc ctc cat gca aac tat cta tct aaa tgt aat taa tta --- --- --- --- --- --- --- --- tga ttt --- --- --- -tg atg ata atg tta -cc ata c-a tt- --- -at atc act act tgg tta gtg tgt att att tag tat gga aga cct att aat aa- --- --- -tt act tat ctt ttg acg atc ttg tt- -at aat tat --- --- --- --- aat ata aaa --a tac tta tga --- --- cat agt aac tca t-- --- --- --- aat tgc tg- acg cga taa att cgt aat aat ctg ttt tgt tca aat ttt tat aag ga- --- --a tct aca ggc ata aaa at- aaa aat ata atc tat aat ata c-- tct tac a-- acg cca tca tga at- --a gca gtg aat taa ttg ctg tta tt- atg gat tta gaa ata gtg gac gat ttt gtg ata tta ata tag tta tta atg atg aaa gga taa acg cgc ata gac tca tcc tat ctg gag cct ccg aat att ttt tcc att ctg ttt tcc aat aat ttt atc gat tct aat gaa tac gaa gtt aat cta agt cat tta gat tat caa agt gtt aac tat ttg atc gat tac att tat ggg ata cct ttg agc cta act aac aat aac gtg aaa tat att ctt tca acc gct ga- -tt ttt tta caa att gga tct gtc att act gag tgc gaa aaa tac ata ctt aaa aat ctt tgt ttt aga aac tgt atc gat ttc --- --- tac ata tac gct gat aaa tat aat aac aag aaa ata gaa tta gca tcg ttt aac aca ata tta cga aat att ttg aga ctc atc aac aat gaa aac ttt aaa tac tta aca gag gaa tca atg ata aaa att tta agc gat gat atg tta tat ata aaa aat gag gat ttc acc cca ctg att ctc att aaa tgg tta gag agt aca c-- caa cca tgt acc gtc gag tta ctt aga tgc ctc aga ata tca -tt tct ttc ccc aca agt tat aaa atc act tta tag tca tcg act ggt tag ttc aat cta cga atg tat aac att ctt aaa caa tat agc att ctt gga taa atc att tcc tag ata cca tat cat cga gtt gat atc tat cgg tat aag taa ttc aca tga taa gat ttc cat aaa ctg cta caa tca t-- aaa aaa att cat ggg aaa tga tat ctt cac gta gat ata ggt gta gtt tcg cag tga ccg tcc tgg ata ata tta tct ata tga tgg gtg gat atg atc agt ccc tgt ata gaa gtt caa agg tta tag cgt aca ata cat gta cta att ctt gga tat atg ata tac cag agc taa aaa tat cat cgt tct aat tgc gga gga gtt gcc aat gac gaa tac att tat tgt ata ggc ggt ata cgc gat cag gag tca tcg ttg ata tct agt atc gat aga tgg aag cca tca aaa cca tat tgg cag aag tat gct aaa atg tgc gaa cca aaa tgt gat atg gg- gtt gcg att tta aac gga tta ata tat gtc ata ggt gga gtc gtt aaa ggt gac aca cat ata cca acg cac tag aga gtt tat cag aag atg gat gga tga atc atc aac gtc ttc caa taa aaa tgt cca ata tgt cga cga ttg ttc atg ctg gaa aga ttt ata tat cta gag gtt aca aca ata gta gtg tag tta atg taa tat cga atc t-- agt cct tag cta taa tcc gat ata tga tga atg gac caa att atc atc att aaa tat tcc tag aat taa tcc tgc tct a

>ON649720.1:156353-158616_Monkeypox_virus_isolate_Monkeypox/PT0024/2022_complete_genome

tta ctg act aca --- tgt a-c tat tt- -ta cat tac tac att ggc tac ggc ata tat acc tat ttc gtc act tcc aca cgc tcc ggt aaa cgg gtg tca tgt gac gag gga ga- -at ctt gat aag agg cat aat caa tgt tgt aat ccg atg tcc acc tgg aga att tgc caa ggt cag atg tag agt tgg tag tga taa cac aaa atg tga aca ctg ccc acc tca tac ata tac cgc aat ccc caa tta ttc t-a ata gat gtc atc aat gta gaa aat gcc caa cag gat cat ttg ata agg taa agt gta ccg gaa cac aga ac- --a aat -gt tcg tgt cat cct ggt tgg ta- tac gct act gat tct tca cag act gaa gat tgt cga gat ttg tgt acc aaa aaa gga gat gtc cat gcg gat act ttg gtg gaa tag atg aa- --g gaa atc cta ttt gta aat cgt gtt g-- -tg ttg gtg aat att gcg act acc tac gta att ata gac ttg atc cat ttc ctc cat gca aac tat cta tct aaa tgt aat taa tta --- --- --- --- --- --- --- --- tga ttt --- --- --- -tg atg ata atg tta -cc ata c-a tt- --- -at atc act act tgg tta gtg tgt att att tag tat gga aga cct att aat aa- --- --- -tt act tat ctt ttg acg atc ttg tt- -at aat tat --- --- --- --- aat ata aaa --a tac tta tga --- --- cat agt aac tca t-- --- --- --- aat tgc tg- acg cga taa att cgt aat aat ctg ttt tgt tca aat ttt tat aag ga- --- --a tct aca ggc ata aaa at- aaa aat ata atc tat aat ata c-- tct tac a-- acg cca tca tga at- --a gca gtg aat taa ttg ctg tta tt- atg gat tta gaa ata gtg gac gat ttt gtg ata tta ata tag tta tta atg atg aaa gga taa acg cgc ata gac tca tcc tat ctg gag cct ccg aat att ttt tcc att ctg ttt tcc aat aat ttt atc gat tct aat gaa tac gaa gtt aat cta agt cat tta gat tat caa agt gtt aac tat ttg atc gat tac att tat ggg ata cct ttg agc cta act aac aat aac gtg aaa tat att ctt tca acc gct ga- -tt ttt tta caa att gga tct gtc att act gag tgc gaa aaa tac ata ctt aaa aat ctt tgt ttt aga aac tgt atc gat ttc --- --- tac ata tac gct gat aaa tat aat aac aag aaa ata gaa tta gca tcg ttt aac aca ata tta cga aat att ttg aga ctc atc aac aat gaa aac ttt aaa tac tta aca gag gaa tca atg ata aaa att tta agc gat gat atg tta tat ata aaa aat gag gat ttc acc cca ctg att ctc att aaa tgg tta gag agt aca c-- caa cca tgt acc gtc gag tta ctt aga tgc ctc aga ata tca -tt tct ttc ccc aca agt tat aaa atc act tta tag tca tcg act ggt tag ttc aat cta cga atg tat aac att ctt aaa caa tat agc att ctt gga taa atc att tcc tag ata cca tat cat cga gtt gat atc tat cgg tat aag taa ttc aca tga taa gat ttc cat aaa ctg cta caa tca t-- aaa aaa att cat ggg aaa tga tat ctt cac gta gat ata ggt gta gtt tcg cag tga ccg tcc tgg ata ata tta tct ata tga tgg gtg gat atg atc agt ccc tgt ata gaa gtt caa agg tta tag cgt aca ata cat gta cta att ctt gga tat atg ata tac cag agc taa aaa tat cat cgt tct aat tgc gga gga gtt gcc aat gac gaa tac att tat tgt ata ggc ggt ata cgc gat cag gag tca tcg ttg ata tct agt atc gat aga tgg aag cca tca aaa cca tat tgg cag aag tat gct aaa atg tgc gaa cca aaa tgt gat atg gg- gtt gcg att tta aac gga tta ata tat gtc ata ggt gga gtc gtt aaa ggt gac aca cat ata cca acg cac tag aga gtt tat cag aag atg gat gga tga atc atc aac gtc ttc caa taa aaa tgt cca ata tgt cga cga ttg ttc atg ctg gaa aga ttt ata tat cta gag gtt aca aca ata gta gtg tag tta atg taa tat cga atc t-- agt cct tag cta taa tcc gat ata tga tga atg gac caa att atc atc att aaa tat tcc tag aat taa tcc tgc tct a

>ON649719.1:156353-158616_Monkeypox_virus_isolate_Monkeypox/PT0012/2022_complete_genome

tta ctg act aca --- tgt a-c tat tt- -ta cat tac tac att ggc tac ggc ata tat acc tat ttc gtc act tcc aca cgc tcc ggt aaa cgg gtg tca tgt gac gag gga ga- -at ctt gat aag agg cat aat caa tgt tgt aat ccg atg tcc acc tgg aga att tgc caa ggt cag atg tag agt tgg tag tga taa cac aaa atg tga aca ctg ccc acc tca tac ata tac cgc aat ccc caa tta ttc t-a ata gat gtc atc aat gta gaa aat gcc caa cag gat cat ttg ata agg taa agt gta ccg gaa cac aga ac- --a aat -gt tcg tgt cat cct ggt tgg ta- tac gct act gat tct tca cag act gaa gat tgt cga gat ttg tgt acc aaa aaa gga gat gtc cat gcg gat act ttg gtg gaa tag atg aa- --g gaa atc cta ttt gta aat cgt gtt g-- -tg ttg gtg aat att gcg act acc tac gta att ata gac ttg atc cat ttc ctc cat gca aac tat cta tct aaa tgt aat taa tta --- --- --- --- --- --- --- --- tga ttt --- --- --- -tg atg ata atg tta -cc ata c-a tt- --- -at atc act act tgg tta gtg tgt att att tag tat gga aga cct att aat aa- --- --- -tt act tat ctt ttg acg atc ttg tt- -at aat tat --- --- --- --- aat ata aaa --a tac tta tga --- --- cat agt aac tca t-- --- --- --- aat tgc tg- acg cga taa att cgt aat aat ctg ttt tgt tca aat ttt tat aag ga- --- --a tct aca ggc ata aaa at- aaa aat ata atc tat aat ata c-- tct tac a-- acg cca tca tga at- --a gca gtg aat taa ttg ctg tta tt- atg gat tta gaa ata gtg gac gat ttt gtg ata tta ata tag tta tta atg atg aaa gga taa acg cgc ata gac tca tcc tat ctg gag cct ccg aat att ttt tcc att ctg ttt tcc aat aat ttt atc gat tct aat gaa tac gaa gtt aat cta agt cat tta gat tat caa agt gtt aac tat ttg atc gat tac att tat ggg ata cct ttg agc cta act aac aat aac gtg aaa tat att ctt tca acc gct ga- -tt ttt tta caa att gga tct gtc att act gag tgc gaa aaa tac ata ctt aaa aat ctt tgt ttt aga aac tgt atc gat ttc --- --- tac ata tac gct gat aaa tat aat aac aag aaa ata gaa tta gca tcg ttt aac aca ata tta cga aat att ttg aga ctc atc aac aat gaa aac ttt aaa tac tta aca gag gaa tca atg ata aaa att tta agc gat gat atg tta tat ata aaa aat gag gat ttc acc cca ctg att ctc att aaa tgg tta gag agt aca c-- caa cca tgt acc gtc gag tta ctt aga tgc ctc aga ata tca -tt tct ttc ccc aca agt tat aaa atc act tta tag tca tcg act ggt tag ttc aat cta cga atg tat aac att ctt aaa caa tat agc att ctt gga taa atc att tcc tag ata cca tat cat cga gtt gat atc tat cgg tat aag taa ttc aca tga taa gat ttc cat aaa ctg cta caa tca t-- aaa aaa att cat ggg aaa tga tat ctt cac gta gat ata ggt gta gtt tcg cag tga ccg tcc tgg ata ata tta tct ata tga tgg gtg gat atg atc agt ccc tgt ata gaa gtt caa agg tta tag cgt aca ata cat gta cta att ctt gga tat atg ata tac cag agc taa aaa tat cat cgt tct aat tgc gga gga gtt gcc aat gac gaa tac att tat tgt ata ggc ggt ata cgc gat cag gag tca tcg ttg ata tct agt atc gat aga tgg aag cca tca aaa cca tat tgg cag aag tat gct aaa atg tgc gaa cca aaa tgt gat atg gg- gtt gcg att tta aac gga tta ata tat gtc ata ggt gga gtc gtt aaa ggt gac aca cat ata cca acg cac tag aga gtt tat cag aag atg gat gga tga atc atc aac gtc ttc caa taa aaa tgt cca ata tgt cga cga ttg ttc atg ctg gaa aga ttt ata tat cta gag gtt aca aca ata gta gtg tag tta atg taa tat cga atc t-- agt cct tag cta taa tcc gat ata tga tga atg gac caa att atc atc att aaa tat tcc tag aat taa tcc tgc tct a

>ON649718.1:156353-158616_Monkeypox_virus_isolate_Monkeypox/PT0018/2022_complete_genome

tta ctg act aca --- tgt a-c tat tt- -ta cat tac tac att ggc tac ggc ata tat acc tat ttc gtc act tcc aca cgc tcc ggt aaa cgg gtg tca tgt gac gag gga ga- -at ctt gat aag agg cat aat caa tgt tgt aat ccg atg tcc acc tgg aga att tgc caa ggt cag atg tag agt tgg tag tga taa cac aaa atg tga aca ctg ccc acc tca tac ata tac cgc aat ccc caa tta ttc t-a ata gat gtc atc aat gta gaa aat gcc caa cag gat cat ttg ata agg taa agt gta ccg gaa cac aga ac- --a aat -gt tcg tgt cat cct ggt tgg ta- tac gct act gat tct tca cag act gaa gat tgt cga gat ttg tgt acc aaa aaa gga gat gtc cat gcg gat act ttg gtg gaa tag atg aa- --g gaa atc cta ttt gta aat cgt gtt g-- -tg ttg gtg aat att gcg act acc tac gta att ata gac ttg atc cat ttc ctc cat gca aac tat cta tct aaa tgt aat taa tta --- --- --- --- --- --- --- --- tga ttt --- --- --- -tg atg ata atg tta -cc ata c-a tt- --- -at atc act act tgg tta gtg tgt att att tag tat gga aga cct att aat aa- --- --- -tt act tat ctt ttg acg atc ttg tt- -at aat tat --- --- --- --- aat ata aaa --a tac tta tga --- --- cat agt aac tca t-- --- --- --- aat tgc tg- acg cga taa att cgt aat aat ctg ttt tgt tca aat ttt tat aag ga- --- --a tct aca ggc ata aaa at- aaa aat ata atc tat aat ata c-- tct tac a-- acg cca tca tga at- --a gca gtg aat taa ttg ctg tta tt- atg gat tta gaa ata gtg gac gat ttt gtg ata tta ata tag tta tta atg atg aaa gga taa acg cgc ata gac tca tcc tat ctg gag cct ccg aat att ttt tcc att ctg ttt tcc aat aat ttt atc gat tct aat gaa tac gaa gtt aat cta agt cat tta gat tat caa agt gtt aac tat ttg atc gat tac att tat ggg ata cct ttg agc cta act aac aat aac gtg aaa tat att ctt tca acc gct ga- -tt ttt tta caa att gga tct gtc att act gag tgc gaa aaa tac ata ctt aaa aat ctt tgt ttt aga aac tgt atc gat ttc --- --- tac ata tac gct gat aaa tat aat aac aag aaa ata gaa tta gca tcg ttt aac aca ata tta cga aat att ttg aga ctc atc aac aat gaa aac ttt aaa tac tta aca gag gaa tca atg ata aaa att tta agc gat gat atg tta tat ata aaa aat gag gat ttc acc cca ctg att ctc att aaa tgg tta gag agt aca c-- caa cca tgt acc gtc gag tta ctt aga tgc ctc aga ata tca -tt tct ttc ccc aca agt tat aaa atc act tta tag tca tcg act ggt tag ttc aat cta cga atg tat aac att ctt aaa caa tat agc att ctt gga taa atc att tcc tag ata cca tat cat cga gtt gat atc tat cgg tat aag taa ttc aca tga taa gat ttc cat aaa ctg cta caa tca t-- aaa aaa att cat ggg aaa tga tat ctt cac gta gat ata ggt gta gtt tcg cag tga ccg tcc tgg ata ata tta tct ata tga tgg gtg gat atg atc agt ccc tgt ata gaa gtt caa agg tta tag cgt aca ata cat gta cta att ctt gga tat atg ata tac cag agc taa aaa tat cat cgt tct aat tgc gga gga gtt gcc aat gac gaa tac att tat tgt ata ggc ggt ata cgc gat cag gag tca tcg ttg ata tct agt atc gat aga tgg aag cca tca aaa cca tat tgg cag aag tat gct aaa atg tgc gaa cca aaa tgt gat atg gg- gtt gcg att tta aac gga tta ata tat gtc ata ggt gga gtc gtt aaa ggt gac aca cat ata cca acg cac tag aga gtt tat cag aag atg gat gga tga atc atc aac gtc ttc caa taa aaa tgt cca ata tgt cga cga ttg ttc atg ctg gaa aga ttt ata tat cta gag gtt aca aca ata gta gtg tag tta atg taa tat cga atc t-- agt cct tag cta taa tcc gat ata tga tga atg gac caa att atc atc att aaa tat tcc tag aat taa tcc tgc tct a

>ON649717.1:156347-158610_Monkeypox_virus_isolate_Monkeypox/PT0019/2022_partial_genome

tta ctg act aca --- tgt a-c tat tt- -ta cat tac tac att ggc tac ggc ata tat acc tat ttc gtc act tcc aca cgc tcc ggt aaa cgg gtg tca tgt gac gag gga ga- -at ctt gat aag agg cat aat caa tgt tgt aat ccg atg tcc acc tgg aga att tgc caa ggt cag atg tag agt tgg tag tga taa cac aaa atg tga aca ctg ccc acc tca tac ata tac cgc aat ccc caa tta ttc t-a ata gat gtc atc aat gta gaa aat gcc caa cag gat cat ttg ata agg taa agt gta ccg gaa cac aga ac- --a aat -gt tcg tgt cat cct ggt tgg ta- tac gct act gat tct tca cag act gaa gat tgt cga gat ttg tgt acc aaa aaa gga gat gtc cat gcg gat act ttg gtg gaa tag atg aa- --g gaa atc cta ttt gta aat cgt gtt g-- -tg ttg gtg aat att gcg act acc tac gta att ata gac ttg atc cat ttc ctc cat gca aac tat cta tct aaa tgt aat taa tta --- --- --- --- --- --- --- --- tga ttt --- --- --- -tg atg ata atg tta -cc ata c-a tt- --- -at atc act act tgg tta gtg tgt att att tag tat gga aga cct att aat aa- --- --- -tt act tat ctt ttg acg atc ttg tt- -at aat tat --- --- --- --- aat ata aaa --a tac tta tga --- --- cat agt aac tca t-- --- --- --- aat tgc tg- acg cga taa att cgt aat aat ctg ttt tgt tca aat ttt tat aag ga- --- --a tct aca ggc ata aaa at- aaa aat ata atc tat aat ata c-- tct tac a-- acg cca tca tga at- --a gca gtg aat taa ttg ctg tta tt- atg gat tta gaa ata gtg gac gat ttt gtg ata tta ata tag tta tta atg atg aaa gga taa acg cgc ata gac tca tcc tat ctg gag cct ccg aat att ttt tcc att ctg ttt tcc aat aat ttt atc gat tct aat gaa tac gaa gtt aat cta agt cat tta gat tat caa agt gtt aac tat ttg atc gat tac att tat ggg ata cct ttg agc cta act aac aat aac gtg aaa tat att ctt tca acc gct ga- -tt ttt tta caa att gga tct gtc att act gag tgc gaa aaa tac ata ctt aaa aat ctt tgt ttt aga aac tgt atc gat ttc --- --- tac ata tac gct gat aaa tat aat aac aag aaa ata gaa tta gca tcg ttt aac aca ata tta cga aat att ttg aga ctc atc aac aat gaa aac ttt aaa tac tta aca gag gaa tca atg ata aaa att tta agc gat gat atg tta tat ata aaa aat gag gat ttc acc cca ctg att ctc att aaa tgg tta gag agt aca c-- caa cca tgt acc gtc gag tta ctt aga tgc ctc aga ata tca -tt tct ttc ccc aca agt tat aaa atc act tta tag tca tcg act ggt tag ttc aat cta cga atg tat aac att ctt aaa caa tat agc att ctt gga taa atc att tcc tag ata cca tat cat cga gtt gat atc tat cgg tat aag taa ttc aca tga taa gat ttc cat aaa ctg cta caa tca t-- aaa aaa att cat ggg aaa tga tat ctt cac gta gat ata ggt gta gtt tcg cag tga ccg tcc tgg ata ata tta tct ata tga tgg gtg gat atg atc agt ccc tgt ata gaa gtt caa agg tta tag cgt aca ata cat gta cta att ctt gga tat atg ata tac cag agc taa aaa tat cat cgt tct aat tgc gga gga gtt gcc aat gac gaa tac att tat tgt ata ggc ggt ata cgc gat cag gag tca tcg ttg ata tct agt atc gat aga tgg aag cca tca aaa cca tat tgg cag aag tat gct aaa atg tgc gaa cca aaa tgt gat atg gg- gtt gcg att tta aac gga tta ata tat gtc ata ggt gga gtc gtt aaa ggt gac aca cat ata cca acg cac tag aga gtt tat cag aag atg gat gga tga atc atc aac gtc ttc caa taa aaa tgt cca ata tgt cga cga ttg ttc atg ctg gaa aga ttt ata tat cta gag gtt aca aca ata gta gtg tag tta atg taa tat cga atc t-- agt cct tag cta taa tcc gat ata tga tga atg gac caa att atc atc att aaa tat tcc tag aat taa tcc tgc tct a

>ON649713.1:155440-157703_Monkeypox_virus_isolate_Monkeypox/PT0020/2022_complete_genome

tta ctg act aca --- tgt a-c tat tt- -ta cat tac tac att ggc tac ggc ata tat acc tat ttc gtc act tcc aca cgc tcc ggt aaa cgg gtg tca tgt gac gag gga ga- -at ctt gat aag agg cat aat caa tgt tgt aat ccg atg tcc acc tgg aga att tgc caa ggt cag atg tag agt tgg tag tga taa cac aaa atg tga aca ctg ccc acc tca tac ata tac cgc aat ccc caa tta ttc t-a ata gat gtc atc aat gta gaa aat gcc caa cag gat cat ttg ata agg taa agt gta ccg gaa cac aga ac- --a aat -gt tcg tgt cat cct ggt tgg ta- tac gct act gat tct tca cag act gaa gat tgt cga gat ttg tgt acc aaa aaa gga gat gtc cat gcg gat act ttg gtg gaa tag atg aa- --g gaa atc cta ttt gta aat cgt gtt g-- -tg ttg gtg aat att gcg act acc tac gta att ata gac ttg atc cat ttc ctc cat gca aac tat cta tct aaa tgt aat taa tta --- --- --- --- --- --- --- --- tga ttt --- --- --- -tg atg ata atg tta -cc ata c-a tt- --- -at atc act act tgg tta gtg tgt att att tag tat gga aga cct att aat aa- --- --- -tt act tat ctt ttg acg atc ttg tt- -at aat tat --- --- --- --- aat ata aaa --a tac tta tga --- --- cat agt aac tca t-- --- --- --- aat tgc tg- acg cga taa att cgt aat aat ctg ttt tgt tca aat ttt tat aag ga- --- --a tct aca ggc ata aaa at- aaa aat ata atc tat aat ata c-- tct tac a-- acg cca tca tga at- --a gca gtg aat taa ttg ctg tta tt- atg gat tta gaa ata gtg gac gat ttt gtg ata tta ata tag tta tta atg atg aaa gga taa acg cgc ata gac tca tcc tat ctg gag cct ccg aat att ttt tcc att ctg ttt tcc aat aat ttt atc gat tct aat gaa tac gaa gtt aat cta agt cat tta gat tat caa agt gtt aac tat ttg atc gat tac att tat ggg ata cct ttg agc cta act aac aat aac gtg aaa tat att ctt tca acc gct ga- -tt ttt tta caa att gga tct gtc att act gag tgc gaa aaa tac ata ctt aaa aat ctt tgt ttt aga aac tgt atc gat ttc --- --- tac ata tac gct gat aaa tat aat aac aag aaa ata gaa tta gca tcg ttt aac aca ata tta cga aat att ttg aga ctc atc aac aat gaa aac ttt aaa tac tta aca gag gaa tca atg ata aaa att tta agc gat gat atg tta tat ata aaa aat gag gat ttc acc cca ctg att ctc att aaa tgg tta gag agt aca c-- caa cca tgt acc gtc gag tta ctt aga tgc ctc aga ata tca -tt tct ttc ccc aca agt tat aaa atc act tta tag tca tcg act ggt tag ttc aat cta cga atg tat aac att ctt aaa caa tat agc att ctt gga taa atc att tcc tag ata cca tat cat cga gtt gat atc tat cgg tat aag taa ttc aca tga taa gat ttc cat aaa ctg cta caa tca t-- aaa aaa att cat ggg aaa tga tat ctt cac gta gat ata ggt gta gtt tcg cag tga ccg tcc tgg ata ata tta tct ata tga tgg gtg gat atg atc agt ccc tgt ata gaa gtt caa agg tta tag cgt aca ata cat gta cta att ctt gga tat atg ata tac cag agc taa aaa tat cat cgt tct aat tgc gga gga gtt gcc aat gac gaa tac att tat tgt ata ggc ggt ata cgc gat cag gag tca tcg ttg ata tct agt atc gat aga tgg aag cca tca aaa cca tat tgg cag aag tat gct aaa atg tgc gaa cca aaa tgt gat atg gg- gtt gcg att tta aac gga tta ata tat gtc ata ggt gga gtc gtt aaa ggt gac aca cat ata cca acg cac tag aga gtt tat cag aag atg gat gga tga atc atc aac gtc ttc caa taa aaa tgt cca ata tgt cga cga ttg ttc atg ctg gaa aga ttt ata tat cta gag gtt aca aca ata gta gtg tag tta atg taa tat cga atc t-- agt cct tag cta taa tcc gat ata tga tga atg gac caa att atc atc att aaa tat tcc tag aat taa tcc tgc tct a

>ON649712.1:156347-158610_Monkeypox_virus_isolate_Monkeypox/PT0025/2022_partial_genome

tta ctg act aca --- tgt a-c tat tt- -ta cat tac tac att ggc tac ggc ata tat acc tat ttc gtc act tcc aca cgc tcc ggt aaa cgg gtg tca tgt gac gag gga ga- -at ctt gat aag agg cat aat caa tgt tgt aat ccg atg tcc acc tgg aga att tgc caa ggt cag atg tag agt tgg tag tga taa cac aaa atg tga aca ctg ccc acc tca tac ata tac cgc aat ccc caa tta ttc t-a ata gat gtc atc aat gta gaa aat gcc caa cag gat cat ttg ata agg taa agt gta ccg gaa cac aga ac- --a aat -gt tcg tgt cat cct ggt tgg ta- tac gct act gat tct tca cag act gaa gat tgt cga gat ttg tgt acc aaa aaa gga gat gtc cat gcg gat act ttg gtg gaa tag atg aa- --g gaa atc cta ttt gta aat cgt gtt g-- -tg ttg gtg aat att gcg act acc tac gta att ata gac ttg atc cat ttc ctc cat gca aac tat cta tct aaa tgt aat taa tta --- --- --- --- --- --- --- --- tga ttt --- --- --- -tg atg ata atg tta -cc ata c-a tt- --- -at atc act act tgg tta gtg tgt att att tag tat gga aga cct att aat aa- --- --- -tt act tat ctt ttg acg atc ttg tt- -at aat tat --- --- --- --- aat ata aaa --a tac tta tga --- --- cat agt aac tca t-- --- --- --- aat tgc tg- acg cga taa att cgt aat aat ctg ttt tgt tca aat ttt tat aag ga- --- --a tct aca ggc ata aaa at- aaa aat ata atc tat aat ata c-- tct tac a-- acg cca tca tga at- --a gca gtg aat taa ttg ctg tta tt- atg gat tta gaa ata gtg gac gat ttt gtg ata tta ata tag tta tta atg atg aaa gga taa acg cgc ata gac tca tcc tat ctg gag cct ccg aat att ttt tcc att ctg ttt tcc aat aat ttt atc gat tct aat gaa tac gaa gtt aat cta agt cat tta gat tat caa agt gtt aac tat ttg atc gat tac att tat ggg ata cct ttg agc cta act aac aat aac gtg aaa tat att ctt tca acc gct ga- -tt ttt tta caa att gga tct gtc att act gag tgc gaa aaa tac ata ctt aaa aat ctt tgt ttt aga aac tgt atc gat ttc --- --- tac ata tac gct gat aaa tat aat aac aag aaa ata gaa tta gca tcg ttt aac aca ata tta cga aat att ttg aga ctc atc aac aat gaa aac ttt aaa tac tta aca gag gaa tca atg ata aaa att tta agc gat gat atg tta tat ata aaa aat gag gat ttc acc cca ctg att ctc att aaa tgg tta gag agt aca c-- caa cca tgt acc gtc gag tta ctt aga tgc ctc aga ata tca -tt tct ttc ccc aca agt tat aaa atc act tta tag tca tcg act ggt tag ttc aat cta cga atg tat aac att ctt aaa caa tat agc att ctt gga taa atc att tcc tag ata cca tat cat cga gtt gat atc tat cgg tat aag taa ttc aca tga taa gat ttc cat aaa ctg cta caa tca t-- aaa aaa att cat ggg aaa tga tat ctt cac gta gat ata ggt gta gtt tcg cag tga ccg tcc tgg ata ata tta tct ata tga tgg gtg gat atg atc agt ccc tgt ata gaa gtt caa agg tta tag cgt aca ata cat gta cta att ctt gga tat atg ata tac cag agc taa aaa tat cat cgt tct aat tgc gga gga gtt gcc aat gac gaa tac att tat tgt ata ggc ggt ata cgc gat cag gag tca tcg ttg ata tct agt atc gat aga tgg aag cca tca aaa cca tat tgg cag aag tat gct aaa atg tgc gaa cca aaa tgt gat atg gg- gtt gcg att tta aac gga tta ata tat gtc ata ggt gga gtc gtt aaa ggt gac aca cat ata cca acg cac tag aga gtt tat cag aag atg gat gga tga atc atc aac gtc ttc caa taa aaa tgt cca ata tgt cga cga ttg ttc atg ctg gaa aga ttt ata tat cta gag gtt aca aca ata gta gtg tag tta atg taa tat cga atc t-- agt cct tag cta taa tcc gat ata tga tga atg gac caa att atc atc att aaa tat tcc tag aat taa tcc tgc tct a

>ON649711.1:156334-158597_Monkeypox_virus_isolate_Monkeypox/PT0026/2022_partial_genome

tta ctg act aca --- tgt a-c tat tt- -ta cat tac tac att ggc tac ggc ata tat acc tat ttc gtc act tcc aca cgc tcc ggt aaa cgg gtg tca tgt gac gag gga ga- -at ctt gat aag agg cat aat caa tgt tgt aat ccg atg tcc acc tgg aga att tgc caa ggt cag atg tag agt tgg tag tga taa cac aaa atg tga aca ctg ccc acc tca tac ata tac cgc aat ccc caa tta ttc t-a ata gat gtc atc aat gta gaa aat gcc caa cag gat cat ttg ata agg taa agt gta ccg gaa cac aga ac- --a aat -gt tcg tgt cat cct ggt tgg ta- tac gct act gat tct tca cag act gaa gat tgt cga gat ttg tgt acc aaa aaa gga gat gtc cat gcg gat act ttg gtg gaa tag atg aa- --g gaa atc cta ttt gta aat cgt gtt g-- -tg ttg gtg aat att gcg act acc tac gta att ata gac ttg atc cat ttc ctc cat gca aac tat cta tct aaa tgt aat taa tta --- --- --- --- --- --- --- --- tga ttt --- --- --- -tg atg ata atg tta -cc ata c-a tt- --- -at atc act act tgg tta gtg tgt att att tag tat gga aga cct att aat aa- --- --- -tt act tat ctt ttg acg atc ttg tt- -at aat tat --- --- --- --- aat ata aaa --a tac tta tga --- --- cat agt aac tca t-- --- --- --- aat tgc tg- acg cga taa att cgt aat aat ctg ttt tgt tca aat ttt tat aag ga- --- --a tct aca ggc ata aaa at- aaa aat ata atc tat aat ata c-- tct tac a-- acg cca tca tga at- --a gca gtg aat taa ttg ctg tta tt- atg gat tta gaa ata gtg gac gat ttt gtg ata tta ata tag tta tta atg atg aaa gga taa acg cgc ata gac tca tcc tat ctg gag cct ccg aat att ttt tcc att ctg ttt tcc aat aat ttt atc gat tct aat gaa tac gaa gtt aat cta agt cat tta gat tat caa agt gtt aac tat ttg atc gat tac att tat ggg ata cct ttg agc cta act aac aat aac gtg aaa tat att ctt tca acc gct ga- -tt ttt tta caa att gga tct gtc att act gag tgc gaa aaa tac ata ctt aaa aat ctt tgt ttt aga aac tgt atc gat ttc --- --- tac ata tac gct gat aaa tat aat aac aag aaa ata gaa tta gca tcg ttt aac aca ata tta cga aat att ttg aga ctc atc aac aat gaa aac ttt aaa tac tta aca gag gaa tca atg ata aaa att tta agc gat gat atg tta tat ata aaa aat gag gat ttc acc cca ctg att ctc att aaa tgg tta gag agt aca c-- caa cca tgt acc gtc gag tta ctt aga tgc ctc aga ata tca -tt tct ttc ccc aca agt tat aaa atc act tta tag tca tcg act ggt tag ttc aat cta cga atg tat aac att ctt aaa caa tat agc att ctt gga taa atc att tcc tag ata cca tat cat cga gtt gat atc tat cgg tat aag taa ttc aca tga taa gat ttc cat aaa ctg cta caa tca t-- aaa aaa att cat ggg aaa tga tat ctt cac gta gat ata ggt gta gtt tcg cag tga ccg tcc tgg ata ata tta tct ata tga tgg gtg gat atg atc agt ccc tgt ata gaa gtt caa agg tta tag cgt aca ata cat gta cta att ctt gga tat atg ata tac cag agc taa aaa tat cat cgt tct aat tgc gga gga gtt gcc aat gac gaa tac att tat tgt ata ggc ggt ata cgc gat cag gag tca tcg ttg ata tct agt atc gat aga tgg aag cca tca aaa cca tat tgg cag aag tat gct aaa atg tgc gaa cca aaa tgt gat atg gg- gtt gcg att tta aac gga tta ata tat gtc ata ggt gga gtc gtt aaa ggt gac aca cat ata cca acg cac tag aga gtt tat cag aag atg gat gga tga atc atc aac gtc ttc caa taa aaa tgt cca ata tgt cga cga ttg ttc atg ctg gaa aga ttt ata tat cta gag gtt aca aca ata gta gtg tag tta atg taa tat cga atc t-- agt cct tag cta taa tcc gat ata tga tga atg gac caa att atc atc att aaa tat tcc tag aat taa tcc tgc tct a

>ON649709.1:156353-158616_Monkeypox_virus_isolate_Monkeypox/PT0028/2022_complete_genome

tta ctg act aca --- tgt a-c tat tt- -ta cat tac tac att ggc tac ggc ata tat acc tat ttc gtc act tcc aca cgc tcc ggt aaa cgg gtg tca tgt gac gag gga ga- -at ctt gat aag agg cat aat caa tgt tgt aat ccg atg tcc acc tgg aga att tgc caa ggt cag atg tag agt tgg tag tga taa cac aaa atg tga aca ctg ccc acc tca tac ata tac cgc aat ccc caa tta ttc t-a ata gat gtc atc aat gta gaa aat gcc caa cag gat cat ttg ata agg taa agt gta ccg gaa cac aga ac- --a aat -gt tcg tgt cat cct ggt tgg ta- tac gct act gat tct tca cag act gaa gat tgt cga gat ttg tgt acc aaa aaa gga gat gtc cat gcg gat act ttg gtg gaa tag atg aa- --g gaa atc cta ttt gta aat cgt gtt g-- -tg ttg gtg aat att gcg act acc tac gta att ata gac ttg atc cat ttc ctc cat gca aac tat cta tct aaa tgt aat taa tta --- --- --- --- --- --- --- --- tga ttt --- --- --- -tg atg ata atg tta -cc ata c-a tt- --- -at atc act act tgg tta gtg tgt att att tag tat gga aga cct att aat aa- --- --- -tt act tat ctt ttg acg atc ttg tt- -at aat tat --- --- --- --- aat ata aaa --a tac tta tga --- --- cat agt aac tca t-- --- --- --- aat tgc tg- acg cga taa att cgt aat aat ctg ttt tgt tca aat ttt tat aag ga- --- --a tct aca ggc ata aaa at- aaa aat ata atc tat aat ata c-- tct tac a-- acg cca tca tga at- --a gca gtg aat taa ttg ctg tta tt- atg gat tta gaa ata gtg gac gat ttt gtg ata tta ata tag tta tta atg atg aaa gga taa acg cgc ata gac tca tcc tat ctg gag cct ccg aat att ttt tcc att ctg ttt tcc aat aat ttt atc gat tct aat gaa tac gaa gtt aat cta agt cat tta gat tat caa agt gtt aac tat ttg atc gat tac att tat ggg ata cct ttg agc cta act aac aat aac gtg aaa tat att ctt tca acc gct ga- -tt ttt tta caa att gga tct gtc att act gag tgc gaa aaa tac ata ctt aaa aat ctt tgt ttt aga aac tgt atc gat ttc --- --- tac ata tac gct gat aaa tat aat aac aag aaa ata gaa tta gca tcg ttt aac aca ata tta cga aat att ttg aga ctc atc aac aat gaa aac ttt aaa tac tta aca gag gaa tca atg ata aaa att tta agc gat gat atg tta tat ata aaa aat gag gat ttc acc cca ctg att ctc att aaa tgg tta gag agt aca c-- caa cca tgt acc gtc gag tta ctt aga tgc ctc aga ata tca -tt tct ttc ccc aca agt tat aaa atc act tta tag tca tcg act ggt tag ttc aat cta cga atg tat aac att ctt aaa caa tat agc att ctt gga taa atc att tcc tag ata cca tat cat cga gtt gat atc tat cgg tat aag taa ttc aca tga taa gat ttc cat aaa ctg cta caa tca t-- aaa aaa att cat ggg aaa tga tat ctt cac gta gat ata ggt gta gtt tcg cag tga ccg tcc tgg ata ata tta tct ata tga tgg gtg gat atg atc agt ccc tgt ata gaa gtt caa agg tta tag cgt aca ata cat gta cta att ctt gga tat atg ata tac cag agc taa aaa tat cat cgt tct aat tgc gga gga gtt gcc aat gac gaa tac att tat tgt ata ggc ggt ata cgc gat cag gag tca tcg ttg ata tct agt atc gat aga tgg aag cca tca aaa cca tat tgg cag aag tat gct aaa atg tgc gaa cca aaa tgt gat atg gg- gtt gcg att tta aac gga tta ata tat gtc ata ggt gga gtc gtt aaa ggt gac aca cat ata cca acg cac tag aga gtt tat cag aag atg gat gga tga atc atc aac gtc ttc caa taa aaa tgt cca ata tgt cga cga ttg ttc atg ctg gaa aga ttt ata tat cta gag gtt aca aca ata gta gtg tag tta atg taa tat cga atc t-- agt cct tag cta taa tcc gat ata tga tga atg gac caa att atc atc att aaa tat tcc tag aat taa tcc tgc tct a

>ON649708.1:156347-158610_Monkeypox_virus_isolate_Monkeypox/PT0023/2022_partial_genome

tta ctg act aca --- tgt a-c tat tt- -ta cat tac tac att ggc tac ggc ata tat acc tat ttc gtc act tcc aca cgc tcc ggt aaa cgg gtg tca tgt gac gag gga ga- -at ctt gat aag agg cat aat caa tgt tgt aat ccg atg tcc acc tgg aga att tgc caa ggt cag atg tag agt tgg tag tga taa cac aaa atg tga aca ctg ccc acc tca tac ata tac cgc aat ccc caa tta ttc t-a ata gat gtc atc aat gta gaa aat gcc caa cag gat cat ttg ata agg taa agt gta ccg gaa cac aga ac- --a aat -gt tcg tgt cat cct ggt tgg ta- tac gct act gat tct tca cag act gaa gat tgt cga gat ttg tgt acc aaa aaa gga gat gtc cat gcg gat act ttg gtg gaa tag atg aa- --g gaa atc cta ttt gta aat cgt gtt g-- -tg ttg gtg aat att gcg act acc tac gta att ata gac ttg atc cat ttc ctc cat gca aac tat cta tct aaa tgt aat taa tta --- --- --- --- --- --- --- --- tga ttt --- --- --- -tg atg ata atg tta -cc ata c-a tt- --- -at atc act act tgg tta gtg tgt att att tag tat gga aga cct att aat aa- --- --- -tt act tat ctt ttg acg atc ttg tt- -at aat tat --- --- --- --- aat ata aaa --a tac tta tga --- --- cat agt aac tca t-- --- --- --- aat tgc tg- acg cga taa att cgt aat aat ctg ttt tgt tca aat ttt tat aag ga- --- --a tct aca ggc ata aaa at- aaa aat ata atc tat aat ata c-- tct tac a-- acg cca tca tga at- --a gca gtg aat taa ttg ctg tta tt- atg gat tta gaa ata gtg gac gat ttt gtg ata tta ata tag tta tta atg atg aaa gga taa acg cgc ata gac tca tcc tat ctg gag cct ccg aat att ttt tcc att ctg ttt tcc aat aat ttt atc gat tct aat gaa tac gaa gtt aat cta agt cat tta gat tat caa agt gtt aac tat ttg atc gat tac att tat ggg ata cct ttg agc cta act aac aat aac gtg aaa tat att ctt tca acc gct ga- -tt ttt tta caa att gga tct gtc att act gag tgc gaa aaa tac ata ctt aaa aat ctt tgt ttt aga aac tgt atc gat ttc --- --- tac ata tac gct gat aaa tat aat aac aag aaa ata gaa tta gca tcg ttt aac aca ata tta cga aat att ttg aga ctc atc aac aat gaa aac ttt aaa tac tta aca gag gaa tca atg ata aaa att tta agc gat gat atg tta tat ata aaa aat gag gat ttc acc cca ctg att ctc att aaa tgg tta gag agt aca c-- caa cca tgt acc gtc gag tta ctt aga tgc ctc aga ata tca -tt tct ttc ccc aca agt tat aaa atc act tta tag tca tcg act ggt tag ttc aat cta cga atg tat aac att ctt aaa caa tat agc att ctt gga taa atc att tcc tag ata cca tat cat cga gtt gat atc tat cgg tat aag taa ttc aca tga taa gat ttc cat aaa ctg cta caa tca t-- aaa aaa att cat ggg aaa tga tat ctt cac gta gat ata ggt gta gtt tcg cag tga ccg tcc tgg ata ata tta tct ata tga tgg gtg gat atg atc agt ccc tgt ata gaa gtt caa agg tta tag cgt aca ata cat gta cta att ctt gga tat atg ata tac cag agc taa aaa tat cat cgt tct aat tgc gga gga gtt gcc aat gac gaa tac att tat tgt ata ggc ggt ata cgc gat cag gag tca tcg ttg ata tct agt atc gat aga tgg aag cca tca aaa cca tat tgg cag aag tat gct aaa atg tgc gaa cca aaa tgt gat atg gg- gtt gcg att tta aac gga tta ata tat gtc ata ggt gga gtc gtt aaa ggt gac aca cat ata cca acg cac tag aga gtt tat cag aag atg gat gga tga atc atc aac gtc ttc caa taa aaa tgt cca ata tgt cga cga ttg ttc atg ctg gaa aga ttt ata tat cta gag gtt aca aca ata gta gtg tag tta atg taa tat cga atc t-- agt cct tag cta taa tcc gat ata tga tga atg gac caa att atc atc att aaa tat tcc tag aat taa tcc tgc tct a

>ON622722.2:156363-158626_Monkeypox_virus_isolate_MPXV_FR_HCL0001_2022_complete_genome

tta ctg act aca --- tgt a-c tat tt- -ta cat tac tac att ggc tac ggc ata tat acc tat ttc gtc act tcc aca cgc tcc ggt aaa cgg gtg tca tgt gac gag gga ga- -at ctt gat aag agg cat aat caa tgt tgt aat ccg atg tcc acc tgg aga att tgc caa ggt cag atg tag agt tgg tag tga taa cac aaa atg tga aca ctg ccc acc tca tac ata tac cgc aat ccc caa tta ttc t-a ata gat gtc atc aat gta gaa aat gcc caa cag gat cat ttg ata agg taa agt gta ccg gaa cac aga ac- --a aat -gt tcg tgt cat cct ggt tgg ta- tac gct act gat tct tca cag act gaa gat tgt cga gat ttg tgt acc aaa aaa gga gat gtc cat gcg gat act ttg gtg gaa tag atg aa- --g gaa atc cta ttt gta aat cgt gtt g-- -tg ttg gtg aat att gcg act acc tac gta att ata gac ttg atc cat ttc ctc cat gca aac tat cta tct aaa tgt aat taa tta --- --- --- --- --- --- --- --- tga ttt --- --- --- -tg atg ata atg tta -cc ata c-a tt- --- -at atc act act tgg tta gtg tgt att att tag tat gga aga cct att aat aa- --- --- -tt act tat ctt ttg acg atc ttg tt- -at aat tat --- --- --- --- aat ata aaa --a tac tta tga --- --- cat agt aac tca t-- --- --- --- aat tgc tg- acg cga taa att cgt aat aat ctg ttt tgt tca aat ttt tat aag ga- --- --a tct aca ggc ata aaa at- aaa aat ata atc tat aat ata c-- tct tac a-- acg cca tca tga at- --a gca gtg aat taa ttg ctg tta tt- atg gat tta gaa ata gtg gac gat ttt gtg ata tta ata tag tta tta atg atg aaa gga taa acg cgc ata gac tca tcc tat ctg gag cct ccg aat att ttt tcc att ctg ttt tcc aat aat ttt atc gat tct aat gaa tac gaa gtt aat cta agt cat tta gat tat caa agt gtt aac tat ttg atc gat tac att tat ggg ata cct ttg agc cta act aac aat aac gtg aaa tat att ctt tca acc gct ga- -tt ttt tta caa att gga tct gtc att act gag tgc gaa aaa tac ata ctt aaa aat ctt tgt ttt aga aac tgt atc gat ttc --- --- tac ata tac gct gat aaa tat aat aac aag aaa ata gaa tta gca tcg ttt aac aca ata tta cga aat att ttg aga ctc atc aac aat gaa aac ttt aaa tac tta aca gag gaa tca atg ata aaa att tta agc gat gat atg tta tat ata aaa aat gag gat ttc acc cca ctg att ctc att aaa tgg tta gag agt aca c-- caa cca tgt acc gtc gag tta ctt aga tgc ctc aga ata tca -tt tct ttc ccc aca agt tat aaa atc act tta tag tca tcg act ggt tag ttc aat cta cga atg tat aac att ctt aaa caa tat agc att ctt gga taa atc att tcc tag ata cca tat cat cga gtt gat atc tat cgg tat aag taa ttc aca tga taa gat ttc cat aaa ctg cta caa tca t-- aaa aaa att cat ggg aaa tga tat ctt cac gta gat ata ggt gta gtt tcg cag tga ccg tcc tgg ata ata tta tct ata tga tgg gtg gat atg atc agt ccc tgt ata gaa gtt caa agg tta tag cgt aca ata cat gta cta att ctt gga tat atg ata tac cag agc taa aaa tat cat cgt tct aat tgc gga gga gtt gcc aat gac gaa tac att tat tgt ata ggc ggt ata cgc gat cag gag tca tcg ttg ata tct agt atc gat aga tgg aag cca tca aaa cca tat tgg cag aag tat gct aaa atg tgc gaa cca aaa tgt gat atg gg- gtt gcg att tta aac gga tta ata tat gtc ata ggt gga gtc gtt aaa ggt gac aca cat ata cca acg cac tag aga gtt tat cag aag atg gat gga tga atc atc aac gtc ttc caa taa aaa tgt cca ata tgt cga cga ttg ttc atg ctg gaa aga ttt ata tat cta gag gtt aca aca ata gta gtg tag tta atg taa tat cga atc t-- agt cct tag cta taa tcc gat ata tga tga atg gac caa att atc atc att aaa tat tcc tag aat taa tcc tgc tct a

>ON644344.1:156584-158847_Monkeypox_virus_isolate_MPXV_FVG-ITA_01_2022_complete_genome

tta ctg act aca --- tgt a-c tat tt- -ta cat tac tac att ggc tac ggc ata tat acc tat ttc gtc act tcc aca cgc tcc ggt aaa cgg gtg tca tgt gac gag gga ga- -at ctt gat aag agg cat aat caa tgt tgt aat ccg atg tcc acc tgg aga att tgc caa ggt cag atg tag agt tgg tag tga taa cac aaa atg tga aca ctg ccc acc tca tac ata tac cgc aat ccc caa tta ttc t-a ata gat gtc atc aat gta gaa aat gcc caa cag gat cat ttg ata agg taa agt gta ccg gaa cac aga ac- --a aat -gt tcg tgt cat cct ggt tgg ta- tac gct act gat tct tca cag act gaa gat tgt cga gat ttg tgt acc aaa aaa gga gat gtc cat gcg gat act ttg gtg gaa tag atg aa- --g gaa atc cta ttt gta aat cgt gtt g-- -tg ttg gtg aat att gcg act acc tac gta att ata gac ttg atc cat ttc ctc cat gca aac tat cta tct aaa tgt aat taa tta --- --- --- --- --- --- --- --- tga ttt --- --- --- -tg atg ata atg tta -cc ata c-a tt- --- -at atc act act tgg tta gtg tgt att att tag tat gga aga cct att aat aa- --- --- -tt act tat ctt ttg acg atc ttg tt- -at aat tat --- --- --- --- aat ata aaa --a tac tta tga --- --- cat agt aac tca t-- --- --- --- aat tgc tg- acg cga taa att cgt aat aat ctg ttt tgt tca aat ttt tat aag ga- --- --a tct aca ggc ata aaa at- aaa aat ata atc tat aat ata c-- tct tac a-- acg cca tca tga at- --a gca gtg aat taa ttg ctg tta tt- atg gat tta gaa ata gtg gac gat ttt gtg ata tta ata tag tta tta atg atg aaa gga taa acg cgc ata gac tca tcc tat ctg gag cct ccg aat att ttt tcc att ctg ttt tcc aat aat ttt atc gat tct aat gaa tac gaa gtt aat cta agt cat tta gat tat caa agt gtt aac tat ttg atc gat tac att tat ggg ata cct ttg agc cta act aac aat aac gtg aaa tat att ctt tca acc gct ga- -tt ttt tta caa att gga tct gtc att act gag tgc gaa aaa tac ata ctt aaa aat ctt tgt ttt aga aac tgt atc gat ttc --- --- tac ata tac gct gat aaa tat aat aac aag aaa ata gaa tta gca tcg ttt aac aca ata tta cga aat att ttg aga ctc atc aac aat gaa aac ttt aaa tac tta aca gag gaa tca atg ata aaa att tta agc gat gat atg tta tat ata aaa aat gag gat ttc acc cca ctg att ctc att aaa tgg tta gag agt aca c-- caa cca tgt acc gtc gag tta ctt aga tgc ctc aga ata tca -tt tct ttc ccc aca agt tat aaa atc act tta tag tca tcg act ggt tag ttc aat cta cga atg tat aac att ctt aaa caa tat agc att ctt gga taa atc att tcc tag ata cca tat cat cga gtt gat atc tat cgg tat aag taa ttc aca tga taa gat ttc cat aaa ctg cta caa tca t-- aaa aaa att cat ggg aaa tga tat ctt cac gta gat ata ggt gta gtt tcg cag tga ccg tcc tgg ata ata tta tct ata tga tgg gtg gat atg atc agt ccc tgt ata gaa gtt caa agg tta tag cgt aca ata cat gta cta att ctt gga tat atg ata tac cag agc taa aaa tat cat cgt tct aat tgc gga gga gtt gcc aat gac gaa tac att tat tgt ata ggc ggt ata cgc gat cag gag tca tcg ttg ata tct agt atc gat aga tgg aag cca tca aaa cca tat tgg cag aag tat gct aaa atg tgc gaa cca aaa tgt gat atg gg- gtt gcg att tta aac gga tta ata tat gtc ata ggt gga gtc gtt aaa ggt gac aca cat ata cca acg cac tag aga gtt tat cag aag atg gat gga tga atc atc aac gtc ttc caa taa aaa tgt cca ata tgt cga cga ttg ttc atg ctg gaa aga ttt ata tat cta gag gtt aca aca ata gta gtg tag tta atg taa tat cga atc t-- agt cct tag cta taa tcc gat ata tga tga atg gac caa att atc atc att aaa tat tcc tag aat taa tcc tgc tct a

>ON595760.2:156270-158533_Monkeypox_virus_isolate_MPXV-CH-38134631/2022_partial_genome

tta ctg act aca --- tgt a-c tat tt- -ta cat tac tac att ggc tac ggc ata tat acc tat ttc gtc act tcc aca cgc tcc ggt aaa cgg gtg tca tgt gac gag gga ga- -at ctt gat aag agg cat aat caa tgt tgt aat ccg atg tcc acc tgg aga att tgc caa ggt cag atg tag agt tgg tag tga taa cac aaa atg tga aca ctg ccc acc tca tac ata tac cgc aat ccc caa tta ttc t-a ata gat gtc atc aat gta gaa aat gcc caa cag gat cat ttg ata agg taa agt gta ccg gaa cac aga ac- --a aat -gt tcg tgt cat cct ggt tgg ta- tac gct act gat tct tca cag act gaa gat tgt cga gat ttg tgt acc aaa aaa gga gat gtc cat gcg gat act ttg gtg gaa tag atg aa- --g gaa atc cta ttt gta aat cgt gtt g-- -tg ttg gtg aat att gcg act acc tac gta att ata gac ttg atc cat ttc ctc cat gca aac tat cta tct aaa tgt aat taa tta --- --- --- --- --- --- --- --- tga ttt --- --- --- -tg atg ata atg tta -cc ata c-a tt- --- -at atc act act tgg tta gtg tgt att att tag tat gga aga cct att aat aa- --- --- -tt act tat ctt ttg acg atc ttg tt- -at aat tat --- --- --- --- aat ata aaa --a tac tta tga --- --- cat agt aac tca t-- --- --- --- aat tgc tg- acg cga taa att cgt aat aat ctg ttt tgt tca aat ttt tat aag ga- --- --a tct aca ggc ata aaa at- aaa aat ata atc tat aat ata c-- tct tac a-- acg cca tca tga at- --a gca gtg aat taa ttg ctg tta tt- atg gat tta gaa ata gtg gac gat ttt gtg ata tta ata tag tta tta atg atg aaa gga taa acg cgc ata gac tca tcc tat ctg gag cct ccg aat att ttt tcc att ctg ttt tcc aat aat ttt atc gat tct aat gaa tac gaa gtt aat cta agt cat tta gat tat caa agt gtt aac tat ttg atc gat tac att tat ggg ata cct ttg agc cta act aac aat aac gtg aaa tat att ctt tca acc gct ga- -tt ttt tta caa att gga tct gtc att act gag tgc gaa aaa tac ata ctt aaa aat ctt tgt ttt aga aac tgt atc gat ttc --- --- tac ata tac gct gat aaa tat aat aac aag aaa ata gaa tta gca tcg ttt aac aca ata tta cga aat att ttg aga ctc atc aac aat gaa aac ttt aaa tac tta aca gag gaa tca atg ata aaa att tta agc gat gat atg tta tat ata aaa aat gag gat ttc acc cca ctg att ctc att aaa tgg tta gag agt aca c-- caa cca tgt acc gtc gag tta ctt aga tgc ctc aga ata tca -tt tct ttc ccc aca agt tat aaa atc act tta tag tca tcg act ggt tag ttc aat cta cga atg tat aac att ctt aaa caa tat agc att ctt gga taa atc att tcc tag ata cca tat cat cga gtt gat atc tat cgg tat aag taa ttc aca tga taa gat ttc cat aaa ctg cta caa tca t-- aaa aaa att cat ggg aaa tga tat ctt cac gta gat ata ggt gta gtt tcg cag tga ccg tcc tgg ata ata tta tct ata tga tgg gtg gat atg atc agt ccc tgt ata gaa gtt caa agg tta tag cgt aca ata cat gta cta att ctt gga tat atg ata tac cag agc taa aaa tat cat cgt tct aat tgc gga gga gtt gcc aat gac gaa tac att tat tgt ata ggc ggt ata cgc gat cag gag tca tcg ttg ata tct agt atc gat aga tgg aag cca tca aaa cca tat tgg cag aag tat gct aaa atg tgc gaa cca aaa tgt gat atg gg- gtt gcg att tta aac gga tta ata tat gtc ata ggt gga gtc gtt aaa ggt gac aca cat ata cca acg cac tag aga gtt tat cag aag atg gat gga tga atc atc aac gtc ttc caa taa aaa tgt cca ata tgt cga cga ttg ttc atg ctg gaa aga ttt ata tat cta gag gtt aca aca ata gta gtg tag tta atg taa tat cga atc t-- agt cct tag cta taa tcc gat ata tga tga atg gac caa att atc atc att aaa tat tcc tag aat taa tcc tgc tct a

>ON637939.1:156427-158690_Monkeypox_virus_isolate_MPXV/Germany/2022/RKI02_complete_genome

tta ctg act aca --- tgt a-c tat tt- -ta cat tac tac att ggc tac ggc ata tat acc tat ttc gtc act tcc aca cgc tcc ggt aaa cgg gtg tca tgt gac gag gga ga- -at ctt gat aag agg cat aat caa tgt tgt aat ccg atg tcc acc tgg aga att tgc caa ggt cag atg tag agt tgg tag tga taa cac aaa atg tga aca ctg ccc acc tca tac ata tac cgc aat ccc caa tta ttc t-a ata gat gtc atc aat gta gaa aat gcc caa cag gat cat ttg ata agg taa agt gta ccg gaa cac aga ac- --a aat -gt tcg tgt cat cct ggt tgg ta- tac gct act gat tct tca cag act gaa gat tgt cga gat ttg tgt acc aaa aaa gga gat gtc cat gcg gat act ttg gtg gaa tag atg aa- --g gaa atc cta ttt gta aat cgt gtt g-- -tg ttg gtg aat att gcg act acc tac gta att ata gac ttg atc cat ttc ctc cat gca aac tat cta tct aaa tgt aat taa tta --- --- --- --- --- --- --- --- tga ttt --- --- --- -tg atg ata atg tta -cc ata c-a tt- --- -at atc act act tgg tta gtg tgt att att tag tat gga aga cct att aat aa- --- --- -tt act tat ctt ttg acg atc ttg tt- -at aat tat --- --- --- --- aat ata aaa --a tac tta tga --- --- cat agt aac tca t-- --- --- --- aat tgc tg- acg cga taa att cgt aat aat ctg ttt tgt tca aat ttt tat aag ga- --- --a tct aca ggc ata aaa at- aaa aat ata atc tat aat ata c-- tct tac a-- acg cca tca tga at- --a gca gtg aat taa ttg ctg tta tt- atg gat tta gaa ata gtg gac gat ttt gtg ata tta ata tag tta tta atg atg aaa gga taa acg cgc ata gac tca tcc tat ctg gag cct ccg aat att ttt tcc att ctg ttt tcc aat aat ttt atc gat tct aat gaa tac gaa gtt aat cta agt cat tta gat tat caa agt gtt aac tat ttg atc gat tac att tat ggg ata cct ttg agc cta act aac aat aac gtg aaa tat att ctt tca acc gct ga- -tt ttt tta caa att gga tct gtc att act gag tgc gaa aaa tac ata ctt aaa aat ctt tgt ttt aga aac tgt atc gat ttc --- --- tac ata tac gct gat aaa tat aat aac aag aaa ata gaa tta gca tcg ttt aac aca ata tta cga aat att ttg aga ctc atc aac aat gaa aac ttt aaa tac tta aca gag gaa tca atg ata aaa att tta agc gat gat atg tta tat ata aaa aat gag gat ttc acc cca ctg att ctc att aaa tgg tta gag agt aca c-- caa cca tgt acc gtc gag tta ctt aga tgc ctc aga ata tca -tt tct ttc ccc aca agt tat aaa atc act tta tag tca tcg act ggt tag ttc aat cta cga atg tat aac att ctt aaa caa tat agc att ctt gga taa atc att tcc tag ata cca tat cat cga gtt gat atc tat cgg tat aag taa ttc aca tga taa gat ttc cat aaa ctg cta caa tca t-- aaa aaa att cat ggg aaa tga tat ctt cac gta gat ata ggt gta gtt tcg cag tga ccg tcc tgg ata ata tta tct ata tga tgg gtg gat atg atc agt ccc tgt ata gaa gtt caa agg tta tag cgt aca ata cat gta cta att ctt gga tat atg ata tac cag agc taa aaa tat cat cgt tct aat tgc gga gga gtt gcc aat gac gaa tac att tat tgt ata ggc ggt ata cgc gat cag gag tca tcg ttg ata tct agt atc gat aga tgg aag cca tca aaa cca tat tgg cag aag tat gct aaa atg tgc gaa cca aaa tgt gat atg gg- gtt gcg att tta aac gga tta ata tat gtc ata ggt gga gtc gtt aaa ggt gac aca cat ata cca acg cac tag aga gtt tat cag aag atg gat gga tga atc atc aac gtc ttc caa taa aaa tgt cca ata tgt cga cga ttg ttc atg ctg gaa aga ttt ata tat cta gag gtt aca aca ata gta gtg tag tta atg taa tat cga atc t-- agt cct tag cta taa tcc gat ata tga tga atg gac caa att atc atc att aaa tat tcc tag aat taa tcc tgc tct a

>ON637938.1:156427-158690_Monkeypox_virus_isolate_MPXV/Germany/2022/RKI01_complete_genome

tta ctg act aca --- tgt a-c tat tt- -ta cat tac tac att ggc tac ggc ata tat acc tat ttc gtc act tcc aca cgc tcc ggt aaa cgg gtg tca tgt gac gag gga ga- -at ctt gat aag agg cat aat caa tgt tgt aat ccg atg tcc acc tgg aga att tgc caa ggt cag atg tag agt tgg tag tga taa cac aaa atg tga aca ctg ccc acc tca tac ata tac cgc aat ccc caa tta ttc t-a ata gat gtc atc aat gta gaa aat gcc caa cag gat cat ttg ata agg taa agt gta ccg gaa cac aga ac- --a aat -gt tcg tgt cat cct ggt tgg ta- tac gct act gat tct tca cag act gaa gat tgt cga gat ttg tgt acc aaa aaa gga gat gtc cat gcg gat act ttg gtg gaa tag atg aa- --g gaa atc cta ttt gta aat cgt gtt g-- -tg ttg gtg aat att gcg act acc tac gta att ata gac ttg atc cat ttc ctc cat gca aac tat cta tct aaa tgt aat taa tta --- --- --- --- --- --- --- --- tga ttt --- --- --- -tg atg ata atg tta -cc ata c-a tt- --- -at atc act act tgg tta gtg tgt att att tag tat gga aga cct att aat aa- --- --- -tt act tat ctt ttg acg atc ttg tt- -at aat tat --- --- --- --- aat ata aaa --a tac tta tga --- --- cat agt aac tca t-- --- --- --- aat tgc tg- acg cga taa att cgt aat aat ctg ttt tgt tca aat ttt tat aag ga- --- --a tct aca ggc ata aaa at- aaa aat ata atc tat aat ata c-- tct tac a-- acg cca tca tga at- --a gca gtg aat taa ttg ctg tta tt- atg gat tta gaa ata gtg gac gat ttt gtg ata tta ata tag tta tta atg atg aaa gga taa acg cgc ata gac tca tcc tat ctg gag cct ccg aat att ttt tcc att ctg ttt tcc aat aat ttt atc gat tct aat gaa tac gaa gtt aat cta agt cat tta gat tat caa agt gtt aac tat ttg atc gat tac att tat ggg ata cct ttg agc cta act aac aat aac gtg aaa tat att ctt tca acc gct ga- -tt ttt tta caa att gga tct gtc att act gag tgc gaa aaa tac ata ctt aaa aat ctt tgt ttt aga aac tgt atc gat ttc --- --- tac ata tac gct gat aaa tat aat aac aag aaa ata gaa tta gca tcg ttt aac aca ata tta cga aat att ttg aga ctc atc aac aat gaa aac ttt aaa tac tta aca gag gaa tca atg ata aaa att tta agc gat gat atg tta tat ata aaa aat gag gat ttc acc cca ctg att ctc att aaa tgg tta gag agt aca c-- caa cca tgt acc gtc gag tta ctt aga tgc ctc aga ata tca -tt tct ttc ccc aca agt tat aaa atc act tta tag tca tcg act ggt tag ttc aat cta cga atg tat aac att ctt aaa caa tat agc att ctt gga taa atc att tcc tag ata cca tat cat cga gtt gat atc tat cgg tat aag taa ttc aca tga taa gat ttc cat aaa ctg cta caa tca t-- aaa aaa att cat ggg aaa tga tat ctt cac gta gat ata ggt gta gtt tcg cag tga ccg tcc tgg ata ata tta tct ata tga tgg gtg gat atg atc agt ccc tgt ata gaa gtt caa agg tta tag cgt aca ata cat gta cta att ctt gga tat atg ata tac cag agc taa aaa tat cat cgt tct aat tgc gga gga gtt gcc aat gac gaa tac att tat tgt ata ggc ggt ata cgc gat cag gag tca tcg ttg ata tct agt atc gat aga tgg aag cca tca aaa cca tat tgg cag aag tat gct aaa atg tgc gaa cca aaa tgt gat atg gg- gtt gcg att tta aac gga tta ata tat gtc ata ggt gga gtc gtt aaa ggt gac aca cat ata cca acg cac tag aga gtt tat cag aag atg gat gga tga atc atc aac gtc ttc caa taa aaa tgt cca ata tgt cga cga ttg ttc atg ctg gaa aga ttt ata tat cta gag gtt aca aca ata gta gtg tag tta atg taa tat cga atc t-- agt cct tag cta taa tcc gat ata tga tga atg gac caa att atc atc att aaa tat tcc tag aat taa tcc tgc tct a

>ON631963.1:156372-158635_Monkeypox_virus_isolate_MPxV/VIDRL01/2022_complete_genome

tta ctg act aca --- tgt a-c tat tt- -ta cat tac tac att ggc tac ggc ata tat acc tat ttc gtc act tcc aca cgc tcc ggt aaa cgg gtg tca tgt gac gag gga ga- -at ctt gat aag agg cat aat caa tgt tgt aat ccg atg tcc acc tgg aga att tgc caa ggt cag atg tag agt tgg tag tga taa cac aaa atg tga aca ctg ccc acc tca tac ata tac cgc aat ccc caa tta ttc t-a ata gat gtc atc aat gta gaa aat gcc caa cag gat cat ttg ata agg taa agt gta ccg gaa cac aga ac- --a aat -gt tcg tgt cat cct ggt tgg ta- tac gct act gat tct tca cag act gaa gat tgt cga gat ttg tgt acc aaa aaa gga gat gtc cat gcg gat act ttg gtg gaa tag atg aa- --g gaa atc cta ttt gta aat cgt gtt g-- -tg ttg gtg aat att gcg act acc tac gta att ata gac ttg atc cat ttc ctc cat gca aac tat cta tct aaa tgt aat taa tta --- --- --- --- --- --- --- --- tga ttt --- --- --- -tg atg ata atg tta -cc ata c-a tt- --- -at atc act act tgg tta gtg tgt att att tag tat gga aga cct att aat aa- --- --- -tt act tat ctt ttg acg atc ttg tt- -at aat tat --- --- --- --- aat ata aaa --a tac tta tga --- --- cat agt aac tca t-- --- --- --- aat tgc tg- acg cga taa att cgt aat aat ctg ttt tgt tca aat ttt tat aag ga- --- --a tct aca ggc ata aaa at- aaa aat ata atc tat aat ata c-- tct tac a-- acg cca tca tga at- --a gca gtg aat taa ttg ctg tta tt- atg gat tta gaa ata gtg gac gat ttt gtg ata tta ata tag tta tta atg atg aaa gga taa acg cgc ata gac tca tcc tat ctg gag cct ccg aat att ttt tcc att ctg ttt tcc aat aat ttt atc gat tct aat gaa tac gaa gtt aat cta agt cat tta gat tat caa agt gtt aac tat ttg atc gat tac att tat ggg ata cct ttg agc cta act aac aat aac gtg aaa tat att ctt tca acc gct ga- -tt ttt tta caa att gga tct gtc att act gag tgc gaa aaa tac ata ctt aaa aat ctt tgt ttt aga aac tgt atc gat ttc --- --- tac ata tac gct gat aaa tat aat aac aag aaa ata gaa tta gca tcg ttt aac aca ata tta cga aat att ttg aga ctc atc aac aat gaa aac ttt aaa tac tta aca gag gaa tca atg ata aaa att tta agc gat gat atg tta tat ata aaa aat gag gat ttc acc cca ctg att ctc att aaa tgg tta gag agt aca c-- caa cca tgt acc gtc gag tta ctt aga tgc ctc aga ata tca -tt tct ttc ccc aca agt tat aaa atc act tta tag tca tcg act ggt tag ttc aat cta cga atg tat aac att ctt aaa caa tat agc att ctt gga taa atc att tcc tag ata cca tat cat cga gtt gat atc tat cgg tat aag taa ttc aca tga taa gat ttc cat aaa ctg cta caa tca t-- aaa aaa att cat ggg aaa tga tat ctt cac gta gat ata ggt gta gtt tcg cag tga ccg tcc tgg ata ata tta tct ata tga tgg gtg gat atg atc agt ccc tgt ata gaa gtt caa agg tta tag cgt aca ata cat gta cta att ctt gga tat atg ata tac cag agc taa aaa tat cat cgt tct aat tgc gga gga gtt gcc aat gac gaa tac att tat tgt ata ggc ggt ata cgc gat cag gag tca tcg ttg ata tct agt atc gat aga tgg aag cca tca aaa cca tat tgg cag aag tat gct aaa atg tgc gaa cca aaa tgt gat atg gg- gtt gcg att tta aac gga tta ata tat gtc ata ggt gga gtc gtt aaa ggt gac aca cat ata cca acg cac tag aga gtt tat cag aag atg gat gga tga atc atc aac gtc ttc caa taa aaa tgt cca ata tgt cga cga ttg ttc atg ctg gaa aga ttt ata tat cta gag gtt aca aca ata gta gtg tag tta atg taa tat cga atc t-- agt cct tag cta taa tcc gat ata tga tga atg gac caa att atc atc att aaa tat tcc tag aat taa tcc tgc tct a

>ON609725.2:156398-158661_Monkeypox_virus_isolate_SLO_complete_genome

tta ctg act aca --- tgt a-c tat tt- -ta cat tac tac att ggc tac ggc ata tat acc tat ttc gtc act tcc aca cgc tcc ggt aaa cgg gtg tca tgt gac gag gga ga- -at ctt gat aag agg cat aat caa tgt tgt aat ccg atg tcc acc tgg aga att tgc caa ggt cag atg tag agt tgg tag tga taa cac aaa atg tga aca ctg ccc acc tca tac ata tac cgc aat ccc caa tta ttc t-a ata gat gtc atc aat gta gaa aat gcc caa cag gat cat ttg ata agg taa agt gta ccg gaa cac aga ac- --a aat -gt tcg tgt cat cct ggt tgg ta- tac gct act gat tct tca cag act gaa gat tgt cga gat ttg tgt acc aaa aaa gga gat gtc cat gcg gat act ttg gtg gaa tag atg aa- --g gaa atc cta ttt gta aat cgt gtt g-- -tg ttg gtg aat att gcg act acc tac gta att ata gac ttg atc cat ttc ctc cat gca aac tat cta tct aaa tgt aat taa tta --- --- --- --- --- --- --- --- tga ttt --- --- --- -tg atg ata atg tta -cc ata c-a tt- --- -at atc act act tgg tta gtg tgt att att tag tat gga aga cct att aat aa- --- --- -tt act tat ctt ttg acg atc ttg tt- -at aat tat --- --- --- --- aat ata aaa --a tac tta tga --- --- cat agt aac tca t-- --- --- --- aat tgc tg- acg cga taa att cgt aat aat ctg ttt tgt tca aat ttt tat aag ga- --- --a tct aca ggc ata aaa at- aaa aat ata atc tat aat ata c-- tct tac a-- acg cca tca tga at- --a gca gtg aat taa ttg ctg tta tt- atg gat tta gaa ata gtg gac gat ttt gtg ata tta ata tag tta tta atg atg aaa gga taa acg cgc ata gac tca tcc tat ctg gag cct ccg aat att ttt tcc att ctg ttt tcc aat aat ttt atc gat tct aat gaa tac gaa gtt aat cta agt cat tta gat tat caa agt gtt aac tat ttg atc gat tac att tat ggg ata cct ttg agc cta act aac aat aac gtg aaa tat att ctt tca acc gct ga- -tt ttt tta caa att gga tct gtc att act gag tgc gaa aaa tac ata ctt aaa aat ctt tgt ttt aga aac tgt atc gat ttc --- --- tac ata tac gct gat aaa tat aat aac aag aaa ata gaa tta gca tcg ttt aac aca ata tta cga aat att ttg aga ctc atc aac aat gaa aac ttt aaa tac tta aca gag gaa tca atg ata aaa att tta agc gat gat atg tta tat ata aaa aat gag gat ttc acc cca ctg att ctc att aaa tgg tta gag agt aca c-- caa cca tgt acc gtc gag tta ctt aga tgc ctc aga ata tca -tt tct ttc ccc aca agt tat aaa atc act tta tag tca tcg act ggt tag ttc aat cta cga atg tat aac att ctt aaa caa tat agc att ctt gga taa atc att tcc tag ata cca tat cat cga gtt gat atc tat cgg tat aag taa ttc aca tga taa gat ttc cat aaa ctg cta caa tca t-- aaa aaa att cat ggg aaa tga tat ctt cac gta gat ata ggt gta gtt tcg cag tga ccg tcc tgg ata ata tta tct ata tga tgg gtg gat atg atc agt ccc tgt ata gaa gtt caa agg tta tag cgt aca ata cat gta cta att ctt gga tat atg ata tac cag agc taa aaa tat cat cgt tct aat tgc gga gga gtt gcc aat gac gaa tac att tat tgt ata ggc ggt ata cgc gat cag gag tca tcg ttg ata tct agt atc gat aga tgg aag cca tca aaa cca tat tgg cag aag tat gct aaa atg tgc gaa cca aaa tgt gat atg gg- gtt gcg att tta aac gga tta ata tat gtc ata ggt gga gtc gtt aaa ggt gac aca cat ata cca acg cac tag aga gtt tat cag aag atg gat gga tga atc atc aac gtc ttc caa taa aaa tgt cca ata tgt cga cga ttg ttc atg ctg gaa aga ttt ata tat cta gag gtt aca aca ata gta gtg tag tta atg taa tat cga atc t-- agt cct tag cta taa tcc gat ata tga tga atg gac caa att atc atc att aaa tat tcc tag aat taa tcc tgc tct a

>ON631241.1:156398-158661_Monkeypox_virus_isolate_2022/2_SLO_complete_genome

tta ctg act aca --- tgt a-c tat tt- -ta cat tac tac att ggc tac ggc ata tat acc tat ttc gtc act tcc aca cgc tcc ggt aaa cgg gtg tca tgt gac gag gga ga- -at ctt gat aag agg cat aat caa tgt tgt aat ccg atg tcc acc tgg aga att tgc caa ggt cag atg tag agt tgg tag tga taa cac aaa atg tga aca ctg ccc acc tca tac ata tac cgc aat ccc caa tta ttc t-a ata gat gtc atc aat gta gaa aat gcc caa cag gat cat ttg ata agg taa agt gta ccg gaa cac aga ac- --a aat -gt tcg tgt cat cct ggt tgg ta- tac gct act gat tct tca cag act gaa gat tgt cga gat ttg tgt acc aaa aaa gga gat gtc cat gcg gat act ttg gtg gaa tag atg aa- --g gaa atc cta ttt gta aat cgt gtt g-- -tg ttg gtg aat att gcg act acc tac gta att ata gac ttg atc cat ttc ctc cat gca aac tat cta tct aaa tgt aat taa tta --- --- --- --- --- --- --- --- tga ttt --- --- --- -tg atg ata atg tta -cc ata c-a tt- --- -at atc act act tgg tta gtg tgt att att tag tat gga aga cct att aat aa- --- --- -tt act tat ctt ttg acg atc ttg tt- -at aat tat --- --- --- --- aat ata aaa --a tac tta tga --- --- cat agt aac tca t-- --- --- --- aat tgc tg- acg cga taa att cgt aat aat ctg ttt tgt tca aat ttt tat aag ga- --- --a tct aca ggc ata aaa at- aaa aat ata atc tat aat ata c-- tct tac a-- acg cca tca tga at- --a gca gtg aat taa ttg ctg tta tt- atg gat tta gaa ata gtg gac gat ttt gtg ata tta ata tag tta tta atg atg aaa gga taa acg cgc ata gac tca tcc tat ctg gag cct ccg aat att ttt tcc att ctg ttt tcc aat aat ttt atc gat tct aat gaa tac gaa gtt aat cta agt cat tta gat tat caa agt gtt aac tat ttg atc gat tac att tat ggg ata cct ttg agc cta act aac aat aac gtg aaa tat att ctt tca acc gct ga- -tt ttt tta caa att gga tct gtc att act gag tgc gaa aaa tac ata ctt aaa aat ctt tgt ttt aga aac tgt atc gat ttc --- --- tac ata tac gct gat aaa tat aat aac aag aaa ata gaa tta gca tcg ttt aac aca ata tta cga aat att ttg aga ctc atc aac aat gaa aac ttt aaa tac tta aca gag gaa tca atg ata aaa att tta agc gat gat atg tta tat ata aaa aat gag gat ttc acc cca ctg att ctc att aaa tgg tta gag agt aca c-- caa cca tgt acc gtc gag tta ctt aga tgc ctc aga ata tca -tt tct ttc ccc aca agt tat aaa atc act tta tag tca tcg act ggt tag ttc aat cta cga atg tat aac att ctt aaa caa tat agc att ctt gga taa atc att tcc tag ata cca tat cat cga gtt gat atc tat cgg tat aag taa ttc aca tga taa gat ttc cat aaa ctg cta caa tca t-- aaa aaa att cat ggg aaa tga tat ctt cac gta gat ata ggt gta gtt tcg cag tga ccg tcc tgg ata ata tta tct ata tga tgg gtg gat atg atc agt ccc tgt ata gaa gtt caa agg tta tag cgt aca ata cat gta cta att ctt gga tat atg ata tac cag agc taa aaa tat cat cgt tct aat tgc gga gga gtt gcc aat gac gaa tac att tat tgt ata ggc ggt ata cgc gat cag gag tca tcg ttg ata tct agt atc gat aga tgg aag cca tca aaa cca tat tgg cag aag tat gct aaa atg tgc gaa cca aaa tgt gat atg gg- gtt gcg att tta aac gga tta ata tat gtc ata ggt gga gtc gtt aaa ggt gac aca cat ata cca acg cac tag aga gtt tat cag aag atg gat gga tga atc atc aac gtc ttc caa taa aaa tgt cca ata tgt cga cga ttg ttc atg ctg gaa aga ttt ata tat cta gag gtt aca aca ata gta gtg tag tta atg taa tat cga atc t-- agt cct tag cta taa tcc gat ata tga tga atg gac caa att atc atc att aaa tat tcc tag aat taa tcc tgc tct a

>ON563414.3:156363-158626_Monkeypox_virus_isolate_MPXV_USA_2022_MA001_complete_genome

tta ctg act aca --- tgt a-c tat tt- -ta cat tac tac att ggc tac ggc ata tat acc tat ttc gtc act tcc aca cgc tcc ggt aaa cgg gtg tca tgt gac gag gga ga- -at ctt gat aag agg cat aat caa tgt tgt aat ccg atg tcc acc tgg aga att tgc caa ggt cag atg tag agt tgg tag tga taa cac aaa atg tga aca ctg ccc acc tca tac ata tac cgc aat ccc caa tta ttc t-a ata gat gtc atc aat gta gaa aat gcc caa cag gat cat ttg ata agg taa agt gta ccg gaa cac aga ac- --a aat -gt tcg tgt cat cct ggt tgg ta- tac gct act gat tct tca cag act gaa gat tgt cga gat ttg tgt acc aaa aaa gga gat gtc cat gcg gat act ttg gtg gaa tag atg aa- --g gaa atc cta ttt gta aat cgt gtt g-- -tg ttg gtg aat att gcg act acc tac gta att ata gac ttg atc cat ttc ctc cat gca aac tat cta tct aaa tgt aat taa tta --- --- --- --- --- --- --- --- tga ttt --- --- --- -tg atg ata atg tta -cc ata c-a tt- --- -at atc act act tgg tta gtg tgt att att tag tat gga aga cct att aat aa- --- --- -tt act tat ctt ttg acg atc ttg tt- -at aat tat --- --- --- --- aat ata aaa --a tac tta tga --- --- cat agt aac tca t-- --- --- --- aat tgc tg- acg cga taa att cgt aat aat ctg ttt tgt tca aat ttt tat aag ga- --- --a tct aca ggc ata aaa at- aaa aat ata atc tat aat ata c-- tct tac a-- acg cca tca tga at- --a gca gtg aat taa ttg ctg tta tt- atg gat tta gaa ata gtg gac gat ttt gtg ata tta ata tag tta tta atg atg aaa gga taa acg cgc ata gac tca tcc tat ctg gag cct ccg aat att ttt tcc att ctg ttt tcc aat aat ttt atc gat tct aat gaa tac gaa gtt aat cta agt cat tta gat tat caa agt gtt aac tat ttg atc gat tac att tat ggg ata cct ttg agc cta act aac aat aac gtg aaa tat att ctt tca acc gct ga- -tt ttt tta caa att gga tct gtc att act gag tgc gaa aaa tac ata ctt aaa aat ctt tgt ttt aga aac tgt atc gat ttc --- --- tac ata tac gct gat aaa tat aat aac aag aaa ata gaa tta gca tcg ttt aac aca ata tta cga aat att ttg aga ctc atc aac aat gaa aac ttt aaa tac tta aca gag gaa tca atg ata aaa att tta agc gat gat atg tta tat ata aaa aat gag gat ttc acc cca ctg att ctc att aaa tgg tta gag agt aca c-- caa cca tgt acc gtc gag tta ctt aga tgc ctc aga ata tca -tt tct ttc ccc aca agt tat aaa atc act tta tag tca tcg act ggt tag ttc aat cta cga atg tat aac att ctt aaa caa tat agc att ctt gga taa atc att tcc tag ata cca tat cat cga gtt gat atc tat cgg tat aag taa ttc aca tga taa gat ttc cat aaa ctg cta caa tca t-- aaa aaa att cat ggg aaa tga tat ctt cac gta gat ata ggt gta gtt tcg cag tga ccg tcc tgg ata ata tta tct ata tga tgg gtg gat atg atc agt ccc tgt ata gaa gtt caa agg tta tag cgt aca ata cat gta cta att ctt gga tat atg ata tac cag agc taa aaa tat cat cgt tct aat tgc gga gga gtt gcc aat gac gaa tac att tat tgt ata ggc ggt ata cgc gat cag gag tca tcg ttg ata tct agt atc gat aga tgg aag cca tca aaa cca tat tgg cag aag tat gct aaa atg tgc gaa cca aaa tgt gat atg gg- gtt gcg att tta aac gga tta ata tat gtc ata ggt gga gtc gtt aaa ggt gac aca cat ata cca acg cac tag aga gtt tat cag aag atg gat gga tga atc atc aac gtc ttc caa taa aaa tgt cca ata tgt cga cga ttg ttc atg ctg gaa aga ttt ata tat cta gag gtt aca aca ata gta gtg tag tta atg taa tat cga atc t-- agt cct tag cta taa tcc gat ata tga tga atg gac caa att atc atc att aaa tat tcc tag aat taa tcc tgc tct a

>ON627808.1:156367-158630_Monkeypox_virus_isolate_MPX/human/USA/UT-UPHL-82200022/2022_complete_genome

tta ctg act aca --- tgt a-c tat tt- -ta cat tac tac att ggc tac ggc ata tat acc tat ttc gtc act tcc aca cgc tcc ggt aaa cgg gtg tca tgt gac gag gga ga- -at ctt gat aag agg cat aat caa tgt tgt aat ccg atg tcc acc tgg aga att tgc caa ggt cag atg tag agt tgg tag tga taa cac aaa atg tga aca ctg ccc acc tca tac ata tac cgc aat ccc caa tta ttc t-a ata gat gtc atc aat gta gaa aat gcc caa cag gat cat ttg ata agg taa agt gta ccg gaa cac aga ac- --a aat -gt tcg tgt cat cct ggt tgg ta- tac gct act gat tct tca cag act gaa gat tgt cga gat ttg tgt acc aaa aaa gga gat gtc cat gcg gat act ttg gtg gaa tag atg aa- --g gaa atc cta ttt gta aat cgt gtt g-- -tg ttg gtg aat att gcg act acc tac gta att ata gac ttg atc cat ttc ctc cat gca aac tat cta tct aaa tgt aat taa tta --- --- --- --- --- --- --- --- tga ttt --- --- --- -tg atg ata atg tta -cc ata c-a tt- --- -at atc act act tgg tta gtg tgt att att tag tat gga aga cct att aat aa- --- --- -tt act tat ctt ttg acg atc ttg tt- -at aat tat --- --- --- --- aat ata aaa --a tac tta tga --- --- cat agt aac tca t-- --- --- --- aat tgc tg- acg cga taa att cgt aat aat ctg ttt tgt tca aat ttt tat aag ga- --- --a tct aca ggc ata aaa at- aaa aat ata atc tat aat ata c-- tct tac a-- acg cca tca tga at- --a gca gtg aat taa ttg ctg tta tt- atg gat tta gaa ata gtg gac gat ttt gtg ata tta ata tag tta tta atg atg aaa gga taa acg cgc ata gac tca tcc tat ctg gag cct ccg aat att ttt tcc att ctg ttt tcc aat aat ttt atc gat tct aat gaa tac gaa gtt aat cta agt cat tta gat tat caa agt gtt aac tat ttg atc gat tac att tat ggg ata cct ttg agc cta act aac aat aac gtg aaa tat att ctt tca acc gct ga- -tt ttt tta caa att gga tct gtc att act gag tgc gaa aaa tac ata ctt aaa aat ctt tgt ttt aga aac tgt atc gat ttc --- --- tac ata tac gct gat aaa tat aat aac aag aaa ata gaa tta gca tcg ttt aac aca ata tta cga aat att ttg aga ctc atc aac aat gaa aac ttt aaa tac tta aca gag gaa tca atg ata aaa att tta agc gat gat atg tta tat ata aaa aat gag gat ttc acc cca ctg att ctc att aaa tgg tta gag agt aca c-- caa cca tgt acc gtc gag tta ctt aga tgc ctc aga ata tca -tt tct ttc ccc aca agt tat aaa atc act tta tag tca tcg act ggt tag ttc aat cta cga atg tat aac att ctt aaa caa tat agc att ctt gga taa atc att tcc tag ata cca tat cat cga gtt gat atc tat cgg tat aag taa ttc aca tga taa gat ttc cat aaa ctg cta caa tca t-- aaa aaa att cat ggg aaa tga tat ctt cac gta gat ata ggt gta gtt tcg cag tga ccg tcc tgg ata ata tta tct ata tga tgg gtg gat atg atc agt ccc tgt ata gaa gtt caa agg tta tag cgt aca ata cat gta cta att ctt gga tat atg ata tac cag agc taa aaa tat cat cgt tct aat tgc gga gga gtt gcc aat gac gaa tac att tat tgt ata ggc ggt ata cgc gat cag gag tca tcg ttg ata tct agt atc gat aga tgg aag cca tca aaa cca tat tgg cag aag tat gct aaa atg tgc gaa cca aaa tgt gat atg gg- gtt gcg att tta aac gga tta ata tat gtc ata ggt gga gtc gtt aaa ggt gac aca cat ata cca acg cac tag aga gtt tat cag aag atg gat gga tga atc atc aac gtc ttc caa taa aaa tgt cca ata tgt cga cga ttg ttc atg ctg gaa aga ttt ata tat cta gag gtt aca aca ata gta gtg tag tta atg taa tat cga atc t-- agt cct tag cta taa tcc gat ata tga tga atg gac caa att atc atc att aaa tat tcc tag aat taa tcc tgc tct a

>ON622713.1:156575-158838_Monkeypox_virus_isolate_MPX/UZ_REGA_2/Belgium/2022_complete_genome

tta ctg act aca --- tgt a-c tat tt- -ta cat tac tac att ggc tac ggc ata tat acc tat ttc gtc act tcc aca cgc tcc ggt aaa cgg gtg tca tgt gac gag gga ga- -at ctt gat aag agg cat aat caa tgt tgt aat ccg atg tcc acc tgg aga att tgc caa ggt cag atg tag agt tgg tag tga taa cac aaa atg tga aca ctg ccc acc tca tac ata tac cgc aat ccc caa tta ttc t-a ata gat gtc atc aat gta gaa aat gcc caa cag gat cat ttg ata agg taa agt gta ccg gaa cac aga ac- --a aat -gt tcg tgt cat cct ggt tgg ta- tac gct act gat tct tca cag act gaa gat tgt cga gat ttg tgt acc aaa aaa gga gat gtc cat gcg gat act ttg gtg gaa tag atg aa- --g gaa atc cta ttt gta aat cgt gtt g-- -tg ttg gtg aat att gcg act acc tac gta att ata gac ttg atc cat ttc ctc cat gca aac tat cta tct aaa tgt aat taa tta --- --- --- --- --- --- --- --- tga ttt --- --- --- -tg atg ata atg tta -cc ata c-a tt- --- -at atc act act tgg tta gtg tgt att att tag tat gga aga cct att aat aa- --- --- -tt act tat ctt ttg acg atc ttg tt- -at aat tat --- --- --- --- aat ata aaa --a tac tta tga --- --- cat agt aac tca t-- --- --- --- aat tgc tg- acg cga taa att cgt aat aat ctg ttt tgt tca aat ttt tat aag ga- --- --a tct aca ggc ata aaa at- aaa aat ata atc tat aat ata c-- tct tac a-- acg cca tca tga at- --a gca gtg aat taa ttg ctg tta tt- atg gat tta gaa ata gtg gac gat ttt gtg ata tta ata tag tta tta atg atg aaa gga taa acg cgc ata gac tca tcc tat ctg gag cct ccg aat att ttt tcc att ctg ttt tcc aat aat ttt atc gat tct aat gaa tac gaa gtt aat cta agt cat tta gat tat caa agt gtt aac tat ttg atc gat tac att tat ggg ata cct ttg agc cta act aac aat aac gtg aaa tat att ctt tca acc gct ga- -tt ttt tta caa att gga tct gtc att act gag tgc gaa aaa tac ata ctt aaa aat ctt tgt ttt aga aac tgt atc gat ttc --- --- tac ata tac gct gat aaa tat aat aac aag aaa ata gaa tta gca tcg ttt aac aca ata tta cga aat att ttg aga ctc atc aac aat gaa aac ttt aaa tac tta aca gag gaa tca atg ata aaa att tta agc gat gat atg tta tat ata aaa aat gag gat ttc acc cca ctg att ctc att aaa tgg tta gag agt aca c-- caa cca tgt acc gtc gag tta ctt aga tgc ctc aga ata tca -tt tct ttc ccc aca agt tat aaa atc act tta tag tca tcg act ggt tag ttc aat cta cga atg tat aac att ctt aaa caa tat agc att ctt gga taa atc att tcc tag ata cca tat cat cga gtt gat atc tat cgg tat aag taa ttc aca tga taa gat ttc cat aaa ctg cta caa tca t-- aaa aaa att cat ggg aaa tga tat ctt cac gta gat ata ggt gta gtt tcg cag tga ccg tcc tgg ata ata tta tct ata tga tgg gtg gat atg atc agt ccc tgt ata gaa gtt caa agg tta tag cgt aca ata cat gta cta att ctt gga tat atg ata tac cag agc taa aaa tat cat cgt tct aat tgc gga gga gtt gcc aat gac gaa tac att tat tgt ata ggc ggt ata cgc gat cag gag tca tcg ttg ata tct agt atc gat aga tgg aag cca tca aaa cca tat tgg cag aag tat gct aaa atg tgc gaa cca aaa tgt gat atg gg- gtt gcg att tta aac gga tta ata tat gtc ata ggt gga gtc gtt aaa ggt gac aca cat ata cca acg cac tag aga gtt tat cag aag atg gat gga tga atc atc aac gtc ttc caa taa aaa tgt cca ata tgt cga cga ttg ttc atg ctg gaa aga ttt ata tat cta gag gtt aca aca ata gta gtg tag tta atg taa tat cga atc t-- agt cct tag cta taa tcc gat ata tga tga atg gac caa att atc atc att aaa tat tcc tag aat taa tcc tgc tct a

>ON622712.1:156575-158838_Monkeypox_virus_isolate_MPX/UZ_REGA_1/Belgium/2022_complete_genome

tta ctg act aca --- tgt a-c tat tt- -ta cat tac tac att ggc tac ggc ata tat acc tat ttc gtc act tcc aca cgc tcc ggt aaa cgg gtg tca tgt gac gag gga ga- -at ctt gat aag agg cat aat caa tgt tgt aat ccg atg tcc acc tgg aga att tgc caa ggt cag atg tag agt tgg tag tga taa cac aaa atg tga aca ctg ccc acc tca tac ata tac cgc aat ccc caa tta ttc t-a ata gat gtc atc aat gta gaa aat gcc caa cag gat cat ttg ata agg taa agt gta ccg gaa cac aga ac- --a aat -gt tcg tgt cat cct ggt tgg ta- tac gct act gat tct tca cag act gaa gat tgt cga gat ttg tgt acc aaa aaa gga gat gtc cat gcg gat act ttg gtg gaa tag atg aa- --g gaa atc cta ttt gta aat cgt gtt g-- -tg ttg gtg aat att gcg act acc tac gta att ata gac ttg atc cat ttc ctc cat gca aac tat cta tct aaa tgt aat taa tta --- --- --- --- --- --- --- --- tga ttt --- --- --- -tg atg ata atg tta -cc ata c-a tt- --- -at atc act act tgg tta gtg tgt att att tag tat gga aga cct att aat aa- --- --- -tt act tat ctt ttg acg atc ttg tt- -at aat tat --- --- --- --- aat ata aaa --a tac tta tga --- --- cat agt aac tca t-- --- --- --- aat tgc tg- acg cga taa att cgt aat aat ctg ttt tgt tca aat ttt tat aag ga- --- --a tct aca ggc ata aaa at- aaa aat ata atc tat aat ata c-- tct tac a-- acg cca tca tga at- --a gca gtg aat taa ttg ctg tta tt- atg gat tta gaa ata gtg gac gat ttt gtg ata tta ata tag tta tta atg atg aaa gga taa acg cgc ata gac tca tcc tat ctg gag cct ccg aat att ttt tcc att ctg ttt tcc aat aat ttt atc gat tct aat gaa tac gaa gtt aat cta agt cat tta gat tat caa agt gtt aac tat ttg atc gat tac att tat ggg ata cct ttg agc cta act aac aat aac gtg aaa tat att ctt tca acc gct ga- -tt ttt tta caa att gga tct gtc att act gag tgc gaa aaa tac ata ctt aaa aat ctt tgt ttt aga aac tgt atc gat ttc --- --- tac ata tac gct gat aaa tat aat aac aag aaa ata gaa tta gca tcg ttt aac aca ata tta cga aat att ttg aga ctc atc aac aat gaa aac ttt aaa tac tta aca gag gaa tca atg ata aaa att tta agc gat gat atg tta tat ata aaa aat gag gat ttc acc cca ctg att ctc att aaa tgg tta gag agt aca c-- caa cca tgt acc gtc gag tta ctt aga tgc ctc aga ata tca -tt tct ttc ccc aca agt tat aaa atc act tta tag tca tcg act ggt tag ttc aat cta cga atg tat aac att ctt aaa caa tat agc att ctt gga taa atc att tcc tag ata cca tat cat cga gtt gat atc tat cgg tat aag taa ttc aca tga taa gat ttc cat aaa ctg cta caa tca t-- aaa aaa att cat ggg aaa tga tat ctt cac gta gat ata ggt gta gtt tcg cag tga ccg tcc tgg ata ata tta tct ata tga tgg gtg gat atg atc agt ccc tgt ata gaa gtt caa agg tta tag cgt aca ata cat gta cta att ctt gga tat atg ata tac cag agc taa aaa tat cat cgt tct aat tgc gga gga gtt gcc aat gac gaa tac att tat tgt ata ggc ggt ata cgc gat cag gag tca tcg ttg ata tct agt atc gat aga tgg aag cca tca aaa cca tat tgg cag aag tat gct aaa atg tgc gaa cca aaa tgt gat atg gg- gtt gcg att tta aac gga tta ata tat gtc ata ggt gga gtc gtt aaa ggt gac aca cat ata cca acg cac tag aga gtt tat cag aag atg gat gga tga atc atc aac gtc ttc caa taa aaa tgt cca ata tgt cga cga ttg ttc atg ctg gaa aga ttt ata tat cta gag gtt aca aca ata gta gtg tag tta atg taa tat cga atc t-- agt cct tag cta taa tcc gat ata tga tga atg gac caa att atc atc att aaa tat tcc tag aat taa tcc tgc tct a

>ON619838.1:156372-158635_Monkeypox_virus_isolate_MPXV_UK_2022_4_complete_genome

tta ctg act aca --- tgt a-c tat tt- -ta cat tac tac att ggc tac ggc ata tat acc tat ttc gtc act tcc aca cgc tcc ggt aaa cgg gtg tca tgt gac gag gga ga- -at ctt gat aag agg cat aat caa tgt tgt aat ccg atg tcc acc tgg aga att tgc caa ggt cag atg tag agt tgg tag tga taa cac aaa atg tga aca ctg ccc acc tca tac ata tac cgc aat ccc caa tta ttc t-a ata gat gtc atc aat gta gaa aat gcc caa cag gat cat ttg ata agg taa agt gta ccg gaa cac aga ac- --a aat -gt tcg tgt cat cct ggt tgg ta- tac gct act gat tct tca cag act gaa gat tgt cga gat ttg tgt acc aaa aaa gga gat gtc cat gcg gat act ttg gtg gaa tag atg aa- --g gaa atc cta ttt gta aat cgt gtt g-- -tg ttg gtg aat att gcg act acc tac gta att ata gac ttg atc cat ttc ctc cat gca aac tat cta tct aaa tgt aat taa tta --- --- --- --- --- --- --- --- tga ttt --- --- --- -tg atg ata atg tta -cc ata c-a tt- --- -at atc act act tgg tta gtg tgt att att tag tat gga aga cct att aat aa- --- --- -tt act tat ctt ttg acg atc ttg tt- -at aat tat --- --- --- --- aat ata aaa --a tac tta tga --- --- cat agt aac tca t-- --- --- --- aat tgc tg- acg cga taa att cgt aat aat ctg ttt tgt tca aat ttt tat aag ga- --- --a tct aca ggc ata aaa at- aaa aat ata atc tat aat ata c-- tct tac a-- acg cca tca tga at- --a gca gtg aat taa ttg ctg tta tt- atg gat tta gaa ata gtg gac gat ttt gtg ata tta ata tag tta tta atg atg aaa gga taa acg cgc ata gac tca tcc tat ctg gag cct ccg aat att ttt tcc att ctg ttt tcc aat aat ttt atc gat tct aat gaa tac gaa gtt aat cta agt cat tta gat tat caa agt gtt aac tat ttg atc gat tac att tat ggg ata cct ttg agc cta act aac aat aac gtg aaa tat att ctt tca acc gct ga- -tt ttt tta caa att gga tct gtc att act gag tgc gaa aaa tac ata ctt aaa aat ctt tgt ttt aga aac tgt atc gat ttc --- --- tac ata tac gct gat aaa tat aat aac aag aaa ata gaa tta gca tcg ttt aac aca ata tta cga aat att ttg aga ctc atc aac aat gaa aac ttt aaa tac tta aca gag gaa tca atg ata aaa att tta agc gat gat atg tta tat ata aaa aat gag gat ttc acc cca ctg att ctc att aaa tgg tta gag agt aca c-- caa cca tgt acc gtc gag tta ctt aga tgc ctc aga ata tca -tt tct ttc ccc aca agt tat aaa atc act tta tag tca tcg act ggt tag ttc aat cta cga atg tat aac att ctt aaa caa tat agc att ctt gga taa atc att tcc tag ata cca tat cat cga gtt gat atc tat cgg tat aag taa ttc aca tga taa gat ttc cat aaa ctg cta caa tca t-- aaa aaa att cat ggg aaa tga tat ctt cac gta gat ata ggt gta gtt tcg cag tga ccg tcc tgg ata ata tta tct ata tga tgg gtg gat atg atc agt ccc tgt ata gaa gtt caa agg tta tag cgt aca ata cat gta cta att ctt gga tat atg ata tac cag agc taa aaa tat cat cgt tct aat tgc gga gga gtt gcc aat gac gaa tac att tat tgt ata ggc ggt ata cgc gat cag gag tca tcg ttg ata tct agt atc gat aga tgg aag cca tca aaa cca tat tgg cag aag tat gct aaa atg tgc gaa cca aaa tgt gat atg gg- gtt gcg att tta aac gga tta ata tat gtc ata ggt gga gtc gtt aaa ggt gac aca cat ata cca acg cac tag aga gtt tat cag aag atg gat gga tga atc atc aac gtc ttc caa taa aaa tgt cca ata tgt cga cga ttg ttc atg ctg gaa aga ttt ata tat cta gag gtt aca aca ata gta gtg tag tta atg taa tat cga atc t-- agt cct tag cta taa tcc gat ata tga tga atg gac caa att atc atc att aaa tat tcc tag aat taa tcc tgc tct a

>ON619837.1:156371-158634_Monkeypox_virus_isolate_MPXV_UK_2022_3_complete_genome

tta ctg act aca --- tgt a-c tat tt- -ta cat tac tac att ggc tac ggc ata tat acc tat ttc gtc act tcc aca cgc tcc ggt aaa cgg gtg tca tgt gac gag gga ga- -at ctt gat aag agg cat aat caa tgt tgt aat ccg atg tcc acc tgg aga att tgc caa ggt cag atg tag agt tgg tag tga taa cac aaa atg tga aca ctg ccc acc tca tac ata tac cgc aat ccc caa tta ttc t-a ata gat gtc atc aat gta gaa aat gcc caa cag gat cat ttg ata agg taa agt gta ccg gaa cac aga ac- --a aat -gt tcg tgt cat cct ggt tgg ta- tac gct act gat tct tca cag act gaa gat tgt cga gat ttg tgt acc aaa aaa gga gat gtc cat gcg gat act ttg gtg gaa tag atg aa- --g gaa atc cta ttt gta aat cgt gtt g-- -tg ttg gtg aat att gcg act acc tac gta att ata gac ttg atc cat ttc ctc cat gca aac tat cta tct aaa tgt aat taa tta --- --- --- --- --- --- --- --- tga ttt --- --- --- -tg atg ata atg tta -cc ata c-a tt- --- -at atc act act tgg tta gtg tgt att att tag tat gga aga cct att aat aa- --- --- -tt act tat ctt ttg acg atc ttg tt- -at aat tat --- --- --- --- aat ata aaa --a tac tta tga --- --- cat agt aac tca t-- --- --- --- aat tgc tg- acg cga taa att cgt aat aat ctg ttt tgt tca aat ttt tat aag ga- --- --a tct aca ggc ata aaa at- aaa aat ata atc tat aat ata c-- tct tac a-- acg cca tca tga at- --a gca gtg aat taa ttg ctg tta tt- atg gat tta gaa ata gtg gac gat ttt gtg ata tta ata tag tta tta atg atg aaa gga taa acg cgc ata gac tca tcc tat ctg gag cct ccg aat att ttt tcc att ctg ttt tcc aat aat ttt atc gat tct aat gaa tac gaa gtt aat cta agt cat tta gat tat caa agt gtt aac tat ttg atc gat tac att tat ggg ata cct ttg agc cta act aac aat aac gtg aaa tat att ctt tca acc gct ga- -tt ttt tta caa att gga tct gtc att act gag tgc gaa aaa tac ata ctt aaa aat ctt tgt ttt aga aac tgt atc gat ttc --- --- tac ata tac gct gat aaa tat aat aac aag aaa ata gaa tta gca tcg ttt aac aca ata tta cga aat att ttg aga ctc atc aac aat gaa aac ttt aaa tac tta aca gag gaa tca atg ata aaa att tta agc gat gat atg tta tat ata aaa aat gag gat ttc acc cca ctg att ctc att aaa tgg tta gag agt aca c-- caa cca tgt acc gtc gag tta ctt aga tgc ctc aga ata tca -tt tct ttc ccc aca agt tat aaa atc act tta tag tca tcg act ggt tag ttc aat cta cga atg tat aac att ctt aaa caa tat agc att ctt gga taa atc att tcc tag ata cca tat cat cga gtt gat atc tat cgg tat aag taa ttc aca tga taa gat ttc cat aaa ctg cta caa tca t-- aaa aaa att cat ggg aaa tga tat ctt cac gta gat ata ggt gta gtt tcg cag tga ccg tcc tgg ata ata tta tct ata tga tgg gtg gat atg atc agt ccc tgt ata gaa gtt caa agg tta tag cgt aca ata cat gta cta att ctt gga tat atg ata tac cag agc taa aaa tat cat cgt tct aat tgc gga gga gtt gcc aat gac gaa tac att tat tgt ata ggc ggt ata cgc gat cag gag tca tcg ttg ata tct agt atc gat aga tgg aag cca tca aaa cca tat tgg cag aag tat gct aaa atg tgc gaa cca aaa tgt gat atg gg- gtt gcg att tta aac gga tta ata tat gtc ata ggt gga gtc gtt aaa ggt gac aca cat ata cca acg cac tag aga gtt tat cag aag atg gat gga tga atc atc aac gtc ttc caa taa aaa tgt cca ata tgt cga cga ttg ttc atg ctg gaa aga ttt ata tat cta gag gtt aca aca ata gta gtg tag tta atg taa tat cga atc t-- agt cct tag cta taa tcc gat ata tga tga atg gac caa att atc atc att aaa tat tcc tag aat taa tcc tgc tct a

>ON619836.1:156378-158641_Monkeypox_virus_isolate_MPXV_UK_2022_2_complete_genome

tta ctg act aca --- tgt a-c tat tt- -ta cat tac tac att ggc tac ggc ata tat acc tat ttc gtc act tcc aca cgc tcc ggt aaa cgg gtg tca tgt gac gag gga ga- -at ctt gat aag agg cat aat caa tgt tgt aat ccg atg tcc acc tgg aga att tgc caa ggt cag atg tag agt tgg tag tga taa cac aaa atg tga aca ctg ccc acc tca tac ata tac cgc aat ccc caa tta ttc t-a ata gat gtc atc aat gta gaa aat gcc caa cag gat cat ttg ata agg taa agt gta ccg gaa cac aga ac- --a aat -gt tcg tgt cat cct ggt tgg ta- tac gct act gat tct tca cag act gaa gat tgt cga gat ttg tgt acc aaa aaa gga gat gtc cat gcg gat act ttg gtg gaa tag atg aa- --g gaa atc cta ttt gta aat cgt gtt g-- -tg ttg gtg aat att gcg act acc tac gta att ata gac ttg atc cat ttc ctc cat gca aac tat cta tct aaa tgt aat taa tta --- --- --- --- --- --- --- --- tga ttt --- --- --- -tg atg ata atg tta -cc ata c-a tt- --- -at atc act act tgg tta gtg tgt att att tag tat gga aga cct att aat aa- --- --- -tt act tat ctt ttg acg atc ttg tt- -at aat tat --- --- --- --- aat ata aaa --a tac tta tga --- --- cat agt aac tca t-- --- --- --- aat tgc tg- acg cga taa att cgt aat aat ctg ttt tgt tca aat ttt tat aag ga- --- --a tct aca ggc ata aaa at- aaa aat ata atc tat aat ata c-- tct tac a-- acg cca tca tga at- --a gca gtg aat taa ttg ctg tta tt- atg gat tta gaa ata gtg gac gat ttt gtg ata tta ata tag tta tta atg atg aaa gga taa acg cgc ata gac tca tcc tat ctg gag cct ccg aat att ttt tcc att ctg ttt tcc aat aat ttt atc gat tct aat gaa tac gaa gtt aat cta agt cat tta gat tat caa agt gtt aac tat ttg atc gat tac att tat ggg ata cct ttg agc cta act aac aat aac gtg aaa tat att ctt tca acc gct ga- -tt ttt tta caa att gga tct gtc att act gag tgc gaa aaa tac ata ctt aaa aat ctt tgt ttt aga aac tgt atc gat ttc --- --- tac ata tac gct gat aaa tat aat aac aag aaa ata gaa tta gca tcg ttt aac aca ata tta cga aat att ttg aga ctc atc aac aat gaa aac ttt aaa tac tta aca gag gaa tca atg ata aaa att tta agc gat gat atg tta tat ata aaa aat gag gat ttc acc cca ctg att ctc att aaa tgg tta gag agt aca c-- caa cca tgt acc gtc gag tta ctt aga tgc ctc aga ata tca -tt tct ttc ccc aca agt tat aaa atc act tta tag tca tcg act ggt tag ttc aat cta cga atg tat aac att ctt aaa caa tat agc att ctt gga taa atc att tcc tag ata cca tat cat cga gtt gat atc tat cgg tat aag taa ttc aca tga taa gat ttc cat aaa ctg cta caa tca t-- aaa aaa att cat ggg aaa tga tat ctt cac gta gat ata ggt gta gtt tcg cag tga ccg tcc tgg ata ata tta tct ata tga tgg gtg gat atg atc agt ccc tgt ata gaa gtt caa agg tta tag cgt aca ata cat gta cta att ctt gga tat atg ata tac cag agc taa aaa tat cat cgt tct aat tgc gga gga gtt gcc aat gac gaa tac att tat tgt ata ggc ggt ata cgc gat cag gag tca tcg ttg ata tct agt atc gat aga tgg aag cca tca aaa cca tat tgg cag aag tat gct aaa atg tgc gaa cca aaa tgt gat atg gg- gtt gcg att tta aac gga tta ata tat gtc ata ggt gga gtc gtt aaa ggt gac aca cat ata cca acg cac tag aga gtt tat cag aag atg gat gga tga atc atc aac gtc ttc caa taa aaa tgt cca ata tgt cga cga ttg ttc atg ctg gaa aga ttt ata tat cta gag gtt aca aca ata gta gtg tag tta atg taa tat cga atc t-- agt cct tag cta taa tcc gat ata tga tga atg gac caa att atc atc att aaa tat tcc tag aat taa tcc tgc tct a

>ON619835.1:156372-158635_Monkeypox_virus_isolate_MPXV_UK_2022_1_complete_genome

tta ctg act aca --- tgt a-c tat tt- -ta cat tac tac att ggc tac ggc ata tat acc tat ttc gtc act tcc aca cgc tcc ggt aaa cgg gtg tca tgt gac gag gga ga- -at ctt gat aag agg cat aat caa tgt tgt aat ccg atg tcc acc tgg aga att tgc caa ggt cag atg tag agt tgg tag tga taa cac aaa atg tga aca ctg ccc acc tca tac ata tac cgc aat ccc caa tta ttc t-a ata gat gtc atc aat gta gaa aat gcc caa cag gat cat ttg ata agg taa agt gta ccg gaa cac aga ac- --a aat -gt tcg tgt cat cct ggt tgg ta- tac gct act gat tct tca cag act gaa gat tgt cga gat ttg tgt acc aaa aaa gga gat gtc cat gcg gat act ttg gtg gaa tag atg aa- --g gaa atc cta ttt gta aat cgt gtt g-- -tg ttg gtg aat att gcg act acc tac gta att ata gac ttg atc cat ttc ctc cat gca aac tat cta tct aaa tgt aat taa tta --- --- --- --- --- --- --- --- tga ttt --- --- --- -tg atg ata atg tta -cc ata c-a tt- --- -at atc act act tgg tta gtg tgt att att tag tat gga aga cct att aat aa- --- --- -tt act tat ctt ttg acg atc ttg tt- -at aat tat --- --- --- --- aat ata aaa --a tac tta tga --- --- cat agt aac tca t-- --- --- --- aat tgc tg- acg cga taa att cgt aat aat ctg ttt tgt tca aat ttt tat aag ga- --- --a tct aca ggc ata aaa at- aaa aat ata atc tat aat ata c-- tct tac a-- acg cca tca tga at- --a gca gtg aat taa ttg ctg tta tt- atg gat tta gaa ata gtg gac gat ttt gtg ata tta ata tag tta tta atg atg aaa gga taa acg cgc ata gac tca tcc tat ctg gag cct ccg aat att ttt tcc att ctg ttt tcc aat aat ttt atc gat tct aat gaa tac gaa gtt aat cta agt cat tta gat tat caa agt gtt aac tat ttg atc gat tac att tat ggg ata cct ttg agc cta act aac aat aac gtg aaa tat att ctt tca acc gct ga- -tt ttt tta caa att gga tct gtc att act gag tgc gaa aaa tac ata ctt aaa aat ctt tgt ttt aga aac tgt atc gat ttc --- --- tac ata tac gct gat aaa tat aat aac aag aaa ata gaa tta gca tcg ttt aac aca ata tta cga aat att ttg aga ctc atc aac aat gaa aac ttt aaa tac tta aca gag gaa tca atg ata aaa att tta agc gat gat atg tta tat ata aaa aat gag gat ttc acc cca ctg att ctc att aaa tgg tta gag agt aca c-- caa cca tgt acc gtc gag tta ctt aga tgc ctc aga ata tca -tt tct ttc ccc aca agt tat aaa atc act tta tag tca tcg act ggt tag ttc aat cta cga atg tat aac att ctt aaa caa tat agc att ctt gga taa atc att tcc tag ata cca tat cat cga gtt gat atc tat cgg tat aag taa ttc aca tga taa gat ttc cat aaa ctg cta caa tca t-- aaa aaa att cat ggg aaa tga tat ctt cac gta gat ata ggt gta gtt tcg cag tga ccg tcc tgg ata ata tta tct ata tga tgg gtg gat atg atc agt ccc tgt ata gaa gtt caa agg tta tag cgt aca ata cat gta cta att ctt gga tat atg ata tac cag agc taa aaa tat cat cgt tct aat tgc gga gga gtt gcc aat gac gaa tac att tat tgt ata ggc ggt ata cgc gat cag gag tca tcg ttg ata tct agt atc gat aga tgg aag cca tca aaa cca tat tgg cag aag tat gct aaa atg tgc gaa cca aaa tgt gat atg gg- gtt gcg att tta aac gga tta ata tat gtc ata ggt gga gtc gtt aaa ggt gac aca cat ata cca acg cac tag aga gtt tat cag aag atg gat gga tga atc atc aac gtc ttc caa taa aaa tgt cca ata tgt cga cga ttg ttc atg ctg gaa aga ttt ata tat cta gag gtt aca aca ata gta gtg tag tta atg taa tat cga atc t-- agt cct tag cta taa tcc gat ata tga tga atg gac caa att atc atc att aaa tat tcc tag aat taa tcc tgc tct a

>ON615424.1:156356-158619_Monkeypox_virus_isolate_MPXV_2022_NL001_partial_genome

tta ctg act aca --- tgt a-c tat tt- -ta cat tac tac att ggc tac ggc ata tat acc tat ttc gtc act tcc aca cgc tcc ggt aaa cgg gtg tca tgt gac gag gga ga- -at ctt gat aag agg cat aat caa tgt tgt aat ccg atg tcc acc tgg aga att tgc caa ggt cag atg tag agt tgg tag tga taa cac aaa atg tga aca ctg ccc acc tca tac ata tac cgc aat ccc caa tta ttc t-a ata gat gtc atc aat gta gaa aat gcc caa cag gat cat ttg ata agg taa agt gta ccg gaa cac aga ac- --a aat -gt tcg tgt cat cct ggt tgg ta- tac gct act gat tct tca cag act gaa gat tgt cga gat ttg tgt acc aaa aaa gga gat gtc cat gcg gat act ttg gtg gaa tag atg aa- --g gaa atc cta ttt gta aat cgt gtt g-- -tg ttg gtg aat att gcg act acc tac gta att ata gac ttg atc cat ttc ctc cat gca aac tat cta tct aaa tgt aat taa tta --- --- --- --- --- --- --- --- tga ttt --- --- --- -tg atg ata atg tta -cc ata c-a tt- --- -at atc act act tgg tta gtg tgt att att tag tat gga aga cct att aat aa- --- --- -tt act tat ctt ttg acg atc ttg tt- -at aat tat --- --- --- --- aat ata aaa --a tac tta tga --- --- cat agt aac tca t-- --- --- --- aat tgc tg- acg cga taa att cgt aat aat ctg ttt tgt tca aat ttt tat aag ga- --- --a tct aca ggc ata aaa at- aaa aat ata atc tat aat ata c-- tct tac a-- acg cca tca tga at- --a gca gtg aat taa ttg ctg tta tt- atg gat tta gaa ata gtg gac gat ttt gtg ata tta ata tag tta tta atg atg aaa gga taa acg cgc ata gac tca tcc tat ctg gag cct ccg aat att ttt tcc att ctg ttt tcc aat aat ttt atc gat tct aat gaa tac gaa gtt aat cta agt cat tta gat tat caa agt gtt aac tat ttg atc gat tac att tat ggg ata cct ttg agc cta act aac aat aac gtg aaa tat att ctt tca acc gct ga- -tt ttt tta caa att gga tct gtc att act gag tgc gaa aaa tac ata ctt aaa aat ctt tgt ttt aga aac tgt atc gat ttc --- --- tac ata tac gct gat aaa tat aat aac aag aaa ata gaa tta gca tcg ttt aac aca ata tta cga aat att ttg aga ctc atc aac aat gaa aac ttt aaa tac tta aca gag gaa tca atg ata aaa att tta agc gat gat atg tta tat ata aaa aat gag gat ttc acc cca ctg att ctc att aaa tgg tta gag agt aca c-- caa cca tgt acc gtc gag tta ctt aga tgc ctc aga ata tca -tt tct ttc ccc aca agt tat aaa atc act tta tag tca tcg act ggt tag ttc aat cta cga atg tat aac att ctt aaa caa tat agc att ctt gga taa atc att tcc tag ata cca tat cat cga gtt gat atc tat cgg tat aag taa ttc aca tga taa gat ttc cat aaa ctg cta caa tca t-- aaa aaa att cat ggg aaa tga tat ctt cac gta gat ata ggt gta gtt tcg cag tga ccg tcc tgg ata ata tta tct ata tga tgg gtg gat atg atc agt ccc tgt ata gaa gtt caa agg tta tag cgt aca ata cat gta cta att ctt gga tat atg ata tac cag agc taa aaa tat cat cgt tct aat tgc gga gga gtt gcc aat gac gaa tac att tat tgt ata ggc ggt ata cgc gat cag gag tca tcg ttg ata tct agt atc gat aga tgg aag cca tca aaa cca tat tgg cag aag tat gct aaa atg tgc gaa cca aaa tgt gat atg gg- gtt gcg att tta aac gga tta ata tat gtc ata ggt gga gtc gtt aaa ggt gac aca cat ata cca acg cac tag aga gtt tat cag aag atg gat gga tga atc atc aac gtc ttc caa taa aaa tgt cca ata tgt cga cga ttg ttc atg ctg gaa aga ttt ata tat cta gag gtt aca aca ata gta gtg tag tta atg taa tat cga atc t-- agt cct tag cta taa tcc gat ata tga tga atg gac caa att atc atc att aaa tat tcc tag aat taa tcc tgc tct a

>ON614676.1:156345-158608_Monkeypox_virus_isolate_INMI-Pt1_partial_genome

tta ctg act aca --- tgt a-c tat tt- -ta cat tac tac att ggc tac ggc ata tat acc tat ttc gtc act tcc aca cgc tcc ggt aaa cgg gtg tca tgt gac gag gga ga- -at ctt gat aag agg cat aat caa tgt tgt aat ccg atg tcc acc tgg aga att tgc caa ggt cag atg tag agt tgg tag tga taa cac aaa atg tga aca ctg ccc acc tca tac ata tac cgc aat ccc caa tta ttc t-a ata gat gtc atc aat gta gaa aat gcc caa cag gat cat ttg ata agg taa agt gta ccg gaa cac aga ac- --a aat -gt tcg tgt cat cct ggt tgg ta- tac gct act gat tct tca cag act gaa gat tgt cga gat ttg tgt acc aaa aaa gga gat gtc cat gcg gat act ttg gtg gaa tag atg aa- --g gaa atc cta ttt gta aat cgt gtt g-- -tg ttg gtg aat att gcg act acc tac gta att ata gac ttg atc cat ttc ctc cat gca aac tat cta tct aaa tgt aat taa tta --- --- --- --- --- --- --- --- tga ttt --- --- --- -tg atg ata atg tta -cc ata c-a tt- --- -at atc act act tgg tta gtg tgt att att tag tat gga aga cct att aat aa- --- --- -tt act tat ctt ttg acg atc ttg tt- -at aat tat --- --- --- --- aat ata aaa --a tac tta tga --- --- cat agt aac tca t-- --- --- --- aat tgc tg- acg cga taa att cgt aat aat ctg ttt tgt tca aat ttt tat aag ga- --- --a tct aca ggc ata aaa at- aaa aat ata atc tat aat ata c-- tct tac a-- acg cca tca tga at- --a gca gtg aat taa ttg ctg tta tt- atg gat tta gaa ata gtg gac gat ttt gtg ata tta ata tag tta tta atg atg aaa gga taa acg cgc ata gac tca tcc tat ctg gag cct ccg aat att ttt tcc att ctg ttt tcc aat aat ttt atc gat tct aat gaa tac gaa gtt aat cta agt cat tta gat tat caa agt gtt aac tat ttg atc gat tac att tat ggg ata cct ttg agc cta act aac aat aac gtg aaa tat att ctt tca acc gct ga- -tt ttt tta caa att gga tct gtc att act gag tgc gaa aaa tac ata ctt aaa aat ctt tgt ttt aga aac tgt atc gat ttc --- --- tac ata tac gct gat aaa tat aat aac aag aaa ata gaa tta gca tcg ttt aac aca ata tta cga aat att ttg aga ctc atc aac aat gaa aac ttt aaa tac tta aca gag gaa tca atg ata aaa att tta agc gat gat atg tta tat ata aaa aat gag gat ttc acc cca ctg att ctc att aaa tgg tta gag agt aca c-- caa cca tgt acc gtc gag tta ctt aga tgc ctc aga ata tca -tt tct ttc ccc aca agt tat aaa atc act tta tag tca tcg act ggt tag ttc aat cta cga atg tat aac att ctt aaa caa tat agc att ctt gga taa atc att tcc tag ata cca tat cat cga gtt gat atc tat cgg tat aag taa ttc aca tga taa gat ttc cat aaa ctg cta caa tca t-- aaa aaa att cat ggg aaa tga tat ctt cac gta gat ata ggt gta gtt tcg cag tga ccg tcc tgg ata ata tta tct ata tga tgg gtg gat atg atc agt ccc tgt ata gaa gtt caa agg tta tag cgt aca ata cat gta cta att ctt gga tat atg ata tac cag agc taa aaa tat cat cgt tct aat tgc gga gga gtt gcc aat gac gaa tac att tat tgt ata ggc ggt ata cgc gat cag gag tca tcg ttg ata tct agt atc gat aga tgg aag cca tca aaa cca tat tgg cag aag tat gct aaa atg tgc gaa cca aaa tgt gat atg gg- gtt gcg att tta aac gga tta ata tat gtc ata ggt gga gtc gtt aaa ggt gac aca cat ata cca acg cac tag aga gtt tat cag aag atg gat gga tga atc atc aac gtc ttc caa taa aaa tgt cca ata tgt cga cga ttg ttc atg ctg gaa aga ttt ata tat cta gag gtt aca aca ata gta gtg tag tta atg taa tat cga atc t-- agt cct tag cta taa tcc gat ata tga tga atg gac caa att atc atc att aaa tat tcc tag aat taa tcc tgc tct a

>ON585038.1:155456-157719_Monkeypox_virus_isolate_Monkeypox/PT0008/2022_complete_genome

tta ctg act aca --- tgt a-c tat tt- -ta cat tac tac att ggc tac ggc ata tat acc tat ttc gtc act tcc aca cgc tcc ggt aaa cgg gtg tca tgt gac gag gga ga- -at ctt gat aag agg cat aat caa tgt tgt aat ccg atg tcc acc tgg aga att tgc caa ggt cag atg tag agt tgg tag tga taa cac aaa atg tga aca ctg ccc acc tca tac ata tac cgc aat ccc caa tta ttc t-a ata gat gtc atc aat gta gaa aat gcc caa cag gat cat ttg ata agg taa agt gta ccg gaa cac aga ac- --a aat -gt tcg tgt cat cct ggt tgg ta- tac gct act gat tct tca cag act gaa gat tgt cga gat ttg tgt acc aaa aaa gga gat gtc cat gcg gat act ttg gtg gaa tag atg aa- --g gaa atc cta ttt gta aat cgt gtt g-- -tg ttg gtg aat att gcg act acc tac gta att ata gac ttg atc cat ttc ctc cat gca aac tat cta tct aaa tgt aat taa tta --- --- --- --- --- --- --- --- tga ttt --- --- --- -tg atg ata atg tta -cc ata c-a tt- --- -at atc act act tgg tta gtg tgt att att tag tat gga aga cct att aat aa- --- --- -tt act tat ctt ttg acg atc ttg tt- -at aat tat --- --- --- --- aat ata aaa --a tac tta tga --- --- cat agt aac tca t-- --- --- --- aat tgc tg- acg cga taa att cgt aat aat ctg ttt tgt tca aat ttt tat aag ga- --- --a tct aca ggc ata aaa at- aaa aat ata atc tat aat ata c-- tct tac a-- acg cca tca tga at- --a gca gtg aat taa ttg ctg tta tt- atg gat tta gaa ata gtg gac gat ttt gtg ata tta ata tag tta tta atg atg aaa gga taa acg cgc ata gac tca tcc tat ctg gag cct ccg aat att ttt tcc att ctg ttt tcc aat aat ttt atc gat tct aat gaa tac gaa gtt aat cta agt cat tta gat tat caa agt gtt aac tat ttg atc gat tac att tat ggg ata cct ttg agc cta act aac aat aac gtg aaa tat att ctt tca acc gct ga- -tt ttt tta caa att gga tct gtc att act gag tgc gaa aaa tac ata ctt aaa aat ctt tgt ttt aga aac tgt atc gat ttc --- --- tac ata tac gct gat aaa tat aat aac aag aaa ata gaa tta gca tcg ttt aac aca ata tta cga aat att ttg aga ctc atc aac aat gaa aac ttt aaa tac tta aca gag gaa tca atg ata aaa att tta agc gat gat atg tta tat ata aaa aat gag gat ttc acc cca ctg att ctc att aaa tgg tta gag agt aca c-- caa cca tgt acc gtc gag tta ctt aga tgc ctc aga ata tca -tt tct ttc ccc aca agt tat aaa atc act tta tag tca tcg act ggt tag ttc aat cta cga atg tat aac att ctt aaa caa tat agc att ctt gga taa atc att tcc tag ata cca tat cat cga gtt gat atc tat cgg tat aag taa ttc aca tga taa gat ttc cat aaa ctg cta caa tca t-- aaa aaa att cat ggg aaa tga tat ctt cac gta gat ata ggt gta gtt tcg cag tga ccg tcc tgg ata ata tta tct ata tga tgg gtg gat atg atc agt ccc tgt ata gaa gtt caa agg tta tag cgt aca ata cat gta cta att ctt gga tat atg ata tac cag agc taa aaa tat cat cgt tct aat tgc gga gga gtt gcc aat gac gaa tac att tat tgt ata ggc ggt ata cgc gat cag gag tca tcg ttg ata tct agt atc gat aga tgg aag cca tca aaa cca tat tgg cag aag tat gct aaa atg tgc gaa cca aaa tgt gat atg gg- gtt gcg att tta aac gga tta ata tat gtc ata ggt gga gtc gtt aaa ggt gac aca cat ata cca acg cac tag aga gtt tat cag aag atg gat gga tga atc atc aac gtc ttc caa taa aaa tgt cca ata tgt cga cga ttg ttc atg ctg gaa aga ttt ata tat cta gag gtt aca aca ata gta gtg tag tta atg taa tat cga atc t-- agt cct tag cta taa tcc gat ata tga tga atg gac caa att atc atc att aaa tat tcc tag aat taa tcc tgc tct a

>ON585037.1:155451-157714_Monkeypox_virus_isolate_Monkeypox/PT0005/2022_complete_genome

tta ctg act aca --- tgt a-c tat tt- -ta cat tac tac att ggc tac ggc ata tat acc tat ttc gtc act tcc aca cgc tcc ggt aaa cgg gtg tca tgt gac gag gga ga- -at ctt gat aag agg cat aat caa tgt tgt aat ccg atg tcc acc tgg aga att tgc caa ggt cag atg tag agt tgg tag tga taa cac aaa atg tga aca ctg ccc acc tca tac ata tac cgc aat ccc caa tta ttc t-a ata gat gtc atc aat gta gaa aat gcc caa cag gat cat ttg ata agg taa agt gta ccg gaa cac aga ac- --a aat -gt tcg tgt cat cct ggt tgg ta- tac gct act gat tct tca cag act gaa gat tgt cga gat ttg tgt acc aaa aaa gga gat gtc cat gcg gat act ttg gtg gaa tag atg aa- --g gaa atc cta ttt gta aat cgt gtt g-- -tg ttg gtg aat att gcg act acc tac gta att ata gac ttg atc cat ttc ctc cat gca aac tat cta tct aaa tgt aat taa tta --- --- --- --- --- --- --- --- tga ttt --- --- --- -tg atg ata atg tta -cc ata c-a tt- --- -at atc act act tgg tta gtg tgt att att tag tat gga aga cct att aat aa- --- --- -tt act tat ctt ttg acg atc ttg tt- -at aat tat --- --- --- --- aat ata aaa --a tac tta tga --- --- cat agt aac tca t-- --- --- --- aat tgc tg- acg cga taa att cgt aat aat ctg ttt tgt tca aat ttt tat aag ga- --- --a tct aca ggc ata aaa at- aaa aat ata atc tat aat ata c-- tct tac a-- acg cca tca tga at- --a gca gtg aat taa ttg ctg tta tt- atg gat tta gaa ata gtg gac gat ttt gtg ata tta ata tag tta tta atg atg aaa gga taa acg cgc ata gac tca tcc tat ctg gag cct ccg aat att ttt tcc att ctg ttt tcc aat aat ttt atc gat tct aat gaa tac gaa gtt aat cta agt cat tta gat tat caa agt gtt aac tat ttg atc gat tac att tat ggg ata cct ttg agc cta act aac aat aac gtg aaa tat att ctt tca acc gct ga- -tt ttt tta caa att gga tct gtc att act gag tgc gaa aaa tac ata ctt aaa aat ctt tgt ttt aga aac tgt atc gat ttc --- --- tac ata tac gct gat aaa tat aat aac aag aaa ata gaa tta gca tcg ttt aac aca ata tta cga aat att ttg aga ctc atc aac aat gaa aac ttt aaa tac tta aca gag gaa tca atg ata aaa att tta agc gat gat atg tta tat ata aaa aat gag gat ttc acc cca ctg att ctc att aaa tgg tta gag agt aca c-- caa cca tgt acc gtc gag tta ctt aga tgc ctc aga ata tca -tt tct ttc ccc aca agt tat aaa atc act tta tag tca tcg act ggt tag ttc aat cta cga atg tat aac att ctt aaa caa tat agc att ctt gga taa atc att tcc tag ata cca tat cat cga gtt gat atc tat cgg tat aag taa ttc aca tga taa gat ttc cat aaa ctg cta caa tca t-- aaa aaa att cat ggg aaa tga tat ctt cac gta gat ata ggt gta gtt tcg cag tga ccg tcc tgg ata ata tta tct ata tga tgg gtg gat atg atc agt ccc tgt ata gaa gtt caa agg tta tag cgt aca ata cat gta cta att ctt gga tat atg ata tac cag agc taa aaa tat cat cgt tct aat tgc gga gga gtt gcc aat gac gaa tac att tat tgt ata ggc ggt ata cgc gat cag gag tca tcg ttg ata tct agt atc gat aga tgg aag cca tca aaa cca tat tgg cag aag tat gct aaa atg tgc gaa cca aaa tgt gat atg gg- gtt gcg att tta aac gga tta ata tat gtc ata ggt gga gtc gtt aaa ggt gac aca cat ata cca acg cac tag aga gtt tat cag aag atg gat gga tga atc atc aac gtc ttc caa taa aaa tgt cca ata tgt cga cga ttg ttc atg ctg gaa aga ttt ata tat cta gag gtt aca aca ata gta gtg tag tta atg taa tat cga atc t-- agt cct tag cta taa tcc gat ata tga tga atg gac caa att atc atc att aaa tat tcc tag aat taa tcc tgc tct a

>ON585035.1:156370-158633_Monkeypox_virus_isolate_Monkeypox/PT0009/2022_complete_genome

tta ctg act aca --- tgt a-c tat tt- -ta cat tac tac att ggc tac ggc ata tat acc tat ttc gtc act tcc aca cgc tcc ggt aaa cgg gtg tca tgt gac gag gga ga- -at ctt gat aag agg cat aat caa tgt tgt aat ccg atg tcc acc tgg aga att tgc caa ggt cag atg tag agt tgg tag tga taa cac aaa atg tga aca ctg ccc acc tca tac ata tac cgc aat ccc caa tta ttc t-a ata gat gtc atc aat gta gaa aat gcc caa cag gat cat ttg ata agg taa agt gta ccg gaa cac aga ac- --a aat -gt tcg tgt cat cct ggt tgg ta- tac gct act gat tct tca cag act gaa gat tgt cga gat ttg tgt acc aaa aaa gga gat gtc cat gcg gat act ttg gtg gaa tag atg aa- --g gaa atc cta ttt gta aat cgt gtt g-- -tg ttg gtg aat att gcg act acc tac gta att ata gac ttg atc cat ttc ctc cat gca aac tat cta tct aaa tgt aat taa tta --- --- --- --- --- --- --- --- tga ttt --- --- --- -tg atg ata atg tta -cc ata c-a tt- --- -at atc act act tgg tta gtg tgt att att tag tat gga aga cct att aat aa- --- --- -tt act tat ctt ttg acg atc ttg tt- -at aat tat --- --- --- --- aat ata aaa --a tac tta tga --- --- cat agt aac tca t-- --- --- --- aat tgc tg- acg cga taa att cgt aat aat ctg ttt tgt tca aat ttt tat aag ga- --- --a tct aca ggc ata aaa at- aaa aat ata atc tat aat ata c-- tct tac a-- acg cca tca tga at- --a gca gtg aat taa ttg ctg tta tt- atg gat tta gaa ata gtg gac gat ttt gtg ata tta ata tag tta tta atg atg aaa gga taa acg cgc ata gac tca tcc tat ctg gag cct ccg aat att ttt tcc att ctg ttt tcc aat aat ttt atc gat tct aat gaa tac gaa gtt aat cta agt cat tta gat tat caa agt gtt aac tat ttg atc gat tac att tat ggg ata cct ttg agc cta act aac aat aac gtg aaa tat att ctt tca acc gct ga- -tt ttt tta caa att gga tct gtc att act gag tgc gaa aaa tac ata ctt aaa aat ctt tgt ttt aga aac tgt atc gat ttc --- --- tac ata tac gct gat aaa tat aat aac aag aaa ata gaa tta gca tcg ttt aac aca ata tta cga aat att ttg aga ctc atc aac aat gaa aac ttt aaa tac tta aca gag gaa tca atg ata aaa att tta agc gat gat atg tta tat ata aaa aat gag gat ttc acc cca ctg att ctc att aaa tgg tta gag agt aca c-- caa cca tgt acc gtc gag tta ctt aga tgc ctc aga ata tca -tt tct ttc ccc aca agt tat aaa atc act tta tag tca tcg act ggt tag ttc aat cta cga atg tat aac att ctt aaa caa tat agc att ctt gga taa atc att tcc tag ata cca tat cat cga gtt gat atc tat cgg tat aag taa ttc aca tga taa gat ttc cat aaa ctg cta caa tca t-- aaa aaa att cat ggg aaa tga tat ctt cac gta gat ata ggt gta gtt tcg cag tga ccg tcc tgg ata ata tta tct ata tga tgg gtg gat atg atc agt ccc tgt ata gaa gtt caa agg tta tag cgt aca ata cat gta cta att ctt gga tat atg ata tac cag agc taa aaa tat cat cgt tct aat tgc gga gga gtt gcc aat gac gaa tac att tat tgt ata ggc ggt ata cgc gat cag gag tca tcg ttg ata tct agt atc gat aga tgg aag cca tca aaa cca tat tgg cag aag tat gct aaa atg tgc gaa cca aaa tgt gat atg gg- gtt gcg att tta aac gga tta ata tat gtc ata ggt gga gtc gtt aaa ggt gac aca cat ata cca acg cac tag aga gtt tat cag aag atg gat gga tga atc atc aac gtc ttc caa taa aaa tgt cca ata tgt cga cga ttg ttc atg ctg gaa aga ttt ata tat cta gag gtt aca aca ata gta gtg tag tta atg taa tat cga atc t-- agt cct tag cta taa tcc gat ata tga tga atg gac caa att atc atc att aaa tat tcc tag aat taa tcc tgc tct a

>ON585034.1:156364-158627_Monkeypox_virus_isolate_Monkeypox/PT0007/2022_complete_genome

tta ctg act aca --- tgt a-c tat tt- -ta cat tac tac att ggc tac ggc ata tat acc tat ttc gtc act tcc aca cgc tcc ggt aaa cgg gtg tca tgt gac gag gga ga- -at ctt gat aag agg cat aat caa tgt tgt aat ccg atg tcc acc tgg aga att tgc caa ggt cag atg tag agt tgg tag tga taa cac aaa atg tga aca ctg ccc acc tca tac ata tac cgc aat ccc caa tta ttc t-a ata gat gtc atc aat gta gaa aat gcc caa cag gat cat ttg ata agg taa agt gta ccg gaa cac aga ac- --a aat -gt tcg tgt cat cct ggt tgg ta- tac gct act gat tct tca cag act gaa gat tgt cga gat ttg tgt acc aaa aaa gga gat gtc cat gcg gat act ttg gtg gaa tag atg aa- --g gaa atc cta ttt gta aat cgt gtt g-- -tg ttg gtg aat att gcg act acc tac gta att ata gac ttg atc cat ttc ctc cat gca aac tat cta tct aaa tgt aat taa tta --- --- --- --- --- --- --- --- tga ttt --- --- --- -tg atg ata atg tta -cc ata c-a tt- --- -at atc act act tgg tta gtg tgt att att tag tat gga aga cct att aat aa- --- --- -tt act tat ctt ttg acg atc ttg tt- -at aat tat --- --- --- --- aat ata aaa --a tac tta tga --- --- cat agt aac tca t-- --- --- --- aat tgc tg- acg cga taa att cgt aat aat ctg ttt tgt tca aat ttt tat aag ga- --- --a tct aca ggc ata aaa at- aaa aat ata atc tat aat ata c-- tct tac a-- acg cca tca tga at- --a gca gtg aat taa ttg ctg tta tt- atg gat tta gaa ata gtg gac gat ttt gtg ata tta ata tag tta tta atg atg aaa gga taa acg cgc ata gac tca tcc tat ctg gag cct ccg aat att ttt tcc att ctg ttt tcc aat aat ttt atc gat tct aat gaa tac gaa gtt aat cta agt cat tta gat tat caa agt gtt aac tat ttg atc gat tac att tat ggg ata cct ttg agc cta act aac aat aac gtg aaa tat att ctt tca acc gct ga- -tt ttt tta caa att gga tct gtc att act gag tgc gaa aaa tac ata ctt aaa aat ctt tgt ttt aga aac tgt atc gat ttc --- --- tac ata tac gct gat aaa tat aat aac aag aaa ata gaa tta gca tcg ttt aac aca ata tta cga aat att ttg aga ctc atc aac aat gaa aac ttt aaa tac tta aca gag gaa tca atg ata aaa att tta agc gat gat atg tta tat ata aaa aat gag gat ttc acc cca ctg att ctc att aaa tgg tta gag agt aca c-- caa cca tgt acc gtc gag tta ctt aga tgc ctc aga ata tca -tt tct ttc ccc aca agt tat aaa atc act tta tag tca tcg act ggt tag ttc aat cta cga atg tat aac att ctt aaa caa tat agc att ctt gga taa atc att tcc tag ata cca tat cat cga gtt gat atc tat cgg tat aag taa ttc aca tga taa gat ttc cat aaa ctg cta caa tca t-- aaa aaa att cat ggg aaa tga tat ctt cac gta gat ata ggt gta gtt tcg cag tga ccg tcc tgg ata ata tta tct ata tga tgg gtg gat atg atc agt ccc tgt ata gaa gtt caa agg tta tag cgt aca ata cat gta cta att ctt gga tat atg ata tac cag agc taa aaa tat cat cgt tct aat tgc gga gga gtt gcc aat gac gaa tac att tat tgt ata ggc ggt ata cgc gat cag gag tca tcg ttg ata tct agt atc gat aga tgg aag cca tca aaa cca tat tgg cag aag tat gct aaa atg tgc gaa cca aaa tgt gat atg gg- gtt gcg att tta aac gga tta ata tat gtc ata ggt gga gtc gtt aaa ggt gac aca cat ata cca acg cac tag aga gtt tat cag aag atg gat gga tga atc atc aac gtc ttc caa taa aaa tgt cca ata tgt cga cga ttg ttc atg ctg gaa aga ttt ata tat cta gag gtt aca aca ata gta gtg tag tta atg taa tat cga atc t-- agt cct tag cta taa tcc gat ata tga tga atg gac caa att atc atc att aaa tat tcc tag aat taa tcc tgc tct a

>ON585033.1:156370-158633_Monkeypox_virus_isolate_Monkeypox/PT0006/2022_complete_genome

tta ctg act aca --- tgt a-c tat tt- -ta cat tac tac att ggc tac ggc ata tat acc tat ttc gtc act tcc aca cgc tcc ggt aaa cgg gtg tca tgt gac gag gga ga- -at ctt gat aag agg cat aat caa tgt tgt aat ccg atg tcc acc tgg aga att tgc caa ggt cag atg tag agt tgg tag tga taa cac aaa atg tga aca ctg ccc acc tca tac ata tac cgc aat ccc caa tta ttc t-a ata gat gtc atc aat gta gaa aat gcc caa cag gat cat ttg ata agg taa agt gta ccg gaa cac aga ac- --a aat -gt tcg tgt cat cct ggt tgg ta- tac gct act gat tct tca cag act gaa gat tgt cga gat ttg tgt acc aaa aaa gga gat gtc cat gcg gat act ttg gtg gaa tag atg aa- --g gaa atc cta ttt gta aat cgt gtt g-- -tg ttg gtg aat att gcg act acc tac gta att ata gac ttg atc cat ttc ctc cat gca aac tat cta tct aaa tgt aat taa tta --- --- --- --- --- --- --- --- tga ttt --- --- --- -tg atg ata atg tta -cc ata c-a tt- --- -at atc act act tgg tta gtg tgt att att tag tat gga aga cct att aat aa- --- --- -tt act tat ctt ttg acg atc ttg tt- -at aat tat --- --- --- --- aat ata aaa --a tac tta tga --- --- cat agt aac tca t-- --- --- --- aat tgc tg- acg cga taa att cgt aat aat ctg ttt tgt tca aat ttt tat aag ga- --- --a tct aca ggc ata aaa at- aaa aat ata atc tat aat ata c-- tct tac a-- acg cca tca tga at- --a gca gtg aat taa ttg ctg tta tt- atg gat tta gaa ata gtg gac gat ttt gtg ata tta ata tag tta tta atg atg aaa gga taa acg cgc ata gac tca tcc tat ctg gag cct ccg aat att ttt tcc att ctg ttt tcc aat aat ttt atc gat tct aat gaa tac gaa gtt aat cta agt cat tta gat tat caa agt gtt aac tat ttg atc gat tac att tat ggg ata cct ttg agc cta act aac aat aac gtg aaa tat att ctt tca acc gct ga- -tt ttt tta caa att gga tct gtc att act gag tgc gaa aaa tac ata ctt aaa aat ctt tgt ttt aga aac tgt atc gat ttc --- --- tac ata tac gct gat aaa tat aat aac aag aaa ata gaa tta gca tcg ttt aac aca ata tta cga aat att ttg aga ctc atc aac aat gaa aac ttt aaa tac tta aca gag gaa tca atg ata aaa att tta agc gat gat atg tta tat ata aaa aat gag gat ttc acc cca ctg att ctc att aaa tgg tta gag agt aca c-- caa cca tgt acc gtc gag tta ctt aga tgc ctc aga ata tca -tt tct ttc ccc aca agt tat aaa atc act tta tag tca tcg act ggt tag ttc aat cta cga atg tat aac att ctt aaa caa tat agc att ctt gga taa atc att tcc tag ata cca tat cat cga gtt gat atc tat cgg tat aag taa ttc aca tga taa gat ttc cat aaa ctg cta caa tca t-- aaa aaa att cat ggg aaa tga tat ctt cac gta gat ata ggt gta gtt tcg cag tga ccg tcc tgg ata ata tta tct ata tga tgg gtg gat atg atc agt ccc tgt ata gaa gtt caa agg tta tag cgt aca ata cat gta cta att ctt gga tat atg ata tac cag agc taa aaa tat cat cgt tct aat tgc gga gga gtt gcc aat gac gaa tac att tat tgt ata ggc ggt ata cgc gat cag gag tca tcg ttg ata tct agt atc gat aga tgg aag cca tca aaa cca tat tgg cag aag tat gct aaa atg tgc gaa cca aaa tgt gat atg gg- gtt gcg att tta aac gga tta ata tat gtc ata ggt gga gtc gtt aaa ggt gac aca cat ata cca acg cac tag aga gtt tat cag aag atg gat gga tga atc atc aac gtc ttc caa taa aaa tgt cca ata tgt cga cga ttg ttc atg ctg gaa aga ttt ata tat cta gag gtt aca aca ata gta gtg tag tta atg taa tat cga atc t-- agt cct tag cta taa tcc gat ata tga tga atg gac caa att atc atc att aaa tat tcc tag aat taa tcc tgc tct a

>ON585032.1:156367-158630_Monkeypox_virus_isolate_Monkeypox/PT0004/2022_complete_genome

tta ctg act aca --- tgt a-c tat tt- -ta cat tac tac att ggc tac ggc ata tat acc tat ttc gtc act tcc aca cgc tcc ggt aaa cgg gtg tca tgt gac gag gga ga- -at ctt gat aag agg cat aat caa tgt tgt aat ccg atg tcc acc tgg aga att tgc caa ggt cag atg tag agt tgg tag tga taa cac aaa atg tga aca ctg ccc acc tca tac ata tac cgc aat ccc caa tta ttc t-a ata gat gtc atc aat gta gaa aat gcc caa cag gat cat ttg ata agg taa agt gta ccg gaa cac aga ac- --a aat -gt tcg tgt cat cct ggt tgg ta- tac gct act gat tct tca cag act gaa gat tgt cga gat ttg tgt acc aaa aaa gga gat gtc cat gcg gat act ttg gtg gaa tag atg aa- --g gaa atc cta ttt gta aat cgt gtt g-- -tg ttg gtg aat att gcg act acc tac gta att ata gac ttg atc cat ttc ctc cat gca aac tat cta tct aaa tgt aat taa tta --- --- --- --- --- --- --- --- tga ttt --- --- --- -tg atg ata atg tta -cc ata c-a tt- --- -at atc act act tgg tta gtg tgt att att tag tat gga aga cct att aat aa- --- --- -tt act tat ctt ttg acg atc ttg tt- -at aat tat --- --- --- --- aat ata aaa --a tac tta tga --- --- cat agt aac tca t-- --- --- --- aat tgc tg- acg cga taa att cgt aat aat ctg ttt tgt tca aat ttt tat aag ga- --- --a tct aca ggc ata aaa at- aaa aat ata atc tat aat ata c-- tct tac a-- acg cca tca tga at- --a gca gtg aat taa ttg ctg tta tt- atg gat tta gaa ata gtg gac gat ttt gtg ata tta ata tag tta tta atg atg aaa gga taa acg cgc ata gac tca tcc tat ctg gag cct ccg aat att ttt tcc att ctg ttt tcc aat aat ttt atc gat tct aat gaa tac gaa gtt aat cta agt cat tta gat tat caa agt gtt aac tat ttg atc gat tac att tat ggg ata cct ttg agc cta act aac aat aac gtg aaa tat att ctt tca acc gct ga- -tt ttt tta caa att gga tct gtc att act gag tgc gaa aaa tac ata ctt aaa aat ctt tgt ttt aga aac tgt atc gat ttc --- --- tac ata tac gct gat aaa tat aat aac aag aaa ata gaa tta gca tcg ttt aac aca ata tta cga aat att ttg aga ctc atc aac aat gaa aac ttt aaa tac tta aca gag gaa tca atg ata aaa att tta agc gat gat atg tta tat ata aaa aat gag gat ttc acc cca ctg att ctc att aaa tgg tta gag agt aca c-- caa cca tgt acc gtc gag tta ctt aga tgc ctc aga ata tca -tt tct ttc ccc aca agt tat aaa atc act tta tag tca tcg act ggt tag ttc aat cta cga atg tat aac att ctt aaa caa tat agc att ctt gga taa atc att tcc tag ata cca tat cat cga gtt gat atc tat cgg tat aag taa ttc aca tga taa gat ttc cat aaa ctg cta caa tca t-- aaa aaa att cat ggg aaa tga tat ctt cac gta gat ata ggt gta gtt tcg cag tga ccg tcc tgg ata ata tta tct ata tga tgg gtg gat atg atc agt ccc tgt ata gaa gtt caa agg tta tag cgt aca ata cat gta cta att ctt gga tat atg ata tac cag agc taa aaa tat cat cgt tct aat tgc gga gga gtt gcc aat gac gaa tac att tat tgt ata ggc ggt ata cgc gat cag gag tca tcg ttg ata tct agt atc gat aga tgg aag cca tca aaa cca tat tgg cag aag tat gct aaa atg tgc gaa cca aaa tgt gat atg gg- gtt gcg att tta aac gga tta ata tat gtc ata ggt gga gtc gtt aaa ggt gac aca cat ata cca acg cac tag aga gtt tat cag aag atg gat gga tga atc atc aac gtc ttc caa taa aaa tgt cca ata tgt cga cga ttg ttc atg ctg gaa aga ttt ata tat cta gag gtt aca aca ata gta gtg tag tta atg taa tat cga atc t-- agt cct tag cta taa tcc gat ata tga tga atg gac caa att atc atc att aaa tat tcc tag aat taa tcc tgc tct a

>ON585031.1:156364-158627_Monkeypox_virus_isolate_Monkeypox/PT0003/2022_complete_genome

tta ctg act aca --- tgt a-c tat tt- -ta cat tac tac att ggc tac ggc ata tat acc tat ttc gtc act tcc aca cgc tcc ggt aaa cgg gtg tca tgt gac gag gga ga- -at ctt gat aag agg cat aat caa tgt tgt aat ccg atg tcc acc tgg aga att tgc caa ggt cag atg tag agt tgg tag tga taa cac aaa atg tga aca ctg ccc acc tca tac ata tac cgc aat ccc caa tta ttc t-a ata gat gtc atc aat gta gaa aat gcc caa cag gat cat ttg ata agg taa agt gta ccg gaa cac aga ac- --a aat -gt tcg tgt cat cct ggt tgg ta- tac gct act gat tct tca cag act gaa gat tgt cga gat ttg tgt acc aaa aaa gga gat gtc cat gcg gat act ttg gtg gaa tag atg aa- --g gaa atc cta ttt gta aat cgt gtt g-- -tg ttg gtg aat att gcg act acc tac gta att ata gac ttg atc cat ttc ctc cat gca aac tat cta tct aaa tgt aat taa tta --- --- --- --- --- --- --- --- tga ttt --- --- --- -tg atg ata atg tta -cc ata c-a tt- --- -at atc act act tgg tta gtg tgt att att tag tat gga aga cct att aat aa- --- --- -tt act tat ctt ttg acg atc ttg tt- -at aat tat --- --- --- --- aat ata aaa --a tac tta tga --- --- cat agt aac tca t-- --- --- --- aat tgc tg- acg cga taa att cgt aat aat ctg ttt tgt tca aat ttt tat aag ga- --- --a tct aca ggc ata aaa at- aaa aat ata atc tat aat ata c-- tct tac a-- acg cca tca tga at- --a gca gtg aat taa ttg ctg tta tt- atg gat tta gaa ata gtg gac gat ttt gtg ata tta ata tag tta tta atg atg aaa gga taa acg cgc ata gac tca tcc tat ctg gag cct ccg aat att ttt tcc att ctg ttt tcc aat aat ttt atc gat tct aat gaa tac gaa gtt aat cta agt cat tta gat tat caa agt gtt aac tat ttg atc gat tac att tat ggg ata cct ttg agc cta act aac aat aac gtg aaa tat att ctt tca acc gct ga- -tt ttt tta caa att gga tct gtc att act gag tgc gaa aaa tac ata ctt aaa aat ctt tgt ttt aga aac tgt atc gat ttc --- --- tac ata tac gct gat aaa tat aat aac aag aaa ata gaa tta gca tcg ttt aac aca ata tta cga aat att ttg aga ctc atc aac aat gaa aac ttt aaa tac tta aca gag gaa tca atg ata aaa att tta agc gat gat atg tta tat ata aaa aat gag gat ttc acc cca ctg att ctc att aaa tgg tta gag agt aca c-- caa cca tgt acc gtc gag tta ctt aga tgc ctc aga ata tca -tt tct ttc ccc aca agt tat aaa atc act tta tag tca tcg act ggt tag ttc aat cta cga atg tat aac att ctt aaa caa tat agc att ctt gga taa atc att tcc tag ata cca tat cat cga gtt gat atc tat cgg tat aag taa ttc aca tga taa gat ttc cat aaa ctg cta caa tca t-- aaa aaa att cat ggg aaa tga tat ctt cac gta gat ata ggt gta gtt tcg cag tga ccg tcc tgg ata ata tta tct ata tga tgg gtg gat atg atc agt ccc tgt ata gaa gtt caa agg tta tag cgt aca ata cat gta cta att ctt gga tat atg ata tac cag agc taa aaa tat cat cgt tct aat tgc gga gga gtt gcc aat gac gaa tac att tat tgt ata ggc ggt ata cgc gat cag gag tca tcg ttg ata tct agt atc gat aga tgg aag cca tca aaa cca tat tgg cag aag tat gct aaa atg tgc gaa cca aaa tgt gat atg gg- gtt gcg att tta aac gga tta ata tat gtc ata ggt gga gtc gtt aaa ggt gac aca cat ata cca acg cac tag aga gtt tat cag aag atg gat gga tga atc atc aac gtc ttc caa taa aaa tgt cca ata tgt cga cga ttg ttc atg ctg gaa aga ttt ata tat cta gag gtt aca aca ata gta gtg tag tta atg taa tat cga atc t-- agt cct tag cta taa tcc gat ata tga tga atg gac caa att atc atc att aaa tat tcc tag aat taa tcc tgc tct a

>ON568298.1:156511-158774_Monkeypox_virus_isolate_MPXV-BY-IMB25241_complete_genome

tta ctg act aca --- tgt a-c tat tt- -ta cat tac tac att ggc tac ggc ata tat acc tat ttc gtc act tcc aca cgc tcc ggt aaa cgg gtg tca tgt gac gag gga ga- -at ctt gat aag agg cat aat caa tgt tgt aat ccg atg tcc acc tgg aga att tgc caa ggt cag atg tag agt tgg tag tga taa cac aaa atg tga aca ctg ccc acc tca tac ata tac cgc aat ccc caa tta ttc t-a ata gat gtc atc aat gta gaa aat gcc caa cag gat cat ttg ata agg taa agt gta ccg gaa cac aga ac- --a aat -gt tcg tgt cat cct ggt tgg ta- tac gct act gat tct tca cag act gaa gat tgt cga gat ttg tgt acc aaa aaa gga gat gtc cat gcg gat act ttg gtg gaa tag atg aa- --g gaa atc cta ttt gta aat cgt gtt g-- -tg ttg gtg aat att gcg act acc tac gta att ata gac ttg atc cat ttc ctc cat gca aac tat cta tct aaa tgt aat taa tta --- --- --- --- --- --- --- --- tga ttt --- --- --- -tg atg ata atg tta -cc ata c-a tt- --- -at atc act act tgg tta gtg tgt att att tag tat gga aga cct att aat aa- --- --- -tt act tat ctt ttg acg atc ttg tt- -at aat tat --- --- --- --- aat ata aaa --a tac tta tga --- --- cat agt aac tca t-- --- --- --- aat tgc tg- acg cga taa att cgt aat aat ctg ttt tgt tca aat ttt tat aag ga- --- --a tct aca ggc ata aaa at- aaa aat ata atc tat aat ata c-- tct tac a-- acg cca tca tga at- --a gca gtg aat taa ttg ctg tta tt- atg gat tta gaa ata gtg gac gat ttt gtg ata tta ata tag tta tta atg atg aaa gga taa acg cgc ata gac tca tcc tat ctg gag cct ccg aat att ttt tcc att ctg ttt tcc aat aat ttt atc gat tct aat gaa tac gaa gtt aat cta agt cat tta gat tat caa agt gtt aac tat ttg atc gat tac att tat ggg ata cct ttg agc cta act aac aat aac gtg aaa tat att ctt tca acc gct ga- -tt ttt tta caa att gga tct gtc att act gag tgc gaa aaa tac ata ctt aaa aat ctt tgt ttt aga aac tgt atc gat ttc --- --- tac ata tac gct gat aaa tat aat aac aag aaa ata gaa tta gca tcg ttt aac aca ata tta cga aat att ttg aga ctc atc aac aat gaa aac ttt aaa tac tta aca gag gaa tca atg ata aaa att tta agc gat gat atg tta tat ata aaa aat gag gat ttc acc cca ctg att ctc att aaa tgg tta gag agt aca c-- caa cca tgt acc gtc gag tta ctt aga tgc ctc aga ata tca -tt tct ttc ccc aca agt tat aaa atc act tta tag tca tcg act ggt tag ttc aat cta cga atg tat aac att ctt aaa caa tat agc att ctt gga taa atc att tcc tag ata cca tat cat cga gtt gat atc tat cgg tat aag taa ttc aca tga taa gat ttc cat aaa ctg cta caa tca t-- aaa aaa att cat ggg aaa tga tat ctt cac gta gat ata ggt gta gtt tcg cag tga ccg tcc tgg ata ata tta tct ata tga tgg gtg gat atg atc agt ccc tgt ata gaa gtt caa agg tta tag cgt aca ata cat gta cta att ctt gga tat atg ata tac cag agc taa aaa tat cat cgt tct aat tgc gga gga gtt gcc aat gac gaa tac att tat tgt ata ggc ggt ata cgc gat cag gag tca tcg ttg ata tct agt atc gat aga tgg aag cca tca aaa cca tat tgg cag aag tat gct aaa atg tgc gaa cca aaa tgt gat atg gg- gtt gcg att tta aac gga tta ata tat gtc ata ggt gga gtc gtt aaa ggt gac aca cat ata cca acg cac tag aga gtt tat cag aag atg gat gga tga atc atc aac gtc ttc caa taa aaa tgt cca ata tgt cga cga ttg ttc atg ctg gaa aga ttt ata tat cta gag gtt aca aca ata gta gtg tag tta atg taa tat cga atc t-- agt cct tag cta taa tcc gat ata tga tga atg gac caa att atc atc att aaa tat tcc tag aat taa tcc tgc tct a

>MT903345.1:156395-158658_Monkeypox_virus_isolate_MPXV-UK_P3

tta ctg act aca --- tgt a-c tat tt- -ta cat tac tac att ggc tac ggc ata tat acc tat ttc gtc act tcc aca cgc tcc ggt aaa cgg gtg tca tgt gac gag gga ga- -at ctt gat aag agg cat aat caa tgt tgt aat ccg atg tcc acc tgg aga att tgc caa ggt cag atg tag agt tgg tag tga taa cac aaa atg tga aca ctg ccc acc tca tac ata tac cgc aat ccc caa tta ttc t-a ata gat gtc atc aat gta gaa aat gcc caa cag gat cat ttg ata agg taa agt gta ccg gaa cac aga ac- --a aat -gt tcg tgt cat cct ggt tgg ta- tac gct act gat tct tca cag act gaa gat tgt cga gat ttg tgt acc aaa aaa gga gat gtc cat gcg gat act ttg gtg gaa tag atg aa- --g gaa atc cta ttt gta aat cgt gtt g-- -tg ttg gtg aat att gcg act acc tac gta att ata gac ttg atc cat ttc ctc cat gca aac tat cta tct aaa tgt aat taa tta --- --- --- --- --- --- --- --- tga ttt --- --- --- -tg atg ata atg tta -cc ata c-a tt- --- -at atc act act tgg tta gtg tgt att att tag tat gga aga cct att aat aa- --- --- -tt act tat ctt ttg acg atc ttg tt- -at aat tat --- --- --- --- aat ata aaa --a tac tta tga --- --- cat agt aac tca t-- --- --- --- aat tgc tg- acg cga taa att cgt aat aat ctg ttt tgt tca aat ttt tat aag ga- --- --a tct aca ggc ata aaa at- aaa aat ata atc tat aat ata c-- tct tac a-- acg cca tca tga at- --a gca gtg aat taa ttg ctg tta tt- atg gat tta gaa ata gtg gac gat ttt gtg ata tta ata tag tta tta atg atg aaa gga taa acg cgc ata gac tca tcc tat ctg gag cct ccg aat att ttt tcc att ctg ttt tcc aat aat ttt atc gat tct aat gaa tac gaa gtt aat cta agt cat tta gat tat caa agt gtt aac tat ttg atc gat tac att tat ggg ata cct ttg agc cta act aac aat aac gtg aaa tat att ctt tca acc gct ga- -tt ttt tta caa att gga tct gtc att act gag tgc gaa aaa tac ata ctt aaa aat ctt tgt ttt aga aac tgt atc gat ttc --- --- tac ata tac gct gat aaa tat aat aac aag aaa ata gaa tta gca tcg ttt aac aca ata tta cga aat att ttg aga ctc atc aac aat gaa aac ttt aaa tac tta aca gag gaa tca atg ata aaa att tta agc gat gat atg tta tat ata aaa aat gag gat ttc acc cca ctg att ctc att aaa tgg tta gag agt aca c-- caa cca tgt acc gtc gag tta ctt aga tgc ctc aga ata tca -tt tct ttc ccc aca agt tat aaa atc act tta tag tca tcg act ggt tag ttc aat cta cga atg tat aac att ctt aaa caa tat agc att ctt gga taa atc att tcc tag ata cca tat cat cga gtt gat atc tat cgg tat aag taa ttc aca tga taa gat ttc cat aaa ctg cta caa tca t-- aaa aaa att cat ggg aaa tga tat ctt cac gta gat ata ggt gta gtt tcg cag tga ccg tcc tgg ata ata tta tct ata tga tgg gtg gat atg atc agt ccc tgt ata gaa gtt caa agg tta tag cgt aca ata cat gta cta att ctt gga tat atg ata tac cag agc taa aaa tat cat cgt tct aat tgc gga gga gtt gcc aat gac gaa tac att tat tgt ata ggc ggt ata cgc gat cag gag tca tcg ttg ata tct agt atc gat aga tgg aag cca tca aaa cca tat tgg cag aag tat gct aaa atg tgc gaa cca aaa tgt gat atg gg- gtt gcg att tta aac gga tta ata tat gtc ata ggt gga gtc gtt aaa ggt gac aca cat ata cca acg cac tag aga gtt tat cag aag atg gat gga tga atc atc aac gtc ttc caa taa aaa tgt cca ata tgt cga cga ttg ttc atg ctg gaa aga ttt ata tat cta gag gtt aca aca ata gta gtg tag tta atg taa tat cga atc t-- agt cct tag cta taa tcc gat ata tga tga atg gac caa att atc atc att aaa tat tcc tag aat taa tcc tgc tct a

>MT903344.1:156381-158644_Monkeypox_virus_isolate_MPXV-UK_P2

tta ctg act aca --- tgt a-c tat tt- -ta cat tac tac att ggc tac ggc ata tat acc tat ttc gtc act tcc aca cgc tcc ggt aaa cgg gtg tca tgt gac gag gga ga- -at ctt gat aag agg cat aat caa tgt tgt aat ccg atg tcc acc tgg aga att tgc caa ggt cag atg tag agt tgg tag tga taa cac aaa atg tga aca ctg ccc acc tca tac ata tac cgc aat ccc caa tta ttc t-a ata gat gtc atc aat gta gaa aat gcc caa cag gat cat ttg ata agg taa agt gta ccg gaa cac aga ac- --a aat -gt tcg tgt cat cct ggt tgg ta- tac gct act gat tct tca cag act gaa gat tgt cga gat ttg tgt acc aaa aaa gga gat gtc cat gcg gat act ttg gtg gaa tag atg aa- --g gaa atc cta ttt gta aat cgt gtt g-- -tg ttg gtg aat att gcg act acc tac gta att ata gac ttg atc cat ttc ctc cat gca aac tat cta tct aaa tgt aat taa tta --- --- --- --- --- --- --- --- tga ttt --- --- --- -tg atg ata atg tta -cc ata c-a tt- --- -at atc act act tgg tta gtg tgt att att tag tat gga aga cct att aat aa- --- --- -tt act tat ctt ttg acg atc ttg tt- -at aat tat --- --- --- --- aat ata aaa --a tac tta tga --- --- cat agt aac tca t-- --- --- --- aat tgc tg- acg cga taa att cgt aat aat ctg ttt tgt tca aat ttt tat aag ga- --- --a tct aca ggc ata aaa at- aaa aat ata atc tat aat ata c-- tct tac a-- acg cca tca tga at- --a gca gtg aat taa ttg ctg tta tt- atg gat tta gaa ata gtg gac gat ttt gtg ata tta ata tag tta tta atg atg aaa gga taa acg cgc ata gac tca tcc tat ctg gag cct ccg aat att ttt tcc att ctg ttt tcc aat aat ttt atc gat tct aat gaa tac gaa gtt aat cta agt cat tta gat tat caa agt gtt aac tat ttg atc gat tac att tat ggg ata cct ttg agc cta act aac aat aac gtg aaa tat att ctt tca acc gct ga- -tt ttt tta caa att gga tct gtc att act gag tgc gaa aaa tac ata ctt aaa aat ctt tgt ttt aga aac tgt atc gat ttc --- --- tac ata tac gct gat aaa tat aat aac aag aaa ata gaa tta gca tcg ttt aac aca ata tta cga aat att ttg aga ctc atc aac aat gaa aac ttt aaa tac tta aca gag gaa tca atg ata aaa att tta agc gat gat atg tta tat ata aaa aat gag gat ttc acc cca ctg att ctc att aaa tgg tta gag agt aca c-- caa cca tgt acc gtc gag tta ctt aga tgc ctc aga ata tca -tt tct ttc ccc aca agt tat aaa atc act tta tag tca tcg act ggt tag ttc aat cta cga atg tat aac att ctt aaa caa tat agc att ctt gga taa atc att tcc tag ata cca tat cat cga gtt gat atc tat cgg tat aag taa ttc aca tga taa gat ttc cat aaa ctg cta caa tca t-- aaa aaa att cat ggg aaa tga tat ctt cac gta gat ata ggt gta gtt tcg cag tga ccg tcc tgg ata ata tta tct ata tga tgg gtg gat atg atc agt ccc tgt ata gaa gtt caa agg tta tag cgt aca ata cat gta cta att ctt gga tat atg ata tac cag agc taa aaa tat cat cgt tct aat tgc gga gga gtt gcc aat gac gaa tac att tat tgt ata ggc ggt ata cgc gat cag gag tca tcg ttg ata tct agt atc gat aga tgg aag cca tca aaa cca tat tgg cag aag tat gct aaa atg tgc gaa cca aaa tgt gat atg gg- gtt gcg att tta aac gga tta ata tat gtc ata ggt gga gtc gtt aaa ggt gac aca cat ata cca acg cac tag aga gtt tat cag aag atg gat gga tga atc atc aac gtc ttc caa taa aaa tgt cca ata tgt cga cga ttg ttc atg ctg gaa aga ttt ata tat cta gag gtt aca aca ata gta gtg tag tta atg taa tat cga atc t-- agt cct tag cta taa tcc gat ata tga tga atg gac caa att atc atc att aaa tat tcc tag aat taa tcc tgc tct a

>MT903343.1:156375-158638_Monkeypox_virus_isolate_MPXV-UK_P1

tta ctg act aca --- tgt a-c tat tt- -ta cat tac tac att ggc tac ggc ata tat acc tat ttc gtc act tcc aca cgc tcc ggt aaa cgg gtg tca tgt gac gag gga ga- -at ctt gat aag agg cat aat caa tgt tgt aat ccg atg tcc acc tgg aga att tgc caa ggt cag atg tag agt tgg tag tga taa cac aaa atg tga aca ctg ccc acc tca tac ata tac cgc aat ccc caa tta ttc t-a ata gat gtc atc aat gta gaa aat gcc caa cag gat cat ttg ata agg taa agt gta ccg gaa cac aga ac- --a aat -gt tcg tgt cat cct ggt tgg ta- tac gct act gat tct tca cag act gaa gat tgt cga gat ttg tgt acc aaa aaa gga gat gtc cat gcg gat act ttg gtg gaa tag atg aa- --g gaa atc cta ttt gta aat cgt gtt g-- -tg ttg gtg aat att gcg act acc tac gta att ata gac ttg atc cat ttc ctc cat gca aac tat cta tct aaa tgt aat taa tta --- --- --- --- --- --- --- --- tga ttt --- --- --- -tg atg ata atg tta -cc ata c-a tt- --- -at atc act act tgg tta gtg tgt att att tag tat gga aga cct att aat aa- --- --- -tt act tat ctt ttg acg atc ttg tt- -at aat tat --- --- --- --- aat ata aaa --a tac tta tga --- --- cat agt aac tca t-- --- --- --- aat tgc tg- acg cga taa att cgt aat aat ctg ttt tgt tca aat ttt tat aag ga- --- --a tct aca ggc ata aaa at- aaa aat ata atc tat aat ata c-- tct tac a-- acg cca tca tga at- --a gca gtg aat taa ttg ctg tta tt- atg gat tta gaa ata gtg gac gat ttt gtg ata tta ata tag tta tta atg atg aaa gga taa acg cgc ata gac tca tcc tat ctg gag cct ccg aat att ttt tcc att ctg ttt tcc aat aat ttt atc gat tct aat gaa tac gaa gtt aat cta agt cat tta gat tat caa agt gtt aac tat ttg atc gat tac att tat ggg ata cct ttg agc cta act aac aat aac gtg aaa tat att ctt tca acc gct ga- -tt ttt tta caa att gga tct gtc att act gag tgc gaa aaa tac ata ctt aaa aat ctt tgt ttt aga aac tgt atc gat ttc --- --- tac ata tac gct gat aaa tat aat aac aag aaa ata gaa tta gca tcg ttt aac aca ata tta cga aat att ttg aga ctc atc aac aat gaa aac ttt aaa tac tta aca gag gaa tca atg ata aaa att tta agc gat gat atg tta tat ata aaa aat gag gat ttc acc cca ctg att ctc att aaa tgg tta gag agt aca c-- caa cca tgt acc gtc gag tta ctt aga tgc ctc aga ata tca -tt tct ttc ccc aca agt tat aaa atc act tta tag tca tcg act ggt tag ttc aat cta cga atg tat aac att ctt aaa caa tat agc att ctt gga taa atc att tcc tag ata cca tat cat cga gtt gat atc tat cgg tat aag taa ttc aca tga taa gat ttc cat aaa ctg cta caa tca t-- aaa aaa att cat ggg aaa tga tat ctt cac gta gat ata ggt gta gtt tcg cag tga ccg tcc tgg ata ata tta tct ata tga tgg gtg gat atg atc agt ccc tgt ata gaa gtt caa agg tta tag cgt aca ata cat gta cta att ctt gga tat atg ata tac cag agc taa aaa tat cat cgt tct aat tgc gga gga gtt gcc aat gac gaa tac att tat tgt ata ggc ggt ata cgc gat cag gag tca tcg ttg ata tct agt atc gat aga tgg aag cca tca aaa cca tat tgg cag aag tat gct aaa atg tgc gaa cca aaa tgt gat atg gg- gtt gcg att tta aac gga tta ata tat gtc ata ggt gga gtc gtt aaa ggt gac aca cat ata cca acg cac tag aga gtt tat cag aag atg gat gga tga atc atc aac gtc ttc caa taa aaa tgt cca ata tgt cga cga ttg ttc atg ctg gaa aga ttt ata tat cta gag gtt aca aca ata gta gtg tag tta atg taa tat cga atc t-- agt cct tag cta taa tcc gat ata tga tga atg gac caa att atc atc att aaa tat tcc tag aat taa tcc tgc tct a

>MT903342.1:156407-158670_Monkeypox_virus_isolate_MPXV-Singapore

tta ctg act aca --- tgt a-c tat tt- -ta cat tac tac att ggc tac ggc ata tat acc tat ttc gtc act tcc aca cgc tcc ggt aaa cgg gtg tca tgt gac gag gga ga- -at ctt gat aag agg cat aat caa tgt tgt aat ccg atg tcc acc tgg aga att tgc caa ggt cag atg tag agt tgg tag tga taa cac aaa atg tga aca ctg ccc acc tca tac ata tac cgc aat ccc caa tta ttc t-a ata gat gtc atc aat gta gaa aat gcc caa cag gat cat ttg ata agg taa agt gta ccg gaa cac aga ac- --a aat -gt tcg tgt cat cct ggt tgg ta- tac gct act gat tct tca cag act gaa gat tgt cga gat ttg tgt acc aaa aaa gga gat gtc cat gcg gat act ttg gtg gaa tag atg aa- --g gaa atc cta ttt gta aat cgt gtt g-- -tg ttg gtg aat att gcg act acc tac gta att ata gac ttg atc cat ttc ctc cat gca aac tat cta tct aaa tgt aat taa tta --- --- --- --- --- --- --- --- tga ttt --- --- --- -tg atg ata atg tta -cc ata c-a tt- --- -at atc act act tgg tta gtg tgt att att tag tat gga aga cct att aat aa- --- --- -tt act tat ctt ttg acg atc ttg tt- -at aat tat --- --- --- --- aat ata aaa --a tac tta tga --- --- cat agt aac tca t-- --- --- --- aat tgc tg- acg cga taa att cgt aat aat ctg ttt tgt tca aat ttt tat aag ga- --- --a tct aca ggc ata aaa at- aaa aat ata atc tat aat ata c-- tct tac a-- acg cca tca tga at- --a gca gtg aat taa ttg ctg tta tt- atg gat tta gaa ata gtg gac gat ttt gtg ata tta ata tag tta tta atg atg aaa gga taa acg cgc ata gac tca tcc tat ctg gag cct ccg aat att ttt tcc att ctg ttt tcc aat aat ttt atc gat tct aat gaa tac gaa gtt aat cta agt cat tta gat tat caa agt gtt aac tat ttg atc gat tac att tat ggg ata cct ttg agc cta act aac aat aac gtg aaa tat att ctt tca acc gct ga- -tt ttt tta caa att gga tct gtc att act gag tgc gaa aaa tac ata ctt aaa aat ctt tgt ttt aga aac tgt atc gat ttc --- --- tac ata tac gct gat aaa tat aat aac aag aaa ata gaa tta gca tcg ttt aac aca ata tta cga aat att ttg aga ctc atc aac aat gaa aac ttt aaa tac tta aca gag gaa tca atg ata aaa att tta agc gat gat atg tta tat ata aaa aat gag gat ttc acc cca ctg att ctc att aaa tgg tta gag agt aca c-- caa cca tgt acc gtc gag tta ctt aga tgc ctc aga ata tca -tt tct ttc ccc aca agt tat aaa atc act tta tag tca tcg act ggt tag ttc aat cta cga atg tat aac att ctt aaa caa tat agc att ctt gga taa atc att tcc tag ata cca tat cat cga gtt gat atc tat cgg tat aag taa ttc aca tga taa gat ttc cat aaa ctg cta caa tca t-- aaa aaa att cat ggg aaa tga tat ctt cac gta gat ata ggt gta gtt tcg cag tga ccg tcc tgg ata ata tta tct ata tga tgg gtg gat atg atc agt ccc tgt ata gaa gtt caa agg tta tag cgt aca ata cat gta cta att ctt gga tat atg ata tac cag agc taa aaa tat cat cgt tct aat tgc gga gga gtt gcc aat gac gaa tac att tat tgt ata ggc ggt ata cgc gat cag gag tca tcg ttg ata tct agt atc gat aga tgg aag cca tca aaa cca tat tgg cag aag tat gct aaa atg tgc gaa cca aaa tgt gat atg gg- gtt gcg att tta aac gga tta ata tat gtc ata ggt gga gtc gtt aaa ggt gac aca cat ata cca acg cac tag aga gtt tat cag aag atg gat gga tga atc atc aac gtc ttc caa taa aaa tgt cca ata tgt cga cga ttg ttc atg ctg gaa aga ttt ata tat cta gag gtt aca aca ata gta gtg tag tta atg taa tat cga atc t-- agt cct tag cta taa tcc gat ata tga tga atg gac caa att atc atc att aaa tat tcc tag aat taa tcc tgc tct a

>MT903341.1:147911-150174_Monkeypox_virus_isolate_MPXV-M5320_M15_Bayelsa

tta ctg act aca --- tgt a-c tat tt- -ta cat tac tac att ggc tac ggc ata tat acc tat ttc gtc act tcc aca cgc tcc ggt aaa cgg gtg tca tgt gac gag gga ga- -at ctt gat aag agg cat aat caa tgt tgt aat ccg atg tcc acc tgg aga att tgc caa ggt cag atg tag agt tgg tag tga taa cac aaa atg tga aca ctg ccc acc tca tac ata tac cgc aat ccc caa tta ttc t-a ata gat gtc atc aat gta gaa aat gcc caa cag gat cat ttg ata agg taa agt gta ccg gaa cac aga ac- --a aat -gt tcg tgt cat cct ggt tgg ta- tac gct act gat tct tca cag act gaa gat tgt cga gat ttg tgt acc aaa aaa gga gat gtc cat gcg gat act ttg gtg gaa tag atg aa- --g gaa atc cta ttt gta aat cgt gtt g-- -tg ttg gtg aat att gcg act acc tac gta att ata gac ttg atc cat ttc ctc cat gca aac tat cta tct aaa tgt aat taa tta --- --- --- --- --- --- --- --- tga ttt --- --- --- -tg atg ata atg tta -cc ata c-a tt- --- -at atc act act tgg tta gtg tgt att att tag tat gga aga cct att aat aa- --- --- -tt act tat ctt ttg acg atc ttg tt- -at aat tat --- --- --- --- aat ata aaa --a tac tta tga --- --- cat agt aac tca t-- --- --- --- aat tgc tg- acg cga taa att cgt aat aat ctg ttt tgt tca aat ttt tat aag ga- --- --a tct aca ggc ata aaa at- aaa aat ata atc tat aat ata c-- tct tac a-- acg cca tca tga at- --a gca gtg aat taa ttg ctg tta tt- atg gat tta gaa ata gtg gac gat ttt gtg ata tta ata tag tta tta atg atg aaa gga taa acg cgc ata gac tca tcc tat ctg gag cct ccg aat att ttt tcc att ctg ttt tcc aat aat ttt atc gat tct aat gaa tac gaa gtt aat cta agt cat tta gat tat caa agt gtt aac tat ttg atc gat tac att tat ggg ata cct ttg agc cta act aac aat aac gtg aaa tat att ctt tca acc gct ga- -tt ttt tta caa att gga tct gtc att act gag tgc gaa aaa tac ata ctt aaa aat ctt tgt ttt aga aac tgt atc gat ttc --- --- tac ata tac gct gat aaa tat aat aac aag aaa ata gaa tta gca tcg ttt aac aca ata tta cga aat att ttg aga ctc atc aac aat gaa aac ttt aaa tac tta aca gag gaa tca atg ata aaa att tta agc gat gat atg tta tat ata aaa aat gag gat ttc acc cca ctg att ctc att aaa tgg tta gag agt aca c-- caa cca tgt acc gtc gag tta ctt aga tgc ctc aga ata tca -tt tct ttc ccc aca agt tat aaa atc act tta tag tca tcg act ggt tag ttc aat cta cga atg tat aac att ctt aaa caa tat agc att ctt gga taa atc att tcc tag ata cca tat cat cga gtt gat atc tat cgg tat aag taa ttc aca tga taa gat ttc cat aaa ctg cta caa tca t-- aaa aaa att cat ggg aaa tga tat ctt cac gta gat ata ggt gta gtt tcg cag tga ccg tcc tgg ata ata tta tct ata tga tgg gtg gat atg atc agt ccc tgt ata gaa gtt caa agg tta tag cgt aca ata cat gta cta att ctt gga tat atg ata tac cag agc taa aaa tat cat cgt tct aat tgc gga gga gtt gcc aat gac gaa tac att tat tgt ata ggc ggt ata cgc gat cag gag tca tcg ttg ata tct agt atc gat aga tgg aag cca tca aaa cca tat tgg cag aag tat gct aaa atg tgc gaa cca aaa tgt gat atg gg- gtt gcg att tta aac gga tta ata tat gtc ata ggt gga gtc gtt aaa ggt gac aca cat ata cca acg cac tag aga gtt tat cag aag atg gat gga tga atc atc aac gtc ttc caa taa aaa tgt cca ata tgt cga cga ttg ttc atg ctg gaa aga ttt ata tat cta gag gtt aca aca ata gta gtg tag tta atg taa tat cga atc t-- agt cct tag cta taa tcc gat ata tga tga atg gac caa att atc atc att aaa tat tcc tag aat taa tcc tgc tct a

>NC_063383.1:156370-158633_Monkeypox_virus_complete_genome

tta ctg act aca --- tgt a-c tat tt- -ta cat tac tac att ggc tac ggc ata tat acc tat ttc gtc act tcc aca cgc tcc ggt aaa cgg gtg tca tgt gac gag gga ga- -at ctt gat aag agg cat aat caa tgt tgt aat ccg atg tcc acc tgg aga att tgc caa ggt cag atg tag agt tgg tag tga taa cac aaa atg tga aca ctg ccc acc tca tac ata tac cgc aat ccc caa tta ttc t-a ata gat gtc atc aat gta gaa aat gcc caa cag gat cat ttg ata agg taa agt gta ccg gaa cac aga ac- --a aat -gt tcg tgt cat cct ggt tgg ta- tac gct act gat tct tca cag act gaa gat tgt cga gat ttg tgt acc aaa aaa gga gat gtc cat gcg gat act ttg gtg gaa tag atg aa- --g gaa atc cta ttt gta aat cgt gtt g-- -tg ttg gtg aat att gcg act acc tac gta att ata gac ttg atc cat ttc ctc cat gca aac tat cta tct aaa tgt aat taa tta --- --- --- --- --- --- --- --- tga ttt --- --- --- -tg atg ata atg tta -cc ata c-a tt- --- -at atc act act tgg tta gtg tgt att att tag tat gga aga cct att aat aa- --- --- -tt act tat ctt ttg acg atc ttg tt- -at aat tat --- --- --- --- aat ata aaa --a tac tta tga --- --- cat agt aac tca t-- --- --- --- aat tgc tg- acg cga taa att cgt aat aat ctg ttt tgt tca aat ttt tat aag ga- --- --a tct aca ggc ata aaa at- aaa aat ata atc tat aat ata c-- tct tac a-- acg cca tca tga at- --a gca gtg aat taa ttg ctg tta tt- atg gat tta gaa ata gtg gac gat ttt gtg ata tta ata tag tta tta atg atg aaa gga taa acg cgc ata gac tca tcc tat ctg gag cct ccg aat att ttt tcc att ctg ttt tcc aat aat ttt atc gat tct aat gaa tac gaa gtt aat cta agt cat tta gat tat caa agt gtt aac tat ttg atc gat tac att tat ggg ata cct ttg agc cta act aac aat aac gtg aaa tat att ctt tca acc gct ga- -tt ttt tta caa att gga tct gtc att act gag tgc gaa aaa tac ata ctt aaa aat ctt tgt ttt aga aac tgt atc gat ttc --- --- tac ata tac gct gat aaa tat aat aac aag aaa ata gaa tta gca tcg ttt aac aca ata tta cga aat att ttg aga ctc atc aac aat gaa aac ttt aaa tac tta aca gag gaa tca atg ata aaa att tta agc gat gat atg tta tat ata aaa aat gag gat ttc acc cca ctg att ctc att aaa tgg tta gag agt aca c-- caa cca tgt acc gtc gag tta ctt aga tgc ctc aga ata tca -tt tct ttc ccc aca agt tat aaa atc act tta tag tca tcg act ggt tag ttc aat cta cga atg tat aac att ctt aaa caa tat agc att ctt gga taa atc att tcc tag ata cca tat cat cga gtt gat atc tat cgg tat aag taa ttc aca tga taa gat ttc cat aaa ctg cta caa tca t-- aaa aaa att cat ggg aaa tga tat ctt cac gta gat ata ggt gta gtt tcg cag tga ccg tcc tgg ata ata tta tct ata tga tgg gtg gat atg atc agt ccc tgt ata gaa gtt caa agg tta tag cgt aca ata cat gta cta att ctt gga tat atg ata tac cag agc taa aaa tat cat cgt tct aat tgc gga gga gtt gcc aat gac gaa tac att tat tgt ata ggc ggt ata cgc gat cag gag tca tcg ttg ata tct agt atc gat aga tgg aag cca tca aaa cca tat tgg cag aag tat gct aaa atg tgc gaa cca aaa tgt gat atg gg- gtt gcg att tta aac gga tta ata tat gtc ata ggt gga gtc gtt aaa ggt gac aca cat ata cca acg cac tag aga gtt tat cag aag atg gat gga tga atc atc aac gtc ttc caa taa aaa tgt cca ata tgt cga cga ttg ttc atg ctg gaa aga ttt ata tat cta gag gtt aca aca ata gta gtg tag tta atg taa tat cga atc t-- agt cct tag cta taa tcc gat ata tga tga atg gac caa att atc atc att aaa tat tcc tag aat taa tcc tgc tct a

>MT903339.1:156518-158781_Monkeypox_virus_isolate_MPXV-M3021_Delta

tta ctg act aca --- tgt a-c tat tt- -ta cat tac tac att ggc tac ggc ata tat acc tat ttc gtc act tcc aca cgc tcc ggt aaa cgg gtg tca tgt gac gag gga ga- -at ctt gat aag agg cat aat caa tgt tgt aat ccg atg tcc acc tgg aga att tgc caa ggt cag atg tag agt tgg tag tga taa cac aaa atg tga aca ctg ccc acc tca tac ata tac cgc aat ccc caa tta ttc t-a ata gat gtc atc aat gta gaa aat gcc caa cag gat cat ttg ata agg taa agt gta ccg gaa cac aga ac- --a aat -gt tcg tgt cat cct ggt tgg ta- tac gct act gat tct tca cag act gaa gat tgt cga gat ttg tgt acc aaa aaa gga gat gtc cat gcg gat act ttg gtg gaa tag atg aa- --g gaa atc cta ttt gta aat cgt gtt g-- -tg ttg gtg aat att gcg act acc tac gta att ata gac ttg atc cat ttc ctc cat gca aac tat cta tct aaa tgt aat taa tta --- --- --- --- --- --- --- --- tga ttt --- --- --- -tg atg ata atg tta -cc ata c-a tt- --- -at atc act act tgg tta gtg tgt att att tag tat gga aga cct att aat aa- --- --- -tt act tat ctt ttg acg atc ttg tt- -at aat tat --- --- --- --- aat ata aaa --a tac tta tga --- --- cat agt aac tca t-- --- --- --- aat tgc tg- acg cga taa att cgt aat aat ctg ttt tgt tca aat ttt tat aag ga- --- --a tct aca ggc ata aaa at- aaa aat ata atc tat aat ata c-- tct tac a-- acg cca tca tga at- --a gca gtg aat taa ttg ctg tta tt- atg gat tta gaa ata gtg gac gat ttt gtg ata tta ata tag tta tta atg atg aaa gga taa acg cgc ata gac tca tcc tat ctg gag cct ccg aat att ttt tcc att ctg ttt tcc aat aat ttt atc gat tct aat gaa tac gaa gtt aat cta agt cat tta gat tat caa agt gtt aac tat ttg atc gat tac att tat ggg ata cct ttg agc cta act aac aat aac gtg aaa tat att ctt tca acc gct ga- -tt ttt tta caa att gga tct gtc att act gag tgc gaa aaa tac ata ctt aaa aat ctt tgt ttt aga aac tgt atc gat ttc --- --- tac ata tac gct gat aaa tat aat aac aag aaa ata gaa tta gca tcg ttt aac aca ata tta cga aat att ttg aga ctc atc aac aat gaa aac ttt aaa tac tta aca gag gaa tca atg ata aaa att tta agc gat gat atg tta tat ata aaa aat gag gat ttc acc cca ctg att ctc att aaa tgg tta gag agt aca c-- caa cca tgt acc gtc gag tta ctt aga tgc ctc aga ata tca -tt tct ttc ccc aca agt tat aaa atc act tta tag tca tcg act ggt tag ttc aat cta cga atg tat aac att ctt aaa caa tat agc att ctt gga taa atc att tcc tag ata cca tat cat cga gtt gat atc tat cgg tat aag taa ttc aca tga taa gat ttc cat aaa ctg cta caa tca t-- aaa aaa att cat ggg aaa tga tat ctt cac gta gat ata ggt gta gtt tcg cag tga ccg tcc tgg ata ata tta tct ata tga tgg gtg gat atg atc agt ccc tgt ata gaa gtt caa agg tta tag cgt aca ata cat gta cta att ctt gga tat atg ata tac cag agc taa aaa tat cat cgt tct aat tgc gga gga gtt gcc aat gac gaa tac att tat tgt ata ggc ggt ata cgc gat cag gag tca tcg ttg ata tct agt atc gat aga tgg aag cca tca aaa cca tat tgg cag aag tat gct aaa atg tgc gaa cca aaa tgt gat atg gg- gtt gcg att tta aac gga tta ata tat gtc ata ggt gga gtc gtt aaa ggt gac aca cat ata cca acg cac tag aga gtt tat cag aag atg gat gga tga atc atc aac gtc ttc caa taa aaa tgt cca ata tgt cga cga ttg ttc atg ctg gaa aga ttt ata tat cta gag gtt aca aca ata gta gtg tag tta atg taa tat cga atc t-- agt cct tag cta taa tcc gat ata tga tga atg gac caa att atc atc att aaa tat tcc tag aat taa tcc tgc tct a

>MT903338.1:156517-158780_Monkeypox_virus_isolate_MPXV-M2957_Lagos

tta ctg act aca --- tgt a-c tat tt- -ta cat tac tac att ggc tac ggc ata tat acc tat ttc gtc act tcc aca cgc tcc ggt aaa cgg gtg tca tgt gac gag gga ga- -at ctt gat aag agg cat aat caa tgt tgt aat ccg atg tcc acc tgg aga att tgc caa ggt cag atg tag agt tgg tag tga taa cac aaa atg tga aca ctg ccc acc tca tac ata tac cgc aat ccc caa tta ttc t-a ata gat gtc atc aat gta gaa aat gcc caa cag gat cat ttg ata agg taa agt gta ccg gaa cac aga ac- --a aat -gt tcg tgt cat cct ggt tgg ta- tac gct act gat tct tca cag act gaa gat tgt cga gat ttg tgt acc aaa aaa gga gat gtc cat gcg gat act ttg gtg gaa tag atg aa- --g gaa atc cta ttt gta aat cgt gtt g-- -tg ttg gtg aat att gcg act acc tac gta att ata gac ttg atc cat ttc ctc cat gca aac tat cta tct aaa tgt aat taa tta --- --- --- --- --- --- --- --- tga ttt --- --- --- -tg atg ata atg tta -cc ata c-a tt- --- -at atc act act tgg tta gtg tgt att att tag tat gga aga cct att aat aa- --- --- -tt act tat ctt ttg acg atc ttg tt- -at aat tat --- --- --- --- aat ata aaa --a tac tta tga --- --- cat agt aac tca t-- --- --- --- aat tgc tg- acg cga taa att cgt aat aat ctg ttt tgt tca aat ttt tat aag ga- --- --a tct aca ggc ata aaa at- aaa aat ata atc tat aat ata c-- tct tac a-- acg cca tca tga at- --a gca gtg aat taa ttg ctg tta tt- atg gat tta gaa ata gtg gac gat ttt gtg ata tta ata tag tta tta atg atg aaa gga taa acg cgc ata gac tca tcc tat ctg gag cct ccg aat att ttt tcc att ctg ttt tcc aat aat ttt atc gat tct aat gaa tac gaa gtt aat cta agt cat tta gat tat caa agt gtt aac tat ttg atc gat tac att tat ggg ata cct ttg agc cta act aac aat aac gtg aaa tat att ctt tca acc gct ga- -tt ttt tta caa att gga tct gtc att act gag tgc gaa aaa tac ata ctt aaa aat ctt tgt ttt aga aac tgt atc gat ttc --- --- tac ata tac gct gat aaa tat aat aac aag aaa ata gaa tta gca tcg ttt aac aca ata tta cga aat att ttg aga ctc atc aac aat gaa aac ttt aaa tac tta aca gag gaa tca atg ata aaa att tta agc gat gat atg tta tat ata aaa aat gag gat ttc acc cca ctg att ctc att aaa tgg tta gag agt aca c-- caa cca tgt acc gtc gag tta ctt aga tgc ctc aga ata tca -tt tct ttc ccc aca agt tat aaa atc act tta tag tca tcg act ggt tag ttc aat cta cga atg tat aac att ctt aaa caa tat agc att ctt gga taa atc att tcc tag ata cca tat cat cga gtt gat atc tat cgg tat aag taa ttc aca tga taa gat ttc cat aaa ctg cta caa tca t-- aaa aaa att cat ggg aaa tga tat ctt cac gta gat ata ggt gta gtt tcg cag tga ccg tcc tgg ata ata tta tct ata tga tgg gtg gat atg atc agt ccc tgt ata gaa gtt caa agg tta tag cgt aca ata cat gta cta att ctt gga tat atg ata tac cag agc taa aaa tat cat cgt tct aat tgc gga gga gtt gcc aat gac gaa tac att tat tgt ata ggc ggt ata cgc gat cag gag tca tcg ttg ata tct agt atc gat aga tgg aag cca tca aaa cca tat tgg cag aag tat gct aaa atg tgc gaa cca aaa tgt gat atg gg- gtt gcg att tta aac gga tta ata tat gtc ata ggt gga gtc gtt aaa ggt gac aca cat ata cca acg cac tag aga gtt tat cag aag atg gat gga tga atc atc aac gtc ttc caa taa aaa tgt cca ata tgt cga cga ttg ttc atg ctg gaa aga ttt ata tat cta gag gtt aca aca ata gta gtg tag tta atg taa tat cga atc t-- agt cct tag cta taa tcc gat ata tga tga atg gac caa att atc atc att aaa tat tcc tag aat taa tcc tgc tct a

>MT903337.1:156516-158779_Monkeypox_virus_isolate_MPXV-M2940_FCT

tta ctg act aca --- tgt a-c tat tt- -ta cat tac tac att ggc tac ggc ata tat acc tat ttc gtc act tcc aca cgc tcc ggt aaa cgg gtg tca tgt gac gag gga ga- -at ctt gat aag agg cat aat caa tgt tgt aat ccg atg tcc acc tgg aga att tgc caa ggt cag atg tag agt tgg tag tga taa cac aaa atg tga aca ctg ccc acc tca tac ata tac cgc aat ccc caa tta ttc t-a ata gat gtc atc aat gta gaa aat gcc caa cag gat cat ttg ata agg taa agt gta ccg gaa cac aga ac- --a aat -gt tcg tgt cat cct ggt tgg ta- tac gct act gat tct tca cag act gaa gat tgt cga gat ttg tgt acc aaa aaa gga gat gtc cat gcg gat act ttg gtg gaa tag atg aa- --g gaa atc cta ttt gta aat cgt gtt g-- -tg ttg gtg aat att gcg act acc tac gta att ata gac ttg atc cat ttc ctc cat gca aac tat cta tct aaa tgt aat taa tta --- --- --- --- --- --- --- --- tga ttt --- --- --- -tg atg ata atg tta -cc ata c-a tt- --- -at atc act act tgg tta gtg tgt att att tag tat gga aga cct att aat aa- --- --- -tt act tat ctt ttg acg atc ttg tt- -at aat tat --- --- --- --- aat ata aaa --a tac tta tga --- --- cat agt aac tca t-- --- --- --- aat tgc tg- acg cga taa att cgt aat aat ctg ttt tgt tca aat ttt tat aag ga- --- --a tct aca ggc ata aaa at- aaa aat ata atc tat aat ata c-- tct tac a-- acg cca tca tga at- --a gca gtg aat taa ttg ctg tta tt- atg gat tta gaa ata gtg gac gat ttt gtg ata tta ata tag tta tta atg atg aaa gga taa acg cgc ata gac tca tcc tat ctg gag cct ccg aat att ttt tcc att ctg ttt tcc aat aat ttt atc gat tct aat gaa tac gaa gtt aat cta agt cat tta gat tat caa agt gtt aac tat ttg atc gat tac att tat ggg ata cct ttg agc cta act aac aat aac gtg aaa tat att ctt tca acc gct ga- -tt ttt tta caa att gga tct gtc att act gag tgc gaa aaa tac ata ctt aaa aat ctt tgt ttt aga aac tgt atc gat ttc --- --- tac ata tac gct gat aaa tat aat aac aag aaa ata gaa tta gca tcg ttt aac aca ata tta cga aat att ttg aga ctc atc aac aat gaa aac ttt aaa tac tta aca gag gaa tca atg ata aaa att tta agc gat gat atg tta tat ata aaa aat gag gat ttc acc cca ctg att ctc att aaa tgg tta gag agt aca c-- caa cca tgt acc gtc gag tta ctt aga tgc ctc aga ata tca -tt tct ttc ccc aca agt tat aaa atc act tta tag tca tcg act ggt tag ttc aat cta cga atg tat aac att ctt aaa caa tat agc att ctt gga taa atc att tcc tag ata cca tat cat cga gtt gat atc tat cgg tat aag taa ttc aca tga taa gat ttc cat aaa ctg cta caa tca t-- aaa aaa att cat ggg aaa tga tat ctt cac gta gat ata ggt gta gtt tcg cag tga ccg tcc tgg ata ata tta tct ata tga tgg gtg gat atg atc agt ccc tgt ata gaa gtt caa agg tta tag cgt aca ata cat gta cta att ctt gga tat atg ata tac cag agc taa aaa tat cat cgt tct aat tgc gga gga gtt gcc aat gac gaa tac att tat tgt ata ggc ggt ata cgc gat cag gag tca tcg ttg ata tct agt atc gat aga tgg aag cca tca aaa cca tat tgg cag aag tat gct aaa atg tgc gaa cca aaa tgt gat atg gg- gtt gcg att tta aac gga tta ata tat gtc ata ggt gga gtc gtt aaa ggt gac aca cat ata cca acg cac tag aga gtt tat cag aag atg gat gga tga atc atc aac gtc ttc caa taa aaa tgt cca ata tgt cga cga ttg ttc atg ctg gaa aga ttt ata tat cta gag gtt aca aca ata gta gtg tag tta atg taa tat cga atc t-- agt cct tag cta taa tcc gat ata tga tga atg gac caa att atc atc att aaa tat tcc tag aat taa tcc tgc tct a

>ON649716.1:156342-158605_Monkeypox_virus_isolate_Monkeypox/PT0016/2022_partial_genome

tta ctg act aca --- tgt a-c tat tt- -ta cat tac tac att ggc tac ggc ata tat acc tat ttc gtc act tcc aca cgc tcc ggt aaa cgg gtg tca tgt gac gag gga ga- -at ctt gat aag agg cat aat caa tgt tgt aat ccg atg tcc acc tgg aga att tgc caa ggt cag atg tag agt tgg tag tga taa cac aaa atg tga aca ctg ccc acc tca tac ata tac cgc aat ccc caa tta ttc t-a ata gat gtc atc aat gta gaa aat gcc caa cag gat cat ttg ata agg taa agt gta ccg gaa cac aga ac- --a aat -gt tcg tgt cat cct ggt tgg ta- tac gct act gat tct tca cag act gaa gat tgt cga gat ttg tgt acc aaa aaa gga gat gtc cat gcg gat act ttg gtg gaa tag atg aa- --g gaa atc cta ttt gta aat cgt gtt g-- -tg ttg gtg aat att gcg act acc tac gta att ata gac ttg atc cat ttc ctc cat gca aac tat cta tct aaa tgt aat taa tta --- --- --- --- --- --- --- --- tga ttt --- --- --- -tg atg ata atg tta -cc ata c-a tt- --- -at atc act act tgg tta gtg tgt att att tag tat gga aga cct att aat aa- --- --- -tt act tat ctt ttg acg atc ttg tt- -at aat tat --- --- --- --- aat ata aaa --a tac tta tga --- --- cat agt aac tca t-- --- --- --- aat tgc tg- acg cga taa att cgt aat aat ctg ttt tgt tca aat ttt tat aag ga- --- --a tct aca ggc ata aaa at- aaa aat ata atc tat aat ata c-- tct tac a-- acg cca tca tga at- --a gca gtg aat taa ttg ctg tta tt- atg gat tta gaa ata gtg gac gat ttt gtg ata tta ata tag tta tta atg atg aaa gga taa acg cgc ata gac tca tcc tat ctg gag cct ccg aat att ttt tcc att ctg ttt tcc aat aat ttt atc gat tct aat gaa tac gaa gtt aat cta agt cat tta gat tat caa agt gtt aac tat ttg atc gat tac att tat ggg ata cct ttg agc cta act aac aat aac gtg aaa tat att ctt tca acc gct ga- -tt ttt tta caa att gga tct gtc att act gag tgc gaa aaa tac ata ctt aaa aat ctt tgt ttt aga aac tgt atc gat ttc --- --- tac ata tac gct gat aaa tat aat aac aag aaa ata gaa tta gca tcg ttt aac aca ata tta cga ?at att ttg aga ctc atc aac aat gaa aac ttt aaa tac tta aca gag gaa tca atg ata aaa att tta agc gat gat atg tta tat ata aaa aat gag gat ttc acc cca ctg att ctc att aaa tgg tta gag agt aca c-- caa cca tgt acc gtc gag tta ctt aga tgc ctc aga ata tca -tt tct ttc ccc aca agt tat aaa atc act tta tag tca tcg act ggt tag ttc aat cta cga atg tat aac att ctt aaa caa tat agc att ctt gga taa atc att tcc tag ata cca tat cat cga gtt gat atc tat cgg tat aag taa ttc aca tga taa gat ttc cat aaa ctg cta caa tca t-- aaa aaa att cat ggg aaa tga tat ctt cac gta gat ata ggt gta gtt tcg cag tga ccg tcc tgg ata ata tta tct ata tga tgg gtg gat atg atc agt ccc tgt ata gaa gtt caa agg tta tag cgt aca ata cat gta cta att ctt gga tat atg ata tac cag agc taa aaa tat cat cgt tct aat tgc gga gga gtt gcc aat gac gaa tac att tat tgt ata ggc ggt ata cgc gat cag gag tca tcg ttg ata tct agt atc gat aga tgg aag cca tca aaa cca tat tgg cag aag tat gct aaa atg tgc gaa cca aaa tgt gat atg gg- gtt gcg att tta aac gga tta ata tat gtc ata ggt gga gtc gtt aaa ggt gac aca cat ata cca acg cac tag aga gtt tat cag aag atg gat gga tga atc atc aac gtc ttc caa taa aaa tgt cca ata tgt cga cga ttg ttc atg ctg gaa aga ttt ata tat cta gag gtt aca aca ata gta gtg tag tta atg taa tat cga atc t-- agt cct tag cta taa tcc gat ata tga tga atg gac caa att atc atc att aaa tat tcc tag aat taa tcc tgc tct a

>ON675438.1:156352-158615_Monkeypox_virus_isolate_MPXV_USA_2022_VA001_complete_genome

tta ctg act aca --- tgt a-c tat tt- -ta cat tac tac att ggc tac ggc ata tat acc tat ttt gtc act tcc aca cgc tcc ggt aaa cgg gtg tca tgt gac gag gga ga- -at ctt gat aag agg cat aat caa tgt tgt aat ccg atg tcc acc tgg aga att tgc caa ggt cag atg tag agt tgg tag tga taa cac aaa atg tga aca ctg ccc acc tca tac ata tac cgc aat ccc caa tta ttc t-a ata gat gtc atc aat gta gaa aat gcc caa cag gat cat ttg ata agg taa agt gta ccg gaa cac aga ac- --a aat -gt tcg tgt cat cct ggt tgg ta- tac gct act gat tct tca cag act gaa gat tgt cga gat ttg tgt acc aaa aaa gga gat gtc cat gcg gat act ttg gtg gaa tag atg aa- --g gaa atc cta ttt gta aat cgt gtt g-- -tg ttg gtg aat att gcg act acc tac gta att ata gac ttg atc cat ttc ctc cat gca aac tat cta tct aaa tgt aat taa tta --- --- --- --- --- --- --- --- tga ttt --- --- --- -tg atg ata atg tta -cc ata c-a tt- --- -at atc act act tgg tta gtg tgt att att tag tat gga aga cct att aat aa- --- --- -tt act tat ctt ttg acg atc ttg tt- -at aat tat --- --- --- --- aat ata aaa --a tac tta tga --- --- cat agt aac tca t-- --- --- --- aat tgc tg- acg cga taa att cgt aat aat ctg ttt tgt tca aat ttt tat aag ga- --- --a tct aca ggc ata aaa at- aaa aat ata atc tat aat ata c-- tct tac a-- acg cca tca tga at- --a gca gtg aat taa ttg ctg tta tt- atg gat tta gaa ata gtg gac gat ttt gtg ata tta ata tag tta tta atg atg aaa gga taa acg cgc ata gac tca tcc tat ctg gag cct ccg aat att ttt tcc att ctg ttt tcc aat aat ttt atc gat tct aat gaa tac gaa gtt aat cta agt cat tta gat tat caa agt gtt aac tat ttg atc gat tac att tat ggg ata cct ttg agc cta act aac aat aac gtg aaa tat att ctt tca acc gct ga- -tt ttt tta caa att gga tct gtc att act gag tgc gaa aaa tac ata ctt aaa aat ctt tgt ttt aga aac tgt atc gat ttc --- --- tac ata tac gct gat aaa tat aat aac aag aaa ata gaa tta gca tcg ttt aac aca ata tta cga aat att ttg aga ctc atc aac aat gaa aac ttt aaa tac tta aca gag gaa tca atg ata aaa att tta agc gat gat atg tta tat ata aaa aat gag gat ttc acc cca ctg att ctc att aaa tgg tta gag agt aca c-- caa cca tgt acc gtc gag tta ctt aga tgc ctc aga ata tca -tt tct ttc ccc aca agt tat aaa atc act tta tag tca tcg act ggt tag ttc aat cta cga atg tat aac att ctt aaa caa tat agc att ctt gga taa atc att tcc tag ata cca tat cat cga gtt gat atc tat cgg tat aag taa ttc aca tga taa gat ttc cat aaa ctg cta caa tca t-- aaa aaa att cat ggg aaa tga tat ctt cac gta gat ata ggt gta gtt tcg cag tga ccg tcc tgg ata ata tta tct ata tga tgg gtg gat atg atc agt ccc tgt ata gaa gtt caa agg tta tag cgt aca ata cat gta cta att ctt gga tat atg ata tac cag agc taa aaa tat cat cgt tct aat tgc gga gga gtt gcc aat gac gaa tac att tat tgt ata ggc ggt ata cgc gat cag gag tca tcg ttg ata tct agt atc gat aga tgg aag cca tca aaa cca tat tgg cag aag tat gct aaa atg tgc gaa cca aaa tgt gat atg gg- gtt gcg att tta aac gga tta ata tat gtc ata ggt gga gtc gtt aaa ggt gac aca cat ata cca acg cac tag aga gtt tat cag aag atg gat gga tga atc atc aac gtc ttc caa taa aaa tgt cca ata tgt cga cga ttg ttc atg ctg gaa aga ttt ata tat cta gag gtt aca aca ata gta gtg tag tta atg taa tat cga atc t-- agt cct tag cta taa tcc gat ata tga tga atg gac caa att atc atc att aaa tat tcc tag aat taa tcc tgc tct a

>MT250197.1:156524-158786_Monkeypox_virus_strain_Singapore_2019

tta ctg act aca --- tgt a-c tat tt- -ta cat tac tac att ggc tac ggc ata tat acc tat ttc gtc act tcc aca cgc tcc ggt aaa cgg gtg tca tgt gac gag gga ga- -at ctt gat aag agg cat aat caa tgt tgt aat ccg atg tcc acc tgg aga att tgc caa ggt cag atg tag agt tgg tag tga taa cac aaa atg tga aca ctg ccc acc tca tac ata tac cgc aat ccc caa tta ttc t-a ata gat gtc atc aat gta gaa aat gcc caa cag gat cat ttg ata agg taa agt gta ccg gaa cac aga ac- --a aat -gt tcg tgt cat cct ggt tgg ta- tac gct act gat tct tca cag act gaa gat tgt cga gat ttg tgt acc aaa aaa gga gat gtc cat gcg gat act ttg gtg gaa tag atg aa- --g gaa atc cta ttt gta aat cgt gtt g-- -tg ttg gtg aat att gcg act acc tac gta att ata gac ttg atc cat ttc ctc cat gca aac tat cta tct aaa tgt aat taa tta --- --- --- --- --- --- --- --- tga ttt --- --- --- -tg atg ata atg tta -cc ata c-a tt- --- -at atc act act tgg tta gtg tgt att att tag tat gga aga cct att aat aa- --- --- -tt act tat ctt ttg acg atc ttg tt- -at aat tat --- --- --- --- aat ata aaa --a tac tta tga --- --- cat agt aac tca t-- --- --- --- aat tgc tg- acg cga taa att cgt aat aat ctg ttt tgt tca aat ttt tat aag ga- --- --a tct aca ggc ata aaa at- aaa aat ata atc tat aat ata c-- tct tac a-- acg cca tca tga at- --a gca gtg aat taa ttg ctg tta tt- atg gat tta gaa ata gtg gac gat ttt gtg ata tta ata tag tta tta atg atg aaa gga taa acg cgc ata gac tca tcc tat ctg gag cct ccg aat att ttt tcc att ctg ttt tcc aat aat ttt atc gat tct aat gaa tac gaa gtt aat cta agt cat tta gat tat caa agt gtt aac tat ttg atc gat tac att tat ggg ata cct ttg agc cta act aac aat aac gtg aaa tat att ctt tca acc gct ga- -tt ttt tta caa att gga tct gtc att act gag tgc gaa aaa tac ata ctt aaa aat ctt tgt ttt aga aac tgt atc gat ttc --- --- tac ata tac gct gat aaa tat aat aac aag aaa ata gaa tta gca tcg ttt aac aca ata tta cga aat att ttg aga ctc atc aac aat gaa aac ttt aaa tac tta aca gag gaa tca atg ata aaa att tta agc gat gat atg tta tat ata aaa aat gag gat ttc acc cca ctg att ctc att aaa tgg tta gag agt aca c-- caa cca tgt acc gtc gag tta ctt aga tgc ctc aga ata tca -tt tct ttc ccc aca agt tat aaa atc act tta tag tca tcg act ggt tag ttc aat cta cga atg tat aac att ctt aaa caa tat agc att ctt gga taa atc att tcc tag ata cca tat cat cga gtt gat atc tat cgg tat aag taa ttc aca tga taa gat ttc cat aaa ctg cta caa tca t-- aaa aaa att cat ggg aaa tga tat ctt cac gta gat ata ggt gta gtt tcg cag tga ccg tcc tgg ata ata tta tct ata tga tgg gtg gat atg atc agt ccc tgt ata gaa gtt caa agg tta tag cgt aca ata cat gta cta att ctt gga tat atg ata tac cag agc t-a aaa tat cat cgt tct aat tgc gga gga gtt gcc aat gac gaa tac att tat tgt ata ggc ggt ata cgc gat cag gag tca tcg ttg ata tct agt atc gat aga tgg aag cca tca aaa cca tat tgg cag aag tat gct aaa atg tgc gaa cca aaa tgt gat atg gg- gtt gcg att tta aac gga tta ata tat gtc ata ggt gga gtc gtt aaa ggt gac aca cat ata cca acg cac tag aga gtt tat cag aag atg gat gga tga atc atc aac gtc ttc caa taa aaa tgt cca ata tgt cga cga ttg ttc atg ctg gaa aga ttt ata tat cta gag gtt aca aca ata gta gtg tag tta atg taa tat cga atc t-- agt cct tag cta taa tcc gat ata tga tga atg gac caa att atc atc att aaa tat tcc tag aat taa tcc tgc tct a

>KJ642617.1:156524-158786_Monkeypox_virus_strain_Nigeria-SE-1971_complete_genome

tta ctg act aca --- tgt a-c tat tt- -ta cat tac tac att ggc tac ggc ata tat acc tat ttc gtc act tcc aca cgc tcc ggt aaa cgg gtg tca tgt gac gag gga ga- -at ctt gat aag agg cat aat caa tgt tgt aat ccg atg tcc acc tgg aga att tgc caa ggt cag atg tag agt tgg tag tga taa cac aaa atg tga aca ctg ccc acc tca tac ata tac cgc aat ccc caa tta ttc t-a ata gat gtc atc aat gta gaa aat gcc caa cag gat cat ttg ata agg taa agt gta ccg gaa cac aga ac- --a aat -gt tcg tgt cat cct ggt tgg ta- tac gct act gat tct tca cag act gaa gat tgt cga gat ttg tgt acc aaa aaa gga gat gtc cat gcg gat act ttg gtg gaa tag atg aa- --g gaa atc cta ttt gta aat cgt gtt g-- -tg ttg gtg aat att gcg act acc tac gta att ata gac ttg atc cat ttc ctc cat gca aac tat cta tct aaa tgt aat taa tta --- --- --- --- --- --- --- --- tga ttt --- --- --- -tg atg ata atg tta -cc ata c-a tt- --- -at atc act act tgg tta gtg tgt att att tag tat gga aga cct att aat aa- --- --- -tt act tat ctt ttg acg atc ttg tt- -at aat tat --- --- --- --- aat ata aaa --a tac tta tga --- --- cat agt aac tca t-- --- --- --- aat tgc tg- acg cga taa att cgt aat aat ctg ttt tgt tca aat ttt tat aag ga- --- --a tct aca ggc ata aaa at- aaa aat ata atc tat aat ata c-- tct tac a-- acg cca tca tga at- --a gca gtg aat taa ttg ctg tta tt- atg gat tta gaa ata gtg gac gat ttt gtg ata tta ata tag tta tta atg atg aaa gga taa acg cgc ata gac tca tcc tat ctg gag cct ccg aat att ttt tcc att ctg ttt tcc aat aat ttt atc gat tct aat gaa tac gaa gtt aat cta agt cat tta gat tat caa agt gtt aac tat ttg atc gat tac att tat ggg ata cct ttg agc cta act aac aat aac gtg aaa tat att ctt tca acc gct ga- -tt ttt tta caa att gga tct gtc att act gag tgc gaa aaa tac ata ctt aaa aat ctt tgt ttt aga aac tgt atc gat ttc --- --- tac ata tac gct gat aaa tat aat aac aag aaa ata gaa tta gca tcg ttt aac aca ata tta cga aat att ttg aga ctc atc aac aat gaa aac ttt aaa tac tta aca gag gaa tca atg ata aaa att tta agc gat gat atg tta tat ata aaa aat gag gat ttc acc cca ctg att ctc att aaa tgg tta gag agt aca c-- caa cca tgt acc gtc gag tta ctt aga tgc ctc aga ata tca -tt tct ttc ccc aca agt tat aaa atc act tta tag tca tcg act ggt tag ttc aat cta cga atg tat aac att ctt aaa caa tat agc att ctt gga taa atc att tcc tag ata cca tat cat cga gtt gat atc tat cgg tat aag taa ttc aca tga taa gat ttc cat aaa ctg cta caa tca t-- aaa aaa att cat ggg aaa tga tat ctt cac gta gat ata ggt gta gtt tcg cag tga ccg tcc tgg ata ata tta tct ata tga tgg gtg gat atg atc agt ccc tgt ata gaa gtt caa agg tta tag cgt aca ata cat gta cta att ctt gga tat atg ata tac cag agc t-a aaa tat cat cgt tct aat tgc gga gga gtt gcc aat gac gaa tac att tat tgt ata ggc ggt ata cgc gat cag gag tca tcg ttg ata tct agt atc gat aga tgg aag cca tca aaa cca tat tgg cag aag tat gct aaa atg tgc gaa cca aaa tgt gat atg gg- gtt gcg att tta aac gga tta ata tat gtc ata ggt gga gtc gtt aaa ggt gac aca cat ata cca acg cac tag aga gtt tat cag aag atg gat gga tga atc atc aac gtc ttc caa taa aaa tgt cca ata tgt cga cga ttg ttc atg ctg gaa aga ttt ata tat cta gag gtt aca aca ata gta gtg tag tta atg taa tat cga atc t-- agt cct tag cta taa tcc gat ata tga tga atg gac caa att atc atc att aaa tat tcc tag aat taa tcc tgc tct a

>ON585029.1:156574-158837_Monkeypox_virus_isolate_Monkeypox/PT0001/2022_partial_genome

tta ctg act aca --- tgt a-c tat tt- -ta cat tac tac att ggc tac ggc ata tat acc tat ttc gtc act tcc aca cgc tcc ggt aaa cgg gtg tca tgt gac gag gga ga- -at ctt gat aag agg cat aat caa tgt tgt aat ccg atg tcc acc tgg aga att tgc caa ggt cag atg tag agt tgg tag tga taa cac aaa atg tga aca ctg ccc acc tca tac ata tac cgc aat ccc caa tta ttc t-a ata gat gtc atc aat gta gaa aat gcc caa cag gat cat ttg ata agg taa agt gta ccg gaa cac aga ac- --a aat -gt tcg tgt cat cct ggt tgg ta- tac gct act gat tct tca cag act gaa gat tgt cga gat ttg tgt acc aaa aaa gga gat gtc cat gcg gat act ttg gtg gaa tag atg aa- --g gaa atc cta ttt gta aat cgt gtt g-- -tg ttg gtg aat att gcg act acc tac gta att ata gac ttg atc cat ttc ctc cat gca aac tat cta tct aaa tgt aat taa tta --- --- --- --- --- --- --- --- tga ttt --- --- --- -tg atg ata atg tta -cc ata c-a tt- --- -at atc act act tgg tta gtg tgt att att tag tat gga aga cct att aat aa- --- --- -tt act tat ctt ttg acg atc ttg tt- -at aat tat --- --- --- --- aat ata aaa --a tac tta tga --- --- cat agt aac tca t-- --- --- --- aat tgc tg- acg cga taa att cgt aat aat ctg ttt tgt tca aat ttt tat aag ga- --- --a tct aca ggc ata aaa at- aaa aat ata atc tat aat ata c-- tct tac a-- acg cca tca tga at- --a gca gtg aat taa ttg ctg tta tt- atg gat tta gaa ata gtg gac gat ttt gtg ata tta ata tag tta tta atg atg aaa gga taa acg cgc ata gac tca tcc tat ctg gag cct ccg aat att ttt tcc att ctg ttt tcc aat aat ttt atc gat tct aat gaa tac gaa gtt aat cta agt cat tta gat tat caa agt gtt aac tat ttg atc gat tac att tat ggg ata cct ttg agc cta act aac aat aac gtg aaa tat att ctt tca acc gct ga- -tt ttt tta caa att gga tct gtc att act gag tgc gaa aaa tac ata ctt aaa aat ctt tgt ttt aga aac tgt atc gat ttc --- --- tac ata tac gct gat aaa tat aat aac aag aaa ata gaa tta gca tcg ttt aac aca ata tta cga aat att ttg aga ctc atc aac aat gaa aac ttt aaa tac tta aca gag gaa tca atg ata aaa att tta agc gat gat atg tta tat ata aaa aat gag gat ttc acc cca ctg att ctc att aaa tgg tta gag agt aca c-- caa cca tgt acc gtc gag tta ctt aga tgc ctc aga ata tca -tt tct ttc ccc aca agt tat aaa atc act tta tag tca tcg act ggt tag ttc aat cta cga atg tat aac att ctt aaa caa tat agc att ctt gga taa atc att tcc tag ata cca tat cat cga gtt gat atc tat cgg tat aag taa ttc aca tga taa gat ttc cat aaa ctg cta caa tca t-- aaa aaa att cat ggg aaa tga tat ctt cac gta gat ata ggt gta gtt tcg cag tga ccg tcc tgg ata ata tta tct ata tga tgg gtg gat atg atc agt ccc tgt ata gaa gtt caa agg tta tag cgt aca ata cat gta cta att ctt gga tat atg ata tac cag agc taa aaa tat cat cgt tct aat tgc gga gga gtt gcc aat gac gaa tac att tat tgt ata ggc ggt ata cgc gat cag gag tca tcg ttg ata tct agt atc gat aga tgg aag cca tca aaa cca tat tgg cag aag tat gct aaa atg tgc gaa cca aaa tgt gat atg gg- gtt gcg att tta aac gga tta ata tat gtc ata ggt gga gtc gtt aaa ggt gac aca cat ata cca acg cac tag aga gtt tat cag aag atg gat gga tga atc atc aac gtc ttc caa t?a aaa tgt ?ca ata tgt cga cga ttg ttc atg ctg gaa aga ttt ata tat cta gag gtt aca aca ata gta gtg tag tta atg taa tat cga atc t-- agt cct tag cta taa tcc gat ata tga tga atg gac caa att atc atc att aaa tat tcc tag aat taa tcc tgc tct a

>ON622720.1:156359-158622_Monkeypox_virus_isolate_MPXV-CH-38156923/2022_partial_genome

tta ctg act aca --- tgt a-c tat tt- -ta cat tac tac att ggc tac ggc ata tat acc tat ttc gtc act tcc aca cgc tcc ggt aaa cgg gtg tca tgt gac ga? gga ga- -at ctt gat aag agg cat aat caa tgt tgt aat ccg atg tcc acc tgg aga att tgc caa ggt cag atg tag agt tgg tag tga taa cac aaa atg tga aca ctg ccc acc tca tac ata tac cgc aat ccc caa tta ttc t-a ata gat gtc atc aat gta gaa aat gcc caa cag gat cat ttg ata agg taa agt gta ccg gaa cac aga ac- --a aat -gt tcg tgt cat cct ggt tgg ta- tac gct act gat tct tca cag act gaa gat tgt cga gat ttg tgt acc ?aa aaa gga gat gtc cat gcg gat act ttg gtg gaa tag atg aa- --g gaa atc cta ttt gta aat cgt gtt g-- -tg ttg gtg aat att gcg act acc tac gta att ata gac ttg atc cat ttc ctc cat gca aac tat cta tct aaa tgt aat taa tta --- --- --- --- --- --- --- --- tga ttt --- --- --- -tg atg ata atg tta -cc ata c-a tt- --- -at atc act act tgg tta gtg tgt att att tag tat gga aga cct att aat aa- --- --- -tt act tat ctt ttg acg atc ttg tt- -at aat tat --- --- --- --- aat ata aaa --a tac tta tga --- --- cat agt aac tca t-- --- --- --- aat tgc tg- acg cga taa att cgt aat aat ctg ttt tgt tca aat ttt tat aag ga- --- --a tct aca ggc ata aaa at- aaa aat ata atc tat aat ata c-- tct tac a-- acg cca tca tga at- --a gca gtg aat taa ttg ctg tta tt- atg gat tta gaa ata gtg gac gat ttt gtg ata tta ata tag tta tta atg atg aaa gga taa acg cgc ata gac tca tcc tat ctg gag cct ccg aat att ttt tcc att ctg ttt tcc aat aat ttt atc gat tct aat gaa tac gaa gtt aat cta agt cat tta gat tat caa agt gtt aac tat ttg atc gat tac att tat ggg ata cct ttg agc cta act aac aat aac gtg aaa tat att ctt tca acc gct ga- -?t ttt tta caa att gga tct gtc att act gag tgc gaa aaa tac ata ctt aaa aat ctt tgt ttt aga aac tgt atc gat ttc --- --- tac ata tac gct gat aaa tat aat aac aag aaa ata gaa tta gca tcg ttt aac aca ata tta cga aat att ttg aga ctc atc aac aat gaa aac ttt aaa tac tta aca gag gaa tca atg ata aaa att tta agc gat gat atg tta tat ata aaa aat gag gat ttc acc cca ctg att ctc att aaa tgg tta gag agt aca c-- caa cca tgt acc gtc gag tta ctt aga tgc ctc aga ata tca -tt tct ttc ccc aca agt tat aaa atc act tta tag tca tcg act ggt tag ttc aat cta cga atg tat aac att ctt aaa caa tat agc att ctt gga taa atc att tcc tag ata cca tat cat cga gtt gat atc tat cgg tat aag taa ttc aca tga taa gat ttc cat aaa ctg cta caa tca t-- ?aa aaa att cat ggg aaa tga tat ctt cac gta gat ata ggt gta gtt tcg cag tga ccg tcc tgg ata ata tta tct ata tga tgg gtg gat atg atc agt ccc tgt ata gaa gtt caa agg tta tag cgt aca ata cat gta cta att ctt gga tat atg ata tac cag agc taa aaa tat cat cgt tct aat tgc gga gga gtt gcc aat gac gaa tac att tat tgt ata ggc ggt ata cgc gat cag gag tca tcg ttg ata tct agt atc gat aga tgg aag cca tca aaa cca tat tgg cag aag tat gct aaa atg tgc gaa cca aaa tgt gat atg gg- gtt gcg att tta aac gga tta ata tat gtc ata ggt gga gtc gtt aaa ggt gac aca cat ata cca acg cac tag aga gtt tat cag aag atg gat gga tga atc atc aac gtc ttc caa taa aaa tgt cca ata tgt cga cga ttg ttc atg ctg gaa aga ttt ata tat cta gag gtt aca aca ata gta gtg tag tta atg taa tat cga atc t-- agt cct tag cta taa tcc gat ata tga tga atg gac caa att atc atc att aaa tat tcc tag aat taa tcc tgc tct a

>ON602722.1:156211-158472_Monkeypox_virus_isolate_MPXV_FRA_2022_TLS67_complete_genome

tta ctg act aca --- tgt a-c tat tt- -ta cat tac tac att ggc tac ggc ata tat acc tat ttc gtc act tcc aca cgc tcc ggt aaa cgg gtg tca tgt gac gag gga ga- -at ctt gat aag agg cat aat caa tgt tgt aat ccg atg tcc acc tgg aga att tgc caa ggt cag atg tag agt tgg tag tga taa cac aaa atg tga aca ctg ccc acc tca tac ata tac cgc aat ccc caa tta ttc t-a ata gat gtc atc aat gta gaa aat gcc caa cag gat cat ttg ata agg taa agt gta ccg gaa cac aga ac- --a aat -gt tcg tgt cat cct ggt tgg ta- tac gct act gat tct tca cag act gaa gat tgt cga gat ttg tgt acc aaa aag g-a gat gtc cat gcg gat act ttg gtg gaa tag atg aa- --g gaa atc cta ttt gta aat cgt gtt g-- -tg ttg gtg aat att gcg act acc tac gta att ata gac ttg atc cat ttc ctc cat gca aac tat cta tct aaa tgt aat taa tta --- --- --- --- --- --- --- --- tga ttt --- --- --- -tg atg ata atg tta -cc ata c-a tt- --- -at atc act act tgg tta gtg tgt att att tag tat gga aga cct att aat aa- --- --- -tt act tat ctt ttg acg atc ttg tt- -at aat tat --- --- --- --- aat ata aaa --a tac tta tga --- --- cat agt aac tca t-- --- --- --- aat tgc tg- acg cga taa att cgt aat aat ctg ttt tgt tca aat ttt tat aag ga- --- --a tct aca ggc ata aaa at- aaa aat ata atc tat aat ata c-- tct tac a-- acg cca tca tga at- --a gca gtg aat taa ttg ctg tta tt- atg gat tta gaa ata gtg gac gat ttt gtg ata tta ata tag tta tta atg atg aaa gga taa acg cgc ata gac tca tcc tat ctg gag cct ccg aat att ttt tcc att ctg ttt tcc aat aat ttt atc gat tct aat gaa tac gaa gtt aat cta agt cat tta gat tat caa agt gtt aac tat ttg atc gat tac att tat ggg ata cct ttg agc cta act aac aat aac gtg aaa tat att ctt tca acc gct ga- -tt ttt tta caa att gga tct gtc att act gag tgc gaa aaa tac ata ctt aaa aat ctt tgt ttt aga aac tgt atc gat ttc --- --- tac ata tac gct gat aaa tat aat aac aag aaa ata gaa tta gca tcg ttt aac aca ata tta cga aat att ttg aga ctc atc aac aat gaa aac ttt aaa tac tta aca gag gaa tca atg ata aaa att tta agc gat gat atg tta tat ata aaa aat gag gat ttc acc cca ctg att ctc att aaa tgg tta gag agt aca c-- caa cca tgt acc gtc gag tta ctt aga tgc ctc aga ata tca -tt tct ttc ccc aca agt tat aaa atc act tta tag tca tcg act ggt tag ttc aat cta cga atg tat aac att ctt aaa caa tat agc att ctt gga taa atc att tcc tag ata cca tat cat cga gtt gat atc tat cgg tat aag taa ttc aca tga taa gat ttc cat aaa ctg cta caa tca t-- -aa aaa att cat ggg aaa tga tat ctt cac gta gat ata ggt gta gtt tcg cag tga ccg tcc tgg ata ata tta tct ata tga tgg gtg gat atg atc agt ccc tgt ata gaa gtt caa agg tta tag cgt aca ata cat gta cta att ctt gga tat atg ata tac cag agc taa aaa tat cat cgt tct aat tgc gga gga gtt gcc aat gac gaa tac att tat tgt ata ggc ggt ata cgc gat ca? gag tca tcg ttg ata tct agt atc gat aga tgg aag cca tca aaa cca tat tgg cag aag tat gct aaa atg tgc gaa cca aaa tgt gat atg gg- gtt gcg att tta aac gga tta ata tat gtc ata ggt gga gtc gtt aaa ggt gac aca cat ata cca acg cac tag aga gtt tat cag aag atg gat gga tga atc atc aac gtc ttc caa taa aaa tgt cca ata tgt cga cga ttg ttc atg ctg gaa aga ttt ata tat cta gag gtt aca aca ata gta gtg tag tta atg taa tat cga atc t-- agt cct tag cta taa tcc gat ata tga tga atg gac caa att atc atc att aaa tat tcc tag aat taa tcc tgc tct a

>ON649710.1:156347-158610_Monkeypox_virus_isolate_Monkeypox/PT0027/2022_partial_genome

tta ctg act aca --- tgt a-c tat tt- -ta cat tac tac att ggc tac ggc ata tat acc tat ttc gtc act tcc aca cgc tcc ggt aaa cgg gtg tca tgt gac gag gga ga- -at ctt gat aag agg cat aat caa tgt tgt aat ccg atg tcc acc tgg aga att tgc caa ggt cag atg tag agt tgg tag tga taa cac aaa atg tga aca ctg ccc acc tca tac ata tac cgc aat ccc caa tta ttc t-a ata gat gtc atc aat gta gaa aat gcc caa cag gat cat ttg ata agg taa agt gta ccg gaa cac aga ac- --a aat -gt tcg tgt cat cct ggt tgg ta- tac gct act gat tct tca cag act gaa gat tgt cga gat ttg tgt acc aaa aaa gga gat gtc cat gcg gat act ttg gtg gaa tag atg aa- --g gaa atc cta ttt gta aat cgt gtt g-- -tg ttg gtg aat att gcg act acc tac gta att ata gac ttg atc cat ttc ctc cat gca aac tat cta tct aaa tgt aat taa tta --- --- --- --- --- --- --- --- tga ttt --- --- --- -tg atg ata atg tta -cc ata c-a tt- --- -at atc act act tgg tta gtg tgt att att tag tat gga aga cct att aat aa- --- --- -tt act tat ctt tt? ??? ??c ttg t?- -a? ??? ??? --- --- --- --- ??? ?ta aaa --a tac tta tga --- --- cat agt aac tca t-- --- --- --- aat tgc tg- acg cga taa att cgt aat aat ctg ttt tgt tca aat ttt tat aag ga- --- --a tct aca ggc ata aaa at- aaa aat ata atc tat aat ata c-- tct tac a-- acg cca tca tga at- --a gca gtg aat taa ttg ctg tta tt- atg gat tta gaa ata gtg gac gat ttt gtg ata tta ata tag tta tta atg atg aaa gga taa acg cgc ata gac tca tcc tat ctg gag cct ccg aat att ttt tcc att ctg ttt tcc aat aat ttt atc gat tct aat gaa tac gaa gtt aat cta agt cat tta gat tat caa agt gtt aac tat ttg atc gat tac att tat ggg ata cct ttg agc cta act aac aat aac gtg aaa tat att ctt tca acc gct ga- -tt ttt tta caa att gga tct gtc att act gag tgc gaa aaa tac ata ctt aaa aat ctt tgt ttt aga aac tgt atc gat ttc --- --- tac ata tac gct gat aaa tat aat aac aag aaa ata gaa tta gca tcg ttt aac aca ata tta cga aat att ttg aga ctc atc aac aat gaa aac ttt aaa tac tta aca gag gaa tca atg ata aaa att tta agc gat gat atg tta tat ata aaa aat gag gat ttc acc cca ctg att ctc att aaa tgg tta gag agt aca c-- caa cca tgt acc gtc gag tta ctt aga tgc ctc aga ata tca -tt tct ttc ccc aca agt tat aaa atc act tta tag tca tcg act ggt tag ttc aat cta cga atg tat aac att ctt aaa caa tat agc att ctt gga taa atc att tcc tag ata cca tat cat cga gtt gat atc tat cgg tat aag taa ttc aca tga taa gat ttc cat aaa ctg cta caa tca t-- aaa aaa att cat ggg aaa tga tat ctt cac gta gat ata ggt gta gtt tcg cag tga ccg tcc tgg ata ata tta tct ata tga tgg gtg gat atg atc agt ccc tgt ata gaa gtt caa agg tta tag cgt aca ata cat gta cta att ctt gga tat atg ata tac cag agc taa aaa tat cat cgt tct aat tgc gga gga gtt gcc aat gac gaa tac att tat tgt ata ggc ggt ata cgc gat cag gag tca tcg ttg ata tct agt atc gat aga tgg aag cca tca aaa cca tat tgg cag aag tat gct aaa atg tgc gaa cca aaa tgt gat atg gg- gtt gcg att tta aac gga tta ata tat gtc ata ggt gga gtc gtt aaa ggt gac aca cat ata cca acg cac tag aga gtt tat cag aag atg gat gga tga atc atc aac gtc ttc caa taa aaa tgt cca ata tgt cga cga ttg ttc atg ctg gaa aga ttt ata tat cta gag gtt aca aca ata gta gtg tag tta atg taa tat cga atc t-- agt cct tag cta taa tcc gat ata tga tga atg gac caa att atc atc att aaa tat tcc tag aat taa tcc tgc tct a

>ON622718.1:155852-158115_Monkeypox_virus_isolate_MPXV/ES0001/HUGTiP/2022_partial_genome

tta ctg act aca --- tgt a-c tat tt- -ta cat tac tac att ggc tac ggc ata tat acc tat ttc gtc act tcc aca cgc tcc ggt aaa cgg gtg tca tgt gac gag gga ga- -at ctt gat aag agg cat aat caa tgt tgt aat ccg atg tcc acc tgg aga att tgc caa ggt cag atg tag agt tgg tag tga taa cac aaa atg tga aca ctg ccc acc tca tac ata tac cgc aat ccc caa tta ttc t-a ata gat gtc atc aat gta gaa aat gcc caa cag gat cat ttg ata agg taa agt gta ccg gaa cac aga ac- --a aat -gt tcg tgt cat cct ggt tgg ta- tac gct act gat tct tca cag act gaa gat tgt cga gat ttg tgt acc aaa aaa gga gat gtc cat gcg gat act ttg gtg gaa tag atg aa- --g gaa atc cta ttt gta aat cgt gtt g-- -tg ttg gtg aat att gcg act acc tac gta att ata gac ttg atc cat ttc ctc cat gca aac tat cta tct aaa tgt aat taa tta --- --- --- --- --- --- --- --- tga ttt --- --- --- -tg atg ata atg tta -cc ata c-a tt- --- -at atc act act tgg tta gtg tgt att att tag tat gga aga cct att aat aa- --- --- -tt act tat ctt ttg acg atc ttg tt- -at aat tat --- --- --- --- aat ata aaa --a tac tta tga --- --- cat agt aac tca t-- --- --- --- aat tgc tg- acg cga taa att cgt aat aat ctg ttt tg? ??? ??? ??? ??? ??? ??- --- --a tct aca ggc ata aaa at- aaa aat ata atc tat aat ata c-- tct tac a-- acg cca tca tga at- --a gca gtg aat taa ttg ctg tta tt- atg gat tta gaa ata gtg gac gat ttt gtg ata tta ata tag tta tta atg atg aaa gga taa acg cgc ata gac tca tcc tat ctg gag cct ccg aat att ttt tcc att ctg ttt tcc aat aat ttt atc gat tct aat gaa tac gaa gtt aat cta agt cat tta gat tat caa agt gtt aac tat ttg atc gat tac att tat ggg ata cct ttg agc cta act aac aat aac gtg aaa tat att ctt tca acc gct ga- -tt ttt tta caa att gga tct gtc att act gag tgc gaa aaa tac ata ctt aaa aat ctt tgt ttt aga aac tgt atc gat ttc --- --- tac ata tac gct gat aaa tat aat aac aag aaa ata gaa tta gca tcg ttt aac aca ata tta cga aat att ttg aga ctc atc aac aat gaa aac ttt aaa tac tta aca gag gaa tca atg ata aaa att tta agc gat gat atg tta tat ata aaa aat gag gat ttc acc cca ctg att ctc att aaa tgg tta gag agt aca c-- caa cca tgt acc gtc gag tta ctt aga tgc ctc aga ata tca -tt tct ttc ccc aca agt tat aaa atc act tta tag tca tcg act ggt tag ttc aat cta cga atg tat aac att ctt aaa caa tat agc att ctt gga taa atc att tcc tag ata cca tat cat cga gtt gat atc tat cgg tat aag taa ttc aca tga taa gat ttc cat aaa ctg cta caa tca t-- aaa aaa att cat ggg aaa tga tat ctt cac gta gat ata ggt gta gtt tcg cag tga ccg tcc tgg ata ata tta tct ata tga tgg gtg gat atg atc agt ccc tgt ata gaa gtt caa agg tta tag cgt aca ata cat gta cta att ctt gga tat atg ata tac cag agc taa aaa tat cat cgt tct aat tgc gga gga gtt gcc aat gac gaa tac att tat tgt ata ggc ggt ata cgc gat cag gag tca tcg ttg ata tct agt atc gat aga tgg aag cca tca aaa cca tat tgg cag aag tat gct aaa atg tgc gaa cca aaa tgt gat atg gg- gtt gcg att tta aac gga tta ata tat gtc ata ggt gga gtc gtt aaa ggt gac aca cat ata cca acg cac tag aga gtt tat cag aag atg gat gga tga atc atc aac gtc ttc caa taa aaa tgt cca ata tgt cga cga ttg ttc atg ctg gaa aga ttt ata tat cta gag gtt aca aca ata gta gtg tag tta atg taa tat cga atc t-- agt cct tag cta taa tcc gat ata tga tga atg gac caa att atc atc att aaa tat tcc tag aat taa tcc tgc tct a

>AY753185.1:158504-160748_Monkeypox_virus_strain_COP-58_complete_genome

tta ctg act aca --- tgt a-c tat tt- -ta cat tac tac att ggc tac ggt --a tat acc tac ttc gtc act tcc aca cgc tcc agt aaa cgg gtg tca tgt gac gag gga ga- -at ctt gat aag agg cat aat caa tgt tgt aat ccg gtg tcc acc tgg aga att tgc caa ggt cag atg tag --t tgg tag tga taa cac aaa atg tga aca ctg ccc acc tca tac ata tac cgc aat ccc caa tta ttc t-a ata gat gtc atc aat gta gaa aat gcc caa cag gat cat ttg ata agg taa agt gta ccg gaa cac aga ac- --a aat -gt tcg tgt cat cct ggt tgg ta- tac gct act gat tct tca cag act gaa gat tgt cga gat ttg tgt acc aaa aaa gga gat gtc cat gcg gat act ttg gtg gaa tag atg aa- --g gaa atc cta ttt gta aat cgt gtt g-- -tg ttg gtg aat att gcg act acc tac gta att ata gac ttg atc cat ttc ctc cat gca acc tat cta tct aaa tgt aat taa tta --- --- --- --- --- --- --- --- tga ttt --- --- --- -tg atg ata atg tta -cc ata c-a tt- --- -at atc gct act tgg tta gtg tgt att att cag tat gga aga cct att aat aa- --- --- -tt act tat ctt ttg acg atc ttg tt- -at aat tat --- --- --- --- aat ata aaa --a tac tta tga cat agt aac agt aac tca t-- --- --- --- aat tgc tg- acg cga taa att cgt aat aat ctg ttt tgt tca aat ttt tat aag ga- --- --a tct aca ggc ata aaa at- aaa aat ata atc tat aat ata c-- tct tac a-- acg cca tca tga at- --a gca gtg aat taa ttg ctg tta tt- atg gat tta gaa ata gtg gac gat ttt gtg ata tta ata tag tta tta atg atg aaa gga taa acg ctc ata gat tca tcc tat ctg gag cct ccg aat att ttt tcc att ctg ttt tcc aat aat ttt atc gat tct aat gaa tac gaa gtt aat cta agt cat tta gat tat caa agt gtt aac tat ttg atc gat tac att tat ggg ata cct ttg agc cta act aac gat aac gtg aaa tat att ctt tca acc gct ga- -tt ttt tta caa att gga tct gcc att act gag tgc gaa aaa tac ata ctt aaa aat ctt tgt ttt aga aac tgt atc gat ttc --- --- tac ata tac gct gat aaa tat aat aac aag aaa ata gaa tta gca tcg ttt aac aca ata tta cga aat att ttg aga ctc atc aac aat gaa aac ttt aaa tac tta aca gag gaa tca --- --- --- --- --- --- -at gat atg tta tat ata aaa aat gag gat ttc acc cca ctg att ctc att aaa tgg tta gag agt aca c-- caa cca tgt acc gtc gag tta ctt aga tgc ctc aga ata tca -tt tct ttc ccc aca agt tat aaa atc act tta tag tca tcg act ggt tag ttc aat cta cga atg tat aac att ctt aaa caa tat agc att ctt gga taa atc att tcc tag ata cca tat cat cga gtt gat atc tat cgg tat aag taa ttc aca tga taa gat ttc cat aaa ctg cta caa tca t-- aaa aaa att cat ggg aaa tga tat ctt cac gta gat --- ggt gta gtt tcg cag tga ccg tcc tgg ata ata tta tct ata tga tgg gtg gat atg atc agt ccc tgt ata gaa gtt caa agg tta tag cgt aca ata cat gta cta att ctt gga tat atg ata tac cag agc t-a aaa tat cct cgt tct aat tgc gga gga gtt gcc aat gac gaa tac att tat tgt ata ggc ggt ata cgc gat cag gag tca tcg ttg ata tct agt atc gat aga tgg aag cca tca aaa cca tat tgg cag aag tat gct aaa atg tgc gaa cca aaa tgt gat atg gg- gtt gcg att tta aac gga tta ata tat gtc ata ggt gga gtc gtt aaa ggt gac aca cat gta ccg acg cac tag aga gtt tat cag aag atg gat gga tga atc atc aac gtc ttc caa taa aaa tgt cca ata tgt cga cga ttg ttc atg ctg gaa aga ttt ata tat ctg gag gtt aca aca ata gta gtg tag tta atg taa tat cga atc tag agt cct tag cta taa tcc gat ata tga tga atg gac caa att atc atc att aaa tat tcc tag aat taa tcc tgc tct a

>AY603973.1:158249-160493_Monkeypox_virus_strain_MPXV-WRAIR7-61_complete_genome

tta ctg act aca --- tgt a-c tat tt- -ta cat tac tac att ggc tac ggt --a tat acc tac ttc gtc act tcc aca cgc tcc agt aaa cgg gtg tca tgt gac gag gga ga- -at ctt gat aag agg cat aat caa tgt tgt aat ccg gtg tcc acc tgg aga att tgc caa ggt cag atg tag --t tgg tag tga taa cac aaa atg tga aca ctg ccc acc tca tac ata tac cgc aat ccc caa tta ttc t-a ata gat gtc atc aat gta gaa aat gcc caa cag gat cat ttg ata agg taa agt gta ccg gaa cac aga ac- --a aat -gt tcg tgt cat cct ggt tgg ta- tac gct act gat tct tca cag act gaa gat tgt cga gat ttg tgt acc aaa aaa gga gat gtc cat gcg gat act ttg gtg gaa tag atg aa- --g gaa atc cta ttt gta aat cgt gtt g-- -tg ttg gtg aat att gcg act acc tac gta att ata gac ttg atc cat ttc ctc cat gca acc tat cta tct aaa tgt aat taa tta --- --- --- --- --- --- --- --- tga ttt --- --- --- -tg atg ata atg tta -cc ata c-a tt- --- -at atc gct act tgg tta gtg tgt att att cag tat gga aga cct att aat aa- --- --- -tt act tat ctt ttg acg atc ttg tt- -at aat tat --- --- --- --- aat ata aaa --a tac tta tga cat agt aac agt aac tca t-- --- --- --- aat tgc tg- acg cga taa att cgt aat aat ctg ttt tgt tca aat ttt tat aag ga- --- --a tct aca ggc ata aaa at- aaa aat ata atc tat aat ata c-- tct tac a-- acg cca tca tga at- --a gca gtg aat taa ttg ctg tta tt- atg gat tta gaa ata gtg gac gat ttt gtg ata tta ata tag tta tta atg atg aaa gga taa acg ctc ata gat tca tcc tat ctg gag cct ccg aat att ttt tcc att ctg ttt tcc aat aat ttt atc gat tct aat gaa tac gaa gtt aat cta agt cat tta gat tat caa agt gtt aac tat ttg atc gat tac att tat ggg ata cct ttg agc cta act aac gat aac gtg aaa tat att ctt tca acc gct ga- -tt ttt tta caa att gga tct gcc att act gag tgc gaa aaa tac ata ctt aaa aat ctt tgt ttt aga aac tgt atc gat ttc --- --- tac ata tac gct gat aaa tat aat aac aag aaa ata gaa tta gca tcg ttt aac aca ata tta cga aat att ttg aga ctc atc aac aat gaa aac ttt aaa tac tta aca gag gaa tca --- --- --- --- --- --- -at gat atg tta tat ata aaa aat gag gat ttc acc cca ctg att ctc att aaa tgg tta gag agt aca c-- caa cca tgt acc gtc gag tta ctt aga tgc ctc aga ata tca -tt tct ttc ccc aca agt tat aaa atc act tta tag tca tcg act ggt tag ttc aat cta cga atg tat aac att ctt aaa caa tat agc att ctt gga taa atc att tcc tag ata cca tat cat cga gtt gat atc tat cgg tat aag taa ttc aca tga taa gat ttc cat aaa ctg cta caa tca t-- aaa aaa att cat ggg aaa tga tat ctt cac gta gat --- ggt gta gtt tcg cag tga ccg tcc tgg ata ata tta tct ata tga tgg gtg gat atg atc agt ccc tgt ata gaa gtt caa agg tta tag cgt aca ata cat gta cta att ctt gga tat atg ata tac cag agc t-a aaa tat cct cgt tct aat tgc gga gga gtt gcc aat gac gaa tac att tat tgt ata ggc ggt ata cgc gat cag gag tca tcg ttg ata tct agt atc gat aga tgg aag cca tca aaa cca tat tgg cag aag tat gct aaa atg tgc gaa cca aaa tgt gat atg gg- gtt gcg att tta aac gga tta ata tat gtc ata ggt gga gtc gtt aaa ggt gac aca cat gta ccg acg cac tag aga gtt tat cag aag atg gat gga tga atc atc aac gtc ttc caa taa aaa tgt cca ata tgt cga cga ttg ttc atg ctg gaa aga ttt ata tat ctg gag gtt aca aca ata gta gtg tag tta atg taa tat cga atc tag agt cct tag cta taa tcc gat ata tga tga atg gac caa att atc atc att aaa tat tcc tag aat taa tcc tgc tct a

>AY741551.1:158320-160566_Monkeypox_virus_isolate_Sierra_Leone_complete_genome

tta ctg act aca --- tgt a-c tat tt- -ta cat tac tac att ggc tac ggt --a tat acc tac ttc gtc act tcc aca cgc tcc agt aaa cgg gtg tca tgt gac gag gga ga- -at ctt gat aag agg cat aat caa tgt tgt aat ccg gtg tcc acc tgg aga att tgc caa ggt cag atg tag --t tgg tag tga taa cac aaa atg tga aca ctg ccc acc tca tac ata tac cgc aat ccc caa tta ttc t-a ata gat gtc atc aat gta gaa aat gcc caa cag gat cat ttg ata agg taa agt gta ccg gaa cac aga ac- --a aat ggt tcg tgt cat cct ggt tgg ta- tac gct act gat tct tca cag act gaa gat tgt cga gat ttg tgt acc aaa aaa gga gat gtc cat gcg gat act ttg gtg gaa tag atg aa- --g gaa atc cta ttt gta aat cgt gtt g-- -tg ttg gtg aat att gcg act acc tac gta att ata gac ttg atc cat ttc ctc cat gca aac tat cta tct aaa tgt aat taa tta --- --- --- --- --- --- --- --- tga ttt --- --- --- -tg atg ata atg tta -cc ata c-a tt- --- -at atc gct act tgg tta gtg tgt att att cag tat gga aga cct att aat aa- --- --- -tt act tat ctt ttg acg atc ttg tt- -at aat tat --- --- --- --- aat ata aaa --a tac tta tga cat agt aac agt aac tca t-- --- --- --- aat tgc tg- acg cga taa att cgt aat aat ctg ttt tgt tca aat ttt tat aag ga- --- --a tct aca ggc ata aaa at- aaa aat ata atc tat aat ata c-- tct tac a-- acg cca tca tga at- --a gca gtg aat taa ttg ctg tta tt- atg gat tta gaa ata gtg gac gat ttt gtg ata tta ata tag tta tta atg atg aaa gga taa acg ctc ata gat tca tcc tat ctg gag cct ccg aat att ttt tcc att ctg ttt tcc aat aat ttt atc gat tct aat gaa tac gaa gtt aat cta agt cat tta gat tat caa agt gtt aac tat ttg atc gat tac att tat ggg ata cct ttg agc cta act aac gat aac gtg aaa tat att ctt tca acc gct ga- -tt ttt tta caa att gga tct gcc att act gag tgc gaa aaa tac ata ctt aaa aat ctt tgt ttt aga aac tgt atc gat ttc --- --- tac ata tac gct gat aaa tat aat aac aag aaa ata gaa tta gca tcg ttt aac aca ata tta cga aat att ttg aga ctc atc aac aat gaa aac ttt aaa tac tta aca gag gaa tca --- --- --- --- --- --- -at gat atg tta tat ata aaa aat gag gat ttc acc cca ctg att ctc att aaa tgg tta gag agt aca c-- caa cca tgt acc gtc gag tta ctt aga tgc ctc aga ata tca ttt tct ttc ccc aca agt tat aaa atc act tta tag tca tcg act ggt tag ttc aat cta cga atg tat aac att ctt aaa caa tat agc att ctt gga taa atc att tcc tag ata cca tat cat cga gtt gat atc tat cgg tat aag taa ttc aca tga taa gat ttc cat aaa ctg cta caa tca t-- aaa aaa att cat ggg aaa tga tat ctt cac gta gat --- ggt gta gtt tcg cag tga ccg tcc tgg ata ata tta tct ata tga tgg gtg gat atg atc agt ccc tgt ata gaa gtt caa agg tta tag cgt aca ata cat gta cta att ctt gga tat atg ata tac cag agc t-a aaa tat cct cgt tct aat tgc gga gga gtt gcc aat gac gaa tac att tat tgt ata ggc ggt ata cgc gat cag gag tca tcg ttg ata tct agt atc gat aga tgg aag cca tca aaa cca tat tgg cag aag tat gct aaa atg tgc gaa cca aaa tgt gat atg gg- gtt gcg att tta aac gga tta ata tat gtc ata ggt gga gtc gtt aaa ggt gac aca cat gta ccg acg cac tag aga gtt tat cag aag atg gat gga tga atc atc aac gtc ttc caa taa aaa tgt cca ata tgt cga cga ttg ttc atg ctg gaa aga ttt ata tat ctg gag gtt aca aca ata gta gtg tag tta atg taa tat cga atc tag agt cct tag cta taa tcc gat ata tga tga atg gac caa att atc atc att aaa tat tcc tag aat taa tcc tgc tct a

>KP849470.1:158947-161191_Monkeypox_virus_isolate_Cote_dIvoire_1971_complete_genome

tta ctg act aca --- tgt a-c tat tt- -ta cat tac tac att ggc tac ggt --a tat acc tac ttc gtc act tcc aca cgc tcc aga aaa cgg gtg tca tgt gac gag gga ga- -at ctt gat aag agg cat aat caa tgt tgt aat ccg gtg tcc acc tgg aga att tgc caa ggt cag atg tag --t tgg tag tga taa cac aaa atg tga aca ctg ccc acc tca tac ata tac cgc aat ccc caa tta ttc t-a ata gat gtc atc aat gta taa aat gcc caa cag gat cat ttg ata agg taa agt gta ccg gaa cac aga ac- --a aat -gt tcg tgt cat cct ggt tgg ta- tac gct act gat tct tca cag act gaa gat tgt cga gat ttg tgt acc aaa aaa gga gat gtc cat gcg gat act ttg gtg gaa tag atg aa- --g gaa atc cta ttt gta aat cgt gtt g-- -tg ttg gtg aat att gcg act acc tac gta att ata gac ttg atc cat ttc ctc cat gca aac tat cta tct aaa tgt aat taa tta --- --- --- --- --- --- --- --- tga ttt --- --- --- -tg atg ata atg tta -cc ata c-a tt- --- -at atc gct act tgg tta gtg tgt att att cag tat gga aga cct att aat aa- --- --- -tt act tat ctt ttg acg atc ttg tt- -at aat tat --- --- --- --- aat ata aaa --a tac tta tga cat agt aac agt aac tca t-- --- --- --- aat tgc tg- acg cga taa att cgt aat aat ctg ttt tgt tca aat ttt tat aag ga- --- --a tct aca ggc ata aaa at- aaa aat ata atc tat aat ata c-- tct tac a-- acg cca tca tga at- --a gca gtg aat taa ttg ctg tta tt- atg gat tta gaa ata gtg gac gat ttt gtg ata tta ata tag tta tta atg atg aaa gga taa acg ctc ata gat tca tcc tat ctg gag cct ccg aat att ttt tcc att ctg ttt tcc aat aat ttt atc gat tct aat gaa tac gaa gtt aat cta agt cat tta gat tat caa agt gtt aac tat ttg atc gat tac att tat ggg ata cct ttg agc cta act aac gat aac gtg aaa tat att ctt tca acc gct ga- -tt ttt tta caa att gga tct gcc att act gag tgc gaa aaa tac ata ctt aaa aat ctt tgt ttt aga aac tgt atc gat ttc --- --- tac ata tac gct gat aaa tat aat aac aag aaa ata gaa tta gca tcg ttt aac aca ata tta cga aat att ttg aga ctc atc aac aat gaa aac ttt aaa tac tta aca gag gaa tca --- --- --- --- --- --- -at gat atg tta tat ata aaa aat gag gat ttc acc cca ctg att ctc att aaa tgg tta gag agt aca c-- caa cca tgt acc gtc gag tta ctt aga tgc ctc aga ata tca -tt tct ttc ccc aca agt tat aaa atc act tta tag tca tcg act ggt tag ttc aat cta cga atg tat aac att ctt aaa caa tat agc att ctt gga taa atc att tcc tag ata cca tat cat cga gtt gat atc tat cgg tat aag taa ttc aca tga taa gat ttc cat aaa ctg cta caa tca t-- aaa aaa att cat ggg aaa tga tat ctt cac gta gat --- ggt gta gtt tcg cag tga ccg tcc tgg ata ata tta tct ata tga tgg gtg gat atg atc agt ccc tgt ata gaa gtt caa agg tta tag cgt aca ata cat gta cta att ctt gga tat atg ata tac cag agc t-a aaa tat cct cgt tct aat tgc gga gga gtt gcc aat gac gaa tac att tat tgt ata ggc ggt ata cgc gat cag gag tca tcg ttg ata tct agt atc gat aga tgg aag cca tca aaa cca tat tgg cag aag tat gct aaa atg tgc gaa cca aaa tgt gat atg gg- gtt gcg att tta aac gga tta ata tat gtc ata ggt gga gtc gtt aaa ggt gac aca cat gta ccg acg cac tag aga gtt tat cag aag atg gat gga tga atc atc aac gtc ttc caa taa aaa tgt cca ata tgt cga cga ttg ttc atg ctg gaa aga ttt ata tat ctg gag gtt aca aca ata gta gtg tag tta atg taa tat cga atc tag agt cct tag cta taa tcc gat ata tga tga atg gac caa att atc atc att aaa tat tcc tag aat taa tcc tgc tct a

>KJ642614.1:149563-151807_Monkeypox_virus_strain_UTC_complete_genome

tta ctg act aca --- tgt a-c tat tt- -ta cat tac tac att ggc tgc ggt --a tat acc tac ttc gtc act tcc aca cgc tcc agt aaa cgg gtg tca tgt gac gag gga ga- -at ctt gat aag agg cat aat caa tgt tgt aat ccg gtg tcc acc tgg aga att tgc caa ggt cag atg tag --t tgg tag tga taa cac aaa atg tga aca ctg ccc acc tca tac ata tac cgc aat ccc caa tta ttc t-a ata gat gtc atc aat gta gaa aat gcc caa cag gat cat ttg ata agg taa agt gta ccg gaa cac aga ac- --a aat -gt tcg tgt cat cct ggt tgg ta- tac gct act gat tct tca cag act gaa gat tgt cga gat ttg tgt acc aaa aaa gga gat gtc cat gcg gat act ttg gtg gaa tag atg aa- --g gaa atc cta ttt gta aat cgt gtt g-- -tg ttg gtg aat att gcg act acc tac gta att ata gac ttg atc cat ttc ctc cat gca acc tat cta tct aaa tgt aat taa tta --- --- --- --- --- --- --- --- tga ttt --- --- --- -tg atg ata atg tta -cc ata c-a tt- --- -at atc gct act tgg tta gtg tgt att att cag tat gga aga cct att aat aa- --- --- -tt act tat ctt ttg acg atc ttg tt- -at aat tat --- --- --- --- aat ata aaa --a tac tta tga cat agt aac agt aac tca t-- --- --- --- aat tgc tg- acg cga taa att cgt aat aat ctg ttt tgt tca aat ttt tat aag ga- --- --a tct aca ggc ata aaa at- aaa aat ata atc tat aat ata c-- tct tac a-- acg cca tca tga at- --a gca gtg aat taa ttg ctg tta tt- atg gat tta gaa ata gtg gac gat ttt gtg ata tta ata tag tta tta atg atg aaa gga taa acg ctc ata gat tca tcc tat ctg gag cct ccg aat att ttt tcc att ctg ttt tcc aat aat ttt atc gat tct aat gaa tac gaa gtt aat cta agt cat tta gat tat caa agt gtt aac tat ttg atc gat tac att tat ggg ata cct ttg agc cta act aac gat aac gtg aaa tat att ctt tca acc gct ga- -tt ttt tta caa att gga tct gcc att act gag tgc gaa aaa tac ata ctt aaa aat ctt tgt ttt aga aac tgt atc gat ttc --- --- tac ata tac gct gat aaa tat aat aac aag aaa ata gaa tta gca tcg ttt aac aca ata tta cga aat att ttg aga ctc atc aac aat gaa aac ttt aaa tac tta aca gag gaa tca --- --- --- --- --- --- -at gat atg tta tat ata aaa aat gag gat ttc acc cca ctg att ctc att aaa tgg tta gag agt aca c-- caa cca tgt acc gtc gag tta ctt aga tgc ctc aga ata tca -tt tct ttc ccc aca agt tat aaa atc act tta tag tca tcg act ggt tag ttc aat cta cga atg tat aac att ctt aaa caa tat agc att ctt gga taa atc att tcc tag ata cca tat cat cga gtt gat atc tat cgg tat aag taa ttc aca tga taa gat ttc cat aaa ctg cta caa tca t-- aaa aaa att cat ggg aaa tga tat ctt cac gta gat --- ggt gta gtt tcg cag tga ccg tcc tgg ata ata tta tct ata tga tgg gtg gat atg atc agt ccc tgt ata gaa gtt caa agg tta tag cgt aca ata cat gta cta att ctt gga tat atg ata tac cag agc t-a aaa tat cct cgt tct aat tgc gga gga gtt gcc aat gac gaa tac att tat tgt ata ggc ggt ata cgc gat cag gag tca tcg ttg ata tct agt atc gat aga tgg aag cca tca aaa cca tat tgg cag aag tat gct aaa atg tgc gaa cca aaa tgt gat atg gg- gtt gcg att tta aac gga tta ata tat gtc ata ggt gga gtc gtt aaa ggt gac aca cat gta ccg acg cac tag aga gtt tat cag aag atg gat gga tga atc atc aac gtc ttc caa taa aaa tgt cca ata tgt cga cga ttg ttc atg ctg gaa aga ttt ata tat ctg gag gtt aca aca ata gta gtg tag tta atg taa tat cga atc tag agt cct tag cta taa tcc gat ata tga tga atg gac caa att atc atc att aaa tat tcc tag aat taa tcc tgc tct a

>DQ011156.1:158783-161027_Monkeypox_virus_strain_Liberia_1970_184_complete_genome

tta ctg act aca --- tgt a-c tat tt- -ta cat tac tac att ggc tac ggt --a tat acc tac ttc gtc act tcc aca cgc tcc aga aaa cgg gtg tca tgt gac gag gga ga- -at ctt gat aag agg cat aat caa tgt tgt aat ccg gtg tcc acc tgg aga att tgc caa ggt cag atg tag --t tgg tag tga taa cac aaa atg tga aca ctg ccc acc tca tac ata tac cgc aat ccc caa tta ttc t-a ata gat gtc atc aat gta taa aat gcc caa cag gat cat ttg ata agg taa agt gta ccg gaa cac aga ac- --a aat -gt tcg tgt cat cct ggt tgg ta- tac gct act gat tct tca cag act gaa gat tgt cga gat ttg tgt acc aaa aaa gga gat gtc cat gcg gat act ttg gtg gaa tag atg aa- --g gaa atc cta ttt gta aat cgt gtt g-- -tg ttg gtg aat att gcg act acc tac gta att ata gac ttg atc cat ttc ctc cat gca aac tat cta tct aaa tgt aat taa tta --- --- --- --- --- --- --- --- tga ttt --- --- --- -tg atg ata atg tta -cc ata c-a tt- --- -at atc gct act tgg tta gtg tgt att att cag tat gga aga cct att aat aa- --- --- -tt act tat ctt ttg acg atc ttg tt- -at aat tat --- --- --- --- aat ata aaa --a tac tta tga cat agt aac agt aac tca t-- --- --- --- aat tgc tg- acg cga taa att cgt aat aat ctg ttt tgt tca aat ttt tat aag ga- --- --a tct aca ggc ata aaa at- aaa aat ata atc tat aat ata c-- tct tac a-- acg cca tca tga at- --a gca gtg aat taa ttg ctg tta tt- atg gat tta gaa ata gtg gac gat ttt gtg ata tta ata tag tta tta atg atg aaa gga taa acg ctc ata gat tca tcc tat ctg gag cct ccg aat att ttt tcc att ctg ttt tcc aat aat ttt atc gat tct aat gaa tac gaa gtt aat cta agt cat tta gat tat caa agt gtt aac tat ttg atc gat tac att tat ggg ata cct ttg agc cta act aac gat aac gtg aaa tat att ctt tca acc gct ga- -tt ttt tta caa att gga tct gcc att act gag tgc gaa aaa tac ata ctt aaa aat ctt tgt ttt aga aac tgt atc gat ttc --- --- tac ata tac gct gat aaa tat aat aac aag aaa ata gaa tta gca tcg ttt aac aca ata tta cga aat att ttg aga ctc atc aac aat gaa aac ttt aaa tac tta aca gag gaa tca --- --- --- --- --- --- -at gat atg tta tat ata aaa aat gag gat ttc acc cca ctg att ctc att aaa tgg tta gag agt aca c-- caa cca tgt acc gtc gag tta ctt aga tgc ctc aga ata tca -tt tct ttc ccc aca agt tat aaa atc act tta tag tca tcg act ggt tag ttc aat cta cga atg tat aac att ctt aaa caa tat agc att ctt gga taa atc att tcc tag ata cca tat cat cga gtt gat atc tat cgg tat aag taa ttc aca tga taa gat ttc cat aaa ctg cta caa tca t-- aaa aaa att cat ggg aaa tga tat ctt cac gta gat --- ggt gta gtt tcg cag tga ccg tcc tgg ata ata tta tct ata tga tgg gtg gat atg atc agt ccc tgt ata gaa gtt caa agg tta tag cgt aca ata cat gta cta att ctt gga tat atg ata tac cag agc t-a aaa tat cct cgt tct aat tgc gga gga gtt gcc aat gac gaa tac att tat tgt ata ggc ggt ata cgc gat cag gag tca tcg ttg ata tct agt atc gat aga tgg aag cca tca aaa cca tat tgg cag aag tat gct aaa atg tgc gaa cca aaa tgt gat atg gg- gtt gcg att tta aac gga tta ata tat gtc ata ggt gga gtc gtt aaa ggt gac aca cat gta ccg acg cac tag aga gtt tat cag aag atg gat gga tga atc atc aac gtc ttc caa taa aaa tgt cca ata tgt cga cga ttg ttc atg ctg gaa aga ttt ata tat ctg gag gtt aca aca ata gta gtg tag tta atg taa tat cga atc tag agt cct tag cta taa tcc gat ata tga tga atg gac caa att atc atc att aaa tat tcc tag aat taa tcc tgc tct a

>KJ642616.1:157970-160214_Monkeypox_virus_strain_PCH_complete_genome

tta ctg act aca --- tgt a-c tat tt- -ta cat tac tac att ggc tac ggt --a tat acc tac ttc gtc act tca aca cgc tcc agt aaa cgg gtg tca tgt gac gag gga ga- -at ctt gat aag agg cat aat caa tgt tgt aat ccg gtg tcc acc tgg aga att tgc caa ggt cag atg tag --t tgg tag tga taa cac aaa atg tga aca ctg ccc acc tca tac ata tac cgc aat ccc caa tta ttc t-a ata gat gtc atc aat gta gaa aat gcc caa cag gat cat ttg ata agg taa agt gta ccg gaa cac aga ac- --a aat -gt tcg tgt cat cct ggt tgg ta- tac gct act gat tct tca cag act gaa gat tgt cga gat ttg tgt acc aaa aaa gga gat gtc cat gcg gat act ttg gtg gaa tag atg aa- --g gaa atc cta ttt gta aat cgt gtt g-- -tg ttg gtg aat att gcg act acc tac gta att ata gac ttg atc cat ttc ctc cat gca acc tat cta tct aaa tgt aat taa tta --- --- --- --- --- --- --- --- tga ttt --- --- --- -tg atg ata atg tta -cc ata c-a tt- --- -at atc gct act tgg tta gtg tgt att att cag tat gga aga cct att aat aa- --- --- -tt act tat ctt ttg acg atc ttg tt- -at aat tat --- --- --- --- aat ata aaa --a tac tta tga cat agt aac agt aac tca t-- --- --- --- aat tgc tg- acg cga taa att cgt aat aat ctg ttt tgt tca aat ttt tat aag ga- --- --a tct aca ggc ata aaa at- aaa aat ata atc tat aat ata c-- tct tac a-- acg cca tca tga at- --a gca gtg aat taa ttg ctg tta tt- atg gat tta gaa ata gtg gac gat ttt gtg ata tta ata tag tta tta atg atg aaa gga taa acg ctc ata gat tca tcc tat ctg gag cct ccg aat att ttt tcc att ctg ttt tcc aat aat ttt atc gat tct aat gaa tac gaa gtt aat cta agt cat tta gat tat caa agt gtt aac tat ttg atc gat tac att tat ggg ata cct ttg agc cta act aac gat aac gtg aaa tat att ctt tca acc gct ga- -tt ttt tta caa att gga tct gcc att act gag tgc gaa aaa tac ata ctt aaa aat ctt tgt ttt aga aac tgt atc gat ttc --- --- tac ata tac gct gat aaa tat aat aac aag aaa ata gaa tta gca tcg ttt aac aca cta tta cga aat att ttg aga ctc atc aac aat gaa aac ttt aaa tac tta aca gag gaa tca --- --- --- --- --- --- -at gat atg tta tat ata aaa aat gag gat ttc acc cca ctg att ctc att aaa tgg tta gag agt aca c-- caa cca tgt acc gtc gag tta ctt aga tgc ctc aga ata tca -tt tct ttc ccc aca agt tat aaa atc act tta tag tca tcg act ggt tag ttc aat cta cga atg tat aac att ctt aaa caa tat agc att ctt gga taa atc att tcc tag ata cca tat cat cga gtt gat atc tat cgg tat aag taa ttc aca tga taa gat ttc cat aaa ctg cta caa tca t-- aaa aaa att cat ggg aaa tga tat ctt cac gta gat --- ggt gta gtt tcg cag tga ccg tcc tgg ata ata tta tct ata tga tgg gtg gat atg atc agt ccc tgt ata gaa gtt caa agg tta tag cgt aca ata cat gta cta att ctt gga tat atg ata tac cag agc t-a aaa tat cct cgt tct aat tgc gga gga gtt gcc aat gac gaa tac att tat tgt ata ggc ggt ata cgc gat cag gag tca tcg ttg ata tct agt atc gat aga tgg aag cca tca aaa cca tat tgg cag aag tat gct aaa atg tgc gaa cca aaa tgt gat atg gg- gtt gcg att tta aac gga tta ata tat gtc ata ggt gga gtt gtt aaa ggt gac aca cat gta ccg acg cac tag aga gtt tat cag aag atg gat gga tga atc atc aac gtc ttc caa taa aaa tgt cca ata tgt cga cga ttg ttc atg ctg gaa aga ttt ata tat ctg gag gtt aca aca ata gta gtg tag tta atg taa tat cga atc tag agt cct tag cta taa tcc gat ata tga tga atg gac caa att atc atc att aaa tat tcc tag aat taa tcc tgc tct a

>MT903346.1:158312-160561_Monkeypox_virus_isolate_MPXV-USA2003_099_Gambian_Rat

tta ctg act aca --- tgt a-c tat tt- -ta cat tac tac att ggc tac ggt --a tat acc tac ttc gtc act tcc aca cgc tcc agt aaa cgg gtg tca tgt gac gag gga ga- -at ctt gat aag agg cat aat caa tgt tgt aat ccg gtg tcc acc tgg aga att tgc caa ggt cag atg tag --t tgg tag tga taa cac aaa atg tga aca ctg ccc acc tca tac ata tac cgc aat ccc caa tta ttc t-a ata gat gtc atc aat gta gaa aat gcc caa cag gat cat ttg ata agg taa agt gta ccg gaa cac aga ac- --a aat -gt tcg tgt cat cct ggt tgg ta- tac gct ac- gat tct tca cag act gaa gat tgt cga gat ttg tgt acc aaa aaa gga gat gtc cat gcg gat act ttg gtg gaa tag atg aa- --g gaa atc cta ttt gta aat cgt gtt g-- -tg ttg gtg aat att gcg act acc tac gta att ata gac ttg atc cat ttc ctc cat gca aac tat cta tct aaa tgt aat taa tta --- --- --- --- --- --- --- --- tga ttt --- --- --- -tg atg ata atg tta -cc ata c-a tt- --- -at atc gct act tgg tta gtg tgt att att cag tat gga aga cct att aat aa- --- --- -tt act tat ctt ttg acg atc ttg tt- -at aat tat --- --- --- --- aat ata aaa --a tac tta tga cat agt aac agt aac tca t-- --- --- --- aat tgc tg- acg cga taa att cgt aat aat ctg ttt tgt tca aat ttt tat aag ga- --- --a tct aca ggc ata aaa at- aaa aat ata atc tat aat ata c-- tct tac a-- acg cca tca tga at- --a gca gtg aat taa ttg ctg tta tt- atg gat tta gaa ata gtg gac gat ttt gtg ata tta ata tag tta tta atg atg aaa gga taa acg ctc ata gat tca tcc tat ctg gag cct ccg aat att ttt tcc att ctg ttt tcc aat aat ttt atc gat tct aat gaa tac gaa gtt aat cta agt cat tta gat tat caa agt gtt aac tat ttg atc gat tac att tat ggg ata cct ttg agc cta act aac gat aac gtg aaa tat att ctt tca acc gct ga- -tt ttt tta caa att gga tct gcc att act gag tgc gaa aaa tac ata ctt aaa aat ctt tgt ttt aga aac tgt atc gat ttc tac ata tac ata tac gct gat aaa tat aat aac aag aaa ata gaa tta gca tcg ttt aac aca ata tta cga aat att ttg aga ctc atc aac aat gaa aac ttt aaa tac tta aca gag gaa tca --- --- --- --- --- --- -at gat atg tta tat ata aaa aat gag gat ttc acc cca ctg att ctc att aaa tgg tta gag agt aca c-- caa cca tgt acc gtc gag tta ctt aga tgc ctc aga ata tca -tt tct ttc ccc aca agt tat aaa atc act tta tag tca tcg act ggt tag ttc aat cta cga atg tat aac att ctt aaa caa tat agc att ctt gga taa atc att tcc tag ata cca tat cat cga gtt gat atc tat cgg tat aag taa ttc aca tga taa gat ttc cat aaa ctg cta caa tca t-- aaa aaa att cat ggg aaa tga tat ctt cac gta gat --- ggt gta gtt tcg cag tga ccg tcc tgg ata ata tta tct ata tga tgg gtg gat atg atc agt ccc tgt ata gaa gtt caa agg tta tag cgt aca ata cat gta cta att ctt gga tat atg ata tac cag agc t-a aaa tat cct cgt tct aat tgc gga gga gtt gcc aat gac gaa tac att tat tgt ata ggc ggt ata cgc gat cag gag tca tcg ttg ata tct agt atc gat aga tgg aag cca tca aaa cca tat tgg cag aag tat gct aaa atg tgc gaa cca aaa tgt gat atg gg- gtt gcg att tta aac gga tta ata tat gtc ata ggt gga gtc gtt aaa ggt gac aca cat gta ccg acg cac tag aga gtt tat cag aag atg gat gga tga atc atc aac gtc ttc caa taa aaa tgt cca ata tgt cga cga ttg ttc atg ctg gaa aga ttt ata tat ctg gag gtt aca aca ata gta gtg tag tta atg taa tat cga atc tag agt cct tag cta taa tcc gat ata tga tga atg gac caa att atc atc att aaa tat tcc tag aat taa tcc tgc tct a

>DQ011157.1:158313-160562_Monkeypox_virus_strain_USA_2003_039_complete_genome

tta ctg act aca --- tgt a-c tat tt- -ta cat tac tac att ggc tac ggt --a tat acc tac ttc gtc act tcc aca cgc tcc agt aaa cgg gtg tca tgt gac gag gga ga- -at ctt gat aag agg cat aat caa tgt tgt aat ccg gtg tcc acc tgg aga att tgc caa ggt cag atg tag --t tgg tag tga taa cac aaa atg tga aca ctg ccc acc tca tac ata tac cgc aat ccc caa tta ttc t-a ata gat gtc atc aat gta gaa aat gcc caa cag gat cat ttg ata agg taa agt gta ccg gaa cac aga ac- --a aat -gt tcg tgt cat cct ggt tgg ta- tac gct ac- gat tct tca cag act gaa gat tgt cga gat ttg tgt acc aaa aaa gga gat gtc cat gcg gat act ttg gtg gaa tag atg aa- --g gaa atc cta ttt gta aat cgt gtt g-- -tg ttg gtg aat att gcg act acc tac gta att ata gac ttg atc cat ttc ctc cat gca aac tat cta tct aaa tgt aat taa tta --- --- --- --- --- --- --- --- tga ttt --- --- --- -tg atg ata atg tta -cc ata c-a tt- --- -at atc gct act tgg tta gtg tgt att att cag tat gga aga cct att aat aa- --- --- -tt act tat ctt ttg acg atc ttg tt- -at aat tat --- --- --- --- aat ata aaa --a tac tta tga cat agt aac agt aac tca t-- --- --- --- aat tgc tg- acg cga taa att cgt aat aat ctg ttt tgt tca aat ttt tat aag ga- --- --a tct aca ggc ata aaa at- aaa aat ata atc tat aat ata c-- tct tac a-- acg cca tca tga at- --a gca gtg aat taa ttg ctg tta tt- atg gat tta gaa ata gtg gac gat ttt gtg ata tta ata tag tta tta atg atg aaa gga taa acg ctc ata gat tca tcc tat ctg gag cct ccg aat att ttt tcc att ctg ttt tcc aat aat ttt atc gat tct aat gaa tac gaa gtt aat cta agt cat tta gat tat caa agt gtt aac tat ttg atc gat tac att tat ggg ata cct ttg agc cta act aac gat aac gtg aaa tat att ctt tca acc gct ga- -tt ttt tta caa att gga tct gcc att act gag tgc gaa aaa tac ata ctt aaa aat ctt tgt ttt aga aac tgt atc gat ttc tac ata tac ata tac gct gat aaa tat aat aac aag aaa ata gaa tta gca tcg ttt aac aca ata tta cga aat att ttg aga ctc atc aac aat gaa aac ttt aaa tac tta aca gag gaa tca --- --- --- --- --- --- -at gat atg tta tat ata aaa aat gag gat ttc acc cca ctg att ctc att aaa tgg tta gag agt aca c-- caa cca tgt acc gtc gag tta ctt aga tgc ctc aga ata tca -tt tct ttc ccc aca agt tat aaa atc act tta tag tca tcg act ggt tag ttc aat cta cga atg tat aac att ctt aaa caa tat agc att ctt gga taa atc att tcc tag ata cca tat cat cga gtt gat atc tat cgg tat aag taa ttc aca tga taa gat ttc cat aaa ctg cta caa tca t-- aaa aaa att cat ggg aaa tga tat ctt cac gta gat --- ggt gta gtt tcg cag tga ccg tcc tgg ata ata tta tct ata tga tgg gtg gat atg atc agt ccc tgt ata gaa gtt caa agg tta tag cgt aca ata cat gta cta att ctt gga tat atg ata tac cag agc t-a aaa tat cct cgt tct aat tgc gga gga gtt gcc aat gac gaa tac att tat tgt ata ggc ggt ata cgc gat cag gag tca tcg ttg ata tct agt atc gat aga tgg aag cca tca aaa cca tat tgg cag aag tat gct aaa atg tgc gaa cca aaa tgt gat atg gg- gtt gcg att tta aac gga tta ata tat gtc ata ggt gga gtc gtt aaa ggt gac aca cat gta ccg acg cac tag aga gtt tat cag aag atg gat gga tga atc atc aac gtc ttc caa taa aaa tgt cca ata tgt cga cga ttg ttc atg ctg gaa aga ttt ata tat ctg gag gtt aca aca ata gta gtg tag tta atg taa tat cga atc tag agt cct tag cta taa tcc gat ata tga tga atg gac caa att atc atc att aaa tat tcc tag aat taa tcc tgc tct a

>DQ011153.1:158313-160562_Monkeypox_virus_strain_USA_2003_044_complete_genome

tta ctg act aca --- tgt a-c tat tt- -ta cat tac tac att ggc tac ggt --a tat acc tac ttc gtc act tcc aca cgc tcc agt aaa cgg gtg tca tgt gac gag gga ga- -at ctt gat aag agg cat aat caa tgt tgt aat ccg gtg tcc acc tgg aga att tgc caa ggt cag atg tag --t tgg tag tga taa cac aaa atg tga aca ctg ccc acc tca tac ata tac cgc aat ccc caa tta ttc t-a ata gat gtc atc aat gta gaa aat gcc caa cag gat cat ttg ata agg taa agt gta ccg gaa cac aga ac- --a aat -gt tcg tgt cat cct ggt tgg ta- tac gct ac- gat tct tca cag act gaa gat tgt cga gat ttg tgt acc aaa aaa gga gat gtc cat gcg gat act ttg gtg gaa tag atg aa- --g gaa atc cta ttt gta aat cgt gtt g-- -tg ttg gtg aat att gcg act acc tac gta att ata gac ttg atc cat ttc ctc cat gca aac tat cta tct aaa tgt aat taa tta --- --- --- --- --- --- --- --- tga ttt --- --- --- -tg atg ata atg tta -cc ata c-a tt- --- -at atc gct act tgg tta gtg tgt att att cag tat gga aga cct att aat aa- --- --- -tt act tat ctt ttg acg atc ttg tt- -at aat tat --- --- --- --- aat ata aaa --a tac tta tga cat agt aac agt aac tca t-- --- --- --- aat tgc tg- acg cga taa att cgt aat aat ctg ttt tgt tca aat ttt tat aag ga- --- --a tct aca ggc ata aaa at- aaa aat ata atc tat aat ata c-- tct tac a-- acg cca tca tga at- --a gca gtg aat taa ttg ctg tta tt- atg gat tta gaa ata gtg gac gat ttt gtg ata tta ata tag tta tta atg atg aaa gga taa acg ctc ata gat tca tcc tat ctg gag cct ccg aat att ttt tcc att ctg ttt tcc aat aat ttt atc gat tct aat gaa tac gaa gtt aat cta agt cat tta gat tat caa agt gtt aac tat ttg atc gat tac att tat ggg ata cct ttg agc cta act aac gat aac gtg aaa tat att ctt tca acc gct ga- -tt ttt tta caa att gga tct gcc att act gag tgc gaa aaa tac ata ctt aaa aat ctt tgt ttt aga aac tgt atc gat ttc tac ata tac ata tac gct gat aaa tat aat aac aag aaa ata gaa tta gca tcg ttt aac aca ata tta cga aat att ttg aga ctc atc aac aat gaa aac ttt aaa tac tta aca gag gaa tca --- --- --- --- --- --- -at gat atg tta tat ata aaa aat gag gat ttc acc cca ctg att ctc att aaa tgg tta gag agt aca c-- caa cca tgt acc gtc gag tta ctt aga tgc ctc aga ata tca -tt tct ttc ccc aca agt tat aaa atc act tta tag tca tcg act ggt tag ttc aat cta cga atg tat aac att ctt aaa caa tat agc att ctt gga taa atc att tcc tag ata cca tat cat cga gtt gat atc tat cgg tat aag taa ttc aca tga taa gat ttc cat aaa ctg cta caa tca t-- aaa aaa att cat ggg aaa tga tat ctt cac gta gat --- ggt gta gtt tcg cag tga ccg tcc tgg ata ata tta tct ata tga tgg gtg gat atg atc agt ccc tgt ata gaa gtt caa agg tta tag cgt aca ata cat gta cta att ctt gga tat atg ata tac cag agc t-a aaa tat cct cgt tct aat tgc gga gga gtt gcc aat gac gaa tac att tat tgt ata ggc ggt ata cgc gat cag gag tca tcg ttg ata tct agt atc gat aga tgg aag cca tca aaa cca tat tgg cag aag tat gct aaa atg tgc gaa cca aaa tgt gat atg gg- gtt gcg att tta aac gga tta ata tat gtc ata ggt gga gtc gtt aaa ggt gac aca cat gta ccg acg cac tag aga gtt tat cag aag atg gat gga tga atc atc aac gtc ttc caa taa aaa tgt cca ata tgt cga cga ttg ttc atg ctg gaa aga ttt ata tat ctg gag gtt aca aca ata gta gtg tag tta atg taa tat cga atc tag agt cct tag cta taa tcc gat ata tga tga atg gac caa att atc atc att aaa tat tcc tag aat taa tcc tgc tct a

>KJ642615.1:156680-158929_Monkeypox_virus_strain_W-Nigeria_complete_genome

tta ctg act aca tgt tgt a-c tat tt- -ta cat tac tac att ggc tac ggt --a tat acc tac ttc gtc act tcc aca cgc tcc agt aaa cgg gtg tca tgt gac gag gga ga- -at ctt gat aag agg cat aat caa tgt tgt aat ccg gtg tcc acc tgg aga att tgc caa ggt cat atg tag --t tgg tag tga taa cac aaa atg tga aca ctg tcc acc tca tac ata tac cgc aat ccc caa tta ttc t-a ata gat gtc atc aat gta gaa aat gcc taa cag gat cat ttg ata agg taa agt gta ccg gaa cac aga ac- --a aat -gt tcg tgt cat cct ggt tgg ta- tac gct act gat tct tca cag act gaa gat tgt cga gat ttg tgt acc aaa aaa gga gat gtc cat gcg gat act ttg gtg gaa tag atg aa- --g gaa atc cta ttt gta aat cgt gtt g-- -tg ttg gtg aat att gcg act acc tac gta att ata gac ttg atc cat ttc ctc cat gca aac tat cta tct aaa tgt aat taa tta --- --- --- --- --- --- --- --- tga ttt --- --- --- -tg atg ata atg tta -cc ata c-a tt- --- -at atc gct act tgg tta gtg tgt att att cag tat gga aga cct att aat aa- --- --- -tt act tat ctt ttg acg atc ttg tt- -at aat tat --- --- --- --- aat ata aaa --a tac tta tga cat agt aac agt aac tca t-- --- --- --- aat tgc tg- acg cga taa att cgt aat aat ctg ttt tgt tca aat ttt tat aag ga- --- --a tct aca ggc ata aaa at- aaa aat ata atc tat aat ata c-- tct tac a-- acg cca tca tga at- --a gca gtg aat taa ttg ctg tta tt- atg gat tta gaa ata gtg gac gat ttt gtg ata tta ata tag tta tta atg atg aaa gga taa acg ctc ata gat tca tcc tat ctg gag cct ccg aat att ttt tcc att ctg ttt tcc aat aat ttt atc gat tct aat gaa tac gaa gtt aat cta agt cat tta gat tat caa agt gtt aac tat ttg atc gat tac att tat ggg ata cct ttg agc cta act aac gat aac gtg aaa tat att ctt tca acc gct gat ttt ttt tta caa att gga tct gcc att act gag tgc gaa aaa tac ata ctt aaa aat ctt tgt ttt aga aac tgt atc gat ttc --- --- tac ata tac gct gat aaa tat aat aac aag aaa ata gaa tta gca tcg ttt aac aca ata tta cga aat att ttg aga ctc atc aac aat gaa aac ttt aaa tac tta aca gag gaa tca --- --- --- --- --- --- -at gat atg tta tat ata aaa aat gag gat ttc acc cca ctg att ctc att aaa tgg tta gag agt aca c-- caa cca tgt acc gtc gag tta ctt aga tgc ctc aga ata tca -tt tct ttc ccc aca agt tat aaa atc act tta tag tca tcg act ggt tag ttc aat cta cga atg tat aac att ctt aaa caa tat agc att ctg gga taa atc att tcc tag ata cca tat cat cga gtt gat atc tat cgg tat aag taa ttc aca tga taa gat ttc cat aaa ctg cta caa tca t-- aaa aaa att cat ggg aaa tga tat ctt cac gta gat --- ggt gta gtt tcg cag tga ccg tcc tgg ata ata tta tct ata tga tgg gtg gat atg atc agt ccc tgt ata gaa gtt caa agg tta tag cgt aca ata cat gta cta att ctt gga tat atg ata tac cag agc t-a aaa tat cct cgt tct aat tgc gga gga gtt gcc aat gac gaa tac att tat tgt ata ggc ggt ata cgc gat cag gag tca tcg ttg ata tct agt atc gat aga tgg aag cca tca aaa cca tat tgg cag aag tat gct aaa atg tgc gaa cca aaa tgt gat atg gg- gtt gcg att tta aac gga tta ata tat gtc ata ggt gga gtc gtt aaa ggt gac aca cat gta ccg acg cac tag aga gtt tat cag aag atg gat gga tga atc atc aac gtc ttc caa taa aaa tgt cca ata tgt cga cga ttg ttc atg ctg gaa aga ttt ata tat ctg gag gtt aca aca ata gta gtg tag tta atg taa tat cga atc tag agt cct tag cta taa tcc gat ata tga tga atg gac caa att atc atc att aaa tat tcc tag aat taa tcc tgc tct a

>HQ407377.1:174485-176745_Cowpox_virus_strain_Austria_1999_complete_genome

ttg ctg act gta --- tgt a-c tat ttt tta cat tac tac att ggc tac ggc aga tat acc tac ttc gtc act gcc aca tgc tcc ggt aaa cgg --g tca tgt gac gag gga gaa tat ctt gat aag agg cat aat caa tgt tgt aat -ca gtg tcc acc tgg aga att tgc caa ggt cag atg ta- --g tgg tag tga taa cac aaa atg tga acg ctg ccc acc tca tac ata tac cgc aat ccc caa tta ttc t-a atg gat gtc atc aat gta gaa aat gcc caa cag gat cat ttg ata agg taa agt gta ccg gaa cac aga aca gta aat -gt tcg tgt ctt cct ggt tgg tat tgc gct act gat tct tca cag act gaa gat tgt cga gat -tg tgt acc aaa aag g-a gat gtc cat gcg gat act ttg gtg gaa tag atg aac aag gaa atc cta ttt gta aat cgt gtt g-- -tg ttg gtg aat att gcg act acc tac gta att ata gac ttg atc cat ttc ctc cat gca aa- --- cta tct aaa tgt aat taa tta --- --- --- --- --- --- --- --- tga ttt --- --- --- -tg atg ata atg tta -cc ata c-a tt- --- -at atc gct --- -gg tta --g tgt att att cag tat g-a aga cct att aat aa- --- --- -tt act tat ctt ttg acg atc ttg tt- -at aat tat --- --- --- --- aat ata aaa --a tac tta tgg --- --- cat agt aac tca t-- --- --- --- aat tgc tg- aca cga taa att cgt aat aat ctg ttt tgt tca aat ttt tat aag gaa t-- --a tct aca ggc ata aaa at- aaa aat ata att tat aat ata c-- tct tac a-- gcg cca tca tga ata aca gca gtg aat taa ttg ctg tta tta atg gat tta gaa ata gtg gac gat ttt gtg ata tta ata tag tta tta atg atg aaa gga taa acg ctc ata gac tca tcc tat ctg gag cct ccg aat a-t ttt tcc att ctg ttt tcc aat aat ttt atc gat tct aat gaa tac gaa gtt aat cta agt cat tta gat tat caa agt gtt aac gat ttg atc gat tac att tat ggg ata cct ttg agc cta act aac gat aac gtg aaa tat att ctt tca acc gct ga- --t ttt tta caa att gga tct gcc att act gag tgc gaa aaa tac ata ctt aaa aat ctt tgt tct aga aac tgt atc gat ttc --- --- tac ata tac gct gat aaa tat aat aac aag aaa ata gaa tca gcg tcg ttt aac aca ata tta cga aat att ttg aga ctc att aac gat gaa aac ttt aaa tac tta aca gag gaa tca atg ata aaa att tta agc gat gat atg tta aat ata aaa aat gag gat ttt gca cca cta att ctc att aaa tgg tta gag agt act caa caa cca tgc acc gtc gag tta ctt aga tgc ctc aga ata tca -tt gct ttc ccc aca agt tat aaa atc act tta tag tca tca act ggt tag ttc aat cta cga atg tat aac att ctt aaa caa tat agc att ctt gga tga atc att tcc tag ata cca tag cat cga gtt gat atc tat cgg tat aag taa ttc gca tga taa gat ttc cat aaa ctg cta caa tca t-a aaa aaa ata cat ggg aaa tga tat ctt cac gta gat ata ggt gta gtt tcg cag tgg ccg tcc tgg ata ata tta tct ata tga tgg gtg gat acg atc agt ccc cat ata gaa gtt caa agg tta tag cgt aca ata cat gta cta att ctt gga tat atg ata tac cag agc t-a aaa tat cct cgt tct aat tgc gga gga gtt gcc gat gac gaa tac att tat tgt ata ggc ggc ata cgc gat cag gat tca tcg ttg aca tct agt att gat aga tgg aag cca tca aaa cca tat tgg cag aag tat gct aaa atg cgc gaa cca aaa tgt gat atg ggg gtt gcg atg tta aac gga tta ata tat gtc ata ggt gga atc gtt aaa ggt g-- aca cgt gta ccg acg cac tag aga gtt tat cag aag atg gat gga tga agc atc aac gtc ttc caa taa aaa tat cca ata tgt cga cga ttg ttc atg atg gca aga ttt ata tat ctg gag gtt aca aca ata gta gtg tag tta atg taa tat cga atc t-- agt cct tag cta taa tcc gat ata tga tga atg gac caa att atc atc att aaa cat tcc tag aat taa tcc cgc tct a

>HQ420893.1:175367-177632_Cowpox_virus_strain_Finland_2000_MAN_complete_genome

ttg ctg act gta --- tgt a-c tat ttt tta cat tac tac att ggc tac ggc aga tat acc tac ttc gtc act gcc aca cgc tcc ggt aaa ctg --g gca tgt gac gag gga gaa tat ctt gat aag agg cat aat caa tgt tgt aat -ca gtg tcc acc tgg aga att tgc caa ggt cag atg ta- --g tgg tag tga taa cac aaa atg tga acg ctg ccc acc tca tac ata tac cgc aat ccc caa tta ctc t-a atg gat gtc atc aat gta gaa aat gcc caa cag gat cat ttg ata agg taa agt gta ccg gaa cac aga aca gta aat -gt tcg tgt ctt cct ggt tgg tat tgt gct act gat tct tca cag act gaa gat tgt cga gat -tg tat acc aaa aag g-a gat gtc cat gcg gat act ttg gtg gaa tag atg aac aag gaa atc cta ttt gta aat cgt gtt g-- -tg ttg gtg aat att gcg act acc tac gta att ata gac ttg atc cat ttc ctc cat gca aa- --- cta tct aaa tgt aat taa tta --- --- --- --- --- --- --- --- tga ttt --- --- --- -tg atg ata atg tta -cc ata c-a tt- --- -at atc gct act tgg tta --g tgt att att cag tat g-a aga cct att aat aat --- --- -tt act tat ctt ttg acg atc ttg tt- -at aat tat --- --- --- --- aat ata aaa --a tac tta tgg --- --- cat agt aac tca t-- --- --- --- aat tgc tg- acg cga taa att cgt aat aat ctg ttt tgt tca aat ttt tat aag ga- --- --a tct aca ggc ata aaa at- aaa aat ata att tat aat ata c-- tct tac agt gcg cca tca tga ata aca gca gtg aat tga ttg ctg tta tta atg gat tta gaa ata gtg gac gat ttt gtg ata tta gta tag tta tta atg atg aaa gga taa acg ctc aca gac tca tcc tat ctg gag cct ccg aat a-t ttt tcc att ctg ttt tcc aat aat ttt atc gat tct aat gaa tac gaa gtt aat cta agt cat tta gat tat caa agt gtt aac gat ttg atc gat tac att tat ggg ata cct ttg agc cta act aac gat aac gtg aaa tat att ctt tca acc gct ga- --t ttt tta caa att gga tct gcc att act gag tgc gaa aaa tac ata ctt aaa aat ctt tgt tct aga aac tgt atc gat ttc --- --- tac ata tac gct gat aaa tat aat aac aag aaa ata gaa tca gcg tcg ttt aac aca ata tta cga aat att ttg aga ctc atc aac gat gaa aac ttt aaa tac tta aca gag gaa tca atg ata aaa att tta agc gat gat atg tta aat ata aaa aat gag gat ttt gca cca cta att ctc att aaa tgg tta gag agt act caa caa tca tgc acc gtc gag tta ctt aga tgc ctc aga ata tca -tt gct ttc ccc aca agt tat aaa atc act tta tag tca tca act ggt tag ttc aat cta cga atg tat aac att ctt aaa caa tat agc att ctt gga tga atc att tcc tag ata cca tag cat cga gtt gat atc tat cgg tat aag taa ttc gca tga taa gat ttc cat aaa ctg cta caa tca t-a aaa aaa ata cat ggg aaa tga tat ctt cac gta gat ata ggt gta gtt tcg cag tgg ctg tcc tgg ata ata tta tct ata tga tgg gtg gat atg atc agt ccc cgt ata gaa gtt caa agg tta tag cgt aca ata cat gta caa att ctt gga tat atg ata tac cag agc t-a aaa tat cct cgt tct aat tgt ggg gga ctg gct gat gac gaa tac att tat tgt ata ggc ggc ata cgc gat cag gat tca tcg ttg aca tct agt att gat aga tgg aag cca tca aaa cca tat tgg cag aag tat gct aaa atg cgc gaa cca aaa tgt gat atg ggg gtt gcg atg tta aac gga tta ata tat gtc ata ggt gga ttc gtt aaa ggt g-- aca cgt gta ccg acg cac tag aga gtt tat cag aag atg gat gga tga agc atc aac gtc ttc caa taa aaa tgt cca ata tgt cga cga ttg ttc atg atg gca aga ttt ata tat ctg gag gtt aca aca ata gta gtg tag tta atg taa tat cga atc t-- agt cct tag cta taa tcc gat ata tga tga atg gac caa att atc atc att aaa cat tcc tag aat taa tcc cgc tct a

>X94355.2:176241-178492_Cowpox_virus_strain_GRI-90_complete_genome

--- --- --- --a --- tgt a-c tat ttt gta cat cac tac att ggt tac ggc aga tat acc tac tcc gtt acc gcc aca cgc tcc ggt aaa cgg --g tca tgt gac gag gga gaa tat ctt gat aag agg cat aat caa tgt tgt aat -ca gtg tcc acc tgg aga att tgc caa ggt cag atg ta- --g tgg tag tga taa cac aaa atg tga acg ctg ccc acc tca tac ata tac cgc aat ccc caa tta ttc t-a acg gat gcc atc aat gta gaa aat gcc caa cag gat cat ttg ata agg taa agt gta ccg gaa cac aga aca gta aat -gt tcg tgt ctt cct ggt tgg tat tgt gct act gat tct tca cag act gaa gat tgt cga gat -tg tgt acc aaa aag t-a gat gtc cat gcg gat act ttg gtg gaa tag atg aac aag gaa atc cta ttt gta aat cgt gtt g-- -tg ttg gtg aat att gcg act acc tac gta att ata gac ttg atc cat ttc ctc cat gca aa- --- cta tct aaa tgt aat taa tta --- --- --- --- --- --- --- --- tga ttt --- --- --- -tg atg ata atg tta -cc ata c-a tt- --- -at atc gct act tgg tta --g tgt att att cag tat g-a aga cct att aat aa- --- --- -tt act tat ctt ttg acg atc ttg tt- -at aat tat --- --- --- --- aat ata aaa --a tac tta tgg --- --- cat agt aac tca t-- --- --- --- aat tgc tg- acg cga taa att cgt aat aat ctg ttt tgt tca aat ttt tat aag ga- --- --a tct aca ggc ata aaa at- aaa aat ata att tat aat ata c-- tct tac a-- gcg cca tca tga ata aca gca gtg aat taa ttg ctg tta tta atg gat tta gaa ata gtg gac gat ttt gtg ata tta ata tag tta tta atg atg aaa gga taa acg ctc ata gac tca tcc tat ctg gag cct ccg aat a-t ttt tcc att ctg ttt tcc aat aat ttt atc gat tct aat gaa tac gaa gtt aat cta agt cat tta gat tat caa agt gtt aac gat ttg atc gat tac att tat ggg ata cct ttg agc cta act aac gat aac gtg aaa tat att ctt tca acc gct ga- --t ttt tta caa att gga tct gcc att act gag tgc gaa aaa tac ata ctt aaa aat ctt tgt tct aga aac tgt atc gat ttc --- --- tac ata tat gct gat aaa tat aat aac aag aaa ata gaa tca gcg tcg ttt aac aca ata tta cga aat att ttg aga ctc atc aac gat gaa aac ttt aaa tac tta aca gag gaa tca atg ata aaa att tta agc gat gat atg tta aat ata aaa aat gag gat ttt gca cca cta att ctc att aaa tgg tta gag agt act caa caa tca tgc acc gtc gag tta ctt aga tgc ctc aga ata tca -tt act ttc ccc aca agt tat aaa atc act tta tag tca tca act ggt tag ttc aat cta cga atg tat aac att ctt aaa caa tat agc att ctt gga tga atc att tcc tag ata cca tag cat cga gtt gat atc tat cgg tat aag taa ttc gca tga taa gat ttc cat aaa ctg cta caa tca t-a aaa aaa ata cat ggg aaa tga tat ctt cac gta gat ata ggt gta gtt tcg cag tgg ccg tcc tgg ata ata tta tct ata tga tgg gtg gat atg atc agt ccc cgt ata gaa gtt caa agg tta tag cgt aca ata cat gta caa att ctt gga tat atg ata tac cag agc t-a aaa tat cct cgt tct aat tgt ggg gga ctg gct gat gac gaa tac att tat tgt ata ggc ggc ata cgc gat cag gat tca tcg ttg aca tct agt att gat aga tgg aag cca tca aaa cca tat tgg cag aag tat gct aaa atg cgc gaa cca aaa tgt gat atg ggg gtt gcg atg tta aac gga tta ata tat gtc ata ggt gga atc gtt aaa ggt g-- aca cgt gta ccg acg cac tag aga gtt tat cag aag atg gat gga tga agc atc aac gtc ttc caa taa aaa tgt cca ata tgt cga cga ttg ttc atg atg gca aga ttt ata tat ctg gag gtt aca aca ata gta gtg tag tta atg taa tat cga atc t-- agt cct tag cta taa tcc gat ata tga tga atg gac caa att atc atc att aaa cat tcc tag aat taa tcc cgc tct a

>EU410304.1:169436-171700_Synthetic_Vaccinia_virus_clone_GLV-1h68

ttg ctg act gca --- tgt a-c tat ttt tta cat tac tac att ggc tac ggc aga tat acc tac ttc gtc act gcc aca cgc tcc ggt aaa cgg --g gca tgt gac gag gga gaa tat ctt gat aag agg cat aat caa tgt tgt aat -ca gtg tcc acc tgg aga att tgc caa ggt tag atg ta- --a tgg taa cga taa cac aaa atg tga acg ctg ccc acc tca tac ata tac cgc aat ccc caa tta ctc t-a atg gat gtc atc aat gta gaa aat gcc caa cag gat cat ttg ata agg taa agt gta ccg gaa cac aga aca gta aat -gt tcg tgt ctt cct ggt tgg tat tgc gct act gat tct tca cag act gaa gat tgt cga gat -tg tat acc aaa aag g-a gat gtc cat gcg gat act ttg gtg gaa tag atg aac aag gaa atc cta ttt gta aat cgt gtt g-- -tg ttg gtg aat att gcg act acc tac gta att ata gac ttg atc cat ttc ctc cat gca aa- --- cta tct aaa tgt aat taa tta --- --- --- --- --- --- --- --- tga ttt --- --- --- -tg atg ata atg tta -cc ata c-a tt- --- -at atc gct act tgg tta --g tgt att att cag tat g-a aga cct att aat aa- --- --- -tt act tat ctt ttg acg atc ttg tt- -at aat tat --- --- --- --- aat ata aaa --a tac tta tgg --- --- cat agt aac tca t-- --- --- --- aat tgc tg- acg cga taa att cgt aat aat ctg ttt tgt tca aat ttt tat aag ga- --- --a tct aca ggc ata aaa at- aaa aat ata att tat aat ata c-- tct tac agc gcg cca tca tga ata aca gca gtg aat tga ttg ctg tta tta atg gat tta gaa ata gtg gac gat ttt gtg ata tta ata tag tta tta atg atg aaa gga taa acg ctc aca gac tca tcc tat ctg gag cct ccg aat a-t ttt tcc att ctg ttt tcc aat aat ttt atc gat tct aat gaa tac gaa gtt aat cta agt cat tta gat tat caa agt gtt aac gat ttg atc gat tac att tat ggg ata cct ttg agc cta act aac gat aac gtg aaa tat att ctt tca acc gct ga- --t ttt tta caa att gga tct gct att acg gag tgt gaa aat tac ata ctt aaa aat ctt tgt tct aga aac tgt atc gat ttc --- --- tac ata tac gct gat aaa tat aat aac aag aaa ata gaa tca gcg tcg ttt aac aca ata tta caa aat att ttg aga ctc atc aac gat gaa aac ttt aaa tac tta aca gag gaa tca atg ata aaa att tta agc gat gat atg tta aat ata aaa aat gag gat ttc gcc cca cta att ctc att aaa tgg tta gag agt act caa caa cca tgc acc gtc gag tta ctt aaa tgc ctc aga ata tca -tt gct ttc ccc aca agt tat aaa atc act tta tag tca tcg act ggt tag ttc aat cta cga atg tat aac att ctt aaa caa tat agc att ctt gga tga atc att tcc tag ata cca tag cat cga gtt gat atc tat cgg tat aag taa ttc gca tga taa gat ttc cat aaa ctg cta caa tca t-a aaa aaa ata cat ggg aaa tga tat ctt cac gta gat ata ggt gta gtt tcg cag tgg ccg tcc tgg ata ata tta tct ata tga tgg gtg gat atg atc agt ccc cgt ata gaa gtt caa agg tta tag cgt aca ata cat gta caa att ctt gga tat atg ata tac cag agc t-a aaa tat cct cgt tct aat tgt ggg gga ctg gct gat gac gaa tac att tat tgt ata ggc ggc ata cgc gat cag gat tca tcg ttg aca tct agt att gat aga tgg aag cca tca aaa cca tat tgg cag aag tat gct aaa atg cgc gaa cca aaa tgt gat atg ggg gtt gcg atg tta aac gga tta ata tat gtt ata ggt gga atc gtt aaa ggt g-- aca cgt gta ccg acg cac tag aga gtt tat cag aag atg gat gga tga agc atc aac gtc ttc caa taa aaa tgt cca ata tgt cga cga ttg ttc atg atg gca aga ttt ata tat ctg gag gtt aca aca ata gta gtg tag tta atg taa tat cga atc t-- agt cct tag cta taa ttc gat ata tga tga atg gac caa att atc atc att aaa cat tcc tag aat taa tcc cgc tct a

>KY369926.1:176107-178360_Cowpox_virus_strain_Kostroma_2015

--- --- --- --a --- tgt a-c tat ttt gta cat cac tac att ggt tac ggc aga tat acc tac tcc gtt gcc gcc aca cgc tcc ggt aaa cgg --g tca tgt gac gag gga gaa tat ctt gat aag agg cat aat caa tgt tgt aat -ca gtg tcc acc tgg aga att tgc caa ggt cag atg ta- --g tgg tag tga taa cac aaa atg tga acg ctg ccc acc tca tac ata tac cgc aat ccc caa tta ttc t-a acg gat gcc atc aat gta gaa aat gtc caa cag gat cat ttg ata agg taa agt gta ccg gaa cac aga aca gta aat -gt tcg tgt ctt cct ggt tgg tat tgt gct act gat tct tca cag act gaa gat tgt cga gat -tg tgt acc aaa aag t-a gat gtc cat gcg gat act ttg gtg gaa tag atg aac aag gaa atc cta ttt gta aat cgt gtt g-- -tg ttg gtg aat att gcg act acc tac gta att ata gac ttg atc cat ttc ctc cat gca aa- --- cta tct aaa tgt aat taa tta --- --- --- --- --- --- --- --- tga ttt --- --- --- -tg atg ata atg tta -cc ata c-a tt- --- -at atc gct act tgg tta --g tgt att att cag tat g-a aga cct att aat aa- --- --- -tt act tat ctt ttg acg atc ttg tt- -at aat tat --- --- --- --- aat ata aaa --a tac tta tgg --- --- cat agt aac tca t-- --- --- --- aat tgc tg- acg cga taa att cgt aat aat ctg ttt tgt tca aat ttt tat aag ga- --- --a tct aca ggc ata aaa at- aaa aat ata att tat aat ata c-- tct tac agc gcg cca tca tga ata aca gca gtg aat tga ttg ctg tta tta atg gat tta gaa ata gtg gac gat ttt gtg ata tta ata tag tta tta atg atg aaa gga taa acg ctc ata gac tca tcc tat ctg gag cct ccg aat a-t ttt tcc att ctg ttt tcc aat aat ttt atc gat tct aat gaa tac gaa gtt aat cta agt cat tta gat tat caa agt gtt aac gat ttg atc gat tac att tat ggg ata cct ttg agc cta act aac gat aac gtg aaa tat att ctt tca acc gct ga- --t ttt tta caa att gga tct gcc att act gag tgc gaa aaa tac ata ctt aaa aat ctt tgt tct aga aac tgt atc gat ttc --- --- tac ata tac gct gat aaa tat aat aac aag aaa ata gaa tca gcg tcg ttt aac aca ata tta cga aat att ttg aga ctc atc aac gat gaa aac ttt aaa tac tta aca gag gaa tca atg ata aaa att tta agc gat gat atg tta aat ata aaa aat gag gat ttc gcc cca ctg att ctc att aaa tgg tta gag agt act caa caa tca tgc acc gtc gag tta ctt aga tgc ctc aga ata tca -tt gct ttc ccc aca agt tat aaa atc act tta tag tca tca act ggt tag ttc aat cta cga atg tat aac att ctt aaa caa tat agc att ctt gga tga atc att tcc tag ata cca tag cat cga gtt gat atc tat cgg tat aag taa ttc gca tga taa gat ttc cat aaa ctg cta taa tca t-a aaa aaa ata cat ggg aaa tga tat ctt cac gta gat ata ggt gta gtt tcg cag tgg ccg tcc tgg ata ata tta tct ata tga tgg gtg gat atg atc agt ccc cgt ata gaa gtt caa agg tta tag cgt aca ata cat gta caa att ctt gga tat atg ata tac cag agc t-a aaa tat cct cgt tct aat tgt ggg gga ctg gct gat gac gaa tac att tat tgt ata ggc ggc ata tgc gat cag gat tca tcg ttg aca tct agt att gat aga tgg aag cca tca aaa cca tat tgg cag aag tat gct aaa atg cgc gaa cca aaa tgt gat atg ggg gtt gcg atg tta aac gga tta ata tat gtc ata ggt gga atc gtt aaa ggt g-- aca cgt gta ccg acg cac tag aga gtt tat cag aag atg gat gga tga agc atc aac gtc ttc caa taa aaa tgt cca ata tgt cga cga ttg ttc atg atg gca aga ttt ata tat ctg gag gtt aca aca ata gta gtg tag tta atg taa tat cga atc t-- agt cct tag cta taa tcc gat ata tga tga atg gac caa att atc atc att aaa cat tcc tag aat taa tcc cgc tct a

>JX489137.1:159078-161342_Vaccinia_virus_strain_Tiantan_clone_TT10_complete_genome

ttg ctg act gca --- tgt a-c tat ttt tta cat tac tac att ggc tac ggc aga tat acc tac ttc gtc act gcc aca cgc tcc ggt aaa cgg --g gca tgt gac gag gga gaa tat ctt gat aag agg cat aat caa tgt tgt aat -ca gtg tcc acc tgg aga att tgc caa ggt tag atg ta- --a tgg taa cga taa cac aaa atg tga acg ctg ccc acc tca tac ata tac cgc aat ccc caa tta ctc t-a atg gat gtc atc aat gta gaa aat gcc caa ccg gat cat ttg ata agg taa agt gta ccg gaa cac aga aca gta aat -gt tcg tgt ctt cct ggt tgg tat tgc gct act gat tct tca cag act gaa gat tgt cga gat -tg tat acc aaa aag g-a gat gtc cat gcg gat act ttg gtg gaa tag atg aac aag gaa atc cta ttt gta aat cgt gtt g-- -tg ttg gtg aat att gcg act acc tac gta att ata gac ttg atc cat ttc ctc cat gca aa- --- cta tct aaa tgt aat taa tta --- --- --- --- --- --- --- --- tga ttt --- --- --- -tg atg ata atg tta -cc ata c-a tt- --- -at atc gct act tgg tta --g tgt att att cag tat g-a aga cct att aat aa- --- --- -tt act tat ctt ttg acg atc ttg tt- -at aat tat --- --- --- --- aat ata aaa --a tac tta tgg --- --- cat agt aac tca t-- --- --- --- aat tgc tg- acg cga taa att cgt aat aat ctg ttt tgt tca aat ttt tat aag ga- --- --a tct aca ggc ata aaa at- aaa aat ata att tat aat ata c-- tct tac agc gcg cca tca tga ata aca gca gtg aat tga ttg ctg tta tta atg gat tta gaa ata gtg gac gat ttt gtg ata tta ata tag tta tta atg atg aaa gga taa acg ctc ata aac tca tcc tat ctg gag cct ccg aat a-t ttt tcc att ctg ttt tcc aat aat ttt atc gat tct aat gaa tac gaa gtt aat cta agt cat tta gat tat caa agc gta aac gat ttg atc gat tac att tat ggg ata cct ttg agc cta act aac gat aac gtg aaa tat att ctt tca acc gct ga- --t ttt tta caa att gga tct gct att acg gag tgt gaa aaa tac ata ctt aaa aat ctt tgt tct aaa aac tgt atc gat ttc --- --- tac ata tac gct gat aaa tat aat aac aag aaa ata gaa tca gcg tcg ttt aac aca ata tta cga aat att ttg aga ctc atc aac gat gaa aac ttt aaa tac tta aca gag gaa tca atg ata aaa att tta agc gat gat atg tta aat ata aaa aat gag gat ttc gcc cca cta att ctc att aaa tgg tta gag agt act caa caa cca tgc acc gtc gag tta ctt aaa tgc ctc aga ata tca -tt gct ttc ccc aca agt tat aaa atc act tta tag tca tcg act ggt tag ttc aat cta cga atg tat aac att ctt aaa caa tat agc att ctt gga tga atc att tcc tag ata cca tag cat cga gtt gat atc tat cgg tat aag taa ttc gcg tga taa aat ttc cat aaa ctg cta caa tca t-a aaa aaa ata cat ggg aaa tga tat ctt cac gta gat ata ggt gta gtt tcg cag tgg ccg tcc tgg ata ata tta tct ata tga tgg gtg gat atg atc agt ccc cgt ata gaa gtt caa agg tta tag cgt aca ata cat gta caa att ctt gga tat atg ata tac cag agc t-a aaa tat cct cgt tct aat tgt ggg gga ctg gct gat gac gaa tac att tat tgt ata ggc ggc ata cgc gat cag gat tca tcg ttg aca tct agt att gat aga tgg aag cca tca aaa cca tat tgg cag aag tat gct aaa atg cgc gaa cca aaa tgt gat atg ggg gtt gcg atg tta aac gga tta ata tat gtc atg ggt gga atc gtt aaa ggt g-- aca cgt gta ccg acg cac tag aga gtt tat cag aag atg gat gga tga agc atc aac gtc ttc caa taa aaa tgt cca ata tgt cga cga ttg ttc atg atg gca aga ttt ata tat ctg gag gtt aca aca ata gta gtg tag tta atg taa tat cga atc t-- agt cct tag cta taa tcc gat ata tga tga atg gac caa att atc atc att aaa tat tcc tag aat taa tcc cgc tct a

>AY678276.1:162195-164454_Vaccinia_virus_strain_Lister_complete_genome

ttg ctg act gca --- tgt a-c tat ttt tta cat tac tac att ggc tac ggc aga tat acc tac ttc gtc act gcc aca cgc tcc ggt aaa cgg --g gca tgt gac gag gga gaa tat ctt gat aag agg cat aat caa tgt tgt aat -cg gtg tcc acc tgg aga att tgc caa ggt tag atg ta- --a tgg taa cga taa cac aaa atg tga acg ctg ccc acc tca tac ata tac cgc aat ccc caa tta ctc t-a atg gat gtc atc aat gta gaa aat gcc caa cag gat cat ttg ata agg taa agt gta ccg gaa cac aga aca gta aat -gt tcg tgt ctt cct ggt tgg tat tgc gct act gat tct tca cag act gaa gat tgt cga gat -tg tat acc aaa aag g-a gat gtc cat gcg gat act ttg gtg gaa tag atg aac aag gaa atc cta ttt gta aat cgt gct g-- -ta ttg gtg aat att gcg act acc tac gta att ata gac ttg atc cat ttc ctc cat gca aa- --- cta tct aaa tgt aat taa tta --- --- --- --- --- --- --- --- tga ttt --- --- --- -tg atg ata atg tta -cc ata c-a tt- --- -at atc gct act tgg tta --g tgt att att cag tat g-a aga cct att aat aa- --- --- -tt act tat ctt ttg acg atc ttg tt- -at aat tat --- --- --- --- aat ata aaa --- -ac tta tgg --- --- cat agt aac tca t-- --- --- --- aat tgc tg- acg cga taa att cgt aat aat ctg ttt tgt tca aat ttt tat aag ga- --- --a tct aca ggc ata aaa at- aaa aat ata att tat aat a-a c-- tct tac a-- acg cca tca tga ata aca gca gtg aat tga ttg ctg tta tta atg gat tta gaa ata gtg gac gat ttt gtg ata tta gta tag tta tta atg atg aaa gga taa acg ctc aca gac tca tcc tat ctg gag cct ccg aat a-t ttt tcc att ctg ttt tcc aat aat ttt atc gat tct aat gaa tac gaa gtt aat cta agt cat tta gat tat caa agc gta aac gat ttg atc gat tac att tat ggg ata cct ttg agc cta act aac gat aac gtg aaa tat att ctt tca acc gct ga- --t ttt tta caa att gga tct gct att act gag tgc gaa aaa tac ata ctt aaa aat ctt tgt tct aga aac tgt atc gat ttc --- --- tac ata tac gct gat aaa tat aat aac aag aaa ata gaa tca gcg tcg ttt aac aca ata tta caa aat att ttg aga ctc atc aac gat gaa aac ttt aaa tac tta aca gag gaa tca atg ata aaa att tta agc gat gat atg tta aat ata aaa aat gag gat ttc gcc cca cta att ctc att aaa tgg tta gag agt act caa caa cca tgc acc gtc gag tta ctt aaa tgc ctc aga ata tca -tt gct ttc ccc aca agt tat aaa atc act tta tag tca tcg act ggt tag ttc aat cta cga atg tat aac att ctt aaa caa tat agc att ctt gga tga atc att tcc tag ata cca tag cat cga gtt gat atc tat cgg tat aag taa ttc gca tga taa gat ttc cat aaa ctg cta caa tca t-a aaa aaa ata cat ggg aaa tga tat ctt cac gta gat ata ggt gta gtt tcg cag tgg ccg tcc tgg ata ata tta tct ata tga tgg gtg gat atg atc agt ccc cgt ata gaa gtt caa agg tta tag cgt aca ata cat gta caa att ctt gga tat atg ata tac cag agc t-a aaa tat cct cgt tct aat tgt ggg gga ctg gct gat gac gaa tac att tat tgt ata ggc ggc ata cgc gat cag gat tca tcg ttg aca tct agt att gat aga tgg aag cca tca aaa cca tat tgg cag aag tat gct aaa atg cgc gaa cca aaa tgt gat atg ggg gtt gcg atg tta aac gga tta ata tat gtt ata ggt gga atc gtt aaa ggt g-- aca cgt gta ccg acg cac tag aga gtt tat cag aag atg gat gga tga agc atc aac gtc ttc caa taa aaa tgt cca ata tgt cga cga ttg ttc atg atg gca aga ttt ata tat ctg gag gtt aca aca ata gta gtg tag tta atg taa tat cga atc t-- agt cct tag cta taa ttc gat ata tga tga atg gac caa att atc atc att aaa cat tcc tag aat taa tcc cgc tct a

>MK336431.1:140243-142507_Vaccinia_virus_isolate_KVAC103

ttg ctg act gta --- tgt a-c tat ttt tta cat tac tac att ggc tac ggc aga tat acc tac tcc gcc act gcc aca cgc tcc ggt aaa cgg --g gca tgt gac gag gga gaa tat ctt gat aag agg cat aat caa tgt tgt aat -ca gtg tcc acc tgg aga att tgc caa ggt tag atg ta- --a tgg taa cga taa cac aaa atg tga acg ctg ccc acc tca tac ata tac cgc aat ccc caa tta ctc t-a atg gat gtc atc aat gta gaa aat gcc caa cag gat cat ttg ata agg taa agt gta ccg gaa cac aga aca gta aat -gt tcg tgt ctt cct ggt tgg tat tgc gct act gat tct tca cag act gaa gat tgt cga gat -tg tat acc aaa aag g-a gat gtc cat gcg gat act ttg gtg gaa tag atg aac aag gaa atc cta ttt gta aat cgt gtt g-- -tg ttg gtg aat att gcg act acc tac gta att ata gac ttg atc cat ttc ctc cat gca aa- --- cta tct aaa tgt aat taa tta --- --- --- --- --- --- --- --- tga ttt --- --- --- -tg atg ata atg tta -cc ata c-a tt- --- -at atc gct act tgg tta --g tgt att att cag tat g-a aga cct att aat aa- --- --- -tt act tat ctt ttg acg atc ttg tt- -at aat tat --- --- --- --- aat ata aaa --a tac tta tgg --- --- cat agt aac tca t-- --- --- --- aat tgc tg- acg cga taa att cgt aat aat ctg ttt tgt tca aat ttt tat aag ga- --- --a tct aca ggc ata aaa at- aaa aat ata att tat aat ata c-- tct tac agc gcg cca tca tga ata aca gca gta aat taa ttg ctg tta tta atg gat tta gaa ata gtg gac gat ttt gtg ata tta gta tag tta tta atg atg aaa tga taa acg ctc aca gac tca tcc tat ctg gag cct ccg aat a-t ttt tcc att ctg ttt tcc aat aat ttt atc gat tct aat gaa tac gaa gtt aat cta agt cat tta gat tat caa agt gtt aac gat ttg atc gat tac att tat ggg ata cct ttg agc cta act aac gat aac gtg aaa tat att ctt tca acc gct ga- --t ttt tta caa att gga tct gct att acg gag tgc gaa aaa tac ata ctt aaa aat ctt tgt tct aga aac tgt atc gat ttc --- --- tac ata tac gct gat aaa tat aat aac aag aaa ata gaa tca gcg tcg ttt aac aca ata tta caa aat att ttg aga ctc atc aac gat gaa aac ttt aaa tac tta aca gag gaa tca atg ata aaa att tta agc gat gat atg tta aat ata aaa aat gag gat ttc gcc cca cta att ctc att aaa tgg tta gag agt act caa caa cca tgc acc gtc gag tta ctt aaa tgc ctc aga ata tca -tt gct ttc ccc aca agt tat aaa atc act tta tag tca tcg act ggt tag ttc aat cta cga atg tat aac att ctt aaa caa tat agc att ctt gga tga atc att tcc tag ata cca tag cat cga gtt gat atc tat cgg tat aag taa ttc gca tga taa gat ttc cat aaa ctg cta caa tca t-a aaa aaa ata cat ggg aaa tga tat ctt cac gta gat ata ggt gta gtt tcg cag tgg ccg tcc tgg ata ata tta tct ata tga tgg gtg gat atg atc agt ccc cgt ata gaa gtt caa agg tta tag cgt aca ata cat gta caa att ctt gga tat atg ata tac cag agc t-a aaa tat cct cgt tct aat tgt ggg gga ctg gct gat gac gaa tac att tat tgt ata ggc ggc ata cgc gat cag gat tca tcg ttg aca tct agt att gat aaa tgg aag cca tca aaa cca tat tgg cag aag tat gct aaa atg cgc gaa cca aaa tgt gat atg ggg gtt gcg atg tta aac gga tta ata tat gtc atg ggt gga atc gtt aaa ggt g-- aca cgt gta ccg acg cac tag aga gtt tat cag aag atg gat gga tga agc atc aac gtc ttc caa taa aaa tgt cca ata tgt cga cga ttg ttc atg atg gca aga ttt ata tat ctg gag gtt aca aca ata gta gtg tag tta atg taa tat cga atc t-- agt cct tag cta taa ttc gat ata tga tga atg gac caa att atc atc att aaa cat tcc tag aat taa tcc cgc tct a

>MH341447.1:162490-164743_Vaccinia_virus_strain_L-IVP_oncoQ

ttg ctg act gca --- tgt a-c tat ttt tta cat tac tac att ggc tac ggc aga tat acc tac ttc gtc act gcc aca cgc tcc ggt aaa cgg --g gca tgt gac gag gga gaa tat ctt gat aag agg cat aat caa tgt tgt aat -cg gtg tcc acc tgg aga att tgc caa ggt tag atg ta- --a tgg taa cga taa cac aaa atg tga acg ctg ccc acc tca tac ata tac cgc aat ccc caa tta ctc t-a atg gat gtc atc aat gta gaa aat gcc caa cag gat cat ttg ata agg taa agt gta ccg gaa cac aga aca gta aat -gt tcg tgt ctt cct ggt tgg tat tgt gct act gat tct tca cag act gaa gat tgt cga gat -tg tat acc aaa aag g-a gat gtc cat gcg gat act ttg gtg gaa tag atg aac aag gaa atc cta ttt gta aat cgt gtt g-- -tg ttg gtg aat att gcg act acc tac gta att ata gac ttg atc cat ttc ctc cat gca aa- --- cta tct aaa tgt aat taa tta --- --- --- --- --- --- --- --- tga ttt --- --- --- -tg atg ata atg tta -cc ata c-a tt- --- -at atc gct act tgg tta --g tgt att att cag tat g-a aga cct att aat aa- --- --- -tt act tat ctt ttg acg atc ttg tt- -at aat tat --- --- --- --- aat ata aaa --a tac tta tgg --- --- cat agt aac tca t-- --- --- --- aat tgc tg- acg cga taa att cgt aat aat ctg ttt tgt tca --- --- --- aag ga- --- --a tct aca ggc ata aaa at- aaa aat ata att tat aat ata c-- tct tac a-- acg cca tca tga ata gca gca gta aat taa ttg ctg tta tta atg gat tta gaa ata gtg gac gat ttt gtg ata tta ata tag tta tta atg atg aaa gga taa acg ctc ata aac tca tcc tat ctg gag cct ccg aat a-t ttt tcc att ctg ttt tcc aat aat ttt atc gat tct aat gaa tac gaa gtt aat cta agt cat tta gat tat caa agt gtt aac gat ttg atc gat tac att tat ggg ata cct ttg agc cta act aac gat aac gtg aaa tat att ctt tca acc gct ga- --t ttt tta caa att gga tct gcc att act gag tgc gaa aaa tac ata ctt aaa aat ctt tgt tct aga aac tgt atc gat ttc --- --- tac ata tac gct gat aaa tat aat aac aag aaa ata gaa tca gcg tcg ttt aac aca ata tta caa aat att ttg aga ctc atc aac gat gaa aac ttt aaa tac tta aca gag gaa tca atg ata aaa att tta agc gat gat atg tta aat ata aaa aat gag gat ttc gcc cca cta att ctc att aaa tgg tta gag agt act caa caa tca tgc acc gtc gag tta ctt aga tgc ctc aga ata tca -tt gct ttc ccc aca agt tat aaa atc act tta tag tca tca act ggt tag ttc aat cta cga atg tat aac att ctt aaa caa tat agc att ctt gga tga atc att tcc tag ata cca tag cat cga gtt gat atc tat cgg tat aag taa ttc gca tga taa gat ttc cat aaa ctg cta caa tca t-a aaa aaa ata cat ggg aaa tga tat ctt cac gta gat ata ggt gta gtt tcg cag tgg ccg tcc tgg ata ata tta tct ata tga tgg gtg gat atg atc agt ccc cgt ata gaa gtt caa agg tta tag cgt aca ata cat gta caa att ctt gga tat atg ata tac cag agc t-a aaa tat cct cgt tct aat tgt ggg gga ctg gct gat gac gaa tac att tat tgt ata ggc ggc ata cgc gat cag gat tca tcg ttg aca tct agt att gat aga tgg aag cca tca aaa cca tat tgg cag aag tat gct aaa atg cgc gaa cca aaa tgt gat atg ggg gtt gcg atg tta aac gga tta ata tat gtt ata ggt gga atc gtt aaa ggt g-- aca cgt gta ccg acg cac tag aga gtt tat cag aag atg gat gga tga agc atc aac gtc ttc caa taa aaa tgt cca ata tgt cga cga ttg ttc atg atg gca aga ttt ata tat ctg gag gtt aca aca ata gta gtg tag tta atg taa tat cga atc t-- agt cct tag cta taa ttc gat ata tga tga atg gac caa att atc atc att aaa tat tcc tag aat taa tcc cgc tct a

>MH341446.1:162356-164609_Vaccinia_virus_strain_L-IVP_oncoM

ttg ctg act gca --- tgt a-c tat ttt tta cat tac tac att ggc tac ggc aga tat acc tac ttc gtc act gcc aca cgc tcc ggt aaa cgg --g gca tgt gac gag gga gaa tat ctt gat aag agg cat aat caa tgt tgt aat -cg gtg tcc acc tgg aga att tgc caa ggt tag atg ta- --a tgg taa cga taa cac aaa atg tga acg ctg ccc acc tca tac ata tac cgc aat ccc caa tta ctc t-a atg gat gtc atc aat gta gaa aat gcc caa cag gat cat ttg ata agg taa agt gta ccg gaa cac aga aca gta aat -gt tcg tgt ctt cct ggt tgg tat tgt gct act gat tct tca cag act gaa gat tgt cga gat -tg tat acc aaa aag g-a gat gtc cat gcg gat act ttg gtg gaa tag atg aac aag gaa atc cta ttt gta aat cgt gtt g-- -tg ttg gtg aat att gcg act acc tac gta att ata gac ttg atc cat ttc ctc cat gca aa- --- cta tct aaa tgt aat taa tta --- --- --- --- --- --- --- --- tga ttt --- --- --- -tg atg ata atg tta -cc ata c-a tt- --- -at atc gct act tgg tta --g tgt att att cag tat g-a aga cct att aat aa- --- --- -tt act tat ctt ttg acg atc ttg tt- -at aat tat --- --- --- --- aat ata aaa --a tac tta tgg --- --- cat agt aac tca t-- --- --- --- aat tgc tg- acg cga taa att cgt aat aat ctg ttt tgt tca --- --- --- aag ga- --- --a tct aca ggc ata aaa at- aaa aat ata att tat aat ata c-- tct tac a-- acg cca tca tga ata gca gca gta aat taa ttg ctg tta tta atg gat tta gaa ata gtg gac gat ttt gtg ata tta ata tag tta tta atg atg aaa gga taa acg ctc ata aac tca tcc tat ctg gag cct ccg aat a-t ttt tcc att ctg ttt tcc aat aat ttt atc gat tct aat gaa tac gaa gtt aat cta agt cat tta gat tat caa agt gtt aac gat ttg atc gat tac att tat ggg ata cct ttg agc cta act aac gat aac gtg aaa tat att ctt tca acc gct ga- --t ttt tta caa att gga tct gcc att act gag tgc gaa aaa tac ata ctt aaa aat ctt tgt tct aga aac tgt atc gat ttc --- --- tac ata tac gct gat aaa tat aat aac aag aaa ata gaa tca gcg tcg ttt aac aca ata tta caa aat att ttg aga ctc atc aac gat gaa aac ttt aaa tac tta aca gag gaa tca atg ata aaa att tta agc gat gat atg tta aat ata aaa aat gag gat ttc gcc cca cta att ctc att aaa tgg tta gag agt act caa caa tca tgc acc gtc gag tta ctt aga tgc ctc aga ata tca -tt gct ttc ccc aca agt tat aaa atc act tta tag tca tca act ggt tag ttc aat cta cga atg tat aac att ctt aaa caa tat agc att ctt gga tga atc att tcc tag ata cca tag cat cga gtt gat atc tat cgg tat aag taa ttc gca tga taa gat ttc cat aaa ctg cta caa tca t-a aaa aaa ata cat ggg aaa tga tat ctt cac gta gat ata ggt gta gtt tcg cag tgg ccg tcc tgg ata ata tta tct ata tga tgg gtg gat atg atc agt ccc cgt ata gaa gtt caa agg tta tag cgt aca ata cat gta caa att ctt gga tat atg ata tac cag agc t-a aaa tat cct cgt tct aat tgt ggg gga ctg gct gat gac gaa tac att tat tgt ata ggc ggc ata cgc gat cag gat tca tcg ttg aca tct agt att gat aga tgg aag cca tca aaa cca tat tgg cag aag tat gct aaa atg cgc gaa cca aaa tgt gat atg ggg gtt gcg atg tta aac gga tta ata tat gtt ata ggt gga atc gtt aaa ggt g-- aca cgt gta ccg acg cac tag aga gtt tat cag aag atg gat gga tga agc atc aac gtc ttc caa taa aaa tgt cca ata tgt cga cga ttg ttc atg atg gca aga ttt ata tat ctg gag gtt aca aca ata gta gtg tag tta atg taa tat cga atc t-- agt cct tag cta taa ttc gat ata tga tga atg gac caa att atc atc att aaa tat tcc tag aat taa tcc cgc tct a

>MH341445.1:162296-164549_Vaccinia_virus_strain_L-IVP_oncoB

ttg ctg act gca --- tgt a-c tat ttt tta cat tac tac att ggc tac ggc aga tat acc tac ttc gtc act gcc aca cgc tcc ggt aaa cgg --g gca tgt gac gag gga gaa tat ctt gat aag agg cat aat caa tgt tgt aat -cg gtg tcc acc tgg aga att tgc caa ggt tag atg ta- --a tgg taa cga taa cac aaa atg tga acg ctg ccc acc tca tac ata tac cgc aat ccc caa tta ctc t-a atg gat gtc atc aat gta gaa aat gcc caa cag gat cat ttg ata agg taa agt gta ccg gaa cac aga aca gta aat -gt tcg tgt ctt cct ggt tgg tat tgt gct act gat tct tca cag act gaa gat tgt cga gat -tg tat acc aaa aag g-a gat gtc cat gcg gat act ttg gtg gaa tag atg aac aag gaa atc cta ttt gta aat cgt gtt g-- -tg ttg gtg aat att gcg act acc tac gta att ata gac ttg atc cat ttc ctc cat gca aa- --- cta tct aaa tgt aat taa tta --- --- --- --- --- --- --- --- tga ttt --- --- --- -tg atg ata atg tta -cc ata c-a tt- --- -at atc gct act tgg tta --g tgt att att cag tat g-a aga cct att aat aa- --- --- -tt act tat ctt ttg acg atc ttg tt- -at aat tat --- --- --- --- aat ata aaa --a tac tta tgg --- --- cat agt aac tca t-- --- --- --- aat tgc tg- acg cga taa att cgt aat aat ctg ttt tgt tca --- --- --- aag ga- --- --a tct aca ggc ata aaa at- aaa aat ata att tat aat ata c-- tct tac a-- acg cca tca tga ata gca gca gta aat taa ttg ctg tta tta atg gat tta gaa ata gtg gac gat ttt gtg ata tta ata tag tta tta atg atg aaa gga taa acg ctc ata aac tca tcc tat ctg gag cct ccg aat a-t ttt tcc att ctg ttt tcc aat aat ttt atc gat tct aat gaa tac gaa gtt aat cta agt cat tta gat tat caa agt gtt aac gat ttg atc gat tac att tat ggg ata cct ttg agc cta act aac gat aac gtg aaa tat att ctt tca acc gct ga- --t ttt tta caa att gga tct gcc att act gag tgc gaa aaa tac ata ctt aaa aat ctt tgt tct aga aac tgt atc gat ttc --- --- tac ata tac gct gat aaa tat aat aac aag aaa ata gaa tca gcg tcg ttt aac aca ata tta caa aat att ttg aga ctc atc aac gat gaa aac ttt aaa tac tta aca gag gaa tca atg ata aaa att tta agc gat gat atg tta aat ata aaa aat gag gat ttc gcc cca cta att ctc att aaa tgg tta gag agt act caa caa tca tgc acc gtc gag tta ctt aga tgc ctc aga ata tca -tt gct ttc ccc aca agt tat aaa atc act tta tag tca tca act ggt tag ttc aat cta cga atg tat aac att ctt aaa caa tat agc att ctt gga tga atc att tcc tag ata cca tag cat cga gtt gat atc tat cgg tat aag taa ttc gca tga taa gat ttc cat aaa ctg cta caa tca t-a aaa aaa ata cat ggg aaa tga tat ctt cac gta gat ata ggt gta gtt tcg cag tgg ccg tcc tgg ata ata tta tct ata tga tgg gtg gat atg atc agt ccc cgt ata gaa gtt caa agg tta tag cgt aca ata cat gta caa att ctt gga tat atg ata tac cag agc t-a aaa tat cct cgt tct aat tgt ggg gga ctg gct gat gac gaa tac att tat tgt ata ggc ggc ata cgc gat cag gat tca tcg ttg aca tct agt att gat aga tgg aag cca tca aaa cca tat tgg cag aag tat gct aaa atg cgc gaa cca aaa tgt gat atg ggg gtt gcg atg tta aac gga tta ata tat gtt ata ggt gga atc gtt aaa ggt g-- aca cgt gta ccg acg cac tag aga gtt tat cag aag atg gat gga tga agc atc aac gtc ttc caa taa aaa tgt cca ata tgt cga cga ttg ttc atg atg gca aga ttt ata tat ctg gag gtt aca aca ata gta gtg tag tta atg taa tat cga atc t-- agt cct tag cta taa ttc gat ata tga tga atg gac caa att atc atc att aaa tat tcc tag aat taa tcc cgc tct a

>JX489136.1:159086-161350_Vaccinia_virus_strain_Tiantan_clone_TT9_complete_genome

ttg ctg act gca --- tgt a-c tat ttt tta cat tac tac att ggc tac ggc aga tat acc tac ttc gtc act gcc aca cgc tcc ggt aaa cgg --g gca tgt gac gag gga gaa tat ctt gat aag agg cat aat caa tgt tgt aat -ca gtg tcc acc tgg aga att tgc caa ggt tag atg ta- --a tgg taa cga taa cac aaa atg tga acg ctg ccc acc tca tac ata tac cgc aat ccc caa tta ctc t-a atg gat gtc atc aat gta gaa aat gcc caa ccg gat cat ttg ata agg taa agt gta ccg gaa cac aga aca gta aat -gt tcg tgt ctt cct ggt tgg tat tgc gct act gat tct tca cag act gaa gat tgt tga aat -tg tgt acc aaa aag g-a gat gtc cat gcg gat act ttg gtg gaa tag atg aac aag gaa atc cta ttt gta aat cgt gtt g-- -tg ttg gtg aat att gcg act acc tac gta att ata gac ttg atc cat ttc ctc cat gca aa- --- cta tct aaa tgt aat taa tta --- --- --- --- --- --- --- --- tga ttt --- --- --- -tg atg ata atg tta -cc ata c-a tt- --- -at atc gct act tgg tta --g tgt att att cag tat g-a aga cct att aat aa- --- --- -tt act tat ctt ttg acg atc ttg tt- -at aat tat --- --- --- --- aat ata aaa --a tac tta tgg --- --- cat agt aac tca t-- --- --- --- aat tgc tg- acg cga taa att cgt aat aat ctg ttt tgt tca aat ttt tat aag ga- --- --a tct aca ggc ata aaa at- aaa aat ata att tat aat ata c-- tct tac agc gcg cca tca tga ata aca gca gtg aat tga ttg ctg tta tta atg gat tta gaa ata gtg gac gat ttt gtg ata tta ata tag tta tta atg atg aaa gga taa acg ctc ata aac tca tcc tat ctg gag cct ccg aat a-t ttt tcc att ctg ttt tcc aat aat ttt atc gat tct aat gaa tac gaa gtt aat cta agt cat tta gat tat caa agc gta aac gat ttg atc gat tat att tat ggg ata cct ttg agc cta act aac gat aac gtg aaa tat att ctt tca acc gct ga- --t ttt tta caa att gga tct gct att acg gag tgt gaa aaa tac ata ctt aaa aat ctt tgt tct aga aac tgt atc gat ttc --- --- tac ata tac gct gat aaa tat aat aac aag aaa ata gaa tca gcg tcg ttt aac aca ata tta cga aat att ttg aga ctc atc aac gat gaa aac ttt aaa tac tta aca gag gaa tca atg ata aaa att tta agc gat gat atg tta aat ata aaa aat gag gat ttc gcc cca cta att ctc att aaa tgg tta gag agt act caa caa tca tgc acc gtc gag tta ctt aaa tgc ctc aga ata tca -tt gct ttc ccc aca agt tat aaa atc act tta tag tca tcg act ggt tag ttc aat cta cga atg tat aac att ctt aaa caa tat agc att ctt gga tga atc att tcc tag ata cca tag cat cga gtt gat atc tat cgg tat aag taa ttc gcg tga taa aat ttc cat aaa ctg cta caa tca t-a aaa aaa ata cat ggg aaa tga tat ctt cac gta gat ata ggt gta gtt tcg cag tgg ccg tcc tgg ata ata tta ttt ata tga tgg gtg gat atg atc agt ccc cgt ata gaa gtt caa agg tta tag cgt aca ata cat gta caa att ctt gga tat atg ata tac cag agc t-a aaa tat cct cgt tct aat tgt ggg gga ctg gct gat gac gaa tac att tat tgt ata ggc ggc ata cgc gat cag gat tca tcg ttg aca tct agt att gat aga tgg aag cca tca aaa cca tat tgg cag aag tat gct aaa atg cgc gaa cca aaa tgt gat atg ggg gtt gcg atg tta aac gga tta ata tat gtc atg ggt gga atc gtt aaa ggt g-- aca cgt gta ccg acg cac tag aga gtt tat cag aag atg gat gga tga agc atc aac gtc ttc caa taa aaa tgt cca ata tgt cga cga ttg ttc atg atg gca aga ttt ata tat ctg gag gtt aca aca ata gta gtg tag tta atg taa tat cga atc t-- agt cct tag cta taa tcc gat ata tga tga atg gac caa att atc atc att aaa tat tcc tag aat taa tcc cgc tct a

>KX061501.1:161607-163871_Vaccinia_virus_strain_Lister_clone_Lister_Butantan

ttg ctg act gca --- tgt a-c tat ttt tta cat tac tac att ggc tac ggc aga tat acc tac ttc gtc act gcc aca cgc tcc ggt aaa cgg --g gca tgt gac gag gga gaa tat ctt gat aag agg cat aat caa tgt tgt aat -cg gtg tcc acc tgg aga att tgc caa ggt tag atg ta- --a tgg taa cga taa cac aaa atg tga acg ctg ccc acc tca tac ata tac cgc aat ccc caa tta ctc t-a atg gat gtc atc aat gta gaa aat gcc caa cag gat cat ttg ata agg taa agt gta ccg gaa cac aga aca gta aat -gt tcg tgt ctt cct ggt tgg tat tgc gct act gat tct tca cag act gaa gat tgt cga gat -tg tat acc aaa aag g-a gat gtc cat gcg gat act ttg gtg gaa tag atg aac aag gaa atc cta ttt gta aat cgt gct g-- -ta ttg gtg aat att gca act acc tac gta att ata gac ttg atc cat ttc ctc cat gca aa- --- cta tct aaa tgt aat taa tta --- --- --- --- --- --- --- --- tga ttt --- --- --- -tg atg ata atg tta -cc ata c-a tt- --- -at atc gct act tgg tta --g tgt att att cag tat g-a aga cct att aat aa- --- --- -tt act tat ctt ttg acg atc ttg tt- -at aat tat --- --- --- --- aat ata aaa --a tac tta tgg --- --- cat agt aac tca t-- --- --- --- aat tgc tg- acg cga taa att cgt aat aat ctg ttt tgt tca aat ttt tat aag ga- --- --a tct aca ggc ata aaa at- aaa aat ata att tat aat ata c-- tct tac agc gcg cca tca tga ata aca gca gtg aat tga ttg ctg tta tta atg gat tta gaa ata gtg gac gat ttt gtg ata tta gta tag tta tta atg atg aaa gga taa acg ctc ata aac tca tcc tat ctg gag cct ccg aat a-t ttt tcc att ctg ttt tcc aat aat ttt atc gat tct aat gaa tac gaa gtt aat cta agt cat tta gat tat caa agc gta aac gat ttg atc gat tac att tat ggg ata cct ttg agc cta act aac gat aac gtg aaa tat att ctt tca acc gct ga- --t ttt tta caa att gga tct gct att acg gag tgt gaa aat tac ata ctt aaa aat ctt tgt tct aga aac tgt atc gat ttc --- --- tac ata tac gct gat aaa tat aat aac aag aaa ata gaa tca gcg tcg ttt aac aca ata tta caa aat att ttg aga ctc atc aac gat gaa aac ttt aaa tac tta aca gag gaa tca atg ata aaa att tta agc gat gat atg tta aat ata aaa aat gag gat ttc gcc cca cta att ctc att aaa tgg tta gag agt act caa caa tca tgc acc gtc gag tta ctt aga tgc ctc aga ata tca -tt gct ttc ccc aca agt tat aaa atc act tta tag tca tcg act ggt tag ttc aat cta cga atg tat aac att ctt aaa caa tat agc att ctt gga tga atc att tcc tag ata cca tag cat cga gtt gat atc tat cgg tat aag taa ttc gca tga taa gat ttc cat aaa ctg cta caa tca t-a aaa aaa ata cat ggg aaa tga tat ctt cac gta gat ata ggt gta gtt tcg cag tgg ccg tcc tgg ata ata tta ttt ata tga tgg gtg gat atg atc agt ccc cgt ata gaa gtt caa agg tta tag cgt aca ata cat gta caa att ctt gga tat atg ata tac cag agc t-a aaa tat cct cgt tct aat tgt ggg gga ctg gct gat gac gaa tac att tat tgt ata ggc ggc ata cgc gat cag gat tca tcg ttg aca tct agt att gat aga tgg aag cca tca aaa cca tat tgg cag aag tat gct aaa atg cgc gaa cca aaa tgt gat atg ggg gtt gcg atg tta aac gga tta ata tat gtt ata ggt gga atc gtt aaa ggt g-- aca cgt gta ccg acg cac tag aga gtt tat cag aag atg gat gga tga agc atc aac gtc ttc caa taa aaa tgt cca ata tgt cga cga ttg ttc atg atg gca aga ttt ata tat ctg gag gtt aca aca ata gta gtg tag tta atg taa tat cga atc t-- agt cct tag cta taa ttc gat ata tga tga atg gac caa att atc atc att aaa cat tcc tag aat taa tcc cgc tct a

>JN654978.1:162098-164358_Vaccinia_virus_strain_Dryvax_clone_DPP11_complete_genome

ttg ctg act gca --- tgt a-c tat ttt tta cat tac tac att ggc tac ggc aga tat acc tac ttc gtc act gcc aca cgc tcc ggt aaa cgg --g gca tgt gac gag gga gaa tat ctt gat aag agg cat aat caa tgt tgt aat -ca gtg tcc acc tgg aga att tgc caa ggt cag atg ta- --g tgg tag tga taa cac aaa atg tga acg ctg ccc acc tca tac ata tac cgc aat ccc caa tta ctc t-a atg gat gtc atc aat gta gaa aat gcc caa cag gat cat ttg ata agg taa agt gta ccg gaa cac aga aca gta aat -gt tcg tgt ctt cct ggt tgg tat tgt gct act gat tct tca cag act gaa gat tgt cga gat -tg tat acc aaa aag g-a gat gtc cat gcg gat act ttg gtg gaa tag atg aac aag gaa atc cta ttt gta aat cgt gct g-- -ta ttg gtg aat att gca act acc tac gta att ata gac ttg atc cat ttt ctc cat gca aa- --- cta tct aaa tgt aat taa tta --- --- --- --- --- --- --- --- tga ttt --- --- --- -tg atg ata atg tta -cc ata c-a tt- --- -at atc gct act tgg tta --g tgt att att cag tat g-a aga cct att aat aa- --- --- -tt act tat ctt ttg acg atc ttg tt- -at aat tat --- --- --- --- aat ata aaa --- -ac tta tgg --- --- cat agt aac tta t-- --- --- --- aat tgc ta- aca cga taa att cgt aat aat ctg ttt tgt tca aat ttt tat aag ga- --- --a tct aca ggc ata aaa at- aaa aat ata atc tat aat ata c-- tct tac a-- acg cca tca tga ata gca gca gta aat taa ttg ctg tta tta atg gat tta gaa ata gtg gac gat ttt gtg ata tta ata tag tta tta atg atg aaa gga taa acg ctc ata aac tca tcc tat ctg gag cct ccg aat a-t ttt tcc att ctg ttt tcc aat aat ttt atc gat tct aat gaa tac gaa gtt aat cta agt cat tta gat tat caa agc gta aac gat ttg atc gat tat att tat ggg ata cct ttg agc cta act aac gat aac gtg aaa tat att ctt tca acc gct ga- --t ttt tta caa att gga tct gct att acg gag tgt gaa aat tac ata ctt aaa aat ctt tgt tct aaa aac tgt atc gat ttc --- --- tac ata tac gct gat aaa tat aat aac aag aaa ata gaa tca gcg tcg ttt aac aca ata tta caa aat att ttg aga ctc atc aac gat gaa aac ttt aaa tac tta aca gag gaa tca atg ata aaa att tta agc gat gat atg tta aat ata aaa aat gag gat ttt gca cca cta att ctc att aaa tgg tta gag agt act caa caa tca tgc acc gtc gag tta ctt aaa tgc ctc aga ata tca -tt gct ttc ccc aca agt tat aaa atc act tta tag tca tcg act ggt tag ttc aat cta cga atg tat aac att ctt aaa caa tat agc att ctt gga tga atc att tcc tag ata cca tag cat cga gtt gat atc tat cgg tat aag taa ttc gca tga taa gat ttc cat aaa ctg cta caa tca t-a aaa aaa ata cat ggg aaa tga tat ctt cac gta gat ata ggt gta gtt tcg cag tgg ccg tcc tgg ata ata tta tct ata tga tgg gtg gat atg atc agt ccc cgt ata gaa gtt caa agg tta tag cgt aca ata cat gta caa att ctt gga tat atg ata tac cag agc t-a aaa tat cct cgt tct aat tgt ggg gga ctg gct gat gac gaa tac att tat tgt ata ggc ggc ata cgc gat cag gat tca tcg ttg aca tct agt att gat aga tgg aag cca tca aaa cca tat tgg cag aag tat gct aaa atg cgc gaa cca aaa tgt gat atg ggg gtt gcg atg tta aac gga tta ata tat gtc atg ggt gga atc gtt aaa ggt g-- aca cgt gta ccg acg cac tag aga gtt tat cag aag atg gat gga tga agc atc aac gtc ttc caa taa aaa tgt cca ata tgt cga cga ttg ttc atg atg gca aga ttt ata tat ctg gag gtt aca aca ata gta gtg tag tta atg taa tat cga atc t-- agt cct tag cta taa ttc gat ata tga tga atg gac caa att atc atc att aaa tat tcc tag aat taa tcc cgc tct a

>AY313848.1:162476-164736_Vaccinia_virus_strain_Acambis_clone_3_complete_genome

ttg ctg act gta --- tgt a-c tat ttt tta cat tac tac att ggc tac ggc aga tat acc tac ttc gtc act gcc aca cgc tcc ggt aaa cgg --g gca tgt gac gag gga gaa tat ctt gat aag agg cat aat caa tgt tgt aat -cg gtg tcc acc tgg aga att tgc caa ggt cag atg ta- --g tgg tag tga taa cac aaa atg tga acg ctg ccc acc tca tac ata tac cgc aat ccc caa tta ctc t-a atg gat gtc atc aat gta gaa aat gcc caa cag gat cat ttg ata agg taa agt gta ccg gaa cac aga aca gta aat -gt tcg tgt ctt cct ggt tgg tat tgt gct act gat tct tca cag act gaa gat tgt cga gat -tg tat acc aaa aag g-a gat gtc cat gcg gat act ttg gtg gaa tag atg aac aag gaa atc cta ttt gta aat cgt gct g-- -ta ttg gtg aat att gca act acc tac gta att ata gac ttg atc cat ttt ctc cat gca aa- --- cta tct aaa tgt aat taa tta --- --- --- --- --- --- --- --- tga ttt --- --- --- -tg atg ata atg tta -cc ata c-a tt- --- -at atc gct act tgg tta --g tgt att att cag tat g-a aga cct att aat aa- --- --- -tt act tat ctt ttg acg atc ttg tt- -at aat tat --- --- --- --- aat ata aaa --- -ac tta tgg --- --- cat agt aac tca t-- --- --- --- aat tgc ta- aca cga taa att cgt aat aat ctg ttt tgt tca aat ttt tat aag ga- --- --a tct aca ggc ata aaa at- aaa aat ata att tat aat ata c-- tct tac a-- acg cca tca tga ata gca gca gta aat taa ttg ctg tta tta atg gat tta gaa ata gtg gac gat ttt gtg ata tta ata tag tta tta atg atg aaa gga taa acg ctc ata aac tca tcc tat ctg gag cct ccg aat a-t ttt tcc att ctg ttt tcc aat aat ttt atc gat tct aat gaa tac gaa gtt aat cta agt cat tta gat tat caa agc gta aac gat ttg atc gat tat att tat ggg ata cct ttg agc cta act aac gat aac gtg aaa tat att ctt tca acc gct ga- --t ttt tta caa att gga tct gct att acg gag tgt gaa aat tac ata ctt aaa aat ctt tgt tct aaa aac tgt atc gat ttc --- --- tac ata tac gct gat aaa tat aat aac aag aaa ata gaa tca gcg tcg ttt aac aca ata tta caa aat att ttg aga ctc atc aac gat gaa aac ttt aaa tac tta aca gag gaa tca atg ata aaa att tta agc gat gat atg tta aat ata aaa aat gag gat ttc gcc cca cta att ctc att aaa tgg tta gag agt act caa caa tca tgc acc gtc gag tta ctt aaa tgc ctc aga ata tca -tt gct ttc ccc aca agt tat aaa atc act tta tag tca tca act ggt tag ttc aat cta cga atg tat aac att ctt aaa caa tat agc att ctt gga tga atc att tcc tag ata cca tag cat cga gtt gat atc tat cgg tat aag taa ttc gca tga taa gat ttc cat aaa ctg cta caa tca t-a aaa aaa ata cat ggg aaa tga tat ctt cac gta gat ata ggt gta gtt tcg cag tgg ccg tcc tgg ata ata tta tct ata tga tgg gtg gat atg atc agt ccc cgt ata gaa gtt caa agg tta tag cgt aca ata cat gta caa att ctt gga tat atg ata tac cag agc t-a aaa tat cct cgt tct aat tgt ggg gga ctg gct gat gac gaa tac att tat tgt ata ggc ggc ata cgc gat cag gat tca tcg ttg aca tct agt att gat aga tgg aag cca tca aaa cca tat tgg cag aag tat gct aaa atg cgc gaa cca aaa tgt gat atg ggg gtt gcg atg tta aac gga tta ata tat gtc atg ggt gga atc gtt aaa ggt g-- aca cgt gta ccg acg cac tag aga gtt tat cag aag atg gat gga tga agc atc aac gtc ttc caa taa aaa tgt cca ata tgt cga cga ttg ttc atg atg gca aga ttt ata tat ctg gag gtt aca aca ata gta gtg tag tta atg taa tat cga atc t-- agt cct tag cta taa ttc gat ata tga tga atg gac caa att atc atc att aaa tat tcc tag aat taa tcc cgc tct a

>JX489138.1:159039-161292_Vaccinia_virus_strain_Tiantan_clone_TT11_complete_genome

ttg ctg act gta --- tgt a-c tat ttt tta cat tac tac att ggc tac ggc aga tat acc tac ttc gtc act gcc aca cgc tcc ggt aaa cgg --g gca tgt gac gag gga gaa tat ctt gat aag agg cat aat caa tgt tgt aat -cg gtg tcc acc tgg aga att tgc caa ggt tag atg ta- --a tgg taa cga taa cac aaa atg tga acg ctg ccc acc tca tac ata tac cgc aat ccc caa tta ctc t-a atg gat gtc atc aat gta gaa aat gcc caa ccg gat cat ttg ata agg taa agt gta ccg gaa cac aga aca gta aat -gt tcg tgt ctt cct ggt tgg tat tgc gct act gat tct tca cag act gaa gat tgt cga gat -tg tat acc aaa aag g-a gat gtc cat gcg gat act ttg gtg gaa tag atg aac aag gaa atc cta ttt gta aat cgt gtt g-- -tg ttg gtg aat att gcg act acc tac gta att ata gac ttg atc cat ttc ctc cat gca aa- --- cta tct aaa tgt aat taa tta --- --- --- --- --- --- --- --- tga ttt --- --- --- -tg atg ata atg tta -cc ata c-a tt- --- -at atc gct act tgg tta --g tgt att att cag tat g-a aga cct att aat aa- --- --- -tt act tat ctt ttg acg atc ttg tt- -at aat tat --- --- --- --- aat ata aaa --a tac tta tgg --- --- cat agt aac tca t-- --- --- --- aat tgc tg- acg cga taa att cgt aat aat ctg ttt tgt tca --- --- --- aag ga- --- --a tct aca ggc ata aaa at- aaa aat ata att tat aat ata c-- tct tac a-- acg cca tca tga ata gca gca gta aat tga ttg ctg tta tta atg gat tta gaa ata gtg gac gat ttt gtg ata tta ata tag tta tta atg atg aaa gga taa acg ctc ata aac tca tcc tat ctg gag cct ccg aat a-t ttt tcc att ctg ttt tcc aat aat ttt atc gat tct aat gaa tac gaa gtt aat cta agt cat tta gat tat caa agc gta aac gat ttg atc gat tac att tat ggg ata cct ttg agc cta act aac gat aac gtg aaa tat att ctt tca acc gct ga- --t ttt tta caa att gga tct gct att acg gag tgt gaa aaa tac ata ctt aaa aat ctt tgt tct aga aac tgt atc gat ttc --- --- tac ata tac gct gat aaa tat aat aac aag aaa ata gaa tca gcg tcg ttt aac aca ata tta cga aat att ttg aga ctc atc aac gat gaa aac ttt aaa tac tta aca gag gaa tca atg ata aaa att tta agc gat gat atg tta aat ata aaa aat gag gat ttc gcc cca cta att ctc att aaa tgg tta gag agt act caa caa cca tgc acc gtc gag tta ctt aaa tgc ctc aga ata tca -tt gct ttc ccc aca agt tat aaa atc act tta tag tca tcg act ggt tag ttc aat cta cga atg tat aac att ctt aaa caa tat agc att ctt gga tga atc att tcc tag ata cca tag cat cga gtt gat atc tat cgg tat aag taa ttc gcg tga taa aat ttc cat aaa ctg cta caa tca t-a aaa aaa ata cat ggg aaa tga tat ctt cac gta gat ata ggt gta gtt tcg cag tgg ccg tcc tgg ata ata tta tct ata tga tgg gtg gat atg atc agt ccc cgt ata gaa gtt caa agg tta tag cgt aca ata cat gta caa att ctt gga tat atg ata tac cag agc t-a aaa tat cct cgt tct aat tgt ggg gga ctg gct gat gac gaa tac att tat tgt ata ggc ggc ata cgc gat cag gat tca tcg ttg aca tct agt att gat aga tgg aag cca tca aaa cca tat tgg cag aag tat gct aaa atg cgc gaa cca aaa tgt gat atg ggg gtt gcg atg tta aac gga tta ata tat gtc atg ggt gga atc gtt aaa ggt g-- aca cgt gta ccg acg cac tag aga gtt tat cag aag atg gat gga tga agc atc aac gtc ttc caa taa aaa tgt cca ata tgt cga cga ttg ttc atg atg gca aga ttt ata tat ctg gag gtt aca aca ata gta gtg tag tta atg taa tat cga atc t-- agt cct tag cta taa tcc gat ata tga tga atg gac caa att atc atc att aaa cat tcc tag aat taa tcc cgc tct a

>JN654977.1:162046-164305_Vaccinia_virus_strain_Dryvax_clone_DPP10_complete_genome

ttg ctg act gta --- tgt a-c tat ttt tta cat tac tac att ggc tac ggc aga tat acc tac ttc gtc act gcc aca cgc tcc ggt aaa cgg --g gca tgt gac gag gga gaa tat ctt gat aag agg cat aat caa tgt tgt aat -ca gtg tcc acc tgg aga att tgc caa ggt tag atg ta- --g tgg tag tga taa cac aaa atg tga acg ctg ccc acc tca tac ata tac cac aat -cc caa tta ttc t-a atg gat gtc atc aat gta gaa aat gcc caa cag gat cat ttg ata agg taa agt gta ccg gaa cac aga aca gta aat -gt tcg tgt ctt cct ggt tgg tat tgc gct act gat tct tca cag act gaa gat tgt tga aat -tg tgt acc aaa aag g-a gat gtc cat gcg gat act ttg gtg gaa tag atg aac aag gaa atc cta ttt gta aat cgt gct g-- -ta ttg gtg aat att gca act acc tac gta att ata gac ttg atc cat ttt ctc cat gca aa- --- cta tct aaa tgt aat taa tta --- --- --- --- --- --- --- --- tga ttt --- --- --- -tg atg ata atg tta -cc ata c-a tt- --- -at atc gct act tgg tta --g tgt att att cag tat g-a aga cct att aat aa- --- --- -tt act tat ctt ttg acg atc ttg tt- -at aat tat --- --- --- --- aat ata aaa --- -ac tta tgg --- --- cat agt aac tta t-- --- --- --- aat tgc ta- aca cga taa att cgt aat aat ctg ttt tgt tca aat ttt tat aag ga- --- --a tct aca ggc ata aaa at- aaa aat ata atc tat aat ata c-- tct tac a-- acg cca tca tga ata gca gca gta aat taa ttg ctg tta tta atg gat tta gaa ata gtg gac gat ttt gtg ata tta ata tag tta tta atg atg aaa gga taa acg ctc ata aac tca tcc tat ctg gag cct ccg aat a-t ttt tcc att ctg ttt tcc aat aat ttt atc gat tct aat gaa tac gaa gtt aat cta agt cat tta gat tat caa agc gta aac gat ttg atc gat tat att tat ggg ata cct ttg agc cta act aac gat aac gtg aaa tat att ctt tca acc gct ga- --t ttt tta caa att gga tct gct att acg gag tgt gaa aat tac ata ctt aaa aat ctt tgt tct aaa aac tgt atc gat ttc --- --- tac ata tac gct gat aaa tat aat aac aag aaa ata gaa tca gcg tcg ttt aac aca ata tta caa aat att ttg aga ctc atc aac gat gaa aac ttt aaa tac tta aca gag gaa tca atg ata aaa att tta agc gat gat atg tta aat ata aaa aat gag gat ttc gcc cca cta att ctc att aaa tgg tta gag agt act caa caa tca tgc acc gtc gag tta ctt aga tgc ctc aga ata tca -tt gct ttc ccc aca agt tat aaa atc act tta tag tca tcg act ggt tag ttc aat cta cga atg tat aac att ctt aaa caa tat agc att ctt gga tga atc att tcc tag ata cca tag cat cga gtt gat atc tat cgg tat aag taa ttc gcg tga taa gat ttc cat aaa ctg cta caa tca t-a aaa aaa ata cat ggg aaa tga tat ctt cac gta gat ata ggt gta gtt tcg cag tgg ccg tcc tgg ata ata tta tct ata tga tgg gtg gat atg atc agt ccc cgt ata gaa gtt caa agg tta tag cgt aca ata cat gta caa att ctt gga tat atg ata tac cag agc t-a aaa tat cct cgt tct aat tgt ggg gga ctg gct gat gac gaa tac att tat tgt ata ggc ggc ata cgc gat cag gat tca tcg ttg aca tct agt att gat aaa tgg aag cca tca aaa cca tat tgg cag aag tat gct aaa atg cgc gaa cca aaa tgt gat atg ggg gtt gcg atg tta aac gga tta ata tat gtt ata ggt gga atc gtt aaa ggt g-- aca cgt gta ccg acg cac tag aga gtt tat cag aag atg gat gga tga agc atc aac gtc ttc caa taa aaa tgt cca ata tgt cga cga ttg ttc atg atg gca aga ttt ata tat ctg gag gtt aca aca ata gta gtg tag tta atg taa tat cga atc t-- agt cct tag cta taa ttc gat ata tga tga atg gac caa att atc atc att aaa tat tcc tag aat taa tcc cgc tct a

>MT227314.1:162428-164688_Vaccinia_virus_strain_Acambis/2019

ttg ctg act gta --- tgt a-c tat ttt tta cat tac tac att ggc tac ggc aga tat acc tac ttc gtc act gcc aca cgc tcc ggt aaa cgg --g gca tgt gac gag gga gaa tat ctt gat aag agg cat aat caa tgt tgt aat -ca gtg tcc acc tgg aga att tgc caa ggt cag atg ta- --g tgg tag tga taa cac aaa atg tga acg ctg ccc acc tca tac ata tac cgc aat ccc caa tta ctc t-a atg gat gtc atc aat gta gaa aat gcc caa cag gat cat ttg ata agg taa agt gta ccg gaa cac aga aca gta aat -gt tcg tgt ctt cct ggt tgg tat tgc gct act gat tct tca cag act gaa gat tgt tga aat -tg tgt acc aaa aag g-a gat gtc cat gcg gat act ttg gtg gaa tag atg aac aag gaa atc cta ttt gta aat cgt gct g-- -ta ttg gtg aat att gca act acc tac gta att ata gac ttg atc cat ttt ctc cat gca aa- --- cta tct aaa tgt aat taa tta --- --- --- --- --- --- --- --- tga ttt --- --- --- -tg atg ata atg tta -cc ata c-a tt- --- -at atc gct act tgg tta --g tgt att att cag tat g-a aga cct att aat aa- --- --- -tt act tat ctt ttg acg atc ttg tt- -at aat tat --- --- --- --- aat ata aaa --- -ac tta tgg --- --- cat agt aac tta t-- --- --- --- aat tgc ta- aca cga taa att cgt aat aat ctg ttt tgt tca aat ttt tat aag ga- --- --a tct aca ggc ata aaa at- aaa aat ata att tat aat ata c-- tct tac a-- acg cca tca tga ata gca gca gta aat taa ttg ctg tta tta atg gat tta gaa ata gtg gac gat ttt gtg ata tta ata tag tta tta atg atg aaa gga taa acg ctc ata aac tca tcc tat ctg gag cct ccg aat a-t ttt tcc att ctg ttt tcc aat aat ttt atc gat tct aat gaa tac gaa gtt aat cta agt cat tta gat tat caa agc gta aac gat ttg atc gat tat att tat ggg ata cct ttg agc cta act aac gat aac gtg aaa tat att ctt tca acc gct ga- --t ttt tta caa att gga tct gct att acg gag tgt gaa aat tac ata ctt aaa aat ctt tgt tct aaa aac tgt atc gat ttc --- --- tac ata tac gct gat aaa tat aat aac aag aaa ata gaa tca gcg tcg ttt aac aca ata tta caa aat att ttg aga ctc atc aac gat gaa aac ttt aaa tac tta aca gag gaa tca atg ata aaa att tta agc gat gat atg tta aat ata aaa aat gag gat ttc gcc cca cta att ctc att aaa tgg tta gag agt act caa caa tca tgc acc gtc gag tta ctt aaa tgc ctc aga ata tca -tt gct ttc ccc aca agt tat aaa atc act tta tag tca tca act ggt tag ttc aat cta cga atg tat aac att ctt aaa caa tat agc att ctt gga tga atc att tcc tag ata cca tag cat cga gtt gat atc tat cgg tat aag taa ttc gcg tga taa aat ttc cat aaa ctg cta caa tca t-a aaa aaa ata cat ggg aaa tga tat ctt cac gta gat ata ggt gta gtt tcg cag tgg ccg tcc tgg ata ata tta tct ata tga tgg gtg gat atg atc agt ccc cgt ata gaa gtt caa agg tta tag cgt aca ata cat gta caa att ctt gga tat atg ata tac cag agc t-a aaa tat cct cgt tct aat tgt ggg gga ctg gct gat gac gaa tac att tat tgt ata ggc ggc ata cgc gat cag gat tca tcg ttg aca tct agt att gat aga tgg aag cca tca aaa cca tat tgg cag aag tat gct aaa atg cgc gaa cca aaa tgt gat atg ggg gtt gcg atg tta aac gga tta ata tat gtc atg ggt gga atc gtt aaa ggt g-- aca cgt gta ccg acg cac tag aga gtt tat cag aag atg gat gga tga agc atc aac gtc ttc caa taa aaa tgt cca ata tgt cga cga ttg ttc atg atg gca aga ttt ata tat ctg gag gtt aca aca ata gta gtg tag tta atg taa tat cga atc t-- agt cct tag cta taa ttc gat ata tga tga atg gac caa att atc atc att aaa tat tcc tag aat taa tcc cgc tct a

>MN974381.1:162719-164979_Vaccinia_virus_strain_TNX-1200

ttg ctg act gta --- tgt a-c tat ttt tta cat tac tac att ggc tac ggc aga tat acc tac ttc gtc act gcc aca cgc tcc ggt aaa cgg --g gca tgt gac gag gga gaa tat ctt gat aag agg cat aat caa tgt tgt aat -ca gtg tcc acc tgg aga att tgc caa ggt cag atg ta- --g tgg tag tga taa cac aaa atg tga acg ctg ccc acc tca tac ata tac cgc aat ccc caa tta ctc t-a atg gat gtc atc aat gta gaa aat gcc caa cag gat cat ttg ata agg taa agt gta ccg gaa cac aga aca gta aat -gt tcg tgt ctt cct ggt tgg tat tgc gct act gat tct tca cag act gaa gat tgt tga aat -tg tgt acc aaa aag g-a gat gtc cat gcg gat act ttg gtg gaa tag atg aac aag gaa atc cta ttt gta aat cgt gct g-- -ta ttg gtg aat att gca act acc tac gta att ata gac ttg atc cat ttt ctc cat gca aa- --- cta tct aaa tgt aat taa tta --- --- --- --- --- --- --- --- tga ttt --- --- --- -tg atg ata atg tta -cc ata c-a tt- --- -at atc gct act tgg tta --g tgt att att cag tat g-a aga cct att aat aa- --- --- -tt act tat ctt ttg acg atc ttg tt- -at aat tat --- --- --- --- aat ata aaa --- -ac tta tgg --- --- cat agt aac tta t-- --- --- --- aat tgc ta- aca cga taa att cgt aat aat ctg ttt tgt tca aat ttt tat aag ga- --- --a tct aca ggc ata aaa at- aaa aat ata att tat aat ata c-- tct tac a-- acg cca tca tga ata gca gca gta aat taa ttg ctg tta tta atg gat tta gaa ata gtg gac gat ttt gtg ata tta ata tag tta tta atg atg aaa gga taa acg ctc ata aac tca tcc tat ctg gag cct ccg aat a-t ttt tcc att ctg ttt tcc aat aat ttt atc gat tct aat gaa tac gaa gtt aat cta agt cat tta gat tat caa agc gta aac gat ttg atc gat tat att tat ggg ata cct ttg agc cta act aac gat aac gtg aaa tat att ctt tca acc gct ga- --t ttt tta caa att gga tct gct att acg gag tgt gaa aat tac ata ctt aaa aat ctt tgt tct aaa aac tgt atc gat ttc --- --- tac ata tac gct gat aaa tat aat aac aag aaa ata gaa tca gcg tcg ttt aac aca ata tta caa aat att ttg aga ctc atc aac gat gaa aac ttt aaa tac tta aca gag gaa tca atg ata aaa att tta agc gat gat atg tta aat ata aaa aat gag gat ttc gcc cca cta att ctc att aaa tgg tta gag agt act caa caa tca tgc acc gtc gag tta ctt aaa tgc ctc aga ata tca -tt gct ttc ccc aca agt tat aaa atc act tta tag tca tca act ggt tag ttc aat cta cga atg tat aac att ctt aaa caa tat agc att ctt gga tga atc att tcc tag ata cca tag cat cga gtt gat atc tat cgg tat aag taa ttc gcg tga taa aat ttc cat aaa ctg cta caa tca t-a aaa aaa ata cat ggg aaa tga tat ctt cac gta gat ata ggt gta gtt tcg cag tgg ccg tcc tgg ata ata tta tct ata tga tgg gtg gat atg atc agt ccc cgt ata gaa gtt caa agg tta tag cgt aca ata cat gta caa att ctt gga tat atg ata tac cag agc t-a aaa tat cct cgt tct aat tgt ggg gga ctg gct gat gac gaa tac att tat tgt ata ggc ggc ata cgc gat cag gat tca tcg ttg aca tct agt att gat aga tgg aag cca tca aaa cca tat tgg cag aag tat gct aaa atg cgc gaa cca aaa tgt gat atg ggg gtt gcg atg tta aac gga tta ata tat gtc atg ggt gga atc gtt aaa ggt g-- aca cgt gta ccg acg cac tag aga gtt tat cag aag atg gat gga tga agc atc aac gtc ttc caa taa aaa tgt cca ata tgt cga cga ttg ttc atg atg gca aga ttt ata tat ctg gag gtt aca aca ata gta gtg tag tta atg taa tat cga atc t-- agt cct tag cta taa ttc gat ata tga tga atg gac caa att atc atc att aaa tat tcc tag aat taa tcc cgc tct a

>MN974380.1:162665-164925_Vaccinia_virus_strain_VACV_li

ttg ctg act gta --- tgt a-c tat ttt tta cat tac tac att ggc tac ggc aga tat acc tac ttc gtc act gcc aca cgc tcc ggt aaa cgg --g gca tgt gac gag gga gaa tat ctt gat aag agg cat aat caa tgt tgt aat -ca gtg tcc acc tgg aga att tgc caa ggt cag atg ta- --g tgg tag tga taa cac aaa atg tga acg ctg ccc acc tca tac ata tac cgc aat ccc caa tta ctc t-a atg gat gtc atc aat gta gaa aat gcc caa cag gat cat ttg ata agg taa agt gta ccg gaa cac aga aca gta aat -gt tcg tgt ctt cct ggt tgg tat tgc gct act gat tct tca cag act gaa gat tgt tga aat -tg tgt acc aaa aag g-a gat gtc cat gcg gat act ttg gtg gaa tag atg aac aag gaa atc cta ttt gta aat cgt gct g-- -ta ttg gtg aat att gca act acc tac gta att ata gac ttg atc cat ttt ctc cat gca aa- --- cta tct aaa tgt aat taa tta --- --- --- --- --- --- --- --- tga ttt --- --- --- -tg atg ata atg tta -cc ata c-a tt- --- -at atc gct act tgg tta --g tgt att att cag tat g-a aga cct att aat aa- --- --- -tt act tat ctt ttg acg atc ttg tt- -at aat tat --- --- --- --- aat ata aaa --- -ac tta tgg --- --- cat agt aac tta t-- --- --- --- aat tgc ta- aca cga taa att cgt aat aat ctg ttt tgt tca aat ttt tat aag ga- --- --a tct aca ggc ata aaa at- aaa aat ata att tat aat ata c-- tct tac a-- acg cca tca tga ata gca gca gta aat taa ttg ctg tta tta atg gat tta gaa ata gtg gac gat ttt gtg ata tta ata tag tta tta atg atg aaa gga taa acg ctc ata aac tca tcc tat ctg gag cct ccg aat a-t ttt tcc att ctg ttt tcc aat aat ttt atc gat tct aat gaa tac gaa gtt aat cta agt cat tta gat tat caa agc gta aac gat ttg atc gat tat att tat ggg ata cct ttg agc cta act aac gat aac gtg aaa tat att ctt tca acc gct ga- --t ttt tta caa att gga tct gct att acg gag tgt gaa aat tac ata ctt aaa aat ctt tgt tct aaa aac tgt atc gat ttc --- --- tac ata tac gct gat aaa tat aat aac aag aaa ata gaa tca gcg tcg ttt aac aca ata tta caa aat att ttg aga ctc atc aac gat gaa aac ttt aaa tac tta aca gag gaa tca atg ata aaa att tta agc gat gat atg tta aat ata aaa aat gag gat ttc gcc cca cta att ctc att aaa tgg tta gag agt act caa caa tca tgc acc gtc gag tta ctt aaa tgc ctc aga ata tca -tt gct ttc ccc aca agt tat aaa atc act tta tag tca tca act ggt tag ttc aat cta cga atg tat aac att ctt aaa caa tat agc att ctt gga tga atc att tcc tag ata cca tag cat cga gtt gat atc tat cgg tat aag taa ttc gcg tga taa aat ttc cat aaa ctg cta caa tca t-a aaa aaa ata cat ggg aaa tga tat ctt cac gta gat ata ggt gta gtt tcg cag tgg ccg tcc tgg ata ata tta tct ata tga tgg gtg gat atg atc agt ccc cgt ata gaa gtt caa agg tta tag cgt aca ata cat gta caa att ctt gga tat atg ata tac cag agc t-a aaa tat cct cgt tct aat tgt ggg gga ctg gct gat gac gaa tac att tat tgt ata ggc ggc ata cgc gat cag gat tca tcg ttg aca tct agt att gat aga tgg aag cca tca aaa cca tat tgg cag aag tat gct aaa atg cgc gaa cca aaa tgt gat atg ggg gtt gcg atg tta aac gga tta ata tat gtc atg ggt gga atc gtt aaa ggt g-- aca cgt gta ccg acg cac tag aga gtt tat cag aag atg gat gga tga agc atc aac gtc ttc caa taa aaa tgt cca ata tgt cga cga ttg ttc atg atg gca aga ttt ata tat ctg gag gtt aca aca ata gta gtg tag tta atg taa tat cga atc t-- agt cct tag cta taa ttc gat ata tga tga atg gac caa att atc atc att aaa tat tcc tag aat taa tcc cgc tct a

>KX781953.1:161670-163923_Vaccinia_virus_strain_L-IVP_clone_14

ttg ctg act gta --- tgt a-c tat ttt tta cat tac tac att ggc tac ggc aga tat acc tac tcc gcc act gcc aca cgc tcc ggt aaa cgg --g gca tgt gac gag gga gaa tat ctt gat aag agg cat aat caa tgt tgt aat -ca gtg tcc acc tgg aga att tgc caa ggt cag atg ta- --g tgg tag tga taa cac aaa atg tga acg ctg ccc acc tca tac ata tac cgc aat ccc caa tta ctc t-a atg gat gtc atc aat gta gaa aat gcc caa ccg gat cat ttg ata agg taa agt gta ccg gaa cac aga aca gta aat -gt tcg tgt ctt cct ggt tgg tat tgc gct act gat tct tca cag act gaa gat tgt cga gat -tg tat acc aaa aag gaa gat gtc cat gcg gat act ttg gtg gaa tag atg aac aag gaa atc cta ttt gta aat cgt gtt g-- -tg ttg gtg aat att gcg act acc tac gta att ata gac ttg atc cat ttc ctc cat gca aa- --- cta tct aaa tgt aat taa tta --- --- --- --- --- --- --- --- tga ttt --- --- --- -tg atg ata atg tta -cc ata c-a tt- --- -at atc gct act tgg tta --g tgt att att cag tat g-a aga cct att aat aa- --- --- -tt act tat ctt ttg acg atc ttg tt- -at aat tat --- --- --- --- aat ata aaa --a tac tta tgg --- --- cat agt aac tca t-- --- --- --- aat tgc tg- acg cga taa att cgt aat aat ctg ttt tgt tca --- --- --- aag ga- --- --a tct aca ggc ata aaa at- aaa aat ata att tat aat a-a c-- tct tac a-- acg cca tca tga ata aca gca gtg aat tga ttg ctg tta tta atg gat tta gaa ata gtg gac gat ttt gtg ata tta gta tag tta tta atg atg aaa gga taa acg ctc ata aac tca tcc tat ctg gag cct ccg aat a-t ttt tcc att ctg ttt tcc aat aat ttt atc gat tct aat gaa tac gaa gtt aat cta agt cat tta gat tat caa agc gta aac gat ttg atc gat tat att tat ggg ata cct ttg agc cta act aac gat aac gtg aaa tat att ctt tca acc gct ga- --t ttt tta caa att gga tct gcc att act gag tgc gaa aaa tac ata ctt aaa aat ctt tgt tct aga aac tgt atc gat ttc --- --- tac ata tac gct gat aaa tat aat aac aag aaa ata gaa tca gcg tcg ttt aac aca ata tta caa aat att ttg aga ctc atc aac gat gaa aac ttt aaa tac tta aca gag gaa tca atg ata aaa att tta agc gat gat atg tta aat ata aaa aat gag gat ttc gcc cca cta att ctc att aaa tgg tta gag agt act caa caa cca tgc acc gtc gag tta ctt aaa tgc ctc aga ata tca -tt gct ttc ccc aca agt tat aaa atc act tta tag tca tcg act ggt tag ttc aat cta cga atg tat aac att ctt aaa caa tat agc att ctt gga tga atc att tcc tag ata cca tag cat cga gtt gat atc tat cgg tat aag taa ttc gca tga taa aat ttc cat aaa ctg cta caa tca t-a aaa aaa ata cat ggg aaa tga tat ctt cac gta gat ata ggt gta gtt tcg cag tgg ccg tcc tgg ata ata tta ttt ata tga tgg gtg gat atg atc agt ccc cgt ata gaa gtt caa agg tta tag cgt aca ata cat gta caa att ctt gga tat atg ata tac cag agc t-a aaa tat cct cgt tct aat tgt ggg gga ctg gct gat gac gaa tac att tat tgt ata ggc ggc ata cgc gat cag gat tca tcg ttg aca tct agt att gat aga tgg aag cca tca aaa cca tat tgg cag aag tat gct aaa atg cgc gaa cca aaa tgt gat atg ggg gtt gcg atg tta aac gga tta ata tat gtt ata ggt gga atc gtt aaa ggt g-- aca cgt gta ccg acg cac tag aga gtt tat cag aag atg gat gga tga agc atc aac gtc ttc caa taa aaa tgt cca ata tgt cga cga ttg ttc atg atg gca aga ttt ata tat ctg gag gtt aca aca ata gta gtg tag tta atg taa tat cga atc t-- agt cct tag cta taa ttc gat ata tga tga atg gac caa att atc atc att aaa cat tcc tag aat taa tcc cgc tct a

>MW960424.1:163277-165537_Vector_synVACV-Delta1Delta3-6_complete_sequence

ttg ctg act gta --- tgt a-c tat ttt tta cat tac tac att ggc tac ggc aga tat acc tac ttc gtc act gcc aca cgc tcc ggt aaa cgg --g gca tgt gac gag gga gaa tat ctt gat aag agg cat aat caa tgt tgt aat -ca gtg tcc acc tgg aga att tgc caa ggt cag atg ta- --g tgg tag tga taa cac aaa atg tga acg ctg ccc acc tca tac ata tac cgc aat ccc caa tta ctc t-a atg gat gtc atc aat gta gaa aat gcc caa cag gat cat ttg ata agg taa agt gta ccg gaa cac aga aca gta aat -gt tcg tgt ctt cct ggt tgg tat tgc gct act gat tct tca cag act gaa gat tgt tga aat -tg tgt acc aaa aag g-a gat gtc cat gcg gat act ttg gtg gaa tag atg aac aag gaa atc cta ttt gta aat cgt gct g-- -ta ttg gtg aat att gca act acc tac gta att ata gac ttg atc cat ttt ctc cat gca aa- --- cta tct aaa tgt aat taa tta --- --- --- --- --- --- --- --- tga ttt --- --- --- -tg atg ata atg tta -cc ata c-a tt- --- -at atc gct act tgg tta --g tgt att att cag tat g-a aga cct att aat aa- --- --- -tt act tat ctt ttg acg atc ttg tt- -at aat tat --- --- --- --- aat ata aaa --- -ac tta tgg --- --- cat agt aac tta t-- --- --- --- aat tgc ta- aca cga taa att cgt aat aat ctg ttt tgt tca aat ttt tat aag ga- --- --a tct aca ggc ata aaa at- aaa aat ata att tat aat ata c-- tct tac a-- acg cca tca tga ata gca gca gta aat taa ttg ctg tta tta atg gat tta gaa ata gtg gac gat ttt gtg ata tta ata tag tta tta atg atg aaa gga taa acg ctc ata aac tca tcc tat ctg gag cct ccg aat a-t ttt tcc att ctg ttt tcc aat aat ttt atc gat tct aat gaa tac gaa gtt aat cta agt cat tta gat tat caa agc gta aac gat ttg atc gat tat att tat ggg ata cct ttg agc cta act aac gat aac gtg aaa tat att ctt tca acc gct ga- --t ttt tta caa att gga tct gct att acg gag tgt gaa aat tac ata ctt aaa aat ctt tgt tct aaa aac tgt atc gat ttc --- --- tac ata tac gct gat aaa tat aat aac aag aaa ata gaa tca gcg tcg ttt aac aca ata tta caa aat att ttg aga ctc atc aac gat gaa aac ttt aaa tac tta aca gag gaa tca atg ata aaa att tta agc gat gat atg tta aat ata aaa aat gag gat ttc gcc cca cta att ctc att aaa tgg tta gag agt act caa caa tca tgc acc gtc gag tta ctt aaa tgc ctc aga ata tca -tt gct ttc ccc aca agt tat aaa atc act tta tag tca tca act ggt tag ttc aat cta cga atg tat aac att ctt aaa caa tat agc att ctt gga tga atc att tcc tag ata cca tag cat cga gtt gat atc tat cgg tat aag taa ttc gcg tga taa aat ttc cat aaa ctg cta caa tca t-a aaa aaa ata cat ggg aaa tga tat ctt cac gta gat ata ggt gta gtt tcg cag tgg ccg tcc tgg ata ata tta tct ata tga tgg gtg gat atg atc agt ccc cgt ata gaa gtt caa agg tta tag cgt aca ata cat gta caa att ctt gga tat atg ata tac cag agc t-a aaa tat cct cgt tct aat tgt ggg gga ctg gct gat gac gaa tac att tat tgt ata ggc ggc ata cgc gat cag gat tca tcg ttg aca tct agt att gat aga tgg aag cca tca aaa cca tat tgg cag aag tat gct aaa atg cgc gaa cca aaa tgt gat atg ggg gtt gcg atg tta aac gga tta ata tat gtc atg ggt gga atc gtt aaa ggt g-- aca cgt gta ccg acg cac tag aga gtt tat cag aag atg gat gga tga agc atc aac gtc ttc caa taa aaa tgt cca ata tgt cga cga ttg ttc atg atg gca aga ttt ata tat ctg gag gtt aca aca ata gta gtg tag tta atg taa tat cga atc t-- agt cct tag cta taa ttc gat ata tga tga atg gac caa att atc atc att aaa tat tcc tag aat taa tcc cgc tct a

>MW960423.1:163278-165538_Vector_synVACV-Delta3-6_complete_sequence

ttg ctg act gta --- tgt a-c tat ttt tta cat tac tac att ggc tac ggc aga tat acc tac ttc gtc act gcc aca cgc tcc ggt aaa cgg --g gca tgt gac gag gga gaa tat ctt gat aag agg cat aat caa tgt tgt aat -ca gtg tcc acc tgg aga att tgc caa ggt cag atg ta- --g tgg tag tga taa cac aaa atg tga acg ctg ccc acc tca tac ata tac cgc aat ccc caa tta ctc t-a atg gat gtc atc aat gta gaa aat gcc caa cag gat cat ttg ata agg taa agt gta ccg gaa cac aga aca gta aat -gt tcg tgt ctt cct ggt tgg tat tgc gct act gat tct tca cag act gaa gat tgt tga aat -tg tgt acc aaa aag g-a gat gtc cat gcg gat act ttg gtg gaa tag atg aac aag gaa atc cta ttt gta aat cgt gct g-- -ta ttg gtg aat att gca act acc tac gta att ata gac ttg atc cat ttt ctc cat gca aa- --- cta tct aaa tgt aat taa tta --- --- --- --- --- --- --- --- tga ttt --- --- --- -tg atg ata atg tta -cc ata c-a tt- --- -at atc gct act tgg tta --g tgt att att cag tat g-a aga cct att aat aa- --- --- -tt act tat ctt ttg acg atc ttg tt- -at aat tat --- --- --- --- aat ata aaa --- -ac tta tgg --- --- cat agt aac tta t-- --- --- --- aat tgc ta- aca cga taa att cgt aat aat ctg ttt tgt tca aat ttt tat aag ga- --- --a tct aca ggc ata aaa at- aaa aat ata att tat aat ata c-- tct tac a-- acg cca tca tga ata gca gca gta aat taa ttg ctg tta tta atg gat tta gaa ata gtg gac gat ttt gtg ata tta ata tag tta tta atg atg aaa gga taa acg ctc ata aac tca tcc tat ctg gag cct ccg aat a-t ttt tcc att ctg ttt tcc aat aat ttt atc gat tct aat gaa tac gaa gtt aat cta agt cat tta gat tat caa agc gta aac gat ttg atc gat tat att tat ggg ata cct ttg agc cta act aac gat aac gtg aaa tat att ctt tca acc gct ga- --t ttt tta caa att gga tct gct att acg gag tgt gaa aat tac ata ctt aaa aat ctt tgt tct aaa aac tgt atc gat ttc --- --- tac ata tac gct gat aaa tat aat aac aag aaa ata gaa tca gcg tcg ttt aac aca ata tta caa aat att ttg aga ctc atc aac gat gaa aac ttt aaa tac tta aca gag gaa tca atg ata aaa att tta agc gat gat atg tta aat ata aaa aat gag gat ttc gcc cca cta att ctc att aaa tgg tta gag agt act caa caa tca tgc acc gtc gag tta ctt aaa tgc ctc aga ata tca -tt gct ttc ccc aca agt tat aaa atc act tta tag tca tca act ggt tag ttc aat cta cga atg tat aac att ctt aaa caa tat agc att ctt gga tga atc att tcc tag ata cca tag cat cga gtt gat atc tat cgg tat aag taa ttc gcg tga taa aat ttc cat aaa ctg cta caa tca t-a aaa aaa ata cat ggg aaa tga tat ctt cac gta gat ata ggt gta gtt tcg cag tgg ccg tcc tgg ata ata tta tct ata tga tgg gtg gat atg atc agt ccc cgt ata gaa gtt caa agg tta tag cgt aca ata cat gta caa att ctt gga tat atg ata tac cag agc t-a aaa tat cct cgt tct aat tgt ggg gga ctg gct gat gac gaa tac att tat tgt ata ggc ggc ata cgc gat cag gat tca tcg ttg aca tct agt att gat aga tgg aag cca tca aaa cca tat tgg cag aag tat gct aaa atg cgc gaa cca aaa tgt gat atg ggg gtt gcg atg tta aac gga tta ata tat gtc atg ggt gga atc gtt aaa ggt g-- aca cgt gta ccg acg cac tag aga gtt tat cag aag atg gat gga tga agc atc aac gtc ttc caa taa aaa tgt cca ata tgt cga cga ttg ttc atg atg gca aga ttt ata tat ctg gag gtt aca aca ata gta gtg tag tta atg taa tat cga atc t-- agt cct tag cta taa ttc gat ata tga tga atg gac caa att atc atc att aaa tat tcc tag aat taa tcc cgc tct a

>MW960422.1:163292-165552_Vector_synVACV-Delta1-3_complete_sequence

ttg ctg act gta --- tgt a-c tat ttt tta cat tac tac att ggc tac ggc aga tat acc tac ttc gtc act gcc aca cgc tcc ggt aaa cgg --g gca tgt gac gag gga gaa tat ctt gat aag agg cat aat caa tgt tgt aat -ca gtg tcc acc tgg aga att tgc caa ggt cag atg ta- --g tgg tag tga taa cac aaa atg tga acg ctg ccc acc tca tac ata tac cgc aat ccc caa tta ctc t-a atg gat gtc atc aat gta gaa aat gcc caa cag gat cat ttg ata agg taa agt gta ccg gaa cac aga aca gta aat -gt tcg tgt ctt cct ggt tgg tat tgc gct act gat tct tca cag act gaa gat tgt tga aat -tg tgt acc aaa aag g-a gat gtc cat gcg gat act ttg gtg gaa tag atg aac aag gaa atc cta ttt gta aat cgt gct g-- -ta ttg gtg aat att gca act acc tac gta att ata gac ttg atc cat ttt ctc cat gca aa- --- cta tct aaa tgt aat taa tta --- --- --- --- --- --- --- --- tga ttt --- --- --- -tg atg ata atg tta -cc ata c-a tt- --- -at atc gct act tgg tta --g tgt att att cag tat g-a aga cct att aat aa- --- --- -tt act tat ctt ttg acg atc ttg tt- -at aat tat --- --- --- --- aat ata aaa --- -ac tta tgg --- --- cat agt aac tta t-- --- --- --- aat tgc ta- aca cga taa att cgt aat aat ctg ttt tgt tca aat ttt tat aag ga- --- --a tct aca ggc ata aaa at- aaa aat ata att tat aat ata c-- tct tac a-- acg cca tca tga ata gca gca gta aat taa ttg ctg tta tta atg gat tta gaa ata gtg gac gat ttt gtg ata tta ata tag tta tta atg atg aaa gga taa acg ctc ata aac tca tcc tat ctg gag cct ccg aat a-t ttt tcc att ctg ttt tcc aat aat ttt atc gat tct aat gaa tac gaa gtt aat cta agt cat tta gat tat caa agc gta aac gat ttg atc gat tat att tat ggg ata cct ttg agc cta act aac gat aac gtg aaa tat att ctt tca acc gct ga- --t ttt tta caa att gga tct gct att acg gag tgt gaa aat tac ata ctt aaa aat ctt tgt tct aaa aac tgt atc gat ttc --- --- tac ata tac gct gat aaa tat aat aac aag aaa ata gaa tca gcg tcg ttt aac aca ata tta caa aat att ttg aga ctc atc aac gat gaa aac ttt aaa tac tta aca gag gaa tca atg ata aaa att tta agc gat gat atg tta aat ata aaa aat gag gat ttc gcc cca cta att ctc att aaa tgg tta gag agt act caa caa tca tgc acc gtc gag tta ctt aaa tgc ctc aga ata tca -tt gct ttc ccc aca agt tat aaa atc act tta tag tca tca act ggt tag ttc aat cta cga atg tat aac att ctt aaa caa tat agc att ctt gga tga atc att tcc tag ata cca tag cat cga gtt gat atc tat cgg tat aag taa ttc gcg tga taa aat ttc cat aaa ctg cta caa tca t-a aaa aaa ata cat ggg aaa tga tat ctt cac gta gat ata ggt gta gtt tcg cag tgg ccg tcc tgg ata ata tta tct ata tga tgg gtg gat atg atc agt ccc cgt ata gaa gtt caa agg tta tag cgt aca ata cat gta caa att ctt gga tat atg ata tac cag agc t-a aaa tat cct cgt tct aat tgt ggg gga ctg gct gat gac gaa tac att tat tgt ata ggc ggc ata cgc gat cag gat tca tcg ttg aca tct agt att gat aga tgg aag cca tca aaa cca tat tgg cag aag tat gct aaa atg cgc gaa cca aaa tgt gat atg ggg gtt gcg atg tta aac gga tta ata tat gtc atg ggt gga atc gtt aaa ggt g-- aca cgt gta ccg acg cac tag aga gtt tat cag aag atg gat gga tga agc atc aac gtc ttc caa taa aaa tgt cca ata tgt cga cga ttg ttc atg atg gca aga ttt ata tat ctg gag gtt aca aca ata gta gtg tag tta atg taa tat cga atc t-- agt cct tag cta taa ttc gat ata tga tga atg gac caa att atc atc att aaa tat tcc tag aat taa tcc cgc tct a

>MW960421.1:163285-165545_Vector_synVACV-Delta5-6_complete_sequence

ttg ctg act gta --- tgt a-c tat ttt tta cat tac tac att ggc tac ggc aga tat acc tac ttc gtc act gcc aca cgc tcc ggt aaa cgg --g gca tgt gac gag gga gaa tat ctt gat aag agg cat aat caa tgt tgt aat -ca gtg tcc acc tgg aga att tgc caa ggt cag atg ta- --g tgg tag tga taa cac aaa atg tga acg ctg ccc acc tca tac ata tac cgc aat ccc caa tta ctc t-a atg gat gtc atc aat gta gaa aat gcc caa cag gat cat ttg ata agg taa agt gta ccg gaa cac aga aca gta aat -gt tcg tgt ctt cct ggt tgg tat tgc gct act gat tct tca cag act gaa gat tgt tga aat -tg tgt acc aaa aag g-a gat gtc cat gcg gat act ttg gtg gaa tag atg aac aag gaa atc cta ttt gta aat cgt gct g-- -ta ttg gtg aat att gca act acc tac gta att ata gac ttg atc cat ttt ctc cat gca aa- --- cta tct aaa tgt aat taa tta --- --- --- --- --- --- --- --- tga ttt --- --- --- -tg atg ata atg tta -cc ata c-a tt- --- -at atc gct act tgg tta --g tgt att att cag tat g-a aga cct att aat aa- --- --- -tt act tat ctt ttg acg atc ttg tt- -at aat tat --- --- --- --- aat ata aaa --- -ac tta tgg --- --- cat agt aac tta t-- --- --- --- aat tgc ta- aca cga taa att cgt aat aat ctg ttt tgt tca aat ttt tat aag ga- --- --a tct aca ggc ata aaa at- aaa aat ata att tat aat ata c-- tct tac a-- acg cca tca tga ata gca gca gta aat taa ttg ctg tta tta atg gat tta gaa ata gtg gac gat ttt gtg ata tta ata tag tta tta atg atg aaa gga taa acg ctc ata aac tca tcc tat ctg gag cct ccg aat a-t ttt tcc att ctg ttt tcc aat aat ttt atc gat tct aat gaa tac gaa gtt aat cta agt cat tta gat tat caa agc gta aac gat ttg atc gat tat att tat ggg ata cct ttg agc cta act aac gat aac gtg aaa tat att ctt tca acc gct ga- --t ttt tta caa att gga tct gct att acg gag tgt gaa aat tac ata ctt aaa aat ctt tgt tct aaa aac tgt atc gat ttc --- --- tac ata tac gct gat aaa tat aat aac aag aaa ata gaa tca gcg tcg ttt aac aca ata tta caa aat att ttg aga ctc atc aac gat gaa aac ttt aaa tac tta aca gag gaa tca atg ata aaa att tta agc gat gat atg tta aat ata aaa aat gag gat ttc gcc cca cta att ctc att aaa tgg tta gag agt act caa caa tca tgc acc gtc gag tta ctt aaa tgc ctc aga ata tca -tt gct ttc ccc aca agt tat aaa atc act tta tag tca tca act ggt tag ttc aat cta cga atg tat aac att ctt aaa caa tat agc att ctt gga tga atc att tcc tag ata cca tag cat cga gtt gat atc tat cgg tat aag taa ttc gcg tga taa aat ttc cat aaa ctg cta caa tca t-a aaa aaa ata cat ggg aaa tga tat ctt cac gta gat ata ggt gta gtt tcg cag tgg ccg tcc tgg ata ata tta tct ata tga tgg gtg gat atg atc agt ccc cgt ata gaa gtt caa agg tta tag cgt aca ata cat gta caa att ctt gga tat atg ata tac cag agc t-a aaa tat cct cgt tct aat tgt ggg gga ctg gct gat gac gaa tac att tat tgt ata ggc ggc ata cgc gat cag gat tca tcg ttg aca tct agt att gat aga tgg aag cca tca aaa cca tat tgg cag aag tat gct aaa atg cgc gaa cca aaa tgt gat atg ggg gtt gcg atg tta aac gga tta ata tat gtc atg ggt gga atc gtt aaa ggt g-- aca cgt gta ccg acg cac tag aga gtt tat cag aag atg gat gga tga agc atc aac gtc ttc caa taa aaa tgt cca ata tgt cga cga ttg ttc atg atg gca aga ttt ata tat ctg gag gtt aca aca ata gta gtg tag tta atg taa tat cga atc t-- agt cct tag cta taa ttc gat ata tga tga atg gac caa att atc atc att aaa tat tcc tag aat taa tcc cgc tct a

>MW960420.1:163293-165553_Vector_synVACV-Delta6_complete_sequence

ttg ctg act gta --- tgt a-c tat ttt tta cat tac tac att ggc tac ggc aga tat acc tac ttc gtc act gcc aca cgc tcc ggt aaa cgg --g gca tgt gac gag gga gaa tat ctt gat aag agg cat aat caa tgt tgt aat -ca gtg tcc acc tgg aga att tgc caa ggt cag atg ta- --g tgg tag tga taa cac aaa atg tga acg ctg ccc acc tca tac ata tac cgc aat ccc caa tta ctc t-a atg gat gtc atc aat gta gaa aat gcc caa cag gat cat ttg ata agg taa agt gta ccg gaa cac aga aca gta aat -gt tcg tgt ctt cct ggt tgg tat tgc gct act gat tct tca cag act gaa gat tgt tga aat -tg tgt acc aaa aag g-a gat gtc cat gcg gat act ttg gtg gaa tag atg aac aag gaa atc cta ttt gta aat cgt gct g-- -ta ttg gtg aat att gca act acc tac gta att ata gac ttg atc cat ttt ctc cat gca aa- --- cta tct aaa tgt aat taa tta --- --- --- --- --- --- --- --- tga ttt --- --- --- -tg atg ata atg tta -cc ata c-a tt- --- -at atc gct act tgg tta --g tgt att att cag tat g-a aga cct att aat aa- --- --- -tt act tat ctt ttg acg atc ttg tt- -at aat tat --- --- --- --- aat ata aaa --- -ac tta tgg --- --- cat agt aac tta t-- --- --- --- aat tgc ta- aca cga taa att cgt aat aat ctg ttt tgt tca aat ttt tat aag ga- --- --a tct aca ggc ata aaa at- aaa aat ata att tat aat ata c-- tct tac a-- acg cca tca tga ata gca gca gta aat taa ttg ctg tta tta atg gat tta gaa ata gtg gac gat ttt gtg ata tta ata tag tta tta atg atg aaa gga taa acg ctc ata aac tca tcc tat ctg gag cct ccg aat a-t ttt tcc att ctg ttt tcc aat aat ttt atc gat tct aat gaa tac gaa gtt aat cta agt cat tta gat tat caa agc gta aac gat ttg atc gat tat att tat ggg ata cct ttg agc cta act aac gat aac gtg aaa tat att ctt tca acc gct ga- --t ttt tta caa att gga tct gct att acg gag tgt gaa aat tac ata ctt aaa aat ctt tgt tct aaa aac tgt atc gat ttc --- --- tac ata tac gct gat aaa tat aat aac aag aaa ata gaa tca gcg tcg ttt aac aca ata tta caa aat att ttg aga ctc atc aac gat gaa aac ttt aaa tac tta aca gag gaa tca atg ata aaa att tta agc gat gat atg tta aat ata aaa aat gag gat ttc gcc cca cta att ctc att aaa tgg tta gag agt act caa caa tca tgc acc gtc gag tta ctt aaa tgc ctc aga ata tca -tt gct ttc ccc aca agt tat aaa atc act tta tag tca tca act ggt tag ttc aat cta cga atg tat aac att ctt aaa caa tat agc att ctt gga tga atc att tcc tag ata cca tag cat cga gtt gat atc tat cgg tat aag taa ttc gcg tga taa aat ttc cat aaa ctg cta caa tca t-a aaa aaa ata cat ggg aaa tga tat ctt cac gta gat ata ggt gta gtt tcg cag tgg ccg tcc tgg ata ata tta tct ata tga tgg gtg gat atg atc agt ccc cgt ata gaa gtt caa agg tta tag cgt aca ata cat gta caa att ctt gga tat atg ata tac cag agc t-a aaa tat cct cgt tct aat tgt ggg gga ctg gct gat gac gaa tac att tat tgt ata ggc ggc ata cgc gat cag gat tca tcg ttg aca tct agt att gat aga tgg aag cca tca aaa cca tat tgg cag aag tat gct aaa atg cgc gaa cca aaa tgt gat atg ggg gtt gcg atg tta aac gga tta ata tat gtc atg ggt gga atc gtt aaa ggt g-- aca cgt gta ccg acg cac tag aga gtt tat cag aag atg gat gga tga agc atc aac gtc ttc caa taa aaa tgt cca ata tgt cga cga ttg ttc atg atg gca aga ttt ata tat ctg gag gtt aca aca ata gta gtg tag tta atg taa tat cga atc t-- agt cct tag cta taa ttc gat ata tga tga atg gac caa att atc atc att aaa tat tcc tag aat taa tcc cgc tct a

>MW960419.1:163281-165541_Vector_synVACV-SFV_complete_sequence

ttg ctg act gta --- tgt a-c tat ttt tta cat tac tac att ggc tac ggc aga tat acc tac ttc gtc act gcc aca cgc tcc ggt aaa cgg --g gca tgt gac gag gga gaa tat ctt gat aag agg cat aat caa tgt tgt aat -ca gtg tcc acc tgg aga att tgc caa ggt cag atg ta- --g tgg tag tga taa cac aaa atg tga acg ctg ccc acc tca tac ata tac cgc aat ccc caa tta ctc t-a atg gat gtc atc aat gta gaa aat gcc caa cag gat cat ttg ata agg taa agt gta ccg gaa cac aga aca gta aat -gt tcg tgt ctt cct ggt tgg tat tgc gct act gat tct tca cag act gaa gat tgt tga aat -tg tgt acc aaa aag g-a gat gtc cat gcg gat act ttg gtg gaa tag atg aac aag gaa atc cta ttt gta aat cgt gct g-- -ta ttg gtg aat att gca act acc tac gta att ata gac ttg atc cat ttt ctc cat gca aa- --- cta tct aaa tgt aat taa tta --- --- --- --- --- --- --- --- tga ttt --- --- --- -tg atg ata atg tta -cc ata c-a tt- --- -at atc gct act tgg tta --g tgt att att cag tat g-a aga cct att aat aa- --- --- -tt act tat ctt ttg acg atc ttg tt- -at aat tat --- --- --- --- aat ata aaa --- -ac tta tgg --- --- cat agt aac tta t-- --- --- --- aat tgc ta- aca cga taa att cgt aat aat ctg ttt tgt tca aat ttt tat aag ga- --- --a tct aca ggc ata aaa at- aaa aat ata att tat aat ata c-- tct tac a-- acg cca tca tga ata gca gca gta aat taa ttg ctg tta tta atg gat tta gaa ata gtg gac gat ttt gtg ata tta ata tag tta tta atg atg aaa gga taa acg ctc ata aac tca tcc tat ctg gag cct ccg aat a-t ttt tcc att ctg ttt tcc aat aat ttt atc gat tct aat gaa tac gaa gtt aat cta agt cat tta gat tat caa agc gta aac gat ttg atc gat tat att tat ggg ata cct ttg agc cta act aac gat aac gtg aaa tat att ctt tca acc gct ga- --t ttt tta caa att gga tct gct att acg gag tgt gaa aat tac ata ctt aaa aat ctt tgt tct aaa aac tgt atc gat ttc --- --- tac ata tac gct gat aaa tat aat aac aag aaa ata gaa tca gcg tcg ttt aac aca ata tta caa aat att ttg aga ctc atc aac gat gaa aac ttt aaa tac tta aca gag gaa tca atg ata aaa att tta agc gat gat atg tta aat ata aaa aat gag gat ttc gcc cca cta att ctc att aaa tgg tta gag agt act caa caa tca tgc acc gtc gag tta ctt aaa tgc ctc aga ata tca -tt gct ttc ccc aca agt tat aaa atc act tta tag tca tca act ggt tag ttc aat cta cga atg tat aac att ctt aaa caa tat agc att ctt gga tga atc att tcc tag ata cca tag cat cga gtt gat atc tat cgg tat aag taa ttc gcg tga taa aat ttc cat aaa ctg cta caa tca t-a aaa aaa ata cat ggg aaa tga tat ctt cac gta gat ata ggt gta gtt tcg cag tgg ccg tcc tgg ata ata tta tct ata tga tgg gtg gat atg atc agt ccc cgt ata gaa gtt caa agg tta tag cgt aca ata cat gta caa att ctt gga tat atg ata tac cag agc t-a aaa tat cct cgt tct aat tgt ggg gga ctg gct gat gac gaa tac att tat tgt ata ggc ggc ata cgc gat cag gat tca tcg ttg aca tct agt att gat aga tgg aag cca tca aaa cca tat tgg cag aag tat gct aaa atg cgc gaa cca aaa tgt gat atg ggg gtt gcg atg tta aac gga tta ata tat gtc atg ggt gga atc gtt aaa ggt g-- aca cgt gta ccg acg cac tag aga gtt tat cag aag atg gat gga tga agc atc aac gtc ttc caa taa aaa tgt cca ata tgt cga cga ttg ttc atg atg gca aga ttt ata tat ctg gag gtt aca aca ata gta gtg tag tta atg taa tat cga atc t-- agt cct tag cta taa ttc gat ata tga tga atg gac caa att atc atc att aaa tat tcc tag aat taa tcc cgc tct a

>MW960418.1:163299-165559_Vector_synVACV-wt_complete_sequence

ttg ctg act gta --- tgt a-c tat ttt tta cat tac tac att ggc tac ggc aga tat acc tac ttc gtc act gcc aca cgc tcc ggt aaa cgg --g gca tgt gac gag gga gaa tat ctt gat aag agg cat aat caa tgt tgt aat -ca gtg tcc acc tgg aga att tgc caa ggt cag atg ta- --g tgg tag tga taa cac aaa atg tga acg ctg ccc acc tca tac ata tac cgc aat ccc caa tta ctc t-a atg gat gtc atc aat gta gaa aat gcc caa cag gat cat ttg ata agg taa agt gta ccg gaa cac aga aca gta aat -gt tcg tgt ctt cct ggt tgg tat tgc gct act gat tct tca cag act gaa gat tgt tga aat -tg tgt acc aaa aag g-a gat gtc cat gcg gat act ttg gtg gaa tag atg aac aag gaa atc cta ttt gta aat cgt gct g-- -ta ttg gtg aat att gca act acc tac gta att ata gac ttg atc cat ttt ctc cat gca aa- --- cta tct aaa tgt aat taa tta --- --- --- --- --- --- --- --- tga ttt --- --- --- -tg atg ata atg tta -cc ata c-a tt- --- -at atc gct act tgg tta --g tgt att att cag tat g-a aga cct att aat aa- --- --- -tt act tat ctt ttg acg atc ttg tt- -at aat tat --- --- --- --- aat ata aaa --- -ac tta tgg --- --- cat agt aac tta t-- --- --- --- aat tgc ta- aca cga taa att cgt aat aat ctg ttt tgt tca aat ttt tat aag ga- --- --a tct aca ggc ata aaa at- aaa aat ata att tat aat ata c-- tct tac a-- acg cca tca tga ata gca gca gta aat taa ttg ctg tta tta atg gat tta gaa ata gtg gac gat ttt gtg ata tta ata tag tta tta atg atg aaa gga taa acg ctc ata aac tca tcc tat ctg gag cct ccg aat a-t ttt tcc att ctg ttt tcc aat aat ttt atc gat tct aat gaa tac gaa gtt aat cta agt cat tta gat tat caa agc gta aac gat ttg atc gat tat att tat ggg ata cct ttg agc cta act aac gat aac gtg aaa tat att ctt tca acc gct ga- --t ttt tta caa att gga tct gct att acg gag tgt gaa aat tac ata ctt aaa aat ctt tgt tct aaa aac tgt atc gat ttc --- --- tac ata tac gct gat aaa tat aat aac aag aaa ata gaa tca gcg tcg ttt aac aca ata tta caa aat att ttg aga ctc atc aac gat gaa aac ttt aaa tac tta aca gag gaa tca atg ata aaa att tta agc gat gat atg tta aat ata aaa aat gag gat ttc gcc cca cta att ctc att aaa tgg tta gag agt act caa caa tca tgc acc gtc gag tta ctt aaa tgc ctc aga ata tca -tt gct ttc ccc aca agt tat aaa atc act tta tag tca tca act ggt tag ttc aat cta cga atg tat aac att ctt aaa caa tat agc att ctt gga tga atc att tcc tag ata cca tag cat cga gtt gat atc tat cgg tat aag taa ttc gcg tga taa aat ttc cat aaa ctg cta caa tca t-a aaa aaa ata cat ggg aaa tga tat ctt cac gta gat ata ggt gta gtt tcg cag tgg ccg tcc tgg ata ata tta tct ata tga tgg gtg gat atg atc agt ccc cgt ata gaa gtt caa agg tta tag cgt aca ata cat gta caa att ctt gga tat atg ata tac cag agc t-a aaa tat cct cgt tct aat tgt ggg gga ctg gct gat gac gaa tac att tat tgt ata ggc ggc ata cgc gat cag gat tca tcg ttg aca tct agt att gat aga tgg aag cca tca aaa cca tat tgg cag aag tat gct aaa atg cgc gaa cca aaa tgt gat atg ggg gtt gcg atg tta aac gga tta ata tat gtc atg ggt gga atc gtt aaa ggt g-- aca cgt gta ccg acg cac tag aga gtt tat cag aag atg gat gga tga agc atc aac gtc ttc caa taa aaa tgt cca ata tgt cga cga ttg ttc atg atg gca aga ttt ata tat ctg gag gtt aca aca ata gta gtg tag tta atg taa tat cga atc t-- agt cct tag cta taa ttc gat ata tga tga atg gac caa att atc atc att aaa tat tcc tag aat taa tcc cgc tct a

>KC207810.1:156359-158612_Vaccinia_virus_strain_TianTan_clone_TP3_complete_genome

ttg ctg act gca --- tgt a-c tat ttt tta cat tac tac att ggc tac ggc aga tat acc tac ttc gtc act gcc aca cgc tcc ggt aaa cgg --g gca tgt gac gag gga gaa tat ctt gat aag agg cat aat caa tgt tgt aat -cg gtg tcc acc tgg aga att tgc caa ggt tag atg ta- --a tgg taa cga taa cac aaa atg tga acg ctg ccc acc tca tac ata tac cgc aat ccc caa tta ctc t-a atg gat gtc atc aat gta gaa aat gcc caa ccg gat cat ttg ata agg taa agt gta ccg gaa cac aga aca gta aat -gt tcg tgt ctt cct ggt tgg tat tgc gct act gat tct tca cag act gaa gat tgt cga gat -tg tat acc aaa aag g-a gat gtc cat gcg gat act ttg gtg gaa tag atg aac aag gaa atc cta ttt gta aat cgt gtt g-- -tg ttg gtg aat att gcg act act tac gta att ata gac ttg atc cat ttc ctc cat gca aa- --- cta tct aaa tgt aat taa tta --- --- --- --- --- --- --- --- tga ttt --- --- --- -tg atg ata atg tta -cc ata c-a tt- --- -at atc gct act tgg tta --g tgt att att cag tat g-a aga cct att aat aa- --- --- -tt act tat ctt ttg acg atc ttg tt- -at aat tat --- --- --- --- aat ata aaa --a tac tta tgg --- --- cat agt aac tca t-- --- --- --- aat tgc tg- acg cga taa att cgt aat aat ctg ttt tgt tca --- --- --- aag ga- --- --a tct aca ggc ata aaa at- aaa aat ata att tat aat ata c-- tct tac a-- acg cca tca tga ata gca gca gta aat taa ttg ctg tta tta atg gat tta gaa ata gtg gac gat ttt gtg ata tta ata tag tta tta atg atg aaa gga taa acg ctc ata aac tca tcc tat ctg gag cct ccg aat a-t ttt tcc att ctg ttt tcc aat aat ttt atc gat tct aat gaa tac gaa gtt aat cta agt cat tta gat tat caa agc gta aac gat ttg atc gat tac att tat ggg ata cct ttg agc cta act aac gat aac gtg aaa tat att ctt tca acc gct ga- --t ttt tta caa att gga tct gct att acg gag tgt gaa aaa tac ata ctt aaa aat ctt tgt tct aga aac tgt atc gat ttc --- --- tac ata tac gct gat aaa tat aat aac aag aaa ata gaa tca gcg tcg ttt aac aca ata tta caa aat att ttg aga ctc atc aac gat gaa aac ttt aaa tac tta aca gag gaa tca atg ata aaa att tta agc gat gat atg tta aat ata aaa aat gag gat ttc gcc cca cta att ctc att aaa tgg tta gag agt act caa caa cca tgc acc gtc gag tta ctt aaa tgc ctc aga ata tca -tt gct ttc ccc aca agt tat aaa atc act tta tag tca tcg act ggt tag ttc aat cta cga atg tat aac att ctt aaa caa tat agc att ctt gga tga atc att tcc tag ata cca tag cat cga gtt gat atc tat cgg tat aag taa ttc gcg tga taa aat ttc cat aaa ctg cta caa tca t-a aaa aaa ata cat ggg aaa tga tat ctt cac gta gat ata ggt gta gtt tcg cag tgg ccg tcc tgg ata ata tta tct ata tga tgg gtg gat atg atc agt ccc cgt ata gaa gtt caa agg tta tag cgt aca ata cat gta caa att ctt gga tat atg ata tac cag agc t-a aaa tat cct cgt tct aat tgt ggg gga ctg gct gat gac gaa tac att tat tgt ata ggc ggc ata cgc gat cag gat tca tcg ttg aca tct agt att gat aaa tgg aag cca tca aaa cca tat tgg cag aag tat gct aaa atg cgc gaa cca aaa tgt gat atg ggg gtt gcg atg tta aac gga tta ata tat gtc atg ggt gga atc gtt aaa ggt g-- aca cgt gta ccg acg cac tag aga gtt tat cag aag atg gat gga tga agc atc aac gtc ttc caa taa aaa tgt cca ata tgt cga cga ttg ttc atg atg gca aga ttt ata tat ctg gag gtt aca aca ata gta gtg tag tta atg taa tat cga atc t-- agt cct tag cta taa tcc gat ata tga tga atg gac caa att atc atc att aaa cat tcc tag aat taa tcc cgc tct a

>JN654986.1:162119-164379_Vaccinia_virus_strain_Dryvax_clone_DPP21_complete_genome

ttg ctg act gta --- tgt a-c tat ttt tta cat tac tac att ggc tac ggc aga tat acc tac ttc gtc act gcc aca cgc tcc ggt aaa cgg --g gca tgt gac gag gga gaa tat ctt gat aag agg cat aat caa tgt tgt aat -ca gtg tcc acc tgg aga att tgc caa ggt cag atg ta- --a tgg taa cga taa cac aaa atg tga acg ctg ccc acc tca tac ata tac cgc aat ccc caa tta ctc t-a atg gat gtc atc aat gta gaa aat gcc caa cag gat cat ttg ata agg taa agt gta ccg gaa cac aga aca gta aat -gt tcg tgt ctt cct ggt tgg tat tgt gct act gat tct tca cag act gaa gat tgt tga aat -tg tgt acc aaa aag g-a gat gtc cat gcg gat act ttg gtg gaa tag atg aac aag gaa atc cta ttt gta aat cgt gct g-- -ta ttg gtg aat att gca act acc tac gta att ata gac ttg atc cat ttt ctc cat gca aa- --- cta tct aaa tgt aat taa tta --- --- --- --- --- --- --- --- tga ttt --- --- --- -tg atg ata atg tta -cc ata c-a tt- --- -at atc gct act tgg tta --g tgt att att cag tat g-a aga cct att aat aa- --- --- -tt act tat ctt ttg acg atc ttg tt- -at aat tat --- --- --- --- aat ata aaa --- -ac tta tgg --- --- cat agt aac tca t-- --- --- --- aat tgc tg- acg cga taa att cgt aat aat ctg ttt tgt tca aat ttt tat aag ga- --- --a tct aca ggc ata aaa at- aaa aat ata att tat aat ata c-- tct tac a-- acg cca tca tga ata gca gca gta aat taa ttg ctg tta tta atg gat tta gaa ata gtg gac gat ttt gtg ata tta ata tag tta tta atg atg aaa gga taa acg ctc ata aac tca tcc tat ctg gag cct ccg aat a-t ttt tcc att ctg ttt tcc aat aat ttt atc gat tct aat gaa tac gaa gtt aat cta agt cat tta gat tat caa agc gta aac gat ttg atc gat tat att tat ggg ata cct ttg agc cta act aac gat aac gtg aaa tat att ctt tca acc gct ga- --t ttt tta caa att gga tct gct att acg gag tgt gaa aat tac ata ctt aaa aat ctt tgt tct aaa aac tgt atc gat ttc --- --- tac ata tac gct gat aaa tat aat aac aag aaa ata gaa tca gcg tcg ttt aac aca ata tta caa aat att ttg aga ctc atc aac gat gaa aac ttt aaa tac tta aca gag gaa tca atg ata aaa att tta agc gat gat atg tta aat ata aaa aat gag gat ttc gcc cca cta att ctc att aaa tgg tta gag agt act caa caa tca tgc acc gtc gag tta ctt aaa tgc ctc aga ata tca -tt gct ttc ccc aca agt tat aaa atc act tta tag tca tcg act ggt tag ttc aat cta cga atg tat aac att ctt aaa caa tat agc att ctt gga tga atc att tcc tag ata cca tag cat cga gtt gat atc tat cgg tat aag taa ttc gcg tga taa aat ttc cat aaa ctg cta caa tca t-a aaa aaa ata cat ggg aaa tga tat ctt cac gta gat ata ggt gta gtt tcg cag tgg ccg tcc tgg ata ata tta ttt ata tga tgg gtg gat atg atc agt ccc cgt ata gaa gtt caa agg tta tag cgt aca ata cat gta caa att ctt gga tat atg ata tac cag agc t-a aaa tat cct cgt tct aat tgt ggg gga ctg gct gat gac gaa tac att tat tgt ata ggc ggc ata cgc gat cag gat tca tcg ttg aca tct agt att gat aga tgg aag cca tca aaa cca tat tgg cag aag tat gct aaa atg cgc gaa cca aaa tgt gat atg ggg gtt gcg atg tta aac gga tta ata tat gtt ata ggt gga atc gtt aaa ggt g-- aca cgt gta ccg acg cac tag aga gtt tat cag aag atg gat gga tga agc atc aac gtc ttc caa taa aaa tgt cca ata tgt cga cga ttg ttc atg atg gca aga ttt ata tat ctg gag gtt aca aca ata gta gtg tag tta atg taa tat cga atc t-- agt cct tag cta taa ttc gat ata tga tga atg gac caa att atc atc att aaa tat tcc tag aat taa tcc cgc tct a

>AY313847.1:162447-164707_Vaccinia_virus_strain_Acambis_clone_2000_complete_genome

ttg ctg act gta --- tgt a-c tat ttt tta cat tac tac att ggc tac ggc aga tat acc tac ttc gtc act gcc aca cgc tcc ggt aaa cgg --g gca tgt gac gag gga gaa tat ctt gat aag agg cat aat caa tgt tgt aat -ca gtg tcc acc tgg aga att tgc caa ggt cag atg ta- --g tgg tag tga taa cac aaa atg tga acg ctg ccc acc tca tac ata tac cgc aat ccc caa tta ctc t-a atg gat gtc atc aat gta gaa aat gcc caa cag gat cat ttg ata agg taa agt gta ccg gaa cac aga aca gta aat -gt tcg tgt ctt cct ggt tgg tat tgc gct act gat tct tca cag act gaa gat tgt tga aat -tg tgt acc aaa aag g-a gat gtc cat gcg gat act ttg gtg gaa tag atg aac aag gaa atc cta ttt gta aat cgt gct g-- -ta ttg gtg aat att gca act acc tac gta att ata gac ttg atc cat ttt ctc cat gca aa- --- cta tct aaa tgt aat taa tta --- --- --- --- --- --- --- --- tga ttt --- --- --- -tg atg ata atg tta -cc ata c-a tt- --- -at atc gct act tgg tta --g tgt att att cag tat g-a aga cct att aat aa- --- --- -tt act tat ctt ttg acg atc ttg tt- -at aat tat --- --- --- --- aat ata aaa --- -ac tta tgg --- --- cat agt aac tta t-- --- --- --- aat tgc ta- aca cga taa att cgt aat aat ctg ttt tgt tca aat ttt tat aag ga- --- --a tct aca ggc ata aaa at- aaa aat ata att tat aat ata c-- tct tac a-- acg cca tca tga ata gca gca gta aat taa ttg ctg tta tta atg gat tta gaa ata gtg gac gat ttt gtg ata tta ata tag tta tta atg atg aaa gga taa acg ctc ata aac tca tcc tat ctg gag cct ccg aat a-t ttt tcc att ctg ttt tcc aat aat ttt atc gat tct aat gaa tac gaa gtt aat cta agt cat tta gat tat caa agc gta aac gat ttg atc gat tat att tat ggg ata cct ttg agc cta act aac gat aac gtg aaa tat att ctt tca acc gct ga- --t ttt tta caa att gga tct gct att acg gag tgt gaa aat tac ata ctt aaa aat ctt tgt tct aaa aac tgt atc gat ttc --- --- tac ata tac gct gat aaa tat aat aac aag aaa ata gaa tca gcg tcg ttt aac aca ata tta caa aat att ttg aga ctc atc aac gat gaa aac ttt aaa tac tta aca gag gaa tca atg ata aaa att tta agc gat gat atg tta aat ata aaa aat gag gat ttc gcc cca cta att ctc att aaa tgg tta gag agt act caa caa tca tgc acc gtc gag tta ctt aaa tgc ctc aga ata tca -tt gct ttc ccc aca agt tat aaa atc act tta tag tca tca act ggt tag ttc aat cta cga atg tat aac att ctt aaa caa tat agc att ctt gga tga atc att tcc tag ata cca tag cat cga gtt gat atc tat cgg tat aag taa ttc gcg tga taa aat ttc cat aaa ctg cta caa tca t-a aaa aaa ata cat ggg aaa tga tat ctt cac gta gat ata ggt gta gtt tcg cag tgg ccg tcc tgg ata ata tta tct ata tga tgg gtg gat atg atc agt ccc cgt ata gaa gtt caa agg tta tag cgt aca ata cat gta caa att ctt gga tat atg ata tac cag agc t-a aaa tat cct cgt tct aat tgt ggg gga ctg gct gat gac gaa tac att tat tgt ata ggc ggc ata cgc gat cag gat tca tcg ttg aca tct agt att gat aga tgg aag cca tca aaa cca tat tgg cag aag tat gct aaa atg cgc gaa cca aaa tgt gat atg ggg gtt gcg atg tta aac gga tta ata tat gtc atg ggt gga atc gtt aaa ggt g-- aca cgt gta ccg acg cac tag aga gtt tat cag aag atg gat gga tga agc atc aac gtc ttc caa taa aaa tgt cca ata tgt cga cga ttg ttc atg atg gca aga ttt ata tat ctg gag gtt aca aca ata gta gtg tag tta atg taa tat cga atc t-- agt cct tag cta taa ttc gat ata tga tga atg gac caa att atc atc att aaa tat tcc tag aat taa tcc cgc tct a

>KC207811.1:162067-164319_Vaccinia_virus_strain_TianTan_clone_TP5_complete_genome

ttg ctg act gca --- tgt a-c tat ttt tta cat tac tac att ggc tac ggc aga tat acc tac ttc gtc act gcc aca cgc tcc ggt aaa cgg --g gca tgt gac gag gga gaa tat ctt gat aag agg cat aat caa tgt tgt aat -cg gtg tcc acc tgg aga att tgc caa ggt tag atg ta- --a tgg taa cga taa cac aaa atg tga acg ctg ccc acc tca tac ata tac cgc aat ccc caa tta ctc t-a atg gat gtc atc aat gta gaa aat gcc caa ccg gat cat ttg ata agg taa agt gta ccg gaa cac aga aca gta aat -gt tcg tgt ctt cct ggt tgg tat tgc gct act gat tct tca cag act gaa gat tgt cga gat -tg tat acc aaa aag g-a gat gtc cat gcg gat act ttg gtg gaa tag atg aac aag gaa atc cta ttt gta aat cgt gtt g-- -tg ttg gtg aat att gcg act act tac gta att ata gac ttg atc cat ttc ctc cat gca aa- --- cta tct aaa tgt aat taa tta --- --- --- --- --- --- --- --- tga ttt --- --- --- -tg atg ata atg tta -cc ata c-a tt- --- -at atc gct act tgg tta --g tgt att att cag tat g-a aga cct att aat aa- --- --- -tt act tat ctt ttg acg atc ttg tt- -at aat tat --- --- --- --- aat ata aaa --a tac tta tgg --- --- cat agt aac tca t-- --- --- --- aat tgc tg- acg cga taa att cgt aat aat ctg ttt tgt tca --- --- --- aag ga- --- --a tct aca ggc ata aaa at- -aa aat ata att tat aat ata c-- tct tac a-- acg cca tca tga ata gca gca gta aat taa ttg ctg tta tta atg gat tta gaa ata gtg gac gat ttt gtg ata tta ata tag tta tta atg atg aaa gga taa acg ctc ata aac tca tcc tat ctg gag cct ccg aat a-t ttt tcc att ctg ttt tcc aat aat ttt atc gat tct aat gaa tac gaa gtt aat cta agt cat tta gat tat caa agc gta aac gat ttg atc gat tac att tat ggg ata cct ttg agc cta act aac gat aac gtg aaa tat att ctt tca acc gct ga- --t ttt tta caa att gga tct gct att acg gag tgt gaa aaa tac ata ctt aaa aat ctt tgt tct aga aac tgt atc gat ttc --- --- tac ata tac gct gat aaa tat aat aac aag aaa ata gaa tca gcg tcg ttt aac aca ata tta caa aat att ttg aga ctc atc aac gat gaa aac ttt aaa tac tta aca gag gaa tca atg ata aaa att tta agc gat gat atg tta aat ata aaa aat gag gat ttc gcc cca cta att ctc att aaa tgg tta gag agt act caa caa cca tgc acc gtc gag tta ctt aaa tgc ctc aga ata tca -tt gct ttc ccc aca agt tat aaa atc act tta tag tca tcg act ggt tag ttc aat cta cga atg tat aac att ctt aaa caa tat agc att ctt gga tga atc att tcc tag ata cca tag cat cga gtt gat atc tat cgg tat aag taa ttc gcg tga taa aat ttc cat aaa ctg cta caa tca t-a aaa aaa ata cat ggg aaa tga tat ctt cac gta gat ata ggt gta gtt tcg cag tgg ccg tcc tgg ata ata tta tct ata tga tgg gtg gat atg atc agt ccc cgt ata gaa gtt caa agg tta tag cgt aca ata cat gta caa att ctt gga tat atg ata tac cag agc t-a aaa tat cct cgt tct aat tgt ggg gga ctg gct gat gac gaa tac att tat tgt ata ggc ggc ata cgc gat cag gat tca tcg ttg aca tct agt att gat aaa tgg aag cca tca aaa cca tat tgg cag aag tat gct aaa atg cgc gaa cca aaa tgt gat atg ggg gtt gcg atg tta aac gga tta ata tat gtc atg ggt gga atc gtt aaa ggt g-- aca cgt gta ccg acg cac tag aga gtt tat cag aag atg gat gga tga agc atc aac gtc ttc caa taa aaa tgt cca ata tgt cga cga ttg ttc atg atg gca aga ttt ata tat ctg gag gtt aca aca ata gta gtg tag tta atg taa tat cga atc t-- agt cct tag cta taa tcc gat ata tga tga atg gac caa att atc atc att aaa cat tcc tag aat taa tcc cgc tct a

>AF095689.1:159334-161586_Vaccinia_virus_(strain_Tian_Tan)_complete_genome

ttg ctg act gca --- tgt a-c tat ttt tta cat tac tac att ggc tac ggc aga tat acc tac ttc gtc act gcc aca cgc tcc ggt aaa c-g --g gca tgt gac gag gga gaa tat ctt gat aag agg cat aat caa tgt tgt aat -cg gtg tcc acc tgg aga att tgc caa ggt tag atg ta- --a tgg taa cga taa cac aaa atg tga acg ctg ccc acc tca tac ata tac cgc aat ccc caa tta ctc t-a atg gat gtc atc aat gta gaa aat gcc caa ccg gat cat ttg ata agg taa agt gta ccg gaa cac aga aca gta aat -gt tcg tgt ctt cct ggt tgg tat tgc gct act gat tct tca cag act gaa gat tgt tga aat -tg tgt acc aaa aag g-a gat gtc cat gcg gat act ttg gtg gaa tag atg aac aag gaa atc cta ttt gta aat cgt gtt g-- -tg ttg gtg aat att gcg act act tac gta att ata gac ttg atc cat ttc ctc cat gca aa- --- cta tct aaa tgt aat taa tta --- --- --- --- --- --- --- --- tga tta --- --- --- -tg atg ata atg tta -cc ata c-a tt- --- -at atc gct act tgg tta --g tgt att att cag tat g-a aga cct att aat aa- --- --- -tt act tat ctt ttg acg atc ttg tt- -at aat tat --- --- --- --- aat ata aaa --a tac tta tgg --- --- cat agt aac tca t-- --- --- --- aat tgc tg- acg cga taa att cgt aat aat ctg ttt tgt tca --- --- --- aag ga- --- --a tct aca ggc ata aaa at- aaa aat ata att tat aat ata c-- tct tac a-- acg cca tca tga ata gca gca gta aat taa ttg ctg tta tta atg gat tta gaa ata gtg gac gat ttt gtg ata tta ata tag tta tta atg atg aaa gga taa acg ctc ata aac tca tcc tat ctg gag cct ccg aat a-t ttt tcc att ctg ttt tcc aat aat ttt atc gat tct aat gaa tac gaa gtt aat cta agt cat tta gat tat caa agt gtt aac gat ttg atc gat tac att tat ggg ata cct ttg agc cta act aac gat aac gtg aaa tat att ctt tca acc gct ga- --t ttt tta caa att gga tct gct att acg gag tgt gaa aaa tac ata ctt aaa aat ctt tgt tct aga aac tgt atc gat ttc --- --- tac ata tac gct gat aaa tat aat aac aag aaa ata gaa tca gcg tcg ttt aac aca ata tta caa aat att ttg aga ctc atc aac gat gaa aac ttt aaa tac tta aca gag gaa tca atg ata aaa att tta agc gat gat atg tta aat ata aaa aat gag gat ttc gcc cca cta att ctc att aaa tgg tta gag agt act caa caa cca tgc acc gtc gag tta ctt aaa tgc ctc aga ata tca -tt gct ttc ccc aca agt tat aaa atc act tta tag tca tcg act ggt tag ttc aat cta cga atg tat aac att ctt aaa caa tat agc att ctt gga tga atc att tcc tag ata cca tag cat cga gtt gat atc tat cgg tat aag taa ttc gcg tga taa aat ttc cat aaa ctg cta caa tca t-a aaa aaa ata cat ggg aaa tga tat ctt cac gta gat ata ggt gta gtt tcg cag tgg ccg tcc tgg ata ata tta tct ata tga tgg gtg gat atg atc agt ccc cgt ata gaa gtt caa agg tta tag cgt aca ata cat gta caa att ctt gga tat atg ata tac cag agc t-a aaa tat cct cgt tct aat tgt ggg gga ctg gct gat gac gaa tac att tat tgt ata ggc ggc ata cgc gat cag gat tca tcg ttg aca tct agt att gat aaa tgg aag cca tca aaa cca tat tgg cag aag tat gct aaa atg cgc gaa cca aaa tgt gat atg ggg gtt gcg atg tta aac gga tta ata tat gtc atg ggt gga atc gtt aaa ggt g-- aca cgt gta ccg acg cac tag aga gtt tat cag aag atg gat gga tga agc atc aac gtc ttc caa taa aaa tgt cca ata tgt cga cga ttg ttc atg atg gca aga ttt ata tat ctg gag gtt aca aca ata gta gtg tag tta atg taa tat cga atc t-- agt cct tag cta taa tcc gat ata tga tga atg gac caa att atc atc att aaa cat tcc tag aat taa tcc cgc tct a

>KF866253.1:161515-163768_Vaccinia_virus_WAU86/88-1_complete_genome

ttg ctg act gta --- tgt a-c tat ttt tta cat tac tac att ggc tac ggc aga tat acc tac ttc gtc act gcc aca cgc tcc ggt aaa cgg --g gca tgt gac gag gga gaa tat ctt gat aag agg cat aat caa tgt tgt aat -cg gtg tcc acc tgg aga att tgc caa ggt tag atg ta- --a tgg taa cga taa cac aaa atg tga acg ctg ccc acc tca tac ata tac cgc aat ccc caa tta ctc t-a atg gat gtc atc aat gta gaa aat gcc caa ccg gat cat ttg ata agg taa agt gta ccg gaa cac aga aca gta aat -gt tcg tgt ctt cct ggt tgg tat tgc gct act gat tct tca cag act gaa gat tgt cga gat -tg tat acc aaa aag gaa gat gtc cat gcg gat act ttg gtg gaa tag atg aac aag gaa atc cta ttt gta aat cgt gtt g-- -tg ttg gtg aat att gcg act acc tac gta att ata gac ttg atc cat ttc ctc cat gca aa- --- cta tct aaa tgt aat taa tta --- --- --- --- --- --- --- --- tga ttt --- --- --- -tg atg ata atg tta -cc ata c-a tt- --- -at atc gct act tgg tta --g tgt att att cag tat g-a aga cct att aat aa- --- --- -tt act tat ctt ttg acg atc ttg tt- -at aat tat --- --- --- --- aat ata aaa --a tac tta tgg --- --- cat agt aac tca t-- --- --- --- aat tgc tg- acg cga taa att cgt aat aat ctg ttt tgt tca --- --- --- aag ga- --- --a tct aca ggc ata aaa at- aaa aat ata att tat aat a-a c-- tct tac a-- acg cca tca tga ata aca gca gtg aat tga ttg ctg tta tta atg gat tta gaa ata gtg gac gat ttt gtg ata tta gta tag tta tta atg atg aaa gga taa acg ctc aca gac tca tcc tat ctg gag cct ccg aat a-t ttt tcc att ctg ttt tcc aat aat ttt atc gat tct aat gaa tac gaa gtt aat cta agt cat tta gat tat caa agc gta aac gat ttg atc gat tac att tat ggg ata cct ttg agc cta act aac gat aac gtg aaa tat att ctt tca acc gct ga- --t ttt tta caa att gga tct gct att acg gag tgt gaa aat tac ata ctt aaa aat ctt tgt tct aga aac tgt atc gat ttc --- --- tac ata tac gct gat aaa tat aat aac aag aaa ata gaa tca gcg tcg ttt aac aca ata tta caa aat att ttg aga ctc atc aac gat gaa aac ttt aaa tac tta aca gag gaa tca atg ata aaa att tta agc gat gat atg tta aat ata aaa aat gag gat ttc gcc cca cta att ctc att aaa tgg tta gag agt act caa caa cca tgc acc gtc gag tta ctt aaa tgc ctc aga ata tca -tt gct ttc ccc aca agt tat aaa atc act tta tag tca tcg act ggt tag ttc aat cta cga atg tat aac att ctt aaa caa tat agc att ctt gga tga atc att tcc tag ata cca tag cat cga gtt gat atc tat cgg tat aag taa ttc gca tga taa gat ttc cat aaa ctg cta caa tca t-a aaa aaa ata cat ggg aaa tga tat ctt cac gta gat ata ggt gta gtt tcg cag tgg ccg tcc tgg ata ata tta tct ata tga tgg gtg gat atg atc agt ccc cgt ata gaa gtt caa agg tta tag cgt aca ata cat gta caa att ctt gga tat atg ata tac cag agc t-a aaa tat cct cgt tct aat tgt ggg gga ctg gct gat gac gaa tac att tat tgt ata ggc ggc ata cgc gat cag gat tca tcg ttg aca tct agt att gat aga tgg aag cca tca aaa cca tat tgg cag aag tat gct aaa atg cgc gaa cca aaa tgt gat atg ggg gtt gcg atg tta aac gga tta ata tat gtt ata ggt gga atc gtt aaa ggt g-- aca cgt gta ccg acg cac tag aga gtt tat cag aag atg gat gga tga agc atc aac gtc ttc caa taa aaa tgt cca ata tgt cga cga ttg ttc atg atg gca aga ttt ata tat ctg gag gtt aca aca ata gta gtg tag tta atg taa tat cga atc t-- agt cct tag cta taa ttc gat ata tga tga atg gac caa att atc atc att aaa cat tcc tag aat taa tcc cgc tct a

>JN654984.1:162173-164433_Vaccinia_virus_strain_Dryvax_clone_DPP19_complete_genome

ttg ctg act gta --- tgt a-c tat ttt tta cat tac tac att ggc tac ggc aga tat acc tac ttc gtc act gcc aca cgc tcc ggt aaa cgg --g gca tgt gac gag gga gaa tat ctt gat aag agg cat aat caa tgt tgt aat -ca gtg tcc acc tgg aga att tgc caa ggt cag atg ta- --g tgg tag tga taa cac aaa atg tga acg ctg ccc acc tca tac ata tac cgc aat ccc caa tta ctc t-a atg gat gtc atc aat gta gaa aat gcc caa cag gat cat ttg ata agg taa agt gta ccg gaa cac aga aca gta aat -gt tcg tgt ctt cct ggt tgg tat tgt gct act gat tct tca cag act gaa gat tgt cga gat -tg tat acc aaa aag g-a gat gtc cat gcg gat act ttg gtg gaa tag atg aac aag gaa atc cta ttt gta aat cgt gct g-- -ta ttg gtg aat att gca act acc tac gta att ata gac ttg atc cat ttt ctc cat gca aa- --- cta tct aaa tgt aat taa tta --- --- --- --- --- --- --- --- tga ttt --- --- --- -tg atg ata atg tta -cc ata c-a tt- --- -at atc gct act tgg tta --g tgt att att cag tat g-a aga cct att aat aa- --- --- -tt act tat ctt ttg acg atc ttg tt- -at aat tat --- --- --- --- aat ata aaa --- -ac tta tgg --- --- cat agt aac tta t-- --- --- --- aat tgc ta- aca cga taa att cgt aat aat ctg ttt tgt tca aat ttt tat aag ga- --- --a tct aca ggc ata aaa at- aaa aat ata atc tat aat ata c-- tct tac a-- acg cca tca tga ata gca gca gta aat taa ttg ctg tta tta atg gat tta gaa ata gtg gac gat ttt gtg ata tta ata tag tta tta atg atg aaa gga taa acg ctc ata aac tca tcc tat ctg gag cct ccg aat a-t ttt tcc att ctg ttt tcc aat aat ttt atc gat tct aat gaa tac gaa gtt aat cta agt cat tta gat tat caa agc gta aac gat ttg atc gat tat att tat ggg ata cct ttg agc cta act aac gat aac gtg aaa tat att ctt tca acc gct ga- --t ttt tta caa att gga tct gct att acg gag tgt gaa aat tac ata ctt aaa aat ctt tgt tct aaa aac tgt atc gat ttc --- --- tac ata tac gct gat aaa tat aat aac aag aaa ata gaa tca gcg tcg ttt aac aca ata tta caa aat att ttg aga ctc atc aac gat gaa aac ttt aaa tac tta aca gag gaa tca atg ata aaa att tta agc gat gat atg tta aat ata aaa aat gag gat ttt gca cca cta att ctc att aaa tgg tta gag agt act caa caa tca tgc acc gtc gag tta ctt aaa tgc ctc aga ata tca -tt gct ttc ccc aca agt tat aaa atc act tta tag tca tca act ggt tag ttc aat cta cga atg tat aac att ctt aaa caa tat agc att ctt gga tga atc att tcc tag ata cca tag cat cga gtt gat atc tat cgg tat aag taa ttc gcg tga taa aat ttc cat aaa ctg cta caa tca t-a aaa aaa ata cat ggg aaa tga tat ctt cac gta gat ata ggt gta gtt tcg cag tgg ccg tcc tgg ata ata tta ttt ata tga tgg gtg gat atg atc agt ccc cgt ata gaa gtt caa agg tta tag cgt aca ata cat gta caa att ctt gga tat atg ata tac cag agc t-a aaa tat cct cgt tct aat tgt ggg gga ctg gct gat gac gaa tac att tat tgt ata ggc ggc ata cgc gat cag gat tca tcg ttg aca tct agt att gat aaa tgg aag cca tca aaa cca tat tgg cag aag tat gct aaa atg cgc gaa cca aaa tgt gat atg ggg gtt gcg atg tta aac gga tta ata tat gtc atg ggt gga atc gtt aaa ggt g-- aca cgt gta ccg acg cac tag aga gtt tat cag aag atg gat gga tga agc atc aac gtc ttc caa taa aaa tgt cca ata tgt cga cga ttg ttc atg atg gca aga ttt ata tat ctg gag gtt aca aca ata gta gtg tag tta atg taa tat cga atc t-- agt cct tag cta taa ttc gat ata tga tga atg gac caa att atc atc att aaa tat tcc tag aat taa tcc cgc tct a

>DQ377945.1:164970-167230_Vaccinia_virus_strain_3737_complete_genome

ttg ctg act gta --- tgt a-c tat ttt tta cat tac tac att ggc tac ggc aga tat acc tac ttc gtc act gcc aca cgc tcc ggt aaa cgg --g gca tgt gac gag gga gaa tat ctt gat aag agg cat aat caa tgt tgt aat -ca gtg tcc acc tgg aga att tgc caa ggt cag atg ta- --g tgg tag tga taa cac aaa atg tga acg ctg ccc acc tca tac ata tac cgc aat ccc caa tta ctc t-a atg gat gtc atc aat gta gaa aat gcc caa cag gat cat ttg ata agg taa agt gta ccg gaa cac aga aca gta aat -gt tcg tgt ctt cct ggt tgg tat tgt gct act gat tct tca cag act gaa gat tgt cga gat -tg tat acc aaa aag g-a gat gtc cat gcg gat act ttg gtg gaa tag atg aac aag gaa atc cta ttt gta aat cgt gct g-- -ta ttg gtg aat att gca act acc tac gta att ata gac ttg atc cat ttc ctc cat gca aa- --- cta tct aaa tgt aat taa tta --- --- --- --- --- --- --- --- tga ttt --- --- --- -tg atg ata atg tta -cc ata c-a tt- --- -at atc gct act tgg tta --g tgt att att cag tat g-a aga cct att aat aa- --- --- -tt act tat ctt ttg acg atc ttg tt- -at aat tat --- --- --- --- aat ata aaa --- -ac tta tgg --- --- cat agt aac tta t-- --- --- --- aat tgc ta- aca cga taa att cgt aat aat ctg ttt tgt tca aat ttt tat aag ga- --- --a tct aca ggc ata aaa at- aaa aat ata att tat aat ata c-- tct tac a-- acg cca tca tga ata gca gca gta aat taa ttg ctg tta tta atg gat tta gaa ata gtg gac gat ttt gtg ata tta ata tag tta tta atg atg aaa gga taa acg ctc ata aac tca tcc tat ctg gag cct ccg aat a-t ttt tcc att ctg ttt tcc aat aat ttt atc gat tct aat gaa tac gaa gtt aat cta agt cat tta gat tat caa agc gta aac gat ttg atc gat tat att tat ggg ata cct ttg agc cta act aac gat aac gtg aaa tat att ctt tca acc gct ga- --t ttt tta caa att gga tct gct att acg gag tgt gaa aat tac ata ctt aaa aat ctt tgt tct aaa aac tgt atc gat ttc --- --- tac ata tac gct gat aaa tat aat aac aag aaa ata gaa tca gcg tcg ttt aac aca ata tta caa aat att ttg aga ctc atc aac gat gaa aac ttt aaa tac tta aca gag gaa tca atg ata aaa att tta agc gat gat atg tta aat ata aaa aat gag gat ttt gca cca cta att ctc att aaa tgg tta gag agt act caa caa tca tgc acc gtc gag tta ctt aaa tgc ctc aga ata tca -tt gct ttc ccc aca agt tat aaa atc act tta tag tca tca act ggt tag ttc aat cta cga atg tat aac att ctt aaa caa tat agc att ctt gga tga atc att tcc tag ata cca tag cat cga gtt gat atc tat cgg tat aag taa ttc gcg tga taa aat ttc cat aaa ctg cta caa tca t-a aaa aaa ata cat ggg aaa tga tat ctt cac gta gat ata ggt gta gtt tcg cag tgg ccg tcc tgg ata ata tta ttt ata tga tgg gtg gat atg atc agt ccc cgt ata gaa gtt caa agg tta tag cgt aca ata cat gta caa att ctt gga tat atg ata tac cag agc t-a aaa tat cct cgt tct aat tgt ggg gga ctg gct gat gac gaa tac att tat tgt ata ggc ggc ata cgc gat cag gat tca tcg ttg aca tct agt att gat aaa tgg aag cca tca aaa cca tat tgg cag aag tat gct aaa atg cgc gaa cca aaa tgt gat atg ggg gtt gcg atg tta aac gga tta ata tat gtt ata ggt gga atc gtt aaa ggt g-- aca cgt gta ccg acg cac tag aga gtt tat cag aag atg gat gga tga agc atc aac gtc ttc caa taa aaa tgt cca ata tgt cga cga ttg ttc atg atg gca aga ttt ata tat ctg gag gtt aca aca ata gta gtg tag tta atg taa tat cga atc t-- agt cct tag cta taa ttc gat ata tga tga atg gac caa att atc atc att aaa tat tcc tag aat taa tcc cgc tct a

>JN654979.1:162286-164543_Vaccinia_virus_strain_Dryvax_clone_DPP12_complete_genome

ttg ctg act gca --- tgt a-c tat ttt tta cat tac tac att ggc tac ggc aga tat acc tac tcc gcc acc --- aca cgc tcc ggt aaa cgg --g gca tgt gac gag gga gaa tat ctt gat aag agg cat aat caa tgt tgt aat -ca gtg tcc acc tgg aga att tgc caa ggt cag atg ta- --g tgg tag tga taa cac aaa atg tga acg ctg ccc acc tca tac ata tac cgc aat ccc caa tta ctc t-a atg gat gtc atc aat gta gaa aat gcc caa cag gat cat ttg ata agg taa agt gta ccg gaa cac aga aca gta aat -gt tcg tgt ctt cct ggt tgg tat tgc gct act gat tct tca cag act gaa gat tgt tga aat -tg tgt acc aaa aag g-a gat gtc cat gcg gat act ttg gtg gaa tag atg aac aag gaa atc cta ttt gta aat cgt gct g-- -ta ttg gtg aat att gca act acc tac gta att ata gac ttg atc cat ttt ctc cat gca aa- --- cta tct aaa tgt aat taa tta --- --- --- --- --- --- --- --- tga ttt --- --- --- -tg atg ata atg tta -cc ata c-a tt- --- -at atc gct act tgg tta --g tgt att att cag tat g-a aga cct att aat aa- --- --- -tt act tat ctt ttg acg atc ttg tt- -at aat tat --- --- --- --- aat ata aaa --- -ac tta tgg --- --- cat agt aac tta t-- --- --- --- aat tgc ta- aca cga taa att cgt aat aat ctg ttt tgt tca aat ttt tat aag ga- --- --a tct aca ggc ata aaa at- aaa aat ata att tat aat ata c-- tct tac a-- acg cca tca tga ata gca gca gta aat taa ttg ctg tta tta atg gat tta gaa ata gtg gac gat ttt gtg ata tta ata tag tta tta atg atg aaa gga taa acg ctc ata aac tca tcc tat ctg gag cct ccg aat a-t ttt tcc att ctg ttt tcc aat aat ttt atc gat tct aat gaa tac gaa gtt aat cta agt cat tta gat tat caa agc gta aac gat ttg atc gat tat att tat ggg ata cct ttg agc cta act aac gat aac gtg aaa tat att ctt tca acc gct ga- --t ttt tta caa att gga tct gct att acg gag tgt gaa aat tac ata ctt aaa aat ctt tgt tct aaa aac tgt atc gat ttc --- --- tac ata tac gct gat aaa tat aat aac aag aaa ata gaa tca gcg tcg ttt aac aca ata tta caa aat att ttg aga ctc atc aac gat gaa aac ttt aaa tac tta aca gag gaa tca atg ata aaa att tta agc gat gat atg tta aat ata aaa aat gag gat ttt gca cca cta att ctc att aaa tgg tta gag agt act caa caa tca tgc acc gtc gag tta ctt aaa tgc ctc aga ata tca -tt gct ttc ccc aca agt tat aaa atc act tta tag tca tcg act ggt tag ttc aat cta cga atg tat aac att ctt aaa caa tat agc att ctt gga tga atc att tcc tag ata cca tag cat cga gtt gat atc tat cgg tat aag taa ttc gca tga taa gat ttc cat aaa ctg cta caa tca t-a aaa aaa ata cat ggg aaa tga tat ctt cac gta gat ata ggt gta gtt tcg cag tgg ccg tcc tgg ata ata tta tct ata tga tgg gtg gat atg atc agt ccc cgt ata gaa gtt caa agg tta tag cgt aca ata cat gta caa att ctt gga tat atg ata tac cag agc t-a aaa tat cct cgt tct aat tgt ggg gga ctg gct gat gac gaa tac att tat tgt ata ggc ggc ata cgc gat cag gat tca tcg ttg aca tct agt att gat aaa tgg aag cca tca aaa cca tat tgg cag aag tat gct aaa atg cgc gaa cca aaa tgt gat atg ggg gtt gcg atg tta aac gga tta ata tat gtt ata ggt gga atc gtt aaa ggt g-- aca cgt gta ccg acg cac tag aga gtt tat cag aag atg gat gga tga agc atc aac gtc ttc caa taa aaa tgt cca ata tgt cga cga ttg ttc atg atg gca aga ttt ata tat ctg gag gtt aca aca ata gta gtg tag tta atg taa tat cga atc t-- agt cct tag cta taa ttc gat ata tga tga atg gac caa att atc atc att aaa tat tcc tag aat taa tcc cgc tct a

>DQ439815.1:164731-166988_Vaccinia_virus_strain_DUKE_complete_genome

ttg ctg act gca --- tgt a-c tat ttt tta cat tac tac att ggc tac ggc aga tat acc tac tcc gcc acc --- aca cgc tcc ggt aaa cgg --g tca tgt gac gag gga gaa tat ctt gat aag agg cat aat caa tgt tgt aat -ca gtg tcc acc tgg aga att tgc caa ggt cag atg ta- --g tgg tag tga taa cac aaa atg tga acg ctg ccc acc tca tac ata tac cgc aat ccc caa tta ctc t-a atg gat gtc atc aat gta gaa aat gcc caa cag gat cat ttg ata agg taa agt gta ccg gaa cac aga aca gta aat -gt tcg tgt ctt cct ggt tgg tat tgt gct act gat tct tca cag act gaa gat tgt cga gat -tg tat acc aaa aag g-a gat gtc cat gcg gat act ttg gtg gaa tag atg aac aag gaa atc cta ttt gta aat cgt gct g-- -ta ttg gtg aat att gca act acc tac gta att ata gac ttg atc cat ttt ctc cat gca aa- --- cta tct aaa tgt aat taa tta --- --- --- --- --- --- --- --- tga ttt --- --- --- -tg atg ata atg tta -cc ata c-a tt- --- -at atc gct act tgg tta --g tgt att att cag tat g-a aga cct att aat aa- --- --- -tt act tat ctt ttg acg atc ttg tt- -at aat tat --- --- --- --- aat ata aaa --- -ac tta tgg --- --- cat agt aac tta t-- --- --- --- aat tgc ta- aca cga taa att cgt aat aat ctg ttt tgt tca aat ttt tat aag ga- --- --a tct aca ggc ata aaa at- aaa aat ata att tat aat ata c-- tct tac a-- acg cca tca tga ata gca gca gta aat taa ttg ctg tta tta atg gat tta gaa ata gtg gac gat ttt gtg ata tta ata tag tta tta atg atg aaa gga taa acg ctc ata aac tca tcc tat ctg gag cct ccg aat a-t ttt tcc att ctg ttt tcc aat aat ttt atc gat tct aat gaa tac gaa gtt aat cta agt cat tta gat tat caa agc gta aac gat ttg atc gat tat att tat ggg ata cct ttg agc cta act aac gat aac gtg aaa tat att ctt tca acc gct ga- --t ttt tta caa att gga tct gct att acg gag tgt gaa aat tac ata ctt aaa aat ctt tgt tct aaa aac tgt atc gat ttc --- --- tac ata tac gct gat aaa tat aat aac aag aaa ata gaa tca gcg tcg ttt aac aca ata tta caa aat att ttg aga ctc atc aac gat gaa aac ttt aaa tac tta aca gag gaa tca atg ata aaa att tta agc gat gat atg tta aat ata aaa aat gag gat ttt gca cca cta att ctc att aaa tgg tta gag agt act caa caa tca tgc acc gtc gag tta ctt aaa tgc ctc aga ata tca -tt gct ttc ccc aca agt tat aaa atc act tta tag tca tca act ggt tag ttc aat cta cga atg tat aac att ctt aaa caa tat agc att ctt gga tga atc att tcc tag ata cca tag cat cga gtt gat atc tat cgg tat aag taa ttc gca tga taa gat ttc cat aaa ctg cta caa tca t-a aaa aaa ata cat ggg aaa tga tat ctt cac gta gat ata ggt gta gtt tcg cag tgg ccg tcc tgg ata ata tta tct ata tga tgg gtg gat atg atc agt ccc cgt ata gaa gtt caa agg tta tag cgt aca ata cat gta caa att ctt gga tat atg ata tac cag agc t-a aaa tat cct cgt tct aat tgt ggg gga ctg gct gat gac gaa tac att tat tgt ata ggc ggc ata cgc gat cag gat tca tcg ttg aca tct agt att gat aaa tgg aag cca tca aaa cca tat tgg cag aag tat gct aaa atg cgc gaa cca aaa tgt gat atg ggg gtt gcg atg tta aac gga tta ata tat gtt ata ggt gga atc gtt aaa ggt g-- aca cgt gta ccg acg cac tag aga gtt tat cag aag atg gat gga tga agc atc aac gtc ttc caa taa aaa tgt cca ata tgt cga cga ttg ttc atg atg gca aga ttt ata tat ctg gag gtt aca aca ata gta gtg tag tta atg taa tat cga atc t-- agt cct tag cta taa tcc gat ata tga tga atg gac caa att atc atc att aaa tat tcc tag aat taa tcc cgc tct a

>JN654985.1:162218-164477_Vaccinia_virus_strain_Dryvax_clone_DPP20_complete_genome

ttg ctg act gta --- tgt a-c tat ttt tta cat tac tac att ggc tac ggc aga tat acc tac ttc gtc act gcc aca cgc tcc ggt aaa cgg --g gca tgt gac gag gga gaa tat ctt gat aag agg cat aat caa tgt tgt aat -ca gtg tcc acc tgg aga att tgc caa ggt tag atg ta- --g tgg tag tga taa cac aaa atg tga acg ctg ccc acc tca tac ata tac cac aat -cc caa tta ttc t-a atg gat gtc atc aat gta gaa aat gcc caa ccg gat cat ttg ata agg taa agt gta ccg gaa cac aga aca gta aat -gt tcg tgt ctt cct ggt tgg tat tgc gct act gat tct tca cag act gaa gat tgt tga aat -tg tgt acc aaa aag g-a gat gtc cat gcg gat act ttg gtg gaa tag atg aac aag gaa atc cta ttt gta aat cgt gct g-- -ta ttg gtg aat att gca act acc tac gta att ata aac ttg atc cat ttc ctc cat gca aa- --- cta tct aaa tgt aat taa tta --- --- --- --- --- --- --- --- tga ttt --- --- --- -tg atg ata atg tta -cc ata c-a tt- --- -at atc gct act tgg tta --g tgt att att cag tat g-a aga cct att aat aa- --- --- -tt act tat ctt ttg acg atc ttg tt- -at aat tat --- --- --- --- aat ata aaa --- -ac tta tgg --- --- cat agt aac tta t-- --- --- --- aat tgc ta- aca cga taa att cgt aat aat ctg ttt tgt tca aat ttt tat aag ga- --- --a tct aca ggc ata aaa at- aaa aat ata att tat aat ata c-- tct tac a-- acg cca tca tga ata gca gca gta aat taa ttg ctg tta tta atg gat tta gaa ata gtg gac gat ttt gtg ata tta ata tag tta tta atg atg aaa gga taa acg ctc ata aac tca tcc tat ctg gag cct ccg aat a-t ttt tcc att ctg ttt tcc aat aat ttt atc gat tct aat gaa tac gaa gtt aat cta agt cat tta gat tat caa agc gta aac gat ttg atc gat tat att tat ggg ata cct ttg agc cta act aac gat aac gtg aaa tat att ctt tca acc gct ga- --t ttt tta caa att gga tct gct att acg gag tgt gaa aat tac ata ctt aaa aat ctt tgt tct aaa aac tgt atc gat ttc --- --- tac ata tac gct gat aaa tat aat aac aag aaa ata gaa tca gcg tcg ttt aac aca ata tta caa aat att ttg aga ctc atc aac gat gaa aac ttt aaa tac tta aca gag gaa tca atg ata aaa att tta agc gat gat atg tta aat ata aaa aat gag gat ttt gca cca cta att ctc att aaa tgg tta gag agt act caa caa tca tgc acc gtc gag tta ctt aaa tgc ctc aga ata tca -tt gct ttc ccc aca agt tat aaa atc act tta tag tca tca act ggt tag ttc aat cta cga atg tat aac att ctt aaa caa tat agc att ctt gga tga atc att tcc tag ata cca tag cat cga gtt gat atc tat cgg tat aag taa ttc gca tga taa gat ttc cat aaa ctg cta caa tca t-a aaa aaa ata cat ggg aaa tga tat ctt cac gta gat ata ggt gta gtt tcg cag tgg ccg tcc tgg ata ata tta tct ata tga tgg gtg gat atg atc agt ccc cgt ata gaa gtt caa agg tta tag cgt aca ata cat gta caa att ctt gga tat atg ata tac cag agc t-a aaa tat cct cgt tct aat tgt ggg gga ctg gct gat gac gaa tac att tat tgt ata ggc ggc ata cgc gat cag gat tca tcg ttg aca tct agt att gat aaa tgg aag cca tca aaa cca tat tgg cag aag tat gct aaa atg cgc gaa cca aaa tgt gat atg ggg gtt gcg atg tta aac gga tta ata tat gtc atg ggt gga atc gtt aaa ggt g-- aca cgt gta ccg acg cac tag aga gtt tat cag aag atg gat gga tga agc atc aac gtc ttc caa taa aaa tgt cca ata tgt cga cga ttg ttc atg atg gca aga ttt ata tat ctg gag gtt aca aca ata gta gtg tag tta atg taa tat cga atc t-- agt cct tag cta taa ttc gat ata tga tga atg gac caa att atc atc att aaa tat tcc tag aat taa tcc cgc tct a

>JN654983.1:162269-164529_Vaccinia_virus_strain_Dryvax_clone_DPP17_complete_genome

ttg ctg act gta --- tgt a-c tat ttt tta cat tac tac att ggc tac ggc aga tat acc tac ttc gtc act gcc aca cgc tcc ggt aaa cgg --g gca tgt gac gag gga gaa tat ctt gat aag agg cat aat caa tgt tgt aat -cg gtg tcc acc tgg aga att tgc caa ggt tag atg ta- --a tgg taa cga taa cac aaa atg tga acg ctg ccc acc tca tac ata tac cgc aat ccc caa tta ctc t-a atg gat gtc atc aat gta gaa aat gcc caa cag gat cat ttg ata agg taa agt gta ccg gaa cac aga aca gta aat -gt tcg tgt ctt cct ggt tgg tat tgt gct act gat tct tca cag act gaa gat tgt cga gat -tg tat acc aaa aag g-a gat gtc cat gcg gat act ttg gtg gaa tag atg aac aag gaa atc cta ttt gta aat cgt gct g-- -ta ttg gtg aat att gca act acc tac gta att ata gac ttg atc cat ttt ctc cat gca aa- --- cta tct aaa tgt aat taa tta --- --- --- --- --- --- --- --- tga ttt --- --- --- -tg atg ata atg tta -cc ata c-a tt- --- -at atc gct act tgg tta --g tgt att att cag tat g-a aga cct att aat aa- --- --- -tt act tat ctt ttg acg atc ttg tt- -at aat tat --- --- --- --- aat ata aaa --- -ac tta tgg --- --- cat agt aac tta t-- --- --- --- aat tgc ta- aca cga taa att cgt aat aat ctg ttt tgt tca aat ttt tat aag ga- --- --a tct aca ggc ata aaa at- aaa aat ata att tat aat ata c-- tct tac a-- acg cca tca tga ata gca gca gta aat taa ttg ctg tta tta atg gat tta gaa ata gtg gac gat ttt gtg ata tta ata tag tta tta atg atg aaa gga taa acg ctc ata aac tca tcc tat ctg gag cct ccg aat a-t ttt tcc att ctg ttt tcc aat aat ttt atc gat tct aat gaa tac gaa gtt aat cta agt cat tta gat tat caa agc gta aac gat ttg atc gat tat att tat ggg ata cct ttg agc cta act aac gat aac gtg aaa tat att ctt tca acc gct ga- --t ttt tta caa att gga tct gct att acg gag tgt gaa aat tac ata ctt aaa aat ctt tgt tct aaa aac tgt atc gat ttc --- --- tac ata tac gct gat aaa tat aat aac aag aaa ata gaa tca gcg tcg ttt aac aca ata tta caa aat att ttg aga ctc atc aac gat gaa aac ttt aaa tac tta aca gag gaa tca atg ata aaa att tta agc gat gat atg tta aat ata aaa aat gag gat ttt gca cca cta att ctc att aaa tgg tta gag agt act caa caa tca tgc acc gtc gag tta ctt aaa tgc ctc aga ata tca -tt gct ttc ccc aca agt tat aaa atc act tta tag tca tca act ggt tag ttc aat cta cga atg tat aac att ctt aaa caa tat agc att ctt gga tga atc att tcc tag ata cca tag cat cga gtt gat atc tat cgg tat aag taa ttc gca tga taa gat ttc cat aaa ctg cta caa tca t-a aaa aaa ata cat ggg aaa tga tat ctt cac gta gat ata ggt gta gtt tcg cag tgg ccg tcc tgg ata ata tta tct ata tga tgg gtg gat atg atc agt ccc cgt ata gaa gtt caa agg tta tag cgt aca ata cat gta caa att ctt gga tat atg ata tac cag agc t-a aaa tat cct cgt tct aat tgt ggg gga ctg gct gat gac gaa tac att tat tgt ata ggc ggc ata cgc gat cag gat tca tcg ttg aca tct agt att gat aaa tgg aag cca tca aaa cca tat tgg cag aag tat gct aaa atg cgc gaa cca aaa tgt gat atg ggg gtt gcg atg tta aac gga tta ata tat gtc atg ggt gga atc gtt aaa ggt g-- aca cgt gta ccg acg cac tag aga gtt tat cag aag atg gat gga tga agc atc aac gtc ttc caa taa aaa tgt cca ata tgt cga cga ttg ttc atg atg gca aga ttt ata tat ctg gag gtt aca aca ata gta gtg tag tta atg taa tat cga atc t-- agt cct tag cta taa ttc gat ata tga tga atg gac caa att atc atc att aaa tat tcc tag aat taa tcc cgc tct a

>JN654981.1:162094-164354_Vaccinia_virus_strain_Dryvax_clone_DPP15_complete_genome

ttg ctg act gca --- tgt a-c tat ttt tta cat tac tac att ggc tac ggc aga tat acc tac ttc gtc act gcc aca cgc tcc ggt aaa cgg --g gca tgt gac gag gga gaa tat ctt gat aag agg cat aat caa tgt tgt aat -cg gtg tcc acc tgg aga att tgc caa ggt tag atg ta- --a tgg taa cga taa cac aaa atg tga acg ctg ccc acc tca tac ata tac cgc aat ccc caa tta ctc t-a atg gat gtc atc aat gta gaa aat gcc caa cag gat cat ttg ata agg taa agt gta ccg gaa cac aga aca gta aat -gt tcg tgt ctt cct ggt tgg tat tgc gct act gat tct tca cag act gaa gat tgt tga aat -tg tgt acc aaa aag g-a gat gtc cat gcg gat act ttg gtg gaa tag atg aac aag gaa atc cta ttt gta aat cgt gct g-- -ta ttg gtg aat att gca act acc tac gta att ata gac ttg atc cat ttt ctc cat gca aa- --- cta tct aaa tgt aat taa tta --- --- --- --- --- --- --- --- tga ttt --- --- --- -tg atg ata atg tta -cc ata c-a tt- --- -at atc gct act tgg tta --g tgt att att cag tat g-a aga cct att aat aa- --- --- -tt act tat ctt ttg acg atc ttg tt- -at aat tat --- --- --- --- aat ata aaa --- -ac tta tgg --- --- cat agt aac tta t-- --- --- --- aat tgc ta- aca cga taa att cgt aat aat ctg ttt tgt tca aat ttt tat aag ga- --- --a tct aca ggc ata aaa at- aaa aat ata atc tat aat ata c-- tct tac a-- acg cca tca tga ata gca gca gta aat taa ttg ctg tta tta atg gat tta gaa ata gtg gac gat ttt gtg ata tta ata tag tta tta atg atg aaa gga taa acg ctc ata aac tca tcc tat ctg gag cct ccg aat a-t ttt tcc att ctg ttt tcc aat aat ttt atc gat tct aat gaa tac gaa gtt aat cta agt cat tta gat tat caa agc gta aac gat ttg atc gat tat att tat ggg ata cct ttg agc cta act aac gat aac gtg aaa tat att ctt tca acc gct ga- --t ttt tta caa att gga tct gct att acg gag tgt gaa aat tac ata ctt aaa aat ctt tgt tct aaa aac tgt atc gat ttc --- --- tac ata tac gct gat aaa tat aat aac aag aaa ata gaa tca gcg tcg ttt aac aca ata tta caa aat att ttg aga ctc atc aac gat gaa aac ttt aaa tac tta aca gag gaa tca atg ata aaa att tta agc gat gat atg tta aat ata aaa aat gag gat ttt gca cca cta att ctc att aaa tgg tta gag agt act caa caa tca tgc acc gtc gag tta ctt aaa tgc ctc aga ata tca -tt gct ttc ccc aca agt tat aaa atc act tta tag tca tca act ggt tag ttc aat cta cga atg tat aac att ctt aaa caa tat agc att ctt gga tga atc att tcc tag ata cca tag cat cga gtt gat atc tat cgg tat aag taa ttc gcg tga taa aat ttc cat aaa ctg cta caa tca t-a aaa aaa ata cat ggg aaa tga tat ctt cac gta gat ata ggt gta gtt tcg cag tgg ccg tcc tgg ata ata tta tct ata tga tgg gtg gat atg atc agt ccc cgt ata gaa gtt caa agg tta tag cgt aca ata cat gta caa att ctt gga tat atg ata tac cag agc t-a aaa tat cct cgt tct aat tgt ggg gga ctg gct gat gac gaa tac att tat tgt ata ggc ggc ata cgc gat cag gat tca tcg ttg aca tct agt att gat aaa tgg aag cca tca aaa cca tat tgg cag aag tat gct aaa atg cgc gaa cca aaa tgt gat atg ggg gtt gcg atg tta aac gga tta ata tat gtt ata ggt gga atc gtt aaa ggt g-- aca cgt gta ccg acg cac tag aga gtt tat cag aag atg gat gga tga agc atc aac gtc ttc caa taa aaa tgt cca ata tgt cga cga ttg ttc atg atg gca aga ttt ata tat ctg gag gtt aca aca ata gta gtg tag tta atg taa tat cga atc t-- agt cct tag cta taa tcc gat ata tga tga atg gac caa att atc atc att aaa cat tcc tag aat taa tcc cgc tct a

>JN654976.1:162090-164349_Vaccinia_virus_strain_Dryvax_clone_DPP9_complete_genome

ttg ctg act gta --- tgt a-c tat ttt tta cat tac tac att ggc tac ggc aga tat acc tac ttc gtc act gcc aca cgc tcc ggt aaa cgg --g gca tgt gac gag gga gaa tat ctt gat aag agg cat aat caa tgt tgt aat -cg gtg tcc acc tgg aga att tgc caa ggt tag atg ta- --a tgg taa cga taa cac aaa atg tga acg ctg ccc acc tca tac ata tac cac aat -cc caa tta ttc t-a atg gat gtc atc aat gta gaa aat gcc caa ccg gat cat ttg ata agg taa agt gta ccg gaa cac aga aca gta aat -gt tcg tgt ctt cct ggt tgg tat tgc gct act gat tct tca cag act gaa gat tgt tga aat -tg tgt acc aaa aag g-a gat gtc cat gcg gat act ttg gtg gaa tag atg aac aag gaa atc cta ttt gta aat cgt gct g-- -ta ttg gtg aat att gca act acc tac gta att ata gac ttg atc cat ttt ctc cat gca aa- --- cta tct aaa tgt aat taa tta --- --- --- --- --- --- --- --- tga ttt --- --- --- -tg atg ata atg tta -cc ata c-a tt- --- -at atc gct act tgg tta --g tgt att att cag tat g-a aga cct att aat aa- --- --- -tt act tat ctt ttg acg atc ttg tt- -at aat tat --- --- --- --- aat ata aaa --- -ac tta tgg --- --- cat agt aac tta t-- --- --- --- aat tgc ta- aca cga taa att cgt aat aat ctg ttt tgt tca aat ttt tat aag ga- --- --a tct aca ggc ata aaa at- aaa aat ata att tat aat ata c-- tct tac a-- acg cca tca tga ata gca gca gta aat taa ttg ctg tta tta atg gat tta gaa ata gtg gac gat ttt gtg ata tta ata tag tta tta atg atg aaa gga taa acg ctc ata aac tca tcc tat ctg gag cct ccg aat a-t ttt tcc att ctg ttt tcc aat aat ttt atc gat tct aat gaa tac gaa gtt aat cta agt cat tta gat tat caa agc gta aac gat ttg atc gat tat att tat ggg ata cct ttg agc cta act aac gat aac gtg aaa tat att ctt tca acc gct ga- --t ttt tta caa att gga tct gct att acg gag tgt gaa aat tac ata ctt aaa aat ctt tgt tct aaa aac tgt atc gat ttc --- --- tac ata tac gct gat aaa tat aat aac aag aaa ata gaa tca gcg tcg ttt aac aca ata tta caa aat att ttg aga ctc atc aac gat gaa aac ttt aaa tac tta aca gag gaa tca atg ata aaa att tta agc gat gat atg tta aat ata aaa aat gag gat ttt gca cca cta att ctc att aaa tgg tta gag agt act caa caa cca tgc acc gtc gag tta ctt aaa tgc ctc aga ata tca -tt gct ttc ccc aca agt tat aaa atc act tta tag tca tcg act ggt tag ttc aat cta cga atg tat aac att ctt aaa caa tat agc att ctt gga tga atc att tcc tag ata cca tag cat cga gtt gat atc tat cgg tat aag taa ttc gcg tga taa aat ttc cat aaa ctg cta caa tca t-a aaa aaa ata cat ggg aaa tga tat ctt cac gta gat ata ggt gta gtt tcg cag tgg ccg tcc tgg ata ata tta ttt ata tga tgg gtg gat atg atc agt ccc cgt ata gaa gtt caa agg tta tag cgt aca ata cat gta caa att ctt gga tat atg ata tac cag agc t-a aaa tat cct cgt tct aat tgt ggg gga ctg gct gat gac gaa tac att tat tgt ata ggc ggc ata cgc gat cag gat tca tcg ttg aca tct agt att gat aga tgg aag cca tca aaa cca tat tgg cag aag tat gct aaa atg cgc gaa cca aaa tgt gat atg ggg gtt gcg atg tta aac gga tta ata tat gtc atg ggt gga atc gtt aaa ggt g-- aca cgt gta ccg acg cac tag aga gtt tat cag aag atg gat gga tga agc atc aac gtc ttc caa taa aaa tgt cca ata tgt cga cga ttg ttc atg atg gca aga ttt ata tat ctg gag gtt aca aca ata gta gtg tag tta atg taa tat cga atc t-- agt cct tag cta taa ttc gat ata tga tga atg gac caa att atc atc att aaa tat tcc tag aat taa tcc cgc tct a

>DQ121394.1:162349-164602_Vaccinia_virus_strain_Lister_clone_VACV107_complete_genome

ttg ctg act gta --- tgt a-c tat ttt tta cat tac tac att ggc tac ggc aga tat acc tac ttc gtc act gcc aca cgc tcc ggt aaa cgg --g gca tgt gac gag gga gaa tat ctt gat aag agg cat aat caa tgt tgt aat -cg gtg tcc acc tgg aga att tgc caa ggt tag atg ta- --a tgg taa cga taa cac aaa atg tga acg ctg ccc acc tca tac ata tac cgc aat ccc caa tta ctc t-a atg gat gtc atc aat gta gaa aat gcc caa cag gat cat ttg ata agg taa agt gta ccg gaa cac aga aca gta aat -gt tcg tgt ctt cct ggt tgg tat tgc gct act gat tct tca cag act gaa gat tgt cga gat -tg tat acc aaa aag g-a gat gtc cat gcg gat act ttg gtg gaa tag atg aac aag gaa atc cta ttt gta aat cgt gtt g-- -tg ttg gtg aat att gcg act acc tac gta att ata gac ttg atc cat ttc ctc cat gca aa- --- cta tct aaa tgt aat taa tta --- --- --- --- --- --- --- --- tga ttt --- --- --- -tg atg ata atg tta -cc ata c-a tt- --- -at atc gct act tgg tta --g tgt att att cag tat g-a aga cct att aat aa- --- --- -tt act tat ctt ttg acg atc ttg tt- -at aat tat --- --- --- --- aat ata aaa --- -ac tta tgg --- --- cat agt aac tta t-- --- --- --- aat tgc tg- acg cga taa att cgt aat aat ctg ttt tgt tca --- --- --- aag ga- --- --a tct aca ggc ata aaa at- aaa aat ata att tat aat ata c-- tct tac agc gcg cca tca tga ata aca gca gtg aat tga ttg ctg tta tta atg gat tta gaa ata gtg gac gat ttt gtg ata tta gta tag tta tta atg atg aaa gga taa acg ctc ata aac tca tcc tat ctg gag cct ccg aat a-t ttt tcc att ctg ttt tcc aat aat ttt atc gat tct aat gaa tac gaa gtt aat cta agt cat tta gat tat caa agt gtt aac gat ttg atc gat tat att tat ggg ata cct ttg agc cta act aac gat aac gtg aaa tat att ctt tca acc gct ga- --t ttt tta caa att gga tct gct att acg gag tgt gaa aat tac ata ctt aaa aat ctt tgt tct aga aac tgt atc gat ttc --- --- tac ata tac gct gat aaa tat aat aac aag aaa ata gaa tca gcg tcg ttt aac aca ata tta caa aat att ttg aga ctc atc aac gat gaa aac ttt aaa tac tta aca gag gaa tca atg ata aaa att tta agc gat gat atg tta aat ata aaa aat gag gat ttc gcc cca cta att ctc att aaa tgg tta gag agt act caa caa cca tgc acc gtc gag tta ctt aaa tgc ctc aga ata tca -tt gct ttc ccc aca agt tat aaa atc act tta tag tca tcg act ggt tag ttc aat cta cga atg tat aac att ctt aaa caa tat agc att ctt gga tga atc att tcc tag ata cca tag cat cga gtt gat atc tat cgg tat aag taa ttc gca tga taa gat ttc cat aaa ctg cta caa tca t-a aaa aaa ata cat ggg aaa tga tat ctt cac gta gat ata ggt gta gtt tcg cag tgg ccg tcc tgg ata ata tta ttt ata tga tgg gtg gat atg atc agt ccc cgt ata gaa gtt caa agg tta tag cgt aca ata cat gta caa att ctt gga tat atg ata tac cag agc t-a aaa tat cct cgt tct aat tgt ggg gga ctg gct gat gac gaa tac att tat tgt ata ggc ggc ata cgc gat cag gat tca tcg ttg aca tct agt att gat aaa tgg aag cca tca aaa cca tat tgg cag aag tat gct aaa atg cgc gaa cca aaa tgt gat atg ggg gtt gcg atg tta aac gga tta ata tat gtt ata ggt gga atc gtt aaa ggt g-- aca cgt gta ccg acg cac tag aga gtt tat cag aag atg gat gga tga agc atc aac gtc ttc caa taa aaa tgt cca ata tgt cga cga ttg ttc atg atg gca aga ttt ata tat ctg gag gtt aca aca ata gta gtg tag tta atg taa tat cga atc t-- agt cct tag cta taa ttc gat ata tga tga atg gac caa att atc atc att aaa cat tcc tag aat taa tcc cgc tct a

>KP233807.1:159976-162224_Vaccinia_virus_strain_L-IVP_complete_genome

ttg ctg act gta --- tgt a-c tat ttt tta cat tac tac att ggc tac ggc aga tat acc tac ttc gtc acc --- --- --- --- --- -aa cgg --g tca tgt gac gag gga gaa tat ctt gat aag agg cat aat caa tgt tgt aat -cg gtg tcc acc tgg aga att tgc caa ggt tag atg ta- --a tgg taa cga taa cac aaa atg tga acg ctg ccc acc tca tac ata tac cgc aat ccc caa tta ctc t-a atg gat gtc atc aat gta gaa aat gcc caa cag gat cat ttg ata agg taa agt gta ccg gaa cac aga aca gta aat -gt tcg tgt ctt cct ggt tgg tat tgc gct act gat tct tca cag act gaa gat tgt cga gat -tg tat acc aaa aag g-a gat gtc cat gcg gat act ttg gtg gaa tag atg aac aag gaa atc cta ttt gta aat cgt gtt g-- -tg ttg gtg aat att gcg act acc tac gta att ata gac ttg atc cat ttc ctc cat gca aa- --- cta tct aaa tgt aat taa tta --- --- --- --- --- --- --- --- tga ttt --- --- --- -tg atg ata atg tta -cc ata c-a tt- --- -at atc gct act tgg tta --g tgt att att cag tat g-a aga cct att aat aa- --- --- -tt act tat ctt ttg acg atc ttg tt- -at aat tat --- --- --- --- aat ata aaa --a tac tta tgg --- --- cat agt aac tca t-- --- --- --- aat tgc tg- acg cga taa att cgt aat aat ctg ttt tgt tca aat ttt tat aag ga- --- --a tct aca ggc ata aaa at- aaa aat ata att tat aat ata c-- tct tac agc gcg cca tca tga ata aca gca gtg aat tga ttg ctg tta tta atg gat tta gaa ata gtg gac gat ttt gtg ata tta gta tag tta tta atg atg aaa tga taa acg ctc aca gac tca tcc tat ctg gag cct ccg aat a-t ttt tcc att ctg ttt tcc aat aat ttt atc gat tct aat gaa tac gaa gtt aat cta agt cat tta gat tat caa agt gtt aac gat ttg atc gat tac att tat ggg ata cct ttg agc cta act aac gat aac gtg aaa tat att ctt tca acc gct ga- --t ttt tta caa att gga tct gct att acg gag tgt gaa aat tac ata ctt aaa aat ctt tgt tct aga aac tgt atc gat ttc --- --- tac ata tac gct gat aaa tat aat aac aag aaa ata gaa tca gcg tcg ttt aac aca ata tta caa aat att ttg aga ctc atc aac gat gaa aac ttt aaa tac tta aca gag gaa tca atg ata aaa att tta agc gat gat atg tta aat ata aaa aat gag gat ttc gcc cca cta att ctc att aaa tgg tta gag agt act caa caa cca tgc acc gtc gag tta ctt aaa tgc ctc aga ata tca -tt gct ttc ccc aca agt tat aaa atc act tta tag tca tcg act ggt tag ttc aat cta cga atg tat aac att ctt aaa caa tat agc att ctt gga tga atc att tcc tag ata cca tag cat cga gtt gat atc tat cgg tat aag taa ttc gca tga taa gat ttc cat aaa ctg cta caa tca t-a aaa aaa ata cat ggg aaa tga tat ctt cac gta gat ata ggt gta gtt tcg cag tgg ccg tcc tgg ata ata tta tct ata tga tgg gtg gat atg atc agt ccc cgt ata gaa gtt caa agg tta tag cgt aca ata cat gta caa att ctt gga tat atg ata tac cag agc t-a aaa tat cct cgt tct aat tgt ggg gga ctg gct gat gac gaa tac att tat tgt ata ggc ggc ata cgc gat cag gat tca tcg ttg aca tct agt att gat aga tgg aag cca tca aaa cca tat tgg cag aag tat gct aaa atg cgc gaa cca aaa tgt gat atg ggg gtt gcg atg tta aac gga tta ata tat gtt ata ggt gga atc gtt aaa ggt g-- aca cgt gta ccg acg cac tag aga gtt tat cag aag atg gat gga tga agc atc aac gtc ttc caa taa aaa tgt cca ata tgt cga cga ttg ttc atg atg gca aga ttt ata tat ctg gag gtt aca aca ata gta gtg tag tta atg taa tat cga atc t-- agt cct tag cta taa ttc gat ata tga tga atg gac caa att atc atc att aaa cat tcc tag aat taa tcc cgc tct a

>MN369532.1:147235-149497_Vaccinia_virus_isolate_VK01

ttg ctg act gca --- tgt a-c tat ttt tta cat tac tac att ggc tac ggc aga tat acc tac tcc gcc acc --- aca cgc tcc ggt aaa cgg --g tca tgt gac aag gga gaa tat ctt gat aag agg cat aat caa tgt tgt aat -cg gtg tcc acc tgg aga att tgc caa ggt tag atg ta- --a tgg taa cga taa cac aaa atg tga acg ctg ccc acc tca tac ata tac cgc aat -cc caa tta ttc t-a atg gat gtc atc aat gta gaa aat gcc caa ccg gat cat ttg ata agg taa agt gta ccg gaa cac aga aca gta aat -gt tcg tgt ctt cct ggt tgg tat tgc gct act gat tct tca cag act gaa gat tgt tga aat -tg tgt acc aaa aag g-a gat gtc cat gcg gat act ttg gtg gaa tag atg aac aag gaa atc cta ttt gta aat cgt gct g-- -ta ttg gtg aat att gca act acc tac gta att ata gac ttg atc cat ttt ctc cat gca aa- --- cta tct aaa tgt aat taa tta --- --- --- --- --- --- tga ttt tga ttt --- --- --- -tg atg ata atg tta -cc ata c-a tt- --- -at atc gct act tgg tta --g tgt att att cag tat g-a aga cct att aat aa- --- --- -tt act tat ctt ttg acg atc ttg tt- -at aat tat --- --- --- --- aat ata aaa --- -ac tta tgg --- --- cat agt aac tta t-- --- --- --- aat tgc ta- aca cga taa att cgt aat aat ctg ttt tgt tca aat ttt tat aag ga- --- --a tct aca ggc ata aaa at- aaa aat ata atc tat aat ata c-- tct tac a-- acg cca tca tga ata gca gca gta aat taa ttg ctg tta tta atg gat tta gaa ata gtg gac gat ttt gtg ata tta ata tag tta tta atg atg aaa gga taa acg ctc ata aac tca tcc tat ctg gag cct ccg aat a-t ttt tcc att ctg ttt tcc aat aat ttt atc gat tct aat gaa tac gaa gtt aat cta agt cat tta gat tat caa agc gta aac gat ttg atc gat tat att tat ggg ata cct ttg agc cta act aac gat aac gtg aaa tat att ctt tca acc gct ga- --t ttt tta caa att gga tct gct att acg gag tgt gaa aat tac ata ctt aaa aat ctt tgt tct aaa aac tgt atc gat ttc --- --- tac ata tac gct gat aaa tat aat aac aag aaa ata gaa tca gcg tcg ttt aac aca ata tta cga aat att ttg aga ctc atc aac gat gaa aac ttt aaa tac tta aca gag gaa tca atg ata aaa att tta agc gat gat atg tta aat ata aaa aat gag gat ttc gcc cca cta att ctc att aaa tgg tta gag agt act caa caa cca tgc acc gtc gag tta ctt aaa tgc ctc aga ata tca -tt gct ttc ccc aca agt tat aaa atc act tta tag tca tcg act ggt tag ttc aat cta cga atg tat aac att ctt aaa caa tat agc att ctt gga tga atc att tcc tag ata cca tag cat cga gtt gat atc tat cgg tat aag taa ttc gcg tga taa aat ttc cat aaa ctg cta caa tca t-a aaa aaa ata cat ggg aaa tga tat ctt cac gta gat ata ggt gta gtt tcg cag tgg ccg tcc tgg ata ata tta ttt ata tga tgg gtg gat atg atc agt ccc cgt ata gaa gtt caa agg tta tag cgt aca ata cat gta caa att ctt gga tat atg ata tac cag agc t-a aaa tat cct cgt tct aat tgt ggg gga ctg gct gat gac gaa tac att tat tgt ata ggc ggc ata cgc gat cag gat tca tcg ttg aca tct agt att gat aaa tgg aag cca tca aaa cca tat tgg cag aag tat gct aaa atg cgc gaa cca aaa tgt gat atg ggg gtt gcg atg tta aac gga tta ata tat gtt ata ggt gga atc gtt aaa ggt g-- aca cgt gta ccg acg cac tag aga gtt tat cag aag atg gat gga tga agc atc aac gtc ttc caa taa aaa tgt cca ata tgt cga cga ttg ttc atg atg gca aga ttt ata tat ctg gag gtt aca aca ata gta gtg tag tta atg taa tat cga atc t-- agt cct tag cta taa ttc gat ata tga tga atg gac caa att atc atc att aaa tat tcc tag aat taa tcc cgc tct a

>BK013339.1:176991-179253_TPA_asm:_Vaccinia_virus_isolate_VK01

ttg ctg act gca --- tgt a-c tat ttt tta cat tac tac att ggc tac ggc aga tat acc tac tcc gcc acc --- aca cgc tcc ggt aaa cgg --g tca tgt gac aag gga gaa tat ctt gat aag agg cat aat caa tgt tgt aat -cg gtg tcc acc tgg aga att tgc caa ggt tag atg ta- --a tgg taa cga taa cac aaa atg tga acg ctg ccc acc tca tac ata tac cgc aat -cc caa tta ttc t-a atg gat gtc atc aat gta gaa aat gcc caa ccg gat cat ttg ata agg taa agt gta ccg gaa cac aga aca gta aat -gt tcg tgt ctt cct ggt tgg tat tgc gct act gat tct tca cag act gaa gat tgt tga aat -tg tgt acc aaa aag g-a gat gtc cat gcg gat act ttg gtg gaa tag atg aac aag gaa atc cta ttt gta aat cgt gct g-- -ta ttg gtg aat att gca act acc tac gta att ata gac ttg atc cat ttt ctc cat gca aa- --- cta tct aaa tgt aat taa tta --- --- --- --- --- --- tga ttt tga ttt --- --- --- -tg atg ata atg tta -cc ata c-a tt- --- -at atc gct act tgg tta --g tgt att att cag tat g-a aga cct att aat aa- --- --- -tt act tat ctt ttg acg atc ttg tt- -at aat tat --- --- --- --- aat ata aaa --- -ac tta tgg --- --- cat agt aac tta t-- --- --- --- aat tgc ta- aca cga taa att cgt aat aat ctg ttt tgt tca aat ttt tat aag ga- --- --a tct aca ggc ata aaa at- aaa aat ata atc tat aat ata c-- tct tac a-- acg cca tca tga ata gca gca gta aat taa ttg ctg tta tta atg gat tta gaa ata gtg gac gat ttt gtg ata tta ata tag tta tta atg atg aaa gga taa acg ctc ata aac tca tcc tat ctg gag cct ccg aat a-t ttt tcc att ctg ttt tcc aat aat ttt atc gat tct aat gaa tac gaa gtt aat cta agt cat tta gat tat caa agc gta aac gat ttg atc gat tat att tat ggg ata cct ttg agc cta act aac gat aac gtg aaa tat att ctt tca acc gct ga- --t ttt tta caa att gga tct gct att acg gag tgt gaa aat tac ata ctt aaa aat ctt tgt tct aaa aac tgt atc gat ttc --- --- tac ata tac gct gat aaa tat aat aac aag aaa ata gaa tca gcg tcg ttt aac aca ata tta cga aat att ttg aga ctc atc aac gat gaa aac ttt aaa tac tta aca gag gaa tca atg ata aaa att tta agc gat gat atg tta aat ata aaa aat gag gat ttc gcc cca cta att ctc att aaa tgg tta gag agt act caa caa cca tgc acc gtc gag tta ctt aaa tgc ctc aga ata tca -tt gct ttc ccc aca agt tat aaa atc act tta tag tca tcg act ggt tag ttc aat cta cga atg tat aac att ctt aaa caa tat agc att ctt gga tga atc att tcc tag ata cca tag cat cga gtt gat atc tat cgg tat aag taa ttc gcg tga taa aat ttc cat aaa ctg cta caa tca t-a aaa aaa ata cat ggg aaa tga tat ctt cac gta gat ata ggt gta gtt tcg cag tgg ccg tcc tgg ata ata tta ttt ata tga tgg gtg gat atg atc agt ccc cgt ata gaa gtt caa agg tta tag cgt aca ata cat gta caa att ctt gga tat atg ata tac cag agc t-a aaa tat cct cgt tct aat tgt ggg gga ctg gct gat gac gaa tac att tat tgt ata ggc ggc ata cgc gat cag gat tca tcg ttg aca tct agt att gat aaa tgg aag cca tca aaa cca tat tgg cag aag tat gct aaa atg cgc gaa cca aaa tgt gat atg ggg gtt gcg atg tta aac gga tta ata tat gtt ata ggt gga atc gtt aaa ggt g-- aca cgt gta ccg acg cac tag aga gtt tat cag aag atg gat gga tga agc atc aac gtc ttc caa taa aaa tgt cca ata tgt cga cga ttg ttc atg atg gca aga ttt ata tat ctg gag gtt aca aca ata gta gtg tag tta atg taa tat cga atc t-- agt cct tag cta taa ttc gat ata tga tga atg gac caa att atc atc att aaa tat tcc tag aat taa tcc cgc tct a

>KM044310.1:152164-154412_Vaccinia_virus_strain_Tashkent_clone_TKT4_complete_genome

ttg ctg act gca --- tgt a-c tat ttt tta cat tac tac att ggc tac ggc aga tat acc tac tcc gcc acc --- --- --- --- --- -aa cgg --g gca tgt gac gag gga gaa tat ctt gat aag agg cat aat caa tgt tgt aat -cg gtg tcc acc tgg aga att tgc caa ggt tag atg ta- --a tgg taa cga taa cac aaa atg tga acg ctg ccc acc tca tac ata tac cgc aat ccc caa tta ttc t-a atg gat gtc atc aat gta gaa aat gcc caa cag gat cat ttg ata agg taa agt gta ccg gaa cac aga aca gta aat -gt tcg tgt ctt cct ggt tgg tat tgc gct act gat tct tca cag act gaa gat tgt cga gat -tg tat acc aaa aag g-a gat gtc cat gcg gat act ttg gtg gaa tag atg aac aag gaa atc cta ttt gta aat cgt gtt g-- -tg ttg gtg aat att gcg act acc tac gta att ata gac ttg atc cat ttc ctc cat gca aa- --- cta tct aaa tgt aat taa tta --- --- --- --- --- --- --- --- tga ttt --- --- --- -tg atg ata atg tta -cc ata c-a tt- --- -at atc gct act tgg tta --g tgt att att cag tat g-a aga cct att aat aa- --- --- -tt act tat ctt ttg acg atc ttg tt- -at aat tat --- --- --- --- aat ata aaa --a tac tta tgg --- --- cat agt aac tca t-- --- --- --- aat tgc tg- acg cga taa att cgt aat aat ctg ttt tgt tca aat ttt tat aag ga- --- --a tct aca ggc ata aaa at- aaa aat ata att tat aat ata c-- tct tac agc gcg cca tca tga ata aca gca gtg aat tga ttg ctg tta tta atg gat tta gaa ata gtg gac gat ttt gtg ata tta gta tag tta tta atg atg aaa tga taa acg ctc aca gac tca tcc tat ctg gag cct cca aat a-t ttt tcc att ctg ttt tcc aat aat ttt atc gat tct aat gaa tac gaa gtt aat cta agt cat tta gat tat caa agc gtt aac gat ttg atc gat tac att tat ggg ata cct ttg agc cta act aac gat aac gtg aaa tat att ctt tca acc gct ga- --t ttt tta caa att gga tct gct att acg gag tgc gaa aaa tac ata ctt aaa aat ctt tgt tct aaa aac tgt atc gat ttc --- --- tac ata tac gct gat aaa tat aat aac aag aaa ata gaa tca gcg tcg ttt aac aca ata tta cga aat att ttg aga ctc atc aac gat gaa aac ttt aaa tac tta aca gag gaa tca atg ata aaa att tta agc gat gat atg tta aat ata aaa aat gag gat ttt gca cca cta att ctc att aaa tgg tta gag agt act caa caa tca tgc acc gtc gag tta ctt aga tgc ctc aga ata tca -tt gct ttc ccc aca agt tat aaa atc act tta tag tca tcg act ggt tag ttc aat cta cga atg tat aac att ctt aaa caa tat agc att ctt gga tga atc att tcc tag ata cca tag cat cga gtt gat atc tat cgg tat aag taa ttc gca tga taa aat ttc cat aaa ctg cta caa tca t-a aaa aaa ata cat ggg aaa tga tat ctt cac gta gat ata ggt gta gtt tcg cag tgg ccg tcc tgg ata ata tta ttt ata tga tgg gtg gat atg atc agt ccc cgt ata gaa gtt caa agg tta tag cgt aca ata cat gta caa att ctt gga tat atg ata tac cag agc t-a aaa tat cct cgt tct aat tgt ggg gga ctg gct gat gac gaa tac att tat tgt ata ggc ggc ata cgc gat cag gat tca tcg ttg aca tct agt att gat aaa tgg aag cca tca aaa cca tat tgg cag aag tat gct aaa atg cgc gaa cca aaa tgt gat atg ggg gtt gcg atg tta aac gga tta ata tat gtc atg ggt gga atc gtt aaa ggt g-- aca cgt gta ccg acg cac tag aga gtt tat cag aag atg gat gga tga agc atc aac gtc ttc caa taa aaa tgt cca ata tgt cga cga ttg ttc atg atg gca aga ttt ata tat ctg gag gtt aca aca ata gta gtg tag tta atg taa tat cga atc t-- agt cct tag cta taa tcc gat ata tga tga atg gac caa att atc atc att aaa cat tcc tag aat taa tcc cgc tct a

>KM044309.1:152595-154843_Vaccinia_virus_strain_Tashkent_clone_TKT3_complete_genome

ttg ctg act gca --- tgt a-c tat ttt tta cat tac tac att ggc tac ggc aga tat acc tac tcc gcc acc --- --- --- --- --- -aa cgg --g gca tgt gac gag gga gaa tat ctt gat aag agg cat aat caa tgt tgt aat -cg gtg tcc acc tgg aga att tgc caa ggt tag atg ta- --a tgg taa cga taa cac aaa atg tga acg ctg ccc acc tca tac ata tac cgc aat ccc caa tta ttc t-a atg gat gtc atc aat gta gaa aat gcc caa cag gat cat ttg ata agg taa agt gta ccg gaa cac aga aca gta aat -gt tcg tgt ctt cct ggt tgg tat tgc gct act gat tct tca cag act gaa gat tgt cga gat -tg tat acc aaa aag g-a gat gtc cat gcg gat act ttg gtg gaa tag atg aac aag gaa atc cta ttt gta aat cgt gtt g-- -tg ttg gtg aat att gcg act acc tac gta att ata gac ttg atc cat ttc ctc cat gca aa- --- cta tct aaa tgt aat taa tta --- --- --- --- --- --- --- --- tga ttt --- --- --- -tg atg ata atg tta -cc ata c-a tt- --- -at atc gct act tgg tta --g tgt att att cag tat g-a aga cct att aat aa- --- --- -tt act tat ctt ttg acg atc ttg tt- -at aat tat --- --- --- --- aat ata aaa --a tac tta tgg --- --- cat agt aac tca t-- --- --- --- aat tgc tg- acg cga taa att cgt aat aat ctg ttt tgt tca aat ttt tat aag ga- --- --a tct aca ggc ata aaa at- aaa aat ata att tat aat ata c-- tct tac agc gcg cca tca tga ata aca gca gtg aat tga ttg ctg tta tta atg gat tta gaa ata gtg gac gat ttt gtg ata tta gta tag tta tta atg atg aaa tga taa acg ctc aca gac tca tcc tat ctg gag cct cca aat a-t ttt tcc att ctg ttt tcc aat aat ttt atc gat tct aat gaa tac gaa gtt aat cta agt cat tta gat tat caa agc gtt aac gat ttg atc gat tac att tat ggg ata cct ttg agc cta act aac gat aac gtg aaa tat att ctt tca acc gct ga- --t ttt tta caa att gga tct gct att acg gag tgc gaa aaa tac ata ctt aaa aat ctt tgt tct aaa aac tgt atc gat ttc --- --- tac ata tac gct gat aaa tat aat aac aag aaa ata gaa tca gcg tcg ttt aac aca ata tta cga aat att ttg aga ctc atc aac gat gaa aac ttt aaa tac tta aca gag gaa tca atg ata aaa att tta agc gat gat atg tta aat ata aaa aat gag gat ttt gca cca cta att ctc att aaa tgg tta gag agt act caa caa tca tgc acc gtc gag tta ctt aga tgc ctc aga ata tca -tt gct ttc ccc aca agt tat aaa atc act tta tag tca tcg act ggt tag ttc aat cta cga atg tat aac att ctt aaa caa tat agc att ctt gga tga atc att tcc tag ata cca tag cat cga gtt gat atc tat cgg tat aag taa ttc gca tga taa aat ttc cat aaa ctg cta caa tca t-a aaa aaa ata cat ggg aaa tga tat ctt cac gta gat ata ggt gta gtt tcg cag tgg ccg tcc tgg ata ata tta ttt ata tga tgg gtg gat atg atc agt ccc cgt ata gaa gtt caa agg tta tag cgt aca ata cat gta caa att ctt gga tat atg ata tac cag agc t-a aaa tat cct cgt tct aat tgt ggg gga ctg gct gat gac gaa tac att tat tgt ata ggc ggc ata cgc gat cag gat tca tcg ttg aca tct agt att gat aaa tgg aag cca tca aaa cca tat tgg cag aag tat gct aaa atg cgc gaa cca aaa tgt gat atg ggg gtt gcg atg tta aac gga tta ata tat gtc atg ggt gga atc gtt aaa ggt g-- aca cgt gta ccg acg cac tag aga gtt tat cag aag atg gat gga tga agc atc aac gtc ttc caa taa aaa tgt cca ata tgt cga cga ttg ttc atg atg gca aga ttt ata tat ctg gag gtt aca aca ata gta gtg tag tta atg taa tat cga atc t-- agt cct tag cta taa tcc gat ata tga tga atg gac caa att atc atc att aaa cat tcc tag aat taa tcc cgc tct a

>BK013342.1:167114-169376_TPA_asm:_Vaccinia_virus_isolate_VK08

ttg ctg act gca --- tgt a-c tat ttt tta cat tac tac att ggc tac ggc aga tat acc tac tcc gcc acc --- aca cgc tcc ggt aaa cgg --g tca tgt gac aag gga gaa tat ctt gat aag agg cat aat caa tgt tgt aat -cg gtg tcc acc tgg aga att tgc caa ggt tag atg ta- --a tgg taa cga taa cac aaa atg tga acg ctg ccc acc tca tac ata tac cgc aat -cc caa tta ttc t-a atg gat gtc atc aat gta gaa aat gcc caa ccg gat cat ttg ata agg taa agt gta ccg gaa cac aga aca gta aat -gt tcg tgt ctt cct ggt tgg tat tgc gct act gat tct tca cag act gaa gat tgt tga aat -tg tgt acc aaa aag g-a gat gtc cat gcg gat act ttg gtg gaa tag atg aac aag gaa atc cta ttt gta aat cgt gct g-- -ta ttg gtg aat att gca act acc tac gta att ata gac ttg atc cat ttt ctc cat gca aa- --- cta tct aaa tgt aat taa tta --- --- --- --- --- --- tga ttt tga ttt --- --- --- -tg atg ata atg tta -cc ata c-a tt- --- -at atc gct act tgg tta --g tgt att att cag tat g-a aga cct att aat aa- --- --- -tt act tat ctt ttg acg atc ttg tt- -at aat tat --- --- --- --- aat ata aaa --- -ac tta tgg --- --- cat agt aac tta t-- --- --- --- aat tgc ta- aca cga taa att cgt aat aat ctg ttt tgt tca aat ttt tat aag ga- --- --a tct aca ggc ata aaa at- aaa aat ata atc tat aat ata c-- tct tac a-- acg cca tca tga ata gca gca gta aat taa ttg ctg tta tta atg gat tta gaa ata gtg gac gat ttt gtg ata tta ata tag tta tta atg atg aaa gga taa acg ctc ata aac tca tcc tat ctg gag cct ccg aat a-t ttt tcc att ctg ttt tcc aat aat ttt atc gat tct aat gaa tac gaa gtt aat cta agt cat tta gat tat caa agc gta aac gat ttg atc gat tat att tat ggg ata cct ttg agc cta act aac gat aac gtg aaa tat att ctt tca acc gct ga- --t ttt tta caa att gga tct gct att acg gag tgt gaa aat tac ata ctt aaa aat ctt tgt tct aaa aac tgt atc gat ttc --- --- tac ata tac gct gat aaa tat aat aac aag aaa ata gaa tca gcg tcg ttt aac aca ata tta cga aat att ttg aga ctc atc aac gat gaa aac ttt aaa tac tta aca gag gaa tca atg ata aaa att tta agc gat gat atg tta aat ata aaa aat gag gat ttc gcc cca cta att ctc att aaa tgg tta gag agt act caa caa cca tgc acc gtc gag tta ctt aaa tgc ctc aga ata tca -tt gct ttc ccc aca agt tat aaa atc act tta tag tca tcg act ggt tag ttc aat cta cga atg tat aac att ctt aaa caa tat agc att ctt gga tga atc att tcc tag ata cca tag cat cga gtt gat atc tat cgg tat aag taa ttc gcg tga taa aat ttc cat aaa ctg cta caa tca t-a aaa aaa ata cat ggg aaa tga tat ctt cac gta gat ata ggt gta gtt tcg cag tgg ccg tcc tgg ata ata tta ttt ata tga tgg gtg gat atg atc agt ccc cgt ata gaa gtt caa agg tta tag cgt aca ata cat gta caa att ctt gga tat atg ata tac cag agc t-a aaa tat cct cgt tct aat tgt ggg gga ctg gyt gat gac gaa tac att tat tgt ata ggc ggc ata cgc gat cag gat tca tcg ttg aca tct agt att gat aaa tgg aag cca tca aaa cca tat tgg cag aag tat gct aaa atg cgc gaa cca aaa tgt gat atg ggg gtt gcg atg tta aac gga tta ata tat gtt ata ggt gga atc gtt aaa ggt g-- aca cgt gta ccg acg cac tag aga gtt tat cag aag atg gat gga tga agc atc aac gtc ttc caa taa aaa tgt cca ata tgt cga cga ttg ttc atg atg gca aga ttt ata tat ctg gag gtt aca aca ata gta gtg tag tta atg taa tat cga atc t-- agt cct tag cta taa ttc gat ata tga tga atg gac caa att atc atc att aaa tat tcc tag aat taa tcc cgc tct a

>KJ125439.1:162896-165142_Vaccinia_virus_strain_IHD-W_clone_IHDW1_complete_genome

ttg ctg act gca --- tgt a-c tat ttt tta cat tac tac att ggc tac ggc aga tat acc tac tcc gcc acc --- --- --- --- --- -aa cgg --g tca tgt gac aag gga gaa tat ctt gat aag agg cat aat caa tgt tgt aat -cg gtg tcc acc tgg aga att tgc caa ggt tag atg ta- --a tgg taa cga taa cac aaa atg tga acg ctg ccc acc tca tac ata tac cac aat ccc caa tta ttc t-a atg gat gtc atc aat gta gaa aat gcc caa ccg gat cat ttg ata agg taa agt gta ccg gaa cac aga aca gta aat -gt tcg tgt ctt cct ggt tgg tat tgc gct act gat tct tca cag act gaa gat tgt tga aat -tg tgt acc aaa aag g-a gat gtc cat gcg gat act ttg gtg gaa tag atg aac aag gaa atc cta ttt gta aat cgt gct g-- -ta ttg gtg aat att gca act acc tac gta att ata gac ttg atc cat ttt ctc cat gca aa- --- cta tct aaa tgt aat taa tta --- --- --- --- --- --- --- --- tga ttt --- --- --- -tg atg ata atg tta -cc ata c-a tt- --- -at atc gct act tgg tta --g tgt att att cag tat g-a aga cct att aat aa- --- --- -tt act tat ctt ttg acg atc ttg tt- -at aat tat --- --- --- --- aat ata aaa --- -ac tta tgg --- --- cat agt aac tca t-- --- --- --- aat tgc tg- acg cga taa att cgt aat aat ctg ttt tgt tca aat ttt tat aag ga- --- --a tct aca ggc ata aaa at- aaa aat ata att tat aat ata c-- tct tac agc gcg cca tca tga ata gca gca gta aat taa ttg ctg tta tta atg gat tta gaa ata gtg gac gat ttt gtg ata tta ata tag tta tta atg atg aaa gga taa acg ctc aca gac tca tcc tat ctg gag cct ccg aat a-t ttt tcc att ctg ttt tcc aat aat ttt atc gat tct aat gaa tac gaa gtt aat cta agt cat tta gat tat caa agt gtt aac gat ttg atc gat tac att tat ggg ata cct ttg agc cta act aac gat aac gtg aaa tat att ctt tca acc gct ga- --t ttt tta caa att gga tct gcc att act gag tgc gaa aaa tac ata ctt aaa aat ctt tgt tct aga aac tgt atc gat ttc --- --- tac ata tac gct gat aaa tat aat aac aag aaa ata gaa tca gcg tcg ttt aac aca ata tta cga aat att ttg aga ctc atc aac gat gaa aac ttt aaa tac tta aca gag gaa tca atg ata aaa att tta agc gat gat atg tta aat ata aaa aat gag gat ttt gca cca cta att ctc att aaa tgg tta gag agt act caa caa tca tgc acc gtc gag tta ctt aga tgc ctc aga ata tca -tt gct ttc ccc aca agt tat aaa atc act tta tag tca tca act ggt tag ttc aat cta cga atg tat aac att ctt aaa caa tat agc att ctt gga tga atc att tcc tag ata cca tag cat cga gtt gat atc tat cgg tat aag taa ttc gca tga taa gat ttc cat aaa ctg cta caa tca t-a aaa aaa ata cat ggg aaa tga tat ctt cac gta gat ata ggt gta gtt tcg cag tgg ccg tcc tgg ata ata tta tct ata tga tgg gtg gat atg atc agt ccc cgt ata gaa gtt caa agg tta tag cgt aca ata cat gta caa att ctt gga tat atg ata tac cag agc t-a aaa tat cct cgt tct aat tgt ggg gga ctg gct gat gac gaa tac att tat tgt ata ggc ggc ata cgc gat cag gat tca tcg ttg aca tct agt att gat aga tgg aag cca tca aaa cca tat tgg cag aag tat gct aaa atg cgc gaa cca aaa tgt gat atg ggg gtt gcg atg tta aac gga tta ata tat gtc atg ggt gga atc gtt aaa ggt g-- aca cgt gta ccg acg cac tag aga gtt tat cag aag atg gat gga tga agc atc aac gtc ttc caa taa aaa tgt cca ata tgt cga cga ttg ttc atg atg gca aga ttt ata tat ctg gag gtt aca aca ata gta gtg tag tta atg taa tat cga atc t-- agt cct tag cta taa tcc gat ata tga tga atg gac caa att atc atc att aaa cat tcc tag aat taa tcc cgc tct a

>KC201194.1:158224-160470_Vaccinia_virus_strain_IHD-W_partial_sequence

ttg ctg act gca --- tgt a-c tat ttt tta cat tac tac att ggc tac ggc aga tat acc tac tcc gcc acc --- --- --- --- --- -aa cgg --g tca tgt gac aag gga gaa tat ctt gat aag agg cat aat caa tgt tgt aat -cg gtg tcc acc tgg aga att tgc caa ggt tag atg ta- --a tgg taa cga taa cac aaa atg tga acg ctg ccc acc tca tac ata tac cac aat ccc caa tta ttc t-a atg gat gtc atc aat gta gaa aat gcc caa ccg gat cat ttg ata agg taa agt gta ccg gaa cac aga aca gta aat -gt tcg tgt ctt cct ggt tgg tat tgc gct act gat tct tca cag act gaa gat tgt tga aat -tg tgt acc aaa aag g-a gat gtc cat gcg gat act ttg gtg gaa tag atg aac aag gaa atc cta ttt gta aat cgt gct g-- -ta ttg gtg aat att gca act acc tac gta att ata gac ttg atc cat ttt ctc cat gca aa- --- cta tct aaa tgt aat taa tta --- --- --- --- --- --- --- --- tga ttt --- --- --- -tg atg ata atg tta -cc ata c-a tt- --- -at atc gct act tgg tta --g tgt att att cag tat g-a aga cct att aat aa- --- --- -tt act tat ctt ttg acg atc ttg tt- -at aat tat --- --- --- --- aat ata aaa --- -ac tta tgg --- --- cat agt aac tca t-- --- --- --- aat tgc tg- acg cga taa att cgt aat aat ctg ttt tgt tca aat ttt tat aag ga- --- --a tct aca ggc ata aaa at- aaa aat ata att tat aat ata c-- tct tac agc gcg cca tca tga ata gca gca gta aat taa ttg ctg tta tta atg gat tta gaa ata gtg gac gat ttt gtg ata tta ata tag tta tta atg atg aaa gga taa acg ctc aca gac tca tcc tat ctg gag cct ccg aat a-t ttt tcc att ctg ttt tcc aat aat ttt atc gat tct aat gaa tac gaa gtt aat cta agt cat tta gat tat caa agt gtt aac gat ttg atc gat tac att tat ggg ata cct ttg agc cta act aac gat aac gtg aaa tat att ctt tca acc gct ga- --t ttt tta caa att gga tct gcc att act gag tgc gaa aaa tac ata ctt aaa aat ctt tgt tct aga aac tgt atc gat ttc --- --- tac ata tac gct gat aaa tat aat aac aag aaa ata gaa tca gcg tcg ttt aac aca ata tta cga aat att ttg aga ctc atc aac gat gaa aac ttt aaa tac tta aca gag gaa tca atg ata aaa att tta agc gat gat atg tta aat ata aaa aat gag gat ttt gca cca cta att ctc att aaa tgg tta gag agt act caa caa tca tgc acc gtc gag tta ctt aga tgc ctc aga ata tca -tt gct ttc ccc aca agt tat aaa atc act tta tag tca tca act ggt tag ttc aat cta cga atg tat aac att ctt aaa caa tat agc att ctt gga tga atc att tcc tag ata cca tag cat cga gtt gat atc tat cgg tat aag taa ttc gca tga taa gat ttc cat aaa ctg cta caa tca t-a aaa aaa ata cat ggg aaa tga tat ctt cac gta gat ata ggt gta gtt tcg cag tgg ccg tcc tgg ata ata tta tct ata tga tgg gtg gat atg atc agt ccc cgt ata gaa gtt caa agg tta tag cgt aca ata cat gta caa att ctt gga tat atg ata tac cag agc t-a aaa tat cct cgt tct aat tgt ggg gga ctg gct gat gac gaa tac att tat tgt ata ggc ggc ata cgc gat cag gat tca tcg ttg aca tct agt att gat aga tgg aag cca tca aaa cca tat tgg cag aag tat gct aaa atg cgc gaa cca aaa tgt gat atg ggg gtt gcg atg tta aac gga tta ata tat gtc atg ggt gga atc gtt aaa ggt g-- aca cgt gta ccg acg cac tag aga gtt tat cag aag atg gat gga tga agc atc aac gtc ttc caa taa aaa tgt cca ata tgt cga cga ttg ttc atg atg gca aga ttt ata tat ctg gag gtt aca aca ata gta gtg tag tta atg taa tat cga atc t-- agt cct tag cta taa tcc gat ata tga tga atg gac caa att atc atc att aaa cat tcc tag aat taa tcc cgc tct a

>AY484669.1:164533-166782_Rabbitpox_virus_complete_genome

ttg ctg act gca --- tgt a-c tat ttt tta cat tac tac att ggc tac ggc aga tat acc tac tcc gcc acc --- aca cgc tcc ggt aaa cgg --g tca tgt gac aag gga gaa tat ctt gat aag agg cat aat caa tgt tgt aat -cg gtg tcc acc tgg aga att tgc caa ggt tag atg ta- --g tgg tag tga taa cac aaa atg tga acg ctg ccc acc tca tac ata tac cac aat -cc caa tta ttc t-a atg gat gtc atc aat gta gaa aat gcc caa cag gat cat ttg ata agg taa agt gta ccg gaa cac aga aca gta aat -gt tcg tgt ctt cct ggt tgg tat tgc gct act gat tct tca cag act gaa gat tgt tga aat -tg tgt acc aaa aag g-a gat gtc cat gcg gat act ttg gtg gaa tag atg aac aag gaa atc cta ttt gta aat cgt gct g-- -ta ttg gtg aat att gca act acc tac gta att ata gac ttg atc cat ttt ctc cat gca aa- --- cta tct aaa tgt aat taa tta --- --- --- --- --- --- --- --- tga ttt --- --- --- -tg atg ata atg tta -cc ata c-a tt- --- -at atc gct act tgg tta --g tgt att att cag tat g-a aga cct att aat aa- --- --- -tt act tat ctt ttg acg atc ttg tt- -at aat tat --- --- --- --- aat ata aaa --- -ac tta tgg --- --- cat agt aac tta t-- --- --- --- aat tgc ta- aca cga taa att cgt aat aat ctg ttt tgt tca --- --- --- aag ga- --- --a tct aca ggc ata aaa at- aaa aat ata att tat aat ata c-- tct tac agc gcg cca tca tga ata aca gca gtg aat tga ttg ctg tta tta atg gat tta gaa ata gtg gac gat ttt gtg ata tta gta tag tta tta atg atg aaa tga taa acg ctc aca gac tca tcc tat ctg gag cct ccg aat a-t ttt tcc att ctg ttt tcc aat aat ttt atc gat tct aat gaa tac gaa gtt aat cta agt cat tta gat tat caa agt gtt aac gat ttg atc gat tac att tat ggg ata cct ttg agc cta act aac gat aac gtg aaa tat att ctt tca acc gct ga- --t ttt tta caa att gga tct gcc att act gag tgc gaa aaa tac ata ctt aaa aat ctt tgt tct aga aac tgt atc gat ttc --- --- tac ata tac gct gat aaa tat aat aac aag aaa ata gaa tca gcg tcg ttt aac aca ata tta caa aat att ttg aaa ctc atc aac gat gaa aac ttt aaa tac tta aca gag gaa tca atg ata aaa att tta agc gat gat atg tta aat ata aaa aat gag gat ttc gcc cca cta att ctc att aaa tgg tta gag agt act caa caa tca tgc acc gtc gag tta ctt aga tgc ctc aga ata tca -tt gct ttc ccc aca agt tat aaa atc act tta tag tca tca act ggt tag ttc aat cta cga atg tat aac att ctt aaa caa tat agc att ctt gga tga atc att tcc tag ata cca tag cat cga gtt gat atc tat cgg tat aag taa ttc gca tga taa gat ttc cat aaa ctg cta caa tca t-a aaa aaa ata cat ggg aaa tga tat ctt cac gta gat ata ggt gta gtt tcg cag tgg ccg tcc tgg ata ata tta ttt ata tga tgg gtg gat atg atc agt ccc cgt ata gaa gtt caa agg tta tag cgt aca ata cat gta caa att ctt gga tat atg ata tac cag agc t-a aaa tat cct cgt tct aat tgt ggg gga ctg gct gat gac gaa tac att tat tgt ata ggc ggc ata cgc gat cag gat tca tcg ttg aca tct agt att gat aaa tgg aag cca tca aaa cca tat tgg cag aag tat gct aaa atg cgc gaa cca aaa tgt gat atg ggg gtt gcg atg tta aac gga tta ata tat gtc atg ggt gga atc gtt aaa ggt g-- aca cgt gta ccg acg cac tag aga gtt tat cag aag atg gat gga tga agc atc aac gtc ttc caa taa aaa tgt cca ata tgt cga cga ttg ttc atg atg gca aga ttt ata tat ctg gag gtt aca aca ata gta gtg tag tta atg taa tat cga atc t-- agt cct tag cta taa tcc gat ata tga tga atg gac caa att atc atc att aaa cat tcc tag aat taa tcc cgc tct a

>BK013343.1:183954-186222_TPA_asm:_Vaccinia_virus_isolate_VK12

ttg ctg act gca --- tgt a-c tat ttt tta cat tac tac att ggc tac ggc aga tat acc tac tcc gcc acc --- aca cgc tcc ggt aaa cgg --g tca tgt gac aag gga gaa tat ctt gat aag agg cat aat caa tgt tgt aat -cg gtg tcc acc tgg aga att tgc caa ggt tag atg ta- --a tgg taa cga taa cac aaa atg tga acg ctg ccc acc tca tac ata tac cac aat -cc caa tta ttc t-a atg gat gtc atc aat gta gaa aat gcc caa ccg gat cat ttg ata agg taa agt gta ccg gaa cac aga aca gta aat -gt tcg tgt ctt cct ggt tgg tat tgc gct act gat tct tca cag act gaa gat tgt tga aat -tg tgt acc aaa aag g-a gat gtc cat gcg gat act ttg gtg gaa tag atg aac aag gaa atc cta ttt gta aat cgt gct g-- -ta ttg gtg aat att gca act acc tac gta att ata gac ttg atc cat ttt ctc cat gca aa- --- cta tct aaa tgt aat taa tta tga tta --- --- --- --- tga ttt tga ttt --- --- --- -tg atg ata atg tta -cc ata c-a tt- --- -at atc gct act tgg tta --g tgt att att cag tat g-a aga cct att aat aa- --- --- -tt act tat ctt ttg acg atc ttg tt- -at aat tat --- --- --- --- aat ata aaa --- -ac tta tgg --- --- cat agt aac tta t-- --- --- --- aat tgc ta- aca cga taa att cgt aat aat ctg ttt tgt tca aat ttt tat aag ga- --- --a tct aca ggc ata aaa at- aaa aat ata atc tat aat ata c-- tct tac a-- acg cca tca tga ata gca gca gta aat taa ttg ctg tta tta atg gat tta gaa ata gtg gac gat ttt gtg ata tta ata tag tta tta atg atg aaa gga taa acg ctc ata aac tca tcc tat ctg gag cct ccg aat a-t ttt tcc att ctg ttt tcc aat aat ttt atc gat tct aat gaa tac gaa gtt aat cta agt cat tta gat tat caa agc gta aac gat ttg atc gat tat att tat ggg ata cct ttg agc cta act aac gat aac gtg aaa tat att ctt tca acc gct ga- --t ttt tta caa att gga tct gct att acg gag tgt gaa aat tac ata ctt aaa aat ctt tgt tct aaa aac tgt atc gat ttc --- --- tac ata tac gct gat aaa tat aat aac aag aaa ata gaa tca gcg tcg ttt aac aca ata tta cga aat att ttg aga ctc atc aac gat gaa aac ttt aaa tac tta aca gag gaa tca atg ata aaa att tta agc gat gat atg tta aat ata aaa aat gag gat ttc gcc cca cta att ctc att aaa tgg tta gag agt act caa caa cca tgc acc gtc gag tta ctt aaa tgc ctc aga ata tca -tt gct ttc ccc aca agt tat aaa atc act tta tag tca tcg act ggt tag ttc aat cta cga atg tat aac att ctt aaa caa tat agc att ctt gga tga atc att tcc tag ata cca tag cat cga gtt gat atc tat cgg tat aag taa ttc gcg tga taa aat ttc cat aaa ctg cta caa tca t-a aaa aaa ata cat ggg aaa tga tat ctt cac gta gat ata ggt gta gtt tcg cag tgg ccg tcc tgg ata ata tta ttt ata tga tgg gtg gat atg atc agt ccc cgt ata gaa gtt caa agg tta tag cgt aca ata cat gta caa att ctt gga tat atg ata tac cag agc t-a aaa tat cct cgt tct aat tgt ggg gga ctg gct gat gac gaa tac att tat tgt ata ggc ggc ata cgc gat cag gat tca tcg ttg aca tct agt att gat aaa tgg aag cca tca aaa cca tat tgg cag aag tat gct aaa atg cgc gaa cca aaa tgt gat atg ggg gtt gca atg tta aac gga tta ata tat gtt ata ggt gga atc gtt aaa ggt g-- aca cgt gta ccg acg cac tag aga gtt tat cag aag atg gat gga tga agc atc aac gtc ttc caa taa aaa tgt cca ata tgt cga cga ttg ttc atg atg gca aga ttt ata tat ctg gag gtt aca aca ata gta gtg tag tta atg taa tat cga atc t-- agt cct tag cta taa ttc gat ata tga tga atg gac caa att atc atc att aaa tat tcc tag aat taa tcc cgc tct a

>BK013340.1:156257-158523_TPA_asm:_Vaccinia_virus_isolate_VK02

ttg ctg act gca --- tgt a-c tat ttt tta cat tac tac att ggc tac ggc aga tat acc tac tcc gcc acc --- aca cgc tcc ggt aaa cgg --g tca tgt gac aag gga gaa tat ctt gat aag agg cat aat caa tgt tgt aat -cg gtg tcc acc tgg aga att tgc caa ggt tag atg ta- --a tgg taa cga taa cac aaa atg tga acg ctg ccc acc tca tac ata tac cgc aat -cc caa tta ttc t-a atg gat gtc atc aat gta gaa aat gcc caa ccg gat cat ttg ata agg taa agt gta ccg gaa cac aga aca gta aat -gt tcg tgt ctt cct ggt tgg tat tgc gct act gat tct tca cag act gaa gat tgt tga aat -tg tgt acc aaa aag g-a gat gtc cat gcg gat act ttg gtg gaa tag atg aac aag gaa atc cta ttt gta aat cgt gct g-- -ta ttg gtg aat att gca act acc tac gta att ata gac ttg atc cat ttt ctc cat gca aa- --- cta tct aaa tgt aat taa tta tga tta --- --- --- --- tga ttt tga ttt --- --- --- -tg atg ata atg tta -cc ata c-a tt- --- -at atc gct act tgg tta --g tgt att att cag tat g-a aga cct att aat aa- --- --- -tt act tat ctt ttg acg atc ttg tt- -at aat tat --- --- --- --- aat ata aaa --- -ac tta tgg --- --- cat agt aac tta t-- --- --- --- aat tgc ta- aca cga taa att cgt aat aat ctg ttt tgt tca aat ttt tat aag ga- --- --a tct aca ggc ata aaa at- aaa aat ata atc tat aat ata c-- tct tac a-- acg cca tca tga ata gca gca gta aat taa ttg ctg tta tta atg gat tta gaa ata gtg gac gat ttt gtg ata tta ata tag tta tta atg atg aaa gga taa acg ctc ata aac tca tcc tat ctg gag cct ccg aat a-t ttt tcc att ctg ttt tcc aat aat ttt atc gat tct aat gaa tac gaa gtt aat cta agt cat tta gat tat caa agc gta aac gat ttg atc gat tat att tat ggg ata cct ttg agc cta act aac gat aac gtg aaa tat att ctt tca acc gct ga- --t ttt tta caa att gga tct gct att acg gag tgt gaa aat tac ata ctt aaa aat ctt tgt tct aaa aac tgt atc gat ttc --- --- tac ata tac gct gat aaa tat aat aac aag aaa ata gaa tca gcg tcg ttt aac aca ata tta cga aat att ttg aga ctc atc aac gat gaa aac ttt aaa tac tta aca gag gaa tca atg ata aaa att tta agc gat gat atg tta aat ata aaa aat gag gat ttc gcc cca cta att ctc att aaa tgg tta gag agt act caa caa cca tgc acc gtc gag tta ctt aaa tgc ctc aga ata tca -tt gct ttc -cc aca agt tat aaa atc act tta tag tca tcg act ggt tag ttc aat cta cga atg tat aac att ctt aaa caa tat agc att ctt gga tga atc att tcc tag ata cca tag cat cga gtt gat atc tat cgg tat aag taa ttc gcg tga taa aat ttc cat aaa ctg cta caa tca t-- aaa aaa ata cat ggg aaa tga tat ctt cac gta gat ata ggt gta gtt tcg cag tgg ccg tcc tgg ata ata tta ttt ata tga tgg gtg gat atg atc agt ccc cgt ata gaa gtt caa agg tta tag cgt aca ata cat gta caa att ctt gga tat atg ata tac cag agc t-a aaa tat cct cgt tct aat tgt ggg gga ctg gct gat gac gaa tac att tat tgt ata ggc ggc ata cgc gat cag gat tca tcg ttg aca tct agt att gat aaa tgg aag cca tca aaa cca tat tgg cag aag tat gct aaa atg cgc gaa cca aaa tgt gat atg ggg gtt gcg atg tta aac gga tta ata tat gtt ata ggt gga atc gtt aaa ggt g-- aca cgt gta ccg acg cac tag aga gtt tat cag aag atg gat gga tga agc atc aac gtc ttc caa taa aaa tgt cca ata tgt tga cga ttg ttc atg atg gca aga ttt ata tat ctg gag gtt aca aca ata gta gtg tag tta atg taa tat caa atc t-- agt cct tag cta taa ttc gat ata tga tga atg gac caa att atc atc att aaa tat tcc tag aat taa tcc cgc tct a

>LC315596.1:166866-169103_BAC_cloning_vector_pLC16m8.8S-BAC_DNA_complete_sequence

ttg ctg act gta --- tgt a-c tat ttt tta cat tac tac att ggc tac ggc aga tat acc tac tcc gcc acc --- --- --- --- --- -aa cgg --g gca tgt gac gag gga gaa tat ctt gat aag agg cat aat caa tgt tgt aat -cg gtg tcc acc tgg aga att tgc caa ggt cag atg ta- --g tgg tag tga taa cac aaa atg tga acg ctg ccc acc tca tac ata tac cgc aat ccc caa tta ctc t-a atg gat gtc atc aat gta gaa aat gcc caa cag gat cat ttg ata agg taa agt gta ccg gaa cac aga aca gta aat -gt tcg tgt ctt cct ggt tgg tat tgc gct act gat tct tca cag act gaa gat tgt cga gat -tg tat acc aaa aag g-a gat gtc cat gcg gat act ttg gtg gaa tag atg aac aag gaa atc cta ttt gta aat cgt gtt g-- -tg ttg gtg aat att gcg act acc tac gta att ata gac ttg atc cat ttc ctc cat gca aa- --- cta tct aaa tgt aat taa tta --- --- --- --- --- --- --- --- tga ttt --- --- --- -tg atg ata atg tta -cc ata c-a tt- --- -at atc gct act tgg tta --g tgt att att cag tat g-a aga cct att aat aa- --- --- -tt act tat ctt ttg acg atc ttg tt- -at aat tat --- --- --- --- aat ata aaa --- -ac tta tgg --- --- cat agt aac tta t-- --- --- --- aat tgc tg- acg cga taa att cgt aat aat ctg ttt tgt tca --- --- --- aag ga- --- --a tct aca ggc ata aaa at- aaa aat ata att tat aat ata c-- tct tac agc gcg cca tca tga ata aca gca gtg aat tga ttg ctg tta tta atg gat tta gaa ata gtg gac gat ttt gtg ata tta gta tag tta tta atg atg aaa gga taa acg ctc ata aac tca tcc tat ctg gag cct ccg aat a-t ttt tcc att ctg ttt tcc aat aat ttt atc gat tct aat gaa tac gaa gtt aat cta agt cat tta gat tat caa agt gtt aac gat ttg atc gat tat att tat ggg ata cct ttg agc cta act aac gat aac gtg aaa tat att ctt tca acc gct ga- --t ttt tta caa att gga tct gcc att act gag tgc gaa aaa tac ata ctt aaa aat ctt tgt tct aga aac tgt atc gat ttc --- --- tac ata tac gct gat aaa tat aat aac aag aaa ata gaa tca gcg tcg ttt aac aca ata tta caa aat att ttg aga ctc atc aac gat gaa aac ttt aaa tac tta aca gag gaa tca atg ata aaa att tta agc gat gat atg tta aat ata aaa aat gag gat ttc gcc cca cta att ctc att aaa tgg tta gag agt act caa caa tca tgc acc gtc gag tta ctt aga tgc ctc aga ata tca -tt gct ttc ccc aca agt tat aaa atc act tta tag tca tcg act ggt tag ttc aat cta cga atg tat aac att ctt aaa caa tat agc att ctt gga tga atc att tcc tag ata cca tag cat cga gtt gat atc tat cgg tat aag taa ttc gca tga taa gat ttc cat aaa ctg cta caa tca t-a aaa aaa ata cat ggg aaa tga tat ctt cac gta gat ata ggt gta gtt tcg cag tgg ccg tcc tgg ata ata tta ttt ata tga tgg gtg gat atg atc agt ccc cgt ata gaa gtt caa agg tta tag cgt aca ata cat gta caa att ctt gga tat atg ata tac cag agc t-a aaa tat cct cgt tct aat tgt ggg gga ctg gct gat gac gaa tac att tat tgt ata ggc ggc ata cgc gat cag gat tca tcg ttg aca tct agt att gat aga tgg aag cca tca aaa cca tat tgg cag aag tat gct aaa atg cgc gaa cca aaa tgt gat atg ggg gtt gcg atg tta aac gga tta ata tat gtc atg ggt gga atc gtt aaa ggt g-- aca cgt gta ccg acg cac tag aga gtt tat cag aag atg gat gga tga agc atc aac gtc ttc caa taa aaa tgt cca ata tgt cga cga ttg ttc atg atg gca aga ttt ata tat ctg gag gtt aca aca ata gta gtg tag tta atg taa tat cga atc t-- agt cct tag cta taa ttc gat ata tga tga atg gac caa att atc atc att aaa cat tcc tag aat taa tcc cgc tct a

>AY678277.1:162304-164541_Vaccinia_virus_strain_LC16mO_complete_genome

ttg ctg act gta --- tgt a-c tat ttt tta cat tac tac att ggc tac ggc aga tat acc tac tcc gcc acc --- --- --- --- --- -aa cgg --g gca tgt gac gag gga gaa tat ctt gat aag agg cat aat caa tgt tgt aat -cg gtg tcc acc tgg aga att tgc caa ggt cag atg ta- --g tgg tag tga taa cac aaa atg tga acg ctg ccc acc tca tac ata tac cgc aat ccc caa tta ctc t-a atg gat gtc atc aat gta gaa aat gcc caa cag gat cat ttg ata agg taa agt gta ccg gaa cac aga aca gta aat -gt tcg tgt ctt cct ggt tgg tat tgc gct act gat tct tca cag act gaa gat tgt cga gat -tg tat acc aaa aag g-a gat gtc cat gcg gat act ttg gtg gaa tag atg aac aag gaa atc cta ttt gta aat cgt gtt g-- -tg ttg gtg aat att gcg act acc tac gta att ata gac ttg atc cat ttc ctc cat gca aa- --- cta tct aaa tgt aat taa tta --- --- --- --- --- --- --- --- tga ttt --- --- --- -tg atg ata atg tta -cc ata c-a tt- --- -at atc gct act tgg tta --g tgt att att cag tat g-a aga cct att aat aa- --- --- -tt act tat ctt ttg acg atc ttg tt- -at aat tat --- --- --- --- aat ata aaa --- -ac tta tgg --- --- cat agt aac tta t-- --- --- --- aat tgc tg- acg cga taa att cgt aat aat ctg ttt tgt tca --- --- --- aag ga- --- --a tct aca ggc ata aaa at- aaa aat ata att tat aat ata c-- tct tac agc gcg cca tca tga ata aca gca gtg aat tga ttg ctg tta tta atg gat tta gaa ata gtg gac gat ttt gtg ata tta gta tag tta tta atg atg aaa gga taa acg ctc ata aac tca tcc tat ctg gag cct ccg aat a-t ttt tcc att ctg ttt tcc aat aat ttt atc gat tct aat gaa tac gaa gtt aat cta agt cat tta gat tat caa agt gtt aac gat ttg atc gat tat att tat ggg ata cct ttg agc cta act aac gat aac gtg aaa tat att ctt tca acc gct ga- --t ttt tta caa att gga tct gcc att act gag tgc gaa aaa tac ata ctt aaa aat ctt tgt tct aga aac tgt atc gat ttc --- --- tac ata tac gct gat aaa tat aat aac aag aaa ata gaa tca gcg tcg ttt aac aca ata tta caa aat att ttg aga ctc atc aac gat gaa aac ttt aaa tac tta aca gag gaa tca atg ata aaa att tta agc gat gat atg tta aat ata aaa aat gag gat ttc gcc cca cta att ctc att aaa tgg tta gag agt act caa caa tca tgc acc gtc gag tta ctt aga tgc ctc aga ata tca -tt gct ttc ccc aca agt tat aaa atc act tta tag tca tcg act ggt tag ttc aat cta cga atg tat aac att ctt aaa caa tat agc att ctt gga tga atc att tcc tag ata cca tag cat cga gtt gat atc tat cgg tat aag taa ttc gca tga taa gat ttc cat aaa ctg cta caa tca t-a aaa aaa ata cat ggg aaa tga tat ctt cac gta gat ata ggt gta gtt tcg cag tgg ccg tcc tgg ata ata tta ttt ata tga tgg gtg gat atg atc agt ccc cgt ata gaa gtt caa agg tta tag cgt aca ata cat gta caa att ctt gga tat atg ata tac cag agc t-a aaa tat cct cgt tct aat tgt ggg gga ctg gct gat gac gaa tac att tat tgt ata ggc ggc ata cgc gat cag gat tca tcg ttg aca tct agt att gat aga tgg aag cca tca aaa cca tat tgg cag aag tat gct aaa atg cgc gaa cca aaa tgt gat atg ggg gtt gcg atg tta aac gga tta ata tat gtc atg ggt gga atc gtt aaa ggt g-- aca cgt gta ccg acg cac tag aga gtt tat cag aag atg gat gga tga agc atc aac gtc ttc caa taa aaa tgt cca ata tgt cga cga ttg ttc atg atg gca aga ttt ata tat ctg gag gtt aca aca ata gta gtg tag tta atg taa tat cga atc t-- agt cct tag cta taa ttc gat ata tga tga atg gac caa att atc atc att aaa cat tcc tag aat taa tcc cgc tct a

>LT966077.1:159778-162025_Vaccinia_virus_WR_strain_WR_-_ATCC_VR-2035_genome_assembly_complete_genome:_monopartite

ttg ctg act gca --- tgt a-c tat ttt tta cat tac tac att ggc tac ggc aga tat acc tac tcc gcc acc --- --- --- --- --- -aa cgg --g tca tgt gac aag gga gaa tat ctt gat aag agg cat aat caa tgt tgt aat -cg gtg tcc acc tgg aga att tgc caa ggt tag atg ta- --a tgg taa cga taa cac aaa atg tga acg ctg ccc acc tca tac ata tac cac aat -cc caa tta ttc t-a atg gat gtc atc aat gta gaa aat gcc caa ccg gat cat ttg ata agg taa agt gta ccg gaa cac aga aca gta aat -gt tcg tgt ctt cct ggt tgg tat tgt gct act gat tct tca cag act gaa gat tgt tga aat -tg tgt acc aaa aag g-a gat gtc cat gcg gat act ttg gtg gaa tag atg aac aag gaa atc cta ttt gta aat cgt gtt g-- -tg ttg gtg aat att gcg act acc tac gta att ata gac ttg atc cat ttc ctc cat gca aa- --- cta tct aaa tgt aat taa tta --- --- --- --- --- --- --- --- tga ttt --- --- --- -tg atg ata atg tta -cc ata c-a tt- --- -at atc gct act tgg tta --g tgt att att cag tat g-a aga cct att aat aa- --- --- -tt act tat ctt ttg acg atc ttg tt- -at aat tat --- --- --- --- aat ata aaa --a tac tta tgg --- --- cat agt aac tca t-- --- --- --- aat tgc tg- acg cga taa att cgt aat aat ctg ttt tgt tca aat ttt tat aag ga- --- --a tct aca ggc ata aaa at- aaa aat ata att tat aat ata c-- tct tac agc gcg cca tca tga ata aca gca gtg aat tga ttg ctg tta tta atg gat tta gaa ata gtg gac gat ttt gtg ata tta gta tag tta tta atg atg aaa gga taa acg ctc ata aac tca tcc tat ctg gag cct ccg aat a-t ttt tcc att ctg ttt tcc aat aat ttt atc gat tct aat gaa tac gaa gtt aat cta agt cat tta gat tat caa agc gtt aac gat ttg atc gat tac att tat ggg ata cct ttg agc cta act aac gat aac gtg aaa tat att ctt tca acc gct ga- --t ttt tta caa att gga tct gct att acg gag tgt gaa aat tac ata ctt aaa aat ctt tgt tct aaa aac tgt atc gat ttc --- --- tac ata tac gct gat aaa tat aat aac aag aaa ata gaa tca gcg tcg ttt aac aca ata tta caa aat att ttg aga ctc atc aac gat gaa aac ttt aaa tac tta aca gag gaa tca atg ata aaa att tta agc gat gat atg tta aat ata aaa aat gag gat ttt gca cca cta att ctc att aaa tgg tta gag agt act caa caa tca tgc acc gtc gag tta ctt aga tgc ctc aga ata tca -tt gct ttc ccc aca agt tat aaa atc act tta tag tca tca act ggt tag ttc aat cta cga atg tat aac att ctt aaa caa tat agc att ctt gga tga atc att tcc tag ata cca tag cat cga gtt gat atc tat cgg tat aag taa ttc gca tga taa gat ttc cat aaa ctg cta caa tca t-a aaa aaa ata cat ggg aaa tga tat ctt cac gta gat ata ggt gta gtt tcg cag tgg ccg tcc tgg ata ata tta tct ata tga tgg gtg gat atg atc agt ccc cgt ata gaa gtt caa agg tta tag cgt aca ata cat gta caa att ctt gga tat atg ata tac cag agc t-a aaa tat cct cgt tct aat tgt ggg gga ctg gct gat gac gaa tac att tat tgt ata ggc ggc ata cgc gat cag gat tca tcg ttg aca tct agt att gat aaa tgg aag cca tca aaa cca tat tgg cag aag tat gct aaa atg cgc gaa cca aaa tgt gat atg ggg gtt gcg atg tta aac gga tta ata tat gtc atg ggt gga atc gtt aaa ggt g-- aca cgt gta ccg acg cac tag aga gtt tat cag aag atg gat gga tga agc atc aac gtc ttc caa taa aaa tgt cca ata tgt cga cga ttg ttc atg atg gca aga ttt ata tat ctg gag gtt aca aca ata gta gtg tag tta atg taa tat cga atc t-- agt cct tag cta taa tcc gat ata tga tga atg gac caa att atc atc att aaa cat tcc tag aat taa tcc cgc tct a

>AY243312.1:159652-161899_Vaccinia_virus_WR_complete_genome

ttg ctg act gca --- tgt a-c tat ttt tta cat tac tac att ggc tac ggc aga tat acc tac tcc gcc acc --- --- --- --- --- -aa cgg --g tca tgt gac aag gga gaa tat ctt gat aag agg cat aat caa tgt tgt aat -cg gtg tcc acc tgg aga att tgc caa ggt tag atg ta- --a tgg taa cga taa cac aaa atg tga acg ctg ccc acc tca tac ata tac cac aat -cc caa tta ttc t-a atg gat gtc atc aat gta gaa aat gcc caa ccg gat cat ttg ata agg taa agt gta ccg gaa cac aga aca gta aat -gt tcg tgt ctt cct ggt tgg tat tgt gct act gat tct tca cag act gaa gat tgt tga aat -tg tgt acc aaa aag g-a gat gtc cat gcg gat act ttg gtg gaa tag atg aac aag gaa atc cta ttt gta aat cgt gtt g-- -tg ttg gtg aat att gcg act acc tac gta att ata gac ttg atc cat ttc ctc cat gca aa- --- cta tct aaa tgt aat taa tta --- --- --- --- --- --- --- --- tga ttt --- --- --- -tg atg ata atg tta -cc ata c-a tt- --- -at atc gct act tgg tta --g tgt att att cag tat g-a aga cct att aat aa- --- --- -tt act tat ctt ttg acg atc ttg tt- -at aat tat --- --- --- --- aat ata aaa --a tac tta tgg --- --- cat agt aac tca t-- --- --- --- aat tgc tg- acg cga taa att cgt aat aat ctg ttt tgt tca aat ttt tat aag ga- --- --a tct aca ggc ata aaa at- aaa aat ata att tat aat ata c-- tct tac agc gcg cca tca tga ata aca gca gtg aat tga ttg ctg tta tta atg gat tta gaa ata gtg gac gat ttt gtg ata tta gta tag tta tta atg atg aaa gga taa acg ctc ata aac tca tcc tat ctg gag cct ccg aat a-t ttt tcc att ctg ttt tcc aat aat ttt atc gat tct aat gaa tac gaa gtt aat cta agt cat tta gat tat caa agc gtt aac gat ttg atc gat tac att tat ggg ata cct ttg agc cta act aac gat aac gtg aaa tat att ctt tca acc gct ga- --t ttt tta caa att gga tct gct att acg gag tgt gaa aat tac ata ctt aaa aat ctt tgt tct aaa aac tgt atc gat ttc --- --- tac ata tac gct gat aaa tat aat aac aag aaa ata gaa tca gcg tcg ttt aac aca ata tta caa aat att ttg aga ctc atc aac gat gaa aac ttt aaa tac tta aca gag gaa tca atg ata aaa att tta agc gat gat atg tta aat ata aaa aat gag gat ttt gca cca cta att ctc att aaa tgg tta gag agt act caa caa tca tgc acc gtc gag tta ctt aga tgc ctc aga ata tca -tt gct ttc ccc aca agt tat aaa atc act tta tag tca tca act ggt tag ttc aat cta cga atg tat aac att ctt aaa caa tat agc att ctt gga tga atc att tcc tag ata cca tag cat cga gtt gat atc tat cgg tat aag taa ttc gca tga taa gat ttc cat aaa ctg cta caa tca t-a aaa aaa ata cat ggg aaa tga tat ctt cac gta gat ata ggt gta gtt tcg cag tgg ccg tcc tgg ata ata tta tct ata tga tgg gtg gat atg atc agt ccc cgt ata gaa gtt caa agg tta tag cgt aca ata cat gta caa att ctt gga tat atg ata tac cag agc t-a aaa tat cct cgt tct aat tgt ggg gga ctg gct gat gac gaa tac att tat tgt ata ggc ggc ata cgc gat cag gat tca tcg ttg aca tct agt att gat aaa tgg aag cca tca aaa cca tat tgg cag aag tat gct aaa atg cgc gaa cca aaa tgt gat atg ggg gtt gcg atg tta aac gga tta ata tat gtc atg ggt gga atc gtt aaa ggt g-- aca cgt gta ccg acg cac tag aga gtt tat cag aag atg gat gga tga agc atc aac gtc ttc caa taa aaa tgt cca ata tgt cga cga ttg ttc atg atg gca aga ttt ata tat ctg gag gtt aca aca ata gta gtg tag tta atg taa tat cga atc t-- agt cct tag cta taa tcc gat ata tga tga atg gac caa att atc atc att aaa cat tcc tag aat taa tcc cgc tct a

>M58054.1:167-2414_Vaccinia_virus_SALF19R_and_SALF21R_genes_complete_cds

ttg ctg act gca --- tgt a-c tat ttt tta cat tac tac att ggc tac ggc aga tat acc tac tcc gcc acc --- --- --- --- --- -aa cgg --g tca tgt gac aag gga gaa tat ctt gat aag agg cat aat caa tgt tgt aat -cg gtg tcc acc tgg aga att tgc caa ggt tag atg ta- --a tgg taa cga taa cac aaa atg tga acg ctg ccc acc tca tac ata tac cac aat -cc caa tta ttc t-a atg gat gtc atc aat gta gaa aat gcc caa ccg gat cat ttg ata agg taa agt gta ccg gaa cac aga aca gta aat -gt tcg tgt ctt cct ggt tgg tat tgt gct act gat tct tca cag act gaa gat tgt tga aat -tg tgt acc aaa aag g-a gat gtc cat gcg gat act ttg gtg gaa tag atg aac aag gaa atc cta ttt gta aat cgt gtt g-- -tg ttg gtg aat att gcg act acc tac gta att ata gac ttg atc cat ttc ctc cat gca aa- --- cta tct aaa tgt aat taa tta --- --- --- --- --- --- --- --- tga ttt --- --- --- -tg atg ata atg tta -cc ata c-a tt- --- -at atc gct act tgg tta --g tgt att att cag tat g-a aga cct att aat aa- --- --- -tt act tat ctt ttg acg atc ttg tt- -at aat tat --- --- --- --- aat ata aaa --a tac tta tgg --- --- cat agt aac tca t-- --- --- --- aat tgc tg- acg cga taa att cgt aat aat ctg ttt tgt tca aat ttt tat aag ga- --- --a tct aca ggc ata aaa at- aaa aat ata att tat aat ata c-- tct tac agc gcg cca tca tga ata aca gca gtg aat tga ttg ctg tta tta atg gat tta gaa ata gtg gac gat ttt gtg ata tta gta tag tta tta atg atg aaa gga taa acg ctc ata aac tca tcc tat ctg gag cct ccg aat a-t ttt tcc att ctg ttt tcc aat aat ttt atc gat tct aat gaa tac gaa gtt aat cta agt cat tta gat tat caa agc gtt aac gat ttg atc gat tac att tat ggg ata cct ttg agc cta act aac gat aac gtg aaa tat att ctt tca acc gct ga- --t ttt tta caa att gga tct gct att acg gag tgt gaa aat tac ata ctt aaa aat ctt tgt tct aaa aac tgt atc gat ttc --- --- tac ata tac gct gat aaa tat aat aac aag aaa ata gaa tca gcg tcg ttt aac aca ata tta caa aat att ttg aga ctc atc aac gat gaa aac ttt aaa tac tta aca gag gaa tca atg ata aaa att tta agc gat gat atg tta aat ata aaa aat gag gat ttt gca cca cta att ctc att aaa tgg tta gag agt act caa caa tca tgc acc gtc gag tta ctt aga tgc ctc aga ata tca -tt gct ttc ccc aca agt tat aaa atc act tta tag tca tca act ggt tag ttc aat cta cga atg tat aac att ctt aaa caa tat agc att ctt gga tga atc att tcc tag ata cca tag cat cga gtt gat atc tat cgg tat aag taa ttc gca tga taa gat ttc cat aaa ctg cta caa tca t-a aaa aaa ata cat ggg aaa tga tat ctt cac gta gat ata ggt gta gtt tcg cag tgg ccg tcc tgg ata ata tta tct ata tga tgg gtg gat atg atc agt ccc cgt ata gaa gtt caa agg tta tag cgt aca ata cat gta caa att ctt gga tat atg ata tac cag agc t-a aaa tat cct cgt tct aat tgt ggg gga ctg gct gat gac gaa tac att tat tgt ata ggc ggc ata cgc gat cag gat tca tcg ttg aca tct agt att gat aaa tgg aag cca tca aaa cca tat tgg cag aag tat gct aaa atg cgc gaa cca aaa tgt gat atg ggg gtt gcg atg tta aac gga tta ata tat gtc atg ggt gga atc gtt aaa ggt g-- aca cgt gta ccg acg cac tag aga gtt tat cag aag atg gat gga tga agc atc aac gtc ttc caa taa aaa tgt cca ata tgt cga cga ttg ttc atg atg gca aga ttt ata tat ctg gag gtt aca aca ata gta gtg tag tta atg taa tat cga atc t-- agt cct tag cta taa tcc gat ata tga tga atg gac caa att atc atc att aaa cat tcc tag aat taa tcc cgc tct a

>D11079.1:17636-19883_Vaccinia_virus_genomic_DNA_42kbp_from_near_the_right_ITR

ttg ctg act gca --- tgt a-c tat ttt tta cat tac tac att ggc tac ggc aga tat acc tac tcc gcc acc --- --- --- --- --- -aa cgg --g tca tgt gac aag gga gaa tat ctt gat aag agg cat aat caa tgt tgt aat -cg gtg tcc acc tgg aga att tgc caa ggt tag atg ta- --a tgg taa cga taa cac aaa atg tga acg ctg ccc acc tca tac ata tac cac aat -cc caa tta ttc t-a atg gat gtc atc aat gta gaa aat gcc caa ccg gat cat ttg ata agg taa agt gta ccg gaa cac aga aca gta aat -gt tcg tgt ctt cct ggt tgg tat tgt gct act gat tct tca cag act gaa gat tgt tga aat -tg tgt acc aaa aag g-a gat gtc cat gcg gat act ttg gtg gaa tag atg aac aag gaa atc cta ttt gta aat cgt gtt g-- -tg ttg gtg aat att gcg act acc tac gta att ata gac ttg atc cat ttc ctc cat gca aa- --- cta tct aaa tgt aat taa tta --- --- --- --- --- --- --- --- tga ttt --- --- --- -tg atg ata atg tta -cc ata c-a tt- --- -at atc gct act tgg tta --g tgt att att cag tat g-a aga cct att aat aa- --- --- -tt act tat ctt ttg acg atc ttg tt- -at aat tat --- --- --- --- aat ata aaa --a tac tta tgg --- --- cat agt aac tca t-- --- --- --- aat tgc tg- acg cga taa att cgt aat aat ctg ttt tgt tca aat ttt tat aag ga- --- --a tct aca ggc ata aaa at- aaa aat ata att tat aat ata c-- tct tac agc gcg cca tca tga ata aca gca gtg aat tga ttg ctg tta tta atg gat tta gaa ata gtg gac gat ttt gtg ata tta gta tag tta tta atg atg aaa gga taa acg ctc ata aac tca tcc tat ctg gag cct ccg aat a-t ttt tcc att ctg ttt tcc aat aat ttt atc gat tct aat gaa tac gaa gtt aat cta agt cat tta gat tat caa agc gtt aac gat ttg atc gat tac att tat ggg ata cct ttg agc cta act aac gat aac gtg aaa tat att ctt tca acc gct ga- --t ttt tta caa att gga tct gct att acg gag tgt gaa aat tac ata ctt aaa aat ctt tgt tct aaa aac tgt atc gat ttc --- --- tac ata tac gct gat aaa tat aat aac aag aaa ata gaa tca gcg tcg ttt aac aca ata tta caa aat att ttg aga ctc atc aac gat gaa aac ttt aaa tac tta aca gag gaa tca atg ata aaa att tta agc gat gat atg tta aat ata aaa aat gag gat ttt gca cca cta att ctc att aaa tgg tta gag agt act caa caa tca tgc acc gtc gag tta ctt aga tgc ctc aga ata tca -tt gct ttc ccc aca agt tat aaa atc act tta tag tca tca act ggt tag ttc aat cta cga atg tat aac att ctt aaa caa tat agc att ctt gga tga atc att tcc tag ata cca tag cat cga gtt gat atc tat cgg tat aag taa ttc gca tga taa gat ttc cat aaa ctg cta caa tca t-a aaa aaa ata cat ggg aaa tga tat ctt cac gta gat ata ggt gta gtt tcg cag tgg ccg tcc tgg ata ata tta tct ata tga tgg gtg gat atg atc agt ccc cgt ata gaa gtt caa agg tta tag cgt aca ata cat gta caa att ctt gga tat atg ata tac cag agc t-a aaa tat cct cgt tct aat tgt ggg gga ctg gct gat gac gaa tac att tat tgt ata ggc ggc ata cgc gat cag gat tca tcg ttg aca tct agt att gat aaa tgg aag cca tca aaa cca tat tgg cag aag tat gct aaa atg cgc gaa cca aaa tgt gat atg ggg gtt gcg atg tta aac gga tta ata tat gtc atg ggt gga atc gtt aaa ggt g-- aca cgt gta ccg acg cac tag aga gtt tat cag aag atg gat gga tga agc atc aac gtc ttc caa taa aaa tgt cca ata tgt cga cga ttg ttc atg atg gca aga ttt ata tat ctg gag gtt aca aca ata gta gtg tag tta atg taa tat cga atc t-- agt cct tag cta taa tcc gat ata tga tga atg gac caa att atc atc att aaa cat tcc tag aat taa tcc cgc tct a

>AY678275.1:162306-164543_Vaccinia_virus_strain_LC16m8_complete_genome

ttg ctg act gta --- tgt t-c tat ttt tta cat tac tac att ggc tac ggc aga tat acc tac tcc gcc acc --- --- --- --- --- -aa cgg --g gca tgt gac gag gga gaa tat ctt gat aag agg cat aat caa tgt tgt aat -cg gtg tcc acc tgg aga att tgc caa ggt cag atg ta- --g tgg tag tga taa cac aaa atg tga acg ctg ccc acc tca tac ata tac cgc aat ccc caa tta ctc t-a atg gat gtc atc aat gta gaa aat gcc caa cag gat cat ttg ata agg taa agt gta ccg gaa cac aga aca gta aat -gt tcg tgt ctt cct ggt tgg tat tgc gct act gat tct tca cag act gaa gat tgt cga gat -tg tat acc aaa aag g-a gat gtc cat gcg gat act ttg gtg gaa tag atg aac aag gaa atc cta ttt gta aat cgt gtt g-- -tg ttg gtg aat att gcg act acc tac gta att ata gac ttg atc cat ttc ctc cat gca aa- --- cta tct aaa tgt aat taa tta --- --- --- --- --- --- --- --- tga ttt --- --- --- -tg atg ata atg tta -cc ata c-a tt- --- -at atc gct act tgg tta --g tgt att att cag tat g-a aga cct att aat aa- --- --- -tt act tat ctt ttg acg atc ttg tt- -at aat tat --- --- --- --- aat ata aaa --- -ac tta tgg --- --- cat agt aac tta t-- --- --- --- aat tgc tg- acg cga taa att cgt aat aat ctg ttt tgt tca --- --- --- aag ga- --- --a tct aca ggc ata aaa at- aaa aat ata att tat aat ata c-- tct tac agc gcg cca tca tga ata aca gca gtg aat tga ttg ctg tta tta atg gat tta gaa ata gtg gac gat ttt gtg ata tta gta tag tta tta atg atg aaa gga taa acg ctc ata aac tca tcc tat ctg gag cct ccg aat a-t ttt tcc att ctg ttt tcc aat aat ttt atc gat tct aat gaa tac gaa gtt aat cta agt cat tta gat tat caa agt gtt aac gat ttg atc gat tat att tat ggg ata cct ttg agc cta act aac gat aac gtg aaa tat att ctt tca acc gct ga- --t ttt tta caa att gga tct gcc att act gag tgc gaa aaa tac ata ctt aaa aat ctt tgt tct aga aac tgt atc gat ttc --- --- tac ata tac gct gat aaa tat aat aac aag aaa ata gaa tca gcg tcg ttt aac aca ata tta caa aat att ttg aga ctc atc aac gat gaa aac ttt aaa tac tta aca gag gaa tca atg ata aaa att tta agc gat gat atg tta aat ata aaa aat gag gat ttc gcc cca cta att ctc att aaa tgg tta gag agt act caa caa tca tgc acc gtc gag tta ctt aga tgc ctc aga ata tca -tt gct ttc ccc aca agt tat aaa atc act tta tag tca tcg act ggt tag ttc aat cta cga atg tat aac att ctt aaa caa tat agc att ctt gga tga atc att tcc tag ata cca tag cat cga gtt gat atc tat cgg tat aag taa ttc gca tga taa gat ttc cat aaa ctg cta caa tca t-a aaa aaa ata cat ggg aaa tga tat ctt cac gta gat ata ggt gta gtt tcg cag tgg ccg tcc tgg ata ata tta ttt ata tga tgg gtg gat atg atc agt ccc cgt ata gaa gtt caa agg tta tag cgt aca ata cat gta caa att ctt gga tat atg ata tac cag agc t-a aaa tat cct cgt tct aat tgt ggg gga ctg gct gat gac gaa tac att tat tgt ata ggc ggc ata cgc gat cag gat tca tcg ttg aca tct agt att gat aga tgg aag cca tca aaa cca tat tgg cag aag tat gct aaa atg cgc gaa cca aaa tgt gat atg ggg gtt gcg atg tta aac gga tta ata tat gtc atg ggt gga atc gtt aaa ggt g-- aca cgt gta ccg acg cac tag aga gtt tat cag aag atg gat gga tga agc atc aac gtc ttc caa taa aaa tgt cca ata tgt cga cga ttg ttc atg atg gca aga ttt ata tat ctg gag gtt aca aca ata gta gtg tag tta atg taa tat cga atc t-- agt cct tag cta taa ttc gat ata tga tga atg gac caa att atc atc att aaa cat tcc tag aat taa tcc cgc tct a

>JX489139.1:159236-161475_Vaccinia_virus_strain_Tiantan_clone_TT12_complete_genome

ttg ctg act gca --- tgt a-c tat ttt tta cat tac tac att ggc tac ggc aga tat acc tac tcc gcc acc --- --- --- --- --- -aa cgg --g tca tgt gac aag gga gaa tat ctt gat aag agg cat aat caa tgt tgt aat -cg gtg tcc acc tgg aga att tgc caa ggt tag atg ta- --a tgg taa cga taa cac aaa atg tga acg ctg ccc acc tca tac ata tac cgc aat ccc caa tta ctc t-a atg gat gtc atc aat gta gaa aat gcc caa ccg gat cat ttg ata agg taa agt gta ccg gaa cac aga aca gta aat -gt tcg tgt ctt cct ggt tgg tat tgc gct act gat tct tca cag act gaa gat tgt tga aat -tg tgt acc aaa aag g-a gat gtc cat gcg gat act ttg gtg gaa tag atg aac aag gaa atc cta ttt gta aat cgt gtt g-- -tg ttg gtg aat att gcg act acc tac gta att ata gac ttg atc cat ttc ctc cat gca aa- --- cta tct aaa tgt aat taa tta --- --- --- --- --- --- --- --- tga ttt --- --- --- -tg atg ata atg tta -cc ata c-a tt- --- -at atc gct act tgg tta --g tgt att att cag tat g-a aga cct att aat aa- --- --- -tt act tat ctt ttg acg atc ttg tt- -at aat tat --- --- --- --- aat ata aaa --a tac tta tgg --- --- cat agt aac tta t-- --- --- --- aat tgc tg- acg cga taa att cgt aat aat ctg ttt tgt tca --- --- --- aag ga- --- --a tct aca ggc ata aaa at- aaa aat ata att tat aat ata c-- tct tac aac gcg cca tca tga ata aca gca gtg aat tga ttg ctg tta tta atg gat tta gaa ata gtg gac gat ttt gtg ata tta ata tag tta tta atg atg aaa gga taa acg ctc ata aac tca tcc tat ctg gag cct ccg aat a-t ttt tcc att ctg ttt tcc aat aat ttt atc gat tct aat gaa tac gaa gtt aat cta agt cat tta gat tat caa agc gta aac gat ttg atc gat tac att tat ggg ata cct ttg agc cta act aac gat aac gtg aaa tat att ctt tca acc gct ga- --t ttt tta caa att gga tct gct att acg gag tgt gaa aaa tac ata ctt aaa aat ctt tgt tct aga aac tgt atc gat ttc --- --- tac ata tac gct gat aaa tat aat aac aag aaa ata gaa tca gcg tcg ttt aac aca ata tta cga aat att ttg aga ctc atc aac gat gaa aac ttt aaa tac tta aca gag gaa tca atg ata aaa att tta agc gat gat atg tta aat ata aaa aat gag gat ttc gcc cca cta att ctc att aaa tgg tta gag agt act caa caa cca tgc acc gtc gag tta ctt aaa tgc ctc aga ata tca -tt gct ttc ccc aca agt tat aaa atc act tta tag tca tcg act ggt tag ttc aat cta cga atg tat aac att ctt aaa caa tat agc att ctt gga tga atc att tcc tag ata cca tag cat cga gtt gat atc tat cgg tat aag taa ttc gcg tga taa aat ttc cat aaa ctg cta caa tca t-a aaa aaa ata cat ggg aaa tga tat ctt cac gta gat ata ggt gta gtt tcg cag tgg ccg tcc tgg ata ata tta tct ata tga tgg gtg gat atg atc agt ccc cgt ata gaa gtt caa agg tta tag cgt aca ata cat gta caa att ctt gga tat atg ata tac cag agc t-a aaa tat cct cgt tct aat tgt ggg gga ctg gct gat gac gaa tac att tat tgt ata ggc ggc ata cgc gat cag gat tca tcg ttg aca tct agt att gat aga tgg aag cca tca aaa cca tat tgg cag aag tat gct aaa atg cgc gaa cca aaa tgt gat atg ggg gtt gcg atg tta aac gga tta ata tat gtc atg ggt gga atc gtt aaa ggt g-- aca cgt gta ccg acg cac tag aga gtt tat cag aag atg gat gga tga agc atc aac gtc ttc caa taa aaa tgt cca ata tgt cga cga ttg ttc atg atg gca aga ttt ata tat ctg gag gtt aca aca ata gta gtg tag tta atg taa tat cga atc t-- agt cct tag cta taa tcc gat ata tga tga atg gac caa att atc atc att aaa tat tcc tag aat taa tcc cgc tct a

>JN654982.1:162293-164537_Vaccinia_virus_strain_Dryvax_clone_DPP16_complete_genome

ttg ctg act gca --- tgt a-c tat ttt tta cat tac tac att ggc tac ggc aga tat acc tac tcc gcc acc --- --- --- --- --- -aa cgg --g tca tgt gac aag gga gaa tat ctt gat aag agg cat aat caa tgt tgt aat -ca gtg tcc acc tgg aga att tgc caa ggt cag atg ta- --g tgg tag tga taa cac aaa atg tga acg ctg ccc acc tca tac ata tac cgc aat ccc caa tta ctc t-a atg gat gtc atc aat gta gaa aat gcc caa ccg gat cat ttg ata agg taa agt gta ccg gaa cac aga aca gta aat -gt tcg tgt ctt cct ggt tgg tat tgt gct act gat tct tca cag act gaa gat tgt tga aat -tg tgt acc aaa aag g-a gat gtc cat gcg gat act ttg gtg gaa tag atg aac aag gaa atc cta ttt gta aat cgt gct g-- -ta ttg gtg aat att gca act acc tac gta att ata gac ttg atc cat ttt ctc cat gca aa- --- cta tct aaa tgt aat taa tta --- --- --- --- --- --- --- --- tga ttt --- --- --- -tg atg ata atg tta -cc ata c-a tt- --- -at atc gct act tgg tta --g tgt att att cag tat g-a aga cct att aat aa- --- --- -tt act tat ctt ttg acg atc ttg tt- -at aat tat --- --- --- --- aat ata aaa --- -ac tta tgg --- --- cat agt aac tta t-- --- --- --- aat tgc ta- aca cga taa att cgt aat aat ctg ttt tgt tca aat ttt tat aag ga- --- --a tct aca ggc ata aaa at- aaa aat ata atc tat aat ata c-- tct tac a-- acg cca tca tga ata gca gca gta aat taa ttg ctg tta tta atg gat tta gaa ata gtg gac gat ttt gtg ata tta ata tag tta tta atg atg aaa gga taa acg ctc ata aac tca tcc tat ctg gag cct ccg aat a-t ttt tcc att ctg ttt tcc aat aat ttt atc gat tct aat gaa tac gaa gtt aat cta agt cat tta gat tat caa agc gta aac gat ttg atc gat tat att tat ggg ata cct ttg agc cta act aac gat aac gtg aaa tat att ctt tca acc gct ga- --t ttt tta caa att gga tct gct att acg gag tgt gaa aat tac ata ctt aaa aat ctt tgt tct aaa aac tgt atc gat ttc --- --- tac ata tac gct gat aaa tat aat aac aag aaa ata gaa tca gcg tcg ttt aac aca ata tta caa aat att ttg aga ctc atc aac gat gaa aac ttt aaa tac tta aca gag gaa tca atg ata aaa att tta agc gat gat atg tta aat ata aaa aat gag gat ttt gca cca cta att ctc att aaa tgg tta gag agt act caa caa tca tgc acc gtc gag tta ctt aaa tgc ctc aga ata tca -tt gct ttc ccc aca agt tat aaa atc act tta tag tca tcg act ggt tag ttc aat cta cga atg tat aac att ctt aaa caa tat agc att ctt gga tga atc att tcc tag ata cca tag cat cga gtt gat atc tat cgg tat aag taa ttc gcg tga taa aat ttc cat aaa ctg cta caa tca t-a aaa aaa ata cat ggg aaa tga tat ctt cac gta gat ata ggt gta gtt tcg cag tgg ccg tcc tgg ata ata tta tct ata tga tgg gtg gat atg atc agt ccc cgt ata gaa gtt caa agg tta tag cgt aca ata cat gta caa att ctt gga tat atg ata tac cag agc t-a aaa tat cct cgt tct aat tgt ggg gga ctg gct gat gac gaa tac att tat tgt ata ggc ggc ata cgc gat cag gat tca tcg ttg aca tct agt att gat aga tgg aag cca tca aaa cca tat tgg cag aag tat gct aaa atg cgc gaa cca aaa tgt gat atg ggg gtt gcg atg tta aac gga tta ata tat gtc atg ggt gga atc gtt aaa ggt g-- aca cgt gta ccg acg cac tag aga gtt tat cag aag atg gat gga tga agc atc aac gtc ttc caa taa aaa tgt cca ata tgt cga cga ttg ttc atg atg gca aga ttt ata tat ctg gag gtt aca aca ata gta gtg tag tta atg taa tat cga atc t-- agt cct tag cta taa ttc gat ata tga tga atg gac caa att atc atc att aaa tat tcc tag aat taa tcc cgc tct a

>JN654980.1:162076-164319_Vaccinia_virus_strain_Dryvax_clone_DPP13_complete_genome

ttg ctg act gta --- tgt a-c tat ttt tta cat tac tac att ggc tac ggc aga tat acc tac tcc gcc acc --- --- --- --- --- -aa cgg --g tca tgt gac gag gga gaa tat ctt gat aag agg cat aat caa tgt tgt aat -ca gtg tcc acc tgg aga att tgc caa ggt tag atg ta- --g tgg tag tga taa cac aaa atg tga acg ctg ccc acc tca tac ata tac cac aat -cc caa tta ttc t-a atg gat gtc atc aat gta gaa aat gcc caa cag gat cat ttg ata agg taa agt gta ccg gaa cac aga aca gta aat -gt tcg tgt ctt cct ggt tgg tat tgc gct act gat tct tca cag act gaa gat tgt tga aat -tg tgt acc aaa aag g-a gat gtc cat gcg gat act ttg gtg gaa tag atg aac aag gaa atc cta ttt gta aat cgt gct g-- -ta ttg gtg aat att gca act acc tac gta att ata gac ttg atc cat ttt ctc cat gca aa- --- cta tct aaa tgt aat taa tta --- --- --- --- --- --- --- --- tga ttt --- --- --- -tg atg ata atg tta -cc ata c-a tt- --- -at atc gct act tgg tta --g tgt att att cag tat g-a aga cct att aat aa- --- --- -tt act tat ctt ttg acg atc ttg tt- -at aat tat --- --- --- --- aat ata aaa --- -ac tta tgg --- --- cat agt aac tca t-- --- --- --- aat tgc tg- acg cga taa att cgt aat aat ctg ttt tgt tca aat ttt tat aag ga- --- --a tct aca ggc ata aaa at- aaa aat ata atc tat aat ata c-- tct tac a-- acg cca tca tga ata gca gca gta aat taa ttg ctg tta tta atg gat tta gaa ata gtg gac gat ttt gtg ata tta ata tag tta tta atg atg aaa gga taa acg ctc ata aac tca tcc tat ctg gag cct ccg aat a-t ttt tcc att ctg ttt tcc aat aat ttt atc gat tct aat gaa tac gaa gtt aat cta agt cat tta gat tat caa agc gta aac gat ttg atc gat tat att tat ggg ata cct ttg agc cta act aac gat aac gtg aaa tat att ctt tca acc gct ga- --t ttt tta caa att gga tct gct att acg gag tgt gaa aat tac ata ctt aaa aat ctt tgt tct aaa aac tgt atc gat ttc --- --- tac ata tac gct gat aaa tat aat aac aag aaa ata gaa tca gcg tcg ttt aac aca ata tta caa aat att ttg aga ctc atc aac gat gaa aac ttt aaa tac tta aca gag gaa tca atg ata aaa att tta agc gat gat atg tta aat ata aaa aat gag gat ttt gca cca cta att ctc att aaa tgg tta gag agt act caa caa tca tgc acc gtc gag tta ctt aaa tgc ctc aga ata tca -tt gct ttc ccc aca agt tat aaa atc act tta tag tca tcg act ggt tag ttc aat cta cga atg tat aac att ctt aaa caa tat agc att ctt gga tga atc att tcc tag ata cca tag cat cga gtt gat atc tat cgg tat aag taa ttc gcg tga taa aat ttc cat aaa ctg cta caa tca t-a aaa aaa ata cat ggg aaa tga tat ctt tac gta gat ata ggt gta gtt tcg cag tgg ccg tcc tgg ata ata tta ttt ata tga tgg gtg gat atg atc agt ccc cgt ata gaa gtt caa agg tta tag cgt aca ata cat gta caa att ctt gga tat atg ata tac cag agc t-a aaa tat cct cgt tct aat tgt ggg gga ctg gct gat gac gaa tac att tat tgt ata ggc ggc ata cgc gat cag gat tca tcg ttg aca tct agt att gat aaa tgg aag cca tca aaa cca tat tgg cag aag tat gct aaa atg cgc gaa cca aaa tgt gat atg ggg gtt gcg atg tta aac gga tta ata tat gtc atg ggt gga atc gtt aaa ggt g-- aca cgt gta ccg acg cac tag aga gtt tat cag aag atg gat gga tga agc atc aac gtc ttc caa taa aaa tgt cca ata tgt cga cga ttg ttc atg atg gca aga ttt ata tat ctg gag gtt aca aca ata gta gtg tag tta atg taa tat cga atc t-- agt cct tag cta taa ttc gat ata tga tga atg gac caa att atc atc att aaa tat tcc tag aat taa tcc cgc tct a

>KY349117.1:171238-173518_Horsepox_virus_strain_MNR_clone_2-1

ttg ctg act gca --- tgt a-c tat ttt tta cat tac tac att ggc tac ggc aga tat acc tac tcc gcc acc --- aca cgc tcc ggt aaa cgg --g tca tgt gac aag gga gaa tat ctt gat aag agg cat aat caa tgt tgt aat -cg gtg tcc acc tgg aga att tgc caa ggt tag atg ta- --a tgg taa cga taa cac aaa atg tga acg ctg ccc acc tca tac ata tac cgc aat -cc caa tta ttc t-a atg gat gtc atc aat gta gaa aat gcc caa ccg gat cat ttg ata agg taa agt gta ccg gaa cac aga aca gta aat -gt tcg tgt ctt cct ggt tgg tat tgc gct act gat tct tca cag act gaa gat tgt tga aat -tg tgt acc aaa aag g-a gat gtc cat gcg gat act ttg gtg gaa tag atg aac aag gaa atc cta ttt gta aat cgt gct g-- -ta ttg gtg aat att gca act acc tac gta att ata gac ttg atc cat ttt ctc cat gca aa- --- cta tct aaa tgt aat taa tta tga tta tga tta tga ttt tga ttt tga ttt --- --- --- -tg atg ata atg tta -cc ata c-a tt- --- -at atc gct act tgg tta --g tgt att att cag tat g-a aga cct att aat aa- --- --- -tt act tat ctt ttg acg atc ttg tt- -at aat tat --- --- --- --- aat ata aaa --- -ac tta tgg --- --- cat agt aac tta t-- --- --- --- aat tgc ta- aca cga taa att cgt aat aat ctg ttt tgt tca aat ttt tat aag ga- --- --a tct aca ggc ata aaa at- aaa aat ata atc tat aat ata c-- tct tac a-- acg cca tca tga ata gca gca gta aat taa ttg ctg tta tta atg gat tta gaa ata gtg gac gat ttt gtg ata tta ata tag tta tta atg atg aaa gga taa acg ctc ata aac tca tcc tat ctg gag cct ccg aat a-t ttt tcc att ctg ttt tcc aat aat ttt atc gat tct aat gaa tac gaa gtt aat cta agt cat tta gat tat caa agc gta aac gat ttg atc gat tat att tat ggg ata cct ttg agc cta act aac gat aac gtg aaa tat att ctt tca acc gct ga- --t ttt tta caa att gga tct gct att acg gag tgt gaa aat tac ata ctt aaa aat ctt tgt tct aaa aac tgt atc gat ttc --- --- tac ata tac gct gat aaa tat aat aac aag aaa ata gaa tca gcg tcg ttt aac aca ata tta caa aat att ttg aga ctc atc aac gat gaa aac ttt aaa tac tta aca gag gaa tca atg ata aaa att tta agc gat gat atg tta aat ata aaa aat gag gat ttc gcc cca cta att ctc att aaa tgg tta gag agt act caa caa cca tgc acc gtc gag tta ctt aaa tgc ctc aga ata tca -tt gct ttc ccc aca agt tat aaa atc act tta tag tca tcg act ggt tag ttc aat cta cga atg tat aac att ctt aaa caa tat agc att ctt gga tga atc att tcc tag ata cca tag cat cga gtt gat atc tat cgg tat aag taa ttc gcg tga taa aat ttc cat aaa ctg cta caa tca t-a aaa aaa ata cat ggg aaa tga tat ctt cac gta gat ata ggt gta gtt tcg cag tgg ccg tcc tgg ata ata tta ttt ata tga tgg gtg gat atg atc agt ccc cgt ata gaa gtt caa agg tta tag cgt aca ata cat gta caa att ctt gga tat atg ata tac cag agc t-a aaa tat cct cgt tct aat tgt ggg gga ctg gct gat gac gaa tac att tat tgt ata ggc ggc ata cgc gat cag gat tca tcg ttg aca tct agt att gat aaa tgg aag cca tca aaa cca tat tgg cag aag tat gct aaa atg cgc gaa cca aaa tgt gat atg ggg gtt gcg atg tta aac gga tta ata tat gtt ata ggt gga atc gtt aaa ggt g-- aca cgt gta ccg acg cac tag aga gtt tat cag aag atg gat gga tga agc atc aac gtc ttc caa taa aaa tgt cca ata tgt cga cga ttg ttc atg atg gca aga ttt ata tat ctg gag gtt aca aca ata gta gtg tag tta atg taa tat cga atc t-- agt cct tag cta taa ttc gat ata tga tga atg gac caa att atc atc att aaa tat tcc tag aat taa tcc cgc tct a

>DQ792504.1:171149-173429_Horsepox_virus_isolate_MNR-76_complete_genome

ttg ctg act gca --- tgt a-c tat ttt tta cat tac tac att ggc tac ggc aga tat acc tac tcc gcc acc --- aca cgc tcc ggt aaa cgg --g tca tgt gac aag gga gaa tat ctt gat aag agg cat aat caa tgt tgt aat -cg gtg tcc acc tgg aga att tgc caa ggt tag atg ta- --a tgg taa cga taa cac aaa atg tga acg ctg ccc acc tca tac ata tac cgc aat -cc caa tta ttc t-a atg gat gtc atc aat gta gaa aat gcc caa ccg gat cat ttg ata agg taa agt gta ccg gaa cac aga aca gta aat -gt tcg tgt ctt cct ggt tgg tat tgc gct act gat tct tca cag act gaa gat tgt tga aat -tg tgt acc aaa aag g-a gat gtc cat gcg gat act ttg gtg gaa tag atg aac aag gaa atc cta ttt gta aat cgt gct g-- -ta ttg gtg aat att gca act acc tac gta att ata gac ttg atc cat ttt ctc cat gca aa- --- cta tct aaa tgt aat taa tta tga tta tga tta tga ttt tga ttt tga ttt --- --- --- -tg atg ata atg tta -cc ata c-a tt- --- -at atc gct act tgg tta --g tgt att att cag tat g-a aga cct att aat aa- --- --- -tt act tat ctt ttg acg atc ttg tt- -at aat tat --- --- --- --- aat ata aaa --- -ac tta tgg --- --- cat agt aac tta t-- --- --- --- aat tgc ta- aca cga taa att cgt aat aat ctg ttt tgt tca aat ttt tat aag ga- --- --a tct aca ggc ata aaa at- aaa aat ata atc tat aat ata c-- tct tac a-- acg cca tca tga ata gca gca gta aat taa ttg ctg tta tta atg gat tta gaa ata gtg gac gat ttt gtg ata tta ata tag tta tta atg atg aaa gga taa acg ctc ata aac tca tcc tat ctg gag cct ccg aat a-t ttt tcc att ctg ttt tcc aat aat ttt atc gat tct aat gaa tac gaa gtt aat cta agt cat tta gat tat caa agc gta aac gat ttg atc gat tat att tat ggg ata cct ttg agc cta act aac gat aac gtg aaa tat att ctt tca acc gct ga- --t ttt tta caa att gga tct gct att acg gag tgt gaa aat tac ata ctt aaa aat ctt tgt tct aaa aac tgt atc gat ttc --- --- tac ata tac gct gat aaa tat aat aac aag aaa ata gaa tca gcg tcg ttt aac aca ata tta caa aat att ttg aga ctc atc aac gat gaa aac ttt aaa tac tta aca gag gaa tca atg ata aaa att tta agc gat gat atg tta aat ata aaa aat gag gat ttc gcc cca cta att ctc att aaa tgg tta gag agt act caa caa cca tgc acc gtc gag tta ctt aaa tgc ctc aga ata tca -tt gct ttc ccc aca agt tat aaa atc act tta tag tca tcg act ggt tag ttc aat cta cga atg tat aac att ctt aaa caa tat agc att ctt gga tga atc att tcc tag ata cca tag cat cga gtt gat atc tat cgg tat aag taa ttc gcg tga taa aat ttc cat aaa ctg cta caa tca t-a aaa aaa ata cat ggg aaa tga tat ctt cac gta gat ata ggt gta gtt tcg cag tgg ccg tcc tgg ata ata tta ttt ata tga tgg gtg gat atg atc agt ccc cgt ata gaa gtt caa agg tta tag cgt aca ata cat gta caa att ctt gga tat atg ata tac cag agc t-a aaa tat cct cgt tct aat tgt ggg gga ctg gct gat gac gaa tac att tat tgt ata ggc ggc ata cgc gat cag gat tca tcg ttg aca tct agt att gat aaa tgg aag cca tca aaa cca tat tgg cag aag tat gct aaa atg cgc gaa cca aaa tgt gat atg ggg gtt gcg atg tta aac gga tta ata tat gtt ata ggt gga atc gtt aaa ggt g-- aca cgt gta ccg acg cac tag aga gtt tat cag aag atg gat gga tga agc atc aac gtc ttc caa taa aaa tgt cca ata tgt cga cga ttg ttc atg atg gca aga ttt ata tat ctg gag gtt aca aca ata gta gtg tag tta atg taa tat cga atc t-- agt cct tag cta taa ttc gat ata tga tga atg gac caa att atc atc att aaa tat tcc tag aat taa tcc cgc tct a

>M35027.1:158666-160902_Vaccinia_virus_Copenhagen_complete_genome

ttg ctg act gca --- tgt a-c tat ttt tta cat tac tac att ggc tac ggc aga tat acc tac tcc gcc acc --- --- --- --- --- -aa cgg --g tca tgt gac aag gga gaa tat ctt gat aag agg cat aat caa tgt tgt aat -cg gtg tcc acc tgg aga att tgc caa ggt tag atg ta- --a tgg taa cga taa cac aaa atg tga acg ctg ccc acc tca tac ata tac cac aat -cc caa tta ttc t-a atg gat gtc atc aat gta gaa aat gcc caa cag gat cat ttg ata agg taa agt gta ccg gaa cac aga aca gta aat -gt tcg tgt ctt cct ggt tgg tat tgc gct act gat tct tca cag act gaa gat tgt tga aat -tg tat acc aaa aag g-a gat gtc cat gcg gat act ttg gtg gaa tag atg aac aag gaa atc cta ttt gta aat cgt gtt g-- -tg ttg gtg aat att gcg act acc tac gta att ata gac ttg atc cat ttc ctc cat gca aa- --- cta tct aaa tgt aat taa tta --- --- --- --- --- --- --- --- tga ttt --- --- --- -tg atg ata atg tta -cc ata c-a tt- --- -at atc gct act tgg tta --g tgt att att cag tat g-a aga cct att aat aa- --- --- -tt act tat ctt ttg acg atc ttg tt- -at aat tat --- --- --- --- aat ata aaa --a tac tta tgg --- --- cat agt aac tca t-- --- --- --- aat tgc tg- acg cga taa att cgt aat aat ctg ttt tgt tca --- --- --- aag ga- --- --a tct aca ggc ata aaa at- aaa aat ata att tat aat ata c-- tct tac a-- acg cca tca tga ata aca gca gtg aat tga ttg ctg tta tta atg gat tta gaa ata gtg gac gat ttt gtg ata tta ata tag tta tta atg atg aaa gga taa acg ctc ata aac tca tcc tat ctg gag cct ccg aat a-t ttt tcc att ctg ttt tcc aat aat ttt atc gat tct aat gaa tac gaa gtt aat cta agt cat tta gat tat caa agt gtt aac gat ttg atc gat tac att tat ggg ata cct ttg agc cta act aac gat aac gtg aaa tat att ctt tca acc gct ga- --t ttt tta caa att gga tct gcc att acg gag tgt gaa aat tac ata ctt aaa aat ctt tgt tct aaa aac tgt atc gat ttc --- --- tac ata tac gct gat aaa tat aat aac aag aaa ata gaa tca gcg tcg ttt aac aca ata tta caa aat att ttg aga ctc atc aac gat gaa aac ttt aaa tac tta aca gag gaa tca atg ata aaa att tta agc gat gat atg tta aat ata aaa aat gag gat ttc gcc cca cta att ctc att aaa tgg tta gag agt act caa caa tca tgc acc gtc gag tta ctt aaa tgc ctc aga ata tca -tt gct ttc ccc aca agt tat aaa atc act tta tag tca tcg act ggt tag ttc aat cta cga atg tat aac att ctt aaa caa tat agc att ctt gga tga atc att tcc tag ata cca tag cat cga gtt gat atc tat cgg tat aag taa ttc gcg tga taa aat ttc cat aaa ctg cta caa tca t-a aaa aaa ata cat ggg aaa tga tat ctt cac gta gat ata ggt gta gtt tcg cag tgg ccg tcc tgg ata ata tta ttt ata tga tgg gtg gat atg atc agt ccc cgt ata gaa gtt caa agg tta tag cgt aca ata cat gta caa att ctt gga tat atg ata tac cag agc t-a aaa tat cct cgt tct aat tgt ggg gga ctg gct gat gac gaa tac att tat tgt ata ggc ggc ata cgc gat cag gat tca tcg ttg aca tct agt att gat aaa tgg aag cca tca aaa cca tat tgg cag aag tat gct aaa atg cgc gaa cca aaa tgt gat atg ggg gtt gcg atg tta aac gga tta ata tat gtt ata ggt gga atc gtt aaa ggt g-- aca cgt gta ccg acg cac tag aga gtt tat cag aag atg gat gga tga agc atc aac gtc ttc caa taa aaa tgt cca ata tgt cga cga ttg ttc atg atg gca aga ttt ata tat ctg gag gtt aca aca ata gta gtg tag tta atg taa tat cga atc t-- agt cct tag cta taa tcc gat ata tga tga atg gac caa att atc atc att aaa cat tcc tag aat taa tcc cgc tct a

>AM501482.1:159967-162204_Vaccinia_virus_Ankara_strain_chorioallantois_vaccinia_virus_Ankara_(CVA)_complete_coding_genome

ttg ctg act gca --- tgt a-c tat ttt tta cat tac tac att ggc tac ggc aga tat acc tac tcc gcc acc --- --- --- --- --- -aa cgg --g gca tgt gac gag gga gaa tat ctt gat aag agg cat aat caa tgt tgt aat -ca gtg tcc acc tgg aga att tgc caa ggt cag atg ta- --g tgg tag tga taa cac aaa atg tga acg ctg ccc acc tca tac ata tac cgc aat ccc caa tta ctc t-a atg gat gtc atc aat gta gaa aat gcc caa cag gat cat ttg ata agg taa agt gta ccg gaa cac aga aca gta aat -gt tcg tgt ctt cct ggt tgg tat tgt gct act gat tct tca cag act gaa gat tgt cga gat -tg tat acc aaa aag g-a gat gtc cat gcg gat act ttg gtg gaa tag atg aac aag gaa atc cta ttt gta aat cgt gct g-- -ta ttg gtg aat att gcg act acc tac gta att ata gac ttg atc cat ttc ctc cat gca aa- --- cta tct aaa tgt aat taa tta --- --- --- --- --- --- --- --- tga ttt --- --- --- -tg atg ata atg tta -cc ata c-a tt- --- -at atc gct act tgg tta --g tgt att att cag tat g-a aga cct att aat aa- --- --- -tt act tat ctt ttg acg atc ttg tt- -at aat tat --- --- --- --- aat ata aaa --- -ac tta tgg --- --- cat agt aac tta t-- --- --- --- aat tgc ta- aca cga taa att cgt aat aat ctg ttt tgt tca --- --- --- aag ga- --- --a tct aca ggc ata aaa at- aaa aat ata att tat aat ata c-- tct tac agc gcg cca tca tga ata aca gca gtg aat tga ttg ctg tta tta atg gat tta gaa ata gtg gac gat ttt gtg ata tta gta tag tta tta atg atg aaa tga taa acg ctc aca gac tca tcc tat ctg gag cct ccg aat a-t ttt tcc att ctg ttt tcc aat aat ttt atc gat tct aat gaa tac gaa gtt aat cta agt cat tta gat tat caa agt gtt aac gat ttg atc gat tac att tat ggg ata cct ttg agc cta act aac gat aac gtg aaa tat att ctt tca acc gct ga- --t ttt tta caa att gga tct gcc att act gag tgc gaa aaa tac ata ctt aaa aat ctt tgt tct aga aac tgt atc gat ttc --- --- tac ata tac gct gat aaa tat aat aac aag aaa ata gaa tca gcg tcg ttt aac aca ata tta cga aat att ttg aga ctc atc aac gat gaa aac ttt aaa tac tta aca gag gaa tca atg ata aaa att tta agc gat gat atg tta aat ata aaa aat gag gat ttt gca cca cta att ctc att aaa tgg tta gag agt act caa caa tca tgc acc gtc gag tta ctt aga tgc ctc aga ata tca -tt gct ttc ccc aca agt tat aaa atc act tta tag tca tca act ggt tag ttc aat cta cga atg tat aac att ctt aaa caa tat agc att ctt gga tga atc att tcc tag ata cca tag cat cga gtt gat atc tat cgg tat aag taa ttc gca tga taa gat ttc cat aaa ctg cta caa tca t-a aaa aaa ata cat ggg aaa tga tat ctt cac gta gat ata ggt gta gtt tcg cag tgg ccg tcc tgg ata ata tta tct ata tga tgg gtg gat atg atc agt ccc cgt ata gaa gtt caa agg tta tag cgt aca ata cat gta caa att ctt gga tat atg ata tac cag agc t-a aaa tat cct cgt tct aat tgt ggg gga ctg gct gat gac gaa tac att tat tgt ata ggc ggc ata cgc gat cag gat tca tcg ttg aca tct agt att gat aga tgg aag cca tca aaa cca tat tgg tag aag tat gct aaa atg cgc gaa cca aaa tgt gat atg ggg gtt gcg atg tta aac gga tta ata tat gtc atg ggt gga atc gtt aaa ggt g-- aca cgt gta ccg acg cac tag aga gtt tat cag aag atg gat gga tga agc atc gac gtc ttc caa taa aaa tgt cca ata tgt cga cga ttg ttc atg atg gca aga ttt ata tat ctg gag gtt aca aca ata gta gtg tag tta atg taa tat cga atc t-- agt cct tag cta taa tcc gat ata tga tga atg gac caa att atc atc att aaa cat tcc tag aat taa tcc cgc tct a

>JX489135.1:158668-160907_Vaccinia_virus_strain_Tiantan_clone_TT8_complete_genome

ttg ctg act gca --- tgt a-c tat ttt tta cat tac tac att ggc tac ggc aga tat acc tac tcc gcc acc --- --- --- --- --- -aa cgg --g tca tgt gac gag gga gaa tat ctt gat aag agg cat aat caa tgt tgt aat -ca gtg tcc acc tgg aga att tgc caa ggt tag atg ta- --a tgg taa cga taa cac aaa atg tga acg ctg ccc acc tca tac ata tac cgc aat ccc caa tta ctc t-a atg gat gtc atc aat gta gaa aat gcc caa ccg gat cat ttg ata agg taa agt gta ccg gaa cac aga aca gta aat -gt tcg tgt ctt cct ggt tgg tat tgc gct act gat tct tca cag act gaa gat tgt tga aat -tg tgt acc aaa aag g-a gat gtc cat gcg gat act ttg gtg gaa tag atg aac aag gaa atc cta ttt gta aat cgt gtt g-- -tg ttg gtg aat att gcg act act tac gta att ata gac ttg atc cat ttc ctc cat gca aa- --- cta tct aaa tgt aat taa tta --- --- --- --- --- --- --- --- tga ttt --- --- --- -tg atg ata atg tta -cc ata c-a tt- --- -at atc gct act tgg tta --g tgt att att cag tat g-a aga cct att aat aa- --- --- -tt act tat ctt ttg acg atc ttg tt- -at aat tat --- --- --- --- aat ata aaa --a tac tta tgg --- --- cat agt aac tca t-- --- --- --- aat tgc tg- acg cga taa att cgt aat aat ctg ttt tgt tca --- --- --- aag ga- --- --a tct aca ggc ata aaa at- aaa aat ata att tat aat ata c-- tct tac agc gcg cca tca tga ata aca gca gtg aat tga ttg ctg tta tta atg gat tta gaa ata gtg gac gat ttt gtg ata tta gta tag tta tta atg atg aaa tga taa acg ctc aca gac tca tcc tat ctg gag cct ccg aat a-t ttt tcc att ctg ttt tcc aat aat ttt atc gat tct aat gaa tac gaa gtt aat cta agt cat tta gat tat caa agc gta aac gat ttg atc gat tac att tat ggg ata cct ttg agc cta act aac gat aac gtg aaa tat att ctt tca acc gct ga- --t ttt tta caa att gga tct gct att acg gag tgt gaa aaa tac ata ctt aaa aat ctt tgt tct aaa aac tgt atc gat ttc --- --- tac ata tac gct gat aaa tat aat aac aag aaa ata gaa tca gcg tcg ttt aac aca ata tta cga aat att ttg aga ctc atc aac gat gaa aac ttt aaa tac tta aca aag gaa tca atg ata aaa att tta agc gat gat atg tta aat ata aaa aat gag gat ttc gcc cca cta att ctc att aaa tgg tta gag agt act caa caa tca tgc acc gtc gag tta ctt aaa tgc ctc aga ata tca -tt gct ttc ccc aca agt tat aaa atc act tta tag tca tcg act ggt tag ttc aat cta cga atg tat aac att ctt aaa caa tat agc att ctt gga tga atc att tcc tag ata cca tag cat cga gtt gat atc tat cgg tat aag taa ttc gcg tga taa aat ttc cat aaa ctg cta caa tca t-a aaa aaa ata cat ggg aaa tga tat ctt cac gta gat ata ggt gta gtt tcg cag tgg ccg tcc tgg ata ata tta tct ata tga tgg gtg gat atg atc agt ccc cgt ata gaa gtt caa agg tta tag cgt aca ata cat gta caa att ctt gga tat atg ata tac cag agc t-a aaa tat cct cgt tct aat tgt ggg gga ctg gct gat gac gaa tac att tat tgt ata ggc ggc ata cgc gat cag gat tca tcg ttg aca tct agt att gat aga tgg aag cca tca aaa cca tat tgg cag aag tat gct aaa atg cgc gaa cca aaa tgt gat atg ggg gtt gcg atg tta aac gga tta ata tat gtc atg ggt gga atc gtt aaa ggt g-- aca cgt gta ccg acg cac tag aga gtt tat cag aag atg gat gga tga agc atc aac gtc ttc caa taa aaa tgt cca ata tgt cga cga ttg ttc atg atg gca aga ttt ata tat ctg gag gtt aca aca ata gta gtg tag tta atg taa tat cga atc t-- agt cct tag cta taa tcc gat ata tga tga atg gac caa att atc atc att aaa tat tcc tag aat taa tcc cgc tct a

>BK013341.1:171175-173455_TPA_asm:_Horsepox_virus_isolate_VK05

ttg ctg act aca --- tgt a-c tat ttt tta cat tac tac att ggc tac ggc aga tat acc tac tcc gcc acc --- aca cgc tcc ggt aaa cgg --g tca tgt gac aag gga gaa tat ctt gat aag agt cat aat caa tgt tgt aat -cg gtg tcc acc tgg aga att tgc caa ggt tag atg ta- --a tgg taa cga taa cac aaa atg tga acg ctg ccc acc tca tac ata tac cgc aat -tc caa tta ttc t-a atg gat gtc atc aat gta gaa aat gcc caa ccg gat cat ttg ata agg taa agt gta ccg gaa cac aga aca gta aat -gt tcg tgt ctt cct ggt tgg tat tgc gct act gat tct tca cag act gaa gat tgt tga aat -tg tgt acc aaa aag g-a gat gtc cat gcg gat act ttg gtg gaa tag atg aac aag gaa atc cta ttt gta aat cgt gct g-- -ta ttg gtg aat att gca act acc tac gta att ata gac ttg atc cat ttt ctc cat gca aa- --- cta tct aaa tgt aat taa tta tga tta tga ttt tga tta tga ttt tga ttt --- --- --- -tg atg ata atg tta -cc ata c-a tt- --- -at atc gct act tgg tta --g tgt att att cag tat g-a aga cct att aat aa- --- --- -tt act tat ctt ttg acg atc ttg tt- -at aat tat --- --- --- --- aat ata aaa --- -ac tta tgg --- --- cat agt aac tta t-- --- --- --- aat tgc ta- aca cga taa att cgt aat aat ctg ttt tgt tca aat ttt tat aag ga- --- --a tct aca ggc ata aaa at- aaa aat ata att tat aat ata c-- tct tac a-- acg cca tca tga ata gca gca gta aat taa ttg ctg tta tta atg gat tta gaa ata gtg gac gat ttt gtg ata tta ata tag tta tta atg atg aaa gga taa acg ctc ata aac tca tcc tat ctg gag cct ccg aat a-t ttt tcc att ctg ttt tcc aat aat ttt atc gat tct aat gaa tac gaa gtt aat cta agt cat tta gat tat caa agc gta aac gat ttg atc gat tat att tat ggg ata cct ttg agc cta act aac gat aac gtg aaa tat att ctt tca acc gct ga- --t ttt tta caa att gga tct gct att acg gag tgt gaa aat tac ata ctt aaa aat ctt tgt tct aaa aac tgt atc gat ttc --- --- tac ata tac gct gat aaa tat aat aac aag aaa ata gaa tca gcg tcg ttt aac aca ata tta caa aat att ttg aga ctc atc aac gat gaa aac ttt aaa tac tta aca gag gaa tca atg ata aaa att tta agc gat gat atg tta aat ata aaa aat gag gat ttc gcc cca cta att ctc att aaa tgg tta gag agt act caa caa cca tgc acc gtc gag tta ctt aaa tgc ctc aga ata tca -tt gct ttc ccc aca agt tat aaa atc act tta tag tca tcg act ggt tag ttc aat cta cga atg tat aac att ctt aaa caa tat agc att ctt gga tga atc att tcc tag ata cca tag cat cga gtt gat atc tat cgg tat aag taa ttc gcg tga taa aat ttc cat aaa ctg cta caa tca t-a aaa aaa ata cat ggg aaa tga tat ctt cac gta gat ata ggt gta gtt tcg cag tgg ccg tcc tgg ata ata tta ttt ata tga tgg gtg gat atg atc agt ccc cgt ata gaa gtt caa agg tta tag cgt aca ata cat gta caa att ctt gga tat atg ata tac cag agc t-a aaa tat cct cgt tct aat tgt ggg gga ctg gct gat gac gaa tac att tat tgt ata ggc ggc ata cgc gat cag gat tca tcg ttg aca tct agt att gat aaa tgg aag cca tca aaa cca tat tgg cag aag tat gct aaa atg cgc gaa cca aaa tgt gat atg ggg gtt gcg atg tta aac gga tta ata tat gtt ata ggt gga atc gtt aaa ggt g-- aca cgt gta ccg acg cac tag aga gtt tat cag aag atg gat gga tga agc atc aac gtc ttc caa taa aaa tgt cca ata tgt cga cga ttg ttc atg atg gca aga ttt ata tat ctg gag gtt aca aca ata gta gtg tag tta atg taa tat cga atc t-- agt cct tag cta taa ttc gat ata tga tga atg gac caa att atc atc att aaa tat tcc tag aat taa tcc cgc tct a

>KJ125438.1:162126-164369_Vaccinia_virus_strain_Dryvax_clone_DPP25_complete_genome

ttg ctg act gta --- tgt a-c tat ttt tta cat tac tac att ggc tac ggc aga tat acc tac tcc gcc acc --- --- --- --- --- -aa cgg --g tca tgt gac aag gga gaa tat ctt gat aag agg cat aat caa tgt tgt aat -cg gtg tcc acc tgg aga att tgc caa ggt tag atg ta- --a tgg taa cga taa cac aaa atg tga acg ctg ccc acc tca tac ata tac cac aat -cc caa tta ttc t-a atg gat gtc atc aat gta gaa aat gcc caa ccg gat cat ttg ata agg taa agt gta ccg gaa cac aga aca gta aat -gt tcg tgt ctt cct ggt tgg tat tgt gct act gat tct tca cag act gaa gat tgt cga gat -tg tat acc aaa aag g-a gat gtc cat gcg gat act ttg gtg gaa tag atg aac aag gaa atc cta ttt gta aat cgt gct g-- -ta ttg gtg aat att gca act acc tac gta att ata gac ttg atc cat ttt ctc cat gca aa- --- cta tct aaa tgt aat taa tta --- --- --- --- --- --- --- --- tga ttt --- --- --- -tg atg ata atg tta -cc ata c-a tt- --- -at atc gct act tgg tta --g tgt att att cag tat g-a aga cct att aat aa- --- --- -tt act tat ctt ttg acg atc ttg tt- -at aat tat --- --- --- --- aat ata aaa --- -ac tta tgg --- --- cat agt aac tta t-- --- --- --- aat tgc ta- aca cga taa att cgt aat aat ctg ttt tgt tca aat ttt tat aag ga- --- --a tct aca ggc ata aaa at- aaa aat ata att tat aat ata c-- tct tac a-- acg cca tca tga ata gca gca gta aat taa ttg ctg tta tta atg gat tta gaa ata gtg gac gat ttt gtg ata tta ata tag tta tta atg atg aaa gga taa acg ctc ata aac tca tcc tat ctg gag cct ccg aat a-t ttt tcc att ctg ttt tcc aat aat ttt atc gat tct aat gaa tac gaa gtt aat cta agt cat tta gat tat caa agc gta aac gat ttg atc gat tat att tat ggg ata cct ttg agc cta act aac gat aac gtg aaa tat att ctt tca acc gct ga- --t ttt tta caa att gga tct gct att acg gag tgt gaa aat tac ata ctt aaa aat ctt tgt tct aga aac tgt atc gat ttc --- --- tac ata tac gct gat aaa tat aat aac aag aaa ata gaa tca gcg tcg ttt aac aca ata tta caa aat att ttg aga ctc atc aac gat gaa aac ttt aaa tac tta aca gag gaa tca atg ata aaa att tta agc gat gat atg tta aat ata aaa aat gag gat ttt gca cca cta att ctc att aaa tgg tta gag agt act caa caa tca tgc acc gtc gag tta ctt aaa tgc ctc aga ata tca -tt gct ttc ccc aca agt tat aaa atc act tta tag tca tca act ggt tag ttc aat cta cga atg tat aac att ctt aaa caa tat agc att ctt gga tga atc att tcc tag ata cca tag cat cga gtt gat atc tat cgg tat aag taa ttc gcg tga taa aat ttc cat aaa ctg cta caa tca t-a aaa aaa ata cat ggg aaa tga tat ctt cac gta gat ata ggt gta gtt tcg cag tgg ccg tcc tgg ata ata tta tct ata tga tgg gtg gat atg atc agt ccc cgt ata gaa gtt caa agg tta tag cgt aca ata cat gta caa att ctt gga tat atg ata tac cag agc t-a aaa tat cct cgt tct aat tgt ggg gga ctg gct gat gac gaa tac att tat tgt ata ggc ggc ata cgc gat cag gat tca tcg ttg aca tct agt att gat aga tgg aag cca tca aaa cca tat tgg cag aag tat gct aaa atg cgc gaa cca aaa tgt gat atg ggg gtt gcg atg tta aac gga tta ata tat gtc atg ggt gga atc gtt aaa ggt g-- aca cgt gta ccg acg cac tag aga gtt tat cag aag atg gat gga tga agc atc aac gtc ttc caa taa aaa tgt cca ata tgt cga cga ttg ttc atg atg gca aga ttt ata tat ctg gag gtt aca aca ata gta gtg tag tta atg taa tat cga atc t-- agt cct tag cta taa ttc gat ata tga tga atg gac caa att atc atc att aaa tat tcc tag aat taa tcc cgc tct a

>MF477237.2:162776-165025_Vaccinia_virus_strain_Mulford_1902_complete_genome

ttg ctg act gca --- tgt a-c tat ttt tta cat tac tac att ggc tac ggc aga tat acc tac tcc gcc acc --- --- --- --- --- -aa cgg --g tca tgt gac aag gga gaa tat ctt gat aag agg cat aat caa tgt tgt aat -cg gtg tcc acc tgg aga att tgc caa ggt tag atg ta- --a tgg taa cga taa cac aaa atg tga acg ctg ccc acc tca tac ata tac cac aat -cc caa tta ttc t-a atg gat gtc atc aat gta gaa aat gcc caa ccg gat cat ttg ata agg taa agt gta ccg gaa cac aga aca gta aat -gt tcg tgt ctt cct ggt tgg tat tgc gct act gat tct tca cag act gaa gat tgt tga aat -tg tgt acc aaa aag g-a gat gtc cat gcg gat act ttg gtg gaa tag atg aac aag gaa atc cta ttt gta aat cgt gct g-- -ta ttg gtg aat att gca act acc tac gta att ata gac ttg atc cat ttt ctc cat gca aa- --- cta tct aaa tgt aat taa tta --- --- --- --- --- --- tga tta tga ttt --- --- --- -tg atg ata atg tta -cc ata c-a tt- --- -at atc gct act tgg tta --g tgt att att cag tat g-a aga cct att aat aa- --- --- -tt act tat ctt ttg acg atc ttg tt- -at aat tat --- --- --- --- aat ata aaa --- -ac tta tgg --- --- cat agt aac tta t-- --- --- --- aat tgc ta- aca cga taa att cgt aat aat ctg ttt tgt tca aat ttt tat aag ga- --- --a tct aca ggc ata aaa at- aaa aat ata att tat aat ata c-- tct tac a-- acg cca tca tga ata gca gca gta aat taa ttg ctg tta tta atg gat tta gaa ata gtg gac gat ttt gtg ata tta ata tag tta tta atg atg aaa gga taa acg ctc ata aac tca tcc tat ctg gag cct ccg aat a-t ttt tcc att ctg ttt tcc aat aat ttt atc gat tct aat gaa tac gaa gtt aat cta agt cat tta gat tat caa agc gta aac gat ttg atc gat tat att tat ggg ata cct ttg agc cta act aac gat aac gtg aaa tat att ctt tca acc gct ga- --t ttt tta caa att gga tct gct att acg gag tgt gaa aat tac ata ctt aaa aat ctt tgt tct aaa aac tgt atc gat ttc --- --- tac ata tac gct gat aaa tat aat aac aag aaa ata gaa tca gcg tcg ttt aac aca ata tta caa aat att ttg aga ctc atc aac gat gaa aac ttt aaa tac tta aca gag gaa tca atg ata aaa att tta agc gat gat atg tta aat ata aaa aat gag gat ttc gcc cca cta att ctc att aaa tgg tta gag agt act caa caa cca tgc acc gtc gag tta ctt aaa tgc ctc aga ata tca -tt gct ttc ccc aca agt tat aaa atc act tta tag tca tcg act ggt tag ttc aat cta cga atg tat aac att ctt aaa caa tat agc att ctt gga tga atc att tcc tag ata cca tag cat cga gtt gat atc tat cgg tat aag taa ttc gcg tga taa aat ttc cat aaa ctg cta caa tca t-a aaa aaa ata cat ggg aaa tga tat ctt cac gta gat ata ggt gta gtt tcg cag tgg ccg tcc tgg ata ata tta ttt ata tga tgg gtg gat atg atc agt ccc cgt ata gaa gtt caa agg tta tag cgt aca ata cat gta caa att ctt gga tat atg ata tac cag agc t-a aaa tat cct cgt tct aat tgt ggg gga ctg gct gat gac gaa tac att tat tgt ata ggc ggc ata cgc gat cag gat tca tcg ttg aca tct agt att gat aaa tgg aag cca tca aaa cca tat tgg cag aag tat gct aaa atg cgc gaa cca aaa tgt gat atg ggg gtt gcg atg tta aac gga tta ata tat gtt ata ggt gga atc gtt aaa ggt g-- aca cgt gta ccg acg cac tag aga gtt tat cag aag atg gat gga tga agc atc aac gtc ttc caa taa aaa tgt cca ata tgt cga cga ttg ttc atg atg gca aga ttt ata tat ctg gag gtt aca aca ata gta gtg tag tta atg taa tat cga atc t-- agt cct tag cta taa ttc gat ata tga tga atg gac caa att atc atc att aaa tat tcc tag aat taa tcc cgc tct a

>NC_055231.1:168766-171032_Orthopoxvirus_Abatino_complete_genome

ttg ctg act gta --- agt a-c tat ttt tta cat tac tac att ggc tac ggc aga tat acc tac ttc gtc act gcc aca cgc tcc ggt aaa cgg --g tca tgt gac gag gga gaa tat ctt gat aag agg tat aat caa tgt tgt aat -ca gtg tcc gcc tgg aga att tgc caa ggt cag atg ta- --g tgg tag cga taa cac aaa atg tga acg ctg ccc acc tca tac ata tac cgc aat ccc caa tta ttc t-a atg gat gtc atc aat gta gaa aat gtc caa cag gat cat ttg ata agg taa agt gta ccg gaa cac aga aca gta aat -gt tcg tgt ctt cct ggt tgg tat tgc gct act gat tct tca cag act gaa gat tgt cga gat -tg tat acc aaa aag g-a gat gtc cgt gcg gat act ttg gtg gaa tag atg aac aag gaa aac cta ttt gta aat cgt gtt g-- -ta ttg gtg aac att gcg act acc tac ata att ata gag ttg gtc ctt ttc ctc cat gta aa- --- cta tct aaa tgt aat taa tta --- --- --- --- --- --- --- --- tga tct tga tat aat atg atg atc atg tta -cc ata c-a tt- --- -at ata gct act ttg gta --g tgt att att cag tat g-a aga cat att aat aa- --- --- -tt act tat c-t ttg aag atc --g tt- -at aat cat --- --- --- --- -aa ata aaa --a tac tta tgg --- -g- cat agt aac tca t-- --- --- --- aat tgc tg- aca cga taa att cgt aat aat ctg ttt tgt tca aat ttt tat aag ga- --- --a tct --a ggc ata aaa ata aaa aac ata att tat aat ac- --- act tac a-- gca cca tca tga ata aca gca gtg act taa ttg ctg tta tta atg gat tta gaa ata gtg gac gat ttt gtg ata tta gta tag tta tta atg atg aaa gga taa atg ctc ata gac tca tcc tat ctg gag cat ccg aat a-t ttt tcc att cta ttt tct agt gat ttt atc gat tct aat gaa tac gaa gtt aat cta agt cat tta gat tat caa agt gtg aac gat ttg atc gat tac att tat ggg ata ccg ttg agc cta act aac gat agc gtg aaa tat att ctt tca act gct ga- --t ttt tta caa att gga tct gct att acg gag tgt gaa aat tac ata ctt aac aat ctc tgt tct aga aac tgt atc gat ttc --- --- tac ata tac gct gat aaa tat aat aac aag aaa ata gaa aca gcg tcg ttt aac aca ata tta cga aat att ttg aga ctc att aac gat gaa aac ttt aaa tac ttg aca gag gaa tca atg ata aaa att tta agc gat gat atg tta aat ata aaa aat gag gat ttc gca cca cta att ctc att aaa tgg tta gag agt aca caa caa cca tgc acc gtc gag tta ctt aga tgc ctc aga ata tca -tt gct ttc ccc aca agt tat aaa atc act tta tag tca tcg act ggt tag ttc aat cta cga atg tat aac att ctt aaa caa tat agc att ctt gga tga atc att tcc tag ata caa tag cat cga gtt gat atc tat cgg tat aag taa ttc aca tga taa gat ttc cat aaa ctg cta caa tcg t-a aaa aaa ata cat ggg aaa tga tat ctt cac gta gat ata ggt gta gtt tcg cag tgg ccg tca tgg ata ata tta tct ata tga tgg gtg gat acg atc agt ccc cgt ata gaa gtt caa agg tta tag cat aca ata cat gta cta att ctt gga tat atg ata tac cag agc t-a aaa tat ccg cgt tct aat tgc gga gga gtt gcc gat gac gaa tac att tat tgt ata ggc ggc ata cgc gat cag gat tca tcg ttg ata tct agt att gat aga tgg aag cca tca aaa cca tat tgg cag acg tat gct aaa atg cga gag cca aaa tgt gat atg ggg gtt gcg atg tta aac gga tta ata tat gtc ata ggt gga gtc gtt aaa ggt g-- aca cat gta ccg aca cac tag aga gtt tat cag aag atg gtt gga tga tgc atc gac gcc ttc caa taa aaa tgt cca ata tgt cga cga ttg ttc atg ctg gaa aga ttt ata tat ctg gag gtt aca cca ata gta ata tag tta atg aaa tat cga atc t-- agt cct tag cta taa tcc gat ata tga tga atg gac caa att atc atc att aaa tat tcc tag aat taa tcc tgc tct a

>KT013210.1:153506-155733_Vaccinia_virus_strain_Cantagalo_isolate_Cantagalo_virus_CM01

ttg ctg act gta --- tgt a-c tat ttt tta cat tac tac att ggc tac ggc aga tat acc tac tcc gcc acc --- --- --- --- --- -aa cgg --g tca tgt gac aag gga gaa tat ctt gat aag agg cat aat c-- --- --- -at -tg gtg tcc acc tgg aga att tgc caa ggt tag atg ta- --a tgg taa cga taa cac aaa atg tga acg ctg ccc acc tca tac ata tac cac aat -cc caa tta ttc t-a atg gat gtc atc aat gta gaa aat gcc caa ccg gat cat ttg ata agg taa agt gta ccg gaa cac aga aca gta aat -gt tcg tgt ctt cct ggt tgg tat tgc gct act gat tct tca cag act gaa gat tgt tga aat -tg tgt acc aaa aag g-a gat gtc cat gcg gat act ttg gtg gaa tag atg aac aag gaa atc cta ttt gta aat cgt gct g-- -ta ttg gtg aat att gca act acc tac gta att ata gac ttg atc cat ttt ctc cat gca aa- --- cta tct aaa tgt aat taa tta --- --- --- --- --- --- --- --- tga ttt --- --- --- -tg atg ata atg tta -cc ata c-a tt- --- -at atc gct act tgg tta --g tgt att att cag tat g-a aga cct att aat aa- --- --- -tt act tat ctt ttg acg atc ttg tt- -at aat tat --- --- --- --- aat ata aaa --- -ac tta tgg --- --- cat agt aac tta t-- --- --- --- aat tgc ta- aca cga taa att cgt aat aat ctg ttt tgt tca --- --- --- aag ga- --- --a tct aca ggc ata aaa at- aaa aat ata att tat aat ata c-- tct tac agc gcg cca tca tga ata aca gca gtg aat tga ttg ctg tta tta atg gat tta gaa ata gtg gac gat ttt gtg ata tta gta tag tta tta atg atg aaa tga taa acg ctc aca gac tca tcc tat ctg gag cct ccg aat a-t ttt tcc att ctg ttt tcc aat aat ttt atc gat tct aat gaa tac gaa gtt aat cta agt cat tta gat tat caa agt gtt aac gat ttg atc gat tac att tat ggg ata cct ttg agc cta act aac gat aac gtg aaa tat att ctt tca acc gct ga- --t ttt tta caa att gga tct gct att acg gag tgt gaa aat tac ata ctt aaa aat ctt tgt tct aaa aac tgt atc gat ttc --- --- tac ata tac gct gat aaa tat aat aac aag aaa ata gaa tca gcg tcg ttt aac aca ata tta caa aat att ttg aga ctc atc aac gat gaa aac ttt aaa tac tta aca gag gaa tca atg ata aaa att tta agc gat gat atg tta aat ata aaa aat gag gat ttc gcc cca cta att ctc att aaa tgg tta gag agt act caa caa tca tgc acc gtc gag tta ctt aga tgc ctc aga ata tca -tt gct ttc ccc aca agt tat aaa atc act tta tag tca tca act agt tag ttc aat cta cga atg tat aac att ctt aaa caa tat agc att ctt gga tga atc att tcc tag ata cca tag cat cga gtt gat atc tat cgg tat aag taa ttt gcg tga taa aat ttc cat aaa ctg cta caa tca t-a aaa aaa ata cat ggg aaa tga tat ctt cac gta gat ata ggt gta gtt tcg cag tgg ccg tcc tgg ata ata tta tct ata tga tgg gtg gat atg atc agt ccc cgt ata gaa gtt caa aag tta tag cgt aca ata cat gta caa att ctt gga tat atg ata tac cag agc t-a aaa tat cct cgt tct aat tgt ggg gga ctg gct gat gac gaa tac att tat tgt ata ggc ggc ata cgc gat cag gat tca tcg ttg aca tct agt att gat aaa tgg aag cca tca aaa cca tat tgg cag aag tat gct aaa atg cgc gaa cca aaa tgt gat atg ggg gtt gcg atg tta aac gga tta ata tat gtt ata ggt gga atc gtt aaa ggt g-- aca cgt gta ccg acg cac tag aga gtt tat cag aag atg gat gga tga agc atc aac gtc ttc caa taa aaa tgt cca ata tgt cga cga ttg ttc atg atg gca aga ttt ata tat ctg gag gtt aca aca ata gta gtg tag tta atg taa tat cga atc t-- agt cct tag cta taa tcc gat ata tga tga atg gac caa att atc atc att aaa cat tcc tag aat taa tcc cgc tct a

>KC813493.1:168221-170466_Cowpox_virus_strain_HumLit08/1_complete_genome

ttg ctg act gta --- tgt a-c tat ttt tta cat tac tac att ggc tac ggc aga tat acc tac ttc gtc act gcc aca cgc tcc ggt aaa cgg --g tca tgt gac gag gga gaa tat ctt gat aag act cat aat cga tgt tgt aat -cg gtg tcc acc tgg aga att tgc caa ggt tag atg ta- --g tgg tag cta taa cac aaa atg tga acg ctg tcc aca tca tac ata tac agc aat tcc caa tta ttc t-a atg gat gcc atc aat gta gaa aat gcc caa agg gat cat ttg ata agg taa agt gta ccg gaa cac aga aca gta aat -gt gcg tgt ctt cct ggt tgg tat tgc gct acc gat tct tca cag act gaa gac tgt cga gat -tg tgt acc aaa aag g-a gct gtc cat gcg gat act ttg gtg gaa tag ata aat tag gaa atc cta ttt gta aat cgt gtt g-- -tg ttg gta aat att gtg acg aca tac gta atc ata gag ttg gtc ctt ttc ctc cat gca aa- --- cta tct aaa tgt aat tag tca --- --- --- --- --- --- --- --- --- --- --- --- --- --- -tg atc atg gta -cc gta caa tt- --- -at att gct --- --g gta --g tat att att cag tat g-a aga cat att aat ac- --- --- -tt atc --- --t ttg aca atc ttg ct- -at aat cat --- --- --- --- aat ata aaa --a tac tta tgg --- -a- cat agt cac tca t-- --- --- --- aat tgc tg- acg cga taa att cgt aat aat ctg ttt tgt tca aat ttt tat aag ga- --- --a tct aca ggc ata aaa at- aaa aat ata att tat aat ac- --- act tac agc gcg cca tca tga ata aca gca gtg aat taa ttg ctg tta tta atg gat tta gaa ata gtg gac gat ttt gtg ata tta ata tag tta tta atg atg aaa gga taa atg ctc ata gac tca tcc tat ctg gag cct ccg aat a-t ttt tcc att ctg ttt tcc aat aat ttt atc gat tct aat gaa tac gaa gtt aat cta agt cat tta gat tat caa agt gtt aac gat ttg atc gat tac att tat ggg ata cct ttg agc cta act aac gat agc gtg aaa tat att ctt tca act gct ga- --t ttt tta caa att gga tct gcc att act gag tgc gaa aat tac ata ctt aaa aat ctg tgt tct aga aac tgt atc gat ttc --- --- tac ata tac gct gat aaa tat aat aac aag aaa ata gaa tca gcg tcg ttt aac aca ata tta cga aat atc ttg aga ctc atc aac gat gaa aac ttt aaa tac tta aca gag gaa tca atg ata aaa att tta agc gat gat atg tta aat ata aaa aat gag gat ttt gca cca cta att ctc att aaa tgg tta gag agt act caa caa tca tgc acc gtc gag tta ctt aga tgc ctc aga ata tca -tt gct ttc ccc aca agt tat aaa atc act tta tag tca tca act ggt tag ttc aat cta cga atg tat aac att ctt aaa caa tat agc att ctt gga tga atc att tcc tag ata cca tag cat cga gtt gat atc tat cgg tat aag taa ttc gca tga taa gat ttc cat aaa ctg cta caa tca t-a aaa aaa ata cat ggg aaa tga tat ctt cac gta gat ata ggt gta gtt tcg cag tgg ccg tca tgg ata ata tta tct ata tga tgg gtg gat atg atc agt ccc cgt ata gga gtt caa agg tta tag cgt aca ata cct gta cta att ctt gga tat atg ata tac cag agc t-a aaa tat cct cgt tct aat tgt gga gga gtt gct gat gat gaa tac att tat tgc ata ggc ggc ata cgc gat cag gat tca tcg ttg ata tct agt att gat aga tgg aag cca tca aaa cca tat tgg caa acg tat gct aaa atg cgc gaa cca aaa tgt gat atg ggg gtt gcg atg tta aac gga tta ata tat gtc atg ggt gga atc gtt aaa ggt g-- aca cgt gta ccg acg cac tag aga gtt tat cag aag atg gat gga tga agc atc aac gtc ttc caa taa aaa tgt cca ata tgt cga cga ttg ttc atg atg gca aga ttt ata tat ctg gag gtt aca aca ata gta gtg tag tta atg taa tat cga atc t-- agt cct tag cta taa tcc gat ata tga tga atg gac caa att atc atc att aaa cat tcc tag aat taa tcc cgc tct a

>MW018153.1:153189-155416_Vaccinia_virus_strain_Cantagalo_isolate_Cantagalo_MI233

ttg ctg act gta --- tgt a-c tat ttt tta cat tac tac att ggc tac ggc aga tat acc tac tcc gcc acc --- --- --- --- --- -aa cgg --g tca tgt gac aag gga gaa tat ctt gat aag agg cat aat c-- --- --- -at -tg gtg tcc acc tgg aga att tgc caa ggt tag atg ta- --a tgg taa cga taa cac aaa atg tga acg ctg ccc acc tca tac ata tac cac aat -cc caa tta ttc t-a atg gat gtc atc aat gta gaa aat gcc caa ccg gat cat ttg ata agg taa agt gta ccg gaa cac aga aca gta aat -gt tcg tgt ctt cct ggt tgg tat tgc gct act gat tct tca cag act gaa gat tgt tga aat -tg tgt acc aaa aag g-a gat gtc cat gcg gat act ttg gtg gaa tag atg aac aag gaa atc cta ttt gta aat cgt gct g-- -ta ttg gtg aat att gca act acc tac gta att ata gac ttg atc cat ttt ctc cat gca aa- --- cta tct aaa tgt aat taa tta --- --- --- --- --- --- --- --- tga ttt --- --- --- -tg atg ata atg tta -cc ata c-a tt- --- -at atc gct act tgg tta --g tgt att att cag tat g-a aga cct att aat aa- --- --- -tt act tat ctt ttg acg atc ttg tt- -at aat tat --- --- --- --- aat ata aaa --- -ac tta tgg --- --- cat agt aac tta t-- --- --- --- aat tgc ta- aca cga taa att cgt aat aat ctg ttt tgt tca --- --- --- aag ga- --- --a tct aca ggc ata aaa at- aaa aat ata att tat aat ata c-- tct tac agc gcg cca tca tga ata aca gca gtg aat tga ttg ctg tta tta atg gat tta gaa ata gtg gac gat ttt gtg ata tta gta tag tta tta atg atg aaa tga taa acg ctc aca gac tca tcc tat ctg gag cct ccg aat a-t ttt tcc att ctg ttt tcc aat aat ttt atc gat tct aat gaa tac gaa gtt aat cta agt cat tta gat tat caa agt gtt aac gat ttg atc gat tac att tat ggg ata cct ttg agc cta act aac gat aac gtg aaa tat att ctt tca acc gct ga- --t ttt tta caa att gga tct gct att acg gag tgt gaa aat tac ata ctt aaa aat ctt tgt tct aaa aac tgt atc gat ttc --- --- tac ata tac gct gat aaa tat aat aac aag aaa ata gaa tca gcg tcg ttt aac aca ata tta caa aat att ttg aga ctc atc aac gat gaa aac ttt aaa tac tta aca gag gaa tca atg ata aaa att tta agc gat gat atg tta aat ata aaa aat gag gat ttc gcc cca cta att ctc att aaa tgg tta gag agt act caa caa tca tgc acc gtc gag tta ctt aga tgc ctc aga ata tca -tt gct ttc ccc aca agt tat aaa atc act tta tag tca tca act agt tag ttc aat cta cga atg tat aac att ctt aaa caa tat agc att ctt gga tga atc att tcc tag ata cca tag cat cga gtt gat atc tat cgg tat aag taa ttt gcg tga taa aat ttc cat aaa ctg cta caa tca t-a aaa aaa ata cat ggg aaa tga tat ctt cac gta gat ata ggt gta gtt tcg cag tgg ccg tcc tgg ata ata tta tct ata tga tgg gtg gat atg atc agt ccc cgt ata gaa gtt caa aag tta tag cgt aca ata cat gta caa att ctt gga tat atg ata tac cag agc t-a aaa tat cct cgt tct aat tgt ggg gga ctg gct gat gac gaa tac att tat tgt ata ggc ggc ata cgc gat cag gat tca tcg ttg aca tct agt att gat aaa tgg aag cca tca aaa cca tat tgg cag aag tat gct aaa atg cgc gaa cca aaa tgt gat atg ggg gtt gcg atg tta aac gga tta ata tat gtt ata ggt gga atc gtt aaa ggt g-- aca cgt gta ccg acg cac tag aga gtt tat cag aag atg gat gga tga agc atc aac gtc ttc caa taa aaa tgt cca ata tgt cga cga ttg ttc atg atg gca aga ttt ata tat ctg gag gtt aca aca ata gta gtg tag tta atg taa tat cga atc t-- agt cct tag cta taa tcc gat ata tga tga atg gac caa att atc atc att aaa cat tcc tag aat taa tcc cgc tct a

>KF179385.1:156623-158850_Vaccinia_virus_complete_genome

ttg ctg act gta --- tgt a-c tat ttt tta cat tac tac att ggc tac ggc aga tat acc tac tcc gcc acc --- --- --- --- --- -aa cgg --g tca tgt gac aag gga gaa tat ctt gat aag agg cat aat c-- --- --- -at -tg gtg tcc acc tgg aga att tgc caa ggt tag atg ta- --a tgg taa cga taa cac aaa atg tga acg ctg ccc acc tca tac ata tac cac aat -cc caa tta ttc t-a atg gat gtc atc aat gta gaa aat gcc caa ccg gat cat ttg ata agg taa agt gta ccg gaa cac aga aca gta aat -gt tcg tgt ctt cct ggt tgg tat tgc gct act gat tct tca cag act gaa gat tgt tga aat -tg tgt acc aaa aag g-a gat gtc cat gcg gat act ttg gtg gaa tag atg aac aag gaa atc cta ttt gta aat cgt gct g-- -ta ttg gtg aat att gca act acc tac gta att ata gac ttg atc cat ttt ctc cat gca aa- --- cta tct aaa tgt aat taa tta --- --- --- --- --- --- --- --- tga ttt --- --- --- -tg atg ata atg tta -cc ata c-a tt- --- -at atc gct act tgg tta --g tgt att att cag tat g-a aga cct att aat aa- --- --- -tt act tat ctt ttg acg atc ttg tt- -at aat tat --- --- --- --- aat ata aaa --- -ac tta tgg --- --- cat agt aac tta t-- --- --- --- aat tgc ta- aca cga taa att cgt aat aat ctg ttt tgt tca --- --- --- aag ga- --- --a tct aca ggc ata aaa at- aaa aat ata att tat aat ata c-- tct tac agc gcg cca tca tga ata aca gca gtg aat tga ttg ctg tta tta atg gat tta gaa ata gtg gac gat ttt gtg ata tta gta tag tta tta atg atg aaa tga taa acg ctc aca gac tta tcc tat ctg gag cct ccg aat a-t ttt tcc att ctg ttt tcc aat aat ttt atc gat tct aat gaa tac gaa gtt aat cta agt cat tta gat tat caa agt gtt aac gat ttg atc gat tac att tat ggg ata cct ttg agc cta act aac gat aac gtg aaa tat att ctt tca acc gct ga- --t ttt tta caa att gga tct gct att acg gag tgt gaa aat tac ata ctt aaa aat ctt tgt tct aaa aac tgt atc gat ttc --- --- tac ata tac gct gat aaa tat aat aac aag aaa ata gaa tca gcg tcg ttt aac aca ata tta caa aat att ttg aga ctc atc aac gat gaa aac ttt aaa tac tta aca gag gaa tca atg ata aaa att tta agc gat gat atg tta aat ata aaa aat gag gat ttc gcc cca cta att ctc att aaa tgg tta gag agt act caa caa tca tgc acc gtc gag tta ctt aga tgc ctc aga ata tca -tt gct ttc ccc aca agt tat aaa atc act tta tag tca tca act agt tag ttc aat cta cga atg tat aac att ctt aaa caa tat agc att ctt gga tga atc att tcc tag ata cca tag cat cga gtt gat atc tat cgg tat aag taa ttt gcg tga taa aat ttc cat aaa ctg cta caa tca t-a aaa aaa ata cat ggg aaa tga tat ctt cac gta gat ata ggt gta gtt tcg cag tgg ccg tcc tgg ata ata tta tct ata tga tgg gtg gat atg atc agt ccc cgt ata gaa gtt caa aag tta tag cgt aca ata cat gta caa att ctt gga tat atg ata tac cag agc t-a aaa tat cct cgt tct aat tgt ggg gga ctg gct gat gac gaa tac att tat tgt ata ggc ggc ata cgc gat cag gat tca tcg ttg aca tct agt att gat aaa tgg aag cca tca aaa cca tat tgg cag aag tat gct aaa atg cgc gaa cca aaa tgt gat atg ggg gtt gcg atg tta aac gga tta ata tat gtt ata ggt gga atc gtt aaa ggt g-- aca cgt gta ccg acg cac tag aga gtt tat cag aag atg gat gga tga agc atc aac gtc ttc caa taa aaa tgt cca ata tgt cga cga ttg ttc atg atg gca aga ttt ata tat ctg gag gtt aca aca ata gta gtg tag tta atg taa tat cga atc t-- agt cct tag cta taa tcc gat ata tga tga atg gac caa att atc atc att aaa cat tcc tag aat taa tcc cgc tct a

>MW018156.1:156968-159195_Vaccinia_virus_strain_Cantagalo_isolate_Cantagalo_ALEH2

ttg ctg act gta --- tgt a-c tat ttt tta cat tac tac att ggc tac agc aga tat acc tac tct gcc acc --- --- --- --- --- -aa cgg --g tca tgt gac aag gga gaa tat ctt gat aag agg cat aat c-- --- --- -at -tg gtg tcc acc tgg aga att tgc caa ggt tag atg ta- --a tgg taa cga taa cac aaa atg tga acg ctg ccc acc tca tac ata tac cac aat -cc caa tta ttc t-a atg gat gtc atc aat gta gaa aat gcc caa ccg gat cat ttg ata agg taa agt gta ccg gaa cac aga aca gta aat -gt tcg tgt ctt cct ggt tgg tat tgc gct act gat tct tca cag act gaa gat tgt tga aat -tg tgt acc aaa aag g-a gat gtc cat gcg gat act ttg gtg gaa tag atg aac aag gaa atc cta ttt gta aat cgt gct g-- -ta ttg gtg aat att gca act acc tac gta att ata gac ttg atc cat ttt ctc cat gca aa- --- cta tct aaa tgt aat taa tta --- --- --- --- --- --- --- --- tga ttt --- --- --- -tg atg ata atg tta -cc ata c-a tt- --- -at atc gct act tgg tta --g tgt att att cag tat g-a aga cct att aat aa- --- --- -tt act tat ctt ttg acg atc ttg tt- -at aat tat --- --- --- --- aat ata aaa --- -ac tta tgg --- --- cat agt aac tta t-- --- --- --- aat tgc ta- aca cga taa att cgt aat aat ctg ttt tgt tca --- --- --- aag ga- --- --a tct aca ggc ata aaa at- aaa aat ata att tat aat ata c-- tct tac agc gcg cca tca tga ata aca gca gtg aat tga ttg ctg tta tta atg gat tta gaa ata gtg gac gat ttt gtg ata tta gta tag tta tta atg atg aaa tga taa acg ctc aca gac tca tcc tat ctg gag cct ccg aat a-t ttt tcc att ctg ttt tcc aat aat ttt atc gat tct aat gaa tac gaa gtt aat cta agt cat tta gat tat caa agt gtt aac gat ttg atc gat tac att tat ggg ata cct ttg agc cta act aac gat aac gtg aaa tat att ctt tca acc gct ga- --t ttt tta caa att gga tct gct att acg gag tgt gaa aat tac ata ctt aaa aat ctt tgt tct aaa aac tgt atc gat ttc --- --- tac ata tac gct gat aaa tat aat aac aag aaa ata gaa tca gcg tcg ttt aac aca ata tta caa aat att ttg aga ctc atc aac gat gaa aac ttt aaa tac tta aca gag gaa tca atg ata aaa att tta agc gat gat atg tta aat ata aaa aat gag gat ttc gcc cca cta att ctc att aaa tgg tta gag agt act caa caa tca tgc acc gtc gag tta ctt aga tgc ctc aga ata tca -tt gct ttc ccc aca agt tat aaa atc act tta tag tca tca act agt tag ttc aat cta cga atg tat aac att ctt aaa caa tat agc att ctt gga tga atc att tcc tag ata cca tag cat cga gtt gat atc tat cgg tat aag taa ttt gcg tga taa aat ttc cat aaa ctg cta caa tca t-a aaa aaa ata cat ggg aaa tga tat ctt cac gta gat ata ggt gta gtt tcg cag tgg ccg tcc tgg ata ata tta tct ata tga tgg gtg gat atg atc agt ccc cgt ata gaa gtt caa aag tta tag cgt aca ata cat gta caa att ctt gga tat atg ata tac cag agc t-a aaa tat cct cgt tct aat tgt ggg gga ctg gct gat gac gaa tac att tat tgt ata ggc ggc ata cgc gat cag gat tca tcg ttg aca tct agt att gat aaa tgg aag cca tca aaa cca tat tgg cag aag tat gct aaa atg cgc gaa cca aaa tgt gat atg ggg gtt gcg atg tta aac gga tta ata tat gtt ata ggt gga atc gtt aaa ggt g-- aca cgt gta ccg acg cac tag aga gtt tat cag aag atg gat gga tga agc atc aac gtc ttc caa taa aaa tgt cca ata tgt cga cga ttg ttc atg atg gca aga ttt ata tat ctg gag gtt aca aca ata gta gtg tag tta atg taa tat cga atc t-- agt cct tag cta taa tcc gat ata tga tga atg gac caa att atc atc att aaa cat tcc tag aat taa tcc cgc tct a

>MW018155.1:153113-155340_Vaccinia_virus_strain_Cantagalo_isolate_Cantagalo_CG04

ttg ctg act gta --- tgt a-c tat ttt tta cat tac tac att ggc tac ggc aga tat acc tac tcc gcc acc --- --- --- --- --- -aa cgg --g tca tgt gac aag gga gaa tat ctt gat aag agg cat aat c-- --- --- -at -tg gtg tcc acc tgg aga att tgc caa ggt tag atg ta- --a tgg taa cga taa cac aaa atg tga acg ctg ccc acc tca tac ata tac cac aat -cc caa tta ttc t-a atg gat gtc atc aat gta gaa aat gcc caa ccg gat cat ttg ata agg taa agt gta ccg gaa cac aga aca gta aat -gt tcg tgt ctt cct ggt tgg tat tgc gct act gat tct tca cag act gaa gat tgt tga aat -tg tgt acc aaa aag g-a gat gtc cat gcg gat act ttg gtg gaa tag atg aac aag gaa atc cta ttt gta aat cgt gct g-- -ta ttg gtg aat att gca act acc tac gta att ata gac ttg atc cat ttt ctc cat gca aa- --- cta tct aaa tgt aat taa tta --- --- --- --- --- --- --- --- tga ttt --- --- --- -tg atg ata atg tta -cc ata c-a tt- --- -at atc gct act tgg tta --g tgt att att cag tat g-a aga cct att aat aa- --- --- -tt act tat ctt ttg acg atc ttg tt- -at aat tat --- --- --- --- aat ata aaa --- -ac tta tgg --- --- cat agt aac tta t-- --- --- --- aat tgc ta- aca cga taa att cgt aat aat ctg ttt tgt tca --- --- --- aag ga- --- --a tct aca ggc ata aaa at- aaa aat ata att tat aat ata c-- tct tac agc gcg cca tca tga ata aca gca gtg aat tga ttg ctg tta tta atg gat tta gaa ata gtg gac gat ttt gtg ata tta gta tag tta tta atg atg aaa tga taa acg ctc aca gac tca tcc tat ctg gag cct ccg aat a-t ttt tcc att ctg ttt tcc aat aat ttt atc aat tct aat gaa tac gaa gtt aat cta agt cat tta gat tat caa agt gtt aac gat ttg atc gat tac att tat ggg ata cct ttg agc cta act aac gat aac gtg aaa tat att ctt tca acc gct ga- --t ttt tta caa att gga tct gct att acg gag tgt gaa aat tac ata ctt aaa aat ctt tgt tct aaa aac tgt atc gat ttc --- --- tac ata tac gct gat aaa tat aat aac aag aaa ata gaa tca gcg tcg ttt aac aca ata tta caa aat att ttg aga ctc atc aac gat gaa aac ttt aaa tac tta aca gag gaa tca atg ata aaa att tta agc gat gat atg tta aat ata aaa aat gag gat ttc gcc cca cta att ctc att aaa tgg tta gag agt act caa caa tca tgc acc gtc gag tta ctt aga tgc ctc aga ata tca -tt gct ttc ccc aca agt tat aaa atc act tta tag tca tca act agt tag ttc aat cta cga atg tat aac att ctt aaa caa tat agc att ctt gga tga atc att tcc tag ata cca tag cat cga gtt gat atc tat cgg tat aag taa ttt gcg tga taa aat ttc cat aaa ctg cta caa tca t-a aaa aaa ata cat ggg aaa tga tat ctt cac gta gat ata ggt gta gtt tcg cag tgg ccg tcc tgg ata ata tta tct ata tga tgg gtg gat atg atc agt ccc cgt ata gaa gtt caa aag tta tag cgt aca ata cat gta caa att ctt gga tat atg ata tac cag agc t-a aaa tat cct cgt tct aat tgt ggg gga ctg gct gat gac gaa tac att tat tgt ata ggc ggc ata cgc gat cag gat tca tcg ttg aca tct agt att gat aaa tgg aag cca tca aaa cca tat tgg cag aag tat gct aaa atg cgc gaa cca aaa tgt gat atg ggg gtt gcg atg tta aac gga tta ata tat gtt ata ggt gga atc gtt aaa ggt g-- aca cgt gta ccg acg cac tag aga gtt tat cag aag atg gat gga tga agc atc aac gtc ttc caa taa aaa tgt cca ata tgt cga cga ttg ttc atg atg gca aga ttt ata tat ctg gag gtt aca aca ata gta gtg tag tta atg taa tat cga atc t-- agt cct tag cta taa tcc gat ata tga tga atg gac caa att atc atc att aaa cat tcc tag aat taa tcc cgc tct a

>OU343156.1:166726-169002_Ectromelia_virus_strain_Mill-Hill_genome_assembly_chromosome:_1

ttg ctg act gta --- agt a-c tat ttt -ta cat tac tac att ggc tac ggc aga tat acc tac ttc gtc act gcc aca cgc tcc ggt aaa cgg --g tca tgt gac gag gga gaa tat ctt gat aa- agg cat aat caa tgt tgt aat -ca gtg tcc acc tgg aga att tgc caa ggt cag atg ta- --g tgg cag tga taa cac aaa atg tga acg ctg cct acc tca tac ata tac cac aat ccc caa tta ttc taa atg gat gtc atc aat gta gaa aat gcc caa cag gat cat ttg ata agg taa agt gta ccg gaa cac ata aca gta aat -gt tcg tgt ctt cct ggt tgg tat tgc gct act gat tct tca cag act gaa gat tgt cga gat -tg tat acc aaa aag g-a gat gtc cgt gcg gat att ttg gtg gaa tag atg aac aag gaa atc cta ttt gta aat cgt gat g-- -ta ttg gtg aac att gcg act acc tac ata att ata gag ttg gtc ctt ttc ctc cat gta aa- --- cta tct aaa tgt aat taa tta --- --- --- --- --- --- --- --- tga tct tga tat aat atg att atc atg tta -cc ata c-a tt- --- -at ata gct act ttg gta --g tgt att att cag tat g-a aga cat att aat aa- --- --- -tt act tat c-t ttg aag atc ttg tt- -at aat cat aaa --- --t ata aaa ata aaa --a tac tta tgg --- -gg cat agt aac tca t-- --- --- --- aat tgc tg- aca cga taa att cgt aat aat ctg ttt tgt tca aat ttt tat aag ga- --- --a tct --a ggc ata aaa at- aaa aac ata att tat aat ac- --- act tac aac ac- cca tca tga ata aca gca gtg aat taa ttg ctg cga tta atg gat tta gaa ata gtg gac gat ttt gtg ata tta ata tag tta tta atg atg aaa gga taa atg ctc ata gac tca tcc tat ctg gag cat ccg aat a-t ttt tcc att cta ttt tct agt gat ttt atc gat tct aat gaa tac gaa gtt aat cta agt cat tta gat tat caa agt gtg aac gat ttg atc gat tac att tat ggg ata cct ttg agc cta act aac gat agc gtg aaa tat att ctt tca act gct ga- --t ttt tta caa att gga tct gct att acg gag tgt gaa aat tac ata ctt aaa aat ctc tgt tct aga aac tgt atc gat ttc --- --- tac ata tac gct gat aaa tat aat aac aag aaa ata gaa aca gcg tcg ttt aac aca ata tta cga aat att ttg aga ctc att aac gat gaa aac ttt aaa tac ttg aca gag gaa tca atg ata aaa att tta agc gat gat atg tta aat ata aaa aat gag gat ttc gca cca cta att ctc att aaa tgg tta gag agt aca caa caa cca tgc acc gtc gag tta ctt aga tgc ctc aga ata tca -tt gct ttc ccc aca agt tat aaa atc act tta tag tca tcg act ggt tag ttc aat cta cga atg tat aac att ctt aaa caa tat agc att ctt gga tga atc att tcc tag ata caa tag cat cga gtt gat atc tat cgg tat aag taa ttc acg tga taa gat ttc cat aaa ctg cta caa tcg t-a aaa aaa ata cat ggg aaa tga tat ctt cac gta gat ata ggt gta gtt tcg cag tgg ctg tca tgg ata ata tta tct ata tga tgg gtg gat acg atc agt ccc cgt ata gaa gtt caa agg tta tag cat aca ata cat gta cta att ctt gga tat atg ata tac cag agc t-a aaa tat ccg cgt tct aat tgc gga gga gtt gtc gat gac gaa tac att tat tgt ata ggt ggc ata cgc gat cag gat tca tcg ttg ata tct gat att gat aga tgg aag cca tca aaa cca tat tgg cag acg tat gct aaa atg cga gag cca aaa tgt gat atg ggg gtt gcg atg tta aac gga tta ata tat gtc ata ggt gga gtc gtt aaa ggt g-- gca cat gta ccg aca cac tag aga gtt tat cag aag atg gat gga tga tgc atc gac gcc ttc caa taa aaa tgt cca ata tgt cga cga ttg ttc atg ctg gaa aga ttt ata tat ctg gag gtt aca cca ata gta gta tag tta atg aaa tat cta atc t-- agt cct tag cta taa tcc gat ata tga tga atg gac caa att atc atc att aaa tat tcc tag aat taa tcc tgc tct a

>OU343112.1:166407-168683_Ectromelia_virus_strain_Hampstead_Egg_genome_assembly_chromosome:_1

ttg ctg act gta --- agt a-c tat ttt -ta cat tac tac att ggc tac ggc aga tat acc tac ttc gtc act gcc aca cgc tcc ggt aaa cgg --g tca tgt gac gag gga gaa tat ctt gat aa- agg cat aat caa tgt tgt aat -ca gtg tcc acc tgg aga att tgc caa ggt cag atg ta- --g tgg cag tga taa cac aaa atg tga acg ctg cct acc tca tac ata tac cac aat ccc caa tta ttc taa atg gat gtc atc aat gta gaa aat gcc caa cag gat cat ttg ata agg taa agt gta ccg gaa cac ata aca gta aat -gt tcg tgt ctt cct ggt tgg tat tgc gct act gat tct tca cag act gaa gat tgt cga gat -tg tat acc aaa aag g-a gat gtc cgt gcg gat att ttg gtg gaa tag atg aac aag gaa atc cta ttt gta aat cgt gat g-- -ta ttg gtg aac att gcg act acc tac ata att ata gag ttg gtc ctt ttc ctc cat gta aa- --- cta tct aaa tgt aat taa tta --- --- --- --- --- --- --- --- tga tct tga tat aat atg att atc atg tta -cc ata c-a tt- --- -at ata gct act ttg gta --g tgt att att cag tat g-a aga cat att aat aa- --- --- -tt act tat c-t ttg aag atc ttg tt- -at aat cat aaa --- --t ata aaa ata aaa --a tac tta tgg --- -gg cat agt aac tca t-- --- --- --- aat tgc tg- aca cga taa att cgt aat aat ctg ttt tgt tca aat ttt tat aag ga- --- --a tct --a ggc ata aaa at- aaa aac ata att tat aat ac- --- act tac aac ac- cca tca tga ata aca gca gtg aat taa ttg ctg cga tta atg gat tta gaa ata gtg gac gat ttt gtg ata tta ata tag tta tta atg atg aaa gga taa atg ctc ata gac tca tcc tat ctg gag cat ccg aat a-t ttt tcc att cta ttt tct agt gat ttt atc gat tct aat gaa tac gaa gtt aat cta agt cat tta gat tat caa agt gtg aac gat ttg atc gat tac att tat ggg ata cct ttg agc cta act aac gat agc gtg aaa tat att ctt tca act gct ga- --t ttt tta caa att gga tct gct att acg gag tgt gaa aat tac ata ctt aaa aat ctc tgt tct aga aac tgt atc gat ttc --- --- tac ata tac gct gat aaa tat aat aac aag aaa ata gaa aca gcg tcg ttt aac aca ata tta cga aat att ttg aga ctc att aac gat gaa aac ttt aaa tac ttg aca gag gaa tca atg ata aaa att tta agc gat gat atg tta aat ata aaa aat gag gat ttc gca cca cta att ctc att aaa tgg tta gag agt aca caa caa cca tgc acc gtc gag tta ctt aga tgc ctc aga ata tca -tt gct ttc ccc aca agt tat aaa atc act tta tag tca tcg act ggt tag ttc aat cta cga atg tat aac att ctt aaa caa tat agc att ctt gga tga atc att tcc tag ata caa tag cat cga gtt gat atc tat cgg tat aag taa ttc acg tga taa gat ttc cat aaa ctg cta caa tcg t-a aaa aaa ata cat ggg aaa tga tat ctt cac gta gat ata ggt gta gtt tcg cag tgg ctg tca tgg ata ata tta tct ata tga tgg gtg gat acg atc agt ccc cgt ata gaa gtt caa agg tta tag cat aca ata cat gta cta att ctt gga tat atg ata tac cag agc t-a aaa tat ccg cgt tct aat tgc gga gga gtt gtc gat gac gaa tac att tat tgt ata ggt ggc ata cgc gat cag gat tca tcg ttg ata tct gat att gat aga tgg aag cca tca aaa cca tat tgg cag acg tat gct aaa atg cga gag cca aaa tgt gat atg ggg gtt gcg atg tta aac gga tta ata tat gtc ata ggt gga gtc gtt aaa ggt g-- gca cat gta ccg aca cac tag aga gtt tat cag aag atg gat gga tga tgc atc gac gcc ttc caa taa aaa tgt cca ata tgt cga cga ttg ttc atg ctg gaa aga ttt ata tat ctg gag gtt aca cca ata gta gta tag tta atg aaa tat cta atc t-- agt cct tag cta taa tcc gat ata tga tga atg gac caa att atc atc att aaa tat tcc tag aat taa tcc tgc tct a

>MW018154.1:153527-155754_Vaccinia_virus_strain_Cantagalo_isolate_Cantagalo_VI04

ttg ctg act gta --- tgt a-c tat ttt tta cat tac tac att ggc tac ggc aga tat acc tac tcc gcc acc --- --- --- --- --- -aa cgg --g tca tgt gac aag gga gaa tat ctt gat aag agg cat aat c-- --- --- -at -tg gtg tcc acc tgg aga att tgc caa ggt tag atg ta- --a tgg taa cga taa cac aaa atg tga acg ctg ccc acc tca tac ata tac cac aat -cc caa tta ttc t-a atg gat gtc atc aat gta gaa aat gcc caa ccg gat cat ttg ata agg taa agt gta ccg gaa cac aga aca gta aat -gt tcg tgt ctt cct ggt tgg tat tgc gct act gat tct tca cag act gaa gat tgt tga aat -tg tgt acc aaa aag g-a gat gtc cat gcg gat act ttg gtg gaa tag atg aac aag gaa atc cta ttt gta aat cgt gct g-- -ta ttg gtg aat att gca act acc tac gta att ata gac ttg atc cat ttt ctc cat gca aa- --- cta tct aaa tgt aat taa tta --- --- --- --- --- --- --- --- tga ttt --- --- --- -tg atg ata atg tta -cc ata c-a tt- --- -at atc gct act tgg tta --g tgt att att cag tat g-a aga cct att aat aa- --- --- -tt act tat ctt ttg acg atc ttg tt- -at aat tat --- --- --- --- aat ata aaa --- -ac tta tgg --- --- cat agt aac tta t-- --- --- --- aat tgc ta- aca cga taa att cgt aat aat ctg ttt tgt tca --- --- --- aag ga- --- --a tct aca ggc ata aaa at- aaa aat ata att tat aat ata c-- tct tac agc gcg cca tca tga ata aca gca gtg aat tga ttg ctg tta tta atg gat tta gaa ata gtg gac gat ttt gtg ata tta gta tag tta tta atg atg aaa tga taa acg ctc aca gac tca tcc tat ctg gag cct ccg aat a-t ttt tcc att ctg ttt tcc aat aat ttt atc gat tct aat gaa tac gaa gtt aat cta agt cat tta gat tat caa agt gtt aac gat ttg atc gat tac att tat ggg ata cct ttg agc cta act aac gat aac gtg aaa tat att ctt tca acc gct ga- --t ttt tta caa att gga tct gct att acg gag tgt gaa aat tac ata ctt aaa aat ctt tgt tct aaa aac tgt atc gat ttc --- --- tac ata tac gct gat aaa tat aat aac aag aaa ata gaa tca gcg tcg ttt aac aca ata tta caa aat att ttg aga ctc atc aac gat gaa aac ttt aaa tac tta aca gag gaa tca atg ata aaa att tta agc gat gat atg tta aat ata aaa aat gag gat ttc gcc cca cta att ctc att aaa tgg tta gag agt act caa caa tca tgc acc gtc gag tta ctt aga tgc ctc aga ata tca -tt gct ttc ccc aca agt tat aaa atc act tta tag tca tca act agt tag ttc aat cta cga atg tat aac att ctt aaa caa tat agc att ctt gga tga atc att tcc tag ata cca tag cat cga gtt gat atc tat cgg tat aag taa ttt gcg tga taa aat ttc cat aaa ctg cta caa tca t-a aaa aaa ata cat ggg aaa tga tat ctt cac gta gat ata ggt gta gtt tcg tag tgg ccg tcc tgg ata ata tta tct ata tga tgg gtg gat atg atc agt ccc cgt ata gaa gtt caa aag tta tag cgt aca ata cat gta caa att ctt aga tat atg ata tac cag agc t-a aaa tat cct cgt tct aat tgt ggg gga ctg gct gat gac gaa tac att tat tgt ata ggc ggc ata cgc gat cag gat tca tcg ttg aca tct agt att gat aaa tgg aag cca tca aaa cca tat tgg cag aag tat gct aaa atg cgc gaa cca aaa tgt gat atg ggg gtt gcg atg tta aac gga tta ata tat gtt ata ggt gga atc gtt aaa ggt g-- aca cgt gta ccg acg cac tag aga gtt tat cag aag atg gat gga tga agc atc aac gtc ttc caa taa aaa tgt cca ata tgt cga cga ttg ttc atg atg gca aga ttt ata tat ctg gag gtt aca aca ata gta gtg tag tta atg taa tat cga atc t-- agt cct tag cta taa tcc gat ata tga tga atg gac caa att atc atc att aaa cat tcc tag aat taa tcc cgc tct a

>KY554976.1:166471-168747_Ectromelia_virus_isolate_Hampstead

ttg ctg act gta --- agt a-c tat ttt -ta cat tac tac att ggc tac ggc aga tat acc tac ttc gtc act gcc aca cgc tcc ggt aaa cgg --g tca tgt gac gag gga gaa tat ctt gat aa- agg cat aat caa tgt tgt aat -ca gtg tcc acc tgg aga att tgc caa ggt cag atg ta- --g tgg cag tga taa cac aaa atg tga acg ctg cct acc tca tac ata tac cac aat ccc caa tta ttc taa atg gat gtc atc aat gta gaa aat gcc caa cag gat cat ttg ata agg taa agt gta ccg gaa cac ata aca gta aat -gt tcg tgt ctt cct ggt tgg tat tgc gct act gat tct tca cag act gaa gat tgt cga gat -tg tat acc aaa aag g-a gat gtc cgt gcg gat att ttg gtg gaa tag atg aac aag gaa atc cta ttt gta aat cgt gat g-- -ta ttg gtg aac att gcg act acc tac ata att ata gag ttg gtc ctt ttc ctc cat gta aa- --- cta tct aaa tgt aat taa tta --- --- --- --- --- --- --- --- tga tct tga tat aat atg att atc atg tta -cc ata c-a tt- --- -at ata gct act ttg gta --g tgt att att cag tat g-a aga cat att aat aa- --- --- -tt act tat c-t ttg aag atc ttg tt- -at aat cat aaa --- --t ata aaa ata aaa --a tac tta tgg --- -gg cat agt aac tca t-- --- --- --- aat tgc tg- aca cga taa att cgt aat aat ctg ttt tgt tca aat ttt tat aag ga- --- --a tct --a ggc ata aaa at- aaa aac ata att tat aat ac- --- act tac aac ac- cca tca tga ata aca gca gtg aat taa ttg ctg cga tta atg gat tta gaa ata gtg gac gat ttt gtg ata tta ata tag tta tta atg atg aaa gga taa atg ctc ata gac tca tcc tat ctg gag cat ccg aat a-t ttt tcc att cta ttt tct agt gat ttt atc gat tct aat gaa tac gaa gtt aat cta agt cat tta gat tat caa agt gtg aac gat ttg atc gat tac att tat ggg ata cct ttg agc cta act aac gat agc gtg aaa tat att ctt tca act gct ga- --t ttt tta caa att gga tct gct att acg gag tgt gaa aat tac ata ctt aaa aat ctc tgt tct aga aac tgt atc gat ttc --- --- tac ata tac gct gat aaa tat aat aac aag aaa ata gaa aca gcg tcg ttt aac aca ata tta cga aat att ttg aga ctc att aac gat gaa aac ttt aaa tac ttg aca gag gaa tca atg ata aaa att tta agc gat gat atg tta aat ata aaa aat gag gat ttc gca cca cta att ctc att aaa tgg tta gag agt aca caa caa cca tgc acc gtc gag tta ctt aga tgc ctc aga ata tca -tt gct ttc ccc aca agt tat aaa atc act tta tag tca tcg act ggt tag ttc aat cta cga atg tat aac att ctt aaa caa tat agc att ctt gga tga atc att tcc tag ata caa tag cat cga gtt gat atc tat cgg tat aag taa ttc acg tga taa gat ttc cat aaa ctg cta caa tcg t-a aaa aaa ata cat ggg aaa tga tat ctt cac gtg gat ata ggt gta gtt tcg cag tgg ctg tca tgg ata ata tta tct ata tga tgg gtg gat acg atc agt ccc cgt ata gaa gtt caa agg tta tag cat aca ata cat gta cta att ctt gga tat atg ata tac cag agc t-a aaa tat ccg cgt tct aat tgc gga gga gtt gtc gat gac gaa tac att tat tgt ata ggt ggc ata cgc gat cag gat tca tcg ttg ata tct gat att gat aga tgg aag cca tca aaa cca tat tgg cag acg tat gct aaa atg cga gag cca aaa tgt gat atg ggg gtt gcg atg tta aac gga tta ata tat gtc ata ggt gga gtc gtt aaa ggt g-- gca cat gta ccg aca cac tag aga gtt tat cag aag atg gat gga tga tgc atc gac gcc ttc caa taa aaa tgt cca ata tgt cga cga ttg ttc atg ctg gaa aga ttt ata tat ctg gag gtt aca cca ata gta gta tag tta atg aaa tat cta atc t-- agt cct tag cta taa tcc gat ata tga tga atg gac caa att atc atc att aaa tat tcc tag aat taa tcc tgc tct a

>OU343159.1:166852-169128_Ectromelia_virus_strain_MouKre_genome_assembly_chromosome:_1

ttg ctg act gta --- agt a-c tat ttt -ta cat tac tac att ggc tac ggc aga tat acc tac ttc gtc act gcc aca cgc tcc ggt aaa cgg --g tca tgt gac gag gga gaa tat ctt gat aa- agg cat aat caa tgt tgt aat -ca gtg tcc acc tgg aga att tgc caa ggt cag atg ta- --g tgg cag tga taa cac aaa atg tga acg ctg cct acc tca tac ata tac cac aat ccc caa tta ttc taa atg gat gtc atc aat gta gaa aat gcc caa cag gat cat ttg ata agg taa agt gta ccg gaa cac ata aca gta aat -gt tcg tgt ctt cct ggt tgg tat tgc gct act gat tct tca cag act gaa gat tgt cga gat -tg tat acc aaa aag g-a gat gtc cgt gcg gat att ttg gtg gaa tag atg aac aag gaa atc cta ttt gta aat cgt gat g-- -ta ttg gtg aac att gcg act acc tac ata att ata gag ttg gtc ctt ttc ctc cat gta aa- --- cta tct aaa tgt aat taa tta --- --- --- --- --- --- --- --- tga tct tga tat aat atg att atc atg tta -cc ata c-a tt- --- -at ata gct act ttg gta --g tgt att att cag tat g-a aga cat att aat aa- --- --- -tt act tat c-t ttg aag atc ttg tt- -at aat cat aaa --- --t ata aaa ata aaa --a tac tta tgg --- -gg cat agt aac tca t-- --- --- --- aat tgc tg- aca cga taa att cgt aat aat ctg ttt tgt tca aat ttt tat aag ga- --- --a tct --a ggc ata aaa at- aaa aac ata att tat aat ac- --- act tac aac ac- cca tca tga ata aca gca gtg aat taa ttg ctg cga tta atg gat tta gaa ata gtg gac gat ttt gtg ata tta ata tag tta tta atg atg aaa gga taa atg ctc ata gac tca tcc tat ctg gag cat ccg aat a-t ttt tcc att cta ttt tct agt gat ttt atc gat tct aat gaa tac gaa gtt aat cta agt cat tta gat tat caa agt gtg aac gat ttg atc gat tac att tat ggg ata cct ttg agc cta act aac gat agc gtg aaa tat att ctt tca act gct ga- --t ttt tta caa att gga tct gct att acg gag tgt gaa aat tac ata ctt aaa aat ctc tgt tct aga aac tgt atc gat ttc --- --- tac ata tac gct gat aaa tat aat aac aag aaa ata gaa aca gcg tcg ttt aac aca ata tta cga aat att ttg aga ctc att aac gat gaa aac ttt aaa tac ttg aca gag gaa tca atg ata aaa att tta agc gat gat atg tta aat ata aaa aat gag gat ttc gca cca cta att ctc att aaa tgg tta gag agt aca caa caa cca tgc acc gtc gag tta ctt aga tgc ctc aga ata tca -tt gct ttc ccc aca agt tat aaa atc act tta tag tca tcg act ggt tag ttc aat cta cga atg tat aac att ctt aaa caa tat agc att ctt gga tga atc att tcc tag ata caa tag cat cga gtt gat atc tat cgg tat aag taa ttc acg tga taa gat ttc cat aaa ctg cta caa tcg t-a aaa aaa ata cat ggg aaa tga tat ctt cac gtg gat ata ggt gta gtt tcg cag tgg ctg tca tgg ata ata tta tct ata tga tgg gtg gat acg atc agt ccc cgt ata gaa gtt caa agg tta tag cat aca ata cat gta cta att ctt gga tat atg ata tac cag agc t-a aaa tat ccg cgt tct aat tgc gga gga gtt gtc gat gac gaa tac att tat tgt ata ggt ggc ata cgc gat cag gat tca tcg ttg ata tct gat att gat aga tgg aag cca tca aaa cca tat tgg cag acg tat gct aaa atg cga gag cca aaa tgt gat atg ggg gtt gcg atg tta aac gga tta ata tat gtc ata ggt gga gtc gtt aaa ggt g-- gca cat gta ccg aca cac tag aga gtt tat cag aag atg gat gga tga tgc atc gac gcc ttc caa taa aaa tgt cca ata tgt cga cga ttg ttc atg ctg gaa aga ttt ata tat ctg gag gtt aca cca ata gta gta tag tta atg aaa tat cta atc t-- agt cct tag cta taa tcc gat ata tga tga atg gac caa att atc atc att aaa tat tcc tag aat taa tcc tgc tct a

>OU343158.1:166535-168811_Ectromelia_virus_strain_Mp1_genome_assembly_chromosome:_1

ttg ctg act gta --- agt a-c tat ttt -ta cat tac tac att ggc tac ggc aga tat acc tac ttc gtc act gcc aca cgc tcc ggt aaa cgg --g tca tgt gac gag gga gaa tat ctt gat aa- agg cat aat caa tgt tgt aat -ca gtg tcc acc tgg aga att tgc caa ggt cag atg ta- --g tgg cag tga taa cac aaa atg tga acg ctg cct acc tca tac ata tac cac aat ccc caa tta ttc taa atg gat gtc atc aat gta gaa aat gcc caa cag gat cat ttg ata agg taa agt gta ccg gaa cac ata aca gta aat -gt tcg tgt ctt cct ggt tgg tat tgc gct act gat tct tca cag act gaa gat tgt cga gat -tg tat acc aaa aag g-a gat gtc cgt gcg gat att ttg gtg gaa tag atg aac aag gaa atc cta ttt gta aat cgt gat g-- -ta ttg gtg aac att gcg act acc tac ata att ata gag ttg gtc ctt ttc ctc cat gta aa- --- cta tct aaa tgt aat taa tta --- --- --- --- --- --- --- --- tga tct tga tat aat atg att atc atg tta -cc ata c-a tt- --- -at ata gct act ttg gta --g tgt att att cag tat g-a aga cat att aat aa- --- --- -tt act tat c-t ttg aag atc ttg tt- -at aat cat aaa --- --t ata aaa ata aaa --a tac tta tgg --- -gg cat agt aac tca t-- --- --- --- aat tgc tg- aca cga taa att cgt aat aat ctg ttt tgt tca aat ttt tat aag ga- --- --a tct --a ggc ata aaa at- aaa aac ata att tat aat ac- --- act tac aac ac- cca tca tga ata aca gca gtg aat taa ttg ctg cga tta atg gat tta gaa ata gtg gac gat ttt gtg ata tta ata tag tta tta atg atg aaa gga taa atg ctc ata gac tca tcc tat ctg gag cat ccg aat a-t ttt tcc att cta ttt tct agt gat ttt atc gat tct aat gaa tac gaa gtt aat cta agt cat tta gat tat caa agt gtg aac gat ttg atc gat tac att tat ggg ata cct ttg agc cta act aac gat agc gtg aaa tat att ctt tca act gct ga- --t ttt tta caa att gga tct gct att acg gag tgt gaa aat tac ata ctt aaa aat ctc tgt tct aga aac tgt atc gat ttc --- --- tac ata tac gct gat aaa tat aat aac aag aaa ata gaa aca gcg tcg ttt aac aca ata tta cga aat att ttg aga ctc att aac gat gaa aac ttt aaa tac ttg aca gag gaa tca atg ata aaa att tta agc gat gat atg tta aat ata aaa aat gag gat ttc gca cca cta att ctc att aaa tgg tta gag agt aca caa caa cca tgc acc gtc gag tta ctt aga tgc ctc aga ata tca -tt gct ttc ccc aca agt tat aaa atc act tta tag tca tcg act ggt tag ttc aat cta cga atg tat aac att ctt aaa caa tat agc att ctt gga tga atc att tcc tag ata caa tag cat cga gtt gat atc tat cgg tat aag taa ttc acg tga taa gat ttc cat aaa ctg cta caa tcg t-a aaa aaa ata cat ggg aaa tga tat ctt cac gtg gat ata ggt gta gtt tcg cag tgg ctg tca tgg ata ata tta tct ata tga tgg gtg gat acg atc agt ccc cgt ata gaa gtt caa agg tta tag cat aca ata cat gta cta att ctt gga tat atg ata tac cag agc t-a aaa tat ccg cgt tct aat tgc gga gga gtt gtc gat gac gaa tac att tat tgt ata ggt ggc ata cgc gat cag gat tca tcg ttg ata tct gat att gat aga tgg aag cca tca aaa cca tat tgg cag acg tat gct aaa atg cga gag cca aaa tgt gat atg ggg gtt gcg atg tta aac gga tta ata tat gtc ata ggt gga gtc gtt aaa ggt g-- gca cat gta ccg aca cac tag aga gtt tat cag aag atg gat gga tga tgc atc gac gcc ttc caa taa aaa tgt cca ata tgt cga cga ttg ttc atg ctg gaa aga ttt ata tat ctg gag gtt aca cca ata gta gta tag tta atg aaa tat cta atc t-- agt cct tag cta taa tcc gat ata tga tga atg gac caa att atc atc att aaa tat tcc tag aat taa tcc tgc tct a

>OU343157.1:167155-169431_Ectromelia_virus_strain_Mp4_genome_assembly_chromosome:_1

ttg ctg act gta --- agt a-c tat ttt -ta cat tac tac att ggc tac ggc aga tat acc tac ttc gtc act gcc aca cgc tcc ggt aaa cgg --g tca tgt gac gag gga gaa tat ctt gat aa- agg cat aat caa tgt tgt aat -ca gtg tcc acc tgg aga att tgc caa ggt cag atg ta- --g tgg cag tga taa cac aaa atg tga acg ctg cct acc tca tac ata tac cac aat ccc caa tta ttc taa atg gat gtc atc aat gta gaa aat gcc caa cag gat cat ttg ata agg taa agt gta ccg gaa cac ata aca gta aat -gt tcg tgt ctt cct ggt tgg tat tgc gct act gat tct tca cag act gaa gat tgt cga gat -tg tat acc aaa aag g-a gat gtc cgt gcg gat att ttg gtg gaa tag atg aac aag gaa atc cta ttt gta aat cgt gat g-- -ta ttg gtg aac att gcg act acc tac ata att ata gag ttg gtc ctt ttc ctc cat gta aa- --- cta tct aaa tgt aat taa tta --- --- --- --- --- --- --- --- tga tct tga tat aat atg att atc atg tta -cc ata c-a tt- --- -at ata gct act ttg gta --g tgt att att cag tat g-a aga cat att aat aa- --- --- -tt act tat c-t ttg aag atc ttg tt- -at aat cat aaa --- --t ata aaa ata aaa --a tac tta tgg --- -gg cat agt aac tca t-- --- --- --- aat tgc tg- aca cga taa att cgt aat aat ctg ttt tgt tca aat ttt tat aag ga- --- --a tct --a ggc ata aaa at- aaa aac ata att tat aat ac- --- act tac aac ac- cca tca tga ata aca gca gtg aat taa ttg ctg cga tta atg gat tta gaa ata gtg gac gat ttt gtg ata tta ata tag tta tta atg atg aaa gga taa atg ctc ata gac tca tcc tat ctg gag cat ccg aat a-t ttt tcc att cta ttt tct agt gat ttt atc gat tct aat gaa tac gaa gtt aat cta agt cat tta gat tat caa agt gtg aac gat ttg atc gat tac att tat ggg ata cct ttg agc cta act aac gat agc gtg aaa tat att ctt tca act gct ga- --t ttt tta caa att gga tct gct att acg gag tgt gaa aat tac ata ctt aaa aat ctc tgt tct aga aac tgt atc gat ttc --- --- tac ata tac gct gat aaa tat aat aac aag aaa ata gaa aca gcg tcg ttt aac aca ata tta cga aat att ttg aga ctc att aac gat gaa aac ttt aaa tac ttg aca gag gaa tca atg ata aaa att tta agc gat gat atg tta aat ata aaa aat gag gat ttc gca cca cta att ctc att aaa tgg tta gag agt aca caa caa cca tgc acc gtc gag tta ctt aga tgc ctc aga ata tca -tt gct ttc ccc aca agt tat aaa atc act tta tag tca tcg act ggt tag ttc aat cta cga atg tat aac att ctt aaa caa tat agc att ctt gga tga atc att tcc tag ata caa tag cat cga gtt gat atc tat cgg tat aag taa ttc acg tga taa gat ttc cat aaa ctg cta caa tcg t-a aaa aaa ata cat ggg aaa tga tat ctt cac gtg gat ata ggt gta gtt tcg cag tgg ctg tca tgg ata ata tta tct ata tga tgg gtg gat acg atc agt ccc cgt ata gaa gtt caa agg tta tag cat aca ata cat gta cta att ctt gga tat atg ata tac cag agc t-a aaa tat ccg cgt tct aat tgc gga gga gtt gtc gat gac gaa tac att tat tgt ata ggt ggc ata cgc gat cag gat tca tcg ttg ata tct gat att gat aga tgg aag cca tca aaa cca tat tgg cag acg tat gct aaa atg cga gag cca aaa tgt gat atg ggg gtt gcg atg tta aac gga tta ata tat gtc ata ggt gga gtc gtt aaa ggt g-- gca cat gta ccg aca cac tag aga gtt tat cag aag atg gat gga tga tgc atc gac gcc ttc caa taa aaa tgt cca ata tgt cga cga ttg ttc atg ctg gaa aga ttt ata tat ctg gag gtt aca cca ata gta gta tag tta atg aaa tat cta atc t-- agt cct tag cta taa tcc gat ata tga tga atg gac caa att atc atc att aaa tat tcc tag aat taa tcc tgc tct a

>OU343155.1:166734-169010_Ectromelia_virus_strain_Mp5_genome_assembly_chromosome:_1

ttg ctg act gta --- agt a-c tat ttt -ta cat tac tac att ggc tac ggc aga tat acc tac ttc gtc act gcc aca cgc tcc ggt aaa cgg --g tca tgt gac gag gga gaa tat ctt gat aa- agg cat aat caa tgt tgt aat -ca gtg tcc acc tgg aga att tgc caa ggt cag atg ta- --g tgg cag tga taa cac aaa atg tga acg ctg cct acc tca tac ata tac cac aat ccc caa tta ttc taa atg gat gtc atc aat gta gaa aat gcc caa cag gat cat ttg ata agg taa agt gta ccg gaa cac ata aca gta aat -gt tcg tgt ctt cct ggt tgg tat tgc gct act gat tct tca cag act gaa gat tgt cga gat -tg tat acc aaa aag g-a gat gtc cgt gcg gat att ttg gtg gaa tag atg aac aag gaa atc cta ttt gta aat cgt gat g-- -ta ttg gtg aac att gcg act acc tac ata att ata gag ttg gtc ctt ttc ctc cat gta aa- --- cta tct aaa tgt aat taa tta --- --- --- --- --- --- --- --- tga tct tga tat aat atg att atc atg tta -cc ata c-a tt- --- -at ata gct act ttg gta --g tgt att att cag tat g-a aga cat att aat aa- --- --- -tt act tat c-t ttg aag atc ttg tt- -at aat cat aaa --- --t ata aaa ata aaa --a tac tta tgg --- -gg cat agt aac tca t-- --- --- --- aat tgc tg- aca cga taa att cgt aat aat ctg ttt tgt tca aat ttt tat aag ga- --- --a tct --a ggc ata aaa at- aaa aac ata att tat aat ac- --- act tac aac ac- cca tca tga ata aca gca gtg aat taa ttg ctg cga tta atg gat tta gaa ata gtg gac gat ttt gtg ata tta ata tag tta tta atg atg aaa gga taa atg ctc ata gac tca tcc tat ctg gag cat ccg aat a-t ttt tcc att cta ttt tct agt gat ttt atc gat tct aat gaa tac gaa gtt aat cta agt cat tta gat tat caa agt gtg aac gat ttg atc gat tac att tat ggg ata cct ttg agc cta act aac gat agc gtg aaa tat att ctt tca act gct ga- --t ttt tta caa att gga tct gct att acg gag tgt gaa aat tac ata ctt aaa aat ctc tgt tct aga aac tgt atc gat ttc --- --- tac ata tac gct gat aaa tat aat aac aag aaa ata gaa aca gcg tcg ttt aac aca ata tta cga aat att ttg aga ctc att aac gat gaa aac ttt aaa tac ttg aca gag gaa tca atg ata aaa att tta agc gat gat atg tta aat ata aaa aat gag gat ttc gca cca cta att ctc att aaa tgg tta gag agt aca caa caa cca tgc acc gtc gag tta ctt aga tgc ctc aga ata tca -tt gct ttc ccc aca agt tat aaa atc act tta tag tca tcg act ggt tag ttc aat cta cga atg tat aac att ctt aaa caa tat agc att ctt gga tga atc att tcc tag ata caa tag cat cga gtt gat atc tat cgg tat aag taa ttc acg tga taa gat ttc cat aaa ctg cta caa tcg t-a aaa aaa ata cat ggg aaa tga tat ctt cac gtg gat ata ggt gta gtt tcg cag tgg ctg tca tgg ata ata tta tct ata tga tgg gtg gat acg atc agt ccc cgt ata gaa gtt caa agg tta tag cat aca ata cat gta cta att ctt gga tat atg ata tac cag agc t-a aaa tat ccg cgt tct aat tgc gga gga gtt gtc gat gac gaa tac att tat tgt ata ggt ggc ata cgc gat cag gat tca tcg ttg ata tct gat att gat aga tgg aag cca tca aaa cca tat tgg cag acg tat gct aaa atg cga gag cca aaa tgt gat atg ggg gtt gcg atg tta aac gga tta ata tat gtc ata ggt gga gtc gtt aaa ggt g-- gca cat gta ccg aca cac tag aga gtt tat cag aag atg gat gga tga tgc atc gac gcc ttc caa taa aaa tgt cca ata tgt cga cga ttg ttc atg ctg gaa aga ttt ata tat ctg gag gtt aca cca ata gta gta tag tta atg aaa tat cta atc t-- agt cct tag cta taa tcc gat ata tga tga atg gac caa att atc atc att aaa tat tcc tag aat taa tcc tgc tct a

>OU343113.1:167053-169329_Ectromelia_virus_strain_Ishibashi_genome_assembly_chromosome:_1

ttg ctg act gta --- agt a-c tat ttt -ta cat tac tac att ggc tac ggc aga tat acc tac ttc gtc act gcc aca cgc tcc ggt aaa cgg --g tca tgt gac gag gga gaa tat ctt gat aa- agg cat aat caa tgt tgt aat -ca gtg tcc acc tgg aga att tgc caa ggt cag atg ta- --g tgg cag tga taa cac aaa atg tga acg ctg cct acc tca tac ata tac cac aat ccc caa tta ttc taa atg gat gtc atc aat gta gaa aat gcc caa cag gat cat ttg ata agg taa agt gta ccg gaa cac ata aca gta aat -gt tcg tgt ctt cct ggt tgg tat tgc gct act gat tct tca cag act gaa gat tgt cga gat -tg tat acc aaa aag g-a gat gtc cgt gcg gat att ttg gtg gaa tag atg aac aag gaa atc cta ttt gta aat cgt gat g-- -ta ttg gtg aac att gcg act acc tac ata att ata gag ttg gtc ctt ttc ctc cat gta aa- --- cta tct aaa tgt aat taa tta --- --- --- --- --- --- --- --- tga tct tga tat aat atg att atc atg tta -cc ata c-a tt- --- -at ata gct act ttg gta --g tgt att att cag tat g-a aga cat att aat aa- --- --- -tt act tat c-t ttg aag atc ttg tt- -at aat cat aaa --- --t ata aaa ata aaa --a tac tta tgg --- -gg cat agt aac tca t-- --- --- --- aat tgc tg- aca cga taa att cgt aat aat ctg ttt tgt tca aat ttt tat aag ga- --- --a tct --a ggc ata aaa at- aaa aac ata att tat aat ac- --- act tac aac ac- cca tca tga ata aca gca gtg aat taa ttg ctg cga tta atg gat tta gaa ata gtg gac gat ttt gtg ata tta ata tag tta tta atg atg aaa gga taa atg ctc ata gac tca tcc tat ctg gag cat ccg aat a-t ttt tcc att cta ttt tct agt gat ttt atc gat tct aat gaa tac gaa gtt aat cta agt cat tta gat tat caa agt gtg aac gat ttg atc gat tac att tat ggg ata cct ttg agc cta act aac gat agc gtg aaa tat att ctt tca act gct ga- --t ttt tta caa att gga tct gct att acg gag tgt gaa aat tac ata ctt aaa aat ctc tgt tct aga aac tgt atc gat ttc --- --- tac ata tac gct gat aaa tat aat aac aag aaa ata gaa aca gcg tcg ttt aac aca ata tta cga aat att ttg aga ctc att aac gat gaa aac ttt aaa tac ttg aca gag gaa tca atg ata aaa att tta agc gat gat atg tta aat ata aaa aat gag gat ttc gca cca cta att ctc att aaa tgg tta gag agt aca caa caa cca tgc acc gtc gag tta ctt aga tgc ctc aga ata tca -tt gct ttc ccc aca agt tat aaa atc act tta tag tca tcg act ggt tag ttc aat cta cga atg tat aac att ctt aaa caa tat agc att ctt gga tga atc att tcc tag ata caa tag cat cga gtt gat atc tat cgg tat aag taa ttc acg tga taa gat ttc cat aaa ctg cta caa tcg t-a aaa aaa ata cat ggg aaa tga tat ctt cac gtg gat ata ggt gta gtt tcg cag tgg ctg tca tgg ata ata tta tct ata tga tgg gtg gat acg atc agt ccc cgt ata gaa gtt caa agg tta tag cat aca ata cat gta cta att ctt gga tat atg ata tac cag agc t-a aaa tat ccg cgt tct aat tgc gga gga gtt gtc gat gac gaa tac att tat tgt ata ggt ggc ata cgc gat cag gat tca tcg ttg ata tct gat att gat aga tgg aag cca tca aaa cca tat tgg cag acg tat gct aaa atg cga gag cca aaa tgt gat atg ggg gtt gcg atg tta aac gga tta ata tat gtc ata ggt gga gtc gtt aaa ggt g-- gca cat gta ccg aca cac tag aga gtt tat cag aag atg gat gga tga tgc atc gac gcc ttc caa taa aaa tgt cca ata tgt cga cga ttg ttc atg ctg gaa aga ttt ata tat ctg gag gtt aca cca ata gta gta tag tta atg aaa tat cta atc t-- agt cct tag cta taa tcc gat ata tga tga atg gac caa att atc atc att aaa tat tcc tag aat taa tcc tgc tct a

>MN912466.1:161255-163531_Ectromelia_virus_WH

ttg ctg act gta --- agt a-c tat ttt -ta cat tac tac att ggc tac ggc aga tat acc tac ttc gtc act gcc aca cgc tcc ggt aaa cgg --g tca tgt gac gag gga gaa tat ctt gat aa- agg cat aat caa tgt tgt aat -ca gtg tcc acc tgg aga att tgc caa ggt cag atg ta- --g tgg cag tga taa cac aaa atg tga acg ctg cct acc tca tac ata tac cac aat ccc caa tta ttc taa atg gat gtc atc aat gta gaa aat gcc caa cag gat cat ttg ata agg taa agt gta ccg gaa cac ata aca gta aat -gt tcg tgt ctt cct ggt tgg tat tgc gct act gat tct tca cag act gaa gat tgt cga gat -tg tat acc aaa aag g-a gat gtc cgt gcg gat att ttg gtg gaa tag atg aac aag gaa atc cta ttt gta aat cgt gat g-- -ta ttg gtg aac att gcg act acc tac ata att ata gag ttg gtc ctt ttc ctc cat gta aa- --- cta tct aaa tgt aat taa tta --- --- --- --- --- --- --- --- tga tct tga tat aat atg att atc atg tta -cc ata c-a tt- --- -at ata gct act ttg gta --g tgt att att cag tat g-a aga cat att aat aa- --- --- -tt act tat c-t ttg aag atc ttg tt- -at aat cat aaa --- --t ata aaa ata aaa --a tac tta tgg --- -gg cat agt aac tca t-- --- --- --- aat tgc tg- aca cga taa att cgt aat aat ctg ttt tgt tca aat ttt tat aag ga- --- --a tct --a ggc ata aaa at- aaa aac ata att tat aat ac- --- act tac aac ac- cca tca tga ata aca gca gtg aat taa ttg ctg cga tta atg gat tta gaa ata gtg gac gat ttt gtg ata tta ata tag tta tta atg atg aaa gga taa atg ctc ata gac tca tcc tat ctg gag cat ccg aat a-t ttt tcc att cta ttt tct agt gat ttt atc gat tct aat gaa tac gaa gtt aat cta agt cat tta gat tat caa agt gtg aac gat ttg atc gat tac att tat ggg ata cct ttg agc cta act aac gat agc gtg aaa tat att ctt tca act gct ga- --t ttt tta caa att gga tct gct att acg gag tgt gaa aat tac ata ctt aaa aat ctc tgt tct aga aac tgt atc gat ttc --- --- tac ata tac gct gat aaa tat aat aac aag aaa ata gaa aca gcg tcg ttt aac aca ata tta cga aat att ttg aga ctc att aac gat gaa aac ttt aaa tac ttg aca gag gaa tca atg ata aaa att tta agc gat gat atg tta aat ata aaa aat gag gat ttc gca cca cta att ctc att aaa tgg tta gag agt aca caa caa cca tgc acc gtc gag tta ctt aga tgc ctc aga ata tca -tt gct ttc ccc aca agt tat aaa atc act tta tag tca tcg act ggt tag ttc aat cta cga atg tat aac att ctt aaa caa tat agc att ctt gga tga atc att tcc tag ata caa tag cat cga gtt gat atc tat cgg tat aag taa ttc acg tga taa gat ttc cat aaa ctg cta caa tcg t-a aaa aaa ata cat ggg aaa tga tat ctt cac gtg gat ata ggt gta gtt tcg cag tgg ctg tca tgg ata ata tta tct ata tga tgg gtg gat acg atc agt ccc cgt ata gaa gtt caa agg tta tag cat aca ata cat gta cta att ctt gga tat atg ata tac cag agc t-a aaa tat ccg cgt tct aat tgc gga gga gtt gtc gat gac gaa tac att tat tgt ata ggt ggc ata cgc gat cag gat tca tcg ttg ata tct gat att gat aga tgg aag cca tca aaa cca tat tgg cag acg tat gct aaa atg cga gag cca aaa tgt gat atg ggg gtt gcg atg tta aac gga tta ata tat gtc ata ggt gga gtc gtt aaa ggt g-- gca cat gta ccg aca cac tag aga gtt tat cag aag atg gat gga tga tgc atc gac gcc ttc caa taa aaa tgt cca ata tgt cga cga ttg ttc atg ctg gaa aga ttt ata tat ctg gag gtt aca cca ata gta gta tag tta atg aaa tat cta atc t-- agt cct tag cta taa tcc gat ata tga tga atg gac caa att atc atc att aaa tat tcc tag aat taa tcc tgc tct a

>JQ410350.1:165087-167363_Ectromelia_virus_ERPV_culture-collection_ATCC:VR-1431_complete_genome

ttg ctg act gta --- agt a-c tat ttt -ta cat tac tac att ggc tac ggc aga tat acc tac ttc gtc act gcc aca cgc tcc ggt aaa cgg --g tca tgt gac gag gga gaa tat ctt gat aa- agg cat aat caa tgt tgt aat -ca gtg tcc acc tgg aga att tgc caa ggt cag atg ta- --g tgg cag tga taa cac aaa atg tga acg ctg cct acc tca tac ata tac cac aat ccc caa tta ttc taa atg gat gtc atc aat gta gaa aat gcc caa cag gat cat ttg ata agg taa agt gta ccg gaa cac ata aca gta aat -gt tcg tgt ctt cct ggt tgg tat tgc gct act gat tct tca cag act gaa gat tgt cga gat -tg tat acc aaa aag g-a gat gtc cgt gcg gat att ttg gtg gaa tag atg aac aag gaa atc cta ttt gta aat cgt gat g-- -ta ttg gtg aac att gcg act acc tac ata att ata gag ttg gtc ctt ttc ctc cat gta aa- --- cta tct aaa tgt aat taa tta --- --- --- --- --- --- --- --- tga tct tga tat aat atg att atc atg tta -cc ata c-a tt- --- -at ata gct act ttg gta --g tgt att att cag tat g-a aga cat att aat aa- --- --- -tt act tat c-t ttg aag atc ttg tt- -at aat cat aaa --- --t ata aaa ata aaa --a tac tta tgg --- -gg cat agt aac tca t-- --- --- --- aat tgc tg- aca cga taa att cgt aat aat ctg ttt tgt tca aat ttt tat aag ga- --- --a tct --a ggc ata aaa at- aaa aac ata att tat aat ac- --- act tac aac ac- cca tca tga ata aca gca gtg aat taa ttg ctg cga tta atg gat tta gaa ata gtg gac gat ttt gtg ata tta ata tag tta tta atg atg aaa gga taa atg ctc ata gac tca tcc tat ctg gag cat ccg aat a-t ttt tcc att cta ttt tct agt gat ttt atc gat tct aat gaa tac gaa gtt aat cta agt cat tta gat tat caa agt gtg aac gat ttg atc gat tac att tat ggg ata cct ttg agc cta act aac gat agc gtg aaa tat att ctt tca act gct ga- --t ttt tta caa att gga tct gct att acg gag tgt gaa aat tac ata ctt aaa aat ctc tgt tct aga aac tgt atc gat ttc --- --- tac ata tac gct gat aaa tat aat aac aag aaa ata gaa aca gcg tcg ttt aac aca ata tta cga aat att ttg aga ctc att aac gat gaa aac ttt aaa tac ttg aca gag gaa tca atg ata aaa att tta agc gat gat atg tta aat ata aaa aat gag gat ttc gca cca cta att ctc att aaa tgg tta gag agt aca caa caa cca tgc acc gtc gag tta ctt aga tgc ctc aga ata tca -tt gct ttc ccc aca agt tat aaa atc act tta tag tca tcg act ggt tag ttc aat cta cga atg tat aac att ctt aaa caa tat agc att ctt gga tga atc att tcc tag ata caa tag cat cga gtt gat atc tat cgg tat aag taa ttc acg tga taa gat ttc cat aaa ctg cta caa tcg t-a aaa aaa ata cat ggg aaa tga tat ctt cac gtg gat ata ggt gta gtt tcg cag tgg ctg tca tgg ata ata tta tct ata tga tgg gtg gat acg atc agt ccc cgt ata gaa gtt caa agg tta tag cat aca ata cat gta cta att ctt gga tat atg ata tac cag agc t-a aaa tat ccg cgt tct aat tgc gga gga gtt gtc gat gac gaa tac att tat tgt ata ggt ggc ata cgc gat cag gat tca tcg ttg ata tct gat att gat aga tgg aag cca tca aaa cca tat tgg cag acg tat gct aaa atg cga gag cca aaa tgt gat atg ggg gtt gcg atg tta aac gga tta ata tat gtc ata ggt gga gtc gtt aaa ggt g-- gca cat gta ccg aca cac tag aga gtt tat cag aag atg gat gga tga tgc atc gac gcc ttc caa taa aaa tgt cca ata tgt cga cga ttg ttc atg ctg gaa aga ttt ata tat ctg gag gtt aca cca ata gta gta tag tta atg aaa tat cta atc t-- agt cct tag cta taa tcc gat ata tga tga atg gac caa att atc atc att aaa tat tcc tag aat taa tcc tgc tct a

>AF012825.2:168113-170389_Ectromelia_virus_strain_Moscow_complete_genome

ttg ctg act gta --- agt a-c tat ttt -ta cat tac tac att ggc tac ggc aga tat acc tac ttc gtc act gcc aca cgc tcc ggt aaa cgg --g tca tgt gac gag gga gaa tat ctt gat aa- agg cat aat caa tgt tgt aat -ca gtg tcc acc tgg aga att tgc caa ggt cag atg ta- --g tgg cag tga taa cac aaa atg tga acg ctg cct acc tca tac ata tac cac aat ccc caa tta ttc taa atg gat gtc atc aat gta gaa aat gcc caa cag gat cat ttg ata agg taa agt gta ccg gaa cac ata aca gta aat -gt tcg tgt ctt cct ggt tgg tat tgc gct act gat tct tca cag act gaa gat tgt cga gat -tg tat acc aaa aag g-a gat gtc cgt gcg gat att ttg gtg gaa tag atg aac aag gaa atc cta ttt gta aat cgt gat g-- -ta ttg gtg aac att gcg act acc tac ata att ata gag ttg gtc ctt ttc ctc cat gta aa- --- cta tct aaa tgt aat taa tta --- --- --- --- --- --- --- --- tga tct tga tat aat atg att atc atg tta -cc ata c-a tt- --- -at ata gct act ttg gta --g tgt att att cag tat g-a aga cat att aat aa- --- --- -tt act tat c-t ttg aag atc ttg tt- -at aat cat aaa --- --t ata aaa ata aaa --a tac tta tgg --- -gg cat agt aac tca t-- --- --- --- aat tgc tg- aca cga taa att cgt aat aat ctg ttt tgt tca aat ttt tat aag ga- --- --a tct --a ggc ata aaa at- aaa aac ata att tat aat ac- --- act tac aac ac- cca tca tga ata aca gca gtg aat taa ttg ctg cga tta atg gat tta gaa ata gtg gac gat ttt gtg ata tta ata tag tta tta atg atg aaa gga taa atg ctc ata gac tca tcc tat ctg gag cat ccg aat a-t ttt tcc att cta ttt tct agt gat ttt atc gat tct aat gaa tac gaa gtt aat cta agt cat tta gat tat caa agt gtg aac gat ttg atc gat tac att tat ggg ata cct ttg agc cta act aac gat agc gtg aaa tat att ctt tca act gct ga- --t ttt tta caa att gga tct gct att acg gag tgt gaa aat tac ata ctt aaa aat ctc tgt tct aga aac tgt atc gat ttc --- --- tac ata tac gct gat aaa tat aat aac aag aaa ata gaa aca gcg tcg ttt aac aca ata tta cga aat att ttg aga ctc att aac gat gaa aac ttt aaa tac ttg aca gag gaa tca atg ata aaa att tta agc gat gat atg tta aat ata aaa aat gag gat ttc gca cca cta att ctc att aaa tgg tta gag agt aca caa caa cca tgc acc gtc gag tta ctt aga tgc ctc aga ata tca -tt gct ttc ccc aca agt tat aaa atc act tta tag tca tcg act ggt tag ttc aat cta cga atg tat aac att ctt aaa caa tat agc att ctt gga tga atc att tcc tag ata caa tag cat cga gtt gat atc tat cgg tat aag taa ttc acg tga taa gat ttc cat aaa ctg cta caa tcg t-a aaa aaa ata cat ggg aaa tga tat ctt cac gtg gat ata ggt gta gtt tcg cag tgg ctg tca tgg ata ata tta tct ata tga tgg gtg gat acg atc agt ccc cgt ata gaa gtt caa agg tta tag cat aca ata cat gta cta att ctt gga tat atg ata tac cag agc t-a aaa tat ccg cgt tct aat tgc gga gga gtt gtc gat gac gaa tac att tat tgt ata ggt ggc ata cgc gat cag gat tca tcg ttg ata tct gat att gat aga tgg aag cca tca aaa cca tat tgg cag acg tat gct aaa atg cga gag cca aaa tgt gat atg ggg gtt gcg atg tta aac gga tta ata tat gtc ata ggt gga gtc gtt aaa ggt g-- gca cat gta ccg aca cac tag aga gtt tat cag aag atg gat gga tga tgc atc gac gcc ttc caa taa aaa tgt cca ata tgt cga cga ttg ttc atg ctg gaa aga ttt ata tat ctg gag gtt aca cca ata gta gta tag tta atg aaa tat cta atc t-- agt cct tag cta taa tcc gat ata tga tga atg gac caa att atc atc att aaa tat tcc tag aat taa tcc tgc tct a

>KJ563295.1:165690-167966_Ectromelia_virus_Naval_complete_genome

ttg ctg act gta --- agt a-c tat ttt -ta cat tac tac att ggc tac ggc aga tat acc tac ttc gtc act gcc aca cgc tcc ggt aaa cgg --g tca tgt gac gag gga gaa tat ctt gat aa- agg cat aat caa tgt tgt aat -ca gtg tcc acc tgg aga att tgc caa ggt cag atg ta- --g tgg cag tga taa cac aaa atg tga acg ctg cct acc tca tac ata tac cac aat ccc caa tta ttc taa atg gat gtc atc aat gta gaa aat gcc caa cag gat cat ttg ata agg taa agt gta ccg gaa cac ata aca gta aat -gt tcg tgt ctt cct ggt tgg tat tgc gct act gat tct tca cag act gaa gat tgt cga gat -tg tat acc aaa aag g-a gat gtc cgt gcg gat att ttg gtg gaa tag atg aac aag gaa atc cta ttt gta aat cgt gat g-- -ta ttg gtg aac att gcg act acc tac ata att ata gag ttg gtc ctt ttc ctc cat gta aa- --- cta tct aaa tgt aat taa tta --- --- --- --- --- --- --- --- tga tct tga tat aat atg att atc atg tta -cc ata c-a tt- --- -at ata gct act ttg gta --g tgt att att cag tat g-a aga cat att aat aa- --- --- -tt act tat c-t ttg aag atc ttg tt- -at aat cat aaa --- --t ata aaa ata aaa --a tac tta tgg --- -gg cat agt aac tca t-- --- --- --- aat tgc tg- aca cga taa att cgt aat aat ctg ttt tgt tca aat ttt tat aag ga- --- --a tct --a ggc ata aaa at- aaa aac ata att tat aat ac- --- act tac aac ac- cca tca tga ata aca gca gtg aat taa ttg ctg cga tta atg gat tta gaa ata gtg gac gat ttt gtg ata tta ata tag tta tta atg atg aaa gga taa atg ctc ata gac tca tcc tat ctg gag cat ccg aat a-t ttt tcc att cta ttt tct agt gat ttt atc gat tct aat gaa tac gaa gtt aat cta agt cat tta gat tat caa agt gtg aac gat ttg atc gat tac att tat ggg ata cct ttg agc cta act aac gat agc gtg aaa tat att ctt tca act gct ga- --t ttt tta caa att gga tct gct att acg gag tgt gaa aat tac ata ctt aaa aat ctc tgt tct aga aac tgt atc gat ttc --- --- tac ata tac gct gat aaa tat aat aac aag aaa ata gaa aca gcg tcg ttt aac aca ata tta cga aat att ttg aga ctc att aac gat gaa aac ttt aaa tac ttg aca gag gaa tca atg ata aaa att tta agc gat gat atg tta aat ata aaa aat gag gat ttc gca cca cta att ctc att aaa tgg tta gag agt aca caa caa cca tgc acc gtc gag tta ctt aga tgc ctc aga ata tca -tt gct ttc ccc aca agt tat aaa atc act tta tag tca tcg act ggt tag ttc aat cta cga atg tat aac att ctt aaa caa tat agc att ctt gga tga atc att tcc tag ata caa tag cat cga gtt gat atc tat cgg tat aag taa ttc acg tga taa gat ttc cat aaa ctg cta caa tcg t-a aaa aaa ata cat ggg aaa tga tat ctt cac gtg gat ata ggt gta gtt tcg cag tgg ctg tca tgg ata ata tta tct ata tga tgg gtg gat acg atc agt ccc cgt ata gaa gtt caa agg tta tag cat aca ata cat gta cta att ctt gga tat atg ata tac cag agc t-a aaa tat ccg cgt tct aat tgc gga gga gtt gtc gat gac gaa tac att tat tgt ata ggt ggc ata cgc gat cag aat tca tcg ttg ata tct gat att gat aga tgg aag cca tca aaa cca tat tgg cag acg tat gct aaa atg cga gag cca aaa tgt gat atg ggg gtt gcg atg tta aac gga tta ata tat gtc ata ggt gga gtc gtt aaa ggt g-- gca cat gta ccg aca cac tag aga gtt tat cag aag atg gat gga tga tgc atc gac gcc ttc caa taa aaa tgt cca ata tgt cga cga ttg ttc atg ctg gaa aga ttt ata tat ctg gag gtt aca cca ata gta gta tag tta atg aaa tat cta atc t-- agt cct tag cta taa tcc gat ata tga tga atg gac caa att atc atc att aaa tat tcc tag aat taa tcc tgc tct a

>MK035758.1:176459-178704_Cowpox_virus_strain_CPXV/Moritz_2015/3a

ttg ctg act gca --- tgt a-c tat ttt tta cat tac tac att ggc tac ggc aga tat acc tac ttc gtc act gcc aca cgc tcc ggt aaa cgg --g gca tgt gac gag gga gaa tat ctt gat aag agg cat aat caa tgt tgt aat -cg gtg tcc acc tgg aga att tgc caa ggt tag atg ta- --g tgg tag cta taa cac aaa atg tga acg ctg tcc aca tca tac ata tac agc aat tcc caa tta ttc t-a atg gat gcc atc aat gta gaa aat gcc caa agg gat cat ttg ata agg taa agt gta ccg gaa cac aga aca gta aat -gt gcg tgt ctt cct ggt tgg tat tgc gct acc gat tct tca cag act gaa gac tgt cga gat -tg tgt acc aaa aag g-a gct gtc cat gcg gat act ttg gtg gaa tag ata aat tag gaa atc cta ttt gta aat cgt gtt g-- -tg ttg gta aat att gtg acg aca tac gta atc ata gag ttg gtc ctt ttc ctc cat gca aa- --- cta tct aaa tgt aat tag tca --- --- --- --- --- --- --- --- --- --- --- --- --- --- -tg atc atg gta -cc gta caa tt- --- -at att gct --- --g gta --g tat att att cag tat g-a aga cat att aat ac- --- --- -tt atc --- --t ttg aca atc ttg ct- -at aat cat --- --- --- --- aat ata aaa --a tac tta tgg --- -a- cat agt cac tca t-- --- --- --- aat tgc tg- acg cga taa att cgt aat aat ctg ttt tgt tca aat ttt tat aag ga- --- --a tct aca ggc ata aaa at- aaa aat ata att tat aat ac- --- act tac agc gcg cca tca tga ata aca gca gtg aat taa ttg ctg tta tta atg gat tta gaa ata gtg gac gat ttt gtg ata tta ata tag tta tta atg atg aaa gga taa atg ctc ata gac tca tcc tat ctg gag cat ccg aat a-t ttt tcc att cta ttt tct agt gat ttt atc gat tct aat gaa tac gaa gtt aat cta agt cat tta gat tat caa agt gtg aac gat ttg atc gat tac att tat ggg ata cct ttg agc cta act aac gat aac gtg aaa tat att ctt tca acc gct ga- --t ttt tta caa att gga tct gcc att act gag tgc gaa aaa tac ata ctt aaa aat ctg tgt tct aga aat tgt atc gat ttc --- --- tac ata tac gct gat aaa tat aat aac aag aaa ata gaa aca gcg tcg ttt aac aca ata tta cga aat att ttg aga ctc att aac gat gaa aac ttt aaa tac ttg aca gag gaa tca atg ata aaa att tta agc gat gat atg tta aat ata aaa aat gag gat ttt gca cca cta att ctc att aaa tgg tta gag agt aca caa caa cca tgc aca gtc gag tta ctt aga tgc ctc aga ata tca -tt gct ttc ccc aca agt tat aaa atc act tta tag tca tcg att ggt tgg ttc aat ata cga atg tat aac att ctt aaa caa tat atc att ctt gga tga atc att tcc tag ata cca tag cat cga gtt gat atc tat cgg tat aag taa ttc aca tga taa gat ttc cat aaa ctg cta taa tcg t-a aaa aaa ata cgt ggg aaa taa tat ctt cac gga gat ata ggt gta gtt ttg cag tgg ccg tca tga ata ata tta tct ata tga tgg gtg gat atg atc agt ccc cgt ata gga gtt caa agg tta tag cgt aca ata cct gta cta att ctt gga tat atg ata tac cag agc t-a aaa tat cct cgt tct aat tgt gga gga gtt gct gat gat gaa tac att tat tgc ata ggc ggc ata cgc gat cag gat tca tcg ttg ata tct agt att gat aga tgg aag cca tca aaa cca tat tgg caa acg tat gct aaa atg cga gag cca aaa tgt gat atg ggt gta gcg atg tta aac gga tta ata tat gtc ata ggc gga gtt gtt aaa ggt g-- aca cgt gta ccg aca cac ttg aga gtt tat cag aag atg gat gga tga tgc atc aac gtc ttc caa taa aaa tgt cca ata tgt cga cga ttg ttc atg ccg gca aaa ttt ata tat ctg gag gct aca aca ata gta gtg tag tta atg gaa tat cga atc t-- agt cct tag cta taa tcc gat ata tga tga atg gac caa att atc atc att aaa tat tcc tag aat taa tcc cgc tct a

>MK035756.1:176238-178483_Cowpox_virus_strain_CPXV/Leo_2015/5

ttg ctg act gca --- tgt a-c tat ttt tta cat tac tac att ggc tac ggc aga tat acc tac ttc gtc act gcc aca cgc tcc ggt aaa cgg --g gca tgt gac gag gga gaa tat ctt gat aag agg cat aat caa tgt tgt aat -cg gtg tcc acc tgg aga att tgc caa ggt tag atg ta- --g tgg tag cta taa cac aaa atg tga acg ctg tcc aca tca tac ata tac agc aat tcc caa tta ttc t-a atg gat gcc atc aat gta gaa aat gcc caa agg gat cat ttg ata agg taa agt gta ccg gaa cac aga aca gta aat -gt gcg tgt ctt cct ggt tgg tat tgc gct acc gat tct tca cag act gaa gac tgt cga gat -tg tgt acc aaa aag g-a gct gtc cat gcg gat act ttg gtg gaa tag ata aat tag gaa atc cta ttt gta aat cgt gtt g-- -tg ttg gta aat att gtg acg aca tac gta atc ata gag ttg gtc ctt ttc ctc cat gca aa- --- cta tct aaa tgt aat tag tca --- --- --- --- --- --- --- --- --- --- --- --- --- --- -tg atc atg gta -cc gta caa tt- --- -at att gct --- --g gta --g tat att att cag tat g-a aga cat att aat ac- --- --- -tt atc --- --t ttg aca atc ttg ct- -at aat cat --- --- --- --- aat ata aaa --a tac tta tgg --- -a- cat agt cac tca t-- --- --- --- aat tgc tg- acg cga taa att cgt aat aat ctg ttt tgt tca aat ttt tat aag ga- --- --a tct aca ggc ata aaa at- aaa aat ata att tat aat ac- --- act tac agc gcg cca tca tga ata aca gca gtg aat taa ttg ctg tta tta atg gat tta gaa ata gtg gac gat ttt gtg ata tta ata tag tta tta atg atg aaa gga taa atg ctc ata gac tca tcc tat ctg gag cat ccg aat a-t ttt tcc att cta ttt tct agt gat ttt atc gat tct aat gaa tac gaa gtt aat cta agt cat tta gat tat caa agt gtg aac gat ttg atc gat tac att tat ggg ata cct ttg agc cta act aac gat aac gtg aaa tat att ctt tca acc gct ga- --t ttt tta caa att gga tct gcc att act gag tgc gaa aaa tac ata ctt aaa aat ctg tgt tct aga aat tgt atc gat ttc --- --- tac ata tac gct gat aaa tat aat aac aag aaa ata gaa aca gcg tcg ttt aac aca ata tta cga aat att ttg aga ctc att aac gat gaa aac ttt aaa tac ttg aca gag gaa tca atg ata aaa att tta agc gat gat atg tta aat ata aaa aat gag gat ttt gca cca cta att ctc att aaa tgg tta gag agt aca caa caa cca tgc aca gtc gag tta ctt aga tgc ctc aga ata tca -tt gct ttc ccc aca agt tat aaa atc act tta tag tca tcg att ggt tgg ttc aat ata cga atg tat aac att ctt aaa caa tat atc att ctt gga tga atc att tcc tag ata cca tag cat cga gtt gat atc tat cgg tat aag taa ttc aca tga taa gat ttc cat aaa ctg cta taa tcg t-a aaa aaa ata cgt ggg aaa taa tat ctt cac gga gat ata ggt gta gtt ttg cag tgg ccg tca tga ata ata tta tct ata tga tgg gtg gat atg atc agt ccc cgt ata gga gtt caa agg tta tag cgt aca ata cct gta cta att ctt gga tat atg ata tac cag agc t-a aaa tat cct cgt tct aat tgt gga gga gtt gct gat gat gaa tac att tat tgc ata ggc ggc ata cgc gat cag gat tca tcg ttg ata tct agt att gat aga tgg aag cca tca aaa cca tat tgg caa acg tat gct aaa atg cga gag cca aaa tgt gat atg ggt gta gcg atg tta aac gga tta ata tat gtc ata ggc gga gtt gtt aaa ggt g-- aca cgt gta ccg aca cac ttg aga gtt tat cag aag atg gat gga tga tgc atc aac gtc ttc caa taa aaa tgt cca ata tgt cga cga ttg ttc atg ccg gca aaa ttt ata tat ctg gag gct aca aca ata gta gtg tag tta atg gaa tat cga atc t-- agt cct tag cta taa tcc gat ata tga tga atg gac caa att atc atc att aaa tat tcc tag aat taa tcc cgc tct a

>MK035754.1:176661-178906_Cowpox_virus_strain_CPXV/Cookie_S963_15

ttg ctg act gca --- tgt a-c tat ttt tta cat tac tac att ggc tac ggc aga tat acc tac ttc gtc act gcc aca cgc tcc ggt aaa cgg --g gca tgt gac gag gga gaa tat ctt gat aag agg cat aat caa tgt tgt aat -cg gtg tcc acc tgg aga att tgc caa ggt tag atg ta- --g tgg tag cta taa cac aaa atg tga acg ctg tcc aca tca tac ata tac agc aat tcc caa tta ttc t-a atg gat gcc atc aat gta gaa aat gcc caa agg gat cat ttg ata agg taa agt gta ccg gaa cac aga aca gta aat -gt gcg tgt ctt cct ggt tgg tat tgc gct acc gat tct tca cag act gaa gac tgt cga gat -tg tgt acc aaa aag g-a gct gtc cat gcg gat act ttg gtg gaa tag ata aat tag gaa atc cta ttt gta aat cgt gtt g-- -tg ttg gta aat att gtg acg aca tac gta atc ata gag ttg gtc ctt ttc ctc cat gca aa- --- cta tct aaa tgt aat tag tca --- --- --- --- --- --- --- --- --- --- --- --- --- --- -tg atc atg gta -cc gta caa tt- --- -at att gct --- --g gta --g tat att att cag tat g-a aga cat att aat ac- --- --- -tt atc --- --t ttg aca atc ttg ct- -at aat cat --- --- --- --- aat ata aaa --a tac tta tgg --- -a- cat agt cac tca t-- --- --- --- aat tgc tg- acg cga taa att cgt aat aat ctg ttt tgt tca aat ttt tat aag ga- --- --a tct aca ggc ata aaa at- aaa aat ata att tat aat ac- --- act tac agc gcg cca tca tga ata aca gca gtg aat taa ttg ctg tta tta atg gat tta gaa ata gtg gac gat ttt gtg ata tta ata tag tta tta atg atg aaa gga taa atg ctc ata gac tca tcc tat ctg gag cat ccg aat a-t ttt tcc att cta ttt tct agt gat ttt atc gat tct aat gaa tac gaa gtt aat cta agt cat tta gat tat caa agt gtg aac gat ttg atc gat tac att tat ggg ata cct ttg agc cta act aac gat aac gtg aaa tat att ctt tca acc gct ga- --t ttt tta caa att gga tct gcc att act gag tgc gaa aaa tac ata ctt aaa aat ctg tgt tct aga aat tgt atc gat ttc --- --- tac ata tac gct gat aaa tat aat aac aag aaa ata gaa aca gcg tcg ttt aac aca ata tta cga aat att ttg aga ctc att aac gat gaa aac ttt aaa tac ttg aca gag gaa tca atg ata aaa att tta agc gat gat atg tta aat ata aaa aat gag gat ttt gca cca cta att ctc att aaa tgg tta gag agt aca caa caa cca tgc aca gtc gag tta ctt aga tgc ctc aga ata tca -tt gct ttc ccc aca agt tat aaa atc act tta tag tca tcg att ggt tgg ttc aat ata cga atg tat aac att ctt aaa caa tat atc att ctt gga tga atc att tcc tag ata cca tag cat cga gtt gat atc tat cgg tat aag taa ttc aca tga taa gat ttc cat aaa ctg cta taa tcg t-a aaa aaa ata cgt ggg aaa taa tat ctt cac gga gat ata ggt gta gtt ttg cag tgg ccg tca tga ata ata tta tct ata tga tgg gtg gat atg atc agt ccc cgt ata gga gtt caa agg tta tag cgt aca ata cct gta cta att ctt gga tat atg ata tac cag agc t-a aaa tat cct cgt tct aat tgt gga gga gtt gct gat gat gaa tac att tat tgc ata ggc ggc ata cgc gat cag gat tca tcg ttg ata tct agt att gat aga tgg aag cca tca aaa cca tat tgg caa acg tat gct aaa atg cga gag cca aaa tgt gat atg ggt gta gcg atg tta aac gga tta ata tat gtc ata ggc gga gtt gtt aaa ggt g-- aca cgt gta ccg aca cac ttg aga gtt tat cag aag atg gat gga tga tgc atc aac gtc ttc caa taa aaa tgt cca ata tgt cga cga ttg ttc atg ccg gca aaa ttt ata tat ctg gag gct aca aca ata gta gtg tag tta atg gaa tat cga atc t-- agt cct tag cta taa tcc gat ata tga tga atg gac caa att atc atc att aaa tat tcc tag aat taa tcc cgc tct a

>MK035752.1:176461-178706_Cowpox_virus_strain_CPXV/Sammy_2015/4a

ttg ctg act gca --- tgt a-c tat ttt tta cat tac tac att ggc tac ggc aga tat acc tac ttc gtc act gcc aca cgc tcc ggt aaa cgg --g gca tgt gac gag gga gaa tat ctt gat aag agg cat aat caa tgt tgt aat -cg gtg tcc acc tgg aga att tgc caa ggt tag atg ta- --g tgg tag cta taa cac aaa atg tga acg ctg tcc aca tca tac ata tac agc aat tcc caa tta ttc t-a atg gat gcc atc aat gta gaa aat gcc caa agg gat cat ttg ata agg taa agt gta ccg gaa cac aga aca gta aat -gt gcg tgt ctt cct ggt tgg tat tgc gct acc gat tct tca cag act gaa gac tgt cga gat -tg tgt acc aaa aag g-a gct gtc cat gcg gat act ttg gtg gaa tag ata aat tag gaa atc cta ttt gta aat cgt gtt g-- -tg ttg gta aat att gtg acg aca tac gta atc ata gag ttg gtc ctt ttc ctc cat gca aa- --- cta tct aaa tgt aat tag tca --- --- --- --- --- --- --- --- --- --- --- --- --- --- -tg atc atg gta -cc gta caa tt- --- -at att gct --- --g gta --g tat att att cag tat g-a aga cat att aat ac- --- --- -tt atc --- --t ttg aca atc ttg ct- -at aat cat --- --- --- --- aat ata aaa --a tac tta tgg --- -a- cat agt cac tca t-- --- --- --- aat tgc tg- acg cga taa att cgt aat aat ctg ttt tgt tca aat ttt tat aag ga- --- --a tct aca ggc ata aaa at- aaa aat ata att tat aat ac- --- act tac agc gcg cca tca tga ata aca gca gtg aat taa ttg ctg tta tta atg gat tta gaa ata gtg gac gat ttt gtg ata tta ata tag tta tta atg atg aaa gga taa atg ctc ata gac tca tcc tat ctg gag cat ccg aat a-t ttt tcc att cta ttt tct agt gat ttt atc gat tct aat gaa tac gaa gtt aat cta agt cat tta gat tat caa agt gtg aac gat ttg atc gat tac att tat ggg ata cct ttg agc cta act aac gat aac gtg aaa tat att ctt tca acc gct ga- --t ttt tta caa att gga tct gcc att act gag tgc gaa aaa tac ata ctt aaa aat ctg tgt tct aga aat tgt atc gat ttc --- --- tac ata tac gct gat aaa tat aat aac aag aaa ata gaa aca gcg tcg ttt aac aca ata tta cga aat att ttg aga ctc att aac gat gaa aac ttt aaa tac ttg aca gag gaa tca atg ata aaa att tta agc gat gat atg tta aat ata aaa aat gag gat ttt gca cca cta att ctc att aaa tgg tta gag agt aca caa caa cca tgc aca gtc gag tta ctt aga tgc ctc aga ata tca -tt gct ttc ccc aca agt tat aaa atc act tta tag tca tcg att ggt tgg ttc aat ata cga atg tat aac att ctt aaa caa tat atc att ctt gga tga atc att tcc tag ata cca tag cat cga gtt gat atc tat cgg tat aag taa ttc aca tga taa gat ttc cat aaa ctg cta taa tcg t-a aaa aaa ata cgt ggg aaa taa tat ctt cac gga gat ata ggt gta gtt ttg cag tgg ccg tca tga ata ata tta tct ata tga tgg gtg gat atg atc agt ccc cgt ata gga gtt caa agg tta tag cgt aca ata cct gta cta att ctt gga tat atg ata tac cag agc t-a aaa tat cct cgt tct aat tgt gga gga gtt gct gat gat gaa tac att tat tgc ata ggc ggc ata cgc gat cag gat tca tcg ttg ata tct agt att gat aga tgg aag cca tca aaa cca tat tgg caa acg tat gct aaa atg cga gag cca aaa tgt gat atg ggt gta gcg atg tta aac gga tta ata tat gtc ata ggc gga gtt gtt aaa ggt g-- aca cgt gta ccg aca cac ttg aga gtt tat cag aag atg gat gga tga tgc atc aac gtc ttc caa taa aaa tgt cca ata tgt cga cga ttg ttc atg ccg gca aaa ttt ata tat ctg gag gct aca aca ata gta gtg tag tta atg gaa tat cga atc t-- agt cct tag cta taa tcc gat ata tga tga atg gac caa att atc atc att aaa tat tcc tag aat taa tcc cgc tct a

>LT896728.1:176254-178499_Cowpox_virus_isolate_Ger/2010/Rat_genome_assembly_complete_genome:_monopartite

ttg ctg act gca --- tgt a-c tat ttt tta cat tac tac att ggc tac ggc aga tat acc tac ttc gtc act gcc aca cgc tcc ggt aaa cgg --g gca tgt gac gag gga gaa tat ctt gat aag agg cat aat caa tgt tgt aat -cg gtg tcc acc tgg aga att tgc caa ggt tag atg ta- --g tgg tag cta taa cac aaa atg tga acg ctg tcc aca tca tac ata tac agc aat tcc caa tta ttc t-a atg gat gcc atc aat gta gaa aat gcc caa agg gat cat ttg ata agg taa agt gta ccg gaa cac aga aca gta aat -gt gcg tgt ctt cct ggt tgg tat tgc gct acc gat tct tca cag act gaa gac tgt cga gat -tg tgt acc aaa aag g-a gct gtc cat gcg gat act ttg gtg gaa tag ata aat tag gaa atc cta ttt gta aat cgt gtt g-- -tg ttg gta aat att gtg acg aca tac gta atc ata gag ttg gtc ctt ttc ctc cat gca aa- --- cta tct aaa tgt aat tag tca --- --- --- --- --- --- --- --- --- --- --- --- --- --- -tg atc atg gta -cc gta caa tt- --- -at att gct --- --g gta --g tat att att cag tat g-a aga cat att aat ac- --- --- -tt atc --- --t ttg aca atc ttg ct- -at aat cat --- --- --- --- aat ata aaa --a tac tta tgg --- -a- cat agt cac tca t-- --- --- --- aat tgc tg- acg cga taa att cgt aat aat ctg ttt tgt tca aat ttt tat aag ga- --- --a tct aca ggc ata aaa at- aaa aat ata att tat aat ac- --- act tac agc gcg cca tca tga ata aca gca gtg aat taa ttg ctg tta tta atg gat tta gaa ata gtg gac gat ttt gtg ata tta ata tag tta tta atg atg aaa gga taa atg ctc ata gac tca tcc tat ctg gag cat ccg aat a-t ttt tcc att cta ttt tct agt gat ttt atc gat tct aat gaa tac gaa gtt aat cta agt cat tta gat tat caa agt gtg aac gat ttg atc gat tac att tat ggg ata cct ttg agc cta act aac gat aac gtg aaa tat att ctt tca acc gct ga- --t ttt tta caa att gga tct gcc att act gag tgc gaa aaa tac ata ctt aaa aat ctg tgt tct aga aat tgt atc gat ttc --- --- tac ata tac gct gat aaa tat aat aac aag aaa ata gaa aca gcg tcg ttt aac aca ata tta cga aat att ttg aga ctc att aac gat gaa aac ttt aaa tac ttg aca gag gaa tca atg ata aaa att tta agc gat gat atg tta aat ata aaa aat gag gat ttt gca cca cta att ctc att aaa tgg tta gag agt aca caa caa cca tgc aca gtc gag tta ctt aga tgc ctc aga ata tca -tt gct ttc ccc aca agt tat aaa atc act tta tag tca tcg att ggt tgg ttc aat ata cga atg tat aac att ctt aaa caa tat atc att ctt gga tga atc att tcc tag ata cca tag cat cga gtt gat atc tat cgg tat aag taa ttc aca tga taa gat ttc cat aaa ctg cta taa tcg t-a aaa aaa ata cgt ggg aaa taa tat ctt cac gga gat ata ggt gta gtt ttg cag tgg ccg tca tga ata ata tta tct ata tga tgg gtg gat atg atc agt ccc cgt ata gga gtt caa agg tta tag cgt aca ata cct gta cta att ctt gga tat atg ata tac cag agc t-a aaa tat cct cgt tct aat tgt gga gga gtt gct gat gat gaa tac att tat tgc ata ggc ggc ata cgc gat cag gat tca tcg ttg ata tct agt att gat aga tgg aag cca tca aaa cca tat tgg caa acg tat gct aaa atg cga gag cca aaa tgt gat atg ggt gta gcg atg tta aac gga tta ata tat gtc ata ggc gga gtt gtt aaa ggt g-- aca cgt gta ccg aca cac ttg aga gtt tat cag aag atg gat gga tga tgc atc aac gtc ttc caa taa aaa tgt cca ata tgt cga cga ttg ttc atg ccg gca aaa ttt ata tat ctg gag gct aca aca ata gta gtg tag tta atg gaa tat cga atc t-- agt cct tag cta taa tcc gat ata tga tga atg gac caa att atc atc att aaa tat tcc tag aat taa tcc cgc tct a

>HQ420895.1:177301-179546_Cowpox_virus_strain_Germany_1980_EP4_complete_genome

ttg ctg act gca --- tgt a-c tat ttt tta cat tac tac att ggc tac ggc aga tat acc tac ttc gtc act gcc aca cgc tcc ggt aaa cgg --g gca tgt gac gag gga gaa tat ctt gat aag agg cat aat caa tgt tgt aat -cg gtg tcc acc tgg aga att tgc caa ggt tag atg ta- --g tgg tag cta taa cac aaa atg tga acg ctg tcc aca tca tac ata tac agc aat tcc caa tta ttc t-a atg gat gcc atc aat gta gaa aat gcc caa agg gat cat ttg ata agg taa agt gta ccg gaa cac aga aca gta aat -gt gcg tgt ctt cct ggt tgg tat tgc gct acc gat tct tca cag act gaa gac tgt cga gat -tg tgt acc aaa aag g-a gct gtc cat gcg gat act ttg gtg gaa tag ata aat tag gaa atc cta ttt gta aat cgt gtt g-- -tg ttg gta aat att gtg acg aca tac gta atc ata gag ttg gtc ctt ttc ctc cat gca aa- --- cta tct aaa tgt aat tag tca --- --- --- --- --- --- --- --- --- --- --- --- --- --- -tg atc atg gta -cc gta caa tt- --- -at att gct --- --g gta --g tat att att cag tat g-a aga cat att aat ac- --- --- -tt atc --- --t ttg aca atc ttg ct- -at aat cat --- --- --- --- aat ata aaa --a tac tta tgg --- -a- cat agt cac tca t-- --- --- --- aat tgc tg- acg cga taa att cgt aat aat ctg ttt tgt tca aat ttt tat aag ga- --- --a tct aca ggc ata aaa at- aaa aat ata att tat aat ac- --- act tac agc gcg cca tca tga ata aca gca gtg aat taa ttg ctg tta tta atg gat tta gaa ata gtg gac gat ttt gtg ata tta ata tag tta tta atg atg aaa gga taa atg ctc ata gac tca tcc tat ctg gag cat ccg aat a-t ttt tcc att cta ttt tct agt gat ttt atc gat tct aat gaa tac gaa gtt aat cta agt cat tta gat tat caa agt gtg aac gat ttg atc gat tac att tat ggg ata cct ttg agc cta act aac gat aac gtg aaa tat att ctt tca acc gct ga- --t ttt tta caa att gga tct gcc att act gag tgc gaa aaa tac ata ctt aaa aat ctg tgt tct aga aat tgt atc gat ttc --- --- tac ata tac gct gat aaa tat aat aac aag aaa ata gaa aca gcg tcg ttt aac aca ata tta cga aat att ttg aga ctc att aac gat gaa aac ttt aaa tac ttg aca gag gaa tca atg ata aaa att tta agc gat gat atg tta aat ata aaa aat gag gat ttt gca cca cta att ctc att aaa tgg tta gag agt aca caa caa cca tgc aca gtc gag tta ctt aga tgc ctc aga ata tca -tt gct ttc ccc aca agt tat aaa atc act tta tag tca tcg att ggt tgg ttc aat ata cga atg tat aac att ctt aaa caa tat atc att ctt gga tga atc att tcc tag ata cca tag cat cga gtt gat atc tat cgg tat aag taa ttc aca tga taa gat ttc cat aaa ctg cta taa tcg t-a aaa aaa ata cgt ggg aaa taa tat ctt cac gga gat ata ggt gta gtt ttg cag tgg ccg tca tga ata ata tta tct ata tga tgg gtg gat atg atc agt ccc cgt ata gga gtt caa agg tta tag cgt aca ata cct gta cta att ctt gga tat atg ata tac cag agc t-a aaa tat cct cgt tct aat tgt gga gga gtt gct gat gat gaa tac att tat tgc ata ggc ggc ata cgc gat cag gat tca tcg ttg ata tct agt att gat aga tgg aag cca tca aaa cca tat tgg caa acg tat gct aaa atg cga gag cca aaa tgt gat atg ggt gta gcg atg tta aac gga tta ata tat gtc ata ggc gga gtt gtt aaa ggt g-- aca cgt gta ccg aca cac ttg aga gtt tat cag aag atg gat gga tga tgc atc aac gtc ttc caa taa aaa tgt cca ata tgt cga cga ttg ttc atg ccg gca aaa ttt ata tat ctg gag gct aca aca ata gta gtg tag tta atg gaa tat cga atc t-- agt cct tag cta taa tcc gat ata tga tga atg gac caa att atc atc att aaa tat tcc tag aat taa tcc cgc tct a

>MK035755.1:176676-178921_Cowpox_virus_strain_CPXV/75/01

ttg ctg act gca --- tgt a-c tat ttt tta cat tac tac att ggc tac ggc aga tat acc tac ttc gtc act gcc aca cgc tcc ggt aaa cgg --g gca tgt gac gag gga gaa tat ctt gat aag agg cat aat caa tgt tgt aat -cg gtg tcc acc tgg aga att tgc caa ggt tag atg ta- --g tgg tag cta taa cac aaa atg tga acg ctg tcc aca tca tac ata tac agc aat tcc caa tta ttc t-a atg gat gcc atc aat gta gaa aat gcc caa agg gat cat ttg ata agg taa agt gta ccg gaa cac aga aca gta aat -gt gcg tgt ctt cct ggt tgg tat tgc gct acc gat tct tca cag act gaa gac tgt cga gat -tg tgt acc aaa aag g-a gct gtc cat gcg gat act ttg gtg gaa tag ata aat tag gaa atc cta ttt gta aat cgt gtt g-- -tg ttg gta aat att gtg acg aca tac gta atc ata gag ttg gtc ctt ttc ctc cat gca aa- --- cta tct aaa tgt aat tag tca --- --- --- --- --- --- --- --- --- --- --- --- --- --- -tg atc atg gta -cc gta caa tt- --- -at att gct --- --g gta --g tat att att cag tat g-a aga cat att aat ac- --- --- -tt atc --- --t ttg aca atc ttg ct- -at aat cat --- --- --- --- aat ata aaa --a tac tta tgg --- -a- cat agt cac tca t-- --- --- --- aat tgc tg- acg cga taa att cgt aat aat ctg ttt tgt tca aat ttt tat aag ga- --- --a tct aca ggc ata aaa at- aaa aat ata att tat aat ac- --- act tac agc gcg cca tca tga ata aca gca gtg aat taa ttg ctg tta tta atg gat tta gaa ata gtg gac gat ttt gtg ata tta ata tag tta tta atg atg aaa gga taa atg ctc ata gac tca tcc tat ctg gag cat ccg aat a-t ttt tcc att cta ttt tct agt gat ttt atc gat tct aat gaa tac gaa gtt aat cta agt cat tta gat tat caa agt gtg aac gat ttg atc gat tac att tat ggg ata cct ttg agc cta act aac gat aac gtg aaa tat att ctt tca acc gct ga- --t ttt tta caa att gga tct gcc att act gag tgc gaa aaa tac ata ctt aaa aat ctg tgt tct aga aat tgt atc gat ttc --- --- tac ata tac gct gat aaa tat aat aac aag aaa ata gaa aca gcg tcg ttt aac aca ata tta cga aat att ttg aga ctc att aac gat gaa aac ttt aaa tac ttg aca gag gaa tca atg ata aaa att tta agc gat gat atg tta aat ata aaa aat gag gat ttt gca cca cta att ctc att aaa tgg tta gag agt aca caa caa cca tgc aca gtc gag tta ctt aga tgc ctc aga ata tca -tt gct ttc ccc aca agt tat aaa atc act tta tag tca tcg att ggt tgg ttc aat ata cga atg tat aac att ctt aaa caa tat atc att ctt gga tga atc att tcc tag ata cca tag cat cga gtt gat atc tat cgg tat aag taa ttc aca tga taa gat ttc cat aaa ctg cta taa tcg t-a aaa aaa ata cgt ggg aaa taa tat ctt cac gga gat ata ggt gta gtt ttg cag tgg ccg tca tga ata ata tta tct ata tga tgg gtg gat atg atc agt ccc cgt ata gga gtt caa agg tta tag cgt aca ata cct gta cta att ctt gga tat atg ata tac cag agc t-a aaa tat cct cgt tct aat tgt gga gga gtt gct gat gat gaa tac att tat tgc ata ggc ggc ata cgc gat cag gat tca tcg ttg ata tct agt att gat aga tgg aag cca tca aaa cca tat tgg caa acg tat gct aaa atg cga gag cca aaa tgt gat atg ggt gta gcg atg tta aac gga tta ata tat gtc ata ggc gga gtt gtt aaa ggt g-- aca tgt gta ccg aca cac ttg aga gtt tat cag aag atg gat gga tga tgc atc aac gtc ttc caa taa aaa tgt cca ata tgt cga cga ttg ttc atg ccg gca aaa ttt ata tat ctg gag gct aca aca ata gta gtg tag tta atg gaa tat cga atc t-- agt cct tag cta taa tcc gat ata tga tga atg gac caa att atc atc att aaa tat tcc tag aat taa tcc cgc tct a

>MK035751.1:177075-179320_Cowpox_virus_strain_CPXV/S2216/04

ttg ctg act gca --- tgt a-c tat ttt tta cat tac tac att ggc tac ggc aga tat acc tac ttc gtc act gcc aca tgc tcc ggt aaa cgg --g gca tgt gac gag gga gaa tat ctt gat aag agg cat aat caa tgt tgt aat -cg gtg tcc acc tgg aga att tgc caa ggt tag atg ta- --g tgg tag cta taa cac aaa atg tga acg ctg tcc aca tca tac ata tac agc aat tcc caa tta ttc t-a atg gat gcc atc aat gta gaa aat gcc caa agg gat cat ttg ata agg taa agt gta ccg gaa cac aga aca gta aat -gt gcg tgt ctt cct ggt tgg tat tgc gct acc gat tct tca cag act gaa gac tgt cga gat -tg tgt acc aaa aag g-a gct gtc cat gcg gat act ttg gtg gaa tag ata aat tag gaa atc cta ttt gta aat cgt gtt g-- -tg ttg gta aat att gtg acg aca tac gta atc ata gag ttg gtc ctt ttc ctc cat gca aa- --- cta tct aaa tgt aat tag tca --- --- --- --- --- --- --- --- --- --- --- --- --- --- -tg atc atg gta -cc gta caa tt- --- -at att gct --- --g gta --g tat att att cag tat g-a aga cat att aat ac- --- --- -tt atc --- --t ttg aca atc ttg ct- -at aat cat --- --- --- --- aat ata aaa --a tac tta tgg --- -a- cat agt cac tca t-- --- --- --- aat tgc tg- acg cga taa att cgt aat aat ctg ttt tgt tca aat ttt tat aag ga- --- --a tct aca ggc ata aaa at- aaa aat ata att tat aat ac- --- act tac agc gcg cca tca tga ata aca gca gtg aat taa ttg ctg tta tta atg gat tta gaa ata gtg gac gat ttt gtg ata tta ata tag tta tta atg atg aaa gga taa atg ctc ata gac tca tcc tat ctg gag cat ccg aat a-t ttt tcc att cta ttt tct agt gat ttt atc gat tct aat gaa tac gaa gtt aat cta agt cat tta gat tat caa agt gtg aac gat ttg atc gat tac att tat ggg ata cct ttg agc cta act aac gat aac gtg aaa tat att ctt tca acc gct ga- --t ttt tta caa att gga tct gcc att act gag tgc gaa aaa tac ata ctt aaa aat ctg tgt tct aga aat tgt atc gat ttc --- --- tac ata tac gct gat aaa tat aat aac aag aaa ata gaa aca gcg tcg ttt aac aca ata tta cga aat att ttg aga ctc att aac gat gaa aac ttt aaa tac ttg aca gag gaa tca atg ata aaa att tta agc gat gat atg tta aat ata aaa aat gag gat ttt gca cca cta att ctc att aaa tgg tta gag agt aca caa caa cca tgc aca gtc gag tta ctt aga tgc ctc aga ata tca -tt gct ttc ccc aca agt tat aaa atc act tta tag tca tcg att ggt tgg ttc aat ata cga atg tat aac att ctt aaa caa tat atc att ctt gga tga atc att tcc tag ata cca tag cat cga gtt gat atc tat cgg tat aag taa ttc aca tga taa gat ttc cat aaa ctg cta taa tcg t-a aaa aaa ata cgt ggg aaa taa tat ctt cac gga gat ata ggt gta gtt ttg cag tgg ccg tca tga ata ata tta tct ata tga tgg gtg gat atg atc agt ccc cgt ata gga gtt caa agg tta tag cgt aca ata cct gta cta att ctt gga tat atg ata tac cag agc t-a aaa tat cct cgt tct aat tgt gga gga gtt gct gat gat gaa tac att tat tgc ata ggc ggc ata cgc gat cag gat tca tcg ttg ata tct agt att gat aga tgg aag cca tca aaa cca tat tgg caa acg tat gct aaa atg cga gag cca aaa tgt gat atg ggt gta gcg atg tta aac gga tta ata tat gtc ata ggc gga gtt gtt aaa ggt g-- aca cgt gta ccg aca cac ttg aga gtt tat cag aag atg gat gga tga tgc atc aac gtc ttc caa taa aaa tgt cca ata tgt cga cga ttg ttc atg ccg gca aaa ttt ata tat ctg gag gct aca aca ata gta gtg tag tta atg gaa tat cga atc t-- agt cct tag cta taa tcc gat ata tga tga atg gac caa att atc atc att aaa tat tcc tag aat taa tcc cgc tct a

>LT896729.1:175828-178073_Cowpox_virus_isolate_Ger/2010/Cat_genome_assembly_complete_genome:_monopartite

ttg ctg act gca --- tgt a-c tat ttt tta cat tac tac att ggc tac ggc aga tat acc tac ttc gtc act gcc aca cgc tcc ggt aaa cgg --g gca tgt gac gag gga gaa tat ctt gat aag act cat aat cga tgt tgt aat -cg gtg tcc acc tgg aga att tgc caa ggt tag atg ta- --g tgg tag cta taa cac aaa atg tga acg ctg tcc aca tca tac ata tac agc aat tcc caa tta ttc t-a atg gat gcc atc aat gta gaa aat gcc caa agg gat cat ttg ata agg taa agt gta cag gaa cac aga aca gta aat -gt gcg tgt ctt cct ggt tgg tat tgc gct acc gat tct tca cag act gaa gac tgt cga gat -tg tgt acc aaa aag g-a gct gtc cat gcg gat act ttg gtg gaa tag ata aat tag gaa atc cta ttt gta aat cgt gtt g-- -tg ttg gta aat att gtg acg aca tac gta atc ata gag ttg gtc ctt ttc ctc cat gca aa- --- cta tct aaa tgt aat tag tca --- --- --- --- --- --- --- --- --- --- --- --- --- --- -tg atc atg gta -cc gta caa tt- --- -at att gct --- --g gta --g tat att att cag tat g-a aga cat att aat ac- --- --- -tt atc --- --t ttg aca atc ttg ct- -at aat cat --- --- --- --- aat ata aaa --a tac tta tgg --- -a- cat agt cac tca t-- --- --- --- aat tgc tg- acg cga taa att cgt aat aat ctg ttt tgt tca aat ttt tat aag ga- --- --a tct aca ggc ata aaa at- aaa aat ata att tat aat ac- --- act tac agc gcg cca tca tga ata aca gca gtg aat tga ttg ctg tta tta atg gat tta gaa ata gtg gac gat ttt gtg ata tta gta tag tta tta atg atg aaa gga taa acg ctc aca gac tca tcc tat ctg gag cct ccg aat a-t ttt tcc att ctg ttt tcc aat aat ttt atc gat tct aat gaa tac gaa gtt aat cta agt cat tta gat tat caa agt gtt aac gat ttg atc gat tac att tat ggg ata cct ttg agc cta act aac gat aac gtg aaa tat att ctt tca acc gct ga- --t ttt tta caa att gga tct gcc att act gag tgc gaa aaa tac ata ctt aaa aat ctg tgt tct aga aat tgt atc gat ttc --- --- tac ata tac gct gat aaa tat aat aac gag aaa ata gaa aca gcg tcg ttt aac aca ata tta cga aat att ttg aga ctc att aac gat gaa aac ttt aaa tac ttg aca gag gaa tca atg ata aaa att tta agc gat gat atg tta aat ata aaa aat gag gat ttt gca cca cta att ctc att aaa tgg tta gag agt aca caa caa cca tgc aca gtc gag tta ctt aga tgc ctc aga ata tca -tt gct ttc ccc aca agt tat aaa atc act tta tag tca tcg att ggt tgg ttc aat ata cga atg tat aac att ctt aaa caa tat atc att ctt gga tga atc att tcc tag ata cca tag cat cga gtt gat atc tat cgg tat aag taa ttc aca tga taa gat ttc cat aaa ctg cta taa tcg t-a aaa aaa ata cgt ggg aga taa tat ctt cac gga gat ata ggt gta gtt ttg cag tgg ccg tca tgg ata ata tta tct ata tga tgg gtg gat atg atc agt ccc cgt ata gga gtt caa agg tta tag cgt aca ata cct gta cta att ctt gga tat atg ata tac cag agc t-a aaa tat cct cgt tct aat tgt gga gga gtt gct gat gat gaa tac att tat tgc ata ggc ggc ata cgc gat cag gat tca tcg ttg ata tct agt att gat aga tgg aag cca tca aaa cca tat tgg caa acg tat gct aaa atg cga gag cca aaa tgt gat atg ggt gta gcg atg tta aac gga tta ata tat gtc ata ggc gga gtt gtt aaa ggt g-- aca cgt gta ccg aca cac ttg aga gtt tat cag aag atg gat gga tga tgc atc aac gtc ttc caa taa aaa tgt cca ata tgt cga cga ttg ttc atg ccg gca aaa ttt ata tat ctg gag gct aca aca ata gta gtg tag tta atg gaa tat cga atc t-- agt cct tag cta taa tcc gat ata tga tga atg gac caa att atc atc att aaa tat tcc tag aat taa tcc cgc tct a

>KY549147.1:168551-170796_Cowpox_virus_isolate_CPXV_K780

ttg ctg act gca --- tgt a-c tat ttt tta cat tac tac att ggc tac ggc aga tat acc tac ttc gtc act gcc aca cgc tcc ggt aaa cgg --g gca tgt gac gag gga gaa tat ctt gat aag agg cat aat caa tgt tgt aat -cg gtg tcc acc tgg aga att tgc caa ggt tag atg ta- --g tgg tag cta taa cac aaa atg tga acg ctg tcc aca tca tac ata tac agc aat tcc caa tta ttc t-a atg gat gcc atc aat gta gaa aat gcc caa agg gat cat ttg ata agg taa agt gta ccg gaa cac aga aca gta aat -gt gcg tgt ctt cct ggt tgg tat tgc gct acc gat tct tca cag act gaa gac tgt cga gat -tg tgt acc aaa aag g-a gct gtc cat gcg gat act ttg gtg gaa tag ata aat tag gaa atc cta ttt gta aat cgt gtt g-- -tg ttg gta aat att gtg acg aca tac gta atc ata gag ttg gtc ctt ttc ctc cat gca aa- --- cta tct aaa tgt aat tag tca --- --- --- --- --- --- --- --- --- --- --- --- --- --- -tg atc atg gta -cc gta caa tt- --- -at att gct --- --g gta --g tat att att cag tat g-a aga cat att aat ac- --- --- -tt atc --- --t ttg aca atc ttg ct- -at aat cat --- --- --- --- aat ata aaa --a tac tta tgg --- -a- cat agt cac tca t-- --- --- --- aat tgc tg- acg cga taa att cgt aat aat ctg ttt tgt tca aat ttt tat aag ga- --- --a tct aca ggc ata aaa at- aaa aat ata att tat aat ac- --- act tac agc gcg cca tca tga ata aca gca gtg aat taa ttg ctg tta tta atg gat tta gaa ata gtg gac gat ttt gtg ata tta ata tag tta tta atg atg aaa gga taa atg ctc ata gac tca tcc tat ctg gag cat ccg aat a-t ttt tcc att cta ttt tct agt gat ttt atc gat tct aat gaa tac gaa gtt aat cta agt cat tta gat tat caa agt gtg aac gat ttg atc gat tac att tat ggg ata cct ttg agc cta act aac gat aac gtg aaa tat att ctt tca acc gct ga- --t ttt tta caa att gga tct gcc att act gag tgc gaa aaa tac ata ctt aaa aat ctg tgt tct aga aat tgt atc gat ttc --- --- tac ata tac gct gat aaa tat aat aac aag aaa ata gaa aca gcg tcg ttt aac aca ata tta cga aat att ttg aga ctc att aac gat gaa aac ttt aaa tac ttg aca gag gaa tca atg ata aaa att tta agc gat gat atg tta aat ata aaa aat gag gat ttt gca cca cta att ctc att aaa tgg tta gag agt aca caa caa cca tgc aca gtc gag tta ctt aga tgc ctc aga ata tca -tt gct ttc ccc aca agt tat aaa atc act tta tag tca tcg att ggt tgg ttc aat ata cga atg tat aac att ctt aaa caa tat atc att ctt gga tga atc att tcc tag ata cca tag cat cga gtt gat atc tat cgg tat aag taa ttc aca tga taa gat ttc cat aaa ctg cta taa tcg t-a aaa aaa ata cgt ggg aaa taa tat ctt cac gga gat ata ggt gta gtt ttg cag tgg ccg tca tga ata ata tta tct ata tga tgg gtg gat atg atc agt ccc cgt ata gga gtt caa agg tta tag cgt aca ata cct gta cta att ctt gga tat atg ata tac cag agc t-a aaa tat cct cgt tct aat tgt gga gga gtt gct gat gat gaa tac att tat tgc ata ggc ggc ata cgc gat cag gat tca tcg ttg ata tct agt att gat aga tgg aag cca tca aaa cca tat tgg caa acg tat gct aaa atg cga gag cca aaa tgt gat atg ggt gta gcg atg tta aac gga tta ata tat gtc ata ggc gga gtt gtt aaa ggt g-- aca tgt gta ccg aca cac ttg aga gtt tat cag aag atg gat gga tga tgc atc aac gtc ttc caa taa aaa tgt cca ata tgt cga cga ttg ttc atg ccg gca aaa ttt ata tat ctg gag gct aca aca ata gta gtg tag tta atg gaa tat cga atc t-- agt cct tag cta taa tcc gat ata tga tga atg gac caa att atc atc att aaa tat tcc tag aat taa tcc cgc tct a

>KY549146.1:168833-171078_Cowpox_virus_isolate_CPXV_K779

ttg ctg act gca --- tgt a-c tat ttt tta cat tac tac att ggc tac ggc aga tat acc tac ttc gtc act gcc aca cgc tcc ggt aaa cgg --g gca tgt gac gag gga gaa tat ctt gat aag agg cat aat caa tgt tgt aat -cg gtg tcc acc tgg aga att tgc caa ggt tag atg ta- --g tgg tag cta taa cac aaa atg tga acg ctg tcc aca tca tac ata tac agc aat tcc caa tta ttc t-a atg gat gcc atc aat gta gaa aat gcc caa agg gat cat ttg ata agg taa agt gta ccg gaa cac aga aca gta aat -gt gcg tgt ctt cct ggt tgg tat tgc gct acc gat tct tca cag act gaa gac tgt cga gat -tg tgt acc aaa aag g-a gct gtc cat gcg gat act ttg gtg gaa tag ata aat tag gaa atc cta ttt gta aat cgt gtt g-- -tg ttg gta aat att gtg acg aca tac gta atc ata gag ttg gtc ctt ttc ctc cat gca aa- --- cta tct aaa tgt aat tag tca --- --- --- --- --- --- --- --- --- --- --- --- --- --- -tg atc atg gta -cc gta caa tt- --- -at att gct --- --g gta --g tat att att cag tat g-a aga cat att aat ac- --- --- -tt atc --- --t ttg aca atc ttg ct- -at aat cat --- --- --- --- aat ata aaa --a tac tta tgg --- -a- cat agt cac tca t-- --- --- --- aat tgc tg- acg cga taa att cgt aat aat ctg ttt tgt tca aat ttt tat aag ga- --- --a tct aca ggc ata aaa at- aaa aat ata att tat aat ac- --- act tac agc gcg cca tca tga ata aca gca gtg aat taa ttg ctg tta tta atg gat tta gaa ata gtg gac gat ttt gtg ata tta ata tag tta tta atg atg aaa gga taa atg ctc ata gac tca tcc tat ctg gag cat ccg aat a-t ttt tcc att cta ttt tct agt gat ttt atc gat tct aat gaa tac gaa gtt aat cta agt cat tta gat tat caa agt gtg aac gat ttg atc gat tac att tat ggg ata cct ttg agc cta act aac gat aac gtg aaa tat att ctt tca acc gct ga- --t ttt tta caa att gga tct gcc att act gag tgc gaa aaa tac ata ctt aaa aat ctg tgt tct aga aat tgt atc gat ttc --- --- tac ata tac gct gat aaa tat aat aac aag aaa ata gaa aca gcg tcg ttt aac aca ata tta cga aat att ttg aga ctc att aac gat gaa aac ttt aaa tac ttg aca gag gaa tca atg ata aaa att tta agc gat gat atg tta aat ata aaa aat gag gat ttt gca cca cta att ctc att aaa tgg tta gag agt aca caa caa cca tgc aca gtc gag tta ctt aga tgc ctc aga ata tca -tt gct ttc ccc aca agt tat aaa atc act tta tag tca tcg att ggt tgg ttc aat ata cga atg tat aac att ctt aaa caa tat atc att ctt gga tga atc att tcc tag ata cca tag cat cga gtt gat atc tat cgg tat aag taa ttc aca tga taa gat ttc cat aaa ctg cta taa tcg t-a aaa aaa ata cgt ggg aaa taa tat ctt cac gga gat ata ggt gta gtt ttg cag tgg ccg tca tga ata ata tta tct ata tga tgg gtg gat atg atc agt ccc cgt ata gga gtt caa agg tta tag cgt aca ata cct gta cta att ctt gga tat atg ata tac cag agc t-a aaa tat cct cgt tct aat tgt gga gga gtt gct gat gat gaa tac att tat tgc ata ggc ggc ata cgc gat cag gat tca tcg ttg ata tct agt att gat aga tgg aag cca tca aaa cca tat tgg caa acg tat gct aaa atg cga gag cca aaa tgt gat atg ggt gta gcg atg tta aac gga tta ata tat gtc ata ggc gga gtt gtt aaa ggt g-- aca tgt gta ccg aca cac ttg aga gtt tat cag aag atg gat gga tga tgc atc aac gtc ttc caa taa aaa tgt cca ata tgt cga cga ttg ttc atg ccg gca aaa ttt ata tat ctg gag gct aca aca ata gta gtg tag tta atg gaa tat cga atc t-- agt cct tag cta taa tcc gat ata tga tga atg gac caa att atc atc att aaa tat tcc tag aat taa tcc cgc tct a

>KC813496.1:163890-166135_Cowpox_virus_strain_HumPad07/1_complete_genome

ttg ctg act gca --- tgt a-c tat ttt tta cat tac tac att ggc tac ggc aga tat acc tac ttc gtc act gcc aca cgc tcc ggt aaa cgg --g gca tgt gac gag gga gaa tat ctt gat aag agg cat aat caa tgt tgt aat -cg gtg tcc acc tgg aga att tgc caa ggt tag atg ta- --g tgg tag cta taa cac aaa atg tga acg ctg tcc aca tca tac ata tac agc aat tcc caa tta ttc t-a atg gat gcc atc aat gta gaa aat gcc caa agg gat cat ttg ata agg taa agt gta ccg gaa cac aga aca gta aat -gt gcg tgt ctt cct ggt tgg tat tgc gct acc gat tct tca cag act gaa gac tgt cga gat -tg tgt acc aaa aag g-a gct gtc cat gcg gat act ttg gtg gaa tag ata aat tag gaa atc cta ttt gta aat cgt gtt g-- -tg ttg gta aat att gtg acg aca tac gta atc ata gag ttg gtc ctt ttc ctc cat gca aa- --- cta tct aaa tgt aat tag tca --- --- --- --- --- --- --- --- --- --- --- --- --- --- -tg atc atg gta -cc gta caa tt- --- -at att gct --- --g gta --g tat att att cag tat g-a aga cat att aat ac- --- --- -tt atc --- --t ttg aca atc ttg ct- -at aat cat --- --- --- --- aat ata aaa --a tac tta tgg --- -a- cat agt cac tca t-- --- --- --- aat tgc tg- acg cga taa att cgt aat aat ctg ttt tgt tca aat ttt tat aag ga- --- --a tct aca ggc ata aaa at- aaa aat ata att tat aat ac- --- act tac agc gcg cca tca tga ata aca gca gtg aat taa ttg ctg tta tta atg gat tta gaa ata gtg gac gat ttt gtg ata tta ata tag tta tta atg atg aaa gga taa atg ctc ata gac tca tcc tat ctg gag cat ccg aat a-t ttt tcc att cta ttt tct agt gat ttt atc gat tct aat gaa tac gaa gtt aat cta agt cat tta gat tat caa agt gtg aac gat ttg atc gat tac att tat ggg ata cct ttg agc cta act aac gat aac gtg aaa tat att ctt tca acc gct ga- --t ttt tta caa att gga tct gcc att act gag tgc gaa aaa tac ata ctt aaa aat ctg tgt tct aga aat tgt atc gat ttc --- --- tac ata tac gct gat aaa tat aat aac aag aaa ata gaa aca gcg tcg ttt aac aca ata tta cga aat att ttg aga ctc att aac gat gaa aac ttt aaa tac ttg aca gag gaa tca atg ata aaa att tta agc gat gat atg tta aat ata aaa aat gag gat ttt gca cca cta att ctc att aaa tgg tta gag agt aca caa caa cca tgc aca gtc gag tta ctt aga tgc ctc aga ata tca -tt gct ttc ccc aca agt tat aaa atc act tta tag tca tcg att ggt tgg ttc aat ata cga atg tat aac att ctt aaa caa tat atc att ctt gga tga atc att tcc tag ata cca tag cat cga gtt gat atc tat cgg tat aag taa ttc aca tga taa gat ttc cat aaa ctg cta taa tcg t-a aaa aaa ata cgt ggg aaa taa tat ctt cac gga gat ata ggt gta gtt ttg cag tgg ccg tca tga ata ata tta tct ata tga tgg gtg gat atg atc agt ccc cgt ata gga gtt caa agg tta tag cgt aca ata cct gta cta att ctt gga tat atg ata tac cag agc t-a aaa tat cct cgt tct aat tgt gga gga gtt gct gat gat gaa tac att tat tgc ata ggc ggc ata cgc gat cag gat tca tcg ttg ata tct agt att gat aga tgg aag cca tca aaa cca tat tgg caa acg tat gct aaa atg cga gag cca aaa tgt gat atg ggt gta gcg atg tta aac gga tta ata tat gtc ata ggc gga gtt gtt aaa ggt g-- aca tgt gta ccg aca cac ttg aga gtt tat cag aag atg gat gga tga tgc atc aac gtc ttc caa taa aaa tgt cca ata tgt cga cga ttg ttc atg ccg gca aaa ttt ata tat ctg gag gct aca aca ata gta gtg tag tta atg gaa tat cga atc t-- agt cct tag cta taa tcc gat ata tga tga atg gac caa att atc atc att aaa tat tcc tag aat taa tcc cgc tct a

>LN879483.1:175773-178018_Cowpox_virus_strain_CPXV_Amadeus_2015_complete_genome

ttg ctg act gta --- tgt a-c tat ttt gta cat cag tac att ggt tac agc aga tat acc tac ttc gtc act gcc aca cgc tcc ggt aaa cgg --g tca tgt gac gaa gga gaa tat ctt gat aag acg cat aat cga tgt tgt aat -cg gtg tcc acc tgg aga att tgc caa ggt tag atg ta- --g tgg tag cta taa cac aaa atg tga acg ctg tcc aca tca tac ata tac agc aat tcc caa tta ttc t-a atg gat gcc atc aat gta gaa aat gcc caa agg gat cat ttg ata agg taa agt gta ccg gaa cac aga aca gta aat -gt gcg tgt ctt cct ggt tgg tat tgc gct acc gat tct tca cag act gaa gac tgt cga gat -tg tgt acc aaa aag g-a gct gtc cat gcg gat act ttg gtg gaa tag ata aat tag gaa atc cta ttt gta aat cgt gtt g-- -tg ttg gta aat att gtg acg aca tac gta atc ata gag ttg gtc ctt ttc ctc cat gca aa- --- cta tct aaa tgt aat tag tca --- --- --- --- --- --- --- --- --- --- --- --- --- --- -tg atc atg gta -cc gta caa tt- --- -at att gct --- --g gta --g tat att att cag tat g-a aga cat att aat ac- --- --- -tt atc --- --t ttg aca atc ttg ct- -at aat cat --- --- --- --- aat ata aaa --a tac tta tgg --- -a- cat agt cac tca t-- --- --- --- aat tgc tg- acg cga taa att cgt aat aat ctg ttt tgt tca aat ttt tat aag ga- --- --a tct aca ggc ata aaa at- aaa aat ata att tat aat ac- --- act tac agc gcg cca tca tga ata aca gca gtg aat taa ttg ctg tta tta atg gat tta gaa ata gtg gac gat ttt gtg ata tta ata tag tta tta atg atg aaa gga taa atg ctc ata gac tca tcc tat ctg gag cat ccg aat a-t ttt tcc att cta ttt tct agt gat ttt atc gat tct aat gaa tac gaa gtt aat cta agt cat tta gat tat caa agt gtg aac gat ttg atc gat tac att tat ggg ata cct ttg agc cta act aac gat aac gtg aaa tat att ctt tca acc gct ga- --t ttt tta caa att gga tct gcc att act gag tgc gaa aaa tac ata ctt aaa aat ctg tgt tct aga aat tgt atc gat ttc --- --- tac ata tac gct gat aaa tat aat aac aag aaa ata gaa aca gcg tcg ttt aac aca ata tta cga aat att ttg aga ctc att aac gat gaa aac ttt aaa tac ttg aca gag gaa tca atg ata aaa att tta agc gat gat atg tta aat ata aaa aat gag gat ttt gca cca cta att ctc att aaa tgg tta gag agt aca caa caa cca tgc aca gtc gag tta ctt aga tgc ctc aga ata tca -tt gct ttc ccc aca agt tat aaa atc act tta tag tca tcg act ggt tgg ttc aat cta cga atg tat aac att ctt aaa caa tat atc att ctt tga tga atc att tcc tag ata cca tag cat cga gtt gat atc tat cgg tat aag taa ttc aca tga taa gat ttc cat aaa ctg cta taa tcg t-a aaa aaa ata cgt ggg aaa taa tat ctt cac gga gat ata ggt gta gtt ttg cag tgg ccg tca tga ata ata tta tct ata tga tgg gtg gat atg atc agt ccc cgt ata gga gtt caa agg tta tag cgt aca ata cct gta cta att ctt gga tat atg ata tac cag agc t-a aaa tat cct cgt tct aat tgt gga gga gtt gct gat gat gaa tac att tat tgc ata ggc ggc ata cgc gat cag gat tca tcg ttg ata tct aat att gat aga tgg aag cca tca aaa cca tat tgg cag acg tat gct aaa atg cgc gag cca aaa tgt gat atg ggg gtt gcg atg tta aac gga tta ata tat gtc ata ggt gga atc gtt aaa ggt g-- aca cat gta ccg acg cac tag aaa gtt tat cag aag atg gat gga tga agc atc aac gtc ttc caa taa aaa tgt cca ata tgt cga cga ttg ttc atg ccg gca aaa ttt ata tat ctg gag gct aca aca ata gta gtg tag tta atg gaa tat cga atc t-- agt cct tag cta taa tcc gat ata tga tga atg gac caa att atc atc att gaa tat tcc tag aat taa tcc cgc tct a

>KC813509.1:163774-166019_Cowpox_virus_strain_HumBer07/1_complete_genome

ttg ctg act gta --- tgt a-c tat ttt gta cat cag tac att ggt tac agc aga tat acc tac ttc gtc act gcc aca cgc tcc ggt aaa cgg --g tca tgt gac gaa gga gaa tat ctt gat aag acg cat aat cga tgt tgt aat -cg gtg tcc acc tgg aga att tgc caa ggt tag atg ta- --g tgg tag cta taa cac aaa atg tga acg ctg tcc aca tca tac ata tac agc aat tcc caa tta ttc t-a atg gat gcc atc aat gta gaa aat gcc caa agg gat cat ttg ata agg taa agt gta ccg gaa cac aga aca gta aat -gt gcg tgt ctt cct ggt tgg tat tgc gct acc gat tct tca cag act gaa gac tgt cga gat -tg tgt acc aaa aag g-a gct gtc cat gcg gat act ttg gtg gaa tag ata aat tag gaa atc cta ttt gta aat cgt gtt g-- -tg ttg gta aat att gtg acg aca tac gta atc ata gag ttg gtc ctt ttc ctc cat gca aa- --- cta tct aaa tgt aat tag tca --- --- --- --- --- --- --- --- --- --- --- --- --- --- -tg atc atg gta -cc gta caa tt- --- -at att gct --- --g gta --g tat att att cag tat g-a aga cat att aat ac- --- --- -tt atc --- --t ttg aca atc ttg ct- -at aat cat --- --- --- --- aat ata aaa --a tac tta tgg --- -a- cat agt cac tca t-- --- --- --- aat tgc tg- acg cga taa att cgt aat aat ctg ttt tgt tca aat ttt tat aag ga- --- --a tct aca ggc ata aaa at- aaa aat ata att tat aat ac- --- act tac agc gcg cca tca tga ata aca gca gtg aat taa ttg ctg tta tta atg gat tta gaa ata gtg gac gat ttt gtg ata tta ata tag tta tta atg atg aaa gga taa atg ctc ata gac tca tcc tat ctg gag cat ccg aat a-t ttt tcc att cta ttt tct agt gat ttt atc gat tct aat gaa tac gaa gtt aat cta agt cat tta gat tat caa agt gtg aac gat ttg atc gat tac att tat ggg ata cct ttg agc cta act aac gat aac gtg aaa tat att ctt tca acc gct ga- --t ttt tta caa att gga tct gcc att act gag tgc gaa aaa tac ata ctt aaa aat ctg tgt tct aga aat tgt atc gat ttc --- --- tac ata tac gct gat aaa tat aat aac aag aaa ata gaa aca gcg tcg ttt aac aca ata tta cga aat att ttg aga ctc att aac gat gaa aac ttt aaa tac ttg aca gag gaa tca atg ata aaa att tta agc gat gat atg tta aat ata aaa aat gag gat ttt gca cca cta att ctc att aaa tgg tta gag agt aca caa caa cca tgc aca gtc gag tta ctt aga tgc ctc aga ata tca -tt gct ttc ccc aca agt tat aaa atc act tta tag tca tcg act ggt tgg ttc aat cta cga atg tat aac att ctt aaa caa tat atc att ctt tga tga atc att tcc tag ata cca tag cat cga gtt gat atc tat cgg tat aag taa ttc aca tga taa gat ttc cat aaa ctg cta taa tcg t-a aaa aaa ata cgt ggg aaa taa tat ctt cac gga gat ata ggt gta gtt ttg cag tgg ccg tca tga ata ata tta tct ata tga tgg gtg gat atg atc agt ccc cgt ata gga gtt caa agg tta tag cgt aca ata cct gta cta att ctt gga tat atg ata tac cag agc t-a aaa tat cct cgt tct aat tgt gga gga gtt gct gat gat gaa tac att tat tgc ata ggc ggc ata cgc gat cag gat tca tcg ttg ata tct aat att gat aga tgg aag cca tca aaa cca tat tgg cag acg tat gct aaa atg cgc gag cca aaa tgt gat atg ggg gtt gcg atg tta aac gga tta ata tat gtc ata ggt gga atc gtt aaa ggt g-- aca cat gta ccg acg cac tag aaa gtt tat cag aag atg gat gga tga agc atc aac gtc ttc caa taa aaa tgt cca ata tgt cga cga ttg ttc atg ccg gca aaa ttt ata tat ctg gag gct aca aca ata gta gtg tag tta atg gaa tat cga atc t-- agt cct tag cta taa tcc gat ata tga tga atg gac caa att atc atc att gaa tat tcc tag aat taa tcc cgc tct a

>KC813502.1:168963-171208_Cowpox_virus_strain_CatBer07/1_complete_genome

ttg ctg act gta --- tgt a-c tat ttt gta cat cag tac att ggt tac agc aga tat acc tac ttc gtc act gcc aca cgc tcc ggt aaa cgg --g tca tgt gac gaa gga gaa tat ctt gat aag acg cat aat cga tgt tgt aat -cg gtg tcc acc tgg aga att tgc caa ggt tag atg ta- --g tgg tag cta taa cac aaa atg tga acg ctg tcc aca tca tac ata tac agc aat tcc caa tta ttc t-a atg gat gcc atc aat gta gaa aat gcc caa agg gat cat ttg ata agg taa agt gta ccg gaa cac aga aca gta aat -gt gcg tgt ctt cct ggt tgg tat tgc gct acc gat tct tca cag act gaa gac tgt cga gat -tg tgt acc aaa aag g-a gct gtc cat gcg gat act ttg gtg gaa tag ata aat tag gaa atc cta ttt gta aat cgt gtt g-- -tg ttg gta aat att gtg acg aca tac gta atc ata gag ttg gtc ctt ttc ctc cat gca aa- --- cta tct aaa tgt aat tag tca --- --- --- --- --- --- --- --- --- --- --- --- --- --- -tg atc atg gta -cc gta caa tt- --- -at att gct --- --g gta --g tat att att cag tat g-a aga cat att aat ac- --- --- -tt atc --- --t ttg aca atc ttg ct- -at aat cat --- --- --- --- aat ata aaa --a tac tta tgg --- -a- cat agt cac tca t-- --- --- --- aat tgc tg- acg cga taa att cgt aat aat ctg ttt tgt tca aat ttt tat aag ga- --- --a tct aca ggc ata aaa at- aaa aat ata att tat aat ac- --- act tac agc gcg cca tca tga ata aca gca gtg aat taa ttg ctg tta tta atg gat tta gaa ata gtg gac gat ttt gtg ata tta ata tag tta tta atg atg aaa gga taa atg ctc ata gac tca tcc tat ctg gag cat ccg aat a-t ttt tcc att cta ttt tct agt gat ttt atc gat tct aat gaa tac gaa gtt aat cta agt cat tta gat tat caa agt gtg aac gat ttg atc gat tac att tat ggg ata cct ttg agc cta act aac gat aac gtg aaa tat att ctt tca acc gct ga- --t ttt tta caa att gga tct gcc att act gag tgc gaa aaa tac ata ctt aaa aat ctg tgt tct aga aat tgt atc gat ttc --- --- tac ata tac gct gat aaa tat aat aac aag aaa ata gaa aca gcg tcg ttt aac aca ata tta cga aat att ttg aga ctc att aac gat gaa aac ttt aaa tac ttg aca gag gaa tca atg ata aaa att tta agc gat gat atg tta aat ata aaa aat gag gat ttt gca cca cta att ctc att aaa tgg tta gag agt aca caa caa cca tgc aca gtc gag tta ctt aga tgc ctc aga ata tca -tt gct ttc ccc aca agt tat aaa atc act tta tag tca tcg act ggt tgg ttc aat cta cga atg tat aac att ctt aaa caa tat atc att ctt tga tga atc att tcc tag ata cca tag cat cga gtt gat atc tat cgg tat aag taa ttc aca tga taa gat ttc cat aaa ctg cta taa tcg t-a aaa aaa ata cgt ggg aaa taa tat ctt cac gga gat ata ggt gta gtt ttg cag tgg ccg tca tga ata ata tta tct ata tga tgg gtg gat atg atc agt ccc cgt ata gga gtt caa agg tta tag cgt aca ata cct gta cta att ctt gga tat atg ata tac cag agc t-a aaa tat cct cgt tct aat tgt gga gga gtt gct gat gat gaa tac att tat tgc ata ggc ggc ata cgc gat cag gat tca tcg ttg ata tct aat att gat aga tgg aag cca tca aaa cca tat tgg cag acg tat gct aaa atg cgc gag cca aaa tgt gat atg ggg gtt gcg atg tta aac gga tta ata tat gtc ata ggt gga atc gtt aaa ggt g-- aca cat gta ccg acg cac tag aaa gtt tat cag aag atg gat gga tga agc atc aac gtc ttc caa taa aaa tgt cca ata tgt cga cga ttg ttc atg ccg gca aaa ttt ata tat ctg gag gct aca aca ata gta gtg tag tta atg gaa tat cga atc t-- agt cct tag cta taa tcc gat ata tga tga atg gac caa att atc atc att gaa tat tcc tag aat taa tcc cgc tct a

>KC813495.1:169869-172114_Cowpox_virus_strain_HumMag07/1_complete_genome

ttg ctg act gta --- tgt a-c tat ttt gta cat cag tac att ggt tac agc aga tat acc tac ttc gtc act gcc aca cgc tcc ggt aaa cgg --g tca tgt gac gaa gga gaa tat ctt gat aag acg cat aat cga tgt tgt aat -cg gtg tcc acc tgg aga att tgc caa ggt tag atg ta- --g tgg tag cta taa cac aaa atg tga acg ctg tcc aca tca tac ata tac agc aat tcc caa tta ttc t-a atg gat gcc atc aat gta gaa aat gcc caa agg gat cat ttg ata agg taa agt gta ccg gaa cac aga aca gta aat -gt gcg tgt ctt cct ggt tgg tat tgc gct acc gat tct tca cag act gaa gac tgt cga gat -tg tgt acc aaa aag g-a gct gtc cat gcg gat act ttg gtg gaa tag ata aat tag gaa atc cta ttt gta aat cgt gtt g-- -tg ttg gta aat att gtg acg aca tac gta atc ata gag ttg gtc ctt ttc ctc cat gca aa- --- cta tct aaa tgt aat tag tca --- --- --- --- --- --- --- --- --- --- --- --- --- --- -tg atc atg gta -cc gta caa tt- --- -at att gct --- --g gta --g tat att att cag tat g-a aga cat att aat ac- --- --- -tt atc --- --t ttg aca atc ttg ct- -at aat cat --- --- --- --- aat ata aaa --a tac tta tgg --- -a- cat agt cac tca t-- --- --- --- aat tgc tg- acg cga taa att cgt aat aat ctg ttt tgt tca aat ttt tat aag ga- --- --a tct aca ggc ata aaa at- aaa aat ata att tat aat ac- --- act tac agc gcg cca tca tga ata aca gca gtg aat taa ttg ctg tta tta atg gat tta gaa ata gtg gac gat ttt gtg ata tta ata tag tta tta atg atg aaa gga taa atg ctc ata gac tca tcc tat ctg gag cat ccg aat a-t ttt tcc att cta ttt tct agt gat ttt atc gat tct aat gaa tac gaa gtt aat cta agt cat tta gat tat caa agt gtg aac gat ttg atc gat tac att tat ggg ata cct ttg agc cta act aac gat aac gtg aaa tat att ctt tca acc gct ga- --t ttt tta caa att gga tct gcc att act gag tgc gaa aaa tac ata ctt aaa aat ctg tgt tct aga aat tgt atc gat ttc --- --- tac ata tac gct gat aaa tat aat aac aag aaa ata gaa aca gcg tcg ttt aac aca ata tta cga aat att ttg aga ctc att aac gat gaa aac ttt aaa tac ttg aca gag gaa tca atg ata aaa att tta agc gat gat atg tta aat ata aaa aat gag gat ttt gca cca cta att ctc att aaa tgg tta gag agt aca caa caa cca tgc aca gtc gag tta ctt aga tgc ctc aga ata tca -tt gct ttc ccc aca agt tat aaa atc act tta tag tca tcg act ggt tgg ttc aat cta cga atg tat aac att ctt aaa caa tat atc att ctt tga tga atc att tcc tag ata cca tag cat cga gtt gat atc tat cgg tat aag taa ttc aca tga taa gat ttc cat aaa ctg cta taa tcg t-a aaa aaa ata cgt ggg aaa taa tat ctt cac gga gat ata ggt gta gtt ttg cag tgg ccg tca tga ata ata tta tct ata tga tgg gtg gat atg atc agt ccc cgt ata gga gtt caa agg tta tag cgt aca ata cct gta cta att ctt gga tat atg ata tac cag agc t-a aaa tat cct cgt tct aat tgt gga gga gtt gct gat gat gaa tac att tat tgc ata ggc ggc ata cgc gat cag gat tca tcg ttg ata tct aat att gat aga tgg aag cca tca aaa cca tat tgg cag acg tat gct aaa atg cgc gag cca aaa tgt gat atg ggg gtt gcg atg tta aac gga tta ata tat gtc ata ggt gga atc gtt aaa ggt g-- aca cat gta ccg acg cac tag aaa gtt tat cag aag atg gat gga tga agc atc aac gtc ttc caa taa aaa tgt cca ata tgt cga cga ttg ttc atg ccg gca aaa ttt ata tat ctg gag gct aca aca ata gta gtg tag tta atg gaa tat cga atc t-- agt cct tag cta taa tcc gat ata tga tga atg gac caa att atc atc att gaa tat tcc tag aat taa tcc cgc tct a

>LT896732.2:176746-178991_Cowpox_virus_isolate_Ger/2017/Alpaca2_genome_assembly_complete_genome:_monopartite

ttg ctg act gca --- tgt a-c tat ttt tta cat tac tac att ggc tac ggc aga tat acc tac ttc gtc act gcc aca cgc tcc ggt aaa cgg --g gca tgt gac gag gga gaa tat ctt gat aag act cat aat cga tgt tgt aat -cg gtg tcc acc tgg aga att tgc caa ggt tag atg ta- --g tgg tag cta taa cac aaa atg tga acg ctg tcc aca tca tac ata tac agc aat tcc caa tta ttc t-a atg gat gcc atc aat gta gaa aat gcc caa agg gat cat ttg ata agg taa agt gta ccg gaa cac aga aca gta aat -gt gcg tgt ctt cct ggt tgg tat tgc gct acc gat tct tca cag act gaa gac tgt cga gat -tg tgt acc aaa aag g-a gct gtc cat gcg gat act ttg gtg gaa tag ata aat tag gaa atc cta ttt gta aat cgt gtt g-- -tg ttg gta aat att gtg acg aca tac gta atc ata gag ttg gtc ctt ttc ctc cat gca aa- --- cta tct aaa tgt aat tag tca --- --- --- --- --- --- --- --- --- --- --- --- --- --- -tg atc atg gta -cc gta caa tt- --- -at att gct --- --g gta --g tat att att cag tat g-a aga cat att aat ac- --- --- -tt atc --- --t ttg aca atc ttg ct- -at aat cat --- --- --- --- aat ata aaa --a tac tta tgg --- -a- cat agt cac tca t-- --- --- --- aat tgc tg- acg cga taa att cgt aat aat ctg ttt tgt tca aat ttt tat aag ga- --- --a tct aca ggc ata aaa at- aaa aat ata att tat aat ac- --- act tac agc gcg cca tca tga ata aca gca gtg aat taa ttg ctg tta tta atg gat tta gaa ata gtg gac gat ttt gtg ata tta ata tag tta tta atg atg aaa gga taa atg ctc ata gac tca tcc tat ctg gag cat ccg aat a-t ttt tcc att cta ttt tct agt gat ttt atc gat tct aat gaa tac gaa gtt aat cta agt cat tta gat tat caa agt gtg aac gat ttg atc gat tac att tat ggg ata cct ttg agc cta act aac gat aac gtg aaa tat att ctt tca acc gct ga- --t ttt tta caa att gga tct gcc att act gag tgc gaa aaa tac ata ctt aaa aat ctg tgt tct aga aat tgt atc gat ttc --- --- tac ata tac gct gat aaa tat aat aac aag aaa ata gaa aca gcg tcg ttt aac aca ata tta cga aat att ttg aga ctc att aac gat gaa aac ttt aaa tac ttg aca gag gaa tca atg ata aaa att tta agc gat gat atg tta aat ata aaa aat gag gat ttt gca cca cta att ctc att aaa tgg tta gag agt aca caa caa cca tgc aca gtc gag tta ctt aga tgc ctc aga ata tca -tt gct ttc ccc aca agt tat aaa atc act tta tag tca tcg att ggt tgg ttc aat ata cga atg tat aac att ctt aaa caa tat atc att ctt gga tga atc att tcc tag ata cca tag cat cga gtt gat atc tat cgg tat aag taa ttc aca tga taa gat ttc cat aaa ctg cta taa tcg t-a aaa aaa ata cgt ggg aaa taa tat ctt cac gga gat ata ggt gta gtt ttg cag tgg ccg tca tga ata ata tta tct ata tga tgg gtg gat atg atc agt ccc cgt ata gga gtt caa agg tta tag cgt aca ata cat gta cta att ctt gga tat atg ata tac cag agc t-a aaa tat cct cgt tct aat tgt gga gga gtt gct gat gat gaa tac att tat tgc ata ggc ggc ata cgc gat cag gat tca tcg ttg ata tct agt att gat aaa tgg aag cca tca aaa cca tat tgg caa acg tat gct aaa atg cga gag cca aaa tgt gat atg ggt gta gcg atg tta aac gga tta ata tat gtc ata ggc gga gtt gtt aaa ggt g-- aca cgt gta ccg aca cac ttg aga gtt tat cag aag atg gat gga tga tgc atc aac gtc ttc caa taa aaa tgt cca ata tgt cga cga ttg ttc atg ccg gca aaa ttt ata tat ctg gag gct aca aca ata gta gtg tag tta atg gaa tat cga atc t-- agt cct tag cta taa tcc gat ata tga tga atg gac caa att atc atc att aaa tat tcc tag aat taa tcc cgc tct a

>KC813511.1:169742-171987_Cowpox_virus_strain_HumGri07/1_complete_genome

ttg ctg act gca --- tgt a-c tat ttt tta cat tac tac att ggc tac ggc aga tat acc tac ttc gtc act gcc aca cgc tcc ggt aaa cgg --g gca tgt gac gag gga gaa tat ctt gat aag act cat aat cga tgt tgt aat -cg gtg tcc acc tgg aga att tgc caa ggt tag atg ta- --g tgg tag cta taa cac aaa atg tga acg ctg tcc aca tca tac ata tac agc aat tcc caa tta ttc t-a atg gat gcc atc aat gta gaa aat gcc caa agg gat cat ttg ata agg taa agt gta ccg gaa cac aga aca gta aat -gt gcg tgt ctt cct ggt tgg tat tgc gct acc gat tct tca cag act gaa gac tgt cga gat -tg tgt acc aaa aag g-a gct gtc cat gcg gat act ttg gtg gaa tag ata aat tag gaa atc cta ttt gta aat cgt gtt g-- -tg ttg gta aat att gtg acg aca tac gta atc ata gag ttg gtc ctt ttc ctc cat gca aa- --- cta tct aaa tgt aat tag tca --- --- --- --- --- --- --- --- --- --- --- --- --- --- -tg atc atg gta -cc gta caa tt- --- -at att gct --- --g gta --g tat att att cag tat g-a aga cat att aat ac- --- --- -tt atc --- --t ttg aca atc ttg ct- -at aat cat --- --- --- --- aat ata aaa --a tac tta tgg --- -a- cat agt cac tca t-- --- --- --- aat tgc tg- acg cga taa att cgt aat aat ctg ttt tgt tca aat ttt tat aag ga- --- --a tct aca ggc ata aaa at- aaa aat ata att tat aat ac- --- act tac agc gcg cca tca tga ata aca gca gtg aat taa ttg ctg tta tta atg gat tta gaa ata gtg gac gat ttt gtg ata tta ata tag tta tta atg atg aaa gga taa atg ctc ata gac tca tcc tat ctg gag cat ccg aat a-t ttt tcc att cta ttt tct agt gat ttt atc gat tct aat gaa tac gaa gtt aat cta agt cat tta gat tat caa agt gtg aac gat ttg atc gat tac att tat ggg ata cct ttg agc cta act aac gat aac gtg aaa tat att ctt tca acc gct ga- --t ttt tta caa att gga tct gcc att act gag tgc gaa aaa tac ata ctt aaa aat ctg tgt tct aga aat tgt atc gat ttc --- --- tac ata tac gct gat aaa tat aat aac aag aaa ata gaa aca gcg tcg ttt aac aca ata tta cga aat att ttg aga ctc att aac gat gaa aac ttt aaa tac ttg aca gag gaa tca atg ata aaa att tta agc gat gat atg tta aat ata aaa aat gag gat ttt gca cca cta att ctc att aaa tgg tta gag agt aca caa caa cca tgc aca gtc gag tta ctt aga tgc ctc aga ata tca -tt gct ttc ccc aca agt tat aaa atc act tta tag tca tcg att ggt tgg ttc aat ata cga atg tat aac att ctt aaa caa tat atc att ctt gga tga atc att tcc tag ata cca tag cat cga gtt gat atc tat cgg tat aag taa ttc aca tga taa gat ttc cat aaa ctg cta taa tcg t-a aaa aaa ata cgt ggg aaa taa tat ctt cac gga gat ata ggt gta gtt ttg cag tgg ccg tca tga ata ata tta tct ata tga tgg gtg gat atg atc agt ccc cgt ata gga gtt caa agg tta tag cgt aca ata cct gta cta att ctt gga tat atg ata tac cag agc t-a aaa tat cct cgt tct aat tgt gga gga gtt gct gat gat gaa tac att tat tgc ata ggc ggc ata cgc gat cag gat tca tcg ttg ata tct agt att gat aga tgg aag cca tca aaa cca tat tgg caa acg tat gct aaa atg cga gag cca aaa tgt gat atg ggt gta gcg atg tta aac gga tta ata tat gtc ata ggc gga gtt gtt aaa ggt g-- aca cgt gta ccg aca cac ttg aga gtt tat cag aag atg gat gga tga tgc atc aac gtc ttc caa taa aaa tgt cca ata tgt cga cga ttg ttc atg ccg gca aaa ttt ata tat ctg gag gct aca aca ata gta gtg tag tta atg gaa tat cga atc t-- agt cct tag cta taa tcc gat ata tga tga atg gac caa att atc atc att aaa tat tcc tag aat taa tcc cgc tct a

>KC813507.1:158221-160466_Cowpox_virus_strain_EleGri07/1_complete_genome

ttg ctg act gca --- tgt a-c tat ttt tta cat tac tac att ggc tac ggc aga tat acc tac ttc gtc act gcc aca cgc tcc ggt aaa cgg --g gca tgt gac gag gga gaa tat ctt gat aag act cat aat cga tgt tgt aat -cg gtg tcc acc tgg aga att tgc caa ggt tag atg ta- --g tgg tag cta taa cac aaa atg tga acg ctg tcc aca tca tac ata tac agc aat tcc caa tta ttc t-a atg gat gcc atc aat gta gaa aat gcc caa agg gat cat ttg ata agg taa agt gta ccg gaa cac aga aca gta aat -gt gcg tgt ctt cct ggt tgg tat tgc gct acc gat tct tca cag act gaa gac tgt cga gat -tg tgt acc aaa aag g-a gct gtc cat gcg gat act ttg gtg gaa tag ata aat tag gaa atc cta ttt gta aat cgt gtt g-- -tg ttg gta aat att gtg acg aca tac gta atc ata gag ttg gtc ctt ttc ctc cat gca aa- --- cta tct aaa tgt aat tag tca --- --- --- --- --- --- --- --- --- --- --- --- --- --- -tg atc atg gta -cc gta caa tt- --- -at att gct --- --g gta --g tat att att cag tat g-a aga cat att aat ac- --- --- -tt atc --- --t ttg aca atc ttg ct- -at aat cat --- --- --- --- aat ata aaa --a tac tta tgg --- -a- cat agt cac tca t-- --- --- --- aat tgc tg- acg cga taa att cgt aat aat ctg ttt tgt tca aat ttt tat aag ga- --- --a tct aca ggc ata aaa at- aaa aat ata att tat aat ac- --- act tac agc gcg cca tca tga ata aca gca gtg aat taa ttg ctg tta tta atg gat tta gaa ata gtg gac gat ttt gtg ata tta ata tag tta tta atg atg aaa gga taa atg ctc ata gac tca tcc tat ctg gag cat ccg aat a-t ttt tcc att cta ttt tct agt gat ttt atc gat tct aat gaa tac gaa gtt aat cta agt cat tta gat tat caa agt gtg aac gat ttg atc gat tac att tat ggg ata cct ttg agc cta act aac gat aac gtg aaa tat att ctt tca acc gct ga- --t ttt tta caa att gga tct gcc att act gag tgc gaa aaa tac ata ctt aaa aat ctg tgt tct aga aat tgt atc gat ttc --- --- tac ata tac gct gat aaa tat aat aac aag aaa ata gaa aca gcg tcg ttt aac aca ata tta cga aat att ttg aga ctc att aac gat gaa aac ttt aaa tac ttg aca gag gaa tca atg ata aaa att tta agc gat gat atg tta aat ata aaa aat gag gat ttt gca cca cta att ctc att aaa tgg tta gag agt aca caa caa cca tgc aca gtc gag tta ctt aga tgc ctc aga ata tca -tt gct ttc ccc aca agt tat aaa atc act tta tag tca tcg att ggt tgg ttc aat ata cga atg tat aac att ctt aaa caa tat atc att ctt gga tga atc att tcc tag ata cca tag cat cga gtt gat atc tat cgg tat aag taa ttc aca tga taa gat ttc cat aaa ctg cta taa tcg t-a aaa aaa ata cgt ggg aaa taa tat ctt cac gga gat ata ggt gta gtt ttg cag tgg ccg tca tga ata ata tta tct ata tga tgg gtg gat atg atc agt ccc cgt ata gga gtt caa agg tta tag cgt aca ata cct gta cta att ctt gga tat atg ata tac cag agc t-a aaa tat cct cgt tct aat tgt gga gga gtt gct gat gat gaa tac att tat tgc ata ggc ggc ata cgc gat cag gat tca tcg ttg ata tct agt att gat aga tgg aag cca tca aaa cca tat tgg caa acg tat gct aaa atg cga gag cca aaa tgt gat atg ggt gta gcg atg tta aac gga tta ata tat gtc ata ggc gga gtt gtt aaa ggt g-- aca cgt gta ccg aca cac ttg aga gtt tat cag aag atg gat gga tga tgc atc aac gtc ttc caa taa aaa tgt cca ata tgt cga cga ttg ttc atg ccg gca aaa ttt ata tat ctg gag gct aca aca ata gta gtg tag tta atg gaa tat cga atc t-- agt cct tag cta taa tcc gat ata tga tga atg gac caa att atc atc att aaa tat tcc tag aat taa tcc cgc tct a

>KC813491.1:169357-171602_Cowpox_virus_strain_BeaBer04/1_complete_genome

ttg ctg act gta --- tgt a-c tat ttt gta cat cag tac att ggt tac agc aga tat acc tac ttc gtc act gcc aca cgc tcc ggt aaa cgg --g tca tgt gac gaa gga gaa tat ctt gat aag acg cat aat cga tgt tgt aat -cg gtg tcc acc tgg aga att tgc caa ggt tag atg ta- --g tgg tag cta taa cac aaa atg tga acg ctg tcc aca tca tac ata tac agc aat tcc caa tta ttc t-a atg gat gcc atc aat gta gaa aat gcc caa agg gat cat ttg ata agg taa agt gta ccg gaa cac aga aca gta aat -gt gcg tgt ctt cct ggt tgg tat tgc gct acc gat tct tca cag act gaa gac tgt cga gat -tg tgt acc aaa aag g-a gct gtc cat gcg gat act ttg gtg gaa tag ata aat tag gaa atc cta ttt gta aat cgt gtt g-- -tg ttg gta aat att gtg acg aca tac gta atc ata gag ttg gtc ctt ttc ctc cat gca aa- --- cta tct aaa tgt aat tag tca --- --- --- --- --- --- --- --- --- --- --- --- --- --- -tg atc atg gta -cc gta caa tt- --- -at att gct --- --g gta --g tat att att cag tat g-a aga cat att aat ac- --- --- -tt atc --- --t ttg aca atc ttg ct- -at aat cat --- --- --- --- aat ata aaa --a tac tta tgg --- -a- cat agt cac tca t-- --- --- --- aat tgc tg- acg cga taa att cgt aat aat ctg ttt tgt tca aat ttt tat aag ga- --- --a tct aca ggc ata aaa at- aaa aat ata att tat aat ac- --- act tac agc gcg cca tca tga ata aca gca gtg aat taa ttg ctg tta tta atg gat tta gaa ata gtg gac gat ttt gtg ata tta ata tag tta tta atg atg aaa gga taa atg ctc ata gac tca tcc tat ctg gag cat ccg aat a-t ttt tcc att cta ttt tct agt gat ttt atc gat tct aat gaa tac gaa gtt aat cta agt cat tta gat tat caa agt gtg aac gat ttg atc gat tac att tat ggg ata cct ttg agc cta act aac gat aac gtg aaa tat att ctt tca acc gct ga- --t ttt tta caa att gga tct gcc att act gag tgc gaa aaa tac ata ctt aaa aat ctg tgt tct aga aat tgt atc gat ttc --- --- tac ata tac gct gat aaa tat aat aac aag aaa ata gaa aca gcg tcg ttt aac aca ata tta cga aat att ttg aga ctc att aac gat gaa aac ttt aaa tac ttg aca gag gaa tca atg ata aaa att tta agc gat gat atg tta aat ata aaa aat gag gat ttt gca cca cta att ctc att aaa tgg tta gag agt aca caa caa cca tgc aca gtc gag tta ctt aga tgc ctc aga ata tca -tt gct ttc ccc aca agt tat aaa atc act tta tag tca tcg att ggt tgg ttc aat ata cga atg tat aac att ctt aaa caa tat atc att ctt gga tga atc att tcc tag ata cca tag cat cga gtt gat atc tat cgg tat aag taa ttc aca tga taa gat ttc cat aaa ctg cta taa tcg t-a aaa aaa ata cgt ggg aaa taa tat ctt cac gga gat ata ggt gta gtt ttg cag tgg ccg tca tga ata ata tta tct ata tga tgg gtg gat atg atc agt ccc cgt ata gga gtt caa agg tta tag cgt aca ata cct gta cta att ctt gga tat atg ata tac cag agc t-a aaa tat cct cgt tct aat tgt gga gga gtt gct gat gat gaa tac att tat tgc ata ggc ggc ata cgc gat cag gat tca tcg ttg ata tct aat att gat aga tgg aag cca tca aaa cca tat tgg cag acg tat gct aaa atg cgc gag cca aaa tgt gat atg ggg gtt gcg atg tta aac gga tta ata tat gtc ata ggt gga atc gtt aaa ggt g-- aca cat gta ccg acg cac tag aaa gtt tat cag aag atg gat gga tga agc atc aac gtc ttc caa taa aaa tgt cca ata tgt cga cga ttg ttc atg ccg gca aaa ttt ata tat ctg gag gct aca aca ata gta gtg tag tta atg gaa tat cga atc t-- agt cct tag cta taa tcc gat ata tga tga atg gac caa att atc atc att gaa tat tcc tag aat taa tcc cgc tct a

>LT993230.1:175800-178042_Cowpox_virus_isolate_Ger/2017/Alpaca1_genome_assembly_complete_genome:_monopartite

ttg ctg act gca --- tgt a-c tat ttt tta cat tac tac att ggc tac ggc aga tat acc tac ttc gtc act gcc aca cgc tcc ggt aaa cgg --g gca tgt gac gag gga gaa tat ctt gat aag act cat aat cga tgt tgt aat -cg gtg tcc acc tgg aga att tgc caa ggt tag atg ta- --g tgg tag cta taa cac aaa atg tga acg ctg tcc aca tca tac ata tac agc aat tcc caa tta ttc t-a atg gat gcc atc aat gta gaa aat gcc caa agg gat cat ttg ata agg taa agt gta ccg gaa cac aga aca gta aat -gt gcg tgt ctt cct ggt tgg tat tgc gct acc gat tct tca cag act gaa gac tgt cga gat -tg tgt acc aaa aag g-a gct gtc cat gcg gat act ttg gtg gaa tag ata aat tag gaa atc cta ttt gta aat cgt gtt g-- -tg ttg gta aat att gtg acg aca tac gta atc ata gag ttg gtc ctt ttc ctc cat gca aa- --- cta tct aaa tgt aat tag tca --- --- --- --- --- --- --- --- --- --- --- --- --- --- -tg atc atg gta -cc gta caa tt- --- -at att gct --- --g gta --g tat att att cag tat g-a aga cat att aat ac- --- --- -tt atc --- --t ttg aca atc ttg ct- -at aat cat --- --- --- --- aat ata aaa --a tac tta tgg --- -a- cat agt cac tca t-- --- --- --- aat tgc tg- acg cga taa att cgt aat aat ctg ttt tgt tca aat ttt tat aag ga- --- --a tct aca ggc ata aaa at- aaa aat ata att tat aat ac- --- act tac agc gcg cca tca tga ata aca gca gtg aat taa ttg ctg tta tta atg gat tta gaa ata gtg gac gat ttt gtg ata tta ata tag tta tta atg atg aaa gga taa atg ctc ata gac tca tcc tat ctg gag cat ccg aat a-t ttt tcc att cta ttt tct agt gat ttt atc gat tct aat gaa tac gaa gtt aat cta agt cat tta gat tat caa agt gtg aac gat ttg atc gat tac att tat ggg ata cct ttg agc cta act aac gat aac gtg aaa tat att ctt tca acc gct ga- --t ttt tta caa att gga tct gcc att act gag tgc gaa aaa tac ata ctt aaa aat ctg tgt tct aga aat tgt atc gat ttc --- --- tac ata tac gct gat aaa tat aat aac gag aaa ata gaa aca gcg tcg ttt aac aca ata tta cga aat att ttg aga ctc att aac gat gaa aac ttt aaa tac ttg aca gag gaa tca atg ata aaa att tta agc gat gat atg tta aat ata aaa aat gag gat ttt gca cca cta att ctc att aaa tgg tta gag agt aca caa caa cca tgc aca gtc gag tta ctt aga tgc ctc aga ata tca -tt gct ttc ccc aca agt tat aaa atc act tta tag tca tcg att ggt tgg ttc aat ata cga atg tat aac att ctt aaa caa tat atc att ctt gga tga atc att tcc tag ata cca tag cat cga gtt gat atc tat cgg tat aag taa ttc aca tga taa gat ttc cat aaa ctg cta taa tcg t-a aaa aaa ata cgt ggg aaa taa tat ctt cac gga gat ata ggt gta gtt ttg cag tgg ccg tca tgg ata ata tta tct ata tga tgg gtg gat atg atc agt ccc cgt ata gga gtt caa agg tta tag cgt aca ata cct gta cta att ctt gga tat atg ata tac cag agc t-a aaa tat cct cgt tct aat tgt gga gga gtt gct gat gat gaa tac att tat tgc ata ggc ggc ata cgc gat cag gat tca tcg ttg ata tct agt att gat aga tgg aag cca tca aaa cca tat tgg caa acg tat gct aaa atg cga gag cca aaa tgt gat atg ggg gtt gcg atg tta aac gga tta ata tat gtc ata ggt gga atc gtt aaa ggt g-- aca cat gta ccg acg cac tag aaa gtt tat cag aag atg gat gga tga agc atc aac gtc ttc caa taa aaa tgt cca ata tgt cga cga ttg ttc atg ctg gta aga ttt ata tat ctg gag gtt ata aca ata gta gtg ccg t-- -tg gac cat cga atc t-- agt cct tag cta taa tcc gat ata tga tga atg gac caa att atc atc att aaa tat tcc tag aat taa tcc cgc tct a

>LT993228.1:175937-178179_Cowpox_virus_isolate_Ger/2017/common_vole_FMEimka_genome_assembly_complete_genome:_monopartite

ttg ctg act gca --- tgt a-c tat ttt tta cat tac tac att ggc tac ggc aga tat acc tac ttc gtc act gcc aca cgc tcc ggt aaa cgg --g gca tgt gac gag gga gaa tat ctt gat aag act cat aat cga tgt tgt aat -cg gtg tcc acc tgg aga att tgc caa ggt tag atg ta- --g tgg tag cta taa cac aaa atg tga acg ctg tcc aca tca tac ata tac agc aat tcc caa tta ttc t-a atg gat gcc atc aat gta gaa aat gcc caa agg gat cat ttg ata agg taa agt gta ccg gaa cac aga aca gta aat -gt gcg tgt ctt cct ggt tgg tat tgc gct acc gat tct tca cag act gaa gac tgt cga gat -tg tgt acc aaa aag g-a gct gtc cat gcg gat act ttg gtg gaa tag ata aat tag gaa atc cta ttt gta aat cgt gtt g-- -tg ttg gta aat att gtg acg aca tac gta atc ata gag ttg gtc ctt ttc ctc cat gca aa- --- cta tct aaa tgt aat tag tca --- --- --- --- --- --- --- --- --- --- --- --- --- --- -tg atc atg gta -cc gta caa tt- --- -at att gct --- --g gta --g tat att att cag tat g-a aga cat att aat ac- --- --- -tt atc --- --t ttg aca atc ttg ct- -at aat cat --- --- --- --- aat ata aaa --a tac tta tgg --- -a- cat agt cac tca t-- --- --- --- aat tgc tg- acg cga taa att cgt aat aat ctg ttt tgt tca aat ttt tat aag ga- --- --a tct aca ggc ata aaa at- aaa aat ata att tat aat ac- --- act tac agc gcg cca tca tga ata aca gca gtg aat taa ttg ctg tta tta atg gat tta gaa ata gtg gac gat ttt gtg ata tta ata tag tta tta atg atg aaa gga taa atg ctc ata gac tca tcc tat ctg gag cat ccg aat a-t ttt tcc att cta ttt tct agt gat ttt atc gat tct aat gaa tac gaa gtt aat cta agt cat tta gat tat caa agt gtg aac gat ttg atc gat tac att tat ggg ata cct ttg agc cta act aac gat aac gtg aaa tat att ctt tca acc gct ga- --t ttt tta caa att gga tct gcc att act gag tgc gaa aaa tac ata ctt aaa aat ctg tgt tct aga aat tgt atc gat ttc --- --- tac ata tac gct gat aaa tat aat aac gag aaa ata gaa aca gcg tcg ttt aac aca ata tta cga aat att ttg aga ctc att aac gat gaa aac ttt aaa tac ttg aca gag gaa tca atg ata aaa att tta agc gat gat atg tta aat ata aaa aat gag gat ttt gca cca cta att ctc att aaa tgg tta gag agt aca caa caa cca tgc aca gtc gag tta ctt aga tgc ctc aga ata tca -tt gct ttc ccc aca agt tat aaa atc act tta tag tca tcg att ggt tgg ttc aat ata cga atg tat aac att ctt aaa caa tat atc att ctt gga tga atc att tcc tag ata cca tag cat cga gtt gat atc tat cgg tat aag taa ttc aca tga taa gat ttc cat aaa ctg cta taa tcg t-a aaa aaa ata cgt ggg aaa taa tat ctt cac gga gat ata ggt gta gtt ttg cag tgg ccg tca tgg ata ata tta tct ata tga tgg gtg gat atg atc agt ccc cgt ata gga gtt caa agg tta tag cgt aca ata cct gta cta att ctt gga tat atg ata tac cag agc t-a aaa tat cct cgt tct aat tgt gga gga gtt gct gat gat gaa tac att tat tgc ata ggc ggc ata cgc gat cag gat tca tcg ttg ata tct agt att gat aga tgg aag cca tca aaa cca tat tgg caa acg tat gct aaa atg cga gag cca aaa tgt gat atg ggg gtt gcg atg tta aac gga tta ata tat gtc ata ggt gga atc gtt aaa ggt g-- aca cat gta ccg acg cac tag aaa gtt tat cag aag atg gat gga tga agc atc aac gtc ttc caa taa aaa tgt cca ata tgt cga cga ttg ttc atg ctg gta aga ttt ata tat ctg gag gtt ata aca ata gta gtg ccg t-- -tg gac cat cga atc t-- agt cct tag cta taa tcc gat ata tga tga atg gac caa att atc atc att aaa tat tcc tag aat taa tcc cgc tct a

>LT896723.1:176491-178736_Cowpox_virus_isolate_Ger/2014/Cat1_genome_assembly_complete_genome:_monopartite

ttg ctg act gca --- tgt a-c tat ttt tta cat tac tac att ggc tac ggc aga tat acc tac ttc gtc act gcc aca cgc tcc ggt aaa cgg --g gca tgt gac gag gga gaa tat ctt gat aag act cat aat cga tgt tgt aat -cg gtg tcc acc tgg aga att tgc caa ggt tag atg ta- --g tgg tag cta taa cac aaa atg tga acg ctg tcc aca tca tac ata tac agc aat tcc caa tta ttc t-a atg gat gcc atc aat gta gaa aat gcc caa agg gat cat ttg ata agg taa agt gta ccg gaa cac aga aca gta aat -gt gcg tgt ctt cct ggt tgg tat tgc gct acc gat tct tca cag act gaa gac tgt cga gat -tg tgt acc aaa aag g-a gct gtc cat gcg gat act ttg gtg gaa tag ata aat tag gaa atc cta ttt gta aat cgt gtt g-- -tg ttg gta aat att gtg acg aca tac gta atc ata gag ttg gtc ctt ttc ctc cat gca aa- --- cta tct aaa tgt aat tag tca --- --- --- --- --- --- --- --- --- --- --- --- --- --- -tg atc atg gta -cc gta caa tt- --- -at att gct --- --g gta --g tat att att cag tat g-a aga cat att aat ac- --- --- -tt atc --- --t ttg aca atc ttg ct- -at aat cat --- --- --- --- aat ata aaa --a tac tta tgg --- -a- cat agt cac tca t-- --- --- --- aat tgc tg- acg cga taa att cgt aat aat ctg ttt tgt tca aat ttt tat aag ga- --- --a tct aca ggc ata aaa at- aaa aat ata att tat aat ac- --- act tac agc gcg cca tca tga ata aca gca gtg aat taa ttg ctg tta tta atg gat tta gaa ata gtg gac gat ttt gtg ata tta ata tag tta tta atg atg aaa gga taa atg ctc ata gac tca tcc tat ctg gag cat ccg aat a-t ttt tcc att cta ttt tct agt gat ttt atc gat tct aat gaa tac gaa gtt aat cta agt cat tta gat tat caa agt gtg aac gat ttg atc gat tac att tat ggg ata cct ttg agc cta act aac gat agc gtg aaa tat att ctt tca acc gct ga- --t ttt tta caa att gga tct gcc att act gag tgc gaa aaa tac ata ctt aaa aat ctg tgt tct aga aat tgt atc gat ttc --- --- tac ata tac gct gat aaa tat aat aac aag aaa ata gaa aca gcg tcg ttt aac aca ata tta cga aat att ttg aga ctc att aac gat gaa aac ttt aaa tac ttg aca gag gaa tca atg ata aaa att tta agc gat gat atg tta aat ata aaa aat gag gat ttt gca cca cta att ctc att aaa tgg tta gag agt aca caa caa cca tgc aca gtc gag tta ctt aga tgc ctc aga ata tca -tt gct ttc ccc aca agt tat aaa atc act tta tag tca tcg att ggt tgg ttc aat ata cga atg tat aac att ctt aaa caa tat atc att ctt gga tga atc att tcc tag ata cca tag cat cga gtt gat atc tat cgg tat aag taa ttc aca tga taa gat ttc cat aaa ctg cta taa tcg t-a aaa aaa ata cgt ggg aaa taa tat ctt cac gga gat ata ggt gta gtt ttg cag tgg ccg tca tga ata ata tta tct ata tga tgg gtg gat atg atc agt ccc cgt ata gga gtt caa agg tta tag cgt aca ata cct gta cta att ctt gga tat atg ata tac cag agc t-a aaa tat cct cgt tct aat tgt gga gga gtt gct gat gat gaa tac att tat tgc ata ggc ggc ata cgc gat cag gat tca tcg ttg ata tct agt att gat aga tgg aag cca tca aaa cca tat tgg caa acg tat gct aaa atg cga gag cca aaa tgt gat atg ggt gta gcg atg tta aac gga tta ata tat gtc ata ggc gga gtt gtt aaa ggt g-- aca cgt gta ccg aca cac ttg aga gtt tat cag aag atg gat gga tga tgc atc aac gtc ttc caa taa aaa tgt cca ata tgt cga cga ttg ttc atg ccg gca aaa ttt ata tat ctg gag gct aca aca ata gta gtg tag tta atg gaa tat cga atc t-- agt cct tag cta taa tcc gat ata tga tga atg gac caa att atc atc att aaa tat tcc tag aat taa tcc cgc tct a

>HQ420898.1:176040-178285_Cowpox_virus_strain_Germany_2002_MKY_complete_genome

ttg ctg act gca --- tgt a-c tat ttt tta cat tac tac att ggt tac ggc aga tat aca tac tcc gtt acc gcc aca cgc tcc ggt aaa cgg --g gca tgt gac gag gga gaa tat ctt gat aag agg cat aat caa tgt tgt aat -cg gtg tcc acc tgg aga att tgc caa ggt tag atg ta- --g tgg tag cta taa cac aaa atg tga acg ctg tcc aca tca tac ata tac agc aat tcc caa tta ttc t-a atg gat gcc atc aat gta gaa aat gcc caa agg gat cat ttg ata agg taa agt gta ccg gaa cac aga aca gta aat -gt gcg tgt ctt cct ggt tgg tat tgc gct acc gat tct tca cag act gaa gac tgt cga gat -tg tgt acc aaa aag g-a gct gtc cat gcg gat act ttg gtg gaa tag ata aat tag gaa atc cta ttt gta aat cgt gtt g-- -tg ttg gta aat att gtg acg aca tac gta atc ata gag ttg gtc ctt ttc ctc cat gca aa- --- cta tct aaa tgt aat tag tca --- --- --- --- --- --- --- --- --- --- --- --- --- --- -tg atc atg gta -cc gta caa tt- --- -at att gct --- --g gta --g tat att att cag tat g-a aga cat att aat ac- --- --- -tt atc --- --t ttg aca atc ttg ct- -at aat cat --- --- --- --- aat ata aaa --a tac tta tgg --- -a- cat agt cac tca t-- --- --- --- aat tgc tg- acg cga taa att cgt aat aat ctg ttt tgt tca aat ttt tat aag ga- --- --a tct aca ggc ata aaa at- aaa aat ata att tat aat ac- --- act tac agc gcg cca tca tga ata aca gca gtg aat taa ttg ctg tta tta atg gat tta gaa ata gtg gac gat ttt gtg ata tta ata tag tta tta atg atg aaa gga taa atg ctc ata gac tca tcc tat ctg gag cat ccg aat a-t ttt tcc att cta ttt tct agt gat ttt atc gat tct aat gaa tac gaa gtt aat cta agt cat tta gat tat caa agt gtg aac gat ttg atc gat tac att tat ggg ata cct ttg agc cta act aac gat aac gtg aaa tat att ctt tca acc gct ga- --t ttt tta caa att gga tct gcc att act gag tgc gaa aaa tac ata ctt aaa aat ctg tgt tct aga aat tgt atc gat ttc --- --- tac ata tac gct gat aaa tat aat aac aag aaa ata gaa aca gcg tcg ttt aac aca ata tta cga aat att ttg aga ctc att aac gat gaa aac ttt aaa tac ttg aca gag gaa tca atg ata aaa att tta agc gat gat atg tta aat ata aaa aat gag gat ttt gca cca cta att ctc att aaa tgg tta gag agt aca caa caa cca tgc aca gtc gag tta ctt aga tgc ctc aga ata tca -tt gct ttc ccc aca agt tat aaa atc act tta tag tca tcg att ggt tgg ttc aat ata cga atg tat aac att ctt aaa caa tat atc att ctt gga tga atc att tcc tag ata cca tag cat cga gtt gat atc tat cgg tat aag taa ttc aca tga taa gat ttc cat aaa ctg cta taa tcg t-a aaa aaa ata cgt ggg aaa taa tat ctt cac gga gat ata ggt gta gtt ttg cag tgg ccg tca tga ata ata tta tct ata tga tgg gtg gat atg atc agt ccc cgt ata gga gtt caa agg tta tag cgt aca ata cct gta cta att ctt gga tat atg ata tac cag agc t-a aaa tat cct cgt tct aat tgt gga gga gtt gct gat gat gaa tac att tat tgc ata ggc ggc ata cgc gat cag gat tca tcg ttg ata tct agt att gat aga tgg aag cca tca aaa cca tat tgg caa acg tat gct aaa atg cga gag cca aaa tgt gat atg ggt gta gcg atg tta aac gga tta ata tat gtc ata ggc gga gtt gtt aaa ggt g-- aca cgt gta ccg aca cac ttg aga gtt tat cag aag atg gat gga tga tgc atc aac gtc ttc caa taa aaa tgt cca ata tgt cga cga ttg ttc atg ccg gca aaa ttt ata tat ctg gag gct aca aca ata gta gtg tag tta atg gaa tat cga atc t-- agt cct tag cta taa tcc gat ata tga tga atg gac caa att atc atc att aaa tat tcc tag aat taa tcc cgc tct a

>LT896719.1:175594-177839_Cowpox_virus_isolate_Ger/2013/Alpaca_genome_assembly_complete_genome:_monopartite

ttg ctg act gta --- tgt a-c tat ttt gta cat cag tac att ggt tac agc aga tat acc tac ttc gtc act gcc aca cgc tcc ggt aaa cgg --g tca tgt gac gaa gga gaa tat ctt gat aag acg cat aat cga tgt tgt aat -cg gtg tcc acc tgg aga att tgc caa ggt tag atg ta- --g tgg tag cta taa cac aaa atg tga acg ctg tcc aca tca tac ata tac agc aat tcc caa tta ttc t-a atg gat gcc atc aat gta gaa aat gcc caa agg gat cat ttg ata agg taa agt gta ccg gaa cac aga aca gta aat -gt gcg tgt ctt cct ggt tgg tat tgc gct acc gat tct tca cag act gaa gac tgt cga gat -tg tgt acc aaa aag g-a gct gtc cat gcg gat act ttg gtg gaa tag ata aat tag gaa atc cta ttt gta aat cgt gtt g-- -tg ttg gta aat att gtg acg aca tac gta atc ata gag ttg gtc ctt ttc ctc cat gca aa- --- cta tct aaa tgt aat tag tca --- --- --- --- --- --- --- --- --- --- --- --- --- --- -tg atc atg gta -cc gta caa tt- --- -at att gct --- --g gta --g tat att att cag tat g-a aga cat att aat ac- --- --- -tt atc --- --t ttg aca atc ttg ct- -at aat cat --- --- --- --- aat ata aaa --a tac tta tgg --- -a- cat agt cac tca t-- --- --- --- aat tgc tg- acg cga taa att cgt aat aat ctg ttt tgt tca aat ttt tat aag ga- --- --a tct aca ggc ata aaa at- aaa aat ata att tat aat ac- --- act tac agc gcg cca tca tga ata aca gca gtg aat taa ttg ctg tta tta atg gat tta gaa ata gtg gac gat ttt gtg ata tta ata tag tta tta atg atg aaa gga taa atg ctc ata gac tca tcc tat ctg gag cat ccg aat a-t ttt tcc att cta ttt tct agt gat ttt atc gat tct aat gaa tac gaa gtt aat cta agt cat tta gat tat caa agt gtg aac gat ttg atc gat tac att tat ggg ata cct ttg agc cta act aac gat aac gtg aaa tat att ctt tca acc gct ga- --t ttt tta caa att gga tct gcc att act gag tgc gaa aaa tac ata ctt aaa aat ctg tgt tct aga aat tgt atc gat ttc --- --- tac ata tac gct gat aaa tat aat aac aag aaa ata gaa aca gcg tcg ttt aac aca ata tta cga aat att ttg aga ctc att aac gat gaa aac ttt aaa tac ttg aca gag gaa tca atg ata aaa att tta agc gat gat atg tta aat ata aaa aat gag gat ttt gca cca cta att ctc att aaa tgg tta gag agt aca caa caa cca tgc aca gtc gag tta ctt aga tgc ctc aga ata tca -tt gct ttc ccc aca agt tat aaa atc act tta tag tca tcg att ggt tgg ttc aat ata cga atg tat aac att ctt aaa caa tat atc att ctt gga tga atc att tcc tag ata cca tag cat cga gtt gat atc tat cgg tat aag taa ttc aca tga taa gat ttc cat aaa ctg cta taa tcg t-a aaa aaa ata cgt ggg aaa taa tat ctt cac gga gat ata ggt gta gtt ttg cag tgg ccg tca tga ata ata tta tct ata tga tgg gtg gat atg atc agt ccc cgt ata gga gtt caa agg tta tag cgt aca ata cct gta cta att ctt gga tat atg ata tac cag agc t-a aaa tat cct cgt tct aat tgt gga gga gtt gct gat gat gaa tac att tat tgc ata ggg ggc ata cgc gat cag gat tca tcg ttg ata tct aat att gat aga tgg aag cca tca aaa cca tat tgg cag acg tat gct aaa atg cgc gag cca aaa tgt gat atg ggt gta gcg atg tta aac gga tta ata tat gtc ata ggc gga gtt gtt aaa ggt g-- aca cgt gta ccg aca cac ttg aga gtt tat cag aag atg gat gga tga tgc atc aac gtc ttc caa taa aaa tgt cca ata tgt cga cga ttg ttc atg ccg gca aaa ttt ata tat ctg gag gct aca aca ata gta gtg tag tta atg gaa tat cga atc t-- agt cct tag cta taa tcc gat ata tga tga atg gac caa att atc atc att aaa tat tcc tag aat taa tcc cgc tct a

>KC813506.1:176961-179205_Cowpox_virus_strain_CatPot07/1_complete_genome

ttg ctg act gta --- tgt a-c tat ttt gta cat cag tac att ggt tac agc aga tat acc tac ttc gtc act gcc aca cgc tcc ggt aaa cgg --g tca tgt gac gaa gga gaa tat ctt gat aag acg cat aat cga tgt tgt aat -cg gtg tcc acc tgg aga att tgc caa ggt tag atg ta- --g tgg tag cta taa cac aaa atg tga acg ctg tcc aca tca tac ata tac agc aat tcc caa tta ttc t-a atg gat gcc atc aat gta gaa aat gcc caa agg gat cat ttg ata agg taa agt gta ccg gaa cac aga aca gta aat -gt gcg tgt ctt cct ggt tgg tat tgc gct acc gat tct tca cag act gaa gac tgt cga gat -tg tgt acc aaa aag g-a gct gtc cat gcg gat act ttg gtg gaa tag ata aat tag gaa atc cta ttt gta aat cgt gtt g-- -tg ttg gta aat att gtg acg aca tac gta atc ata gag ttg gtc ctt ttc ctc cat gca aa- --- cta tct aaa tgt aat tag tca --- --- --- --- --- --- --- --- --- --- --- --- --- --- -tg atc atg gta -cc gta caa tt- --- -at att gct --- --g gta --g tat att att cag tat g-a aga cat att aat ac- --- --- -tt atc --- --t ttg aca atc ttg ct- -at aat cat --- --- --- --- aat ata aaa --a tac tta tgg --- -a- cat agt cac tca t-- --- --- --- aat tgc tg- acg cga taa att cgt aat aat ctg ttt tgt tca aat ttt tat aag ga- --- --a tct aca ggc ata aaa -t- aaa aat ata att tat aat ac- --- act tac agc gcg cca tca tga ata aca gca gtg aat taa ttg ctg tta tta atg gat tta gaa ata gtg gac gat ttt gtg ata tta ata tag tta tta atg atg aaa gga taa atg ctc ata gac tca tcc tat ctg gag cat ccg aat a-t ttt tcc att cta ttt tct agt gat ttt atc gat tct aat gaa tac gaa gtt aat cta agt cat tta gat tat caa agt gtg aac gat ttg atc gat tac att tat ggg ata cct ttg agc cta act aac gat aac gtg aaa tat att ctt tca acc gct ga- --t ttt tta caa att gga tct gcc att act gag tgc gaa aaa tac ata ctt aaa aat ctg tgt tct aga aat tgt atc gat ttc --- --- tac ata tac gct gat aaa tat aat aac aag aaa ata gaa aca gcg tcg ttt aac aca ata tta cga aat att ttg aga ctc att aac gat gaa aac ttt aaa tac ttg aca gag gaa tca atg ata aaa att tta agc gat gat atg tta aat ata aaa aat gag gat ttt gca cca cta att ctc att aaa tgg tta gag agt aca caa caa cca tgc aca gtc gag tta ctt aga tgc ctc aga ata tca -tt gct ttc ccc aca agt tat aaa atc act tta tag tca tcg att ggt tgg ttc aat ata cga atg tat aac att ctt aaa caa tat atc att ctt gga tga atc att tcc tag ata cca tag cat cga gtt gat atc tat cgg tat aag taa ttc aca tga taa gat ttc cat aaa ctg cta taa tcg t-a aaa aaa ata cgt ggg aaa taa tat ctt cac gga gat ata ggt gta gtt ttg cag tgg ccg tca tga ata ata tta tct ata tga tgg gtg gat atg atc agt ccc cgt ata gga gtt caa agg tta tag cgt aca ata cct gta cta att ctt gga tat atg ata tac cag agc t-a aaa tat cct cgt tct aat tgt gga gga gtt gct gat gat gaa tac att tat tgc ata ggc ggc ata cgc gat cag gat tca tcg ttg ata tct agt att gat aga tgg aag cca tca aaa cca tat tgg caa acg tat gct aaa atg cga gag cca aaa tgt gat atg ggt gta gcg atg tta aac gga tta ata tat gtc ata ggt gga atc gtt aaa ggt g-- aca cgt gta ccg aca cac ttg aga gtt tat cag aag atg gat gga tga agc atc aac gtc ttc caa taa aaa tgt cca ata tgt cga cga ttg ttc atg ccg gca aaa ttt ata tat ctg gag gct aca aca ata gta gtg tag tta atg gaa tat cga atc t-- agt cct tag cta taa tcc gat ata tga tga atg gac caa att atc atc att gaa tat tcc tag aat taa tcc cgc tct a

>KY463519.1:167145-169379_Cowpox_virus_strain_Germany_1971_EP1

--- --- --- --a --- tgt a-c tat ttt ata cat tac tac att ggt tac ggc aga tat aca tac tcc gtt acc gcc aca cgc ttc ggt aaa cgg --g tca tgt gcc gag gga gaa tat ctt gat aag act cat aat cga tgt tgt aat -cg gtg tcc acc tgg aga att tgc caa ggt tag atg ta- --g tgg tag cta taa cac aaa atg tga acg ctg tcc aca tca tac ata tac agc aat tcc caa tta ttc t-a atg gat gcc atc aat gta gaa aat gcc caa agg gat cat ttg ata agg taa agt gta ccg gaa cac aga aca gta aat -gt gcg tgt ctt cct ggt tgg tat tgc gct acc gat tct tca cag act gaa gac tgt cga gat -tg tgt acc aaa aag g-a gct gtc cat gcg gat act ttg gtg gaa tag ata aat tag gaa atc cta ttt gta aat cgt gtt g-- -tg ttg gta aat att gtg acg aca tac gta atc ata gag ttg gtc ctt ttc ctc cat gca aa- --- cta tct aaa tgt aat tag tca --- --- --- --- --- --- --- --- --- --- --- --- --- --- -tg atc atg gta -cc gta caa tt- --- -at att gct --- --g gta --g tat att att cag tat g-a aga cat att aat ac- --- --- -tt atc --- --t ttg aca atc ttg ct- -at aat cat --- --- --- --- aat ata aaa --a tac tta tgg --- -a- cat agt cac tca t-- --- --- --- aat tgc tg- acg cga taa att cgt aat aat ctg ttt tgt tca aat ttt tat aag ga- --- --a tct aca ggc ata aaa at- aaa aat ata att tat aat ac- --- act tac agc gcg cca tca tga ata aca gca gtg aat taa ttg ctg tta tta atg gat tta gaa ata gtg gac gat ttt gtg ata tta ata tag tta tta atg atg aaa gga taa atg ctc ata gac tca tcc tat ctg gag cat ccg aat a-t ttt tcc att cta ttt tct agt gat ttt atc gat tct aat gaa tac gaa gtt aat cta agt cat tta gat tat caa agt gtg aac gat ttg atc gat tac att tat ggg ata cct ttg agc cta act aac gat aac gtg aaa tat att ctt tca acc gct ga- --t ttt tta caa att gga tct gcc att act gag tgc gaa aaa tac ata ctt aaa aat ctg tgt tct aga aat tgt atc gat ttc --- --- tac ata tac gct gat aaa tat aat aac aag aaa ata gaa aca gcg tcg ttt aac aca ata tta cga aat att ttg aga ctc att aac gat gaa aac ttt aaa tac ttg aca gag gaa tca atg ata aaa att tta agc gat gat atg tta aat ata aaa aat gag gat ttt gca cca cta att ctc att aaa tgg tta gag agt aca caa caa cca tgc aca gtc gag tta ctt aga tgc ctc aga ata tca -tt gct ttc ccc aca agt tat aaa atc act tta tag tca tcg act ggt tag ttc aat cta cga atg tat aac att ctt aaa caa tat atc att ctt gga tga atc att tcc tag ata cca tag cat cga gtt gat atc tat cgg tat aag taa ttc aca tga taa gat ttc cat aaa ctg cta taa tcg t-a aaa aaa ata cgt ggg aaa taa tat ctt cac gga gat ata ggt gta gtt ttg cag tgg ccg tca tga ata ata tta tct ata tga tgg gtg gat atg atc agt ccc cgt ata gga gtt caa agg tta tag cgt aca ata cct gta cta att ctt gga tat atg ata tac cag agc t-a aaa tat cct cgt tct aat tgt gga gga gtt gct gat gat gaa tac att tat tgc ata ggc ggc ata cga gat cag gat tca tcg ttg ata tct agt att gat aga tgg aag cca tca aaa cca tat tgg caa acg tat gct aaa atg cga gag cca aaa tgt gat atg ggt gta gcg atg tta aac gga tta ata tat gtc ata ggc gga gtt gtt aaa ggt g-- aca cgt gta ccg aca cac ttg aga gtt tat cag aag atg gat gga tga tgc atc aac gtc ttc caa taa aaa tgt cca ata tgt cga cga ttg ttc atg ccg gca aaa ttt ata tat ctg gag gct aca aca ata gta gtg tag tta atg gaa tat cga atc t-- agt cct tag cta taa tcc gat ata tga tga atg gac caa att atc atc att aaa tat tcc tag aat taa tcc cgc tct a

>LT993231.1:176600-178834_Cowpox_virus_isolate_Ger/2015/Prairie-dog_genome_assembly_complete_genome:_monopartite

--- --- --- --a --- tgt a-c tat ttt ata cat tac tac att ggt tac ggc aga tat aca tac tcc gtt acc gcc aca cgc ttc ggt aaa cgg --g tca tgt gcc gag gga gaa tat ctt gat aag act cat aat cga tgt tgt aat -cg gtg tcc acc tgg aga att tgc caa ggt tag atg ta- --g tgg tag cta taa cac aaa atg tga acg ctg tcc aca tca tac ata tac agc aat tcc caa tta ttc t-a atg gat gcc atc aat gta gaa aat gcc caa agg gat cat ttg ata agg taa agt gta ccg gaa cac aga aca gta aat -gt gcg tgt ctt cct ggt tgg tat tgc gct acc gat tct tca cag act gaa gac tgt cga gat -tg tgt acc aaa aag g-a gct gtc cat gcg gat act ttg gtg gaa tag ata aat tag gaa atc cta ttt gta aat cgt gtt g-- -tg ttg gta aat att gtg acg aca tac gta atc ata gag ttg gtc ctt ttc ctc cat gca aa- --- cta tct aaa tgt aat tag tca --- --- --- --- --- --- --- --- --- --- --- --- --- --- -tg atc atg gta -cc gta caa tt- --- -at att gct --- --g gta --g tat att att cag tat g-a aga cat att aat ac- --- --- -tt atc --- --t ttg aca atc ttg ct- -at aat cat --- --- --- --- aat ata aaa --a tac tta tgg --- -a- cat agt cac tca t-- --- --- --- aat tgc tg- acg cga taa att cgt aat aat ctg ttt tgt tca aat ttt tat aag ga- --- --a tct aca ggc ata aaa at- aaa aat ata att tat aat ac- --- act tac agc gcg cca tca tga ata aca gca gtg aat taa ttg ctg tta tta atg gat tta gaa ata gtg gac gat ttt gtg ata tta ata tag tta tta atg atg aaa gga taa atg ctc ata gac tca tcc tat ctg gag cat ccg aat a-t ttt tcc att cta ttt tct agt gat ttt atc gat tct aat gaa tac gaa gtt aat cta agt cat tta gat tat caa agt gtg aac gat ttg atc gat tac att tat ggg ata cct ttg agc cta act aac gat aac gtg aaa tat att ctt tca acc gct ga- --t ttt tta caa att gga tct gcc att act gag tgc gaa aaa tac ata ctt aaa aat ctg tgt tct aga aat tgt atc gat ttc --- --- tac ata tac gct gat aaa tat aat aac aag aaa ata gaa aca gcg tcg ttt aac aca ata tta cga aat att ttg aga ctc att aac gat gaa aac ttt aaa tac ttg aca gag gaa tca atg ata aaa att tta agc gat gat atg tta aat ata aaa aat gag gat ttt gca cca cta att ctc att aaa tgg tta gag agt aca caa caa cca tgc aca gtc gag tta ctt aga tgc ctc aga ata tca -tt gct ttc ccc aca agt tat aaa atc act tta tag tca tcg att ggt tgg ttc aat ata cga atg tat aac att ctt aaa caa tat atc att ctt gga tga atc att tcc tag ata cca tag cat cga gtt gat atc tat cgg tat aag taa ttc aca tga taa gat ttc cat aaa ctg cta taa tcg t-a aaa aaa ata cgt ggg aaa taa tat ctt cac gga gat ata ggt gta gtt ttg cag tgg ccg tca tga ata ata tta tct ata tga tgg gtg gat atg atc agt ccc cgt ata gga gtt caa agg tta tag cgt aca ata cct gta cta att ctt gga tat atg ata tac cag agc t-a aaa tat cct cgt tct aat tgt gga gga gtt gct gat gat gaa tac att tat tgc ata ggc ggc ata cgc gat cag gat tca tcg ttg ata tct agt att gat aaa tgg aag cca tca aaa cca tat tgg caa acg tat gct aaa atg cga gag cca aaa tgt gat atg ggt gta gcg atg tta aac gga tta ata tat gtc ata ggc gga gtt gtt aaa ggt g-- aca cgt gta ccg aca cac ttg aga gtt tat cag aag atg gat gga tga tgc atc aac gtc ttc caa taa aaa tgt cca ata tgt cga cga ttg ttc atg ccg gca aaa ttt ata tat ctg gag gct aca aca ata gta gtg tag tta atg gaa tat cga atc t-- agt cct tag cta taa tcc gat ata tga tga atg gac caa att atc atc att aaa tat tcc tag aat taa tcc cgc tct a

>LT896733.1:175953-178187_Cowpox_virus_isolate_Ger/2015/Cat3_genome_assembly_complete_genome:_monopartite

--- --- --- --a --- tgt a-c tat ttt ata cat tac tac att ggt tac ggc aga tat aca tac tcc gtt acc gcc aca cgc ttc ggt aaa cgg --g tca tgt gcc gag gga gaa tat ctt gat aag act cat aat cga tgt tgt aat -cg gtg tcc acc tgg aga att tgc caa ggt tag atg ta- --g tgg tag cta taa cac aaa atg tga acg ctg tcc aca tca tac ata tac agc aat tcc caa tta ttc t-a atg gat gcc atc aat gta gaa aat gcc caa agg gat cat ttg ata agg taa agt gta ccg gaa cac aga aca gta aat -gt gcg tgt ctt cct ggt tgg tat tgc gct acc gat tct tca cag act gaa gac tgt cga gat -tg tgt acc aaa aag g-a gct gtc cat gcg gat act ttg gtg gaa tag ata aat tag gaa atc cta ttt gta aat cgt gtt g-- -tg ttg gta aat att gtg acg aca tac gta atc ata gag ttg gtc ctt ttc ctc cat gca aa- --- cta tct aaa tgt aat tag tca --- --- --- --- --- --- --- --- --- --- --- --- --- --- -tg atc atg gta -cc gta caa tt- --- -at att gct --- --g gta --g tat att att cag tat g-a aga cat att aat ac- --- --- -tt atc --- --t ttg aca atc ttg ct- -at aat cat --- --- --- --- aat ata aaa --a tac tta tgg --- -a- cat agt cac tca t-- --- --- --- aat tgc tg- acg cga taa att cgt aat aat ctg ttt tgt tca aat ttt tat aag ga- --- --a tct aca ggc ata aaa at- aaa aat ata att tat aat ac- --- act tac agc gcg cca tca tga ata aca gca gtg aat taa ttg ctg tta tta atg gat tta gaa ata gtg gac gat ttt gtg ata tta ata tag tta tta atg atg aaa gga taa atg ctc ata gac tca tcc tat ctg gag cat ccg aat a-t ttt tcc att cta ttt tct agt gat ttt atc gat tct aat gaa tac gaa gtt aat cta agt cat tta gat tat caa agt gtg aac gat ttg atc gat tac att tat ggg ata cct ttg agc cta act aac gat aac gtg aaa tat att ctt tca acc gct ga- --t ttt tta caa att gga tct gcc att act gag tgc gaa aaa tac ata ctt aaa aat ctg tgt tct aga aat tgt atc gat ttc --- --- tac ata tac gct gat aaa tat aat aac aag aaa ata gaa aca gcg tcg ttt aac aca ata tta cga aat att ttg aga ctc att aac gat gaa aac ttt aaa tac ttg aca gag gaa tca atg ata aaa att tta agc gat gat atg tta aat ata aaa aat gag gat ttt gca cca cta att ctc att aaa tgg tta gag agt aca caa caa cca tgc aca gtc gag tta ctt aga tgc ctc aga ata tca -tt gct ttc ccc aca agt tat aaa atc act tta tag tca tcg att ggt tgg ttc aat ata cga atg tat aac att ctt aaa caa tat atc att ctt gga tga atc att tcc tag ata cca tag cat cga gtt gat atc tat cgg tat aag taa ttc aca tga taa gat ttc cat aaa ctg cta taa tcg t-a aaa aaa ata cgt ggg aaa taa tat ctt cac gga gat ata ggt gta gtt ttg cag tgg ccg tca tga ata ata tta tct ata tga tgg gtg gat atg atc agt ccc cgt ata gga gtt caa agg tta tag cgt aca ata cct gta cta att ctt gga tat atg ata tac cag agc t-a aaa tat cct cgt tct aat tgt gga gga gtt gct gat gat gaa tac att tat tgc ata ggc ggc ata cgc gat cag gat tca tcg ttg ata tct agt att gat aaa tgg aag cca tca aaa cca tat tgg caa acg tat gct aaa atg cga gag cca aaa tgt gat atg ggt gta gcg atg tta aac gga tta ata tat gtc ata ggc gga gtt gtt aaa ggt g-- aca cgt gta ccg aca cac ttg aga gtt tat cag aag atg gat gga tga tgc atc aac gtc ttc caa taa aaa tgt cca ata tgt cga cga ttg ttc atg ccg gca aaa ttt ata tat ctg gag gct aca aca ata gta gtg tag tta atg gaa tat cga atc t-- agt cct tag cta taa tcc gat ata tga tga atg gac caa att atc atc att aaa tat tcc tag aat taa tcc cgc tct a

>LT896730.1:175812-178057_Cowpox_virus_isolate_Ger/2010/Racoon_genome_assembly_complete_genome:_monopartite

ttg ctg act gca --- tgt a-c tat ttt tta cat tac tac att ggc tac ggc aga tat acc tac ttc gtc act gcc aca cgc tcc ggt aaa cgg --g gca tgt gac gag gga gaa tat ctt gat aag act cat aat cga tgt tgt aat -cg gtg tcc acc tgg aga att tgc caa ggt tag atg ta- --g tgg tag cta taa cac aaa atg tga acg ctg tcc aca tca tac ata tac agc aat tcc caa tta ttc t-a atg gat gcc atc aat gta gaa aat gcc caa agg gat cat ttg ata agg taa agt gta cag gaa cac aga aca gta aat -gt gcg tgt ctt cct ggt tgg tat tgc gct acc gat tct tca cag act gaa gac tgt cga gat -tg tgt acc aaa aag g-a gct gtc cat gcg gat act ttg gtg gaa tag ata aat tag gaa atc cta ttt gta aat cgt gtt g-- -tg ttg gta aat att gtg acg aca tac gta atc ata gag ttg gtc ctt ttc ctc cat gca aa- --- cta tct aaa tgt aat tag tca --- --- --- --- --- --- --- --- --- --- --- --- --- --- -tg atc atg gta -cc gta caa tt- --- -at att gct --- --g gta --g tat att att cag tat g-a aga cat att aat ac- --- --- -tt atc --- --t ttg aca atc ttg ct- -at aat cat --- --- --- --- aat ata aaa --a tac tta tgg --- -a- cat agt cac tca t-- --- --- --- aat tgc tg- acg cga taa att cgt aat aat ctg ttt tgt tca aat ttt tat aag ga- --- --a tct aca ggc ata aaa at- aaa aat ata att tat aat ac- --- act tac agc gcg cca tca tga ata aca gca gtg aat taa ttg ctg tta tta atg gat tta gaa ata gtg gac gat ttt gtg ata tta ata tag tta tta atg atg aaa gga taa atg ctc ata gac tca tcc tat ctg gag cat ccg aat a-t ttt tcc att cta ttt tct agt gat ttt atc gat tct aat gaa tac gaa gtt aat cta agt cat tta gat tat caa agt gtg aac gat ttg atc gat tac att tat ggg ata cct ttg agc cta act aac gat agc gtg aaa tat att ctt tca act gct ga- --t ttt tta caa att gga tat gct att acc gaa tgt gaa aat tac ata ctt aaa aat ctg tgt tct aga aat tgt atc gat ttc --- --- tac ata tac gct gat aaa tat aat aac gag aaa ata gaa aca gcg tcg ttt aac aca ata tta cga aat att ttg aga ctc att aac gat gaa aac ttt aaa tac ttg aca gag gaa tca atg ata aaa att tta agc gat gat atg tta aat ata aaa aat gag gat ttt gca cca cta att ctc att aaa tgg tta gag agt aca caa caa cca tgc aca gtc gag tta ctt aga tgc ctc aga ata tca -tt gct ttc ccc aca agt tat aaa atc act tta tag tca tcg att ggt tgg ttc aat ata cga atg tat aac att ctt aaa caa tat atc att ctt gga tga atc att tcc tag ata cca tag cat cga gtt gat atc tat cgg tat aag taa ttc aca tga taa gat ttc cat aaa ctg cta taa tcg t-a aaa aaa ata cgt ggg aga taa tat ctt cac gga gat ata ggt gta gtt ttg cag tgg ccg tca tgg ata ata tta tct ata tga tgg gtg gat atg atc agt ccc cgt ata gga gtt caa agg tta tag cgt aca ata cct gta cta att ctt gga tat atg ata tac cag agc t-a aaa tat cct cgt tct aat tgt gga gga gtt gct gat gat gaa tac att tat tgc ata ggc ggc ata cgc gat cag gat tca tcg ttg ata tct agt att gat aga tgg aag cca tca aaa cca tat tgg caa acg tat gct aaa atg cga gag cca aaa tgt gat atg ggt gta gcg atg tta aac gga tta ata tat gtc ata ggc gga gtt gtt aaa ggt g-- aca cgt gta ccg aca cac ttg aga gtt tat cag aag atg gat gga tga tgc atc aac gtc ttc caa taa aaa tgt cca ata tgt cga cga ttg ttc atg ccg gca aaa ttt ata tat ctg gag gct aca aca ata gta gtg tag tta atg gaa tat cga atc t-- agt cct tag cta taa tcc gat ata tga tga atg gac caa att atc atc att aaa tat tcc tag aat taa tcc cgc tct a

>LT896718.1:176372-178606_Cowpox_virus_isolate_Ger/2010/Alpaca_genome_assembly_complete_genome:_monopartite

--- --- --- --a --- tgt a-c tat ttt ata cat tac tac att ggt tac ggc aga tat aca tac tcc gtt acc gcc aca cgc ttc ggt aaa cgg --g tca tgt gcc gag gga gaa tat ctt gat aag act cat aat cga tgt tgt aat -cg gtg tcc acc tgg aga att tgc caa ggt tag atg ta- --g tgg tag cta taa cac aaa atg tga acg ctg tcc aca tca tac ata tac agc aat tcc caa tta ttc t-a atg gat gcc atc aat gta gaa aat gcc caa agg gat cat ttg ata agg taa agt gta ccg gaa cac aga aca gta aat -gt gcg tgt ctt cct ggt tgg tat tgc gct acc gat tct tca cag act gaa gac tgt cga gat -tg tgt acc aaa aag g-c gct gtc cat gcg gat act ttg gtg gaa tag ata aat tag gaa atc cta ttt gta aat cgt gtt g-- -tg ttg gta aat att gtg acg aca tac gta atc ata gag ttg gtc ctt ttc ctc cat gca aa- --- cta tct aaa tgt aat tag tca --- --- --- --- --- --- --- --- --- --- --- --- --- --- -tg atc atg gta -cc gta caa tt- --- -at att gct --- --g gta --g tat att att cag tat g-a aga cat att aat ac- --- --- -tt atc --- --t ttg aca atc ttg ct- -at aat cat --- --- --- --- aat ata aaa --a tac tta tgg --- -a- cat agt cac tca t-- --- --- --- aat tgc tg- acg cga taa att cgt aat aat ctg ttt tgt tca aat ttt tat aag ga- --- --a tct aca ggc ata aaa at- aaa aat ata att tat aat ac- --- act tac agc gcg cca tca tga ata aca gca gtg aat taa ttg ctg tta tta atg gat tta gaa ata gtg gac gat ttt gtg ata tta ata tag tta tta atg atg aaa gga taa atg ctc ata gac tca tcc tat ctg gag cat ccg aat a-t ttt tcc att cta ttt tct agt gat ttt atc gat tct aat gaa tac gaa gtt aat cta agt cat tta gat tat caa agt gtg aac gat ttg atc gat tac att tat ggg ata cct ttg agc cta act aac gat aac gtg aaa tat att ctt tca acc gct ga- --t ttt tta caa att gga tct gcc att act gag tgc gaa aaa tac ata ctt aaa aat ctg tgt tct aga aat tgt atc gat ttc --- --- tac ata tac gct gat aaa tat aat aac aag aaa ata gaa aca gcg tcg ttt aac aca ata tta cga aat att ttg aga ctc att aac gat gaa aac ttt aaa tac ttg aca gag gaa tca atg ata aaa att tta agc gat gat atg tta aat ata aaa aat gag gat ttt gca cca cta att ctc att aaa tgg tta gag agt aca caa caa cca tgc aca gtc gag tta ctt aga tgc ctc aga ata tca -tt gct ttc ccc aca agt tat aaa atc act tta tag tca tcg att tgt tgg ttc aat ata cga atg tat aac att ctt aaa caa tat atc att ctt gga tga atc att tcc tag ata cca tag cat cga gtt gat atc tat cgg tat aag taa ttc aca tga taa gat ttc cat aaa ctg cta taa tcg t-a aaa aaa ata cgt ggg aaa taa tat ctt cac gga gat ata ggt gta gtt ttg cag tgg ccg tca tga ata ata tta tct ata tga tgg gtg gat atg atc agt ccc cgt ata gga gtt caa agg tta tag cgt aca ata cct gta cta att ctt gga tat atg ata tac cag agc t-a aaa tat cct cgt tct aat tgt gga gga gtt gct gat gat gaa tac att tat tgc ata ggc ggc ata cgc gat cag gat tca tcg ttg ata tct agt att gat aga tgg aag cca tca aaa cca tat tgg caa acg tat gct aaa atg cga gag cca aaa tgt gat atg ggt gta gcg atg tta aac gga tta ata tat gtc ata ggc gga gtt gtt aaa ggt g-- aca cgt gta ccg aca cac ttg aga gtt tat cag aag atg gat gga tga tgc atc aac gtc ttc caa taa aaa tgt cca ata tgt cga cga ttg ttc atg ccg gca aaa ttt ata tat ctg gag gct aca aca ata gta gtg tag tta atg gaa tat cga atc t-- agt cct tag cta taa tcc gat ata tga tga atg gac caa att atc atc att aaa tat tcc tag aat taa tcc cgc tct a

>AF482758.2:176786-179054_Cowpox_virus_strain_Brighton_Red_complete_genome

ttg ctg act gta --- tgt a-c tat ttt gta cat cag tac att ggt tac agc aga tat acc tac ttc gtc act gcc aca cgc tcc ggt aaa cgg --g tca tgt gac gac gga gaa tat ctt gat aag acg cat aat caa tgt tgt aat -cg gtg tcc acc tgg aga att tgc caa gat cag atg ta- --g cgg tag cga taa cac aaa atg tga acg ctg ccc acc tca tac ata tac cac agt acc caa tta ttc t-a atg gat gtc atc aat gta gga aat gcc caa cag gat cat ttg ata agg taa agt gta ccg gaa cac aga aca gta aat -gt tcg tgt ctt cct ggt tgg ttt tgc gct act gat tct tcg aag act gaa gat tgt cga gat -tg tat acc aaa aag a-a aat gtc cat gtg gat act ttg gtg gaa tag atg aac tcg gaa atc ctc ttt gta aat cgt gtt g-- -tg ttg gtg aat att gcg acg aca tac gta atc ata gag ttg gtc ctt ttc ctc cat gca aa- --- cta tct aaa tgt aat tag cca --- --- --- --- --- --- --- --- --- --- --- --- --- --- -tg atc atg tta -cc ata c-a tc- --- -at atc gct act tgg t-a --g cgt att att cag tat g-a aga tct att aat aat aat tat ctt att tat ttc tt- --g --- --- --- -at att aat cgg tgt cct aat aat ata aaa --- tac tta tgg --- --- --a aat aaa ttg t-a aac aag atg aat taa t-- aca caa tac atc ctt -at act cag ctt -gt tca aa- ttt tat aaa gaa tca tat tca tta ggc ata taa ta- aaa aat ata att tat aat ac- --- act tac a-- gcg tca tca tgc ata aca gca gtg aat taa ttg ctg tta tta atg gat tta gaa ata gtg gac gat ttt gtg ata tta ata tag tta tta atg atg aaa gga taa acg ctc ata gac tca tgc tat ctg gag cat ccg aat a-t ttt tcc att cta ttt tct agt gat ttt atc gat tct aat gac tac gaa gtt aat cta agt cat tta gat tat caa agt gtg aac gat ttg atc gat tac att tat ggg ata cct ttg agc cta act aac gat aac gtg aaa tat att ctt tca acc gct ga- --t ttt tta caa att gga tct gcc att act gag tgc gaa aaa tac ata ctt aaa aat ctt tgt tct aga aac tgt atc gat ttc --- --- tac ata tac gct gat aaa tat aat aac aag aaa ata gaa tca gcg tcg ttt aac aca ata tta cga aat att ttg aga ctc att aac gat gaa aac ttt aaa tac ttg aca gag gaa tca atg ata aaa att tta agc gat gat atg tta aat ata aaa aat gag gat ttc gca cca cta att ctc att aaa tgg tta gag agt aca caa caa cca tgc acc gtc gag tta ctt aga tgc ctc aga ata tca -tt gct ttc ccc aca agt tat aaa atc act tta tag tca tcg act ggt tag ttc aat cta cga atg tat aac att ctt aaa caa tat agc att ctt gga tga atc att tcc tag ata cca tag cat cga gtt gat atc tat cgg tat aag taa ttc gca tga taa gat ttc cat aaa ctg cta caa tcg t-a aaa aaa ata cat ggg aga tga tat ctt cac gta gat ata ggt gta gtt tcg cag tgg ccg tcc tgg ata ata tta tct ata tga tgg gtg gat atg atc agt ccc cgt ata gaa gtt caa agg tta tag cgt aca ata cat gta caa att ctt gga tat atg ata tac cag agc t-a aaa tat cct cgt tct aat tgc gga gga gtt gcc gat gac gaa tac att tat tgt ata ggc ggc ata cgc gat cag gat tca tcg ttg ata tct agt att gat aga tgg aag cca tca aaa cca tat tgg cag acg tat gct aaa atg cgc gaa cca aaa tgt gat atg ggg gtt gcg atg tta aac gga tta ata tat gtc ata ggt gga atc gtt aaa ggt g-- aca cat gta ccg acg cac tag aga gtt tat cag aag atg gat gga tga agc atc aac gtc ttc caa taa aaa tgt cca ata tgt cga cga ttg ttc atg ctg gta aga ttt ata tat ctg gag gtt ata aca ata gta gtg ccg tta atg gac cat cga atc t-- agt cct tag cta taa tcc gat ata tga tga atg gac caa att atc atc att aaa tat tcc tag aat taa tcc cgc tct a

>LT896720.1:174538-176769_Cowpox_virus_isolate_Ger/2015/Human1_genome_assembly_complete_genome:_monopartite

--- --- --- --a --- tgt a-c tat ttt gta cat cac tac att ggt tac agc aga tat acc tac tcc gcc a-- -cc aca tgc tcc ggt aaa cgg --g tca tgt gaa gag gga gaa tat ctt gat agg acg cat aat caa tgt tgt aat -cg gtg tcc tcc tgg aga att tgc caa ggt tag atg ta- --g tgg tag cta taa cac aaa atg tga acg ctg tcc aca tca tac ata tac agc aat tcc caa tta ttc t-a atg gat gcc atc aat gta gaa aat gcc caa agg gat cat ttg ata agg taa agt gta cag gaa cac aga aca gta aat -gt gcg tgt ctt cct ggt tgg tat tgc gct acc gat tct tca cag act gaa gac tgt cga gat -tg tgt acc aaa aag g-a gct gtc cat gcg gat act ttg gtg gaa tag ata aat tag gaa atc cta ttt gta aat cgt gtt g-- -tg ttg gta aat att gtg acg aca tac gta atc ata gag ttg gtc ctt ttc ctc cat gca aa- --- cta tct aaa tgt aat tag tca --- --- --- --- --- --- --- --- --- --- --- --- --- --- -tg atc atg gta -cc gta caa tt- --- -at att gct --- --g gta --g tat att att cag tat g-a aga cat att aat ac- --- --- -tt atc --- --t ttg aca atc ttg ct- -at aat cat --- --- --- --- aat ata aaa --a tac tta tgg --- -a- cat agt cac tca t-- --- --- --- aat tgc tg- acg cga taa att cgt aat aat ctg ttt tgt tca aat ttt tat aag ga- --- --a tct aca ggc ata aaa at- aaa aat ata att tat aat ac- --- act tac agc gcg cca tca tga ata aca gca gtg aat taa ttg ctg tta tta atg gat tta gaa ata gtg gac gat ttt gtg ata tta ata tag tta tta atg atg aaa gga taa atg ctc ata gac tca tcc tat ctg gag cat ccg aat a-t ttt tcc att cta ttt tct agt gat ttt atc gat tct aat gaa tac gaa gtt aat cta agt cat tta gat tat caa agt gtg aac gat ttg atc gat tac att tat ggg ata cct ttg agc cta act aac gat aac gtg aaa tat att ctt tca acc gct ga- --t ttt tta caa att gga tct gcc att act gag tgc gaa aaa tac ata ctt aaa aat ctg tgt tct aga aat tgt atc gat ttc --- --- tac ata tac gct gat aaa tat aat aac aag aaa ata gaa aca gcg tcg ttt aac aca ata tta cga aat att ttg aga ctc att aac gat gaa aac ttt aaa tac ttg aca gag gaa tca atg ata aaa att tta agc gat gat atg tta aat ata aaa aat gag gat ttt gca cca cta att ctc att aaa tgg tta gag agt aca caa caa cca tgc aca gtc gag tta ctt aga tgc ctc aga ata tca -tt gct ttc ccc aca agt tat aaa atc act tta tag tca tcg att ggt tgg ttc aat ata cga atg tat aac att ctt aaa caa tat atc att ctt gga tga atc att tcc tag ata cca tag cat cga gtt gat atc tat cgg tat aag taa ttc aca tga taa gat ttc cat aaa ctg cta taa tcg t-a aaa aaa ata cgt ggg aaa taa tat ctt cac gga gat ata ggt gta gtt ttg cag tgg ccg tca tga ata ata tta tct ata tga tgg gtg gat atg atc agt ccc cgt ata gga gtt caa agg tta tag cgt aca ata cct gta cta att ctt gga tat atg ata tac cag agc t-a aaa tat cct cgt tct aat tgt gga gga gtt gct gat gat gaa tac att tat tgc ata ggg ggc ata cgc gat cag gat tca tcg ttg ata tct aat att gat aga tgg aag cca tca aaa cca tat tgg cag acg tat gct aaa atg cga gag cca aaa tgt gat atg ggt gta gcg atg tta aac gga tta ata tat gtc ata ggc gga gtt gtt aaa ggt g-- aca cgt gta ccg aca cac ttg aga gtt tat cag aag atg gat gga tga tgc atc aac gtc ttc caa taa aaa tgt cca ata tgt cga cga ttg ttc atg ccg gca aaa ttt ata tat ctg gag gct aca aca ata gta gtg tag tta atg gaa tat cga atc t-- agt cct tag cta taa tcc gat ata tga tga atg gac caa att atc atc att aaa tat tcc tag aat taa tcc cgc tct a

>LT896722.1:175911-178145_Cowpox_virus_isolate_Ger/2007/Vole_genome_assembly_complete_genome:_monopartite

--- --- --- --a --- tgt a-c tat ttt ata cat tac tac att ggt tac ggc aga tat aca tac tcc gtt acc gcc aca cgc ttc ggt aaa cgg --g tca tgt gcc gag gga gaa tat ctt gat aag act cat aat cga tgt tgt aat -cg gtg tcc acc tgg aga att tgc caa ggt tag atg ta- --g tgg tag cta taa cac aaa atg tga acg ctg tcc aca tca tac ata tac agc aat tcc caa tta ttc t-a atg gat gcc atc aat gta gaa aat gcc caa agg gat cat ttg ata agg taa agt gta ccg gaa cac aga aca gta aat -gt gcg tgt ctt cct ggt tgg tat tgc gct acc gat tct tca cag act gaa gac tgt cga gat -tg tgt acc aaa aag g-a gct gtc cat gcg gat act ttg gtg gaa tag ata aat tag gaa atc cta ttt gta aat cgt gtt g-- -tg ttg gta aat att gtg acg aca tac gta atc ata gag ttg gtc ctt ttc ctc cat gca aa- --- cta tct aaa tgt aat tag tca --- --- --- --- --- --- --- --- --- --- --- --- --- --- -tg atc atg gta -cc gta caa tt- --- -at att gct --- --g gta --g tat att att cag tat g-a aga cat att aat ac- --- --- -tt atc --- --t ttg aca atc ttg ct- -at aat cat --- --- --- --- aat ata aaa --a tac tta tgg --- -a- cat agt cac tca t-- --- --- --- aat tgc tg- acg cga taa att cgt aat aat ctg ttt tgt tca aat ttt tat aag ga- --- --a tct aca ggc ata aaa at- aaa aat ata att tat aat ac- --- act tac agc gcg cca tca tga ata aca gca gtg aat taa ttg ctg tta tta atg gat tta gaa ata gtg gac gat ttt gtg ata tta gta tag tta tta atg atg aaa gga taa acg ctc ata gac tca tcc tat ctg gag cat ccg aat a-t ttt tcc att cta ttt tct agt gat ttt atc gat tct aat gaa tac gaa gtt aat cta agt cat tta gat tat caa agt gtg aac gat ttg atc gat tac att tat ggg ata cct ttg agc cta act aac gat agc gtg aaa tat att ctt tca act gct ga- --t ttt tta caa att gga tct gct att acc gaa tgt gaa aat tac ata ctt aaa aat ctg tgt tct aga aat tgt atc gat ttc --- --- tac ata tac gct gat aaa tat aat aac aag aaa ata gaa aca gcg tcg ttt aac aca ata tta cga aat att ttg aga ctc att aac gat gaa aac ttt aaa tac ttg aca gag gaa tca atg ata aaa att tta agc gat gat atg tta aat ata aaa aat gag gat ttt gca cca cta att ctc att aaa tgg tta gag agt aca caa caa cca tgc aca gtc gag tta ctt aga tgc ctc aga ata tca -tt gct ttc ccc aca agt tat aaa atc act tta tag tca tcg att ggt tgg ttc aat ata cga atg tat aac att ctt aaa caa tat atc att ctt gga tga atc att tcc tag ata cca tag cat cga gtt gat atc tat cgg tat aag taa ttc aca tga taa gat ttc cat aaa ctg cta taa tcg t-a aaa aaa ata cgt ggg aga taa tat ctt cac gga gat ata ggt gta gtt ttg cag tgg ccg tca tgg ata ata tta tct ata tga tgg gtg gat atg atc agt ccc cgt ata gga gtt caa agg tta tag cgt aca ata cct gta cta att ctt gga tat atg ata tac cag agc t-a aaa tat cct cgt tct aat tgt gga gga gtt gct gat gat gaa tac att tat tgc ata ggc ggc ata cgc gat cag gat tca tcg ttg ata tct agt att gat aga tgg aag cca tca aaa cca tat tgg caa acg tat gct aaa atg cga gag cca aaa tgt gat atg ggt gta gcg atg tta aac gga tta ata tat gtc ata ggc gga gtt gtt aaa ggt g-- aca cgt gta ccg aca cac ttg aga gtt tat cag aag atg gat gga tga tgc atc aac gtc ttc caa taa aaa tgt cca ata tgt cga cga ttg ttc atg ccg gca aaa ttt ata tat ctg gag gct aca aca ata gta gtg tag tta atg gaa tat cga atc t-- agt cct tag cta taa tcc gat ata tga tga atg gac caa att atc atc att aaa tat tcc tag aat taa tcc cgc tct a

>KY549144.1:177159-179431_Cowpox_virus_isolate_CPXV_Catpox5wv1

ttg ctg act gta --- tgt a-c tat ttt gta cat cag tac att ggt tac agc aga tat acc tac ttc gtc act gcc aca cgc tcc ggt aaa cgg --g tca tgt gac gac gga gaa tat ctt gat aag acg cat aat caa tgt tgt aat -cg gtg tcc acc tgg aga att tgc caa gat cag atg ta- --g cgg tag cga taa cac aaa atg tga acg ctg ccc acc tca tac ata tac cac agt acc caa tta ttc t-a atg gat gcc atc aat gta gaa aat gcc caa cag gat cat ttg ata agg taa agt gta ccg gaa cac aga aca gta aat -gt tcg tgt ctt cct ggt tgg ttt tgc gct act gat tct tcg aag act gaa gat tgt cga gat -tg tat acc aaa aag a-a aat gtc cat gcg gat act ttg gtg gaa tag atg aac tcg gaa atc ctc ttt gta aat cgt gtt g-- -tg ttg gtg aat att gcg acg aca tac gta atc ata gag ttg gtc ctt ttc ctc cat gca aa- --- cta tct aaa tgt aat tag cca --- --- --- --- --- --- --- --- --- --- --- --- --- --- -tg atc atg tta ccc ata c-a tc- --- -at atc gct act tgg t-a --g cgt att att cag tat g-a aga tct att aat aat aat tat ctt att tat ttc ttg acg --- --- --- -at att aat cgg tgt cct aat aat ata aaa --- tac tta tgg --- --- --a aat aaa ttg t-a aac aag atg aat taa t-- aca caa tac atc ctt -at act cag ctt -gt tca aa- ttt tat aac gaa tca tat tca tta ggc ata taa ta- aaa aat ata att tat aat ac- --- act tac a-- gcg tca tca tgc ata aca gca gtg aat taa ttg ctg tta tta atg gat tta gaa ata gtg gac gat ttt gtg ata tta ata tag tta tta atg atg aaa gga taa acg ctc ata gac tca tgc tat ctg gag cat ccg aat a-t ttt tcc att cta ttt tct agt gat ttt atc gat tct aat gac tac gaa gtt aat cta agt cat tta gat tat caa agt gtg aac gat ttg atc gat tac att tat ggg ata cct ttg agc cta act aac gat aac gtg aaa tat att ctt tca acc gct ga- --t ttt tta caa att gga tct gcc att act gag tgc gaa aaa tac ata ctt aaa aat ctt tgt tct aga aac tgt atc gat ttc --- --- tac ata tac gct gat aaa tat aat aac aag aaa ata gaa tca gcg tcg ttt aac aca ata tta cga aat att ttg aga ctc att aac gat gaa aac ttt aaa tac ttg aca gag gaa tca atg ata aaa att tta agc gat gat atg tta aat ata aaa aat gag gat ttc gca cca cta att ctc att aaa tgg tta gag agt aca caa caa cca tgc acc gtc gag tta ctt aga tgc ctc aga ata tca -tt gct ttc ccc aca agt tat aaa atc act tta tag tca tcg act ggt tag ttc aat cta cga atg tat aac att ctt aaa caa tat agc att ctt gga tga atc att tcc tag ata cca tag cat cga gtt gat atc tat cgg tat aag taa ttc aca tga taa gat ttc cat aaa ctg cta caa tcg t-a aaa aaa ata cat ggg aaa tga tat ctt cac gta gat ata ggt gta gtt tcg cag tgg ccg tcc tgg ata ata tta tct ata tga tgg gtg gat atg atc agt ccc cgt ata gaa gtt caa agg tta tag cgt aca ata cat gta caa att ctt gga tat atg ata tac cag agc t-a aaa tat cct cgt tct aat tgc gga gga gtt gcc gat gac gaa tac att tat tgt ata ggc ggc ata cgc gat cag gat tca tcg ttg ata tct aat att gat aga tgg aag cca tca aaa cca tat tgg cag acg tat gct aaa atg cgc gag cca aaa tgt gat atg ggg gtt gcg atg tta aac gga tta ata tat gtc ata ggt gga atc gtt aaa ggt g-- aca cat gta ccg acg cac tag aaa gtt tat cag aag atg gat gga tga agc atc agc gcc ttc caa taa aaa tgt cca ata tgt cga cga ttg ttc atg ctg gta aga ttt ata tat ctg gag gtt ata aca ata gta gtg ccg tta atg gac cat cga atc t-- agt cct tag cta taa tcc gat ata tga tga atg gac caa att atc atc att aaa tat tcc tag aat taa tcc cgc tct a

>KC813510.1:175815-178049_Cowpox_virus_strain_HumGra07/1_complete_genome

--- --- --- --a --- tgt a-c tat ttt ata cat tac tac att ggt tac ggc aga tat aca tac tcc gtt acc gcc aca cgc tcc ggt aaa cgg --g tca tgt gcc gag gga gaa tat ctt gat aag act cat aat cga tgt tgt aat -cg gtg tcc acc tgg aga att tgc caa ggt tag atg ta- --g tgg tag cta taa cac aaa atg tga acg ctg tcc aca tca tac ata tac agc aat tcc caa tta ttc t-a atg gat gcc atc aat gta gaa aat gcc caa agg gat cat ttg ata agg taa agt gta ccg gaa aac aga aca gta aat -gt gcg tgt ctt cct ggt tgg tat tgc gct acc gat tct tca cag act gaa gac tgt cga gat -tg tgt acc aaa aag g-a gct gtc cat gcg gat act ttg gtg gaa tag ata aat tag gaa atc cta ttt gta aat cgt gtt g-- -tg ttg gta aat att gtg acg aca tac gta atc ata gag ttg gtc ctt ttc ctc cat gca aa- --- cta tct aaa tgt aat tag tca --- --- --- --- --- --- --- --- --- --- --- --- --- --- -tg atc atg gta -cc gta caa tt- --- -at att gct --- --g gta --g tat att att cag tat g-a aga cat att aat ac- --- --- -tt atc --- --t ttg aca atc ttg ct- -at aat cat --- --- --- --- aat ata aaa --a tac tta tgg --- -a- cat agt cac tca t-- --- --- --- aat tgc tg- acg cga taa att cgt aat aat ctg ttt tgt tca aat ttt tat aag ga- --- --a tct aca ggc ata aaa at- aaa aat ata att tat aat ac- --- act tac agc gcg cca tca tga ata aca gca gtg aat taa ttg ctg tta tta atg gat tta gaa ata gtg gac gat ttt gtg ata tta ata tag tta tta atg atg aaa gga taa atg ctc ata gac tca tcc tat ctg gag cat ccg aat a-t ttt tcc att cta ttt tct agt gat ttt atc gat tct aat gaa tac gaa gtt aat cta agt cat tta gat tat caa agt gtg aac gat ttg atc gat tac att tat ggg ata cct ttg agc cta act aac gat agc gtg aaa tat att ctt tca act gct ga- --t ttt tta caa att gga tct gct att acc gaa tgt gaa aat tac ata ctt aaa aat ctg tgt tct aga aat tgt atc gat ttc --- --- tac ata tac gct gat aaa tat aat aac aag aaa ata gaa aca gcg tcg ttt aac aca ata tta cga aat att ttg aga ctc att aac gat gaa aac ttt aaa tac ttg aca gag gaa tca atg ata aaa att tta agc gat gat atg tta aat ata aaa aat gag gat ttt gca cca cta att ctc att aaa tgg tta gag agt aca caa caa cca tgc aca gtc gag tta ctt aga tgc ctc aga ata tca -tt gct ttc ccc aca agt tat aaa atc act tta tag tca tcg att ggt tgg ttc aat ata cga atg tat aac att ctt aaa caa tat atc att ctt gga tga atc att tcc tag ata cca tag cat cga gtt gat atc tat cgg tat aag taa ttc aca tga taa gat ttc cat aaa ctg cta taa tcg t-a aaa aaa ata cgt ggg aga taa tat ctt cac gga gat ata ggt gta gtt ttg cag tgg ccg tcc tgg ata ata tta tct ata tga tgg gtg gat atg atc agt ccc cgt ata gaa gtt caa agg tta tag cgt aca ata cat gta caa att ctt gga tat atg ata tac cag aac t-a aaa tat cct cgt tct aat tgt gga gga gtt gct gat gat gaa tac att tat tgc ata ggc ggc ata cgc gat cag gat tca tcg ttg ata tct agt att gat aga tgg aag cca tca aaa cca tat tgg caa acg tat gct aaa atg cga gag cca aaa tgt gat atg ggt gta acg atg tta aac gga tta ata tat gtc ata ggc gga gtt gtt aaa ggt g-- aca cgt gta ccg aca cac ttg aga gtt tat cag aag atg gat gga tga tgc atc aac gtc ttc caa taa aaa tgt cca ata tgt cga cga ttg ttc atg ccg gca aaa ttt ata tat ctg gag gct aca aca ata gta gtg tag tta atg gaa tat cga atc t-- agt cct tag cta taa tcc gat ata tga tga atg gac caa att atc atc att aaa tat tcc tag aat taa tcc cgc tct a

>HQ420897.1:176953-179187_Cowpox_virus_strain_Germany_1998_2_complete_genome

--- --- --- --a --- tgt a-c tat ttt ata cat tac tac att ggt tac ggc aga tat aca tac tcc gtt acc gcc aca cgc ttc ggt aaa cgg --g tca tgt gcc gag gga gaa tat ctt gat aag act cat aat cga tgt tgt aat -cg gtg tcc acc tgg aga att tgc caa ggt tag atg ta- --g tgg tag cta taa cac aaa atg tga acg ctg tcc aca tca tac ata tac agc aat tcc caa tta ttc t-a atg gat gcc atc aat gta gaa aat gcc caa agg gat cat ttg ata agg taa agt gta ccg gaa cac aga aca gta aat -gt gcg tgt ctt cct ggt tgg tat tgc gct acc gat tct tca cag act gaa gac tgt cga gat -tg tgt acc aaa aag g-a gct gtc cat gcg gat act ttg gtg gaa tag ata aat tag gaa atc cta ttt gta aat cgt gtt g-- -tg ttg gta aat att gtg acg aca tac gta atc ata gag ttg gtc ctt ttc ctc cat gca aa- --- cta tct aaa tgt aat tag tca --- --- --- --- --- --- --- --- --- --- --- --- --- --- -tg atc atg gta -cc gta caa tt- --- -at att gct --- --g gta --g tat att att cag tat g-a aga cat att aat ac- --- --- -tt atc --- --t ttg aca atc ttg ct- -at aat cat --- --- --- --- aat ata aaa --a tac tta tgg --- -a- cat agt cac tca t-- --- --- --- aat tgc tg- acg cga taa att cgt aat aat ctg ttt tgt tca aat ttt tat aag ga- --- --a tct aca ggc ata aaa at- aaa aat ata att tat aat ac- --- act tac agc gcg cca tca tga ata aca gca gtg aat taa ttg ctg tta tta atg gat tta gaa ata gtg gac gat ttt gtg ata tta gta tag tta tta atg atg aaa gga taa acg ctc ata gac tca tcc tat ctg gag cat ccg aat a-t ttt tcc att cta ttt tct agt gat ttt atc gat tct aat gaa tac gaa gtt aat cta agt cat tta gat tat caa agt gtg aac gat ttg atc gat tac att tat ggg ata cct ttg agc cta act aac gat agc gtg aaa tat att ctt tca act gct ga- --t ttt tta caa att gga tct gct att acc gaa tgt gaa aat tac ata ctt aaa aat ctg tgt tct aga aat tgt atc gat ttc --- --- tac ata tac gct gat aaa tat aat aac aag aaa ata gaa aca gcg tcg ttt aac aca ata tta cga aat att ttg aga ctc att aac gat gaa aac ttt aaa tac ttg aca gag gaa tca atg ata aaa att tta agc gat gat atg tta aat ata aaa aat gag gat ttt gca cca cta att ctc att aaa tgg tta gag agt aca caa caa cca tgc aca gtc gag tta ctt aga tgc ctc aga ata tca -tt gct ttc ccc aca agt tat aaa atc act tta tag tca tcg att ggt tgg ttc aat ata cga atg tat aac att ctt aaa caa tat atc att ctt gga tga atc att tcc tag ata cca tag cat cga gtt gat atc tat cgg tat aag taa ttc aca tga taa gat ttc cat aaa ctg cta taa tcg t-a aaa aaa ata cgt ggg aga taa tat ctt cac gga gat ata ggt gta gtt ttg cag tgg ccg tca tgg ata ata tta tct ata tga tgg gtg gat atg atc agt ccc cgt ata gga gtt caa agg tta tag cgt aca ata cct gta cta att ctt gga tat atg ata tac cag agc t-a aaa tat cct cgt tct aat tgt gga gga gtt gct gat gat gaa tac att tat tgc ata ggc ggc ata cgc gat cag gat tca tcg ttg ata tct agt att gat aga tgg aag cca tca aaa cca tat tgg caa acg tat gct aaa atg cga gag cca aaa tgt gat atg ggt gta gcg atg tta aac gga tta ata tat gtc ata ggc gga gtt gtt aaa ggt g-- aca cgt gta ccg aca cac ttg aga gtt tat cag aag atg gat gga tga tgc atc aac gtc ttc caa taa aaa tgt cca ata tgt cga cga ttg ttc atg ccg gca aaa ttt ata tat ctg gag gct aca aca ata gta gtg tag tta atg gaa tat cga atc t-- agt cct tag cta taa tcc gat ata tga tga atg gac caa att atc atc att aaa tat tcc tag aat taa tcc cgc tct a

>KT184690.1:160309-162494_Vaccinia_virus_strain_IOC_clone_B141

ttg ctg act gca --- tgt a-c tat ttt tta cat tac tac att ggc tac ggc aga tat acc tac tcc gcc acc --- --- --- --- --- -aa cgg --g tca tgt gac aag gga gaa tat ctt gat aag agg cat aat c-- --- --- -at -cg gtg tcc acc tgg aga att tgc caa ggt tag atg ta- --a tgg taa cga taa cac aaa atg tga acg ctg ccc acc tca tac ata tac cac aat -cc caa tta ttc t-a atg gat gtc atc aat gta gaa aat gcc caa ccg gat cat ttg ata agg taa agt gta ccg gaa cac aga aca gta aat -gt tcg tgt ctt cct ggt tgg tat tgc gct act gat tct tca cag act gaa gat tgt tga aat -tg tgt acc aaa aag g-a gat gtc cat gcg gat act ttg gtg gaa tag atg aac aag gaa atc cta ttt gta aat cgt gct g-- -ta ttg gtg aat att gca act acc tac gta att ata gac ttg atc cat ttt ctc cat gca aa- --- cta tct aaa tgt aat taa tta --- --- --- --- --- --- --- --- tga ttt --- --- --- -tg atg ata atg tta -cc ata c-a tt- --- -at atc gct act tgg tta --g tgt att att cag tat g-a aga cct att aat aa- --- --- -tt act tat ctt ttg acg atc ttg tt- -at aat tat --- --- --- --- aat ata aaa --a tac tta tgg --- --- cat agt aac tca t-- --- --- --- aat tgc tg- acg cga taa att cgt aat aat ctg ttt tgt tca --- --- --- aag ga- --- --a tct aca ggc ata aaa at- aaa aat ata att tat aat a-a c-- tct tac a-- acg cca tca tga ata gca gca gta aat taa ttg ctg tta tta atg gat tta gaa ata gtg gac gat ttt gtg ata tta ata tag tta tta atg atg aaa gga taa acg ctc ata aac tca tcc tat ctg gag cct ccg aat a-t ttt tcc att ctg ttt tcc aat aat ttt atc gat tct aat gaa tac gaa gtt aat cta agt cat tta gat tat caa agc gta aac gat ttg atc gat tat att tat ggg at- --- --- -gc cta act aac gat aac gtg aaa tat att ctt tca acc gct ga- --t ttt tta caa att gga tct gct att acg gag tgt gaa aat tac ata ctt aaa aat ctt tgt tct aaa aac tgt atc gat ttc --- --- tac ata tac gct gat aaa tat aat aac aag aaa ata gaa tca gcg tcg ttt aac aca ata tta caa aat att ttg aga ctc atc aac gat gaa aac ttt aaa tac tta aca gag gaa tca atg ata aaa att tta agc gat gat atg tta aat ata aaa aat gag gat ttt gca cca cta att ctc att aaa tgg tta gag agt act caa caa tca tgc acc gtc gag tta ctt aga tgc ctc aga ata tca -tt gct ttc ccc aca agt tat aaa atc act tta tag tca tcg act ggt tag ttc aat cta cga atg tat aa- --- --- --- --- --- --c att ctt gga tga atc att tcc tag ata cca tag cat cga gtt gat atc tat cgg tat aag taa ttc gca tga taa gat ttc cat aaa ctg cta caa tca t-a aaa aaa ata cat ggg aaa tga tat ctt cac gta gat ata ggt gta gtt tcg cag tgg ccg tcc tgg ata ata tta ttt ata tga tgg gtg gat atg atc agt ccc cgt ata gaa gtt caa agg tta tag cgt aca ata cat gta caa att ctt gga tat atg ata tac cag agc t-a aaa tat cct cgt tct aat tgt ggg gga ctg gct gat gac gaa tac att tat tgt ata ggc ggc ata cgc gat cag gat tca tcg ttg aca tct agt att gat aaa tgg aag cca tca aaa cca tat tgg cag aag tat gct aaa atg cgc gaa cca aaa tgt gat atg ggg gtt gcg atg tta aac gga tta ata tat gtt at- --- --- --- --- --a ggt g-- aca cgt gta ccg acg cac tag aga gtt tat cag aag atg gat gga tga agc atc aac gtc ttc caa taa aaa tgt cca ata tgt cga cga ttg ttc atg atg gca aga ttt ata tat ctg gag gtt aca aca ata gta gtg tag tta atg taa tat cga atc t-- agt cct tag cta taa tcc gat ata tga tga atg gac caa att atc atc att aaa cat tcc tag aat taa tcc cgc tct a

>KT184691.1:159867-162050_Vaccinia_virus_strain_IOC_clone_B388

ttg ctg act gca --- tgt a-c tat ttt tta cat tac tac att ggc tac ggc aga tat acc tac tcc gcc acc --- --- --- --- --- -aa cgg --g tca tgt gac aag gga gaa tat ctt gat aag agg cat aat c-- --- --- -at -cg gtg tcc acc tgg aga att tgc caa ggt tag atg ta- --a tgg taa cga taa cac aaa atg tga acg ctg ccc acc tca tac ata tac cac aat -cc caa tta ttc t-a atg gat gtc atc aat gta gaa aat gcc caa ccg gat cat ttg ata agg taa agt gta ccg gaa cac aga aca gta aat -gt tcg tgt ctt cct ggt tgg tat tgc gct act gat tct tca cag act gaa gat tgt cga gat -tg tat acc aaa aag g-a gat gtc cat gcg gat act ttg gtg gaa tag atg aac aag gaa atc cta ttt gta aat cgt gtt g-- -tg ttg gtg aat att gcg act acc tac gta att ata gac ttg atc cat ttc ctc cat gca aa- --- cta tct aaa tgt aat taa tta --- --- --- --- --- --- --- --- tga ttt --- --- --- -tg atg ata atg tta -cc ata c-a tt- --- -at atc gct act tgg tta --g tgt att att cag tat g-a aga cct att aat aa- --- --- -tt act tat ctt ttg acg atc ttg tt- -at aat tat --- --- --- --- aat ata aaa --- -ac tta tgg --- --- cat agt aac tta t-- --- --- --- aat tgc ta- aca cga taa att cgt aat aat ctg ttt tgt tca --- --- --- aag ga- --- --a tct aca ggc ata aaa at- aaa aat ata att tat aat a-a c-- tct tac a-- acg cca tca tga ata gca gca gta aat taa ttg ctg tta tta atg gat tta gaa ata gtg gac gat ttt gtg ata tta ata tag tta tta atg atg aaa gga taa acg ctc ata aac tca tcc tat ctg gag cct ccg aat a-t ttt tcc att ctg ttt tcc aat aat ttt atc gat tct aat gaa tac gaa gtt aat cta agt cat tta gat tat caa agc gta aac gat ttg atc gat tat att tat ggg at- --- --- -gc cta act aac gat aac gtg aaa tat att ctt tca acc gct ga- --t ttt tta caa att gga tct gct att acg gag tgt gaa aat tac ata ctt aaa aat ctt tgt tct aaa aac tgt atc gat ttc --- --- tac ata tac gct gat aaa tat aat aac aag aaa ata gaa tca gcg tcg ttt aac aca ata tta caa aat att ttg aga ctc atc aac gat gaa aac ttt aaa tac tta aca gag gaa tca atg ata aaa att tta agc gat gat atg tta aat ata aaa aat gag gat ttt gca cca cta att ctc att aaa tgg tta gag agt act caa caa tca tgc acc gtc gag tta ctt aga tgc ctc aga ata tca -tt gct ttc ccc aca agt tat aaa atc act tta tag tca tcg act ggt tag ttc aat cta cga atg tat aa- --- --- --- --- --- --c att ctt gga tga atc att tcc tag ata cca tag cat cga gtt gat atc tat cgg tat aag taa ttc gca tga taa gat ttc cat aaa ctg cta caa tca t-a aaa aaa ata cat ggg aaa tga tat ctt cac gta gat ata ggt gta gtt tcg cag tgg ccg tcc tgg ata ata tta ttt ata tga tgg gtg gat atg atc agt ccc cgt ata gaa gtt caa agg tta tag cgt aca ata cat gta caa att ctt gga tat atg ata tac cag agc t-a aaa tat cct cgt tct aat tgt ggg gga ctg gct gat gac gaa tac att tat tgt ata ggc ggc ata cgc gat cag gat tca tcg ttg aca tct agt att gat aaa tgg aag cca tca aaa cca tat tgg cag aag tat gct aaa atg cgc gaa cca aaa tgt gat atg ggg gtt gcg atg tta aac gga tta ata tat gtt at- --- --- --- --- --a ggt g-- aca cgt gta ccg acg cac tag aga gtt tat cag aag atg gat gga tga agc atc aac gtc ttc caa taa aaa tgt cca ata tgt cga cga ttg ttc atg atg gca aga ttt ata tat ctg gag gtt aca aca ata gta gtg tag tta atg taa tat cga atc t-- agt cct tag cta taa tcc gat ata tga tga atg gac caa att atc atc att aaa cat tcc tag aat taa tcc cgc tct a

>MK035759.1:175725-177959_Cowpox_virus_strain_CPXV/Cepad_332

--- --- --- --a --- tgt a-c tat ttt ata cat tac tac att ggt tac ggc aga tat aca tac tcc gtt acc gcc aca cgc tcc ggt aaa cgg --g tca tgt gcc gag gga gaa tat ctt gat aag act cat aat cga tgt tgt aat -cg gtg tcc acc tgg aga att tgc caa ggt tag atg ta- --g tgg tag cta taa cac aaa atg tga acg ctg tcc aca tca tac ata tac agc aat tcc caa tta ttc t-a atg gat gcc atc aat gta gaa aat gcc caa agg gat cat ttg ata agg taa agt gta ccg gaa aac aga aca gta aat -gt gcg tgt ctt cct ggt tgg tat tgc gct acc gat tct tca cag act gaa gac tgt cga gat -tg tgt acc aaa aag g-a gct gtc cat gcg gat act ttg gtg gaa tag ata aat tag gaa atc cta ttt gta aat cgt gtt g-- -tg ttg gta aat att gtg acg aca tac gta atc ata gag ttg gtc ctt ttc ctc cat gca aa- --- cta tct aaa tgt aat tag tca --- --- --- --- --- --- --- --- --- --- --- --- --- --- -tg atc atg gta -cc gta caa tt- --- -at att gct --- --g gta --g tat att att cag tat g-a aga cat att aat ac- --- --- -tt atc --- --t ttg aca atc ttg ct- -at aat cat --- --- --- --- aat ata aaa --a tac tta tgg --- -a- cat agt cac tca t-- --- --- --- aat tgc tg- acg cga taa att cgt aat aat ctg ttt tgt tca aat ttt tat aag ga- --- --a tct aca ggc ata aaa at- aaa aat ata att tat aat ac- --- act tac agc gcg cca tca tga ata aca gca gtg aat taa ttg ctg tta tta atg gat tta gaa ata gtg gac gat ttt gtg ata tta ata tag tta tta atg atg aaa gga taa atg ctc ata gac tca tcc tat ctg gag cat ccg aat a-t ttt tcc att cta ttt tct agt gat ttt atc gat tct aat gaa tac gaa gtt aat cta agt cat tta gat tat caa agt gtg aac gat ttg atc gat tac att tat ggg ata cct ttg agc cta act aac gat agc gtg aaa tat att ctt tca act gct ga- --t ttt tta caa att gga tct gct att acc gaa tgt gaa aat tac ata ctt aaa aat ctg tgt tct aga aat tgt atc gat ttc --- --- tac ata tac gct gat aaa tat aat aac aag aaa ata gaa aca gcg tcg ttt aac aca ata tta cga aat att ttg aga ctc att aac gat gaa aac ttt aaa tac ttg aca gag gaa tca atg ata aaa att tta agc gat gat atg tta aat ata aaa aat gag gat ttt gca cca cta att ctc att aaa tgg tta gag agt aca caa caa cca tgc aca gtc gag tta ctt aga tgc ctc aga ata tca -tt gct ttc ccc aca agt tat aaa atc act tta tag tca tcg att ggt tgg ttc aat ata cga atg tat aac att ctt aaa caa tat atc att ctt gga tga atc att tcc tag ata cca tag cat cga gtt gat atc tat cgg tat aag taa ttc aca tga taa gat ttc cat aaa ctg cta taa tcg t-a aaa aaa ata cgt ggg aga taa tat ctt cac gga gat ata ggt gta gtt ttg cag tgg ccg tcc tgg ata ata tta tct ata tga tgg gtg gat atg atc agt ccc cgt ata gaa gtt caa agg tta tag cgt aca ata cat gta caa att ctt gga tat atg ata tac cag aac t-a aaa tat cct cgt tct aat tgt gga gga gtt gct gat gat gaa tac att tat tgc ata ggc ggc ata cgc gat cag gat tca tcg ttg ata tct agt att gat aga tgg aag cca tca aaa cca tat tgg caa acg tat gct aaa atg cga gag cca aaa tgt gat atg ggt gta gcg atg tta aac gga tta ata tat gtc ata ggc gga gtt gtt aaa ggt g-- aca cgt gta ccg aca cac ttg aga gtt tat cag aag atg gat gga tga tgc atc aac gtc ttc caa taa aaa tgt cca ata tgt cga cga ctg ttc atg ccg gca aaa ttt ata tat ctg gag gct aca aca ata gta gtg tag tta atg gaa tat cga atc t-- agt cct tag cta taa tcc gat ata tga tga atg gac caa att atc atc att aaa cat tcc tag aat taa tcc cgc tct a

>MK035757.1:176020-178254_Cowpox_virus_strain_CPXV/Rat_Koelle

--- --- --- --a --- tgt a-c tat ttt ata cat tac tac att ggt tac ggc aga tat aca tac tcc gtt acc gcc aca cgc tcc ggt aaa cgg --g tca tgt gcc gag gga gaa tat ctt gat aag act cat aat cga tgt tgt aat -cg gtg tcc acc tgg aga att tgc caa ggt tag atg ta- --g tgg tag cta taa cac aaa atg tga acg ctg tcc aca tca tac ata tac agc aat tcc caa tta ttc t-a atg gat gcc atc aat gta gaa aat gcc caa agg gat cat ttg ata agg taa agt gta ccg gaa aac aga aca gta aat -gt gcg tgt ctt cct ggt tgg tat tgc gct acc gat tct tca cag act gaa gac tgt cga gat -tg tgt acc aaa aag g-a gct gtc cat gcg gat act ttg gtg gaa tag ata aat tag gaa atc cta ttt gta aat cgt gtt g-- -tg ttg gta aat att gtg acg aca tac gta atc ata gag ttg gtc ctt ttc ctc cat gca aa- --- cta tct aaa tgt aat tag tca --- --- --- --- --- --- --- --- --- --- --- --- --- --- -tg atc atg gta -cc gta caa tt- --- -at att gct --- --g gta --g tat att att cag tat g-a aga cat att aat ac- --- --- -tt atc --- --t ttg aca atc ttg ct- -at aat cat --- --- --- --- aat ata aaa --a tac tta tgg --- -a- cat agt cac tca t-- --- --- --- aat tgc tg- acg cga taa att cgt aat aat ctg ttt tgt tca aat ttt tat aag ga- --- --a tct aca ggc ata aaa at- aaa aat ata att tat aat ac- --- act tac agc gcg cca tca tga ata aca gca gtg aat taa ttg ctg tta tta atg gat tta gaa ata gtg gac gat ttt gtg ata tta ata tag tta tta atg atg aaa gga taa atg ctc ata gac tca tcc tat ctg gag cat ccg aat a-t ttt tcc att cta ttt tct agt gat ttt atc gat tct aat gaa tac gaa gtt aat cta agt cat tta gat tat caa agt gtg aac gat ttg atc gat tac att tat ggg ata cct ttg agc cta act aac gat agc gtg aaa tat att ctt tca act gct ga- --t ttt tta caa att gga tct gct att acc gaa tgt gaa aat tac ata ctt aaa aat ctg tgt tct aga aat tgt atc gat ttc --- --- tac ata tac gct gat aaa tat aat aac aag aaa ata gaa aca gcg tcg ttt aac aca ata tta cga aat att ttg aga ctc att aac gat gaa aac ttt aaa tac ttg aca gag gaa tca atg ata aaa att tta agc gat gat atg tta aat ata aaa aat gag gat ttt gca cca cta att ctc att aaa tgg tta gag agt aca caa caa cca tgc aca gtc gag tta ctt aga tgc ctc aga ata tca -tt gct ttc ccc aca agt tat aaa atc act tta tag tca tcg att ggt tgg ttc aat ata cga atg tat aac att ctt aaa caa tat atc att ctt gga tga atc att tcc tag ata cca tag cat cga gtt gat atc tat cgg tat aag taa ttc aca tga taa gat ttc cat aaa ctg cta taa tcg t-a aaa aaa ata cgt ggg aga taa tat ctt cac gga gat ata ggt gta gtt ttg cag tgg ccg tcc tgg ata ata tta tct ata tga tgg gtg gat atg atc agt ccc cgt ata gaa gtt caa agg tta tag cgt aca ata cat gta caa att ctt gga tat atg ata tac cag aac t-a aaa tat cct cgt tct aat tgt gga gga gtt gct gat gat gaa tac att tat tgc ata ggc ggc ata cgc gat cag gat tca tcg ttg ata tct agt att gat aga tgg aag cca tca aaa cca tat tgg caa acg tat gct aaa atg cga gag cca aaa tgt gat atg ggt gta gcg atg tta aac gga tta ata tat gtc ata ggc gga gtt gtt aaa ggt g-- aca cgt gta ccg aca cac ttg aga gtt tat cag aag atg gat gga tga tgc atc aac gtc ttc caa taa aaa tgt cca ata tgt cga cga ctg ttc atg ccg gca aaa ttt ata tat ctg gag gct aca aca ata gta gtg tag tta atg gaa tat cga atc t-- agt cct tag cta taa tcc gat ata tga tga atg gac caa att atc atc att aaa cat tcc tag aat taa tcc cgc tct a

>MK035753.1:175727-177961_Cowpox_virus_strain_CPXV/Cepad_333

--- --- --- --a --- tgt a-c tat ttt ata cat tac tac att ggt tac ggc aga tat aca tac tcc gtt acc gcc aca cgc tcc ggt aaa cgg --g tca tgt gcc gag gga gaa tat ctt gat aag act cat aat cga tgt tgt aat -cg gtg tcc acc tgg aga att tgc caa ggt tag atg ta- --g tgg tag cta taa cac aaa atg tga acg ctg tcc aca tca tac ata tac agc aat tcc caa tta ttc t-a atg gat gcc atc aat gta gaa aat gcc caa agg gat cat ttg ata agg taa agt gta ccg gaa aac aga aca gta aat -gt gcg tgt ctt cct ggt tgg tat tgc gct acc gat tct tca cag act gaa gac tgt cga gat -tg tgt acc aaa aag g-a gct gtc cat gcg gat act ttg gtg gaa tag ata aat tag gaa atc cta ttt gta aat cgt gtt g-- -tg ttg gta aat att gtg acg aca tac gta atc ata gag ttg gtc ctt ttc ctc cat gca aa- --- cta tct aaa tgt aat tag tca --- --- --- --- --- --- --- --- --- --- --- --- --- --- -tg atc atg gta -cc gta caa tt- --- -at att gct --- --g gta --g tat att att cag tat g-a aga cat att aat ac- --- --- -tt atc --- --t ttg aca atc ttg ct- -at aat cat --- --- --- --- aat ata aaa --a tac tta tgg --- -a- cat agt cac tca t-- --- --- --- aat tgc tg- acg cga taa att cgt aat aat ctg ttt tgt tca aat ttt tat aag ga- --- --a tct aca ggc ata aaa at- aaa aat ata att tat aat ac- --- act tac agc gcg cca tca tga ata aca gca gtg aat taa ttg ctg tta tta atg gat tta gaa ata gtg gac gat ttt gtg ata tta ata tag tta tta atg atg aaa gga taa atg ctc ata gac tca tcc tat ctg gag cat ccg aat a-t ttt tcc att cta ttt tct agt gat ttt atc gat tct aat gaa tac gaa gtt aat cta agt cat tta gat tat caa agt gtg aac gat ttg atc gat tac att tat ggg ata cct ttg agc cta act aac gat agc gtg aaa tat att ctt tca act gct ga- --t ttt tta caa att gga tct gct att acc gaa tgt gaa aat tac ata ctt aaa aat ctg tgt tct aga aat tgt atc gat ttc --- --- tac ata tac gct gat aaa tat aat aac aag aaa ata gaa aca gcg tcg ttt aac aca ata tta cga aat att ttg aga ctc att aac gat gaa aac ttt aaa tac ttg aca gag gaa tca atg ata aaa att tta agc gat gat atg tta aat ata aaa aat gag gat ttt gca cca cta att ctc att aaa tgg tta gag agt aca caa caa cca tgc aca gtc gag tta ctt aga tgc ctc aga ata tca -tt gct ttc ccc aca agt tat aaa atc act tta tag tca tcg att ggt tgg ttc aat ata cga atg tat aac att ctt aaa caa tat atc att ctt gga tga atc att tcc tag ata cca tag cat cga gtt gat atc tat cgg tat aag taa ttc aca tga taa gat ttc cat aaa ctg cta taa tcg t-a aaa aaa ata cgt ggg aga taa tat ctt cac gga gat ata ggt gta gtt ttg cag tgg ccg tcc tgg ata ata tta tct ata tga tgg gtg gat atg atc agt ccc cgt ata gaa gtt caa agg tta tag cgt aca ata cat gta caa att ctt gga tat atg ata tac cag aac t-a aaa tat cct cgt tct aat tgt gga gga gtt gct gat gat gaa tac att tat tgc ata ggc ggc ata cgc gat cag gat tca tcg ttg ata tct agt att gat aga tgg aag cca tca aaa cca tat tgg caa acg tat gct aaa atg cga gag cca aaa tgt gat atg ggt gta gcg atg tta aac gga tta ata tat gtc ata ggc gga gtt gtt aaa ggt g-- aca cgt gta ccg aca cac ttg aga gtt tat cag aag atg gat gga tga tgc atc aac gtc ttc caa taa aaa tgt cca ata tgt cga cga ctg ttc atg ccg gca aaa ttt ata tat ctg gag gct aca aca ata gta gtg tag tta atg gaa tat cga atc t-- agt cct tag cta taa tcc gat ata tga tga atg gac caa att atc atc att aaa cat tcc tag aat taa tcc cgc tct a

>MK035749.1:161232-163466_Cowpox_virus_strain_CPXV/Rat_Biederstein

--- --- --- --a --- tgt a-c tat ttt ata cat tac tac att ggt tac ggc aga tat aca tac tcc gtt acc gcc aca cgc tcc ggt aaa cgg --g tca tgt gcc gag gga gaa tat ctt gat aag act cat aat cga tgt tgt aat -cg gtg tcc acc tgg aga att tgc caa ggt tag atg ta- --g tgg tag cta taa cac aaa atg tga acg ctg tcc aca tca tac ata tac agc aat tcc caa tta ttc t-a atg gat gcc atc aat gta gaa aat gcc caa agg gat cat ttg ata agg taa agt gta ccg gaa aac aga aca gta aat -gt gcg tgt ctt cct ggt tgg tat tgc gct acc gat tct tca cag act gaa gac tgt cga gat -tg tgt acc aaa aag g-a gct gtc cat gcg gat act ttg gtg gaa tag ata aat tag gaa atc cta ttt gta aat cgt gtt g-- -tg ttg gta aat att gtg acg aca tac gta atc ata gag ttg gtc ctt ttc ctc cat gca aa- --- cta tct aaa tgt aat tag tca --- --- --- --- --- --- --- --- --- --- --- --- --- --- -tg atc atg gta -cc gta caa tt- --- -at att gct --- --g gta --g tat att att cag tat g-a aga cat att aat ac- --- --- -tt atc --- --t ttg aca atc ttg ct- -at aat cat --- --- --- --- aat ata aaa --a tac tta tgg --- -a- cat agt cac tca t-- --- --- --- aat tgc tg- acg cga taa att cgt aat aat ctg ttt tgt tca aat ttt tat aag ga- --- --a tct aca ggc ata aaa at- aaa aat ata att tat aat ac- --- act tac agc gcg cca tca tga ata aca gca gtg aat taa ttg ctg tta tta atg gat tta gaa ata gtg gac gat ttt gtg ata tta ata tag tta tta atg atg aaa gga taa atg ctc ata gac tca tcc tat ctg gag cat ccg aat a-t ttt tcc att cta ttt tct agt gat ttt atc gat tct aat gaa tac gaa gtt aat cta agt cat tta gat tat caa agt gtg aac gat ttg atc gat tac att tat ggg ata cct ttg agc cta act aac gat agc gtg aaa tat att ctt tca act gct ga- --t ttt tta caa att gga tct gct att acc gaa tgt gaa aat tac ata ctt aaa aat ctg tgt tct aga aat tgt atc gat ttc --- --- tac ata tac gct gat aaa tat aat aac aag aaa ata gaa aca gcg tcg ttt aac aca ata tta cga aat att ttg aga ctc att aac gat gaa aac ttt aaa tac ttg aca gag gaa tca atg ata aaa att tta agc gat gat atg tta aat ata aaa aat gag gat ttt gca cca cta att ctc att aaa tgg tta gag agt aca caa caa cca tgc aca gtc gag tta ctt aga tgc ctc aga ata tca -tt gct ttc ccc aca agt tat aaa atc act tta tag tca tcg att ggt tgg ttc aat ata cga atg tat aac att ctt aaa caa tat atc att ctt gga tga atc att tcc tag ata cca tag cat cga gtt gat atc tat cgg tat aag taa ttc aca tga taa gat ttc cat aaa ctg cta taa tcg t-a aaa aaa ata cgt ggg aga taa tat ctt cac gga gat ata ggt gta gtt ttg cag tgg ccg tcc tgg ata ata tta tct ata tga tgg gtg gat atg atc agt ccc cgt ata gaa gtt caa agg tta tag cgt aca ata cat gta caa att ctt gga tat atg ata tac cag aac t-a aaa tat cct cgt tct aat tgt gga gga gtt gct gat gat gaa tac att tat tgc ata ggc ggc ata cgc gat cag gat tca tcg ttg ata tct agt att gat aga tgg aag cca tca aaa cca tat tgg caa acg tat gct aaa atg cga gag cca aaa tgt gat atg ggt gta gcg atg tta aac gga tta ata tat gtc ata ggc gga gtt gtt aaa ggt g-- aca cgt gta ccg aca cac ttg aga gtt tat cag aag atg gat gga tga tgc atc aac gtc ttc caa taa aaa tgt cca ata tgt cga cga ctg ttc atg ccg gca aaa ttt ata tat ctg gag gct aca aca ata gta gtg tag tta atg gaa tat cga atc t-- agt cct tag cta taa tcc gat ata tga tga atg gac caa att atc atc att aaa cat tcc tag aat taa tcc cgc tct a

>MK035748.1:175637-177871_Cowpox_virus_strain_CPXV/Boa_Marl

--- --- --- --a --- tgt a-c tat ttt ata cat tac tac att ggt tac ggc aga tat aca tac tcc gtt acc gcc aca cgc tcc ggt aaa cgg --g tca tgt gcc gag gga gaa tat ctt gat aag act cat aat cga tgt tgt aat -cg gtg tcc acc tgg aga att tgc caa ggt tag atg ta- --g tgg tag cta taa cac aaa atg tga acg ctg tcc aca tca tac ata tac agc aat tcc caa tta ttc t-a atg gat gcc atc aat gta gaa aat gcc caa agg gat cat ttg ata agg taa agt gta ccg gaa aac aga aca gta aat -gt gcg tgt ctt cct ggt tgg tat tgc gct acc gat tct tca cag act gaa gac tgt cga gat -tg tgt acc aaa aag g-a gct gtc cat gcg gat act ttg gtg gaa tag ata aat tag gaa atc cta ttt gta aat cgt gtt g-- -tg ttg gta aat att gtg acg aca tac gta atc ata gag ttg gtc ctt ttc ctc cat gca aa- --- cta tct aaa tgt aat tag tca --- --- --- --- --- --- --- --- --- --- --- --- --- --- -tg atc atg gta -cc gta caa tt- --- -at att gct --- --g gta --g tat att att cag tat g-a aga cat att aat ac- --- --- -tt atc --- --t ttg aca atc ttg ct- -at aat cat --- --- --- --- aat ata aaa --a tac tta tgg --- -a- cat agt cac tca t-- --- --- --- aat tgc tg- acg cga taa att cgt aat aat ctg ttt tgt tca aat ttt tat aag ga- --- --a tct aca ggc ata aaa at- aaa aat ata att tat aat ac- --- act tac agc gcg cca tca tga ata aca gca gtg aat taa ttg ctg tta tta atg gat tta gaa ata gtg gac gat ttt gtg ata tta ata tag tta tta atg atg aaa gga taa atg ctc ata gac tca tcc tat ctg gag cat ccg aat a-t ttt tcc att cta ttt tct agt gat ttt atc gat tct aat gaa tac gaa gtt aat cta agt cat tta gat tat caa agt gtg aac gat ttg atc gat tac att tat ggg ata cct ttg agc cta act aac gat agc gtg aaa tat att ctt tca act gct ga- --t ttt tta caa att gga tct gct att acc gaa tgt gaa aat tac ata ctt aaa aat ctg tgt tct aga aat tgt atc gat ttc --- --- tac ata tac gct gat aaa tat aat aac aag aaa ata gaa aca gcg tcg ttt aac aca ata tta cga aat att ttg aga ctc att aac gat gaa aac ttt aaa tac ttg aca gag gaa tca atg ata aaa att tta agc gat gat atg tta aat ata aaa aat gag gat ttt gca cca cta att ctc att aaa tgg tta gag agt aca caa caa cca tgc aca gtc gag tta ctt aga tgc ctc aga ata tca -tt gct ttc ccc aca agt tat aaa atc act tta tag tca tcg att ggt tgg ttc aat ata cga atg tat aac att ctt aaa caa tat atc att ctt gga tga atc att tcc tag ata cca tag cat cga gtt gat atc tat cgg tat aag taa ttc aca tga taa gat ttc cat aaa ctg cta taa tcg t-a aaa aaa ata cgt ggg aga taa tat ctt cac gga gat ata ggt gta gtt ttg cag tgg ccg tcc tgg ata ata tta tct ata tga tgg gtg gat atg atc agt ccc cgt ata gaa gtt caa agg tta tag cgt aca ata cat gta caa att ctt gga tat atg ata tac cag aac t-a aaa tat cct cgt tct aat tgt gga gga gtt gct gat gat gaa tac att tat tgc ata ggc ggc ata cgc gat cag gat tca tcg ttg ata tct agt att gat aga tgg aag cca tca aaa cca tat tgg caa acg tat gct aaa atg cga gag cca aaa tgt gat atg ggt gta gcg atg tta aac gga tta ata tat gtc ata ggc gga gtt gtt aaa ggt g-- aca cgt gta ccg aca cac ttg aga gtt tat cag aag atg gat gga tga tgc atc aac gtc ttc caa taa aaa tgt cca ata tgt cga cga ctg ttc atg ccg gca aaa ttt ata tat ctg gag gct aca aca ata gta gtg tag tta atg gaa tat cga atc t-- agt cct tag cta taa tcc gat ata tga tga atg gac caa att atc atc att aaa cat tcc tag aat taa tcc cgc tct a

>MK035747.1:175639-177873_Cowpox_virus_strain_CPXV/Rat_Marl

--- --- --- --a --- tgt a-c tat ttt ata cat tac tac att ggt tac ggc aga tat aca tac tcc gtt acc gcc aca cgc tcc ggt aaa cgg --g tca tgt gcc gag gga gaa tat ctt gat aag act cat aat cga tgt tgt aat -cg gtg tcc acc tgg aga att tgc caa ggt tag atg ta- --g tgg tag cta taa cac aaa atg tga acg ctg tcc aca tca tac ata tac agc aat tcc caa tta ttc t-a atg gat gcc atc aat gta gaa aat gcc caa agg gat cat ttg ata agg taa agt gta ccg gaa aac aga aca gta aat -gt gcg tgt ctt cct ggt tgg tat tgc gct acc gat tct tca cag act gaa gac tgt cga gat -tg tgt acc aaa aag g-a gct gtc cat gcg gat act ttg gtg gaa tag ata aat tag gaa atc cta ttt gta aat cgt gtt g-- -tg ttg gta aat att gtg acg aca tac gta atc ata gag ttg gtc ctt ttc ctc cat gca aa- --- cta tct aaa tgt aat tag tca --- --- --- --- --- --- --- --- --- --- --- --- --- --- -tg atc atg gta -cc gta caa tt- --- -at att gct --- --g gta --g tat att att cag tat g-a aga cat att aat ac- --- --- -tt atc --- --t ttg aca atc ttg ct- -at aat cat --- --- --- --- aat ata aaa --a tac tta tgg --- -a- cat agt cac tca t-- --- --- --- aat tgc tg- acg cga taa att cgt aat aat ctg ttt tgt tca aat ttt tat aag ga- --- --a tct aca ggc ata aaa at- aaa aat ata att tat aat ac- --- act tac agc gcg cca tca tga ata aca gca gtg aat taa ttg ctg tta tta atg gat tta gaa ata gtg gac gat ttt gtg ata tta ata tag tta tta atg atg aaa gga taa atg ctc ata gac tca tcc tat ctg gag cat ccg aat a-t ttt tcc att cta ttt tct agt gat ttt atc gat tct aat gaa tac gaa gtt aat cta agt cat tta gat tat caa agt gtg aac gat ttg atc gat tac att tat ggg ata cct ttg agc cta act aac gat agc gtg aaa tat att ctt tca act gct ga- --t ttt tta caa att gga tct gct att acc gaa tgt gaa aat tac ata ctt aaa aat ctg tgt tct aga aat tgt atc gat ttc --- --- tac ata tac gct gat aaa tat aat aac aag aaa ata gaa aca gcg tcg ttt aac aca ata tta cga aat att ttg aga ctc att aac gat gaa aac ttt aaa tac ttg aca gag gaa tca atg ata aaa att tta agc gat gat atg tta aat ata aaa aat gag gat ttt gca cca cta att ctc att aaa tgg tta gag agt aca caa caa cca tgc aca gtc gag tta ctt aga tgc ctc aga ata tca -tt gct ttc ccc aca agt tat aaa atc act tta tag tca tcg att ggt tgg ttc aat ata cga atg tat aac att ctt aaa caa tat atc att ctt gga tga atc att tcc tag ata cca tag cat cga gtt gat atc tat cgg tat aag taa ttc aca tga taa gat ttc cat aaa ctg cta taa tcg t-a aaa aaa ata cgt ggg aga taa tat ctt cac gga gat ata ggt gta gtt ttg cag tgg ccg tcc tgg ata ata tta tct ata tga tgg gtg gat atg atc agt ccc cgt ata gaa gtt caa agg tta tag cgt aca ata cat gta caa att ctt gga tat atg ata tac cag aac t-a aaa tat cct cgt tct aat tgt gga gga gtt gct gat gat gaa tac att tat tgc ata ggc ggc ata cgc gat cag gat tca tcg ttg ata tct agt att gat aga tgg aag cca tca aaa cca tat tgg caa acg tat gct aaa atg cga gag cca aaa tgt gat atg ggt gta gcg atg tta aac gga tta ata tat gtc ata ggc gga gtt gtt aaa ggt g-- aca cgt gta ccg aca cac ttg aga gtt tat cag aag atg gat gga tga tgc atc aac gtc ttc caa taa aaa tgt cca ata tgt cga cga ctg ttc atg ccg gca aaa ttt ata tat ctg gag gct aca aca ata gta gtg tag tta atg gaa tat cga atc t-- agt cct tag cta taa tcc gat ata tga tga atg gac caa att atc atc att aaa cat tcc tag aat taa tcc cgc tct a

>MK035746.1:161686-163920_Cowpox_virus_strain_CPXV/Boy_Biederstein

--- --- --- --a --- tgt a-c tat ttt ata cat tac tac att ggt tac ggc aga tat aca tac tcc gtt acc gcc aca cgc tcc ggt aaa cgg --g tca tgt gcc gag gga gaa tat ctt gat aag act cat aat cga tgt tgt aat -cg gtg tcc acc tgg aga att tgc caa ggt tag atg ta- --g tgg tag cta taa cac aaa atg tga acg ctg tcc aca tca tac ata tac agc aat tcc caa tta ttc t-a atg gat gcc atc aat gta gaa aat gcc caa agg gat cat ttg ata agg taa agt gta ccg gaa aac aga aca gta aat -gt gcg tgt ctt cct ggt tgg tat tgc gct acc gat tct tca cag act gaa gac tgt cga gat -tg tgt acc aaa aag g-a gct gtc cat gcg gat act ttg gtg gaa tag ata aat tag gaa atc cta ttt gta aat cgt gtt g-- -tg ttg gta aat att gtg acg aca tac gta atc ata gag ttg gtc ctt ttc ctc cat gca aa- --- cta tct aaa tgt aat tag tca --- --- --- --- --- --- --- --- --- --- --- --- --- --- -tg atc atg gta -cc gta caa tt- --- -at att gct --- --g gta --g tat att att cag tat g-a aga cat att aat ac- --- --- -tt atc --- --t ttg aca atc ttg ct- -at aat cat --- --- --- --- aat ata aaa --a tac tta tgg --- -a- cat agt cac tca t-- --- --- --- aat tgc tg- acg cga taa att cgt aat aat ctg ttt tgt tca aat ttt tat aag ga- --- --a tct aca ggc ata aaa at- aaa aat ata att tat aat ac- --- act tac agc gcg cca tca tga ata aca gca gtg aat taa ttg ctg tta tta atg gat tta gaa ata gtg gac gat ttt gtg ata tta ata tag tta tta atg atg aaa gga taa atg ctc ata gac tca tcc tat ctg gag cat ccg aat a-t ttt tcc att cta ttt tct agt gat ttt atc gat tct aat gaa tac gaa gtt aat cta agt cat tta gat tat caa agt gtg aac gat ttg atc gat tac att tat ggg ata cct ttg agc cta act aac gat agc gtg aaa tat att ctt tca act gct ga- --t ttt tta caa att gga tct gct att acc gaa tgt gaa aat tac ata ctt aaa aat ctg tgt tct aga aat tgt atc gat ttc --- --- tac ata tac gct gat aaa tat aat aac aag aaa ata gaa aca gcg tcg ttt aac aca ata tta cga aat att ttg aga ctc att aac gat gaa aac ttt aaa tac ttg aca gag gaa tca atg ata aaa att tta agc gat gat atg tta aat ata aaa aat gag gat ttt gca cca cta att ctc att aaa tgg tta gag agt aca caa caa cca tgc aca gtc gag tta ctt aga tgc ctc aga ata tca -tt gct ttc ccc aca agt tat aaa atc act tta tag tca tcg att ggt tgg ttc aat ata cga atg tat aac att ctt aaa caa tat atc att ctt gga tga atc att tcc tag ata cca tag cat cga gtt gat atc tat cgg tat aag taa ttc aca tga taa gat ttc cat aaa ctg cta taa tcg t-a aaa aaa ata cgt ggg aga taa tat ctt cac gga gat ata ggt gta gtt ttg cag tgg ccg tcc tgg ata ata tta tct ata tga tgg gtg gat atg atc agt ccc cgt ata gaa gtt caa agg tta tag cgt aca ata cat gta caa att ctt gga tat atg ata tac cag aac t-a aaa tat cct cgt tct aat tgt gga gga gtt gct gat gat gaa tac att tat tgc ata ggc ggc ata cgc gat cag gat tca tcg ttg ata tct agt att gat aga tgg aag cca tca aaa cca tat tgg caa acg tat gct aaa atg cga gag cca aaa tgt gat atg ggt gta gcg atg tta aac gga tta ata tat gtc ata ggc gga gtt gtt aaa ggt g-- aca cgt gta ccg aca cac ttg aga gtt tat cag aag atg gat gga tga tgc atc aac gtc ttc caa taa aaa tgt cca ata tgt cga cga ctg ttc atg ccg gca aaa ttt ata tat ctg gag gct aca aca ata gta gtg tag tta atg gaa tat cga atc t-- agt cct tag cta taa tcc gat ata tga tga atg gac caa att atc atc att aaa cat tcc tag aat taa tcc cgc tct a

>LN864565.1:175540-177774_Cowpox_virus_complete_genome_strain_Ratpox09

--- --- --- --a --- tgt a-c tat ttt ata cat tac tac att ggt tac ggc aga tat aca tac tcc gtt acc gcc aca cgc tcc ggt aaa cgg --g tca tgt gcc gag gga gaa tat ctt gat aag act cat aat cga tgt tgt aat -cg gtg tcc acc tgg aga att tgc caa ggt tag atg ta- --g tgg tag cta taa cac aaa atg tga acg ctg tcc aca tca tac ata tac agc aat tcc caa tta ttc t-a atg gat gcc atc aat gta gaa aat gcc caa agg gat cat ttg ata agg taa agt gta ccg gaa aac aga aca gta aat -gt gcg tgt ctt cct ggt tgg tat tgc gct acc gat tct tca cag act gaa gac tgt cga gat -tg tgt acc aaa aag g-a gct gtc cat gcg gat act ttg gtg gaa tag ata aat tag gaa atc cta ttt gta aat cgt gtt g-- -tg ttg gta aat att gtg acg aca tac gta atc ata gag ttg gtc ctt ttc ctc cat gca aa- --- cta tct aaa tgt aat tag tca --- --- --- --- --- --- --- --- --- --- --- --- --- --- -tg atc atg gta -cc gta caa tt- --- -at att gct --- --g gta --g tat att att cag tat g-a aga cat att aat ac- --- --- -tt atc --- --t ttg aca atc ttg ct- -at aat cat --- --- --- --- aat ata aaa --a tac tta tgg --- -a- cat agt cac tca t-- --- --- --- aat tgc tg- acg cga taa att cgt aat aat ctg ttt tgt tca aat ttt tat aag ga- --- --a tct aca ggc ata aaa at- aaa aat ata att tat aat ac- --- act tac agc gcg cca tca tga ata aca gca gtg aat taa ttg ctg tta tta atg gat tta gaa ata gtg gac gat ttt gtg ata tta ata tag tta tta atg atg aaa gga taa atg ctc ata gac tca tcc tat ctg gag cat ccg aat a-t ttt tcc att cta ttt tct agt gat ttt atc gat tct aat gaa tac gaa gtt aat cta agt cat tta gat tat caa agt gtg aac gat ttg atc gat tac att tat ggg ata cct ttg agc cta act aac gat agc gtg aaa tat att ctt tca act gct ga- --t ttt tta caa att gga tct gct att acc gaa tgt gaa aat tac ata ctt aaa aat ctg tgt tct aga aat tgt atc gat ttc --- --- tac ata tac gct gat aaa tat aat aac aag aaa ata gaa aca gcg tcg ttt aac aca ata tta cga aat att ttg aga ctc att aac gat gaa aac ttt aaa tac ttg aca gag gaa tca atg ata aaa att tta agc gat gat atg tta aat ata aaa aat gag gat ttt gca cca cta att ctc att aaa tgg tta gag agt aca caa caa cca tgc aca gtc gag tta ctt aga tgc ctc aga ata tca -tt gct ttc ccc aca agt tat aaa atc act tta tag tca tcg att ggt tgg ttc aat ata cga atg tat aac att ctt aaa caa tat atc att ctt gga tga atc att tcc tag ata cca tag cat cga gtt gat atc tat cgg tat aag taa ttc aca tga taa gat ttc cat aaa ctg cta taa tcg t-a aaa aaa ata cgt ggg aga taa tat ctt cac gga gat ata ggt gta gtt ttg cag tgg ccg tcc tgg ata ata tta tct ata tga tgg gtg gat atg atc agt ccc cgt ata gaa gtt caa agg tta tag cgt aca ata cat gta caa att ctt gga tat atg ata tac cag aac t-a aaa tat cct cgt tct aat tgt gga gga gtt gct gat gat gaa tac att tat tgc ata ggc ggc ata cgc gat cag gat tca tcg ttg ata tct agt att gat aga tgg aag cca tca aaa cca tat tgg caa acg tat gct aaa atg cga gag cca aaa tgt gat atg ggt gta gcg atg tta aac gga tta ata tat gtc ata ggc gga gtt gtt aaa ggt g-- aca cgt gta ccg aca cac ttg aga gtt tat cag aag atg gat gga tga tgc atc aac gtc ttc caa taa aaa tgt cca ata tgt cga cga ctg ttc atg ccg gca aaa ttt ata tat ctg gag gct aca aca ata gta gtg tag tta atg gaa tat cga atc t-- agt cct tag cta taa tcc gat ata tga tga atg gac caa att atc atc att aaa cat tcc tag aat taa tcc cgc tct a

>KC813512.1:167858-170092_Cowpox_virus_strain_HumKre08/1_complete_genome

--- --- --- --a --- tgt a-c tat ttt ata cat tac tac att ggt tac ggc aga tat aca tac tcc gtt acc gcc aca cgc tcc ggt aaa cgg --g tca tgt gcc gag gga gaa tat ctt gat aag act cat aat cga tgt tgt aat -cg gtg tcc acc tgg aga att tgc caa ggt tag atg ta- --g tgg tag cta taa cac aaa atg tga acg ctg tcc aca tca tac ata tac agc aat tcc caa tta ttc t-a atg gat gcc atc aat gta gaa aat gcc caa agg gat cat ttg ata agg taa agt gta ccg gaa aac aga aca gta aat -gt gcg tgt ctt cct ggt tgg tat tgc gct acc gat tct tca cag act gaa gac tgt cga gat -tg tgt acc aaa aag g-a gct gtc cat gcg gat act ttg gtg gaa tag ata aat tag gaa atc cta ttt gta aat cgt gtt g-- -tg ttg gta aat att gtg acg aca tac gta atc ata gag ttg gtc ctt ttc ctc cat gca aa- --- cta tct aaa tgt aat tag tca --- --- --- --- --- --- --- --- --- --- --- --- --- --- -tg atc atg gta -cc gta caa tt- --- -at att gct --- --g gta --g tat att att cag tat g-a aga cat att aat ac- --- --- -tt atc --- --t ttg aca atc ttg ct- -at aat cat --- --- --- --- aat ata aaa --a tac tta tgg --- -a- cat agt cac tca t-- --- --- --- aat tgc tg- acg cga taa att cgt aat aat ctg ttt tgt tca aat ttt tat aag ga- --- --a tct aca ggc ata aaa at- aaa aat ata att tat aat ac- --- act tac agc gcg cca tca tga ata aca gca gtg aat taa ttg ctg tta tta atg gat tta gaa ata gtg gac gat ttt gtg ata tta ata tag tta tta atg atg aaa gga taa atg ctc ata gac tca tcc tat ctg gag cat ccg aat a-t ttt tcc att cta ttt tct agt gat ttt atc gat tct aat gaa tac gaa gtt aat cta agt cat tta gat tat caa agt gtg aac gat ttg atc gat tac att tat ggg ata cct ttg agc cta act aac gat agc gtg aaa tat att ctt tca act gct ga- --t ttt tta caa att gga tct gct att acc gaa tgt gaa aat tac ata ctt aaa aat ctg tgt tct aga aat tgt atc gat ttc --- --- tac ata tac gct gat aaa tat aat aac aag aaa ata gaa aca gcg tcg ttt aac aca ata tta cga aat att ttg aga ctc att aac gat gaa aac ttt aaa tac ttg aca gag gaa tca atg ata aaa att tta agc gat gat atg tta aat ata aaa aat gag gat ttt gca cca cta att ctc att aaa tgg tta gag agt aca caa caa cca tgc aca gtc gag tta ctt aga tgc ctc aga ata tca -tt gct ttc ccc aca agt tat aaa atc act tta tag tca tcg att ggt tgg ttc aat ata cga atg tat aac att ctt aaa caa tat atc att ctt gga tga atc att tcc tag ata cca tag cat cga gtt gat atc tat cgg tat aag taa ttc aca tga taa gat ttc cat aaa ctg cta taa tcg t-a aaa aaa ata cgt ggg aga taa tat ctt cac gga gat ata ggt gta gtt ttg cag tgg ccg tcc tgg ata ata tta tct ata tga tgg gtg gat atg atc agt ccc cgt ata gaa gtt caa agg tta tag cgt aca ata cat gta caa att ctt gga tat atg ata tac cag aac t-a aaa tat cct cgt tct aat tgt gga gga gtt gct gat gat gaa tac att tat tgc ata ggc ggc ata cgc gat cag gat tca tcg ttg ata tct agt att gat aga tgg aag cca tca aaa cca tat tgg caa acg tat gct aaa atg cga gag cca aaa tgt gat atg ggt gta gcg atg tta aac gga tta ata tat gtc ata ggc gga gtt gtt aaa ggt g-- aca cgt gta ccg aca cac ttg aga gtt tat cag aag atg gat gga tga tgc atc aac gtc ttc caa taa aaa tgt cca ata tgt cga cga ctg ttc atg ccg gca aaa ttt ata tat ctg gag gct aca aca ata gta gtg tag tta atg gaa tat cga atc t-- agt cct tag cta taa tcc gat ata tga tga atg gac caa att atc atc att aaa cat tcc tag aat taa tcc cgc tct a

>KC813508.1:168183-170417_Cowpox_virus_strain_HumAac09/1_complete_genome

--- --- --- --a --- tgt a-c tat ttt ata cat tac tac att ggt tac ggc aga tat aca tac tcc gtt acc gcc aca cgc tcc ggt aaa cgg --g tca tgt gcc gag gga gaa tat ctt gat aag act cat aat cga tgt tgt aat -cg gtg tcc acc tgg aga att tgc caa ggt tag atg ta- --g tgg tag cta taa cac aaa atg tga acg ctg tcc aca tca tac ata tac agc aat tcc caa tta ttc t-a atg gat gcc atc aat gta gaa aat gcc caa agg gat cat ttg ata agg taa agt gta ccg gaa aac aga aca gta aat -gt gcg tgt ctt cct ggt tgg tat tgc gct acc gat tct tca cag act gaa gac tgt cga gat -tg tgt acc aaa aag g-a gct gtc cat gcg gat act ttg gtg gaa tag ata aat tag gaa atc cta ttt gta aat cgt gtt g-- -tg ttg gta aat att gtg acg aca tac gta atc ata gag ttg gtc ctt ttc ctc cat gca aa- --- cta tct aaa tgt aat tag tca --- --- --- --- --- --- --- --- --- --- --- --- --- --- -tg atc atg gta -cc gta caa tt- --- -at att gct --- --g gta --g tat att att cag tat g-a aga cat att aat ac- --- --- -tt atc --- --t ttg aca atc ttg ct- -at aat cat --- --- --- --- aat ata aaa --a tac tta tgg --- -a- cat agt cac tca t-- --- --- --- aat tgc tg- acg cga taa att cgt aat aat ctg ttt tgt tca aat ttt tat aag ga- --- --a tct aca ggc ata aaa at- aaa aat ata att tat aat ac- --- act tac agc gcg cca tca tga ata aca gca gtg aat taa ttg ctg tta tta atg gat tta gaa ata gtg gac gat ttt gtg ata tta ata tag tta tta atg atg aaa gga taa atg ctc ata gac tca tcc tat ctg gag cat ccg aat a-t ttt tcc att cta ttt tct agt gat ttt atc gat tct aat gaa tac gaa gtt aat cta agt cat tta gat tat caa agt gtg aac gat ttg atc gat tac att tat ggg ata cct ttg agc cta act aac gat agc gtg aaa tat att ctt tca act gct ga- --t ttt tta caa att gga tct gct att acc gaa tgt gaa aat tac ata ctt aaa aat ctg tgt tct aga aat tgt atc gat ttc --- --- tac ata tac gct gat aaa tat aat aac aag aaa ata gaa aca gcg tcg ttt aac aca ata tta cga aat att ttg aga ctc att aac gat gaa aac ttt aaa tac ttg aca gag gaa tca atg ata aaa att tta agc gat gat atg tta aat ata aaa aat gag gat ttt gca cca cta att ctc att aaa tgg tta gag agt aca caa caa cca tgc aca gtc gag tta ctt aga tgc ctc aga ata tca -tt gct ttc ccc aca agt tat aaa atc act tta tag tca tcg att ggt tgg ttc aat ata cga atg tat aac att ctt aaa caa tat atc att ctt gga tga atc att tcc tag ata cca tag cat cga gtt gat atc tat cgg tat aag taa ttc aca tga taa gat ttc cat aaa ctg cta taa tcg t-a aaa aaa ata cgt ggg aga taa tat ctt cac gga gat ata ggt gta gtt ttg cag tgg ccg tcc tgg ata ata tta tct ata tga tgg gtg gat atg atc agt ccc cgt ata gaa gtt caa agg tta tag cgt aca ata cat gta caa att ctt gga tat atg ata tac cag aac t-a aaa tat cct cgt tct aat tgt gga gga gtt gct gat gat gaa tac att tat tgc ata ggc ggc ata cgc gat cag gat tca tcg ttg ata tct agt att gat aga tgg aag cca tca aaa cca tat tgg caa acg tat gct aaa atg cga gag cca aaa tgt gat atg ggt gta gcg atg tta aac gga tta ata tat gtc ata ggc gga gtt gtt aaa ggt g-- aca cgt gta ccg aca cac ttg aga gtt tat cag aag atg gat gga tga tgc atc aac gtc ttc caa taa aaa tgt cca ata tgt cga cga ctg ttc atg ccg gca aaa ttt ata tat ctg gag gct aca aca ata gta gtg tag tta atg gaa tat cga atc t-- agt cct tag cta taa tcc gat ata tga tga atg gac caa att atc atc att aaa cat tcc tag aat taa tcc cgc tct a

>KC813505.1:168104-170338_Cowpox_virus_strain_RatKre08/2_complete_genome

--- --- --- --a --- tgt a-c tat ttt ata cat tac tac att ggt tac ggc aga tat aca tac tcc gtt acc gcc aca cgc tcc ggt aaa cgg --g tca tgt gcc gag gga gaa tat ctt gat aag act cat aat cga tgt tgt aat -cg gtg tcc acc tgg aga att tgc caa ggt tag atg ta- --g tgg tag cta taa cac aaa atg tga acg ctg tcc aca tca tac ata tac agc aat tcc caa tta ttc t-a atg gat gcc atc aat gta gaa aat gcc caa agg gat cat ttg ata agg taa agt gta ccg gaa aac aga aca gta aat -gt gcg tgt ctt cct ggt tgg tat tgc gct acc gat tct tca cag act gaa gac tgt cga gat -tg tgt acc aaa aag g-a gct gtc cat gcg gat act ttg gtg gaa tag ata aat tag gaa atc cta ttt gta aat cgt gtt g-- -tg ttg gta aat att gtg acg aca tac gta atc ata gag ttg gtc ctt ttc ctc cat gca aa- --- cta tct aaa tgt aat tag tca --- --- --- --- --- --- --- --- --- --- --- --- --- --- -tg atc atg gta -cc gta caa tt- --- -at att gct --- --g gta --g tat att att cag tat g-a aga cat att aat ac- --- --- -tt atc --- --t ttg aca atc ttg ct- -at aat cat --- --- --- --- aat ata aaa --a tac tta tgg --- -a- cat agt cac tca t-- --- --- --- aat tgc tg- acg cga taa att cgt aat aat ctg ttt tgt tca aat ttt tat aag ga- --- --a tct aca ggc ata aaa at- aaa aat ata att tat aat ac- --- act tac agc gcg cca tca tga ata aca gca gtg aat taa ttg ctg tta tta atg gat tta gaa ata gtg gac gat ttt gtg ata tta ata tag tta tta atg atg aaa gga taa atg ctc ata gac tca tcc tat ctg gag cat ccg aat a-t ttt tcc att cta ttt tct agt gat ttt atc gat tct aat gaa tac gaa gtt aat cta agt cat tta gat tat caa agt gtg aac gat ttg atc gat tac att tat ggg ata cct ttg agc cta act aac gat agc gtg aaa tat att ctt tca act gct ga- --t ttt tta caa att gga tct gct att acc gaa tgt gaa aat tac ata ctt aaa aat ctg tgt tct aga aat tgt atc gat ttc --- --- tac ata tac gct gat aaa tat aat aac aag aaa ata gaa aca gcg tcg ttt aac aca ata tta cga aat att ttg aga ctc att aac gat gaa aac ttt aaa tac ttg aca gag gaa tca atg ata aaa att tta agc gat gat atg tta aat ata aaa aat gag gat ttt gca cca cta att ctc att aaa tgg tta gag agt aca caa caa cca tgc aca gtc gag tta ctt aga tgc ctc aga ata tca -tt gct ttc ccc aca agt tat aaa atc act tta tag tca tcg att ggt tgg ttc aat ata cga atg tat aac att ctt aaa caa tat atc att ctt gga tga atc att tcc tag ata cca tag cat cga gtt gat atc tat cgg tat aag taa ttc aca tga taa gat ttc cat aaa ctg cta taa tcg t-a aaa aaa ata cgt ggg aga taa tat ctt cac gga gat ata ggt gta gtt ttg cag tgg ccg tcc tgg ata ata tta tct ata tga tgg gtg gat atg atc agt ccc cgt ata gaa gtt caa agg tta tag cgt aca ata cat gta caa att ctt gga tat atg ata tac cag aac t-a aaa tat cct cgt tct aat tgt gga gga gtt gct gat gat gaa tac att tat tgc ata ggc ggc ata cgc gat cag gat tca tcg ttg ata tct agt att gat aga tgg aag cca tca aaa cca tat tgg caa acg tat gct aaa atg cga gag cca aaa tgt gat atg ggt gta gcg atg tta aac gga tta ata tat gtc ata ggc gga gtt gtt aaa ggt g-- aca cgt gta ccg aca cac ttg aga gtt tat cag aag atg gat gga tga tgc atc aac gtc ttc caa taa aaa tgt cca ata tgt cga cga ctg ttc atg ccg gca aaa ttt ata tat ctg gag gct aca aca ata gta gtg tag tta atg gaa tat cga atc t-- agt cct tag cta taa tcc gat ata tga tga atg gac caa att atc atc att aaa cat tcc tag aat taa tcc cgc tct a

>KC813503.1:175881-178115_Cowpox_virus_strain_RatGer09/1_complete_genome

--- --- --- --a --- tgt a-c tat ttt ata cat tac tac att ggt tac ggc aga tat aca tac tcc gtt acc gcc aca cgc tcc ggt aaa cgg --g tca tgt gcc gag gga gaa tat ctt gat aag act cat aat cga tgt tgt aat -cg gtg tcc acc tgg aga att tgc caa ggt tag atg ta- --g tgg tag cta taa cac aaa atg tga acg ctg tcc aca tca tac ata tac agc aat tcc caa tta ttc t-a atg gat gcc atc aat gta gaa aat gcc caa agg gat cat ttg ata agg taa agt gta ccg gaa aac aga aca gta aat -gt gcg tgt ctt cct ggt tgg tat tgc gct acc gat tct tca cag act gaa gac tgt cga gat -tg tgt acc aaa aag g-a gct gtc cat gcg gat act ttg gtg gaa tag ata aat tag gaa atc cta ttt gta aat cgt gtt g-- -tg ttg gta aat att gtg acg aca tac gta atc ata gag ttg gtc ctt ttc ctc cat gca aa- --- cta tct aaa tgt aat tag tca --- --- --- --- --- --- --- --- --- --- --- --- --- --- -tg atc atg gta -cc gta caa tt- --- -at att gct --- --g gta --g tat att att cag tat g-a aga cat att aat ac- --- --- -tt atc --- --t ttg aca atc ttg ct- -at aat cat --- --- --- --- aat ata aaa --a tac tta tgg --- -a- cat agt cac tca t-- --- --- --- aat tgc tg- acg cga taa att cgt aat aat ctg ttt tgt tca aat ttt tat aag ga- --- --a tct aca ggc ata aaa at- aaa aat ata att tat aat ac- --- act tac agc gcg cca tca tga ata aca gca gtg aat taa ttg ctg tta tta atg gat tta gaa ata gtg gac gat ttt gtg ata tta ata tag tta tta atg atg aaa gga taa atg ctc ata gac tca tcc tat ctg gag cat ccg aat a-t ttt tcc att cta ttt tct agt gat ttt atc gat tct aat gaa tac gaa gtt aat cta agt cat tta gat tat caa agt gtg aac gat ttg atc gat tac att tat ggg ata cct ttg agc cta act aac gat agc gtg aaa tat att ctt tca act gct ga- --t ttt tta caa att gga tct gct att acc gaa tgt gaa aat tac ata ctt aaa aat ctg tgt tct aga aat tgt atc gat ttc --- --- tac ata tac gct gat aaa tat aat aac aag aaa ata gaa aca gcg tcg ttt aac aca ata tta cga aat att ttg aga ctc att aac gat gaa aac ttt aaa tac ttg aca gag gaa tca atg ata aaa att tta agc gat gat atg tta aat ata aaa aat gag gat ttt gca cca cta att ctc att aaa tgg tta gag agt aca caa caa cca tgc aca gtc gag tta ctt aga tgc ctc aga ata tca -tt gct ttc ccc aca agt tat aaa atc act tta tag tca tcg att ggt tgg ttc aat ata cga atg tat aac att ctt aaa caa tat atc att ctt gga tga atc att tcc tag ata cca tag cat cga gtt gat atc tat cgg tat aag taa ttc aca tga taa gat ttc cat aaa ctg cta taa tcg t-a aaa aaa ata cgt ggg aga taa tat ctt cac gga gat ata ggt gta gtt ttg cag tgg ccg tcc tgg ata ata tta tct ata tga tgg gtg gat atg atc agt ccc cgt ata gaa gtt caa agg tta tag cgt aca ata cat gta caa att ctt gga tat atg ata tac cag aac t-a aaa tat cct cgt tct aat tgt gga gga gtt gct gat gat gaa tac att tat tgc ata ggc ggc ata cgc gat cag gat tca tcg ttg ata tct agt att gat aga tgg aag cca tca aaa cca tat tgg caa acg tat gct aaa atg cga gag cca aaa tgt gat atg ggt gta gcg atg tta aac gga tta ata tat gtc ata ggc gga gtt gtt aaa ggt g-- aca cgt gta ccg aca cac ttg aga gtt tat cag aag atg gat gga tga tgc atc aac gtc ttc caa taa aaa tgt cca ata tgt cga cga ctg ttc atg ccg gca aaa ttt ata tat ctg gag gct aca aca ata gta gtg tag tta atg gaa tat cga atc t-- agt cct tag cta taa tcc gat ata tga tga atg gac caa att atc atc att aaa cat tcc tag aat taa tcc cgc tct a

>KC813501.1:167862-170096_Cowpox_virus_strain_RatAac09/1_complete_genome

--- --- --- --a --- tgt a-c tat ttt ata cat tac tac att ggt tac ggc aga tat aca tac tcc gtt acc gcc aca cgc tcc ggt aaa cgg --g tca tgt gcc gag gga gaa tat ctt gat aag act cat aat cga tgt tgt aat -cg gtg tcc acc tgg aga att tgc caa ggt tag atg ta- --g tgg tag cta taa cac aaa atg tga acg ctg tcc aca tca tac ata tac agc aat tcc caa tta ttc t-a atg gat gcc atc aat gta gaa aat gcc caa agg gat cat ttg ata agg taa agt gta ccg gaa aac aga aca gta aat -gt gcg tgt ctt cct ggt tgg tat tgc gct acc gat tct tca cag act gaa gac tgt cga gat -tg tgt acc aaa aag g-a gct gtc cat gcg gat act ttg gtg gaa tag ata aat tag gaa atc cta ttt gta aat cgt gtt g-- -tg ttg gta aat att gtg acg aca tac gta atc ata gag ttg gtc ctt ttc ctc cat gca aa- --- cta tct aaa tgt aat tag tca --- --- --- --- --- --- --- --- --- --- --- --- --- --- -tg atc atg gta -cc gta caa tt- --- -at att gct --- --g gta --g tat att att cag tat g-a aga cat att aat ac- --- --- -tt atc --- --t ttg aca atc ttg ct- -at aat cat --- --- --- --- aat ata aaa --a tac tta tgg --- -a- cat agt cac tca t-- --- --- --- aat tgc tg- acg cga taa att cgt aat aat ctg ttt tgt tca aat ttt tat aag ga- --- --a tct aca ggc ata aaa at- aaa aat ata att tat aat ac- --- act tac agc gcg cca tca tga ata aca gca gtg aat taa ttg ctg tta tta atg gat tta gaa ata gtg gac gat ttt gtg ata tta ata tag tta tta atg atg aaa gga taa atg ctc ata gac tca tcc tat ctg gag cat ccg aat a-t ttt tcc att cta ttt tct agt gat ttt atc gat tct aat gaa tac gaa gtt aat cta agt cat tta gat tat caa agt gtg aac gat ttg atc gat tac att tat ggg ata cct ttg agc cta act aac gat agc gtg aaa tat att ctt tca act gct ga- --t ttt tta caa att gga tct gct att acc gaa tgt gaa aat tac ata ctt aaa aat ctg tgt tct aga aat tgt atc gat ttc --- --- tac ata tac gct gat aaa tat aat aac aag aaa ata gaa aca gcg tcg ttt aac aca ata tta cga aat att ttg aga ctc att aac gat gaa aac ttt aaa tac ttg aca gag gaa tca atg ata aaa att tta agc gat gat atg tta aat ata aaa aat gag gat ttt gca cca cta att ctc att aaa tgg tta gag agt aca caa caa cca tgc aca gtc gag tta ctt aga tgc ctc aga ata tca -tt gct ttc ccc aca agt tat aaa atc act tta tag tca tcg att ggt tgg ttc aat ata cga atg tat aac att ctt aaa caa tat atc att ctt gga tga atc att tcc tag ata cca tag cat cga gtt gat atc tat cgg tat aag taa ttc aca tga taa gat ttc cat aaa ctg cta taa tcg t-a aaa aaa ata cgt ggg aga taa tat ctt cac gga gat ata ggt gta gtt ttg cag tgg ccg tcc tgg ata ata tta tct ata tga tgg gtg gat atg atc agt ccc cgt ata gaa gtt caa agg tta tag cgt aca ata cat gta caa att ctt gga tat atg ata tac cag aac t-a aaa tat cct cgt tct aat tgt gga gga gtt gct gat gat gaa tac att tat tgc ata ggc ggc ata cgc gat cag gat tca tcg ttg ata tct agt att gat aga tgg aag cca tca aaa cca tat tgg caa acg tat gct aaa atg cga gag cca aaa tgt gat atg ggt gta gcg atg tta aac gga tta ata tat gtc ata ggc gga gtt gtt aaa ggt g-- aca cgt gta ccg aca cac ttg aga gtt tat cag aag atg gat gga tga tgc atc aac gtc ttc caa taa aaa tgt cca ata tgt cga cga ctg ttc atg ccg gca aaa ttt ata tat ctg gag gct aca aca ata gta gtg tag tta atg gaa tat cga atc t-- agt cct tag cta taa tcc gat ata tga tga atg gac caa att atc atc att aaa cat tcc tag aat taa tcc cgc tct a

>DQ437593.1:175920-178154_Cowpox_virus_strain_Germany_91-3_complete_genome

--- --- --- --a --- tgt a-c tat ttt ata cat tac tac att ggt tac ggc aga tat aca tac tcc gtt acc gcc aca cgc ttc ggt aaa cgg --g tca tgt gcc gag gga gaa tat ctt gat aag act cat aat cga tgt tgt aat -cg gtg tcc acc tgg aga att tgc caa ggt tag atg ta- --g tgg tag cta taa cac aaa atg tga acg ctg tcc aca tca tac ata tac agc aat tcc caa tta ttc t-a atg gat gcc atc aat gta gaa aat gcc caa agg gat cat ttg ata agg taa agt gta ccg gaa cac aga aca gta aat -gt gcg tgt ctt cct ggt tgg tat tgc gct acc gat tct tca cag act gaa gac tgt cga gat -tg tgt acc aaa aag g-a gct gtc cat gcg gat act ttg gtg gaa tag ata aat tag gaa atc cta ttt gta aat cgt gtt g-- -tg ttg gta aat att gtg acg aca tac gta atc ata gag ttg gtc ctt ttc ctc cat gca aa- --- cta tct aaa tgt aat tag tca --- --- --- --- --- --- --- --- --- --- --- --- --- --- -tg atc atg gta -cc gta caa tt- --- -at att gct --- --g gta --g tat att att cag tat g-a aga cat att aat ac- --- --- -tt atc --- --t ttg aca atc ttg ct- -at aat cat --- --- --- --- aat ata aaa --a tac tta tgg --- -a- cat agt cac tca t-- --- --- --- aat tgc tg- acg cga taa att cgt aat aat ctg ttt tgt tca aat ttt tat aag ga- --- --a tct aca ggc ata aaa at- aaa aat ata att tat aat ac- --- act tac agc gcg cca tca tga ata aca gca gtg aat taa ttg ctg tta tta atg gat tta gaa ata gtg gac gat ttt gtg ata tta gta tag tta tta atg atg aaa gga taa acg ctc ata gac tca tcc tat ctg gag cat ccg aat a-t ttt tcc att cta ttt tct agt gat ttt atc gat tct aat gaa tac gaa gtt aat cta agt cat tta gat tat caa agt gtg aac gat ttg atc gat tac att tat ggg ata cct ttg agc cta act aac gat agc gtg aaa tat att ctt tca act gct ga- --t ttt tta caa att gga tct gct att acc gaa tgt gaa aat tac ata ctt aaa aat ctg tgt tct aga aat tgt atc gat ttc --- --- tac ata tac gct gat aaa tat aat aac aag aaa ata gaa aca gcg tcg ttt aac aca ata tta cga aat att ttg aga ctc att aac gat gaa aac ttt aaa tac ttg aca gag gaa tca atg ata aaa att tta agc gat gat atg tta aat ata aaa aat gag gat ttt gca cca cta att ctc att aaa tgg tta gag agt aca caa caa cca tgc aca gtc gag tta ctt aga tgc ctc aga ata tca -tt gct ttc ccc aca agt tat aaa atc act tta tag tca tcg att ggt tgg ttc aat ata cga atg tat aac att ctt aaa caa tat atc att ctt gga tga atc att tcc tag ata cca tag cat cga gtt gat atc tat cgg tat aag taa ttc aca tga taa gat ttc cat aaa ctg cta taa tcg t-a aaa aaa ata cgt ggg aga taa tat ctt cac gga gat ata ggt gta gtt ttg cag tgg ccg tca tgg ata ata tta tct ata tga tgg gtg gat atg atc agt ccc cgt ata gga gtt caa agg tta tag cgt aca ata cct gta cta att ctt gga tat atg ata tac cag agc t-a aaa tat cct cgt tct aat tgt gga gga gtt gct gat gat gaa tac att tat tgc ata ggc ggc ata cgc gat cag gat tca tcg ttg ata tct agt att gat aga tgg aag cca tca aaa cca tat tgg caa acg tat gct aaa atg cga gag cca aaa tgt gat atg ggt gta gcg atg tta aac gga tta ata tat gtc ata ggc gga gtt gtt aaa ggc g-- aca cgt gta ccg aca cac ttg aga gtt tat cag aag atg gat gga tga tgc atc aac gtc ttc caa taa aaa tgt cca ata tgt cga cga ttg ttc atg ccg gca aaa ttt ata tat ctg gag gct aca aca ata gta gtg tag tta atg gaa tat cga atc t-- agt cct tag cta taa tcc gat ata tga tga atg gac caa att atc atc att aaa tat tcc tag aat taa tcc cgc tct a

>LN864566.1:175649-177882_Cowpox_virus_complete_genome_strain_FM2292

--- --- --- --a --- tgt a-c tat ttt ata cat tac tac att ggt tac ggc aga tat aca tac tcc gtt acc gcc aca cgc ttc ggt aaa cgg --g tca tgt gcc gag gga gaa tat ctt gat aag act cat aat cga tgt tgt aat -cg gtg tcc acc tgg aga att tgc caa ggt tag atg ta- --g tgg tag cta taa cac aaa atg tga acg ctg tcc aca tca tac ata tac agc aat tcc caa tta ttc t-a atg gat gcc atc aat gta gaa aat gcc caa agg gat cat ttg ata agg taa agt gta ccg gaa cac aga aca gta aat -gt gcg tgt ctt cct ggt tgg tat tgc gct acc gat tct tca cag act gaa gac tgt cga gat -tg tgt acc aaa aag g-a gct gtc cat gcg gat act ttg gtg gaa tag ata aat tag gaa atc cta ttt gta aat cgt gtt g-- -tg ttg gta aat att gtg acg aca tac gta atc ata gag ttg gtc ctt ttc ctc cat gca aa- --- cta tct aaa tgt aat tag tca --- --- --- --- --- --- --- --- --- --- --- --- --- --- -tg atc atg gta -cc gta caa tt- --- -at att gct --- --g gta --g tat att att cag tat g-- aga cat att aat ac- --- --- -tt atc --- --t ttg aca atc ttg ct- -at aat cat --- --- --- --- aat ata aaa --a tac tta tgg --- -a- cat agt cac tca t-- --- --- --- aat tgc tg- acg cga taa att cgt aat aat ctg ttt tgt tca aat ttt tat aag ga- --- --a tct aca ggc ata aaa at- aaa aat ata att tat aat ac- --- act tac agc gcg cca tca tga ata aca gca gtg aat taa ttg ctg tta tta atg gat tta gaa ata gtg gac gat ttt gtg ata tta gta tag tta tta atg atg aaa gga taa acg ctc ata gac tca tcc tat ctg gag cat ccg aat a-t ttt tcc att cta ttt tct agt gat ttt atc gat tct aat gaa tac gaa gtt aat cta agt cat tta gat tat caa agt gtg aac gat ttg atc gat tac att tat ggg ata cct ttg agc cta act aac gat agc gtg aaa tat att ctt tca act gct ga- --t ttt tta caa att gga tct gct att acc gaa tgt gaa aat tac ata ctt aaa aat ctg tgt tct aga aat tgt atc gat ttc --- --- tac ata tac gct gat aaa tat aat aac aag aaa ata gaa aca gcg tcg ttt aac aca ata tta cga aat att ttg aga ctc att aac gat gaa aac ttt aaa tac ttg aca gag gaa tca atg ata aaa att tta agc gat gat atg tta aat ata aaa aat gag gat ttt gca cca cta att ctc att aaa tgg tta gag agt aca caa caa cca tgc aca gtc gag tta ctt aga tgc ctc aga ata tca -tt gct ttc ccc aca agt tat aaa atc act tta tag tca tcg att ggt tgg ttc aat ata cga atg tat aac att ctt aaa caa tat atc att ctt gga tga atc att tcc tag ata cca tag cat cga gtt gat atc tat cgg tat aag taa ttc aca tga taa gat ttc cat aaa ctg cta taa tcg t-a aaa aaa ata cgt ggg aga taa tat ctt cac gga gat ata ggt gta gtt ttg cag tgg ccg tca tgg ata ata tta tct ata tga tgg gtg gat atg atc agt ccc cgt ata gga gtt caa agg tta tag cgt aca ata cct gta cta att ctt gga tat atg ata tac cag agc t-a aaa tat cct cgt tct aat tgt gga gga gtt gct gat gat gaa tac att tat tgc ata ggc ggc ata cgc gat cag gat tca tcg ttg ata tct agt att gat aga tgg aag cca tca aaa cca tat tgg caa acg tat gct aaa atg cga gag cca aaa tgt gat atg ggt gta gcg atg tta aac gga tta ata tat gtc ata ggc gga gtt gtt aaa ggt g-- aca cgt gta ccg aca cac ttg aga gtt tat cag aag atg gat gga tga tgc atc aac gtc ttc caa taa aaa tgt cca ata tgt cga cga ttg ttc atg ccg gca aaa ttt ata tat ctg gag gct aca aca ata gta gtg tag tta atg gaa tat cga atc t-- agt cct tag cta taa tcc gat ata tga tga atg gac caa att atc atc att aaa tat tcc tag aat taa tcc cgc tct a

>KY569022.1:165853-168127_Cowpox_virus_strain_CPXV_CheTopCut_DK_2011

ttg ctg act gta --- tgt a-c tat ttt gta cat cag tac att ggt tac agc aga tat acc tac ttc gtc act gcc aca cac tcc ggt aaa cgg --g tca tgt gaa gag gga gaa tat ctt gat aag acg cat aat caa tgt tgt aat -cg gtg tcc acc tgg aga att tgc caa gat cag atg ta- --g cgg tag cga taa cac aaa atg tga acg ctg ccc acc tca tac ata tac cgc aat ccc caa tta ttc t-a atg gat gcc atc aat gta gaa aat gcc caa cag gat cat ttg ata agg taa agt gta ccg gaa cac aga aca gta aat -gt tcg tgt att cct ggt tgg tat tgc gct act gat tct tca tgg acc gaa gac tgt att gtg -tg cgt acc aaa aaa g-a gat gtc cgt gcg gat act ttg gtg gaa tag atg aac tcg gaa atc ctc ttt gta aat cgt gtt g-- -tg ttg gtg aat att gcg acg aca tac gta atc ata gag ttg gtc ctt ttc ctc cat gca aa- --- cta tct aaa tgt aat tag cca --- --- --- --- --- --- --- --- --- --- --- --- --- --- -tt atc atg tta ccc ata c-a tca cat cat atc gct act tgg t-a --g cgt att att cag tat g-a aga tct att aat aat aat tat ctt att tat ttc ttg acg --- --- --- -at att aat cgg agt cct aat aat ata aaa --- tac tta tgg --- --- --a cat aaa ttg t-a aac aag atg a-- taa t-- aca cga t-- atc ctt -at aat cag ctt -gt tca aat ttt tat aaa gaa tta tat tca tta ggc ata taa ta- aaa aat ata att tat aat ac- --- act tac a-- gcg tca tca tgc ata aca gca gtg aat taa ttg ctg tta tta atg gat tta gaa ata gtg gac gat ttt gtg ata tta ata tag tta tta atg atg aaa gga taa acg ctc ata gac tca tcc tat ctg gag cat cag aat a-t ttt tcc att ctg ttt tcc aat aat ttt atc gat tct aat gac tac gaa gtt aat cta agt cat tta gat tat caa agt gtt aac gat ttg atc gat tac att tat ggg ata cct ttg agc cta act aac gat aac gtg aaa tat att ctt tca aca gct ga- --t ttt tta caa att gga tct gcc att act gag tgc gaa aaa tac ata ctt aaa aat ctc tgt tct aga aac tgt atc gat ttc --- --- tac ata tac gct gat aaa tat aat aac aag aaa ata gaa tca gcg tcg ttt aac aca ata tta cga aat att ttg aga ctc att aac gat gaa aac ttt aaa tac ttg aca gag gaa tca atg ata aaa att tta agc gat gat atg tta aat ata aaa aat gag gat ttc gca cca cta att ctc att aaa tgg tta gag agt aca caa caa cca tgc acc gtc gag tta ctt aga tgc ctc aga ata tca -tt gct ttc ccc aca agt tat aaa atc act tta tag tca tcg act ggt tag ttc aat cta cga atg tat aac att ctt aaa caa tat agc att ctt gga tga atc att tcc tag ata caa tag cat cga gtt gat atc tat cgg tat aag taa ttc gca tga taa gat ttc cat aaa ctg cta caa tcg t-a aaa aaa ata cat ggg aaa tga tat ctt cac gta gat ata ggt gta gtt tcg cag tgg ccg tcc tgg ata ata tta tct ata tga tgg gtg gat atg atc agt ccc cgt ata gaa gtt caa agg tta tag cgt aca ata cat gta caa att ctt gga tat atg ata tac cag agc t-a aaa tat cct cgt tct aat tgc gga gga gtt gcc gat gac gaa tac att tat tgt ata ggc ggc ata cgc gat cag gat tca tcg ttg ata tct aat att gat aga tgg aag cca tca aaa cca tat tgg cag acg tat gct aaa atg cgc gag cca aaa tgt gat atg ggg gtt gcg atg tta aac gga tta ata tat gtc ata ggt gga atc gtt aaa ggt g-- aca cat gta ccg acg cac tag aga gtt tat cag aag atg gat gga tga agc atc agc gcc ttc caa taa aaa tgt cca ata tgt cga cga ttg ttc atg ctg gta aga ttt ata tat ctg gag gtt ata aca ata gta gtg ccg tta atg gac cat cga atc t-- agt cct tag cta taa tcc gat ata tga tga atg gac caa att atc atc att aaa tat tcc tag aat taa tcc cgc tct a

>KY569020.1:165769-168043_Cowpox_virus_strain_CPXV_CheNuru_DK_2012

ttg ctg act gta --- tgt a-c tat ttt gta cat cag tac att ggt tac agc aga tat acc tac ttc gtc act gcc aca cac tcc ggt aaa cgg --g tca tgt gaa gag gga gaa tat ctt gat aag acg cat aat caa tgt tgt aat -cg gtg tcc acc tgg aga att tgc caa gat cag atg ta- --g cgg tag cga taa cac aaa atg tga acg ctg ccc acc tca tac ata tac cgc aat ccc caa tta ttc t-a atg gat gcc atc aat gta gaa aat gcc caa cag gat cat ttg ata agg taa agt gta ccg gaa cac aga aca gta aat -gt tcg tgt att cct ggt tgg tat tgc gct act gat tct tca tgg acc gaa gac tgt att gtg -tg cgt acc aaa aaa g-a gat gtc cgt gcg gat act ttg gtg gaa tag atg aac tcg gaa atc ctc ttt gta aat cgt gtt g-- -tg ttg gtg aat att gcg acg aca tac gta atc ata gag ttg gtc ctt ttc ctc cat gca aa- --- cta tct aaa tgt aat tag cca --- --- --- --- --- --- --- --- --- --- --- --- --- --- -tt atc atg tta ccc ata c-a tca cat cat atc gct act tgg t-a --g cgt att att cag tat g-a aga tct att aat aat aat tat ctt att tat ttc ttg acg --- --- --- -at att aat cgg agt cct aat aat ata aaa --- tac tta tgg --- --- --a cat aaa ttg t-a aac aag atg a-- taa t-- aca cga t-- atc ctt -at aat cag ctt -gt tca aat ttt tat aaa gaa tta tat tca tta ggc ata taa ta- aaa aat ata att tat aat ac- --- act tac a-- gcg tca tca tgc ata aca gca gtg aat taa ttg ctg tta tta atg gat tta gaa ata gtg gac gat ttt gtg ata tta ata tag tta tta atg atg aaa gga taa acg ctc ata gac tca tcc tat ctg gag cat cag aat a-t ttt tcc att ctg ttt tcc aat aat ttt atc gat tct aat gac tac gaa gtt aat cta agt cat tta gat tat caa agt gtt aac gat ttg atc gat tac att tat ggg ata cct ttg agc cta act aac gat aac gtg aaa tat att ctt tca aca gct ga- --t ttt tta caa att gga tct gcc att act gag tgc gaa aaa tac ata ctt aaa aat ctc tgt tct aga aac tgt atc gat ttc --- --- tac ata tac gct gat aaa tat aat aac aag aaa ata gaa tca gcg tcg ttt aac aca ata tta cga aat att ttg aga ctc att aac gat gaa aac ttt aaa tac ttg aca gag gaa tca atg ata aaa att tta agc gat gat atg tta aat ata aaa aat gag gat ttc gca cca cta att ctc att aaa tgg tta gag agt aca caa caa cca tgc acc gtc gag tta ctt aga tgc ctc aga ata tca -tt gct ttc ccc aca agt tat aaa atc act tta tag tca tcg act ggt tag ttc aat cta cga atg tat aac att ctt aaa caa tat agc att ctt gga tga atc att tcc tag ata caa tag cat cga gtt gat atc tat cgg tat aag taa ttc gca tga taa gat ttc cat aaa ctg cta caa tcg t-a aaa aaa ata cat ggg aaa tga tat ctt cac gta gat ata ggt gta gtt tcg cag tgg ccg tcc tgg ata ata tta tct ata tga tgg gtg gat atg atc agt ccc cgt ata gaa gtt caa agg tta tag cgt aca ata cat gta caa att ctt gga tat atg ata tac cag agc t-a aaa tat cct cgt tct aat tgc gga gga gtt gcc gat gac gaa tac att tat tgt ata ggc ggc ata cgc gat cag gat tca tcg ttg ata tct aat att gat aga tgg aag cca tca aaa cca tat tgg cag acg tat gct aaa atg cgc gag cca aaa tgt gat atg ggg gtt gcg atg tta aac gga tta ata tat gtc ata ggt gga atc gtt aaa ggt g-- aca cat gta ccg acg cac tag aga gtt tat cag aag atg gat gga tga agc atc agc gcc ttc caa taa aaa tgt cca ata tgt cga cga ttg ttc atg ctg gta aga ttt ata tat ctg gag gtt ata aca ata gta gtg ccg tta atg gac cat cga atc t-- agt cct tag cta taa tcc gat ata tga tga atg gac caa att atc atc att aaa tat tcc tag aat taa tcc cgc tct a

>KY569019.1:174717-176991_Cowpox_virus_strain_CPXV_CheNova_DK_2014

ttg ctg act gta --- tgt a-c tat ttt gta cat cag tac att ggt tac agc aga tat acc tac ttc gtc act gcc aca cac tcc ggt aaa cgg --g tca tgt gaa gag gga gaa tat ctt gat aag acg cat aat caa tgt tgt aat -cg gtg tcc acc tgg aga att tgc caa gat cag atg ta- --g cgg tag cga taa cac aaa atg tga acg ctg ccc acc tca tac ata tac cgc aat ccc caa tta ttc t-a atg gat gcc atc aat gta gaa aat gcc caa cag gat cat ttg ata agg taa agt gta ccg gaa cac aga aca gta aat -gt tcg tgt att cct ggt tgg tat tgc gct act gat tct tca tgg acc gaa gac tgt att gtg -tg cgt acc aaa aaa g-a gat gtc cgt gcg gat act ttg gtg gaa tag atg aac tcg gaa atc ctc ttt gta aat cgt gtt g-- -tg ttg gtg aat att gcg acg aca tac gta atc ata gag ttg gtc ctt ttc ctc cat gca aa- --- cta tct aaa tgt aat tag cca --- --- --- --- --- --- --- --- --- --- --- --- --- --- -tt atc atg tta ccc ata c-a tca cat cat atc gct act tgg t-a --g cgt att att cag tat g-a aga tct att aat aat aat tat ctt att tat ttc ttg acg --- --- --- -at att aat cgg agt cct aat aat ata aaa --- tac tta tgg --- --- --a cat aaa ttg t-a aac aag atg a-- taa t-- aca cga t-- atc ctt -at aat cag ctt -gt tca aat ttt tat aaa gaa tta tat tca tta ggc ata taa ta- aaa aat ata att tat aat ac- --- act tac a-- gcg tca tca tgc ata aca gca gtg aat taa ttg ctg tta tta atg gat tta gaa ata gtg gac gat ttt gtg ata tta ata tag tta tta atg atg aaa gga taa acg ctc ata gac tca tcc tat ctg gag cat cag aat a-t ttt tcc att ctg ttt tcc aat aat ttt atc gat tct aat gac tac gaa gtt aat cta agt cat tta gat tat caa agt gtt aac gat ttg atc gat tac att tat ggg ata cct ttg agc cta act aac gat aac gtg aaa tat att ctt tca aca gct ga- --t ttt tta caa att gga tct gcc att act gag tgc gaa aaa tac ata ctt aaa aat ctc tgt tct aga aac tgt atc gat ttc --- --- tac ata tac gct gat aaa tat aat aac aag aaa ata gaa tca gcg tcg ttt aac aca ata tta cga aat att ttg aga ctc att aac gat gaa aac ttt aaa tac ttg aca gag gaa tca atg ata aaa att tta agc gat gat atg tta aat ata aaa aat gag gat ttc gca cca cta att ctc att aaa tgg tta gag agt aca caa caa cca tgc acc gtc gag tta ctt aga tgc ctc aga ata tca -tt gct ttc ccc aca agt tat aaa atc act tta tag tca tcg act ggt tag ttc aat cta cga atg tat aac att ctt aaa caa tat agc att ctt gga tga atc att tcc tag ata caa tag cat cga gtt gat atc tat cgg tat aag taa ttc gca tga taa gat ttc cat aaa ctg cta caa tcg t-a aaa aaa ata cat ggg aaa tga tat ctt cac gta gat ata ggt gta gtt tcg cag tgg ccg tcc tgg ata ata tta tct ata tga tgg gtg gat atg atc agt ccc cgt ata gaa gtt caa agg tta tag cgt aca ata cat gta caa att ctt gga tat atg ata tac cag agc t-a aaa tat cct cgt tct aat tgc gga gga gtt gcc gat gac gaa tac att tat tgt ata ggc ggc ata cgc gat cag gat tca tcg ttg ata tct aat att gat aga tgg aag cca tca aaa cca tat tgg cag acg tat gct aaa atg cgc gag cca aaa tgt gat atg ggg gtt gcg atg tta aac gga tta ata tat gtc ata ggt gga atc gtt aaa ggt g-- aca cat gta ccg acg cac tag aga gtt tat cag aag atg gat gga tga agc atc agc gcc ttc caa taa aaa tgt cca ata tgt cga cga ttg ttc atg ctg gta aga ttt ata tat ctg gag gtt ata aca ata gta gtg ccg tta atg gac cat cga atc t-- agt cct tag cta taa tcc gat ata tga tga atg gac caa att atc atc att aaa tat tcc tag aat taa tcc cgc tct a

>KY569018.1:166064-168338_Cowpox_virus_strain_CPXV_CheHurley_DK_2012

ttg ctg act gta --- tgt a-c tat ttt gta cat cag tac att ggt tac agc aga tat acc tac ttc gtc act gcc aca cac tcc ggt aaa cgg --g tca tgt gaa gag gga gaa tat ctt gat aag acg cat aat caa tgt tgt aat -cg gtg tcc acc tgg aga att tgc caa gat cag atg ta- --g cgg tag cga taa cac aaa atg tga acg ctg ccc acc tca tac ata tac cgc aat ccc caa tta ttc t-a atg gat gcc atc aat gta gaa aat gcc caa cag gat cat ttg ata agg taa agt gta ccg gaa cac aga aca gta aat -gt tcg tgt att cct ggt tgg tat tgc gct act gat tct tca tgg acc gaa gac tgt att gtg -tg cgt acc aaa aaa g-a gat gtc cgt gcg gat act ttg gtg gaa tag atg aac tcg gaa atc ctc ttt gta aat cgt gtt g-- -tg ttg gtg aat att gcg acg aca tac gta atc ata gag ttg gtc ctt ttc ctc cat gca aa- --- cta tct aaa tgt aat tag cca --- --- --- --- --- --- --- --- --- --- --- --- --- --- -tt atc atg tta ccc ata c-a tca cat cat atc gct act tgg t-a --g cgt att att cag tat g-a aga tct att aat aat aat tat ctt att tat ttc ttg acg --- --- --- -at att aat cgg agt cct aat aat ata aaa --- tac tta tgg --- --- --a cat aaa ttg t-a aac aag atg a-- taa t-- aca cga t-- atc ctt -at aat cag ctt -gt tca aat ttt tat aaa gaa tta tat tca tta ggc ata taa ta- aaa aat ata att tat aat ac- --- act tac a-- gcg tca tca tgc ata aca gca gtg aat taa ttg ctg tta tta atg gat tta gaa ata gtg gac gat ttt gtg ata tta ata tag tta tta atg atg aaa gga taa acg ctc ata gac tca tcc tat ctg gag cat cag aat a-t ttt tcc att ctg ttt tcc aat aat ttt atc gat tct aat gac tac gaa gtt aat cta agt cat tta gat tat caa agt gtt aac gat ttg atc gat tac att tat ggg ata cct ttg agc cta act aac gat aac gtg aaa tat att ctt tca aca gct ga- --t ttt tta caa att gga tct gcc att act gag tgc gaa aaa tac ata ctt aaa aat ctc tgt tct aga aac tgt atc gat ttc --- --- tac ata tac gct gat aaa tat aat aac aag aaa ata gaa tca gcg tcg ttt aac aca ata tta cga aat att ttg aga ctc att aac gat gaa aac ttt aaa tac ttg aca gag gaa tca atg ata aaa att tta agc gat gat atg tta aat ata aaa aat gag gat ttc gca cca cta att ctc att aaa tgg tta gag agt aca caa caa cca tgc acc gtc gag tta ctt aga tgc ctc aga ata tca -tt gct ttc ccc aca agt tat aaa atc act tta tag tca tcg act ggt tag ttc aat cta cga atg tat aac att ctt aaa caa tat agc att ctt gga tga atc att tcc tag ata caa tag cat cga gtt gat atc tat cgg tat aag taa ttc gca tga taa gat ttc cat aaa ctg cta caa tcg t-a aaa aaa ata cat ggg aaa tga tat ctt cac gta gat ata ggt gta gtt tcg cag tgg ccg tcc tgg ata ata tta tct ata tga tgg gtg gat atg atc agt ccc cgt ata gaa gtt caa agg tta tag cgt aca ata cat gta caa att ctt gga tat atg ata tac cag agc t-a aaa tat cct cgt tct aat tgc gga gga gtt gcc gat gac gaa tac att tat tgt ata ggc ggc ata cgc gat cag gat tca tcg ttg ata tct aat att gat aga tgg aag cca tca aaa cca tat tgg cag acg tat gct aaa atg cgc gag cca aaa tgt gat atg ggg gtt gcg atg tta aac gga tta ata tat gtc ata ggt gga atc gtt aaa ggt g-- aca cat gta ccg acg cac tag aga gtt tat cag aag atg gat gga tga agc atc agc gcc ttc caa taa aaa tgt cca ata tgt cga cga ttg ttc atg ctg gta aga ttt ata tat ctg gag gtt ata aca ata gta gtg ccg tta atg gac cat cga atc t-- agt cct tag cta taa tcc gat ata tga tga atg gac caa att atc atc att aaa tat tcc tag aat taa tcc cgc tct a

>MK035750.1:165520-167755_Cowpox_virus_strain_CPXV/Compiegne_2009

--- --- --- --a --- tgt a-c tat ttt ata cat tac tac att ggt tac ggc aga tat aca tac tcc gtt acc gcc aca cgc tcc ggt aaa cgg --g tca tgt gcc gag gga gaa tat ctt gat aag act cat aat cga tgt tgt aat -cg gtg tcc acc tgg aga att tgc caa ggt tag atg ta- --g tgg tag cta taa cac aaa atg tga acg ctg tcc aca tca tac ata tac agc aat tcc caa tta ttc t-a atg gat gcc atc aat gta gaa aat gcc caa agg gat cat ttg ata agg taa agt gta ccg gaa aac aga aca gta aat -gt gcg tgt ctt cct ggt tgg tat tgc gct acc gat tct tca cag act gaa gac tgt cga gat -tg tgt acc aaa aag g-a gct gtc cat gcg gat act ttg gtg gaa tag ata aat tag gaa atc cta ttt gta aat cgt gtt g-- -tg ttg gta aat att gtg acg aca tac gta atc ata gag ttg gtc ctt ttc ctc cat gca aa- --- cta tct aaa tgt aat tag tca --- --- --- --- --- --- --- --- --- --- --- --- --- --- -tg atc atg gta -cc gta caa tt- --- -at att gct --- --g gta --g tat att att cag tat g-a aga cat att aat ac- --- --- -tt atc --- --t ttg aca atc ttg ct- -at aat cat --- --- --- --- aat ata aaa --a tac tta tgg --- -a- cat agt cac tca t-- --- --- --- aat tgc tg- acg cga taa att cgt aat aat ctg ttt tgt tca aat ttt tat aag ga- --- --a tct aca ggc ata aaa at- aaa aat ata att tat aat ac- --- act tac agc gcg cca tca tga ata aca gca gtg aat taa ttg ctg tta tta atg gat tta gaa ata gtg gac gat ttt gtg ata tta ata tag tta tta atg atg aaa gga taa atg ctc ata gac tca tcc tat ctg gag cat ccg aat a-t ttt tcc att cta ttt tct agt gat ttt atc gat tct aat gaa tac gaa gtt aat cta agt cat tta gat tat caa agt gtg aac gat ttg atc gat tac att tat ggg ata cct ttg agc cta act aac gat agc gtg aaa tat att ctt tca act gct ga- --t ttt tta caa att gga tct gct att acc gaa tgt gaa aat tac ata ctt aaa aat ctg tgt tct aga aat tgt atc gat ttc --- --- tac ata tac gct gat aaa tat aat aac aag aaa ata gaa aca gcg tcg ttt aac aca ata tta cga aat att ttg aga ctc att aac gat gaa aac ttt aaa tac ttg aca gag gaa tca atg ata aaa att tta agc gat gat atg tta aat ata aaa aat gag gat ttt gca cca cta att ctc att aaa tgg tta gag agt aca caa caa cca tgc aca gtc gag tta ctt aga tgc ctc aga ata tca -tt gct ttc ccc aca agt tat aaa atc act tta tag tca tcg att ggt tgg ttc aat ata cga atg tat aac att ctt aaa caa tat atc att ctt gga tga atc att tcc tag ata cca tag cat cga gtt gat atc tat cgg tat aag taa ttc aca tga taa gat ttc cat aaa ctg cta taa tcg taa aaa aaa ata cgt ggg aga taa tat ctt cac gga gat ata ggt gta gtt ttg cag tgg ccg tcc tgg ata ata tta tct ata tga tgg gtg gat atg atc agt ccc cgt ata gaa gtt caa agg tta tag cgt aca ata cat gta caa att ctt gga tat atg ata tac cag aac t-a aaa tat cct cgt tct aat tgt gga gga gtt gct gat gat gaa tac att tat tgc ata ggc ggc ata cgc gat cag gat tca tcg ttg ata tct agt att gat aga tgg aag cca tca aaa cca tat tgg caa acg tat gct aaa atg cga gag cca aaa tgt gat atg ggt gta gcg atg tta aac gga tta ata tat gtc ata ggc gga gtt gtt aaa ggt g-- aca cgt gta ccg aca cac ttg aga gtt tat cag aag atg gat gga tga tgc atc aac gtc ttc caa taa aaa tgt cca ata tgt cga cga ctg ttc atg ccg gca aaa ttt ata tat ctg gag gct aca aca ata gta gtg tag tta atg gaa tat cga atc t-- agt cct tag cta taa tcc gat ata tga tga atg gac caa att atc atc att aaa cat tcc tag aat taa tcc cgc tct a

>KY569021.1:166123-168397_Cowpox_virus_strain_CPXV_CheGrey_DK_2010

ttg ctg act gta --- tgt a-c tat ttt gta cat cag tac att ggt tac agc aga tat acc tac ttc gtc act gcc aca cac tcc ggt aaa cgg --g tca tgt gaa gag gga gaa tat ctt gat aag acg cat aat caa tgt tgt aat -cg gtg tcc acc tgg aga att tgc caa gat cag atg ta- --g cgg tag cga taa cac aaa atg tga acg ctg ccc acc tca tac ata tac cgc aat ccc caa tta ttc t-a atg gat gcc atc aat gta gaa aat gcc caa cag gat cat ttg ata agg taa agt gta ccg gaa cac aga aca gta aat -gt tcg tgt att cct ggt tgg tat tgc gct act gat tct tca tgg acc gaa gac tgt att gtg -tg cgt acc aaa aaa g-a gat gtc cgt gcg gat act ttg gtg gaa tag atg aac tcg gaa atc ctc ttt gta aat cgt gtt g-- -tg ttg gtg aat att gcg acg aca tac gta atc ata gag ttg gtc ctt ttc ctc cat gca aa- --- cta tct aaa tgt aat tag cca --- --- --- --- --- --- --- --- --- --- --- --- --- --- -tt atc atg tta ccc ata c-a tca cat cat atc gct act tgg t-a --g cgt att att cag tat g-a aga tct att aat aat aat tat ctt att tat ttc ttg acg --- --- --- -at att aat cgg agt cct aat aat ata aaa --- tac tta tgg --- --- --a cat aaa ttg t-a aac aag atg a-- taa t-- aca cga t-- atc ctt -at aat cag ctt -gt tca aat ttt tat aaa gaa tta tat tca tta ggc ata taa ta- aaa aat ata att tat aat ac- --- act tac a-- gcg tca tca tgc ata aca gca gtg aat taa ttg ctg tta tta atg gat tta gaa ata gtg gac gat ttt gtg ata tta ata tag tta tta atg atg aaa gga taa acg ctc ata gac tca tcc tat ctg gag cat cag aat a-t ttt tcc att ctg ttt tcc aat aat ttt atc gat tct aat gac tac gaa gtt aat cta agt cat tta gat tat caa agt gtt aac gat ttg atc gat tac att tat ggg ata cct ttg agc cta act aac gat aac gtg aaa tat att ctt tca aca gct ga- --t ttt tta caa att gga tct gcc att act gag tgc gaa aaa tac ata ctt aaa aat ctc tgt tct aga aac tgt atc gat ttc --- --- tac ata tac gct gat aaa tat aat aac aag aaa ata gaa tca gcg tcg ttt aac aca ata tta cga aat att ttg aga ctc att aac gat gaa aac ttt aaa tac ttg aca gag gaa tca atg ata aaa att tta agc gat gat atg tta aat ata aaa aat gag gat ttc gca cca cta att ctc att aaa tgg tta gag agt aca caa caa cca tgc acc gtc gag tta ctt aga tgc ctc aga ata tca -tt gct ttc ccc aca agt tat aaa atc act tta tag tca tcg act ggt tag ttc aat cta cga atg tat aac att ctt aaa caa tat agc att ctt gga tga atc att tcc tag ata caa tag cat cga gtt gat atc tat cgg tat aag taa ttc gca tga taa gat ttc cat aaa ctg cta caa tcg t-a aaa aaa ata cat ggg aaa tga tat ctt cac gta gat ata ggt gta gtt tcg cag tgg ccg tcc tgg ata ata tta tct ata tga tgg gtg gat atg atc agt ccc cgt ata gaa gtt caa agg tta tag cgt aca ata cat gta caa att ctt gga tat atg ata tac cag agc t-a aaa tat cct cgt tct aat tgt gga gga gtt gcc gat gac gaa tac att tat tgt ata ggc ggc ata cgc gat cag gat tca tcg ttg ata tct aat att gat aga tgg aag cca tca aaa cca tat tgg cag acg tat gct aaa atg cgc gag cca aaa tgt gat atg ggg gtt gcg atg tta aac gga tta ata tat gtc ata ggt gga atc gtt aaa ggt g-- aca cat gta ccg acg cac tag aga gtt tat cag aag atg gat gga tga agc atc agc gcc ttc caa taa aaa tgt cca ata tgt cga cga ttg ttc atg ctg gta aga ttt ata tat ctg gag gtt ata aca ata gta gtg ccg tta atg gac cat cga atc t-- agt cct tag cta taa tcc gat ata tga tga atg gac caa att atc atc att aaa tat tcc tag aat taa tcc cgc tct a

>KY549145.1:177538-179810_Cowpox_virus_isolate_CPXV_K428

ttg ctg act gta --- tgt a-c tat ttt gta cat cag tac att ggt tac agc aga tat acc tac ttc gtc act gcc aca tgc tcc ggt aaa cgg --g tca tgt gac gaa gga gaa tat ctt gat aag acg cat aat caa tgt tgt aat -cg gtg tcc acc tgg aga att tgc caa gat cag atg ta- --g cgg tag cga taa cac aaa atg tga acg ctg ccc acc tca tac ata tac cac agt acc caa tta ttc t-a atg gat gtc atc aat gta gga aat gcc caa cag gat cat ttg ata agg taa agt gta ccg gaa cac aga aca gta aat -gt tcg tgt ctt cct ggt tgg ttt tgc gct act gat tct tcg aag act gaa gat tgt cga gat -tg tat acc aaa aag a-a aat gtc cat gcg gat act ttg gtg gaa tag atg aac tcg gaa atc ctc ttt gta aat cgt gtt g-- -tg ttg gtg aat att gcg acg aca tac gta atc ata gag ttg gtc ctt ttc ctc cat gca aa- --- cta tct aaa tgt aat tag cca --- --- --- --- --- --- --- --- --- --- --- --- --- --- -tg atc atg tta ccc ata c-a tc- --- -at atc gct act tgg t-a --g cgt att att cag tat g-a aga tct att aat aat aat tat ctt att tat ttc ttg acg --- --- --- -at att aat cgg tgt cct aat aat ata aaa --- tac tta tgg --- --- --a aat aaa ttg t-a aac aag atg aat taa t-- aca caa tac atc ctt -at act cag ctt -gt tca aa- ttt tat aaa gaa tca tat tca tta ggc ata taa ta- aaa aat ata att tat aat ac- --- act tac a-- gcg tca tca tgc ata aca gca gtg aat taa ttg ctg tta tta atg gat tta gaa ata gtg gac gat ttt gtg ata tta ata tag tta tta atg atg aaa gga taa acg ctc ata gac tca tgc tat ctg gag cat ccg aat a-t ttt tcc att cta ttt tct agt gat ttt atc gat tct aat gac tac gaa gtt aat cta agt cat tta gat tat caa agt gtt aac gat ttg atc gat tac att tat ggg ata cct ttg agc cta act aac gat aac gtg aaa tat att ctt tca acc gct ga- --t ttt tta caa att gga tct gcc att act gag tgc gaa aaa tac ata ctt aaa aat ctt tgt tct aga aac tgt atc gat ttc --- --- tac ata tac gct gat aaa tat aat aac aag aaa ata gaa tca gcg tcg ttt aac aca ata tta cga aat att ttg aga ctc att aac gat gaa aac ttt aaa tac ttg aca gag gaa tca atg ata aaa att tta agc gat gat atg tta aat ata aaa aat gag gat ttc gca cca cta att ctc att aaa tgg tta gag agt aca caa caa cca tgc acc gtc gag tta ctt aga tgc ctc aga ata tca -tt gct ttc ccc aca agt tat aaa atc act tta tag tca tcg act ggt tag ttc aat cta cga atg tat aac att ctt aaa caa tat agc att ctt gga tga atc att tcc tag ata cca tag cat cga gtt gat atc tat cgg tat aag taa ttc gca tga taa gat ttc cat aaa ctg cta caa tcg t-a aaa aaa ata cat ggg aaa tga tat ctt cac gta gat ata ggt gta gtt tcg cag tgg ccg tcc tgg ata ata tta tct ata tga tgg gtg gat atg atc agt ccc cgt ata gaa gtt caa agg tta tag cgt aca aca cat gta caa att ctt gga tat atg ata tac cag agc t-a aaa tat cct cgt tct aat tgc gga gga gtt gcc gat gac gaa tac att tat tgt ata ggc ggc ata cgc gat cag gat tca tcg ttg ata tct agt att gat aga tgg aag cca tca aaa cca tat tgg cag acg tat gct aaa atg cgc gaa cca aaa tgt gat atg ggg gtt gcg atg tta aac gga tta ata tat gtc ata ggt gga atc gtt aaa ggt g-- atg cgt gta ccg acg cac tag aaa gtt tat cag aag atg gat gga tga agc atc agc gcc ttc caa taa aaa tgt cca ata tgt cga cga ttg ttc atg ctg gta aga ttt ata tat ctg gag gtt ata aca ata gta gtg ccg tta atg gac cat cga atc t-- agt cct tag cta taa tcc gat ata tga tga atg gac caa att atc atc att aaa tat tcc tag aat taa tcc cgc tct a

>HQ420900.1:175746-178018_Cowpox_virus_strain_UK2000_K2984_complete_genome

ttg ctg act gta --- tgt a-c tat ttt gta cat cag tac att ggt tac agc aga tat acc tac ttc gtc act gcc aca tgc tcc ggt aaa cgg --g tca tgt gac gaa gga gaa tat ctt gat aag acg cat aat caa tgt tgt aat -cg gtg tcc acc tgg aga att tgc caa gat cag atg ta- --g cgg tag cta taa cac aaa atg tga acg ctg tcc acc tca tac att tac cac agt acc caa tta ttc t-a atg gat gtc atc aat gta gga aat gcc caa cag gat cat ttg ata agg taa agt gta ccg gaa cac aga aca gta aat -gt tcg tgt ctt cct ggt tgg ttt tgc gct act gat tct tcg aag act gaa gat tgt cga gat -tg tat acc aaa aag a-a aat gtc cat gcg gat act ttg gtg gaa tag atg aac tcg gaa atc ctc ttt gta aat cgt gtt g-- -tg ttg gtg aat att gcg acg aca tac gta atc ata gag ttg gtc ctt ttc ctc cat gca aa- --- cta tct aaa tgt aat tag cca --- --- --- --- --- --- --- --- --- --- --- --- --- --- -tg atc atg tta ccc ata c-a tc- --- -at atc gct act tgg t-a --g cgt att att cag tat g-a aga tct att aat aat aat tat ctt att tat ttc ttg acg --- --- --- -at att aat cgg tgt cct aat aat ata aaa --- tac tta tgg --- --- --a aat aaa ttg t-a aac aag atg aat taa t-- aca caa tac atc ctt -at act cag ctt -gt tca aa- ttt tat aac gaa tca tat tca tta ggc ata taa ta- aaa aat ata att tat aat ac- --- act tac a-- gcg tca tca tgc ata aca gca gtg aat taa ttg ctg tta tta atg gat tta gaa ata gtg gac gat ttt gtg ata tta ata tag tta tta atg atg aaa gga taa acg ctc ata gac tca tgc tat ctg gag cat ccg aat a-t ttt tcc att cta ttt tct agt gat ttt atc gat tct aat gac tac gaa gtt aat cta agt cat tta gat tat caa agt gtg aac gat ttg atc gat tac att tat ggg ata cct ttg agc cta act aac gat aac gtg aaa tat att ctt tca acc gct ga- --t ttt tta caa att gga tct gcc att act gag tgc gaa aaa tac ata ctt aaa aat ctt tgt tct aga aac tgt atc gat ttc --- --- tac ata tac gct gat aaa tat aat aac aag aaa ata gaa tca gcg tcg ttt aac aca ata tta cga aat att ttg aga ctc att aac gat gaa aac ttt aaa tac ttg aca gag gaa tca atg ata aaa att tta agc gat gat atg tta aat ata aaa aat gag gat ttc gca cca cta att ctc att aaa tgg tta gag agt aca caa caa cca tgc acc gtc gag tta ctt aga tgc ctc aga ata tca -tt gct ttc ccc aca agt tat aaa atc act tta tag tca tcg act ggt tag ttc aat cta cga atg tat aac att ctt aaa caa tat agc att ctt gga tga atc att tcc tag ata cca tag cat cga gtt gat atc tat cgg tat aag taa ttc aca tga taa gat ttc cat aaa ctg cta caa tcg t-a aaa aaa ata cat ggg aaa tga tat ctt cac gta gat ata ggt gta gtt tcg cag tgg ccg tcc tgg ata ata tta tct ata tga tgg gtg gat atg atc agt ccc cgt ata gaa gtt caa agg tta tag cgt aca ata cat gta caa att ctt gga tat atg ata tac cag agc t-a aaa tat cct cgt tct aat tgc gga gga gtt gcc gat gac gaa tac att tat tgt ata ggc ggc ata cgc gat cag gat tca tcg ttg ata tct agt att gat aga tgg aag cca tca aaa cca tat tgg cag acg tat gct aaa atg cgc gag cca aaa tgt gat atg ggg gtt gcg atg tta aac gga tta ata tat gtc ata ggt gga atc gtt aaa ggt g-- aca cat gta ccg acg cac tag aaa gtt tat cag aag atg gat gga tga agc atc agc gcc ttc caa taa aaa tgt cca ata tgt cga cga ttg ttc atg ctg gta aga ttt ata tat ctg gag gtt ata aca ata gta gtg ccg tta atg gac cat cga atc t-- agt cct tag cta taa tcc gat ata tga tga atg gac caa att atc atc att aaa tat tcc tag aat taa tcc cgc tct a

>KY549150.2:176701-178969_Cowpox_virus_isolate_CPXV_K4207

ttg ctg act gta --- tgt a-c tat ttt gta cat cag tac att ggt tac agc aga tat acc tac ttc gtc act gcc aca tgc tcc ggt aaa cgg --g aca tgt gaa gag gga gaa tat ctt gat aag acg cat aat caa tgt tgt aat -cg gtg tcc tcc tgg aga att tgc caa ggt tag atg ta- --a tgg taa cga taa cac aaa atg tga acg ctg tcc acc tca tac ata tac cac agt acc caa tta ttc t-a atg gat gcc atc aat gta gaa aat gcc caa cag gat cat ttg ata agg taa agt gta ccg gaa cac aga aca gta aat -gt tcg tgt ctt cct ggt tgg ttt tgc gct act gat tct tcg aag act gaa gat tgt cga gat -tg tat acc aaa aag g-a aat gtc cat gcg gat act ttg gtg gaa tag atg aac tcg gaa atc ctc ttt gta aat cgt gtt g-- -tg ttg gtg aat att gcg acg aca tac gta atc ata gag ttg gtc ctt ttc ctc cat gca aa- --- cta tct aaa tgt aat tag cca --- --- --- --- --- --- --- --- --- --- --- --- --- --- -tg atc atg tta -cc ata c-a tc- --- -at atc gct act tgg t-a --g cgt att att cag tat g-a aga tct att aat aat aat tat ctt att tat ttc tt- --g --- --- --- -at att aat cgg tgt cct aat aat ata aaa --- tac tta tgg --- --- --a aat aaa ttg t-a aac aag atg aat taa t-- aca caa tac atc ctt -at act cag ctt -gt tca aa- ttt tat aaa gaa tca tat tca tta ggc ata taa ta- aaa aat ata att tat aat ac- --- act tac a-- gcg tca tca tgc ata aca gca gtg aat taa ttg ctg tta tta atg gat tta gaa ata gtg gac gat ttt gtg ata tta ata tag tta tta atg atg aaa gga taa acg ctc ata gac tca tgc tat ctg gag cat ccg aat a-t ttt tcc att cta ttt tct agt gat ttt atc gat tct aat gac tac gaa gtt aat cta agt cat tta gat tat caa agt gtg aac gat ttg atc gat tac att tat ggg ata cct ttg agc cta act aac gat aac gtg aaa tat att ctt tca acc gct ga- --t ttt tta caa att gga tct gcc att act gag tgc gaa aaa tac ata ctt aaa aat ctt tgt tct aga aac tgt atc gat ttc --- --- tac ata tac gct gat aaa tat aat aac aag aaa ata gaa tca gcg tcg ttt aac aca ata tta cga aat att ttg aga ctc att aac gat gaa aac ttt aaa tac ttg aca gag gaa tca atg ata aaa att tta agc gat gat atg tta aat ata aaa aat gag gat ttc gca cca cta att ctc att aaa tgg tta gag agt aca caa caa cca tgc acc gtc gag tta ctt aga tgc ctc aga ata tca -tt gct ttc ccc aca agt tat aaa atc act tta tag tca tcg act ggt tag ttc aat cta cga atg tat aac att ctt aaa caa tat agc att ctt gga tga atc att tcc tag ata cca tag cat cga gtt gat atc tat cgg tat aag taa ttc gca tga taa gat ttc cat aaa ctg cta caa tcg t-a aaa aaa ata cat ggg aga tga tat ctt cac gta gat ata ggt gta gtt tcg cag tgg ccg tcc tgg ata ata tta tct ata tga tgg gtg gat atg atc agt ccc cgt ata gaa gtt caa agg tta tag cgt aca aca cat gta caa att ctt gga tat atg ata tac cag agc t-a aaa tat cct cgt tct aat tgc gga gga gtt gcc gat gac gaa tac att tat tgt ata ggc ggc ata cgc gat cag gat tca tcg ttg ata tct agt att gat aga tgg aag cca tca aaa cca tat tgg cag acg tat gct aaa atg cgc gag cca aaa tgt gat atg ggg gtt gcg atg tta aac gga tta ata tat gtc ata ggt gga atc gtt aaa ggt g-- atg cgt gta ccg acg cac tag aga gtt tat cag aag atg gat gga tga agc atc agc gcc ttc caa taa aaa tgt cca ata tgt cga cga ttg ttc atg ctg gta aga ttt ata tat ctg gag gtt ata aca ata gta gtg ccg tta atg gac cat cga atc t-- agt cct tag cta taa tcc gat ata tga tga atg gac caa att atc atc att aaa tat tcc tag aat taa tcc cgc tct a

>LT993232.1:174533-176764_Cowpox_virus_isolate_Ger/2015/Human2_genome_assembly_complete_genome:_monopartite

--- --- --- --a --- tgt a-c tat ttt gta cat cac tac att ggt tac agc aga tat acc tac tcc gcc a-- -cc aca tgc tcc ggt aaa cgg --g tca tgt gaa gag gga gaa tat ctt gat agg acg cat aat caa tgt tgt aat -cg gtg tcc tcc tgg aga att tgc caa ggt tag atg ta- --g tgg tag cta taa cac aaa atg tga acg ctg tcc aca tca tac ata tac agc aat tcc caa tta ttc t-a atg gat gcc atc aat gta gaa aat gcc caa agg gat cat ttg ata agg taa agt gta cag gaa cac aga aca gta aat -gt gcg tgt ctt cct ggt tgg tat tgc gct acc gat tct tca cag act gaa gac tgt cga gat -tg tgt acc aaa aag g-a gct gtc cat gcg gat act ttg gtg gaa tag ata aat tag gaa atc cta ttt gta aat cgt gtt g-- -tg ttg gta aat att gtg acg aca tac gta atc ata gag ttg gtc ctt ttc ctc cat gca aa- --- cta tct aaa tgt aat tag tca --- --- --- --- --- --- --- --- --- --- --- --- --- --- -tg atc atg gta -cc gta caa tt- --- -at att gct --- --g gta --g tat att att cag tat g-a aga cat att aat ac- --- --- -tt atc --- --t ttg aca atc ttg ct- -at aat cat --- --- --- --- aat ata aaa --a tac tta tgg --- -a- cat agt cac tca t-- --- --- --- aat tgc tg- acg cga taa att cgt aat aat ctg ttt tgt tca aat ttt tat aag ga- --- --a tct aca ggc ata aaa at- aaa aat ata att tat aat ac- --- act tac agc gcg cca tca tga ata aca gca gtg aat taa ttg ctg tta tta atg gat tta gaa ata gtg gac gat ttt gtg ata tta ata tag tta tta atg atg aaa gga taa atg ctc ata gac tca tcc tat ctg gag cat ccg aat a-t ttt tcc att cta ttt tct agt gat ttt atc gat tct aat gaa tac gaa gtt aat cta agt cat tta gat tat caa agt gtg aac gat ttg atc gat tac att tat ggg ata cct ttg agc cta act aac gat agc gtg aaa tat att ctt tca act gct ga- --t ttt tta caa att gga tat gct att acc gaa tgt gaa aat tac ata ctt aaa aat ctg tgt tct aga aat tgt atc gat ttc --- --- tac ata tac gct gat aaa tat aat aac gag aaa ata gaa aca gcg tcg ttt aac aca ata tta cga aat att ttg aga ctc att aac gat gaa aac ttt aaa tac ttg aca gag gaa tca atg ata aaa att tta agc gat gat atg tta aat ata aaa aat gag gat ttt gca cca cta att ctc att aaa tgg tta gag agt aca caa caa cca tgc aca gtc gag tta ctt aga tgc ctc aga ata tca -tt gct ttc ccc aca agt tat aaa atc act tta tag tca tcg att ggt tgg ttc aat ata cga atg tat aac att ctt aaa caa tat atc att ctt gga tga atc att tcc tag ata cca tag cat cga gtt gat atc tat cgg tat aag taa ttc aca tga taa gat ttc cat aaa ctg cta taa tcg t-a aaa aaa ata cgt ggg aga taa tat ctt cac gga gat ata ggt gta gtt ttg cag tgg ccg tca tgg ata ata tta tct ata tga tgg gtg gat atg atc agt ccc cgt ata gga gtt caa agg tta tag cgt aca ata cct gta cta att ctt gga tat atg ata tac cag agc t-a aaa tat cct cgt tct aat tgt gga gga gtt gct gat gat gaa tac att tat tgc ata ggc ggc ata cgc gat cag gat tca tcg ttg ata tct agt att gat aga tgg aag cca tca aaa cca tat tgg caa acg tat gct aaa atg cga gag cca aaa tgt gat atg ggt gta gcg atg tta aac gga tta ata tat gtc ata ggc gga gtt gtt aaa ggt g-- aca cgt gta ccg aca cac ttg aga gtt tat cag aag atg gat gga tga tgc atc aac gtc ttc caa taa aaa tgt cca ata tgt cga cga ttg ttc atg ccg gca aaa ttt ata tat ctg gag gct aca aca ata gta gtg tag tta atg gaa tat cga atc t-- agt cct tag cta taa tcc gat ata tga tga atg gac caa att atc atc att aaa tat tcc tag aat taa tcc cgc tct a

>LT896731.1:175906-178137_Cowpox_virus_isolate_Ger/2015/Cat4_genome_assembly_complete_genome:_monopartite

--- --- --- --a --- tgt a-c tat ttt gta cat cac tac att ggt tac agc aga tat acc tac tcc gcc a-- -cc aca tgc tcc ggt aaa cgg --g tca tgt gaa gag gga gaa tat ctt gat agg acg cat aat caa tgt tgt aat -cg gtg tcc tcc tgg aga att tgc caa ggt tag atg ta- --g tgg tag cta taa cac aaa atg tga acg ctg tcc aca tca tac ata tac agc aat tcc caa tta ttc t-a atg gat gcc atc aat gta gaa aat gcc caa agg gat cat ttg ata agg taa agt gta cag gaa cac aga aca gta aat -gt gcg tgt ctt cct ggt tgg tat tgc gct acc gat tct tca cag act gaa gac tgt cga gat -tg tgt acc aaa aag g-a gct gtc cat gcg gat act ttg gtg gaa tag ata aat tag gaa atc cta ttt gta aat cgt gtt g-- -tg ttg gta aat att gtg acg aca tac gta atc ata gag ttg gtc ctt ttc ctc cat gca aa- --- cta tct aaa tgt aat tag tca --- --- --- --- --- --- --- --- --- --- --- --- --- --- -tg atc atg gta -cc gta caa tt- --- -at att gct --- --g gta --g tat att att cag tat g-a aga cat att aat ac- --- --- -tt atc --- --t ttg aca atc ttg ct- -at aat cat --- --- --- --- aat ata aaa --a tac tta tgg --- -a- cat agt cac tca t-- --- --- --- aat tgc tg- acg cga taa att cgt aat aat ctg ttt tgt tca aat ttt tat aag ga- --- --a tct aca ggc ata aaa at- aaa aat ata att tat aat ac- --- act tac agc gcg cca tca tga ata aca gca gtg aat taa ttg ctg tta tta atg gat tta gaa ata gtg gac gat ttt gtg ata tta ata tag tta tta atg atg aaa gga taa atg ctc ata gac tca tcc tat ctg gag cat ccg aat a-t ttt tcc att cta ttt tct agt gat ttt atc gat tct aat gaa tac gaa gtt aat cta agt cat tta gat tat caa agt gtg aac gat ttg atc gat tac att tat ggg ata cct ttg agc cta act aac gat agc gtg aaa tat att ctt tca act gct ga- --t ttt tta caa att gga tat gct att acc gaa tgt gaa aat tac ata ctt aaa aat ctg tgt tct aga aat tgt atc gat ttc --- --- tac ata tac gct gat aaa tat aat aac gag aaa ata gaa aca gcg tcg ttt aac aca ata tta cga aat att ttg aga ctc att aac gat gaa aac ttt aaa tac ttg aca gag gaa tca atg ata aaa att tta agc gat gat atg tta aat ata aaa aat gag gat ttt gca cca cta att ctc att aaa tgg tta gag agt aca caa caa cca tgc aca gtc gag tta ctt aga tgc ctc aga ata tca -tt gct ttc ccc aca agt tat aaa atc act tta tag tca tcg att ggt tgg ttc aat ata cga atg tat aac att ctt aaa caa tat atc att ctt gga tga atc att tcc tag ata cca tag cat cga gtt gat atc tat cgg tat aag taa ttc aca tga taa gat ttc cat aaa ctg cta taa tcg t-a aaa aaa ata cgt ggg aga taa tat ctt cac gga gat ata ggt gta gtt ttg cag tgg ccg tca tgg ata ata tta tct ata tga tgg gtg gat atg atc agt ccc cgt ata gga gtt caa agg tta tag cgt aca ata cct gta cta att ctt gga tat atg ata tac cag agc t-a aaa tat cct cgt tct aat tgt gga gga gtt gct gat gat gaa tac att tat tgc ata ggc ggc ata cgc gat cag gat tca tcg ttg ata tct agt att gat aga tgg aag cca tca aaa cca tat tgg caa acg tat gct aaa atg cga gag cca aaa tgt gat atg ggt gta gcg atg tta aac gga tta ata tat gtc ata ggc gga gtt gtt aaa ggt g-- aca cgt gta ccg aca cac ttg aga gtt tat cag aag atg gat gga tga tgc atc aac gtc ttc caa taa aaa tgt cca ata tgt cga cga ttg ttc atg ccg gca aaa ttt ata tat ctg gag gct aca aca ata gta gtg tag tta atg gaa tat cga atc t-- agt cct tag cta taa tcc gat ata tga tga atg gac caa att atc atc att aaa tat tcc tag aat taa tcc cgc tct a

>LT896726.1:175753-177984_Cowpox_virus_isolate_Ger/2012/Alpaca_genome_assembly_complete_genome:_monopartite

--- --- --- --a --- tgt a-c tat ttt gta cat cac tac att ggt tac agc aga tat acc tac tcc gcc a-- -cc aca tgc tcc ggt aaa cgg --g tca tgt gaa gag gga gaa tat ctt gat agg acg cat aat caa tgt tgt aat -cg gtg tcc tcc tgg aga att tgc caa ggt tag atg ta- --g tgg tag cta taa cac aaa atg tga acg ctg tcc aca tca tac ata tac agc aat tcc caa tta ttc t-a atg gat gcc atc aat gta gaa aat gcc caa agg gat cat ttg ata agg taa agt gta cag gaa cac aga aca gta aat -gt gcg tgt ctt cct ggt tgg tat tgc gct acc gat tct tca cag act gaa gac tgt cga gat -tg tgt acc aaa aag g-a gct gtc cat gcg gat act ttg gtg gaa tag ata aat tag gaa atc cta ttt gta aat cgt gtt g-- -tg ttg gta aat att gtg acg aca tac gta atc ata gag ttg gtc ctt ttc ctc cat gca aa- --- cta tct aaa tgt aat tag tca --- --- --- --- --- --- --- --- --- --- --- --- --- --- -tg atc atg gta -cc gta caa tt- --- -at att gct --- --g gta --g tat att att cag tat g-a aga cat att aat ac- --- --- -tt atc --- --t ttg aca atc ttg ct- -at aat cat --- --- --- --- aat ata aaa --a tac tta tgg --- -a- cat agt cac tca t-- --- --- --- aat tgc tg- acg cga taa att cgt aat aat ctg ttt tgt tca aat ttt tat aag ga- --- --a tct aca ggc ata aaa at- aaa aat ata att tat aat ac- --- act tac agc gcg cca tca tga ata aca gca gtg aat taa ttg ctg tta tta atg gat tta gaa ata gtg gac gat ttt gtg ata tta ata tag tta tta atg atg aaa gga taa atg ctc ata gac tca tcc tat ctg gag cat ccg aat a-t ttt tcc att cta ttt tct agt gat ttt atc gat tct aat gaa tac gaa gtt aat cta agt cat tta gat tat caa agt gtg aac gat ttg atc gat tac att tat ggg ata cct ttg agc cta act aac gat agc gtg aaa tat att ctt tca act gct ga- --t ttt tta caa att gga tat gct att acc gaa tgt gaa aat tac ata ctt aaa aat ctg tgt tct aga aat tgt atc gat ttc --- --- tac ata tac gct gat aaa tat aat aac gag aaa ata gaa aca gcg tcg ttt aac aca ata tta cga aat att ttg aga ctc att aac gat gaa aac ttt aaa tac ttg aca gag gaa tca atg ata aaa att tta agc gat gat atg tta aat ata aaa aat gag gat ttt gca cca cta att ctc att aaa tgg tta gag agt aca caa caa cca tgc aca gtc gag tta ctt aga tgc ctc aga ata tca -tt gct ttc ccc aca agt tat aaa atc act tta tag tca tcg att ggt tgg ttc aat ata cga atg tat aac att ctt aaa caa tat atc att ctt gga tga atc att tcc tag ata cca tag cat cga gtt gat atc tat cgg tat aag taa ttc aca tga taa gat ttc cat aaa ctg cta taa tcg t-a aaa aaa ata cgt ggg aga taa tat ctt cac gga gat ata ggt gta gtt ttg cag tgg ccg tca tgg ata ata tta tct ata tga tgg gtg gat atg atc agt ccc cgt ata gga gtt caa agg tta tag cgt aca ata cct gta cta att ctt gga tat atg ata tac cag agc t-a aaa tat cct cgt tct aat tgt gga gga gtt gct gat gat gaa tac att tat tgc ata ggc ggc ata cgc gat cag gat tca tcg ttg ata tct agt att gat aga tgg aag cca tca aaa cca tat tgg caa acg tat gct aaa atg cga gag cca aaa tgt gat atg ggt gta gcg atg tta aac gga tta ata tat gtc ata ggc gga gtt gtt aaa ggt g-- aca cgt gta ccg aca cac ttg aga gtt tat cag aag atg gat gga tga tgc atc aac gtc ttc caa taa aaa tgt cca ata tgt cga cga ttg ttc atg ccg gca aaa ttt ata tat ctg gag gct aca aca ata gta gtg tag tta atg gaa tat cga atc t-- agt cct tag cta taa tcc gat ata tga tga atg gac caa att atc atc att aaa tat tcc tag aat taa tcc cgc tct a

>LT896725.1:177051-179282_Cowpox_virus_isolate_Ger/2014/Cat2_genome_assembly_complete_genome:_monopartite

--- --- --- --a --- tgt a-c tat ttt gta cat cac tac att ggt tac agc aga tat acc tac tcc gcc a-- -cc aca tgc tcc ggt aaa cgg --g tca tgt gaa gag gga gaa tat ctt gat agg acg cat aat caa tgt tgt aat -cg gtg tcc tcc tgg aga att tgc caa ggt tag atg ta- --g tgg tag cta taa cac aaa atg tga acg ctg tcc aca tca tac ata tac agc aat tcc caa tta ttc t-a atg gat gcc atc aat gta gaa aat gcc caa agg gat cat ttg ata agg taa agt gta cag gaa cac aga aca gta aat -gt gcg tgt ctt cct ggt tgg tat tgc gct acc gat tct tca cag act gaa gac tgt cga gat -tg tgt acc aaa aag g-a gct gtc cat gcg gat act ttg gtg gaa tag ata aat tag gaa atc cta ttt gta aat cgt gtt g-- -tg ttg gta aat att gtg acg aca tac gta atc ata gag ttg gtc ctt ttc ctc cat gca aa- --- cta tct aaa tgt aat tag tca --- --- --- --- --- --- --- --- --- --- --- --- --- --- -tg atc atg gta -cc gta caa tt- --- -at att gct --- --g gta --g tat att att cag tat g-a aga cat att aat ac- --- --- -tt atc --- --t ttg aca atc ttg ct- -at aat cat --- --- --- --- aat ata aaa --a tac tta tgg --- -a- cat agt cac tca t-- --- --- --- aat tgc tg- acg cga taa att cgt aat aat ctg ttt tgt tca aat ttt tat aag ga- --- --a tct aca ggc ata aaa at- aaa aat ata att tat aat ac- --- act tac agc gcg cca tca tga ata aca gca gtg aat taa ttg ctg tta tta atg gat tta gaa ata gtg gac gat ttt gtg ata tta ata tag tta tta atg atg aaa gga taa atg ctc ata gac tca tcc tat ctg gag cat ccg aat a-t ttt tcc att cta ttt tct agt gat ttt atc gat tct aat gaa tac gaa gtt aat cta agt cat tta gat tat caa agt gtg aac gat ttg atc gat tac att tat ggg ata cct ttg agc cta act aac gat agc gtg aaa tat att ctt tca act gct ga- --t ttt tta caa att gga tat gct att acc gaa tgt gaa aat tac ata ctt aaa aat ctg tgt tct aga aat tgt atc gat ttc --- --- tac ata tac gct gat aaa tat aat aac gag aaa ata gaa aca gcg tcg ttt aac aca ata tta cga aat att ttg aga ctc att aac gat gaa aac ttt aaa tac ttg aca gag gaa tca atg ata aaa att tta agc gat gat atg tta aat ata aaa aat gag gat ttt gca cca cta att ctc att aaa tgg tta gag agt aca caa caa cca tgc aca gtc gag tta ctt aga tgc ctc aga ata tca -tt gct ttc ccc aca agt tat aaa atc act tta tag tca tcg att ggt tgg ttc aat ata cga atg tat aac att ctt aaa caa tat atc att ctt gga tga atc att tcc tag ata cca tag cat cga gtt gat atc tat cgg tat aag taa ttc aca tga taa gat ttc cat aaa ctg cta taa tcg t-a aaa aaa ata cgt ggg aga taa tat ctt cac gga gat ata ggt gta gtt ttg cag tgg ccg tca tgg ata ata tta tct ata tga tgg gtg gat atg atc agt ccc cgt ata gga gtt caa agg tta tag cgt aca ata cct gta cta att ctt gga tat atg ata tac cag agc t-a aaa tat cct cgt tct aat tgt gga gga gtt gct gat gat gaa tac att tat tgc ata ggc ggc ata cgc gat cag gat tca tcg ttg ata tct agt att gat aga tgg aag cca tca aaa cca tat tgg caa acg tat gct aaa atg cga gag cca aaa tgt gat atg ggt gta gcg atg tta aac gga tta ata tat gtc ata ggc gga gtt gtt aaa ggt g-- aca cgt gta ccg aca cac ttg aga gtt tat cag aag atg gat gga tga tgc atc aac gtc ttc caa taa aaa tgt cca ata tgt cga cga ttg ttc atg ccg gca aaa ttt ata tat ctg gag gct aca aca ata gta gtg tag tta atg gaa tat cga atc t-- agt cct tag cta taa tcc gat ata tga tga atg gac caa att atc atc att aaa tat tcc tag aat taa tcc cgc tct a

>KY549148.1:177111-179383_Cowpox_virus_isolate_CPXV_1639

ttg ctg act gta --- tgt a-c tat ttt gta cat cag tac att ggt tac agc aga tat acc tac ttc gtc act gcc aca tgc tcc ggt aaa cgg --g aca tgt gaa gag gga gaa tat ctt gat aag acg cat aat caa tgt tgt aat -cg gtg tcc tcc tgg aga att tgc caa gtt tag atg ta- --a tgg taa cga taa cac aaa atg tga acg ctg ccc acc tca tac ata tac cac agt acc caa tta ttc t-a atg gat gtc atc aat gta gga aat gcc caa cag gat cat ttg ata agg taa agt gta ccg gaa cac aga aca gta aat -gt tcg tgt ctt cct ggt tgg ttt tgc gct act gat tct tcg aag act gaa gat tgt cga gat -tg tat acc aaa aag a-a aat gtc cat gcg gat act ttg gtg gaa tag atg aac tcg gaa atc ctc ttt gta aat cgt gtt g-- -tg ttg gtg aat att gcg acg aca tac gta atc ata gag ttg gtc ctt ttc ctc cat gca aa- --- cta tct aaa tgt aat tag cca --- --- --- --- --- --- --- --- --- --- --- --- --- --- -tg atc atg tta ccc ata c-a tc- --- -at atc gct act tgg t-a --g cgt att att cag tat g-a aga tct att aat aat aat tat ctt att tat ttc ttg acg --- --- --- -at att aat cgg tgt cct aat aat ata aaa --- tac tta tgg --- --- --a aat aaa ttg t-a aac aag atg aat taa t-- aca caa tac atc ctt -at act cag ctt -gt tca aa- ttt tat aaa gaa tca tat tca tta ggc ata taa ta- aaa aat ata att tat aat ac- --- act tac a-- gcg tca tca tgc ata aca gca gtg aat taa ttg ctg tta tta atg gat tta gaa ata gtg gac gat ttt gtg ata tta gta tag tta tta atg atg aaa gga taa acg ctc ata gac tca tgc tat ctg gag cat ccg aat a-t ttt tcc att cta ttt tct agt gat ttt atc gat tct aat gac tac gaa gtt aat cta agt cat tta gat tat caa agt gtg aac gat ttg atc gat tac att tat ggg ata cct ttg agc cta act aac gat aac gtg aaa tat att ctt tca acc gct ga- --t ttt tta caa att gga tct gcc att act gag tgc gaa aaa tac ata ctt aaa aat ctt tgt tct aga aac tgt atc gat ttc --- --- tac ata tac gct gat aaa tat aat aac aag aaa ata gaa tca gcg tcg ttt aac aca ata tta cga aat att ttg aga ctc att aac gat gaa aac ttt aaa tac ttg aca gag gaa tca atg ata aaa att tta agc gat gat atg tta aat ata aaa aat gag gat ttc gca cca cta att ctc att aaa tgg tta gag agt aca caa caa cca tgc acc gtc gag tta ctt aga tgc ctc aga ata tca -tt gct ttc ccc aca agt tat aaa atc act tta tag tca tcg act ggt tag ttc aat cta cga atg tat aac att ctt aaa caa tat agc att ctt gga tga atc att tcc tag ata cca tag cat cga gtt gat atc tat cgg tat aag taa ttc gca tga taa gat ttc cat aaa ctg cta caa tcg t-a aaa aaa ata cat ggg aaa tga tat ctt cac gta gat ata ggt gta gtt tcg cag tgg ccg tcc tgg ata ata tta tct ata tga tgg gtg gat atg atc agt ccc cgt ata gaa gtt caa agg tta tag cgt aca aca cat gta caa att ctt gga tat atg ata tac cag agc t-a aaa tat cct cgt tct aat tgc gga gga gtt gcc gat gac gaa tac att tat tgt ata ggc ggc ata cgc gat cag gat tca tcg ttg ata tct agt att gat aga tgg aag cca tca aaa cca tat tgg cag acg tat gct aaa atg cgc gag cca aaa tgt gat atg ggg gtt gcg atg tta aac gga tta ata tat gtc ata ggt gga atc gtt aaa ggt g-- aca cat gta ccg acg cac tag aaa gtt tat cag aag atg gat gga tga agc atc agc gcc ttc caa taa aaa tgt cca ata tgt cga cga ttg ttc atg ctg gta aga ttt ata tat ctg gag gtt ata aca ata gta gtg ccg tta atg gac cat cga atc t-- agt cct tag cta taa tcc gat ata tga tga atg gac caa att atc atc att aaa tat tcc tag aat taa tcc cgc tct a

>LT896727.1:176642-178873_Cowpox_virus_isolate_Ger/2015/Cat2_genome_assembly_complete_genome:_monopartite

--- --- --- --a --- tgt a-c tat ttt gta cat cac tac att ggt tac agc aga tat acc tac tcc gcc a-- -cc aca tgc tcc ggt aaa cgg --g aca tgt gaa gag gga gaa tat ctt gat agg acg cat aat caa tgt tgt aat -cg gtg tcc tcc tgg aga att tgc caa ggt tag atg ta- --a tgg taa cga taa cac aaa atg tga acg ctg tcc acc tca tac att tac cac agt acc caa tta ttc t-a atg gat gtc atc aat gta gaa aat gcc caa agg gat cat ttg ata agg taa agt gta cag gaa cac aga aca gta aat -gt tcg tgt ctt cct ggt tgg ttt tgt gct act gat tct tcg aag act gaa gat tgt cga gat -tg tgt acc aaa aag g-a gct gtc cat gcg gat act ttg gtg gaa tag ata aat tag gaa atc cta ttt gta aat cgt gtt g-- -tg ttg gta aat att gtg acg aca tac gta atc ata gag ttg gtc ctt ttc ctc cat gca aa- --- cta tct aaa tgt aat tag tca --- --- --- --- --- --- --- --- --- --- --- --- --- --- -tg atc atg gta -cc gta caa tt- --- -at att gct --- --g gta --g tat att att cag tat g-a aga cat att aat ac- --- --- -tt atc --- --t ttg aca atc ttg ct- -at aat cat --- --- --- --- aat ata aaa --a tac tta tgg --- -a- cat agt cac tca t-- --- --- --- aat tgc tg- acg cga taa att cgt aat aat ctg ttt tgt tca aat ttt tat aag ga- --- --a tct aca ggc ata aaa at- aaa aat ata att tat aat ac- --- act tac agc gcg cca tca tga ata aca gca gtg aat taa ttg ctg tta tta atg gat tta gaa ata gtg gac gat ttt gtg ata tta ata tag tta tta atg atg aaa gga taa atg ctc ata gac tca tcc tat ctg gag cat ccg aat a-t ttt tcc att cta ttt tct agt gat ttt atc gat tct aat gaa tac gaa gtt aat cta agt cat tta gat tat caa agt gtg aac gat ttg atc gat tac att tat ggg ata cct ttg agc cta act aac gat agc gtg aaa tat att ctt tca act gct ga- --t ttt tta caa att gga tat gct att acc gaa tgt gaa aat tac ata ctt aaa aat ctg tgt tct aga aat tgt atc gat ttc --- --- tac ata tac gct gat aaa tat aat aac gag aaa ata gaa aca gcg tcg ttt aac aca ata tta cga aat att ttg aga ctc att aac gat gaa aac ttt aaa tac ttg aca gag gaa tca atg ata aaa att tta agc gat gat atg tta aat ata aaa aat gag gat ttt gca cca cta att ctc att aaa tgg tta gag agt aca caa caa cca tgc aca gtc gag tta ctt aga tgc ctc aga ata tca -tt gct ttc ccc aca agt tat aaa atc act tta tag tca tcg att ggt tgg ttc aat ata cga atg tat aac att ctt aaa caa tat atc att ctt gga tga atc att tcc tag ata cca tag cat cga gtt gat atc tat cgg tat aag taa ttc aca tga taa gat ttc cat aaa ctg cta taa tcg t-a aaa aaa ata cgt ggg aga taa tat ctt cac gga gat ata ggt gta gtt ttg cag tgg ccg tca tgg ata ata tta tct ata tga tgg gtg gat atg atc agt ccc cgt ata gga gtt caa agg tta tag cgt aca ata cct gta cta att ctt gga tat atg ata tac cag agc t-a aaa tat cct cgt tct aat tgt gga gga gtt gct gat gat gaa tac att tat tgc ata ggc ggc ata cgc gat cag gat tca tcg ttg ata tct agt att gat aga tgg aag cca tca aaa cca tat tgg caa acg tat gct aaa atg cga gag cca aaa tgt gat atg ggt gta gcg atg tta aac gga tta ata tat gtc ata ggc gga gtt gtt aaa ggt g-- aca cgt gta ccg aca cac ttg aga gtt tat cag aag atg gat gga tga tgc atc aac gtc ttc caa taa aaa tgt cca ata tgt cga cga ttg ttc atg ccg gca aaa ttt ata tat ctg gag gct aca aca ata gta gtg tag tta atg gaa tat cga atc t-- agt cct tag cta taa tcc gat ata tga tga atg gac caa att atc atc att aaa tat tcc tag aat taa tcc cgc tct a

>KY549149.1:177612-179885_Cowpox_virus_isolate_CPXV_K2739

ttg ctg act gta --- tgt a-c tat ttt gta cat cag tac att ggt tac agc aga tat acc tac ttc gtc act gcc aca tgc tcc ggt aaa cgg --g tca tgt gac gaa gga gaa tat ctt gat aag acg cat aat caa tgt tgt aat -cg gtg tcc tcc tgg aga att tgc caa gat cag atg ta- --g cgg tag cga taa cac aaa atg tga acg ctg tcc acc tca tac ata tac cac agt acc caa tta ttc t-a atg gat gcc atc aat gta gaa aat gcc caa cag gat cat ttg ata agg taa agt gta ccg gaa cac aga aca gta aat -gt tcg tgt ctt cct ggt tgg ttt tgt gct act gat tct tcg aag act gaa gat tgt cga gat -tg tat acc aaa aag g-a aat gtc cat gcg gat act ttg gtg gaa tag atg aac tcg gaa atc ctc ttt gta aat cgt gtt g-- -tg ttg gtg aat att gcg acg aca tac gta atc ata gag ttg gtc ctt ttc ctc cat gca aa- --- cta tct aaa tgt aat tag cca --- --- --- --- --- --- --- --- --- --- --- --- --- --- -tg atc atg tta -cc ata c-a tc- --- -at ata gct act tgg t-a --g cgt att att cag tat g-a aga tct att aat aat aat tat ctt att tat ttc ttg acg --- --- --- -at att aat cgg tgt cct aat aat ata aaa --- tac tta tgg --- --- --a aat aaa ttg t-a aac aag atg aat taa tac aca caa tac atc ctt -at act cag ctt -gt tca aa- ttt tat aaa gaa tca tat tca tta ggc ata taa ta- aaa aat ata att tat aat ac- --- act tac a-- gcg tca tca tgc ata aca gca gtg aat taa ttg ctg tta tta atg gat tta gaa ata gtg gac gat ttt gtg ata tta gta tag tta tta atg atg aaa gga taa acg ctc ata gac tca tgc tat ctg gag cat cag aat a-t ttt tcc att cta ttt tct agt gat ttt atc gat tct aat gac tac gaa gtt aat cta agt cat tta gat tat caa agt gtt aat gat ttg atc gat tac att tat ggg ata cct ttg agc cta act aac gat aac gtg aaa tat att ctt tca acc gct ga- --t ttt tta caa att gga tct gcc att act gag tgc gaa aaa tac ata ctt aaa aat ctt tgt tct aga aac tgt atc gat ttc --- --- tac ata tac gct gat aaa tat aat aac aag aaa ata gaa tca gcg tcg ttt aac aca ata tta cga aat att ttg aga ctc att aac gat gaa aac ttt aaa tac ttg aca gag gaa tca atg ata aaa att tta agc gat gat atg tta aat ata aaa aat gag gat ttc gca cca cta att ctc att aaa tgg tta gag agt aca caa caa cca tgc acc gtc gag tta ctt aga tgc ctc aga ata tca -tt gct ttc ccc aca agt tat aaa atc act tta tag tca tcg act ggt tag ttc aat cta cga atg tat aac att ctt aaa caa tat agc att ctt gga tga atc att tcc tag ata cca tag cat cga gtt gat atc tat cgg tat aag taa ttc aca tga taa gat ttc cat aaa ctg cta caa tcg t-a aaa aaa ata cat ggg aga tga tat ctt cac gta gat ata ggt gta gtt tcg cag tgg ccg tcc tgg ata ata tta tct ata tga tgg gtg gat atg atc agt ccc cgt ata gaa gtt caa agg tta tag cgt aca ata cat gta caa att ctt gga tat atg ata tac cag agc t-a aaa tat cct cgt tct aat tgc gga gga gtt gcc gat gac gaa tac att tat tgt ata ggc ggc ata cgc gat cag gat tca tcg ttg ata tct agt att gat aga tgg aag cca tca aaa cca tat tgg cag acg tat gct aaa atg cgc gaa cca aaa tgt gat atg ggg gtt gcg atg tta aac gga tta ata tat gtc ata ggt gga atc gtt aaa ggt g-- atg cgt gta ccg acg cac tag aaa gtt tat cag aag atg gat gga tga agc atc agc gcc ttc caa taa aaa tgt cca ata tgt cga cga ttg ttc atg ctg gta aga ttt ata tat ctg gag gtt ata aca ata gta gtg ccg tta atg gac cat cga atc t-- agt cct tag cta taa tcc gat ata tga tga atg gac caa att atc atc att aaa tat tcc tag aat taa tcc cgc tct a

>KC813504.1:169254-171485_Cowpox_virus_strain_RatHei09/1_complete_genome

--- --- --- --a --- tgt a-c tat ttt gta cat cac tac att ggt tac agc aga tat acc tac tcc gcc a-- -cc aca tgc tcc ggt aaa cgg --g aca tgt gaa gag gga gaa tat ctt gat agg acg cat aat caa tgt tgt aat -cg gtg tcc tcc tgg aga att tgc caa ggt tag atg ta- --a tgg taa cga taa cac aaa atg tga acg ctg tcc acc tca tac att tac cac agt acc caa tta ttc t-a atg gat gtc atc aat gta gaa aat gcc caa agg gat cat ttg ata agg taa agt gta cag gaa cac aga aca gta aat -gt tcg tgt ctt cct ggt tgg ttt tgt gct act gat tct tcg aag act gaa gat tgt cga gat -tg tgt acc aaa aag g-a gct gtc cat gcg gat act ttg gtg gaa tag ata aat tag gaa atc cta ttt gta aat cgt gtt g-- -tg ttg gta aat att gtg acg aca tac gta atc ata gag ttg gtc ctt ttc ctc cat gca aa- --- cta tct aaa tgt aat tag tca --- --- --- --- --- --- --- --- --- --- --- --- --- --- -tg atc atg gta -cc gta caa tt- --- -at att gct --- --g gta --g tat att att cag tat g-a aga cat att aat ac- --- --- -tt atc --- --t ttg aca atc ttg ct- -at aat cat --- --- --- --- aat ata aaa --a tac tta tgg --- -a- cat agt cac tca t-- --- --- --- aat tgc tg- acg cga taa att cgt aat aat ctg ttt tgt tca aat ttt tat aag ga- --- --a tct aca ggc ata aaa at- aaa aat ata att tat aat ac- --- act tac agc gcg cca tca tga ata aca gca gtg aat taa ttg ctg tta tta atg gat tta gaa ata gtg gac gat ttt gtg ata tta ata tag tta tta atg atg aaa gga taa atg ctc ata gac tca tcc tat ctg gag cat ccg aat a-t ttt tcc att cta ttt tct agt gat ttt atc gat tct aat gaa tac gaa gtt aat cta agt cat tta gat tat caa agt gtg aac gat ttg atc gat tac att tat ggg ata cct ttg agc cta act aac gat agc gtg aaa tat att ctt tca act gct ga- --t ttt tta caa att gga tat gct att acc gaa tgt gaa aat tac ata ctt aaa aat ctg tgt tct aga aat tgt atc gat ttc --- --- tac ata tac gct gat aaa tat aat aac gag aaa ata gaa aca gcg tcg ttt aac aca ata tta cga aat att ttg aga ctc att aac gat gaa aac ttt aaa tac ttg aca gag gaa tca atg ata aaa att tta agc gat gat atg tta aat ata aaa aat gag gat ttt gca cca cta att ctc att aaa tgg tta gag agt aca caa caa cca tgc aca gtc gag tta ctt aga tgc ctc aga ata tca -tt gct ttc ccc aca agt tat aaa atc act tta tag tca tcg att ggt tgg ttc aat ata cga atg tat aac att ctt aaa caa tat atc att ctt gga tga atc att tcc tag ata cca tag cat cga gtt gat atc tat cgg tat aag taa ttc aca tga taa gat ttc cat aaa ctg cta taa tcg t-a aaa aaa ata cgt ggg aga taa tat ctt cac gga gat ata ggt gta gtt ttg cag tgg ccg tca tgg ata ata tta tct ata tga tgg gtg gat atg atc agt ccc cgt ata gga gtt caa agg tta tag cgt aca ata cct gta cta att ctt gga tat atg ata tac cag agc t-a aaa tat cct cgt tct aat tgt gga gga gtt gct gat gat gaa tac att tat tgc ata ggc ggc ata cgc gat cag gat tca tcg ttg ata tct agt att gat aga tgg aag cca tca aaa cca tat tgg caa acg tat gct aaa atg cga gag cca aaa tgt gat atg ggt gta gcg atg tta aac gga tta ata tat gtc ata ggc gga gtt gtt aaa ggt g-- aca cgt gta ccg aca cac ttg aga gtt tat cag aag atg gat gga tga tgc atc aac gtc ttc caa taa aaa tgt cca ata tgt cga cga ttg ttc atg ccg gca aaa ttt ata tat ctg gag gct aca aca ata gta gtg tag tta atg gaa tat cga atc t-- agt cct tag cta taa tcc gat ata tga tga atg gac caa att atc atc att aaa tat tcc tag aat taa tcc cgc tct a

>KC813500.1:158321-160552_Cowpox_virus_strain_MonKre08/4_complete_genome

--- --- --- --a --- tgt a-c tat ttt gta cat cac tac att ggt tac agc aga tat acc tac tcc gcc a-- -cc aca tgc tcc ggt aaa cgg --g aca tgt gaa gag gga gaa tat ctt gat agg acg cat aat caa tgt tgt aat -cg gtg tcc tcc tgg aga att tgc caa ggt tag atg ta- --a tgg taa cga taa cac aaa atg tga acg ctg tcc acc tca tac att tac cac agt acc caa tta ttc t-a atg gat gtc atc aat gta gaa aat gcc caa agg gat cat ttg ata agg taa agt gta cag gaa cac aga aca gta aat -gt tcg tgt ctt cct ggt tgg ttt tgt gct act gat tct tcg aag act gaa gat tgt cga gat -tg tgt acc aaa aag g-a gct gtc cat gcg gat act ttg gtg gaa tag ata aat tag gaa atc cta ttt gta aat cgt gtt g-- -tg ttg gta aat att gtg acg aca tac gta atc ata gag ttg gtc ctt ttc ctc cat gca aa- --- cta tct aaa tgt aat tag tca --- --- --- --- --- --- --- --- --- --- --- --- --- --- -tg atc atg gta -cc gta caa tt- --- -at att gct --- --g gta --g tat att att cag tat g-a aga cat att aat ac- --- --- -tt atc --- --t ttg aca atc ttg ct- -at aat cat --- --- --- --- aat ata aaa --a tac tta tgg --- -a- cat agt cac tca t-- --- --- --- aat tgc tg- acg cga taa att cgt aat aat ctg ttt tgt tca aat ttt tat aag ga- --- --a tct aca ggc ata aaa at- aaa aat ata att tat aat ac- --- act tac agc gcg cca tca tga ata aca gca gtg aat taa ttg ctg tta tta atg gat tta gaa ata gtg gac gat ttt gtg ata tta ata tag tta tta atg atg aaa gga taa atg ctc ata gac tca tcc tat ctg gag cat ccg aat a-t ttt tcc att cta ttt tct agt gat ttt atc gat tct aat gaa tac gaa gtt aat cta agt cat tta gat tat caa agt gtg aac gat ttg atc gat tac att tat ggg ata cct ttg agc cta act aac gat agc gtg aaa tat att ctt tca act gct ga- --t ttt tta caa att gga tat gct att acc gaa tgt gaa aat tac ata ctt aaa aat ctg tgt tct aga aat tgt atc gat ttc --- --- tac ata tac gct gat aaa tat aat aac gag aaa ata gaa aca gcg tcg ttt aac aca ata tta cga aat att ttg aga ctc att aac gat gaa aac ttt aaa tac ttg aca gag gaa tca atg ata aaa att tta agc gat gat atg tta aat ata aaa aat gag gat ttt gca cca cta att ctc att aaa tgg tta gag agt aca caa caa cca tgc aca gtc gag tta ctt aga tgc ctc aga ata tca -tt gct ttc ccc aca agt tat aaa atc act tta tag tca tcg att ggt tgg ttc aat ata cga atg tat aac att ctt aaa caa tat atc att ctt gga tga atc att tcc tag ata cca tag cat cga gtt gat atc tat cgg tat aag taa ttc aca tga taa gat ttc cat aaa ctg cta taa tcg t-a aaa aaa ata cgt ggg aga taa tat ctt cac gga gat ata ggt gta gtt ttg cag tgg ccg tca tgg ata ata tta tct ata tga tgg gtg gat atg atc agt ccc cgt ata gga gtt caa agg tta tag cgt aca ata cct gta cta att ctt gga tat atg ata tac cag agc t-a aaa tat cct cgt tct aat tgt gga gga gtt gct gat gat gaa tac att tat tgc ata ggc ggc ata cgc gat cag gat tca tcg ttg ata tct agt att gat aga tgg aag cca tca aaa cca tat tgg caa acg tat gct aaa atg cga gag cca aaa tgt gat atg ggt gta gcg atg tta aac gga tta ata tat gtc ata ggc gga gtt gtt aaa ggt g-- aca cgt gta ccg aca cac ttg aga gtt tat cag aag atg gat gga tga tgc atc aac gtc ttc caa taa aaa tgt cca ata tgt cga cga ttg ttc atg ccg gca aaa ttt ata tat ctg gag gct aca aca ata gta gtg tag tta atg gaa tat cga atc t-- agt cct tag cta taa tcc gat ata tga tga atg gac caa att atc atc att aaa tat tcc tag aat taa tcc cgc tct a

>KC813498.1:177299-179530_Cowpox_virus_strain_JagKre08/2_complete_genome

--- --- --- --a --- tgt a-c tat ttt gta cat cac tac att ggt tac agc aga tat acc tac tcc gcc a-- -cc aca tgc tcc ggt aaa cgg --g aca tgt gaa gag gga gaa tat ctt gat agg acg cat aat caa tgt tgt aat -cg gtg tcc tcc tgg aga att tgc caa ggt tag atg ta- --a tgg taa cga taa cac aaa atg tga acg ctg tcc acc tca tac att tac cac agt acc caa tta ttc t-a atg gat gtc atc aat gta gaa aat gcc caa agg gat cat ttg ata agg taa agt gta cag gaa cac aga aca gta aat -gt tcg tgt ctt cct ggt tgg ttt tgt gct act gat tct tcg aag act gaa gat tgt cga gat -tg tgt acc aaa aag g-a gct gtc cat gcg gat act ttg gtg gaa tag ata aat tag gaa atc cta ttt gta aat cgt gtt g-- -tg ttg gta aat att gtg acg aca tac gta atc ata gag ttg gtc ctt ttc ctc cat gca aa- --- cta tct aaa tgt aat tag tca --- --- --- --- --- --- --- --- --- --- --- --- --- --- -tg atc atg gta -cc gta caa tt- --- -at att gct --- --g gta --g tat att att cag tat g-a aga cat att aat ac- --- --- -tt atc --- --t ttg aca atc ttg ct- -at aat cat --- --- --- --- aat ata aaa --a tac tta tgg --- -a- cat agt cac tca t-- --- --- --- aat tgc tg- acg cga taa att cgt aat aat ctg ttt tgt tca aat ttt tat aag ga- --- --a tct aca ggc ata aaa at- aaa aat ata att tat aat ac- --- act tac agc gcg cca tca tga ata aca gca gtg aat taa ttg ctg tta tta atg gat tta gaa ata gtg gac gat ttt gtg ata tta ata tag tta tta atg atg aaa gga taa atg ctc ata gac tca tcc tat ctg gag cat ccg aat a-t ttt tcc att cta ttt tct agt gat ttt atc gat tct aat gaa tac gaa gtt aat cta agt cat tta gat tat caa agt gtg aac gat ttg atc gat tac att tat ggg ata cct ttg agc cta act aac gat agc gtg aaa tat att ctt tca act gct ga- --t ttt tta caa att gga tat gct att acc gaa tgt gaa aat tac ata ctt aaa aat ctg tgt tct aga aat tgt atc gat ttc --- --- tac ata tac gct gat aaa tat aat aac gag aaa ata gaa aca gcg tcg ttt aac aca ata tta cga aat att ttg aga ctc att aac gat gaa aac ttt aaa tac ttg aca gag gaa tca atg ata aaa att tta agc gat gat atg tta aat ata aaa aat gag gat ttt gca cca cta att ctc att aaa tgg tta gag agt aca caa caa cca tgc aca gtc gag tta ctt aga tgc ctc aga ata tca -tt gct ttc ccc aca agt tat aaa atc act tta tag tca tcg att ggt tgg ttc aat ata cga atg tat aac att ctt aaa caa tat atc att ctt gga tga atc att tcc tag ata cca tag cat cga gtt gat atc tat cgg tat aag taa ttc aca tga taa gat ttc cat aaa ctg cta taa tcg t-a aaa aaa ata cgt ggg aga taa tat ctt cac gga gat ata ggt gta gtt ttg cag tgg ccg tca tgg ata ata tta tct ata tga tgg gtg gat atg atc agt ccc cgt ata gga gtt caa agg tta tag cgt aca ata cct gta cta att ctt gga tat atg ata tac cag agc t-a aaa tat cct cgt tct aat tgt gga gga gtt gct gat gat gaa tac att tat tgc ata ggc ggc ata cgc gat cag gat tca tcg ttg ata tct agt att gat aga tgg aag cca tca aaa cca tat tgg caa acg tat gct aaa atg cga gag cca aaa tgt gat atg ggt gta gcg atg tta aac gga tta ata tat gtc ata ggc gga gtt gtt aaa ggt g-- aca cgt gta ccg aca cac ttg aga gtt tat cag aag atg gat gga tga tgc atc aac gtc ttc caa taa aaa tgt cca ata tgt cga cga ttg ttc atg ccg gca aaa ttt ata tat ctg gag gct aca aca ata gta gtg tag tta atg gaa tat cga atc t-- agt cct tag cta taa tcc gat ata tga tga atg gac caa att atc atc att aaa tat tcc tag aat taa tcc cgc tct a

>KC813497.1:176983-179214_Cowpox_virus_strain_JagKre08/1_complete_genome

--- --- --- --a --- tgt a-c tat ttt gta cat cac tac att ggt tac agc aga tat acc tac tcc gcc a-- -cc aca tgc tcc ggt aaa cgg --g aca tgt gaa gag gga gaa tat ctt gat agg acg cat aat caa tgt tgt aat -cg gtg tcc tcc tgg aga att tgc caa ggt tag atg ta- --a tgg taa cga taa cac aaa atg tga acg ctg tcc acc tca tac att tac cac agt acc caa tta ttc t-a atg gat gtc atc aat gta gaa aat gcc caa agg gat cat ttg ata agg taa agt gta cag gaa cac aga aca gta aat -gt tcg tgt ctt cct ggt tgg ttt tgt gct act gat tct tcg aag act gaa gat tgt cga gat -tg tgt acc aaa aag g-a gct gtc cat gcg gat act ttg gtg gaa tag ata aat tag gaa atc cta ttt gta aat cgt gtt g-- -tg ttg gta aat att gtg acg aca tac gta atc ata gag ttg gtc ctt ttc ctc cat gca aa- --- cta tct aaa tgt aat tag tca --- --- --- --- --- --- --- --- --- --- --- --- --- --- -tg atc atg gta -cc gta caa tt- --- -at att gct --- --g gta --g tat att att cag tat g-a aga cat att aat ac- --- --- -tt atc --- --t ttg aca atc ttg ct- -at aat cat --- --- --- --- aat ata aaa --a tac tta tgg --- -a- cat agt cac tca t-- --- --- --- aat tgc tg- acg cga taa att cgt aat aat ctg ttt tgt tca aat ttt tat aag ga- --- --a tct aca ggc ata aaa at- aaa aat ata att tat aat ac- --- act tac agc gcg cca tca tga ata aca gca gtg aat taa ttg ctg tta tta atg gat tta gaa ata gtg gac gat ttt gtg ata tta ata tag tta tta atg atg aaa gga taa atg ctc ata gac tca tcc tat ctg gag cat ccg aat a-t ttt tcc att cta ttt tct agt gat ttt atc gat tct aat gaa tac gaa gtt aat cta agt cat tta gat tat caa agt gtg aac gat ttg atc gat tac att tat ggg ata cct ttg agc cta act aac gat agc gtg aaa tat att ctt tca act gct ga- --t ttt tta caa att gga tat gct att acc gaa tgt gaa aat tac ata ctt aaa aat ctg tgt tct aga aat tgt atc gat ttc --- --- tac ata tac gct gat aaa tat aat aac gag aaa ata gaa aca gcg tcg ttt aac aca ata tta cga aat att ttg aga ctc att aac gat gaa aac ttt aaa tac ttg aca gag gaa tca atg ata aaa att tta agc gat gat atg tta aat ata aaa aat gag gat ttt gca cca cta att ctc att aaa tgg tta gag agt aca caa caa cca tgc aca gtc gag tta ctt aga tgc ctc aga ata tca -tt gct ttc ccc aca agt tat aaa atc act tta tag tca tcg att ggt tgg ttc aat ata cga atg tat aac att ctt aaa caa tat atc att ctt gga tga atc att tcc tag ata cca tag cat cga gtt gat atc tat cgg tat aag taa ttc aca tga taa gat ttc cat aaa ctg cta taa tcg t-a aaa aaa ata cgt ggg aga taa tat ctt cac gga gat ata ggt gta gtt ttg cag tgg ccg tca tgg ata ata tta tct ata tga tgg gtg gat atg atc agt ccc cgt ata gga gtt caa agg tta tag cgt aca ata cct gta cta att ctt gga tat atg ata tac cag agc t-a aaa tat cct cgt tct aat tgt gga gga gtt gct gat gat gaa tac att tat tgc ata ggc ggc ata cgc gat cag gat tca tcg ttg ata tct agt att gat aga tgg aag cca tca aaa cca tat tgg caa acg tat gct aaa atg cga gag cca aaa tgt gat atg ggt gta gcg atg tta aac gga tta ata tat gtc ata ggc gga gtt gtt aaa ggt g-- aca cgt gta ccg aca cac ttg aga gtt tat cag aag atg gat gga tga tgc atc aac gtc ttc caa taa aaa tgt cca ata tgt cga cga ttg ttc atg ccg gca aaa ttt ata tat ctg gag gct aca aca ata gta gtg tag tta atg gaa tat cga atc t-- agt cct tag cta taa tcc gat ata tga tga atg gac caa att atc atc att aaa tat tcc tag aat taa tcc cgc tct a

>KC813492.1:169106-171337_Cowpox_virus_strain_HumLan08/1_complete_genome

--- --- --- --a --- tgt a-c tat ttt gta cat cac tac att ggt tac agc aga tat acc tac tcc gcc a-- -cc aca tgc tcc ggt aaa cgg --g aca tgt gaa gag gga gaa tat ctt gat agg acg cat aat caa tgt tgt aat -cg gtg tcc tcc tgg aga att tgc caa ggt tag atg ta- --a tgg taa cga taa cac aaa atg tga acg ctg tcc acc tca tac att tac cac agt acc caa tta ttc t-a atg gat gtc atc aat gta gaa aat gcc caa agg gat cat ttg ata agg taa agt gta cag gaa cac aga aca gta aat -gt tcg tgt ctt cct ggt tgg ttt tgt gct act gat tct tcg aag act gaa gat tgt cga gat -tg tgt acc aaa aag g-a gct gtc cat gcg gat act ttg gtg gaa tag ata aat tag gaa atc cta ttt gta aat cgt gtt g-- -tg ttg gta aat att gtg acg aca tac gta atc ata gag ttg gtc ctt ttc ctc cat gca aa- --- cta tct aaa tgt aat tag tca --- --- --- --- --- --- --- --- --- --- --- --- --- --- -tg atc atg gta -cc gta caa tt- --- -at att gct --- --g gta --g tat att att cag tat g-a aga cat att aat ac- --- --- -tt atc --- --t ttg aca atc ttg ct- -at aat cat --- --- --- --- aat ata aaa --a tac tta tgg --- -a- cat agt cac tca t-- --- --- --- aat tgc tg- acg cga taa att cgt aat aat ctg ttt tgt tca aat ttt tat aag ga- --- --a tct aca ggc ata aaa at- aaa aat ata att tat aat ac- --- act tac agc gcg cca tca tga ata aca gca gtg aat taa ttg ctg tta tta atg gat tta gaa ata gtg gac gat ttt gtg ata tta ata tag tta tta atg atg aaa gga taa atg ctc ata gac tca tcc tat ctg gag cat ccg aat a-t ttt tcc att cta ttt tct agt gat ttt atc gat tct aat gaa tac gaa gtt aat cta agt cat tta gat tat caa agt gtg aac gat ttg atc gat tac att tat ggg ata cct ttg agc cta act aac gat agc gtg aaa tat att ctt tca act gct ga- --t ttt tta caa att gga tat gct att acc gaa tgt gaa aat tac ata ctt aaa aat ctg tgt tct aga aat tgt atc gat ttc --- --- tac ata tac gct gat aaa tat aat aac gag aaa ata gaa aca gcg tcg ttt aac aca ata tta cga aat att ttg aga ctc att aac gat gaa aac ttt aaa tac ttg aca gag gaa tca atg ata aaa att tta agc gat gat atg tta aat ata aaa aat gag gat ttt gca cca cta att ctc att aaa tgg tta gag agt aca caa caa cca tgc aca gtc gag tta ctt aga tgc ctc aga ata tca -tt gct ttc ccc aca agt tat aaa atc act tta tag tca tcg att ggt tgg ttc aat ata cga atg tat aac att ctt aaa caa tat atc att ctt gga tga atc att tcc tag ata cca tag cat cga gtt gat atc tat cgg tat aag taa ttc aca tga taa gat ttc cat aaa ctg cta taa tcg t-a aaa aaa ata cgt ggg aga taa tat ctt cac gga gat ata ggt gta gtt ttg cag tgg ccg tca tgg ata ata tta tct ata tga tgg gtg gat atg atc agt ccc cgt ata gga gtt caa agg tta tag cgt aca ata cct gta cta att ctt gga tat atg ata tac cag agc t-a aaa tat cct cgt tct aat tgt gga gga gtt gct gat gat gaa tac att tat tgc ata ggc ggc ata cgc gat cag gat tca tcg ttg ata tct agt att gat aga tgg aag cca tca aaa cca tat tgg caa acg tat gct aaa atg cga gag cca aaa tgt gat atg ggt gta gcg atg tta aac gga tta ata tat gtc ata ggc gga gtt gtt aaa ggt g-- aca cgt gta ccg aca cac ttg aga gtt tat cag aag atg gat gga tga tgc atc aac gtc ttc caa taa aaa tgt cca ata tgt cga cga ttg ttc atg ccg gca aaa ttt ata tat ctg gag gct aca aca ata gta gtg tag tta atg gaa tat cga atc t-- agt cct tag cta taa tcc gat ata tga tga atg gac caa att atc atc att aaa tat tcc tag aat taa tcc cgc tct a

>MG012796.1:159720-161899_Vaccinia_virus_isolate_CEyV2

--- --- --- --- --- --- --- --- --- --- --- --- --- --- --- --- --- --- --- --- --- --- --- --- --- --- --- --- --- -aa cgg --g taa tgt gac aag gga gaa tat ctt gat aag agg cat aat caa tgt tgt aat -cg gtg tcc acc tgg aga att tgc caa ggt tag atg ta- --a cgg taa cga tga cac aaa atg tga acg ctg ccc acc tca tac ata tac cac aat -cc aaa tta ttc t-a acg gat gtc atc aat gta gaa aat gcc caa ccg gac cat ttg ata agg taa agt gta ccg gaa cac aga aca gta aat -gt tgg tgt ctt cct ggt tgg tat ttt gct act gat tct tca cag act gaa gat att tga aat -tg tct acc aaa aag g-a gat ttc cat gcg gat act ttg gtg gaa tag atg aac acg gaa atc cta ttt gta aat cgt att g-- -tg ttg gtg aat att gcg act acc tac gaa att ata gac ttg atc cat ttc ctc cat gac aa- --- cta tct aaa tgt aat taa tta --- --- --- --- --- --- --- --- tga ttt --- --- --- -tg atg ata atg ttc -cc ata c-a tt- --- -at atc gct act tgg tta --g tgc att att cag tat t-a aga cct atg aat aa- --- --- -tt act tat ctt ttg acg atc ttg tt- -at aat tat --- --- --- --- aat aca aaa --a tac tta tag --- --- cat agt aac tca t-- --- --- --- aat tgc tg- acg cga taa att cgt aat aat ctc ttt tgt tca aat atg tat aag ga- --- --a tct aca gct ata aaa at- aaa aat ata att tat aat ata c-- tct tac agc gcg cca tca tga ata aca gca gtg cat tga ttg ctg tta tta atg gat tta gaa aca gtg gac gat tgt atg cta tta gta tag tta tta atg atg aaa gga taa acg ctc ata aac tca tcc tat ctg gag cct ccg aat a-t ttt tcc att ctg ttt tgc aat aat ttt atc gat tct aat gaa tac gaa gtt aat cta agt cgt tta gat tat caa agc gtt aac gat ttg atc gat tac att tat ggg ata cct ttg agc cta act aac aat aac gtg aaa tat att ctt tca acc cct ga- --t ttt tta caa att gga tct gct att acg gtg tgt gac aat tac cta ctt aaa aat cct tgt tct atg aac tgt atc gat ttc --- --- tac ata tac gct gtt aaa tat aat aac aac aaa att gaa tca tcg tcg ttt aac ata ata tta caa aat atc ttg aga ctc atc aac gaa gaa aac ttt aaa tac tta aca gag gaa tca atg ata aaa att tta agc gat gat atg tta aat ata aaa aat gag gat ttt tca cca cta att gtc att aaa tgg tta gag aat acg caa caa tca tgc acc gtc gag tta ctt aga tgc ctc aga ata tca -tt gct ttc gcc aca agt tat aaa atc act tta tag tcc tca act ggt tag ttg aat cta cga atg tat aac att ctt aaa caa tat agc att ctt gga tga ctc att tcc tag ata cca cag cat cga gtt gat agc tat cgg tat aag taa ttc gca tga taa gat ttc cat aaa ctg cta caa tcg t-a aaa aaa ata cat ggg aaa tga tat ctt cac gta ggt ata ggt gta gtt tcg cag tgg ccg tcc tgg ata ata tta tct ata tga tgg gtg gat atg ttc agt acc cgt ata gaa gtt caa agg tta tag cgt aca ata cat gta caa att ctt ggc tat atg aaa tac cag agc t-a aaa tat cct cgt tct att tgt gcg gga ctg gct gat gac gaa tac att tat tgt ata ggc ggc ata cgc gat cag gat tca tcg ttg acc tct agt att gat aaa tgg aat cca tca aaa cca tat tgg cag aag tgt gct aaa gtg cgc gaa cca aaa tgt gat atg ggg gtt gcg atg tta aac gga tta ata gat gtc atg ggt gga atc gtt aaa ggt g-- aca cgt gta ccg acg cac tag aga gtt tat cag aat atg gat gga tga agc atc aac gtc ttc cac taa aaa tgt cca ata tgt cga cga ctg ttc atg atg gca aca att ata tat ctg gag gtt aca aca ata gta gtt tag tta atg taa tat cga atc t-- agt cct tag cta tta cac gat ata tga tga atg gac caa att atc atc att aaa cat tcc tag aat taa tcc cgc tct a

>LR800245.1:154734-156956_Variola_virus_genome_assembly_complete_genome:_monopartite

--- --- --- --a --- tgt a-c tat ttt ata cat tac tac att ggt tac ggc aga tat acc tac tcc gcc a-- -cc aca tgc tcc ggt aaa cgg --g tca tgt gtc gag gga gaa tat ctt gat aag act cat aat cga tgt tgt aat -cg gtg tcc acc tgg aga att tgc caa ggt tag atg ta- --g tgg tag cta taa cac aaa atg tga acg ctg tcc aca tca tac ata tac agc aat tcc caa tta ttc t-a atg gat gcc atc aat gta gaa aat gcc caa a-g gat cat ttg ata agg taa agt gta ccg gaa cac aga aca gta aa- --t gtg tgt ctt cct ggt tgg tat tgc gct acc gat tct tca cag act gaa gac tgt cga gat -tg tgt acc aaa aag g-a gct gtc cat gcg gat act ttg gtg gaa tag ata aat tag gaa atc cta ttt gta aat cgt gtt g-- -tg ttg gta aat att gtg aca aca tac gta atc ata gag ttg gtc ctt ttc ctc cat gca aa- --- cta tct aaa tgt aat tag tca --- --- --- --- --- --- --- --- --- --- --- --- --- --- -tg atc atg gta -cc gta caa tt- --- -at att gct --- --g gta --g tat att att cag tat g-a aga cat att aat ac- --- --- -tt atc --- --t ttg aca atc ttg ct- -at aat cat --- --- --- --- aat ata aaa --a tac tta tgg --- -a- cat agt cac tca t-- --- --- --- aat tgc tg- acg cga taa att cgt aat aat ctg ttt tgt tca aat ttt tat aag ga- --- --a tct aca ggc at- --- --- aaa aat ata att tat aat ac- --- act tac agc gcg cca tca tga ata aca gca gtg aat taa ttg ctg tta tta atg gat tta gaa ata gtg gac gat ttt gtg ata tta ata tag tta tta atg atg aaa gga taa atg ctc ata gac tca tcc tat ctg gag cat ccg aat a-t ttt tcc att cta ttt tct agt gat ttt atc gat tct aat gaa tac gaa gtt aat cta agt cat tta gat tat caa agt gtg aac gat ttg atc gat tac att tat ggg ata cct ttg agc cta act aac gat agc gtg aaa tat att ctt tca act gct ga- --t ttt tta caa att gga tct gct att acc gaa tgt gaa aat tac ata ctt aaa aat ctg tgt tct aga aat tgt atc gat ttc --- --- tac ata tac gct gat aaa tat aat aac aag aaa ata gaa aca gcg tcg ttt aac aca ata tta cga aat att tta aga ctc att aac gat gaa aac ttt aaa tac ttg aca gag gaa tca atg ata aaa att tta agt gat gat atg tta aat ata aaa aat gag gat ttt gca cca cta att ctc att aaa tgg tta gag agt aca caa caa cca tgc aca gtc gag tta ctt aga tgc ctc aga ata tca -tt gct ttc ccc aca agt tat aaa atc act tta tag tca tcg att ggt tgg ttc aat ata cga atg tat aac att ctt aaa caa tat atc att ctt gga tga atc att tcc tag ata cca tag cat cga gtt gat atc tat cgg tat aag taa ttc aca tga taa gat ttc cat aaa ctg cta taa tcg t-a aaa aaa ata cgt ggg aga taa tat ctt cac gga gat ata ggt gta gtt ttg cag tgg ccg tcc tgg ata ata tta tct ata tga tgg gtg gat atg atc agt ccc cgt ata gaa gtt caa agg tta tag cgt aca ata cat gta caa att ctt gga tat atg ata tac cgg agc t-a aaa tat cct cgt tct aat tgt gga gga gtt gct gat gat gaa tac att tat tgc ata ggc ggc ata cgc gat cag gat tca tcg ttg ata tct agt att gat aga tgg aag cca tca aaa cca tat tgg caa acg tat gct aaa atg cga gag cca aaa tgt gat att ggt gta gcg atg tta aac gga tta ata tat gtc ata ggc gga gtt gtt aaa ggt g-- aca cgt gta ccg aca cac ttg aga gtt tat cag aag atg gat gga tga tgc atc aac gtc ttc caa taa aaa tgt cca ata tgt cga cga ttg ttc atg ccg gca aaa ttt ata tat ctg gag gct aca aca ata gta gtg tag tta atg gaa tat caa atc t-- agt cct tag cta taa tcc gat ata tga tga atg gac caa att atc atc att aaa tat tcc tag aat taa tcc cgc tct a

>KC813494.1:167300-169570_Cowpox_virus_strain_HumLue09/1_complete_genome

ttg ctg act gtg --- tgt a-c tat ttt gta cat cag tac att ggt tac agc aga tat acc tac ttc gtc act gcc aca cgc tcc ggt aaa cgg --g tca tgt gac gaa gga gaa tat ctt gat aag acg cat aat caa tgt tgt aat -cg gtg tcc acc tgg aga att tgc caa gat cag atg ta- --g cgg tag cga taa cac aaa atg tga acg ctg ccc acc tca tac ata tac cgc agt acc caa tta ttc t-a atg gat gtc atc aat gta gga aat gcc caa agg gat cat ttg ata agg taa agt gta cca gaa cac aga aca gta aat -gt tcg tgt att cct ggt tgg tat tgc gct act gat tct tca tgg acc gaa gac tgt att gtg -tg cgt acc aaa aaa g-a gat gtc cat gcg gat act ttg gtg gaa tag atg aac tcg gaa atc ctc ttt gta aat cgt gtt g-- -tg ttg gtg aat att gcg acg aca tac gta atc ata gag ttg gtc ctt ttc ctc cat gca aa- --- cta tct aaa tgt aat tag cca --- --- --- --- --- --- --- --- --- --- --- --- --- --- -tg atc atg tta ccc ata c-a tc- --- -at atc gct act tgg t-a --g cgt att att cag tat g-a aga tct att aat aat aat tat ctt att tat ttc ttg acg --- --- --- -at att aat cgg tgt cct aat aat ata aaa --- tac tta tgg --- --- --a aat aaa ttg t-a aac aag atg aat taa t-- aca caa tac atc ct- --t aat tag ctt -gt tca aa- ttt tat aaa gaa tta tat tca tta ggc ata taa ta- aaa aat ata att tat aat ac- --- act tac a-- gcg tca tca tgc ata aca gca gtg aat taa ttg ctg tta tta atg gat tta gaa ata gtg gac gat ttt gtg ata tta ata tag tta tta atg atg aaa gga taa acg ctc ata gac tca tgc tat ctg gag cat ccg aat a-t ttt tcc att cta ttt tct agt gat ttt atc gat tct aat gac tac gaa gtt aat cta agt cat tta gat tat caa agt gtt aac gat ttg atc gat tac att tat ggg ata cct ttg agc cta act aac gat aac gtg aaa tat att ctt tca aca gct ga- --t ttt tta caa att gga tct gcc att act gag tgc gaa aaa tac ata ctt aaa aat ctc tgt tct aga aac tgt atc gat ttc --- --- tac ata tac gct gat aaa tat aat aac aag aaa ata gaa tca gcg tcg ttt aac aca ata tta cga aat att ttg aga ctc att aac gat gaa aac ttt aaa tac ttg aca gag gaa tca atg ata aaa att tta agc gat gat atg tta aat ata aaa aat gag gat ttc gca cca cta att ctc att aaa tgg tta gag agt aca caa caa cca tgc acc gtc gag tta ctt aga tgc ctc aga ata tca -tt gct ttc ccc aca agt tat aaa atc act tta tag tca tcg act ggt tag ttc aat cta cga atg tat aac att ctt aaa caa tat agc att ctt gga tga atc att tcc tag ata cca tag cat cga gtt gat atc tat cgg tat aag taa ttc gca tga taa gat ttc cat aaa ctg cta caa tcg t-a aaa aaa ata cat ggg aga tga tat ctt cac gta gat ata ggt gta gtt tcg cag tgg ccg ttc tgg ata ata tta tct ata tga tgg gtg gat atg atc agt ccc cgt ata gaa gtt caa agg tta tag cgt aca ata cat gta caa att ctt gga tat atg ata tac cag agc t-a aaa tat cct cgt tct aat tgc gga gga gtg gct gat gac gaa tac att tat tgt ata ggc ggc ata cgc gat cag gat tca tcg ttg ata tct aat att gat aga tgg aag cca tca aaa cca tat tgg cag acg tat gct aaa atg cgc gag cca aaa tgt gat atg ggg gtt gcg atg tta aac gga tta ata tat gtc ata ggt gga atc gtt aaa ggt g-- aca cat gta ccg acg cac tag aaa gtt tat cag aag atg gat gga tga agc atc aac gtc ttc caa taa aaa tgt cca ata tgt cga cga ttg ttc atg ctg gta aga ttt ata tat ctg gag gtt ata aca ata gta gtg ccg tta atg gac cat cga atc t-- agt cct tag cta taa tcc gat ata tga tga atg gac caa att atc atc att aaa tat tcc tag aat taa tcc cgc tct a

>HQ420899.1:175341-177611_Cowpox_virus_strain_Norway_1994_MAN_complete_genome

ttg ctg act gta --- tgt a-c tat ttt gta cat cag tac att ggt tac agc aga tat acc tac ttc gtc act gcc aca cgc tcc ggt aaa cgg --g tca tgt gac gaa gga gaa tat ctt gat aag acg cat aat caa tgt tgt aat -cg gtg tcc acc tgg aga att tgc caa gat cag atg ta- --g cgg tag cga taa cac aaa atg tga acg ctg ccc acc tca tac ata tac cgc aat ccc caa tta ttc t-a atg gat gcc atc aat gta gaa aat gcc caa cag gat cat ttg ata agg taa agt gta ccg gaa cac aga aca gta aat -gt tcg tgt att cct ggt tgg tat tgc gct act gat tct tca tgg acc gaa gac tgt att acg -tg cgt acc aaa aaa g-a gat gtc cat gcg gat act ttg gtg gaa tag atg aac tcg gaa atc ctc ttt gta aat cgt gtt g-- -tg ttg gtg aat att gcg acg aca tac gta atc ata gag ttg gtc ctt ttc ctc cat gca aa- --- cta tct aaa tgt aat tag cca --- --- --- --- --- --- --- --- --- --- --- --- --- --- -tg atc atg tta ccc ata c-a tc- --- -at atc gct act tgg t-a --g cgt att att cag tat g-a aga tct att aat aat aat tat ctt att tat ttc ttg acg --- --- --- -at att aat cgg tgt cct aat aat ata aaa --- tac tta tgg --- --- --a aat aaa ttg taa aac aag atg aat taa t-- aca caa t-c atc ct- --t aat tag ctt -gt tca aa- ttt tat aaa gaa tca tat tca tta ggc ata aca ta- aaa aat ata att tat aat ac- --- act tac a-- gcg tca tca tga ata aca gca gtg act taa ttg ctg tta tta atg gat tta gaa ata gtg gac gat ttt gtg ata tta gta tag tta tta atg atg aaa gga taa atg ctc ata gac tca tgc tat ctg gag cat ccg aat a-t ttt tcc att cta ttt tct agt gat ttt atc gat tct aat gac tac gaa gtt aat cta agt cat tta gat tat caa agt gtt aac gat ttg atc gat tac att tat ggg ata cct ttg agc cta act aac gat aac gtg aaa tat att ctt tca acc gct ga- --t ttt tta caa att gga tct gcc att act gag tgc gaa aaa tac ata ctt aaa aat ctc tgt tct aga aac tgt atc gat ttc --- --- tac ata tac gct gat aaa tat aat aac aag aaa ata gaa tca gcg tcg ttt aac aca ata tta cga aat att ttg aga ctc att aac gat gaa aac ttt aaa tac ttg aca gag gaa tca atg ata aaa att tta agc gat gat atg tta aat ata aaa aat gag gat ttt gca cca cta att ctc att aaa tgg tta gag agt aca caa caa cca tgc acc gtc gag tta ctt aga tgc ctc aga ata tca -tt gct ttc ccc aca agt tat aaa atc act tta tag tca tca act ggt tag ttc aat cta cga atg tat aac att ctt aaa caa tat agc att ctt gga tga atc att tcc tag ata cca tag cat cga gtt gat atc tat cgg tat aag taa ttc gca tga taa gat ttc cat aaa ctg cta caa tcg t-a aaa aaa ata cat ggg aga tga tat ctt cac gta gat ata ggt gta gtt tcg cag tgt cag tcc tgg ata ata tta tat ata tga tgg gtg gat atg atc aat ccc cgt ata gga gtt caa agg tta tag cgt aca aca cat gta caa att ctt gga tat atg ata tac cag agc t-a aaa tat cct cgt tct aat tgt gga gga gtt gcc gat gac gaa tac att tat tgt ata ggc ggc ata cgc gat cag gat tca tcg ttg ata tct aat att gat aga tgg aag cca tca aaa cca tat tgg cag acg tat gct aaa atg cgc gaa cca aaa tgt gat atg ggg gtt gcg atg tta aac gga tta ata tat gtc ata ggt gga atc gtt aaa ggt g-- aca cat gta ccg acg cac tag aga gtt tat cag aag atg gat gga tga agc atc aac gtc ttc caa taa aaa tgt cca ata tgt cga cga ttg ttc atg ctg gta aga ttt ata tat ctg gag gtt ata aca ata gta gtg ccg tta atg gac cat cga atc t-- agt cct tag cta taa tcc gat ata tga tga atg gac caa att atc atc att aaa tat tcc tag aat taa tcc cgc tct a

>LT896724.1:175302-177570_Cowpox_virus_isolate_Ger/2015/Cat1_genome_assembly_complete_genome:_monopartite

ttg ctg act gta --- tgt a-c tat ttt gta cat cag tac att ggt tac agc aga tat acc tac ttc gtc act gcc aca cgc tcc ggt aaa cgg --g tca tgt gac gaa gga gaa tat ctt gat aag acg cat aat caa tgt tgt aat -cg gtg tcc acc tgg aga att tgc caa gat cag atg ta- --a tgg taa cga taa cac aaa atg tga acg ctg tcc acc tca tac att tac cac agt acc caa tta ttc t-a atg gat gtc atc aat gta gga aat gcc caa agg gat cat ttg ata agg taa agt gta ccg gaa cac aga aca gta aat -gt tcg tgt ctt cct ggt tgg ttt tgc gct act gat tct tca tgg acc gaa gac tgt att gcg -tg cgt acc aaa aaa g-a gat gtc cat gcg gat act ttg gtg gaa tag atg aac tcg gaa atc ctc ttt gta aat cgt gtt g-- -tg ttg gtg aat att gcg acg aca tac gta atc ata gag ttg gtc ctt ttc ctc cat gta aa- --- cta tct aaa tgt aat tag cca --- --- --- --- --- --- --- --- --- --- --- --- --- --- -tg atc atg tta ccc ata c-a tc- --- -at atc gct act tgg t-a --g cgt att att cag tat g-a aga tct att aat aat aat tat ctt att tat ttc ttg acg --- --- --- -at att aat cg- -gt cct aat aat ata aaa --- tac tta tgg --- --- --a aat aaa ttg t-a aac aag atg aat taa t-- aca caa tac atc ct- --t aat tag ctt -gt tca aa- ttt tat aaa gaa tca tat tca tta ggc ata taa ta- aaa aat ata att tat aat ac- --- act tac a-- gcg tca tca tgc ata aca gca gtg aat taa ttg ctg tta tta atg gat tta gaa ata gtg gac gat ttt gtg ata tta ata tag tta tta atg atg aaa gga taa acg ctc ata gac tca tgc tat ctg gag cat ccg aat a-t ttt tcc att cta ttt tct agt gat ttt atc gat tct aat gac tac gaa gtt aat cta agt cat tta gat tat caa agt gtt aac gat ttg atc gat tac att tat ggg ata cct ttg agc cta act aac gat aac gtg aaa tat att ctt tca acc gct ga- --t ttt tta caa att gga tct gcc att act gag tgc gaa aaa tac ata ctt aaa aat ctt tgt tct aga aac tgt atc gat ttc --- --- tac ata tac gct gat aaa tat aat aac aag aaa ata gaa tca gcg tcg ttt aac aca ata tta cga aat att ttg aga ctc att aac gat gaa aac ttt aaa tac ttg aca gag gaa tca atg ata aaa att tta agc gat gat atg tta aat ata aaa aat gag gat ttc gca cca cta att ctc att aaa tgg tta gag agt aca caa caa caa tgt acc gtc gag tta ctt aga tgc ctc aga ata tca -tt gct ttc ccc aca agt tat aaa atc act tta tag tca tcg cct ggt tag ttc aat cta cga atg tat aac att ctt aaa caa tat agc att ctt gga tga atc att tcc tag ata cca tag cat cga gtt gat atc tat cgg tat aag taa ttc gca tga taa gat ttc cat aaa ctg cta caa tcg t-a aaa aaa ata cat ggg aga tga tat ctt cac gta gat ata ggt gta gtt tcg cag tgt cag tcc tgg ata ata tta tct ata tga tgg gtg gat atg atc aat ccc cgt ata gga gtt caa agg tta tag cgt aca ata cat gta caa att ctt gga tat atg ata tac cag agc t-a aaa tat cct cgt tct aat tgc gga gga gtt gct gat gac gaa tac att tat tgt ata ggc ggc ata cgc gat cag gat tca tcg ttg ata tct aat att gat aga tgg aag cca tca aaa cca tat tgg cag acg tat gct aaa atg cgc gaa cca aaa tgt gat atg ggg gtt gcg atg tta aac gga tta ata tat gtc ata ggt gga atc gtt aaa ggt g-- aca cat gta ccg acg cac tag aaa gtt tat cag aag atg gat gga tga agc atc aac gtc ttc caa taa aaa tgt cca ata tgt cga cga ttg ttc atg ctg gta aga ttt ata tat ctg gag gtt ata aca ata gta gtg ccg tta atg gac cat cga atc t-- agt cct tag cta taa tcc gat ata tga tga atg gac caa att atc atc att aaa tat tcc tag aat taa tcc cgc tct a

>LT896721.1:176067-178331_Cowpox_virus_isolate_Ger_2010_MKY_genome_assembly_complete_genome:_monopartite

--- --- --- --a --- tgt a-c tat ttt gta cat cac tac att ggt tac agc aga tat acc tac tcc gcc a-- -cc aca tgc tcc ggt aaa cgg --g tca tgt gaa gag gga gaa tat ctt gat agg acg cat aat caa tgt tgt aat -cg gtg tcc tcc tgg aga att tgc caa ggt tag atg ta- --a tgg taa cga taa cac aaa atg tga acg ctg tcc acc tca tac att tac cgc att acc caa tta ttc t-a atg gat gtc atc aat gta gaa aat gcc caa agg gat cat ttg ata agg taa agt gta ccg gaa cac aga aca gta aat -gt tcg tgt ctt cct ggt tgg ttt tgc gct act gat tct tcg aag act gaa gat tgt cga gat -tg tat acc aaa aag g-a gat gtc cat gcg gat act ttg gtg gaa tag atg aac gag gaa atc ctc ttt gta aat cgt gtt g-- -tg ttg gtg aat att gtg acc aca tac gta atc ata tag ttg gtc ctt ttc ctt cat gca aa- --- cta tct aaa tgc aat tag tca --- --- --- --- --- --- --- --- --- --- --- --- --- --- -tg gcc gtg tta -cc atc a-a tc- --- -at ata gct act tgg t-a --g tgt att att cag tat g-a aga cct att aat aat aat tat ctt att tat ttc ttg acg atc ttc taa aat gtt att aag agt cat aat att ata aaa --- tac tta tgg --- --- --a cat aaa ttg t-a aac aag atg a-- taa t-- aca cga tat cct t-- -at aat cag ctt -gt tca aat ttt tat aaa gaa tta tat tca tta ggc ata aca ta- aaa aat ata att tat aat ac- --- act tac a-- gca cca tca tga ata aca gca gtg act taa ttg ctg tta tta atg gat tta gaa ata gtg gac gat ttt gtg ata tta gta tag tta tta atg atg aaa gga taa atg ctc ata gac tca tcc tat ctg gag cat cag aat a-t ttt tcc att ctg ttt tcc aat aat ttt atc gat tct aat gaa tac gaa gtt aat cta agt cat tta gat tat caa agt gtg aac gat ttg atc gat tac att tat ggg ata cct ttg agc cta act aac gat agc gtg aaa tat att ctt tca acc gct ga- --t ttt tta caa att gga tct gcc att act gag tgc gaa aaa tac ata ctt aaa aat ctt tgt tct aga aac tgt atc gat ttc --- --- tac ata tac gct gat aaa tat aat aac aag aaa ata gaa tca gcg tcg ttt aac aca ata tta cga aat att ttg aga ctc att aac gat gaa aac ttt aaa tac ttg aca gag gaa tca atg ata aaa att tta agc gat gat atg tta aat ata aaa aat gag gat ttt gcc cca ctg att ctc att aaa tgg tta gag agt act caa caa cca tgc acc gtc gag tta ctt aga tgc ctc aga ata tca -tt gct ttc ccc aca agt tat aaa atc act tta tag tca tca act ggt tag ttc aat cta cga atg tat aac att ctt aaa caa tat agc att att gga tga atc att tcc tag ata cca tag cat cga gtt gat atc tat cgg tat aag taa ttc gca tga taa gat ttc cat aaa ctg cta caa tca t-a aaa aaa ata cat ggg aaa tga tat ctt cac gta gat ata ggt gta gtt tcg cag tgg ccg tcc tgg ata ata tta tct ata tga tgg gtg gat atg atc agt ccc cgt ata gaa gtt caa agg tta tag cgt aca ata cat gta caa att ctt gga tat atg ata tac cag agc t-a aaa tat cct cgt tct aat tgt ggg gga ctg acc gat gac gaa tac att tat tgt ata ggc ggc ata cgc gat cag gat tca tcg ttg ata tct aat att gat aga tgg aag cca tca aaa cca tat tgg cag acg tat gct aaa atg cgc gaa cca aaa tgt gat atg ggg gtt gcg atg tta aac gga tta ata tat gtc ata ggt gga atc gtt aaa ggt g-- atg cgt ata ccg acg cac tag aga gtt tat cag aag atg gat gga tga agc atc aac gtc ttc caa taa aaa tgt cca ata tgt cga cga ttg ttc atg ctg gaa aga ttt ata tat ctg gag gtt aca cca ata gta gta tag tta atg gaa tat cta atc t-- att cct tag cta taa tcc gat ata tga cga atg gac caa att atc atc gtt aaa tat tcc tag aat taa tcc tgc tct a

>LR812035.1:175918-178181_Cowpox_virus_isolate_GerMygEK_938/17_genome_assembly_complete_genome:_monopartite

--- --- --- --a --- tgt a-c tat ttt gta cat cac tac att ggt tac agc aga tat acc tac tcc gcc a-- -cc aca tgc tcc ggt aaa cgg --g tca tgt gaa gag gga gaa tat ctt gat agg acg cat aat caa tgt tgt aat -cg gtg tcc tcc tgg aga att tgc caa ggt tag atg ta- --a tgg taa cga taa cac aaa atg tga acg ctg tcc acc tca tac att tac cgc att acc caa tta ttc t-a atg gat gtc atc aat gta gaa aat gcc caa agg gat cat ttg ata agg taa agt gta ccg gaa cac aga aca gta aat -gt tcg tgt ctt cct ggt tgg ttt tgc gct act gat tct tcg aag act gaa gat tgt cga gat -tg tat acc aaa aag g-a aat gtc cat gcg gat act ttg gtg gaa tag atg aac gag gaa atc ctc ttt gta aat cgt gtt g-- -tg ttg gtg aat att gtg acc aca tac gta atc ata tag ttg gtc ctt ttc ctc cat gca aa- --- cta tct aaa tgc aat tag tca --- --- --- --- --- --- --- --- --- --- --- --- --- --- -tg gtc gtg tta --c atc a-a tca --- tat ata gct act tgg t-a --g tgt att att cag tat g-a aga cct att aat aat aat tat ctt att tat ttc ttg acg atc ttc taa aat gtt att aag agt cat aat att ata aaa --- tac tt- -gg --- --- --a cat aaa ttg t-a aac aag atg a-- taa t-- aca cga tat cct t-- -at aat cag ctt -gt tca aat ttt tat aaa gaa tta tat tca tta ggc ata aca ta- aaa aat ata att tat aat ac- --- act tac a-- gca cca tca tga ata aca gca gtg act taa ttg ctg tta tta atg gat tta gaa ata gtg gac gat ttt gtg ata tta gta tag tta tta atg atg aaa gga taa atg ctc ata gac tca tcc tat ctg gag cat cag aat a-t ttt tcc att ctg ttt tcc aat aat ttt atc gat tct aat gaa tac gaa gtt aat cta agt cat tta gat tat caa agt gtt aac gat ttg atc gat tac att tat ggg ata cct ttg agc cta act aac gat agc gtg aaa tat att ctt tca acc gct ga- --t ttt tta caa att gga tct gcc att act gag tgc gaa aaa tac ata ctt aaa aat ctt tgt tct aga aac tgt atc gat ttc --- --- tac ata tac gct gat aaa tat aat aac aag aaa ata gaa tca gcg tca ttt aac aca ata tta cga aat att ttg aga ctc att aac gat gaa aac ttt aaa tac ttg aca gag gaa tca atg ata aaa att tta agc gat gat atg tta aat ata aaa aat gag gat ttc gcc cca ctg att ctc att aaa tgg tta gag agt act caa caa cca tgc acc gtc gag tta ctt aga tgc ctc aga ata tca -tt gct ttc ccc aca agt tat aaa atc act tta tag tca tca act ggt tag ttc aat cta cga atg tat aac att ctt aaa caa tat agc att att gga tga atc att tcc tag ata cca tag cat cga gtt gat atc tat cgg tat aag taa ttc gca tga taa gat ttc cat aaa ctg cta caa tca t-a aaa aaa ata cat ggg aaa tga tat ctt cac gta gat ata ggt gta gtt tcg cag tgg ccg tcc tgg ata ata tta tct ata tga tgg gtg gat atg atc agt ccc cgt ata gaa gtt caa agg tta tag cgt aca ata cat gta caa att ctt gga tat atg ata tac cag agc t-a aaa tat cct cgt tct aat tgt ggg gga ctg acc gat gac gaa tac att tat tgt ata ggc ggc ata cgc gat cag gat tca tcg ttg ata tct aat att gat aga tgg aag cca tca aaa cca tat tgg cag acg tat gct aaa atg cgc gaa cca aaa tgt gat atg ggg gtt gcg atg tta aac gga tta ata tat gtc ata ggt gga atc gtt aaa ggt g-- atg cgt ata ccg acg cac tag aga gtt tat cag aag atg gat gga tga agc atc aac gtc ttc caa taa aaa tgt cca ata tgt cga cga ttg ttc atg ctg gaa aga ttt ata tat ctg gag gtt aca cca ata gta gta tag tta atg gaa tat cta atc t-- att cct tag cta taa tcc gat ata tga cga atg gac caa att atc atc gtt aaa tat tcc tag aat taa tcc tgc tct a

>LT993226.1:175432-177701_Cowpox_virus_isolate_Ger/2014/Human_genome_assembly_complete_genome:_monopartite

ttg ctg act gta --- tgt a-c tat ttt gta cat cag tac att ggt tac agc aga tat acc tac ttc gtc act gcc aca cgc tcc ggt aaa cgg --a tca tgt gac gaa gga gaa tat ctt gat aag acg cat aat caa tgt tgt aat -cg gtg tcc acc tgg aga att tgc caa gat cag atg ta- --g tgg tag cga taa cac aaa atg tga acg ctg tcc acc tca tac att tac cac agt acc caa tta ttc t-a atg gat gtc atc aat gta gga aat gcc caa agg gat cat ttg ata agg taa agt gta ccg gaa cac aga aca gta aat -gt tcg tgt ctt cct ggt tgg ttt tgc gct acc gat tct tca ttg acc gaa gac tgt att gcg -tg cgt acc aaa aaa g-a gat gtc cat gcg gat act ttg gtg gaa tag atg aac tcg gaa atc ctc ttt gta aat cgt gtt g-- -tg ttg gtg aat att gcg acg aca tac gta atc ata gag ttg gtc ctt ttc ctc cat gca aa- --- cta tct aaa tgt aat tag cca --- --- --- --- --- --- --- --- --- --- --- --- --- --- -tg atc atg tta ccc ata c-a tc- --- -at atc gct act tgg t-a --g cgt att att -ag tat g-a aga tct att aat aat aat tat ctt att tat ttc ttg acg --- --- --- -at att aat cgg tgt cct aat aat ata aaa --- tac tta tgg --- --- --a aat aaa ttg t-a aac aag atg aat taa t-- aca caa tac atc ct- --t aat tag ctt -gt tca aa- ttt tat aaa gaa tca tat tca tta ggc ata taa ta- aaa aat ata att tat aat ac- --- act tac a-- gcg tca tca tgc ata aca gca gtg aat taa ttg ctg tta tta atg gat tta gaa ata gtg gac gat ttt gtg ata tta gta tag tta tta atg atg aaa gga taa acg ctc ata gac tca tcc tat ctg gag cat cag aat a-t ttt tcc att cta ttt tct agt gat ttt atc gat tct aat gac tac gaa gtt aat cta agt cat tta gat tat caa agt gtt aac gat ttg atc gat tac att tat ggg ata cct ttg agc cta act aac gat aac gtg aaa tat att ctt tca acc gct ga- --t ttt tta caa att gga tct gcc att act gag tgc gaa aaa tac ata ctt aaa aat ctt tgt tct aga aac tgt atc gat ttc --- --- tac ata tac gct gat aaa tat aat aac aag aaa ata gaa tca gcg tcg ttt aac aca ata tta cga aat att ttg aga ctc att aac gat gaa aac ttt aaa tac ttg aca gag gaa tca atg ata aaa att tta agc gat gat atg tta aat ata aaa aat gag gat ttc gca cca cta att ctc att aaa tgg tta gag agt aca cga caa cca tgc acc gtc gag tta ctt aga tgc ctc aga ata tca -tt gct ttc ccc aca agt tat aaa gtc act tta tag tca tcg act ggt tag ttc aat cta cga atg tat aac att ctt aaa caa tat agc att ctt gga tga atc att tcc tag ata cca tag cat cga gtt gat atc tat cgg tat aag taa ttc gca tga taa gat ttc cat aaa ctg cta caa tcg t-a aaa aaa ata cat ggg aga tga tat ctt cac gta gat ata ggt gta gtt tcg cag tgg ccg tcc tgg ata ata tta tct ata tga tgg gtg gat atg atc aat ccc cgt ata gga gtt caa agg tta tag cgt aca ata cat gta caa att ctt gga tat atg ata tac cag agc t-a aaa tat cct cgt tct aat tgt gga gga gtg gct gat gac gaa tac att tat tgt ata ggc ggc ata cgc gat cag gat tca tcg ttg ata tct agt att gat aga tgg aag cca tca aaa cca tat tgg cag acg tat gct aaa atg cgc gaa cca aaa tgt gat atg ggg gtt gcg atg tta aac gga tta ata tat gtc ata ggt gga atc gtt aaa ggt g-- atg cgt gta ccg acg cac tag aga gtt tat cag aag atg gat gga tga agc atc aac gtc ttc caa taa aaa tgt caa ata tgt cga cga ttg ttc atg ctg gaa aga ttt ata tat ctg gag gtt aca cca ata gta gta tag tta atg gaa tat cta atc t-- agt cct tag cta taa tcc gat ata tga tga atg gac caa att atc atc att aaa tat ttc tag aat taa tcc cgc tct a

>KY549151.1:172119-174387_Cowpox_virus_isolate_Norwayfeline

ttg ctg act gta --- tgt acc tat ttt gta cat cag tac att ggt tac agc aga tat acc tac ttc gtc act gcc aca cgc tcc ggt aaa cgg --g tca tgt gac gaa gga gaa tat ctt gat aag acg cat aat caa tgt tgt aat -cg gtg tcc acc tgg aga att tgc caa gat cag atg ta- --g cgg tag cga taa cac aaa atg tga acg ctg ccc acc tca tac ata tac cgc aat ccc caa tta ttc t-a atg gat gcc atc aat gta gaa aat gcc caa cag gat cat ttg ata agg taa agt gta ccg gaa cac aga aca gta aat -gt tcg tgt att cct ggt tgg tat tgc gct act gat tct tca tgg acc gaa gac tgt att acg -tg cgt acc aaa aaa g-a gat gtc cat gcg gat act ttg gtg gaa tag atg aac tcg gaa atc ctc ttt gta aat cgt gtt g-- -tg ttg gtg aat att gcg acg aca tac gta atc ata gag ttg gtc ctt ttc ctc cat gca aa- --- cta tct aaa tgt aat tag cca --- --- --- --- --- --- --- --- --- --- --- --- --- --- -tg atc atg tta ccc ata c-a tc- --- -at atc gct act tgg t-a --g cgt att att cag tat g-a aga tct att aat aat aat tat ctt att tat ttc ttg acg --- --- --- -at att aat cgg tgt cct aat aat ata aaa --- tac tta tgg --- --- --a aat aaa ttg taa aac aag atg aat taa t-- aca caa t-c atc c-- --- -at tag ctt -gt tca aa- ttt tat aaa gaa tca tat tca tta ggc ata aca ta- aaa aat ata att tat aat ac- --- act tac a-- gcg tca tca tga ata aca gca gtg act taa ttg ctg tta tta atg gat tta gaa ata gtg gac gat ttt gtg ata tta gta tag tta tta atg atg aaa gga taa atg ctc ata gac tca tcc tat ctg gag cat ccg aat a-t ttt tcc att cta ttt tct agt gat ttt atc gat tct aat gac tac gaa gtt aat cta agt cat tta gat tat caa agt gtt aac gat ttg atc gat tac att tat ggg ata cct ttg agc cta act aac gat aac gtg aaa tat att ctt tca acc gct ga- --t ttt tta caa att gga tct gcc att act gag tgc gaa aaa tac ata ctt aaa aat ctt tgt tct aga aac tgt atc gat ttc --- --- tac ata tac gct gat aaa tat aat aac aag aaa ata gaa tca gcg tcg ttt aac aca ata tta cga aat att ttg aga ctc att aac gat gaa aac ttt aaa tac ttg aca gag gaa tca atg ata aaa att tta agc gat gat atg tta aat ata aaa aat gag gat ttc gca cca cta att ctc att aaa tgg tta gag agt aca caa caa cca tgc acc gtc gag tta ctt aga tgc ctc aga ata tca -tt gct ttc ccc aca agt tat aaa atc act tta tag tca tca act ggt tag ttc aat cta cga atg tat aac att ctt aaa caa tat agc att ctt gga tga atc att tcc tag ata cca tag cat cga gtt gat atc tat cgg tat aag taa ttc gca tga taa gat ttc cat aaa ctg cta caa tcg t-a aaa aaa ata cat ggg aga tga tat ctt cac gta gat ata ggt gta gtt tcg cag tgt cag tcc tgg ata ata tta tat ata tga tgg gtg gat atg atc aat ccc cgt ata gga gtt caa agg tta tag cgt aca ata cat gta caa att ctt gga tat atg ata tac cag agc t-a aaa tat cct cgt tct aat tgt ggg gga ctg gct gat gac gaa tac att tat tgt ata ggc gga ata cgc gat cag gat tca tcg ttg ata tct agt att gat aga tgg aag cca tca aaa cca tat tgg cag acg tat gct aaa atg cgc gag cca aaa tgt gat atg ggg gtt gcg atg tta aac gga tta ata tat gtc ata ggt gga atc gtt aaa ggt g-- aca cat gta ccg acg cac tag aaa gtt tat cag aag atg gat gga tga tgc atc agc gcc ttc caa taa aaa tgt cca ata tgt cga cga ttg ttc atg ctg gta aga ttt ata tat ctg gag gtt ata aca ata gta gtg ccg tta atg gac cat cga atc t-- agt cct tag cta taa tcc gat ata tga tga atg gac caa att atc atc att aaa tat tcc tag aat taa tcc cgc tct a

>KP768318.1:158980-161215_Camelpox_virus_strain_0408151v_complete_genome

--- --- --- --a --- tgt a-c tat ttt ata cat tac tac att ggt tac ggc aga tat acc tac tcc gcc a-- -cc aca tgc tcc ggt aaa cgg --g tca tgt gcc gag gga gaa tat ctt gat aag act aat aat cga tgt tgt aat -cg gtg tcc acc tgg aga att tgc caa ggt tag atg tag -tg tgg tag ata taa cac aaa atg tga acg ctg tca aaa tca tac ata tac agc aat tcc caa tta tta t-a atg gat gcc acc aat gta gaa aat acc caa agg gat cat ttg ata agg taa agt gta ccg gaa cac aga aca gta aat -gt gcg tgt ctt cct ggt tgg tat tgc gct acc gat tct tca cag act gaa gac tgt cga gat -tg tgt acc aaa aag g-a gct gtt cat gcg gat act ttg gtg gaa tag ata aat tag gaa atc cta ttt gta aat cgt gtt g-- -tg ttg gta aat att gtg acg aca tac gta atc ata gag ttg gtc ctt ttc ctc cat gca aa- --- cta tct aaa tgt aat tag tca --- --- --- --- --- --- --- --- --- --- --- --- --- --- -tg atc atg gta -cc gta taa tt- --- -at att gct --- --g g-a --g tat att att cag tat g-a aga cat att aat ac- --- --- -tt a-- --- --- ttg aca atc ttg ct- -at aat cat --- --- --- --- aat ata aaa aaa tac tta tgg --- -a- cat agt cac tca t-- --- --- --- aat tgc tg- acg cga taa att cgt aat aat ctg ttt tgt tca aat ttt tat aag ga- --- --a tct aca tgc ata aaa at- aaa aat ata att tat aat aca ctt act tac agt gcg cca tca tga ata aca gca gtg aat taa ttg ctg tta tta atg gat tta gaa ata gtg gac gat ttt gtg ata ttg ata tag tta tta atg atg aaa gga taa atg ctc ata gac tca tcc tat ctg gag cat ccg aat a-t ttt tcc att cta ttt tct agt gat ttt atc gat tct aat aaa tac gaa gtt aat cta agt cat tta gat tat caa agt gtg aac gat ttg atc gat tac att tat ggg ata cca ttg agc cta act aac gat atc gtg aaa tat att ctt tca act gct ga- --t ttt tta caa att gga tct gct att acc gaa tgt gaa aat tac ata ctt aaa aat ctg tgt tct aga aat tgt atc gat ttc --- --- tac ata tac gct gat aaa tat aat aac aag aaa ata gaa aca gcg tcg ttt aac aca ata tta tta aat att ttg aga ctc att aac gat gaa aac ttt aaa tac ttg aca gag gaa tca atg ata aaa ttt tta agc gat gat atg tta aat ata aaa aat gag gat ttt gca cca cta att ctc att aaa tgg tta gag agt aca caa caa cca tgc aca gtc gag tta ctt aga tgc ctc aga ata tca -tt gct ttc ccc aca agt tat aaa atc act tta tag tca tcg att ggt tgg ttc aat ata cga atg tat aac att ctt aaa caa tat atc att ctt gga tga atc att tcc tag ata cca tag cat cga gtt gat atc tat cgg tat aag taa ttc aca tga taa gat ttc cat aaa ctg cta taa tcg t-a aaa aaa ata cgt ggg ata taa tat ctt cac gga gat ata ggt gta gtt ttg cag tgg ccg tcc tgg ata ata tta tct ata tga tgg gtg gat atg atc agt ccc cgt ata gaa gtt caa agg tta tag cgt aca ata cat gta caa att ctt gga tat atg ata tac cag agc t-a aaa tat cct cgt tct aat tgt gga gga gtt gct gat gat gaa tac att tat tgc ata ggc ggt ata cgc gat cag gat tca tcg ttg ata tct agt att gat aga tgg aag cca tca aaa cca tat tgg caa acg tat gct aaa ata cga gag cca aaa tgt gat atg ggt gta gcg atg tta aac gga tta ata tat gtc ata ggc gga gtt gtt aaa ggt g-- aca cgt gta ccg aca cac ttg aga gtt tat cag aag atg gat gga tga tgc atc aac gtc ttc caa taa aaa tgt cca ata tgt cga cga ttg ttc atg ccg gca aaa ttt ata tat ctg gag gct aca aca ata gta gtg tag tta atg gaa tat cga atc t-- agt cct tag cta taa tcc gat ata tga tga atg gac caa att atc atc att aaa tat tcc tag aat taa tcc cgc tct a

>MK910851.1:158739-160970_Camelpox_virus_strain_Negev2016

--- --- --- --a --- tgt a-c tat ttt ata cat tac tac att ggt tac ggc aga tat acc tac tcc gcc a-- -cc aca tgc tcc ggt aaa cgg --g tca tgt gcc gag gga gaa tat ctt gat aag act aat aat cga tgt tgt aat -cg gtg tcc acc tgg aga att tgc caa ggt tag atg tag -tg tgg tag cta taa cac aaa atg tga acg ctg tca aca tca tac ata tac agc aat tcc caa tta tta t-a atg gat gcc acc aat gta gaa aat acc caa agg gat cat ttg ata agg taa agt gta ccg gaa cac aga aca gta aat -gt gcg tgt ctt cct ggt tgg tat tgc gct acc gat tct tca cag act gaa gac tgt cga gat -tg tgt acc aaa aag g-a gct gtt cat gcg gat act ttg gtg gaa tag ata aat tag gaa atc cta ttt gta aat cgt gtt g-- -tg ttg gta aat att gtg acg aca tac gta atc ata gag ttg gtc ctt ttc ctc cat gca aa- --- cta tct aaa tgt aat tag tca --- --- --- --- --- --- --- --- --- --- --- --- --- --- -tg atc atg gta -cc gta taa tt- --- -at att gct --- --g g-a --g tat att att cag tat g-a aga cat att aat ac- --- --- -tt a-- --- --- ttg aca atc ttg ct- -at aat cat --- --- --- --- aat ata aaa aaa tac tta tgg --- -a- cat agt cac tca t-- --- --- --- aat tgc tg- acg cga taa att cgt aat aat ctg ttt tgt tca aat ttt tat aag ga- --- --a tct aca tgc ata aaa at- aaa aat ata att tat aat ac- --- act tac agt gcg cca tca tga ata aca gca gtg aat taa ttg ctg tta tta atg gat tta gaa ata gtg gac gat ttt gtg ata ttg ata tag tta tta atg atg aaa gga taa atg ctc ata gac tca tcc tat ctg gag cat ccg aat a-t ttt tcc att cta ttt tct agt gat ttt atc gat tct aat aaa tac gaa gtt aat cta agt cat tta gat tat caa agt gtg aac gat ttg atc aat tac att tat ggg ata cca ttg agc cta act aac gat atc gtg aaa tat att ctt tca act gct ga- --t ttt tta caa att gga tct gct att acc gaa tgt gaa aat tac ata ctt aaa aat ctg tgt tct aga aat tgt atc gat ttc --- --- tac ata tac gct gat aaa tat aat aac aag aaa ata gaa aca gcg tcg ttt aac aca ata tta tta aat att ttg aga ctc att aac gat gaa aac ttt aaa tac ttg aca gag gaa tca atg ata aaa ttt tta agc gat gat atg tta aat ata aaa aat gag gat ttt gca cca cta att ctc att aaa tgg tta gag agt aca caa caa cca tgc aca gtc gag tta ctt aga tgc ctc aga ata tca -tt gct ttc ccc aca agt tat aaa atc act tta tag tca tcg att ggt tgg ttc aat ata cga atg tat aac att ctt aaa caa tat atc att ctt gga tga atc att tcc tag ata cca tag cat cga gtt gat atc tat cgg tat aag taa ttc aca tga taa gat ttc cat aaa ctg cta taa tcg t-a aaa aaa ata cgt ggg aga taa tat ctt cac gga gat ata ggt gta gtt ttg cag tgg ccg tcc tgg ata ata tta tct ata tga tgg gtg gat atg atc agt ccc cgt ata gaa gtt caa agg tta tag cgt aca ata cat gta caa att ctt gga tat atg ata tac cag agc t-a aaa tat cct cgt tct aat tgt gga gga gtt gct gat gat gaa tac att tat tgc ata ggc ggt ata cgc gat cag gat tca tcg ttg ata tct agt att gat aga tgg aag cca tca aaa cca tat tgg caa acg tat gct aaa ata cga gag cca aaa tgt gat atg ggt gta gcg atg tta aac gga tta ata tat gtc ata ggc gga gtt gtt aaa ggt g-- aca cgt gta ccg aca cac ttg aga gtt tat cag aag atg gat gga tga tgc atc aac gtc ttc caa taa aaa tgt cca ata tgt cga cga ttg ttc atg ccg gca aaa ttt ata tat ctg gag gct aca aca ata gta gtg tag tta atg gaa tat cga atc t-- agt cct tag cta taa tcc gat ata tga tga atg gac caa att atc atc att aaa tat tcc tag aat taa tcc cgc tct a

>MZ300860.1:158656-160887_Camelpox_virus_strain_D1621/20_complete_genome

--- --- --- --a --- tgt a-c tat ttt ata cat tac tac att ggt tac ggc aga tat acc tac tcc gcc a-- -cc aca tgc tcc ggt aaa cgg --g tca tgt gcc gag gga gaa tat ctt gat aag act aat aat cga tgt tgt aat -cg gtg tcc acc tgg aga att tgc caa ggt tag atg tag -tg tgg tag cta taa cac aaa atg tga acg ctg tca aca tca tac ata tac agc aat tcc caa tta tta t-a atg gat gcc acc aat gta gaa aat acc caa agg gat cat ttg ata agg taa agt gta ccg gaa cac aga aca gta aat -gt gcg tgt ctt cct ggt tgg tat tgc gct acc gat tct tca cag act gaa gac tgt cga gat -tg tgt acc aaa aag g-a gct gtt cat gcg gat act ttg gtg gaa tag ata aat tag gaa atc cta ttt gta aat cgt gtt g-- -tg ttg gta aat att gtg acg aca tac gta atc ata gag ttg gtc ctt ttc ctc cat gca aa- --- cta tct aaa tgt aat tag tca --- --- --- --- --- --- --- --- --- --- --- --- --- --- -tg atc atg gta -cc gta taa tt- --- -at att gct --- --g g-a --g tat att att cag tat g-a aga cat att aat ac- --- --- -tt a-- --- --- ttg aca atc ttg ct- -at aat cat --- --- --- --- aat ata aaa aaa tac tta tgg --- -a- cat agt cac tca t-- --- --- --- aat tgc tg- acg cga taa att cgt aat aat ctg ttt tgt tca aat ttt tat aag ga- --- --a tct aca tgc ata aaa at- aaa aat ata att tat aat ac- --- act tac agt gcg cca tca tga ata aca gca gtg aat taa ttg ctg tta tta atg gat tta gaa ata gtg gac gat ttt gtg ata ttg ata tag tta tta atg atg aaa gga taa atg ctc ata gac tca tcc tat ctg gag cat ccg aat a-t ttt tcc att cta ttt tct agt gat ttt atc gat tct aat aaa tac gaa gtt aat cta agt cat tta gat tat caa agt gtg aac gat ttg atc aat tac att tat ggg ata cca ttg agc cta act aac gat atc gtg aaa tat att ctt tca act gct ga- --t ttt tta caa att gga tct gct att acc gaa tgt gaa aat tac ata ctt aaa aat ctg tgt tct aga aat tgt atc gat ttc --- --- tac ata tac gct gat aaa tat aat aac aag aaa ata gaa aca gcg tcg ttt aac aca ata tta tta aat att ttg aga ctc att aac gat gaa aac ttt aaa tac ttg aca gag gaa tca atg ata aaa ttt tta agc gat gat atg tta aat ata aaa aat gag gat ttt gca cca cta att ctc att aaa tgg tta gag agt aca caa caa cca tgc aca gtc gag tta ctt aga tgc ctc aga ata tca -tt gct ttc ccc aca agt tat aaa atc act tta tag tca tcg att ggt tgg ttc aat ata cga atg tat aac att ctt aaa caa tat atc att ctt gga tga atc att tcc tag ata cca tag cat cga gtt gat atc tat cgg tat aag taa ttc aca tga taa gat ttc cat aaa ctg cta taa tcg t-a aaa aaa ata cgt ggg aga taa tat ctt cac gga gat ata ggt gta gtt ttg cag tgg ccg tcc tgg ata ata tta tct ata tga tgg gtg gat atg atc agt ccc cgt ata gaa gtt caa agg tta tag cgt aca ata cat gta caa att ctt gga tat atg ata tac cag agc t-a aaa tat cct cgt tct aat tgt gga gga gtt gct gat gat gaa tac att tat tgc ata ggc ggt ata cgc gat cag gat tca tcg ttg ata tct agt att gat aga tgg aag cca tca aaa cca tat tgg caa acg tat gct aaa ata cga gag cca aaa tgt gat atg ggt gta gcg atg tta aac gga tta ata tat gtc ata ggc gga gtt gtt aaa ggt g-- aca cgt gta ccg aca cac ttg aga gtt tat cag aag atg gat gga tga tgc atc aac gtc ttc caa taa aaa tgt cca ata tgt cga cga ttg ttc atg ccg gca aaa ttt ata tat ctg gag gct aca aca ata gta gtg tag tta atg gaa tat cga atc t-- agt cct tag cta taa tcc gat ata tga tga atg gac caa att atc atc att aaa tat tcc tag aat taa tcc cgc tct a

>MZ300859.1:158657-160888_Camelpox_virus_strain_D1734/20_complete_genome

--- --- --- --a --- tgt a-c tat ttt ata cat tac tac att ggt tac ggc aga tat acc tac tcc gcc a-- -cc aca tgc tcc ggt aaa cgg --g tca tgt gcc gag gga gaa tat ctt gat aag act aat aat cga tgt tgt aat -cg gtg tcc acc tgg aga att tgc caa ggt tag atg tag -tg tgg tag cta taa cac aaa atg tga acg ctg tca aca tca tac ata tac agc aat tcc caa tta tta t-a atg gat gcc acc aat gta gaa aat acc caa agg gat cat ttg ata agg taa agt gta ccg gaa cac aga aca gta aat -gt gcg tgt ctt cct ggt tgg tat tgc gct acc gat tct tca cag act gaa gac tgt cga gat -tg tgt acc aaa aag g-a gct gtt cat gcg gat act ttg gtg gaa tag ata aat tag gaa atc cta ttt gta aat cgt gtt g-- -tg ttg gta aat att gtg acg aca tac gta atc ata gag ttg gtc ctt ttc ctc cat gca aa- --- cta tct aaa tgt aat tag tca --- --- --- --- --- --- --- --- --- --- --- --- --- --- -tg atc atg gta -cc gta taa tt- --- -at att gct --- --g g-a --g tat att att cag tat g-a aga cat att aat ac- --- --- -tt a-- --- --- ttg aca atc ttg ct- -at aat cat --- --- --- --- aat ata aaa aaa tac tta tgg --- -a- cat agt cac tca t-- --- --- --- aat tgc tg- acg cga taa att cgt aat aat ctg ttt tgt tca aat ttt tat aag ga- --- --a tct aca tgc ata aaa at- aaa aat ata att tat aat ac- --- act tac agt gcg cca tca tga ata aca gca gtg aat taa ttg ctg tta tta atg gat tta gaa ata gtg gac gat ttt gtg ata ttg ata tag tta tta atg atg aaa gga taa atg ctc ata gac tca tcc tat ctg gag cat ccg aat a-t ttt tcc att cta ttt tct agt gat ttt atc gat tct aat aaa tac gaa gtt aat cta agt cat tta gat tat caa agt gtg aac gat ttg atc aat tac att tat ggg ata cca ttg agc cta act aac gat atc gtg aaa tat att ctt tca act gct ga- --t ttt tta caa att gga tct gct att acc gaa tgt gaa aat tac ata ctt aaa aat ctg tgt tct aga aat tgt atc gat ttc --- --- tac ata tac gct gat aaa tat aat aac aag aaa ata gaa aca gcg tcg ttt aac aca ata tta tta aat att ttg aga ctc att aac gat gaa aac ttt aaa tac ttg aca gag gaa tca atg ata aaa ttt tta agc gat gat atg tta aat ata aaa aat gag gat ttt gca cca cta att ctc att aaa tgg tta gag agt aca caa caa cca tgc aca gtc gag tta ctt aga tgc ctc aga ata tca -tt gct ttc ccc aca agt tat aaa atc act tta tag tca tcg att ggt tgg ttc aat ata cga atg tat aac att ctt aaa caa tat atc att ctt gga tga atc att tcc tag ata cca tag cat cga gtt gat atc tat cgg tat aag taa ttc aca tga taa gat ttc cat aaa ctg cta taa tcg t-a aaa aaa ata cgt ggg aga taa tat ctt cac gga gat ata ggt gta gtt ttg cag tgg ccg tcc tgg ata ata tta tct ata tga tgg gtg gat atg atc agt ccc cgt ata gaa gtt caa agg tta tag cgt aca ata cat gta caa att ctt gga tat atg ata tac cag agc t-a aaa tat cct cgt tct aat tgt gga gga gtt gct gat gat gaa tac att tat tgc ata ggc ggt ata cgc gat cag gat tca tcg ttg ata tct agt att gat aga tgg aag cca tca aaa cca tat tgg caa acg tat gct aaa ata cga gag cca aaa tgt gat atg ggt gta gcg atg tta aac gga tta ata tat gtc ata ggc gga gtt gtt aaa ggt g-- aca cgt gta ccg aca cac ttg aga gtt tat cag aag atg gat gga tga tgc atc aac gtc ttc caa taa aaa tgt cca ata tgt cga cga ttg ttc atg ccg gca aaa ttt ata tat ctg gag gct aca aca ata gta gtg tag tta atg gaa tat cga atc t-- agt cct tag cta taa tcc gat ata tga tga atg gac caa att atc atc att aaa tat tcc tag aat taa tcc cgc tct a

>MZ300858.1:158657-160888_Camelpox_virus_strain_D1795/20_complete_genome

--- --- --- --a --- tgt a-c tat ttt ata cat tac tac att ggt tac ggc aga tat acc tac tcc gcc a-- -cc aca tgc tcc ggt aaa cgg --g tca tgt gcc gag gga gaa tat ctt gat aag act aat aat cga tgt tgt aat -cg gtg tcc acc tgg aga att tgc caa ggt tag atg tag -tg tgg tag cta taa cac aaa atg tga acg ctg tca aca tca tac ata tac agc aat tcc caa tta tta t-a atg gat gcc acc aat gta gaa aat acc caa agg gat cat ttg ata agg taa agt gta ccg gaa cac aga aca gta aat -gt gcg tgt ctt cct ggt tgg tat tgc gct acc gat tct tca cag act gaa gac tgt cga gat -tg tgt acc aaa aag g-a gct gtt cat gcg gat act ttg gtg gaa tag ata aat tag gaa atc cta ttt gta aat cgt gtt g-- -tg ttg gta aat att gtg acg aca tac gta atc ata gag ttg gtc ctt ttc ctc cat gca aa- --- cta tct aaa tgt aat tag tca --- --- --- --- --- --- --- --- --- --- --- --- --- --- -tg atc atg gta -cc gta taa tt- --- -at att gct --- --g g-a --g tat att att cag tat g-a aga cat att aat ac- --- --- -tt a-- --- --- ttg aca atc ttg ct- -at aat cat --- --- --- --- aat ata aaa aaa tac tta tgg --- -a- cat agt cac tca t-- --- --- --- aat tgc tg- acg cga taa att cgt aat aat ctg ttt tgt tca aat ttt tat aag ga- --- --a tct aca tgc ata aaa at- aaa aat ata att tat aat ac- --- act tac agt gcg cca tca tga ata aca gca gtg aat taa ttg ctg tta tta atg gat tta gaa ata gtg gac gat ttt gtg ata ttg ata tag tta tta atg atg aaa gga taa atg ctc ata gac tca tcc tat ctg gag cat ccg aat a-t ttt tcc att cta ttt tct agt gat ttt atc gat tct aat aaa tac gaa gtt aat cta agt cat tta gat tat caa agt gtg aac gat ttg atc aat tac att tat ggg ata cca ttg agc cta act aac gat atc gtg aaa tat att ctt tca act gct ga- --t ttt tta caa att gga tct gct att acc gaa tgt gaa aat tac ata ctt aaa aat ctg tgt tct aga aat tgt atc gat ttc --- --- tac ata tac gct gat aaa tat aat aac aag aaa ata gaa aca gcg tcg ttt aac aca ata tta tta aat att ttg aga ctc att aac gat gaa aac ttt aaa tac ttg aca gag gaa tca atg ata aaa ttt tta agc gat gat atg tta aat ata aaa aat gag gat ttt gca cca cta att ctc att aaa tgg tta gag agt aca caa caa cca tgc aca gtc gag tta ctt aga tgc ctc aga ata tca -tt gct ttc ccc aca agt tat aaa atc act tta tag tca tcg att ggt tgg ttc aat ata cga atg tat aac att ctt aaa caa tat atc att ctt gga tga atc att tcc tag ata cca tag cat cga gtt gat atc tat cgg tat aag taa ttc aca tga taa gat ttc cat aaa ctg cta taa tcg t-a aaa aaa ata cgt ggg aga taa tat ctt cac gga gat ata ggt gta gtt ttg cag tgg ccg tcc tgg ata ata tta tct ata tga tgg gtg gat atg atc agt ccc cgt ata gaa gtt caa agg tta tag cgt aca ata cat gta caa att ctt gga tat atg ata tac cag agc t-a aaa tat cct cgt tct aat tgt gga gga gtt gct gat gat gaa tac att tat tgc ata ggc ggt ata cgc gat cag gat tca tcg ttg ata tct agt att gat aga tgg aag cca tca aaa cca tat tgg caa acg tat gct aaa ata cga gag cca aaa tgt gat atg ggt gta gcg atg tta aac gga tta ata tat gtc ata ggc gga gtt gtt aaa ggt g-- aca cgt gta ccg aca cac ttg aga gtt tat cag aag atg gat gga tga tgc atc aac gtc ttc caa taa aaa tgt cca ata tgt cga cga ttg ttc atg ccg gca aaa ttt ata tat ctg gag gct aca aca ata gta gtg tag tta atg gaa tat cga atc t-- agt cct tag cta taa tcc gat ata tga tga atg gac caa att atc atc att aaa tat tcc tag aat taa tcc cgc tct a

>MZ300857.1:158656-160887_Camelpox_virus_strain_D1804/20_complete_genome

--- --- --- --a --- tgt a-c tat ttt ata cat tac tac att ggt tac ggc aga tat acc tac tcc gcc a-- -cc aca tgc tcc ggt aaa cgg --g tca tgt gcc gag gga gaa tat ctt gat aag act aat aat cga tgt tgt aat -cg gtg tcc acc tgg aga att tgc caa ggt tag atg tag -tg tgg tag cta taa cac aaa atg tga acg ctg tca aca tca tac ata tac agc aat tcc caa tta tta t-a atg gat gcc acc aat gta gaa aat acc caa agg gat cat ttg ata agg taa agt gta ccg gaa cac aga aca gta aat -gt gcg tgt ctt cct ggt tgg tat tgc gct acc gat tct tca cag act gaa gac tgt cga gat -tg tgt acc aaa aag g-a gct gtt cat gcg gat act ttg gtg gaa tag ata aat tag gaa atc cta ttt gta aat cgt gtt g-- -tg ttg gta aat att gtg acg aca tac gta atc ata gag ttg gtc ctt ttc ctc cat gca aa- --- cta tct aaa tgt aat tag tca --- --- --- --- --- --- --- --- --- --- --- --- --- --- -tg atc atg gta -cc gta taa tt- --- -at att gct --- --g g-a --g tat att att cag tat g-a aga cat att aat ac- --- --- -tt a-- --- --- ttg aca atc ttg ct- -at aat cat --- --- --- --- aat ata aaa aaa tac tta tgg --- -a- cat agt cac tca t-- --- --- --- aat tgc tg- acg cga taa att cgt aat aat ctg ttt tgt tca aat ttt tat aag ga- --- --a tct aca tgc ata aaa at- aaa aat ata att tat aat ac- --- act tac agt gcg cca tca tga ata aca gca gtg aat taa ttg ctg tta tta atg gat tta gaa ata gtg gac gat ttt gtg ata ttg ata tag tta tta atg atg aaa gga taa atg ctc ata gac tca tcc tat ctg gag cat ccg aat a-t ttt tcc att cta ttt tct agt gat ttt atc gat tct aat aaa tac gaa gtt aat cta agt cat tta gat tat caa agt gtg aac gat ttg atc aat tac att tat ggg ata cca ttg agc cta act aac gat atc gtg aaa tat att ctt tca act gct ga- --t ttt tta caa att gga tct gct att acc gaa tgt gaa aat tac ata ctt aaa aat ctg tgt tct aga aat tgt atc gat ttc --- --- tac ata tac gct gat aaa tat aat aac aag aaa ata gaa aca gcg tcg ttt aac aca ata tta tta aat att ttg aga ctc att aac gat gaa aac ttt aaa tac ttg aca gag gaa tca atg ata aaa ttt tta agc gat gat atg tta aat ata aaa aat gag gat ttt gca cca cta att ctc att aaa tgg tta gag agt aca caa caa cca tgc aca gtc gag tta ctt aga tgc ctc aga ata tca -tt gct ttc ccc aca agt tat aaa atc act tta tag tca tcg att ggt tgg ttc aat ata cga atg tat aac att ctt aaa caa tat atc att ctt gga tga atc att tcc tag ata cca tag cat cga gtt gat atc tat cgg tat aag taa ttc aca tga taa gat ttc cat aaa ctg cta taa tcg t-a aaa aaa ata cgt ggg aga taa tat ctt cac gga gat ata ggt gta gtt ttg cag tgg ccg tcc tgg ata ata tta tct ata tga tgg gtg gat atg atc agt ccc cgt ata gaa gtt caa agg tta tag cgt aca ata cat gta caa att ctt gga tat atg ata tac cag agc t-a aaa tat cct cgt tct aat tgt gga gga gtt gct gat gat gaa tac att tat tgc ata ggc ggt ata cgc gat cag gat tca tcg ttg ata tct agt att gat aga tgg aag cca tca aaa cca tat tgg caa acg tat gct aaa ata cga gag cca aaa tgt gat atg ggt gta gcg atg tta aac gga tta ata tat gtc ata ggc gga gtt gtt aaa ggt g-- aca cgt gta ccg aca cac ttg aga gtt tat cag aag atg gat gga tga tgc atc aac gtc ttc caa taa aaa tgt cca ata tgt cga cga ttg ttc atg ccg gca aaa ttt ata tat ctg gag gct aca aca ata gta gtg tag tta atg gaa tat cga atc t-- agt cct tag cta taa tcc gat ata tga tga atg gac caa att atc atc att aaa tat tcc tag aat taa tcc cgc tct a

>MZ300856.1:158593-160824_Camelpox_virus_strain_D1865/20_complete_genome

--- --- --- --a --- tgt a-c tat ttt ata cat tac tac att ggt tac ggc aga tat acc tac tcc gcc a-- -cc aca tgc tcc ggt aaa cgg --g tca tgt gcc gag gga gaa tat ctt gat aag act aat aat cga tgt tgt aat -cg gtg tcc acc tgg aga att tgc caa ggt tag atg tag -tg tgg tag cta taa cac aaa atg tga acg ctg tca aca tca tac ata tac agc aat tcc caa tta tta t-a atg gat gcc acc aat gta gaa aat acc caa agg gat cat ttg ata agg taa agt gta ccg gaa cac aga aca gta aat -gt gcg tgt ctt cct ggt tgg tat tgc gct acc gat tct tca cag act gaa gac tgt cga gat -tg tgt acc aaa aag g-a gct gtt cat gcg gat act ttg gtg gaa tag ata aat tag gaa atc cta ttt gta aat cgt gtt g-- -tg ttg gta aat att gtg acg aca tac gta atc ata gag ttg gtc ctt ttc ctc cat gca aa- --- cta tct aaa tgt aat tag tca --- --- --- --- --- --- --- --- --- --- --- --- --- --- -tg atc atg gta -cc gta taa tt- --- -at att gct --- --g g-a --g tat att att cag tat g-a aga cat att aat ac- --- --- -tt a-- --- --- ttg aca atc ttg ct- -at aat cat --- --- --- --- aat ata aaa aaa tac tta tgg --- -a- cat agt cac tca t-- --- --- --- aat tgc tg- acg cga taa att cgt aat aat ctg ttt tgt tca aat ttt tat aag ga- --- --a tct aca tgc ata aaa at- aaa aat ata att tat aat ac- --- act tac agt gcg cca tca tga ata aca gca gtg aat taa ttg ctg tta tta atg gat tta gaa ata gtg gac gat ttt gtg ata ttg ata tag tta tta atg atg aaa gga taa atg ctc ata gac tca tcc tat ctg gag cat ccg aat a-t ttt tcc att cta ttt tct agt gat ttt atc gat tct aat aaa tac gaa gtt aat cta agt cat tta gat tat caa agt gtg aac gat ttg atc aat tac att tat ggg ata cca ttg agc cta act aac gat atc gtg aaa tat att ctt tca act gct ga- --t ttt tta caa att gga tct gct att acc gaa tgt gaa aat tac ata ctt aaa aat ctg tgt tct aga aat tgt atc gat ttc --- --- tac ata tac gct gat aaa tat aat aac aag aaa ata gaa aca gcg tcg ttt aac aca ata tta tta aat att ttg aga ctc att aac gat gaa aac ttt aaa tac ttg aca gag gaa tca atg ata aaa ttt tta agc gat gat atg tta aat ata aaa aat gag gat ttt gca cca cta att ctc att aaa tgg tta gag agt aca caa caa cca tgc aca gtc gag tta ctt aga tgc ctc aga ata tca -tt gct ttc ccc aca agt tat aaa atc act tta tag tca tcg att ggt tgg ttc aat ata cga atg tat aac att ctt aaa caa tat atc att ctt gga tga atc att tcc tag ata cca tag cat cga gtt gat atc tat cgg tat aag taa ttc aca tga taa gat ttc cat aaa ctg cta taa tcg t-a aaa aaa ata cgt ggg aga taa tat ctt cac gga gat ata ggt gta gtt ttg cag tgg ccg tcc tgg ata ata tta tct ata tga tgg gtg gat atg atc agt ccc cgt ata gaa gtt caa agg tta tag cgt aca ata cat gta caa att ctt gga tat atg ata tac cag agc t-a aaa tat cct cgt tct aat tgt gga gga gtt gct gat gat gaa tac att tat tgc ata ggc ggt ata cgc gat cag gat tca tcg ttg ata tct agt att gat aga tgg aag cca tca aaa cca tat tgg caa acg tat gct aaa ata cga gag cca aaa tgt gat atg ggt gta gcg atg tta aac gga tta ata tat gtc ata ggc gga gtt gtt aaa ggt g-- aca cgt gta ccg aca cac ttg aga gtt tat cag aag atg gat gga tga tgc atc aac gtc ttc caa taa aaa tgt cca ata tgt cga cga ttg ttc atg ccg gca aaa ttt ata tat ctg gag gct aca aca ata gta gtg tag tta atg gaa tat cga atc t-- agt cct tag cta taa tcc gat ata tga tga atg gac caa att atc atc att aaa tat tcc tag aat taa tcc cgc tct a

>AF438165.1:160821-163057_Camelpox_virus_M-96_from_Kazakhstan_complete_genome

--- --- --- --a --- tgt a-c tat ttt ata cat tac tac att ggt tac ggc aga tat acc tac tcc gcc a-- -cc aca tgc tcc ggt aaa cgg --g tca tgt gcc gag gga gaa tat ctt gat aag act aat aat cga tgt tgt aat -cg gtg tcc acc tgg aga att tgc caa ggt tag atg tag -tg tgg tag ata taa cac aaa atg tga acg ctg tca aaa tca tac ata tac agc aat tcc caa tta tta t-a atg gat gcc acc aat gta gaa aat acc caa agg gat cat ttg ata agg taa agt gta ccg gaa cac aga aca gta aat -gt gcg tgt ctt cct ggt tgg tat tgc gct acc gat tct tca cag act gaa gac tgt cga gat -tg tgt acc aaa aag g-a gct gtt cat gcg gat act ttg gtg gaa tag ata aat tag gaa atc cta ttt gta aat cgt gtt g-- -tg ttg gta aat att gtg acg aca tac gta atc ata gag ttg gtc ctt ttc ctc cat gca aa- --- cta tct aaa tgt aat tag tca --- --- --- --- --- --- --- --- --- --- --- --- --- --- -tg atc atg gta -cc gta taa tt- --- -at att gct --- --g g-a --g tat att att cag tat g-a aga cat att aat ac- --- --- -tt a-- --- --- ttg aca atc ttg ct- -at aat cat --- --- --- --- aat ata aaa aaa tac tta tgg --- -a- cat agt cac tca t-- --- --- --- aat tgc tg- acg cga taa att cgt aat aat ctg ttt tgt tca aat ttt tat aag ga- --- --a tct aca tgc ata aaa at- aaa aat ata att tat aat aca ctt act tac agt gcg cca tca tga ata aca gca gtg aat taa ttg ctg tta tta atg gat tta gaa ata gtg gac gat ttt gtg ata ttg ata tag tta tta atg atg aaa gga taa atg ctc ata gac tca tcc tat ctg gag cat ccg aat a-t ttt tcc att cta ttt tct agt gat ttt atc gat tct aat aaa tac gaa gtt aat cta agt cat tta gat tat caa agt gtg aac gat ttg atc gat tac att tat ggg ata cca ttg agc cta act aac gat atc gtg aaa tat att ctt tca act gct ga- --t ttt tta caa att gga tct gct att acc gaa tgt gaa aat tac ata ctt aaa aat ctg tgt tct aga aat tgt atc gat ttc --- --- tac ata tac gct gat aaa tat aat aac aag aaa ata gaa aca gcg tcg ttt aac aca ata tta tta aat att ttg aga ctc att aac gat gaa aac ttt aaa tac ttg aca gag gaa tca atg ata aaa ttt tta agc gat gat atg tta aat ata aaa aat gag gat ttt gca cca cta att ctc att aaa tgg tta gag agt aca caa caa cca tgc aca gtc gag tta ctt aga tgc ctc aga ata tca -tt gct ttc ccc aca agt tat aaa atc act tta tag tca tcg att ggt tgg ttc aat ata cga atg tat aac att ctt aaa caa tat atc att ctt gga tga atc att tcc tag ata cca tag cat cga gtt gat atc tat cgg tat aag taa ttc aca tga taa gat ttc cat aaa ctg cta taa tcg taa aaa aaa ata cgt ggg ata taa tat ctt cac gga gat ata ggt gta gtt ttg cag tgg ccg tcc tgg ata ata tta tct ata tga tgg gtg gat atg atc agt ccc cgt ata gaa gtt caa agg tta tag cgt aca ata cat gta caa att ctt gga tat atg ata tac cag agc t-a aaa tat cct cgt tct aat tgt gga gga gtt gct gat gat gaa tac att tat tgc ata ggc ggt ata cgc gat cag gat tca tcg ttg ata tct agt att gat aga tgg aag cca tca aaa cca tat tgg caa acg tat gct aaa ata cga gag cca aaa tgt gat atg ggt gta gcg atg tta aac gga tta ata tat gtc ata ggc gga gtt gtt aaa ggt g-- aca cgt gta ccg aca cac ttg aga gtt tat cag aag atg gat gga tga tgc atc aac gtc ttc caa taa aaa tgt cca ata tgt cga cga ttg ttc atg ccg gca aaa ttt ata tat ctg gag gct aca aca ata gta gtg tag tta atg gaa tat cga atc t-- agt cct tag cta taa tcc gat ata tga tga atg gac caa att atc atc att aaa tat tcc tag aat taa tcc cgc tct a

>AY009089.1:158937-161172_Camelpox_virus_CMS_complete_genome

--- --- --- --a --- tgt a-c tat ttt ata cat tac tac att ggt tac ggc aga tat acc tac tcc gcc a-- -cc aca tgc tcc ggt aaa cgg --g tca tgt gcc gag gga gaa tat ctt gat aag act aat aat cga tgt tgt aat -cg gtg tcc acc tgg aga att tgc caa ggt tag atg tag -tg tgg tag ata taa cac aaa atg tga acg ctg tca aaa tca tac ata tac agc aat tcc caa tta tta t-a atg gat gcc acc aat gta gaa aat acc caa agg gat cat ttg ata agg taa agt gta ccg gaa cac aga aca gta aat -gt gcg tgt ctt cct ggt tgg tat tgc gct acc gat tct tca cag act gaa gac tgt cga gat -tg tgt acc aaa aag g-a gct gtt cat gcg gat act ttg gtg gaa tag ata aat tag gaa atc cta ttt gta aat cgt gtt g-- -tg ttg gta aat att gtg acg aca tac gta atc ata gag ttg gtc ctt ttc ctc cat gca aa- --- cta tct aaa tgt aat tag tca --- --- --- --- --- --- --- --- --- --- --- --- --- --- -tg atc atg gta -cc gta taa tt- --- -at att gct --- --g g-a --g tat att att cag tat g-a aga cat att aat ac- --- --- -tt a-- --- --- ttg aca atc ttg ct- -at aat cat --- --- --- --- aat ata aaa aaa tac tta tgg --- -a- cat agt cac tca t-- --- --- --- aat tgc tg- acg cga taa att cgt aat aat ctg ttt tgt tca aat ttt tat aag ga- --- --a tct aca tgc ata aaa at- aaa aat ata att tat aat aca ctt act tac agt gcg cca tca tga ata aca gca gtg aat taa ttg ctg tta tta atg gat tta gaa ata gtg gac gat ttt gtg ata ttg ata tag tta tta atg atg aaa gga taa atg ctc ata gac tca tcc tat ctg gag cat ccg aat a-t ttt tcc att cta ttt tct agt gat ttt atc gat tct aat aaa tac gaa gtt aat cta agt cat tta gat tat caa agt gtg aac gat ttg atc gat tac att tat ggg ata cca ttg agc cta act aac gat atc gtg aaa tat att ctt tca act gct ga- --t ttt tta caa att gga tct gct att acc gaa tgt gaa aat tac ata ctt aaa aat ctg tgt tct aga aat tgt atc gat ttc --- --- tac ata tac gct gat aaa tat aat aac aag aaa ata gaa aca gcg tcg ttt aac aca ata tta tta aat att ttg aga ctc att aac gat gaa aac ttt aaa tac ttg aca gag gaa tca atg ata aaa ttt tta agc gat gat atg tta aat ata aaa aat gag gat ttt gca cca cta att ctc att aaa tgg tta gag agt aca caa caa cca tgc aca gtc gag tta ctt aga tgc ctc aga ata tca -tt gct ttc ccc aca agt tat aaa atc act tta tag tca tcg att ggt tgg ttc aat ata cga atg tat aac att ctt aaa caa tat atc att ctt gga tga atc att tcc tag ata cca tag cat cga gtt gat atc tat cgg tat aag taa ttc aca tga taa gat ttc cat aaa ctg cta taa tcg t-a aaa aaa ata cgt ggg ata taa tat ctt cac gga gat ata ggt gta gtt ttg cag tgg ccg tcc tgg ata ata tta tct ata tga tgg gtg gat atg atc agt ccc cgt ata gaa gtt caa agg tta tag cgt aca ata cat gta caa att ctt gga tat atg ata tac cag agc t-a aaa tat cct cgt tct aat tgt gga gga gtt gct gat gat gaa tac att tat tgc ata ggc ggt ata cgc gat cag gat tca tcg ttg ata tct agt att gat aga tgg aag cca tca aaa cca tat tgg caa acg tat gct aaa ata cga gag cca aaa tgt gat atg ggt gta gcg atg tta aac gga tta ata tat gtc ata ggc gga gtt gtt aaa ggt g-- aca cgt gta ccg aca cac ttg aga gtt tat cac aag atg gat gga tga tgc atc aac gtc ttc caa taa aaa tgt cca ata tgt cga cga ttg ttc atg ccg gca aaa ttt ata tat ctg gag gct aca aca ata gta gtg tag tta atg gaa tat cga atc t-- agt cct tag cta taa tcc gat ata tga tga atg gac caa att atc atc att aaa tat tcc tag aat taa tcc cgc tct a

>MG012795.1:147873-150098_Vaccinia_virus_isolate_CEyV1

ttg ctg act gta --- tgt a-c tat ttt tta cat tac tac ttt ggc tac ggc aga tat acc tac tcc gcc acc --- --- --- --- --- -aa cgg --g tca tgt gac aag gga gaa tat ctt gat aag acg cat aat c-- --- --- -at -tg gtg tcc acc tgg aga att tgc caa ggt tag atg ta- --a tgg taa cga taa cac aaa atg tga acg ctg ccc acc tca tac ata tac cac aat -cc caa tta ttc t-a atg gat gtc atc aat gta gaa aat gcg caa ccg tat aat ttg aaa agg taa agt gta ccg gaa cac aga aca gta aat -gt tcg tga ctt cct ggt tgg tat tgc gct act gat tct tca cag act gaa gat tgt tga aat -tg tgt acc aaa aag g-a gat gtc cat gcg gat act ttg gtg aaa tag atg aat aag gaa atc cta tta gta aat cgt gct g-- -ta ttg gtg aat att gca act acc tcc gta att ata gac ttg atc cat ttt ctc cct gca aa- --- cta tct aaa tgt aat taa tta --- --- --- --- --- --- --- --- tga ttt --- --- --- -tg atg ata atg tta -cc ata c-a gt- --- -at atc gct act tgg tta --g tgt att att cag tat g-a aga cct att aat aa- --- --- -tt aca tat ctt ttg acg gtc tta tt- -at aat tat --- --- --- --- aat ata aaa --- -ac tta tgg --- --- cat agt aac tta t-- --- --- --- tat tgc ta- aca cga taa att cgt aat aat ctg ttt tgt tca --- --- --- aag ga- --- --a tct gca ggc gta aaa ac- aaa aat ata att tat aat ata c-- tct tac agc gcg cca tcg tga ata aca gca gtg aat tga ttg ctg tta tta atg gaa tta gaa ata gtg gac gat ttt gtg ata tta gta tag taa tta att atg taa tga tga acg ctc aca gac tta tcc tat ctg cag cct ccg aat a-t ttt tcc att cag ttt tcc aat aat ttt atc gat tct aat gaa tac aaa gtt aat cta agt cat tta gat tat caa agt gat aac gat ttg atc gat tac att tat ggg ata cct ttg agc cta act aac gat agc gtg aaa tat att ctt tca acc gct ga- --t tta tta caa att gga tcc gct att aca gag tgt gaa aat tac gta ctt aaa aat ctt tgt tct aaa aac tgt atc gat ttc --- --- tac ata tac gct gat aaa tat aat aac acg aaa ata gaa tca gcg tcg ttt aac aca ata tta caa aat att ttg aga ctc atc aac gat gaa aac ttt aaa tcc tta aca gag gaa tca atg ata aaa att tta agc gat gat atg tta aat ata aaa aat gag gat ttc gcc cca cga att ctc att aaa tgg tta gcg agt act caa caa tca tgc acc gtc gag tta ctt aga tgc ctc aga ata tca -tt gct ttc ccc aca agt tat aaa atc act tta tag tca tca act agt tag ttc aat cta cga atg tat aac att ctt aaa taa tat agc att ctt gga tga atc att tcc tag ata cct tag cat cga ttt gat atc tat cgg tat aag taa ttt gcg tga taa aat ttc cat aaa ctg cta caa tca t-a aaa aaa ata cat ggg aaa tga tat ctt cac gta gat ata ggt gta gtt tcg cag tgg ccg gcc tgg ata ata tta tct ata tga tgg gtg gct atg atc agt ccc cgt ata gaa gtt caa aag tta tag cgt aca ata cat gta cga atg ctt gga tat atg ata tac cag agc t-a gaa tat cct cgt tct aat tgt ggg gga ctg tct cat gac gat tac att tag tgt ata ggt ggc ata cgc gat cag gat tca tcg ttg aca tct agt att gat aaa tgg aag cca tca aaa cca tat tgg cag aag tat gct aaa atg cgc gaa cca aaa tgt gat atg ggg gtt gcg atg tta aac gga tta ata tat gtt ata ggc gga atc gtt aaa ggt a-- aca cgt gta ccg acg cac tag aga ttt tat cag aag atg gat ggg tga agc atc aac gtc ttc caa caa aaa tgt cca ata tgt cga cga ttg ttc atg atg gca aga ttt ata tat ctg gag gtt aca aca ata gta gtg tag tta atg taa tat cga atc t-- agt cct tag cta taa tcc gat ata tga tga atg gac caa att atc atc att aaa cac tcc tag aat caa tcc cgc tc- -

>OM460002.1:175594-177860_Cowpox_virus_isolate_No-H2_complete_genome

--- --- --- --a --- tgt a-c tat ttt gta cat cac tac att ggt tac agc aga tat acc tac tcc gcc a-- -cc aca tgc tcc ggt aaa cgg --g aca tgt gaa gag gga gaa tat ctt gat agg acg cat aat caa tgt tgt aat -cg gtg tcc tcc tgg aga att tgc caa ggt tag atg ta- --a tgg taa cga taa cac aaa atg tga acg ctg tcc acc tca tac att tac cgc agt acc caa tta ttc t-a atg gat gtc atc aat gta gaa aat gcc caa agg gat cat ttg ata agg taa agt gta ccg gaa cac aga aca gta aat -gt tcg tgt ctt cct ggt tgg ttt tgc gct act gat tct tcg aag act gaa gat tgt cga gat -tg tat acc aaa aag g-a gat gtc cat gcg gat act ttg gtg gaa tag atg aac aag gaa atc ctc ttt gta aat cgt gtt gtg ctg ttg gtg gat att gcg acc tcg tac gcg gtc ata tag tcg gtg --- ttc atc cat gca aa- --- cta tct aaa tgt aat tag tca --- --- --- --- --- --- --- --- --- --- --- --- --- --- -tg gcc gtg tta -cc atc a-a tc- --- -at ata gct act tgg t-a --g tgt att att cag tat g-a aga cct att aat aat aat tat ctt att tat ttc ttg acg atc ttc taa aat gtt att aag agt cat aat att ata aaa --- tac tta tgg --- --- --a cat aaa ttg t-a aac aag atg a-- taa t-- aca cga tat cct ctt -at aat cag ctt -gt tca aat ttt tat aaa gaa tta tat tca tta ggc ata aca ta- aaa aat ata att tat aat ac- --- act tac a-- gca cca tca tga ata aca gca gtg act taa ttg ctg tta tta atg gat tta gaa ata gtg gac gat ttt gtg ata tta gta tag tta tta atg atg aaa gga taa atg ctc ata gac tca tcc tat ctg gag cct ccg aat a-t ttt tcc att ctg ttt tcc aat aat ttt atc gat tct aat gaa tac gaa gtt aat cta agt cat tta gat tat caa agt gtg aac gat ttg atc gat tac att tat ggg ata cct ttg agc cta act aac gat agc gtg aaa tat att ctt tca acc gct ga- --t ttt tta caa att gga tct gcc att act gag tgc gaa aaa tac ata ctt aaa aat ctt tgt tct aga aac tgt atc gat ttc --- --- tac ata tac gct gat aaa tat aat aac aag aaa ata gaa tca gcg tcg ttt aac aca ata tta cga aat att ttg aga ctc att aac gat gaa aac ttt aaa tac ttg aca gag gaa tca atg ata aaa att tta agc gat gat atg tta aat ata aaa aat gag gat ttc gca cca cta att ctc att aaa tgg tta gag agt aca caa caa cca tgc acc gtc gag tta ctt aga tgc ctc aga ata tca -tt gct ttc cac aca agt tat aaa atc act tta tag tca tcg act ggt tag ttc aat cta cga atg tat aac att ctt aaa caa tat agc att ctt gga tga atc att tcc tag ata caa tag cat cga gtt gat atc tat cgg tat aag taa ttc aca tga taa gat ttc cat aaa ctg cta caa tca t-a aaa aaa ata cat ggg aaa tga tat ctt cac gta gat ata ggt gta gtt tcg cag tgg ccg tcc tgg ata ata tta tct ata tga tgg gtg gat atg atc agt ccc cgt ata gaa gtt caa agg tta tag cgt aca ata cat gta caa att ctt gga tat atg ata tac cag agc t-a aaa tat cct cgt tct aat tgt ggg gga ctg gct gat gac gaa tac att tat tgt ata ggc ggc ata cgc gat cag gat tca tcg ttg ata tct aat att gat aga tgg aag cca tca aaa caa tat tgg cag acg tat gct aaa atg cgc gaa cca aaa tgt gat atg gga gtt gcg atg tta aac gga tta ata tat gtc ata ggt gga atc gtt aaa ggt g-- aca cgt gta ccg acg cac tag aga gtt tat cag aag atg gat gga tga agc atc aac gtc ttc caa taa aaa tgt cca ata tgt cga cga ttg ttc atg ctg gaa aga ttt ata tat ctg gag gtt aca cca ata gta gta tag tta atg gaa tat cta atc t-- agt cct tag cta taa tcc gat ata tga tga atg gac caa att atc atc att aaa tat tcc tag aat taa tcc tgc tct a

>DQ437594.1:158345-160564_Taterapox_virus_strain_Dahomey_1968_complete_genome

--- --- --- --a --- tgt a-c tat ttt ata cat tac tac att ggt tac ggc aga tat acc tac tcc gcc a-- -cc aca tgc tcc ggt aaa cgg --g tca tgt acc gag gga gaa tat ctt gat aag act cat aat tga tgt tgt aat -cg gtg tcc acc tgg aga att tgc caa ggt tag atg ta- --g tgg tag cta taa cac aaa atg tga acg ctg tcc aca tca tac ata tac agc aat tcc caa ttc ttc t-a atg gat gcc atc aat gta gaa aat gcc caa agg gat cat ttg ata agg taa agt gta ccg gaa cac aga aca gta aat -gt gcg tgt ctt cct ggt tgg tat tgc gct acc gat tct tca cag act aaa gac tgt cga gat -tg tgt acc aaa aag g-a gat gtc cat gcg gat act ttg gtg gaa tag ata aat tag gaa atc cta ttt gta aat cgt gtt g-- -tg ttg gta aat att gtg acg aca tac gta atc ata gag ttg gtc ctt ttc ctc cat gca aa- --- cta tct aaa t-- aat tag tca --- --- --- --- --- --- --- --- --- --- --- --- --- --- -tg atc atg gta -cc gta caa tt- --- -at att gct --- --g gta --g tat att att cag tat g-a aga cat att aat ac- --- --- -tt atc --- --- ttg aca atc ttg ct- -at aat cat --- --- --- --- aat ata aaa --a tac tta tgg --- -a- cat agt cac tca t-- --- --- --- aat tgc tg- acg cga taa att cgt aat aat ctg ttt tgt tca aat ttt tat aag ga- --- --a tct aca ggc ata aaa at- -aa aat ata att tat aat ac- --- a-- tac agc gcg cca tta tga ata aca gca gtg aat taa ttg ctg tta tta atg gat tta gaa ata gtg gac gat ttt gtg ata tta ata tag tta tta atg atg aaa gga taa atg ctc ata gac tca tcc tat ctg gag cat ccg aat a-t ttt tcc att cta ttt tct agt gat ttt atc gat tct aat gaa tac gaa gtt aat cta agt cat tta gat tat caa agt atg aac gat ttg atc gat tac att tat ggg ata cct ttg agc cta act aac gat agc gtg aaa tat att ctt tca act gct ga- --t ttt tta caa att gga tct gct att acc gaa tgt gaa aat tac ata ctt aaa aat ctg tgt tct aga aat tgt atc gat ttc --- --- tac ata tac gct gat aaa tat aat aac aag aaa ata gaa aca gcg tcg ttt aac aca ata tta cga aat att ttg aga ctc att aac gat gaa aac ttt aaa tac ttg aca gag gaa tca atg ata aaa att tta agc gat gat atg tta aat ata aaa aat gag gat ttt gca cca cta att ctc att aaa tgg tta gag agt aca caa caa cca tgc aca gtc gag tt- --- -ga tgc ctc aga ata tca -tt gct ttc ccc aca agt tat aaa atc act tta tag tca tcg att ggt tgg ttc aat ata cga atg tat aac att ctt aaa caa tat atc att ctt gga tga atc att tcc tag ata cca tag cat cga gtt gat atc tat cgg tat aag taa ttc aca tga taa gat ttc cat aaa ctg cta taa tcg t-- aaa aaa ata cgt ggg aga taa tat ctt cac gga gat ata ggt gta gtt ttg cag tgg ccg ttc tgg ata ata tta tct ata tga tgg gtg gat atg atc agt ccc cgt ata gaa gtt caa agg tta tag cgt aca ata cat gta caa att ctt gga tat atg ata tac cag agc t-a aaa tat cct cgt tct aat tgt gga gga gtt gct gat gat gaa tac att tat tgc ata ggc ggc ata cgc aat cag gat tca tcg ttg ata tct aat att aat aga tgg aag cca tca aaa cca tat tgg caa acg tat gct aaa atg cga gag cca aaa tgt gat atg ggt gta gcg atg tta aac gga tta ata tat gtc ata ggc gga gtt gtt aaa ggt g-- aca cgt gta ccg aca cac ttg aga gtt tat cag aag atg gat gga tga tgc atc aac gtc ttc caa taa aaa tgt aca ata tgt cga cga ttg ttc atg ccg gca aaa ttt ata tat ctg gag gct aca aca ata gta gtg tag tta atg gaa tat cga atc t-- agt cct tag cta taa tcc gat ata tga tga atg gac caa att atc atc att aaa tat tcc tag aat taa tcc cgc tct a

>LR800247.1:153348-155571_Variola_virus_genome_assembly_complete_genome:_monopartite

--- --- --- --a --- tgt a-c tat ttt ata cat tac tac att ggt tac ggc aga tat acc tac tcc gcc a-- -cc aca tgc tcc ggt aaa cgg --g tca tgt gtc aag gga gaa tat ctt gat aag act cat aat cga tgt tgt aat -ca gtg tcc -cc tgg aga att tgc caa ggt tag atg ta- --g tgg tag cta taa cac aaa atg tga acg ctg tcc aca tca tac ata tac agc aat tcc caa tta ttt t-a atg gat gcc atc aat gta gaa aat gcc caa a-g gat cat ttg ata agg taa agt gta ccg gaa cac aaa aca gta aat -gt gtg tgt ctt cct ggt tgg tat tgc gct acc aat tct tca cag act gaa gac tgt cga gat -tg tgt acc aaa aag g-a gct gtc cat gcg gat act ttg gtg gaa tag ata aat tag gaa atc cta ttt gta aat cgt gtt g-- -tg ttg gta aat att gtg aca aca tac gta atc ata gag ttg gtc ctt ttc ctc cat gca aa- --- cta tct aaa tgt aat tag tca --- --- --- --- --- --- --- --- --- --- --- --- --- --- -tg atc atg gta -cc gta caa tt- --- -at att gct --- --g gta --g tat att att cag tat g-a aga cat att aat ac- --- --- -tt atc --- --t ttg aca atc ttg ct- -at aat cat --- --- --- --- aat ata aaa --a tac tta tgg --- -a- cat agt cac tca t-- --- --- --- aat tgc tg- acg cga taa att cgt aat aat ctg ttt tgt tca aat ttt tat aag ga- --- --a tct aca ggc at- --- --- aaa aat ata att tat aat ac- --- act tac agc gcg cca tca tga ata aca gca gtg aat taa ttg ctg tta tta atg gat tta gaa ata gtg gac gat ttt gtg ata tta ata tag tta tta atg atg aaa gga taa atg ctc ata gac tca tcc tat ctg gag cat ccg aat a-t ttt tcc att cta ttt tct agt gat ttt atc gat tct aat gaa tac aaa gtt aat cta agt cat tta gat tat caa agt gtg aac gat ttg atc gat tac att tat ggg ata cct ttg agc cta act aac gat agc gtg aaa tat att ctt tca act gct ga- --t ttt tta caa att gga tct gct att acc gaa tgt gaa aat tac ata ctt aaa aat ctg tgt tct aga aa? ??? ?tc gat ttc --- --- tac ata tac gct gat aaa tat aat aac aag aaa ata gaa aca gcg tcg ttt aac aca ata tta cga aat att ttg aga ctc att aac gat gaa aac ttt aaa tac ttg aca gag gaa tca atg ata aaa att tta agt gat gat atg tta aat ata aaa aat gag gat ttt gca cca cta att ctc att aaa tgg tta gag agt aca caa caa cca tgc aca gtc gag tta ctt aga tgc ctc aga ata tca -tt gct ttc ccc aca agt tat aaa atc act tta tag tca ttg att ggt tgg ttc aat ata cga atg tat aac att ctt aaa caa tat atc att ctt gga tga atc att tcc tag ata cca tag cat cga gtt gat atc tat cgg tat aag taa ttc aca taa taa gat ttc cat aaa ctg cta taa tcg t-a aaa aaa ata cgt ggg aga taa tat ctt cac gga gat ata ggt gta gtt ttg cag tgg ccg tcc tgg ata ata tta tct ata tga tgg gtg gat atg atc agt ccc cgt ata gaa gtt caa agg tta tag cgt aca ata cat gta caa att ctt gga tat atg ata tac cag agc t-a aaa tat cct cgt tct aat tgt gga gga gtt gct gat gat gaa tac att tat tgc ata ggc ggc ata cgc gat cag gat tca tcg ttg ata tct agt att gat aga tgg aag cca tca aaa cca tat tgg caa acg tat gct aaa atg cga gag cca aaa tgt gat att ggt gta gcg atg tta aac gaa tta ata tat gtc ata ggc gga gtt gtt aaa ggt g-- aca cgt gta ccg aca cac ttg aga gtt tat cag aag atg gat gga tga tgc atc aac gtc ttc caa taa aaa tgt cca ata tgg cga cga ttg ttc atg ccg gca aaa ttt ata tat ctg gag gct aca aca ata gta gtg tag tta atg gaa tat caa atc t-- agt cct tag cta taa tcc gat ata tga tga atg gac caa att atc atc att aaa tat tcc tag aat taa tcc cgc tct a

>HQ420896.1:175002-177271_Cowpox_virus_strain_Germany_1990_2_complete_genome

ttg ctg act gta --- tgt a-c tat ttt gta cat cag tac att ggt tac agc aga tat acc tac ttc gtc act gcc aca cgc tcc ggt aaa cgg --g tca tgt gac gaa gga gaa tat ctt gat aag acg cat aat caa tgt tgt aat -cg gtg tcc acc tgg aga att tgc caa ggt tag atg ta- --a tgg caa cga taa cac aaa atg tga acg ctg tcc acc tca tac att tac cac agt acc caa tta ttc t-a atg gat gtc atc aat gta tga aat gcc caa agg gat cat ttg ata agg tac gat gtg atg gaa cac aga aca gta aat -gt tcg tgt ctt ccg ggt tgg ttt tgc gct act gat tct tcg aag act gaa gat tgt cga gat -tg tat acc aaa aag g-a gat gtc cat gcg gat act ttg gtg gaa tag ata aat tag gaa atc ctc ttt gta aat cgt gtt g-- -tg ttg gtg aat att gcg acg aca tac gta atc ata gag ttg gtc ctt ttc ctc cat gca aa- --- cta tct aaa tgt aat tag cca --- --- --- --- --- --- --- --- --- --- --- --- --- --- -tg atc atg tta ccc ata c-a tc- --- -at atc gct act tgg taa --g cgt att att cag tat g-a aga tct att aat aat aat tat ctt att tat ttc ttg acg --- --- --- -at att aat cg- -gt cct aat aat ata aaa --- tac tta tgg --- --- --a aat aaa ttg t-a aac aag atg aat taa t-- aca caa tac atc ct- --t aat tag ctt -gt tca aa- ttt tat aaa gaa tca tat tca tta ggc ata taa ta- aaa aat ata att tat aat ac- --- act tac a-- gcg tca tca tgc ata aca gca gtg aat taa ttg ctg tta tta atg gat tta gaa ata gtg gaa gat ttt gtg ata tta gta tag tta tta atg atg aaa gga taa atg ctc ata gac tca tcc tat ctg gag cat cag aat a-t ttt tcc att ctg ttt tcc aat aat ttt atc gat tct aat gaa tac gaa gtt aat cta agt cat tta gat tat caa agt gtg aac gat ttg atc gat tac att tat ggg ata cct ttg agc cta act aac gat agc gtg aaa tat att ctt tca acc gct ga- --t ttt tta caa att gga tct gcc att act gag tgc gaa aaa tac ata ctt aaa aat ctt tgt tct aga aac tgt atc gat ttc --- --- tac ata tac gct gat aaa tat aat aac aag aaa ata gaa tca gcg tcg ttt aac aca ata tta cga aat att ttg aga ctc att aat gat gaa aac ttt aaa tac ttg aca gag gaa tca atg ata aaa att tta agc gat gat atg tta aat ata aaa aat gag gat ttt gca cca cta att ctc att aaa tgg tta gag agt aca caa caa cca tgt acc gtc gag tta ctt aga tgc ctc aga ata tca -tt gct ttc ccc aca agt tat aaa atc act tta tag tca tcg act ggt tag ttc aat cta cga atg tat aac att ctt aaa caa tat agt att ctt gga tga atc att tcc tag ata caa tag cat cga gtt gat atc tat cgg tat aag taa ttc gca tga taa gat ttc cat aaa ctg cta caa tcg t-a aaa aaa ata cat ggg aga tga tat ctt cac gta gat ata ggt gta gtt tcg cag tgg ccg ttc tgg ata ata tta tct ata tga tgg gtg gat atg atc aat ccc cgt ata gga gtt caa agg tta tag cgt aca ata cat gta caa att ctt gga tat atg ata tac cag agc t-a aaa tat cct cgt tct aat tgt gga gga gtt gcc gat gac gaa tac att tat tgt ata ggc ggc ata cgc gat cag gat tca tcg ttg ata tct aat att gat aga tgg aag cca tca aaa cca tat tgg cag acg tat gct aaa atg cgc gaa cca aaa tgt gat atg ggg gtt gcg atg tta aac gga tta ata tat gtc ata ggt gga atc gtt aaa ggt g-- aca cat gta ccg acg cac tag aaa gtt tat cag aag atg gat gga tga tgc atc agc gcc ttc caa taa aaa tgt cca ata tgg cta cga ttg ttc atg ctg gta aga ttt ata tat ctg gag gtt ata aca ata gta gtg ccg tta atg gac cat cga atc t-- agt cct tag cta taa tcc gat ata tga tga atg gac caa att atc atc att aaa tat tcc tag aat taa tcc cgc tct a

>KC813499.1:175125-177395_Cowpox_virus_strain_MarLei07/1_complete_genome

ttg ctg act gta --- tgt a-c tat ttt gta cat cag tac att ggt tac agc aga tat acc tac ttc gtc act gcc aca cgc tcc ggt aaa cgg --a tca tgt gac gaa gga gaa tat ctt gat aag acg cat aat caa tgt tgt aat -cg gtg tcc acc tgg aga att tgc caa ggt tag atg ta- --a tgg taa cga taa cac aaa atg tga acg ctg tcc acc tca tac att tac cac agt acc caa tta ttc t-a atg gat gtc atc aat gta tga aat gcc caa agg gat cat ttg ata agg tac gat gtg atg gaa cac aga aca gta aat -gt tcg tgt ctt ccg ggt tgg ttt tgc gct act gat tct tcg aag act gaa gat tgt cga gat -tg tat acc aaa aag g-a gat gtc cat gcg gat act ttg gcg gaa tag ata aat tag gaa atc ctc ttt gta aat cgt gtt g-- -tg ttg gtg aat att gcg acg aca tac gta atc ata gag ttg gtc ctt ttc ctc cat gca aa- --- cta tct aaa tgt aat tag cca --- --- --- --- --- --- --- --- --- --- --- --- --- --- -tg atc atg tta ccc ata c-a tc- --- -at atc gct act tgg t-a --g cgt att att cag tat g-a aga tct att aat aat aat tat ctt att tat ttc ttg acg --- --- --- -at att aat cgg tgt cct aat aat ata aaa --- tac tta tgg --- --- --a aat aaa ttg t-a aac aag atg aat taa t-- aca caa tac atc ct- --t aat tag ctt -gt tca aa- ttt tat aaa gaa tca tat tca tta ggc ata taa ta- aaa aat ata att tat aat ac- --- act tac a-- gcg tca tca tgc ata aca gca gtg aat taa ttg ctg tta tta atg gat tta gaa ata gtg gac gat ttt gtg ata tta ata tag tta tta atg atg aaa gga tac acg ctc ata gac tca tcc tat ctg gag cat cag aat a-t ttt tcc att ctg ttt tcc aat aat ttt atc gat tct aat gaa tac gaa gtt aat cta agt cat tta gat tat caa agt gtg aac gat ttg atc gat tac att tat ggg ata cct ttg agc cta act aac gat agc gtg aaa tat att ctt tca aca gct ga- --t ttt tta caa att gaa tct gcc att acg gag tgt gaa aat tac ata ctt aaa aat ctt tgt tct aga aac tgt atc gat ttc --- --- tac ata tac gct gat aaa tat aat aac aag aaa ata gaa tca gcg tcg ttt aac aca ata tta cga aat att ttg aga ctc att aac gat gaa aac ttt aaa tac ttg aca gag gaa tca atg ata aaa att tta agc gat gat atg tta aat ata aaa aat gag gat ttc gca cca cta att ctc att aaa tgg tta gag agt aca caa caa cca tgc acc ctc gag tta ctt aga tgc ctc aga ata tca -tt gct ttc ccc aca agt tat aaa atc act tta tag tca tca act ggt tag ttc aat cta cga atg tat aac att ctt aaa caa tat agc att ctt gga tga atc att tcc tag ata cca tag cat cga gtt gat atc tat cgg cat aag taa ttc gca tga taa gat ttc cat aaa ctg cta caa tcg t-a aaa aaa ata cat ggg aga tga tat ctt cac gta gat ata ggt gta gtt tcg cag tgg ccg tcc tgg ata ata tta tct ata tga tgg gtg gat atg atc aat ccc cgt ata gga gtt caa agg tta tag cgt aca aca cat gta caa att ctt gga tat atg ata tac cag agc t-a aaa tat cct cgt tct aat tgt gga gga gtg gtc gat gac gaa tac att tat tgt ata ggc ggc ata cgc gat cag gat tca tcg ttg ata tct agt att gat aga tgg aag cca tca aaa cca tat tgg cag acg tat gct aaa atg cgc gaa cca aaa tgt gat atg ggg gtt gcg atg tta aac gga tta ata tat gtc ata ggt gga atc gtt aaa ggt g-- aca cat gta ccg acg cac tag aga gtt tat cag aag atg gat gga tga tgc atc agc gcc ttc caa taa aaa tgt cca ata tgg cta cga ttg ttc atg ctg gta aga ttt ata tat ctg gag gtt ata aca ata gta gtg ccg tta atg gaa cat cga atc t-- agt cct tag cta taa tcc gat ata tga tga atg gac caa att atc atc att aaa tat tcc tag aat taa tcc cgc tct a

>LT883663.1:174218-176488_Cowpox_virus_strain_France_Amiens_2016_genome_assembly_complete_genome:_monopartite

ttg ctg act gta --- tgt a-c tat ttt gta cat cag tac att ggt tac agc aga tat acc tac ttc gtc act gcc aca cgc tcc ggt aaa cgg --g tca tgt gac gaa gga gaa tat ctt gat aag acg cat aat caa tgt tgt aat -cg gtg tcc acc tgg aga att tgc caa ggt tag atg ta- --a tgg taa cga taa cac aaa atg tga acg ctg tcc acc tca tac att tac cac agt acc caa tta ttc t-a atg gat gtc atc aat gta tga aat gcc caa agg gat cat ttg ata agg tac gat gtg atg gaa cac aga aca gta aat -gt tcg tgt ctt ccg ggt tgg ttt tgc gct act gat tct tcg aag act gaa gat tgt cga gat -tg tat acc aaa aag g-a gat gtc cat gcg gat act ttg gcg gaa tag ata aat tag gaa atc ctc ttt gta aat cgt gtt g-- -tg ttg gtg aat att gcg acg aca tac gta atc ata gag ttg gtc ctt ttc ctc cat gca aa- --- cta tct aaa tgt aat tag cca --- --- --- --- --- --- --- --- --- --- --- --- --- --- -tg atc atg tta ccc ata c-a tc- --- -at atc gct act tgg t-a --g cgt att att cag tat g-a aga tct att aat aat aat tat ctt att tat ttc ttg acg --- --- --- -at att aat cgg tgt cct aat aat ata aaa --- tac tta tgg --- --- --a aat aaa ttg t-a aac aag atg aat taa t-- aca caa tac atc ct- --t aat tag ctt -gt tca aa- ttt tat aaa gaa tca tat tca tta ggc ata taa ta- aaa aat ata att tat aat ac- --- act tac a-- gcg tca tca tgc ata aca gca gtg aat taa ttg ctg tta tca atg gat tta gaa ata gtg gac gat ttt gtg ata tta ata tag tta tta atg atg aaa ggg tac acg ctc ata gac tca tcc tat ctg gag cat cag aat a-t ttt tcc att ctg ttt tcc aat aat ttt atc gat tct aat gaa tac gaa gtt aat cta agt cat tta gat tat caa agt gtg aac gat ttg atc gat tac att tat ggg ata cct ttg agc cta act aac gat agc gtg aaa tat att ctt tca aca gct ga- --t ttt tta caa att gaa tct gcc att acg gag tgt gaa aat tac ata ctt aaa aat ctt tgt tct aga aac tgt atc gat ttc --- --- tac ata tac gct gat aaa tat aat aac aag aaa ata gaa tca gcg tcg ttt aac aca ata tta cga aat att ttg aga ctc att aac gat gaa aac ttt aaa tac ttg aca gag gaa tca atg ata aaa att tta agc gat gat atg tta aat ata aaa aat gag gat ttc gca cca cta att ctc att aaa tgg tta gag agt aca caa caa cca tgc acc ctc gag tta ctt aga tgc ctc aga ata tca -tt gct ttc ccc aca agt tat aaa atc act tta tag tca tca act ggt tag tcc aat cta cga atg tat aac att ctt aaa caa tat agc att ctt gga tga atc att tcc tag ata cca tag cat cga gtt gat atc tat cgg cat aag taa ttc gca tga taa gat ttc cat aaa ctg cta caa tcg t-a aaa aaa ata cat ggg aga tga tat ctt cac gta gat ata ggt gta gtt tcg cag tgg ccg tcc tgg ata ata tta tct ata tga tgg gtg gat atg atc aat ccc cgt ata gga gtt caa agg tta tag cgt aca ata cat gta caa att ctt gga tat atg ata tac cag agc t-a aaa tat cct cgt tct aat tgt ggg gga ctg act gat gac gaa tac att tat tgt ata ggc ggc ata cgc gat cag gat tca tcg ttg ata tct agt att gat aga tgg aag cca tca aaa cca tat tgg cag acg tat gct aaa atg cgc gaa cca aaa tgt gat atg ggg gtt gcg atg tta aac gga tta ata tat gtc ata ggt gga atc gtt aaa ggt g-- atg cgt gta ccg acg cac tag aga gtt tat cag aag atg gat gga tga agc atc aac gtc ttc caa taa aaa tgt cca ata tgt cga cga ttg ttc atg ctg gta aga ttt ata tat ctg gag gtt ata aca ata gta gtg ccg tta atg gac cat cga atc t-- agt cct tag cta taa tcc gat ata tga tga atg gac caa att atc atc att aaa tat tcc tag aat taa tcc cgc tct a

>HQ420894.1:174955-177225_Cowpox_virus_strain_France_2001_Nancy_complete_genome

ttg ctg act gta --- tgt a-c tat ttt gta cat cag tac att ggt tac agc aga tat acc tac ttc gtc act gcc aca cgc tcc ggt aaa cgg --g tca tgt gac gaa gga gaa tat ctt gat aag acg cat aat caa tgt tgt aat -cg gtg tcc acc tgg aga att tgc caa ggt tag atg ta- --a tgg taa cga taa cac aaa atg tga acg ctg tcc acc tca tac att tac cac agt acc caa tta ttc t-a atg gat gtc atc aat gta tga aat gcc caa agg gat cat ttg ata agg tac gat gtg atg gaa cac aga aca gta aat -gt tcg tgt ctt ccg ggt tgg ttt tgc gct act gat tct tcg aag act gaa gat tgt cga gat -tg tat acc aaa aag g-a gat gtc cat gcg gat act ttg gcg gaa tag ata aat tag gaa atc ctc ttt gta aat cgt gtt g-- -tg ttg gtg aat att gcg acg aca tac gta atc ata gag ttg gtc ctt ttc ctc cat gca aa- --- cta tct aaa tgt aat tag cca --- --- --- --- --- --- --- --- --- --- --- --- --- --- -tg atc atg tta ccc ata c-a tc- --- -at atc gct act tgg t-a --g cgt att att cag tat g-a aga tct att aat aat aat tat ctt att tat ttc ttg acg --- --- --- -at att aat cgg tgt cct aat aat ata aaa --- tac tta tgg --- --- --a aat aaa ttg t-a aac aag atg aat taa t-- aca caa tac atc ct- --t aat tag ctt -gt tca aa- ttt tat aaa gaa tca tat tca tta ggc ata taa ta- aaa aat ata att tat aat ac- --- act tac a-- gcg tca tca tgc ata aca gca gtg aat taa ttg ctg tta tca atg gat tta gaa ata gtg gac gat ttt gtg ata tta ata tag tta tta atg atg aaa gga tac acg ctc ata gac tca tcc tat ctg gag cat cag aat a-t ttt tcc att ctg ttt tcc aat aat ttt atc gat tct aat gaa tac gaa gtt aat cta agt cat tta gat tat caa agt gtg aac gat ttg atc gat tac att tat ggg ata cct ttg agc cta act aac gat agc gtg aaa tat att ctt tca aca gct ga- --t ttt tta caa att gaa tct gcc att acg gag tgt gaa aat tac ata ctt aaa aat ctt tgt tct aga aac tgt atc gat ttc --- --- tac ata tac gct gat aaa tat aat aac aag aaa ata gaa tca gcg tcg ttt aac aca ata tta cga aat att ttg aga ctc att aac gat gaa aac ttt aaa tac ttg aca gag gaa tca atg ata aaa att tta agc gat gat atg tta aat ata aaa aat gag gat ttc gca cca cta att ctc att aaa tgg tta gag agt aca caa caa cca tgc acc ctc gag tta ctt aga tgc ctc aga ata tca -tt gct ttc ccc aca agt tat aaa atc act tta tag tca tca act ggt tag tcc aat cta cga atg tat aac att ctt aaa caa tat agc att ctt gga tga atc att tcc tag ata cca tag cat cga gtt gat atc tat cgg cat aag taa ttc gca tga taa gat ttc cat aaa ctg cta caa tcg t-a aaa aaa ata cat ggg aga tga tat ctt cac gta gat ata ggt gta gtt tcg cag tgg ccg tcc tgg ata ata tta tct ata tga tgg gtg gat atg atc aat ccc cgt ata gga gtt caa agg tta tag cgt aca ata cat gta caa att ctt gga tat atg ata tac cag agc t-a aaa tat cct cgt tct aat tgt ggg gga ctg act gat gac gaa tac att tat tgt ata ggc ggc ata cgc gat cag gat tca tcg ttg ata tct agt att gat aga tgg aag cca tta aaa cca tat tgg cag acg tat gct aaa atg cgc gaa cca aaa tgt gat atg ggg gtt gcg atg tta aac gga tta ata tat gtc ata ggt gga atc gtt aaa ggt g-- atg cgt gta ccg acg cac tag aga gtt tat cag aag atg gat gga tga agc atc aac gtc ttc caa taa aaa tgt cca ata tgt cga cga ttg ttc atg ctg gta aga ttt ata tat ctg gag gtt ata aca ata gta gtg ccg tta atg gac cat cga atc t-- agt cct tag cta taa tcc gat ata tga tga atg gac caa att atc atc att aaa tat tcc tag aat taa tcc cgc tct a

>MN244298.1:174274-176510_Akhmeta_virus_strain_A40_complete_genome

--- -tg act ata --- tgt a-c tat ttt gta cat tgg tac aat ggt tac agc aga tac tga tca tcc acc a-- --- -ca tcc tcc tgt aga cag --a tca tgt aat gag gga gaa tat ctt gat aga aag act aat caa tgt tgc aaa -ct gtg tcc tcc tgg aga gtt cgc cga ggt tcg atg ta- --g tgg cgg caa taa tac aaa atg tga aaa atg tcc aca cga tac ata tac agc agt acc caa tta ttc t-a acg gat gtc atc aat gta gaa aat gct cag tgg gat cat tta ata agg taa agt gta ctg gag cac ata acc gcg aat -gt tcg tgt ctt cct ggt tgg ttt tgt gct acc gat tct tca cag act gaa aat tgt cga gat -tg tat acc aaa aag g-a gat gtc cat gcg gat act ttg gtg gaa tag ata att tag gaa atc ctc ttt gta aat cgt gtt g-- -tg atg gtg aat att gta act acc tac gta atc ata gag ttg gta ttt ttc ctc cat gca aa- --- cta tct aaa tgt aat tag tca --- --- --- --- --- --- --- --- --- --- --- --- --- --- -tg atc atg -ta -cc at- --a tt- --- -at atc gct act t-g gta --g tat att att cag tat g-a ata cct att aat aa- --- --- -tt a-- -at ctt ttg acg atc ttg tt- -at aat cat --- --- --- --- aat ata aaa --a tac tta tgg --- -g- aat agt cac tca t-- --- --- --- aat tac tg- acg cga taa att cat aat aat cag ctt gtt taa aat ttt tat aag ga- --- --a tct aca ggc ata aaa at- aaa aat ata att tat aat ac- --- act tat agc gc- cca tca tga ata aca gca gtg aat taa ttg ctg tta tta atg gat tta gaa ata gtg gac gat ttt gtg ata tta ata tag tta tta atg atg aaa gga taa acg ctc ata gac tca tcc tat ctg gag cct ccg aat a-t ttt tcc att ctg ttt tcc gat aat ttt atc gat tct aat gaa tac gaa gtt aat cta agt cat tta gat tat caa agc gta aac gat ttg atc gat tac att tat ggg ata cct ttg agc cta act aac gat agc gtg aaa tat att ctt tca aca gct ga- --t ttt tta caa att aga tct gcc att act gag tgt gaa aat tac ata ctt aaa aat ctg tgt tct aga aat tgt atc gat ttt --- --- tac ata tac gcc gat aaa tat aat aac aag aaa ata aaa aca gcg gca ttt aac aca ata tta cga aat att ttg aga ctc atc aac gat gaa aac ttt aaa tac tta aca gag gaa tca atg ata aaa att tta agc gat gat atg tta aat ata aaa aat gag gat ttt gca cca cta att ctc att aaa tgg tta gag agt aca caa caa cca tgc acg gtc gag tta ctt aga tgc ctc aga ata tca -tt gct ttc ccc tca agt tat aaa atc act tta tag tca tcg aca ggt cag ttc aat ata tga atg tat aac att att aaa taa tat atc att ctt gga tga atc att tcc tag ata cca tag cat cga gtt gat atc tat cgg tat aag taa ttc aca tga taa aat ttc cat aaa ctg cta caa tcg t-a aaa aaa ata cgt ggg aga tga tat ctt cac gga gat ata ggt gta gtt ttg cag tgg ccg tca tgg aca ata tta tct ata tga tgg gtg gat atg atc ggt ccc cgt ata aga gtt caa agg tta tag cgt aca ata cat gta cta att ctt gga tat atg ata tac cag agt t-a aaa tat cct cgt tct aat tgt gga gga gtt gct gat gat gaa tac att tat tgc ata ggc ggc ata cgc gat cag gat tta tcg ttg ata tct agt att gat aga tgg aag cca tca aaa tca tat tgg caa acg tat gct aaa atg aga gag cca aaa tgt gat atg ggt gta gcg atg tta aac gga tta ata tat gtc ata ggc gga gtt gtt aaa ggt g-- aca cgt gta ccg aca cac ttg aga gtt tat cag aag atg gat gga tga tgc atc aac gcc ttc caa taa aaa tgt cca ata tgt cga cga ttg ttc acg ccg gaa aaa ttt ata tat ctg gag gtt aca aca ata gta gtg tag tta atg gaa tat cga atc t-- agt cct tag cta cga tcc gat ata tga tga atg gac caa att atc atc att aaa cat tcc tag aat taa tcc tgc tct a

>MN244297.1:174147-176383_Akhmeta_virus_strain_A39_complete_genome

--- -tg act ata --- tgt a-c tat ttt gta cat tgg tac aat ggt tac agc aga tac tga tca tcc acc a-- --- -ca tcc tcc tgt aga cag --a tca tgt aat gag gga gaa tat ctt gat aga aag act aat caa tgt tgc aaa -ct gtg tcc tcc tgg aga gtt cgc cga ggt tcg atg ta- --g tgg cgg caa taa tac aaa atg tga aaa atg tcc aca cga tac ata tac agc agt acc caa tta ttc t-a acg gat gtc atc aat gta gaa aat gct cag tgg gat cat tta ata agg taa agt gta ctg gag cac ata acc gcg aat -gt tcg tgt ctt cct ggt tgg ttt tgt gct acc gat tct tca cag act gaa aat tgt cga gat -tg tat acc aaa aag g-a gat gtc cat gcg gat act ttg gtg gaa tag ata att tag gaa atc ctc ttt gta aat cgt gtt g-- -tg atg gtg aat att gta act acc tac gta atc ata gag ttg gta ttt ttc ctc cat gca aa- --- cta tct aaa tgt aat tag tca --- --- --- --- --- --- --- --- --- --- --- --- --- --- -tg atc atg -ta -cc at- --a tt- --- -at atc gct act t-g gta --g tat att att cag tat g-a ata cct att aat aa- --- --- -tt a-- -at ctt ttg acg atc ttg tt- -at aat cat --- --- --- --- aat ata aaa --a tac tta tgg --- -g- aat agt cac tca t-- --- --- --- aat tac tg- acg cga taa att cat aat aat cag ctt gtt taa aat ttt tat aag ga- --- --a tct aca ggc ata aaa at- aaa aat ata att tat aat ac- --- act tat agc gc- cca tca tga ata aca gca gtg aat taa ttg ctg tta tta atg gat tta gaa ata gtg gac gat ttt gtg ata tta ata tag tta tta atg atg aaa gga taa acg ctc ata gac tca tcc tat ctg gag cct ccg aat a-t ttt tcc att ctg ttt tcc gat aat ttt atc gat tct aat gaa tac gaa gtt aat cta agt cat tta gat tat caa agc gta aac gat ttg atc gat tac att tat ggg ata cct ttg agc cta act aac gat agc gtg aaa tat att ctt tca aca gct ga- --t ttt tta caa att aga tct gcc att act gag tgt gaa aat tac ata ctt aaa aat ctg tgt tct aga aat tgt atc gat ttt --- --- tac ata tac gcc gat aaa tat aat aac aag aaa ata aaa aca gcg gca ttt aac aca ata tta cga aat att ttg aga ctc atc aac gat gaa aac ttt aaa tac tta aca gag gaa tca atg ata aaa att tta agc gat gat atg tta aat ata aaa aat gag gat ttt gca cca cta att ctc att aaa tgg tta gag agt aca caa caa cca tgc acg gtc gag tta ctt aga tgc ctc aga ata tca -tt gct ttc ccc tca agt tat aaa atc act tta tag tca tcg aca ggt cag ttc aat ata tga atg tat aac att att aaa taa tat atc att ctt gga tga atc att tcc tag ata cca tag cat cga gtt gat atc tat cgg tat aag taa ttc aca tga taa aat ttc cat aaa ctg cta caa tcg t-a aaa aaa ata cgt ggg aga tga tat ctt cac gga gat ata ggt gta gtt ttg cag tgg ccg tca tgg aca ata tta tct ata tga tgg gtg gat atg atc ggt ccc cgt ata aga gtt caa agg tta tag cgt aca ata cat gta cta att ctt gga tat atg ata tac cag agt t-a aaa tat cct cgt tct aat tgt gga gga gtt gct gat gat gaa tac att tat tgc ata ggc ggc ata cgc gat cag gat tta tcg ttg ata tct agt att gat aga tgg aag cca tca aaa tca tat tgg caa acg tat gct aaa atg aga gag cca aaa tgt gat atg ggt gta gcg atg tta aac gga tta ata tat gtc ata ggc gga gtt gtt aaa ggt g-- aca cgt gta ccg aca cac ttg aga gtt tat cag aag atg gat gga tga tgc atc aac gcc ttc caa taa aaa tgt cca ata tgt cga cga ttg ttc acg ccg gaa aaa ttt ata tat ctg gag gtt aca aca ata gta gtg tag tta atg gaa tat cga atc t-- agt cct tag cta cga tcc gat ata tga tga atg gac caa att atc atc att aaa cat tcc tag aat taa tcc tgc tct a

>MN244296.1:174985-177141_Akhmeta_virus_strain_G66_complete_genome

--- --- --- --- --- --- --- --- --- --- --- --- --- --- --- --- --- --- --- --- --- --- --- --- --- --- --- --- --- --- --- --- tca tgc aat gag gga gaa tat ctt gat aga acg act aat caa tgt tgc aaa -ct gtg tcc tcc tgg aga gtt cgc cga ggt tcg atg ta- --g tgg cgg caa taa tac aaa atg tga aaa atg tcc aca cga tac ata tac agc agt acc caa tta ttc t-a acg gat gtc atc aat gta gaa aat gct cag tgg gat cat tta ata agg taa agt gta ctg gag cac ata acc gcg aat -gt tcg tgt ctt cct ggt tgg ttt tgt gct acc gat tct tca cag act gaa aat tgt cga gat -tg tat acc aaa aag g-a gat gtc cat gcg gat act ttg gtg gaa tag ata att tag gaa atc ctc ttt gta aat cgt gtt g-- -cg atg gtg aat att gta act acc tac gta atc ata gag ttg gta ttt ttc ctc cat gca aa- --- cta tct aaa tgt aat tag tca --- --- --- --- --- --- --- --- --- --- --- --- --- --- -tg atc atg -ta -cc at- --a tt- --- -at atc gct act t-g gta --g tat att att cag tat g-a aga cct att aat aa- --- --- -tt a-- -at ctt ttg acg atc ttg tt- -at aat cat --- --- --- --- aat ata aaa --a tac tta tgg --- -g- aat agt cac tca t-- --- --- --- aat tgc tg- acg cga taa att cat aat aat cag ctt gtt taa aat ttt tat aag ga- --- --a tct aca ggc ata aaa at- aaa aat ata att tat aat ac- --- act tat agc ac- cca tca tga ata aca gca gtg aat taa ttg ctg tta tta atg gat tta gaa ata gtg gac gat ttt gtg ata tta ata tag tta tta atg atg aaa gga taa acg ctc ata gac tca tcc tat ctg gag cct ccg aat a-t ttt tcc att ctg ttt tcc gat aat ttt atc gat tct aat gaa tac gaa gtt aat cta agt cat tta gat tat caa agc gta aac gat ttg atc gat tac att tat ggg ata cct ttg agc cta act aac gat agc gtg aaa tat att ctt tca aca gcc ga- --t ttt tta caa att aga tcg gcc att act gag tgt gaa aat tac ata ctt aaa aat ctg tgt tct aga aat tgt atc gat ttt --- --- tac ata tac gcc gat aaa tat aat aac aag aaa ata aaa aca gcg gca ttt aac aca ata tta cga aat att ttg aga ctc atc aac gat gaa aac ttt aaa tac tta aca gag gaa tca atg ata aaa att tta agc gat gat atg tta aat ata aaa aat gag gat ttt gca cca cta att ctc att aaa tgg tta gag agt aca caa caa cca tgc aca gtc gag tta ctt aga tgc ctc aga ata tca -tt gct ttc ccc tca agt tat aaa atc act tta tag tca tcg aca ggt cag ttc aat ata tga atg tat aac att att aaa taa tat atc att ctt gga tga atc att tcc tag ata cca tag cat cga gtt gat atc tat cgg tat aag taa ttc aca tga taa aat ttc cat aaa ctg cta caa tcg t-a aaa aaa ata cgt ggg aga tga tat ctt cac gga gat ata ggt gta gtt ttg cag tgg ccg tca tgg aca ata tta tct ata tga tgg gtg gat atg atc ggt ccc cgt ata aga gtt caa agg tta tag cgt aca ata cat gta cta att ctt gga tat atg ata tac cag agt t-a aaa tat cct cgt tct aat tgt gga gga gtt gct gat gat gaa tac att tat tgc ata ggc ggc ata cgc gat cag gat tta tcg ttg ata tct agt att gat aga tgg aag cca tca aaa tca tat tgg caa acg tat gct aaa atg cga gag cca aaa tgt gat atg ggt gta gcg atg tta aac gga tta ata tat gtc ata ggc gga gtt gtt aaa ggt g-- aca cgt gta ccg aca cac ttg aga gtt tat cag aag atg gat gga tga tgc atc aac gcc ttc caa taa aaa tgt cca ata tgt cga cga ttg ttc atg ccg gaa aaa ttt ata tat ctg gag gtt aca aca ata gta gtg tag tta atg gaa tat cga atc t-- agt cct tag cta cga tcc gat ata tga tga atg gac caa att atc atc att aaa cat tcc tag aat taa tcc tgc tct a

>MH607142.1:175006-177162_Akhmeta_virus_isolate_Akhmeta_2013-85_complete_genome

--- --- --- --- --- --- --- --- --- --- --- --- --- --- --- --- --- --- --- --- --- --- --- --- --- --- --- --- --- --- --- --- tca tgc aat gag gga gaa tat ctt gat aga acg act aat caa tgt tgc aaa -ct gtg tcc tcc tgg aga gtt cgc cga ggt tcg atg ta- --g tgg cgg caa taa tac aaa atg tga aaa atg tcc aca cga tac ata tac agc agt acc caa tta ttc t-a acg gat gtc atc aat gta gaa aat gct cag tgg gat cat tta ata agg taa agt gta ctg gag cac ata acc gcg aat -gt tcg tgt ctt cct ggt tgg ttt tgt gct acc gat tct tca cag act gaa aat tgt cga gat -tg tat acc aaa aag g-a gat gtc cat gcg gat act ttg gtg gaa tag ata att tag gaa atc ctc ttt gta aat cgt gtt g-- -cg atg gtg aat att gta act acc tac gta atc ata gag ttg gta ttt ttc ctc cat gca aa- --- cta tct aaa tgt aat tag tca --- --- --- --- --- --- --- --- --- --- --- --- --- --- -tg atc atg -ta -cc at- --a tt- --- -at atc gct act t-g gta --g tat att att cag tat g-a aga cct att aat aa- --- --- -tt a-- -at ctt ttg acg atc ttg tt- -at aat cat --- --- --- --- aat ata aaa --a tac tta tgg --- -g- aat agt cac tca t-- --- --- --- aat tgc tg- acg cga taa att cat aat aat cag ctt gtt taa aat ttt tat aag ga- --- --a tct aca ggc ata aaa at- aaa aat ata att tat aat ac- --- act tat agc ac- cca tca tga ata aca gca gtg aat taa ttg ctg tta tta atg gat tta gaa ata gtg gac gat ttt gtg ata tta ata tag tta tta atg atg aaa gga taa acg ctc ata gac tca tcc tat ctg gag cct ccg aat a-t ttt tcc att ctg ttt tcc gat aat ttt atc gat tct aat gaa tac gaa gtt aat cta agt cat tta gat tat caa agc gta aac gat ttg atc gat tac att tat ggg ata cct ttg agc cta act aac gat agc gtg aaa tat att ctt tca aca gcc ga- --t ttt tta caa att aga tcg gcc att act gag tgt gaa aat tac ata ctt aaa aat ctg tgt tct aga aat tgt atc gat ttt --- --- tac ata tac gcc gat aaa tat aat aac aag aaa ata aaa aca gcg gca ttt aac aca ata tta cga aat att ttg aga ctc atc aac gat gaa aac ttt aaa tac tta aca gag gaa tca atg ata aaa att tta agc gat gat atg tta aat ata aaa aat gag gat ttt gca cca cta att ctc att aaa tgg tta gag agt aca caa caa cca tgc aca gtc gag tta ctt aga tgc ctc aga ata tca -tt gct ttc ccc tca agt tat aaa atc act tta tag tca tcg aca ggt cag ttc aat ata tga atg tat aac att att aaa taa tat atc att ctt gga tga atc att tcc tag ata cca tag cat cga gtt gat atc tat cgg tat aag taa ttc aca tga taa aat ttc cat aaa ctg cta caa tcg t-a aaa aaa ata cgt ggg aga tga tat ctt cac gga gat ata ggt gta gtt ttg cag tgg ccg tca tgg aca ata tta tct ata tga tgg gtg gat atg atc ggt ccc cgt ata aga gtt caa agg tta tag cgt aca ata cat gta cta att ctt gga tat atg ata tac cag agt t-a aaa tat cct cgt tct aat tgt gga gga gtt gct gat gat gaa tac att tat tgc ata ggc ggc ata cgc gat cag gat tta tcg ttg ata tct agt att gat aga tgg aag cca tca aaa tca tat tgg caa acg tat gct aaa atg cga gag cca aaa tgt gat atg ggt gta gcg atg tta aac gga tta ata tat gtc ata ggc gga gtt gtt aaa ggt g-- aca cgt gta ccg aca cac ttg aga gtt tat cag aag atg gat gga tga tgc atc aac gcc ttc caa taa aaa tgt cca ata tgt cga cga ttg ttc atg ccg gaa aaa ttt ata tat ctg gag gtt aca aca ata gta gtg tag tta atg gaa tat cga atc t-- agt cct tag cta cga tcc gat ata tga tga atg gac caa att atc atc att aaa cat tcc tag aat taa tcc tgc tct a

>NC_055230.1:175012-177168_Akhmeta_virus_isolate_Akhmeta_2013-88_complete_genome

--- --- --- --- --- --- --- --- --- --- --- --- --- --- --- --- --- --- --- --- --- --- --- --- --- --- --- --- --- --- --- --- tca tgc aat gag gga gaa tat ctt gat aga acg act aat caa tgt tgc aaa -ct gtg tcc tcc tgg aga gtt cgc cga ggt tcg atg ta- --g tgg cgg caa taa tac aaa atg tga aaa atg tcc aca cga tac ata tac agc agt acc caa tta ttc t-a acg gat gtc atc aat gta gaa aat gct cag tgg gat cat tta ata agg taa agt gta ctg gag cac ata acc gcg aat -gt tcg tgt ctt cct ggt tgg ttt tgt gct acc gat tct tca cag act gaa aat tgt cga gat -tg tat acc aaa aag g-a gat gtc cat gcg gat act ttg gtg gaa tag ata att tag gaa atc ctc ttt gta aat cgt gtt g-- -cg atg gtg aat att gta act acc tac gta atc ata gag ttg gta ttt ttc ctc cat gca aa- --- cta tct aaa tgt aat tag tca --- --- --- --- --- --- --- --- --- --- --- --- --- --- -tg atc atg -ta -cc at- --a tt- --- -at atc gct act t-g gta --g tat att att cag tat g-a aga cct att aat aa- --- --- -tt a-- -at ctt ttg acg atc ttg tt- -at aat cat --- --- --- --- aat ata aaa --a tac tta tgg --- -g- aat agt cac tca t-- --- --- --- aat tgc tg- acg cga taa att cat aat aat cag ctt gtt taa aat ttt tat aag ga- --- --a tct aca ggc ata aaa at- aaa aat ata att tat aat ac- --- act tat agc ac- cca tca tga ata aca gca gtg aat taa ttg ctg tta tta atg gat tta gaa ata gtg gac gat ttt gtg ata tta ata tag tta tta atg atg aaa gga taa acg ctc ata gac tca tcc tat ctg gag cct ccg aat a-t ttt tcc att ctg ttt tcc gat aat ttt atc gat tct aat gaa tac gaa gtt aat cta agt cat tta gat tat caa agc gta aac gat ttg atc gat tac att tat ggg ata cct ttg agc cta act aac gat agc gtg aaa tat att ctt tca aca gcc ga- --t ttt tta caa att aga tcg gcc att act gag tgt gaa aat tac ata ctt aaa aat ctg tgt tct aga aat tgt atc gat ttt --- --- tac ata tac gcc gat aaa tat aat aac aag aaa ata aaa aca gcg gca ttt aac aca ata tta cga aat att ttg aga ctc atc aac gat gaa aac ttt aaa tac tta aca gag gaa tca atg ata aaa att tta agc gat gat atg tta aat ata aaa aat gag gat ttt gca cca cta att ctc att aaa tgg tta gag agt aca caa caa cca tgc aca gtc gag tta ctt aga tgc ctc aga ata tca -tt gct ttc ccc tca agt tat aaa atc act tta tag tca tcg aca ggt cag ttc aat ata tga atg tat aac att att aaa taa tat atc att ctt gga tga atc att tcc tag ata cca tag cat cga gtt gat atc tat cgg tat aag taa ttc aca tga taa aat ttc cat aaa ctg cta caa tcg t-a aaa aaa ata cgt ggg aga tga tat ctt cac gga gat ata ggt gta gtt ttg cag tgg ccg tca tgg aca ata tta tct ata tga tgg gtg gat atg atc ggt ccc cgt ata aga gtt caa agg tta tag cgt aca ata cat gta cta att ctt gga tat atg ata tac cag agt t-a aaa tat cct cgt tct aat tgt gga gga gtt gct gat gat gaa tac att tat tgc ata ggc ggc ata cgc gat cag gat tta tcg ttg ata tct agt att gat aga tgg aag cca tca aaa tca tat tgg caa acg tat gct aaa atg cga gag cca aaa tgt gat atg ggt gta gcg atg tta aac gga tta ata tat gtc ata ggc gga gtt gtt aaa ggt g-- aca cgt gta ccg aca cac ttg aga gtt tat cag aag atg gat gga tga tgc atc aac gcc ttc caa taa aaa tgt cca ata tgt cga cga ttg ttc atg ccg gaa aaa ttt ata tat ctg gag gtt aca aca ata gta gtg tag tta atg gaa tat cga atc t-- agt cct tag cta cga tcc gat ata tga tga atg gac caa att atc atc att aaa cat tcc tag aat taa tcc tgc tct a

>MH607143.1:172434-174670_Akhmeta_virus_isolate_Vani_2010_complete_genome

--- -tg act ata --- tgt a-c tat ttt gta cat tgg cac aat ggt tac agc aaa tac tga tca tcc acc a-- --- -ca tcc tcc tgt aga cag --a tca tgt aat gag gga gaa tat ctt gat aga aat act aat caa tgt tgc aaa -ct gtg tcc tcc tgg aga gtt cgc cga ggt tcg atg ta- --g tgg cgg caa taa tac aaa atg tga aaa atg tcc aca cga tac ata tac agc agt acc caa tta ttc t-a acg gat gtc atc aat gta gaa aat gcc cag tgg gat cat tta ata agg taa agt gta ctg gag cac ata acc gcg aat -gt tcg tgt ctt cct ggt tgg ttt tgt gct acc gat tct tca cag act gaa aat tgt cga gat -tg tat acc aaa aag g-a gat gtc cat gcg gat act ttg gtg gaa tag ata att tag gaa atc ctc ttt gta aat cgt gtt g-- -cg atg gtg aat att gta act acc tac gta atc ata gag ttg gta ttt ttc ctc cat gca aa- --- cta tct aaa tgt aat tag tca --- --- --- --- --- --- --- --- --- --- --- --- --- --- -tg atc atg -ta -cc at- --a tt- --- -at atc gct act t-g gta --g tat att att cag tat g-a ata cct att aat aa- --- --- -tt a-- -at ctt ttg acg atc ttg tt- -at aat cat --- --- --- --- aat ata aaa --a tac tta tgg --- -g- aat agt cac tca t-- --- --- --- aat tac tg- acg cga taa att cat aat aat cag ctt gtt taa aat ttt tat aag ga- --- --a tct aca ggc ata aaa at- aaa aat ata att tat aat ac- --- act tat agc gc- cca tca tga ata aca gca gtg aat taa ttg ctg tta tta atg gat tta gaa ata gtg gac gat ttt gtg ata tta ata tag tta tta atg atg aaa gga taa acg ctc ata gac tca tcc tat ctg gag cct ccg aat a-t ttt tcc att ctg ttt tcc gat aat ttt atc gat tct aat gaa tac gaa gtt aat cta agt cat tta gat tat caa agc gta aac gat ttg atc gat tac att tat ggg ata cct ttg agc cta act aac gat agc gtg aaa tat att ctt tca aca gct ga- --t ttt tta caa att aga tcg gcc att act gag tgt gaa aat tac ata ctt aaa aat ctg tgt tct aga aat tgt atc gat ttt --- --- tac ata tac gcc gat aaa tat aat aac aag aaa ata aaa aca gcg gca ttt aac aca ata tta cga aat att ttg aga ctc atc aac gat gaa aac ttt aaa tac tta aca gag gaa tca atg ata aaa att tta agc gat gat atg tta aat ata aaa aat gag gat ttt gca cca cta att ctc att aaa tgg tta gag agt aca caa caa cca tgc aca gtc gag tta ctt aga tgc ctc aga ata tca -tt gct ttc ccc tca agt tat aaa atc act tta tag tca tcg aca ggt cag ttc aat ata tga atg tat aac att att aaa taa tat atc att ctt gga tga atc att tcc tag ata cca tag cat cga gtt gat atc tat cgg tat aag taa ttc aca tga taa aat ttc cat aaa ctg cta caa tcg t-a aaa aaa ata cgt ggg aga tga tat ctt cac gga gat ata ggt gta gtt ttg cag tgg ccg tca tgg aca ata tta tct ata tga tgg gtg gat atg atc ggt ccc cgt ata aga gtt caa agg tta tag cgt aca ata cat gta cta att ctt gga tat atg ata tac cag agt t-a aaa tat cct cgt tct aat tgt gga gga gtt gct gat gat gaa tac att tat tgc ata ggc ggc ata cgc gat cag gat tta tcg ttg ata tct agt att gat aga tgg aag cca tca aaa tca tat tgg caa acg tat gct aaa atg aga gag cca aaa tgt gat atg ggt gta gcg atg tta aac gga tta ata tat gtc ata ggc gga gtt gtt aaa ggt g-- aca cgt gta ccg aca cgc ttg aga gtt tat cag aag atg gat gga tga tgc atc aac gcc ttc caa taa aaa tgt cca ata tgt cga cga ttg ttc acg ccg gaa aaa ttt ata tat ctg gag gtt aca aca ata gta gtg tag tta atg gaa tat cga atc t-- agt cct tag cta cga tcc gat ata tga tga atg gac caa att atc atc att aaa cat tcc tag aat taa tcc tgc tct a
