## Supplementary figures and images for "Genomic regions insertion and deletion in Monkeypox virus causing multi-country outbreak-2022"

### Supplementary Figure 1

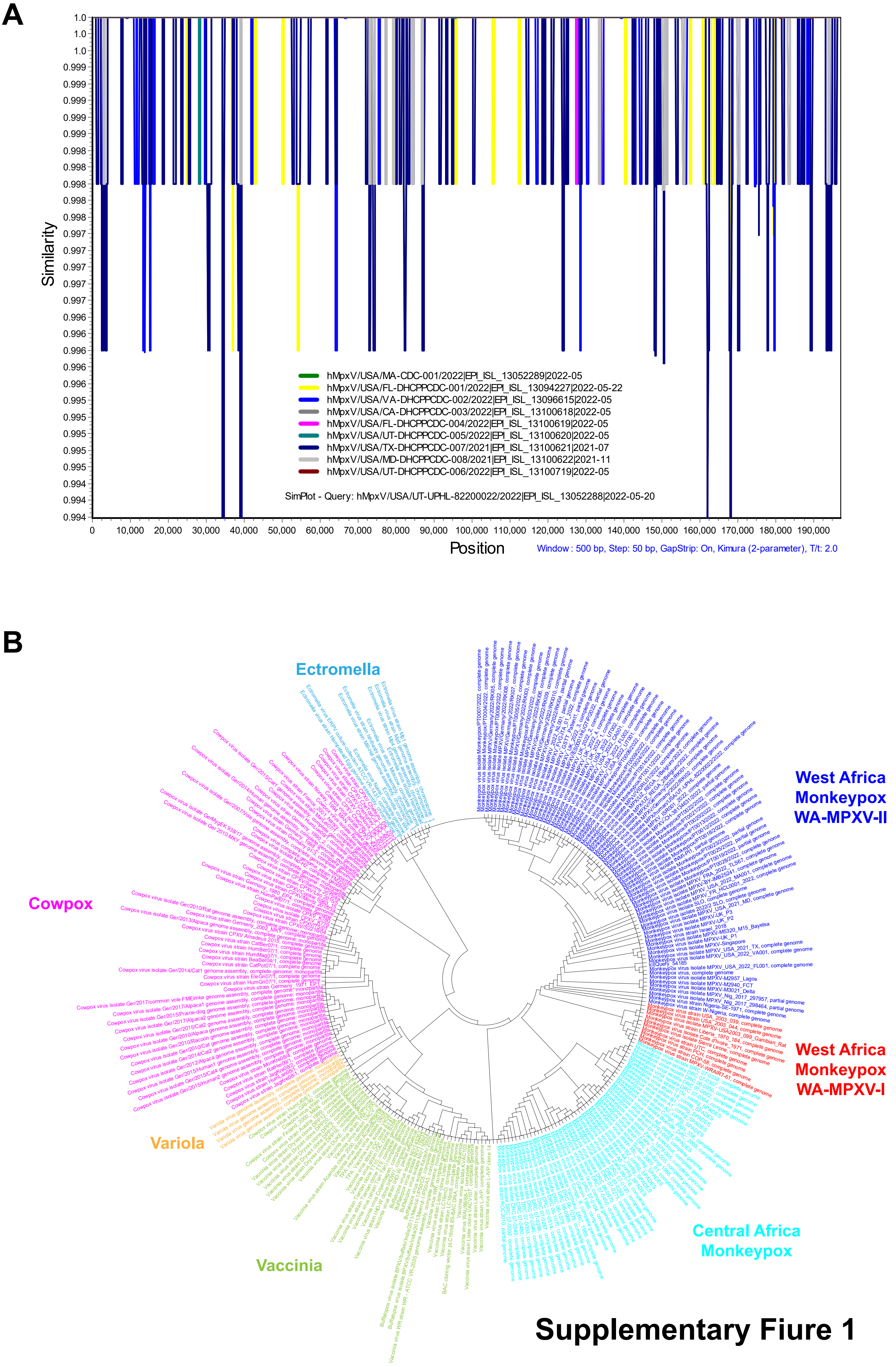


Supplementary Fiure 1
